# Phyllosphere-inhabiting endophytic bacteria feature a stationary phase-like lifestyle

**DOI:** 10.1101/2021.05.10.443510

**Authors:** André C. Velásquez, José C. Huguet-Tapia, Sheng Yang He

## Abstract

Plants are in constant association with a variety of microbes, and although much is known about how symbiotic and pathogenic microbes interact with plants, less is known about the population dynamics, adaptive traits, and transcriptional features of endophytic commensal microbes that live inside leaves. In this study, we evaluated the long-term population and transcriptional dynamics of two bacterial microbiota endophytes and compared them to those of a commensal-simulating non-pathogenic mutant of the bacterial pathogen *Pseudomonas syringae*. We found that population densities of all three endophytic phyllosphere bacteria remained static over a long period of time, which was caused by a continual equilibrium between bacterial multiplication and death, as evidenced by treatment of plants with antibiotics that only targeted dividing bacteria or by *in planta* visualization of bacteria carrying a fluorescent division reporter. Population stasis could not be explained by a lack of resources, as Arabidopsis leaves could support population densities up to 100 times higher than the normal microbiota populations, nor was population stasis reversed by significantly quenching PAMP-triggered immunity. Long-term temporal *in planta* transcriptomic analysis of these three bacterial endophytes revealed up-regulation of protein translation, the generation of energy, and the response to stress, and interestingly, for the microbiota strains, the longer the bacteria remained inside plants, the greater the up-regulation of some of these processes. Further transcriptomic analysis of *in planta* populations of commensal-simulating *Pseudomonas syringae* revealed a remarkable resemblance to those of *in vitro* bacteria in stationary phase, a metabolically active physiological state in which the production of secondary metabolites and stress responses are induced. This study provides novel insight into how endophytic bacteria survive and thrive inside plant leaves, and reshapes our current understanding of what it means to be part of the endophytic microbiota in the phyllosphere.

## INTRODUCTION

Plants are exposed to a multitude of microbes during their life cycles. Interactions of plants with microbes may range from causing no observable effect (commensal interactions) to forming intricate symbiotic relationships with specialized organs for nutrient acquisition (Hassani *et al*. 2018). Of the myriad extant microbes, only very few are capable of causing disease in a particular host: probably only tens of microbes will cause disease in a particular host, while most biotrophic pathogens have a very narrow host range (Morris and Moury 2019). Due to their great impact on crop production and natural ecosystems, symbiotic and pathogenic microbes have received great attention in the past. By contrast, insight into the lives of the vast number of commensal microbes is lagging behind.

Of commensal bacteria that live on the phyllosphere (above-ground parts) of plants, most live on the surface as epiphytes, probably because the plant interior (specifically the apoplast) exerts a strong selection pressure on the type of microbes that can grow and multiply. Not only are bacterial epiphytes more than 100 times more abundant than endophytes, but also the microbiota composition is different (Chen *et al*. 2020b). Commensal bacterial endophyte population numbers are low; in Arabidopsis (*Arabidopsis thaliana*) the number of culturable endophytes appears to be fewer than 10^4^ CFU cm^-2^ (Chen *et al*. 2020b, Xin *et al*. 2016). By contrast, the population density of pathogenic endophytic bacteria, such as *Pseudomonas syringae*, can increase to almost the carrying capacity of the plant (Xin *et al*. 2018), unless the plant surveillance systems mounts effector-triggered immunity (ETI) upon effector recognition (Deslandes *et al*. 2002, Macho *et al*. 2012). Interestingly, in the absence of virulence-promoting effector proteins and toxins, non-pathogenic mutants of *P. syringae*, such as the *ΔhrcCΔCFA* mutant (defective in the type III secretion system and coronatine production), are unable to multiply to high levels, resembling commensal bacteria that normally reside in the leaf apoplast (Worley et al. 2013).

The transcriptomes of phyllosphere-inhabiting pathogenic bacteria have been analyzed in the past few years to obtain clues into the processes that influence plant colonization. A microarray-based approach revealed differences in gene expression between epiphytic and apoplastic pathogenic *Pseudomonas syringae*, from which it was inferred that bacteria residing in the plant apoplast experience a more taxing osmotic stress than those that live on the surface of the leaves (at 48–72 hours post-inoculation; Yu *et al*. 2013). A more recent transcriptomics study of the early responses of *Pseudomonas syringae* (6 hours after inoculation) showed a strong correlation between bacterial genes responsive to the plant immune system and future bacterial densities (Nobori *et al*. 2018). However, to date, the nature of the long-term bacterial population homeostasis and associated transcriptomic dynamics of non-pathogenic endophytic commensal phyllosphere bacteria has not been evaluated, leaving a significant gap in the understanding of population and molecular features that are important for long-term adaptation and survival of commensal microbiota to the apoplastic environment of the phyllosphere.

There are several non-mutually exclusive plausible explanations for the inability of a microbe to survive and/or multiply inside the plant apoplast: (a) plant surface barriers do not allow for microbe invasion into plants (Ellinger *et al*. 2013, Lee *et al*. 2019). (b) a nutritional deficiency in the plant’s apoplast that the microbe cannot overcome (Bezrutczyk *et al*. 2018). (c) the concentration of free water in the apoplast is too low for the microbe to multiply (Wright and Beattie 2004). (d) the microbe cannot counteract plant immune defenses (Cunnac *et al*. 2011, Stoitsova *et al*. 2008, Fan *et al*. 2011). (e) host inducers of microbial virulence and/or survival mechanisms are not present in plants, or are actively sequestered (Anderson *et al*. 2014, Wang *et al*. 2020); and (f) for biotrophic microbes that cause plant cell death, niche destruction does not allow the microbe to grow (Balint-Kurti 2019).

We conducted a detailed study to understand the population dynamics of two common bacterial commensal endophytes as well as a commensal-simulating mutant of *P. syringae* pv. *tomato* DC3000 in Arabidopsis leaves. We then used multi-time-point transcriptomic analysis to infer the biological processes important for a long-term commensal lifestyle in the leaf apoplast. Our results point to a previously unrecognized stationary phase-like lifestyle for phyllosphere-inhabiting endophytic bacteria. This population stasis was caused by equilibrium in the rates of multiplication and death of phyllosphere-inhabiting endophytes. This finding has significant implications in understanding commensal plant-endophytic microbiota interactions and in guiding the application of endophytic commensal microbiota in agricultural and natural ecosystem settings.

## RESULTS

### Non-pathogenic phyllosphere bacterial population densities are static

A naïve assumption for the outcome after phyllosphere bacterial strains lacking virulence mechanisms are inoculated into plant leaves would be that the plant immune system would eventually clear out those invading microbes by producing antibacterial compounds. We tested this possibility by inoculating a disarmed (no longer pathogenic) strain, *Pseudomonas syringae* pv. *tomato* (*Pst*) DC3000 Δ*hrcC*Δ*CFA*, into Arabidopsis leaves and evaluating the population densities over the course of four weeks. *Pst* Δ*hrcC*Δ*CFA* has been rendered essentially non-pathogenic by eliminating its two main virulence determinants: the ability to form a type III secretion system (*hrcC*) and the production of the phytotoxin coronatine (*CFA*) (Worley et al. 2013). As such, *Pst* Δ*hrcC*Δ*CFA* is unable to cause disease. As can be observed in Figure 1A, the bacterial population densities remained unchanged over the course of 4 weeks, suggesting that for this particular plant–microbe interaction, antimicrobial compounds are not significantly altering the bacterial population densities.

**Figure 1.**
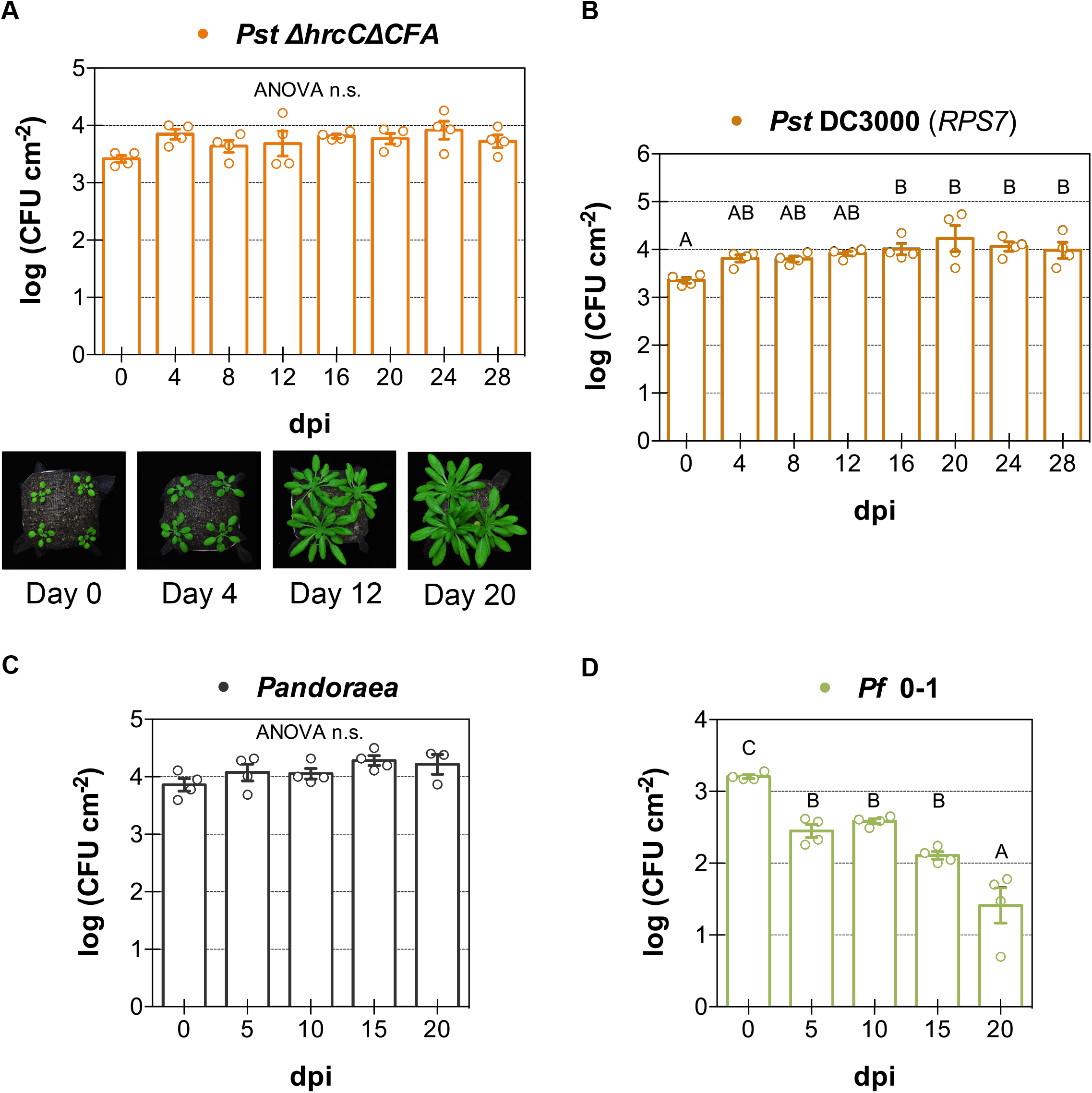
Non-pathogenic phyllosphere bacteria experience *in planta* bacterial population stasis. **(A)** Bacterial population density numbers of non-pathogenic *P. syringae* pv. *tomato* (*Pst*) *ΔhrcCΔCFA* (2 × 10^6^ CFU mL^-1^), a mutant of *Pst* DC3000 that does not produce coronatine or a functional type III secretion system, over the course of 28 days after infection of Arabidopsis accession Col-0. Photographs of representative infected plants are shown beneath the graph. **(B)** Bacterial population density numbers of *Pst* DC3000 (2 × 10^6^ CFU mL^-1^) over the course of 28 days after infection of Arabidopsis accession Bu-22. Bu-22 carries *RPS7*, a gene that recognizes the effector AvrPto, and triggers effector-triggered immunity. **(C)** Bacterial population density numbers of endophytic microbiota strain *Pandoraea* sp. Col-0-28 (10^6^ CFU mL^-1^) over the course of 20 days after infection of Col-0 plants. **(D)** Bacterial population density numbers of non-pathogenic *P. fluorescens* (*Pf*) 0-1 (2 × 10^5^ CFU mL^-1^), a bacterium isolated from soil, over the course of 20 days after infection of Col-0 plants. There is a decrease in population density observed over time. Individual biological repetitions for each treatment are shown as open circles. Error bars indicate the standard error of the mean. When appropriate, different letters indicate differences in means, as judged by a Tukey HSD test (*p* < 0.05). dpi indicates days post-inoculation.

Phyllosphere-inhabiting bacteria also include pathogenic strains that trigger ETI in incompatible plants, and are therefore unable to cause disease. Bu-22 is an Arabidopsis accession in which the *Pst* virulence-promoting effector AvrPto is recognized in order to mount ETI (Velásquez *et al*. 2017). Bu-22 plants carry a single locus responsible for AvrPto recognition that we named *RPS7* (*<u>R</u>esistance to <u>Ps</u>eudomonas syringae <u>7</u>*). When we inoculated *Pst* DC3000 into the Arabidopsis accession Bu-22, there was only a slight increase in growth very early in the interaction (before 4 days post-inoculation [dpi]), after which the bacterial population densities remained static (Figure 1B). Taken together, these results suggest that for phyllosphere microbes that trigger ETI and for non-pathogenic bacteria, the plant immune system maintains the microbial population numbers without fully eliminating bacteria from the plant apoplast.

We wanted to know if the population stasis phenomenon was widespread and if it also applied to normal non-pathogenic leaf microbiota strains. We inoculated into Arabidopsis plants three bacterial strains that had been previously isolated from Arabidopsis leaves: the β-Proteobacteria *Achromobacter xylosoxidans* Col-0-50 and *Pandoraea* sp. Col-0-28 (Tao *et al*. 2020), and the gram-positive Actinobacteria *Rhodococcus* sp. 964 (Bodenhausen *et al*. 2014) into Arabidopsis plants. These microbiota strains were chosen because they did not cause any disease-like symptoms when inoculated into leaves at very high population densities (Bodenhausen *et al*. 2014, Tao *et al*. 2020). After three weeks, population densities did not change for either Proteobacteria strain (Figure 1C and Figure S1A), while there was a very small initial increase in growth for *Rhodococcus* (Supplementary Figure S1A). We also tested additional *R* gene–effector pair interactions that caused ETI in Arabidopsis. For three additional interactions— recognition of AvrRpt2, AvrRps4, and AvrPphB by RSP2, RPS4 and RPS5, respectively— population stasis of *Pst* was observed (other than the small initial population density increase; Supplementary Figure S1B), similar to the effect observed during the interaction between RPS7 and AvrPto. The only bacterial strain for which we did not observe population stasis was *Pseudomonas fluorescens* 0-1, a bacterium originally isolated from soil. For this strain, the population density numbers decreased over time, but never reached zero, not even after almost 3 weeks after inoculation (Figure 1D). Overall, these results suggest that population stasis occurs for most phyllosphere non-pathogenic bacteria (8 out of 9 strains) once they are inside plants, and that plant defenses, including ETI, keep bacterial population densities constant.

### Bacterial population stasis occurs irrespective of bacterial population density or a functional PTI system

It is possible that the population stasis phenomenon is only observed at a specific population density, one at which the carrying capacity of the system has been reached. To test this possibility, we inoculated *Pst ΔhrcCΔCFA* into Arabidopsis leaves at four different initial population densities (covering 3 orders of magnitude) and evaluated the resulting population densities over 2 weeks. Irrespective of the bacterial inoculum, the population density numbers did not change over time (Figure 2A). At the highest initial population density (5 x 10^8^ CFU mL^-1^), inoculated leaves did not continue to grow over the course of the experiment, in accordance with the well-known dichotomy between growth and defense, in which only one process at a time is prioritized for resources by plants (Supplementary Figure S2A, Huot *et al*. 2014). Not only that, an earlier onset-senescence phenotype was also observed for leaves infiltrated at the highest inoculum. This result suggests that the growth–defense tradeoffs might not be important for plant–commensal interactions at the natural endophyte population density that exists in nature. Additionally, the lack of differences in the overall population stasis phenotype at different initial population densities suggests that a lack of resources cannot explain why phyllosphere non-pathogenic bacteria are unable to multiply to achieve high population densities, as populations of up to 10^6^ CFU cm^-2^ were able to be maintained inside leaves.

**Figure 2.**
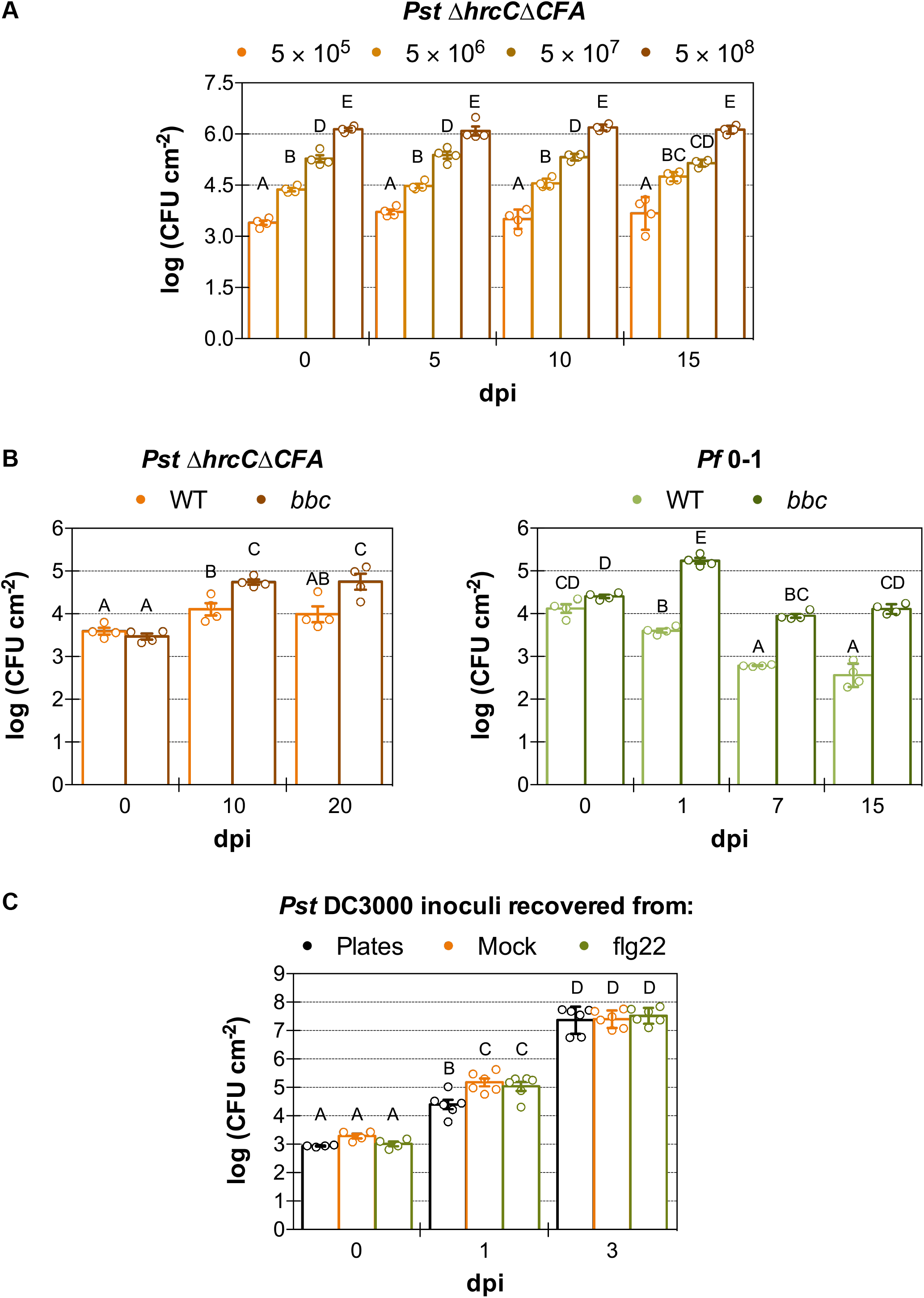
Bacterial population stasis is a reversible physiological state independent of bacterial density or PAMP-triggered immunity. **(A)** Bacterial population density in Col-0 plants after infiltration with different inoculum concentrations of *Pst ΔhrcCΔCFA* (in CFU mL^-1^). Bacterial population density remains static for over two weeks, irrespective of the starting bacterial inoculum concentration. **(B)** Compromising PTI is not sufficient to have sustained growth of non-pathogens. Bacterial population density in Col-0 and PTI-compromised triple mutant *bak1 bkk1 cerk1* (*bbc*) in *Pst ΔhrcCΔCFA* and *P. fluorescens* 0-1 (2 × 10^6^ CFU mL^-1^) over the course of up to 20 days. **(C)** Bacteria recovered from tissue experiencing flagellin-induced population stasis are as capable as causing disease as those from naïve plants. Plants were inoculated with bacteria (2 × 10^5^ CFU mL^-1^) recovered from mock- or PTI-primed plants (with DMSO or flg22, an elicitor from bacterial flagellin that induces PTI). As an inoculum control, bacteria grown *in vitro* on LM plates were used. Individual biological repetitions for each treatment are shown as open circles. Error bars indicate the standard error of the mean. When appropriate, different letters indicate differences in means, as judged by a Tukey HSD test (*p* < 0.05). dpi indicates days post-inoculation.

Suppression of PTI alone does not seem to confer a dramatic increase in bacterial population density *in planta* (less than a 10-fold increase, Xin *et al*. 2016), at least not in the few days after inoculation when the samples were collected. We reasoned, however, that perhaps evaluating population densities over the course of weeks rather than days would show a small but continuous increase in bacterial population densities in PTI-compromised mutant plants. However, this was not the case when we inoculated a PTI mutant deficient in three PTI co-receptors (*bak1 bkk1 cerk1*, involved in the recognition of multiple microbe-associated molecular patterns [MAMPs]; Xin *et al*. 2016) with multiple bacterial strains and compared the population densities to that of wild-type plants (Figure 2B and Supplementary Figure S2B). Even though population densities of *Pst ΔhrcCΔCFA* and *Rhodococcus* were larger in the PTI triple mutant than in the wild type, these populations did not continue to increase over time and remained static after an initial growth (Figure 2B and Supplementary Figure S2B). Additionally, for *P. fluorescens*, there was a decrease in population density over time, as had been observed for wild-type plants (Figure 1D), even though population density initially increased in the PTI triple mutant (Figure 2B). The plant immune defense is composed of many layers, and we can infer from these results that the failure of non-pathogenic microbes to continuously increase their population densities is not exclusively determined by PTI.

### Reversibility of bacterial stasis after PTI priming

Virulent bacteria may be rendered non-pathogenic in plants in which PTI priming has occurred, such as when a PTI inducer is added hours before bacterial inoculation, allowing for plant defenses to be primed and in high alert before pathogen arrival (Zipfel *et al*. 2004). We wanted to test whether PTI-mediated defenses render bacteria in a physiological state that could easily be reversed if the conditions became favorable. To do this, we recovered bacteria from plants pre-treated or not with an epitope of the PTI inducer flagellin, flg22. Before recovery, mock-treated virulent *Pst* DC3000 bacteria were able to multiply to the plant’s carrying capacity, while bacteria in PTI-primed plants did not increase in population density (Supplementary Figure S2C).

We used the recovered bacteria to inoculate naïve plants and found that there were no differences in population densities from bacteria recovered from PTI-primed or mock-treated plants (Figure 2C). The only difference observed was the higher earlier population densities of bacteria recovered from plants (at 1 dpi), compared to those that used an *in vitro* grown inoculum. This difference is likely reflective of the very low expression of genes involved in virulence mechanisms in bacteria grown in rich media (Nobori *et al*. 2018). The inability of bacteria to grow in PTI-primed plant tissue disappeared over time, further supporting the conclusion that this induced, static bacterial physiological state is reversible (Supplementary Figure S2D).

### Equilibrium between bacterial multiplication and death explains static population densities

Population density stability after the colonization of non-pathogenic phyllosphere microbes might be reflective of bacterial cells that completely cease to divide, or of an equilibrium between the rates of death and multiplication of such cells. To try to differentiate between these two possibilities, we used β-lactam antibiotics, which only target cells that are actively dividing (Spoering and Lewis 2001). We confirmed the *in vitro* cell-multiplication inhibitory effect of the bacterial cell-wall biosynthesis-targeting β-lactam antibiotic carbenicillin on *Pst ΔhrcCΔCFA* (Figure 3A). As expected, cells in the logarithmic phase were killed by carbenicillin as soon as they tried to divide (Figure 3A). However, when the cells had reached stationary phase, adding a β-lactam antibiotic had no effect on them. The reduction in population density for both the mock and carbenicillin treatments for stationary cells at later time points was likely caused by nutrient depletion in the medium (Navarro Llorens *et al*. 2010).

**Figure 3.**
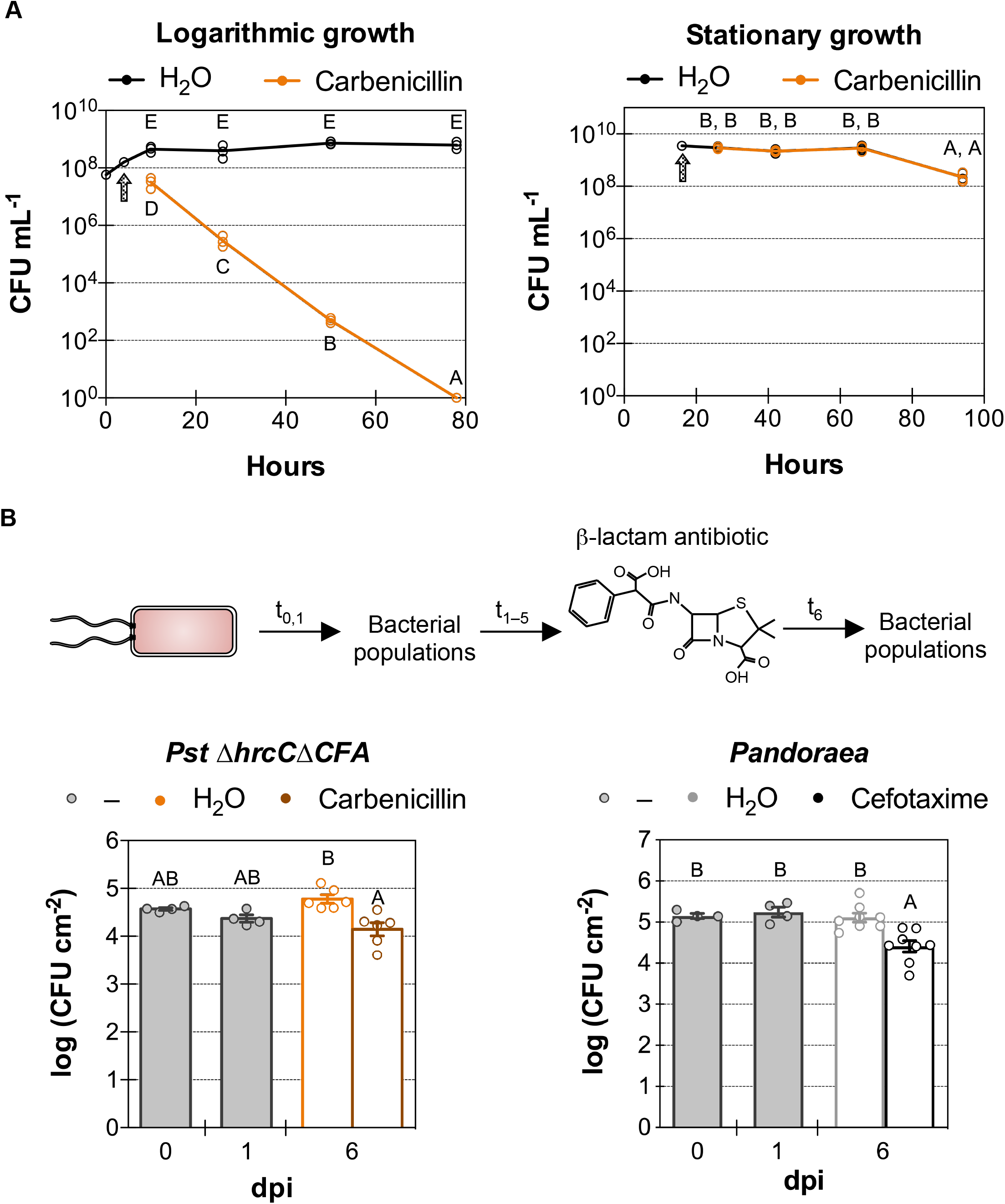
Apparent bacterial population stasis is caused by an equilibrium between individual cell death and multiplication. **(A)** Stationary culture density is unaffected by the antibiotic carbenicillin, which only targets actively dividing bacteria. *In vitro* growth (in LM media) of *P. syringae* pv. *tomato (Pst) ΔhrcCΔCFA* cultures in logarithmic or stationary phase after the addition of 400 µg mL^-1^ of carbenicillin or H_2_O control. The arrow indicates the time at which H_2_O or the antibiotic was added to the culture. The y-axis is a logarithmic scale. **(B)** *In planta* bacterial population density of *Pst ΔhrcCΔCFA* (10^7^ CFU mL^-1^) or *Pandoraea* sp. Col-0-28 (5 × 10^7^ CFU mL^-1^) in Col-0 plants after the addition of 400 µg mL^-1^ of carbenicillin or cefotaxime, β-lactam antibiotics that target cell wall biosynthesis and only kill dividing bacteria. Over 70% of the population is killed over the course of 5 days. Individual biological repetitions for each treatment are shown as open circles. Error bars indicate the standard error of the mean. Different letters indicate differences in means, as judged by a Tukey HSD test (*p* < 0.05). dpi indicates days post-inoculation. t_x_ indicates the number of days post inoculation.

To know if we can use carbenicillin to kill dividing bacteria inside plants, we inoculated Arabidopsis leaves with *Pst* DC3000, followed with infiltration of carbenicillin 1 and 2 days after inoculation. *Pst* DC3000 is virulent in Arabidopsis Col-0 plants and is expected to rapidly multiply once inside the apoplast. A greater than 100,000-fold increase in population density was observed between days 1 and 3 for the H_2_O control, which corresponds on average to approximately eleven cell divisions per inoculated bacterial cell (Supplementary Figure S3A). By contrast, in leaves infiltrated with carbenicillin, bacterial populations numbers were even lower at day 3 than they were before treatment (at day 1). More than 99.99% of cells attempted to divide and were killed by the β-lactam antibiotic. We can infer from this result that carbenicillin is very effective in killing dividing *Pst* DC3000 cells within plants.

We used carbenicillin to evaluate whether non-pathogenic *Pst ΔhrcCΔCFA* was able to multiply inside plants, or if cells had been rendered truly in stasis. Population densities of *Pst ΔhrcCΔCFA* treated with the antibiotic were reduced after 5 consecutive days of treatment (at 6 days post-infiltration) when compared to the mock H_2_O control (Figure 3B). Approximately 75% of all bacterial cells attempted to divide over 5 days and were killed by the antibiotic. When using conditions that favored even more bacterial multiplication (by increasing the relative humidity to over 99%), more than 98% of dividing bacterial cells were killed (Supplementary Figure S3B).

A similar antibiotic treatment experiment using microbiota strains *Pandoraea* and *Rhodococcus* yielded similar results. Five days after β-lactam antibiotic treatment, reduction in bacterial population densities was approximately 80% for *Pandoraea* (this experiment used a different β-lactam antibiotic, cefotaxime, as *Pandoraea* is resistant to carbenicillin; Figure 3B), while for *Rhodococcus* this reduction was almost 90% (Supplementary Figure S3C). Overall, these results suggest that bacterial endophytes are actively multiplying and dying inside plants, and that both rates are equivalent. This equilibrium would cause microbiota population densities to effectively remain static over time.

### Visualization of population stasis of commensal bacteria

In order to differentiate multiplying from static bacteria inside plants, we generated a bacterial system whereby fluorescence signal was used as a reporter of bacterial stasis. We integrated into the *Pst* genome a reporter that expressed two fluorescent proteins: mCerluean3 (Markwardt *et al*. 2011), expressed constitutively, and mCitrine (Griesbeck *et al*. 2001), expressed under the control of an inducible tetracycline promoter (Bertram and Hillen 2008; Figure 4A). If we grow the inoculum under the presence of a tetracycline analogue (anhydrotetracycline), bacteria would be doubly fluorescent for mCerulean3 and mCitrine. When inoculating such bacteria into plants, the mCitrine signal would become diluted if the bacteria divide, as anhydrotetracycline would no longer be available to induce fluorescent protein expression. If multiple cell divisions occur, the mCitrine signal would become diluted past the point of detection, and the cells would only be detected by the mCerulean3 signal (Figure 4A). Therefore, static cells that do not multiply would be fluorescent for both fluorescent proteins, while newly divided cells would not express mCitrine and after several divisions would be fluorescent only for mCerulean3.

**Figure 4.**
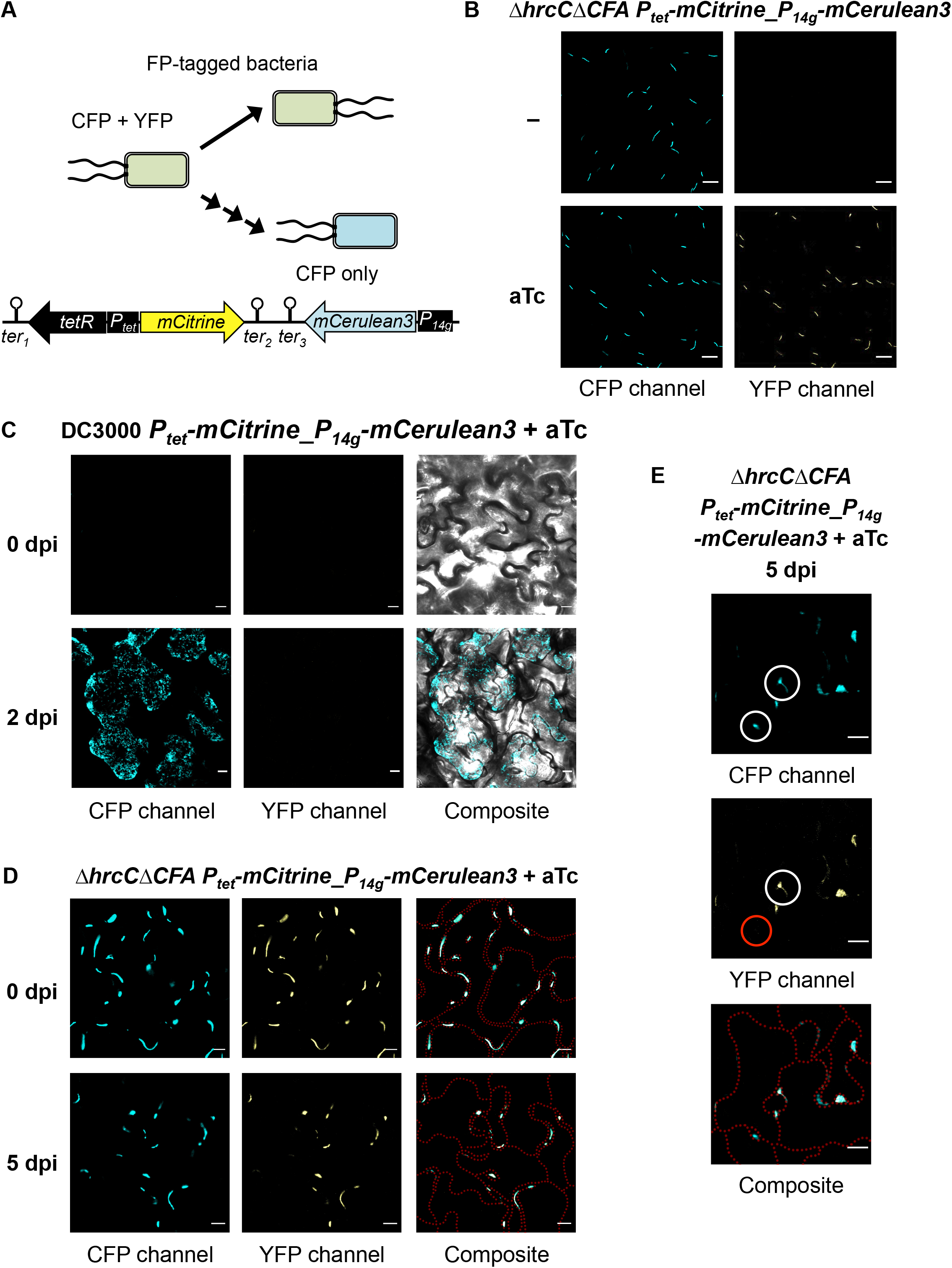
Visualization of *in planta* population stasis of bacterial endophytes. **(A)** Construct integrated into the *P. syringae* pv. *tomato* (*Pst*) genome. The tetracycline promoter (*P*_*tet*_) controls the bidirectional expression of the *tetR* transcriptional regulator and of fluorescent protein mCitrine. The strong constitutive *P*_*14g*_ promoter controls the expression of fluorescent protein mCerulean3. There are 3 different transcriptional terminators (*ter*) in the construct. Above the construct is a schematic representation of the dilution of fluorescent protein (FP) signal when a bacterium divides. mCitrine signal is conditionally expressed only in the presence of anhydrotetracycline (aTc); if the inducer is absent and bacteria multiply (represented by multiple consecutive arrows), only the constitutive mCerulean3 signal would be observed. **(B)** Confocal images of *Pst ΔhrcCΔCFA* carrying the fluorescent division reporter *P*_*tet*_*-mCitrine_P*_*14g*_*-mCerulean3* after *in vitro* growth with or without the addition of 67 ng µL^-1^ of aTc to induce mCitrine expression. **(C)** Confocal images of virulent *Pst* DC3000 carrying the fluorescent division reporter (5 × 10^7^ CFU mL^-1^; inoculum grown with 80 ng mL^-1^ aTc) 2 hours (0 dpi) and 2 days after inoculation of Col-0 plants. Fluorescent signal from mCerulean3 is only observed at the high population densities achieved at 2 dpi, while no inducible mCitrine signal is observed in the samples. **(D)** Confocal images of non-pathogenic *Pst ΔhrcCΔCFA* carrying the fluorescent division reporter (10^9^ CFU mL^-1^; inoculum grown with 80 ng mL^-1^ aTc) 3 hours (0 dpi) and 5 days after inoculation of Col-0 plants. Signal of both FPs mostly overlap. **(E)** Close-up of two bacterial aggregates, one with and one without mCitrine signal, indicating a static sub-population (white circle overlaid on the YFP-channel image) and one that arose from in planta bacterial multiplication (red circle), respectively. Cyan represents signal from mCerulean3. Yellow represents signal from mCitrine. Composite image of both FPs is also shown; in Figure 4C, the transmitted light image is overlaid, while in Figures 4D and 4E, the cell outlines are overlaid as red dotted lines. White bars represent 10 µm. dpi indicates days post-inoculation.

We first tested the inducibility of our reporter system *in vitro*. Indeed, *Pst ΔhrcCΔCFA* cells carrying the reporter were detected by only mCerulen3 signal in the absence of anhydrotetracycline, and by both mCerulean3 and mCitrine signals when anhydrotetracycline was added to the *in vitro* medium (Figure 4B). Note that the inducible mCitrine signal was overall lower, on average, than that of the constitutive mCerulean3 fluorescent protein.

We then tested the function of our reporter *in planta*. Virulent *Pst* DC3000 carrying the reporter and originally grown *in vitro* in the presence of the anhydrotetracycline inducer was inoculated into Arabidopsis Col-0 plants at a low density that was well below the detection level of our confocal microscopy set-up (day 0; Figure 4C). After 2 days of growth *in planta*, only mCerulean3 signal became detectable because bacteria multiplied to population densities well past the detection minimum, while mCitrine signal never became detectable, as mCitrine expression was absent inside plants, and the original mCitrine signal from the inoculum was diluted. At this point of the infection with *Pst* DC3000, symptoms were starting to appear on the leaves and the entirety of multiple plant cells had been colonized; the leaf colonization seemed to have reached beyond the apoplastic space. Most likely, these plant cells were dead due to bacterial infection, as at later time points *Pst* DC3000 causes wilting and necrosis in Arabidopsis (Velásquez *et al*. 2017). At the leaf surface, it was easier to discern individual bacterial cells and colonization of intercellular spaces by *Pst* DC3000 (Supplementary Figure S4).

To visualize a phyllosphere-inhabiting microbial endophyte *in planta*, we inoculated *Pst ΔhrcCΔCFA* carrying the cell division reporter into Arabidopsis leaves. On day 0, *Pst ΔhrcCΔCFA* had colonized multiple non-continuous regions of the apoplast of plant cells. Both mCerulean3 and mCitrine signals were observed, and their signals overlapped (Figure 4D). These bacterium-colonized apoplast areas are most likely those in which a liquid environment is present, while those that are not colonized are likely full of pockets of air. It might be that only in pathogenic organisms, the virulence mechanisms would allow access of other areas by using effectors that promote the water soaking of the entire leaf apoplast (Xin *et al*. 2016; Figure 4C and Supplementary Figure S4). On day 5, signal from both fluorescent proteins *in Pst ΔhrcCΔCFA* was still observed (Figure 4D). However, at this point, while some bacterial aggregates had signal from both fluorescent proteins, others had signal only from mCerulean3 (Figures 4D and 4E). This indicates that within a leaf, some of the bacterial cells of the population remain static (at least until 5 days post-inoculation), while other cells are multiplying and give rise to new bacterial aggregates (those devoid of mCitrine signal). These results seem to agree with the results obtained with the β-lactam antibiotic treatment (Figure 3B and Supplementary Figure S3B), in which some of the cells in the population divided while others remained in stasis.

### *In planta* transcriptomics analysis of phyllosphere-inhabiting bacteria

To try to understand which biological processes are affected during long-term survival of non-pathogenic bacteria inside plants and to gain a glimpse into the causes of population stasis, we performed RNA-Sequencing (RNA-Seq) from Arabidopsis leaves inoculated with bacteria. We focused on two microbiota strains and a disarmed mutant of *Pseudomonas*: *A. xylosoxidans, Pandoraea* sp., and *Pst ΔhrcCΔCFA*, respectively. We evaluated the transcriptomes of the inocula before, and at 6, 24, and 168 hours after inoculation of these three bacterial strains into plants. For *A. xylosoxidans*, we sequenced only the endophytic bacteria by eliminating the surface-inhabiting bacteria with sodium hypochlorite. For *Pst ΔhrcCΔCFA*, we also included *in vitro* populations of cells that were in logarithmic or stationary phase of growth. The bacterial population densities of the samples used for sequencing are shown in Supplementary Figures S5A, S5B, and S5C. At the very high inoculum needed for performing RNA-Seq (2 × 10^9^ CFU mL^-1^; otherwise the sequencing would be overwhelmed by mostly plant RNAs), there was a slight decrease in population densities between 6 and 24 hours. This was most likely caused by a strong initial PTI response due to the overabundance of MAMPs in the inoculum.

For performing RNA-Sequencing, we used a pipeline that we originally described in Nobori *et al*. (2018), in which bacterial mRNA transcripts are enriched using custom-designed probes during the process of preparing the libraries. To be thorough in calculating differential gene expression (DGE), we used three different methods (for a comparison of the number of differentially expressed genes identified by each of the three methods, see Supplementary Figure S5D): (1) alignment of reads with HISAT2 (Kim *et al*. 2019), calculation of transcript frequency with StringTie (Kovaka *et al*. 2019), and DGE with DESeq2 (Love *et al*. 2014); (2) alignment of reads with HISAT2 and DGE with Cufflinks (Trapnell *et al*. 2013), and (3) pseudo-alignment of reads with Salmon (Patro *et al*. 2017), and DGE with DESeq2. Differentially expressed genes present in at least one of the analyses were used for subsequent analysis. A principal component analysis (PCA) indicated that all the samples of every treatment clustered together separately from samples from other treatments (Supplementary Figures S5E and S5F).

*In planta* gene expression patterns over time of select biological pathways that were enriched in differentially expressed genes in at least one of the different treatment comparisons are presented in Figures 5A (for *Pst ΔhrcCΔCFA*; also see Supplementary Figure S6), 5B (for *A. xylosoxidans*) and 5C (for *Pandoraea* sp.). A complete list of biological pathways enriched in differentially expressed up or down-regulated genes is shown in Supplementary Table S1. We also performed an analysis for biological pathways enriched in different expression profiles over time to identify sets of genes whose expression increased or decreased once phyllosphere bacteria were inside plants. The expression profiles were calculated using STEM (Ernst and Bar-Joseph 2006) and are shown in Supplementary Table S2. Finally, an analysis to find biological pathways enriched in orthologous genes whose differential expression followed the same pattern of up- or down-regulation in all three strains was performed.

**Figure 5.**
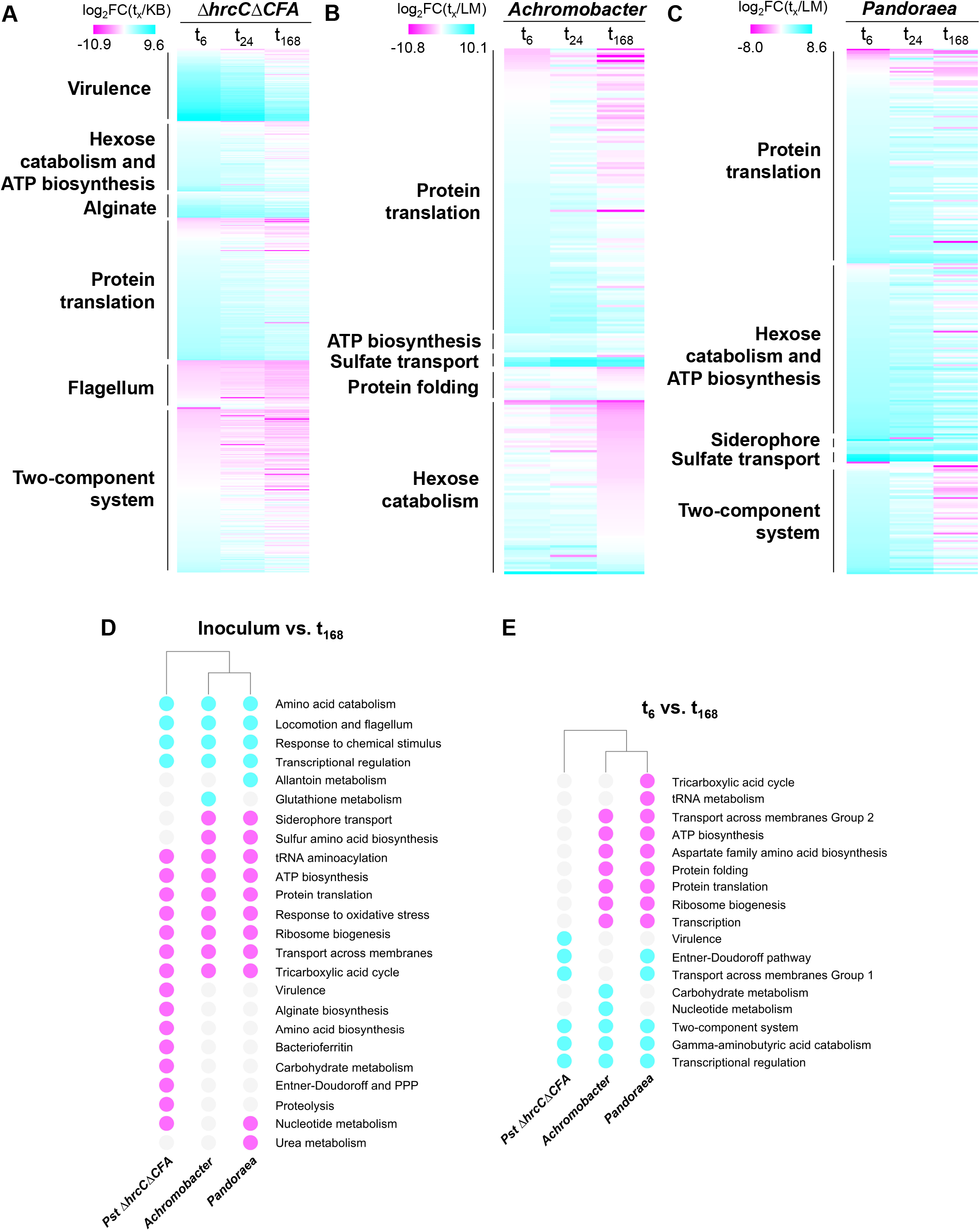
Transcriptome analysis of three phyllosphere-inhabiting bacteria. **(A)** Long-term *in planta* transcriptome heatmap of gene expression of select categories in non-pathogenic *P. syringae* pv. *tomato* (*Pst*) *ΔhrcCΔCFA*. **(B)** Long-term *in planta* transcriptome heatmap of select categories of bacterial microbiota member *Achromobacter xylosoxidans* Col-0-50. Only endophytic bacteria were used for the analysis. **(C)** Long-term *in planta* transcriptome heatmap of select categories of bacterial microbiota member *Pandoraea* sp. Col-0-28. (**D**) Up- or down-regulation of expression of enriched biological processes of differentially expressed groups of orthologous genes shared between *Pst ΔhrcCΔCFA, A. xylosoxidans*, and *Pandoraea* sp., shared between two strains, or unique to each strain, when comparing the inoculum to 168 hours post-inoculation or **(E)** 6 to 168 hours post-inoculation into Col-0 plants. For the transcriptome heatmaps, gene expression values of select categories were graphed in ascending order of their log_2_ fold change (FC) of the ratio between 6 (t_6_), 24 (t_24_) or 168 (t_168_) hpi and the expression in the inoculum (KB or LM). FC was color coded for repression (magenta), no change (white), and induction (cyan), with the same color scheme used for up-regulation or down-regulation of biological processes. PPP refers to the Pentose phosphate pathway, while KB and LM refer to the bacterial inoculi, which were grown in King’s B medium or modified LB, respectively.

As expected, expression of alginate biosynthesis genes and virulence-promoting mechanisms—including genes for coronatine biosynthesis and type III secretion system structural components and effectors—was up-regulated inside plants for *Pst ΔhrcCΔCFA* (Nobori *et al*. 2018). Note that this strain is non-pathogenic, and as such, virulence-promoting mechanisms should not play an essential role in bacterial survival. Primary metabolic pathways for energy and intermediary metabolite generation involved in ATP biosynthesis and hexose catabolism, including the Entner-Doudoroff pathway, Tricarboxylic acid cycle, and the Pentose phosphate pathway (see Supplementary Figure S6), were also up-regulated inside plants. These same primary metabolic pathways were up-regulated *in planta* when compared to the inoculum for the microbiota strains *Achromobacter* and *Pandoraea* (Figure 5D and 5E); however, only in the microbiota endophytes was expression of ATP biosynthesis genes up-regulated the longer the bacteria stayed inside the leaves. The response to oxidative stress was also up-regulated *in planta* for all three strains, perhaps as an adaptation to the harsher environment encountered by microbes in the apoplast.

Expression of protein translation genes, especially those involved in ribosome biogenesis, was also up-regulated inside plants when compared to the inoculum (Figures 5A, 5B, 5C, and 5D), the only exception being tRNA-coding genes whose expression was in general lower inside plants than in the inoculum irrespective of the time point analyzed (Supplementary Figure S6). Expression of protein translation was higher in both microbiota endophytes after 7 days post-inoculation (t_168_) compared to earlier inoculation times, while no difference was observed for *Pst ΔhrcCΔCFA* (Figure 5E). Transcriptional regulation processes seemed to be reduced in all three strains at 168 hours post-inoculation (hpi), and so did two-component system expression (Supplementary Tables S2A, S2B, S2C and Figure 5D and 5E), perhaps because processes involved in sensing and responding to the plant environment are more important during the initial interaction of the microbe with its host. Only in the microbiota endophytes were genes involved directly in transcription, such as the different RNA polymerase subunits, expressed higher after 168 hpi. For both *Pst ΔhrcCΔCFA* and *Achromobacter*, flagellum biosynthesis was lower in plants when compared to the inoculum, and these genes were down-regulated even further as time inside plants went by (Supplementary Tables S2A and S2B and Supplementary Figure S6). Flagellum biosynthesis repression may be reflective of the strong pressure that flagellin detection by plants has over microbiota adaptation (Colaianni *et al*. 2021).

As secondary metabolites can be responsible for regulating microbe–microbe and plant– microbe interactions, we used antiSMASH (Blin *et al*. 2019) to identify potential secondary metabolite biosynthesis clusters in all three phyllosphere-inhabiting bacteria (Supplementary Table S3). The analysis confirmed the presence of known secondary metabolite clusters in *Pseudomonas syringae*, such as those for the biosynthesis of the phytotoxin coronatine and the siderophores pyoverdin and yersiniabactin. Other than those three clusters, a region involved in the biosynthesis of the dipeptide N-acetylglutaminylglutamine amide (NAGGN) showed a clear and stable up-regulation inside plants (Supplementary Table S3A). NAGGN might function in osmoregulation in *Pseudomonas*, as has already been shown in other bacteria (Sagot *et al*. 2010), and protect *Pst ΔhrcCΔCFA* from osmotic stress. No differentially expressed biosynthetic clusters were observed in *Pandoraea*, while in *A. xylosoxidans*, expression of a cluster involved in the biosynthesis of the osmoprotectant ectoine was up-regulated in plants, especially early during plant colonization (Supplementary Table S3B). This result independently confirmed the findings of the biological pathway enrichment analysis done previously (Supplementary Tables S1C and S2B). A secondary metabolite cluster in *A. xylosoxidans* for the production of resorcinol, which could potentially have antimicrobial activity (Schöner *et al*. 2015), was induced early in the interaction of *A. xylosoxidans* with plants, perhaps because this metabolite confers competition advantages for growth for this microbiota strain. Finally, expression of genes predicted to be involved in the biosynthesis of a desferrioxamine-like siderophore was down-regulated inside plants in *A. xylosoxidans*, which suggests that this endophyte does not experience iron limitation, similar to what has been observed during pathogenic infections of *Pst* DC3000 inside plants (Jones and Wildermuth 2011).

### Non-pathogenic endophytes resemble more closely the physiological status of stationary-phase bacteria

As the density of populations inside plants seems to not change for non-pathogenic bacterial endophytes (see Figure 1A, 1C and Supplementary Figure S1A), we hypothesized that bacteria could have entered a physiological status that closely resembles that of bacteria grown *in vitro* when they reach stationary phase. We observed that overall gene expression *in vitro* was more similar between each *in vitro* population (the inoculum, logarithmic and stationary populations) than with the overall gene expression observed inside plants (see PCAs in Supplementary Figures S5E and S5F). However, when we compared the bacterial processes of *Pst ΔhrcCΔCFA* that were enriched for up- and down-regulated genes inside plants with populations of cells that were grown *in vitro* to logarithmic and stationary phase (Supplementary Tables S1A and S1B), bacterial populations that were actively dividing and had reached logarithmic phase were different from populations in stationary phase or from bacterial populations that had been inside plants for 168 hours (7 days; Figures 6A and 6B). Not only that, but the majority of different biological processes between logarithmic and stationary populations were the same as those observed between logarithmic populations and at 168 hpi. Protein translation, and generation of metabolite precursors and energy processes were down-regulated in stationary phase and after 168 hpi, while virulence-promoting mechanisms, flagellum biosynthesis, and two-component systems were up-regulated when compared to logarithmic phase populations. This reinforces the idea that bacteria inside plants more closely resemble *in vitro* bacteria that have reached stationary phase, and correlates with their inability to increase in population density inside plants.

**Figure 6.**
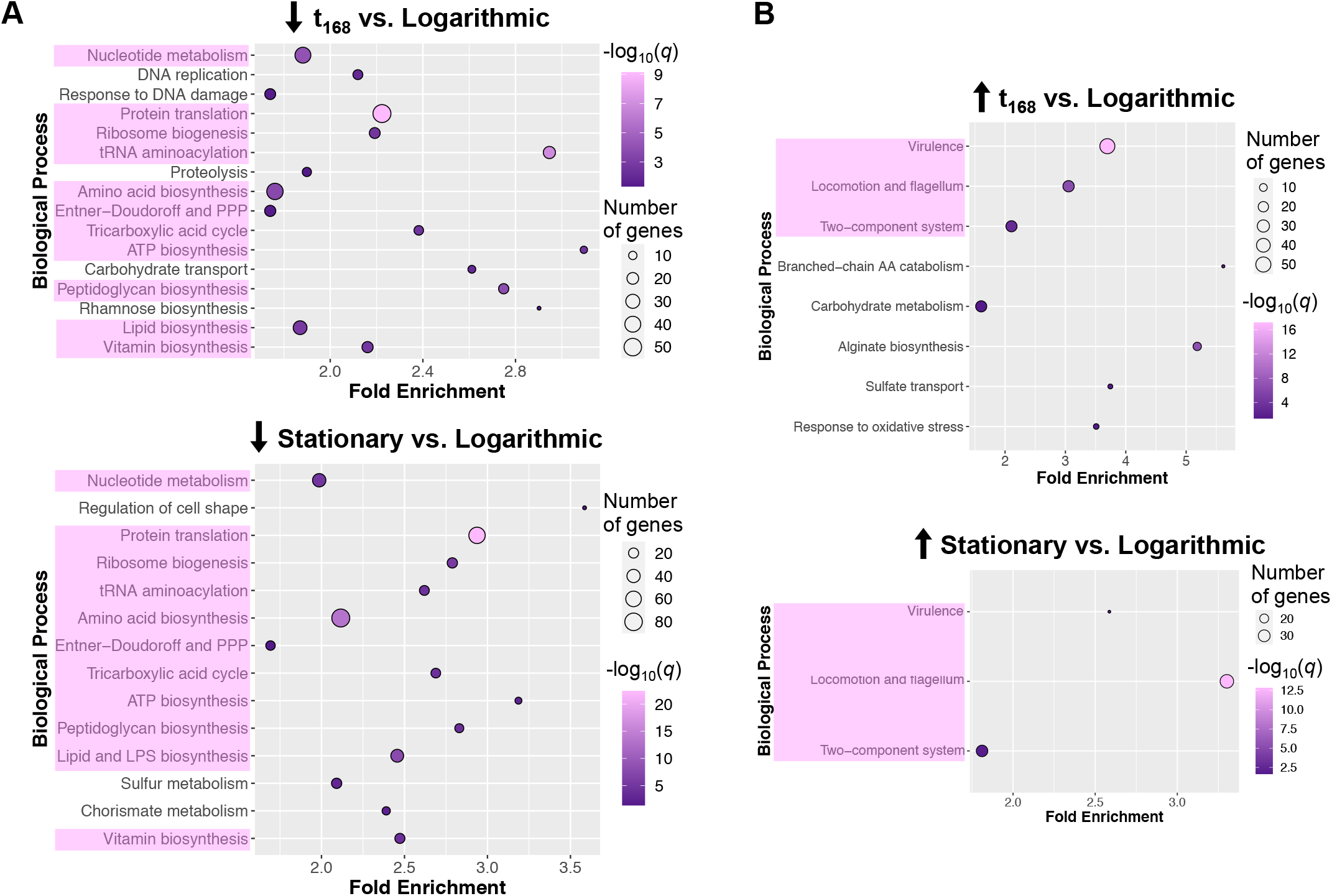
Static phyllosphere-inhabiting bacterial populations show similar transcriptional responses to populations in stationary phase. **(A)** Bubble graph of enriched biological processes in differentially expressed genes (DEGs) down-regulated at 168 hours post-inoculation (t_168_) or in *in vitro* stationary-phase populations, when compared to *in vitro* logarithmic-phase populations. Processes found in both comparisons are highlighted in pink. **(B)** Bubble graph of the same comparisons as in D, but using the enriched biological processes from up-regulated DEGs. For the bubble graphs, the fold-enrichment, number of significant DEGs, and the log_10_ of the adjusted *p*-value (*q*) is shown for each biological process. PPP refers to the Pentose phosphate pathway.

### Bacterial gene expression of select biological processes during long-term adaptation of a non-pathogen inside plants

We decided to corroborate the RNA-Seq results by performing qRT-PCR of select genes. We selected genes involved in virulence (the sigma factor master regulator of virulence *hrpL*; Lam *et al*. 2014, and the type III effector *avrPto*; Nguyen *et al*. 2010), transcription (the housekeeping RNA polymerase sigma factor *rpoD*; Feklístov *et al*. 2014), and bacterial cell division (*ftsZ*; McQuillen and Xiao 2020). For *avrPto, hrpL*, and *rpoD*, gene expression was extremely low in the inoculum, increased exponentially by 6 hours after inoculation into plants, and then decreased over time, with the lowest level by 168 hours (Figure 7A and Supplementary Figure S8A). For example, the expression of *rpoD* at 6 hours post-inoculation was almost 200 times greater than in the inoculum, and 6 times greater than the expression at 168 hours. *rpoD* has been used in the past as a qPCR reference gene (Smith *et al*. 2018); however, its expression, at least under certain conditions, is too variable for this purpose, as observed in our experiments. *ftsZ* showed the same trend of early increased expression and subsequent decrease *in planta*; however, the relative expression in the inoculum was more variable (Figure 7A and Supplementary Figure S8B). Overall, qRT-PCR results were in agreement with the transcriptome results for the above-mentioned three biological processes.

**Figure 7.**
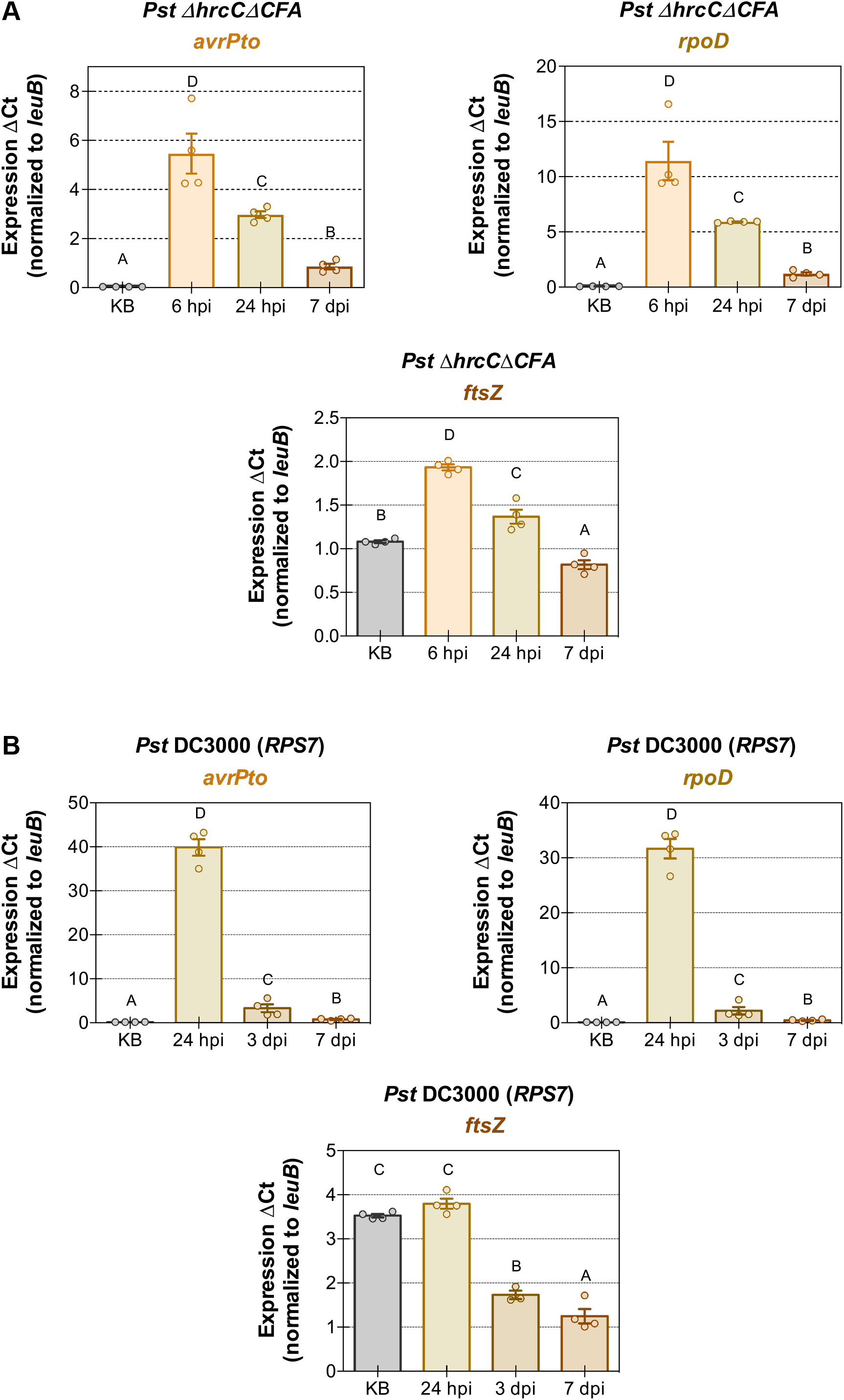
Bacteria experiencing effector-triggered immunity have a transcriptional response that mimics non-pathogenic endophytic bacteria. **(A)** *In planta* gene expression of a type III effector (*avrPto*), the housekeeping RNA polymerase sigma factor (*rpoD*), and a gene involved in bacterial cell division (*ftsZ*) after inoculation with *Pst ΔhrcCΔCFA* into Col-0 plants. **(B)** *In planta* gene expression of the same genes as in 4D after inoculation with *Pst* DC3000 into Bu-22 plants, an interaction that triggers effector-triggered immunity. For quantitative reverse-transcriptase PCR, gene expression was evaluated using *leuB* as the reference gene and the ΔCt method. Error bars indicate the standard error of the mean. Different letters indicate differences in means, as judged by a Tukey HSD test (*p* < 0.05), which was performed between the ΔCt values. hpi indicates hours post-inoculation, dpi indicates days post-inoculation, and KB refers to the bacterial inoculum, which was grown in King’s B medium.

### Bacterial gene expression in ETI-causing interactions is similar to that observed for non-pathogens

As both non-pathogens and ETI-causing bacterial strains had the same apparent population stasis in plants (see Figures 1A and 1B), we evaluated gene expression for *Pst* DC3000 after AvrPto recognition (via *RPS7* in Bu-22 plants) for the same genes described in the previous section to try to understand if gene expression was similarly affected. Initially, we tried using a high inoculum for this experiment, similar to was used for *Pst ΔhrcCΔCFA*. However, even though a faster hypersensitive response is observed at a high inoculum in Bu-22 plants (Velásquez *et al*. 2017), ETI restriction of bacterial multiplication is compromised (Supplementary Figure S8C). As such, we decided to use a lower inoculum for evaluating gene expression in an incompatible interaction (Supplementary Figure S8D). We also started measuring gene expression 1 day after inoculation and not at 6 hours, as in ETI interactions, there is usually a small initial growth of the microbe before population stasis (Figure 1B and Supplementary Figure S1B).

The same gene expression pattern as that observed with *Pst ΔhrcCΔCFA* was observed under ETI conditions with *Pst* DC3000 for *avrPto, hrpL, rpoD*, and *ftsZ* (Figure 7B and Supplementary Figures S8B and S8E), with the highest expression for each of these genes observed at 24 hours after inoculation into Bu-22 plants, and then decreasing afterwards. As we were unable to perform RNA-Sequencing of ETI-inducing strains to characterize population stasis using a high inoculum (see the explanation above), we evaluated the expression of a few additional genes under ETI conditions: an additional gene involved in virulence (the phytotoxin biosynthesis gene coronafacate ligase *cfl*; Worley *et al*. 2013, which is deleted in *Pst ΔhrcCΔCFA*), a gene involved in last step of the Entner-Doudoroff pathway (the pyruvate kinase *pyk*; Schormann *et al*. 2019), and a gene involved in protein translation (tRNA^fMet^-formyl transferase *fmt*, which formylates the initiator tRNA^fMet^ that carries the first amino acid used in protein translation in Eubacteria; Laursen *et al*. 2005). Expression of these three genes showed a similar pattern to that observed previously; gene expression was very low in the inoculum, increased by 24 hours after inoculation, and decreased at later time points (Supplementary Figure S8E).

## DISCUSSION

In this study, we characterized the population dynamics behind bacterial endophyte survival and multiplication, and used transcriptomics to infer the biological processes important for microbial multiplication. We observed that long-term population densities of phyllosphere-inhabiting bacteria seemed to remain static over time. This bacterial population stasis lasted for several weeks and was observed for both ETI-inducing and non-pathogenic microbes. We decided to study this bacterial population stasis phenomenon by using a combination of fluorescent reporters, antibiotics targeting dividing bacteria, and *in planta* transcriptomics. The apparent prevalent population stasis was caused by equilibrium in the rates of multiplication and death of the microbe, and likely not by lack of sufficient resources or of an appropriate niche for microbial growth. *In planta* transcriptomes of bacterial endophytes suggested that their physiology more closely resembles the physiology that bacteria experience during stationary phase, a condition in which there is selective transcription of certain biological processes and in which stress responses are more active.

We observed that microbiota population density numbers were in apparent stasis, with the immune system unable to eliminate (or perhaps sometimes even detect) bacterial endophytes. Even if phyllosphere endophytes are in a stationary-like state, this does not mean that bacteria are not metabolically active, irrespective of the lack changes in their population density numbers. Bacteria in stationary phase are not inert; for example, during stationary phase protein expression is constant (Gefen *et al*. 2014). Also, it is during such phase that secondary metabolites, such as osmoprotectants and competition-facilitator antibiotics, are synthesized (Navarro Llorens *et al*. 2010). Bacteria are usually in a nutrient-deprived stationary state in nature, as abundant resources are a rare occurrence. Population stasis at high bacterial densities suggests that it is not a lack of resources that prevents non-pathogenic endophytes at low densities from multiplying and increasing their numbers once they are inside leaves (Figure 2A). Commensal phyllosphere bacteria might have evolved to maintain low population numbers via a slow multiplication rate that matches the death rate, in order to not alert the immune system of their presence, as would occur during the colonization of a pathogen—which normally would carry immune-dampening mechanisms. Most likely, long-term evaluation of phyllosphere bacteria, in contrast to previous reports that focused on the early events after infections, are more likely reflective of the commensal experience in nature.

Even four weeks after the initial inoculation, and after a significant growth of the inoculated plants had occurred (see Figure 1A), the endophytic population densities remained stable, almost as if the plants were not affected by the presence of such microbes. This makes us wonder if there is a lower population density threshold (<10^3^ CFU cm^-2^, less than the culturable density of endophytes recovered from leaves; Chen *et al*. 2020b, Xin *et al*. 2016) that would allow for unrestricted growth of microbiota needed for seedling colonization by microbes in nature. Regarding an upper population density in which the immune system becomes activated to eliminate the invading endophytes, no such threshold was observed (Figure 2A), except at inoculum densities in excess of 10^6^ CFU cm^-2^, which is at least two orders of magnitude higher than naturally observed endophyte densities (and even then, the population density was reduced to only about 10^6^ CFU cm^-2^, after which static population densities were observed; Supplementary Figures S5A, S5B, and S5C). The only bacterial strain for which apparent population stasis was not observed was *P. fluorescens* 0-1. This bacterium was originally isolated from soil (Silby *et al*. 2009), and perhaps lacks the adaptation mechanisms to survive as a leaf endophyte. Another possibility would be for *P. fluorescens* 0-1 to carry a MAMP that would trigger such a strong PTI response in Arabidopsis that the population density would be reduced (at an inoculum of 10^8^ CFU mL^-1^, inoculation of *P. fluorescens* into Arabidopsis Col-0 causes cell death; A. Velásquez, unpublished).

During ETI interactions, apparent population stasis is also observed, phenocopying the response observed for microbiota bacteria lacking virulence mechanisms. Population density stasis was observed after activation by both NLRs (NOD-like receptors) of the coiled-coil and TIR classes of Resistance proteins (RPS2 and RPS5 are of the coiled-coil class, while RPS4 is a TIR NLR; Cui *et al*. 2015), suggesting that this phenomenon is widespread. The initial growth observed for ETI-inducing strains in the first day after inoculation implies that the ETI-mediated population restriction mechanisms have not kicked in. In the host–pathogen battle experienced by ETI-inducing bacteria, the initial gains by bacterial virulence mechanisms seem to be outdone by the plant immune system at later time points. Additionally, niche destruction caused by the hypersensitive response of the cells undergoing effector recognition is not enough to eliminate bacteria. Having potential pathogenic populations always present in plants undergoing ETI suggests that these plants could serve as potential reservoirs for the infection of nearby plants not carrying the corresponding *R* genes. This could have implications for disease management during agricultural production, assuming that bacteria could be disseminated from plants undergoing ETI.

Gene expression in *Pst* DC3000 after plants undergo ETI showed a pattern reminiscent of the one observed for non-pathogenic *Pst ΔhrcCΔCFA* (Figures 7A and 7B). Under ETI conditions, there was a dramatic reduction of expression over time inside plants of genes involved in virulence: for the type III effector *avrPto*, the coronatine biosynthesis gene *cfl*, and the master regulator of virulence *hrpL* (Figures 7B and Supplementary Figure S8E). The down-regulation of *avrPto* expression, which in *RPS7*-carrying plants betrays *Pseudomonas syringae* by alerting plants of its presence, might be an adaptation to quench the strong immune response experienced by bacteria during ETI, and “transform” *Pst* DC3000 into a non-pathogenic persisting microbe, or it might occur due to a lack of type III secretion system-inducing compounds present in the plant apoplast during ETI.

Transcriptomes of all three phyllosphere-inhabiting bacteria, *Achromobacter, Pandoraea* and *Pst ΔhrcCΔCFA*, showed that the expression of the machinery for protein translation and for the generation of energy and intermediate metabolites was up-regulated inside plants when compared to the inoculum, which was grown *in vitro*. Once bacteria were in the apoplast, the above-mentioned processes had a higher expression for the two microbiota endophytes the longer bacteria resided in the apoplast, perhaps suggesting that they have a better adaptation than *Pst ΔhrcCΔCFA* for long-term survival inside plants, especially as this non-pathogen has been disarmed for its main adaptation mechanisms for apoplast survival: the virulence-promoting type III effectors and coronatine. The phyllosphere bacteria have adapted to colonize the apoplast, and as such, there was an induced expression of mechanisms involved in the response to stress, while the expression of genes putatively involved in the biosynthesis of secondary metabolites was up-regulated, including two clusters that synthesize osmoprotectants: N-acetylglutaminylglutamine for *Pst ΔhrcCΔCFA*, and ectoine for *A. xylosoxidans*.

The apparent population stasis in non-pathogenic microbiota was caused by bacterial multiplication and death at equivalent rates, as demonstrated by the killing of any bacteria that attempted to divide under β-lactam antibiotic treatment (Figure 3B and Supplementary Figures S3B and S3C). This was further confirmed using a division fluorescent reporter in *Pst ΔhrcCΔCFA* (Figure 4D and 4E). In immune-activated tissue, there is a transient non-lethal effect against virulent bacteria; this physiological state can easily be reversed if the proper environmental conditions are met (Figure 2C and Supplementary Figure S2D). It was also apparent that there is phenotypic heterogeneity of the clonal population of microbiota: some bacteria are physiologically able to multiply (depending on the environmental conditions, up to 98% of the population), while others are in a quiescent state, almost equivalent to being in stationary phase. Understanding the cues for multiplication and stasis of *in planta* bacterial populations could be used to manipulate pathogenic microbes into entering a stationary state and not causing disease. This might require adaptation and optimization of current techniques for single-cell bacterial RNA-Seq (Blattman *et al*. 2020, Kuchina *et al*. 2021) for use in future *in planta* transcriptomics aimed to better understand phenotypic heterogeneity.

Both the transcriptome of *Pst ΔhrcCΔCFA* and their *in planta* bacterial population density numbers (Figures 1A and 6) implied that bacteria were experiencing a physiological state that closely resembled the one that bacteria would experience in stationary phase. Additionally, it could well be that the physiology of pathogenic microbes, at least before reaching the plant’s carrying capacity, resembles that of bacteria in logarithmic phase—except for the activation of virulence mechanisms, which occurs only inside the host. Expression of the RNA polymerase sigma factor involved in adaptation to stationary phase physiological conditions, *rpoS*, showed almost no change at 168 hpi or stationary phase, compared to the expression observed in logarithmic-phase bacterial populations (Supplementary Table S4A). This could reflect a more important role of post-transcriptional regulation in modifying *rpoS* activity in *P. syringae* (Jaishankar and Srivastava 2017). By contrast, the mRNA expression of two other regulators of stationary phase, *relA* and *spoT*, involved in the bacterial stringent response (by synthesizing and degrading guanosine tetra and pentaphosphate [(p)ppGpp)]; Jaishankar and Srivastava 2017), was down-regulated in both stationary phase and *in planta* after 168 hpi compared to the expression in logarithmic phase (Supplementary Table S4A). This result reinforces the idea that long-term bacteria inside plants resemble stationary-phase populations of bacteria.

Microbiota endophytes seem to be in a stationary-like phase physiological state, one in which their population density numbers are in equilibrium, possibly to not alert the plant immune system of their presence. By no means does this imply that bacteria are in a quiescent state; while the population density might not change, the bacteria are still metabolically active and adapting to their surrounding environment. A better understanding of the nature of the population dynamics and transcriptomic features associated with a commensal lifestyle sets a foundation for future engineering of commensal microbiota in agricultural settings.

## MATERIALS AND METHODS

All experiments reported in this study were done at least thrice, except for the RNA-Sequencing experiments.

### *In vitro* bacterial growth and antibiotics used

Supplementary Table S5 lists the bacterial strains used in this study. *Escherichia coli* strains were grown in LB (Lennox) medium at 37 °C, while all other strains were grown in either a modified LB medium (LM: 10 g L^- 1^ tryptone, 6 g L^-1^, yeast extract, 1.5 g L^-1^, KH_2_PO_4_, 0.6 g NaCl, and 0.4 g MgSO_4_?7H_2_O) or King’s B medium at 30 °C.

Antibiotics and derivatives were used at the following concentrations: 100 µg mL^-1^ ampicillin, 67–80 ng mL^-1^ anhydrotetracycline, 400 µg mL^-1^ carbenicillin, 400 µg mL^-1^ cefotaxime, 75 µg mL^-1^ cycloheximide (to prevent fungal growth), 1–15 µg mL^-1^ gentamycin, 50 µg mL^-1^ kanamycin, 100 µg mL^-1^ rifampicin, and 50 µg mL^-1^ spectinomycin. Diaminopimelic acid (DAP) was used at a concentration between 200–400 µg mL^-1^. flg22, an epitope of bacterial flagellin and inducer of PAMP-triggered immunity (PTI), was used at a concentration of 500 nM.

To evaluate the response of logarithmic- and stationary-phase grown *Pseudomonas syringae* pv. *tomato* (*Pst*) *ΔhrcCΔCFA* cultures to the antibiotic carbenicillin, bacteria were grown in liquid LM overnight at 30 °C. For logarithmic-phase samples, overnight bacterial suspensions were diluted with LM to 10^8^ CFU mL^- 1^, and grown for 4 hours before antibiotic treatment. Stationary-phase samples were not diluted. H_2_O control or 400 µg mL^-1^ carbenicillin were added to the cultures and grown for over 3 days. Before plating serial dilutions of the samples to determine population densities, cultures were washed twice with 0.25 mM MgCl_2_ to remove any trace of the antibiotic from the bacterial suspension.

### Plant growth conditions

Arabidopsis plants were grown in a growth chamber with the following conditions: 12-hour day length, 80 µmoles m^-2^ s^-1^ of photon flux, a temperature of 24.5 °C during the day and 23.0 °C during the night, and a relative humidity between 65% and 75%. Supplementary Table S6 lists the plant material used for this study. For Supplementary Figure 3B, and for the *in planta* RNA-Seq samples of *Achromobacter* and *Pandoraea* (Supplementary Figures 5B and 5C), the relative humidity was increased to 99% after bacterial inoculation. Seeds were stratified for 2 to 5 days before sowing. Plants were watered with one-half strength Hoagland’s solution when needed. All plants were grown under a partially covered transparent plastic dome.

### Bacterial population density quantification assays

Bacterial inoculum suspensions were prepared from cultures grown in plates to stationary phase. Bacteria were resuspended in 0.25 mM MgCl_2_ to the appropriate OD_600_, after which they were infiltrated using a needleless syringe into the abaxial side of Arabidopsis leaves. To determine the *in planta* population density numbers, leaf-disc punches were collected from plants and ground in 250 µL of 10 mM MgCl_2_ using the TissueLyser II (QIAGEN; 2 cycles of 30 seconds at 25 Hz) and 3-mm zirconium oxide beads (Glen Mills Inc.). Serial dilutions of the ground tissue were spotted onto plates with the appropriate antibiotics and grown overnight. Colony forming units (CFUs) per cm^2^ were determined for each sample, while CFUs per mL were determined for each inoculum.

To determine endophytic bacterial populations, leaves were placed in a 0.825% sodium hypochlorite solution for 1 minute, and then washed twice in distilled H_2_O for a minute. Leaves were blotted dry, after which leaf discs were collected as described above.

To recover bacteria from leaves to use as inoculum, leaf discs were collected and ground in 0.25 mM MgCl_2_. The plant debris was filtered through a 70-µM-pore nylon filter (Corning Inc.), and bacterial suspensions were washed twice in 0.25 mM MgCl_2_ at 4,000 *g* for 5 minutes. For plant inoculation, DMSO-primed recovered bacteria were diluted 100 times, while flg22-primed recovered bacteria were diluted 5 times, to achieve equivalent inoculum numbers.

To prime plant defenses with flg22, 20 to 22 hours before bacterial inoculation, leaves were infiltrated with mock 0.005% DMSO or with 500 nM of the PTI inducer flg22.

For *in planta* antibiotic treatments, 400 µg mL^-1^ of one of two β-lactam antibiotics, carbenicillin or cefotaxime, was infiltrated into plants every day until the end of the experiment starting at 1 day post-inoculation with the respective bacterial endophyte. H_2_O was used as a mock control.

### Cloning

The dual fluorescent division reporter pUC18-mini-Tn7T-Gm::*tetR(BD)-P*_*tet*_*-mCitrine_mCerulean3-BCD2-P*_*14g*_ was created in two steps. In the first step, a DNA fragment containing a strong transcriptional terminator, TpheA-1 (Chen *et al*. 2013), repressor *tetR(BD)* [sequence is the fusion of two alleles of *tetR*, carrying 50 codons of *tetR(B)* and the last 158 codons of *tetR(D)* (Kamionka *et al*. 2006)], the tetracycline promoter from *E. coli* Tn10 (which controls expression of both *tetR(BD)* and *mCitrine*; GenBank accession number AF162223), fluorescent protein mCitrine (Griesbeck *et al*. 2001), and two strong transcriptional terminators in opposite orientation, ECK125109870 and L3S2P56 (Chen *et al*. 2013), was synthesized by GenScript®. This DNA fragment was cloned into pUC18-mini-Tn7T-Gm (Choi & Schweizer 2006; cut with NsiI and StuI [New England Biolabs®, Inc.]) using Gibson assembly.

In the second step, two DNA fragments were PCR amplified using primer pairs AVL003 and AVL004, and AVL005 and AVL006 (Supplementary Table S7): one amplicon included the promoter P14g and translational coupler BCD2 from plasmid pBG42 (Zobel et *al*. 2015), while the other amplicon included the gene coding for the fluorescent protein mCerulean3 (Markwardt *et al*. 2011). These fragments were introduced into the previously generated intermediate plasmid (after cutting the plasmid with SpeI [New England Biolabs®, Inc.]) by Gibson assembly to generate the division reporter pUC18-mini-Tn7T-Gm::*tetR(BD)-P*_*tet*_*-mCitrine_mCerulean3-BCD2-P*_*14g*_.

### Gene integration into the *Pseudomonas* genome

Integration into the *Pseudomonas* genome was performed by site-specific transposition by Tn7 into the unique attTn7 site of *Pst* DC3000. For this, triparental mating was set up between the *Pseudomonas* recipient strain, the strain containing the transposase helper plasmid pTNS3 (Choi *et al*. 2008), and the *E. coli* RHO5 donor strain (Kvitko *et al*. 2012; carrying plasmids pBG42_mod or pUC18-mini-Tn7T-Gm with the genes to be integrated). Three to six days after conjugation and selection in plates containing 1.5 µg mL^-1^ of gentamycin, colony PCRs were set up to identify the putative transconjugants using three primer pairs: AVL007 and AVL008, AVL009 and AVL010, and AVL007 and AVL009 (Supplementary Table S7).

### Confocal microscopy

Images were taken using the Nikon A1Rsi confocal microscope with a 20X objective (numerical aperture of 0.7 and a pinhole of 1.2 airy units). Both the CFP and YFP channels are detected using gallium arsenide phosphide photomultiplier (PMT) detectors. For the CFP channel, an excitation of 443.6 nm and an emission between 467 to 502 nm were used. For the YFP channel, an excitation of 513.9 nm and an emission of 530 to 600 nm were used. Images were acquired at a gain and offset at which the negative control (Arabidopsis plants infected with non-fluorescent *Pst* DC3000 or *Pst ΔhrcCΔCFA*) had no signal being detected.

### RNA extraction

Frozen plant tissue was ground using the TissueLyser II (QIAGEN; 2 cycles of 30 seconds at 27 Hz) and 3-mm zirconium oxide beads (Glen Mills Inc.). RNA was extracted using TRIzol™ (Thermo Fisher Scientific) and the Direct-zol RNA miniprep kit (Zymo Research), using a 30-minute on-column treatment with 12 U of DNase I. A second DNase I treatment to remove all residual genomic DNA was performed using 30 U of recombinant DNase I (Roche) and 10 U of Protector RNase inhibitor (Roche) for 1 hour at 37 °C. RNA was purified a second time using TRIzol™ and the Direct-zol RNA miniprep kit.

To prepare logarithmic- and stationary-phase grown *Pst ΔhrcCΔCFA* cultures for RNA extraction, bacteria were grown in liquid KB overnight at 30 °C. For logarithmic-phase samples, overnight bacterial suspensions were diluted with KB to 10^8^ CFU mL^-1^ and grown for 6 hours. Stationary-phase samples were not diluted and grown for 6 additional hours.

To extract RNA from bacterial cultures grown *in vitro*, 10^9^ CFUs of bacteria were resuspended in 150 µL of buffer TE (10 mM Tris-HCl, 1 mM EDTA, pH 8.0) with 5 mg mL^-1^ of lysozyme (Roche) for 5 minutes at room temperature. For the *in vitro* liquid-grown logarithmic- and stationary-phase bacteria, an extra step that used the RNA stabilizer RNAprotect Bacteria Reagent (QIAGEN), was used before the addition of lysozyme. After incubation with lysozyme, 400 µL of buffer RLT (QIAGEN) were added to the suspension, which was then applied through QIA shredder columns (QIAGEN). The eluate was used for RNA purification using the RNeasy mini kit (QIAGEN). The only modification to the protocol of the kit was the extension of the on column DNase I (QIAGEN) digestion to 30 minutes. After RNA purification, a second DNase I treatment and purification using the Direct-zol RNA miniprep kit was performed as described above.

To check for the presence of genomic DNA (gDNA) contamination in the RNA samples, a PCR using 50–100 ng of RNA was performed using primers AVL011 and AVL012 (to check for *Pst* gDNA contamination), AVL013 and AVL014 (to check for *Pandoraea* sp. Col-0-28 gDNA contamination), AVL015 and AVL016 (to check for *Achromobacter xylosoxidans* sp. Col-0-50 gDNA contamination), and AVL017 and AVL018 (to check for *A. thaliana* gDNA contamination) (Supplementary Table S7).

### cDNA preparation and Quantitative Reverse transcriptase-PCR (qRT-PCR)

Three-hundred ng to two µg of RNA were used to prepare cDNA using 2.5 µM of random hexamer primers (Integrated DNA Technologies, Inc.), 40 U of Protector RNase inhibitor and 200 U of M-MLV reverse transcriptase (Invitrogen™), following the protocol described by the manufacturer. Between 0.4 and 1 µL of cDNA was used with appropriate primers (Supplementary Table S7) and SYBR® Green PCR master mix (Applied Biosystems) for qRT-PCR. Two technical replicates were done per sample. PCR was performed on the 7500 Fast Real-Time PCR System (Applied Biosystems(tm)) using automatic C_T_ threshold detection. Bacterial transcripts were normalized to the *leuB* (*PSPTO_2175*) reference gene using the ΔC_T_ method (Livak and Schmittgen 2001). In previous RNA-Seq experiments, *leuB* showed very little variability in expression under multiple conditions. The efficiency of all primer pairs was validated to be between 90 to 110%. Amplicon melting curves had a single predominant peak.

### RNA-Sequencing and bioinformatic analysis

RNA integrity was evaluated using the 2100 BioAnalyzer (Agilent Technologies, Inc.) and RNA concentration was determined with the Qubit™ RNA HS assay kit (Thermo Fisher Scientific). Libraries for RNA-Seq were prepared using the protocol described in Nobori *et al*. (2018), but using the newest iteration of the library preparation kit from NuGEN® (Tecan Genomics, Inc.), the Universal RNA-Seq with NuQuant® kit, and skipping the use of the MICROBEnrich kit. This protocol allows the removal of plant and bacterial rRNAs and of the most abundant Arabidopsis transcripts. Libraries were pooled and sequenced in the Illumina® HiSeq 4000 using HiSeq 4000 SBS reagents to obtain single 50-bp reads. Base calling was done by Illumina Real Time Analysis (RTA version 2.7.7).

The read quality was evaluated with FASTQC (version 0.11.7; Andrews 2010), while adapters and low quality sequences were trimmed with Cutadapt (version 2.9; Martin 2011). Three different pipelines were used to calculate differential gene expression (DGE), as DGE varies depending on the models used for transcript estimation (Trapnell *et al*. 2013). In the first pipeline, reads were aligned to the reference genomes using HISAT2 (version 2.1.0; Kim *et al*. 2019). Aligned reads were processed using SAMTools (version 1.9; Li *et al*. 2009), after which StringTie (version 2.1.3; Kovaka *et al*. 2019) was used to calculate transcript frequency (average expression per treatment is shown on Supplementary Tables S4A, S4B, and S4C), to finally estimate DGE using DESeq2 (Love *et al*. 2014). On the second pipeline, reads were also aligned with HISAT2, but this time DGE was calculated directly using Cuffdiff (version 2.2.1; Trapnell *et al*. 2013). In the third pipeline, reads were pseudo-aligned using Salmon (version 0.11.3 for *Pst ΔhrcCΔCFA* or 1.2.1 for the other two endophytes; Patro *et al*. 2017), after which DESeq2 was performed for DGE.

Principal component analysis (PCA) of transcript reads after DESeq2 analysis used the rlog and plotPCA functions of DESeq2 in Rstudio (version 1.3.1093). Significantly up- or down-regulated genes were identified as those in any of the three pipelines in which the absolute value of the logarithm (in base 2) of the fold change was greater than 0.58, and the adjusted *p* value of the statistical test was lesser than 0.05. GO (gene ontology) analysis used the Cytoscape application BINGO (version 3.0.4; Maere *et al*. 2005); these GO results were summarized with REVIGO (Supek *et al*. 2011), after which manual curation of the enriched pathways was done. PaintOmics 3 (version 0.4.5; Hernández-de-Diego *et al*. 2018) was used for initial visualization of *Pst ΔhrcCΔCFA* gene expression in biological pathways. We used antiSMASH (version 6.0.0; Blin *et al*. 2019) to identify clusters of genes predicted to be involved in the production of secondary metabolites.

STEM (Short-Time series Expression Miner; version 1.3.13; Ernst and Bar-Joseph 2006) analysis was used to look for enriched patterns of expression over time between the samples collected for the inoculum, 6, 24, and 168 hours post-inoculation (hpi) into Col-0 plants. Analysis used the logarithmically (in base 2) normalized expression data obtained from StringTie. GO pathway enrichment analysis used the BINGO application, and summarization of the results was done as described above.

Orthologous gene groups between the three bacterial strains were identified using OrthoMCL (Li *et al*. 2003). Each orthologous group may have more than one gene from each strain. Genes that had the same pattern of differential up- or down-regulation for a comparison between two treatments in one, two or all three strains were identified. We only compared the inoculum or 6 hpi with 168 hpi, as we expected these two comparisons to be the most significant for the long-term adaptation of phyllosphere microbiota to plants. Biological process enrichment analysis of orthologous groups used the *Pst ΔhrcCΔCFA* gene belonging to such group for the analysis when available.

The reference genomes used for analysis are those from *Pst* DC3000: GenBank accession AE016853.1 for the genomic DNA, and GenBank accessions AE016855.1 and AE016854.1 for the two plasmids; for *Achromobacter xylosoxidans* Col-0-50 and *Pandoraea* sp. Col-0-28, the genome sequences are under the BioProject PRJNA555902 in the Sequence Read Archive data, sample runs SRR9732406 and SRR9732397, respectively. Reference databases for GO analysis for *Achromobacter* and *Pandoraea* were created using InterProScan (version 5.47-82.0).

Graphs and statistical analysis were performed using Prism (version 6.0b) or Rstudio (Bubble graphs and Venn diagrams). Upset plots and the heatmaps for Figures 5D and 5E were graphed using TBTools (version 1.0; Chen *et al*. 2020a).

## ACKNOWLEDGEMENTS

We would like to thank Dr. Melinda Frame for her help using the confocal microscope; Cody Keilen and James Klug for their help maintaining the growth chambers; Dr. Kevin Childs, Dr. Jie Wang, Dr. James Kremer, and OE Biotechnology (China) for their bioinformatics expertise; Dr. Brian Kvitko for suggesting using carbenicillin for killing dividing bacteria; the undergraduate students at the He laboratory for preparing laboratory and growth chamber materials; and Dr. Kyaw Aung, Dr. Jonghum Kim, and Dr. Anne Rea for critically reviewing the manuscript.

Funding for this study was provided by the United States National Institute of Health AI155441 grant (to SYH).

## SUPPLEMENTARY INFORMATION

**Supplementary References**

**Supplementary Figure S1. Endophytic bacterial populations experience stasis inside plants**.

**Supplementary Figure S2. Bacterial population stasis after priming disappears over time and is not determined by PAMP-triggered immunity**.

**Supplementary Figure S3. Equilibrium in bacterial multiplication and death causes apparent bacterial population stasis**.

**Supplementary Figure S4. Visualization of pathogenic *Pseudomonas syringae* (*Pst*) carrying the fluorescent division reporter on the surface of Arabidopsis leaves**.

**Supplementary Figure S5. Transcriptome and population analysis of three phyllosphere-inhabiting bacteria**.

**Supplementary Figure S6. Expression of genes from select biological pathways for *Pseudomonas syringae* pv. *tomato ΔhrcCΔCFA***.

**Supplementary Figure S7. Number of shared and unique differentially expressed genes between the three analyzed phyllosphere-inhabiting endophytes**.

**Supplementary Figure S8. *In planta* gene expression for non-pathogenic and effector triggered immunity-inducing bacteria show a similar response**

**Supplementary Table S1. Enriched biological processes of differentially expressed genes (DEG) in bacterial endophytes**.

**Supplementary Table S2. Enriched biological processes in significantly different expression profiles after STEM (Short-Time series Expression Miner) analysis**

**Supplementary Table S3. Secondary metabolite biosynthetic clusters whose expression is differentially regulated inside plants**.

**Supplementary Table S4. Average gene expression in bacterial endophytes**.

**Supplementary Table S5. Bacterial strains used in this study**.

**Supplementary Table S6. Plant genotypes used in this study**.

**Supplementary Table S7. Primer sequences used in this study**.

