## Supplementary Information for "Phyllosphere-inhabiting endophytic bacteria feature a stationary phase-like lifestyle"

**Supplementary Figure S1**

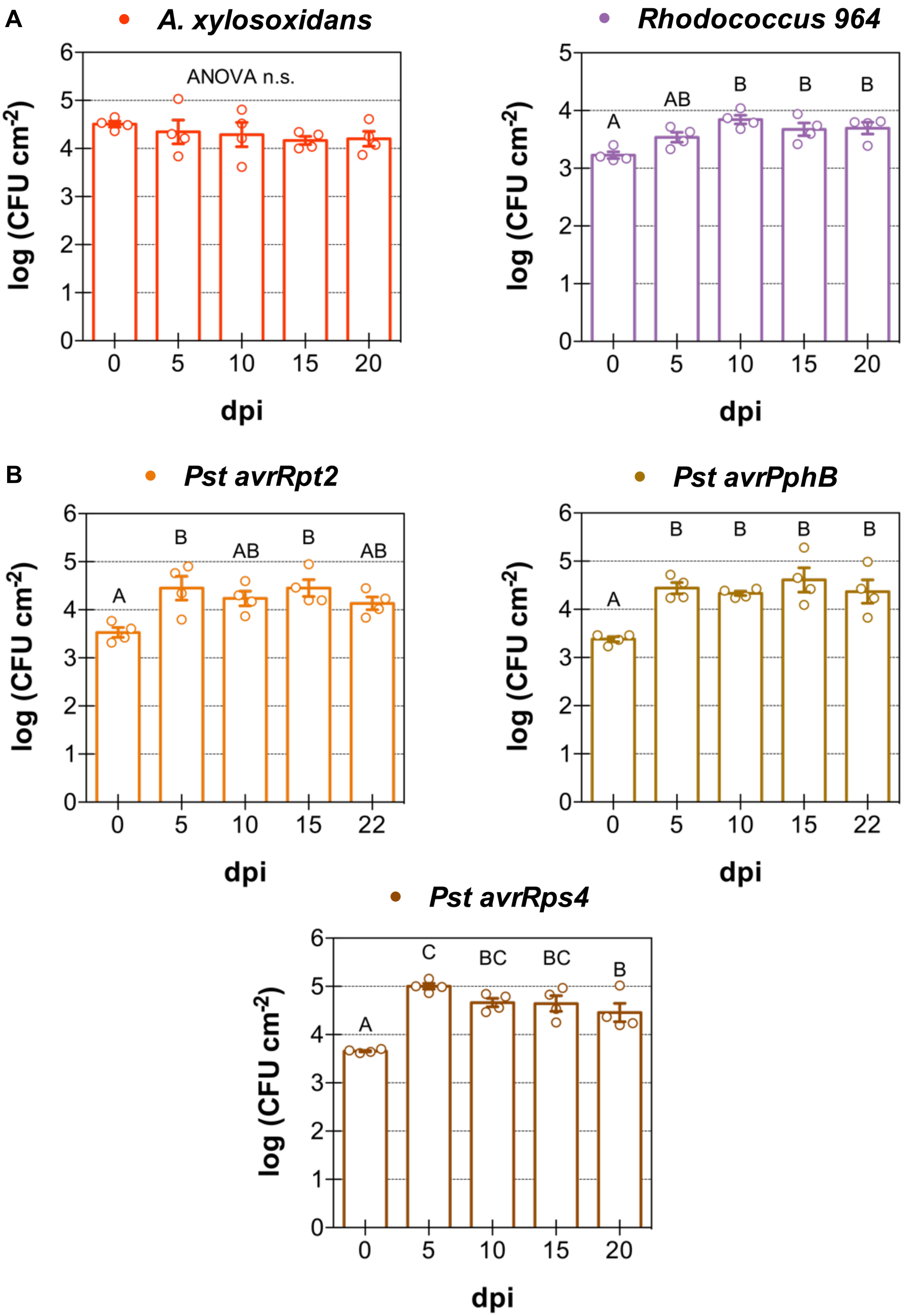

**Supplementary Figure S1. Endophytic bacterial populations experience stasis inside plants. (A)** Bacterial population density of gram-negative and gram-positive microbiota members *Achromobacter xylosoxidans* Col-0-50 ($10^7$ CFU $mL^{-1}$) and *Rhodococcus* sp. 964 ($10^6$ CFU $mL^{-1}$) over the course of 20 days after infection of Col-0 plants. **(B)** Bacterial population density of three effector-triggered immunity-inducing strains ($5 \times 10^5$ – $2 \times 10^6$ CFU $mL^{-1}$) over the course of 3 weeks after infection of Col-0 plants. Effector AvrRpt2 is recognized by RPS2, AvrPphB is recognized by RPS5, while AvrRps4 is recognized by RPS4 in Col-0. The first two R proteins are of the coiled coil class of NLRs, while RPS4 is of the TIR class of NLRs.

Individual biological repetitions for each treatment are shown as open circles. Error bars indicate the standard error of the mean. When appropriate, different letters indicate differences in means, as judged by a Tukey HSD test ($p < 0.05$). dpi indicates days post-inoculation.

**Supplementary Figure S2**

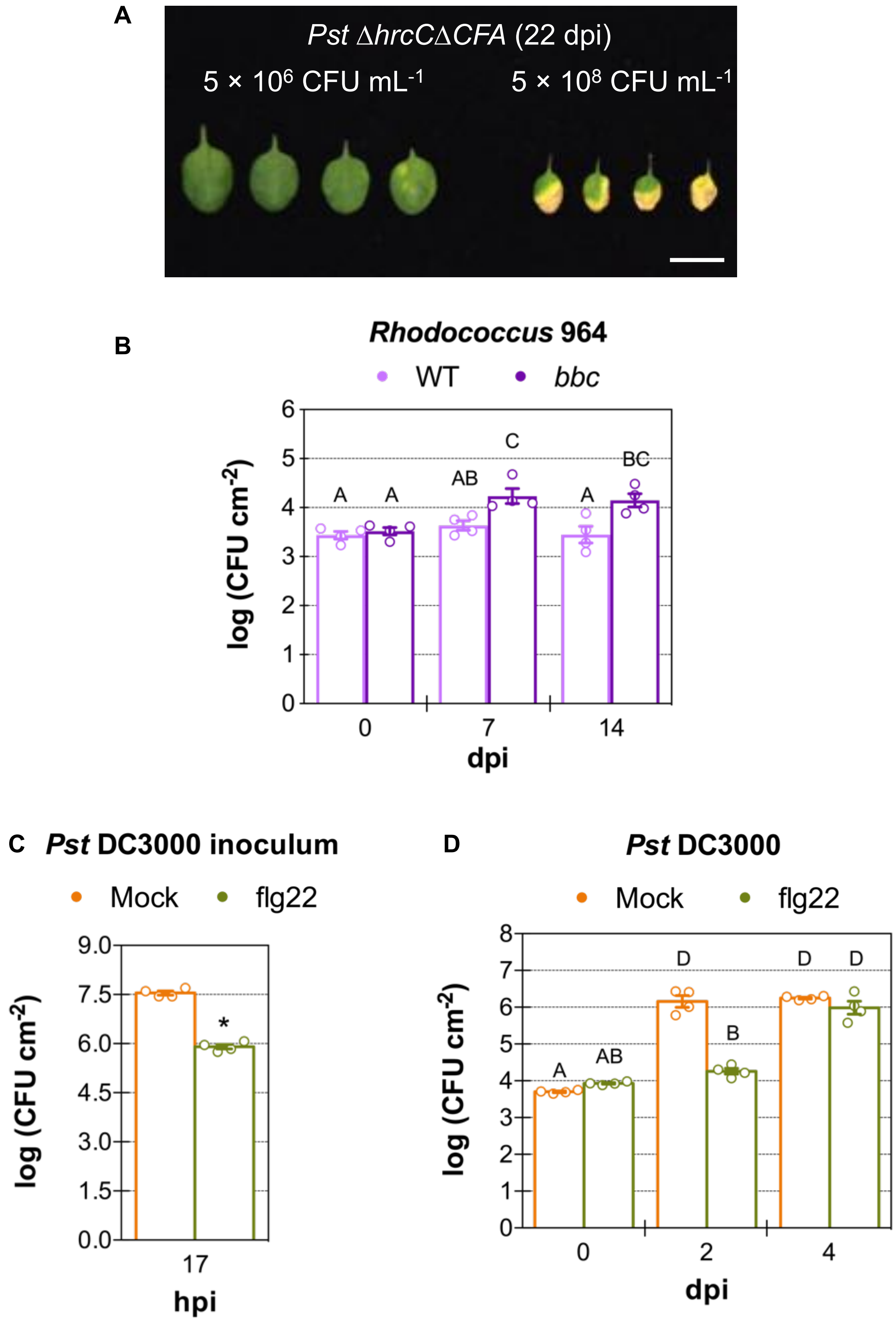

**Supplementary Figure S2. Bacterial population stasis after priming disappears over time and is not determined by PAMP-triggered immunity. (A)** Photographs of Col-0 leaves inoculated with non-pathogenic *Pseudomonas syringae* pv. *tomato* (*Pst*) Δ*hrcC*Δ*CFA* at two different inoculum levels 22 days after infiltration. Notice how the leaves infiltrated at the higher inoculum are chlorotic and have barely grown in the course of 22 days. The white bar represents 1 cm. **(B)** Bacterial population density of *Rhodococcus* sp. 964 ($5 \times 10^5$ CFU $mL^{-1}$) over the course of 2 weeks after infiltration of Col-0 and PTI-compromised triple mutant *bak1 bkk1 cerk1* (*bbc*). **(C)** Bacterial population density of plants from which bacteria were recovered and used to inoculate the plants shown in Figure 2C. Plants were pre-treated for 22 hours with 0.005% DMSO or 500 nM flg22, after which leaves were inoculated with *Pst* DC3000 ($10^9$ CFU $mL^{-1}$). Samples were collected 17 hpi with *Pst* DC3000. **(D)** Bacterial population density of plants inoculated with *Pst* DC3000 ($10^6$ CFU $mL^{-1}$) in which PTI had been primed previously for 21 hours by infiltrating plants with 0.005% DMSO or 500 nM flg22. Four days after inoculation, the protective effect of PTI disappears and bacterial densities are equivalent in both DMSO- and flg22-treated leaves.

Individual biological repetitions for each treatment are shown as open circles. Error bars indicate the standard error of the mean. When appropriate, different letters indicate differences in means, as judged by a Tukey HSD test ($p < 0.05$). An asterisk indicates a significant Student's *t*-test ($p < 0.05$). dpi indicates days post-inoculation, while hpi indicates hours post-inoculation.

Supplementary Figure S3

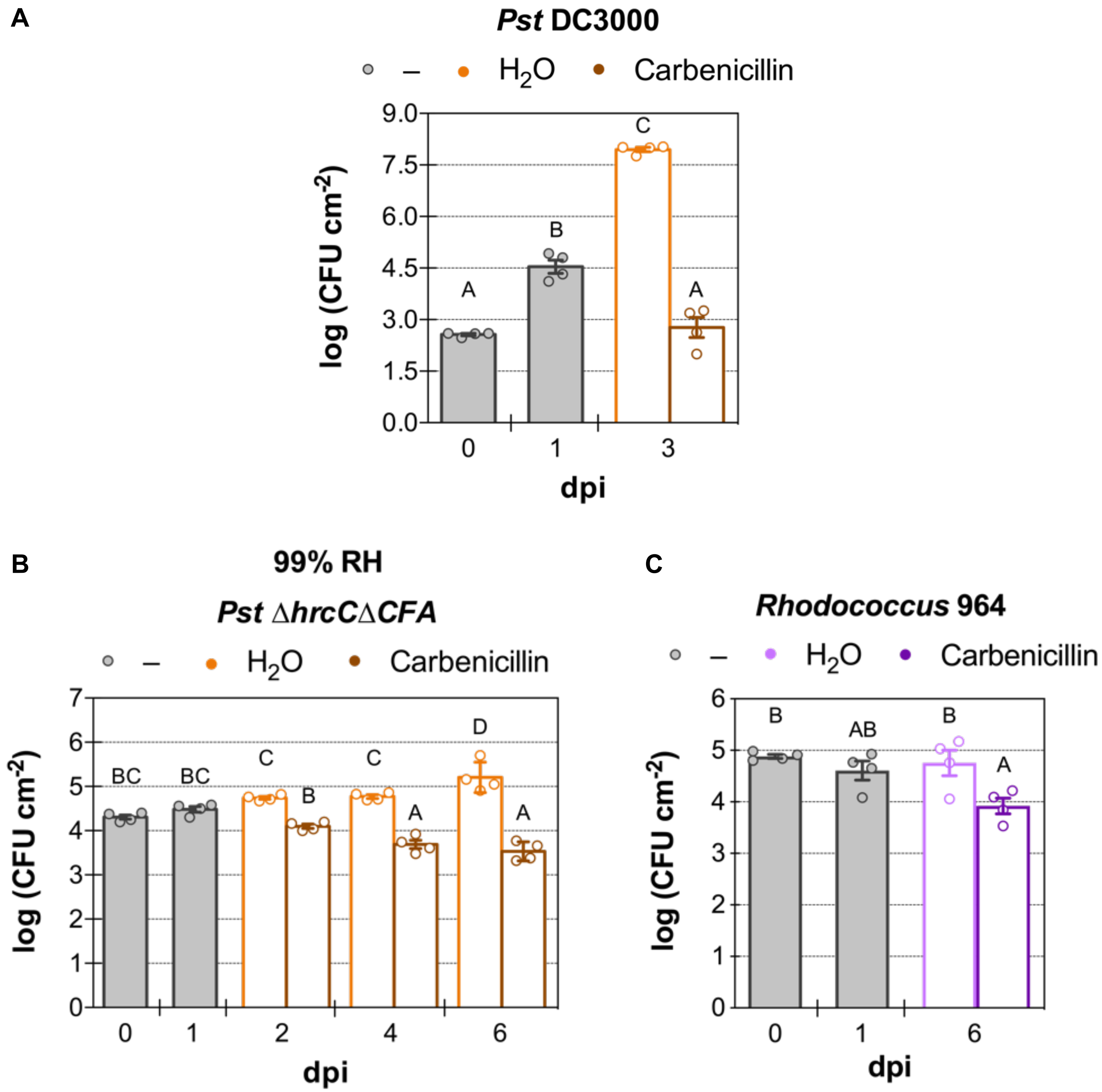

**Supplementary Figure S3. Equilibrium in bacterial multiplication and death causes apparent bacterial population stasis (A)** The antibiotic carbenicillin is effective at killing dividing bacteria in plants. Bacterial population density of virulent *Pseudomonas syringae* pv. *tomato* (*Pst*) DC3000 ($10^5$ CFU $mL^{-1}$) after infiltration with 400 µg $mL^{-1}$ of carbenicillin, an antibiotic that only kills dividing bacteria. Notice the dramatic difference in bacterial populations between the $H_2O$ control and carbenicillin, with carbenicillin having 0.001% of the population of the water control. **(B)** Same experiment as in Figure 3B, but, using a relative humidity of over 99% throughout the experiment. *Pst ∆hrcC∆CFA* was inoculated at $10^7$ CFU $mL^{-1}$, and 400 µg $mL^{-1}$ of carbenicillin was infiltrated daily starting from day 1 post-inoculation. Over 98% of the population is killed over the course of 5 days. **(C)** *In planta* population density of *Rhodococcus* sp. 964 ($2 \times 10^7$ CFU $mL^{-1}$) after addition of 400 µg $mL^{-1}$ of carbenicillin. Over 87% of the population is killed after 5 days.

Individual biological repetitions for each treatment are shown as open circles. Error bars indicate the standard error of the mean. Different letters indicate differences in means, as judged by a Tukey HSD test ($p < 0.05$). dpi indicates days post-inoculation.

**Supplementary Figure S4**

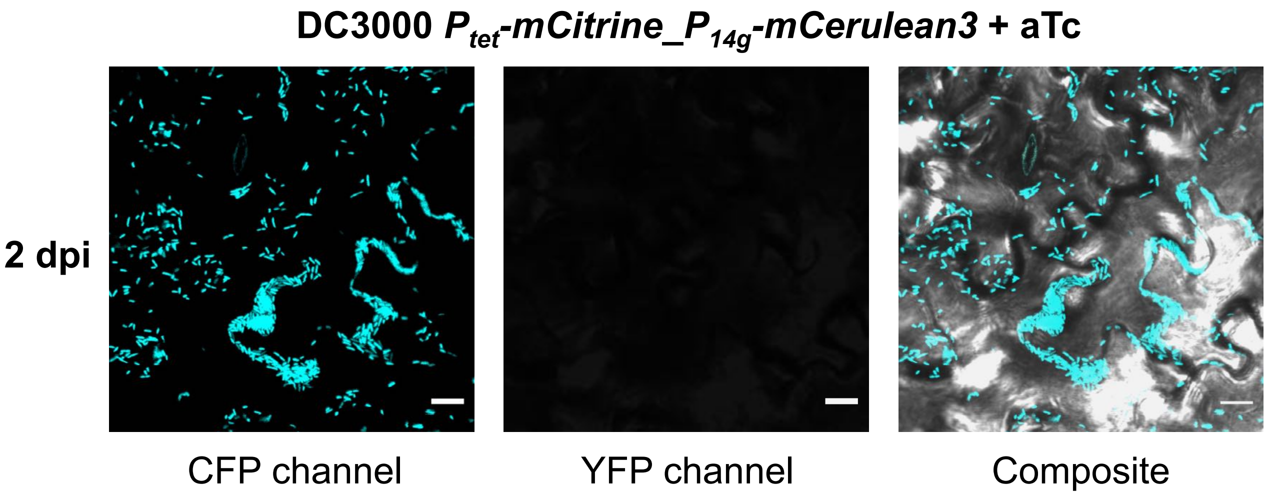

**Supplementary Figure S4. Visualization of pathogenic *Pseudomonas syringae* (*Pst*) carrying the fluorescent division reporter on the surface of Arabidopsis leaves.** Confocal images of dual-labeled virulent *Pst* DC3000 ($5 \times 10^7$ CFU $mL^{-1}$; inoculum grown with 80 ng $mL^{-1}$ anhydrotetracycline [aTc]) 2 days after inoculation of Col-0 plants. Images were taken at the leaf surface, where individual fluorescent bacteria can be observed. No inducible mCitrine signal is observed in the samples, as it was diluted during *in planta* bacterial multiplication.

Cyan represents signal from mCerulean3. Yellow represents signal from mCitrine. Composite image of both fluorescent proteins and the transmitted light image is shown. White bars represent 10 µm.

### Supplementary Figure S5

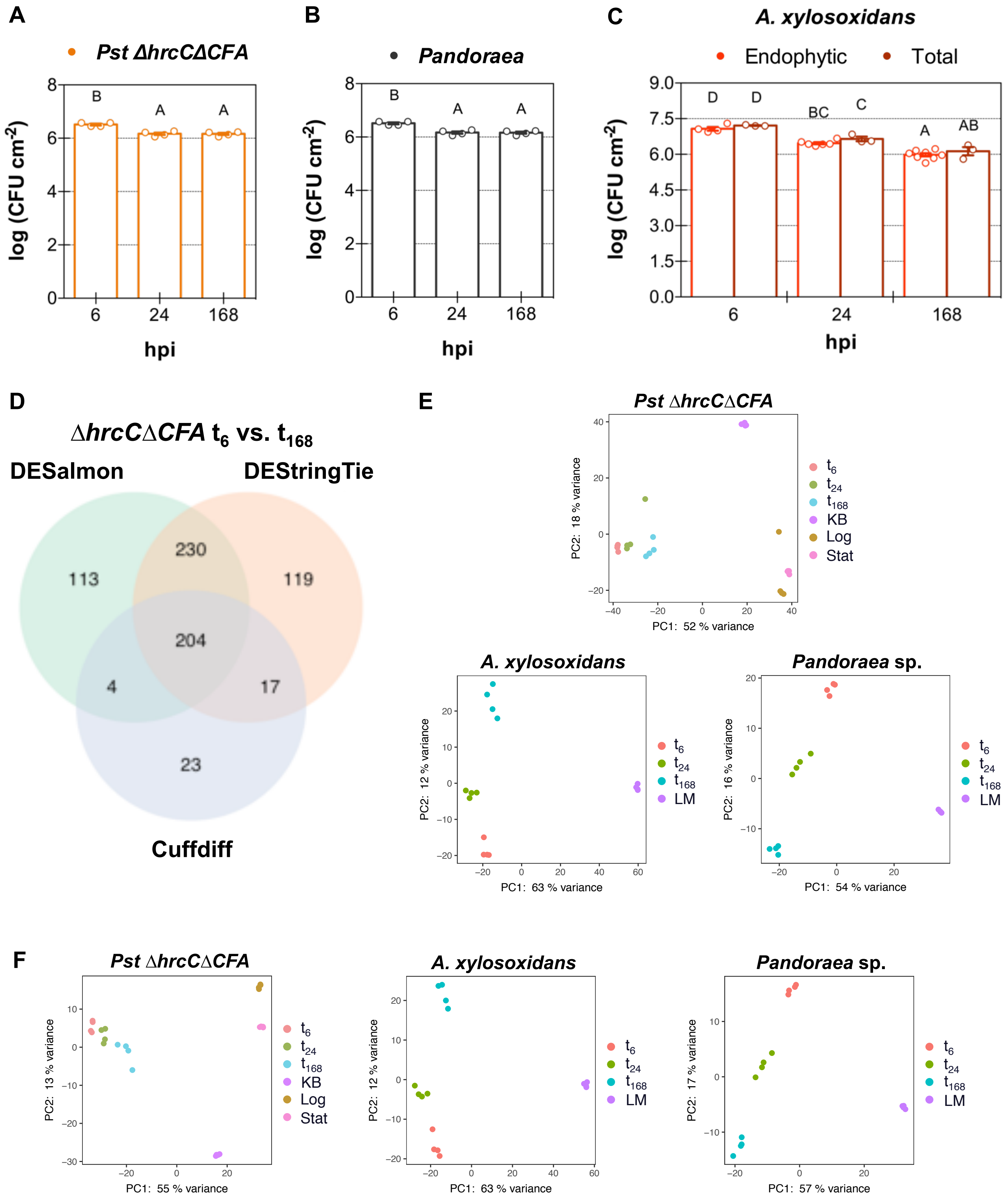

**Supplementary Figure S5. Transcriptome and population analysis of three phyllosphere-inhabiting bacteria. (A)** Bacterial population density of non-pathogenic *Pseudomonas syringae* pv. *tomato* (*Pst*) Δ*hrcC*Δ*CFA* ($2 \times 10^9$ CFU $mL^{-1}$) over the course of 7 days in Col-0 plants. These samples were used for the bacterial RNA-Seq analysis presented in Figure 5A. **(B)** Bacterial population density of bacterial microbiota *Pandoraea* sp. Col-0-28 ($2 \times 10^9$ CFU $mL^{-1}$) over the course of 7 days in Col-0 plants. These samples were used for the bacterial RNA-Seq analysis presented in Figure 5B. **(C)** Total and endophytic bacterial population density of *Achromobacter xylosoxidans* Col-0-50 ($2 \times 10^9$ CFU $mL^{-1}$) over the course of 7 days in Col-0 plants. Endophytic samples were used for the bacterial RNA-Seq analysis presented in Figure 5C. Leaves for endophytic samples were surface sterilized with sodium hypochlorite. **(D)** Venn diagram showing the number of differentially expressed genes between 6 ($t_6$) and 168 ($t_{168}$) hpi with *Pst* Δ*hrcC*Δ*CFA* into Arabidopsis leaves after using one of three methods: DESeq2 after pseudo-alignment with Salmon (DESalmon), DESeq2 after alignment with HISAT2 and analysis with StringTie (DEStringTie), and Cuffdiff after alignment with HISAT2 (Cuffdiff). **(E)** Principal component analysis (PCA) of transcript reads pseudo-aligned with Salmon, and then analyzed with DESeq2 with the rlog and plotPCA functions in R. There were three experiments: Col-0 plants inoculated with *Pst* Δ*hrcC*Δ*CFA*, *Achromobacter xylosoxidans* Col-0-50, or *Pandoraea* sp. Col-0-28 at 6 ($t_6$), 24 ($t_{24}$), and 168 ($t_{168}$, 7 days) hpi, and their respective inoculi recovered from King's (KB) or modified LB (LM) medium. *In vitro* logarithmic (Log) or stationary (Stat) phase treatments were also included for *Pst* Δ*hrcC*Δ*CFA*. **(F)** PCA of transcript reads aligned with HISAT2, and then analyzed with StringTie and DESeq2. Same three experiments as in Supplementary Figure 5E were analyzed.

Individual biological repetitions for each treatment are shown as filled circles. Error bars indicate the standard error of the mean. When appropriate, different letters indicate differences in means, as judged by a Tukey HSD test ($p < 0.05$). hpi indicates hours post-inoculation. dpi indicates days post-inoculation.

**Supplementary Figure S6**

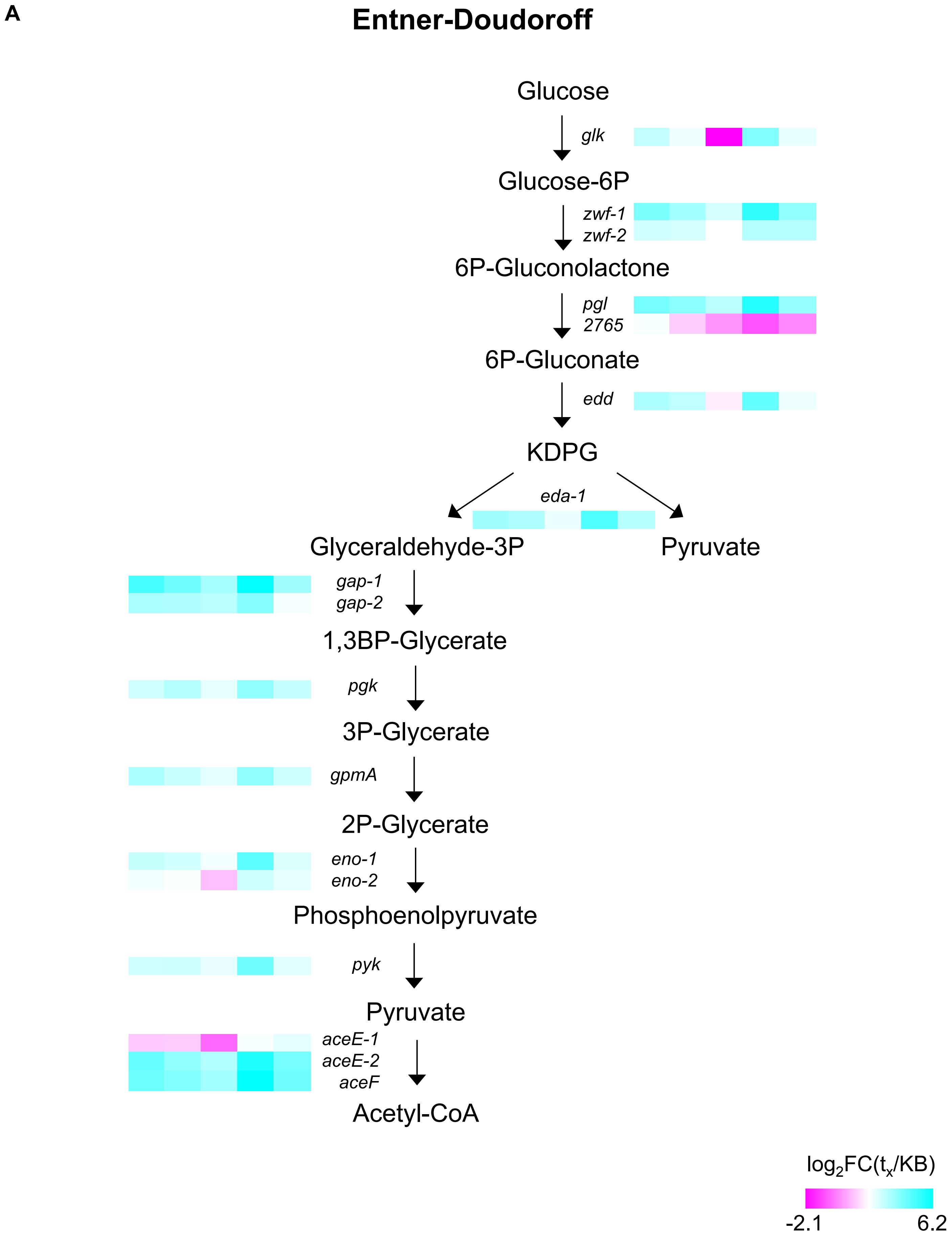

**Supplementary Figure S6**

**B**

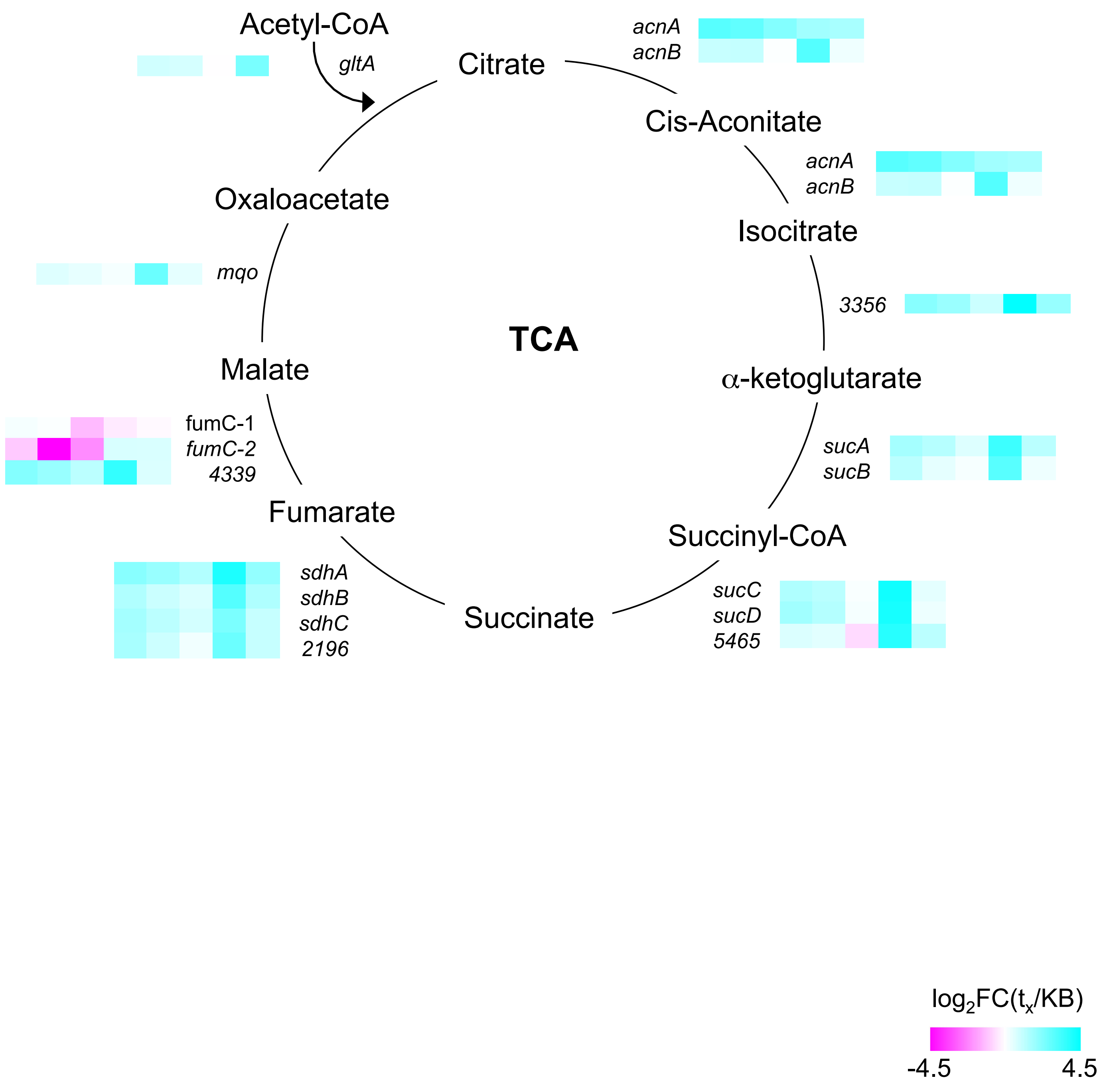

**Supplementary Figure S6**

**C**

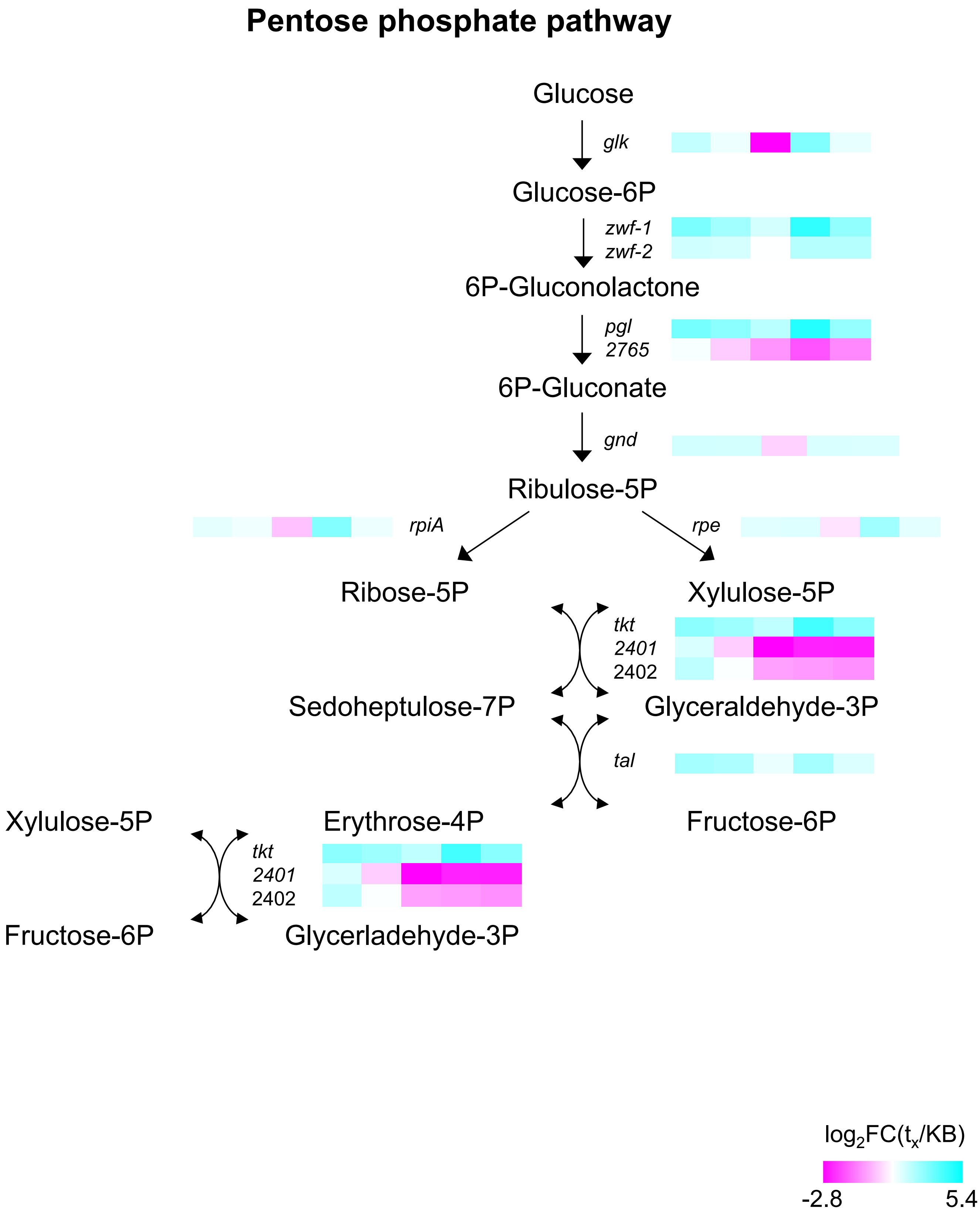

**Supplementary Figure S6**

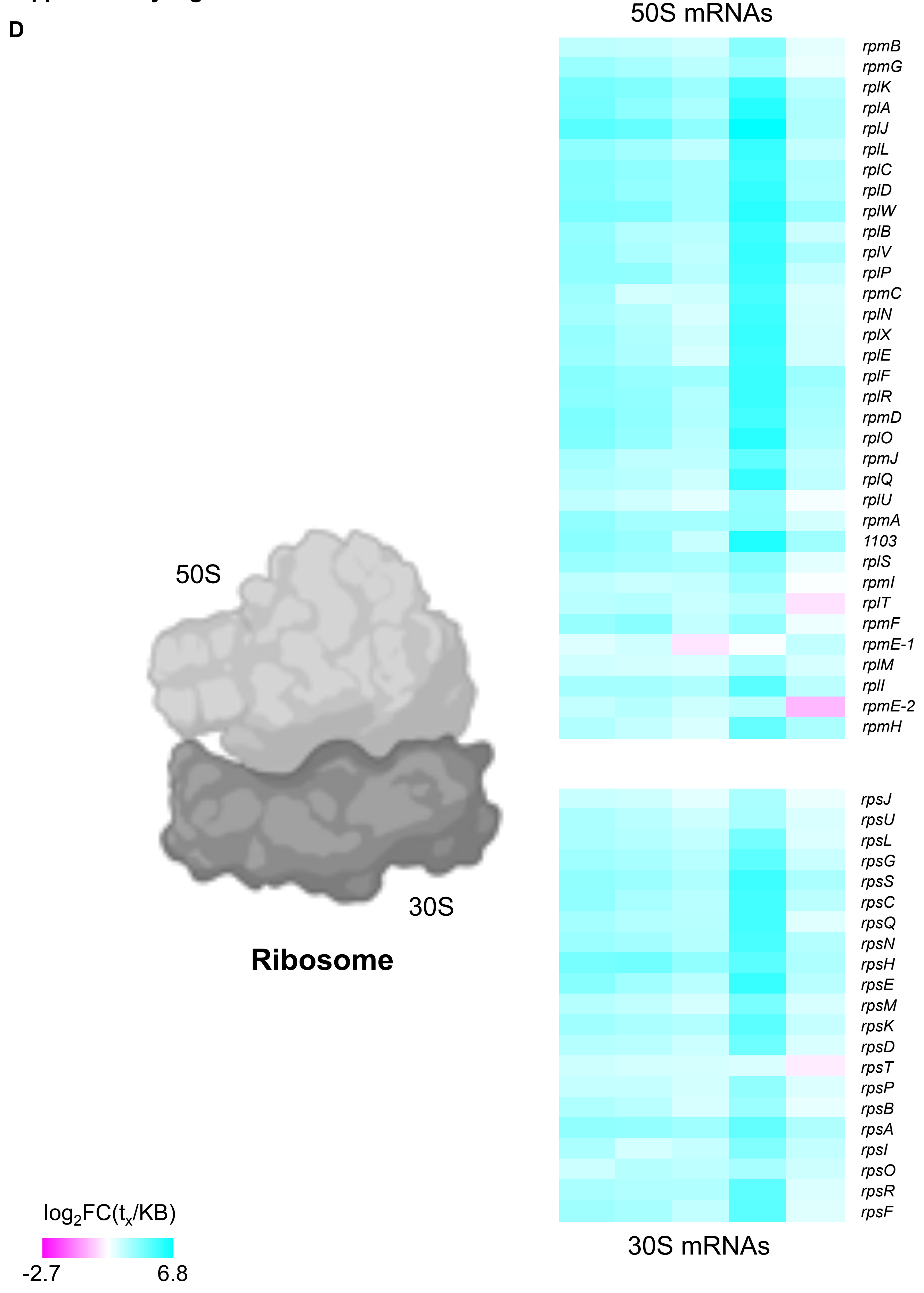

Supplementary Figure S6

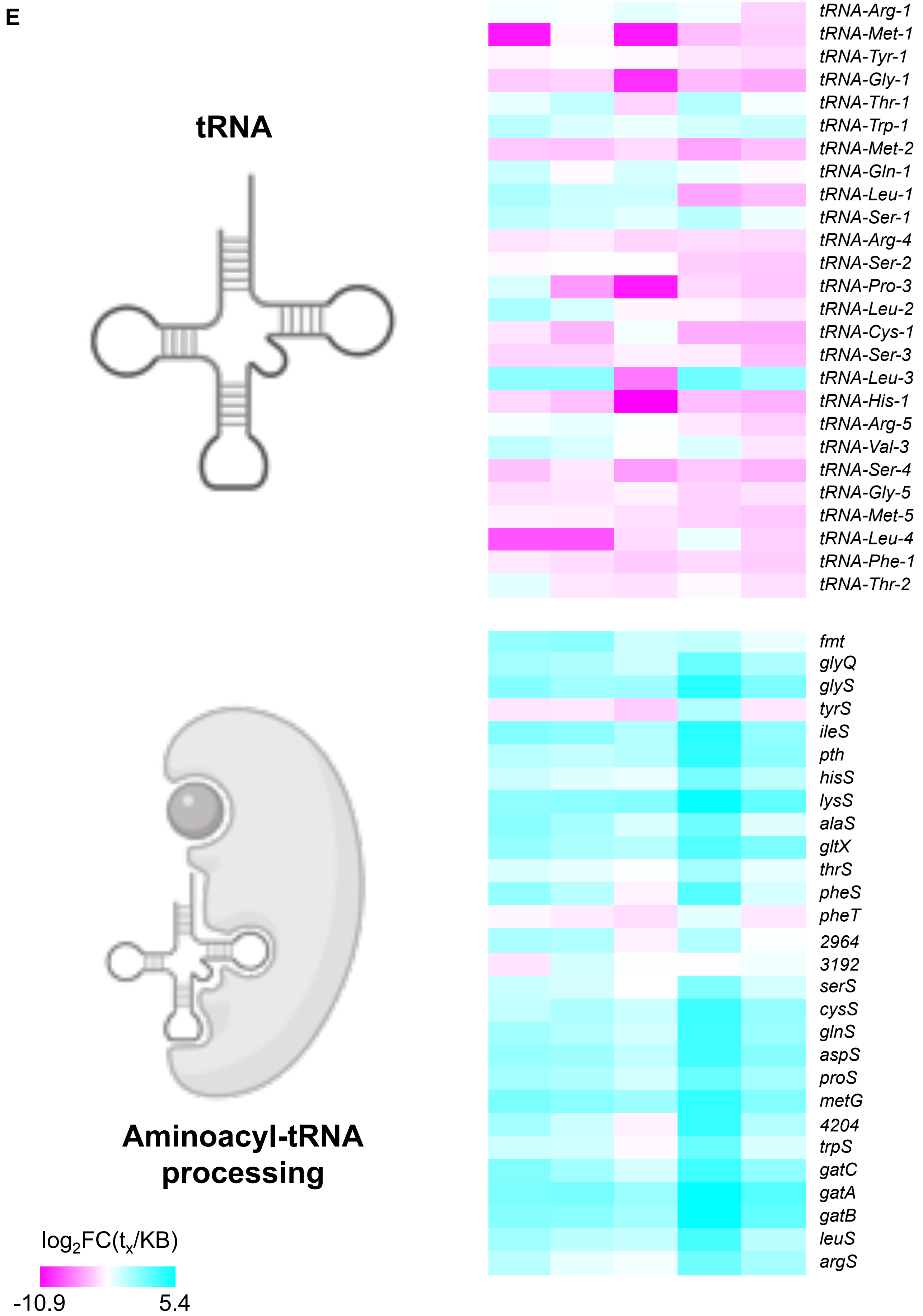

**Supplementary Figure S6**

**F**

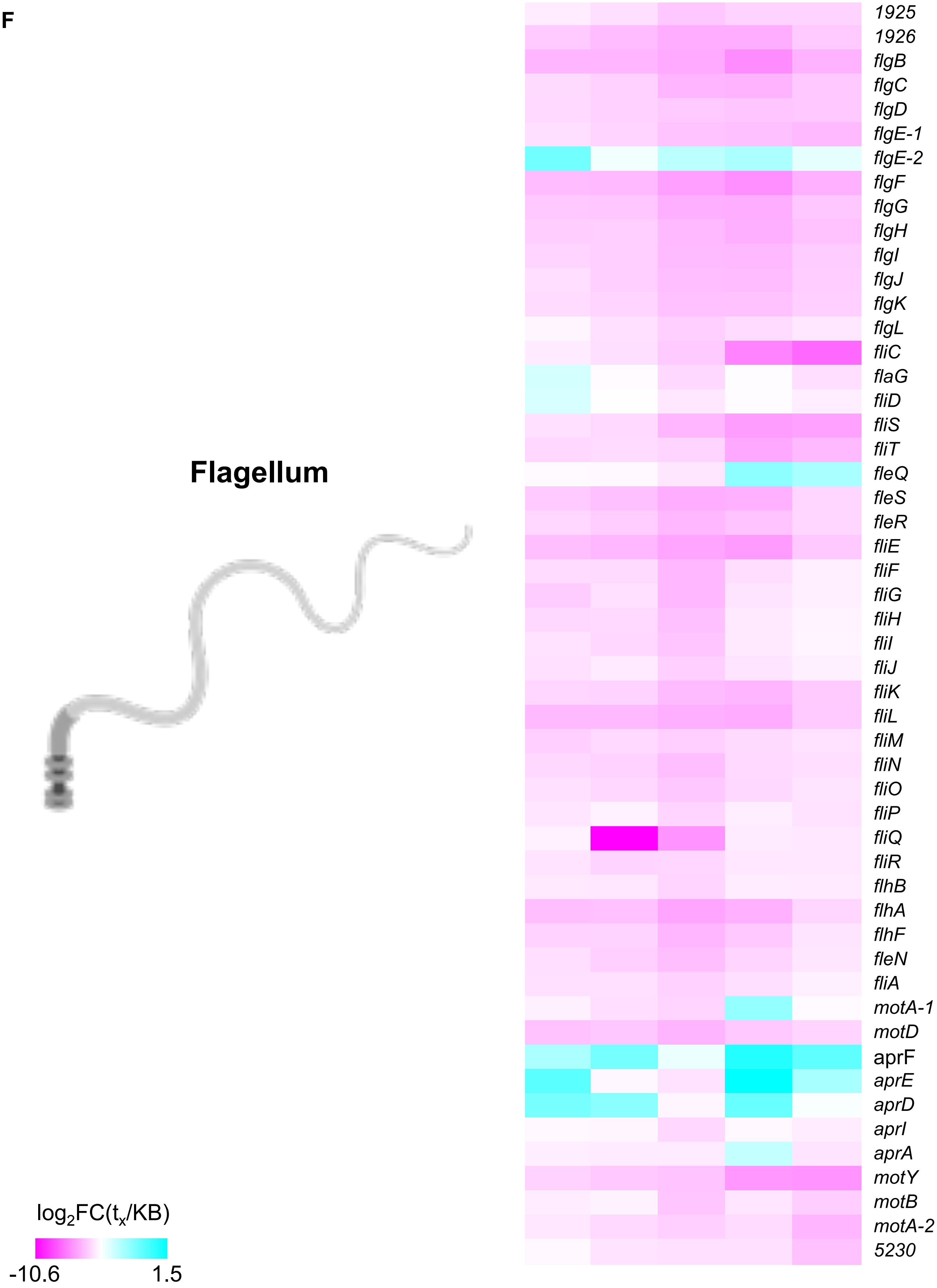

Supplementary Figure S6

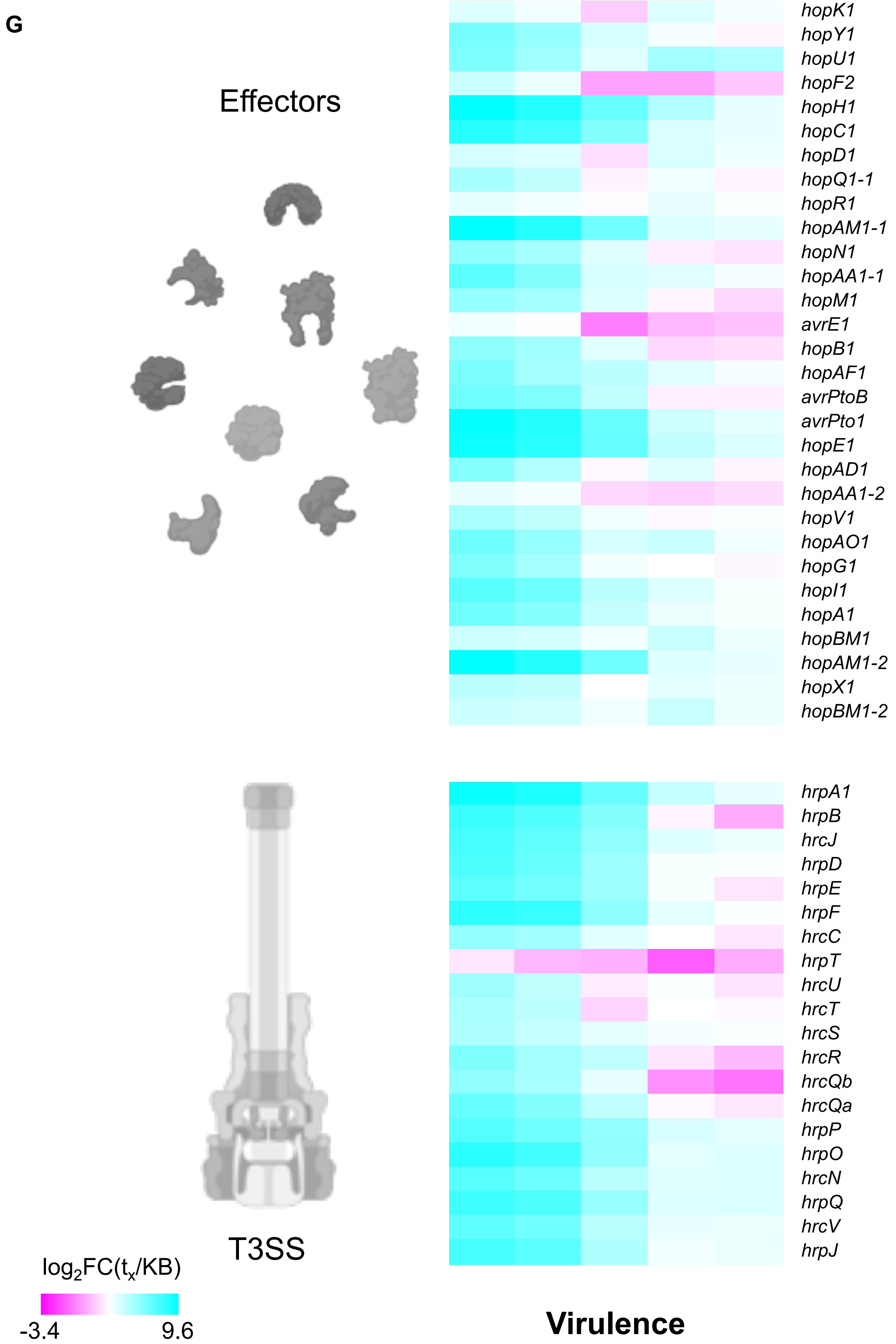

**Supplementary Figure S6**

**H**

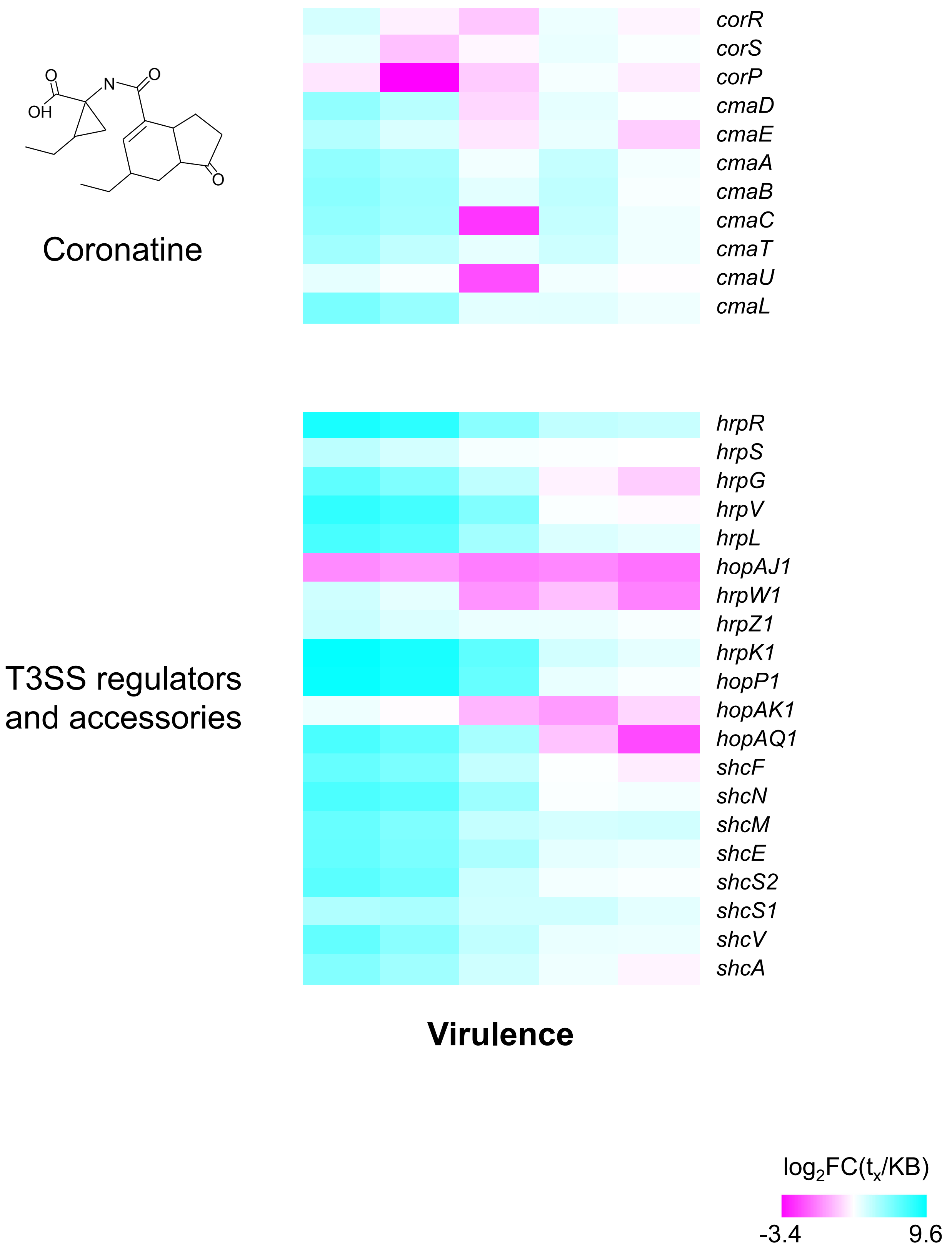

**Supplementary Figure S6. Expression of genes from select biological pathways for *Pseudomonas syringae pv. tomato* ∆*hrcC*∆*CFA*.** Pathways represented were **(A)** the Entner-Doudoroff pathway (with the addition in the pathway of the conversion of pyruvate to acetyl-CoA), **(B)** the Tricarboxylic acid cycle (TCA), **(C)** the Pentose Phosphate pathway, **(D)** Ribosome biogenesis genes, **(E)** tRNAs and genes involved in tRNA aminoacylation, **(F)** Flagellar biosynthesis genes, and **(G)** and **(H)** Virulence-promoting genes: structural and regulatory components of the Type III secretion system (T3SS), effectors, and coronatine biosynthesis genes.

Expression of each gene was calculated as the average $\log_2$ fold change (FC) of the ratio between the treatment and the inoculum. Expression of each gene is represented by five rectangles corresponding to each treatment in the following order: $t_6$, $t_{24}$, $t_{168}$, Log, and Stat. FC was color coded for repression (magenta), no change (white), and induction (cyan).

### Supplementary Figure S7

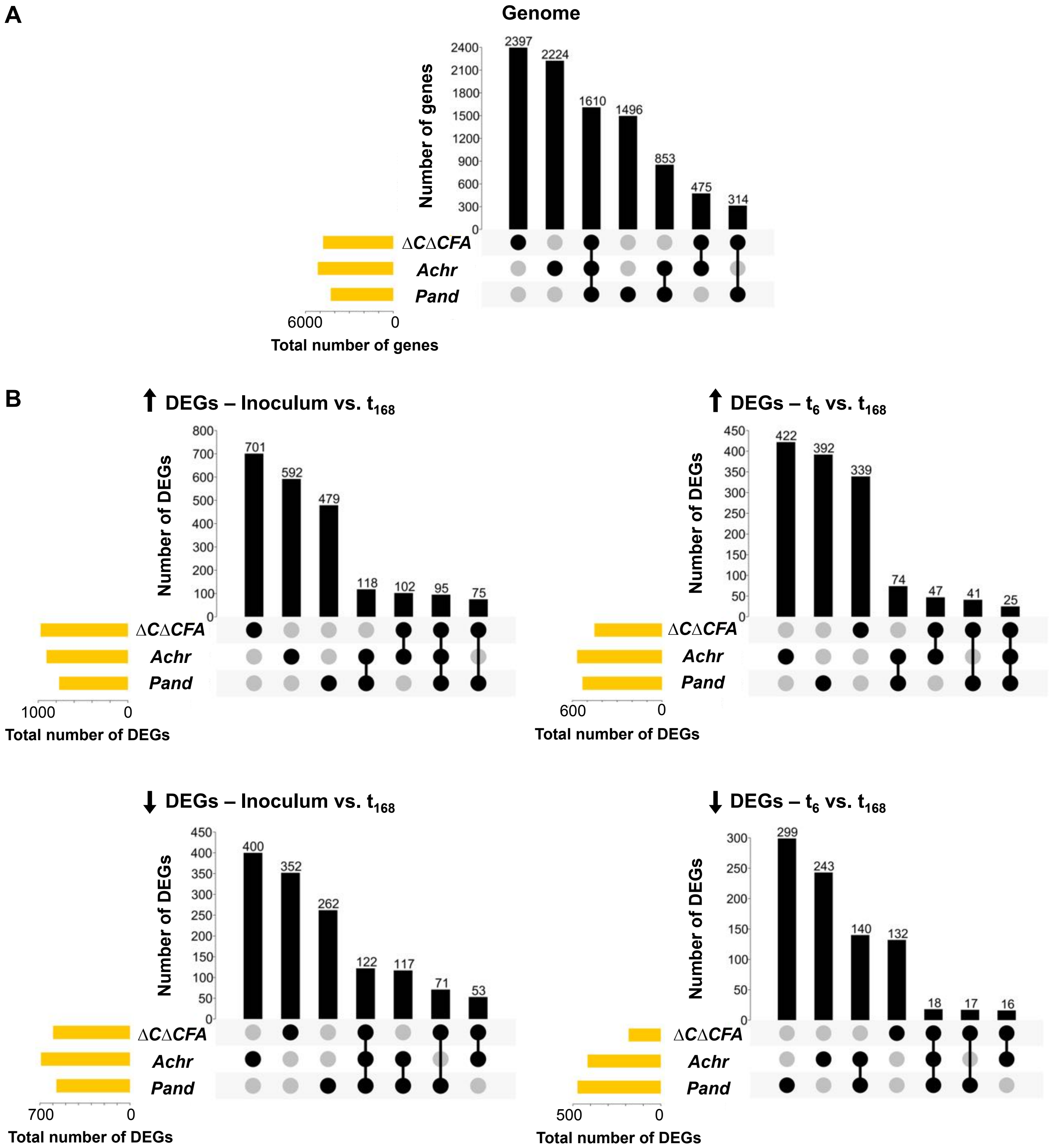

**Supplementary Figure S7. Number of shared and unique differentially expressed genes between the three analyzed phyllosphere-inhabiting endophytes. (A)** Upset graph showing the number of unique and shared orthologous gene groups in the genomes of *Pst* ∆*hrcC*∆*CFA* (∆*C*∆*CFA*), *Achromobacter xylosoxidans* Col-0-50 (*Achr*), and *Pandoraea* sp. Col-0-28 (*Pand*). The core genome encompasses 1,610 gene groups shared by all three bacterial strains. **(B)** Upset graphs showing the number of unique and shared orthologous up-regulated and down-regulated differentially expressed gene (DEG) groups in *Pst* ∆*hrcC*∆*CFA*, *A. xylosoxidans* Col-0-50, and *Pandoraea* sp. Col-0-28 in the inoculum or 6 hpi ($t_6$) of Col-0 plants, when compared to 168 hpi ($t_{168}$).

### Supplementary Figure S8

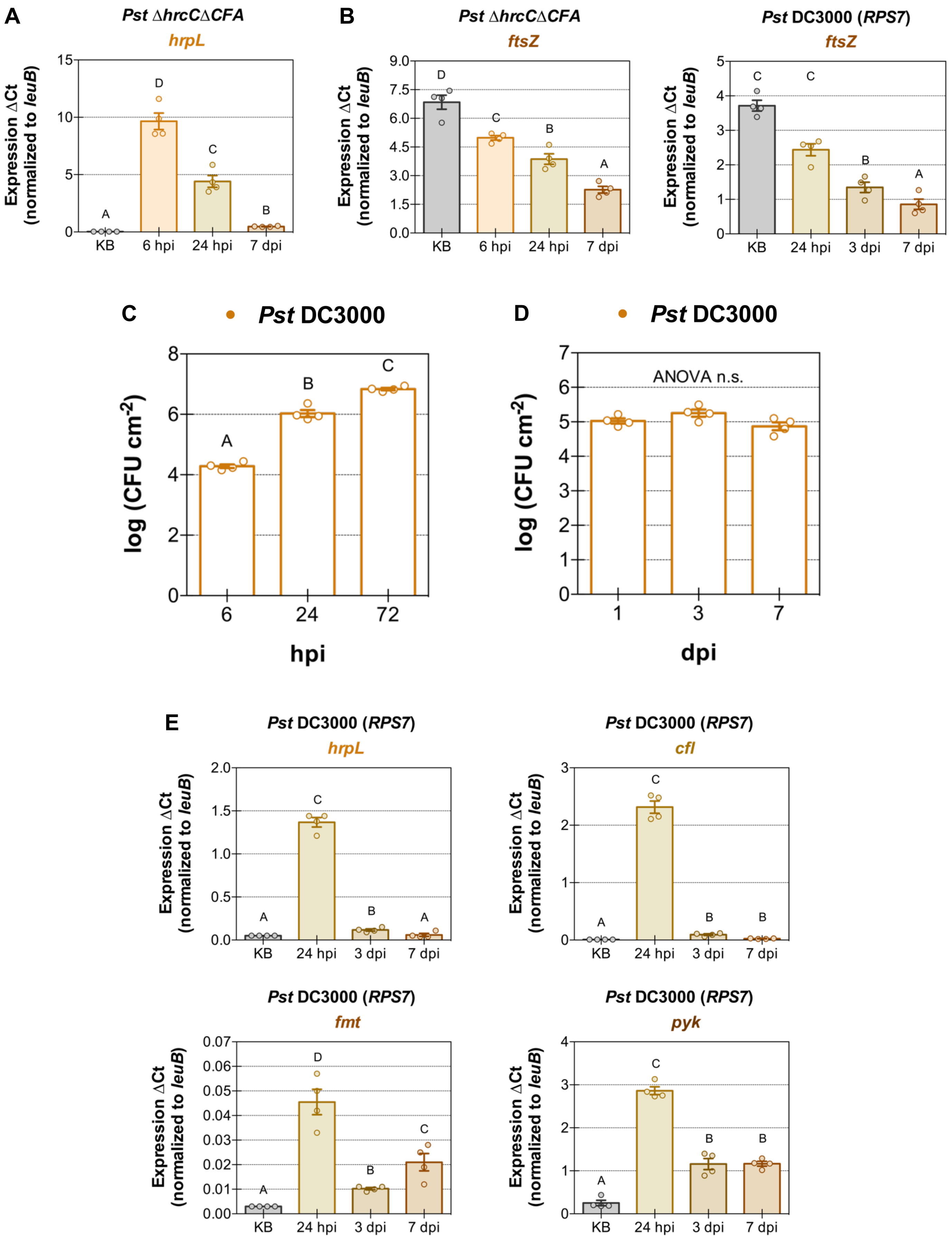

**Supplementary Figure S8. *In planta* gene expression for non-pathogenic and effector triggered immunity-inducing bacteria show a similar response. (A)** *In planta* gene expression of the RNA polymerase sigma factor controlling virulence in *Pseudomonas syringae* (*hrpL*) after inoculation of *Pst* Δ*hrcC*Δ*CFA* into Col-0 plants. **(B)** An experimental repeat of *ftsZ* expression after *Pst* Δ*hrcC*Δ*CFA* inoculation into Col-0 plants or *Pst* DC3000 inoculation into Bu-22 plants. Related to Figures 7A and 7B. **(C)** Bacterial population density after high inoculum infiltration of *Pst* DC3000 ($10^7$ CFU $mL^{-1}$) into Arabidopsis Bu-22 plants. Notice how the containment of bacterial growth is lost at high inoculum. **(D)** Bacterial population density after low inoculum infiltration of *Pst* DC3000 ($4 \times 10^5$ CFU $mL^{-1}$) into Arabidopsis Bu-22 plants. The same plants were also used to collect samples for qPCR. **(E)** *In planta* gene expression of the virulence master regulator *hrpL*, a gene involved in the biosynthesis of the phytotoxin coronatine (coronafacate ligase, *cfl*), a gene involved in mRNA translation ($tRNA^{fMet}$-formyl transferase, *fmt*), and a gene involved in the final step of the Entner-Doudoroff pathway (pyruvate kinase, *pyk*) after inoculation of *Pst* DC3000 into Bu-22 plants.

Individual biological repetitions for each treatment are shown as filled circles. Error bars indicate the standard error of the mean. When appropriate, different letters indicate differences in means, as judged by a Tukey HSD test ($p < 0.05$). hpi indicates hours post-inoculation. dpi indicates days post-inoculation.

**Supplementary Table S1. Enriched biological processes of differentially expressed genes (DEG) in bacterial endophytes. (A)** Up-regulated processes in *Pseudomonas syringae* pv. *tomato* (Pst) Δ*hrcC*Δ*CFA*. Comparisons were done between the inoculum, 6 ($t_6$), 24 ($t_{24}$) and 168 ($t_{168}$) hours post-inoculation, and *in vitro* populations grown to logarithmic (Log) or stationary (Stat) phase. An asterisk denotes that a biological process had less than 5 DEGs.

| **Up-regulated processes in *Pst* Δ*hrcC*Δ*CFA*** | | | | | | | |
|---|---|---|---|---|---|---|---|
| **$t_6$ vs. $t_{24}$** | **$t_6$ vs. $t_{168}$** | **$t_6$ vs. KB** | **$t_6$ vs. Stat** | **$t_{24}$ vs. $t_{168}$** | **$t_{24}$ vs. KB** | **$t_{24}$ vs. Stat** | **$t_{168}$ vs. KB** |
| Virulence<br>Deoxyribonucleotide metabolism* | Virulence<br>Transcriptional regulation<br>Transport across membranes and ATP<br>Response to chemical stimulus | Virulence<br>Primary metabolism<br>Nucleotide metabolism<br>Protein translation<br>Ribosome biogenesis<br>tRNA aminoacylation<br>Amino acid biosynthesis<br>Carbohydrate metabolism<br>Entner-Doudoroff and Pentose phosphate pathways<br>Tricarboxylic acid cycle<br>ATP biosynthesis<br>Lipid and LPS biosynthesis<br>Peptidoglycan biosynthesis<br>Alginate biosynthesis<br>Transport across membranes<br>Sulfur metabolism<br>Response to oxidative stress | Virulence<br>Protein translation<br>Valine biosynthesis*<br>Carbohydrate metabolism<br>Alginate biosynthesis<br>Transport across membranes<br>Sulfate transport*<br>Bacterioferritin*<br>Response to chemical stimulus<br>Response to oxidative stress | Virulence<br>Regulation of transcription<br>Transport across membranes | Virulence<br>Primary metabolism<br>Nucleotide metabolism<br>Transcription<br>Protein translation<br>Ribosome biogenesis<br>tRNA aminoacylation<br>Amino acid biosynthesis<br>Carbohydrate metabolism<br>Entner-Doudoroff and Pentose phosphate pathways<br>Acetyl-CoA metabolism and Tricarboxylic acid cycle<br>ATP biosynthesis<br>Alginate biosynthesis<br>Transport across membranes<br>Response to oxidative stress | Virulence<br>Protein translation<br>Valine biosynthesis*<br>Carbohydrate metabolism<br>Alginate biosynthesis<br>Transport across membranes<br>Phosphate transport<br>Response to chemical stimulus<br>Response to oxidative stress<br>Bacterioferritin*<br>3,4-dihydroxybenzoate metabolism* | Virulence<br>Primary metabolism<br>Nucleotide metabolism<br>Transcription<br>Protein translation<br>Ribosome biogenesis<br>tRNA aminoacylation<br>Amino acid metabolism<br>Carbohydrate metabolism<br>Entner-Doudoroff and Pentose phosphate pathways<br>Tricarboxylic acid cycle<br>Alginate biosynthesis<br>Transport across membranes and ATP<br>Response to oxidative stress |

| **Up-regulated processes in *Pst* Δ*hrcC*Δ*CFA*** | | | | | | |
|---|---|---|---|---|---|---|
| **$t_{168}$ vs. Stat** | **KB vs. Stat** | **Log vs. $t_6$** | **Log vs. $t_{24}$** | **Log vs. $t_{168}$** | **Log vs. KB** | **Log vs. Stat** |
| Virulence<br>Protein translation<br>Valine biosynthesis<br>Carbohydrate metabolism<br>Alginate biosynthesis<br>Transport across membranes<br>Sulfate transport<br>Bacterioferritin*<br>3,4-dihydroxybenzoate metabolism* | Locomotion and flagellum | Primary metabolism<br>Nucleotide metabolism<br>DNA replication<br>DNA repair<br>Regulation of cell shape*<br>Protein translation<br>Ribosome biogenesis<br>tRNA aminoacylation<br>Protein folding*<br>Amino acid biosynthesis<br>Carbohydrate metabolism<br>Entner-Doudoroff and Pentose phosphate pathways<br>Tricarboxylic acid cycle<br>ATP biosynthesis<br>Fermentation<br>Lipid biosynthesis<br>Peptidoglycan biosynthesis<br>Rhamnose biosynthesis<br>Vitamin biosynthesis<br>Cobalamin biosynthesis<br>Siderophore biosynthesis | Primary metabolism<br>Nucleotide metabolism<br>DNA replication<br>DNA repair<br>Protein translation<br>Ribosome biogenesis<br>tRNA aminoacylation<br>Amino acid biosynthesis<br>Carbohydrate metabolism<br>Entner-Doudoroff and Pentose phosphate pathways<br>Tricarboxylic acid cycle<br>ATP biosynthesis<br>Lipid and LPS biosynthesis<br>Peptidoglycan biosynthesis<br>Vitamin biosynthesis<br>Riboflavin biosynthesis | Primary metabolism<br>Nucleotide metabolism<br>DNA replication<br>Response to DNA damage<br>Protein translation<br>Ribosome biogenesis<br>tRNA aminoacylation<br>Proteolysis<br>Amino acid biosynthesis<br>Carbohydrate metabolism<br>Entner-Doudoroff and Pentose phosphate pathways<br>Tricarboxylic acid cycle<br>ATP biosynthesis<br>Carbohydrate transport<br>Peptidoglycan biosynthesis<br>Rhamnose biosynthesis<br>Lipid biosynthesis<br>Vitamin biosynthesis | Primary metabolism<br>Nucleotide metabolism<br>DNA replication<br>DNA repair<br>Cytokinesis<br>Regulation of cell shape*<br>Transcription<br>Protein translation<br>Ribosome biogenesis<br>tRNA aminoacylation<br>Proteolysis<br>Amino acid biosynthesis<br>Carbohydrate metabolism<br>Entner-Doudoroff and Pentose phosphate pathways<br>Tricarboxylic acid cycle<br>ATP biosynthesis<br>One-carbon metabolism<br>Peptidoglycan biosynthesis<br>Rhamnose biosynthesis<br>Lipid and LPS biosynthesis<br>Vitamin biosynthesis | Primary metabolism<br>Nucleotide metabolism<br>Regulation of cell shape*<br>Protein translation<br>Ribosome biogenesis<br>tRNA aminoacylation<br>Amino acid biosynthesis<br>Carbohydrate metabolism<br>Entner-Doudoroff and Pentose phosphate pathways<br>Tricarboxylic acid cycle<br>ATP biosynthesis<br>Lipid and LPS biosynthesis<br>Peptidoglycan biosynthesis<br>Sulfur metabolism<br>Chorismate metabolism<br>Vitamin biosynthesis |

**Supplementary Table S1. Enriched biological processes of differentially expressed genes (DEG) in bacterial endophytes. (B)** Down-regulated processes in *Pst ΔhrcCΔCFA*.

Comparisons were done between the inoculum, 6 ($t_6$), 24 ($t_{24}$) and 168 ($t_{168}$) hours post-inoculation, and *in vitro* populations grown to logarithmic (Log) or stationary (Stat) phase. An asterisk denotes that a biological process had less than 5 DEGs.

| **Down-regulated processes in *Pst ΔhrcCΔCFA*** | | | | | | | |
|---|---|---|---|---|---|---|---|
| **$t_6$ vs. KB** | **$t_6$ vs. $t_{168}$** | **$t_6$ vs. KB** | **$t_6$ vs. Stat** | **$t_{24}$ vs. $t_{168}$** | **$t_{24}$ vs. KB** | **$t_{24}$ vs. Stat** | **$t_{168}$ vs. KB** |
| Locomotion and flagellum | – | – | – | – | Locomotion and flagellum<br>Fatty acid biosynthesis | Locomotion and flagellum<br>LPS biosynthesis*<br>Siderophore biosynthesis | Locomotion and flagellum |

| **Down-regulated processes in *Pst ΔhrcCΔCFA*** | | | | | | |
|---|---|---|---|---|---|---|
| **$t_{168}$ vs. Stat** | **KB vs. Stat** | **Log vs. $t_6$** | **Log vs. $t_{24}$** | **Log vs. $t_{168}$** | **Log vs. KB** | **Log vs. Stat** |
| Peptidoglycan biosynthesis<br>Siderophore biosynthesis | Primary metabolism<br>Nucleotide metabolism<br>DNA repair<br>Protein translation<br>Ribosome biogenesis<br>tRNA aminoacylation<br>Protein secretion<br>Amino acid biosynthesis<br>Carbohydrate metabolism<br>ATP biosynthesis<br>One-carbon metabolism<br>Peptidoglycan biosynthesis<br>Alginate biosynthesis<br>Vitamin biosynthesis<br>Siderophore biosynthesis | Virulence<br>Locomotion and flagellum<br>Two-component system<br>Alginate biosynthesis<br>Response to oxidative stress<br>Bacterioferritin* | Virulence<br>Locomotion and flagellum<br>Two-component system<br>Branched-chain amino acid catabolism*<br>Carbohydrate metabolism<br>Alginate biosynthesis<br>Response to oxidative stress<br>Bacterioferritin* | Virulence<br>Locomotion and flagellum<br>Two-component system<br>Branched-chain amino acid catabolism*<br>Carbohydrate metabolism<br>Alginate biosynthesis<br>Sulfate transport*<br>Response to oxidative stress | Virulence<br>Locomotion and flagellum<br>Two-component system | Virulence<br>Locomotion and flagellum<br>Two-component system |

**Supplementary Table S1. Enriched biological processes of differentially expressed genes (DEG) in bacterial endophytes. (C)** Up-regulated processes in *Achromobacter xylosoxidans* Col-0-50.
Comparisons were done between the inoculum, 6 ($t_6$), 24 ($t_{24}$) and 168 ($t_{168}$) hours post-inoculation. An asterisk denotes that a biological process had less than 5 DEGs.

| **Up-regulated processes in *Achromobacter xylosoxidans* Col-0-50** | | | | | |
|---|---|---|---|---|---|
| **$t_6$ vs. $t_{24}$** | **$t_6$ vs. $t_{168}$** | **$t_6$ vs. LM** | **$t_{24}$ vs. $t_{168}$** | **$t_{24}$ vs. LM** | **$t_{168}$ vs. LM** |
| – | – | Primary metabolism<br>Protein translation | Osmolyte (ectoine) biosynthesis* | Primary metabolism<br>RNA primer synthesis for DNA replication*<br>Protein translation<br>Carbohydrate metabolism<br>ATP biosynthesis<br>Histidine biosynthesis<br>Osmolyte (ectoine) biosynthesis* | Primary metabolism<br>Protein translation<br>Protein folding<br>Ion transport<br>ATP biosynthesis<br>Osmolyte (ectoine) biosynthesis* |

**Supplementary Table S1. Enriched biological processes of differentially expressed genes (DEG) in bacterial endophytes. (D)** Down-regulated processes in *Achromobacter xylosoxidans* Col-0-50.
Comparisons were done between the inoculum, 6 ($t_6$), 24 ($t_{24}$) and 168 ($t_{168}$) hours post-inoculation. An asterisk denotes that a biological process had less than 5 DEGs.

| **Down-regulated processes in *Achromobacter xylosoxidans* Col-0-50** | | | | | |
|---|---|---|---|---|---|
| **$t_6$ vs. $t_{24}$** | **$t_6$ vs. $t_{168}$** | **$t_6$ vs. LM** | **$t_{24}$ vs. $t_{168}$** | **$t_{24}$ vs. LM** | **$t_{169}$ vs. LM** |
| Protein folding<br>Ion transport<br>Sulfate transport | Primary metabolism<br>Ion transport<br>ATP biosynthesis<br>Protein translation<br>Protein folding<br>Serine family amino acid catabolism*<br>Sulfate transport | Amino acid metabolism<br>Glutathione metabolism | Protein translation<br>Protein folding<br>ATP biosynthesis* | Locomotion and flagellum | – |

**Supplementary Table S1. Enriched biological processes of differentially expressed genes (DEG) in bacterial endophytes. (E)** Up-regulated processes in *Pandoraea* sp. Col-0-28.
Comparisons were done between the inoculum, 6 ($t_6$), 24 ($t_{24}$) and 168 ($t_{168}$) hours post-inoculation. An asterisk denotes that a biological process had less than 5 DEGs.

| **Up-regulated processes in *Pandoraea* sp. Col-0-28** | | | | | |
|---|---|---|---|---|---|
| **$t_6$ vs. $t_{24}$** | **$t_6$ vs. $t_{168}$** | **$t_6$ vs. LM** | **$t_{24}$ vs. $t_{168}$** | **$t_{24}$ vs. LM** | **$t_{168}$ vs. LM** |
| Cell division<br>Peptidoglycan biosynthesis*<br>Sulfate transport* | Two-component system | – | Two-component system | Nucleotide metabolism<br>Tricarboxylic acid cycle<br>Aerobic respiration<br>Sulfur compound biosynthesis | Primary metabolism<br>Nucleotide metabolism<br>Transcription termination*<br>Protein translation<br>tRNA aminoacylation<br>Protein folding<br>Amino acid biosynthesis<br>Urea metabolism*<br>Tricarboxylic acid cycle<br>Aerobic respiration<br>ATP biosynthesis<br>Sulfur compound biosynthesis<br>Response to oxidative stress<br>Siderophore transport* |

**Supplementary Table S1. Enriched biological processes of differentially expressed genes (DEG) in bacterial endophytes. (F)** Down-regulated processes in *Pandoraea* sp. Col-0-28.
Comparisons were done between the inoculum, 6 ($t_6$), 24 ($t_{24}$) and 168 ($t_{168}$) hours post-inoculation. An asterisk denotes that a biological process had less than 5 DEGs.

| **Down-regulated processes in *Pandoraea* sp. Col-0-28** | | | | | |
|---|---|---|---|---|---|
| **$t_6$ vs. $t_{24}$** | **$t_6$ vs. $t_{168}$** | **$t_6$ vs. LM** | **$t_{24}$ vs. $t_{168}$** | **$t_{24}$ vs. LM** | **$t_{168}$ vs. LM** |
| Protein translation<br>Tricarboxylic acid cycle*<br>Trehalose metabolism*<br>Nitrate transport | Primary metabolism<br>Nucleotide biosynthesis<br>Transcription<br>Protein translation<br>Ribosome biogenesis<br>Methionine biosynthesis*<br>Tricarboxylic acid cycle<br>ATP biosynthesis<br>Siderophore transport* | Primary metabolism<br>Protein translation<br>Branched-chain amino acid biosynthesis<br>Allantoin metabolism<br>Lipid metabolism | Primary metabolism<br>Protein translation<br>Tricarboxylic acid cycle<br>ATP biosynthesis<br>Siderophore transport* | Allantoin metabolism | – |

**Supplementary Table S2. Enriched biological processes in significantly different expression profiles after STEM (Short-Time series Expression Miner) analysis. (A)** Enriched processes in *Pseudomonas syringae* pv. *tomato* (*Pst*) ∆*hrcC*∆*CFA*.

An image of the expression profile is shown below the list of processes. The time points used for the analysis were the inoculum, 6, 24, and 168 hours post-inoculation. An asterisk denotes that a biological process had less than 5 genes.

| **Enriched *Pst* ∆*hrcC*∆*CFA* biological processes in Profile:** | | | | | | |
|---|---|---|---|---|---|---|
| **1** | **11** | **26** | **39** | **44** | **45** | **47** |
| Flagellum and locomotion | Flagellum and locomotion<br>Nucleoside transport | Transcriptional regulation | | Virulence<br>Glutamine family amino acid metabolism<br>Lysine metabolism*<br>Carbohydrate transport*<br>Vitamin biosynthesis | Primary metabolism<br>Purine ribonucleotide biosynthesis<br>Protein translation | Virulence<br>Primary metabolism<br>Nucleotide metabolism<br>Transcription<br>Protein translation<br>Ribosome biogenesis<br>tRNA aminoacylation<br>Amino acid biosynthesis<br>Carbohydrate metabolism<br>Entner-Doudoroff and Pentose phosphate<br>Tricarboxylic acid cycle<br>Alginate biosynthesis<br>Transport across membranes<br>Response to oxidative stress |

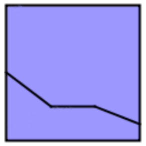
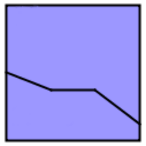
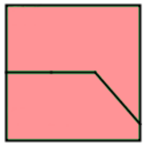
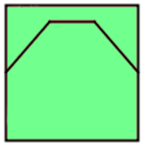
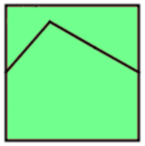
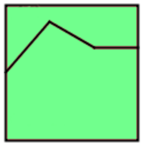
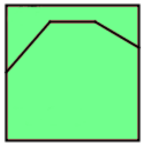

**Supplementary Table S2. Enriched biological processes in significantly different expression profiles after STEM (Short-Time series Expression Miner) analysis. (B)** Enriched processes in *Achromobacter xylosoxidans* Col-0-50.

An image of the expression profile is shown below the list of processes. The time points used for the analysis were the inoculum, 6, 24, and 168 hours post-inoculation. An asterisk denotes that a biological process had less than 5 genes.

| **Enriched *Achromobacter xylosoxidans* Col-0-50 biological processes in Profile:** | | | | |
|---|---|---|---|---|
| **9** | **26** | **34** | **39** | **47** |
| Flagellum and locomotion | Transcriptional regulation | Primary metabolism<br>Nucleotide metabolism<br>Vitamin biosynthesis | Primary metabolism<br>Histidine biosynthesis<br>Lipid biosynthesis | Primary metabolism<br>Protein translation<br>ATP biosynthesis<br>Osmolyte (Ectoine) biosynthesis* |

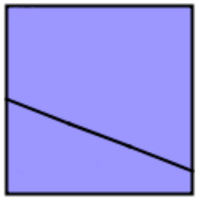
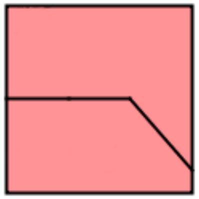
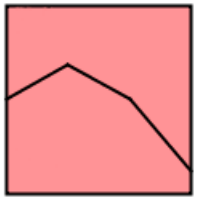
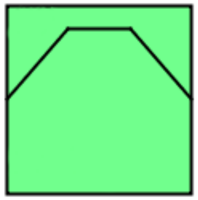
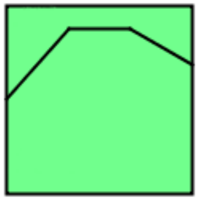

**Supplementary Table S2. Enriched biological processes in significantly different expression profiles after STEM (Short-Time series Expression Miner) analysis. (C)** Enriched processes in *Pandoraea* sp. Col-0-28.

An image of the expression profile is shown below the list of processes. The time points used for the analysis were the inoculum, 6, 24, and 168 hours post-inoculation. An asterisk denotes that a biological process had less than 5 genes.

| **Enriched *Pandoraea* sp. Col-0-28 biological processes in Profile:** | | | |
|---|---|---|---|
| **39** | **44** | **47** | **49** |
| Transcriptional regulation | Two-component system<br>Transposition | Primary metabolism<br>Nucleotide metabolism<br>tRNA aminoacylation<br>RNA processing<br>Amino acid biosynthesis<br>Vitamin biosynthesis<br>Ubiquinone biosynthesis | Primary metabolism<br>Protein translation<br>Tricarboxylic acid cycle<br>ATP biosynthesis<br>Inositol phosphate dephosphorylation* |

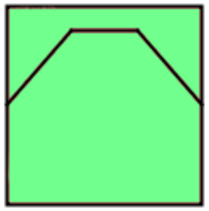
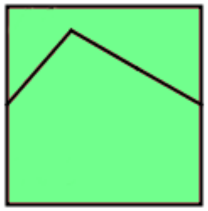
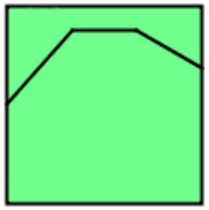
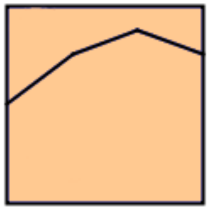

**Supplementary Table S3. Secondary metabolite biosynthetic clusters whose expression is differentially regulated inside plants. (A)** Secondary metabolite biosynthesis clusters in *Pseudomonas syringae* pv. *tomato* Δ*hrcC*Δ*CFA*. The yersiniabactin cluster did not show differential expression, but is included as it is cluster already known to be associated with secondary metabolite biosynthesis.

Gene expression used the Transcripts per million (TPM) values calculated by StringTie in the inoculum (KB), 6 ($t_6$) and 168 ($t_{168}$) hours post-inoculation. Relative expression was calculated using the logarithm in base 2 of the fold change between two treatments (an arbitrary addition of a value of 0.01 to each TPM was used to avoid values whose result would be undetermined). Genes that are up-regulated are highlighted in cyan, while those that are down-regulated are highlighted in magenta. A column indicating if a gene was differentially regulated, as determined by DESeq2 (adjusted $p$ < 0.05) is also shown.

| Locus name | Gene name | $log_2(t_6/KB)$ | < 0.05 | $log_2(t_{168}/KB)$ | < 0.05 | $log_2(t_6/t_{168})$ | < 0.05 | Annotation description |
|---|---|---|---|---|---|---|---|---|
| | | | | | | | | **N-acetylglutaminylglutamine amide** |
| *PSPTO_1627* | *PSPTO_1627* | -0.11 | | -1.64 | yes | 1.53 | | Short chain dehydrogenase |
| *PSPTO_1631* | *PSPTO_1631* | 4.20 | yes | 3.59 | yes | 0.61 | | |
| *PSPTO_1632* | *PSPTO_1632* | 4.13 | yes | 2.31 | yes | 1.81 | yes | GNAT family acetyltransferase |
| *PSPTO_1633* | *asnB* | 3.41 | yes | 2.29 | yes | 1.12 | | Asparagine synthetase |
| | | | | | | | | **Coronatine** |
| *PSPTO_4671* | | 0.30 | yes | -1.13 | yes | 1.43 | | Alkaline D-peptidase |
| *PSPTO_4680* | *cfl* | | | | | | | Coronafacic acid synthetase, ligase component; deleted in *Pst* Δ*hrcC*Δ*CFA*. |
| *PSPTO_4681* | *cfa1* | | | | | | | Coronafacic acid synthetase, acyl carrier protein component; deleted in *Pst* Δ*hrcC*Δ*CFA*. |
| *PSPTO_4682* | *cfa2* | | | | | | | Coronafacic acid synthetase, dehydratase component; deleted in *Pst* Δ*hrcC*Δ*CFA*. |
| *PSPTO_4683* | *cfa3* | | | | | | | Coronafacic acid beta-ketoacyl synthetase component; deleted in *Pst* Δ*hrcC*Δ*CFA*. |
| *PSPTO_4684* | *cfa4* | | | | | | | Coronafacic acid synthetase component; deleted in *Pst* Δ*hrcC*Δ*CFA*. |
| *PSPTO_4685* | *cfa5* | | | | | | | Coronafacic acid synthetase, ligase component; deleted in *Pst* Δ*hrcC*Δ*CFA*. |
| *PSPTO_4686* | *cfa6* | | | | | | | Coronafacic acid polyketide synthase I; deleted in *Pst* Δ*hrcC*Δ*CFA*. |
| *PSPTO_4687* | *cfa7* | | | | | | | Coronafacic acid polyketide synthetase II; deleted in *Pst* Δ*hrcC*Δ*CFA*. |
| *PSPTO_4689* | *cfa8* | | | | | | | Crotonyl-CoA reductase; deleted in *Pst* Δ*hrcC*Δ*CFA*. |
| *PSPTO_4690* | *cfa9* | | | | | | | CFA synthetase, thioesterase component; deleted in *Pst* Δ*hrcC*Δ*CFA*. |
| *PSPTO_4699* | | 0.06 | yes | -1.11 | yes | 1.17 | | Non-ribosomal peptide synthetase, terminal component |
| *PSPTO_4704* | *corR* | 1.63 | | -0.75 | | 2.39 | | DNA-binding response regulator CorR |
| *PSPTO_4705* | *corS* | 0.89 | | -0.11 | yes | 1.00 | | Sensor histidine kinase CorS |
| *PSPTO_4706* | *corP* | -0.32 | | -0.68 | yes | 0.36 | | Response regulator CorP |
| *PSPTO_4707* | *cmaD* | 4.17 | yes | -0.52 | | 4.68 | yes | Coronamic acid synthetase CmaD |
| *PSPTO_4708* | *cmaE* | 2.84 | yes | -0.32 | | 3.16 | yes | Coronamic acid synthetase CmaE |
| *PSPTO_4709* | *cmaA* | 4.13 | yes | 0.50 | | 3.63 | yes | Coronamic acid synthetase CmaA |
| *PSPTO_4710* | *cmaB* | 4.43 | yes | 1.08 | yes | 3.34 | yes | Coronamic acid synthetase CmaB |
| *PSPTO_4711* | *cmaC* | 4.12 | yes | -2.69 | yes | 6.81 | yes | Coronamic acid synthetase CmaC |
| *PSPTO_4712* | *cmaT* | 3.55 | yes | 0.91 | yes | 2.64 | yes | Coronamic acid synthetase, thioesterase component |
| *PSPTO_4714* | *cmaU* | 0.95 | | -2.35 | yes | 3.31 | yes | CmaU protein |
| *PSPTO_4723* | *cmaL* | 5.04 | yes | 1.05 | yes | 3.98 | yes | |

| Locus name | Gene name | $log_2(t_6/KB)$ | < 0.05 | $log_2(t_{168}/KB)$ | < 0.05 | $log_2(t_6/t_{168})$ | < 0.05 | Annotation description |
|---|---|---|---|---|---|---|---|---|
| | | | | **Pyoverdin** | | | | |
| *PSPTO_2120* | *PSPTO_2120* | -0.19 | yes | -2.89 | yes | 2.70 | yes | Peptidase, M20/M25/M40 family |
| *PSPTO_2123* | *PSPTO_2123* | 0.00 | yes | 0.65 | | -0.65 | yes | Sensor histidine kinase |
| *PSPTO_2128* | *PSPTO_2128* | 3.55 | yes | 1.05 | yes | 2.50 | yes | Response regulator |
| *PSPTO_2130* | *PSPTO_2130* | 3.80 | yes | 2.27 | yes | 1.53 | | LuxR family DNA-binding response regulator |
| *PSPTO_2131* | *PSPTO_2131* | 3.02 | yes | 0.60 | yes | 2.42 | yes | Sensor histidine kinase |
| *PSPTO_2133* | *sigX* | 0.34 | yes | -0.61 | yes | 0.95 | yes | RNA polymerase sigma-70 family protein |
| *PSPTO_2134* | *pvdG* | -6.22 | yes | -6.22 | | 0.00 | | Pyoverdine synthetase, thioesterase component |
| *PSPTO_2135* | *pvsA, pvdL* | -1.36 | yes | -1.08 | yes | -0.28 | yes | Pyoverdine chromophore precursor synthetase |
| *PSPTO_2136* | *daT, pvdH* | 0.18 | | 0.22 | | -0.04 | | 2,4-diaminobutyrate 4-transaminase |
| *PSPTO_2137* | *PSPTO_2137* | 0.45 | | -1.86 | yes | 2.31 | | MbtH-like protein |
| *PSPTO_2140* | *PSPTO_2140* | -0.51 | | 0.44 | | -0.96 | | Cation ABC transporter ATP-binding protein |
| *PSPTO_2141* | *PSPTO_2141* | -0.56 | yes | -1.81 | yes | 1.25 | | Cation ABC transporter substrate-binding protein |
| *PSPTO_2146* | *PSPTO_2146* | 0.18 | | -8.51 | yes | 8.69 | yes | Pyoverdine biosynthesis regulatory gene |
| *PSPTO_2147* | *PSPTO_2147* | 0.71 | yes | -0.15 | yes | 0.86 | | Pyoverdine sidechain peptide synthetase I, epsilon-Lys module |
| *PSPTO_2148* | *PSPTO_2148* | 0.17 | | -0.24 | | 0.41 | | Pyoverdine sidechain peptide synthetase II, D-Asp-L-Thr component |
| *PSPTO_2149* | *pvdD* | 0.40 | | 0.42 | | -0.02 | | Pyoverdine sidechain peptide synthetase III, L-Thr-L-Ser component |
| *PSPTO_2150* | *PSPTO_2150* | 0.08 | yes | -0.22 | | 0.30 | | Pyoverdine sidechain peptide synthetase IV, D-Asp-L-Ser component |
| *PSPTO_2151* | *fpvA* | 1.88 | yes | -0.61 | | 2.49 | yes | TonB-dependent siderophore receptor |
| *PSPTO_2152* | *PSPTO_2152* | 0.08 | | -0.57 | | 0.65 | | TonB-dependent siderophore receptor |
| *PSPTO_2153* | *pvdE* | 0.62 | yes | -0.38 | yes | 1.00 | | Pyoverdine ABC transporter, ATP-binding/permease protein |
| *PSPTO_2154* | *pvdO* | 0.81 | yes | -1.57 | | 2.38 | yes | |
| *PSPTO_2155* | *pvdN* | -1.14 | yes | 0.13 | yes | -1.27 | yes | Class V aminotransferase |
| *PSPTO_2156* | *pvdM* | 1.40 | | -3.75 | yes | 5.15 | yes | Renal dipeptidase family protein |
| *PSPTO_2158* | *PSPTO_2158* | 0.94 | | 0.34 | | 0.59 | | Outer membrane efflux protein |
| *PSPTO_2159* | *PSPTO_2159* | -0.13 | yes | -1.56 | yes | 1.42 | | Macrolide ABC efflux protein |
| *PSPTO_2160* | *PSPTO_2160* | -2.32 | yes | 0.20 | yes | -2.51 | | RND family efflux transporter MFP subunit |
| *PSPTO_2161* | *pvdQ* | -0.23 | yes | -0.85 | yes | 0.63 | | Penicillin amidase family protein |
| | | | | **Yersiniabactin** | | | | |
| *PSPTO_2583* | | -0.78 | | -6.57 | | 5.79 | | Glutamine amidotransferase, class-II protein |
| *PSPTO_2590* | | 2.50 | yes | 2.69 | yes | -0.19 | yes | Luciferase family protein |
| *PSPTO_2591* | | 0.22 | yes | 0.33 | | -0.11 | | Diguanylate cyclase |
| *PSPTO_2592* | *saxG* | 0.97 | | 0.39 | | 0.58 | | Aliphatic isothiocyanate resistance protein SaxG; AcrB/AcrD/AcrF family |
| *PSPTO_2593* | | 1.68 | yes | -3.55 | yes | 5.23 | yes | Multidrug resistance protein, AcrA/AcrE family |
| *PSPTO_2595* | *pchA* | 0.44 | | -0.03 | | 0.46 | | Isochorismate synthase |
| *PSPTO_2596* | *pchB* | 0.25 | yes | -1.26 | yes | 1.51 | | Isochorismate pyruvate-lyase |
| *PSPTO_2597* | *irp5, ybtE* | 1.16 | | -0.28 | | 1.44 | | Yersiniabactin synthetase, salycilate ligase component |
| *PSPTO_2598* | *irp4, ybtT* | 0.34 | | -0.53 | | 0.88 | | Yersiniabactin synthetase, thioesterase component |
| *PSPTO_2599* | *irp3, ybtU* | -0.45 | | -0.67 | | 0.23 | | Yersiniabactin synthetase, thiazolinyl reductase component |
| *PSPTO_2600* | *irp1, hmwp1* | 0.26 | yes | -0.53 | yes | 0.79 | | Yersiniabactin polyketide/non-ribosomal peptide synthetase |
| *PSPTO_2601* | *ybtX* | 0.81 | | -1.78 | yes | 2.59 | yes | Membrane protein |
| *PSPTO_2602* | *hmwp2* | 0.33 | yes | -1.12 | yes | 1.45 | | Yersiniabactin non-ribosomal peptide synthetase |
| *PSPTO_2603* | *ybtQ* | -0.18 | | -1.22 | | 1.04 | | ABC transporter ATP-binding protein/permease |
| *PSPTO_2604* | *ybtP* | 0.04 | yes | -1.64 | yes | 1.68 | | ABC transporter ATP-binding protein/permease |
| *PSPTO_2605* | *fyuA* | -0.09 | yes | -0.22 | | 0.13 | yes | TonB-dependent siderophore receptor |
| *PSPTO_2606* | *ybtA* | 0.03 | yes | -1.52 | yes | 1.56 | | AraC family transcriptional regulator |
| *PSPTO_2607* | | 0.06 | | 0.06 | | 0.00 | | TonB-dependent siderophore receptor |
| *PSPTO_2612* | | -0.02 | | -1.48 | | 1.46 | | GNAT family acetyltransferase |
| *PSPTO_2614* | | 2.34 | yes | -0.54 | | 2.87 | yes | Dioxygenase, TauD/TfdA family |
| *PSPTO_2617* | | 1.76 | | 0.85 | | 0.91 | | LysR family transcriptional regulator |
| *PSPTO_2618* | *argD-2* | 0.47 | | -1.98 | yes | 2.45 | | Acetylornithine aminotransferase |

**Supplementary Table S3. Secondary metabolite biosynthetic clusters whose expression is differentially regulated inside plants. (B)** Secondary metabolite biosynthesis clusters in *Achromobacter xylosoxidans* Col-0-50.

Gene expression used the Transcripts per million (TPM) values calculated by StringTie in the inoculum (LM), 6 ($t_6$) and 168 ($t_{168}$) hours post-inoculation. Relative expression was calculated using the logarithm in base 2 of the fold change between two treatments (an arbitrary addition of a value of 0.01 to each TPM was used to avoid values whose result would be undetermined). Genes that are up-regulated are highlighted in cyan, while those that are down-regulated are highlighted in magenta. A column indicating if a gene was differentially regulated, as determined by DESeq2 (adjusted $p$ < 0.05) is also shown.

| Locus name | Gene name | $\log_2(t_6/KB)$ | < 0.05 | $\log_2(t_{168}/KB)$ | < 0.05 | $\log_2(t_6/t_{168})$ | < 0.05 | Annotation description |
|---|---|---|---|---|---|---|---|---|
| **Siderophore, similar to desferrioxamine** | | | | | | | | |
| *FLPKDCKG_00714* | *bcr_1* | -3.54 | yes | -4.17 | yes | 0.63 | | Bicyclomycin resistance protein |
| *FLPKDCKG_00715* | *pchR_3* | -3.40 | yes | -3.24 | yes | -0.16 | yes | Regulatory protein PchR |
| *FLPKDCKG_00718* | *iucC* | -4.35 | yes | -3.88 | yes | -0.47 | | Aerobactin synthase |
| *FLPKDCKG_00719* | *iucB* | -3.11 | yes | -4.56 | yes | 1.45 | | N(6)-hydroxylysine O-acetyltransferase |
| *FLPKDCKG_00720* | *iucD* | -4.45 | yes | -3.83 | yes | -0.61 | | L-lysine N6-monooxygenase |
| *FLPKDCKG_00721* | *fetA* | 0.51 | | -1.88 | | 2.38 | | Putative iron export ATP-binding protein FetA |
| *FLPKDCKG_00722* | *fetB* | 0.09 | | -7.42 | yes | 7.51 | | Putative iron export permease protein FetB. |
| **Ectoine** | | | | | | | | |
| *FLPKDCKG_01732* | *ectD_1* | 7.08 | yes | 3.59 | yes | 3.48 | yes | Ectoine dioxygenase |
| *FLPKDCKG_01733* | *ectC* | 7.77 | yes | 4.29 | yes | 3.48 | yes | L-ectoine synthase |
| *FLPKDCKG_01734* | *ectB* | 8.38 | yes | 4.55 | yes | 3.83 | yes | Diaminobutyrate--2-oxoglutarate transaminase |
| *FLPKDCKG_01735* | *ectA* | 6.60 | yes | 2.10 | yes | 4.50 | yes | L-2,4-diaminobutyric acid acetyltransferase |
| *FLPKDCKG_01736* | *yusO* | 3.49 | yes | 0.21 | yes | 3.28 | yes | Putative HTH-type transcriptional regulator YusO |
| ***Non-ribosomal peptide synthase-like*** | | | | | | | | |
| *FLPKDCKG_01927* | *livH_6* | -0.03 | | -0.28 | | 0.25 | | High-affinity branched-chain amino acid transport system permease protein LivH |
| *FLPKDCKG_01928* | *livF_6* | 1.76 | | 1.23 | | 0.53 | | High-affinity branched-chain amino acid transport ATP-binding protein LivF |
| *FLPKDCKG_01929* | *lptB_3* | -0.11 | | 3.07 | | -3.17 | | Lipopolysaccharide export system ATP-binding protein LptB |
| *FLPKDCKG_01930* | *gno_1* | 0.86 | yes | 1.93 | | -1.07 | yes | Gluconate 5-dehydrogenase |
| *FLPKDCKG_01932* | - | -0.23 | | 0.88 | | -1.11 | | |
| *FLPKDCKG_01933* | *fabG2* | 0.70 | | -1.05 | | 1.76 | | Putative oxidoreductase |
| *FLPKDCKG_01934* | *kipR_2* | 0.58 | | -1.23 | yes | 1.81 | | HTH-type transcriptional regulator KipR |
| *FLPKDCKG_01947* | *dhbF* | 0.21 | | -0.99 | | 1.20 | yes | Dimodular nonribosomal peptide synthase |
| *FLPKDCKG_01952* | *tctD_1* | 0.42 | | -0.90 | | 1.32 | | Transcriptional regulatory protein tctD |
| *FLPKDCKG_01953* | *qseC_2* | 1.01 | | 0.34 | yes | 0.67 | | Sensor protein QseC |
| *FLPKDCKG_01954* | *proP_3* | 3.32 | yes | 1.89 | yes | 1.43 | | Proline/betaine transporter |
| *FLPKDCKG_01957* | *dosC_2* | 1.20 | | -2.40 | yes | 3.59 | yes | Diguanylate cyclase DosC |
| *FLPKDCKG_01958* | *pyrB_1* | -0.60 | yes | -1.66 | | 1.06 | yes | Aspartate carbamoyltransferase catalytic subunit |
| **Resorcinol** | | | | | | | | |
| *FLPKDCKG_03746* | *yajR_2* | 1.88 | | -1.74 | | 3.62 | | Inner membrane transport protein YajR |
| *FLPKDCKG_03748* | *bioC_1* | 1.45 | | -1.47 | | 2.92 | | Malonyl-[acyl-carrier protein] O-methyltransferase |
| *FLPKDCKG_03751* | *acuR* | 0.80 | | -1.33 | | 2.13 | | Transcriptional regulator AcuR |
| *FLPKDCKG_03754* | - | -1.57 | yes | -3.91 | yes | 2.33 | | |
| *FLPKDCKG_03756* | *glcC_2* | 1.33 | | -1.77 | | 3.10 | | Glc operon transcriptional activator |
| *FLPKDCKG_03766* | *fabH2* | 1.39 | | -1.18 | | 2.57 | | 3-oxoacyl-[acyl-carrier-protein] synthase 3 protein 2 |
| *FLPKDCKG_03774* | *yijE_8* | 1.88 | yes | -1.01 | | 2.90 | | Putative cystine transporter YijE |
| *FLPKDCKG_03775* | *bmr3_1* | 1.77 | yes | -0.45 | | 2.22 | | Multidrug resistance protein 3 |
| *FLPKDCKG_03777* | *rkpK_2* | 1.44 | yes | -0.75 | | 2.18 | | UDP-glucose 6-dehydrogenase |
| *FLPKDCKG_03778* | - | 1.60 | yes | -0.52 | | 2.12 | | UDP-glucose 4-epimerase |
| *FLPKDCKG_03779* | *pseB* | 1.35 | | -0.81 | | 2.16 | | UDP-N-acetylglucosamine 4,6-dehydratase (inverting) |
| *FLPKDCKG_03780* | *pseC* | 1.84 | yes | -0.53 | | 2.37 | | UDP-4-amino-4,6-dideoxy-N-acetyl-beta-L-altrosamine transaminase |

**Supplementary Table S4. Average gene expression in bacterial endophytes. (A)** Gene expression in *Pseudomonas syringae* pv. *tomato* Δ*hrcC*Δ*CFA*.

Gene expression was calculated as Transcripts per million (TPM) using StringTie in the inoculum, 6 ($t_6$), 24 ($t_{24}$) and 168 ($t_{168}$) hours post-inoculation, and *in vitro* populations grown to logarithmic or stationary phase. Each treatment had four biological replicates. When available, gene name for each locus is shown. DNA template strand, coordinates for the start and end of each gene, and a brief annotation are also shown.

| Locus name | Gene name | Strand | Gene start (bp) | Gene end (bp) | Average TPM Inoculum | Average TPM $t_6$ | Average TPM $t_{24}$ | Average TPM $t_{168}$ | Average TPM Logarithmic | Average TPM Stationary | Annotation description |
|---|---|---|---|---|---|---|---|---|---|---|---|
| *PSPTO_0001* | *dnaA* | + | 339 | 1874 | 9.30 | 31.17 | 26.35 | 21.30 | 107.60 | 29.64 | Chromosomal replication initiation protein |
| *PSPTO_0002* | *dnaN* | + | 1913 | 3016 | 9.17 | 29.63 | 31.39 | 12.28 | 117.46 | 27.91 | DNA polymerase III subunit beta |
| *PSPTO_0003* | *recF* | + | 3039 | 4142 | 17.87 | 47.00 | 29.28 | 20.94 | 126.30 | 30.55 | DNA replication and repair protein Recife |
| *PSPTO_0004* | *gyrB* | + | 4147 | 6564 | 13.74 | 48.53 | 29.87 | 21.18 | 155.97 | 42.92 | DNA gyrase subunit B |
| *PSPTO_0005* | | + | 7092 | 8798 | 25.32 | 36.61 | 29.83 | 18.09 | 68.67 | 30.28 | Type I restriction-modification system, M subunit |
| *PSPTO_0006* | | + | 8861 | 10168 | 9.64 | 12.71 | 15.42 | 5.92 | 33.35 | 9.94 | Type I restriction-modification system, S subunit, Eco family |
| *PSPTO_0007* | | + | 10179 | 11063 | 4.05 | 3.96 | 8.27 | 1.24 | 20.33 | 4.74 | Hypothetical protein |
| *PSPTO_0008* | | + | 11067 | 14366 | 9.31 | 8.37 | 7.31 | 3.77 | 22.02 | 7.98 | HsdR family type I site-specific deoxyribonuclease |
| *PSPTO_0009* | | - | 14512 | 15741 | 31.38 | 27.31 | 16.91 | 12.30 | 94.97 | 23.17 | ISPsy3, transposase |
| *PSPTO_0011* | | + | 16743 | 17453 | 26.77 | 44.40 | 9.17 | 18.76 | 70.58 | 36.89 | Hypothetical protein |
| *PSPTO_0012* | | + | 17514 | 18017 | 12.80 | 14.91 | 18.13 | 7.71 | 37.92 | 21.84 | Hypothetical protein |
| *PSPTO_0013* | | + | 17992 | 18561 | 8.38 | 2.72 | 3.49 | 4.03 | 12.78 | 7.20 | Hypothetical protein |
| *PSPTO_0014* | | - | 18573 | 18863 | 5.91 | 4.57 | 7.33 | 0.25 | 5.40 | 6.54 | Hypothetical protein |
| *PSPTO_0015* | | + | 19321 | 19932 | 4.76 | 4.42 | 5.16 | 4.00 | 18.09 | 6.54 | Hypothetical protein |
| *PSPTO_0016* | | + | 19968 | 20435 | 9.79 | 6.11 | 5.52 | 2.61 | 20.76 | 7.21 | Hypothetical protein |
| *PSPTO_0017* | | - | 20738 | 23143 | 7.48 | 5.25 | 7.15 | 3.64 | 10.53 | 7.88 | Helicase/SNF2 family domain protein |
| *PSPTO_0018* | | - | 23290 | 24099 | 0.00 | 0.00 | 0.00 | 0.00 | 0.00 | 0.00 | ISPsy4, transposition helper protein |
| *PSPTO_0019* | | - | 24096 | 25118 | 0.00 | 0.00 | 0.00 | 0.00 | 0.00 | 0.00 | ISPsy4, transposase |
| *PSPTO_0020* | | - | 25187 | 25537 | 6.32 | 0.06 | 8.44 | 1.45 | 7.78 | 4.94 | Hypothetical protein |
| *PSPTO_0021* | | - | 25556 | 27796 | 2.97 | 4.72 | 2.31 | 3.49 | 9.91 | 4.92 | Hypothetical protein |
| *PSPTO_0022* | | + | 28478 | 29323 | 72.47 | 78.96 | 59.67 | 43.95 | 59.91 | 40.76 | DNA-binding protein |
| *PSPTO_0023* | | + | 29320 | 29919 | 19.15 | 33.02 | 22.88 | 13.06 | 30.62 | 20.55 | Hypothetical protein |
| *PSPTO_0024* | | + | 30850 | 30987 | 19.20 | 9.42 | 7.21 | 10.11 | 12.18 | 13.67 | Hypothetical protein |
| *PSPTO_0025* | | + | 31338 | 31967 | 21.08 | 27.86 | 33.22 | 21.02 | 38.94 | 30.50 | Haloacid dehalogenase-like family hydrolase |
| *PSPTO_0026* | | - | 32022 | 33068 | 6.72 | 8.32 | 10.51 | 4.10 | 12.63 | 6.24 | Hypothetical protein |
| *PSPTO_0027* | | + | 33254 | 33976 | 2.11 | 9.70 | 4.98 | 3.72 | 8.17 | 2.90 | Hypothetical protein |
| *PSPTO_0028* | | + | 33978 | 35933 | 0.68 | 1.46 | 1.92 | 0.00 | 4.64 | 1.73 | Transposase |
| *PSPTO_0029* | | + | 35933 | 36868 | 1.23 | 5.39 | 0.83 | 3.84 | 9.92 | 2.86 | Transposition helper protein |
| *PSPTO_0030* | | + | 36881 | 38767 | 4.23 | 3.98 | 2.22 | 5.15 | 12.41 | 7.39 | Hypothetical protein |
| *PSPTO_0031* | | + | 39345 | 40082 | 5.89 | 5.98 | 3.36 | 2.21 | 9.19 | 4.54 | Ser/Thr protein phosphatase family protein |
| *PSPTO_0032* | | - | 40884 | 41852 | 3.37 | 5.73 | 3.81 | 0.38 | 5.76 | 4.45 | Hypothetical protein |
| *PSPTO_0033* | | - | 41852 | 42757 | 1.03 | 3.96 | 4.99 | 2.82 | 5.03 | 2.55 | ParB family protein |
| *PSPTO_0034* | | - | 42754 | 44310 | 8.63 | 9.93 | 9.41 | 7.40 | 17.16 | 10.42 | Recombinase |
| *PSPTO_0035* | | - | 45183 | 46736 | 0.00 | 0.00 | 0.00 | 0.00 | 0.00 | 0.00 | ISPsy5, transposase |
| *PSPTO_0036* | | - | 46754 | 47113 | 0.00 | 0.00 | 0.00 | 0.00 | 0.00 | 0.00 | ISPsy5, Orf1 |
| *PSPTO_0037* | | + | 47448 | 52358 | 5.73 | 11.48 | 9.38 | 5.91 | 79.96 | 18.86 | Helicase domain-containing protein |
| *PSPTO_0038* | | + | 52403 | 54493 | 3.75 | 2.62 | 3.89 | 3.85 | 22.52 | 8.75 | Hypothetical protein |
| *PSPTO_0039* | | - | 54971 | 56524 | 0.00 | 0.00 | 0.00 | 0.00 | 0.00 | 0.00 | ISPsy5, transposase |
| *PSPTO_0040* | | - | 56542 | 56901 | 0.00 | 0.00 | 0.00 | 0.00 | 0.00 | 0.00 | ISPsy5, Orf1 |
| *PSPTO_0041* | | + | 56961 | 57347 | 18.32 | 13.52 | 13.09 | 13.60 | 51.80 | 19.08 | Hypothetical protein |
| *PSPTO_0042* | | + | 57351 | 58214 | - | - | - | - | - | - | |
| *PSPTO_0043* | | + | 58344 | 59906 | 92.49 | 76.61 | 49.17 | 35.22 | 75.51 | 100.38 | Cytidine/deoxycytidylate deaminase family protein |
| *PSPTO_0044* | *hopK1* | - | 60407 | 61423 | 49.32 | 121.97 | 70.33 | 30.95 | 121.44 | 62.41 | Type III effector HopK1 |
| *PSPTO_0045* | | + | 61526 | 61684 | 17.98 | 35.12 | 36.65 | 9.84 | 116.98 | 63.21 | Hypothetical protein |
| *PSPTO_0046* | | + | 62078 | 64063 | 14.96 | 10.95 | 12.20 | 9.18 | 20.93 | 19.09 | Hypothetical protein |
| *PSPTO_0047* | | + | 64066 | 65970 | 12.64 | 7.73 | 11.24 | 11.42 | 28.87 | 16.58 | UvrD/REP helicase family protein |
| *PSPTO_0048* | | + | 67445 | 68611 | 1.73 | 1.12 | 7.01 | 0.11 | 17.15 | 4.26 | Hypothetical protein |
| *PSPTO_0049* | | + | 68863 | 69606 | 19.00 | 28.15 | 25.74 | 10.11 | 45.88 | 20.10 | Hypothetical protein |
| *PSPTO_0050* | | + | 69623 | 70954 | 9.97 | 0.91 | 2.25 | 3.37 | 15.33 | 13.12 | Hypothetical protein |
| *PSPTO_0051* | | + | 71041 | 71484 | 7.44 | 4.83 | 5.26 | 2.90 | 39.69 | 19.06 | Hypothetical protein |
| *PSPTO_0052* | | + | 71711 | 73480 | 22.75 | 11.82 | 12.23 | 4.87 | 94.30 | 23.10 | Hypothetical protein |
| *PSPTO_0053* | | - | 73641 | 75086 | 7.32 | 7.48 | 7.13 | 9.45 | 35.08 | 17.31 | Hypothetical protein |
| *PSPTO_0054* | | - | 75073 | 75684 | 8.26 | 10.50 | 4.00 | 6.52 | 22.85 | 11.96 | Hypothetical protein |
| *PSPTO_0055* | | + | 75949 | 76971 | 0.00 | 0.00 | 0.00 | 0.00 | 0.00 | 0.00 | ISPsy4, transposase |
| *PSPTO_0056* | | + | 76968 | 77777 | 0.00 | 0.00 | 0.00 | 0.00 | 0.00 | 0.00 | ISPsy4, transposition helper protein |
| *PSPTO_0057* | | - | 78120 | 79010 | 11.35 | 5.81 | 11.09 | 4.19 | 17.15 | 11.58 | Transposition helper protein |
| *PSPTO_0058* | | - | 79007 | 80794 | 13.74 | 13.37 | 11.65 | 12.04 | 35.72 | 24.63 | Transposase |
| *PSPTO_0059* | | - | 80922 | 81542 | 18.21 | 9.03 | 16.40 | 9.74 | 35.87 | 26.50 | Hypothetical protein |
| *PSPTO_0060* | | + | 81816 | 82382 | - | - | - | - | - | - | |
| *PSPTO_0061* | *hopY1* | + | 82500 | 83363 | 4.43 | 156.59 | 68.45 | 13.59 | 5.82 | 4.10 | Type III effector HopY1 |
| *PSPTO_0062* | | - | 83552 | 84859 | 7.74 | 21.05 | 23.04 | 4.97 | 24.25 | 7.59 | Citrate transporter |
| *PSPTO_0063* | | + | 85159 | 85557 | 33.64 | 37.00 | 38.02 | 7.80 | 45.38 | 46.44 | Hypothetical protein |
| *PSPTO_0064* | | - | 85784 | 87181 | 68.18 | 93.84 | 78.37 | 37.74 | 164.53 | 86.62 | Hypothetical protein |
| *PSPTO_0065* | *recG* | - | 87268 | 89343 | 7.80 | 12.18 | 8.33 | 4.89 | 31.32 | 7.38 | ATP-dependent DNA helicase RecG |
| *PSPTO_0066* | *oxyR* | - | 89355 | 90278 | 26.02 | 20.53 | 28.18 | 3.67 | 76.11 | 11.72 | Oxidative stress regulatory protein OxyR |
| *PSPTO_0067* | *tonB-1* | - | 90371 | 91117 | 14.54 | 43.07 | 29.71 | 14.85 | 224.17 | 130.41 | TonB protein |
| *PSPTO_0068* | *exbD-1* | - | 91114 | 91539 | 4.20 | 14.24 | 11.98 | 9.16 | 138.71 | 110.57 | TonB system transport protein ExbD |
| *PSPTO_0069* | *exbB-1* | - | 91543 | 92499 | 39.21 | 69.50 | 61.02 | 32.91 | 299.82 | 245.71 | TonB system transport protein ExbB |
| *PSPTO_0070* | | + | 92710 | 93558 | 13.13 | 24.16 | 23.22 | 16.88 | 26.71 | 22.65 | Hypothetical protein |
| *PSPTO_0071* | | - | 93607 | 94344 | 8.18 | 28.44 | 30.12 | 11.95 | 88.24 | 32.05 | Lipoprotein |
| *PSPTO_0072* | | - | 94399 | 94779 | 51.46 | 34.11 | 19.77 | 14.27 | 89.75 | 34.22 | Endoribonuclease L-PSP |
| *PSPTO_0073* | *spoT* | - | 94805 | 96910 | 13.45 | 31.96 | 14.82 | 15.99 | 100.97 | 39.49 | Guanosine-3',5'-bis(diphosphate) 3'-pyrophosphohydrolase |
| *PSPTO_0074* | *rpoZ* | - | 96971 | 97234 | 212.33 | 616.85 | 485.77 | 324.28 | 895.11 | 651.57 | DNA-directed RNA polymerase subunit omega |
| *PSPTO_0075* | *gmk* | - | 97382 | 98026 | 12.04 | 34.31 | 25.44 | 17.35 | 76.00 | 22.82 | Guanylate kinase |
| *PSPTO_0076* | | - | 98040 | 98903 | 12.43 | 25.71 | 22.11 | 6.56 | 49.62 | 19.23 | Hypothetical protein |

| Locus name | Gene name | Strand | Gene start (bp) | Gene end (bp) | Average TPM Inoculum | Average TPM $t_6$ | Average TPM $t_{24}$ | Average TPM $t_{168}$ | Average TPM Logarithmic | Average TPM Stationary | Annotation description |
|---|---|---|---|---|---|---|---|---|---|---|---|
| *PSPTO_0077* | *rph* | + | 99223 | 99945 | 119.03 | 145.83 | 120.82 | 62.82 | 263.71 | 86.59 | Ribonuclease PH |
| *PSPTO_0078* | | + | 99962 | 100333 | 10.04 | 17.54 | 10.69 | 18.55 | 30.07 | 15.71 | Hypothetical protein |
| *PSPTO_0079* | | - | 100403 | 101182 | 43.87 | 75.25 | 64.67 | 34.13 | 111.00 | 41.51 | Exodeoxyribonuclease III |
| *PSPTO_0080* | *pyrE* | + | 101265 | 101906 | 30.10 | 63.12 | 57.20 | 29.47 | 188.89 | 67.77 | Orotate phosphoribosyltransferase |
| *PSPTO_0081* | | + | 101917 | 102549 | 10.39 | 45.87 | 22.08 | 12.12 | 195.90 | 64.86 | Hypothetical protein |
| *PSPTO_0082* | *argB* | - | 102801 | 103706 | 28.36 | 72.50 | 61.30 | 25.50 | 161.91 | 54.25 | Acetylglutamate kinase |
| *PSPTO_0083* | *algC* | - | 103735 | 105132 | 17.63 | 108.47 | 88.21 | 43.76 | 170.44 | 66.86 | Phosphomannomutase |
| *PSPTO_0084* | *dut* | - | 106375 | 106830 | 7.62 | 18.87 | 17.94 | 7.94 | 73.90 | 17.52 | Deoxyuridine 5'triphosphate nucleotidohydrolase |
| *PSPTO_0085* | *coaBC* | - | 106837 | 108084 | 12.16 | 38.49 | 40.27 | 16.58 | 85.06 | 19.73 | Phosphopantothenoylcysteine decarboxylase/phosphopantothenate--cysteine ligase |
| *PSPTO_0086* | *radC* | + | 108181 | 108855 | 9.36 | 23.65 | 15.08 | 5.92 | 28.26 | 16.95 | DNA repair protein RadC |
| *PSPTO_0087* | | - | 108910 | 110496 | 18.45 | 28.94 | 28.17 | 16.97 | 90.38 | 36.33 | Family 5 extracellular solute-binding protein |
| *PSPTO_0088* | | - | 110557 | 110691 | 18.98 | 8.07 | 18.17 | 16.55 | 33.20 | 13.55 | Hypothetical protein |
| *PSPTO_0089* | *rpmB* | + | 110860 | 111093 | 415.55 | 1401.00 | 1310.43 | 1030.31 | 3742.77 | 669.86 | 50S ribosomal protein L28 |
| *PSPTO_0090* | *rpmG* | + | 111105 | 111260 | 44.93 | 294.67 | 223.33 | 158.29 | 282.96 | 64.58 | 50S ribosomal protein L33 |
| *PSPTO_0091* | | - | 111368 | 111733 | 85.13 | 75.26 | 45.83 | 8.07 | 117.91 | 48.74 | Hypothetical protein |
| *PSPTO_0092* | *aldA* | + | 111972 | 113465 | 8.49 | 17.66 | 21.52 | 13.12 | 98.50 | 16.75 | Aldehyde dehydrogenase family protein. Conversion of indole-3-acetaldehyde to indole-3-acetic acid (an auxin). |
| *PSPTO_0093* | | - | 113528 | 113779 | 4.59 | 18.24 | 0.78 | 4.10 | 21.68 | 6.56 | Hypothetical protein |
| *PSPTO_0094* | | + | 113892 | 115040 | 5.79 | 5.59 | 5.64 | 3.23 | 17.53 | 14.25 | Membrane protein |
| *PSPTO_0095* | | - | 115037 | 116665 | 17.67 | 9.38 | 9.27 | 5.01 | 18.26 | 14.88 | Phospholipase D |
| *PSPTO_0096* | | - | 116690 | 118111 | 58.73 | 62.12 | 50.96 | 23.96 | 45.86 | 63.89 | GntR family transcriptional regulator/aminotransferase, class I |
| *PSPTO_0097* | | + | 118233 | 118538 | 60.87 | 8.72 | 35.68 | 19.31 | 18.94 | 51.32 | Hypothetical protein |
| *PSPTO_0098* | | - | 118544 | 119857 | 13.50 | 23.24 | 25.17 | 9.74 | 40.45 | 19.91 | Hypothetical protein |
| *PSPTO_0099* | | + | 119909 | 120316 | 16.70 | 32.18 | 30.86 | 8.40 | 68.88 | 21.33 | Hypothetical protein |
| *PSPTO_0100* | *lrp* | - | 120331 | 120819 | 7.63 | 2.49 | 17.92 | 4.53 | 33.57 | 10.38 | Leucine-responsive regulatory protein |
| *PSPTO_0101* | *dadA* | + | 120976 | 122277 | 127.83 | 64.29 | 73.02 | 32.66 | 619.25 | 424.77 | D-amino acid dehydrogenase small subunit |
| *PSPTO_0102* | | + | 122249 | 122602 | 65.37 | 73.09 | 60.86 | 31.52 | 435.32 | 315.94 | Endoribonuclease L-PSP family protein |
| *PSPTO_0103* | *dadX* | + | 122682 | 123755 | 14.13 | 9.41 | 12.81 | 3.51 | 182.13 | 111.55 | Alanine racemase |
| *PSPTO_0104* | | - | 123740 | 123844 | 7.64 | 0.00 | 27.40 | 0.00 | 80.48 | 77.24 | Hypothetical protein |
| *PSPTO_0105* | | + | 123831 | 124430 | 9.31 | 10.47 | 10.76 | 3.69 | 91.58 | 54.66 | DNA-binding protein |
| *PSPTO_0106* | *cycB* | + | 124455 | 124778 | 4.97 | 3.87 | 2.84 | 2.41 | 36.35 | 16.83 | Cytochrome c5 |
| *PSPTO_0107* | | + | 124875 | 126806 | 13.71 | 15.97 | 13.71 | 10.29 | 39.96 | 39.07 | Hypothetical protein |
| *PSPTO_0108* | | - | 126885 | 127703 | 2.93 | 3.78 | 9.62 | 1.34 | 12.94 | 61.57 | Membrane protein |
| *PSPTO_0109* | | - | 127696 | 128949 | 10.08 | 5.55 | 11.86 | 10.39 | 42.32 | 180.53 | Hypothetical protein |
| *PSPTO_0110* | | - | 128977 | 129861 | 28.17 | 9.82 | 15.30 | 8.95 | 150.53 | 367.87 | Metallo-beta-lactamase superfamily protein |
| *PSPTO_0111* | | + | 130125 | 131444 | 7.53 | 6.11 | 6.49 | 5.19 | 14.09 | 3.23 | Sigma-54 dependent transcriptional regulator |
| *PSPTO_0112* | *xpt* | - | 131808 | 132377 | 2.97 | 2.68 | 0.56 | 1.20 | 31.65 | 5.66 | Xanthine phosphoribosyltransferase |
| *PSPTO_0113* | *rep* | - | 132432 | 134441 | 8.94 | 27.01 | 20.28 | 10.95 | 68.50 | 19.45 | ATP-dependent DNA helicase Rep |
| *PSPTO_0114* | | + | 134647 | 136311 | 6.30 | 2.34 | 3.31 | 2.61 | 23.41 | 28.24 | GGDEF domain/EAL domain protein |
| *PSPTO_0115* | *norA* | - | 136326 | 137708 | 30.78 | 15.06 | 13.67 | 7.47 | 57.28 | 159.24 | Multidrug efflux protein NorA |
| *PSPTO_0116* | | + | 137865 | 138785 | 7.71 | 9.83 | 3.36 | 3.61 | 14.66 | 6.45 | LysR family transcriptional regulator |
| *PSPTO_0117* | | + | 138969 | 140945 | 39.04 | 13.89 | 13.29 | 10.91 | 16.76 | 8.31 | Methyl-accepting chemotaxis protein |
| *PSPTO_0118* | | + | 141265 | 142917 | 6.33 | 7.40 | 3.13 | 1.40 | 14.90 | 7.60 | Na/Pi cotransporter family protein |
| *PSPTO_0119* | | + | 142975 | 144354 | 16.98 | 22.63 | 19.57 | 11.95 | 21.02 | 28.14 | Peptidase, M16 family |
| *PSPTO_0120* | | + | 144451 | 145638 | 8.27 | 16.53 | 16.94 | 5.55 | 31.04 | 18.35 | Membrane protein |
| *PSPTO_0121* | | - | 145643 | 146884 | 5.73 | 14.24 | 8.75 | 4.43 | 25.24 | 10.45 | Membrane protein |
| *PSPTO_0123* | | + | 147445 | 147960 | 14.71 | 19.80 | 18.32 | 13.60 | 38.00 | 18.89 | ISPsy8, transposase OrfA |
| *PSPTO_0124* | | + | 148017 | 148796 | 16.66 | 17.21 | 12.20 | 9.73 | 25.38 | 16.94 | ISPsy8, transposase OrfB |
| *PSPTO_0125* | *argH* | - | 148983 | 150377 | 11.41 | 43.23 | 35.16 | 13.70 | 114.39 | 21.16 | Argininosuccinate lyase |
| *PSPTO_0126* | *algZ* | + | 150512 | 151594 | 90.78 | 41.06 | 45.70 | 22.87 | 48.22 | 51.94 | Alginate biosynthesis protein AlgZ/FimS |
| *PSPTO_0127* | *algR* | + | 151591 | 152337 | 52.54 | 47.80 | 43.86 | 19.95 | 52.91 | 48.60 | Alginate biosynthesis regulatory protein AlgR |
| *PSPTO_0128* | *hemC* | + | 152560 | 153501 | 8.17 | 31.27 | 28.00 | 14.35 | 54.90 | 20.33 | Porphobilinogen deaminase |
| *PSPTO_0129* | *hemD* | + | 153498 | 154274 | 29.23 | 89.65 | 72.35 | 20.59 | 106.76 | 63.62 | Uroporphyrinogen-III synthase |
| *PSPTO_0130* | | + | 154316 | 155446 | 18.11 | 83.12 | 69.43 | 26.75 | 121.62 | 60.45 | Uroporphyrin-III C-methyltransferase |
| *PSPTO_0131* | | + | 155443 | 156687 | 7.44 | 28.46 | 16.64 | 13.53 | 50.75 | 21.51 | HemY protein |
| *PSPTO_0132* | | + | 156902 | 157429 | 12.57 | 8.29 | 6.82 | 4.97 | 20.31 | 9.62 | DsbB family protein |
| *PSPTO_0133* | | + | 157434 | 157586 | 664.25 | 1085.61 | 965.91 | 770.34 | 972.33 | 416.24 | Hypothetical protein |
| *PSPTO_0134* | *algQ* | + | 157604 | 158077 | 59.26 | 167.29 | 157.62 | 108.63 | 106.98 | 69.40 | Transcriptional regulator AlgQ |
| *PSPTO_0135* | | - | 158162 | 158836 | 27.99 | 22.54 | 19.85 | 23.85 | 59.56 | 26.72 | FKBP-type peptidyl-prolyl cis-trans isomerase |
| *PSPTO_0136* | *algR3* | + | 159053 | 160009 | 107.31 | 259.74 | 259.79 | 132.84 | 643.75 | 386.19 | Alginate regulatory protein AlgR3 |
| *PSPTO_0137* | | - | 160006 | 160494 | 21.21 | 9.70 | 30.09 | 11.13 | 45.28 | 20.87 | Hypothetical protein |
| *PSPTO_0138* | | + | 160496 | 162451 | 5.92 | 18.27 | 21.65 | 8.78 | 47.44 | 12.13 | ABC transporter ATP-binding protein |
| *PSPTO_0139* | | + | 162468 | 163061 | 5.97 | 15.52 | 7.59 | 4.93 | 22.77 | 7.50 | Hypothetical protein |
| *PSPTO_0140* | | + | 163128 | 163760 | 17.79 | 22.72 | 19.79 | 11.42 | 51.24 | 19.21 | Homoserine/homoserine lactone efflux protein |
| *PSPTO_0141* | | + | 163941 | 165839 | 7.79 | 7.42 | 13.80 | 7.95 | 16.45 | 10.70 | Lead uptake protein |
| *PSPTO_0142* | | + | 166054 | 166284 | 45.79 | 79.20 | 84.75 | 96.70 | 23.65 | 24.53 | Hypothetical protein |
| *PSPTO_0143* | | - | 166389 | 166676 | 15.93 | 38.31 | 21.95 | 15.44 | 42.32 | 34.04 | Hypothetical protein |
| *PSPTO_0144* | *prlC* | - | 166673 | 168724 | 14.51 | 64.89 | 52.95 | 22.50 | 89.00 | 33.97 | Oligopeptidase A |
| *PSPTO_0145* | | + | 168837 | 169382 | 20.88 | 36.28 | 18.68 | 17.64 | 75.82 | 24.78 | Hypothetical protein |
| *PSPTO_0146* | | - | 169523 | 170578 | 2.41 | 3.62 | 3.73 | 0.00 | 5.61 | 3.49 | Hypothetical protein |
| *PSPTO_0147* | | + | 170766 | 171197 | 10.82 | 7.56 | 17.47 | 6.64 | 37.65 | 15.30 | Hypothetical protein |
| *PSPTO_0148* | | + | 171406 | 173142 | 4.41 | 2.44 | 3.01 | 1.56 | 13.90 | 5.77 | Oligopeptide transporter |
| *PSPTO_0149* | | + | 173456 | 174133 | 19.25 | 22.24 | 22.34 | 13.42 | 51.87 | 31.34 | Hypothetical protein |
| *PSPTO_0150* | | + | 174152 | 174586 | 8.46 | 15.11 | 5.89 | 7.87 | 34.30 | 14.04 | Lipoprotein |
| *PSPTO_0151* | | - | 174587 | 174820 | 11.22 | 56.02 | 13.31 | 14.50 | 24.44 | 14.70 | Hypothetical protein |
| *PSPTO_0152* | *osmC* | - | 174938 | 175369 | 55.87 | 280.87 | 198.10 | 91.17 | 60.95 | 110.61 | Osmotically inducible protein |
| *PSPTO_0153* | | + | 175580 | 176581 | 15.70 | 21.54 | 13.23 | 6.41 | 24.43 | 6.48 | Luciferase family protein |
| *PSPTO_0154* | | - | 176728 | 176952 | 18.75 | 58.00 | 91.87 | 33.21 | 15.13 | 20.62 | Hypothetical protein |
| *PSPTO_0155* | | - | 177102 | 177317 | 19.05 | 138.89 | 110.84 | 62.18 | 34.03 | 46.47 | Hypothetical protein |
| *PSPTO_0156* | | + | 177485 | 177811 | 217.29 | 729.00 | 551.52 | 196.66 | 462.40 | 308.20 | Hypothetical protein |

| Locus name | Gene name | Strand | Gene start (bp) | Gene end (bp) | Average TPM Inoculum | Average TPM $t_6$ | Average TPM $t_{24}$ | Average TPM $t_{168}$ | Average TPM Logarithmic | Average TPM Stationary | Annotation description |
|---|---|---|---|---|---|---|---|---|---|---|---|
| *PSPTO_0157* | *trpI* | - | 177812 | 178720 | 4.04 | 6.13 | 7.58 | 4.94 | 12.62 | 6.68 | trpBA operon transcriptional activator |
| *PSPTO_0158* | *trpB* | + | 178848 | 180077 | 12.98 | 38.09 | 34.87 | 17.76 | 113.47 | 32.44 | Tryptophan synthase subunit beta |
| *PSPTO_0159* | *trpA* | + | 180074 | 180886 | 8.66 | 20.98 | 22.30 | 6.30 | 50.23 | 23.99 | Tryptophan synthase subunit alpha |
| *PSPTO_0161* | | + | 181013 | 181528 | 113.29 | 94.52 | 66.73 | 36.65 | 43.39 | 25.43 | Hypothetical protein |
| *PSPTO_0162* | | - | 181714 | 182538 | 20.73 | 69.98 | 54.71 | 37.53 | 147.90 | 37.10 | 3-oxoadipate enol-lactone hydrolase family protein |
| *PSPTO_0163* | | - | 182751 | 183083 | 5.08 | 9.01 | 3.52 | 6.51 | 17.73 | 8.05 | Hypothetical protein |
| *PSPTO_0164* | | - | 183857 | 184807 | 16.14 | 16.63 | 13.49 | 9.76 | 33.53 | 21.27 | LysR family transcriptional regulator |
| *PSPTO_0165* | | + | 184922 | 186427 | 14.25 | 20.50 | 19.98 | 6.37 | 94.93 | 22.90 | Sulfatase family protein |
| *PSPTO_0166* | | + | 186464 | 187390 | 25.22 | 49.01 | 31.05 | 19.05 | 109.16 | 32.90 | Glycine/betaine family, ABC transporter, substrate-binding protein |
| *PSPTO_0167* | | + | 187452 | 187874 | 146.45 | 293.59 | 211.55 | 115.02 | 258.85 | 138.99 | Hypothetical protein |
| *PSPTO_0168* | | + | 187946 | 189514 | 4.82 | 7.81 | 8.57 | 2.12 | 20.82 | 8.22 | Sulfate transporter family protein |
| *PSPTO_0169* | *aroE* | - | 189618 | 190442 | 6.30 | 17.62 | 8.13 | 12.66 | 26.10 | 8.46 | Shikimate 5-dehydrogenase |
| *PSPTO_0170* | *hemF* | - | 190442 | 191356 | 10.76 | 19.97 | 20.41 | 9.37 | 37.98 | 9.86 | Coproporphyrinogen III oxidase |
| *PSPTO_0171* | | + | 192549 | 193526 | 0.00 | 0.00 | 0.00 | 0.00 | 0.00 | 0.00 | ISPssy, transposase |
| *PSPTO_0172* | | + | 194114 | 194407 | 3.55 | 9.11 | 13.05 | 2.22 | 19.52 | 6.38 | Hypothetical protein |
| *PSPTO_0173* | *qor* | + | 194503 | 195480 | 67.91 | 97.68 | 70.76 | 50.47 | 156.00 | 106.35 | Quinone oxidoreductase |
| *PSPTO_0174* | | - | 195488 | 196045 | 19.67 | 52.47 | 39.49 | 13.31 | 85.02 | 27.77 | SUA5/yciO/yrdC family domain protein |
| *PSPTO_0175* | | - | 196084 | 197268 | 2.98 | 4.58 | 0.92 | 2.74 | 20.46 | 6.05 | DNA processing protein DprA |
| *PSPTO_0176* | | - | 197323 | 198348 | 17.02 | 17.97 | 18.29 | 7.06 | 71.55 | 17.63 | LysM domain-containing protein |
| *PSPTO_0177* | *def-1* | + | 198474 | 198980 | 28.94 | 31.29 | 49.37 | 17.57 | 95.72 | 30.77 | Polypeptide deformylase |
| *PSPTO_0178* | *fmt* | + | 199041 | 199985 | 2.67 | 10.22 | 5.80 | 1.77 | 50.48 | 8.25 | Methionyl-tRNA formyltransferase |
| *PSPTO_0179* | *sun* | + | 199982 | 201328 | 3.33 | 6.55 | 9.76 | 3.78 | 73.13 | 8.28 | Sun protein |
| *PSPTO_0180* | *trkA* | + | 201325 | 202698 | 8.24 | 11.59 | 10.66 | 6.28 | 57.26 | 10.65 | Trk system potassium uptake protein TrkA |
| *PSPTO_0181* | | + | 202781 | 205195 | 4.52 | 6.91 | 4.20 | 4.53 | 9.34 | 5.60 | Molybdopterin oxidoreductase subunit alpha |
| *PSPTO_0182* | | - | 205278 | 206165 | 7.80 | 29.19 | 33.16 | 12.65 | 63.57 | 23.30 | Lipid A biosynthesis lauroyl acyltransferase |
| *PSPTO_0183* | *tag* | - | 206203 | 206754 | 15.77 | 20.40 | 16.64 | 16.19 | 33.72 | 25.09 | DNA-3-methyladenine glycosidase I |
| *PSPTO_0184* | *glyQ* | + | 206834 | 207781 | 15.36 | 59.19 | 48.91 | 31.85 | 140.85 | 50.66 | Glycyl-tRNA synthetase subunit alpha |
| *PSPTO_0185* | *glyS* | + | 207778 | 209832 | 5.19 | 31.40 | 20.59 | 21.24 | 121.58 | 35.34 | Glycyl-tRNA synthetase subunit beta |
| *PSPTO_0186* | | + | 209846 | 210397 | 6.53 | 16.85 | 14.47 | 4.69 | 41.84 | 13.63 | D,D-heptose 1,7-bisphosphate phosphatase |
| *PSPTO_0187* | *hdtS* | + | ` | 211225 | 3.61 | 5.43 | 2.23 | 0.00 | 39.42 | 9.88 | HdtS protein |
| *PSPTO_0188* | | - | 211395 | 212333 | - | - | - | - | - | - | |
| *PSPTO_0189* | | + | 212468 | 213511 | 2.82 | 7.23 | 2.64 | 0.00 | 3.93 | 2.77 | Nitrilase |
| *PSPTO_0190* | | + | 213513 | 214433 | - | - | - | - | - | - | |
| *PSPTO_0191* | | + | 214448 | 214963 | - | - | - | - | - | - | |
| *PSPTO_0192* | | + | 215807 | 217363 | 5.26 | 9.28 | 10.48 | 7.67 | 12.96 | 6.98 | Recombinase |
| *PSPTO_0193* | | - | 217450 | 218259 | 0.00 | 0.00 | 0.00 | 0.00 | 0.00 | 0.00 | ISPsy4, transposition helper protein |
| *PSPTO_0194* | | - | 218256 | 219278 | 0.00 | 0.00 | 0.00 | 0.00 | 0.00 | 0.00 | ISPsy4, transposase |
| *PSPTO_0196* | | - | 219554 | 221107 | 0.00 | 0.00 | 0.00 | 0.00 | 0.00 | 0.00 | ISPsy5, transposase |
| *PSPTO_0197* | | - | 221125 | 221484 | 0.00 | 0.00 | 0.00 | 0.00 | 0.00 | 0.00 | ISPsy5, Orf1 |
| *PSPTO_0200* | | - | 222528 | 223745 | 3.01 | 4.95 | 2.12 | 4.05 | 24.26 | 3.19 | Hypothetical protein |
| *PSPTO_0201* | | - | 224484 | 225383 | 2.34 | 9.69 | 6.99 | 5.52 | 64.20 | 5.14 | Hypothetical protein |
| *PSPTO_0202* | | - | 225386 | 226549 | 4.46 | 17.26 | 10.40 | 5.47 | 67.64 | 13.23 | Membrane protein |
| *PSPTO_0203* | | - | 226554 | 227672 | 1.83 | 5.47 | 2.04 | 4.82 | 39.52 | 3.04 | Cysteine synthase |
| *PSPTO_0205* | | - | 231080 | 232075 | 2.66 | 9.30 | 11.88 | 6.56 | 27.97 | 24.68 | Hypothetical protein |
| *PSPTO_0206* | | - | 232469 | 232945 | 234.39 | 224.57 | 172.05 | 74.45 | 152.10 | 202.38 | Hypothetical protein |
| *PSPTO_0207* | | - | 233124 | 233427 | - | - | - | - | - | - | |
| *PSPTO_0208* | | - | 233476 | 234099 | 13.80 | 14.02 | 28.42 | 5.89 | 54.36 | 16.83 | Hypothetical protein |
| *PSPTO_0209* | *lsyA-1* | - | 234287 | 235525 | 2.11 | 1.37 | 2.25 | 2.28 | 5.25 | 2.95 | Diaminopimelate decarboxylase |
| *PSPTO_0210* | | - | 235527 | 236402 | 1.49 | 0.00 | 3.84 | 0.21 | 4.21 | 2.47 | Membrane protein |
| *PSPTO_0211* | *iucD* | - | 236402 | 237685 | 6.22 | 4.09 | 0.13 | 0.83 | 4.84 | 3.12 | L-lysine 6-monooxygenase |
| *PSPTO_0212* | *lysR* | + | 237840 | 238775 | 3.60 | 8.94 | 15.25 | 6.53 | 11.87 | 5.08 | DNA-binding transcriptional regulator LysR |
| *PSPTO_0213* | | + | 239026 | 239304 | 115.71 | 85.43 | 54.27 | 44.25 | 64.88 | 44.49 | Proteic killer protein |
| *PSPTO_0214* | | + | 239314 | 239619 | 30.18 | 30.52 | 15.08 | 13.95 | 35.38 | 20.82 | Virulence-associated protein |
| *PSPTO_0215* | | - | 239653 | 239934 | 42.03 | 91.93 | 111.65 | 2.60 | 86.02 | 36.96 | Hypothetical protein |
| *PSPTO_0216* | | + | 240015 | 240155 | 134.44 | 50.65 | 29.38 | 30.88 | 100.01 | 76.21 | Hypothetical protein |
| *PSPTO_0217* | *glnK* | + | 240354 | 240692 | 139.02 | 331.17 | 214.17 | 75.60 | 227.14 | 86.90 | Nitrogen regulatory protein P-II |
| *PSPTO_0218* | *amt-1* | + | 240728 | 242065 | 5.12 | 32.73 | 25.00 | 11.34 | 16.11 | 7.29 | Ammonium transporter |
| *PSPTO_0219* | | + | 242307 | 242732 | 169.21 | 113.55 | 117.04 | 88.97 | 251.50 | 184.56 | Hypothetical protein |
| *PSPTO_0220* | | + | 242831 | 243160 | 456.80 | 174.75 | 190.55 | 94.31 | 457.53 | 239.25 | Hypothetical protein |
| *PSPTO_0221* | | - | 243235 | 243927 | 14.18 | 16.87 | 8.19 | 8.19 | 21.09 | 21.03 | HAD-superfamily hydrolase |
| *PSPTO_0222* | *xerC* | - | 243920 | 244819 | 16.11 | 21.82 | 13.92 | 17.37 | 40.40 | 22.13 | integrase/recombinase XerC |
| *PSPTO_0223* | | - | 244822 | 245556 | 7.74 | 19.57 | 10.79 | 9.51 | 76.05 | 20.66 | Hypothetical protein |
| *PSPTO_0224* | *dapF* | - | 245570 | 246400 | 15.73 | 30.24 | 26.22 | 11.89 | 108.35 | 29.31 | Diaminopimelate epimerase |
| *PSPTO_0225* | *lsyA-2* | - | 246405 | 247652 | 11.98 | 51.27 | 28.48 | 15.17 | 146.83 | 39.86 | Diaminopimelate decarboxylase |
| *PSPTO_0226* | *lppL* | - | 247662 | 247808 | 86.79 | 178.22 | 83.18 | 27.01 | 201.39 | 69.80 | Lipoprotein LppL |
| *PSPTO_0227* | *cyaY* | + | 248015 | 248347 | 12.20 | 14.58 | 23.64 | 21.38 | 33.56 | 15.13 | CyaY protein |
| *PSPTO_0228* | | + | 248350 | 248586 | 5.31 | 19.83 | 15.48 | 0.00 | 8.22 | 4.54 | Hypothetical protein |
| *PSPTO_0229* | *rnk* | + | 248724 | 249131 | 3.45 | 10.28 | 1.49 | 8.30 | 17.22 | 3.02 | Regulator of nucleoside diphosphate kinase |
| *PSPTO_0230* | *cyaA* | - | 249596 | 252442 | 4.96 | 4.42 | 5.37 | 1.84 | 6.63 | 4.22 | Adenylate cyclase |
| *PSPTO_0231* | | - | 252625 | 252873 | 37.11 | 51.22 | 51.85 | 13.54 | 39.42 | 35.73 | Hypothetical protein |
| *PSPTO_0232* | | - | 253090 | 253365 | 23.22 | 108.88 | 74.76 | 61.73 | 226.87 | 50.90 | Hypothetical protein |
| *PSPTO_0233* | *gltP* | - | 253459 | 254790 | 5.28 | 5.45 | 5.63 | 3.35 | 16.57 | 5.67 | Proton/glutamate symporter |
| *PSPTO_0234* | *rimK* | - | 255243 | 256148 | 8.74 | 20.75 | 21.83 | 10.39 | 55.15 | 40.32 | Ribosomal protein S6 modification protein |
| *PSPTO_0235* | | - | 256145 | 256621 | 6.09 | 13.18 | 12.59 | 9.11 | 32.18 | 11.59 | Hypothetical protein |
| *PSPTO_0236* | | - | 256638 | 257420 | 28.24 | 46.37 | 27.43 | 15.69 | 40.80 | 23.26 | EAL domain-containing protein |
| *PSPTO_0237* | *hslR* | + | 257588 | 257995 | 7.14 | 12.92 | 15.78 | 9.20 | 39.72 | 8.58 | Heat shock protein 15 |
| *PSPTO_0238* | *hslO* | + | 258156 | 259058 | 17.86 | 26.33 | 23.96 | 21.03 | 109.70 | 46.65 | Chaperonin, 33 kDa |
| *PSPTO_0239* | *pckA* | + | 259238 | 260782 | 26.25 | 65.22 | 74.73 | 51.26 | 206.10 | 39.42 | Phosphoenolpyruvate carboxykinase |
| *PSPTO_0240* | | + | 261079 | 261330 | 57.52 | 185.91 | 163.45 | 34.71 | 93.48 | 41.60 | Prevent-host-death family protein |

| Locus name | Gene name | Strand | Gene start (bp) | Gene end (bp) | Average TPM Inoculum | Average TPM $t_6$ | Average TPM $t_{24}$ | Average TPM $t_{168}$ | Average TPM Logarithmic | Average TPM Stationary | Annotation description |
|---|---|---|---|---|---|---|---|---|---|---|---|
| *PSPTO_0241* | | + | 261343 | 261693 | 3.74 | 41.52 | 50.37 | 19.85 | 20.81 | 8.16 | Hypothetical protein |
| *PSPTO_0242* | | - | 262839 | 263411 | 1.56 | 3.05 | 1.49 | 0.00 | 1.47 | 1.01 | Hypothetical protein |
| *PSPTO_0243* | | - | 263550 | 264371 | 4.23 | 8.69 | 3.81 | 0.22 | 2.09 | 1.12 | Hypothetical protein |
| *PSPTO_0244* | | - | 265284 | 265997 | 9.87 | 19.90 | 25.27 | 0.52 | 33.63 | 10.46 | Hypothetical protein |
| *PSPTO_0245* | | - | 266158 | 267015 | 7.57 | 14.50 | 17.60 | 2.76 | 16.19 | 6.87 | GNAT family acetyltransferase |
| *PSPTO_0246* | | - | 267161 | 267574 | 17.46 | 13.95 | 60.43 | 2.96 | 35.01 | 10.52 | Hypothetical protein |
| *PSPTO_0247* | *dehII* | + | 267549 | 268217 | 49.50 | 89.44 | 110.35 | 20.78 | 76.71 | 21.48 | Haloacid dehalogenase |
| *PSPTO_0248* | | + | 268284 | 269558 | 13.15 | 14.44 | 17.53 | 7.80 | 15.18 | 6.30 | Oxidoreductase, FAD-binding protein |
| *PSPTO_0249* | | + | 269650 | 271287 | 7.20 | 16.09 | 18.96 | 3.24 | 9.68 | 6.70 | Peptide ABC transporter periplasmic peptide-binding protein |
| *PSPTO_0250* | | + | 271319 | 272275 | 3.62 | 4.75 | 6.33 | 0.41 | 3.14 | 3.27 | Peptide ABC transporter permease |
| *PSPTO_0251* | | + | 272272 | 273150 | 2.92 | 11.02 | 11.37 | 2.92 | 3.08 | 3.61 | Peptide ABC transporter permease |
| *PSPTO_0252* | | + | 273152 | 274993 | 4.88 | 11.05 | 13.43 | 4.97 | 6.23 | 5.28 | Peptide ABC transporter ATP-binding protein |
| *PSPTO_0253* | | + | 274990 | 275301 | 7.96 | 13.85 | 6.30 | 5.48 | 7.44 | 7.02 | Hypothetical protein |
| *PSPTO_0254* | | + | 275280 | 276689 | 5.47 | 6.46 | 15.60 | 0.02 | 5.45 | 4.05 | Pyridine nucleotide-disulfide oxidoreductase family protein |
| *PSPTO_0255* | | + | 276686 | 277840 | 9.31 | 11.58 | 17.92 | 3.38 | 15.05 | 10.11 | Oxidoreductase, FAD-binding protein |
| *PSPTO_0256* | *argE-1* | + | 277837 | 278994 | 6.09 | 7.92 | 7.19 | 4.68 | 8.71 | 7.70 | Acetylornithine deacetylase |
| *PSPTO_0257* | *gabD-1* | + | 279109 | 280566 | 7.19 | 21.48 | 38.49 | 4.59 | 17.78 | 13.55 | Succinate-semialdehyde dehydrogenase |
| *PSPTO_0258* | | - | 280829 | 281794 | 3.80 | 10.47 | 5.01 | 2.01 | 6.60 | 4.17 | D-isomer specific 2-hydroxyacid dehydrogenase family protein |
| *PSPTO_0259* | *gabT-1* | + | 282043 | 283347 | 4.09 | 20.29 | 28.12 | 4.93 | 12.28 | 4.09 | 4-aminobutyrate aminotransferase |
| *PSPTO_0260* | | + | 283423 | 284079 | 8.01 | 11.18 | 20.58 | 2.58 | 4.57 | 3.55 | Hypothetical protein |
| *PSPTO_0261* | | + | 284127 | 284615 | 34.47 | 54.51 | 40.38 | 18.37 | 60.21 | 16.73 | AsnC family transcriptional regulator |
| *PSPTO_0262* | | - | 284644 | 285102 | 4.73 | 7.07 | 6.49 | 3.97 | 10.69 | 7.89 | Hypothetical protein |
| *PSPTO_0263* | | + | 285536 | 287032 | 2.88 | 5.98 | 3.72 | 1.83 | 7.12 | 2.90 | Methyl-accepting chemotaxis protein |
| *PSPTO_0264* | | - | 287061 | 288644 | 0.92 | 1.35 | 1.28 | 0.46 | 2.24 | 2.17 | Peptide ABC transporter ATP-binding protein |
| *PSPTO_0265* | | - | 288646 | 289668 | 1.65 | 0.29 | 2.47 | 0.00 | 1.79 | 1.45 | Peptide ABC transporter permease |
| *PSPTO_0266* | | - | 289668 | 290735 | 1.58 | 1.47 | 0.61 | 0.00 | 1.12 | 1.35 | Peptide ABC transporter permease |
| *PSPTO_0267* | | - | 290737 | 292653 | 1.58 | 0.72 | 0.13 | 1.40 | 1.75 | 1.57 | Family 5 extracellular solute-binding protein |
| *PSPTO_0268* | | - | 292765 | 295398 | 2.49 | 3.42 | 2.17 | 1.38 | 3.64 | 2.37 | TonB system transport protein |
| *PSPTO_0269* | | - | 295690 | 296091 | 0.85 | 6.09 | 7.16 | 0.00 | 1.75 | 0.66 | Biopolymer transport protein ExbD |
| *PSPTO_0270* | | - | 296091 | 296798 | 1.16 | 3.45 | 0.00 | 0.37 | 1.66 | 1.61 | Biopolymer transport protein ExbB |
| *PSPTO_0271* | | - | 296849 | 297670 | 1.45 | 4.73 | 1.00 | 4.26 | 3.15 | 2.01 | TonB protein |
| *PSPTO_0272* | *iscS-1* | - | 298850 | 300016 | 26.80 | 33.66 | 64.10 | 16.89 | 41.42 | 24.84 | Cysteine desulfurase |
| *PSPTO_0273* | | + | 300484 | 300954 | 12.82 | 5.98 | 8.62 | 3.70 | 41.32 | 8.83 | AsnC family transcriptional regulator |
| *PSPTO_0274* | | + | 301038 | 301292 | 6.56 | 31.31 | 23.93 | 7.42 | 13.18 | 10.05 | Hypothetical protein |
| *PSPTO_0275* | | - | 301329 | 302462 | 5.35 | 9.10 | 7.97 | 8.77 | 30.55 | 10.67 | DNA-binding protein |
| *PSPTO_0276* | | - | 302698 | 303681 | 11.38 | 22.10 | 15.19 | 12.27 | 21.03 | 10.46 | Haloacid dehalogenase-like family hydrolase |
| *PSPTO_0277* | | + | 303927 | 304415 | 2.92 | 5.23 | 5.64 | 3.47 | 16.05 | 5.63 | Hypothetical protein |
| *PSPTO_0278* | | - | 304528 | 304968 | 3.91 | 14.63 | 11.52 | 0.00 | 17.05 | 16.95 | Hypothetical protein |
| *PSPTO_0279* | | - | 304985 | 305341 | 5.88 | 10.47 | 20.01 | 2.59 | 38.48 | 11.05 | Hypothetical protein |
| *PSPTO_0280* | | - | 305562 | 306266 | 9.17 | 7.15 | 6.57 | 6.00 | 37.87 | 17.41 | Methionine aminopeptidase |
| *PSPTO_0281* | | - | 306711 | 307076 | 17.88 | 46.07 | 45.95 | 36.76 | 27.01 | 19.37 | Hypothetical protein |
| *PSPTO_0282* | | - | 307260 | 307940 | 2.45 | 2.66 | 5.09 | 3.53 | 6.28 | 3.75 | Membrane protein |
| *PSPTO_0283* | | - | 308181 | 308624 | 9.74 | 2.07 | 0.28 | 5.62 | 48.19 | 3.17 | Hypothetical protein |
| *PSPTO_0284* | | + | 308896 | 309108 | 7.89 | 9.69 | 4.62 | 7.27 | 5.11 | 5.02 | Hypothetical protein |
| *PSPTO_0285* | | - | 309736 | 310818 | 2.63 | 10.27 | 4.60 | 8.24 | 55.13 | 12.06 | Hypothetical protein |
| *PSPTO_0286* | | - | 310815 | 312821 | - | - | - | - | - | - | |
| *PSPTO_0287* | | + | 314157 | 315395 | 20.82 | 20.85 | 20.81 | 2.32 | 41.92 | 25.86 | Site-specific recombinase, phage integrase family |
| *PSPTO_0288* | | + | 316071 | 316271 | 7.59 | 10.45 | 13.95 | 6.47 | 53.93 | 5.45 | Excisionase |
| *PSPTO_0289* | | + | 316268 | 316423 | 15.51 | 14.33 | 9.82 | 12.53 | 132.57 | 9.34 | Hypothetical protein |
| *PSPTO_0290* | | + | 316546 | 316782 | 10.86 | 8.69 | 0.00 | 11.00 | 34.75 | 6.37 | Hypothetical protein |
| *PSPTO_0291* | | - | 316980 | 317789 | 0.00 | 0.00 | 0.00 | 0.00 | 0.00 | 0.00 | ISPsy4, transposition helper protein |
| *PSPTO_0292* | | - | 317786 | 318808 | 0.00 | 0.00 | 0.00 | 0.00 | 0.00 | 0.00 | ISPsy4, transposase |
| *PSPTO_0293* | | + | 319381 | 319689 | 20.22 | 17.28 | 24.71 | 6.80 | 25.97 | 15.97 | Hypothetical protein |
| *PSPTO_0294* | | + | 319703 | 319903 | 37.27 | 48.30 | 45.07 | 17.48 | 40.55 | 36.55 | Hypothetical protein |
| *PSPTO_0295* | | - | 320707 | 320928 | 24.69 | 27.47 | 54.67 | 0.00 | 37.51 | 61.97 | Hypothetical protein |
| *PSPTO_0296* | | + | 322507 | 322830 | 167.00 | 276.84 | 172.25 | 144.36 | 796.36 | 203.21 | Hypothetical protein |
| *PSPTO_0297* | | + | 322827 | 323117 | 14.92 | 29.18 | 1.35 | 3.16 | 77.74 | 19.96 | Hypothetical protein |
| *PSPTO_0298* | | + | 323239 | 323856 | 63.55 | 26.85 | 51.98 | 15.80 | 63.87 | 13.13 | Resolvase |
| *PSPTO_0299* | | + | 324439 | 325494 | 5.34 | 10.28 | 7.14 | 2.53 | 12.81 | 5.37 | Monooxygenase |
| *PSPTO_0300* | *gabD-2* | + | 325765 | 327207 | 19.97 | 37.66 | 45.07 | 31.13 | 122.71 | 23.15 | Succinate-semialdehyde dehydrogenase |
| *PSPTO_0301* | *gabT-2* | + | 327391 | 328671 | 13.72 | 27.37 | 29.26 | 17.32 | 103.81 | 19.44 | 4-aminobutyrate aminotransferase |
| *PSPTO_0302* | | + | 328701 | 328877 | 2.53 | 4.26 | 3.04 | 0.74 | 9.65 | 2.36 | Hypothetical protein |
| *PSPTO_0303* | | + | 328831 | 330219 | 16.09 | 26.20 | 19.92 | 14.97 | 66.62 | 20.23 | Response regulator |
| *PSPTO_0304* | | - | 330437 | 331435 | 13.82 | 33.91 | 35.87 | 16.77 | 27.48 | 93.04 | Sensory box/GGDEF domain protein |
| *PSPTO_0305* | | - | 331569 | 332807 | 60.59 | 171.31 | 126.49 | 33.84 | 232.07 | 42.84 | Fatty acid desaturase |
| *PSPTO_0306* | | + | 332910 | 334841 | 6.49 | 13.85 | 6.15 | 8.70 | 34.61 | 16.56 | Sensory box/GGDEF domain/EAL domain-containing protein |
| *PSPTO_0307* | | + | 335032 | 335214 | 8.23 | 649.24 | 529.09 | 367.52 | 39.29 | 15.84 | Hypothetical protein |
| *PSPTO_0308* | *sbp* | + | 335361 | 336377 | 4.36 | 115.00 | 103.27 | 96.60 | 16.64 | 4.21 | Sulfate-binding protein |
| *PSPTO_0309* | *cysT* | + | 336472 | 337293 | 2.20 | 11.32 | 5.23 | 6.64 | 10.40 | 2.97 | Sulfate ABC transporter permease CysT |
| *PSPTO_0310* | *cysW* | + | 337307 | 338179 | 4.53 | 10.97 | 12.44 | 7.79 | 13.25 | 7.61 | Sulfate ABC transporter permease CysW |
| *PSPTO_0311* | *cysA* | + | 338182 | 339162 | 2.27 | 4.24 | 4.00 | 4.49 | 8.77 | 2.43 | Sulfate ABC transporter, ATP-binding protein CysA |
| *PSPTO_0312* | | - | 339425 | 339949 | 14.47 | 27.02 | 16.39 | 16.34 | 15.06 | 9.55 | Hypothetical protein |
| *PSPTO_0313* | | + | 340034 | 340738 | 33.33 | 7.88 | 28.49 | 16.53 | 21.70 | 17.47 | Cyclic nucleotide-binding protein |
| *PSPTO_0314* | | + | 340857 | 341858 | 158.63 | 88.79 | 75.02 | 35.71 | 191.00 | 119.18 | Iron ABC transporter periplasmic iron-binding protein |
| *PSPTO_0315* | | + | 342066 | 343682 | 5.32 | 3.62 | 8.38 | 2.12 | 37.64 | 9.02 | Iron ABC transporter permease |
| *PSPTO_0316* | *gcvT-1* | + | 343836 | 344918 | 30.47 | 81.27 | 69.65 | 47.46 | 160.83 | 56.75 | Glycine cleavage system T protein |
| *PSPTO_0317* | *gcvH-1* | + | 344962 | 345348 | 11.34 | 30.99 | 39.30 | 19.56 | 51.86 | 16.85 | Glycine cleavage system H protein |
| *PSPTO_0318* | | - | 345431 | 345751 | 418.56 | 922.28 | 778.68 | 236.78 | 232.72 | 251.25 | Hypothetical protein |
| *PSPTO_0319* | | - | 345891 | 347675 | 22.81 | 18.71 | 17.92 | 8.99 | 16.61 | 15.75 | Type IV pilus biogenesis protein |

| Locus name | Gene name | Strand | Gene start (bp) | Gene end (bp) | Average TPM Inoculum | Average TPM $t_6$ | Average TPM $t_{24}$ | Average TPM $t_{168}$ | Average TPM Logarithmic | Average TPM Stationary | Annotation description |
|---|---|---|---|---|---|---|---|---|---|---|---|
| *PSPTO_0320* | | - | 347786 | 348133 | 6.19 | 2.30 | 0.70 | 0.00 | 10.92 | 3.68 | Hypothetical protein |
| *PSPTO_0321* | | + | 348259 | 348738 | 7.49 | 8.17 | 8.86 | 18.45 | 15.68 | 5.61 | AsnC family transcriptional regulator |
| *PSPTO_0322* | | - | 349816 | 351231 | 10.27 | 47.11 | 24.16 | 15.38 | 86.68 | 23.69 | Adenylate cyclase |
| *PSPTO_0323* | *argE-2* | + | 351346 | 352494 | 27.82 | 52.90 | 43.26 | 18.61 | 74.72 | 35.21 | Acetylornithine deacetylase |
| *PSPTO_0324* | *argA* | + | 352570 | 353916 | 6.33 | 15.11 | 12.88 | 6.99 | 26.28 | 13.05 | N-acetylglutamate synthase |
| *PSPTO_0325* | *gshA* | - | 354040 | 355629 | 16.48 | 32.64 | 24.76 | 18.77 | 100.32 | 32.07 | Glutamate-cysteine ligase |
| *PSPTO_0326* | | - | 355873 | 356256 | 2.27 | 10.74 | 13.70 | 0.00 | 47.57 | 5.77 | Hypothetical protein |
| *PSPTO_0327* | | - | 356256 | 358577 | 5.67 | 13.01 | 9.95 | 12.32 | 83.90 | 12.44 | S1 RNA binding domain-containing protein |
| *PSPTO_0328* | *ompR* | + | 358877 | 359617 | 30.08 | 144.04 | 104.25 | 13.99 | 149.50 | 23.49 | DNA-binding response regulator OmpR |
| *PSPTO_0329* | *envZ* | + | 359769 | 361091 | 16.44 | 40.28 | 20.24 | 14.21 | 88.55 | 30.81 | Osmolarity sensor protein EnvZ |
| *PSPTO_0330* | | - | 361341 | 361808 | 9.56 | 80.01 | 35.59 | 32.00 | 28.43 | 10.05 | Hypothetical protein |
| *PSPTO_0331* | | + | 361787 | 362011 | 16.89 | 250.71 | 136.47 | 52.89 | 39.29 | 16.60 | Hypothetical protein |
| *PSPTO_0332* | | - | 362035 | 362322 | 32.56 | 509.70 | 282.35 | 163.77 | 24.48 | 15.81 | Hypothetical protein |
| *PSPTO_0333* | | - | 362552 | 363391 | 16.93 | 52.50 | 52.19 | 20.17 | 33.83 | 39.95 | Hypothetical protein |
| *PSPTO_0334* | *algB* | + | 363589 | 364935 | 27.79 | 36.82 | 28.57 | 20.64 | 94.54 | 45.78 | Alginate biosynthesis transcriptional regulator AlgB |
| *PSPTO_0335* | | + | 364946 | 366733 | 15.34 | 24.76 | 19.92 | 10.21 | 67.17 | 34.24 | Sensory box histidine kinase |
| *PSPTO_0336* | | - | 366822 | 367070 | 26.33 | 41.70 | 26.22 | 0.00 | 45.42 | 29.17 | Hypothetical protein |
| *PSPTO_0337* | | - | 367096 | 367374 | 118.42 | 67.37 | 65.78 | 18.71 | 76.63 | 43.46 | Prevent-host-death family protein |
| *PSPTO_0338* | | - | 367634 | 368413 | 8.18 | 47.06 | 46.76 | 15.86 | 81.12 | 44.16 | N-acetylmuramoyl-L-alanine amidase family protein |
| *PSPTO_0339* | | - | 368509 | 370542 | 7.90 | 21.59 | 14.21 | 5.43 | 62.28 | 19.76 | Diguanylate cyclase |
| *PSPTO_0340* | | - | 370539 | 371414 | 16.96 | 43.86 | 48.67 | 27.48 | 240.53 | 63.22 | Endonuclease/exonuclease/phosphatase family protein |
| *PSPTO_0341* | *dsbA* | - | 371424 | 372068 | 63.54 | 65.23 | 79.55 | 43.48 | 312.40 | 131.39 | Thiol:disulfide interchange protein DsbA |
| *PSPTO_0342* | | - | 372072 | 372212 | 182.83 | 176.50 | 112.51 | 83.93 | 561.04 | 229.05 | Hypothetical protein |
| *PSPTO_0343* | *engB* | + | 372362 | 372997 | 15.12 | 19.85 | 9.52 | 4.61 | 70.59 | 18.84 | GTP-binding protein EngB |
| *PSPTO_0344* | *polI* | - | 373243 | 376020 | 5.14 | 15.14 | 17.02 | 10.85 | 83.31 | 26.33 | DNA polymerase I |
| *PSPTO_0345* | | + | 376083 | 376370 | 27.01 | 32.32 | 71.09 | 34.65 | 91.66 | 58.06 | Hypothetical protein |
| *PSPTO_0346* | *thrB* | + | 376407 | 377357 | 6.05 | 12.36 | 11.71 | 10.70 | 43.12 | 24.70 | Homoserine kinase |
| *PSPTO_0347* | | - | 377461 | 378507 | 3.66 | 5.98 | 9.82 | 1.23 | 11.89 | 5.49 | LacI family transcriptional regulator |
| *PSPTO_0348* | | + | 378728 | 380987 | - | - | - | - | - | - | |
| *PSPTO_0349* | | + | 381023 | 382348 | 1.67 | 3.43 | 2.90 | 1.28 | 5.08 | 2.51 | Xenobiotic compound monooxygenase, DszA family, A subunit |
| *PSPTO_0350* | | + | 382479 | 383414 | 4.86 | 13.45 | 12.95 | 10.80 | 39.61 | 11.01 | Alkanesulfonate transporter substrate-binding subunit |
| *PSPTO_0351* | | + | 383497 | 384534 | 5.18 | 37.67 | 22.09 | 10.25 | 69.27 | 13.18 | Oxidoreductase, aldo/keto reductase family |
| *PSPTO_0352* | *ntrC* | - | 384711 | 386147 | 22.43 | 35.05 | 21.22 | 12.34 | 23.90 | 7.40 | Nitrogen regulation protein NtrC |
| *PSPTO_0353* | *ntrB* | - | 386144 | 387229 | 33.65 | 40.71 | 28.91 | 12.55 | 20.12 | 6.76 | Nitrogen regulation protein NtrB |
| *PSPTO_0354* | | - | 387189 | 387347 | 92.61 | 42.49 | 50.43 | 43.19 | 12.84 | 5.58 | Hypothetical protein |
| *PSPTO_0355* | | - | 387513 | 388145 | 4.96 | 22.56 | 7.75 | 1.24 | 26.31 | 12.55 | Hypothetical protein |
| *PSPTO_0356* | | - | 388142 | 388684 | 29.32 | 34.27 | 38.77 | 11.14 | 59.93 | 21.93 | Hypothetical protein |
| *PSPTO_0357* | *aroH* | - | 388763 | 389320 | 9.59 | 43.72 | 55.88 | 26.75 | 91.75 | 19.28 | Chorismate mutase |
| *PSPTO_0358* | | - | 389541 | 391001 | 0.43 | 0.18 | 0.28 | 0.00 | 0.44 | 0.43 | ISPsy6, transposase |
| *PSPTO_0359* | *glnA-1* | - | 391090 | 392496 | 104.55 | 326.49 | 208.70 | 64.12 | 385.42 | 77.80 | Glutamine synthetase, type I |
| *PSPTO_0360* | *thiI* | + | 392832 | 394286 | 33.04 | 70.33 | 52.82 | 36.39 | 160.00 | 44.00 | Thiamin biosynthesis protein ThiI |
| *PSPTO_0361* | *typA* | + | 394429 | 396249 | 8.01 | 61.64 | 37.02 | 33.26 | 252.23 | 28.12 | GTP-binding protein TypA |
| *PSPTO_0362* | | - | 396350 | 397132 | 6.75 | 14.34 | 12.23 | 6.95 | 17.23 | 8.64 | DeoR family transcriptional regulator |
| *PSPTO_0363* | *polS* | - | 397298 | 398071 | 3.47 | 6.80 | 3.84 | 0.00 | 4.31 | 3.91 | Sorbitol dehydrogenase |
| *PSPTO_0364* | | - | 398111 | 399427 | 1.03 | 2.38 | 2.05 | 0.00 | 2.35 | 0.98 | Sugar ABC transporter periplasmic sugar-binding protein |
| *PSPTO_0365* | | + | 399709 | 400608 | 6.55 | 14.23 | 12.29 | 8.35 | 24.47 | 7.96 | AraC family transcriptional regulator |
| *PSPTO_0366* | *nhaP* | - | 400609 | 401868 | 3.94 | 6.64 | 7.35 | 2.51 | 9.84 | 5.00 | Na+/H+ antiporter NhaP |
| *PSPTO_0367* | | - | 402145 | 402474 | 50.16 | 11596.65 | 5717.79 | 980.31 | 234.91 | 31.12 | Hypothetical protein |
| *PSPTO_0368* | | - | 402612 | 402947 | 17.25 | 3438.73 | 2634.17 | 449.55 | 66.38 | 13.63 | Hypothetical protein |
| *PSPTO_0369* | | + | 403338 | 404663 | 6.52 | 38.01 | 28.00 | 7.49 | 41.94 | 4.98 | Outer membrane porin, OprD family |
| *PSPTO_0370* | | + | 404865 | 406241 | 3.21 | 15.14 | 7.45 | 1.75 | 6.63 | 5.43 | MATE efflux family protein |
| *PSPTO_0371* | *iaaL* | + | 406323 | 407510 | 4.49 | 121.27 | 59.24 | 14.97 | 15.12 | 8.14 | Indoleacetate-lysine ligase |
| *PSPTO_0372* | | - | 407868 | 408890 | 20.09 | 26.89 | 20.70 | 14.34 | 36.67 | 25.76 | ISPsy7, transposase |
| *PSPTO_0373* | | - | 408969 | 411536 | 7.67 | 11.89 | 12.21 | 11.30 | 27.20 | 20.64 | Rhs family protein |
| *PSPTO_0374* | | - | 411649 | 412236 | 43.39 | 19.38 | 41.14 | 12.60 | 47.95 | 38.82 | Hypothetical protein |
| *PSPTO_0375* | | - | 412540 | 415701 | 1.21 | 8.63 | 6.14 | 3.30 | 13.95 | 1.14 | Cation efflux family protein |
| *PSPTO_0376* | | - | 415726 | 416994 | 2.80 | 16.96 | 24.42 | 10.47 | 16.25 | 1.89 | Cation efflux family protein |
| *PSPTO_0377* | | - | 417062 | 418288 | 4.12 | 154.22 | 137.07 | 28.89 | 15.88 | 0.87 | Metal ion efflux outer membrane protein |
| *PSPTO_0378* | | + | 418542 | 419216 | 36.93 | 213.35 | 196.12 | 36.81 | 77.92 | 23.88 | DNA-binding heavy metal response regulator |
| *PSPTO_0379* | | + | 419213 | 420604 | 10.02 | 28.30 | 18.75 | 11.54 | 24.09 | 10.13 | Heavy metal sensor histidine kinase |
| *PSPTO_0380* | | + | 420719 | 421120 | 34.98 | 74.22 | 41.79 | 35.66 | 50.29 | 42.59 | Hypothetical protein |
| *PSPTO_0381* | | - | 421174 | 421584 | 24.64 | 40.59 | 46.84 | 15.90 | 87.25 | 36.92 | Hypothetical protein |
| *PSPTO_0382* | *ligB* | - | 421625 | 423310 | 10.37 | 7.19 | 1.47 | 2.41 | 12.39 | 9.18 | NAD-dependent DNA ligase LigB |
| *PSPTO_0383* | *metK* | - | 423533 | 424723 | 16.99 | 47.67 | 52.43 | 18.86 | 168.64 | 16.16 | S-adenosylmethionine synthetase |
| *PSPTO_0384* | | - | 424744 | 425739 | 38.22 | 45.20 | 44.65 | 17.37 | 88.57 | 20.08 | ArsR family transcriptional regulator |
| *PSPTO_0385* | *tkt* | + | 425999 | 427996 | 15.42 | 83.05 | 63.72 | 39.21 | 241.90 | 86.39 | Transketolase |
| *PSPTO_0386* | *epd* | + | 428140 | 429183 | 13.22 | 19.89 | 26.81 | 9.95 | 60.74 | 25.06 | D-erythrose 4-phosphate dehydrogenase |
| *PSPTO_0387* | *pgk* | + | 429206 | 430369 | 7.22 | 16.10 | 26.44 | 11.10 | 47.74 | 19.64 | Phosphoglycerate kinase |
| *PSPTO_0388* | | + | 430397 | 430630 | 22.16 | 107.71 | 77.24 | 32.99 | 188.45 | 77.42 | Lipoprotein |
| *PSPTO_0389* | | + | 430717 | 431043 | 37.17 | 160.23 | 146.03 | 119.44 | 455.40 | 172.70 | Lipoprotein |
| *PSPTO_0390* | *fba* | + | 431157 | 432221 | 16.49 | 74.22 | 60.64 | 29.38 | 204.61 | 55.90 | Fructose-bisphosphate aldolase |
| *PSPTO_0391* | | - | 432379 | 432798 | 50.11 | 58.02 | 71.73 | 15.89 | 149.06 | 61.70 | Hypothetical protein |
| *PSPTO_0393* | | - | 432854 | 433264 | 34.97 | 71.99 | 61.03 | 30.10 | 101.77 | 38.04 | Hypothetical protein |
| *PSPTO_0394* | | + | 433493 | 434470 | 1.94 | 3.86 | 16.29 | 10.90 | 8.18 | 1.27 | Renal dipeptidase family protein |
| *PSPTO_0395* | | + | 434554 | 435084 | 1.47 | 0.00 | 10.33 | 5.27 | 5.24 | 1.96 | Hypothetical protein |
| *PSPTO_0396* | | + | 435100 | 437160 | 2.56 | 4.13 | 10.54 | 6.04 | 6.32 | 2.10 | NADH:flavin oxidoreductase/NADH oxidase family protein |
| *PSPTO_0397* | | + | 437235 | 439244 | 2.03 | 2.18 | 3.30 | 1.28 | 3.86 | 2.05 | Iron-sulfur cluster-binding protein |
| *PSPTO_0398* | *etfA-1* | + | 439478 | 440464 | 3.49 | 1.77 | 7.46 | 1.13 | 3.28 | 2.01 | Electron transfer flavoprotein subunit alpha |
| *PSPTO_0399* | *etfB-1* | + | 440454 | 441251 | 1.42 | 1.56 | 2.57 | 0.00 | 2.73 | 2.04 | Electron transfer flavoprotein subunit beta |

| Locus name | Gene name | Strand | Gene start (bp) | Gene end (bp) | Average TPM Inoculum | Average TPM $t_6$ | Average TPM $t_{24}$ | Average TPM $t_{168}$ | Average TPM Logarithmic | Average TPM Stationary | Annotation description |
|---|---|---|---|---|---|---|---|---|---|---|---|
| *PSPTO_0400* | | - | 441859 | 443655 | 23.50 | 14.32 | 20.82 | 12.30 | 22.69 | 22.71 | Chitinase |
| *PSPTO_0401* | | - | 443858 | 445090 | 3.70 | 3.09 | 10.12 | 10.98 | 8.19 | 3.81 | Iron-sulfur cluster-binding protein, Rieske family |
| *PSPTO_0402* | | + | 445454 | 446554 | 7.60 | 4.99 | 12.35 | 13.49 | 31.05 | 12.09 | Iron-sulfur cluster-binding protein |
| *PSPTO_0403* | | - | 446683 | 446949 | 51.26 | 24.94 | 50.03 | 17.28 | 60.74 | 20.72 | Hypothetical protein |
| *PSPTO_0404* | | + | 447686 | 448522 | 14.15 | 48.32 | 32.30 | 17.54 | 132.15 | 70.35 | Hypothetical protein |
| *PSPTO_0405* | *msrA* | - | 448752 | 449399 | 54.27 | 87.06 | 51.92 | 49.36 | 176.83 | 103.27 | Peptide methionine sulfoxide reductase |
| *PSPTO_0406* | | - | 449550 | 452243 | 21.79 | 30.76 | 31.08 | 13.36 | 64.27 | 25.47 | Sensory box/GGDEF domain/EAL domain-containing protein |
| *PSPTO_0407* | | - | 452492 | 453880 | 3.53 | 6.35 | 2.91 | 3.85 | 9.40 | 4.60 | Drug resistance transporter, EmrB/QacA family |
| *PSPTO_0408* | | - | 453914 | 455239 | 20.19 | 8.78 | 11.71 | 2.09 | 38.90 | 9.68 | Hypothetical protein |
| *PSPTO_0409* | | - | 455239 | 456681 | 25.31 | 10.11 | 11.13 | 5.74 | 41.80 | 8.09 | TldD/PmbA family protein |
| *PSPTO_0410* | | - | 456805 | 457845 | 10.92 | 21.15 | 14.25 | 5.32 | 47.79 | 14.03 | Threonine aldolase, low-specificity |
| *PSPTO_0411* | | + | 457946 | 458692 | 12.59 | 6.05 | 7.37 | 2.33 | 19.16 | 9.50 | Membrane protein |
| *PSPTO_0412* | | - | 458865 | 460061 | 6.12 | 22.58 | 22.92 | 10.89 | 54.51 | 16.98 | Hypothetical protein |
| *PSPTO_0413* | | + | 460408 | 461172 | 19.18 | 46.31 | 26.05 | 21.08 | 91.44 | 33.76 | Hypothetical protein |
| *PSPTO_0414* | *mutM* | + | 461218 | 462030 | 7.65 | 13.83 | 19.56 | 12.46 | 58.64 | 23.34 | Formamidopyrimidine-DNA glycosylase |
| *PSPTO_0415* | | + | 462120 | 462458 | 31.52 | 51.13 | 63.16 | 25.93 | 76.47 | 61.84 | Hypothetical protein |
| *PSPTO_0416* | | - | 462467 | 462718 | 52.95 | 67.43 | 59.02 | 30.23 | 90.65 | 70.21 | Ferredoxin |
| *PSPTO_0417* | | - | 462809 | 463288 | 37.18 | 70.60 | 68.15 | 24.26 | 48.42 | 30.89 | Pantetheine-phosphate adenylyltransferase |
| *PSPTO_0418* | | - | 463456 | 464883 | 4.29 | 9.58 | 4.63 | 4.93 | 13.74 | 12.28 | Coniferyl aldehyde dehydrogenase |
| *PSPTO_0419* | | + | 465161 | 465823 | 42.76 | 44.48 | 33.77 | 19.63 | 95.20 | 42.96 | TetR family transcriptional regulator |
| *PSPTO_0420* | | + | 465877 | 466734 | 17.11 | 29.60 | 26.30 | 12.70 | 58.28 | 41.72 | Rhodanese-like domain-containing protein |
| *PSPTO_0421* | | - | 466742 | 467755 | 13.71 | 17.35 | 15.65 | 8.39 | 54.10 | 25.33 | Alpha/beta fold family hydrolase |
| *PSPTO_0422* | | - | 467890 | 468183 | 29.55 | 37.16 | 19.81 | 4.27 | 137.75 | 39.42 | Virulence-associated protein |
| *PSPTO_0423* | | - | 468196 | 468474 | 25.32 | 24.36 | 27.11 | 8.58 | 66.27 | 17.67 | Proteic killer protein |
| *PSPTO_0424* | | - | 468737 | 469321 | 4.20 | 9.19 | 8.80 | 5.76 | 68.65 | 11.48 | Methyltransferase |
| *PSPTO_0425* | | - | 469321 | 470814 | 5.28 | 16.46 | 24.67 | 6.66 | 58.47 | 14.43 | Hypothetical protein |
| *PSPTO_0426* | | - | 470811 | 472163 | 6.13 | 14.77 | 17.62 | 6.83 | 69.89 | 16.49 | Peptidase, M16 family |
| *PSPTO_0427* | *ftsY* | + | 472419 | 473960 | 49.25 | 36.83 | 29.08 | 20.29 | 83.21 | 41.69 | Signal recognition particle-docking protein FtsY |
| *PSPTO_0428* | *ftsE* | + | 473957 | 474628 | 17.31 | 34.33 | 29.88 | 25.78 | 91.24 | 68.45 | Cell division ATP-binding protein FtsE |
| *PSPTO_0429* | *ftsX* | + | 474625 | 475659 | 92.71 | 91.98 | 66.08 | 41.69 | 113.42 | 83.44 | Putative protein insertion permease FtsX |
| *PSPTO_0430* | *rpoH* | + | 475778 | 476632 | 413.81 | 517.55 | 480.66 | 223.26 | 457.87 | 407.31 | RNA polymerase sigma-32 factor |
| *PSPTO_0431* | *mtgA* | - | 476853 | 477563 | 7.51 | 29.50 | 14.55 | 14.07 | 45.46 | 16.89 | Monofunctional biosynthetic peptidoglycan transglycosylase |
| *PSPTO_0432* | | + | 477644 | 478024 | 12.15 | 13.28 | 16.88 | 8.18 | 37.22 | 16.98 | Hypothetical protein |
| *PSPTO_0433* | *thiS* | + | 478133 | 478333 | 36.09 | 57.25 | 30.62 | 51.30 | 187.16 | 57.35 | Thiamine biosynthesis protein ThiS |
| *PSPTO_0434* | *thiG* | + | 478385 | 479179 | 10.17 | 11.71 | 35.94 | 25.93 | 104.41 | 39.58 | Thiazole biosynthesis protein ThiG |
| *PSPTO_0435* | | + | 479190 | 479924 | 17.36 | 26.76 | 27.89 | 20.15 | 76.07 | 34.43 | tRNA (guanine-N(7)-)-methyltransferase |
| *PSPTO_0436* | | - | 479967 | 481025 | 2.93 | 9.14 | 6.01 | 6.71 | 42.08 | 8.81 | Hypothetical protein |
| *PSPTO_0437* | *folA* | - | 481216 | 481728 | 20.13 | 20.55 | 11.98 | 2.69 | 24.67 | 8.74 | Dihydrofolate reductase |
| *PSPTO_0438* | | + | 481854 | 483233 | 5.03 | 7.76 | 1.77 | 1.04 | 14.16 | 4.43 | Hypothetical protein |
| *PSPTO_0439* | | + | 483226 | 484614 | 4.49 | 7.17 | 9.48 | 1.22 | 22.16 | 7.87 | Hypothetical protein |
| *PSPTO_0440* | *betI* | + | 484957 | 485550 | 16.09 | 11.61 | 14.57 | 3.52 | 367.70 | 8.46 | Regulatory protein BetI |
| *PSPTO_0441* | *betB* | + | 485655 | 487127 | 7.69 | 21.77 | 20.00 | 5.50 | 268.84 | 11.95 | Betaine aldehyde dehydrogenase BADH |
| *PSPTO_0442* | | + | 487139 | 487240 | 11.93 | 46.71 | 94.38 | 8.04 | 628.38 | 42.42 | Hypothetical protein |
| *PSPTO_0443* | *betA* | + | 487268 | 488974 | 3.35 | 15.21 | 19.91 | 4.61 | 138.87 | 7.68 | Choline dehydrogenase |
| *PSPTO_0444* | | + | 489138 | 489656 | 0.54 | 0.00 | 0.93 | 0.00 | 5.90 | 5.66 | RNA polymerase sigma-70 family protein |
| *PSPTO_0445* | | + | 489653 | 490645 | 1.19 | 0.00 | 1.94 | 0.39 | 6.76 | 5.97 | Regulatory protein |
| *PSPTO_0446* | | + | 490766 | 491899 | 1.29 | 0.18 | 0.98 | 0.00 | 2.68 | 2.03 | Membrane protein |
| *PSPTO_0447* | | + | 491896 | 492078 | 2.12 | 3.14 | 0.00 | 0.00 | 1.29 | 1.82 | Hypothetical protein |
| *PSPTO_0448* | | + | 492163 | 492777 | 11.31 | 27.89 | 24.94 | 10.02 | 53.88 | 20.14 | Glutathione S-transferase |
| *PSPTO_0449* | | + | 493002 | 493517 | 14.71 | 19.80 | 18.32 | 13.60 | 38.00 | 18.87 | ISPsy8, transposase OrfA |
| *PSPTO_0450* | | + | 493574 | 494353 | 16.66 | 17.37 | 12.15 | 9.73 | 25.35 | 16.93 | ISPsy8, transposase OrfB |
| *PSPTO_0451* | | - | 494461 | 494964 | 150.54 | 53.91 | 60.22 | 25.14 | 46.86 | 44.59 | Hypothetical protein |
| *PSPTO_0452* | | + | 495158 | 496057 | 11.33 | 15.49 | 26.28 | 9.02 | 43.17 | 27.36 | Putative acyltransferase |
| *PSPTO_0453* | | - | 496037 | 496756 | 266.19 | 126.18 | 87.48 | 49.26 | 98.70 | 120.19 | Hypothetical protein |
| *PSPTO_0454* | | + | 496831 | 497829 | 15.72 | 23.45 | 20.78 | 21.43 | 23.51 | 19.43 | Hypothetical protein |
| *PSPTO_0455* | *fdhA-1* | - | 497858 | 499057 | 15.73 | 16.49 | 43.73 | 30.23 | 15.40 | 19.61 | Glutathione-independent formaldehyde dehydrogenase |
| *PSPTO_0456* | *purU-1* | - | 499114 | 499971 | 39.49 | 15.69 | 33.82 | 21.46 | 15.80 | 19.73 | Formyltetrahydrofolate deformylase |
| *PSPTO_0457* | *soxG-1* | - | 500142 | 500774 | 1.89 | 3.17 | 8.89 | 0.00 | 4.65 | 1.90 | Sarcosine oxidase, gamma subunit |
| *PSPTO_0458* | *soxA-1* | - | 500811 | 503831 | 2.18 | 2.93 | 7.13 | 6.79 | 7.31 | 2.51 | Sarcosine oxidase, alpha subunit |
| *PSPTO_0459* | *soxD-1* | - | 503828 | 504124 | 4.94 | 3.66 | 1.16 | 7.96 | 11.60 | 5.46 | Sarcosine oxidase subunit delta |
| *PSPTO_0460* | *soxB-1* | - | 504140 | 505390 | 2.36 | 3.27 | 8.24 | 2.58 | 8.71 | 4.96 | Sarcosine oxidase, beta subunit |
| *PSPTO_0461* | *glyA-1* | - | 505408 | 506661 | 1.27 | 0.69 | 7.72 | 7.99 | 7.32 | 1.74 | Serine hydroxymethyltransferase |
| *PSPTO_0462* | *cbcV* | - | 506918 | 508096 | 2.41 | 5.45 | 8.27 | 4.60 | 11.22 | 5.66 | Glycine/betaine/L-proline ABC transporter, ATP-binding subunit |
| *PSPTO_0463* | *cbcW* | - | 508093 | 508938 | 3.20 | 5.67 | 4.51 | 6.93 | 12.40 | 7.02 | Glycine/betaine/L-proline ABC transporter, permease protein |
| *PSPTO_0464* | *cbcX* | - | 509006 | 509953 | 8.39 | 24.92 | 26.96 | 34.29 | 31.53 | 32.81 | Glycine betaine/L-proline ABC transporter, periplasmic substrate-binding protein |
| *PSPTO_0465* | | + | 510737 | 511840 | 56.47 | 32.71 | 45.28 | 41.13 | 12.80 | 3.79 | AraC family transcriptional regulator |
| *PSPTO_0466* | | - | 512022 | 513647 | 5.90 | 4.32 | 6.76 | 6.58 | 9.74 | 6.83 | Methyl-accepting chemotaxis protein |
| *PSPTO_0467* | | - | 513782 | 513937 | 12.61 | 9.74 | 23.11 | 0.90 | 28.52 | 4.15 | Hypothetical protein |
| *PSPTO_0468* | | + | 514044 | 514301 | 53.21 | 98.16 | 94.55 | 94.59 | 127.17 | 117.50 | Hypothetical protein |
| *PSPTO_0469* | | + | 514298 | 514600 | 8.52 | 34.11 | 25.38 | 19.86 | 21.54 | 24.10 | Hypothetical protein |
| *PSPTO_0470* | | - | 514680 | 515393 | 14.12 | 32.95 | 16.33 | 14.20 | 59.28 | 8.89 | Exonuclease |
| *PSPTO_0471* | | - | 515390 | 517258 | 20.80 | 44.66 | 43.89 | 13.61 | 112.19 | 9.04 | Nucleotidyltransferase |
| *PSPTO_0472* | | + | 517608 | 518000 | 32.05 | 42.59 | 34.88 | 47.72 | 26.09 | 30.85 | Response regulator |
| *PSPTO_0473* | *hopAS1* | - | 518079 | 521327 | - | - | - | - | - | - | |
| *PSPTO_0474* | | - | 521328 | 522167 | 1.31 | 0.00 | 2.54 | 0.66 | 1.34 | 1.31 | Truncated type III effector HopAS1; premature stop codon |
| *PSPTO_0475* | | - | 522512 | 522943 | 14.04 | 6.25 | 6.36 | 1.81 | 23.72 | 15.87 | Hypothetical protein |
| *PSPTO_0476* | | - | 522940 | 523386 | 4.95 | 3.91 | 0.00 | 3.79 | 8.05 | 4.38 | Hypothetical protein |
| *PSPTO_0477* | | - | 523383 | 523826 | 1.42 | 3.34 | 3.69 | 2.50 | 6.18 | 2.60 | Hypothetical protein |
| *PSPTO_0478* | | - | 523823 | 524749 | 3.86 | 3.39 | 3.59 | 0.59 | 8.17 | 7.08 | Hypothetical protein |

| Locus name | Gene name | Strand | Gene start (bp) | Gene end (bp) | Average TPM Inoculum | Average TPM $t_{6}$ | Average TPM $t_{24}$ | Average TPM $t_{168}$ | Average TPM Logarithmic | Average TPM Stationary | Annotation description |
|---|---|---|---|---|---|---|---|---|---|---|---|
| *PSPTO_0479* | | - | 524756 | 525322 | 0.75 | 2.02 | 1.38 | 0.00 | 3.08 | 2.56 | Hypothetical protein |
| *PSPTO_0480* | *glcB-1* | - | 525681 | 527858 | 22.02 | 47.19 | 54.39 | 32.96 | 46.69 | 10.96 | Malate synthase G |
| *PSPTO_0481* | | + | 528181 | 528699 | 41.90 | 39.53 | 53.56 | 8.31 | 25.35 | 19.88 | ACT domain-containing protein |
| *PSPTO_0482* | | - | 528719 | 529069 | 30.21 | 26.84 | 28.35 | 0.15 | 30.05 | 12.67 | Hypothetical protein |
| *PSPTO_0483* | *rarD-1* | - | 529381 | 530268 | 2.91 | 10.25 | 8.95 | 5.19 | 16.12 | 4.62 | RarD protein |
| *PSPTO_0484* | | - | 530394 | 531368 | 5.90 | 17.28 | 13.68 | 14.86 | 61.73 | 20.80 | Hypothetical protein |
| *PSPTO_0485* | | + | 531564 | 532583 | 0.75 | 0.98 | 1.16 | 0.00 | 2.33 | 1.69 | LacI family transcriptional regulator |
| *PSPTO_0486* | | + | 532567 | 533568 | 0.66 | 2.07 | 0.82 | 1.27 | 2.82 | 1.26 | ABC transporter substrate-binding protein |
| *PSPTO_0487* | | + | 533568 | 534437 | 1.83 | 0.00 | 0.00 | 0.00 | 1.32 | 0.99 | ABC transporter permease |
| *PSPTO_0488* | | + | 534434 | 535228 | 1.95 | 0.00 | 2.48 | 0.00 | 2.94 | 2.52 | ABC transporter permease |
| *PSPTO_0489* | | + | 535225 | 536268 | 5.38 | 2.97 | 1.46 | 6.00 | 9.09 | 7.54 | ABC transporter ATP-binding protein |
| *PSPTO_0490* | | + | 536293 | 537126 | 1.52 | 2.81 | 2.05 | 1.31 | 3.22 | 1.97 | Ser/Thr protein phosphatase family protein |
| *PSPTO_0491* | | - | 537253 | 539169 | 4.36 | 7.76 | 11.02 | 2.07 | 7.44 | 3.36 | Tat (twin-arginine translocation) pathway signal sequence domain-containing protein |
| *PSPTO_0492* | *modE* | - | 539251 | 540015 | 11.59 | 14.96 | 18.14 | 2.09 | 22.28 | 12.10 | Molybdate transport regulator ModE |
| *PSPTO_0493* | *comF* | - | 540207 | 540941 | 3.98 | 3.37 | 0.89 | 1.52 | 4.65 | 3.96 | Competence protein ComF |
| *PSPTO_0494* | *bioB* | + | 541032 | 542090 | 59.50 | 127.44 | 112.52 | 43.30 | 246.91 | 20.45 | Biotin synthetase |
| *PSPTO_0495* | *bioF* | + | 542154 | 543344 | 6.25 | 8.25 | 14.00 | 9.39 | 68.53 | 8.56 | 8-amino-7-oxononanoate synthase |
| *PSPTO_0496* | *bioH* | + | 543337 | 544068 | 6.90 | 3.57 | 4.46 | 1.97 | 23.82 | 7.71 | BioH protein |
| *PSPTO_0497* | *bioC* | + | 544061 | 544870 | 3.19 | 0.33 | 8.91 | 3.18 | 30.93 | 3.96 | Biotin synthesis protein BioC |
| *PSPTO_0498* | *bioD* | + | 544902 | 545582 | 8.38 | 15.65 | 10.13 | 3.80 | 32.33 | 8.32 | Dethiobiotin synthetase |
| *PSPTO_0499* | | + | 545737 | 546033 | 109.77 | 292.44 | 263.94 | 142.88 | 94.00 | 174.98 | Hypothetical protein |
| *PSPTO_0500* | | + | 546382 | 548187 | 31.04 | 30.74 | 26.54 | 12.83 | 39.53 | 13.60 | Acyl-CoA dehydrogenase family protein |
| *PSPTO_0501* | *hopU1* | - | 548376 | 549170 | 0.32 | 9.45 | 3.98 | 0.70 | 3.65 | 2.57 | Type III effector HopU1 |
| *PSPTO_0502* | *hopF2* | - | 549423 | 550037 | 15.58 | 60.99 | 25.95 | 6.57 | 6.70 | 9.28 | Type III effector HopF2 |
| *PSPTO_0503* | *shcF* | - | 550174 | 550569 | 0.84 | 357.57 | 210.75 | 18.25 | 4.30 | 3.57 | Type III chaperone protein ShcF |
| *PSPTO_0505* | | + | 552012 | 553307 | 7.69 | 3.16 | 6.16 | 4.94 | 5.86 | 4.76 | Diguanylate cyclase |
| *PSPTO_0506* | | + | 553477 | 555255 | 16.73 | 30.99 | 34.06 | 16.84 | 116.03 | 16.47 | Acyl-CoA dehydrogenase family protein |
| *PSPTO_0507* | | + | 555671 | 555832 | 146.38 | 701.29 | 523.56 | 198.83 | 261.63 | 165.71 | Hypothetical protein |
| *PSPTO_0508* | | - | 555873 | 557705 | 3.80 | 14.37 | 14.92 | 7.09 | 23.60 | 10.48 | Hypothetical protein |
| *PSPTO_0509* | *pqqE* | - | 557668 | 558837 | 5.90 | 14.94 | 17.81 | 6.37 | 36.79 | 15.09 | Coenzyme PQQ synthesis protein E |
| *PSPTO_0510* | *pqqD* | - | 558830 | 559126 | 52.92 | 53.71 | 62.39 | 39.66 | 94.25 | 66.38 | Coenzyme PQQ synthesis protein D |
| *PSPTO_0511* | *pqqC* | - | 559123 | 559878 | 23.36 | 35.85 | 24.70 | 10.26 | 73.13 | 36.20 | Coenzyme PQQ synthesis protein C |
| *PSPTO_0512* | *pqqB* | - | 559887 | 560798 | 21.28 | 56.00 | 24.55 | 3.52 | 145.42 | 74.52 | Coenzyme PQQ synthesis protein B |
| *PSPTO_0513* | *pqqA* | - | 560896 | 560970 | 16.02 | 182.41 | 117.65 | 14.01 | 31.79 | 52.24 | Coenzyme PQQ synthesis protein A |
| *PSPTO_0514* | *pqqF* | - | 561106 | 563445 | 4.52 | 10.95 | 6.30 | 4.85 | 31.93 | 12.23 | Coenzyme PQQ synthesis protein F |
| *PSPTO_0515* | | + | 563660 | 564175 | 14.78 | 19.80 | 18.32 | 13.60 | 38.14 | 18.93 | ISPsy8, transposase OrfA |
| *PSPTO_0516* | | + | 564232 | 565011 | 16.64 | 17.21 | 12.20 | 9.73 | 25.42 | 17.02 | ISPsy8, transposase OrfB |
| *PSPTO_0517* | | - | 565108 | 565914 | 13.64 | 7.92 | 8.34 | 4.72 | 85.42 | 266.43 | Hydrolase |
| *PSPTO_0518* | | - | 565932 | 567566 | 28.36 | 8.09 | 5.36 | 6.73 | 170.27 | 351.67 | Tryptophan 2-monooxygenase |
| *PSPTO_0519* | | - | 567763 | 568203 | 2.42 | 5.55 | 6.24 | 3.37 | 20.31 | 4.18 | AsnC family transcriptional regulator |
| *PSPTO_0520* | | - | 568236 | 569312 | 15.17 | 15.60 | 12.64 | 7.21 | 48.38 | 13.73 | Membrane protein |
| *PSPTO_0521* | | + | 569661 | 569945 | 15.87 | 17.05 | 6.72 | 21.94 | 43.28 | 14.38 | DNA-binding protein |
| *PSPTO_0522* | | + | 569945 | 571171 | 6.25 | 11.24 | 11.14 | 2.71 | 41.14 | 17.95 | Hypothetical protein |
| *PSPTO_0523* | | - | 571196 | 572326 | 7.37 | 10.61 | 4.96 | 4.23 | 25.31 | 9.50 | Fic family protein |
| *PSPTO_0524* | | + | 572573 | 573871 | 16.83 | 30.14 | 24.72 | 6.14 | 38.94 | 16.17 | Peptidase, M20/M25/M40 family |
| *PSPTO_0525* | | - | 574042 | 574635 | 3.20 | 6.88 | 1.44 | 0.00 | 11.51 | 3.55 | Lipoprotein |
| *PSPTO_0526* | | - | 574681 | 574893 | 5.65 | 10.31 | 4.95 | 0.00 | 4.66 | 3.83 | Hypothetical protein |
| *PSPTO_0527* | | - | 575375 | 577327 | 7.76 | 16.69 | 15.03 | 9.69 | 43.95 | 27.32 | Hypothetical protein |
| *PSPTO_0528* | | - | 577324 | 578211 | 1.29 | 7.18 | 3.38 | 3.39 | 26.39 | 7.57 | Hypothetical protein |
| *PSPTO_0529* | | - | 578208 | 579008 | 5.43 | 9.09 | 10.32 | 4.40 | 36.93 | 7.38 | Hypothetical protein |
| *PSPTO_0530* | | - | 579598 | 580119 | 4.66 | 23.04 | 7.09 | 5.34 | 6.57 | 5.56 | Hypothetical protein |
| *PSPTO_0531* | | + | 580669 | 581433 | 4.83 | 7.78 | 6.28 | 4.12 | 2.48 | 6.68 | Type IV secretion system protein |
| *PSPTO_0532* | | + | 581917 | 582171 | 0.27 | 0.00 | 0.00 | 2.87 | 3.03 | 3.04 | Lipoprotein |
| *PSPTO_0533* | | + | 582174 | 583040 | - | - | - | - | - | - | |
| *PSPTO_0534* | | + | 583081 | 584058 | 0.00 | 0.00 | 0.00 | 0.00 | 0.00 | 0.00 | ISPssy, transposase |
| *PSPTO_0535* | | - | 584780 | 586021 | 3.70 | 8.53 | 6.46 | 2.96 | 10.58 | 3.89 | Site-specific recombinase, phage integrase family |
| *PSPTO_0536* | | - | 586100 | 588589 | 19.18 | 20.75 | 22.82 | 7.86 | 95.95 | 24.32 | Sensory box/GGDEF domain/EAL domain-containing protein |
| *PSPTO_0537* | *rpoD* | - | 588846 | 590696 | 20.96 | 91.39 | 72.10 | 48.11 | 174.18 | 93.70 | RNA polymerase sigma-70 factor |
| *PSPTO_0538* | *dnaG* | - | 590764 | 592722 | 30.48 | 35.43 | 23.64 | 24.18 | 85.76 | 34.65 | DNA primase |
| *PSPTO_0539* | *rpsU* | - | 593192 | 593407 | 89.54 | 417.98 | 321.28 | 224.32 | 437.47 | 179.05 | 30S ribosomal protein S21 |
| *PSPTO_0540* | *gcp* | + | 593605 | 594630 | 4.61 | 5.68 | 12.92 | 1.02 | 14.93 | 7.65 | O-sialoglycoprotein endopeptidase |
| *PSPTO_0541* | | - | 594683 | 595261 | 44.29 | 79.79 | 51.97 | 27.44 | 180.80 | 44.25 | Membrane protein |
| *PSPTO_0542* | *folB-1* | + | 595327 | 595680 | 5.14 | 10.59 | 6.85 | 6.29 | 38.01 | 6.81 | Dihydroneopterin aldolase |
| *PSPTO_0543* | *folK-1* | + | 595671 | 596195 | 2.69 | 6.49 | 3.48 | 9.78 | 38.83 | 5.26 | 2-amino-4-hydroxy-6-hydroxymethyldihydropteridine pyrophosphokinase |
| *PSPTO_0544* | *cca* | - | 596465 | 597694 | 5.36 | 25.73 | 15.41 | 9.68 | 31.08 | 15.64 | tRNA nucleotidyltransferase |
| *PSPTO_0545* | | - | 597792 | 599354 | 57.98 | 154.57 | 145.76 | 95.79 | 20.65 | 149.86 | SpoVR like family protein |
| *PSPTO_0546* | | - | 599351 | 600622 | 95.70 | 231.49 | 197.39 | 138.32 | 40.47 | 219.95 | Hypothetical protein |
| *PSPTO_0547* | | - | 600744 | 602666 | 204.29 | 295.34 | 295.82 | 169.26 | 86.83 | 352.67 | Hypothetical protein |
| *PSPTO_0548* | | - | 602952 | 603272 | 20.31 | 37.14 | 53.53 | 18.27 | 86.60 | 25.44 | Thiosulfate sulfurtransferase |
| *PSPTO_0549* | *apaH* | - | 603269 | 604171 | 22.01 | 32.75 | 26.22 | 14.18 | 68.79 | 21.30 | Bis(5'-nucleosyl)-tetraphosphatase, symmetrical |
| *PSPTO_0550* | *apaG* | - | 604171 | 604551 | 60.86 | 36.02 | 64.72 | 28.21 | 102.66 | 40.80 | ApaG protein |
| *PSPTO_0551* | *ksgA* | - | 604665 | 605471 | 3.92 | 24.38 | 5.38 | 9.96 | 82.70 | 9.42 | Dimethyladenosine transferase |
| *PSPTO_0552* | *pdxA* | - | 605468 | 606457 | 10.08 | 36.32 | 21.00 | 13.93 | 153.70 | 18.67 | Pyridoxal phosphate biosynthetic protein PdxA |
| *PSPTO_0553* | *surA* | - | 606454 | 607740 | 6.96 | 24.05 | 22.94 | 17.43 | 146.32 | 23.38 | Peptidyl-prolyl cis-trans isomerase SurA |
| *PSPTO_0554* | | - | 607757 | 610474 | 9.50 | 55.30 | 55.59 | 45.37 | 207.54 | 48.47 | Organic solvent tolerance protein |
| *PSPTO_0555* | | + | 610670 | 611695 | 49.77 | 24.88 | 20.13 | 20.65 | 51.70 | 24.13 | Hypothetical protein |
| *PSPTO_0556* | | + | 611814 | 612446 | 17.87 | 36.11 | 16.52 | 13.64 | 45.62 | 28.73 | Nucleotidyltransferase family protein |
| *PSPTO_0557* | *djlA* | + | 612447 | 613214 | 23.54 | 24.95 | 22.03 | 15.89 | 69.07 | 43.64 | DnaJ-like protein DjlA |
| *PSPTO_0558* | | - | 613245 | 614243 | 21.10 | 19.48 | 11.17 | 5.80 | 33.17 | 10.52 | Hypothetical protein |

| Locus name | Gene name | Strand | Gene start (bp) | Gene end (bp) | Average TPM Inoculum | Average TPM $t_{6}$ | Average TPM $t_{24}$ | Average TPM $t_{168}$ | Average TPM Logarithmic | Average TPM Stationary | Annotation description |
|---|---|---|---|---|---|---|---|---|---|---|---|
| *PSPTO_0559* | | + | 614343 | 616358 | 6.93 | 4.60 | 4.55 | 2.60 | 7.47 | 4.67 | Sensory box histidine kinase |
| *PSPTO_0560* | | + | 616355 | 616984 | 2.74 | 6.40 | 2.51 | 1.45 | 14.06 | 7.34 | LuxR family DNA-binding response regulator |
| *PSPTO_0561* | | + | 617232 | 617405 | 0.85 | 5.02 | 0.00 | 0.00 | 1.94 | 2.56 | Hypothetical protein |
| *PSPTO_0562* | | + | 617525 | 618649 | 20.36 | 3.57 | 3.68 | 1.50 | 5.21 | 1.56 | Polyamine ABC transporter ATP-binding protein |
| *PSPTO_0563* | | + | 618732 | 619766 | 18.28 | 2.54 | 3.92 | 5.37 | 9.94 | 1.59 | Polyamine ABC transporter periplasmic polyamine-binding protein |
| *PSPTO_0564* | | + | 619844 | 621091 | 9.76 | 5.30 | 4.99 | 5.11 | 6.84 | 2.98 | Polyamine ABC transporter permease |
| *PSPTO_0565* | | + | 621103 | 621924 | 5.13 | 6.74 | 3.42 | 0.00 | 8.93 | 3.85 | Polyamine ABC transporter permease |
| *PSPTO_0566* | *rpe* | + | 622080 | 622754 | 40.88 | 62.59 | 66.47 | 33.36 | 166.25 | 62.13 | Ribulose-phosphate 3-epimerase |
| *PSPTO_0567* | *gph-1* | + | 622769 | 623569 | 15.95 | 25.18 | 19.74 | 16.48 | 63.65 | 16.86 | Phosphoglycolate phosphatase |
| *PSPTO_0568* | *trpE* | + | 623639 | 625120 | 14.94 | 25.29 | 25.51 | 5.27 | 71.20 | 15.96 | Anthranilate synthase, component I |
| *PSPTO_0569* | | - | 625342 | 627264 | 9.56 | 44.16 | 38.34 | 19.87 | 40.47 | 8.68 | Autotransporting lipase, GDSL family |
| *PSPTO_0570* | *prtN* | - | 627371 | 627637 | 6.64 | 24.19 | 33.88 | 13.42 | 32.28 | 14.55 | Transcriptional regulator PrtN |
| *PSPTO_0571* | | - | 628136 | 628783 | 18.92 | 32.98 | 15.52 | 10.05 | 41.08 | 20.45 | Repressor protein c2 |
| *PSPTO_0572* | | + | 629847 | 630677 | 11.54 | 19.68 | 22.60 | 6.59 | 47.62 | 17.17 | Bacteriocin |
| *PSPTO_0573* | | + | 631257 | 631727 | 2.58 | 0.00 | 2.67 | 0.00 | 3.74 | 2.60 | Hypothetical protein |
| *PSPTO_0574* | | + | 631705 | 632046 | 4.80 | 1.71 | 0.00 | 0.00 | 8.10 | 4.31 | Lipoprotein |
| *PSPTO_0575* | | + | 632092 | 632682 | 3.40 | 5.09 | 26.40 | 5.43 | 34.62 | 13.08 | Hypothetical protein |
| *PSPTO_0576* | | + | 632679 | 632867 | 8.42 | 43.77 | 65.57 | 31.72 | 112.15 | 34.51 | Hypothetical protein |
| *PSPTO_0577* | | + | 632886 | 634382 | 2.32 | 8.94 | 10.08 | 3.35 | 13.16 | 5.73 | Tail sheath protein |
| *PSPTO_0578* | | + | 634443 | 634790 | 1.97 | 19.72 | 16.24 | 7.54 | 34.28 | 13.73 | Hypothetical protein |
| *PSPTO_0579* | | + | 634787 | 635083 | 2.02 | 6.34 | 19.94 | 10.57 | 15.34 | 6.51 | Phage protein (sigma factor domain protein) |
| *PSPTO_0580* | | + | 635214 | 637358 | 3.03 | 5.12 | 8.38 | 4.76 | 11.05 | 6.71 | Tail tape measure protein |
| *PSPTO_0581* | | + | 637355 | 639178 | 5.08 | 6.85 | 9.14 | 9.62 | 9.64 | 6.95 | DNA circulation protein |
| *PSPTO_0582* | | + | 639401 | 640042 | 2.98 | 4.10 | 4.65 | 3.86 | 8.44 | 6.68 | Hypothetical protein |
| *PSPTO_0583* | | - | 640284 | 640973 | 3.68 | 11.85 | 12.90 | 3.97 | 32.69 | 13.36 | Hypothetical protein |
| *PSPTO_0584* | | - | 641322 | 642014 | 2.66 | 3.94 | 10.90 | 1.86 | 8.74 | 5.58 | Sir2 family transcriptional regulator |
| *PSPTO_0585* | | - | 642156 | 642623 | 3.64 | 0.32 | 5.54 | 0.00 | 7.24 | 7.57 | Hypothetical protein |
| *PSPTO_0586* | | + | 644311 | 645321 | 9.66 | 10.09 | 5.80 | 7.78 | 8.97 | 9.42 | Site-specific recombinase, phage integrase family |
| *PSPTO_0587* | | + | 645345 | 647486 | 14.48 | 18.79 | 14.46 | 14.70 | 27.55 | 17.52 | Site-specific recombinase, phage integrase family |
| *PSPTO_0588* | *hopH1* | - | 647588 | 648244 | 0.90 | 697.11 | 276.53 | 48.57 | 7.06 | 1.69 | Type III effector HopH1 |
| *PSPTO_0589* | *hopC1* | - | 648896 | 649705 | 1.77 | 499.02 | 244.89 | 45.70 | 4.80 | 3.36 | Type III effector HopC1 |
| *PSPTO_0591* | | + | 649880 | 650980 | 17.62 | 10.07 | 11.26 | 3.93 | 12.14 | 15.49 | ImpB/mucB/samB family protein |
| *PSPTO_0592* | *trpG* | + | 651383 | 651982 | 21.16 | 34.30 | 26.78 | 9.65 | 89.78 | 19.42 | Anthranilate synthase component II |
| *PSPTO_0593* | *trpD* | + | 652004 | 653053 | 11.38 | 28.10 | 15.38 | 19.03 | 76.64 | 15.26 | Anthranilate phosphoribosyltransferase |
| *PSPTO_0594* | *trpC* | + | 653050 | 653886 | 12.35 | 34.18 | 33.66 | 12.17 | 87.29 | 15.53 | Indole-3-glycerol phosphate synthase |
| *PSPTO_0595* | | + | 653921 | 654616 | 11.33 | 27.18 | 17.64 | 15.09 | 87.45 | 19.06 | Hypothetical protein |
| *PSPTO_0596* | *crp* | - | 654628 | 655272 | 26.64 | 11.78 | 6.13 | 6.59 | 8.37 | 8.70 | Catabolite gene activator Crp |
| *PSPTO_0597* | | + | 655482 | 655904 | 16.77 | 117.25 | 85.47 | 37.29 | 78.76 | 27.79 | OsmC/Ohr family protein |
| *PSPTO_0598* | *speD* | + | 656147 | 656941 | 10.82 | 17.64 | 19.30 | 10.85 | 35.31 | 10.92 | S-adenosylmethionine decarboxylase |
| *PSPTO_0599* | | - | 657002 | 657649 | 62.49 | 98.02 | 96.28 | 56.21 | 213.67 | 96.52 | Hypothetical protein |
| *PSPTO_0600* | | - | 657771 | 658109 | 16.39 | 29.28 | 17.09 | 22.35 | 89.64 | 24.17 | HIT family protein |
| *PSPTO_0601* | | - | 658212 | 658988 | 9.09 | 8.46 | 10.60 | 6.73 | 45.44 | 14.47 | Short chain dehydrogenase/reductase family oxidoreductase |
| *PSPTO_0602* | | - | 659042 | 659974 | 7.20 | 13.91 | 12.48 | 2.19 | 24.97 | 9.82 | Hypothetical protein |
| *PSPTO_0603* | | - | 660073 | 660501 | 8.68 | 17.12 | 16.50 | 11.51 | 22.96 | 8.36 | Hypothetical protein |
| *PSPTO_0604* | *argC* | + | 660635 | 661669 | 12.02 | 88.16 | 66.55 | 33.89 | 166.10 | 43.66 | N-acetyl-gamma-glutamyl-phosphate reductase |
| *PSPTO_0605* | | + | 661872 | 662222 | 203.36 | 219.33 | 280.14 | 181.58 | 245.09 | 56.35 | HesB/YadR/YfhF family protein |
| *PSPTO_0606* | | - | 662286 | 663356 | 16.60 | 56.19 | 40.20 | 15.26 | 80.01 | 40.42 | Hypothetical protein |
| *PSPTO_0607* | | - | 663384 | 664808 | 122.01 | 136.89 | 119.02 | 91.05 | 268.87 | 161.08 | Peptidase, M23/M37 family |
| *PSPTO_0608* | | - | 664805 | 664906 | 297.15 | 510.12 | 417.33 | 259.87 | 418.17 | 483.62 | Hypothetical protein |
| *PSPTO_0609* | *tyrS* | + | 665010 | 666221 | 43.97 | 21.45 | 23.43 | 9.95 | 138.87 | 22.84 | Tyrosyl-tRNA synthetase |
| *PSPTO_0610* | *birA* | + | 672113 | 673072 | 10.98 | 14.06 | 13.92 | 10.13 | 35.68 | 17.29 | BirA bifunctional protein |
| *PSPTO_0611* | | + | 673062 | 673811 | 11.31 | 14.79 | 17.11 | 2.06 | 22.37 | 15.05 | Pantothenate kinase |
| *PSPTO_0612* | | + | 673819 | 674247 | 17.18 | 22.01 | 22.06 | 10.38 | 35.16 | 28.70 | Hypothetical protein |
| *PSPTO_0613* | *secE* | + | 674963 | 675331 | 27.50 | 101.87 | 78.24 | 70.32 | 158.88 | 97.33 | Preprotein translocase subunit SecE |
| *PSPTO_0614* | *nusG* | + | 675341 | 675874 | 15.42 | 59.80 | 75.77 | 37.67 | 112.85 | 65.87 | Transcription antitermination protein NusG |
| *PSPTO_0615* | *rplK* | + | 675993 | 676424 | 13.61 | 167.07 | 138.55 | 82.46 | 427.17 | 48.93 | 50S ribosomal protein L11 |
| *PSPTO_0616* | *rplA* | + | 676424 | 677119 | 26.08 | 345.04 | 207.85 | 122.14 | 1445.17 | 116.10 | 50S ribosomal protein L1 |
| *PSPTO_0617* | *rplJ* | + | 677322 | 677822 | 19.10 | 411.35 | 320.74 | 150.41 | 2063.88 | 82.30 | 50S ribosomal protein L10 |
| *PSPTO_0618* | *rplL* | + | 677902 | 678267 | 17.54 | 137.60 | 99.96 | 56.84 | 688.80 | 52.72 | 50S ribosomal protein L7/L12 |
| *PSPTO_0619* | *rpoB* | + | 678489 | 682562 | 19.43 | 70.65 | 60.48 | 39.93 | 346.21 | 40.90 | DNA-directed RNA polymerase subunit beta |
| *PSPTO_0620* | *rpoC* | + | 682627 | 686826 | 37.57 | 120.37 | 88.18 | 51.61 | 651.38 | 63.42 | DNA-directed RNA polymerase subunit beta' |
| *PSPTO_0621* | *rpsL* | + | 687024 | 687395 | 124.67 | 578.80 | 500.49 | 387.56 | 1547.92 | 240.03 | 30S ribosomal protein S12 |
| *PSPTO_0622* | *rpsG* | + | 687518 | 687988 | 51.72 | 301.03 | 259.41 | 192.69 | 1018.54 | 136.96 | 30S ribosomal protein S7 |
| *PSPTO_0623* | *fusA* | + | 688019 | 690124 | 62.66 | 332.73 | 282.36 | 203.62 | 1681.21 | 179.04 | Translation elongation factor G |
| *PSPTO_0624* | *tuf* | + | 690156 | 691349 | 61.30 | 433.91 | 339.42 | 243.76 | 1677.18 | 191.37 | Translation elongation factor Tu |
| *PSPTO_0625* | *rpsJ* | + | 691491 | 691802 | 323.59 | 893.16 | 797.05 | 540.71 | 1569.35 | 460.78 | 30S ribosomal protein S10 |
| *PSPTO_0626* | *rplC* | + | 691884 | 692519 | 67.75 | 699.03 | 537.76 | 377.49 | 2330.71 | 317.87 | 50S ribosomal protein L3 |
| *PSPTO_0627* | *rplD* | + | 692532 | 693134 | 37.12 | 377.65 | 267.78 | 208.22 | 1575.67 | 164.40 | 50S ribosomal protein L4 |
| *PSPTO_0628* | *rplW* | + | 693131 | 693430 | 41.93 | 499.99 | 449.11 | 239.34 | 2101.77 | 290.60 | 50S ribosomal protein L23 |
| *PSPTO_0629* | *rplB* | + | 693445 | 694269 | 85.42 | 621.16 | 347.98 | 307.28 | 3033.47 | 225.19 | 50S ribosomal protein L2 |
| *PSPTO_0630* | *rpsS* | + | 694286 | 694561 | 244.88 | 1814.08 | 1544.20 | 1059.40 | 8612.21 | 1129.85 | 30S ribosomal protein S19 |
| *PSPTO_0631* | *rplV* | + | 694575 | 694907 | 84.84 | 652.01 | 391.37 | 276.83 | 3426.72 | 396.49 | 50S ribosomal protein L22 |
| *PSPTO_0632* | *rpsC* | + | 694921 | 695607 | 34.42 | 261.41 | 168.10 | 128.29 | 1075.48 | 119.01 | 30S ribosomal protein S3 |
| *PSPTO_0633* | *rplP* | + | 695620 | 696033 | 33.07 | 257.69 | 248.93 | 119.89 | 1206.68 | 97.30 | 50S ribosomal protein L16 |
| *PSPTO_0634* | *rpmC* | + | 696033 | 696224 | 27.49 | 161.90 | 60.66 | 68.01 | 798.02 | 57.22 | 50S ribosomal protein L29 |
| *PSPTO_0635* | *rpsQ* | + | 696227 | 696493 | 35.50 | 189.78 | 144.88 | 133.66 | 1123.74 | 63.55 | 30S ribosomal protein S17 |
| *PSPTO_0636* | *rplN* | + | 696517 | 696885 | 46.11 | 241.72 | 181.30 | 96.51 | 1641.45 | 102.14 | 50S ribosomal protein L14 |
| *PSPTO_0637* | *rplX* | + | 696897 | 697211 | 50.70 | 354.59 | 220.91 | 128.51 | 1965.45 | 119.50 | 50S ribosomal protein L24 |
| *PSPTO_0638* | *rplE* | + | 697232 | 697771 | 45.05 | 279.08 | 206.73 | 96.06 | 1599.69 | 105.54 | 50S ribosomal protein L5 |

| Locus name | Gene name | Strand | Gene start (bp) | Gene end (bp) | Average TPM Inoculum | Average TPM $t_6$ | Average TPM $t_{24}$ | Average TPM $t_{168}$ | Average TPM Logarithmic | Average TPM Stationary | Annotation description |
|---|---|---|---|---|---|---|---|---|---|---|---|
| *PSPTO_0639* | *rpsN* | + | 697785 | 698090 | 9.41 | 58.94 | 51.21 | 37.23 | 269.99 | 37.87 | 30S ribosomal protein S14 |
| *PSPTO_0640* | *rpsH* | + | 698305 | 698697 | 53.07 | 655.28 | 716.52 | 392.79 | 1070.75 | 236.79 | 30S ribosomal protein S8 |
| *PSPTO_0641* | *rplF* | + | 698710 | 699243 | 69.02 | 627.44 | 469.57 | 414.24 | 2586.89 | 426.05 | 50S ribosomal protein L6 |
| *PSPTO_0642* | *rplR* | + | 699254 | 699604 | 53.76 | 460.04 | 409.99 | 220.74 | 2068.83 | 282.24 | 50S ribosomal protein L18 |
| *PSPTO_0643* | *rpsE* | + | 699608 | 700108 | 28.87 | 261.12 | 163.32 | 102.72 | 1130.21 | 103.38 | 30S ribosomal protein S5 |
| *PSPTO_0644* | *rpmD* | + | 700111 | 700287 | 39.61 | 437.67 | 307.37 | 167.73 | 1260.76 | 186.01 | 50S ribosomal protein L30 |
| *PSPTO_0645* | *rplO* | + | 700291 | 700725 | 43.88 | 454.43 | 324.33 | 158.50 | 2227.51 | 182.11 | 50S ribosomal protein L15 |
| *PSPTO_0646* | *secY* | + | 700726 | 702060 | 53.49 | 329.59 | 248.64 | 208.71 | 1431.08 | 213.60 | Preprotein translocase subunit SecY |
| *PSPTO_0647* | *rpmJ* | + | 702089 | 702205 | 158.91 | 788.69 | 517.46 | 525.25 | 3000.56 | 484.62 | 50S ribosomal protein L36 |
| *PSPTO_0648* | *rpsM* | + | 702336 | 702692 | 113.01 | 438.31 | 356.40 | 247.44 | 1340.59 | 239.44 | 30S ribosomal protein S13 |
| *PSPTO_0649* | *rpsK* | + | 702723 | 703112 | 61.69 | 359.50 | 289.75 | 253.05 | 1259.29 | 180.28 | 30S ribosomal protein S11 |
| *PSPTO_0650* | *rpsD* | + | 703131 | 703751 | 37.86 | 146.36 | 138.95 | 97.45 | 546.28 | 76.33 | 30S ribosomal protein S4 |
| *PSPTO_0651* | *rpoA* | + | 703774 | 704775 | 46.35 | 200.11 | 173.22 | 129.18 | 1019.45 | 122.22 | DNA-directed RNA polymerase subunit alpha |
| *PSPTO_0652* | *rplQ* | + | 704817 | 705203 | 29.50 | 123.87 | 110.64 | 73.92 | 1213.88 | 96.90 | 50S ribosomal protein L17 |
| *PSPTO_0653* | | + | 705466 | 705930 | 53.96 | 208.72 | 148.78 | 91.91 | 61.12 | 12.79 | Bacterioferritin |
| *PSPTO_0654* | *uvrA* | - | 706024 | 709002 | 17.30 | 24.09 | 22.17 | 11.90 | 46.20 | 22.90 | Excinuclease ABC subunit A |
| *PSPTO_0655* | | + | 708988 | 710382 | 61.80 | 113.41 | 84.46 | 56.29 | 225.14 | 96.51 | Major facilitator family transporter |
| *PSPTO_0656* | *ssB* | + | 710392 | 710961 | 43.99 | 127.70 | 114.31 | 68.61 | 292.49 | 106.56 | Single-stranded DNA-binding protein |
| *PSPTO_0657* | | - | 711177 | 711767 | 25.60 | 19.50 | 18.26 | 10.20 | 27.77 | 13.16 | Hypothetical protein |
| *PSPTO_0658* | | + | 711853 | 712398 | 30.04 | 28.78 | 28.56 | 22.03 | 59.99 | 35.95 | Hypothetical protein |
| *PSPTO_0659* | | - | 712538 | 713281 | 13.95 | 32.87 | 30.77 | 10.77 | 52.61 | 18.55 | Inositol monophosphatase family protein |
| *PSPTO_0660* | | - | 713278 | 714039 | 19.74 | 48.28 | 20.89 | 11.00 | 42.35 | 13.92 | Glycerophosphoryl diester phosphodiesterase |
| *PSPTO_0661* | *ugpC* | - | 714046 | 715152 | 1.72 | 2.92 | 1.85 | 0.33 | 3.68 | 1.64 | Glycerol-3-phosphate ABC transporter, ATP-binding protein |
| *PSPTO_0662* | *ugpE* | - | 715149 | 716042 | 2.21 | 1.52 | 4.18 | 0.00 | 3.23 | 2.14 | Glycerol-3-phosphate ABC transporter permease |
| *PSPTO_0663* | *ugpA* | - | 716039 | 716932 | 6.71 | 6.47 | 9.22 | 7.42 | 9.83 | 4.30 | Glycerol-3-phosphate ABC transporter permease |
| *PSPTO_0664* | *ugpB* | - | 716929 | 718365 | 1.72 | 1.82 | 3.04 | 0.00 | 2.83 | 1.72 | Glycerol-3-phosphate ABC transporter substrate-binding protein |
| *PSPTO_0665* | | + | 718531 | 719661 | 7.08 | 9.39 | 4.79 | 1.30 | 23.49 | 7.68 | ROK family protein |
| *PSPTO_0666* | | - | 719689 | 720210 | 15.28 | 15.10 | 11.15 | 11.00 | 47.74 | 16.55 | Hypothetical protein |
| *PSPTO_0667* | | - | 720816 | 721586 | 8.92 | 10.69 | 9.35 | 9.11 | 16.13 | 19.42 | Short-chain dehydrogenase/reductase family oxidoreductase |
| *PSPTO_0668* | | - | 721761 | 722429 | 19.03 | 16.79 | 12.19 | 11.00 | 12.96 | 16.18 | Hypothetical protein |
| *PSPTO_0669* | | + | 723766 | 724125 | 0.00 | 0.00 | 0.00 | 0.00 | 0.00 | 0.00 | ISPsy5, Orf1 |
| *PSPTO_0670* | | + | 724143 | 725696 | 0.00 | 0.00 | 0.00 | 0.00 | 0.00 | 0.00 | ISPsy5, transposase |
| *PSPTO_0671* | | - | 726055 | 728361 | 5.99 | 4.41 | 4.76 | 2.13 | 9.67 | 9.44 | TonB-dependent receptor |
| *PSPTO_0672* | *exbD-2* | - | 728432 | 728836 | 3.51 | 4.78 | 0.00 | 5.59 | 4.57 | 4.64 | TonB system transport protein ExbD |
| *PSPTO_0673* | *exbB-2* | - | 728838 | 729530 | 1.62 | 1.49 | 0.45 | 2.90 | 4.40 | 4.19 | TonB system transport protein ExbB |
| *PSPTO_0674* | | - | 729541 | 730344 | 1.31 | 0.94 | 1.67 | 0.00 | 2.92 | 1.93 | TonB domain protein |
| *PSPTO_0675* | *cpoF* | - | 730513 | 731331 | 2.96 | 32.86 | 18.10 | 4.01 | 6.05 | 8.52 | Arylesterase |
| *PSPTO_0676* | | + | 731587 | 733890 | 21.82 | 39.69 | 44.47 | 18.85 | 22.98 | 28.32 | Mechanosensitive ion channel family protein |
| *PSPTO_0677* | | + | 734058 | 734936 | 8.19 | 28.01 | 33.73 | 15.13 | 63.76 | 16.80 | Lipopolysaccharide biosynthesis protein |
| *PSPTO_0678* | | + | 734929 | 735858 | 4.98 | 24.12 | 13.42 | 7.32 | 73.70 | 17.53 | NAD-dependent epimerase/dehydratase family protein |
| *PSPTO_0679* | *ompW* | - | 736029 | 736715 | 15.75 | 1006.99 | 661.46 | 112.83 | 395.90 | 2177.54 | Outer membrane protein OmpW |
| *PSPTO_0680* | | - | 736819 | 737343 | 11.41 | 601.23 | 521.57 | 93.19 | 22.88 | 53.63 | Lipoprotein |
| *PSPTO_0681* | | - | 737352 | 738617 | 5.73 | 222.59 | 154.78 | 30.90 | 11.41 | 9.45 | Permease |
| *PSPTO_0682* | | - | 738619 | 739329 | 2.88 | 7.25 | 14.90 | 4.77 | 10.17 | 5.17 | ABC transporter ATP-binding protein |
| *PSPTO_0683* | | - | 739399 | 739986 | 84.62 | 188.26 | 219.31 | 88.62 | 140.81 | 36.17 | Hypothetical protein |
| *PSPTO_0684* | | + | 740161 | 740712 | 61.59 | 33.25 | 20.12 | 22.67 | 16.03 | 26.92 | Hypothetical protein |
| *PSPTO_0685* | | + | 740788 | 741144 | 17.08 | 51.65 | 31.40 | 7.39 | 60.05 | 32.92 | Hypothetical protein |
| *PSPTO_0686* | *trx-1* | - | 741199 | 742071 | 13.60 | 31.52 | 14.60 | 11.97 | 54.10 | 15.75 | Thioredoxin |
| *PSPTO_0687* | | - | 742244 | 742897 | 3.76 | 3.09 | 4.83 | 1.42 | 15.76 | 6.81 | Hypothetical protein |
| *PSPTO_0688* | | - | 742894 | 743343 | 10.53 | 9.61 | 31.06 | 17.07 | 57.19 | 14.50 | Lipoprotein |
| *PSPTO_0689* | | + | 743520 | 743984 | 20.86 | 27.75 | 31.55 | 7.18 | 76.31 | 22.35 | Hypothetical protein |
| *PSPTO_0690* | *ribD* | + | 743981 | 745126 | 8.60 | 25.38 | 18.64 | 10.82 | 65.60 | 19.39 | Riboflavin biosynthesis protein RibD |
| *PSPTO_0691* | *ribE* | + | 745174 | 745836 | 12.78 | 30.35 | 31.04 | 2.50 | 163.33 | 34.67 | Riboflavin synthase subunit alpha |
| *PSPTO_0692* | *ribBA-1* | + | 745854 | 746945 | 8.80 | 30.33 | 18.91 | 9.69 | 130.47 | 27.61 | Bifunctional 3,4-dihydroxy-2-butanone 4-phosphate synthase/GTP cyclohydrolase II-like protein |
| *PSPTO_0693* | *ribH-1* | + | 747074 | 747550 | 16.48 | 41.19 | 30.91 | 18.57 | 182.19 | 59.58 | 6,7-dimethyl-8-ribityllumazine synthase |
| *PSPTO_0694* | *nusB* | + | 747547 | 748044 | 10.00 | 29.30 | 23.85 | 13.26 | 118.59 | 49.44 | N utilization substance protein B |
| *PSPTO_0695* | *thiL* | + | 748061 | 749026 | 3.21 | 8.06 | 4.56 | 2.20 | 27.73 | 7.94 | Thiamine monophosphate kinase |
| *PSPTO_0696* | *ribA* | + | 749113 | 749778 | 16.99 | 65.33 | 31.07 | 39.24 | 110.64 | 39.11 | GTP cyclohydrolase II |
| *PSPTO_0697* | | + | 749775 | 750191 | 25.97 | 28.99 | 17.00 | 9.28 | 34.89 | 14.46 | Hypothetical protein |
| *PSPTO_0698* | *dxs* | - | 750560 | 752455 | 10.09 | 31.53 | 34.85 | 10.44 | 75.11 | 17.20 | Deoxyxylulose-5-phosphate synthase |
| *PSPTO_0699* | *ispA* | - | 752619 | 753506 | 3.56 | 9.82 | 17.04 | 1.09 | 41.18 | 9.66 | Geranyltranstransferase |
| *PSPTO_0700* | *xseB* | - | 753503 | 753748 | 13.14 | 69.51 | 40.36 | 26.73 | 79.85 | 21.43 | Exodeoxyribonuclease VII small subunit |
| *PSPTO_0701* | *metC-1* | - | 753852 | 755018 | 10.19 | 11.57 | 12.38 | 4.37 | 24.71 | 8.42 | Cystathionine beta-lyase |
| *PSPTO_0702* | | - | 755088 | 756107 | 7.15 | 12.16 | 11.14 | 2.91 | 26.53 | 9.53 | Hypothetical protein |
| *PSPTO_0703* | | - | 756309 | 757223 | 11.59 | 19.01 | 37.28 | 15.45 | 63.06 | 18.56 | Putative S-transferase |
| *PSPTO_0704* | | + | 757479 | 758255 | 4.58 | 5.67 | 3.97 | 2.29 | 13.39 | 10.09 | O-antigen ABC transporter, permease protein |
| *PSPTO_0705* | | + | 758252 | 758956 | 0.05 | 2.15 | 0.00 | 0.00 | 2.24 | 1.62 | O-antigen ABC transporter, ATP-binding protein |
| *PSPTO_0706* | | + | 759038 | 759799 | 12.16 | 1.43 | 1.45 | 3.34 | 3.53 | 2.28 | Lipopolysaccharide biosynthesis protein |
| *PSPTO_0707* | | + | 759801 | 760415 | 4.96 | 2.87 | 2.56 | 1.48 | 5.53 | 3.55 | Hypothetical protein |
| *PSPTO_0708* | | + | 760423 | 761061 | 1.55 | 0.39 | 2.39 | 2.29 | 5.89 | 2.72 | HAD-superfamily hydrolase |
| *PSPTO_0709* | | + | 761067 | 761789 | 3.73 | 0.00 | 0.00 | 3.31 | 7.60 | 5.57 | Hypothetical protein |
| *PSPTO_0710* | | + | 761810 | 762106 | 10.60 | 1.21 | 12.64 | 8.96 | 12.33 | 12.61 | Hypothetical protein |
| *PSPTO_0711* | | + | 762110 | 763885 | 8.46 | 6.60 | 7.24 | 2.58 | 12.70 | 9.48 | Hypothetical protein |
| *PSPTO_0712* | | + | 763882 | 764487 | 3.48 | 3.26 | 1.62 | 2.38 | 5.59 | 4.56 | Hypothetical protein |
| *PSPTO_0713* | | + | 764541 | 765584 | 12.30 | 11.65 | 9.81 | 8.58 | 13.81 | 15.64 | Hypothetical protein |
| *PSPTO_0714* | | + | 765763 | 768054 | 8.46 | 10.97 | 9.13 | 9.58 | 18.70 | 11.23 | Autotransporter |
| *PSPTO_0715* | | + | 768108 | 768905 | 5.47 | 3.98 | 8.15 | 13.95 | 12.34 | 7.74 | Hypothetical protein |
| *PSPTO_0716* | | - | 768928 | 769200 | 8.43 | 26.34 | 14.75 | 2.04 | 13.98 | 14.33 | DNA-binding protein |
| *PSPTO_0717* | | - | 769224 | 769466 | 47.04 | 81.96 | 72.67 | 22.49 | 91.32 | 51.52 | DNA-binding protein |

| Locus name | Gene name | Strand | Gene start (bp) | Gene end (bp) | Average TPM Inoculum | Average TPM $t_6$ | Average TPM $t_{24}$ | Average TPM $t_{168}$ | Average TPM Logarithmic | Average TPM Stationary | Annotation description |
|---|---|---|---|---|---|---|---|---|---|---|---|
| *PSPTO_0718* | | + | 769587 | 769985 | 13.60 | 15.70 | 16.38 | 7.51 | 30.50 | 20.00 | Hypothetical protein |
| *PSPTO_0719* | | + | 769966 | 770142 | 47.36 | 37.61 | 38.45 | 29.55 | 104.55 | 34.99 | Hypothetical protein |
| *PSPTO_0720* | | + | 770144 | 770413 | 42.25 | 36.26 | 13.22 | 10.65 | 35.57 | 24.17 | Hypothetical protein |
| *PSPTO_0721* | | - | 770709 | 771440 | 6.80 | 10.27 | 8.91 | 6.18 | 14.17 | 6.53 | Transcriptional regulator |
| *PSPTO_0722* | *ppa-1* | - | 771799 | 772326 | 82.34 | 101.39 | 97.24 | 26.71 | 342.97 | 66.76 | Inorganic pyrophosphatase |
| *PSPTO_0723* | | - | 772437 | 773246 | 5.68 | 9.92 | 10.96 | 11.81 | 53.25 | 18.61 | Hypothetical protein |
| *PSPTO_0724* | | - | 773250 | 773897 | 5.82 | 14.92 | 4.69 | 2.37 | 22.20 | 8.12 | DedA family protein |
| *PSPTO_0725* | | - | 774090 | 774560 | 10.54 | 14.77 | 22.56 | 2.13 | 21.33 | 9.04 | GNAT family acetyltransferase |
| *PSPTO_0726* | *eutC* | - | 774632 | 775495 | 6.88 | 20.90 | 11.00 | 5.75 | 16.03 | 11.26 | Ethanolamine ammonia-lyase, light subunit |
| *PSPTO_0727* | *eutB* | - | 775492 | 776886 | 22.43 | 66.37 | 49.83 | 15.57 | 42.55 | 23.57 | Ethanolamine ammonia-lyase, heavy subunit |
| *PSPTO_0728* | | - | 777182 | 778702 | 43.38 | 111.11 | 124.32 | 63.76 | 285.70 | 24.84 | Aldehyde dehydrogenase family protein |
| *PSPTO_0729* | *mpl* | + | 778928 | 780277 | 8.53 | 31.89 | 19.14 | 17.94 | 66.05 | 19.88 | UDP-N-acetylmuramate:L-alanyl-gamma-D-glutamyl-meso-diaminopimelate ligase |
| *PSPTO_0730* | | + | 780274 | 780903 | 6.18 | 31.33 | 14.29 | 5.37 | 46.25 | 17.17 | Aromatic acid decarboxylase |
| *PSPTO_0731* | | + | 780884 | 781180 | 6.35 | 21.70 | 16.29 | 10.25 | 39.77 | 19.44 | Lipoprotein |
| *PSPTO_0732* | | - | 781219 | 781860 | 32.00 | 24.48 | 31.89 | 18.83 | 37.87 | 36.15 | Hypothetical protein |
| *PSPTO_0733* | | - | 781888 | 783615 | 46.68 | 37.02 | 23.21 | 18.34 | 53.44 | 32.67 | MORN repeat-containing protein |
| *PSPTO_0734* | | - | 783756 | 784211 | 27.22 | 19.88 | 16.70 | 17.10 | 16.68 | 17.80 | MaoC-like domain protein |
| *PSPTO_0735* | | + | 784409 | 784771 | 9.86 | 11.28 | 10.36 | 0.00 | 17.74 | 10.61 | LrgA family protein |
| *PSPTO_0736* | | + | 784768 | 785484 | 22.08 | 28.32 | 21.61 | 26.29 | 34.56 | 27.91 | Membrane protein |
| *PSPTO_0737* | | + | 785498 | 786088 | 19.74 | 24.91 | 28.14 | 24.99 | 65.02 | 47.47 | ATP-dependent protease La domain-containing protein |
| *PSPTO_0738* | | - | 786218 | 786781 | 13.04 | 18.11 | 41.16 | 28.74 | 34.14 | 45.28 | Lipoprotein |
| *PSPTO_0739* | | - | 786811 | 787449 | 15.99 | 21.02 | 29.60 | 21.03 | 27.18 | 30.81 | Lipoprotein |
| *PSPTO_0740* | *smtA* | - | 787469 | 788134 | 25.52 | 37.11 | 19.28 | 16.63 | 49.47 | 35.02 | SmtA protein |
| *PSPTO_0741* | | - | 788215 | 788520 | 31.06 | 40.16 | 42.84 | 18.28 | 78.20 | 44.90 | Hypothetical protein |
| *PSPTO_0742* | | - | 788635 | 789492 | 4.34 | 40.60 | 32.25 | 16.45 | 14.22 | 21.01 | MaoC-like domain protein |
| *PSPTO_0743* | | - | 789523 | 790878 | 12.52 | 55.59 | 49.64 | 20.62 | 30.59 | 35.08 | 3-ketoacyl-(acyl-carrier-protein) reductase |
| *PSPTO_0744* | | + | 790987 | 792264 | 10.84 | 14.67 | 12.38 | 9.32 | 11.76 | 16.07 | Acetyl-CoA acetyltransferase |
| *PSPTO_0745* | | + | 792445 | 792675 | 25.88 | 24.69 | 35.64 | 4.55 | 74.64 | 19.60 | Hypothetical protein |
| *PSPTO_0746* | | + | 792708 | 794105 | 3.89 | 3.41 | 7.40 | 3.60 | 23.86 | 7.07 | Hypothetical protein |
| *PSPTO_0747* | | - | 794113 | 795141 | 5.61 | 12.00 | 1.35 | 6.73 | 23.82 | 9.64 | Transporter |
| *PSPTO_0748* | | + | 795585 | 796481 | 60.27 | 102.10 | 95.87 | 45.84 | 145.48 | 40.24 | RIO1 family protein |
| *PSPTO_0749* | | - | 796601 | 797002 | 29.39 | 22.64 | 11.43 | 6.29 | 37.32 | 17.19 | Heavy metal-dependent transcriptional regulator |
| *PSPTO_0750* | | - | 796999 | 799197 | 8.04 | 18.17 | 15.02 | 4.80 | 20.09 | 10.82 | Copper-translocating P-type ATPase |
| *PSPTO_0751* | | - | 799267 | 799647 | 48.93 | 57.05 | 87.31 | 16.80 | 91.00 | 35.00 | Hypothetical protein |
| *PSPTO_0752* | | + | 799794 | 799991 | 14.87 | 50.67 | 18.82 | 3.61 | 15.30 | 16.70 | CopZ protein |
| *PSPTO_0753* | | + | 800138 | 801334 | 5.67 | 10.01 | 9.75 | 8.08 | 9.45 | 7.44 | Bcr/CflA family multidrug resistance transporter |
| *PSPTO_0754* | | - | 801383 | 802432 | 6.07 | 19.69 | 17.68 | 3.41 | 11.79 | 7.22 | Alcohol dehydrogenase |
| *PSPTO_0755* | | + | 802509 | 803432 | 2.57 | 3.06 | 4.01 | 3.59 | 3.87 | 1.94 | LysR family transcriptional regulator |
| *PSPTO_0756* | | + | 803869 | 804561 | 7.53 | 7.38 | 14.38 | 8.06 | 18.61 | 15.95 | Transcriptional regulator |
| *PSPTO_0757* | *add* | - | 804854 | 805807 | 15.87 | 40.28 | 23.90 | 7.15 | 71.79 | 16.14 | Adenosine deaminase |
| *PSPTO_0758* | | - | 805886 | 806851 | 16.52 | 9.33 | 11.85 | 5.97 | 12.02 | 6.41 | 2OG-Fe(II) oxygenase family oxidoreductase |
| *PSPTO_0759* | | - | 806929 | 808023 | 96.62 | 28.69 | 17.77 | 12.95 | 16.85 | 9.06 | Bmp family protein |
| *PSPTO_0760* | | - | 808253 | 809044 | 1.14 | 0.00 | 0.60 | 0.00 | 14.13 | 2.94 | Iron(III) dicitrate transport system, ATP-binding protein FecE |
| *PSPTO_0761* | *fecD* | - | 809048 | 810016 | 2.86 | 2.45 | 1.60 | 1.35 | 10.49 | 5.63 | Iron(III) dicitrate transport system, permease protein FecD |
| *PSPTO_0762* | *fecC* | - | 810013 | 810987 | 3.48 | 7.59 | 2.77 | 1.32 | 7.50 | 5.55 | Iron-dicitrate transporter permease subunit |
| *PSPTO_0763* | *fecB* | - | 811003 | 811923 | 0.97 | 4.09 | 0.13 | 1.70 | 9.68 | 4.95 | Iron(III) dicitrate transport system, periplasmic iron-binding protein FecB |
| *PSPTO_0764* | | + | 812170 | 813267 | 3.65 | 3.66 | 1.00 | 1.31 | 3.88 | 4.45 | Calcium/proton antiporter |
| *PSPTO_0765* | *atzB* | - | 813434 | 814789 | 2.31 | 4.55 | 4.45 | 0.81 | 7.17 | 6.73 | Hydroxydechloroatrazine ethylaminohydrolase |
| *PSPTO_0766* | | - | 814913 | 815638 | 11.71 | 26.72 | 40.94 | 5.77 | 47.22 | 16.16 | Short chain dehydrogenase |
| *PSPTO_0767* | | - | 815889 | 816815 | 2.97 | 6.62 | 1.40 | 0.00 | 1.89 | 1.85 | ABC transporter permease |
| *PSPTO_0768* | | - | 816815 | 817915 | 5.17 | 2.75 | 0.60 | 0.00 | 3.35 | 2.66 | ABC transporter permease |
| *PSPTO_0769* | | - | 817927 | 819480 | 18.68 | 5.33 | 2.66 | 0.83 | 2.34 | 2.59 | ABC transporter ATP-binding protein |
| *PSPTO_0770* | | - | 820004 | 820597 | 2.07 | 2.34 | 5.02 | 5.99 | 22.89 | 8.55 | Hypothetical protein |
| *PSPTO_0771* | | - | 820630 | 821286 | 14.31 | 6.82 | 10.93 | 0.84 | 31.90 | 9.39 | TetR family transcriptional regulator |
| *PSPTO_0772* | | - | 821385 | 822782 | 13.75 | 11.73 | 7.98 | 3.52 | 11.39 | 7.12 | Xanthine/uracil permease family protein |
| *PSPTO_0773* | | - | 822968 | 823888 | 6.01 | 5.62 | 15.22 | 1.55 | 17.13 | 10.60 | LysR family transcriptional regulator |
| *PSPTO_0774* | | + | 824016 | 825362 | 12.52 | 50.09 | 53.83 | 15.58 | 12.18 | 11.65 | Beta-alanine--pyruvate aminotransferase |
| *PSPTO_0775* | *mmsA-1* | + | 825456 | 826949 | 8.14 | 28.38 | 30.58 | 16.37 | 12.96 | 11.48 | Methylmalonate-semialdehyde dehydrogenase |
| *PSPTO_0776* | *recC* | + | 832977 | 836453 | 5.06 | 4.36 | 7.64 | 5.70 | 18.71 | 7.26 | Exodeoxyribonuclease V subunit gamma |
| *PSPTO_0777* | *recB* | + | 836450 | 840142 | 3.90 | 5.83 | 3.43 | 1.78 | 18.88 | 7.03 | Exodeoxyribonuclease V subunit beta |
| *PSPTO_0778* | *recD* | + | 840139 | 842232 | 2.69 | 3.65 | 7.23 | 7.73 | 14.55 | 6.19 | Exodeoxyribonuclease V subunit alpha |
| *PSPTO_0779* | | - | 842459 | 843613 | 1.28 | 1.66 | 0.54 | 6.66 | 1.80 | 1.60 | Hypothetical protein |
| *PSPTO_0780* | | + | 843955 | 844932 | 0.00 | 0.00 | 0.00 | 0.00 | 0.00 | 0.00 | ISPssy, transposase |
| *PSPTO_0781* | | - | 845447 | 846391 | 2.63 | 6.69 | 3.51 | 0.91 | 16.41 | 5.03 | LysR family transcriptional regulator |
| *PSPTO_0782* | *mmsA-2* | + | 846549 | 848075 | 89.47 | 6.43 | 4.39 | 6.29 | 5.42 | 111.75 | Methylmalonate-semialdehyde dehydrogenase |
| *PSPTO_0783* | *mmsB* | + | 848110 | 849000 | 57.90 | 6.62 | 8.83 | 10.92 | 4.53 | 84.82 | 3-hydroxyisobutyrate dehydrogenase |
| *PSPTO_0784* | | + | 849115 | 851085 | 4.21 | 2.68 | 5.25 | 0.28 | 6.53 | 6.26 | TonB-dependent siderophore receptor |
| *PSPTO_0785* | | - | 851370 | 852716 | 2.50 | 1.40 | 2.48 | 2.98 | 11.41 | 7.61 | Sensor histidine kinase |
| *PSPTO_0786* | | - | 852746 | 853510 | 1.58 | 1.37 | 2.54 | 2.87 | 16.50 | 14.56 | DNA-binding response regulator |
| *PSPTO_0787* | *ggt* | + | 853803 | 855530 | 6.12 | 11.34 | 10.99 | 5.67 | 16.01 | 4.64 | Gamma-glutamyltranspeptidase |
| *PSPTO_0788* | | - | 855610 | 856377 | 1.89 | 3.67 | 2.32 | 3.05 | 5.57 | 2.43 | Phosphonate ABC transporter, permease protein |
| *PSPTO_0789* | | - | 856374 | 857201 | 2.66 | 5.45 | 4.10 | 2.20 | 4.95 | 1.66 | Phosphonate ABC transporter, permease protein |
| *PSPTO_0790* | | - | 857195 | 857992 | 2.07 | 3.31 | 0.35 | 4.96 | 6.37 | 2.22 | Phosphonate ABC transporter, ATP-binding protein |
| *PSPTO_0791* | | - | 857989 | 858840 | 9.67 | 23.85 | 4.99 | 10.37 | 27.84 | 15.18 | Phosphonate ABC transporter periplasmic phosphonate-binding protein |
| *PSPTO_0792* | | + | 858886 | 859020 | 4.00 | 0.00 | 0.00 | 2.86 | 12.83 | 8.16 | Hypothetical protein |
| *PSPTO_0793* | *selD* | + | 859077 | 860111 | 3.65 | 12.19 | 9.02 | 9.87 | 20.92 | 6.60 | Selenide, water dikinase |
| *PSPTO_0794* | | + | 860111 | 861211 | 2.28 | 5.58 | 4.62 | 7.17 | 10.20 | 3.66 | Hypothetical protein |
| *PSPTO_0795* | | + | 861246 | 862010 | 2.73 | 3.98 | 9.80 | 8.24 | 17.07 | 6.87 | Membrane protein |
| *PSPTO_0796* | *ispB* | - | 862083 | 863051 | 39.36 | 42.80 | 39.10 | 14.51 | 146.59 | 50.13 | Octylprenyl diphosphate synthase |

| Locus name | Gene name | Strand | Gene start (bp) | Gene end (bp) | Average TPM Inoculum | Average TPM $t_6$ | Average TPM $t_{24}$ | Average TPM $t_{168}$ | Average TPM Logarithmic | Average TPM Stationary | Annotation description |
|---|---|---|---|---|---|---|---|---|---|---|---|
| *PSPTO_0797* | *rplU* | + | 863291 | 863602 | 177.59 | 570.07 | 425.27 | 296.15 | 1306.89 | 213.43 | 50S ribosomal protein L21 |
| *PSPTO_0798* | *rpmA* | + | 863635 | 863910 | 16.78 | 128.59 | 90.09 | 86.55 | 125.27 | 37.20 | 50S ribosomal protein L27 |
| *PSPTO_0799* | | + | 864127 | 865350 | 10.94 | 48.40 | 32.11 | 17.80 | 75.52 | 21.53 | GTP-binding protein, GTP1/Obg family |
| *PSPTO_0800* | *proB* | + | 865441 | 866559 | 71.18 | 66.39 | 57.22 | 27.87 | 104.97 | 54.83 | Glutamate 5-kinase |
| *PSPTO_0801* | *creA* | + | 866585 | 867055 | 18.86 | 29.43 | 23.67 | 13.46 | 56.89 | 29.65 | CreA protein |
| *PSPTO_0802* | *rpsT* | - | 867260 | 867538 | 32.47 | 82.16 | 73.37 | 70.67 | 65.29 | 28.59 | 30S ribosomal protein S20 |
| *PSPTO_0803* | | + | 867642 | 867740 | 792.94 | 1147.94 | 774.86 | 657.40 | 1925.67 | 349.64 | Hypothetical protein |
| *PSPTO_0804* | | + | 867789 | 869327 | 6.05 | 9.46 | 6.34 | 3.46 | 13.21 | 7.26 | Membrane protein, MviN family |
| *PSPTO_0805* | *ribF* | + | 869460 | 870386 | 20.64 | 39.90 | 42.35 | 28.70 | 123.90 | 56.60 | Riboflavin biosynthesis protein RibF |
| *PSPTO_0806* | *ileS* | + | 870407 | 873238 | 5.15 | 31.24 | 27.68 | 15.79 | 121.53 | 26.11 | Isoleucyl-tRNA synthetase |
| *PSPTO_0807* | *lspA* | + | 873231 | 873752 | 17.77 | 45.00 | 40.15 | 52.85 | 257.40 | 39.24 | Lipoprotein signal peptidase |
| *PSPTO_0808* | | + | 873745 | 874221 | 11.72 | 30.22 | 24.24 | 23.38 | 120.80 | 23.89 | FKBP-type peptidyl-prolyl cis-trans isomerase |
| *PSPTO_0809* | *ispH* | + | 874305 | 875252 | 6.11 | 21.99 | 21.23 | 13.36 | 113.06 | 25.46 | Hydroxymethylbutenyl pyrophosphate reductase |
| *PSPTO_0810* | | - | 875259 | 875759 | 5.67 | 5.09 | 24.59 | 7.65 | 31.19 | 11.91 | Type IV pilus biogenesis protein |
| *PSPTO_0811* | | + | 875891 | 876364 | 106.09 | 113.62 | 91.70 | 52.59 | 231.84 | 42.61 | Pillin |
| *PSPTO_0812* | | + | 876364 | 876879 | 17.91 | 18.05 | 12.41 | 2.40 | 32.08 | 7.07 | Hypothetical protein |
| *PSPTO_0813* | | + | 876876 | 877583 | 17.92 | 19.21 | 6.26 | 5.37 | 55.00 | 11.70 | Hypothetical protein |
| *PSPTO_0814* | | + | 877595 | 878101 | 15.39 | 16.56 | 23.00 | 3.64 | 38.34 | 15.91 | Hypothetical protein |
| *PSPTO_0815* | | + | 878088 | 881291 | 21.92 | 23.93 | 21.41 | 11.18 | 64.68 | 22.76 | Type IV pilus-associated protein |
| *PSPTO_0816* | | + | 881303 | 881704 | 11.35 | 26.45 | 13.98 | 8.18 | 76.68 | 20.68 | Type IV pilus biogenesis protein |
| *PSPTO_0817* | | + | 881733 | 882836 | 6.90 | 13.34 | 7.61 | 5.42 | 50.86 | 14.02 | Oxidoreductase, FAD-binding protein |
| *PSPTO_0818* | | + | 882967 | 883410 | 4.77 | 10.31 | 3.87 | 0.00 | 17.24 | 5.41 | MarR family transcriptional regulator |
| *PSPTO_0819* | | + | 883449 | 884195 | 5.31 | 19.89 | 8.22 | 4.99 | 18.20 | 6.99 | Hypothetical protein |
| *PSPTO_0820* | | - | 884192 | 887302 | 2.42 | 2.19 | 3.43 | 2.00 | 5.79 | 2.84 | AcrB/AcrD/AcrF family protein |
| *PSPTO_0821* | | - | 887299 | 888414 | 1.82 | 2.68 | 0.44 | 0.00 | 4.47 | 1.16 | RND family efflux transporter MFP subunit |
| *PSPTO_0822* | | - | 888496 | 889113 | 8.10 | 12.50 | 5.06 | 6.90 | 31.15 | 9.61 | TetR family transcriptional regulator |
| *PSPTO_0823* | *pilR* | - | 889587 | 890927 | 5.69 | 8.16 | 5.50 | 9.12 | 27.10 | 14.47 | Type 4 fimbriae expression regulatory protein pilR |
| *PSPTO_0824* | *pilS* | - | 890930 | 892525 | 7.73 | 6.65 | 3.22 | 5.12 | 18.75 | 14.35 | Sensor protein PilS |
| *PSPTO_0825* | | - | 892515 | 892841 | 22.07 | 18.30 | 12.77 | 0.00 | 37.82 | 16.76 | Hypothetical protein |
| *PSPTO_0826* | *comL* | - | 892888 | 893904 | 20.93 | 51.65 | 42.92 | 20.97 | 134.74 | 40.14 | Competence lipoprotein ComL |
| *PSPTO_0827* | *rluD* | + | 894060 | 895022 | 5.80 | 22.87 | 20.93 | 14.20 | 49.70 | 15.02 | Ribosomal large subunit pseudouridine synthase D |
| *PSPTO_0828* | | + | 895019 | 895747 | 3.39 | 5.07 | 2.82 | 7.55 | 11.67 | 5.23 | Hypothetical protein |
| *PSPTO_0829* | *clpB* | + | 895919 | 898483 | 36.25 | 69.39 | 49.18 | 21.24 | 137.65 | 109.95 | ClpB protein |
| *PSPTO_0830* | | - | 899534 | 900991 | - | - | - | - | - | - | |
| *PSPTO_0831* | | - | 900988 | 902916 | 17.04 | 10.90 | 9.00 | 8.47 | 18.70 | 13.70 | Hypothetical protein |
| *PSPTO_0832* | | - | 903025 | 903834 | 0.00 | 0.00 | 0.00 | 0.00 | 0.00 | 0.00 | ISPsy4, transposition helper protein |
| *PSPTO_0833* | | - | 903831 | 904853 | 0.00 | 0.00 | 0.00 | 0.00 | 0.00 | 0.00 | ISPsy4, transposase |
| *PSPTO_0834* | | + | 905575 | 906534 | 4.19 | 60.45 | 38.32 | 10.42 | 8.92 | 4.53 | Alcohol dehydrogenase |
| *PSPTO_0835* | | + | 906604 | 907278 | 0.86 | 50.72 | 28.35 | 7.91 | 5.95 | 1.98 | RibD C-terminal domain protein |
| *PSPTO_0836* | | + | 907344 | 907931 | 1.66 | 18.75 | 6.33 | 0.00 | 3.50 | 0.60 | Hypothetical protein |
| *PSPTO_0837* | | + | 907933 | 908535 | 0.43 | 35.02 | 11.94 | 1.22 | 4.83 | 0.79 | Hypothetical protein |
| *PSPTO_0838* | | + | 908556 | 909749 | 7.43 | 25.19 | 21.09 | 16.33 | 49.36 | 18.99 | Major facilitator family transporter |
| *PSPTO_0840* | | + | 910795 | 911211 | 20.67 | 55.49 | 62.50 | 58.57 | 89.81 | 72.37 | Hypothetical protein |
| *PSPTO_0841* | | - | 911253 | 911717 | - | - | - | - | - | - | |
| *PSPTO_0842* | | + | 911777 | 912136 | 0.00 | 0.00 | 0.00 | 0.00 | 0.00 | 0.00 | ISPsy5, Orf1 |
| *PSPTO_0844* | | - | 912397 | 913374 | 0.00 | 0.00 | 0.00 | 0.00 | 0.00 | 0.00 | ISPssy, transposase |
| *PSPTO_0846* | | - | 914941 | 916050 | - | - | - | - | - | - | Hypothetical protein |
| *PSPTO_0847* | | - | 916047 | 916415 | 12.64 | 16.90 | 16.15 | 16.70 | 19.37 | 11.92 | |
| *PSPTO_0848* | | - | 916412 | 917785 | 9.52 | 14.06 | 8.97 | 6.82 | 23.32 | 11.06 | Hypothetical protein |
| *PSPTO_0849* | | - | 917806 | 918744 | 13.17 | 21.87 | 13.23 | 14.28 | 24.45 | 14.34 | Hypothetical protein |
| *PSPTO_0850* | | - | 918744 | 919160 | 8.91 | 5.06 | 0.76 | 0.00 | 7.90 | 4.89 | Hypothetical protein |
| *PSPTO_0851* | | - | 919701 | 920189 | 21.88 | 44.34 | 45.29 | 11.71 | 49.58 | 26.05 | Hypothetical protein |
| *PSPTO_0852* | *hopAJ1* | - | 920451 | 921692 | 33.49 | 193.15 | 107.68 | 42.67 | 37.96 | 33.73 | Type III helper protein HopAJ1 |
| *PSPTO_0853* | | + | 923620 | 924033 | 40.20 | 29.83 | 35.37 | 35.22 | 86.18 | 143.53 | DnaK suppressor protein |
| *PSPTO_0854* | | + | 924071 | 924292 | 12.50 | 19.93 | 17.42 | 14.43 | 46.71 | 67.99 | Hypothetical protein |
| *PSPTO_0855* | | + | 925982 | 927079 | 0.94 | 1.97 | 2.79 | 2.48 | 22.08 | 4.90 | ParA family protein |
| *PSPTO_0856* | | + | 927082 | 927900 | 8.30 | 9.43 | 9.45 | 10.18 | 31.87 | 12.47 | Hypothetical protein |
| *PSPTO_0857* | | - | 928074 | 928778 | 10.57 | 6.00 | 15.59 | 2.03 | 44.24 | 19.80 | Hypothetical protein |
| *PSPTO_0858* | | - | 928775 | 929074 | 16.55 | 14.71 | 8.20 | 0.36 | 32.96 | 14.39 | Hypothetical protein |
| *PSPTO_0859* | | - | 929071 | 931740 | 12.83 | 13.96 | 10.17 | 5.89 | 26.18 | 13.60 | Hypothetical protein |
| *PSPTO_0860* | | - | 932028 | 932483 | 11.25 | 5.07 | 6.92 | 2.85 | 16.15 | 14.01 | Hypothetical protein |
| *PSPTO_0861* | | - | 932461 | 933957 | 2.16 | 1.65 | 2.66 | 2.93 | 4.75 | 3.60 | Hypothetical protein |
| *PSPTO_0862* | | - | 933947 | 934870 | 2.89 | 3.39 | 1.68 | 0.00 | 3.27 | 1.98 | Hypothetical protein |
| *PSPTO_0863* | | - | 934867 | 935535 | 2.15 | 2.14 | 2.27 | 0.00 | 3.04 | 2.57 | Hypothetical protein |
| *PSPTO_0864* | | - | 935655 | 936464 | 0.00 | 0.00 | 0.00 | 0.00 | 0.00 | 0.00 | ISPsy4, transposition helper protein |
| *PSPTO_0865* | | - | 936461 | 937483 | 0.00 | 0.00 | 0.00 | 0.00 | 0.00 | 0.00 | ISPsy4, transposase |
| *PSPTO_0866* | | - | 937527 | 937892 | 1.97 | 4.49 | 0.00 | 0.68 | 5.76 | 3.47 | Hypothetical protein |
| *PSPTO_0867* | | - | 937908 | 938276 | 1.31 | 5.15 | 0.00 | 0.00 | 8.76 | 4.02 | Hypothetical protein |
| *PSPTO_0868* | | - | 938296 | 938445 | 4.10 | 0.00 | 14.21 | 0.00 | 9.93 | 3.04 | Hypothetical protein |
| *PSPTO_0869* | | - | 938532 | 938897 | 2.46 | 1.09 | 3.40 | 0.00 | 5.45 | 2.15 | Hypothetical protein |
| *PSPTO_0871* | | + | 939884 | 941068 | 19.35 | 36.87 | 22.03 | 14.73 | 27.96 | 27.83 | Macrolide efflux protein |
| *PSPTO_0873* | | + | 942019 | 943119 | 2.87 | 152.98 | 50.96 | 10.08 | 32.29 | 3.25 | Amidinotransferase family protein |
| *PSPTO_0874* | | + | 943206 | 944459 | 0.83 | 57.36 | 18.55 | 5.14 | 11.45 | 2.82 | Nikkomycin biosynthesis domain protein |
| *PSPTO_0875* | | + | 944512 | 945756 | 0.67 | 42.28 | 12.97 | 1.47 | 8.50 | 1.58 | Hypothetical protein |
| *PSPTO_0876* | *hopD1* | + | 946222 | 948339 | 16.38 | 49.78 | 40.10 | 12.25 | 43.60 | 24.62 | Type III effector HopD1 |
| *PSPTO_0877* | *hopQ1-1* | - | 948455 | 949798 | 19.28 | 190.66 | 95.08 | 17.29 | 27.74 | 17.55 | Type III effector HopQ1-1 |
| *PSPTO_0878* | | + | 949832 | 950149 | 21.69 | 12.90 | 7.28 | 1.34 | 29.11 | 15.92 | Hypothetical protein |
| *PSPTO_0879* | | - | 950387 | 951844 | 28.50 | 14.82 | 16.02 | 8.87 | 26.02 | 21.19 | Hypothetical protein |
| *PSPTO_0880* | | - | 951854 | 952603 | 3.98 | 6.86 | 8.75 | 3.18 | 13.92 | 5.63 | Hypothetical protein |

| Locus name | Gene name | Strand | Gene start (bp) | Gene end (bp) | Average TPM Inoculum | Average TPM $t_6$ | Average TPM $t_{24}$ | Average TPM $t_{168}$ | Average TPM Logarithmic | Average TPM Stationary | Annotation description |
|---|---|---|---|---|---|---|---|---|---|---|---|
| *PSPTO_0881* | | - | 952643 | 953524 | 6.87 | 6.52 | 6.17 | 8.02 | 19.05 | 11.82 | Hypothetical protein |
| *PSPTO_0882* | | - | 953552 | 954085 | 18.19 | 10.40 | 7.07 | 12.86 | 38.51 | 19.25 | Hypothetical protein |
| *PSPTO_0883* | *hopR1* | + | 954407 | 960280 | 7.05 | 13.88 | 9.54 | 6.89 | 13.28 | 8.06 | Type III effector HopR1 |
| *PSPTO_0884* | | - | 960352 | 961344 | 9.65 | 18.80 | 8.51 | 7.81 | 24.63 | 9.20 | LacI family transcriptional regulator |
| *PSPTO_0885* | | - | 961425 | 962906 | 2.89 | 22.67 | 8.70 | 2.92 | 8.22 | 2.67 | Sucrose-6-phosphate hydrolase |
| *PSPTO_0886* | *malK* | - | 962924 | 964069 | 4.54 | 25.35 | 12.24 | 4.38 | 11.26 | 4.02 | Maltose/maltodextrin ABC transporter ATP-binding protein |
| *PSPTO_0887* | | - | 964097 | 964939 | 5.72 | 10.69 | 9.64 | 5.60 | 8.95 | 5.43 | Sugar ABC transporter permease |
| *PSPTO_0888* | | - | 964936 | 965898 | 9.20 | 32.70 | 12.05 | 2.94 | 36.66 | 27.47 | Sugar ABC transporter permease |
| *PSPTO_0889* | | - | 965920 | 967194 | 2.46 | 31.85 | 11.13 | 4.40 | 8.39 | 3.26 | Sugar ABC transporter periplasmic sugar-binding protein |
| *PSPTO_0890* | *scrY* | + | 967317 | 968939 | 4.62 | 27.32 | 16.76 | 5.17 | 9.96 | 5.03 | Sucrose porin |
| *PSPTO_0891* | | + | 969035 | 969265 | 72.71 | 75.58 | 55.49 | 19.98 | 33.23 | 24.99 | Hypothetical protein |
| *PSPTO_0892* | | - | 969369 | 972419 | 10.78 | 21.10 | 18.69 | 15.74 | 23.44 | 15.01 | Autotransporter |
| *PSPTO_0893* | | + | 972764 | 974032 | 8.33 | 14.91 | 26.12 | 31.92 | 27.82 | 10.16 | Outer membrane protein P1 |
| *PSPTO_0894* | | + | 974212 | 974712 | 1.40 | 19.48 | 27.52 | 37.12 | 42.40 | 10.51 | Lipoprotein |
| *PSPTO_0895* | | - | 974837 | 975343 | 2.49 | 21.43 | 39.83 | 36.53 | 31.55 | 5.56 | Hypothetical protein |
| *PSPTO_0896* | | - | 975585 | 977240 | 5.15 | 9.93 | 9.39 | 8.75 | 21.89 | 16.15 | Sensor histidine kinase/response regulator |
| *PSPTO_0897* | | - | 977617 | 978240 | 14.97 | 34.33 | 28.23 | 32.52 | 23.43 | 38.13 | LuxR family DNA-binding response regulator |
| *PSPTO_0898* | | + | 978514 | 980127 | 6.85 | 11.15 | 14.94 | 14.16 | 15.53 | 15.42 | Sensor histidine kinase/response regulator |
| *PSPTO_0899* | | - | 980143 | 980610 | 7.18 | 3.01 | 4.77 | 5.07 | 5.39 | 6.82 | Hypothetical protein |
| *PSPTO_0900* | | + | 980624 | 980905 | 2.66 | 0.00 | 2.46 | 0.00 | 3.01 | 6.08 | Hypothetical protein |
| *PSPTO_0901* | *hopAG1* | + | 981500 | 981958 | 6.33 | 41.55 | 19.96 | 7.92 | 5.00 | 7.72 | Disrupted non-functional type III effector HopAG1; has an insertion element disruption. |
| *PSPTO_0902* | | - | 982088 | 983065 | 0.00 | 0.00 | 0.00 | 0.00 | 0.00 | 0.00 | ISPssy, transposase |
| *PSPTO_0903* | | - | 983280 | 984257 | 0.00 | 0.00 | 0.00 | 0.00 | 0.00 | 0.00 | ISPssy, transposase |
| *PSPTO_0904* | | + | 984490 | 986004 | - | - | - | - | - | - | |
| *PSPTO_0905* | *hopAH1* | + | 986113 | 987381 | 6.60 | 5.43 | 10.70 | 0.69 | 17.22 | 18.13 | Harpin-like protein |
| *PSPTO_0906* | *hopAI1* | + | 987569 | 988354 | 2.15 | 3.42 | 7.68 | 6.06 | 33.31 | 6.95 | Most likely not a type III effector, no hrp-box present. |
| *PSPTO_0907* | | + | 988523 | 989734 | 3.30 | 4.09 | 8.62 | 7.24 | 40.53 | 12.83 | Hypothetical protein |
| *PSPTO_0908* | *cheB-1* | - | 989768 | 990844 | 4.65 | 4.01 | 5.67 | 1.46 | 13.84 | 8.97 | Protein-glutamate methylesterase CheB |
| *PSPTO_0909* | *cheD* | - | 990834 | 991340 | 6.06 | 3.14 | 1.93 | 0.00 | 9.81 | 5.41 | Chemoreceptor glutamine deamidase CheD |
| *PSPTO_0910* | *cheR-1* | - | 991337 | 992146 | 2.82 | 1.29 | 2.87 | 0.80 | 5.71 | 3.42 | Chemotaxis protein methyltransferase CheR |
| *PSPTO_0911* | *cheW-1* | - | 992156 | 992695 | 2.79 | 0.69 | 0.00 | 2.68 | 3.32 | 2.67 | Chemotaxis protein CheW |
| *PSPTO_0912* | | - | 992732 | 994387 | 7.59 | 4.71 | 2.78 | 2.65 | 7.87 | 5.84 | Methyl-accepting chemotaxis protein |
| *PSPTO_0913* | *cheA-1* | - | 994416 | 996518 | 3.05 | 3.37 | 3.10 | 1.05 | 3.87 | 3.40 | Chemotaxis sensor histidine kinase CheA |
| *PSPTO_0914* | | - | 996565 | 996855 | 5.11 | 5.81 | 2.09 | 1.34 | 3.78 | 6.35 | STAS domain-containing protein |
| *PSPTO_0915* | *cheY-1* | - | 996882 | 997250 | 8.52 | 4.41 | 0.00 | 5.04 | 7.08 | 9.28 | Chemotaxis protein CheY |
| *PSPTO_0916* | | - | 997260 | 998474 | 10.31 | 3.86 | 6.19 | 3.35 | 7.27 | 8.73 | Methyl-accepting chemotaxis protein |
| *PSPTO_0917* | *ndH* | - | 998903 | 1000201 | 13.31 | 16.52 | 11.76 | 4.59 | 29.84 | 22.42 | NADH dehydrogenase |
| *PSPTO_0918* | | - | 1000297 | 1000404 | 9.56 | 0.00 | 47.54 | 13.98 | 12.60 | 5.81 | Hypothetical protein |
| *PSPTO_0919* | | - | 1000418 | 1000942 | 14.26 | 17.23 | 12.76 | 12.50 | 27.82 | 13.82 | MOSC domain-containing protein |
| *PSPTO_0920* | | - | 1000948 | 1001577 | 30.77 | 36.56 | 36.29 | 20.86 | 45.73 | 21.04 | Hypothetical protein |
| *PSPTO_0921* | | - | 1001589 | 1001720 | 2.19 | 10.43 | 0.00 | 5.55 | 10.02 | 2.25 | Hypothetical protein |
| *PSPTO_0922* | | - | 1002367 | 1002576 | 8.55 | 24.29 | 18.61 | 23.58 | 127.17 | 36.10 | Hypothetical protein |
| *PSPTO_0923* | *coaE* | - | 1002573 | 1003196 | 11.14 | 40.21 | 27.08 | 18.39 | 94.50 | 31.46 | Dephospho-CoA kinase |
| *PSPTO_0924* | *pilD* | - | 1003193 | 1004065 | 8.18 | 10.15 | 9.93 | 5.38 | 27.44 | 9.09 | Type IV pilus prepilin peptidase PilD |
| *PSPTO_0925* | *pilC* | - | 1004067 | 1005284 | 15.92 | 22.52 | 22.65 | 10.07 | 44.67 | 20.45 | Type IV pilus biogenesis protein PilC |
| *PSPTO_0926* | *pilB* | - | 1005287 | 1006981 | 33.39 | 55.56 | 42.42 | 27.73 | 73.64 | 33.74 | Type IV pilus biogenesis protein PilB |
| *PSPTO_0927* | | + | 1007206 | 1007634 | 3526.89 | 1083.91 | 914.34 | 585.08 | 306.09 | 437.47 | Type IV pilus biogenesis protein |
| *PSPTO_0928* | | - | 1007830 | 1008198 | 57.55 | 85.62 | 62.06 | 38.48 | 125.48 | 73.49 | Hypothetical protein |
| *PSPTO_0929* | *gsT* | + | 1008356 | 1008976 | 38.65 | 82.41 | 49.83 | 10.54 | 152.33 | 17.86 | Glutathione S-transferase |
| *PSPTO_0930* | | + | 1009217 | 1009909 | 25.93 | 24.85 | 24.20 | 22.93 | 35.44 | 22.47 | Hypothetical protein |
| *PSPTO_0931* | | + | 1009906 | 1010313 | 21.88 | 39.75 | 34.37 | 12.67 | 47.90 | 39.50 | Hypothetical protein |
| *PSPTO_0932* | | - | 1010358 | 1010903 | 13.69 | 57.68 | 27.31 | 10.94 | 61.33 | 17.20 | Hypothetical protein |
| *PSPTO_0933* | | + | 1011024 | 1012022 | 13.62 | 18.62 | 18.60 | 9.61 | 33.72 | 20.22 | Hypothetical protein |
| *PSPTO_0934* | | + | 1012077 | 1013225 | 10.09 | 13.98 | 10.84 | 5.51 | 53.07 | 19.19 | Trypsin domain-containing protein |
| *PSPTO_0935* | | + | 1013292 | 1014425 | 4.33 | 5.51 | 8.60 | 1.34 | 29.50 | 7.36 | Hypothetical protein |
| *PSPTO_0936* | | + | 1014418 | 1015176 | 18.70 | 11.41 | 5.74 | 5.61 | 42.56 | 9.97 | Hypothetical protein |
| *PSPTO_0937* | | + | 1015169 | 1016263 | 4.42 | 6.76 | 4.46 | 1.72 | 18.17 | 5.75 | Hypothetical protein |
| *PSPTO_0938* | | + | 1016263 | 1017213 | 13.94 | 23.41 | 19.97 | 10.66 | 39.37 | 14.54 | Hypothetical protein |
| *PSPTO_0939* | | + | 1017283 | 1017960 | 39.60 | 71.17 | 49.49 | 12.33 | 160.30 | 53.28 | HAD-superfamily hydrolase |
| *PSPTO_0940* | *terZ* | + | 1018008 | 1018601 | 27.34 | 228.08 | 156.84 | 32.98 | 242.12 | 40.01 | Tellurium resistance protein TerZ |
| *PSPTO_0941* | *terA* | + | 1018598 | 1019803 | 44.83 | 82.87 | 68.05 | 25.00 | 150.87 | 39.75 | Tellurium resistance protein TerA |
| *PSPTO_0942* | *terB* | + | 1019855 | 1020304 | 31.25 | 53.95 | 48.03 | 25.33 | 88.35 | 31.82 | Tellurium resistance protein TerB |
| *PSPTO_0943* | *terC* | + | 1020331 | 1021365 | 20.14 | 57.85 | 56.50 | 19.45 | 167.01 | 37.38 | Tellurium resistance protein TerC |
| *PSPTO_0944* | *terD* | + | 1021393 | 1021968 | 22.70 | 98.18 | 65.57 | 26.34 | 237.74 | 40.88 | Tellurium resistance protein TerD |
| *PSPTO_0945* | *terE* | + | 1022006 | 1022584 | 19.66 | 110.91 | 116.93 | 45.08 | 203.92 | 43.06 | Tellurium resistance protein TerE |
| *PSPTO_0946* | | - | 1022855 | 1024093 | 14.81 | 21.78 | 25.20 | 15.78 | 57.42 | 34.57 | Tellurium resistance protein |
| *PSPTO_0947* | | - | 1024154 | 1024888 | 10.27 | 15.72 | 18.78 | 13.47 | 38.18 | 22.51 | Hypothetical protein |
| *PSPTO_0948* | *nadC* | - | 1025270 | 1026118 | 53.73 | 61.38 | 68.51 | 26.70 | 118.09 | 44.90 | Nicotinate-nucleotide pyrophosphorylase |
| *PSPTO_0949* | | + | 1026472 | 1028727 | 36.34 | 46.97 | 36.76 | 22.26 | 75.82 | 77.05 | Hypothetical protein |
| *PSPTO_0950* | *ampD* | + | 1029038 | 1029586 | 47.79 | 30.13 | 23.86 | 8.42 | 39.08 | 13.40 | N-acetyl-anhydromuranmyl-L-alanine amidase |
| *PSPTO_0951* | | + | 1029602 | 1030438 | 17.77 | 10.87 | 11.20 | 5.44 | 20.05 | 8.25 | AmpE protein |
| *PSPTO_0952* | | - | 1030586 | 1031284 | 9.82 | 7.59 | 11.05 | 4.71 | 19.75 | 9.73 | TatD family hydrolase |
| *PSPTO_0953* | *fruR* | - | 1031386 | 1032381 | 5.81 | 12.98 | 14.83 | 4.67 | 15.74 | 6.91 | DNA-binding transcriptional regulator FruR |
| *PSPTO_0954* | | + | 1032736 | 1035603 | 7.88 | 25.26 | 18.01 | 7.02 | 19.03 | 12.49 | Phosphoenolpyruvate-protein phosphotransferase,EI/HPr/EIIA components |
| *PSPTO_0955* | *fruK* | + | 1035603 | 1036544 | 4.39 | 26.33 | 17.19 | 16.70 | 20.27 | 13.36 | 1-phosphofructokinase |
| *PSPTO_0956* | | + | 1036557 | 1038311 | 9.61 | 29.05 | 21.20 | 14.96 | 27.16 | 11.91 | Phosphotransferase system, fructose-specific IIBC component |
| *PSPTO_0957* | *phbA-1* | - | 1038333 | 1039511 | 8.17 | 24.49 | 24.32 | 7.60 | 42.86 | 13.23 | Acetyl-CoA acetyltransferase |
| *PSPTO_0958* | | - | 1039650 | 1040516 | 2.97 | 8.69 | 2.63 | 1.20 | 19.82 | 6.31 | Hypothetical protein |
| *PSPTO_0959* | *pgi* | - | 1040763 | 1042427 | 35.35 | 71.18 | 57.95 | 31.08 | 351.03 | 64.79 | Glucose-6-phosphate isomerase |

| Locus name | Gene name | Strand | Gene start (bp) | Gene end (bp) | Average TPM Inoculum | Average TPM $t_6$ | Average TPM $t_{24}$ | Average TPM $t_{168}$ | Average TPM Logarithmic | Average TPM Stationary | Annotation description |
|---|---|---|---|---|---|---|---|---|---|---|---|
| *PSPTO_0960* | *panC* | - | 1042670 | 1043521 | 3.99 | 17.96 | 19.07 | 17.57 | 102.28 | 17.05 | Pantoate--beta-alanine ligase |
| *PSPTO_0961* | *panB* | - | 1043518 | 1044318 | 16.27 | 24.34 | 33.07 | 9.49 | 89.74 | 23.17 | 3-methyl-2-oxobutanoate hydroxymethyltransferase |
| *PSPTO_0962* | *folK-2* | - | 1044547 | 1045053 | 2.33 | 6.30 | 1.45 | 9.56 | 43.21 | 10.70 | 2-amino-4-hydroxy-6-hydroxymethyldihydropteridine pyrophosphokinase |
| *PSPTO_0963* | *pcnB* | - | 1045046 | 1046449 | 18.17 | 47.06 | 38.03 | 33.20 | 182.44 | 53.67 | Poly(A) polymerase |
| *PSPTO_0964* | | - | 1047150 | 1048586 | 12.77 | 26.98 | 25.08 | 16.91 | 82.18 | 36.70 | Sigma-54 dependent transcriptional regulator/response regulator |
| *PSPTO_0965* | | - | 1048609 | 1051557 | 6.88 | 11.63 | 8.54 | 4.04 | 22.49 | 12.63 | Sensor histidine kinase |
| *PSPTO_0966* | | - | 1051547 | 1051723 | 15.42 | 27.17 | 0.00 | 8.58 | 10.18 | 7.77 | Hypothetical protein |
| *PSPTO_0967* | | - | 1051822 | 1052709 | 5.60 | 16.38 | 14.59 | 19.39 | 35.92 | 13.71 | Glutamyl-Q tRNA(Asp) synthetase |
| *PSPTO_0968* | | - | 1053296 | 1054756 | 0.30 | 0.38 | 0.04 | 0.36 | 0.41 | 0.34 | ISPsy6, transposase |
| *PSPTO_0969* | *dksA* | - | 1054836 | 1055276 | 15.87 | 43.14 | 29.71 | 29.55 | 115.08 | 43.60 | DnaK suppressor protein |
| *PSPTO_0970* | | + | 1055626 | 1056798 | 6.81 | 9.66 | 17.79 | 5.94 | 22.37 | 12.84 | Class I and II aminotransferase |
| *PSPTO_0971* | *sfsA* | + | 1056798 | 1057511 | 5.04 | 16.18 | 18.78 | 2.92 | 23.00 | 15.12 | Sugar fermentation stimulation protein |
| *PSPTO_0972* | | - | 1057534 | 1057866 | 27.30 | 29.88 | 24.63 | 8.90 | 32.52 | 21.45 | Iron-sulfur cluster-binding protein, Rieske family |
| *PSPTO_0973* | *deoC* | + | 1058084 | 1058863 | 3.35 | 7.28 | 0.00 | 2.34 | 7.55 | 9.66 | Deoxyribose-phosphate aldolase |
| *PSPTO_0974* | | + | 1059152 | 1059259 | 5.02 | 0.00 | 0.00 | 0.00 | 10.81 | 3.60 | Hypothetical protein |
| *PSPTO_0975* | | - | 1059244 | 1059588 | 28.15 | 49.40 | 46.71 | 27.90 | 112.21 | 51.38 | Pentapeptide repeat-containing protein |
| *PSPTO_0976* | | - | 1059690 | 1061228 | 40.12 | 50.69 | 30.02 | 23.74 | 239.81 | 68.62 | Hypothetical protein |
| *PSPTO_0977* | | + | 1061359 | 1063680 | 10.71 | 16.75 | 13.28 | 8.74 | 44.30 | 9.97 | Penicillin-binding protein |
| *PSPTO_0978* | | + | 1063668 | 1064438 | 19.27 | 34.64 | 21.35 | 23.04 | 80.07 | 22.01 | Hypothetical protein |
| *PSPTO_0979* | | + | 1064438 | 1064767 | 8.42 | 21.62 | 22.54 | 10.56 | 34.15 | 12.29 | Hypothetical protein |
| *PSPTO_0980* | | - | 1064777 | 1065220 | 27.49 | 56.52 | 58.22 | 15.17 | 54.98 | 20.63 | Hypothetical protein |
| *PSPTO_0981* | *ilvB* | + | 1065591 | 1067315 | 10.33 | 73.33 | 48.04 | 37.75 | 82.71 | 19.70 | Acetolactate synthase large subunit |
| *PSPTO_0982* | *ilvN* | + | 1067318 | 1067809 | 27.07 | 270.35 | 152.74 | 146.82 | 464.54 | 69.18 | Acetolactate synthase small subunit |
| *PSPTO_0983* | *ilvC* | + | 1067857 | 1068873 | 10.29 | 108.82 | 70.29 | 37.79 | 159.12 | 18.38 | Ketol-acid reductoisomerase |
| *PSPTO_0984* | *pssA-1* | + | 1069018 | 1069869 | 7.93 | 36.46 | 19.17 | 14.17 | 53.75 | 19.43 | CDP-diacylglycerol--serine O-phosphatidyltransferase |
| *PSPTO_0985* | | + | 1069934 | 1070947 | 29.22 | 39.91 | 39.84 | 22.30 | 24.69 | 23.85 | Hypothetical protein |
| *PSPTO_0986* | | + | 1070947 | 1071579 | 12.98 | 22.53 | 29.75 | 14.10 | 22.68 | 23.30 | Membrane protein |
| *PSPTO_0987* | | + | 1077352 | 1077582 | 6.55 | 8.63 | 5.51 | 2.41 | 4.50 | 2.42 | Hypothetical protein |
| *PSPTO_0988* | | - | 1077641 | 1080460 | 3.19 | 5.11 | 9.91 | 4.40 | 63.47 | 12.66 | Hypothetical protein |
| *PSPTO_0989* | | - | 1080457 | 1081164 | 1.76 | 3.70 | 1.89 | 5.64 | 56.73 | 11.80 | Hypothetical protein |
| *PSPTO_0990* | | - | 1081154 | 1082431 | 3.77 | 11.35 | 13.49 | 1.97 | 49.70 | 13.62 | Hypothetical protein |
| *PSPTO_0991* | | + | 1082577 | 1083380 | 21.36 | 27.23 | 18.55 | 12.14 | 60.34 | 25.53 | Hypothetical protein |
| *PSPTO_0992* | *rimI* | + | 1083373 | 1083828 | 6.41 | 32.56 | 21.32 | 14.46 | 35.89 | 16.15 | Ribosomal-protein-alanine acetyltransferase |
| *PSPTO_0993* | | - | 1083838 | 1084245 | 37.49 | 194.42 | 128.17 | 67.95 | 54.01 | 44.52 | Hypothetical protein |
| *PSPTO_0994* | | + | 1084494 | 1085132 | 17.72 | 10.70 | 12.34 | 7.84 | 21.80 | 6.50 | Carbonic anhydrase |
| *PSPTO_0995* | | - | 1085145 | 1086818 | 4.97 | 6.12 | 3.94 | 0.58 | 8.76 | 3.62 | Methyl-accepting chemotaxis protein |
| *PSPTO_0996* | | - | 1086926 | 1087585 | 12.04 | 20.93 | 15.92 | 9.97 | 29.21 | 9.42 | HD domain-containing protein |
| *PSPTO_0997* | | - | 1087661 | 1089664 | 10.89 | 2.77 | 3.22 | 1.97 | 5.44 | 4.90 | Acyltransferase family protein |
| *PSPTO_0998* | *phnA* | + | 1089883 | 1090257 | 21.40 | 19.52 | 10.62 | 12.95 | 57.17 | 34.84 | Alkylphosphonate utilization operon protein PhnA |
| *PSPTO_0999* | | + | 1090603 | 1091826 | 5.02 | 14.07 | 8.50 | 4.31 | 23.35 | 30.46 | Major facilitator family transporter |
| *PSPTO_1000* | | - | 1091889 | 1092608 | 42.65 | 41.16 | 37.90 | 31.64 | 87.22 | 68.17 | CDP-alcohol phosphatidyltransferase |
| *PSPTO_1001* | | - | 1092695 | 1094710 | 34.07 | 54.91 | 48.84 | 29.97 | 86.42 | 61.02 | Hypothetical protein |
| *PSPTO_1002* | | + | 1094884 | 1096449 | - | - | - | - | - | - | |
| *PSPTO_1003* | | - | 1096637 | 1097074 | 19.94 | 9.42 | 25.17 | 11.53 | 31.57 | 13.56 | GNAT family acetyltransferase |
| *PSPTO_1004* | | - | 1097150 | 1098391 | 6.15 | 4.89 | 9.48 | 6.61 | 9.73 | 7.62 | Diguanylate cyclase |
| *PSPTO_1005* | *gmd* | + | 1098690 | 1099736 | 35.86 | 35.37 | 33.00 | 12.79 | 33.68 | 31.84 | GDP-mannose 4,6-dehydratase |
| *PSPTO_1006* | | + | 1099737 | 1099838 | 5.49 | 6.41 | 0.00 | 22.49 | 8.10 | 3.45 | Hypothetical protein |
| *PSPTO_1007* | | - | 1099814 | 1100773 | 13.29 | 20.81 | 11.75 | 8.47 | 20.61 | 13.03 | GDP-6-deoxy-D-lyxo-4-hexulose reductase |
| *PSPTO_1008* | | + | 1100918 | 1102318 | 53.87 | 12.56 | 15.74 | 10.10 | 10.80 | 13.46 | Methyl-accepting chemotaxis protein |
| *PSPTO_1009* | | - | 1102902 | 1103528 | 4.55 | 4.83 | 1.68 | 1.66 | 14.49 | 5.02 | Isochorismatase family protein |
| *PSPTO_1010* | *phoD* | - | 1103683 | 1105254 | 5.13 | 3.72 | 2.20 | 2.47 | 6.81 | 3.98 | Alkaline phosphatase D |
| *PSPTO_1011* | | - | 1105251 | 1105814 | 6.26 | 0.00 | 1.30 | 0.46 | 4.85 | 2.61 | GNAT family acetyltransferase |
| *PSPTO_1012* | | + | 1106014 | 1106421 | 2.35 | 0.00 | 2.38 | 0.00 | 2.07 | 1.88 | TonB system transport protein |
| *PSPTO_1013* | | + | 1106421 | 1107206 | 6.11 | 6.81 | 3.19 | 1.16 | 3.64 | 2.98 | TonB domain protein |
| *PSPTO_1014* | | + | 1107227 | 1109509 | 6.04 | 7.44 | 2.60 | 6.51 | 12.70 | 10.42 | TonB-dependent receptor |
| *PSPTO_1015* | | + | 1109641 | 1110651 | 12.87 | 11.43 | 5.28 | 5.71 | 10.40 | 9.47 | Integrase/recombinase XerC |
| *PSPTO_1016* | | + | 1110675 | 1112279 | 17.22 | 18.87 | 14.97 | 14.43 | 33.37 | 20.79 | Hypothetical protein |
| *PSPTO_1017* | | + | 1112173 | 1112532 | 0.00 | 0.00 | 0.00 | 0.00 | 0.00 | 0.00 | ISPsy5, Orf1 |
| *PSPTO_1018* | | + | 1112550 | 1112735 | - | - | - | - | - | - | |
| *PSPTO_1019* | | + | 1112778 | 1113137 | 0.00 | 0.00 | 0.00 | 0.00 | 0.00 | 0.00 | ISPsy5, Orf1 |
| *PSPTO_1020* | | + | 1113155 | 1114708 | 0.00 | 0.00 | 0.00 | 0.00 | 0.00 | 0.00 | ISPsy5, transposase |
| *PSPTO_1021* | | + | 1114754 | 1115488 | - | - | - | - | - | - | |
| *PSPTO_1022* | *hopAM1-1* | - | 1115514 | 1116344 | 0.52 | 393.68 | 163.64 | 22.67 | 1.28 | 1.00 | Type III effector HopAM1-1 |
| *PSPTO_1023* | | + | 1116554 | 1117102 | 3.83 | 0.00 | 2.37 | 0.00 | 0.44 | 0.60 | MotA/TolQ/ExbB proton channel family protein |
| *PSPTO_1024* | | + | 1117166 | 1118434 | 7.43 | 2.17 | 1.43 | 0.70 | 2.28 | 1.98 | Phospholipase D |
| *PSPTO_1025* | | + | 1118505 | 1119704 | 4.55 | 17.82 | 14.57 | 4.93 | 25.22 | 7.99 | Transglycosylase |
| *PSPTO_1026* | *wssA* | + | 1119987 | 1121132 | 5.19 | 16.03 | 19.96 | 3.21 | 11.53 | 7.82 | Cell morphology protein |
| *PSPTO_1027* | *wssB* | + | 1121129 | 1123348 | 7.46 | 3.20 | 4.73 | 3.46 | 9.08 | 4.39 | Cellulose synthase, catalytic subunit |
| *PSPTO_1028* | | + | 1123363 | 1125618 | 2.36 | 0.68 | 4.02 | 2.85 | 7.78 | 2.45 | Cellulose synthase regulator protein |
| *PSPTO_1029* | *wssD* | + | 1125622 | 1126833 | 0.81 | 1.89 | 1.32 | 1.41 | 9.11 | 0.90 | Endoglucanase |
| *PSPTO_1030* | *wssE* | + | 1127013 | 1130705 | 1.89 | 1.24 | 1.82 | 0.82 | 7.59 | 1.93 | Cellulose synthase operon protein C |
| *PSPTO_1031* | *wssF* | + | 1130721 | 1131386 | 3.63 | 0.75 | 3.57 | 1.10 | 15.91 | 3.32 | Cell morphology protein |
| *PSPTO_1032* | *wssG* | + | 1131383 | 1132057 | 1.97 | 4.01 | 1.44 | 0.00 | 7.27 | 2.75 | Cell morphology protein |
| *PSPTO_1033* | *wssH* | + | 1132076 | 1133491 | 5.15 | 4.17 | 3.56 | 6.58 | 8.47 | 5.24 | Cellulose acetylase, subunit WssH |
| *PSPTO_1034* | *wssI* | + | 1133503 | 1134654 | 1.18 | 4.60 | 3.45 | 5.12 | 6.31 | 1.91 | Cell morphology protein |
| *PSPTO_1035* | | - | 1135027 | 1136406 | 21.04 | 29.23 | 34.03 | 32.46 | 37.32 | 25.08 | Colicin/pyosin nuclease family protein |
| *PSPTO_1036* | | - | 1136613 | 1137074 | 1.21 | 1.45 | 1.02 | 0.56 | 1.93 | 1.56 | Hypothetical protein |
| *PSPTO_1037* | | - | 1137307 | 1137768 | 7.42 | 4.18 | 7.70 | 1.98 | 45.14 | 54.58 | Hypothetical protein |
| *PSPTO_1038* | | - | 1137765 | 1138214 | 43.93 | 23.29 | 20.77 | 14.79 | 88.57 | 142.39 | Hypothetical protein |

| Locus name | Gene name | Strand | Gene start (bp) | Gene end (bp) | Average TPM Inoculum | Average TPM $t_6$ | Average TPM $t_{24}$ | Average TPM $t_{168}$ | Average TPM Logarithmic | Average TPM Stationary | Annotation description |
|---|---|---|---|---|---|---|---|---|---|---|---|
| *PSPTO_1039* | | + | 1138505 | 1139509 | 43.82 | 28.45 | 22.56 | 8.13 | 25.06 | 12.09 | Response regulator |
| *PSPTO_1040* | | + | 1139513 | 1139995 | 17.76 | 24.36 | 6.91 | 7.22 | 16.24 | 7.79 | Sensory box protein |
| *PSPTO_1041* | *aroA* | + | 1140091 | 1141347 | 5.78 | 29.61 | 29.80 | 8.17 | 52.14 | 16.63 | 3-phosphoshikimate 1-carboxyvinyltransferase |
| *PSPTO_1042* | *chrR* | - | 1141370 | 1142032 | 7.11 | 20.93 | 18.28 | 4.46 | 21.92 | 12.58 | Transcriptional activator ChrR |
| *PSPTO_1043* | | - | 1142032 | 1142640 | 21.90 | 36.32 | 35.52 | 14.53 | 33.59 | 21.99 | RNA polymerase sigma factor |
| *PSPTO_1044* | | - | 1143054 | 1143233 | 13.34 | 7.60 | 4.56 | 12.32 | 8.88 | 6.26 | Hypothetical protein |
| *PSPTO_1045* | | - | 1143628 | 1144032 | 6.79 | 8.74 | 5.59 | 0.00 | 20.07 | 5.65 | Virulence-associated protein |
| *PSPTO_1046* | | - | 1144032 | 1144262 | 42.50 | 61.88 | 58.24 | 34.09 | 60.67 | 21.59 | Virulence-associated protein |
| *PSPTO_1047* | | - | 1144377 | 1145258 | 5.81 | 3.82 | 1.61 | 3.42 | 9.89 | 3.85 | Hypothetical protein |
| *PSPTO_1048* | | - | 1145233 | 1146204 | 3.29 | 0.15 | 4.84 | 1.51 | 4.79 | 3.82 | Membrane protein |
| *PSPTO_1049* | *dctM* | - | 1146288 | 1147568 | 8.82 | 5.89 | 2.74 | 1.81 | 7.49 | 4.61 | TRAP dicarboxylate transporter subunit DctM |
| *PSPTO_1050* | *dctQ* | - | 1147569 | 1148096 | 1.88 | 6.57 | 4.37 | 3.48 | 3.49 | 2.66 | TRAP dicarboxylate transporter, DctQ subunit |
| *PSPTO_1051* | *dctP* | - | 1148160 | 1149185 | 3.97 | 10.50 | 3.81 | 3.44 | 6.75 | 2.42 | TRAP dicarboxylate transporter subunit DctP |
| *PSPTO_1052* | | - | 1149242 | 1150126 | 4.58 | 6.91 | 4.07 | 8.30 | 9.10 | 4.84 | Senescence marker protein-30 family protein |
| *PSPTO_1053* | | - | 1150183 | 1151010 | 10.25 | 14.79 | 8.89 | 10.30 | 12.63 | 6.56 | Hypothetical protein |
| *PSPTO_1054* | | - | 1151189 | 1152457 | 3.96 | 2.97 | 6.15 | 4.87 | 4.90 | 3.40 | Outer membrane porin, OprD family |
| *PSPTO_1055* | | + | 1152451 | 1152669 | 17.52 | 31.04 | 22.95 | 18.85 | 7.52 | 5.51 | Hypothetical protein |
| *PSPTO_1056* | | + | 1152922 | 1153686 | 18.21 | 27.61 | 22.49 | 8.38 | 18.15 | 16.29 | Short chain dehydrogenase/reductase family oxidoreductase |
| *PSPTO_1057* | | - | 1153738 | 1154148 | 11.62 | 21.54 | 10.31 | 14.66 | 19.78 | 5.60 | Hypothetical protein |
| *PSPTO_1058* | | - | 1154148 | 1154387 | 101.96 | 214.69 | 105.95 | 74.43 | 191.03 | 26.74 | Hypothetical protein |
| *PSPTO_1059* | | - | 1154539 | 1156428 | 7.32 | 3.56 | 1.82 | 3.69 | 6.66 | 4.81 | Methyl-accepting chemotaxis protein |
| *PSPTO_1060* | | + | 1156430 | 1156573 | 27.32 | 3.47 | 5.81 | 8.11 | 2.77 | 1.14 | Hypothetical protein |
| *PSPTO_1061* | | + | 1156819 | 1158699 | 6.43 | 10.45 | 8.94 | 6.87 | 11.20 | 7.73 | Methyl-accepting chemotaxis protein |
| *PSPTO_1062* | | - | 1158828 | 1159091 | 640.45 | 768.16 | 753.40 | 796.82 | 152.91 | 705.24 | Hypothetical protein |
| *PSPTO_1064* | | - | 1161594 | 1162319 | - | - | - | - | - | - | |
| *PSPTO_1065* | | - | 1162781 | 1163902 | 20.32 | 25.15 | 28.76 | 21.83 | 66.02 | 24.04 | DnaJ domain-containing protein |
| *PSPTO_1066* | | + | 1164246 | 1166303 | 127.91 | 32.59 | 31.76 | 14.00 | 18.78 | 30.36 | Methyl-accepting chemotaxis protein |
| *PSPTO_1067* | | + | 1166480 | 1167493 | 13.66 | 29.16 | 24.40 | 21.93 | 56.94 | 31.24 | Glycosyl transferase family protein |
| *PSPTO_1068* | | - | 1167577 | 1169136 | 3.05 | 8.59 | 8.91 | 5.11 | 18.85 | 8.92 | Membrane protein |
| *PSPTO_1069* | | - | 1169280 | 1169678 | 0.55 | 0.00 | 14.68 | 2.61 | 25.90 | 3.58 | Membrane protein |
| *PSPTO_1070* | | - | 1169689 | 1170639 | 9.55 | 26.11 | 35.34 | 18.53 | 91.04 | 28.50 | Lipopolysaccharide biosynthesis protein |
| *PSPTO_1071* | | - | 1170626 | 1171576 | 18.21 | 28.63 | 29.01 | 15.91 | 103.61 | 35.32 | Glycosyl transferase family protein |
| *PSPTO_1072* | | - | 1171573 | 1172688 | 22.58 | 44.15 | 28.01 | 31.95 | 163.40 | 47.26 | DegT/DnrJ/EryC1/StrS family aminotransferase |
| *PSPTO_1073* | | - | 1172691 | 1174586 | 9.45 | 13.04 | 9.49 | 11.52 | 53.37 | 14.98 | Membrane protein |
| *PSPTO_1074* | | - | 1174632 | 1179317 | 9.70 | 26.47 | 19.26 | 14.47 | 96.83 | 15.66 | Glycosyl transferase family protein |
| *PSPTO_1075* | | - | 1179310 | 1180674 | 6.32 | 16.70 | 12.47 | 11.50 | 59.86 | 9.14 | O-antigen ABC transporter, ATP-binding protein |
| *PSPTO_1076* | | - | 1180664 | 1181491 | 9.52 | 24.02 | 10.39 | 9.14 | 47.88 | 12.17 | O-antigen ABC transporter, permease protein |
| *PSPTO_1077* | *rfbC-1* | - | 1181491 | 1182036 | 12.63 | 16.31 | 24.12 | 6.96 | 49.84 | 12.63 | DTDP-4-dehydrorhamnose 3,5-epimerase |
| *PSPTO_1078* | | - | 1182707 | 1183636 | 6.50 | 16.22 | 8.60 | 8.42 | 37.95 | 21.20 | Lipopolysaccharide biosynthesis protein |
| *PSPTO_1079* | *rfbA* | - | 1183678 | 1184568 | 15.97 | 33.38 | 16.78 | 20.73 | 83.22 | 36.66 | Glucose-1-phosphate thymidylyltransferase |
| *PSPTO_1080* | *rfbD* | - | 1184565 | 1185470 | 51.25 | 68.07 | 31.22 | 38.55 | 208.85 | 69.54 | DTDP-4-dehydrorhamnose reductase |
| *PSPTO_1081* | *rfbB-1* | - | 1185470 | 1186543 | 13.28 | 48.96 | 42.73 | 25.83 | 224.00 | 81.40 | DTDP-glucose 4,6-dehydratase |
| *PSPTO_1082* | | + | 1187114 | 1187332 | 7.14 | 18.87 | 16.12 | 10.78 | 32.41 | 56.74 | Hypothetical protein |
| *PSPTO_1083* | | - | 1188596 | 1188997 | 66.00 | 45.74 | 33.38 | 18.00 | 29.72 | 36.43 | Peptidase, S24 family |
| *PSPTO_1084* | | - | 1190000 | 1190890 | 10.16 | 25.44 | 20.76 | 12.96 | 30.41 | 19.38 | Integrase/recombinase XerD |
| *PSPTO_1085* | | - | 1190965 | 1191342 | 53.24 | 44.32 | 60.81 | 18.82 | 39.55 | 36.25 | Hypothetical protein |
| *PSPTO_1086* | *hsdM* | + | 1191539 | 1193269 | 9.36 | 18.91 | 17.04 | 8.22 | 48.33 | 17.93 | Type I restriction-modification system, M subunit |
| *PSPTO_1087* | | + | 1193259 | 1194527 | 10.35 | 15.44 | 20.28 | 4.47 | 49.73 | 18.95 | Type I restriction-modification system subunit S |
| *PSPTO_1088* | | + | 1194524 | 1195576 | 9.02 | 11.12 | 9.54 | 6.17 | 45.96 | 19.47 | Hypothetical protein |
| *PSPTO_1089* | | + | 1195573 | 1198695 | 8.66 | 14.64 | 8.96 | 5.11 | 43.14 | 16.79 | Type I restriction-modification enzyme, R subunit |
| *PSPTO_1090* | | - | 1198881 | 1199858 | 2.74 | 13.77 | 3.87 | 2.51 | 3.23 | 11.47 | Hypothetical protein |
| *PSPTO_1091* | | - | 1199858 | 1200274 | 10.96 | 56.85 | 31.37 | 7.66 | 8.52 | 63.32 | Hypothetical protein |
| *PSPTO_1092* | *mobB-1* | + | 1201192 | 1201506 | 1.43 | 5.34 | 0.00 | 2.91 | 3.59 | 3.48 | Mobilization protein MobB |
| *PSPTO_1093* | | + | 1201509 | 1203926 | 10.83 | 8.46 | 7.85 | 10.31 | 14.42 | 8.50 | Relaxase/mobilization nuclease domain protein |
| *PSPTO_1094* | | + | 1203937 | 1204314 | 49.40 | 41.90 | 22.15 | 13.96 | 63.86 | 41.19 | Hypothetical protein |
| *PSPTO_1095* | | + | 1205057 | 1206079 | 0.00 | 0.00 | 0.00 | 0.00 | 0.00 | 0.00 | ISPsy4, transposase |
| *PSPTO_1096* | | + | 1206076 | 1206885 | 0.00 | 0.00 | 0.00 | 0.00 | 0.00 | 0.00 | ISPsy4, transposition helper protein |
| *PSPTO_1097* | | + | 1207129 | 1209657 | 1.51 | 4.99 | 4.11 | 3.22 | 11.30 | 6.16 | Membrane protein |
| *PSPTO_1098* | | - | 1211249 | 1212802 | 0.00 | 0.00 | 0.00 | 0.00 | 0.00 | 0.00 | ISPsy5, transposase |
| *PSPTO_1099* | | - | 1212820 | 1213179 | 0.00 | 0.00 | 0.00 | 0.00 | 0.00 | 0.00 | ISPsy5, Orf1 |
| *PSPTO_1100* | | - | 1214486 | 1215673 | 91.79 | 65.29 | 38.59 | 19.31 | 151.94 | 82.21 | Site-specific recombinase, phage integrase family |
| *PSPTO_1101* | *ychF* | - | 1215889 | 1216989 | 6.61 | 22.37 | 7.21 | 6.81 | 68.29 | 17.48 | GTP-binding protein YchF |
| *PSPTO_1102* | *pth* | - | 1217011 | 1217595 | 6.75 | 19.13 | 16.15 | 19.44 | 142.95 | 36.64 | Peptidyl-tRNA hydrolase |
| *PSPTO_1103* | | - | 1217760 | 1218371 | 7.97 | 69.34 | 51.49 | 22.23 | 496.15 | 46.87 | Ribosomal 5S rRNA E-loop binding protein Ctc/L25/TL5 |
| *PSPTO_1104* | *prsA* | - | 1218509 | 1219450 | 18.15 | 32.22 | 38.99 | 37.65 | 116.90 | 69.36 | Ribose-phosphate pyrophosphokinase |
| *PSPTO_1105* | *ispE* | - | 1219609 | 1220457 | 330.75 | 291.38 | 270.42 | 218.59 | 198.65 | 203.28 | 4-diphosphocytidyl-2-C-methyl-D-erythritol kinase |
| *PSPTO_1106* | *lolB* | - | 1220460 | 1221077 | 3.46 | 16.86 | 13.65 | 4.65 | 37.85 | 11.79 | Outer membrane lipoprotein LolB |
| *PSPTO_1107* | | - | 1221082 | 1222809 | 19.16 | 38.40 | 35.88 | 23.11 | 109.06 | 43.14 | TPR domain-containing protein |
| *PSPTO_1108* | *hemA* | + | 1223036 | 1224313 | 16.44 | 24.49 | 25.65 | 10.65 | 58.84 | 36.21 | Glutamyl-tRNA reductase |
| *PSPTO_1109* | *prfA* | + | 1224310 | 1225392 | 13.31 | 25.64 | 20.82 | 12.40 | 85.87 | 44.94 | Peptide chain release factor 1 |
| *PSPTO_1110* | *hemK* | + | 1225393 | 1226226 | 14.46 | 19.99 | 16.53 | 4.44 | 66.30 | 35.62 | HemK protein |
| *PSPTO_1111* | *moeB* | + | 1226211 | 1226990 | 3.90 | 16.24 | 8.53 | 4.57 | 44.93 | 19.58 | Molybdopterin biosynthesis protein MoeB |
| *PSPTO_1112* | *murI* | + | 1226983 | 1227810 | 13.94 | 17.45 | 14.00 | 12.91 | 29.06 | 18.28 | Glutamate racemase |
| *PSPTO_1114* | | - | 1228770 | 1228994 | 4.86 | 12.90 | 5.31 | 2.47 | 10.96 | 11.59 | Hypothetical protein |
| *PSPTO_1115* | | - | 1229099 | 1229602 | 13.09 | 30.18 | 34.00 | 21.32 | 36.90 | 36.79 | Membrane protein |
| *PSPTO_1116* | *cfa* | - | 1229595 | 1230863 | 5.76 | 19.45 | 21.86 | 9.26 | 21.29 | 10.62 | Cyclopropane-fatty-acyl-phospholipid synthase |
| *PSPTO_1117* | | - | 1230850 | 1231533 | 6.82 | 19.12 | 22.10 | 1.29 | 22.07 | 7.38 | Hypothetical protein |
| *PSPTO_1118* | | - | 1231647 | 1232894 | 5.62 | 16.70 | 10.71 | 9.05 | 21.68 | 10.70 | Amine oxidase, flavin-containing protein |
| *PSPTO_1119* | | - | 1232891 | 1233679 | 8.32 | 24.80 | 22.27 | 10.94 | 21.09 | 11.47 | Short chain dehydrogenase/reductase family oxidoreductase |

| Locus name | Gene name | Strand | Gene start (bp) | Gene end (bp) | Average TPM Inoculum | Average TPM $t_6$ | Average TPM $t_{24}$ | Average TPM $t_{168}$ | Average TPM Logarithmic | Average TPM Stationary | Annotation description |
|---|---|---|---|---|---|---|---|---|---|---|---|
| *PSPTO_1120* | | - | 1233676 | 1234098 | 7.36 | 10.34 | 13.66 | 5.07 | 14.63 | 10.37 | Hypothetical protein |
| *PSPTO_1121* | *phrB* | - | 1234091 | 1235539 | 6.34 | 23.51 | 16.32 | 9.82 | 23.85 | 24.61 | Deoxyribodipyrimidine photolyase |
| *PSPTO_1122* | | - | 1235549 | 1236457 | 9.48 | 26.14 | 22.33 | 9.85 | 29.96 | 15.12 | MerR family transcriptional regulator |
| *PSPTO_1123* | | - | 1236463 | 1237419 | 19.63 | 36.03 | 44.71 | 15.00 | 50.50 | 21.25 | Hypothetical protein |
| *PSPTO_1124* | | - | 1237618 | 1237740 | 9.55 | 3.80 | 0.00 | 1.21 | 15.26 | 6.50 | Hypothetical protein |
| *PSPTO_1125* | | - | 1237715 | 1237930 | 25.66 | 33.56 | 27.63 | 22.57 | 47.14 | 24.33 | Hypothetical protein |
| *PSPTO_1126* | | - | 1237927 | 1238913 | 27.35 | 39.37 | 31.86 | 12.67 | 95.20 | 50.19 | Amine oxidase, flavin-containing protein |
| *PSPTO_1127* | | + | 1239201 | 1240103 | 24.37 | 43.74 | 33.19 | 15.17 | 35.53 | 27.27 | Hypothetical protein |
| *PSPTO_1128* | *hemH* | + | 1240127 | 1241149 | 13.35 | 29.86 | 20.62 | 7.15 | 44.91 | 27.85 | Ferrochelatase |
| *PSPTO_1129* | *uraA* | - | 1241259 | 1242479 | 7.47 | 19.17 | 14.67 | 17.73 | 59.60 | 15.07 | Uracil transporter |
| *PSPTO_1130* | *upp* | - | 1242541 | 1243179 | 31.24 | 24.67 | 32.70 | 14.87 | 75.21 | 26.30 | Uracil phosphoribosyltransferase |
| *PSPTO_1131* | *hpt* | + | 1243387 | 1243944 | 8.02 | 12.10 | 7.64 | 10.52 | 32.52 | 10.61 | Hypoxanthine phosphoribosyltransferase |
| *PSPTO_1132* | | + | 1244010 | 1244504 | 12.64 | 16.26 | 6.86 | 17.83 | 62.98 | 18.01 | Hypothetical protein |
| *PSPTO_1133* | | - | 1245042 | 1245773 | 9.52 | 26.43 | 13.55 | 19.37 | 54.09 | 25.73 | Amino acid ABC transporter permease |
| *PSPTO_1134* | | - | 1245872 | 1246693 | 16.07 | 74.10 | 36.47 | 53.66 | 56.99 | 10.92 | Amino acid ABC transporter substrate-binding protein |
| *PSPTO_1135* | | + | 1246804 | 1247700 | 10.01 | 36.07 | 24.14 | 21.43 | 39.68 | 10.80 | Homocysteine S-methyltransferase family protein |
| *PSPTO_1136* | *mqo* | - | 1247783 | 1249291 | 26.93 | 40.77 | 36.26 | 30.49 | 169.59 | 37.47 | Malate:quinone oxidoreductase |
| *PSPTO_1137* | | + | 1249976 | 1250086 | 29.00 | 32.79 | 10.07 | 16.51 | 28.47 | 24.28 | Hypothetical protein |
| *PSPTO_1138* | | + | 1250473 | 1251054 | 19.97 | 29.42 | 25.42 | 7.42 | 71.75 | 16.93 | Lipoprotein |
| *PSPTO_1139* | | - | 1251135 | 1251386 | 7.39 | 30.74 | 25.39 | 11.39 | 7.20 | 3.48 | Hypothetical protein |
| *PSPTO_1140* | | - | 1251599 | 1252762 | 12.83 | 29.36 | 21.82 | 5.50 | 64.51 | 24.72 | Hypothetical protein |
| *PSPTO_1141* | | + | 1252869 | 1253051 | 61.83 | 49.78 | 39.71 | 19.20 | 30.76 | 22.00 | Hypothetical protein |
| *PSPTO_1142* | | + | 1253051 | 1253311 | 22.64 | 16.03 | 35.22 | 0.70 | 44.65 | 19.31 | Hypothetical protein |
| *PSPTO_1143* | | + | 1253340 | 1253969 | 7.48 | 6.95 | 15.22 | 3.74 | 34.57 | 14.01 | Hypothetical protein |
| *PSPTO_1144* | | + | 1254153 | 1254971 | 15.24 | 29.16 | 24.68 | 14.59 | 86.68 | 15.72 | Lipoprotein |
| *PSPTO_1145* | | - | 1255028 | 1255612 | 8.18 | 28.49 | 11.08 | 8.48 | 16.44 | 6.97 | Hypothetical protein |
| *PSPTO_1146* | *rsmC* | - | 1255859 | 1256857 | 2.39 | 5.42 | 4.37 | 5.65 | 11.21 | 3.66 | Ribosomal RNA small subunit methyltransferase C |
| *PSPTO_1147* | | + | 1257094 | 1257726 | 7.74 | 11.86 | 7.13 | 1.36 | 24.22 | 12.45 | LysE family transporter |
| *PSPTO_1148* | *alkK* | + | 1257917 | 1259599 | 12.24 | 24.26 | 34.31 | 15.04 | 71.64 | 24.31 | Medium-chain-fatty-acid--CoA ligase |
| *PSPTO_1149* | | + | 1259909 | 1261846 | 20.30 | 18.47 | 36.19 | 24.40 | 46.09 | 14.38 | Hypothetical protein |
| *PSPTO_1150* | | + | 1261851 | 1263239 | 22.60 | 34.07 | 35.99 | 25.36 | 69.08 | 14.68 | Hypothetical protein |
| *PSPTO_1151* | | + | 1263491 | 1266223 | 9.32 | 20.09 | 17.31 | 8.26 | 20.80 | 15.95 | LuxR family transcriptional regulator |
| *PSPTO_1152* | | - | 1266241 | 1267428 | 4.05 | 1.05 | 2.67 | 1.25 | 5.14 | 4.47 | HDIG domain protein |
| *PSPTO_1153* | | - | 1267728 | 1268270 | 7.26 | 1.84 | 6.16 | 3.17 | 5.76 | 5.55 | NAD(P)H-flavin oxidoreductase |
| *PSPTO_1154* | | - | 1268283 | 1269062 | 8.90 | 19.60 | 20.37 | 2.26 | 12.03 | 8.02 | Alpha/beta fold family hydrolase |
| *PSPTO_1155* | | - | 1269120 | 1269503 | 5.32 | 34.79 | 21.74 | 5.77 | 13.24 | 6.24 | Endoribonuclease L-PSP family protein |
| *PSPTO_1156* | | - | 1269514 | 1270254 | 4.47 | 12.16 | 6.59 | 1.73 | 6.10 | 3.41 | Isochorismatase family protein |
| *PSPTO_1157* | | - | 1270254 | 1271405 | 16.49 | 14.22 | 18.43 | 3.94 | 5.17 | 4.56 | Luciferase family protein |
| *PSPTO_1158* | | + | 1271668 | 1272324 | 3.43 | 11.31 | 20.15 | 0.33 | 11.23 | 5.05 | TetR family transcriptional regulator |
| *PSPTO_1159* | | + | 1272558 | 1273649 | 27.05 | 10.74 | 10.79 | 4.21 | 3.02 | 3.92 | Bmp family protein |
| *PSPTO_1160* | | + | 1273646 | 1275157 | 5.83 | 6.29 | 3.94 | 0.00 | 2.27 | 3.15 | ABC transporter ATP-binding protein |
| *PSPTO_1161* | | + | 1275150 | 1275344 | 4.90 | 0.00 | 8.79 | 0.00 | 1.86 | 1.90 | Hypothetical protein |
| *PSPTO_1162* | | + | 1275360 | 1276270 | - | - | - | - | - | - | |
| *PSPTO_1163* | | + | 1276270 | 1277196 | - | - | - | - | - | - | |
| *PSPTO_1164* | | + | 1277310 | 1277999 | 256.65 | 324.20 | 326.30 | 110.86 | 522.12 | 169.29 | OmpA family protein |
| *PSPTO_1165* | | + | 1278328 | 1279428 | 28.42 | 43.32 | 41.36 | 18.77 | 90.57 | 48.42 | Hypothetical protein |
| *PSPTO_1166* | | + | 1279469 | 1280446 | 0.00 | 0.00 | 0.00 | 0.00 | 0.00 | 0.00 | ISPssy, transposase |
| *PSPTO_1167* | | + | 1281800 | 1282393 | 0.57 | 4.72 | 0.79 | 0.00 | 13.44 | 2.61 | Hypothetical protein |
| *PSPTO_1168* | | - | 1282914 | 1283837 | 6.16 | 10.01 | 9.38 | 0.99 | 22.74 | 7.56 | Putative acyltransferase |
| *PSPTO_1169* | *pta* | - | 1283902 | 1285992 | 7.81 | 6.48 | 11.35 | 4.99 | 34.68 | 10.71 | Phosphate acetyltransferase |
| *PSPTO_1170* | | - | 1286070 | 1286297 | 31.75 | 18.90 | 13.56 | 5.77 | 19.41 | 10.98 | Hypothetical protein |
| *PSPTO_1171* | | + | 1286390 | 1286875 | 32.67 | 102.56 | 66.93 | 53.22 | 107.64 | 53.19 | FKBP-type peptidyl-prolyl cis-trans isomerase |
| *PSPTO_1172* | | + | 1287005 | 1287493 | 18.81 | 50.77 | 48.41 | 63.92 | 176.91 | 70.10 | Glutathione peroxidase |
| *PSPTO_1173* | | + | 1287676 | 1288779 | 8.78 | 9.32 | 10.46 | 8.45 | 19.99 | 7.70 | Oxidoreductase FAD/FMN-binding protein |
| *PSPTO_1174* | | - | 1288748 | 1289959 | 2.71 | 2.82 | 3.32 | 0.78 | 2.96 | 1.64 | Glycoside hydrolase family protein |
| *PSPTO_1175* | | + | 1290659 | 1291255 | 0.28 | 11.08 | 1.59 | 1.53 | 2.55 | 2.30 | Membrane protein |
| *PSPTO_1176* | | - | 1291353 | 1291697 | 1.72 | 0.24 | 7.94 | 0.12 | 2.32 | 1.28 | Hypothetical protein |
| *PSPTO_1177* | *cysZ* | - | 1291883 | 1292644 | 16.25 | 27.10 | 36.09 | 11.33 | 43.11 | 17.96 | CysZ protein |
| *PSPTO_1178* | *trxB* | + | 1293002 | 1293964 | 25.30 | 236.00 | 218.57 | 49.05 | 100.89 | 37.70 | Thioredoxin reductase |
| *PSPTO_1179* | *hopJ1* | - | 1294195 | 1294536 | 11.88 | 56.19 | 43.29 | 27.81 | 63.17 | 31.14 | HopJ1 protein |
| *PSPTO_1180* | | - | 1294529 | 1294825 | 28.13 | 96.39 | 65.77 | 40.79 | 170.12 | 59.08 | Hypothetical protein |
| *PSPTO_1181* | *folB-2* | - | 1294869 | 1295240 | 18.86 | 58.01 | 64.02 | 36.78 | 57.67 | 37.69 | Dihydroneopterin aldolase |
| *PSPTO_1182* | *folE-1* | - | 1295240 | 1295803 | 14.90 | 99.09 | 98.94 | 44.52 | 91.77 | 42.03 | GTP cyclohydrolase I |
| *PSPTO_1183* | | - | 1295825 | 1296535 | 25.43 | 166.47 | 153.08 | 61.20 | 185.13 | 85.45 | Dihydromonapterin reductase |
| *PSPTO_1184* | | - | 1296770 | 1297342 | 102.12 | 51.17 | 40.05 | 30.58 | 47.27 | 59.50 | Hypothetical protein |
| *PSPTO_1185* | | - | 1297494 | 1298099 | 80.61 | 100.27 | 95.52 | 46.51 | 61.99 | 131.14 | Hypothetical protein |
| *PSPTO_1186* | | + | 1298287 | 1298739 | 39.25 | 20.09 | 43.73 | 13.59 | 74.10 | 32.80 | Flavodoxin |
| *PSPTO_1187* | | + | 1298852 | 1299772 | 20.80 | 21.87 | 23.92 | 9.50 | 56.23 | 26.58 | Short chain dehydrogenase/reductase family oxidoreductase |
| *PSPTO_1188* | | + | 1299927 | 1300904 | 0.00 | 0.00 | 0.00 | 0.00 | 0.00 | 0.00 | ISPssy, transposase |
| *PSPTO_1189* | | - | 1301655 | 1303208 | 0.00 | 0.00 | 0.00 | 0.00 | 0.00 | 0.00 | ISPsy5, transposase |
| *PSPTO_1190* | | - | 1303226 | 1303585 | 0.00 | 0.00 | 0.00 | 0.00 | 0.00 | 0.00 | ISPsy5, Orf1 |
| *PSPTO_1191* | | - | 1303986 | 1304789 | 36.66 | 57.61 | 50.61 | 30.35 | 70.99 | 55.94 | Hypothetical protein |
| *PSPTO_1192* | | - | 1304872 | 1305615 | 3.38 | 1.61 | 2.89 | 3.91 | 2.81 | 4.00 | Hypothetical protein |
| *PSPTO_1193* | | - | 1305776 | 1307740 | 5.14 | 12.98 | 2.91 | 0.00 | 4.52 | 6.88 | Hypothetical protein |
| *PSPTO_1194* | *nfrB* | - | 1307737 | 1309857 | 19.57 | 27.66 | 21.43 | 18.04 | 14.34 | 18.13 | Bacteriophage N4 adsorption protein B |
| *PSPTO_1195* | | - | 1309870 | 1312011 | 43.23 | 44.06 | 47.29 | 21.12 | 45.20 | 77.23 | Hypothetical protein |
| *PSPTO_1196* | | - | 1312251 | 1313501 | 3.96 | 19.19 | 21.99 | 5.58 | 5.24 | 5.85 | UDP-N-acetylglucosamine 2-epimerase |
| *PSPTO_1197* | | - | 1314205 | 1314471 | 5.92 | 25.84 | 16.56 | 1.67 | 15.77 | 8.40 | Hypothetical protein |
| *PSPTO_1198* | | - | 1314528 | 1314650 | 10.91 | 149.94 | 51.50 | 14.59 | 119.97 | 40.14 | Hypothetical protein |

| Locus name | Gene name | Strand | Gene start (bp) | Gene end (bp) | Average TPM Inoculum | Average TPM $t_6$ | Average TPM $t_{24}$ | Average TPM $t_{168}$ | Average TPM Logarithmic | Average TPM Stationary | Annotation description |
|---|---|---|---|---|---|---|---|---|---|---|---|
| *PSPTO_1199* | | - | 1314638 | 1314775 | 15.91 | 24.68 | 4.88 | 3.99 | 26.34 | 13.08 | Hypothetical protein |
| *PSPTO_1200* | | + | 1314899 | 1315387 | 37.99 | 14.68 | 25.41 | 13.94 | 34.31 | 43.97 | Hypothetical protein |
| *PSPTO_1201* | | - | 1315456 | 1315788 | 66.04 | 23.43 | 29.89 | 3.73 | 41.08 | 35.68 | Lipoprotein |
| *PSPTO_1202* | | - | 1316033 | 1316365 | 223.45 | 399.02 | 383.42 | 171.04 | 37.65 | 129.14 | Hypothetical protein |
| *PSPTO_1203* | | + | 1316530 | 1317072 | 310.98 | 74.54 | 52.85 | 38.99 | 92.96 | 92.87 | RNA polymerase sigma-70 family protein |
| *PSPTO_1204* | | + | 1317069 | 1318031 | 7.77 | 5.41 | 13.72 | 4.02 | 32.54 | 37.97 | Regulatory protein |
| *PSPTO_1205* | | - | 1318123 | 1318551 | 197.93 | 492.47 | 477.21 | 185.91 | 1530.90 | 223.70 | Ribosomal subunit interface protein |
| *PSPTO_1206* | | + | 1318800 | 1321301 | 2.94 | 2.02 | 3.59 | 0.52 | 41.87 | 94.20 | TonB-dependent siderophore receptor |
| *PSPTO_1207* | *fecA* | - | 1321452 | 1323800 | 7.83 | 9.61 | 4.42 | 1.96 | 51.79 | 25.35 | Iron(III) dicitrate transport protein fecA |
| *PSPTO_1208* | | - | 1324328 | 1325305 | 2.13 | 4.41 | 1.17 | 0.76 | 23.60 | 18.48 | Regulatory protein |
| *PSPTO_1209* | | - | 1325302 | 1325808 | 1.51 | 0.00 | 0.00 | 1.45 | 45.05 | 36.48 | RNA polymerase sigma-70 family protein |
| *PSPTO_1210* | | + | 1326007 | 1326351 | 0.39 | 3.55 | 6.65 | 3.19 | 14.16 | 22.24 | Hypothetical protein |
| *PSPTO_1211* | | + | 1326348 | 1327913 | 1.94 | 2.83 | 2.19 | 3.68 | 16.87 | 14.97 | Membrane protein |
| *PSPTO_1212* | | + | 1327913 | 1328245 | 2.56 | 4.16 | 1.14 | 3.73 | 21.47 | 20.41 | Hypothetical protein |
| *PSPTO_1213* | | - | 1328234 | 1329109 | 17.13 | 18.02 | 22.34 | 4.87 | 41.49 | 27.79 | AraC family transcriptional regulator |
| *PSPTO_1214* | | + | 1329246 | 1330148 | 2.10 | 1.11 | 2.55 | 0.58 | 5.76 | 2.09 | Membrane protein |
| *PSPTO_1215* | | - | 1330200 | 1331174 | 9.27 | 25.91 | 20.80 | 11.32 | 75.77 | 14.98 | D-isomer specific 2-hydroxyacid dehydrogenase family protein |
| *PSPTO_1216* | | - | 1331262 | 1332209 | 51.74 | 25.52 | 20.36 | 11.68 | 87.91 | 44.16 | LysR family transcriptional regulator |
| *PSPTO_1217* | | + | 1332394 | 1333890 | 2.79 | 2.95 | 3.01 | 2.03 | 5.15 | 8.86 | Outer membrane efflux protein |
| *PSPTO_1218* | | + | 1333887 | 1336073 | 2.47 | 1.86 | 2.22 | 2.77 | 3.52 | 3.21 | Fusaric acid resistance protein |
| *PSPTO_1219* | | + | 1336058 | 1336258 | 2.42 | 1.98 | 4.26 | 0.00 | 2.45 | 2.49 | Hypothetical protein |
| *PSPTO_1220* | | + | 1336284 | 1337144 | 5.08 | 2.91 | 1.14 | 1.27 | 5.11 | 3.68 | Fusaric acid resistance protein |
| *PSPTO_1221* | | + | 1338122 | 1338751 | 1.90 | 8.44 | 11.37 | 23.04 | 43.38 | 8.68 | LysE family transporter |
| *PSPTO_1222* | | + | 1340118 | 1340312 | 1.23 | 0.00 | 0.40 | 0.00 | 1.63 | 0.97 | Hypothetical protein |
| *PSPTO_1223* | | - | 1340384 | 1341202 | 17.00 | 9.48 | 10.68 | 6.07 | 12.61 | 16.76 | Short chain dehydrogenase |
| *PSPTO_1224* | | - | 1341219 | 1341683 | 22.90 | 37.29 | 22.10 | 6.88 | 49.47 | 22.25 | Hypothetical protein |
| *PSPTO_1225* | | + | 1341834 | 1343702 | - | - | - | - | - | - | |
| *PSPTO_1226* | | + | 1343596 | 1343955 | 0.00 | 0.00 | 0.00 | 0.00 | 0.00 | 0.00 | ISPsy5, Orf1 |
| *PSPTO_1227* | | + | 1343973 | 1345526 | 0.00 | 0.00 | 0.00 | 0.00 | 0.00 | 0.00 | ISPsy5, transposase |
| *PSPTO_1228* | | + | 1345791 | 1346540 | - | - | - | - | - | - | |
| *PSPTO_1229* | | + | 1346595 | 1347572 | 0.00 | 0.00 | 0.00 | 0.00 | 0.00 | 0.00 | ISPssy, transposase |
| *PSPTO_1230* | | - | 1347973 | 1348083 | 2.31 | 13.60 | 1.10 | 1.98 | 8.73 | 6.26 | Hypothetical protein |
| *PSPTO_1231* | | + | 1348208 | 1350976 | 6.25 | 4.66 | 3.41 | 2.21 | 10.32 | 4.99 | Insecticidal toxin protein |
| *PSPTO_1232* | *algA* | - | 1351082 | 1352533 | 8.79 | 185.62 | 115.98 | 60.79 | 54.35 | 96.49 | Alginate biosynthesis protein AlgA |
| *PSPTO_1233* | *algF* | - | 1352670 | 1353338 | 7.36 | 268.51 | 208.24 | 123.43 | 78.00 | 75.94 | Alginate biosynthesis protein AlgF |
| *PSPTO_1234* | *algJ* | - | 1353351 | 1354526 | 4.11 | 78.63 | 63.29 | 46.71 | 26.34 | 18.75 | Alginate biosynthesis protein AlgJ |
| *PSPTO_1235* | *algI* | - | 1354537 | 1356093 | 2.43 | 37.44 | 24.82 | 12.94 | 10.84 | 7.32 | Alginate biosynthesis protein AlgI |
| *PSPTO_1236* | *algL* | - | 1356280 | 1357416 | 2.54 | 46.60 | 30.39 | 29.39 | 10.40 | 7.41 | Alginate lyase |
| *PSPTO_1237* | *algX* | - | 1357425 | 1358864 | 4.15 | 42.94 | 40.21 | 26.93 | 16.93 | 10.71 | Alginate biosynthesis protein AlgX |
| *PSPTO_1238* | *algG* | - | 1358879 | 1360489 | 3.75 | 50.88 | 35.18 | 26.43 | 12.19 | 8.62 | Alginate biosynthesis protein AlgG |
| *PSPTO_1239* | *algE* | - | 1360516 | 1361997 | 2.55 | 46.72 | 42.02 | 29.25 | 10.68 | 7.13 | Alginate biosynthesis protein AlgE |
| *PSPTO_1240* | *algK* | - | 1361994 | 1363406 | 2.17 | 32.43 | 16.78 | 19.05 | 8.46 | 7.52 | Alginate biosynthesis protein AlgK |
| *PSPTO_1241* | *alg44* | - | 1363409 | 1364581 | 3.07 | 52.28 | 39.20 | 25.89 | 9.89 | 10.76 | Alginate biosynthesis protein Alg44 |
| *PSPTO_1242* | *alg8* | - | 1364622 | 1366103 | 5.92 | 32.87 | 21.59 | 25.74 | 6.23 | 9.37 | Alginate biosynthesis protein Alg8 |
| *PSPTO_1243* | *algD* | - | 1366206 | 1367522 | 37.45 | 158.73 | 103.77 | 65.42 | 22.19 | 44.78 | GDP-mannose 6-dehydrogenase AlgD |
| *PSPTO_1244* | | - | 1368333 | 1369112 | 11.94 | 45.34 | 40.84 | 16.86 | 86.89 | 13.21 | Hypothetical protein |
| *PSPTO_1245* | | + | 1369648 | 1370481 | 5.54 | 11.80 | 3.76 | 3.63 | 15.64 | 7.87 | Polysaccharide deacetylase family protein |
| *PSPTO_1246* | | + | 1370776 | 1372170 | 77.92 | 123.29 | 111.78 | 26.75 | 81.40 | 22.03 | PhoH-like protein |
| *PSPTO_1247* | *moaC* | + | 1372312 | 1372797 | 32.46 | 26.90 | 29.60 | 7.59 | 35.26 | 14.74 | Molybdenum cofactor biosynthesis protein C |
| *PSPTO_1248* | *moaD* | + | 1372794 | 1373039 | 13.23 | 17.84 | 9.22 | 8.94 | 18.95 | 10.52 | Molybdenum cofactor biosynthesis protein D |
| *PSPTO_1249* | *moaE* | + | 1373042 | 1373488 | 7.39 | 2.25 | 3.34 | 2.07 | 9.22 | 4.33 | Molybdenum cofactor biosynthesis protein E |
| *PSPTO_1250* | | + | 1373711 | 1374394 | 9.41 | 6.09 | 1.37 | 7.85 | 17.74 | 4.86 | CsgG family protein |
| *PSPTO_1251* | | + | 1374416 | 1374793 | 2.88 | 0.87 | 9.73 | 1.45 | 11.77 | 3.27 | Hypothetical protein |
| *PSPTO_1252* | | + | 1374790 | 1375452 | 1.47 | 1.80 | 3.30 | 0.00 | 12.15 | 1.74 | Lipoprotein |
| *PSPTO_1253* | *rhlB* | - | 1375533 | 1376726 | 28.76 | 23.00 | 35.22 | 24.00 | 67.01 | 13.67 | ATP-dependent RNA helicase RhlB |
| *PSPTO_1254* | *estA* | - | 1377109 | 1377768 | 33.92 | 46.47 | 18.85 | 21.91 | 76.53 | 29.96 | Carboxylesterase |
| *PSPTO_1255* | | + | 1378253 | 1379281 | 134.56 | 140.34 | 95.52 | 44.56 | 215.32 | 15.10 | Amino acid ABC transporter substrate-binding protein |
| *PSPTO_1256* | | + | 1379352 | 1380533 | 20.29 | 24.81 | 23.40 | 9.29 | 64.61 | 7.57 | Amino acid ABC transporter permease |
| *PSPTO_1257* | | + | 1380544 | 1381641 | 13.34 | 19.79 | 12.80 | 1.02 | 35.27 | 6.44 | Amino acid ABC transporter permease |
| *PSPTO_1258* | | + | 1381652 | 1382416 | 20.21 | 44.47 | 18.70 | 12.94 | 110.33 | 6.33 | Amino acid ABC transporter ATP-binding protein |
| *PSPTO_1259* | | - | 1382406 | 1382582 | 2.54 | 0.00 | 0.00 | 6.45 | 7.74 | 0.81 | Hypothetical protein |
| *PSPTO_1260* | | + | 1382572 | 1383801 | 2.96 | 8.26 | 1.80 | 5.47 | 6.48 | 3.40 | Cyanate MFS transporter |
| *PSPTO_1261* | | + | 1383842 | 1384366 | 7.99 | 25.26 | 21.59 | 11.59 | 23.38 | 9.20 | Hypothetical protein |
| *PSPTO_1262* | | + | 1384359 | 1384676 | 1.91 | 3.43 | 16.53 | 10.49 | 11.12 | 6.56 | GIY-YIG nuclease superfamily protein |
| *PSPTO_1263* | | + | 1384741 | 1385673 | 4.69 | 11.54 | 8.04 | 3.24 | 16.73 | 7.56 | Hypothetical protein |
| *PSPTO_1264* | *nrdH* | - | 1385856 | 1386068 | 0.98 | 9.97 | 5.26 | 0.00 | 16.07 | 5.24 | Glutaredoxin NrdH |
| *PSPTO_1265* | | - | 1386213 | 1387214 | 6.27 | 20.81 | 17.17 | 10.13 | 53.14 | 21.82 | Nucleoid-associated family protein |
| *PSPTO_1266* | | + | 1387607 | 1387888 | 92.15 | 33.14 | 83.48 | 27.32 | 334.72 | 21.91 | HU family DNA-binding protein |
| *PSPTO_1267* | | - | 1388286 | 1389311 | 10.63 | 24.35 | 22.30 | 11.25 | 43.29 | 14.24 | Hypothetical protein |
| *PSPTO_1268* | *valS* | - | 1389375 | 1392221 | 9.66 | 31.71 | 38.96 | 24.63 | 134.67 | 77.20 | Valyl-tRNA synthetase |
| *PSPTO_1269* | | - | 1392581 | 1393006 | 14.39 | 39.22 | 50.80 | 29.97 | 140.25 | 55.14 | Hypothetical protein |
| *PSPTO_1270* | *holC* | - | 1393014 | 1393442 | 3.51 | 40.69 | 23.73 | 11.86 | 144.93 | 34.94 | DNA polymerase III subunit chi |
| *PSPTO_1271* | *pepA* | - | 1393501 | 1394991 | 27.02 | 49.83 | 50.62 | 26.03 | 183.52 | 71.72 | Cytosol aminopeptidase |
| *PSPTO_1272* | | + | 1395310 | 1396437 | 6.91 | 28.92 | 24.69 | 7.84 | 50.78 | 18.52 | Membrane protein |
| *PSPTO_1273* | | + | 1396424 | 1397485 | 8.26 | 20.37 | 25.56 | 17.12 | 55.11 | 34.25 | Membrane protein |
| *PSPTO_1274* | | - | 1397612 | 1397824 | 226.51 | 793.53 | 783.97 | 626.24 | 646.34 | 172.21 | Cold shock domain family protein |
| *PSPTO_1275* | *gcvT-2* | - | 1398228 | 1399457 | 10.70 | 7.34 | 5.53 | 3.49 | 623.59 | 70.64 | Glycine cleavage system T protein |
| *PSPTO_1276* | *gcvP* | - | 1399420 | 1402284 | 16.92 | 8.26 | 10.89 | 4.32 | 1099.88 | 197.19 | Glycine dehydrogenase |
| *PSPTO_1277* | *gcvH-2* | - | 1402295 | 1402678 | 54.13 | 20.61 | 34.28 | 8.31 | 2153.47 | 527.27 | Glycine cleavage system H protein |

| Locus name | Gene name | Strand | Gene start (bp) | Gene end (bp) | Average TPM Inoculum | Average TPM $t_6$ | Average TPM $t_{24}$ | Average TPM $t_{168}$ | Average TPM Logarithmic | Average TPM Stationary | Annotation description |
|---|---|---|---|---|---|---|---|---|---|---|---|
| *PSPTO_1278* | | - | 1402983 | 1405100 | 25.13 | 13.29 | 11.02 | 6.37 | 14.56 | 5.70 | Sensory box protein/response regulator |
| *PSPTO_1279* | | - | 1405110 | 1407274 | - | - | - | - | - | - | |
| *PSPTO_1280* | *tyrR* | - | 1407395 | 1408903 | 4.48 | 13.49 | 9.27 | 2.29 | 22.16 | 10.43 | Transcriptional regulator TyrR |
| *PSPTO_1281* | | - | 1409228 | 1409584 | 232.91 | 152.75 | 125.38 | 49.31 | 100.32 | 132.52 | Hypothetical protein |
| *PSPTO_1282* | | - | 1409689 | 1410147 | 10.46 | 22.73 | 23.75 | 13.17 | 43.35 | 20.14 | Hypothetical protein |
| *PSPTO_1283* | | - | 1410144 | 1410908 | 3.91 | 9.39 | 5.81 | 0.24 | 11.12 | 11.30 | Heme oxygenase |
| *PSPTO_1284* | | - | 1410889 | 1413462 | 5.75 | 7.09 | 6.71 | 4.33 | 15.25 | 13.68 | TonB-dependent outer membrane heme receptor |
| *PSPTO_1285* | | - | 1413583 | 1414488 | 4.84 | 9.30 | 8.06 | 7.25 | 16.03 | 10.03 | Regulatory protein |
| *PSPTO_1286* | | - | 1414616 | 1415158 | 3.90 | 1.98 | 6.36 | 4.66 | 36.10 | 17.45 | RNA polymerase sigma-70 family protein |
| *PSPTO_1287* | *gap-1* | - | 1415258 | 1416259 | 16.13 | 370.88 | 182.84 | 75.49 | 1191.20 | 82.67 | Glyceraldehyde 3-phosphate dehydrogenase, type I |
| *PSPTO_1288* | *edd* | + | 1416479 | 1418305 | 14.62 | 58.70 | 42.61 | 13.26 | 202.83 | 20.02 | 6-phosphogluconate dehydratase |
| *PSPTO_1289* | *glk* | + | 1418302 | 1419267 | 9.99 | 27.95 | 13.50 | 2.27 | 85.11 | 14.78 | Glucokinase |
| *PSPTO_1290* | | + | 1419318 | 1420055 | 9.22 | 22.90 | 25.59 | 8.93 | 58.33 | 12.93 | DNA-binding response regulator |
| *PSPTO_1291* | | + | 1420117 | 1421523 | 2.73 | 10.25 | 2.94 | 3.49 | 22.27 | 3.60 | Sensor histidine kinase |
| *PSPTO_1292* | | + | 1421695 | 1422981 | 80.70 | 140.31 | 98.88 | 85.41 | 87.39 | 15.84 | Glucose ABC transporter, periplasmic glucose-binding protein |
| *PSPTO_1293* | | + | 1423086 | 1423994 | 8.42 | 21.89 | 6.94 | 13.58 | 24.38 | 4.64 | Glucose ABC transporter permease |
| *PSPTO_1294* | | + | 1423987 | 1424832 | 17.18 | 38.75 | 22.79 | 8.99 | 46.97 | 5.85 | Glucose ABC transporter permease |
| *PSPTO_1295* | *gltK* | + | 1424835 | 1425995 | 50.96 | 92.41 | 63.12 | 52.68 | 184.47 | 20.87 | Glucose ABC transporter ATP-binding protein |
| *PSPTO_1296* | *oprB* | + | 1426129 | 1427490 | 51.53 | 135.99 | 123.45 | 103.81 | 134.75 | 19.07 | Porin B |
| *PSPTO_1297* | | + | 1427498 | 1427632 | 0.74 | 8.36 | 0.00 | 0.00 | 6.21 | 0.98 | Hypothetical protein |
| *PSPTO_1298* | | + | 1427604 | 1428479 | 1.98 | 17.67 | 14.92 | 9.75 | 40.99 | 6.33 | Hypothetical protein |
| *PSPTO_1299* | *hexR* | - | 1428566 | 1429432 | 7.45 | 22.32 | 8.87 | 6.46 | 36.88 | 12.49 | Hex regulon repressor |
| *PSPTO_1300* | *zwf-1* | + | 1429651 | 1431120 | 16.61 | 154.62 | 81.17 | 35.24 | 555.98 | 105.87 | Glucose-6-phosphate 1-dehydrogenase |
| *PSPTO_1301* | *pgl* | + | 1431107 | 1431820 | 11.11 | 115.56 | 79.95 | 36.61 | 461.80 | 67.55 | 6-phosphogluconolactonase |
| *PSPTO_1302* | *eda-1* | + | 1431849 | 1432523 | 7.48 | 38.42 | 28.72 | 10.83 | 146.72 | 25.47 | Keto-hydroxyglutarate-aldolase/keto-deoxy-phosphogluconate aldolase |
| *PSPTO_1303* | | - | 1432654 | 1432893 | 56.99 | 160.10 | 133.48 | 67.37 | 173.51 | 80.48 | Hypothetical protein |
| *PSPTO_1304* | | + | 1433113 | 1433583 | 29.36 | 474.49 | 249.88 | 96.23 | 21.66 | 4.88 | Hypothetical protein |
| *PSPTO_1305* | | - | 1433678 | 1435186 | 5.53 | 9.05 | 9.56 | 9.15 | 15.99 | 28.62 | Diguanylate cyclase |
| *PSPTO_1306* | | - | 1435390 | 1436754 | 3.96 | 5.09 | 4.44 | 11.03 | 12.57 | 7.95 | Heavy metal sensor histidine kinase |
| *PSPTO_1307* | | - | 1436744 | 1437421 | 1.71 | 1.26 | 3.58 | 0.00 | 9.27 | 5.20 | DNA-binding heavy metal response regulator |
| *PSPTO_1308* | | - | 1437432 | 1440536 | 5.07 | 6.39 | 6.46 | 2.19 | 10.96 | 6.91 | AcrB/AcrD/AcrF family protein |
| *PSPTO_1309* | | - | 1440533 | 1441699 | 5.98 | 7.72 | 6.08 | 8.87 | 18.23 | 8.07 | HlyD family secretion protein |
| *PSPTO_1310* | | + | 1441968 | 1442945 | 0.00 | 0.00 | 0.00 | 0.00 | 0.00 | 0.00 | ISPssy, transposase |
| *PSPTO_1311* | | + | 1443185 | 1443562 | 32.42 | 16.86 | 24.17 | 24.14 | 18.61 | 59.66 | Hypothetical protein |
| *PSPTO_1312* | | + | 1443875 | 1444459 | 2.22 | 4.16 | 3.58 | 6.38 | 7.77 | 5.11 | Fimbral biosynthesis protein |
| *PSPTO_1313* | | + | 1444523 | 1445275 | 7.49 | 6.04 | 10.33 | 4.34 | 12.32 | 10.17 | Pili assembly chaperone |
| *PSPTO_1314* | | + | 1445452 | 1446216 | - | - | - | - | - | - | |
| *PSPTO_1315* | | - | 1446346 | 1447323 | 0.00 | 0.00 | 0.00 | 0.00 | 0.00 | 0.00 | ISPssy, transposase |
| *PSPTO_1316* | | + | 1447484 | 1449169 | - | - | - | - | - | - | |
| *PSPTO_1317* | | + | 1449166 | 1449948 | 2.75 | 2.54 | 6.91 | 1.64 | 2.45 | 3.95 | Fimbral chaperone protein |
| *PSPTO_1318* | | + | 1449950 | 1450948 | 15.29 | 5.60 | 5.31 | 8.95 | 6.05 | 12.23 | Hypothetical protein |
| *PSPTO_1319* | | + | 1450996 | 1451283 | 56.88 | 10.17 | 25.96 | 19.05 | 17.50 | 21.16 | Hypothetical protein |
| *PSPTO_1320* | | - | 1451301 | 1452053 | 470.18 | 55.12 | 49.51 | 31.72 | 32.83 | 95.00 | Hypothetical protein |
| *PSPTO_1321* | | + | 1452046 | 1452201 | 2136.29 | 149.66 | 103.55 | 107.74 | 85.63 | 327.78 | Hypothetical protein |
| *PSPTO_1322* | | - | 1452168 | 1452641 | 3.72 | 1.89 | 7.46 | 0.00 | 4.15 | 7.75 | GNAT family acetyltransferase |
| *PSPTO_1323* | | - | 1452704 | 1453648 | 118.00 | 35.61 | 29.70 | 20.45 | 28.09 | 40.59 | Chemotaxis protein CheV |
| *PSPTO_1324* | *dsbB* | + | 1454243 | 1454752 | 1091.80 | 943.38 | 1158.72 | 1082.28 | 341.40 | 6373.72 | Disulfide oxidoreductase |
| *PSPTO_1325* | *cyoA* | + | 1455218 | 1456159 | 34.66 | 112.83 | 103.63 | 51.18 | 246.60 | 212.91 | Cytochrome o ubiquinol oxidase subunit II |
| *PSPTO_1326* | *cyoB* | + | 1456163 | 1458175 | 26.76 | 96.38 | 90.62 | 44.43 | 364.92 | 208.21 | Cytochrome o ubiquinol oxidase subunit I |
| *PSPTO_1327* | *cyoC* | + | 1458179 | 1458811 | 22.01 | 55.42 | 53.55 | 29.15 | 221.89 | 84.49 | Cytochrome o ubiquinol oxidase subunit III |
| *PSPTO_1328* | *cyoD* | + | 1458811 | 1459143 | 34.27 | 48.67 | 50.67 | 45.32 | 240.59 | 131.19 | Cytochrome o ubiquinol oxidase subunit IV |
| *PSPTO_1329* | *cyoE* | + | 1459154 | 1460041 | 20.70 | 44.55 | 46.13 | 20.94 | 209.47 | 72.03 | Protoheme IX farnesyltransferase |
| *PSPTO_1330* | | + | 1460299 | 1461183 | 35.06 | 59.90 | 71.19 | 44.81 | 74.40 | 37.18 | Glycosyl transferase family protein |
| *PSPTO_1331* | | + | 1461281 | 1461868 | 8.81 | 21.16 | 20.24 | 11.93 | 50.33 | 26.89 | Acetyltransferase |
| *PSPTO_1332* | *ilvE* | + | 1462008 | 1463027 | 84.80 | 84.45 | 62.82 | 38.61 | 249.85 | 195.36 | Branched-chain amino acid aminotransferase |
| *PSPTO_1333* | | - | 1463091 | 1464086 | 31.85 | 23.82 | 22.11 | 16.13 | 97.18 | 27.73 | Hypothetical protein |
| *PSPTO_1334* | | - | 1464236 | 1465888 | 59.78 | 25.36 | 22.15 | 9.98 | 11.72 | 7.78 | Methyl-accepting chemotaxis protein |
| *PSPTO_1335* | | - | 1466008 | 1466799 | 27.19 | 9.84 | 13.54 | 5.36 | 28.36 | 11.86 | Cyclohexadienyl dehydratase |
| *PSPTO_1336* | | + | 1466973 | 1467269 | 30.01 | 17.61 | 19.91 | 5.75 | 20.22 | 10.92 | ArsR family transcriptional regulator |
| *PSPTO_1337* | | + | 1467326 | 1468336 | 2.53 | 3.67 | 4.41 | 4.01 | 37.47 | 9.54 | Oxidoreductase zinc-binding protein |
| *PSPTO_1338* | *sodC* | + | 1468578 | 1469096 | 11.50 | 35.57 | 46.09 | 15.25 | 13.01 | 19.73 | Superoxide dismutase, Cu-Zn |
| *PSPTO_1339* | | - | 1469164 | 1470351 | 3.29 | 6.41 | 2.30 | 2.49 | 5.12 | 5.20 | Membrane protein |
| *PSPTO_1340* | | - | 1470549 | 1471325 | 64.62 | 614.83 | 412.80 | 96.42 | 479.86 | 45.65 | Carbonic anhydrase |
| *PSPTO_1341* | | - | 1471584 | 1472552 | 19.04 | 15.34 | 10.78 | 10.71 | 38.15 | 12.69 | Hypothetical protein |
| *PSPTO_1342* | | - | 1472620 | 1473678 | 16.64 | 18.23 | 25.30 | 21.31 | 53.90 | 20.35 | Hypothetical protein |
| *PSPTO_1343* | | + | 1473898 | 1475637 | 3.08 | 8.21 | 4.12 | 2.09 | 52.41 | 11.22 | Hypothetical protein |
| *PSPTO_1344* | | + | 1475743 | 1476315 | 6.92 | 5.58 | 8.48 | 0.00 | 47.88 | 11.60 | Hypothetical protein |
| *PSPTO_1345* | *gnl* | + | 1476333 | 1477349 | 23.47 | 34.52 | 23.77 | 17.44 | 61.37 | 39.02 | Gluconolactonase |
| *PSPTO_1346* | | - | 1477371 | 1478648 | 9.69 | 22.96 | 14.40 | 7.50 | 23.81 | 14.33 | Sensory box/GGDEF domain protein |
| *PSPTO_1347* | | + | 1479036 | 1480382 | 8.23 | 7.62 | 10.17 | 4.59 | 12.17 | 10.70 | Gluconate transporter family protein |
| *PSPTO_1348* | | - | 1480410 | 1482701 | 3.03 | 7.32 | 4.91 | 1.73 | 8.14 | 4.41 | Sensory box/GGDEF domain/EAL domain-containing protein |
| *PSPTO_1349* | | - | 1482921 | 1488389 | 2.88 | 7.40 | 6.02 | 2.97 | 28.86 | 6.12 | Hypothetical protein |
| *PSPTO_1350* | | - | 1488377 | 1489579 | 15.35 | 12.49 | 19.29 | 6.89 | 43.63 | 14.07 | Hypothetical protein |
| *PSPTO_1351* | | - | 1489605 | 1489901 | 15.34 | 20.85 | 24.35 | 6.39 | 62.42 | 18.73 | Hypothetical protein |
| *PSPTO_1352* | | - | 1490163 | 1490849 | 11.79 | 26.62 | 11.53 | 4.47 | 47.54 | 14.61 | GntR family transcriptional regulator |
| *PSPTO_1353* | | - | 1490846 | 1491241 | 2.87 | 21.96 | 14.27 | 7.15 | 9.51 | 5.88 | Hypothetical protein |
| *PSPTO_1354* | | - | 1491241 | 1491918 | 2.17 | 14.87 | 3.14 | 3.68 | 4.22 | 2.86 | Isochorismatase family protein |
| *PSPTO_1355* | | - | 1491915 | 1493450 | 2.66 | 13.93 | 5.15 | 4.73 | 4.45 | 3.72 | ABC transporter ATP-binding protein |
| *PSPTO_1356* | | - | 1493453 | 1494139 | 2.60 | 23.41 | 16.69 | 0.76 | 4.79 | 2.72 | Isochorismatase family protein |

| Locus name | Gene name | Strand | Gene start (bp) | Gene end (bp) | Average TPM Inoculum | Average TPM $t_6$ | Average TPM $t_{24}$ | Average TPM $t_{168}$ | Average TPM Logarithmic | Average TPM Stationary | Annotation description |
|---|---|---|---|---|---|---|---|---|---|---|---|
| *PSPTO_1357* | *amiE* | - | 1494145 | 1495161 | 6.52 | 29.17 | 22.92 | 6.32 | 5.03 | 2.67 | Aliphatic amidase |
| *PSPTO_1358* | | - | 1495186 | 1496106 | 6.61 | 7.42 | 5.56 | 0.61 | 1.65 | 1.66 | ABC transporter permease |
| *PSPTO_1359* | | - | 1496099 | 1497211 | 1.95 | 5.44 | 1.70 | 0.00 | 2.84 | 1.26 | ABC transporter permease |
| *PSPTO_1360* | | - | 1497221 | 1498330 | 9.40 | 18.49 | 8.88 | 7.48 | 2.22 | 1.37 | Bmp family protein |
| *PSPTO_1361* | | - | 1498345 | 1500150 | 8.06 | 13.25 | 6.68 | 5.28 | 1.49 | 1.46 | Allophanate hydrolase |
| *PSPTO_1362* | | - | 1500375 | 1500770 | 5.51 | 9.22 | 7.93 | 1.85 | 11.05 | 7.36 | Hypothetical protein |
| *PSPTO_1363* | | - | 1500812 | 1502233 | 9.97 | 13.85 | 9.27 | 11.34 | 18.28 | 12.46 | Sugar transporter family protein |
| *PSPTO_1364* | | + | 1502395 | 1502811 | 15.87 | 41.38 | 34.98 | 9.36 | 42.92 | 18.35 | Hypothetical protein |
| *PSPTO_1365* | *gstA* | + | 1502901 | 1503527 | 14.78 | 45.35 | 24.26 | 17.78 | 28.44 | 22.71 | Glutathione S-transferase |
| *PSPTO_1366* | | - | 1503611 | 1503772 | 33.63 | 39.24 | 17.02 | 8.11 | 57.40 | 22.03 | Hypothetical protein |
| *PSPTO_1367* | | + | 1503935 | 1504381 | 7.35 | 11.13 | 6.30 | 4.06 | 12.88 | 9.43 | GNAT family acetyltransferase |
| *PSPTO_1368* | | - | 1504444 | 1504731 | 20.79 | 23.32 | 25.62 | 9.34 | 29.52 | 24.31 | Lipoprotein |
| *PSPTO_1369* | *shcN* | + | 1504948 | 1505427 | 7.05 | 462.03 | 197.92 | 37.70 | 6.28 | 4.45 | Type III chaperone protein ShcN |
| *PSPTO_1370* | *hopN1* | + | 1505441 | 1506493 | 8.78 | 163.45 | 91.32 | 20.63 | 7.52 | 6.81 | Type III effector HopN1 |
| *PSPTO_1371* | | - | 1506530 | 1507474 | 20.43 | 94.59 | 134.00 | 34.28 | 14.04 | 7.51 | Effector locus protein |
| *PSPTO_1372* | *hopAA1-1* | + | 1507846 | 1509306 | 1.58 | 114.78 | 39.72 | 4.41 | 3.47 | 2.04 | Type III effector HopAA1-1 |
| *PSPTO_1373* | *hrpW1* | - | 1509390 | 1510664 | 4.95 | 1098.74 | 649.96 | 133.38 | 5.73 | 4.75 | Type III helper protein HrpW1 |
| *PSPTO_1374* | *shcM* | + | 1510970 | 1511464 | 1.27 | 152.15 | 101.56 | 14.24 | 3.28 | 2.39 | Type III chaperone ShcM |
| *PSPTO_1375* | *hopM1* | + | 1511461 | 1513599 | 5.33 | 82.40 | 56.45 | 13.69 | 4.88 | 3.73 | Type III effector HopM1 |
| *PSPTO_1376* | *shcE* | + | 1513672 | 1514067 | 55.44 | 19.00 | 22.64 | 16.80 | 18.36 | 15.10 | Type III chaperone ShcE |
| *PSPTO_1377* | *avrE1* | - | 1514116 | 1519503 | 27.26 | 40.25 | 27.03 | 8.33 | 14.25 | 15.64 | Type III effector AvrE |
| *PSPTO_1378* | *hrpH* | + | 1519724 | 1521211 | 6.22 | 78.14 | 35.14 | 8.43 | 9.59 | 6.48 | Membrane-bound lytic murein transglycosylase D |
| *PSPTO_1379* | *hrpR* | + | 1522289 | 1523233 | 22.95 | 81.22 | 45.84 | 8.52 | 12.84 | 7.25 | Type III transcriptional regulator HrpR |
| *PSPTO_1380* | *hrpS* | + | 1523279 | 1524187 | 8.88 | 36.28 | 23.20 | 15.33 | 14.59 | 10.72 | Type III transcriptional regulator HrpS |
| *PSPTO_1381* | *hrpA1* | + | 1524281 | 1524622 | 11.39 | 6843.30 | 4372.96 | 627.28 | 50.56 | 21.47 | Type III helper protein HrpA1 |
| *PSPTO_1382* | *hrpZ1* | + | 1524678 | 1525790 | 3.61 | 2636.39 | 1505.92 | 242.41 | 11.71 | 6.98 | Type III restriction system endonuclease |
| *PSPTO_1383* | *hrpB* | + | 1525818 | 1526192 | 3.66 | 595.85 | 333.23 | 87.32 | 3.33 | 1.66 | Type III secretion protein HrpB |
| *PSPTO_1384* | *hrcJ* | + | 1526201 | 1527007 | 2.39 | 301.53 | 145.05 | 41.85 | 5.69 | 3.73 | Type III secretion protein HrcJ |
| *PSPTO_1385* | *hrpD* | + | 1527004 | 1527534 | 2.91 | 306.23 | 155.19 | 38.26 | 3.52 | 3.35 | Type III secretion protein HrpD |
| *PSPTO_1386* | *hrpE* | + | 1527570 | 1528151 | 2.10 | 138.04 | 78.89 | 26.54 | 2.61 | 1.66 | Type III secretion protein HrpE |
| *PSPTO_1387* | *hrpF* | + | 1528246 | 1528470 | 1.99 | 472.03 | 359.71 | 39.16 | 4.15 | 2.31 | Type III secretion protein HrpF |
| *PSPTO_1388* | *hrpG* | + | 1528457 | 1528888 | 1.10 | 732.91 | 423.14 | 60.70 | 1.99 | 1.36 | Type III secretion protein HrpG |
| *PSPTO_1389* | *hrcC* | + | 1528890 | 1530989 | 1.13 | 18.38 | 12.90 | 2.48 | 1.15 | 0.91 | Outer-membrane type III secretion protein HrcC |
| *PSPTO_1390* | *hrpT* | + | 1530986 | 1531189 | 233.22 | 188.63 | 120.58 | 113.19 | 52.01 | 108.20 | Type III secretion protein |
| *PSPTO_1391* | *hrpV* | + | 1531192 | 1531551 | 69.65 | 110.97 | 68.04 | 35.26 | 27.87 | 47.97 | Negative regulator of hrp expression HrpV |
| *PSPTO_1392* | *hrcU* | - | 1532093 | 1533172 | 4.22 | 52.63 | 21.33 | 3.56 | 4.96 | 3.31 | Type III secretion protein HrcU |
| *PSPTO_1393* | *hrcT* | - | 1533169 | 1533963 | 2.39 | 21.12 | 13.48 | 1.61 | 2.47 | 2.28 | Type III secretion protein HrcT |
| *PSPTO_1394* | *hrcS* | - | 1533964 | 1534230 | 3.07 | 25.80 | 14.20 | 6.18 | 4.08 | 3.64 | Type III secretion protein HrcS |
| *PSPTO_1395* | *hrcR* | - | 1534238 | 1534840 | 2.89 | 80.30 | 31.13 | 14.94 | 2.29 | 1.52 | Type III secretion protein HrcR |
| *PSPTO_1396* | *hrcQb* | - | 1534888 | 1535301 | 8.61 | 154.85 | 85.29 | 16.14 | 3.07 | 2.37 | Type III secretion protein HrcQb |
| *PSPTO_1397* | *hrcQa* | - | 1535291 | 1536007 | 4.32 | 243.99 | 108.06 | 22.77 | 4.11 | 3.49 | Type III secretion protein HrcQa |
| *PSPTO_1398* | *hrpP* | - | 1536004 | 1536573 | 1.61 | 134.92 | 65.23 | 27.11 | 4.48 | 3.37 | Type III secretion protein HrpP |
| *PSPTO_1399* | *hrpO* | - | 1536580 | 1537026 | 3.14 | 818.20 | 421.31 | 55.92 | 6.64 | 7.83 | Type III secretion protein HrpO |
| *PSPTO_1400* | *hrcN* | - | 1537029 | 1538378 | 1.52 | 124.94 | 67.69 | 9.73 | 3.53 | 3.78 | Type III secretion cytoplasmic ATPase HrcN |
| *PSPTO_1401* | *hrpQ* | - | 1538375 | 1539367 | 1.51 | 220.19 | 153.20 | 23.37 | 3.62 | 3.82 | Type III secretion protein HrpQ |
| *PSPTO_1402* | *hrcV* | - | 1539401 | 1541488 | 1.98 | 125.32 | 81.86 | 12.17 | 3.53 | 3.39 | Type III secretion protein HrcV |
| *PSPTO_1403* | *hrpJ* | - | 1541488 | 1542594 | 2.34 | 291.64 | 173.45 | 19.06 | 3.18 | 3.94 | Type III secretion protein HrpJ |
| *PSPTO_1404* | *hrpL* | + | 1542838 | 1543392 | 6.50 | 740.76 | 377.55 | 64.75 | 3.74 | 1.21 | RNA polymerase sigma factor HrpL |
| *PSPTO_1405* | *hrpK1* | + | 1543479 | 1545821 | 7.60 | 418.39 | 241.48 | 35.73 | 8.23 | 6.45 | Type III helper protein HrpK1 |
| *PSPTO_1406* | *hopB1* | + | 1545896 | 1547296 | 13.22 | 254.40 | 153.85 | 29.07 | 9.30 | 10.09 | Type III effector HopB1 |
| *PSPTO_1407* | | - | 1547542 | 1548381 | 1.77 | 1.84 | 0.96 | 0.00 | 2.97 | 2.98 | ISPssy transposase or derivative |
| *PSPTO_1408* | | + | 1548919 | 1549104 | 6.27 | 246.09 | 124.55 | 69.95 | 13.87 | 9.30 | Hypothetical protein |
| *PSPTO_1409* | | + | 1549288 | 1550133 | 3.51 | 57.78 | 38.92 | 10.90 | 7.18 | 8.18 | Hypothetical protein |
| *PSPTO_1410* | | + | 1550627 | 1551043 | 879.11 | 431.41 | 429.47 | 210.68 | 69.92 | 325.34 | Exchangeable effector locus protein |
| *PSPTO_1411* | | - | 1551198 | 1551773 | 98.77 | 56.47 | 63.15 | 49.25 | 79.84 | 39.32 | Hypothetical protein |
| *PSPTO_1412* | *queA* | + | 1551979 | 1553043 | 8.51 | 25.31 | 28.08 | 11.57 | 56.12 | 17.82 | S-adenosylmethionine--tRNA ribosyltransferase-isomerase |
| *PSPTO_1413* | *tgt* | + | 1553058 | 1554173 | 7.22 | 18.37 | 19.44 | 10.08 | 63.08 | 13.03 | Queuine tRNA-ribosyltransferase |
| *PSPTO_1414* | *yajC* | + | 1554227 | 1554562 | 19.49 | 90.80 | 56.51 | 33.30 | 294.31 | 49.48 | Preprotein translocase subunit YajC |
| *PSPTO_1415* | *secD* | + | 1554625 | 1556493 | 3.20 | 19.70 | 10.92 | 12.10 | 117.91 | 9.36 | Protein-export membrane protein SecD |
| *PSPTO_1416* | *secF* | + | 1556503 | 1557414 | 5.40 | 20.19 | 17.03 | 6.09 | 82.66 | 7.63 | Protein-export membrane protein SecF |
| *PSPTO_1417* | | + | 1557523 | 1558062 | 1528.71 | 1723.28 | 1607.79 | 907.45 | 503.45 | 1731.15 | Hypothetical protein |
| *PSPTO_1418* | | - | 1558205 | 1559665 | 0.22 | 0.42 | 0.10 | 0.38 | 0.45 | 0.37 | ISPsy6, transposase |
| *PSPTO_1419* | *suhB* | - | 1559737 | 1560552 | 39.05 | 66.87 | 60.39 | 43.81 | 104.60 | 77.84 | Inositol-1-monophosphatase |
| *PSPTO_1420* | | + | 1560768 | 1561541 | 5.92 | 22.41 | 22.71 | 12.72 | 29.85 | 14.03 | RNA methyltransferase, TrmH family, group 1 |
| *PSPTO_1421* | *cysE* | + | 1561541 | 1562317 | 3.80 | 12.77 | 8.75 | 17.34 | 33.02 | 12.68 | Serine O-acetyltransferase |
| *PSPTO_1422* | | + | 1562477 | 1562968 | 267.61 | 164.41 | 171.13 | 95.23 | 80.78 | 18.23 | Rrf2 family protein |
| *PSPTO_1423* | *iscS-2* | + | 1563013 | 1564227 | 135.29 | 125.11 | 164.82 | 106.91 | 191.21 | 24.39 | Cysteine desulfurase |
| *PSPTO_1424* | *iscU* | + | 1564310 | 1564696 | 144.15 | 128.74 | 150.90 | 83.86 | 197.85 | 32.09 | Iron-binding protein IscU |
| *PSPTO_1425* | *iscA* | + | 1564739 | 1565062 | 57.46 | 52.61 | 63.58 | 68.99 | 148.47 | 19.76 | Iron-binding protein IscA |
| *PSPTO_1426* | *hscB* | + | 1565074 | 1565595 | 53.00 | 38.00 | 58.20 | 37.23 | 122.91 | 19.51 | Co-chaperone Hsc20 |
| *PSPTO_1427* | *hscA* | + | 1565640 | 1567502 | 14.93 | 21.71 | 14.43 | 8.21 | 70.50 | 5.20 | Chaperone protein HscA |
| *PSPTO_1428* | | + | 1567506 | 1567847 | 16.34 | 14.45 | 12.74 | 24.00 | 74.90 | 7.66 | Ferredoxin, 2Fe-2S |
| *PSPTO_1429* | | + | 1567856 | 1568062 | 16.68 | 8.32 | 0.00 | 16.92 | 36.65 | 3.90 | Hypothetical protein |
| *PSPTO_1430* | *ndk* | + | 1568159 | 1568584 | 59.89 | 185.14 | 170.83 | 119.38 | 1003.42 | 145.47 | Nucleoside diphosphate kinase |
| *PSPTO_1431* | | + | 1568618 | 1569766 | 11.01 | 46.43 | 31.31 | 27.76 | 74.96 | 44.61 | Radical SAM enzyme, Cfr family |
| *PSPTO_1432* | *pilF* | + | 1569793 | 1570551 | 22.38 | 56.51 | 53.65 | 44.48 | 163.19 | 89.42 | Type IV pilus biogenesis protein PilF |
| *PSPTO_1433* | | + | 1570551 | 1571546 | 17.48 | 42.86 | 43.87 | 21.05 | 151.62 | 50.38 | Hypothetical protein |
| *PSPTO_1434* | *ispG* | + | 1571550 | 1572659 | 16.81 | 38.99 | 27.18 | 29.79 | 130.25 | 34.97 | 1-hydroxy-2-methyl-2-(E)-butenyl 4-diphosphate synthase |
| *PSPTO_1435* | *hisS* | + | 1572682 | 1573971 | 16.94 | 36.56 | 27.62 | 23.68 | 125.86 | 44.62 | Histidyl-tRNA synthetase |

| Locus name | Gene name | Strand | Gene start (bp) | Gene end (bp) | Average TPM Inoculum | Average TPM $t_6$ | Average TPM $t_{24}$ | Average TPM $t_{168}$ | Average TPM Logarithmic | Average TPM Stationary | Annotation description |
|---|---|---|---|---|---|---|---|---|---|---|---|
| *PSPTO_1436* | | + | 1574010 | 1574651 | 30.73 | 80.90 | 74.60 | 43.40 | 267.30 | 73.82 | Hypothetical protein |
| *PSPTO_1437* | | + | 1574644 | 1575798 | 10.05 | 27.79 | 11.81 | 16.75 | 77.95 | 20.40 | PQQ enzyme repeat domain-containing protein |
| *PSPTO_1438* | *engA* | + | 1575936 | 1577405 | 5.07 | 10.07 | 11.54 | 1.26 | 38.18 | 15.48 | GTP-binding protein EngA |
| *PSPTO_1439* | | - | 1577530 | 1578990 | 0.29 | 0.30 | 0.14 | 0.32 | 0.41 | 0.36 | ISPsy6, transposase |
| *PSPTO_1440* | | + | 1579125 | 1580273 | 15.27 | 57.15 | 53.27 | 35.70 | 162.61 | 79.01 | Class I and II aminotransferase |
| *PSPTO_1441* | | + | 1580261 | 1581052 | 5.05 | 32.67 | 16.41 | 10.57 | 54.12 | 31.93 | Hydrolase |
| *PSPTO_1443* | | - | 1581671 | 1582540 | 39.15 | 22.85 | 23.65 | 18.45 | 67.64 | 42.59 | Carboxyphosphonoenolpyruvate phosphonomutase |
| *PSPTO_1444* | *leuA* | - | 1582912 | 1584669 | 48.60 | 184.60 | 162.68 | 113.64 | 403.87 | 164.80 | 2-isopropylmalate synthase |
| *PSPTO_1445* | | - | 1584963 | 1585787 | 6.77 | 12.56 | 16.21 | 11.19 | 42.51 | 15.25 | Peptidase, M23/M37 family |
| *PSPTO_1446* | *xseA* | - | 1585794 | 1587185 | 12.47 | 15.00 | 14.75 | 7.70 | 31.74 | 13.82 | Exodeoxyribonuclease VII large subunit |
| *PSPTO_1447* | | + | 1587200 | 1587388 | 6.77 | 7.26 | 11.03 | 3.68 | 3.82 | 5.15 | Hypothetical protein |
| *PSPTO_1448* | | + | 1587464 | 1588012 | 9.21 | 18.90 | 24.48 | 8.29 | 40.53 | 14.21 | Hypothetical protein |
| *PSPTO_1449* | *guaB* | + | 1588107 | 1589576 | 31.03 | 58.16 | 70.99 | 55.60 | 155.25 | 95.09 | Inosine 5'-monophosphate dehydrogenase |
| *PSPTO_1450* | *guaA* | + | 1589700 | 1591277 | 4.91 | 23.65 | 16.95 | 17.35 | 128.84 | 41.05 | GMP synthase |
| *PSPTO_1451* | | - | 1592228 | 1593409 | 9.17 | 28.34 | 29.49 | 16.47 | 74.14 | 40.72 | Hypothetical protein |
| *PSPTO_1452* | | - | 1593498 | 1593614 | 31.46 | 7.23 | 11.51 | 34.08 | 121.70 | 51.31 | Hypothetical protein |
| *PSPTO_1453* | *lsc-1* | - | 1593629 | 1594924 | 13.05 | 15.36 | 18.13 | 17.20 | 56.26 | 26.32 | Levansucrase |
| *PSPTO_1454* | | - | 1595545 | 1596534 | 18.07 | 18.77 | 8.86 | 7.04 | 20.50 | 14.17 | Hypothetical protein |
| *PSPTO_1455* | | + | 1598062 | 1600338 | 1.47 | 2.54 | 1.95 | 4.14 | 14.38 | 4.02 | Hypothetical protein |
| *PSPTO_1456* | *cumA* | + | 1600668 | 1602041 | 4.52 | 11.60 | 11.60 | 12.84 | 33.74 | 20.35 | Multicopper oxidase |
| *PSPTO_1457* | | + | 1602041 | 1602550 | 25.93 | 20.42 | 32.34 | 37.53 | 68.42 | 47.86 | Cytidine/deoxycytidylate deaminase family protein |
| *PSPTO_1458* | | - | 1602547 | 1603917 | 12.41 | 44.64 | 34.80 | 21.10 | 75.80 | 35.59 | Periplasmic binding domain/transglycosylase SLT domain fusion protein |
| *PSPTO_1459* | *purL* | + | 1604253 | 1608149 | 11.37 | 34.78 | 24.64 | 20.12 | 129.60 | 62.29 | Phosphoribosylformylglycinamidine synthase |
| *PSPTO_1460* | | + | 1608155 | 1608466 | 4.20 | 21.42 | 19.76 | 8.67 | 42.29 | 29.41 | Hypothetical protein |
| *PSPTO_1461* | | - | 1608605 | 1609231 | 29.49 | 27.69 | 34.31 | 18.58 | 76.51 | 44.95 | Hypothetical protein |
| *PSPTO_1462* | | - | 1609228 | 1609779 | 44.68 | 93.05 | 53.28 | 28.44 | 129.87 | 48.92 | MutT/nudix family protein |
| *PSPTO_1463* | | + | 1609794 | 1609940 | 12.67 | 21.39 | 7.68 | 17.14 | 36.08 | 17.75 | Hypothetical protein |
| *PSPTO_1464* | | + | 1609930 | 1610526 | 44.40 | 78.22 | 56.91 | 28.27 | 138.17 | 66.42 | MutT/nudix family protein |
| *PSPTO_1465* | | + | 1610585 | 1611109 | 32.33 | 57.29 | 39.33 | 32.29 | 67.72 | 50.22 | Transferase, hexapeptide repeat protein |
| *PSPTO_1466* | | + | 1611109 | 1611327 | 18.57 | 19.11 | 15.75 | 0.85 | 24.03 | 17.39 | Hypothetical protein |
| *PSPTO_1467* | | + | 1611959 | 1612426 | 5.95 | 20.49 | 20.84 | 6.69 | 25.33 | 10.72 | Hypothetical protein |
| *PSPTO_1468* | *purT* | + | 1612530 | 1613711 | 7.28 | 12.27 | 16.34 | 10.61 | 46.28 | 14.38 | Phosphoribosylglycinamide formyltransferase 2 |
| *PSPTO_1469* | | - | 1613824 | 1615140 | 2.96 | 6.94 | 0.96 | 2.18 | 14.67 | 3.28 | Major facilitator family transporter |
| *PSPTO_1470* | | - | 1615271 | 1616518 | 4.73 | 11.30 | 7.66 | 6.70 | 62.26 | 16.99 | Hypothetical protein |
| *PSPTO_1471* | | - | 1616531 | 1617343 | 9.11 | 20.66 | 14.67 | 9.81 | 77.31 | 28.22 | Membrane protein |
| *PSPTO_1472* | *ffh* | + | 1617559 | 1618935 | 14.40 | 37.95 | 39.80 | 38.05 | 124.90 | 48.84 | Signal recognition particle protein Ffh |
| *PSPTO_1473* | *rpsP* | + | 1619184 | 1619441 | 95.95 | 278.64 | 278.08 | 211.50 | 714.44 | 185.28 | 30S ribosomal protein S16 |
| *PSPTO_1474* | *rimM* | + | 1619447 | 1619986 | 32.59 | 164.27 | 148.41 | 117.38 | 471.53 | 95.93 | 16S rRNA-processing protein RimM |
| *PSPTO_1475* | *trmD* | + | 1619992 | 1620744 | 76.22 | 461.06 | 314.81 | 248.19 | 1891.73 | 253.96 | tRNA (guanine-N1)-methyltransferase |
| *PSPTO_1476* | *rplS* | + | 1620789 | 1621139 | 14.35 | 92.06 | 78.94 | 68.95 | 132.41 | 24.10 | 50S ribosomal protein L19 |
| *PSPTO_1477* | | - | 1621273 | 1622733 | 0.24 | 0.32 | 0.33 | 0.07 | 0.44 | 0.31 | ISPsy6, transposase |
| *PSPTO_1478* | *xerD* | + | 1622850 | 1623746 | 2.92 | 5.85 | 3.29 | 2.21 | 7.91 | 4.70 | Integrase/recombinase XerD |
| *PSPTO_1479* | *dsbC* | + | 1623892 | 1624632 | 13.07 | 18.35 | 21.18 | 9.20 | 45.18 | 16.80 | Thiol:disulfide interchange protein DsbC |
| *PSPTO_1480* | *hom* | + | 1624779 | 1626083 | 12.92 | 18.77 | 27.02 | 17.49 | 122.18 | 28.51 | Homoserine dehydrogenase |
| *PSPTO_1481* | *thrC* | + | 1626188 | 1627597 | 7.61 | 18.54 | 17.81 | 7.22 | 91.59 | 18.07 | Threonine synthase |
| *PSPTO_1482* | | + | 1627718 | 1631359 | 4.44 | 8.65 | 9.07 | 4.10 | 13.26 | 4.31 | Sensory box histidine kinase/response regulator |
| *PSPTO_1483* | | + | 1631483 | 1632772 | 10.68 | 8.85 | 11.17 | 3.79 | 7.79 | 7.76 | Response regulator/EAL domain protein |
| *PSPTO_1484* | | - | 1632745 | 1633371 | 4.33 | 2.91 | 1.76 | 1.89 | 6.37 | 5.80 | LuxR family DNA-binding response regulator |
| *PSPTO_1485* | | + | 1633695 | 1633991 | 268.15 | 858.44 | 744.18 | 426.37 | 287.51 | 310.39 | Hypothetical protein |
| *PSPTO_1486* | | - | 1634164 | 1634937 | 10.59 | 4.71 | 11.05 | 6.19 | 19.27 | 14.82 | Hypothetical protein |
| *PSPTO_1487* | | + | 1635337 | 1635879 | 6.26 | 19.78 | 11.99 | 5.05 | 60.64 | 27.55 | Hypothetical protein |
| *PSPTO_1488* | *recJ* | + | 1635953 | 1637662 | 2.87 | 11.48 | 8.40 | 6.97 | 27.40 | 10.14 | Single-stranded-DNA-specific exonuclease RecJ |
| *PSPTO_1489* | | + | 1637777 | 1638883 | 4.29 | 61.98 | 40.46 | 13.14 | 12.89 | 14.53 | Xenobiotic reductase |
| *PSPTO_1490* | | - | 1638947 | 1641034 | 8.88 | 7.69 | 9.02 | 5.09 | 7.19 | 10.70 | Sensory box histidine kinase/response regulator |
| *PSPTO_1491* | | + | 1641328 | 1642074 | 40.19 | 262.17 | 225.06 | 25.52 | 77.37 | 111.43 | Hypothetical protein |
| *PSPTO_1492* | | - | 1642132 | 1648086 | 2.90 | 5.41 | 3.46 | 1.58 | 4.73 | 7.55 | Hypothetical protein |
| *PSPTO_1493* | | + | 1648356 | 1649978 | 61.14 | 25.15 | 26.77 | 12.69 | 22.19 | 21.38 | Methyl-accepting chemotaxis protein |
| *PSPTO_1494* | | + | 1649986 | 1650522 | 20.58 | 11.06 | 16.26 | 13.19 | 14.60 | 9.54 | Chemotaxis protein CheW |
| *PSPTO_1495* | | + | 1650519 | 1651799 | 13.40 | 7.98 | 12.65 | 3.29 | 13.22 | 8.88 | Chemotaxis protein methyltransferase CheR |
| *PSPTO_1496* | | + | 1651799 | 1652488 | 10.95 | 11.17 | 5.07 | 6.75 | 15.92 | 6.31 | Chemotaxis protein CheW |
| *PSPTO_1497* | | + | 1652485 | 1654866 | 10.44 | 11.88 | 10.61 | 4.64 | 17.01 | 8.03 | Sensor histidine kinase/response regulator |
| *PSPTO_1498* | | + | 1654863 | 1655873 | 34.45 | 43.85 | 46.44 | 20.06 | 63.62 | 37.93 | Protein-glutamate methylesterase CheB |
| *PSPTO_1499* | | + | 1655919 | 1656923 | 19.04 | 26.88 | 26.46 | 26.56 | 65.50 | 37.11 | Response regulator/GGDEF domain-containing protein |
| *PSPTO_1500* | *prfB* | + | 1657245 | 1658126 | 9.30 | 61.15 | 73.91 | 44.29 | 236.93 | 97.97 | Hypothetical protein |
| *PSPTO_1501* | *lysS* | + | 1658196 | 1659698 | 3.04 | 15.22 | 16.76 | 18.15 | 114.52 | 31.14 | Lysyl-tRNA synthetase |
| *PSPTO_1502* | | + | 1659777 | 1660502 | 11.44 | 19.37 | 10.35 | 14.90 | 35.86 | 12.21 | TetR family transcriptional regulator |
| *PSPTO_1503* | | + | 1660644 | 1661921 | 57.41 | 54.46 | 48.43 | 21.50 | 83.12 | 59.06 | Hypothetical protein |
| *PSPTO_1504* | | + | 1662172 | 1663068 | 5.46 | 6.27 | 4.90 | 3.68 | 15.77 | 12.79 | Lipoprotein |
| *PSPTO_1505* | | + | 1663065 | 1663385 | 4.76 | 8.77 | 4.63 | 5.53 | 13.23 | 6.21 | Lipoprotein |
| *PSPTO_1506* | | - | 1663462 | 1664244 | 23.82 | 348.50 | 293.67 | 95.61 | 129.97 | 118.97 | OmpA family protein |
| *PSPTO_1507* | | - | 1664303 | 1664716 | 64.46 | 614.32 | 396.91 | 118.66 | 214.10 | 169.09 | Hypothetical protein |
| *PSPTO_1508* | *ppc* | + | 1665095 | 1667731 | 9.66 | 36.98 | 34.92 | 17.04 | 36.39 | 14.61 | Phosphoenolpyruvate carboxylase |
| *PSPTO_1509* | *adk* | + | 1668011 | 1668658 | 16.66 | 42.69 | 29.45 | 26.08 | 129.38 | 26.99 | Adenylate kinase |
| *PSPTO_1510* | | + | 1668746 | 1669420 | 15.12 | 22.50 | 25.67 | 1.35 | 29.59 | 10.44 | Hypothetical protein |
| *PSPTO_1511* | | + | 1669644 | 1669955 | 109.59 | 130.63 | 86.91 | 90.98 | 74.56 | 212.11 | Hypothetical protein |
| *PSPTO_1512* | | + | 1669952 | 1670821 | 4.71 | 8.94 | 8.06 | 6.27 | 17.38 | 14.70 | Hypothetical protein |
| *PSPTO_1513* | | + | 1670963 | 1671652 | 3.94 | 4.08 | 3.04 | 0.00 | 4.53 | 5.46 | Hypothetical protein |
| *PSPTO_1514* | | - | 1671744 | 1672397 | 195.11 | 39.05 | 28.35 | 19.58 | 7.95 | 6.09 | Hypothetical protein |
| *PSPTO_1515* | | + | 1672410 | 1673273 | 22.49 | 17.38 | 18.78 | 16.08 | 26.08 | 19.57 | Hypothetical protein |

| Locus name | Gene name | Strand | Gene start (bp) | Gene end (bp) | Average TPM Inoculum | Average TPM $t_6$ | Average TPM $t_{24}$ | Average TPM $t_{168}$ | Average TPM Logarithmic | Average TPM Stationary | Annotation description |
|---|---|---|---|---|---|---|---|---|---|---|---|
| *PSPTO_1516* | | - | 1673275 | 1673403 | 2.29 | 0.00 | 11.44 | 0.00 | 11.22 | 4.75 | Hypothetical protein |
| *PSPTO_1517* | | + | 1673427 | 1673654 | 6.56 | 3.15 | 7.24 | 8.58 | 19.74 | 9.92 | Hypothetical protein |
| *PSPTO_1518* | | - | 1673702 | 1674391 | 17.07 | 27.81 | 19.27 | 13.72 | 48.81 | 14.80 | Hypothetical protein |
| *PSPTO_1519* | | - | 1674599 | 1674949 | 4.54 | 3.51 | 5.96 | 0.00 | 6.06 | 3.81 | Hypothetical protein |
| *PSPTO_1520* | *plsB* | - | 1675061 | 1677562 | 6.12 | 7.39 | 5.40 | 4.98 | 28.53 | 6.87 | Glycerol-3-phosphate acyltransferase |
| *PSPTO_1521* | | - | 1677754 | 1678131 | 24.62 | 20.07 | 21.23 | 8.49 | 68.52 | 14.10 | Hypothetical protein |
| *PSPTO_1522* | | - | 1678151 | 1678960 | 7.93 | 14.53 | 13.39 | 5.54 | 54.09 | 9.97 | RRNA large subunit methyltransferase A |
| *PSPTO_1523* | *dapE* | - | 1678960 | 1680111 | 5.99 | 20.71 | 14.52 | 12.37 | 69.62 | 12.37 | Succinyl-diaminopimelate desuccinylase |
| *PSPTO_1524* | | - | 1680211 | 1682739 | 10.33 | 11.71 | 12.29 | 9.12 | 25.35 | 17.96 | Glycosyl transferase family protein |
| *PSPTO_1525* | | + | 1682938 | 1683768 | 17.12 | 21.05 | 20.44 | 18.19 | 32.91 | 21.71 | ThiF family protein |
| *PSPTO_1526* | *sufE* | - | 1683759 | 1684166 | 1.79 | 5.65 | 14.34 | 3.83 | 21.95 | 8.29 | SufE protein |
| *PSPTO_1527* | *csdA* | - | 1684163 | 1685368 | 3.80 | 12.06 | 10.95 | 7.77 | 32.56 | 13.36 | Cysteine sulfinate desulfinase |
| *PSPTO_1528* | | - | 1685429 | 1686463 | 8.95 | 40.15 | 24.23 | 12.08 | 154.06 | 65.15 | Tetrahydrodipicolinate succinylase |
| *PSPTO_1529* | | - | 1686511 | 1686858 | 21.42 | 44.06 | 47.56 | 29.31 | 106.53 | 77.03 | ArsC family protein |
| *PSPTO_1530* | | - | 1687418 | 1688266 | 4.83 | 17.48 | 17.83 | 6.09 | 7.58 | 7.57 | Hypothetical protein |
| *PSPTO_1531* | | - | 1688403 | 1689596 | 8.64 | 25.72 | 22.47 | 11.67 | 68.87 | 30.13 | Class I and II aminotransferase |
| *PSPTO_1532* | *glnD* | - | 1689699 | 1692395 | 5.85 | 25.14 | 24.47 | 10.81 | 76.85 | 42.86 | [protein-pII] uridylyltransferase |
| *PSPTO_1533* | *map-1* | - | 1692462 | 1693244 | 9.87 | 38.07 | 35.18 | 27.30 | 121.48 | 53.35 | Methionine aminopeptidase |
| *PSPTO_1534* | *rpsB* | + | 1693557 | 1694300 | 64.93 | 286.10 | 235.27 | 137.61 | 411.55 | 102.27 | 30S ribosomal protein S2 |
| *PSPTO_1535* | *tsf* | + | 1694491 | 1695354 | 26.84 | 122.63 | 156.92 | 88.30 | 577.70 | 74.89 | Translation elongation factor Ts |
| *PSPTO_1536* | *pyrH* | + | 1695560 | 1696303 | 37.55 | 60.69 | 61.22 | 31.28 | 180.64 | 47.49 | Uridylate kinase |
| *PSPTO_1537* | *frr* | + | 1696300 | 1696857 | 20.65 | 45.59 | 37.87 | 49.66 | 197.30 | 53.84 | Ribosome recycling factor |
| *PSPTO_1538* | *uppS* | + | 1696874 | 1697629 | 13.31 | 30.53 | 25.71 | 15.78 | 125.69 | 26.45 | Undecaprenyl diphosphate synthase |
| *PSPTO_1539* | *cdsA-1* | + | 1697629 | 1698444 | 39.03 | 24.08 | 21.37 | 6.10 | 59.73 | 18.67 | Phosphatidate cytidylyltransferase |
| *PSPTO_1540* | *dxr* | + | 1698441 | 1699631 | 28.84 | 33.33 | 28.75 | 14.44 | 78.39 | 19.15 | 1-deoxy-D-xylulose 5-phosphate reductoisomerase |
| *PSPTO_1541* | | + | 1699749 | 1701101 | 27.67 | 65.15 | 55.73 | 48.15 | 88.68 | 33.89 | Membrane-associated zinc metalloprotease |
| *PSPTO_1542* | | + | 1701189 | 1703576 | 10.74 | 38.67 | 41.70 | 26.00 | 147.79 | 27.12 | Outer membrane protein |
| *PSPTO_1543* | *ompH* | + | 1703622 | 1704125 | 9.05 | 37.01 | 21.43 | 14.38 | 153.72 | 13.86 | Outer membrane protein OmpH |
| *PSPTO_1544* | *lpxD* | + | 1704128 | 1705183 | 5.05 | 24.38 | 11.88 | 7.15 | 130.11 | 10.30 | UDP-3-O-[3-hydroxymyristoyl] glucosamine N-acyltransferase |
| *PSPTO_1545* | *fabZ* | + | 1705293 | 1705733 | 16.47 | 10.93 | 11.15 | 5.18 | 71.17 | 13.81 | (3R)-hydroxymyristoyl-ACP dehydratase |
| *PSPTO_1546* | *lpxA* | + | 1705730 | 1706506 | 7.70 | 13.10 | 14.72 | 3.27 | 86.32 | 9.78 | UDP-N-acetylglucosamine acyltransferase |
| *PSPTO_1547* | *lpxB* | + | 1706509 | 1707651 | 3.56 | 15.94 | 8.23 | 3.53 | 56.69 | 3.33 | Lipid A disaccharide synthase |
| *PSPTO_1548* | *rnhB* | + | 1707663 | 1708316 | 9.12 | 16.33 | 14.56 | 7.25 | 68.15 | 6.64 | Ribonuclease HII |
| *PSPTO_1549* | *dnaE* | + | 1708384 | 1711905 | 11.25 | 24.70 | 22.49 | 9.45 | 93.95 | 16.94 | DNA polymerase III subunit alpha |
| *PSPTO_1550* | *accA* | + | 1712034 | 1712981 | 42.69 | 29.18 | 28.51 | 13.53 | 153.21 | 29.76 | Acetyl-CoA carboxylase carboxyltransferase subunit alpha |
| *PSPTO_1551* | | + | 1713103 | 1714470 | 6.97 | 8.24 | 9.31 | 6.98 | 26.85 | 6.82 | Cell cycle protein mesJ |
| *PSPTO_1552* | *pyrG* | + | 1714734 | 1716365 | 16.30 | 54.18 | 49.17 | 30.45 | 151.91 | 58.97 | CTP synthase |
| *PSPTO_1553* | *kdsA* | + | 1716380 | 1717225 | 7.91 | 11.71 | 16.02 | 13.51 | 67.54 | 13.71 | 2-dehydro-3-deoxyphosphooctonate aldolase |
| *PSPTO_1554* | *eno-1* | + | 1717364 | 1718650 | 37.71 | 97.84 | 82.33 | 47.77 | 601.70 | 70.34 | Enolase |
| *PSPTO_1555* | | + | 1718840 | 1719118 | 2.18 | 9.62 | 8.90 | 0.66 | 23.25 | 4.10 | Hypothetical protein |
| *PSPTO_1556* | *ispD* | + | 1719132 | 1719842 | 4.02 | 5.76 | 7.09 | 2.28 | 26.16 | 5.53 | 4-diphosphocytidyl-2C-methyl-D-erythritol synthase |
| *PSPTO_1557* | | - | 1719849 | 1720745 | 4.99 | 13.55 | 6.80 | 7.52 | 26.05 | 6.26 | LysR family transcriptional regulator |
| *PSPTO_1558* | *adhC* | + | 1720850 | 1721962 | 24.05 | 17.04 | 17.22 | 7.72 | 125.90 | 28.62 | Alcohol dehydrogenase, class III |
| *PSPTO_1559* | | + | 1722062 | 1722907 | 10.42 | 22.03 | 19.79 | 13.24 | 132.11 | 38.16 | Esterase |
| *PSPTO_1560* | *ispF* | + | 1722961 | 1723434 | 100.24 | 105.29 | 114.31 | 61.11 | 130.97 | 54.60 | 2-C-methyl-D-erythritol 2,4-cyclodiphosphate synthase |
| *PSPTO_1561* | | + | 1723431 | 1724489 | 5.47 | 19.86 | 12.08 | 6.95 | 31.44 | 9.84 | Hypothetical protein |
| *PSPTO_1562* | *surE* | + | 1724477 | 1725226 | 10.01 | 24.01 | 9.73 | 11.52 | 60.65 | 14.70 | Stationary-phase survival protein SurE |
| *PSPTO_1563* | *pcm* | + | 1725268 | 1725903 | 8.80 | 19.01 | 15.31 | 8.28 | 54.77 | 20.13 | Protein-L-isoaspartate O-methyltransferase |
| *PSPTO_1564* | *nlpD* | + | 1726109 | 1726975 | 517.64 | 603.95 | 535.55 | 304.40 | 223.43 | 216.64 | Lipoprotein NlpD |
| *PSPTO_1565* | *rpoS* | + | 1727080 | 1728087 | 62.37 | 79.09 | 93.68 | 67.19 | 50.23 | 65.22 | RNA polymerase sigma-38 factor |
| *PSPTO_1566* | | - | 1728164 | 1728262 | 3.26 | 16.84 | 0.00 | 0.00 | 1.61 | 0.37 | Hypothetical protein |
| *PSPTO_1567* | | + | 1728650 | 1730110 | 16.28 | 21.67 | 21.01 | 12.52 | 34.54 | 23.14 | ISPsy6, transposase |
| *PSPTO_1568* | *hopAF1* | - | 1730311 | 1731165 | 0.59 | 18.93 | 6.14 | 3.53 | 1.30 | 0.78 | Type III effector HopAF1 |
| *PSPTO_1570* | | + | 1731720 | 1731875 | 3.29 | 10.39 | 2.35 | 14.32 | 22.95 | 6.22 | Hypothetical protein |
| *PSPTO_1571* | | + | 1731922 | 1733493 | 7.42 | 16.99 | 20.56 | 11.03 | 25.39 | 15.63 | Hypothetical protein |
| *PSPTO_1572* | | + | 1733797 | 1734111 | 3.08 | 7.25 | 4.69 | 0.00 | 6.28 | 3.90 | Hypothetical protein |
| *PSPTO_1573* | | - | 1734488 | 1734670 | 7.61 | 0.00 | 6.18 | 7.01 | 3.73 | 4.60 | Hypothetical protein |
| *PSPTO_1574* | | - | 1734733 | 1736043 | 3.73 | 7.56 | 4.68 | 2.52 | 11.65 | 6.20 | Mechanosensitive ion channel family protein |
| *PSPTO_1575* | | + | 1736307 | 1736618 | 2.57 | 0.11 | 11.89 | 0.41 | 3.57 | 3.40 | Hypothetical protein |
| *PSPTO_1576* | | + | 1736641 | 1736997 | 3.35 | 12.33 | 17.25 | 2.57 | 8.76 | 4.51 | Hypothetical protein |
| *PSPTO_1577* | | + | 1737114 | 1738226 | 13.72 | 15.64 | 15.34 | 10.20 | 34.14 | 15.04 | Hypothetical protein |
| *PSPTO_1578* | | - | 1738236 | 1739129 | 18.48 | 12.57 | 7.37 | 2.64 | 32.37 | 11.32 | LysR family transcriptional regulator |
| *PSPTO_1579* | | + | 1739236 | 1740444 | 3.68 | 6.18 | 3.11 | 0.00 | 6.25 | 5.86 | Major facilitator family transporter |
| *PSPTO_1580* | | + | 1740493 | 1741521 | 6.21 | 7.79 | 2.89 | 3.00 | 9.97 | 6.57 | Hypothetical protein |
| *PSPTO_1581* | | - | 1741579 | 1742217 | 12.81 | 7.24 | 5.94 | 1.43 | 12.91 | 9.37 | Hypothetical protein |
| *PSPTO_1582* | | + | 1742565 | 1743326 | 8.83 | 9.35 | 9.56 | 3.81 | 13.48 | 8.30 | Short-chain dehydrogenase/reductase family oxidoreductase |
| *PSPTO_1583* | | - | 1743431 | 1744135 | 11.10 | 15.34 | 15.66 | 9.08 | 24.35 | 14.92 | Hypothetical protein |
| *PSPTO_1584* | | - | 1744138 | 1745070 | 9.18 | 6.58 | 1.85 | 9.96 | 9.37 | 5.16 | Alcohol dehydrogenase |
| *PSPTO_1585* | | - | 1745195 | 1745941 | 24.73 | 10.19 | 9.30 | 10.36 | 13.09 | 16.70 | Hypothetical protein |
| *PSPTO_1586* | | - | 1745971 | 1746342 | 8.67 | 5.29 | 4.08 | 1.29 | 3.77 | 4.10 | Hypothetical protein |
| *PSPTO_1587* | *srmB* | + | 1746321 | 1747682 | 16.98 | 29.35 | 32.10 | 23.11 | 89.75 | 38.15 | ATP-dependent RNA helicase SrmB |
| *PSPTO_1588* | | + | 1747822 | 1748181 | 38.52 | 73.02 | 57.66 | 26.35 | 18.27 | 27.64 | Hypothetical protein |
| *PSPTO_1589* | | - | 1748298 | 1749161 | 13.34 | 12.64 | 10.87 | 2.37 | 14.74 | 14.51 | Lipoprotein |
| *PSPTO_1590* | | - | 1749231 | 1749959 | 6.03 | 18.10 | 17.74 | 8.52 | 17.73 | 20.57 | Hypothetical protein |
| *PSPTO_1591* | | + | 1749986 | 1750108 | 10.71 | 136.48 | 81.02 | 43.06 | 64.07 | 66.38 | Hypothetical protein |
| *PSPTO_1592* | | - | 1750102 | 1750671 | 46.26 | 51.38 | 57.41 | 13.43 | 95.98 | 37.02 | Hypothetical protein |
| *PSPTO_1593* | | + | 1751280 | 1752116 | 34.64 | 46.74 | 53.49 | 14.05 | 70.46 | 22.80 | Hypothetical protein |
| *PSPTO_1594* | | + | 1752749 | 1753153 | 12.16 | 34.17 | 9.25 | 10.74 | 23.36 | 15.16 | Hypothetical protein |
| *PSPTO_1595* | | + | 1753172 | 1753849 | 122.86 | 387.61 | 323.28 | 164.56 | 112.73 | 69.39 | Hypothetical protein |

| Locus name | Gene name | Strand | Gene start (bp) | Gene end (bp) | Average TPM Inoculum | Average TPM $t_6$ | Average TPM $t_{24}$ | Average TPM $t_{168}$ | Average TPM Logarithmic | Average TPM Stationary | Annotation description |
|---|---|---|---|---|---|---|---|---|---|---|---|
| *PSPTO_1596* | | + | 1753846 | 1754037 | 135.75 | 586.36 | 550.50 | 285.28 | 77.72 | 75.67 | Hypothetical protein |
| *PSPTO_1597* | | + | 1754264 | 1755235 | 30.94 | 41.61 | 35.12 | 15.16 | 44.84 | 44.89 | Ribonuclease BN |
| *PSPTO_1598* | *def-2* | + | 1755282 | 1755821 | 10.49 | 29.34 | 26.16 | 18.68 | 42.12 | 20.93 | Polypeptide deformylase |
| *PSPTO_1599* | | - | 1755846 | 1756310 | 8.68 | 10.99 | 13.59 | 12.78 | 15.12 | 13.53 | GNAT family acetyltransferase |
| *PSPTO_1600* | | + | 1756386 | 1757399 | 18.35 | 50.71 | 35.63 | 12.03 | 23.04 | 27.69 | ABC transporter periplasmic substrate-binding protein |
| *PSPTO_1601* | | + | 1757534 | 1758034 | 67.60 | 88.02 | 72.52 | 90.50 | 106.43 | 52.91 | Hypothetical protein |
| *PSPTO_1602* | *cdd* | + | 1758068 | 1758505 | 14.23 | 39.27 | 32.63 | 7.02 | 26.19 | 19.39 | Cytosine deaminase |
| *PSPTO_1603* | | + | 1758542 | 1758661 | 13.68 | 7.74 | 13.03 | 18.59 | 30.77 | 19.02 | Hypothetical protein |
| *PSPTO_1604* | | + | 1758790 | 1759611 | 22.68 | 22.55 | 17.34 | 10.77 | 38.94 | 16.03 | Hydrolase |
| *PSPTO_1605* | | + | 1759601 | 1760776 | 12.45 | 19.25 | 10.39 | 8.98 | 25.89 | 12.61 | Sensory box histidine kinase |
| *PSPTO_1606* | | + | 1760910 | 1762121 | 14.61 | 21.43 | 15.57 | 12.48 | 25.95 | 13.60 | Sensor histidine kinase |
| *PSPTO_1607* | | + | 1762336 | 1762656 | 250.71 | 332.65 | 303.76 | 74.09 | 232.14 | 321.97 | Hypothetical protein |
| *PSPTO_1608* | | + | 1762728 | 1763891 | 39.09 | 94.36 | 81.46 | 33.52 | 104.69 | 83.73 | Hypothetical protein |
| *PSPTO_1609* | | - | 1763976 | 1764284 | 41.82 | 754.51 | 876.28 | 264.86 | 103.67 | 46.71 | Hypothetical protein |
| *PSPTO_1610* | | + | 1764582 | 1766756 | 3.45 | 3.50 | 5.14 | 2.81 | 9.76 | 5.36 | TonB-dependent siderophore receptor |
| *PSPTO_1611* | | - | 1766753 | 1766866 | 1.71 | 0.00 | 2.76 | 0.00 | 6.30 | 3.33 | Hypothetical protein |
| *PSPTO_1612* | | + | 1766931 | 1768247 | 12.49 | 7.29 | 4.86 | 4.46 | 6.50 | 5.92 | Major facilitator family transporter |
| *PSPTO_1613* | | + | 1768630 | 1769010 | 4.02 | 5.70 | 0.00 | 5.29 | 4.94 | 2.23 | LuxR family transcriptional regulator |
| *PSPTO_1614* | | + | 1769379 | 1769918 | 0.34 | 2.72 | 0.61 | 0.00 | 5.47 | 0.68 | Hypothetical protein |
| *PSPTO_1615* | | + | 1770750 | 1771622 | 3.75 | 1.09 | 1.09 | 0.00 | 8.66 | 5.20 | Hypothetical protein |
| *PSPTO_1616* | | - | 1771627 | 1772703 | 6.10 | 6.40 | 7.36 | 13.02 | 11.82 | 8.00 | Diguanylate cyclase |
| *PSPTO_1617* | | - | 1773000 | 1774217 | 3.09 | 6.40 | 6.69 | 4.73 | 5.21 | 2.93 | Major facilitator family transporter |
| *PSPTO_1618* | | + | 1774328 | 1775236 | 32.89 | 23.94 | 21.61 | 4.36 | 41.46 | 10.41 | LysR family transcriptional regulator |
| *PSPTO_1619* | | - | 1775263 | 1775943 | 25.20 | 20.00 | 22.16 | 6.32 | 25.78 | 14.63 | Hypothetical protein |
| *PSPTO_1620* | | - | 1775996 | 1776808 | 5.00 | 10.54 | 10.40 | 2.42 | 22.62 | 8.14 | Hypothetical protein |
| *PSPTO_1621* | | - | 1776801 | 1778099 | 31.09 | 32.59 | 19.99 | 13.89 | 64.52 | 29.97 | Hypothetical protein |
| *PSPTO_1622* | | + | 1779033 | 1780019 | 6.25 | 7.85 | 14.39 | 9.43 | 6.61 | 5.23 | Glycine betaine-binding protein |
| *PSPTO_1623* | | + | 1780267 | 1780578 | 162.45 | 239.88 | 204.86 | 94.62 | 111.46 | 33.63 | Hypothetical protein |
| *PSPTO_1624* | | + | 1780575 | 1782233 | 20.73 | 82.52 | 84.81 | 24.59 | 55.53 | 6.91 | Sodium:solute symporter family protein |
| *PSPTO_1625* | | - | 1782364 | 1782522 | 5.12 | 16.00 | 0.00 | 0.00 | 1.64 | 0.47 | Hypothetical protein |
| *PSPTO_1626* | | + | 1782846 | 1783223 | 3.31 | 18.22 | 4.20 | 11.70 | 4.03 | 2.61 | Hypothetical protein |
| *PSPTO_1627* | | + | 1783328 | 1784143 | 11.35 | 10.53 | 12.87 | 3.63 | 6.98 | 4.04 | Short chain dehydrogenase |
| *PSPTO_1628* | | + | 1784377 | 1784709 | 10.77 | 12.37 | 21.54 | 3.13 | 7.27 | 7.08 | Hypothetical protein |
| *PSPTO_1629* | *csrA-1* | + | 1784879 | 1785073 | 6.77 | 7.00 | 9.56 | 0.00 | 15.10 | 7.36 | Carbon storage regulator |
| *PSPTO_1630* | | - | 1785076 | 1785303 | 35.99 | 42.59 | 36.43 | 5.87 | 68.07 | 26.71 | Hypothetical protein |
| *PSPTO_1631* | *ggnC* | - | 1785376 | 1786572 | 1.91 | 35.20 | 39.35 | 23.08 | 7.89 | 13.28 | Hypothetical protein |
| *PSPTO_1632* | *ggnB* | - | 1786655 | 1788403 | 4.44 | 77.69 | 53.62 | 22.12 | 16.11 | 23.99 | GNAT family acetyltransferase |
| *PSPTO_1633* | *asnB* | - | 1788407 | 1790179 | 7.97 | 85.01 | 73.96 | 39.10 | 21.31 | 24.35 | Asparagine synthetase |
| *PSPTO_1634* | | + | 1790464 | 1790646 | 108.33 | 179.85 | 150.39 | 24.92 | 260.95 | 222.76 | Hypothetical protein |
| *PSPTO_1635* | | + | 1790682 | 1791086 | 26.54 | 29.50 | 42.28 | 21.30 | 47.56 | 44.63 | Hypothetical protein |
| *PSPTO_1636* | | - | 1791102 | 1792163 | 8.57 | 12.73 | 15.52 | 2.98 | 12.59 | 16.46 | Hypothetical protein |
| *PSPTO_1637* | | - | 1792171 | 1794648 | 3.35 | 7.38 | 6.19 | 3.67 | 10.41 | 9.29 | ABC transporter permease |
| *PSPTO_1638* | | - | 1794648 | 1795316 | 23.97 | 28.42 | 10.14 | 7.88 | 25.86 | 29.12 | ABC transporter ATP-binding protein |
| *PSPTO_1639* | | - | 1795541 | 1797523 | 7.05 | 9.02 | 6.54 | 10.55 | 19.11 | 12.35 | Bifunctional tRNA (mnm(5)s(2)U34)-methyltransferase/FAD-dependent cmnm(5)s(2)U34 oxidoreductase |
| *PSPTO_1640* | | + | 1797671 | 1799167 | 8.28 | 19.18 | 8.92 | 6.90 | 12.08 | 13.28 | Hypothetical protein |
| *PSPTO_1641* | | - | 1799219 | 1799500 | 7.83 | 4.43 | 8.47 | 0.00 | 7.29 | 6.82 | Hypothetical protein |
| *PSPTO_1642* | | - | 1799627 | 1800028 | 3.50 | 6.48 | 7.01 | 0.36 | 10.98 | 4.70 | Hypothetical protein |
| *PSPTO_1643* | | - | 1800039 | 1800626 | 88.77 | 136.98 | 187.55 | 128.20 | 286.35 | 112.19 | Hypothetical protein |
| *PSPTO_1644* | *recQ* | + | 1800803 | 1802932 | 3.77 | 41.07 | 24.06 | 11.96 | 37.91 | 19.48 | ATP-dependent DNA helicase RecQ |
| *PSPTO_1645* | | + | 1803026 | 1803460 | 15.20 | 53.49 | 39.90 | 6.06 | 48.03 | 24.06 | MarR family transcriptional regulator |
| *PSPTO_1646* | | + | 1803599 | 1805752 | 14.04 | 27.34 | 22.73 | 9.49 | 52.31 | 25.32 | LysM domain-containing protein |
| *PSPTO_1647* | | + | 1805874 | 1806056 | 4.43 | 1.97 | 6.26 | 7.31 | 19.09 | 13.72 | Lipoprotein |
| *PSPTO_1648* | *aer-1* | + | 1806405 | 1807970 | 14.58 | 11.32 | 4.98 | 8.20 | 10.62 | 12.82 | Aerotaxis receptor |
| *PSPTO_1649* | | + | 1808975 | 1811980 | 1.52 | 0.72 | 1.37 | 0.00 | 2.07 | 1.27 | Autotransporter |
| *PSPTO_1650* | | + | 1812034 | 1815201 | 1.36 | 2.15 | 0.39 | 0.00 | 1.70 | 1.74 | Autotransporter |
| *PSPTO_1651* | | + | 1815561 | 1815851 | 5.44 | 12.21 | 9.87 | 2.25 | 17.92 | 3.43 | Hypothetical protein |
| *PSPTO_1652* | | - | 1815904 | 1816479 | 2.08 | 0.00 | 6.60 | 7.11 | 3.68 | 2.29 | Hypothetical protein |
| *PSPTO_1653* | | + | 1816911 | 1818464 | 3.31 | 8.59 | 7.42 | 8.65 | 45.68 | 7.70 | Hypothetical protein |
| *PSPTO_1654* | | + | 1818461 | 1819735 | 2.92 | 5.78 | 4.42 | 3.38 | 26.19 | 3.91 | Hypothetical protein |
| *PSPTO_1655* | | - | 1820002 | 1820478 | 1.73 | 10.53 | 3.84 | 10.57 | 13.40 | 7.89 | Hypothetical protein |
| *PSPTO_1656* | | - | 1820528 | 1822417 | 10.15 | 9.88 | 9.93 | 9.46 | 35.79 | 16.02 | Hypothetical protein |
| *PSPTO_1657* | | + | 1822740 | 1824410 | 15.56 | 14.72 | 14.64 | 9.01 | 22.05 | 16.12 | Hypothetical protein |
| *PSPTO_1658* | | - | 1824436 | 1825245 | 0.00 | 0.00 | 0.00 | 0.00 | 0.00 | 0.00 | ISPsy4, transposition helper protein |
| *PSPTO_1659* | | - | 1825242 | 1826264 | 0.00 | 0.00 | 0.00 | 0.00 | 0.00 | 0.00 | ISPsy4, transposase |
| *PSPTO_1660* | | + | 1827359 | 1829776 | 16.80 | 19.60 | 12.43 | 9.38 | 37.38 | 30.37 | Helicase/SNF2 family domain protein |
| *PSPTO_1661* | *nrdB* | - | 1830031 | 1831278 | 16.17 | 51.19 | 57.68 | 27.16 | 146.55 | 78.47 | Ribonucleotide-diphosphate reductase subunit beta |
| *PSPTO_1662* | | + | 1831336 | 1831533 | 277.19 | 240.28 | 220.09 | 217.34 | 224.85 | 218.26 | Hypothetical protein |
| *PSPTO_1663* | | - | 1831947 | 1832516 | 10.48 | 8.35 | 14.96 | 5.77 | 13.67 | 11.30 | GNAT family acetyltransferase |
| *PSPTO_1664* | | - | 1832622 | 1833269 | 3.72 | 1.26 | 4.11 | 0.00 | 2.98 | 6.84 | Hypothetical protein |
| *PSPTO_1665* | | - | 1833391 | 1833840 | 2.48 | 1.45 | 0.00 | 0.00 | 1.56 | 1.43 | Hypothetical protein |
| *PSPTO_1666* | | - | 1833875 | 1834627 | 8.70 | 6.49 | 6.16 | 5.58 | 4.27 | 5.60 | Crp/Fnr family transcriptional regulator |
| *PSPTO_1667* | | + | 1834878 | 1835768 | 5.77 | 6.42 | 2.15 | 1.61 | 9.07 | 42.47 | Universal stress protein family |
| *PSPTO_1668* | | - | 1835852 | 1836100 | 2.86 | 3.50 | 0.00 | 5.41 | 13.99 | 72.24 | Hypothetical protein |
| *PSPTO_1669* | *tdcD* | + | 1836085 | 1837308 | 2.07 | 0.27 | 5.46 | 0.40 | 5.41 | 29.35 | Propionate kinase |
| *PSPTO_1670* | *xfp* | + | 1837381 | 1839888 | 2.63 | 4.17 | 2.04 | 2.10 | 3.82 | 4.81 | Xylulose-5-phosphate/fructose-6-phosphate phosphoketolase |
| *PSPTO_1671* | *nrdA* | - | 1840004 | 1842916 | 31.61 | 63.03 | 46.63 | 26.36 | 281.60 | 155.68 | Ribonucleotide-diphosphate reductase subunit alpha |
| *PSPTO_1672* | | + | 1843614 | 1844357 | 8.47 | 4.18 | 5.44 | 5.16 | 6.20 | 5.98 | DNA-binding response regulator |
| *PSPTO_1673* | | + | 1844578 | 1845972 | 3.01 | 4.23 | 3.07 | 1.21 | 4.21 | 4.07 | Sensor histidine kinase |
| *PSPTO_1674* | *fhuF* | + | 1845930 | 1846676 | 2.92 | 2.16 | 3.02 | 2.68 | 7.32 | 4.56 | Ferric iron reductase protein FhuF |

| Locus name | Gene name | Strand | Gene start (bp) | Gene end (bp) | Average TPM Inoculum | Average TPM $t_6$ | Average TPM $t_{24}$ | Average TPM $t_{168}$ | Average TPM Logarithmic | Average TPM Stationary | Annotation description |
|---|---|---|---|---|---|---|---|---|---|---|---|
| *PSPTO_1675* | | - | 1846677 | 1847321 | 19.34 | 31.15 | 24.03 | 16.40 | 55.84 | 36.10 | Siderophore biosynthesis protein |
| *PSPTO_1676* | | + | 1847662 | 1848135 | 7.24 | 12.49 | 7.99 | 5.00 | 16.44 | 7.37 | Hypothetical protein |
| *PSPTO_1677* | | - | 1848378 | 1849166 | 45.25 | 32.38 | 41.13 | 12.78 | 44.70 | 41.77 | Dienelactone hydrolase family protein |
| *PSPTO_1678* | | + | 1849346 | 1849696 | 80.72 | 50.80 | 24.04 | 15.73 | 178.36 | 50.49 | Hypothetical protein |
| *PSPTO_1679* | *phoP* | + | 1849724 | 1850401 | 43.02 | 29.41 | 19.99 | 19.06 | 125.84 | 39.29 | Transcriptional regulatory protein PhoP |
| *PSPTO_1680* | *phoQ* | + | 1850398 | 1851744 | 12.72 | 7.40 | 11.04 | 7.86 | 58.66 | 16.28 | Sensor protein PhoQ |
| *PSPTO_1681* | | - | 1851726 | 1852010 | 27.49 | 8.83 | 6.39 | 3.51 | 18.51 | 8.90 | Hypothetical protein |
| *PSPTO_1682* | *dctA-1* | + | 1852071 | 1853453 | 61.17 | 47.72 | 24.17 | 11.55 | 22.82 | 10.59 | C4-dicarboxylate transport protein |
| *PSPTO_1683* | | - | 1853519 | 1854013 | 11.72 | 33.71 | 13.65 | 11.05 | 160.76 | 24.89 | Peptidase, SprT family |
| *PSPTO_1684* | | + | 1854218 | 1855042 | 32.82 | 32.16 | 39.14 | 31.18 | 66.88 | 45.79 | Hypothetical protein |
| *PSPTO_1685* | | - | 1855049 | 1855720 | 18.49 | 17.78 | 19.77 | 8.79 | 33.65 | 18.32 | Hypothetical protein |
| *PSPTO_1686* | *wrbA* | - | 1855816 | 1856421 | 7.64 | 23.06 | 20.21 | 5.79 | 102.08 | 24.89 | Trp repressor binding protein |
| *PSPTO_1687* | *arsC* | - | 1856418 | 1856771 | 23.08 | 90.91 | 35.86 | 10.63 | 232.05 | 49.90 | Arsenate reductase |
| *PSPTO_1688* | | + | 1856901 | 1857368 | 74.71 | 68.35 | 71.52 | 54.47 | 80.00 | 61.17 | Thioredoxin |
| *PSPTO_1689* | | - | 1857514 | 1857921 | 17.21 | 29.44 | 19.90 | 26.17 | 23.56 | 13.87 | Lipoprotein |
| *PSPTO_1690* | | - | 1857918 | 1858355 | 15.16 | 63.18 | 24.50 | 6.18 | 49.11 | 13.41 | Hypothetical protein |
| *PSPTO_1691* | *gacS* | - | 1858439 | 1861192 | 3.78 | 7.38 | 8.37 | 8.65 | 16.20 | 8.70 | Sensor histidine kinase/response regulator GacS |
| *PSPTO_1692* | *cysM* | + | 1861465 | 1862367 | 22.71 | 53.62 | 50.56 | 27.87 | 64.98 | 19.44 | Cysteine synthase B |
| *PSPTO_1693* | | + | 1862367 | 1863722 | 50.12 | 64.56 | 45.10 | 36.26 | 62.60 | 18.34 | 23S rRNA m(5)U1939 methyltransferase |
| *PSPTO_1694* | *relA* | + | 1863888 | 1866122 | 29.72 | 59.72 | 60.53 | 25.05 | 116.85 | 26.19 | GTP pyrophosphokinase |
| *PSPTO_1695* | | + | 1866236 | 1867069 | 32.11 | 24.57 | 27.60 | 10.20 | 78.89 | 18.88 | Nucleoside triphosphate pyrophosphohydrolase |
| *PSPTO_1696* | | + | 1867100 | 1867639 | 51.72 | 37.32 | 33.16 | 32.60 | 98.12 | 26.34 | Hypothetical protein |
| *PSPTO_1697* | | - | 1867739 | 1867918 | 36.22 | 2495.42 | 932.36 | 140.71 | 139.36 | 11.92 | Hypothetical protein |
| *PSPTO_1698* | | - | 1868418 | 1869131 | 137.81 | 123.57 | 133.54 | 91.69 | 479.80 | 243.60 | Hypothetical protein |
| *PSPTO_1699* | *purN* | - | 1869140 | 1869790 | 103.59 | 38.79 | 31.69 | 16.18 | 129.19 | 66.85 | Phosphoribosylglycinamide formyltransferase |
| *PSPTO_1700* | *purM* | - | 1869790 | 1870848 | 14.69 | 43.11 | 40.27 | 15.67 | 97.99 | 39.59 | Phosphoribosylformylglycinamidine cyclo-ligase |
| *PSPTO_1701* | | + | 1871179 | 1872279 | 23.07 | 15.13 | 13.77 | 8.51 | 31.04 | 15.79 | Hypothetical protein |
| *PSPTO_1702* | | + | 1872319 | 1873392 | 20.92 | 11.81 | 17.64 | 9.64 | 26.71 | 12.60 | Permease |
| *PSPTO_1703* | | + | 1873622 | 1874326 | 10.15 | 28.25 | 6.96 | 23.34 | 39.37 | 17.54 | DNA replication initiation factor |
| *PSPTO_1704* | | + | 1874338 | 1875084 | 11.27 | 20.65 | 18.07 | 13.99 | 47.13 | 23.53 | NAD-dependent deacetylase |
| *PSPTO_1705* | | + | 1875154 | 1875882 | 8.67 | 268.34 | 221.14 | 106.75 | 197.58 | 86.03 | NLP/P60 family protein |
| *PSPTO_1706* | | + | 1875977 | 1876522 | 23.12 | 24.03 | 29.20 | 9.27 | 32.99 | 33.27 | Lipoprotein |
| *PSPTO_1707* | | - | 1876688 | 1878016 | 18.48 | 50.17 | 27.41 | 11.99 | 18.52 | 22.77 | L-sorbosone dehydrogenase |
| *PSPTO_1708* | *cobO* | + | 1878441 | 1879052 | 3.94 | 7.03 | 7.09 | 7.52 | 30.08 | 10.59 | Cob(I)alamin adenosyltransferase |
| *PSPTO_1709* | | + | 1879092 | 1880387 | 2.02 | 7.53 | 9.87 | 0.99 | 35.37 | 7.87 | Cobyrinic acid a,c-diamide synthase |
| *PSPTO_1710* | | + | 1880384 | 1881034 | 2.87 | 10.16 | 2.18 | 4.06 | 48.13 | 11.14 | Nitroreductase family protein |
| *PSPTO_1711* | *cobD* | + | 1881031 | 1881939 | 4.92 | 13.55 | 9.74 | 2.48 | 54.47 | 10.22 | Cobalamin biosynthesis protein CobD |
| *PSPTO_1712* | *cobC* | + | 1881932 | 1882936 | 2.92 | 5.04 | 3.00 | 2.45 | 42.82 | 6.96 | Cobalamin biosynthesis protein CobC |
| *PSPTO_1713* | *cobQ* | + | 1882933 | 1884393 | 8.76 | 11.27 | 8.56 | 3.08 | 88.20 | 12.81 | Cobyric acid synthase |
| *PSPTO_1714* | *cobP* | + | 1884394 | 1884915 | 2.61 | 6.35 | 3.05 | 0.00 | 52.98 | 5.36 | Cobinamide kinase/cobinamide phosphate guanylyltransferase |
| *PSPTO_1715* | *cobT* | + | 1884912 | 1885964 | 2.73 | 3.94 | 8.22 | 2.43 | 43.69 | 6.17 | Nicotinate-nucleotide--dimethylbenzimidazole phosphoribosyltransferase |
| *PSPTO_1716* | | + | 1885961 | 1886533 | 5.53 | 3.45 | 6.30 | 3.12 | 24.08 | 5.84 | Alpha-ribazole-5'-phosphate phosphatase |
| *PSPTO_1717* | *cobS* | + | 1886539 | 1887270 | 3.65 | 15.39 | 1.57 | 3.78 | 36.86 | 8.89 | Cobalamin (5'-phosphate) synthase |
| *PSPTO_1718* | | + | 1887368 | 1888576 | 11.64 | 31.01 | 15.74 | 6.59 | 24.64 | 18.73 | Transporter |
| *PSPTO_1719* | | + | 1888757 | 1889308 | 21.95 | 39.92 | 47.71 | 7.56 | 36.45 | 15.68 | Glutathione peroxidase |
| *PSPTO_1720* | | - | 1889959 | 1891233 | 25.96 | 76.84 | 70.38 | 51.46 | 231.14 | 24.89 | Outer membrane protein |
| *PSPTO_1721* | | - | 1891475 | 1891954 | 53.98 | 359.56 | 255.27 | 107.09 | 181.41 | 132.97 | Hypothetical protein |
| *PSPTO_1722* | | - | 1892174 | 1892479 | 36.75 | 23.88 | 36.76 | 30.22 | 31.55 | 46.29 | Hypothetical protein |
| *PSPTO_1723* | | + | 1892463 | 1892606 | 68.58 | 30.47 | 44.69 | 12.15 | 39.08 | 47.48 | Hypothetical protein |
| *PSPTO_1724* | | + | 1893124 | 1894602 | 4.27 | 8.79 | 10.43 | 9.96 | 10.13 | 6.37 | Hypothetical protein |
| *PSPTO_1725* | | + | 1894737 | 1894919 | 617.07 | 741.94 | 536.63 | 179.97 | 209.87 | 274.95 | Hypothetical protein |
| *PSPTO_1726* | | - | 1895010 | 1895324 | 30.48 | 38.62 | 74.99 | 16.66 | 29.79 | 35.46 | Hypothetical protein |
| *PSPTO_1727* | | + | 1895371 | 1895523 | 25.75 | 35.72 | 37.14 | 14.37 | 31.60 | 26.80 | Hypothetical protein |
| *PSPTO_1728* | | + | 1895553 | 1896152 | 2.97 | 7.84 | 3.52 | 10.67 | 8.10 | 6.40 | Hypothetical protein |
| *PSPTO_1729* | | + | 1896253 | 1897215 | 4.12 | 4.40 | 7.32 | 6.73 | 15.65 | 9.97 | Membrane protein |
| *PSPTO_1730* | | - | 1897240 | 1897854 | 23.72 | 52.71 | 57.89 | 17.87 | 45.84 | 51.40 | NAD(P)H dehydrogenase, quinone family |
| *PSPTO_1731* | *fumC-1* | - | 1897847 | 1899241 | 32.31 | 36.96 | 34.00 | 14.06 | 25.18 | 30.33 | Class II fumarate hydratase |
| *PSPTO_1732* | | + | 1899411 | 1899938 | 73.96 | 426.61 | 436.32 | 276.15 | 1011.16 | 503.16 | Hypothetical protein |
| *PSPTO_1733* | *bolA* | + | 1899960 | 1900247 | 3.87 | 35.55 | 25.25 | 0.92 | 73.97 | 41.46 | BolA protein |
| *PSPTO_1734* | | + | 1900419 | 1901372 | 22.82 | 35.72 | 22.97 | 26.22 | 37.01 | 33.11 | Putative rhodanese-related sulfurtransferase |
| *PSPTO_1735* | | + | 1901376 | 1902008 | 4.53 | 9.31 | 9.70 | 5.63 | 20.67 | 16.32 | Hypothetical protein |
| *PSPTO_1736* | | + | 1902001 | 1903827 | 5.14 | 9.26 | 7.35 | 6.00 | 33.58 | 19.42 | ABC transporter ATP-binding protein/permease |
| *PSPTO_1737* | | + | 1903951 | 1906407 | 45.15 | 31.76 | 26.95 | 21.85 | 19.17 | 20.72 | Sensory box/GGDEF domain/EAL domain-containing protein |
| *PSPTO_1738* | | + | 1906438 | 1907223 | 87.31 | 92.57 | 87.89 | 46.78 | 46.37 | 46.81 | Hypothetical protein |
| *PSPTO_1739* | | - | 1907254 | 1908180 | 57.59 | 75.71 | 61.06 | 24.33 | 38.41 | 67.52 | Diguanylate cyclase |
| *PSPTO_1740* | | - | 1908427 | 1909167 | 8.21 | 21.72 | 17.79 | 12.95 | 67.07 | 30.61 | Putative oxoacyl-(acyl carrier protein) reductase |
| *PSPTO_1741* | *gph-2* | - | 1909195 | 1909866 | 25.14 | 55.67 | 54.08 | 42.68 | 194.39 | 96.54 | Phosphoglycolate phosphatase |
| *PSPTO_1742* | *ubiG* | - | 1909870 | 1910589 | 48.56 | 61.52 | 55.08 | 31.12 | 120.17 | 57.62 | 3-demethylubiquinone-9 3-methyltransferase |
| *PSPTO_1743* | | - | 1910654 | 1911985 | 20.05 | 39.17 | 27.41 | 28.88 | 66.17 | 33.91 | N-ethylammeline chlorohydrolase |
| *PSPTO_1744* | | + | 1912105 | 1913181 | 7.24 | 16.27 | 16.22 | 5.24 | 27.43 | 18.89 | Initiation factor 2 subunit family |
| *PSPTO_1745* | *gyrA* | + | 1913484 | 1916270 | 9.48 | 30.62 | 29.96 | 18.44 | 201.78 | 80.02 | DNA gyrase subunit A |
| *PSPTO_1746* | *serC* | + | 1916456 | 1917541 | 5.37 | 21.34 | 8.16 | 16.98 | 91.64 | 38.03 | Phosphoserine aminotransferase |
| *PSPTO_1747* | *pheA* | + | 1917541 | 1918635 | 7.00 | 5.86 | 10.79 | 5.48 | 79.44 | 30.16 | Chorismate mutase/prephenate dehydratase |
| *PSPTO_1748* | | + | 1918702 | 1920309 | 7.74 | 13.34 | 10.80 | 3.74 | 65.02 | 19.18 | Prephenate dehydrogenase/3-phosphoshikimate 1-carboxyvinyltransferase family protein |
| *PSPTO_1749* | *cmk* | + | 1920306 | 1920995 | 306.11 | 1036.33 | 816.45 | 598.08 | 2068.15 | 751.92 | Cytidylate kinase |
| *PSPTO_1750* | *rpsA* | + | 1921117 | 1922808 | 53.09 | 397.09 | 387.78 | 306.36 | 940.70 | 234.74 | 30S ribosomal protein S1 |
| *PSPTO_1751* | *ihfB* | + | 1923124 | 1923420 | 171.14 | 474.37 | 351.57 | 140.93 | 425.75 | 99.45 | Integration host factor subunit beta |
| *PSPTO_1752* | | + | 1923445 | 1923687 | 14.32 | 19.76 | 27.95 | 27.93 | 40.76 | 11.90 | Hypothetical protein |
| *PSPTO_1753* | | + | 1924100 | 1925542 | 4.30 | 4.85 | 9.28 | 3.71 | 16.16 | 6.36 | Hypothetical protein |

| Locus name | Gene name | Strand | Gene start (bp) | Gene end (bp) | Average TPM Inoculum | Average TPM $t_6$ | Average TPM $t_{24}$ | Average TPM $t_{168}$ | Average TPM Logarithmic | Average TPM Stationary | Annotation description |
|---|---|---|---|---|---|---|---|---|---|---|---|
| *PSPTO_1754* | | + | 1925715 | 1926695 | 25.88 | 19.22 | 14.62 | 12.90 | 39.86 | 27.39 | UDP-glucose 4-epimerase |
| *PSPTO_1755* | | + | 1926688 | 1927704 | 37.27 | 35.04 | 15.75 | 17.85 | 52.69 | 25.94 | Glycoside hydrolase family protein |
| *PSPTO_1756* | *wbpM* | + | 1927805 | 1929799 | 20.55 | 27.49 | 21.74 | 16.73 | 33.99 | 24.25 | Nucleotide sugar epimerase/dehydratase WbpM |
| *PSPTO_1757* | | + | 1929959 | 1930288 | 5.46 | 11.70 | 7.80 | 10.21 | 20.93 | 9.32 | Competence protein |
| *PSPTO_1758* | | - | 1930396 | 1930965 | 85.50 | 256.91 | 198.73 | 73.45 | 137.68 | 79.66 | TetR family transcriptional regulator |
| *PSPTO_1759* | | - | 1931039 | 1931878 | 1.85 | 2.82 | 10.79 | 2.30 | 5.66 | 2.66 | Hypothetical protein |
| *PSPTO_1760* | | + | 1932296 | 1933573 | 11.60 | 30.90 | 22.41 | 11.29 | 57.63 | 17.98 | Threonine/serine transporter |
| *PSPTO_1761* | | - | 1933649 | 1933855 | 128.40 | 173.43 | 160.31 | 50.26 | 139.12 | 89.68 | Hypothetical protein |
| *PSPTO_1762* | | - | 1933979 | 1934461 | 19.52 | 25.16 | 9.78 | 7.12 | 50.49 | 29.56 | Hypothetical protein |
| *PSPTO_1763* | | - | 1934465 | 1934869 | 110.28 | 88.53 | 72.82 | 29.14 | 231.69 | 154.16 | Hypothetical protein |
| *PSPTO_1764* | | + | 1935065 | 1936201 | 9.50 | 10.11 | 5.34 | 14.76 | 19.94 | 11.75 | Hypothetical protein |
| *PSPTO_1765* | *efp* | + | 1936301 | 1936870 | 52.45 | 126.73 | 102.38 | 58.78 | 330.29 | 42.48 | Translation elongation factor P |
| *PSPTO_1766* | | - | 1936962 | 1937981 | 2.31 | 7.55 | 2.20 | 1.26 | 8.51 | 2.44 | Lipase |
| *PSPTO_1767* | *ohr* | - | 1938022 | 1938441 | 1.85 | 7.75 | 3.33 | 0.00 | 16.24 | 1.64 | Organic hydroperoxide resistance protein |
| *PSPTO_1768* | | - | 1938658 | 1939131 | 17.89 | 16.10 | 26.21 | 9.00 | 39.40 | 12.88 | MarR family transcriptional regulator |
| *PSPTO_1769* | | + | 1939317 | 1940063 | 4.37 | 10.05 | 9.53 | 5.92 | 17.56 | 7.88 | Membrane protein |
| *PSPTO_1770* | | - | 1940053 | 1940907 | 4.15 | 5.28 | 5.56 | 7.00 | 16.44 | 4.56 | LysR family transcriptional regulator |
| *PSPTO_1771* | | + | 1940924 | 1941235 | 1.75 | 2.30 | 3.51 | 4.70 | 2.61 | 1.34 | Hypothetical protein |
| *PSPTO_1772* | | - | 1941263 | 1942609 | 5.89 | 12.29 | 11.09 | 10.88 | 30.67 | 10.41 | Phospho-2-dehydro-3-deoxyheptonate aldolase |
| *PSPTO_1773* | | + | 1942781 | 1943542 | 5.71 | 17.63 | 9.60 | 6.66 | 17.54 | 15.90 | Hypothetical protein |
| *PSPTO_1774* | | + | 1943679 | 1943858 | 106.69 | 307.91 | 159.58 | 199.54 | 338.76 | 332.92 | Hypothetical protein |
| *PSPTO_1775* | | + | 1943997 | 1945685 | 8.14 | 63.11 | 41.39 | 32.68 | 109.08 | 58.80 | ATP-dependent RNA helicase, DEAD box family |
| *PSPTO_1776* | | - | 1945772 | 1946539 | 10.38 | 14.37 | 17.93 | 2.27 | 16.96 | 8.83 | Hypothetical protein |
| *PSPTO_1777* | *tpm* | + | 1946642 | 1947304 | 10.75 | 13.27 | 5.37 | 2.92 | 12.37 | 7.69 | Thiopurine s-methyltransferase |
| *PSPTO_1778* | *htpX* | - | 1947589 | 1948476 | 94.04 | 350.49 | 250.14 | 65.38 | 192.68 | 81.75 | Heat shock protein HtpX |
| *PSPTO_1779* | | - | 1948745 | 1949956 | 33.19 | 31.12 | 30.71 | 15.83 | 170.35 | 60.41 | Class I and II aminotransferase |
| *PSPTO_1780* | | + | 1950147 | 1950542 | 31.39 | 63.51 | 70.70 | 25.24 | 69.94 | 55.21 | Methionine sulfoxide reductase B |
| *PSPTO_1781* | | + | 1950687 | 1951172 | 19.93 | 52.03 | 32.46 | 5.60 | 33.11 | 32.56 | Glutathione peroxidase |
| *PSPTO_1782* | | - | 1951178 | 1953529 | 19.60 | 16.17 | 12.88 | 5.80 | 34.83 | 21.59 | Sensor histidine kinase/response regulator |
| *PSPTO_1783* | | + | 1953610 | 1954500 | 9.49 | 17.73 | 17.77 | 7.67 | 50.18 | 13.72 | Hypothetical protein |
| *PSPTO_1784* | | + | 1954497 | 1954979 | 11.42 | 10.75 | 6.83 | 8.41 | 66.05 | 24.57 | GAF domain-containing protein |
| *PSPTO_1785* | | - | 1955032 | 1955802 | 241.83 | 146.99 | 159.27 | 81.29 | 124.47 | 81.59 | Hypothetical protein |
| *PSPTO_1786* | | - | 1956613 | 1957335 | 3.55 | 11.95 | 8.68 | 1.73 | 4.73 | 4.62 | Hypothetical protein |
| *PSPTO_1787* | | + | 1957524 | 1958429 | 8.29 | 18.08 | 8.68 | 3.99 | 14.58 | 5.81 | LysR family transcriptional regulator |
| *PSPTO_1788* | | + | 1958574 | 1959545 | 4.54 | 10.29 | 7.60 | 1.13 | 8.79 | 5.18 | ABC transporter, periplasmic substrate-binding protein, aliphatic sulfonates family |
| *PSPTO_1789* | | + | 1959574 | 1961844 | 5.81 | 9.04 | 8.28 | 2.81 | 7.64 | 5.91 | Dimethylsulfoxide reductase |
| *PSPTO_1790* | | - | 1961852 | 1963045 | 1.57 | 4.65 | 2.36 | 0.87 | 2.65 | 1.59 | Acyl-CoA dehydrogenase family protein |
| *PSPTO_1791* | | - | 1963078 | 1964067 | 2.76 | 7.67 | 5.65 | 5.41 | 6.16 | 3.73 | ABC transporter, periplasmic substrate-binding protein, aliphatic sulfonates family |
| *PSPTO_1792* | | - | 1964256 | 1965839 | 2.22 | 18.28 | 11.91 | 8.83 | 3.60 | 2.34 | Rhodanese-like domain-containing protein |
| *PSPTO_1793* | | - | 1965836 | 1966450 | 6.03 | 45.66 | 40.57 | 28.50 | 3.80 | 2.40 | Hypothetical protein |
| *PSPTO_1794* | | + | 1966670 | 1967575 | 6.28 | 6.23 | 6.67 | 8.29 | 8.69 | 4.75 | LysR family transcriptional regulator |
| *PSPTO_1795* | | - | 1967637 | 1968725 | 14.64 | 17.85 | 15.83 | 8.95 | 7.66 | 5.49 | Alkanesulfonate monooxygenase |
| *PSPTO_1796* | | + | 1968984 | 1969841 | 1.15 | 3.85 | 6.24 | 3.66 | 3.44 | 1.89 | Sulfonate ABC transporter permease |
| *PSPTO_1797* | | + | 1969844 | 1970656 | 1.67 | 6.41 | 2.60 | 0.16 | 2.20 | 1.56 | Sulfonate ABC transporter ATP-binding protein |
| *PSPTO_1798* | | + | 1970708 | 1971010 | 4.29 | 27.47 | 12.31 | 2.15 | 25.73 | 15.28 | Hypothetical protein |
| *PSPTO_1799* | | + | 1971103 | 1971924 | 4.12 | 6.24 | 8.32 | 4.57 | 11.13 | 7.65 | Ser/Thr protein phosphatase family protein |
| *PSPTO_1800* | | - | 1971940 | 1972251 | 71.23 | 71.98 | 56.83 | 26.88 | 82.06 | 60.61 | Hypothetical protein |
| *PSPTO_1801* | *trkH* | - | 1972879 | 1974333 | 5.01 | 6.84 | 6.01 | 4.92 | 13.03 | 8.68 | Potassium uptake protein TrkH |
| *PSPTO_1802* | | + | 1974508 | 1975068 | 5.23 | 21.10 | 18.91 | 0.23 | 25.56 | 11.80 | Nitroreductase family protein |
| *PSPTO_1803* | | - | 1975082 | 1976434 | 3.49 | 5.91 | 7.16 | 5.76 | 9.73 | 8.20 | Sensor histidine kinase |
| *PSPTO_1804* | | + | 1976425 | 1976517 | 1.18 | 7.23 | 13.48 | 0.00 | 6.45 | 8.51 | Hypothetical protein |
| *PSPTO_1805* | | - | 1976521 | 1976985 | 57.66 | 851.84 | 1569.25 | 211.07 | 60.17 | 50.31 | Hypothetical protein |
| *PSPTO_1806* | | - | 1977171 | 1977848 | 48.36 | 65.47 | 48.84 | 33.77 | 80.34 | 52.18 | DNA-binding response regulator |
| *PSPTO_1807* | | - | 1977856 | 1978233 | 76.00 | 41.96 | 50.21 | 16.16 | 36.43 | 27.10 | Hypothetical protein |
| *PSPTO_1808* | | - | 1978400 | 1978699 | 25.96 | 81.52 | 111.87 | 53.27 | 139.50 | 39.07 | Hypothetical protein |
| *PSPTO_1809* | *ispZ* | - | 1978701 | 1979297 | 50.61 | 154.63 | 114.44 | 76.88 | 267.80 | 73.86 | Intracellular septation protein A |
| *PSPTO_1810* | | + | 1979363 | 1980226 | 41.66 | 62.65 | 48.00 | 32.86 | 92.11 | 61.26 | PHP domain-containing protein |
| *PSPTO_1811* | | + | 1980240 | 1980869 | 20.91 | 31.89 | 25.50 | 37.49 | 105.82 | 45.56 | Sua5/YciO/YrdC/YwlC family protein |
| *PSPTO_1812* | | + | 1980869 | 1981687 | 20.67 | 38.47 | 29.41 | 18.50 | 121.85 | 55.12 | Hypothetical protein |
| *PSPTO_1813* | | + | 1981703 | 1982476 | 2.79 | 13.29 | 2.79 | 8.74 | 21.09 | 5.96 | Hypothetical protein |
| *PSPTO_1814* | | + | 1982538 | 1982840 | 184.44 | 51.47 | 46.76 | 28.34 | 32.61 | 55.98 | Hypothetical protein |
| *PSPTO_1815* | *rluB* | + | 1983025 | 1984251 | 13.43 | 44.01 | 35.18 | 28.79 | 38.09 | 29.53 | Ribosomal large subunit pseudouridine synthase B |
| *PSPTO_1816* | | + | 1984314 | 1984526 | 49.23 | 50.91 | 23.29 | 36.50 | 62.99 | 31.99 | Hypothetical protein |
| *PSPTO_1817* | | + | 1984523 | 1985932 | 4.15 | 11.05 | 8.93 | 6.04 | 16.90 | 6.71 | Aromatic amino acid permease |
| *PSPTO_1818* | | + | 1985964 | 1986377 | 12.14 | 10.00 | 22.68 | 12.91 | 51.38 | 23.09 | Hypothetical protein |
| *PSPTO_1819* | | - | 1986679 | 1987878 | 4.05 | 4.96 | 3.94 | 3.50 | 14.90 | 5.59 | Major facilitator superfamily transporter |
| *PSPTO_1820* | | - | 1988016 | 1988195 | 32.22 | 28.53 | 43.35 | 9.05 | 58.04 | 38.32 | Hypothetical protein |
| *PSPTO_1821* | *phhB* | - | 1988647 | 1989003 | 327.08 | 106.50 | 147.86 | 40.20 | 225.28 | 232.38 | Pterin-4-alpha-carbinolamine dehydratase |
| *PSPTO_1822* | *phhA* | - | 1989038 | 1989835 | 366.20 | 63.02 | 56.58 | 25.47 | 72.20 | 139.73 | Phenylalanine-4-hydroxylase |
| *PSPTO_1823* | *phhR* | + | 1990110 | 1991675 | 4.56 | 6.37 | 5.98 | 1.20 | 14.05 | 11.26 | Phenylalanine hydroxylase transcriptional activator PhhR |
| *PSPTO_1824* | | - | 1991772 | 1992038 | 6.34 | 8.67 | 5.50 | 0.00 | 6.77 | 36.84 | Hypothetical protein |
| *PSPTO_1825* | *acs* | + | 1992395 | 1994350 | 100.66 | 265.60 | 206.53 | 61.06 | 308.72 | 27.00 | Acetyl-CoA synthetase |
| *PSPTO_1826* | | + | 1994884 | 1995660 | 40.32 | 29.84 | 37.45 | 28.63 | 225.21 | 83.19 | Arginine/ornithine ABC transporter, periplasmic arginine/ornithine-binding protein |
| *PSPTO_1827* | | + | 1995745 | 1996434 | 2.73 | 2.23 | 5.29 | 3.14 | 22.71 | 11.89 | Arginine/ornithine ABC transporter, permease protein |
| *PSPTO_1828* | | + | 1996431 | 1997129 | 2.15 | 3.08 | 8.08 | 2.63 | 33.50 | 15.74 | Arginine/ornithine ABC transporter, permease protein |
| *PSPTO_1829* | | + | 1997131 | 1998243 | 3.76 | 7.89 | 6.83 | 2.68 | 43.97 | 13.65 | Hypothetical protein |
| *PSPTO_1830* | *hisP* | + | 1998266 | 1999030 | 8.15 | 5.07 | 14.01 | 4.51 | 55.41 | 24.07 | Histidine ABC transporter ATP-binding protein |
| *PSPTO_1831* | *argR* | + | 1999103 | 2000158 | 3.34 | 6.25 | 3.05 | 3.00 | 28.82 | 11.27 | Transcriptional regulator ArgR |
| *PSPTO_1832* | *argD-1* | + | 2000395 | 2001612 | 6.70 | 24.74 | 38.82 | 13.96 | 210.38 | 72.86 | Acetylornithine delta-aminotransferase |

| Locus name | Gene name | Strand | Gene start (bp) | Gene end (bp) | Average TPM Inoculum | Average TPM $t_6$ | Average TPM $t_{24}$ | Average TPM $t_{168}$ | Average TPM Logarithmic | Average TPM Stationary | Annotation description |
|---|---|---|---|---|---|---|---|---|---|---|---|
| *PSPTO_1833* | *aruF* | + | 2001746 | 2002777 | 3.16 | 3.26 | 13.03 | 2.29 | 66.96 | 33.64 | Arginine N-succinyltransferase subunit alpha |
| *PSPTO_1834* | *aruG* | + | 2002841 | 2003866 | 3.53 | 9.15 | 5.99 | 3.44 | 71.66 | 29.30 | Arginine N-succinyltransferase subunit beta |
| *PSPTO_1835* | *astD* | + | 2003866 | 2005332 | 3.76 | 6.70 | 6.29 | 4.91 | 47.51 | 17.53 | Succinylglutamic semialdehyde dehydrogenase |
| *PSPTO_1836* | *astB* | + | 2005413 | 2006759 | 7.83 | 15.83 | 9.00 | 4.15 | 79.17 | 16.48 | Succinylarginine dihydrolase |
| *PSPTO_1838* | *astE* | + | 2007120 | 2008127 | 7.22 | 20.89 | 18.69 | 7.60 | 73.34 | 13.51 | Succinylglutamate desuccinylase |
| *PSPTO_1839* | *ribH-2* | - | 2008266 | 2008778 | 13.47 | 183.22 | 147.01 | 190.06 | 48.50 | 21.30 | 6,7-dimethyl-8-ribityllumazine synthase |
| *PSPTO_1840* | | - | 2008800 | 2008958 | 60.50 | 351.06 | 289.99 | 330.63 | 104.76 | 50.49 | Hypothetical protein |
| *PSPTO_1841* | | + | 2009291 | 2010295 | 32.95 | 56.52 | 28.79 | 24.08 | 106.90 | 41.84 | Threonine aldolase, low-specificity |
| *PSPTO_1842* | *alaS* | + | 2010440 | 2013064 | 9.54 | 53.02 | 35.33 | 16.55 | 78.74 | 15.31 | Alanyl-tRNA synthetase |
| *PSPTO_1843* | | + | 2013155 | 2014387 | 16.74 | 84.52 | 68.11 | 29.54 | 234.76 | 20.78 | Aspartate kinase |
| *PSPTO_1844* | *csrA-2* | + | 2014559 | 2014747 | 152.77 | 409.85 | 233.08 | 111.62 | 591.71 | 97.53 | Carbon storage regulator |
| *PSPTO_1845* | | - | 2015294 | 2015785 | 4.48 | 11.03 | 7.08 | 2.91 | 5.24 | 6.84 | Hypothetical protein |
| *PSPTO_1846* | *mgtE* | + | 2016109 | 2017551 | 9.41 | 15.20 | 6.63 | 8.52 | 11.85 | 9.64 | Magnesium transporter |
| *PSPTO_1847* | | - | 2017756 | 2018082 | 387.37 | 460.48 | 393.69 | 215.28 | 403.03 | 313.78 | Transcriptional repressor |
| *PSPTO_1848* | | - | 2018527 | 2018760 | 26.81 | 72.26 | 30.01 | 16.65 | 12.56 | 12.81 | Hypothetical protein |
| *PSPTO_1849* | | - | 2018748 | 2019263 | 39.14 | 97.55 | 42.26 | 22.62 | 5.63 | 8.91 | Hypothetical protein |
| *PSPTO_1850* | | - | 2019792 | 2021390 | 10.54 | 23.69 | 16.17 | 10.47 | 44.33 | 18.45 | Hypothetical protein |
| *PSPTO_1851* | | - | 2021357 | 2023519 | 17.22 | 24.00 | 25.95 | 16.82 | 73.66 | 24.36 | Hypothetical protein |
| *PSPTO_1852* | | - | 2023987 | 2026401 | 7.07 | 12.97 | 14.67 | 9.40 | 30.14 | 19.23 | Mechanosensitive ion channel family protein |
| *PSPTO_1853* | | - | 2026720 | 2026827 | 28.37 | 10.76 | 8.98 | 9.00 | 12.78 | 12.02 | Hypothetical protein |
| *PSPTO_1854* | | + | 2026946 | 2027335 | 8.12 | 17.56 | 12.73 | 5.57 | 11.88 | 11.47 | Hypothetical protein |
| *PSPTO_1855* | | + | 2027444 | 2029525 | 6.93 | 12.11 | 10.95 | 11.13 | 14.43 | 17.23 | TonB-dependent receptor |
| *PSPTO_1856* | | + | 2030104 | 2031066 | 3.06 | 5.51 | 1.34 | 0.00 | 8.70 | 3.37 | AraC family transcriptional regulator |
| *PSPTO_1857* | *saxC* | - | 2031166 | 2032194 | 12.05 | 8.84 | 11.69 | 5.06 | 17.84 | 17.24 | AraC family transcriptional regulator |
| *PSPTO_1858* | *saxA* | + | 2032383 | 2033264 | 6.36 | 36.46 | 23.04 | 1.45 | 11.19 | 5.32 | Aliphatic isothiocyanate resistance protein SaxA |
| *PSPTO_1859* | *saxB* | + | 2033359 | 2033904 | 24.37 | 37.08 | 36.26 | 20.94 | 39.79 | 28.46 | Isothiocyanate resistance protein SaxB; isochorismatase family |
| *PSPTO_1860* | | - | 2034000 | 2034491 | 14.14 | 26.04 | 27.09 | 2.85 | 16.32 | 9.38 | Hypothetical protein |
| *PSPTO_1861* | | - | 2034565 | 2035377 | 3.72 | 8.60 | 6.59 | 2.51 | 4.55 | 3.34 | Short chain dehydrogenase |
| *PSPTO_1862* | | - | 2035455 | 2036456 | 6.45 | 5.97 | 7.23 | 8.12 | 7.56 | 5.23 | Oxidoreductase zinc-binding protein |
| *PSPTO_1863* | | - | 2036536 | 2037396 | 7.77 | 13.45 | 13.54 | 1.09 | 9.17 | 8.02 | Alpha/beta fold family hydrolase |
| *PSPTO_1864* | | - | 2037413 | 2038201 | 5.16 | 18.36 | 15.78 | 6.86 | 6.92 | 5.32 | Short chain dehydrogenase/reductase family oxidoreductase |
| *PSPTO_1865* | | - | 2038243 | 2039325 | 10.74 | 98.93 | 72.32 | 15.32 | 11.15 | 7.38 | NADH:flavin oxidoreductase/NADH oxidase family protein |
| *PSPTO_1866* | | - | 2039589 | 2040188 | 4.34 | 2.96 | 9.55 | 0.00 | 6.14 | 4.63 | TetR family transcriptional regulator |
| *PSPTO_1867* | | + | 2040296 | 2041211 | - | - | - | - | - | - | |
| *PSPTO_1868* | *hss* | - | 2041244 | 2042656 | 7.31 | 13.13 | 10.81 | 7.71 | 15.88 | 14.52 | Homospermidine synthase |
| *PSPTO_1869* | | - | 2043020 | 2043310 | 5.32 | 15.44 | 7.83 | 11.76 | 4.79 | 20.88 | Hypothetical protein |
| *PSPTO_1870* | | - | 2043345 | 2045420 | 27.22 | 38.85 | 36.79 | 18.74 | 10.44 | 27.23 | Sensory box histidine kinase/response regulator |
| *PSPTO_1871* | | - | 2045768 | 2047219 | 14.83 | 9.02 | 7.93 | 6.44 | 3.34 | 3.79 | Cytosine/purines uracil thiamine allantoin permease |
| *PSPTO_1872* | | - | 2047315 | 2047410 | 13.31 | 0.00 | 10.32 | 2.66 | 2.90 | 7.14 | Hypothetical protein |
| *PSPTO_1873* | | + | 2047587 | 2048282 | 22.99 | 11.39 | 19.48 | 10.34 | 11.81 | 15.61 | GntR family transcriptional regulator |
| *PSPTO_1874* | *hyuE* | + | 2048293 | 2049024 | 10.34 | 10.25 | 9.57 | 12.92 | 15.28 | 15.34 | Hydantoin racemase |
| *PSPTO_1875* | *fdH* | - | 2049167 | 2050402 | 6.37 | 24.68 | 15.64 | 6.67 | 8.33 | 9.48 | Glutathione-dependent formaldehyde dehydrogenase |
| *PSPTO_1876* | | + | 2050480 | 2050683 | 1.90 | 10.36 | 2.90 | 1.43 | 7.22 | 3.29 | Hypothetical protein |
| *PSPTO_1877* | | + | 2050724 | 2051125 | 79.62 | 92.13 | 70.43 | 35.16 | 65.51 | 83.84 | ISPsy11, transposase OrfA |
| *PSPTO_1878* | | + | 2051139 | 2051957 | 6.97 | 8.78 | 5.55 | 4.34 | 8.67 | 7.83 | ISPsy11, transposase OrfB |
| *PSPTO_1879* | | - | 2051888 | 2052739 | 49.50 | 81.07 | 75.77 | 29.13 | 104.22 | 75.32 | LysR family transcriptional regulator |
| *PSPTO_1880* | | + | 2052948 | 2053790 | 6.65 | 10.55 | 6.44 | 5.94 | 18.69 | 8.19 | 3-hydroxyisobutyrate dehydrogenase |
| *PSPTO_1881* | | + | 2053794 | 2055311 | 4.21 | 3.69 | 4.11 | 0.12 | 14.13 | 5.34 | Aldehyde dehydrogenase family protein |
| *PSPTO_1882* | | + | 2055316 | 2056473 | 6.90 | 7.07 | 2.91 | 1.26 | 16.49 | 7.18 | Peptidase, M20/M25/M40 family |
| *PSPTO_1883* | | - | 2056905 | 2057831 | 24.56 | 29.06 | 21.90 | 8.91 | 36.11 | 18.00 | LysR family transcriptional regulator |
| *PSPTO_1884* | | + | 2058001 | 2058765 | 2.32 | 3.43 | 6.74 | 6.34 | 8.60 | 4.97 | Hypothetical protein |
| *PSPTO_1885* | | + | 2058772 | 2059413 | 32.56 | 20.19 | 24.89 | 11.42 | 113.67 | 22.34 | Efflux protein, LysE family |
| *PSPTO_1886* | *sdaA* | - | 2059427 | 2060803 | 57.65 | 30.76 | 41.51 | 32.70 | 366.11 | 33.09 | L-serine dehydratase 1 |
| *PSPTO_1887* | | + | 2060932 | 2061843 | 6.45 | 9.70 | 10.72 | 7.76 | 15.84 | 8.69 | LysR family transcriptional regulator |
| *PSPTO_1888* | | - | 2061856 | 2063133 | 3.40 | 5.72 | 8.12 | 6.67 | 8.15 | 7.77 | Membrane protein |
| *PSPTO_1889* | | - | 2063891 | 2065276 | 2.79 | 3.25 | 6.96 | 2.40 | 6.71 | 4.61 | Hypothetical protein |
| *PSPTO_1890* | *gabT-3* | - | 2065300 | 2066547 | 4.15 | 2.68 | 5.22 | 3.83 | 5.11 | 3.69 | 4-aminobutyrate aminotransferase |
| *PSPTO_1891* | *pcD* | + | 2066812 | 2068302 | 4.94 | 1.97 | 1.83 | 2.03 | 4.70 | 5.88 | Piperideine-6-carboxylate dehydrogenase |
| *PSPTO_1892* | | + | 2068346 | 2069635 | 4.64 | 1.55 | 3.74 | 4.01 | 3.71 | 6.49 | Hypothetical protein |
| *PSPTO_1893* | | - | 2069648 | 2071021 | 1.21 | 7.93 | 7.19 | 3.55 | 11.17 | 4.57 | Sensor histidine kinase |
| *PSPTO_1894* | | - | 2071018 | 2071680 | 2.56 | 3.41 | 6.64 | 1.59 | 11.36 | 6.67 | DNA-binding response regulator |
| *PSPTO_1895* | | + | 2071848 | 2072111 | 8.29 | 1180.50 | 945.54 | 145.65 | 12.68 | 10.17 | Hypothetical protein |
| *PSPTO_1896* | | + | 2072111 | 2072656 | 7.14 | 137.27 | 160.09 | 43.54 | 13.35 | 10.59 | Cytochrome b561 family protein |
| *PSPTO_1897* | | + | 2072695 | 2073993 | 6.20 | 23.49 | 26.34 | 5.83 | 6.58 | 4.95 | BNR/Asp-box repeat-containing protein |
| *PSPTO_1898* | | + | 2073990 | 2074610 | 3.55 | 59.20 | 35.39 | 11.79 | 7.55 | 6.10 | Membrane protein |
| *PSPTO_1899* | | + | 2074621 | 2076654 | 4.20 | 28.94 | 19.97 | 9.31 | 7.38 | 6.38 | Sulfatase family protein |
| *PSPTO_1900* | | + | 2076844 | 2076939 | 63.17 | 13.05 | 38.28 | 19.81 | 51.81 | 44.07 | Hypothetical protein |
| *PSPTO_1901* | *bphO* | + | 2077011 | 2077625 | 25.81 | 23.35 | 28.15 | 27.74 | 39.55 | 18.80 | Bacteriophytochrome heme oxygenase BphO |
| *PSPTO_1902* | *bphP* | + | 2077622 | 2079859 | 11.91 | 12.09 | 13.41 | 11.11 | 26.14 | 16.73 | Bacteriophytochrome histidine kinase |
| *PSPTO_1903* | | - | 2079912 | 2080598 | 2.54 | 1.14 | 3.73 | 3.45 | 2.70 | 3.58 | Hypothetical protein |
| *PSPTO_1904* | | - | 2080595 | 2084023 | 4.97 | 2.98 | 3.12 | 2.54 | 2.97 | 2.76 | Membrane protein |
| *PSPTO_1905* | | + | 2084126 | 2084365 | 4.36 | 5.38 | 5.16 | 0.00 | 2.94 | 1.55 | Hypothetical protein |
| *PSPTO_1906* | | - | 2084332 | 2085657 | 3.38 | 7.18 | 5.45 | 3.09 | 4.66 | 3.16 | Major facilitator family transporter |
| *PSPTO_1907* | *pobA* | - | 2085974 | 2087158 | 3.09 | 4.76 | 3.84 | 1.99 | 4.85 | 2.53 | P-hydroxybenzoate hydroxylase |
| *PSPTO_1908* | *pobR* | + | 2087309 | 2088178 | 8.13 | 17.45 | 11.99 | 6.37 | 13.57 | 3.25 | Transcriptional regulator PobR |
| *PSPTO_1909* | | - | 2088230 | 2089603 | 20.03 | 39.69 | 40.47 | 17.14 | 161.99 | 22.06 | Decarboxylase family protein |
| *PSPTO_1910* | | + | 2089979 | 2090728 | 23.06 | 47.16 | 34.81 | 32.50 | 45.59 | 49.89 | Hypothetical protein |
| *PSPTO_1911* | | + | 2091127 | 2092728 | 22.12 | 10.63 | 6.68 | 2.37 | 8.14 | 5.96 | Response regulator/TPR domain protein |
| *PSPTO_1912* | | + | 2092725 | 2093420 | 16.05 | 12.57 | 15.10 | 6.30 | 15.40 | 12.15 | Sensor histidine kinase |

| Locus name | Gene name | Strand | Gene start (bp) | Gene end (bp) | Average TPM Inoculum | Average TPM $t_6$ | Average TPM $t_{24}$ | Average TPM $t_{168}$ | Average TPM Logarithmic | Average TPM Stationary | Annotation description |
|---|---|---|---|---|---|---|---|---|---|---|---|
| *PSPTO_1913* | | + | 2093596 | 2093892 | 2886.21 | 399.84 | 410.38 | 158.12 | 162.45 | 366.29 | Hypothetical protein |
| *PSPTO_1914* | | + | 2093986 | 2094708 | 339.93 | 68.23 | 60.26 | 24.94 | 52.79 | 68.50 | Short chain dehydrogenase/reductase family oxidoreductase |
| *PSPTO_1915* | | + | 2094705 | 2095346 | 226.66 | 63.47 | 50.58 | 20.54 | 47.90 | 54.89 | Transferase, hexapeptide repeat protein |
| *PSPTO_1916* | | + | 2095352 | 2096368 | 136.87 | 48.18 | 27.27 | 18.98 | 30.83 | 42.53 | 3-oxoacyl-(acyl-carrier-protein) synthase III |
| *PSPTO_1917* | | - | 2096418 | 2097665 | 97.66 | 26.51 | 23.61 | 19.21 | 21.52 | 33.88 | Major facilitator family transporter |
| *PSPTO_1918* | | - | 2097934 | 2101455 | 38.49 | 9.28 | 9.86 | 3.86 | 15.84 | 14.81 | Glycosyl transferase family protein |
| *PSPTO_1919* | | - | 2101530 | 2102255 | 52.81 | 15.15 | 11.09 | 7.45 | 20.92 | 26.90 | Hypothetical protein |
| *PSPTO_1920* | | - | 2102255 | 2103373 | 267.46 | 52.55 | 56.00 | 30.50 | 68.44 | 138.61 | DegT/DnrJ/EryC1/StrS family aminotransferase |
| *PSPTO_1921* | | + | 2103671 | 2105014 | 16.60 | 4.20 | 6.88 | 1.59 | 10.16 | 6.95 | Glutamine synthetase |
| *PSPTO_1922* | | + | 2105069 | 2106466 | 34.86 | 10.59 | 13.01 | 7.91 | 25.84 | 21.05 | Major facilitator family transporter |
| *PSPTO_1923* | | - | 2106563 | 2107309 | 48.12 | 43.01 | 42.84 | 27.21 | 70.47 | 52.20 | Hypothetical protein |
| *PSPTO_1924* | | - | 2107392 | 2107859 | 114.53 | 77.67 | 57.25 | 31.40 | 70.26 | 55.79 | Hypothetical protein |
| *PSPTO_1925* | *flgM* | - | 2107909 | 2108223 | 847.83 | 491.50 | 339.07 | 171.04 | 244.87 | 244.75 | Negative regulator of flagellin synthesis FlgM |
| *PSPTO_1926* | *flgA* | - | 2108384 | 2109130 | 862.83 | 190.24 | 125.65 | 83.52 | 84.01 | 192.01 | Flagellar basal body P-ring biosynthesis protein FlgA |
| *PSPTO_1927* | | + | 2109221 | 2110147 | 100.42 | 70.88 | 52.69 | 19.55 | 47.52 | 42.87 | Chemotaxis protein CheV |
| *PSPTO_1928* | *cheR-2* | + | 2110184 | 2111026 | 22.42 | 19.56 | 21.44 | 20.20 | 28.01 | 28.10 | Chemotaxis protein methyltransferase CheR |
| *PSPTO_1929* | | + | 2111107 | 2112567 | 16.27 | 21.61 | 21.07 | 12.54 | 34.53 | 23.17 | ISPsy6, transposase |
| *PSPTO_1930* | *cynS* | + | 2112744 | 2113211 | 31.79 | 89.59 | 34.57 | 23.26 | 69.66 | 22.03 | Cyanate lyase |
| *PSPTO_1931* | | + | 2113222 | 2113695 | 8.07 | 23.21 | 14.90 | 8.18 | 31.99 | 9.85 | Hypothetical protein |
| *PSPTO_1932* | | + | 2113745 | 2113903 | 1.20 | 3.43 | 0.95 | 0.00 | 5.41 | 3.31 | Hypothetical protein |
| *PSPTO_1933* | *flgB* | + | 2113970 | 2114377 | 4660.91 | 558.09 | 554.77 | 397.40 | 171.82 | 521.17 | Flagellar basal-body rod protein FlgB |
| *PSPTO_1934* | *flgC* | + | 2114381 | 2114821 | 520.26 | 182.18 | 142.33 | 62.95 | 59.31 | 112.15 | Flagellar basal body rod protein FlgC |
| *PSPTO_1935* | *flgD* | + | 2114837 | 2115526 | 202.46 | 69.60 | 53.94 | 44.42 | 38.29 | 43.20 | Basal-body rod modification protein FlgD |
| *PSPTO_1936* | *flgE-1* | + | 2115552 | 2116877 | 371.98 | 149.57 | 109.60 | 66.73 | 62.38 | 50.39 | Flagellar hook protein FlgE |
| *PSPTO_1937* | *flgE-2* | + | 2117067 | 2118827 | 4.01 | 7.26 | 4.25 | 5.30 | 5.69 | 4.47 | Flagellar hook protein FlgE |
| *PSPTO_1938* | | + | 2118936 | 2119079 | 53.17 | 31.43 | 8.66 | 3.62 | 35.64 | 26.21 | Hypothetical protein |
| *PSPTO_1939* | *flgF* | + | 2119303 | 2120043 | 1805.17 | 264.85 | 242.54 | 116.98 | 72.16 | 186.73 | Flagellar basal body rod protein FlgF |
| *PSPTO_1940* | *flgG* | + | 2120090 | 2120878 | 954.54 | 207.35 | 185.61 | 96.98 | 94.63 | 193.42 | Flagellar basal-body rod protein FlgG |
| *PSPTO_1941* | *flgH* | + | 2120963 | 2121676 | 317.57 | 77.63 | 80.26 | 44.45 | 32.54 | 55.92 | Flagellar L-ring protein FlgH |
| *PSPTO_1942* | *flgI* | + | 2121690 | 2122799 | 135.35 | 41.07 | 34.12 | 19.38 | 18.20 | 31.47 | Flagellar P-ring protein FlgI |
| *PSPTO_1943* | *flgJ* | + | 2122810 | 2124060 | 182.69 | 71.02 | 46.35 | 28.25 | 27.11 | 43.92 | Peptidoglycan hydrolase FlgJ |
| *PSPTO_1944* | *flgK* | + | 2124072 | 2126132 | 169.18 | 61.75 | 49.16 | 28.72 | 30.03 | 43.06 | Flagellar hook-associated protein FlgK |
| *PSPTO_1945* | *flgL* | + | 2126146 | 2127738 | 97.51 | 74.66 | 41.88 | 24.96 | 36.06 | 50.20 | Flagellar hook-associated protein FlgL |
| *PSPTO_1946* | | + | 2127827 | 2131402 | 8.43 | 11.73 | 7.25 | 6.27 | 15.30 | 9.04 | Glycosyl transferase family protein |
| *PSPTO_1947* | | + | 2131534 | 2134440 | 8.00 | 12.28 | 6.24 | 2.71 | 17.78 | 10.73 | Glycosyl transferase family protein |
| *PSPTO_1948* | | + | 2134726 | 2135652 | 17.38 | 12.97 | 14.07 | 14.22 | 52.49 | 18.41 | 3-oxoacyl-(acyl-carrier-protein) synthase III |
| *PSPTO_1949* | *fliC* | + | 2135981 | 2136829 | 1320.29 | 748.49 | 537.16 | 288.03 | 37.73 | 17.20 | Flagellin |
| *PSPTO_1950* | *flaG* | + | 2136896 | 2137294 | 432.52 | 520.25 | 393.77 | 144.66 | 402.20 | 172.03 | Flagellin FlaG |
| *PSPTO_1951* | *fliD* | + | 2137378 | 2138856 | 214.08 | 252.08 | 208.37 | 112.61 | 200.70 | 135.07 | Flagellar hook-associated protein FliD |
| *PSPTO_1952* | *fliS* | + | 2138982 | 2139371 | 981.43 | 412.97 | 337.59 | 122.82 | 58.55 | 67.63 | Flagellar protein FliS |
| *PSPTO_1953* | *fliT* | + | 2139397 | 2139693 | 161.33 | 51.84 | 58.57 | 48.13 | 13.20 | 22.53 | Hypothetical protein |
| *PSPTO_1954* | *fleQ* | + | 2139870 | 2141345 | 55.40 | 49.19 | 46.70 | 27.49 | 88.64 | 79.30 | Transcriptional regulator FleQ |
| *PSPTO_1955* | *fleS* | + | 2141576 | 2142682 | 438.54 | 95.91 | 71.90 | 43.31 | 47.60 | 137.03 | Sensor histidine kinase FleS |
| *PSPTO_1956* | *fleR* | + | 2142682 | 2144097 | 200.84 | 64.50 | 48.55 | 26.45 | 36.81 | 62.62 | Sigma-54 dependent transcriptional regulator/response regulator FleR |
| *PSPTO_1957* | *fliE* | + | 2144312 | 2144668 | 1927.88 | 305.77 | 254.85 | 144.18 | 106.88 | 397.90 | Flagellar hook-basal body complex protein FliE |
| *PSPTO_1958* | *fliF* | + | 2144684 | 2146468 | 140.87 | 50.64 | 48.69 | 18.06 | 53.73 | 89.26 | Flagellar M-ring protein FliF |
| *PSPTO_1959* | *fliG* | + | 2146476 | 2147477 | 132.95 | 31.05 | 54.88 | 17.50 | 62.48 | 86.21 | Flagellar motor switch protein FliG |
| *PSPTO_1960* | *fliH* | + | 2147493 | 2148311 | 88.34 | 29.19 | 30.31 | 14.49 | 47.96 | 62.59 | Flagellar assembly protein H |
| *PSPTO_1961* | *fliI* | + | 2148301 | 2149659 | 44.19 | 19.65 | 14.10 | 8.40 | 23.66 | 32.57 | Flagellum-specific ATP synthase FliI |
| *PSPTO_1962* | *fliJ* | + | 2149666 | 2150115 | 19.58 | 8.46 | 10.81 | 5.07 | 9.15 | 13.03 | Flagellar biosynthesis chaperone |
| *PSPTO_1963* | | + | 2150198 | 2150500 | 133.44 | 63.08 | 72.74 | 64.36 | 36.53 | 104.35 | STAS domain-containing protein |
| *PSPTO_1964* | | + | 2150505 | 2152211 | 8.71 | 10.85 | 8.30 | 8.35 | 13.54 | 12.67 | Response regulator |
| *PSPTO_1965* | | + | 2152245 | 2152601 | 29.72 | 66.36 | 67.03 | 41.10 | 66.82 | 63.62 | Hpt domain-containing protein |
| *PSPTO_1966* | *fliK* | + | 2152700 | 2154124 | 520.70 | 163.61 | 150.36 | 74.17 | 62.05 | 118.45 | Flagellar hook-length control protein FliK |
| *PSPTO_1967* | | - | 2154255 | 2154434 | 17.94 | 6.09 | 10.92 | 5.09 | 8.09 | 13.44 | Hypothetical protein |
| *PSPTO_1968* | *fliL* | + | 2154679 | 2155182 | 868.72 | 122.04 | 119.58 | 87.76 | 79.11 | 191.63 | Flagellar basal body-associated protein FliL |
| *PSPTO_1969* | *fliM* | + | 2155192 | 2156160 | 155.69 | 40.59 | 52.38 | 41.37 | 54.84 | 68.50 | Flagellar motor switch protein FliM |
| *PSPTO_1970* | *fliN* | + | 2156222 | 2156680 | 279.18 | 90.93 | 78.43 | 43.89 | 95.95 | 108.85 | Flagellar motor switch protein FliN |
| *PSPTO_1971* | *fliO* | + | 2156681 | 2157121 | 161.36 | 67.08 | 52.46 | 32.65 | 53.63 | 74.93 | Flagellar protein FliO |
| *PSPTO_1972* | *fliP* | + | 2157121 | 2157873 | 32.84 | 15.99 | 22.30 | 9.56 | 20.21 | 14.61 | Flagellar biosynthetic protein FliP |
| *PSPTO_1973* | *fliQ* | + | 2157899 | 2158168 | 15.36 | 10.39 | 0.00 | 0.68 | 8.68 | 8.01 | Flagellar biosynthetic protein FliQ |
| *PSPTO_1974* | *fliR* | + | 2158184 | 2158960 | 44.25 | 19.91 | 12.50 | 13.71 | 22.23 | 22.85 | Flagellar biosynthetic protein FliR |
| *PSPTO_1975* | *flhB* | + | 2158962 | 2160098 | 16.63 | 8.90 | 8.72 | 4.99 | 9.82 | 9.10 | Flagellar biosynthetic protein FlhB |
| *PSPTO_1976* | *flhA* | + | 2160485 | 2162614 | 832.88 | 132.24 | 140.15 | 63.23 | 90.28 | 258.77 | Flagellar biosynthesis protein FlhA |
| *PSPTO_1977* | *flhF* | + | 2162626 | 2163954 | 339.35 | 94.70 | 96.27 | 42.48 | 70.55 | 157.88 | Flagellar biosynthesis protein FlhF |
| *PSPTO_1978* | *fleN* | + | 2164079 | 2164903 | 175.37 | 68.67 | 44.23 | 28.19 | 53.79 | 86.71 | Flagellar synthesis regulator FleN |
| *PSPTO_1979* | *fliA* | + | 2164900 | 2165640 | 413.81 | 175.10 | 173.01 | 113.25 | 168.58 | 273.21 | Motility sigma factor FliA |
| *PSPTO_1980* | *cheY-2* | + | 2165893 | 2166267 | 232.13 | 146.26 | 127.48 | 71.48 | 181.75 | 189.82 | Chemotaxis protein CheY |
| *PSPTO_1981* | *cheZ* | + | 2166386 | 2167072 | 79.31 | 37.80 | 48.68 | 22.81 | 47.30 | 58.15 | Chemotaxis protein CheZ |
| *PSPTO_1982* | *cheA-2* | + | 2167106 | 2169352 | 81.02 | 45.49 | 22.10 | 27.18 | 32.50 | 29.54 | Chemotaxis sensor histidine kinase CheA |
| *PSPTO_1983* | *cheB-2* | + | 2169403 | 2170575 | 25.55 | 9.85 | 17.44 | 5.18 | 21.73 | 15.17 | Protein-glutamate methylesterase CheB |
| *PSPTO_1984* | *motA-1* | + | 2170575 | 2171315 | 12.44 | 8.20 | 4.83 | 3.72 | 19.44 | 10.92 | Flagellar motor protein |
| *PSPTO_1985* | *motD* | + | 2171322 | 2172209 | 105.07 | 18.56 | 21.13 | 12.48 | 22.83 | 30.97 | MotD protein |
| *PSPTO_1986* | | + | 2172255 | 2173043 | 139.40 | 22.78 | 16.26 | 14.49 | 19.47 | 30.93 | ParA family protein |
| *PSPTO_1987* | | + | 2173119 | 2174006 | 246.10 | 60.86 | 44.38 | 28.76 | 26.96 | 25.75 | Chemotaxis protein CheW |
| *PSPTO_1988* | *cheW-2* | + | 2174064 | 2174543 | 183.93 | 88.81 | 72.40 | 49.54 | 40.17 | 42.85 | Chemotaxis protein CheW |
| *PSPTO_1989* | | + | 2174586 | 2174945 | 66.24 | 46.14 | 26.67 | 19.13 | 21.94 | 21.71 | Hypothetical protein |
| *PSPTO_1990* | *apt* | - | 2175076 | 2175624 | 31.30 | 34.25 | 50.67 | 21.15 | 63.59 | 30.48 | Adenine phosphoribosyltransferase |
| *PSPTO_1991* | *anr* | - | 2175686 | 2176420 | 5.43 | 23.92 | 11.31 | 0.48 | 44.57 | 28.08 | Transcriptional activator Anr |

| Locus name | Gene name | Strand | Gene start (bp) | Gene end (bp) | Average TPM Inoculum | Average TPM $t_6$ | Average TPM $t_{24}$ | Average TPM $t_{168}$ | Average TPM Logarithmic | Average TPM Stationary | Annotation description |
|---|---|---|---|---|---|---|---|---|---|---|---|
| *PSPTO_1992* | | + | 2176562 | 2177041 | 124.97 | 26.59 | 19.66 | 15.53 | 2.73 | 13.23 | Hypothetical protein |
| *PSPTO_1993* | *hemN* | - | 2177061 | 2178443 | 6.86 | 6.69 | 9.92 | 6.19 | 32.23 | 154.00 | Oxygen-independent coproporphyrinogen III oxidase |
| *PSPTO_1994* | | - | 2178539 | 2179222 | 2.27 | 2.94 | 6.99 | 0.00 | 4.75 | 3.77 | Membrane protein |
| *PSPTO_1995* | *ccoS* | - | 2179215 | 2179418 | 5.77 | 18.17 | 4.61 | 0.00 | 11.67 | 7.02 | Cbb3-type cytochrome oxidase maturation protein |
| *PSPTO_1996* | | - | 2179397 | 2181868 | 6.42 | 8.31 | 5.89 | 2.72 | 13.72 | 5.86 | Copper-translocating P-type ATPase |
| *PSPTO_1997* | | - | 2181871 | 2182407 | 2.66 | 10.56 | 1.22 | 0.00 | 22.69 | 8.01 | Hypothetical protein |
| *PSPTO_1998* | | - | 2182418 | 2183830 | 6.34 | 12.15 | 8.04 | 7.32 | 14.51 | 11.60 | FixG-related protein |
| *PSPTO_1999* | | - | 2183895 | 2184347 | 10.18 | 30.23 | 47.82 | 12.78 | 50.85 | 31.95 | PIN domain-containing protein |
| *PSPTO_2000* | | - | 2184347 | 2184598 | 28.51 | 69.54 | 84.53 | 0.00 | 50.26 | 38.79 | Prevent-host-death family protein |
| *PSPTO_2001* | *ccoP* | - | 2184736 | 2185713 | 3.48 | 8.57 | 4.88 | 2.95 | 20.66 | 7.14 | Cytochrome c oxidase, cbb3-type subunit III |
| *PSPTO_2002* | *ccoQ* | - | 2185710 | 2185901 | 2.17 | 6.99 | 20.16 | 3.71 | 11.19 | 4.91 | Cytochrome c oxidase, cbb3-type, CcoQ subunit |
| *PSPTO_2003* | *ccoO* | - | 2185907 | 2186515 | 5.57 | 14.56 | 6.01 | 3.35 | 28.38 | 11.94 | Cbb3-type cytochrome c oxidase subunit II |
| *PSPTO_2004* | *ccoN* | - | 2186515 | 2187957 | 5.54 | 10.22 | 9.16 | 6.10 | 18.33 | 8.54 | Cbb3-type cytochrome c oxidase subunit I |
| *PSPTO_2005* | | + | 2187950 | 2188942 | 9.73 | 25.11 | 21.89 | 6.60 | 28.91 | 17.20 | Hypothetical protein |
| *PSPTO_2006* | | - | 2188944 | 2189417 | 5.79 | 2.07 | 2.77 | 0.65 | 6.69 | 4.00 | Hypothetical protein |
| *PSPTO_2007* | | - | 2189494 | 2191350 | 7.06 | 7.60 | 4.19 | 3.40 | 8.37 | 5.60 | Hypothetical protein |
| *PSPTO_2008* | | + | 2191370 | 2192347 | 0.00 | 0.00 | 0.00 | 0.00 | 0.00 | 0.00 | ISPssy, transposase |
| *PSPTO_2009* | | - | 2192645 | 2194264 | 1.12 | 4.95 | 3.67 | 4.93 | 8.02 | 5.93 | Hypothetical protein |
| *PSPTO_2010* | | - | 2194390 | 2195367 | 0.00 | 0.00 | 0.00 | 0.00 | 0.00 | 0.00 | ISPssy, transposase |
| *PSPTO_2011* | | + | 2196049 | 2198358 | 3.68 | 8.12 | 3.67 | 0.79 | 10.49 | 9.25 | Autotransporter |
| *PSPTO_2012* | | - | 2198423 | 2198959 | 0.86 | 4.23 | 0.00 | 0.00 | 8.68 | 1.01 | Hypothetical protein |
| *PSPTO_2013* | | - | 2199398 | 2200060 | 1.08 | 0.00 | 0.49 | 2.80 | 1.84 | 0.60 | Hypothetical protein |
| *PSPTO_2014* | *aer-2* | - | 2200378 | 2201943 | 12.44 | 8.06 | 5.95 | 1.85 | 6.53 | 4.30 | Aerotaxis receptor |
| *PSPTO_2015* | | - | 2202040 | 2202840 | 6.51 | 11.97 | 13.30 | 0.00 | 10.25 | 6.06 | CAAX amino terminal protease family protein |
| *PSPTO_2016* | *acnA* | - | 2202961 | 2205705 | 6.42 | 50.88 | 45.18 | 29.68 | 20.43 | 18.89 | Aconitate hydratase 1 |
| *PSPTO_2017* | | + | 2205885 | 2206958 | 3.85 | 8.69 | 10.30 | 1.11 | 11.35 | 4.82 | Hypothetical protein |
| *PSPTO_2018* | *sirA* | + | 2207025 | 2207279 | 46.61 | 31.53 | 60.82 | 11.15 | 115.16 | 55.08 | Sulfur transfer protein SirA |
| *PSPTO_2019* | *pdxB* | + | 2207368 | 2208510 | 4.47 | 9.92 | 4.96 | 2.25 | 16.40 | 6.77 | Erythronate-4-phosphate dehydrogenase |
| *PSPTO_2020* | | - | 2208732 | 2209064 | 682.41 | 1405.93 | 838.88 | 582.46 | 248.39 | 884.33 | Hypothetical protein |
| *PSPTO_2021* | | + | 2209190 | 2210959 | 6.69 | 9.95 | 9.60 | 8.39 | 18.34 | 12.03 | ABC transporter ATP-binding protein/permease |
| *PSPTO_2022* | | - | 2211013 | 2211294 | 27.14 | 128.23 | 108.21 | 61.25 | 498.23 | 56.77 | Hypothetical protein |
| *PSPTO_2023* | | - | 2211408 | 2211950 | 20.51 | 13.68 | 13.72 | 13.97 | 11.62 | 11.72 | Exonuclease |
| *PSPTO_2024* | | + | 2212207 | 2213046 | 17.54 | 29.01 | 22.32 | 13.19 | 45.35 | 20.37 | Hypothetical protein |
| *PSPTO_2025* | | - | 2213108 | 2213275 | 204.54 | 275.07 | 179.01 | 109.51 | 195.43 | 149.00 | Hypothetical protein |
| *PSPTO_2026* | | - | 2213388 | 2214752 | 20.49 | 13.51 | 17.20 | 5.13 | 11.14 | 9.01 | Ethanolamine permease family protein |
| *PSPTO_2027* | | - | 2214849 | 2214965 | 3.93 | 9.16 | 5.93 | 0.00 | 20.85 | 5.01 | Hypothetical protein |
| *PSPTO_2028* | *pyrF* | + | 2215104 | 2215742 | 10.00 | 14.00 | 13.63 | 5.68 | 29.56 | 14.48 | Orotidine 5-phosphate decarboxylase |
| *PSPTO_2029* | | - | 2215916 | 2217076 | 15.52 | 32.88 | 20.76 | 14.29 | 17.66 | 16.54 | Oxidoreductase zinc-binding protein |
| *PSPTO_2030* | | + | 2217088 | 2218089 | 14.40 | 22.85 | 24.96 | 12.04 | 28.08 | 21.75 | Muramoyltetrapeptide carboxypeptidase |
| *PSPTO_2031* | | - | 2218103 | 2218594 | 3.24 | 4.17 | 4.30 | 3.29 | 8.66 | 6.98 | Benzoate transport C-terminal domain protein |
| *PSPTO_2032* | | + | 2218643 | 2219245 | 33.39 | 30.75 | 35.34 | 20.57 | 50.58 | 32.83 | Glutathione S-transferase family protein |
| *PSPTO_2033* | | + | 2219277 | 2219663 | 8.93 | 15.76 | 10.66 | 5.48 | 31.25 | 25.07 | Lipoprotein |
| *PSPTO_2034* | | - | 2219685 | 2220062 | 10.14 | 26.76 | 29.42 | 14.25 | 64.20 | 23.18 | Hypothetical protein |
| *PSPTO_2035* | *folE-2* | - | 2220137 | 2220682 | 11.61 | 17.87 | 23.57 | 7.60 | 58.28 | 19.39 | GTP cyclohydrolase I |
| *PSPTO_2036* | | + | 2220919 | 2221095 | 14.38 | 4.33 | 0.00 | 0.00 | 7.84 | 8.61 | Lipoprotein |
| *PSPTO_2037* | | - | 2221165 | 2221722 | 6.78 | 17.84 | 19.96 | 7.99 | 29.51 | 16.43 | Smr domain protein |
| *PSPTO_2038* | | - | 2221769 | 2222086 | 25.07 | 6.39 | 20.64 | 3.69 | 8.87 | 10.42 | Hypothetical protein |
| *PSPTO_2039* | | - | 2222263 | 2222853 | 114.04 | 134.00 | 96.54 | 78.91 | 215.10 | 76.72 | Isochorismatase family protein |
| *PSPTO_2040* | | + | 2223115 | 2224023 | 11.52 | 28.30 | 27.37 | 17.87 | 59.63 | 25.63 | HemK family modification methylase |
| *PSPTO_2041* | | - | 2224103 | 2224906 | 80.83 | 26.04 | 30.09 | 11.82 | 6.12 | 3.48 | Hypothetical protein |
| *PSPTO_2042* | | - | 2225003 | 2226034 | 18.47 | 20.10 | 11.37 | 8.10 | 19.63 | 21.37 | Hypothetical protein |
| *PSPTO_2043* | *aroC* | + | 2226176 | 2227267 | 13.48 | 48.77 | 55.85 | 37.34 | 101.80 | 41.42 | Chorismate synthase |
| *PSPTO_2044* | | + | 2227299 | 2228468 | 2.58 | 4.28 | 2.07 | 5.67 | 14.78 | 4.56 | Transporter |
| *PSPTO_2045* | | + | 2228465 | 2229079 | 6.74 | 9.97 | 5.41 | 6.89 | 41.19 | 5.92 | Methylthioribulose-1-phosphate dehydratase |
| *PSPTO_2046* | | + | 2229161 | 2229706 | 4.94 | 16.15 | 17.17 | 10.70 | 64.34 | 7.00 | ARD/ARD' family protein |
| *PSPTO_2047* | | + | 2229723 | 2230406 | 3.49 | 2.84 | 16.60 | 4.40 | 31.57 | 3.87 | HAD-superfamily hydrolase |
| *PSPTO_2048* | | + | 2230681 | 2231403 | 264.25 | 168.78 | 134.64 | 66.67 | 57.18 | 119.48 | LuxR family transcriptional regulator |
| *PSPTO_2049* | | - | 2231544 | 2231822 | 117.73 | 1292.62 | 804.73 | 316.61 | 61.02 | 62.54 | Hypothetical protein |
| *PSPTO_2050* | | + | 2231957 | 2232181 | 42.00 | 30.46 | 33.64 | 21.77 | 27.98 | 43.23 | Lipoprotein |
| *PSPTO_2051* | | + | 2232185 | 2232730 | 25.32 | 21.86 | 43.92 | 11.28 | 14.21 | 23.12 | Ankyrin domain protein |
| *PSPTO_2052* | | + | 2232829 | 2233347 | 12.63 | 12.62 | 31.51 | 11.64 | 38.38 | 24.05 | Hypothetical protein |
| *PSPTO_2053* | | - | 2233344 | 2234261 | 13.73 | 20.71 | 12.05 | 9.67 | 14.24 | 9.41 | Membrane protein |
| *PSPTO_2054* | | + | 2234416 | 2234607 | 105.11 | 94.60 | 52.58 | 69.20 | 57.17 | 59.95 | Hypothetical protein |
| *PSPTO_2055* | *speE* | + | 2234806 | 2235666 | 6.34 | 19.60 | 13.92 | 8.71 | 58.99 | 13.31 | Spermidine synthase |
| *PSPTO_2056* | | + | 2235931 | 2238018 | 4.97 | 18.38 | 11.59 | 10.11 | 92.29 | 9.43 | Membrane protein |
| *PSPTO_2057* | | + | 2238692 | 2239033 | 485.99 | 687.82 | 389.86 | 174.39 | 36.69 | 112.89 | Hypothetical protein |
| *PSPTO_2058* | | + | 2239033 | 2239221 | 27.88 | 16.58 | 22.34 | 16.75 | 5.52 | 13.82 | Hypothetical protein |
| *PSPTO_2059* | | + | 2239542 | 2240951 | 13.33 | 49.10 | 35.25 | 13.77 | 16.42 | 17.61 | Hypothetical protein |
| *PSPTO_2060* | | + | 2240964 | 2241923 | 16.31 | 38.31 | 24.59 | 15.69 | 15.09 | 14.06 | Hypothetical protein |
| *PSPTO_2061* | | + | 2241920 | 2242750 | 28.40 | 62.53 | 40.76 | 26.91 | 19.96 | 25.91 | Transglutaminase-like domain protein |
| *PSPTO_2062* | | - | 2242794 | 2243165 | 45.08 | 47.73 | 87.32 | 60.70 | 60.85 | 49.35 | Hypothetical protein |
| *PSPTO_2063* | | + | 2243781 | 2244509 | 23.89 | 67.63 | 54.60 | 21.07 | 82.25 | 32.58 | Hypothetical protein |
| *PSPTO_2064* | | - | 2244569 | 2245540 | 36.49 | 14.60 | 22.60 | 4.85 | 22.91 | 18.03 | Magnesium and cobalt transport protein |
| *PSPTO_2065* | | - | 2245666 | 2246355 | 17.07 | 37.30 | 19.90 | 10.04 | 71.76 | 21.40 | Enoyl-CoA hydratase |
| *PSPTO_2066* | | - | 2246481 | 2246774 | 17.61 | 15.38 | 13.25 | 6.06 | 28.26 | 17.39 | Hypothetical protein |
| *PSPTO_2067* | | + | 2246765 | 2246884 | 23.41 | 22.23 | 14.15 | 0.00 | 45.88 | 26.63 | Hypothetical protein |
| *PSPTO_2068* | | + | 2246940 | 2247269 | 41.65 | 31.83 | 19.63 | 33.57 | 45.49 | 14.93 | Hypothetical protein |
| *PSPTO_2069* | | + | 2247841 | 2248026 | 52.08 | 84.03 | 61.30 | 36.16 | 11.93 | 18.46 | Hypothetical protein |
| *PSPTO_2070* | | - | 2248078 | 2248446 | 43.65 | 24.24 | 43.61 | 25.97 | 22.94 | 20.77 | Cytochrome c2 |

| Locus name | Gene name | Strand | Gene start (bp) | Gene end (bp) | Average TPM Inoculum | Average TPM $t_6$ | Average TPM $t_{24}$ | Average TPM $t_{168}$ | Average TPM Logarithmic | Average TPM Stationary | Annotation description |
|---|---|---|---|---|---|---|---|---|---|---|---|
| *PSPTO_2071* | | - | 2248514 | 2249044 | 33.23 | 21.11 | 13.59 | 1.04 | 13.08 | 15.48 | Hypothetical protein |
| *PSPTO_2072* | | - | 2249264 | 2249767 | 19.28 | 21.50 | 14.80 | 8.07 | 19.25 | 20.49 | Auxin-binding protein |
| *PSPTO_2073* | | - | 2249888 | 2250436 | 38.36 | 25.63 | 36.74 | 15.56 | 41.40 | 27.65 | Hypothetical protein |
| *PSPTO_2074* | | - | 2250531 | 2250824 | 5.18 | 4.14 | 3.21 | 2.44 | 6.49 | 5.28 | Hypothetical protein |
| *PSPTO_2075* | | + | 2251135 | 2251380 | 1.24 | 2.37 | 0.00 | 0.00 | 1.35 | 2.53 | Hypothetical protein |
| *PSPTO_2076* | | - | 2251753 | 2252214 | 0.77 | 0.00 | 2.36 | 0.00 | 0.82 | 0.95 | Hypothetical protein |
| *PSPTO_2077* | | + | 2252628 | 2252993 | 11.39 | 0.00 | 6.41 | 2.87 | 6.55 | 11.56 | Hypothetical protein |
| *PSPTO_2078* | | + | 2253004 | 2253381 | 6.77 | 2.56 | 3.97 | 11.84 | 7.77 | 12.36 | Hypothetical protein |
| *PSPTO_2079* | | + | 2253384 | 2253746 | 33.33 | 19.62 | 38.93 | 11.49 | 13.29 | 20.10 | Hypothetical protein |
| *PSPTO_2080* | | + | 2253746 | 2254126 | 24.83 | 8.94 | 8.34 | 13.43 | 13.57 | 18.44 | Hypothetical protein |
| *PSPTO_2081* | | + | 2254123 | 2254575 | 6.20 | 7.06 | 1.99 | 2.59 | 7.87 | 6.26 | Hypothetical protein |
| *PSPTO_2082* | | - | 2254666 | 2255499 | 5.62 | 2.63 | 6.52 | 6.65 | 3.48 | 3.86 | Tail length tape measure protein, internal deletion |
| *PSPTO_2083* | | - | 2255556 | 2256083 | 20.36 | 35.02 | 21.44 | 3.76 | 14.87 | 12.62 | Hypothetical protein |
| *PSPTO_2084* | | - | 2256299 | 2256550 | 3.32 | 12.57 | 3.01 | 0.00 | 4.96 | 3.17 | Hypothetical protein |
| *PSPTO_2085* | | - | 2256568 | 2256963 | 5.09 | 3.75 | 2.62 | 0.00 | 6.88 | 4.58 | Hypothetical protein |
| *PSPTO_2086* | | - | 2257112 | 2257294 | 29.99 | 23.57 | 26.48 | 35.26 | 23.41 | 35.78 | Hypothetical protein |
| *PSPTO_2087* | | + | 2257405 | 2257713 | 1.10 | 0.00 | 0.20 | 1.50 | 2.39 | 0.86 | Hypothetical protein |
| *PSPTO_2088* | | + | 2257716 | 2258114 | 0.00 | 0.00 | 0.00 | 2.33 | 0.27 | 0.38 | Hypothetical protein |
| *PSPTO_2089* | | + | 2258114 | 2261395 | 3.55 | 3.18 | 2.21 | 1.19 | 2.71 | 2.53 | Host specificity protein J, internal deletion |
| *PSPTO_2090* | | + | 2261392 | 2261760 | 1.13 | 2.21 | 1.97 | 0.00 | 3.90 | 2.81 | Hypothetical protein |
| *PSPTO_2091* | | + | 2261757 | 2262419 | 1.25 | 1.73 | 2.23 | 5.09 | 2.41 | 2.55 | Hypothetical protein |
| *PSPTO_2092* | | + | 2262447 | 2263874 | 20.02 | 11.35 | 9.85 | 4.17 | 7.01 | 13.35 | Tail fiber domain protein |
| *PSPTO_2093* | | + | 2263954 | 2264487 | 15.37 | 37.89 | 29.10 | 29.08 | 43.98 | 33.09 | Lysozyme |
| *PSPTO_2094* | | + | 2264484 | 2264999 | 8.79 | 16.05 | 6.60 | 8.45 | 23.82 | 14.91 | Hypothetical protein |
| *PSPTO_2095* | | + | 2265012 | 2265224 | 3.25 | 9.60 | 3.90 | 0.00 | 18.81 | 9.72 | Hypothetical protein |
| *PSPTO_2096* | | + | 2265221 | 2266465 | 9.93 | 15.88 | 6.86 | 4.38 | 10.69 | 9.24 | Hypothetical protein |
| *PSPTO_2097* | | - | 2266931 | 2267680 | 7.57 | 3.82 | 6.86 | 0.49 | 10.58 | 7.20 | 3-ketoacyl-(acyl-carrier-protein) reductase |
| *PSPTO_2098* | | + | 2267792 | 2268358 | 6.53 | 11.19 | 23.48 | 3.33 | 25.90 | 6.40 | Isochorismatase family protein |
| *PSPTO_2099* | | - | 2268366 | 2271122 | 5.67 | 13.22 | 9.50 | 2.68 | 60.80 | 9.20 | Helicase/SNF2 family domain protein |
| *PSPTO_2100* | | - | 2271197 | 2271721 | 10.80 | 28.85 | 21.29 | 17.04 | 77.83 | 25.26 | Hypothetical protein |
| *PSPTO_2101* | *mfd* | - | 2271733 | 2275185 | 6.48 | 18.03 | 15.87 | 11.70 | 90.65 | 20.42 | Transcription-repair coupling factor |
| *PSPTO_2102* | *gap-2* | + | 2275390 | 2276838 | 37.46 | 158.65 | 147.05 | 124.83 | 291.36 | 44.47 | Glyceraldehyde 3-phosphate dehydrogenase, type I |
| *PSPTO_2103* | | + | 2276999 | 2278261 | 7.88 | 1.63 | 0.62 | 1.40 | 7.34 | 4.97 | Membrane protein |
| *PSPTO_2104* | | - | 2278340 | 2279638 | 6.40 | 21.35 | 27.79 | 10.76 | 13.58 | 7.30 | Major facilitator family transporter |
| *PSPTO_2105* | | + | 2280026 | 2281057 | 11.04 | 60.18 | 33.82 | 7.93 | 7.62 | 6.17 | Thiamine biosynthesis lipoprotein |
| *PSPTO_2106* | *sthA* | + | 2281200 | 2282594 | 39.91 | 59.97 | 38.14 | 25.31 | 187.45 | 36.68 | Soluble pyridine nucleotide transhydrogenase |
| *PSPTO_2107* | | - | 2282637 | 2283359 | 10.16 | 27.51 | 20.03 | 27.33 | 40.47 | 19.01 | Glycerophosphoryl diester phosphodiesterase |
| *PSPTO_2108* | | - | 2283419 | 2284006 | 26.16 | 41.16 | 32.05 | 32.64 | 86.56 | 35.02 | Hypothetical protein |
| *PSPTO_2109* | *lolC* | + | 2284109 | 2285359 | 6.66 | 12.73 | 7.41 | 4.19 | 21.85 | 8.88 | Lipoprotein releasing system transmembrane protein LolC |
| *PSPTO_2110* | *lolD* | + | 2285367 | 2286050 | 4.96 | 10.43 | 3.27 | 4.12 | 31.79 | 5.82 | Lipoprotein releasing system, ATP-binding protein LolD |
| *PSPTO_2111* | *lolE* | + | 2286086 | 2287369 | 7.51 | 10.18 | 8.94 | 4.31 | 46.31 | 11.69 | Lipoprotein releasing system transmembrane protein LolE |
| *PSPTO_2112* | | + | 2287479 | 2288309 | 16.45 | 42.92 | 32.24 | 21.50 | 80.63 | 36.89 | 7-cyano-7-deazaguanine reductase |
| *PSPTO_2113* | | - | 2288651 | 2288920 | 124.84 | 139.08 | 109.18 | 70.47 | 81.10 | 71.01 | Hypothetical protein |
| *PSPTO_2114* | | - | 2289022 | 2289612 | 12.27 | 22.64 | 14.99 | 1.17 | 4.87 | 3.37 | Hypothetical protein |
| *PSPTO_2115* | *vacJ* | + | 2289830 | 2290531 | 22.12 | 52.57 | 48.02 | 28.06 | 117.05 | 40.38 | VacJ lipoprotein |
| *PSPTO_2116* | | - | 2290619 | 2290918 | 80.40 | 187.22 | 171.35 | 56.25 | 762.23 | 94.02 | Hypothetical protein |
| *PSPTO_2117* | | + | 2291187 | 2292371 | 118.13 | 101.12 | 72.05 | 63.89 | 79.29 | 78.81 | Response regulator |
| *PSPTO_2118* | | + | 2292368 | 2292853 | 94.12 | 73.15 | 76.37 | 74.15 | 55.49 | 65.97 | Anti-anti-sigma factor |
| *PSPTO_2119* | *tal* | - | 2292928 | 2293857 | 45.64 | 170.78 | 151.92 | 62.31 | 169.02 | 78.27 | Transaldolase |
| *PSPTO_2120* | | + | 2294146 | 2295393 | 11.24 | 9.85 | 7.40 | 1.51 | 14.37 | 7.89 | Peptidase, M20/M25/M40 family |
| *PSPTO_2121* | | + | 2295503 | 2295946 | 12.60 | 43.12 | 17.85 | 10.10 | 23.64 | 12.97 | RulA protein |
| *PSPTO_2122* | | + | 2295980 | 2296273 | 31.01 | 59.03 | 53.70 | 26.23 | 95.89 | 28.46 | Hypothetical protein |
| *PSPTO_2123* | | + | 2296562 | 2297482 | 10.66 | 10.64 | 16.42 | 16.69 | 21.78 | 15.39 | Sensor histidine kinase |
| *PSPTO_2124* | | + | 2298115 | 2298372 | 200.95 | 169.32 | 150.15 | 62.28 | 41.03 | 73.07 | Hypothetical protein |
| *PSPTO_2125* | | - | 2298408 | 2298773 | 20.90 | 35.07 | 28.59 | 29.98 | 11.57 | 16.24 | Hypothetical protein |
| *PSPTO_2126* | | + | 2299104 | 2300114 | 5.32 | 17.67 | 15.06 | 9.13 | 28.65 | 15.82 | YjbN family TIM-barrel protein |
| *PSPTO_2127* | | + | 2300229 | 2300660 | 56.85 | 52.11 | 57.59 | 46.99 | 62.27 | 71.25 | Universal stress protein family |
| *PSPTO_2128* | | - | 2300687 | 2301151 | 12.43 | 145.75 | 110.87 | 25.82 | 31.89 | 35.28 | Response regulator |
| *PSPTO_2129* | | - | 2301148 | 2303388 | 7.12 | 55.29 | 32.90 | 12.69 | 14.42 | 12.86 | Sensory box histidine kinase/response regulator |
| *PSPTO_2130* | | - | 2303472 | 2304140 | 6.06 | 84.47 | 66.41 | 29.23 | 10.52 | 6.30 | LuxR family DNA-binding response regulator |
| *PSPTO_2131* | | - | 2304137 | 2305321 | 13.69 | 110.83 | 74.90 | 20.73 | 19.24 | 11.25 | Sensor histidine kinase |
| *PSPTO_2132* | | + | 2305545 | 2306069 | 8.72 | 3318.72 | 2777.22 | 519.94 | 9.83 | 20.30 | Hypothetical protein |
| *PSPTO_2133* | *pvdS* | - | 2306144 | 2306680 | 2.52 | 26.05 | 17.94 | 7.42 | 58.28 | 102.89 | RNA polymerase sigma-70 family protein |
| *PSPTO_2134* | *pvdG* | + | 2307009 | 2307755 | 0.74 | 0.00 | 0.00 | 0.00 | 14.32 | 21.42 | Pyoverdine synthetase, thioesterase component |
| *PSPTO_2135* | *pvsA, pvdL* | + | 2307860 | 2320870 | 2.17 | 0.84 | 1.48 | 1.02 | 5.12 | 11.87 | Pyoverdine chromophore precursor synthetase |
| *PSPTO_2136* | *daT, pvdH* | + | 2320980 | 2322446 | 1.60 | 1.82 | 1.08 | 1.87 | 16.59 | 42.36 | 2,4-diaminobutyrate 4-transaminase |
| *PSPTO_2137* | | + | 2322528 | 2322755 | 4.93 | 6.74 | 0.00 | 1.35 | 60.80 | 83.14 | MbtH-like protein |
| *PSPTO_2138* | | - | 2322875 | 2323804 | 2.29 | 0.86 | 2.23 | 0.00 | 21.12 | 16.80 | ABC transporter periplasmic substrate-binding protein |
| *PSPTO_2139* | | - | 2323805 | 2324704 | 0.61 | 0.00 | 0.00 | 0.00 | 7.68 | 4.74 | Cation ABC transporter permease |
| *PSPTO_2140* | | - | 2324704 | 2325456 | 1.42 | 0.99 | 4.08 | 1.94 | 12.70 | 12.20 | Cation ABC transporter ATP-binding protein |
| *PSPTO_2141* | | - | 2325453 | 2326406 | 4.26 | 2.88 | 4.18 | 1.21 | 44.08 | 35.98 | Cation ABC transporter substrate-binding protein |
| *PSPTO_2142* | | - | 2326421 | 2326993 | 1.03 | 0.82 | 5.62 | 4.36 | 33.97 | 26.41 | Hypothetical protein |
| *PSPTO_2143* | | - | 2326990 | 2327322 | 2.68 | 0.00 | 0.00 | 1.10 | 41.56 | 50.42 | Hypothetical protein |
| *PSPTO_2144* | | - | 2327319 | 2327861 | 1.34 | 1.49 | 1.45 | 0.00 | 36.51 | 57.04 | Hypothetical protein |
| *PSPTO_2145* | | - | 2327858 | 2329093 | 0.78 | 1.46 | 0.66 | 0.00 | 17.01 | 39.59 | Iron-regulated membrane protein |
| *PSPTO_2146* | | + | 2329386 | 2330363 | 3.64 | 4.12 | 2.68 | 0.00 | 20.35 | 25.96 | Pyoverdine biosynthesis regulatory gene |
| *PSPTO_2147* | | + | 2330649 | 2333963 | 2.82 | 4.62 | 4.85 | 2.55 | 14.95 | 22.80 | Pyoverdine sidechain peptide synthetase I, epsilon-Lys module |
| *PSPTO_2148* | | + | 2333960 | 2341837 | 1.37 | 1.54 | 1.78 | 1.16 | 5.98 | 13.74 | Pyoverdine sidechain peptide synthetase II, D-Asp-L-Thr component |
| *PSPTO_2149* | *pvdD* | + | 2341852 | 2348307 | 1.06 | 1.40 | 1.02 | 1.42 | 8.97 | 17.08 | Pyoverdine sidechain peptide synthetase III, L-Thr-L-Ser component |

| Locus name | Gene name | Strand | Gene start (bp) | Gene end (bp) | Average TPM Inoculum | Average TPM $t_6$ | Average TPM $t_{24}$ | Average TPM $t_{168}$ | Average TPM Logarithmic | Average TPM Stationary | Annotation description |
|---|---|---|---|---|---|---|---|---|---|---|---|
| *PSPTO_2150* | | + | 2348309 | 2356936 | 1.27 | 1.35 | 1.07 | 1.09 | 6.07 | 16.42 | Pyoverdine sidechain peptide synthetase IV, D-Asp-L-Ser component |
| *PSPTO_2151* | *fpvA* | - | 2357035 | 2359512 | 3.52 | 12.97 | 7.32 | 2.30 | 22.68 | 6.85 | TonB-dependent siderophore receptor |
| *PSPTO_2152* | | - | 2359772 | 2362003 | 3.17 | 3.36 | 2.41 | 2.13 | 42.45 | 65.23 | TonB-dependent siderophore receptor |
| *PSPTO_2153* | *pvdE* | - | 2362311 | 2363975 | 2.37 | 3.63 | 6.00 | 1.82 | 10.85 | 16.90 | Pyoverdine ABC transporter, ATP-binding/permease protein |
| *PSPTO_2154* | *pvdO* | - | 2364041 | 2364889 | 3.66 | 6.42 | 1.73 | 1.23 | 14.63 | 19.52 | Hypothetical protein |
| *PSPTO_2155* | *pvdN* | - | 2364968 | 2366251 | 1.71 | 0.77 | 0.80 | 1.87 | 19.33 | 24.59 | Class V aminotransferase |
| *PSPTO_2156* | *pvdM* | - | 2366294 | 2367637 | 1.96 | 5.18 | 2.68 | 0.14 | 25.87 | 33.06 | Renal dipeptidase family protein |
| *PSPTO_2158* | | - | 2369567 | 2370970 | 0.94 | 1.80 | 1.44 | 1.19 | 3.80 | 5.79 | Outer membrane efflux protein |
| *PSPTO_2159* | | - | 2370963 | 2372933 | 0.80 | 0.73 | 0.64 | 0.27 | 4.33 | 5.79 | Macrolide ABC efflux protein |
| *PSPTO_2160* | | - | 2372934 | 2374106 | 0.97 | 0.19 | 0.00 | 1.11 | 4.28 | 5.46 | RND family efflux transporter MFP subunit |
| *PSPTO_2161* | *pvdQ* | + | 2374310 | 2376631 | 1.92 | 1.64 | 3.12 | 1.06 | 7.23 | 11.46 | Penicillin amidase family protein |
| *PSPTO_2162* | | - | 2376685 | 2377179 | 13.98 | 135.94 | 88.87 | 75.07 | 59.30 | 73.26 | Hypothetical protein |
| *PSPTO_2163* | *aspC* | - | 2377145 | 2378341 | 27.07 | 31.42 | 19.67 | 12.15 | 80.86 | 33.05 | Aspartate aminotransferase |
| *PSPTO_2164* | *uvrB* | + | 2378720 | 2380735 | 6.30 | 21.44 | 18.86 | 6.93 | 35.59 | 27.41 | Excinuclease ABC subunit B |
| *PSPTO_2165* | | - | 2381162 | 2382529 | 3.20 | 5.46 | 3.04 | 4.74 | 3.31 | 3.42 | Group II intron, maturase |
| *PSPTO_2166* | *gltX* | + | 2383120 | 2384601 | 9.09 | 43.41 | 30.33 | 26.78 | 113.52 | 63.18 | Glutamyl-tRNA synthetase |
| *PSPTO_2167* | | + | 2385750 | 2386772 | 20.89 | 27.94 | 22.80 | 14.86 | 37.53 | 26.76 | ISPsy7, transposase |
| *PSPTO_2168* | | + | 2387378 | 2387839 | 23.58 | 16.20 | 18.20 | 14.60 | 20.40 | 14.61 | 4-hydroxybenzoyl-CoA thioesterase |
| *PSPTO_2169* | | + | 2387868 | 2388821 | 5.02 | 7.61 | 8.01 | 6.45 | 11.54 | 7.08 | Dihydrouridine synthase family protein |
| *PSPTO_2170* | | + | 2388952 | 2389395 | 718.64 | 786.08 | 408.13 | 468.98 | 821.02 | 993.49 | Heat shock protein, Hsp20 family |
| *PSPTO_2171* | | - | 2389452 | 2391644 | 41.15 | 19.70 | 23.29 | 12.53 | 18.03 | 22.03 | Sensory box/GGDEF domain/EAL domain-containing protein |
| *PSPTO_2172* | | - | 2391713 | 2392585 | 40.58 | 66.05 | 47.42 | 34.95 | 75.66 | 50.02 | LysR family transcriptional regulator |
| *PSPTO_2173* | *leuC* | + | 2392730 | 2394157 | 358.59 | 640.24 | 680.97 | 360.40 | 773.76 | 438.23 | Isopropylmalate isomerase large subunit |
| *PSPTO_2174* | *leuD* | + | 2394154 | 2394795 | 173.42 | 503.92 | 511.60 | 261.93 | 529.27 | 343.07 | Isopropylmalate isomerase small subunit |
| *PSPTO_2175* | *leuB* | + | 2394942 | 2396024 | 85.39 | 299.75 | 282.84 | 161.82 | 709.20 | 370.75 | 3-isopropylmalate dehydrogenase |
| *PSPTO_2176* | *asd-1* | + | 2396095 | 2397207 | 55.91 | 204.07 | 195.02 | 132.78 | 453.49 | 277.61 | Aspartate-semialdehyde dehydrogenase |
| *PSPTO_2177* | *dgoK* | + | 2397373 | 2398377 | 16.72 | 436.10 | 267.69 | 62.55 | 11.81 | 75.83 | 2-dehydro-3-deoxygalactonokinase |
| *PSPTO_2178* | *eda-2* | + | 2398379 | 2399014 | 12.50 | 278.55 | 181.98 | 30.53 | 15.82 | 21.58 | 2-dehydro-3-deoxy-6-phosphogalactonate aldolase |
| *PSPTO_2179* | | + | 2399075 | 2400223 | 9.19 | 264.35 | 186.17 | 33.40 | 35.85 | 17.77 | Galactonate dehydratase |
| *PSPTO_2180* | | + | 2400319 | 2401755 | 6.22 | 75.48 | 38.90 | 15.18 | 10.73 | 4.77 | MFS transporter, phthalate permease family |
| *PSPTO_2181* | | + | 2401837 | 2402646 | 12.04 | 42.18 | 39.71 | 10.48 | 32.17 | 19.42 | IclR family transcriptional regulator |
| *PSPTO_2182* | | - | 2402658 | 2403710 | 7.18 | 46.61 | 34.00 | 20.31 | 39.08 | 27.72 | Oxidoreductase zinc-binding protein |
| *PSPTO_2183* | | + | 2403892 | 2404785 | 16.88 | 14.19 | 14.80 | 11.29 | 31.42 | 22.74 | AraC family transcriptional regulator |
| *PSPTO_2184* | | - | 2404839 | 2406479 | - | - | - | - | - | - | |
| *PSPTO_2185* | *etfB-2* | + | 2407018 | 2407767 | 38.91 | 85.90 | 87.24 | 29.15 | 226.50 | 77.51 | Electron transfer flavoprotein subunit beta |
| *PSPTO_2186* | *etfA-2* | + | 2407768 | 2408697 | 13.64 | 27.12 | 40.32 | 21.71 | 99.93 | 34.83 | Electron transfer flavoprotein subunit alpha |
| *PSPTO_2187* | | + | 2408818 | 2409645 | 12.59 | 12.99 | 19.21 | 5.03 | 31.73 | 14.35 | Hypothetical protein |
| *PSPTO_2188* | | + | 2409656 | 2410009 | 8.25 | 4.25 | 8.55 | 0.31 | 29.74 | 11.81 | Lipoprotein |
| *PSPTO_2189* | | + | 2410006 | 2410821 | 6.49 | 13.05 | 16.53 | 12.04 | 31.58 | 11.27 | OmpA family protein |
| *PSPTO_2190* | | + | 2411039 | 2412478 | 13.77 | 18.63 | 17.44 | 4.02 | 30.95 | 15.26 | GntR family transcriptional regulator |
| *PSPTO_2191* | | + | 2412513 | 2413247 | 12.78 | 29.91 | 17.96 | 17.31 | 28.79 | 22.09 | Hypothetical protein |
| *PSPTO_2192* | | + | 2413342 | 2413599 | 22.54 | 34.54 | 19.53 | 42.22 | 30.72 | 42.07 | Hypothetical protein |
| *PSPTO_2193* | | + | 2413657 | 2414262 | 23.09 | 32.05 | 27.22 | 21.28 | 31.95 | 40.44 | Hypothetical protein |
| *PSPTO_2194* | *gltA* | - | 2414332 | 2415621 | 133.37 | 239.17 | 224.12 | 132.33 | 698.41 | 134.18 | Citrate synthase I |
| *PSPTO_2195* | *sdhC* | + | 2415985 | 2416359 | 58.33 | 179.26 | 134.96 | 101.34 | 295.99 | 118.76 | Succinate dehydrogenase, cytochrome b556 subunit |
| *PSPTO_2196* | | + | 2416353 | 2416721 | 61.57 | 180.41 | 116.28 | 73.86 | 383.26 | 125.23 | Succinate dehydrogenase, hydrophobic membrane anchor protein |
| *PSPTO_2197* | *sdhA* | + | 2416725 | 2418497 | 26.54 | 115.89 | 94.87 | 68.53 | 434.36 | 100.48 | Succinate dehydrogenase, flavoprotein subunit |
| *PSPTO_2198* | *sdhB* | + | 2418510 | 2419214 | 71.76 | 190.87 | 137.76 | 112.72 | 586.86 | 193.17 | Succinate dehydrogenase, iron-sulfur protein |
| *PSPTO_2199* | *sucA* | + | 2419474 | 2422305 | 32.35 | 98.03 | 79.78 | 49.31 | 353.36 | 75.55 | 2-oxoglutarate dehydrogenase E1 component |
| *PSPTO_2200* | *sucB* | + | 2422347 | 2423567 | 43.65 | 101.63 | 60.39 | 49.19 | 341.18 | 53.89 | Dihydrolipoamide succinyltransferase |
| *PSPTO_2201* | *lpdA* | + | 2423740 | 2425176 | 51.11 | 139.00 | 95.58 | 58.38 | 600.81 | 82.09 | Dihydrolipoamide dehydrogenase |
| *PSPTO_2202* | *sucC* | + | 2425385 | 2426551 | 54.68 | 147.40 | 128.17 | 59.32 | 1035.15 | 76.76 | Succinyl-CoA synthetase subunit beta |
| *PSPTO_2203* | *sucD* | + | 2426551 | 2427432 | 39.85 | 126.97 | 100.85 | 45.16 | 705.45 | 49.73 | Succinyl-CoA synthetase subunit alpha |
| *PSPTO_2204* | | - | 2427573 | 2429033 | 0.25 | 0.25 | 0.12 | 0.36 | 0.46 | 0.39 | ISPsy6, transposase |
| *PSPTO_2205* | | - | 2429130 | 2429711 | 1.56 | 0.00 | 2.37 | 1.12 | 2.01 | 0.68 | Hypothetical protein |
| *PSPTO_2206* | *brnQ* | + | 2430007 | 2431320 | 8.51 | 7.65 | 7.21 | 5.75 | 13.76 | 10.64 | Branched-chain amino acid transport system II carrier protein |
| *PSPTO_2207* | | - | 2431418 | 2431678 | 1352.80 | 1107.84 | 842.96 | 505.90 | 1141.81 | 1200.41 | Lipoprotein |
| *PSPTO_2208* | *htpG* | + | 2431992 | 2433899 | 71.22 | 108.89 | 82.92 | 37.80 | 651.34 | 113.32 | Heat shock protein HtpG |
| *PSPTO_2209* | | + | 2433990 | 2434718 | 11.42 | 47.51 | 33.96 | 14.27 | 31.38 | 16.79 | Dienelactone hydrolase family protein |
| *PSPTO_2210* | *fabB* | - | 2434906 | 2436129 | 61.31 | 58.26 | 45.97 | 24.07 | 231.84 | 35.69 | 3-oxoacyl-(acyl carrier protein) synthase I |
| *PSPTO_2211* | *fabA* | - | 2436140 | 2436655 | 144.19 | 75.84 | 74.25 | 36.66 | 225.20 | 39.33 | 3-hydroxydecanoyl-(acyl carrier protein) dehydratase |
| *PSPTO_2212* | | - | 2436876 | 2438801 | 10.05 | 4.40 | 5.22 | 5.97 | 10.72 | 6.79 | Sensor histidine kinase/response regulator |
| *PSPTO_2213* | *gpsA* | + | 2439122 | 2440147 | 37.34 | 51.31 | 41.43 | 14.82 | 50.59 | 34.23 | Glycerol-3-phosphate dehydrogenase (NAD(P)+) |
| *PSPTO_2214* | | + | 2440187 | 2440531 | 31.79 | 96.54 | 60.07 | 10.60 | 75.89 | 56.26 | Hypothetical protein |
| *PSPTO_2215* | *sixA* | + | 2440528 | 2440983 | 16.83 | 22.72 | 50.26 | 10.30 | 42.68 | 28.94 | Phosphohistidine phosphatase SixA |
| *PSPTO_2216* | | + | 2441051 | 2441494 | 6.53 | 11.11 | 14.29 | 12.77 | 23.08 | 11.54 | 4-hydroxybenzoyl-CoA thioesterase |
| *PSPTO_2217* | | + | 2441614 | 2442423 | 15.02 | 17.50 | 21.56 | 10.43 | 30.59 | 21.42 | Hypothetical protein |
| *PSPTO_2218* | | + | 2442420 | 2443274 | 13.26 | 10.71 | 8.09 | 1.07 | 16.14 | 12.27 | Alpha/beta fold family hydrolase |
| *PSPTO_2219* | | + | 2443350 | 2443514 | 13.31 | 20.23 | 46.55 | 24.11 | 50.71 | 16.18 | Hypothetical protein |
| *PSPTO_2220* | | + | 2443525 | 2444244 | 3.32 | 10.63 | 7.73 | 4.37 | 12.94 | 3.60 | Hypothetical protein |
| *PSPTO_2221* | | + | 2444345 | 2445394 | 22.12 | 19.45 | 15.56 | 10.29 | 31.02 | 32.17 | Lipoprotein |
| *PSPTO_2222* | | - | 2445595 | 2446629 | 16.92 | 38.01 | 25.38 | 9.67 | 28.37 | 25.00 | Sensor histidine kinase |
| *PSPTO_2223* | | - | 2446626 | 2447318 | 13.43 | 15.00 | 24.62 | 9.92 | 12.69 | 14.91 | DNA-binding response regulator |
| *PSPTO_2224* | | + | 2447757 | 2449760 | 12.56 | 14.15 | 8.02 | 6.04 | 9.72 | 16.53 | Hypothetical protein |
| *PSPTO_2225* | | + | 2450057 | 2452378 | 4.89 | 6.38 | 4.77 | 3.55 | 6.03 | 7.79 | Autotransporter |
| *PSPTO_2226* | | + | 2452514 | 2454061 | 15.09 | 10.06 | 8.62 | 5.05 | 11.36 | 7.94 | Hypothetical protein |
| *PSPTO_2227* | | - | 2454099 | 2455211 | 20.12 | 21.83 | 25.45 | 21.31 | 32.10 | 23.38 | Fimbrial protein |
| *PSPTO_2228* | *fimD* | - | 2455258 | 2457771 | 9.38 | 8.56 | 6.25 | 3.81 | 9.27 | 10.77 | Outer membrane usher protein fimD |
| *PSPTO_2229* | *papD* | - | 2457811 | 2458566 | 5.32 | 3.87 | 1.74 | 5.57 | 6.63 | 8.02 | Chaperone protein PapD |

| Locus name | Gene name | Strand | Gene start (bp) | Gene end (bp) | Average TPM Inoculum | Average TPM $t_6$ | Average TPM $t_{24}$ | Average TPM $t_{168}$ | Average TPM Logarithmic | Average TPM Stationary | Annotation description |
|---|---|---|---|---|---|---|---|---|---|---|---|
| *PSPTO_2230* | *fimA* | - | 2458675 | 2459226 | 8.96 | 7.00 | 26.99 | 5.00 | 14.31 | 11.21 | Type I pilus biogensis protein FimA |
| *PSPTO_2231* | | - | 2459655 | 2462603 | 16.68 | 12.40 | 11.40 | 11.53 | 19.74 | 14.23 | YD repeat protein |
| *PSPTO_2232* | | + | 2464986 | 2466809 | 10.71 | 8.44 | 5.97 | 2.98 | 10.09 | 12.75 | Hypothetical protein |
| *PSPTO_2233* | | + | 2466929 | 2468260 | 18.49 | 21.48 | 15.59 | 17.52 | 21.12 | 19.81 | Hypothetical protein |
| *PSPTO_2234* | | - | 2468395 | 2469372 | 0.00 | 0.00 | 0.00 | 0.00 | 0.00 | 0.00 | ISPssy, transposase |
| *PSPTO_2235* | | + | 2469471 | 2470577 | 3.69 | 5.17 | 5.54 | 3.24 | 7.53 | 6.52 | Hypothetical protein |
| *PSPTO_2236* | | + | 2470632 | 2472905 | 3.72 | 1.37 | 4.77 | 2.14 | 2.82 | 2.89 | Hypothetical protein |
| *PSPTO_2237* | | + | 2472944 | 2475934 | 25.25 | 19.01 | 10.08 | 6.86 | 14.66 | 19.24 | Hypothetical protein |
| *PSPTO_2238* | | + | 2475931 | 2476416 | 1.50 | 10.31 | 3.86 | 0.19 | 3.35 | 2.50 | Lipoprotein |
| *PSPTO_2239* | | + | 2476507 | 2481195 | 16.70 | 15.03 | 15.42 | 12.91 | 25.95 | 22.88 | YD repeat protein |
| *PSPTO_2240* | | - | 2481297 | 2481560 | 3.01 | 9.62 | 11.36 | 0.00 | 3.72 | 2.64 | LuxR family transcriptional regulator |
| *PSPTO_2241* | | + | 2481789 | 2481899 | 8.97 | 2.41 | 6.61 | 0.00 | 6.60 | 6.66 | Hypothetical protein |
| *PSPTO_2242* | *kdpA* | + | 2481902 | 2483596 | 0.87 | 0.00 | 1.84 | 0.66 | 1.38 | 1.29 | Potassium-transporting ATPase subunit A |
| *PSPTO_2243* | *kdpB* | + | 2483607 | 2485685 | 1.11 | 3.44 | 2.99 | 1.77 | 1.82 | 0.94 | Potassium-transporting ATPase subunit B |
| *PSPTO_2244* | *kdpC* | + | 2485740 | 2486312 | 1.55 | 8.45 | 0.78 | 9.33 | 1.95 | 1.09 | Potassium-transporting ATPase subunit C |
| *PSPTO_2245* | *kdpD* | + | 2486379 | 2489042 | 8.17 | 8.50 | 7.80 | 6.23 | 12.40 | 11.92 | Sensor protein KdpD |
| *PSPTO_2246* | *kdpE* | + | 2489039 | 2489731 | 18.85 | 37.90 | 38.56 | 16.92 | 34.37 | 25.94 | KDP operon transcriptional regulatory protein kdpE |
| *PSPTO_2247* | | + | 2489838 | 2490116 | 504.85 | 119.66 | 165.49 | 97.48 | 131.81 | 175.62 | Lipoprotein |
| *PSPTO_2248* | | + | 2490454 | 2491371 | 12.36 | 12.28 | 8.56 | 14.07 | 19.09 | 12.10 | MoxR protein |
| *PSPTO_2249* | | + | 2491341 | 2492324 | 5.89 | 10.20 | 4.25 | 0.00 | 6.45 | 3.73 | Hypothetical protein |
| *PSPTO_2250* | | + | 2492420 | 2494345 | 6.33 | 9.23 | 10.57 | 5.16 | 8.56 | 7.69 | Transglutaminase-like domain protein |
| *PSPTO_2251* | | + | 2494370 | 2495131 | 13.50 | 31.36 | 28.13 | 8.40 | 24.13 | 26.61 | Hypothetical protein |
| *PSPTO_2252* | | + | 2495289 | 2496086 | - | - | - | - | - | - | |
| *PSPTO_2253* | | - | 2496096 | 2496341 | 8.88 | 108.78 | 94.88 | 14.67 | 6.32 | 13.21 | Hypothetical protein |
| *PSPTO_2254* | | - | 2496461 | 2497936 | 2.17 | 3.34 | 3.19 | 2.02 | 13.45 | 28.88 | Methyl-accepting chemotaxis protein |
| *PSPTO_2255* | | - | 2498021 | 2498857 | 36.96 | 25.71 | 27.36 | 5.29 | 24.24 | 19.83 | TatD family hydrolase |
| *PSPTO_2256* | | + | 2499048 | 2500469 | 17.33 | 19.07 | 20.99 | 10.65 | 13.43 | 17.48 | Transglycosylase |
| *PSPTO_2257* | | + | 2500576 | 2501010 | 12.26 | 21.37 | 16.84 | 19.88 | 27.82 | 9.88 | DoxD-like family protein |
| *PSPTO_2258* | | + | 2501132 | 2501576 | - | - | - | - | - | - | |
| *PSPTO_2259* | | + | 2501881 | 2504172 | 13.35 | 10.90 | 9.43 | 8.87 | 10.35 | 11.16 | Sigma-54 dependent transcriptional regulator |
| *PSPTO_2260* | | + | 2504194 | 2505423 | 8.78 | 8.19 | 8.85 | 0.93 | 11.67 | 9.41 | Hypothetical protein |
| *PSPTO_2261* | | + | 2505420 | 2506136 | 27.91 | 22.14 | 21.00 | 10.81 | 22.56 | 21.79 | Racemase |
| *PSPTO_2262* | | - | 2506204 | 2507247 | 8.57 | 9.86 | 18.54 | 10.94 | 6.69 | 8.93 | Hypothetical protein |
| *PSPTO_2263* | | + | 2507215 | 2507652 | 2.88 | 4.98 | 4.99 | 4.42 | 3.56 | 4.17 | Hypothetical protein |
| *PSPTO_2264* | | + | 2508087 | 2508416 | 38.22 | 100.60 | 82.64 | 36.84 | 131.59 | 69.96 | Hypothetical protein |
| *PSPTO_2265* | *greB-1* | + | 2508463 | 2508936 | 22.27 | 47.62 | 43.43 | 36.91 | 44.39 | 43.50 | Transcription elongation factor GreB |
| *PSPTO_2266* | | - | 2509214 | 2511730 | 3.73 | 12.47 | 9.55 | 4.96 | 26.84 | 6.97 | Permease |
| *PSPTO_2267* | | - | 2511730 | 2512413 | 15.47 | 28.79 | 31.05 | 12.10 | 43.13 | 11.22 | ABC transporter ATP-binding protein |
| *PSPTO_2268* | | + | 2512424 | 2513029 | 13.04 | 40.15 | 28.80 | 10.07 | 63.28 | 17.59 | Acyl-CoA thioesterase I |
| *PSPTO_2269* | | + | 2513086 | 2513367 | 15.31 | 15.23 | 26.37 | 6.77 | 25.25 | 13.69 | Hypothetical protein |
| *PSPTO_2270* | | + | 2513476 | 2514441 | 29.63 | 48.29 | 46.63 | 25.39 | 142.55 | 68.03 | ErfK/YbiS/YcfS/YnhG family protein |
| *PSPTO_2271* | | - | 2514550 | 2514663 | 8.83 | 1.59 | 8.66 | 2.14 | 6.94 | 3.33 | Hypothetical protein |
| *PSPTO_2272* | *oprI* | - | 2514998 | 2515249 | 154.83 | 992.44 | 791.02 | 388.30 | 3687.69 | 394.85 | Outer membrane lipoprotein OprI |
| *PSPTO_2273* | | + | 2515482 | 2515655 | 118.23 | 113.16 | 82.21 | 101.28 | 88.80 | 88.80 | Hypothetical protein |
| *PSPTO_2274* | | + | 2515722 | 2516009 | 54.72 | 145.97 | 86.90 | 61.59 | 85.46 | 28.16 | Hypothetical protein |
| *PSPTO_2275* | *aroF-1* | - | 2516096 | 2517172 | 5.99 | 34.09 | 23.06 | 21.67 | 76.95 | 18.08 | Phospho-2-dehydro-3-deoxyheptonate aldolase |
| *PSPTO_2276* | *panE-1* | + | 2517416 | 2518387 | 5.90 | 15.01 | 8.54 | 19.96 | 17.61 | 7.69 | 2-dehydropantoate 2-reductase |
| *PSPTO_2277* | | + | 2518391 | 2519311 | 3.00 | 11.72 | 7.38 | 4.62 | 14.78 | 4.80 | 5'-nucleotidase |
| *PSPTO_2278* | | + | 2519390 | 2519893 | 52.04 | 39.23 | 36.47 | 21.22 | 16.30 | 32.44 | Universal stress protein family |
| *PSPTO_2279* | *cysB* | + | 2520059 | 2521033 | 15.48 | 35.37 | 30.62 | 10.76 | 74.46 | 38.73 | Cys regulon transcriptional activator |
| *PSPTO_2280* | *cysH* | - | 2521178 | 2521912 | 27.68 | 77.18 | 56.71 | 25.31 | 75.78 | 37.39 | Phosphoadenosine phosphosulfate reductase |
| *PSPTO_2281* | | + | 2521957 | 2522670 | 9.52 | 21.89 | 13.90 | 16.89 | 27.24 | 11.09 | Phosphoserine phosphatase |
| *PSPTO_2282* | *pabB* | - | 2522822 | 2524165 | 4.11 | 8.55 | 5.28 | 6.45 | 17.63 | 6.76 | Para-aminobenzoate synthase, component I |
| *PSPTO_2283* | | - | 2524255 | 2525235 | 7.53 | 11.30 | 5.94 | 2.53 | 33.16 | 15.02 | Hypothetical protein |
| *PSPTO_2284* | | - | 2525232 | 2526773 | 6.25 | 9.57 | 6.12 | 2.26 | 34.07 | 16.97 | Membrane protein |
| *PSPTO_2285* | | - | 2526778 | 2527329 | 34.46 | 44.29 | 22.21 | 17.77 | 66.96 | 49.58 | Hypothetical protein |
| *PSPTO_2286* | | + | 2527650 | 2528348 | 367.03 | 52.54 | 49.17 | 16.96 | 366.97 | 16.97 | GntR family transcriptional regulator |
| *PSPTO_2287* | *prpB* | + | 2528345 | 2529238 | 93.45 | 15.30 | 22.90 | 7.52 | 211.80 | 8.73 | Methylisocitrate lyase |
| *PSPTO_2288* | *prpC* | + | 2529613 | 2530740 | 89.10 | 25.74 | 37.18 | 21.48 | 316.49 | 11.34 | 2-methylcitrate synthase |
| *PSPTO_2289* | | + | 2530833 | 2533424 | 18.23 | 4.33 | 9.38 | 4.05 | 132.95 | 3.82 | Aconitate hydratase |
| *PSPTO_2290* | | + | 2533545 | 2534735 | 7.50 | 7.93 | 4.70 | 1.79 | 78.36 | 6.08 | Hypothetical protein |
| *PSPTO_2291* | | - | 2534741 | 2535559 | 42.21 | 23.12 | 36.03 | 14.72 | 50.60 | 27.58 | Hypothetical protein |
| *PSPTO_2292* | *ppsA* | + | 2535753 | 2538128 | 44.21 | 59.40 | 47.23 | 28.24 | 130.30 | 81.74 | Phosphoenolpyruvate synthase |
| *PSPTO_2293* | | + | 2538321 | 2539313 | 3.56 | 1.05 | 3.55 | 1.12 | 18.21 | 7.37 | Esterase/lipase/thioesterase family protein |
| *PSPTO_2294* | *rraA* | + | 2539402 | 2539890 | 29.37 | 64.98 | 41.61 | 18.13 | 68.63 | 31.77 | Ribonuclease activity regulator protein RraA |
| *PSPTO_2295* | *cmaX* | + | 2539902 | 2540897 | 33.01 | 49.78 | 50.75 | 28.62 | 66.80 | 52.09 | CmaX protein |
| *PSPTO_2296* | *crfX* | + | 2540983 | 2541231 | 283.22 | 73.90 | 61.18 | 70.81 | 78.40 | 84.36 | CrfX protein |
| *PSPTO_2297* | | + | 2541234 | 2542058 | 175.64 | 162.22 | 127.02 | 94.68 | 258.28 | 168.21 | Membrane protein |
| *PSPTO_2298* | *sigX* | + | 2542234 | 2542731 | 2034.01 | 2572.54 | 2185.11 | 1329.43 | 2336.89 | 1033.23 | RNA polymerase sigma factor SigX |
| *PSPTO_2299* | *oprF* | + | 2542838 | 2543872 | 1225.77 | 3344.61 | 2545.61 | 1394.69 | 7844.81 | 1405.71 | Outer membrane porin OprF |
| *PSPTO_2300* | *cobA* | - | 2543960 | 2544721 | 19.14 | 23.36 | 8.81 | 3.03 | 2.51 | 2.51 | Uroporphyrin-III C-methyltransferase |
| *PSPTO_2301* | *nasA* | - | 2544732 | 2547458 | 1.67 | 14.29 | 10.24 | 7.87 | 0.87 | 1.11 | Nitrate reductase |
| *PSPTO_2302* | | - | 2547449 | 2548678 | 3.21 | 11.03 | 6.24 | 7.03 | 1.82 | 2.37 | Assimilatory nitrate reductase electron transfer subunit domain protein |
| *PSPTO_2303* | | - | 2548961 | 2550550 | 1.66 | 11.51 | 7.91 | 6.49 | 1.11 | 1.29 | Serine/threonine protein kinase |
| *PSPTO_2304* | | - | 2550630 | 2551841 | 5.24 | 29.57 | 19.27 | 12.50 | 1.99 | 2.21 | Nitrate transporter |
| *PSPTO_2305* | *lsc-2* | + | 2552817 | 2554064 | 3.21 | 9.97 | 15.02 | 18.86 | 4.79 | 4.78 | Levansucrase |
| *PSPTO_2306* | *nasT* | - | 2554122 | 2554697 | 5.39 | 8.89 | 11.97 | 0.23 | 3.92 | 3.30 | Response regulator NasT |
| *PSPTO_2307* | | - | 2554712 | 2555923 | 9.83 | 22.43 | 13.69 | 10.72 | 8.30 | 3.97 | Hypothetical protein |
| *PSPTO_2308* | | - | 2555920 | 2556117 | 20.69 | 8.94 | 26.80 | 0.00 | 12.57 | 5.34 | Hypothetical protein |

| Locus name | Gene name | Strand | Gene start (bp) | Gene end (bp) | Average TPM Inoculum | Average TPM $t_6$ | Average TPM $t_{24}$ | Average TPM $t_{168}$ | Average TPM Logarithmic | Average TPM Stationary | Annotation description |
|---|---|---|---|---|---|---|---|---|---|---|---|
| *PSPTO_2309* | *pyrD* | + | 2556168 | 2557199 | 16.78 | 22.20 | 25.05 | 26.68 | 52.50 | 25.34 | Dihydroorotate dehydrogenase |
| *PSPTO_2310* | *rmf* | - | 2557345 | 2557560 | 1706.10 | 6211.12 | 6053.82 | 4282.51 | 369.40 | 1460.00 | Ribosome modulation factor-related protein |
| *PSPTO_2311* | | + | 2558146 | 2560434 | 5.75 | 15.34 | 17.68 | 9.66 | 26.37 | 10.31 | 23S rRNA m(2)G2445 methyltransferase |
| *PSPTO_2312* | | + | 2560473 | 2561996 | 6.52 | 9.66 | 6.47 | 6.63 | 13.25 | 5.31 | CHASE domain/PAS domain protein |
| *PSPTO_2313* | | + | 2561986 | 2562957 | 4.92 | 8.61 | 4.85 | 3.90 | 13.20 | 4.03 | Sensory box/GGDEF domain protein |
| *PSPTO_2314* | *dacB* | - | 2563001 | 2564467 | 25.47 | 23.59 | 32.70 | 16.81 | 37.28 | 28.35 | D-alanyl-D-alanine carboxypeptidase |
| *PSPTO_2315* | | + | 2564794 | 2565138 | 22.73 | 47.40 | 65.81 | 42.07 | 147.02 | 18.28 | Hypothetical protein |
| *PSPTO_2316* | | - | 2565244 | 2566677 | 6.25 | 8.25 | 8.29 | 2.77 | 4.61 | 7.77 | Sensor histidine kinase/response regulator |
| *PSPTO_2317* | | + | 2566716 | 2568320 | 5.92 | 21.64 | 18.98 | 5.65 | 15.29 | 14.64 | Hypothetical protein |
| *PSPTO_2318* | | + | 2568386 | 2568700 | 3.15 | 20.02 | 24.12 | 16.60 | 40.03 | 23.86 | Hypothetical protein |
| *PSPTO_2319* | | - | 2569753 | 2570166 | 14.71 | 0.00 | 4.57 | 4.91 | 5.88 | 4.03 | Resolvase |
| *PSPTO_2320* | | - | 2571887 | 2573140 | 0.54 | 0.70 | 2.35 | 1.61 | 2.06 | 1.30 | Hypothetical protein |
| *PSPTO_2321* | | + | 2573654 | 2574676 | 0.00 | 0.00 | 0.00 | 0.00 | 0.00 | 0.00 | ISPsy4, transposase |
| *PSPTO_2322* | | + | 2574673 | 2575482 | 0.00 | 0.00 | 0.00 | 0.00 | 0.00 | 0.00 | ISPsy4, transposition helper protein |
| *PSPTO_2323* | | - | 2575936 | 2577726 | 3.93 | 4.93 | 4.66 | 2.46 | 4.17 | 4.43 | Hypothetical protein |
| *PSPTO_2324* | | - | 2577833 | 2578810 | 0.00 | 0.00 | 0.00 | 0.00 | 0.00 | 0.00 | ISPssy, transposase |
| *PSPTO_2325* | | - | 2580097 | 2580357 | 1.91 | 3.28 | 4.52 | 0.00 | 9.85 | 3.10 | Hypothetical protein |
| *PSPTO_2326* | | - | 2581120 | 2583375 | 5.53 | 5.16 | 5.83 | 3.49 | 24.93 | 16.05 | Extracellular solute-binding protein/sensory box protein |
| *PSPTO_2327* | | - | 2583384 | 2583497 | - | - | - | - | - | - | |
| *PSPTO_2328* | | - | 2583651 | 2584628 | 0.00 | 0.00 | 0.00 | 0.00 | 0.00 | 0.00 | ISPssy, transposase |
| *PSPTO_2329* | | - | 2584669 | 2585202 | - | - | - | - | - | - | |
| *PSPTO_2330* | | - | 2585606 | 2585998 | 9.47 | 12.26 | 18.66 | 13.78 | 9.33 | 9.75 | Response regulator |
| *PSPTO_2331* | | - | 2586250 | 2587581 | 20.18 | 52.80 | 40.14 | 17.12 | 76.68 | 28.92 | Deoxyguanosinetriphosphate triphosphohydrolase-like protein |
| *PSPTO_2332* | | + | 2587685 | 2588152 | 14.15 | 70.35 | 75.24 | 33.00 | 47.42 | 26.14 | Hypothetical protein |
| *PSPTO_2333* | | - | 2588101 | 2588493 | 24.46 | 117.76 | 126.68 | 61.43 | 219.86 | 58.57 | Hypothetical protein |
| *PSPTO_2334* | | - | 2588498 | 2588812 | 127.66 | 515.65 | 376.67 | 166.36 | 630.00 | 176.05 | Hypothetical protein |
| *PSPTO_2335* | *exoZ* | + | 2589210 | 2590280 | 12.32 | 8.57 | 6.97 | 5.92 | 8.50 | 8.24 | Exopolysaccharide production protein ExoZ |
| *PSPTO_2336* | | + | 2590311 | 2590601 | 87.00 | 126.04 | 83.08 | 62.47 | 149.47 | 156.18 | Hypothetical protein |
| *PSPTO_2337* | *pcaQ* | + | 2590640 | 2591569 | 11.43 | 11.86 | 13.44 | 9.05 | 20.53 | 19.37 | Pca operon transcriptional activator PcaQ |
| *PSPTO_2338* | *pcaH* | + | 2591697 | 2592416 | 25.40 | 38.53 | 46.26 | 21.40 | 52.21 | 9.66 | Protocatechuate 3,4-dioxygenase subunit beta |
| *PSPTO_2339* | *pcaG* | + | 2592428 | 2593030 | 18.84 | 43.59 | 51.25 | 20.70 | 59.20 | 10.49 | Protocatechuate 3,4-dioxygenase subunit alpha |
| *PSPTO_2340* | *pcaK* | + | 2593098 | 2594447 | 9.85 | 16.10 | 11.86 | 11.47 | 25.32 | 5.05 | 4-hydroxybenzoate transporter |
| *PSPTO_2341* | *pcaB* | + | 2594472 | 2595833 | 8.57 | 24.04 | 11.99 | 10.20 | 34.32 | 6.20 | 3-carboxy-cis,cis-muconate cycloisomerase |
| *PSPTO_2342* | *pcaC-1* | + | 2595830 | 2596228 | 12.97 | 22.58 | 22.74 | 24.33 | 35.89 | 5.19 | 4-carboxymuconolactone decarboxylase |
| *PSPTO_2343* | | - | 2596721 | 2597977 | 20.60 | 17.22 | 7.98 | 2.55 | 13.14 | 7.84 | Outer membrane porin, OprD family |
| *PSPTO_2344* | | + | 2598221 | 2599087 | 6.50 | 10.71 | 6.15 | 1.16 | 15.32 | 6.52 | Membrane protein |
| *PSPTO_2345* | | - | 2599195 | 2600514 | 8.22 | 11.75 | 8.98 | 4.20 | 9.38 | 7.59 | Transporter |
| *PSPTO_2346* | | + | 2601132 | 2603039 | 8.21 | 15.15 | 16.17 | 4.69 | 9.53 | 12.39 | 4-hydroxyphenylpyruvate dioxygenase |
| *PSPTO_2347* | | + | 2603144 | 2603815 | 8.70 | 9.67 | 9.53 | 7.27 | 21.08 | 9.57 | TetR family transcriptional regulator |
| *PSPTO_2348* | | - | 2603819 | 2604673 | 3.83 | 4.88 | 5.49 | 0.66 | 13.48 | 5.00 | Shikimate 5-dehydrogenase |
| *PSPTO_2349* | | + | 2604859 | 2605749 | 12.24 | 13.90 | 17.42 | 8.94 | 19.54 | 13.57 | RNA pseudouridine synthase family protein |
| *PSPTO_2350* | | - | 2605826 | 2606050 | 1467.64 | 1012.34 | 792.26 | 319.11 | 532.05 | 478.79 | Lipoprotein |
| *PSPTO_2351* | *mobA* | - | 2606146 | 2606751 | 53.74 | 168.62 | 152.01 | 82.15 | 554.29 | 53.61 | Molybdopterin-guanine dinucleotide biosynthesis protein MobA |
| *PSPTO_2352* | *moaB* | + | 2606827 | 2607366 | 11.61 | 40.80 | 26.82 | 14.54 | 52.01 | 27.04 | Molybdenum cofactor biosynthesis protein B |
| *PSPTO_2353* | *moeA* | + | 2607373 | 2608584 | 3.15 | 24.73 | 14.99 | 7.70 | 30.32 | 11.10 | Molybdopterin biosynthesis protein MoeA |
| *PSPTO_2354* | | + | 2608674 | 2609600 | 20.52 | 23.43 | 18.39 | 10.50 | 17.48 | 21.50 | Glycosyl transferase family protein |
| *PSPTO_2355* | | - | 2609648 | 2611456 | 3.00 | 4.71 | 6.76 | 3.66 | 9.10 | 6.18 | Glutathione-regulated potassium-efflux system protein |
| *PSPTO_2356* | | - | 2611521 | 2611904 | 48.02 | 49.67 | 53.90 | 31.52 | 88.01 | 56.23 | Hypothetical protein |
| *PSPTO_2358* | | - | 2612511 | 2613539 | 34.86 | 45.63 | 48.68 | 20.76 | 44.59 | 50.57 | Hypothetical protein |
| *PSPTO_2359* | *allD* | + | 2613769 | 2614800 | 2.23 | 3.61 | 7.56 | 2.85 | 7.41 | 7.39 | Ureidoglycolate dehydrogenase |
| *PSPTO_2360* | | + | 2614830 | 2615099 | 29.64 | 95.05 | 66.83 | 29.00 | 27.97 | 22.61 | Hypothetical protein |
| *PSPTO_2361* | | + | 2615212 | 2615442 | 38.29 | 104.22 | 77.94 | 11.04 | 78.12 | 77.87 | Hypothetical protein |
| *PSPTO_2362* | *map-2* | + | 2615439 | 2616230 | 14.04 | 35.77 | 26.66 | 8.38 | 49.22 | 46.30 | Methionine aminopeptidase |
| *PSPTO_2363* | | - | 2616307 | 2616546 | 10.66 | 8.15 | 22.00 | 0.00 | 15.51 | 7.22 | Hypothetical protein |
| *PSPTO_2364* | *endA* | - | 2616752 | 2617453 | 1.48 | 9.36 | 6.03 | 0.72 | 31.11 | 6.22 | Endonuclease I |
| *PSPTO_2365* | | - | 2617458 | 2617709 | 18.00 | 47.83 | 67.34 | 33.85 | 147.71 | 31.58 | Hypothetical protein |
| *PSPTO_2366* | | + | 2617846 | 2618070 | 145.51 | 233.51 | 186.76 | 95.71 | 315.36 | 149.29 | Lipoprotein |
| *PSPTO_2367* | *rbsB-1* | + | 2618254 | 2619213 | 18.82 | 47.02 | 66.72 | 40.39 | 73.44 | 21.66 | Ribose ABC transporter periplasmic ribose-binding protein |
| *PSPTO_2368* | *rbsA-1* | + | 2619267 | 2620820 | 2.52 | 3.56 | 9.97 | 4.97 | 62.79 | 1.86 | Ribose ABC transporter ATP-binding protein |
| *PSPTO_2369* | *rbsC-1* | + | 2620817 | 2621809 | 2.20 | 5.32 | 3.13 | 0.06 | 82.22 | 1.92 | Ribose ABC transporter permease |
| *PSPTO_2370* | *rbsR* | + | 2621811 | 2622827 | 2.24 | 9.30 | 6.75 | 2.97 | 123.10 | 3.07 | Ribose operon repressor |
| *PSPTO_2371* | *rbsK* | + | 2622858 | 2623778 | 17.96 | 26.43 | 10.20 | 11.77 | 114.68 | 9.91 | Ribokinase |
| *PSPTO_2372* | | + | 2623775 | 2624179 | 24.51 | 43.48 | 29.32 | 13.79 | 175.33 | 19.75 | Ribose ABC transporter protein |
| *PSPTO_2373* | | + | 2624219 | 2625244 | 23.73 | 49.23 | 41.48 | 18.22 | 163.03 | 27.66 | Inosine/uridine-preferring nucleoside hydrolase |
| *PSPTO_2374* | | - | 2625268 | 2625723 | 31.88 | 78.96 | 48.08 | 8.69 | 23.52 | 8.69 | Hypothetical protein |
| *PSPTO_2375* | | + | 2625780 | 2626115 | 26.94 | 97.98 | 68.52 | 29.92 | 66.03 | 29.83 | Lipoprotein |
| *PSPTO_2376* | | - | 2626181 | 2626393 | 181.01 | 435.08 | 291.03 | 232.28 | 1140.94 | 34.84 | Cold shock domain family protein |
| *PSPTO_2377* | | - | 2626699 | 2627004 | 191.45 | 166.04 | 123.54 | 67.16 | 158.27 | 203.77 | Hypothetical protein |
| *PSPTO_2378* | *thrS* | + | 2627367 | 2629289 | 208.17 | 362.88 | 307.25 | 223.39 | 777.55 | 293.27 | Threonyl-tRNA synthetase |
| *PSPTO_2379* | *infC* | + | 2629307 | 2629840 | 625.00 | 1678.40 | 1517.70 | 1032.90 | 2819.49 | 778.96 | Translation initiation factor IF-3 |
| *PSPTO_2380* | *rpmI* | + | 2629901 | 2630095 | 97.66 | 311.28 | 267.71 | 289.34 | 597.05 | 108.23 | 50S ribosomal protein L35 |
| *PSPTO_2381* | *rplT* | + | 2630125 | 2630481 | 28.69 | 104.37 | 115.19 | 78.36 | 111.07 | 23.19 | 50S ribosomal protein L20 |
| *PSPTO_2382* | *pheS* | + | 2630585 | 2631601 | 7.54 | 37.04 | 21.37 | 5.38 | 88.10 | 13.72 | Phenylalanyl-tRNA synthetase subunit alpha |
| *PSPTO_2383* | *pheT* | + | 2631630 | 2634008 | 57.10 | 45.18 | 31.00 | 21.07 | 87.02 | 29.29 | Phenylalanyl-tRNA synthetase subunit beta |
| *PSPTO_2384* | *ihfA* | + | 2634012 | 2634314 | 333.46 | 282.92 | 234.91 | 90.91 | 200.27 | 137.08 | Integration host factor subunit alpha |
| *PSPTO_2385* | | + | 2634295 | 2634651 | 210.90 | 147.82 | 158.25 | 68.65 | 115.83 | 66.46 | Transcriptional regulator |
| *PSPTO_2386* | | - | 2635219 | 2636106 | 8.34 | 0.97 | 3.52 | 2.54 | 6.58 | 15.41 | C-5 cytosine-specific DNA methylase family protein |
| *PSPTO_2387* | | + | 2636500 | 2637522 | 0.00 | 0.00 | 0.00 | 0.00 | 0.00 | 0.00 | ISPsy4, transposase |
| *PSPTO_2388* | | + | 2637519 | 2638328 | 0.00 | 0.00 | 0.00 | 0.00 | 0.00 | 0.00 | ISPsy4, transposition helper protein |

| Locus name | Gene name | Strand | Gene start (bp) | Gene end (bp) | Average TPM Inoculum | Average TPM $t_{6}$ | Average TPM $t_{24}$ | Average TPM $t_{168}$ | Average TPM Logarithmic | Average TPM Stationary | Annotation description |
|---|---|---|---|---|---|---|---|---|---|---|---|
| *PSPTO_2389* | | - | 2638723 | 2639505 | 0.91 | 5.17 | 2.91 | 1.64 | 3.45 | 2.18 | Hypothetical protein |
| *PSPTO_2390* | | + | 2639556 | 2640326 | 10.13 | 23.08 | 12.63 | 10.36 | 39.39 | 17.29 | Hypothetical protein |
| *PSPTO_2391* | | - | 2640619 | 2640939 | 5.19 | 10.28 | 5.57 | 0.00 | 6.78 | 7.27 | Hypothetical protein |
| *PSPTO_2392* | | + | 2640996 | 2641355 | 3.74 | 3.60 | 3.92 | 3.22 | 2.30 | 3.18 | ISPsy5, Orf1 |
| *PSPTO_2394* | | - | 2643148 | 2644068 | 13.69 | 34.82 | 31.10 | 27.38 | 38.02 | 40.46 | Radical SAM domain-containing protein |
| *PSPTO_2395* | | - | 2644361 | 2645314 | 12.18 | 26.99 | 24.14 | 9.47 | 36.84 | 19.35 | Transcriptional regulator |
| *PSPTO_2396* | | - | 2645396 | 2646196 | 31.93 | 42.87 | 32.12 | 8.51 | 10.58 | 12.11 | Short chain dehydrogenase/reductase family oxidoreductase |
| *PSPTO_2397* | | - | 2646227 | 2646991 | 6.98 | 39.51 | 29.31 | 4.55 | 4.23 | 2.08 | Short chain dehydrogenase |
| *PSPTO_2398* | *rbsA-2* | - | 2647023 | 2648552 | 6.49 | 23.16 | 18.20 | 5.40 | 1.85 | 1.31 | Ribose ABC transporter ATP-binding protein |
| *PSPTO_2399* | *rbsB-2* | - | 2648568 | 2649500 | 28.05 | 114.15 | 53.48 | 10.48 | 3.37 | 2.70 | Ribose ABC transporter periplasmic ribose-binding protein |
| *PSPTO_2400* | *rbsC-2* | - | 2649567 | 2650571 | 20.61 | 42.93 | 27.83 | 3.42 | 6.60 | 4.69 | Ribose ABC transporter permease |
| *PSPTO_2401* | | + | 2650935 | 2651771 | 42.98 | 73.98 | 29.54 | 6.33 | 8.17 | 8.05 | Transketolase |
| *PSPTO_2402* | | + | 2651761 | 2652693 | 12.43 | 32.18 | 13.25 | 6.12 | 5.74 | 5.32 | Transketolase, C-terminal subunit |
| *PSPTO_2403* | | + | 2652724 | 2654199 | 6.19 | 21.73 | 14.66 | 8.99 | 3.66 | 4.22 | Glycerol kinase |
| *PSPTO_2404* | | - | 2654285 | 2654584 | 119.99 | 154.49 | 97.80 | 35.56 | 102.59 | 52.59 | ArsR family transcriptional regulator |
| *PSPTO_2405* | *xenA* | + | 2654680 | 2655771 | 7.67 | 21.63 | 16.03 | 2.50 | 32.22 | 12.81 | Xenobiotic reductase A |
| *PSPTO_2406* | | + | 2655936 | 2656382 | 6.19 | 34.88 | 43.21 | 6.90 | 22.38 | 24.96 | Hypothetical protein |
| *PSPTO_2407* | | + | 2656519 | 2657163 | 19.54 | 15.52 | 21.33 | 7.16 | 17.39 | 9.73 | Hypothetical protein |
| *PSPTO_2408* | *ureD-1* | - | 2657199 | 2657969 | 9.15 | 7.92 | 5.27 | 14.49 | 5.18 | 8.15 | Urease accessory protein UreD |
| *PSPTO_2409* | *ureG-1* | - | 2658053 | 2658715 | 3.21 | 4.98 | 4.50 | 2.36 | 2.61 | 2.06 | Urease accessory protein UreG |
| *PSPTO_2410* | | - | 2658743 | 2660455 | - | - | - | - | - | - | |
| *PSPTO_2411* | | - | 2660459 | 2661154 | 2.04 | 6.34 | 1.76 | 5.78 | 4.24 | 5.76 | Urease, beta/gamma subunit |
| *PSPTO_2412* | *ureF-1* | - | 2661170 | 2661901 | 2.47 | 5.20 | 0.00 | 1.75 | 3.20 | 4.57 | Urease accessory protein UreF |
| *PSPTO_2413* | | - | 2661898 | 2662338 | 1.44 | 0.00 | 0.00 | 0.00 | 2.36 | 1.97 | Urease accessory protein UreE |
| *PSPTO_2414* | | - | 2662361 | 2663050 | 1.69 | 1.28 | 4.40 | 0.00 | 3.54 | 3.50 | Branched-chain amino acid ABC transporter ATP-binding protein |
| *PSPTO_2415* | | - | 2663053 | 2663790 | 8.02 | 9.62 | 8.95 | 5.75 | 9.42 | 11.34 | Branched-chain amino acid ABC transporter ATP-binding protein |
| *PSPTO_2416* | | - | 2663787 | 2664929 | 1.66 | 1.43 | 1.90 | 0.27 | 2.26 | 3.57 | Branched-chain amino acid ABC transporter permease |
| *PSPTO_2417* | | - | 2664931 | 2665812 | 3.24 | 3.99 | 3.35 | 7.62 | 2.75 | 5.27 | Branched-chain amino acid ABC transporter permease |
| *PSPTO_2418* | | - | 2665900 | 2667081 | 1.92 | 3.58 | 1.06 | 2.67 | 2.05 | 2.46 | Branched-chain amino acid ABC transporter substrate-binding protein |
| *PSPTO_2419* | | + | 2667858 | 2669006 | 1.90 | 1.60 | 3.76 | 2.39 | 2.27 | 3.69 | Regulatory protein |
| *PSPTO_2420* | | + | 2669003 | 2669635 | 0.85 | 0.99 | 1.53 | 4.34 | 0.91 | 0.92 | Transcription antitermination protein |
| *PSPTO_2421* | *amdA* | + | 2669857 | 2671392 | 2.32 | 1.12 | 0.96 | 4.40 | 2.43 | 3.03 | Amidase |
| *PSPTO_2422* | | - | 2671508 | 2673427 | 6.38 | 7.51 | 11.98 | 4.38 | 19.87 | 7.42 | Hypothetical protein |
| *PSPTO_2423* | | - | 2673558 | 2674532 | 8.24 | 7.42 | 9.05 | 2.42 | 3.67 | 4.32 | ABC transporter, periplasmic substrate-binding protein, aliphatic sulfonates family |
| *PSPTO_2424* | | - | 2674609 | 2675568 | 4.40 | 4.93 | 2.29 | 0.56 | 2.50 | 2.85 | D-isomer specific 2-hydroxyacid dehydrogenase family protein |
| *PSPTO_2425* | | - | 2675565 | 2676911 | 4.94 | 31.80 | 28.29 | 14.15 | 4.46 | 4.24 | Monooxygenase, NtaA/SnaA/SoxA family |
| *PSPTO_2426* | | + | 2677987 | 2678964 | 0.00 | 0.00 | 0.00 | 0.00 | 0.00 | 0.00 | ISPssy, transposase |
| *PSPTO_2427* | | + | 2679142 | 2680236 | 8.52 | 31.88 | 29.74 | 25.56 | 38.60 | 27.22 | Serine hydroxymethyltransferase |
| *PSPTO_2428* | | + | 2680315 | 2681655 | 0.94 | 3.51 | 5.28 | 1.86 | 8.37 | 2.17 | Multidrug resistance protein NorM |
| *PSPTO_2429* | | + | 2681681 | 2683111 | 0.72 | 5.63 | 1.00 | 2.10 | 15.38 | 1.91 | CapK domain protein |
| *PSPTO_2430* | | + | 2683108 | 2684091 | 0.61 | 0.00 | 0.00 | 0.00 | 5.78 | 0.95 | Pyridoxal-phosphate dependent enzyme family protein |
| *PSPTO_2431* | | + | 2684150 | 2685067 | 0.70 | 0.18 | 1.19 | 2.42 | 3.12 | 1.30 | Hypothetical protein |
| *PSPTO_2432* | | - | 2685197 | 2686174 | 0.00 | 0.00 | 0.00 | 0.00 | 0.00 | 0.00 | ISPssy, transposase |
| *PSPTO_2433* | | - | 2686388 | 2686519 | 0.00 | 0.00 | 0.00 | 0.00 | 1.43 | 0.17 | Hypothetical protein |
| *PSPTO_2434* | | + | 2686542 | 2687483 | 0.20 | 1.97 | 0.00 | 2.95 | 1.64 | 1.07 | D-isomer specific 2-hydroxyacid dehydrogenase family protein |
| *PSPTO_2435* | | + | 2687499 | 2688914 | 1.84 | 3.71 | 2.80 | 2.68 | 2.66 | 2.45 | Oxidoreductase, FAD-binding |
| *PSPTO_2436* | | + | 2689146 | 2689505 | 0.00 | 0.00 | 0.00 | 0.00 | 0.00 | 0.00 | ISPsy5, Orf1 |
| *PSPTO_2437* | | + | 2689523 | 2691076 | 0.00 | 0.00 | 0.00 | 0.00 | 0.00 | 0.00 | ISPsy5, transposase |
| *PSPTO_2438* | | + | 2691134 | 2692657 | 7.43 | 12.27 | 11.84 | 7.69 | 27.78 | 20.33 | Amine oxidase, flavin-containing protein |
| *PSPTO_2439* | | + | 2692713 | 2693924 | 12.94 | 13.03 | 14.94 | 6.52 | 19.20 | 12.81 | Acyl-CoA dehydrogenase family protein |
| *PSPTO_2440* | | + | 2693952 | 2695139 | 9.17 | 6.00 | 6.98 | 4.46 | 16.88 | 13.70 | Acyl-CoA dehydrogenase family protein |
| *PSPTO_2441* | | + | 2695933 | 2698065 | 26.13 | 21.60 | 24.22 | 12.80 | 14.85 | 19.33 | Methyl-accepting chemotaxis protein |
| *PSPTO_2442* | | + | 2698089 | 2699552 | 6.38 | 6.41 | 8.17 | 4.91 | 3.75 | 7.37 | Chemotaxis protein CheW |
| *PSPTO_2443* | | + | 2699607 | 2699732 | 20.90 | 32.46 | 69.14 | 24.42 | 6.59 | 26.44 | Hypothetical protein |
| *PSPTO_2444* | | - | 2699806 | 2700291 | 8.05 | 10.47 | 8.59 | 4.91 | 11.42 | 6.38 | GNAT family acetyltransferase |
| *PSPTO_2445* | | - | 2700352 | 2700789 | 11.54 | 10.69 | 1.80 | 8.20 | 8.64 | 6.74 | Hypothetical protein |
| *PSPTO_2446* | | - | 2700804 | 2701694 | 105.96 | 62.18 | 48.86 | 42.42 | 77.09 | 89.09 | Metallo-beta-lactamase superfamily protein |
| *PSPTO_2447* | | + | 2701802 | 2702686 | 11.23 | 8.15 | 11.82 | 7.37 | 14.41 | 12.17 | LysR family transcriptional regulator |
| *PSPTO_2448* | | - | 2702812 | 2704752 | 8.32 | 4.93 | 7.32 | 0.30 | 3.53 | 7.40 | Methyl-accepting chemotaxis protein |
| *PSPTO_2449* | *soxG-2* | - | 2705019 | 2705591 | 4.49 | 4.59 | 6.11 | 0.00 | 6.29 | 15.59 | Sarcosine oxidase, gamma subunit |
| *PSPTO_2450* | *soxA-2* | - | 2705595 | 2708501 | 3.44 | 2.20 | 2.07 | 1.45 | 5.39 | 14.44 | Sarcosine oxidase, alpha subunit |
| *PSPTO_2451* | *soxD-2* | - | 2708498 | 2708803 | 3.04 | 0.00 | 0.80 | 0.00 | 8.66 | 29.04 | Sarcosine oxidase subunit delta |
| *PSPTO_2452* | *soxB-2* | - | 2708817 | 2710058 | 3.82 | 3.08 | 3.51 | 1.05 | 8.30 | 22.26 | Sarcosine oxidase, beta subunit |
| *PSPTO_2453* | *folD-1* | - | 2710040 | 2710942 | 1.54 | 1.56 | 1.45 | 0.00 | 5.44 | 13.85 | Methylenetetrahydrofolate dehydrogenase/methenyltetrahydrofolate cyclohydrolase |
| *PSPTO_2454* | *purU-2* | - | 2710939 | 2711805 | 4.41 | 1.33 | 2.64 | 2.10 | 8.17 | 17.12 | Formyltetrahydrofolate deformylase |
| *PSPTO_2455* | | + | 2712390 | 2712626 | 21.37 | 3.97 | 10.28 | 14.88 | 7.61 | 11.13 | Hypothetical protein |
| *PSPTO_2456* | | - | 2712652 | 2712768 | 5.61 | 11.94 | 10.67 | 17.72 | 9.94 | 13.96 | Hypothetical protein |
| *PSPTO_2457* | | + | 2712992 | 2714518 | 29.57 | 24.28 | 28.74 | 14.74 | 146.69 | 41.07 | Hypothetical protein |
| *PSPTO_2458* | | + | 2714515 | 2715129 | 28.92 | 17.67 | 24.84 | 18.79 | 60.59 | 30.51 | Hypothetical protein |
| *PSPTO_2459* | | - | 2715226 | 2715516 | - | - | - | - | - | - | |
| *PSPTO_2460* | | - | 2715565 | 2717118 | 0.00 | 0.00 | 0.00 | 0.00 | 0.00 | 0.00 | ISPsy5, transposase |
| *PSPTO_2461* | | - | 2717136 | 2717495 | 0.00 | 0.00 | 0.00 | 0.00 | 0.00 | 0.00 | ISPsy5, Orf1 |
| *PSPTO_2462* | | - | 2717389 | 2718021 | - | - | - | - | - | - | |
| *PSPTO_2463* | | - | 2718060 | 2720171 | 16.82 | 6.01 | 12.71 | 4.53 | 3.78 | 2.98 | TonB-dependent siderophore receptor |
| *PSPTO_2464* | | - | 2720385 | 2721689 | 9.92 | 9.74 | 11.57 | 7.17 | 8.38 | 11.52 | NRAMP family Mn2+/Fe2+ transporter |
| *PSPTO_2465* | | + | 2722266 | 2723288 | 20.04 | 24.97 | 19.11 | 13.11 | 33.26 | 25.11 | ISPsy7, transposase |
| *PSPTO_2466* | | + | 2723961 | 2724710 | 9.93 | 22.60 | 17.44 | 8.21 | 13.63 | 14.24 | Hydrolase |
| *PSPTO_2467* | | - | 2724717 | 2725970 | 13.29 | 6.70 | 7.92 | 6.93 | 8.21 | 10.53 | Major facilitator family transporter |
| *PSPTO_2468* | | - | 2726077 | 2726259 | 2.15 | 0.73 | 0.00 | 0.00 | 2.89 | 1.68 | Hypothetical protein |

| Locus name | Gene name | Strand | Gene start (bp) | Gene end (bp) | Average TPM Inoculum | Average TPM $t_6$ | Average TPM $t_{24}$ | Average TPM $t_{168}$ | Average TPM Logarithmic | Average TPM Stationary | Annotation description |
|---|---|---|---|---|---|---|---|---|---|---|---|
| *PSPTO_2469* | | - | 2726261 | 2726632 | 9.62 | 23.17 | 10.36 | 5.45 | 16.42 | 9.77 | Hypothetical protein |
| *PSPTO_2470* | | + | 2726819 | 2727910 | 7.84 | 10.68 | 10.34 | 8.98 | 8.01 | 8.51 | Senescence marker protein-30 family protein |
| *PSPTO_2471* | | + | 2728020 | 2728853 | 7.71 | 14.20 | 13.91 | 18.24 | 6.56 | 11.11 | Membrane protein |
| *PSPTO_2472* | | + | 2729060 | 2731045 | 41.61 | 18.53 | 18.79 | 3.77 | 3.53 | 3.18 | Methyl-accepting chemotaxis protein |
| *PSPTO_2473* | | + | 2731062 | 2731982 | 36.42 | 20.36 | 11.75 | 15.85 | 5.94 | 6.88 | Periplasmic substrate-binding protein |
| *PSPTO_2474* | | - | 2732375 | 2733325 | 9.02 | 8.96 | 8.36 | 5.12 | 8.44 | 8.26 | DNA/RNA non-specific endonuclease |
| *PSPTO_2475* | | + | 2733477 | 2735273 | 59.73 | 15.25 | 15.01 | 11.31 | 19.80 | 14.24 | Methyl-accepting chemotaxis protein |
| *PSPTO_2476* | | + | 2735382 | 2735693 | 46.39 | 84.91 | 78.17 | 52.92 | 13.66 | 16.49 | Hypothetical protein |
| *PSPTO_2477* | | - | 2735702 | 2736100 | 12.33 | 19.70 | 21.78 | 6.30 | 17.55 | 9.52 | Endoribonuclease L-PSP family protein |
| *PSPTO_2478* | *topB* | - | 2736166 | 2738154 | 2.89 | 5.76 | 2.73 | 2.78 | 5.61 | 3.33 | DNA topoisomerase III |
| *PSPTO_2479* | | - | 2738331 | 2739158 | 31.49 | 18.49 | 21.00 | 20.22 | 3.52 | 25.56 | Hypothetical protein |
| *PSPTO_2480* | | + | 2739439 | 2741328 | 8.96 | 8.62 | 3.16 | 0.39 | 3.47 | 3.00 | Methyl-accepting chemotaxis protein |
| *PSPTO_2481* | *tonB-2* | + | 2741606 | 2742427 | 3.10 | 26.72 | 20.98 | 16.79 | 2.25 | 2.34 | TonB protein |
| *PSPTO_2482* | *exbB-3* | + | 2742562 | 2743290 | 1.99 | 2.14 | 2.89 | 3.40 | 1.61 | 5.58 | TonB system transport protein ExbB |
| *PSPTO_2483* | *exbD-3* | + | 2743322 | 2743693 | 3.98 | 13.60 | 9.68 | 5.44 | 2.73 | 13.29 | TonB system transport protein ExbD |
| *PSPTO_2484* | | + | 2743976 | 2746378 | 4.47 | 3.97 | 3.88 | 4.45 | 4.25 | 4.22 | TonB-dependent siderophore receptor |
| *PSPTO_2485* | | + | 2746748 | 2747770 | 20.11 | 25.87 | 23.69 | 14.79 | 35.48 | 25.29 | ISPsy7, transposase |
| *PSPTO_2486* | | - | 2748125 | 2749162 | 2.93 | 16.18 | 10.98 | 5.33 | 7.00 | 4.04 | ABC transporter periplasmic substrate-binding protein |
| *PSPTO_2487* | | + | 2749395 | 2750210 | 10.65 | 53.90 | 46.94 | 23.90 | 6.48 | 12.16 | Aldolase II superfamily protein |
| *PSPTO_2488* | | + | 2750281 | 2751189 | 9.87 | 3.07 | 11.84 | 6.05 | 5.67 | 4.07 | Glutaminase |
| *PSPTO_2489* | | + | 2751293 | 2752369 | 9.73 | 9.65 | 14.06 | 10.34 | 12.24 | 11.02 | Hypothetical protein |
| *PSPTO_2490* | | + | 2752511 | 2754322 | 3.75 | 2.70 | 4.68 | 5.52 | 15.02 | 5.70 | ABC transporter ATP-binding protein/permease |
| *PSPTO_2491* | | + | 2754410 | 2755447 | 29.37 | 85.06 | 67.61 | 59.15 | 149.22 | 41.13 | Oxidoreductase, aldo/keto reductase family |
| *PSPTO_2492* | | + | 2755531 | 2756331 | 8.93 | 68.99 | 30.66 | 12.09 | 17.29 | 15.68 | Short-chain dehydrogenase/reductase family oxidoreductase |
| *PSPTO_2493* | | + | 2756324 | 2758150 | 2.43 | 16.12 | 15.64 | 6.52 | 5.48 | 6.77 | Glycosyl hydrolase family protein |
| *PSPTO_2494* | | - | 2758195 | 2759103 | 4.64 | 21.06 | 7.77 | 6.99 | 9.02 | 4.73 | LysR family transcriptional regulator |
| *PSPTO_2495* | | + | 2759406 | 2760287 | 4.36 | 7.44 | 1.14 | 5.84 | 6.07 | 5.42 | Alpha/beta fold family hydrolase |
| *PSPTO_2496* | | - | 2760925 | 2762343 | 13.25 | 36.81 | 32.20 | 15.80 | 12.22 | 30.53 | Major facilitator family transporter |
| *PSPTO_2497* | | - | 2762532 | 2762657 | 4.67 | 10.10 | 12.20 | 0.00 | 9.94 | 10.07 | Hypothetical protein |
| *PSPTO_2498* | | - | 2762711 | 2762833 | 4.10 | 0.00 | 0.00 | 0.00 | 1.83 | 1.51 | Hypothetical protein |
| *PSPTO_2499* | *mntH* | + | 2762823 | 2764055 | 4.79 | 4.18 | 2.58 | 4.98 | 6.19 | 4.25 | Manganese transport protein MntH |
| *PSPTO_2500* | | + | 2764117 | 2765568 | 7.91 | 13.02 | 12.50 | 9.30 | 6.64 | 12.76 | Major facilitator family transporter |
| *PSPTO_2501* | | - | 2765573 | 2765842 | 265.69 | 49.25 | 59.35 | 19.48 | 20.49 | 15.05 | Hypothetical protein |
| *PSPTO_2502* | | - | 2765910 | 2766113 | 78.02 | 92.30 | 89.10 | 46.01 | 97.25 | 56.04 | Hypothetical protein |
| *PSPTO_2503* | | - | 2766177 | 2766746 | 30.54 | 36.43 | 38.78 | 21.23 | 44.62 | 23.58 | Hypothetical protein |
| *PSPTO_2504* | | + | 2766894 | 2767460 | 5.12 | 1.95 | 2.16 | 0.66 | 5.65 | 9.24 | Efflux protein, LysE family |
| *PSPTO_2505* | | - | 2767470 | 2768303 | 33.64 | 69.59 | 50.15 | 27.86 | 53.30 | 58.33 | AraC family transcriptional regulator |
| *PSPTO_2506* | | - | 2768415 | 2768723 | 118.76 | 190.30 | 142.19 | 51.25 | 44.35 | 66.80 | Lipoprotein |
| *PSPTO_2507* | | - | 2768816 | 2769673 | 9.31 | 215.97 | 166.95 | 44.04 | 10.55 | 11.49 | Hypothetical protein |
| *PSPTO_2508* | | + | 2769761 | 2770189 | 8.78 | 21.60 | 16.30 | 0.00 | 10.13 | 10.09 | Hypothetical protein |
| *PSPTO_2509* | | + | 2770328 | 2772217 | 3.67 | 11.46 | 8.69 | 5.02 | 8.42 | 6.45 | Penicillin-binding protein |
| *PSPTO_2510* | *poxB* | + | 2772382 | 2774100 | 3.51 | 30.33 | 21.73 | 14.38 | 14.80 | 15.71 | Pyruvate dehydrogenase |
| *PSPTO_2511* | | + | 2774360 | 2776321 | 12.41 | 7.79 | 6.92 | 4.73 | 6.51 | 7.32 | Methyl-accepting chemotaxis protein |
| *PSPTO_2513* | | + | 2777452 | 2777559 | 148.81 | 82.00 | 100.57 | 66.33 | 88.70 | 51.10 | Hypothetical protein |
| *PSPTO_2514* | | + | 2777569 | 2778087 | 76.40 | 89.41 | 109.15 | 36.66 | 109.74 | 103.06 | Hypothetical protein |
| *PSPTO_2515* | | + | 2778105 | 2778638 | 81.90 | 102.37 | 76.87 | 43.37 | 96.65 | 93.55 | Lipoprotein |
| *PSPTO_2516* | *moaA* | - | 2778665 | 2779663 | 56.04 | 44.06 | 42.62 | 24.93 | 46.27 | 46.37 | Molybdenum cofactor biosynthesis protein A |
| *PSPTO_2517* | | - | 2779927 | 2781276 | 3.59 | 4.50 | 4.58 | 2.07 | 3.44 | 5.12 | Sodium/hydrogen exchanger family protein |
| *PSPTO_2518* | | - | 2781389 | 2782441 | 3.84 | 5.71 | 5.78 | 2.15 | 4.55 | 3.82 | Extracellular solute-binding domain protein |
| *PSPTO_2519* | | - | 2782438 | 2783622 | 2.75 | 2.91 | 3.34 | 3.52 | 3.36 | 2.68 | Oxidoreductase, FAD-binding protein |
| *PSPTO_2520* | | - | 2783619 | 2785010 | 3.64 | 2.99 | 4.92 | 1.97 | 3.73 | 2.94 | Pyridine nucleotide-disulfide oxidoreductase family protein |
| *PSPTO_2521* | | - | 2785007 | 2785297 | 2.15 | 4.13 | 3.23 | 0.00 | 2.41 | 1.54 | Hypothetical protein |
| *PSPTO_2522* | | - | 2785312 | 2786106 | 2.79 | 1.88 | 10.75 | 2.95 | 2.87 | 2.35 | ABC transporter permease |
| *PSPTO_2523* | | - | 2786110 | 2786982 | 3.95 | 3.43 | 2.89 | 1.90 | 2.15 | 2.43 | ABC transporter permease |
| *PSPTO_2524* | | - | 2786979 | 2788007 | 4.94 | 4.40 | 6.22 | 5.56 | 2.03 | 1.78 | ABC transporter ATP-binding protein |
| *PSPTO_2525* | | + | 2788249 | 2789088 | 8.55 | 13.63 | 12.09 | 3.99 | 9.89 | 4.35 | IclR family transcriptional regulator |
| *PSPTO_2526* | | + | 2789232 | 2790911 | 5.97 | 4.80 | 2.68 | 0.00 | 1.90 | 1.72 | Methyl-accepting chemotaxis protein |
| *PSPTO_2527* | | - | 2790980 | 2791942 | 6.23 | 15.07 | 5.12 | 7.60 | 20.80 | 8.93 | Hypothetical protein |
| *PSPTO_2528* | | - | 2792008 | 2792637 | 27.15 | 13.21 | 17.37 | 11.45 | 28.90 | 25.21 | Membrane protein |
| *PSPTO_2529* | | - | 2793112 | 2793453 | 6.30 | 6.98 | 2.59 | 12.69 | 19.58 | 10.91 | FKBP-type peptidyl-prolyl cis-trans isomerase |
| *PSPTO_2530* | | - | 2793450 | 2793578 | 17.93 | 0.00 | 0.00 | 3.63 | 26.43 | 49.27 | Hypothetical protein |
| *PSPTO_2531* | | - | 2793591 | 2793875 | 28.39 | 29.56 | 23.90 | 8.61 | 15.70 | 85.38 | Hypothetical protein |
| *PSPTO_2532* | | + | 2794541 | 2796109 | 9.10 | 25.75 | 36.17 | 4.58 | 50.46 | 17.59 | Membrane protein, TerC family |
| *PSPTO_2533* | | - | 2796179 | 2796754 | 4.98 | 8.66 | 2.35 | 3.71 | 13.53 | 4.83 | GNAT family acetyltransferase |
| *PSPTO_2534* | | - | 2796816 | 2797496 | 3.95 | 5.95 | 5.17 | 2.96 | 6.33 | 4.07 | Hypothetical protein |
| *PSPTO_2535* | | - | 2797493 | 2800933 | 4.65 | 4.33 | 3.61 | 1.20 | 4.34 | 3.18 | Hypothetical protein |
| *PSPTO_2536* | | - | 2800881 | 2801324 | 4.68 | 5.67 | 0.83 | 0.00 | 6.84 | 7.60 | Hypothetical protein |
| *PSPTO_2537* | | - | 2801296 | 2801763 | 7.97 | 4.13 | 2.10 | 0.00 | 5.94 | 8.49 | Hypothetical protein |
| *PSPTO_2538* | | - | 2801756 | 2803798 | 3.66 | 2.79 | 3.15 | 0.82 | 3.61 | 2.86 | Rhs element Vgr protein |
| *PSPTO_2539* | *hcp-1* | - | 2803874 | 2804389 | 5.05 | 4.62 | 1.13 | 1.19 | 2.75 | 1.47 | Secreted protein Hcp |
| *PSPTO_2540* | | + | 2804890 | 2805876 | - | - | - | - | - | - | |
| *PSPTO_2541* | | - | 2806011 | 2806988 | 0.00 | 0.00 | 0.00 | 0.00 | 0.00 | 0.00 | ISPssy, transposase |
| *PSPTO_2542* | | + | 2807021 | 2807629 | - | - | - | - | - | - | |
| *PSPTO_2543* | | + | 2807665 | 2808165 | 0.70 | 2.86 | 2.82 | 0.00 | 0.73 | 0.67 | Hypothetical protein |
| *PSPTO_2544* | | + | 2808195 | 2809670 | 1.23 | 3.05 | 4.57 | 0.95 | 2.62 | 4.11 | Hypothetical protein |
| *PSPTO_2545* | | + | 2809686 | 2810093 | 0.58 | 1.19 | 0.00 | 2.70 | 3.20 | 5.09 | Hypothetical protein |
| *PSPTO_2546* | | + | 2810191 | 2811978 | 0.53 | 2.01 | 1.71 | 0.93 | 2.53 | 2.79 | Hypothetical protein |
| *PSPTO_2547* | | + | 2811942 | 2812949 | 2.48 | 0.07 | 4.94 | 4.29 | 3.47 | 3.50 | Hypothetical protein |
| *PSPTO_2548* | | + | 2812959 | 2815562 | 4.04 | 4.32 | 3.23 | 0.55 | 4.52 | 4.22 | ClpB protein |

| Locus name | Gene name | Strand | Gene start (bp) | Gene end (bp) | Average TPM Inoculum | Average TPM $t_{6}$ | Average TPM $t_{24}$ | Average TPM $t_{168}$ | Average TPM Logarithmic | Average TPM Stationary | Annotation description |
|---|---|---|---|---|---|---|---|---|---|---|---|
| *PSPTO_2549* | | + | 2815564 | 2817093 | 3.21 | 3.05 | 6.47 | 1.11 | 3.77 | 2.85 | Sigma-54 dependent transcriptional regulator |
| *PSPTO_2550* | | + | 2817143 | 2817283 | 4.29 | 5.42 | 0.00 | 0.00 | 5.31 | 4.11 | Hypothetical protein |
| *PSPTO_2551* | | + | 2817337 | 2817828 | 2.75 | 0.44 | 4.83 | 0.00 | 5.30 | 4.05 | Hypothetical protein |
| *PSPTO_2552* | | + | 2817858 | 2819156 | 5.06 | 6.07 | 1.33 | 1.71 | 4.30 | 3.07 | Hypothetical protein |
| *PSPTO_2553* | | + | 2819156 | 2820028 | 2.31 | 0.66 | 0.00 | 4.44 | 3.08 | 2.26 | Hypothetical protein |
| *PSPTO_2554* | | + | 2820042 | 2823548 | 5.70 | 3.64 | 2.59 | 1.53 | 6.36 | 3.45 | Hypothetical protein |
| *PSPTO_2555* | *phnC* | + | 2823826 | 2824659 | 2.24 | 1.70 | 0.00 | 0.00 | 1.07 | 1.71 | Phosphonates ABC transporter ATP-binding protein |
| *PSPTO_2556* | *phnD* | + | 2824697 | 2825707 | 3.57 | 3.13 | 4.25 | 3.51 | 3.10 | 2.61 | Phosphonates ABC transporter periplasmic phosphonates-binding protein |
| *PSPTO_2557* | *phnE* | + | 2825735 | 2826523 | 21.91 | 6.57 | 7.97 | 5.34 | 4.48 | 7.39 | Phosphonates ABC transporter permease |
| *PSPTO_2558* | *phnF* | + | 2826562 | 2827281 | 5.91 | 8.13 | 5.31 | 2.33 | 5.48 | 10.52 | Transcriptional regulator PhnF |
| *PSPTO_2559* | *phnG* | + | 2827296 | 2827760 | 3.16 | 15.96 | 3.79 | 0.00 | 4.04 | 5.57 | Phosphonate metabolism protein PhnG |
| *PSPTO_2560* | *phnH* | + | 2827760 | 2828356 | 1.54 | 5.90 | 4.73 | 8.19 | 3.09 | 3.84 | PhnH protein |
| *PSPTO_2561* | *phnI* | + | 2828356 | 2829435 | 4.05 | 4.63 | 2.87 | 1.02 | 4.48 | 5.42 | Phosphonate metabolism protein PhnI |
| *PSPTO_2562* | *phnJ* | + | 2829432 | 2830319 | 5.55 | 4.24 | 3.80 | 1.03 | 6.51 | 6.71 | Phosphonate metabolism protein PhnJ |
| *PSPTO_2563* | *phnK* | + | 2830316 | 2831137 | 2.05 | 3.43 | 3.33 | 0.00 | 3.10 | 2.83 | Phosphonate C-P lyase system protein PhnK |
| *PSPTO_2564* | | + | 2831170 | 2831886 | 3.17 | 3.18 | 1.37 | 1.50 | 3.39 | 2.93 | Phosphonate ABC transporter ATPase |
| *PSPTO_2565* | *phnM* | + | 2831880 | 2833025 | 10.60 | 26.14 | 22.57 | 7.40 | 19.62 | 12.33 | Phosphonate metabolism protein PhnM |
| *PSPTO_2566* | *phnN* | + | 2833025 | 2833591 | 4.61 | 20.77 | 15.85 | 0.00 | 10.03 | 5.83 | ATP-binding protein PhnN |
| *PSPTO_2567* | *phnP* | + | 2833582 | 2834331 | 3.48 | 7.45 | 10.88 | 12.13 | 14.72 | 7.21 | Phosphonate metabolism protein, PhnP |
| *PSPTO_2568* | *gcd-1* | + | 2834486 | 2836852 | 2.14 | 1.74 | 3.46 | 0.85 | 5.95 | 3.25 | Glucose dehydrogenase |
| *PSPTO_2569* | | - | 2836915 | 2838438 | 7.01 | 11.04 | 15.51 | 10.66 | 8.41 | 15.38 | Amidase family protein |
| *PSPTO_2570* | | - | 2838609 | 2839760 | 6.67 | 10.29 | 6.80 | 5.75 | 5.60 | 7.63 | Peptidase, M20/M25/M40 family |
| *PSPTO_2571* | | - | 2839757 | 2841184 | 4.39 | 3.57 | 3.42 | 4.44 | 3.50 | 3.04 | Peptidase, M20/M25/M40 family |
| *PSPTO_2572* | | - | 2841181 | 2842194 | 0.64 | 2.34 | 1.24 | 0.00 | 1.02 | 0.79 | Peptide ABC transporter ATP-binding protein |
| *PSPTO_2573* | | - | 2842191 | 2843192 | 1.58 | 0.00 | 5.23 | 5.21 | 1.81 | 1.87 | Peptide ABC transporter ATP-binding protein |
| *PSPTO_2574* | | - | 2843209 | 2844117 | 0.96 | 1.65 | 1.21 | 6.31 | 1.28 | 1.35 | Peptide ABC transporter permease |
| *PSPTO_2575* | | - | 2844126 | 2845043 | 4.58 | 18.08 | 26.12 | 3.25 | 1.41 | 1.31 | Peptide ABC transporter permease |
| *PSPTO_2576* | | - | 2845146 | 2846666 | 1.03 | 1.86 | 2.20 | 7.86 | 1.20 | 1.29 | Peptide ABC transporter periplasmic peptide-binding protein |
| *PSPTO_2577* | | + | 2846878 | 2847771 | 2.58 | 3.20 | 4.05 | 0.34 | 6.68 | 3.25 | LysR family transcriptional regulator |
| *PSPTO_2578* | | + | 2847956 | 2848861 | 3.46 | 11.68 | 16.63 | 5.13 | 1.60 | 1.37 | Hypothetical protein |
| *PSPTO_2579* | | + | 2848873 | 2849433 | 43.16 | 25.81 | 14.70 | 4.72 | 3.15 | 5.28 | Carbonic anhydrase-like protein |
| *PSPTO_2580* | | + | 2849485 | 2850651 | 5.19 | 7.72 | 7.67 | 7.34 | 6.08 | 3.81 | Membrane protein |
| *PSPTO_2581* | | - | 2850700 | 2851329 | 13.25 | 13.03 | 12.04 | 9.89 | 19.00 | 9.51 | DNA-binding protein |
| *PSPTO_2582* | | + | 2851604 | 2852937 | - | - | - | - | - | - | |
| *PSPTO_2583* | | + | 2853098 | 2854018 | 0.94 | 0.54 | 0.00 | 0.00 | 1.05 | 0.89 | Glutamine amidotransferase, class-II protein |
| *PSPTO_2584* | | + | 2854015 | 2854698 | 1.01 | 1.61 | 1.42 | 0.80 | 1.85 | 0.93 | Hypothetical protein |
| *PSPTO_2585* | | + | 2854714 | 2856054 | 1.43 | 0.76 | 2.94 | 1.05 | 2.18 | 1.62 | Glutamate synthase family protein |
| *PSPTO_2586* | *amt-2* | + | 2856588 | 2857913 | 2.04 | 2.16 | 2.63 | 1.09 | 1.88 | 3.53 | Ammonium transporter |
| *PSPTO_2587* | | + | 2857952 | 2858074 | 15.20 | 0.00 | 14.87 | 0.00 | 7.03 | 7.80 | Hypothetical protein |
| *PSPTO_2588* | | - | 2858488 | 2859525 | 6.71 | 4.89 | 5.38 | 8.70 | 9.29 | 4.63 | Luciferase family protein |
| *PSPTO_2589* | | - | 2859734 | 2860069 | 141.12 | 43.70 | 50.65 | 10.66 | 21.34 | 21.68 | Hypothetical protein |
| *PSPTO_2590* | | + | 2860393 | 2861493 | 11.68 | 66.19 | 38.46 | 75.47 | 3.45 | 4.78 | Luciferase family protein |
| *PSPTO_2591* | | + | 2861816 | 2863402 | 6.04 | 7.03 | 9.86 | 7.59 | 6.70 | 5.01 | Diguanylate cyclase |
| *PSPTO_2592* | *saxG* | - | 2863432 | 2866575 | 2.03 | 3.99 | 5.56 | 2.66 | 14.99 | 3.26 | Aliphatic isothiocyanate resistance protein SaxG; AcrB/AcrD/AcrF family |
| *PSPTO_2593* | | - | 2866609 | 2867769 | 6.69 | 21.41 | 29.84 | 0.56 | 19.16 | 13.87 | Multidrug resistance protein, AcrA/AcrE family |
| *PSPTO_2594* | *nfxB* | + | 2867950 | 2868504 | 27.42 | 33.93 | 24.61 | 4.95 | 34.29 | 33.94 | Transcriptional regulator NfxB |
| *PSPTO_2595* | *pchA* | - | 2868541 | 2869950 | 5.60 | 7.57 | 7.10 | 5.50 | 96.46 | 148.07 | Isochorismate synthase |
| *PSPTO_2596* | *pchB* | - | 2869947 | 2870267 | 5.04 | 6.01 | 3.05 | 2.10 | 107.55 | 150.93 | Isochorismate pyruvate-lyase |
| *PSPTO_2597* | *irp5, ybtE* | - | 2870283 | 2871851 | 4.30 | 9.63 | 4.79 | 3.55 | 142.37 | 163.51 | Yersiniabactin synthetase, salycilate ligase component |
| *PSPTO_2598* | *irp4, ybtT* | - | 2871864 | 2872679 | 7.24 | 9.19 | 13.78 | 5.01 | 212.23 | 200.12 | Yersiniabactin synthetase, thioesterase component |
| *PSPTO_2599* | *irp3, ybtU* | - | 2872651 | 2873733 | 3.81 | 2.79 | 2.79 | 2.38 | 139.94 | 150.27 | Yersiniabactin synthetase, thiazolinyl reductase component |
| *PSPTO_2600* | *irp1, hmwp1* | - | 2873730 | 2883251 | 7.90 | 9.48 | 11.62 | 5.47 | 184.66 | 230.81 | Yersiniabactin polyketide/non-ribosomal peptide synthetase |
| *PSPTO_2601* | *ybtX* | - | 2883238 | 2884464 | 6.62 | 11.64 | 13.07 | 1.93 | 242.06 | 247.48 | Membrane protein |
| *PSPTO_2602* | *hmwp2* | - | 2884461 | 2890634 | 7.50 | 9.43 | 7.92 | 3.45 | 176.22 | 205.70 | Yersiniabactin non-ribosomal peptide synthetase |
| *PSPTO_2603* | *ybtQ* | - | 2890695 | 2892440 | 4.93 | 4.35 | 6.83 | 2.11 | 119.64 | 128.07 | ABC transporter ATP-binding protein/permease |
| *PSPTO_2604* | *ybtP* | - | 2892437 | 2894218 | 6.25 | 6.42 | 4.94 | 2.00 | 126.89 | 167.40 | ABC transporter ATP-binding protein/permease |
| *PSPTO_2605* | *fyuA* | - | 2894373 | 2896430 | 51.74 | 48.63 | 37.61 | 44.50 | 433.72 | 908.79 | TonB-dependent siderophore receptor |
| *PSPTO_2606* | *ybtA* | + | 2896523 | 2897461 | 14.79 | 15.13 | 13.74 | 5.14 | 14.35 | 29.08 | AraC family transcriptional regulator |
| *PSPTO_2607* | | - | 2897458 | 2899956 | 1.78 | 1.86 | 1.18 | 1.86 | 2.34 | 3.45 | TonB-dependent siderophore receptor |
| *PSPTO_2608* | | + | 2900350 | 2900955 | 10.59 | 33.66 | 25.26 | 12.99 | 41.31 | 175.24 | Sco1/SenC family protein |
| *PSPTO_2609* | | + | 2900955 | 2901443 | 7.03 | 32.36 | 24.82 | 3.42 | 50.12 | 107.21 | Hypothetical protein |
| *PSPTO_2610* | | - | 2901682 | 2902140 | 2.73 | 3.83 | 12.37 | 3.99 | 7.79 | 19.20 | Phosphoesterase |
| *PSPTO_2611* | | - | 2902216 | 2902797 | 8.26 | 11.62 | 10.94 | 3.63 | 7.74 | 9.70 | tRNA synthetase class II family protein |
| *PSPTO_2612* | | - | 2903125 | 2903628 | 10.25 | 10.11 | 16.56 | 3.68 | 22.98 | 8.33 | GNAT family acetyltransferase |
| *PSPTO_2613* | | - | 2903711 | 2904286 | 28.41 | 42.19 | 58.49 | 13.76 | 59.48 | 23.53 | Hypothetical protein |
| *PSPTO_2614* | | + | 2904461 | 2905300 | 14.42 | 72.92 | 74.57 | 9.93 | 34.95 | 42.30 | Dioxygenase, TauD/TfdA family |
| *PSPTO_2615* | | - | 2905681 | 2906178 | 12.43 | 21.52 | 20.18 | 15.23 | 21.35 | 9.82 | GAF domain-containing protein |
| *PSPTO_2616* | | + | 2906392 | 2908293 | 29.41 | 18.12 | 15.72 | 10.85 | 5.71 | 4.25 | Methyl-accepting chemotaxis protein |
| *PSPTO_2617* | | - | 2908370 | 2909275 | 3.94 | 13.42 | 7.73 | 7.14 | 14.79 | 7.23 | LysR family transcriptional regulator |
| *PSPTO_2618* | *argD-2* | + | 2909379 | 2910581 | 3.63 | 5.02 | 5.09 | 0.91 | 3.08 | 2.96 | Acetylornithine aminotransferase |
| *PSPTO_2620* | | - | 2911293 | 2911733 | 156.81 | 438.54 | 358.26 | 70.61 | 27.60 | 28.55 | Hypothetical protein |
| *PSPTO_2621* | *polB* | + | 2911958 | 2914321 | 4.67 | 5.33 | 7.80 | 3.97 | 9.53 | 8.80 | DNA polymerase II |
| *PSPTO_2622* | | - | 2914318 | 2915163 | 7.64 | 9.59 | 9.54 | 3.14 | 4.86 | 8.23 | Sir2 family transcriptional regulator |
| *PSPTO_2623* | | + | 2915330 | 2916172 | 3.18 | 14.91 | 14.20 | 8.31 | 47.57 | 20.32 | Phenazine biosynthesis protein, PhzF family |
| *PSPTO_2624* | | - | 2916314 | 2917153 | 2.13 | 3.27 | 7.45 | 0.78 | 6.33 | 4.57 | Hypothetical protein |
| *PSPTO_2625* | | - | 2917150 | 2918368 | - | - | - | - | - | - | |
| *PSPTO_2626* | | - | 2918440 | 2919312 | 10.28 | 9.76 | 14.38 | 3.53 | 9.39 | 6.62 | LysR family transcriptional regulator |
| *PSPTO_2627* | | + | 2919555 | 2920397 | 1.89 | 21.03 | 12.96 | 4.32 | 4.20 | 3.47 | Amino acid ABC transporter substrate-binding protein |
| *PSPTO_2628* | | + | 2920441 | 2921175 | 0.47 | 0.00 | 1.28 | 0.00 | 0.42 | 0.39 | Amino acid ABC transporter permease |

| Locus name | Gene name | Strand | Gene start (bp) | Gene end (bp) | Average TPM Inoculum | Average TPM $t_6$ | Average TPM $t_{24}$ | Average TPM $t_{168}$ | Average TPM Logarithmic | Average TPM Stationary | Annotation description |
|---|---|---|---|---|---|---|---|---|---|---|---|
| *PSPTO_2629* | | + | 2921165 | 2921953 | 1.74 | 1.83 | 3.83 | 3.64 | 1.53 | 2.18 | Amino acid ABC transporter permease |
| *PSPTO_2630* | | + | 2921940 | 2922776 | 2.18 | 6.66 | 3.45 | 0.00 | 4.47 | 1.78 | Amino acid ABC transporter ATP-binding protein |
| *PSPTO_2631* | | + | 2922804 | 2923397 | 1.32 | 2.90 | 1.44 | 0.00 | 4.95 | 2.89 | Hypothetical protein |
| *PSPTO_2632* | | + | 2923595 | 2926390 | 6.81 | 13.56 | 16.97 | 8.39 | 10.20 | 10.08 | Rhodanese domain protein/cystathionine beta-lyase |
| *PSPTO_2633* | | + | 2926602 | 2927351 | 3.02 | 7.55 | 15.98 | 4.87 | 1.04 | 1.44 | Cation ABC transporter ATP-binding protein |
| *PSPTO_2634* | | + | 2927348 | 2928217 | 3.22 | 17.69 | 12.55 | 2.08 | 2.12 | 2.22 | Cation ABC transporter permease |
| *PSPTO_2635* | | + | 2928229 | 2929125 | 5.08 | 55.99 | 77.40 | 18.29 | 3.11 | 3.89 | Cation ABC transporter substrate-binding protein |
| *PSPTO_2636* | *galM* | - | 2929175 | 2930323 | 7.21 | 196.78 | 228.33 | 87.67 | 15.58 | 8.61 | Aldose 1-epimerase |
| *PSPTO_2637* | | + | 2930623 | 2931495 | 13.50 | 53.47 | 54.35 | 27.84 | 23.20 | 24.29 | Senescence marker protein-30 family protein |
| *PSPTO_2638* | *abP* | + | 2931657 | 2932670 | 21.08 | 245.11 | 183.19 | 62.65 | 25.50 | 17.12 | L-arabinose ABC transporter periplasmic L-arabinose-binding protein |
| *PSPTO_2639* | | + | 2932726 | 2934249 | 6.65 | 56.46 | 36.42 | 20.20 | 7.67 | 4.26 | L-arabinose transporter ATP-binding protein |
| *PSPTO_2640* | *araH* | + | 2934266 | 2935255 | 5.81 | 52.15 | 29.78 | 19.52 | 7.96 | 4.90 | L-arabinose transporter permease protein |
| *PSPTO_2641* | | + | 2935266 | 2935373 | 8.14 | 55.08 | 26.98 | 25.72 | 12.16 | 7.41 | Lipoprotein |
| *PSPTO_2642* | | - | 2935352 | 2936602 | 3.15 | 3.97 | 2.69 | 3.84 | 5.88 | 2.97 | Sensor histidine kinase |
| *PSPTO_2643* | | - | 2936674 | 2937375 | 1.47 | 1.80 | 0.22 | 2.25 | 1.33 | 2.33 | DNA-binding response regulator |
| *PSPTO_2644* | | - | 2937426 | 2938205 | 6.69 | 4.51 | 3.67 | 1.00 | 3.63 | 3.09 | MltA-interacting protein |
| *PSPTO_2645* | | - | 2938265 | 2939854 | 3.24 | 4.59 | 3.63 | 0.41 | 2.41 | 3.16 | Hypothetical protein |
| *PSPTO_2646* | | + | 2940161 | 2940871 | 62.16 | 55.24 | 55.05 | 34.25 | 75.68 | 78.40 | TetR family transcriptional regulator |
| *PSPTO_2647* | | + | 2940935 | 2941675 | 36.72 | 30.78 | 30.21 | 23.01 | 97.97 | 72.63 | Hypothetical protein |
| *PSPTO_2648* | | + | 2941675 | 2942142 | 16.00 | 10.75 | 17.78 | 12.68 | 51.82 | 31.77 | Hypothetical protein |
| *PSPTO_2649* | | + | 2942151 | 2943113 | 42.07 | 20.07 | 13.58 | 22.89 | 27.09 | 58.43 | Hypothetical protein |
| *PSPTO_2650* | | + | 2943186 | 2943347 | 3.31 | 2.58 | 10.00 | 3.16 | 8.77 | 18.00 | Membrane protein |
| *PSPTO_2651* | | - | 2943337 | 2943714 | 2.93 | 6.47 | 7.23 | 5.90 | 2.26 | 5.26 | Response regulator |
| *PSPTO_2652* | | - | 2943711 | 2946692 | 5.76 | 3.43 | 4.99 | 0.90 | 2.99 | 4.95 | Bacteriophytochrome |
| *PSPTO_2653* | | - | 2947054 | 2948691 | - | - | - | - | - | - | |
| *PSPTO_2654* | | + | 2949573 | 2950250 | 2.47 | 15.80 | 14.88 | 5.15 | 4.99 | 18.97 | 4Fe-4S binding protein |
| *PSPTO_2655* | | + | 2950250 | 2951233 | 2.47 | 8.55 | 6.26 | 4.33 | 4.22 | 7.26 | Oxidoreductase, molybdopterin-binding subunit |
| *PSPTO_2656* | | + | 2951420 | 2953459 | 2.93 | 5.56 | 6.52 | 6.05 | 6.16 | 7.70 | Aldehyde oxidase and xanthine dehydrogenase family protein |
| *PSPTO_2657* | | + | 2953462 | 2954430 | 2.62 | 6.29 | 4.64 | 6.45 | 3.43 | 7.62 | Xanthine dehydrogenase accessory factor XdhC |
| *PSPTO_2658* | | - | 2954411 | 2955004 | 4.13 | 2.05 | 4.69 | 4.37 | 6.53 | 5.87 | Hypothetical protein |
| *PSPTO_2659* | | + | 2955177 | 2955761 | 20.11 | 17.31 | 10.20 | 6.57 | 10.89 | 17.72 | NAD(P)H-flavin oxidoreductase |
| *PSPTO_2660* | | - | 2956044 | 2957435 | 2.92 | 2.54 | 3.44 | 2.09 | 3.04 | 2.57 | Aldehyde dehydrogenase family protein |
| *PSPTO_2661* | | + | 2957538 | 2958416 | 4.25 | 4.97 | 4.84 | 3.75 | 7.27 | 4.96 | LysR family transcriptional regulator |
| *PSPTO_2662* | | - | 2958422 | 2959516 | 4.25 | 4.54 | 6.76 | 2.77 | 4.22 | 9.45 | Tartrate dehydrogenase |
| *PSPTO_2663* | | + | 2959629 | 2960534 | 3.78 | 5.25 | 4.51 | 5.94 | 5.75 | 3.28 | LysR family transcriptional regulator |
| *PSPTO_2664* | | - | 2960581 | 2961735 | 4.12 | 4.08 | 3.16 | 6.64 | 6.50 | 4.44 | Spermidine/putrescine ABC transporter ATP-binding protein |
| *PSPTO_2665* | | - | 2961732 | 2962547 | 6.05 | 3.41 | 11.35 | 0.00 | 5.17 | 4.85 | Spermidine/putrescine ABC transporter permease |
| *PSPTO_2666* | | - | 2962554 | 2963408 | 7.56 | 3.24 | 2.76 | 3.64 | 6.33 | 5.68 | Spermidine/putrescine ABC transporter permease |
| *PSPTO_2667* | | - | 2963495 | 2964520 | 5.00 | 7.26 | 4.19 | 3.45 | 5.69 | 2.99 | Spermidine/putrescine ABC transporter periplasmic spermidine/putrescine-binding protein |
| *PSPTO_2668* | *ribBA-2* | - | 2964810 | 2965919 | 4.76 | 7.30 | 7.27 | 0.00 | 8.92 | 7.36 | Bifunctional 3,4-dihydroxy-2-butanone 4-phosphate synthase/GTP cyclohydrolase II-like protein |
| *PSPTO_2669* | | + | 2966014 | 2966304 | 11.37 | 13.21 | 11.86 | 3.78 | 20.64 | 12.52 | Hypothetical protein |
| *PSPTO_2670* | | - | 2966404 | 2966967 | 15.70 | 13.19 | 20.26 | 14.86 | 22.29 | 15.49 | MutT/nudix family protein |
| *PSPTO_2671* | | - | 2966969 | 2967721 | 24.15 | 31.96 | 19.22 | 7.24 | 20.46 | 19.93 | GntR family transcriptional regulator |
| *PSPTO_2672* | | - | 2967912 | 2968397 | 32.93 | 14.56 | 17.69 | 10.25 | 22.11 | 14.43 | NAD(P)H-flavin oxidoreductase |
| *PSPTO_2673* | *aldB* | - | 2968411 | 2969892 | 6.55 | 8.24 | 9.88 | 5.54 | 8.82 | 7.97 | Aldehyde dehydrogenase family protein. Conversion of indole-3-acetaldehyde to indole-3-acetic acid (an auxin). |
| *PSPTO_2674* | | - | 2969889 | 2970737 | 3.76 | 6.78 | 12.72 | 0.43 | 6.02 | 4.81 | 3-oxoadipate enol-lactone hydrolase family protein |
| *PSPTO_2675* | | - | 2970772 | 2971362 | 4.77 | 5.32 | 4.81 | 2.67 | 6.43 | 2.97 | Hypothetical protein |
| *PSPTO_2676* | *fdhA-2* | + | 2971649 | 2972788 | 4.75 | 31.47 | 30.83 | 6.81 | 3.39 | 5.21 | Glutathione-independent formaldehyde dehydrogenase |
| *PSPTO_2677* | | + | 2972883 | 2973743 | 4.67 | 16.29 | 24.67 | 4.11 | 5.81 | 8.11 | Short chain dehydrogenase/reductase family oxidoreductase |
| *PSPTO_2678* | *hopP1* | + | 2973945 | 2974919 | 3.62 | 384.73 | 269.12 | 46.67 | 4.14 | 5.04 | Type III helper protein HopP1 |
| *PSPTO_2679* | | + | 2974894 | 2975016 | 0.31 | 28.81 | 47.32 | 4.44 | 0.88 | 0.24 | Hypothetical protein |
| *PSPTO_2680* | *gabD-3* | + | 2975356 | 2976786 | 8.24 | 26.92 | 29.85 | 11.67 | 67.52 | 34.81 | Succinate-semialdehyde dehydrogenase |
| *PSPTO_2681* | *pbpG* | - | 2976862 | 2977797 | 12.14 | 15.21 | 17.52 | 5.12 | 97.54 | 22.82 | Penicillin-binding protein 7 |
| *PSPTO_2683* | | - | 2978728 | 2978910 | 45.75 | 373.32 | 369.29 | 101.21 | 13.12 | 50.77 | Hypothetical protein |
| *PSPTO_2684* | | - | 2979083 | 2979439 | 1518.63 | 874.31 | 955.72 | 394.78 | 277.06 | 452.01 | Hypothetical protein |
| *PSPTO_2685* | | + | 2979641 | 2980417 | 8.07 | 17.90 | 13.11 | 1.87 | 17.33 | 8.67 | Shikimate 5-dehydrogenase |
| *PSPTO_2686* | | - | 2980482 | 2981147 | 15.83 | 19.53 | 18.03 | 16.48 | 24.57 | 15.94 | Hypothetical protein |
| *PSPTO_2687* | | + | 2981249 | 2981812 | 52.97 | 59.11 | 99.69 | 56.52 | 135.19 | 66.33 | Hypothetical protein |
| *PSPTO_2688* | | + | 2982076 | 2983230 | 5.05 | 4.47 | 5.18 | 0.65 | 16.41 | 5.08 | Major facilitator family transporter |
| *PSPTO_2689* | | - | 2983227 | 2984129 | 9.07 | 8.09 | 6.77 | 1.88 | 26.25 | 8.15 | LysR family transcriptional regulator |
| *PSPTO_2690* | | + | 2984262 | 2984504 | 22.36 | 1458.07 | 1265.18 | 276.82 | 21.31 | 9.57 | Lipoprotein |
| *PSPTO_2691* | | + | 2984539 | 2985300 | 30.21 | 251.85 | 244.50 | 72.57 | 15.42 | 13.80 | Membrane protein, TerC family |
| *PSPTO_2692* | | - | 2985402 | 2985581 | 123.05 | 207.13 | 274.45 | 158.50 | 19.92 | 154.42 | Hypothetical protein |
| *PSPTO_2693* | | + | 2985801 | 2986496 | 4.95 | 14.28 | 4.13 | 0.82 | 10.92 | 5.38 | Hypothetical protein |
| *PSPTO_2694* | | - | 2986568 | 2987851 | 4.53 | 8.95 | 14.44 | 3.07 | 28.55 | 5.65 | Hypothetical protein |
| *PSPTO_2695* | | - | 2988478 | 2989539 | 28.86 | 74.26 | 62.69 | 16.93 | 23.63 | 26.65 | Hypothetical protein |
| *PSPTO_2696* | | - | 2989651 | 2990037 | 86.88 | 63.72 | 35.00 | 33.05 | 20.21 | 30.86 | MutT/nudix family protein |
| *PSPTO_2697* | | - | 2990361 | 2991437 | 29.67 | 70.85 | 63.49 | 30.56 | 25.46 | 46.83 | Oxidoreductase zinc-binding protein |
| *PSPTO_2698* | | + | 2991704 | 2992018 | 61.56 | 27.78 | 16.98 | 26.62 | 18.97 | 22.54 | Hypothetical protein |
| *PSPTO_2699* | | + | 2992163 | 2994622 | 6.36 | 6.50 | 4.85 | 2.03 | 5.80 | 9.04 | GAF domain/GGDEF domain/EAL domain protein |
| *PSPTO_2700* | | + | 2994802 | 2995356 | 5265.58 | 1243.87 | 911.12 | 443.98 | 144.23 | 687.61 | Hypothetical protein |
| *PSPTO_2701* | | - | 2995844 | 2996782 | 5.84 | 11.28 | 10.52 | 8.79 | 82.44 | 9.48 | Fructokinase |
| *PSPTO_2702* | *xylB* | - | 2996807 | 2998288 | 7.77 | 13.52 | 9.74 | 9.91 | 78.29 | 5.51 | Xylulokinase |
| *PSPTO_2703* | *uxuB* | - | 2998299 | 2999780 | 2.70 | 20.70 | 19.34 | 9.03 | 143.04 | 5.39 | D-mannonate oxidoreductase |
| *PSPTO_2704* | | - | 2999799 | 3000912 | - | - | - | - | - | - | |
| *PSPTO_2705* | | - | 3000944 | 3001777 | 5.27 | 13.41 | 13.65 | 10.56 | 80.66 | 6.65 | Mannitol ABC transporter permease |
| *PSPTO_2706* | | - | 3001788 | 3002720 | 3.66 | 14.60 | 17.00 | 5.66 | 71.08 | 7.00 | Mannitol ABC transporter permease |
| *PSPTO_2707* | | - | 3002808 | 3004130 | 9.23 | 51.70 | 46.08 | 23.29 | 135.79 | 54.95 | Mannitol ABC transporter, periplasmic mannitol-binding protein |
| *PSPTO_2708* | *mltR* | - | 3004371 | 3005276 | 7.87 | 19.79 | 17.45 | 8.60 | 27.47 | 12.72 | Transcriptional activator MltR |

| Locus name | Gene name | Strand | Gene start (bp) | Gene end (bp) | Average TPM Inoculum | Average TPM $t_6$ | Average TPM $t_{24}$ | Average TPM $t_{168}$ | Average TPM Logarithmic | Average TPM Stationary | Annotation description |
|---|---|---|---|---|---|---|---|---|---|---|---|
| *PSPTO_2709* | | + | 3005404 | 3006189 | 18.15 | 15.89 | 19.52 | 14.94 | 29.15 | 20.41 | Precorrin 6A synthase |
| *PSPTO_2710* | | - | 3006174 | 3007382 | 5.34 | 12.88 | 7.09 | 3.34 | 25.37 | 8.29 | Hypothetical protein |
| *PSPTO_2711* | | - | 3007489 | 3007848 | 61.57 | 68.93 | 86.24 | 47.16 | 82.33 | 38.90 | Response regulator |
| *PSPTO_2712* | | + | 3008195 | 3011707 | 4.07 | 14.47 | 12.63 | 5.09 | 21.93 | 8.81 | Sensor histidine kinase/response regulator |
| *PSPTO_2713* | | + | 3011736 | 3012602 | 5.12 | 9.06 | 6.84 | 5.16 | 19.41 | 5.42 | Chemotaxis protein methyltransferase CheR |
| *PSPTO_2714* | | + | 3012599 | 3013213 | 10.51 | 15.45 | 8.69 | 12.01 | 19.85 | 13.29 | Protein-glutamate methylesterase |
| *PSPTO_2715* | | + | 3013207 | 3014388 | 4.30 | 10.99 | 9.79 | 4.74 | 23.05 | 8.58 | Sensor histidine kinase/response regulator |
| *PSPTO_2716* | | + | 3014480 | 3014857 | 245.76 | 88.45 | 85.37 | 42.48 | 70.24 | 138.98 | Response regulator |
| *PSPTO_2717* | | - | 3014885 | 3016915 | 6.49 | 18.12 | 20.63 | 7.41 | 20.84 | 16.80 | Sensory box histidine kinase/response regulator |
| *PSPTO_2718* | | - | 3016912 | 3018396 | 18.83 | 34.48 | 31.90 | 18.98 | 24.02 | 20.56 | Hypothetical protein |
| *PSPTO_2719* | | + | 3018617 | 3019429 | 18.03 | 48.84 | 44.62 | 24.20 | 46.61 | 20.03 | Enoyl-CoA hydratase |
| *PSPTO_2720* | | + | 3019482 | 3020318 | 15.96 | 35.43 | 39.79 | 13.35 | 45.26 | 34.65 | MutT/nudix family protein |
| *PSPTO_2721* | | - | 3020286 | 3021065 | 6.95 | 15.04 | 16.00 | 3.51 | 24.42 | 16.55 | Membrane protein |
| *PSPTO_2722* | | - | 3021171 | 3022319 | 2.38 | 40.66 | 30.17 | 4.12 | 11.13 | 2.40 | Oxidoreductase, Gfo/Idh/MocA family |
| *PSPTO_2723* | | - | 3022420 | 3023187 | 3.92 | 3.77 | 10.32 | 4.34 | 10.08 | 5.47 | Short chain dehydrogenase |
| *PSPTO_2724* | | - | 3023215 | 3024282 | 3.16 | 10.30 | 5.39 | 4.00 | 9.06 | 4.36 | Hypothetical protein |
| *PSPTO_2725* | | - | 3024485 | 3024802 | 17.92 | 26.87 | 27.40 | 3.44 | 70.84 | 6.17 | SCP-2 sterol transfer family protein |
| *PSPTO_2726* | | - | 3024858 | 3025568 | 54.10 | 49.85 | 55.74 | 24.15 | 76.19 | 30.40 | Hypothetical protein |
| *PSPTO_2727* | | - | 3025629 | 3025724 | 10.07 | 7.68 | 15.88 | 0.88 | 3.03 | 4.66 | Hypothetical protein |
| *PSPTO_2728* | | + | 3025774 | 3026796 | 28.38 | 35.42 | 28.05 | 25.70 | 41.65 | 21.04 | Peptidase, U7 family |
| *PSPTO_2729* | | - | 3027056 | 3027550 | 9.47 | 36.57 | 26.37 | 38.86 | 28.13 | 17.22 | Hypothetical protein |
| *PSPTO_2730* | *cysI* | - | 3027534 | 3029192 | 8.33 | 86.55 | 51.61 | 48.06 | 40.68 | 50.27 | Sulfite reductase |
| *PSPTO_2731* | | + | 3029769 | 3030842 | 24.69 | 39.72 | 30.88 | 23.14 | 14.19 | 30.76 | Periplasmic sugar-binding domain protein |
| *PSPTO_2732* | *metH* | - | 3030852 | 3034571 | 7.98 | 17.03 | 18.58 | 7.43 | 88.16 | 26.74 | 5-methyltetrahydrofolate--homocysteine methyltransferase |
| *PSPTO_2733* | *cti* | + | 3034729 | 3037041 | 4.52 | 6.84 | 6.20 | 6.56 | 8.41 | 4.35 | Esterified fatty acid cis/trans isomerase |
| *PSPTO_2734* | | + | 3037217 | 3038239 | 20.85 | 55.10 | 38.60 | 23.23 | 29.01 | 92.50 | Acetyltransferase family protein |
| *PSPTO_2735* | *yhgI* | + | 3038310 | 3038903 | 363.88 | 353.69 | 353.29 | 207.29 | 221.87 | 170.66 | YhgI protein |
| *PSPTO_2736* | | - | 3039164 | 3041113 | 12.70 | 2.26 | 3.24 | 3.77 | 83.16 | 9.98 | Biotin carboxylase |
| *PSPTO_2737* | | - | 3041110 | 3041958 | 16.44 | 3.87 | 4.79 | 12.87 | 178.37 | 18.86 | Gamma-carboxygeranoyl-CoA hydratase |
| *PSPTO_2738* | | - | 3041973 | 3043580 | 17.76 | 6.52 | 14.57 | 8.65 | 158.94 | 14.02 | Carboxyl transferase domain protein |
| *PSPTO_2739* | | - | 3043594 | 3044937 | 26.40 | 4.80 | 7.13 | 7.71 | 211.02 | 18.55 | Acyl-CoA dehydrogenase |
| *PSPTO_2740* | | + | 3045260 | 3046282 | 0.00 | 0.00 | 0.00 | 0.00 | 0.00 | 0.00 | ISPsy4, transposase |
| *PSPTO_2741* | | + | 3046279 | 3047088 | 0.00 | 0.00 | 0.00 | 0.00 | 0.00 | 0.00 | ISPsy4, transposition helper protein |
| *PSPTO_2742* | *mvaB* | - | 3047462 | 3048361 | 11.43 | 15.98 | 20.02 | 10.76 | 123.23 | 27.73 | Hydroxymethylglutaryl-CoA lyase |
| *PSPTO_2743* | | - | 3048392 | 3048784 | 12.98 | 13.28 | 11.94 | 2.41 | 278.11 | 25.15 | MerR family transcriptional regulator |
| *PSPTO_2744* | | - | 3048952 | 3049881 | 11.67 | 26.02 | 15.58 | 22.88 | 31.73 | 15.20 | LysR family transcriptional regulator |
| *PSPTO_2745* | | + | 3050066 | 3051295 | 10.31 | 19.72 | 20.64 | 3.32 | 14.92 | 12.96 | Acyl-CoA dehydrogenase family protein |
| *PSPTO_2746* | | - | 3051325 | 3052680 | 7.21 | 9.46 | 8.84 | 2.88 | 3.89 | 5.41 | OmpA family protein |
| *PSPTO_2747* | *xthA* | - | 3052800 | 3053612 | 7.98 | 32.50 | 22.88 | 16.39 | 62.27 | 19.93 | Exodeoxyribonuclease III |
| *PSPTO_2748* | | + | 3053803 | 3054474 | 10.49 | 16.82 | 23.68 | 10.93 | 18.48 | 17.56 | Hypothetical protein |
| *PSPTO_2749* | | + | 3054534 | 3056258 | 4.35 | 13.50 | 12.52 | 2.91 | 23.74 | 7.78 | Hypothetical protein |
| *PSPTO_2750* | | + | 3056255 | 3059944 | 9.19 | 10.65 | 16.42 | 5.59 | 21.44 | 9.04 | Hypothetical protein |
| *PSPTO_2751* | | + | 3060050 | 3060967 | 14.70 | 14.77 | 14.33 | 7.33 | 30.74 | 26.28 | LysR family transcriptional regulator |
| *PSPTO_2752* | *tpx* | + | 3061058 | 3061558 | 65.25 | 134.96 | 109.71 | 71.09 | 282.51 | 39.90 | Thiol peroxidase |
| *PSPTO_2753* | | + | 3061677 | 3063011 | 9.06 | 16.15 | 27.39 | 17.23 | 34.10 | 14.98 | HlyD family secretion protein |
| *PSPTO_2754* | | + | 3063008 | 3066119 | - | - | - | - | - | - | |
| *PSPTO_2755* | | + | 3066116 | 3069223 | 5.10 | 3.94 | 2.55 | 2.69 | 12.53 | 3.51 | AcrB/AcrD/AcrF family protein |
| *PSPTO_2756* | | + | 3069220 | 3070692 | 4.20 | 4.98 | 6.21 | 0.83 | 19.85 | 4.63 | Outer membrane efflux protein |
| *PSPTO_2757* | | + | 3070860 | 3072971 | 5.07 | 8.71 | 8.66 | 5.98 | 6.07 | 6.04 | GGDEF domain/EAL domain protein |
| *PSPTO_2758* | | + | 3073084 | 3073830 | 25.17 | 33.20 | 27.74 | 24.73 | 41.15 | 25.70 | Short-chain dehydrogenase/reductase family oxidoreductase |
| *PSPTO_2759* | | - | 3073877 | 3074059 | 122.38 | 407.07 | 363.59 | 188.20 | 66.23 | 319.76 | Hypothetical protein |
| *PSPTO_2760* | | + | 3074330 | 3076348 | 9.23 | 52.63 | 51.58 | 21.24 | 13.22 | 21.96 | Alpha-amylase family protein |
| *PSPTO_2761* | | + | 3076368 | 3079694 | 7.37 | 33.98 | 28.29 | 17.65 | 11.88 | 16.18 | Alpha-amylase family protein |
| *PSPTO_2762* | *glgB* | + | 3079691 | 3081916 | 13.41 | 22.46 | 21.56 | 13.48 | 10.30 | 12.89 | 1,4-alpha-glucan-branching protein |
| *PSPTO_2763* | | - | 3081957 | 3084740 | 10.98 | 7.25 | 9.04 | 8.22 | 8.44 | 10.82 | Autotransporter |
| *PSPTO_2764* | | - | 3084908 | 3086140 | 4.70 | 2.85 | 3.76 | 1.49 | 6.39 | 4.23 | Major facilitator superfamily transporter |
| *PSPTO_2765* | | + | 3086383 | 3087462 | 15.73 | 18.20 | 11.73 | 8.49 | 5.82 | 7.93 | Hypothetical protein |
| *PSPTO_2766* | | - | 3087525 | 3089090 | 9.84 | 24.43 | 21.37 | 16.14 | 61.51 | 11.95 | ABC transporter ATP-binding protein |
| *PSPTO_2767* | | - | 3089486 | 3090349 | 10.64 | 51.90 | 56.62 | 41.49 | 44.02 | 21.63 | Lipopolysaccharide core biosynthesis domain protein |
| *PSPTO_2768* | | - | 3090795 | 3090971 | 11.62 | 58.31 | 34.59 | 10.31 | 1.72 | 1.93 | Hypothetical protein |
| *PSPTO_2769* | | + | 3091241 | 3091942 | 41.71 | 20.23 | 22.59 | 11.27 | 20.26 | 24.39 | Lipoprotein |
| *PSPTO_2770* | | - | 3091981 | 3092088 | 5.97 | 22.40 | 20.60 | 6.79 | 13.45 | 10.98 | Hypothetical protein |
| *PSPTO_2771* | | + | 3092048 | 3092578 | 2.53 | 10.47 | 3.58 | 0.00 | 4.74 | 4.51 | GNAT family acetyltransferase |
| *PSPTO_2772* | | + | 3092691 | 3093116 | 9.97 | 26.93 | 30.85 | 7.26 | 42.57 | 18.04 | Hypothetical protein |
| *PSPTO_2773* | | + | 3093232 | 3093675 | 30.85 | 74.24 | 40.46 | 24.86 | 33.38 | 24.14 | Hypothetical protein |
| *PSPTO_2774* | | + | 3093688 | 3094083 | 10.47 | 40.95 | 30.08 | 15.21 | 54.91 | 31.08 | Hypothetical protein |
| *PSPTO_2775* | | + | 3094410 | 3095261 | 83.84 | 84.25 | 83.50 | 56.14 | 9.14 | 12.39 | Amino acid ABC transporter substrate-binding protein |
| *PSPTO_2776* | | + | 3095325 | 3095987 | 15.43 | 14.26 | 27.38 | 18.15 | 6.46 | 5.55 | Amino acid ABC transporter permease |
| *PSPTO_2777* | | + | 3095997 | 3096659 | 7.72 | 11.30 | 19.20 | 12.66 | 6.43 | 4.61 | Amino acid ABC transporter permease |
| *PSPTO_2778* | | + | 3096656 | 3097447 | 13.23 | 21.78 | 24.74 | 19.03 | 17.55 | 11.17 | Amino acid ABC transporter ATP-binding protein |
| *PSPTO_2779* | | + | 3097484 | 3097870 | 13.43 | 17.02 | 12.43 | 41.45 | 39.34 | 17.28 | Endoribonuclease L-PSP family protein |
| *PSPTO_2780* | | + | 3098025 | 3098789 | 12.76 | 17.51 | 20.06 | 17.39 | 35.81 | 13.28 | IclR family transcriptional regulator |
| *PSPTO_2781* | | + | 3098807 | 3100042 | 6.55 | 5.87 | 9.61 | 12.85 | 21.94 | 8.01 | Hypothetical protein |
| *PSPTO_2782* | *kdgK* | + | 3100071 | 3100994 | 14.62 | 17.61 | 14.04 | 9.16 | 22.61 | 13.18 | 2-dehydro-3-deoxygluconokinase |
| *PSPTO_2783* | | + | 3101370 | 3103616 | 7.63 | 16.16 | 9.36 | 5.13 | 14.21 | 10.45 | UvrD/REP helicase family protein |
| *PSPTO_2784* | | - | 3103627 | 3103893 | 72.45 | 26.56 | 37.47 | 13.15 | 22.46 | 59.68 | Hypothetical protein |
| *PSPTO_2785* | | + | 3104282 | 3105361 | 22.55 | 20.42 | 19.35 | 23.14 | 26.38 | 19.62 | Putrescine ABC transporter periplasmic putrescine-binding protein |
| *PSPTO_2786* | | - | 3105372 | 3106070 | 5.55 | 7.61 | 5.93 | 23.85 | 5.09 | 4.19 | ABC transporter ATP-binding protein |
| *PSPTO_2787* | | - | 3106089 | 3106937 | 1.73 | 0.16 | 3.66 | 4.20 | 1.48 | 2.63 | ABC transporter permease |

| Locus name | Gene name | Strand | Gene start (bp) | Gene end (bp) | Average TPM Inoculum | Average TPM $t_6$ | Average TPM $t_{24}$ | Average TPM $t_{168}$ | Average TPM Logarithmic | Average TPM Stationary | Annotation description |
|---|---|---|---|---|---|---|---|---|---|---|---|
| *PSPTO_2788* | | - | 3106934 | 3107944 | 3.86 | 15.28 | 4.17 | 3.31 | 3.53 | 3.25 | ABC transporter substrate-binding protein |
| *PSPTO_2789* | | + | 3108133 | 3108279 | 3.50 | 5.94 | 0.00 | 4.93 | 3.11 | 3.07 | Hypothetical protein |
| *PSPTO_2790* | | + | 3108311 | 3109129 | 16.34 | 15.58 | 12.91 | 12.19 | 11.73 | 18.69 | Hypothetical protein |
| *PSPTO_2791* | | - | 3109136 | 3110131 | 7.55 | 37.32 | 40.30 | 21.99 | 21.19 | 11.09 | Oxidoreductase, aldo/keto reductase family |
| *PSPTO_2792* | *lexA-1* | + | 3110567 | 3111175 | 16.87 | 39.64 | 32.53 | 17.34 | 20.04 | 8.10 | LexA repressor |
| *PSPTO_2793* | | + | 3111175 | 3111795 | 1.60 | 3.81 | 6.92 | 0.60 | 5.64 | 2.95 | Hypothetical protein |
| *PSPTO_2794* | | + | 3111755 | 3113218 | 2.69 | 0.81 | 1.35 | 0.92 | 2.88 | 2.60 | Hypothetical protein |
| *PSPTO_2795* | | + | 3113215 | 3116310 | 2.55 | 4.08 | 4.36 | 2.13 | 6.40 | 3.92 | Error-prone DNA polymerase |
| *PSPTO_2796* | | + | 3116381 | 3117232 | 32.16 | 14.67 | 14.09 | 13.54 | 3.27 | 3.90 | Hypothetical protein |
| *PSPTO_2797* | | + | 3119828 | 3119998 | 6.23 | 16.88 | 17.46 | 0.00 | 2.78 | 3.46 | Hypothetical protein |
| *PSPTO_2798* | | - | 3120003 | 3120638 | 5.53 | 6.06 | 4.06 | 0.00 | 2.50 | 5.69 | Transporter |
| *PSPTO_2799* | | + | 3120730 | 3121314 | 13.67 | 26.27 | 6.94 | 9.46 | 9.64 | 17.53 | Hypothetical protein |
| *PSPTO_2800* | | + | 3121311 | 3122255 | 21.85 | 16.76 | 10.52 | 4.90 | 20.87 | 18.54 | Polysaccharide deacetylase family protein |
| *PSPTO_2801* | | - | 3122263 | 3123522 | 1.71 | 3.33 | 1.28 | 5.40 | 2.94 | 3.23 | Prolidase |
| *PSPTO_2802* | *proP* | - | 3123525 | 3124841 | 2.14 | 2.04 | 1.38 | 0.70 | 1.71 | 1.92 | Proline/betaine transporter |
| *PSPTO_2803* | | - | 3125032 | 3125928 | 1.97 | 4.16 | 2.70 | 0.00 | 6.74 | 3.64 | LysR family transcriptional regulator |
| *PSPTO_2804* | | - | 3125953 | 3126858 | 6.65 | 12.03 | 11.40 | 4.50 | 17.05 | 7.84 | LysR family transcriptional regulator |
| *PSPTO_2805* | | + | 3127013 | 3130060 | 6.21 | 16.31 | 17.73 | 9.22 | 19.02 | 27.73 | Oxidoreductase, FAD-binding |
| *PSPTO_2806* | | - | 3130139 | 3131026 | 25.86 | 46.73 | 47.60 | 32.83 | 37.90 | 27.19 | Dienelactone hydrolase family protein |
| *PSPTO_2807* | | + | 3131142 | 3132299 | 10.77 | 9.38 | 3.34 | 3.52 | 4.21 | 5.71 | Diguanylate cyclase |
| *PSPTO_2808* | | + | 3132407 | 3133357 | 12.33 | 9.45 | 6.72 | 4.13 | 7.76 | 5.90 | LysR family transcriptional regulator |
| *PSPTO_2809* | | + | 3133704 | 3135164 | 4.37 | 9.89 | 9.25 | 2.87 | 5.94 | 2.74 | Hypothetical protein |
| *PSPTO_2810* | | + | 3135161 | 3136135 | 1.00 | 1.82 | 0.00 | 0.40 | 1.28 | 0.67 | Peptide ABC transporter permease |
| *PSPTO_2811* | | + | 3136132 | 3137052 | 1.52 | 4.23 | 2.71 | 3.00 | 3.55 | 2.17 | Peptide ABC transporter permease |
| *PSPTO_2812* | | + | 3137049 | 3138062 | 1.41 | 9.64 | 3.20 | 1.35 | 3.93 | 2.00 | Peptide ABC transporter ATP-binding protein |
| *PSPTO_2813* | | + | 3138052 | 3139065 | - | - | - | - | - | - | |
| *PSPTO_2814* | | + | 3139118 | 3140683 | 2.31 | 3.97 | 1.95 | 3.28 | 4.25 | 2.66 | Peptide ABC transporter periplasmic peptide-binding protein |
| *PSPTO_2815* | | + | 3140755 | 3141276 | 1.12 | 0.00 | 0.88 | 0.00 | 4.76 | 2.31 | Hypothetical protein |
| *PSPTO_2816* | | + | 3141276 | 3141866 | 3.25 | 0.60 | 4.49 | 0.00 | 2.88 | 2.77 | Alkylhydroperoxidase AhpD domain-containing protein |
| *PSPTO_2817* | | + | 3141929 | 3142114 | 1.11 | 0.00 | 3.38 | 0.00 | 2.30 | 1.48 | Hypothetical protein |
| *PSPTO_2818* | | - | 3142091 | 3143482 | 1.94 | 4.35 | 4.81 | 6.07 | 9.01 | 5.22 | Fatty acid desaturase |
| *PSPTO_2819* | | + | 3143666 | 3144454 | 16.54 | 23.29 | 31.46 | 6.72 | 39.41 | 18.57 | Hypothetical protein |
| *PSPTO_2820* | | + | 3144448 | 3145563 | 3.26 | 4.00 | 3.66 | 0.26 | 10.70 | 4.85 | Hypothetical protein |
| *PSPTO_2821* | | - | 3145551 | 3146459 | 3.11 | 4.11 | 7.75 | 2.64 | 9.52 | 5.58 | Hypothetical protein |
| *PSPTO_2822* | | - | 3146484 | 3147287 | 3.30 | 10.08 | 6.89 | 5.86 | 11.43 | 4.90 | Hypothetical protein |
| *PSPTO_2823* | | + | 3147654 | 3148403 | 4.02 | 9.65 | 5.89 | 7.24 | 18.80 | 13.14 | Hypothetical protein |
| *PSPTO_2824* | | + | 3148532 | 3149995 | 1.33 | 6.32 | 5.30 | 2.29 | 14.39 | 6.20 | Auxin-responsive GH3-related protein |
| *PSPTO_2825* | | + | 3149980 | 3151056 | 2.01 | 3.97 | 1.60 | 0.12 | 6.62 | 2.56 | Iron-sulfur cluster-binding protein, Rieske family |
| *PSPTO_2826* | | - | 3151069 | 3151368 | 1.99 | 2.23 | 0.00 | 0.00 | 2.55 | 2.06 | Hypothetical protein |
| *PSPTO_2827* | | + | 3151403 | 3151645 | 13.74 | 34.52 | 20.87 | 17.62 | 10.33 | 16.38 | Hypothetical protein |
| *PSPTO_2828* | *syrR* | - | 3151815 | 3152588 | 103.34 | 33.12 | 29.05 | 73.32 | 28.00 | 19.26 | Transcriptional regulator SyrR |
| *PSPTO_2829* | *syfA* | + | 3152879 | 3162385 | 2.25 | 1.93 | 1.28 | 1.87 | 1.41 | 1.70 | Non-ribosomal peptide synthetase SyfA |
| *PSPTO_2830* | *syfB* | + | 3162382 | 3180171 | 2.97 | 2.41 | 3.27 | 2.33 | 2.19 | 2.91 | Non-ribosomal peptide synthetase SyfB |
| *PSPTO_2831* | *syfC* | + | 3180168 | 3181319 | 4.92 | 5.57 | 4.65 | 1.93 | 3.94 | 4.79 | Syringafactin efflux protein SyfC |
| *PSPTO_2832* | *syfD* | + | 3181322 | 3183328 | 5.15 | 4.25 | 7.33 | 3.47 | 7.78 | 5.47 | Syringafactin efflux protein SyfD |
| *PSPTO_2833* | | - | 3183334 | 3184002 | 19.76 | 5.46 | 10.39 | 10.66 | 38.46 | 11.36 | LuxR family transcriptional regulator |
| *PSPTO_2834* | | + | 3184331 | 3185293 | 12.68 | 17.98 | 24.28 | 14.42 | 22.49 | 20.37 | Beta-lactamase |
| *PSPTO_2835* | | - | 3185763 | 3185915 | 46.36 | 54.17 | 76.08 | 11.53 | 54.29 | 71.58 | Hypothetical protein |
| *PSPTO_2836* | | + | 3186079 | 3187056 | 0.45 | 0.07 | 0.24 | 0.59 | 0.99 | 0.61 | ISPssy, transposase |
| *PSPTO_2837* | | + | 3187252 | 3187752 | - | - | - | - | - | - | |
| *PSPTO_2838* | | - | 3188801 | 3189223 | 1.20 | 0.00 | 0.98 | 3.52 | 0.99 | 1.61 | Hypothetical protein |
| *PSPTO_2839* | | + | 3189278 | 3189637 | 0.00 | 0.00 | 0.00 | 0.00 | 0.00 | 0.00 | ISPsy5, Orf1 |
| *PSPTO_2840* | | + | 3189655 | 3191208 | 0.00 | 0.00 | 0.00 | 0.00 | 0.00 | 0.00 | ISPsy5, transposase |
| *PSPTO_2841* | | + | 3191955 | 3193469 | 1.70 | 2.09 | 2.35 | 1.28 | 2.39 | 3.32 | ISPsy14, transposase |
| *PSPTO_2842* | | + | 3193490 | 3194245 | 7.00 | 2.32 | 4.12 | 2.46 | 4.24 | 4.64 | ISPsy14, transposition helper protein |
| *PSPTO_2843* | | + | 3194491 | 3194874 | 126.62 | 35.84 | 35.52 | 22.27 | 38.99 | 40.54 | Hypothetical protein |
| *PSPTO_2844* | | + | 3195189 | 3195731 | 7.88 | 8.62 | 15.55 | 1.04 | 11.21 | 13.23 | Hypothetical protein |
| *PSPTO_2845* | | + | 3195668 | 3196132 | 2.89 | 8.72 | 6.38 | 4.41 | 13.41 | 4.52 | Hypothetical protein |
| *PSPTO_2846* | | + | 3196253 | 3198439 | 12.18 | 9.49 | 10.09 | 4.60 | 16.55 | 9.23 | TonB-dependent siderophore receptor |
| *PSPTO_2847* | | + | 3198450 | 3199280 | 3.49 | 3.77 | 2.64 | 1.29 | 9.47 | 4.52 | ThiJ/PfpI family protein |
| *PSPTO_2848* | | - | 3199419 | 3200018 | - | - | - | - | - | - | |
| *PSPTO_2849* | | - | 3200389 | 3200676 | 0.00 | 1.68 | 1.00 | 0.00 | 0.37 | 15.43 | Cytochrome c family protein |
| *PSPTO_2850* | | - | 3200905 | 3201393 | 0.93 | 2.81 | 3.02 | 1.50 | 1.51 | 75.51 | Hypothetical protein |
| *PSPTO_2851* | | - | 3201393 | 3201959 | 0.48 | 2.41 | 2.52 | 0.65 | 1.90 | 76.49 | SCO1/SenC family protein |
| *PSPTO_2852* | | + | 3203321 | 3203902 | 18.50 | 9.33 | 2.91 | 6.54 | 7.27 | 17.38 | Hypothetical protein |
| *PSPTO_2853* | | + | 3204648 | 3207002 | 8.11 | 6.08 | 5.07 | 4.64 | 6.93 | 54.19 | TonB-dependent receptor |
| *PSPTO_2855* | | + | 3208312 | 3208635 | 3.71 | 3.60 | 2.99 | 2.82 | 3.15 | 4.63 | DNA-binding protein |
| *PSPTO_2856* | | - | 3209245 | 3210549 | 10.06 | 67.63 | 49.25 | 41.44 | 142.62 | 100.81 | Site-specific recombinase, phage integrase family |
| *PSPTO_2857* | | + | 3213297 | 3214631 | 2.79 | 3.81 | 3.45 | 3.98 | 9.85 | 4.94 | Site-specific recombinase, phage integrase family |
| *PSPTO_2858* | | + | 3215113 | 3215469 | 4.80 | 7.86 | 10.51 | 0.00 | 3.47 | 7.47 | Hypothetical protein |
| *PSPTO_2859* | | + | 3215716 | 3218316 | 6.94 | 4.03 | 4.49 | 2.70 | 7.88 | 6.46 | Hypothetical protein |
| *PSPTO_2860* | | + | 3218240 | 3219619 | 7.01 | 3.60 | 1.56 | 2.04 | 4.05 | 4.27 | Helicase domain-containing protein |
| *PSPTO_2861* | | + | 3219607 | 3219978 | 3.95 | 13.01 | 9.27 | 12.81 | 5.47 | 11.10 | 4Fe-4S binding protein |
| *PSPTO_2862* | | + | 3219975 | 3220970 | 30.41 | 16.11 | 17.26 | 11.55 | 23.33 | 27.73 | Oxidoreductase, molybdopterin-binding subunit |
| *PSPTO_2863* | | + | 3220976 | 3223186 | 5.69 | 4.81 | 3.05 | 2.26 | 3.90 | 4.46 | Aldehyde oxidase and xanthine dehydrogenase family protein |
| *PSPTO_2864* | | + | 3223226 | 3224140 | 70.73 | 15.20 | 14.63 | 14.99 | 19.26 | 40.08 | LysR family transcriptional regulator |
| *PSPTO_2865* | | - | 3224146 | 3224934 | 3.79 | 0.06 | 6.80 | 0.00 | 5.50 | 6.43 | 4-carboxymuconolactone decarboxylase |
| *PSPTO_2866* | | - | 3224931 | 3225332 | 4.53 | 10.12 | 0.34 | 0.00 | 4.86 | 6.47 | Cupin family protein |
| *PSPTO_2867* | | + | 3225935 | 3226375 | 21.27 | 8.57 | 12.90 | 10.30 | 9.49 | 22.97 | TspO/MBR family protein |

| Locus name | Gene name | Strand | Gene start (bp) | Gene end (bp) | Average TPM Inoculum | Average TPM $t_6$ | Average TPM $t_{24}$ | Average TPM $t_{168}$ | Average TPM Logarithmic | Average TPM Stationary | Annotation description |
|---|---|---|---|---|---|---|---|---|---|---|---|
| *PSPTO_2868* | | - | 3226393 | 3227571 | 24.43 | 27.05 | 16.91 | 15.10 | 10.16 | 32.07 | Hypothetical protein |
| *PSPTO_2869* | | + | 3227547 | 3227669 | 46.67 | 10.44 | 22.22 | 0.00 | 14.03 | 39.90 | Hypothetical protein |
| *PSPTO_2870* | | + | 3227747 | 3229276 | 8.36 | 12.42 | 13.74 | 5.27 | 24.37 | 10.28 | Hypothetical protein |
| *PSPTO_2871* | | + | 3229288 | 3232422 | 1.80 | 1.73 | 1.68 | 1.00 | 16.64 | 2.43 | Hypothetical protein |
| *PSPTO_2872* | | + | 3232419 | 3235118 | 1.80 | 1.59 | 0.64 | 0.04 | 14.86 | 1.86 | HopL1 protein |
| *PSPTO_2873* | | + | 3235122 | 3236159 | 3.11 | 4.70 | 1.20 | 1.07 | 30.84 | 4.21 | Hypothetical protein |
| *PSPTO_2874* | | + | 3236186 | 3238147 | 5.37 | 4.07 | 3.45 | 0.56 | 17.55 | 3.64 | PpkA-related protein |
| *PSPTO_2875* | | + | 3238149 | 3238841 | 2.46 | 0.00 | 6.78 | 0.00 | 8.69 | 3.21 | ABC transporter ATP-binding protein |
| *PSPTO_2876* | | + | 3238825 | 3240048 | 2.30 | 0.82 | 1.24 | 0.75 | 6.08 | 13.81 | ABC transporter permease |
| *PSPTO_2877* | | + | 3240077 | 3241585 | 1.40 | 2.14 | 0.73 | 0.49 | 15.54 | 1.38 | Hypothetical protein |
| *PSPTO_2878* | | + | 3241618 | 3242358 | 2.89 | 0.51 | 3.54 | 1.73 | 24.18 | 1.90 | Lipoprotein |
| *PSPTO_2879* | | + | 3242366 | 3242866 | 2.11 | 2.76 | 0.00 | 0.00 | 13.00 | 1.22 | Lipoprotein |
| *PSPTO_2880* | | + | 3242922 | 3244193 | 12.34 | 4.01 | 5.18 | 0.63 | 4.24 | 4.57 | Hypothetical protein |
| *PSPTO_2881* | | - | 3244444 | 3244542 | 2.40 | 0.00 | 0.00 | 0.00 | 4.90 | 10.51 | Hypothetical protein |
| *PSPTO_2882* | | - | 3244586 | 3245884 | - | - | - | - | - | - | |
| *PSPTO_2883* | | - | 3246195 | 3247556 | 25.99 | 4.82 | 6.68 | 3.08 | 1.57 | 2.82 | Methyl-accepting chemotaxis protein |
| *PSPTO_2885* | | - | 3248015 | 3248374 | 0.10 | 0.00 | 0.00 | 1.83 | 0.13 | 0.00 | ISPsy5, Orf1 |
| *PSPTO_2886* | | - | 3249237 | 3249740 | - | - | - | - | - | - | |
| *PSPTO_2887* | | + | 3249839 | 3250816 | 0.00 | 0.00 | 0.00 | 0.00 | 0.00 | 0.00 | ISPssy, transposase |
| *PSPTO_2888* | | + | 3250923 | 3251069 | 1.93 | 4.45 | 0.00 | 0.00 | 2.47 | 1.34 | Hypothetical protein |
| *PSPTO_2889* | | + | 3251328 | 3251441 | 10.91 | 11.48 | 0.00 | 0.00 | 7.01 | 10.26 | Hypothetical protein |
| *PSPTO_2890* | | + | 3251438 | 3251683 | 3.24 | 6.41 | 1.05 | 0.00 | 1.82 | 4.79 | Hypothetical protein |
| *PSPTO_2891* | | + | 3251661 | 3253055 | 4.82 | 3.09 | 3.25 | 0.26 | 2.47 | 3.60 | UDP-glucose 6-dehydrogenase |
| *PSPTO_2892* | | + | 3253146 | 3253244 | 0.49 | 0.00 | 0.00 | 3.31 | 0.47 | 1.53 | Hypothetical protein |
| *PSPTO_2893* | | + | 3253364 | 3254194 | 12.37 | 24.76 | 22.13 | 9.80 | 23.22 | 11.89 | PAP2 superfamily protein |
| *PSPTO_2894* | | + | 3254445 | 3256850 | 6.02 | 4.26 | 4.00 | 3.53 | 5.90 | 7.81 | Lectin repeat domain protein |
| *PSPTO_2895* | | - | 3257076 | 3258095 | 8.09 | 38.54 | 24.73 | 32.45 | 23.88 | 31.03 | Hypothetical protein |
| *PSPTO_2896* | | - | 3258416 | 3260020 | 21.78 | 29.27 | 35.97 | 19.42 | 24.24 | 33.08 | Sensory box histidine kinase/response regulator |
| *PSPTO_2897* | | - | 3260162 | 3261304 | 3.43 | 5.23 | 1.15 | 2.93 | 2.25 | 3.54 | Hypothetical protein |
| *PSPTO_2898* | *glf* | - | 3261307 | 3262488 | 4.19 | 7.59 | 3.26 | 1.04 | 3.49 | 5.10 | UDP-galactopyranose mutase |
| *PSPTO_2899* | | - | 3262476 | 3263660 | 2.46 | 3.56 | 3.55 | 0.88 | 1.51 | 2.38 | Hypothetical protein |
| *PSPTO_2900* | *galE* | + | 3264060 | 3265088 | 8.38 | 8.29 | 9.67 | 4.13 | 6.57 | 7.35 | UDP-glucose 4-epimerase |
| *PSPTO_2901* | | - | 3265109 | 3265657 | 12.90 | 23.34 | 19.47 | 23.62 | 31.45 | 22.66 | DNA-binding protein |
| *PSPTO_2902* | | - | 3265770 | 3266534 | 4.33 | 7.06 | 3.08 | 2.43 | 2.57 | 2.83 | Glutamine amidotransferase |
| *PSPTO_2903* | | + | 3266662 | 3267564 | 3.19 | 12.23 | 9.21 | 0.00 | 14.33 | 4.47 | LysR family transcriptional regulator |
| *PSPTO_2904* | *vanA* | + | 3267712 | 3268776 | 6.47 | 84.21 | 62.48 | 19.21 | 33.25 | 7.07 | Vanillate O-demethylase, oxygenase subunit |
| *PSPTO_2905* | *vanB* | + | 3268846 | 3269796 | 6.21 | 24.31 | 29.70 | 15.14 | 12.73 | 6.41 | Vanillate O-demethylase oxidoreductase |
| *PSPTO_2906* | *vanR* | - | 3269797 | 3270507 | 11.54 | 21.55 | 7.96 | 10.05 | 10.54 | 9.59 | Transcriptional regulator VanR |
| *PSPTO_2907* | | - | 3270583 | 3272553 | 20.87 | 9.26 | 4.64 | 3.31 | 3.63 | 6.06 | GGDEF domain/EAL domain protein |
| *PSPTO_2908* | | - | 3272857 | 3273549 | 88.25 | 67.11 | 81.67 | 35.05 | 6.44 | 29.87 | Hypothetical protein |
| *PSPTO_2909* | | - | 3273689 | 3274588 | 4.76 | 9.13 | 3.45 | 1.01 | 7.18 | 5.55 | LysR family transcriptional regulator |
| *PSPTO_2910* | | + | 3274733 | 3276076 | 6.41 | 13.69 | 18.35 | 7.67 | 6.44 | 6.88 | Tartrate transporter |
| *PSPTO_2911* | | + | 3276096 | 3277370 | 3.60 | 2.30 | 21.98 | 6.34 | 3.03 | 1.46 | Mandelate racemase/muconate lactonizing protein |
| *PSPTO_2912* | | - | 3277425 | 3278258 | 2.61 | 4.27 | 3.91 | 4.14 | 3.73 | 2.77 | Glutamine ABC transporter, periplasmic amino acid-binding protein |
| *PSPTO_2913* | | - | 3278322 | 3279059 | 2.04 | 1.01 | 2.80 | 6.97 | 3.17 | 1.89 | Glutamine ABC transporter ATP-binding protein |
| *PSPTO_2914* | | - | 3279070 | 3279810 | 2.95 | 3.81 | 6.56 | 3.72 | 5.17 | 3.54 | Glutamine ABC transporter, permease protein |
| *PSPTO_2915* | | - | 3279817 | 3280518 | 1.95 | 2.65 | 4.18 | 1.30 | 1.61 | 2.17 | Glutamine ABC transporter, permease protein |
| *PSPTO_2916* | | - | 3280640 | 3281392 | 7.41 | 3.47 | 6.40 | 6.32 | 5.44 | 4.80 | Short chain dehydrogenase/reductase family oxidoreductase |
| *PSPTO_2917* | | - | 3281523 | 3282293 | 2.43 | 2.74 | 2.55 | 0.68 | 3.19 | 2.83 | LamB/YcsF family protein |
| *PSPTO_2918* | | - | 3282298 | 3282705 | 1.34 | 0.00 | 6.62 | 0.00 | 1.80 | 1.82 | Hypothetical protein |
| *PSPTO_2919* | | - | 3282705 | 3283676 | 1.39 | 4.04 | 2.65 | 0.00 | 1.94 | 1.18 | Urea amidolyase-related protein |
| *PSPTO_2920* | | - | 3283673 | 3284344 | 2.01 | 1.72 | 1.66 | 1.76 | 1.23 | 3.75 | Hypothetical protein |
| *PSPTO_2921* | *accC-1* | - | 3284341 | 3285693 | 1.06 | 2.30 | 1.98 | 1.01 | 1.44 | 1.77 | Acetyl-CoA carboxylase, biotin carboxylase |
| *PSPTO_2922* | | - | 3285734 | 3286171 | 1.36 | 5.94 | 2.87 | 0.30 | 1.06 | 1.52 | Acetyl-CoA carboxylase, biotin carboxyl carrier protein |
| *PSPTO_2923* | | - | 3286291 | 3287199 | 8.07 | 11.00 | 13.22 | 6.97 | 4.86 | 5.51 | Nitrogen assimilation transcriptional regulator |
| *PSPTO_2924* | | + | 3287122 | 3287283 | 7.42 | 3.13 | 4.79 | 0.00 | 2.44 | 1.26 | Hypothetical protein |
| *PSPTO_2925* | *aspB* | + | 3287487 | 3288695 | 2.41 | 2.27 | 3.63 | 1.30 | 1.81 | 2.76 | Aspartate aminotransferase |
| *PSPTO_2926* | | + | 3288706 | 3290265 | 3.79 | 6.59 | 4.27 | 1.57 | 3.19 | 6.15 | Multidrug transporter |
| *PSPTO_2927* | *oprE* | + | 3290338 | 3291669 | 6.08 | 8.24 | 7.72 | 6.27 | 3.11 | 7.43 | Outer membrane porin OprE |
| *PSPTO_2928* | | + | 3291838 | 3293511 | 18.37 | 13.58 | 14.92 | 9.95 | 16.01 | 18.52 | DnaJ domain-containing protein |
| *PSPTO_2929* | | + | 3293501 | 3294463 | 53.54 | 47.66 | 43.92 | 25.13 | 62.05 | 53.93 | Hypothetical protein |
| *PSPTO_2930* | | - | 3294625 | 3294933 | 6.48 | 0.57 | 9.08 | 0.00 | 8.20 | 7.66 | Hypothetical protein |
| *PSPTO_2931* | | - | 3294960 | 3296144 | 12.09 | 19.05 | 15.81 | 6.62 | 25.14 | 17.60 | Hypothetical protein |
| *PSPTO_2932* | | - | 3296385 | 3297392 | 14.96 | 78.85 | 68.44 | 17.95 | 4.72 | 6.00 | Oxidoreductase, aldo/keto reductase family |
| *PSPTO_2933* | | + | 3297568 | 3298149 | 93.31 | 172.33 | 158.69 | 40.40 | 62.79 | 49.12 | TetR family transcriptional regulator |
| *PSPTO_2934* | | - | 3298155 | 3299129 | 5.11 | 5.75 | 18.15 | 5.58 | 6.81 | 8.91 | Urea amidolyase-related protein |
| *PSPTO_2935* | | - | 3299126 | 3299998 | 1.66 | 9.05 | 8.62 | 4.39 | 2.45 | 3.25 | Hypothetical protein |
| *PSPTO_2936* | | - | 3300087 | 3300842 | 2.35 | 15.77 | 12.47 | 6.41 | 2.34 | 2.70 | LamB/YcsF family protein |
| *PSPTO_2937* | | + | 3300969 | 3301886 | 2.24 | 9.85 | 4.26 | 0.00 | 7.34 | 3.62 | LysR family transcriptional regulator |
| *PSPTO_2938* | | - | 3301897 | 3302214 | 1.50 | 10.05 | 2.77 | 5.76 | 7.09 | 3.04 | Hypothetical protein |
| *PSPTO_2939* | | - | 3302230 | 3302742 | 13.08 | 20.83 | 8.43 | 11.81 | 11.41 | 10.29 | Hypothetical protein |
| *PSPTO_2940* | | - | 3302739 | 3303980 | 2.25 | 0.00 | 1.17 | 0.15 | 1.25 | 1.90 | Thiolase family protein |
| *PSPTO_2941* | *fcs* | - | 3303977 | 3305746 | 4.45 | 5.26 | 6.37 | 3.63 | 5.90 | 6.83 | Feruloyl-CoA synthase |
| *PSPTO_2942* | | - | 3305774 | 3305878 | 3.75 | 14.67 | 0.00 | 0.00 | 2.15 | 1.80 | Hypothetical protein |
| *PSPTO_2943* | *vdh* | - | 3306019 | 3307467 | 5.65 | 8.32 | 5.73 | 2.45 | 8.05 | 5.06 | Vanillin dehydrogenase |
| *PSPTO_2944* | | - | 3307573 | 3308403 | 6.72 | 9.12 | 5.11 | 5.75 | 12.90 | 7.02 | P-hydroxycinnamoyl CoA hydratase/lyase |
| *PSPTO_2945* | | + | 3308615 | 3309103 | 4.14 | 13.71 | 11.87 | 6.00 | 11.65 | 10.86 | MarR family transcriptional regulator |
| *PSPTO_2946* | | - | 3309146 | 3310468 | 3.63 | 0.68 | 2.60 | 3.36 | 2.64 | 2.79 | Outer membrane porin, OprD family |
| *PSPTO_2947* | | + | 3310822 | 3312036 | 3.46 | 7.92 | 3.55 | 9.29 | 3.60 | 3.14 | Major facilitator family transporter |

| Locus name | Gene name | Strand | Gene start (bp) | Gene end (bp) | Average TPM Inoculum | Average TPM $t_6$ | Average TPM $t_{24}$ | Average TPM $t_{168}$ | Average TPM Logarithmic | Average TPM Stationary | Annotation description |
|---|---|---|---|---|---|---|---|---|---|---|---|
| *PSPTO_2948* | | - | 3312511 | 3313457 | - | - | - | - | - | - | |
| *PSPTO_2949* | | - | 3313545 | 3314312 | 6.62 | 26.28 | 16.31 | 7.37 | 3.95 | 3.59 | 3-alpha-hydroxysteroid dehydrogenase |
| *PSPTO_2950* | *xylcC* | - | 3314398 | 3315879 | 5.73 | 33.83 | 19.92 | 5.82 | 3.23 | 3.22 | Benzaldehyde dehydrogenase |
| *PSPTO_2951* | | + | 3316244 | 3317929 | 3.50 | 3.32 | 6.63 | 2.26 | 8.06 | 3.03 | Sigma-54 dependent transcriptional regulator |
| *PSPTO_2952* | | - | 3317979 | 3320042 | 11.67 | 34.85 | 40.08 | 11.47 | 11.16 | 15.85 | Glycosidase |
| *PSPTO_2953* | | - | 3320793 | 3321053 | 5.60 | 5.50 | 6.21 | 0.00 | 4.44 | 4.10 | LysR family transcriptional regulator |
| *PSPTO_2954* | | - | 3321297 | 3321902 | 4.97 | 4.31 | 4.04 | 5.48 | 6.09 | 5.36 | ThiJ/PfpI family protein |
| *PSPTO_2955* | *ftrA* | + | 3322024 | 3322977 | 14.03 | 18.32 | 29.04 | 15.96 | 37.09 | 20.03 | Transcriptional activator FtrA |
| *PSPTO_2956* | *estB* | - | 3322937 | 3324091 | 3.66 | 1.39 | 2.99 | 6.22 | 4.58 | 4.92 | Esterase |
| *PSPTO_2957* | *medH* | - | 3324119 | 3325273 | 2.19 | 1.90 | 2.01 | 2.06 | 1.50 | 1.72 | Methanol dehydrogenase, NAD-dependent |
| *PSPTO_2958* | *rhtC* | - | 3325310 | 3325954 | 2.08 | 2.46 | 2.90 | 0.00 | 3.61 | 1.85 | Threonine efflux protein |
| *PSPTO_2959* | | - | 3326074 | 3327729 | 0.61 | 0.68 | 1.83 | 1.87 | 1.37 | 0.89 | Hypothetical protein |
| *PSPTO_2960* | | - | 3327729 | 3329852 | 1.51 | 0.99 | 0.00 | 0.00 | 2.03 | 1.86 | Pyridoxal-phosphate dependent enzyme family/ornithine cyclodeaminase family protein |
| *PSPTO_2961* | *pvdA* | - | 3329849 | 3331171 | 1.01 | 1.30 | 1.65 | 0.69 | 1.97 | 1.23 | L-lysine 6-monooxygenase |
| *PSPTO_2962* | | - | 3331226 | 3332179 | 0.88 | 0.92 | 0.62 | 2.79 | 2.12 | 2.52 | Hypothetical protein |
| *PSPTO_2963* | | - | 3332197 | 3333336 | 0.90 | 1.80 | 1.22 | 1.66 | 2.08 | 3.00 | Oxidoreductase, FAD-binding protein |
| *PSPTO_2964* | | - | 3333375 | 3334859 | 1.70 | 0.73 | 3.27 | 1.75 | 1.53 | 2.10 | Hypothetical protein |
| *PSPTO_2965* | | - | 3334867 | 3335259 | 2.64 | 8.84 | 6.26 | 10.35 | 6.73 | 4.11 | Hypothetical protein |
| *PSPTO_2966* | | - | 3335609 | 3336574 | 4.24 | 11.68 | 13.14 | 1.90 | 13.98 | 6.04 | Site-specific recombinase, phage integrase family |
| *PSPTO_2967* | | + | 3336803 | 3337246 | 1.78 | 11.45 | 14.25 | 0.00 | 4.30 | 3.78 | Hypothetical protein |
| *PSPTO_2968* | | - | 3337272 | 3338255 | 3.97 | 7.83 | 0.12 | 3.95 | 5.64 | 6.60 | Alcohol dehydrogenase |
| *PSPTO_2969* | | - | 3338318 | 3338806 | 2.38 | 9.30 | 1.70 | 2.25 | 5.23 | 4.50 | Hypothetical protein |
| *PSPTO_2970* | | + | 3338861 | 3339220 | 0.00 | 0.00 | 0.00 | 0.00 | 0.00 | 0.00 | ISPsy5, Orf1 |
| *PSPTO_2971* | | + | 3339238 | 3340791 | 0.00 | 0.00 | 0.00 | 0.00 | 0.00 | 0.00 | ISPsy5, transposase |
| *PSPTO_2972* | | + | 3341158 | 3342237 | 11.80 | 21.72 | 12.06 | 23.66 | 43.26 | 28.30 | Oxidoreductase, 2-nitropropane dioxygenase family |
| *PSPTO_2973* | *modA* | + | 3342393 | 3343106 | 33.89 | 21.15 | 32.04 | 16.96 | 79.73 | 31.76 | Molybdate ABC transporter periplasmic molybdate-binding protein |
| *PSPTO_2974* | *modB* | + | 3343107 | 3343787 | 8.56 | 7.37 | 11.25 | 2.36 | 25.42 | 11.98 | Molybdate ABC transporter permease |
| *PSPTO_2975* | *modC* | + | 3343791 | 3344879 | 7.20 | 4.97 | 5.46 | 3.21 | 23.83 | 9.36 | Molybdate ABC transporter ATP-binding protein |
| *PSPTO_2976* | | + | 3344968 | 3346023 | 6.31 | 28.88 | 16.76 | 6.59 | 9.84 | 12.24 | DNA topoisomerase, type I |
| *PSPTO_2977* | | - | 3346580 | 3347215 | 10.06 | 14.88 | 9.86 | 7.41 | 24.25 | 17.98 | Hypothetical protein |
| *PSPTO_2978* | | - | 3348163 | 3348912 | 12.56 | 9.87 | 15.77 | 5.30 | 14.21 | 8.98 | Chitin-binding protein |
| *PSPTO_2979* | | - | 3349079 | 3349465 | 7.03 | 6.81 | 1.27 | 0.00 | 3.12 | 3.47 | Hypothetical protein |
| *PSPTO_2980* | | - | 3349467 | 3350480 | 11.37 | 2.10 | 13.88 | 2.69 | 2.68 | 3.77 | 2OG-Fe(II) oxygenase family oxidoreductase |
| *PSPTO_2981* | | - | 3350706 | 3351389 | 55.90 | 24.31 | 12.72 | 7.23 | 6.34 | 12.87 | Isochorismatase family protein |
| *PSPTO_2982* | | - | 3351821 | 3353551 | 6.22 | 5.06 | 8.93 | 2.09 | 9.20 | 9.23 | Hypothetical protein |
| *PSPTO_2983* | | + | 3353925 | 3354110 | 7.65 | 3.09 | 1.52 | 2.80 | 1.12 | 1.03 | Histidine kinase phosphoacceptor domain protein |
| *PSPTO_2984* | | - | 3354211 | 3355611 | 3.96 | 4.47 | 10.62 | 5.84 | 2.62 | 3.09 | Xenobiotic compound monooxygenase, DszA family |
| *PSPTO_2985* | | - | 3356072 | 3357352 | 5.11 | 4.84 | 7.87 | 11.71 | 1.27 | 1.28 | Acyl-CoA dehydrogenase family protein |
| *PSPTO_2986* | | - | 3357472 | 3358248 | 1.24 | 0.89 | 1.41 | 0.72 | 1.41 | 2.05 | Branched-chain amino acid ABC transporter ATP-binding protein |
| *PSPTO_2987* | | - | 3358253 | 3359584 | 2.67 | 2.59 | 3.95 | 0.00 | 2.96 | 2.16 | Hypothetical protein |
| *PSPTO_2988* | | - | 3359635 | 3360702 | 3.91 | 2.88 | 3.64 | 2.68 | 1.44 | 3.67 | Branched-chain amino acid ABC transporter permease |
| *PSPTO_2989* | | - | 3360726 | 3361607 | 0.57 | 0.85 | 0.00 | 0.60 | 0.47 | 1.30 | Branched-chain amino acid ABC transporter permease |
| *PSPTO_2990* | | - | 3361611 | 3362405 | 2.91 | 0.83 | 2.10 | 1.97 | 0.49 | 1.97 | Branched-chain amino acid ABC transporter ATP-binding protein |
| *PSPTO_2991* | | - | 3362402 | 3363760 | 3.81 | 1.95 | 1.80 | 0.26 | 1.43 | 2.53 | Hypothetical protein |
| *PSPTO_2992* | | + | 3363774 | 3365012 | 2.63 | 3.69 | 2.12 | 5.71 | 1.44 | 2.38 | Acyl-CoA dehydrogenase family protein |
| *PSPTO_2993* | | - | 3365025 | 3366011 | 3.62 | 3.56 | 3.32 | 2.39 | 2.57 | 2.48 | Ribose ABC transporter permease |
| *PSPTO_2994* | | - | 3366008 | 3367012 | 3.54 | 4.61 | 3.70 | 3.65 | 2.51 | 2.74 | Ribose ABC transporter permease |
| *PSPTO_2995* | | - | 3366999 | 3368567 | 2.16 | 2.58 | 3.29 | 1.90 | 0.97 | 1.49 | Ribose ABC transporter, ATP-binding protein |
| *PSPTO_2996* | | - | 3368584 | 3369588 | 15.17 | 4.51 | 10.51 | 8.69 | 1.78 | 2.49 | Ribose ABC transporter, periplasmic ribose-binding protein |
| *PSPTO_2997* | | - | 3369813 | 3371441 | 12.56 | 6.79 | 5.68 | 1.95 | 3.87 | 5.15 | Methyl-accepting chemotaxis protein |
| *PSPTO_2998* | | + | 3371648 | 3372076 | 33.94 | 60.95 | 50.73 | 24.61 | 141.28 | 31.37 | Hypothetical protein |
| *PSPTO_2999* | | + | 3372037 | 3372789 | 20.20 | 33.29 | 31.99 | 9.39 | 16.79 | 6.85 | Hypothetical protein |
| *PSPTO_3000* | | + | 3373117 | 3373884 | 2.30 | 81.79 | 133.29 | 5.95 | 2.13 | 3.03 | Papain cysteine protease family protein |
| *PSPTO_3001* | *xylR* | - | 3373902 | 3375089 | 9.70 | 21.31 | 24.50 | 3.85 | 6.96 | 5.55 | Xylose operon regluatory protein |
| *PSPTO_3002* | *xylA* | + | 3375268 | 3376584 | 11.91 | 29.36 | 25.84 | 15.76 | 7.38 | 11.51 | Xylose isomerase |
| *PSPTO_3003* | *xylF* | + | 3376725 | 3377726 | 6.70 | 25.72 | 21.70 | 14.41 | 2.15 | 2.29 | D-xylose ABC transporter, periplasmic-D xylose binding protein |
| *PSPTO_3004* | *xylG* | + | 3377835 | 3379421 | 6.42 | 11.20 | 7.23 | 6.11 | 4.70 | 4.28 | Xylose transporter ATP-binding subunit |
| *PSPTO_3005* | *xylH* | + | 3379422 | 3380558 | 2.35 | 5.86 | 5.46 | 4.65 | 3.40 | 3.25 | D-xylose ABC transporter permease |
| *PSPTO_3006* | *amnE* | - | 3380581 | 3381015 | 3.13 | 2.06 | 1.65 | 10.00 | 5.18 | 4.69 | Deaminase AmnE |
| *PSPTO_3007* | | - | 3381057 | 3382640 | 2.46 | 4.08 | 2.08 | 1.74 | 2.72 | 2.91 | Amine oxidase, flavin-containing protein |
| *PSPTO_3008* | | - | 3382663 | 3383592 | 2.55 | 3.18 | 2.71 | 0.00 | 3.71 | 3.98 | Amino acid ABC transporter substrate-binding protein |
| *PSPTO_3009* | | - | 3383621 | 3384406 | 1.04 | 1.11 | 1.25 | 1.87 | 2.52 | 1.86 | Amino acid ABC transporter ATP-binding protein |
| *PSPTO_3010* | | - | 3384399 | 3385373 | 3.24 | 0.56 | 0.96 | 1.06 | 3.04 | 2.89 | Amino acid ABC transporter permease |
| *PSPTO_3011* | | - | 3385417 | 3387882 | 1.85 | 1.77 | 0.38 | 2.18 | 0.90 | 2.18 | TonB-dependent sideophore receptor |
| *PSPTO_3012* | | - | 3388401 | 3388955 | - | - | - | - | - | - | |
| *PSPTO_3013* | | + | 3389035 | 3390057 | 0.00 | 0.00 | 0.00 | 0.00 | 0.00 | 0.00 | ISPsy4, transposase |
| *PSPTO_3014* | | + | 3390054 | 3390863 | 0.00 | 0.00 | 0.00 | 0.00 | 0.00 | 0.00 | ISPsy4, transposition helper protein |
| *PSPTO_3015* | | - | 3390873 | 3391037 | - | - | - | - | - | - | |
| *PSPTO_3016* | | + | 3391425 | 3391778 | 15.32 | 13.81 | 7.25 | 2.59 | 4.51 | 6.47 | Hypothetical protein |
| *PSPTO_3017* | | - | 3391855 | 3392907 | 38.53 | 17.43 | 10.77 | 4.54 | 44.61 | 18.76 | LacI family transcriptional regulator |
| *PSPTO_3018* | | + | 3393046 | 3393969 | 6.85 | 3.76 | 2.89 | 1.81 | 4.48 | 3.02 | Hypothetical protein |
| *PSPTO_3019* | | + | 3393989 | 3395158 | 5.09 | 5.28 | 8.32 | 6.76 | 5.96 | 4.84 | Oxidoreductase, Gfo/Idh/MocA family |
| *PSPTO_3020* | | - | 3395538 | 3396407 | 6.28 | 4.72 | 10.16 | 5.16 | 16.13 | 7.07 | Hypothetical protein |
| *PSPTO_3021* | | + | 3396692 | 3397567 | 9.93 | 37.07 | 28.94 | 14.66 | 70.32 | 11.78 | LysR family transcriptional regulator |
| *PSPTO_3022* | *pgsA* | - | 3397895 | 3398461 | 28.16 | 23.29 | 24.85 | 14.21 | 49.68 | 30.63 | CDP-diacylglycerol--glycerol-3-phosphate 3-phosphatidyltransferase |
| *PSPTO_3023* | *uvrC* | - | 3398488 | 3400311 | 7.99 | 17.97 | 18.43 | 7.26 | 40.16 | 14.78 | Excinuclease ABC subunit C |
| *PSPTO_3024* | *gacA* | - | 3400314 | 3400958 | 32.01 | 82.78 | 78.65 | 27.14 | 100.23 | 34.25 | DNA-binding response regulator GacA |
| *PSPTO_3025* | | - | 3401126 | 3401488 | 29.08 | 37.50 | 43.68 | 18.96 | 19.55 | 28.26 | DNA-binding protein |
| *PSPTO_3026* | | + | 3401570 | 3401815 | 428.27 | 370.95 | 239.50 | 125.60 | 133.51 | 105.18 | Hypothetical protein |

| Locus name | Gene name | Strand | Gene start (bp) | Gene end (bp) | Average TPM Inoculum | Average TPM $t_6$ | Average TPM $t_{24}$ | Average TPM $t_{168}$ | Average TPM Logarithmic | Average TPM Stationary | Annotation description |
|---|---|---|---|---|---|---|---|---|---|---|---|
| *PSPTO_3027* | *aroF-2* | + | 3402517 | 3403599 | 17.20 | 22.20 | 8.78 | 10.04 | 30.07 | 21.92 | Phospho-2-dehydro-3-deoxyheptonate aldolase |
| *PSPTO_3028* | *ppiC-1* | + | 3403680 | 3403961 | 59.42 | 127.69 | 132.37 | 43.86 | 171.56 | 46.90 | Peptidyl-prolyl cis-trans isomerase C |
| *PSPTO_3029* | | + | 3404189 | 3406057 | 3.52 | 9.41 | 6.32 | 1.44 | 11.84 | 5.12 | Peptide ABC transporter periplasmic peptide-binding protein |
| *PSPTO_3030* | | + | 3406058 | 3407122 | 2.97 | 7.45 | 4.25 | 1.22 | 7.67 | 5.39 | Peptide ABC transporter permease |
| *PSPTO_3031* | | + | 3407122 | 3408144 | 2.35 | 1.77 | 3.67 | 2.80 | 7.06 | 3.07 | Peptide ABC transporter permease |
| *PSPTO_3032* | | + | 3408146 | 3409723 | 2.28 | 1.43 | 4.07 | 0.70 | 8.65 | 4.56 | Peptide ABC transporter ATP-binding protein |
| *PSPTO_3033* | | - | 3409902 | 3411077 | 8.53 | 18.13 | 16.86 | 6.43 | 16.21 | 21.07 | Twitching motility protein |
| *PSPTO_3034* | *ppiC-2* | + | 3411226 | 3411501 | 24.69 | 22.45 | 17.06 | 13.14 | 16.38 | 19.63 | Peptidyl-prolyl cis-trans isomerase C |
| *PSPTO_3035* | *pgm* | - | 3411577 | 3413223 | 23.29 | 34.72 | 37.64 | 18.71 | 35.64 | 38.86 | Phosphoglucomutase, alpha-D-glucose phosphate-specific |
| *PSPTO_3036* | | - | 3413324 | 3415609 | 3.76 | 3.07 | 5.17 | 4.17 | 6.25 | 4.65 | Hypothetical protein |
| *PSPTO_3037* | | - | 3415757 | 3417163 | 2.07 | 3.44 | 5.26 | 5.83 | 3.63 | 4.44 | Transcriptional regulator |
| *PSPTO_3038* | | - | 3417166 | 3418668 | 1.77 | 2.29 | 0.16 | 0.00 | 1.36 | 1.95 | Hypothetical protein |
| *PSPTO_3039* | | - | 3419246 | 3420268 | 37.81 | 196.98 | 192.09 | 51.49 | 43.04 | 64.24 | Alcohol dehydrogenase |
| *PSPTO_3040* | | + | 3420579 | 3421493 | 34.86 | 35.30 | 24.12 | 26.46 | 32.56 | 34.60 | LysR family transcriptional regulator |
| *PSPTO_3041* | | - | 3421941 | 3422894 | 0.78 | 1.77 | 0.15 | 0.00 | 1.61 | 1.05 | Membrane protein |
| *PSPTO_3042* | | - | 3423125 | 3423613 | - | - | - | - | - | - | |
| *PSPTO_3043* | | - | 3423692 | 3424648 | 3.29 | 4.88 | 3.08 | 5.44 | 4.31 | 6.30 | 5'-nucleotidase |
| *PSPTO_3044* | | - | 3424661 | 3425506 | 5.31 | 7.27 | 7.26 | 0.44 | 5.67 | 8.19 | ISPsy12, transposase OrfB |
| *PSPTO_3045* | | - | 3425515 | 3425703 | 1.49 | 4.36 | 8.52 | 0.00 | 3.37 | 3.91 | ISPsy12, transposase OrfA |
| *PSPTO_3046* | | - | 3425886 | 3427889 | 24.78 | 14.01 | 7.64 | 2.14 | 27.30 | 6.05 | Sigma-54 dependent transcriptional regulator |
| *PSPTO_3047* | | + | 3428075 | 3429097 | 11.10 | 4.77 | 11.12 | 6.02 | 5.51 | 4.45 | Myo-inositol 2-dehydrogenase |
| *PSPTO_3048* | | + | 3429094 | 3429882 | 7.26 | 7.73 | 8.67 | 6.58 | 8.77 | 9.50 | Hypothetical protein |
| *PSPTO_3049* | | + | 3429981 | 3430907 | 11.92 | 7.41 | 8.21 | 5.83 | 7.18 | 8.96 | Sugar ABC transporter, periplasmic sugar-binding |
| *PSPTO_3050* | | - | 3431400 | 3432371 | 16.63 | 12.91 | 12.09 | 27.07 | 9.59 | 8.93 | AraC family transcriptional regulator |
| *PSPTO_3051* | | + | 3432613 | 3433146 | 143.29 | 56.55 | 70.75 | 37.06 | 39.62 | 74.72 | 2,4'-dihydroxyacetophenone dioxygenase |
| *PSPTO_3052* | | + | 3433170 | 3434411 | 7.84 | 13.20 | 12.27 | 7.92 | 7.45 | 9.39 | Major facilitator family transporter |
| *PSPTO_3053* | | + | 3434473 | 3435234 | 4.83 | 9.24 | 0.00 | 3.87 | 6.06 | 4.61 | Hypothetical protein |
| *PSPTO_3054* | | + | 3435263 | 3435994 | 3.91 | 6.71 | 4.18 | 3.38 | 3.80 | 2.93 | Short chain dehydrogenase/reductase family oxidoreductase |
| *PSPTO_3055* | | - | 3436113 | 3436262 | 0.35 | 0.00 | 0.00 | 0.00 | 0.96 | 1.07 | Hypothetical protein |
| *PSPTO_3056* | | - | 3436246 | 3437280 | 6.80 | 7.19 | 5.17 | 7.71 | 7.01 | 8.21 | AraC family transcriptional regulator |
| *PSPTO_3057* | | - | 3437310 | 3438662 | 3.81 | 0.89 | 4.36 | 4.50 | 3.72 | 3.11 | MmgE/PrpD family protein |
| *PSPTO_3058* | | - | 3438739 | 3439716 | 5.94 | 12.40 | 10.13 | 4.27 | 6.51 | 6.08 | Glycine betaine transporter periplasmic subunit |
| *PSPTO_3059* | | - | 3439826 | 3440791 | 3.67 | 6.77 | 5.41 | 3.05 | 2.54 | 3.56 | Glycine betaine/L-proline ABC transporter permease |
| *PSPTO_3060* | | - | 3440841 | 3441878 | 5.00 | 1.14 | 8.09 | 2.19 | 1.92 | 6.48 | Glycine betaine/L-proline ABC transporter ATP-binding protein |
| *PSPTO_3061* | | + | 3442333 | 3443256 | 2.85 | 4.97 | 1.70 | 2.41 | 5.32 | 3.61 | LysR family transcriptional regulator |
| *PSPTO_3062* | | + | 3443659 | 3444909 | 6.34 | 10.51 | 5.84 | 6.69 | 8.15 | 4.65 | Iron-sulfur cluster-binding protein, rieske family |
| *PSPTO_3063* | | + | 3445128 | 3446039 | 1.05 | 1.12 | 3.03 | 0.00 | 3.43 | 2.15 | Dihydrodipicolinate synthase |
| *PSPTO_3064* | | + | 3446032 | 3447549 | 2.04 | 3.20 | 4.50 | 1.20 | 6.32 | 3.67 | Aldehyde dehydrogenase family protein |
| *PSPTO_3065* | | + | 3447546 | 3448892 | 12.87 | 9.27 | 8.06 | 4.68 | 10.55 | 7.41 | Hypothetical protein |
| *PSPTO_3066* | | + | 3448873 | 3449457 | 4.56 | 5.48 | 8.39 | 1.25 | 4.07 | 4.81 | Membrane protein |
| *PSPTO_3067* | | + | 3449676 | 3450368 | 7.06 | 7.91 | 8.72 | 4.90 | 56.33 | 6.30 | Hypothetical protein |
| *PSPTO_3068* | | - | 3450465 | 3451592 | 5.21 | 3.76 | 8.79 | 3.17 | 11.17 | 7.80 | Major facilitator family transporter |
| *PSPTO_3069* | | + | 3451778 | 3452404 | 34.97 | 30.12 | 32.64 | 6.93 | 27.46 | 19.54 | TetR family transcriptional regulator |
| *PSPTO_3070* | | + | 3452473 | 3452928 | 11.46 | 19.54 | 15.73 | 2.41 | 13.24 | 6.33 | GNAT family acetyltransferase |
| *PSPTO_3071* | | + | 3453003 | 3454862 | 8.10 | 11.49 | 10.57 | 5.39 | 20.13 | 8.96 | ABC transporter ATP-binding protein/permease |
| *PSPTO_3072* | | - | 3454877 | 3455776 | 7.50 | 8.63 | 5.03 | 6.56 | 7.40 | 5.61 | AraC family transcriptional regulator |
| *PSPTO_3073* | | + | 3455936 | 3456694 | 2.09 | 5.96 | 3.36 | 0.00 | 3.05 | 2.94 | Short chain dehydrogenase/reductase family oxidoreductase |
| *PSPTO_3074* | | - | 3456718 | 3457659 | 13.96 | 7.07 | 1.64 | 6.71 | 11.25 | 9.68 | AraC family transcriptional regulator |
| *PSPTO_3075* | | + | 3458021 | 3459073 | 9.00 | 6.87 | 9.64 | 1.51 | 7.26 | 7.49 | Oxidoreductase, aldo/keto reductase family |
| *PSPTO_3076* | | + | 3459207 | 3459788 | 52.01 | 116.17 | 94.63 | 55.09 | 20.98 | 34.00 | TetR family transcriptional regulator |
| *PSPTO_3077* | | + | 3459847 | 3460950 | 1.75 | 9.03 | 11.26 | 1.59 | 3.73 | 1.44 | NADH:flavin oxidoreductase / NADH oxidase family protein |
| *PSPTO_3078* | | + | 3461005 | 3461391 | 13.16 | 20.74 | 12.76 | 24.21 | 14.71 | 8.43 | Hypothetical protein |
| *PSPTO_3079* | | + | 3461451 | 3462107 | 6.80 | 12.60 | 20.55 | 4.47 | 11.45 | 10.62 | Hypothetical protein |
| *PSPTO_3080* | | - | 3462124 | 3463221 | 11.70 | 9.32 | 6.98 | 2.39 | 13.08 | 9.23 | NADH:flavin oxidoreductase / NADH oxidase family |
| *PSPTO_3081* | | + | 3463360 | 3464283 | 4.30 | 4.12 | 4.78 | 0.40 | 5.22 | 4.07 | LysR family transcriptional regulator |
| *PSPTO_3082* | | - | 3464325 | 3465197 | 3.57 | 8.02 | 4.09 | 6.15 | 6.57 | 4.96 | Glutamine/glutamate ABC transporter periplasmic glutamine/glutamate-binding protein |
| *PSPTO_3083* | | - | 3465194 | 3465955 | 4.27 | 8.18 | 10.40 | 3.79 | 7.80 | 6.37 | Glutamine/glutamate ABC transporter, ATP-binding protein |
| *PSPTO_3084* | | - | 3465952 | 3466611 | 3.01 | 5.65 | 4.90 | 2.30 | 2.34 | 3.08 | Glutamine/glutamate ABC transporter, permease protein |
| *PSPTO_3085* | | - | 3466592 | 3467257 | 3.33 | 5.03 | 9.33 | 6.43 | 5.59 | 3.71 | Glutamine/glutamate ABC transporter, permease protein |
| *PSPTO_3086* | | + | 3467721 | 3468359 | 5.08 | 88.95 | 62.25 | 31.77 | 5.33 | 4.81 | Transcriptional regulator |
| *PSPTO_3087* | *hopAB2, avrPtoB* | - | 3468470 | 3470131 | 10.68 | 445.50 | 284.51 | 52.92 | 9.23 | 9.26 | Type III effector HopAB2 (AvrPtoB) |
| *PSPTO_3088* | | + | 3470485 | 3472113 | 11.34 | 7.13 | 2.91 | 4.74 | 10.17 | 6.80 | Nickel ABC transporter, periplasmic nickel-binding protein |
| *PSPTO_3089* | *nikB* | + | 3472110 | 3473060 | 3.00 | 4.81 | 6.42 | 1.63 | 4.90 | 4.03 | Nickel ABC transporter, permease protein |
| *PSPTO_3090* | | + | 3473065 | 3473982 | 1.46 | 0.62 | 2.50 | 0.00 | 2.93 | 1.69 | Nickel ABC transporter, permease protein |
| *PSPTO_3091* | | + | 3473979 | 3475625 | 4.35 | 8.45 | 4.23 | 0.68 | 5.15 | 6.69 | Nickel ABC transporter, ATP-binding protein |
| *PSPTO_3092* | | + | 3475664 | 3476500 | 5.89 | 3.04 | 1.75 | 1.72 | 8.52 | 13.42 | Hypothetical protein |
| *PSPTO_3094* | | + | 3477551 | 3478012 | 7.87 | 29.38 | 20.86 | 10.12 | 10.42 | 7.02 | Lipoprotein |
| *PSPTO_3095* | | - | 3479336 | 3479625 | - | - | - | - | - | - | |
| *PSPTO_3096* | | + | 3480059 | 3480703 | 3.32 | 20.57 | 4.16 | 9.04 | 8.35 | 9.53 | Hypothetical protein |
| *PSPTO_3097* | | + | 3480696 | 3480854 | - | - | - | - | - | - | |
| *PSPTO_3098* | | - | 3480906 | 3482834 | 8.69 | 10.02 | 6.18 | 3.35 | 5.21 | 5.30 | Methyl-accepting chemotaxis protein |
| *PSPTO_3099* | *mexE* | + | 3483136 | 3484380 | 17.29 | 60.86 | 67.63 | 11.02 | 16.18 | 6.26 | Multidrug efflux RND membrane fusion protein MexE |
| *PSPTO_3100* | *saxF* | + | 3484393 | 3487584 | 4.48 | 18.72 | 9.77 | 2.42 | 9.80 | 4.74 | Aliphatic isothiocyanate resistance protein SaxF; AcrB/AcrD/AcrF family |
| *PSPTO_3101* | | + | 3487584 | 3488981 | 1.79 | 7.32 | 7.26 | 3.10 | 7.06 | 2.07 | Outer membrane efflux protein |
| *PSPTO_3102* | | + | 3489064 | 3490233 | 17.96 | 43.19 | 21.78 | 16.77 | 32.11 | 21.80 | TPR domain-containing protein |
| *PSPTO_3103* | | + | 3490721 | 3491083 | 36.05 | 105.59 | 143.38 | 44.00 | 265.52 | 51.05 | Transcriptional regulator |
| *PSPTO_3104* | | + | 3491068 | 3491214 | 1.40 | 24.61 | 16.27 | 12.91 | 134.12 | 22.89 | Hypothetical protein |
| *PSPTO_3105* | | - | 3491560 | 3492276 | 170.03 | 183.80 | 140.84 | 58.71 | 115.22 | 130.36 | Hypothetical protein |
| *PSPTO_3106* | *gloA* | + | 3492265 | 3492786 | 44.86 | 97.75 | 91.31 | 44.55 | 120.70 | 83.47 | Lactoylglutathione lyase |

| Locus name | Gene name | Strand | Gene start (bp) | Gene end (bp) | Average TPM Inoculum | Average TPM $t_6$ | Average TPM $t_{24}$ | Average TPM $t_{168}$ | Average TPM Logarithmic | Average TPM Stationary | Annotation description |
|---|---|---|---|---|---|---|---|---|---|---|---|
| *PSPTO_3107* | *ahpF* | - | 3492875 | 3494437 | 6.16 | 147.59 | 155.82 | 21.94 | 25.67 | 8.54 | Alkyl hydroperoxide reductase subunit F |
| *PSPTO_3108* | *ahpC* | - | 3494562 | 3495125 | 68.35 | 1517.24 | 1328.22 | 247.09 | 679.82 | 112.89 | Alkyl hydroperoxide reductase |
| *PSPTO_3109* | | - | 3495236 | 3496195 | 101.13 | 141.92 | 95.40 | 59.71 | 343.91 | 71.13 | Cointegrate resolution protein S |
| *PSPTO_3110* | | + | 3496381 | 3497391 | 72.79 | 18.34 | 14.55 | 1.63 | 22.71 | 6.94 | Cointegrate resolution protein T |
| *PSPTO_3111* | | - | 3497402 | 3498673 | 16.89 | 16.95 | 10.39 | 8.28 | 12.93 | 16.46 | Sensory box histidine kinase |
| *PSPTO_3112* | *gor-1* | - | 3498764 | 3500122 | 51.46 | 43.41 | 39.20 | 21.85 | 37.07 | 28.33 | Glutathione reductase |
| *PSPTO_3113* | *galU* | - | 3500226 | 3501065 | 96.32 | 170.29 | 155.18 | 60.56 | 165.92 | 100.12 | UTP-glucose-1-phosphate uridylyltransferase |
| *PSPTO_3114* | | + | 3501396 | 3501677 | 60.69 | 39.72 | 30.73 | 6.02 | 50.29 | 55.07 | Lipoprotein |
| *PSPTO_3115* | | - | 3501750 | 3502208 | 41.68 | 89.05 | 54.47 | 26.14 | 67.90 | 67.98 | Hypothetical protein |
| *PSPTO_3116* | | + | 3502271 | 3502576 | 29.41 | 73.29 | 66.40 | 19.62 | 220.52 | 44.73 | Hypothetical protein |
| *PSPTO_3117* | | + | 3502591 | 3502773 | 12.01 | 37.17 | 18.22 | 0.00 | 86.39 | 15.16 | Hypothetical protein |
| *PSPTO_3118* | *alkB* | + | 3502831 | 3503517 | 8.79 | 4.35 | 5.40 | 9.07 | 28.53 | 12.05 | DNA alkylation damage repair protein AlkB |
| *PSPTO_3119* | | - | 3503526 | 3505343 | 24.80 | 25.19 | 25.63 | 13.88 | 40.58 | 32.62 | Glycosyl hydrolase family protein |
| *PSPTO_3120* | | - | 3505396 | 3506214 | 17.22 | 59.28 | 44.63 | 24.61 | 58.46 | 49.75 | Cof-like hydrolase family protein |
| *PSPTO_3121* | *zwf-2* | - | 3506207 | 3507739 | 18.50 | 41.17 | 37.82 | 18.96 | 65.81 | 64.07 | Glucose-6-phosphate 1-dehydrogenase |
| *PSPTO_3122* | *gnd* | - | 3507739 | 3508713 | 39.00 | 75.35 | 74.75 | 27.71 | 68.87 | 66.35 | 6-phosphogluconate dehydrogenase |
| *PSPTO_3123* | | + | 3509027 | 3509200 | 1740.69 | 955.16 | 811.17 | 595.98 | 251.58 | 113.50 | Hypothetical protein |
| *PSPTO_3124* | | - | 3509337 | 3509828 | 47.77 | 42.06 | 33.55 | 27.67 | 85.65 | 67.83 | GNAT family acetyltransferase |
| *PSPTO_3125* | *glgA* | + | 3510581 | 3512035 | 52.26 | 82.08 | 57.99 | 46.04 | 32.35 | 55.79 | Glycogen synthase |
| *PSPTO_3126* | | + | 3512055 | 3513869 | 11.97 | 39.44 | 27.72 | 18.21 | 13.62 | 33.14 | Alpha-amylase family protein |
| *PSPTO_3127* | *malQ* | + | 3513859 | 3515937 | 9.38 | 26.50 | 18.92 | 8.03 | 9.75 | 21.55 | 4-alpha-glucanotransferase |
| *PSPTO_3128* | | + | 3515934 | 3518717 | 62.77 | 65.31 | 52.60 | 36.28 | 25.63 | 45.81 | Glycosyl hydrolase family protein |
| *PSPTO_3129* | | + | 3518735 | 3519148 | 60.43 | 269.11 | 188.76 | 83.19 | 58.58 | 98.88 | Hypothetical protein |
| *PSPTO_3130* | *glgX* | + | 3519308 | 3521491 | 7.79 | 40.41 | 26.06 | 20.57 | 14.81 | 24.40 | Glycogen operon protein GlgX |
| *PSPTO_3131* | | + | 3521488 | 3522501 | 8.72 | 9.81 | 12.32 | 4.71 | 6.73 | 10.76 | Hypothetical protein |
| *PSPTO_3132* | | + | 3522528 | 3523301 | 4.72 | 5.51 | 14.49 | 5.76 | 5.76 | 6.54 | Hypothetical protein |
| *PSPTO_3133* | | + | 3523298 | 3523924 | 3.71 | 6.93 | 5.33 | 5.05 | 3.61 | 6.75 | Methyltransferase |
| *PSPTO_3134* | | + | 3523893 | 3524546 | 14.05 | 14.41 | 8.69 | 3.40 | 6.50 | 11.15 | Glycosyl transferase family protein |
| *PSPTO_3135* | | - | 3524581 | 3525600 | 66.06 | 36.12 | 29.04 | 16.11 | 29.76 | 70.31 | Hypothetical protein |
| *PSPTO_3136* | | - | 3525969 | 3526577 | 8.77 | 93.48 | 67.81 | 12.54 | 10.52 | 4.40 | Azoreductase |
| *PSPTO_3137* | | - | 3526652 | 3527611 | 9.27 | 16.89 | 4.35 | 9.12 | 12.78 | 9.35 | LysR family transcriptional regulator |
| *PSPTO_3138* | | - | 3527691 | 3528482 | 4.71 | 12.99 | 5.87 | 6.72 | 12.94 | 10.20 | 3-oxoadipate enol-lactone hydrolase |
| *PSPTO_3139* | | + | 3528610 | 3528954 | 6.25 | 17.27 | 7.33 | 4.11 | 12.20 | 7.74 | Hypothetical protein |
| *PSPTO_3140* | | - | 3528949 | 3529419 | 4.20 | 3.38 | 6.75 | 0.00 | 15.08 | 6.30 | Hypothetical protein |
| *PSPTO_3141* | | - | 3529416 | 3530246 | 8.19 | 13.16 | 8.28 | 3.56 | 19.12 | 8.49 | Undecaprenyl pyrophosphate phosphatase |
| *PSPTO_3142* | *pnuC* | + | 3530526 | 3531089 | 9.30 | 17.00 | 10.30 | 7.53 | 13.31 | 11.46 | Nicotinamide mononucleotide transporter PnuC |
| *PSPTO_3143* | | + | 3531086 | 3531622 | 4.25 | 5.79 | 13.36 | 9.33 | 11.59 | 8.18 | Nicotinamide-nucleotide adenylyltransferase |
| *PSPTO_3144* | | - | 3531672 | 3532976 | 5.22 | 7.48 | 8.66 | 4.22 | 4.09 | 5.42 | Sigma-54 dependent transcriptional regulator |
| *PSPTO_3145* | | + | 3533269 | 3533571 | 712.71 | 173.22 | 275.28 | 154.21 | 293.80 | 68.80 | Hypothetical protein |
| *PSPTO_3146* | | - | 3534233 | 3535255 | 20.60 | 26.18 | 21.77 | 15.35 | 36.71 | 26.27 | ISPsy7, transposase |
| *PSPTO_3147* | | - | 3535672 | 3536283 | 3.11 | 10.89 | 0.63 | 0.00 | 21.94 | 7.18 | Hypothetical protein |
| *PSPTO_3148* | | - | 3536337 | 3537365 | 2.69 | 14.70 | 7.82 | 0.90 | 40.27 | 7.87 | Magnesium chelatase, subunit ChlI |
| *PSPTO_3149* | *cobN* | - | 3537365 | 3541129 | 3.24 | 13.52 | 10.94 | 2.64 | 41.45 | 10.24 | Cobaltochelatase subunit CobN |
| *PSPTO_3150* | | - | 3541133 | 3542197 | 5.25 | 27.49 | 25.94 | 7.95 | 63.38 | 17.26 | Cobalamin synthesis protein/P47K family protein |
| *PSPTO_3151* | | + | 3542904 | 3543089 | 15.19 | 41.86 | 18.11 | 13.64 | 112.23 | 34.22 | Hypothetical protein |
| *PSPTO_3152* | | + | 3543110 | 3543832 | 20.76 | 35.47 | 28.00 | 9.83 | 118.09 | 34.06 | Membrane protein |
| *PSPTO_3153* | | + | 3543795 | 3544274 | - | - | - | - | - | - | |
| *PSPTO_3154* | *cobM* | + | 3544307 | 3545056 | 74.03 | 16.50 | 26.41 | 19.89 | 45.22 | 39.15 | Precorrin-4 C(11)-methyltransferase |
| *PSPTO_3155* | | - | 3545226 | 3548708 | 303.33 | 60.38 | 82.88 | 43.17 | 140.91 | 97.83 | Indolepyruvate ferredoxin oxidoreductase |
| *PSPTO_3156* | | + | 3548894 | 3549361 | 8.93 | 21.37 | 20.01 | 5.80 | 19.46 | 12.18 | AsnC family transcriptional regulator |
| *PSPTO_3157* | | + | 3549774 | 3550406 | 101.44 | 34.74 | 35.11 | 11.27 | 158.62 | 558.66 | Hypothetical protein |
| *PSPTO_3158* | | + | 3550528 | 3551664 | - | - | - | - | - | - | |
| *PSPTO_3159* | | - | 3551698 | 3552942 | 21.68 | 51.55 | 40.69 | 32.38 | 18.64 | 38.86 | Mutlidrug resistance protein |
| *PSPTO_3160* | | + | 3553049 | 3553594 | 4.30 | 19.05 | 18.25 | 0.00 | 11.66 | 4.77 | Transcriptional regulator |
| *PSPTO_3161* | | + | 3553635 | 3554975 | 3.82 | 10.84 | 8.50 | 10.23 | 19.22 | 7.30 | Protein kinase |
| *PSPTO_3162* | | - | 3554994 | 3555698 | 19.20 | 16.73 | 15.47 | 9.38 | 38.53 | 16.38 | Hypothetical protein |
| *PSPTO_3163* | | - | 3555900 | 3556673 | 9.52 | 6.43 | 4.32 | 8.27 | 14.56 | 3.32 | Enoyl-CoA hydratase/isomerase family protein |
| *PSPTO_3164* | *phbA-2* | - | 3556756 | 3557964 | 29.87 | 26.15 | 31.08 | 23.40 | 30.22 | 12.63 | Acetyl-CoA acetyltransferase |
| *PSPTO_3165* | | + | 3558140 | 3558619 | 30.47 | 19.02 | 27.89 | 9.50 | 11.51 | 22.75 | Lipoprotein |
| *PSPTO_3166* | | + | 3558705 | 3559667 | 10.56 | 8.33 | 4.31 | 3.81 | 4.35 | 6.33 | Hypothetical protein |
| *PSPTO_3167* | | - | 3559693 | 3559998 | 5.45 | 17.68 | 11.50 | 0.00 | 4.37 | 3.45 | Hypothetical protein |
| *PSPTO_3168* | | + | 3560098 | 3560238 | 21.86 | 128.45 | 98.49 | 77.54 | 18.86 | 16.03 | Hypothetical protein |
| *PSPTO_3169* | | + | 3560546 | 3560809 | 2.57 | 7.88 | 6.30 | 7.25 | 7.09 | 1.88 | Hypothetical protein |
| *PSPTO_3170* | | + | 3560854 | 3563547 | 2.93 | 1.61 | 5.52 | 2.06 | 8.22 | 3.06 | Glycosyl transferase family protein |
| *PSPTO_3171* | | - | 3563563 | 3564213 | 3.41 | 7.50 | 6.03 | 3.41 | 11.19 | 9.49 | TetR family transcriptional regulator |
| *PSPTO_3172* | | - | 3564236 | 3565597 | 9.57 | 17.05 | 12.85 | 10.90 | 19.14 | 13.83 | Membrane protein |
| *PSPTO_3173* | | - | 3565701 | 3565955 | 76.90 | 99.64 | 89.43 | 26.28 | 135.68 | 91.45 | Hypothetical protein |
| *PSPTO_3174* | | + | 3566109 | 3566426 | 52.83 | 129.67 | 158.55 | 32.87 | 38.83 | 38.28 | Hypothetical protein |
| *PSPTO_3175* | | - | 3566539 | 3566868 | 21.04 | 61.91 | 24.24 | 14.25 | 30.44 | 22.70 | Hypothetical protein |
| *PSPTO_3176* | *lasB* | + | 3567159 | 3568229 | 25.44 | 59.76 | 37.33 | 24.30 | 17.45 | 35.93 | Thermolysin metallopeptidase |
| *PSPTO_3177* | | + | 3568261 | 3568560 | 36.20 | 45.90 | 33.09 | 17.66 | 16.77 | 33.60 | Hypothetical protein |
| *PSPTO_3178* | | + | 3568814 | 3570063 | - | - | - | - | - | - | |
| *PSPTO_3179* | | + | 3570056 | 3571735 | 12.87 | 19.68 | 16.31 | 11.98 | 47.72 | 19.40 | PqiB family protein |
| *PSPTO_3180* | | + | 3571732 | 3572355 | 11.11 | 34.69 | 24.09 | 11.32 | 35.18 | 16.32 | Lipoprotein |
| *PSPTO_3181* | | + | 3572634 | 3573083 | 11.14 | 37.70 | 23.02 | 13.60 | 17.00 | 14.83 | Transcriptional regulator |
| *PSPTO_3182* | | - | 3573111 | 3574022 | 12.25 | 18.23 | 19.19 | 4.60 | 26.59 | 9.98 | LysR family transcriptional regulator |
| *PSPTO_3183* | | + | 3574134 | 3574970 | 3.49 | 11.99 | 13.86 | 2.42 | 4.84 | 3.87 | Hypothetical protein |
| *PSPTO_3184* | | - | 3574983 | 3575846 | 7.55 | 14.06 | 10.28 | 2.98 | 6.23 | 8.29 | Putative oxidoreductase |
| *PSPTO_3185* | | + | 3575974 | 3576898 | - | - | - | - | - | - | |

| Locus name | Gene name | Strand | Gene start (bp) | Gene end (bp) | Average TPM Inoculum | Average TPM $t_6$ | Average TPM $t_{24}$ | Average TPM $t_{168}$ | Average TPM Logarithmic | Average TPM Stationary | Annotation description |
|---|---|---|---|---|---|---|---|---|---|---|---|
| *PSPTO_3186* | | - | 3576922 | 3578451 | 3.76 | 6.00 | 10.66 | 2.59 | 5.26 | 5.32 | Sodium:solute symporter family protein |
| *PSPTO_3187* | | - | 3578448 | 3578708 | 0.99 | 0.00 | 0.00 | 0.00 | 1.23 | 1.48 | Hypothetical protein |
| *PSPTO_3188* | | - | 3578832 | 3579767 | 5.31 | 17.18 | 20.00 | 6.28 | 20.41 | 7.51 | Choline/ethanolamine kinase |
| *PSPTO_3189* | | - | 3579764 | 3580717 | 4.61 | 20.34 | 23.81 | 5.99 | 19.30 | 6.74 | Hypothetical protein |
| *PSPTO_3190* | | - | 3580727 | 3582007 | 12.15 | 21.77 | 20.46 | 13.36 | 16.03 | 13.11 | Peptidase, M20/M25/M40 family |
| *PSPTO_3191* | | - | 3581985 | 3582743 | 3.42 | 6.40 | 2.94 | 0.13 | 14.08 | 4.32 | DeoR family transcriptional regulator |
| *PSPTO_3192* | | - | 3582812 | 3584239 | 6.71 | 23.77 | 21.32 | 4.56 | 21.25 | 7.14 | Amidase family protein |
| *PSPTO_3193* | | - | 3584534 | 3586141 | 17.46 | 51.25 | 41.78 | 47.67 | 45.97 | 29.35 | Metalloprotease |
| *PSPTO_3194* | | + | 3586369 | 3587346 | 0.00 | 0.00 | 0.00 | 0.00 | 0.00 | 0.00 | ISPssy, transposase |
| *PSPTO_3195* | | + | 3587832 | 3589148 | 15.66 | 55.81 | 40.52 | 23.87 | 114.35 | 55.60 | Sensor histidine kinase |
| *PSPTO_3196* | | - | 3589345 | 3590259 | 13.63 | 10.47 | 12.93 | 8.69 | 11.37 | 10.06 | LysR family transcriptional regulator |
| *PSPTO_3197* | | + | 3590420 | 3591976 | 2.23 | 1.89 | 3.20 | 2.74 | 2.05 | 2.46 | Membrane protein |
| *PSPTO_3198* | | + | 3591973 | 3592959 | 8.35 | 13.53 | 6.48 | 2.81 | 5.57 | 7.36 | Luciferase family protein |
| *PSPTO_3199* | | + | 3593053 | 3594246 | 10.63 | 10.71 | 4.32 | 9.16 | 11.50 | 9.55 | Luciferase family protein |
| *PSPTO_3200* | | - | 3594296 | 3595168 | 2.61 | 2.07 | 4.05 | 0.63 | 1.99 | 1.33 | Hypothetical protein |
| *PSPTO_3201* | | - | 3595165 | 3595944 | 1.81 | 2.60 | 1.26 | 0.00 | 3.72 | 2.06 | Hypothetical protein |
| *PSPTO_3202* | | - | 3596031 | 3597002 | 3.23 | 4.50 | 3.05 | 0.00 | 4.78 | 3.22 | Oxidoreductase, 2-nitropropane dioxygenase family |
| *PSPTO_3203* | | - | 3597083 | 3597547 | 3.36 | 0.47 | 0.00 | 10.46 | 7.53 | 4.83 | Hypothetical protein |
| *PSPTO_3204* | | - | 3597525 | 3598238 | 6.24 | 5.65 | 0.53 | 0.78 | 7.56 | 6.26 | Hypothetical protein |
| *PSPTO_3205* | | - | 3598222 | 3599412 | 8.04 | 9.17 | 7.73 | 0.00 | 6.81 | 5.39 | Isocitrate/isopropylmalate family dehydrogenase |
| *PSPTO_3206* | | - | 3599409 | 3600495 | - | - | - | - | - | - | |
| *PSPTO_3207* | | - | 3600713 | 3601621 | 4.75 | 1.32 | 1.07 | 2.62 | 2.71 | 3.55 | LysR family transcriptional regulator |
| *PSPTO_3208* | | + | 3601794 | 3602810 | 0.00 | 0.00 | 0.00 | 0.00 | 0.01 | 0.02 | ISPssy, transposase |
| *PSPTO_3210* | | - | 3604010 | 3606985 | 2.08 | 2.37 | 1.52 | 1.78 | 1.80 | 2.69 | Filamentous hemagglutinin family protein |
| *PSPTO_3211* | | - | 3608136 | 3608651 | 0.81 | 5.29 | 0.00 | 3.90 | 1.48 | 1.07 | Hypothetical protein |
| *PSPTO_3212* | | + | 3609072 | 3609431 | 0.00 | 0.00 | 0.00 | 0.00 | 0.00 | 0.00 | ISPsy5, Orf1 |
| *PSPTO_3213* | | + | 3609449 | 3611002 | 0.00 | 0.00 | 0.00 | 0.00 | 0.00 | 0.00 | ISPsy5, transposase |
| *PSPTO_3214* | | - | 3614640 | 3616358 | - | - | - | - | - | - | |
| *PSPTO_3215* | | + | 3616413 | 3616772 | 0.00 | 0.00 | 0.00 | 0.00 | 0.00 | 0.00 | ISPsy5, Orf1 |
| *PSPTO_3216* | | + | 3616790 | 3618343 | 0.00 | 0.00 | 0.00 | 0.00 | 0.00 | 0.00 | ISPsy5, transposase |
| *PSPTO_3217* | | - | 3619682 | 3619849 | 2.19 | 0.65 | 0.00 | 0.00 | 17.70 | 7.82 | Hypothetical protein |
| *PSPTO_3218* | | - | 3620468 | 3621664 | 37.37 | 38.05 | 26.14 | 21.11 | 57.69 | 49.05 | Hypothetical protein |
| *PSPTO_3219* | | - | 3621773 | 3621898 | 5.09 | 7.66 | 9.37 | 0.00 | 9.94 | 12.72 | Hypothetical protein |
| *PSPTO_3220* | | - | 3621993 | 3623504 | 18.78 | 10.49 | 9.22 | 4.20 | 21.69 | 10.85 | ISPsy5, transposase |
| *PSPTO_3221* | | - | 3623523 | 3623882 | 1.10 | 0.00 | 3.85 | 0.00 | 2.03 | 3.20 | ISPsy5, Orf1 |
| *PSPTO_3222* | | - | 3623952 | 3624167 | 11.32 | 2.31 | 1.79 | 0.00 | 9.40 | 9.40 | Hypothetical protein |
| *PSPTO_3223* | | + | 3624316 | 3625293 | 17.73 | 22.38 | 19.78 | 15.97 | 36.15 | 28.75 | IS52, transposase |
| *PSPTO_3224* | | + | 3625392 | 3626414 | 0.00 | 0.00 | 0.00 | 0.00 | 0.00 | 0.00 | ISPsy4, transposase |
| *PSPTO_3225* | | + | 3626411 | 3627220 | 0.00 | 0.00 | 0.00 | 0.00 | 0.00 | 0.00 | ISPsy4, transposition helper protein |
| *PSPTO_3226* | | - | 3627715 | 3628134 | 0.81 | 14.44 | 0.58 | 3.10 | 3.40 | 2.48 | Hypothetical protein |
| *PSPTO_3227* | | - | 3628302 | 3628430 | 0.00 | 1.76 | 0.00 | 0.00 | 0.53 | 0.96 | Hypothetical protein |
| *PSPTO_3228* | | - | 3628880 | 3629050 | 3.90 | 9.50 | 14.88 | 7.67 | 27.89 | 12.87 | Hypothetical protein |
| *PSPTO_3229* | | - | 3629677 | 3648501 | 6.49 | 6.08 | 5.82 | 4.43 | 12.59 | 5.34 | Filamentous hemagglutinin, intein-containing |
| *PSPTO_3230* | | - | 3648541 | 3650250 | 5.74 | 5.24 | 3.47 | 2.26 | 6.39 | 4.53 | Hemolysin activator protein, HlyB family |
| *PSPTO_3231* | | + | 3650472 | 3651347 | 21.78 | 9.11 | 8.74 | 4.87 | 15.85 | 14.50 | LysR family transcriptional regulator |
| *PSPTO_3232* | | - | 3651385 | 3651810 | 5.92 | 12.42 | 3.28 | 14.04 | 21.95 | 11.42 | Hypothetical protein |
| *PSPTO_3233* | | - | 3652016 | 3653257 | 15.50 | 28.57 | 21.49 | 9.17 | 90.17 | 24.21 | Peptidase, M20/M25/M40 family |
| *PSPTO_3234* | | - | 3653342 | 3654829 | 3.92 | 10.83 | 5.76 | 3.82 | 12.50 | 5.29 | Sulfate permease family protein |
| *PSPTO_3235* | | + | 3655252 | 3656660 | - | - | - | - | - | - | |
| *PSPTO_3236* | | + | 3656657 | 3657052 | 321.22 | 34.74 | 91.52 | 8.91 | 112.39 | 73.84 | Hypothetical protein |
| *PSPTO_3237* | | + | 3657206 | 3659521 | 129.32 | 23.99 | 17.75 | 12.08 | 5.39 | 4.41 | Methyl-accepting chemotaxis protein |
| *PSPTO_3238* | | - | 3659577 | 3660401 | 5.53 | 4.33 | 2.08 | 3.09 | 4.41 | 3.08 | TonB protein |
| *PSPTO_3239* | *tolR-1* | - | 3660407 | 3660853 | 1.70 | 0.00 | 0.00 | 0.23 | 4.00 | 4.42 | TolR protein |
| *PSPTO_3240* | *tolQ-1* | - | 3660863 | 3661573 | 0.81 | 1.84 | 0.44 | 0.71 | 1.90 | 1.30 | TolQ protein |
| *PSPTO_3241* | | - | 3661628 | 3663550 | 1.90 | 2.11 | 0.34 | 1.55 | 1.14 | 1.57 | Phytase domain-containing protein |
| *PSPTO_3242* | | - | 3663587 | 3666103 | 7.08 | 3.55 | 4.47 | 3.96 | 2.80 | 5.01 | TonB-dependent receptor |
| *PSPTO_3243* | | + | 3666583 | 3667497 | 1.94 | 3.86 | 0.36 | 1.13 | 4.73 | 6.65 | Diguanylate cyclase |
| *PSPTO_3244* | | + | 3667507 | 3668544 | 2.15 | 2.51 | 3.81 | 3.16 | 2.02 | 2.10 | AraC family transcriptional regulator |
| *PSPTO_3245* | | - | 3668552 | 3669226 | 5.98 | 10.11 | 14.07 | 4.57 | 6.30 | 3.44 | DNA-binding response regulator |
| *PSPTO_3246* | | - | 3669338 | 3670036 | 3.01 | 5.32 | 11.16 | 4.90 | 3.13 | 1.85 | Hypothetical protein |
| *PSPTO_3247* | | + | 3670593 | 3671759 | 2.24 | 7.31 | 13.89 | 6.04 | 4.26 | 2.72 | Mandelate racemase/muconate lactonizing protein |
| *PSPTO_3248* | | + | 3671845 | 3673467 | 1.59 | 1.63 | 3.65 | 3.44 | 2.28 | 1.92 | Dipeptide ABC transporter, periplasmic dipeptide-binding protein |
| *PSPTO_3249* | *dppB* | + | 3673508 | 3674545 | 6.67 | 2.96 | 3.73 | 4.22 | 3.96 | 2.36 | Dipeptide ABC transporter, permease protein DppB |
| *PSPTO_3250* | *dppC* | + | 3674545 | 3675450 | 2.44 | 6.46 | 12.23 | 2.46 | 8.26 | 3.59 | Dipeptide ABC transporter, permease protein DppC |
| *PSPTO_3251* | | + | 3675513 | 3676352 | 2.37 | 6.41 | 7.05 | 3.27 | 4.89 | 1.69 | Dipeptide ABC transporter ATP-binding protein |
| *PSPTO_3252* | | + | 3676363 | 3677127 | 2.80 | 7.28 | 12.81 | 1.20 | 4.65 | 1.99 | Dipeptide ABC transporter ATP-binding protein |
| *PSPTO_3253* | | + | 3677263 | 3677985 | 8.48 | 11.93 | 9.45 | 1.26 | 9.85 | 6.77 | Hypothetical protein |
| *PSPTO_3254* | | - | 3678347 | 3679081 | 12.20 | 16.73 | 9.13 | 6.89 | 21.79 | 15.38 | GntR family transcriptional regulator |
| *PSPTO_3255* | | + | 3679190 | 3679905 | - | - | - | - | - | - | |
| *PSPTO_3256* | | + | 3680085 | 3681266 | 4.70 | 7.65 | 10.95 | 1.71 | 53.15 | 13.79 | Iron ABC transporter, periplasmic iron-binding protein |
| *PSPTO_3257* | | + | 3681275 | 3682285 | 7.36 | 4.05 | 2.55 | 2.17 | 22.70 | 6.09 | Iron ABC transporter, permease protein |
| *PSPTO_3258* | | + | 3682282 | 3683073 | 5.19 | 4.34 | 8.20 | 7.24 | 34.66 | 5.66 | Iron ABC transporter ATP-binding protein |
| *PSPTO_3259* | | + | 3683180 | 3685186 | 5.43 | 25.07 | 18.43 | 11.60 | 36.90 | 16.79 | Peptidase, U32 family |
| *PSPTO_3260* | | - | 3685233 | 3685862 | 3.20 | 8.17 | 6.59 | 1.36 | 8.75 | 7.03 | GNAT family acetyltransferase |
| *PSPTO_3261* | | - | 3685988 | 3686242 | 13.35 | 31.95 | 15.43 | 12.13 | 8.69 | 9.49 | Hypothetical protein |
| *PSPTO_3262* | *nirB* | + | 3686459 | 3689020 | 2.33 | 32.91 | 17.02 | 27.28 | 2.54 | 2.49 | Nitrite reductase [NAD(P)H], large subunit |
| *PSPTO_3263* | *nirD* | + | 3689020 | 3689406 | 7.39 | 35.18 | 21.47 | 8.73 | 5.36 | 7.11 | Nitrite reductase [NAD(P)H], small subunit |
| *PSPTO_3264* | | - | 3689464 | 3690462 | 5.52 | 33.33 | 34.20 | 6.73 | 12.06 | 9.64 | Alpha/beta fold family hydrolase |
| *PSPTO_3265* | | + | 3690850 | 3691779 | 17.86 | 21.66 | 20.81 | 11.08 | 20.56 | 10.44 | LysR family transcriptional regulator |

| Locus name | Gene name | Strand | Gene start (bp) | Gene end (bp) | Average TPM Inoculum | Average TPM $t_6$ | Average TPM $t_{24}$ | Average TPM $t_{168}$ | Average TPM Logarithmic | Average TPM Stationary | Annotation description |
|---|---|---|---|---|---|---|---|---|---|---|---|
| *PSPTO_3266* | *pstB-1* | - | 3691780 | 3692559 | 18.75 | 29.16 | 21.65 | 17.44 | 43.04 | 13.93 | Phosphate transporter ATP-binding protein |
| *PSPTO_3267* | *pstA* | - | 3692559 | 3693431 | 7.24 | 15.30 | 9.15 | 5.20 | 22.70 | 6.02 | Phosphate ABC transporter permease |
| *PSPTO_3268* | *pstC* | - | 3693440 | 3694435 | 5.08 | 18.49 | 16.08 | 7.60 | 11.40 | 3.34 | Phosphate ABC transporter permease |
| *PSPTO_3269* | *pstS* | - | 3694517 | 3695542 | 60.35 | 86.77 | 87.25 | 28.47 | 31.56 | 3.68 | Phosphate ABC transporter substrate-binding protein |
| *PSPTO_3270* | | + | 3696039 | 3696323 | 242.34 | 1226.42 | 858.71 | 414.72 | 150.04 | 634.88 | Hypothetical protein |
| *PSPTO_3271* | | + | 3696409 | 3696741 | 10.38 | 106.67 | 117.15 | 11.72 | 32.84 | 45.74 | Hypothetical protein |
| *PSPTO_3272* | | - | 3696717 | 3698447 | 4.51 | 15.87 | 13.93 | 12.11 | 15.51 | 12.09 | Hypothetical protein |
| *PSPTO_3273* | | + | 3698579 | 3699079 | 15.33 | 16.16 | 23.34 | 7.51 | 23.39 | 24.15 | Hypothetical protein |
| *PSPTO_3274* | | - | 3699860 | 3700882 | 20.18 | 26.50 | 20.62 | 13.68 | 34.75 | 25.68 | ISPsy7, transposase |
| *PSPTO_3275* | | - | 3701313 | 3702215 | 4.89 | 3.18 | 2.00 | 0.49 | 8.24 | 2.68 | LysR family transcriptional regulator |
| *PSPTO_3276* | | - | 3702234 | 3702353 | 17.45 | 26.67 | 5.20 | 2.48 | 6.66 | 10.90 | Hypothetical protein |
| *PSPTO_3277* | | + | 3702391 | 3703407 | 2.85 | 4.11 | 3.84 | 2.69 | 6.74 | 6.91 | Membrane protein |
| *PSPTO_3278* | | - | 3703435 | 3704679 | 18.36 | 20.89 | 20.80 | 17.39 | 27.23 | 16.85 | Hypothetical protein |
| *PSPTO_3279* | | - | 3704803 | 3706428 | 3.25 | 3.45 | 2.23 | 3.36 | 4.59 | 3.68 | Methyl-accepting chemotaxis protein |
| *PSPTO_3280* | *glxK* | - | 3706609 | 3707748 | 6.48 | 7.55 | 10.13 | 4.52 | 46.67 | 7.67 | Glycerate kinase |
| *PSPTO_3281* | | - | 3707859 | 3708950 | 4.80 | 5.75 | 6.29 | 10.08 | 11.02 | 5.90 | Carbohydrate diacid regulator |
| *PSPTO_3282* | | + | 3709119 | 3710270 | 3.90 | 18.15 | 18.72 | 4.17 | 33.43 | 7.91 | Lipoprotein |
| *PSPTO_3283* | | - | 3710282 | 3710995 | 3.74 | 9.49 | 9.56 | 9.64 | 10.21 | 5.43 | GntR family transcriptional regulator |
| *PSPTO_3284* | | + | 3711465 | 3712790 | 2.56 | 8.43 | 11.96 | 4.72 | 3.04 | 2.61 | Major facilitator family transporter |
| *PSPTO_3285* | *gudD* | + | 3712842 | 3714194 | 1.63 | 14.91 | 10.23 | 3.78 | 2.40 | 1.17 | Glucarate dehydratase |
| *PSPTO_3286* | | + | 3714339 | 3714992 | 17.67 | 11.58 | 10.24 | 3.05 | 14.38 | 5.17 | Hypothetical protein |
| *PSPTO_3287* | | + | 3715029 | 3715985 | 6.90 | 4.60 | 9.28 | 5.55 | 6.86 | 3.72 | D-isomer specific 2-hydroxyacid dehydrogenase family protein |
| *PSPTO_3288* | | - | 3716180 | 3716509 | 5.07 | 428.64 | 267.91 | 39.87 | 6.60 | 2.14 | Hypothetical protein |
| *PSPTO_3289* | | + | 3716755 | 3716973 | 1.51 | 3713.10 | 2177.19 | 359.15 | 2.84 | 0.37 | Hypothetical protein |
| *PSPTO_3290* | | + | 3717304 | 3718629 | 3.95 | 41.19 | 28.84 | 5.86 | 4.65 | 2.31 | Outer membrane porin, OprD family |
| *PSPTO_3291* | | - | 3718924 | 3720552 | 5.07 | 2.34 | 1.80 | 2.27 | 22.08 | 6.35 | Methyl-accepting chemotaxis protein |
| *PSPTO_3292* | *hopAH2-1* | + | 3721153 | 3722367 | 2.98 | 4.15 | 2.77 | 2.69 | 15.59 | 3.68 | Not an effector. |
| *PSPTO_3293* | *hopAH2-2* | + | 3722818 | 3724068 | 24.67 | 48.15 | 46.82 | 29.12 | 46.01 | 25.79 | Not an effector. |
| *PSPTO_3294* | | - | 3724173 | 3726308 | 5.04 | 2.79 | 2.05 | 1.03 | 45.93 | 10.97 | TonB-dependent siderophore receptor |
| *PSPTO_3295* | *pncA* | + | 3726505 | 3727155 | 11.82 | 19.41 | 12.63 | 1.49 | 19.90 | 11.21 | Pyrazinamidase/nicotinamidase |
| *PSPTO_3296* | | + | 3727290 | 3728225 | 47.76 | 32.18 | 34.84 | 17.44 | 44.67 | 43.78 | Transporter |
| *PSPTO_3297* | | + | 3728312 | 3728995 | 53.38 | 299.80 | 235.93 | 67.59 | 134.20 | 105.33 | DNA-binding heavy metal response regulator |
| *PSPTO_3298* | | + | 3728973 | 3730370 | 5.07 | 14.16 | 16.65 | 4.85 | 8.97 | 6.97 | Sensor histidine kinase |
| *PSPTO_3299* | *phaG-1* | + | 3730941 | 3731717 | 14.62 | 10.17 | 11.29 | 5.42 | 4.88 | 12.42 | 3-hydroxyacyl-CoA-acyl carrier protein transferase |
| *PSPTO_3300* | | + | 3731816 | 3733003 | 3.56 | 1.23 | 3.22 | 0.95 | 3.00 | 2.98 | RND family efflux transporter MFP subunit |
| *PSPTO_3301* | | + | 3733000 | 3734100 | 1.88 | 1.96 | 2.06 | 4.40 | 1.61 | 2.08 | RND family efflux transporter MFP subunit |
| *PSPTO_3302* | | + | 3734097 | 3737132 | 4.87 | 8.42 | 8.76 | 3.74 | 4.55 | 4.68 | AcrB/AcrD/AcrF family protein |
| *PSPTO_3303* | | + | 3737129 | 3737317 | 106.97 | 150.98 | 97.80 | 36.27 | 85.28 | 59.48 | Hypothetical protein |
| *PSPTO_3304* | | + | 3737479 | 3738675 | 7.88 | 12.75 | 5.45 | 4.83 | 8.33 | 11.38 | Major facilitator family transporter |
| *PSPTO_3305* | | + | 3738804 | 3739646 | 26.50 | 23.17 | 23.25 | 8.13 | 20.69 | 18.93 | Hypothetical protein |
| *PSPTO_3306* | | - | 3739697 | 3740419 | 7.13 | 13.08 | 9.75 | 5.14 | 13.47 | 10.84 | Phospholipase/carboxylesterase family protein |
| *PSPTO_3307* | *gspD* | - | 3740495 | 3742786 | 5.69 | 12.79 | 8.53 | 5.63 | 16.66 | 11.74 | General secretion pathway protein D |
| *PSPTO_3308* | *gspN* | - | 3742834 | 3743361 | 3.90 | 20.71 | 19.35 | 12.54 | 21.12 | 15.00 | General secretion pathway protein N |
| *PSPTO_3309* | *gspM* | - | 3743358 | 3743969 | 2.30 | 2.64 | 8.51 | 1.62 | 3.93 | 8.07 | General secretion pathway protein M |
| *PSPTO_3310* | *gspL* | - | 3743950 | 3745014 | 3.20 | 9.81 | 5.24 | 3.31 | 6.02 | 9.31 | General secretion pathway protein L |
| *PSPTO_3311* | *gspK* | - | 3745011 | 3745745 | 23.97 | 37.81 | 19.35 | 21.00 | 12.81 | 31.03 | General secretion pathway protein K |
| *PSPTO_3312* | *gspJ* | - | 3745742 | 3746374 | 3.63 | 16.74 | 4.96 | 2.69 | 6.75 | 15.51 | General secretion pathway protein J |
| *PSPTO_3313* | *gspI* | - | 3746371 | 3746754 | 3.41 | 12.49 | 6.17 | 14.88 | 7.99 | 19.19 | General secretion pathway protein I |
| *PSPTO_3314* | *gspH* | - | 3746754 | 3747188 | 4.45 | 8.82 | 3.66 | 6.87 | 11.46 | 21.30 | General secretion pathway protein H |
| *PSPTO_3315* | *gspG* | - | 3747197 | 3747628 | 9.71 | 19.52 | 15.33 | 35.64 | 18.55 | 30.61 | General secretion pathway protein G |
| *PSPTO_3316* | *gspF* | - | 3747654 | 3748856 | 18.03 | 16.06 | 12.53 | 7.29 | 13.59 | 24.79 | General secretion pathway protein F |
| *PSPTO_3317* | *gspE* | - | 3748853 | 3750610 | 5.71 | 10.47 | 8.81 | 1.64 | 13.01 | 10.22 | General secretion pathway protein E |
| *PSPTO_3318* | | + | 3751384 | 3754125 | 4.23 | 3.95 | 2.93 | 0.53 | 5.69 | 4.01 | Beta-glucosidase |
| *PSPTO_3319* | | - | 3754165 | 3754809 | 15.41 | 19.21 | 10.95 | 11.12 | 26.56 | 16.18 | TetR family transcriptional regulator |
| *PSPTO_3320* | | - | 3754868 | 3755323 | 5.03 | 3.93 | 5.51 | 1.82 | 6.28 | 8.84 | Membrane protein |
| *PSPTO_3321* | *bar* | - | 3755364 | 3755903 | 2.16 | 3.41 | 2.72 | 0.41 | 3.97 | 2.80 | Phosphinothricin N-acetyltransferase |
| *PSPTO_3322* | | + | 3756009 | 3756899 | 2.04 | 1.90 | 0.33 | 1.47 | 7.20 | 3.64 | LysR family transcriptional regulator |
| *PSPTO_3323* | | + | 3757178 | 3758761 | 4.79 | 26.88 | 38.75 | 23.93 | 28.35 | 13.90 | Aldehyde dehydrogenase family protein |
| *PSPTO_3324* | | + | 3758993 | 3759100 | 1.02 | 3.93 | 0.00 | 0.00 | 2.35 | 1.67 | Hypothetical protein |
| *PSPTO_3325* | | + | 3759367 | 3760389 | 21.02 | 27.69 | 21.96 | 14.35 | 37.49 | 27.01 | ISPsy7, transposase |
| *PSPTO_3326* | | - | 3760741 | 3762108 | 3.20 | 5.46 | 3.04 | 4.74 | 3.31 | 3.42 | Group II intron, maturase |
| *PSPTO_3327* | | + | 3762732 | 3763907 | 5.70 | 19.23 | 18.79 | 18.75 | 9.67 | 8.80 | Sarcosine oxidase |
| *PSPTO_3328* | *tliF, aprF* | - | 3763908 | 3765278 | 12.61 | 17.92 | 22.38 | 13.68 | 31.61 | 24.52 | ABC transporter permease; aprF. |
| *PSPTO_3329* | *tliE, aprE* | - | 3765280 | 3766533 | 6.53 | 13.02 | 5.21 | 2.84 | 18.88 | 9.39 | ABC transporter substrate-binding protein; aprE. |
| *PSPTO_3330* | *tliD, aprD* | - | 3766614 | 3768413 | 9.29 | 16.36 | 15.06 | 7.00 | 17.54 | 9.53 | ABC transporter ATP-binding protein; aprD. |
| *PSPTO_3331* | *aprI* | - | 3768494 | 3768877 | 41.26 | 34.33 | 31.61 | 13.26 | 34.86 | 24.81 | Protease inhibitor Inh; aprI; hypothetically of AprA. |
| *PSPTO_3332* | *aprA* | - | 3768937 | 3770364 | 11.61 | 7.27 | 6.70 | 6.55 | 14.91 | 5.37 | Alkaline metalloendoprotease; aprA; degrades flagellin |
| *PSPTO_3333* | | - | 3770841 | 3771512 | 204.56 | 645.79 | 465.06 | 128.26 | 223.62 | 130.69 | Membrane protein |
| *PSPTO_3334* | | + | 3771878 | 3772786 | 7.14 | 18.26 | 14.04 | 3.06 | 10.45 | 5.65 | LysR family transcriptional regulator |
| *PSPTO_3335* | | - | 3772772 | 3773551 | 2.84 | 2.49 | 6.16 | 2.11 | 3.23 | 2.67 | ABC transporter periplasmic substrate-binding protein |
| *PSPTO_3336* | | - | 3773567 | 3774868 | 4.92 | 4.36 | 6.25 | 0.71 | 4.43 | 6.42 | Citrate transporter |
| *PSPTO_3337* | | + | 3775152 | 3775739 | 21.85 | 109.67 | 91.06 | 26.80 | 81.61 | 20.38 | Hypothetical protein |
| *PSPTO_3338* | | + | 3775862 | 3776254 | 15.72 | 19.26 | 25.03 | 5.26 | 28.89 | 18.92 | DsrE family protein |
| *PSPTO_3339* | | + | 3776256 | 3776618 | 12.71 | 30.51 | 24.06 | 13.85 | 43.93 | 24.80 | DsrF family protein |
| *PSPTO_3340* | | + | 3776618 | 3776917 | 17.46 | 28.43 | 42.46 | 10.69 | 70.22 | 26.78 | DsrH family protein |
| *PSPTO_3341* | | + | 3776914 | 3777249 | 10.96 | 23.36 | 19.77 | 24.07 | 67.12 | 28.45 | Hypothetical protein |
| *PSPTO_3342* | | + | 3777246 | 3778247 | 6.59 | 14.98 | 14.55 | 13.91 | 46.45 | 23.04 | Hypothetical protein |
| *PSPTO_3343* | | + | 3778356 | 3779354 | 23.60 | 35.49 | 40.14 | 16.24 | 20.53 | 30.27 | Glutathione S-transferase domain protein |
| *PSPTO_3344* | *cysG* | - | 3779406 | 3780800 | 9.53 | 16.51 | 13.08 | 10.98 | 64.10 | 19.40 | Siroheme synthase |

| Locus name | Gene name | Strand | Gene start (bp) | Gene end (bp) | Average TPM Inoculum | Average TPM $t_6$ | Average TPM $t_{24}$ | Average TPM $t_{168}$ | Average TPM Logarithmic | Average TPM Stationary | Annotation description |
|---|---|---|---|---|---|---|---|---|---|---|---|
| *PSPTO_3345* | *serS* | - | 3780801 | 3782081 | 12.14 | 26.60 | 22.65 | 12.21 | 78.73 | 23.04 | Seryl-tRNA synthetase |
| *PSPTO_3346* | | - | 3782098 | 3782472 | 20.81 | 49.88 | 33.04 | 32.94 | 84.60 | 32.92 | CrcB family protein |
| *PSPTO_3347* | *rarA* | - | 3782472 | 3783794 | 9.34 | 18.16 | 14.81 | 11.78 | 45.46 | 24.87 | Recombination factor protein RarA |
| *PSPTO_3348* | *lolA* | - | 3783812 | 3784435 | 7.88 | 21.29 | 18.36 | 21.30 | 49.04 | 24.72 | Outer membrane lipoprotein carrier protein |
| *PSPTO_3349* | *ftsK* | - | 3784512 | 3786917 | 13.52 | 17.58 | 16.09 | 7.08 | 50.96 | 26.12 | Cell division protein FtsK |
| *PSPTO_3350* | *aat* | + | 3787395 | 3788084 | 4.84 | 8.35 | 4.20 | 5.69 | 20.49 | 7.25 | Leucyl/phenylalanyl-tRNA--protein transferase |
| *PSPTO_3351* | | + | 3788145 | 3788852 | 68.27 | 48.53 | 55.84 | 31.39 | 68.87 | 57.16 | Hypothetical protein |
| *PSPTO_3352* | *infA* | + | 3788954 | 3789172 | 69.35 | 147.72 | 172.32 | 97.13 | 208.81 | 47.66 | Translation initiation factor IF-1 |
| *PSPTO_3353* | *clpA* | - | 3789289 | 3791562 | 75.22 | 215.23 | 185.64 | 125.55 | 118.75 | 99.17 | ATP-dependent clp protease, ATP-binding subunit ClpA |
| *PSPTO_3354* | | - | 3791593 | 3791955 | 79.03 | 154.20 | 92.98 | 37.35 | 81.19 | 43.68 | Hypothetical protein |
| *PSPTO_3355* | | + | 3792176 | 3792445 | 131.32 | 130.81 | 94.20 | 28.75 | 33.57 | 76.64 | Cold shock domain family protein |
| *PSPTO_3356* | | + | 3793079 | 3795310 | 43.72 | 190.42 | 151.57 | 82.88 | 1002.16 | 157.85 | Isocitrate dehydrogenase |
| *PSPTO_3357* | | + | 3795436 | 3795882 | 2.51 | 3.73 | 6.48 | 3.21 | 20.65 | 5.79 | MutT/nudix family protein |
| *PSPTO_3358* | *trmU* | + | 3795943 | 3797073 | 6.07 | 25.19 | 19.16 | 12.10 | 51.19 | 18.27 | tRNA (5-methyl aminomethyl-2-thiouridylate)-methyltransferase |
| *PSPTO_3359* | | + | 3797073 | 3797693 | 3.32 | 21.14 | 13.84 | 0.00 | 55.43 | 10.24 | Hypothetical protein |
| *PSPTO_3360* | *purB* | + | 3797825 | 3799195 | 16.25 | 59.38 | 60.65 | 38.33 | 127.37 | 40.89 | Adenylosuccinate lyase |
| *PSPTO_3361* | | + | 3799256 | 3800422 | 13.97 | 25.42 | 22.77 | 13.72 | 84.92 | 37.02 | Hypothetical protein |
| *PSPTO_3362* | | + | 3800415 | 3800846 | 9.38 | 22.42 | 12.15 | 21.45 | 35.30 | 16.31 | GNAT family acetyltransferase |
| *PSPTO_3363* | | + | 3801024 | 3801770 | 11.31 | 25.12 | 12.65 | 8.93 | 22.78 | 13.72 | Type II and III secretion system family protein |
| *PSPTO_3364* | *aceA* | + | 3802047 | 3803372 | 59.72 | 145.96 | 173.49 | 129.24 | 29.53 | 25.95 | Isocitrate lyase |
| *PSPTO_3365* | *nuoA* | + | 3803847 | 3804260 | 148.83 | 237.37 | 169.98 | 143.35 | 649.06 | 229.45 | NADH:ubiquinone oxidoreductase subunit A |
| *PSPTO_3366* | *nuoB* | + | 3804271 | 3804945 | 56.81 | 141.80 | 118.89 | 70.73 | 391.03 | 114.47 | NADH dehydrogenase subunit B |
| *PSPTO_3367* | *nuoCD* | + | 3805020 | 3806801 | 13.84 | 53.90 | 54.03 | 23.29 | 192.65 | 33.70 | Bifunctional NADH:ubiquinone oxidoreductase subunit C/D |
| *PSPTO_3368* | *nuoE* | + | 3806804 | 3807301 | 19.61 | 53.37 | 72.24 | 27.11 | 252.36 | 39.81 | NADH dehydrogenase subunit E |
| *PSPTO_3369* | *nuoF* | + | 3807298 | 3808656 | 20.42 | 65.97 | 79.30 | 26.98 | 210.55 | 31.47 | NADH dehydrogenase I subunit F |
| *PSPTO_3370* | *nuoG* | + | 3808794 | 3811511 | 11.80 | 41.75 | 46.75 | 17.06 | 188.86 | 24.41 | NADH dehydrogenase subunit G |
| *PSPTO_3371* | *nuoH* | + | 3811508 | 3812515 | 14.91 | 37.53 | 35.91 | 7.24 | 101.83 | 14.31 | NADH:ubiquinone oxidoreductase subunit H |
| *PSPTO_3372* | *nuoI* | + | 3812527 | 3813075 | 13.47 | 34.16 | 53.75 | 11.70 | 176.07 | 21.49 | NADH dehydrogenase subunit I |
| *PSPTO_3373* | *nuoJ* | + | 3813086 | 3813595 | 6.36 | 26.04 | 18.24 | 5.16 | 88.48 | 10.33 | NADH:ubiquinone oxidoreductase subunit J |
| *PSPTO_3374* | *nouK* | + | 3813592 | 3813900 | 13.06 | 35.30 | 13.87 | 5.94 | 84.46 | 11.56 | NADH:ubiquinone oxidoreductase subunit K |
| *PSPTO_3375* | *nuoL* | + | 3813897 | 3815750 | 7.53 | 17.82 | 22.00 | 11.63 | 79.35 | 7.83 | NADH:ubiquinone oxidoreductase subunit L |
| *PSPTO_3376* | *nuoM* | + | 3815790 | 3817322 | 10.26 | 18.94 | 16.80 | 11.29 | 82.59 | 8.64 | NADH:ubiquinone oxidoreductase subunit M |
| *PSPTO_3377* | *nuoN* | + | 3817330 | 3818799 | 6.84 | 12.81 | 18.04 | 5.16 | 82.84 | 7.45 | NADH:ubiquinone oxidoreductase subunit N |
| *PSPTO_3378* | | - | 3818856 | 3819920 | 5.69 | 6.80 | 7.25 | 3.53 | 6.05 | 4.16 | Membrane protein |
| *PSPTO_3379* | | - | 3820066 | 3821745 | 49.69 | 14.79 | 14.79 | 9.89 | 9.37 | 9.48 | Methyl-accepting chemotaxis protein |
| *PSPTO_3380* | | - | 3821872 | 3823191 | 7.19 | 20.08 | 17.78 | 8.61 | 16.64 | 10.19 | Sensor histidine kinase |
| *PSPTO_3381* | | - | 3823188 | 3823856 | 18.38 | 55.79 | 87.00 | 24.57 | 32.76 | 23.97 | DNA-binding response regulator |
| *PSPTO_3382* | | - | 3823856 | 3824161 | 59.21 | 119.93 | 110.95 | 46.95 | 44.44 | 40.13 | Lipoprotein |
| *PSPTO_3383* | | - | 3824161 | 3824469 | 55.77 | 147.03 | 197.77 | 61.67 | 52.90 | 38.87 | Hypothetical protein |
| *PSPTO_3384* | | - | 3824533 | 3824739 | 13.33 | 6.53 | 4.91 | 19.28 | 9.86 | 11.14 | Hypothetical protein |
| *PSPTO_3385* | | + | 3824867 | 3826042 | 14.53 | 16.88 | 20.83 | 6.00 | 15.66 | 11.31 | Site-specific recombinase, phage integrase family |
| *PSPTO_3386* | | - | 3826104 | 3828458 | 2.61 | 12.28 | 10.96 | 8.68 | 17.56 | 7.19 | Hypothetical protein |
| *PSPTO_3387* | | - | 3829451 | 3829780 | 1770.98 | 4103.90 | 3583.16 | 1546.73 | 582.60 | 1223.69 | Hypothetical protein |
| *PSPTO_3388* | | - | 3830042 | 3830836 | 5.69 | 10.92 | 7.66 | 0.00 | 11.46 | 15.05 | DNA adenine methylase |
| *PSPTO_3389* | | - | 3831038 | 3831583 | 2.89 | 4.75 | 5.23 | 4.53 | 15.99 | 3.24 | Lysozyme |
| *PSPTO_3390* | | - | 3831622 | 3832665 | 4.22 | 2.85 | 3.89 | 1.78 | 5.86 | 2.99 | Tail protein D |
| *PSPTO_3391* | | - | 3832757 | 3832963 | 4.59 | 0.69 | 6.37 | 0.00 | 1.50 | 1.07 | Tail protein X |
| *PSPTO_3392* | | - | 3832938 | 3833702 | 3.87 | 11.16 | 4.37 | 0.00 | 12.82 | 5.20 | Hypothetical protein |
| *PSPTO_3393* | | - | 3833793 | 3836543 | 3.45 | 6.78 | 4.43 | 4.49 | 8.94 | 4.17 | Tail tape measure protein |
| *PSPTO_3394* | | - | 3836682 | 3836984 | 0.39 | 4.93 | 10.49 | 0.00 | 3.70 | 0.83 | Hypothetical protein |
| *PSPTO_3395* | | - | 3837000 | 3837509 | 3.71 | 5.96 | 4.47 | 4.74 | 5.05 | 2.35 | Major tail tube protein |
| *PSPTO_3396* | | - | 3837522 | 3838685 | 4.08 | 6.21 | 8.19 | 2.19 | 7.72 | 3.80 | Major tail sheath protein |
| *PSPTO_3397* | | - | 3838779 | 3839321 | 9.01 | 5.41 | 2.11 | 4.10 | 4.62 | 4.65 | Tail fiber assembly domain protein |
| *PSPTO_3398* | | - | 3839330 | 3841399 | 5.91 | 5.14 | 3.89 | 1.62 | 9.21 | 5.34 | Tail fiber protein H |
| *PSPTO_3399* | | - | 3841396 | 3842001 | 2.23 | 2.64 | 3.29 | 0.00 | 4.09 | 1.10 | Tail protein I |
| *PSPTO_3400* | | - | 3842003 | 3842884 | 7.33 | 3.02 | 3.65 | 2.63 | 5.87 | 3.50 | Baseplate assembly protein J |
| *PSPTO_3401* | | - | 3842881 | 3843207 | 10.80 | 15.74 | 4.51 | 8.64 | 9.51 | 4.92 | Baseplate assembly protein W |
| *PSPTO_3402* | | - | 3843211 | 3843489 | 1.91 | 10.70 | 3.60 | 1.76 | 11.00 | 2.86 | Hypothetical protein |
| *PSPTO_3403* | | - | 3843532 | 3844095 | 2.38 | 6.63 | 7.19 | 0.00 | 7.77 | 3.33 | Baseplate assembly protein V |
| *PSPTO_3404* | | - | 3844095 | 3844622 | 4.18 | 5.97 | 3.20 | 12.27 | 14.04 | 9.34 | Hypothetical protein |
| *PSPTO_3405* | | - | 3844615 | 3845271 | 1.90 | 4.12 | 1.92 | 4.20 | 6.38 | 1.79 | Hypothetical protein |
| *PSPTO_3406* | | - | 3845268 | 3845582 | 0.96 | 4.48 | 0.00 | 1.66 | 1.80 | 0.86 | Hypothetical protein |
| *PSPTO_3407* | | - | 3845585 | 3846580 | 4.38 | 9.74 | 4.75 | 1.45 | 8.66 | 2.04 | Hypothetical protein |
| *PSPTO_3408* | | - | 3846666 | 3847010 | 3.05 | 13.47 | 13.55 | 6.03 | 9.73 | 3.79 | Hypothetical protein |
| *PSPTO_3409* | *clpP-1* | - | 3847007 | 3848152 | 3.74 | 6.67 | 5.44 | 5.22 | 10.74 | 4.46 | ATP-dependent Clp protease, proteolytic subunit ClpP |
| *PSPTO_3410* | | - | 3848149 | 3849630 | 1.61 | 4.08 | 6.32 | 0.71 | 4.66 | 2.23 | Portal protein |
| *PSPTO_3411* | | - | 3849627 | 3849833 | 1.62 | 4.30 | 0.00 | 0.00 | 5.64 | 2.06 | Hypothetical protein |
| *PSPTO_3412* | | - | 3849835 | 3851859 | 3.04 | 4.61 | 5.34 | 1.65 | 6.55 | 3.57 | Terminase large subunit |
| *PSPTO_3413* | | - | 3851864 | 3852433 | 1.98 | 7.78 | 1.73 | 2.28 | 7.13 | 2.18 | Hypothetical protein |
| *PSPTO_3414* | | - | 3852658 | 3852804 | 0.80 | 1.19 | 0.00 | 9.74 | 10.33 | 2.65 | Hypothetical protein |
| *PSPTO_3415* | | - | 3852806 | 3853192 | 1.48 | 9.88 | 4.11 | 0.73 | 3.49 | 1.14 | Hypothetical protein |
| *PSPTO_3416* | | - | 3853189 | 3853512 | 1.20 | 0.00 | 0.49 | 3.62 | 3.00 | 1.04 | Holin |
| *PSPTO_3417* | | - | 3854040 | 3854399 | 7.66 | 1.21 | 10.16 | 0.00 | 4.82 | 3.45 | Hypothetical protein |
| *PSPTO_3418* | | - | 3854392 | 3856611 | 4.40 | 4.32 | 4.18 | 3.81 | 4.97 | 4.51 | DNA primase domain protein |
| *PSPTO_3419* | | - | 3856601 | 3856828 | 2.20 | 5.26 | 6.47 | 0.00 | 3.32 | 3.51 | C4-type zinc finger protein, DksA/TraR family |
| *PSPTO_3420* | | - | 3856821 | 3857420 | 8.04 | 11.88 | 22.77 | 11.69 | 18.05 | 17.16 | Hypothetical protein |
| *PSPTO_3421* | | + | 3857990 | 3858721 | 33.21 | 65.88 | 45.56 | 13.86 | 113.57 | 41.70 | Repressor protein c2 |
| *PSPTO_3422* | | + | 3859929 | 3860504 | 5.33 | 8.99 | 19.30 | 3.65 | 13.68 | 10.79 | Hypothetical protein |
| *PSPTO_3423* | | + | 3860513 | 3860797 | 2.17 | 12.84 | 8.41 | 0.00 | 8.45 | 3.12 | Hypothetical protein |

| Locus name | Gene name | Strand | Gene start (bp) | Gene end (bp) | Average TPM Inoculum | Average TPM $t_6$ | Average TPM $t_{24}$ | Average TPM $t_{168}$ | Average TPM Logarithmic | Average TPM Stationary | Annotation description |
|---|---|---|---|---|---|---|---|---|---|---|---|
| *PSPTO_3424* | | + | 3860938 | 3861312 | 3.19 | 3.33 | 14.71 | 0.49 | 12.18 | 3.16 | Hypothetical protein |
| *PSPTO_3425* | | + | 3861309 | 3861632 | 4.07 | 20.45 | 6.94 | 8.44 | 10.54 | 4.79 | Transcriptional regulator |
| *PSPTO_3426* | | + | 3861683 | 3862105 | 5.67 | 6.91 | 6.36 | 5.64 | 6.94 | 3.36 | Hypothetical protein |
| *PSPTO_3427* | | + | 3862800 | 3864584 | 3.61 | 3.42 | 4.30 | 0.72 | 5.30 | 3.24 | C-5 cytosine-specific DNA methylase family protein |
| *PSPTO_3428* | | + | 3864581 | 3865039 | 5.00 | 3.35 | 3.71 | 2.87 | 7.25 | 3.92 | Hypothetical protein |
| *PSPTO_3429* | | + | 3865036 | 3865278 | 30.55 | 26.91 | 20.97 | 25.71 | 26.04 | 19.64 | DNA-binding protein |
| *PSPTO_3430* | | + | 3865354 | 3865710 | 22.88 | 49.16 | 42.13 | 18.16 | 109.47 | 37.40 | 6-pyruvoyl tetrahydrobiopterin synthase |
| *PSPTO_3432* | | - | 3866884 | 3867693 | 0.00 | 0.00 | 0.00 | 0.00 | 0.00 | 0.00 | ISPsy4, transposition helper protein |
| *PSPTO_3433* | | - | 3867690 | 3868712 | 0.00 | 0.00 | 0.00 | 0.00 | 0.00 | 0.00 | ISPsy4, transposase |
| *PSPTO_3435* | | + | 3875338 | 3876225 | - | - | - | - | - | - | |
| *PSPTO_3436* | | + | 3876391 | 3876705 | 17.79 | 25.26 | 11.92 | 8.32 | 31.03 | 14.69 | Lipoprotein |
| *PSPTO_3437* | | - | 3876731 | 3878539 | 22.37 | 31.27 | 35.05 | 20.11 | 47.14 | 23.10 | Peptidase, M24 family protein |
| *PSPTO_3438* | *iscS-3* | - | 3878597 | 3879778 | 6.48 | 62.59 | 55.02 | 27.09 | 9.50 | 11.62 | Cysteine desulfurase |
| *PSPTO_3439* | | + | 3880359 | 3881480 | 2.53 | 3.66 | 3.92 | 4.28 | 4.77 | 4.12 | DegT/DnrJ/EryC1/StrS family aminotransferase |
| *PSPTO_3440* | | + | 3881486 | 3883000 | 2.71 | 3.99 | 8.42 | 3.88 | 4.77 | 5.51 | Polysaccharide biosynthesis protein |
| *PSPTO_3441* | | + | 3883004 | 3883939 | 2.73 | 0.61 | 5.43 | 0.00 | 4.18 | 2.16 | Glycosyl transferase family protein |
| *PSPTO_3442* | | + | 3883929 | 3884396 | 1.90 | 3.22 | 5.39 | 0.40 | 4.85 | 3.95 | Low molecular weight phosphotyrosine protein phosphatase family protein |
| *PSPTO_3443* | | + | 3884397 | 3886658 | 1.33 | 1.70 | 1.05 | 3.72 | 2.51 | 1.98 | Tyrosine-protein kinase |
| *PSPTO_3444* | *rfaH* | + | 3886710 | 3887222 | 2.07 | 7.66 | 0.00 | 0.00 | 2.32 | 2.65 | Transcriptional activator RfaH |
| *PSPTO_3445* | | + | 3887311 | 3888312 | 3.47 | 2.18 | 3.75 | 1.43 | 3.79 | 3.78 | Glycosyl transferase family protein |
| *PSPTO_3446* | | + | 3888309 | 3889547 | 3.85 | 1.43 | 5.63 | 0.44 | 3.24 | 3.74 | Glycoside hydrolase family protein |
| *PSPTO_3447* | | + | 3889623 | 3890732 | 3.73 | 8.92 | 7.46 | 5.38 | 13.86 | 9.96 | Glycoside hydrolase family protein |
| *PSPTO_3448* | | + | 3890803 | 3892032 | 2.45 | 1.44 | 1.71 | 0.76 | 2.97 | 2.18 | Glycoside hydrolase family protein |
| *PSPTO_3449* | | + | 3892045 | 3893454 | 2.04 | 3.10 | 3.08 | 0.71 | 1.98 | 1.85 | Capsular polysaccharide biosynthesis protein |
| *PSPTO_3450* | | + | 3893560 | 3894654 | 2.33 | 0.61 | 4.26 | 0.00 | 2.66 | 3.20 | Capsular polysaccharide biosynthesis protein |
| *PSPTO_3451* | *ssuE* | + | 3894902 | 3895483 | 43.79 | 56.98 | 63.04 | 59.79 | 20.67 | 15.30 | FMN reductase, NADH-dependent |
| *PSPTO_3452* | | - | 3895494 | 3896102 | 19.69 | 64.75 | 41.56 | 40.38 | 59.94 | 16.37 | LysE family transporter |
| *PSPTO_3453* | | - | 3896226 | 3896501 | 27.88 | 178.27 | 231.71 | 90.34 | 27.03 | 28.05 | Lipoprotein |
| *PSPTO_3454* | | - | 3896590 | 3897480 | 2.50 | 5.33 | 10.31 | 5.58 | 7.75 | 4.87 | LysR family transcriptional regulator |
| *PSPTO_3455* | | + | 3897610 | 3898311 | 126.06 | 16.52 | 21.00 | 11.53 | 112.05 | 83.61 | 3-oxoacid CoA-transferase, subunit A family |
| *PSPTO_3456* | | + | 3898357 | 3899013 | 65.15 | 14.33 | 23.27 | 4.34 | 71.73 | 38.62 | 3-oxoacid CoA-transferase, subunit B family |
| *PSPTO_3457* | | + | 3899457 | 3900875 | 6.41 | 1.78 | 2.69 | 0.29 | 10.13 | 3.41 | Short-chain fatty acid transporter |
| *PSPTO_3458* | | - | 3900878 | 3901207 | 6.20 | 7.85 | 11.34 | 1.46 | 10.36 | 5.78 | Hypothetical protein |
| *PSPTO_3459* | | - | 3901204 | 3901938 | 8.63 | 5.01 | 10.04 | 1.52 | 12.44 | 10.79 | Hypothetical protein |
| *PSPTO_3460* | | - | 3902047 | 3902979 | 7.14 | 22.82 | 28.64 | 17.95 | 50.09 | 30.65 | D-isomer specific 2-hydroxyacid dehydrogenase family protein |
| *PSPTO_3461* | *glpQ* | + | 3903238 | 3904356 | 12.48 | 20.98 | 15.01 | 11.54 | 21.86 | 11.73 | Glycerophosphoryl diester phosphodiesterase |
| *PSPTO_3462* | | - | 3904401 | 3906587 | 3.11 | 1.98 | 2.89 | 1.84 | 21.67 | 17.74 | TonB-dependent siderophore receptor |
| *PSPTO_3463* | *pssA-2* | + | 3906829 | 3908172 | 11.14 | 43.68 | 33.04 | 3.84 | 85.94 | 26.74 | CDP-diacylglycerol--serine O-phosphatidyltransferase |
| *PSPTO_3464* | | - | 3908246 | 3910801 | 4.71 | 13.57 | 8.99 | 8.52 | 5.94 | 6.57 | ATP-dependent DNA ligase |
| *PSPTO_3465* | | - | 3910838 | 3911737 | 11.79 | 27.19 | 7.70 | 14.67 | 6.79 | 7.85 | KU domain protein |
| *PSPTO_3466* | *ssuD* | + | 3912138 | 3913277 | 4.47 | 183.00 | 165.70 | 100.07 | 5.66 | 4.64 | Alkanesulfonate monooxygenase |
| *PSPTO_3467* | | + | 3913277 | 3914383 | 3.24 | 64.31 | 37.67 | 46.92 | 3.83 | 4.71 | Sigma-54 dependent transcriptional regulator |
| *PSPTO_3468* | | - | 3914690 | 3916093 | 6.22 | 10.51 | 13.83 | 4.44 | 15.97 | 11.55 | Hypothetical protein |
| *PSPTO_3469* | | - | 3916142 | 3916711 | 18.62 | 31.84 | 49.08 | 13.76 | 33.75 | 24.57 | DNA-binding protein |
| *PSPTO_3470* | | + | 3916833 | 3917771 | 10.85 | 11.67 | 10.39 | 1.10 | 22.60 | 18.50 | Membrane protein |
| *PSPTO_3471* | | + | 3917931 | 3920849 | 7.25 | 20.01 | 14.21 | 6.99 | 24.48 | 10.79 | Putative monovalent cation/H+ antiporter subunit A |
| *PSPTO_3472* | | + | 3920849 | 3921193 | 6.76 | 25.59 | 23.21 | 1.51 | 19.48 | 15.05 | Potassium efflux system protein PhaC |
| *PSPTO_3473* | | + | 3921199 | 3922875 | 9.41 | 13.30 | 18.00 | 10.62 | 20.17 | 13.35 | Monovalent cation/proton antiporter, MnhD/PhaD subunit |
| *PSPTO_3474* | | + | 3922872 | 3923360 | 14.55 | 8.33 | 10.96 | 7.52 | 23.35 | 17.36 | Putative monovalent cation/H+ antiporter subunit E |
| *PSPTO_3475* | | + | 3923357 | 3923635 | 4.35 | 22.66 | 7.53 | 11.66 | 29.22 | 15.09 | Potassium efflux system protein PhaF |
| *PSPTO_3476* | | + | 3923632 | 3924012 | 8.56 | 20.24 | 10.64 | 4.88 | 42.88 | 19.70 | Potassium efflux system protein PhaG |
| *PSPTO_3477* | | - | 3924034 | 3924501 | 4.92 | 26.61 | 17.59 | 14.07 | 43.38 | 21.57 | Hypothetical protein |
| *PSPTO_3478* | | + | 3924946 | 3925185 | 18.47 | 88.90 | 79.49 | 58.14 | 15.65 | 18.11 | Hypothetical protein |
| *PSPTO_3479* | | - | 3925206 | 3926774 | 4.48 | 3.10 | 5.18 | 4.76 | 3.80 | 5.35 | Hypothetical protein |
| *PSPTO_3480* | | - | 3926984 | 3928585 | 28.61 | 21.47 | 24.29 | 13.37 | 30.60 | 15.86 | Methyl-accepting chemotaxis protein |
| *PSPTO_3481* | | + | 3929300 | 3929530 | 33.44 | 12.54 | 5.41 | 14.34 | 6.97 | 13.31 | Hypothetical protein |
| *PSPTO_3482* | | + | 3929574 | 3931601 | 9.73 | 9.78 | 12.42 | 9.97 | 6.27 | 9.77 | Rhs element Vgr protein |
| *PSPTO_3483* | | + | 3931598 | 3932446 | 226.44 | 257.56 | 168.09 | 167.94 | 104.88 | 169.48 | Hypothetical protein |
| *PSPTO_3484* | | + | 3932472 | 3933272 | 7.58 | 21.22 | 19.78 | 20.00 | 27.94 | 20.93 | Lipoprotein |
| *PSPTO_3485* | | + | 3933269 | 3935545 | 6.99 | 15.59 | 12.65 | 15.92 | 25.47 | 14.98 | Lipase family protein |
| *PSPTO_3486* | | + | 3935770 | 3935886 | 4.90 | 15.78 | 16.09 | 0.00 | 8.90 | 10.82 | Hypothetical protein |
| *PSPTO_3487* | | - | 3936017 | 3936421 | 18.19 | 47.37 | 49.90 | 34.36 | 41.14 | 29.38 | DnaK suppressor protein |
| *PSPTO_3488* | | - | 3936725 | 3937750 | 6.99 | 87.77 | 50.47 | 22.13 | 12.96 | 6.23 | Sugar ABC transporter permease |
| *PSPTO_3489* | | - | 3937795 | 3939372 | 5.09 | 114.36 | 77.85 | 24.42 | 8.62 | 4.13 | Sugar ABC transporter ATP-binding protein |
| *PSPTO_3490* | | - | 3939442 | 3940371 | 7.57 | 230.21 | 166.22 | 39.12 | 8.06 | 3.47 | Sugar ABC transporter periplasmic sugar-binding protein |
| *PSPTO_3491* | | + | 3941852 | 3942220 | 17.19 | 26.23 | 13.58 | 12.48 | 11.44 | 8.94 | Hypothetical protein |
| *PSPTO_3492* | | - | 3942270 | 3943289 | 12.42 | 40.61 | 40.07 | 23.02 | 17.45 | 11.92 | Oxidoreductase, Gfo/Idh/MocA family |
| *PSPTO_3493* | | - | 3943355 | 3944215 | 33.11 | 135.48 | 104.43 | 49.72 | 35.94 | 23.97 | IolH protein |
| *PSPTO_3494* | *iolG* | - | 3944268 | 3945278 | 7.71 | 101.51 | 88.77 | 17.85 | 18.16 | 5.41 | Myo-inositol 2-dehydrogenase |
| *PSPTO_3495* | | - | 3945362 | 3947299 | 5.01 | 81.33 | 58.45 | 28.62 | 19.67 | 8.05 | IolD protein |
| *PSPTO_3496* | *iolI* | - | 3947363 | 3948193 | 3.46 | 40.01 | 26.53 | 11.88 | 11.11 | 2.62 | IolI protein |
| *PSPTO_3497* | *mmsA-3* | - | 3948230 | 3949732 | 7.41 | 77.67 | 51.84 | 18.25 | 23.06 | 8.09 | Methylmalonate-semialdehyde dehydrogenase |
| *PSPTO_3498* | | - | 3949760 | 3950569 | 5.15 | 107.13 | 72.41 | 28.45 | 18.67 | 4.96 | IolB protein |
| *PSPTO_3499* | | - | 3950599 | 3951504 | 3.12 | 73.58 | 54.05 | 17.48 | 16.24 | 4.78 | IolE protein |
| *PSPTO_3500* | | - | 3951574 | 3953511 | 4.69 | 65.00 | 46.08 | 14.77 | 10.35 | 3.08 | IolC protein |
| *PSPTO_3501* | | + | 3953783 | 3954691 | 20.35 | 38.53 | 40.85 | 18.02 | 49.70 | 20.43 | RpiR family transcriptional regulator |
| *PSPTO_3502* | | - | 3954745 | 3955743 | 4.38 | 15.01 | 24.66 | 6.97 | 12.33 | 12.61 | Membrane protein |
| *PSPTO_3503* | | - | 3955740 | 3956990 | 5.89 | 18.84 | 7.29 | 3.45 | 8.49 | 13.13 | Cardiolipin synthetase 2 |
| *PSPTO_3504* | | - | 3956987 | 3957700 | 39.94 | 32.74 | 18.45 | 21.18 | 16.09 | 17.12 | Endonuclease/exonuclease/phosphatase family protein |

| Locus name | Gene name | Strand | Gene start (bp) | Gene end (bp) | Average TPM Inoculum | Average TPM $t_6$ | Average TPM $t_{24}$ | Average TPM $t_{168}$ | Average TPM Logarithmic | Average TPM Stationary | Annotation description |
|---|---|---|---|---|---|---|---|---|---|---|---|
| *PSPTO_3505* | | + | 3957995 | 3958534 | 55.63 | 26.12 | 40.04 | 15.43 | 38.86 | 24.07 | Cytochrome b561 |
| *PSPTO_3506* | | - | 3958536 | 3959276 | 17.75 | 27.93 | 39.18 | 8.47 | 44.73 | 22.24 | Phosphorylase, family 2 |
| *PSPTO_3507* | *nagZ* | - | 3959286 | 3960296 | 7.62 | 14.54 | 15.73 | 14.18 | 33.46 | 12.28 | Beta-hexosaminidase |
| *PSPTO_3508* | *psrA* | - | 3960420 | 3961136 | 25.35 | 26.19 | 26.84 | 8.99 | 29.07 | 15.25 | Transcriptional regulator PsrA |
| *PSPTO_3509* | | - | 3961269 | 3961391 | 7.54 | 0.00 | 0.00 | 18.62 | 21.93 | 10.44 | Hypothetical protein |
| *PSPTO_3510* | *lexA-2* | + | 3961393 | 3962001 | 19.60 | 109.81 | 75.09 | 22.33 | 67.03 | 30.84 | LexA repressor |
| *PSPTO_3511* | | + | 3962011 | 3962481 | 7.46 | 23.49 | 32.26 | 7.74 | 35.10 | 21.06 | Cell division inhibitor |
| *PSPTO_3512* | | - | 3962536 | 3962799 | 92.92 | 63.74 | 76.95 | 57.91 | 83.80 | 78.10 | Hypothetical protein |
| *PSPTO_3513* | | - | 3962834 | 3963355 | 125.29 | 60.47 | 56.80 | 25.40 | 36.37 | 41.96 | Hypothetical protein |
| *PSPTO_3514* | *topA* | - | 3963478 | 3966087 | 24.69 | 52.47 | 46.39 | 28.69 | 81.04 | 30.46 | DNA topoisomerase I |
| *PSPTO_3515* | | - | 3966201 | 3966464 | 39.81 | 36.25 | 30.58 | 8.96 | 43.63 | 19.97 | Hypothetical protein |
| *PSPTO_3516* | *fadA* | - | 3966542 | 3967717 | 17.50 | 31.17 | 34.33 | 15.70 | 51.54 | 9.41 | 3-ketoacyl-CoA thiolase |
| *PSPTO_3517* | *fadB* | - | 3967722 | 3969887 | 50.05 | 53.46 | 56.51 | 17.54 | 95.21 | 65.57 | Multifunctional fatty acid oxidation complex subunit alpha |
| *PSPTO_3518* | | + | 3970942 | 3971067 | 198.14 | 54.62 | 60.16 | 30.91 | 14.89 | 12.46 | Hypothetical protein |
| *PSPTO_3519* | | - | 3971084 | 3971521 | 10.73 | 28.96 | 40.82 | 9.23 | 32.89 | 17.16 | Hypothetical protein |
| *PSPTO_3520* | | + | 3971652 | 3972113 | 128.87 | 238.64 | 260.32 | 80.71 | 354.65 | 398.15 | Universal stress protein family |
| *PSPTO_3521* | | - | 3972182 | 3974092 | 14.12 | 34.47 | 26.02 | 21.17 | 48.15 | 19.23 | ABC transporter ATP-binding protein |
| *PSPTO_3522* | | + | 3974297 | 3976225 | 8.68 | 16.17 | 7.97 | 6.29 | 55.88 | 14.90 | Lytic murein transglycosylase |
| *PSPTO_3523* | | - | 3976273 | 3977082 | 11.83 | 19.09 | 18.20 | 12.79 | 32.67 | 32.97 | MOSC domain-containing protein |
| *PSPTO_3524* | | - | 3977175 | 3978101 | 113.20 | 31.62 | 37.10 | 11.42 | 5.49 | 4.17 | Chemotaxis protein CheV |
| *PSPTO_3525* | | + | 3978552 | 3978983 | 28.87 | 49.63 | 24.48 | 13.45 | 19.62 | 25.44 | Hypothetical protein |
| *PSPTO_3526* | | + | 3979129 | 3979791 | 27.95 | 22.15 | 51.48 | 15.59 | 39.32 | 36.70 | LuxR family DNA-binding response regulator |
| *PSPTO_3527* | | - | 3979775 | 3980692 | 85.01 | 57.35 | 33.58 | 16.99 | 36.90 | 67.71 | Hypothetical protein |
| *PSPTO_3528* | | - | 3981164 | 3982141 | 18.11 | 22.63 | 20.30 | 17.15 | 36.66 | 29.31 | IS52, transposase |
| *PSPTO_3529* | *pslA* | + | 3982432 | 3983868 | 10.06 | 10.35 | 7.32 | 2.69 | 9.63 | 6.53 | Capsular polysaccharide biosynthesis protein PslA |
| *PSPTO_3530* | *pslB* | + | 3983868 | 3985325 | 6.27 | 5.53 | 3.57 | 1.64 | 7.09 | 3.51 | Mannose-1-phosphate guanylyltransferase/mannose-6-phosphate isomerase PslB |
| *PSPTO_3531* | *pslD* | + | 3985471 | 3986247 | 3.71 | 1.41 | 1.65 | 0.00 | 6.13 | 3.49 | Lipoprotein PslD |
| *PSPTO_3532* | *pslE* | + | 3986265 | 3988256 | 2.32 | 1.76 | 1.24 | 2.01 | 5.56 | 2.05 | Exopolysaccharide biosynthesis protein PslE |
| *PSPTO_3533* | *pslF* | + | 3988256 | 3989434 | 3.90 | 8.49 | 6.43 | 4.43 | 9.32 | 4.29 | Glycosyl transferase, group 1 family protein PslF |
| *PSPTO_3534* | *pslG* | + | 3989436 | 3990749 | 5.60 | 10.51 | 5.39 | 4.62 | 20.10 | 9.15 | Glycosyl hydrolase, family 5 PslG |
| *PSPTO_3535* | *pslH* | + | 3990753 | 3991970 | 4.18 | 1.63 | 3.01 | 1.04 | 6.76 | 3.96 | Glycosyl transferase, group 1 family protein PslH |
| *PSPTO_3536* | *pslI* | + | 3991961 | 3993061 | 4.98 | 8.29 | 9.47 | 5.55 | 11.29 | 8.65 | Glycosyl transferase, group 1 family protein PslI |
| *PSPTO_3537* | *pslJ* | + | 3993061 | 3994476 | 4.46 | 7.31 | 2.79 | 3.53 | 11.08 | 4.13 | Membrane protein PslJ |
| *PSPTO_3538* | | + | 3994473 | 3995294 | 3.37 | 3.92 | 3.96 | 0.00 | 9.89 | 4.96 | Transferase, hexapeptide repeat protein |
| *PSPTO_3539* | *pslK* | + | 3995310 | 3996725 | 1.80 | 2.03 | 4.42 | 0.00 | 2.75 | 2.41 | Membrane protein PslK |
| *PSPTO_3540* | | - | 3996659 | 3997612 | 1.54 | 0.93 | 3.36 | 3.96 | 2.24 | 2.59 | LysR family transcriptional regulator |
| *PSPTO_3541* | | + | 3997736 | 3998896 | 2.82 | 1.71 | 1.19 | 0.00 | 3.76 | 3.89 | C4-dicarboxylate transporter/malic acid transport protein |
| *PSPTO_3542* | | + | 3998961 | 3999878 | 2.72 | 5.63 | 2.21 | 0.87 | 2.92 | 2.89 | Hypothetical protein |
| *PSPTO_3543* | | + | 3999972 | 4000283 | 203.43 | 77.48 | 67.21 | 26.72 | 33.72 | 36.98 | Hypothetical protein |
| *PSPTO_3544* | | + | 4000561 | 4001706 | 21.77 | 104.51 | 84.98 | 34.82 | 65.39 | 7.45 | Hypothetical protein |
| *PSPTO_3545* | | + | 4001872 | 4002549 | 13.01 | 93.15 | 84.30 | 45.38 | 27.33 | 40.71 | ThiJ/PfpI family protein |
| *PSPTO_3546* | | + | 4002668 | 4002811 | 30.45 | 30.44 | 30.19 | 16.55 | 6.09 | 8.64 | Hypothetical protein |
| *PSPTO_3547* | | - | 4002957 | 4003892 | 6.45 | 13.57 | 12.46 | 2.57 | 9.37 | 8.41 | LysR family transcriptional regulator |
| *PSPTO_3548* | | - | 4003986 | 4005086 | 31.24 | 113.26 | 87.32 | 47.95 | 67.35 | 40.45 | Hypothetical protein |
| *PSPTO_3549* | *aefR* | + | 4005294 | 4005896 | 40.11 | 69.29 | 72.79 | 47.76 | 54.37 | 35.11 | Transcriptional regulator AefR |
| *PSPTO_3550* | *fahA* | - | 4005909 | 4007204 | 120.91 | 315.09 | 271.42 | 185.03 | 271.56 | 110.87 | Fumarylacetoacetase |
| *PSPTO_3551* | *hmgA* | - | 4007204 | 4008508 | 146.00 | 366.22 | 304.00 | 191.75 | 232.39 | 120.19 | Homogentisate 1,2-dioxygenase |
| *PSPTO_3552* | | + | 4008637 | 4009428 | 3.78 | 5.23 | 2.93 | 0.00 | 7.94 | 5.35 | IclR family transcriptional regulator |
| *PSPTO_3553* | *hppD* | - | 4009715 | 4010806 | 42.89 | 96.96 | 110.34 | 60.55 | 383.56 | 121.35 | 4-hydroxyphenylpyruvate dioxygenase |
| *PSPTO_3554* | *maiA* | - | 4010949 | 4011584 | 70.84 | 176.79 | 157.33 | 126.57 | 278.29 | 74.52 | Maleylacetoacetate isomerase |
| *PSPTO_3555* | *glcD* | + | 4011746 | 4013245 | 9.27 | 14.53 | 19.34 | 7.96 | 11.78 | 5.32 | Glycolate oxidase subunit GlcD |
| *PSPTO_3556* | *glcE* | + | 4013245 | 4014303 | 3.99 | 11.44 | 15.16 | 12.46 | 9.18 | 2.69 | Glycolate oxidase FAD binding subunit |
| *PSPTO_3557* | *glcF* | + | 4014307 | 4015540 | - | - | - | - | - | - | |
| *PSPTO_3558* | *glcG* | + | 4015576 | 4015983 | 22.74 | 60.28 | 58.89 | 36.60 | 39.32 | 25.75 | GlcG protein |
| *PSPTO_3559* | *glcB-2* | + | 4016048 | 4018225 | 23.27 | 40.98 | 36.28 | 16.61 | 25.22 | 19.32 | Malate synthase G |
| *PSPTO_3560* | | - | 4018366 | 4019574 | 9.83 | 13.60 | 19.03 | 12.21 | 16.68 | 19.97 | GDA1/CD39 family protein |
| *PSPTO_3561* | | - | 4019732 | 4020703 | 2.26 | 1.28 | 3.17 | 0.13 | 4.11 | 5.16 | Hypothetical protein |
| *PSPTO_3562* | | - | 4020895 | 4020996 | 9.25 | 10.89 | 0.00 | 0.00 | 3.29 | 3.24 | Hypothetical protein |
| *PSPTO_3563* | *gntR* | - | 4021034 | 4022050 | 6.18 | 9.05 | 6.14 | 2.60 | 9.22 | 8.92 | Gluconate utilization system GNT-I transcriptional repressor |
| *PSPTO_3564* | *gntV* | + | 4022334 | 4022822 | 12.27 | 36.98 | 32.07 | 28.56 | 23.15 | 31.01 | Gluconokinase |
| *PSPTO_3565* | | + | 4022920 | 4024272 | 6.67 | 10.57 | 16.33 | 11.60 | 11.93 | 13.60 | Gluconate permease |
| *PSPTO_3566* | *csrA-3* | + | 4024829 | 4025017 | 74.73 | 157.62 | 112.34 | 46.93 | 63.84 | 52.48 | Carbon storage regulator |
| *PSPTO_3567* | | - | 4025135 | 4026535 | 6.95 | 11.36 | 8.21 | 3.15 | 5.36 | 8.51 | Hypothetical protein |
| *PSPTO_3568* | | - | 4027084 | 4027209 | 1.07 | 0.00 | 0.00 | 0.00 | 0.66 | 8.19 | Hypothetical protein |
| *PSPTO_3569* | | - | 4027298 | 4027558 | 108.39 | 47.56 | 105.89 | 56.10 | 19.22 | 54.76 | Hypothetical protein |
| *PSPTO_3570* | | - | 4027965 | 4028558 | 52.76 | 205.83 | 150.19 | 66.94 | 245.74 | 82.24 | GNAT family acetyltransferase |
| *PSPTO_3571* | | - | 4028602 | 4028718 | 48.35 | 13.46 | 46.26 | 9.33 | 11.33 | 12.49 | Hypothetical protein |
| *PSPTO_3572* | | + | 4028816 | 4029760 | 47.61 | 30.94 | 28.28 | 10.78 | 33.23 | 19.28 | Hypothetical protein |
| *PSPTO_3573* | | + | 4029815 | 4030279 | 4.61 | 4.11 | 6.14 | 4.95 | 12.99 | 5.14 | Putative acetyltransferase |
| *PSPTO_3574* | | + | 4030399 | 4032519 | 11.31 | 8.00 | 12.67 | 6.83 | 125.43 | 98.74 | TonB-dependent siderophore receptor |
| *PSPTO_3575* | | - | 4032615 | 4033136 | 41.37 | 44.52 | 52.67 | 11.62 | 19.91 | 25.84 | Hypothetical protein |
| *PSPTO_3576* | *tvrR* | + | 4033365 | 4033988 | 20.18 | 30.75 | 29.71 | 7.27 | 43.35 | 27.09 | TetR-like virulence regulator |
| *PSPTO_3577* | | + | 4034169 | 4035797 | 34.98 | 20.77 | 6.07 | 5.45 | 8.71 | 7.05 | Methyl-accepting chemotaxis protein |
| *PSPTO_3578* | | + | 4035879 | 4036367 | 15.34 | 22.61 | 35.20 | 29.69 | 17.34 | 24.79 | Hypothetical protein |
| *PSPTO_3579* | | - | 4036406 | 4037287 | 24.14 | 40.62 | 23.89 | 19.16 | 53.89 | 25.18 | Hypothetical protein |
| *PSPTO_3580* | | - | 4037428 | 4039068 | 8.48 | 11.05 | 9.44 | 5.24 | 9.01 | 8.83 | Methyl-accepting chemotaxis protein |
| *PSPTO_3581* | | - | 4039263 | 4040078 | 18.61 | 27.95 | 15.22 | 12.34 | 21.97 | 10.64 | GNAT family acetyltransferase |
| *PSPTO_3582* | *katB* | + | 4040557 | 4042089 | 59.13 | 745.85 | 680.55 | 150.50 | 18.10 | 8.53 | Catalase |
| *PSPTO_3583* | | + | 4042152 | 4042706 | 5.47 | 100.77 | 98.04 | 35.36 | 5.46 | 3.96 | Ankyrin domain protein |

| Locus name | Gene name | Strand | Gene start (bp) | Gene end (bp) | Average TPM Inoculum | Average TPM $t_6$ | Average TPM $t_{24}$ | Average TPM $t_{168}$ | Average TPM Logarithmic | Average TPM Stationary | Annotation description |
|---|---|---|---|---|---|---|---|---|---|---|---|
| *PSPTO_3584* | | - | 4042719 | 4044659 | 6.70 | 11.62 | 7.88 | 5.87 | 10.80 | 6.37 | Sensory box histidine kinase/response regulator |
| *PSPTO_3585* | | - | 4044986 | 4045195 | 40.23 | 56.10 | 38.30 | 28.13 | 28.97 | 25.06 | Hypothetical protein |
| *PSPTO_3586* | | - | 4045502 | 4046386 | 12.41 | 8.39 | 19.06 | 6.73 | 18.74 | 11.76 | LysR family transcriptional regulator |
| *PSPTO_3587* | | + | 4046458 | 4046967 | 7.38 | 380.86 | 157.99 | 48.46 | 6.56 | 4.69 | Hypothetical protein |
| *PSPTO_3588* | | + | 4047092 | 4047925 | 6.39 | 95.55 | 50.55 | 13.36 | 13.98 | 11.92 | Phenazine biosynthesis family protein |
| *PSPTO_3589* | | - | 4047932 | 4049083 | 5.08 | 21.53 | 19.38 | 2.97 | 17.65 | 10.69 | Transporter |
| *PSPTO_3590* | | + | 4049295 | 4049951 | 13.28 | 14.71 | 17.97 | 11.13 | 11.72 | 12.94 | HAD-superfamily hydrolase |
| *PSPTO_3591* | *rarD-2* | + | 4050049 | 4050972 | 13.44 | 12.96 | 7.58 | 9.38 | 16.72 | 15.21 | RarD protein |
| *PSPTO_3592* | | - | 4050976 | 4052091 | 1.48 | 5.53 | 6.90 | 4.94 | 11.20 | 4.80 | ATPase |
| *PSPTO_3593* | | - | 4052158 | 4053603 | 3.90 | 9.04 | 7.57 | 3.27 | 12.78 | 7.72 | Sulfate permease family protein |
| *PSPTO_3594* | *ampC* | + | 4053747 | 4054898 | 6.16 | 7.55 | 3.10 | 6.64 | 17.42 | 8.73 | Beta-lactamase |
| *PSPTO_3595* | *ada* | - | 4054891 | 4055997 | 7.87 | 11.30 | 6.17 | 3.31 | 12.39 | 10.01 | Ada regulatory protein |
| *PSPTO_3596* | | + | 4056266 | 4057120 | 11.92 | 1.71 | 7.90 | 1.71 | 56.28 | 133.50 | Membrane protein |
| *PSPTO_3597* | | + | 4057129 | 4058337 | 11.91 | 8.16 | 6.06 | 7.64 | 163.81 | 151.29 | Hypothetical protein |
| *PSPTO_3598* | | + | 4058362 | 4059684 | 15.39 | 6.61 | 6.95 | 3.27 | 269.74 | 163.00 | Dyp-type peroxidase family protein |
| *PSPTO_3599* | | + | 4059714 | 4060538 | 11.73 | 10.59 | 4.67 | 3.46 | 233.05 | 91.73 | Hypothetical protein |
| *PSPTO_3600* | | - | 4060605 | 4061252 | 7.15 | 8.23 | 7.30 | 4.86 | 21.92 | 10.87 | Oxidoreductase, molybdopterin-binding protein |
| *PSPTO_3601* | | - | 4061372 | 4061992 | 6.51 | 11.93 | 12.99 | 4.62 | 4.18 | 6.71 | Hypothetical protein |
| *PSPTO_3602* | | - | 4062065 | 4062670 | 35.18 | 58.45 | 56.49 | 26.74 | 10.86 | 31.48 | Hypothetical protein |
| *PSPTO_3603* | | + | 4062841 | 4063530 | 4.26 | 12.00 | 6.93 | 1.35 | 5.09 | 3.96 | DNA-binding heavy metal response regulator |
| *PSPTO_3604* | | + | 4063509 | 4064897 | 13.90 | 10.10 | 6.44 | 5.59 | 4.97 | 7.57 | Heavy metal sensor histidine kinase |
| *PSPTO_3605* | | - | 4064916 | 4065584 | 438.98 | 173.75 | 129.15 | 84.13 | 26.23 | 65.17 | Lyase |
| *PSPTO_3606* | | + | 4065632 | 4065790 | 293.74 | 64.57 | 46.09 | 42.26 | 16.92 | 29.94 | Hypothetical protein |
| *PSPTO_3607* | | - | 4066177 | 4066986 | 0.00 | 0.00 | 0.00 | 0.00 | 0.00 | 0.00 | ISPsy4, transposition helper protein |
| *PSPTO_3608* | | - | 4066983 | 4068005 | 0.00 | 0.00 | 0.00 | 0.00 | 0.00 | 0.00 | ISPsy4, transposase |
| *PSPTO_3609* | | + | 4068085 | 4069107 | 0.97 | 3.89 | 1.59 | 3.47 | 1.87 | 0.84 | Site-specific recombinase, phage integrase family |
| *PSPTO_3610* | | + | 4069104 | 4070783 | 0.78 | 1.61 | 0.29 | 0.65 | 0.73 | 1.20 | Hypothetical protein |
| *PSPTO_3611* | | + | 4071121 | 4071154 | - | - | - | - | - | - | |
| *PSPTO_3612* | | + | 4071237 | 4071596 | 0.00 | 0.00 | 0.00 | 0.00 | 0.00 | 0.00 | ISPsy5, Orf1 |
| *PSPTO_3613* | | + | 4071614 | 4073167 | 0.46 | 0.00 | 0.00 | 0.00 | 0.89 | 0.56 | ISPsy5, transposase |
| *PSPTO_3614* | | + | 4073215 | 4074165 | - | - | - | - | - | - | |
| *PSPTO_3615* | | + | 4074346 | 4075947 | 0.70 | 3.02 | 2.17 | 0.11 | 1.90 | 1.13 | Site-specific recombinase, phage integrase family |
| *PSPTO_3616* | | - | 4077186 | 4078172 | 1.84 | 6.11 | 4.74 | 2.25 | 13.54 | 3.93 | Hypothetical protein |
| *PSPTO_3617* | | + | 4079962 | 4080414 | 23.42 | 9.96 | 3.93 | 0.00 | 6.51 | 5.68 | MarR family transcriptional regulator |
| *PSPTO_3618* | | + | 4080432 | 4082498 | 2.96 | 4.38 | 4.04 | 1.00 | 5.23 | 3.16 | Membrane protein |
| *PSPTO_3619* | | + | 4082495 | 4082704 | 0.93 | 11.02 | 0.00 | 0.00 | 3.86 | 1.05 | Membrane protein |
| *PSPTO_3620* | | + | 4082701 | 4083585 | 4.29 | 3.88 | 3.82 | 2.90 | 7.62 | 6.26 | HlyD family secretion protein |
| *PSPTO_3621* | | + | 4083582 | 4085015 | 3.65 | 9.20 | 5.49 | 0.10 | 7.82 | 6.08 | Outer membrane efflux protein |
| *PSPTO_3624* | | + | 4085993 | 4086826 | 20.95 | 23.83 | 25.81 | 18.93 | 24.32 | 17.28 | Ion transport protein |
| *PSPTO_3625* | | + | 4086925 | 4087923 | 5.91 | 30.04 | 19.20 | 25.56 | 20.65 | 9.61 | Sulfate ABC transporter substrate-binding protein |
| *PSPTO_3626* | | - | 4087984 | 4088391 | 43.65 | 41.09 | 39.41 | 15.50 | 18.65 | 17.79 | Lipoprotein |
| *PSPTO_3627* | *cycH* | - | 4088403 | 4089584 | - | - | - | - | - | - | |
| *PSPTO_3628* | *cycL* | - | 4089581 | 4090051 | 2.77 | 5.81 | 3.60 | 0.00 | 7.85 | 4.02 | Cytochrome c-type biogenesis protein CycL |
| *PSPTO_3629* | *dsbE* | - | 4090048 | 4090584 | 5.53 | 9.07 | 18.64 | 8.53 | 15.96 | 7.10 | Thiol:disulfide interchange protein DsbE |
| *PSPTO_3630* | *ccmF* | - | 4090581 | 4092554 | 2.14 | 3.28 | 4.65 | 0.93 | 8.25 | 3.81 | Cytochrome c-type biogenesis protein CcmF |
| *PSPTO_3631* | *ccmE* | - | 4092551 | 4093024 | 5.21 | 10.59 | 13.54 | 3.60 | 20.50 | 10.03 | Cytochrome c-type biogenesis protein CcmE |
| *PSPTO_3632* | *ccmD* | - | 4093021 | 4093197 | 1.76 | 0.91 | 8.00 | 0.00 | 8.59 | 4.14 | Heme exporter protein CcmD |
| *PSPTO_3633* | *ccmC* | - | 4093194 | 4093997 | 4.47 | 9.70 | 3.30 | 2.11 | 18.65 | 14.13 | Heme exporter protein CcmC |
| *PSPTO_3634* | *ccmB* | - | 4094053 | 4094724 | 0.86 | 5.86 | 9.73 | 0.82 | 7.25 | 8.54 | Heme exporter protein CcmB |
| *PSPTO_3635* | *ccmA* | - | 4094721 | 4095359 | 1.68 | 3.29 | 1.23 | 2.59 | 5.85 | 2.89 | Heme exporter protein CcmA |
| *PSPTO_3636* | | + | 4096486 | 4097199 | 16.60 | 34.93 | 20.90 | 6.02 | 15.79 | 7.55 | Hypothetical protein |
| *PSPTO_3637* | | + | 4097199 | 4097522 | 8.23 | 23.59 | 14.26 | 8.28 | 24.95 | 11.90 | FlhB domain-containing protein |
| *PSPTO_3638* | *recR* | - | 4097544 | 4098146 | 9.26 | 14.72 | 16.58 | 11.61 | 27.11 | 12.67 | Recombination protein RecR |
| *PSPTO_3639* | | - | 4098257 | 4099141 | 4.82 | 10.09 | 2.57 | 8.96 | 11.51 | 5.65 | LysR family transcriptional regulator |
| *PSPTO_3640* | | + | 4099238 | 4100374 | 8.16 | 10.07 | 13.84 | 0.16 | 7.66 | 11.40 | Hypothetical protein |
| *PSPTO_3641* | | + | 4100364 | 4100756 | 11.51 | 4.76 | 14.00 | 0.00 | 8.32 | 10.35 | Endoribonuclease L-PSP family protein |
| *PSPTO_3642* | | + | 4100776 | 4101960 | - | - | - | - | - | - | |
| *PSPTO_3643* | | + | 4101990 | 4102847 | 1.99 | 0.17 | 5.20 | 0.00 | 2.49 | 2.36 | GNAT family acetyltransferase |
| *PSPTO_3644* | *aldC* | + | 4103003 | 4104478 | 8.62 | 6.08 | 13.32 | 7.58 | 11.01 | 6.72 | Aldehyde dehydrogenase family protein. |
| *PSPTO_3645* | | - | 4104596 | 4104922 | 78.97 | 151.63 | 119.56 | 35.29 | 373.59 | 179.10 | Hypothetical protein |
| *PSPTO_3646* | *dnaZX* | - | 4104984 | 4107215 | 5.40 | 31.32 | 33.74 | 21.66 | 101.73 | 41.63 | DNA polymerase III subunits gamma and tau |
| *PSPTO_3647* | | + | 4107316 | 4108116 | 12.17 | 8.21 | 10.43 | 2.07 | 5.79 | 9.65 | Hypothetical protein |
| *PSPTO_3648* | *plcA1* | + | 4108208 | 4109890 | 5.30 | 6.61 | 6.18 | 4.82 | 3.62 | 3.62 | Acid phosphatase |
| *PSPTO_3649* | | - | 4109900 | 4111024 | 2.78 | 3.46 | 3.27 | 3.23 | 2.54 | 3.31 | Mechanosensitive ion channel family protein |
| *PSPTO_3650* | | + | 4111338 | 4112618 | 2.52 | 2.31 | 1.72 | 8.02 | 12.94 | 3.03 | Hypothetical protein |
| *PSPTO_3651* | | - | 4112598 | 4114151 | 0.00 | 0.00 | 0.00 | 0.00 | 0.00 | 0.00 | ISPsy5, transposase |
| *PSPTO_3652* | | - | 4114169 | 4114528 | 0.00 | 0.00 | 0.00 | 0.00 | 0.00 | 0.00 | ISPsy5, Orf1 |
| *PSPTO_3653* | | + | 4114588 | 4114797 | - | - | - | - | - | - | |
| *PSPTO_3654* | | + | 4116337 | 4116825 | 3.13 | 6.21 | 1.32 | 7.15 | 9.50 | 10.48 | GNAT family acetyltransferase |
| *PSPTO_3655* | | - | 4116878 | 4118044 | 6.37 | 12.00 | 11.51 | 7.51 | 8.18 | 7.19 | Hypothetical protein |
| *PSPTO_3656* | *ligA* | - | 4118486 | 4120849 | 2.75 | 9.33 | 8.15 | 3.73 | 24.92 | 9.09 | NAD-dependent DNA ligase LigA |
| *PSPTO_3657* | *zipA* | - | 4120977 | 4121843 | 64.31 | 165.57 | 124.29 | 42.42 | 168.74 | 69.39 | Cell division protein ZipA |
| *PSPTO_3658* | | - | 4122095 | 4125583 | 8.82 | 17.57 | 17.37 | 5.74 | 37.14 | 13.07 | Chromosome segregation SMC protein |
| *PSPTO_3659* | | - | 4125587 | 4126246 | 10.21 | 10.35 | 16.13 | 4.57 | 23.49 | 8.74 | GntR family transcriptional regulator |
| *PSPTO_3660* | | + | 4126521 | 4127975 | 42.43 | 22.87 | 19.25 | 5.48 | 115.43 | 31.43 | Xanthine dehydrogenase, N-terminal subunit |
| *PSPTO_3661* | | + | 4127968 | 4130346 | 24.91 | 22.69 | 23.25 | 5.17 | 73.24 | 34.05 | Xanthine dehydrogenase, C-terminal subunit |
| *PSPTO_3662* | | + | 4130333 | 4131208 | 14.20 | 8.09 | 11.47 | 2.66 | 22.44 | 24.11 | Xanthine dehydrogenase accessory factor XdhC |
| *PSPTO_3663* | *guaD* | + | 4131205 | 4132518 | 12.41 | 16.64 | 8.00 | 3.32 | 26.62 | 24.38 | Guanine aminohydrolase |
| *PSPTO_3664* | | - | 4132974 | 4133723 | 10.89 | 13.65 | 8.07 | 9.28 | 34.04 | 29.03 | GntR family transcriptional regulator |

| Locus name | Gene name | Strand | Gene start (bp) | Gene end (bp) | Average TPM Inoculum | Average TPM $t_6$ | Average TPM $t_{24}$ | Average TPM $t_{168}$ | Average TPM Logarithmic | Average TPM Stationary | Annotation description |
|---|---|---|---|---|---|---|---|---|---|---|---|
| *PSPTO_3665* | | - | 4133845 | 4134216 | 13.74 | 12.88 | 13.29 | 2.00 | 26.34 | 7.22 | Hypothetical protein |
| *PSPTO_3666* | | + | 4134568 | 4135497 | 67.95 | 48.65 | 48.43 | 49.68 | 61.61 | 27.73 | Polysaccharide deacetylase family protein |
| *PSPTO_3667* | | + | 4135494 | 4136024 | 31.68 | 38.23 | 52.11 | 22.39 | 66.53 | 18.87 | Hypothetical protein |
| *PSPTO_3668* | | + | 4136051 | 4137046 | 20.25 | 31.07 | 35.73 | 20.73 | 61.59 | 18.74 | Allantoicase |
| *PSPTO_3669* | *allA* | + | 4137185 | 4137697 | 12.44 | 23.97 | 7.64 | 24.02 | 28.99 | 11.33 | Ureidoglycolate hydrolase |
| *PSPTO_3670* | | + | 4137715 | 4138980 | 9.45 | 14.35 | 18.65 | 12.95 | 25.35 | 13.37 | Membrane protein |
| *PSPTO_3671* | | + | 4139347 | 4140132 | 44.42 | 19.83 | 15.83 | 2.21 | 12.34 | 6.84 | Hypothetical protein |
| *PSPTO_3672* | | - | 4140274 | 4141182 | 13.63 | 25.59 | 15.55 | 6.58 | 21.44 | 11.63 | Membrane protein |
| *PSPTO_3673* | | + | 4141223 | 4141345 | 90.82 | 3.04 | 17.80 | 10.76 | 26.29 | 6.79 | Hypothetical protein |
| *PSPTO_3674* | | - | 4141330 | 4141848 | 7.37 | 25.68 | 14.29 | 0.75 | 19.83 | 2.37 | AsnC family transcriptional regulator |
| *PSPTO_3675* | | + | 4142002 | 4143018 | 4.81 | 13.54 | 8.18 | 4.04 | 19.58 | 4.63 | 1-aminocyclopropane-1-carboxylate deaminase |
| *PSPTO_3676* | | + | 4143187 | 4143426 | 131.97 | 79.05 | 68.78 | 5.58 | 79.20 | 31.17 | Hypothetical protein |
| *PSPTO_3677* | | + | 4143416 | 4143700 | 30.63 | 23.86 | 33.46 | 12.21 | 33.69 | 9.04 | Hypothetical protein |
| *PSPTO_3678* | | - | 4143736 | 4144590 | 15.61 | 9.07 | 15.09 | 7.31 | 13.43 | 7.76 | LysR family transcriptional regulator |
| *PSPTO_3679* | | + | 4144665 | 4145597 | 8.25 | 9.38 | 9.81 | 4.55 | 7.21 | 6.87 | Membrane protein |
| *PSPTO_3680* | | - | 4145626 | 4147656 | 9.03 | 12.59 | 13.66 | 5.29 | 8.54 | 10.01 | Methyl-accepting chemotaxis protein |
| *PSPTO_3681* | | - | 4147806 | 4148291 | 22.79 | 26.77 | 14.19 | 9.86 | 31.74 | 25.90 | Hypothetical protein |
| *PSPTO_3682* | | - | 4148294 | 4149421 | 5.69 | 2.77 | 5.28 | 7.00 | 8.50 | 7.34 | Hypothetical protein |
| *PSPTO_3683* | | + | 4149604 | 4149789 | 20.39 | 5.38 | 6.17 | 12.43 | 15.19 | 8.03 | Hypothetical protein |
| *PSPTO_3684* | | + | 4149987 | 4150388 | 2.84 | 6.01 | 6.44 | 2.27 | 3.89 | 1.46 | Hypothetical protein |
| *PSPTO_3685* | | + | 4150495 | 4152135 | 4.31 | 6.69 | 5.98 | 4.31 | 8.74 | 3.69 | Methyl-accepting chemotaxis protein |
| *PSPTO_3686* | | - | 4152149 | 4153348 | 2.97 | 5.42 | 2.39 | 1.33 | 6.51 | 4.61 | Carbohydrate transporter |
| *PSPTO_3687* | | - | 4153613 | 4154905 | 13.49 | 43.52 | 40.92 | 4.91 | 16.69 | 13.89 | L-sorbosone dehydrogenase |
| *PSPTO_3688* | | - | 4154902 | 4155330 | 13.80 | 82.24 | 44.32 | 10.20 | 21.05 | 17.17 | Membrane protein |
| *PSPTO_3689* | | - | 4155617 | 4157155 | 8.42 | 11.29 | 14.94 | 5.17 | 18.44 | 7.83 | Hypothetical protein |
| *PSPTO_3690* | | - | 4157261 | 4158121 | 4.48 | 3.04 | 3.93 | 0.00 | 25.30 | 5.51 | Hypothetical protein |
| *PSPTO_3691* | | - | 4158118 | 4159659 | 5.83 | 7.33 | 11.12 | 5.94 | 52.04 | 12.40 | Ea59 protein |
| *PSPTO_3692* | | + | 4160122 | 4162290 | 18.38 | 4.61 | 7.08 | 2.32 | 14.97 | 35.09 | TonB-dependent siderophore receptor |
| *PSPTO_3693* | | + | 4162721 | 4163008 | 24.54 | 33.78 | 11.52 | 7.81 | 57.75 | 13.70 | Hypothetical protein |
| *PSPTO_3694* | | + | 4163440 | 4165758 | 4.25 | 10.31 | 7.89 | 5.67 | 12.08 | 8.31 | Glycosyl transferase family protein |
| *PSPTO_3695* | | - | 4165817 | 4166473 | 6.60 | 13.51 | 5.87 | 7.22 | 12.90 | 12.96 | TetR family transcriptional regulator |
| *PSPTO_3696* | | + | 4166962 | 4169445 | 9.26 | 9.06 | 12.35 | 3.87 | 7.87 | 4.05 | Sensory box histidine kinase/response regulator |
| *PSPTO_3697* | | + | 4169483 | 4169599 | 5.80 | 3.90 | 1.66 | 1.09 | 8.46 | 6.67 | Hypothetical protein |
| *PSPTO_3698* | | + | 4169607 | 4170215 | 6.72 | 9.91 | 16.19 | 3.01 | 19.67 | 10.18 | TetR family transcriptional regulator |
| *PSPTO_3699* | | + | 4170590 | 4172221 | 42.24 | 24.29 | 11.91 | 7.74 | 5.85 | 4.60 | Methyl-accepting chemotaxis protein |
| *PSPTO_3700* | | + | 4172325 | 4173146 | 72.45 | 48.82 | 47.39 | 14.17 | 23.64 | 67.45 | Oxidoreductase, aldo/keto reductase family |
| *PSPTO_3701* | | + | 4173317 | 4174336 | 5.25 | 5.33 | 3.15 | 2.73 | 1.90 | 2.27 | Cation transporter |
| *PSPTO_3702* | | + | 4174615 | 4176069 | 5.08 | 7.13 | 6.19 | 5.51 | 5.51 | 4.06 | Phosphate transporter family protein |
| *PSPTO_3703* | | - | 4176175 | 4177338 | 4.89 | 7.70 | 2.45 | 2.13 | 6.08 | 7.19 | HlyD family secretion protein |
| *PSPTO_3704* | | - | 4177335 | 4178963 | 8.64 | 11.87 | 5.81 | 3.79 | 9.43 | 9.27 | Transporter |
| *PSPTO_3705* | | + | 4179326 | 4180423 | 78.84 | 26.09 | 51.05 | 21.30 | 53.28 | 103.83 | Enoly-CoA hydratase/isomerase family protein |
| *PSPTO_3706* | *acdA* | + | 4180427 | 4181578 | 28.27 | 10.83 | 16.04 | 14.32 | 27.18 | 43.24 | Acyl-CoA dehydrogenase |
| *PSPTO_3707* | | - | 4181623 | 4186854 | 8.15 | 5.86 | 6.31 | 2.82 | 8.74 | 8.71 | Hypothetical protein |
| *PSPTO_3708* | | - | 4186991 | 4187575 | 45.65 | 110.41 | 124.66 | 41.80 | 25.41 | 52.98 | FMN reductase, NADPH-dependent |
| *PSPTO_3709* | | + | 4187546 | 4187986 | 2503.59 | 783.24 | 615.33 | 328.30 | 550.53 | 558.00 | Hypothetical protein |
| *PSPTO_3710* | | - | 4188036 | 4188590 | 11.60 | 30.45 | 35.71 | 11.03 | 31.57 | 27.75 | GNAT family acetyltransferase |
| *PSPTO_3711* | *dnaQ* | - | 4188612 | 4189385 | 65.71 | 101.99 | 84.12 | 29.28 | 133.67 | 61.18 | DNA polymerase III subunit epsilon |
| *PSPTO_3712* | *rnhA* | - | 4189457 | 4189909 | 16.16 | 66.21 | 60.13 | 22.15 | 78.99 | 31.24 | Ribonuclease HI |
| *PSPTO_3713* | | - | 4189956 | 4190717 | 10.71 | 38.66 | 43.83 | 13.40 | 38.41 | 15.88 | Hypothetical protein |
| *PSPTO_3714* | | + | 4190799 | 4191578 | 40.73 | 104.89 | 90.11 | 46.63 | 128.16 | 51.36 | Hydroxyacylglutathione hydrolase |
| *PSPTO_3715* | *mltD* | + | 4191678 | 4193270 | 16.28 | 101.92 | 93.18 | 75.88 | 232.65 | 62.03 | Membrane-bound lytic murein transglycosylase D |
| *PSPTO_3716* | | + | 4193408 | 4195237 | 21.64 | 44.50 | 25.25 | 15.95 | 52.54 | 22.47 | ABC transporter substrate-binding protein |
| *PSPTO_3717* | | + | 4195234 | 4197090 | 4.55 | 14.74 | 18.26 | 7.08 | 31.62 | 10.99 | ABC transporter substrate-binding protein |
| *PSPTO_3718* | | + | 4197092 | 4198165 | 4.18 | 6.73 | 7.32 | 2.93 | 19.31 | 6.43 | ABC transporter permease |
| *PSPTO_3719* | | + | 4198167 | 4199186 | 5.74 | 17.28 | 5.56 | 4.07 | 34.06 | 10.61 | ABC transporter permease |
| *PSPTO_3720* | | + | 4199188 | 4200798 | 41.14 | 30.77 | 23.87 | 17.07 | 73.02 | 20.66 | ABC transporter ATP-binding protein |
| *PSPTO_3721* | *fabI* | + | 4200823 | 4201617 | 28.11 | 36.34 | 36.61 | 9.30 | 126.35 | 11.61 | Enoyl-(acyl-carrier-protein) reductase |
| *PSPTO_3722* | | - | 4201727 | 4203610 | 4.32 | 16.44 | 10.18 | 6.25 | 84.47 | 7.61 | Peptidyl-prolyl cis-trans isomerase D |
| *PSPTO_3723* | *hupB* | - | 4203816 | 4204088 | 52.71 | 137.92 | 142.78 | 68.92 | 358.85 | 72.70 | DNA-binding protein HU-beta |
| *PSPTO_3724* | *lon-1* | - | 4204235 | 4206631 | 109.09 | 193.95 | 162.65 | 82.64 | 368.50 | 103.37 | ATP-dependent protease La |
| *PSPTO_3725* | *clpX* | - | 4206842 | 4208125 | 115.06 | 206.61 | 184.62 | 100.45 | 502.91 | 119.33 | ATP-dependent Clp protease ATP-binding subunit ClpX |
| *PSPTO_3726* | *clpP-2* | - | 4208232 | 4208882 | 109.33 | 186.59 | 193.98 | 83.60 | 479.35 | 89.31 | ATP-dependent Clp protease, proteolytic subunit ClpP |
| *PSPTO_3727* | *tig* | - | 4208965 | 4210275 | 41.95 | 88.50 | 78.95 | 37.98 | 277.53 | 50.89 | Trigger factor |
| *PSPTO_3728* | | - | 4210718 | 4210834 | 5.49 | 0.00 | 0.00 | 19.75 | 5.29 | 11.89 | Hypothetical protein |
| *PSPTO_3729* | | + | 4211413 | 4211892 | 7.18 | 33.88 | 9.20 | 26.72 | 130.21 | 17.36 | Lipoprotein |
| *PSPTO_3730* | | - | 4212288 | 4214144 | 0.93 | 2.02 | 0.00 | 0.21 | 1.49 | 1.22 | Membrane protein |
| *PSPTO_3731* | | + | 4214279 | 4215256 | 0.00 | 0.00 | 0.00 | 0.00 | 0.00 | 0.00 | ISPssy, transposase |
| *PSPTO_3732* | | - | 4215455 | 4215694 | 0.36 | 2.23 | 10.37 | 0.00 | 4.35 | 1.82 | Hypothetical protein |
| *PSPTO_3733* | *folD-2* | + | 4216904 | 4217758 | 17.61 | 53.39 | 41.01 | 28.61 | 81.62 | 29.92 | Methylenetetrahydrofolate dehydrogenase/methenyltetrahydrofolate cyclohydrolase |
| *PSPTO_3734* | | - | 4217893 | 4219353 | 0.40 | 0.07 | 0.21 | 0.00 | 0.40 | 0.44 | ISPsy6, transposase |
| *PSPTO_3735* | | - | 4219432 | 4221204 | 18.19 | 178.83 | 66.22 | 66.07 | 8.73 | 8.83 | ABC transporter periplasmic substrate-binding protein |
| *PSPTO_3736* | | - | 4221290 | 4221562 | 16.85 | 264.69 | 66.75 | 45.59 | 11.39 | 10.36 | Hypothetical protein |
| *PSPTO_3737* | | - | 4221575 | 4222375 | 9.58 | 60.62 | 15.10 | 22.42 | 3.82 | 5.78 | ABC transporter permease |
| *PSPTO_3738* | | - | 4222386 | 4223252 | 13.08 | 82.65 | 20.42 | 15.66 | 6.62 | 9.58 | ABC transporter permease |
| *PSPTO_3739* | | - | 4223245 | 4224360 | 7.96 | 56.34 | 18.54 | 28.48 | 1.59 | 1.91 | ABC transporter ATP-binding protein |
| *PSPTO_3740* | | - | 4224360 | 4225454 | 13.25 | 80.03 | 25.64 | 16.06 | 1.72 | 1.15 | ABC transporter ATP-binding protein |
| *PSPTO_3741* | | + | 4225753 | 4227606 | 15.72 | 30.31 | 27.73 | 17.95 | 13.18 | 4.97 | Sigma-54 dependent transcriptional regulator |
| *PSPTO_3742* | *cysS* | - | 4227791 | 4229173 | 11.23 | 26.80 | 37.68 | 25.05 | 192.94 | 55.15 | Cysteinyl-tRNA synthetase |
| *PSPTO_3743* | *glnS* | - | 4229193 | 4230902 | 10.96 | 42.87 | 33.60 | 21.63 | 177.25 | 47.29 | Glutaminyl-tRNA synthetase |

| Locus name | Gene name | Strand | Gene start (bp) | Gene end (bp) | Average TPM Inoculum | Average TPM $t_6$ | Average TPM $t_{24}$ | Average TPM $t_{168}$ | Average TPM Logarithmic | Average TPM Stationary | Annotation description |
|---|---|---|---|---|---|---|---|---|---|---|---|
| *PSPTO_3744* | *ppiB* | + | 4231095 | 4231598 | 84.67 | 144.87 | 180.16 | 75.01 | 803.82 | 91.64 | Peptidyl-prolyl cis-trans isomerase B |
| *PSPTO_3745* | *lpxH* | + | 4231595 | 4232341 | 6.40 | 19.00 | 16.56 | 3.75 | 56.47 | 15.60 | UDP-2,3-diacylglucosamine hydrolase |
| *PSPTO_3746* | | - | 4232799 | 4233821 | 20.12 | 27.03 | 21.56 | 14.50 | 36.79 | 25.50 | ISPsy7, transposase |
| *PSPTO_3747* | | - | 4234161 | 4235699 | 6.37 | 23.92 | 12.62 | 2.41 | 9.63 | 5.79 | Drug resistance transporter, EmrB/QacA family |
| *PSPTO_3748* | | - | 4235726 | 4236934 | 4.59 | 28.58 | 13.93 | 4.01 | 8.58 | 11.03 | Multidrug resistance protein |
| *PSPTO_3749* | | - | 4237025 | 4237513 | 13.93 | 72.56 | 45.66 | 7.54 | 11.48 | 49.96 | Multiple antibiotic resistance protein MarR |
| *PSPTO_3750* | | - | 4237660 | 4238283 | 5.73 | 8.34 | 6.29 | 5.51 | 19.81 | 8.41 | Hypothetical protein |
| *PSPTO_3751* | | - | 4238367 | 4238837 | 9.50 | 5.66 | 27.35 | 8.26 | 26.46 | 11.46 | Hypothetical protein |
| *PSPTO_3752* | *acnB* | + | 4239241 | 4241841 | 81.09 | 161.86 | 167.74 | 84.51 | 667.62 | 99.55 | Aconitate hydratase 2 |
| *PSPTO_3753* | | - | 4242087 | 4243355 | 37.29 | 14.87 | 15.31 | 7.02 | 24.23 | 21.19 | Methyl-accepting chemotaxis protein |
| *PSPTO_3754* | | - | 4243461 | 4244354 | 6.64 | 8.20 | 4.25 | 1.29 | 8.99 | 8.73 | Hypothetical protein |
| *PSPTO_3755* | | + | 4245060 | 4245632 | 18.53 | 29.32 | 25.77 | 14.76 | 26.69 | 20.30 | Membrane protein |
| *PSPTO_3756* | | + | 4245936 | 4247096 | 15.47 | 12.22 | 6.38 | 3.13 | 13.81 | 10.57 | Hypothetical protein |
| *PSPTO_3757* | | - | 4249716 | 4254572 | 17.02 | 32.74 | 29.20 | 10.17 | 351.36 | 28.50 | Hypothetical protein |
| *PSPTO_3758* | | + | 4255345 | 4256094 | 2.75 | 18.93 | 18.15 | 11.18 | 25.91 | 9.00 | Hypothetical protein |
| *PSPTO_3759* | | + | 4256300 | 4257259 | 8.40 | 22.25 | 19.34 | 7.46 | 46.72 | 17.98 | MoxR protein |
| *PSPTO_3760* | | + | 4257264 | 4258208 | 2.57 | 5.83 | 7.22 | 0.00 | 12.34 | 3.43 | Hypothetical protein |
| *PSPTO_3761* | | + | 4258205 | 4258699 | 0.53 | 4.02 | 1.66 | 0.00 | 9.68 | 1.87 | Hypothetical protein |
| *PSPTO_3762* | | + | 4258692 | 4259750 | 2.98 | 2.63 | 1.73 | 2.24 | 9.62 | 4.39 | Von Willebrand factor type A domain-containing protein |
| *PSPTO_3763* | | + | 4259747 | 4261465 | 4.02 | 8.48 | 8.48 | 1.39 | 14.41 | 4.23 | TPR domain-containing protein |
| *PSPTO_3764* | | + | 4261462 | 4263117 | 2.72 | 3.94 | 3.93 | 3.13 | 16.62 | 3.82 | Hypothetical protein |
| *PSPTO_3765* | *sbcD* | + | 4263160 | 4264404 | 3.53 | 2.40 | 3.63 | 3.40 | 18.22 | 3.89 | Exonuclease SbcD |
| *PSPTO_3766* | *sbcC* | + | 4264401 | 4268045 | 5.41 | 9.52 | 10.83 | 3.11 | 34.41 | 7.75 | Exonuclease SbcC |
| *PSPTO_3767* | | + | 4268271 | 4268531 | 102.07 | 1884.17 | 2089.65 | 193.45 | 95.72 | 198.69 | Hypothetical protein |
| *PSPTO_3768* | | - | 4268669 | 4269841 | 19.22 | 74.68 | 92.58 | 41.84 | 112.26 | 39.50 | Hypothetical protein |
| *PSPTO_3769* | | - | 4269921 | 4270319 | 11.41 | 63.64 | 72.68 | 27.12 | 102.03 | 21.79 | Hypothetical protein |
| *PSPTO_3770* | | - | 4270412 | 4271404 | 7.47 | 154.95 | 119.31 | 34.69 | 13.22 | 7.92 | Hypothetical protein |
| *PSPTO_3771* | | - | 4271484 | 4272791 | 2.63 | 53.56 | 49.53 | 13.05 | 5.18 | 2.80 | Major facilitator family transporter |
| *PSPTO_3772* | | + | 4273005 | 4274747 | 2.71 | 47.58 | 38.21 | 9.95 | 5.55 | 2.75 | Dihydroxy-acid dehydratase |
| *PSPTO_3773* | | + | 4274862 | 4275575 | 31.04 | 64.70 | 42.88 | 10.45 | 83.45 | 30.31 | GntR family transcriptional regulator |
| *PSPTO_3774* | | + | 4275912 | 4276496 | 0.35 | 1.72 | 2.81 | 2.73 | 1.86 | 0.34 | Hypothetical protein |
| *PSPTO_3775* | | + | 4276503 | 4276787 | 2.97 | 3.06 | 3.68 | 5.33 | 4.20 | 4.11 | Hypothetical protein |
| *PSPTO_3776* | | + | 4277420 | 4277866 | 6.37 | 10.65 | 14.27 | 7.56 | 85.83 | 12.17 | Hypothetical protein |
| *PSPTO_3777* | | - | 4278166 | 4280556 | 4.91 | 8.43 | 6.40 | 4.32 | 13.05 | 7.33 | Membrane protein |
| *PSPTO_3778* | | - | 4280584 | 4281606 | 9.32 | 8.58 | 13.55 | 9.18 | 23.84 | 11.01 | BNR/Asp-box repeat-containing protein |
| *PSPTO_3779* | | - | 4281838 | 4283193 | 6.67 | 6.28 | 7.25 | 10.28 | 10.29 | 9.89 | Hypothetical protein |
| *PSPTO_3780* | | - | 4283714 | 4284871 | 7.37 | 4.40 | 8.38 | 4.04 | 36.31 | 10.86 | Glutathionylspermidine synthase |
| *PSPTO_3781* | | - | 4284883 | 4285605 | 5.27 | 9.36 | 6.44 | 4.75 | 49.70 | 11.86 | Hypothetical protein |
| *PSPTO_3782* | | - | 4285657 | 4286064 | 5.07 | 6.16 | 9.58 | 0.00 | 28.45 | 6.89 | Hypothetical protein |
| *PSPTO_3783* | | - | 4286096 | 4286752 | 13.26 | 4.18 | 13.38 | 11.91 | 43.86 | 13.25 | Hypothetical protein |
| *PSPTO_3784* | | - | 4286749 | 4287774 | 7.96 | 6.35 | 5.73 | 7.62 | 34.67 | 12.81 | Potassium channel protein |
| *PSPTO_3785* | | - | 4287862 | 4288419 | 7.04 | 17.35 | 22.56 | 9.02 | 30.09 | 11.72 | Rhomboid family protein |
| *PSPTO_3786* | | - | 4288450 | 4289157 | 18.93 | 107.92 | 147.20 | 29.56 | 84.41 | 25.77 | PspA/IM30 family protein |
| *PSPTO_3787* | | - | 4289184 | 4289837 | 45.50 | 217.78 | 185.82 | 48.28 | 140.34 | 29.55 | Hypothetical protein |
| *PSPTO_3788* | *pepN* | - | 4290096 | 4292762 | 11.19 | 30.00 | 27.72 | 18.34 | 74.23 | 29.29 | Aminopeptidase N |
| *PSPTO_3789* | | - | 4292775 | 4293575 | 36.58 | 45.15 | 35.46 | 32.88 | 95.23 | 40.03 | Hypothetical protein |
| *PSPTO_3790* | | - | 4293605 | 4293865 | 82.98 | 210.34 | 146.08 | 103.82 | 283.40 | 197.59 | Hypothetical protein |
| *PSPTO_3791* | | - | 4293894 | 4294748 | 7.43 | 13.32 | 23.38 | 10.39 | 49.22 | 19.64 | Rhomboid family protein |
| *PSPTO_3792* | | - | 4294763 | 4295734 | 5.96 | 19.23 | 22.89 | 12.00 | 51.08 | 30.76 | Serine/threonine protein phosphatase |
| *PSPTO_3793* | *ppnK* | - | 4295734 | 4296624 | 9.33 | 14.98 | 24.19 | 17.36 | 45.16 | 26.81 | Inorganic polyphosphate/ATP-NAD kinase |
| *PSPTO_3794* | | + | 4296666 | 4297079 | 3.09 | 0.00 | 5.76 | 11.10 | 6.77 | 6.19 | Hypothetical protein |
| *PSPTO_3795* | | + | 4297265 | 4297558 | 23.96 | 46.15 | 37.86 | 66.36 | 105.19 | 105.33 | Hypothetical protein |
| *PSPTO_3796* | | - | 4297651 | 4298718 | 2.65 | 6.70 | 8.99 | 15.66 | 24.68 | 17.53 | Diguanylate cyclase |
| *PSPTO_3797* | | + | 4299116 | 4300099 | 9.59 | 14.86 | 7.80 | 7.62 | 21.23 | 9.83 | Hypothetical protein |
| *PSPTO_3798* | *arsB* | - | 4300065 | 4301345 | 2.56 | 5.26 | 7.23 | 2.08 | 9.75 | 5.33 | Arsenical pump membrane protein |
| *PSPTO_3799* | *fsr* | - | 4301345 | 4302553 | 3.47 | 6.43 | 7.54 | 6.48 | 15.08 | 7.25 | Fosmidomycin resistance protein |
| *PSPTO_3800* | | - | 4303185 | 4304207 | 20.02 | 25.12 | 21.17 | 16.38 | 36.07 | 25.49 | ISPsy7, transposase |
| *PSPTO_3801* | | - | 4304466 | 4305803 | 3.72 | 5.72 | 2.72 | 1.25 | 4.29 | 4.24 | Major facilitator family transporter |
| *PSPTO_3802* | | - | 4305951 | 4306895 | 8.53 | 15.59 | 13.45 | 5.54 | 15.35 | 12.77 | 1-aminocyclopropane-1-carboxylate deaminase |
| *PSPTO_3803* | *fadH* | - | 4306921 | 4308960 | 6.11 | 13.41 | 9.23 | 9.35 | 13.80 | 5.72 | 2,4-dienoyl-CoA reductase |
| *PSPTO_3804* | | + | 4309166 | 4310299 | 16.39 | 18.35 | 20.91 | 8.26 | 26.81 | 13.81 | Hydrolase |
| *PSPTO_3805* | | + | 4310302 | 4311342 | 13.59 | 17.18 | 19.70 | 4.63 | 34.09 | 27.63 | AraC family transcriptional regulator |
| *PSPTO_3806* | *ilvA-1* | + | 4311900 | 4313423 | 5.27 | 8.88 | 12.27 | 4.20 | 29.51 | 7.38 | Threonine dehydratase |
| *PSPTO_3807* | | - | 4313633 | 4314457 | 13.19 | 18.06 | 20.50 | 13.83 | 39.47 | 15.67 | Lipoprotein |
| *PSPTO_3808* | | - | 4314740 | 4316200 | 0.33 | 0.42 | 0.29 | 0.05 | 0.48 | 0.32 | ISPsy6, transposase |
| *PSPTO_3809* | | - | 4316872 | 4317597 | 6.82 | 16.14 | 6.38 | 3.05 | 54.87 | 14.48 | Oxidoreductase |
| *PSPTO_3810* | *metZ* | - | 4317687 | 4318898 | 12.95 | 29.02 | 34.16 | 13.90 | 124.08 | 40.49 | O-succinylhomoserine sulfhydrylase |
| *PSPTO_3811* | *purF* | - | 4318919 | 4320427 | 13.28 | 36.06 | 24.75 | 7.95 | 90.42 | 28.79 | Amidophosphoribosyltransferase |
| *PSPTO_3812* | | - | 4320440 | 4321027 | 90.94 | 58.13 | 22.78 | 29.41 | 56.62 | 40.54 | CvpA family protein |
| *PSPTO_3813* | | - | 4321146 | 4321796 | 25.97 | 22.94 | 35.01 | 15.44 | 50.77 | 25.98 | Hypothetical protein |
| *PSPTO_3814* | *folC* | - | 4321780 | 4323084 | 17.83 | 17.36 | 12.16 | 14.70 | 37.08 | 13.12 | Bifunctional folylpolyglutamate synthase/dihydrofolate synthase |
| *PSPTO_3815* | *accD* | - | 4323081 | 4324001 | 33.86 | 26.81 | 31.30 | 26.56 | 86.49 | 26.64 | Acetyl-CoA carboxylase subunit beta |
| *PSPTO_3816* | *trpF* | - | 4324254 | 4324874 | 18.02 | 81.92 | 72.14 | 40.43 | 99.38 | 48.00 | N-(5'-phosphoribosyl)anthranilate isomerase |
| *PSPTO_3817* | *truA* | - | 4324935 | 4325759 | 6.57 | 9.27 | 11.49 | 2.89 | 12.28 | 7.97 | tRNA pseudouridine synthase A |
| *PSPTO_3818* | | - | 4325920 | 4328763 | 31.20 | 35.89 | 33.60 | 18.44 | 60.83 | 34.98 | Hypothetical protein |
| *PSPTO_3819* | *asd-2* | - | 4328873 | 4329886 | 85.11 | 47.03 | 30.17 | 17.84 | 35.14 | 40.32 | Aspartate-semialdehyde dehydrogenase |
| *PSPTO_3820* | | - | 4329972 | 4330532 | 51.71 | 20.64 | 14.16 | 12.19 | 29.69 | 27.16 | Hypothetical protein |
| *PSPTO_3821* | | - | 4330552 | 4331154 | 319.54 | 68.69 | 78.70 | 41.25 | 42.10 | 105.12 | Lipoprotein |
| *PSPTO_3822* | | - | 4331281 | 4331940 | 8.73 | 26.48 | 19.96 | 14.30 | 40.74 | 19.16 | TetR family transcriptional regulator |

| Locus name | Gene name | Strand | Gene start (bp) | Gene end (bp) | Average TPM Inoculum | Average TPM $t_6$ | Average TPM $t_{24}$ | Average TPM $t_{168}$ | Average TPM Logarithmic | Average TPM Stationary | Annotation description |
|---|---|---|---|---|---|---|---|---|---|---|---|
| *PSPTO_3823* | | - | 4332060 | 4332722 | 3.63 | 8.52 | 8.77 | 5.55 | 17.90 | 7.95 | Membrane protein |
| *PSPTO_3824* | | - | 4333052 | 4333837 | 57.16 | 28.32 | 35.93 | 31.07 | 62.59 | 46.42 | TatD family deoxyribonuclease |
| *PSPTO_3825* | *pilZ* | - | 4333852 | 4334208 | 39.85 | 65.01 | 41.48 | 20.57 | 87.01 | 45.76 | Type IV pilus biogenesis protein PilZ |
| *PSPTO_3826* | *holB* | - | 4334243 | 4335229 | 12.00 | 15.98 | 9.85 | 7.25 | 32.23 | 10.35 | DNA polymerase III subunit delta' |
| *PSPTO_3827* | *tmk* | - | 4335222 | 4335854 | 26.27 | 18.12 | 33.45 | 7.02 | 65.51 | 28.16 | Thymidylate kinase |
| *PSPTO_3828* | | - | 4335874 | 4337013 | 8.99 | 21.24 | 20.42 | 10.15 | 47.21 | 19.83 | Hypothetical protein |
| *PSPTO_3829* | *pabC* | - | 4337017 | 4337832 | 5.94 | 10.59 | 10.28 | 0.26 | 25.53 | 11.38 | 4-amino-4-deoxychorismate lyase |
| *PSPTO_3830* | *fabF* | - | 4337832 | 4339076 | 9.65 | 22.54 | 17.28 | 14.63 | 72.41 | 40.55 | 3-oxoacyl-(acyl carrier protein) synthase II |
| *PSPTO_3831* | *acpP* | - | 4339190 | 4339426 | 132.64 | 297.46 | 276.36 | 146.67 | 340.20 | 68.50 | Acyl carrier protein |
| *PSPTO_3832* | *fabG* | - | 4339619 | 4340362 | 105.93 | 126.40 | 114.53 | 66.05 | 366.33 | 133.22 | 3-ketoacyl-(acyl-carrier-protein) reductase |
| *PSPTO_3833* | *fabD* | - | 4340378 | 4341316 | 44.87 | 43.21 | 46.23 | 25.40 | 144.78 | 35.04 | Malonyl CoA-acyl carrier protein transacylase |
| *PSPTO_3834* | *plsX* | - | 4341433 | 4342425 | 15.18 | 43.04 | 36.11 | 29.46 | 162.69 | 37.97 | Fatty acid/phospholipid synthesis protein PlsX |
| *PSPTO_3835* | *rpmF* | - | 4342460 | 4342642 | 26.26 | 175.19 | 228.38 | 79.54 | 180.80 | 36.09 | 50S ribosomal protein L32 |
| *PSPTO_3836* | | - | 4342656 | 4343183 | 177.62 | 308.31 | 281.85 | 222.89 | 489.09 | 163.62 | Hypothetical protein |
| *PSPTO_3837* | *maf-1* | + | 4343290 | 4343868 | 31.63 | 26.11 | 26.86 | 13.52 | 106.41 | 54.13 | Maf protein |
| *PSPTO_3838* | *sppA* | - | 4343921 | 4344919 | 8.76 | 14.54 | 13.23 | 13.69 | 52.54 | 13.60 | Signal peptide peptidase SppA, 36K type |
| *PSPTO_3839* | | - | 4344943 | 4345605 | 11.06 | 23.60 | 15.36 | 16.38 | 47.81 | 15.20 | HAD-superfamily hydrolase |
| *PSPTO_3840* | *rluC* | - | 4345595 | 4346551 | 11.78 | 18.86 | 22.88 | 11.60 | 53.27 | 11.95 | Ribosomal large subunit pseudouridine synthase C |
| *PSPTO_3841* | | + | 4347221 | 4350607 | 11.66 | 60.86 | 53.99 | 34.13 | 215.24 | 49.91 | Ribonuclease, Rne/Rng family protein |
| *PSPTO_3842* | *murB* | - | 4350890 | 4351909 | 1.96 | 4.19 | 6.63 | 2.51 | 20.56 | 6.21 | UDP-N-acetylenolpyruvoylglucosamine reductase |
| *PSPTO_3843* | *kdsB* | - | 4351944 | 4352708 | 8.49 | 35.45 | 11.92 | 17.62 | 70.67 | 20.45 | 3-deoxy-D-manno-octulosonate cytidylyltransferase |
| *PSPTO_3844* | | - | 4352705 | 4352890 | 17.47 | 38.81 | 32.56 | 9.95 | 97.58 | 29.04 | Hypothetical protein |
| *PSPTO_3845* | *lpxK* | - | 4352908 | 4353903 | 7.90 | 26.85 | 12.04 | 20.73 | 46.02 | 13.59 | Tetraacyldisaccharide 4'-kinase |
| *PSPTO_3846* | | - | 4353903 | 4354334 | 9.73 | 23.99 | 12.54 | 0.85 | 65.74 | 18.08 | Biopolymer transport protein ExbD/TolR family |
| *PSPTO_3847* | | - | 4354331 | 4354975 | 12.58 | 24.16 | 24.65 | 8.91 | 31.43 | 9.38 | MotA/TolQ/ExbB proton channel family protein |
| *PSPTO_3848* | | - | 4355091 | 4357307 | 4.28 | 4.11 | 9.45 | 4.00 | 6.61 | 5.40 | DNA internalization-related competence protein ComEC/Rec2 |
| *PSPTO_3849* | | + | 4357545 | 4359572 | 2.51 | 4.76 | 4.42 | 0.63 | 4.02 | 2.63 | Rhs element Vgr protein |
| *PSPTO_3850* | | + | 4359569 | 4360420 | 3.41 | 2.35 | 3.91 | 2.15 | 4.92 | 3.54 | Hypothetical protein |
| *PSPTO_3851* | | + | 4360417 | 4363731 | 3.65 | 4.70 | 3.27 | 1.59 | 5.71 | 4.00 | Hypothetical protein |
| *PSPTO_3852* | | + | 4363695 | 4364615 | 7.84 | 5.92 | 3.70 | 3.86 | 7.06 | 5.21 | Hypothetical protein |
| *PSPTO_3853* | | + | 4364612 | 4365511 | 5.36 | 11.75 | 10.20 | 3.10 | 10.71 | 5.01 | Hypothetical protein |
| *PSPTO_3854* | | - | 4365545 | 4366333 | 5.52 | 12.73 | 15.01 | 4.41 | 15.16 | 4.84 | Inner membrane transport permease |
| *PSPTO_3855* | | - | 4366330 | 4367262 | 10.86 | 22.75 | 22.36 | 14.05 | 29.05 | 11.87 | ABC transporter ATP-binding protein |
| *PSPTO_3856* | | - | 4367430 | 4368053 | 10.89 | 14.69 | 14.63 | 5.61 | 20.85 | 9.94 | Glutathione S-transferase family protein |
| *PSPTO_3857* | | + | 4368229 | 4370676 | 15.72 | 29.92 | 31.90 | 13.54 | 18.44 | 22.74 | Acyl-CoA dehydrogenase |
| *PSPTO_3858* | | + | 4370753 | 4371163 | 54.16 | 98.21 | 75.36 | 46.64 | 72.64 | 35.50 | Hypothetical protein |
| *PSPTO_3859* | | + | 4371357 | 4372211 | 9.41 | 8.85 | 8.65 | 7.29 | 11.38 | 10.89 | Hypothetical protein |
| *PSPTO_3860* | *aceE-1* | - | 4372314 | 4374983 | 10.88 | 7.87 | 8.06 | 4.55 | 12.64 | 17.72 | Pyruvate dehydrogenase, E1 component |
| *PSPTO_3861* | | + | 4375128 | 4375580 | 27.37 | 37.24 | 42.99 | 9.73 | 40.39 | 41.93 | MarR family transcriptional regulator |
| *PSPTO_3862* | | - | 4375583 | 4376686 | 15.64 | 9.96 | 10.40 | 2.55 | 13.19 | 34.01 | Hypothetical protein |
| *PSPTO_3863* | *psyR* | + | 4376977 | 4377720 | 71.26 | 16.52 | 11.07 | 9.03 | 19.39 | 124.47 | Transcriptional regulator PsyR |
| *PSPTO_3864* | *psyI* | - | 4377652 | 4378386 | 238.84 | 37.32 | 37.72 | 27.21 | 36.80 | 151.83 | Autoinducer synthesis protein PsyI |
| *PSPTO_3865* | | + | 4378606 | 4379802 | 7.94 | 7.03 | 7.75 | 2.38 | 6.35 | 4.71 | Hypothetical protein |
| *PSPTO_3866* | | + | 4379795 | 4380328 | 1.54 | 2.05 | 0.04 | 5.48 | 2.87 | 1.74 | Hypothetical protein |
| *PSPTO_3867* | | + | 4380318 | 4381115 | 2.37 | 2.86 | 1.03 | 2.12 | 5.07 | 2.68 | Dihydrodipicolinate synthetase family protein |
| *PSPTO_3868* | | + | 4381115 | 4381972 | 3.61 | 3.95 | 5.09 | 2.60 | 5.26 | 3.69 | Hypothetical protein |
| *PSPTO_3869* | | + | 4381956 | 4382192 | 3.61 | 0.00 | 2.07 | 0.77 | 4.25 | 3.46 | Hypothetical protein |
| *PSPTO_3870* | | - | 4382255 | 4383436 | 3.61 | 10.39 | 5.34 | 5.00 | 13.87 | 4.70 | Patatin family protein |
| *PSPTO_3871* | *htrB* | - | 4383469 | 4384401 | 7.15 | 28.05 | 20.80 | 10.04 | 29.12 | 10.83 | Lipid A biosynthesis lauroyl acyltransferase |
| *PSPTO_3872* | *minC* | + | 4384522 | 4385262 | 86.73 | 75.14 | 60.71 | 49.15 | 120.80 | 83.68 | Septum site-determining protein MinC |
| *PSPTO_3873* | *minD* | + | 4385492 | 4386304 | 37.52 | 97.04 | 88.35 | 61.42 | 185.51 | 90.30 | Septum site-determining protein MinD |
| *PSPTO_3874* | *minE* | + | 4386304 | 4386558 | 5.47 | 14.35 | 22.67 | 17.96 | 20.39 | 9.04 | Cell division topological specificity factor MinE |
| *PSPTO_3875* | *rluA* | + | 4386649 | 4387284 | 4.87 | 23.11 | 9.50 | 19.79 | 50.55 | 25.49 | Ribosomal large subunit pseudouridine synthase A |
| *PSPTO_3876* | | + | 4387656 | 4388945 | 14.36 | 24.55 | 22.78 | 9.10 | 83.78 | 22.84 | Aspartyl aminopeptidase |
| *PSPTO_3877* | | - | 4389003 | 4389722 | 26.46 | 9.09 | 8.50 | 8.89 | 16.07 | 12.42 | GntR family transcriptional regulator |
| *PSPTO_3878* | | + | 4390080 | 4391147 | 82.10 | 78.62 | 47.53 | 20.42 | 14.72 | 31.14 | ABC transporter substrate-binding protein |
| *PSPTO_3879* | | + | 4391168 | 4392031 | 14.26 | 9.22 | 13.29 | 9.20 | 8.56 | 9.77 | Hypothetical protein |
| *PSPTO_3880* | | + | 4392028 | 4392870 | 3.99 | 12.13 | 9.45 | 3.03 | 5.63 | 4.54 | Polyamine ABC transporter permease |
| *PSPTO_3881* | | + | 4392857 | 4393654 | 7.54 | 7.42 | 13.85 | 2.55 | 7.95 | 5.91 | Polyamine ABC transporter permease |
| *PSPTO_3882* | | + | 4393656 | 4394645 | 15.42 | 18.50 | 17.46 | 4.20 | 30.47 | 20.22 | Polyamine ABC transporter, ATP-binding protein |
| *PSPTO_3883* | | + | 4394799 | 4395380 | 16.01 | 19.87 | 20.21 | 7.21 | 21.03 | 14.01 | HAD-superfamily hydrolase |
| *PSPTO_3884* | | - | 4395399 | 4396361 | 18.12 | 50.82 | 50.89 | 21.52 | 77.34 | 26.72 | Oxidoreductase zinc-binding protein |
| *PSPTO_3885* | *prc* | + | 4396500 | 4398581 | 33.44 | 69.67 | 66.58 | 24.38 | 163.86 | 70.50 | Tail-specific protease |
| *PSPTO_3886* | | + | 4398715 | 4400487 | 2.71 | 4.46 | 2.26 | 4.98 | 2.82 | 3.82 | EAL domain/GGDEF domain protein |
| *PSPTO_3887* | | - | 4400679 | 4401044 | 11.93 | 20.97 | 34.79 | 16.83 | 22.15 | 20.19 | Hypothetical protein |
| *PSPTO_3888* | | - | 4401047 | 4401388 | 20.96 | 36.84 | 55.28 | 8.26 | 28.04 | 22.35 | Hypothetical protein |
| *PSPTO_3889* | | - | 4401489 | 4401890 | 28.93 | 65.96 | 55.70 | 23.58 | 34.22 | 45.25 | Hypothetical protein |
| *PSPTO_3890* | | - | 4402330 | 4403103 | 127.24 | 146.63 | 116.14 | 79.60 | 346.01 | 134.02 | FKBP-type peptidyl-prolyl cis-trans isomerase |
| *PSPTO_3891* | | - | 4403310 | 4404725 | 4.88 | 6.70 | 3.79 | 0.55 | 4.30 | 4.49 | Hypothetical protein |
| *PSPTO_3892* | | + | 4404888 | 4405505 | 6.91 | 252.98 | 153.32 | 36.29 | 32.78 | 11.80 | Hypothetical protein |
| *PSPTO_3893* | | - | 4405578 | 4405967 | 24.42 | 79.99 | 49.28 | 22.62 | 53.63 | 33.30 | Glyoxalase |
| *PSPTO_3894* | | - | 4406024 | 4407214 | 13.66 | 28.23 | 24.76 | 15.17 | 20.19 | 15.21 | CAIB/BAIF family protein |
| *PSPTO_3895* | | - | 4407337 | 4407732 | 5.58 | 2.48 | 13.67 | 5.63 | 5.05 | 2.94 | Hypothetical protein |
| *PSPTO_3896* | | + | 4407959 | 4408084 | 0.49 | 2.28 | 5.83 | 0.00 | 0.63 | 0.00 | Hypothetical protein |
| *PSPTO_3897* | *rdgC* | + | 4408438 | 4409358 | 7.37 | 16.92 | 20.18 | 4.39 | 119.91 | 8.25 | Recombination associated protein rdgC |
| *PSPTO_3898* | | + | 4409447 | 4410394 | 3.47 | 1.98 | 1.89 | 0.00 | 7.47 | 3.63 | Sodium/bile acid symporter family protein |
| *PSPTO_3899* | | - | 4410922 | 4412796 | 8.65 | 9.08 | 13.10 | 7.21 | 21.56 | 10.91 | Acyltransferase family protein |
| *PSPTO_3900* | | + | 4413040 | 4416513 | 6.08 | 6.94 | 12.00 | 2.70 | 14.26 | 8.19 | Sensory box histidine kinase/response regulator |
| *PSPTO_3901* | | - | 4416569 | 4417507 | 14.56 | 33.75 | 29.18 | 23.13 | 67.94 | 40.03 | Hypothetical protein |

| Locus name | Gene name | Strand | Gene start (bp) | Gene end (bp) | Average TPM Inoculum | Average TPM $t_6$ | Average TPM $t_{24}$ | Average TPM $t_{168}$ | Average TPM Logarithmic | Average TPM Stationary | Annotation description |
|---|---|---|---|---|---|---|---|---|---|---|---|
| *PSPTO_3902* | *cysK* | - | 4417685 | 4418659 | 23.71 | 64.59 | 46.56 | 33.23 | 92.61 | 38.05 | Cysteine synthase A |
| *PSPTO_3903* | | + | 4418787 | 4419857 | 30.06 | 53.20 | 27.09 | 17.55 | 25.79 | 28.36 | Hypothetical protein |
| *PSPTO_3904* | | - | 4419919 | 4420071 | 210.34 | 99.13 | 75.88 | 52.39 | 88.54 | 83.85 | Hypothetical protein |
| *PSPTO_3905* | | + | 4420195 | 4421040 | 12.54 | 21.89 | 18.55 | 7.45 | 39.13 | 17.87 | AAA ATPase |
| *PSPTO_3906* | | + | 4421080 | 4422258 | 7.26 | 10.71 | 9.66 | 9.40 | 20.62 | 10.48 | Hypothetical protein |
| *PSPTO_3907* | | + | 4422424 | 4423176 | 3.59 | 8.31 | 7.39 | 7.99 | 10.17 | 6.83 | Hypothetical protein |
| *PSPTO_3908* | | - | 4423209 | 4423733 | 8.88 | 42.32 | 18.69 | 16.40 | 43.12 | 17.45 | Hypothetical protein |
| *PSPTO_3909* | | - | 4423780 | 4424568 | 23.87 | 20.20 | 26.69 | 12.65 | 41.31 | 26.51 | Hypothetical protein |
| *PSPTO_3910* | | - | 4424645 | 4425121 | 23.04 | 46.02 | 45.26 | 35.50 | 64.37 | 28.24 | Hypothetical protein |
| *PSPTO_3911* | | + | 4425254 | 4427311 | 4.42 | 12.15 | 11.62 | 6.84 | 38.95 | 13.68 | Protease II |
| *PSPTO_3912* | | + | 4427438 | 4427872 | 34.37 | 53.88 | 22.90 | 1.26 | 89.35 | 37.44 | Cyclic nucleotide-binding protein |
| *PSPTO_3913* | | + | 4427938 | 4428312 | 297.04 | 256.70 | 212.53 | 136.35 | 605.50 | 283.78 | Hypothetical protein |
| *PSPTO_3914* | *copA* | + | 4428608 | 4430377 | 7.05 | 15.58 | 9.82 | 8.11 | 50.41 | 20.63 | Copper resistance protein A |
| *PSPTO_3915* | | + | 4430482 | 4431258 | 3.85 | 5.89 | 6.83 | 2.37 | 31.96 | 9.02 | Copper resistance protein B |
| *PSPTO_3916* | | + | 4431331 | 4431828 | 10.94 | 18.63 | 10.44 | 4.11 | 28.94 | 15.24 | Hypothetical protein |
| *PSPTO_3917* | | + | 4431937 | 4432317 | 33.56 | 160.67 | 120.37 | 36.18 | 84.76 | 65.78 | Hypothetical protein |
| *PSPTO_3918* | | - | 4432322 | 4432771 | 10.50 | 47.26 | 15.94 | 8.22 | 20.46 | 23.49 | Hypothetical protein |
| *PSPTO_3919* | | + | 4432983 | 4433576 | 6.13 | 31.33 | 21.13 | 8.36 | 62.52 | 17.31 | Nitroreductase family protein |
| *PSPTO_3920* | | - | 4433652 | 4434584 | 2.54 | 12.70 | 4.16 | 7.21 | 27.88 | 8.85 | D-isomer specific 2-hydroxyacid dehydrogenase family protein |
| *PSPTO_3921* | | - | 4434655 | 4434948 | 4.08 | 8.71 | 22.55 | 6.21 | 26.23 | 10.30 | Hypothetical protein |
| *PSPTO_3922* | *rnd* | - | 4434945 | 4436078 | 20.03 | 23.62 | 30.47 | 11.75 | 47.69 | 17.72 | Ribonuclease D |
| *PSPTO_3923* | | - | 4436285 | 4437928 | 4.72 | 14.87 | 9.17 | 7.50 | 16.04 | 6.17 | Membrane protein |
| *PSPTO_3924* | *sfcA* | - | 4438161 | 4439882 | 9.19 | 38.22 | 34.40 | 26.94 | 137.97 | 10.00 | Malate dehydrogenase |
| *PSPTO_3925* | | - | 4440165 | 4440647 | 545.65 | 323.03 | 293.62 | 172.68 | 102.13 | 440.85 | Gas vesicle protein |
| *PSPTO_3926* | | - | 4440888 | 4441297 | - | - | - | - | - | - | |
| *PSPTO_3927* | | - | 4441402 | 4442067 | 22.53 | 52.55 | 53.08 | 19.98 | 36.51 | 34.35 | Hypothetical protein |
| *PSPTO_3929* | | - | 4442900 | 4443118 | 0.00 | 1.74 | 3.88 | 4.13 | 5.11 | 1.82 | Cold shock domain family protein |
| *PSPTO_3930* | | - | 4443121 | 4444302 | 1.77 | 5.34 | 3.53 | 1.10 | 13.12 | 3.25 | Retron reverse transcriptase |
| *PSPTO_3931* | | + | 4445395 | 4445538 | 1.08 | 1.04 | 0.00 | 0.00 | 1.94 | 0.13 | Hypothetical protein |
| *PSPTO_3932* | | - | 4445527 | 4446045 | 1.28 | 0.00 | 0.00 | 0.00 | 1.38 | 0.89 | Hypothetical protein |
| *PSPTO_3933* | | - | 4446042 | 4446575 | 1.88 | 5.92 | 6.70 | 2.28 | 4.21 | 3.95 | Lysozyme |
| *PSPTO_3934* | | - | 4446636 | 4446914 | 2.17 | 0.52 | 4.18 | 0.00 | 1.77 | 1.51 | Tail fiber assembly domain protein |
| *PSPTO_3935* | | - | 4446907 | 4447263 | 1.29 | 14.08 | 1.83 | 0.00 | 1.09 | 1.35 | Hypothetical protein |
| *PSPTO_3936* | | - | 4447263 | 4448261 | 3.40 | 6.42 | 3.66 | 2.64 | 2.62 | 5.00 | Tail fiber domain protein |
| *PSPTO_3937* | | - | 4448289 | 4448951 | 1.22 | 1.73 | 2.00 | 4.63 | 2.35 | 2.44 | Hypothetical protein |
| *PSPTO_3938* | | - | 4448948 | 4449316 | 1.00 | 1.86 | 1.97 | 0.00 | 3.90 | 2.22 | Hypothetical protein |
| *PSPTO_3939* | | - | 4449313 | 4452045 | - | - | - | - | - | - | |
| *PSPTO_3940* | | - | 4452042 | 4452668 | - | - | - | - | - | - | |
| *PSPTO_3941* | | - | 4452671 | 4453456 | - | - | - | - | - | - | |
| *PSPTO_3942* | | + | 4453493 | 4454095 | 1.11 | 0.37 | 0.26 | 0.00 | 0.90 | 0.79 | Hypothetical protein |
| *PSPTO_3943* | | + | 4454139 | 4454333 | 5.46 | 0.00 | 5.02 | 0.00 | 2.04 | 2.29 | Carbon storage regulator |
| *PSPTO_3944* | | + | 4454323 | 4455000 | 2.29 | 0.00 | 4.18 | 0.82 | 4.22 | 4.36 | Hypothetical protein |
| *PSPTO_3945* | | + | 4454997 | 4455197 | 0.81 | 2.48 | 5.41 | 0.00 | 0.38 | 0.24 | Hypothetical protein |
| *PSPTO_3946* | | + | 4455286 | 4455753 | 1.96 | 0.29 | 2.09 | 2.87 | 2.12 | 1.33 | Hypothetical protein |
| *PSPTO_3947* | | + | 4455743 | 4456000 | 3.33 | 4.66 | 0.00 | 0.00 | 5.70 | 3.79 | Hypothetical protein |
| *PSPTO_3948* | | + | 4455993 | 4456376 | 3.72 | 8.13 | 0.00 | 0.00 | 4.65 | 2.16 | Hypothetical protein |
| *PSPTO_3949* | | - | 4457531 | 4458574 | 86.65 | 84.00 | 72.22 | 44.95 | 151.17 | 108.22 | Site-specific recombinase, phage integrase family |
| *PSPTO_3950* | *purC* | - | 4458800 | 4459510 | 14.96 | 77.48 | 51.59 | 35.44 | 254.91 | 59.41 | Phosphoribosylaminoimidazolesuccinocarboxamide synthase |
| *PSPTO_3951* | | - | 4459539 | 4460297 | 20.93 | 57.58 | 63.63 | 39.87 | 226.33 | 56.29 | Metallo-beta-lactamase superfamily protein |
| *PSPTO_3952* | | - | 4460298 | 4461416 | 12.08 | 44.05 | 34.60 | 25.20 | 157.22 | 44.31 | Lipoprotein |
| *PSPTO_3953* | *dapA* | - | 4461435 | 4462313 | 68.89 | 178.62 | 135.38 | 77.73 | 279.92 | 90.24 | Dihydrodipicolinate synthase |
| *PSPTO_3954* | | + | 4462639 | 4463202 | 18.80 | 71.34 | 67.23 | 26.52 | 236.26 | 51.81 | Hypothetical protein |
| *PSPTO_3955* | | + | 4463212 | 4463685 | 5.58 | 37.64 | 41.51 | 28.48 | 118.52 | 17.70 | AhpC/Tsa family protein |
| *PSPTO_3956* | | - | 4463752 | 4464822 | 2.97 | 9.96 | 10.25 | 4.23 | 10.79 | 4.80 | Membrane protein, PerM family |
| *PSPTO_3957* | | - | 4464855 | 4465106 | 17.30 | 12.63 | 7.12 | 3.62 | 17.00 | 9.94 | Hypothetical protein |
| *PSPTO_3958* | | + | 4465247 | 4466680 | 9.59 | 35.81 | 25.22 | 12.07 | 88.64 | 21.60 | Hypothetical protein |
| *PSPTO_3959* | *nadA* | - | 4466782 | 4467840 | 18.12 | 22.55 | 20.56 | 11.46 | 27.38 | 15.37 | Quinolinate synthetase |
| *PSPTO_3960* | *pglA* | + | 4468247 | 4469845 | 8.10 | 9.14 | 6.86 | 8.15 | 10.30 | 7.46 | Polygalacturonase |
| *PSPTO_3961* | | + | 4470140 | 4470340 | 5.20 | 3.15 | 18.24 | 7.85 | 9.48 | 5.12 | Hypothetical protein |
| *PSPTO_3962* | | + | 4470547 | 4470651 | 0.64 | 0.21 | 0.00 | 8.85 | 5.43 | 2.42 | Hypothetical protein |
| *PSPTO_3963* | | - | 4470781 | 4471758 | 0.00 | 0.00 | 0.00 | 0.00 | 0.00 | 0.00 | ISPssy, transposase |
| *PSPTO_3965* | | + | 4472738 | 4474291 | 8.44 | 11.56 | 8.36 | 7.77 | 7.92 | 9.56 | Hypothetical protein |
| *PSPTO_3966* | | + | 4474543 | 4474902 | 13.71 | 9.25 | 11.85 | 2.13 | 8.66 | 11.20 | ISPsy5, Orf1 |
| *PSPTO_3968* | *exsB* | - | 4476573 | 4477253 | 16.99 | 20.91 | 17.51 | 11.42 | 32.91 | 22.56 | ExsB protein |
| *PSPTO_3969* | | - | 4477264 | 4477911 | 5.75 | 12.45 | 12.77 | 6.48 | 13.50 | 10.12 | Radical SAM domain-containing protein |
| *PSPTO_3970* | | - | 4478083 | 4478901 | 40.27 | 97.24 | 78.43 | 78.86 | 212.38 | 89.15 | Hypothetical protein |
| *PSPTO_3971* | *pal* | - | 4478908 | 4479408 | 187.57 | 343.30 | 325.39 | 149.41 | 1382.47 | 178.09 | Peptidoglycan-associated lipoprotein |
| *PSPTO_3972* | *tolB* | - | 4479461 | 4480762 | 284.17 | 254.86 | 214.83 | 152.96 | 886.60 | 185.14 | TolB protein |
| *PSPTO_3973* | *tolA* | - | 4480759 | 4481829 | 121.20 | 104.86 | 71.80 | 47.62 | 124.69 | 75.93 | TolA protein |
| *PSPTO_3974* | *tolR-2* | - | 4481829 | 4482293 | 16.14 | 31.86 | 43.67 | 39.80 | 126.63 | 28.44 | TolR protein |
| *PSPTO_3975* | *tolQ-2* | - | 4482308 | 4483003 | 28.47 | 62.74 | 46.26 | 34.71 | 187.96 | 48.73 | TolQ protein |
| *PSPTO_3976* | | - | 4482993 | 4483460 | 52.67 | 72.01 | 80.61 | 31.38 | 151.92 | 60.74 | Hypothetical protein |
| *PSPTO_3977* | *ruvB* | - | 4483595 | 4484533 | 13.62 | 27.16 | 12.48 | 9.20 | 45.84 | 19.08 | Holliday junction DNA helicase RuvB |
| *PSPTO_3978* | *ruvA* | - | 4484657 | 4485265 | 21.98 | 21.42 | 30.95 | 16.45 | 69.53 | 30.46 | Holliday junction DNA helicase RuvA |
| *PSPTO_3979* | *ruvC* | - | 4485374 | 4485904 | 54.40 | 41.43 | 44.73 | 26.66 | 125.11 | 67.89 | Crossover junction endodeoxyribonuclease RuvC |
| *PSPTO_3980* | | - | 4486035 | 4486781 | 6.07 | 9.38 | 15.33 | 16.36 | 73.93 | 36.48 | Hypothetical protein |
| *PSPTO_3981* | *aspS* | - | 4486896 | 4488671 | 11.85 | 55.76 | 49.46 | 29.20 | 188.06 | 70.20 | Aspartyl-tRNA synthetase |
| *PSPTO_3982* | | - | 4488956 | 4489213 | 11.05 | 10.50 | 4.45 | 0.00 | 2.96 | 1.90 | Hypothetical protein |
| *PSPTO_3983* | | - | 4489660 | 4490133 | 19.55 | 124.80 | 124.40 | 55.74 | 239.62 | 51.01 | Dps family protein |

| Locus name | Gene name | Strand | Gene start (bp) | Gene end (bp) | Average TPM Inoculum | Average TPM $t_6$ | Average TPM $t_{24}$ | Average TPM $t_{168}$ | Average TPM Logarithmic | Average TPM Stationary | Annotation description |
|---|---|---|---|---|---|---|---|---|---|---|---|
| *PSPTO_3984* | | + | 4490988 | 4491209 | 21.00 | 33.49 | 18.52 | 32.45 | 55.72 | 28.88 | Cold shock domain family protein |
| *PSPTO_3985* | | - | 4491291 | 4491497 | 13.62 | 28.94 | 33.48 | 1.89 | 24.23 | 17.64 | Hypothetical protein |
| *PSPTO_3986* | | - | 4491494 | 4491925 | 76.00 | 63.68 | 63.89 | 54.26 | 106.64 | 91.67 | Hypothetical protein |
| *PSPTO_3987* | *oprD* | + | 4492830 | 4494176 | 232.56 | 295.81 | 227.40 | 255.86 | 263.55 | 29.66 | Porin D |
| *PSPTO_3988* | *proS* | + | 4494353 | 4496068 | 18.02 | 67.92 | 57.55 | 34.82 | 166.59 | 67.75 | Prolyl-tRNA synthetase |
| *PSPTO_3989* | | + | 4496135 | 4497091 | 2.94 | 9.39 | 9.43 | 3.66 | 50.46 | 12.87 | Lipoprotein |
| *PSPTO_3990* | *dinP* | - | 4497127 | 4498188 | 5.40 | 14.43 | 15.12 | 8.73 | 15.42 | 7.56 | DNA polymerase IV |
| *PSPTO_3991* | | + | 4499143 | 4501785 | 8.91 | 12.79 | 18.15 | 7.69 | 41.04 | 12.99 | Membrane protein |
| *PSPTO_3992* | | + | 4501785 | 4503119 | 8.82 | 19.22 | 12.06 | 6.93 | 60.31 | 18.42 | Hypothetical protein |
| *PSPTO_3993* | *kup* | - | 4503214 | 4505109 | 10.05 | 10.85 | 7.37 | 3.48 | 6.70 | 5.39 | Potassium uptake protein |
| *PSPTO_3994* | | + | 4505431 | 4506519 | 10.98 | 18.55 | 10.36 | 4.64 | 13.32 | 12.50 | Hypothetical protein |
| *PSPTO_3995* | | + | 4507088 | 4508254 | 56.54 | 67.08 | 64.16 | 19.72 | 186.49 | 26.88 | Abortive infection protein, internal deletion |
| *PSPTO_3996* | | - | 4509044 | 4510597 | 0.47 | 0.00 | 0.00 | 0.00 | 0.75 | 0.49 | ISPsy5, transposase |
| *PSPTO_3997* | | - | 4510615 | 4510974 | 0.00 | 0.00 | 0.00 | 0.00 | 0.00 | 0.00 | ISPsy5, Orf1 |
| *PSPTO_3998* | | + | 4511447 | 4511806 | 0.00 | 0.00 | 0.00 | 0.00 | 0.00 | 0.00 | ISPsy5, Orf1 |
| *PSPTO_3999* | | + | 4511824 | 4513377 | 0.00 | 0.00 | 0.00 | 0.00 | 0.00 | 0.00 | ISPsy5, transposase |
| *PSPTO_4001* | *avrPto1* | - | 4514766 | 4515260 | 5.20 | 3457.08 | 1618.49 | 278.43 | 19.18 | 10.15 | Type III effector protein AvrPto1 |
| *PSPTO_4002* | | - | 4515377 | 4517031 | - | - | - | - | - | - | |
| *PSPTO_4003* | | - | 4517028 | 4517564 | 0.52 | 0.00 | 0.00 | 0.97 | 0.91 | 1.17 | Hypothetical protein |
| *PSPTO_4004* | | - | 4517561 | 4518493 | 1.27 | 1.84 | 1.58 | 0.00 | 1.93 | 0.96 | Hypothetical protein |
| *PSPTO_4005* | | + | 4518582 | 4519010 | 3.73 | 4.10 | 1.31 | 0.00 | 3.34 | 2.40 | Hypothetical protein |
| *PSPTO_4006* | | - | 4518980 | 4521406 | 3.00 | 2.36 | 2.74 | 1.26 | 3.50 | 2.72 | Tail fiber protein H |
| *PSPTO_4007* | | - | 4521415 | 4521939 | 1.90 | 5.69 | 3.23 | 3.75 | 2.50 | 1.34 | Hypothetical protein |
| *PSPTO_4008* | | - | 4521997 | 4522404 | 2.29 | 4.82 | 8.13 | 5.33 | 3.35 | 2.15 | Hypothetical protein |
| *PSPTO_4009* | *cro* | - | 4522401 | 4522631 | 10.19 | 43.03 | 46.33 | 2.41 | 8.31 | 4.56 | Regulatory protein Cro |
| *PSPTO_4010* | | + | 4522782 | 4523486 | 121.96 | 172.78 | 132.49 | 67.27 | 305.87 | 188.69 | Repressor protein cI |
| *PSPTO_4011* | | - | 4524096 | 4524380 | 6.83 | 11.57 | 15.03 | 3.09 | 30.37 | 16.95 | Hypothetical protein |
| *PSPTO_4012* | | - | 4524602 | 4525363 | 15.99 | 10.40 | 8.97 | 5.76 | 15.56 | 10.66 | Hypothetical protein |
| *PSPTO_4013* | | + | 4525463 | 4526485 | 0.00 | 0.00 | 0.00 | 0.00 | 0.00 | 0.00 | ISPsy4, transposase |
| *PSPTO_4014* | | + | 4526482 | 4527291 | 0.00 | 0.00 | 0.00 | 0.00 | 0.00 | 0.00 | ISPsy4, transposition helper protein |
| *PSPTO_4015* | | - | 4527317 | 4527565 | 5.40 | 8.19 | 1.26 | 0.00 | 3.95 | 3.82 | Hypothetical protein |
| *PSPTO_4016* | | - | 4527558 | 4527764 | 0.70 | 0.00 | 0.00 | 0.00 | 1.21 | 2.17 | Hypothetical protein |
| *PSPTO_4017* | | - | 4527761 | 4528300 | 5.01 | 5.59 | 3.55 | 3.43 | 5.94 | 4.93 | Hypothetical protein |
| *PSPTO_4018* | | - | 4528754 | 4530373 | 17.99 | 23.00 | 23.96 | 11.47 | 36.05 | 20.12 | Hypothetical protein |
| *PSPTO_4019* | | + | 4530408 | 4531739 | 13.17 | 13.15 | 14.13 | 6.57 | 23.39 | 12.84 | Ribosomal protein S12 methylthiotransferase |
| *PSPTO_4020* | | + | 4531842 | 4532282 | 11.29 | 8.74 | 14.89 | 6.78 | 6.81 | 12.15 | GNAT family acetyltransferase |
| *PSPTO_4021* | | + | 4532376 | 4533086 | 27.68 | 79.21 | 64.42 | 22.97 | 193.53 | 78.99 | S4 domain-containing protein |
| *PSPTO_4022* | | + | 4533168 | 4533629 | 1.86 | 11.31 | 6.96 | 3.17 | 41.81 | 9.10 | Hypothetical protein |
| *PSPTO_4023* | | + | 4533616 | 4534335 | 3.27 | 3.73 | 6.30 | 8.62 | 25.72 | 5.10 | Hypothetical protein |
| *PSPTO_4024* | *fnr-1* | - | 4534415 | 4535194 | 32.31 | 135.03 | 114.12 | 89.45 | 72.92 | 139.06 | Ferredoxin-NADP reductase |
| *PSPTO_4025* | | + | 4535318 | 4536244 | 7.51 | 5.49 | 13.21 | 4.79 | 18.11 | 5.04 | LysR family transcriptional regulator |
| *PSPTO_4026* | *dgkA* | - | 4536246 | 4536611 | 1.54 | 9.45 | 8.19 | 4.20 | 17.10 | 3.60 | Diacylglycerol kinase |
| *PSPTO_4027* | | - | 4536695 | 4537342 | 6.67 | 8.39 | 13.94 | 1.61 | 25.10 | 8.77 | LuxR family DNA-binding response regulator |
| *PSPTO_4028* | | - | 4537422 | 4537547 | 24.26 | 87.14 | 32.87 | 32.70 | 74.18 | 53.06 | Hypothetical protein |
| *PSPTO_4029* | | - | 4537594 | 4538304 | 8.37 | 7.65 | 4.19 | 6.00 | 25.69 | 7.26 | Hypothetical protein |
| *PSPTO_4030* | | + | 4538509 | 4538925 | 289.68 | 91.21 | 107.64 | 55.01 | 175.06 | 131.93 | Hypothetical protein |
| *PSPTO_4031* | | + | 4539117 | 4540238 | 20.05 | 46.41 | 58.09 | 21.62 | 95.44 | 31.02 | Decarboxylase family protein |
| *PSPTO_4032* | *recX* | - | 4540243 | 4540701 | 3.83 | 7.44 | 8.56 | 0.00 | 24.46 | 7.38 | RecX protein |
| *PSPTO_4033* | *recA* | - | 4540716 | 4541780 | 23.62 | 62.81 | 45.01 | 14.35 | 65.74 | 22.24 | RecA protein |
| *PSPTO_4034* | | - | 4541874 | 4542374 | 33.79 | 57.55 | 62.20 | 24.62 | 34.02 | 25.84 | CinA domain protein |
| *PSPTO_4035* | | + | 4543232 | 4543858 | 7.54 | 9.03 | 11.20 | 4.78 | 7.76 | 6.54 | Hypothetical protein |
| *PSPTO_4036* | | - | 4543840 | 4544319 | 6.23 | 11.15 | 8.73 | 6.89 | 11.91 | 10.25 | Hypothetical protein |
| *PSPTO_4037* | | - | 4544346 | 4544879 | 14.50 | 37.29 | 27.24 | 28.63 | 42.63 | 32.40 | Lysozyme |
| *PSPTO_4038* | | - | 4544959 | 4546386 | 19.31 | 10.69 | 9.30 | 3.95 | 6.99 | 13.05 | Tail fiber domain protein |
| *PSPTO_4039* | | - | 4546414 | 4547076 | 1.22 | 1.73 | 2.00 | 4.63 | 2.35 | 2.46 | Hypothetical protein |
| *PSPTO_4040* | | - | 4547073 | 4547441 | 0.99 | 1.86 | 1.97 | 0.00 | 3.81 | 2.72 | Hypothetical protein |
| *PSPTO_4041* | | - | 4547438 | 4550719 | 5.85 | 2.15 | 2.06 | 1.30 | 3.29 | 2.64 | Host specificity protein J, internal deletion |
| *PSPTO_4042* | | - | 4550719 | 4551117 | 0.13 | 0.00 | 2.08 | 0.00 | 0.29 | 0.48 | Hypothetical protein |
| *PSPTO_4043* | | - | 4551120 | 4551722 | 5.05 | 1.38 | 4.14 | 6.08 | 7.08 | 3.18 | Hypothetical protein |
| *PSPTO_4044* | | - | 4551722 | 4552306 | 4.57 | 6.70 | 1.40 | 0.00 | 5.45 | 2.33 | Hypothetical protein |
| *PSPTO_4045* | | - | 4552335 | 4552634 | 9.44 | 1.25 | 5.35 | 0.55 | 15.11 | 6.47 | Hypothetical protein |
| *PSPTO_4046* | | - | 4552631 | 4556299 | 8.53 | 5.13 | 3.65 | 2.83 | 6.18 | 3.89 | Tail tape measure protein |
| *PSPTO_4047* | | - | 4556356 | 4556718 | 126.28 | 100.98 | 116.42 | 76.76 | 91.00 | 61.45 | Hypothetical protein |
| *PSPTO_4048* | | - | 4556782 | 4557033 | 6.74 | 0.99 | 19.20 | 0.00 | 8.27 | 6.88 | Hypothetical protein |
| *PSPTO_4049* | | - | 4557051 | 4557449 | 2.14 | 3.13 | 0.00 | 3.61 | 2.72 | 1.76 | Hypothetical protein |
| *PSPTO_4050* | | - | 4557596 | 4558348 | 3.96 | 2.13 | 0.65 | 0.00 | 3.54 | 2.88 | Hypothetical protein |
| *PSPTO_4051* | | + | 4558522 | 4559097 | 4.18 | 6.60 | 2.47 | 3.64 | 4.81 | 5.64 | Hypothetical protein |
| *PSPTO_4052* | | + | 4559094 | 4559960 | 10.66 | 5.03 | 8.31 | 3.80 | 9.80 | 5.79 | Hypothetical protein |
| *PSPTO_4053* | | - | 4560148 | 4560246 | 12.51 | 16.38 | 18.36 | 0.00 | 9.07 | 3.15 | Hypothetical protein |
| *PSPTO_4054* | | + | 4560424 | 4561236 | 3.95 | 4.35 | 3.54 | 1.76 | 3.91 | 2.95 | Hypothetical protein |
| *PSPTO_4055* | | + | 4561445 | 4561639 | 0.36 | 0.00 | 0.00 | 0.00 | 1.84 | 1.00 | Hypothetical protein |
| *PSPTO_4056* | | + | 4561688 | 4562275 | 2.67 | 1.71 | 0.54 | 0.32 | 3.48 | 1.74 | Hypothetical protein |
| *PSPTO_4057* | | + | 4562518 | 4563495 | 12.91 | 7.98 | 9.34 | 4.64 | 13.71 | 10.14 | Site-specific recombinase, phage integrase family |
| *PSPTO_4058* | *mutS* | + | 4563546 | 4566113 | 16.60 | 18.02 | 20.57 | 9.07 | 123.38 | 30.60 | DNA mismatch repair protein MutS |
| *PSPTO_4059* | *fdxA* | + | 4566360 | 4566683 | 11.29 | 31.01 | 3.83 | 5.47 | 33.50 | 7.92 | Ferredoxin |
| *PSPTO_4060* | | - | 4566810 | 4568270 | 0.21 | 0.28 | 0.34 | 0.04 | 0.45 | 0.34 | ISPsy6, transposase |
| *PSPTO_4061* | | - | 4568345 | 4569220 | 5.97 | 22.43 | 7.70 | 9.61 | 17.71 | 4.65 | LysR family transcriptional regulator |
| *PSPTO_4062* | | + | 4569320 | 4570489 | 4.95 | 1.77 | 3.28 | 4.11 | 3.78 | 2.54 | Mandelate racemase/muconate lactonizing protein |
| *PSPTO_4063* | *dctA-2* | + | 4570592 | 4571899 | 4.62 | 7.15 | 2.87 | 1.69 | 4.48 | 4.09 | C4-dicarboxylate transport protein |

| Locus name | Gene name | Strand | Gene start (bp) | Gene end (bp) | Average TPM Inoculum | Average TPM $t_6$ | Average TPM $t_{24}$ | Average TPM $t_{168}$ | Average TPM Logarithmic | Average TPM Stationary | Annotation description |
|---|---|---|---|---|---|---|---|---|---|---|---|
| *PSPTO_4064* | | - | 4571945 | 4572535 | 8.94 | 8.71 | 12.53 | 10.49 | 20.66 | 8.66 | TetR family transcriptional regulator |
| *PSPTO_4065* | | + | 4572655 | 4573413 | 8.52 | 11.38 | 11.51 | 5.63 | 38.21 | 5.04 | Short-chain dehydrogenase/reductase family oxidoreductase |
| *PSPTO_4066* | | - | 4573483 | 4573776 | 1.03 | 7.36 | 3.05 | 0.00 | 8.51 | 4.45 | Hypothetical protein |
| *PSPTO_4067* | | - | 4573831 | 4574727 | 5.57 | 6.34 | 9.11 | 9.60 | 8.63 | 6.03 | Short-chain dehydrogenase/reductase family oxidoreductase |
| *PSPTO_4068* | | + | 4574899 | 4575966 | 9.11 | 10.84 | 6.34 | 5.20 | 30.29 | 21.52 | AraC family transcriptional regulator |
| *PSPTO_4069* | | + | 4576000 | 4576164 | 1.37 | 5.77 | 1.56 | 3.16 | 12.55 | 11.55 | Hypothetical protein |
| *PSPTO_4070* | | + | 4576250 | 4576687 | 1.65 | 2.20 | 6.59 | 0.00 | 6.03 | 5.41 | Hypothetical protein |
| *PSPTO_4071* | | - | 4576746 | 4577321 | 11.88 | 23.67 | 7.05 | 7.07 | 20.75 | 11.09 | Hypothetical protein |
| *PSPTO_4072* | | - | 4577419 | 4577766 | 2.65 | 17.83 | 1.19 | 5.25 | 6.22 | 5.02 | Regulatory protein |
| *PSPTO_4073* | *fmdA* | - | 4577872 | 4579104 | 2.54 | 36.13 | 8.47 | 0.11 | 5.91 | 2.54 | Formamidase |
| *PSPTO_4074* | | - | 4579229 | 4579918 | 2.16 | 10.86 | 4.54 | 0.00 | 1.53 | 1.23 | Branched-chain amino acid ABC transporter ATP-binding protein |
| *PSPTO_4075* | | - | 4579929 | 4580690 | 2.84 | 10.03 | 2.48 | 2.05 | 2.60 | 3.05 | Branched-chain amino acid ABC transporter ATP-binding protein |
| *PSPTO_4076* | | - | 4580687 | 4581862 | 0.55 | 7.59 | 0.00 | 0.33 | 0.72 | 0.99 | Branched-chain amino acid ABC transporter permease |
| *PSPTO_4077* | | - | 4581873 | 4582754 | 1.34 | 10.20 | 1.25 | 2.10 | 1.54 | 1.28 | Branched-chain amino acid ABC transporter permease |
| *PSPTO_4078* | | - | 4582872 | 4584116 | 1.73 | 42.10 | 2.22 | 0.00 | 1.84 | 1.51 | Branched-chain amino acid ABC transporter substrate-binding protein |
| *PSPTO_4079* | | + | 4584274 | 4587759 | 30.96 | 10.38 | 6.32 | 6.48 | 4.62 | 4.54 | Sensor histidine kinase/response regulator |
| *PSPTO_4080* | | + | 4587746 | 4588414 | 22.98 | 12.28 | 22.51 | 8.73 | 23.11 | 13.69 | LuxR family DNA-binding response regulator |
| *PSPTO_4081* | | - | 4588320 | 4591742 | 17.31 | 25.09 | 19.29 | 12.98 | 60.42 | 25.19 | Rhs family protein |
| *PSPTO_4082* | | + | 4592224 | 4592958 | 17.18 | 28.31 | 29.47 | 10.48 | 41.27 | 13.53 | Short chain dehydrogenase/reductase family oxidoreductase |
| *PSPTO_4083* | | - | 4592960 | 4593838 | 4.32 | 4.47 | 1.63 | 3.13 | 6.43 | 5.48 | Membrane protein |
| *PSPTO_4084* | | + | 4594116 | 4598948 | 8.32 | 6.60 | 8.64 | 5.04 | 7.09 | 4.61 | Mannuronan C-5-epimerase |
| *PSPTO_4087* | | - | 4599541 | 4599825 | 98.21 | 51.81 | 52.10 | 42.14 | 71.52 | 84.90 | Hypothetical protein |
| *PSPTO_4088* | | + | 4599923 | 4600845 | - | - | - | - | - | - | |
| *PSPTO_4089* | | + | 4600859 | 4601647 | 3.02 | 5.34 | 1.08 | 0.00 | 4.03 | 5.70 | Alpha/beta fold family hydrolase |
| *PSPTO_4090* | *ppiA* | + | 4601797 | 4602360 | 27.12 | 77.52 | 54.54 | 37.89 | 302.28 | 40.32 | Peptidyl-prolyl cis-trans isomerase A |
| *PSPTO_4091* | | + | 4602462 | 4604294 | 3.46 | 10.70 | 6.37 | 5.71 | 34.90 | 7.68 | ABC transporter ATP-binding protein/permease |
| *PSPTO_4092* | | - | 4604386 | 4605513 | 11.02 | 16.87 | 13.50 | 8.06 | 26.22 | 17.65 | Hypothetical protein |
| *PSPTO_4093* | | - | 4605553 | 4610457 | 7.12 | 7.31 | 5.45 | 4.00 | 9.20 | 6.40 | Leucine rich repeat domain protein |
| *PSPTO_4094* | | - | 4610578 | 4611699 | 7.42 | 38.70 | 30.49 | 15.39 | 57.06 | 25.44 | 3-oxoacyl-(acyl carrier protein) synthase III |
| *PSPTO_4095* | *hrpA* | - | 4611940 | 4615851 | 3.35 | 8.98 | 7.97 | 7.51 | 43.71 | 11.29 | ATP-dependent helicase HrpA |
| *PSPTO_4096* | | + | 4616010 | 4616150 | 3.91 | 6.58 | 6.88 | 1.07 | 26.44 | 5.40 | Hypothetical protein |
| *PSPTO_4097* | *fadD-1* | - | 4616449 | 4618140 | 3.40 | 10.09 | 8.79 | 4.02 | 27.09 | 4.88 | Long-chain-fatty-acid--CoA ligase |
| *PSPTO_4098* | *fadD-2* | - | 4618496 | 4620184 | 10.64 | 16.44 | 17.74 | 7.18 | 32.18 | 16.00 | Long-chain-fatty-acid--CoA ligase |
| *PSPTO_4099* | | - | 4620249 | 4620398 | 92.26 | 91.11 | 67.24 | 36.52 | 262.54 | 134.03 | Hypothetical protein |
| *PSPTO_4100* | | + | 4620529 | 4621050 | 20.51 | 28.10 | 35.73 | 18.35 | 68.50 | 26.00 | MaoC-like domain protein |
| *PSPTO_4101* | *hopAK1* | + | 4621223 | 4622890 | 6.37 | 337.82 | 182.88 | 29.49 | 19.50 | 21.40 | Type III helper protein HopAK1 |
| *PSPTO_4102* | | + | 4623165 | 4623362 | 35.53 | 90.86 | 125.41 | 130.76 | 96.05 | 45.78 | Hypothetical protein |
| *PSPTO_4103* | | - | 4623412 | 4623762 | 278.76 | 262.38 | 177.81 | 63.14 | 58.18 | 85.76 | Hypothetical protein |
| *PSPTO_4104* | *hepA* | + | 4623994 | 4626840 | 4.87 | 18.03 | 16.41 | 8.13 | 65.91 | 16.73 | ATP-dependent helicase HepA |
| *PSPTO_4105* | | - | 4626944 | 4628383 | 16.05 | 8.51 | 11.19 | 3.97 | 28.80 | 10.09 | Major facilitator family transporter |
| *PSPTO_4106* | | + | 4628608 | 4630878 | 11.60 | 11.82 | 7.05 | 4.41 | 8.26 | 9.21 | GGDEF domain/EAL domain protein |
| *PSPTO_4107* | | - | 4631003 | 4631317 | 11.24 | 16.14 | 6.97 | 19.47 | 40.08 | 14.20 | Hypothetical protein |
| *PSPTO_4108* | *livK* | + | 4631636 | 4632760 | 67.48 | 78.40 | 81.57 | 28.59 | 151.37 | 20.28 | Leucine ABC transporter subunit substrate-binding protein LivK |
| *PSPTO_4109* | | + | 4632986 | 4633909 | 2.97 | 6.07 | 4.20 | 0.00 | 22.92 | 3.29 | High-affinity branched-chain amino acid ABC transporter permease |
| *PSPTO_4110* | | + | 4633906 | 4635171 | 4.41 | 7.43 | 5.25 | 8.14 | 26.30 | 3.15 | Leucine/isoleucine/valine transporter permease subunit |
| *PSPTO_4111* | | + | 4635168 | 4635935 | 6.97 | 21.64 | 4.27 | 1.60 | 70.10 | 4.10 | Leucine/isoleucine/valine transporter ATP-binding subunit |
| *PSPTO_4112* | | + | 4635936 | 4636637 | 9.53 | 8.25 | 14.89 | 0.00 | 31.76 | 6.01 | High-affinity amino acid ABC transporter, ATP-binding protein |
| *PSPTO_4113* | | + | 4636922 | 4637602 | 18.66 | 16.40 | 18.65 | 17.81 | 50.68 | 20.27 | Lipoprotein |
| *PSPTO_4114* | | + | 4637699 | 4639312 | 6.35 | 17.65 | 18.59 | 9.69 | 63.48 | 24.50 | Hypothetical protein |
| *PSPTO_4115* | *slyB* | - | 4639663 | 4640127 | 122.30 | 76.73 | 102.81 | 63.58 | 93.62 | 85.00 | Lipoprotein SlyB |
| *PSPTO_4116* | *pdxH* | - | 4640275 | 4640922 | 3.84 | 4.59 | 9.74 | 1.38 | 23.36 | 7.06 | Pyridoxamine 5'-phosphate oxidase |
| *PSPTO_4117* | | - | 4641196 | 4642341 | 6.02 | 28.62 | 23.23 | 11.49 | 73.09 | 14.96 | OmpA family protein |
| *PSPTO_4118* | | + | 4642344 | 4642460 | 17.16 | 0.00 | 7.50 | 0.00 | 23.95 | 11.09 | Hypothetical protein |
| *PSPTO_4119* | *estC* | - | 4642495 | 4643643 | 17.27 | 24.11 | 23.87 | 5.91 | 47.21 | 20.42 | Carboxylesterase |
| *PSPTO_4120* | | - | 4643754 | 4645898 | 2.89 | 8.90 | 8.50 | 3.20 | 21.38 | 3.95 | DinG family ATP-dependent helicase |
| *PSPTO_4121* | | + | 4646006 | 4646470 | 12.59 | 9.78 | 16.12 | 4.43 | 14.20 | 7.82 | Hypothetical protein |
| *PSPTO_4122* | *tsk* | + | 4646738 | 4647652 | 14.10 | 6.61 | 3.98 | 0.94 | 9.00 | 3.49 | Nucleoside-specific channel-forming protein |
| *PSPTO_4123* | | + | 4647679 | 4648710 | 15.22 | 9.28 | 26.40 | 2.70 | 30.28 | 10.42 | Purine nucleoside permease |
| *PSPTO_4124* | | - | 4648789 | 4649547 | 39.24 | 34.80 | 28.62 | 28.48 | 64.04 | 45.41 | Hypothetical protein |
| *PSPTO_4125* | | - | 4649540 | 4649815 | 49.49 | 12.13 | 15.47 | 28.00 | 28.23 | 27.02 | Hypothetical protein |
| *PSPTO_4126* | | + | 4650033 | 4650530 | 49.33 | 66.28 | 48.95 | 50.78 | 135.49 | 93.21 | Hypothetical protein |
| *PSPTO_4127* | | + | 4650534 | 4651007 | 8.83 | 13.85 | 15.71 | 7.20 | 28.16 | 19.87 | Hypothetical protein |
| *PSPTO_4128* | | - | 4651186 | 4653621 | 4.03 | 3.42 | 3.34 | 1.66 | 8.45 | 7.11 | TonB-dependent siderophore receptor |
| *PSPTO_4129* | | - | 4653734 | 4654285 | 38.46 | 43.57 | 25.13 | 7.93 | 54.31 | 53.48 | Hypothetical protein |
| *PSPTO_4130* | | + | 4654399 | 4654893 | 19.47 | 46.25 | 32.19 | 33.62 | 40.91 | 21.14 | Hypothetical protein |
| *PSPTO_4131* | | - | 4655506 | 4655757 | 33.02 | 66.28 | 40.47 | 5.74 | 12.78 | 24.11 | Hypothetical protein |
| *PSPTO_4132* | | - | 4655885 | 4656538 | 2.87 | 5.03 | 8.27 | 3.39 | 16.16 | 10.04 | Repressor protein c2 |
| *PSPTO_4133* | | + | 4657254 | 4659728 | 8.25 | 18.06 | 14.04 | 16.01 | 56.18 | 16.90 | Penicillin amidase family protein |
| *PSPTO_4134* | | + | 4660133 | 4661179 | 11.26 | 33.42 | 21.53 | 8.53 | 13.52 | 15.27 | Hypothetical protein |
| *PSPTO_4135* | | + | 4661176 | 4662891 | 3.16 | 16.34 | 11.01 | 3.40 | 5.47 | 9.78 | ATP-dependent DNA ligase |
| *PSPTO_4136* | | + | 4663235 | 4664017 | 39.45 | 47.65 | 32.26 | 22.13 | 26.05 | 5.79 | Amino acid ABC transporter substrate-binding protein |
| *PSPTO_4137* | | + | 4664114 | 4664827 | 3.11 | 5.89 | 1.55 | 0.00 | 6.01 | 2.23 | Amino acid ABC transporter permease |
| *PSPTO_4138* | *hisM* | + | 4664824 | 4665534 | 4.60 | 15.61 | 4.83 | 1.29 | 10.12 | 1.71 | Histidine/lysine/arginine/ornithine ABC transporter permease HisM |
| *PSPTO_4139* | | + | 4665547 | 4666665 | 5.50 | 7.18 | 5.12 | 9.14 | 16.33 | 5.05 | Hypothetical protein |
| *PSPTO_4140* | | + | 4666675 | 4667439 | 4.43 | 12.40 | 12.63 | 6.29 | 16.01 | 3.56 | Amino acid ABC transporter ATP-binding protein |
| *PSPTO_4141* | | + | 4667817 | 4670327 | 10.39 | 9.26 | 9.59 | 5.46 | 19.02 | 11.94 | Helicase domain-containing protein |
| *PSPTO_4142* | | + | 4670324 | 4670980 | 12.64 | 20.64 | 52.07 | 22.49 | 21.62 | 29.04 | Hypothetical protein |
| *PSPTO_4143* | | - | 4671062 | 4671280 | 537.38 | 476.98 | 469.01 | 155.78 | 155.88 | 232.66 | Hypothetical protein |
| *PSPTO_4144* | | - | 4671393 | 4671959 | 24.31 | 127.46 | 94.06 | 54.51 | 314.01 | 61.12 | Deoxycytidine triphosphate deaminase |

| Locus name | Gene name | Strand | Gene start (bp) | Gene end (bp) | Average TPM Inoculum | Average TPM $t_6$ | Average TPM $t_{24}$ | Average TPM $t_{168}$ | Average TPM Logarithmic | Average TPM Stationary | Annotation description |
|---|---|---|---|---|---|---|---|---|---|---|---|
| *PSPTO_4145* | *capB* | + | 4672279 | 4672488 | 20.24 | 108.41 | 153.11 | 13.61 | 90.02 | 13.48 | Cold shock protein CapB |
| *PSPTO_4146* | | - | 4672629 | 4673723 | 6.17 | 24.43 | 24.99 | 10.96 | 41.55 | 11.88 | ParA family protein |
| *PSPTO_4147* | *metG* | + | 4673910 | 4675958 | 8.44 | 59.42 | 47.38 | 34.75 | 167.77 | 53.96 | Methionyl-tRNA synthetase |
| *PSPTO_4148* | | + | 4676089 | 4676964 | 17.59 | 24.39 | 20.23 | 13.56 | 122.08 | 43.98 | Iron-sulfur cluster-binding protein |
| *PSPTO_4149* | *nth* | + | 4676961 | 4677599 | 7.98 | 9.79 | 7.41 | 4.77 | 36.11 | 16.33 | Endonuclease III |
| *PSPTO_4150* | | - | 4677643 | 4678005 | 2589.16 | 841.01 | 810.22 | 581.59 | 325.13 | 488.69 | Hypothetical protein |
| *PSPTO_4151* | | - | 4677998 | 4678624 | 14.59 | 22.58 | 30.29 | 23.14 | 13.96 | 19.16 | LuxR family DNA-binding response regulator |
| *PSPTO_4152* | | + | 4679391 | 4680368 | 0.00 | 0.00 | 0.00 | 0.00 | 0.00 | 0.00 | ISPssy, transposase |
| *PSPTO_4153* | | + | 4681237 | 4681635 | 2.52 | 4.94 | 3.82 | 10.22 | 3.23 | 4.65 | Hypothetical protein |
| *PSPTO_4154* | | + | 4681977 | 4682252 | 3.47 | 2.01 | 0.00 | 3.98 | 2.74 | 3.27 | Hypothetical protein |
| *PSPTO_4155* | *argG* | - | 4682334 | 4683551 | 20.84 | 88.96 | 68.59 | 36.30 | 105.41 | 27.46 | Argininosuccinate synthase |
| *PSPTO_4156* | *motY* | - | 4683672 | 4684577 | 90.21 | 25.42 | 19.38 | 16.26 | 4.59 | 4.07 | Sodium-type flagellar protein MotY |
| *PSPTO_4157* | | + | 4684726 | 4685769 | 59.09 | 105.26 | 109.69 | 55.35 | 575.58 | 76.04 | Dihydroorotase, homodimeric type |
| *PSPTO_4158* | *rnt* | + | 4685766 | 4686443 | 4.61 | 19.97 | 10.75 | 1.54 | 64.04 | 8.68 | Ribonuclease T |
| *PSPTO_4159* | | + | 4686627 | 4686845 | 7.02 | 20.57 | 8.08 | 13.20 | 168.25 | 207.00 | Bacterioferritin-associated ferredoxin |
| *PSPTO_4160* | *bfr* | + | 4687046 | 4687516 | 398.36 | 672.63 | 472.32 | 185.00 | 164.20 | 8.48 | Bacterioferritin |
| *PSPTO_4161* | | - | 4687604 | 4687930 | 63.94 | 169.83 | 138.96 | 82.38 | 397.75 | 70.02 | Glutaredoxin-like protein |
| *PSPTO_4162* | | - | 4688019 | 4690217 | 8.08 | 5.70 | 6.65 | 4.71 | 6.75 | 4.73 | Oxidoreductase, molybdopterin-binding protein |
| *PSPTO_4163* | | - | 4690272 | 4690565 | 76.22 | 63.68 | 53.73 | 17.48 | 98.24 | 57.54 | Hypothetical protein |
| *PSPTO_4164* | *argF* | + | 4691015 | 4691935 | 8.27 | 31.22 | 36.61 | 13.35 | 44.24 | 23.81 | Ornithine carbamoyltransferase |
| *PSPTO_4165* | | + | 4691941 | 4693002 | 5.48 | 12.21 | 9.24 | 4.36 | 28.80 | 11.18 | ABC transporter ATP-binding protein |
| *PSPTO_4166* | | - | 4693011 | 4693481 | 12.68 | 24.56 | 6.64 | 3.80 | 21.12 | 14.84 | YbaK/ebsC protein |
| *PSPTO_4167* | *glpF* | + | 4693833 | 4694690 | 16.76 | 78.44 | 32.98 | 19.98 | 371.37 | 131.93 | Glycerol uptake facilitator protein |
| *PSPTO_4168* | *glpK* | + | 4694720 | 4696225 | 7.38 | 58.29 | 24.73 | 16.99 | 301.60 | 68.00 | Glycerol kinase |
| *PSPTO_4169* | *glpR* | + | 4696336 | 4697091 | 16.80 | 42.18 | 17.94 | 19.70 | 61.10 | 21.01 | Glycerol-3-phosphate regulon repressor |
| *PSPTO_4170* | *glpD* | + | 4697683 | 4699221 | 35.74 | 142.79 | 73.23 | 42.44 | 549.41 | 822.55 | Glycerol-3-phosphate dehydrogenase |
| *PSPTO_4171* | | + | 4699577 | 4700503 | 14.71 | 31.98 | 51.26 | 31.64 | 136.30 | 16.11 | Amino acid ABC transporter substrate-binding protein |
| *PSPTO_4172* | | + | 4700634 | 4701380 | 2.84 | 11.55 | 13.42 | 11.30 | 58.74 | 5.27 | Amino acid ABC transporter permease |
| *PSPTO_4173* | | + | 4701380 | 4702045 | 3.87 | 13.23 | 8.88 | 9.48 | 80.06 | 4.79 | Amino acid ABC transporter permease |
| *PSPTO_4174* | | + | 4702042 | 4702776 | 7.04 | 15.95 | 10.03 | 1.91 | 107.74 | 6.28 | Amino acid ABC transporter ATP-binding protein |
| *PSPTO_4175* | | + | 4702835 | 4704766 | 7.88 | 7.35 | 8.60 | 2.44 | 31.84 | 7.48 | Sensor histidine kinase |
| *PSPTO_4176* | | + | 4704777 | 4706105 | 5.19 | 3.50 | 1.37 | 5.39 | 28.47 | 5.15 | Sigma-54 dependent transcriptional regulator/response regulator |
| *PSPTO_4177* | | - | 4706341 | 4706988 | 38.38 | 92.19 | 71.87 | 23.47 | 70.36 | 37.54 | 2-hydroxychromene-2-carboxylate isomerase |
| *PSPTO_4178* | | - | 4707042 | 4708034 | 3.01 | 4.09 | 2.63 | 3.48 | 81.55 | 3.65 | Alpha/beta fold family hydrolase |
| *PSPTO_4179* | *metE* | - | 4708220 | 4710520 | 8.04 | 11.49 | 14.86 | 3.54 | 755.37 | 7.83 | 5-methyltetrahydropteroyltriglutamate/homocysteine S-methyltransferase |
| *PSPTO_4180* | *metR* | + | 4710627 | 4711544 | 3.29 | 4.06 | 5.11 | 0.00 | 71.95 | 19.31 | Transcriptional activator MetR |
| *PSPTO_4181* | | - | 4711599 | 4712009 | 4.50 | 3.08 | 12.96 | 4.95 | 34.26 | 9.08 | MutT/nudix family protein |
| *PSPTO_4182* | | - | 4712006 | 4713097 | 9.80 | 19.48 | 20.18 | 13.06 | 53.99 | 19.20 | D-alanine--D-alanine ligase |
| *PSPTO_4183* | *rpmE-1* | - | 4713249 | 4713530 | 272.35 | 499.71 | 659.43 | 224.99 | 308.74 | 841.09 | 50S ribosomal protein L31 type B |
| *PSPTO_4184* | | + | 4713603 | 4713803 | 14.03 | 20.37 | 13.99 | 0.00 | 10.86 | 7.92 | Hypothetical protein |
| *PSPTO_4185* | | + | 4713825 | 4714955 | 1.92 | 5.15 | 3.12 | 3.43 | 3.64 | 4.79 | Membrane protein |
| *PSPTO_4186* | | + | 4715051 | 4715731 | 0.49 | 0.63 | 1.08 | 1.39 | 0.66 | 0.57 | Hypothetical protein |
| *PSPTO_4187* | | - | 4716078 | 4716731 | 0.22 | 0.76 | 0.00 | 0.00 | 0.14 | 0.15 | Hypothetical protein |
| *PSPTO_4188* | | + | 4716949 | 4717662 | 0.67 | 0.00 | 0.00 | 0.91 | 0.77 | 1.36 | Hypothetical protein |
| *PSPTO_4189* | | + | 4717841 | 4718530 | 0.22 | 0.00 | 0.46 | 0.00 | 1.05 | 0.37 | Hypothetical protein |
| *PSPTO_4190* | | - | 4718581 | 4719450 | 0.47 | 1.07 | 1.14 | 1.71 | 1.04 | 1.06 | Hypothetical protein |
| *PSPTO_4191* | | + | 4719830 | 4722967 | 14.41 | 7.49 | 12.40 | 5.71 | 4.09 | 12.79 | Hypothetical protein |
| *PSPTO_4192* | | + | 4723046 | 4723645 | 16.73 | 14.34 | 23.10 | 2.57 | 38.98 | 25.84 | Hypothetical protein |
| *PSPTO_4193* | | + | 4723642 | 4724409 | 9.35 | 10.52 | 12.02 | 16.64 | 23.17 | 20.02 | Iron utilization protein |
| *PSPTO_4194* | | - | 4724431 | 4725090 | 4.14 | 10.66 | 13.84 | 13.28 | 9.47 | 12.46 | Hypothetical protein |
| *PSPTO_4195* | | - | 4725161 | 4725529 | 12.90 | 29.08 | 14.16 | 9.04 | 44.28 | 57.97 | Hypothetical protein |
| *PSPTO_4196* | *gcd* | + | 4726006 | 4728249 | 7.05 | 42.58 | 38.03 | 15.45 | 178.55 | 34.24 | Glucose dehydrogenase |
| *PSPTO_4197* | | - | 4729025 | 4730221 | 3.18 | 5.05 | 7.47 | 4.35 | 13.43 | 6.70 | Hypothetical protein |
| *PSPTO_4198* | | - | 4730218 | 4731225 | 2.77 | 7.16 | 4.26 | 5.50 | 12.91 | 4.03 | Cobalamin synthesis protein/P47K family protein |
| *PSPTO_4199* | | - | 4731222 | 4731521 | 1.31 | 6.86 | 5.13 | 0.00 | 7.24 | 2.10 | Hypothetical protein |
| *PSPTO_4200* | | - | 4731544 | 4732293 | 3.72 | 5.46 | 5.81 | 3.50 | 11.14 | 4.25 | Short chain dehydrogenase/reductase family oxidoreductase |
| *PSPTO_4201* | | - | 4732325 | 4733086 | 4.40 | 39.70 | 36.68 | 10.95 | 35.11 | 8.34 | Short chain dehydrogenase/reductase family oxidoreductase |
| *PSPTO_4202* | | - | 4733097 | 4734437 | 3.55 | 12.14 | 9.42 | 3.45 | 11.66 | 4.66 | Major facilitator family transporter |
| *PSPTO_4203* | | - | 4734465 | 4735346 | 5.04 | 36.39 | 16.59 | 10.29 | 22.37 | 4.52 | Polysaccharide deacetylase family protein |
| *PSPTO_4204* | | - | 4735471 | 4736799 | 2.75 | 13.24 | 15.66 | 5.66 | 6.79 | 3.72 | Hypothetical protein |
| *PSPTO_4205* | | + | 4737182 | 4738183 | 8.34 | 10.29 | 14.97 | 10.26 | 15.95 | 10.66 | GntR family transcriptional regulator |
| *PSPTO_4206* | | - | 4738525 | 4738947 | 13.02 | 14.40 | 15.46 | 4.37 | 12.03 | 10.12 | Globin family protein |
| *PSPTO_4207* | | - | 4738947 | 4739822 | 15.91 | 16.50 | 11.30 | 6.35 | 21.34 | 12.61 | Hypothetical protein |
| *PSPTO_4208* | | - | 4739826 | 4742180 | 8.18 | 3.84 | 3.78 | 3.35 | 4.78 | 3.79 | HAMP domain/GGDEF domain/EAL domain protein |
| *PSPTO_4209* | | - | 4742177 | 4742848 | 49.52 | 26.97 | 18.49 | 12.10 | 16.60 | 13.63 | Hypothetical protein |
| *PSPTO_4210* | *lon-2* | + | 4743030 | 4745447 | 44.26 | 39.06 | 35.54 | 15.96 | 126.44 | 55.54 | ATP-dependent protease Lon2 |
| *PSPTO_4211* | | + | 4745655 | 4746047 | 25.87 | 85.00 | 59.01 | 37.35 | 142.35 | 49.26 | Lipoprotein |
| *PSPTO_4212* | | + | 4746148 | 4746891 | 6.12 | 15.38 | 8.23 | 7.52 | 39.04 | 12.56 | Methyltransferase |
| *PSPTO_4213* | | + | 4746888 | 4747847 | 5.13 | 11.62 | 14.56 | 3.05 | 47.26 | 11.48 | tRNA mo(5)U34 methyltransferase |
| *PSPTO_4214* | *pdxJ* | - | 4747856 | 4748596 | 5.52 | 18.21 | 15.49 | 6.46 | 40.34 | 8.56 | Pyridoxal phosphate biosynthetic protein PdxJ |
| *PSPTO_4215* | *recO* | - | 4748659 | 4749342 | 1.03 | 3.75 | 3.74 | 0.00 | 14.03 | 4.70 | DNA repair protein RecO |
| *PSPTO_4216* | *era* | - | 4749403 | 4750305 | 10.41 | 15.63 | 10.55 | 14.49 | 82.86 | 15.03 | GTP-binding protein Era |
| *PSPTO_4217* | *rnc* | - | 4750298 | 4750987 | 9.65 | 30.78 | 20.21 | 12.11 | 118.24 | 19.80 | Ribonuclease III |
| *PSPTO_4218* | | - | 4750984 | 4751361 | 42.71 | 40.60 | 23.38 | 32.05 | 114.16 | 35.56 | Hypothetical protein |
| *PSPTO_4219* | *lepB* | - | 4751478 | 4752332 | 15.89 | 40.09 | 38.31 | 24.86 | 226.97 | 37.45 | Signal peptidase I |
| *PSPTO_4220* | *lepA* | - | 4752342 | 4754129 | 11.11 | 33.44 | 23.29 | 25.00 | 74.02 | 23.22 | GTP-binding protein LepA |
| *PSPTO_4221* | *mucD* | - | 4754310 | 4755749 | 13.94 | 49.65 | 48.01 | 14.45 | 96.08 | 38.89 | Serine protease, MucD |
| *PSPTO_4222* | *mucB* | - | 4756035 | 4756991 | 37.68 | 89.29 | 60.80 | 39.10 | 72.13 | 96.83 | Sigma factor algU regulatory protein MucB |
| *PSPTO_4223* | *mucA* | - | 4757000 | 4757590 | 513.12 | 963.94 | 880.69 | 420.42 | 268.14 | 653.81 | Sigma factor algU negative regulatory protein MucA |

| Locus name | Gene name | Strand | Gene start (bp) | Gene end (bp) | Average TPM Inoculum | Average TPM $t_{6}$ | Average TPM $t_{24}$ | Average TPM $t_{168}$ | Average TPM Logarithmic | Average TPM Stationary | Annotation description |
|---|---|---|---|---|---|---|---|---|---|---|---|
| *PSPTO_4224* | *rpoE, algU* | - | 4757624 | 4758205 | 3039.75 | 2880.02 | 2207.88 | 1108.15 | 1050.25 | 2008.47 | RNA polymerase sigma-24 factor |
| *PSPTO_4225* | *nadB* | + | 4758704 | 4760320 | 17.10 | 27.51 | 22.50 | 9.53 | 26.83 | 17.71 | L-aspartate oxidase |
| *PSPTO_4226* | | - | 4760289 | 4760753 | 7.22 | 15.32 | 24.24 | 17.86 | 28.19 | 19.16 | Hypothetical protein |
| *PSPTO_4227* | | - | 4760737 | 4760991 | 18.53 | 35.39 | 38.29 | 17.60 | 56.96 | 20.28 | Hypothetical protein |
| *PSPTO_4228* | | + | 4761138 | 4762085 | 47.92 | 45.40 | 43.53 | 30.56 | 92.78 | 33.47 | Hypothetical protein |
| *PSPTO_4229* | | + | 4762199 | 4763020 | 30.36 | 22.60 | 27.28 | 22.88 | 48.04 | 35.04 | Hypothetical protein |
| *PSPTO_4230* | *tctE* | - | 4763026 | 4764408 | 10.04 | 23.15 | 14.69 | 14.06 | 24.56 | 19.11 | Sensor histidine kinase TctE |
| *PSPTO_4231* | *tctD* | - | 4764401 | 4765072 | 7.96 | 15.95 | 2.98 | 1.80 | 14.34 | 11.19 | DNA-binding response regulator TctD |
| *PSPTO_4232* | *tctC* | + | 4765571 | 4766548 | 6.25 | 14.28 | 11.19 | 7.18 | 9.96 | 7.29 | TctC protein |
| *PSPTO_4233* | *tctB* | + | 4766611 | 4767069 | 14.03 | 7.35 | 13.93 | 2.43 | 8.96 | 6.76 | Membrane protein TctB |
| *PSPTO_4234* | *tctA* | + | 4767071 | 4768585 | 4.69 | 5.52 | 9.47 | 6.72 | 7.29 | 4.14 | Membrane protein TctA |
| *PSPTO_4235* | | + | 4768578 | 4769612 | 1.37 | 3.89 | 3.42 | 1.27 | 12.43 | 8.33 | Membrane protein |
| *PSPTO_4236* | *ung* | - | 4769631 | 4770323 | 27.22 | 55.36 | 46.07 | 20.94 | 70.17 | 57.54 | Uracil-DNA glycosylase |
| *PSPTO_4237* | | - | 4770504 | 4770962 | 359.13 | 216.79 | 217.41 | 103.48 | 136.49 | 400.80 | Lipoprotein |
| *PSPTO_4238* | | + | 4771535 | 4772602 | 8.64 | 310.37 | 160.48 | 68.68 | 2.35 | 2.67 | ABC transporter substrate-binding protein |
| *PSPTO_4239* | | + | 4772630 | 4773445 | 2.48 | 89.84 | 59.99 | 21.76 | 3.47 | 1.85 | ABC transporter permease |
| *PSPTO_4240* | | + | 4773442 | 4774224 | 1.09 | 140.43 | 56.02 | 19.30 | 0.71 | 1.27 | ABC transporter ATP-binding protein |
| *PSPTO_4241* | | + | 4774221 | 4774946 | 2.20 | 122.26 | 55.32 | 26.48 | 2.81 | 1.49 | Hypothetical protein |
| *PSPTO_4242* | | + | 4774958 | 4775575 | 2.62 | 144.60 | 51.25 | 22.00 | 5.66 | 4.64 | Hypothetical protein |
| *PSPTO_4243* | | + | 4775685 | 4779365 | 2.22 | 4.85 | 4.52 | 2.53 | 3.34 | 2.97 | Urea amidolyase-related protein |
| *PSPTO_4244* | | + | 4779401 | 4781215 | 3.86 | 5.81 | 3.38 | 0.22 | 4.04 | 2.97 | Allophanate hydrolase |
| *PSPTO_4245* | | + | 4781326 | 4784661 | 5.54 | 4.59 | 5.48 | 3.01 | 5.16 | 5.38 | Hypothetical protein |
| *PSPTO_4246* | | + | 4784654 | 4784902 | 22.85 | 9.34 | 12.53 | 10.51 | 31.10 | 24.93 | Hypothetical protein |
| *PSPTO_4247* | *rsuA* | + | 4784962 | 4785654 | 5.03 | 7.02 | 6.79 | 3.99 | 13.28 | 10.55 | Ribosomal small subunit pseudouridine synthase A |
| *PSPTO_4248* | *phaG-2* | + | 4785951 | 4786835 | 3.73 | 7.66 | 4.68 | 2.09 | 3.42 | 10.54 | 3-hydroxyacyl-CoA-acyl carrier protein transferase |
| *PSPTO_4249* | | - | 4787793 | 4788511 | - | - | - | - | - | - | |
| *PSPTO_4250* | | + | 4788681 | 4789817 | 14.12 | 23.88 | 18.42 | 11.40 | 35.71 | 24.32 | DNA-binding protein |
| *PSPTO_4251* | | - | 4790778 | 4792331 | 0.00 | 0.00 | 0.00 | 0.00 | 0.00 | 0.00 | ISPsy5, transposase |
| *PSPTO_4252* | | - | 4792349 | 4792708 | 0.00 | 0.00 | 0.00 | 0.00 | 0.00 | 0.00 | ISPsy5, Orf1 |
| *PSPTO_4253* | | - | 4793542 | 4794519 | 0.00 | 0.00 | 0.00 | 0.00 | 0.00 | 0.00 | ISPssy, transposase |
| *PSPTO_4254* | *gor-2* | + | 4794539 | 4795939 | 9.01 | 8.65 | 7.81 | 8.64 | 10.02 | 8.90 | Glutathione reductase |
| *PSPTO_4255* | | + | 4796392 | 4796724 | 0.71 | 57.32 | 20.61 | 7.42 | 3.46 | 0.47 | Carboxymuconolactone decarboxylase family protein |
| *PSPTO_4256* | | + | 4796769 | 4797875 | 4.50 | 10.43 | 6.55 | 1.66 | 5.13 | 3.03 | NADH:flavin oxidoreductase/NADH oxidase family protein |
| *PSPTO_4257* | | + | 4797904 | 4798122 | 0.31 | 11.65 | 5.03 | 0.00 | 2.18 | 0.71 | 4-oxalocrotonate tautomerase |
| *PSPTO_4258* | | + | 4798233 | 4798754 | 0.25 | 0.00 | 0.00 | 2.50 | 0.51 | 0.42 | NAD(P)H dehydrogenase, quinone family |
| *PSPTO_4259* | *kefC* | + | 4798755 | 4800578 | 3.38 | 2.21 | 2.12 | 3.24 | 3.41 | 3.46 | Glutathione-regulated potassium-efflux system protein |
| *PSPTO_4260* | *trxA* | - | 4800613 | 4800942 | 1.36 | 2.16 | 4.55 | 4.30 | 3.76 | 4.76 | Thioredoxin |
| *PSPTO_4261* | | - | 4801007 | 4801705 | 2.10 | 5.02 | 2.78 | 3.90 | 2.78 | 2.95 | Isomerase |
| *PSPTO_4262* | | - | 4801709 | 4802299 | 2.40 | 4.67 | 0.77 | 5.09 | 1.36 | 2.02 | TetR family transcriptional regulator |
| *PSPTO_4263* | | - | 4802393 | 4803166 | 1.38 | 10.90 | 7.21 | 0.00 | 1.48 | 0.93 | Short chain dehydrogenase/reductase family oxidoreductase |
| *PSPTO_4264* | | - | 4803256 | 4803810 | 0.51 | 74.54 | 43.94 | 2.08 | 5.26 | 1.35 | Hypothetical protein |
| *PSPTO_4265* | *metC-2* | + | 4804230 | 4805399 | 1.84 | 2.15 | 1.15 | 2.92 | 1.76 | 1.45 | Cystathionine beta-lyase |
| *PSPTO_4266* | | + | 4805477 | 4806364 | 0.61 | 0.00 | 0.70 | 1.26 | 0.57 | 1.41 | Membrane protein |
| *PSPTO_4267* | | - | 4806458 | 4807027 | 6.72 | 22.81 | 23.58 | 11.81 | 40.87 | 18.74 | TetR family transcriptional regulator |
| *PSPTO_4268* | | + | 4807420 | 4807878 | - | - | - | - | - | - | |
| *PSPTO_4269* | | + | 4807947 | 4808969 | 0.00 | 0.00 | 0.00 | 0.00 | 0.00 | 0.00 | ISPsy4, transposase |
| *PSPTO_4270* | | + | 4808966 | 4809775 | 0.00 | 0.00 | 0.00 | 0.00 | 0.00 | 0.00 | ISPsy4, transposition helper protein |
| *PSPTO_4271* | | + | 4809832 | 4810875 | - | - | - | - | - | - | |
| *PSPTO_4272* | | - | 4811663 | 4813147 | 1.25 | 4.14 | 0.51 | 3.97 | 7.38 | 3.98 | Hypothetical protein |
| *PSPTO_4273* | | + | 4813818 | 4814905 | - | - | - | - | - | - | |
| *PSPTO_4274* | | - | 4814851 | 4815513 | 8.80 | 18.13 | 10.73 | 3.87 | 6.21 | 5.83 | TetR family transcriptional regulator |
| *PSPTO_4275* | | + | 4815638 | 4816423 | 49.32 | 10.50 | 13.36 | 10.73 | 6.06 | 4.75 | Short-chain dehydrogenase/reductase family oxidoreductase |
| *PSPTO_4276* | | - | 4817118 | 4818041 | 13.09 | 16.48 | 14.99 | 4.46 | 16.28 | 10.70 | LysR family transcriptional regulator |
| *PSPTO_4277* | | + | 4818257 | 4819090 | 11.71 | 6.63 | 2.04 | 4.07 | 12.48 | 10.08 | Esterase/lipase/thioesterase family protein |
| *PSPTO_4278* | | + | 4819268 | 4820440 | 8.31 | 3.08 | 2.68 | 4.95 | 9.64 | 6.48 | Major facilitator family transporter |
| *PSPTO_4279* | | + | 4820633 | 4820926 | 5.09 | 5.80 | 2.54 | 1.12 | 11.87 | 5.03 | Hypothetical protein |
| *PSPTO_4280* | | + | 4820934 | 4821812 | 3.75 | 1.96 | 5.41 | 3.35 | 9.91 | 4.10 | Membrane protein |
| *PSPTO_4281* | | + | 4821822 | 4822262 | 8.41 | 9.73 | 15.47 | 13.57 | 20.17 | 11.83 | Hypothetical protein |
| *PSPTO_4282* | | + | 4822314 | 4822454 | 13.59 | 0.00 | 16.74 | 8.18 | 29.36 | 19.26 | Hypothetical protein |
| *PSPTO_4283* | *pnlA* | - | 4822443 | 4823687 | 2.81 | 1.30 | 2.34 | 2.63 | 13.16 | 19.14 | Pectin lyase |
| *PSPTO_4284* | | - | 4823844 | 4824290 | 10.30 | 6.19 | 2.74 | 0.42 | 59.51 | 133.61 | GNAT family acetyltransferase |
| *PSPTO_4285* | *adhB* | - | 4824315 | 4825463 | 9.38 | 11.24 | 5.27 | 9.02 | 135.93 | 587.62 | Alcohol dehydrogenase II |
| *PSPTO_4286* | | - | 4825598 | 4826818 | 18.41 | 20.01 | 18.56 | 9.80 | 47.59 | 29.28 | Hypothetical protein |
| *PSPTO_4287* | | + | 4828342 | 4829352 | 19.85 | 13.40 | 14.42 | 21.84 | 19.57 | 23.52 | Binary cytotoxin component |
| *PSPTO_4288* | | - | 4829410 | 4830027 | 6.21 | 16.68 | 6.35 | 5.38 | 26.09 | 13.41 | Hypothetical protein |
| *PSPTO_4289* | | - | 4830062 | 4830679 | 26.03 | 51.32 | 32.45 | 7.80 | 58.51 | 25.74 | Hypothetical protein |
| *PSPTO_4290* | *bglX* | + | 4831075 | 4833372 | 4.56 | 24.40 | 10.57 | 10.56 | 64.61 | 12.69 | Beta-glucosidase |
| *PSPTO_4291* | | + | 4833443 | 4835200 | 2.08 | 2.42 | 5.54 | 3.60 | 19.17 | 6.87 | Sensor histidine kinase |
| *PSPTO_4292* | | + | 4835207 | 4836541 | 2.38 | 1.49 | 2.68 | 4.23 | 17.65 | 3.05 | Sigma-54 dependent transcriptional regulator/response regulator |
| *PSPTO_4293* | | + | 4836767 | 4839148 | 9.80 | 11.14 | 9.49 | 4.14 | 17.36 | 12.59 | Sensory box histidine kinase/response regulator |
| *PSPTO_4294* | *hscC* | + | 4839316 | 4841010 | 30.36 | 17.80 | 21.29 | 14.63 | 60.60 | 34.10 | Chaperone protein HscC |
| *PSPTO_4295* | | + | 4841016 | 4842656 | 32.53 | 18.11 | 17.56 | 11.90 | 38.33 | 37.14 | DnaJ domain-containing protein |
| *PSPTO_4296* | | + | 4842839 | 4844164 | 4.27 | 12.01 | 5.49 | 3.90 | 16.66 | 3.14 | Metabolite-proton symporter |
| *PSPTO_4297* | | + | 4844667 | 4845029 | 50.34 | 862.59 | 970.77 | 153.13 | 24.85 | 88.87 | Hypothetical protein |
| *PSPTO_4298* | | + | 4845168 | 4845791 | 15.34 | 27.98 | 25.54 | 14.43 | 48.87 | 18.34 | Hypothetical protein |
| *PSPTO_4299* | | - | 4845792 | 4846031 | 16.19 | 23.56 | 21.58 | 21.32 | 72.54 | 33.34 | Hypothetical protein |
| *PSPTO_4300* | | + | 4846462 | 4847976 | 13.19 | 26.93 | 17.68 | 13.11 | 47.69 | 31.02 | Drug resistance transporter, EmrB/QacA family |
| *PSPTO_4301* | | - | 4847990 | 4849123 | 5.89 | 5.00 | 2.69 | 2.42 | 15.80 | 10.27 | Hypothetical protein |
| *PSPTO_4302* | | - | 4849586 | 4850257 | 47.41 | 50.81 | 46.16 | 24.33 | 48.15 | 33.02 | TetR family transcriptional regulator |

| Locus name | Gene name | Strand | Gene start (bp) | Gene end (bp) | Average TPM Inoculum | Average TPM $t_6$ | Average TPM $t_{24}$ | Average TPM $t_{168}$ | Average TPM Logarithmic | Average TPM Stationary | Annotation description |
|---|---|---|---|---|---|---|---|---|---|---|---|
| *PSPTO_4303* | | + | 4850512 | 4851672 | 27.34 | 92.29 | 57.00 | 22.14 | 164.02 | 28.24 | RND family efflux transporter MFP subunit |
| *PSPTO_4304* | *saxB* | + | 4851676 | 4854810 | 13.83 | 40.24 | 23.57 | 9.29 | 105.13 | 13.03 | Isothiocyanate resistance protein SaxB; isochorismatase family |
| *PSPTO_4305* | | + | 4854807 | 4856267 | 19.54 | 45.12 | 33.14 | 18.99 | 130.92 | 20.21 | Outer membrane efflux protein |
| *PSPTO_4306* | *pcaT* | - | 4856343 | 4857644 | 5.19 | 13.94 | 12.11 | 11.82 | 29.10 | 10.77 | Dicarboxylic acid transport protein |
| *PSPTO_4307* | *catF* | - | 4857791 | 4858990 | 4.82 | 6.94 | 4.25 | 0.92 | 15.51 | 18.01 | 3-oxoadipyl-CoA thiolase |
| *PSPTO_4308* | *catJ* | - | 4858990 | 4859769 | 3.00 | 5.92 | 5.02 | 3.21 | 14.09 | 10.63 | 3-oxoadipate:succinyl-CoA transferase subunit B |
| *PSPTO_4309* | *catI* | - | 4859769 | 4860626 | 4.44 | 2.98 | 9.76 | 2.26 | 25.47 | 14.34 | 3-oxoadipate:succinyl-CoA transferase subunit A |
| *PSPTO_4310* | *pcaR* | - | 4860784 | 4861626 | 6.93 | 11.18 | 8.09 | 4.35 | 20.18 | 11.17 | Pca regulon regulatory protein PcaR |
| *PSPTO_4312* | | + | 4862886 | 4864358 | 7.24 | 18.38 | 15.91 | 8.54 | 67.13 | 29.19 | Phosphate transporter family protein |
| *PSPTO_4313* | | - | 4864414 | 4864623 | 10.83 | 12.05 | 3.46 | 0.89 | 32.84 | 14.27 | Hypothetical protein |
| *PSPTO_4314* | *purU-3* | - | 4864702 | 4865553 | 15.80 | 36.57 | 26.04 | 27.96 | 113.51 | 29.06 | Formyltetrahydrofolate deformylase |
| *PSPTO_4315* | | + | 4865912 | 4866292 | 81.67 | 412.28 | 429.73 | 327.09 | 1222.63 | 143.42 | Transcriptional regulator |
| *PSPTO_4316* | *sbcB* | - | 4866365 | 4867795 | 11.67 | 19.49 | 15.04 | 8.89 | 40.76 | 15.46 | Exodeoxyribonuclease I |
| *PSPTO_4317* | | + | 4868002 | 4868709 | 7.47 | 11.59 | 3.83 | 2.76 | 28.95 | 11.51 | Hypothetical protein |
| *PSPTO_4318* | | + | 4868706 | 4869686 | 7.19 | 4.78 | 1.98 | 0.80 | 13.74 | 5.90 | Integral membrane protein |
| *PSPTO_4319* | | + | 4869673 | 4871265 | 1.48 | 4.82 | 0.99 | 0.25 | 7.93 | 1.51 | Hypothetical protein |
| *PSPTO_4320* | | + | 4871262 | 4872482 | 3.82 | 1.43 | 1.20 | 0.00 | 12.33 | 3.45 | Hypothetical protein |
| *PSPTO_4321* | | + | 4872780 | 4873772 | 50.92 | 46.21 | 24.88 | 24.11 | 120.32 | 51.53 | Hypothetical protein |
| *PSPTO_4322* | | + | 4873769 | 4875589 | 70.68 | 62.69 | 58.26 | 31.36 | 188.37 | 63.84 | Hypothetical protein |
| *PSPTO_4323* | | + | 4875594 | 4876073 | 30.28 | 29.57 | 29.17 | 24.48 | 67.96 | 34.01 | Hypothetical protein |
| *PSPTO_4324* | | + | 4876653 | 4876949 | 31.86 | 10.84 | 6.56 | 4.32 | 17.92 | 9.46 | Hypothetical protein |
| *PSPTO_4325* | | + | 4876949 | 4878457 | 42.80 | 26.28 | 22.40 | 15.77 | 22.05 | 18.40 | Hypothetical protein |
| *PSPTO_4326* | | + | 4878641 | 4879582 | 4.42 | 7.04 | 6.96 | 9.99 | 7.39 | 5.88 | Hypothetical protein |
| *PSPTO_4327* | | + | 4879585 | 4879914 | 7.71 | 2.08 | 1.43 | 3.81 | 6.19 | 7.63 | Hypothetical protein |
| *PSPTO_4328* | | - | 4879876 | 4880313 | 7.67 | 3.13 | 1.69 | 1.55 | 10.14 | 13.50 | ISPs1, transposase OrfB |
| *PSPTO_4329* | | - | 4880361 | 4880729 | 11.11 | 11.16 | 40.32 | 11.72 | 23.10 | 19.69 | ISPs1, transposase OrfA |
| *PSPTO_4330* | | + | 4880833 | 4881072 | 18.27 | 7.37 | 15.95 | 6.69 | 14.25 | 12.86 | Hypothetical protein |
| *PSPTO_4331* | *hopE1* | + | 4881167 | 4881802 | 1.27 | 704.91 | 328.53 | 71.98 | 6.46 | 3.21 | Type III effector HopE1 |
| *PSPTO_4332* | | + | 4882082 | 4883365 | 3.11 | 19.93 | 14.37 | 5.44 | 6.18 | 4.05 | Hypothetical protein |
| *PSPTO_4333* | | + | 4883603 | 4884652 | 3.16 | 9.54 | 6.86 | 2.52 | 14.20 | 4.45 | MoxR protein |
| *PSPTO_4334* | | + | 4884649 | 4885992 | 4.38 | 10.35 | 10.91 | 3.86 | 12.59 | 7.06 | Hypothetical protein |
| *PSPTO_4335* | | - | 4886033 | 4886392 | 40.19 | 25.36 | 23.50 | 15.07 | 9.73 | 27.38 | Hypothetical protein |
| *PSPTO_4336* | | - | 4886581 | 4887033 | 189.81 | 38.60 | 36.44 | 21.00 | 66.81 | 16.76 | Lipoprotein |
| *PSPTO_4337* | *pyk* | + | 4887159 | 4888610 | 37.27 | 83.68 | 86.98 | 54.53 | 434.20 | 64.04 | Pyruvate kinase |
| *PSPTO_4338* | | - | 4888692 | 4889630 | 12.61 | 19.21 | 24.76 | 6.36 | 28.71 | 20.06 | Hypothetical protein |
| *PSPTO_4339* | | + | 4889810 | 4891333 | 13.61 | 61.85 | 48.45 | 31.66 | 168.18 | 21.19 | Fumarate hydratase, class I |
| *PSPTO_4340* | | - | 4891381 | 4894203 | 18.24 | 13.59 | 14.94 | 12.26 | 19.59 | 16.30 | Insecticidal toxin protein |
| *PSPTO_4341* | | + | 4894462 | 4897403 | - | - | - | - | - | - | |
| *PSPTO_4342* | | + | 4897445 | 4902292 | 6.39 | 5.04 | 6.35 | 2.57 | 9.31 | 5.78 | Insecticidal toxin protein |
| *PSPTO_4343* | | + | 4902312 | 4906652 | 3.66 | 1.65 | 2.81 | 1.85 | 5.79 | 4.15 | Insecticidal toxin protein |
| *PSPTO_4344* | | + | 4906645 | 4909305 | 5.75 | 3.41 | 2.90 | 1.38 | 6.51 | 4.41 | Insecticidal toxin protein |
| *PSPTO_4345* | | + | 4909571 | 4909786 | 0.09 | 2.58 | 3.40 | 0.00 | 0.33 | 0.20 | Hypothetical protein |
| *PSPTO_4346* | | + | 4910339 | 4910683 | 2.47 | 12.00 | 19.08 | 6.37 | 9.32 | 5.99 | Hypothetical protein |
| *PSPTO_4347* | | + | 4910862 | 4911839 | 0.00 | 0.00 | 0.00 | 0.00 | 0.00 | 0.00 | ISPssy, transposase |
| *PSPTO_4348* | | - | 4912008 | 4912493 | 10.59 | 9.27 | 10.04 | 5.12 | 16.03 | 11.40 | Hypothetical protein |
| *PSPTO_4349* | | - | 4912598 | 4913755 | 5.91 | 7.94 | 10.05 | 1.63 | 16.49 | 13.87 | Phospholipase D |
| *PSPTO_4350* | | - | 4913757 | 4914338 | 33.09 | 13.63 | 29.08 | 8.28 | 29.68 | 31.26 | Hypothetical protein |
| *PSPTO_4351* | | - | 4914419 | 4914757 | 2.96 | 1.18 | 3.19 | 0.00 | 12.21 | 5.65 | Hypothetical protein |
| *PSPTO_4352* | | - | 4914754 | 4915860 | 5.11 | 17.30 | 3.68 | 7.54 | 27.31 | 8.19 | Hydrolase |
| *PSPTO_4353* | | + | 4916238 | 4916993 | 1.67 | 0.00 | 0.90 | 0.00 | 1.94 | 1.67 | Hypothetical protein |
| *PSPTO_4354* | | + | 4916993 | 4917799 | 2.18 | 2.29 | 8.60 | 4.07 | 3.37 | 3.63 | Acyltransferase |
| *PSPTO_4355* | | - | 4917757 | 4918356 | 7.73 | 12.76 | 4.35 | 10.64 | 14.52 | 6.09 | Hypothetical protein |
| *PSPTO_4356* | | + | 4918521 | 4918835 | 39.81 | 155.05 | 150.31 | 22.58 | 15.76 | 4.90 | ArsR family transcriptional regulator |
| *PSPTO_4357* | | + | 4918868 | 4920034 | 3.96 | 16.35 | 15.83 | 4.54 | 7.78 | 3.79 | Major facilitator family transporter |
| *PSPTO_4358* | | + | 4920056 | 4921105 | 5.47 | 21.46 | 25.28 | 8.27 | 33.57 | 10.67 | Oxidoreductase FAD/FMN-binding protein |
| *PSPTO_4359* | | - | 4921196 | 4922065 | 40.53 | 57.26 | 52.30 | 29.40 | 142.83 | 48.28 | Hypothetical protein |
| *PSPTO_4360* | | - | 4922149 | 4923069 | 31.16 | 13.91 | 19.44 | 13.12 | 44.90 | 17.57 | Dehydrogenase |
| *PSPTO_4361* | *iciA* | - | 4923213 | 4924112 | 7.26 | 10.23 | 6.92 | 1.46 | 14.90 | 6.16 | Chromosome initiation inhibitor |
| *PSPTO_4362* | | + | 4924208 | 4924810 | 3.16 | 0.36 | 4.14 | 0.00 | 4.94 | 8.09 | LysE family transporter |
| *PSPTO_4363* | *sodB* | + | 4925003 | 4925584 | 89.07 | 310.62 | 245.53 | 129.41 | 335.13 | 21.26 | Superoxide dismutase, Fe |
| *PSPTO_4364* | | - | 4925639 | 4925737 | 8.64 | 6.74 | 0.00 | 7.40 | 2.86 | 6.45 | Hypothetical protein |
| *PSPTO_4365* | | + | 4925971 | 4928022 | 7.51 | 11.45 | 7.66 | 6.14 | 30.11 | 7.65 | GGDEF domain/EAL domain protein |
| *PSPTO_4366* | | + | 4928198 | 4929556 | 8.39 | 39.04 | 34.01 | 12.39 | 208.64 | 43.04 | Iron-regulated protein A |
| *PSPTO_4367* | | + | 4929853 | 4931280 | 4.09 | 6.58 | 10.81 | 5.91 | 31.03 | 14.01 | Lipoprotein |
| *PSPTO_4368* | | + | 4931341 | 4932405 | 9.50 | 26.07 | 14.88 | 5.30 | 95.68 | 18.08 | Lipoprotein |
| *PSPTO_4369* | | + | 4932422 | 4933519 | 11.38 | 17.42 | 13.63 | 13.73 | 51.12 | 15.54 | Lipoprotein |
| *PSPTO_4370* | | - | 4933544 | 4933771 | 207.99 | 192.43 | 110.10 | 104.71 | 31.19 | 70.03 | Hypothetical protein |
| *PSPTO_4371* | *inaA* | - | 4933768 | 4934490 | 56.84 | 81.68 | 72.98 | 88.28 | 22.20 | 14.14 | InaA protein |
| *PSPTO_4372* | | - | 4934495 | 4935175 | 101.92 | 149.15 | 183.49 | 202.21 | 16.06 | 4.49 | Hypothetical protein |
| *PSPTO_4373* | *colS* | - | 4935376 | 4936665 | 4.69 | 9.68 | 10.18 | 1.58 | 17.98 | 8.22 | Sensor histidine kinase ColS |
| *PSPTO_4374* | *colR* | - | 4936655 | 4937338 | 14.27 | 9.90 | 16.98 | 10.09 | 12.70 | 8.20 | DNA-binding response regulator ColR |
| *PSPTO_4375* | | + | 4937515 | 4938258 | 9.64 | 11.06 | 6.06 | 4.53 | 7.57 | 4.39 | PAP2 superfamily protein |
| *PSPTO_4376* | *groEL* | - | 4938343 | 4939986 | 104.17 | 278.44 | 165.46 | 104.00 | 2689.34 | 328.86 | Chaperonin, 60 kDa |
| *PSPTO_4377* | *groES* | - | 4940036 | 4940329 | 211.36 | 406.27 | 247.60 | 132.91 | 1826.11 | 187.54 | Chaperonin, 10 kDa |
| *PSPTO_4378* | | - | 4940443 | 4940553 | 182.20 | 189.48 | 111.70 | 48.76 | 115.35 | 49.75 | Hypothetical protein |
| *PSPTO_4379* | | - | 4940585 | 4941079 | 45.40 | 126.07 | 92.86 | 23.08 | 118.69 | 54.98 | Hypothetical protein |
| *PSPTO_4380* | | + | 4941398 | 4942156 | 14.22 | 5.88 | 30.89 | 32.66 | 111.16 | 35.30 | 3-ketoacyl-(acyl-carrier-protein) reductase |
| *PSPTO_4381* | | - | 4942205 | 4943212 | 14.47 | 26.06 | 27.76 | 12.13 | 77.21 | 85.90 | Hypothetical protein |
| *PSPTO_4382* | | + | 4943287 | 4943733 | 9.36 | 19.31 | 10.96 | 1.09 | 17.51 | 8.07 | MGMT family protein |

| Locus name | Gene name | Strand | Gene start (bp) | Gene end (bp) | Average TPM Inoculum | Average TPM $t_6$ | Average TPM $t_{24}$ | Average TPM $t_{168}$ | Average TPM Logarithmic | Average TPM Stationary | Annotation description |
|---|---|---|---|---|---|---|---|---|---|---|---|
| *PSPTO_4383* | | + | 4943788 | 4945338 | 3.82 | 8.74 | 8.65 | 5.70 | 25.19 | 12.96 | AmpG protein |
| *PSPTO_4384* | | + | 4945371 | 4945694 | 0.69 | 1.95 | 0.00 | 4.42 | 24.05 | 4.70 | Hypothetical protein |
| *PSPTO_4385* | | + | 4946342 | 4948633 | 7.32 | 12.03 | 9.30 | 8.16 | 38.15 | 14.95 | Rhs element Vgr protein |
| *PSPTO_4386* | | + | 4948630 | 4950201 | 8.20 | 3.12 | 3.38 | 1.78 | 23.16 | 6.22 | Hypothetical protein |
| *PSPTO_4387* | | + | 4951238 | 4952605 | 5.15 | 5.24 | 2.41 | 4.62 | 10.06 | 5.77 | Hypothetical protein |
| *PSPTO_4388* | | + | 4953597 | 4953755 | 4.56 | 3.02 | 10.84 | 1.21 | 7.34 | 3.46 | Hypothetical protein |
| *PSPTO_4389* | | - | 4953854 | 4955407 | 0.00 | 0.00 | 0.00 | 0.00 | 0.00 | 0.00 | ISPsy5, transposase |
| *PSPTO_4390* | | - | 4955425 | 4955784 | 0.00 | 0.00 | 0.00 | 0.00 | 0.00 | 0.00 | ISPsy5, Orf1 |
| *PSPTO_4392* | | - | 4956360 | 4957190 | 11.32 | 18.30 | 15.60 | 8.92 | 73.82 | 25.91 | Mechanosensitive ion channel family protein |
| *PSPTO_4393* | | - | 4957313 | 4957792 | 35.65 | 102.75 | 102.02 | 61.10 | 345.74 | 132.69 | Hypothetical protein |
| *PSPTO_4394* | *panE-2* | + | 4957962 | 4958882 | 4.59 | 24.61 | 5.78 | 6.61 | 26.88 | 11.37 | 2-dehydropantoate 2-reductase |
| *PSPTO_4395* | | + | 4958927 | 4960963 | 7.92 | 17.47 | 14.38 | 9.57 | 26.11 | 10.57 | Sensory box histidine kinase |
| *PSPTO_4396* | | + | 4960979 | 4961557 | 12.26 | 29.50 | 8.54 | 12.42 | 49.00 | 20.80 | ATP:cob(I)alamin adenosyltransferase |
| *PSPTO_4397* | | - | 4961699 | 4962649 | 6.92 | 18.60 | 15.95 | 9.99 | 54.26 | 15.65 | MutT/nudix family protein |
| *PSPTO_4398* | | - | 4962492 | 4963274 | 4.67 | 16.63 | 14.31 | 16.06 | 90.37 | 22.32 | Glutathione S-transferase family protein |
| *PSPTO_4399* | *argJ* | - | 4963332 | 4964549 | 4.66 | 13.43 | 11.87 | 11.92 | 101.11 | 17.23 | Glutamate N-acetyltransferase/amino-acid acetyltransferase |
| *PSPTO_4400* | *secA* | - | 4964661 | 4967402 | 14.66 | 66.99 | 48.23 | 34.16 | 238.92 | 72.35 | Preprotein translocase subunit SecA |
| *PSPTO_4401* | | + | 4967701 | 4968162 | 11.54 | 33.63 | 39.10 | 25.62 | 80.73 | 48.25 | Hypothetical protein |
| *PSPTO_4402* | *lpxC* | - | 4968298 | 4969209 | 18.14 | 78.74 | 57.56 | 31.27 | 130.49 | 38.03 | UDP-3-0-acyl N-acetylglucosamine deacetylase |
| *PSPTO_4403* | *ftsZ* | - | 4969322 | 4970509 | 27.99 | 159.27 | 92.50 | 49.61 | 503.06 | 177.64 | Cell division protein FtsZ |
| *PSPTO_4404* | *ftsA* | - | 4970597 | 4971853 | 10.00 | 69.66 | 32.39 | 8.33 | 178.49 | 42.45 | Cell division protein FtsA |
| *PSPTO_4405* | *ftsQ* | - | 4971868 | 4972737 | 13.47 | 53.69 | 33.03 | 11.27 | 157.90 | 63.09 | Cell division protein FtsQ |
| *PSPTO_4406* | | - | 4972741 | 4973700 | 9.88 | 30.46 | 14.59 | 7.01 | 82.79 | 34.08 | D-alanine--D-alanine ligase |
| *PSPTO_4407* | *murC* | - | 4973697 | 4975157 | 4.67 | 14.18 | 8.30 | 6.45 | 73.62 | 20.18 | UDP-N-acetylmuramate--L-alanine ligase |
| *PSPTO_4408* | *murG* | - | 4975150 | 4976220 | 4.64 | 24.40 | 7.68 | 0.42 | 87.71 | 18.91 | UDP-N-acetylglucosamine-N-acetylmuramyl- (pentapeptide) pyrophosphoryl-undecaprenol N- acetylglucosamine transferase |
| *PSPTO_4409* | *ftsW* | - | 4976210 | 4977424 | 4.82 | 26.05 | 14.45 | 4.65 | 70.52 | 20.89 | Cell division protein FtsW |
| *PSPTO_4410* | *murD* | - | 4977421 | 4978767 | 4.11 | 17.79 | 10.98 | 6.24 | 75.29 | 20.84 | UDP-N-acetylmuramoylalanine--D-glutamate ligase |
| *PSPTO_4411* | *mraY* | - | 4978773 | 4979855 | 3.70 | 12.64 | 4.23 | 0.96 | 42.23 | 9.09 | Phospho-N-acetylmuramoyl-pentapeptide-transferase |
| *PSPTO_4412* | *murF* | - | 4979855 | 4981225 | 6.40 | 22.05 | 19.99 | 4.07 | 96.55 | 32.03 | UDP-N-acetylmuramoylalanyl-D-glutamyl-2,6-diaminopimelate--D-alanyl-D-alanyl ligase |
| *PSPTO_4413* | *murE* | - | 4981218 | 4982681 | 5.90 | 24.24 | 10.49 | 5.82 | 63.91 | 21.47 | UDP-N-acetylmuramoylalanyl-D-glutamate--2,6-diaminopimelate ligase |
| *PSPTO_4414* | | - | 4982681 | 4984408 | 7.96 | 42.97 | 22.86 | 8.39 | 102.14 | 37.69 | Penicillin-binding protein |
| *PSPTO_4415* | *ftsL* | - | 4984408 | 4984701 | 33.38 | 74.59 | 49.10 | 29.18 | 77.63 | 34.31 | Cell division protein FtsL |
| *PSPTO_4416* | *mraW* | - | 4984698 | 4985639 | 30.21 | 86.32 | 59.60 | 10.30 | 140.84 | 65.16 | S-adenosyl-methyltransferase MraW |
| *PSPTO_4417* | | - | 4985642 | 4986097 | 32.71 | 84.49 | 57.31 | 18.54 | 176.56 | 66.41 | MarZ family protein |
| *PSPTO_4418* | | - | 4986848 | 4987717 | 28.27 | 39.28 | 45.89 | 15.08 | 68.79 | 38.58 | Tetrapyrrole methylase family protein |
| *PSPTO_4419* | | + | 4987997 | 4989811 | 16.74 | 49.06 | 37.46 | 20.51 | 114.50 | 41.41 | Lipoprotein |
| *PSPTO_4420* | | + | 4989808 | 4990194 | 10.80 | 68.46 | 20.49 | 9.94 | 65.62 | 20.18 | Hypothetical protein |
| *PSPTO_4421* | *gmhA* | + | 4990370 | 4990957 | 81.68 | 75.30 | 62.42 | 33.15 | 116.52 | 78.27 | Phosphoheptose isomerase |
| *PSPTO_4422* | | + | 4990954 | 4991532 | 15.37 | 28.18 | 38.09 | 27.10 | 50.81 | 28.99 | Lipoprotein |
| *PSPTO_4423* | *sspB* | - | 4991593 | 4992006 | 11.04 | 39.12 | 19.11 | 13.55 | 25.60 | 12.59 | Stringent starvation protein B |
| *PSPTO_4424* | *sspA* | - | 4992024 | 4992641 | 13.47 | 20.96 | 4.41 | 5.13 | 15.92 | 7.25 | Stringent starvation protein A |
| *PSPTO_4425* | *rpsI* | - | 4992922 | 4993314 | 12.04 | 55.33 | 26.75 | 34.63 | 123.24 | 36.53 | 30S ribosomal protein S9 |
| *PSPTO_4426* | *rplM* | - | 4993329 | 4993757 | 151.59 | 367.50 | 336.37 | 300.04 | 726.56 | 323.18 | 50S ribosomal protein L13 |
| *PSPTO_4427* | | + | 4994153 | 4995163 | 2.99 | 15.54 | 13.30 | 4.86 | 25.70 | 2.90 | AraC family transcriptional regulator |
| *PSPTO_4428* | | - | 4995238 | 4996332 | 5.08 | 18.75 | 11.77 | 5.63 | 75.30 | 9.78 | ATPase |
| *PSPTO_4429* | *trpS* | - | 4996395 | 4997744 | 7.22 | 14.97 | 14.57 | 6.33 | 63.35 | 12.62 | Tryptophanyl-tRNA synthetase |
| *PSPTO_4430* | | - | 4997784 | 4998413 | 12.52 | 38.30 | 20.21 | 17.46 | 90.40 | 18.68 | Hypothetical protein |
| *PSPTO_4431* | | + | 4998559 | 4999002 | 233.05 | 231.02 | 238.33 | 124.88 | 836.11 | 98.15 | Hypothetical protein |
| *PSPTO_4432* | *cysN/C* | - | 4999132 | 5001030 | 4.76 | 57.29 | 43.40 | 24.96 | 109.72 | 21.42 | Bifunctional sulfate adenylyltransferase subunit 1/adenylylsulfate kinase protein |
| *PSPTO_4433* | *cysD* | - | 5001041 | 5001958 | 13.11 | 135.22 | 115.97 | 84.64 | 110.25 | 33.20 | Sulfate adenylyltransferase subunit 2 |
| *PSPTO_4434* | | - | 5002112 | 5002870 | 13.96 | 23.02 | 13.60 | 11.23 | 46.83 | 17.27 | Hypothetical protein |
| *PSPTO_4435* | | + | 5002982 | 5004142 | 10.82 | 19.31 | 16.86 | 12.97 | 42.31 | 24.20 | Trypsin domain-containing protein |
| *PSPTO_4436* | | - | 5004167 | 5004619 | 9.25 | 14.10 | 14.74 | 6.55 | 18.73 | 13.79 | Hypothetical protein |
| *PSPTO_4437* | *hisC* | - | 5005376 | 5006428 | 8.93 | 40.99 | 22.56 | 19.53 | 166.91 | 25.78 | Histidinol-phosphate aminotransferase |
| *PSPTO_4438* | *hisD* | - | 5006428 | 5007774 | 5.95 | 47.05 | 23.58 | 6.30 | 115.50 | 19.73 | Histidinol dehydrogenase |
| *PSPTO_4439* | *hisG* | - | 5007899 | 5008534 | 7.48 | 35.89 | 30.68 | 12.12 | 134.50 | 29.23 | ATP phosphoribosyltransferase |
| *PSPTO_4440* | | + | 5008564 | 5008692 | 1.55 | 9.76 | 19.48 | 0.00 | 61.18 | 8.27 | Hypothetical protein |
| *PSPTO_4441* | *murA* | - | 5008689 | 5009954 | 9.86 | 44.07 | 53.94 | 21.84 | 116.99 | 24.69 | UDP-N-acetylglucosamine 1-carboxyvinyltransferase |
| *PSPTO_4442* | | - | 5009977 | 5010216 | 54.35 | 117.59 | 106.02 | 105.14 | 284.82 | 68.99 | Toluene tolerance protein |
| *PSPTO_4443* | | - | 5010331 | 5010636 | 8.58 | 37.67 | 15.70 | 13.23 | 52.15 | 15.76 | STAS domain-containing protein |
| *PSPTO_4444* | | - | 5010633 | 5011283 | 22.09 | 39.96 | 24.47 | 6.76 | 78.39 | 26.63 | Toluene tolerance protein |
| *PSPTO_4445* | | - | 5011313 | 5011780 | 72.63 | 105.78 | 72.52 | 55.62 | 153.07 | 69.06 | Mce-like protein |
| *PSPTO_4446* | | - | 5011780 | 5012577 | 58.68 | 75.12 | 53.98 | 22.23 | 85.59 | 39.29 | Membrane protein |
| *PSPTO_4447* | | - | 5012577 | 5013386 | 12.20 | 24.39 | 25.81 | 9.27 | 36.92 | 14.27 | Toluene tolerance ABC transporter, ATP-binding protein |
| *PSPTO_4448* | | + | 5013691 | 5014665 | 18.21 | 43.88 | 39.76 | 28.39 | 112.67 | 42.06 | Sugar isomerase |
| *PSPTO_4449* | | + | 5014665 | 5015204 | 14.42 | 36.62 | 20.01 | 19.42 | 65.77 | 19.07 | Phosphatase, YrbI family |
| *PSPTO_4450* | | + | 5015213 | 5015785 | 84.43 | 109.66 | 84.99 | 73.22 | 264.89 | 74.62 | Hypothetical protein |
| *PSPTO_4451* | | + | 5015772 | 5016320 | 79.05 | 142.52 | 139.70 | 68.15 | 346.32 | 119.63 | OstA family protein |
| *PSPTO_4452* | | + | 5016321 | 5017046 | 32.26 | 73.93 | 58.91 | 42.74 | 317.92 | 82.10 | ABC transporter ATP-binding protein |
| *PSPTO_4453* | *rpoN* | + | 5017229 | 5018722 | 51.68 | 42.58 | 52.08 | 23.14 | 122.76 | 40.36 | RNA polymerase sigma-54 factor |
| *PSPTO_4454* | *yfiA* | + | 5018797 | 5019105 | 434.54 | 653.20 | 447.35 | 214.87 | 363.39 | 150.20 | Ribosomal subunit interface protein |
| *PSPTO_4455* | *ptsN* | + | 5019117 | 5019581 | 52.18 | 98.81 | 70.56 | 24.97 | 232.67 | 45.07 | PTS transporter subunit IIA-like nitrogen-regulatory protein PtsN |
| *PSPTO_4456* | | + | 5019584 | 5020441 | 35.67 | 42.48 | 37.79 | 23.57 | 183.26 | 31.50 | Hypothetical protein |
| *PSPTO_4457* | *ptsH* | + | 5020458 | 5020730 | 10.43 | 51.72 | 56.60 | 4.40 | 179.65 | 27.14 | Phosphocarrier protein HPr |
| *PSPTO_4458* | | - | 5020745 | 5020873 | 7.67 | 12.59 | 1.76 | 0.78 | 29.47 | 5.05 | Hypothetical protein |
| *PSPTO_4459* | *sodA* | - | 5020901 | 5021512 | 2.40 | 1.56 | 14.31 | 5.83 | 17.83 | 10.58 | Superoxide dismutase |
| *PSPTO_4460* | | - | 5021513 | 5021974 | 5.48 | 13.11 | 3.71 | 1.21 | 33.12 | 30.98 | Hypothetical protein |
| *PSPTO_4461* | *fumC-2* | - | 5022007 | 5023383 | 8.05 | 4.20 | 0.36 | 1.98 | 13.09 | 12.91 | Class II fumarate hydratase |
| *PSPTO_4462* | | - | 5023376 | 5023771 | 13.84 | 1.15 | 2.10 | 0.00 | 3.33 | 9.51 | Hypothetical protein |

| Locus name | Gene name | Strand | Gene start (bp) | Gene end (bp) | Average TPM Inoculum | Average TPM $t_6$ | Average TPM $t_{24}$ | Average TPM $t_{168}$ | Average TPM Logarithmic | Average TPM Stationary | Annotation description |
|---|---|---|---|---|---|---|---|---|---|---|---|
| *PSPTO_4463* | *pmbA* | - | 5024076 | 5025422 | 16.51 | 27.99 | 25.12 | 18.22 | 79.13 | 35.09 | PmbA protein |
| *PSPTO_4464* | | + | 5025530 | 5026051 | 46.19 | 89.57 | 94.69 | 35.73 | 216.79 | 57.05 | Hypothetical protein |
| *PSPTO_4465* | *tldD* | - | 5026100 | 5027539 | 19.28 | 41.65 | 38.89 | 17.47 | 93.10 | 37.12 | TldD protein |
| *PSPTO_4466* | | - | 5027542 | 5028387 | 46.44 | 70.07 | 70.22 | 32.59 | 101.97 | 47.90 | Hydrolase |
| *PSPTO_4467* | | - | 5028436 | 5032251 | 2.35 | 6.57 | 3.02 | 3.17 | 18.41 | 3.26 | Hypothetical protein |
| *PSPTO_4468* | *cafA* | - | 5032433 | 5033890 | 10.14 | 16.97 | 14.47 | 5.52 | 105.49 | 12.64 | Ribonuclease G |
| *PSPTO_4469* | *maf-2* | - | 5033887 | 5034489 | 7.81 | 7.21 | 13.10 | 1.81 | 36.13 | 3.72 | Maf protein |
| *PSPTO_4470* | *mreD* | - | 5034557 | 5035048 | 4.00 | 4.98 | 9.05 | 1.86 | 26.79 | 3.80 | Rod shape-determining protein MreD |
| *PSPTO_4471* | *mreC* | - | 5035041 | 5036120 | 3.31 | 12.39 | 7.74 | 8.86 | 84.33 | 12.15 | Rod shape-determining protein MreC |
| *PSPTO_4472* | *mreB* | - | 5036351 | 5037388 | 20.11 | 59.49 | 46.92 | 37.98 | 144.71 | 41.77 | Rod shape-determining protein MreB |
| *PSPTO_4473* | *gatC* | + | 5037595 | 5037882 | 26.74 | 169.37 | 105.75 | 50.16 | 449.81 | 138.62 | Aspartyl/glutamyl-tRNA amidotransferase subunit C |
| *PSPTO_4474* | *gatA* | + | 5037897 | 5039348 | 5.45 | 38.10 | 40.41 | 23.97 | 228.51 | 68.62 | Aspartyl/glutamyl-tRNA amidotransferase subunit A |
| *PSPTO_4475* | *gatB* | + | 5039359 | 5040804 | 8.00 | 45.78 | 45.23 | 30.52 | 317.48 | 80.12 | Aspartyl/glutamyl-tRNA amidotransferase subunit B |
| *PSPTO_4476* | | + | 5040903 | 5041271 | 2.90 | 8.75 | 6.84 | 1.01 | 102.88 | 33.74 | RplA family protein |
| *PSPTO_4477* | | + | 5041338 | 5042369 | 7.02 | 7.39 | 12.75 | 9.36 | 39.25 | 9.97 | K+-dependent Na+/Ca+ exchanger-like protein |
| *PSPTO_4478* | *pcaC-2* | + | 5042401 | 5042784 | 4.28 | 7.48 | 3.11 | 0.00 | 42.66 | 12.71 | 4-carboxymuconolactone decarboxylase |
| *PSPTO_4479* | | - | 5042771 | 5043721 | 10.31 | 10.45 | 16.90 | 9.03 | 24.09 | 11.06 | Membrane protein |
| *PSPTO_4480* | | - | 5043826 | 5045409 | 4.65 | 4.49 | 3.34 | 1.13 | 4.34 | 4.11 | Tat (twin-arginine translocation) pathway signal sequence domain-containing protein |
| *PSPTO_4481* | | + | 5045558 | 5046574 | 3.04 | 12.46 | 9.76 | 3.93 | 17.17 | 8.68 | Hypothetical protein |
| *PSPTO_4482* | | + | 5046654 | 5049662 | 9.11 | 15.20 | 8.50 | 4.61 | 34.43 | 20.76 | Hypothetical protein |
| *PSPTO_4483* | | + | 5049655 | 5049996 | 5.20 | 15.26 | 14.63 | 8.41 | 32.35 | 9.69 | Hypothetical protein |
| *PSPTO_4484* | | - | 5050151 | 5050504 | 45.97 | 701.27 | 529.96 | 208.62 | 127.79 | 9.46 | Hypothetical protein |
| *PSPTO_4485* | | - | 5050863 | 5052323 | 0.30 | 0.33 | 0.29 | 0.04 | 0.44 | 0.36 | ISPsy6, transposase |
| *PSPTO_4486* | *pnp* | - | 5052402 | 5054507 | 15.41 | 63.81 | 52.15 | 22.01 | 216.58 | 16.30 | Polyribonucleotide nucleotidyltransferase |
| *PSPTO_4487* | *rpsO* | - | 5054683 | 5054952 | 29.69 | 76.50 | 114.49 | 98.18 | 150.01 | 73.68 | 30S ribosomal protein S15 |
| *PSPTO_4488* | *truB* | - | 5055111 | 5056028 | 25.08 | 40.51 | 24.93 | 20.59 | 87.39 | 10.79 | tRNA pseudouridine synthase B |
| *PSPTO_4489* | *rbfA* | - | 5056032 | 5056448 | 8.83 | 35.20 | 16.27 | 14.62 | 160.16 | 13.23 | Ribosome-binding factor A |
| *PSPTO_4490* | *infB* | - | 5056608 | 5059133 | 10.72 | 38.85 | 30.77 | 23.06 | 240.46 | 18.57 | Translation initiation factor IF-2 |
| *PSPTO_4491* | *nusA* | - | 5059161 | 5060642 | 10.93 | 55.75 | 43.03 | 38.33 | 266.90 | 25.57 | N utilization substance protein A |
| *PSPTO_4492* | | - | 5060702 | 5061178 | 16.94 | 69.16 | 50.08 | 23.61 | 81.23 | 30.26 | Hypothetical protein |
| *PSPTO_4493* | *secG* | - | 5061595 | 5061975 | 198.84 | 185.99 | 144.50 | 125.54 | 258.56 | 54.48 | Preprotein translocase subunit SecG |
| *PSPTO_4494* | *tpiA* | - | 5061980 | 5062735 | 197.40 | 247.27 | 243.48 | 178.37 | 265.95 | 113.31 | Triosephosphate isomerase |
| *PSPTO_4495* | *glmM* | - | 5062802 | 5064145 | 10.15 | 28.78 | 18.73 | 7.84 | 129.69 | 23.96 | Phosphoglucosamine mutase |
| *PSPTO_4496* | *folP* | - | 5064162 | 5065013 | 24.22 | 37.49 | 49.70 | 15.57 | 125.18 | 39.28 | Dihydropteroate synthase |
| *PSPTO_4497* | *ftsH* | - | 5065022 | 5066926 | 61.47 | 193.34 | 177.51 | 72.87 | 229.80 | 129.64 | Cell division protein FtsH |
| *PSPTO_4498* | | - | 5067120 | 5067770 | 62.35 | 59.70 | 44.91 | 49.95 | 85.86 | 52.82 | FtsJ cell division protein |
| *PSPTO_4499* | | + | 5067874 | 5068185 | 35.02 | 147.89 | 149.51 | 110.29 | 377.52 | 49.90 | Hypothetical protein |
| *PSPTO_4500* | *greA* | - | 5068265 | 5068741 | 20.71 | 28.05 | 32.46 | 16.13 | 111.71 | 17.95 | Transcription elongation factor GreA |
| *PSPTO_4501* | *carB* | - | 5068738 | 5071959 | 18.19 | 35.46 | 19.74 | 14.41 | 187.93 | 27.26 | Carbamoyl-phosphate synthase large subunit |
| *PSPTO_4502* | *carA* | - | 5072095 | 5073231 | 45.63 | 51.10 | 33.33 | 23.73 | 153.21 | 44.20 | Carbamoyl-phosphate synthase small subunit |
| *PSPTO_4503* | *dapB* | - | 5073447 | 5074250 | 14.24 | 16.75 | 12.44 | 12.56 | 145.03 | 33.88 | Dihydrodipicolinate reductase |
| *PSPTO_4504* | *dnaJ* | - | 5074273 | 5075415 | 26.80 | 24.77 | 19.99 | 18.48 | 156.06 | 47.02 | DnaJ protein |
| *PSPTO_4505* | *dnaK* | - | 5075741 | 5077657 | 108.21 | 151.01 | 109.26 | 60.86 | 633.92 | 259.24 | Molecular chaperone DnaK |
| *PSPTO_4506* | *grpE* | - | 5077768 | 5078331 | 252.50 | 180.41 | 157.19 | 74.60 | 449.84 | 153.52 | Heat shock protein GrpE |
| *PSPTO_4507* | *recN* | + | 5078600 | 5080273 | 16.37 | 74.15 | 54.35 | 23.13 | 46.17 | 33.05 | DNA repair protein RecN |
| *PSPTO_4508* | *fur* | - | 5080338 | 5080745 | 73.44 | 148.96 | 114.42 | 53.89 | 157.48 | 48.72 | Ferric uptake regulation protein |
| *PSPTO_4509* | *omlA* | + | 5080844 | 5081365 | 148.63 | 155.48 | 97.59 | 72.28 | 311.58 | 128.68 | Outer membrane lipoprotein OmlA |
| *PSPTO_4510* | | + | 5081470 | 5081871 | 79.61 | 92.13 | 70.41 | 35.02 | 65.42 | 83.81 | ISPsy11, transposase OrfA |
| *PSPTO_4511* | | + | 5081895 | 5082692 | 8.89 | 10.85 | 7.19 | 4.46 | 10.42 | 9.79 | ISPsy11, transposase OrfB |
| *PSPTO_4512* | | - | 5082748 | 5083062 | 7.34 | 19.94 | 3.64 | 2.47 | 17.97 | 6.80 | Hypothetical protein |
| *PSPTO_4513* | | - | 5083055 | 5083528 | 8.72 | 6.96 | 5.10 | 4.74 | 21.06 | 11.42 | Hypothetical protein |
| *PSPTO_4514* | | - | 5083497 | 5084900 | 7.65 | 14.43 | 15.72 | 4.17 | 29.55 | 14.58 | Sodium-dependent transporter |
| *PSPTO_4515* | *smpB* | + | 5085014 | 5085496 | 46.10 | 67.11 | 65.59 | 34.64 | 156.34 | 47.13 | SsrA-binding protein |
| *PSPTO_4516* | | + | 5085788 | 5086072 | 123.79 | 544.53 | 449.99 | 151.84 | 150.02 | 147.92 | Hypothetical protein |
| *PSPTO_4517* | | + | 5087432 | 5088382 | 3.42 | 0.97 | 5.32 | 2.31 | 9.12 | 4.25 | Fatty acid desaturase |
| *PSPTO_4518* | | + | 5088445 | 5093706 | 5.63 | 4.39 | 4.09 | 1.33 | 9.01 | 5.66 | Non-ribosomal peptide synthetase, initiating component |
| *PSPTO_4519* | | + | 5093815 | 5104113 | 5.79 | 3.14 | 2.76 | 1.93 | 7.81 | 4.24 | Non-ribosomal peptide synthetase, terminal component |
| *PSPTO_4520* | | + | 5104150 | 5106501 | 9.37 | 6.55 | 7.10 | 2.77 | 15.90 | 8.63 | Methyltransferase domain protein |
| *PSPTO_4521* | | + | 5106533 | 5106739 | 4.52 | 0.00 | 0.00 | 0.00 | 6.13 | 6.66 | Hypothetical protein |
| *PSPTO_4522* | | + | 5106886 | 5108169 | 5.04 | 4.04 | 5.49 | 5.63 | 10.04 | 5.88 | Membrane protein |
| *PSPTO_4523* | | + | 5108505 | 5108936 | 4.21 | 10.01 | 18.67 | 14.55 | 25.24 | 12.31 | AsnC family transcriptional regulator |
| *PSPTO_4524* | | + | 5109010 | 5109819 | 3.40 | 21.78 | 14.26 | 6.24 | 24.42 | 8.63 | Hypothetical protein |
| *PSPTO_4525* | | + | 5110540 | 5110707 | 1.64 | 0.00 | 0.00 | 0.22 | 1.83 | 3.12 | Hypothetical protein |
| *PSPTO_4526* | | - | 5111194 | 5112639 | 8.46 | 17.81 | 8.82 | 8.19 | 9.69 | 6.16 | 6-aminohexanoate-cyclic-dimer hydrolase |
| *PSPTO_4527* | | - | 5112686 | 5113618 | 6.71 | 6.37 | 2.96 | 8.18 | 7.86 | 6.49 | Hypothetical protein |
| *PSPTO_4528* | | - | 5113687 | 5114808 | 53.10 | 85.07 | 90.14 | 129.77 | 36.46 | 16.83 | AraC family transcriptional regulator |
| *PSPTO_4529* | | + | 5114872 | 5115360 | 46.32 | 54.38 | 38.22 | 43.94 | 304.24 | 80.94 | Hypothetical protein |
| *PSPTO_4530* | *katG* | + | 5115931 | 5118201 | 58.95 | 539.99 | 491.90 | 133.75 | 186.65 | 51.56 | Catalase/peroxidase HPI |
| *PSPTO_4531* | | - | 5118280 | 5119584 | 26.78 | 6.55 | 5.16 | 3.46 | 8.26 | 2.82 | Methyl-accepting chemotaxis protein |
| *PSPTO_4532* | | - | 5119679 | 5119987 | 25.31 | 20.90 | 16.31 | 3.23 | 67.01 | 16.49 | Hypothetical protein |
| *PSPTO_4533* | *nhaA-1* | - | 5120036 | 5121214 | 7.72 | 3.05 | 5.67 | 2.45 | 23.31 | 6.63 | Sodium-proton antiporter NhaA |
| *PSPTO_4534* | | - | 5121372 | 5122886 | 6.26 | 6.71 | 7.46 | 2.26 | 12.92 | 6.41 | Peptide ABC transporter substrate-binding protein |
| *PSPTO_4535* | | - | 5122936 | 5123916 | 2.85 | 4.58 | 3.08 | 5.07 | 6.39 | 3.92 | Peptide ABC transporter ATP-binding protein |
| *PSPTO_4536* | | - | 5123913 | 5124917 | 1.92 | 2.67 | 1.46 | 3.13 | 5.06 | 2.42 | Peptide ABC transporter ATP-binding protein |
| *PSPTO_4537* | | - | 5124914 | 5125777 | 9.60 | 6.32 | 7.21 | 2.80 | 8.66 | 6.59 | Peptide ABC transporter permease |
| *PSPTO_4538* | | - | 5125774 | 5126721 | 3.59 | 0.00 | 4.79 | 0.00 | 4.35 | 3.48 | Peptide ABC transporter permease |
| *PSPTO_4539* | | + | 5127060 | 5127824 | 60.96 | 22.54 | 31.57 | 6.35 | 23.56 | 16.84 | LuxR family transcriptional regulator, autoinducer-regulated |
| *PSPTO_4540* | | + | 5127919 | 5128869 | 15.53 | 11.97 | 12.58 | 14.51 | 14.73 | 12.77 | Proline iminopeptidase |
| *PSPTO_4541* | | + | 5129126 | 5130751 | 43.78 | 12.04 | 7.76 | 10.75 | 8.10 | 7.01 | Methyl-accepting chemotaxis protein |

| Locus name | Gene name | Strand | Gene start (bp) | Gene end (bp) | Average TPM Inoculum | Average TPM $t_6$ | Average TPM $t_{24}$ | Average TPM $t_{168}$ | Average TPM Logarithmic | Average TPM Stationary | Annotation description |
|---|---|---|---|---|---|---|---|---|---|---|---|
| *PSPTO_4542* | | + | 5130995 | 5131495 | 56.86 | 119.35 | 58.71 | 38.09 | 195.92 | 57.03 | GAF domain-containing protein |
| *PSPTO_4543* | | + | 5131860 | 5133683 | 15.40 | 14.85 | 8.57 | 3.57 | 12.27 | 6.80 | GAF domain/GGDEF domain/EAL domain protein |
| *PSPTO_4544* | | + | 5133760 | 5134560 | 3.83 | 9.20 | 7.65 | 2.78 | 4.83 | 3.76 | Membrane protein |
| *PSPTO_4545* | | + | 5134785 | 5134985 | 117.78 | 196.92 | 148.85 | 117.11 | 19.95 | 41.40 | Hypothetical protein |
| *PSPTO_4546* | | - | 5135614 | 5135853 | 50.15 | 279.83 | 210.17 | 148.86 | 121.39 | 91.14 | Hypothetical protein |
| *PSPTO_4547* | *garD* | - | 5136020 | 5137573 | 9.07 | 21.68 | 22.40 | 9.69 | 23.99 | 8.70 | D-galactarate dehydratase |
| *PSPTO_4548* | | - | 5137584 | 5138969 | 5.54 | 11.56 | 15.41 | 8.73 | 17.83 | 9.05 | MFS transporter, phthalate permease family |
| *PSPTO_4549* | | - | 5139135 | 5140046 | 2.72 | 20.62 | 34.10 | 9.13 | 15.40 | 3.43 | 5-dehydro-4-deoxyglucarate dehydratase |
| *PSPTO_4550* | | - | 5140337 | 5141086 | 42.64 | 81.90 | 75.40 | 26.74 | 139.14 | 68.50 | GntR family transcriptional regulator |
| *PSPTO_4551* | | - | 5141184 | 5142917 | 4.03 | 29.92 | 16.74 | 2.63 | 5.95 | 5.80 | ABC transporter ATP-binding protein |
| *PSPTO_4552* | | + | 5142907 | 5143017 | 3.77 | 30.29 | 0.00 | 0.00 | 8.26 | 11.83 | Hypothetical protein |
| *PSPTO_4553* | | - | 5143007 | 5143570 | 16.96 | 35.06 | 26.03 | 21.49 | 44.49 | 75.81 | DNA-binding response regulator |
| *PSPTO_4554* | | - | 5143567 | 5144826 | 8.55 | 10.58 | 5.70 | 4.23 | 32.02 | 40.73 | Sensor histidine kinase |
| *PSPTO_4555* | | + | 5145020 | 5145733 | 18.27 | 32.89 | 31.33 | 8.51 | 57.09 | 15.66 | Hypothetical protein |
| *PSPTO_4556* | *phcA* | + | 5145818 | 5147497 | 4.62 | 6.08 | 4.77 | 3.46 | 13.07 | 7.60 | PhcA |
| *PSPTO_4557* | | + | 5147594 | 5149201 | 6.30 | 21.26 | 18.84 | 6.99 | 79.45 | 13.24 | Peptide ABC transporter periplasmic peptide-binding protein |
| *PSPTO_4558* | | + | 5149459 | 5151081 | 48.77 | 67.22 | 65.63 | 19.19 | 394.28 | 20.26 | Dipeptide ABC transporter, periplasmic dipeptide-binding protein |
| *PSPTO_4559* | | + | 5151411 | 5153003 | 7.37 | 13.80 | 9.71 | 2.84 | 80.94 | 5.57 | Dipeptide ABC transporter substrate-binding protein |
| *PSPTO_4560* | | + | 5153159 | 5154538 | 24.85 | 20.82 | 26.35 | 8.90 | 274.92 | 7.12 | Outer membrane porin, OprD family |
| *PSPTO_4561* | *dppA* | + | 5154604 | 5156199 | 27.46 | 34.09 | 40.45 | 19.73 | 388.16 | 19.59 | Dipeptide ABC transporter substrate-binding protein |
| *PSPTO_4562* | | + | 5156274 | 5157284 | 6.00 | 13.75 | 14.16 | 10.69 | 101.09 | 5.79 | Dipeptide ABC transporter permease |
| *PSPTO_4563* | *dppC* | + | 5157295 | 5158206 | 3.78 | 8.04 | 9.51 | 5.46 | 58.13 | 4.01 | Dipeptide ABC transporter permease |
| *PSPTO_4564* | | + | 5158217 | 5159185 | 7.12 | 24.95 | 20.05 | 5.78 | 123.45 | 7.03 | Dipeptide ABC transporter ATP-binding protein |
| *PSPTO_4565* | | + | 5159185 | 5159541 | - | - | - | - | - | - | |
| *PSPTO_4566* | | + | 5159584 | 5159943 | 0.00 | 0.00 | 0.00 | 0.00 | 0.00 | 0.00 | ISPsy5, Orf1 |
| *PSPTO_4567* | | + | 5159961 | 5161514 | 0.00 | 0.00 | 0.00 | 0.00 | 0.00 | 0.00 | ISPsy5, transposase |
| *PSPTO_4568* | | + | 5161531 | 5162226 | - | - | - | - | - | - | |
| *PSPTO_4569* | | - | 5162342 | 5164810 | 4.61 | 3.06 | 3.96 | 1.72 | 11.41 | 6.17 | Iron-uptake factor |
| *PSPTO_4570* | | - | 5165012 | 5166025 | 6.67 | 9.75 | 7.70 | 3.17 | 10.36 | 7.94 | Binary cytotoxin component |
| *PSPTO_4571* | | - | 5166061 | 5167260 | 0.88 | 0.32 | 0.00 | 0.46 | 2.78 | 1.53 | Binary cytotoxin component |
| *PSPTO_4572* | | - | 5167536 | 5168615 | 31.90 | 48.26 | 50.79 | 33.58 | 63.33 | 20.04 | Pyridoxal-dependent decarboxylase, pyridoxal binding domain protein |
| *PSPTO_4573* | | - | 5168635 | 5168745 | 68.57 | 77.72 | 72.82 | 56.34 | 68.67 | 34.27 | Hypothetical protein |
| *PSPTO_4574* | | + | 5169133 | 5171616 | 7.84 | 6.27 | 8.32 | 5.89 | 18.88 | 11.73 | Hypothetical protein |
| *PSPTO_4575* | *opuCA* | - | 5171664 | 5172821 | 2.51 | 15.55 | 11.68 | 5.04 | 14.95 | 9.84 | Glycine betaine/choline OpuC ABC transporter, ATP-binding protein |
| *PSPTO_4576* | *opuCV* | - | 5172818 | 5173471 | 7.92 | 10.86 | 12.24 | 7.55 | 19.97 | 10.11 | Glycine betaine/choline OpuC ABC transporter, permease protein |
| *PSPTO_4577* | *opuCC* | - | 5173485 | 5174378 | 2.02 | 10.59 | 8.47 | 8.30 | 16.03 | 9.10 | Glycine betaine/choline OpuC ABC transporter, periplasmic substrate-binding protein |
| *PSPTO_4578* | *opuCD* | - | 5174393 | 5175106 | 1.92 | 10.09 | 6.29 | 7.80 | 10.33 | 6.62 | Glycine betaine/carnitine/choline ABC transporter, permease protein |
| *PSPTO_4579* | *prfC* | + | 5175451 | 5177034 | 11.40 | 36.50 | 40.33 | 25.47 | 107.40 | 29.44 | Peptide chain release factor 3 |
| *PSPTO_4580* | | + | 5177384 | 5177626 | 11.97 | 1.89 | 0.00 | 1.51 | 18.01 | 227.05 | Hypothetical protein |
| *PSPTO_4581* | | - | 5177708 | 5178382 | 5.14 | 16.80 | 14.62 | 11.38 | 70.20 | 37.40 | FKBP-type peptidyl-prolyl cis-trans isomerase |
| *PSPTO_4582* | | - | 5178480 | 5178788 | 10.99 | 42.90 | 21.33 | 26.55 | 21.13 | 27.21 | Hypothetical protein |
| *PSPTO_4583* | | - | 5178900 | 5179388 | 7.73 | 11.26 | 11.24 | 6.16 | 20.65 | 27.52 | Hypothetical protein |
| *PSPTO_4584* | | - | 5179514 | 5179843 | 61.00 | 58.49 | 37.59 | 17.65 | 12.79 | 28.60 | Hypothetical protein |
| *PSPTO_4585* | | - | 5180018 | 5182453 | 9.72 | 16.58 | 16.24 | 6.83 | 63.98 | 17.74 | ATP-dependent protease |
| *PSPTO_4586* | | + | 5182661 | 5183218 | 224.37 | 1896.06 | 1559.73 | 742.71 | 699.47 | 532.52 | Hypothetical protein |
| *PSPTO_4587* | | + | 5183281 | 5185134 | 2.61 | 10.88 | 6.39 | 3.08 | 32.34 | 7.10 | Hypothetical protein |
| *PSPTO_4588* | *hopS2* | - | 5185073 | 5185606 | 1.67 | 33.55 | 52.92 | 4.81 | 2.90 | 1.85 | |
| *PSPTO_4589* | *shcS2* | - | 5185621 | 5186094 | 3.37 | 191.56 | 110.24 | 29.84 | 6.77 | 5.40 | Type III chaperone ShcS2 |
| *PSPTO_4590* | *hopT2* | - | 5186263 | 5186646 | 15.21 | 16.54 | 20.02 | 15.78 | 24.63 | 16.25 | Premature stop codon in this non-functional type III effector |
| *PSPTO_4591* | *hopO1-3* | - | 5186668 | 5186958 | - | - | - | - | - | - | Possibly a pseudogene |
| *PSPTO_4592* | *hopO1-3* | - | 5187067 | 5187573 | 7.62 | 5.21 | 12.27 | 3.63 | 22.63 | 18.48 | Possibly a pseudogene |
| *PSPTO_4593* | *hopT1-2* | - | 5187920 | 5189089 | 13.23 | 13.68 | 7.26 | 13.16 | 22.75 | 18.88 | Probably not a type III effector |
| *PSPTO_4594* | *hopO1-2* | - | 5189095 | 5189991 | 12.31 | 11.05 | 15.39 | 8.50 | 29.88 | 18.19 | Probably not a type III effector |
| *PSPTO_4595* | | - | 5190048 | 5190437 | 16.89 | 12.35 | 11.00 | 19.48 | 40.39 | 25.04 | Hypothetical protein |
| *PSPTO_4596* | | + | 5190629 | 5191606 | 0.00 | 0.00 | 0.00 | 0.00 | 0.00 | 0.00 | ISPssy, transposase |
| *PSPTO_4597* | *hopS1* | - | 5191741 | 5192097 | 4.16 | 179.53 | 78.70 | 17.42 | 12.32 | 12.13 | Possibly a pseudogene |
| *PSPTO_4599* | *shcS1* | - | 5192110 | 5192583 | 9.66 | 715.19 | 417.61 | 37.43 | 13.27 | 11.38 | Type III chaperone ShcS1 |
| *PSPTO_4600* | | + | 5192782 | 5193135 | 65.86 | 23.64 | 14.91 | 24.38 | 10.60 | 29.95 | Lipoprotein |
| *PSPTO_4601* | *greB-2* | + | 5193212 | 5193709 | 16.41 | 12.61 | 22.14 | 12.28 | 7.53 | 10.77 | Transcription elongation factor GreB |
| *PSPTO_4602* | | - | 5193762 | 5195453 | 17.18 | 20.50 | 27.15 | 14.12 | 182.28 | 29.36 | Putative ABC transporter ATP-binding protein |
| *PSPTO_4603* | | + | 5195601 | 5196866 | 17.17 | 20.66 | 16.55 | 7.14 | 34.50 | 34.52 | Site-specific recombinase, phage integrase family |
| *PSPTO_4604* | | + | 5196863 | 5198458 | 14.58 | 17.09 | 15.33 | 4.49 | 24.39 | 32.46 | Site-specific recombinase, phage integrase family |
| *PSPTO_4605* | | + | 5198455 | 5200692 | 14.26 | 17.49 | 13.16 | 7.24 | 32.34 | 35.23 | Hypothetical protein |
| *PSPTO_4606* | | + | 5200695 | 5201093 | 7.74 | 17.07 | 13.28 | 1.19 | 29.54 | 22.70 | Hypothetical protein |
| *PSPTO_4607* | | + | 5201523 | 5203982 | 4.24 | 12.47 | 12.24 | 15.57 | 54.17 | 18.85 | Hypothetical protein |
| *PSPTO_4608* | | + | 5204349 | 5205311 | 8.99 | 12.27 | 3.90 | 5.86 | 14.09 | 9.73 | Site-specific recombinase, phage integrase family |
| *PSPTO_4609* | | - | 5205285 | 5205530 | 10.48 | 7.44 | 14.67 | 16.00 | 10.22 | 5.34 | Hypothetical protein |
| *PSPTO_4610* | | - | 5206026 | 5206406 | 77.51 | 375.18 | 248.65 | 326.56 | 657.88 | 201.54 | Hypothetical protein |
| *PSPTO_4611* | | - | 5206737 | 5207777 | 17.45 | 77.22 | 30.42 | 22.96 | 50.61 | 23.45 | Hypothetical protein |
| *PSPTO_4612* | | - | 5207887 | 5208783 | 11.64 | 9.22 | 11.45 | 1.69 | 29.07 | 7.67 | LysR family transcriptional regulator |
| *PSPTO_4613* | | + | 5208901 | 5210070 | 42.06 | 37.73 | 33.56 | 17.53 | 38.69 | 40.30 | C4-dicarboxylate transporter/malic acid transport protein |
| *PSPTO_4614* | | - | 5210079 | 5210681 | 13.83 | 13.06 | 10.67 | 9.20 | 19.91 | 16.20 | Hypothetical protein |
| *PSPTO_4615* | *ppa-2* | - | 5210681 | 5211226 | 2.70 | 14.40 | 0.58 | 12.05 | 10.13 | 10.99 | Inorganic pyrophosphatase |
| *PSPTO_4616* | *eno-2* | - | 5211342 | 5212625 | 8.11 | 10.20 | 8.89 | 5.60 | 19.17 | 12.44 | Enolase |
| *PSPTO_4617* | | + | 5212667 | 5212909 | 6.20 | 0.51 | 0.00 | 12.94 | 9.03 | 5.01 | Hypothetical protein |
| *PSPTO_4618* | | - | 5213444 | 5213767 | 10.61 | 8.42 | 8.12 | 11.22 | 12.02 | 6.24 | Hypothetical protein |
| *PSPTO_4619* | | - | 5214210 | 5215568 | 3.81 | 4.20 | 1.99 | 3.80 | 6.56 | 3.78 | Voltage-gated chloride channel family protein |
| *PSPTO_4620* | | - | 5215805 | 5216128 | 6.10 | 5.78 | 3.87 | 0.00 | 9.75 | 9.25 | Hypothetical protein |
| *PSPTO_4621* | | - | 5216139 | 5216882 | 11.22 | 7.83 | 7.37 | 0.71 | 13.62 | 13.16 | Transporter |

| Locus name | Gene name | Strand | Gene start (bp) | Gene end (bp) | Average TPM Inoculum | Average TPM $t_6$ | Average TPM $t_{24}$ | Average TPM $t_{168}$ | Average TPM Logarithmic | Average TPM Stationary | Annotation description |
|---|---|---|---|---|---|---|---|---|---|---|---|
| *PSPTO_4622* | | + | 5216974 | 5217408 | 129.76 | 52.88 | 48.01 | 42.42 | 78.92 | 86.86 | MerR family transcriptional regulator |
| *PSPTO_4623* | | + | 5218247 | 5218624 | 12.89 | 19.10 | 19.76 | 8.61 | 37.93 | 23.04 | Hypothetical protein |
| *PSPTO_4624* | | - | 5218750 | 5220369 | 13.69 | 3.83 | 4.13 | 3.05 | 4.31 | 2.97 | Methyl-accepting chemotaxis protein |
| *PSPTO_4625* | | + | 5220631 | 5221653 | 0.00 | 0.00 | 0.00 | 0.00 | 0.00 | 0.00 | ISPsy4, transposase |
| *PSPTO_4626* | | + | 5221650 | 5222459 | 0.00 | 0.00 | 0.00 | 0.00 | 0.00 | 0.00 | ISPsy4, transposition helper protein |
| *PSPTO_4627* | | + | 5224073 | 5226526 | 3.41 | 5.32 | 5.14 | 2.13 | 16.06 | 7.11 | Hypothetical protein |
| *PSPTO_4628* | | + | 5226849 | 5227055 | 12.08 | 21.64 | 10.14 | 18.07 | 17.64 | 10.14 | Hypothetical protein |
| *PSPTO_4629* | | + | 5228357 | 5228464 | 9.98 | 9.74 | 0.00 | 0.00 | 6.90 | 10.24 | Hypothetical protein |
| *PSPTO_4630* | | - | 5228484 | 5228768 | 38.26 | 12.20 | 3.40 | 0.00 | 24.19 | 30.76 | PbsX family transcriptional regulator |
| *PSPTO_4631* | | + | 5230653 | 5234489 | 5.13 | 6.65 | 9.89 | 9.21 | 17.98 | 12.17 | Sensory box/GGDEF domain/EAL domain-containing protein |
| *PSPTO_4632* | *glyA-2* | + | 5234654 | 5235907 | 71.81 | 150.44 | 128.96 | 57.55 | 1837.06 | 137.46 | Serine hydroxymethyltransferase |
| *PSPTO_4633* | | + | 5236578 | 5237111 | 72.72 | 83.53 | 82.82 | 51.54 | 235.13 | 85.10 | Hypothetical protein |
| *PSPTO_4634* | | + | 5237118 | 5237831 | 15.20 | 17.25 | 11.91 | 5.68 | 101.08 | 18.27 | N-acetylmuramoyl-L-alanine amidase |
| *PSPTO_4635* | | + | 5238570 | 5239139 | 2.58 | 3.62 | 3.65 | 1.93 | 9.23 | 3.60 | Hypothetical protein |
| *PSPTO_4636* | | - | 5239167 | 5240144 | 7.24 | 2.68 | 9.91 | 6.09 | 39.05 | 8.12 | Cobalamin synthesis protein/P47K family protein |
| *PSPTO_4637* | | - | 5240240 | 5240437 | 18.67 | 38.85 | 36.25 | 32.66 | 162.38 | 18.51 | Hypothetical protein |
| *PSPTO_4638* | *cstA* | - | 5240465 | 5242540 | 16.52 | 33.35 | 23.32 | 11.85 | 153.23 | 15.92 | Carbon starvation protein CstA |
| *PSPTO_4639* | | + | 5242826 | 5243194 | 52.66 | 35.93 | 59.49 | 18.19 | 41.74 | 30.21 | Hypothetical protein |
| *PSPTO_4640* | *radA* | - | 5243215 | 5244582 | 9.35 | 24.36 | 20.11 | 15.12 | 65.88 | 48.40 | DNA repair protein RadA |
| *PSPTO_4641* | *mscL* | + | 5244777 | 5245223 | 123.49 | 110.02 | 105.69 | 81.33 | 85.12 | 134.41 | Large conductance mechanosensitive channel protein |
| *PSPTO_4642* | *fnr-2* | - | 5245423 | 5246199 | 27.53 | 31.30 | 33.06 | 29.14 | 70.27 | 45.29 | Ferredoxin--NADP reductase |
| *PSPTO_4643* | | - | 5246633 | 5247610 | 0.00 | 0.00 | 0.00 | 0.00 | 0.00 | 0.00 | ISPssy, transposase |
| *PSPTO_4644* | | + | 5247729 | 5247962 | 7.51 | 15.19 | 1.97 | 0.00 | 6.61 | 7.31 | LuxR family transcriptional regulator |
| *PSPTO_4645* | | + | 5248092 | 5249216 | 5.55 | 8.14 | 11.88 | 5.79 | 33.79 | 13.91 | Methyltransferase |
| *PSPTO_4646* | | - | 5249304 | 5249471 | 14.37 | 10.91 | 12.43 | 18.44 | 27.19 | 113.34 | Hypothetical protein |
| *PSPTO_4647* | *cioB* | - | 5249493 | 5250500 | 9.04 | 36.22 | 33.74 | 23.79 | 13.39 | 74.73 | Cyanide-insensitive terminal oxidase CioB |
| *PSPTO_4648* | *cioA* | - | 5250504 | 5251943 | 7.75 | 23.40 | 16.65 | 17.89 | 8.31 | 70.31 | Cyanide-insensitive terminal oxidase CioA |
| *PSPTO_4649* | | + | 5251947 | 5252120 | 64.43 | 63.06 | 31.65 | 32.57 | 18.90 | 356.91 | Hypothetical protein |
| *PSPTO_4650* | | - | 5252337 | 5254538 | 4.23 | 18.74 | 14.45 | 8.35 | 8.51 | 7.13 | Aldehyde oxidase and xanthine dehydrogenase family protein |
| *PSPTO_4651* | | - | 5254535 | 5255524 | 2.76 | 10.62 | 10.91 | 3.95 | 5.24 | 4.26 | Oxidoreductase, molybdopterin-binding subunit |
| *PSPTO_4652* | | - | 5255521 | 5256051 | 4.08 | 16.99 | 20.02 | 11.72 | 5.85 | 5.37 | Iron-sulfur cluster-binding protein |
| *PSPTO_4653* | | + | 5256316 | 5257611 | 10.87 | 8.76 | 11.13 | 10.36 | 13.05 | 8.82 | Xanthine/uracil permease family protein |
| *PSPTO_4654* | *trmA* | + | 5257608 | 5258693 | 2.50 | 3.86 | 8.08 | 1.94 | 31.57 | 6.48 | tRNA (uracil-5-)-methyltransferase |
| *PSPTO_4656* | | + | 5259642 | 5260925 | 31.69 | 10.01 | 11.01 | 10.59 | 26.79 | 18.01 | Membrane protein |
| *PSPTO_4657* | | + | 5261062 | 5261952 | 61.41 | 45.13 | 39.15 | 20.08 | 59.66 | 32.14 | Zinc metallopeptidase |
| *PSPTO_4658* | | + | 5261957 | 5262946 | 19.33 | 36.28 | 39.01 | 8.88 | 64.16 | 27.99 | Hypothetical protein |
| *PSPTO_4659* | | - | 5263123 | 5263677 | 6.98 | 15.54 | 7.55 | 7.99 | 16.90 | 9.81 | HAD-superfamily hydrolase |
| *PSPTO_4660* | *tesB* | - | 5263710 | 5264579 | 9.50 | 8.95 | 6.16 | 12.96 | 24.06 | 13.21 | Acyl-CoA thioesterase II |
| *PSPTO_4661* | | - | 5264717 | 5265286 | 17.76 | 20.26 | 9.71 | 11.58 | 11.78 | 12.75 | GNAT family acetyltransferase |
| *PSPTO_4662* | | + | 5265421 | 5266338 | 11.43 | 24.96 | 13.03 | 7.82 | 32.28 | 13.48 | Histone deacetylase family protein |
| *PSPTO_4663* | | + | 5266428 | 5267666 | 2.25 | 4.01 | 2.97 | 1.20 | 13.19 | 4.24 | Hypothetical protein |
| *PSPTO_4664* | | - | 5267751 | 5269079 | 7.54 | 16.49 | 21.86 | 6.67 | 50.67 | 16.49 | ATP-dependent RNA helicase rhlE |
| *PSPTO_4665* | | - | 5269194 | 5269808 | 11.14 | 18.54 | 16.07 | 6.56 | 41.95 | 14.47 | Exonuclease |
| *PSPTO_4666* | | - | 5269827 | 5270303 | 31.69 | 55.29 | 41.10 | 22.00 | 73.37 | 30.85 | Hypothetical protein |
| *PSPTO_4667* | | - | 5270574 | 5271371 | 70.80 | 140.25 | 69.69 | 18.54 | 95.07 | 64.37 | Hypothetical protein |
| *PSPTO_4668* | | + | 5271474 | 5272136 | 3.57 | 13.94 | 10.97 | 4.07 | 23.73 | 7.45 | Membrane protein |
| *PSPTO_4669* | | + | 5272223 | 5273314 | 7.86 | 17.46 | 15.57 | 9.87 | 47.49 | 16.95 | Endonuclease/exonuclease/phosphatase family protein |
| *PSPTO_4670* | | - | 5273330 | 5274127 | 4.45 | 7.22 | 2.28 | 6.95 | 19.32 | 9.20 | Hypothetical protein |
| *PSPTO_4671* | | - | 5274267 | 5275259 | 12.41 | 15.32 | 9.82 | 5.67 | 27.76 | 19.21 | Alkaline D-peptidase |
| *PSPTO_4672* | | + | 5275303 | 5275476 | 2.99 | 0.00 | 1.85 | 6.77 | 3.77 | 4.39 | Hypothetical protein |
| *PSPTO_4673* | | + | 5275543 | 5275791 | 51.06 | 95.66 | 69.03 | 85.52 | 13.27 | 35.37 | Hypothetical protein |
| *PSPTO_4674* | | - | 5276101 | 5276277 | 4.35 | 8.01 | 2.07 | 4.14 | 6.01 | 7.70 | Hypothetical protein |
| *PSPTO_4675* | | - | 5277517 | 5278287 | 6.98 | 23.49 | 15.58 | 11.22 | 31.76 | 14.98 | Sir2 family transcriptional regulator |
| *PSPTO_4676* | | + | 5279840 | 5280049 | 35.75 | 18.60 | 35.05 | 19.44 | 67.11 | 41.79 | Hypothetical protein |
| *PSPTO_4677* | | + | 5280438 | 5281079 | 3.16 | 4.71 | 0.76 | 0.00 | 4.30 | 3.67 | Hypothetical protein |
| *PSPTO_4678* | | - | 5280963 | 5281343 | - | - | - | - | - | - | |
| *PSPTO_4679* | | - | 5281410 | 5281888 | - | - | - | - | - | - | |
| *PSPTO_4680* | *cfl* | + | 5282597 | 5284126 | 0.00 | 0.12 | 0.00 | 0.00 | 0.01 | 0.00 | Coronafacic acid synthetase, ligase component |
| *PSPTO_4681* | *cfa1* | + | 5284213 | 5284464 | 0.00 | 0.00 | 0.00 | 0.00 | 0.00 | 0.00 | Coronafacic acid synthetase, acyl carrier protein component |
| *PSPTO_4682* | *cfa2* | + | 5284476 | 5284973 | 0.00 | 0.00 | 0.00 | 0.00 | 0.00 | 0.00 | Coronafacic acid synthetase, dehydratase component |
| *PSPTO_4683* | *cfa3* | + | 5284967 | 5286112 | 0.00 | 0.00 | 0.00 | 0.00 | 0.00 | 0.00 | Coronafacic acid beta-ketoacyl synthetase component |
| *PSPTO_4684* | *cfa4* | + | 5286109 | 5286642 | 0.00 | 0.00 | 0.00 | 0.00 | 0.00 | 0.00 | Coronafacic acid synthetase component |
| *PSPTO_4685* | *cfa5* | + | 5286639 | 5288057 | 0.00 | 0.00 | 0.00 | 0.00 | 0.00 | 0.00 | Coronafacic acid synthetase, ligase component |
| *PSPTO_4686* | *cfa6* | + | 5287945 | 5296104 | 0.00 | 0.00 | 0.00 | 0.00 | 0.00 | 0.00 | Coronafacic acid polyketide synthase I |
| *PSPTO_4687* | *cfa7* | + | 5296110 | 5302310 | 0.00 | 0.00 | 0.00 | 0.00 | 0.00 | 0.00 | Coronafacic acid polyketide synthetase II |
| *PSPTO_4688* | | + | 5302307 | 5302750 | 0.00 | 0.00 | 0.00 | 0.00 | 0.00 | 0.00 | Hypothetical protein |
| *PSPTO_4689* | *cfa8* | + | 5302800 | 5304005 | 0.00 | 0.00 | 0.00 | 0.00 | 0.00 | 0.00 | Crotonyl-CoA reductase |
| *PSPTO_4690* | *cfa9* | + | 5304061 | 5304804 | 0.00 | 0.11 | 0.00 | 0.00 | 0.00 | 0.00 | CFA synthetase, thioesterase component |
| *PSPTO_4691* | *hopAD1* | + | 5305803 | 5307701 | 2.86 | 70.14 | 20.30 | 2.71 | 6.60 | 2.62 | Type III effector HopAD1 |
| *PSPTO_4692* | | - | 5307739 | 5308098 | - | - | - | - | - | - | |
| *PSPTO_4693* | | - | 5308353 | 5309906 | 0.00 | 0.00 | 0.00 | 0.00 | 0.00 | 0.00 | ISPsy5, transposase |
| *PSPTO_4694* | | - | 5309924 | 5310283 | 0.00 | 0.00 | 0.00 | 0.00 | 0.00 | 0.00 | ISPsy5, Orf1 |
| *PSPTO_4695* | | + | 5310495 | 5310746 | 10.43 | 9.80 | 9.20 | 4.31 | 16.55 | 15.72 | ISPsy13, transposase OrfA |
| *PSPTO_4696* | | + | 5310778 | 5312067 | 9.21 | 22.75 | 11.48 | 16.20 | 22.68 | 20.80 | ISPsy13, transposase OrfB |
| *PSPTO_4697* | | - | 5312676 | 5313014 | 3.08 | 17.53 | 9.50 | 5.30 | 15.74 | 15.12 | ISPs1, transposase OrfB |
| *PSPTO_4699* | | - | 5313602 | 5327197 | 12.19 | 12.71 | 8.59 | 5.65 | 17.76 | 11.06 | Non-ribosomal peptide synthetase, terminal component |
| *PSPTO_4702* | | + | 5329495 | 5330472 | 0.00 | 0.00 | 0.00 | 0.00 | 0.00 | 0.00 | ISPssy, transposase |
| *PSPTO_4703* | *hopAQ1* | + | 5330802 | 5331053 | 6.11 | 48.95 | 56.59 | 21.62 | 21.80 | 12.61 | Type III effector HopAQ1 |
| *PSPTO_4704* | *corR* | + | 5331204 | 5331803 | 6.53 | 20.23 | 5.70 | 3.86 | 10.70 | 5.94 | DNA-binding response regulator CorR |

| Locus name | Gene name | Strand | Gene start (bp) | Gene end (bp) | Average TPM Inoculum | Average TPM $t_6$ | Average TPM $t_{24}$ | Average TPM $t_{168}$ | Average TPM Logarithmic | Average TPM Stationary | Annotation description |
|---|---|---|---|---|---|---|---|---|---|---|---|
| *PSPTO_4705* | *corS* | + | 5331800 | 5333275 | 9.61 | 17.79 | 5.42 | 8.88 | 17.02 | 10.98 | Sensor histidine kinase CorS |
| *PSPTO_4706* | *corP* | - | 5333538 | 5333711 | 19.27 | 15.40 | 1.83 | 12.03 | 25.01 | 16.18 | Response regulator CorP |
| *PSPTO_4707* | *cmaD* | + | 5333913 | 5334143 | 14.01 | 251.54 | 91.14 | 9.77 | 27.18 | 15.39 | Coronamic acid synthetase CmaD |
| *PSPTO_4708* | *cmaE* | + | 5334154 | 5335002 | 32.77 | 234.65 | 89.44 | 26.22 | 56.70 | 20.82 | Coronamic acid synthetase CmaE |
| *PSPTO_4709* | *cmaA* | + | 5334999 | 5336786 | 11.89 | 208.72 | 120.59 | 16.84 | 54.74 | 16.00 | Coronamic acid synthetase CmaA |
| *PSPTO_4710* | *cmaB* | + | 5336773 | 5337702 | 21.05 | 452.47 | 243.74 | 44.66 | 113.65 | 25.68 | Coronamic acid synthetase CmaB |
| *PSPTO_4711* | *cmaC* | + | 5337712 | 5338134 | 5.73 | 99.66 | 62.69 | 0.88 | 26.25 | 8.59 | Coronamic acid synthetase CmaC |
| *PSPTO_4712* | *cmaT* | + | 5338144 | 5338905 | 6.80 | 79.82 | 35.73 | 12.83 | 25.43 | 10.11 | Coronamic acid synthetase, thioesterase component |
| *PSPTO_4713* | | + | 5338911 | 5339549 | 5.40 | 47.87 | 33.20 | 2.91 | 18.53 | 10.96 | Alanyl tRNA synthetase-related protein |
| *PSPTO_4714* | *cmaU* | + | 5339568 | 5340179 | 26.98 | 52.27 | 32.96 | 5.27 | 38.02 | 26.54 | CmaU protein |
| *PSPTO_4716* | | + | 5341362 | 5341859 | 1.18 | 11.25 | 0.00 | 0.00 | 6.17 | 3.99 | Hypothetical protein |
| *PSPTO_4717* | | + | 5341867 | 5342136 | 4.53 | 2.43 | 0.58 | 0.00 | 6.27 | 4.39 | Hypothetical protein |
| *PSPTO_4718* | *hopAA1-2* | - | 5342748 | 5344211 | 64.23 | 122.24 | 86.45 | 45.01 | 42.23 | 47.10 | Type III effector HopAA1-2 |
| *PSPTO_4719* | | + | 5344569 | 5344736 | 7.94 | 0.32 | 0.90 | 0.00 | 2.57 | 7.40 | Hypothetical protein |
| *PSPTO_4720* | *hopV1* | - | 5345081 | 5346253 | 12.77 | 118.11 | 63.83 | 17.55 | 12.22 | 15.24 | Type III effector HopV1 |
| *PSPTO_4721* | *shcV* | - | 5346338 | 5346730 | 3.99 | 237.88 | 85.77 | 20.60 | 7.02 | 6.65 | Type III chaperone ShcV |
| *PSPTO_4722* | *hopAO1* | - | 5347118 | 5348524 | 3.30 | 143.25 | 51.26 | 9.68 | 13.72 | 4.56 | Type III effector HopAO1 |
| *PSPTO_4723* | *cmaL* | - | 5349028 | 5349336 | 13.87 | 456.27 | 211.06 | 28.81 | 30.13 | 20.81 | |
| *PSPTO_4724* | *hopD* | + | 5350102 | 5350647 | 26.12 | 25.88 | 29.35 | 19.09 | 34.71 | 18.90 | Type III effector HopD |
| *PSPTO_4725* | | + | 5350688 | 5351665 | 17.97 | 22.55 | 20.12 | 16.06 | 36.44 | 28.86 | IS52, transposase |
| *PSPTO_4726* | | + | 5351817 | 5353451 | - | - | - | - | - | - | |
| *PSPTO_4727* | *hopG1* | - | 5353685 | 5355166 | 4.79 | 128.98 | 51.87 | 6.47 | 4.83 | 4.46 | Type III effector HopG1 |
| *PSPTO_4729* | | - | 5356558 | 5357367 | 0.00 | 0.00 | 0.00 | 0.00 | 0.00 | 0.00 | ISPsy4, transposition helper protein |
| *PSPTO_4730* | | - | 5357364 | 5358386 | 0.00 | 0.00 | 0.00 | 0.00 | 0.00 | 0.00 | ISPsy4, transposase |
| *PSPTO_4732* | *hopQ1-2* | + | 5359854 | 5360336 | 0.38 | 6.38 | 10.38 | 0.00 | 1.63 | 0.97 | No hrp box controlling this putaive type III effector. |
| *PSPTO_4733* | | - | 5361115 | 5361654 | 1.39 | 48.12 | 28.25 | 2.36 | 18.26 | 7.43 | Hypothetical protein |
| *PSPTO_4734* | | - | 5362019 | 5362246 | 17.88 | 18.93 | 19.60 | 12.41 | 33.35 | 24.10 | Hypothetical protein |
| *PSPTO_4735* | *hrpB* | - | 5362728 | 5365250 | 51.66 | 78.73 | 70.82 | 61.74 | 243.77 | 216.11 | ATP-dependent helicase HrpB |
| *PSPTO_4737* | | - | 5365761 | 5367314 | 0.00 | 0.00 | 0.00 | 0.00 | 0.00 | 0.00 | ISPsy5, transposase |
| *PSPTO_4738* | | - | 5367332 | 5367691 | 0.00 | 0.00 | 0.00 | 0.00 | 0.00 | 0.00 | ISPsy5, Orf1 |
| *PSPTO_4741* | | - | 5368513 | 5368935 | 0.33 | 1.47 | 5.83 | 0.00 | 3.63 | 0.83 | Hypothetical protein |
| *PSPTO_4742* | | - | 5368939 | 5370987 | 0.48 | 2.43 | 2.42 | 0.00 | 15.18 | 0.69 | Site-specific recombinase, phage integrase family |
| *PSPTO_4743* | | - | 5370987 | 5372888 | 0.41 | 2.63 | 1.39 | 0.00 | 17.33 | 0.59 | Hypothetical protein |
| *PSPTO_4744* | | - | 5372875 | 5374362 | 4.88 | 10.71 | 13.60 | 5.08 | 62.63 | 15.12 | Site-specific recombinase, phage integrase family |
| *PSPTO_4745* | | - | 5374442 | 5376511 | 12.77 | 4.30 | 5.67 | 2.20 | 6.43 | 4.12 | ATP-dependent helicase HrpB |
| *PSPTO_4746* | | + | 5378945 | 5381041 | 4.16 | 7.61 | 3.20 | 2.76 | 77.68 | 9.20 | Site-specific recombinase, phage integrase family domain protein |
| *PSPTO_4747* | | + | 5381038 | 5381676 | 4.23 | 5.39 | 7.93 | 3.23 | 46.33 | 6.66 | Hypothetical protein |
| *PSPTO_4748* | | + | 5381679 | 5383103 | 14.80 | 15.28 | 13.14 | 4.68 | 58.44 | 16.60 | Site-specific recombinase, phage integrase family |
| *PSPTO_4749* | | - | 5383175 | 5384062 | - | - | - | - | - | - | |
| *PSPTO_4750* | | - | 5384172 | 5384651 | 5.89 | 12.39 | 4.84 | 7.68 | 16.36 | 9.16 | Hypothetical protein |
| *PSPTO_4751* | | - | 5384766 | 5387489 | 11.90 | 22.30 | 26.41 | 12.34 | 40.88 | 28.55 | UvrD/REP helicase family protein |
| *PSPTO_4752* | | - | 5388074 | 5389039 | 104.64 | 134.81 | 97.02 | 48.13 | 109.31 | 152.02 | Hypothetical protein |
| *PSPTO_4753* | | + | 5389184 | 5390275 | 18.03 | 24.40 | 19.93 | 5.41 | 50.92 | 29.44 | Hypothetical protein |
| *PSPTO_4754* | | - | 5391079 | 5391924 | 19.06 | 20.30 | 21.38 | 14.94 | 63.51 | 22.40 | Hypothetical protein |
| *PSPTO_4755* | | + | 5392110 | 5392484 | 18.41 | 30.91 | 24.65 | 21.47 | 66.45 | 25.31 | Hypothetical protein |
| *PSPTO_4756* | | - | 5393362 | 5393496 | 4.93 | 0.00 | 0.00 | 0.00 | 6.34 | 5.26 | Hypothetical protein |
| *PSPTO_4757* | | + | 5393657 | 5393878 | 0.55 | 0.00 | 0.00 | 0.00 | 4.35 | 1.66 | Hypothetical protein |
| *PSPTO_4758* | | + | 5394944 | 5395078 | 0.00 | 0.00 | 0.00 | 0.00 | 6.36 | 1.51 | Hypothetical protein |
| *PSPTO_4759* | | - | 5395243 | 5395767 | 4.21 | 0.00 | 3.41 | 0.00 | 2.24 | 2.93 | Hypothetical protein |
| *PSPTO_4760* | | + | 5396264 | 5396704 | 23.17 | 71.24 | 43.04 | 46.35 | 92.49 | 63.61 | DNA-binding protein |
| *PSPTO_4761* | | + | 5397049 | 5399070 | 15.75 | 33.48 | 27.25 | 16.93 | 90.86 | 31.31 | Hypothetical protein |
| *PSPTO_4762* | | + | 5399103 | 5399777 | 6.80 | 15.01 | 24.33 | 3.40 | 40.86 | 16.75 | Von Willebrand factor type A domain-containing protein |
| *PSPTO_4763* | | + | 5399781 | 5400563 | 5.71 | 8.87 | 11.05 | 5.34 | 20.19 | 7.71 | Hypothetical protein |
| *PSPTO_4764* | | - | 5402565 | 5404118 | 0.47 | 0.00 | 0.00 | 0.00 | 0.75 | 0.49 | ISPsy5, transposase |
| *PSPTO_4765* | | - | 5404136 | 5404495 | 0.00 | 0.00 | 0.00 | 0.00 | 0.00 | 0.00 | ISPsy5, Orf1 |
| *PSPTO_4766* | | - | 5404538 | 5404846 | 87.37 | 61.14 | 47.06 | 2.94 | 54.58 | 17.49 | Hypothetical protein |
| *PSPTO_4767* | | - | 5405456 | 5406439 | 1.08 | 1.76 | 1.66 | 0.00 | 1.26 | 1.41 | Pentapeptide repeat-containing protein |
| *PSPTO_4768* | | - | 5406651 | 5407460 | 0.00 | 0.00 | 0.00 | 0.00 | 0.00 | 0.00 | ISPsy4, transposition helper protein |
| *PSPTO_4769* | | - | 5407457 | 5408479 | 0.00 | 0.00 | 0.00 | 0.00 | 0.00 | 0.00 | ISPsy4, transposase |
| *PSPTO_4770* | | - | 5409688 | 5410536 | 14.91 | 12.71 | 9.95 | 9.23 | 42.59 | 22.87 | Hypothetical protein |
| *PSPTO_4771* | | + | 5410639 | 5411769 | 13.72 | 11.18 | 12.69 | 4.84 | 15.35 | 13.62 | Hypothetical protein |
| *PSPTO_4772* | | + | 5412009 | 5413049 | 14.00 | 16.31 | 20.97 | 11.34 | 20.81 | 26.43 | Hypothetical protein |
| *PSPTO_4773* | | - | 5413440 | 5413844 | 7.50 | 15.20 | 2.41 | 0.00 | 12.93 | 9.61 | ATP-dependent helicase, N-terminal domain protein |
| *PSPTO_4774* | | - | 5413895 | 5415073 | 14.87 | 16.98 | 24.44 | 7.28 | 37.32 | 18.53 | Membrane protein |
| *PSPTO_4775* | | + | 5415268 | 5416608 | 8.74 | 40.19 | 23.94 | 3.03 | 31.29 | 9.03 | GntR family transcriptional regulator/aminotransferase, class I |
| *PSPTO_4776* | *hopI1* | - | 5416707 | 5418173 | 5.43 | 417.99 | 230.88 | 32.77 | 13.22 | 6.74 | Type III effector HopI1 |
| *PSPTO_4777* | | - | 5418553 | 5419176 | 2.70 | 10.46 | 5.68 | 2.86 | 13.90 | 2.35 | Glutathione S-transferase family protein |
| *PSPTO_4778* | | - | 5419226 | 5419609 | 7.11 | 28.01 | 28.50 | 5.59 | 78.62 | 14.63 | Endoribonuclease L-PSP family protein |
| *PSPTO_4779* | | + | 5419777 | 5420472 | 3.84 | 6.89 | 11.96 | 2.04 | 17.12 | 8.40 | Hypothetical protein |
| *PSPTO_4780* | | + | 5420592 | 5421131 | 39.20 | 66.83 | 32.77 | 24.17 | 33.28 | 43.58 | Hypothetical protein |
| *PSPTO_4781* | | - | 5421139 | 5422152 | 19.00 | 22.61 | 34.20 | 15.69 | 40.23 | 29.71 | Hypothetical protein |
| *PSPTO_4782* | *gal* | - | 5422185 | 5423111 | 16.76 | 20.73 | 32.87 | 14.55 | 33.12 | 19.76 | D-galactose 1-dehydrogenase |
| *PSPTO_4784* | | + | 5424383 | 5425483 | 2.55 | 8.38 | 4.37 | 3.50 | 16.67 | 6.65 | Diguanylate cyclase |
| *PSPTO_4785* | | + | 5425544 | 5425654 | 0.46 | 0.85 | 4.20 | 0.00 | 2.50 | 4.03 | Hypothetical protein |
| *PSPTO_4786* | | + | 5425668 | 5426981 | 5.41 | 1.46 | 3.00 | 2.69 | 5.76 | 6.64 | Methyl-accepting chemotaxis protein |
| *PSPTO_4787* | | + | 5427008 | 5427757 | 7.59 | 9.82 | 3.61 | 3.73 | 9.91 | 7.99 | Sensor histidine kinase |
| *PSPTO_4788* | *kptA* | + | 5427872 | 5428435 | 59.01 | 49.72 | 55.39 | 17.58 | 158.21 | 58.86 | RNA 2'-phosphotransferase |
| *PSPTO_4789* | | + | 5428800 | 5429219 | 56.58 | 65.07 | 48.24 | 44.90 | 99.18 | 47.11 | Hypothetical protein |
| *PSPTO_4790* | | + | 5429235 | 5430128 | 8.05 | 29.02 | 11.22 | 4.52 | 34.98 | 21.26 | Cation efflux family protein |

| Locus name | Gene name | Strand | Gene start (bp) | Gene end (bp) | Average TPM Inoculum | Average TPM $t_6$ | Average TPM $t_{24}$ | Average TPM $t_{168}$ | Average TPM Logarithmic | Average TPM Stationary | Annotation description |
|---|---|---|---|---|---|---|---|---|---|---|---|
| *PSPTO_4791* | | - | 5430135 | 5431466 | 67.17 | 62.48 | 53.96 | 41.96 | 77.56 | 63.98 | Hypothetical protein |
| *PSPTO_4792* | | - | 5431547 | 5433007 | 0.24 | 0.36 | 0.10 | 0.34 | 0.41 | 0.32 | ISPsy6, transposase |
| *PSPTO_4793* | | + | 5433349 | 5433828 | 6.37 | 1.76 | 7.82 | 2.18 | 30.86 | 5.25 | AsnC family transcriptional regulator |
| *PSPTO_4794* | *rluE* | - | 5433842 | 5434474 | 4.31 | 10.06 | 9.45 | 4.99 | 40.77 | 12.66 | Ribosomal large subunit pseudouridine synthase E |
| *PSPTO_4795* | | - | 5434554 | 5436017 | 5.95 | 19.86 | 11.47 | 10.79 | 62.11 | 12.92 | AMP nucleosidase |
| *PSPTO_4796* | | - | 5436176 | 5438182 | 29.63 | 11.93 | 9.99 | 3.38 | 6.24 | 12.72 | Sensor histidine kinase |
| *PSPTO_4797* | | - | 5438118 | 5438336 | 30.93 | 20.23 | 18.76 | 56.70 | 8.12 | 64.23 | Hypothetical protein |
| *PSPTO_4798* | | + | 5438617 | 5439414 | 33.73 | 38.29 | 47.82 | 36.82 | 52.57 | 24.65 | Bifunctional hydroxy-methylpyrimidine kinase/ hydroxy-phosphomethylpyrimidine kinase |
| *PSPTO_4799* | *thiE* | + | 5439438 | 5440055 | 8.57 | 17.26 | 22.15 | 7.24 | 23.73 | 14.07 | Thiamine-phosphate pyrophosphorylase |
| *PSPTO_4800* | *hemL* | + | 5440261 | 5441544 | 5.32 | 21.41 | 15.74 | 17.97 | 63.76 | 17.36 | Glutamate-1-semialdehyde-2,1-aminomutase |
| *PSPTO_4801* | | + | 5441619 | 5441774 | 59.78 | 74.40 | 32.49 | 6.02 | 57.50 | 39.04 | Hypothetical protein |
| *PSPTO_4802* | | + | 5441840 | 5442388 | 39.67 | 42.48 | 43.98 | 28.32 | 71.30 | 45.08 | Hypothetical protein |
| *PSPTO_4803* | | - | 5442431 | 5442757 | 8.73 | 17.71 | 13.15 | 8.53 | 21.07 | 13.21 | Hypothetical protein |
| *PSPTO_4804* | *miaB* | + | 5442888 | 5444216 | 23.85 | 81.20 | 57.20 | 44.81 | 158.56 | 160.55 | tRNA-i(6)A37 modification enzyme MiaB |
| *PSPTO_4805* | | + | 5444394 | 5445416 | 44.79 | 93.81 | 74.26 | 60.05 | 282.29 | 108.26 | PhoH-like protein |
| *PSPTO_4806* | | + | 5445409 | 5445909 | 288.07 | 130.57 | 129.29 | 84.11 | 412.29 | 109.52 | Hypothetical protein |
| *PSPTO_4807* | | + | 5445915 | 5446757 | 59.02 | 53.71 | 38.86 | 18.64 | 123.67 | 35.78 | Metal ion transporter |
| *PSPTO_4808* | *cutE* | + | 5446799 | 5448319 | 5.31 | 6.06 | 6.08 | 1.31 | 36.64 | 8.87 | Apolipoprotein N-acyltransferase |
| *PSPTO_4809* | | - | 5448371 | 5449141 | 24.72 | 23.98 | 24.89 | 16.19 | 28.24 | 19.14 | Hypothetical protein |
| *PSPTO_4810* | | - | 5449212 | 5449562 | 15.94 | 10.40 | 8.36 | 13.04 | 1.94 | 5.39 | Hypothetical protein |
| *PSPTO_4811* | | - | 5449572 | 5449691 | 13.10 | 21.08 | 0.00 | 8.78 | 8.03 | 5.22 | Hypothetical protein |
| *PSPTO_4812* | *leuS* | + | 5449874 | 5452480 | 8.38 | 33.08 | 23.01 | 19.78 | 128.23 | 22.81 | Leucyl-tRNA synthetase |
| *PSPTO_4813* | | + | 5452570 | 5453175 | 19.99 | 72.12 | 48.65 | 41.37 | 418.52 | 70.13 | Lipoprotein |
| *PSPTO_4814* | *holA* | + | 5453217 | 5454254 | 2.28 | 12.45 | 9.76 | 6.21 | 49.19 | 7.99 | DNA polymerase III subunit delta |
| *PSPTO_4815* | | + | 5454259 | 5454465 | 9.65 | 44.14 | 16.90 | 4.05 | 61.53 | 14.21 | Hypothetical protein |
| *PSPTO_4816* | | + | 5454589 | 5454786 | 579.55 | 265.92 | 232.83 | 154.33 | 68.77 | 57.99 | Hypothetical protein |
| *PSPTO_4817* | | + | 5454870 | 5456207 | 21.64 | 40.00 | 41.30 | 23.69 | 90.10 | 33.85 | HopAJ2 protein |
| *PSPTO_4818* | *lipA* | - | 5456472 | 5457428 | 44.56 | 45.85 | 43.54 | 31.59 | 98.10 | 42.01 | Lipoic acid synthetase |
| *PSPTO_4819* | *lipB* | - | 5457472 | 5458128 | 79.06 | 109.18 | 75.30 | 43.34 | 176.49 | 72.08 | Lipoate-protein ligase B |
| *PSPTO_4820* | | - | 5458128 | 5458403 | 13.80 | 15.47 | 35.91 | 23.95 | 83.87 | 20.69 | Hypothetical protein |
| *PSPTO_4821* | | - | 5458494 | 5459654 | 10.90 | 35.28 | 30.11 | 13.80 | 171.14 | 22.87 | D-alanyl-D-alanine carboxypeptidase |
| *PSPTO_4822* | | - | 5459896 | 5460924 | 20.21 | 45.53 | 29.09 | 14.79 | 154.11 | 31.14 | Lipoprotein, rlpA family |
| *PSPTO_4823* | *mltB* | - | 5460924 | 5461931 | 14.16 | 15.81 | 16.24 | 6.49 | 60.86 | 12.65 | Membrane-bound lytic murein transglycosylase B |
| *PSPTO_4824* | *rodA* | - | 5461949 | 5463094 | 6.51 | 8.35 | 6.94 | 11.25 | 34.21 | 4.28 | Rod-shape-determining protein RodA |
| *PSPTO_4825* | | - | 5463084 | 5464979 | 2.31 | 5.76 | 5.08 | 8.44 | 53.82 | 5.87 | Penicillin-binding protein |
| *PSPTO_4826* | | - | 5465009 | 5465476 | 4.35 | 19.79 | 32.80 | 5.01 | 101.22 | 18.05 | Hypothetical protein |
| *PSPTO_4827* | | - | 5465486 | 5465857 | 3.12 | 15.60 | 32.82 | 7.13 | 59.52 | 25.24 | Iojap-like protein |
| *PSPTO_4828* | *nadD* | - | 5465933 | 5466601 | 5.04 | 28.93 | 16.16 | 25.91 | 81.34 | 27.21 | Nicotinate (nicotinamide) nucleotide adenylyltransferase |
| *PSPTO_4829* | *proA* | - | 5466615 | 5467880 | 22.54 | 46.96 | 38.34 | 24.51 | 79.74 | 39.68 | Gamma-glutamyl phosphate reductase |
| *PSPTO_4830* | | + | 5468163 | 5468834 | 16.12 | 61.03 | 62.11 | 8.71 | 108.68 | 26.60 | 3-methyladenine DNA glycosylase |
| *PSPTO_4831* | | + | 5468838 | 5470154 | 5.04 | 12.28 | 9.76 | 3.15 | 17.93 | 8.69 | PAP2 superfamily protein/DedA family protein |
| *PSPTO_4832* | | + | 5470302 | 5472320 | 8.04 | 13.03 | 11.62 | 3.78 | 5.14 | 11.65 | Hypothetical protein |
| *PSPTO_4833* | | - | 5472317 | 5475016 | 5.22 | 2.59 | 5.43 | 2.89 | 9.67 | 9.92 | Sensory box histidine kinase |
| *PSPTO_4834* | | - | 5475075 | 5475530 | 40.48 | 24.01 | 32.08 | 20.29 | 25.12 | 37.31 | Hypothetical protein |
| *PSPTO_4835* | | + | 5475603 | 5476076 | 11.28 | 17.60 | 8.06 | 5.50 | 13.94 | 11.26 | Hypothetical protein |
| *PSPTO_4836* | | + | 5476234 | 5477016 | 9.64 | 15.39 | 13.48 | 14.54 | 30.58 | 18.25 | DNA binding response regulator, LuxR family |
| *PSPTO_4837* | | - | 5477023 | 5478678 | 3.77 | 7.32 | 6.84 | 3.94 | 7.89 | 4.93 | Response regulator |
| *PSPTO_4838* | | + | 5478894 | 5479043 | 487.74 | 299.69 | 114.10 | 139.53 | 181.85 | 202.08 | Hypothetical protein |
| *PSPTO_4839* | | + | 5479006 | 5480085 | 94.41 | 114.61 | 83.19 | 45.08 | 77.41 | 104.16 | Hypothetical protein |
| *PSPTO_4840* | | + | 5480085 | 5480627 | 25.90 | 40.47 | 26.42 | 20.76 | 48.12 | 63.67 | MutT/nudix family protein |
| *PSPTO_4841* | *sui1* | + | 5480837 | 5481208 | 42.98 | 37.77 | 42.22 | 18.59 | 68.48 | 53.69 | Translation initiation factor Sui1 |
| *PSPTO_4842* | *speA* | + | 5481297 | 5483210 | 7.53 | 21.14 | 21.43 | 10.87 | 68.25 | 21.83 | Biosynthetic arginine decarboxylase |
| *PSPTO_4843* | | + | 5483479 | 5484387 | 125.92 | 75.51 | 53.19 | 14.00 | 57.98 | 69.26 | Esterase/lipase/thioesterase family protein |
| *PSPTO_4844* | *dinF* | - | 5484381 | 5485721 | 3.42 | 8.11 | 12.55 | 0.69 | 11.50 | 12.21 | DNA-damage-inducible protein F |
| *PSPTO_4845* | | + | 5485965 | 5490914 | 3.36 | 4.25 | 3.02 | 5.60 | 24.44 | 5.62 | Lipoprotein |
| *PSPTO_4846* | | - | 5491161 | 5491454 | 27.12 | 20.32 | 24.38 | 23.92 | 28.33 | 13.47 | Hypothetical protein |
| *PSPTO_4847* | | + | 5491550 | 5493898 | 2.18 | 4.22 | 1.19 | 0.62 | 11.28 | 3.58 | Penicillin-binding protein |
| *PSPTO_4848* | | - | 5494003 | 5494410 | 124.22 | 109.02 | 100.87 | 83.28 | 194.52 | 90.94 | Response regulator |
| *PSPTO_4849* | | - | 5494451 | 5494657 | 258.88 | 110.25 | 99.07 | 106.60 | 78.93 | 136.92 | Hypothetical protein |
| *PSPTO_4850* | | + | 5494865 | 5495938 | 20.18 | 12.16 | 13.86 | 5.19 | 44.80 | 15.04 | Hypothetical protein |
| *PSPTO_4851* | | + | 5495959 | 5497218 | 3.76 | 1.74 | 4.85 | 3.30 | 17.12 | 4.44 | Type II/III secretion system protein |
| *PSPTO_4852* | | + | 5497215 | 5498405 | 4.65 | 1.84 | 2.36 | 0.92 | 17.22 | 4.29 | Hypothetical protein |
| *PSPTO_4853* | | + | 5498405 | 5499661 | 5.19 | 5.79 | 5.37 | 0.00 | 15.21 | 3.07 | Type II/IV secretion system protein |
| *PSPTO_4854* | | + | 5499680 | 5500567 | 6.23 | 3.57 | 1.09 | 0.00 | 13.20 | 3.17 | Hypothetical protein |
| *PSPTO_4855* | | + | 5500569 | 5501453 | 6.50 | 2.50 | 1.32 | 1.78 | 9.61 | 3.62 | Membrane protein |
| *PSPTO_4856* | | + | 5501495 | 5502247 | 13.22 | 8.39 | 10.25 | 3.46 | 30.12 | 8.84 | TPR domain-containing protein |
| *PSPTO_4857* | | + | 5502244 | 5502516 | 17.90 | 16.22 | 7.42 | 0.00 | 73.16 | 23.11 | Hypothetical protein |
| *PSPTO_4858* | *dsbD* | + | 5502597 | 5504411 | 5.66 | 12.87 | 14.49 | 5.81 | 25.63 | 9.04 | Thiol:disulfide interchange protein DsbD |
| *PSPTO_4859* | *aroQ* | + | 5504541 | 5504993 | 48.11 | 81.19 | 71.76 | 37.57 | 160.05 | 22.29 | 3-dehydroquinate dehydratase |
| *PSPTO_4860* | *accB* | + | 5505070 | 5505525 | 53.84 | 91.02 | 85.37 | 86.25 | 616.35 | 48.92 | Acetyl-CoA carboxylase, biotin carboxyl carrier protein |
| *PSPTO_4861* | *accC-2* | + | 5505543 | 5506886 | 14.23 | 43.04 | 39.22 | 22.33 | 178.50 | 14.08 | Acetyl-CoA carboxylase, biotin carboxylase |
| *PSPTO_4862* | *prmA* | + | 5507022 | 5507900 | 8.43 | 34.83 | 17.11 | 17.84 | 98.94 | 38.92 | Ribosomal protein L11 methyltransferase |
| *PSPTO_4863* | | + | 5507925 | 5509286 | 4.84 | 15.71 | 20.73 | 18.72 | 60.05 | 22.12 | Hypothetical protein |
| *PSPTO_4864* | | + | 5509603 | 5510616 | 24.32 | 64.55 | 46.27 | 53.30 | 84.74 | 52.81 | Hypothetical protein |
| *PSPTO_4865* | *fis* | + | 5510619 | 5510933 | 8.41 | 39.41 | 12.88 | 15.97 | 40.88 | 12.03 | Global DNA-binding transcriptional dual regulator Fis |
| *PSPTO_4866* | *purH* | + | 5511016 | 5512620 | 2.93 | 27.48 | 18.50 | 11.52 | 137.86 | 16.66 | Phosphoribosylaminoimidazolecarboxamide formyltransferase/IMP cyclohydrolase |
| *PSPTO_4867* | *purD* | + | 5512819 | 5514114 | 15.46 | 24.80 | 21.40 | 13.59 | 81.05 | 12.92 | Phosphoribosylamine--glycine ligase |
| *PSPTO_4868* | *retS* | + | 5514202 | 5516991 | 5.26 | 10.37 | 7.21 | 4.86 | 38.01 | 8.12 | Sensor histidine kinase/response regulator RetS |
| *PSPTO_4869* | | + | 5517061 | 5517657 | 7.48 | 6.37 | 4.58 | 5.80 | 16.41 | 9.26 | Membrane protein, MarC family |

| Locus name | Gene name | Strand | Gene start (bp) | Gene end (bp) | Average TPM Inoculum | Average TPM $t_6$ | Average TPM $t_{24}$ | Average TPM $t_{168}$ | Average TPM Logarithmic | Average TPM Stationary | Annotation description |
|---|---|---|---|---|---|---|---|---|---|---|---|
| *PSPTO_4870* | | - | 5517664 | 5518299 | 3.27 | 2.42 | 4.73 | 0.00 | 4.48 | 2.95 | Hypothetical protein |
| *PSPTO_4871* | | - | 5518336 | 5518971 | 6.72 | 6.82 | 5.63 | 0.58 | 8.70 | 5.61 | Hypothetical protein |
| *PSPTO_4872* | | - | 5518964 | 5519662 | 6.81 | 0.80 | 0.94 | 6.19 | 5.87 | 3.92 | Hypothetical protein |
| *PSPTO_4873* | | - | 5519721 | 5520530 | 1.22 | 2.80 | 2.25 | 0.91 | 3.16 | 2.18 | Hypothetical protein |
| *PSPTO_4874* | | - | 5520896 | 5522599 | 4.70 | 10.82 | 5.93 | 3.84 | 28.73 | 8.93 | CbiG protein/precorrin-3B C17-methyltransferase |
| *PSPTO_4875* | *cobI* | - | 5522599 | 5523330 | 6.59 | 12.35 | 22.47 | 12.51 | 61.37 | 15.94 | Precorrin-2 C(20)-methyltransferase |
| *PSPTO_4876* | *cobH* | - | 5523330 | 5523956 | 4.09 | 6.75 | 6.68 | 12.14 | 44.07 | 10.96 | Precorrin-8X methylmutase |
| *PSPTO_4877* | *cobG* | - | 5523953 | 5525383 | 6.31 | 18.11 | 17.15 | 10.24 | 30.13 | 14.02 | Cobalamin biosynthesis protein CobG |
| *PSPTO_4878* | *cobL* | + | 5525424 | 5526626 | 1.27 | 7.73 | 8.19 | 5.90 | 17.21 | 9.03 | Precorrin-6Y C5,15-methyltransferase |
| *PSPTO_4879* | *cbiD* | + | 5526626 | 5527738 | 2.91 | 5.71 | 4.78 | 1.52 | 16.82 | 7.04 | Cobalamin biosynthesis protein CbiD |
| *PSPTO_4880* | | + | 5527735 | 5528475 | 2.64 | 6.77 | 7.75 | 3.60 | 36.90 | 9.00 | Cobalt-precorrin-6x reductase |
| *PSPTO_4881* | | + | 5528813 | 5529274 | 24.08 | 46.42 | 46.16 | 15.03 | 117.55 | 47.46 | Hypothetical protein |
| *PSPTO_4882* | | + | 5529288 | 5529722 | 7.60 | 34.83 | 28.07 | 11.46 | 58.58 | 12.46 | Hypothetical protein |
| *PSPTO_4883* | | + | 5529806 | 5531179 | 7.26 | 34.03 | 22.43 | 11.97 | 37.05 | 4.90 | Membrane protein |
| *PSPTO_4884* | | + | 5532595 | 5534901 | 4.24 | 5.65 | 3.07 | 0.45 | 10.36 | 4.69 | Hypothetical protein |
| *PSPTO_4885* | | + | 5535156 | 5536421 | 51.93 | 63.87 | 29.82 | 11.47 | 7.60 | 4.39 | Branched-chain amino acid ABC transporter substrate-binding protein |
| *PSPTO_4886* | | + | 5536635 | 5538224 | 3.78 | 9.84 | 8.69 | 3.68 | 5.16 | 1.79 | Branched-chain amino acid ABC transporter permease |
| *PSPTO_4887* | | + | 5538224 | 5539303 | 6.78 | 19.82 | 6.27 | 2.80 | 4.41 | 3.40 | Branched-chain amino acid ABC transporter permease |
| *PSPTO_4888* | | + | 5539300 | 5540160 | 3.74 | 16.77 | 8.06 | 0.00 | 4.29 | 1.54 | Branched-chain amino acid ABC transporter ATP-binding protein |
| *PSPTO_4889* | | + | 5540278 | 5540976 | 15.17 | 16.64 | 20.24 | 5.14 | 9.55 | 7.25 | Branched-chain amino acid ABC transporter ATP-binding protein |
| *PSPTO_4890* | *ureD-2* | + | 5541025 | 5541870 | 10.64 | 21.57 | 11.18 | 5.95 | 26.33 | 11.00 | Urease accessory protein UreD |
| *PSPTO_4891* | *ureA* | + | 5541956 | 5542258 | 14.55 | 39.30 | 37.55 | 9.14 | 49.90 | 17.07 | Urease subunit gamma |
| *PSPTO_4892* | | + | 5542269 | 5542811 | 12.86 | 28.30 | 15.07 | 9.91 | 35.52 | 13.90 | Phosphinothricin N-acetyltransferase |
| *PSPTO_4893* | *ttr* | + | 5542822 | 5543355 | 9.72 | 37.97 | 22.33 | 16.17 | 36.73 | 15.12 | Tabtoxin resistance protein |
| *PSPTO_4894* | *ureB* | + | 5543352 | 5543660 | 3.17 | 13.51 | 7.25 | 18.44 | 34.39 | 8.85 | Urease subunit beta |
| *PSPTO_4895* | *ureC* | + | 5543886 | 5545586 | 8.32 | 28.94 | 13.56 | 11.73 | 61.11 | 17.44 | Urease subunit alpha |
| *PSPTO_4896* | | + | 5545779 | 5546909 | 87.15 | 45.80 | 35.65 | 30.97 | 110.36 | 51.54 | Sensor histidine kinase |
| *PSPTO_4897* | *cbpA* | - | 5546945 | 5547889 | 32.30 | 30.46 | 23.31 | 14.56 | 102.95 | 48.03 | Curved-DNA-binding protein |
| *PSPTO_4898* | *yegD* | + | 5548200 | 5549465 | 2.62 | 10.74 | 3.56 | 5.08 | 24.58 | 5.71 | Heat shock protein YegD |
| *PSPTO_4899* | | - | 5549550 | 5550560 | 11.48 | 18.23 | 7.42 | 8.84 | 31.72 | 20.68 | Membrane protein |
| *PSPTO_4900* | *psiF* | - | 5550749 | 5551039 | 242.23 | 111.54 | 120.70 | 67.55 | 9.43 | 81.28 | Phosphate starvation-inducible protein PsiF |
| *PSPTO_4901* | | + | 5551193 | 5552074 | 5.26 | 7.23 | 6.81 | 4.10 | 9.12 | 6.88 | Membrane protein |
| *PSPTO_4902* | | - | 5552071 | 5552304 | 6.47 | 0.74 | 7.45 | 0.00 | 3.86 | 4.56 | Hypothetical protein |
| *PSPTO_4903* | *osmE* | + | 5552577 | 5552915 | 50.00 | 142.45 | 101.81 | 69.72 | 59.34 | 30.82 | DNA-binding transcriptional activator OsmE |
| *PSPTO_4905* | | - | 5553785 | 5554186 | 62.22 | 81.41 | 65.24 | 31.48 | 58.65 | 63.40 | ISPsy11, transposase OrfA |
| *PSPTO_4906* | | + | 5554594 | 5555133 | 36.64 | 382.93 | 338.57 | 236.31 | 29.43 | 28.90 | Bacterioferritin |
| *PSPTO_4907* | | - | 5555300 | 5557366 | 9.28 | 17.15 | 12.89 | 6.57 | 20.19 | 12.57 | Hypothetical protein |
| *PSPTO_4908* | | + | 5557588 | 5558220 | 13.99 | 32.86 | 31.75 | 8.58 | 102.17 | 22.29 | TetR family transcriptional regulator |
| *PSPTO_4909* | *ureE* | + | 5558405 | 5558905 | 16.58 | 25.41 | 14.13 | 0.78 | 8.94 | 3.33 | Urease accessory protein UreE |
| *PSPTO_4910* | *ureF-2* | + | 5558902 | 5559576 | 6.94 | 34.65 | 20.42 | 1.64 | 14.84 | 3.92 | Urease accessory protein UreF |
| *PSPTO_4911* | *ureG-2* | + | 5559597 | 5560214 | 102.05 | 63.29 | 42.08 | 21.20 | 12.30 | 8.20 | Urease accessory protein UreG |
| *PSPTO_4912* | | + | 5560239 | 5560811 | 10.27 | 22.44 | 7.89 | 14.65 | 6.29 | 3.16 | Urease accessory protein |
| *PSPTO_4913* | | + | 5561015 | 5562229 | 12.07 | 16.39 | 15.30 | 6.94 | 40.46 | 12.79 | Hypothetical protein |
| *PSPTO_4914* | | - | 5562260 | 5563024 | 5.67 | 39.82 | 29.22 | 12.47 | 59.95 | 21.17 | Short chain dehydrogenase/reductase family oxidoreductase |
| *PSPTO_4915* | | - | 5563744 | 5564460 | 7.30 | 8.92 | 3.37 | 4.21 | 16.31 | 3.67 | High affinity branched-chain amino acid ABC transporter ATP-binding protein |
| *PSPTO_4916* | | - | 5564457 | 5565332 | 9.05 | 10.94 | 8.38 | 10.05 | 20.95 | 3.75 | High affinity branched-chain amino acid ABC transporter ATP-binding protein |
| *PSPTO_4917* | *braE* | - | 5565329 | 5566642 | 12.95 | 15.57 | 10.57 | 4.38 | 16.88 | 7.25 | High-affinity branched-chain amino acid ABC transporter, permease protein BraE |
| *PSPTO_4918* | *braD* | - | 5566645 | 5567559 | 7.95 | 4.89 | 6.32 | 6.91 | 11.62 | 3.09 | High-affinity branched-chain amino acid ABC transporter, permease protein BraD |
| *PSPTO_4919* | | - | 5567722 | 5568864 | 80.34 | 37.69 | 28.16 | 47.54 | 48.44 | 10.89 | High affinity branched-chain amino acid ABC transporter periplasmic amino acid-binding protein |
| *PSPTO_4920* | | + | 5569357 | 5570262 | 5.17 | 10.13 | 10.11 | 6.51 | 26.95 | 13.74 | LysR family transcriptional regulator |
| *PSPTO_4921* | *pncB* | + | 5570362 | 5571585 | 16.54 | 12.05 | 5.85 | 11.23 | 39.99 | 13.55 | Nicotinate phosphoribosyltransferase |
| *PSPTO_4922* | *nadE* | + | 5571612 | 5572439 | 34.05 | 47.74 | 39.36 | 23.77 | 91.39 | 48.36 | NH(3)-dependent NAD(+) synthetase |
| *PSPTO_4923* | *azu* | - | 5572680 | 5573090 | 41.24 | 85.85 | 45.21 | 27.63 | 123.73 | 82.27 | Azurin |
| *PSPTO_4924* | | - | 5573283 | 5574176 | 7.87 | 8.92 | 1.30 | 3.65 | 24.25 | 8.32 | Transglutaminase-like domain protein |
| *PSPTO_4925* | | - | 5574176 | 5576671 | 9.24 | 5.32 | 4.50 | 3.98 | 14.95 | 3.52 | Hypothetical protein |
| *PSPTO_4926* | | - | 5576759 | 5580037 | 10.85 | 15.81 | 10.21 | 5.85 | 21.99 | 9.04 | Transglutaminase-like superfamily domain protein |
| *PSPTO_4927* | | + | 5580309 | 5583053 | 10.08 | 9.45 | 6.44 | 6.64 | 14.55 | 10.62 | Hypothetical protein |
| *PSPTO_4928* | | - | 5583101 | 5585416 | 28.80 | 53.95 | 58.64 | 57.41 | 85.34 | 32.10 | Hypothetical protein |
| *PSPTO_4929* | *dnaB* | - | 5585537 | 5586931 | 11.20 | 17.77 | 14.62 | 5.88 | 49.30 | 22.81 | Replicative DNA helicase |
| *PSPTO_4930* | *rplI* | - | 5587055 | 5587501 | 23.79 | 123.50 | 123.95 | 104.20 | 499.05 | 83.39 | 50S ribosomal protein L9 |
| *PSPTO_4931* | | - | 5587521 | 5588417 | 20.50 | 72.69 | 82.60 | 39.26 | 531.89 | 35.02 | Membrane protein |
| *PSPTO_4932* | *rpsR* | - | 5588457 | 5588687 | 42.06 | 202.92 | 176.68 | 174.54 | 822.95 | 82.42 | 30S ribosomal protein S18 |
| *PSPTO_4933* | *rpsF* | - | 5588716 | 5589141 | 78.25 | 457.13 | 391.87 | 240.46 | 1564.44 | 142.65 | 30S ribosomal protein S6 |
| *PSPTO_4934* | | - | 5589356 | 5590108 | 17.01 | 23.02 | 39.21 | 28.13 | 59.41 | 29.93 | TrmH family RNA methyltransferase |
| *PSPTO_4935* | *vacB* | - | 5590105 | 5592738 | 27.61 | 89.24 | 62.19 | 34.77 | 171.35 | 51.58 | Ribonuclease R |
| *PSPTO_4936* | | - | 5593402 | 5595336 | 9.27 | 6.31 | 3.28 | 2.98 | 7.05 | 3.58 | Methyl-accepting chemotaxis protein |
| *PSPTO_4937* | *purA* | - | 5595509 | 5596801 | 4.44 | 8.40 | 12.05 | 5.34 | 69.09 | 7.31 | Adenylosuccinate synthetase |
| *PSPTO_4938* | | - | 5596856 | 5598043 | 7.46 | 13.40 | 8.40 | 9.03 | 28.23 | 8.86 | ATP phosphoribosyltransferase regulatory subunit |
| *PSPTO_4939* | *hflC* | - | 5598312 | 5599181 | 19.94 | 53.19 | 40.58 | 23.02 | 128.85 | 48.73 | HflC protein |
| *PSPTO_4940* | *hflK* | - | 5599181 | 5600368 | 24.78 | 93.05 | 67.27 | 33.48 | 194.04 | 62.61 | HflK protein |
| *PSPTO_4941* | *hflX* | - | 5600465 | 5601766 | 34.72 | 56.04 | 57.89 | 35.94 | 202.02 | 93.32 | GTP-binding protein HflX |
| *PSPTO_4942* | *hfq* | - | 5601779 | 5602039 | 291.37 | 322.85 | 288.56 | 176.53 | 858.36 | 409.21 | Hfq protein |
| *PSPTO_4943* | *miaA* | - | 5602132 | 5603103 | 110.90 | 82.60 | 65.76 | 47.57 | 100.90 | 61.43 | TRNA delta(2)-isopentenylpyrophosphate transferase |
| *PSPTO_4944* | *mutL* | - | 5603104 | 5605041 | 7.45 | 19.21 | 13.66 | 8.77 | 61.72 | 23.90 | DNA mismatch repair protein MutL |
| *PSPTO_4945* | | - | 5605041 | 5606456 | 8.36 | 20.76 | 20.49 | 11.74 | 54.19 | 28.66 | N-acetylmuramoyl-L-alanine amidase family protein |
| *PSPTO_4946* | | - | 5606471 | 5606899 | 13.25 | 34.08 | 25.55 | 12.25 | 33.38 | 23.28 | Hypothetical protein |
| *PSPTO_4947* | | - | 5606929 | 5608419 | 10.61 | 23.78 | 11.69 | 7.34 | 37.08 | 16.26 | Hypothetical protein |
| *PSPTO_4948* | | + | 5608483 | 5609571 | 7.81 | 26.36 | 11.72 | 5.63 | 39.27 | 14.23 | Iron-sulfur cluster-binding protein |
| *PSPTO_4949* | | - | 5609673 | 5610284 | 4.31 | 10.63 | 7.51 | 0.00 | 22.19 | 7.36 | Membrane protein |

| Locus name | Gene name | Strand | Gene start (bp) | Gene end (bp) | Average TPM Inoculum | Average TPM $t_6$ | Average TPM $t_{24}$ | Average TPM $t_{168}$ | Average TPM Logarithmic | Average TPM Stationary | Annotation description |
|---|---|---|---|---|---|---|---|---|---|---|---|
| *PSPTO_4950* | *orn* | - | 5610381 | 5610917 | 21.51 | 44.00 | 53.09 | 27.38 | 111.92 | 23.86 | Oligoribonuclease |
| *PSPTO_4951* | | + | 5611026 | 5612057 | 17.55 | 36.89 | 36.00 | 18.04 | 85.80 | 30.84 | Hypothetical protein |
| *PSPTO_4952* | *motB* | - | 5612121 | 5613146 | 23.26 | 13.69 | 16.29 | 4.47 | 11.18 | 5.53 | Flagellar motor protein MotB |
| *PSPTO_4953* | *motA-2* | - | 5613150 | 5614001 | 53.00 | 26.85 | 17.93 | 14.02 | 19.56 | 6.47 | Flagellar motor protein MotA |
| *PSPTO_4954* | | + | 5614208 | 5615743 | 35.73 | 25.62 | 29.33 | 23.40 | 73.76 | 52.33 | Hypothetical protein |
| *PSPTO_4955* | | + | 5615789 | 5617630 | 23.63 | 99.00 | 76.59 | 38.07 | 174.08 | 43.82 | Bifunctional thiosulfate sulfurtransferase/phosphatidylserine decarboxylase |
| *PSPTO_4956* | | + | 5617882 | 5619669 | 9.95 | 36.29 | 31.17 | 13.50 | 77.65 | 31.43 | Hypothetical protein |
| *PSPTO_4957* | *serB* | - | 5619706 | 5620962 | 2.48 | 14.44 | 17.03 | 7.91 | 27.54 | 12.48 | ACT domain protein/phosphoserine phosphatase SerB |
| *PSPTO_4958* | | + | 5621051 | 5622592 | 11.65 | 33.63 | 17.62 | 17.89 | 66.91 | 13.74 | HAMP domain protein |
| *PSPTO_4959* | | - | 5622904 | 5623599 | 60.88 | 81.97 | 71.11 | 50.51 | 189.96 | 112.65 | Lipoprotein |
| *PSPTO_4960* | *parC* | - | 5623783 | 5626038 | 17.45 | 12.01 | 10.27 | 11.55 | 55.22 | 11.99 | DNA topoisomerase IV subunit A |
| *PSPTO_4961* | | - | 5626049 | 5626582 | 3.74 | 0.92 | 4.98 | 3.83 | 28.51 | 6.35 | Hypothetical protein |
| *PSPTO_4962* | | - | 5626579 | 5627571 | 13.86 | 17.52 | 15.08 | 4.99 | 46.19 | 19.20 | Hypothetical protein |
| *PSPTO_4963* | *parE* | - | 5627568 | 5629472 | 11.23 | 26.94 | 17.63 | 15.10 | 51.83 | 21.34 | DNA topoisomerase IV subunit B |
| *PSPTO_4964* | | - | 5629562 | 5630179 | 50.57 | 65.70 | 71.27 | 43.16 | 138.56 | 75.56 | Hypothetical protein |
| *PSPTO_4965* | | - | 5630267 | 5631037 | 39.87 | 30.88 | 24.40 | 22.53 | 52.17 | 32.58 | Ser/Thr protein phosphatase |
| *PSPTO_4966* | | - | 5631161 | 5631547 | 0.88 | 6.36 | 3.90 | 0.00 | 4.80 | 3.61 | Hypothetical protein |
| *PSPTO_4967* | | - | 5631682 | 5632344 | 1.23 | 4.23 | 5.45 | 3.51 | 1.69 | 4.74 | Hypothetical protein |
| *PSPTO_4968* | | - | 5633077 | 5633718 | 1.14 | 1.19 | 0.00 | 0.00 | 1.35 | 1.17 | Hypothetical protein |
| *PSPTO_4969* | | + | 5633916 | 5636768 | 3.02 | 7.81 | 5.47 | 2.74 | 11.16 | 4.38 | Hypothetical protein |
| *PSPTO_4970* | | + | 5636765 | 5641774 | 9.08 | 10.71 | 7.25 | 9.04 | 22.15 | 14.62 | YD repeat protein |
| *PSPTO_4971* | | - | 5641904 | 5642881 | 0.00 | 0.00 | 0.00 | 0.00 | 0.00 | 0.00 | ISPssy, transposase |
| *PSPTO_4972* | | - | 5643184 | 5643636 | 25.92 | 13.79 | 20.34 | 4.01 | 58.74 | 20.83 | Hypothetical protein |
| *PSPTO_4973* | | - | 5643627 | 5644244 | 125.59 | 130.67 | 110.10 | 39.99 | 328.14 | 97.61 | MutT/nudix family protein |
| *PSPTO_4974* | | + | 5644453 | 5645199 | 78.69 | 27.48 | 30.84 | 30.43 | 68.07 | 53.18 | Lipoprotein |
| *PSPTO_4975* | | - | 5645168 | 5646457 | 2.63 | 4.12 | 1.99 | 3.85 | 18.43 | 9.10 | Cytosine/purines uracil thiamine allantoin permease |
| *PSPTO_4976* | *thiC* | - | 5646584 | 5648473 | 3.24 | 6.29 | 5.96 | 5.59 | 75.38 | 11.87 | Thiamin biosynthesis protein ThiC |
| *PSPTO_4977* | | + | 5648897 | 5650336 | 24.64 | 37.20 | 30.35 | 25.09 | 114.36 | 37.49 | Outer membrane efflux protein TolC |
| *PSPTO_4978* | *kdtA* | - | 5650394 | 5651737 | 18.49 | 27.76 | 21.64 | 17.28 | 44.48 | 30.85 | 3-deoxy-D-manno-octulosonic-acid transferase |
| *PSPTO_4979* | | + | 5651806 | 5652138 | 15.94 | 24.84 | 16.74 | 9.97 | 34.48 | 18.37 | Multidrug resistance protein |
| *PSPTO_4980* | | + | 5652187 | 5653362 | 68.27 | 102.46 | 71.34 | 41.92 | 97.53 | 40.76 | Oxidoreductase, FAD-binding protein |
| *PSPTO_4981* | | + | 5653359 | 5654174 | 12.42 | 30.54 | 33.80 | 13.46 | 82.29 | 48.33 | Oxidoreductase, aldo/keto reductase family |
| *PSPTO_4982* | | + | 5654211 | 5655167 | 5.38 | 23.60 | 23.43 | 7.24 | 47.80 | 21.56 | Hypothetical protein |
| *PSPTO_4983* | *rfaE* | - | 5655241 | 5656665 | 10.08 | 50.45 | 38.35 | 17.91 | 71.52 | 29.32 | Lipopolysaccharide biosynthesis protein RfaE |
| *PSPTO_4984* | *msbA* | - | 5656790 | 5658664 | 11.65 | 30.79 | 24.89 | 18.56 | 59.79 | 25.39 | Lipid A ABC transporter, ATP-binding/permease protein |
| *PSPTO_4985* | | + | 5658863 | 5659543 | 11.36 | 16.91 | 16.76 | 14.86 | 33.45 | 22.53 | Toluene tolerance protein |
| *PSPTO_4986* | | - | 5659585 | 5660760 | 3.51 | 11.43 | 6.70 | 2.40 | 38.86 | 7.50 | Membrane protein |
| *PSPTO_4987* | | - | 5660750 | 5661487 | 7.38 | 11.29 | 14.67 | 8.05 | 71.73 | 10.69 | Hypothetical protein |
| *PSPTO_4988* | | - | 5661555 | 5662511 | 3.25 | 10.87 | 6.53 | 11.21 | 52.16 | 6.00 | Hypothetical protein |
| *PSPTO_4989* | | - | 5662542 | 5663978 | 1.75 | 4.21 | 11.53 | 9.56 | 30.06 | 6.83 | Hypothetical protein |
| *PSPTO_4990* | | - | 5663948 | 5664844 | 2.62 | 10.16 | 4.48 | 8.85 | 42.68 | 16.72 | Hypothetical protein |
| *PSPTO_4991* | | - | 5664844 | 5665974 | 5.07 | 11.36 | 9.03 | 5.49 | 42.96 | 19.61 | Glycoside hydrolase family protein |
| *PSPTO_4992* | | - | 5665958 | 5667715 | 3.90 | 34.01 | 27.82 | 15.00 | 73.41 | 23.61 | Carbamoyltransferase family protein |
| *PSPTO_4993* | | + | 5667943 | 5671257 | 1.63 | 2.91 | 2.79 | 1.39 | 3.57 | 2.78 | Hypothetical protein |
| *PSPTO_4994* | | - | 5671299 | 5672852 | 0.00 | 0.00 | 0.00 | 0.00 | 0.00 | 0.00 | ISPsy5, transposase |
| *PSPTO_4995* | | - | 5672870 | 5673229 | 0.00 | 0.00 | 0.00 | 0.00 | 0.00 | 0.00 | ISPsy5, Orf1 |
| *PSPTO_4996* | | + | 5673289 | 5676210 | 5.85 | 6.57 | 7.58 | 4.82 | 11.37 | 7.87 | Hypothetical protein |
| *PSPTO_4997* | | - | 5676226 | 5677671 | 6.18 | 9.96 | 7.79 | 7.12 | 50.87 | 7.84 | Hypothetical protein |
| *PSPTO_4998* | | - | 5677702 | 5678463 | 10.76 | 15.95 | 20.52 | 13.39 | 157.15 | 21.84 | Lipopolysaccharide biosynthesis protein |
| *PSPTO_4999* | | - | 5678460 | 5679194 | 7.58 | 21.20 | 12.22 | 4.33 | 45.80 | 12.07 | Lipopolysaccharide core biosynthesis protein |
| *PSPTO_5000* | *waaP* | - | 5679194 | 5680000 | 4.92 | 15.95 | 16.71 | 10.10 | 61.45 | 14.82 | Lipopolysaccharide core biosynthesis protein WaaP |
| *PSPTO_5001* | *waaG* | - | 5680000 | 5681121 | 12.43 | 16.99 | 15.70 | 5.97 | 79.49 | 19.62 | Lipopolysaccharide core biosynthesis protein WaaG |
| *PSPTO_5002* | *waaC* | - | 5681121 | 5682185 | 6.96 | 14.73 | 10.86 | 0.61 | 27.74 | 10.61 | Lipopolysaccharide heptosyltransferase |
| *PSPTO_5003* | *rfaF* | - | 5682188 | 5683222 | 3.21 | 12.54 | 3.33 | 4.65 | 31.45 | 11.17 | ADP-heptose--LPS heptosyltransferase II |
| *PSPTO_5004* | *glnE* | - | 5683352 | 5686309 | 11.37 | 15.89 | 14.19 | 8.81 | 50.73 | 17.71 | Glutamate-ammonia-ligase adenylyltransferase |
| *PSPTO_5005* | *aceE-2* | + | 5686694 | 5689339 | 9.44 | 128.31 | 60.52 | 35.71 | 417.26 | 98.58 | Pyruvate dehydrogenase, E1 component |
| *PSPTO_5006* | *aceF* | + | 5689482 | 5691128 | 3.87 | 45.36 | 31.44 | 18.87 | 294.09 | 45.05 | Pyruvate dehydrogenase complex, E2 component, dihydrolipoamide acetyltransferase |
| *PSPTO_5007* | *dbpA* | + | 5691491 | 5692870 | 8.07 | 19.59 | 12.00 | 7.06 | 32.26 | 8.08 | ATP-independent RNA helicase DbpA |
| *PSPTO_5008* | | + | 5693047 | 5694225 | 3.33 | 10.35 | 2.72 | 2.42 | 22.08 | 3.80 | Hypothetical protein |
| *PSPTO_5009* | | + | 5694361 | 5696559 | 10.49 | 11.58 | 14.23 | 7.17 | 23.93 | 10.44 | Membrane protein |
| *PSPTO_5010* | | - | 5696522 | 5697277 | 24.66 | 6.89 | 6.69 | 3.00 | 13.54 | 9.21 | Amino acid ABC transporter, periplasmic amino acid-binding protein |
| *PSPTO_5011* | | + | 5697503 | 5697961 | 31.61 | 57.32 | 39.19 | 24.94 | 89.45 | 42.08 | GNAT family acetyltransferase |
| *PSPTO_5012* | | - | 5698019 | 5698375 | 24.82 | 37.28 | 18.18 | 9.77 | 51.76 | 40.53 | Hypothetical protein |
| *PSPTO_5013* | | - | 5698372 | 5698896 | 16.14 | 22.96 | 22.82 | 14.38 | 48.03 | 20.90 | Hypothetical protein |
| *PSPTO_5014* | | - | 5698929 | 5701088 | 16.92 | 20.67 | 15.64 | 11.78 | 9.28 | 11.16 | Response regulator/sensory box/GGDEF domain/EAL domain protein |
| *PSPTO_5015* | | + | 5701382 | 5702053 | 88.37 | 283.60 | 250.20 | 180.09 | 129.75 | 140.94 | Lyase |
| *PSPTO_5016* | | - | 5702150 | 5703637 | 9.50 | 9.91 | 14.65 | 9.96 | 80.93 | 19.35 | Proline permease |
| *PSPTO_5017* | | + | 5704088 | 5708041 | 57.18 | 9.61 | 10.14 | 5.77 | 225.08 | 118.29 | Bifunctional putA protein |
| *PSPTO_5018* | | - | 5708165 | 5709001 | 8.76 | 8.50 | 12.04 | 5.51 | 40.56 | 16.71 | ISPsy9, transposase OrfB |
| *PSPTO_5019* | | - | 5709001 | 5709540 | 10.27 | 14.87 | 6.39 | 5.18 | 31.99 | 10.13 | ISPsy9, transposase OrfA |
| *PSPTO_5020* | | + | 5709754 | 5711556 | 16.53 | 19.32 | 13.47 | 10.90 | 25.51 | 15.01 | Acyl-CoA dehydrogenase family protein |
| *PSPTO_5021* | | - | 5711579 | 5714200 | 2.67 | 2.48 | 3.23 | 2.55 | 3.50 | 3.13 | Hypothetical protein |
| *PSPTO_5022* | | - | 5714337 | 5717465 | 3.92 | 2.20 | 2.51 | 2.91 | 3.90 | 3.30 | Hypothetical protein |
| *PSPTO_5023* | | + | 5717662 | 5719458 | 32.62 | 22.13 | 18.78 | 7.01 | 47.51 | 12.40 | Sodium/hydrogen exchanger family protein |
| *PSPTO_5024* | | + | 5719482 | 5722886 | 17.95 | 28.95 | 25.62 | 15.98 | 92.82 | 38.16 | Hypothetical protein |
| *PSPTO_5025* | | - | 5723444 | 5723815 | 11.31 | 8.20 | 2.20 | 4.10 | 16.36 | 11.58 | Hypothetical protein |
| *PSPTO_5026* | | - | 5724039 | 5724149 | 5.20 | 5.91 | 0.00 | 0.00 | 3.04 | 3.95 | Hypothetical protein |
| *PSPTO_5027* | | + | 5724197 | 5724871 | 13.15 | 20.96 | 14.36 | 7.67 | 40.19 | 17.99 | GntR family transcriptional regulator |
| *PSPTO_5028* | | + | 5725001 | 5726464 | 4.77 | 8.62 | 8.09 | 3.04 | 17.69 | 10.45 | Hypothetical protein |

| Locus name | Gene name | Strand | Gene start (bp) | Gene end (bp) | Average TPM Inoculum | Average TPM $t_6$ | Average TPM $t_{24}$ | Average TPM $t_{168}$ | Average TPM Logarithmic | Average TPM Stationary | Annotation description |
|---|---|---|---|---|---|---|---|---|---|---|---|
| *PSPTO_5029* | | - | 5726735 | 5727205 | 4.30 | 7.16 | 12.49 | 2.49 | 15.10 | 7.53 | Hypothetical protein |
| *PSPTO_5030* | | - | 5727198 | 5733161 | 6.86 | 10.02 | 8.45 | 4.49 | 22.46 | 8.40 | Sensor histidine kinase/response regulator |
| *PSPTO_5031* | *pilJ* | - | 5733209 | 5735251 | 7.29 | 12.93 | 9.29 | 3.45 | 22.88 | 10.01 | Type IV pilus biogenesis protein PilJ |
| *PSPTO_5032* | *pilI* | - | 5735284 | 5735823 | 6.85 | 15.53 | 14.08 | 2.78 | 27.19 | 8.36 | Type IV pilus protein PilI |
| *PSPTO_5033* | *pilH* | - | 5735838 | 5736203 | 27.24 | 39.53 | 19.08 | 20.18 | 77.14 | 34.37 | Type IV pilus response regulator PilH |
| *PSPTO_5034* | *pilG* | - | 5736250 | 5736660 | 48.45 | 97.76 | 68.50 | 36.09 | 209.13 | 75.81 | Type IV pilus response regulator PilG |
| *PSPTO_5035* | *gshB* | + | 5736905 | 5737864 | 22.86 | 39.43 | 42.61 | 23.10 | 72.82 | 20.03 | Glutathione synthetase |
| *PSPTO_5036* | | + | 5738123 | 5739016 | 34.57 | 53.84 | 25.14 | 26.36 | 69.12 | 52.04 | TonB domain protein |
| *PSPTO_5037* | | + | 5739092 | 5739664 | 43.03 | 66.34 | 58.37 | 23.34 | 132.38 | 52.15 | Hypothetical protein |
| *PSPTO_5038* | | + | 5739664 | 5740104 | 30.11 | 50.44 | 42.82 | 9.65 | 189.35 | 55.21 | Hypothetical protein |
| *PSPTO_5039* | *pyrR* | + | 5740347 | 5740859 | 13.26 | 31.08 | 17.47 | 16.10 | 150.74 | 48.48 | Pyrimidine operon regulatory protein PyrR |
| *PSPTO_5040* | *pyrB* | + | 5740872 | 5741876 | 11.10 | 29.91 | 13.62 | 15.93 | 131.09 | 25.27 | Aspartate carbamoyltransferase |
| *PSPTO_5041* | | + | 5741873 | 5743144 | 7.91 | 13.10 | 13.29 | 13.04 | 72.30 | 21.81 | Dihydroorotase, multifunctional complex type |
| *PSPTO_5043* | | - | 5743362 | 5743916 | 25.96 | 50.63 | 58.38 | 29.64 | 106.02 | 25.14 | Hypothetical protein |
| *PSPTO_5044* | | + | 5744017 | 5744697 | 8.91 | 153.11 | 141.91 | 121.91 | 66.34 | 45.48 | NLP/P60 family protein |
| *PSPTO_5045* | | - | 5744794 | 5745828 | 18.61 | 21.47 | 29.12 | 14.37 | 51.46 | 15.62 | Twitching motility protein |
| *PSPTO_5046* | | + | 5745884 | 5746570 | 51.10 | 58.02 | 34.71 | 15.12 | 68.14 | 33.41 | Hypothetical protein |
| *PSPTO_5047* | *proC* | + | 5746623 | 5747441 | 7.59 | 25.29 | 17.21 | 5.56 | 80.17 | 16.39 | Pyrroline-5-carboxylate reductase |
| *PSPTO_5048* | | + | 5747467 | 5748057 | 9.22 | 15.03 | 17.71 | 10.05 | 53.63 | 13.62 | Hypothetical protein |
| *PSPTO_5049* | | + | 5748347 | 5749486 | 18.33 | 28.59 | 21.74 | 28.32 | 113.95 | 33.83 | Homoserine O-acetyltransferase |
| *PSPTO_5050* | *metW* | + | 5749494 | 5750114 | 10.14 | 32.77 | 29.41 | 14.87 | 123.36 | 27.14 | MetW protein |
| *PSPTO_5051* | | + | 5750148 | 5750741 | 2.61 | 14.92 | 5.20 | 4.65 | 63.63 | 10.99 | Ham1 protein |
| *PSPTO_5052* | | + | 5750738 | 5751952 | 5.70 | 49.34 | 25.10 | 5.94 | 32.42 | 8.58 | HemN family oxidoreductase |
| *PSPTO_5053* | | + | 5751962 | 5752285 | 4.28 | 25.23 | 12.61 | 0.00 | 27.76 | 9.78 | Hypothetical protein |
| *PSPTO_5054* | | - | 5752893 | 5754557 | 1.01 | 5.40 | 3.44 | 0.79 | 5.24 | 2.71 | Hypothetical protein |
| *PSPTO_5055* | | - | 5754554 | 5756578 | 7.67 | 9.36 | 7.89 | 10.34 | 15.23 | 10.47 | Hypothetical protein |
| *PSPTO_5056* | | - | 5756578 | 5759135 | - | - | - | - | - | - | |
| *PSPTO_5057* | *ilvD* | - | 5759478 | 5761325 | 10.58 | 21.70 | 26.71 | 23.73 | 70.65 | 32.18 | Dihydroxy-acid dehydratase |
| *PSPTO_5058* | | + | 5761665 | 5762873 | 5.92 | 13.40 | 12.71 | 9.74 | 39.55 | 12.74 | Class I and II aminotransferase |
| *PSPTO_5059* | | - | 5762976 | 5763545 | 12.52 | 14.53 | 16.69 | 13.63 | 24.16 | 10.82 | Hypothetical protein |
| *PSPTO_5060* | | + | 5763742 | 5764647 | 2.25 | 15.68 | 15.20 | 7.49 | 6.41 | 1.89 | Oxidoreductase, acting on the CH-OH group of donors, NAD or NADP as acceptor |
| *PSPTO_5061* | | + | 5764733 | 5765947 | 1.52 | 8.43 | 16.41 | 1.75 | 6.09 | 1.86 | HopAN1 protein |
| *PSPTO_5062* | | + | 5765947 | 5766585 | 1.22 | 2.11 | 7.09 | 3.89 | 5.78 | 1.19 | Putative aldolase |
| *PSPTO_5063* | | + | 5766721 | 5768178 | 2.43 | 6.63 | 3.58 | 2.60 | 7.72 | 2.39 | Gluconate transporter family protein |
| *PSPTO_5064* | | + | 5768201 | 5768986 | 6.76 | 13.16 | 1.20 | 3.25 | 11.10 | 4.46 | Hydroxypyruvate isomerase |
| *PSPTO_5065* | | + | 5769093 | 5769608 | 16.87 | 20.21 | 25.19 | 11.29 | 37.77 | 18.82 | ISPsy8, transposase OrfA |
| *PSPTO_5066* | | + | 5769665 | 5770444 | 16.60 | 15.22 | 11.70 | 11.48 | 25.13 | 16.12 | ISPsy8, transposase OrfB |
| *PSPTO_5067* | | - | 5770690 | 5771085 | 15.21 | 80.30 | 46.13 | 4.41 | 67.73 | 22.24 | 4-hydroxybenzoyl-CoA thioesterase |
| *PSPTO_5068* | *ahcY* | + | 5771439 | 5772848 | 12.80 | 45.56 | 34.29 | 24.04 | 494.92 | 73.49 | Adenosylhomocysteinase |
| *PSPTO_5069* | *metF* | + | 5773038 | 5773883 | 4.61 | 10.63 | 11.68 | 5.55 | 176.79 | 25.36 | 5,10-methylenetetrahydrofolate reductase |
| *PSPTO_5070* | | + | 5774194 | 5776083 | 6.76 | 25.64 | 23.64 | 9.74 | 78.46 | 23.68 | ATP-dependent RNA helicase rhlE |
| *PSPTO_5071* | | - | 5776365 | 5776943 | 32.48 | 167.04 | 98.89 | 47.59 | 545.66 | 29.79 | Hypothetical protein |
| *PSPTO_5072* | | - | 5776971 | 5777525 | 51.79 | 168.03 | 139.56 | 42.87 | 292.14 | 31.85 | Cytochrome b561 |
| *PSPTO_5073* | | - | 5777562 | 5777801 | 24.08 | 187.71 | 56.15 | 7.66 | 143.40 | 12.50 | Hypothetical protein |
| *PSPTO_5074* | | - | 5777855 | 5779732 | 3.04 | 2.71 | 2.48 | 3.05 | 16.30 | 3.46 | Amine oxidase, flavin-containing protein |
| *PSPTO_5075* | *bioA* | + | 5779943 | 5781349 | 3.46 | 13.11 | 13.59 | 6.53 | 48.29 | 9.67 | Adenosylmethionine--8-amino-7-oxononanoate transaminase |
| *PSPTO_5076* | *rsmE* | + | 5781466 | 5782185 | 8.35 | 19.28 | 20.05 | 5.71 | 56.00 | 13.58 | 16S ribosomal RNA methyltransferase RsmE |
| *PSPTO_5077* | | - | 5782307 | 5782921 | 21.50 | 37.42 | 21.05 | 17.65 | 50.46 | 20.40 | Hemolysin III |
| *PSPTO_5078* | | + | 5783044 | 5783970 | 3.52 | 0.86 | 1.92 | 6.05 | 8.00 | 2.89 | Malonate utilization transcriptional regulator |
| *PSPTO_5079* | *madM* | - | 5783981 | 5784745 | 18.51 | 17.46 | 13.56 | 14.17 | 18.68 | 10.64 | Malonate transporter subunit MadM |
| *PSPTO_5080* | *madL* | - | 5784747 | 5785172 | 21.11 | 21.45 | 15.26 | 17.86 | 11.98 | 10.99 | Malonate transporter subunit MadL |
| *PSPTO_5081* | | - | 5785242 | 5786177 | 14.06 | 11.45 | 8.20 | 0.79 | 10.00 | 7.39 | Malonyl CoA-acyl carrier protein transacylase |
| *PSPTO_5082* | | - | 5786174 | 5786809 | 10.34 | 1.68 | 7.53 | 4.35 | 3.48 | 3.34 | Phosphoribosyl-dephospho-CoA transferase |
| *PSPTO_5083* | *mdcE* | - | 5786806 | 5787606 | 16.70 | 6.03 | 2.99 | 3.23 | 5.07 | 2.89 | Malonate decarboxylase subunit gamma |
| *PSPTO_5084* | *mdcD* | - | 5787603 | 5788454 | 31.79 | 9.14 | 17.72 | 7.79 | 6.41 | 2.08 | Malonate decarboxylase subunit beta |
| *PSPTO_5085* | *mdcC* | - | 5788447 | 5788746 | 15.14 | 0.00 | 14.26 | 0.00 | 11.38 | 8.21 | Malonate decarboxylase subunit delta |
| *PSPTO_5086* | *mdcB* | - | 5788748 | 5789623 | 32.19 | 7.99 | 2.40 | 7.98 | 4.52 | 4.56 | 2-(5''-triphosphoribosyl)-3'-dephosphocoenzyme-A synthase |
| *PSPTO_5087* | *mdcA* | - | 5789623 | 5791299 | 254.07 | 20.02 | 18.56 | 30.69 | 9.44 | 41.18 | Malonate decarboxylase subunit alpha |
| *PSPTO_5088* | | - | 5791333 | 5791434 | 4.92 | 0.00 | 11.26 | 0.00 | 6.44 | 7.64 | Hypothetical protein |
| *PSPTO_5089* | | + | 5791486 | 5791944 | 3.44 | 0.91 | 7.46 | 0.00 | 6.75 | 5.35 | Hypothetical protein |
| *PSPTO_5090* | | - | 5792018 | 5792797 | 11.96 | 49.40 | 51.54 | 15.94 | 47.58 | 26.45 | ParA family protein |
| *PSPTO_5091* | | + | 5793103 | 5793822 | 23.34 | 14.32 | 14.05 | 9.29 | 25.14 | 13.71 | Hypothetical protein |
| *PSPTO_5092* | | + | 5793798 | 5794610 | 15.17 | 16.01 | 15.17 | 11.16 | 26.09 | 12.94 | Acyltransferase family protein |
| *PSPTO_5093* | | + | 5794585 | 5794845 | 23.00 | 12.53 | 18.55 | 7.77 | 42.26 | 15.39 | Acyl carrier protein |
| *PSPTO_5094* | | + | 5794849 | 5795109 | 8.70 | 9.48 | 6.47 | 3.51 | 12.69 | 6.10 | Acyl carrier protein |
| *PSPTO_5095* | | + | 5795106 | 5795651 | 9.58 | 17.20 | 4.65 | 1.96 | 40.53 | 8.99 | Membrane protein |
| *PSPTO_5096* | | + | 5795770 | 5797344 | 2.44 | 5.03 | 3.25 | 4.50 | 15.28 | 1.83 | AMP-binding protein |
| *PSPTO_5097* | | + | 5797337 | 5798071 | 5.19 | 5.34 | 4.51 | 4.53 | 13.41 | 4.77 | Glycosyl transferase family protein |
| *PSPTO_5098* | | + | 5798068 | 5799015 | 15.12 | 7.36 | 13.23 | 6.55 | 15.54 | 7.64 | Hypothetical protein |
| *PSPTO_5099* | *hutH-1* | + | 5798996 | 5800543 | 5.49 | 3.87 | 3.29 | 2.54 | 18.90 | 5.52 | Histidine ammonia-lyase |
| *PSPTO_5100* | | + | 5800533 | 5800958 | 8.64 | 7.03 | 2.23 | 9.80 | 20.42 | 4.05 | 4-hydroxybenzoyl-CoA thioesterase |
| *PSPTO_5101* | | + | 5800955 | 5801569 | 3.97 | 6.00 | 2.49 | 1.90 | 17.40 | 4.49 | Hypothetical protein |
| *PSPTO_5102* | | + | 5801541 | 5803916 | 2.98 | 4.55 | 4.23 | 2.88 | 12.46 | 4.06 | Membrane protein |
| *PSPTO_5103* | | + | 5803913 | 5805085 | 1.25 | 2.05 | 1.55 | 2.06 | 7.38 | 2.41 | Membrane protein |
| *PSPTO_5104* | | + | 5805072 | 5806319 | 1.53 | 3.21 | 5.42 | 4.70 | 15.20 | 2.69 | Hypothetical protein |
| *PSPTO_5105* | | + | 5806331 | 5807050 | 2.09 | 2.45 | 1.30 | 1.03 | 11.59 | 3.06 | Hypothetical protein |
| *PSPTO_5106* | | + | 5807047 | 5807529 | 3.20 | 5.34 | 4.53 | 0.00 | 11.21 | 1.80 | Lipoprotein |
| *PSPTO_5107* | | + | 5807526 | 5808722 | 3.21 | 5.50 | 0.76 | 0.00 | 15.60 | 4.56 | 3-oxoacyl-(acyl carrier protein) synthase I |
| *PSPTO_5108* | | + | 5808719 | 5809174 | 5.70 | 12.47 | 7.64 | 0.80 | 33.61 | 11.03 | Hypothetical protein |

| Locus name | Gene name | Strand | Gene start (bp) | Gene end (bp) | Average TPM Inoculum | Average TPM $t_6$ | Average TPM $t_{24}$ | Average TPM $t_{168}$ | Average TPM Logarithmic | Average TPM Stationary | Annotation description |
|---|---|---|---|---|---|---|---|---|---|---|---|
| *PSPTO_5109* | | + | 5809171 | 5809899 | 3.74 | 9.15 | 17.35 | 11.50 | 24.91 | 6.55 | 3-ketoacyl-(acyl-carrier-protein) reductase |
| *PSPTO_5110* | | + | 5809899 | 5811125 | 3.37 | 8.12 | 10.27 | 5.33 | 21.07 | 4.54 | 3-oxoacyl-(acyl carrier protein) synthase II |
| *PSPTO_5111* | | - | 5811163 | 5812056 | 1.52 | 2.05 | 9.76 | 0.00 | 9.12 | 1.58 | LysR family transcriptional regulator |
| *PSPTO_5112* | | + | 5812173 | 5813429 | 7.97 | 8.28 | 4.79 | 4.77 | 18.47 | 20.16 | Major facilitator superfamily protein |
| *PSPTO_5113* | | + | 5813507 | 5814775 | 6.94 | 25.25 | 10.86 | 13.86 | 41.06 | 14.92 | Hypothetical protein |
| *PSPTO_5114* | | + | 5814772 | 5815416 | 3.36 | 6.55 | 11.17 | 0.00 | 30.73 | 7.69 | Hypothetical protein |
| *PSPTO_5115* | | + | 5815715 | 5817064 | 6.07 | 5.24 | 5.24 | 4.94 | 6.81 | 4.15 | Gluconate transporter family protein |
| *PSPTO_5116* | | + | 5817065 | 5817925 | 0.97 | 5.01 | 6.10 | 0.03 | 6.20 | 2.53 | RpiR family transcriptional regulator |
| *PSPTO_5117* | | + | 5817929 | 5819443 | 2.72 | 4.51 | 9.09 | 4.50 | 10.52 | 3.07 | N-acyl-D-amino acid deacylase family protein |
| *PSPTO_5118* | *hemE* | - | 5819471 | 5820535 | 7.90 | 44.23 | 29.48 | 5.31 | 41.10 | 13.25 | Uroporphyrinogen decarboxylase |
| *PSPTO_5119* | | - | 5820666 | 5823896 | 11.89 | 9.94 | 6.38 | 6.92 | 16.84 | 10.91 | Hypothetical protein |
| *PSPTO_5120* | | - | 5824060 | 5825619 | 5.50 | 4.84 | 1.66 | 3.68 | 4.89 | 3.56 | Amidase family protein |
| *PSPTO_5121* | *gltD* | - | 5825777 | 5827195 | 8.80 | 40.08 | 22.48 | 12.65 | 59.87 | 7.20 | Glutamate synthase, small subunit |
| *PSPTO_5122* | | + | 5827210 | 5827308 | 3.89 | 16.72 | 15.97 | 0.00 | 20.27 | 0.30 | Hypothetical protein |
| *PSPTO_5123* | *gltB* | - | 5827318 | 5831763 | 20.14 | 50.29 | 33.90 | 15.08 | 49.93 | 6.26 | Glutamate synthase, large subunit |
| *PSPTO_5124* | | - | 5831970 | 5832068 | 27.11 | 9.77 | 19.02 | 4.23 | 9.58 | 32.43 | Hypothetical protein |
| *PSPTO_5125* | | - | 5832123 | 5833748 | 40.83 | 58.06 | 60.56 | 36.36 | 176.23 | 80.79 | Hypothetical protein |
| *PSPTO_5126* | *aroB* | - | 5833759 | 5834862 | 34.66 | 79.45 | 58.32 | 48.10 | 158.22 | 52.32 | 3-dehydroquinate synthase |
| *PSPTO_5127* | *aroK* | - | 5834981 | 5835499 | 27.05 | 40.44 | 20.23 | 1.51 | 73.91 | 22.10 | Shikimate kinase |
| *PSPTO_5128* | *pilQ* | - | 5835504 | 5837660 | 26.64 | 19.49 | 11.14 | 6.34 | 24.64 | 11.22 | Type IV pilus biogenesis protein PilQ |
| *PSPTO_5129* | *pilP* | - | 5837657 | 5838184 | 53.39 | 13.28 | 23.28 | 15.48 | 25.63 | 15.50 | Type IV pilus biogenesis protein PilP |
| *PSPTO_5130* | *pilO* | - | 5838181 | 5838804 | 31.94 | 20.15 | 12.84 | 5.83 | 22.42 | 11.72 | Type IV pilus biogenesis protein PilO |
| *PSPTO_5131* | *pilN* | - | 5838801 | 5839427 | 27.14 | 10.70 | 12.21 | 10.82 | 26.79 | 14.62 | Type IV pilus biogenesis protein PilN |
| *PSPTO_5132* | *pilM* | - | 5839427 | 5840491 | 41.26 | 16.42 | 24.77 | 8.43 | 29.67 | 23.06 | Type IV pilus assembly protein PilM |
| *PSPTO_5133* | | + | 5840698 | 5843142 | 210.67 | 54.73 | 50.84 | 25.19 | 72.16 | 75.34 | Penicillin-binding protein |
| *PSPTO_5134* | | - | 5843502 | 5844770 | 13.40 | 93.40 | 65.05 | 24.10 | 212.93 | 31.32 | Malic enzyme family protein |
| *PSPTO_5135* | | - | 5844940 | 5845683 | 2.31 | 7.05 | 2.25 | 4.29 | 7.44 | 5.29 | Staphylococcal nuclease-like protein |
| *PSPTO_5136* | *rpmE-2* | - | 5845754 | 5845981 | 63.58 | 194.00 | 248.09 | 156.96 | 221.12 | 37.42 | 50S ribosomal protein L31 |
| *PSPTO_5137* | *priA* | + | 5846150 | 5848369 | 6.49 | 8.16 | 10.99 | 4.07 | 20.21 | 10.28 | Primosomal protein N` |
| *PSPTO_5138* | *argS* | + | 5848467 | 5850203 | 9.52 | 26.56 | 13.79 | 11.31 | 78.05 | 35.69 | Arginyl-tRNA synthetase |
| *PSPTO_5139* | | + | 5850205 | 5850900 | 9.52 | 31.76 | 21.67 | 14.24 | 109.94 | 31.12 | Hypothetical protein |
| *PSPTO_5140* | *hslV* | + | 5851055 | 5851585 | 155.55 | 293.99 | 196.08 | 118.24 | 616.17 | 164.87 | Heat shock protein HslV |
| *PSPTO_5141* | *hslU* | + | 5851651 | 5852988 | 17.29 | 45.74 | 39.34 | 21.01 | 219.94 | 44.57 | Heat shock protein HslVU, ATPase subunit HslU |
| *PSPTO_5142* | | + | 5853065 | 5853451 | 6.30 | 15.60 | 7.58 | 4.75 | 105.03 | 24.93 | Hypothetical protein |
| *PSPTO_5143* | | + | 5853760 | 5855439 | - | - | - | - | - | - | |
| *PSPTO_5144* | *phaB* | + | 5855493 | 5856350 | 7.95 | 7.83 | 13.10 | 5.15 | 6.82 | 9.02 | Poly(3-hydroxyalkanoate) depolymerase |
| *PSPTO_5145* | *phaC* | + | 5856501 | 5858192 | 2.97 | 1.98 | 1.43 | 1.73 | 1.98 | 3.98 | Poly(3-hydroxyalkanoate) polymerase |
| *PSPTO_5146* | *phaD* | + | 5858240 | 5858854 | 1.46 | 2.86 | 6.00 | 0.64 | 2.12 | 1.67 | Transcriptional regulator PhaD |
| *PSPTO_5147* | | - | 5859216 | 5859989 | 141.30 | 141.11 | 139.78 | 74.28 | 251.51 | 139.65 | Polyhydroxyalkanoate granule-associated protein PhaF |
| *PSPTO_5148* | | - | 5860027 | 5860473 | 181.69 | 134.77 | 110.19 | 51.87 | 179.21 | 99.29 | Polyhydroxyalkanoate granule-associated protein PhaI |
| *PSPTO_5149* | | - | 5860577 | 5860855 | 32.59 | 56.13 | 29.40 | 5.50 | 79.42 | 24.04 | Hypothetical protein |
| *PSPTO_5150* | *ubiE* | + | 5861000 | 5861770 | 41.39 | 73.84 | 61.03 | 39.43 | 141.01 | 61.65 | Ubiquinone/menaquinone biosynthesis methyltransferase UbiE |
| *PSPTO_5151* | | + | 5861770 | 5862393 | 4.28 | 12.90 | 13.36 | 14.65 | 42.10 | 15.76 | Hypothetical protein |
| *PSPTO_5152* | *ubiB* | + | 5862390 | 5864009 | 3.52 | 10.76 | 17.00 | 6.91 | 41.45 | 8.92 | Ubiquinone biosynthesis protein UbiB |
| *PSPTO_5153* | *hisI* | + | 5864077 | 5864469 | 33.45 | 103.69 | 93.40 | 80.60 | 169.01 | 70.61 | Phosphoribosyl-AMP cyclohydrolase |
| *PSPTO_5154* | *hisE* | + | 5864472 | 5864804 | 16.15 | 41.34 | 42.29 | 27.22 | 91.00 | 30.49 | Phosphoribosyl-ATP pyrophosphatase |
| *PSPTO_5155* | *tatA* | + | 5864849 | 5865124 | 18.76 | 84.93 | 110.14 | 43.63 | 233.56 | 64.03 | Sec-independent protein translocase TatA |
| *PSPTO_5156* | *tatB* | + | 5865129 | 5865590 | 16.14 | 54.21 | 60.23 | 37.72 | 184.70 | 53.21 | Sec-independent protein translocase TatB |
| *PSPTO_5157* | *tatC* | + | 5865587 | 5866387 | 9.49 | 40.14 | 35.32 | 26.71 | 96.47 | 31.14 | Sec-independent protein translocase TatC |
| *PSPTO_5158* | | + | 5866384 | 5867091 | 3.39 | 3.32 | 6.07 | 7.04 | 31.95 | 12.66 | Hypothetical protein |
| *PSPTO_5159* | | + | 5867181 | 5869097 | 17.00 | 2.79 | 4.31 | 4.03 | 3.23 | 2.92 | Methyl-accepting chemotaxis protein |
| *PSPTO_5160* | | + | 5869300 | 5871243 | 2.31 | 1.37 | 1.23 | 1.28 | 1.89 | 1.60 | Methyl-accepting chemotaxis protein |
| *PSPTO_5161* | *mdoH* | - | 5871526 | 5874105 | 9.60 | 35.92 | 24.46 | 11.79 | 103.68 | 31.50 | Periplasmic glucan biosynthesis protein |
| *PSPTO_5162* | *mdoG* | - | 5874098 | 5876023 | 32.74 | 77.97 | 81.72 | 39.02 | 155.65 | 80.09 | Periplasmic glucan biosynthesis protein |
| *PSPTO_5163* | *dtd* | - | 5876289 | 5876726 | 40.94 | 27.01 | 25.94 | 10.97 | 31.27 | 17.73 | D-tyrosyl-tRNA(Tyr) deacylase |
| *PSPTO_5164* | *pip* | - | 5876768 | 5877739 | 22.73 | 38.21 | 24.51 | 18.85 | 58.26 | 27.15 | Proline iminopeptidase |
| *PSPTO_5165* | *glgP* | + | 5878027 | 5880477 | 22.14 | 26.64 | 20.51 | 14.12 | 20.38 | 26.54 | Glycogen phosphorylase |
| *PSPTO_5166* | | + | 5880710 | 5883529 | 3.79 | 6.98 | 2.89 | 2.78 | 10.41 | 5.25 | Membrane protein |
| *PSPTO_5167* | | + | 5883540 | 5884904 | 3.29 | 14.93 | 6.79 | 1.08 | 22.95 | 6.46 | Hypothetical protein |
| *PSPTO_5168* | *fbp* | + | 5885038 | 5886048 | 75.48 | 136.53 | 106.52 | 47.70 | 186.86 | 75.09 | Fructose-1,6-bisphosphatase |
| *PSPTO_5169* | | + | 5886304 | 5886558 | 202.06 | 63.58 | 29.32 | 25.45 | 77.15 | 81.21 | Lipoprotein |
| *PSPTO_5170* | *blc* | + | 5886562 | 5887140 | 66.25 | 34.07 | 19.66 | 20.45 | 65.20 | 67.70 | Lipoprotein Blc |
| *PSPTO_5171* | | - | 5887357 | 5888721 | 4.56 | 3.62 | 8.91 | 5.49 | 27.34 | 3.62 | Atz/Trz family protein |
| *PSPTO_5172* | *hutC* | + | 5888885 | 5889634 | 11.44 | 20.40 | 21.33 | 14.13 | 49.86 | 13.27 | Histidine utilization repressor |
| *PSPTO_5173* | | + | 5889631 | 5890251 | 15.06 | 5.42 | 8.21 | 8.90 | 35.67 | 11.38 | Hypothetical protein |
| *PSPTO_5174* | *nhaA-2* | + | 5890278 | 5891453 | 2.50 | 3.45 | 0.46 | 1.88 | 9.41 | 2.05 | Sodium-proton antiporter NhaA |
| *PSPTO_5175* | | + | 5891558 | 5892091 | 156.72 | 130.57 | 103.01 | 46.79 | 153.08 | 47.96 | Hypothetical protein |
| *PSPTO_5176* | | + | 5892197 | 5892763 | 63.59 | 65.52 | 57.59 | 24.57 | 75.66 | 46.33 | RNA polymerase sigma-70 family protein |
| *PSPTO_5177* | | + | 5892760 | 5893467 | 22.81 | 36.91 | 36.09 | 20.15 | 46.70 | 34.24 | Hypothetical protein |
| *PSPTO_5178* | | - | 5893481 | 5894434 | 35.54 | 38.17 | 34.49 | 17.02 | 77.29 | 21.56 | Serine O-acetyltransferase |
| *PSPTO_5179* | | + | 5894631 | 5895629 | 13.42 | 32.43 | 26.76 | 14.11 | 56.40 | 12.01 | D-cysteine desulfhydrase |
| *PSPTO_5180* | | + | 5895850 | 5896656 | 20.29 | 60.70 | 29.57 | 34.25 | 124.01 | 31.55 | Cystine transporter subunit |
| *PSPTO_5181* | | + | 5896653 | 5897321 | 5.34 | 11.82 | 9.52 | 6.48 | 44.08 | 9.84 | Amino acid ABC transporter permease |
| *PSPTO_5182* | | + | 5897325 | 5898068 | 8.83 | 19.05 | 8.71 | 6.52 | 35.90 | 8.16 | Putative amino-acid ABC transporter ATP-binding protein YecC |
| *PSPTO_5183* | | + | 5898642 | 5899361 | 1.41 | 1.20 | 3.57 | 0.00 | 2.81 | 2.25 | Hypothetical protein |
| *PSPTO_5184* | | + | 5899755 | 5901077 | 1.22 | 56.46 | 30.09 | 29.92 | 2.19 | 1.87 | Acyl-CoA dehydrogenase family protein |
| *PSPTO_5185* | | + | 5901103 | 5902308 | 0.43 | 11.54 | 6.03 | 0.97 | 1.11 | 0.66 | Acyl-CoA dehydrogenase family protein, Sulphate-starvation-induced SfnB |
| *PSPTO_5186* | | + | 5902308 | 5903666 | 1.40 | 8.19 | 5.37 | 6.03 | 2.37 | 1.44 | DszA family monooxygenase |
| *PSPTO_5187* | *metQ-1* | + | 5903778 | 5904566 | 5.25 | 13.93 | 10.14 | 0.83 | 4.47 | 3.14 | D-methionine-binding lipoprotein MetQ |

| Locus name | Gene name | Strand | Gene start (bp) | Gene end (bp) | Average TPM Inoculum | Average TPM $t_6$ | Average TPM $t_{24}$ | Average TPM $t_{168}$ | Average TPM Logarithmic | Average TPM Stationary | Annotation description |
|---|---|---|---|---|---|---|---|---|---|---|---|
| *PSPTO_5188* | *metN-1* | + | 5904662 | 5905693 | 1.02 | 4.19 | 4.45 | 0.00 | 2.16 | 1.16 | D-methionine ABC transporter ATP-binding protein |
| *PSPTO_5189* | *metI-1* | + | 5905683 | 5906327 | 5.51 | 7.55 | 7.29 | 10.21 | 10.28 | 8.89 | D-methionine ABC transporter permease |
| *PSPTO_5190* | | + | 5906379 | 5907002 | 4.26 | 15.51 | 23.98 | 9.29 | 25.52 | 13.09 | Hypothetical protein |
| *PSPTO_5191* | | - | 5907261 | 5910362 | 3.03 | 10.22 | 4.91 | 7.76 | 33.88 | 7.30 | AcrB/AcrD/AcrF family protein |
| *PSPTO_5192* | | - | 5910359 | 5911429 | 6.53 | 9.31 | 15.51 | 8.39 | 41.64 | 11.95 | RND family efflux transporter MFP subunit |
| *PSPTO_5193* | | - | 5911426 | 5912544 | 11.46 | 16.19 | 13.95 | 5.42 | 36.35 | 14.79 | RND family efflux transporter MFP subunit |
| *PSPTO_5194* | | - | 5912741 | 5913538 | 15.00 | 19.90 | 17.62 | 16.91 | 34.73 | 18.47 | Hypothetical protein |
| *PSPTO_5195* | | - | 5913818 | 5914837 | 2.94 | 17.85 | 7.54 | 15.76 | 9.06 | 3.73 | ABC transporter substrate-binding protein |
| *PSPTO_5196* | | - | 5914876 | 5915742 | 1.58 | 13.59 | 4.70 | 5.33 | 4.87 | 2.71 | ABC transporter permease |
| *PSPTO_5197* | | - | 5915834 | 5916694 | 0.98 | 15.69 | 12.16 | 17.42 | 3.57 | 1.86 | ABC transporter ATP-binding protein |
| *PSPTO_5198* | | - | 5916711 | 5917607 | 2.44 | 66.26 | 59.94 | 46.31 | 7.97 | 3.70 | Dioxygenase, TauD/TfdA family |
| *PSPTO_5199* | | - | 5917797 | 5917961 | 6.04 | 9.18 | 0.00 | 0.00 | 5.17 | 8.12 | Hypothetical protein |
| *PSPTO_5200* | | + | 5917963 | 5920923 | 1.70 | 1.97 | 1.15 | 0.75 | 2.25 | 2.26 | Autotransporter |
| *PSPTO_5201* | | + | 5921091 | 5921993 | 20.03 | 22.86 | 20.63 | 15.35 | 46.91 | 15.81 | Hypothetical protein |
| *PSPTO_5202* | | + | 5922182 | 5923180 | 28.53 | 28.28 | 33.19 | 27.06 | 92.14 | 43.14 | Hypothetical protein |
| *PSPTO_5203* | | + | 5923177 | 5924256 | 14.41 | 16.93 | 17.85 | 17.13 | 72.60 | 23.91 | Hypothetical protein |
| *PSPTO_5204* | | + | 5924253 | 5926277 | 9.42 | 9.99 | 10.28 | 9.09 | 42.01 | 15.54 | EF hand domain-containing protein |
| *PSPTO_5205* | | + | 5926826 | 5927059 | 1.06 | 0.00 | 6.67 | 3.41 | 10.27 | 1.07 | Hypothetical protein |
| *PSPTO_5206* | | + | 5927129 | 5928061 | 0.72 | 2.35 | 1.16 | 1.05 | 4.43 | 1.25 | EF hand domain-containing protein |
| *PSPTO_5207* | | + | 5928071 | 5928466 | 0.75 | 2.17 | 1.99 | 0.00 | 9.28 | 1.54 | Hypothetical protein |
| *PSPTO_5208* | | + | 5928634 | 5928801 | 11.07 | 6.72 | 4.40 | 4.66 | 16.09 | 8.21 | Hypothetical protein |
| *PSPTO_5209* | | + | 5928798 | 5929976 | 6.66 | 8.42 | 12.78 | 3.47 | 67.39 | 12.19 | Hypothetical protein |
| *PSPTO_5210* | | + | 5931087 | 5932019 | 0.61 | 3.79 | 2.46 | 0.59 | 3.62 | 1.03 | EF hand domain-containing protein |
| *PSPTO_5211* | | + | 5932029 | 5932424 | 0.88 | 0.00 | 0.00 | 0.00 | 5.13 | 1.35 | Hypothetical protein |
| *PSPTO_5212* | | - | 5932848 | 5934401 | 0.00 | 0.00 | 0.00 | 0.00 | 0.00 | 0.00 | ISPsy5, transposase |
| *PSPTO_5213* | | - | 5934419 | 5934778 | 0.00 | 0.00 | 0.00 | 0.00 | 0.00 | 0.00 | ISPsy5, Orf1 |
| *PSPTO_5214* | | + | 5936100 | 5936459 | 0.00 | 0.00 | 0.00 | 0.00 | 0.00 | 0.00 | ISPsy5, Orf1 |
| *PSPTO_5215* | | + | 5936477 | 5938030 | 0.00 | 0.00 | 0.00 | 0.00 | 0.00 | 0.00 | ISPsy5, transposase |
| *PSPTO_5216* | | - | 5938115 | 5938897 | 8.58 | 119.73 | 99.46 | 58.89 | 130.09 | 17.52 | Lipoprotein, NLPA family |
| *PSPTO_5217* | | - | 5939068 | 5940018 | 5.44 | 19.59 | 16.06 | 11.02 | 7.52 | 5.51 | Sigma-54-binding protein |
| *PSPTO_5218* | | - | 5940101 | 5940682 | 4.63 | 73.37 | 65.33 | 31.82 | 4.19 | 3.34 | Hypothetical protein |
| *PSPTO_5219* | | - | 5940898 | 5942208 | 3.54 | 7.45 | 8.24 | 3.50 | 3.02 | 2.58 | MFS transporter, phthalate permease family |
| *PSPTO_5220* | | + | 5942813 | 5943805 | 15.92 | 20.13 | 15.78 | 9.40 | 21.10 | 23.36 | AraC family transcriptional regulator |
| *PSPTO_5221* | *ubiF* | - | 5943909 | 5945138 | 3.49 | 10.88 | 13.89 | 8.00 | 20.25 | 7.71 | 2-octaprenyl-3-methyl-6-methoxy-1,4-benzoquinol hydroxylase |
| *PSPTO_5222* | *ubiH* | - | 5945190 | 5946377 | 5.41 | 32.37 | 27.17 | 21.24 | 61.17 | 20.00 | 2-octaprenyl-6-methoxyphenyl hydroxylase |
| *PSPTO_5223* | *pepP* | - | 5946374 | 5947708 | 8.07 | 33.85 | 25.79 | 10.83 | 49.26 | 17.10 | Xaa-Pro aminopeptidase |
| *PSPTO_5224* | | - | 5947784 | 5948338 | 27.70 | 68.21 | 53.19 | 17.55 | 137.95 | 32.98 | Hypothetical protein |
| *PSPTO_5225* | | + | 5948489 | 5948698 | 379.56 | 262.15 | 203.62 | 155.89 | 215.27 | 288.31 | Hypothetical protein |
| *PSPTO_5226* | | + | 5948695 | 5949024 | 98.87 | 170.71 | 125.30 | 91.06 | 161.84 | 174.44 | Hypothetical protein |
| *PSPTO_5227* | | + | 5949328 | 5949939 | 4.81 | 13.51 | 13.83 | 4.93 | 33.90 | 17.12 | 5-formyltetrahydrofolate cyclo-ligase |
| *PSPTO_5228* | | - | 5949986 | 5950138 | 47.99 | 20.99 | 35.53 | 23.61 | 50.30 | 32.13 | Hypothetical protein |
| *PSPTO_5229* | | + | 5950332 | 5950781 | 38.54 | 25.41 | 13.25 | 30.96 | 72.01 | 41.64 | Hypothetical protein |
| *PSPTO_5230* | | - | 5950789 | 5951196 | 153.37 | 131.14 | 69.07 | 64.38 | 65.08 | 26.70 | Flagellar basal body-associated protein FliL-like protein |
| *PSPTO_5231* | | + | 5951327 | 5952304 | 19.67 | 26.94 | 16.96 | 12.83 | 46.04 | 14.81 | Oxidoreductase zinc-binding protein |
| *PSPTO_5232* | | + | 5952403 | 5954370 | 3.64 | 6.60 | 4.66 | 3.25 | 9.95 | 6.10 | Pyocin/colicin protein |
| *PSPTO_5233* | | + | 5954360 | 5954632 | 2.68 | 3.96 | 9.83 | 3.11 | 32.44 | 10.37 | Colicin/pyocin immunity family protein |
| *PSPTO_5234* | | + | 5954680 | 5955255 | 4.59 | 4.44 | 3.51 | 0.00 | 10.03 | 4.78 | Hypothetical protein |
| *PSPTO_5235* | | - | 5955571 | 5955927 | 2.76 | 0.00 | 0.00 | 0.00 | 1.79 | 2.10 | Hypothetical protein |
| *PSPTO_5236* | | - | 5955924 | 5956037 | 1.43 | 0.00 | 0.00 | 0.00 | 2.61 | 0.49 | Hypothetical protein |
| *PSPTO_5237* | | - | 5956131 | 5956589 | 2.20 | 2.07 | 1.07 | 3.12 | 2.88 | 2.76 | TonB domain protein |
| *PSPTO_5238* | *glpT* | + | 5956864 | 5958213 | 5.23 | 9.47 | 15.04 | 3.00 | 54.07 | 11.72 | Glycerol-3-phosphate transporter |
| *PSPTO_5239* | | - | 5958214 | 5958882 | 12.46 | 24.40 | 14.10 | 18.22 | 34.87 | 15.39 | ChaC-related protein |
| *PSPTO_5240* | | - | 5959090 | 5960058 | 6.81 | 67.76 | 54.60 | 31.44 | 63.83 | 23.26 | CDP-6-deoxy-delta-3,4-glucoseen reductase |
| *PSPTO_5241* | *ubiD* | - | 5960058 | 5961524 | 13.05 | 43.37 | 36.93 | 16.25 | 83.81 | 26.10 | 3-octaprenyl-4-hydroxybenzoate carboxy-lyase |
| *PSPTO_5242* | *rho* | - | 5961653 | 5962912 | 8.86 | 57.26 | 46.82 | 30.37 | 114.86 | 27.77 | Transcription termination factor Rho |
| *PSPTO_5243* | *trx-2* | - | 5963144 | 5963473 | 127.90 | 653.70 | 495.37 | 131.92 | 521.42 | 135.72 | Thioredoxin |
| *PSPTO_5244* | | - | 5963595 | 5964299 | 5.44 | 17.97 | 7.49 | 0.55 | 16.38 | 10.00 | GntR family transcriptional regulator |
| *PSPTO_5245* | | + | 5964550 | 5965335 | 29.96 | 50.61 | 23.32 | 7.48 | 9.05 | 7.94 | Amino acid ABC transporter substrate-binding protein |
| *PSPTO_5246* | | + | 5965365 | 5966030 | 2.69 | 7.02 | 4.10 | 5.85 | 2.65 | 3.02 | Amino acid ABC transporter permease |
| *PSPTO_5247* | | + | 5966030 | 5966677 | 6.83 | 10.60 | 3.54 | 2.58 | 4.78 | 2.80 | Amino acid ABC transporter permease |
| *PSPTO_5248* | | + | 5966664 | 5967398 | 4.18 | 24.95 | 5.40 | 0.89 | 4.24 | 2.67 | Amino acid ABC transporter ATP-binding protein |
| *PSPTO_5249* | *ppx* | + | 5967782 | 5969284 | 7.62 | 34.98 | 34.19 | 11.26 | 85.24 | 13.20 | Exopolyphosphatase |
| *PSPTO_5250* | *ppk* | - | 5969271 | 5971481 | 18.18 | 29.23 | 22.63 | 16.93 | 63.35 | 80.75 | Polyphosphate kinase |
| *PSPTO_5251* | *hemB* | - | 5971498 | 5972508 | 42.20 | 51.63 | 43.37 | 10.00 | 102.93 | 103.11 | Delta-aminolevulinic acid dehydratase |
| *PSPTO_5252* | | + | 5972740 | 5973402 | 10.41 | 56.12 | 32.01 | 16.42 | 65.41 | 17.93 | DedA family protein |
| *PSPTO_5253* | *elbB* | + | 5973609 | 5974277 | 33.96 | 56.70 | 58.96 | 26.45 | 85.72 | 40.25 | Enhancing lycopene biosynthesis protein 2 |
| *PSPTO_5254* | | + | 5974390 | 5975448 | 19.94 | 25.52 | 21.21 | 9.06 | 32.98 | 25.69 | Radical SAM domain-containing protein |
| *PSPTO_5255* | *cynT* | + | 5975730 | 5976395 | 20.09 | 208.81 | 266.87 | 71.25 | 111.37 | 33.52 | Carbonic anhydrase |
| *PSPTO_5256* | | + | 5976499 | 5978031 | 5.60 | 29.53 | 43.39 | 14.28 | 18.91 | 8.75 | Sulfate transporter family protein |
| *PSPTO_5257* | | - | 5978140 | 5978790 | 127.12 | 45.95 | 53.31 | 30.07 | 33.66 | 59.88 | Hypothetical protein |
| *PSPTO_5258* | *ctaG* | + | 5979094 | 5979582 | 13.60 | 23.90 | 9.13 | 10.78 | 13.56 | 15.62 | Cytochrome c oxidase assembly protein |
| *PSPTO_5259* | | + | 5979646 | 5980278 | 8.62 | 11.38 | 27.46 | 13.43 | 60.79 | 27.97 | Sco1/SenC family protein |
| *PSPTO_5260* | *metQ-2* | - | 5980418 | 5981191 | 17.11 | 29.95 | 32.34 | 30.37 | 110.91 | 26.44 | D-methionine-binding lipoprotein MetQ |
| *PSPTO_5261* | *metI-2* | - | 5981244 | 5981918 | 15.53 | 12.82 | 8.84 | 9.97 | 45.62 | 12.02 | D-methionine ABC transporter permease |
| *PSPTO_5262* | *metN-2* | - | 5981918 | 5982925 | 3.36 | 11.80 | 15.18 | 22.19 | 30.21 | 7.38 | DL-methionine transporter ATP-binding subunit |
| *PSPTO_5263* | *katE* | + | 5983233 | 5985383 | 3.20 | 29.56 | 23.29 | 12.79 | 9.00 | 6.18 | Catalase |
| *PSPTO_5264* | | - | 5985584 | 5986300 | 17.46 | 32.44 | 16.81 | 27.96 | 33.32 | 30.15 | Lipoprotein |
| *PSPTO_5265* | *znuB* | - | 5986339 | 5987127 | 15.64 | 10.89 | 17.34 | 5.12 | 17.80 | 9.74 | Zinc ABC transporter permease |
| *PSPTO_5266* | *znuC* | - | 5987120 | 5987914 | 7.25 | 10.73 | 11.15 | 6.59 | 13.43 | 6.79 | Zinc ABC transporter ATP-binding protein |

| Locus name | Gene name | Strand | Gene start (bp) | Gene end (bp) | Average TPM Inoculum | Average TPM $t_6$ | Average TPM $t_{24}$ | Average TPM $t_{168}$ | Average TPM Logarithmic | Average TPM Stationary | Annotation description |
|---|---|---|---|---|---|---|---|---|---|---|---|
| *PSPTO_5267* | | - | 5987914 | 5988396 | 23.53 | 37.87 | 32.56 | 18.09 | 31.98 | 19.63 | Zinc uptake regulation protein |
| *PSPTO_5268* | *znuA* | + | 5988464 | 5989399 | 15.82 | 36.34 | 53.58 | 27.03 | 35.50 | 19.53 | Zinc ABC transporter periplasmic zinc-binding protein |
| *PSPTO_5269* | *betT* | - | 5989584 | 5991578 | 6.38 | 10.80 | 13.57 | 5.18 | 60.59 | 13.82 | Choline transporter |
| *PSPTO_5270* | *hutU* | + | 5992065 | 5993888 | 2.95 | 5.14 | 7.32 | 4.79 | 25.35 | 1.91 | Urocanate hydratase |
| *PSPTO_5271* | | + | 5994001 | 5994969 | 6.36 | 6.89 | 12.04 | 7.15 | 17.87 | 4.31 | Histidine transporter, periplasmic histidine-binding protein |
| *PSPTO_5272* | | + | 5995043 | 5995894 | 4.54 | 9.43 | 9.04 | 2.09 | 18.28 | 3.93 | Histidine ABC transporter, permease protein |
| *PSPTO_5273* | | + | 5995891 | 5996721 | 4.05 | 11.96 | 8.50 | 3.48 | 25.58 | 6.91 | Histidine ABC transporter, ATP-binding protein |
| *PSPTO_5274* | *hutH-2* | + | 5996734 | 5998275 | 2.14 | 3.62 | 4.33 | 2.97 | 10.99 | 3.96 | Histidine ammonia-lyase |
| *PSPTO_5275* | *hutH-3* | + | 5998330 | 5999877 | 4.23 | 12.17 | 7.75 | 7.25 | 16.92 | 8.87 | Histidine ammonia-lyase |
| *PSPTO_5276* | *proY* | + | 5999980 | 6001356 | 2.09 | 5.81 | 5.53 | 2.82 | 29.94 | 10.68 | Proline-specific permease ProY |
| *PSPTO_5277* | *hutI* | + | 6001372 | 6002577 | 4.25 | 6.21 | 6.00 | 4.81 | 32.02 | 12.92 | Imidazolonepropionase |
| *PSPTO_5278* | | + | 6002591 | 6003391 | 3.81 | 15.12 | 10.41 | 14.70 | 51.84 | 33.28 | Formimidoylglutamase |
| *PSPTO_5279* | *cadA-1* | - | 6003536 | 6005794 | 30.86 | 28.65 | 21.03 | 11.56 | 181.98 | 110.55 | Cadmium-translocating P-type ATPase |
| *PSPTO_5280* | *cadR* | + | 6005880 | 6006326 | 16.52 | 9.01 | 8.06 | 11.89 | 49.35 | 6.53 | Transcriptional regulator CadR |
| *PSPTO_5281* | | - | 6006348 | 6006584 | 8.05 | 13.14 | 18.33 | 0.00 | 22.83 | 10.90 | Hypothetical protein |
| *PSPTO_5282* | *thyA* | - | 6006968 | 6007939 | 17.05 | 31.65 | 10.31 | 20.53 | 111.25 | 29.15 | Thymidylate synthase |
| *PSPTO_5283* | *lgt* | - | 6007992 | 6008804 | 16.92 | 21.75 | 22.60 | 9.71 | 64.55 | 26.01 | Prolipoprotein diacylglyceryl transferase |
| *PSPTO_5284* | *ptsP* | - | 6009168 | 6011447 | 25.23 | 58.06 | 49.23 | 22.21 | 112.52 | 69.38 | Phosphoenolpyruvate-protein phosphotransferase PtsP |
| *PSPTO_5285* | | - | 6011469 | 6011948 | 60.20 | 169.08 | 145.83 | 54.85 | 249.08 | 103.59 | MutT/nudix family protein |
| *PSPTO_5286* | | + | 6012091 | 6012747 | 26.87 | 78.84 | 55.58 | 18.11 | 174.80 | 94.73 | HAD-superfamily hydrolase |
| *PSPTO_5287* | | - | 6012761 | 6013189 | 13.23 | 19.34 | 23.49 | 3.17 | 34.07 | 15.13 | Hypothetical protein |
| *PSPTO_5288* | *ilvA-2* | - | 6013350 | 6014894 | 4.99 | 9.64 | 11.01 | 4.74 | 38.14 | 9.74 | Threonine dehydratase |
| *PSPTO_5289* | *rpiA* | + | 6015042 | 6015713 | 4.45 | 6.52 | 5.55 | 2.80 | 27.13 | 5.87 | Ribose 5-phosphate isomerase |
| *PSPTO_5290* | | - | 6015759 | 6016697 | 9.62 | 11.04 | 9.76 | 4.66 | 12.85 | 15.62 | Hypothetical protein |
| *PSPTO_5291* | | - | 6016775 | 6017707 | 34.63 | 26.48 | 18.28 | 9.11 | 35.20 | 60.82 | Hypothetical protein |
| *PSPTO_5292* | | - | 6017891 | 6018556 | 14.43 | 27.36 | 29.84 | 16.58 | 72.73 | 27.45 | Fumarylacetoacetate hydrolase family protein |
| *PSPTO_5293* | | - | 6018624 | 6020018 | 13.25 | 35.48 | 34.61 | 17.30 | 68.13 | 35.60 | Oxidoreductase, FAD-binding protein |
| *PSPTO_5294* | *serA* | + | 6020179 | 6021408 | 14.86 | 66.64 | 63.43 | 33.84 | 1314.30 | 56.19 | D-3-phosphoglycerate dehydrogenase |
| *PSPTO_5295* | | - | 6021581 | 6022018 | 16.61 | 84.19 | 73.43 | 39.93 | 31.38 | 23.95 | Hypothetical protein |
| *PSPTO_5296* | | - | 6022156 | 6022890 | 32.43 | 40.21 | 19.37 | 14.34 | 35.35 | 31.35 | 2OG-Fe(II) oxygenase family oxidoreductase |
| *PSPTO_5297* | | + | 6022933 | 6023691 | 17.23 | 37.70 | 39.39 | 24.43 | 92.94 | 38.15 | Hypothetical protein |
| *PSPTO_5298* | | + | 6023742 | 6023933 | 9.73 | 12.71 | 26.33 | 0.00 | 45.93 | 24.67 | Hypothetical protein |
| *PSPTO_5299* | | - | 6023930 | 6025000 | 31.16 | 32.47 | 21.34 | 18.93 | 49.96 | 32.44 | Hypothetical protein |
| *PSPTO_5300* | | + | 6025145 | 6027532 | 7.61 | 21.54 | 14.92 | 11.11 | 52.79 | 21.24 | Penicillin amidase family protein |
| *PSPTO_5301* | *potI* | - | 6027838 | 6028719 | 6.35 | 15.98 | 8.65 | 10.98 | 44.78 | 18.68 | Putrescine ABC transporter permease |
| *PSPTO_5302* | *potH* | - | 6028716 | 6029669 | 3.11 | 5.20 | 3.58 | 0.00 | 21.08 | 5.97 | Putrescine ABC transporter permease |
| *PSPTO_5303* | *potG* | - | 6029633 | 6030775 | 7.35 | 8.50 | 8.72 | 7.70 | 43.85 | 16.68 | Putrescine ABC transporter ATP-binding protein |
| *PSPTO_5304* | | - | 6030902 | 6032455 | 0.85 | 0.28 | 0.54 | 0.32 | 4.49 | 1.20 | ISPsy5, transposase |
| *PSPTO_5305* | | - | 6032473 | 6032832 | 0.00 | 0.00 | 0.00 | 0.00 | 0.00 | 0.00 | ISPsy5, Orf1 |
| *PSPTO_5306* | | - | 6032868 | 6033992 | 9.87 | 22.29 | 19.42 | 15.20 | 87.29 | 20.09 | Putrescine ABC transporter periplasmic putrescine-binding protein |
| *PSPTO_5307* | | - | 6034164 | 6035261 | 51.31 | 65.65 | 62.09 | 35.01 | 236.59 | 40.99 | Putrescine ABC transporter periplasmic putrescine-binding protein |
| *PSPTO_5308* | | - | 6035392 | 6036759 | 30.90 | 16.59 | 18.31 | 13.23 | 34.23 | 11.26 | Class III aminotransferase |
| *PSPTO_5309* | | - | 6036829 | 6038187 | 13.85 | 30.30 | 34.47 | 8.15 | 108.40 | 20.08 | Glutamine synthetase |
| *PSPTO_5310* | | + | 6038783 | 6040159 | 9.25 | 67.79 | 53.82 | 27.20 | 128.73 | 30.16 | Glutamine synthetase |
| *PSPTO_5311* | | + | 6040789 | 6041421 | 10.74 | 5.76 | 7.96 | 8.21 | 32.36 | 13.73 | TetR family transcriptional regulator |
| *PSPTO_5312* | | - | 6041425 | 6042567 | 2.59 | 12.80 | 2.24 | 8.00 | 6.78 | 4.60 | Hypothetical protein |
| *PSPTO_5313* | | - | 6042702 | 6042917 | 1.79 | 69.77 | 71.13 | 37.18 | 3.67 | 2.29 | Molybdenum-pterin binding domain-containing protein |
| *PSPTO_5314* | | - | 6042943 | 6043755 | 1.09 | 87.90 | 39.07 | 78.25 | 7.39 | 2.26 | Aliphatic sulfonates transport ATP-binding subunit |
| *PSPTO_5315* | *ssuC* | - | 6043752 | 6044549 | 1.18 | 85.43 | 49.33 | 57.15 | 5.30 | 2.85 | Aliphatic sulfonates ABC transporter permease |
| *PSPTO_5316* | | - | 6044578 | 6045543 | 1.28 | 145.64 | 75.64 | 76.04 | 5.04 | 1.86 | Sulfonate ABC transporter periplasmic sulfonate-binding protein |
| *PSPTO_5317* | | - | 6045856 | 6046494 | 66.18 | 240.27 | 152.46 | 127.18 | 36.90 | 39.47 | Antioxidant, AhpC/Tsa family |
| *PSPTO_5318* | | - | 6046664 | 6048010 | 25.95 | 21.66 | 19.13 | 16.51 | 56.34 | 25.63 | Outer membrane porin, OprD family |
| *PSPTO_5319* | *tauA* | + | 6048768 | 6049745 | 2.79 | 18.07 | 16.92 | 10.92 | 6.52 | 2.99 | Taurine ABC transporter, periplasmic taurine-binding protein |
| *PSPTO_5320* | *tauB* | + | 6049763 | 6050548 | 3.06 | 14.48 | 6.48 | 11.50 | 6.01 | 3.72 | Taurine transporter ATP-binding subunit |
| *PSPTO_5321* | *tauC* | + | 6050545 | 6051402 | 7.20 | 4.72 | 3.44 | 4.05 | 8.60 | 4.49 | Taurine ABC transporter permease |
| *PSPTO_5322* | | - | 6051749 | 6052189 | 63.26 | 34.90 | 13.63 | 11.97 | 35.88 | 14.98 | Hypothetical protein |
| *PSPTO_5323* | | + | 6052188 | 6052643 | 37.31 | 15.63 | 13.99 | 8.31 | 46.17 | 17.59 | TrmH family RNA methyltransferase |
| *PSPTO_5324* | *secB* | - | 6052712 | 6053206 | 9.82 | 72.51 | 49.67 | 16.57 | 269.17 | 18.03 | Protein-export protein SecB |
| *PSPTO_5325* | | - | 6053247 | 6053498 | 30.86 | 76.20 | 72.17 | 56.60 | 412.93 | 33.15 | Glutaredoxin |
| *PSPTO_5326* | | - | 6053501 | 6053914 | 31.65 | 58.36 | 86.87 | 27.28 | 198.11 | 37.20 | Rhodanese-like domain-containing protein |
| *PSPTO_5327* | *gpmA* | + | 6054071 | 6055603 | 14.20 | 56.91 | 36.23 | 23.08 | 95.04 | 33.96 | Phosphoglycerate mutase |
| *PSPTO_5328* | | + | 6055756 | 6057078 | 30.00 | 40.22 | 33.26 | 21.31 | 125.82 | 43.75 | Peptidase, M23/M37 family |
| *PSPTO_5329* | | + | 6057112 | 6058449 | 15.01 | 29.98 | 24.42 | 14.13 | 99.27 | 38.90 | Carboxyl-terminal protease family protein |
| *PSPTO_5330* | | + | 6058453 | 6059235 | 8.91 | 18.86 | 9.41 | 13.06 | 49.82 | 20.72 | Hypothetical protein |
| *PSPTO_5331* | | + | 6059422 | 6061314 | 45.47 | 28.93 | 28.64 | 19.81 | 44.18 | 32.03 | Hypothetical protein |
| *PSPTO_5332* | | + | 6061497 | 6063029 | 9.75 | 8.72 | 9.72 | 17.17 | 16.92 | 16.54 | Trypsin domain-containing protein |
| *PSPTO_5333* | | - | 6063034 | 6063843 | 6.98 | 13.18 | 12.47 | 2.28 | 49.01 | 17.39 | Hypothetical protein |
| *PSPTO_5334* | *hisF* | - | 6063976 | 6064746 | 2.39 | 10.99 | 3.30 | 7.99 | 75.36 | 9.51 | Imidazole glycerol phosphate synthase cyclase subunit |
| *PSPTO_5335* | *hisA* | - | 6064766 | 6065503 | 3.43 | 19.67 | 19.87 | 11.99 | 124.82 | 15.27 | Phosphoribosylformimino-5-aminoimidazole carboxamide ribotide isomerase |
| *PSPTO_5336* | | - | 6065609 | 6065869 | 3.99 | 16.28 | 21.82 | 12.74 | 68.27 | 8.55 | Hypothetical protein |
| *PSPTO_5337* | *hisH* | - | 6065874 | 6066512 | 5.99 | 20.08 | 14.75 | 13.50 | 66.94 | 12.78 | Amidotransferase |
| *PSPTO_5338* | *hisB* | - | 6066512 | 6067105 | 8.14 | 31.07 | 26.70 | 23.61 | 87.73 | 18.83 | Imidazoleglycerol-phosphate dehydratase |
| *PSPTO_5339* | *tyrB* | - | 6067198 | 6068397 | 6.11 | 19.83 | 24.20 | 9.52 | 67.62 | 13.45 | Aromatic-amino-acid aminotransferase |
| *PSPTO_5340* | | - | 6068635 | 6070296 | 29.66 | 28.66 | 19.18 | 8.49 | 27.60 | 4.30 | MFS permease-like protein |
| *PSPTO_5341* | | + | 6070797 | 6073022 | 7.14 | 18.91 | 24.07 | 12.43 | 78.34 | 51.90 | Hypothetical protein |
| *PSPTO_5342* | *mutY* | + | 6073107 | 6074174 | 9.69 | 25.24 | 20.58 | 19.43 | 97.61 | 58.78 | A/G-specific adenine glycosylase |
| *PSPTO_5343* | | + | 6074171 | 6074443 | 19.20 | 40.31 | 45.03 | 44.04 | 112.18 | 57.11 | Hypothetical protein |
| *PSPTO_5344* | | + | 6074827 | 6076035 | 9.73 | 24.09 | 13.20 | 9.74 | 56.11 | 17.35 | Site-specific recombinase, phage integrase family |
| *PSPTO_5345* | | + | 6076193 | 6077149 | 13.99 | 27.80 | 23.13 | 14.23 | 60.54 | 19.93 | Hypothetical protein |

| Locus name | Gene name | Strand | Gene start (bp) | Gene end (bp) | Average TPM Inoculum | Average TPM $t_6$ | Average TPM $t_{24}$ | Average TPM $t_{168}$ | Average TPM Logarithmic | Average TPM Stationary | Annotation description |
|---|---|---|---|---|---|---|---|---|---|---|---|
| *PSPTO_5346* | | + | 6078314 | 6079507 | 0.84 | 2.17 | 1.52 | 0.00 | 2.40 | 2.00 | Hypothetical protein |
| *PSPTO_5347* | | + | 6079628 | 6079879 | 2.49 | 7.65 | 3.84 | 5.30 | 1.68 | 3.61 | Hypothetical protein |
| *PSPTO_5348* | | - | 6080753 | 6081853 | 18.03 | 15.52 | 10.39 | 6.95 | 31.32 | 11.91 | Hypothetical protein |
| *PSPTO_5349* | | + | 6082956 | 6083183 | 18.61 | 25.28 | 27.89 | 0.00 | 26.12 | 20.64 | Prevent-host-death family protein |
| *PSPTO_5350* | | + | 6083180 | 6083553 | - | - | - | - | - | - | |
| *PSPTO_5351* | | + | 6083709 | 6083987 | 139.40 | 117.64 | 99.12 | 36.98 | 104.68 | 72.26 | Lipoprotein |
| *PSPTO_5352* | | + | 6084300 | 6085646 | 147.57 | 64.31 | 56.64 | 24.42 | 17.57 | 26.66 | Methyl-accepting chemotaxis protein |
| *PSPTO_5353* | *shcA* | + | 6085815 | 6086198 | 16.25 | 411.36 | 197.72 | 57.39 | 24.78 | 14.82 | Type III chaperone protein ShcA |
| *PSPTO_5354* | *hopA1* | + | 6086243 | 6087385 | 8.75 | 375.00 | 213.47 | 39.80 | 14.83 | 10.95 | Type III effector HopA1 |
| *PSPTO_5355* | | + | 6087509 | 6087994 | 4.43 | 124.95 | 64.82 | 7.79 | 40.27 | 10.57 | Hypothetical protein |
| *PSPTO_5356* | *gabP* | + | 6088042 | 6089433 | 5.16 | 55.39 | 25.97 | 10.16 | 23.86 | 5.51 | Gamma-aminobutyrate permease |
| *PSPTO_5357* | | - | 6089958 | 6090731 | 23.84 | 34.04 | 25.34 | 3.51 | 70.48 | 24.71 | Amino acid ABC transporter ATP-binding protein |
| *PSPTO_5358* | | + | 6090746 | 6091510 | 17.59 | 35.55 | 27.72 | 7.53 | 75.97 | 15.34 | Amino acid ABC transporter substrate-binding protein |
| *PSPTO_5359* | | + | 6091577 | 6092272 | 2.39 | 5.50 | 2.46 | 0.00 | 20.45 | 2.54 | Amino acid ABC transporter permease |
| *PSPTO_5360* | | + | 6092269 | 6092958 | 4.26 | 6.34 | 12.72 | 0.00 | 25.12 | 3.69 | Amino acid ABC transporter permease |
| *PSPTO_5361* | | + | 6092960 | 6094231 | 6.12 | 8.28 | 7.21 | 3.67 | 25.93 | 6.64 | Hypothetical protein |
| *PSPTO_5362* | | + | 6094527 | 6094841 | 4.49 | 7.62 | 18.02 | 24.00 | 22.65 | 8.57 | Hypothetical protein |
| *PSPTO_5363* | | - | 6095369 | 6096382 | 30.09 | 8.17 | 12.80 | 3.23 | 6.32 | 3.10 | DnaJ domain-containing protein |
| *PSPTO_5364* | | - | 6096783 | 6097085 | 3.13 | 0.00 | 5.23 | 0.13 | 5.44 | 4.25 | Hypothetical protein |
| *PSPTO_5365* | | + | 6098758 | 6099795 | 12.66 | 52.48 | 27.31 | 40.99 | 131.17 | 34.48 | Hypothetical protein |
| *PSPTO_5367* | | + | 6101513 | 6101872 | 0.00 | 0.00 | 0.00 | 0.00 | 0.00 | 0.00 | ISPsy5, Orf1 |
| *PSPTO_5368* | | + | 6101890 | 6103443 | 0.00 | 0.00 | 0.00 | 0.00 | 0.00 | 0.00 | ISPsy5, transposase |
| *PSPTO_5370* | | + | 6104051 | 6104860 | 0.00 | 0.00 | 0.00 | 0.00 | 0.00 | 0.00 | ISPsy4, transposition helper protein |
| *PSPTO_5371* | | + | 6105124 | 6105483 | 0.81 | 2.44 | 0.23 | 0.00 | 2.24 | 1.24 | ISPsy5, Orf1 |
| *PSPTO_5373* | | + | 6105848 | 6106930 | 2.56 | 6.63 | 11.59 | 0.48 | 9.71 | 4.63 | Hypothetical protein |
| *PSPTO_5374* | | + | 6107632 | 6107955 | 104.54 | 104.35 | 69.00 | 14.79 | 118.82 | 41.80 | Hypothetical protein |
| *PSPTO_5375* | | + | 6107952 | 6108296 | 19.68 | 37.80 | 22.00 | 14.78 | 84.25 | 29.24 | Hypothetical protein |
| *PSPTO_5376* | | - | 6108448 | 6109113 | 23.68 | 15.39 | 8.78 | 6.19 | 91.02 | 21.87 | GntR family transcriptional regulator |
| *PSPTO_5377* | | + | 6109369 | 6110601 | 8.84 | 4.28 | 0.40 | 1.21 | 30.02 | 6.04 | Membrane protein |
| *PSPTO_5378* | | + | 6110636 | 6111406 | 6.49 | 10.22 | 9.01 | 6.61 | 48.77 | 8.02 | LamB/YcsF family protein |
| *PSPTO_5379* | | + | 6111403 | 6112209 | 4.86 | 10.58 | 1.54 | 0.00 | 63.18 | 5.62 | Hypothetical protein |
| *PSPTO_5380* | | + | 6112219 | 6113838 | 5.67 | 10.45 | 3.74 | 3.31 | 76.72 | 9.07 | Allophanate hydrolase |
| *PSPTO_5381* | | + | 6113835 | 6115574 | 3.90 | 6.12 | 5.90 | 4.11 | 59.24 | 6.81 | Biotin carboxylase |
| *PSPTO_5382* | | + | 6115993 | 6116622 | 10.27 | 9.93 | 18.81 | 7.76 | 29.89 | 21.32 | ACP phosphodiesterase |
| *PSPTO_5383* | | - | 6116679 | 6117293 | 7.18 | 16.99 | 9.56 | 7.52 | 26.82 | 14.29 | Phosphoglycerate mutase family protein |
| *PSPTO_5384* | | - | 6117371 | 6117712 | 22.15 | 23.57 | 21.11 | 5.21 | 62.48 | 25.41 | Hypothetical protein |
| *PSPTO_5385* | | + | 6118066 | 6118803 | 22.88 | 19.65 | 24.38 | 9.40 | 8.94 | 4.49 | Hypothetical protein |
| *PSPTO_5386* | | - | 6118825 | 6118932 | 1.78 | 0.00 | 7.28 | 0.00 | 3.44 | 2.73 | Hypothetical protein |
| *PSPTO_5387* | | - | 6119030 | 6119320 | 50.99 | 19.60 | 30.72 | 21.87 | 44.15 | 21.21 | Stability cassette protein |
| *PSPTO_5388* | *relB* | - | 6119310 | 6119549 | 52.35 | 39.89 | 14.24 | 2.80 | 62.17 | 12.56 | Bifunctional antitoxin/transcriptional repressor RelB |
| *PSPTO_5389* | | - | 6119724 | 6120185 | 3.66 | 1.65 | 4.36 | 5.71 | 4.28 | 2.59 | GNAT family acetyltransferase |
| *PSPTO_5390* | | + | 6120522 | 6121052 | 15.48 | 18.22 | 9.49 | 6.58 | 17.60 | 12.68 | Hypothetical protein |
| *PSPTO_5391* | | - | 6121092 | 6122366 | 15.73 | 23.23 | 19.84 | 18.73 | 107.76 | 33.44 | Outer membrane porin, OprD family |
| *PSPTO_5392* | | - | 6122596 | 6122823 | 164.88 | 167.54 | 178.70 | 96.31 | 16.98 | 91.24 | Hypothetical protein |
| *PSPTO_5393* | | - | 6123022 | 6124128 | 4.78 | 12.73 | 12.24 | 4.37 | 53.80 | 7.22 | Hypothetical protein |
| *PSPTO_5394* | | - | 6124134 | 6125012 | 10.17 | 36.80 | 32.91 | 16.15 | 140.55 | 20.35 | Carbon-nitrogen hydrolase family protein |
| *PSPTO_5395* | | - | 6125320 | 6128232 | 3.88 | 6.39 | 4.70 | 3.43 | 10.05 | 5.91 | Class III aminotransferase |
| *PSPTO_5396* | *rfbB-2* | + | 6128443 | 6129525 | 10.77 | 34.08 | 37.49 | 20.78 | 92.61 | 19.92 | DTDP-glucose 4,6-dehydratase |
| *PSPTO_5397* | *rfbC-2* | + | 6129606 | 6130151 | 19.62 | 48.71 | 45.32 | 13.69 | 111.82 | 22.06 | DTDP-4-dehydrorhamnose 3,5-epimerase |
| *PSPTO_5398* | | + | 6130322 | 6132130 | 5.14 | 8.95 | 5.97 | 6.90 | 17.29 | 6.98 | Sensor histidine kinase |
| *PSPTO_5399* | | + | 6132127 | 6133512 | 6.16 | 13.74 | 9.37 | 2.92 | 40.45 | 10.89 | Sigma-54 dependent transcriptional regulator/response regulator |
| *PSPTO_5400* | | + | 6133499 | 6133975 | 2.63 | 4.46 | 4.47 | 5.46 | 16.99 | 4.90 | Hypothetical protein |
| *PSPTO_5401* | *cysQ* | - | 6134052 | 6134861 | 3.73 | 19.57 | 6.86 | 8.25 | 47.90 | 17.29 | 3'(2'),5'-bisphosphate nucleotidase |
| *PSPTO_5402* | | - | 6134891 | 6135457 | 11.26 | 39.03 | 57.40 | 27.21 | 105.20 | 36.10 | MutT/nudix family protein |
| *PSPTO_5403* | | + | 6135556 | 6136218 | 15.88 | 24.22 | 12.21 | 16.27 | 34.77 | 22.01 | HAD-superfamily hydrolase |
| *PSPTO_5404* | | - | 6136260 | 6136700 | 36.15 | 56.05 | 57.38 | 45.20 | 29.30 | 58.02 | LysM domain/BON superfamily protein |
| *PSPTO_5405* | | + | 6136852 | 6137733 | 9.33 | 18.71 | 22.55 | 20.60 | 35.50 | 23.04 | LysR family transcriptional regulator |
| *PSPTO_5406* | *fdhD* | + | 6137836 | 6138672 | 77.95 | 46.65 | 40.86 | 32.19 | 19.25 | 46.21 | Formate dehydrogenase accessory protein FdhD |
| *PSPTO_5407* | | + | 6138669 | 6141005 | 31.91 | 29.82 | 31.23 | 19.79 | 13.45 | 25.73 | Oxidoreductase, molybdopterin-binding protein |
| *PSPTO_5408* | | + | 6141173 | 6141679 | 337.98 | 716.14 | 560.76 | 154.63 | 129.22 | 93.36 | Hypothetical protein |
| *PSPTO_5409* | | - | 6141738 | 6143666 | 8.90 | 26.34 | 17.75 | 13.71 | 36.93 | 14.65 | ATP-dependent DNA helicase domain protein |
| *PSPTO_5410* | | + | 6143847 | 6144659 | 6.52 | 4.50 | 3.98 | 0.04 | 12.53 | 6.30 | Hypothetical protein |
| *PSPTO_5411* | | - | 6145421 | 6146974 | 0.00 | 0.00 | 0.00 | 0.00 | 0.00 | 0.00 | ISPsy5, transposase |
| *PSPTO_5412* | | - | 6146992 | 6147351 | 0.00 | 0.00 | 0.00 | 0.00 | 0.00 | 0.00 | ISPsy5, Orf1 |
| *PSPTO_5413* | | - | 6149242 | 6151620 | 13.75 | 24.28 | 21.36 | 10.77 | 115.12 | 24.65 | EF hand domain-containing protein |
| *PSPTO_5414* | | - | 6152531 | 6153547 | 6.28 | 17.40 | 9.41 | 6.31 | 118.59 | 20.57 | Lipoprotein |
| *PSPTO_5415* | | - | 6153553 | 6155589 | 8.08 | 7.20 | 5.58 | 3.94 | 52.09 | 9.87 | Rhs element Vgr protein |
| *PSPTO_5416* | | - | 6155611 | 6156606 | 10.17 | 6.21 | 9.65 | 7.31 | 44.14 | 10.31 | Serine/threonine protein kinase |
| *PSPTO_5417* | | - | 6156615 | 6157343 | 5.12 | 6.59 | 4.77 | 6.96 | 37.81 | 6.95 | Serine/threonine phosphoprotein phosphatase |
| *PSPTO_5418* | | - | 6157343 | 6160921 | 4.13 | 7.07 | 5.81 | 1.35 | 48.10 | 5.30 | Hypothetical protein |
| *PSPTO_5419* | | - | 6160884 | 6161759 | 5.79 | 2.36 | 3.86 | 4.06 | 58.04 | 7.65 | Hypothetical protein |
| *PSPTO_5420* | | - | 6161765 | 6163096 | 3.47 | 2.56 | 7.01 | 3.34 | 49.65 | 4.54 | Hypothetical protein |
| *PSPTO_5421* | | - | 6163099 | 6163599 | 4.60 | 7.48 | 5.78 | 1.46 | 63.52 | 6.55 | Lipoprotein |
| *PSPTO_5422* | | - | 6163605 | 6164801 | 3.71 | 7.46 | 6.45 | 2.47 | 69.95 | 5.61 | FHA domain-containing protein |
| *PSPTO_5423* | | - | 6164819 | 6164983 | 9.36 | 21.97 | 20.55 | 0.00 | 143.53 | 11.64 | Hypothetical protein |
| *PSPTO_5424* | | - | 6165047 | 6166564 | 4.31 | 5.70 | 6.35 | 0.95 | 25.12 | 4.45 | Sigma-54 dependent transcriptional regulator |
| *PSPTO_5425* | | - | 6166575 | 6169223 | 1.83 | 3.77 | 1.75 | 0.00 | 25.22 | 3.24 | ClpB protein |
| *PSPTO_5426* | | - | 6169237 | 6170244 | 2.63 | 4.75 | 2.66 | 3.31 | 22.70 | 3.57 | Hypothetical protein |
| *PSPTO_5427* | | - | 6170208 | 6171761 | 3.26 | 8.28 | 6.51 | 6.40 | 34.72 | 5.88 | Hypothetical protein |

| Locus name | Gene name | Strand | Gene start (bp) | Gene end (bp) | Average TPM Inoculum | Average TPM $t_6$ | Average TPM $t_{24}$ | Average TPM $t_{168}$ | Average TPM Logarithmic | Average TPM Stationary | Annotation description |
|---|---|---|---|---|---|---|---|---|---|---|---|
| *PSPTO_5428* | | - | 6173232 | 6174209 | 0.00 | 0.00 | 0.00 | 0.00 | 0.00 | 0.00 | ISPssy, transposase |
| *PSPTO_5430* | | - | 6175762 | 6176286 | 12.89 | 35.97 | 28.43 | 13.73 | 306.04 | 25.24 | Hypothetical protein |
| *PSPTO_5431* | | - | 6176297 | 6176704 | 8.39 | 9.30 | 5.08 | 15.71 | 154.81 | 13.90 | Hypothetical protein |
| *PSPTO_5432* | | - | 6176718 | 6178196 | 20.67 | 53.87 | 40.17 | 28.14 | 543.63 | 43.22 | Type VI protein secretion system |
| *PSPTO_5433* | | - | 6178225 | 6178731 | 26.35 | 56.80 | 48.34 | 35.82 | 645.04 | 43.40 | Hypothetical protein |
| *PSPTO_5434* | | - | 6178764 | 6180320 | 13.08 | 37.57 | 29.71 | 29.82 | 236.10 | 43.68 | Hypothetical protein |
| *PSPTO_5435* | *hcp-2* | + | 6181106 | 6181624 | 37.67 | 141.29 | 105.14 | 34.26 | 1134.43 | 26.12 | Secreted protein Hcp |
| *PSPTO_5436* | | + | 6181756 | 6183798 | 4.04 | 2.86 | 3.17 | 3.58 | 57.56 | 5.51 | Rhs element Vgr protein |
| *PSPTO_5437* | | + | 6183842 | 6184525 | 4.63 | 2.88 | 5.54 | 0.00 | 36.51 | 4.32 | Hypothetical protein |
| *PSPTO_5438* | | + | 6184525 | 6189243 | 8.40 | 13.55 | 12.24 | 7.50 | 77.05 | 15.08 | Rhs family protein |
| *PSPTO_5439* | | + | 6189796 | 6190467 | - | - | - | - | - | - | |
| *PSPTO_5440* | | - | 6190597 | 6191574 | 0.00 | 0.00 | 0.00 | 0.00 | 0.00 | 0.00 | ISPssy, transposase |
| *PSPTO_5441* | | + | 6192157 | 6193251 | - | - | - | - | - | - | |
| *PSPTO_5442* | | - | 6193709 | 6194068 | 8.68 | 11.97 | 10.97 | 5.22 | 22.31 | 12.57 | Hypothetical protein |
| *PSPTO_5443* | | - | 6194168 | 6195721 | 0.00 | 0.00 | 0.00 | 0.00 | 0.00 | 0.00 | ISPsy5, transposase |
| *PSPTO_5444* | | + | 6195901 | 6196260 | 0.00 | 0.00 | 0.00 | 0.00 | 0.00 | 0.00 | ISPsy5, Orf1 |
| *PSPTO_5445* | | + | 6196278 | 6197831 | 0.00 | 0.00 | 0.00 | 0.00 | 0.00 | 0.00 | ISPsy5, transposase |
| *PSPTO_5446* | | + | 6197883 | 6198005 | 5.95 | 6.52 | 12.97 | 0.00 | 7.29 | 5.90 | Hypothetical protein |
| *PSPTO_5447* | | + | 6198383 | 6198811 | 5.88 | 0.86 | 8.48 | 4.73 | 33.07 | 6.62 | Hypothetical protein |
| *PSPTO_5448* | | - | 6198893 | 6201319 | 4.88 | 3.90 | 2.08 | 3.36 | 16.44 | 4.81 | Hypothetical protein |
| *PSPTO_5449* | | - | 6201300 | 6201800 | 4.33 | 9.49 | 10.38 | 1.76 | 27.49 | 7.84 | Hypothetical protein |
| *PSPTO_5450* | | - | 6201983 | 6202288 | 2.93 | 8.99 | 3.80 | 1.70 | 26.84 | 15.92 | Hypothetical protein |
| *PSPTO_5451* | | - | 6202574 | 6204055 | 13.06 | 18.17 | 20.72 | 6.52 | 64.64 | 33.40 | Hypothetical protein |
| *PSPTO_5452* | | + | 6204261 | 6204899 | 9.65 | 31.31 | 21.16 | 8.74 | 66.69 | 12.26 | HAD-superfamily hydrolase |
| *PSPTO_5453* | | + | 6204920 | 6205927 | 9.64 | 23.99 | 27.55 | 6.91 | 64.11 | 18.05 | Oxidoreductase, aldo/keto reductase family |
| *PSPTO_5454* | | + | 6206247 | 6206984 | 29.48 | 42.09 | 26.16 | 23.39 | 95.54 | 70.89 | GntR family transcriptional regulator |
| *PSPTO_5455* | | + | 6207790 | 6208566 | 2.18 | 0.00 | 0.00 | 0.72 | 4.04 | 2.88 | Hypothetical protein |
| *PSPTO_5456* | | + | 6208627 | 6209556 | 5.51 | 4.66 | 4.30 | 3.56 | 4.84 | 3.85 | Hypothetical protein |
| *PSPTO_5457* | *mgoA* | + | 6209553 | 6213002 | 2.60 | 2.22 | 2.43 | 2.87 | 3.81 | 2.73 | Ornithine acetyl transferase inhibitor |
| *PSPTO_5458* | | + | 6213036 | 6213512 | 2.77 | 7.59 | 13.37 | 4.24 | 5.98 | 2.84 | Hypothetical protein |
| *PSPTO_5459* | | + | 6219079 | 6219174 | 4877.98 | 2967.82 | 2096.28 | 2642.75 | 21669.22 | 4569.63 | |
| *PSPTO_5460* | | - | 6219738 | 6220958 | 45.07 | 5.52 | 3.39 | 3.94 | 43.27 | 88.75 | CAIB/BAIF family protein |
| *PSPTO_5461* | | - | 6220971 | 6222152 | 450.94 | 18.71 | 25.28 | 9.11 | 144.05 | 306.09 | Acyl-CoA dehydrogenase |
| *PSPTO_5462* | | + | 6222308 | 6223228 | 14.91 | 39.33 | 28.09 | 7.98 | 40.12 | 13.01 | LysR family transcriptional regulator |
| *PSPTO_5463* | | + | 6223654 | 6225109 | - | - | - | - | - | - | |
| *PSPTO_5464* | *pntB* | + | 6225106 | 6226560 | 2.33 | 2.23 | 0.65 | 0.77 | 5.57 | 2.10 | NAD(P) transhydrogenase subunit beta |
| *PSPTO_5465* | | + | 6226740 | 6228233 | 10.44 | 16.29 | 14.95 | 6.69 | 150.04 | 24.32 | Acetyl-CoA hydrolase/transferase family protein |
| *PSPTO_5466* | | + | 6228554 | 6228769 | 1199.65 | 235.55 | 240.67 | 89.05 | 1427.39 | 2851.96 | Hypothetical protein |
| *PSPTO_5467* | | + | 6229085 | 6229402 | 24.50 | 103.75 | 61.77 | 43.75 | 316.82 | 96.77 | Hypothetical protein |
| *PSPTO_5468* | | + | 6229490 | 6229795 | 12.50 | 52.82 | 31.26 | 11.08 | 101.70 | 26.90 | Hypothetical protein |
| *PSPTO_5469* | | + | 6230154 | 6232115 | 6.99 | 14.93 | 9.58 | 7.66 | 47.23 | 17.88 | Hypothetical protein |
| *PSPTO_5470* | | - | 6232116 | 6232661 | 22.65 | 1.41 | 7.02 | 2.10 | 11.16 | 10.69 | Hypothetical protein |
| *PSPTO_5471* | | - | 6232666 | 6233094 | 6.99 | 3.34 | 8.30 | 0.00 | 7.68 | 5.83 | Hypothetical protein |
| *PSPTO_5472* | | - | 6233359 | 6233694 | 29.17 | 225.84 | 313.93 | 182.65 | 144.10 | 15.08 | HU family DNA-binding protein |
| *PSPTO_5473* | *rubB* | - | 6233823 | 6234971 | 17.81 | 31.78 | 35.03 | 17.44 | 65.78 | 20.97 | Rubredoxin reductase |
| *PSPTO_5474* | *rubA* | - | 6235020 | 6235187 | 24.57 | 115.61 | 112.66 | 18.64 | 168.67 | 43.22 | Rubredoxin |
| *PSPTO_5475* | | + | 6235375 | 6235935 | 4.17 | 10.19 | 8.99 | 8.27 | 37.05 | 8.92 | Chorismate-pyruvate lyase |
| *PSPTO_5476* | *ubiA* | + | 6235935 | 6236825 | 3.23 | 7.52 | 6.53 | 2.70 | 22.54 | 4.88 | 4-hydroxybenzoate octaprenyltransferase |
| *PSPTO_5477* | *phoB* | + | 6236994 | 6237683 | 43.53 | 56.94 | 37.83 | 20.95 | 30.45 | 10.60 | Phosphate regulon transcriptional regulatory protein PhoB |
| *PSPTO_5478* | *phoR* | + | 6237744 | 6239066 | 5.53 | 8.35 | 11.59 | 8.54 | 15.36 | 5.50 | Phosphate regulon sensor protein phoR |
| *PSPTO_5479* | | + | 6239166 | 6240506 | 6.88 | 17.05 | 12.63 | 6.89 | 50.85 | 13.61 | Transporter |
| *PSPTO_5480* | | - | 6240594 | 6241400 | 4.69 | 8.23 | 6.92 | 4.66 | 19.34 | 9.04 | Peptidase, M23/M37 family |
| *PSPTO_5481* | | - | 6241657 | 6246309 | 9.52 | 5.06 | 7.11 | 4.54 | 11.67 | 9.41 | Hypothetical protein |
| *PSPTO_5482* | | - | 6246533 | 6247426 | 25.20 | 12.28 | 10.83 | 8.38 | 9.56 | 4.38 | Response regulator |
| *PSPTO_5483* | *phoU* | - | 6247629 | 6248390 | 8.18 | 17.46 | 23.50 | 13.10 | 74.02 | 7.48 | Phosphate transport system protein PhoU |
| *PSPTO_5484* | *pstB-2* | - | 6248461 | 6249294 | 27.95 | 38.32 | 34.17 | 11.68 | 116.42 | 27.02 | Phosphate transporter ATP-binding protein |
| *PSPTO_5485* | | - | 6249361 | 6251034 | 11.60 | 14.76 | 10.33 | 4.65 | 27.39 | 7.29 | Phosphate ABC transporter permease |
| *PSPTO_5486* | | - | 6251046 | 6253079 | 9.12 | 13.18 | 6.90 | 4.41 | 17.13 | 6.06 | Phosphate ABC transporter permease |
| *PSPTO_5487* | | - | 6253460 | 6254458 | 40.72 | 33.96 | 41.65 | 14.39 | 12.63 | 5.15 | Phosphate ABC transporter substrate-binding protein |
| *PSPTO_5488* | | - | 6254628 | 6255965 | 18.82 | 25.78 | 18.17 | 14.57 | 35.57 | 17.57 | Major facilitator family transporter |
| *PSPTO_5489* | | + | 6256348 | 6256749 | 23.26 | 19.71 | 29.84 | 9.90 | 32.59 | 10.81 | Cytosolic long-chain acyl-CoA thioester hydrolase family protein |
| *PSPTO_5490* | | + | 6256840 | 6257751 | 53.00 | 51.82 | 29.31 | 20.62 | 133.47 | 41.33 | Aldose 1-epimerase family protein |
| *PSPTO_5491* | | - | 6257777 | 6258325 | 29.24 | 22.90 | 37.21 | 9.21 | 60.60 | 35.28 | Hypothetical protein |
| *PSPTO_5492* | | - | 6258394 | 6258639 | 113.19 | 108.77 | 88.05 | 37.25 | 56.13 | 66.59 | Hypothetical protein |
| *PSPTO_5493* | *purK* | - | 6258927 | 6260009 | 7.27 | 4.57 | 13.19 | 3.17 | 25.97 | 7.30 | Phosphoribosylaminoimidazole carboxylase ATPase subunit |
| *PSPTO_5494* | *purE* | - | 6260019 | 6260525 | 5.24 | 36.67 | 22.58 | 20.73 | 77.20 | 23.86 | Phosphoribosylaminoimidazole carboxylase catalytic subunit |
| *PSPTO_5495* | | + | 6260843 | 6261475 | 4.14 | 4.92 | 1.67 | 1.32 | 7.32 | 4.98 | Hypothetical protein |
| *PSPTO_5496* | | + | 6261517 | 6261672 | 14.29 | 10.33 | 8.82 | 1.64 | 17.34 | 16.18 | Hypothetical protein |
| *PSPTO_5497* | | + | 6261731 | 6262363 | 3.96 | 1.03 | 1.35 | 4.44 | 15.67 | 7.33 | Hypothetical protein |
| *PSPTO_5498* | | - | 6262399 | 6263310 | 13.50 | 7.61 | 6.02 | 5.00 | 20.52 | 9.72 | LysR family transcriptional regulator |
| *PSPTO_5499* | *aspA* | + | 6263605 | 6265029 | 9.56 | 26.41 | 28.91 | 15.51 | 394.12 | 34.27 | Aspartate ammonia-lyase |
| *PSPTO_5500* | | + | 6265312 | 6266763 | 6.25 | 24.68 | 16.44 | 9.42 | 153.93 | 42.10 | Sodium/alanine transporter |
| *PSPTO_5501* | *ansA* | + | 6266793 | 6267797 | 2.99 | 10.89 | 10.72 | 3.14 | 63.20 | 6.34 | L-asparaginase I |
| *PSPTO_5502* | | - | 6267861 | 6268748 | 9.30 | 12.31 | 22.03 | 3.58 | 24.09 | 19.27 | Hypothetical protein |
| *PSPTO_5503* | | - | 6268881 | 6270092 | 3.21 | 7.05 | 2.02 | 4.76 | 14.72 | 7.66 | YeeE/YedE family protein |
| *PSPTO_5504* | | - | 6270147 | 6271709 | 2.66 | 5.11 | 2.83 | 0.47 | 9.13 | 4.39 | GntR family transcriptional regulator |
| *PSPTO_5505* | | + | 6271800 | 6272438 | 30.42 | 15.92 | 10.45 | 1.00 | 69.09 | 89.83 | Transcriptional regulator |
| *PSPTO_5506* | | + | 6272435 | 6272881 | 12.43 | 5.99 | 9.19 | 6.03 | 40.44 | 42.92 | GNAT family acetyltransferase |
| *PSPTO_5507* | | + | 6272897 | 6273565 | 11.28 | 11.64 | 12.32 | 4.81 | 38.03 | 43.86 | GNAT family acetyltransferase |

| Locus name | Gene name | Strand | Gene start (bp) | Gene end (bp) | Average TPM Inoculum | Average TPM $t_6$ | Average TPM $t_{24}$ | Average TPM $t_{168}$ | Average TPM Logarithmic | Average TPM Stationary | Annotation description |
|---|---|---|---|---|---|---|---|---|---|---|---|
| *PSPTO_5508* | | + | 6273896 | 6274411 | 14.74 | 19.80 | 18.32 | 13.60 | 38.07 | 18.96 | ISPsy8, transposase OrfA |
| *PSPTO_5509* | | + | 6274468 | 6275247 | 16.63 | 17.24 | 12.20 | 9.73 | 25.32 | 16.97 | ISPsy8, transposase OrfB |
| *PSPTO_5510* | *oadA* | - | 6275563 | 6277371 | 128.97 | 73.90 | 43.00 | 20.32 | 175.84 | 13.95 | Pyruvate carboxylase subunit B |
| *PSPTO_5511* | | - | 6277474 | 6278889 | 114.18 | 177.11 | 101.75 | 29.10 | 418.46 | 22.70 | Pyruvate carboxylase subunit A |
| *PSPTO_5512* | | + | 6279118 | 6280053 | 1.72 | 2.71 | 1.43 | 5.05 | 5.42 | 2.42 | LysR family transcriptional regulator |
| *PSPTO_5513* | | - | 6280007 | 6280228 | 14.79 | 10.58 | 14.58 | 20.54 | 11.06 | 9.80 | Hypothetical protein |
| *PSPTO_5514* | | + | 6280450 | 6280761 | 21.35 | 40.54 | 25.50 | 46.25 | 32.06 | 22.72 | Hypothetical protein |
| *PSPTO_5515* | | - | 6280855 | 6281721 | 5.24 | 35.41 | 20.60 | 10.55 | 78.13 | 9.97 | RpiR family transcriptional regulator |
| *PSPTO_5516* | *uvrD* | + | 6282042 | 6284225 | 9.58 | 23.12 | 15.45 | 12.17 | 92.38 | 22.13 | DNA helicase II |
| *PSPTO_5517* | | + | 6284450 | 6285292 | 37.73 | 384.47 | 320.72 | 55.53 | 376.42 | 68.00 | Hypothetical protein |
| *PSPTO_5518* | | + | 6285423 | 6285830 | 131.33 | 108.73 | 78.27 | 52.83 | 178.73 | 67.17 | Hypothetical protein |
| *PSPTO_5519* | | - | 6285870 | 6287633 | 6.57 | 16.74 | 8.50 | 2.62 | 16.86 | 8.33 | Sodium/hydrogen exchanger family protein |
| *PSPTO_5520* | | + | 6287926 | 6288411 | 27.61 | 29.92 | 16.94 | 8.83 | 43.23 | 14.09 | Cytosolic long-chain acyl-CoA thioester hydrolase family protein |
| *PSPTO_5521* | *pdxY* | - | 6288458 | 6289324 | 3.78 | 7.64 | 4.88 | 0.00 | 13.25 | 4.96 | Pyridoxal kinase |
| *PSPTO_5522* | | + | 6289443 | 6289835 | 68.64 | 100.55 | 46.05 | 34.17 | 147.31 | 56.02 | Hypothetical protein |
| *PSPTO_5523* | | - | 6289884 | 6290855 | 12.24 | 22.09 | 13.50 | 8.05 | 38.46 | 17.27 | Cobalamin synthesis protein/P47K family protein |
| *PSPTO_5524* | | + | 6290908 | 6291279 | 11.39 | 27.72 | 27.57 | 4.50 | 35.53 | 17.95 | Hypothetical protein |
| *PSPTO_5525* | | - | 6291286 | 6291936 | 9.13 | 25.19 | 30.80 | 20.28 | 12.32 | 6.40 | Hypothetical protein |
| *PSPTO_5526* | | - | 6291936 | 6293147 | 5.77 | 18.58 | 28.01 | 7.34 | 5.72 | 3.27 | Cobalamin synthesis protein/P47K family protein |
| *PSPTO_5527* | | + | 6293289 | 6293687 | 33.20 | 27.26 | 26.14 | 8.28 | 11.99 | 12.98 | Hypothetical protein |
| *PSPTO_5528* | | - | 6293693 | 6294916 | 17.49 | 22.47 | 22.97 | 9.11 | 12.02 | 13.76 | N-acetylmuramoyl-L-alanine amidase family protein |
| *PSPTO_5529* | | - | 6295115 | 6295720 | 64.23 | 65.43 | 37.92 | 21.09 | 39.52 | 41.10 | Hypothetical protein |
| *PSPTO_5530* | *cls* | + | 6295809 | 6297248 | 9.87 | 16.56 | 12.58 | 3.97 | 15.23 | 7.66 | Cardiolipin synthetase |
| *PSPTO_5531* | | + | 6297321 | 6298361 | 12.44 | 19.01 | 14.50 | 15.82 | 26.30 | 21.56 | Hypothetical protein |
| *PSPTO_5532* | *cadA-2* | - | 6298291 | 6300216 | 3.13 | 8.15 | 6.36 | 2.65 | 7.55 | 4.95 | Cadmium-translocating P-type ATPase |
| *PSPTO_5533* | | - | 6300213 | 6301283 | 1.97 | 7.60 | 11.28 | 5.51 | 7.87 | 4.02 | Hypothetical protein |
| *PSPTO_5534* | | - | 6301280 | 6302317 | 2.50 | 7.79 | 3.77 | 0.00 | 6.73 | 3.47 | Hypothetical protein |
| *PSPTO_5535* | | - | 6302329 | 6304275 | 3.35 | 7.67 | 8.26 | 1.14 | 8.75 | 3.28 | Hypothetical protein |
| *PSPTO_5536* | | + | 6304618 | 6305235 | 3.58 | 4.77 | 8.22 | 1.32 | 13.07 | 4.79 | CDP-alcohol phosphatidyltransferase |
| *PSPTO_5537* | | + | 6305268 | 6307025 | 5.48 | 7.07 | 11.57 | 1.04 | 23.89 | 9.26 | Hypothetical protein |
| *PSPTO_5538* | | + | 6307022 | 6308377 | 3.31 | 5.47 | 1.44 | 2.43 | 7.12 | 3.58 | Dual specificity phosphatase, catalytic domain-containing protein |
| *PSPTO_5539* | | + | 6308401 | 6308856 | 1.61 | 1.81 | 3.10 | 3.75 | 4.12 | 2.67 | Hypothetical protein |
| *PSPTO_5540* | | + | 6308865 | 6309485 | 1.69 | 3.74 | 0.00 | 0.00 | 7.09 | 3.35 | Acyltransferase |
| *PSPTO_5541* | *cdsA-2* | + | 6309485 | 6310414 | 5.77 | 4.51 | 1.42 | 3.95 | 16.99 | 5.69 | Phosphatidate cytidylyltransferase |
| *PSPTO_5542* | | + | 6310816 | 6312435 | 7.93 | 74.28 | 72.12 | 15.12 | 135.39 | 41.58 | Glucan biosynthesis protein D |
| *PSPTO_5543* | | - | 6312909 | 6314462 | 0.00 | 0.00 | 0.00 | 0.00 | 0.00 | 0.00 | ISPsy5, transposase |
| *PSPTO_5544* | | - | 6314480 | 6314839 | 0.00 | 0.00 | 0.00 | 0.00 | 0.00 | 0.00 | ISPsy5, Orf1 |
| *PSPTO_5545* | | + | 6315100 | 6316401 | 3.09 | 2.56 | 4.46 | 1.57 | 4.38 | 2.95 | Membrane protein |
| *PSPTO_5546* | | + | 6316403 | 6317749 | 2.94 | 3.45 | 2.61 | 0.00 | 4.59 | 2.35 | Hypothetical protein |
| *PSPTO_5547* | | + | 6317770 | 6319656 | 3.30 | 3.37 | 3.97 | 0.91 | 5.03 | 5.01 | Acyltransferase family protein |
| *PSPTO_5548* | | - | 6319798 | 6320436 | 5.87 | 1.53 | 9.45 | 1.43 | 11.38 | 5.52 | LuxR family DNA-binding response regulator |
| *PSPTO_5549* | | - | 6320436 | 6321503 | 6.08 | 16.34 | 9.95 | 2.89 | 16.80 | 8.32 | Sensor histidine kinase |
| *PSPTO_5550* | | - | 6321553 | 6322857 | 3.94 | 5.43 | 1.36 | 5.05 | 11.16 | 5.79 | ABC transporter ATP-binding protein |
| *PSPTO_5551* | | - | 6322889 | 6324643 | 1.25 | 1.29 | 2.29 | 0.00 | 2.18 | 1.15 | ABC transporter permease |
| *PSPTO_5552* | | + | 6324864 | 6324989 | 27.79 | 90.90 | 12.55 | 9.69 | 45.78 | 25.29 | Hypothetical protein |
| *PSPTO_5553* | | - | 6325129 | 6327051 | 7.98 | 5.71 | 3.44 | 2.92 | 6.75 | 3.71 | Methyl-accepting chemotaxis protein |
| *PSPTO_5554* | | - | 6327228 | 6329258 | 4.23 | 3.54 | 5.05 | 2.55 | 4.69 | 4.17 | Methyl-accepting chemotaxis protein |
| *PSPTO_5555* | | + | 6329502 | 6329735 | 3.77 | 3.97 | 9.81 | 0.00 | 3.31 | 2.08 | Hypothetical protein |
| *PSPTO_5556* | | + | 6330285 | 6331262 | 0.00 | 0.00 | 0.00 | 0.00 | 0.00 | 0.00 | ISPssy, transposase |
| *PSPTO_5557* | *cysC* | - | 6331415 | 6332083 | 13.28 | 12.88 | 16.14 | 6.74 | 40.71 | 18.10 | Adenylylsulfate kinase |
| *PSPTO_5558* | | + | 6332275 | 6333726 | 4.12 | 6.41 | 8.39 | 5.11 | 5.80 | 4.90 | Hypothetical protein |
| *PSPTO_5559* | | + | 6333723 | 6334697 | 1.50 | 5.23 | 6.73 | 7.75 | 5.23 | 3.37 | Hypothetical protein |
| *PSPTO_5560* | | + | 6334735 | 6336954 | 3.49 | 7.22 | 9.49 | 6.51 | 5.60 | 3.61 | TonB-dependent receptor |
| *PSPTO_5561* | | + | 6336954 | 6337733 | 2.46 | 3.17 | 3.36 | 0.47 | 3.26 | 3.19 | Iron-compound ABC transporter, ATP-binding protein |
| *PSPTO_5562* | | + | 6337734 | 6338684 | 8.36 | 9.69 | 5.61 | 5.00 | 9.74 | 12.05 | Iron compound ABC transporter substrate-binding protein |
| *PSPTO_5563* | | + | 6338681 | 6339721 | 4.65 | 5.58 | 5.78 | 3.19 | 7.17 | 3.89 | Hemin ABC transporter, permease protein |
| *PSPTO_5564* | | + | 6339803 | 6340780 | 0.00 | 0.00 | 0.00 | 0.00 | 0.00 | 0.00 | ISPssy, transposase |
| *PSPTO_5565* | | - | 6341287 | 6341928 | 0.85 | 0.83 | 0.95 | 0.00 | 14.00 | 6.91 | Hypothetical protein |
| *PSPTO_5566* | | - | 6342781 | 6344472 | 1.61 | 3.89 | 8.28 | 3.78 | 29.13 | 8.40 | Transposition protein, TnsD-related protein |
| *PSPTO_5567* | | + | 6345505 | 6346263 | 7.92 | 5.31 | 6.81 | 6.06 | 11.25 | 7.77 | Membrane protein |
| *PSPTO_5568* | | - | 6346410 | 6346559 | 2.30 | 8.62 | 0.00 | 0.00 | 8.22 | 5.30 | Hypothetical protein |
| *PSPTO_5569* | | + | 6347028 | 6348653 | 7.89 | 16.76 | 17.20 | 15.93 | 41.39 | 24.67 | Methyl-accepting chemotaxis protein |
| *PSPTO_5570* | | - | 6349026 | 6349463 | - | - | - | - | - | - | |
| *PSPTO_5571* | | + | 6349624 | 6350646 | 20.49 | 27.90 | 22.99 | 15.16 | 37.57 | 26.52 | ISPsy7, transposase |
| *PSPTO_5572* | | - | 6350829 | 6352334 | - | - | - | - | - | - | |
| *PSPTO_5573* | | + | 6352913 | 6354199 | 14.92 | 14.03 | 16.23 | 11.71 | 32.38 | 19.99 | Sensor histidine kinase |
| *PSPTO_5574* | | + | 6354199 | 6354822 | - | - | - | - | - | - | |
| *PSPTO_5575* | | + | 6354877 | 6355899 | 0.00 | 0.00 | 0.00 | 0.00 | 0.00 | 0.00 | ISPsy4, transposase |
| *PSPTO_5576* | | + | 6355896 | 6356705 | 0.00 | 0.00 | 0.00 | 0.00 | 0.00 | 0.00 | ISPsy4, transposition helper protein |
| *PSPTO_5577* | | + | 6356752 | 6356841 | - | - | - | - | - | - | |
| *PSPTO_5578* | | + | 6356934 | 6357614 | 6.42 | 15.12 | 10.46 | 11.52 | 13.45 | 5.82 | Hypothetical protein |
| *PSPTO_5579* | | - | 6357958 | 6358392 | 8.89 | 16.55 | 14.55 | 4.78 | 23.31 | 11.28 | MerR family transcriptional regulator |
| *PSPTO_5580* | | + | 6358486 | 6359448 | 6.03 | 4.32 | 7.20 | 0.84 | 14.85 | 8.04 | Sodium:bile acid symporter family protein |
| *PSPTO_5581* | | - | 6359624 | 6359956 | 4.69 | 8.38 | 11.99 | 0.00 | 25.87 | 11.29 | Hypothetical protein |
| *PSPTO_5582* | | + | 6360062 | 6361039 | 0.00 | 0.00 | 0.00 | 0.00 | 0.00 | 0.00 | ISPssy, transposase |
| *PSPTO_5583* | *ais* | - | 6361325 | 6362011 | 6.91 | 4.43 | 4.24 | 4.21 | 10.80 | 6.18 | Ais protein |
| *PSPTO_5584* | | + | 6362330 | 6363979 | 3.94 | 5.80 | 3.82 | 3.79 | 8.49 | 5.68 | Membrane protein |
| *PSPTO_5585* | | + | 6364181 | 6365509 | 5.34 | 4.85 | 4.22 | 3.60 | 6.50 | 3.33 | UDP-glucose 6-dehydrogenase |
| *PSPTO_5586* | | + | 6365506 | 6366504 | 3.58 | 4.92 | 1.70 | 4.31 | 5.32 | 5.11 | Capsular polysaccharide biosynthesis protein |

| Locus name | Gene name | Strand | Gene start (bp) | Gene end (bp) | Average TPM Inoculum | Average TPM $t_6$ | Average TPM $t_{24}$ | Average TPM $t_{168}$ | Average TPM Logarithmic | Average TPM Stationary | Annotation description |
|---|---|---|---|---|---|---|---|---|---|---|---|
| *PSPTO_5587* | | + | 6366513 | 6367256 | 1.48 | 2.53 | 2.80 | 2.25 | 4.46 | 3.79 | Glycosyl transferase family protein |
| *PSPTO_5588* | | + | 6367333 | 6369063 | 10.30 | 9.13 | 10.13 | 4.08 | 10.50 | 6.44 | Dolichyl-phosphate-mannose-protein mannosyltransferase family protein |
| *PSPTO_5589* | | + | 6369107 | 6370087 | 8.74 | 12.56 | 10.86 | 12.79 | 12.98 | 9.50 | Hypothetical protein |
| *PSPTO_5590* | | + | 6370444 | 6370803 | 0.00 | 0.00 | 0.00 | 0.00 | 0.00 | 0.00 | ISPsy5, Orf1 |
| *PSPTO_5591* | | + | 6370821 | 6372374 | 0.00 | 0.00 | 0.00 | 0.00 | 0.00 | 0.00 | ISPsy5, transposase |
| *PSPTO_5592* | | - | 6373114 | 6373995 | 2.10 | 4.30 | 0.70 | 1.19 | 11.26 | 4.19 | Hypothetical protein |
| *PSPTO_5593* | | - | 6374081 | 6374518 | - | - | - | - | - | - | |
| *PSPTO_5594* | | - | 6374515 | 6376038 | - | - | - | - | - | - | |
| *PSPTO_5595* | *glmS* | - | 6376404 | 6378239 | 9.68 | 30.67 | 36.68 | 18.99 | 74.01 | 11.46 | Glucosamine--fructose-6-phosphate aminotransferase |
| *PSPTO_5596* | | - | 6378249 | 6379016 | 4.49 | 69.71 | 70.04 | 27.82 | 37.97 | 7.35 | DeoR family transcriptional regulator |
| *PSPTO_5597* | *glmU* | - | 6379386 | 6380753 | 11.27 | 40.72 | 37.40 | 17.16 | 87.56 | 52.04 | UDP-N-acetylglucosamine pyrophosphorylase |
| *PSPTO_5598* | *atpC* | - | 6380879 | 6381298 | 15.66 | 64.67 | 36.25 | 22.84 | 526.70 | 58.82 | F0F1 ATP synthase subunit epsilon |
| *PSPTO_5599* | *atpD* | - | 6381349 | 6382728 | 22.39 | 76.80 | 66.75 | 20.51 | 550.11 | 52.22 | F0F1 ATP synthase subunit beta |
| *PSPTO_5600* | *atpG* | - | 6382756 | 6383616 | 35.94 | 140.98 | 105.73 | 38.65 | 992.37 | 78.41 | F0F1 ATP synthase subunit gamma |
| *PSPTO_5601* | *atpA* | - | 6383667 | 6385211 | 25.02 | 104.54 | 73.71 | 38.04 | 915.66 | 77.01 | F0F1 ATP synthase subunit alpha |
| *PSPTO_5602* | *atpH* | - | 6385233 | 6385769 | 50.60 | 187.91 | 185.76 | 141.62 | 1951.74 | 166.35 | F0F1 ATP synthase subunit delta |
| *PSPTO_5603* | *atpF* | - | 6385782 | 6386252 | 38.87 | 165.08 | 176.97 | 68.96 | 1309.61 | 149.77 | F0F1 ATP synthase subunit B |
| *PSPTO_5604* | *atpE* | - | 6386310 | 6386567 | 10.52 | 50.01 | 27.45 | 20.28 | 265.90 | 36.63 | F0F1 ATP synthase subunit C |
| *PSPTO_5605* | *atpB* | - | 6386706 | 6387575 | 12.82 | 32.55 | 47.98 | 31.13 | 146.44 | 36.45 | F0F1 ATP synthase subunit A |
| *PSPTO_5606* | *atpI* | - | 6387592 | 6387999 | 55.86 | 45.74 | 66.29 | 31.51 | 75.35 | 43.40 | F0F1 ATP synthase subunit I |
| *PSPTO_5607* | | - | 6388146 | 6389018 | 3.24 | 14.44 | 14.56 | 6.29 | 76.69 | 16.40 | ParB family protein |
| *PSPTO_5608* | | - | 6389028 | 6389819 | 13.59 | 62.43 | 44.62 | 18.41 | 191.93 | 57.52 | ParA family protein |
| *PSPTO_5609* | *gidB* | - | 6389838 | 6390473 | 7.18 | 21.92 | 21.45 | 7.39 | 112.29 | 46.87 | Glucose-inhibited division protein B |
| *PSPTO_5610* | *gidA* | - | 6390479 | 6392371 | 12.56 | 44.50 | 34.26 | 28.48 | 128.14 | 90.40 | Glucose-inhibited division protein A |
| *PSPTO_5611* | *trmE* | - | 6392951 | 6394321 | 10.03 | 8.31 | 4.03 | 2.03 | 18.70 | 7.76 | tRNA modification GTPase TrmE |
| *PSPTO_5612* | | - | 6394400 | 6396088 | 6.41 | 26.03 | 20.83 | 14.34 | 108.24 | 24.04 | Inner membrane protein, 60 kDa |
| *PSPTO_5613* | | - | 6396091 | 6396336 | 47.87 | 67.59 | 54.16 | 27.78 | 123.97 | 86.98 | Hypothetical protein |
| *PSPTO_5614* | *rnpA* | - | 6396329 | 6396730 | 15.15 | 91.07 | 53.10 | 35.66 | 158.36 | 53.04 | Ribonuclease P protein component |
| *PSPTO_5615* | *rpmH* | - | 6396747 | 6396881 | 296.62 | 1242.23 | 908.70 | 595.73 | 5079.49 | 1384.46 | 50S ribosomal protein L34 |
| *PSPTO_5616* | | - | 522206 | 522316 | 18.79 | 20.02 | 12.48 | 11.72 | 30.44 | 19.45 | Hypothetical protein |
| *PSPTO_5617* | | - | 938980 | 939307 | - | - | - | - | - | - | |
| *PSPTO_5619* | | + | 981249 | 981500 | 2.37 | 80.06 | 47.83 | 4.09 | 2.97 | 3.15 | Hypothetical protein |
| *PSPTO_5620* | | - | 1731272 | 1731397 | 0.00 | 42.26 | 31.20 | 0.00 | 0.00 | 0.50 | |
| *PSPTO_5621* | | + | 923360 | 923584 | 169.06 | 125.87 | 96.72 | 77.73 | 324.43 | 342.53 | Hypothetical protein |
| *PSPTO_5622* | | + | 1548470 | 1548718 | 12.75 | 38998.29 | 26556.75 | 3525.35 | 54.89 | 85.48 | Hypothetical protein |
| *PSPTO_5623* | | + | 5355978 | 5356022 | - | - | - | - | - | - | |
| *PSPTO_5624* | *pvdP* | + | 2367818 | 2369452 | 1.90 | 2.15 | 0.97 | 4.73 | 11.06 | 25.35 | Tat (twin-arginine translocation) pathway signal sequence domain-containing protein |
| *PSPTO_5625* | | + | 4827116 | 4828339 | 1.03 | 3.67 | 9.16 | 1.49 | 10.99 | 2.43 | |
| *PSPTO_5627* | | + | 5358542 | 5359036 | - | - | - | - | - | - | |
| *PSPTO_5628* | | + | 5359162 | 5359782 | - | - | - | - | - | - | |
| *PSPTO_5629* | | - | 5328266 | 5329000 | 32.98 | 24.19 | 19.96 | 9.13 | 24.47 | 18.28 | Insertion sequence |
| *PSPTO_5630* | | + | 191660 | 192508 | 5.76 | 15.33 | 16.16 | 8.64 | 48.43 | 7.64 | Hypothetical protein |
| *PSPTO_5631* | | + | 5423493 | 5424341 | 2.71 | 6.53 | 7.97 | 6.80 | 33.98 | 6.38 | Hypothetical protein |
| *PSPTO_5632* | | + | 5376643 | 5378961 | 25.27 | 10.82 | 12.62 | 5.31 | 74.13 | 14.52 | Hypothetical protein |
| *PSPTO_5633* | *HopBM1* | + | 15948 | 16421 | 115.21 | 441.27 | 370.05 | 170.03 | 499.16 | 184.82 | Type III effector HopBM1 |
| *PSPTO_5634* | | + | 1244571 | 1244858 | 25.58 | 66.44 | 76.45 | 46.60 | 205.75 | 66.11 | Hypothetical protein |
| *PSPTO_5635* | | + | 1581112 | 1581555 | 985.32 | 259.90 | 157.16 | 104.55 | 82.65 | 80.60 | Hypothetical protein |
| *PSPTO_5636* | | + | 1228041 | 1228448 | 27.07 | 23.98 | 19.86 | 9.64 | 39.60 | 50.93 | Hypothetical protein |
| *PSPTO_5637* | | + | 2006821 | 2007108 | 8.53 | 20.97 | 10.26 | 24.26 | 121.99 | 18.55 | Hypothetical protein |
| *PSPTO_5638* | | - | 2611919 | 2612308 | 17.26 | 12.40 | 34.38 | 24.26 | 28.40 | 35.91 | Hypothetical protein |
| *PSPTO_5639* | | + | 2777027 | 2777347 | 18.65 | 26.63 | 29.24 | 57.99 | 4.08 | 3.53 | Hypothetical protein |
| *PSPTO_5640* | | + | 2910705 | 2911238 | 28.42 | 60.26 | 59.93 | 25.49 | 54.44 | 29.42 | Hypothetical protein |
| *PSPTO_5641* | | + | 2978065 | 2978625 | 16.32 | 28.64 | 21.09 | 12.10 | 27.09 | 12.04 | Hypothetical protein |
| *PSPTO_5642* | | - | 3476622 | 3477131 | 10.95 | 55.69 | 44.08 | 1.29 | 4.44 | 9.74 | Hypothetical protein |
| *PSPTO_5643* | | - | 3476773 | 3477240 | 18.33 | 111.76 | 22.52 | 18.48 | 6.10 | 6.20 | Hypothetical protein |
| *PSPTO_5644* | | + | 4862483 | 4862680 | 45.78 | 79.24 | 89.31 | 47.21 | 77.42 | 30.80 | Hypothetical protein |
| *PSPTO_5645* | | - | 6174792 | 6175730 | 7.79 | 32.34 | 24.24 | 12.40 | 275.62 | 24.48 | Hypothetical protein |
| *PSPTO_5646* | | - | 6174382 | 6174792 | 1.60 | 5.68 | 1.40 | 0.18 | 37.62 | 2.18 | Hypothetical protein |
| *PSPTO_5647* | *rsmY* | + | 555344 | 555465 | 6490.44 | 806.34 | 733.16 | 1210.10 | 530.00 | 1780.49 | RsmY |
| *PSPTO_5648* | *P26* | + | 678273 | 678338 | 11.69 | 57.64 | 110.42 | 64.45 | 477.23 | 47.15 | P26 |
| *PSPTO_5649* | *prrF* | - | 1058888 | 1058999 | 27.89 | 25.73 | 36.22 | 34.38 | 53.04 | 758.88 | PrrF |
| *PSPTO_5650* | | - | 1255678 | 1255819 | 145.75 | 249.75 | 232.24 | 150.04 | 180.10 | 96.06 | Hypothetical protein |
| *PSPTO_5651* | *t44* | + | 1693384 | 1693536 | 4896.45 | 6526.56 | 5590.46 | 4437.63 | 15106.55 | 4044.62 | T44 |
| *PSPTO_5652* | *PrrB_RsmZ* | + | 1728435 | 1728566 | 926.66 | 3184.12 | 2782.65 | 3926.20 | 519.63 | 5221.20 | PrrB_RsmZ |
| *PSPTO_5653* | *Cobalamin-1* | + | 1878136 | 1878350 | 26.89 | 95.89 | 30.66 | 11.62 | 293.51 | 89.57 | Cobalamin-1 |
| *PSPTO_5654* | *FMN* | - | 2008912 | 2009078 | 936.49 | 3557.73 | 2639.12 | 2394.97 | 1022.23 | 505.63 | FMN |
| *PSPTO_5655* | *Cobalamin-2* | - | 3542313 | 3542537 | 20.36 | 73.90 | 99.38 | 27.61 | 123.48 | 47.56 | Cobalamin-2 |
| *PSPTO_5656* | *Cobalamin-3* | + | 3542594 | 3542808 | 193.23 | 784.83 | 662.56 | 476.11 | 1197.68 | 384.14 | Cobalamin-3 |
| *PSPTO_5657* | *prrF2* | + | 3549491 | 3549603 | 29.72 | 21.85 | 23.98 | 14.39 | 45.93 | 424.94 | PrrF2 |
| *PSPTO_5658* | *Cobalamin-4* | + | 3679925 | 3680123 | 14.74 | 43.47 | 83.05 | 23.22 | 301.58 | 131.47 | Cobalamin-4 |
| *PSPTO_5659* | | - | 4116013 | 4116114 | 838.40 | 437.86 | 412.11 | 235.20 | 81.96 | 62.40 | |
| *PSPTO_5660* | *P16* | + | 4332812 | 4333007 | 34.06 | 36.08 | 15.13 | 29.63 | 11.23 | 20.12 | P16 |
| *PSPTO_5661* | *RNaseP* | - | 4986420 | 4986712 | 9377.44 | 9169.99 | 14305.31 | 11238.00 | 12364.34 | 15963.15 | RNaseP |
| *PSPTO_5662* | *S15* | - | 5054896 | 5055007 | 17.45 | 51.37 | 62.90 | 70.77 | 146.57 | 53.66 | S15 |
| *PSPTO_5663* | *tmRNA* | + | 5086183 | 5086569 | 157584.77 | 192198.53 | 284679.72 | 366059.77 | 172210.26 | 375315.40 | TmRNA |
| *PSPTO_5664* | *P24* | - | 5433053 | 5433294 | 28.93 | 1097.73 | 897.76 | 691.94 | 22.90 | 9.35 | P24 |
| *PSPTO_5665* | | - | 5648630 | 5648734 | 46.82 | 60.18 | 79.85 | 58.70 | 375.70 | 127.46 | Hypothetical protein |
| *PSPTO_5666* | *6S* | + | 5949055 | 5949233 | 11349.01 | 3203.01 | 3400.85 | 2686.59 | 650.83 | 1018.55 | 6S |
| *PSPTO_5667* | *P1* | + | 6038511 | 6038690 | 24.48 | 108.58 | 70.14 | 33.96 | 114.06 | 30.15 | P1 |

| Locus name | Gene name | Strand | Gene start (bp) | Gene end (bp) | Average TPM Inoculum | Average TPM $t_6$ | Average TPM $t_{24}$ | Average TPM $t_{168}$ | Average TPM Logarithmic | Average TPM Stationary | Annotation description |
|---|---|---|---|---|---|---|---|---|---|---|---|
| *PSPTO_5668* | *Psr1* | - | 1046593 | 1046951 | 51079.52 | 55607.17 | 42830.51 | 58663.69 | 14374.99 | 39471.46 | Psr1 |
| *PSPTO_5669* | *Psr2* | - | 1778383 | 1778751 | 16671.70 | 64608.08 | 65934.42 | 110606.94 | 3896.41 | 2915.59 | Psr2 |
| *PSPTO_5670* | *Spot42* | - | 373014 | 373131 | 14.74 | 14.73 | 2.89 | 4.04 | 25.35 | 10.76 | Spot42 |
| *PSPTO_5671* | *rsmX* | - | 4170293 | 4170371 | 179.48 | 5.45 | 9.96 | 2.32 | 1.30 | 7.20 | RsmX |
| *PSPTO_5672* | *rsmX-1* | - | 3244255 | 3244374 | 65.63 | 0.00 | 5.47 | 0.00 | 1.00 | 0.60 | RsmX-1 |
| *PSPTO_5673* | *rsmX-3* | + | 6144830 | 6144943 | 58.45 | 10.26 | 3.22 | 6.86 | 0.21 | 1.43 | RsmX-3 |
| *PSPTO_5674* | *rsmX-4* | + | 6145122 | 6145235 | 226.63 | 5.86 | 18.04 | 4.82 | 3.80 | 8.31 | RsmX-4 |
| *PSPTO_5675* | *rsmX-5* | - | 6198038 | 6198149 | 49.70 | 3.97 | 2.93 | 0.00 | 28.12 | 37.32 | RsmX-5 |
| *PSPTO_A0001* | *repA* | + | 351 | 1664 | 0.80 | 0.14 | 0.00 | 0.14 | 1.23 | 0.48 | RepA |
| *PSPTO_A0002* | *rulA* | + | 1803 | 2228 | 0.00 | 0.00 | 0.00 | 0.00 | 0.00 | 0.00 | RulA |
| *PSPTO_A0003* | | + | 2221 | 2469 | - | - | - | - | - | - | |
| *PSPTO_A0004* | | + | 2530 | 3540 | 8.47 | 5.55 | 1.39 | 3.11 | 3.91 | 4.20 | Phage integrase family site specific recombinase |
| *PSPTO_A0005* | *hopAM1-2* | - | 5731 | 6561 | 0.52 | 393.51 | 163.44 | 22.67 | 1.28 | 1.00 | HopAM1-2 |
| *PSPTO_A0006* | | + | 6708 | 7748 | - | - | - | - | - | - | |
| *PSPTO_A0007* | | - | 7782 | 8255 | 0.00 | 0.00 | 0.00 | 0.00 | 0.00 | 0.00 | Hypothetical protein |
| *PSPTO_A0008* | | - | 8252 | 9136 | 0.00 | 0.00 | 0.00 | 0.00 | 0.00 | 0.00 | ParB family partitioning protein |
| *PSPTO_A0009* | | - | 9357 | 10484 | 0.00 | 0.00 | 0.00 | 0.00 | 0.00 | 0.00 | ParaA family ATPase |
| *PSPTO_A0010* | | - | 11224 | 14193 | 35.32 | 76.33 | 55.99 | 35.11 | 86.70 | 34.20 | Tn3 family transposase |
| *PSPTO_A0011* | | + | 14315 | 14860 | 259.68 | 108.98 | 81.32 | 46.05 | 121.21 | 84.23 | Resolvase, putative |
| *PSPTO_A0012* | *HopX1* | - | 14914 | 16068 | 48.67 | 281.30 | 229.75 | 49.53 | 101.45 | 79.33 | HopX1 |
| *PSPTO_A0013* | | - | 16332 | 17354 | 0.00 | 0.00 | 0.00 | 0.00 | 0.00 | 0.00 | ISPsy4, transposase |
| *PSPTO_A0014* | | + | 17627 | 18256 | 0.00 | 0.00 | 0.00 | 0.00 | 0.00 | 0.00 | ParaA family ATPase |
| *PSPTO_A0015* | | + | 18277 | 18495 | 0.00 | 0.00 | 0.00 | 0.00 | 0.00 | 0.00 | Plasmid partition protein ParB, truncated |
| *PSPTO_A0016* | | + | 18841 | 19242 | - | - | - | - | - | - | |
| *PSPTO_A0017* | *shc01* | + | 19718 | 20146 | 0.00 | 0.00 | 0.00 | 0.00 | 0.00 | 0.00 | Shc01 |
| *PSPTO_A0018* | *hopO1-1* | + | 20399 | 21250 | 0.00 | 0.00 | 0.00 | 0.00 | 0.00 | 0.00 | HopO1-1 |
| *PSPTO_A0019* | *hopT1-1* | + | 21259 | 22395 | 0.00 | 0.00 | 0.00 | 0.00 | 0.00 | 0.00 | HopT1-1 |
| *PSPTO_A0020* | | - | 22604 | 22771 | - | - | - | - | - | - | |
| *PSPTO_A0021* | | - | 23291 | 23956 | 76.68 | 32.32 | 34.12 | 10.69 | 23.09 | 15.29 | Resolvase, putative |
| *PSPTO_A0022* | | + | 24081 | 24662 | 0.00 | 0.00 | 0.00 | 0.00 | 0.00 | 0.01 | Hypothetical protein |
| *PSPTO_A0023* | | - | 24612 | 24962 | 1.73 | 1.47 | 4.21 | 4.46 | 1.24 | 1.67 | Hypothetical protein |
| *PSPTO_A0024* | | + | 25096 | 25473 | 1.78 | 2.77 | 3.80 | 3.67 | 1.67 | 1.43 | Hypothetical protein |
| *PSPTO_A0025* | | + | 25520 | 26356 | 7.46 | 2.35 | 1.08 | 0.22 | 2.48 | 1.59 | Hypothetical protein |
| *PSPTO_A0026* | | + | 26414 | 26602 | 0.00 | 0.00 | 0.00 | 0.00 | 0.00 | 0.00 | Hypothetical protein |
| *PSPTO_A0027* | *mobC* | - | 26676 | 26954 | 0.00 | 0.00 | 0.00 | 0.00 | 0.00 | 0.00 | Mobilization protein MobC, putative |
| *PSPTO_A0028* | *mobB* | + | 27399 | 27728 | 107.67 | 48.80 | 37.64 | 17.10 | 71.56 | 26.76 | Mobilization protein MobB, putative |
| *PSPTO_A0029* | *mobA* | + | 27739 | 29691 | 15.80 | 6.89 | 4.73 | 1.57 | 19.74 | 7.69 | Relaxase/mobilization nuclease MobA, putative |
| *PSPTO_A0030* | | - | 30218 | 30841 | 0.00 | 0.00 | 0.00 | 0.00 | 0.00 | 0.00 | Glycosyl hydrolase family protein |
| *PSPTO_A0031* | | - | 31647 | 32828 | 0.06 | 0.00 | 0.00 | 0.00 | 0.19 | 0.08 | Hypothetical protein |
| *PSPTO_A0032* | *lsc-3* | - | 32908 | 34203 | 0.66 | 0.51 | 1.50 | 0.43 | 4.75 | 1.76 | Lsc-3 |
| *PSPTO_A0033* | | + | 34296 | 34694 | 0.00 | 0.00 | 0.00 | 0.00 | 0.07 | 0.01 | Hypothetical protein |
| *PSPTO_A0034* | | - | 34863 | 36812 | 0.00 | 0.00 | 0.00 | 0.00 | 0.00 | 0.00 | GGDEF domain/EAL domain-containing protein |
| *PSPTO_A0035* | | + | 37434 | 38135 | 24.43 | 19.07 | 14.21 | 7.65 | 21.87 | 14.63 | GntR family transcriptional regulator |
| *PSPTO_A0036* | | + | 38204 | 39397 | 0.00 | 0.00 | 0.00 | 0.00 | 0.00 | 0.00 | Hydroxyglutarate oxidase |
| *PSPTO_A0037* | | + | 39438 | 40784 | 0.00 | 0.00 | 0.00 | 0.00 | 0.00 | 0.00 | Major facilitator family transporter |
| *PSPTO_A0038* | | + | 40825 | 41010 | 0.00 | 0.00 | 0.00 | 0.00 | 0.00 | 0.00 | Hypothetical protein |
| *PSPTO_A0039* | | - | 41032 | 41367 | 0.00 | 0.00 | 0.00 | 0.00 | 0.00 | 0.00 | PbsX family transcriptional regulator |
| *PSPTO_A0040* | | + | 41631 | 42164 | 9.87 | 8.43 | 3.45 | 7.78 | 18.88 | 11.98 | Hypothetical protein |
| *PSPTO_A0041* | | + | 42480 | 42752 | 1.54 | 0.95 | 3.44 | 1.67 | 2.48 | 1.50 | Putative lipoprotein |
| *PSPTO_A0042* | *pilT* | + | 42755 | 43246 | 3.91 | 4.53 | 4.43 | 0.00 | 6.49 | 4.59 | PilT |
| *PSPTO_A0043* | *traH* | + | 43283 | 43756 | 4.42 | 6.61 | 7.98 | 3.03 | 11.33 | 5.39 | TraH protein, putative |
| *PSPTO_A0044* | *traI* | + | 43753 | 44553 | 2.69 | 0.69 | 2.90 | 3.22 | 4.89 | 3.28 | TraI |
| *PSPTO_A0045* | | + | 44550 | 45731 | 5.26 | 1.55 | 3.57 | 5.47 | 8.03 | 4.21 | Type IV pilus biogenesis protein |
| *PSPTO_A0046* | | + | 46304 | 46513 | 0.27 | 0.00 | 0.00 | 0.00 | 0.51 | 0.11 | Hypothetical protein |
| *PSPTO_A0047* | | + | 46510 | 47061 | 0.38 | 1.50 | 1.13 | 0.00 | 0.93 | 0.48 | Endonuclease |
| *PSPTO_A0048* | *traK-2* | + | 47372 | 47659 | 0.89 | 0.00 | 1.86 | 2.87 | 3.04 | 0.59 | TraK-2 |
| *PSPTO_A0049* | | + | 47811 | 51980 | 4.81 | 4.37 | 3.70 | 1.83 | 10.23 | 6.64 | DNA primase, putative |
| *PSPTO_A0050* | *traL-2* | + | 52006 | 52359 | 7.24 | 7.17 | 5.39 | 4.84 | 8.97 | 4.45 | TraL-2 |
| *PSPTO_A0051* | *traM-2* | + | 52356 | 53063 | 1.34 | 0.51 | 1.39 | 4.74 | 3.03 | 1.52 | TraM-2 |
| *PSPTO_A0052* | *traN-2* | + | 53120 | 54178 | 2.30 | 2.88 | 2.68 | 0.74 | 5.56 | 2.68 | TraN-2 |
| *PSPTO_A0053* | *traO-2* | + | 54184 | 55542 | 2.71 | 3.87 | 3.35 | 0.44 | 8.21 | 8.10 | TraO-2 |
| *PSPTO_A0054* | *traP-2* | + | 55584 | 56249 | 5.40 | 0.75 | 4.37 | 0.97 | 12.15 | 5.18 | TraP-2 |
| *PSPTO_A0055* | *traQ-2* | + | 56258 | 56800 | 4.52 | 4.07 | 1.26 | 0.00 | 5.77 | 3.98 | TraQ-2 |
| *PSPTO_A0056* | *traR-2* | + | 56876 | 57271 | 4.31 | 3.10 | 2.47 | 2.82 | 6.66 | 3.67 | TraR-2 |
| *PSPTO_A0057* | | - | 57315 | 57590 | 0.00 | 0.00 | 0.00 | 0.00 | 0.00 | 0.00 | Hypothetical protein |
| *PSPTO_A0058* | | - | 57568 | 57768 | 0.00 | 0.00 | 0.00 | 0.00 | 0.00 | 0.00 | Hypothetical protein |
| *PSPTO_A0059* | *traT* | + | 57953 | 58612 | 0.99 | 0.09 | 0.00 | 0.99 | 1.47 | 0.89 | TraT |
| *PSPTO_A0060* | *traU-2* | + | 58596 | 61643 | 1.30 | 1.73 | 1.44 | 1.23 | 3.45 | 1.75 | TraU-2 |
| *PSPTO_A0061* | | + | 61667 | 61903 | 0.00 | 0.00 | 0.00 | 0.00 | 0.00 | 0.00 | Hypothetical protein |
| *PSPTO_A0062* | *traW-2* | + | 61946 | 63160 | 1.06 | 0.92 | 0.34 | 1.84 | 3.89 | 1.81 | TraW-2 |
| *PSPTO_A0063* | *traX-2* | + | 63150 | 63752 | 0.71 | 0.00 | 0.00 | 0.15 | 1.09 | 0.53 | TraX-2 |
| *PSPTO_A0064* | *traY-2* | + | 63781 | 65925 | 0.47 | 0.19 | 0.00 | 0.21 | 0.62 | 0.38 | TraY-2 |
| *PSPTO_A0065* | | + | 66269 | 66919 | 1.85 | 1.86 | 1.86 | 0.38 | 1.99 | 1.11 | Exclusion-determining protein, putative |
| *PSPTO_A0066* | *trbA* | + | 67012 | 68211 | 1.12 | 1.00 | 0.32 | 0.00 | 2.21 | 1.65 | TrbA |
| *PSPTO_A0067* | *trbB* | + | 68297 | 69382 | 0.82 | 0.25 | 0.45 | 0.00 | 1.79 | 0.92 | TrbB |
| *PSPTO_A0068* | | + | 69369 | 71657 | 4.23 | 5.50 | 3.61 | 2.00 | 8.23 | 4.37 | TraG/TraD family conjugal transfer protein |
| *PSPTO_A0069* | | + | 71685 | 72107 | 0.00 | 0.00 | 0.00 | 0.00 | 0.00 | 0.00 | Hypothetical protein |
| *PSPTO_A0070* | | + | 72068 | 72691 | 0.00 | 0.00 | 0.00 | 0.00 | 0.00 | 0.00 | Hypothetical protein |
| *PSPTO_A0071* | | + | 72879 | 73307 | 0.00 | 0.00 | 0.00 | 0.00 | 0.00 | 0.00 | GntR family transcriptional regulator |

| Locus name | Gene name | Strand | Gene start (bp) | Gene end (bp) | Average TPM Inoculum | Average TPM $t_6$ | Average TPM $t_{24}$ | Average TPM $t_{168}$ | Average TPM Logarithmic | Average TPM Stationary | Annotation description |
|---|---|---|---|---|---|---|---|---|---|---|---|
| *PSPTO_A0072* | | + | 3564 | 5705 | 3.52 | 13.10 | 8.40 | 2.88 | 8.13 | 5.73 | Phage integrase family site specific recombinase |
| *PSPTO_A0073* | | + | 19220 | 19279 | - | - | - | - | - | - | Similar to ultraviolet light resistance protein RulB, truncation |
| *PSPTO_B0001* | *repA* | + | 350 | 1723 | 10.02 | 10.49 | 7.63 | 6.19 | 22.01 | 6.41 | RepA protein, putative |
| *PSPTO_B0002* | | - | 1841 | 3070 | 31.38 | 27.31 | 16.91 | 12.30 | 94.97 | 23.17 | ISPsy3, transposase |
| *PSPTO_B0003* | *HopBM1* | + | 3277 | 3750 | 115.21 | 441.27 | 370.05 | 170.03 | 499.16 | 184.82 | Type III effector HopBM1 |
| *PSPTO_B0004* | | - | 3798 | 3947 | - | - | - | - | - | - | |
| *PSPTO_B0005* | *plcA2* | - | 4424 | 6106 | 19.26 | 712.52 | 452.04 | 111.30 | 107.99 | 54.37 | PlcA2 |
| *PSPTO_B0006* | | - | 7250 | 7378 | 20.87 | 56.60 | 22.99 | 0.32 | 76.07 | 59.93 | Hypothetical protein |
| *PSPTO_B0007* | *pilM* | + | 7588 | 8028 | 40.93 | 45.93 | 27.43 | 13.36 | 61.18 | 44.93 | PilM protein, putative |
| *PSPTO_B0008* | | - | 8099 | 8356 | 16.08 | 6.73 | 22.73 | 0.25 | 32.90 | 13.12 | ISPsy3, transposase, internal deletion |
| *PSPTO_B0009* | | - | 8450 | 9316 | 46.97 | 51.68 | 58.36 | 23.27 | 31.09 | 42.83 | Hypothetical protein |
| *PSPTO_B0010* | | + | 9517 | 10932 | 225.33 | 401.20 | 325.76 | 156.25 | 870.88 | 306.32 | Major facilitator family transporter |
| *PSPTO_B0011* | | + | 11077 | 11826 | - | - | - | - | - | - | Partial ORF |
| *PSPTO_B0012* | | - | 12597 | 13655 | 89.59 | 62.64 | 57.21 | 40.58 | 88.80 | 60.54 | Hypothetical protein |
| *PSPTO_B0013* | | - | 14664 | 15446 | 7.46 | 6.60 | 7.93 | 14.41 | 19.00 | 5.56 | Hypothetical protein |
| *PSPTO_B0014* | | + | 15706 | 16129 | - | - | - | - | - | - | Truncated IS5 family transposase |
| *PSPTO_B0015* | | + | 16128 | 16424 | 42.80 | 57.01 | 45.69 | 41.76 | 97.70 | 73.66 | Hypothetical protein |
| *PSPTO_B0016* | | + | 16446 | 16799 | 11.63 | 12.99 | 26.32 | 30.51 | 33.46 | 22.85 | Hypothetical protein |
| *PSPTO_B0017* | | + | 17333 | 18283 | - | - | - | - | - | - | |
| *PSPTO_B0018* | *stbB* | - | 18458 | 18877 | 41.14 | 35.20 | 47.79 | 5.69 | 132.80 | 30.41 | StbB |
| *PSPTO_B0019* | *stbC* | - | 18874 | 19167 | 666.03 | 402.91 | 294.84 | 92.35 | 362.19 | 90.45 | StbC |
| *PSPTO_B0020* | | - | 19504 | 20451 | 42.46 | 65.29 | 71.17 | 52.22 | 364.70 | 187.57 | Hypothetical protein |
| *PSPTO_B0021* | | - | 21094 | 21777 | 148.81 | 204.67 | 144.32 | 78.09 | 167.86 | 103.69 | Hypothetical protein |
| *PSPTO_B0022* | | + | 22172 | 22372 | 157.76 | 115.61 | 92.17 | 57.71 | 128.46 | 131.32 | DnaK suppressor domain-containing protein |
| *PSPTO_B0023* | | + | 22410 | 22631 | 230.12 | 491.20 | 308.66 | 191.77 | 587.71 | 271.35 | Hypothetical protein |
| *PSPTO_B0024* | | - | 22964 | 23584 | 91.03 | 44.01 | 45.98 | 14.86 | 38.60 | 26.50 | Resolvase, putative |
| *PSPTO_B0025* | | + | 23792 | 24130 | 1570.59 | 434.71 | 452.80 | 149.55 | 198.12 | 262.36 | Hypothetical protein |
| *PSPTO_B0026* | | + | 24130 | 24417 | 156.01 | 115.68 | 90.23 | 42.63 | 107.70 | 94.14 | Hypothetical protein |
| *PSPTO_B0027* | | + | 24479 | 24895 | 35.35 | 35.86 | 48.32 | 10.95 | 48.55 | 30.97 | Hypothetical protein |
| *PSPTO_B0028* | | - | 24916 | 25191 | 3.74 | 1.69 | 8.77 | 5.67 | 6.13 | 5.05 | Hypothetical protein |
| *PSPTO_B0029* | | + | 25478 | 25732 | 4.92 | 0.95 | 11.70 | 4.90 | 6.87 | 5.84 | Hypothetical protein |
| *PSPTO_B0030* | | + | 25779 | 26615 | 21.09 | 6.26 | 12.92 | 4.63 | 7.42 | 4.52 | Hypothetical protein |
| *PSPTO_B0031* | | + | 26683 | 26883 | 6.09 | 0.54 | 3.26 | 5.92 | 4.28 | 3.05 | Hypothetical protein |
| *PSPTO_B0032* | *mobC* | - | 26950 | 27228 | 352.44 | 329.82 | 315.39 | 153.60 | 669.33 | 79.64 | MobC protein |
| *PSPTO_B0033* | | - | 27272 | 27553 | 304.70 | 214.19 | 204.84 | 141.22 | 226.45 | 108.91 | Hypothetical protein |
| *PSPTO_B0034* | | - | 27526 | 27870 | 351.27 | 78.77 | 109.38 | 58.83 | 103.03 | 60.21 | Hypothetical protein |
| *PSPTO_B0035* | *mobB* | + | 28130 | 28459 | 122.96 | 51.67 | 38.10 | 18.88 | 76.55 | 28.60 | Mobilization protein MobB, putative |
| *PSPTO_B0036* | *mobA* | + | 28470 | 30425 | 48.21 | 17.97 | 14.28 | 10.19 | 63.70 | 31.04 | Relaxase/mobilization nuclease MobA, putative |
| *PSPTO_B0037* | | + | 30519 | 30728 | 13.18 | 8.22 | 19.69 | 15.25 | 19.56 | 13.92 | Hypothetical protein |
| *PSPTO_B0038* | | + | 30760 | 30954 | - | - | - | - | - | - | Partial ORF |
| *PSPTO_B0039* | | - | 31003 | 31812 | 0.00 | 0.00 | 0.00 | 0.00 | 0.00 | 0.00 | Transposase/IS protein |
| *PSPTO_B0040* | | - | 31809 | 32831 | 0.00 | 0.00 | 0.00 | 0.00 | 0.00 | 0.00 | ISPsy4, transposase |
| *PSPTO_B0041* | | + | 32917 | 33429 | - | - | - | - | - | - | Partial ORF |
| *PSPTO_B0042* | | + | 33497 | 34693 | 15.44 | 12.12 | 21.05 | 12.92 | 36.34 | 32.99 | Hydroxyglutarate oxidase |
| *PSPTO_B0043* | | + | 34735 | 36081 | 3.12 | 6.05 | 8.73 | 3.95 | 9.29 | 4.79 | Major facilitator family transporter |
| *PSPTO_B0044* | | + | 36137 | 36322 | 2.78 | 9.24 | 18.22 | 25.64 | 12.99 | 4.55 | Hypothetical protein |
| *PSPTO_B0045* | | - | 36339 | 36692 | 42.42 | 138.25 | 190.61 | 132.60 | 189.99 | 137.97 | PbsX family transcriptional regulator |
| *PSPTO_B0046* | | + | 36937 | 37470 | 17.44 | 17.07 | 16.06 | 16.58 | 39.97 | 21.22 | Hypothetical protein |
| *PSPTO_B0047* | | + | 37460 | 37768 | 2.54 | 4.13 | 0.00 | 0.00 | 4.56 | 1.66 | Hypothetical protein |
| *PSPTO_B0048* | | + | 37786 | 38058 | 1.55 | 0.95 | 3.44 | 1.67 | 2.48 | 1.50 | Putative lipoprotein |
| *PSPTO_B0049* | *pilT* | + | 38100 | 38552 | 3.64 | 4.79 | 4.64 | 0.00 | 6.62 | 4.69 | PilT protein, putative |
| *PSPTO_B0050* | *traH* | + | 38589 | 39062 | 4.41 | 6.61 | 7.98 | 3.03 | 11.32 | 5.39 | TraH protein, putative |
| *PSPTO_B0051* | *traI* | + | 39059 | 39859 | 2.69 | 0.69 | 2.90 | 3.22 | 4.90 | 3.28 | TraI protein |
| *PSPTO_B0052* | | + | 39856 | 41037 | 5.50 | 1.55 | 3.57 | 5.83 | 8.47 | 4.38 | Type IV pilus biogenesis protein |
| *PSPTO_B0053* | | + | 41038 | 41733 | 3.91 | 5.36 | 2.69 | 1.60 | 8.65 | 4.52 | Hypothetical protein |
| *PSPTO_B0054* | *nuc* | + | 41772 | 42281 | 3.15 | 3.75 | 5.51 | 2.16 | 7.16 | 3.38 | Nuc |
| *PSPTO_B0055* | | + | 42318 | 42563 | 25.70 | 7.69 | 3.48 | 0.00 | 8.94 | 4.57 | Hypothetical protein |
| *PSPTO_B0056* | *traK-1* | + | 42592 | 42879 | 9.94 | 13.61 | 10.73 | 10.13 | 18.93 | 6.97 | TraK-1 |
| *PSPTO_B0057* | *traC* | + | 42922 | 47595 | 21.13 | 13.02 | 9.57 | 6.50 | 28.04 | 15.39 | DNA primase TraC |
| *PSPTO_B0058* | *traL-1* | + | 47621 | 47974 | 7.25 | 7.17 | 5.39 | 4.84 | 8.99 | 4.46 | TraL-1 |
| *PSPTO_B0059* | *traM-1* | + | 48001 | 48678 | 1.34 | 0.53 | 1.45 | 4.06 | 3.03 | 1.51 | TraM-1 |
| *PSPTO_B0060* | *traN-1* | + | 48735 | 49793 | 2.30 | 2.88 | 2.68 | 0.74 | 5.56 | 2.68 | TraN-1 |
| *PSPTO_B0061* | *traO-1* | + | 49799 | 51157 | 2.70 | 3.87 | 3.35 | 0.44 | 8.20 | 8.10 | TraO-1 |
| *PSPTO_B0062* | *traP-1* | + | 51154 | 51864 | 5.23 | 0.70 | 4.61 | 0.96 | 12.17 | 5.22 | TraP-1 |
| *PSPTO_B0063* | *traQ-1* | + | 51873 | 52415 | 4.52 | 4.07 | 1.26 | 0.00 | 5.77 | 3.98 | TraQ-1 |
| *PSPTO_B0064* | *traR-1* | + | 52485 | 52886 | 4.52 | 3.06 | 2.60 | 2.77 | 6.84 | 3.87 | TraR-1 |
| *PSPTO_B0065* | | + | 52968 | 53579 | - | - | - | - | - | - | |
| *PSPTO_B0066* | *traU-1* | + | 53554 | 56610 | 6.42 | 4.28 | 3.66 | 4.96 | 11.14 | 6.43 | TraU-1 |
| *PSPTO_B0067* | *traW-1* | + | 56621 | 57835 | 6.69 | 4.61 | 6.06 | 7.11 | 24.34 | 13.24 | TraW-1 |
| *PSPTO_B0068* | *traX-1* | + | 57825 | 58427 | 4.85 | 4.07 | 2.85 | 2.58 | 11.64 | 6.72 | TraX-1 |
| *PSPTO_B0069* | *traY-1* | + | 58458 | 60602 | 19.86 | 15.69 | 13.35 | 16.36 | 29.74 | 19.68 | TraY-1 |
| *PSPTO_B0070* | | + | 60678 | 60950 | 462.61 | 181.12 | 195.78 | 115.49 | 143.24 | 146.73 | Hypothetical protein |
| *PSPTO_B0071* | | + | 60947 | 61594 | 209.11 | 193.33 | 224.93 | 138.64 | 231.87 | 160.58 | Surface exclusion protein, putative |
| *PSPTO_B0072* | | + | 61687 | 62889 | 9.90 | 12.67 | 8.72 | 10.35 | 23.39 | 15.99 | Hypothetical protein |
| *PSPTO_B0073* | | - | 62879 | 63256 | 10.33 | 20.59 | 18.66 | 14.56 | 26.74 | 10.72 | PilT domain-containing protein |
| *PSPTO_B0074* | | - | 63253 | 63453 | 47.99 | 74.34 | 93.07 | 36.00 | 206.78 | 51.92 | Hypothetical protein |
| *PSPTO_B0075* | *trbB* | + | 63546 | 64631 | 5.48 | 2.89 | 1.99 | 1.38 | 7.65 | 3.83 | TrbB protein, putative |
| *PSPTO_B0076* | *trbC* | + | 64618 | 66906 | 6.24 | 6.05 | 5.32 | 3.16 | 13.29 | 6.94 | TrbC |
| *PSPTO_B0077* | | + | 66984 | 67412 | 8.82 | 9.85 | 4.11 | 7.66 | 20.21 | 11.88 | GntR family transcriptional regulator |

| Locus name | Gene name | Strand | Gene start (bp) | Gene end (bp) | Average TPM Inoculum | Average TPM $t_6$ | Average TPM $t_{24}$ | Average TPM $t_{168}$ | Average TPM Logarithmic | Average TPM Stationary | Annotation description |
|---|---|---|---|---|---|---|---|---|---|---|---|
| *PSPTO_B0078* | | - | 6549 | 6839 | 153.91 | 5200.29 | 3452.75 | 603.29 | 241.38 | 218.73 | |
| *PSPTO_r01* | *ps16SA* | + | 666741 | 668258 | 85433.69 | 43904.27 | 42725.62 | 45618.52 | 48367.64 | 57362.31 | Ps16SA |
| *PSPTO_r02* | *ps23SA* | + | 668802 | 671704 | 20690.22 | 9613.30 | 10521.27 | 13571.51 | 12763.05 | 13481.66 | Ps23SA |
| *PSPTO_r03* | *ps5SA* | + | 671852 | 671971 | 2.72 | 19.78 | 26.78 | 40.98 | 1.84 | 1.16 | Ps5SA |
| *PSPTO_r04* | *ps16SB* | + | 827555 | 829072 | 85379.01 | 43863.64 | 42694.65 | 45588.67 | 48325.21 | 57294.16 | Ps16SB |
| *PSPTO_r05* | *ps23SB* | + | 829616 | 832518 | 20048.45 | 9442.54 | 10251.63 | 13509.84 | 12490.81 | 13358.96 | Ps23SB |
| *PSPTO_r06* | *ps5SB* | + | 832666 | 832785 | 3.51 | 20.51 | 26.08 | 40.69 | 1.92 | 0.53 | Ps5SB |
| *PSPTO_r07* | *ps16SC* | + | 1072092 | 1073609 | 85431.37 | 43891.81 | 42714.42 | 45606.89 | 48366.73 | 57361.05 | Ps16SC |
| *PSPTO_r08* | *ps23SC* | + | 1074153 | 1077055 | 20061.37 | 9451.59 | 10260.13 | 13517.58 | 12503.26 | 13375.69 | Ps23SC |
| *PSPTO_r09* | *ps5SC* | + | 1077203 | 1077322 | 3.55 | 20.18 | 25.27 | 39.86 | 2.73 | 2.44 | Ps5SC |
| *PSPTO_r10* | *ps5SD* | - | 3869438 | 3869557 | 2.95 | 18.13 | 27.12 | 38.47 | 3.10 | 2.35 | Ps5SD |
| *PSPTO_r11* | *ps23SD* | - | 3869705 | 3872607 | 19805.98 | 9183.63 | 9996.45 | 13045.52 | 12256.05 | 12991.45 | Ps23SD |
| *PSPTO_r12* | *ps16SD* | - | 3873151 | 3874668 | 84354.65 | 38533.49 | 38908.13 | 40100.04 | 47521.47 | 56095.05 | Ps16SD |
| *PSPTO_r13* | *ps5SE* | - | 6213909 | 6214028 | 2.57 | 18.23 | 24.65 | 40.36 | 2.75 | 2.25 | Ps5SE |
| *PSPTO_r14* | *ps23SE* | - | 6214176 | 6217078 | 20294.51 | 9430.25 | 10275.81 | 13395.73 | 12537.45 | 13277.10 | Ps23SE |
| *PSPTO_r15* | *ps16SE* | - | 6217622 | 6219139 | 84213.29 | 38428.37 | 38831.57 | 40000.91 | 46231.00 | 55883.73 | Ps16SE |
| *PSPTO_t01* | *tRNA-Arg-1* | - | 323982 | 324058 | 11.36 | 13.22 | 12.78 | 16.72 | 14.03 | 3.11 | tRNA-Arg-1 |
| *PSPTO_t02* | *tRNA-Met-1* | - | 586008 | 586084 | 9.72 | 0.00 | 8.17 | 0.00 | 1.39 | 2.19 | tRNA-Met-1 |
| *PSPTO_t03* | *tRNA-Ile-1* | + | 668362 | 668438 | 210.37 | 201.65 | 184.99 | 160.94 | 175.14 | 86.55 | tRNA-Ile-1 |
| *PSPTO_t04* | *tRNA-Ala-1* | + | 668483 | 668558 | 74.79 | 57.27 | 48.08 | 31.38 | 12.72 | 9.63 | tRNA-Ala-1 |
| *PSPTO_t05* | *tRNA-Tyr-1* | + | 674391 | 674475 | 164.23 | 121.15 | 163.33 | 142.70 | 80.27 | 56.62 | tRNA-Tyr-1 |
| *PSPTO_t06* | *tRNA-Gly-1* | + | 674500 | 674573 | 121.70 | 26.37 | 32.19 | 0.25 | 15.38 | 9.94 | tRNA-Gly-1 |
| *PSPTO_t07* | *tRNA-Thr-1* | + | 674600 | 674675 | 8.73 | 12.27 | 21.89 | 2.41 | 26.98 | 9.98 | tRNA-Thr-1 |
| *PSPTO_t08* | *tRNA-Trp-1* | + | 674841 | 674916 | 124.75 | 338.73 | 206.62 | 162.75 | 237.85 | 285.01 | tRNA-Trp-1 |
| *PSPTO_t09* | *tRNA-Ile-2* | + | 829176 | 829252 | 210.35 | 201.49 | 185.05 | 160.94 | 174.81 | 86.36 | tRNA-Ile-2 |
| *PSPTO_t10* | *tRNA-Ala-2* | + | 829297 | 829372 | 74.89 | 57.27 | 48.21 | 31.38 | 12.48 | 9.63 | tRNA-Ala-2 |
| *PSPTO_t11* | *tRNA-Asn-1* | + | 898785 | 898860 | 190.50 | 198.85 | 385.51 | 330.81 | 89.53 | 133.49 | tRNA-Asn-1 |
| *PSPTO_t12* | *tRNA-Pro-1* | + | 898890 | 898966 | 80.42 | 151.02 | 129.07 | 58.97 | 80.12 | 97.21 | tRNA-Pro-1 |
| *PSPTO_t13* | *tRNA-Lys-1* | + | 898972 | 899047 | 160.21 | 171.75 | 220.33 | 107.74 | 181.38 | 220.35 | tRNA-Lys-1 |
| *PSPTO_t14* | *tRNA-Pro-2* | + | 899143 | 899219 | 57.59 | 150.70 | 120.63 | 77.96 | 40.44 | 51.56 | tRNA-Pro-2 |
| *PSPTO_t15* | *tRNA-Ile-3* | + | 1073713 | 1073789 | 210.53 | 201.65 | 185.05 | 160.94 | 175.14 | 86.74 | tRNA-Ile-3 |
| *PSPTO_t16* | *tRNA-Ala-3* | + | 1073834 | 1073909 | 74.63 | 57.27 | 48.21 | 31.38 | 12.59 | 9.63 | tRNA-Ala-3 |
| *PSPTO_t17* | *tRNA-Met-2* | - | 1162588 | 1162664 | 84.20 | 16.79 | 14.30 | 29.27 | 5.93 | 12.80 | tRNA-Met-2 |
| *PSPTO_t18* | *tRNA-Gln-1* | - | 1219496 | 1219570 | 33.91 | 74.53 | 30.10 | 61.13 | 43.98 | 29.22 | tRNA-Gln-1 |
| *PSPTO_t19* | *tRNA-Leu-1* | - | 1551777 | 1551861 | 34.66 | 117.60 | 74.05 | 71.26 | 2.40 | 4.66 | tRNA-Leu-1 |
| *PSPTO_t20* | *tRNA-Glu-1* | + | 1956097 | 1956172 | 279.89 | 112.59 | 151.51 | 116.68 | 69.09 | 79.22 | tRNA-Glu-1 |
| *PSPTO_t21* | *tRNA-Gly-2* | + | 1956175 | 1956250 | 150.17 | 122.85 | 103.94 | 56.89 | 94.76 | 44.48 | tRNA-Gly-2 |
| *PSPTO_t22* | *tRNA-Glu-2* | + | 1956350 | 1956425 | 41.31 | 48.25 | 63.55 | 34.48 | 23.47 | 37.40 | tRNA-Glu-2 |
| *PSPTO_t23* | *tRNA-Gly-3* | + | 1956430 | 1956505 | 113.93 | 73.34 | 64.02 | 35.16 | 41.44 | 19.67 | tRNA-Gly-3 |
| *PSPTO_t24* | *tRNA-Ser-1* | + | 2014810 | 2014900 | 2228.11 | 5867.65 | 4454.21 | 3337.88 | 5944.90 | 2927.70 | tRNA-Ser-1 |
| *PSPTO_t25* | *tRNA-Arg-2* | + | 2015010 | 2015086 | 48.86 | 55.42 | 96.19 | 39.10 | 86.26 | 33.15 | tRNA-Arg-2 |
| *PSPTO_t26* | *tRNA-Arg-3* | + | 2015174 | 2015250 | 34.48 | 44.63 | 53.36 | 12.10 | 35.29 | 12.71 | tRNA-Arg-3 |
| *PSPTO_t27* | *tRNA-Arg-4* | + | 2019304 | 2019380 | 140.98 | 64.95 | 80.52 | 40.58 | 49.35 | 44.77 | tRNA-Arg-4 |
| *PSPTO_t28* | *tRNA-Ser-2* | - | 2266664 | 2266753 | 41.77 | 36.49 | 41.53 | 38.94 | 9.91 | 8.57 | tRNA-Ser-2 |
| *PSPTO_t29* | *tRNA-Ala-4* | + | 2384751 | 2384826 | 287.28 | 235.53 | 281.25 | 115.01 | 174.49 | 95.14 | tRNA-Ala-4 |
| *PSPTO_t30* | *tRNA-Glu-3* | + | 2384916 | 2384991 | 105.55 | 131.62 | 137.59 | 119.67 | 77.67 | 74.83 | tRNA-Glu-3 |
| *PSPTO_t31* | *tRNA-Ala-5* | + | 2385044 | 2385119 | 225.88 | 208.92 | 260.47 | 104.36 | 135.62 | 86.82 | tRNA-Ala-5 |
| *PSPTO_t32* | *tRNA-Glu-4* | + | 2385164 | 2385239 | 33.72 | 26.91 | 48.58 | 18.82 | 14.05 | 31.67 | tRNA-Glu-4 |
| *PSPTO_t33* | *tRNA-Pro-3* | + | 2634736 | 2634812 | 8.93 | 15.47 | 0.43 | 0.00 | 2.86 | 1.67 | tRNA-Pro-3 |
| *PSPTO_t34* | *tRNA-Val-1* | + | 2988034 | 2988110 | 39.34 | 23.16 | 18.67 | 29.42 | 2.50 | 2.97 | tRNA-Val-1 |
| *PSPTO_t35* | *tRNA-Val-2* | + | 2988276 | 2988352 | 38.22 | 23.16 | 28.49 | 29.42 | 2.13 | 1.99 | tRNA-Val-2 |
| *PSPTO_t36* | *tRNA-Leu-2* | + | 3346191 | 3346276 | 18.33 | 63.83 | 33.82 | 14.35 | 12.88 | 8.23 | tRNA-Leu-2 |
| *PSPTO_t37* | *tRNA-Cys-1* | - | 3351545 | 3351618 | 38.99 | 17.75 | 4.46 | 45.12 | 3.49 | 3.28 | tRNA-Cys-1 |
| *PSPTO_t38* | *tRNA-Gly-4* | - | 3397753 | 3397828 | 108.58 | 65.69 | 57.50 | 27.36 | 38.99 | 17.96 | tRNA-Gly-4 |
| *PSPTO_t39* | *tRNA-Asn-2* | - | 3657062 | 3657137 | 116.87 | 49.26 | 89.82 | 55.32 | 21.99 | 18.93 | tRNA-Asn-2 |
| *PSPTO_t40* | *tRNA-Ser-3* | + | 3771634 | 3771721 | 61.43 | 16.61 | 17.16 | 39.94 | 35.38 | 8.70 | tRNA-Ser-3 |
| *PSPTO_t41* | *tRNA-Ala-6* | - | 3872851 | 3872926 | 67.29 | 54.84 | 53.85 | 30.14 | 12.47 | 9.16 | tRNA-Ala-6 |
| *PSPTO_t42* | *tRNA-Ile-4* | - | 3872971 | 3873047 | 204.98 | 192.78 | 182.86 | 158.32 | 169.95 | 84.63 | tRNA-Ile-4 |
| *PSPTO_t43* | *tRNA-Leu-3* | - | 4216266 | 4216350 | 0.54 | 2.94 | 2.88 | 0.00 | 4.55 | 2.42 | tRNA-Leu-3 |
| *PSPTO_t44* | *tRNA-His-1* | - | 4216407 | 4216482 | 19.64 | 6.02 | 2.90 | 0.00 | 2.85 | 1.93 | tRNA-His-1 |
| *PSPTO_t45* | *tRNA-Arg-5* | - | 4216534 | 4216610 | 233.48 | 266.90 | 331.21 | 252.16 | 120.42 | 59.00 | tRNA-Arg-5 |
| *PSPTO_t46* | *tRNA-Asp-1* | - | 4316299 | 4316375 | 34.85 | 34.44 | 49.70 | 11.67 | 23.27 | 20.20 | tRNA-Asp-1 |
| *PSPTO_t47* | *tRNA-Asp-2* | - | 4316490 | 4316566 | 83.74 | 66.76 | 96.81 | 38.04 | 95.72 | 39.56 | tRNA-Asp-2 |
| *PSPTO_t48* | *tRNA-Val-3* | - | 4316635 | 4316710 | 1937.08 | 4835.14 | 3531.02 | 1801.17 | 3245.12 | 905.52 | tRNA-Val-3 |
| *PSPTO_t49* | *tRNA-Asp-3* | - | 4408094 | 4408170 | 31.13 | 13.75 | 31.27 | 13.02 | 16.60 | 11.81 | tRNA-Asp-3 |
| *PSPTO_t50* | *tRNA-Val-4* | - | 4408191 | 4408266 | 1246.95 | 2818.57 | 2254.20 | 1300.46 | 2765.03 | 759.36 | tRNA-Val-4 |
| *PSPTO_t51* | *tRNA-Ser-4* | - | 4458619 | 4458708 | 54.36 | 9.03 | 26.76 | 2.90 | 10.28 | 5.82 | tRNA-Ser-4 |
| *PSPTO_t52* | *tRNA-Lys-2* | - | 4476434 | 4476509 | 59.03 | 63.42 | 91.29 | 65.95 | 55.93 | 69.67 | tRNA-Lys-2 |
| *PSPTO_t53* | *tRNA-Met-3* | + | 4498448 | 4498524 | 59.81 | 22.72 | 21.29 | 12.28 | 5.92 | 4.41 | tRNA-Met-3 |
| *PSPTO_t54* | *tRNA-Met-4* | + | 4498716 | 4498792 | 58.85 | 22.72 | 21.29 | 12.28 | 5.58 | 3.93 | tRNA-Met-4 |
| *PSPTO_t55* | *tRNA-Gly-5* | + | 4787411 | 4787484 | 71.90 | 27.68 | 31.71 | 46.76 | 18.44 | 30.76 | tRNA-Gly-5 |
| *PSPTO_t56* | *tRNA-Met-5* | - | 5061291 | 5061367 | 65.79 | 40.88 | 40.78 | 26.42 | 16.66 | 12.63 | tRNA-Met-5 |
| *PSPTO_t57* | *tRNA-Leu-4* | - | 5061463 | 5061548 | 1.59 | 0.00 | 0.00 | 0.56 | 2.14 | 0.38 | tRNA-Leu-4 |
| *PSPTO_t58* | *tRNA-Leu-5* | + | 5592964 | 5593050 | 53.63 | 173.90 | 119.50 | 92.01 | 98.83 | 34.84 | tRNA-Leu-5 |
| *PSPTO_t59* | *tRNA-Leu-6* | + | 5593211 | 5593297 | 209.67 | 343.76 | 343.42 | 222.34 | 148.29 | 66.34 | tRNA-Leu-6 |
| *PSPTO_t60* | *tRNA-Phe-1* | + | 6074544 | 6074619 | 248.60 | 122.74 | 88.57 | 51.47 | 81.90 | 59.20 | tRNA-Phe-1 |
| *PSPTO_t61* | *tRNA-Thr-2* | + | 6097142 | 6097217 | 150.72 | 227.57 | 74.93 | 64.98 | 129.88 | 58.30 | tRNA-Thr-2 |
| *PSPTO_t62* | *tRNA-Ala-7* | - | 6217322 | 6217397 | 67.27 | 54.84 | 53.85 | 30.14 | 12.47 | 9.16 | tRNA-Ala-7 |
| *PSPTO_t63* | *tRNA-Ile-5* | - | 6217442 | 6217518 | 204.29 | 192.78 | 182.86 | 158.32 | 169.98 | 84.63 | tRNA-Ile-5 |

**Supplementary Table S4. Average gene expression in bacterial endophytes. (B)** Gene expression in *Achromobacter xylosoxidans* Col-0-50.

Gene expression was calculated as Transcripts per million (TPM) using StringTie in the inoculum, 6 ($t_6$), 24 ($t_{24}$) and 168 ($t_{168}$) hours post-inoculation. Each treatment had four biological replicates. When available, gene name for each locus is shown. DNA template strand, coordinates for the start and end of each gene, and a brief annotation are also shown.

| Locus name | Gene name | Strand | Gene start (bp) | Gene end (bp) | Average TPM Inoculum | Average TPM $t_6$ | Average TPM $t_{24}$ | Average TPM $t_{168}$ | Annotation description |
|---|---|---|---|---|---|---|---|---|---|
| *FLPKDCKG_00001* | | - | 365 | 838 | 2.82 | 4.32 | 1.37 | 0.87 | Hypothetical protein |
| *FLPKDCKG_00002* | *csbC* | + | 1056 | 2348 | 0.58 | 1.80 | 0.26 | 0.19 | putative metabolite transport protein CsbC |
| *FLPKDCKG_00003* | *glpK-1* | + | 2527 | 3516 | 0.19 | 0.32 | 1.10 | 0.00 | Glycerol kinase |
| *FLPKDCKG_00004* | *glpK-2* | + | 3461 | 4057 | 0.05 | 0.61 | 0.00 | 0.23 | Glycerol kinase |
| *FLPKDCKG_00005* | *glpD2-1* | + | 4304 | 4900 | 0.77 | 0.41 | 0.41 | 0.07 | Glycerol-3-phosphate dehydrogenase 2 |
| *FLPKDCKG_00006* | *glpD1* | + | 4882 | 5697 | 0.15 | 1.19 | 0.21 | 0.40 | Glycerol-3-phosphate dehydrogenase 1 |
| *FLPKDCKG_00007* | | + | 5725 | 6678 | 0.54 | 0.92 | 0.82 | 0.29 | putative FAD-linked oxidoreductase |
| *FLPKDCKG_00008* | | + | 6624 | 7277 | 1.17 | 0.94 | 0.21 | 0.00 | Hypothetical protein |
| *FLPKDCKG_00009* | *glpP-1* | + | 7279 | 7953 | 0.85 | 3.05 | 3.40 | 0.98 | Glycerol uptake operon antiterminator regulatory protein |
| *FLPKDCKG_00010* | | - | 8010 | 8657 | 30.66 | 12.01 | 4.21 | 2.55 | Hypothetical protein |
| *FLPKDCKG_00011* | | - | 8654 | 9676 | 2.97 | 3.87 | 0.77 | 0.86 | Hypothetical protein |
| *FLPKDCKG_00012* | | - | 9717 | 10931 | 8.33 | 7.94 | 5.86 | 1.78 | Hypothetical protein |
| *FLPKDCKG_00013* | *kdsA-1* | + | 11226 | 12023 | 4.68 | 4.52 | 5.63 | 2.35 | 2-dehydro-3-deoxyphosphooctonate aldolase |
| *FLPKDCKG_00014* | | + | 12144 | 12392 | 3.13 | 5.63 | 5.85 | 2.11 | Hypothetical protein |
| *FLPKDCKG_00015* | | - | 12416 | 12721 | 12.31 | 11.25 | 6.08 | 1.23 | Hypothetical protein |
| *FLPKDCKG_00016* | | - | 12792 | 13187 | 6.14 | 1.90 | 4.12 | 2.96 | Hypothetical protein |
| *FLPKDCKG_00017* | | - | 13439 | 13684 | 0.14 | 0.00 | 1.48 | 0.00 | Hypothetical protein |
| *FLPKDCKG_00018* | | + | 14145 | 15467 | 0.42 | 0.14 | 0.10 | 0.13 | Hypothetical protein |
| *FLPKDCKG_00019* | *rutC-1* | + | 15440 | 15796 | 3.14 | 1.82 | 1.70 | 0.80 | Putative aminoacrylate peracid reductase RutC |
| *FLPKDCKG_00020* | *dmlR-1* | - | 15971 | 16780 | 2.20 | 2.21 | 2.64 | 0.79 | HTH-type transcriptional regulator DmlR |
| *FLPKDCKG_00021* | *tfdR* | - | 16719 | 16892 | 9.97 | 18.10 | 8.92 | 4.15 | HTH-type transcriptional regulator TdfR |
| *FLPKDCKG_00022* | *dthadh* | + | 17039 | 18187 | 1.94 | 6.00 | 4.40 | 0.76 | D-threo-3-hydroxyaspartate dehydratase |
| *FLPKDCKG_00023* | *yabJ-1* | + | 18214 | 18609 | 2.53 | 6.64 | 0.55 | 0.62 | 2-iminobutanoate/2-iminopropanoate deaminase |
| *FLPKDCKG_00024* | | + | 18709 | 18981 | 9.50 | 50.70 | 41.90 | 16.05 | Hypothetical protein |
| *FLPKDCKG_00025* | | - | 19076 | 19363 | 86.91 | 146.51 | 113.64 | 36.71 | Hypothetical protein |
| *FLPKDCKG_00026* | | + | 19486 | 19884 | 7.08 | 12.55 | 7.59 | 3.31 | Hypothetical protein |
| *FLPKDCKG_00027* | | + | 19975 | 20295 | 17.49 | 63.13 | 31.87 | 14.21 | Hypothetical protein |
| *FLPKDCKG_00028* | *cspA-1* | + | 20665 | 20868 | 358.47 | 666.06 | 503.96 | 161.68 | putative cold shock protein A |
| *FLPKDCKG_00029* | | + | 21091 | 21654 | 6.26 | 14.84 | 12.42 | 4.38 | Hypothetical protein |
| *FLPKDCKG_00030* | *cspA-2* | + | 21948 | 22154 | 47.76 | 143.49 | 132.38 | 44.49 | Cold shock protein CspA |
| *FLPKDCKG_00031* | | - | 22253 | 22798 | 17.66 | 34.84 | 34.94 | 5.18 | Hypothetical protein |
| *FLPKDCKG_00032* | *loiP* | + | 23061 | 23816 | 21.59 | 113.14 | 78.82 | 5.10 | Metalloprotease LoiP |
| *FLPKDCKG_00033* | *tig* | + | 23985 | 25295 | 31.80 | 88.61 | 65.81 | 19.47 | Trigger factor |
| *FLPKDCKG_00034* | *clpP-1* | + | 25300 | 25953 | 46.99 | 72.58 | 71.86 | 17.01 | ATP-dependent Clp protease proteolytic subunit |
| *FLPKDCKG_00035* | *clpX* | + | 26055 | 27353 | 39.27 | 82.22 | 49.73 | 18.46 | ATP-dependent Clp protease ATP-binding subunit ClpX |
| *FLPKDCKG_00036* | *lon-1* | + | 27604 | 30054 | 67.51 | 123.83 | 120.97 | 49.86 | Lon protease |
| *FLPKDCKG_00037* | *lexA* | - | 30165 | 30815 | 47.12 | 70.20 | 27.88 | 8.91 | LexA repressor |
| *FLPKDCKG_00038* | *tyrB* | - | 30971 | 32170 | 75.40 | 90.31 | 61.44 | 27.39 | Aromatic-amino-acid aminotransferase |
| *FLPKDCKG_00039* | *uvrB* | + | 32248 | 34368 | 11.83 | 19.99 | 12.91 | 5.01 | UvrABC system protein B |
| *FLPKDCKG_00040* | *yfkJ* | + | 34618 | 35109 | 454.62 | 413.01 | 270.84 | 106.68 | Low molecular weight protein-tyrosine-phosphatase YfkJ |
| *FLPKDCKG_00041* | *iscR* | + | 35302 | 35844 | 168.78 | 450.84 | 336.70 | 81.04 | HTH-type transcriptional regulator IscR |
| *FLPKDCKG_00042* | *iscS-1* | + | 35863 | 37074 | 74.76 | 156.96 | 114.08 | 27.65 | Cysteine desulfurase IscS |
| *FLPKDCKG_00043* | *iscU* | + | 37108 | 37512 | 78.72 | 150.04 | 110.33 | 27.04 | Iron-sulfur cluster assembly scaffold protein IscU |
| *FLPKDCKG_00044* | *iscA* | + | 37515 | 37838 | 33.87 | 154.76 | 125.61 | 22.44 | Iron-binding protein IscA |
| *FLPKDCKG_00045* | *hscB* | + | 37842 | 38354 | 14.54 | 71.24 | 56.51 | 12.62 | Co-chaperone protein HscB |
| *FLPKDCKG_00046* | *hscA* | + | 38419 | 40281 | 4.13 | 13.58 | 13.14 | 4.73 | Chaperone protein HscA |
| *FLPKDCKG_00047* | *fdx-1* | + | 40291 | 40632 | 9.68 | 28.87 | 18.67 | 4.38 | 2Fe-2S ferredoxin |
| *FLPKDCKG_00048* | | + | 40632 | 40826 | 9.93 | 20.27 | 29.18 | 9.40 | Hypothetical protein |
| *FLPKDCKG_00049* | *mgsA* | + | 40930 | 41820 | 37.92 | 31.11 | 19.96 | 10.58 | Methylglyoxal synthase |
| *FLPKDCKG_00050* | *fhuA-1* | + | 42059 | 44179 | 4.19 | 4.32 | 2.95 | 1.90 | Ferrichrome-iron receptor |
| *FLPKDCKG_00051* | *echA8-1* | - | 44195 | 44998 | 10.67 | 4.71 | 7.32 | 1.68 | putative enoyl-CoA hydratase echA8 |
| *FLPKDCKG_00052* | *lptB-1* | + | 45232 | 46053 | 12.54 | 9.53 | 5.78 | 1.46 | Lipopolysaccharide export system ATP-binding protein LptB |
| *FLPKDCKG_00053* | *livH-1* | + | 46053 | 46931 | 13.11 | 6.94 | 0.64 | 0.54 | High-affinity branched-chain amino acid transport system permease protein LivH |
| *FLPKDCKG_00054* | | + | 46934 | 47971 | 8.82 | 5.12 | 3.48 | 0.91 | Hypothetical protein |
| *FLPKDCKG_00055* | *livJ* | + | 48009 | 49190 | 33.67 | 4.11 | 3.32 | 1.84 | Leu/Ile/Val-binding protein |
| *FLPKDCKG_00056* | *livF-1* | + | 49198 | 50025 | 8.24 | 1.94 | 4.46 | 0.15 | High-affinity branched-chain amino acid transport ATP-binding protein LivF |
| *FLPKDCKG_00057* | | + | 50022 | 51875 | 7.23 | 2.27 | 3.29 | 1.19 | Long-chain-fatty-acid--CoA ligase FadD15 |
| *FLPKDCKG_00058* | *pcaF-1* | + | 51872 | 53026 | 4.92 | 2.01 | 1.71 | 1.56 | Beta-ketoadipyl-CoA thiolase |
| *FLPKDCKG_00059* | *kstR2-1* | + | 53023 | 53676 | 4.03 | 3.01 | 1.44 | 0.93 | HTH-type transcriptional repressor KstR2 |
| *FLPKDCKG_00060* | | - | 53688 | 54515 | 3.16 | 5.70 | 1.36 | 2.09 | Hypothetical protein |
| *FLPKDCKG_00061* | *ydcO-1* | - | 54512 | 55765 | 0.91 | 0.51 | 0.34 | 0.30 | Inner membrane protein YdcO |
| *FLPKDCKG_00062* | *gabR-1* | + | 55941 | 57449 | 3.44 | 4.39 | 2.49 | 0.64 | HTH-type transcriptional regulatory protein GabR |
| *FLPKDCKG_00063* | | - | 57506 | 57829 | 1.69 | 2.90 | 0.56 | 0.98 | Hypothetical protein |
| *FLPKDCKG_00064* | *lysS* | - | 57845 | 59368 | 11.65 | 18.68 | 18.82 | 4.34 | Lysine--tRNA ligase |
| *FLPKDCKG_00065* | *yueD-1* | - | 59379 | 59861 | 39.86 | 53.85 | 46.56 | 13.36 | Benzil reductase ((S)-benzoin forming) |

| Locus name | Gene name | Strand | Gene start (bp) | Gene end (bp) | Average TPM Inoculum | Average TPM $t_6$ | Average TPM $t_{24}$ | Average TPM $t_{168}$ | Annotation description |
|---|---|---|---|---|---|---|---|---|---|
| *FLPKDCKG_00066* | *yueD-2* | - | 59879 | 60133 | 7.15 | 13.72 | 15.22 | 5.81 | Benzil reductase ((S)-benzoin forming) |
| *FLPKDCKG_00067* | *prfB-1* | - | 60176 | 61336 | 29.69 | 53.24 | 32.24 | 10.81 | Peptide chain release factor RF2 |
| *FLPKDCKG_00068* | *nicT* | + | 61545 | 62855 | 1.79 | 34.56 | 26.38 | 7.91 | Putative metabolite transport protein NicT |
| *FLPKDCKG_00069* | *recJ* | - | 63111 | 64811 | 3.07 | 6.09 | 4.03 | 1.17 | Single-stranded-DNA-specific exonuclease RecJ |
| *FLPKDCKG_00070* | | - | 64793 | 65731 | 5.09 | 9.62 | 6.67 | 2.82 | Hypothetical protein |
| *FLPKDCKG_00071* | *lolE* | + | 65862 | 67142 | 7.81 | 16.93 | 8.88 | 2.51 | Lipoprotein-releasing system transmembrane protein LolE |
| *FLPKDCKG_00072* | *lolD* | + | 67135 | 67824 | 8.05 | 8.25 | 9.26 | 1.90 | Lipoprotein-releasing system ATP-binding protein LolD |
| *FLPKDCKG_00073* | *yjjV* | + | 67889 | 68704 | 2.14 | 7.58 | 6.71 | 0.64 | putative metal-dependent hydrolase YjjV |
| *FLPKDCKG_00074* | *cpxA-1* | - | 68757 | 70151 | 0.75 | 0.92 | 1.87 | 1.30 | Sensor histidine kinase CpxA |
| *FLPKDCKG_00075* | *cpxR* | - | 70151 | 70846 | 2.70 | 3.01 | 0.42 | 0.57 | Transcriptional regulatory protein CpxR |
| *FLPKDCKG_00076* | *cirA-1* | + | 71071 | 73092 | 1.62 | 1.05 | 0.47 | 0.55 | Colicin I receptor |
| *FLPKDCKG_00077* | *yclQ-1* | + | 73158 | 74120 | 1.14 | 0.12 | 0.56 | 0.00 | putative ABC transporter solute-binding protein YclQ |
| *FLPKDCKG_00078* | *feuB* | + | 74148 | 75146 | 0.08 | 0.38 | 0.00 | 0.00 | Iron-uptake system permease protein FeuB |
| *FLPKDCKG_00079* | *fepD* | + | 75139 | 76122 | 0.23 | 0.00 | 0.00 | 0.26 | Ferric enterobactin transport system permease protein FepD |
| *FLPKDCKG_00080* | *fecE-1* | + | 76119 | 76877 | 0.08 | 0.00 | 0.37 | 0.00 | Fe(3+) dicitrate transport ATP-binding protein FecE |
| *FLPKDCKG_00081* | *cpdB* | - | 76977 | 79082 | 0.13 | 0.23 | 0.00 | 0.40 | 2',3'-cyclic-nucleotide 2'-phosphodiesterase/3'-nucleotidase |
| *FLPKDCKG_00082* | *bfrD-1* | + | 79329 | 81434 | 0.72 | 0.84 | 1.30 | 0.53 | putative TonB-dependent receptor BfrD |
| *FLPKDCKG_00083* | *thiM* | - | 81477 | 82298 | 0.70 | 2.19 | 0.11 | 1.36 | Hydroxyethylthiazole kinase |
| *FLPKDCKG_00084* | *mprA-1* | - | 82618 | 83289 | 10.37 | 1.72 | 0.43 | 0.05 | Transcriptional repressor MprA |
| *FLPKDCKG_00085* | | + | 83479 | 83937 | 0.14 | 0.00 | 0.00 | 0.00 | Hypothetical protein |
| *FLPKDCKG_00086* | | + | 83959 | 84636 | 0.13 | 0.00 | 0.92 | 0.00 | Hypothetical protein |
| *FLPKDCKG_00087* | *yxdL* | + | 84646 | 85260 | 10.82 | 8.38 | 2.73 | 3.48 | ABC transporter ATP-binding protein YxdL |
| *FLPKDCKG_00088* | *fabG-1* | - | 85276 | 86037 | 5.85 | 5.04 | 3.23 | 2.08 | 3-oxoacyl-[acyl-carrier-protein] reductase |
| *FLPKDCKG_00089* | *sdh* | + | 86184 | 86915 | 4.81 | 4.64 | 5.12 | 2.55 | Serine 3-dehydrogenase |
| *FLPKDCKG_00090* | *fdx-2* | - | 86921 | 87259 | 6.15 | 2.19 | 5.55 | 2.66 | 2Fe-2S ferredoxin |
| *FLPKDCKG_00091* | *crtF* | - | 87467 | 88492 | 2.28 | 7.27 | 2.21 | 2.61 | Demethylspheroidene O-methyltransferase |
| *FLPKDCKG_00092* | *leuE-1* | + | 88683 | 89333 | 24.76 | 24.28 | 13.65 | 6.44 | Leucine efflux protein |
| *FLPKDCKG_00093* | *speE-1* | - | 89364 | 91901 | 7.92 | 5.55 | 3.04 | 1.70 | Polyamine aminopropyltransferase |
| *FLPKDCKG_00094* | *atsA* | - | 91977 | 93620 | 163.12 | 46.11 | 24.06 | 15.95 | Arylsulfatase |
| *FLPKDCKG_00095* | | - | 93667 | 94689 | 69.42 | 29.69 | 9.71 | 8.34 | Hypothetical protein |
| *FLPKDCKG_00096* | | - | 94745 | 96064 | 5.66 | 3.11 | 1.49 | 0.86 | Hypothetical protein |
| *FLPKDCKG_00097* | | - | 96061 | 97572 | 7.25 | 4.04 | 1.86 | 1.24 | Hypothetical protein |
| *FLPKDCKG_00098* | | - | 97557 | 98576 | 11.79 | 7.33 | 1.19 | 1.58 | Hypothetical protein |
| *FLPKDCKG_00099* | | - | 98563 | 99063 | 6.65 | 0.54 | 0.44 | 0.03 | Hypothetical protein |
| *FLPKDCKG_00100* | | - | 99060 | 100004 | 16.21 | 8.82 | 2.44 | 1.67 | Hypothetical protein |
| *FLPKDCKG_00101* | *nirQ* | - | 100009 | 100980 | 20.73 | 9.57 | 5.92 | 3.52 | Denitrification regulatory protein NirQ |
| *FLPKDCKG_00102* | *pkn1-1* | - | 101072 | 102100 | 87.17 | 42.58 | 27.30 | 6.17 | Serine/threonine-protein kinase pkn1 |
| *FLPKDCKG_00103* | | + | 102837 | 103604 | 9.40 | 7.83 | 5.19 | 1.51 | Hypothetical protein |
| *FLPKDCKG_00104* | | + | 103664 | 105073 | 18.26 | 6.73 | 11.66 | 10.94 | Hypothetical protein |
| *FLPKDCKG_00105* | *hspA* | - | 105089 | 105532 | 117.28 | 23.24 | 41.97 | 51.32 | Spore protein SP21 |
| *FLPKDCKG_00106* | *groL-1* | - | 105808 | 107430 | 13.44 | 5.40 | 8.37 | 12.72 | 60 kDa chaperonin |
| *FLPKDCKG_00107* | *groS-1* | - | 107462 | 107779 | 18.38 | 9.84 | 11.81 | 1.61 | 10 kDa chaperonin |
| *FLPKDCKG_00108* | | - | 107939 | 108559 | 6.62 | 3.48 | 3.01 | 2.00 | Hypothetical protein |
| *FLPKDCKG_00109* | *kefF* | + | 108764 | 109360 | 7.61 | 12.85 | 6.97 | 3.23 | Glutathione-regulated potassium-efflux system ancillary protein KefF |
| *FLPKDCKG_00110* | *betI-1* | + | 109380 | 109778 | 5.48 | 12.34 | 1.97 | 2.72 | HTH-type transcriptional regulator BetI |
| *FLPKDCKG_00111* | | + | 109736 | 109990 | 1.10 | 4.44 | 0.00 | 2.11 | Hypothetical protein |
| *FLPKDCKG_00112* | | + | 110064 | 110456 | 2.08 | 5.01 | 1.84 | 2.53 | Hypothetical protein |
| *FLPKDCKG_00113* | | + | 110588 | 111856 | 3.46 | 7.29 | 3.16 | 0.90 | Hypothetical protein |
| *FLPKDCKG_00114* | *fliC-1* | - | 112037 | 113584 | 0.13 | 0.95 | 0.76 | 0.72 | Flagellin |
| *FLPKDCKG_00115* | *trpI-1* | - | 113919 | 114794 | 2.89 | 3.88 | 3.32 | 1.02 | HTH-type transcriptional regulator TrpI |
| *FLPKDCKG_00116* | *trpB-1* | + | 114900 | 116108 | 0.20 | 0.76 | 0.00 | 0.64 | Tryptophan synthase beta chain |
| *FLPKDCKG_00117* | *trpA-1* | + | 116105 | 116896 | 0.11 | 0.37 | 0.35 | 0.80 | Tryptophan synthase alpha chain |
| *FLPKDCKG_00118* | *comEC* | + | 117081 | 119516 | 0.17 | 0.23 | 0.18 | 0.49 | ComE operon protein 3 |
| *FLPKDCKG_00119* | *ylmB-1* | - | 119566 | 120876 | 0.31 | 0.63 | 0.02 | 0.80 | N-formyl-4-amino-5-aminomethyl-2-methylpyrimidine deformylase |
| *FLPKDCKG_00120* | *proP-1* | - | 120906 | 122231 | 0.31 | 0.43 | 0.21 | 0.18 | Proline/betaine transporter |
| *FLPKDCKG_00121* | *cysL-1* | + | 122361 | 123338 | 3.69 | 4.77 | 1.15 | 2.01 | HTH-type transcriptional regulator CysL |
| *FLPKDCKG_00122* | | - | 123361 | 123675 | 2.16 | 5.06 | 1.68 | 0.48 | Hypothetical protein |
| *FLPKDCKG_00123* | *cynR-1* | - | 124397 | 125284 | 4.34 | 6.50 | 3.45 | 0.88 | HTH-type transcriptional regulator CynR |
| *FLPKDCKG_00124* | *alkJ-1* | + | 125416 | 127059 | 11.65 | 27.19 | 17.06 | 14.43 | Alcohol dehydrogenase [acceptor] |
| *FLPKDCKG_00125* | *cfa-1* | - | 127198 | 127347 | 5.28 | 14.48 | 7.64 | 0.00 | Cyclopropane-fatty-acyl-phospholipid synthase |
| *FLPKDCKG_00126* | *cfa-2* | - | 127296 | 128375 | 30.74 | 49.45 | 39.79 | 11.17 | Cyclopropane-fatty-acyl-phospholipid synthase |
| *FLPKDCKG_00127* | *aroL* | + | 128829 | 129377 | 3.48 | 13.22 | 7.50 | 1.92 | Shikimate kinase 2 |
| *FLPKDCKG_00128* | | + | 129437 | 130372 | 8.58 | 12.05 | 6.07 | 2.01 | Tropinesterase |
| *FLPKDCKG_00129* | | + | 130542 | 131387 | 22.80 | 31.03 | 29.21 | 6.14 | 3',5'-nucleoside bisphosphate phosphatase |
| *FLPKDCKG_00130* | *greB* | + | 131479 | 132030 | 9.58 | 17.17 | 11.54 | 2.89 | Transcription elongation factor GreB |
| *FLPKDCKG_00131* | | - | 132214 | 133245 | 11.54 | 11.22 | 11.73 | 5.27 | Solute-binding protein |
| *FLPKDCKG_00132* | *dctM-1* | - | 133319 | 134617 | 1.82 | 5.05 | 2.55 | 1.01 | C4-dicarboxylate TRAP transporter large permease protein DctM |
| *FLPKDCKG_00133* | *yiaM-1* | - | 134614 | 135165 | 1.50 | 1.59 | 3.65 | 0.97 | 2,3-diketo-L-gulonate TRAP transporter small permease protein YiaM |
| *FLPKDCKG_00134* | *purL* | + | 135368 | 139579 | 20.02 | 45.47 | 25.57 | 8.95 | Phosphoribosylformylglycinamidine synthase |

| Locus name | Gene name | Strand | Gene start (bp) | Gene end (bp) | Average TPM Inoculum | Average TPM $t_6$ | Average TPM $t_{24}$ | Average TPM $t_{168}$ | Annotation description |
|---|---|---|---|---|---|---|---|---|---|
| *FLPKDCKG_00135* | *trg-1* | + | 139819 | 141444 | 1.59 | 1.56 | 0.43 | 0.06 | Methyl-accepting chemotaxis protein III |
| *FLPKDCKG_00136* | | - | 141478 | 142230 | 0.33 | 0.00 | 1.77 | 0.77 | Hypothetical protein |
| *FLPKDCKG_00137* | *mipA* | - | 142261 | 143022 | 0.56 | 0.54 | 0.73 | 0.28 | MltA-interacting protein |
| *FLPKDCKG_00138* | *qseB-1* | + | 143147 | 143836 | 0.43 | 1.23 | 2.94 | 0.00 | Transcriptional regulatory protein QseB |
| *FLPKDCKG_00139* | *qseC-1* | + | 143833 | 145191 | 0.25 | 0.28 | 0.00 | 1.11 | Sensor protein QseC |
| *FLPKDCKG_00140* | | - | 145246 | 146232 | 0.90 | 1.67 | 1.67 | 1.02 | Hypothetical protein |
| *FLPKDCKG_00141* | | - | 146251 | 147204 | 0.23 | 0.44 | 0.00 | 0.00 | Hypothetical protein |
| *FLPKDCKG_00142* | *yjiE-1* | + | 147318 | 148235 | 0.71 | 0.73 | 1.28 | 0.22 | HTH-type transcriptional regulator YjiE |
| *FLPKDCKG_00143* | | + | 148366 | 149838 | 47.02 | 61.86 | 41.24 | 16.45 | Hypothetical protein |
| *FLPKDCKG_00144* | *crcB-1* | - | 149994 | 150365 | 1.89 | 9.62 | 13.29 | 5.60 | Putative fluoride ion transporter CrcB |
| *FLPKDCKG_00145* | | + | 150696 | 151253 | 6.20 | 10.48 | 16.03 | 4.16 | Hypothetical protein |
| *FLPKDCKG_00146* | | + | 151501 | 151803 | 0.63 | 0.22 | 0.00 | 1.15 | Hypothetical protein |
| *FLPKDCKG_00147* | | + | 151746 | 152492 | 1.71 | 2.28 | 0.37 | 0.21 | Hypothetical protein |
| *FLPKDCKG_00148* | | + | 152502 | 153800 | 1.20 | 1.08 | 1.20 | 1.05 | Hypothetical protein |
| *FLPKDCKG_00149* | | - | 153876 | 154631 | 1.69 | 2.30 | 1.89 | 1.03 | Hypothetical protein |
| *FLPKDCKG_00150* | | - | 154641 | 166340 | 0.16 | 0.59 | 0.24 | 0.15 | Hypothetical protein |
| *FLPKDCKG_00151* | | - | 166334 | 168115 | 0.12 | 0.00 | 0.00 | 0.16 | Hypothetical protein |
| *FLPKDCKG_00152* | | + | 168154 | 169290 | 0.35 | 0.37 | 0.00 | 0.00 | Hypothetical protein |
| *FLPKDCKG_00153* | | - | 169323 | 170216 | 0.05 | 0.08 | 0.00 | 0.39 | Hypothetical protein |
| *FLPKDCKG_00154* | *hxuB-1* | - | 170261 | 171055 | 0.00 | 0.38 | 0.49 | 0.00 | Heme/hemopexin transporter protein HuxB |
| *FLPKDCKG_00155* | *pleD-1* | + | 171368 | 172282 | 0.49 | 0.93 | 1.38 | 0.44 | Response regulator PleD |
| *FLPKDCKG_00156* | *luxQ* | + | 172275 | 174959 | 0.30 | 0.39 | 0.19 | 0.60 | Autoinducer 2 sensor kinase/phosphatase LuxQ |
| *FLPKDCKG_00157* | *dmlR-2* | - | 174968 | 175867 | 0.73 | 5.05 | 1.83 | 0.66 | HTH-type transcriptional regulator DmlR |
| *FLPKDCKG_00158* | *stp-1* | + | 175979 | 177514 | 0.48 | 0.49 | 0.15 | 0.10 | Multidrug resistance protein Stp |
| *FLPKDCKG_00159* | | - | 177533 | 178201 | 6.68 | 3.92 | 3.33 | 0.51 | Putative O-methyltransferase/MSMEI_4947 |
| *FLPKDCKG_00160* | *slmA-1* | + | 178285 | 178947 | 3.16 | 1.41 | 0.00 | 0.74 | Nucleoid occlusion factor SlmA |
| *FLPKDCKG_00161* | *bphD* | + | 179047 | 179880 | 3.38 | 2.40 | 3.41 | 0.66 | 2-hydroxy-6-oxo-6-phenylhexa-2,4-dienoate hydrolase |
| *FLPKDCKG_00162* | *tsaC1* | + | 179892 | 180635 | 2.21 | 3.89 | 3.86 | 1.29 | 4-formylbenzenesulfonate dehydrogenase TsaC1/TsaC2 |
| *FLPKDCKG_00163* | | - | 180655 | 181650 | 6.96 | 6.47 | 2.86 | 1.35 | Hypothetical protein |
| *FLPKDCKG_00164* | | - | 181647 | 182564 | 19.41 | 15.46 | 5.77 | 3.54 | Hypothetical protein |
| *FLPKDCKG_00165* | *pucI* | - | 183151 | 184497 | 0.67 | 9.00 | 10.01 | 2.81 | putative allantoin permease |
| *FLPKDCKG_00166* | *pepIP* | - | 184794 | 185588 | 3.91 | 6.52 | 4.86 | 3.20 | Proline iminopeptidase |
| *FLPKDCKG_00167* | *murA-1* | - | 185664 | 186965 | 8.86 | 16.21 | 9.16 | 4.63 | UDP-N-acetylglucosamine 1-carboxyvinyltransferase |
| *FLPKDCKG_00168* | | - | 187057 | 187533 | 20.90 | 22.61 | 18.96 | 5.09 | Hypothetical protein |
| *FLPKDCKG_00169* | | + | 187595 | 188392 | 2.84 | 3.68 | 2.14 | 1.53 | Hypothetical protein |
| *FLPKDCKG_00170* | | - | 188453 | 189160 | 25.05 | 29.12 | 24.74 | 9.06 | Hypothetical protein |
| *FLPKDCKG_00171* | *aphA-1* | - | 189370 | 190155 | 0.75 | 5.28 | 5.26 | 2.10 | Aminoglycoside 3'-phosphotransferase |
| *FLPKDCKG_00172* | | - | 190152 | 190601 | 6.60 | 12.52 | 18.21 | 6.47 | Hypothetical protein |
| *FLPKDCKG_00173* | | + | 190767 | 191492 | 4.33 | 4.85 | 8.99 | 1.11 | Hypothetical protein |
| *FLPKDCKG_00174* | *nucA-1* | - | 191527 | 192435 | 1.02 | 1.43 | 1.33 | 0.85 | Nuclease |
| *FLPKDCKG_00175* | *pgrR-1* | - | 192584 | 193483 | 0.94 | 3.11 | 1.81 | 0.87 | HTH-type transcriptional regulator PgrR |
| *FLPKDCKG_00176* | | + | 193797 | 194639 | 1.39 | 3.94 | 7.82 | 0.98 | Hypothetical protein |
| *FLPKDCKG_00177* | | - | 194723 | 195538 | 14.01 | 35.92 | 28.01 | 10.12 | Hypothetical protein |
| *FLPKDCKG_00178* | *fhuA-2* | + | 196651 | 197994 | 0.54 | 1.42 | 0.43 | 0.34 | Ferrichrome-iron receptor |
| *FLPKDCKG_00179* | *fhuA-3* | + | 198133 | 199746 | 1.48 | 1.77 | 0.52 | 0.55 | Ferrichrome-iron receptor |
| *FLPKDCKG_00180* | *fhuA-4* | + | 199797 | 200300 | 1.19 | 1.17 | 1.49 | 1.85 | Ferrichrome-iron receptor |
| *FLPKDCKG_00181* | *ycdT-1* | + | 200442 | 201617 | 0.84 | 1.97 | 3.59 | 1.07 | putative diguanylate cyclase YcdT |
| *FLPKDCKG_00182* | | + | 201659 | 202546 | 1.53 | 1.83 | 3.99 | 1.20 | Hypothetical protein |
| *FLPKDCKG_00183* | | - | 202658 | 203080 | 2.24 | 3.73 | 1.90 | 0.75 | Hypothetical protein |
| *FLPKDCKG_00184* | *hmo* | + | ` | 204454 | 3.75 | 3.47 | 8.54 | 2.34 | 4-hydroxymandelate oxidase |
| *FLPKDCKG_00185* | | + | 204611 | 205285 | 6.52 | 11.08 | 12.10 | 3.45 | Hypothetical protein |
| *FLPKDCKG_00186* | | + | 205322 | 205948 | 4.08 | 7.65 | 4.14 | 0.97 | Hypothetical protein |
| *FLPKDCKG_00187* | | + | 206241 | 206873 | 4.92 | 15.20 | 8.67 | 3.34 | Hypothetical protein |
| *FLPKDCKG_00188* | | + | 207038 | 208045 | 0.29 | 2.16 | 1.39 | 0.10 | Hypothetical protein |
| *FLPKDCKG_00189* | | + | 208035 | 208301 | 0.22 | 0.00 | 0.00 | 0.00 | Hypothetical protein |
| *FLPKDCKG_00190* | | + | 208309 | 209073 | 0.25 | 0.00 | 0.00 | 0.00 | Hypothetical protein |
| *FLPKDCKG_00191* | *flhB-1* | + | 209073 | 210152 | 0.17 | 0.14 | 0.47 | 0.00 | Flagellar biosynthetic protein FlhB |
| *FLPKDCKG_00192* | *flhA-1* | + | 210149 | 212227 | 0.19 | 0.63 | 0.72 | 0.26 | Flagellar biosynthesis protein FlhA |
| *FLPKDCKG_00193* | *flgG-1* | + | 212264 | 212992 | 0.28 | 0.83 | 0.00 | 0.12 | Flagellar basal-body rod protein FlgG |
| *FLPKDCKG_00194* | *flgG-2* | + | 213004 | 213786 | 0.15 | 0.07 | 1.27 | 0.41 | Flagellar basal-body rod protein FlgG |
| *FLPKDCKG_00195* | | + | 213795 | 214757 | 0.18 | 0.61 | 0.00 | 0.00 | Hypothetical protein |
| *FLPKDCKG_00196* | *flgH-1* | + | 214754 | 215359 | 0.47 | 1.91 | 2.33 | 0.03 | Flagellar L-ring protein |
| *FLPKDCKG_00197* | *flgI-1* | + | 215359 | 216498 | 0.98 | 3.27 | 1.30 | 1.04 | Flagellar P-ring protein |
| *FLPKDCKG_00198* | *flgB-1* | + | 216520 | 216897 | 0.12 | 0.00 | 0.00 | 0.90 | Flagellar basal body rod protein FlgB |
| *FLPKDCKG_00199* | *flgC-1* | + | 216900 | 217337 | 0.00 | 0.00 | 0.00 | 0.00 | Flagellar basal-body rod protein FlgC |
| *FLPKDCKG_00200* | *fliE-1* | + | 217334 | 217630 | 0.16 | 0.00 | 0.00 | 0.00 | Flagellar hook-basal body complex protein FliE |
| *FLPKDCKG_00201* | *fliF-1* | + | 217648 | 219075 | 0.13 | 0.53 | 1.34 | 0.00 | Flagellar M-ring protein |
| *FLPKDCKG_00202* | | + | 219075 | 219596 | 0.08 | 0.52 | 0.00 | 0.00 | Hypothetical protein |
| *FLPKDCKG_00203* | *yscL-1* | + | 219593 | 220375 | 0.36 | 0.00 | 0.00 | 0.11 | Yop proteins translocation protein L |

| Locus name | Gene name | Strand | Gene start (bp) | Gene end (bp) | Average TPM Inoculum | Average TPM $t_6$ | Average TPM $t_{24}$ | Average TPM $t_{168}$ | Annotation description |
|---|---|---|---|---|---|---|---|---|---|
| *FLPKDCKG_00204* | *fliI-1* | + | 220372 | 221709 | 0.34 | 0.96 | 0.93 | 0.50 | Flagellum-specific ATP synthase |
| *FLPKDCKG_00205* | | + | 221706 | 222170 | 0.17 | 0.00 | 0.00 | 0.00 | Hypothetical protein |
| *FLPKDCKG_00206* | | - | 221995 | 222810 | 0.03 | 0.00 | 0.00 | 0.00 | Hypothetical protein |
| *FLPKDCKG_00207* | | + | 222843 | 223232 | 0.50 | 0.00 | 0.00 | 0.00 | Hypothetical protein |
| *FLPKDCKG_00208* | *flgE-1* | + | 223244 | 224443 | 0.12 | 0.27 | 0.28 | 0.00 | Flagellar hook protein FlgE |
| *FLPKDCKG_00209* | | + | 224443 | 225213 | 0.17 | 1.76 | 3.28 | 0.85 | Hypothetical protein |
| *FLPKDCKG_00210* | *fliN-1* | + | 225256 | 225603 | 0.02 | 0.00 | 0.00 | 0.00 | Flagellar motor switch protein FliN |
| *FLPKDCKG_00211* | | + | 225600 | 225926 | 0.00 | 0.00 | 0.00 | 0.00 | Hypothetical protein |
| *FLPKDCKG_00212* | *fliP-1* | + | 225923 | 226672 | 0.12 | 0.00 | 0.00 | 0.00 | Flagellar biosynthetic protein FliP |
| *FLPKDCKG_00213* | *fliA-1* | + | 226684 | 227460 | 1.16 | 4.59 | 3.56 | 0.72 | RNA polymerase sigma factor FliA |
| *FLPKDCKG_00214* | *bepA-1* | + | 227521 | 228105 | 3.57 | 3.04 | 4.13 | 2.54 | Beta-barrel assembly-enhancing protease |
| *FLPKDCKG_00215* | | + | 228213 | 228638 | 1.89 | 4.25 | 2.73 | 0.00 | Hypothetical protein |
| *FLPKDCKG_00216* | | - | 228663 | 229049 | 2.15 | 8.81 | 3.10 | 1.16 | Hypothetical protein |
| *FLPKDCKG_00217* | | - | 229057 | 229620 | 3.05 | 11.48 | 6.00 | 0.99 | Hypothetical protein |
| *FLPKDCKG_00218* | | - | 229646 | 230764 | 2.22 | 15.14 | 2.91 | 0.23 | Hypothetical protein |
| *FLPKDCKG_00219* | | - | 230761 | 231663 | 0.43 | 1.11 | 1.59 | 0.00 | Hypothetical protein |
| *FLPKDCKG_00220* | | - | 231675 | 233855 | 0.34 | 0.74 | 0.58 | 0.14 | Hypothetical protein |
| *FLPKDCKG_00221* | | - | 233899 | 234033 | 1.65 | 2.80 | 3.61 | 2.63 | Hypothetical protein |
| *FLPKDCKG_00222* | *wapA-1* | + | 234273 | 249455 | 1.23 | 3.48 | 2.09 | 0.41 | tRNA3(Ser)-specific nuclease WapA |
| *FLPKDCKG_00223* | *fliA-2* | + | 251644 | 252432 | 0.50 | 1.60 | 2.69 | 0.53 | RNA polymerase sigma factor FliA |
| *FLPKDCKG_00224* | *wapA-2* | + | 252587 | 266317 | 0.72 | 2.82 | 1.38 | 0.36 | tRNA3(Ser)-specific nuclease WapA |
| *FLPKDCKG_00225* | *fhuE-1* | + | 268791 | 270986 | 2.04 | 1.98 | 1.20 | 0.50 | FhuE receptor |
| *FLPKDCKG_00226* | | - | 271070 | 271672 | 3.18 | 16.69 | 6.77 | 1.35 | Hypothetical protein |
| *FLPKDCKG_00227* | | - | 271730 | 272173 | 2.51 | 14.08 | 15.28 | 4.05 | Hypothetical protein |
| *FLPKDCKG_00228* | *pheA-1* | - | 272344 | 273207 | 2.56 | 0.73 | 1.41 | 1.89 | Prephenate dehydratase |
| *FLPKDCKG_00229* | *cynR-2* | - | 273231 | 274112 | 1.64 | 4.57 | 3.84 | 0.46 | HTH-type transcriptional regulator CynR |
| *FLPKDCKG_00230* | *citE* | + | 274724 | 275044 | 0.00 | 0.00 | 0.00 | 0.00 | Citrate lyase subunit beta-like protein |
| *FLPKDCKG_00231* | *uctC-1* | + | 275079 | 276482 | 0.75 | 1.81 | 0.42 | 0.00 | Acetyl-CoA:oxalate CoA-transferase |
| *FLPKDCKG_00232* | | + | 276564 | 277400 | 0.54 | 1.42 | 1.29 | 0.38 | Hypothetical protein |
| *FLPKDCKG_00233* | | + | 277471 | 278457 | 0.62 | 0.36 | 1.47 | 0.13 | Hypothetical protein |
| *FLPKDCKG_00234* | | + | 278523 | 278975 | 4.98 | 9.14 | 4.74 | 0.49 | Hypothetical protein |
| *FLPKDCKG_00235* | | - | 279697 | 280125 | 8.38 | 10.82 | 5.68 | 0.87 | Hypothetical protein |
| *FLPKDCKG_00236* | *fom3* | + | 280272 | 282185 | 1.36 | 5.36 | 2.50 | 1.16 | 2-hydroxyethylphosphonate methyltransferase |
| *FLPKDCKG_00237* | *pchR-1* | - | 282121 | 283161 | 0.15 | 0.04 | 0.00 | 0.00 | Regulatory protein PchR |
| *FLPKDCKG_00238* | *fhuA-5* | + | 283388 | 285496 | 1.59 | 2.45 | 1.68 | 0.38 | Ferrichrome-iron receptor |
| *FLPKDCKG_00239* | | - | 285604 | 286698 | 3.39 | 15.62 | 8.28 | 2.25 | Hypothetical protein |
| *FLPKDCKG_00240* | | - | 286998 | 287831 | 8.08 | 21.59 | 19.58 | 5.46 | Hypothetical protein |
| *FLPKDCKG_00241* | | - | 287999 | 290011 | 3.35 | 5.34 | 5.67 | 1.60 | Hypothetical protein |
| *FLPKDCKG_00242* | | + | 290276 | 291103 | 6.98 | 5.18 | 6.00 | 1.59 | Hypothetical protein |
| *FLPKDCKG_00243* | | - | 291615 | 291788 | 0.70 | 4.72 | 0.51 | 0.00 | Hypothetical protein |
| *FLPKDCKG_00244* | | - | 292416 | 293354 | 6.33 | 33.44 | 14.72 | 3.24 | Hypothetical protein |
| *FLPKDCKG_00245* | | + | 293838 | 294050 | 62.94 | 117.13 | 95.82 | 18.07 | Hypothetical protein |
| *FLPKDCKG_00246* | | - | 294827 | 295675 | 7.17 | 19.79 | 9.58 | 4.62 | Hypothetical protein |
| *FLPKDCKG_00247* | | - | 295824 | 296618 | 0.29 | 0.17 | 0.00 | 0.66 | Hypothetical protein |
| *FLPKDCKG_00248* | *aiiA* | - | 296752 | 297522 | 0.75 | 0.00 | 0.97 | 0.00 | N-acyl homoserine lactonase |
| *FLPKDCKG_00249* | | - | 297527 | 298498 | 0.80 | 0.24 | 0.83 | 1.25 | Hypothetical protein |
| *FLPKDCKG_00250* | *yjmC-1* | - | 298526 | 299572 | 1.59 | 0.25 | 0.00 | 0.40 | putative oxidoreductase YjmC |
| *FLPKDCKG_00251* | | - | 299607 | 300599 | 0.63 | 0.38 | 0.06 | 0.00 | Hypothetical protein |
| *FLPKDCKG_00252* | *oxyR-1* | + | 300708 | 301607 | 4.49 | 7.23 | 4.01 | 1.64 | Hydrogen peroxide-inducible genes activator |
| *FLPKDCKG_00253* | | - | 301746 | 302915 | 0.44 | 2.11 | 3.71 | 1.35 | Hypothetical protein |
| *FLPKDCKG_00254* | | - | 302940 | 304298 | 0.97 | 3.28 | 7.53 | 1.39 | Hypothetical protein |
| *FLPKDCKG_00255* | *hbpA-1* | - | 304309 | 305880 | 2.19 | 8.52 | 12.01 | 4.17 | Heme-binding protein A |
| *FLPKDCKG_00256* | *oppF-1* | - | 305912 | 306952 | 1.51 | 4.67 | 5.11 | 0.97 | Oligopeptide transport ATP-binding protein OppF |
| *FLPKDCKG_00257* | *oppD-1* | - | 306949 | 307971 | 1.23 | 2.51 | 5.27 | 0.27 | Oligopeptide transport ATP-binding protein OppD |
| *FLPKDCKG_00258* | *ddpC-1* | - | 307968 | 308912 | 1.11 | 6.43 | 4.01 | 0.71 | putative D,D-dipeptide transport system permease protein DdpC |
| *FLPKDCKG_00259* | *dppB-1* | - | 308909 | 309883 | 0.90 | 1.90 | 1.72 | 0.00 | Dipeptide transport system permease protein DppB |
| *FLPKDCKG_00260* | *hpyO* | - | 309880 | 311226 | 0.67 | 4.13 | 2.93 | 0.41 | FAD-dependent urate hydroxylase |
| *FLPKDCKG_00261* | *gltC-1* | - | 311350 | 312315 | 2.64 | 6.47 | 7.18 | 1.41 | HTH-type transcriptional regulator GltC |
| *FLPKDCKG_00262* | | - | 312487 | 313212 | 1.48 | 1.99 | 1.33 | 0.30 | Hypothetical protein |
| *FLPKDCKG_00263* | *sigK-1* | - | 313209 | 313778 | 6.75 | 7.43 | 8.87 | 3.49 | ECF RNA polymerase sigma factor SigK |
| *FLPKDCKG_00264* | | - | 313869 | 314417 | 8.77 | 8.98 | 11.01 | 2.56 | Hypothetical protein |
| *FLPKDCKG_00265* | | + | 314598 | 315299 | 1.16 | 2.58 | 2.17 | 1.52 | Hypothetical protein |
| *FLPKDCKG_00266* | | - | 315315 | 315605 | 0.72 | 2.41 | 2.15 | 2.02 | Hypothetical protein |
| *FLPKDCKG_00267* | *aam-1* | - | 315740 | 317263 | 0.00 | 6.99 | 7.17 | 4.36 | Acylamidase |
| *FLPKDCKG_00268* | | - | 317278 | 318258 | 0.13 | 14.45 | 12.06 | 13.14 | Hypothetical protein |
| *FLPKDCKG_00269* | *gltC-2* | + | 318402 | 319337 | 0.55 | 0.93 | 2.89 | 0.43 | HTH-type transcriptional regulator GltC |
| *FLPKDCKG_00270* | | - | 319334 | 319621 | 10.81 | 12.64 | 14.02 | 6.23 | Hypothetical protein |
| *FLPKDCKG_00271* | | - | 319701 | 319910 | 10.02 | 21.72 | 29.43 | 12.10 | Hypothetical protein |
| *FLPKDCKG_00272* | | + | 319995 | 320234 | 33.76 | 59.01 | 34.08 | 10.06 | Hypothetical protein |

| Locus name | Gene name | Strand | Gene start (bp) | Gene end (bp) | Average TPM Inoculum | Average TPM $t_6$ | Average TPM $t_{24}$ | Average TPM $t_{168}$ | Annotation description |
|---|---|---|---|---|---|---|---|---|---|
| *FLPKDCKG_00273* | | - | 320495 | 321421 | 0.15 | 2.25 | 0.72 | 0.02 | 2-ketogluconate reductase |
| *FLPKDCKG_00274* | | - | 321649 | 322644 | 0.52 | 1.89 | 1.83 | 0.00 | Hypothetical protein |
| *FLPKDCKG_00275* | *hcaR-1* | + | 322820 | 323710 | 1.82 | 3.60 | 3.08 | 0.23 | Hca operon transcriptional activator HcaR |
| *FLPKDCKG_00276* | | - | 323834 | 324418 | 1.41 | 4.18 | 3.92 | 0.07 | Hypothetical protein |
| *FLPKDCKG_00277* | | - | 324415 | 325719 | 2.10 | 4.79 | 5.03 | 0.88 | Hypothetical protein |
| *FLPKDCKG_00278* | | - | 325923 | 326312 | 24.60 | 44.39 | 31.15 | 2.43 | Hypothetical protein |
| *FLPKDCKG_00279* | | - | 326562 | 327227 | 4.21 | 7.29 | 9.15 | 0.56 | Hypothetical protein |
| *FLPKDCKG_00280* | *guaB* | + | 327391 | 328851 | 60.31 | 97.76 | 50.68 | 23.26 | Inosine-5'-monophosphate dehydrogenase |
| *FLPKDCKG_00281* | *guaA-1* | + | 328972 | 330564 | 6.43 | 15.43 | 14.38 | 2.97 | GMP synthase [glutamine-hydrolyzing] |
| *FLPKDCKG_00282* | | - | 330909 | 331313 | 0.21 | 0.06 | 1.24 | 0.67 | Hypothetical protein |
| *FLPKDCKG_00283* | | - | 331300 | 331755 | 0.19 | 0.64 | 0.82 | 2.02 | Hypothetical protein |
| *FLPKDCKG_00284* | *fpvA-1* | - | 331808 | 334021 | 7.41 | 5.36 | 2.97 | 1.07 | Ferripyoverdine receptor |
| *FLPKDCKG_00285* | *nimR-1* | - | 334223 | 335062 | 5.77 | 2.32 | 1.39 | 1.14 | HTH-type transcriptional regulator NimR |
| *FLPKDCKG_00286* | | + | 335402 | 335890 | 9.54 | 8.45 | 6.00 | 0.88 | Hypothetical protein |
| *FLPKDCKG_00287* | *emrB-1* | - | 335767 | 337287 | 1.29 | 3.61 | 2.35 | 0.96 | Multidrug export protein EmrB |
| *FLPKDCKG_00288* | *dmlR-3* | + | 337582 | 338484 | 5.13 | 6.83 | 6.21 | 2.99 | HTH-type transcriptional regulator DmlR |
| *FLPKDCKG_00289* | *xecA1* | - | 339157 | 340185 | 2.20 | 34.61 | 45.50 | 16.57 | 2-hydroxypropyl-CoM lyase |
| *FLPKDCKG_00290* | | - | 340225 | 341208 | 1.66 | 33.80 | 31.63 | 14.52 | Hypothetical protein |
| *FLPKDCKG_00291* | *msuE* | - | 341381 | 341881 | 4.37 | 29.91 | 25.79 | 11.22 | FMN reductase (NADPH) |
| *FLPKDCKG_00292* | *metR-1* | + | 342086 | 343003 | 4.62 | 10.64 | 7.90 | 2.26 | HTH-type transcriptional regulator MetR |
| *FLPKDCKG_00293* | | + | 343060 | 343956 | 0.97 | 8.14 | 4.18 | 2.16 | Hypothetical protein |
| *FLPKDCKG_00294* | | - | 344123 | 345085 | 0.21 | 1.40 | 0.00 | 0.52 | Zinc-type alcohol dehydrogenase-like protein |
| *FLPKDCKG_00295* | | - | 345246 | 345557 | 14.99 | 8.11 | 15.00 | 7.82 | Hypothetical protein |
| *FLPKDCKG_00296* | | + | 345695 | 346075 | 3.67 | 5.05 | 4.09 | 1.28 | HTH-type transcriptional regulator |
| *FLPKDCKG_00297* | | + | 346072 | 346551 | 4.01 | 3.43 | 5.18 | 0.66 | Hypothetical protein |
| *FLPKDCKG_00298* | *gstB-1* | + | 346548 | 347207 | 3.75 | 2.96 | 2.43 | 1.59 | Glutathione S-transferase GstB |
| *FLPKDCKG_00299* | | + | 347318 | 347911 | 4.40 | 8.52 | 2.14 | 2.19 | Hypothetical protein |
| *FLPKDCKG_00300* | | + | 348055 | 349263 | 1.17 | 2.65 | 1.70 | 0.49 | Hypothetical protein |
| *FLPKDCKG_00301* | | + | 349410 | 349973 | 11.98 | 3.46 | 3.84 | 0.20 | Hypothetical protein |
| *FLPKDCKG_00302* | *ctpF* | + | 350065 | 352731 | 2.61 | 2.44 | 1.18 | 0.81 | putative cation-transporting ATPase F |
| *FLPKDCKG_00303* | | + | 352769 | 354328 | 2.09 | 2.78 | 2.42 | 0.00 | Hypothetical protein |
| *FLPKDCKG_00304* | *rclR-1* | - | 354340 | 355278 | 4.61 | 5.09 | 2.85 | 1.55 | RCS-specific HTH-type transcriptional activator RclR |
| *FLPKDCKG_00305* | *fabG-2* | + | 355447 | 356310 | 0.50 | 2.46 | 1.43 | 1.13 | 3-oxoacyl-[acyl-carrier-protein] reductase FabG |
| *FLPKDCKG_00306* | *nimR-2* | - | 356507 | 357340 | 1.08 | 4.77 | 4.50 | 0.87 | HTH-type transcriptional regulator NimR |
| *FLPKDCKG_00307* | | + | 357437 | 358189 | 0.38 | 0.90 | 0.00 | 1.56 | Hypothetical protein |
| *FLPKDCKG_00308* | *yqjH-1* | + | 358195 | 358920 | 0.22 | 1.98 | 0.22 | 0.38 | NADPH-dependent ferric-chelate reductase |
| *FLPKDCKG_00309* | | + | 359085 | 359600 | 0.21 | 1.74 | 0.87 | 0.00 | Hypothetical protein |
| *FLPKDCKG_00310* | *rstB-1* | + | 359737 | 361077 | 12.01 | 12.02 | 7.00 | 0.73 | Sensor protein RstB |
| *FLPKDCKG_00311* | | + | 361549 | 362316 | 4.23 | 3.09 | 3.42 | 1.74 | Hypothetical protein |
| *FLPKDCKG_00312* | *cueR* | + | 362414 | 362824 | 2.00 | 5.71 | 5.21 | 1.94 | HTH-type transcriptional regulator CueR |
| *FLPKDCKG_00313* | *dmlR-4* | - | 362859 | 363839 | 1.43 | 3.76 | 1.62 | 1.63 | HTH-type transcriptional regulator DmlR |
| *FLPKDCKG_00314* | *pbuE-1* | + | 363941 | 365176 | 0.68 | 1.17 | 1.20 | 0.40 | Purine efflux pump PbuE |
| *FLPKDCKG_00315* | *gltC-3* | - | 365191 | 366090 | 3.65 | 8.51 | 6.40 | 2.03 | HTH-type transcriptional regulator GltC |
| *FLPKDCKG_00316* | *argH-1* | + | 366200 | 367720 | 0.46 | 1.39 | 0.23 | 0.17 | Argininosuccinate lyase |
| *FLPKDCKG_00317* | | + | 367770 | 368747 | 2.65 | 3.13 | 1.66 | 1.18 | Hypothetical protein |
| *FLPKDCKG_00318* | *yncA* | - | 368787 | 369278 | 2.90 | 7.05 | 7.18 | 0.00 | L-methionine sulfoximine/L-methionine sulfone acetyltransferase |
| *FLPKDCKG_00319* | *feaR-1* | + | 369700 | 370695 | 1.93 | 5.44 | 5.14 | 3.11 | Transcriptional activator FeaR |
| *FLPKDCKG_00320* | | - | 370815 | 371222 | 3.79 | 7.92 | 6.00 | 1.14 | Hypothetical protein |
| *FLPKDCKG_00321* | *cdhR-1* | + | 371350 | 372312 | 3.21 | 7.24 | 4.29 | 2.77 | HTH-type transcriptional regulator CdhR |
| *FLPKDCKG_00322* | *pcpR-1* | - | 372441 | 373331 | 3.17 | 2.88 | 2.68 | 1.03 | PCP degradation transcriptional activation protein |
| *FLPKDCKG_00323* | *ribZ-1* | + | 373504 | 374904 | 0.41 | 0.57 | 0.39 | 0.00 | Riboflavin transporter RibZ |
| *FLPKDCKG_00324* | *nucA-2* | - | 374925 | 377027 | 0.83 | 0.81 | 0.63 | 0.17 | Nuclease |
| *FLPKDCKG_00325* | *lsrF* | - | 377133 | 377984 | 1.82 | 1.09 | 2.38 | 0.13 | 3-hydroxy-5-phosphonooxypentane-2,4-dione thiolase |
| *FLPKDCKG_00326* | *yhdN-1* | - | 378017 | 378997 | 1.39 | 0.77 | 0.48 | 0.00 | General stress protein 69 |
| *FLPKDCKG_00327* | *rbsC* | - | 378994 | 380025 | 0.72 | 0.09 | 1.11 | 0.00 | Ribose import permease protein RbsC |
| *FLPKDCKG_00328* | *rbsA-1* | - | 380022 | 381545 | 1.06 | 0.59 | 0.44 | 0.32 | Ribose import ATP-binding protein RbsA |
| *FLPKDCKG_00329* | *alsB* | - | 381542 | 382576 | 8.64 | 1.58 | 1.66 | 1.79 | D-allose-binding periplasmic protein |
| *FLPKDCKG_00330* | *lyx* | + | 382767 | 384257 | 1.29 | 1.50 | 3.26 | 0.42 | L-xylulose/3-keto-L-gulonate kinase |
| *FLPKDCKG_00331* | *glpD2-2* | + | 384254 | 385975 | 3.33 | 4.92 | 5.08 | 2.50 | Glycerol-3-phosphate dehydrogenase 2 |
| *FLPKDCKG_00332* | *yurK* | + | 385972 | 386730 | 2.38 | 5.11 | 2.03 | 1.26 | putative HTH-type transcriptional regulator YurK |
| *FLPKDCKG_00333* | | - | 386790 | 387389 | 9.48 | 7.52 | 2.81 | 4.25 | Hypothetical protein |
| *FLPKDCKG_00334* | | + | 387735 | 388448 | 8.68 | 58.38 | 33.37 | 6.69 | Gluconate 2-dehydrogenase subunit 3 |
| *FLPKDCKG_00335* | | + | 388452 | 390227 | 7.75 | 55.23 | 27.58 | 7.37 | Gluconate 2-dehydrogenase flavoprotein |
| *FLPKDCKG_00336* | | + | 390238 | 391554 | 4.91 | 49.64 | 24.24 | 10.40 | Gluconate 2-dehydrogenase cytochrome c subunit |
| *FLPKDCKG_00337* | *comR-1* | - | 391663 | 392301 | 1.74 | 5.53 | 2.46 | 0.00 | HTH-type transcriptional repressor ComR |
| *FLPKDCKG_00338* | *sugE-1* | - | 392298 | 392615 | 8.05 | 11.75 | 5.09 | 3.12 | Quaternary ammonium compound-resistance protein SugE |
| *FLPKDCKG_00339* | *bdcA-1* | - | 392681 | 393418 | 3.15 | 1.28 | 2.79 | 0.79 | Cyclic-di-GMP-binding biofilm dispersal mediator protein |
| *FLPKDCKG_00340* | | + | 393570 | 394004 | 8.75 | 16.97 | 14.99 | 1.79 | Hypothetical protein |
| *FLPKDCKG_00341* | | - | 394052 | 394297 | 16.87 | 20.92 | 11.70 | 4.80 | Hypothetical protein |

| Locus name | Gene name | Strand | Gene start (bp) | Gene end (bp) | Average TPM Inoculum | Average TPM $t_6$ | Average TPM $t_{24}$ | Average TPM $t_{168}$ | Annotation description |
|---|---|---|---|---|---|---|---|---|---|
| *FLPKDCKG_00342* | | - | 394391 | 394906 | 4.48 | 4.62 | 2.74 | 3.96 | putative N-acetyltransferase |
| *FLPKDCKG_00343* | | + | 395150 | 395668 | 5.38 | 1.03 | 4.36 | 0.40 | Hypothetical protein |
| *FLPKDCKG_00344* | *fan1-1* | + | 395735 | 396250 | 0.92 | 4.68 | 4.62 | 0.00 | Fanconi-associated nuclease 1 |
| *FLPKDCKG_00345* | *fan1-2* | + | 396250 | 397125 | 0.92 | 2.93 | 0.22 | 0.67 | Fanconi-associated nuclease 1 |
| *FLPKDCKG_00346* | *fan1-3* | + | 397091 | 397396 | 0.31 | 0.86 | 0.62 | 0.18 | Fanconi-associated nuclease 1 |
| *FLPKDCKG_00347* | | - | 397264 | 398241 | 0.58 | 2.45 | 0.19 | 0.18 | Hypothetical protein |
| *FLPKDCKG_00348* | *dinG-1* | + | 398693 | 399661 | 0.80 | 4.32 | 0.52 | 1.08 | putative ATP-dependent helicase DinG |
| *FLPKDCKG_00349* | *ctb-1* | - | 399731 | 400135 | 0.33 | 1.16 | 0.00 | 0.00 | Group 3 truncated hemoglobin ctb |
| *FLPKDCKG_00350* | | - | 400261 | 400701 | 0.21 | 0.00 | 0.00 | 0.00 | Hypothetical protein |
| *FLPKDCKG_00351* | | + | 401142 | 401606 | 0.07 | 0.00 | 0.00 | 0.00 | Hypothetical protein |
| *FLPKDCKG_00352* | *dmlR-5* | + | 401975 | 402859 | 1.23 | 1.22 | 0.43 | 0.14 | HTH-type transcriptional regulator DmlR |
| *FLPKDCKG_00353* | *fes* | + | 403028 | 404605 | 1.68 | 0.00 | 1.26 | 0.42 | Enterochelin esterase |
| *FLPKDCKG_00354* | | + | 404713 | 405264 | 3.07 | 3.21 | 0.50 | 0.00 | Hypothetical protein |
| *FLPKDCKG_00355* | *aam-2* | + | 405394 | 406866 | 4.19 | 3.81 | 1.52 | 2.77 | Acylamidase |
| *FLPKDCKG_00356* | | - | 406709 | 407425 | 17.63 | 9.99 | 7.14 | 2.78 | Hypothetical protein |
| *FLPKDCKG_00357* | *yoaH* | + | 407467 | 408219 | 22.87 | 10.61 | 7.71 | 3.63 | Putative methyl-accepting chemotaxis protein YoaH |
| *FLPKDCKG_00358* | *mcpA* | + | 408165 | 408752 | 11.20 | 8.33 | 5.53 | 3.24 | Methyl-accepting chemotaxis protein McpA |
| *FLPKDCKG_00359* | *cheW-1* | + | 408757 | 409272 | 9.68 | 5.28 | 2.81 | 3.93 | Chemotaxis protein CheW |
| *FLPKDCKG_00360* | *wspC* | + | 409269 | 410576 | 3.24 | 2.15 | 2.95 | 0.67 | putative biofilm formation methyltransferase WspC |
| *FLPKDCKG_00361* | | + | 410569 | 411258 | 5.01 | 4.08 | 4.21 | 2.72 | Hypothetical protein |
| *FLPKDCKG_00362* | *frzE* | + | 411255 | 413516 | 3.57 | 3.80 | 2.18 | 2.94 | Gliding motility regulatory protein |
| *FLPKDCKG_00363* | *cheB-1* | + | 413513 | 414529 | 8.94 | 16.69 | 8.24 | 1.73 | Chemotaxis response regulator protein-glutamate methylesterase |
| *FLPKDCKG_00364* | *pleD-2* | + | 414566 | 415609 | 7.12 | 11.30 | 8.91 | 3.10 | Response regulator PleD |
| *FLPKDCKG_00365* | *phnV-1* | - | 415652 | 417307 | 0.33 | 1.70 | 1.97 | 0.28 | Putative 2-aminoethylphosphonate transport system permease protein PhnV |
| *FLPKDCKG_00366* | *phnS-1* | - | 417349 | 418320 | 1.11 | 1.61 | 0.00 | 0.18 | Putative 2-aminoethylphosphonate-binding periplasmic protein |
| *FLPKDCKG_00367* | *phnT-1* | - | 418322 | 419035 | 0.07 | 0.15 | 0.53 | 0.00 | Putative 2-aminoethylphosphonate import ATP-binding protein PhnT |
| *FLPKDCKG_00368* | | - | 419032 | 420150 | 0.09 | 0.00 | 0.00 | 0.06 | Outer membrane porin protein |
| *FLPKDCKG_00369* | *decR-1* | - | 420299 | 420811 | 4.66 | 8.84 | 7.94 | 2.78 | DNA-binding transcriptional activator DecR |
| *FLPKDCKG_00370* | *mdeA-1* | + | 420958 | 422244 | 18.79 | 13.47 | 9.86 | 7.24 | L-methionine gamma-lyase |
| *FLPKDCKG_00371* | *nicB-1* | - | 422278 | 423597 | 2.28 | 3.69 | 0.86 | 1.63 | Nicotinate dehydrogenase subunit B |
| *FLPKDCKG_00372* | | + | 423644 | 424618 | 2.21 | 4.15 | 1.58 | 0.66 | Hypothetical protein |
| *FLPKDCKG_00373* | *nicF-1* | - | 424627 | 425280 | 2.07 | 2.04 | 3.48 | 0.94 | Maleamate amidohydrolase |
| *FLPKDCKG_00374* | *nicX* | - | 425277 | 426350 | 1.38 | 1.22 | 0.95 | 1.99 | 2,5-dihydroxypyridine 5,6-dioxygenase |
| *FLPKDCKG_00375* | *nicD* | - | 426387 | 427220 | 0.96 | 1.27 | 2.17 | 1.29 | N-formylmaleamate deformylase |
| *FLPKDCKG_00376* | | + | 427535 | 428692 | 3.27 | 2.75 | 3.32 | 3.36 | 6-hydroxynicotinate 3-monooxygenase |
| *FLPKDCKG_00377* | *nicA* | + | 428737 | 429267 | 0.51 | 0.92 | 0.26 | 1.94 | Nicotinate dehydrogenase subunit A |
| *FLPKDCKG_00378* | *nicB-2* | + | 429264 | 432251 | 0.55 | 1.14 | 1.23 | 0.86 | Nicotinate dehydrogenase subunit B |
| *FLPKDCKG_00379* | *naiP* | + | 432461 | 433777 | 0.70 | 0.77 | 1.58 | 1.73 | Putative niacin/nicotinamide transporter NaiP |
| *FLPKDCKG_00380* | *nicR* | - | 433874 | 434341 | 8.28 | 7.13 | 3.75 | 0.89 | HTH-type transcriptional repressor NicR |
| *FLPKDCKG_00381* | *smvA* | - | 434469 | 436007 | 3.93 | 2.54 | 2.14 | 0.37 | Methyl viologen resistance protein SmvA |
| *FLPKDCKG_00382* | *teaD-1* | - | 437109 | 437555 | 170.56 | 4.62 | 3.62 | 3.49 | TRAP-T-associated universal stress protein TeaD |
| *FLPKDCKG_00383* | *rspR-1* | - | 437689 | 438375 | 6.66 | 9.92 | 9.33 | 3.57 | HTH-type transcriptional repressor RspR |
| *FLPKDCKG_00384* | *acdS-1* | + | 438654 | 439715 | 0.23 | 0.28 | 0.00 | 0.21 | 1-aminocyclopropane-1-carboxylate deaminase |
| *FLPKDCKG_00385* | | - | 439768 | 440754 | 0.94 | 2.28 | 1.53 | 0.55 | Hypothetical protein |
| *FLPKDCKG_00386* | *lcfB-1* | - | 440842 | 442773 | 0.99 | 3.27 | 2.23 | 1.21 | Long-chain-fatty-acid--CoA ligase |
| *FLPKDCKG_00387* | *pigA* | - | 442791 | 443924 | 0.84 | 1.08 | 0.20 | 1.36 | Flavoprotein desaturase PigA |
| *FLPKDCKG_00388* | | - | 443934 | 445130 | 2.76 | 2.88 | 0.88 | 0.28 | Putative acyl-CoA dehydrogenase FadE17 |
| *FLPKDCKG_00389* | *fadA-1* | - | 445142 | 446323 | 2.88 | 2.99 | 2.05 | 0.00 | 3-ketoacyl-CoA thiolase |
| *FLPKDCKG_00390* | *fadJ-1* | - | 446349 | 448469 | 3.01 | 2.89 | 1.59 | 0.56 | Fatty acid oxidation complex subunit alpha |
| *FLPKDCKG_00391* | *dmlR-6* | + | 448752 | 449678 | 4.11 | 12.84 | 5.62 | 1.57 | HTH-type transcriptional regulator DmlR |
| *FLPKDCKG_00392* | | + | 449959 | 451257 | 4.65 | 3.84 | 0.53 | 0.56 | Hypothetical protein |
| *FLPKDCKG_00393* | | + | 451284 | 451847 | 3.84 | 2.99 | 1.05 | 0.00 | Hypothetical protein |
| *FLPKDCKG_00394* | | + | 451933 | 452685 | 3.34 | 6.58 | 4.28 | 1.43 | Hypothetical protein |
| *FLPKDCKG_00395* | | - | 452740 | 453159 | 0.65 | 0.58 | 0.30 | 0.00 | Hypothetical protein |
| *FLPKDCKG_00396* | *eno-1* | - | 453245 | 454525 | 0.26 | 0.57 | 1.16 | 0.62 | Enolase |
| *FLPKDCKG_00397* | *trpI-2* | + | 454737 | 455615 | 1.10 | 1.33 | 1.73 | 0.00 | HTH-type transcriptional regulator TrpI |
| *FLPKDCKG_00398* | | + | 455670 | 457010 | 4.96 | 9.75 | 6.13 | 1.21 | Hypothetical protein |
| *FLPKDCKG_00399* | *sodC* | + | 457202 | 457729 | 48.97 | 107.12 | 73.53 | 25.11 | Superoxide dismutase [Cu-Zn] |
| *FLPKDCKG_00400* | *gstB-2* | - | 457780 | 458421 | 9.01 | 21.41 | 13.22 | 2.95 | Glutathione S-transferase GstB |
| *FLPKDCKG_00401* | | - | 458605 | 459825 | 0.65 | 0.36 | 1.34 | 0.47 | Hypothetical protein |
| *FLPKDCKG_00402* | *map-1* | - | 459926 | 460717 | 6.44 | 18.09 | 15.61 | 3.45 | Methionine aminopeptidase |
| *FLPKDCKG_00403* | | - | 460718 | 460963 | 20.37 | 37.89 | 53.13 | 7.34 | Hypothetical protein |
| *FLPKDCKG_00404* | | - | 461050 | 462210 | 11.68 | 30.63 | 32.49 | 11.30 | Hypothetical protein |
| *FLPKDCKG_00405* | | - | 462247 | 462498 | 33.67 | 75.24 | 73.79 | 33.61 | Hypothetical protein |
| *FLPKDCKG_00406* | | + | 462674 | 463786 | 10.16 | 16.74 | 16.43 | 4.49 | Hypothetical protein |
| *FLPKDCKG_00407* | *arfA* | - | 463876 | 464670 | 1.27 | 12.68 | 14.11 | 4.62 | Peptidoglycan-binding protein ArfA |
| *FLPKDCKG_00408* | | - | 464673 | 469457 | 1.65 | 7.97 | 5.45 | 1.14 | Hypothetical protein |
| *FLPKDCKG_00409* | | - | 469481 | 470884 | 2.39 | 6.32 | 8.67 | 0.63 | Hypothetical protein |
| *FLPKDCKG_00410* | *cwlK* | - | 470955 | 471806 | 12.28 | 47.60 | 38.45 | 6.10 | Peptidoglycan L-alanyl-D-glutamate endopeptidase CwlK |

| Locus name | Gene name | Strand | Gene start (bp) | Gene end (bp) | Average TPM Inoculum | Average TPM $t_6$ | Average TPM $t_{24}$ | Average TPM $t_{168}$ | Annotation description |
|---|---|---|---|---|---|---|---|---|---|
| *FLPKDCKG_00411* | | + | 471998 | 472924 | 19.88 | 73.02 | 51.20 | 14.47 | Hypothetical protein |
| *FLPKDCKG_00412* | *vgrG1-1* | + | 472951 | 475827 | 4.11 | 15.67 | 13.45 | 4.81 | Actin cross-linking toxin VgrG1 |
| *FLPKDCKG_00413* | | + | 475839 | 476816 | 7.20 | 29.14 | 23.13 | 5.16 | Hypothetical protein |
| *FLPKDCKG_00414* | | + | 476809 | 477588 | 3.85 | 25.15 | 15.02 | 5.26 | Hypothetical protein |
| *FLPKDCKG_00415* | | + | 477581 | 479881 | 4.69 | 28.18 | 22.64 | 7.36 | Hypothetical protein |
| *FLPKDCKG_00416* | | + | 479878 | 480666 | 2.15 | 21.31 | 12.42 | 2.57 | Hypothetical protein |
| *FLPKDCKG_00417* | | - | 480679 | 481320 | 1.55 | 4.45 | 14.71 | 2.88 | Hypothetical protein |
| *FLPKDCKG_00418* | *vgrG1-2* | - | 481301 | 486496 | 3.17 | 7.54 | 7.64 | 3.72 | Actin cross-linking toxin VgrG1 |
| *FLPKDCKG_00419* | | - | 486522 | 487301 | 13.16 | 25.61 | 20.48 | 11.81 | Hypothetical protein |
| *FLPKDCKG_00420* | | - | 487360 | 488706 | 11.26 | 32.19 | 20.64 | 8.61 | Hypothetical protein |
| *FLPKDCKG_00421* | | - | 488784 | 489359 | 11.79 | 72.32 | 36.31 | 9.67 | Hypothetical protein |
| *FLPKDCKG_00422* | | + | 489591 | 490838 | 25.75 | 84.13 | 69.62 | 23.59 | Hypothetical protein |
| *FLPKDCKG_00423* | | + | 490831 | 491547 | 25.20 | 69.28 | 58.49 | 22.20 | Hypothetical protein |
| *FLPKDCKG_00424* | | + | 491550 | 492320 | 68.00 | 253.65 | 171.31 | 42.88 | Hypothetical protein |
| *FLPKDCKG_00425* | *hcpA* | + | 492361 | 492864 | 35.87 | 175.94 | 197.21 | 51.19 | Major exported protein |
| *FLPKDCKG_00426* | *iraD* | + | 492935 | 493420 | 8.16 | 92.77 | 80.43 | 22.82 | Anti-adapter protein IraD |
| *FLPKDCKG_00427* | | + | 493470 | 495308 | 1.73 | 12.38 | 11.29 | 3.73 | Hypothetical protein |
| *FLPKDCKG_00428* | | + | 495281 | 496372 | 2.81 | 23.05 | 18.57 | 3.58 | Hypothetical protein |
| *FLPKDCKG_00429* | *clpB-1* | + | 496392 | 499076 | 2.27 | 14.40 | 14.17 | 3.98 | Chaperone protein ClpB |
| *FLPKDCKG_00430* | *vgrG1-3* | + | 499093 | 501786 | 1.65 | 11.08 | 8.47 | 2.37 | Actin cross-linking toxin VgrG1 |
| *FLPKDCKG_00431* | | + | 501779 | 502765 | 0.84 | 4.07 | 2.93 | 1.24 | Hypothetical protein |
| *FLPKDCKG_00432* | | + | 502749 | 505682 | 3.74 | 21.61 | 18.87 | 3.35 | Hypothetical protein |
| *FLPKDCKG_00433* | | + | 505921 | 506988 | 2.55 | 14.58 | 22.24 | 2.33 | Hypothetical protein |
| *FLPKDCKG_00434* | | + | 506996 | 508033 | 1.40 | 11.37 | 7.64 | 0.96 | Hypothetical protein |
| *FLPKDCKG_00435* | | + | 508066 | 509136 | 1.13 | 4.11 | 5.96 | 0.29 | Hypothetical protein |
| *FLPKDCKG_00436* | | + | 509441 | 510325 | 2.51 | 9.63 | 7.77 | 0.48 | Hypothetical protein |
| *FLPKDCKG_00437* | | + | 510430 | 511359 | 0.29 | 0.78 | 0.63 | 0.19 | Hypothetical protein |
| *FLPKDCKG_00438* | | + | 511437 | 512768 | 0.47 | 1.42 | 0.78 | 0.46 | Carotenoid cleavage oxygenase |
| *FLPKDCKG_00439* | | - | 512885 | 513637 | 0.12 | 0.63 | 1.01 | 0.00 | Hypothetical protein |
| *FLPKDCKG_00440* | *yiaD-1* | - | 513627 | 514283 | 0.07 | 0.68 | 2.60 | 0.90 | putative lipoprotein YiaD |
| *FLPKDCKG_00441* | | - | 514280 | 517579 | 0.88 | 2.67 | 2.74 | 0.72 | Hypothetical protein |
| *FLPKDCKG_00442* | | - | 517590 | 518492 | 3.06 | 8.54 | 9.95 | 2.06 | Hypothetical protein |
| *FLPKDCKG_00443* | | - | 518715 | 519395 | 6.18 | 4.07 | 0.00 | 3.00 | Hypothetical protein |
| *FLPKDCKG_00444* | | - | 519397 | 520035 | 2.60 | 1.62 | 1.70 | 0.99 | Hypothetical protein |
| *FLPKDCKG_00445* | | + | 520917 | 521138 | 0.00 | 1.91 | 0.00 | 0.00 | Hypothetical protein |
| *FLPKDCKG_00446* | | + | 521239 | 522465 | 3.11 | 1.08 | 1.92 | 1.62 | Hypothetical protein |
| *FLPKDCKG_00447* | *fldA-1* | - | 522520 | 523959 | 3.74 | 2.95 | 4.51 | 0.59 | Cinnamoyl-CoA:phenyllactate CoA-transferase |
| *FLPKDCKG_00448* | *citA-1* | + | 524143 | 525354 | 1.81 | 3.65 | 1.82 | 1.69 | Putative citrate synthase 2 |
| *FLPKDCKG_00449* | | - | 525426 | 525905 | 5.85 | 9.31 | 6.77 | 3.31 | Hypothetical protein |
| *FLPKDCKG_00450* | *speG-1* | - | 525998 | 527311 | 13.66 | 14.12 | 8.99 | 2.51 | Spermidine N(1)-acetyltransferase |
| *FLPKDCKG_00451* | | + | 527468 | 527980 | 1.14 | 0.52 | 0.44 | 0.00 | putative HTH-type transcriptional regulator/GBAA_1941/BAS1801 |
| *FLPKDCKG_00452* | | + | 527977 | 528747 | 1.14 | 1.07 | 0.08 | 0.00 | Hypothetical protein |
| *FLPKDCKG_00453* | *gstB-3* | - | 528811 | 529416 | 0.30 | 1.64 | 0.00 | 0.72 | Glutathione S-transferase GST-6.0 |
| *FLPKDCKG_00454* | | - | 529548 | 530675 | 1.23 | 2.91 | 2.84 | 1.36 | Hypothetical protein |
| *FLPKDCKG_00455* | *hcaR-2* | + | 530873 | 531748 | 1.64 | 3.64 | 5.08 | 2.54 | Hca operon transcriptional activator HcaR |
| *FLPKDCKG_00456* | *phnO-1* | + | 531829 | 532257 | 2.68 | 3.79 | 5.43 | 0.05 | Aminoalkylphosphonate N-acetyltransferase |
| *FLPKDCKG_00457* | | + | 532343 | 533161 | 6.51 | 3.30 | 1.80 | 3.16 | Hypothetical protein |
| *FLPKDCKG_00458* | *yhaJ-1* | - | 533232 | 534134 | 9.45 | 18.18 | 5.10 | 1.07 | HTH-type transcriptional regulator YhaJ |
| *FLPKDCKG_00459* | *exuT-1* | + | 534242 | 535450 | 3.07 | 14.35 | 9.54 | 4.93 | Hexuronate transporter |
| *FLPKDCKG_00460* | *yieH-1* | + | 535525 | 536193 | 6.72 | 8.22 | 10.83 | 1.47 | 6-phosphogluconate phosphatase |
| *FLPKDCKG_00461* | | - | 536245 | 536628 | 7.15 | 7.52 | 0.73 | 0.08 | Hypothetical protein |
| *FLPKDCKG_00462* | *ytfF-1* | - | 536762 | 537715 | 12.40 | 1.10 | 0.18 | 0.45 | Inner membrane protein YtfF |
| *FLPKDCKG_00463* | *adaA* | - | 537826 | 538716 | 6.85 | 9.06 | 4.10 | 2.35 | Bifunctional transcriptional activator/DNA repair enzyme AdaA |
| *FLPKDCKG_00464* | *sigK-2* | - | 538810 | 540099 | 1.66 | 1.23 | 1.56 | 0.58 | ECF RNA polymerase sigma factor SigK |
| *FLPKDCKG_00465* | | - | 540096 | 540503 | 5.03 | 0.20 | 2.42 | 1.05 | Hypothetical protein |
| *FLPKDCKG_00466* | | - | 540545 | 540958 | 17.77 | 6.09 | 15.20 | 3.82 | Hypothetical protein |
| *FLPKDCKG_00467* | | - | 541116 | 541469 | 7.48 | 4.92 | 5.43 | 8.64 | Hypothetical protein |
| *FLPKDCKG_00468* | *ramB* | - | 541580 | 541771 | 3.96 | 5.93 | 3.21 | 2.18 | HTH-type transcriptional regulator RamB |
| *FLPKDCKG_00469* | | - | 541768 | 542154 | 5.99 | 21.30 | 4.69 | 7.07 | Hypothetical protein |
| *FLPKDCKG_00470* | *motB-1* | - | 542284 | 543225 | 1.44 | 0.75 | 0.92 | 0.00 | Motility protein B |
| *FLPKDCKG_00471* | *pomA* | - | 543278 | 544030 | 6.08 | 2.53 | 1.46 | 0.29 | Chemotaxis protein PomA |
| *FLPKDCKG_00472* | *ydaF-1* | - | 544321 | 544890 | 5.57 | 8.35 | 9.20 | 1.67 | Putative ribosomal N-acetyltransferase YdaF |
| *FLPKDCKG_00473* | | + | 545015 | 546214 | 6.55 | 4.19 | 2.36 | 1.15 | Hypothetical protein |
| *FLPKDCKG_00474* | *nimR-3* | - | 546673 | 547479 | 1.04 | 4.56 | 0.50 | 1.65 | HTH-type transcriptional regulator NimR |
| *FLPKDCKG_00475* | | + | 547610 | 547867 | 0.44 | 1.68 | 0.56 | 0.71 | Hypothetical protein |
| *FLPKDCKG_00476* | | + | 547987 | 548493 | 2.42 | 4.18 | 3.78 | 2.21 | Hypothetical protein |
| *FLPKDCKG_00477* | | + | 548483 | 548809 | 2.69 | 13.89 | 11.63 | 4.30 | Hypothetical protein |
| *FLPKDCKG_00478* | | - | 548921 | 549454 | 62.30 | 177.36 | 103.45 | 37.29 | Hypothetical protein |
| *FLPKDCKG_00479* | | - | 549569 | 550105 | 1.86 | 4.51 | 2.96 | 1.38 | Hypothetical protein |

| Locus name | Gene name | Strand | Gene start (bp) | Gene end (bp) | Average TPM Inoculum | Average TPM $t_6$ | Average TPM $t_{24}$ | Average TPM $t_{168}$ | Annotation description |
|---|---|---|---|---|---|---|---|---|---|
| *FLPKDCKG_00480* | *kstR2-2* | + | 550230 | 550769 | 3.44 | 17.28 | 3.75 | 4.69 | HTH-type transcriptional repressor KstR2 |
| *FLPKDCKG_00481* | | - | 550854 | 551828 | 0.00 | 1.45 | 3.62 | 1.08 | Hypothetical protein |
| *FLPKDCKG_00482* | | + | 551995 | 553053 | 1.90 | 1.99 | 2.56 | 1.51 | Hypothetical protein |
| *FLPKDCKG_00483* | *gpr-1* | + | 553199 | 554209 | 3.66 | 8.44 | 5.13 | 2.45 | L-glyceraldehyde 3-phosphate reductase |
| *FLPKDCKG_00484* | *yhjE-1* | - | 554251 | 555585 | 0.48 | 0.66 | 2.24 | 0.43 | Inner membrane metabolite transport protein YhjE |
| *FLPKDCKG_00485* | *alkJ-2* | - | 555654 | 557306 | 0.46 | 1.51 | 0.02 | 0.00 | Alcohol dehydrogenase [acceptor] |
| *FLPKDCKG_00486* | *kdgR-1* | + | 557512 | 558351 | 5.25 | 4.15 | 1.59 | 1.23 | Transcriptional regulator KdgR |
| *FLPKDCKG_00487* | | - | 558372 | 559673 | 0.93 | 1.22 | 1.87 | 0.46 | Hypothetical protein |
| *FLPKDCKG_00488* | *aziB2-1* | - | 559670 | 560692 | 0.14 | 0.00 | 0.00 | 0.37 | 3-hydroxy-5-methyl-1-naphthoate 3-O-methyltransferase |
| *FLPKDCKG_00489* | *fyuA-1* | - | 560715 | 562772 | 0.07 | 0.00 | 0.17 | 0.00 | Pesticin receptor |
| *FLPKDCKG_00490* | *pchR-2* | + | 562888 | 563904 | 4.71 | 1.43 | 1.91 | 0.29 | Regulatory protein PchR |
| *FLPKDCKG_00491* | | - | 563941 | 565116 | 47.86 | 6.05 | 6.16 | 0.78 | Hypothetical protein |
| *FLPKDCKG_00492* | *stp-2* | - | 565215 | 566660 | 1.90 | 1.85 | 1.40 | 0.56 | Multidrug resistance protein Stp |
| *FLPKDCKG_00493* | | - | 566711 | 567139 | 35.43 | 22.70 | 26.14 | 7.04 | Hypothetical protein |
| *FLPKDCKG_00494* | | + | 567402 | 568292 | 0.70 | 2.74 | 1.96 | 0.67 | N-carbamoyl-D-amino acid hydrolase |
| *FLPKDCKG_00495* | *amiC-1* | + | 568332 | 569498 | 0.52 | 1.17 | 2.02 | 0.69 | Aliphatic amidase expression-regulating protein |
| *FLPKDCKG_00496* | *livH-2* | + | 569554 | 570387 | 0.37 | 0.57 | 0.50 | 0.35 | High-affinity branched-chain amino acid transport system permease protein LivH |
| *FLPKDCKG_00497* | *glnQ-1* | + | 570380 | 572329 | 0.18 | 0.54 | 1.58 | 1.62 | Glutamine transport ATP-binding protein GlnQ |
| *FLPKDCKG_00498* | *livF-2* | + | 572326 | 573036 | 0.39 | 0.38 | 3.03 | 0.23 | High-affinity branched-chain amino acid transport ATP-binding protein LivF |
| *FLPKDCKG_00499* | *amiC-2* | + | 573063 | 574223 | 6.73 | 5.09 | 1.63 | 0.91 | Aliphatic amidase expression-regulating protein |
| *FLPKDCKG_00500* | *amiR* | + | 574226 | 574879 | 6.55 | 8.01 | 2.76 | 1.72 | Aliphatic amidase regulator |
| *FLPKDCKG_00501* | | + | 574876 | 576369 | 4.45 | 3.18 | 2.51 | 1.31 | Hypothetical protein |
| *FLPKDCKG_00502* | *ahpD-1* | + | 576571 | 576978 | 28.34 | 24.07 | 13.13 | 7.36 | Alkyl hydroperoxide reductase AhpD |
| *FLPKDCKG_00503* | | + | 576988 | 577596 | 5.34 | 8.40 | 11.98 | 1.53 | Hypothetical protein |
| *FLPKDCKG_00504* | | + | 577666 | 578487 | 4.31 | 13.89 | 8.97 | 1.35 | Hypothetical protein |
| *FLPKDCKG_00505* | | - | 578595 | 578885 | 8.85 | 39.80 | 14.41 | 7.63 | Hypothetical protein |
| *FLPKDCKG_00506* | *tsr-1* | + | 579186 | 580820 | 3.43 | 6.05 | 4.39 | 2.00 | Methyl-accepting chemotaxis protein I |
| *FLPKDCKG_00507* | *zitB-1* | + | 580961 | 581446 | 5.20 | 14.71 | 12.29 | 5.89 | Zinc transporter ZitB |
| *FLPKDCKG_00508* | | - | 581269 | 581976 | 3.23 | 11.38 | 7.49 | 1.98 | Hypothetical protein |
| *FLPKDCKG_00509* | | + | 582814 | 583590 | 1.49 | 9.22 | 6.49 | 3.42 | putative cation efflux system protein |
| *FLPKDCKG_00510* | | + | 583587 | 583790 | 14.96 | 16.95 | 16.80 | 2.19 | Hypothetical protein |
| *FLPKDCKG_00511* | *yodB* | + | 583816 | 584367 | 14.92 | 14.59 | 10.44 | 5.00 | Cytochrome b561 |
| *FLPKDCKG_00512* | | + | 585167 | 586450 | 11.22 | 18.13 | 8.77 | 4.56 | Hypothetical protein |
| *FLPKDCKG_00513* | | + | 586460 | 588415 | 8.71 | 24.18 | 11.16 | 2.39 | Hypothetical protein |
| *FLPKDCKG_00514* | | - | 588734 | 588958 | 3.72 | 17.15 | 20.17 | 1.54 | Hypothetical protein |
| *FLPKDCKG_00515* | | + | 589719 | 590048 | 12.34 | 11.90 | 9.57 | 2.39 | Hypothetical protein |
| *FLPKDCKG_00516* | *proA-1* | + | 590120 | 590848 | 0.43 | 1.16 | 3.52 | 0.00 | 4-hydroxy-4-methyl-2-oxoglutarate aldolase/4-carboxy-4-hydroxy-2-oxoadipate aldolase |
| *FLPKDCKG_00517* | | + | 590845 | 592104 | 0.38 | 0.45 | 1.50 | 0.45 | Aspartate aminotransferase |
| *FLPKDCKG_00518* | *fliY-1* | + | 592178 | 592999 | 0.07 | 0.40 | 0.55 | 0.53 | L-cystine-binding protein FliY |
| *FLPKDCKG_00519* | *glnM-1* | + | 593016 | 593705 | 0.06 | 0.00 | 1.14 | 0.00 | putative glutamine ABC transporter permease protein GlnM |
| *FLPKDCKG_00520* | *yhdY-1* | + | 593698 | 594387 | 0.13 | 0.55 | 0.00 | 0.35 | Inner membrane amino-acid ABC transporter permease protein YhdY |
| *FLPKDCKG_00521* | *glnQ-2* | + | 594450 | 595184 | 0.25 | 6.99 | 3.62 | 0.34 | Glutamine transport ATP-binding protein GlnQ |
| *FLPKDCKG_00522* | *cmpR-1* | + | 595277 | 596242 | 3.47 | 6.00 | 2.29 | 1.11 | HTH-type transcriptional activator CmpR |
| *FLPKDCKG_00523* | *cysB-1* | + | 597370 | 598323 | 4.95 | 9.26 | 6.37 | 1.23 | HTH-type transcriptional regulator CysB |
| *FLPKDCKG_00524* | | - | 598258 | 598992 | 1.35 | 1.97 | 4.46 | 0.19 | Glucose 1-dehydrogenase |
| *FLPKDCKG_00525* | | - | 599145 | 599882 | 0.76 | 2.65 | 0.00 | 0.00 | Hypothetical protein |
| *FLPKDCKG_00526* | | - | 599896 | 600552 | 13.35 | 10.66 | 12.13 | 3.35 | Hypothetical protein |
| *FLPKDCKG_00527* | | - | 600668 | 602362 | 0.47 | 0.88 | 2.02 | 0.00 | Hypothetical protein |
| *FLPKDCKG_00528* | *prs2* | - | 602385 | 603257 | 0.77 | 1.15 | 0.00 | 0.70 | Ribose-phosphate pyrophosphokinase 2 |
| *FLPKDCKG_00529* | | - | 603348 | 603998 | 6.08 | 7.56 | 7.51 | 2.13 | Hypothetical protein |
| *FLPKDCKG_00530* | *dkgB-1* | - | 604070 | 604942 | 14.32 | 13.47 | 9.31 | 4.92 | 2,5-diketo-D-gluconic acid reductase B |
| *FLPKDCKG_00531* | *pgrR-2* | + | 605051 | 605950 | 2.30 | 2.86 | 0.78 | 0.92 | HTH-type transcriptional regulator PgrR |
| *FLPKDCKG_00532* | *nolA-1* | - | 606035 | 607063 | 2.39 | 4.87 | 4.81 | 1.81 | Nodulation protein NolA |
| *FLPKDCKG_00533* | | - | 607366 | 607746 | 58.72 | 71.84 | 54.44 | 11.17 | Hypothetical protein |
| *FLPKDCKG_00534* | *dnaE2* | - | 607749 | 611126 | 5.39 | 8.16 | 4.70 | 1.05 | Error-prone DNA polymerase |
| *FLPKDCKG_00535* | | - | 611134 | 611292 | 1.40 | 0.00 | 0.79 | 0.00 | Hypothetical protein |
| *FLPKDCKG_00536* | | - | 611292 | 612602 | 1.07 | 0.58 | 0.38 | 0.00 | Hypothetical protein |
| *FLPKDCKG_00537* | | - | 612875 | 613228 | 2.45 | 4.21 | 8.32 | 0.65 | Hypothetical protein |
| *FLPKDCKG_00538* | *pctC* | + | 613553 | 614605 | 0.22 | 10.89 | 11.81 | 2.87 | Methyl-accepting chemotaxis protein PctC |
| *FLPKDCKG_00539* | *lysP-1* | - | 614671 | 616212 | 1.31 | 6.51 | 5.16 | 2.09 | Lysine-specific permease |
| *FLPKDCKG_00540* | | + | 616374 | 617093 | 11.02 | 11.88 | 8.26 | 1.24 | Hypothetical protein |
| *FLPKDCKG_00541* | *iclR-1* | + | 617104 | 617910 | 2.43 | 4.18 | 3.13 | 1.76 | Transcriptional repressor IclR |
| *FLPKDCKG_00542* | *bacC-1* | - | 617941 | 618735 | 0.72 | 0.73 | 1.17 | 0.17 | Dihydroanticapsin 7-dehydrogenase |
| *FLPKDCKG_00543* | | - | 618737 | 619714 | 0.86 | 1.96 | 1.09 | 1.47 | Hypothetical protein |
| *FLPKDCKG_00544* | *ilvB* | - | 619757 | 621490 | 0.49 | 0.74 | 0.79 | 0.57 | Acetolactate synthase large subunit |
| *FLPKDCKG_00545* | | - | 621474 | 622337 | 0.26 | 0.72 | 0.37 | 0.48 | Hypothetical protein |
| *FLPKDCKG_00546* | *srpC-1* | - | 622728 | 624149 | 0.22 | 0.99 | 2.48 | 0.21 | Solvent efflux pump outer membrane protein SrpC |
| *FLPKDCKG_00547* | *aaeA* | - | 624205 | 625128 | 1.18 | 1.70 | 3.37 | 0.85 | p-hydroxybenzoic acid efflux pump subunit AaeA |
| *FLPKDCKG_00548* | *aaeX* | - | 625131 | 625343 | 0.61 | 0.12 | 4.39 | 0.89 | Protein AaeX |

| Locus name | Gene name | Strand | Gene start (bp) | Gene end (bp) | Average TPM Inoculum | Average TPM $t_6$ | Average TPM $t_{24}$ | Average TPM $t_{168}$ | Annotation description |
|---|---|---|---|---|---|---|---|---|---|
| *FLPKDCKG_00549* | *aaeB* | - | 625340 | 627391 | 0.65 | 1.59 | 1.99 | 0.22 | p-hydroxybenzoic acid efflux pump subunit AaeB |
| *FLPKDCKG_00550* | *slyA* | - | 627388 | 627861 | 2.76 | 9.29 | 15.30 | 2.44 | Transcriptional regulator SlyA |
| *FLPKDCKG_00551* | *queD* | - | 628180 | 628623 | 5.84 | 19.44 | 10.09 | 4.72 | 6-carboxy-5,6,7,8-tetrahydropterin synthase |
| *FLPKDCKG_00552* | *queE-1* | - | 628620 | 629252 | 36.14 | 73.06 | 33.50 | 11.94 | 7-carboxy-7-deazaguanine synthase |
| *FLPKDCKG_00553* | *dbpA* | - | 629312 | 630700 | 13.98 | 18.29 | 9.10 | 4.90 | ATP-dependent RNA helicase DbpA |
| *FLPKDCKG_00554* | *iclR-2* | - | 630832 | 631593 | 2.29 | 3.28 | 1.43 | 0.00 | Transcriptional repressor IclR |
| *FLPKDCKG_00555* | *xsc* | + | 631754 | 633745 | 1.85 | 1.11 | 0.39 | 0.79 | Sulfoacetaldehyde acetyltransferase |
| *FLPKDCKG_00556* | *pta-1* | + | 633745 | 634497 | 0.17 | 0.00 | 0.00 | 0.00 | Phosphate acetyltransferase |
| *FLPKDCKG_00557* | *appA* | + | 634629 | 636215 | 1.01 | 0.67 | 0.00 | 0.00 | Oligopeptide-binding protein AppA |
| *FLPKDCKG_00558* | *dppB-2* | + | 636228 | 637196 | 0.47 | 0.00 | 0.00 | 0.00 | Dipeptide transport system permease protein DppB |
| *FLPKDCKG_00559* | *dppC-1* | + | 637193 | 638095 | 0.79 | 0.03 | 0.58 | 0.23 | Dipeptide transport system permease protein DppC |
| *FLPKDCKG_00560* | *gsiA-1* | + | 638095 | 639726 | 2.45 | 0.84 | 0.00 | 0.30 | Glutathione import ATP-binding protein GsiA |
| *FLPKDCKG_00561* | *nfdA-1* | + | 639719 | 641395 | 1.60 | 0.86 | 0.00 | 0.21 | N-substituted formamide deformylase |
| *FLPKDCKG_00562* | *yciC-1* | + | 641397 | 642389 | 0.23 | 0.00 | 1.18 | 0.00 | Putative metal chaperone YciC |
| *FLPKDCKG_00563* | | + | 642538 | 642930 | 0.95 | 0.00 | 0.00 | 0.00 | Hypothetical protein |
| *FLPKDCKG_00564* | *puuB-1* | + | 642970 | 644412 | 0.84 | 7.86 | 5.82 | 2.05 | Gamma-glutamylputrescine oxidoreductase |
| *FLPKDCKG_00565* | *novR-1* | - | 644476 | 645195 | 0.51 | 53.65 | 71.31 | 13.18 | Decarboxylase NovR |
| *FLPKDCKG_00566* | | - | 645224 | 646219 | 0.65 | 75.92 | 107.22 | 19.88 | Hypothetical protein |
| *FLPKDCKG_00567* | *tsh* | - | 646389 | 647402 | 0.70 | 4.01 | 1.55 | 0.44 | Temperature-sensitive hemagglutinin tsh autotransporter |
| *FLPKDCKG_00568* | *pic* | - | 647399 | 651778 | 0.66 | 1.08 | 1.45 | 0.53 | Serine protease pic autotransporter |
| *FLPKDCKG_00569* | *tauR* | - | 652010 | 653533 | 6.68 | 6.67 | 4.95 | 3.31 | HTH-type transcriptional regulator TauR |
| *FLPKDCKG_00570* | | - | 654343 | 654447 | 0.66 | 0.86 | 0.96 | 0.00 | Hypothetical protein |
| *FLPKDCKG_00571* | | + | 655385 | 655933 | 1.49 | 3.85 | 3.36 | 0.18 | Hypothetical protein |
| *FLPKDCKG_00572* | *ssuC-1* | - | 655941 | 656768 | 1.50 | 2.47 | 1.34 | 0.53 | Putative aliphatic sulfonates transport permease protein SsuC |
| *FLPKDCKG_00573* | *tauB-1* | - | 656761 | 657597 | 1.26 | 0.95 | 1.29 | 0.00 | Taurine import ATP-binding protein TauB |
| *FLPKDCKG_00574* | *tauA-1* | - | 657691 | 658719 | 5.82 | 2.46 | 1.62 | 0.43 | Taurine-binding periplasmic protein |
| *FLPKDCKG_00575* | *gltR-1* | - | 658981 | 659832 | 2.48 | 3.28 | 3.11 | 0.96 | HTH-type transcriptional regulator GltR |
| *FLPKDCKG_00576* | *eamA-1* | + | 659923 | 660870 | 0.15 | 2.14 | 0.49 | 0.13 | putative amino-acid metabolite efflux pump |
| *FLPKDCKG_00577* | *folE* | - | 660944 | 661489 | 19.30 | 25.41 | 12.80 | 6.05 | GTP cyclohydrolase 1 |
| *FLPKDCKG_00578* | *nreC-1* | - | 661588 | 662244 | 1.60 | 1.00 | 1.63 | 1.59 | Oxygen regulatory protein NreC |
| *FLPKDCKG_00579* | *degS-1* | - | 662241 | 663659 | 2.75 | 6.91 | 4.13 | 1.58 | Signal transduction histidine-protein kinase/phosphatase DegS |
| *FLPKDCKG_00580* | *cstA* | + | 663976 | 666063 | 2.74 | 17.26 | 18.93 | 10.37 | Carbon starvation protein A |
| *FLPKDCKG_00581* | | + | 666081 | 666293 | 2.18 | 6.53 | 8.99 | 4.89 | Hypothetical protein |
| *FLPKDCKG_00582* | | - | 666476 | 667339 | 0.68 | 0.13 | 1.94 | 0.46 | Hypothetical protein |
| *FLPKDCKG_00583* | *norG-1* | + | 667491 | 668852 | 1.41 | 1.83 | 1.90 | 0.79 | HTH-type transcriptional regulator NorG |
| *FLPKDCKG_00584* | *hcaT* | - | 668901 | 670073 | 0.90 | 0.78 | 0.38 | 0.00 | putative 3-phenylpropionic acid transporter |
| *FLPKDCKG_00585* | | - | 670178 | 670879 | 0.54 | 1.22 | 2.31 | 0.00 | Hypothetical protein |
| *FLPKDCKG_00586* | | + | 670993 | 671661 | 4.64 | 5.47 | 10.01 | 1.28 | Hypothetical protein |
| *FLPKDCKG_00587* | *ybiH* | - | 671658 | 672305 | 1.63 | 3.97 | 2.75 | 2.10 | putative HTH-type transcriptional regulator YbiH |
| *FLPKDCKG_00588* | *pepE-1* | - | 672417 | 673673 | 0.83 | 2.89 | 0.67 | 0.61 | putative dipeptidase PepE |
| *FLPKDCKG_00589* | *dps* | + | 673882 | 674433 | 128.75 | 120.87 | 67.32 | 31.73 | DNA protection during starvation protein |
| *FLPKDCKG_00590* | | - | 674500 | 675000 | 6.69 | 12.32 | 9.09 | 2.77 | Hypothetical protein |
| *FLPKDCKG_00591* | | - | 675164 | 678031 | 0.34 | 4.91 | 2.15 | 0.99 | Extracellular serine protease |
| *FLPKDCKG_00592* | *msrA3* | - | 678461 | 679147 | 0.38 | 68.92 | 39.93 | 8.78 | Peptide methionine sulfoxide reductase MsrA 3 |
| *FLPKDCKG_00593* | *msrB-1* | - | 679181 | 679678 | 0.69 | 125.08 | 78.26 | 21.73 | Peptide methionine sulfoxide reductase MsrB |
| *FLPKDCKG_00594* | | - | 679773 | 680087 | 4.61 | 269.63 | 151.05 | 30.31 | Hypothetical protein |
| *FLPKDCKG_00595* | *msrP-1* | - | 680130 | 680921 | 3.20 | 294.37 | 116.88 | 17.89 | Protein-methionine-sulfoxide reductase catalytic subunit MsrP |
| *FLPKDCKG_00596* | | - | 680932 | 681570 | 1.30 | 25.58 | 10.65 | 1.82 | Hypothetical protein |
| *FLPKDCKG_00597* | *sphR* | + | 681768 | 682499 | 8.63 | 31.94 | 18.46 | 6.09 | Alkaline phosphatase synthesis transcriptional regulatory protein SphR |
| *FLPKDCKG_00598* | *resE* | + | 682496 | 683968 | 2.13 | 9.84 | 7.19 | 0.57 | Sensor histidine kinase ResE |
| *FLPKDCKG_00599* | *pigC* | - | 683997 | 686594 | 0.40 | 1.93 | 1.65 | 0.00 | Prodigiosin synthesizing transferase PigC |
| *FLPKDCKG_00600* | | - | 686591 | 687238 | 0.82 | 1.44 | 0.00 | 0.00 | Hypothetical protein |
| *FLPKDCKG_00601* | | - | 687331 | 688488 | 7.98 | 0.75 | 0.67 | 0.64 | Hypothetical protein |
| *FLPKDCKG_00602* | | - | 688671 | 690059 | 8.26 | 11.53 | 7.76 | 4.85 | Hypothetical protein |
| *FLPKDCKG_00603* | | - | 690056 | 690421 | 0.00 | 0.00 | 0.00 | 0.00 | Hypothetical protein |
| *FLPKDCKG_00604* | *mvaA* | - | 690619 | 691908 | 0.43 | 0.96 | 0.00 | 0.00 | 3-hydroxy-3-methylglutaryl-coenzyme A reductase |
| *FLPKDCKG_00605* | *ytrA* | + | 692049 | 692900 | 2.06 | 4.02 | 1.96 | 0.00 | HTH-type transcriptional repressor YtrA |
| *FLPKDCKG_00606* | *norG-2* | + | 692903 | 693478 | 1.67 | 3.18 | 1.11 | 0.00 | HTH-type transcriptional regulator NorG |
| *FLPKDCKG_00607* | *queE-2* | - | 693469 | 694260 | 1.76 | 1.29 | 0.02 | 0.28 | 7-carboxy-7-deazaguanine synthase |
| *FLPKDCKG_00608* | | - | 694226 | 694408 | 99.58 | 8.14 | 8.94 | 1.18 | Hypothetical protein |
| *FLPKDCKG_00609* | *nrdD* | - | 694473 | 696473 | 12.13 | 1.71 | 1.67 | 0.58 | Anaerobic ribonucleoside-triphosphate reductase |
| *FLPKDCKG_00610* | | - | 696613 | 696756 | 0.62 | 0.00 | 1.30 | 0.00 | Hypothetical protein |
| *FLPKDCKG_00611* | | - | 696766 | 696963 | 2.10 | 0.57 | 1.21 | 0.00 | Hypothetical protein |
| *FLPKDCKG_00612* | | - | 696960 | 698570 | 1.91 | 0.67 | 1.14 | 0.46 | Hypothetical protein |
| *FLPKDCKG_00613* | | - | 698756 | 698983 | 0.45 | 2.22 | 0.89 | 0.00 | Hypothetical protein |
| *FLPKDCKG_00614* | | - | 698923 | 699747 | 0.46 | 2.29 | 2.83 | 0.89 | Hypothetical protein |
| *FLPKDCKG_00615* | *cpg2-1* | - | 699760 | 701229 | 1.75 | 10.28 | 6.28 | 3.13 | Carboxypeptidase G2 |
| *FLPKDCKG_00616* | *gltC-4* | + | 701293 | 702219 | 1.81 | 2.76 | 1.33 | 1.98 | HTH-type transcriptional regulator GltC |
| *FLPKDCKG_00617* | *mdtC-1* | - | 702250 | 702570 | 0.65 | 0.00 | 0.00 | 0.00 | Multidrug resistance protein MdtC |

| Locus name | Gene name | Strand | Gene start (bp) | Gene end (bp) | Average TPM Inoculum | Average TPM $t_6$ | Average TPM $t_{24}$ | Average TPM $t_{168}$ | Annotation description |
|---|---|---|---|---|---|---|---|---|---|
| *FLPKDCKG_00618* | *mdtC-2* | - | 702609 | 705317 | 0.16 | 0.18 | 0.34 | 0.05 | Multidrug resistance protein MdtC |
| *FLPKDCKG_00619* | *mdtB-1* | - | 705314 | 708391 | 0.59 | 0.39 | 0.56 | 0.11 | Multidrug resistance protein MdtB |
| *FLPKDCKG_00620* | *mdtA-1* | - | 708388 | 709560 | 1.60 | 1.02 | 0.42 | 0.35 | Multidrug resistance protein MdtA |
| *FLPKDCKG_00621* | *dmlR-7* | - | 709619 | 710593 | 1.56 | 4.20 | 2.22 | 0.52 | HTH-type transcriptional regulator DmlR |
| *FLPKDCKG_00622* | *hpxO-1* | + | 710720 | 711370 | 0.17 | 0.00 | 0.08 | 0.00 | FAD-dependent urate hydroxylase |
| *FLPKDCKG_00623* | *hpxO-2* | + | 711503 | 711901 | 0.33 | 0.00 | 0.44 | 0.00 | FAD-dependent urate hydroxylase |
| *FLPKDCKG_00624* | | + | 711963 | 712817 | 6.48 | 8.37 | 6.47 | 4.21 | Hypothetical protein |
| *FLPKDCKG_00625* | | - | 712837 | 714144 | 2.06 | 1.63 | 0.80 | 0.25 | Hypothetical protein |
| *FLPKDCKG_00626* | *narT* | + | 714329 | 715588 | 0.80 | 0.36 | 0.00 | 0.05 | putative nitrate transporter NarT |
| *FLPKDCKG_00627* | *narK* | + | 715608 | 716993 | 1.88 | 8.22 | 2.35 | 1.06 | Nitrate/nitrite transporter NarK |
| *FLPKDCKG_00628* | *narG* | + | 717143 | 720898 | 1.07 | 2.31 | 0.82 | 0.30 | Respiratory nitrate reductase 1 alpha chain |
| *FLPKDCKG_00629* | *narH* | + | 720895 | 722457 | 0.99 | 0.70 | 1.51 | 0.29 | Respiratory nitrate reductase 1 beta chain |
| *FLPKDCKG_00630* | *narJ* | + | 722472 | 723119 | 0.62 | 0.00 | 1.30 | 0.59 | Nitrate reductase molybdenum cofactor assembly chaperone NarJ |
| *FLPKDCKG_00631* | *narI* | + | 723137 | 723820 | 0.33 | 1.13 | 1.94 | 0.59 | Respiratory nitrate reductase 1 gamma chain |
| *FLPKDCKG_00632* | | + | 723836 | 724129 | 1.94 | 3.26 | 4.82 | 0.00 | Hypothetical protein |
| *FLPKDCKG_00633* | *ppiC-1* | + | 724117 | 724605 | 0.59 | 2.70 | 2.62 | 0.00 | Peptidyl-prolyl cis-trans isomerase C |
| *FLPKDCKG_00634* | *moaA2* | + | 724602 | 725645 | 0.43 | 0.00 | 0.33 | 0.60 | GTP 3',8-cyclase 2 |
| *FLPKDCKG_00635* | *icfA* | + | 725649 | 726308 | 1.51 | 3.30 | 2.50 | 0.10 | Carbonic anhydrase |
| *FLPKDCKG_00636* | *mobB* | + | 726305 | 726802 | 3.49 | 3.74 | 5.32 | 1.15 | Molybdopterin-guanine dinucleotide biosynthesis adapter protein |
| *FLPKDCKG_00637* | *narX* | + | 726903 | 728840 | 3.38 | 3.91 | 2.99 | 0.97 | Nitrate/nitrite sensor protein NarX |
| *FLPKDCKG_00638* | *degU-1* | + | 728854 | 729525 | 4.19 | 8.63 | 4.46 | 1.24 | Transcriptional regulatory protein DegU |
| *FLPKDCKG_00639* | *yhbU* | + | 729609 | 730625 | 2.33 | 0.71 | 1.00 | 0.68 | putative protease YhbU |
| *FLPKDCKG_00640* | | + | 730616 | 731512 | 1.09 | 0.00 | 0.82 | 0.00 | Hypothetical protein |
| *FLPKDCKG_00641* | | + | 731509 | 731943 | 1.77 | 0.00 | 1.03 | 0.00 | Hypothetical protein |
| *FLPKDCKG_00642* | *mdeA-2* | + | 732025 | 733413 | 8.48 | 26.83 | 96.35 | 25.39 | L-methionine gamma-lyase |
| *FLPKDCKG_00643* | *pytH-1* | + | 733419 | 734273 | 2.83 | 6.11 | 28.07 | 11.17 | Pyrethroid hydrolase |
| *FLPKDCKG_00644* | *rclR-2* | - | 734325 | 735266 | 1.16 | 4.44 | 7.49 | 3.60 | RCS-specific HTH-type transcriptional activator RclR |
| *FLPKDCKG_00645* | | + | 735470 | 736027 | 2.58 | 9.55 | 32.53 | 20.04 | Hypothetical protein |
| *FLPKDCKG_00646* | | + | 736024 | 737136 | 1.50 | 4.44 | 13.75 | 8.56 | putative acyl-CoA dehydrogenase fadE25 |
| *FLPKDCKG_00647* | *cmpC-1* | + | 737133 | 737945 | 0.99 | 2.62 | 14.95 | 3.94 | Bicarbonate transport ATP-binding protein CmpC |
| *FLPKDCKG_00648* | *nrtA-1* | + | 737935 | 739125 | 2.54 | 15.14 | 40.06 | 13.86 | Nitrate transport protein NrtA |
| *FLPKDCKG_00649* | *cmpB-1* | + | 739122 | 739946 | 2.06 | 6.57 | 10.82 | 5.30 | Bicarbonate transport system permease protein CmpB |
| *FLPKDCKG_00650* | | + | 740781 | 742154 | 2.05 | 2.01 | 3.03 | 2.03 | Hypothetical protein |
| *FLPKDCKG_00651* | | + | 742158 | 742613 | 0.83 | 3.44 | 0.57 | 2.13 | Hypothetical protein |
| *FLPKDCKG_00652* | *rspR-2* | - | 742629 | 743318 | 2.57 | 11.69 | 5.68 | 4.26 | HTH-type transcriptional repressor RspR |
| *FLPKDCKG_00653* | | + | 743524 | 745254 | 15.91 | 14.59 | 26.58 | 4.97 | 6-deoxy-6-sulfo-D-gluconate dehydratase |
| *FLPKDCKG_00654* | *proA-2* | + | 745293 | 746012 | 8.08 | 6.34 | 12.49 | 8.21 | 4-hydroxy-4-methyl-2-oxoglutarate aldolase/4-carboxy-4-hydroxy-2-oxoadipate aldolase |
| *FLPKDCKG_00655* | | + | 746077 | 747066 | 3.14 | 4.64 | 6.27 | 4.88 | Hypothetical protein |
| *FLPKDCKG_00656* | | - | 747168 | 748142 | 0.27 | 0.52 | 0.00 | 0.73 | Hypothetical protein |
| *FLPKDCKG_00657* | *dapA-1* | - | 748156 | 749058 | 0.90 | 0.03 | 0.48 | 0.35 | 4-hydroxy-tetrahydrodipicolinate synthase |
| *FLPKDCKG_00658* | *rspR-3* | + | 749336 | 750058 | 3.00 | 2.49 | 4.78 | 0.29 | HTH-type transcriptional repressor RspR |
| *FLPKDCKG_00659* | | - | 750152 | 751138 | 0.74 | 1.77 | 1.19 | 1.35 | Hypothetical protein |
| *FLPKDCKG_00660* | *catB* | - | 751222 | 752346 | 0.93 | 2.97 | 0.28 | 0.19 | Muconate cycloisomerase 1 |
| *FLPKDCKG_00661* | *benM-1* | + | 752453 | 753337 | 1.69 | 3.61 | 0.89 | 0.31 | HTH-type transcriptional regulator BenM |
| *FLPKDCKG_00662* | | + | 753437 | 753934 | 6.99 | 15.62 | 18.77 | 4.52 | Hypothetical protein |
| *FLPKDCKG_00663* | *cmoA* | - | 753954 | 754622 | 3.88 | 8.38 | 8.89 | 2.11 | Carboxy-S-adenosyl-L-methionine synthase |
| *FLPKDCKG_00664* | *tgl* | - | 754724 | 755881 | 3.04 | 17.36 | 10.48 | 4.41 | Protein-glutamine gamma-glutamyltransferase |
| *FLPKDCKG_00665* | | + | 756185 | 756793 | 6.34 | 19.96 | 8.48 | 2.96 | putative S-adenosylmethionine-dependent methyltransferase/MSMEI_2290 |
| *FLPKDCKG_00666* | | - | 756820 | 757395 | 1.87 | 10.50 | 7.62 | 2.17 | Hypothetical protein |
| *FLPKDCKG_00667* | | - | 757392 | 757490 | 7.51 | 41.01 | 43.61 | 9.24 | Hypothetical protein |
| *FLPKDCKG_00668* | *pgrR-3* | - | 757714 | 758652 | 2.27 | 5.39 | 3.65 | 0.33 | HTH-type transcriptional regulator PgrR |
| *FLPKDCKG_00669* | *yjgH-1* | + | 758791 | 759189 | 0.52 | 0.95 | 0.14 | 0.76 | RutC family protein YjgH |
| *FLPKDCKG_00670* | | - | 759271 | 760812 | 14.27 | 5.75 | 5.01 | 0.39 | Hypothetical protein |
| *FLPKDCKG_00671* | | - | 761061 | 761627 | 5.60 | 35.13 | 21.69 | 5.57 | Hypothetical protein |
| *FLPKDCKG_00672* | | + | 761933 | 762766 | 15.44 | 46.93 | 25.19 | 12.22 | Hypothetical protein |
| *FLPKDCKG_00673* | *pcpR-2* | - | 762850 | 763776 | 1.10 | 1.28 | 1.93 | 0.00 | PCP degradation transcriptional activation protein |
| *FLPKDCKG_00674* | *pcp* | + | 763870 | 764517 | 0.40 | 0.00 | 0.31 | 0.00 | Pyrrolidone-carboxylate peptidase |
| *FLPKDCKG_00675* | | + | 764531 | 765418 | 0.15 | 0.00 | 0.44 | 0.00 | Soluble epoxide hydrolase |
| *FLPKDCKG_00676* | | + | 765688 | 766656 | 0.08 | 0.37 | 2.47 | 2.09 | Hypothetical protein |
| *FLPKDCKG_00677* | *glnH-1* | + | 766762 | 767601 | 0.05 | 0.56 | 1.45 | 0.79 | ABC transporter glutamine-binding protein GlnH |
| *FLPKDCKG_00678* | *fliY-2* | + | 767633 | 768493 | 0.10 | 1.03 | 1.82 | 0.26 | L-cystine-binding protein FliY |
| *FLPKDCKG_00679* | *aspC-1* | + | 768555 | 769799 | 0.00 | 0.33 | 1.12 | 0.00 | Aspartate aminotransferase |
| *FLPKDCKG_00680* | *glnM-2* | + | 769796 | 770533 | 0.14 | 0.51 | 0.61 | 0.00 | putative glutamine ABC transporter permease protein GlnM |
| *FLPKDCKG_00681* | *occM-1* | + | 770523 | 771341 | 0.13 | 0.72 | 0.46 | 0.33 | Octopine transport system permease protein OccM |
| *FLPKDCKG_00682* | *tcyC-1* | + | 771328 | 772176 | 0.18 | 1.40 | 2.00 | 0.95 | L-cystine import ATP-binding protein TcyC |
| *FLPKDCKG_00683* | | + | 772203 | 772802 | 0.60 | 2.14 | 1.93 | 1.53 | Hypothetical protein |
| *FLPKDCKG_00684* | *metC-1* | + | 772825 | 774030 | 0.46 | 1.72 | 4.89 | 0.17 | Cystathionine beta-lyase |
| *FLPKDCKG_00685* | | + | 774153 | 774500 | 2.25 | 6.61 | 22.49 | 4.41 | Hypothetical protein |
| *FLPKDCKG_00686* | *cysB-2* | + | 774514 | 775443 | 0.45 | 1.98 | 7.05 | 2.69 | HTH-type transcriptional regulator CysB |

| Locus name | Gene name | Strand | Gene start (bp) | Gene end (bp) | Average TPM Inoculum | Average TPM $t_6$ | Average TPM $t_{24}$ | Average TPM $t_{168}$ | Annotation description |
|---|---|---|---|---|---|---|---|---|---|
| *FLPKDCKG_00687* | *glpE-1* | + | 775622 | 777829 | 0.05 | 0.07 | 1.98 | 1.30 | Thiosulfate sulfurtransferase GlpE |
| *FLPKDCKG_00688* | | + | 778028 | 778954 | 0.61 | 1.34 | 0.80 | 0.69 | Putative thiosulfate sulfurtransferase |
| *FLPKDCKG_00689* | | + | 778955 | 780271 | 0.10 | 0.00 | 0.53 | 1.04 | Hypothetical protein |
| *FLPKDCKG_00690* | | + | 780740 | 781894 | 0.93 | 4.47 | 2.02 | 0.99 | Hypothetical protein |
| *FLPKDCKG_00691* | | - | 782005 | 782493 | 2.70 | 2.64 | 4.01 | 1.37 | Hypothetical protein |
| *FLPKDCKG_00692* | *tmoT-1* | - | 782616 | 783230 | 0.86 | 1.14 | 0.45 | 0.00 | Response regulator protein TmoT |
| *FLPKDCKG_00693* | *fixL-1* | - | 783227 | 784666 | 2.63 | 2.73 | 1.64 | 1.15 | Sensor protein FixL |
| *FLPKDCKG_00694* | | - | 784735 | 785889 | 0.40 | 3.25 | 3.59 | 1.36 | Hypothetical protein |
| *FLPKDCKG_00695* | | - | 785996 | 786448 | 4.27 | 4.55 | 2.48 | 5.11 | Hypothetical protein |
| *FLPKDCKG_00696* | *clcA-1* | + | 786546 | 788330 | 0.47 | 5.18 | 1.71 | 0.17 | H(+)/Cl(-) exchange transporter ClcA |
| *FLPKDCKG_00697* | *cmpR-2* | - | 788404 | 789300 | 2.19 | 1.41 | 0.64 | 1.05 | HTH-type transcriptional activator CmpR |
| *FLPKDCKG_00698* | *argC-1* | + | 789434 | 790378 | 0.20 | 0.00 | 0.49 | 0.00 | N-acetyl-gamma-glutamyl-phosphate reductase |
| *FLPKDCKG_00699* | | - | 790456 | 791199 | 0.36 | 1.08 | 0.64 | 0.17 | Hypothetical protein |
| *FLPKDCKG_00700* | *pcpR-3* | - | 791432 | 792355 | 1.03 | 7.37 | 1.78 | 0.23 | PCP degradation transcriptional activation protein |
| *FLPKDCKG_00701* | *caiC* | + | 792494 | 794077 | 0.12 | 0.59 | 0.00 | 0.14 | putative crotonobetaine/carnitine-CoA ligase |
| *FLPKDCKG_00702* | *proP-2* | + | 794113 | 795426 | 0.69 | 1.06 | 0.21 | 0.05 | Proline/betaine transporter |
| *FLPKDCKG_00703* | | + | 795423 | 796565 | 0.37 | 0.00 | 1.27 | 0.38 | Hypothetical protein |
| *FLPKDCKG_00704* | | + | 796562 | 796903 | 0.64 | 1.03 | 1.10 | 0.00 | Hypothetical protein |
| *FLPKDCKG_00705* | *stoA-1* | - | 796955 | 797779 | 0.45 | 1.41 | 0.58 | 1.92 | Sporulation thiol-disulfide oxidoreductase A |
| *FLPKDCKG_00706* | | + | 797922 | 798995 | 2.88 | 1.94 | 2.30 | 0.62 | Nitronate monooxygenase |
| *FLPKDCKG_00707* | *fliY-3* | + | 799421 | 800170 | 22.71 | 16.91 | 22.56 | 14.87 | L-cystine-binding protein FliY |
| *FLPKDCKG_00708* | *yecS-1* | + | 800287 | 800940 | 2.06 | 4.92 | 4.31 | 1.44 | L-cystine transport system permease protein YecS |
| *FLPKDCKG_00709* | *artM-1* | + | 800921 | 801718 | 2.28 | 2.36 | 7.09 | 1.45 | Arginine transport ATP-binding protein ArtM |
| *FLPKDCKG_00710* | *fepC-1* | - | 801792 | 803648 | 3.17 | 1.71 | 1.32 | 0.97 | Ferric enterobactin transport ATP-binding protein FepC |
| *FLPKDCKG_00711* | *yfiZ-1* | - | 803645 | 804685 | 2.91 | 0.00 | 0.27 | 0.63 | putative siderophore transport system permease protein YfiZ |
| *FLPKDCKG_00712* | *yhfQ* | - | 804685 | 805806 | 9.20 | 1.07 | 0.36 | 0.44 | Putative ABC transporter substrate-binding lipoprotein YhfQ |
| *FLPKDCKG_00713* | *fpvA-2* | - | 805850 | 808066 | 59.01 | 5.90 | 6.01 | 4.04 | Ferripyoverdine receptor |
| *FLPKDCKG_00714* | *bcr-1* | - | 808222 | 809463 | 12.13 | 1.03 | 0.98 | 0.66 | Bicyclomycin resistance protein |
| *FLPKDCKG_00715* | *pchR-3* | - | 809582 | 810547 | 15.61 | 1.47 | 4.88 | 1.64 | Regulatory protein PchR |
| *FLPKDCKG_00716* | *bnzA* | - | 810699 | 811865 | 2.54 | 0.30 | 0.45 | 0.49 | Benzene 1,2-dioxygenase subunit alpha |
| *FLPKDCKG_00717* | | - | 811905 | 812654 | 3.85 | 1.19 | 0.00 | 1.14 | Hypothetical protein |
| *FLPKDCKG_00718* | *iucC* | - | 812651 | 814504 | 25.85 | 1.26 | 1.37 | 1.74 | Aerobactin synthase |
| *FLPKDCKG_00719* | *iucB* | - | 814501 | 815106 | 29.04 | 3.37 | 1.31 | 1.22 | N(6)-hydroxylysine O-acetyltransferase |
| *FLPKDCKG_00720* | *iucD* | - | 815124 | 816521 | 10.95 | 0.49 | 0.50 | 0.76 | L-lysine N6-monooxygenase |
| *FLPKDCKG_00721* | *fetA* | + | 816902 | 817528 | 1.98 | 2.81 | 1.61 | 0.53 | putative iron export ATP-binding protein FetA |
| *FLPKDCKG_00722* | *fetB* | + | 817519 | 818319 | 1.70 | 1.82 | 0.70 | 0.00 | putative iron export permease protein FetB |
| *FLPKDCKG_00723* | | + | 818453 | 819094 | 1.09 | 0.06 | 1.86 | 0.60 | putative membrane protein |
| *FLPKDCKG_00724* | *icd-1* | - | 819240 | 821468 | 4.43 | 47.64 | 41.97 | 12.44 | Isocitrate dehydrogenase [NADP] |
| *FLPKDCKG_00725* | *rstA-1* | + | 821914 | 823455 | 1.46 | 1.57 | 2.15 | 0.85 | Transcriptional regulatory protein RstA |
| *FLPKDCKG_00726* | *ilvA-1* | + | 823651 | 825228 | 5.89 | 2.73 | 2.56 | 1.25 | L-threonine dehydratase biosynthetic IlvA |
| *FLPKDCKG_00727* | *fhuC* | - | 825225 | 826052 | 0.26 | 0.00 | 0.00 | 0.23 | Iron(3+)-hydroxamate import ATP-binding protein FhuC |
| *FLPKDCKG_00728* | | - | 826057 | 826389 | 0.00 | 0.00 | 0.00 | 0.00 | Azurin |
| *FLPKDCKG_00729* | *hmuU-1* | - | 826350 | 827597 | 0.12 | 0.02 | 0.00 | 0.00 | Hemin transport system permease protein HmuU |
| *FLPKDCKG_00730* | *btuF-1* | - | 827610 | 828731 | 0.60 | 0.30 | 1.94 | 0.12 | Vitamin B12-binding protein |
| *FLPKDCKG_00731* | *cusR* | + | 828925 | 829608 | 1.82 | 1.03 | 3.47 | 2.79 | Transcriptional regulatory protein CusR |
| *FLPKDCKG_00732* | *arlS* | + | 829601 | 830971 | 1.28 | 1.00 | 1.58 | 0.60 | Signal transduction histidine-protein kinase ArlS |
| *FLPKDCKG_00733* | | - | 831036 | 832616 | 0.65 | 1.61 | 1.74 | 0.97 | Alpha-ketoglutaric semialdehyde dehydrogenase |
| *FLPKDCKG_00734* | | - | 832676 | 833590 | 2.90 | 8.76 | 6.20 | 1.61 | putative 5-dehydro-4-deoxyglucarate dehydratase |
| *FLPKDCKG_00735* | *garD* | + | 833841 | 835415 | 1.05 | 2.24 | 2.54 | 1.15 | Galactarate dehydratase (L-threo-forming) |
| *FLPKDCKG_00736* | *gnl-1* | + | 835474 | 836523 | 0.96 | 5.61 | 3.21 | 0.65 | Gluconolactonase |
| *FLPKDCKG_00737* | | + | 836548 | 837525 | 0.70 | 1.35 | 3.45 | 0.68 | Hypothetical protein |
| *FLPKDCKG_00738* | *proQ* | - | 837616 | 838317 | 20.87 | 19.27 | 20.43 | 7.71 | RNA chaperone ProQ |
| *FLPKDCKG_00739* | | + | 838361 | 838885 | 7.49 | 7.23 | 6.52 | 2.92 | Hypothetical protein |
| *FLPKDCKG_00740* | *pcoC* | + | 839052 | 839408 | 1.73 | 10.58 | 7.41 | 4.80 | Copper resistance protein C |
| *FLPKDCKG_00741* | *yebZ* | + | 839410 | 840297 | 3.12 | 7.18 | 8.99 | 3.19 | Inner membrane protein YebZ |
| *FLPKDCKG_00742* | | - | 840355 | 841836 | 0.21 | 0.77 | 1.90 | 0.09 | Hypothetical protein |
| *FLPKDCKG_00743* | *guaA-2* | - | 841905 | 842588 | 2.04 | 14.62 | 14.44 | 3.60 | GMP synthase [glutamine-hydrolyzing] |
| *FLPKDCKG_00744* | *ntaA* | - | 842804 | 844180 | 0.12 | 1.03 | 2.58 | 1.51 | Nitrilotriacetate monooxygenase component A |
| *FLPKDCKG_00745* | *ssuB-1* | - | 844239 | 845105 | 0.05 | 0.00 | 0.91 | 0.72 | Aliphatic sulfonates import ATP-binding protein SsuB |
| *FLPKDCKG_00746* | *ssuC-2* | - | 845124 | 845948 | 0.00 | 0.62 | 2.78 | 0.96 | Putative aliphatic sulfonates transport permease protein SsuC |
| *FLPKDCKG_00747* | *ssuC-3* | - | 845945 | 846775 | 0.00 | 0.55 | 5.07 | 1.76 | Putative aliphatic sulfonates transport permease protein SsuC |
| *FLPKDCKG_00748* | *ssuA-1* | - | 846789 | 847883 | 0.12 | 0.66 | 2.73 | 2.90 | Putative aliphatic sulfonates-binding protein |
| *FLPKDCKG_00749* | | - | 847941 | 849461 | 0.28 | 0.56 | 6.49 | 3.24 | Hypothetical protein |
| *FLPKDCKG_00750* | *cpdA-1* | - | 849604 | 851745 | 0.21 | 1.05 | 0.28 | 0.10 | 3',5'-cyclic adenosine monophosphate phosphodiesterase CpdA |
| *FLPKDCKG_00751* | | - | 851866 | 852546 | 2.17 | 2.03 | 8.54 | 1.96 | Hypothetical protein |
| *FLPKDCKG_00752* | *cynR-3* | - | 852676 | 853620 | 2.43 | 3.12 | 1.29 | 2.14 | HTH-type transcriptional regulator CynR |
| *FLPKDCKG_00753* | *csd-1* | + | 853802 | 855007 | 3.94 | 1.48 | 1.80 | 0.42 | putative cysteine desulfurase |
| *FLPKDCKG_00754* | *gsiB-1* | + | 855052 | 856605 | 2.60 | 3.94 | 2.44 | 0.88 | Glutathione-binding protein GsiB |
| *FLPKDCKG_00755* | *gsiC-1* | + | 856616 | 857572 | 0.60 | 0.81 | 1.44 | 0.21 | Glutathione transport system permease protein GsiC |

| Locus name | Gene name | Strand | Gene start (bp) | Gene end (bp) | Average TPM Inoculum | Average TPM $t_6$ | Average TPM $t_{24}$ | Average TPM $t_{168}$ | Annotation description |
|---|---|---|---|---|---|---|---|---|---|
| *FLPKDCKG_00756* | *gsiD-1* | + | 857569 | 858399 | 0.73 | 1.35 | 0.00 | 0.30 | Glutathione transport system permease protein GsiD |
| *FLPKDCKG_00757* | *oppD-2* | + | 858396 | 859262 | 0.77 | 0.71 | 0.88 | 0.00 | Oligopeptide transport ATP-binding protein OppD |
| *FLPKDCKG_00758* | *oppF-2* | + | 859259 | 860098 | 0.48 | 3.21 | 0.53 | 1.08 | Oligopeptide transport ATP-binding protein OppF |
| *FLPKDCKG_00759* | | + | 860221 | 860619 | 0.55 | 3.74 | 2.96 | 1.49 | Hypothetical protein |
| *FLPKDCKG_00760* | *pfeA* | + | 860707 | 862776 | 0.21 | 1.23 | 0.89 | 0.82 | Ferric enterobactin receptor |
| *FLPKDCKG_00761* | *cmpR-3* | - | 862766 | 863680 | 2.81 | 5.27 | 2.19 | 0.90 | HTH-type transcriptional activator CmpR |
| *FLPKDCKG_00762* | *metC-2* | + | 863789 | 864937 | 0.75 | 0.85 | 0.39 | 0.08 | Cystathionine beta-lyase MetC |
| *FLPKDCKG_00763* | *yhdW-1* | + | 865071 | 866192 | 0.22 | 0.00 | 0.00 | 0.00 | Putative amino-acid ABC transporter-binding protein YhdW |
| *FLPKDCKG_00764* | | + | 866212 | 866925 | 0.07 | 0.21 | 1.18 | 0.00 | Hypothetical protein |
| *FLPKDCKG_00765* | *glnP-1* | + | 866877 | 867359 | 0.00 | 0.00 | 1.28 | 0.00 | putative glutamine ABC transporter permease protein GlnP |
| *FLPKDCKG_00766* | *yhdY-2* | + | 867356 | 868453 | 0.15 | 0.46 | 0.16 | 0.00 | Inner membrane amino-acid ABC transporter permease protein YhdY |
| *FLPKDCKG_00767* | *glnQ-3* | + | 868450 | 869205 | 1.02 | 1.60 | 0.34 | 0.00 | Glutamine transport ATP-binding protein GlnQ |
| *FLPKDCKG_00768* | *fecI-1* | + | 869293 | 869796 | 3.16 | 5.83 | 5.28 | 0.90 | putative RNA polymerase sigma factor FecI |
| *FLPKDCKG_00769* | | + | 869783 | 870769 | 0.42 | 0.83 | 1.07 | 0.23 | Hypothetical protein |
| *FLPKDCKG_00770* | *iutA* | + | 870881 | 873313 | 0.84 | 1.46 | 0.48 | 0.31 | Ferric aerobactin receptor |
| *FLPKDCKG_00771* | *gcvA-1* | - | 873338 | 874210 | 2.15 | 2.99 | 1.19 | 1.46 | Glycine cleavage system transcriptional activator |
| *FLPKDCKG_00772* | *aguA* | + | 874340 | 875461 | 0.59 | 0.67 | 0.70 | 0.57 | Agmatine deiminase |
| *FLPKDCKG_00773* | *dadA-1* | + | 875522 | 877195 | 3.30 | 4.33 | 4.85 | 0.91 | D-amino acid dehydrogenase |
| *FLPKDCKG_00774* | *yceI* | + | 877491 | 878081 | 11.37 | 13.40 | 8.13 | 1.84 | Protein YceI |
| *FLPKDCKG_00775* | | - | 878379 | 878642 | 2.67 | 14.33 | 7.21 | 0.35 | Hypothetical protein |
| *FLPKDCKG_00776* | | - | 878698 | 879534 | 21.74 | 59.52 | 30.25 | 11.16 | Hypothetical protein |
| *FLPKDCKG_00777* | *teaD-2* | + | 879818 | 880243 | 26.23 | 14.60 | 6.91 | 4.25 | TRAP-T-associated universal stress protein TeaD |
| *FLPKDCKG_00778* | | + | 880356 | 881099 | 4.26 | 2.77 | 1.54 | 1.72 | Hypothetical protein |
| *FLPKDCKG_00779* | *xpkA* | + | 881096 | 883474 | 3.15 | 3.59 | 1.40 | 0.71 | Xylulose-5-phosphate phosphoketolase |
| *FLPKDCKG_00780* | | - | 883475 | 884848 | 29.35 | 5.99 | 5.03 | 0.99 | Ribonuclease |
| *FLPKDCKG_00781* | *hrp1-1* | + | 885013 | 885498 | 6.34 | 3.20 | 0.23 | 1.30 | Hypoxic response protein 1 |
| *FLPKDCKG_00782* | | + | 885495 | 886016 | 4.57 | 0.34 | 0.00 | 0.32 | Hypothetical protein |
| *FLPKDCKG_00783* | *nreB* | + | 886166 | 887254 | 13.12 | 16.42 | 12.38 | 5.50 | Oxygen sensor histidine kinase NreB |
| *FLPKDCKG_00784* | *uvrY-1* | + | 887251 | 887904 | 4.57 | 9.72 | 6.49 | 1.14 | Response regulator UvrY |
| *FLPKDCKG_00785* | *fnr-1* | + | 888094 | 888822 | 7.20 | 8.05 | 3.23 | 1.45 | Fumarate and nitrate reduction regulatory protein |
| *FLPKDCKG_00786* | | + | 888942 | 889361 | 9.31 | 7.63 | 6.29 | 2.52 | Hypothetical protein |
| *FLPKDCKG_00787* | | - | 889346 | 890578 | 5.25 | 6.87 | 5.48 | 3.18 | Hypothetical protein |
| *FLPKDCKG_00788* | *teaD-3* | + | 890783 | 891241 | 4.83 | 1.07 | 0.00 | 0.00 | TRAP-T-associated universal stress protein TeaD |
| *FLPKDCKG_00789* | *mntH-1* | + | 891461 | 892585 | 0.48 | 0.47 | 1.08 | 0.00 | Divalent metal cation transporter MntH |
| *FLPKDCKG_00790* | | - | 892582 | 893214 | 2.10 | 1.02 | 0.00 | 0.53 | Hypothetical protein |
| *FLPKDCKG_00791* | *nrdZ* | + | 893404 | 894900 | 1.57 | 1.94 | 0.49 | 0.43 | Vitamin B12-dependent ribonucleoside-diphosphate reductase |
| *FLPKDCKG_00792* | | - | 895198 | 896043 | 0.88 | 0.00 | 0.63 | 0.71 | Hypothetical protein |
| *FLPKDCKG_00793* | | - | 896061 | 896246 | 5.97 | 0.00 | 0.00 | 3.90 | Hypothetical protein |
| *FLPKDCKG_00794* | | + | 896486 | 897133 | 1.05 | 0.72 | 0.00 | 1.48 | Hypothetical protein |
| *FLPKDCKG_00795* | *adhA-1* | - | 897218 | 897577 | 0.28 | 1.16 | 0.00 | 0.00 | putative alcohol dehydrogenase AdhA |
| *FLPKDCKG_00796* | *adhA-2* | - | 897602 | 898069 | 0.71 | 0.00 | 0.00 | 0.85 | putative alcohol dehydrogenase AdhA |
| *FLPKDCKG_00797* | *osmY-1* | - | 898223 | 898870 | 5.60 | 1.81 | 0.69 | 2.65 | Osmotically-inducible protein Y |
| *FLPKDCKG_00798* | | - | 898965 | 899390 | 49.10 | 2.04 | 2.40 | 10.00 | 18 kDa heat shock protein |
| *FLPKDCKG_00799* | | + | 899520 | 899876 | 5.43 | 20.16 | 15.28 | 5.08 | Hypothetical protein |
| *FLPKDCKG_00800* | *arsC-1* | + | 899921 | 900421 | 3.55 | 7.17 | 7.50 | 1.77 | Glutaredoxin arsenate reductase |
| *FLPKDCKG_00801* | *acr3* | + | 900425 | 901507 | 1.41 | 3.78 | 2.36 | 0.00 | Arsenical-resistance protein Acr3 |
| *FLPKDCKG_00802* | | + | 902223 | 902708 | 0.59 | 1.39 | 1.02 | 1.14 | Hypothetical protein |
| *FLPKDCKG_00803* | *arsH* | + | 902740 | 903504 | 2.48 | 2.08 | 3.48 | 1.95 | NADPH-dependent FMN reductase ArsH |
| *FLPKDCKG_00804* | | - | 903535 | 903840 | 2.30 | 5.15 | 3.47 | 0.50 | Hypothetical protein |
| *FLPKDCKG_00805* | | - | 903908 | 905134 | 2.67 | 8.79 | 8.19 | 2.08 | Hypothetical protein |
| *FLPKDCKG_00806* | *yfcG-1* | - | 905241 | 905870 | 9.78 | 13.48 | 13.05 | 3.61 | Disulfide-bond oxidoreductase YfcG |
| *FLPKDCKG_00807* | *rutR-1* | - | 905923 | 906540 | 5.48 | 11.90 | 9.38 | 0.62 | HTH-type transcriptional regulator RutR |
| *FLPKDCKG_00808* | | + | 906672 | 907817 | 4.69 | 2.68 | 3.22 | 1.89 | Hypothetical protein |
| *FLPKDCKG_00809* | *mqo* | + | 908206 | 909906 | 0.76 | 3.00 | 2.70 | 1.97 | Malate:quinone oxidoreductase |
| *FLPKDCKG_00810* | | + | 910034 | 910999 | 3.31 | 11.94 | 12.89 | 1.62 | Hypothetical protein |
| *FLPKDCKG_00811* | *yhjE-2* | + | 911249 | 912574 | 1.90 | 4.87 | 1.90 | 1.23 | Inner membrane metabolite transport protein YhjE |
| *FLPKDCKG_00812* | *aphA-2* | + | 912571 | 913590 | 3.62 | 5.15 | 4.44 | 2.91 | Acetylpolyamine aminohydrolase |
| *FLPKDCKG_00813* | *cysL-2* | + | 913655 | 914644 | 3.34 | 1.80 | 2.98 | 0.60 | HTH-type transcriptional regulator CysL |
| *FLPKDCKG_00814* | *gstB-4* | + | 914800 | 915426 | 10.28 | 7.71 | 3.57 | 1.78 | Glutathione S-transferase GST-6.0 |
| *FLPKDCKG_00815* | | + | 915563 | 916093 | 17.08 | 8.97 | 8.90 | 3.88 | Hypothetical protein |
| *FLPKDCKG_00816* | | + | 916206 | 917462 | 0.10 | 0.00 | 1.19 | 0.00 | Hypothetical protein |
| *FLPKDCKG_00817* | | + | 917459 | 917932 | 0.30 | 0.00 | 0.00 | 0.00 | Hypothetical protein |
| *FLPKDCKG_00818* | | - | 918004 | 918786 | 0.06 | 2.21 | 3.31 | 0.00 | Hypothetical protein |
| *FLPKDCKG_00819* | | - | 919228 | 919575 | 2.48 | 7.35 | 4.76 | 0.27 | Hypothetical protein |
| *FLPKDCKG_00820* | *dauA* | + | 919695 | 920840 | 1.97 | 1.84 | 0.47 | 0.30 | FAD-dependent catabolic D-arginine dehydrogenase DauA |
| *FLPKDCKG_00821* | *yabJ-2* | + | 920852 | 921229 | 10.14 | 5.74 | 0.27 | 1.14 | 2-iminobutanoate/2-iminopropanoate deaminase |
| *FLPKDCKG_00822* | | + | 921259 | 922704 | 3.66 | 3.32 | 0.87 | 1.44 | Hypothetical protein |
| *FLPKDCKG_00823* | *gsiB-2* | + | 922755 | 924332 | 1.58 | 1.64 | 2.94 | 0.50 | Glutathione-binding protein GsiB |
| *FLPKDCKG_00824* | *gsiC-2* | + | 924329 | 925375 | 0.64 | 0.48 | 0.40 | 0.00 | Glutathione transport system permease protein GsiC |

| Locus name | Gene name | Strand | Gene start (bp) | Gene end (bp) | Average TPM Inoculum | Average TPM $t_6$ | Average TPM $t_{24}$ | Average TPM $t_{168}$ | Annotation description |
|---|---|---|---|---|---|---|---|---|---|
| *FLPKDCKG_00825* | *gsiD-2* | + | 925389 | 926264 | 0.70 | 1.34 | 0.03 | 0.44 | Glutathione transport system permease protein GsiD |
| *FLPKDCKG_00826* | *oppD-3* | + | 926275 | 927288 | 0.63 | 0.86 | 0.40 | 0.41 | Oligopeptide transport ATP-binding protein OppD |
| *FLPKDCKG_00827* | *gsiA-2* | + | 927285 | 928277 | 1.57 | 1.66 | 1.10 | 0.92 | Glutathione import ATP-binding protein GsiA |
| *FLPKDCKG_00828* | *dan-1* | + | 928309 | 929763 | 1.79 | 2.61 | 2.37 | 1.36 | D-aminoacylase |
| *FLPKDCKG_00829* | | + | 929842 | 930492 | 4.33 | 6.21 | 5.32 | 0.00 | Hypothetical protein |
| *FLPKDCKG_00830* | *steT* | + | 930801 | 932204 | 1.58 | 1.74 | 6.13 | 2.05 | Serine/threonine exchanger SteT |
| *FLPKDCKG_00831* | *ufaA1* | + | 932261 | 933115 | 6.50 | 4.31 | 4.46 | 4.06 | Tuberculostearic acid methyltransferase UfaA1 |
| *FLPKDCKG_00832* | | - | 933144 | 934292 | 1.62 | 6.38 | 4.10 | 1.57 | Hypothetical protein |
| *FLPKDCKG_00833* | *ykfA* | - | 934322 | 935458 | 1.30 | 2.36 | 0.95 | 0.93 | putative murein peptide carboxypeptidase |
| *FLPKDCKG_00834* | *rluB-1* | - | 935670 | 936299 | 2.40 | 5.02 | 2.49 | 0.79 | Ribosomal large subunit pseudouridine synthase B |
| *FLPKDCKG_00835* | | - | 936587 | 937693 | 6.11 | 4.77 | 3.32 | 2.50 | Hypothetical protein |
| *FLPKDCKG_00836* | | - | 937766 | 938491 | 1.93 | 4.44 | 4.37 | 1.53 | Hypothetical protein |
| *FLPKDCKG_00837* | | + | 939216 | 939590 | 6.25 | 8.75 | 4.80 | 7.32 | Hypothetical protein |
| *FLPKDCKG_00838* | | + | 939587 | 940075 | 1.13 | 7.81 | 4.38 | 1.10 | Hypothetical protein |
| *FLPKDCKG_00839* | | + | 940146 | 941546 | 0.83 | 3.14 | 0.37 | 0.41 | putative CtpA-like serine protease |
| *FLPKDCKG_00840* | | + | 941586 | 942212 | 1.26 | 2.77 | 6.04 | 0.86 | Hypothetical protein |
| *FLPKDCKG_00841* | *esiB-1* | + | 942400 | 943773 | 1.09 | 6.43 | 6.23 | 2.02 | Secretory immunoglobulin A-binding protein EsiB |
| *FLPKDCKG_00842* | *cmpR-4* | - | 944989 | 945855 | 2.26 | 2.83 | 4.37 | 0.61 | HTH-type transcriptional activator CmpR |
| *FLPKDCKG_00843* | *argH-2* | + | 946003 | 947223 | 0.66 | 0.29 | 0.80 | 0.23 | Argininosuccinate lyase |
| *FLPKDCKG_00844* | | + | 947177 | 947512 | 1.28 | 0.13 | 0.00 | 0.00 | Hypothetical protein |
| *FLPKDCKG_00845* | | + | 947566 | 948531 | 0.43 | 0.00 | 1.14 | 0.15 | Hypothetical protein |
| *FLPKDCKG_00846* | *psdht* | - | 948559 | 949539 | 1.59 | 3.36 | 3.89 | 1.36 | Phenylserine dehydratase |
| *FLPKDCKG_00847* | *tauA-2* | + | 949757 | 950770 | 0.36 | 0.03 | 1.76 | 0.49 | Taurine-binding periplasmic protein |
| *FLPKDCKG_00848* | *ssuC-4* | + | 950776 | 951549 | 0.06 | 0.24 | 1.46 | 0.33 | Putative aliphatic sulfonates transport permease protein SsuC |
| *FLPKDCKG_00849* | *tauB-2* | + | 951546 | 952355 | 0.12 | 0.80 | 0.00 | 0.32 | Taurine import ATP-binding protein TauB |
| *FLPKDCKG_00850* | | + | 952431 | 953840 | 1.45 | 3.12 | 1.32 | 0.00 | Hypothetical protein |
| *FLPKDCKG_00851* | | - | 953886 | 954020 | 0.00 | 0.00 | 0.00 | 0.00 | Hypothetical protein |
| *FLPKDCKG_00852* | | - | 954005 | 954343 | 0.14 | 0.35 | 0.00 | 0.52 | Hypothetical protein |
| *FLPKDCKG_00853* | | - | 954520 | 954831 | 0.00 | 0.00 | 0.00 | 0.00 | Hypothetical protein |
| *FLPKDCKG_00854* | | - | 954828 | 955622 | 1.51 | 0.00 | 0.74 | 1.02 | Hypothetical protein |
| *FLPKDCKG_00855* | *xerC-1* | + | 955722 | 956939 | 1.12 | 2.60 | 1.13 | 0.73 | Tyrosine recombinase XerC |
| *FLPKDCKG_00856* | *ycdT-2* | + | 957029 | 958582 | 0.38 | 2.11 | 1.36 | 0.49 | putative diguanylate cyclase YcdT |
| *FLPKDCKG_00857* | | - | 958579 | 958986 | 0.64 | 1.47 | 1.22 | 0.02 | Hypothetical protein |
| *FLPKDCKG_00858* | *soxR* | + | 959114 | 959578 | 11.74 | 10.14 | 12.50 | 4.18 | Redox-sensitive transcriptional activator SoxR |
| *FLPKDCKG_00859* | *ydaM-1* | + | 959765 | 961528 | 1.26 | 1.10 | 1.67 | 0.42 | putative diguanylate cyclase YdaM |
| *FLPKDCKG_00860* | | - | 961623 | 961772 | 35.57 | 63.73 | 54.12 | 22.41 | Hypothetical protein |
| *FLPKDCKG_00861* | | + | 962157 | 962333 | 0.91 | 40.79 | 34.67 | 1.21 | Hypothetical protein |
| *FLPKDCKG_00862* | | - | 962334 | 962591 | 0.85 | 70.27 | 38.83 | 3.14 | Hypothetical protein |
| *FLPKDCKG_00863* | *yxaF* | + | 962739 | 963317 | 6.95 | 15.44 | 13.56 | 3.24 | putative HTH-type transcriptional regulator YxaF |
| *FLPKDCKG_00864* | | + | 963398 | 963877 | 2.49 | 4.23 | 10.06 | 1.56 | Hypothetical protein |
| *FLPKDCKG_00865* | | - | 963898 | 964338 | 0.62 | 0.00 | 3.97 | 1.43 | Hypothetical protein |
| *FLPKDCKG_00866* | | - | 964485 | 965147 | 0.07 | 0.00 | 0.00 | 0.00 | Hypothetical protein |
| *FLPKDCKG_00867* | *xcpV-1* | - | 965144 | 965551 | 0.14 | 0.00 | 0.00 | 2.99 | Type II secretion system protein I |
| *FLPKDCKG_00868* | | - | 965580 | 966107 | 0.09 | 0.36 | 0.00 | 0.12 | Hypothetical protein |
| *FLPKDCKG_00869* | | - | 966104 | 966688 | 0.07 | 0.00 | 0.00 | 0.00 | Hypothetical protein |
| *FLPKDCKG_00870* | | - | 966672 | 967295 | 0.00 | 0.00 | 0.22 | 0.64 | Hypothetical protein |
| *FLPKDCKG_00871* | | - | 967401 | 967988 | 0.10 | 0.50 | 0.00 | 0.00 | Hypothetical protein |
| *FLPKDCKG_00872* | | - | 968124 | 969116 | 0.19 | 0.44 | 0.41 | 0.11 | Hypothetical protein |
| *FLPKDCKG_00873* | *fecI-2* | - | 969216 | 969731 | 0.08 | 0.63 | 0.00 | 0.00 | putative RNA polymerase sigma factor FecI |
| *FLPKDCKG_00874* | *pupA-1* | - | 969762 | 970481 | 0.40 | 1.89 | 0.75 | 1.47 | Ferric-pseudobactin 358 receptor |
| *FLPKDCKG_00875* | *hxuA-1* | - | 970607 | 982909 | 0.34 | 0.68 | 0.61 | 0.25 | Heme/hemopexin-binding protein |
| *FLPKDCKG_00876* | *phoA2-1* | - | 983095 | 984309 | 0.12 | 0.53 | 0.00 | 0.00 | Alkaline phosphatase L |
| *FLPKDCKG_00877* | *phoA2-2* | - | 984439 | 985674 | 0.10 | 0.71 | 0.44 | 0.44 | Alkaline phosphatase L |
| *FLPKDCKG_00878* | *epsF-1* | - | 985919 | 987127 | 0.22 | 0.00 | 0.34 | 0.47 | Type II secretion system protein F |
| *FLPKDCKG_00879* | *hxcR-1* | - | 987136 | 988581 | 0.14 | 0.16 | 0.04 | 0.00 | putative type II secretion system protein HxcR |
| *FLPKDCKG_00880* | | - | 988578 | 988754 | 0.25 | 0.00 | 1.64 | 0.00 | Hypothetical protein |
| *FLPKDCKG_00881* | | - | 988759 | 989355 | 0.00 | 0.00 | 0.00 | 0.00 | Hypothetical protein |
| *FLPKDCKG_00882* | | - | 989352 | 990587 | 0.03 | 0.00 | 0.00 | 0.00 | Hypothetical protein |
| *FLPKDCKG_00883* | | - | 990587 | 991582 | 0.00 | 0.04 | 0.55 | 0.30 | Hypothetical protein |
| *FLPKDCKG_00884* | *xcpT-1* | - | 991554 | 992015 | 0.00 | 0.00 | 0.00 | 0.00 | Type II secretion system protein G |
| *FLPKDCKG_00885* | | + | 992394 | 992864 | 0.00 | 0.13 | 0.00 | 0.00 | Hypothetical protein |
| *FLPKDCKG_00886* | | + | 992868 | 994397 | 0.55 | 1.37 | 1.77 | 0.95 | Hypothetical protein |
| *FLPKDCKG_00887* | | - | 994476 | 994808 | 21.87 | 38.52 | 18.39 | 7.12 | Hypothetical protein |
| *FLPKDCKG_00888* | *mdtL-1* | + | 994952 | 996157 | 0.50 | 1.99 | 0.98 | 0.00 | Multidrug resistance protein MdtL |
| *FLPKDCKG_00889* | | - | 996189 | 996662 | 0.00 | 0.00 | 3.20 | 0.00 | Hypothetical protein |
| *FLPKDCKG_00890* | *pcpR-4* | + | 996758 | 997654 | 2.23 | 5.03 | 2.83 | 0.60 | PCP degradation transcriptional activation protein |
| *FLPKDCKG_00891* | *echA8-2* | + | 997725 | 998552 | 12.55 | 21.79 | 5.69 | 3.49 | putative enoyl-CoA hydratase echA8 |
| *FLPKDCKG_00892* | *chrA* | - | 998634 | 999893 | 9.21 | 17.55 | 18.66 | 9.84 | Chromate transport protein |
| *FLPKDCKG_00893* | *rstB-2* | + | 1000152 | 1001465 | 2.06 | 2.72 | 4.16 | 0.91 | Sensor protein RstB |

| Locus name | Gene name | Strand | Gene start (bp) | Gene end (bp) | Average TPM Inoculum | Average TPM $t_6$ | Average TPM $t_{24}$ | Average TPM $t_{168}$ | Annotation description |
|---|---|---|---|---|---|---|---|---|---|
| *FLPKDCKG_00894* | | - | 1001493 | 1002602 | 18.89 | 9.62 | 6.17 | 1.55 | Hypothetical protein |
| *FLPKDCKG_00895* | | - | 1002668 | 1003648 | 57.62 | 16.78 | 14.40 | 4.01 | Hypothetical protein |
| *FLPKDCKG_00896* | *corC-1* | - | 1004164 | 1005540 | 2.54 | 2.27 | 1.73 | 0.58 | Magnesium and cobalt efflux protein CorC |
| *FLPKDCKG_00897* | | + | 1006335 | 1007069 | 35.78 | 101.70 | 52.57 | 14.91 | Gluconate 2-dehydrogenase subunit 3 |
| *FLPKDCKG_00898* | | + | 1007062 | 1008834 | 42.60 | 94.52 | 59.25 | 13.78 | Gluconate 2-dehydrogenase flavoprotein |
| *FLPKDCKG_00899* | | + | 1008831 | 1009733 | 12.20 | 25.84 | 17.43 | 6.08 | Cytochrome c4 |
| *FLPKDCKG_00900* | | + | 1010158 | 1010529 | 2.90 | 4.21 | 6.96 | 3.08 | Hypothetical protein |
| *FLPKDCKG_00901* | | + | 1010550 | 1010657 | 7.39 | 2.41 | 8.04 | 0.60 | Hypothetical protein |
| *FLPKDCKG_00902* | | + | 1010702 | 1011415 | 1.14 | 0.72 | 0.00 | 0.37 | Hypothetical protein |
| *FLPKDCKG_00903* | | + | 1011457 | 1011861 | 47.14 | 71.18 | 68.08 | 21.95 | Hypothetical protein |
| *FLPKDCKG_00904* | *fhuA-6* | + | 1011910 | 1014087 | 4.94 | 22.16 | 26.22 | 10.66 | Ferrichrome-iron receptor |
| *FLPKDCKG_00905* | | - | 1014089 | 1015186 | 1.11 | 30.32 | 19.89 | 2.97 | Hypothetical protein |
| *FLPKDCKG_00906* | | - | 1015469 | 1015651 | 3.00 | 1784.74 | 1552.18 | 189.29 | Hypothetical protein |
| *FLPKDCKG_00907* | *cph2-1* | - | 1015824 | 1016729 | 34.87 | 62.47 | 39.72 | 10.28 | Phytochrome-like protein cph2 |
| *FLPKDCKG_00908* | *yccS-1* | + | 1017085 | 1019196 | 0.31 | 1.46 | 1.28 | 0.51 | Inner membrane protein YccS |
| *FLPKDCKG_00909* | | + | 1019324 | 1020160 | 26.72 | 3.29 | 1.27 | 0.10 | Hypothetical protein |
| *FLPKDCKG_00910* | | - | 1020186 | 1021133 | 2.34 | 0.67 | 1.84 | 0.82 | 2-ketogluconate reductase |
| *FLPKDCKG_00911* | | + | 1021244 | 1022422 | 0.40 | 60.82 | 46.47 | 2.93 | Hypothetical protein |
| *FLPKDCKG_00912* | | - | 1022525 | 1022677 | 0.00 | 57.56 | 34.31 | 1.87 | Hypothetical protein |
| *FLPKDCKG_00913* | | - | 1022719 | 1022883 | 0.00 | 879.15 | 667.01 | 86.26 | Hypothetical protein |
| *FLPKDCKG_00914* | *yraA-1* | - | 1022961 | 1023515 | 0.64 | 722.28 | 352.64 | 32.33 | Putative cysteine protease YraA |
| *FLPKDCKG_00915* | | + | 1023646 | 1024677 | 1.07 | 77.36 | 45.55 | 3.72 | Hypothetical protein |
| *FLPKDCKG_00916* | | - | 1024699 | 1024977 | 102.90 | 116.92 | 127.34 | 36.34 | Hypothetical protein |
| *FLPKDCKG_00917* | | + | 1025238 | 1025576 | 11.49 | 133.09 | 109.58 | 36.57 | Hypothetical protein |
| *FLPKDCKG_00918* | *ydaD* | + | 1025678 | 1026565 | 1.02 | 397.40 | 244.86 | 25.97 | General stress protein 39 |
| *FLPKDCKG_00919* | | - | 1026597 | 1027235 | 0.14 | 91.17 | 51.02 | 4.64 | Hypothetical protein |
| *FLPKDCKG_00920* | | - | 1027263 | 1027856 | 0.06 | 122.20 | 81.41 | 10.22 | Hypothetical protein |
| *FLPKDCKG_00921* | | + | 1028073 | 1028357 | 0.00 | 64.51 | 46.57 | 3.24 | Hypothetical protein |
| *FLPKDCKG_00922* | | - | 1028394 | 1028672 | 0.80 | 462.64 | 247.50 | 43.78 | Hypothetical protein |
| *FLPKDCKG_00923* | *asr* | + | 1028829 | 1029227 | 1.80 | 1098.41 | 485.03 | 38.27 | Acid shock protein |
| *FLPKDCKG_00924* | *zraR-1* | - | 1029262 | 1030632 | 19.97 | 138.61 | 88.63 | 18.39 | Transcriptional regulatory protein ZraR |
| *FLPKDCKG_00925* | *zraR-2* | + | 1030810 | 1031748 | 41.87 | 175.97 | 93.20 | 29.21 | Transcriptional regulatory protein ZraR |
| *FLPKDCKG_00926* | *rpoN-1* | + | 1031951 | 1033402 | 12.41 | 482.48 | 235.43 | 32.84 | RNA polymerase sigma-54 factor |
| *FLPKDCKG_00927* | | - | 1033516 | 1033869 | 1.40 | 2555.87 | 1486.05 | 106.77 | Hypothetical protein |
| *FLPKDCKG_00928* | *cckA* | - | 1034009 | 1037023 | 13.90 | 55.41 | 34.60 | 7.67 | Sensor kinase CckA |
| *FLPKDCKG_00929* | | - | 1037125 | 1037838 | 5.23 | 2.25 | 5.06 | 2.87 | Hypothetical protein |
| *FLPKDCKG_00930* | *flhD-1* | + | 1038139 | 1038342 | 7.60 | 15.89 | 19.22 | 11.02 | Flagellar transcriptional regulator FlhD |
| *FLPKDCKG_00931* | | - | 1038514 | 1038903 | 2.42 | 18.89 | 7.46 | 4.81 | Hypothetical protein |
| *FLPKDCKG_00932* | *ynfM* | - | 1038932 | 1040227 | 1.30 | 3.05 | 2.34 | 1.99 | Inner membrane transport protein YnfM |
| *FLPKDCKG_00933* | *hcaR-3* | + | 1040332 | 1041240 | 1.74 | 6.75 | 2.43 | 1.00 | Hca operon transcriptional activator HcaR |
| *FLPKDCKG_00934* | | + | 1041400 | 1043025 | 25.51 | 40.35 | 34.72 | 12.67 | Hypothetical protein |
| *FLPKDCKG_00935* | *cph2-2* | - | 1043060 | 1044286 | 7.43 | 7.98 | 8.36 | 1.77 | Phytochrome-like protein cph2 |
| *FLPKDCKG_00936* | *aroG-1* | + | 1044489 | 1045610 | 20.12 | 25.71 | 21.69 | 7.76 | Phospho-2-dehydro-3-deoxyheptonate aldolase, Phe-sensitive |
| *FLPKDCKG_00937* | *dadA1-1* | - | 1045623 | 1046867 | 3.04 | 4.99 | 3.70 | 0.70 | D-amino acid dehydrogenase 1 |
| *FLPKDCKG_00938* | *ydaF-2* | + | 1046987 | 1047589 | 4.08 | 5.88 | 2.34 | 2.76 | Putative ribosomal N-acetyltransferase YdaF |
| *FLPKDCKG_00939* | | + | 1047712 | 1048218 | 2.09 | 5.03 | 2.29 | 1.66 | Hypothetical protein |
| *FLPKDCKG_00940* | *phoP-1* | + | 1048596 | 1049315 | 3.54 | 5.31 | 4.64 | 1.68 | Transcriptional regulatory protein PhoP |
| *FLPKDCKG_00941* | *mcl2* | + | 1049367 | 1050209 | 1.74 | 5.15 | 1.81 | 0.76 | (3S)-malyl-CoA thioesterase |
| *FLPKDCKG_00942* | *yheS-1* | + | 1050414 | 1052048 | 1.20 | 4.98 | 2.93 | 0.18 | putative ABC transporter ATP-binding protein YheS |
| *FLPKDCKG_00943* | *psd* | + | 1052199 | 1053065 | 6.38 | 12.69 | 14.56 | 2.80 | Phosphatidylserine decarboxylase proenzyme |
| *FLPKDCKG_00944* | *mdlB-1* | + | 1053210 | 1054418 | 0.58 | 3.66 | 2.64 | 1.81 | (S)-mandelate dehydrogenase |
| *FLPKDCKG_00945* | *cpdA-2* | + | 1054508 | 1055695 | 3.18 | 5.62 | 3.30 | 1.00 | 3',5'-cyclic adenosine monophosphate phosphodiesterase CpdA |
| *FLPKDCKG_00946* | *decR-2* | - | 1055727 | 1056182 | 1.84 | 28.61 | 5.97 | 2.72 | DNA-binding transcriptional activator DecR |
| *FLPKDCKG_00947* | *pbuE-2* | + | 1056275 | 1057486 | 1.16 | 1.99 | 1.55 | 1.31 | Purine efflux pump PbuE |
| *FLPKDCKG_00948* | *yegE-1* | - | 1057494 | 1058465 | 12.38 | 8.34 | 5.18 | 1.29 | putative diguanylate cyclase YegE |
| *FLPKDCKG_00949* | *metQ-1* | - | 1058607 | 1059428 | 12.22 | 56.47 | 32.91 | 3.25 | D-methionine-binding lipoprotein MetQ |
| *FLPKDCKG_00950* | *rocF* | + | 1059583 | 1060521 | 12.88 | 25.08 | 24.20 | 3.26 | Arginase |
| *FLPKDCKG_00951* | *kdgR-2* | - | 1060525 | 1061265 | 2.56 | 5.11 | 7.28 | 0.96 | Pectin degradation repressor protein KdgR |
| *FLPKDCKG_00952* | | + | 1061770 | 1062684 | 12.15 | 6.20 | 3.89 | 1.29 | Hypothetical protein |
| *FLPKDCKG_00953* | *adh* | + | 1062710 | 1063771 | 15.84 | 9.78 | 7.59 | 2.45 | Alcohol dehydrogenase |
| *FLPKDCKG_00954* | | + | 1063891 | 1064268 | 11.24 | 18.67 | 27.71 | 4.20 | Hypothetical protein |
| *FLPKDCKG_00955* | *ribN-1* | + | 1064346 | 1065254 | 2.71 | 10.23 | 7.90 | 3.08 | Riboflavin transporter |
| *FLPKDCKG_00956* | *gstB-5* | - | 1065306 | 1065947 | 13.65 | 20.86 | 13.51 | 5.64 | Glutathione S-transferase GST-6.0 |
| *FLPKDCKG_00957* | *yliI* | - | 1065953 | 1067125 | 24.63 | 17.50 | 12.43 | 4.89 | Aldose sugar dehydrogenase YliI |
| *FLPKDCKG_00958* | *gst-1* | + | 1067245 | 1067859 | 27.58 | 29.49 | 26.02 | 6.68 | Glutathione S-transferase GST-4.5 |
| *FLPKDCKG_00959* | *cmpR-5* | - | 1067950 | 1068903 | 2.43 | 4.06 | 0.30 | 2.05 | HTH-type transcriptional activator CmpR |
| *FLPKDCKG_00960* | | + | 1069023 | 1070003 | 0.43 | 0.52 | 1.44 | 0.67 | Hypothetical protein |
| *FLPKDCKG_00961* | | + | 1070012 | 1071508 | 11.37 | 19.05 | 10.13 | 4.23 | N-acyl-D-aspartate deacylase |
| *FLPKDCKG_00962* | | + | 1071501 | 1072244 | 9.51 | 26.25 | 11.69 | 3.57 | Aspartate racemase |

| Locus name | Gene name | Strand | Gene start (bp) | Gene end (bp) | Average TPM Inoculum | Average TPM $t_6$ | Average TPM $t_{24}$ | Average TPM $t_{168}$ | Annotation description |
|---|---|---|---|---|---|---|---|---|---|
| *FLPKDCKG_00963* | | + | 1072340 | 1072792 | 52.59 | 42.85 | 29.42 | 9.29 | Hypothetical protein |
| *FLPKDCKG_00964* | *livF-3* | - | 1073367 | 1073669 | 16.03 | 28.31 | 21.30 | 5.03 | High-affinity branched-chain amino acid transport ATP-binding protein LivF |
| *FLPKDCKG_00965* | *livF-4* | - | 1073699 | 1074067 | 19.83 | 16.20 | 16.27 | 17.11 | High-affinity branched-chain amino acid transport ATP-binding protein LivF |
| *FLPKDCKG_00966* | *lptB-2* | - | 1074068 | 1074835 | 10.26 | 8.73 | 6.60 | 4.57 | Lipopolysaccharide export system ATP-binding protein LptB |
| *FLPKDCKG_00967* | *livH-3* | - | 1074832 | 1076067 | 5.15 | 6.18 | 6.69 | 3.55 | High-affinity branched-chain amino acid transport system permease protein LivH |
| *FLPKDCKG_00968* | *livH-4* | - | 1076067 | 1076993 | 7.98 | 6.88 | 5.53 | 5.74 | High-affinity branched-chain amino acid transport system permease protein LivH |
| *FLPKDCKG_00969* | *braC-1* | - | 1077184 | 1078299 | 132.37 | 76.82 | 82.60 | 54.57 | Leucine-, isoleucine-, valine-, threonine-, and alanine-binding protein |
| *FLPKDCKG_00970* | *yhhQ* | - | 1078695 | 1079039 | 5.38 | 12.93 | 5.82 | 1.74 | Inner membrane protein YhhQ |
| *FLPKDCKG_00971* | *proC-1* | - | 1079244 | 1080077 | 10.54 | 15.72 | 8.30 | 2.41 | Pyrroline-5-carboxylate reductase |
| *FLPKDCKG_00972* | *ugpB-1* | + | 1080272 | 1081585 | 33.80 | 30.31 | 24.64 | 10.47 | sn-glycerol-3-phosphate-binding periplasmic protein UgpB |
| *FLPKDCKG_00973* | *ugpA-1* | + | 1081820 | 1082701 | 2.12 | 6.99 | 5.84 | 2.23 | sn-glycerol-3-phosphate transport system permease protein UgpA |
| *FLPKDCKG_00974* | *araQ* | + | 1082720 | 1083571 | 3.31 | 4.78 | 3.85 | 1.43 | L-arabinose transport system permease protein AraQ |
| *FLPKDCKG_00975* | *ugpC-1* | + | 1083634 | 1084722 | 1.82 | 1.23 | 4.58 | 0.99 | sn-glycerol-3-phosphate import ATP-binding protein UgpC |
| *FLPKDCKG_00976* | *ompV* | - | 1084802 | 1085704 | 24.28 | 13.61 | 15.75 | 2.94 | Outer membrane protein OmpV |
| *FLPKDCKG_00977* | *pbpX* | - | 1085770 | 1086873 | 10.16 | 15.71 | 12.20 | 1.31 | Putative penicillin-binding protein PbpX |
| *FLPKDCKG_00978* | *rstB-3* | - | 1087022 | 1088314 | 3.15 | 3.94 | 3.03 | 1.22 | Sensor protein RstB |
| *FLPKDCKG_00979* | *rstA-2* | - | 1088323 | 1089006 | 4.39 | 1.64 | 0.91 | 1.28 | Transcriptional regulatory protein RstA |
| *FLPKDCKG_00980* | *braC-2* | - | 1089219 | 1090346 | 126.66 | 77.88 | 59.12 | 21.62 | Leucine-, isoleucine-, valine-, threonine-, and alanine-binding protein |
| *FLPKDCKG_00981* | | + | 1090582 | 1091244 | 1.23 | 13.74 | 23.16 | 7.49 | Hypothetical protein |
| *FLPKDCKG_00982* | *pelA* | - | 1091313 | 1092587 | 1.02 | 16.58 | 15.00 | 2.41 | Pectate trisaccharide-lyase |
| *FLPKDCKG_00983* | *bepA-2* | - | 1092584 | 1094566 | 1.13 | 23.17 | 26.61 | 9.62 | Beta-barrel assembly-enhancing protease |
| *FLPKDCKG_00984* | | - | 1094559 | 1095590 | 0.62 | 15.32 | 21.36 | 3.00 | Hypothetical protein |
| *FLPKDCKG_00985* | *kpsT* | - | 1095712 | 1096425 | 0.18 | 9.99 | 12.71 | 2.15 | Polysialic acid transport ATP-binding protein KpsT |
| *FLPKDCKG_00986* | *kpsM* | - | 1096435 | 1097775 | 0.28 | 29.19 | 28.63 | 6.13 | Polysialic acid transport protein KpsM |
| *FLPKDCKG_00987* | | - | 1097783 | 1098934 | 0.59 | 34.99 | 39.09 | 8.06 | Hypothetical protein |
| *FLPKDCKG_00988* | *pglJ-1* | - | 1098991 | 1101051 | 1.73 | 67.23 | 69.47 | 18.64 | N-acetylgalactosamine-N,N'-diacetylbacillosaminyl-diphospho-undecaprenol 4-alpha-N-acetylgalactosaminyltransferase |
| *FLPKDCKG_00989* | | - | 1101048 | 1103528 | 5.17 | 102.88 | 109.70 | 37.14 | Hypothetical protein |
| *FLPKDCKG_00990* | *wbpA-1* | - | 1103572 | 1104849 | 3.84 | 85.45 | 87.96 | 27.81 | UDP-N-acetyl-D-glucosamine 6-dehydrogenase |
| *FLPKDCKG_00991* | *dps2* | - | 1105632 | 1106132 | 85.52 | 109.25 | 68.73 | 24.01 | DNA protection during starvation protein 2 |
| *FLPKDCKG_00992* | *oxyR-2* | - | 1106250 | 1107200 | 8.68 | 13.66 | 16.86 | 5.81 | Hydrogen peroxide-inducible genes activator |
| *FLPKDCKG_00993* | *recG* | - | 1107289 | 1109379 | 6.22 | 14.95 | 11.88 | 2.98 | ATP-dependent DNA helicase RecG |
| *FLPKDCKG_00994* | | - | 1109416 | 1109802 | 65.56 | 138.19 | 88.20 | 20.22 | RutC family protein |
| *FLPKDCKG_00995* | *gltC-5* | - | 1109937 | 1110839 | 1.24 | 2.92 | 2.30 | 1.37 | HTH-type transcriptional regulator GltC |
| *FLPKDCKG_00996* | | + | 1110901 | 1112289 | 0.49 | 2.08 | 1.40 | 1.98 | Hypothetical protein |
| *FLPKDCKG_00997* | | + | 1112286 | 1112669 | 2.02 | 6.37 | 6.13 | 2.64 | Hypothetical protein |
| *FLPKDCKG_00998* | | + | 1112724 | 1113701 | 1.74 | 2.29 | 2.15 | 1.05 | Hypothetical protein |
| *FLPKDCKG_00999* | *fpr-1* | - | 1113811 | 1114587 | 19.18 | 50.07 | 63.10 | 20.90 | Ferredoxin--NADP reductase |
| *FLPKDCKG_01000* | | + | 1114887 | 1115411 | 57.79 | 16.69 | 18.69 | 19.81 | putative binding protein component of ABC iron transporter |
| *FLPKDCKG_01001* | *futA2* | + | 1115408 | 1115929 | 29.87 | 5.16 | 9.92 | 11.27 | Iron uptake protein A2 |
| *FLPKDCKG_01002* | *phnV-2* | + | 1116093 | 1117763 | 1.39 | 0.55 | 0.36 | 0.66 | Putative 2-aminoethylphosphonate transport system permease protein PhnV |
| *FLPKDCKG_01003* | *fbpC* | + | 1117760 | 1118545 | 2.19 | 0.67 | 0.79 | 0.08 | Fe(3+) ions import ATP-binding protein FbpC |
| *FLPKDCKG_01004* | | - | 1118661 | 1119638 | 1.34 | 3.15 | 0.00 | 0.69 | Hypothetical protein |
| *FLPKDCKG_01005* | *rtcB* | + | 1120055 | 1121191 | 1.09 | 1.72 | 0.93 | 0.42 | RNA-splicing ligase RtcB |
| *FLPKDCKG_01006* | *prfB-2* | + | 1121191 | 1121814 | 1.58 | 3.37 | 3.53 | 4.11 | Peptide chain release factor RF2 |
| *FLPKDCKG_01007* | | - | 1121873 | 1122778 | 12.38 | 9.18 | 15.16 | 4.29 | Putative quercetin 2,3-dioxygenase |
| *FLPKDCKG_01008* | *yhhW-1* | - | 1122929 | 1123627 | 0.74 | 2.34 | 6.24 | 2.43 | Quercetin 2,3-dioxygenase |
| *FLPKDCKG_01009* | *yhaJ-2* | + | 1123816 | 1124733 | 7.98 | 14.54 | 14.12 | 2.50 | HTH-type transcriptional regulator YhaJ |
| *FLPKDCKG_01010* | *glnG-1* | - | 1124805 | 1126307 | 1.74 | 72.38 | 59.32 | 11.27 | Nitrogen regulation protein NR(I) |
| *FLPKDCKG_01011* | *glnL* | - | 1126307 | 1127371 | 2.23 | 164.99 | 111.33 | 19.20 | Nitrogen regulation protein NR(II) |
| *FLPKDCKG_01012* | | - | 1127563 | 1128975 | 18.26 | 636.66 | 443.68 | 93.34 | 3-hydroxylaminophenol mutase |
| *FLPKDCKG_01013* | *hemN-1* | - | 1129160 | 1130383 | 1.74 | 7.23 | 4.50 | 1.73 | Oxygen-independent coproporphyrinogen-III oxidase-like protein YqeR |
| *FLPKDCKG_01014* | | - | 1130414 | 1131037 | 13.08 | 29.44 | 31.86 | 5.54 | Non-canonical purine NTP pyrophosphatase |
| *FLPKDCKG_01015* | *ylaB* | + | 1131164 | 1133020 | 0.29 | 1.40 | 0.29 | 0.01 | Putative cyclic-di-GMP phosphodiesterase YlaB |
| *FLPKDCKG_01016* | *tap-1* | - | 1132889 | 1134598 | 0.17 | 1.26 | 0.81 | 0.14 | Methyl-accepting chemotaxis protein IV |
| *FLPKDCKG_01017* | | + | 1134832 | 1135992 | 5.49 | 9.80 | 8.23 | 3.16 | Cyclohex-1-ene-1-carbonyl-CoA dehydrogenase |
| *FLPKDCKG_01018* | *prpD-1* | + | 1135995 | 1137359 | 2.06 | 3.56 | 1.02 | 0.20 | 2-methylcitrate dehydratase |
| *FLPKDCKG_01019* | *rph-1* | - | 1137376 | 1137636 | 1.30 | 3.56 | 0.62 | 1.85 | Ribonuclease PH |
| *FLPKDCKG_01020* | *rph-2* | - | 1137633 | 1138109 | 4.04 | 9.05 | 3.09 | 0.00 | Ribonuclease PH |
| *FLPKDCKG_01021* | | + | 1138289 | 1139218 | 10.56 | 12.33 | 13.07 | 2.98 | Hypothetical protein |
| *FLPKDCKG_01022* | *lysP-2* | - | 1139299 | 1140882 | 3.54 | 12.29 | 7.50 | 2.19 | Lysine-specific permease |
| *FLPKDCKG_01023* | *amn* | - | 1141041 | 1142540 | 21.95 | 19.38 | 16.68 | 3.32 | AMP nucleosidase |
| *FLPKDCKG_01024* | | + | 1142749 | 1143096 | 0.35 | 4.46 | 11.25 | 11.20 | Hypothetical protein |
| *FLPKDCKG_01025* | | + | 1143351 | 1145243 | 0.39 | 0.52 | 2.52 | 1.95 | putative TonB-dependent receptor |
| *FLPKDCKG_01026* | *mepH-1* | + | 1145456 | 1146049 | 1.44 | 52.11 | 39.70 | 6.59 | Murein DD-endopeptidase MepH |
| *FLPKDCKG_01027* | | - | 1146283 | 1147053 | 20.37 | 71.50 | 38.64 | 12.37 | putative oxidoreductase |
| *FLPKDCKG_01028* | *cc4-1* | + | 1147331 | 1148056 | 24.73 | 18.61 | 10.61 | 2.88 | Cytochrome c4 |
| *FLPKDCKG_01029* | *tsdA-1* | + | 1148053 | 1148979 | 15.89 | 11.56 | 5.13 | 0.95 | Thiosulfate dehydrogenase |
| *FLPKDCKG_01030* | *gmk* | + | 1149160 | 1149792 | 59.03 | 178.41 | 97.41 | 32.97 | Guanylate kinase |
| *FLPKDCKG_01031* | *rpoZ* | + | 1149839 | 1150042 | 91.99 | 180.65 | 168.84 | 48.63 | DNA-directed RNA polymerase subunit omega |

| Locus name | Gene name | Strand | Gene start (bp) | Gene end (bp) | Average TPM Inoculum | Average TPM $t_6$ | Average TPM $t_{24}$ | Average TPM $t_{168}$ | Annotation description |
|---|---|---|---|---|---|---|---|---|---|
| *FLPKDCKG_01032* | | - | 1150143 | 1150228 | 8.20 | 24.94 | 32.21 | 7.22 | tRNA |
| *FLPKDCKG_01033* | *glnQ-4* | - | 1152476 | 1153204 | 10.94 | 20.16 | 20.47 | 6.58 | Glutamine transport ATP-binding protein GlnQ |
| *FLPKDCKG_01034* | *glnP-2* | - | 1153201 | 1153857 | 9.58 | 16.34 | 12.32 | 3.44 | Glutamine transport system permease protein GlnP |
| *FLPKDCKG_01035* | *glnH-2* | - | 1154024 | 1154776 | 91.23 | 80.19 | 58.60 | 22.72 | Glutamine-binding periplasmic protein |
| *FLPKDCKG_01036* | | + | 1154942 | 1156003 | 14.30 | 11.48 | 4.13 | 1.50 | Hypothetical protein |
| *FLPKDCKG_01037* | *apc3-1* | - | 1156034 | 1159657 | 1.18 | 2.87 | 1.16 | 0.39 | Acetophenone carboxylase gamma subunit |
| *FLPKDCKG_01038* | | - | 1159744 | 1160253 | 0.82 | 22.89 | 11.06 | 2.26 | Hypothetical protein |
| *FLPKDCKG_01039* | *nudC-1* | - | 1160412 | 1161182 | 33.56 | 33.49 | 28.67 | 15.58 | NADH pyrophosphatase |
| *FLPKDCKG_01040* | | + | 1161460 | 1162434 | 13.61 | 8.53 | 11.09 | 10.67 | Solute-binding protein |
| *FLPKDCKG_01041* | | + | 1162442 | 1162942 | 4.18 | 6.01 | 8.89 | 6.49 | Hypothetical protein |
| *FLPKDCKG_01042* | *dctM-2* | + | 1162935 | 1164236 | 2.72 | 4.66 | 3.70 | 3.81 | C4-dicarboxylate TRAP transporter large permease protein DctM |
| *FLPKDCKG_01043* | | + | 1164576 | 1165646 | 65.78 | 17.31 | 19.87 | 18.10 | Outer membrane porin protein |
| *FLPKDCKG_01044* | *livF-5* | - | 1165738 | 1166457 | 5.06 | 1.75 | 7.88 | 0.61 | High-affinity branched-chain amino acid transport ATP-binding protein LivF |
| *FLPKDCKG_01045* | *glnQ-5* | - | 1166450 | 1167172 | 3.84 | 2.94 | 1.50 | 2.58 | Glutamine transport ATP-binding protein GlnQ |
| *FLPKDCKG_01046* | | - | 1167153 | 1168142 | 3.69 | 2.80 | 1.14 | 1.60 | Hypothetical protein |
| *FLPKDCKG_01047* | *livH-5* | - | 1168144 | 1168995 | 2.83 | 4.88 | 3.00 | 0.05 | High-affinity branched-chain amino acid transport system permease protein LivH |
| *FLPKDCKG_01048* | | - | 1169041 | 1170291 | 10.58 | 5.15 | 7.76 | 3.86 | Leu/Ile/Val-binding protein |
| *FLPKDCKG_01049* | | - | 1170654 | 1171460 | 16.01 | 87.64 | 46.79 | 11.93 | Hypothetical protein |
| *FLPKDCKG_01050* | *yofA-1* | - | 1171562 | 1172473 | 3.62 | 4.81 | 5.56 | 0.21 | HTH-type transcriptional regulator YofA |
| *FLPKDCKG_01051* | *amiD-1* | - | 1172629 | 1173975 | 3.42 | 5.05 | 3.10 | 1.05 | Putative amidase AmiD |
| *FLPKDCKG_01052* | | - | 1174067 | 1174750 | 0.65 | 2.86 | 0.54 | 0.00 | Hypothetical protein |
| *FLPKDCKG_01053* | *dctM-3* | - | 1174830 | 1176113 | 0.58 | 0.34 | 0.71 | 0.58 | C4-dicarboxylate TRAP transporter large permease protein DctM |
| *FLPKDCKG_01054* | *uehB* | - | 1176110 | 1176730 | 0.51 | 1.70 | 0.76 | 0.00 | Ectoine/5-hydroxyectoine TRAP transporter small permease protein UehB |
| *FLPKDCKG_01055* | | - | 1176817 | 1177518 | 5.79 | 2.29 | 2.25 | 1.35 | Lactate-binding periplasmic protein |
| *FLPKDCKG_01056* | | - | 1177515 | 1177862 | 2.70 | 10.65 | 2.76 | 0.00 | Hypothetical protein |
| *FLPKDCKG_01057* | *yofA-2* | - | 1178301 | 1179209 | 3.06 | 3.31 | 4.14 | 1.51 | HTH-type transcriptional regulator YofA |
| *FLPKDCKG_01058* | *ribN-2* | + | 1179297 | 1180250 | 1.07 | 1.91 | 2.85 | 1.23 | Riboflavin transporter |
| *FLPKDCKG_01059* | *fabV* | + | 1180379 | 1181575 | 1.85 | 2.51 | 2.80 | 1.08 | Enoyl-[acyl-carrier-protein] reductase [NADH] |
| *FLPKDCKG_01060* | *serB-1* | - | 1181639 | 1182361 | 0.20 | 1.13 | 0.69 | 0.17 | Phosphoserine phosphatase |
| *FLPKDCKG_01061* | *gcvA-2* | - | 1182375 | 1183283 | 0.45 | 2.34 | 0.10 | 0.21 | Glycine cleavage system transcriptional activator |
| *FLPKDCKG_01062* | *potA-1* | + | 1183526 | 1184602 | 0.54 | 2.14 | 1.58 | 1.31 | Spermidine/putrescine import ATP-binding protein PotA |
| *FLPKDCKG_01063* | *potH-1* | + | 1184616 | 1186262 | 0.60 | 2.03 | 2.09 | 0.32 | Putrescine transport system permease protein PotH |
| *FLPKDCKG_01064* | | + | 1186285 | 1187268 | 0.83 | 3.14 | 3.43 | 0.79 | putative binding protein component of ABC iron transporter |
| *FLPKDCKG_01065* | *ytrE* | - | 1187369 | 1188040 | 10.61 | 11.12 | 12.22 | 5.65 | ABC transporter ATP-binding protein YtrE |
| *FLPKDCKG_01066* | *tesA* | + | 1188102 | 1188698 | 8.36 | 25.90 | 17.11 | 7.80 | Esterase TesA |
| *FLPKDCKG_01067* | *ppiD* | - | 1188746 | 1190716 | 3.97 | 10.41 | 12.38 | 4.18 | Peptidyl-prolyl cis-trans isomerase D |
| *FLPKDCKG_01068* | *cpg2-2* | + | 1191057 | 1192556 | 49.44 | 28.31 | 23.18 | 3.74 | Carboxypeptidase G2 |
| *FLPKDCKG_01069* | *ugpQ* | + | 1192803 | 1193552 | 13.05 | 14.97 | 13.94 | 4.30 | Glycerophosphodiester phosphodiesterase, cytoplasmic |
| *FLPKDCKG_01070* | *rlmJ* | + | 1193565 | 1194455 | 6.53 | 11.31 | 7.31 | 4.12 | Ribosomal RNA large subunit methyltransferase J |
| *FLPKDCKG_01071* | *pdhD* | - | 1194518 | 1195954 | 0.36 | 9.49 | 27.58 | 5.51 | Dihydrolipoyl dehydrogenase |
| *FLPKDCKG_01072* | | - | 1196159 | 1196887 | 5.17 | 225.44 | 290.47 | 74.49 | Hybrid peroxiredoxin hyPrx5 |
| *FLPKDCKG_01073* | *actP-1* | - | 1197057 | 1197257 | 4.91 | 16.86 | 7.45 | 0.40 | Copper-transporting P-type ATPase |
| *FLPKDCKG_01074* | *actP-2* | + | 1197419 | 1199692 | 2.99 | 8.93 | 4.89 | 2.14 | Copper-transporting P-type ATPase |
| *FLPKDCKG_01075* | *hmrR* | + | 1199719 | 1200111 | 2.79 | 5.56 | 1.16 | 0.00 | HTH-type transcriptional regulator HmrR |
| *FLPKDCKG_01076* | *rlmD* | - | 1200175 | 1201680 | 5.01 | 8.89 | 11.09 | 2.32 | 23S rRNA (uracil(1939)-C(5))-methyltransferase RlmD |
| *FLPKDCKG_01077* | | - | 1201847 | 1202599 | 0.51 | 61.51 | 100.56 | 49.16 | Hypothetical protein |
| *FLPKDCKG_01078* | | - | 1202961 | 1203569 | 3.94 | 314.60 | 158.52 | 46.63 | Hypothetical protein |
| *FLPKDCKG_01079* | | - | 1203737 | 1204516 | 10.33 | 28.64 | 18.75 | 7.08 | Hypothetical protein |
| *FLPKDCKG_01080* | *nlpD-1* | - | 1204513 | 1205373 | 44.85 | 71.34 | 44.96 | 18.30 | Murein hydrolase activator NlpD |
| *FLPKDCKG_01081* | *pcm-1* | - | 1205391 | 1206215 | 8.36 | 27.30 | 16.41 | 4.86 | Protein-L-isoaspartate O-methyltransferase |
| *FLPKDCKG_01082* | *surE* | - | 1206200 | 1206958 | 18.71 | 21.90 | 21.97 | 5.94 | 5'-nucleotidase SurE |
| *FLPKDCKG_01083* | *plsY* | + | 1207147 | 1207788 | 2.19 | 6.45 | 7.66 | 3.80 | putative glycerol-3-phosphate acyltransferase |
| *FLPKDCKG_01084* | *tsaD* | - | 1207858 | 1208889 | 1.91 | 5.02 | 2.59 | 2.90 | tRNA N6-adenosine threonylcarbamoyltransferase |
| *FLPKDCKG_01085* | *adeP* | + | 1208998 | 1210290 | 5.18 | 11.38 | 5.56 | 3.37 | Adenine permease AdeP |
| *FLPKDCKG_01086* | | - | 1210375 | 1210731 | 20.55 | 19.21 | 16.87 | 2.15 | Hypothetical protein |
| *FLPKDCKG_01087* | *limB-1* | + | 1211109 | 1212131 | 16.62 | 45.45 | 45.06 | 16.19 | Limonene 1,2-monooxygenase |
| *FLPKDCKG_01088* | *trxC* | + | 1212329 | 1212703 | 26.03 | 45.25 | 66.43 | 24.46 | Putative thioredoxin 2 |
| *FLPKDCKG_01089* | *yigZ* | - | 1212737 | 1213324 | 11.23 | 55.02 | 53.68 | 19.69 | IMPACT family member YigZ |
| *FLPKDCKG_01090* | | - | 1213397 | 1214212 | 0.38 | 1.71 | 10.32 | 2.89 | Hypothetical protein |
| *FLPKDCKG_01091* | *ssuC-5* | - | 1214221 | 1214994 | 0.36 | 3.07 | 10.31 | 3.56 | Putative aliphatic sulfonates transport permease protein SsuC |
| *FLPKDCKG_01092* | *ssuA-2* | - | 1214991 | 1215989 | 1.33 | 18.08 | 47.72 | 19.91 | Putative aliphatic sulfonates-binding protein |
| *FLPKDCKG_01093* | *tauB-3* | - | 1216062 | 1216832 | 1.79 | 105.18 | 156.63 | 39.13 | Taurine import ATP-binding protein TauB |
| *FLPKDCKG_01094* | | - | 1216945 | 1218162 | 1.21 | 3.05 | 2.18 | 0.19 | Hypothetical protein |
| *FLPKDCKG_01095* | *ybiT* | - | 1218349 | 1219953 | 16.31 | 15.10 | 11.57 | 4.47 | putative ABC transporter ATP-binding protein YbiT |
| *FLPKDCKG_01096* | *ldcA* | - | 1220098 | 1221135 | 1.91 | 9.88 | 8.60 | 2.17 | Murein tetrapeptide carboxypeptidase |
| *FLPKDCKG_01097* | *tadA* | - | 1221132 | 1221623 | 15.55 | 47.48 | 41.50 | 7.93 | tRNA-specific adenosine deaminase |
| *FLPKDCKG_01098* | | - | 1221634 | 1222482 | 27.87 | 93.43 | 57.26 | 15.09 | Hypothetical protein |
| *FLPKDCKG_01099* | | - | 1222595 | 1223092 | 182.62 | 691.83 | 447.05 | 85.13 | Hypothetical protein |
| *FLPKDCKG_01100* | *mrpG* | - | 1223323 | 1223661 | 4.80 | 12.83 | 7.13 | 5.13 | Na(+)/H(+) antiporter subunit G |

| Locus name | Gene name | Strand | Gene start (bp) | Gene end (bp) | Average TPM Inoculum | Average TPM $t_6$ | Average TPM $t_{24}$ | Average TPM $t_{168}$ | Annotation description |
|---|---|---|---|---|---|---|---|---|---|
| *FLPKDCKG_01101* | *mrpF* | - | 1223658 | 1223939 | 2.27 | 9.67 | 1.44 | 0.29 | Na(+)/H(+) antiporter subunit F |
| *FLPKDCKG_01102* | *mrpE* | - | 1223936 | 1224418 | 2.63 | 6.93 | 1.85 | 1.47 | Na(+)/H(+) antiporter subunit E |
| *FLPKDCKG_01103* | *mrpD* | - | 1224418 | 1226049 | 1.10 | 2.24 | 2.57 | 1.17 | Na(+)/H(+) antiporter subunit D |
| *FLPKDCKG_01104* | *mnhC1* | - | 1226046 | 1226390 | 7.31 | 9.89 | 14.83 | 0.79 | Na(+)/H(+) antiporter subunit C1 |
| *FLPKDCKG_01105* | *mrpA* | - | 1226390 | 1229305 | 3.27 | 4.51 | 3.82 | 2.11 | Na(+)/H(+) antiporter subunit A |
| *FLPKDCKG_01106* | | + | 1229857 | 1230843 | 8.51 | 11.33 | 12.95 | 2.85 | Hypothetical protein |
| *FLPKDCKG_01107* | | + | 1231832 | 1232227 | 0.26 | 0.00 | 0.00 | 0.00 | Hypothetical protein |
| *FLPKDCKG_01108* | | - | 1232240 | 1233112 | 0.71 | 0.95 | 1.29 | 0.49 | Hypothetical protein |
| *FLPKDCKG_01109* | *smc-1* | - | 1233550 | 1236177 | 4.06 | 3.48 | 3.24 | 0.70 | Chromosome partition protein Smc |
| *FLPKDCKG_01110* | *sbcD* | - | 1236174 | 1237340 | 11.18 | 11.03 | 11.63 | 4.35 | Nuclease SbcCD subunit D |
| *FLPKDCKG_01111* | *dadA1-2* | - | 1237484 | 1238785 | 3.73 | 105.89 | 49.50 | 14.93 | D-amino acid dehydrogenase 1 |
| *FLPKDCKG_01112* | *lrp-1* | + | 1238921 | 1239388 | 5.08 | 11.60 | 3.62 | 1.21 | Leucine-responsive regulatory protein |
| *FLPKDCKG_01113* | | + | 1239432 | 1240196 | 8.63 | 13.10 | 9.62 | 4.64 | Hypothetical protein |
| *FLPKDCKG_01114* | | + | 1240228 | 1240452 | 2.10 | 5.51 | 3.54 | 0.00 | Hypothetical protein |
| *FLPKDCKG_01115* | | + | 1240449 | 1241024 | 0.39 | 1.38 | 2.33 | 0.00 | Hypothetical protein |
| *FLPKDCKG_01116* | | + | 1241119 | 1241505 | 1.14 | 2.71 | 3.53 | 2.57 | Hypothetical protein |
| *FLPKDCKG_01117* | | + | 1241525 | 1243465 | 0.52 | 3.73 | 2.17 | 0.97 | Hypothetical protein |
| *FLPKDCKG_01118* | *mmgC-1* | - | 1243562 | 1244689 | 12.54 | 9.97 | 6.37 | 1.56 | Acyl-CoA dehydrogenase |
| *FLPKDCKG_01119* | *dmlR-8* | - | 1244822 | 1245727 | 2.54 | 2.68 | 0.28 | 0.00 | HTH-type transcriptional regulator DmlR |
| *FLPKDCKG_01120* | *azoB-1* | + | 1245863 | 1246747 | 0.74 | 0.00 | 0.93 | 0.53 | NAD(P)H azoreductase |
| *FLPKDCKG_01121* | *ywnA* | - | 1246722 | 1247162 | 5.74 | 4.75 | 7.69 | 1.50 | Putative HTH-type transcriptional regulator YwnA |
| *FLPKDCKG_01122* | *trxB-1* | + | 1247256 | 1248215 | 1.45 | 2.39 | 3.57 | 0.69 | Thioredoxin reductase |
| *FLPKDCKG_01123* | | - | 1248283 | 1248918 | 6.27 | 5.53 | 8.86 | 1.13 | Hypothetical protein |
| *FLPKDCKG_01124* | | + | 1249032 | 1249499 | 4.64 | 12.06 | 11.57 | 3.98 | Hypothetical protein |
| *FLPKDCKG_01125* | | - | 1249554 | 1250513 | 1.09 | 4.33 | 3.04 | 1.92 | Hypothetical protein |
| *FLPKDCKG_01126* | | + | 1250712 | 1251866 | 0.40 | 1.35 | 0.91 | 0.19 | Outer membrane porin protein |
| *FLPKDCKG_01127* | | - | 1252051 | 1252461 | 1.51 | 0.00 | 1.55 | 0.00 | Hypothetical protein |
| *FLPKDCKG_01128* | *btr-1* | - | 1252515 | 1252874 | 0.69 | 2.20 | 0.00 | 0.16 | HTH-type transcriptional activator Btr |
| *FLPKDCKG_01129* | | - | 1253001 | 1254245 | 2.04 | 3.30 | 1.46 | 0.18 | Hypothetical protein |
| *FLPKDCKG_01130* | | + | 1254479 | 1254931 | 0.63 | 6.74 | 3.69 | 1.83 | Hypothetical protein |
| *FLPKDCKG_01131* | *cat* | + | 1255050 | 1255700 | 1.26 | 4.37 | 3.95 | 1.06 | Chloramphenicol acetyltransferase |
| *FLPKDCKG_01132* | *copR* | - | 1255675 | 1256037 | 0.53 | 3.54 | 1.99 | 1.26 | Transcriptional activator protein CopR |
| *FLPKDCKG_01133* | | - | 1256067 | 1256372 | 0.44 | 5.99 | 2.01 | 0.00 | Hypothetical protein |
| *FLPKDCKG_01134* | | - | 1256525 | 1258255 | 19.41 | 98.98 | 47.15 | 16.30 | Hypothetical protein |
| *FLPKDCKG_01135* | | - | 1258368 | 1258676 | 21.19 | 214.05 | 80.98 | 14.49 | Hypothetical protein |
| *FLPKDCKG_01136* | *ycjP* | - | 1258693 | 1259505 | 4.37 | 18.48 | 11.71 | 5.66 | Inner membrane ABC transporter permease protein YcjP |
| *FLPKDCKG_01137* | *araP* | - | 1259498 | 1260388 | 4.39 | 16.85 | 5.56 | 1.28 | L-arabinose transport system permease protein AraP |
| *FLPKDCKG_01138* | *ugpC-2* | - | 1260385 | 1261473 | 2.61 | 10.23 | 4.49 | 2.31 | sn-glycerol-3-phosphate import ATP-binding protein UgpC |
| *FLPKDCKG_01139* | *ugpC-3* | - | 1261473 | 1262558 | 4.74 | 20.58 | 6.44 | 0.82 | sn-glycerol-3-phosphate import ATP-binding protein UgpC |
| *FLPKDCKG_01140* | *glpR* | + | 1262788 | 1263579 | 31.17 | 178.95 | 47.74 | 10.87 | Glycerol-3-phosphate regulon repressor |
| *FLPKDCKG_01141* | *glpD2-3* | + | 1263683 | 1265281 | 4.02 | 31.66 | 7.39 | 3.96 | Glycerol-3-phosphate dehydrogenase 2 |
| *FLPKDCKG_01142* | | - | 1265394 | 1265933 | 1.72 | 8.35 | 10.74 | 0.59 | 3'3'-cGAMP-specific phosphodiesterase 2 |
| *FLPKDCKG_01143* | *yfkM* | - | 1266169 | 1266849 | 37.07 | 80.52 | 43.83 | 10.36 | General stress protein 18 |
| *FLPKDCKG_01144* | *curA-1* | - | 1266887 | 1267924 | 29.51 | 47.49 | 21.98 | 12.09 | NADPH-dependent curcumin reductase |
| *FLPKDCKG_01145* | *kstR2-3* | + | 1268045 | 1268821 | 5.94 | 7.21 | 4.19 | 0.72 | HTH-type transcriptional repressor KstR2 |
| *FLPKDCKG_01146* | | + | 1268824 | 1269792 | 2.22 | 1.49 | 0.00 | 0.77 | Hypothetical protein |
| *FLPKDCKG_01147* | *ybhF-1* | + | 1269789 | 1270754 | 3.94 | 2.67 | 2.54 | 0.97 | putative ABC transporter ATP-binding protein YbhF |
| *FLPKDCKG_01148* | *ybhS* | + | 1270751 | 1271896 | 3.27 | 7.23 | 0.83 | 0.33 | Inner membrane transport permease YbhS |
| *FLPKDCKG_01149* | *oprM-1* | + | 1271905 | 1273383 | 1.21 | 3.03 | 3.08 | 1.71 | Outer membrane protein OprM |
| *FLPKDCKG_01150* | *ompR-1* | + | 1273526 | 1274242 | 0.93 | 3.08 | 0.00 | 0.00 | Transcriptional regulatory protein OmpR |
| *FLPKDCKG_01151* | *proS-1* | + | 1274328 | 1274810 | 14.61 | 17.90 | 11.57 | 2.16 | Proline--tRNA ligase |
| *FLPKDCKG_01152* | *yccS-2* | + | 1274828 | 1276810 | 1.06 | 0.51 | 0.45 | 0.09 | Inner membrane protein YccS |
| *FLPKDCKG_01153* | | + | 1276814 | 1277608 | 1.59 | 2.25 | 4.31 | 0.94 | Hypothetical protein |
| *FLPKDCKG_01154* | | + | 1277700 | 1278134 | 69.55 | 91.77 | 108.88 | 46.80 | Hypothetical protein |
| *FLPKDCKG_01155* | | - | 1278209 | 1278388 | 7.85 | 23.18 | 32.43 | 11.14 | Hypothetical protein |
| *FLPKDCKG_01156* | | + | 1278821 | 1279672 | 0.43 | 0.44 | 0.26 | 0.39 | Hypothetical protein |
| *FLPKDCKG_01157* | *nemA* | - | 1279725 | 1280783 | 1.55 | 1.63 | 1.55 | 0.42 | N-ethylmaleimide reductase |
| *FLPKDCKG_01158* | *sdpR-1* | - | 1280829 | 1281170 | 4.77 | 2.71 | 8.95 | 4.32 | Transcriptional repressor SdpR |
| *FLPKDCKG_01159* | *arfB* | - | 1281305 | 1281721 | 12.90 | 13.44 | 24.58 | 8.30 | Peptidyl-tRNA hydrolase ArfB |
| *FLPKDCKG_01160* | *catD-1* | + | 1281925 | 1282332 | 16.71 | 19.72 | 28.34 | 14.55 | Putative oxidoreductase CatD |
| *FLPKDCKG_01161* | | + | 1282578 | 1282736 | 25.07 | 122.96 | 117.30 | 38.90 | Hypothetical protein |
| *FLPKDCKG_01162* | | + | 1282769 | 1284043 | 9.38 | 70.58 | 52.99 | 28.31 | Hypothetical protein |
| *FLPKDCKG_01163* | *prxU* | + | 1284165 | 1284704 | 28.08 | 118.56 | 74.00 | 17.36 | Selenocysteine-containing peroxiredoxin PrxU |
| *FLPKDCKG_01164* | *frc* | - | 1284821 | 1286257 | 39.61 | 158.97 | 103.02 | 39.83 | Formyl-CoA:oxalate CoA-transferase |
| *FLPKDCKG_01165* | *phrB* | + | 1286564 | 1287982 | 1.03 | 3.52 | 1.11 | 0.49 | Deoxyribodipyrimidine photo-lyase |
| *FLPKDCKG_01166* | *aer* | + | 1288144 | 1288623 | 1.26 | 1.92 | 1.15 | 0.26 | Aerotaxis receptor |
| *FLPKDCKG_01167* | *tsr-2* | + | 1288608 | 1289831 | 0.34 | 0.82 | 0.76 | 0.20 | Methyl-accepting chemotaxis protein I |
| *FLPKDCKG_01168* | | - | 1289848 | 1290555 | 0.27 | 1.66 | 0.00 | 0.00 | Hypothetical protein |
| *FLPKDCKG_01169* | *pdxA2-1* | - | 1290568 | 1291620 | 0.42 | 1.84 | 0.47 | 0.00 | 4-hydroxythreonine-4-phosphate dehydrogenase 2 |

| Locus name | Gene name | Strand | Gene start (bp) | Gene end (bp) | Average TPM Inoculum | Average TPM $t_6$ | Average TPM $t_{24}$ | Average TPM $t_{168}$ | Annotation description |
|---|---|---|---|---|---|---|---|---|---|
| *FLPKDCKG_01170* | | - | 1291617 | 1292768 | 0.23 | 0.59 | 0.05 | 0.00 | Hypothetical protein |
| *FLPKDCKG_01171* | *trpI-3* | + | 1292845 | 1293720 | 3.81 | 2.24 | 1.93 | 0.59 | HTH-type transcriptional regulator TrpI |
| *FLPKDCKG_01172* | | - | 1293749 | 1294081 | 29.24 | 17.60 | 12.03 | 8.23 | Hypothetical protein |
| *FLPKDCKG_01173* | *ydhP-1* | + | 1294309 | 1294701 | 1.44 | 3.74 | 5.54 | 0.02 | Inner membrane transport protein YdhP |
| *FLPKDCKG_01174* | *ydhP-2* | + | 1294701 | 1295510 | 1.62 | 2.68 | 3.07 | 1.54 | Inner membrane transport protein YdhP |
| *FLPKDCKG_01175* | *gpx1* | + | 1295526 | 1296020 | 11.63 | 11.36 | 9.65 | 3.33 | Hydroperoxy fatty acid reductase gpx1 |
| *FLPKDCKG_01176* | *lrpC* | - | 1296128 | 1296595 | 6.50 | 17.21 | 9.16 | 0.66 | HTH-type transcriptional regulator LrpC |
| *FLPKDCKG_01177* | *yijE-1* | + | 1296718 | 1297623 | 0.41 | 3.39 | 2.94 | 0.42 | putative cystine transporter YijE |
| *FLPKDCKG_01178* | *yfcG-2* | + | 1297744 | 1298442 | 33.00 | 32.77 | 14.82 | 8.97 | Disulfide-bond oxidoreductase YfcG |
| *FLPKDCKG_01179* | *phnO-2* | + | 1298455 | 1298919 | 30.67 | 35.46 | 29.36 | 9.04 | Aminoalkylphosphonate N-acetyltransferase |
| *FLPKDCKG_01180* | | + | 1298960 | 1299235 | 9.96 | 15.73 | 19.43 | 7.92 | Hypothetical protein |
| *FLPKDCKG_01181* | *lplJ* | + | 1299232 | 1300008 | 5.20 | 10.47 | 10.48 | 4.32 | Lipoate-protein ligase LplJ |
| *FLPKDCKG_01182* | | + | 1300075 | 1300887 | 7.02 | 9.52 | 8.67 | 3.29 | Hypothetical protein |
| *FLPKDCKG_01183* | *gdhA-1* | - | 1301052 | 1302341 | 106.77 | 36.67 | 27.31 | 12.37 | Glutamate dehydrogenase |
| *FLPKDCKG_01184* | *aknOx* | + | 1302832 | 1304505 | 37.32 | 9.28 | 6.14 | 2.43 | Aclacinomycin-N/aclacinomycin-A oxidase |
| *FLPKDCKG_01185* | *hutG-1* | - | 1304585 | 1304827 | 2.23 | 4.63 | 4.08 | 0.00 | Formimidoylglutamase |
| *FLPKDCKG_01186* | *hutG-2* | - | 1304797 | 1305528 | 2.02 | 2.79 | 2.57 | 1.22 | Formimidoylglutamase |
| *FLPKDCKG_01187* | *glnQ-6* | - | 1305525 | 1306307 | 2.80 | 2.55 | 3.27 | 0.38 | Glutamine transport ATP-binding protein GlnQ |
| *FLPKDCKG_01188* | *yecS-2* | - | 1306304 | 1306972 | 2.61 | 1.85 | 2.47 | 0.47 | L-cystine transport system permease protein YecS |
| *FLPKDCKG_01189* | *glnH-3* | - | 1306969 | 1307712 | 11.57 | 17.93 | 8.79 | 3.60 | Glutamine-binding periplasmic protein |
| *FLPKDCKG_01190* | *kdgR-3* | - | 1307835 | 1308572 | 6.17 | 6.86 | 6.47 | 0.56 | Transcriptional regulator KdgR |
| *FLPKDCKG_01191* | *ankX* | - | 1308834 | 1309757 | 3.45 | 5.68 | 6.18 | 3.71 | Phosphocholine transferase AnkX |
| *FLPKDCKG_01192* | *ycfH* | - | 1309797 | 1310567 | 2.75 | 9.05 | 3.97 | 2.94 | putative metal-dependent hydrolase YcfH |
| *FLPKDCKG_01193* | *holB* | - | 1310643 | 1311692 | 4.20 | 9.44 | 5.15 | 1.86 | DNA polymerase III subunit delta' |
| *FLPKDCKG_01194* | *tmk-1* | - | 1311689 | 1312312 | 6.78 | 11.87 | 7.76 | 2.68 | Thymidylate kinase |
| *FLPKDCKG_01195* | *mltG* | - | 1312309 | 1313448 | 8.51 | 10.92 | 7.77 | 3.89 | Endolytic murein transglycosylase |
| *FLPKDCKG_01196* | | + | 1313465 | 1313554 | 2.73 | 3.13 | 15.09 | 4.44 | Hypothetical protein |
| *FLPKDCKG_01197* | *ygfZ* | + | 1313606 | 1314604 | 12.73 | 6.20 | 4.97 | 2.10 | tRNA-modifying protein YgfZ |
| *FLPKDCKG_01198* | | - | 1314611 | 1315300 | 5.48 | 6.77 | 9.74 | 2.18 | Hypothetical protein |
| *FLPKDCKG_01199* | | - | 1315358 | 1315795 | 6.84 | 9.56 | 1.03 | 2.05 | Putative esterase |
| *FLPKDCKG_01200* | *mmsB-1* | - | 1315797 | 1316165 | 14.26 | 11.61 | 4.01 | 1.70 | 3-hydroxyisobutyrate dehydrogenase |
| *FLPKDCKG_01201* | | + | 1316217 | 1316717 | 8.82 | 1.85 | 5.03 | 0.87 | Hypothetical protein |
| *FLPKDCKG_01202* | *crt-1* | - | 1316767 | 1317942 | 14.08 | 9.47 | 12.62 | 4.30 | Short-chain-enoyl-CoA hydratase |
| *FLPKDCKG_01203* | *acrC-1* | - | 1317939 | 1319093 | 9.27 | 12.76 | 8.34 | 5.07 | Acryloyl-CoA reductase (NADH) |
| *FLPKDCKG_01204* | *pasI* | - | 1319304 | 1319651 | 3.61 | 13.66 | 7.52 | 1.42 | Persistence and stress-resistance antitoxin PasI |
| *FLPKDCKG_01205* | *pasT* | - | 1319641 | 1320075 | 20.44 | 53.79 | 33.94 | 9.54 | Persistence and stress-resistance toxin PasT |
| *FLPKDCKG_01206* | *smpB* | + | 1320181 | 1320648 | 41.44 | 43.75 | 26.79 | 10.47 | SsrA-binding protein |
| *FLPKDCKG_01207* | *dosC-1* | - | 1320688 | 1321878 | 2.94 | 5.55 | 5.37 | 1.92 | Diguanylate cyclase DosC |
| *FLPKDCKG_01208* | *hflK-1* | - | 1321905 | 1322831 | 21.88 | 16.16 | 23.22 | 10.27 | Modulator of FtsH protease HflK |
| *FLPKDCKG_01209* | | - | 1322962 | 1323402 | 53.55 | 38.22 | 50.97 | 13.61 | Hypothetical protein |
| *FLPKDCKG_01210* | *guaA-3* | + | 1323570 | 1324307 | 9.62 | 24.07 | 16.81 | 6.49 | GMP synthase [glutamine-hydrolyzing] |
| *FLPKDCKG_01211* | *ada* | + | 1324401 | 1325243 | 5.98 | 2.33 | 3.89 | 0.44 | Bifunctional transcriptional activator/DNA repair enzyme Ada |
| *FLPKDCKG_01212* | *sigE-1* | + | 1325371 | 1325874 | 3.06 | 0.02 | 6.16 | 0.00 | ECF RNA polymerase sigma factor SigE |
| *FLPKDCKG_01213* | | + | 1325878 | 1326792 | 1.36 | 2.82 | 4.32 | 0.38 | Hypothetical protein |
| *FLPKDCKG_01214* | *fhuA-7* | + | 1326925 | 1329390 | 0.30 | 0.97 | 0.62 | 0.00 | Ferrichrome-iron receptor |
| *FLPKDCKG_01215* | *ppsA-1* | - | 1329511 | 1331877 | 24.61 | 39.67 | 35.05 | 15.33 | Phosphoenolpyruvate synthase |
| *FLPKDCKG_01216* | *ppsR* | + | 1332069 | 1332896 | 14.56 | 27.42 | 17.56 | 7.27 | Phosphoenolpyruvate synthase regulatory protein |
| *FLPKDCKG_01217* | | - | 1332932 | 1333369 | 2.64 | 9.42 | 5.99 | 4.30 | Hypothetical protein |
| *FLPKDCKG_01218* | *trmH* | - | 1333452 | 1334240 | 1.14 | 0.78 | 4.11 | 0.27 | tRNA (guanosine(18)-2'-O)-methyltransferase |
| *FLPKDCKG_01219* | *rnhB-1* | - | 1334237 | 1334623 | 1.38 | 5.96 | 4.11 | 0.65 | Ribonuclease HII |
| *FLPKDCKG_01220* | *rnhB-2* | - | 1334587 | 1334844 | 0.44 | 4.82 | 7.81 | 0.72 | Ribonuclease HII |
| *FLPKDCKG_01221* | *lpxB* | - | 1334831 | 1336027 | 1.97 | 5.14 | 6.00 | 1.65 | Lipid-A-disaccharide synthase |
| *FLPKDCKG_01222* | *lpxA-1* | - | 1336024 | 1336818 | 7.99 | 15.95 | 19.04 | 7.44 | Acyl-[acyl-carrier-protein]--UDP-N-acetylglucosamine O-acyltransferase |
| *FLPKDCKG_01223* | *fabZ* | - | 1336819 | 1337223 | 13.31 | 29.87 | 38.44 | 8.62 | 3-hydroxyacyl-[acyl-carrier-protein] dehydratase FabZ |
| *FLPKDCKG_01224* | *lpxD* | - | 1337345 | 1338442 | 19.38 | 47.02 | 36.69 | 10.83 | UDP-3-O-acylglucosamine N-acyltransferase |
| *FLPKDCKG_01225* | *skp* | - | 1338457 | 1339017 | 188.85 | 346.60 | 233.92 | 57.97 | Chaperone protein Skp |
| *FLPKDCKG_01226* | *bamA* | - | 1339125 | 1341470 | 15.46 | 43.16 | 32.22 | 10.22 | Outer membrane protein assembly factor BamA |
| *FLPKDCKG_01227* | *rseP* | - | 1341524 | 1342357 | 11.11 | 26.93 | 17.75 | 8.27 | Regulator of sigma-E protease RseP |
| *FLPKDCKG_01228* | *dxr* | - | 1342920 | 1344119 | 8.36 | 25.21 | 19.56 | 5.92 | 1-deoxy-D-xylulose 5-phosphate reductoisomerase |
| *FLPKDCKG_01229* | *cdsA* | - | 1344116 | 1344970 | 2.93 | 3.63 | 7.28 | 2.46 | Phosphatidate cytidylyltransferase |
| *FLPKDCKG_01230* | *uppS* | - | 1345032 | 1345796 | 4.30 | 7.41 | 12.80 | 6.32 | Ditrans,polycis-undecaprenyl-diphosphate synthase ((2E,6E)-farnesyl-diphosphate specific) |
| *FLPKDCKG_01231* | *frr* | - | 1345966 | 1346526 | 12.51 | 23.76 | 28.06 | 7.45 | Ribosome-recycling factor |
| *FLPKDCKG_01232* | *pyrH* | - | 1346542 | 1347204 | 24.61 | 33.05 | 23.83 | 13.70 | Uridylate kinase |
| *FLPKDCKG_01233* | *tsf* | - | 1347356 | 1348234 | 10.01 | 55.18 | 47.80 | 26.03 | Elongation factor Ts |
| *FLPKDCKG_01234* | *rpsB* | - | 1348417 | 1349166 | 14.23 | 77.72 | 75.28 | 22.37 | 30S ribosomal protein S2 |
| *FLPKDCKG_01235* | *map-2* | + | 1350011 | 1350832 | 27.45 | 57.78 | 29.75 | 16.04 | Methionine aminopeptidase |
| *FLPKDCKG_01236* | *glnD* | + | 1350852 | 1353458 | 4.07 | 8.56 | 7.06 | 2.74 | Bifunctional uridylyltransferase/uridylyl-removing enzyme |
| *FLPKDCKG_01237* | | + | 1353533 | 1354246 | 10.64 | 16.01 | 9.04 | 3.33 | Hypothetical protein |
| *FLPKDCKG_01238* | *dsbB-1* | + | 1354313 | 1354798 | 4.54 | 12.79 | 6.12 | 2.97 | Disulfide bond formation protein B |

| Locus name | Gene name | Strand | Gene start (bp) | Gene end (bp) | Average TPM Inoculum | Average TPM $t_6$ | Average TPM $t_{24}$ | Average TPM $t_{168}$ | Annotation description |
|---|---|---|---|---|---|---|---|---|---|
| *FLPKDCKG_01239* | | + | 1354897 | 1355409 | 0.97 | 3.30 | 1.52 | 1.02 | Hypothetical protein |
| *FLPKDCKG_01240* | *purF* | - | 1355487 | 1357007 | 11.18 | 26.24 | 22.18 | 8.53 | Amidophosphoribosyltransferase |
| *FLPKDCKG_01241* | *cvpA* | - | 1357038 | 1357526 | 16.89 | 46.41 | 34.30 | 5.20 | Colicin V production protein |
| *FLPKDCKG_01242* | *dedD* | - | 1357526 | 1358365 | 20.29 | 33.98 | 22.31 | 7.38 | Cell division protein DedD |
| *FLPKDCKG_01243* | *folC* | - | 1358457 | 1359767 | 16.56 | 25.73 | 15.51 | 5.24 | Dihydrofolate synthase/folylpolyglutamate synthase |
| *FLPKDCKG_01244* | | + | 1359932 | 1360348 | 3.10 | 0.66 | 0.00 | 1.25 | Hypothetical protein |
| *FLPKDCKG_01245* | | + | 1360427 | 1361398 | 1.17 | 0.78 | 1.04 | 0.70 | Hypothetical protein |
| *FLPKDCKG_01246* | | + | 1361395 | 1363791 | 0.59 | 1.36 | 1.07 | 0.49 | Acyl-CoA dehydrogenase fadE12 |
| *FLPKDCKG_01247* | *kdgR-4* | - | 1363829 | 1364566 | 7.00 | 9.53 | 8.04 | 1.24 | Transcriptional regulator KdgR |
| *FLPKDCKG_01248* | | + | 1364710 | 1365087 | 24.79 | 3.66 | 0.99 | 0.35 | Hypothetical protein |
| *FLPKDCKG_01249* | *fliD* | + | 1365232 | 1366623 | 70.85 | 3.46 | 0.95 | 0.42 | Flagellar hook-associated protein 2 |
| *FLPKDCKG_01250* | *fliS* | + | 1366642 | 1367070 | 19.91 | 7.42 | 4.79 | 0.65 | Flagellar protein FliS |
| *FLPKDCKG_01251* | *fliT* | + | 1367119 | 1367472 | 8.79 | 0.58 | 2.93 | 0.00 | Flagellar protein FliT |
| *FLPKDCKG_01252* | | + | 1367469 | 1368827 | 2.79 | 0.29 | 0.06 | 0.25 | Hypothetical protein |
| *FLPKDCKG_01253* | *flhB-2* | + | 1368827 | 1369150 | 4.77 | 1.39 | 3.57 | 1.06 | Flagellar biosynthetic protein FlhB |
| *FLPKDCKG_01254* | *ycgR* | + | 1369162 | 1369959 | 4.92 | 4.84 | 1.75 | 0.52 | Flagellar brake protein YcgR |
| *FLPKDCKG_01255* | *fliE-2* | - | 1370096 | 1370428 | 58.96 | 6.38 | 5.85 | 2.69 | Flagellar hook-basal body complex protein FliE |
| *FLPKDCKG_01256* | *fliF-2* | + | 1370777 | 1372453 | 5.88 | 0.24 | 0.90 | 0.78 | Flagellar M-ring protein |
| *FLPKDCKG_01257* | *fliG* | + | 1372443 | 1373453 | 5.63 | 0.29 | 0.00 | 0.62 | Flagellar motor switch protein FliG |
| *FLPKDCKG_01258* | *fliH* | + | 1373446 | 1374189 | 3.65 | 0.04 | 0.12 | 0.00 | Flagellar assembly protein FliH |
| *FLPKDCKG_01259* | *fliI-2* | + | 1374186 | 1375661 | 2.46 | 0.70 | 0.72 | 0.28 | Flagellum-specific ATP synthase |
| *FLPKDCKG_01260* | *fliJ* | + | 1375654 | 1376103 | 7.70 | 0.18 | 0.00 | 0.45 | Flagellar FliJ protein |
| *FLPKDCKG_01261* | *fliK* | + | 1376124 | 1377494 | 2.56 | 1.31 | 0.15 | 0.46 | Flagellar hook-length control protein |
| *FLPKDCKG_01262* | | + | 1377755 | 1378276 | 11.11 | 3.54 | 1.20 | 0.81 | Hypothetical protein |
| *FLPKDCKG_01263* | *fliM* | + | 1378279 | 1379286 | 13.55 | 5.48 | 2.33 | 0.57 | Flagellar motor switch protein FliM |
| *FLPKDCKG_01264* | *fliN-2* | + | 1379279 | 1379821 | 5.52 | 0.00 | 1.61 | 0.11 | Flagellar motor switch protein FliN |
| *FLPKDCKG_01265* | *fliO* | + | 1379835 | 1380158 | 1.55 | 0.09 | 1.90 | 0.00 | Flagellar protein FliO |
| *FLPKDCKG_01266* | *fliP-2* | + | 1380181 | 1380837 | 2.00 | 3.18 | 1.32 | 0.29 | Flagellar biosynthetic protein FliP |
| *FLPKDCKG_01267* | *fliP-3* | + | 1380837 | 1380962 | 0.57 | 0.00 | 0.00 | 0.00 | Flagellar biosynthetic protein FliP |
| *FLPKDCKG_01268* | | + | 1380972 | 1381241 | 0.33 | 0.00 | 0.00 | 0.00 | Hypothetical protein |
| *FLPKDCKG_01269* | | + | 1381276 | 1382064 | 0.09 | 0.75 | 0.33 | 0.74 | Hypothetical protein |
| *FLPKDCKG_01270* | *tar-1* | - | 1382168 | 1382770 | 1.42 | 3.26 | 0.50 | 0.00 | Methyl-accepting chemotaxis protein II |
| *FLPKDCKG_01271* | *tar-2* | - | 1384088 | 1385818 | 1.79 | 0.33 | 0.52 | 0.41 | Methyl-accepting chemotaxis protein II |
| *FLPKDCKG_01272* | | - | 1385955 | 1386428 | 2.06 | 1.01 | 0.90 | 0.00 | Hypothetical protein |
| *FLPKDCKG_01273* | *trg-2* | - | 1386458 | 1386916 | 0.71 | 0.00 | 0.00 | 0.45 | Methyl-accepting chemotaxis protein III |
| *FLPKDCKG_01274* | *tsr-3* | - | 1386913 | 1388328 | 2.44 | 1.04 | 0.95 | 0.57 | Methyl-accepting chemotaxis protein I |
| *FLPKDCKG_01275* | *tar-3* | - | 1388443 | 1390359 | 2.18 | 0.49 | 0.39 | 0.41 | Methyl-accepting chemotaxis protein II |
| *FLPKDCKG_01276* | *flgL* | - | 1390500 | 1391759 | 14.80 | 2.79 | 0.36 | 0.00 | Flagellar hook-associated protein 3 |
| *FLPKDCKG_01277* | *flgK* | - | 1391808 | 1393484 | 28.63 | 6.19 | 4.43 | 0.82 | Flagellar hook-associated protein 1 |
| *FLPKDCKG_01278* | *flgJ* | - | 1393558 | 1394541 | 4.38 | 0.31 | 2.18 | 0.00 | Peptidoglycan hydrolase FlgJ |
| *FLPKDCKG_01279* | *flgI-2* | - | 1394547 | 1395683 | 4.14 | 0.43 | 0.00 | 0.04 | Flagellar P-ring protein |
| *FLPKDCKG_01280* | *flgH-2* | - | 1395686 | 1396378 | 8.84 | 0.39 | 2.31 | 0.87 | Flagellar L-ring protein |
| *FLPKDCKG_01281* | *flgG-3* | - | 1396378 | 1397163 | 19.09 | 1.50 | 0.22 | 0.37 | Flagellar basal-body rod protein FlgG |
| *FLPKDCKG_01282* | *flgF* | - | 1397233 | 1397994 | 20.64 | 4.45 | 0.69 | 1.40 | Flagellar basal-body rod protein FlgF |
| *FLPKDCKG_01283* | *flgE-2* | - | 1398033 | 1399520 | 37.85 | 2.73 | 1.33 | 1.01 | Flagellar hook protein FlgE |
| *FLPKDCKG_01284* | *flgD* | - | 1399592 | 1400287 | 47.55 | 1.62 | 1.59 | 1.44 | Basal-body rod modification protein FlgD |
| *FLPKDCKG_01285* | *flgC-2* | - | 1400348 | 1400767 | 42.50 | 3.24 | 0.50 | 0.00 | Flagellar basal-body rod protein FlgC |
| *FLPKDCKG_01286* | *flgB-2* | - | 1400790 | 1401197 | 46.83 | 1.06 | 3.26 | 0.00 | Flagellar basal body rod protein FlgB |
| *FLPKDCKG_01287* | *flgA* | + | 1401366 | 1402079 | 16.33 | 16.43 | 4.29 | 2.05 | Flagella basal body P-ring formation protein FlgA |
| *FLPKDCKG_01288* | *flgM* | + | 1402207 | 1402500 | 89.27 | 99.18 | 79.77 | 15.58 | Negative regulator of flagellin synthesis |
| *FLPKDCKG_01289* | | + | 1402537 | 1403013 | 29.83 | 29.56 | 19.85 | 3.78 | Hypothetical protein |
| *FLPKDCKG_01290* | *flhF* | - | 1403094 | 1405907 | 1.85 | 1.21 | 0.92 | 0.50 | Flagellar biosynthesis protein FlhF |
| *FLPKDCKG_01291* | *flhA-2* | - | 1405904 | 1408024 | 2.80 | 0.49 | 0.57 | 0.17 | Flagellar biosynthesis protein FlhA |
| *FLPKDCKG_01292* | *flhB-3* | - | 1408028 | 1409182 | 4.26 | 1.67 | 1.72 | 0.64 | Flagellar biosynthetic protein FlhB |
| *FLPKDCKG_01293* | *mcp4* | - | 1409379 | 1410926 | 13.79 | 5.44 | 3.28 | 2.19 | Methyl-accepting chemotaxis protein 4 |
| *FLPKDCKG_01294* | | - | 1411132 | 1411764 | 4.73 | 6.45 | 2.38 | 0.83 | Hypothetical protein |
| *FLPKDCKG_01295* | *cheY-1* | - | 1411773 | 1412162 | 4.87 | 4.69 | 2.14 | 2.79 | Chemotaxis protein CheY |
| *FLPKDCKG_01296* | *cheB-2* | - | 1412213 | 1413271 | 3.35 | 2.23 | 1.32 | 0.88 | Chemotaxis response regulator protein-glutamate methylesterase |
| *FLPKDCKG_01297* | *cheR-1* | - | 1413317 | 1414153 | 2.62 | 0.63 | 1.97 | 0.33 | Chemotaxis protein methyltransferase |
| *FLPKDCKG_01298* | *tar-4* | - | 1414180 | 1415886 | 2.91 | 1.25 | 0.41 | 0.29 | Methyl-accepting chemotaxis protein II |
| *FLPKDCKG_01299* | *cheW-2* | - | 1416024 | 1416524 | 14.86 | 1.94 | 2.36 | 0.50 | Chemotaxis protein CheW |
| *FLPKDCKG_01300* | *cheA* | - | 1416540 | 1418225 | 8.37 | 0.82 | 2.00 | 0.17 | Chemotaxis protein CheA |
| *FLPKDCKG_01301* | *cheY-2* | - | 1418710 | 1419045 | 18.53 | 5.62 | 1.65 | 0.00 | Chemotaxis protein CheY |
| *FLPKDCKG_01302* | *motB-2* | - | 1419093 | 1420082 | 20.55 | 1.01 | 1.17 | 0.00 | Motility protein B |
| *FLPKDCKG_01303* | *motA* | - | 1420094 | 1420963 | 12.63 | 1.59 | 0.22 | 0.20 | Motility protein A |
| *FLPKDCKG_01304* | *flhC* | - | 1421081 | 1421662 | 21.19 | 2.25 | 3.51 | 0.37 | Flagellar transcriptional regulator FlhC |
| *FLPKDCKG_01305* | *flhD-2* | - | 1421694 | 1422017 | 28.31 | 6.48 | 1.07 | 2.76 | Flagellar transcriptional regulator FlhD |
| *FLPKDCKG_01306* | *fliA-3* | + | 1422449 | 1423210 | 10.15 | 3.52 | 5.74 | 1.74 | RNA polymerase sigma factor FliA |
| *FLPKDCKG_01307* | *cfa-3* | + | 1423418 | 1425016 | 8.09 | 2.50 | 0.13 | 1.07 | Cyclopropane-fatty-acyl-phospholipid synthase |

| Locus name | Gene name | Strand | Gene start (bp) | Gene end (bp) | Average TPM Inoculum | Average TPM $t_6$ | Average TPM $t_{24}$ | Average TPM $t_{168}$ | Annotation description |
|---|---|---|---|---|---|---|---|---|---|
| *FLPKDCKG_01308* | *fliC-2* | + | 1425232 | 1426923 | 551.81 | 20.58 | 12.30 | 6.00 | Flagellin |
| *FLPKDCKG_01309* | | + | 1427321 | 1428193 | 6.36 | 4.26 | 4.38 | 1.21 | Hypothetical protein |
| *FLPKDCKG_01310* | | + | 1428307 | 1428501 | 259.41 | 127.59 | 108.89 | 28.91 | Hypothetical protein |
| *FLPKDCKG_01311* | *pgrR-4* | - | 1428642 | 1429550 | 9.39 | 12.78 | 10.10 | 2.60 | HTH-type transcriptional regulator PgrR |
| *FLPKDCKG_01312* | *fabG-3* | - | 1429722 | 1430495 | 3.21 | 4.69 | 0.60 | 0.71 | 3-oxoacyl-[acyl-carrier-protein] reductase FabG |
| *FLPKDCKG_01313* | *lpdA* | - | 1430762 | 1432534 | 7.23 | 11.04 | 15.48 | 4.77 | Dihydrolipoyl dehydrogenase |
| *FLPKDCKG_01314* | *aceF* | - | 1432547 | 1434214 | 8.75 | 9.84 | 10.24 | 4.89 | Dihydrolipoyllysine-residue acetyltransferase component of pyruvate dehydrogenase complex |
| *FLPKDCKG_01315* | *aceE-1* | - | 1434227 | 1436935 | 18.72 | 41.03 | 25.45 | 13.08 | Pyruvate dehydrogenase E1 component |
| *FLPKDCKG_01316* | *fixL-2* | + | 1437368 | 1439122 | 10.19 | 13.31 | 10.07 | 4.26 | Sensor protein FixL |
| *FLPKDCKG_01317* | *tmoT-2* | + | 1439119 | 1439745 | 28.94 | 30.01 | 29.54 | 14.60 | Response regulator protein TmoT |
| *FLPKDCKG_01318* | *folD* | + | 1439908 | 1440651 | 7.72 | 16.48 | 13.66 | 2.71 | Bifunctional protein FolD protein |
| *FLPKDCKG_01319* | *prlC* | + | 1440872 | 1442938 | 3.78 | 6.94 | 6.98 | 4.18 | Oligopeptidase A |
| *FLPKDCKG_01320* | *ypmQ-1* | + | 1443948 | 1444562 | 10.72 | 25.69 | 16.28 | 5.50 | SCO1 protein |
| *FLPKDCKG_01321* | | + | 1444705 | 1444956 | 115.26 | 202.03 | 138.78 | 44.06 | Hypothetical protein |
| *FLPKDCKG_01322* | *oprM-2* | - | 1445038 | 1446528 | 0.77 | 5.06 | 5.75 | 1.57 | Outer membrane protein OprM |
| *FLPKDCKG_01323* | *mepB* | - | 1446525 | 1449707 | 0.72 | 2.91 | 3.61 | 1.75 | Multidrug/solvent efflux pump membrane transporter MepB |
| *FLPKDCKG_01324* | *mexA* | - | 1449722 | 1450915 | 0.41 | 2.09 | 2.79 | 1.79 | Multidrug resistance protein MexA |
| *FLPKDCKG_01325* | *dmlR-9* | + | 1451213 | 1452136 | 8.11 | 27.60 | 18.29 | 4.23 | HTH-type transcriptional regulator DmlR |
| *FLPKDCKG_01326* | *rluF* | - | 1452226 | 1452930 | 2.48 | 7.81 | 3.22 | 3.24 | 23S rRNA pseudouridine(2604) synthase |
| *FLPKDCKG_01327* | *ubiC* | + | 1453084 | 1453707 | 0.56 | 5.27 | 1.34 | 0.00 | Chorismate pyruvate-lyase |
| *FLPKDCKG_01328* | *emrB-2* | - | 1453790 | 1455214 | 0.25 | 0.59 | 1.28 | 0.00 | Multidrug export protein EmrB |
| *FLPKDCKG_01329* | *polS* | - | 1455271 | 1455969 | 0.19 | 0.00 | 1.19 | 0.11 | Sorbitol dehydrogenase |
| *FLPKDCKG_01330* | | - | 1456095 | 1456694 | 6.37 | 7.45 | 8.66 | 3.44 | putative HTH-type transcriptional regulator |
| *FLPKDCKG_01331* | *curA-2* | - | 1456772 | 1457809 | 0.13 | 0.83 | 0.84 | 0.25 | NADPH-dependent curcumin reductase |
| *FLPKDCKG_01332* | *nemR* | - | 1457900 | 1458511 | 2.54 | 4.06 | 7.71 | 3.65 | HTH-type transcriptional repressor NemR |
| *FLPKDCKG_01333* | | - | 1458556 | 1459092 | 1.97 | 12.90 | 4.42 | 0.68 | Hypothetical protein |
| *FLPKDCKG_01334* | *gsiA-3* | - | 1459215 | 1461083 | 1.22 | 1.44 | 2.20 | 1.35 | Glutathione import ATP-binding protein GsiA |
| *FLPKDCKG_01335* | *gsiD-3* | - | 1461089 | 1461967 | 1.60 | 3.72 | 0.67 | 0.14 | Glutathione transport system permease protein GsiD |
| *FLPKDCKG_01336* | *dppB-3* | - | 1461964 | 1462908 | 1.16 | 4.87 | 1.48 | 0.09 | Dipeptide transport system permease protein DppB |
| *FLPKDCKG_01337* | *dppA-1* | - | 1463038 | 1464552 | 5.99 | 10.51 | 9.03 | 2.99 | Periplasmic dipeptide transport protein |
| *FLPKDCKG_01338* | *ywrD-1* | - | 1464626 | 1466254 | 17.00 | 40.61 | 23.27 | 5.48 | Putative gamma-glutamyltransferase YwrD |
| *FLPKDCKG_01339* | *yofA-3* | + | 1466360 | 1467262 | 1.52 | 0.82 | 1.92 | 0.64 | HTH-type transcriptional regulator YofA |
| *FLPKDCKG_01340* | *plcN-1* | - | 1467366 | 1469546 | 0.24 | 1.00 | 0.22 | 0.00 | Non-hemolytic phospholipase C |
| *FLPKDCKG_01341* | | - | 1469583 | 1469762 | 1.20 | 1.96 | 0.00 | 0.00 | Hypothetical protein |
| *FLPKDCKG_01342* | | - | 1469783 | 1471777 | 0.05 | 0.22 | 0.12 | 0.29 | Hypothetical protein |
| *FLPKDCKG_01343* | | + | 1472112 | 1472468 | 0.24 | 0.00 | 0.00 | 0.00 | Hypothetical protein |
| *FLPKDCKG_01344* | *xcpT-2* | + | 1472544 | 1472972 | 0.09 | 0.00 | 0.08 | 0.15 | Type II secretion system protein G |
| *FLPKDCKG_01345* | *gspH* | + | 1472944 | 1473432 | 0.70 | 1.13 | 0.00 | 0.00 | Putative type II secretion system protein H |
| *FLPKDCKG_01346* | *xcpV-2* | + | 1473419 | 1473823 | 0.00 | 0.00 | 1.80 | 0.00 | Type II secretion system protein I |
| *FLPKDCKG_01347* | *xcpW* | + | 1473820 | 1474434 | 0.97 | 2.04 | 2.84 | 0.38 | Type II secretion system protein J |
| *FLPKDCKG_01348* | | + | 1474431 | 1475375 | 0.36 | 0.58 | 1.40 | 0.00 | Hypothetical protein |
| *FLPKDCKG_01349* | | + | 1475372 | 1476364 | 0.09 | 0.00 | 0.47 | 0.00 | Hypothetical protein |
| *FLPKDCKG_01350* | | + | 1476453 | 1476947 | 0.00 | 0.00 | 0.00 | 0.00 | Hypothetical protein |
| *FLPKDCKG_01351* | | + | 1476947 | 1477702 | 0.00 | 0.75 | 0.00 | 0.00 | Hypothetical protein |
| *FLPKDCKG_01352* | *cirA-2* | + | 1478147 | 1480300 | 0.62 | 0.91 | 1.35 | 0.10 | Colicin I receptor |
| *FLPKDCKG_01353* | *stoA-2* | + | 1480300 | 1480821 | 0.00 | 0.90 | 2.36 | 0.00 | Sporulation thiol-disulfide oxidoreductase A |
| *FLPKDCKG_01354* | *hxcR-2* | - | 1480852 | 1482264 | 0.13 | 0.09 | 0.14 | 0.00 | putative type II secretion system protein HxcR |
| *FLPKDCKG_01355* | *gspD* | - | 1482279 | 1484891 | 0.19 | 0.59 | 0.19 | 0.54 | Putative type II secretion system protein D |
| *FLPKDCKG_01356* | *epsF-2* | + | 1485373 | 1486269 | 0.23 | 2.11 | 0.45 | 0.00 | Type II secretion system protein F |
| *FLPKDCKG_01357* | *glnB* | - | 1486611 | 1486949 | 228.85 | 185.92 | 149.50 | 50.71 | Nitrogen regulatory protein P-II 1 |
| *FLPKDCKG_01358* | *nadE* | - | 1487088 | 1489193 | 9.33 | 14.61 | 10.47 | 2.22 | Glutamine-dependent NAD(+) synthetase |
| *FLPKDCKG_01359* | *ybdG* | - | 1489248 | 1490585 | 5.00 | 6.11 | 10.35 | 3.30 | Miniconductance mechanosensitive channel YbdG |
| *FLPKDCKG_01360* | *argH-3* | + | 1490646 | 1492214 | 21.64 | 25.23 | 20.99 | 10.41 | Argininosuccinate lyase |
| *FLPKDCKG_01361* | *ppsA-2* | + | 1492389 | 1494428 | 2.39 | 2.56 | 5.27 | 0.74 | Phosphoenolpyruvate synthase |
| *FLPKDCKG_01362* | *lpdV* | - | 1494603 | 1496006 | 8.35 | 7.62 | 4.55 | 1.14 | Dihydrolipoyl dehydrogenase |
| *FLPKDCKG_01363* | *bkdB* | - | 1496009 | 1497364 | 13.11 | 10.34 | 6.18 | 2.39 | Lipoamide acyltransferase component of branched-chain alpha-keto acid dehydrogenase complex |
| *FLPKDCKG_01364* | *bkdA2* | - | 1497366 | 1498409 | 18.42 | 13.70 | 5.23 | 1.92 | 2-oxoisovalerate dehydrogenase subunit beta |
| *FLPKDCKG_01365* | *bkdA1* | - | 1498413 | 1499645 | 28.82 | 20.67 | 16.32 | 5.43 | 2-oxoisovalerate dehydrogenase subunit alpha |
| *FLPKDCKG_01366* | *fhuA-8* | - | 1499887 | 1502268 | 0.36 | 1.02 | 0.98 | 0.23 | Ferrichrome-iron receptor |
| *FLPKDCKG_01367* | | - | 1502330 | 1503301 | 0.76 | 0.00 | 0.09 | 0.28 | Hypothetical protein |
| *FLPKDCKG_01368* | *fecI-3* | - | 1503298 | 1503810 | 2.67 | 1.91 | 2.39 | 1.32 | putative RNA polymerase sigma factor FecI |
| *FLPKDCKG_01369* | | + | 1503979 | 1504305 | 5.03 | 14.93 | 10.57 | 6.34 | Hypothetical protein |
| *FLPKDCKG_01370* | *ydjA* | - | 1504429 | 1505022 | 17.92 | 20.32 | 18.20 | 6.13 | Putative NAD(P)H nitroreductase YdjA |
| *FLPKDCKG_01371* | *merR1* | - | 1505148 | 1505582 | 45.83 | 71.04 | 41.26 | 9.98 | Mercuric resistance operon regulatory protein |
| *FLPKDCKG_01372* | | - | 1505686 | 1505916 | 109.80 | 158.85 | 62.79 | 17.36 | Hypothetical protein |
| *FLPKDCKG_01373* | *cadA-1* | + | 1505965 | 1508121 | 7.90 | 12.53 | 5.07 | 2.06 | putative cadmium-transporting ATPase |
| *FLPKDCKG_01374* | *speG-2* | - | 1508292 | 1508876 | 32.24 | 56.32 | 29.72 | 7.56 | Spermidine N(1)-acetyltransferase |
| *FLPKDCKG_01375* | *cmpR-6* | - | 1509104 | 1509748 | 1.03 | 2.23 | 2.75 | 0.00 | HTH-type transcriptional activator CmpR |
| *FLPKDCKG_01376* | *ttuB* | + | 1510239 | 1511501 | 0.91 | 4.86 | 1.03 | 0.32 | Putative tartrate transporter |

| Locus name | Gene name | Strand | Gene start (bp) | Gene end (bp) | Average TPM Inoculum | Average TPM $t_6$ | Average TPM $t_{24}$ | Average TPM $t_{168}$ | Annotation description |
|---|---|---|---|---|---|---|---|---|---|
| *FLPKDCKG_01377* | | + | 1511555 | 1513048 | 1.78 | 4.69 | 2.93 | 1.02 | Hypothetical protein |
| *FLPKDCKG_01378* | *ribC* | - | 1513101 | 1513805 | 18.59 | 25.90 | 24.99 | 9.50 | Riboflavin synthase |
| *FLPKDCKG_01379* | *puuC-1* | - | 1513917 | 1515350 | 49.52 | 70.45 | 54.52 | 26.47 | Aldehyde dehydrogenase PuuC |
| *FLPKDCKG_01380* | *lldR-1* | + | 1515680 | 1516483 | 4.73 | 8.48 | 4.53 | 2.24 | Putative L-lactate dehydrogenase operon regulatory protein |
| *FLPKDCKG_01381* | *ped* | + | 1516590 | 1517366 | 0.44 | 1.80 | 2.40 | 0.75 | (S)-1-Phenylethanol dehydrogenase |
| *FLPKDCKG_01382* | | - | 1517391 | 1518050 | 7.34 | 5.81 | 3.21 | 2.05 | Hypothetical protein |
| *FLPKDCKG_01383* | | - | 1518043 | 1518522 | 4.32 | 3.76 | 0.57 | 0.92 | Hypothetical protein |
| *FLPKDCKG_01384* | *xthA* | + | 1518717 | 1519499 | 24.91 | 20.38 | 10.43 | 3.69 | Exodeoxyribonuclease III |
| *FLPKDCKG_01385* | | + | 1519570 | 1519645 | 43.60 | 128.55 | 106.47 | 30.85 | tRNA |
| *FLPKDCKG_01386* | | - | 1519762 | 1520280 | 5.18 | 21.52 | 14.77 | 3.41 | Hypothetical protein |
| *FLPKDCKG_01387* | | - | 1520255 | 1520638 | 4.42 | 12.03 | 8.75 | 1.42 | Hypothetical protein |
| *FLPKDCKG_01388* | | - | 1520639 | 1521424 | 8.68 | 5.01 | 3.79 | 2.15 | Amino acid--[acyl-carrier-protein] ligase |
| *FLPKDCKG_01389* | | - | 1521482 | 1522438 | 8.54 | 6.33 | 3.24 | 0.85 | Hypothetical protein |
| *FLPKDCKG_01390* | | - | 1522453 | 1523346 | 12.82 | 5.45 | 4.09 | 1.89 | Hypothetical protein |
| *FLPKDCKG_01391* | *dltC-1* | - | 1523362 | 1523610 | 20.84 | 5.16 | 1.05 | 1.60 | D-alanine--poly(phosphoribitol) ligase subunit 2 |
| *FLPKDCKG_01392* | *aziB2-2* | - | 1523642 | 1524631 | 14.88 | 7.45 | 4.68 | 0.67 | 3-hydroxy-5-methyl-1-naphthoate 3-O-methyltransferase |
| *FLPKDCKG_01393* | | + | 1524973 | 1525737 | 4.11 | 3.80 | 2.86 | 0.31 | Hypothetical protein |
| *FLPKDCKG_01394* | | + | 1525861 | 1526562 | 7.17 | 8.83 | 5.62 | 1.37 | Hypothetical protein |
| *FLPKDCKG_01395* | | - | 1526550 | 1526753 | 6.16 | 3.78 | 0.58 | 0.88 | Hypothetical protein |
| *FLPKDCKG_01396* | | - | 1526750 | 1527673 | 3.80 | 9.13 | 3.48 | 0.81 | Hypothetical protein |
| *FLPKDCKG_01397* | | - | 1527796 | 1528494 | 6.62 | 9.99 | 4.80 | 3.08 | Hypothetical protein |
| *FLPKDCKG_01398* | *lprI-1* | + | 1528678 | 1530069 | 2.71 | 2.78 | 2.18 | 1.38 | Lipoprotein LprI |
| *FLPKDCKG_01399* | *amiD-2* | - | 1530077 | 1531465 | 0.66 | 0.49 | 0.62 | 0.39 | Putative amidase AmiD |
| *FLPKDCKG_01400* | *exuT-2* | - | 1531462 | 1532802 | 1.21 | 3.24 | 0.22 | 0.93 | Hexuronate transporter |
| *FLPKDCKG_01401* | *purR* | + | 1532943 | 1533983 | 4.83 | 11.69 | 6.81 | 3.27 | HTH-type transcriptional repressor PurR |
| *FLPKDCKG_01402* | | + | 1534165 | 1534548 | 3.55 | 2.94 | 4.90 | 1.64 | Hypothetical protein |
| *FLPKDCKG_01403* | *comR-2* | + | 1534755 | 1535336 | 8.28 | 4.73 | 9.47 | 0.24 | HTH-type transcriptional repressor ComR |
| *FLPKDCKG_01404* | *nepI-1* | + | 1535434 | 1536642 | 0.51 | 1.67 | 0.54 | 0.36 | Purine ribonucleoside efflux pump NepI |
| *FLPKDCKG_01405* | *nepI-2* | - | 1536701 | 1537894 | 0.53 | 0.27 | 0.95 | 0.35 | Purine ribonucleoside efflux pump NepI |
| *FLPKDCKG_01406* | *dmlR-10* | + | 1538072 | 1539016 | 1.46 | 3.83 | 1.45 | 0.72 | HTH-type transcriptional regulator DmlR |
| *FLPKDCKG_01407* | | + | 1539151 | 1539951 | 8.19 | 16.56 | 7.83 | 4.88 | Hypothetical protein |
| *FLPKDCKG_01408* | *benM-2* | + | 1539972 | 1540985 | 0.87 | 2.68 | 3.23 | 0.14 | HTH-type transcriptional regulator BenM |
| *FLPKDCKG_01409* | *abo-1* | - | 1541025 | 1542212 | 0.61 | 0.97 | 1.28 | 0.51 | 4-methylaminobutanoate oxidase (formaldehyde-forming) |
| *FLPKDCKG_01410* | *gcvA-3* | - | 1542438 | 1543403 | 6.22 | 14.10 | 7.07 | 2.22 | Glycine cleavage system transcriptional activator |
| *FLPKDCKG_01411* | | + | 1543497 | 1545410 | 1.28 | 1.85 | 0.73 | 0.11 | Putative multidrug export ATP-binding/permease protein |
| *FLPKDCKG_01412* | *msbA-1* | + | 1546113 | 1547105 | 1.16 | 0.21 | 1.31 | 0.19 | Lipid A export ATP-binding/permease protein MsbA |
| *FLPKDCKG_01413* | *pac* | - | 1547071 | 1549659 | 1.62 | 1.42 | 1.07 | 0.32 | Penicillin G acylase |
| *FLPKDCKG_01414* | *cynR-4* | + | 1549809 | 1550771 | 3.55 | 3.29 | 1.83 | 1.58 | HTH-type transcriptional regulator CynR |
| *FLPKDCKG_01415* | | - | 1550758 | 1551102 | 1.33 | 5.76 | 1.69 | 1.75 | Hypothetical protein |
| *FLPKDCKG_01416* | | - | 1551177 | 1551569 | 3.39 | 4.30 | 5.26 | 1.07 | Hypothetical protein |
| *FLPKDCKG_01417* | | - | 1551572 | 1552135 | 3.18 | 3.58 | 0.51 | 0.61 | Hypothetical protein |
| *FLPKDCKG_01418* | *bacC-2* | - | 1552194 | 1552958 | 3.29 | 3.66 | 2.67 | 1.66 | Dihydroanticapsin 7-dehydrogenase |
| *FLPKDCKG_01419* | | - | 1553043 | 1554674 | 1.73 | 3.81 | 2.75 | 0.19 | Long-chain-fatty-acid--CoA ligase FadD13 |
| *FLPKDCKG_01420* | | - | 1554671 | 1555843 | 2.55 | 2.22 | 2.47 | 1.27 | putative acetyl-CoA acetyltransferase |
| *FLPKDCKG_01421* | *kipR-1* | - | 1555888 | 1556715 | 5.23 | 11.41 | 7.21 | 4.22 | HTH-type transcriptional regulator KipR |
| *FLPKDCKG_01422* | *siaP* | + | 1556951 | 1557964 | 2.27 | 5.11 | 4.08 | 0.78 | Sialic acid-binding periplasmic protein SiaP |
| *FLPKDCKG_01423* | *siaT-1* | + | 1557961 | 1559853 | 0.94 | 0.64 | 3.47 | 0.59 | Sialic acid TRAP transporter permease protein SiaT |
| *FLPKDCKG_01424* | *atm1-1* | + | 1560250 | 1562154 | 12.55 | 2.97 | 4.34 | 3.72 | ATM1-type heavy metal exporter |
| *FLPKDCKG_01425* | *rhtC-1* | - | 1562242 | 1562871 | 0.78 | 1.64 | 0.00 | 0.44 | Threonine efflux protein |
| *FLPKDCKG_01426* | *lrp-2* | + | 1562959 | 1563486 | 7.98 | 4.82 | 0.66 | 2.20 | Leucine-responsive regulatory protein |
| *FLPKDCKG_01427* | *inhA-1* | - | 1563443 | 1564102 | 2.52 | 3.90 | 4.67 | 2.54 | Isonitrile hydratase |
| *FLPKDCKG_01428* | *inhA-2* | + | 1564142 | 1565113 | 5.28 | 4.65 | 4.80 | 1.38 | Isonitrile hydratase |
| *FLPKDCKG_01429* | | + | 1565205 | 1565591 | 7.69 | 10.63 | 4.19 | 4.58 | Hypothetical protein |
| *FLPKDCKG_01430* | *fabG-4* | - | 1565633 | 1566355 | 0.25 | 0.88 | 2.96 | 0.00 | 3-oxoacyl-[acyl-carrier-protein] reductase FabG |
| *FLPKDCKG_01431* | | - | 1566352 | 1567356 | 0.72 | 0.35 | 3.30 | 0.00 | Hypothetical protein |
| *FLPKDCKG_01432* | | + | 1567515 | 1568549 | 4.65 | 3.90 | 6.76 | 1.96 | Hypothetical protein |
| *FLPKDCKG_01433* | *menE* | + | 1568554 | 1570161 | 1.78 | 3.10 | 3.57 | 0.53 | 2-succinylbenzoate--CoA ligase |
| *FLPKDCKG_01434* | *yydK* | + | 1570173 | 1570916 | 2.77 | 5.81 | 7.63 | 0.75 | putative HTH-type transcriptional regulator YydK |
| *FLPKDCKG_01435* | | + | 1571083 | 1572369 | 1.37 | 1.17 | 1.90 | 0.47 | Hypothetical protein |
| *FLPKDCKG_01436* | *nsaD-1* | - | 1572500 | 1573099 | 1.61 | 2.66 | 0.74 | 2.06 | 2-hydroxychromene-2-carboxylate isomerase |
| *FLPKDCKG_01437* | *yhjE-3* | - | 1573096 | 1574436 | 1.47 | 1.72 | 0.67 | 0.59 | Inner membrane metabolite transport protein YhjE |
| *FLPKDCKG_01438* | *novR-2* | - | 1574480 | 1575265 | 0.27 | 0.06 | 0.60 | 0.00 | Decarboxylase NovR |
| *FLPKDCKG_01439* | *cynR-5* | - | 1575428 | 1576303 | 1.70 | 1.68 | 1.88 | 0.55 | HTH-type transcriptional regulator CynR |
| *FLPKDCKG_01440* | *tetA-1* | - | 1576446 | 1577642 | 0.28 | 0.28 | 0.21 | 0.00 | Tetracycline resistance protein, class C |
| *FLPKDCKG_01441* | *tetR* | + | 1577724 | 1578323 | 13.41 | 2.52 | 7.96 | 1.05 | Tetracycline repressor protein class A from transposon 1721 |
| *FLPKDCKG_01442* | | + | 1578320 | 1578766 | 1.82 | 0.53 | 1.09 | 0.00 | Bifunctional NMN adenylyltransferase/Nudix hydrolase |
| *FLPKDCKG_01443* | | + | 1578814 | 1579623 | 3.40 | 4.81 | 4.33 | 1.18 | Hypothetical protein |
| *FLPKDCKG_01444* | *nimR-4* | - | 1579640 | 1580398 | 1.45 | 0.44 | 1.56 | 1.17 | HTH-type transcriptional regulator NimR |
| *FLPKDCKG_01445* | *fptA-1* | + | 1580537 | 1582696 | 0.15 | 0.20 | 0.40 | 0.14 | Fe(3+)-pyochelin receptor |

| Locus name | Gene name | Strand | Gene start (bp) | Gene end (bp) | Average TPM Inoculum | Average TPM $t_6$ | Average TPM $t_{24}$ | Average TPM $t_{168}$ | Annotation description |
|---|---|---|---|---|---|---|---|---|---|
| *FLPKDCKG_01446* | | + | 1582816 | 1584210 | 1.89 | 7.58 | 2.88 | 1.53 | Hypothetical protein |
| *FLPKDCKG_01447* | *pgrR-5* | - | 1584189 | 1585109 | 1.83 | 2.60 | 3.74 | 0.10 | HTH-type transcriptional regulator PgrR |
| *FLPKDCKG_01448* | *fabG-5* | + | 1585236 | 1585979 | 3.75 | 3.89 | 4.32 | 0.89 | 3-oxoacyl-[acyl-carrier-protein] reductase FabG |
| *FLPKDCKG_01449* | | + | 1586033 | 1586719 | 1.71 | 1.22 | 5.75 | 0.98 | Hypothetical protein |
| *FLPKDCKG_01450* | | - | 1586746 | 1587438 | 1.31 | 1.85 | 0.59 | 0.00 | Hypothetical protein |
| *FLPKDCKG_01451* | | - | 1587440 | 1588366 | 2.00 | 4.76 | 3.09 | 0.00 | Hypothetical protein |
| *FLPKDCKG_01452* | *crt-2* | + | 1588503 | 1589297 | 0.71 | 1.75 | 0.85 | 0.05 | Short-chain-enoyl-CoA hydratase |
| *FLPKDCKG_01453* | *uctC-2* | + | 1589294 | 1591465 | 0.55 | 0.99 | 1.80 | 0.91 | Acetyl-CoA:oxalate CoA-transferase |
| *FLPKDCKG_01454* | | + | 1591642 | 1592625 | 1.30 | 2.55 | 0.80 | 0.15 | Hypothetical protein |
| *FLPKDCKG_01455* | | - | 1592643 | 1592963 | 2.31 | 1.29 | 2.44 | 0.00 | Hypothetical protein |
| *FLPKDCKG_01456* | *bfrD-2* | - | 1593145 | 1595244 | 0.21 | 0.23 | 0.84 | 0.31 | putative TonB-dependent receptor BfrD |
| *FLPKDCKG_01457* | *paaF-1* | + | 1595773 | 1596564 | 1.22 | 4.95 | 5.57 | 2.11 | 2,3-dehydroadipyl-CoA hydratase |
| *FLPKDCKG_01458* | *gmr-1* | - | 1596586 | 1598583 | 56.09 | 79.25 | 46.35 | 26.41 | Cyclic di-GMP phosphodiesterase Gmr |
| *FLPKDCKG_01459* | | - | 1598897 | 1599892 | 1.15 | 0.97 | 0.58 | 0.19 | Hypothetical protein |
| *FLPKDCKG_01460* | *araC-1* | + | 1600113 | 1601888 | 2.14 | 4.34 | 3.38 | 2.30 | L-arabonate dehydratase |
| *FLPKDCKG_01461* | | + | 1601907 | 1602590 | 1.04 | 2.39 | 4.18 | 0.31 | Hypothetical protein |
| *FLPKDCKG_01462* | *gltC-6* | - | 1602610 | 1603539 | 2.40 | 1.87 | 3.38 | 1.36 | HTH-type transcriptional regulator GltC |
| *FLPKDCKG_01463* | *paaH-1* | + | 1603692 | 1604654 | 0.48 | 0.79 | 0.00 | 0.69 | 3-hydroxyadipyl-CoA dehydrogenase |
| *FLPKDCKG_01464* | *kce-1* | + | 1604651 | 1605505 | 1.01 | 0.00 | 0.94 | 0.00 | 3-keto-5-aminohexanoate cleavage enzyme |
| *FLPKDCKG_01465* | *ttrA* | - | 1605578 | 1608766 | 3.77 | 1.41 | 0.18 | 0.14 | Tetrathionate reductase subunit A |
| *FLPKDCKG_01466* | *ttrC* | - | 1608804 | 1609838 | 8.01 | 3.49 | 2.15 | 0.13 | Tetrathionate reductase subunit C |
| *FLPKDCKG_01467* | *ttrB* | - | 1609852 | 1610619 | 31.67 | 6.71 | 1.14 | 0.00 | Tetrathionate reductase subunit B |
| *FLPKDCKG_01468* | | - | 1610747 | 1611295 | 4.58 | 13.58 | 6.03 | 1.56 | Hypothetical protein |
| *FLPKDCKG_01469* | *yhhW-2* | + | 1611445 | 1612308 | 4.49 | 9.32 | 8.80 | 3.92 | Quercetin 2,3-dioxygenase |
| *FLPKDCKG_01470* | *fecI-4* | + | 1612620 | 1613168 | 0.97 | 0.00 | 0.00 | 0.34 | putative RNA polymerase sigma factor FecI |
| *FLPKDCKG_01471* | | + | 1613395 | 1614189 | 0.51 | 1.54 | 0.66 | 0.00 | Hypothetical protein |
| *FLPKDCKG_01472* | *fptA-2* | + | 1614291 | 1616786 | 0.50 | 0.51 | 1.15 | 0.37 | Fe(3+)-pyochelin receptor |
| *FLPKDCKG_01473* | | - | 1616832 | 1617008 | 1.14 | 3.78 | 2.85 | 1.19 | Hypothetical protein |
| *FLPKDCKG_01474* | *mdlB-2* | - | 1617215 | 1618372 | 0.26 | 0.20 | 0.53 | 0.50 | (S)-mandelate dehydrogenase |
| *FLPKDCKG_01475* | *gltR-2* | + | 1618539 | 1619438 | 1.45 | 1.95 | 0.81 | 0.95 | HTH-type transcriptional regulator GltR |
| *FLPKDCKG_01476* | *proA-3* | + | 1619570 | 1620238 | 0.62 | 1.78 | 3.23 | 0.67 | 4-hydroxy-4-methyl-2-oxoglutarate aldolase/4-carboxy-4-hydroxy-2-oxoadipate aldolase |
| *FLPKDCKG_01477* | *slcC-1* | + | 1620235 | 1621278 | 0.34 | 2.32 | 0.99 | 1.45 | (S)-sulfolactate dehydrogenase |
| *FLPKDCKG_01478* | | + | 1621356 | 1622135 | 0.41 | 2.31 | 1.74 | 0.72 | Hypothetical protein |
| *FLPKDCKG_01479* | *sadH* | - | 1622349 | 1623344 | 0.77 | 38.27 | 50.73 | 4.81 | Putative oxidoreductase SadH |
| *FLPKDCKG_01480* | *pchR-4* | + | 1624447 | 1624854 | 3.43 | 1.01 | 4.07 | 2.03 | Regulatory protein PchR |
| *FLPKDCKG_01481* | | - | 1624949 | 1625365 | 2.18 | 9.61 | 5.83 | 0.23 | Hypothetical protein |
| *FLPKDCKG_01482* | | - | 1625379 | 1625753 | 18.02 | 21.09 | 17.09 | 4.45 | Hypothetical protein |
| *FLPKDCKG_01483* | | - | 1625895 | 1626788 | 0.00 | 1.58 | 0.75 | 0.00 | Hypothetical protein |
| *FLPKDCKG_01484* | | + | 1626817 | 1628406 | 0.06 | 0.21 | 0.39 | 0.00 | Hypothetical protein |
| *FLPKDCKG_01485* | *ntaB* | - | 1628593 | 1629093 | 0.01 | 0.27 | 0.00 | 0.15 | FMN reductase (NADH) NtaB |
| *FLPKDCKG_01486* | *styA* | - | 1629093 | 1630334 | 0.10 | 0.03 | 0.00 | 0.21 | Styrene monooxygenase StyA |
| *FLPKDCKG_01487* | *bacC-3* | - | 1630403 | 1631191 | 0.23 | 1.39 | 0.00 | 0.00 | Dihydroanticapsin 7-dehydrogenase |
| *FLPKDCKG_01488* | *tam-1* | - | 1631557 | 1632186 | 10.10 | 13.31 | 9.81 | 2.84 | Trans-aconitate 2-methyltransferase |
| *FLPKDCKG_01489* | *pckG* | - | 1632317 | 1634173 | 24.82 | 43.60 | 36.65 | 15.88 | Phosphoenolpyruvate carboxykinase [GTP] |
| *FLPKDCKG_01490* | *acsA-1* | - | 1634347 | 1635969 | 4.47 | 2.91 | 2.53 | 0.97 | Acetyl-coenzyme A synthetase |
| *FLPKDCKG_01491* | *fadR-1* | + | 1636087 | 1636812 | 8.37 | 7.29 | 11.39 | 1.67 | Fatty acid metabolism regulator protein |
| *FLPKDCKG_01492* | *fadH-1* | - | 1636776 | 1637687 | 0.56 | 1.78 | 0.50 | 0.83 | putative 2,4-dienoyl-CoA reductase |
| *FLPKDCKG_01493* | *echA8-3* | + | 1637827 | 1638618 | 0.75 | 0.45 | 0.37 | 0.45 | putative enoyl-CoA hydratase echA8 |
| *FLPKDCKG_01494* | *mdtK* | + | 1638740 | 1640134 | 0.69 | 1.09 | 0.72 | 0.56 | Multidrug resistance protein MdtK |
| *FLPKDCKG_01495* | *amaB-1* | + | 1640161 | 1641432 | 4.32 | 2.66 | 3.79 | 0.97 | N-carbamoyl-L-amino acid hydrolase |
| *FLPKDCKG_01496* | | + | 1641451 | 1641786 | 0.79 | 2.10 | 1.03 | 0.47 | Hypothetical protein |
| *FLPKDCKG_01497* | *yijE-2* | + | 1641783 | 1642739 | 0.60 | 2.43 | 0.25 | 2.95 | putative cystine transporter YijE |
| *FLPKDCKG_01498* | *yijE-3* | - | 1642836 | 1643720 | 0.73 | 3.70 | 0.00 | 0.81 | putative cystine transporter YijE |
| *FLPKDCKG_01499* | *ydaF-3* | - | 1643711 | 1644445 | 2.53 | 2.51 | 1.61 | 0.70 | Putative ribosomal N-acetyltransferase YdaF |
| *FLPKDCKG_01500* | *gabR-2* | + | 1644597 | 1646123 | 4.12 | 4.10 | 4.22 | 0.48 | HTH-type transcriptional regulatory protein GabR |
| *FLPKDCKG_01501* | *rhtB-1* | - | 1646135 | 1646767 | 7.55 | 4.06 | 5.58 | 2.32 | Homoserine/homoserine lactone efflux protein |
| *FLPKDCKG_01502* | | + | 1647239 | 1648075 | 22.97 | 14.30 | 8.72 | 5.15 | Hypothetical protein |
| *FLPKDCKG_01503* | *garB* | - | 1648155 | 1649513 | 11.92 | 10.30 | 7.95 | 4.60 | Glutathione amide reductase |
| *FLPKDCKG_01504* | *aceE-2* | - | 1649727 | 1652387 | 1.05 | 1.20 | 1.69 | 0.18 | Pyruvate dehydrogenase E1 component |
| *FLPKDCKG_01505* | | - | 1652661 | 1652987 | 1.83 | 0.81 | 0.07 | 0.00 | Cytochrome c-555 |
| *FLPKDCKG_01506* | *nirK* | - | 1653001 | 1654083 | 12.06 | 2.63 | 3.64 | 0.00 | Copper-containing nitrite reductase |
| *FLPKDCKG_01507* | *ntcA* | + | 1654265 | 1655032 | 7.54 | 2.63 | 1.43 | 0.17 | Global nitrogen regulator |
| *FLPKDCKG_01508* | | + | 1655210 | 1655932 | 62.57 | 71.22 | 40.12 | 4.06 | Hypothetical protein |
| *FLPKDCKG_01509* | *pkn1-2* | - | 1656421 | 1657248 | 2.89 | 2.08 | 2.58 | 1.03 | Serine/threonine-protein kinase pkn1 |
| *FLPKDCKG_01510* | *norB* | - | 1657424 | 1659709 | 0.50 | 0.48 | 1.14 | 0.11 | Nitric oxide reductase subunit B |
| *FLPKDCKG_01511* | *mhqR-1* | + | 1659935 | 1660390 | 5.54 | 10.41 | 13.39 | 1.45 | HTH-type transcriptional regulator MhqR |
| *FLPKDCKG_01512* | *racX* | + | 1660412 | 1661140 | 3.43 | 7.58 | 1.88 | 2.40 | putative amino-acid racemase |
| *FLPKDCKG_01513* | *glxR-1* | - | 1661154 | 1661864 | 5.81 | 2.96 | 3.51 | 1.09 | CRP-like cAMP-activated global transcriptional regulator |
| *FLPKDCKG_01514* | | + | 1662060 | 1663697 | 26.25 | 9.42 | 8.29 | 2.58 | Long-chain-fatty-acid--CoA ligase |

| Locus name | Gene name | Strand | Gene start (bp) | Gene end (bp) | Average TPM Inoculum | Average TPM $t_6$ | Average TPM $t_{24}$ | Average TPM $t_{168}$ | Annotation description |
|---|---|---|---|---|---|---|---|---|---|
| *FLPKDCKG_01515* | *dmdC-1* | + | 1663732 | 1665501 | 21.47 | 7.95 | 8.35 | 2.62 | 3-methylmercaptopropionyl-CoA dehydrogenase |
| *FLPKDCKG_01516* | *ssuA-3* | + | 1665719 | 1666732 | 1.49 | 39.93 | 84.36 | 20.11 | Putative aliphatic sulfonates-binding protein |
| *FLPKDCKG_01517* | *ssuC-6* | + | 1666729 | 1667547 | 0.40 | 2.74 | 9.99 | 4.21 | Putative aliphatic sulfonates transport permease protein SsuC |
| *FLPKDCKG_01518* | *msbA-2* | + | 1667535 | 1667789 | 0.00 | 2.96 | 5.35 | 0.00 | Lipid A export ATP-binding/permease protein MsbA |
| *FLPKDCKG_01519* | *tauB-4* | + | 1667789 | 1668274 | 0.72 | 9.54 | 15.37 | 1.70 | Taurine import ATP-binding protein TauB |
| *FLPKDCKG_01520* | | - | 1668296 | 1669477 | 0.12 | 4.32 | 8.33 | 0.50 | putative transporter |
| *FLPKDCKG_01521* | | + | 1669770 | 1670252 | 3.56 | 7.95 | 3.03 | 0.00 | putative HTH-type transcriptional regulator |
| *FLPKDCKG_01522* | | - | 1670325 | 1671368 | 0.85 | 1.46 | 0.50 | 0.55 | Hypothetical protein |
| *FLPKDCKG_01523* | *yfcG-3* | - | 1671399 | 1672028 | 2.42 | 3.22 | 1.48 | 0.95 | Disulfide-bond oxidoreductase YfcG |
| *FLPKDCKG_01524* | *dmlR-11* | - | 1672130 | 1673035 | 1.00 | 1.23 | 2.99 | 0.51 | HTH-type transcriptional regulator DmlR |
| *FLPKDCKG_01525* | *yofA-4* | - | 1673019 | 1673972 | 3.04 | 1.38 | 2.31 | 0.85 | HTH-type transcriptional regulator YofA |
| *FLPKDCKG_01526* | | + | 1674067 | 1675401 | 0.40 | 0.48 | 0.36 | 0.16 | putative MFS-type transporter |
| *FLPKDCKG_01527* | *gcvA-4* | - | 1675439 | 1676323 | 3.28 | 1.89 | 2.03 | 1.08 | Glycine cleavage system transcriptional activator |
| *FLPKDCKG_01528* | *ywqN* | + | 1676449 | 1677111 | 0.32 | 0.22 | 0.00 | 0.31 | Putative NAD(P)H-dependent FMN-containing oxidoreductase YwqN |
| *FLPKDCKG_01529* | | - | 1677020 | 1677958 | 0.42 | 0.16 | 1.12 | 0.45 | Hypothetical protein |
| *FLPKDCKG_01530* | *virF* | - | 1678079 | 1678957 | 5.06 | 4.56 | 5.94 | 1.20 | Virulence regulon transcriptional activator VirF |
| *FLPKDCKG_01531* | *putB* | - | 1679021 | 1679980 | 0.23 | 0.29 | 0.26 | 0.00 | Proline dehydrogenase 2 |
| *FLPKDCKG_01532* | | - | 1680061 | 1680975 | 0.41 | 1.17 | 0.27 | 0.32 | Hypothetical protein |
| *FLPKDCKG_01533* | *dadA-2* | - | 1681117 | 1682364 | 2.83 | 10.31 | 5.14 | 1.05 | D-amino acid dehydrogenase |
| *FLPKDCKG_01534* | *nsrR* | - | 1682546 | 1682983 | 5.13 | 13.22 | 10.60 | 3.74 | HTH-type transcriptional repressor NsrR |
| *FLPKDCKG_01535* | *hmp* | - | 1683003 | 1684220 | 2.49 | 4.25 | 2.19 | 1.89 | Flavohemoprotein |
| *FLPKDCKG_01536* | *sfp* | - | 1684320 | 1685018 | 1.37 | 3.64 | 4.97 | 3.48 | 4'-phosphopantetheinyl transferase sfp |
| *FLPKDCKG_01537* | | + | 1685142 | 1685840 | 11.53 | 30.44 | 18.17 | 8.60 | Hypothetical protein |
| *FLPKDCKG_01538* | | + | 1685837 | 1686721 | 1.35 | 3.63 | 0.44 | 0.64 | Hypothetical protein |
| *FLPKDCKG_01539* | *acpP-1* | + | 1686702 | 1686962 | 2.33 | 11.06 | 5.85 | 9.79 | Acyl carrier protein |
| *FLPKDCKG_01540* | *acpM* | + | 1686979 | 1687239 | 1.76 | 0.79 | 3.64 | 2.02 | Meromycolate extension acyl carrier protein |
| *FLPKDCKG_01541* | *yciB-1* | + | 1687236 | 1687787 | 0.28 | 0.71 | 2.32 | 1.23 | intracellular septation protein A |
| *FLPKDCKG_01542* | *lgrA* | + | 1687791 | 1689485 | 0.29 | 1.67 | 1.88 | 0.77 | Linear gramicidin synthase subunit A |
| *FLPKDCKG_01543* | | + | 1689472 | 1690224 | 0.08 | 1.77 | 1.80 | 0.89 | Undecaprenyl-phosphate mannosyltransferase |
| *FLPKDCKG_01544* | | + | 1690221 | 1691144 | 0.21 | 2.77 | 1.32 | 0.97 | Hypothetical protein |
| *FLPKDCKG_01545* | | + | 1691178 | 1691774 | 0.23 | 3.57 | 1.33 | 0.39 | Hypothetical protein |
| *FLPKDCKG_01546* | | + | 1691758 | 1694097 | 0.52 | 1.04 | 1.16 | 0.29 | Hypothetical protein |
| *FLPKDCKG_01547* | | + | 1694255 | 1694815 | 0.86 | 2.60 | 0.35 | 0.40 | Hypothetical protein |
| *FLPKDCKG_01548* | *kasA* | + | 1694812 | 1696014 | 1.39 | 3.36 | 1.15 | 0.38 | 3-oxoacyl-[acyl-carrier-protein] synthase 1 |
| *FLPKDCKG_01549* | | + | 1696011 | 1696466 | 0.52 | 3.40 | 1.14 | 0.78 | Hypothetical protein |
| *FLPKDCKG_01550* | *fabG-6* | + | 1696463 | 1697194 | 0.75 | 3.60 | 4.50 | 1.74 | 3-oxoacyl-[acyl-carrier-protein] reductase FabG |
| *FLPKDCKG_01551* | *fabF-1* | + | 1697194 | 1698417 | 1.39 | 3.73 | 5.10 | 2.10 | 3-oxoacyl-[acyl-carrier-protein] synthase 2 |
| *FLPKDCKG_01552* | | + | 1698511 | 1698972 | 67.75 | 74.49 | 60.46 | 16.94 | Hypothetical protein |
| *FLPKDCKG_01553* | *dinB* | - | 1698975 | 1700057 | 3.61 | 9.01 | 8.50 | 1.40 | DNA polymerase IV |
| *FLPKDCKG_01554* | | + | 1700228 | 1700653 | 5.19 | 8.99 | 6.81 | 1.17 | Hypothetical protein |
| *FLPKDCKG_01555* | | - | 1700714 | 1701169 | 48.63 | 32.47 | 31.94 | 6.95 | Hypothetical protein |
| *FLPKDCKG_01556* | *dapb3* | + | 1701321 | 1703240 | 32.92 | 43.00 | 26.54 | 5.62 | Dipeptidyl aminopeptidase BIII |
| *FLPKDCKG_01557* | *adrA-1* | - | 1703230 | 1704459 | 3.62 | 4.27 | 3.49 | 1.05 | putative diguanylate cyclase AdrA |
| *FLPKDCKG_01558* | *pgaA* | + | 1705065 | 1707527 | 1.27 | 0.46 | 1.54 | 0.00 | Poly-beta-1,6-N-acetyl-D-glucosamine export protein |
| *FLPKDCKG_01559* | *pgaB* | + | 1707535 | 1709568 | 1.37 | 2.09 | 0.52 | 0.73 | Poly-beta-1,6-N-acetyl-D-glucosamine N-deacetylase |
| *FLPKDCKG_01560* | *pgaC* | + | 1709565 | 1710902 | 0.57 | 1.06 | 1.50 | 0.49 | Poly-beta-1,6-N-acetyl-D-glucosamine synthase |
| *FLPKDCKG_01561* | *pgaD* | + | 1710899 | 1711360 | 0.09 | 0.00 | 0.00 | 0.00 | Biofilm PGA synthesis protein PgaD |
| *FLPKDCKG_01562* | *fsr-1* | - | 1711410 | 1712660 | 1.50 | 2.87 | 1.58 | 0.84 | Fosmidomycin resistance protein |
| *FLPKDCKG_01563* | *nimR-5* | + | 1712826 | 1713632 | 4.17 | 1.10 | 2.08 | 0.96 | HTH-type transcriptional regulator NimR |
| *FLPKDCKG_01564* | *mrcA-1* | + | 1713848 | 1716544 | 5.24 | 20.68 | 15.91 | 6.75 | Penicillin-binding protein 1A |
| *FLPKDCKG_01565* | *pphA-1* | - | 1716663 | 1717382 | 10.79 | 40.65 | 20.14 | 7.65 | Serine/threonine-protein phosphatase 1 |
| *FLPKDCKG_01566* | | + | 1717568 | 1718743 | 8.82 | 5.01 | 11.13 | 5.60 | Hypothetical protein |
| *FLPKDCKG_01567* | | + | 1718740 | 1718940 | 9.54 | 11.80 | 12.14 | 7.18 | Hypothetical protein |
| *FLPKDCKG_01568* | *dkgA* | - | 1719014 | 1719841 | 41.91 | 29.60 | 36.70 | 15.08 | 2,5-diketo-D-gluconic acid reductase A |
| *FLPKDCKG_01569* | *atm1-2* | + | 1720137 | 1721939 | 2.27 | 2.59 | 1.77 | 0.76 | ATM1-type heavy metal exporter |
| *FLPKDCKG_01570* | | + | 1721992 | 1722378 | 2.10 | 1.24 | 5.33 | 1.33 | Hypothetical protein |
| *FLPKDCKG_01571* | | - | 1722383 | 1723369 | 0.24 | 0.38 | 1.03 | 0.42 | Hypothetical protein |
| *FLPKDCKG_01572* | *gltC-7* | + | 1723478 | 1724320 | 1.36 | 2.60 | 2.56 | 0.39 | HTH-type transcriptional regulator GltC |
| *FLPKDCKG_01573* | | - | 1724363 | 1724893 | 0.16 | 0.99 | 1.41 | 1.32 | Hypothetical protein |
| *FLPKDCKG_01574* | | - | 1724971 | 1726461 | 0.03 | 0.43 | 2.50 | 0.38 | Hypothetical protein |
| *FLPKDCKG_01575* | *tauB-5* | - | 1726455 | 1727240 | 0.72 | 0.99 | 3.31 | 2.96 | Taurine import ATP-binding protein TauB |
| *FLPKDCKG_01576* | *cmpB-2* | - | 1727234 | 1728253 | 0.77 | 0.69 | 5.67 | 2.12 | Bicarbonate transport system permease protein CmpB |
| *FLPKDCKG_01577* | *cmpA* | - | 1728240 | 1729271 | 0.22 | 0.42 | 4.35 | 0.22 | Bicarbonate-binding protein CmpA |
| *FLPKDCKG_01578* | | - | 1729421 | 1729741 | 1.88 | 0.86 | 1.32 | 0.79 | Hypothetical protein |
| *FLPKDCKG_01579* | | - | 1729888 | 1730766 | 3.20 | 5.60 | 2.21 | 2.18 | Hypothetical protein |
| *FLPKDCKG_01580* | | - | 1730769 | 1731758 | 3.47 | 4.87 | 2.51 | 0.12 | Hypothetical protein |
| *FLPKDCKG_01581* | *gcvA-5* | + | 1731866 | 1732849 | 4.24 | 4.08 | 2.13 | 1.61 | Glycine cleavage system transcriptional activator |
| *FLPKDCKG_01582* | | - | 1732939 | 1733154 | 76.17 | 33.16 | 19.39 | 2.67 | Hypothetical protein |
| *FLPKDCKG_01583* | | - | 1733412 | 1734362 | 5.65 | 55.68 | 30.49 | 5.21 | Cellobiose 2-epimerase |

| Locus name | Gene name | Strand | Gene start (bp) | Gene end (bp) | Average TPM Inoculum | Average TPM $t_6$ | Average TPM $t_{24}$ | Average TPM $t_{168}$ | Annotation description |
|---|---|---|---|---|---|---|---|---|---|
| *FLPKDCKG_01584* | | - | 1734512 | 1734724 | 90.36 | 410.06 | 334.86 | 70.88 | Hypothetical protein |
| *FLPKDCKG_01585* | | - | 1735091 | 1735315 | 70.98 | 179.18 | 92.03 | 15.42 | Hypothetical protein |
| *FLPKDCKG_01586* | *yijE-4* | - | 1735358 | 1736263 | 0.85 | 1.90 | 2.92 | 0.58 | putative cystine transporter YijE |
| *FLPKDCKG_01587* | *lrp-3* | + | 1736424 | 1736894 | 1.80 | 2.79 | 3.67 | 0.87 | Leucine-responsive regulatory protein |
| *FLPKDCKG_01588* | | + | 1737006 | 1737239 | 142.92 | 102.83 | 54.31 | 8.52 | Hypothetical protein |
| *FLPKDCKG_01589* | | - | 1737324 | 1737917 | 115.11 | 924.30 | 496.76 | 64.63 | Hypothetical protein |
| *FLPKDCKG_01590* | | - | 1738164 | 1738418 | 3.95 | 8.50 | 10.93 | 1.86 | Hypothetical protein |
| *FLPKDCKG_01591* | | - | 1738638 | 1739519 | 0.68 | 3.10 | 4.97 | 0.10 | Hypothetical protein |
| *FLPKDCKG_01592* | | + | 1739833 | 1740651 | 2.66 | 3.05 | 3.13 | 0.91 | Hypothetical protein |
| *FLPKDCKG_01593* | | + | 1740661 | 1741614 | 2.41 | 1.59 | 3.56 | 1.77 | Hypothetical protein |
| *FLPKDCKG_01594* | *ypeA-1* | - | 1741704 | 1742138 | 2.49 | 4.97 | 0.00 | 0.55 | Acetyltransferase YpeA |
| *FLPKDCKG_01595* | *phzF-1* | - | 1742429 | 1743346 | 3.30 | 10.61 | 3.61 | 1.11 | Trans-2,3-dihydro-3-hydroxyanthranilate isomerase |
| *FLPKDCKG_01596* | | - | 1743504 | 1743809 | 1.54 | 7.59 | 4.05 | 2.67 | Hypothetical protein |
| *FLPKDCKG_01597* | | - | 1743879 | 1744181 | 3.77 | 16.01 | 7.38 | 2.52 | Hypothetical protein |
| *FLPKDCKG_01598* | | + | 1744512 | 1744670 | 3.08 | 15.59 | 13.49 | 3.85 | Hypothetical protein |
| *FLPKDCKG_01599* | | + | 1744670 | 1744906 | 2.53 | 5.67 | 6.74 | 3.33 | Hypothetical protein |
| *FLPKDCKG_01600* | | - | 1745028 | 1745360 | 45.29 | 40.15 | 19.32 | 4.50 | Hypothetical protein |
| *FLPKDCKG_01601* | | - | 1745742 | 1745818 | 2.07 | 0.00 | 4.56 | 0.00 | tRNA |
| *FLPKDCKG_01602* | *bluR* | - | 1746110 | 1746541 | 198.74 | 578.65 | 298.96 | 68.50 | HTH-type transcriptional repressor BluR |
| *FLPKDCKG_01603* | *ihfA* | - | 1746599 | 1746925 | 29.06 | 74.96 | 47.99 | 16.71 | Integration host factor subunit alpha |
| *FLPKDCKG_01604* | *pheT* | - | 1746941 | 1749358 | 4.19 | 11.20 | 13.39 | 4.18 | Phenylalanine--tRNA ligase beta subunit |
| *FLPKDCKG_01605* | *pheS* | - | 1749372 | 1750394 | 2.14 | 13.99 | 10.32 | 6.01 | Phenylalanine--tRNA ligase alpha subunit |
| *FLPKDCKG_01606* | *rplT* | - | 1750472 | 1750831 | 4.09 | 26.90 | 32.71 | 7.08 | 50S ribosomal protein L20 |
| *FLPKDCKG_01607* | *rpmI* | - | 1750849 | 1751046 | 35.96 | 194.14 | 263.01 | 97.44 | 50S ribosomal protein L35 |
| *FLPKDCKG_01608* | *comM* | - | 1751491 | 1752669 | 0.94 | 2.07 | 0.53 | 0.31 | Competence protein ComM |
| *FLPKDCKG_01609* | | - | 1753122 | 1753373 | 36.95 | 24.17 | 24.99 | 2.67 | Hypothetical protein |
| *FLPKDCKG_01610* | | - | 1753560 | 1753991 | 102.60 | 85.32 | 94.47 | 31.77 | Hypothetical protein |
| *FLPKDCKG_01611* | *glnK* | + | 1754537 | 1754875 | 3.06 | 689.87 | 467.07 | 130.75 | Nitrogen regulatory protein P-II 2 |
| *FLPKDCKG_01612* | *amtB* | + | 1754907 | 1756145 | 0.89 | 112.62 | 145.92 | 48.49 | Ammonia channel |
| *FLPKDCKG_01613* | *ptsI* | - | 1756372 | 1758147 | 4.28 | 15.90 | 8.65 | 3.74 | Phosphoenolpyruvate-protein phosphotransferase |
| *FLPKDCKG_01614* | *ptsH* | - | 1758191 | 1758460 | 33.70 | 62.33 | 29.94 | 14.41 | Phosphocarrier protein HPr |
| *FLPKDCKG_01615* | *manX* | - | 1758571 | 1758972 | 13.19 | 30.35 | 21.59 | 3.86 | PTS system mannose-specific EIIAB component |
| *FLPKDCKG_01616* | *gshB* | - | 1758990 | 1759967 | 36.23 | 37.81 | 30.40 | 7.48 | Glutathione synthetase |
| *FLPKDCKG_01617* | *infC* | - | 1760131 | 1760631 | 25.24 | 66.22 | 85.27 | 34.77 | Translation initiation factor IF-3 |
| *FLPKDCKG_01618* | *thrS* | - | 1760703 | 1762655 | 74.82 | 99.02 | 72.24 | 22.87 | Threonine--tRNA ligase |
| *FLPKDCKG_01619* | *rfbD* | - | 1762998 | 1763912 | 6.37 | 10.16 | 6.38 | 2.80 | dTDP-4-dehydrorhamnose reductase |
| *FLPKDCKG_01620* | *rcsB-1* | + | 1764206 | 1764940 | 1.69 | 3.16 | 1.45 | 2.29 | Transcriptional regulatory protein RcsB |
| *FLPKDCKG_01621* | *hemN-2* | - | 1764995 | 1766422 | 15.80 | 5.57 | 1.19 | 0.74 | Oxygen-independent coproporphyrinogen III oxidase |
| *FLPKDCKG_01622* | | + | 1766585 | 1766848 | 27.09 | 53.83 | 38.12 | 10.55 | Hypothetical protein |
| *FLPKDCKG_01623* | *rgtC* | - | 1766949 | 1768427 | 0.99 | 3.58 | 2.80 | 0.28 | Lipopolysaccharide core galacturonosyltransferase RgtC |
| *FLPKDCKG_01624* | *lldR-2* | + | 1768732 | 1769466 | 3.73 | 6.76 | 3.94 | 1.67 | Putative L-lactate dehydrogenase operon regulatory protein |
| *FLPKDCKG_01625* | *ilvG-1* | + | 1769564 | 1771279 | 4.08 | 4.73 | 1.87 | 2.03 | Acetolactate synthase isozyme 2 large subunit |
| *FLPKDCKG_01626* | *amaB-2* | + | 1771298 | 1772539 | 9.96 | 3.26 | 3.48 | 2.49 | N-carbamoyl-L-amino acid hydrolase |
| *FLPKDCKG_01627* | *puuC-2* | + | 1772554 | 1773744 | 5.55 | 2.15 | 2.04 | 1.39 | Aldehyde dehydrogenase PuuC |
| *FLPKDCKG_01628* | *puuC-3* | + | 1773741 | 1774052 | 7.93 | 3.94 | 5.48 | 0.98 | Aldehyde dehydrogenase PuuC |
| *FLPKDCKG_01629* | *hbpA-2* | + | 1774248 | 1775822 | 15.56 | 6.63 | 10.07 | 2.70 | Heme-binding protein A |
| *FLPKDCKG_01630* | *dppB-4* | + | 1775916 | 1776923 | 1.95 | 1.82 | 0.92 | 0.94 | Dipeptide transport system permease protein DppB |
| *FLPKDCKG_01631* | *dppC-2* | + | 1776920 | 1777759 | 2.40 | 1.92 | 1.40 | 2.31 | Dipeptide transport system permease protein DppC |
| *FLPKDCKG_01632* | *gsiA-4* | + | 1777769 | 1779403 | 7.72 | 4.67 | 4.32 | 0.60 | Glutathione import ATP-binding protein GsiA |
| *FLPKDCKG_01633* | *bauB* | + | 1779431 | 1779751 | 3.41 | 0.00 | 2.49 | 0.00 | Beta-alanine degradation protein BauB |
| *FLPKDCKG_01634* | *aphA-3* | + | 1779768 | 1780826 | 2.73 | 0.78 | 0.29 | 1.57 | Acetylpolyamine aminohydrolase |
| *FLPKDCKG_01635* | *ribN-3* | + | 1780835 | 1781731 | 1.03 | 2.15 | 0.71 | 0.43 | Riboflavin transporter |
| *FLPKDCKG_01636* | *gntR-1* | - | 1781744 | 1782868 | 4.62 | 12.26 | 5.54 | 4.11 | HTH-type transcriptional regulator GntR |
| *FLPKDCKG_01637* | *yiaM-2* | + | 1782902 | 1783444 | 13.47 | 22.27 | 15.37 | 5.80 | 2,3-diketo-L-gulonate TRAP transporter small permease protein YiaM |
| *FLPKDCKG_01638* | *siaM* | + | 1783481 | 1784761 | 6.63 | 16.18 | 7.74 | 1.64 | Sialic acid TRAP transporter large permease protein SiaM |
| *FLPKDCKG_01639* | *yiaO* | + | 1784821 | 1785837 | 29.83 | 69.57 | 28.91 | 10.06 | 2,3-diketo-L-gulonate-binding periplasmic protein YiaO |
| *FLPKDCKG_01640* | *araC-2* | + | 1785892 | 1787673 | 9.53 | 25.87 | 7.10 | 3.40 | L-arabonate dehydratase |
| *FLPKDCKG_01641* | *ybaN* | - | 1787769 | 1788227 | 2.85 | 12.72 | 3.00 | 2.78 | Inner membrane protein YbaN |
| *FLPKDCKG_01642* | | + | 1788290 | 1788499 | 6.56 | 4.44 | 8.01 | 9.50 | Hypothetical protein |
| *FLPKDCKG_01643* | | + | 1788679 | 1789212 | 0.79 | 3.67 | 2.73 | 0.00 | Hypothetical protein |
| *FLPKDCKG_01644* | | + | 1789406 | 1789720 | 6.11 | 10.41 | 9.48 | 0.88 | Hypothetical protein |
| *FLPKDCKG_01645* | | - | 1789728 | 1789937 | 67.24 | 93.95 | 30.31 | 10.07 | Hypothetical protein |
| *FLPKDCKG_01646* | | - | 1790059 | 1790652 | 3.89 | 31.89 | 18.60 | 4.32 | Hypothetical protein |
| *FLPKDCKG_01647* | *dhmA1* | + | 1790748 | 1791683 | 1.24 | 38.19 | 18.28 | 3.83 | Haloalkane dehalogenase 1 |
| *FLPKDCKG_01648* | | - | 1791649 | 1791867 | 1.58 | 209.84 | 73.16 | 4.23 | Hypothetical protein |
| *FLPKDCKG_01649* | | - | 1792451 | 1792527 | 55.03 | 195.19 | 152.80 | 31.42 | tRNA |
| *FLPKDCKG_01650* | *trpF* | - | 1792783 | 1793436 | 28.07 | 48.99 | 25.44 | 7.66 | N-(5'-phosphoribosyl)anthranilate isomerase |
| *FLPKDCKG_01651* | *takP-1* | - | 1793529 | 1794623 | 6.09 | 11.52 | 10.13 | 1.27 | Alpha-keto acid-binding periplasmic protein TakP |
| *FLPKDCKG_01652* | *truA* | - | 1794660 | 1795466 | 13.22 | 44.72 | 19.25 | 5.46 | tRNA pseudouridine synthase A |

| Locus name | Gene name | Strand | Gene start (bp) | Gene end (bp) | Average TPM Inoculum | Average TPM $t_6$ | Average TPM $t_{24}$ | Average TPM $t_{168}$ | Annotation description |
|---|---|---|---|---|---|---|---|---|---|
| *FLPKDCKG_01653* | | - | 1795485 | 1797353 | 15.87 | 19.42 | 23.81 | 9.05 | Hypothetical protein |
| *FLPKDCKG_01654* | *asd* | - | 1797617 | 1798744 | 36.04 | 50.96 | 40.20 | 22.32 | Aspartate-semialdehyde dehydrogenase |
| *FLPKDCKG_01655* | *leuB* | - | 1798975 | 1800051 | 15.65 | 23.25 | 24.63 | 8.38 | 3-isopropylmalate dehydrogenase |
| *FLPKDCKG_01656* | *leuD-1* | - | 1800089 | 1800739 | 38.83 | 49.90 | 49.61 | 12.80 | 3-isopropylmalate dehydratase small subunit |
| *FLPKDCKG_01657* | *leuC-1* | - | 1800762 | 1802165 | 34.73 | 49.23 | 37.95 | 11.92 | 3-isopropylmalate dehydratase large subunit |
| *FLPKDCKG_01658* | *bepC* | + | 1802680 | 1804194 | 42.32 | 7.44 | 6.24 | 5.98 | Outer membrane efflux protein BepC |
| *FLPKDCKG_01659* | | + | 1804218 | 1804430 | 28.50 | 9.90 | 7.52 | 4.65 | Hypothetical protein |
| *FLPKDCKG_01660* | *apxIB-1* | + | 1804414 | 1806579 | 9.94 | 2.60 | 1.62 | 1.54 | Toxin RTX-I translocation ATP-binding protein |
| *FLPKDCKG_01661* | *prsE-1* | + | 1806587 | 1808023 | 11.84 | 2.19 | 2.26 | 1.74 | Type I secretion system membrane fusion protein PrsE |
| *FLPKDCKG_01662* | *degU-2* | + | 1808043 | 1808699 | 4.48 | 1.53 | 1.95 | 2.05 | Transcriptional regulatory protein DegU |
| *FLPKDCKG_01663* | *rapA2* | + | 1808785 | 1819206 | 118.37 | 73.40 | 40.56 | 7.66 | Calcium-binding lectin RapA2 |
| *FLPKDCKG_01664* | *algE5* | + | 1819343 | 1819795 | 27.28 | 61.44 | 36.46 | 6.27 | Poly(beta-D-mannuronate) C5 epimerase 5 |
| *FLPKDCKG_01665* | | + | 1819930 | 1820613 | 8.73 | 20.29 | 11.83 | 0.71 | Hypothetical protein |
| *FLPKDCKG_01666* | *pdeB* | + | 1820631 | 1822613 | 4.36 | 7.85 | 6.38 | 1.99 | Cyclic di-GMP phosphodiesterase PdeB |
| *FLPKDCKG_01667* | *bepA-3* | - | 1822772 | 1823611 | 14.44 | 13.57 | 17.50 | 5.76 | Beta-barrel assembly-enhancing protease |
| *FLPKDCKG_01668* | *aroC* | - | 1823657 | 1824724 | 30.69 | 37.77 | 31.80 | 6.80 | Chorismate synthase |
| *FLPKDCKG_01669* | | - | 1824760 | 1825896 | 3.33 | 2.19 | 2.24 | 1.28 | putative signaling protein |
| *FLPKDCKG_01670* | *gmr-2* | + | 1826757 | 1828721 | 5.52 | 7.68 | 4.09 | 1.46 | Cyclic di-GMP phosphodiesterase Gmr |
| *FLPKDCKG_01671* | *kdpD-1* | + | 1828820 | 1830256 | 1.95 | 3.54 | 3.64 | 1.23 | Sensor protein KdpD |
| *FLPKDCKG_01672* | *torR* | - | 1830238 | 1830957 | 4.25 | 11.49 | 8.79 | 2.47 | TorCAD operon transcriptional regulatory protein TorR |
| *FLPKDCKG_01673* | | - | 1831172 | 1832659 | 10.48 | 21.47 | 8.23 | 3.83 | Hypothetical protein |
| *FLPKDCKG_01674* | | - | 1832656 | 1833027 | 14.68 | 25.96 | 11.45 | 5.35 | Hypothetical protein |
| *FLPKDCKG_01675* | | + | 1833045 | 1834358 | 23.59 | 34.18 | 18.15 | 5.63 | Hypothetical protein |
| *FLPKDCKG_01676* | *kdsD-1* | + | 1834391 | 1834849 | 138.19 | 136.82 | 88.24 | 28.43 | Arabinose 5-phosphate isomerase KdsD |
| *FLPKDCKG_01677* | *dusA* | - | 1834971 | 1835990 | 12.69 | 29.16 | 16.18 | 6.09 | tRNA-dihydrouridine(20/20a) synthase |
| *FLPKDCKG_01678* | *thrB-1* | + | 1836176 | 1837144 | 4.66 | 13.91 | 7.39 | 2.65 | Homoserine kinase |
| *FLPKDCKG_01679* | | + | 1837209 | 1838012 | 21.83 | 16.59 | 14.96 | 0.75 | Hypothetical protein |
| *FLPKDCKG_01680* | | - | 1838249 | 1839220 | 1.70 | 2.37 | 1.84 | 0.28 | Hypothetical protein |
| *FLPKDCKG_01681* | | - | 1839234 | 1840079 | 0.70 | 0.95 | 0.62 | 0.21 | Hypothetical protein |
| *FLPKDCKG_01682* | | - | 1840076 | 1841734 | 0.17 | 0.00 | 0.62 | 0.12 | Putative conjugal transfer protein |
| *FLPKDCKG_01683* | *outD-1* | - | 1841739 | 1843100 | 0.17 | 0.74 | 0.00 | 0.23 | Type II secretion system protein D |
| *FLPKDCKG_01684* | | - | 1843097 | 1843939 | 0.31 | 0.00 | 0.00 | 0.00 | Hypothetical protein |
| *FLPKDCKG_01685* | | - | 1843936 | 1844688 | 0.39 | 0.00 | 0.00 | 0.00 | Hypothetical protein |
| *FLPKDCKG_01686* | | - | 1845293 | 1846657 | 0.17 | 0.14 | 0.38 | 0.20 | Hypothetical protein |
| *FLPKDCKG_01687* | | - | 1846654 | 1846920 | 0.33 | 0.00 | 0.00 | 0.00 | Hypothetical protein |
| *FLPKDCKG_01688* | *oatA-1* | + | 1847337 | 1848512 | 3.90 | 4.60 | 5.52 | 0.97 | O-acetyltransferase OatA |
| *FLPKDCKG_01689* | *rfaQ-1* | + | 1848643 | 1849563 | 6.78 | 7.21 | 9.32 | 1.24 | Lipopolysaccharide core heptosyltransferase RfaQ |
| *FLPKDCKG_01690* | *mshA-1* | - | 1849671 | 1850813 | 23.50 | 4.12 | 7.45 | 1.92 | D-inositol 3-phosphate glycosyltransferase |
| *FLPKDCKG_01691* | | - | 1851544 | 1851690 | 2.70 | 10.28 | 1.18 | 5.56 | Hypothetical protein |
| *FLPKDCKG_01692* | | - | 1853003 | 1853081 | 4.45 | 0.77 | 7.41 | 4.75 | tRNA |
| *FLPKDCKG_01693* | *rpoD-1* | - | 1853089 | 1855356 | 46.81 | 96.20 | 85.90 | 32.30 | RNA polymerase sigma factor RpoD |
| *FLPKDCKG_01694* | | - | 1856113 | 1856817 | 9.84 | 31.47 | 33.34 | 6.33 | Hypothetical protein |
| *FLPKDCKG_01695* | *dnaG* | - | 1856768 | 1858105 | 2.14 | 18.38 | 12.15 | 4.55 | DNA primase |
| *FLPKDCKG_01696* | *rpsU* | - | 1858194 | 1858406 | 149.42 | 321.73 | 242.28 | 89.47 | 30S ribosomal protein S21 |
| *FLPKDCKG_01697* | *gpt* | - | 1858892 | 1859434 | 11.57 | 13.96 | 16.50 | 7.63 | Xanthine phosphoribosyltransferase |
| *FLPKDCKG_01698* | *purA* | - | 1859518 | 1860813 | 9.02 | 21.40 | 21.89 | 14.87 | Adenylosuccinate synthetase |
| *FLPKDCKG_01699* | *hisZ* | - | 1860890 | 1862047 | 12.24 | 33.15 | 24.01 | 9.28 | ATP phosphoribosyltransferase regulatory subunit |
| *FLPKDCKG_01700* | *hflC-1* | - | 1862307 | 1863209 | 19.41 | 23.78 | 27.26 | 10.27 | Modulator of FtsH protease HflC |
| *FLPKDCKG_01701* | *hflK-2* | - | 1863228 | 1864532 | 24.45 | 44.11 | 32.41 | 5.92 | Modulator of FtsH protease HflK |
| *FLPKDCKG_01702* | *hflX* | - | 1864498 | 1865604 | 8.46 | 28.98 | 17.61 | 6.13 | GTPase HflX |
| *FLPKDCKG_01703* | *hfq* | - | 1865689 | 1865925 | 376.10 | 538.15 | 360.41 | 123.29 | RNA-binding protein Hfq |
| *FLPKDCKG_01704* | *hisC-1* | - | 1866087 | 1867154 | 1.82 | 9.53 | 8.53 | 2.50 | Histidinol-phosphate aminotransferase |
| *FLPKDCKG_01705* | *der-1* | - | 1867158 | 1868513 | 3.58 | 12.60 | 10.29 | 5.46 | GTPase Der |
| *FLPKDCKG_01706* | *bamB* | - | 1868539 | 1869699 | 8.38 | 16.58 | 20.00 | 6.44 | Outer membrane protein assembly factor BamB |
| *FLPKDCKG_01707* | | - | 1869705 | 1870343 | 4.83 | 6.62 | 9.07 | 5.32 | Hypothetical protein |
| *FLPKDCKG_01708* | *hisS* | - | 1870345 | 1871640 | 2.40 | 18.49 | 6.75 | 3.02 | Histidine--tRNA ligase |
| *FLPKDCKG_01709* | *ispG* | - | 1871688 | 1872974 | 3.76 | 9.11 | 12.35 | 1.98 | 4-hydroxy-3-methylbut-2-en-1-yl diphosphate synthase (flavodoxin) |
| *FLPKDCKG_01710* | | - | 1872990 | 1873490 | 9.40 | 11.29 | 14.94 | 4.17 | Hypothetical protein |
| *FLPKDCKG_01711* | *rlmN-1* | - | 1873487 | 1874641 | 9.82 | 24.77 | 19.57 | 8.12 | Dual-specificity RNA methyltransferase RlmN |
| *FLPKDCKG_01712* | *ndk* | - | 1874682 | 1875107 | 20.92 | 75.91 | 44.45 | 27.10 | Nucleoside diphosphate kinase |
| *FLPKDCKG_01713* | *valS* | + | 1875520 | 1878414 | 11.74 | 22.65 | 12.30 | 4.59 | Valine--tRNA ligase |
| *FLPKDCKG_01714* | *puuB-2* | - | 1878488 | 1879777 | 3.75 | 1.03 | 0.80 | 0.63 | Gamma-glutamylputrescine oxidoreductase |
| *FLPKDCKG_01715* | *tcyC-2* | - | 1879787 | 1880518 | 2.40 | 1.81 | 1.61 | 0.96 | L-cystine import ATP-binding protein TcyC |
| *FLPKDCKG_01716* | *yecS-3* | - | 1880515 | 1881168 | 0.87 | 0.37 | 1.40 | 0.00 | L-cystine transport system permease protein YecS |
| *FLPKDCKG_01717* | *artJ* | - | 1881359 | 1882111 | 12.53 | 10.69 | 7.20 | 3.82 | ABC transporter arginine-binding protein 1 |
| *FLPKDCKG_01718* | *puuR-1* | - | 1882202 | 1882879 | 5.67 | 10.18 | 2.67 | 2.46 | HTH-type transcriptional regulator PuuR |
| *FLPKDCKG_01719* | *trmD* | - | 1883214 | 1884032 | 2.29 | 5.10 | 4.80 | 1.70 | tRNA (guanine-N(1)-)-methyltransferase |
| *FLPKDCKG_01720* | *rimM* | - | 1884032 | 1884652 | 5.24 | 25.42 | 10.25 | 5.02 | Ribosome maturation factor RimM |
| *FLPKDCKG_01721* | *rpsP* | - | 1884720 | 1884980 | 70.91 | 231.28 | 211.58 | 68.22 | 30S ribosomal protein S16 |

| Locus name | Gene name | Strand | Gene start (bp) | Gene end (bp) | Average TPM Inoculum | Average TPM $t_6$ | Average TPM $t_{24}$ | Average TPM $t_{168}$ | Annotation description |
|---|---|---|---|---|---|---|---|---|---|
| *FLPKDCKG_01722* | | + | 1885479 | 1886060 | 366.15 | 124.80 | 51.05 | 27.50 | Hypothetical protein |
| *FLPKDCKG_01723* | *tusA* | - | 1886138 | 1886392 | 4.30 | 5.95 | 7.40 | 1.11 | Sulfurtransferase TusA |
| *FLPKDCKG_01724* | *alaS-1* | - | 1886394 | 1889018 | 24.77 | 46.03 | 34.79 | 10.43 | Alanine--tRNA ligase |
| *FLPKDCKG_01725* | *ptlF* | - | 1889052 | 1889339 | 0.95 | 2.77 | 0.72 | 0.06 | Type IV secretion system protein PtlF |
| *FLPKDCKG_01726* | *ugpB-2* | - | 1889734 | 1891074 | 30.66 | 56.25 | 44.32 | 15.68 | sn-glycerol-3-phosphate-binding periplasmic protein UgpB |
| *FLPKDCKG_01727* | *dadA1-3* | - | 1891079 | 1892335 | 15.65 | 31.72 | 20.57 | 5.81 | D-amino acid dehydrogenase 1 |
| *FLPKDCKG_01728* | *ascD-1* | - | 1892529 | 1893575 | 4.27 | 6.40 | 4.60 | 4.05 | CDP-6-deoxy-L-threo-D-glycero-4-hexulose-3-dehydrase reductase |
| *FLPKDCKG_01729* | *panD* | - | 1893651 | 1894019 | 6.38 | 14.87 | 10.64 | 4.47 | Aspartate 1-decarboxylase |
| *FLPKDCKG_01730* | *sugE-2* | + | 1894287 | 1894604 | 0.68 | 5.42 | 6.42 | 2.41 | Quaternary ammonium compound-resistance protein SugE |
| *FLPKDCKG_01731* | | - | 1894653 | 1895525 | 2.90 | 12.04 | 4.48 | 1.53 | Hypothetical protein |
| *FLPKDCKG_01732* | *ectD-1* | - | 1895745 | 1896671 | 1.40 | 190.57 | 152.34 | 17.04 | Ectoine dioxygenase |
| *FLPKDCKG_01733* | *ectC* | - | 1896687 | 1897091 | 1.28 | 282.25 | 250.29 | 25.21 | L-ectoine synthase |
| *FLPKDCKG_01734* | *ectB* | - | 1897088 | 1898395 | 1.10 | 371.92 | 255.60 | 26.17 | Diaminobutyrate--2-oxoglutarate transaminase |
| *FLPKDCKG_01735* | *ectA* | - | 1898516 | 1899076 | 8.45 | 817.89 | 368.32 | 36.19 | L-2,4-diaminobutyric acid acetyltransferase |
| *FLPKDCKG_01736* | *yusO* | - | 1899073 | 1899621 | 40.91 | 459.44 | 283.40 | 47.18 | putative HTH-type transcriptional regulator YusO |
| *FLPKDCKG_01737* | *hexR-1* | + | 1899869 | 1900816 | 4.51 | 5.81 | 2.52 | 1.03 | HTH-type transcriptional regulator HexR |
| *FLPKDCKG_01738* | | + | 1900896 | 1902236 | 4.29 | 5.94 | 3.68 | 2.29 | putative aminotransferase |
| *FLPKDCKG_01739* | | - | 1902244 | 1903515 | 0.47 | 2.66 | 1.19 | 0.41 | Hypothetical protein |
| *FLPKDCKG_01740* | *fadR-2* | - | 1903512 | 1904135 | 2.05 | 6.37 | 3.26 | 1.66 | Fatty acid metabolism regulator protein |
| *FLPKDCKG_01741* | *parC* | - | 1904274 | 1906595 | 1.12 | 3.80 | 3.73 | 0.50 | DNA topoisomerase 4 subunit A |
| *FLPKDCKG_01742* | | - | 1906631 | 1907242 | 1.59 | 8.89 | 7.03 | 0.18 | Hypothetical protein |
| *FLPKDCKG_01743* | *parE* | - | 1907242 | 1909203 | 3.04 | 5.84 | 8.25 | 1.60 | DNA topoisomerase 4 subunit B |
| *FLPKDCKG_01744* | *trxA* | + | 1909533 | 1909859 | 204.75 | 328.95 | 249.73 | 101.99 | Thioredoxin 1 |
| *FLPKDCKG_01745* | *rho* | + | 1909977 | 1911236 | 12.53 | 25.50 | 26.85 | 11.62 | Transcription termination factor Rho |
| *FLPKDCKG_01746* | | - | 1911360 | 1914998 | 5.95 | 11.25 | 7.45 | 1.97 | Hypothetical protein |
| *FLPKDCKG_01747* | *glnE* | + | 1915082 | 1917847 | 6.62 | 10.53 | 6.45 | 2.57 | Bifunctional glutamine synthetase adenylyltransferase/adenylyl-removing enzyme |
| *FLPKDCKG_01748* | | + | 1917947 | 1918621 | 48.77 | 88.71 | 44.71 | 20.62 | Hypothetical protein |
| *FLPKDCKG_01749* | *nphR-1* | - | 1918658 | 1919629 | 17.64 | 16.45 | 17.39 | 8.39 | Transcriptional activator NphR |
| *FLPKDCKG_01750* | *cycA-1* | - | 1919779 | 1920351 | 0.54 | 0.38 | 0.00 | 0.20 | Cytochrome c-552 |
| *FLPKDCKG_01751* | *aauA* | - | 1920348 | 1920863 | 0.82 | 1.03 | 0.28 | 3.46 | Aralkylamine dehydrogenase light chain |
| *FLPKDCKG_01752* | *mauD* | - | 1920847 | 1921485 | 0.78 | 0.09 | 0.10 | 0.01 | Methylamine utilization protein MauD |
| *FLPKDCKG_01753* | | - | 1921506 | 1922063 | 0.79 | 0.00 | 0.00 | 1.21 | Hypothetical protein |
| *FLPKDCKG_01754* | *aauB* | - | 1922060 | 1923250 | 1.46 | 1.77 | 0.96 | 0.41 | Aralkylamine dehydrogenase heavy chain |
| *FLPKDCKG_01755* | | - | 1923282 | 1924163 | 0.83 | 2.35 | 1.09 | 0.00 | Hypothetical protein |
| *FLPKDCKG_01756* | *styD* | - | 1924232 | 1925740 | 2.29 | 3.41 | 2.56 | 0.32 | Phenylacetaldehyde dehydrogenase |
| *FLPKDCKG_01757* | *qorA-1* | - | 1925904 | 1926893 | 10.46 | 11.39 | 18.71 | 6.53 | Quinone oxidoreductase 1 |
| *FLPKDCKG_01758* | | - | 1926957 | 1927937 | 4.82 | 7.20 | 8.25 | 1.37 | Hypothetical protein |
| *FLPKDCKG_01759* | *yijE-5* | + | 1928029 | 1928919 | 1.42 | 2.68 | 5.32 | 0.26 | putative cystine transporter YijE |
| *FLPKDCKG_01760* | *marR-1* | + | 1928950 | 1929450 | 12.36 | 38.11 | 31.08 | 6.40 | Multiple antibiotic resistance protein MarR |
| *FLPKDCKG_01761* | *oprM-3* | + | 1929610 | 1930983 | 5.28 | 21.60 | 21.44 | 1.83 | Outer membrane protein OprM |
| *FLPKDCKG_01762* | *emrA* | + | 1931004 | 1932263 | 3.07 | 15.16 | 15.74 | 2.16 | Multidrug export protein EmrA |
| *FLPKDCKG_01763* | *emrB-3* | + | 1932276 | 1933895 | 0.87 | 6.26 | 4.78 | 0.72 | Multidrug export protein EmrB |
| *FLPKDCKG_01764* | | - | 1933953 | 1934909 | 2.01 | 2.25 | 6.47 | 1.51 | Hypothetical protein |
| *FLPKDCKG_01765* | *dmlA-1* | - | 1934936 | 1936000 | 13.48 | 18.55 | 23.02 | 4.01 | D-malate dehydrogenase [decarboxylating] |
| *FLPKDCKG_01766* | | - | 1936148 | 1937671 | 45.64 | 49.77 | 35.14 | 10.68 | Hypothetical protein |
| *FLPKDCKG_01767* | | + | 1937831 | 1938562 | 64.86 | 82.32 | 52.13 | 14.20 | putative transcriptional regulatory protein |
| *FLPKDCKG_01768* | *purD* | + | 1938742 | 1940049 | 8.23 | 23.96 | 18.22 | 4.55 | Phosphoribosylamine--glycine ligase |
| *FLPKDCKG_01769* | *hemF* | + | 1940062 | 1940973 | 2.34 | 10.01 | 8.65 | 4.07 | Oxygen-dependent coproporphyrinogen-III oxidase |
| *FLPKDCKG_01770* | *nadD* | + | 1940970 | 1941557 | 0.85 | 3.75 | 4.07 | 3.15 | putative nicotinate-nucleotide adenylyltransferase |
| *FLPKDCKG_01771* | *rsfS* | + | 1941631 | 1942035 | 101.52 | 119.83 | 65.59 | 14.06 | Ribosomal silencing factor RsfS |
| *FLPKDCKG_01772* | *rlmH* | + | 1942044 | 1942514 | 10.48 | 26.02 | 20.80 | 4.64 | Ribosomal RNA large subunit methyltransferase H |
| *FLPKDCKG_01773* | *yhdE* | + | 1942564 | 1943175 | 1.45 | 3.28 | 5.23 | 0.29 | Maf-like protein YhdE |
| *FLPKDCKG_01774* | *dctA-1* | - | 1943248 | 1944588 | 0.34 | 14.34 | 4.54 | 1.45 | C4-dicarboxylate transport protein |
| *FLPKDCKG_01775* | *dctB* | + | 1944860 | 1946812 | 3.10 | 5.31 | 4.30 | 0.84 | C4-dicarboxylate transport sensor protein DctB |
| *FLPKDCKG_01776* | *dctD* | + | 1946914 | 1948299 | 1.49 | 4.25 | 4.60 | 0.21 | C4-dicarboxylate transport transcriptional regulatory protein DctD |
| *FLPKDCKG_01777* | | - | 1948425 | 1948536 | 139.79 | 260.49 | 284.66 | 234.78 | rRNA |
| *FLPKDCKG_01778* | | - | 1948737 | 1951615 | 120070.92 | 116961.90 | 154023.16 | 209023.74 | rRNA |
| *FLPKDCKG_01779* | | - | 1951912 | 1951987 | 11.98 | 3.34 | 4.97 | 0.00 | tRNA |
| *FLPKDCKG_01780* | | - | 1952018 | 1952093 | 7.33 | 11.41 | 20.18 | 0.75 | tRNA |
| *FLPKDCKG_01781* | | - | 1952229 | 1953746 | 177440.48 | 142855.16 | 131570.81 | 126345.70 | rRNA |
| *FLPKDCKG_01782* | *rng* | + | 1954987 | 1956468 | 5.48 | 12.64 | 11.66 | 2.73 | Ribonuclease G |
| *FLPKDCKG_01783* | | + | 1956802 | 1957503 | 3.09 | 13.05 | 10.05 | 5.21 | Hypothetical protein |
| *FLPKDCKG_01784* | *cph2-3* | - | 1957592 | 1959064 | 5.62 | 6.96 | 2.82 | 1.86 | Phytochrome-like protein cph2 |
| *FLPKDCKG_01785* | | - | 1959331 | 1959558 | 61.78 | 7.15 | 4.13 | 4.61 | Hypothetical protein |
| *FLPKDCKG_01786* | *kefC-1* | - | 1959704 | 1961521 | 3.23 | 5.40 | 5.95 | 2.28 | Glutathione-regulated potassium-efflux system protein KefC |
| *FLPKDCKG_01787* | *msbA-3* | - | 1961572 | 1963245 | 7.67 | 10.01 | 9.18 | 1.84 | Lipid A export ATP-binding/permease protein MsbA |
| *FLPKDCKG_01788* | *rfaF-1* | + | 1963456 | 1964427 | 5.53 | 9.98 | 5.85 | 4.53 | ADP-heptose--LPS heptosyltransferase 2 |
| *FLPKDCKG_01789* | | + | 1964449 | 1965207 | 1.91 | 4.87 | 4.04 | 0.27 | Hypothetical protein |
| *FLPKDCKG_01790* | | + | 1965288 | 1965482 | 1.65 | 2.53 | 6.38 | 0.89 | Hypothetical protein |

| Locus name | Gene name | Strand | Gene start (bp) | Gene end (bp) | Average TPM Inoculum | Average TPM $t_6$ | Average TPM $t_{24}$ | Average TPM $t_{168}$ | Annotation description |
|---|---|---|---|---|---|---|---|---|---|
| *FLPKDCKG_01791* | *rfaG* | - | 1965533 | 1966681 | 2.89 | 5.78 | 1.93 | 0.64 | Lipopolysaccharide core biosynthesis protein RfaG |
| *FLPKDCKG_01792* | *icaB-1* | - | 1966656 | 1967363 | 30.67 | 23.16 | 22.98 | 2.22 | Poly-beta-1,6-N-acetyl-D-glucosamine N-deacetylase |
| *FLPKDCKG_01793* | *bshA-1* | - | 1967399 | 1968514 | 2.71 | 9.67 | 8.86 | 1.79 | N-acetyl-alpha-D-glucosaminyl L-malate synthase |
| *FLPKDCKG_01794* | *sunS* | - | 1968511 | 1969266 | 4.71 | 25.64 | 6.69 | 4.64 | SPBc2 prophage-derived glycosyltransferase SunS |
| *FLPKDCKG_01795* | *icaB-2* | - | 1969263 | 1970111 | 7.50 | 15.44 | 10.58 | 1.91 | Poly-beta-1,6-N-acetyl-D-glucosamine N-deacetylase |
| *FLPKDCKG_01796* | *epsD* | - | 1970108 | 1971199 | 6.44 | 15.06 | 13.73 | 1.91 | Putative glycosyltransferase EpsD |
| *FLPKDCKG_01797* | *dnaE-1* | - | 1971199 | 1974426 | 12.90 | 21.09 | 9.60 | 3.14 | DNA polymerase III subunit alpha |
| *FLPKDCKG_01798* | *dnaE-2* | - | 1974470 | 1974685 | 22.62 | 51.99 | 37.30 | 12.51 | DNA polymerase III subunit alpha |
| *FLPKDCKG_01799* | | + | 1974815 | 1975714 | 24.63 | 78.01 | 40.47 | 9.09 | Hypothetical protein |
| *FLPKDCKG_01800* | *cmpD-1* | - | 1975786 | 1977081 | 2.29 | 3.60 | 2.48 | 0.38 | Bicarbonate transport ATP-binding protein CmpD |
| *FLPKDCKG_01801* | *cmpB-3* | - | 1977090 | 1978823 | 1.59 | 1.58 | 1.71 | 0.82 | Bicarbonate transport system permease protein CmpB |
| *FLPKDCKG_01802* | | - | 1978994 | 1982935 | 2.28 | 5.11 | 3.96 | 0.74 | Hypothetical protein |
| *FLPKDCKG_01803* | *argA* | + | 1983072 | 1984430 | 8.83 | 11.94 | 7.93 | 2.58 | Amino-acid acetyltransferase |
| *FLPKDCKG_01804* | | + | 1984549 | 1984821 | 84.41 | 124.92 | 62.06 | 19.58 | putative Fe(2+)-trafficking protein |
| *FLPKDCKG_01805* | *plcN-2* | + | 1984957 | 1986909 | 7.47 | 10.22 | 5.32 | 0.76 | Non-hemolytic phospholipase C |
| *FLPKDCKG_01806* | *phoU* | - | 1986845 | 1987888 | 23.83 | 16.81 | 24.08 | 12.67 | Phosphate-specific transport system accessory protein PhoU |
| *FLPKDCKG_01807* | *rpiA* | - | 1988045 | 1988725 | 10.17 | 9.29 | 19.88 | 3.41 | Ribose-5-phosphate isomerase A |
| *FLPKDCKG_01808* | *aidB-1* | - | 1988762 | 1990426 | 11.35 | 5.24 | 6.18 | 2.13 | Putative acyl-CoA dehydrogenase AidB |
| *FLPKDCKG_01809* | *nylA* | - | 1990481 | 1992010 | 21.01 | 11.63 | 11.66 | 3.09 | 6-aminohexanoate-cyclic-dimer hydrolase |
| *FLPKDCKG_01810* | | - | 1992199 | 1993578 | 3.10 | 6.20 | 4.74 | 1.50 | Hypothetical protein |
| *FLPKDCKG_01811* | | + | 1993810 | 1994478 | 28.62 | 45.85 | 19.89 | 7.46 | Hypothetical protein |
| *FLPKDCKG_01812* | | - | 1994517 | 1995242 | 2.94 | 16.12 | 6.94 | 2.43 | Putative 3-methyladenine DNA glycosylase |
| *FLPKDCKG_01813* | *spxA* | - | 1995247 | 1995603 | 7.21 | 14.73 | 3.76 | 1.54 | Regulatory protein Spx |
| *FLPKDCKG_01814* | *ydcV-1* | - | 1995600 | 1996409 | 9.34 | 19.83 | 17.80 | 1.41 | Inner membrane ABC transporter permease protein YdcV |
| *FLPKDCKG_01815* | *potH-2* | - | 1996406 | 1997326 | 1.83 | 4.16 | 3.81 | 1.49 | Putrescine transport system permease protein PotH |
| *FLPKDCKG_01816* | *potA-2* | - | 1997323 | 1998450 | 2.59 | 4.71 | 3.40 | 1.77 | Spermidine/putrescine import ATP-binding protein PotA |
| *FLPKDCKG_01817* | *spuD* | - | 1998456 | 1999559 | 12.72 | 15.43 | 15.74 | 6.72 | Putrescine-binding periplasmic protein SpuD |
| *FLPKDCKG_01818* | *kdkA* | - | 1999710 | 2000375 | 6.26 | 11.16 | 9.93 | 3.61 | 3-deoxy-D-manno-octulosonic acid kinase |
| *FLPKDCKG_01819* | *glbO* | - | 2000431 | 2000853 | 6.24 | 14.71 | 5.10 | 1.83 | Group 2 truncated hemoglobin GlbO |
| *FLPKDCKG_01820* | | - | 2000853 | 2003384 | 4.44 | 5.70 | 4.57 | 3.11 | Hypothetical protein |
| *FLPKDCKG_01821* | | + | 2003510 | 2004484 | 22.25 | 19.23 | 16.42 | 8.75 | Solute-binding protein |
| *FLPKDCKG_01822* | *siaQ* | + | 2004596 | 2005096 | 1.25 | 0.06 | 2.49 | 0.00 | Sialic acid TRAP transporter small permease protein SiaQ |
| *FLPKDCKG_01823* | *dctM-4* | + | 2005130 | 2006407 | 0.83 | 3.43 | 1.83 | 1.45 | C4-dicarboxylate TRAP transporter large permease protein DctM |
| *FLPKDCKG_01824* | | - | 2006529 | 2006723 | 5.20 | 7.27 | 8.16 | 2.67 | Hypothetical protein |
| *FLPKDCKG_01825* | *argD-1* | + | 2006966 | 2008147 | 22.99 | 25.67 | 13.18 | 4.75 | Acetylornithine aminotransferase |
| *FLPKDCKG_01826* | *argF* | + | 2008186 | 2009145 | 22.37 | 48.58 | 35.38 | 12.99 | Ornithine carbamoyltransferase |
| *FLPKDCKG_01827* | | - | 2009213 | 2010100 | 1.82 | 2.17 | 0.33 | 1.04 | 2-(acetamidomethylene)succinate hydrolase |
| *FLPKDCKG_01828* | | - | 2010124 | 2010687 | 7.46 | 2.42 | 4.08 | 1.71 | Hypothetical protein |
| *FLPKDCKG_01829* | | - | 2010700 | 2011230 | 2.82 | 2.26 | 0.00 | 0.90 | Hypothetical protein |
| *FLPKDCKG_01830* | | - | 2011233 | 2012189 | 0.27 | 0.00 | 1.79 | 0.18 | Hypothetical protein |
| *FLPKDCKG_01831* | | - | 2012248 | 2013114 | 0.90 | 4.45 | 3.52 | 0.15 | 4-sulfomuconolactone hydrolase |
| *FLPKDCKG_01832* | *cynR-6* | + | 2013242 | 2014171 | 4.75 | 9.44 | 9.33 | 1.67 | HTH-type transcriptional regulator CynR |
| *FLPKDCKG_01833* | *argG* | + | 2014282 | 2015622 | 49.50 | 68.17 | 50.24 | 17.00 | Argininosuccinate synthase |
| *FLPKDCKG_01834* | *kup* | - | 2016037 | 2017965 | 5.92 | 9.29 | 7.36 | 2.87 | Low affinity potassium transport system protein kup |
| *FLPKDCKG_01835* | *basS* | - | 2017962 | 2019077 | 2.22 | 5.01 | 2.97 | 0.43 | Sensor protein BasS |
| *FLPKDCKG_01836* | *qseB-2* | - | 2019164 | 2019835 | 61.20 | 59.57 | 57.48 | 18.63 | Transcriptional regulatory protein QseB |
| *FLPKDCKG_01837* | *exbD-1* | - | 2019901 | 2020380 | 11.17 | 15.70 | 12.71 | 15.55 | Biopolymer transport protein ExbD |
| *FLPKDCKG_01838* | *exbB-1* | - | 2020380 | 2021369 | 5.40 | 6.94 | 9.37 | 4.71 | Biopolymer transport protein ExbB |
| *FLPKDCKG_01839* | *fhaB* | - | 2021371 | 2022072 | 10.44 | 5.21 | 9.65 | 10.00 | Filamentous hemagglutinin |
| *FLPKDCKG_01840* | | - | 2022588 | 2022663 | 17.63 | 23.99 | 28.46 | 9.48 | tRNA |
| *FLPKDCKG_01841* | *hupB* | - | 2022811 | 2023083 | 153.32 | 195.30 | 212.57 | 125.70 | DNA-binding protein HU-beta |
| *FLPKDCKG_01842* | | - | 2023473 | 2024153 | 3.13 | 3.58 | 2.90 | 1.00 | PKHD-type hydroxylase |
| *FLPKDCKG_01843* | *mgl* | - | 2024208 | 2025452 | 6.80 | 9.13 | 9.83 | 2.38 | L-methionine gamma-lyase |
| *FLPKDCKG_01844* | *rlmB* | - | 2025504 | 2026241 | 7.50 | 14.54 | 14.30 | 4.18 | 23S rRNA (guanosine-2'-O-)-methyltransferase RlmB |
| *FLPKDCKG_01845* | *rnr-1* | - | 2026375 | 2028840 | 12.75 | 29.16 | 18.87 | 4.82 | Ribonuclease R |
| *FLPKDCKG_01846* | | + | 2029037 | 2029121 | 6.76 | 5.64 | 10.85 | 2.92 | tRNA |
| *FLPKDCKG_01847* | | - | 2029401 | 2030012 | 3.13 | 5.82 | 4.44 | 1.84 | Hypothetical protein |
| *FLPKDCKG_01848* | | + | 2030205 | 2030289 | 11.13 | 13.31 | 16.85 | 4.62 | tRNA |
| *FLPKDCKG_01849* | | - | 2030429 | 2030872 | 18.56 | 18.60 | 7.52 | 5.03 | Acetyltransferase |
| *FLPKDCKG_01850* | *eamA-2* | + | 2030991 | 2031899 | 2.61 | 4.26 | 4.73 | 0.44 | putative amino-acid metabolite efflux pump |
| *FLPKDCKG_01851* | | + | 2031931 | 2032875 | 2.05 | 3.88 | 0.75 | 0.03 | Hypothetical protein |
| *FLPKDCKG_01852* | | + | 2033045 | 2034268 | 1.47 | 8.30 | 14.93 | 6.31 | Hypothetical protein |
| *FLPKDCKG_01853* | *gltA-1* | + | 2034455 | 2036125 | 4.60 | 17.66 | 9.70 | 3.47 | Glutamate synthase [NADPH] large chain |
| *FLPKDCKG_01854* | *ywiE* | - | 2036157 | 2037626 | 5.02 | 58.82 | 30.10 | 5.29 | putative cardiolipin synthase YwiE |
| *FLPKDCKG_01855* | | - | 2037638 | 2037970 | 34.40 | 164.67 | 147.43 | 27.75 | Hypothetical protein |
| *FLPKDCKG_01856* | *egtD* | + | 2038349 | 2039326 | 58.94 | 347.79 | 203.65 | 38.16 | Histidine N-alpha-methyltransferase |
| *FLPKDCKG_01857* | *egtB* | + | 2039364 | 2040680 | 9.57 | 71.28 | 59.60 | 12.13 | Hercynine oxygenase |
| *FLPKDCKG_01858* | | + | 2040820 | 2040896 | 33.14 | 101.76 | 86.92 | 24.44 | tRNA |
| *FLPKDCKG_01859* | | - | 2041074 | 2041985 | 2.94 | 289.71 | 199.18 | 18.68 | Hypothetical protein |

| Locus name | Gene name | Strand | Gene start (bp) | Gene end (bp) | Average TPM Inoculum | Average TPM $t_6$ | Average TPM $t_{24}$ | Average TPM $t_{168}$ | Annotation description |
|---|---|---|---|---|---|---|---|---|---|
| *FLPKDCKG_01860* | | - | 2042145 | 2042336 | 4.95 | 938.31 | 445.22 | 51.62 | Hypothetical protein |
| *FLPKDCKG_01861* | | - | 2042466 | 2042720 | 73.44 | 72.84 | 57.95 | 19.77 | Hypothetical protein |
| *FLPKDCKG_01862* | | + | 2042968 | 2043159 | 171.46 | 158.20 | 72.82 | 47.71 | Hypothetical protein |
| *FLPKDCKG_01863* | *yddG* | - | 2043178 | 2044113 | 13.62 | 29.90 | 20.53 | 11.47 | Aromatic amino acid exporter YddG |
| *FLPKDCKG_01864* | | + | 2044258 | 2044518 | 9.24 | 13.33 | 5.60 | 5.20 | Hypothetical protein |
| *FLPKDCKG_01865* | | - | 2044553 | 2046130 | 14.22 | 19.18 | 11.54 | 3.55 | putative signaling protein |
| *FLPKDCKG_01866* | | - | 2046473 | 2046745 | 46.16 | 11.29 | 5.41 | 3.58 | Hypothetical protein |
| *FLPKDCKG_01867* | | - | 2046726 | 2047256 | 19.93 | 8.97 | 6.03 | 1.07 | Hypothetical protein |
| *FLPKDCKG_01868* | | - | 2047246 | 2047707 | 11.53 | 7.60 | 0.00 | 0.00 | Hypothetical protein |
| *FLPKDCKG_01869* | | - | 2047899 | 2048348 | 19.03 | 9.81 | 5.30 | 1.06 | Hypothetical protein |
| *FLPKDCKG_01870* | | - | 2048360 | 2049391 | 25.14 | 11.13 | 8.98 | 1.38 | Hypothetical protein |
| *FLPKDCKG_01871* | | - | 2049406 | 2050071 | 14.19 | 7.21 | 4.58 | 0.48 | Hypothetical protein |
| *FLPKDCKG_01872* | | - | 2050073 | 2051254 | 13.81 | 5.68 | 2.89 | 1.10 | Hypothetical protein |
| *FLPKDCKG_01873* | | - | 2051251 | 2051604 | 24.99 | 12.16 | 11.56 | 0.77 | Hypothetical protein |
| *FLPKDCKG_01874* | | - | 2051613 | 2052356 | 14.38 | 7.86 | 2.33 | 1.10 | Hypothetical protein |
| *FLPKDCKG_01875* | | - | 2052391 | 2053419 | 53.60 | 14.78 | 10.16 | 2.38 | Hypothetical protein |
| *FLPKDCKG_01876* | | - | 2053421 | 2053720 | 40.07 | 7.92 | 7.38 | 2.37 | Hypothetical protein |
| *FLPKDCKG_01877* | | - | 2053730 | 2054329 | 31.18 | 12.47 | 8.02 | 1.70 | Hypothetical protein |
| *FLPKDCKG_01878* | | - | 2054322 | 2055491 | 17.00 | 8.40 | 4.23 | 1.86 | Hypothetical protein |
| *FLPKDCKG_01879* | | - | 2055439 | 2055993 | 12.13 | 8.83 | 3.09 | 0.01 | Hypothetical protein |
| *FLPKDCKG_01880* | | - | 2056171 | 2056620 | 14.84 | 9.33 | 8.03 | 5.80 | Hypothetical protein |
| *FLPKDCKG_01881* | | - | 2056624 | 2057064 | 26.11 | 22.52 | 7.82 | 1.05 | Hypothetical protein |
| *FLPKDCKG_01882* | | - | 2057078 | 2058553 | 24.27 | 11.68 | 6.02 | 1.42 | Hypothetical protein |
| *FLPKDCKG_01883* | | - | 2058564 | 2059157 | 9.92 | 4.28 | 5.51 | 0.79 | Hypothetical protein |
| *FLPKDCKG_01884* | | - | 2059650 | 2059907 | 0.65 | 0.00 | 2.68 | 0.00 | Hypothetical protein |
| *FLPKDCKG_01885* | | + | 2060217 | 2060978 | 23.88 | 5.76 | 4.81 | 3.19 | Hypothetical protein |
| *FLPKDCKG_01886* | | + | 2060983 | 2062800 | 6.84 | 11.99 | 7.19 | 2.71 | Hypothetical protein |
| *FLPKDCKG_01887* | | - | 2062874 | 2063095 | 58.84 | 197.87 | 84.81 | 26.04 | Hypothetical protein |
| *FLPKDCKG_01888* | | - | 2063264 | 2064403 | 12.13 | 15.94 | 12.96 | 6.19 | Hypothetical protein |
| *FLPKDCKG_01889* | | - | 2064543 | 2065247 | 2.40 | 5.89 | 9.11 | 3.54 | Hypothetical protein |
| *FLPKDCKG_01890* | | - | 2065410 | 2065922 | 1.84 | 1.52 | 2.08 | 0.87 | Hypothetical protein |
| *FLPKDCKG_01891* | | + | 2066107 | 2066466 | 6.07 | 21.95 | 7.38 | 3.62 | Hypothetical protein |
| *FLPKDCKG_01892* | | + | 2067286 | 2067567 | 25.73 | 26.69 | 18.77 | 15.12 | Hypothetical protein |
| *FLPKDCKG_01893* | *hyaC* | + | 2067677 | 2068369 | 7.29 | 8.91 | 2.88 | 0.91 | putative Ni/Fe-hydrogenase 1 B-type cytochrome subunit |
| *FLPKDCKG_01894* | | - | 2068498 | 2068956 | 17.92 | 24.53 | 24.50 | 5.94 | Cytochrome c' |
| *FLPKDCKG_01895* | *purB* | - | 2069013 | 2070389 | 14.70 | 35.30 | 17.01 | 5.23 | Adenylosuccinate lyase |
| *FLPKDCKG_01896* | *yibF-1* | + | 2070551 | 2071162 | 14.10 | 38.81 | 27.19 | 6.81 | putative GST-like protein YibF |
| *FLPKDCKG_01897* | | - | 2071224 | 2072042 | 1.83 | 2.86 | 2.12 | 0.10 | Hypothetical protein |
| *FLPKDCKG_01898* | *mnmA* | - | 2072165 | 2073283 | 12.51 | 23.07 | 17.99 | 3.11 | tRNA-specific 2-thiouridylase MnmA |
| *FLPKDCKG_01899* | *gcvA-6* | - | 2073296 | 2074216 | 2.24 | 7.32 | 4.03 | 2.28 | Glycine cleavage system transcriptional activator |
| *FLPKDCKG_01900* | | + | 2074408 | 2075397 | 6.62 | 8.15 | 14.42 | 7.62 | Hypothetical protein |
| *FLPKDCKG_01901* | | + | 2075468 | 2076343 | 0.98 | 4.19 | 4.25 | 0.77 | Hypothetical protein |
| *FLPKDCKG_01902* | *pntAA* | + | 2076529 | 2077647 | 25.44 | 15.73 | 14.25 | 7.58 | NAD(P) transhydrogenase subunit alpha part 1 |
| *FLPKDCKG_01903* | *pntA* | + | 2077652 | 2077969 | 21.09 | 12.35 | 9.61 | 2.39 | NAD(P) transhydrogenase subunit alpha |
| *FLPKDCKG_01904* | *pntB-1* | + | 2077966 | 2078919 | 7.71 | 2.31 | 3.62 | 1.27 | NAD(P) transhydrogenase subunit beta |
| *FLPKDCKG_01905* | *pntB-2* | + | 2078919 | 2079404 | 12.28 | 14.19 | 6.88 | 2.82 | NAD(P) transhydrogenase subunit beta |
| *FLPKDCKG_01906* | *lutA* | - | 2079499 | 2080728 | 5.58 | 3.45 | 7.58 | 4.09 | Lactate utilization protein A |
| *FLPKDCKG_01907* | | - | 2080734 | 2081837 | 3.78 | 7.59 | 9.94 | 2.47 | putative FAD-linked oxidoreductase |
| *FLPKDCKG_01908* | | - | 2081850 | 2082638 | 47.65 | 42.99 | 50.60 | 17.83 | putative FAD-linked oxidoreductase |
| *FLPKDCKG_01909* | | - | 2082660 | 2083349 | 10.05 | 9.91 | 12.95 | 9.20 | putative FAD-linked oxidoreductase |
| *FLPKDCKG_01910* | | - | 2083618 | 2085027 | 35.76 | 36.92 | 20.29 | 6.53 | putative FAD-linked oxidoreductase |
| *FLPKDCKG_01911* | *aroG-2* | - | 2085126 | 2086199 | 44.97 | 7.68 | 15.79 | 3.63 | Phospho-2-dehydro-3-deoxyheptonate aldolase, Phe-sensitive |
| *FLPKDCKG_01912* | *tldD* | - | 2086468 | 2087844 | 28.07 | 59.87 | 34.05 | 11.13 | Metalloprotease TldD |
| *FLPKDCKG_01913* | | + | 2088700 | 2089086 | 303.93 | 611.74 | 362.66 | 114.97 | DNA-binding protein Bv3F |
| *FLPKDCKG_01914* | | - | 2089172 | 2089810 | 51.83 | 81.70 | 77.84 | 18.48 | Hypothetical protein |
| *FLPKDCKG_01915* | *pepN* | + | 2089963 | 2092668 | 21.83 | 42.22 | 25.08 | 8.34 | Aminopeptidase N |
| *FLPKDCKG_01916* | *fbp* | + | 2092679 | 2093698 | 23.95 | 47.56 | 27.97 | 8.91 | Fructose-1,6-bisphosphatase class 1 |
| *FLPKDCKG_01917* | *ygdR* | - | 2093770 | 2093991 | 497.09 | 4915.30 | 1912.73 | 204.93 | putative lipoprotein YgdR |
| *FLPKDCKG_01918* | *thrC-1* | - | 2094146 | 2095555 | 1.34 | 5.46 | 6.41 | 2.19 | Threonine synthase |
| *FLPKDCKG_01919* | *hom* | - | 2095676 | 2096980 | 2.23 | 5.95 | 5.21 | 2.77 | Homoserine dehydrogenase |
| *FLPKDCKG_01920* | *alaC* | - | 2097184 | 2098368 | 9.36 | 20.65 | 17.02 | 7.97 | Glutamate-pyruvate aminotransferase AlaC |
| *FLPKDCKG_01921* | | + | 2098705 | 2099166 | 325.71 | 442.02 | 262.72 | 59.38 | Hypothetical protein |
| *FLPKDCKG_01922* | *bcp* | + | 2099163 | 2099630 | 118.56 | 437.20 | 222.44 | 31.39 | Putative peroxiredoxin bcp |
| *FLPKDCKG_01923* | | + | 2099689 | 2101389 | 43.69 | 140.98 | 84.99 | 16.73 | PhoH-like protein |
| *FLPKDCKG_01924* | | - | 2101515 | 2101646 | 3.05 | 4.82 | 5.86 | 1.22 | Hypothetical protein |
| *FLPKDCKG_01925* | *nagK-1* | - | 2101688 | 2102383 | 27.71 | 40.67 | 38.63 | 19.28 | Fumarylpyruvate hydrolase |
| *FLPKDCKG_01926* | | - | 2102465 | 2102767 | 0.49 | 2.62 | 1.52 | 0.00 | Hypothetical protein |
| *FLPKDCKG_01927* | *livH-6* | - | 2102767 | 2104644 | 0.37 | 0.37 | 0.58 | 0.31 | High-affinity branched-chain amino acid transport system permease protein LivH |
| *FLPKDCKG_01928* | *livF-6* | - | 2104658 | 2105350 | 0.31 | 1.09 | 0.72 | 0.75 | High-affinity branched-chain amino acid transport ATP-binding protein LivF |

| Locus name | Gene name | Strand | Gene start (bp) | Gene end (bp) | Average TPM Inoculum | Average TPM $t_6$ | Average TPM $t_{24}$ | Average TPM $t_{168}$ | Annotation description |
|---|---|---|---|---|---|---|---|---|---|
| *FLPKDCKG_01929* | *lptB-3* | - | 2105350 | 2106111 | 0.12 | 0.11 | 0.00 | 1.04 | Lipopolysaccharide export system ATP-binding protein LptB |
| *FLPKDCKG_01930* | *gno-1* | - | 2106122 | 2106901 | 0.51 | 0.94 | 1.36 | 1.99 | Gluconate 5-dehydrogenase |
| *FLPKDCKG_01931* | | + | 2107118 | 2107516 | 1.96 | 1.94 | 2.44 | 7.05 | Hypothetical protein |
| *FLPKDCKG_01932* | | + | 2107607 | 2108818 | 0.44 | 0.37 | 0.81 | 0.82 | Hypothetical protein |
| *FLPKDCKG_01933* | *fabG2* | + | 2108815 | 2109579 | 2.28 | 3.72 | 3.19 | 1.09 | putative oxidoreductase |
| *FLPKDCKG_01934* | *kipR-2* | + | 2109594 | 2110433 | 2.71 | 4.07 | 3.03 | 1.15 | HTH-type transcriptional regulator KipR |
| *FLPKDCKG_01935* | *ppa* | - | 2110434 | 2110964 | 43.76 | 93.01 | 63.47 | 24.07 | Inorganic pyrophosphatase |
| *FLPKDCKG_01936* | | - | 2110994 | 2112607 | 19.64 | 28.51 | 20.60 | 4.75 | Hypothetical protein |
| *FLPKDCKG_01937* | *hemX* | - | 2112612 | 2113805 | 26.43 | 36.73 | 35.32 | 9.98 | Putative uroporphyrinogen-III C-methyltransferase |
| *FLPKDCKG_01938* | *hemD* | - | 2113843 | 2114655 | 4.16 | 8.17 | 8.74 | 2.59 | Uroporphyrinogen-III synthase |
| *FLPKDCKG_01939* | *hemC* | - | 2114655 | 2115599 | 9.17 | 33.12 | 14.90 | 7.33 | Porphobilinogen deaminase |
| *FLPKDCKG_01940* | | - | 2115609 | 2115731 | 30.35 | 66.83 | 31.33 | 13.59 | Hypothetical protein |
| *FLPKDCKG_01941* | | - | 2115718 | 2116521 | 23.25 | 30.97 | 13.79 | 7.50 | Hypothetical protein |
| *FLPKDCKG_01942* | | - | 2116552 | 2117250 | 9.58 | 12.03 | 16.24 | 8.51 | Hypothetical protein |
| *FLPKDCKG_01943* | *sucD* | - | 2117512 | 2118393 | 30.13 | 73.02 | 74.00 | 34.29 | Succinate--CoA ligase [ADP-forming] subunit alpha |
| *FLPKDCKG_01944* | *sucC* | - | 2118419 | 2119579 | 31.83 | 80.60 | 68.98 | 24.73 | Succinate--CoA ligase [ADP-forming] subunit beta |
| *FLPKDCKG_01945* | | - | 2119590 | 2120192 | 18.40 | 60.63 | 27.91 | 13.35 | Hypothetical protein |
| *FLPKDCKG_01946* | | - | 2120335 | 2121123 | 1.22 | 4.10 | 3.97 | 2.02 | Hypothetical protein |
| *FLPKDCKG_01947* | *dhbF* | - | 2121062 | 2124307 | 6.15 | 7.09 | 6.30 | 3.09 | Dimodular nonribosomal peptide synthase |
| *FLPKDCKG_01948* | *metE* | - | 2124474 | 2126771 | 45.70 | 267.31 | 224.71 | 102.62 | 5-methyltetrahydropteroyltriglutamate--homocysteine methyltransferase |
| *FLPKDCKG_01949* | *metR-2* | + | 2126967 | 2127884 | 8.15 | 19.36 | 18.15 | 3.78 | HTH-type transcriptional regulator MetR |
| *FLPKDCKG_01950* | *recX* | - | 2127931 | 2128668 | 4.19 | 8.35 | 6.33 | 0.34 | Regulatory protein RecX |
| *FLPKDCKG_01951* | *recA* | - | 2128770 | 2129834 | 22.39 | 45.03 | 43.68 | 13.70 | Protein RecA |
| *FLPKDCKG_01952* | *tctD-1* | + | 2130213 | 2130902 | 15.65 | 20.93 | 18.47 | 8.35 | Transcriptional regulatory protein tctD |
| *FLPKDCKG_01953* | *qseC-2* | + | 2130943 | 2132394 | 2.72 | 5.49 | 8.14 | 3.45 | Sensor protein QseC |
| *FLPKDCKG_01954* | *proP-3* | + | 2132706 | 2134382 | 29.99 | 300.35 | 257.16 | 111.23 | Proline/betaine transporter |
| *FLPKDCKG_01955* | | - | 2134813 | 2135022 | 148.31 | 278.08 | 151.96 | 79.96 | Hypothetical protein |
| *FLPKDCKG_01956* | *rdgC* | + | 2135174 | 2136073 | 10.62 | 39.18 | 22.76 | 5.49 | Recombination-associated protein RdgC |
| *FLPKDCKG_01957* | *dosC-2* | - | 2136324 | 2137736 | 0.93 | 2.15 | 0.92 | 0.17 | Diguanylate cyclase DosC |
| *FLPKDCKG_01958* | *pyrB-1* | - | 2137867 | 2139159 | 16.18 | 10.69 | 8.48 | 5.13 | Aspartate carbamoyltransferase catalytic subunit |
| *FLPKDCKG_01959* | | - | 2139260 | 2139673 | 7.34 | 10.11 | 8.48 | 3.29 | Hypothetical protein |
| *FLPKDCKG_01960* | | - | 2139676 | 2140101 | 25.03 | 23.21 | 21.33 | 5.58 | Hypothetical protein |
| *FLPKDCKG_01961* | | - | 2140260 | 2140391 | 1883.93 | 914.25 | 436.89 | 164.56 | Hypothetical protein |
| *FLPKDCKG_01962* | | + | 2140560 | 2140919 | 77.91 | 99.98 | 85.35 | 22.37 | Hypothetical protein |
| *FLPKDCKG_01963* | | + | 2140983 | 2141384 | 47.61 | 77.54 | 60.24 | 30.96 | Hypothetical protein |
| *FLPKDCKG_01964* | *mnmC-1* | - | 2141391 | 2142641 | 2.28 | 5.44 | 5.83 | 1.33 | tRNA 5-methylaminomethyl-2-thiouridine biosynthesis bifunctional protein MnmC |
| *FLPKDCKG_01965* | | - | 2142899 | 2143243 | 124.13 | 75.29 | 32.36 | 16.09 | Hypothetical protein |
| *FLPKDCKG_01966* | | - | 2143466 | 2143542 | 83.20 | 73.94 | 71.69 | 15.92 | tRNA |
| *FLPKDCKG_01967* | | - | 2143563 | 2143943 | 0.84 | 1.42 | 0.00 | 0.00 | Hypothetical protein |
| *FLPKDCKG_01968* | *norG-3* | + | 2144113 | 2145552 | 11.44 | 25.57 | 15.54 | 3.96 | HTH-type transcriptional regulator NorG |
| *FLPKDCKG_01969* | *yohD* | + | 2145549 | 2146118 | 2.16 | 7.27 | 5.74 | 0.89 | Inner membrane protein YohD |
| *FLPKDCKG_01970* | | - | 2146154 | 2146426 | 1.79 | 3.94 | 4.63 | 1.35 | Hypothetical protein |
| *FLPKDCKG_01971* | | - | 2146623 | 2146868 | 6.88 | 9.99 | 6.12 | 9.07 | Hypothetical protein |
| *FLPKDCKG_01972* | | + | 2147007 | 2147612 | 23.85 | 20.78 | 13.31 | 4.48 | Hypothetical protein |
| *FLPKDCKG_01973* | *asnC-1* | + | 2147658 | 2148116 | 12.35 | 31.92 | 21.49 | 5.09 | Regulatory protein AsnC |
| *FLPKDCKG_01974* | | + | 2148242 | 2148571 | 122.75 | 144.50 | 72.31 | 17.11 | Hypothetical protein |
| *FLPKDCKG_01975* | *pdxA2-2* | + | 2148568 | 2149548 | 16.03 | 26.93 | 17.52 | 6.14 | 4-hydroxythreonine-4-phosphate dehydrogenase 2 |
| *FLPKDCKG_01976* | | + | 2149632 | 2150609 | 4.49 | 5.88 | 4.20 | 1.19 | Hypothetical protein |
| *FLPKDCKG_01977* | *dnaB* | - | 2150676 | 2152058 | 11.36 | 16.70 | 17.49 | 2.36 | Replicative DNA helicase |
| *FLPKDCKG_01978* | | - | 2152055 | 2152894 | 4.40 | 17.85 | 7.43 | 3.77 | Hypothetical protein |
| *FLPKDCKG_01979* | *rplI* | - | 2153040 | 2153495 | 12.84 | 67.81 | 65.65 | 45.49 | 50S ribosomal protein L9 |
| *FLPKDCKG_01980* | *rpsR* | - | 2153510 | 2153782 | 32.31 | 133.33 | 102.24 | 45.65 | 30S ribosomal protein S18 |
| *FLPKDCKG_01981* | *priB* | - | 2153889 | 2154212 | 44.05 | 226.95 | 215.00 | 112.67 | Primosomal replication protein N |
| *FLPKDCKG_01982* | *rpsF* | - | 2154264 | 2154638 | 29.12 | 84.48 | 78.55 | 46.07 | 30S ribosomal protein S6 |
| *FLPKDCKG_01983* | | - | 2154873 | 2155787 | 0.99 | 2.36 | 3.28 | 0.95 | Hypothetical protein |
| *FLPKDCKG_01984* | *folE2* | - | 2155795 | 2156592 | 10.20 | 18.63 | 17.12 | 5.97 | GTP cyclohydrolase FolE2 |
| *FLPKDCKG_01985* | *dxs-1* | - | 2156784 | 2158646 | 6.42 | 14.52 | 11.09 | 3.09 | 1-deoxy-D-xylulose-5-phosphate synthase |
| *FLPKDCKG_01986* | *ispA* | - | 2158700 | 2159611 | 21.63 | 37.17 | 32.09 | 9.72 | Farnesyl diphosphate synthase |
| *FLPKDCKG_01987* | *xseB* | - | 2159619 | 2159885 | 14.44 | 25.76 | 19.86 | 4.15 | Exodeoxyribonuclease 7 small subunit |
| *FLPKDCKG_01988* | *tsaQ1-1* | - | 2159968 | 2160822 | 2.32 | 3.11 | 2.41 | 0.30 | HTH-type transcriptional regulator TsaQ1/TsaQ2 |
| *FLPKDCKG_01989* | | + | 2160916 | 2161884 | 1.13 | 3.69 | 2.60 | 1.91 | Hypothetical protein |
| *FLPKDCKG_01990* | *wecH* | + | 2162102 | 2163112 | 2.05 | 36.22 | 24.05 | 5.29 | O-acetyltransferase WecH |
| *FLPKDCKG_01991* | *bphA1* | + | 2163544 | 2164692 | 6.30 | 19.10 | 16.17 | 5.14 | Biphenyl 2,3-dioxygenase subunit alpha |
| *FLPKDCKG_01992* | | - | 2164759 | 2165718 | 0.39 | 0.83 | 0.88 | 0.42 | Hypothetical protein |
| *FLPKDCKG_01993* | *fbpA* | - | 2165975 | 2166997 | 0.13 | 1.40 | 0.03 | 0.75 | Fe(3+)-binding periplasmic protein |
| *FLPKDCKG_01994* | *phnX* | - | 2167112 | 2167933 | 0.56 | 0.05 | 0.38 | 0.59 | Phosphonoacetaldehyde hydrolase |
| *FLPKDCKG_01995* | *potH-3* | - | 2167961 | 2169754 | 0.18 | 0.02 | 0.19 | 0.00 | Putrescine transport system permease protein PotH |
| *FLPKDCKG_01996* | *cysA-1* | - | 2169732 | 2170862 | 0.40 | 1.01 | 1.40 | 0.11 | Sulfate/thiosulfate import ATP-binding protein CysA |
| *FLPKDCKG_01997* | *cmpR-7* | + | 2171023 | 2171886 | 2.69 | 1.77 | 5.45 | 2.47 | HTH-type transcriptional activator CmpR |

| Locus name | Gene name | Strand | Gene start (bp) | Gene end (bp) | Average TPM Inoculum | Average TPM $t_6$ | Average TPM $t_{24}$ | Average TPM $t_{168}$ | Annotation description |
|---|---|---|---|---|---|---|---|---|---|
| *FLPKDCKG_01998* | | + | 2171939 | 2172508 | 1.44 | 3.03 | 3.12 | 3.95 | Hypothetical protein |
| *FLPKDCKG_01999* | *sseB* | - | 2172582 | 2173433 | 10.14 | 12.43 | 10.18 | 5.15 | Putative thiosulfate sulfurtransferase SseB |
| *FLPKDCKG_02000* | *ribN-4* | - | 2173470 | 2174309 | 13.05 | 17.48 | 10.52 | 2.87 | Riboflavin transporter |
| *FLPKDCKG_02001* | *htpX-1* | + | 2174479 | 2175360 | 17.45 | 16.10 | 21.52 | 8.61 | Protease HtpX |
| *FLPKDCKG_02002* | *sdaA* | - | 2175507 | 2176898 | 12.85 | 13.92 | 12.09 | 3.80 | L-serine dehydratase 1 |
| *FLPKDCKG_02003* | | + | 2177074 | 2178735 | 8.56 | 18.60 | 7.84 | 0.38 | Hypothetical protein |
| *FLPKDCKG_02004* | *clcD-1* | - | 2179857 | 2180645 | 18.74 | 51.93 | 39.50 | 13.24 | Carboxymethylenebutenolidase |
| *FLPKDCKG_02005* | *zupT-1* | - | 2180761 | 2181600 | 7.90 | 42.41 | 20.22 | 7.29 | Zinc transporter ZupT |
| *FLPKDCKG_02006* | *polA* | - | 2181751 | 2184468 | 25.33 | 29.61 | 12.88 | 4.22 | DNA polymerase I |
| *FLPKDCKG_02007* | *log* | + | 2184557 | 2185183 | 41.84 | 78.03 | 47.05 | 21.35 | Cytokinin riboside 5'-monophosphate phosphoribohydrolase |
| *FLPKDCKG_02008* | *abgB* | - | 2185251 | 2186675 | 3.89 | 4.46 | 2.63 | 0.82 | p-aminobenzoyl-glutamate hydrolase subunit B |
| *FLPKDCKG_02009* | | - | 2186746 | 2187729 | 0.75 | 2.13 | 1.42 | 0.15 | Hypothetical protein |
| *FLPKDCKG_02010* | *gbpR-1* | + | 2187863 | 2188807 | 2.59 | 3.05 | 1.48 | 0.23 | HTH-type transcriptional regulator GbpR |
| *FLPKDCKG_02011* | *allS* | - | 2188827 | 2189687 | 2.05 | 6.37 | 2.51 | 1.92 | HTH-type transcriptional activator AllS |
| *FLPKDCKG_02012* | | + | 2189794 | 2190186 | 1.37 | 1.72 | 1.51 | 1.58 | Hypothetical protein |
| *FLPKDCKG_02013* | *typA* | - | 2190366 | 2192189 | 4.63 | 10.10 | 11.78 | 6.96 | GTP-binding protein TypA/BipA |
| *FLPKDCKG_02014* | *truB* | - | 2192186 | 2192935 | 5.87 | 23.41 | 19.83 | 6.27 | tRNA pseudouridine synthase B |
| *FLPKDCKG_02015* | *rbfA* | - | 2192975 | 2193376 | 8.13 | 16.67 | 9.69 | 8.27 | 30S ribosome-binding factor |
| *FLPKDCKG_02016* | *infB* | - | 2193400 | 2196453 | 15.62 | 28.24 | 33.80 | 12.77 | Translation initiation factor IF-2 |
| *FLPKDCKG_02017* | *nusA* | - | 2196546 | 2198024 | 20.93 | 44.95 | 31.58 | 11.43 | Transcription termination/antitermination protein NusA |
| *FLPKDCKG_02018* | *rimP* | - | 2198021 | 2198569 | 26.67 | 72.48 | 58.82 | 21.71 | Ribosome maturation factor RimP |
| *FLPKDCKG_02019* | *rluB-2* | - | 2198928 | 2200664 | 3.74 | 10.85 | 10.43 | 6.99 | Ribosomal large subunit pseudouridine synthase B |
| *FLPKDCKG_02020* | *scpB* | - | 2201160 | 2201927 | 8.83 | 24.79 | 19.30 | 7.74 | Segregation and condensation protein B |
| *FLPKDCKG_02021* | | + | 2201997 | 2202073 | 95.68 | 90.43 | 86.23 | 26.94 | tRNA |
| *FLPKDCKG_02022* | *dmlR-12* | - | 2203035 | 2203934 | 36.59 | 49.12 | 39.36 | 10.38 | HTH-type transcriptional regulator DmlR |
| *FLPKDCKG_02023* | *yjiA-1* | - | 2204226 | 2205233 | 3.39 | 4.94 | 3.21 | 4.27 | putative GTP-binding protein YjiA |
| *FLPKDCKG_02024* | *yciC-2* | - | 2205262 | 2206575 | 2.70 | 6.15 | 12.57 | 12.25 | Putative metal chaperone YciC |
| *FLPKDCKG_02025* | | - | 2206761 | 2206836 | 8.01 | 9.13 | 15.26 | 3.30 | tRNA |
| *FLPKDCKG_02026* | | + | 2207048 | 2207123 | 5.97 | 7.49 | 10.87 | 4.23 | tRNA |
| *FLPKDCKG_02027* | *qseB-3* | + | 2207268 | 2207927 | 16.44 | 50.06 | 33.28 | 13.07 | Transcriptional regulatory protein QseB |
| *FLPKDCKG_02028* | *qseC-3* | + | 2207924 | 2209267 | 4.96 | 14.35 | 15.90 | 3.48 | Sensor protein QseC |
| *FLPKDCKG_02029* | *htrA* | + | 2209332 | 2210798 | 2.50 | 5.27 | 6.41 | 3.76 | Putative serine protease HtrA |
| *FLPKDCKG_02030* | *nagL-1* | - | 2210831 | 2211475 | 7.07 | 15.02 | 15.77 | 4.99 | Maleylpyruvate isomerase |
| *FLPKDCKG_02031* | *ispH* | - | 2211526 | 2212509 | 15.33 | 23.78 | 10.55 | 5.86 | 4-hydroxy-3-methylbut-2-enyl diphosphate reductase |
| *FLPKDCKG_02032* | *fkpB* | - | 2212509 | 2212985 | 25.57 | 40.94 | 42.24 | 11.32 | FKBP-type 16 kDa peptidyl-prolyl cis-trans isomerase |
| *FLPKDCKG_02033* | | + | 2213037 | 2213714 | 0.49 | 0.76 | 0.00 | 0.73 | Hypothetical protein |
| *FLPKDCKG_02034* | *kynB-1* | + | 2213820 | 2214449 | 9.42 | 29.60 | 23.84 | 10.96 | Kynurenine formamidase |
| *FLPKDCKG_02035* | *kynA* | + | 2214657 | 2215505 | 1.48 | 2.36 | 1.73 | 1.50 | Tryptophan 2,3-dioxygenase |
| *FLPKDCKG_02036* | *rpmE2* | + | 2215692 | 2215958 | 20.13 | 32.67 | 38.82 | 12.69 | 50S ribosomal protein L31 type B |
| *FLPKDCKG_02037* | | + | 2216029 | 2217813 | 8.30 | 12.15 | 10.27 | 2.85 | Hypothetical protein |
| *FLPKDCKG_02038* | *norM* | + | 2217810 | 2219216 | 1.26 | 5.30 | 2.87 | 0.28 | Multidrug resistance protein NorM |
| *FLPKDCKG_02039* | *paaK* | - | 2219329 | 2220645 | 10.61 | 9.63 | 4.06 | 2.49 | Phenylacetate-coenzyme A ligase |
| *FLPKDCKG_02040* | *paaI* | - | 2220742 | 2221215 | 13.78 | 15.07 | 8.64 | 5.38 | Acyl-coenzyme A thioesterase PaaI |
| *FLPKDCKG_02041* | *paaG-1* | - | 2221215 | 2222003 | 8.61 | 14.02 | 16.24 | 4.44 | 1,2-epoxyphenylacetyl-CoA isomerase |
| *FLPKDCKG_02042* | *amnC-1* | - | 2222019 | 2223683 | 16.20 | 12.37 | 15.41 | 8.75 | 2-aminomuconic 6-semialdehyde dehydrogenase |
| *FLPKDCKG_02043* | *paaE-1* | - | 2223729 | 2224817 | 8.16 | 11.91 | 12.91 | 3.10 | 1,2-phenylacetyl-CoA epoxidase, subunit E |
| *FLPKDCKG_02044* | *paaD* | - | 2224832 | 2225338 | 4.81 | 3.76 | 6.31 | 2.32 | Putative 1,2-phenylacetyl-CoA epoxidase, subunit D |
| *FLPKDCKG_02045* | *paaC* | - | 2225335 | 2226099 | 7.99 | 11.58 | 11.23 | 4.89 | 1,2-phenylacetyl-CoA epoxidase, subunit C |
| *FLPKDCKG_02046* | *paaB* | - | 2226115 | 2226399 | 16.40 | 19.91 | 22.30 | 8.38 | 1,2-phenylacetyl-CoA epoxidase, subunit B |
| *FLPKDCKG_02047* | *paaA* | - | 2226489 | 2227478 | 41.73 | 79.10 | 50.57 | 16.23 | 1,2-phenylacetyl-CoA epoxidase, subunit A |
| *FLPKDCKG_02048* | *paaX* | + | 2227651 | 2228580 | 5.26 | 12.94 | 7.63 | 1.53 | Transcriptional repressor PaaX |
| *FLPKDCKG_02049* | *nnr-1* | - | 2228591 | 2229220 | 1.80 | 9.98 | 6.06 | 2.76 | Bifunctional NAD(P)H-hydrate repair enzyme Nnr |
| *FLPKDCKG_02050* | *orn* | - | 2229264 | 2229809 | 23.50 | 39.37 | 28.27 | 5.97 | Oligoribonuclease |
| *FLPKDCKG_02051* | *htpX-2* | + | 2230071 | 2231135 | 11.67 | 11.44 | 6.23 | 2.35 | Protease HtpX |
| *FLPKDCKG_02052* | *rsgA* | + | 2231132 | 2232094 | 6.77 | 10.29 | 11.20 | 4.04 | Small ribosomal subunit biogenesis GTPase RsgA |
| *FLPKDCKG_02053* | *nudL* | - | 2232126 | 2232857 | 9.30 | 34.16 | 18.20 | 5.88 | putative Nudix hydrolase NudL |
| *FLPKDCKG_02054* | *rplS* | - | 2232977 | 2233357 | 21.14 | 60.96 | 101.11 | 28.85 | 50S ribosomal protein L19 |
| *FLPKDCKG_02055* | *arnE-1* | + | 2233598 | 2234470 | 10.30 | 16.95 | 9.42 | 1.58 | 4-amino-4-deoxy-L-arabinose-phosphoundecaprenol flippase subunit ArnE |
| *FLPKDCKG_02056* | | - | 2234551 | 2234967 | 3.98 | 8.46 | 9.41 | 3.00 | Hypothetical protein |
| *FLPKDCKG_02057* | | - | 2235068 | 2236309 | 4.61 | 11.63 | 7.51 | 2.12 | Hypothetical protein |
| *FLPKDCKG_02058* | *cysNC* | - | 2236694 | 2237992 | 0.32 | 4.99 | 23.68 | 9.00 | Bifunctional enzyme CysN/CysC |
| *FLPKDCKG_02059* | *cysD-1* | - | 2238036 | 2238944 | 0.44 | 19.04 | 46.12 | 21.64 | Sulfate adenylyltransferase subunit 2 |
| *FLPKDCKG_02060* | *cysH* | - | 2238984 | 2239730 | 0.05 | 4.99 | 23.62 | 7.23 | Thioredoxin-dependent 5'-adenylylsulfate reductase |
| *FLPKDCKG_02061* | *cysA-2* | - | 2239727 | 2240791 | 0.10 | 5.82 | 25.60 | 9.13 | Sulfate/thiosulfate import ATP-binding protein CysA |
| *FLPKDCKG_02062* | *cysW-1* | - | 2240817 | 2241716 | 0.24 | 4.36 | 24.57 | 9.47 | Sulfate transport system permease protein CysW |
| *FLPKDCKG_02063* | *cysT* | - | 2241713 | 2242603 | 0.05 | 1.54 | 18.09 | 9.56 | Sulfate transport system permease protein CysT |
| *FLPKDCKG_02064* | *sbp* | - | 2242628 | 2243656 | 0.44 | 26.38 | 109.14 | 38.33 | Sulfate-binding protein |
| *FLPKDCKG_02065* | *ahpC* | - | 2243882 | 2244523 | 125.70 | 363.60 | 540.97 | 117.45 | Alkyl hydroperoxide reductase subunit C |
| *FLPKDCKG_02066* | *dmdC-2* | - | 2244677 | 2246503 | 14.51 | 19.54 | 13.57 | 4.10 | 3-methylmercaptopropionyl-CoA dehydrogenase |

| Locus name | Gene name | Strand | Gene start (bp) | Gene end (bp) | Average TPM Inoculum | Average TPM $t_6$ | Average TPM $t_{24}$ | Average TPM $t_{168}$ | Annotation description |
|---|---|---|---|---|---|---|---|---|---|
| *FLPKDCKG_02067* | *aspT-1* | - | 2246665 | 2248368 | 31.73 | 12.68 | 11.11 | 5.77 | Aspartate/alanine antiporter |
| *FLPKDCKG_02068* | *asD* | - | 2248406 | 2250220 | 160.89 | 29.33 | 22.24 | 9.15 | Bifunctional aspartate aminotransferase and L-aspartate beta-decarboxylase |
| *FLPKDCKG_02069* | *aspT-2* | - | 2250280 | 2251971 | 278.20 | 45.45 | 31.29 | 14.49 | Aspartate/alanine antiporter |
| *FLPKDCKG_02070* | *etfA* | - | 2252167 | 2253096 | 27.51 | 40.80 | 40.67 | 22.75 | Electron transfer flavoprotein subunit alpha |
| *FLPKDCKG_02071* | *etfB* | - | 2253098 | 2253847 | 70.28 | 95.88 | 96.77 | 34.68 | Electron transfer flavoprotein subunit beta |
| *FLPKDCKG_02072* | *hdaH* | - | 2253930 | 2254853 | 18.03 | 24.23 | 17.85 | 3.23 | Histone deacetylase-like amidohydrolase |
| *FLPKDCKG_02073* | | - | 2254882 | 2256075 | 8.21 | 15.20 | 12.94 | 4.39 | Hypothetical protein |
| *FLPKDCKG_02074* | | - | 2256126 | 2257175 | 1.44 | 2.52 | 2.52 | 1.34 | Hypothetical protein |
| *FLPKDCKG_02075* | *iclR-3* | + | 2257244 | 2258020 | 6.43 | 7.13 | 4.77 | 1.37 | Transcriptional repressor IclR |
| *FLPKDCKG_02076* | | + | 2258017 | 2258997 | 4.90 | 6.57 | 4.82 | 3.34 | Hypothetical protein |
| *FLPKDCKG_02077* | *cysM* | - | 2259120 | 2260043 | 38.73 | 35.15 | 25.45 | 9.19 | Cysteine synthase B |
| *FLPKDCKG_02078* | *comEA* | - | 2260192 | 2260734 | 0.36 | 0.95 | 0.00 | 1.07 | ComE operon protein 1 |
| *FLPKDCKG_02079* | *hldD* | - | 2260932 | 2261921 | 10.17 | 20.76 | 15.37 | 3.90 | ADP-L-glycero-D-manno-heptose-6-epimerase |
| *FLPKDCKG_02080* | *rfaE* | - | 2261918 | 2262859 | 13.77 | 25.74 | 16.80 | 4.59 | D-beta-D-heptose 7-phosphate kinase |
| *FLPKDCKG_02081* | *lapB* | - | 2262856 | 2264073 | 11.55 | 18.42 | 11.15 | 2.60 | Lipopolysaccharide assembly protein B |
| *FLPKDCKG_02082* | *lapA* | - | 2264134 | 2264448 | 35.93 | 41.20 | 24.28 | 5.57 | Lipopolysaccharide assembly protein A |
| *FLPKDCKG_02083* | *ihfB* | - | 2264640 | 2264999 | 5.57 | 58.74 | 67.49 | 44.65 | Integration host factor subunit beta |
| *FLPKDCKG_02084* | *rpsA-1* | - | 2265003 | 2266715 | 25.06 | 87.41 | 105.86 | 49.63 | 30S ribosomal protein S1 |
| *FLPKDCKG_02085* | *cmk* | - | 2266891 | 2267580 | 29.95 | 99.68 | 58.48 | 25.06 | Cytidylate kinase |
| *FLPKDCKG_02086* | *aroA* | - | 2267613 | 2268935 | 1.12 | 3.34 | 5.14 | 2.30 | 3-phosphoshikimate 1-carboxyvinyltransferase |
| *FLPKDCKG_02087* | *tyrA-1* | - | 2268942 | 2269844 | 0.70 | 4.15 | 5.02 | 1.09 | Prephenate dehydrogenase |
| *FLPKDCKG_02088* | *hisC-2* | - | 2269841 | 2270962 | 7.26 | 20.66 | 16.41 | 4.66 | Histidinol-phosphate aminotransferase |
| *FLPKDCKG_02089* | *pheA-2* | - | 2270971 | 2272056 | 8.85 | 15.78 | 14.45 | 5.95 | P-protein |
| *FLPKDCKG_02090* | *serC* | - | 2272049 | 2273179 | 11.27 | 31.47 | 23.92 | 7.04 | Phosphoserine aminotransferase |
| *FLPKDCKG_02091* | *gyrA* | - | 2273179 | 2275848 | 20.31 | 41.39 | 32.20 | 9.71 | DNA gyrase subunit A |
| *FLPKDCKG_02092* | *psaB* | + | 2276262 | 2276855 | 382.34 | 369.60 | 338.06 | 134.68 | Photosystem I P700 chlorophyll a apoprotein A2 |
| *FLPKDCKG_02093* | *ubiG-1* | + | 2277003 | 2277731 | 12.03 | 44.47 | 16.16 | 3.82 | Ubiquinone biosynthesis O-methyltransferase |
| *FLPKDCKG_02094* | *cbbZC-1* | + | 2277728 | 2278402 | 8.15 | 26.59 | 8.08 | 5.71 | Phosphoglycolate phosphatase, chromosomal |
| *FLPKDCKG_02095* | *pbuE-3* | + | 2278783 | 2279958 | 0.41 | 1.78 | 0.77 | 1.31 | Purine efflux pump PbuE |
| *FLPKDCKG_02096* | *ssrA* | + | 2280094 | 2280482 | 7087.13 | 18294.28 | 15503.19 | 7908.16 | tmRNA |
| *FLPKDCKG_02097* | *czcD* | - | 2280543 | 2281436 | 10.49 | 13.48 | 4.91 | 2.84 | Metal cation efflux system protein CzcD |
| *FLPKDCKG_02098* | *wecB* | + | 2282060 | 2283085 | 5.80 | 25.80 | 12.76 | 1.49 | UDP-N-acetylglucosamine 2-epimerase |
| *FLPKDCKG_02099* | | + | 2283118 | 2285328 | 1.68 | 5.31 | 3.70 | 1.10 | Hypothetical protein |
| *FLPKDCKG_02100* | | + | 2285347 | 2287044 | 0.97 | 3.52 | 1.89 | 0.50 | Hypothetical protein |
| *FLPKDCKG_02101* | | + | 2287041 | 2289779 | 0.63 | 4.46 | 1.77 | 0.59 | Hypothetical protein |
| *FLPKDCKG_02102* | *algJ-1* | + | 2289817 | 2290815 | 0.61 | 4.65 | 3.12 | 0.43 | putative alginate O-acetylase AlgJ |
| *FLPKDCKG_02103* | *algJ-2* | + | 2290820 | 2291920 | 0.04 | 5.93 | 0.31 | 0.02 | putative alginate O-acetylase AlgJ |
| *FLPKDCKG_02104* | *patA-1* | + | 2291927 | 2293507 | 0.60 | 2.31 | 3.63 | 0.16 | Peptidoglycan O-acetyltransferase |
| *FLPKDCKG_02105* | *dltC-2* | + | 2293533 | 2293772 | 8.00 | 10.46 | 6.23 | 0.00 | D-alanine--poly(phosphoribitol) ligase subunit 2 |
| *FLPKDCKG_02106* | *dltA-1* | + | 2293795 | 2295270 | 6.11 | 9.49 | 5.27 | 1.69 | D-alanine--poly(phosphoribitol) ligase subunit 1 |
| *FLPKDCKG_02107* | | + | 2295267 | 2295947 | 8.21 | 20.01 | 7.60 | 1.97 | Hypothetical protein |
| *FLPKDCKG_02108* | *mmgC-2* | + | 2295957 | 2297087 | 4.11 | 5.53 | 5.12 | 0.56 | Acyl-CoA dehydrogenase |
| *FLPKDCKG_02109* | | - | 2297100 | 2297837 | 3.37 | 12.75 | 5.67 | 1.60 | Hypothetical protein |
| *FLPKDCKG_02110* | | - | 2298109 | 2298522 | 389.53 | 126.71 | 49.54 | 18.74 | Hypothetical protein |
| *FLPKDCKG_02111* | *mltB-1* | - | 2298642 | 2299835 | 1.74 | 4.11 | 3.85 | 1.22 | Membrane-bound lytic murein transglycosylase B |
| *FLPKDCKG_02112* | | - | 2299832 | 2300548 | 4.56 | 9.49 | 3.93 | 2.83 | Hypothetical protein |
| *FLPKDCKG_02113* | | - | 2300553 | 2301395 | 5.89 | 6.93 | 7.89 | 0.37 | Hypothetical protein |
| *FLPKDCKG_02114* | | + | 2301515 | 2302654 | 3.92 | 4.46 | 3.14 | 0.46 | Hypothetical protein |
| *FLPKDCKG_02115* | | - | 2302659 | 2303129 | 2.10 | 3.18 | 1.61 | 2.22 | Hypothetical protein |
| *FLPKDCKG_02116* | | - | 2303144 | 2305429 | 5.81 | 11.78 | 10.21 | 3.29 | putative ABC transporter ATP-binding protein |
| *FLPKDCKG_02117* | *cphA-1* | + | 2305690 | 2308305 | 13.26 | 20.76 | 17.07 | 8.03 | Cyanophycin synthetase |
| *FLPKDCKG_02118* | *cphA-2* | + | 2308368 | 2310938 | 6.14 | 12.73 | 11.45 | 5.54 | Cyanophycin synthetase |
| *FLPKDCKG_02119* | | - | 2311003 | 2311122 | 22.26 | 49.06 | 29.97 | 0.62 | Hypothetical protein |
| *FLPKDCKG_02120* | | + | 2311559 | 2311759 | 278.66 | 5518.09 | 3488.65 | 412.22 | Hypothetical protein |
| *FLPKDCKG_02121* | | + | 2311786 | 2311950 | 315.04 | 7349.25 | 3738.05 | 393.93 | Hypothetical protein |
| *FLPKDCKG_02122* | *osmY-2* | + | 2312028 | 2312366 | 57.97 | 1800.77 | 972.61 | 104.59 | Osmotically-inducible protein Y |
| *FLPKDCKG_02123* | *phzF-2* | - | 2312548 | 2313420 | 2.58 | 8.73 | 4.35 | 1.48 | Trans-2,3-dihydro-3-hydroxyanthranilate isomerase |
| *FLPKDCKG_02124* | | - | 2314165 | 2314905 | 11.18 | 35.97 | 14.42 | 3.89 | Hypothetical protein |
| *FLPKDCKG_02125* | *cdiA-1* | - | 2314902 | 2316992 | 1.97 | 6.22 | 3.49 | 0.48 | 16S rRNA endonuclease CdiA |
| *FLPKDCKG_02126* | *cdiA-2* | - | 2316940 | 2324889 | 0.33 | 0.72 | 0.62 | 0.16 | tRNA nuclease CdiA |
| *FLPKDCKG_02127* | *shlB-1* | - | 2324915 | 2326636 | 1.20 | 1.50 | 1.26 | 0.54 | Hemolysin transporter protein ShlB |
| *FLPKDCKG_02128* | | - | 2326807 | 2328204 | 2.21 | 2.51 | 2.99 | 1.91 | Hypothetical protein |
| *FLPKDCKG_02129* | *efeU* | - | 2328253 | 2329098 | 4.71 | 9.37 | 7.63 | 4.64 | Ferrous iron permease EfeU |
| *FLPKDCKG_02130* | | - | 2329112 | 2329474 | 5.14 | 6.84 | 8.42 | 2.29 | Hypothetical protein |
| *FLPKDCKG_02131* | *tpd* | - | 2329477 | 2330013 | 23.88 | 33.52 | 38.79 | 9.91 | 34 kDa membrane antigen |
| *FLPKDCKG_02132* | | - | 2330237 | 2330806 | 30.56 | 49.57 | 37.88 | 7.41 | Hypothetical protein |
| *FLPKDCKG_02133* | *phaB-1* | - | 2330927 | 2331664 | 76.60 | 106.17 | 63.93 | 13.56 | Acetoacetyl-CoA reductase |
| *FLPKDCKG_02134* | *phaC-1* | - | 2331740 | 2333341 | 33.06 | 58.06 | 42.71 | 11.32 | Poly(3-hydroxyalkanoate) polymerase subunit PhaC |
| *FLPKDCKG_02135* | *yfiH* | - | 2333501 | 2334265 | 8.14 | 28.55 | 26.26 | 3.47 | Laccase domain protein YfiH |

| Locus name | Gene name | Strand | Gene start (bp) | Gene end (bp) | Average TPM Inoculum | Average TPM $t_6$ | Average TPM $t_{24}$ | Average TPM $t_{168}$ | Annotation description |
|---|---|---|---|---|---|---|---|---|---|
| *FLPKDCKG_02136* | *rluD* | - | 2334253 | 2335263 | 9.60 | 23.37 | 24.07 | 7.98 | Ribosomal large subunit pseudouridine synthase D |
| *FLPKDCKG_02137* | *bamD* | + | 2335317 | 2336159 | 138.64 | 123.03 | 98.45 | 43.94 | Outer membrane protein assembly factor BamD |
| *FLPKDCKG_02138* | *dinG-2* | - | 2336240 | 2338327 | 3.45 | 7.04 | 7.69 | 2.80 | putative ATP-dependent helicase DinG |
| *FLPKDCKG_02139* | *rpsA-2* | + | 2338555 | 2340930 | 4.33 | 8.64 | 6.51 | 1.45 | 30S ribosomal protein S1 |
| *FLPKDCKG_02140* | | + | 2341244 | 2341747 | 4.75 | 2.30 | 4.40 | 1.23 | Hypothetical protein |
| *FLPKDCKG_02141* | *trpS* | - | 2341820 | 2343172 | 18.47 | 37.04 | 23.65 | 4.81 | Tryptophan--tRNA ligase |
| *FLPKDCKG_02142* | | - | 2343316 | 2343639 | 6.49 | 1.20 | 3.30 | 0.69 | Hypothetical protein |
| *FLPKDCKG_02143* | | - | 2343646 | 2345358 | 6.27 | 4.54 | 2.25 | 0.54 | Hypothetical protein |
| *FLPKDCKG_02144* | | - | 2345355 | 2345636 | 5.60 | 1.88 | 0.00 | 2.07 | Hypothetical protein |
| *FLPKDCKG_02145* | | - | 2345660 | 2346292 | 18.53 | 4.29 | 7.75 | 2.73 | Hypothetical protein |
| *FLPKDCKG_02146* | *fhuA-9* | - | 2346441 | 2348942 | 0.38 | 0.50 | 0.00 | 0.28 | Ferrichrome-iron receptor |
| *FLPKDCKG_02147* | | - | 2349033 | 2349983 | 0.26 | 0.92 | 1.13 | 0.30 | Hypothetical protein |
| *FLPKDCKG_02148* | *fecI-5* | - | 2349976 | 2350509 | 0.95 | 0.00 | 0.60 | 1.63 | putative RNA polymerase sigma factor FecI |
| *FLPKDCKG_02149* | *zapE-1* | - | 2350630 | 2351502 | 1.94 | 3.43 | 3.65 | 0.22 | Cell division protein ZapE |
| *FLPKDCKG_02150* | *zapE-2* | - | 2351502 | 2351720 | 2.72 | 6.99 | 6.58 | 1.21 | Cell division protein ZapE |
| *FLPKDCKG_02151* | *lpd-1* | - | 2351797 | 2353224 | 23.34 | 13.66 | 15.07 | 7.67 | Dihydrolipoyl dehydrogenase |
| *FLPKDCKG_02152* | *sucB-1* | - | 2353345 | 2354577 | 70.33 | 35.72 | 28.68 | 16.90 | Dihydrolipoyllysine-residue succinyltransferase component of 2-oxoglutarate dehydrogenase complex |
| *FLPKDCKG_02153* | *sucA* | - | 2354681 | 2357548 | 73.31 | 71.33 | 52.96 | 23.98 | 2-oxoglutarate dehydrogenase E1 component |
| *FLPKDCKG_02154* | *lrp-4* | - | 2357878 | 2358348 | 1.05 | 1.36 | 2.97 | 0.64 | Leucine-responsive regulatory protein |
| *FLPKDCKG_02155* | *acdS-2* | + | 2359038 | 2359556 | 1.51 | 82.39 | 45.53 | 18.51 | 1-aminocyclopropane-1-carboxylate deaminase |
| *FLPKDCKG_02156* | *pgdA-1* | + | 2359783 | 2360727 | 5.77 | 30.01 | 24.11 | 6.57 | Peptidoglycan deacetylase |
| *FLPKDCKG_02157* | *amaB-3* | + | 2360739 | 2362433 | 2.93 | 21.48 | 15.83 | 4.10 | N-carbamoyl-L-amino acid hydrolase |
| *FLPKDCKG_02158* | *argE-1* | + | 2362437 | 2363669 | 3.41 | 16.91 | 16.79 | 5.93 | Acetylornithine deacetylase |
| *FLPKDCKG_02159* | | - | 2363722 | 2365773 | 1.87 | 6.07 | 5.18 | 2.53 | Hypothetical protein |
| *FLPKDCKG_02160* | *aceE-3* | + | 2366043 | 2368757 | 29.32 | 16.73 | 6.43 | 1.14 | Pyruvate dehydrogenase E1 component |
| *FLPKDCKG_02161* | *maeB-1* | + | 2369001 | 2371298 | 7.17 | 10.85 | 7.83 | 3.36 | NADP-dependent malic enzyme |
| *FLPKDCKG_02162* | | + | 2371450 | 2371902 | 11.18 | 27.31 | 7.88 | 3.29 | Hypothetical protein |
| *FLPKDCKG_02163* | | - | 2371940 | 2373049 | 2.37 | 3.43 | 1.61 | 0.78 | Hypothetical protein |
| *FLPKDCKG_02164* | *gltA-2* | - | 2373151 | 2374461 | 15.74 | 43.77 | 45.82 | 18.83 | Citrate synthase |
| *FLPKDCKG_02165* | | - | 2374481 | 2374741 | 61.45 | 152.03 | 116.03 | 23.61 | Hypothetical protein |
| *FLPKDCKG_02166* | *sdhB* | - | 2374768 | 2375484 | 19.58 | 42.15 | 49.28 | 14.02 | Succinate dehydrogenase iron-sulfur subunit |
| *FLPKDCKG_02167* | *sdhA-1* | - | 2375503 | 2376168 | 46.12 | 82.44 | 67.60 | 29.22 | Succinate dehydrogenase flavoprotein subunit |
| *FLPKDCKG_02168* | *sdhA-2* | - | 2376162 | 2377280 | 14.65 | 43.52 | 42.53 | 12.80 | Succinate dehydrogenase flavoprotein subunit |
| *FLPKDCKG_02169* | *sdhD* | - | 2377284 | 2377667 | 7.70 | 26.26 | 15.75 | 6.20 | Succinate dehydrogenase hydrophobic membrane anchor subunit |
| *FLPKDCKG_02170* | *sdhC* | - | 2377668 | 2378084 | 24.20 | 63.71 | 46.12 | 15.41 | Succinate dehydrogenase cytochrome b556 subunit |
| *FLPKDCKG_02171* | *mngR* | - | 2378206 | 2378985 | 9.33 | 48.20 | 20.08 | 13.78 | Mannosyl-D-glycerate transport/metabolism system repressor MngR |
| *FLPKDCKG_02172* | *mdh-1* | + | 2379343 | 2380332 | 103.08 | 202.53 | 134.52 | 58.89 | Malate dehydrogenase |
| *FLPKDCKG_02173* | *prpB* | + | 2380598 | 2381488 | 6.90 | 16.01 | 17.12 | 9.57 | 2-methylisocitrate lyase |
| *FLPKDCKG_02174* | *prpC* | + | 2381579 | 2382784 | 16.18 | 24.70 | 31.49 | 17.71 | 2-methylcitrate synthase |
| *FLPKDCKG_02175* | *prpD-2* | + | 2382867 | 2384318 | 12.53 | 10.93 | 10.73 | 3.43 | 2-methylcitrate dehydratase |
| *FLPKDCKG_02176* | | + | 2384720 | 2385148 | 11.58 | 27.31 | 20.77 | 8.33 | Hypothetical protein |
| *FLPKDCKG_02177* | *rlmL* | + | 2386088 | 2386954 | 6.67 | 5.21 | 3.88 | 2.73 | Ribosomal RNA large subunit methyltransferase L |
| *FLPKDCKG_02178* | | + | 2387215 | 2387403 | 505.76 | 97.09 | 49.42 | 18.90 | Hypothetical protein |
| *FLPKDCKG_02179* | | + | 2387558 | 2388073 | 3.11 | 5.00 | 7.84 | 1.54 | Hypothetical protein |
| *FLPKDCKG_02180* | | + | 2388223 | 2389131 | 3.45 | 21.48 | 12.98 | 2.38 | Hypothetical protein |
| *FLPKDCKG_02181* | | + | 2389135 | 2389860 | 0.87 | 7.50 | 3.14 | 0.68 | Hypothetical protein |
| *FLPKDCKG_02182* | *lpxH-1* | + | 2389981 | 2390838 | 0.25 | 0.06 | 0.92 | 0.84 | UDP-2,3-diacylglucosamine hydrolase |
| *FLPKDCKG_02183* | *acsA-2* | + | 2390967 | 2392634 | 46.68 | 44.97 | 33.59 | 8.35 | Acetyl-coenzyme A synthetase |
| *FLPKDCKG_02184* | *dapH* | + | 2392682 | 2393203 | 30.18 | 31.32 | 25.41 | 6.56 | 2,3,4,5-tetrahydropyridine-2,6-dicarboxylate N-acetyltransferase |
| *FLPKDCKG_02185* | *hslO* | + | 2393290 | 2394219 | 24.98 | 36.16 | 21.90 | 7.41 | 33 kDa chaperonin |
| *FLPKDCKG_02186* | | + | 2394264 | 2395001 | 1.80 | 3.43 | 3.22 | 3.99 | Hypothetical protein |
| *FLPKDCKG_02187* | *ftsB* | - | 2394955 | 2395383 | 7.44 | 45.04 | 23.28 | 9.65 | Cell division protein FtsB |
| *FLPKDCKG_02188* | *eno-2* | - | 2395448 | 2396734 | 16.51 | 47.75 | 43.03 | 13.63 | Enolase |
| *FLPKDCKG_02189* | | - | 2396739 | 2397023 | 22.84 | 61.25 | 41.90 | 8.72 | Hypothetical protein |
| *FLPKDCKG_02190* | *pyrG* | - | 2397251 | 2398903 | 37.33 | 96.26 | 58.90 | 23.21 | CTP synthase |
| *FLPKDCKG_02191* | | + | 2399099 | 2400106 | 1.47 | 10.26 | 11.81 | 1.33 | Hypothetical protein |
| *FLPKDCKG_02192* | *dppA-2* | - | 2400187 | 2402886 | 3.38 | 5.22 | 2.83 | 1.70 | D-aminopeptidase |
| *FLPKDCKG_02193* | | - | 2402907 | 2404001 | 0.91 | 3.14 | 2.82 | 0.77 | Beta-peptidyl aminopeptidase BapA |
| *FLPKDCKG_02194* | *gsiD-4* | - | 2404171 | 2405073 | 0.59 | 2.46 | 0.56 | 0.48 | Glutathione transport system permease protein GsiD |
| *FLPKDCKG_02195* | *gsiC-3* | - | 2405081 | 2406001 | 0.81 | 1.07 | 3.75 | 0.84 | Glutathione transport system permease protein GsiC |
| *FLPKDCKG_02196* | *gsiB-3* | - | 2406094 | 2407647 | 2.32 | 9.44 | 12.79 | 4.46 | Glutathione-binding protein GsiB |
| *FLPKDCKG_02197* | *gsiA-5* | - | 2407713 | 2409599 | 0.73 | 2.82 | 2.35 | 1.32 | Glutathione import ATP-binding protein GsiA |
| *FLPKDCKG_02198* | *iaaA* | - | 2409646 | 2410626 | 2.38 | 4.43 | 3.27 | 1.13 | Isoaspartyl peptidase |
| *FLPKDCKG_02199* | *ybbH* | - | 2410623 | 2411498 | 4.53 | 2.28 | 2.39 | 2.99 | putative HTH-type transcriptional regulator YbbH |
| *FLPKDCKG_02200* | | + | 2411687 | 2412145 | 2.76 | 8.72 | 1.06 | 1.38 | Acetyltransferase |
| *FLPKDCKG_02201* | *scoB-1* | - | 2412223 | 2412876 | 27.57 | 19.69 | 27.32 | 7.20 | putative succinyl-CoA:3-ketoacid coenzyme A transferase subunit B |
| *FLPKDCKG_02202* | *scoA-1* | - | 2412889 | 2413587 | 109.96 | 105.97 | 77.71 | 24.96 | putative succinyl-CoA:3-ketoacid coenzyme A transferase subunit A |
| *FLPKDCKG_02203* | | - | 2413885 | 2414334 | 11.59 | 17.50 | 9.20 | 4.31 | Hypothetical protein |
| *FLPKDCKG_02204* | | + | 2414447 | 2416093 | 54.48 | 55.75 | 32.54 | 10.55 | Electron transfer flavoprotein-ubiquinone oxidoreductase |

| Locus name | Gene name | Strand | Gene start (bp) | Gene end (bp) | Average TPM Inoculum | Average TPM $t_6$ | Average TPM $t_{24}$ | Average TPM $t_{168}$ | Annotation description |
|---|---|---|---|---|---|---|---|---|---|
| *FLPKDCKG_02205* | *galE-1* | + | 2416107 | 2416952 | 8.45 | 13.31 | 4.82 | 2.45 | UDP-glucose 4-epimerase |
| *FLPKDCKG_02206* | | + | 2417024 | 2418142 | 0.64 | 1.04 | 0.45 | 0.01 | Putative cytochrome P450 123 |
| *FLPKDCKG_02207* | | + | 2418139 | 2418483 | 1.28 | 3.06 | 3.39 | 0.39 | Hypothetical protein |
| *FLPKDCKG_02208* | *rspR-4* | + | 2418534 | 2419415 | 10.32 | 11.58 | 12.37 | 1.93 | HTH-type transcriptional repressor RspR |
| *FLPKDCKG_02209* | *bauA-1* | + | 2419612 | 2420946 | 0.72 | 2.12 | 1.65 | 1.04 | Beta-alanine--pyruvate aminotransferase |
| *FLPKDCKG_02210* | *ltaE-1* | - | 2420985 | 2421995 | 3.33 | 7.44 | 2.66 | 3.10 | Low specificity L-threonine aldolase |
| *FLPKDCKG_02211* | *yijE-6* | - | 2422038 | 2422925 | 2.26 | 2.28 | 1.74 | 0.59 | putative cystine transporter YijE |
| *FLPKDCKG_02212* | *acs* | - | 2423078 | 2425057 | 21.24 | 86.17 | 60.08 | 20.77 | Acetyl-coenzyme A synthetase |
| *FLPKDCKG_02213* | | - | 2425325 | 2426665 | 1.67 | 4.35 | 5.68 | 1.30 | Hypothetical protein |
| *FLPKDCKG_02214* | | - | 2426662 | 2427069 | 31.29 | 47.64 | 43.08 | 14.01 | Hypothetical protein |
| *FLPKDCKG_02215* | *acsA-3* | - | 2427066 | 2428955 | 23.76 | 35.71 | 26.31 | 10.45 | Acetyl-coenzyme A synthetase |
| *FLPKDCKG_02216* | *mepH-2* | + | 2429270 | 2429929 | 18.68 | 44.14 | 42.68 | 19.17 | Murein DD-endopeptidase MepH |
| *FLPKDCKG_02217* | | - | 2430120 | 2430449 | 169.05 | 97.66 | 79.28 | 49.70 | Hypothetical protein |
| *FLPKDCKG_02218* | *cysB-3* | - | 2430685 | 2431629 | 8.98 | 52.68 | 97.95 | 23.10 | HTH-type transcriptional regulator CysB |
| *FLPKDCKG_02219* | *mmsA* | - | 2431723 | 2433216 | 36.25 | 21.17 | 18.90 | 10.36 | Methylmalonate-semialdehyde dehydrogenase [acylating] |
| *FLPKDCKG_02220* | | - | 2433313 | 2434530 | 13.44 | 12.22 | 12.14 | 4.28 | Hypothetical protein |
| *FLPKDCKG_02221* | *dmlR-13* | + | 2434702 | 2435622 | 2.52 | 8.08 | 5.38 | 0.58 | HTH-type transcriptional regulator DmlR |
| *FLPKDCKG_02222* | *lptF* | - | 2435722 | 2436864 | 7.11 | 14.22 | 10.28 | 0.96 | Lipopolysaccharide export system permease protein LptF |
| *FLPKDCKG_02223* | *pepA* | + | 2436878 | 2438356 | 42.85 | 32.69 | 32.84 | 7.07 | Cytosol aminopeptidase |
| *FLPKDCKG_02224* | *holC* | + | 2438405 | 2438866 | 45.28 | 131.22 | 69.62 | 15.37 | DNA polymerase III subunit chi |
| *FLPKDCKG_02225* | | + | 2438927 | 2439373 | 6.42 | 17.63 | 14.53 | 8.48 | Hypothetical protein |
| *FLPKDCKG_02226* | *yccA* | + | 2439537 | 2440247 | 128.03 | 29.46 | 80.06 | 50.11 | Modulator of FtsH protease YccA |
| *FLPKDCKG_02227* | *dkgB-2* | + | 2440314 | 2441231 | 5.35 | 4.31 | 5.42 | 2.29 | 2,5-diketo-D-gluconic acid reductase B |
| *FLPKDCKG_02228* | | + | 2441236 | 2442522 | 0.60 | 0.43 | 0.81 | 0.37 | Hypothetical protein |
| *FLPKDCKG_02229* | | + | 2442614 | 2443576 | 0.68 | 0.35 | 0.27 | 0.13 | Hypothetical protein |
| *FLPKDCKG_02230* | *doeB* | + | 2443608 | 2444585 | 0.66 | 1.33 | 0.00 | 0.14 | N-alpha-acetyl-L-2,4-diaminobutyric acid deacetylase |
| *FLPKDCKG_02231* | *thrC-2* | + | 2444589 | 2445686 | 0.64 | 1.40 | 0.76 | 0.33 | Threonine synthase |
| *FLPKDCKG_02232* | *cynR-7* | - | 2445758 | 2446675 | 0.98 | 5.55 | 3.22 | 0.99 | HTH-type transcriptional regulator CynR |
| *FLPKDCKG_02233* | *recO* | - | 2446695 | 2447240 | 2.27 | 13.01 | 5.67 | 0.61 | DNA repair protein RecO |
| *FLPKDCKG_02234* | *era* | - | 2447275 | 2448165 | 16.33 | 58.25 | 23.83 | 11.20 | GTPase Era |
| *FLPKDCKG_02235* | *rnc* | - | 2448162 | 2448923 | 16.96 | 32.60 | 20.40 | 9.30 | Ribonuclease 3 |
| *FLPKDCKG_02236* | *lepB* | - | 2448926 | 2449810 | 11.01 | 34.23 | 22.85 | 8.03 | Signal peptidase I |
| *FLPKDCKG_02237* | *lepA* | - | 2449833 | 2451626 | 3.91 | 14.82 | 10.71 | 5.74 | Elongation factor 4 |
| *FLPKDCKG_02238* | *degQ* | - | 2451851 | 2453341 | 17.38 | 71.18 | 68.14 | 24.23 | Periplasmic pH-dependent serine endoprotease DegQ |
| *FLPKDCKG_02239* | *mucB* | - | 2453382 | 2454458 | 11.76 | 73.94 | 64.43 | 15.32 | Sigma factor AlgU regulatory protein MucB |
| *FLPKDCKG_02240* | | - | 2454458 | 2454967 | 45.19 | 119.67 | 101.44 | 37.61 | Hypothetical protein |
| *FLPKDCKG_02241* | *rpoE-1* | - | 2454980 | 2455579 | 71.35 | 219.29 | 126.71 | 33.67 | ECF RNA polymerase sigma-E factor |
| *FLPKDCKG_02242* | | - | 2455576 | 2456070 | 19.60 | 94.97 | 71.73 | 13.48 | Hypothetical protein |
| *FLPKDCKG_02243* | *fabF-2* | - | 2456072 | 2457301 | 21.69 | 47.79 | 47.66 | 17.46 | 3-oxoacyl-[acyl-carrier-protein] synthase 2 |
| *FLPKDCKG_02244* | *acpP-2* | - | 2457491 | 2457730 | 89.73 | 139.35 | 133.29 | 43.78 | Acyl carrier protein |
| *FLPKDCKG_02245* | *fabG-7* | - | 2457947 | 2458699 | 6.19 | 23.73 | 22.12 | 6.55 | 3-oxoacyl-[acyl-carrier-protein] reductase FabG |
| *FLPKDCKG_02246* | *fabD* | - | 2458704 | 2459639 | 5.84 | 16.41 | 19.24 | 6.47 | Malonyl CoA-acyl carrier protein transacylase |
| *FLPKDCKG_02247* | *fabH-1* | - | 2459729 | 2460541 | 5.54 | 12.79 | 12.20 | 4.71 | 3-oxoacyl-[acyl-carrier-protein] synthase 3 |
| *FLPKDCKG_02248* | *fabH-2* | - | 2460538 | 2460714 | 6.61 | 20.22 | 19.18 | 4.12 | 3-oxoacyl-[acyl-carrier-protein] synthase 3 |
| *FLPKDCKG_02249* | *plsX* | - | 2460714 | 2461757 | 4.96 | 23.27 | 21.22 | 9.98 | Phosphate acyltransferase |
| *FLPKDCKG_02250* | *rpmF* | - | 2461841 | 2462023 | 64.37 | 143.86 | 102.84 | 34.68 | 50S ribosomal protein L32 |
| *FLPKDCKG_02251* | *yceD* | - | 2462076 | 2462549 | 32.61 | 87.58 | 101.76 | 32.03 | Large ribosomal RNA subunit accumulation protein YceD |
| *FLPKDCKG_02252* | *yceF* | + | 2462870 | 2463472 | 24.59 | 39.33 | 37.02 | 16.32 | Maf-like protein YceF |
| *FLPKDCKG_02253* | *rsmI-1* | + | 2463469 | 2464179 | 3.78 | 11.76 | 9.96 | 2.26 | Ribosomal RNA small subunit methyltransferase I |
| *FLPKDCKG_02254* | *amiD-3* | + | 2464182 | 2464967 | 1.64 | 5.88 | 4.10 | 3.22 | N-acetylmuramoyl-L-alanine amidase AmiD |
| *FLPKDCKG_02255* | *msrQ* | - | 2465022 | 2465675 | 1.84 | 11.00 | 7.78 | 1.83 | Protein-methionine-sulfoxide reductase heme-binding subunit MsrQ |
| *FLPKDCKG_02256* | *msrP-2* | - | 2465672 | 2466640 | 7.94 | 14.32 | 14.92 | 3.40 | Protein-methionine-sulfoxide reductase catalytic subunit MsrP |
| *FLPKDCKG_02257* | *ppaX* | - | 2466805 | 2467470 | 1.47 | 3.91 | 1.51 | 0.57 | Pyrophosphatase PpaX |
| *FLPKDCKG_02258* | *rluC* | - | 2467472 | 2468398 | 1.88 | 8.66 | 6.62 | 3.08 | Ribosomal large subunit pseudouridine synthase C |
| *FLPKDCKG_02259* | *rne* | + | 2468946 | 2472020 | 4.02 | 21.09 | 22.53 | 13.64 | Ribonuclease E |
| *FLPKDCKG_02260* | *pagL* | - | 2472165 | 2472713 | 77.42 | 30.15 | 38.08 | 18.08 | Lipid A deacylase PagL |
| *FLPKDCKG_02261* | *accD* | - | 2472961 | 2473839 | 12.19 | 29.28 | 25.43 | 12.14 | Acetyl-coenzyme A carboxylase carboxyl transferase subunit beta |
| *FLPKDCKG_02262* | *trpA-2* | - | 2473877 | 2474743 | 14.55 | 21.54 | 18.70 | 4.92 | Tryptophan synthase alpha chain |
| *FLPKDCKG_02263* | *trpB-2* | - | 2474745 | 2475944 | 26.00 | 44.14 | 22.13 | 8.78 | Tryptophan synthase beta chain |
| *FLPKDCKG_02264* | *argE-2* | - | 2476097 | 2477248 | 29.37 | 30.28 | 23.90 | 5.64 | Acetylornithine deacetylase |
| *FLPKDCKG_02265* | *proP-4* | + | 2477383 | 2478696 | 1.61 | 3.70 | 2.29 | 0.74 | Proline/betaine transporter |
| *FLPKDCKG_02266* | *relA* | - | 2478816 | 2481107 | 6.32 | 8.56 | 4.58 | 0.87 | GTP pyrophosphokinase |
| *FLPKDCKG_02267* | *mtfA* | + | 2481165 | 2482049 | 6.63 | 11.44 | 5.33 | 2.19 | Protein MtfA |
| *FLPKDCKG_02268* | *fldA-2* | + | 2482046 | 2482555 | 1.76 | 0.74 | 2.22 | 0.00 | Flavodoxin |
| *FLPKDCKG_02269* | *yabJ-3* | - | 2482720 | 2483073 | 23.60 | 28.78 | 28.82 | 8.20 | 2-iminobutanoate/2-iminopropanoate deaminase |
| *FLPKDCKG_02270* | *pncB2* | - | 2483144 | 2484319 | 22.39 | 47.50 | 32.78 | 6.11 | Nicotinate phosphoribosyltransferase 2 |
| *FLPKDCKG_02271* | | + | 2484476 | 2484799 | 216.23 | 225.54 | 162.67 | 35.90 | Ferredoxin 1 |
| *FLPKDCKG_02272* | *fpr-2* | + | 2484846 | 2485646 | 13.73 | 27.31 | 18.50 | 4.50 | Ferredoxin--NADP reductase |
| *FLPKDCKG_02273* | *phaC-2* | + | 2485883 | 2487109 | 80.33 | 37.65 | 23.47 | 14.71 | Poly(3-hydroxyalkanoate) polymerase subunit PhaC |

| Locus name | Gene name | Strand | Gene start (bp) | Gene end (bp) | Average TPM Inoculum | Average TPM $t_6$ | Average TPM $t_{24}$ | Average TPM $t_{168}$ | Annotation description |
|---|---|---|---|---|---|---|---|---|---|
| *FLPKDCKG_02274* | *rsxB* | + | 2487165 | 2487806 | 6.67 | 8.40 | 8.85 | 3.23 | Electron transport complex subunit RsxB |
| *FLPKDCKG_02275* | *nth* | + | 2487803 | 2488438 | 1.75 | 5.22 | 5.59 | 1.14 | Endonuclease III |
| *FLPKDCKG_02276* | *drrA* | + | 2488805 | 2489578 | 39.32 | 22.05 | 21.30 | 6.74 | Daunorubicin/doxorubicin resistance ATP-binding protein DrrA |
| *FLPKDCKG_02277* | *livF-7* | + | 2489575 | 2490285 | 51.98 | 21.63 | 21.68 | 6.83 | High-affinity branched-chain amino acid transport ATP-binding protein LivF |
| *FLPKDCKG_02278* | | + | 2490347 | 2491561 | 88.38 | 24.47 | 19.70 | 14.02 | Hypothetical protein |
| *FLPKDCKG_02279* | *livH-7* | + | 2491660 | 2492547 | 15.45 | 7.42 | 8.29 | 7.38 | High-affinity branched-chain amino acid transport system permease protein LivH |
| *FLPKDCKG_02280* | | + | 2492544 | 2493527 | 5.41 | 2.40 | 5.16 | 2.15 | Hypothetical protein |
| *FLPKDCKG_02281* | | + | 2493634 | 2493801 | 13.02 | 22.05 | 10.32 | 2.80 | Hypothetical protein |
| *FLPKDCKG_02282* | *proX-1* | + | 2493774 | 2494643 | 13.45 | 12.77 | 10.03 | 2.75 | Glycine betaine/proline betaine-binding periplasmic protein |
| *FLPKDCKG_02283* | *ilvE-1* | + | 2494716 | 2495561 | 25.17 | 25.47 | 17.68 | 5.54 | Branched-chain-amino-acid aminotransferase |
| *FLPKDCKG_02284* | *plcN-3* | + | 2495644 | 2497794 | 2.07 | 3.35 | 2.36 | 0.99 | Non-hemolytic phospholipase C |
| *FLPKDCKG_02285* | | + | 2497776 | 2498198 | 1.27 | 4.56 | 2.66 | 2.32 | Hypothetical protein |
| *FLPKDCKG_02286* | *livF-8* | - | 2498221 | 2498985 | 2.21 | 1.77 | 3.11 | 1.16 | High-affinity branched-chain amino acid transport ATP-binding protein LivF |
| *FLPKDCKG_02287* | *cysA-3* | - | 2498982 | 2500820 | 3.23 | 1.14 | 1.37 | 0.61 | Sulfate/thiosulfate import ATP-binding protein CysA |
| *FLPKDCKG_02288* | *livH-8* | - | 2500826 | 2501863 | 2.44 | 3.84 | 2.18 | 0.24 | High-affinity branched-chain amino acid transport system permease protein LivH |
| *FLPKDCKG_02289* | | - | 2501997 | 2503160 | 48.05 | 31.18 | 21.89 | 11.21 | Leu/Ile/Val-binding protein |
| *FLPKDCKG_02290* | *bioY* | - | 2503408 | 2503965 | 1.14 | 5.93 | 1.62 | 0.45 | Biotin transporter BioY |
| *FLPKDCKG_02291* | *bioN* | - | 2504019 | 2504615 | 0.69 | 2.13 | 1.52 | 1.51 | Energy-coupling factor transporter transmembrane protein BioN |
| *FLPKDCKG_02292* | *bioM* | - | 2504612 | 2505289 | 2.70 | 6.73 | 4.55 | 0.65 | Biotin transport ATP-binding protein BioM |
| *FLPKDCKG_02293* | *msrB-2* | + | 2505342 | 2505740 | 78.37 | 79.20 | 68.43 | 20.81 | Peptide methionine sulfoxide reductase MsrB |
| *FLPKDCKG_02294* | *yciB-2* | + | 2505737 | 2506303 | 24.07 | 51.89 | 39.12 | 9.99 | Intracellular septation protein |
| *FLPKDCKG_02295* | *bolA* | + | 2506380 | 2506649 | 40.91 | 118.70 | 61.36 | 11.80 | DNA-binding transcriptional regulator BolA |
| *FLPKDCKG_02296* | | + | 2506667 | 2507443 | 75.91 | 102.99 | 75.96 | 27.13 | putative parvulin-type peptidyl-prolyl cis-trans isomerase |
| *FLPKDCKG_02297* | *ligA* | - | 2507641 | 2509740 | 2.12 | 6.29 | 5.25 | 1.16 | DNA ligase |
| *FLPKDCKG_02298* | | - | 2509740 | 2509961 | 11.09 | 27.67 | 9.17 | 8.98 | Hypothetical protein |
| *FLPKDCKG_02299* | *zipA* | - | 2509903 | 2510802 | 12.93 | 16.01 | 12.37 | 3.13 | Cell division protein ZipA |
| *FLPKDCKG_02300* | *smc-2* | - | 2510849 | 2514382 | 11.54 | 10.56 | 8.64 | 2.94 | Chromosome partition protein Smc |
| *FLPKDCKG_02301* | *lplT* | + | 2514695 | 2516011 | 3.29 | 7.68 | 7.67 | 2.42 | Lysophospholipid transporter LplT |
| *FLPKDCKG_02302* | | - | 2516076 | 2517005 | 3.45 | 11.70 | 4.11 | 2.40 | Hypothetical protein |
| *FLPKDCKG_02303* | *tsaB* | - | 2517002 | 2518258 | 12.13 | 24.21 | 16.27 | 4.43 | tRNA threonylcarbamoyladenosine biosynthesis protein TsaB |
| *FLPKDCKG_02304* | *ompR-2* | + | 2518390 | 2519124 | 319.08 | 762.03 | 459.35 | 116.10 | Transcriptional regulatory protein OmpR |
| *FLPKDCKG_02305* | *envZ-1* | + | 2519159 | 2520553 | 9.17 | 44.63 | 31.91 | 5.93 | Osmolarity sensor protein EnvZ |
| *FLPKDCKG_02306* | | + | 2520613 | 2521161 | 167.84 | 610.43 | 597.68 | 169.86 | Alkyl hydroperoxide reductase subunit C |
| *FLPKDCKG_02307* | *ahpD-2* | + | 2521235 | 2521753 | 21.86 | 233.90 | 242.35 | 65.18 | Alkyl hydroperoxide reductase AhpD |
| *FLPKDCKG_02308* | | - | 2521867 | 2523951 | 32.78 | 48.06 | 28.53 | 6.57 | Hypothetical protein |
| *FLPKDCKG_02309* | *ispF* | - | 2524225 | 2524710 | 10.56 | 17.49 | 13.45 | 2.22 | 2-C-methyl-D-erythritol 2,4-cyclodiphosphate synthase |
| *FLPKDCKG_02310* | *ispD* | - | 2524707 | 2525390 | 5.36 | 2.10 | 1.52 | 1.12 | 2-C-methyl-D-erythritol 4-phosphate cytidylyltransferase |
| *FLPKDCKG_02311* | *mfd* | + | 2525420 | 2528902 | 3.45 | 6.63 | 4.02 | 1.94 | Transcription-repair-coupling factor |
| *FLPKDCKG_02312* | *serB-2* | + | 2528926 | 2529783 | 4.06 | 7.36 | 4.89 | 1.13 | Phosphoserine phosphatase |
| *FLPKDCKG_02313* | *bfrD-3* | - | 2529845 | 2532064 | 0.72 | 1.51 | 0.48 | 1.04 | putative TonB-dependent receptor BfrD |
| *FLPKDCKG_02314* | | - | 2532297 | 2532626 | 1.13 | 6.00 | 2.88 | 0.88 | Hypothetical protein |
| *FLPKDCKG_02315* | *nuoN* | - | 2532639 | 2534123 | 2.41 | 10.86 | 6.67 | 3.37 | NADH-quinone oxidoreductase subunit N |
| *FLPKDCKG_02316* | *nuoM* | - | 2534133 | 2535635 | 4.47 | 12.92 | 13.07 | 2.94 | NADH-quinone oxidoreductase subunit M |
| *FLPKDCKG_02317* | *nuoL* | - | 2535632 | 2537683 | 4.37 | 12.68 | 9.65 | 4.38 | NADH-quinone oxidoreductase subunit L |
| *FLPKDCKG_02318* | | - | 2537701 | 2538009 | 6.62 | 25.94 | 12.99 | 3.86 | NADH-quinone oxidoreductase subunit 11 |
| *FLPKDCKG_02319* | *nuoJ* | - | 2538006 | 2538656 | 6.33 | 16.15 | 10.16 | 5.40 | NADH-quinone oxidoreductase subunit J |
| *FLPKDCKG_02320* | *nuoI* | - | 2538674 | 2539162 | 6.45 | 15.19 | 23.95 | 3.24 | NADH-quinone oxidoreductase subunit I |
| *FLPKDCKG_02321* | *nuoH* | - | 2539184 | 2540257 | 6.31 | 19.71 | 14.19 | 4.82 | NADH-quinone oxidoreductase subunit H |
| *FLPKDCKG_02322* | *nqo3* | - | 2540257 | 2542584 | 9.06 | 25.64 | 22.13 | 6.13 | NADH-quinone oxidoreductase chain 3 |
| *FLPKDCKG_02323* | *nuoF* | - | 2542600 | 2543967 | 15.96 | 35.60 | 21.06 | 6.46 | NADH-quinone oxidoreductase subunit F |
| *FLPKDCKG_02324* | *nqo2* | - | 2543964 | 2544458 | 24.39 | 38.84 | 39.78 | 3.38 | NADH-quinone oxidoreductase subunit 2 |
| *FLPKDCKG_02325* | *nuoD* | - | 2544505 | 2545761 | 21.12 | 47.50 | 32.29 | 6.61 | NADH-quinone oxidoreductase subunit D |
| *FLPKDCKG_02326* | *nqo5* | - | 2545765 | 2546388 | 53.16 | 78.14 | 47.92 | 11.39 | NADH-quinone oxidoreductase chain 5 |
| *FLPKDCKG_02327* | *nuoB* | - | 2546474 | 2546950 | 26.89 | 61.66 | 46.58 | 9.39 | NADH-quinone oxidoreductase subunit B |
| *FLPKDCKG_02328* | *ndhC* | - | 2546993 | 2547352 | 127.81 | 368.82 | 178.94 | 37.53 | NAD(P)H-quinone oxidoreductase subunit 3 |
| *FLPKDCKG_02329* | | - | 2547770 | 2548933 | 876.37 | 630.51 | 640.82 | 348.89 | Outer membrane porin protein |
| *FLPKDCKG_02330* | *ttgR-1* | + | 2549371 | 2549985 | 1.05 | 1.90 | 3.49 | 3.12 | HTH-type transcriptional regulator TtgR |
| *FLPKDCKG_02331* | | + | 2549982 | 2550428 | 1.02 | 6.25 | 6.76 | 4.95 | Hypothetical protein |
| *FLPKDCKG_02332* | | + | 2550425 | 2550859 | 1.57 | 2.78 | 2.15 | 6.10 | Hypothetical protein |
| *FLPKDCKG_02333* | | - | 2550830 | 2551597 | 1.01 | 6.06 | 3.99 | 1.10 | Hypothetical protein |
| *FLPKDCKG_02334* | | - | 2551594 | 2552052 | 2.32 | 1.85 | 4.56 | 1.20 | Hypothetical protein |
| *FLPKDCKG_02335* | *dagK* | - | 2552425 | 2553417 | 5.38 | 9.74 | 4.69 | 1.45 | Diacylglycerol kinase |
| *FLPKDCKG_02336* | *ywrD-2* | - | 2553476 | 2555278 | 1.71 | 2.68 | 2.24 | 0.73 | Putative gamma-glutamyltransferase YwrD |
| *FLPKDCKG_02337* | *gsiB-4* | - | 2555371 | 2556873 | 6.51 | 4.83 | 3.53 | 1.09 | Glutathione-binding protein GsiB |
| *FLPKDCKG_02338* | *cmpR-8* | - | 2557106 | 2558002 | 2.59 | 4.54 | 3.03 | 1.28 | HTH-type transcriptional activator CmpR |
| *FLPKDCKG_02339* | *ttgR-2* | - | 2558089 | 2558628 | 0.62 | 1.56 | 0.00 | 1.97 | HTH-type transcriptional regulator TtgR |
| *FLPKDCKG_02340* | *ttgW* | - | 2558625 | 2558726 | 2.62 | 6.91 | 3.12 | 1.97 | putative HTH-type transcriptional regulator TtgW |
| *FLPKDCKG_02341* | *acrA-1* | + | 2558969 | 2560144 | 0.89 | 2.18 | 1.37 | 0.96 | Multidrug efflux pump subunit AcrA |
| *FLPKDCKG_02342* | *acrB-1* | + | 2560151 | 2563288 | 0.53 | 0.63 | 0.67 | 0.20 | Multidrug efflux pump subunit AcrB |

| Locus name | Gene name | Strand | Gene start (bp) | Gene end (bp) | Average TPM Inoculum | Average TPM $t_6$ | Average TPM $t_{24}$ | Average TPM $t_{168}$ | Annotation description |
|---|---|---|---|---|---|---|---|---|---|
| *FLPKDCKG_02343* | *oprM-4* | + | 2563285 | 2564676 | 1.05 | 0.56 | 1.24 | 0.15 | Outer membrane protein OprM |
| *FLPKDCKG_02344* | | - | 2564686 | 2565348 | 5.04 | 12.52 | 7.11 | 1.77 | Hypothetical protein |
| *FLPKDCKG_02345* | *srpC-2* | - | 2565628 | 2566803 | 0.51 | 0.00 | 0.27 | 0.41 | Solvent efflux pump outer membrane protein SrpC |
| *FLPKDCKG_02346* | *bepE-1* | - | 2566814 | 2569933 | 0.87 | 1.11 | 1.19 | 0.50 | Efflux pump membrane transporter BepE |
| *FLPKDCKG_02347* | *mdtA-2* | - | 2569950 | 2571080 | 1.10 | 1.87 | 0.62 | 1.79 | Multidrug resistance protein MdtA |
| *FLPKDCKG_02348* | | - | 2571271 | 2573241 | 10.64 | 5.90 | 4.37 | 1.08 | Glycine betaine transporter |
| *FLPKDCKG_02349* | *ttgR-3* | - | 2573384 | 2574007 | 23.71 | 42.71 | 26.42 | 8.34 | HTH-type transcriptional regulator TtgR |
| *FLPKDCKG_02350* | *srpA* | + | 2574174 | 2575361 | 1.04 | 0.60 | 1.11 | 1.34 | Solvent efflux pump periplasmic linker SrpA |
| *FLPKDCKG_02351* | *acrF* | + | 2575358 | 2578531 | 0.81 | 0.57 | 0.33 | 0.13 | Multidrug export protein AcrF |
| *FLPKDCKG_02352* | *mepC* | + | 2578534 | 2580054 | 1.05 | 0.36 | 0.91 | 0.71 | Multidrug/solvent efflux pump outer membrane protein MepC |
| *FLPKDCKG_02353* | | + | 2580300 | 2581436 | 0.27 | 0.23 | 0.43 | 0.45 | Hypothetical protein |
| *FLPKDCKG_02354* | *dppA-3* | + | 2581511 | 2583073 | 0.30 | 0.13 | 0.63 | 0.32 | Periplasmic dipeptide transport protein |
| *FLPKDCKG_02355* | *dppA-4* | - | 2583376 | 2584935 | 1.26 | 2.83 | 0.91 | 0.68 | Periplasmic dipeptide transport protein |
| *FLPKDCKG_02356* | | - | 2585117 | 2585737 | 8.14 | 14.85 | 6.57 | 2.06 | Hypothetical protein |
| *FLPKDCKG_02357* | *gsiA-6* | - | 2585828 | 2587483 | 0.22 | 0.90 | 0.19 | 0.53 | Glutathione import ATP-binding protein GsiA |
| *FLPKDCKG_02358* | *gsiD-5* | - | 2587480 | 2588301 | 0.06 | 0.32 | 0.00 | 0.00 | Glutathione transport system permease protein GsiD |
| *FLPKDCKG_02359* | *gsiC-4* | - | 2588298 | 2589242 | 0.14 | 0.60 | 0.00 | 0.26 | Glutathione transport system permease protein GsiC |
| *FLPKDCKG_02360* | | - | 2589304 | 2590167 | 0.43 | 0.00 | 2.29 | 0.00 | Hypothetical protein |
| *FLPKDCKG_02361* | *asnC-2* | - | 2590758 | 2591183 | 20.82 | 11.83 | 4.29 | 2.87 | Regulatory protein AsnC |
| *FLPKDCKG_02362* | *fecA-1* | + | 2591480 | 2593636 | 0.18 | 0.63 | 0.97 | 0.49 | Fe(3+) dicitrate transport protein FecA |
| *FLPKDCKG_02363* | | - | 2593691 | 2594587 | 1.69 | 0.82 | 1.13 | 1.66 | Beta-lactamase |
| *FLPKDCKG_02364* | *ampR* | + | 2594689 | 2595573 | 1.81 | 4.24 | 2.14 | 0.42 | HTH-type transcriptional activator AmpR |
| *FLPKDCKG_02365* | *yedI* | - | 2595606 | 2596517 | 4.72 | 7.95 | 9.00 | 0.92 | Inner membrane protein YedI |
| *FLPKDCKG_02366* | | + | 2596681 | 2596989 | 3.92 | 5.88 | 2.15 | 2.70 | Hypothetical protein |
| *FLPKDCKG_02367* | *ytfE* | - | 2597087 | 2597767 | 1.66 | 2.76 | 1.35 | 0.00 | Iron-sulfur cluster repair protein YtfE |
| *FLPKDCKG_02368* | *norR2* | + | 2597918 | 2599438 | 2.59 | 6.53 | 2.04 | 0.21 | Nitric oxide reductase transcription regulator NorR2 |
| *FLPKDCKG_02369* | | - | 2599452 | 2599802 | 3.16 | 18.98 | 9.48 | 1.84 | Hypothetical protein |
| *FLPKDCKG_02370* | *bphP-1* | - | 2599887 | 2601017 | 2.53 | 21.60 | 10.89 | 0.61 | Bacteriophytochrome |
| *FLPKDCKG_02371* | | - | 2601055 | 2602584 | 2.31 | 57.35 | 33.27 | 2.81 | Blue-light-activated protein |
| *FLPKDCKG_02372* | *gtaB-1* | - | 2602684 | 2603517 | 0.91 | 122.10 | 78.08 | 4.88 | UTP--glucose-1-phosphate uridylyltransferase |
| *FLPKDCKG_02373* | *pleD-3* | + | 2603851 | 2604351 | 27.65 | 110.31 | 62.34 | 10.94 | Response regulator PleD |
| *FLPKDCKG_02374* | *apaH-1* | - | 2604312 | 2605139 | 0.91 | 13.78 | 15.73 | 1.57 | Bis(5'-nucleosyl)-tetraphosphatase [symmetrical] |
| *FLPKDCKG_02375* | *bacC-4* | + | 2605296 | 2606060 | 2.49 | 10.30 | 4.04 | 2.66 | Dihydroanticapsin 7-dehydrogenase |
| *FLPKDCKG_02376* | *phaB-2* | - | 2606071 | 2606811 | 0.58 | 8.11 | 1.74 | 2.99 | Acetoacetyl-CoA reductase |
| *FLPKDCKG_02377* | *citN* | - | 2606850 | 2608190 | 0.16 | 1.00 | 1.76 | 1.41 | Citrate transporter |
| *FLPKDCKG_02378* | | + | 2608569 | 2612627 | 0.62 | 1.89 | 0.98 | 0.93 | Extracellular serine protease |
| *FLPKDCKG_02379* | *tctD-2* | + | 2612855 | 2613553 | 13.08 | 14.75 | 12.05 | 2.70 | Transcriptional regulatory protein tctD |
| *FLPKDCKG_02380* | *qseC-4* | + | 2613522 | 2614964 | 0.53 | 2.09 | 2.21 | 0.03 | Sensor protein QseC |
| *FLPKDCKG_02381* | | + | 2614954 | 2616132 | 0.26 | 0.58 | 0.00 | 0.07 | Hypothetical protein |
| *FLPKDCKG_02382* | | - | 2616080 | 2619013 | 0.42 | 0.54 | 0.45 | 0.06 | putative TonB-dependent receptor |
| *FLPKDCKG_02383* | | - | 2619135 | 2620154 | 0.72 | 0.99 | 1.80 | 0.81 | Hypothetical protein |
| *FLPKDCKG_02384* | *fecI-6* | - | 2620195 | 2620722 | 0.82 | 1.48 | 2.92 | 0.04 | putative RNA polymerase sigma factor FecI |
| *FLPKDCKG_02385* | | - | 2620900 | 2621955 | 1.07 | 1.88 | 0.50 | 0.45 | Hypothetical protein |
| *FLPKDCKG_02386* | *pupB-1* | - | 2622050 | 2624503 | 0.24 | 0.64 | 0.12 | 0.27 | Ferric-pseudobactin BN7/BN8 receptor |
| *FLPKDCKG_02387* | | - | 2624590 | 2625540 | 0.89 | 0.00 | 0.42 | 0.00 | Hypothetical protein |
| *FLPKDCKG_02388* | *fecI-7* | - | 2625547 | 2626074 | 4.41 | 0.00 | 0.39 | 1.23 | putative RNA polymerase sigma factor FecI |
| *FLPKDCKG_02389* | | - | 2626180 | 2626644 | 0.46 | 2.52 | 0.00 | 0.00 | Hypothetical protein |
| *FLPKDCKG_02390* | *fptA-3* | - | 2626644 | 2628770 | 0.51 | 0.83 | 1.14 | 0.15 | Fe(3+)-pyochelin receptor |
| *FLPKDCKG_02391* | *ssuB-2* | - | 2628867 | 2629550 | 0.15 | 0.00 | 0.00 | 0.00 | Aliphatic sulfonates import ATP-binding protein SsuB |
| *FLPKDCKG_02392* | *ssuC-7* | - | 2629543 | 2630337 | 0.11 | 0.00 | 0.33 | 0.00 | Putative aliphatic sulfonates transport permease protein SsuC |
| *FLPKDCKG_02393* | | - | 2630334 | 2631329 | 0.31 | 0.77 | 0.00 | 0.19 | Hypothetical protein |
| *FLPKDCKG_02394* | *anr-1* | + | 2631391 | 2632110 | 0.56 | 1.70 | 1.33 | 0.71 | Transcriptional activator protein Anr |
| *FLPKDCKG_02395* | *ctb-2* | + | 2632237 | 2632950 | 1.30 | 10.01 | 10.40 | 2.49 | Group 3 truncated hemoglobin ctb |
| *FLPKDCKG_02396* | | - | 2632999 | 2633727 | 32.09 | 109.68 | 63.24 | 8.27 | Hypothetical protein |
| *FLPKDCKG_02397* | *recD* | - | 2634134 | 2640187 | 1.56 | 1.17 | 1.44 | 0.63 | RecBCD enzyme subunit RecD |
| *FLPKDCKG_02398* | | + | 2640374 | 2641159 | 13.49 | 19.21 | 8.08 | 2.21 | Hypothetical protein |
| *FLPKDCKG_02399* | | + | 2641239 | 2642171 | 4.79 | 11.69 | 6.00 | 2.58 | Hypothetical protein |
| *FLPKDCKG_02400* | | - | 2642187 | 2642801 | 0.54 | 2.04 | 1.18 | 1.45 | Hypothetical protein |
| *FLPKDCKG_02401* | | - | 2643021 | 2643620 | 10.08 | 19.83 | 14.65 | 5.91 | Hypothetical protein |
| *FLPKDCKG_02402* | *trg-3* | - | 2643762 | 2644751 | 0.11 | 2.20 | 1.92 | 0.17 | Methyl-accepting chemotaxis protein III |
| *FLPKDCKG_02403* | | - | 2644748 | 2645401 | 0.07 | 3.60 | 4.19 | 0.00 | Hypothetical protein |
| *FLPKDCKG_02404* | | + | 2645721 | 2646023 | 2.10 | 3.07 | 3.09 | 0.67 | Hypothetical protein |
| *FLPKDCKG_02405* | | - | 2646155 | 2648269 | 0.72 | 1.08 | 0.84 | 0.08 | Hypothetical protein |
| *FLPKDCKG_02406* | *pbuE-4* | - | 2648439 | 2649632 | 1.42 | 6.51 | 2.08 | 0.60 | Purine efflux pump PbuE |
| *FLPKDCKG_02407* | | + | 2650382 | 2651224 | 3.72 | 4.89 | 3.17 | 1.64 | Hypothetical protein |
| *FLPKDCKG_02408* | *adhT-1* | + | 2651405 | 2652424 | 2.02 | 2.85 | 2.33 | 1.36 | Alcohol dehydrogenase |
| *FLPKDCKG_02409* | | - | 2652643 | 2652727 | 4.26 | 18.53 | 6.02 | 3.98 | tRNA |
| *FLPKDCKG_02410* | | + | 2653061 | 2653240 | 20.60 | 107.12 | 104.69 | 14.16 | Hypothetical protein |
| *FLPKDCKG_02411* | *secG* | - | 2653328 | 2653831 | 2.91 | 8.67 | 7.56 | 1.77 | Protein-export membrane protein SecG |

| Locus name | Gene name | Strand | Gene start (bp) | Gene end (bp) | Average TPM Inoculum | Average TPM $t_6$ | Average TPM $t_{24}$ | Average TPM $t_{168}$ | Annotation description |
|---|---|---|---|---|---|---|---|---|---|
| *FLPKDCKG_02412* | *tpiA* | - | 2653927 | 2654643 | 8.76 | 18.03 | 19.27 | 6.46 | Triosephosphate isomerase |
| *FLPKDCKG_02413* | | - | 2654724 | 2655701 | 11.88 | 30.67 | 16.88 | 7.45 | 2-haloacrylate reductase |
| *FLPKDCKG_02414* | | + | 2655823 | 2655910 | 9.54 | 27.26 | 24.48 | 7.90 | tRNA |
| *FLPKDCKG_02415* | | + | 2656158 | 2656790 | 4.48 | 3.88 | 4.65 | 4.01 | Hypothetical protein |
| *FLPKDCKG_02416* | | - | 2656827 | 2658089 | 4.42 | 5.65 | 3.14 | 1.43 | Hypothetical protein |
| *FLPKDCKG_02417* | *nepI-3* | - | 2658198 | 2659433 | 0.30 | 0.12 | 0.51 | 0.16 | Purine ribonucleoside efflux pump NepI |
| *FLPKDCKG_02418* | *rclR-3* | + | 2659534 | 2660520 | 3.09 | 8.33 | 5.49 | 1.68 | RCS-specific HTH-type transcriptional activator RclR |
| *FLPKDCKG_02419* | *pcpR-5* | - | 2660461 | 2661402 | 1.68 | 1.40 | 1.42 | 1.14 | PCP degradation transcriptional activation protein |
| *FLPKDCKG_02420* | | + | 2661495 | 2662418 | 1.47 | 84.80 | 47.33 | 18.36 | Nitrilase |
| *FLPKDCKG_02421* | | + | 2662871 | 2663536 | 0.43 | 1.96 | 4.71 | 1.59 | Hypothetical protein |
| *FLPKDCKG_02422* | *tdcB-1* | - | 2663538 | 2664755 | 1.78 | 4.26 | 5.50 | 1.43 | L-threonine ammonia-lyase |
| *FLPKDCKG_02423* | | - | 2664771 | 2665499 | 5.43 | 10.64 | 9.49 | 3.01 | Hypothetical protein |
| *FLPKDCKG_02424* | *pnp* | - | 2665708 | 2667864 | 3.60 | 53.74 | 46.85 | 20.42 | Polyribonucleotide nucleotidyltransferase |
| *FLPKDCKG_02425* | *rpsO* | - | 2667955 | 2668224 | 9.47 | 114.22 | 106.71 | 33.12 | 30S ribosomal protein S15 |
| *FLPKDCKG_02426* | | + | 2668542 | 2669060 | 11.34 | 16.73 | 10.34 | 1.63 | Hypothetical protein |
| *FLPKDCKG_02427* | *pcs* | - | 2669119 | 2669892 | 32.30 | 75.98 | 36.41 | 7.76 | Phosphatidylcholine synthase |
| *FLPKDCKG_02428* | *ilvC* | - | 2670125 | 2671141 | 15.62 | 32.05 | 43.16 | 18.09 | Ketol-acid reductoisomerase (NADP(+)) |
| *FLPKDCKG_02429* | *ilvH* | - | 2671212 | 2671703 | 71.29 | 95.23 | 96.92 | 18.82 | Acetolactate synthase isozyme 3 small subunit |
| *FLPKDCKG_02430* | *ilvI-1* | - | 2671714 | 2673429 | 16.28 | 20.17 | 21.71 | 6.71 | Acetolactate synthase isozyme 3 large subunit |
| *FLPKDCKG_02431* | *yccS-3* | + | 2673924 | 2676146 | 12.04 | 13.60 | 7.15 | 2.76 | Inner membrane protein YccS |
| *FLPKDCKG_02432* | | - | 2676223 | 2676993 | 4.52 | 11.76 | 4.07 | 3.10 | Hypothetical protein |
| *FLPKDCKG_02433* | *bdhA-1* | - | 2677010 | 2677792 | 19.38 | 31.03 | 20.51 | 6.56 | D-beta-hydroxybutyrate dehydrogenase |
| *FLPKDCKG_02434* | *kefC-2* | - | 2677941 | 2679236 | 3.36 | 6.86 | 5.39 | 1.61 | Glutathione-regulated potassium-efflux system protein KefC |
| *FLPKDCKG_02435* | | + | 2679495 | 2679585 | 40.46 | 50.40 | 40.35 | 4.37 | tRNA |
| *FLPKDCKG_02436* | | - | 2679730 | 2680359 | 3.79 | 2.79 | 5.02 | 1.48 | Hypothetical protein |
| *FLPKDCKG_02437* | *irtA-1* | + | 2680493 | 2682286 | 0.10 | 0.35 | 0.50 | 0.03 | Iron import ATP-binding/permease protein IrtA |
| *FLPKDCKG_02438* | *irtB* | + | 2682280 | 2684046 | 0.07 | 1.15 | 0.00 | 0.33 | Iron import ATP-binding/permease protein IrtB |
| *FLPKDCKG_02439* | *hmuU-2* | + | 2684046 | 2684987 | 0.09 | 0.16 | 0.00 | 0.00 | Hemin transport system permease protein HmuU |
| *FLPKDCKG_02440* | *hmuU-3* | + | 2684984 | 2685931 | 0.11 | 0.36 | 0.00 | 0.00 | Hemin transport system permease protein HmuU |
| *FLPKDCKG_02441* | *yusV* | + | 2685928 | 2686704 | 0.24 | 0.18 | 0.00 | 0.00 | putative siderophore transport system ATP-binding protein YusV |
| *FLPKDCKG_02442* | *yclQ-2* | + | 2686701 | 2687681 | 0.18 | 0.04 | 0.09 | 0.00 | putative ABC transporter solute-binding protein YclQ |
| *FLPKDCKG_02443* | *fcuA-1* | + | 2687791 | 2690061 | 0.18 | 0.00 | 0.42 | 0.29 | Ferrichrome receptor FcuA |
| *FLPKDCKG_02444* | | - | 2690077 | 2691054 | 0.76 | 0.63 | 0.42 | 0.79 | Hypothetical protein |
| *FLPKDCKG_02445* | *yagE* | - | 2691173 | 2692081 | 5.43 | 14.64 | 4.95 | 2.17 | putative 2-keto-3-deoxy-galactonate aldolase YagE |
| *FLPKDCKG_02446* | | - | 2692095 | 2693222 | 6.75 | 8.42 | 5.79 | 2.01 | L-talarate/galactarate dehydratase |
| *FLPKDCKG_02447* | *gltR-3* | + | 2693381 | 2694259 | 2.21 | 3.96 | 7.52 | 0.66 | HTH-type transcriptional regulator GltR |
| *FLPKDCKG_02448* | *atzC* | - | 2694274 | 2695506 | 6.93 | 15.39 | 15.37 | 3.39 | N-isopropylammelide isopropyl amidohydrolase |
| *FLPKDCKG_02449* | | - | 2695534 | 2695734 | 232.74 | 429.37 | 299.97 | 63.25 | Hypothetical protein |
| *FLPKDCKG_02450* | | + | 2695926 | 2696426 | 29.28 | 61.09 | 30.65 | 13.22 | Hypothetical protein |
| *FLPKDCKG_02451* | | + | 2696778 | 2697116 | 466.57 | 1056.39 | 528.83 | 185.17 | Hypothetical protein |
| *FLPKDCKG_02452* | | - | 2697352 | 2697600 | 228.14 | 552.78 | 338.52 | 45.30 | Hypothetical protein |
| *FLPKDCKG_02453* | *serS* | - | 2697901 | 2699247 | 10.50 | 15.09 | 15.03 | 4.32 | Serine--tRNA ligase |
| *FLPKDCKG_02454* | *rarA* | - | 2699290 | 2700630 | 12.34 | 21.99 | 15.14 | 6.39 | Replication-associated recombination protein A |
| *FLPKDCKG_02455* | | - | 2700765 | 2701556 | 11.00 | 18.12 | 7.33 | 2.94 | Hydrolase |
| *FLPKDCKG_02456* | | + | 2701730 | 2702998 | 3.03 | 8.32 | 9.95 | 5.82 | D-threonine aldolase |
| *FLPKDCKG_02457* | *gsiB-5* | + | 2703148 | 2704707 | 5.59 | 8.93 | 17.00 | 9.12 | Glutathione-binding protein GsiB |
| *FLPKDCKG_02458* | *gsiC-5* | + | 2704793 | 2705734 | 0.40 | 3.46 | 1.03 | 0.97 | Glutathione transport system permease protein GsiC |
| *FLPKDCKG_02459* | *ddpC-2* | + | 2705762 | 2706631 | 0.62 | 1.78 | 0.43 | 0.78 | putative D,D-dipeptide transport system permease protein DdpC |
| *FLPKDCKG_02460* | *oppD-4* | + | 2706634 | 2707641 | 0.75 | 0.36 | 2.84 | 1.77 | Oligopeptide transport ATP-binding protein OppD |
| *FLPKDCKG_02461* | *oppF-3* | + | 2707638 | 2708639 | 1.65 | 4.66 | 4.94 | 2.97 | Oligopeptide transport ATP-binding protein OppF |
| *FLPKDCKG_02462* | *hexR-2* | + | 2708672 | 2709544 | 1.22 | 0.52 | 4.31 | 1.57 | HTH-type transcriptional regulator HexR |
| *FLPKDCKG_02463* | *dan-2* | + | 2709559 | 2711040 | 1.85 | 3.54 | 5.41 | 2.38 | D-aminoacylase |
| *FLPKDCKG_02464* | *yabJ-4* | + | 2711124 | 2711456 | 1.77 | 4.39 | 4.68 | 3.98 | 2-iminobutanoate/2-iminopropanoate deaminase |
| *FLPKDCKG_02465* | *lolA* | - | 2711965 | 2712558 | 72.91 | 153.91 | 84.29 | 29.33 | Outer-membrane lipoprotein carrier protein |
| *FLPKDCKG_02466* | *ftsK* | - | 2712598 | 2714985 | 13.72 | 27.24 | 12.99 | 5.89 | DNA translocase FtsK |
| *FLPKDCKG_02467* | *trxB-2* | + | 2715151 | 2715978 | 40.37 | 64.12 | 34.73 | 12.85 | Thioredoxin reductase |
| *FLPKDCKG_02468* | *smrA* | + | 2715965 | 2716651 | 2.10 | 11.22 | 8.17 | 2.79 | putative DNA endonuclease SmrA |
| *FLPKDCKG_02469* | *emrE* | + | 2716648 | 2716977 | 2.87 | 15.11 | 4.97 | 1.47 | Multidrug transporter EmrE |
| *FLPKDCKG_02470* | *actP-3* | - | 2717051 | 2719144 | 1.06 | 2.44 | 2.45 | 0.68 | Cation/acetate symporter ActP |
| *FLPKDCKG_02471* | | - | 2719144 | 2719419 | 3.12 | 1.89 | 5.54 | 3.08 | Hypothetical protein |
| *FLPKDCKG_02472* | | + | 2720155 | 2720244 | 152.90 | 5.71 | 0.00 | 6.20 | Hypothetical protein |
| *FLPKDCKG_02473* | *kdpA* | + | 2720244 | 2722040 | 100.88 | 2.73 | 1.37 | 12.98 | Potassium-transporting ATPase potassium-binding subunit |
| *FLPKDCKG_02474* | *kdpB* | + | 2722063 | 2724222 | 15.83 | 1.74 | 1.95 | 7.31 | Potassium-transporting ATPase ATP-binding subunit |
| *FLPKDCKG_02475* | *kdpC* | + | 2724253 | 2724861 | 4.62 | 1.17 | 0.36 | 7.35 | Potassium-transporting ATPase KdpC subunit |
| *FLPKDCKG_02476* | *kdpD-2* | + | 2724913 | 2727699 | 3.10 | 1.83 | 2.44 | 1.38 | Sensor protein KdpD |
| *FLPKDCKG_02477* | *kdpE* | + | 2727709 | 2728401 | 1.13 | 0.94 | 3.87 | 2.88 | KDP operon transcriptional regulatory protein KdpE |
| *FLPKDCKG_02478* | | + | 2728564 | 2729319 | 34.32 | 2.34 | 1.89 | 1.21 | Hypothetical protein |
| *FLPKDCKG_02479* | | + | 2729459 | 2729782 | 305.48 | 0.75 | 0.25 | 4.49 | Hypothetical protein |
| *FLPKDCKG_02480* | *dapA-2* | + | 2729786 | 2730685 | 67.96 | 0.39 | 1.07 | 2.08 | 4-hydroxy-tetrahydrodipicolinate synthase |

| Locus name | Gene name | Strand | Gene start (bp) | Gene end (bp) | Average TPM Inoculum | Average TPM $t_6$ | Average TPM $t_{24}$ | Average TPM $t_{168}$ | Annotation description |
|---|---|---|---|---|---|---|---|---|---|
| *FLPKDCKG_02481* | *tap-2* | - | 2730663 | 2732207 | 5.86 | 0.91 | 1.37 | 0.50 | Methyl-accepting chemotaxis protein IV |
| *FLPKDCKG_02482* | *rfaL* | - | 2732452 | 2733723 | 1.64 | 4.47 | 1.88 | 0.88 | O-antigen ligase |
| *FLPKDCKG_02483* | *rfaQ-2* | - | 2733838 | 2734794 | 15.16 | 22.37 | 20.05 | 3.87 | Lipopolysaccharide core heptosyltransferase RfaQ |
| *FLPKDCKG_02484* | *icd-2* | - | 2735047 | 2736303 | 87.28 | 93.15 | 55.30 | 21.15 | Isocitrate dehydrogenase [NADP] |
| *FLPKDCKG_02485* | | + | 2736509 | 2736934 | 40.89 | 111.87 | 70.67 | 9.82 | Hypothetical protein |
| *FLPKDCKG_02486* | *cc4-2* | - | 2737004 | 2737366 | 11.06 | 18.38 | 9.42 | 5.80 | Cytochrome c4 |
| *FLPKDCKG_02487* | | - | 2737382 | 2737735 | 23.22 | 73.70 | 48.27 | 12.40 | Cytochrome c-554(548) |
| *FLPKDCKG_02488* | *lon-2* | + | 2737914 | 2738795 | 30.89 | 42.19 | 17.04 | 6.71 | Lon protease |
| *FLPKDCKG_02489* | | + | 2738818 | 2739990 | 14.91 | 16.29 | 10.66 | 5.62 | Hypothetical protein |
| *FLPKDCKG_02490* | | - | 2740031 | 2740588 | 6.04 | 18.84 | 8.69 | 3.16 | L-methionine sulfoximine/L-methionine sulfone acetyltransferase |
| *FLPKDCKG_02491* | | - | 2740681 | 2741061 | 44.97 | 64.31 | 49.85 | 26.20 | Hypothetical protein |
| *FLPKDCKG_02492* | | - | 2741373 | 2741972 | 6.98 | 11.32 | 6.11 | 2.66 | Hypothetical protein |
| *FLPKDCKG_02493* | | + | 2742084 | 2744837 | 30.02 | 36.27 | 36.46 | 7.90 | putative zinc protease |
| *FLPKDCKG_02494* | | + | 2745117 | 2745506 | 27.56 | 24.65 | 26.15 | 7.33 | Hypothetical protein |
| *FLPKDCKG_02495* | *dnaJ* | - | 2745866 | 2746996 | 2.95 | 6.01 | 15.40 | 7.62 | Chaperone protein DnaJ |
| *FLPKDCKG_02496* | *dnaK-1* | - | 2747106 | 2749031 | 84.21 | 47.53 | 139.08 | 75.69 | Chaperone protein DnaK |
| *FLPKDCKG_02497* | | - | 2749113 | 2749517 | 18.94 | 23.09 | 50.60 | 15.52 | Hypothetical protein |
| *FLPKDCKG_02498* | *grpE* | - | 2749532 | 2750089 | 51.83 | 35.76 | 70.07 | 40.81 | Protein GrpE |
| *FLPKDCKG_02499* | | - | 2750297 | 2750530 | 59.94 | 105.10 | 78.99 | 18.73 | Hypothetical protein |
| *FLPKDCKG_02500* | *hemH* | - | 2750543 | 2751634 | 50.65 | 65.56 | 54.01 | 16.58 | Ferrochelatase |
| *FLPKDCKG_02501* | *hrcA* | - | 2751680 | 2752684 | 55.25 | 87.78 | 51.71 | 18.25 | Heat-inducible transcription repressor HrcA |
| *FLPKDCKG_02502* | *nadK* | + | 2752814 | 2753683 | 31.37 | 36.24 | 17.32 | 7.91 | NAD kinase |
| *FLPKDCKG_02503* | *recN* | + | 2753726 | 2755387 | 11.76 | 13.00 | 6.94 | 2.72 | DNA repair protein RecN |
| *FLPKDCKG_02504* | *fur-1* | - | 2755527 | 2755946 | 49.24 | 91.41 | 84.38 | 23.24 | Ferric uptake regulation protein |
| *FLPKDCKG_02505* | *bamE* | + | 2756103 | 2756639 | 52.60 | 61.81 | 47.98 | 12.42 | Outer membrane protein assembly factor BamE |
| *FLPKDCKG_02506* | *dapB* | + | 2756656 | 2757447 | 12.42 | 20.52 | 15.72 | 4.10 | 4-hydroxy-tetrahydrodipicolinate reductase |
| *FLPKDCKG_02507* | *murB* | - | 2757512 | 2758549 | 2.56 | 6.35 | 6.13 | 2.67 | UDP-N-acetylenolpyruvoylglucosamine reductase |
| *FLPKDCKG_02508* | | + | 2758871 | 2758946 | 35.46 | 105.01 | 177.20 | 76.35 | tRNA |
| *FLPKDCKG_02509* | *azoR-1* | - | 2759198 | 2759836 | 1.23 | 1.98 | 1.42 | 0.00 | FMN-dependent NADH-azoreductase |
| *FLPKDCKG_02510* | *phnO-3* | - | 2759833 | 2760279 | 1.19 | 1.53 | 1.61 | 2.40 | Aminoalkylphosphonate N-acetyltransferase |
| *FLPKDCKG_02511* | *gabR-3* | + | 2760393 | 2761802 | 4.59 | 7.38 | 5.11 | 1.22 | HTH-type transcriptional regulatory protein GabR |
| *FLPKDCKG_02512* | | - | 2761853 | 2762332 | 1.14 | 3.69 | 2.35 | 0.00 | Hypothetical protein |
| *FLPKDCKG_02513* | | + | 2762561 | 2762636 | 61.92 | 166.96 | 206.12 | 99.93 | tRNA |
| *FLPKDCKG_02514* | | + | 2763113 | 2763188 | 55.00 | 143.94 | 202.49 | 84.97 | tRNA |
| *FLPKDCKG_02515* | | + | 2763243 | 2763316 | 36.98 | 131.55 | 78.12 | 23.73 | tRNA |
| *FLPKDCKG_02516* | | + | 2763386 | 2763754 | 86.02 | 500.26 | 323.27 | 74.18 | Hypothetical protein |
| *FLPKDCKG_02517* | | + | 2763757 | 2764347 | 12.82 | 70.03 | 79.38 | 12.25 | Hypothetical protein |
| *FLPKDCKG_02518* | | - | 2764390 | 2764842 | 2.19 | 6.28 | 5.80 | 0.46 | Hypothetical protein |
| *FLPKDCKG_02519* | *mhqP* | - | 2764939 | 2765340 | 1.60 | 3.07 | 0.83 | 0.31 | Putative oxidoreductase MhqP |
| *FLPKDCKG_02520* | *dmlR-14* | + | 2765466 | 2766398 | 4.30 | 2.82 | 0.53 | 1.45 | HTH-type transcriptional regulator DmlR |
| *FLPKDCKG_02521* | *mutT1* | + | 2766409 | 2766990 | 58.24 | 62.17 | 36.55 | 8.94 | putative 8-oxo-dGTP diphosphatase 1 |
| *FLPKDCKG_02522* | *aat* | + | 2766987 | 2767742 | 4.40 | 8.59 | 6.84 | 0.62 | Leucyl/phenylalanyl-tRNA--protein transferase |
| *FLPKDCKG_02523* | *ate* | + | 2767870 | 2768625 | 92.14 | 79.31 | 40.79 | 11.19 | Putative arginyl-tRNA--protein transferase |
| *FLPKDCKG_02524* | *pyrD* | + | 2768671 | 2769711 | 10.65 | 22.80 | 15.71 | 5.82 | Dihydroorotate dehydrogenase (quinone) |
| *FLPKDCKG_02525* | | - | 2769857 | 2770435 | 55.87 | 331.33 | 460.49 | 219.26 | Hypothetical protein |
| *FLPKDCKG_02526* | | + | 2770819 | 2771796 | 35.27 | 45.61 | 63.38 | 38.50 | Hypothetical protein |
| *FLPKDCKG_02527* | *yiaJ* | - | 2771925 | 2772788 | 20.56 | 27.66 | 15.86 | 13.28 | DNA-binding transcriptional repressor YiaJ |
| *FLPKDCKG_02528* | *cinA* | - | 2772946 | 2773758 | 17.07 | 11.02 | 8.55 | 5.43 | CinA-like protein |
| *FLPKDCKG_02529* | | - | 2773805 | 2774608 | 16.67 | 12.64 | 13.27 | 6.11 | Hypothetical protein |
| *FLPKDCKG_02530* | | + | 2774672 | 2775055 | 42.17 | 82.23 | 45.12 | 14.07 | Hypothetical protein |
| *FLPKDCKG_02531* | *rhlE-1* | - | 2775144 | 2776589 | 5.71 | 29.28 | 29.03 | 7.74 | ATP-dependent RNA helicase RhlE |
| *FLPKDCKG_02532* | *pchR-5* | - | 2776834 | 2777808 | 0.90 | 2.54 | 2.71 | 0.60 | Regulatory protein PchR |
| *FLPKDCKG_02533* | *fyuA-2* | + | 2777886 | 2780003 | 0.33 | 1.57 | 0.48 | 0.07 | Pesticin receptor |
| *FLPKDCKG_02534* | | + | 2780020 | 2781270 | 0.03 | 0.51 | 0.00 | 0.00 | Hypothetical protein |
| *FLPKDCKG_02535* | *pqsH* | + | 2781273 | 2782439 | 1.51 | 4.56 | 3.78 | 0.46 | 2-heptyl-3-hydroxy-4(1H)-quinolone synthase |
| *FLPKDCKG_02536* | | - | 2782446 | 2782781 | 1.06 | 7.51 | 3.12 | 0.00 | Hypothetical protein |
| *FLPKDCKG_02537* | | - | 2782778 | 2783527 | 2.50 | 4.11 | 3.28 | 1.72 | Hypothetical protein |
| *FLPKDCKG_02538* | *ilvE-2* | + | 2783714 | 2784634 | 71.39 | 59.67 | 36.48 | 10.79 | Branched-chain-amino-acid aminotransferase |
| *FLPKDCKG_02539* | | + | 2784714 | 2784926 | 66.68 | 88.32 | 48.86 | 22.09 | Hypothetical protein |
| *FLPKDCKG_02540* | | + | 2784961 | 2785383 | 16.38 | 14.76 | 22.55 | 5.51 | Hypothetical protein |
| *FLPKDCKG_02541* | | + | 2785395 | 2786495 | 4.65 | 14.14 | 7.83 | 3.83 | Hypothetical protein |
| *FLPKDCKG_02542* | *bepA-4* | - | 2786539 | 2788062 | 10.12 | 19.93 | 13.75 | 3.42 | Beta-barrel assembly-enhancing protease |
| *FLPKDCKG_02543* | *gtaB-2* | + | 2788120 | 2788965 | 42.60 | 55.75 | 40.92 | 10.06 | UTP--glucose-1-phosphate uridylyltransferase |
| *FLPKDCKG_02544* | | - | 2788994 | 2789872 | 8.52 | 15.14 | 14.87 | 2.06 | Hypothetical protein |
| *FLPKDCKG_02545* | | - | 2789969 | 2791075 | 2.34 | 3.75 | 4.76 | 1.15 | Outer membrane porin protein |
| *FLPKDCKG_02546* | *mgtA-1* | - | 2791130 | 2793931 | 0.51 | 1.28 | 0.38 | 0.33 | Magnesium-transporting ATPase, P-type 1 |
| *FLPKDCKG_02547* | | - | 2794351 | 2794683 | 43.30 | 16.35 | 27.83 | 20.89 | Hypothetical protein |
| *FLPKDCKG_02548* | | - | 2795128 | 2796132 | 0.96 | 6.22 | 3.17 | 5.54 | Hypothetical protein |
| *FLPKDCKG_02549* | *pepP* | + | 2796508 | 2797851 | 13.81 | 29.36 | 18.63 | 6.37 | Xaa-Pro aminopeptidase |

| Locus name | Gene name | Strand | Gene start (bp) | Gene end (bp) | Average TPM Inoculum | Average TPM $t_6$ | Average TPM $t_{24}$ | Average TPM $t_{168}$ | Annotation description |
|---|---|---|---|---|---|---|---|---|---|
| *FLPKDCKG_02550* | | - | 2797920 | 2798384 | 11.99 | 20.90 | 15.43 | 11.34 | Hypothetical protein |
| *FLPKDCKG_02551* | *ubiH* | + | 2798465 | 2799502 | 7.91 | 14.04 | 9.34 | 2.88 | 2-octaprenyl-6-methoxyphenol hydroxylase |
| *FLPKDCKG_02552* | *dus* | + | 2799716 | 2800741 | 29.06 | 53.87 | 25.56 | 11.06 | putative tRNA-dihydrouridine synthase |
| *FLPKDCKG_02553* | *fis* | + | 2800778 | 2801017 | 23.54 | 73.92 | 55.74 | 19.71 | DNA-binding protein Fis |
| *FLPKDCKG_02554* | *purH* | + | 2801121 | 2802710 | 4.96 | 18.03 | 14.61 | 6.30 | Bifunctional purine biosynthesis protein PurH |
| *FLPKDCKG_02555* | *ruvC* | + | 2802710 | 2803258 | 2.06 | 6.71 | 8.61 | 4.11 | Crossover junction endodeoxyribonuclease RuvC |
| *FLPKDCKG_02556* | *ruvA* | + | 2803358 | 2803930 | 5.42 | 13.04 | 8.95 | 5.15 | Holliday junction ATP-dependent DNA helicase RuvA |
| *FLPKDCKG_02557* | | - | 2804040 | 2804960 | 30.96 | 32.40 | 21.59 | 6.86 | Hypothetical protein |
| *FLPKDCKG_02558* | *tdcB-2* | - | 2805074 | 2806039 | 7.99 | 13.18 | 11.56 | 2.54 | L-threonine dehydratase catabolic TdcB |
| *FLPKDCKG_02559* | *ruvB* | - | 2806134 | 2807207 | 10.80 | 13.29 | 8.87 | 3.82 | Holliday junction ATP-dependent DNA helicase RuvB |
| *FLPKDCKG_02560* | *envZ-2* | - | 2807310 | 2808860 | 2.00 | 11.99 | 7.28 | 3.96 | Osmolarity sensor protein EnvZ |
| *FLPKDCKG_02561* | *ompR-3* | - | 2808860 | 2809612 | 13.38 | 60.41 | 19.66 | 6.08 | Transcriptional regulatory protein OmpR |
| *FLPKDCKG_02562* | | + | 2809856 | 2811088 | 7.50 | 38.17 | 22.38 | 6.64 | Hypothetical protein |
| *FLPKDCKG_02563* | *recQ* | + | 2811182 | 2813011 | 3.67 | 11.31 | 13.01 | 2.25 | ATP-dependent DNA helicase RecQ |
| *FLPKDCKG_02564* | *cydB-1* | - | 2813123 | 2814178 | 5.54 | 10.56 | 7.65 | 1.96 | Cytochrome bd-I ubiquinol oxidase subunit 2 |
| *FLPKDCKG_02565* | *appC-1* | - | 2814175 | 2815518 | 5.14 | 9.29 | 6.04 | 1.64 | Cytochrome bd-II ubiquinol oxidase subunit 1 |
| *FLPKDCKG_02566* | *glnS* | + | 2815655 | 2817421 | 13.84 | 32.94 | 27.36 | 9.93 | Glutamine--tRNA ligase |
| *FLPKDCKG_02567* | *minC* | + | 2817639 | 2818556 | 8.27 | 15.63 | 11.69 | 5.37 | Septum site-determining protein MinC |
| *FLPKDCKG_02568* | *minD* | + | 2818746 | 2819561 | 27.58 | 44.11 | 39.57 | 13.53 | Septum site-determining protein MinD |
| *FLPKDCKG_02569* | *minE* | + | 2819565 | 2819822 | 5.52 | 18.06 | 10.93 | 4.29 | Cell division topological specificity factor |
| *FLPKDCKG_02570* | | - | 2819954 | 2820178 | 98.89 | 875.29 | 514.01 | 139.53 | Hypothetical protein |
| *FLPKDCKG_02571* | | - | 2820369 | 2820902 | 9.27 | 22.91 | 11.62 | 3.73 | Hypothetical protein |
| *FLPKDCKG_02572* | *ettA-1* | - | 2821668 | 2823341 | 30.61 | 73.46 | 35.63 | 19.36 | Energy-dependent translational throttle protein EttA |
| *FLPKDCKG_02573* | *yokD* | + | 2823492 | 2824310 | 4.49 | 7.58 | 1.51 | 2.71 | SPBc2 prophage-derived aminoglycoside N(3')-acetyltransferase-like protein YokD |
| *FLPKDCKG_02574* | | - | 2824323 | 2825249 | 2.58 | 1.89 | 2.36 | 1.81 | Hypothetical protein |
| *FLPKDCKG_02575* | *nimR-6* | + | 2825348 | 2826136 | 8.63 | 8.45 | 8.92 | 2.25 | HTH-type transcriptional regulator NimR |
| *FLPKDCKG_02576* | | + | 2826247 | 2826768 | 20.45 | 35.64 | 14.78 | 4.77 | Hypothetical protein |
| *FLPKDCKG_02577* | *oppF-4* | - | 2827262 | 2828257 | 5.45 | 1.92 | 1.06 | 2.91 | Oligopeptide transport ATP-binding protein OppF |
| *FLPKDCKG_02578* | *oppD-5* | - | 2828254 | 2829234 | 1.14 | 1.22 | 1.13 | 0.72 | Oligopeptide transport ATP-binding protein OppD |
| *FLPKDCKG_02579* | *dppC-3* | - | 2829384 | 2830298 | 1.93 | 0.49 | 1.57 | 0.09 | Dipeptide transport system permease protein DppC |
| *FLPKDCKG_02580* | *gsiC-6* | - | 2830298 | 2831287 | 1.34 | 1.28 | 2.57 | 0.64 | Glutathione transport system permease protein GsiC |
| *FLPKDCKG_02581* | *dppA-5* | - | 2831351 | 2832922 | 56.09 | 5.01 | 7.36 | 9.34 | Periplasmic dipeptide transport protein |
| *FLPKDCKG_02582* | *badR-1* | - | 2833149 | 2833640 | 2.50 | 7.34 | 4.11 | 2.05 | Transcriptional activatory protein BadR |
| *FLPKDCKG_02583* | | + | 2833985 | 2836234 | 1.75 | 4.59 | 5.05 | 3.74 | Hypothetical protein |
| *FLPKDCKG_02584* | | + | 2836246 | 2837790 | 0.88 | 3.07 | 3.22 | 2.08 | Hypothetical protein |
| *FLPKDCKG_02585* | *ophA1* | + | 2837813 | 2838790 | 1.26 | 4.07 | 6.29 | 4.60 | Phthalate dioxygenase reductase |
| *FLPKDCKG_02586* | *antA-1* | + | 2838817 | 2840130 | 1.99 | 3.76 | 4.01 | 3.86 | Anthranilate 1,2-dioxygenase large subunit |
| *FLPKDCKG_02587* | *bnzB-1* | + | 2840127 | 2840618 | 1.60 | 2.60 | 2.26 | 2.46 | Benzene 1,2-dioxygenase subunit beta |
| *FLPKDCKG_02588* | *sauT* | + | 2840628 | 2841254 | 0.65 | 1.07 | 0.79 | 0.90 | putative sulfoacetate--CoA ligase |
| *FLPKDCKG_02589* | *lcfB-2* | + | 2841248 | 2842258 | 0.72 | 2.78 | 2.98 | 1.75 | Long-chain-fatty-acid--CoA ligase |
| *FLPKDCKG_02590* | | + | 2842255 | 2843430 | 0.53 | 1.25 | 1.48 | 1.97 | Hypothetical protein |
| *FLPKDCKG_02591* | | + | 2843427 | 2844668 | 0.93 | 3.11 | 2.83 | 2.18 | putative oxidoreductase |
| *FLPKDCKG_02592* | *kynB-2* | + | 2844700 | 2845485 | 2.49 | 12.31 | 6.53 | 8.14 | Kynurenine formamidase |
| *FLPKDCKG_02593* | *fabG-8* | + | 2845524 | 2846252 | 0.74 | 3.86 | 1.38 | 1.44 | 3-oxoacyl-[acyl-carrier-protein] reductase FabG |
| *FLPKDCKG_02594* | | + | 2846416 | 2847564 | 6.59 | 4.71 | 4.67 | 4.94 | Hypothetical protein |
| *FLPKDCKG_02595* | *livH-9* | + | 2847690 | 2848730 | 0.61 | 0.39 | 1.66 | 0.43 | High-affinity branched-chain amino acid transport system permease protein LivH |
| *FLPKDCKG_02596* | *cysA-4* | + | 2848727 | 2850520 | 0.51 | 0.77 | 1.81 | 0.49 | Sulfate/thiosulfate import ATP-binding protein CysA |
| *FLPKDCKG_02597* | *livF-9* | + | 2850520 | 2851263 | 1.05 | 8.39 | 4.91 | 1.69 | High-affinity branched-chain amino acid transport ATP-binding protein LivF |
| *FLPKDCKG_02598* | *pglI-1* | - | 2851318 | 2852253 | 21.10 | 61.60 | 34.11 | 5.89 | GalNAc(5)-diNAcBac-PP-undecaprenol beta-1,3-glucosyltransferase |
| *FLPKDCKG_02599* | *dmlR-15* | + | 2852482 | 2853441 | 8.19 | 21.72 | 11.63 | 6.98 | HTH-type transcriptional regulator DmlR |
| *FLPKDCKG_02600* | | + | 2854096 | 2854224 | 4.13 | 9.03 | 13.31 | 5.18 | Hypothetical protein |
| *FLPKDCKG_02601* | *kdgK* | - | 2854465 | 2855388 | 7.96 | 3.87 | 5.08 | 1.35 | 2-dehydro-3-deoxygluconokinase |
| *FLPKDCKG_02602* | *kdgA* | - | 2855381 | 2856025 | 16.38 | 10.72 | 7.28 | 5.62 | KHG/KDPG aldolase |
| *FLPKDCKG_02603* | *iclR-4* | - | 2856050 | 2856841 | 6.24 | 9.42 | 5.39 | 1.86 | Transcriptional repressor IclR |
| *FLPKDCKG_02604* | *uvrA-1* | + | 2857080 | 2862839 | 6.13 | 12.24 | 9.93 | 3.93 | UvrABC system protein A |
| *FLPKDCKG_02605* | *pabB* | + | 2862848 | 2863984 | 0.80 | 1.64 | 0.75 | 1.35 | Aminodeoxychorismate synthase component 1 |
| *FLPKDCKG_02606* | *ilvE-3* | + | 2864061 | 2864693 | 20.17 | 32.33 | 18.69 | 7.23 | putative branched-chain-amino-acid aminotransferase |
| *FLPKDCKG_02607* | *pcpR-6* | - | 2864728 | 2865630 | 2.94 | 6.35 | 5.08 | 0.85 | PCP degradation transcriptional activation protein |
| *FLPKDCKG_02608* | *nagG* | + | 2865824 | 2867152 | 1.21 | 221.59 | 264.94 | 41.30 | Salicylate 5-hydroxylase, large oxygenase component |
| *FLPKDCKG_02609* | *nagH* | + | 2867155 | 2867628 | 2.12 | 221.12 | 258.80 | 35.84 | Salicylate 5-hydroxylase, small oxygenase component |
| *FLPKDCKG_02610* | *nagAb* | + | 2867644 | 2867955 | 1.10 | 270.98 | 276.91 | 41.54 | Naphthalene 1,2-dioxygenase/salicylate 5-hydroxylase systems, ferredoxin component |
| *FLPKDCKG_02611* | *tcdA* | - | 2867998 | 2868834 | 10.63 | 41.08 | 47.16 | 10.08 | tRNA threonylcarbamoyladenosine dehydratase |
| *FLPKDCKG_02612* | *mdtA-3* | - | 2868883 | 2870157 | 1.21 | 3.18 | 6.24 | 1.85 | Multidrug resistance protein MdtA |
| *FLPKDCKG_02613* | | - | 2870150 | 2871532 | 0.19 | 1.23 | 0.00 | 0.62 | Hypothetical protein |
| *FLPKDCKG_02614* | *macB-1* | - | 2871624 | 2872814 | 0.57 | 1.23 | 1.03 | 0.00 | Macrolide export ATP-binding/permease protein MacB |
| *FLPKDCKG_02615* | *yknY* | - | 2872833 | 2873537 | 0.55 | 2.97 | 0.00 | 0.57 | putative ABC transporter ATP-binding protein YknY |
| *FLPKDCKG_02616* | | - | 2874379 | 2875218 | 0.51 | 2.29 | 1.51 | 0.62 | Hypothetical protein |
| *FLPKDCKG_02617* | | - | 2876561 | 2877412 | 1.19 | 4.13 | 3.20 | 0.49 | Hypothetical protein |
| *FLPKDCKG_02618* | | - | 2877868 | 2878047 | 4.36 | 3.75 | 9.76 | 3.45 | Hypothetical protein |

| Locus name | Gene name | Strand | Gene start (bp) | Gene end (bp) | Average TPM Inoculum | Average TPM $t_6$ | Average TPM $t_{24}$ | Average TPM $t_{168}$ | Annotation description |
|---|---|---|---|---|---|---|---|---|---|
| *FLPKDCKG_02619* | | - | 2878587 | 2879447 | 1.34 | 2.57 | 2.46 | 0.00 | Hypothetical protein |
| *FLPKDCKG_02620* | *hcaR-4* | - | 2879640 | 2880548 | 1.04 | 1.22 | 0.40 | 0.00 | Hca operon transcriptional activator HcaR |
| *FLPKDCKG_02621* | *tdeA* | - | 2880891 | 2882318 | 1.85 | 4.53 | 3.72 | 0.56 | Toxin and drug export protein A |
| *FLPKDCKG_02622* | *macB-2* | - | 2882324 | 2884282 | 2.44 | 5.49 | 3.80 | 0.72 | Macrolide export ATP-binding/permease protein MacB |
| *FLPKDCKG_02623* | *macA* | - | 2884284 | 2885468 | 4.86 | 12.17 | 13.80 | 1.94 | Macrolide export protein MacA |
| *FLPKDCKG_02624* | | + | 2885884 | 2887092 | 5.39 | 13.61 | 8.09 | 3.65 | Hypothetical protein |
| *FLPKDCKG_02625* | | + | 2887101 | 2887748 | 3.41 | 4.89 | 3.45 | 2.86 | Hypothetical protein |
| *FLPKDCKG_02626* | *pdxH* | + | 2887855 | 2888484 | 68.67 | 84.90 | 46.61 | 9.25 | Pyridoxine/pyridoxamine 5'-phosphate oxidase |
| *FLPKDCKG_02627* | *C1-hpah-1* | + | 2888529 | 2889053 | 4.74 | 16.19 | 12.02 | 0.38 | p-hydroxyphenylacetate 3-hydroxylase, reductase component |
| *FLPKDCKG_02628* | *lhgO* | + | 2889050 | 2890183 | 6.84 | 23.91 | 12.85 | 3.17 | L-2-hydroxyglutarate oxidase LhgO |
| *FLPKDCKG_02629* | *purU* | + | 2890187 | 2891041 | 6.42 | 25.22 | 12.37 | 3.19 | Formyltetrahydrofolate deformylase |
| *FLPKDCKG_02630* | *dmlR-16* | - | 2891070 | 2891975 | 0.82 | 0.90 | 1.07 | 0.44 | HTH-type transcriptional regulator DmlR |
| *FLPKDCKG_02631* | *mdh-2* | + | 2892061 | 2893245 | 0.81 | 0.45 | 0.88 | 0.00 | NAD-dependent methanol dehydrogenase |
| *FLPKDCKG_02632* | *tauD-1* | + | 2893303 | 2894178 | 0.82 | 1.01 | 2.32 | 0.00 | Alpha-ketoglutarate-dependent taurine dioxygenase |
| *FLPKDCKG_02633* | | + | 2894202 | 2895182 | 0.34 | 1.40 | 0.18 | 0.00 | Hypothetical protein |
| *FLPKDCKG_02634* | *mdlB-3* | + | 2895205 | 2896362 | 0.35 | 0.00 | 0.00 | 0.00 | (S)-mandelate dehydrogenase |
| *FLPKDCKG_02635* | | + | 2896359 | 2896781 | 0.28 | 0.58 | 0.00 | 0.00 | Hypothetical protein |
| *FLPKDCKG_02636* | | + | 2896793 | 2897557 | 0.59 | 1.34 | 0.00 | 0.65 | Ureidoglycolate lyase |
| *FLPKDCKG_02637* | *hcaR-5* | - | 2897599 | 2898489 | 0.98 | 0.94 | 0.74 | 1.53 | Hca operon transcriptional activator HcaR |
| *FLPKDCKG_02638* | *aidB-2* | + | 2898632 | 2900332 | 0.10 | 0.52 | 1.04 | 1.75 | Putative acyl-CoA dehydrogenase AidB |
| *FLPKDCKG_02639* | | + | 2900389 | 2901366 | 0.24 | 0.45 | 1.05 | 0.85 | Hypothetical protein |
| *FLPKDCKG_02640* | *yfdE-1* | + | 2901380 | 2902501 | 0.12 | 0.37 | 0.31 | 0.98 | Acetyl-CoA:oxalate CoA-transferase |
| *FLPKDCKG_02641* | *yhdG* | + | 2902822 | 2904225 | 6.71 | 63.28 | 31.80 | 3.95 | putative amino acid permease YhdG |
| *FLPKDCKG_02642* | *ggt-1* | - | 2904306 | 2906063 | 5.19 | 8.47 | 4.37 | 1.12 | Gamma-glutamyltranspeptidase |
| *FLPKDCKG_02643* | *cynR-8* | + | 2906189 | 2907079 | 2.21 | 0.09 | 1.64 | 1.31 | HTH-type transcriptional regulator CynR |
| *FLPKDCKG_02644* | | - | 2907182 | 2908165 | 35.82 | 40.26 | 30.72 | 18.15 | Hypothetical protein |
| *FLPKDCKG_02645* | | + | 2908443 | 2909309 | 3.91 | 10.96 | 1.53 | 2.31 | Hypothetical protein |
| *FLPKDCKG_02646* | *fumC* | - | 2909391 | 2910782 | 55.18 | 70.71 | 56.67 | 17.37 | Fumarate hydratase class II |
| *FLPKDCKG_02647* | *amiC-3* | + | 2911417 | 2912796 | 64.84 | 41.75 | 29.51 | 10.49 | N-acetylmuramoyl-L-alanine amidase AmiC |
| *FLPKDCKG_02648* | | + | 2913112 | 2915547 | 2.03 | 4.71 | 2.46 | 0.56 | putative hemoglobin and hemoglobin-haptoglobin-binding protein 3 |
| *FLPKDCKG_02649* | | - | 2915596 | 2916234 | 2.06 | 3.37 | 7.98 | 1.31 | putative membrane protein |
| *FLPKDCKG_02650* | *mutL-1* | + | 2916332 | 2917036 | 29.28 | 47.19 | 17.20 | 7.98 | DNA mismatch repair protein MutL |
| *FLPKDCKG_02651* | *mutL-2* | + | 2917033 | 2918283 | 8.80 | 16.10 | 9.08 | 3.07 | DNA mismatch repair protein MutL |
| *FLPKDCKG_02652* | *miaA* | + | 2918293 | 2919243 | 3.63 | 9.01 | 7.60 | 0.43 | tRNA dimethylallyltransferase |
| *FLPKDCKG_02653* | | + | 2919328 | 2919822 | 1.60 | 10.07 | 4.57 | 1.53 | Hypothetical protein |
| *FLPKDCKG_02654* | *purM* | - | 2919881 | 2920930 | 15.38 | 27.76 | 17.99 | 5.97 | Phosphoribosylformylglycinamidine cyclo-ligase |
| *FLPKDCKG_02655* | *hda* | + | 2921247 | 2921948 | 10.11 | 14.00 | 11.23 | 3.31 | DnaA regulatory inactivator Hda |
| *FLPKDCKG_02656* | *serB1* | + | 2921945 | 2922643 | 21.63 | 36.42 | 21.03 | 7.18 | Phosphoserine phosphatase SerB1 |
| *FLPKDCKG_02657* | *pcnB* | + | 2922644 | 2924029 | 6.01 | 11.09 | 7.21 | 2.09 | Poly(A) polymerase I |
| *FLPKDCKG_02658* | *yihV* | - | 2924581 | 2925477 | 4.94 | 5.50 | 3.58 | 2.03 | Sulfofructose kinase |
| *FLPKDCKG_02659* | | - | 2925533 | 2926363 | 12.49 | 15.49 | 11.84 | 3.61 | Citrate lyase subunit beta-like protein |
| *FLPKDCKG_02660* | *yfdE-2* | - | 2926363 | 2927556 | 4.38 | 5.72 | 1.89 | 1.16 | Acetyl-CoA:oxalate CoA-transferase |
| *FLPKDCKG_02661* | *meh* | - | 2927553 | 2928383 | 8.41 | 5.59 | 2.16 | 3.37 | Mesaconyl-C(4)-CoA hydratase |
| *FLPKDCKG_02662* | *gltC-8* | + | 2928469 | 2929413 | 10.32 | 15.51 | 13.76 | 3.49 | HTH-type transcriptional regulator GltC |
| *FLPKDCKG_02663* | *paaG-2* | - | 2929509 | 2930309 | 0.67 | 2.58 | 0.39 | 1.76 | 1,2-epoxyphenylacetyl-CoA isomerase |
| *FLPKDCKG_02664* | *mcd* | - | 2930306 | 2931418 | 0.24 | 0.10 | 0.75 | 0.26 | (2S)-methylsuccinyl-CoA dehydrogenase |
| *FLPKDCKG_02665* | | - | 2931448 | 2932638 | 0.03 | 0.00 | 0.24 | 0.00 | Putative acyl-CoA dehydrogenase FadE17 |
| *FLPKDCKG_02666* | | + | 2932833 | 2933231 | 0.00 | 0.00 | 0.00 | 0.00 | Hypothetical protein |
| *FLPKDCKG_02667* | *fadA-2* | + | 2933228 | 2934400 | 0.07 | 0.00 | 0.00 | 0.00 | 3-ketoacyl-CoA thiolase |
| *FLPKDCKG_02668* | *lcfB-3* | + | 2934418 | 2935935 | 0.06 | 0.00 | 0.00 | 0.22 | Long-chain-fatty-acid--CoA ligase |
| *FLPKDCKG_02669* | | + | 2935991 | 2936968 | 0.09 | 0.00 | 0.47 | 0.34 | Hypothetical protein |
| *FLPKDCKG_02670* | | + | 2936988 | 2937941 | 0.47 | 0.20 | 0.18 | 0.22 | Nitronate monooxygenase |
| *FLPKDCKG_02671* | | - | 2938304 | 2939287 | 2.07 | 1.99 | 0.61 | 0.25 | Hypothetical protein |
| *FLPKDCKG_02672* | *cysL-3* | - | 2939606 | 2940505 | 10.19 | 8.03 | 6.33 | 2.49 | HTH-type transcriptional regulator CysL |
| *FLPKDCKG_02673* | | + | 2940654 | 2941568 | 0.44 | 0.99 | 0.41 | 1.27 | Putative short-chain type dehydrogenase/reductase |
| *FLPKDCKG_02674* | | + | 2941576 | 2942436 | 0.23 | 0.00 | 0.67 | 0.20 | Hypothetical protein |
| *FLPKDCKG_02675* | *uctC-3* | + | 2942433 | 2943659 | 0.24 | 1.01 | 0.61 | 0.12 | Acetyl-CoA:oxalate CoA-transferase |
| *FLPKDCKG_02676* | | + | 2943915 | 2946704 | 1.23 | 3.31 | 2.06 | 0.82 | Extracellular serine protease |
| *FLPKDCKG_02677* | *bdlA-1* | + | 2946913 | 2948364 | 0.47 | 0.02 | 0.85 | 0.13 | Biofilm dispersion protein BdlA |
| *FLPKDCKG_02678* | *fumB-1* | - | 2948477 | 2950000 | 4.18 | 6.85 | 8.38 | 4.96 | Fumarate hydratase class I, anaerobic |
| *FLPKDCKG_02679* | *caiD* | - | 2950078 | 2950845 | 19.94 | 28.66 | 27.64 | 10.46 | Carnitinyl-CoA dehydratase |
| *FLPKDCKG_02680* | *glyQ* | + | 2951116 | 2952027 | 12.99 | 19.35 | 16.84 | 4.02 | Glycine--tRNA ligase alpha subunit |
| *FLPKDCKG_02681* | *glyS* | + | 2952486 | 2954162 | 5.35 | 7.55 | 3.09 | 3.99 | Glycine--tRNA ligase beta subunit |
| *FLPKDCKG_02682* | | + | 2954159 | 2954698 | 3.59 | 4.09 | 6.79 | 2.78 | D-glycero-beta-D-manno-heptose-1,7-bisphosphate 7-phosphatase |
| *FLPKDCKG_02683* | *plsC-1* | + | 2954698 | 2955426 | 9.56 | 14.10 | 6.96 | 3.09 | 1-acyl-sn-glycerol-3-phosphate acyltransferase |
| *FLPKDCKG_02684* | | + | 2955413 | 2956396 | 13.45 | 14.02 | 6.80 | 0.97 | Hypothetical protein |
| *FLPKDCKG_02685* | *phnW* | + | 2956449 | 2957585 | 7.37 | 18.02 | 9.91 | 5.17 | 2-aminoethylphosphonate--pyruvate transaminase |
| *FLPKDCKG_02686* | *gloA* | + | 2957597 | 2957992 | 7.35 | 23.84 | 15.81 | 4.91 | Lactoylglutathione lyase |
| *FLPKDCKG_02687* | *def* | + | 2958242 | 2958643 | 3.38 | 12.24 | 12.00 | 2.37 | Peptide deformylase |

| Locus name | Gene name | Strand | Gene start (bp) | Gene end (bp) | Average TPM Inoculum | Average TPM $t_6$ | Average TPM $t_{24}$ | Average TPM $t_{168}$ | Annotation description |
|---|---|---|---|---|---|---|---|---|---|
| *FLPKDCKG_02688* | *fadE-1* | - | 2958631 | 2960904 | 16.35 | 13.82 | 10.20 | 2.42 | Acyl-coenzyme A dehydrogenase |
| *FLPKDCKG_02689* | *spmB* | + | 2961053 | 2962408 | 3.49 | 6.41 | 3.94 | 0.88 | Spore maturation protein B |
| *FLPKDCKG_02690* | *pykA* | - | 2962357 | 2963799 | 13.57 | 13.15 | 8.58 | 1.75 | Pyruvate kinase II |
| *FLPKDCKG_02691* | *glxK* | + | 2963894 | 2965048 | 15.14 | 12.43 | 3.39 | 2.43 | Glycerate 3-kinase |
| *FLPKDCKG_02692* | *rsmA* | - | 2965121 | 2965912 | 1.81 | 5.67 | 7.06 | 3.24 | Ribosomal RNA small subunit methyltransferase A |
| *FLPKDCKG_02693* | *surA-1* | - | 2965931 | 2967073 | 8.39 | 9.70 | 11.85 | 4.31 | Chaperone SurA |
| *FLPKDCKG_02694* | *surA-2* | - | 2967119 | 2967496 | 11.52 | 15.53 | 15.13 | 5.84 | Chaperone SurA |
| *FLPKDCKG_02695* | *lptD* | - | 2967493 | 2969748 | 6.96 | 11.94 | 10.90 | 2.79 | LPS-assembly protein LptD |
| *FLPKDCKG_02696* | *srkA* | + | 2970055 | 2971095 | 11.31 | 16.90 | 5.92 | 2.47 | Stress response kinase A |
| *FLPKDCKG_02697* | *cugP* | + | 2971095 | 2971781 | 4.02 | 12.02 | 9.24 | 3.14 | UTP--glucose-1-phosphate uridylyltransferase |
| *FLPKDCKG_02698* | | + | 2971938 | 2972636 | 15.71 | 64.25 | 57.67 | 19.44 | Hypothetical protein |
| *FLPKDCKG_02699* | *mhpA-1* | + | 2972759 | 2974393 | 1.68 | 2.53 | 1.45 | 1.39 | 3-(3-hydroxy-phenyl)propionate/3-hydroxycinnamic acid hydroxylase |
| *FLPKDCKG_02700* | | + | 2974390 | 2974611 | 2.35 | 7.31 | 6.93 | 0.26 | Hypothetical protein |
| *FLPKDCKG_02701* | | + | 2974608 | 2975063 | 2.12 | 14.91 | 5.38 | 1.29 | putative HTH-type transcriptional regulator |
| *FLPKDCKG_02702* | *phhB* | - | 2975120 | 2975446 | 38.96 | 98.08 | 33.86 | 18.86 | Putative pterin-4-alpha-carbinolamine dehydratase |
| *FLPKDCKG_02703* | *yiaD-2* | - | 2975458 | 2976105 | 119.92 | 140.01 | 89.10 | 39.88 | putative lipoprotein YiaD |
| *FLPKDCKG_02704* | *hadL* | - | 2976190 | 2976873 | 5.07 | 7.68 | 4.37 | 2.46 | (S)-2-haloacid dehalogenase |
| *FLPKDCKG_02705* | *azoR2* | + | 2977216 | 2977839 | 0.63 | 1.66 | 0.40 | 0.00 | FMN-dependent NADH-azoreductase 2 |
| *FLPKDCKG_02706* | *prpE* | - | 2977967 | 2978911 | 13.18 | 14.75 | 9.00 | 1.82 | Bis(5'-nucleosyl)-tetraphosphatase PrpE [asymmetrical] |
| *FLPKDCKG_02707* | | - | 2979234 | 2979309 | 36.57 | 12.65 | 24.59 | 4.26 | tRNA |
| *FLPKDCKG_02708* | *bdcA-2* | - | 2979483 | 2980229 | 1.41 | 3.17 | 2.77 | 0.99 | Cyclic-di-GMP-binding biofilm dispersal mediator protein |
| *FLPKDCKG_02709* | *comR-3* | + | 2980392 | 2980991 | 18.82 | 25.31 | 17.78 | 7.46 | HTH-type transcriptional repressor ComR |
| *FLPKDCKG_02710* | *yedQ* | + | 2981003 | 2982598 | 3.85 | 16.89 | 9.55 | 1.33 | putative diguanylate cyclase YedQ |
| *FLPKDCKG_02711* | | - | 2982605 | 2984677 | 9.86 | 74.16 | 31.55 | 6.75 | Hypothetical protein |
| *FLPKDCKG_02712* | *ltaS2-1* | + | 2984973 | 2986901 | 6.02 | 11.74 | 8.93 | 2.73 | Lipoteichoic acid synthase 2 |
| *FLPKDCKG_02713* | *tag* | - | 2986918 | 2987559 | 10.79 | 9.28 | 12.83 | 1.01 | DNA-3-methyladenine glycosylase 1 |
| *FLPKDCKG_02714* | | + | 2987778 | 2988251 | 11.93 | 35.77 | 16.21 | 11.16 | Hypothetical protein |
| *FLPKDCKG_02715* | *est-1* | - | 2988294 | 2989157 | 40.34 | 56.61 | 33.84 | 11.65 | Carboxylesterase |
| *FLPKDCKG_02716* | | - | 2989154 | 2989312 | 72.54 | 21.23 | 26.49 | 3.37 | Hypothetical protein |
| *FLPKDCKG_02717* | *nimR-7* | - | 2989514 | 2990296 | 8.35 | 5.74 | 6.91 | 1.27 | HTH-type transcriptional regulator NimR |
| *FLPKDCKG_02718* | *fur-2* | + | 2990404 | 2990844 | 15.17 | 13.50 | 8.20 | 4.56 | Ferric uptake regulation protein |
| *FLPKDCKG_02719* | | + | 2990841 | 2991260 | 5.73 | 5.18 | 6.81 | 3.09 | Hypothetical protein |
| *FLPKDCKG_02720* | *cysL-4* | + | 2991712 | 2992623 | 0.67 | 4.24 | 0.62 | 0.76 | HTH-type transcriptional regulator CysL |
| *FLPKDCKG_02721* | *tsr-4* | + | 2992772 | 2994388 | 0.23 | 1.31 | 0.93 | 0.55 | Methyl-accepting chemotaxis protein I |
| *FLPKDCKG_02722* | | + | 2994505 | 2994792 | 0.61 | 3.46 | 3.48 | 0.00 | Hypothetical protein |
| *FLPKDCKG_02723* | *yjgH-2* | + | 2994894 | 2995325 | 3.08 | 3.39 | 6.50 | 1.67 | RutC family protein YjgH |
| *FLPKDCKG_02724* | *fpvA-3* | - | 2995336 | 2997726 | 0.59 | 0.82 | 1.03 | 0.29 | Ferripyoverdine receptor |
| *FLPKDCKG_02725* | | - | 2997830 | 2998819 | 1.17 | 4.04 | 0.80 | 0.02 | Hypothetical protein |
| *FLPKDCKG_02726* | *fecI-8* | - | 2998816 | 2999322 | 3.67 | 5.74 | 0.79 | 0.68 | putative RNA polymerase sigma factor FecI |
| *FLPKDCKG_02727* | | - | 2999549 | 2999624 | 15.08 | 22.69 | 9.13 | 9.50 | tRNA |
| *FLPKDCKG_02728* | | + | 2999690 | 2999911 | 0.00 | 0.00 | 0.00 | 0.00 | Hypothetical protein |
| *FLPKDCKG_02729* | *metQ-2* | - | 2999842 | 3000639 | 2.65 | 16.92 | 26.93 | 6.35 | D-methionine-binding lipoprotein MetQ |
| *FLPKDCKG_02730* | *metI-1* | - | 3000689 | 3001342 | 2.69 | 11.21 | 19.62 | 5.06 | D-methionine transport system permease protein MetI |
| *FLPKDCKG_02731* | *metN-1* | - | 3001299 | 3002387 | 4.96 | 27.29 | 31.40 | 9.90 | Methionine import ATP-binding protein MetN |
| *FLPKDCKG_02732* | *yicI* | - | 3002562 | 3004787 | 17.81 | 40.60 | 28.97 | 8.69 | Alpha-xylosidase |
| *FLPKDCKG_02733* | *mnmC-2* | - | 3005059 | 3006930 | 1.07 | 2.25 | 0.41 | 0.36 | tRNA 5-methylaminomethyl-2-thiouridine biosynthesis bifunctional protein MnmC |
| *FLPKDCKG_02734* | *suhB-1* | + | 3007076 | 3007951 | 242.48 | 122.68 | 106.44 | 31.11 | Inositol-1-monophosphatase |
| *FLPKDCKG_02735* | *fadD-1* | - | 3007971 | 3009671 | 25.10 | 15.95 | 12.20 | 3.82 | Long-chain-fatty-acid--CoA ligase |
| *FLPKDCKG_02736* | *yxeP-1* | + | 3009748 | 3010944 | 40.34 | 55.05 | 42.33 | 8.80 | putative hydrolase YxeP |
| *FLPKDCKG_02737* | *cmpR-9* | - | 3010998 | 3011894 | 1.49 | 9.79 | 6.86 | 0.41 | HTH-type transcriptional activator CmpR |
| *FLPKDCKG_02738* | | + | 3012034 | 3013125 | 7.32 | 5.30 | 3.71 | 1.29 | Hypothetical protein |
| *FLPKDCKG_02739* | | + | 3013229 | 3013639 | 42.11 | 41.50 | 32.16 | 12.15 | putative acyl-CoA thioester hydrolase |
| *FLPKDCKG_02740* | | + | 3013650 | 3015122 | 7.38 | 14.69 | 8.49 | 2.12 | Magnesium transporter MgtE |
| *FLPKDCKG_02741* | *pagP* | + | 3015256 | 3015810 | 13.47 | 27.65 | 18.43 | 3.16 | Lipid A palmitoyltransferase PagP |
| *FLPKDCKG_02742* | | - | 3015848 | 3016459 | 56.47 | 22.16 | 6.92 | 2.19 | Hypothetical protein |
| *FLPKDCKG_02743* | *decR-3* | - | 3016686 | 3017144 | 15.07 | 5.53 | 3.18 | 2.43 | DNA-binding transcriptional activator DecR |
| *FLPKDCKG_02744* | *bacG* | - | 3017235 | 3018014 | 9.58 | 10.22 | 6.42 | 3.42 | NADPH-dependent reductase BacG |
| *FLPKDCKG_02745* | | + | 3018180 | 3018716 | 223.19 | 49.46 | 11.41 | 9.87 | 3-mercaptopropionate dioxygenase |
| *FLPKDCKG_02746* | *ilvI-2* | - | 3018806 | 3020500 | 2.99 | 2.47 | 1.10 | 1.60 | Acetolactate synthase isozyme 3 large subunit |
| *FLPKDCKG_02747* | | - | 3020559 | 3021548 | 5.23 | 1.97 | 3.00 | 1.75 | Hypothetical protein |
| *FLPKDCKG_02748* | *rspR-5* | - | 3021545 | 3022258 | 4.35 | 3.10 | 1.50 | 0.59 | HTH-type transcriptional repressor RspR |
| *FLPKDCKG_02749* | | - | 3022473 | 3023192 | 1.61 | 7.93 | 4.49 | 2.09 | Hypothetical protein |
| *FLPKDCKG_02750* | *secA* | - | 3023205 | 3025940 | 9.44 | 31.62 | 22.21 | 8.20 | Protein translocase subunit SecA |
| *FLPKDCKG_02751* | *mepM-1* | - | 3026116 | 3027099 | 10.99 | 25.09 | 22.53 | 16.15 | Murein DD-endopeptidase MepM |
| *FLPKDCKG_02752* | | + | 3027227 | 3027634 | 15.84 | 51.52 | 37.45 | 9.93 | Hypothetical protein |
| *FLPKDCKG_02753* | *lpxC* | - | 3027834 | 3028757 | 41.58 | 86.50 | 44.22 | 14.20 | UDP-3-O-acyl-N-acetylglucosamine deacetylase |
| *FLPKDCKG_02754* | *ftsZ* | - | 3028997 | 3030181 | 22.60 | 51.09 | 61.70 | 21.70 | Cell division protein FtsZ |
| *FLPKDCKG_02755* | *ftsA* | - | 3030372 | 3031598 | 11.95 | 24.22 | 17.95 | 5.15 | Cell division protein FtsA |
| *FLPKDCKG_02756* | *ftsQ* | - | 3031603 | 3032427 | 8.38 | 17.07 | 12.69 | 3.75 | Cell division protein FtsQ |

| Locus name | Gene name | Strand | Gene start (bp) | Gene end (bp) | Average TPM Inoculum | Average TPM $t_6$ | Average TPM $t_{24}$ | Average TPM $t_{168}$ | Annotation description |
|---|---|---|---|---|---|---|---|---|---|
| *FLPKDCKG_02757* | *ddlB* | - | 3032440 | 3033390 | 18.57 | 19.44 | 15.08 | 4.41 | D-alanine--D-alanine ligase B |
| *FLPKDCKG_02758* | *murC* | - | 3033387 | 3034793 | 7.98 | 8.68 | 10.34 | 3.81 | UDP-N-acetylmuramate--L-alanine ligase |
| *FLPKDCKG_02759* | *murG* | - | 3034790 | 3035452 | 6.03 | 7.94 | 5.53 | 2.96 | UDP-N-acetylglucosamine--N-acetylmuramyl-(pentapeptide) pyrophosphoryl-undecaprenol N-acetylglucosamine transferase |
| *FLPKDCKG_02760* | *ftsW* | - | 3035391 | 3037061 | 6.43 | 6.21 | 4.41 | 3.03 | putative peptidoglycan glycosyltransferase FtsW |
| *FLPKDCKG_02761* | *murD* | - | 3037058 | 3038590 | 7.19 | 10.76 | 6.82 | 5.35 | UDP-N-acetylmuramoylalanine--D-glutamate ligase |
| *FLPKDCKG_02762* | *mraY* | - | 3038587 | 3039756 | 9.85 | 10.95 | 7.52 | 2.14 | Phospho-N-acetylmuramoyl-pentapeptide-transferase |
| *FLPKDCKG_02763* | *murE* | - | 3039746 | 3042610 | 13.66 | 14.83 | 10.82 | 2.95 | UDP-N-acetylmuramoyl-L-alanyl-D-glutamate--2,6-diaminopimelate ligase |
| *FLPKDCKG_02764* | *penA* | - | 3042607 | 3044334 | 11.27 | 13.91 | 9.86 | 2.93 | putative peptidoglycan D,D-transpeptidase PenA |
| *FLPKDCKG_02765* | *ftsL* | - | 3044331 | 3044621 | 6.74 | 7.74 | 8.51 | 2.62 | Cell division protein FtsL |
| *FLPKDCKG_02766* | *rsmH* | - | 3044621 | 3045715 | 120.47 | 151.18 | 97.49 | 29.99 | Ribosomal RNA small subunit methyltransferase H |
| *FLPKDCKG_02767* | *mraZ* | - | 3045723 | 3046175 | 77.38 | 95.78 | 54.12 | 15.12 | Transcriptional regulator MraZ |
| *FLPKDCKG_02768* | *uup* | + | 3046986 | 3048821 | 4.25 | 4.80 | 6.88 | 2.21 | ABC transporter ATP-binding protein uup |
| *FLPKDCKG_02769* | *pyrC-1* | - | 3048869 | 3049933 | 7.80 | 7.24 | 10.80 | 1.77 | Dihydroorotase |
| *FLPKDCKG_02770* | *yxeP-2* | - | 3049945 | 3051138 | 24.21 | 32.62 | 19.62 | 3.36 | putative hydrolase YxeP |
| *FLPKDCKG_02771* | | + | 3051286 | 3051564 | 490.24 | 1682.32 | 1068.35 | 266.57 | Hypothetical protein |
| *FLPKDCKG_02772* | *gcvA-7* | - | 3051877 | 3052692 | 2.67 | 4.38 | 2.69 | 1.87 | Glycine cleavage system transcriptional activator |
| *FLPKDCKG_02773* | *rhtB-2* | + | 3052913 | 3053518 | 1.05 | 1.93 | 3.56 | 0.43 | Homoserine/homoserine lactone efflux protein |
| *FLPKDCKG_02774* | | - | 3053643 | 3054008 | 54.00 | 58.87 | 31.77 | 11.57 | Hypothetical protein |
| *FLPKDCKG_02775* | | - | 3054342 | 3054453 | 139.79 | 260.49 | 284.66 | 234.78 | rRNA |
| *FLPKDCKG_02776* | | - | 3054656 | 3057532 | 116163.96 | 110011.45 | 125620.61 | 134115.04 | rRNA |
| *FLPKDCKG_02777* | | - | 3057831 | 3057906 | 17.92 | 4.30 | 3.95 | 0.00 | tRNA |
| *FLPKDCKG_02778* | | - | 3057937 | 3058012 | 8.51 | 11.41 | 21.13 | 0.75 | tRNA |
| *FLPKDCKG_02779* | | - | 3058148 | 3059665 | 176679.70 | 143357.61 | 131929.56 | 126969.33 | rRNA |
| *FLPKDCKG_02780* | *lrp-5* | - | 3060989 | 3061477 | 3.40 | 8.18 | 8.38 | 4.19 | Leucine-responsive regulatory protein |
| *FLPKDCKG_02781* | *hpd* | + | 3061622 | 3062734 | 23.28 | 26.98 | 16.24 | 10.45 | 4-hydroxyphenylpyruvate dioxygenase |
| *FLPKDCKG_02782* | *msrA2* | - | 3062820 | 3063374 | 84.55 | 86.47 | 46.41 | 25.89 | Peptide methionine sulfoxide reductase MsrA 2 |
| *FLPKDCKG_02783* | *cyt* | - | 3063413 | 3063748 | 7.19 | 15.60 | 10.08 | 2.31 | Cytochrome c-552 |
| *FLPKDCKG_02784* | *ilvD-1* | - | 3063732 | 3065417 | 24.16 | 14.88 | 11.67 | 3.31 | Dihydroxy-acid dehydratase |
| *FLPKDCKG_02785* | *lgt* | + | 3065508 | 3066296 | 8.36 | 21.99 | 22.57 | 7.67 | Prolipoprotein diacylglyceryl transferase |
| *FLPKDCKG_02786* | | - | 3066271 | 3066972 | 7.72 | 54.66 | 34.46 | 12.91 | Hypothetical protein |
| *FLPKDCKG_02787* | *zapA* | - | 3067343 | 3067654 | 38.89 | 105.26 | 48.96 | 18.05 | Cell division protein ZapA |
| *FLPKDCKG_02788* | *smc-3* | - | 3067654 | 3068100 | 47.69 | 140.69 | 153.25 | 44.18 | Chromosome partition protein Smc |
| *FLPKDCKG_02789* | | + | 3068585 | 3069049 | 4.77 | 6.84 | 9.18 | 3.92 | Hypothetical protein |
| *FLPKDCKG_02790* | | + | 3069063 | 3070568 | 7.41 | 12.54 | 14.99 | 5.69 | Hypothetical protein |
| *FLPKDCKG_02791* | *ywrD-3* | + | 3070772 | 3072376 | 12.55 | 17.80 | 8.72 | 2.16 | Putative gamma-glutamyltransferase YwrD |
| *FLPKDCKG_02792* | *glxR-2* | + | 3072429 | 3073334 | 8.70 | 15.08 | 17.54 | 5.78 | 2-hydroxy-3-oxopropionate reductase |
| *FLPKDCKG_02793* | *azoR-2* | - | 3073387 | 3074019 | 15.42 | 86.66 | 48.56 | 11.16 | FMN-dependent NADH-azoreductase |
| *FLPKDCKG_02794* | *dmlR-17* | + | 3074144 | 3075085 | 7.52 | 30.20 | 19.01 | 5.27 | HTH-type transcriptional regulator DmlR |
| *FLPKDCKG_02795* | *ghrA-1* | - | 3075079 | 3075468 | 6.64 | 23.03 | 10.79 | 0.73 | Glyoxylate/hydroxypyruvate reductase A |
| *FLPKDCKG_02796* | *ghrA-2* | - | 3075432 | 3076007 | 7.30 | 13.71 | 6.36 | 0.83 | Glyoxylate/hydroxypyruvate reductase A |
| *FLPKDCKG_02797* | | - | 3076119 | 3077642 | 0.38 | 2.04 | 1.56 | 0.35 | Hypothetical protein |
| *FLPKDCKG_02798* | | - | 3077649 | 3078122 | 0.02 | 1.37 | 2.56 | 0.66 | Hypothetical protein |
| *FLPKDCKG_02799* | *rssA* | - | 3078341 | 3079852 | 2.41 | 7.56 | 3.75 | 1.19 | Swarming motility regulation sensor protein RssA |
| *FLPKDCKG_02800* | *tctD-3* | - | 3079881 | 3080567 | 21.97 | 70.98 | 45.02 | 13.04 | Transcriptional regulatory protein tctD |
| *FLPKDCKG_02801* | *rppH* | - | 3080548 | 3081120 | 68.26 | 172.85 | 84.29 | 31.71 | RNA pyrophosphohydrolase |
| *FLPKDCKG_02802* | *proS-2* | + | 3081186 | 3082916 | 24.94 | 57.64 | 35.46 | 8.64 | Proline--tRNA ligase |
| *FLPKDCKG_02803* | *ybgC* | + | 3082974 | 3083396 | 15.33 | 74.95 | 66.74 | 14.38 | Acyl-CoA thioesterase YbgC |
| *FLPKDCKG_02804* | *exbB-2* | + | 3083399 | 3084079 | 9.59 | 33.02 | 28.21 | 3.47 | Biopolymer transport protein ExbB |
| *FLPKDCKG_02805* | *exbD-2* | + | 3084079 | 3084537 | 19.45 | 48.31 | 41.07 | 17.52 | Biopolymer transport protein ExbD |
| *FLPKDCKG_02806* | *iga* | + | 3084577 | 3085668 | 14.63 | 34.18 | 38.85 | 11.59 | IgA-specific serine endopeptidase autotransporter |
| *FLPKDCKG_02807* | | - | 3085944 | 3086032 | 51.80 | 102.71 | 47.59 | 13.92 | tRNA |
| *FLPKDCKG_02808* | *pal* | + | 3087039 | 3087539 | 211.04 | 369.76 | 288.04 | 107.27 | Outer membrane protein P6 |
| *FLPKDCKG_02809* | *cpoB* | + | 3087625 | 3088317 | 28.62 | 49.65 | 39.81 | 19.73 | Cell division coordinator CpoB |
| *FLPKDCKG_02810* | *hmuU-4* | + | 3088317 | 3089360 | 1.00 | 2.42 | 3.27 | 1.25 | Hemin transport system permease protein HmuU |
| *FLPKDCKG_02811* | *fecE-2* | + | 3089357 | 3090157 | 0.56 | 2.83 | 2.34 | 1.45 | Fe(3+) dicitrate transport ATP-binding protein FecE |
| *FLPKDCKG_02812* | *ybaK-1* | - | 3090211 | 3090708 | 12.76 | 15.95 | 6.55 | 5.07 | Cys-tRNA(Pro)/Cys-tRNA(Cys) deacylase YbaK |
| *FLPKDCKG_02813* | *acoC-1* | + | 3090851 | 3091660 | 1.20 | 0.93 | 1.49 | 1.15 | Dihydrolipoyllysine-residue acetyltransferase component of acetoin cleaving system |
| *FLPKDCKG_02814* | *lrp-6* | - | 3091673 | 3092143 | 3.53 | 7.17 | 5.40 | 2.30 | Leucine-responsive regulatory protein |
| *FLPKDCKG_02815* | | + | 3092444 | 3093343 | 2.02 | 1.98 | 1.10 | 0.20 | S-adenosylmethionine/S-adenosylhomocysteine transporter |
| *FLPKDCKG_02816* | *fusA-1* | - | 3093405 | 3095516 | 47.01 | 32.64 | 28.13 | 10.75 | Elongation factor G |
| *FLPKDCKG_02817* | *teaD-4* | - | 3095796 | 3096260 | 257.03 | 269.85 | 168.58 | 39.98 | TRAP-T-associated universal stress protein TeaD |
| *FLPKDCKG_02818* | *blh* | - | 3096348 | 3096692 | 107.83 | 88.99 | 48.64 | 19.38 | Beta-lactamase hydrolase-like protein |
| *FLPKDCKG_02819* | *lysN-1* | - | 3096750 | 3098183 | 5.15 | 6.90 | 6.29 | 1.30 | 2-aminoadipate transaminase |
| *FLPKDCKG_02820* | | + | 3098320 | 3099246 | 1.08 | 3.28 | 1.29 | 0.31 | Hypothetical protein |
| *FLPKDCKG_02821* | *eamB* | + | 3099243 | 3099887 | 0.97 | 0.20 | 0.00 | 0.13 | Cysteine/O-acetylserine efflux protein |
| *FLPKDCKG_02822* | *yohK* | - | 3099939 | 3100631 | 0.84 | 0.74 | 0.17 | 1.66 | Inner membrane protein YohK |
| *FLPKDCKG_02823* | *lrgA* | - | 3100621 | 3101025 | 0.10 | 0.00 | 0.64 | 0.00 | Antiholin-like protein LrgA |
| *FLPKDCKG_02824* | *gltC-9* | + | 3101121 | 3102008 | 1.43 | 1.00 | 1.53 | 1.01 | HTH-type transcriptional regulator GltC |
| *FLPKDCKG_02825* | *lip3* | + | 3102080 | 3102967 | 10.12 | 24.62 | 14.80 | 3.38 | Lipase 3 |

| Locus name | Gene name | Strand | Gene start (bp) | Gene end (bp) | Average TPM Inoculum | Average TPM $t_6$ | Average TPM $t_{24}$ | Average TPM $t_{168}$ | Annotation description |
|---|---|---|---|---|---|---|---|---|---|
| *FLPKDCKG_02826* | *dapAL* | + | 3103225 | 3104136 | 0.16 | 0.90 | 0.74 | 0.38 | putative DapA-like lyase |
| *FLPKDCKG_02827* | *rbsA-2* | + | 3104213 | 3104968 | 0.06 | 0.62 | 0.00 | 0.28 | Ribose import ATP-binding protein RbsA |
| *FLPKDCKG_02828* | *livF-10* | + | 3104961 | 3105680 | 0.06 | 0.00 | 0.60 | 0.00 | High-affinity branched-chain amino acid transport ATP-binding protein LivF |
| *FLPKDCKG_02829* | *amiC-4* | + | 3105768 | 3106973 | 0.07 | 0.49 | 0.34 | 0.13 | Aliphatic amidase expression-regulating protein |
| *FLPKDCKG_02830* | *livH-10* | + | 3107086 | 3107988 | 0.41 | 0.96 | 0.00 | 0.00 | High-affinity branched-chain amino acid transport system permease protein LivH |
| *FLPKDCKG_02831* | | + | 3107990 | 3108955 | 13.45 | 21.63 | 16.85 | 6.05 | Hypothetical protein |
| *FLPKDCKG_02832* | *rspR-6* | + | 3108955 | 3109605 | 2.74 | 6.87 | 4.48 | 1.95 | HTH-type transcriptional repressor RspR |
| *FLPKDCKG_02833* | | - | 3109580 | 3110071 | 19.33 | 29.65 | 28.88 | 6.58 | Hypothetical protein |
| *FLPKDCKG_02834* | *yhaJ-3* | + | 3110165 | 3111046 | 3.53 | 6.53 | 7.32 | 0.21 | HTH-type transcriptional regulator YhaJ |
| *FLPKDCKG_02835* | *chbG* | - | 3111098 | 3111913 | 1.98 | 20.03 | 7.39 | 0.76 | Chitooligosaccharide deacetylase ChbG |
| *FLPKDCKG_02836* | | + | 3111950 | 3112345 | 6.69 | 33.70 | 24.96 | 7.75 | Hypothetical protein |
| *FLPKDCKG_02837* | *arnT-1* | - | 3112302 | 3113519 | 1.30 | 19.73 | 7.23 | 1.05 | Undecaprenyl phosphate-alpha-4-amino-4-deoxy-L-arabinose arabinosyl transferase |
| *FLPKDCKG_02838* | *arnT-2* | - | 3113586 | 3113891 | 1.39 | 35.98 | 3.96 | 0.00 | Undecaprenyl phosphate-alpha-4-amino-4-deoxy-L-arabinose arabinosyl transferase |
| *FLPKDCKG_02839* | | - | 3113891 | 3114916 | 8.28 | 45.93 | 12.46 | 1.50 | putative glycosyltransferase |
| *FLPKDCKG_02840* | | + | 3115212 | 3115715 | 4.08 | 10.42 | 8.98 | 2.70 | Hypothetical protein |
| *FLPKDCKG_02841* | *gltS* | + | 3115864 | 3117072 | 2.90 | 5.86 | 4.50 | 1.29 | Sodium/glutamate symporter |
| *FLPKDCKG_02842* | | + | 3117075 | 3118571 | 4.91 | 8.31 | 4.64 | 1.36 | Hypothetical protein |
| *FLPKDCKG_02843* | | + | 3118614 | 3118877 | 11.24 | 17.72 | 12.76 | 1.49 | Hypothetical protein |
| *FLPKDCKG_02844* | | + | 3118877 | 3119344 | 3.22 | 5.53 | 15.23 | 6.51 | Hypothetical protein |
| *FLPKDCKG_02845* | | - | 3119372 | 3119539 | 38.46 | 30.28 | 11.30 | 2.75 | Hypothetical protein |
| *FLPKDCKG_02846* | | - | 3119900 | 3119974 | 256.98 | 1569.74 | 851.20 | 379.37 | tRNA |
| *FLPKDCKG_02847* | *dnaQ* | - | 3120035 | 3120757 | 9.40 | 15.78 | 15.85 | 4.74 | DNA polymerase III subunit epsilon |
| *FLPKDCKG_02848* | *mexB* | - | 3120762 | 3123878 | 3.35 | 6.19 | 3.93 | 1.42 | Multidrug resistance protein MexB |
| *FLPKDCKG_02849* | *mdtA-4* | - | 3123908 | 3124996 | 0.67 | 1.98 | 3.09 | 1.48 | Multidrug resistance protein MdtA |
| *FLPKDCKG_02850* | *rnhA* | - | 3125237 | 3125698 | 2.86 | 5.72 | 8.20 | 1.70 | Ribonuclease HI |
| *FLPKDCKG_02851* | *ubiE-1* | - | 3125748 | 3126527 | 13.01 | 26.30 | 18.11 | 6.64 | Ubiquinone/menaquinone biosynthesis C-methyltransferase UbiE |
| *FLPKDCKG_02852* | *gloB-1* | + | 3126548 | 3127369 | 12.71 | 160.31 | 89.45 | 49.43 | Hydroxyacylglutathione hydrolase |
| *FLPKDCKG_02853* | *mltD* | + | 3127377 | 3128795 | 1.73 | 40.18 | 31.25 | 14.33 | Membrane-bound lytic murein transglycosylase D |
| *FLPKDCKG_02854* | *fabI-1* | + | 3128920 | 3129705 | 15.88 | 32.72 | 20.51 | 7.69 | Enoyl-[acyl-carrier-protein] reductase [NADH] FabI |
| *FLPKDCKG_02855* | | - | 3129779 | 3130177 | 24.74 | 55.74 | 52.37 | 13.69 | Hypothetical protein |
| *FLPKDCKG_02856* | *yebE* | + | 3130357 | 3131034 | 27.27 | 22.15 | 43.80 | 29.55 | Inner membrane protein YebE |
| *FLPKDCKG_02857* | | - | 3131110 | 3132096 | 1.63 | 4.09 | 7.54 | 4.26 | Hypothetical protein |
| *FLPKDCKG_02858* | *nadC* | - | 3132125 | 3133048 | 5.28 | 10.09 | 13.90 | 5.02 | putative nicotinate-nucleotide pyrophosphorylase [carboxylating] |
| *FLPKDCKG_02859* | | - | 3133088 | 3134023 | 2.73 | 18.79 | 8.96 | 2.86 | Hypothetical protein |
| *FLPKDCKG_02860* | | + | 3134258 | 3134494 | 167.09 | 592.47 | 345.42 | 49.34 | Hypothetical protein |
| *FLPKDCKG_02861* | *btrV* | - | 3134527 | 3134862 | 3.48 | 26.78 | 19.11 | 3.65 | Putative anti-sigma factor antagonist BtrV |
| *FLPKDCKG_02862* | *btrW* | + | 3135058 | 3135489 | 40.84 | 84.51 | 45.63 | 20.28 | Serine/threonine-protein kinase BtrW |
| *FLPKDCKG_02863* | *dadX-1* | + | 3135620 | 3136765 | 4.87 | 10.28 | 6.18 | 4.92 | Alanine racemase, catabolic |
| *FLPKDCKG_02864* | | - | 3136844 | 3139312 | 1.79 | 5.11 | 5.43 | 1.47 | Hypothetical protein |
| *FLPKDCKG_02865* | *rsbU* | + | 3139422 | 3141425 | 5.48 | 7.32 | 5.65 | 1.82 | Phosphoserine phosphatase RsbU |
| *FLPKDCKG_02866* | | + | 3141422 | 3142618 | 2.01 | 2.59 | 2.98 | 0.41 | Hypothetical protein |
| *FLPKDCKG_02867* | | + | 3142776 | 3143348 | 13.87 | 33.75 | 20.11 | 8.66 | Hypothetical protein |
| *FLPKDCKG_02868* | *sigW* | + | 3143546 | 3144157 | 7.18 | 5.36 | 8.60 | 1.71 | ECF RNA polymerase sigma factor SigW |
| *FLPKDCKG_02869* | *spiA* | - | 3144231 | 3146135 | 0.56 | 0.95 | 0.32 | 0.35 | Type III secretion system outer membrane protein SpiA |
| *FLPKDCKG_02870* | | - | 3146171 | 3146569 | 0.11 | 0.00 | 1.42 | 0.00 | Hypothetical protein |
| *FLPKDCKG_02871* | *yscU* | - | 3146566 | 3147636 | 0.42 | 0.97 | 0.00 | 0.00 | Yop proteins translocation protein U |
| *FLPKDCKG_02872* | | - | 3147621 | 3148421 | 0.00 | 0.00 | 0.00 | 0.00 | Hypothetical protein |
| *FLPKDCKG_02873* | | - | 3148437 | 3148703 | 0.00 | 0.60 | 0.00 | 0.00 | Hypothetical protein |
| *FLPKDCKG_02874* | *fliP-4* | - | 3148711 | 3149379 | 0.27 | 0.44 | 0.00 | 0.32 | Flagellar biosynthetic protein FliP |
| *FLPKDCKG_02875* | *yscQ* | - | 3149376 | 3150455 | 0.18 | 0.00 | 0.00 | 0.26 | Yop proteins translocation protein Q |
| *FLPKDCKG_02876* | | - | 3150468 | 3151028 | 0.19 | 0.00 | 0.80 | 0.26 | Hypothetical protein |
| *FLPKDCKG_02877* | | - | 3151025 | 3151531 | 0.18 | 0.00 | 0.00 | 0.31 | Hypothetical protein |
| *FLPKDCKG_02878* | *yscN* | - | 3151528 | 3152859 | 0.21 | 0.96 | 0.00 | 0.14 | putative ATP synthase YscN |
| *FLPKDCKG_02879* | *yscL-2* | - | 3152862 | 3153512 | 0.19 | 1.24 | 1.23 | 0.00 | Yop proteins translocation protein L |
| *FLPKDCKG_02880* | | - | 3153482 | 3154138 | 0.50 | 1.48 | 0.00 | 0.00 | Hypothetical protein |
| *FLPKDCKG_02881* | *nolT* | - | 3154141 | 3154986 | 0.35 | 0.28 | 0.00 | 0.00 | Nodulation protein NolT |
| *FLPKDCKG_02882* | | - | 3154986 | 3155402 | 0.22 | 0.00 | 0.42 | 0.00 | Hypothetical protein |
| *FLPKDCKG_02883* | | - | 3155408 | 3155821 | 0.41 | 0.00 | 0.00 | 0.00 | Hypothetical protein |
| *FLPKDCKG_02884* | *sicA-1* | - | 3155909 | 3156370 | 1.14 | 1.70 | 1.07 | 0.10 | Chaperone protein SicA |
| *FLPKDCKG_02885* | | - | 3156395 | 3157615 | 1.23 | 0.15 | 0.32 | 0.62 | Hypothetical protein |
| *FLPKDCKG_02886* | | - | 3157658 | 3158578 | 1.93 | 0.89 | 0.57 | 1.79 | Hypothetical protein |
| *FLPKDCKG_02887* | *sicA-2* | - | 3158592 | 3159086 | 1.42 | 0.00 | 0.00 | 0.84 | Chaperone protein SicA |
| *FLPKDCKG_02888* | | - | 3159117 | 3159413 | 1.79 | 0.00 | 0.00 | 0.00 | Hypothetical protein |
| *FLPKDCKG_02889* | | - | 3159443 | 3160057 | 2.33 | 0.61 | 0.45 | 0.07 | Hypothetical protein |
| *FLPKDCKG_02890* | | + | 3160372 | 3161472 | 0.72 | 1.11 | 0.63 | 0.36 | Hypothetical protein |
| *FLPKDCKG_02891* | | + | 3161469 | 3161849 | 0.57 | 0.00 | 0.84 | 0.00 | Hypothetical protein |
| *FLPKDCKG_02892* | | + | 3161872 | 3162279 | 0.64 | 1.57 | 0.00 | 0.00 | Hypothetical protein |
| *FLPKDCKG_02893* | | + | 3162276 | 3162647 | 0.27 | 0.00 | 0.55 | 0.32 | Hypothetical protein |
| *FLPKDCKG_02894* | *ssaV* | + | 3162656 | 3164749 | 0.83 | 0.89 | 0.58 | 0.46 | Secretion system apparatus protein SsaV |

| Locus name | Gene name | Strand | Gene start (bp) | Gene end (bp) | Average TPM Inoculum | Average TPM $t_6$ | Average TPM $t_{24}$ | Average TPM $t_{168}$ | Annotation description |
|---|---|---|---|---|---|---|---|---|---|
| *FLPKDCKG_02895* | | + | 3164775 | 3166136 | 0.49 | 1.09 | 0.00 | 0.09 | Hypothetical protein |
| *FLPKDCKG_02896* | | + | 3166163 | 3166438 | 1.01 | 0.00 | 5.22 | 1.80 | Hypothetical protein |
| *FLPKDCKG_02897* | | + | 3166449 | 3166715 | 1.82 | 0.21 | 4.50 | 1.19 | Hypothetical protein |
| *FLPKDCKG_02898* | | + | 3167351 | 3168268 | 0.29 | 0.27 | 0.00 | 0.00 | Hypothetical protein |
| *FLPKDCKG_02899* | | + | 3168265 | 3170052 | 0.30 | 0.08 | 0.59 | 0.54 | Hypothetical protein |
| *FLPKDCKG_02900* | *egtA* | - | 3170122 | 3171600 | 1.21 | 1.68 | 1.67 | 0.99 | Glutamate--cysteine ligase EgtA |
| *FLPKDCKG_02901* | *cmpR-10* | - | 3171745 | 3172650 | 3.83 | 4.32 | 5.06 | 1.49 | HTH-type transcriptional activator CmpR |
| *FLPKDCKG_02902* | | + | 3172793 | 3173785 | 5.54 | 10.45 | 5.00 | 1.64 | Hypothetical protein |
| *FLPKDCKG_02903* | | + | 3173812 | 3175074 | 0.85 | 2.87 | 0.78 | 1.50 | Hypothetical protein |
| *FLPKDCKG_02904* | | - | 3175123 | 3175773 | 14.88 | 14.16 | 7.68 | 3.48 | Hypothetical protein |
| *FLPKDCKG_02905* | | - | 3175943 | 3176701 | 0.40 | 3.49 | 0.13 | 0.00 | Hypothetical protein |
| *FLPKDCKG_02906* | | - | 3177015 | 3177395 | 2.46 | 8.21 | 3.69 | 0.94 | Hypothetical protein |
| *FLPKDCKG_02907* | *yofA-5* | + | 3177479 | 3178336 | 1.71 | 3.81 | 4.46 | 0.38 | HTH-type transcriptional regulator YofA |
| *FLPKDCKG_02908* | | - | 3178438 | 3178528 | 56.78 | 121.54 | 46.38 | 15.33 | tRNA |
| *FLPKDCKG_02909* | | - | 3178707 | 3180257 | 72.09 | 238.05 | 149.67 | 29.22 | Hypothetical protein |
| *FLPKDCKG_02910* | | - | 3180254 | 3181513 | 164.14 | 533.89 | 304.81 | 59.28 | Hypothetical protein |
| *FLPKDCKG_02911* | | - | 3181526 | 3183451 | 336.59 | 730.56 | 416.39 | 76.36 | Hypothetical protein |
| *FLPKDCKG_02912* | | + | 3184317 | 3185513 | 1.06 | 1.82 | 1.74 | 0.56 | Hypothetical protein |
| *FLPKDCKG_02913* | | + | 3185500 | 3187956 | 1.41 | 3.22 | 2.17 | 0.34 | Hypothetical protein |
| *FLPKDCKG_02914* | *ribD* | - | 3187953 | 3189113 | 5.93 | 5.02 | 9.15 | 1.41 | Riboflavin biosynthesis protein RibD |
| *FLPKDCKG_02915* | *pncA* | + | 3189315 | 3189884 | 11.29 | 4.92 | 3.93 | 1.62 | Nicotinamidase/pyrazinamidase |
| *FLPKDCKG_02916* | *cdhR-2* | + | 3189897 | 3190856 | 3.61 | 4.99 | 2.54 | 1.64 | HTH-type transcriptional regulator CdhR |
| *FLPKDCKG_02917* | *nrdR* | - | 3190904 | 3191383 | 19.50 | 34.58 | 21.96 | 6.14 | Transcriptional repressor NrdR |
| *FLPKDCKG_02918* | *glyA1* | - | 3191588 | 3192838 | 87.10 | 143.64 | 104.40 | 41.02 | Serine hydroxymethyltransferase 1 |
| *FLPKDCKG_02919* | | - | 3193135 | 3193791 | 208.03 | 255.73 | 160.54 | 31.45 | Hypothetical protein |
| *FLPKDCKG_02920* | *tyrS* | - | 3193864 | 3195093 | 11.97 | 13.99 | 10.67 | 4.55 | Tyrosine--tRNA ligase |
| *FLPKDCKG_02921* | *mepM-2* | + | 3195351 | 3196760 | 4.24 | 43.16 | 43.89 | 21.94 | Murein DD-endopeptidase MepM |
| *FLPKDCKG_02922* | *anmK* | + | 3196838 | 3197947 | 1.41 | 3.63 | 7.94 | 2.55 | Anhydro-N-acetylmuramic acid kinase |
| *FLPKDCKG_02923* | *erpA* | - | 3198053 | 3198424 | 290.90 | 256.56 | 147.61 | 65.90 | Iron-sulfur cluster insertion protein ErpA |
| *FLPKDCKG_02924* | | - | 3198510 | 3199256 | 4.90 | 10.07 | 5.20 | 0.98 | Hypothetical protein |
| *FLPKDCKG_02925* | *argC-2* | - | 3199297 | 3200361 | 38.41 | 60.26 | 35.47 | 11.01 | N-acetyl-gamma-glutamyl-phosphate reductase |
| *FLPKDCKG_02926* | *rpsI* | - | 3200550 | 3200942 | 14.22 | 38.41 | 34.35 | 18.86 | 30S ribosomal protein S9 |
| *FLPKDCKG_02927* | *rplM* | - | 3200952 | 3201380 | 51.67 | 123.35 | 93.87 | 45.12 | 50S ribosomal protein L13 |
| *FLPKDCKG_02928* | *livF-11* | - | 3201745 | 3202458 | 1.22 | 1.81 | 2.47 | 1.86 | High-affinity branched-chain amino acid transport ATP-binding protein LivF |
| *FLPKDCKG_02929* | *lptB-4* | - | 3202455 | 3203243 | 2.60 | 7.18 | 5.63 | 1.60 | Lipopolysaccharide export system ATP-binding protein LptB |
| *FLPKDCKG_02930* | | - | 3203236 | 3203451 | 2.02 | 0.00 | 6.06 | 1.48 | Hypothetical protein |
| *FLPKDCKG_02931* | | - | 3203508 | 3204101 | 0.54 | 0.98 | 1.72 | 0.00 | Hypothetical protein |
| *FLPKDCKG_02932* | *livH-11* | - | 3204104 | 3204985 | 2.31 | 3.16 | 2.38 | 0.94 | High-affinity branched-chain amino acid transport system permease protein LivH |
| *FLPKDCKG_02933* | *braC-3* | - | 3205233 | 3206387 | 21.01 | 28.36 | 26.51 | 13.04 | Leucine-, isoleucine-, valine-, threonine-, and alanine-binding protein |
| *FLPKDCKG_02934* | *takP-2* | - | 3206645 | 3207877 | 22.23 | 18.87 | 13.39 | 3.06 | Alpha-keto acid-binding periplasmic protein TakP |
| *FLPKDCKG_02935* | *pmbA* | - | 3208011 | 3209375 | 17.55 | 33.24 | 32.34 | 7.35 | Metalloprotease PmbA |
| *FLPKDCKG_02936* | | + | 3209434 | 3209982 | 54.59 | 105.20 | 48.80 | 18.09 | Hypothetical protein |
| *FLPKDCKG_02937* | *estB* | + | 3209998 | 3210675 | 7.48 | 14.37 | 12.50 | 3.49 | Carboxylesterase 2 |
| *FLPKDCKG_02938* | | + | 3210999 | 3212006 | 3.68 | 3.90 | 1.33 | 0.31 | Hypothetical protein |
| *FLPKDCKG_02939* | *ssuB-3* | + | 3212006 | 3212818 | 1.55 | 0.70 | 0.71 | 0.21 | Aliphatic sulfonates import ATP-binding protein SsuB |
| *FLPKDCKG_02940* | *ribX-1* | + | 3212822 | 3213658 | 0.76 | 1.85 | 0.00 | 0.25 | Riboflavin transport system permease protein RibX |
| *FLPKDCKG_02941* | *rutB* | + | 3213668 | 3214360 | 1.85 | 0.62 | 0.00 | 0.41 | Peroxyureidoacrylate/ureidoacrylate amidohydrolase RutB |
| *FLPKDCKG_02942* | | - | 3214408 | 3216849 | 5.85 | 6.79 | 4.77 | 1.32 | Acetyltransferase Pat |
| *FLPKDCKG_02943* | *gltX* | + | 3216970 | 3218376 | 8.09 | 14.16 | 7.66 | 5.48 | Glutamate--tRNA ligase |
| *FLPKDCKG_02944* | *emrB-4* | - | 3218450 | 3219643 | 1.19 | 2.70 | 1.68 | 0.79 | Multidrug export protein EmrB |
| *FLPKDCKG_02945* | | - | 3219773 | 3220279 | 8.87 | 6.03 | 4.94 | 0.53 | Hypothetical protein |
| *FLPKDCKG_02946* | *cynR-9* | - | 3220468 | 3221367 | 4.51 | 3.21 | 4.01 | 0.90 | HTH-type transcriptional regulator CynR |
| *FLPKDCKG_02947* | *menH-1* | + | 3221442 | 3222293 | 1.28 | 0.31 | 2.69 | 1.27 | 2-succinyl-6-hydroxy-2,4-cyclohexadiene-1-carboxylate synthase |
| *FLPKDCKG_02948* | | + | 3222438 | 3223418 | 1.54 | 5.11 | 4.68 | 1.02 | Hypothetical protein |
| *FLPKDCKG_02949* | *lrp-7* | - | 3223964 | 3224413 | 23.29 | 33.99 | 15.12 | 10.40 | Leucine-responsive regulatory protein |
| *FLPKDCKG_02950* | | + | 3224549 | 3228058 | 1.79 | 38.91 | 20.71 | 10.39 | Hypothetical protein |
| *FLPKDCKG_02951* | *nrdA* | + | 3228362 | 3231292 | 94.65 | 106.62 | 77.20 | 26.72 | Ribonucleoside-diphosphate reductase 1 subunit alpha |
| *FLPKDCKG_02952* | *nrdB* | + | 3231313 | 3232512 | 90.40 | 93.11 | 69.46 | 16.26 | Ribonucleoside-diphosphate reductase subunit beta |
| *FLPKDCKG_02953* | *hctB* | + | 3232858 | 3233505 | 341.87 | 508.78 | 502.07 | 218.47 | Histone H1-like protein HC2 |
| *FLPKDCKG_02954* | | + | 3233685 | 3234245 | 10.70 | 17.38 | 11.08 | 5.00 | Hypothetical protein |
| *FLPKDCKG_02955* | *pigI* | - | 3234310 | 3234519 | 4.20 | 37.23 | 23.45 | 0.50 | putative L-prolyl-AMP ligase PigI |
| *FLPKDCKG_02956* | *dltA-2* | - | 3234485 | 3235756 | 1.34 | 7.66 | 8.42 | 1.59 | D-alanine--poly(phosphoribitol) ligase subunit 1 |
| *FLPKDCKG_02957* | *patA-2* | - | 3235773 | 3236900 | 1.21 | 6.79 | 2.68 | 2.19 | Peptidoglycan O-acetyltransferase |
| *FLPKDCKG_02958* | | - | 3236908 | 3237114 | 2.05 | 19.64 | 12.00 | 0.61 | Hypothetical protein |
| *FLPKDCKG_02959* | | - | 3237111 | 3238289 | 5.87 | 32.00 | 17.89 | 6.14 | Hypothetical protein |
| *FLPKDCKG_02960* | *acpP-3* | - | 3238267 | 3238506 | 15.34 | 89.64 | 74.16 | 31.88 | Acyl carrier protein |
| *FLPKDCKG_02961* | *slyB* | - | 3238921 | 3239424 | 197.03 | 480.01 | 328.33 | 108.19 | Outer membrane lipoprotein SlyB |
| *FLPKDCKG_02962* | *adoK* | - | 3239645 | 3240586 | 3.95 | 6.93 | 6.03 | 2.30 | Adenosine kinase |
| *FLPKDCKG_02963* | | - | 3240599 | 3242011 | 1.58 | 1.05 | 1.85 | 1.47 | Hypothetical protein |

| Locus name | Gene name | Strand | Gene start (bp) | Gene end (bp) | Average TPM Inoculum | Average TPM $t_6$ | Average TPM $t_{24}$ | Average TPM $t_{168}$ | Annotation description |
|---|---|---|---|---|---|---|---|---|---|
| *FLPKDCKG_02964* | *prmA-1* | - | 3242037 | 3242963 | 2.27 | 4.34 | 6.25 | 3.70 | Ribosomal protein L11 methyltransferase |
| *FLPKDCKG_02965* | *accC-1* | - | 3243066 | 3244412 | 5.84 | 20.19 | 18.28 | 6.32 | Biotin carboxylase |
| *FLPKDCKG_02966* | *accB-1* | - | 3244422 | 3244883 | 5.29 | 20.95 | 20.10 | 9.83 | Biotin carboxyl carrier protein of acetyl-CoA carboxylase |
| *FLPKDCKG_02967* | *aroQ1* | - | 3244972 | 3245406 | 38.30 | 131.62 | 113.02 | 33.39 | 3-dehydroquinate dehydratase 1 |
| *FLPKDCKG_02968* | *resA-1* | - | 3245496 | 3246026 | 7.87 | 23.65 | 15.94 | 3.66 | Thiol-disulfide oxidoreductase ResA |
| *FLPKDCKG_02969* | *mpl* | + | 3246123 | 3247493 | 10.94 | 26.41 | 20.15 | 5.74 | UDP-N-acetylmuramate--L-alanyl-gamma-D-glutamyl-meso-2,6-diaminoheptandioate ligase |
| *FLPKDCKG_02970* | | + | 3247543 | 3248145 | 3.69 | 13.23 | 6.17 | 4.79 | Hypothetical protein |
| *FLPKDCKG_02971* | *rnr-2* | + | 3248211 | 3250118 | 3.77 | 12.11 | 7.97 | 3.14 | Ribonuclease R |
| *FLPKDCKG_02972* | *tctD-4* | + | 3251102 | 3251773 | 6.34 | 7.03 | 4.40 | 1.48 | Transcriptional regulatory protein tctD |
| *FLPKDCKG_02973* | *qseC-5* | + | 3251760 | 3253055 | 0.98 | 2.88 | 0.76 | 0.29 | Sensor protein QseC |
| *FLPKDCKG_02974* | | + | 3253037 | 3254245 | 1.56 | 8.33 | 3.54 | 0.50 | Hypothetical protein |
| *FLPKDCKG_02975* | | + | 3254385 | 3255482 | 13.60 | 175.16 | 93.92 | 35.00 | Hypothetical protein |
| *FLPKDCKG_02976* | *cysW-2* | + | 3255631 | 3257190 | 0.85 | 14.24 | 11.60 | 2.55 | Sulfate transport system permease protein CysW |
| *FLPKDCKG_02977* | | + | 3257166 | 3257408 | 2.42 | 25.01 | 6.18 | 2.87 | Hypothetical protein |
| *FLPKDCKG_02978* | *potA-3* | + | 3257401 | 3258489 | 1.78 | 36.54 | 24.45 | 9.57 | Spermidine/putrescine import ATP-binding protein PotA |
| *FLPKDCKG_02979* | | + | 3258520 | 3259425 | 25.05 | 41.60 | 26.67 | 14.51 | Hypothetical protein |
| *FLPKDCKG_02980* | *aroE-1* | + | 3259422 | 3260285 | 5.03 | 8.31 | 6.04 | 1.42 | Shikimate dehydrogenase (NADP(+)) |
| *FLPKDCKG_02981* | *mtgA* | + | 3260290 | 3261024 | 11.33 | 26.21 | 18.18 | 4.65 | Biosynthetic peptidoglycan transglycosylase |
| *FLPKDCKG_02982* | *lptG* | + | 3261116 | 3262294 | 29.26 | 45.25 | 29.69 | 11.63 | Lipopolysaccharide export system permease protein LptG |
| *FLPKDCKG_02983* | *apaH-2* | + | 3262291 | 3263121 | 12.85 | 15.59 | 16.02 | 7.88 | Bis(5'-nucleosyl)-tetraphosphatase, symmetrical |
| *FLPKDCKG_02984* | *glnQ-7* | - | 3264020 | 3264745 | 0.40 | 0.98 | 1.03 | 0.87 | Glutamine transport ATP-binding protein GlnQ |
| *FLPKDCKG_02985* | *glnP-3* | - | 3264742 | 3265458 | 0.48 | 2.35 | 0.86 | 1.11 | putative glutamine ABC transporter permease protein GlnP |
| *FLPKDCKG_02986* | *glnP-4* | - | 3265459 | 3266160 | 0.00 | 0.00 | 0.72 | 0.16 | Glutamine transport system permease protein GlnP |
| *FLPKDCKG_02987* | *glnH-4* | - | 3266189 | 3267016 | 0.18 | 0.00 | 0.38 | 0.00 | ABC transporter glutamine-binding protein GlnH |
| *FLPKDCKG_02988* | | - | 3267085 | 3268302 | 0.41 | 0.47 | 0.00 | 0.19 | Aspartate aminotransferase |
| *FLPKDCKG_02989* | *proA-4* | - | 3268299 | 3268973 | 0.72 | 1.35 | 2.01 | 0.69 | 4-hydroxy-4-methyl-2-oxoglutarate aldolase/4-carboxy-4-hydroxy-2-oxoadipate aldolase |
| *FLPKDCKG_02990* | *gltR-4* | + | 3269140 | 3270036 | 9.15 | 9.43 | 4.87 | 1.72 | HTH-type transcriptional regulator GltR |
| *FLPKDCKG_02991* | *plsC-2* | - | 3271012 | 3271752 | 9.13 | 26.14 | 21.44 | 4.19 | 1-acyl-sn-glycerol-3-phosphate acyltransferase |
| *FLPKDCKG_02992* | *pyrC-2* | - | 3271749 | 3273053 | 1.15 | 2.72 | 2.38 | 0.91 | Dihydroorotase |
| *FLPKDCKG_02993* | *pyrB-2* | - | 3273050 | 3274039 | 1.74 | 4.70 | 7.32 | 2.25 | Aspartate carbamoyltransferase |
| *FLPKDCKG_02994* | *yqgF* | - | 3274059 | 3274460 | 6.25 | 19.93 | 18.12 | 3.46 | Putative pre-16S rRNA nuclease |
| *FLPKDCKG_02995* | | - | 3274453 | 3275040 | 15.83 | 20.72 | 30.32 | 24.09 | Hypothetical protein |
| *FLPKDCKG_02996* | *rubA2* | - | 3275168 | 3275332 | 189.77 | 276.70 | 242.39 | 92.24 | Rubredoxin-2 |
| *FLPKDCKG_02997* | *thiD-1* | + | 3275488 | 3276297 | 9.42 | 32.28 | 20.85 | 5.95 | Hydroxymethylpyrimidine/phosphomethylpyrimidine kinase |
| *FLPKDCKG_02998* | *thiE* | + | 3276294 | 3276959 | 5.21 | 7.19 | 6.54 | 1.17 | Thiamine-phosphate synthase |
| *FLPKDCKG_02999* | *hemL* | + | 3276965 | 3278242 | 3.83 | 10.80 | 5.90 | 1.81 | Glutamate-1-semialdehyde 2,1-aminomutase |
| *FLPKDCKG_03000* | *slt-1* | - | 3278344 | 3279087 | 0.41 | 3.40 | 2.09 | 0.41 | Soluble lytic murein transglycosylase |
| *FLPKDCKG_03001* | | + | 3279395 | 3281122 | 17.41 | 5.65 | 4.51 | 1.66 | Hypothetical protein |
| *FLPKDCKG_03002* | *fabG-9* | - | 3281253 | 3282059 | 1.57 | 27.09 | 15.95 | 7.97 | 3-oxoacyl-[acyl-carrier-protein] reductase FabG |
| *FLPKDCKG_03003* | *yecD* | - | 3282073 | 3282684 | 4.08 | 29.47 | 23.64 | 7.08 | Isochorismatase family protein YecD |
| *FLPKDCKG_03004* | *bnzB-2* | - | 3282681 | 3283214 | 2.75 | 42.91 | 45.16 | 8.79 | Benzene 1,2-dioxygenase subunit beta |
| *FLPKDCKG_03005* | *antA-2* | - | 3283205 | 3284548 | 3.87 | 72.29 | 50.78 | 18.19 | Anthranilate 1,2-dioxygenase large subunit |
| *FLPKDCKG_03006* | *clcD-2* | - | 3284558 | 3285802 | 5.24 | 65.08 | 55.06 | 20.47 | Carboxymethylenebutenolidase |
| *FLPKDCKG_03007* | | - | 3285844 | 3287031 | 7.94 | 89.04 | 77.19 | 21.33 | Leu/Ile/Val-binding protein |
| *FLPKDCKG_03008* | *cdhR-3* | + | 3287294 | 3288280 | 24.84 | 31.06 | 19.10 | 4.84 | HTH-type transcriptional regulator CdhR |
| *FLPKDCKG_03009* | *yabJ-5* | - | 3288752 | 3289156 | 16.06 | 23.52 | 28.56 | 22.40 | 2-iminobutanoate/2-iminopropanoate deaminase |
| *FLPKDCKG_03010* | *fadM* | - | 3289208 | 3289633 | 1.48 | 2.34 | 2.23 | 3.09 | Long-chain acyl-CoA thioesterase FadM |
| *FLPKDCKG_03011* | *bclA-1* | - | 3289657 | 3291297 | 1.50 | 3.78 | 2.53 | 4.10 | Benzoate--CoA ligase |
| *FLPKDCKG_03012* | | - | 3291358 | 3292530 | 4.87 | 4.91 | 9.33 | 6.15 | putative acyl-CoA dehydrogenase fadE25 |
| *FLPKDCKG_03013* | | - | 3292530 | 3293387 | 4.90 | 5.00 | 6.97 | 4.37 | putative enoyl-CoA hydratase echA12 |
| *FLPKDCKG_03014* | *fabG-10* | - | 3293426 | 3294217 | 4.33 | 9.74 | 6.91 | 4.25 | 3-oxoacyl-[acyl-carrier-protein] reductase FabG |
| *FLPKDCKG_03015* | *sdgC* | - | 3294214 | 3296565 | 1.64 | 2.49 | 2.62 | 4.71 | Salicyloyl-CoA 5-hydroxylase |
| *FLPKDCKG_03016* | *dosP-1* | - | 3297684 | 3300260 | 1.47 | 2.37 | 2.65 | 1.52 | Oxygen sensor protein DosP |
| *FLPKDCKG_03017* | *ydaM-2* | - | 3300300 | 3301097 | 0.73 | 2.70 | 2.02 | 2.46 | putative diguanylate cyclase YdaM |
| *FLPKDCKG_03018* | | - | 3301094 | 3301471 | 1.52 | 3.66 | 2.48 | 1.39 | Hypothetical protein |
| *FLPKDCKG_03019* | | - | 3301510 | 3302049 | 2.54 | 1.95 | 2.13 | 3.96 | Hypothetical protein |
| *FLPKDCKG_03020* | *mcpH* | - | 3302073 | 3304067 | 2.25 | 4.26 | 1.93 | 0.97 | Methyl-accepting chemotaxis protein McpH |
| *FLPKDCKG_03021* | *livF-12* | - | 3304285 | 3304989 | 0.50 | 1.34 | 1.93 | 2.99 | High-affinity branched-chain amino acid transport ATP-binding protein LivF |
| *FLPKDCKG_03022* | *lptB-5* | - | 3304989 | 3306752 | 0.93 | 1.84 | 1.12 | 2.90 | Lipopolysaccharide export system ATP-binding protein LptB |
| *FLPKDCKG_03023* | | - | 3306755 | 3307387 | 0.44 | 0.24 | 2.21 | 0.30 | Hypothetical protein |
| *FLPKDCKG_03024* | *livH-12* | - | 3307350 | 3307673 | 0.63 | 4.37 | 0.00 | 3.14 | High-affinity branched-chain amino acid transport system permease protein LivH |
| *FLPKDCKG_03025* | *amiC-5* | - | 3307826 | 3308989 | 10.70 | 15.66 | 21.52 | 14.79 | Aliphatic amidase expression-regulating protein |
| *FLPKDCKG_03026* | *badR-2* | + | 3309250 | 3309744 | 26.90 | 36.40 | 22.76 | 7.33 | Transcriptional activatory protein BadR |
| *FLPKDCKG_03027* | | + | 3309855 | 3310895 | 4.06 | 5.91 | 1.45 | 0.42 | Hypothetical protein |
| *FLPKDCKG_03028* | *argJ* | + | 3311018 | 3312268 | 16.45 | 15.68 | 12.21 | 7.66 | Arginine biosynthesis bifunctional protein ArgJ |
| *FLPKDCKG_03029* | | + | 3312543 | 3313409 | 3.49 | 6.25 | 4.01 | 1.44 | Hypothetical protein |
| *FLPKDCKG_03030* | *nudG* | + | 3313402 | 3314364 | 1.09 | 1.95 | 3.23 | 1.26 | CTP pyrophosphohydrolase |
| *FLPKDCKG_03031* | | + | 3314662 | 3316098 | 5.25 | 1.23 | 1.10 | 0.26 | Hypothetical protein |
| *FLPKDCKG_03032* | *sugC* | + | 3316103 | 3317224 | 2.97 | 0.61 | 0.00 | 0.00 | Trehalose import ATP-binding protein SugC |

| Locus name | Gene name | Strand | Gene start (bp) | Gene end (bp) | Average TPM Inoculum | Average TPM $t_6$ | Average TPM $t_{24}$ | Average TPM $t_{168}$ | Annotation description |
|---|---|---|---|---|---|---|---|---|---|
| *FLPKDCKG_03033* | *ugpA-2* | + | 3317221 | 3318165 | 1.46 | 0.68 | 0.00 | 0.20 | sn-glycerol-3-phosphate transport system permease protein UgpA |
| *FLPKDCKG_03034* | *sugB* | + | 3318155 | 3319054 | 1.31 | 0.80 | 0.00 | 0.33 | Trehalose transport system permease protein SugB |
| *FLPKDCKG_03035* | | + | 3319064 | 3319993 | 3.22 | 3.22 | 2.44 | 0.14 | Hypothetical protein |
| *FLPKDCKG_03036* | *betI-2* | + | 3319980 | 3320645 | 17.59 | 11.87 | 5.90 | 0.53 | HTH-type transcriptional regulator BetI |
| *FLPKDCKG_03037* | *dan-3* | + | 3320653 | 3322152 | 3.99 | 1.79 | 2.15 | 0.33 | D-aminoacylase |
| *FLPKDCKG_03038* | *dadA1-4* | + | 3322203 | 3323456 | 1.15 | 1.01 | 2.17 | 0.03 | D-amino acid dehydrogenase 1 |
| *FLPKDCKG_03039* | | - | 3323530 | 3324528 | 2.96 | 1.13 | 0.81 | 0.25 | Hypothetical protein |
| *FLPKDCKG_03040* | *oprM-5* | - | 3324738 | 3326315 | 1.05 | 5.50 | 1.49 | 0.52 | Outer membrane protein OprM |
| *FLPKDCKG_03041* | *mdtC-3* | - | 3326312 | 3329422 | 1.35 | 5.12 | 2.85 | 0.73 | Multidrug resistance protein MdtC |
| *FLPKDCKG_03042* | *mdtC-4* | - | 3329419 | 3329793 | 1.19 | 7.68 | 1.89 | 0.88 | Multidrug resistance protein MdtC |
| *FLPKDCKG_03043* | *mdtB-2* | - | 3329814 | 3332516 | 1.73 | 5.78 | 3.33 | 0.83 | Multidrug resistance protein MdtB |
| *FLPKDCKG_03044* | | + | 3333569 | 3333850 | 4.12 | 3.64 | 10.22 | 4.74 | Hypothetical protein |
| *FLPKDCKG_03045* | *zapD* | - | 3333942 | 3334712 | 15.09 | 39.33 | 22.09 | 6.63 | Cell division protein ZapD |
| *FLPKDCKG_03046* | *coaE-1* | - | 3334737 | 3335384 | 29.95 | 51.30 | 23.44 | 6.18 | Dephospho-CoA kinase |
| *FLPKDCKG_03047* | *outO* | - | 3335458 | 3336315 | 8.68 | 13.80 | 7.72 | 2.90 | Type 4 prepilin-like proteins leader peptide-processing enzyme |
| *FLPKDCKG_03048* | | + | 3336498 | 3337274 | 275.37 | 154.39 | 98.73 | 26.28 | Hypothetical protein |
| *FLPKDCKG_03049* | *tdiR* | - | 3337408 | 3337806 | 1.38 | 5.18 | 0.07 | 0.32 | Transcriptional regulatory protein TdiR |
| *FLPKDCKG_03050* | *panC* | - | 3337938 | 3338792 | 5.05 | 8.60 | 15.68 | 4.18 | Pantothenate synthetase |
| *FLPKDCKG_03051* | *scpA* | - | 3338838 | 3339728 | 12.82 | 37.08 | 23.04 | 4.49 | Segregation and condensation protein A |
| *FLPKDCKG_03052* | | - | 3339745 | 3339933 | 304.70 | 609.04 | 410.60 | 131.25 | Hypothetical protein |
| *FLPKDCKG_03053* | *hsrA* | + | 3340104 | 3341486 | 14.14 | 46.86 | 31.08 | 6.13 | putative transport protein HsrA |
| *FLPKDCKG_03054* | *glpK-3* | + | 3341458 | 3342996 | 11.06 | 26.57 | 19.08 | 7.78 | Glycerol kinase |
| *FLPKDCKG_03055* | | - | 3343040 | 3343597 | 1.68 | 9.78 | 4.07 | 1.20 | Hypothetical protein |
| *FLPKDCKG_03056* | *gbpR-2* | - | 3343597 | 3344532 | 1.53 | 5.23 | 2.11 | 0.61 | HTH-type transcriptional regulator GbpR |
| *FLPKDCKG_03057* | *sdgD-1* | + | 3344702 | 3345754 | 1.65 | 168.61 | 195.03 | 30.76 | Gentisate 1,2-dioxygenase |
| *FLPKDCKG_03058* | *nagK-2* | + | 3345814 | 3346512 | 0.52 | 56.16 | 93.23 | 23.94 | Fumarylpyruvate hydrolase |
| *FLPKDCKG_03059* | *nagL-2* | + | 3346525 | 3347163 | 0.27 | 93.70 | 94.43 | 16.85 | Maleylpyruvate isomerase |
| *FLPKDCKG_03060* | *pchB* | + | 3347160 | 3347558 | 0.87 | 101.39 | 144.97 | 29.79 | Isochorismate pyruvate lyase |
| *FLPKDCKG_03061* | | + | 3347551 | 3348555 | 0.58 | 110.75 | 134.34 | 40.32 | Hypothetical protein |
| *FLPKDCKG_03062* | *sthA* | + | 3348638 | 3349990 | 0.41 | 51.14 | 67.18 | 19.88 | Soluble pyridine nucleotide transhydrogenase |
| *FLPKDCKG_03063* | *mhbM* | + | 3350010 | 3351230 | 0.40 | 53.88 | 75.19 | 16.21 | 3-hydroxybenzoate 6-hydroxylase |
| *FLPKDCKG_03064* | *braC-4* | + | 3351257 | 3352432 | 0.51 | 76.04 | 91.27 | 20.84 | Leucine-, isoleucine-, valine-, threonine-, and alanine-binding protein |
| *FLPKDCKG_03065* | *livH-13* | + | 3352448 | 3353362 | 0.30 | 17.62 | 27.11 | 6.88 | High-affinity branched-chain amino acid transport system permease protein LivH |
| *FLPKDCKG_03066* | | + | 3353370 | 3354338 | 0.29 | 17.19 | 24.91 | 6.04 | Hypothetical protein |
| *FLPKDCKG_03067* | *phnT-2* | + | 3354331 | 3355086 | 0.11 | 12.04 | 16.97 | 2.97 | Putative 2-aminoethylphosphonate import ATP-binding protein PhnT |
| *FLPKDCKG_03068* | *livF-13* | + | 3355086 | 3355778 | 0.56 | 19.74 | 36.73 | 4.18 | High-affinity branched-chain amino acid transport ATP-binding protein LivF |
| *FLPKDCKG_03069* | *ble* | - | 3355811 | 3356173 | 7.82 | 11.65 | 7.03 | 3.78 | Bleomycin resistance protein |
| *FLPKDCKG_03070* | | + | 3356259 | 3356618 | 0.63 | 7.55 | 7.00 | 3.84 | Hypothetical protein |
| *FLPKDCKG_03071* | | - | 3356753 | 3357769 | 7.07 | 8.01 | 4.43 | 4.91 | Hypothetical protein |
| *FLPKDCKG_03072* | *glnQ-8* | - | 3358033 | 3358773 | 26.02 | 20.78 | 15.16 | 8.49 | Glutamine transport ATP-binding protein GlnQ |
| *FLPKDCKG_03073* | *yhdY-3* | - | 3358834 | 3359937 | 19.05 | 15.48 | 17.00 | 5.15 | Inner membrane amino-acid ABC transporter permease protein YhdY |
| *FLPKDCKG_03074* | *glnM-3* | - | 3359939 | 3361129 | 15.06 | 27.45 | 19.63 | 9.60 | putative glutamine ABC transporter permease protein GlnM |
| *FLPKDCKG_03075* | *yhdW-2* | - | 3361255 | 3362271 | 262.86 | 181.91 | 168.23 | 54.34 | Putative amino-acid ABC transporter-binding protein YhdW |
| *FLPKDCKG_03076* | *ftsE* | + | 3362587 | 3363300 | 28.01 | 75.34 | 44.65 | 11.77 | Cell division ATP-binding protein FtsE |
| *FLPKDCKG_03077* | *ftsX* | + | 3363297 | 3364205 | 4.19 | 6.56 | 9.57 | 1.45 | Cell division protein FtsX |
| *FLPKDCKG_03078* | *cheD* | + | 3364621 | 3365397 | 6.30 | 9.28 | 1.54 | 0.74 | Chemoreceptor glutamine deamidase CheD |
| *FLPKDCKG_03079* | *cdhR-4* | - | 3365410 | 3366381 | 0.92 | 2.78 | 0.51 | 0.65 | HTH-type transcriptional regulator CdhR |
| *FLPKDCKG_03080* | | + | 3366675 | 3367364 | 3.28 | 9.19 | 5.93 | 0.34 | Hypothetical protein |
| *FLPKDCKG_03081* | | - | 3367451 | 3368233 | 19.82 | 16.72 | 11.70 | 6.54 | Hypothetical protein |
| *FLPKDCKG_03082* | | - | 3368230 | 3369546 | 24.04 | 16.03 | 12.10 | 5.63 | Hypothetical protein |
| *FLPKDCKG_03083* | | - | 3370327 | 3371103 | 34.33 | 23.41 | 20.57 | 9.33 | Hypothetical protein |
| *FLPKDCKG_03084* | *mutY* | + | 3371231 | 3372307 | 14.47 | 36.99 | 20.88 | 6.00 | Adenine DNA glycosylase |
| *FLPKDCKG_03085* | | - | 3372506 | 3373324 | 14.13 | 22.15 | 10.84 | 4.42 | Hypothetical protein |
| *FLPKDCKG_03086* | *mdlY* | + | 3373526 | 3374971 | 5.66 | 6.43 | 2.05 | 0.98 | Mandelamide hydrolase |
| *FLPKDCKG_03087* | *por-1* | - | 3375003 | 3376565 | 0.37 | 0.88 | 1.00 | 0.23 | Polyol:NADP oxidoreductase |
| *FLPKDCKG_03088* | | - | 3376553 | 3377764 | 0.66 | 1.50 | 0.72 | 0.67 | D-mannonate dehydratase |
| *FLPKDCKG_03089* | *glnQ-9* | - | 3377792 | 3378643 | 0.29 | 1.31 | 0.49 | 0.83 | Glutamine transport ATP-binding protein GlnQ |
| *FLPKDCKG_03090* | *glnP-5* | - | 3378649 | 3379308 | 0.23 | 0.36 | 0.00 | 1.40 | Glutamine transport system permease protein GlnP |
| *FLPKDCKG_03091* | *artQ* | - | 3379305 | 3379970 | 0.58 | 3.77 | 0.52 | 1.06 | Arginine transport system permease protein ArtQ |
| *FLPKDCKG_03092* | *idnD* | - | 3380002 | 3381039 | 0.25 | 0.76 | 1.46 | 0.28 | L-idonate 5-dehydrogenase (NAD(P)(+)) |
| *FLPKDCKG_03093* | *mltF* | - | 3381052 | 3381885 | 0.99 | 3.09 | 2.10 | 1.47 | Membrane-bound lytic murein transglycosylase F |
| *FLPKDCKG_03094* | *artP-1* | - | 3382016 | 3382882 | 3.52 | 12.40 | 5.92 | 6.17 | Arginine-binding extracellular protein ArtP |
| *FLPKDCKG_03095* | *lutR-1* | - | 3382977 | 3383705 | 13.63 | 36.26 | 25.27 | 20.58 | HTH-type transcriptional regulator LutR |
| *FLPKDCKG_03096* | *metH* | - | 3383885 | 3387688 | 7.37 | 8.92 | 7.35 | 3.35 | Methionine synthase |
| *FLPKDCKG_03097* | *cirA-3* | + | 3388108 | 3390153 | 3.76 | 11.12 | 12.68 | 5.98 | Colicin I receptor |
| *FLPKDCKG_03098* | *btuF-2* | + | 3390208 | 3390603 | 2.57 | 1.01 | 3.33 | 1.67 | Vitamin B12-binding protein |
| *FLPKDCKG_03099* | *btuF-3* | + | 3390603 | 3391139 | 0.36 | 3.64 | 1.10 | 0.00 | Vitamin B12-binding protein |
| *FLPKDCKG_03100* | *thiG* | + | 3391387 | 3392196 | 16.40 | 15.48 | 8.85 | 5.78 | Thiazole synthase |
| *FLPKDCKG_03101* | *thiD-2* | + | 3392193 | 3393035 | 5.08 | 8.54 | 4.82 | 1.17 | Hydroxymethylpyrimidine/phosphomethylpyrimidine kinase |

| Locus name | Gene name | Strand | Gene start (bp) | Gene end (bp) | Average TPM Inoculum | Average TPM $t_6$ | Average TPM $t_{24}$ | Average TPM $t_{168}$ | Annotation description |
|---|---|---|---|---|---|---|---|---|---|
| *FLPKDCKG_03102* | *pcm-2* | + | 3393177 | 3393857 | 76.32 | 87.72 | 57.93 | 18.10 | Protein-L-isoaspartate O-methyltransferase |
| *FLPKDCKG_03103* | *tolC-1* | + | 3393956 | 3395287 | 2.22 | 8.22 | 5.41 | 2.45 | Outer membrane protein TolC |
| *FLPKDCKG_03104* | | - | 3395303 | 3395791 | 10.53 | 10.28 | 11.92 | 5.06 | D-beta-D-heptose 1-phosphate adenylyltransferase |
| *FLPKDCKG_03105* | | - | 3395825 | 3396757 | 1.96 | 8.18 | 5.15 | 2.03 | Hypothetical protein |
| *FLPKDCKG_03106* | *ubiB* | - | 3396939 | 3398492 | 8.02 | 15.83 | 12.18 | 5.43 | putative protein kinase UbiB |
| *FLPKDCKG_03107* | | - | 3398496 | 3399134 | 6.22 | 6.58 | 6.54 | 2.79 | Hypothetical protein |
| *FLPKDCKG_03108* | | - | 3399330 | 3400328 | 68.31 | 11.88 | 33.28 | 25.82 | Hypothetical protein |
| *FLPKDCKG_03109* | *ubiE-2* | - | 3400505 | 3401281 | 29.71 | 44.97 | 26.89 | 7.53 | Ubiquinone/menaquinone biosynthesis C-methyltransferase UbiE |
| *FLPKDCKG_03110* | *phoB* | + | 3401443 | 3402141 | 19.88 | 25.62 | 34.56 | 19.56 | Phosphate regulon transcriptional regulatory protein PhoB |
| *FLPKDCKG_03111* | *phoR-1* | + | 3402154 | 3403461 | 4.57 | 14.21 | 10.03 | 2.65 | Phosphate regulon sensor protein PhoR |
| *FLPKDCKG_03112* | *czcR-1* | + | 3403526 | 3404212 | 7.34 | 6.99 | 7.80 | 2.11 | Transcriptional activator protein CzcR |
| *FLPKDCKG_03113* | *cusS-1* | + | 3404209 | 3405609 | 1.03 | 1.31 | 0.24 | 0.45 | Sensor kinase CusS |
| *FLPKDCKG_03114* | | + | 3405684 | 3406178 | 0.82 | 2.09 | 0.85 | 0.50 | Auracyanin-B |
| *FLPKDCKG_03115* | *copA* | + | 3406423 | 3407988 | 0.23 | 1.51 | 0.46 | 0.22 | Copper resistance protein A |
| *FLPKDCKG_03116* | *copB* | + | 3407992 | 3409077 | 1.77 | 4.92 | 2.74 | 0.50 | Copper resistance protein B |
| *FLPKDCKG_03117* | | + | 3409096 | 3409572 | 5.85 | 3.60 | 7.11 | 4.58 | Hypothetical protein |
| *FLPKDCKG_03118* | *serA* | + | 3409718 | 3410917 | 12.00 | 8.18 | 12.01 | 3.14 | D-3-phosphoglycerate dehydrogenase |
| *FLPKDCKG_03119* | | - | 3411041 | 3411280 | 2.35 | 8.23 | 2.66 | 2.77 | Hypothetical protein |
| *FLPKDCKG_03120* | *yfbR* | - | 3411353 | 3411625 | 17.05 | 28.98 | 22.56 | 6.77 | 5'-deoxynucleotidase YfbR |
| *FLPKDCKG_03121* | *cmpR-11* | - | 3411698 | 3412573 | 3.17 | 4.31 | 4.68 | 0.70 | HTH-type transcriptional activator CmpR |
| *FLPKDCKG_03122* | | + | 3412715 | 3413146 | 46.81 | 78.98 | 59.96 | 24.53 | Hypothetical protein |
| *FLPKDCKG_03123* | *kipI-1* | + | 3413171 | 3413905 | 2.96 | 8.43 | 7.03 | 4.21 | Kinase A inhibitor |
| *FLPKDCKG_03124* | *kipA-1* | + | 3413902 | 3414903 | 4.55 | 3.36 | 6.39 | 0.30 | KipI antagonist |
| *FLPKDCKG_03125* | | + | 3414959 | 3415729 | 3.26 | 4.06 | 5.39 | 1.99 | Hypothetical protein |
| *FLPKDCKG_03126* | *glcC-1* | - | 3415818 | 3416564 | 5.20 | 2.71 | 4.36 | 2.04 | Glc operon transcriptional activator |
| *FLPKDCKG_03127* | | - | 3416630 | 3417268 | 1.52 | 3.23 | 2.03 | 2.00 | Hypothetical protein |
| *FLPKDCKG_03128* | *uspE* | + | 3417528 | 3418397 | 66.61 | 27.21 | 19.66 | 7.00 | Universal stress protein E |
| *FLPKDCKG_03129* | | - | 3418435 | 3419715 | 8.50 | 10.60 | 10.41 | 4.30 | Hypothetical protein |
| *FLPKDCKG_03130* | *rebM* | - | 3419785 | 3420399 | 10.05 | 7.21 | 11.79 | 1.46 | Demethylrebeccamycin-D-glucose O-methyltransferase |
| *FLPKDCKG_03131* | *metX* | - | 3420396 | 3421667 | 10.10 | 4.32 | 6.23 | 2.03 | Homoserine O-acetyltransferase |
| *FLPKDCKG_03132* | *dasR-1* | - | 3421879 | 3422634 | 10.37 | 4.47 | 7.52 | 3.07 | HTH-type transcriptional repressor DasR |
| *FLPKDCKG_03133* | *suyA* | + | 3422817 | 3423098 | 1.32 | 3.36 | 1.95 | 0.00 | (2R)-sulfolactate sulfo-lyase subunit alpha |
| *FLPKDCKG_03134* | *suyB-1* | + | 3423129 | 3424304 | 0.95 | 0.98 | 0.69 | 0.41 | (2R)-sulfolactate sulfo-lyase subunit beta |
| *FLPKDCKG_03135* | | + | 3424556 | 3425569 | 0.10 | 0.64 | 0.27 | 0.35 | Hypothetical protein |
| *FLPKDCKG_03136* | *slcC-2* | + | 3425583 | 3426518 | 0.34 | 1.57 | 2.20 | 0.13 | (S)-sulfolactate dehydrogenase |
| *FLPKDCKG_03137* | *comC* | + | 3426549 | 3427556 | 9.19 | 5.08 | 4.19 | 1.09 | (2R)-3-sulfolactate dehydrogenase (NADP(+)) |
| *FLPKDCKG_03138* | | + | 3427632 | 3428240 | 1.38 | 7.61 | 1.68 | 0.78 | Hypothetical protein |
| *FLPKDCKG_03139* | | - | 3428262 | 3429161 | 1.43 | 7.73 | 2.03 | 0.79 | Hypothetical protein |
| *FLPKDCKG_03140* | *rpaR* | + | 3429280 | 3429531 | 5.89 | 16.72 | 10.74 | 1.53 | HTH-type quorum sensing-dependent transcriptional regulator RpaR |
| *FLPKDCKG_03141* | | - | 3429521 | 3430033 | 7.30 | 4.32 | 8.98 | 4.57 | Hypothetical protein |
| *FLPKDCKG_03142* | | + | 3430122 | 3431015 | 5.56 | 3.50 | 0.88 | 1.23 | Monoacylglycerol lipase |
| *FLPKDCKG_03143* | *gshA* | + | 3431048 | 3432619 | 24.05 | 27.79 | 19.86 | 6.80 | Glutamate--cysteine ligase |
| *FLPKDCKG_03144* | *blc-1* | + | 3432658 | 3433179 | 38.05 | 59.25 | 29.34 | 12.43 | Outer membrane lipoprotein Blc |
| *FLPKDCKG_03145* | *astE-1* | - | 3433304 | 3434248 | 0.59 | 3.83 | 1.25 | 0.55 | Succinylglutamate desuccinylase |
| *FLPKDCKG_03146* | | - | 3434334 | 3435326 | 0.23 | 0.21 | 0.79 | 0.44 | Hypothetical protein |
| *FLPKDCKG_03147* | *yjiE-2* | - | 3435501 | 3436415 | 0.94 | 1.29 | 0.90 | 0.17 | HTH-type transcriptional regulator YjiE |
| *FLPKDCKG_03148* | *nlhH-1* | - | 3436441 | 3437325 | 3.92 | 2.96 | 2.18 | 0.61 | Carboxylesterase NlhH |
| *FLPKDCKG_03149* | *rspR-7* | - | 3437318 | 3438079 | 4.14 | 3.32 | 2.19 | 2.08 | HTH-type transcriptional repressor RspR |
| *FLPKDCKG_03150* | *nitA* | - | 3438164 | 3439201 | 0.72 | 0.43 | 1.53 | 0.64 | Aliphatic nitrilase |
| *FLPKDCKG_03151* | *livF-14* | - | 3439221 | 3439913 | 0.42 | 0.08 | 0.90 | 0.00 | High-affinity branched-chain amino acid transport ATP-binding protein LivF |
| *FLPKDCKG_03152* | *lptB-6* | - | 3439900 | 3440760 | 0.32 | 1.24 | 1.17 | 0.50 | Lipopolysaccharide export system ATP-binding protein LptB |
| *FLPKDCKG_03153* | | - | 3440750 | 3441730 | 0.47 | 0.62 | 0.82 | 0.00 | Hypothetical protein |
| *FLPKDCKG_03154* | *livH-14* | - | 3441733 | 3442647 | 0.39 | 0.25 | 0.00 | 0.87 | High-affinity branched-chain amino acid transport system permease protein LivH |
| *FLPKDCKG_03155* | *amiC-6* | - | 3442644 | 3443813 | 0.50 | 1.52 | 2.16 | 0.85 | Aliphatic amidase expression-regulating protein |
| *FLPKDCKG_03156* | | - | 3443872 | 3444783 | 0.82 | 1.69 | 1.69 | 1.92 | Ureidoglycolate lyase |
| *FLPKDCKG_03157* | *sdgD-2* | - | 3444873 | 3445997 | 3.18 | 3.63 | 6.60 | 3.76 | Gentisate 1,2-dioxygenase |
| *FLPKDCKG_03158* | | + | 3446205 | 3446465 | 8.02 | 24.56 | 8.97 | 7.15 | Hypothetical protein |
| *FLPKDCKG_03159* | | - | 3446708 | 3447856 | 0.29 | 0.95 | 1.47 | 0.27 | Outer membrane porin protein |
| *FLPKDCKG_03160* | *ytnP* | - | 3448117 | 3449052 | 0.84 | 1.17 | 1.40 | 0.00 | putative quorum-quenching lactonase YtnP |
| *FLPKDCKG_03161* | *sigJ* | + | 3449350 | 3450228 | 0.95 | 3.45 | 2.25 | 0.54 | ECF RNA polymerase sigma factor SigJ |
| *FLPKDCKG_03162* | | + | 3450356 | 3450787 | 1.43 | 2.16 | 1.81 | 0.11 | Hypothetical protein |
| *FLPKDCKG_03163* | *pgrR-6* | + | 3450946 | 3451950 | 0.26 | 1.82 | 0.76 | 0.85 | HTH-type transcriptional regulator PgrR |
| *FLPKDCKG_03164* | *ohrB-1* | - | 3451955 | 3452449 | 0.09 | 0.00 | 0.00 | 0.00 | Organic hydroperoxide resistance protein OhrB |
| *FLPKDCKG_03165* | *azoB-2* | - | 3452470 | 3453225 | 1.35 | 1.36 | 0.46 | 0.61 | NAD(P)H azoreductase |
| *FLPKDCKG_03166* | | - | 3453236 | 3453715 | 1.92 | 2.31 | 1.21 | 0.00 | Hypothetical protein |
| *FLPKDCKG_03167* | *pbuE-5* | + | 3454104 | 3455282 | 2.44 | 6.42 | 3.53 | 0.49 | Purine efflux pump PbuE |
| *FLPKDCKG_03168* | | - | 3455303 | 3455377 | 75.80 | 190.27 | 119.04 | 33.68 | tRNA |
| *FLPKDCKG_03169* | | + | 3455607 | 3456512 | 15.37 | 25.05 | 22.88 | 10.46 | Cytochrome c-555 |
| *FLPKDCKG_03170* | *paaH-2* | - | 3456609 | 3458135 | 7.55 | 8.71 | 5.81 | 3.30 | 3-hydroxyadipyl-CoA dehydrogenase |

| Locus name | Gene name | Strand | Gene start (bp) | Gene end (bp) | Average TPM Inoculum | Average TPM $t_6$ | Average TPM $t_{24}$ | Average TPM $t_{168}$ | Annotation description |
|---|---|---|---|---|---|---|---|---|---|
| *FLPKDCKG_03171* | *uctC-4* | - | 3458190 | 3459419 | 24.49 | 18.78 | 15.20 | 3.15 | Acetyl-CoA:oxalate CoA-transferase |
| *FLPKDCKG_03172* | *acrC-2* | - | 3459973 | 3461121 | 3.51 | 2.59 | 4.42 | 1.13 | Acryloyl-CoA reductase (NADH) |
| *FLPKDCKG_03173* | *kstR2-4* | - | 3461157 | 3461780 | 3.78 | 5.95 | 4.37 | 3.97 | HTH-type transcriptional repressor KstR2 |
| *FLPKDCKG_03174* | | + | 3461980 | 3462951 | 13.43 | 26.16 | 8.78 | 4.62 | Nitronate monooxygenase |
| *FLPKDCKG_03175* | *paaF-2* | + | 3462948 | 3463688 | 1.28 | 1.19 | 1.69 | 0.88 | 2,3-dehydroadipyl-CoA hydratase |
| *FLPKDCKG_03176* | *fadB-1* | - | 3463804 | 3464637 | 9.74 | 6.29 | 4.28 | 3.74 | putative enoyl-CoA hydratase |
| *FLPKDCKG_03177* | *nnr-2* | + | 3464728 | 3465597 | 5.49 | 4.56 | 3.79 | 0.64 | Bifunctional NAD(P)H-hydrate repair enzyme Nnr |
| *FLPKDCKG_03178* | | + | 3465719 | 3465952 | 2517.58 | 219.88 | 55.67 | 47.75 | Hypothetical protein |
| *FLPKDCKG_03179* | *hbpA-3* | + | 3466446 | 3468059 | 3.89 | 9.45 | 11.00 | 5.39 | Heme-binding protein A |
| *FLPKDCKG_03180* | *gltC-10* | - | 3468140 | 3469054 | 1.70 | 2.42 | 4.27 | 1.28 | HTH-type transcriptional regulator GltC |
| *FLPKDCKG_03181* | *argE-3* | + | 3469221 | 3470657 | 1.98 | 5.83 | 3.82 | 0.66 | Acetylornithine deacetylase |
| *FLPKDCKG_03182* | *astE-2* | + | 3470711 | 3471682 | 5.03 | 12.27 | 6.58 | 1.98 | Succinylglutamate desuccinylase |
| *FLPKDCKG_03183* | *hbpA-4* | + | 3471747 | 3473387 | 3.16 | 4.42 | 1.55 | 0.31 | Heme-binding protein A |
| *FLPKDCKG_03184* | *dppB-5* | + | 3473438 | 3474415 | 1.50 | 2.34 | 1.22 | 0.95 | Dipeptide transport system permease protein DppB |
| *FLPKDCKG_03185* | *ddpC-3* | + | 3474421 | 3475338 | 1.69 | 2.15 | 1.06 | 0.38 | putative D,D-dipeptide transport system permease protein DdpC |
| *FLPKDCKG_03186* | *oppD-6* | + | 3475469 | 3476452 | 2.26 | 4.73 | 1.67 | 0.00 | Oligopeptide transport ATP-binding protein OppD |
| *FLPKDCKG_03187* | *oppF-5* | + | 3476449 | 3477459 | 2.08 | 2.91 | 0.73 | 1.19 | Oligopeptide transport ATP-binding protein OppF |
| *FLPKDCKG_03188* | | + | 3477465 | 3478370 | 1.72 | 2.25 | 4.84 | 0.44 | Hypothetical protein |
| *FLPKDCKG_03189* | *cpg2-3* | + | 3478404 | 3479534 | 0.88 | 2.24 | 1.13 | 0.97 | Carboxypeptidase G2 |
| *FLPKDCKG_03190* | | - | 3479619 | 3480539 | 0.50 | 1.92 | 1.40 | 0.60 | Hypothetical protein |
| *FLPKDCKG_03191* | *dapD-1* | + | 3480697 | 3481677 | 7.00 | 5.89 | 3.05 | 1.06 | 2,3,4,5-tetrahydropyridine-2,6-dicarboxylate N-succinyltransferase |
| *FLPKDCKG_03192* | *cdhR-5* | - | 3481709 | 3482641 | 3.37 | 4.56 | 3.99 | 1.43 | HTH-type transcriptional regulator CdhR |
| *FLPKDCKG_03193* | *isfD* | + | 3482770 | 3483606 | 0.39 | 1.26 | 0.85 | 1.19 | Sulfoacetaldehyde reductase |
| *FLPKDCKG_03194* | *egtC* | - | 3483573 | 3484412 | 12.35 | 5.85 | 5.69 | 1.27 | Gamma-glutamyl-hercynylcysteine sulfoxide hydrolase |
| *FLPKDCKG_03195* | *ltaE-2* | + | 3484551 | 3485606 | 2.56 | 4.41 | 1.97 | 0.00 | Low specificity L-threonine aldolase |
| *FLPKDCKG_03196* | *tetD-1* | - | 3485631 | 3486458 | 4.99 | 2.15 | 3.61 | 1.72 | Transposon Tn10 TetD protein |
| *FLPKDCKG_03197* | *rep* | - | 3486531 | 3488594 | 1.82 | 3.43 | 3.66 | 1.40 | ATP-dependent DNA helicase Rep |
| *FLPKDCKG_03198* | *priA* | - | 3488650 | 3490755 | 0.77 | 1.81 | 2.16 | 0.63 | Primosomal protein N' |
| *FLPKDCKG_03199* | *hemE* | - | 3491118 | 3492191 | 18.77 | 24.28 | 15.44 | 5.52 | Uroporphyrinogen decarboxylase |
| *FLPKDCKG_03200* | | - | 3492361 | 3493125 | 0.96 | 1.43 | 2.21 | 0.14 | Hypothetical protein |
| *FLPKDCKG_03201* | *atpC* | - | 3493324 | 3493749 | 5.07 | 31.14 | 25.57 | 11.13 | ATP synthase epsilon chain |
| *FLPKDCKG_03202* | *atpD1* | - | 3493759 | 3495159 | 14.11 | 33.90 | 42.37 | 25.18 | ATP synthase subunit beta 1 |
| *FLPKDCKG_03203* | *atpG* | - | 3495203 | 3496108 | 9.05 | 32.45 | 41.82 | 18.45 | ATP synthase gamma chain |
| *FLPKDCKG_03204* | *atpA* | - | 3496208 | 3497749 | 12.11 | 36.25 | 37.59 | 19.68 | ATP synthase subunit alpha |
| *FLPKDCKG_03205* | *atpH* | - | 3497781 | 3498320 | 16.41 | 43.80 | 51.42 | 32.98 | ATP synthase subunit delta |
| *FLPKDCKG_03206* | *atpF* | - | 3498333 | 3498803 | 11.17 | 44.23 | 53.78 | 40.26 | ATP synthase subunit b |
| *FLPKDCKG_03207* | *atpE* | - | 3498946 | 3499188 | 53.43 | 89.94 | 113.38 | 54.84 | ATP synthase subunit c |
| *FLPKDCKG_03208* | *atpB* | - | 3499306 | 3500187 | 15.89 | 40.72 | 31.21 | 12.30 | ATP synthase subunit a |
| *FLPKDCKG_03209* | | - | 3500275 | 3500748 | 54.61 | 146.84 | 125.28 | 56.79 | Hypothetical protein |
| *FLPKDCKG_03210* | | - | 3501231 | 3502787 | 34.97 | 413.95 | 237.04 | 52.93 | Hypothetical protein |
| *FLPKDCKG_03211* | *echA8-4* | - | 3503080 | 3503856 | 62.14 | 84.25 | 59.17 | 14.00 | putative enoyl-CoA hydratase echA8 |
| *FLPKDCKG_03212* | | + | 3503896 | 3505671 | 30.76 | 55.94 | 27.94 | 7.40 | Hypothetical protein |
| *FLPKDCKG_03213* | | - | 3506187 | 3507179 | 2.53 | 10.16 | 7.46 | 2.73 | Hypothetical protein |
| *FLPKDCKG_03214* | *pcaJ-1* | - | 3507217 | 3507921 | 0.37 | 2.13 | 2.70 | 0.00 | 3-oxoadipate CoA-transferase subunit B |
| *FLPKDCKG_03215* | *pcaI-1* | - | 3507923 | 3508588 | 0.49 | 1.39 | 0.50 | 0.23 | 3-oxoadipate CoA-transferase subunit A |
| *FLPKDCKG_03216* | | - | 3508589 | 3509557 | 0.06 | 1.24 | 0.98 | 0.00 | Hypothetical protein |
| *FLPKDCKG_03217* | *gltC-11* | - | 3509691 | 3510572 | 1.45 | 5.69 | 3.31 | 0.68 | HTH-type transcriptional regulator GltC |
| *FLPKDCKG_03218* | *maiA* | + | 3510696 | 3511457 | 0.85 | 3.47 | 2.84 | 0.53 | Maleate isomerase |
| *FLPKDCKG_03219* | *yhaJ-4* | - | 3511513 | 3512439 | 0.82 | 0.15 | 1.10 | 0.23 | HTH-type transcriptional regulator YhaJ |
| *FLPKDCKG_03220* | *catD-2* | + | 3512593 | 3513393 | 1.09 | 2.76 | 1.96 | 0.21 | 3-oxoadipate enol-lactonase 2 |
| *FLPKDCKG_03221* | *trxB-3* | - | 3513472 | 3514374 | 2.93 | 7.86 | 4.21 | 3.06 | Thioredoxin reductase |
| *FLPKDCKG_03222* | *folB* | + | 3514497 | 3514856 | 25.24 | 25.87 | 17.96 | 4.74 | Dihydroneopterin aldolase |
| *FLPKDCKG_03223* | *ttcA* | + | 3514859 | 3515875 | 8.32 | 16.21 | 9.76 | 5.07 | tRNA 2-thiocytidine biosynthesis protein TtcA |
| *FLPKDCKG_03224* | | - | 3515992 | 3516948 | 4.76 | 10.18 | 9.25 | 5.47 | Hydroxypyruvate reductase |
| *FLPKDCKG_03225* | *dsbC* | - | 3517012 | 3517815 | 9.57 | 23.48 | 13.52 | 8.42 | putative thiol:disulfide interchange protein DsbC |
| *FLPKDCKG_03226* | *ubiF* | - | 3517984 | 3519153 | 2.98 | 12.13 | 9.42 | 1.35 | 2-octaprenyl-3-methyl-6-methoxy-1,4-benzoquinol hydroxylase |
| *FLPKDCKG_03227* | *mltA* | - | 3519150 | 3520451 | 5.90 | 10.90 | 5.19 | 3.17 | Membrane-bound lytic murein transglycosylase A |
| *FLPKDCKG_03228* | *apaG* | - | 3520493 | 3520867 | 85.44 | 173.41 | 80.50 | 26.72 | Protein ApaG |
| *FLPKDCKG_03229* | *rpe* | + | 3520937 | 3521632 | 31.85 | 46.94 | 41.42 | 10.12 | Ribulose-phosphate 3-epimerase |
| *FLPKDCKG_03230* | *cbbZC-2* | + | 3521644 | 3522351 | 234.01 | 224.32 | 126.26 | 37.35 | Phosphoglycolate phosphatase, chromosomal |
| *FLPKDCKG_03231* | *trpE-1* | + | 3522625 | 3523668 | 20.87 | 10.64 | 7.83 | 2.64 | Anthranilate synthase component 1 |
| *FLPKDCKG_03232* | *trpE-2* | + | 3523668 | 3524144 | 15.11 | 13.75 | 4.10 | 4.52 | Anthranilate synthase component 1 |
| *FLPKDCKG_03233* | *trpG* | + | 3524162 | 3524770 | 19.77 | 29.12 | 11.33 | 4.56 | Anthranilate synthase component 2 |
| *FLPKDCKG_03234* | *trpD* | + | 3524783 | 3525814 | 11.83 | 21.99 | 13.22 | 6.18 | Anthranilate phosphoribosyltransferase |
| *FLPKDCKG_03235* | *trpC* | + | 3525811 | 3526599 | 21.96 | 25.85 | 21.34 | 8.96 | Indole-3-glycerol phosphate synthase |
| *FLPKDCKG_03236* | *acoR* | - | 3526617 | 3528509 | 7.08 | 6.24 | 2.85 | 1.98 | Acetoin catabolism regulatory protein |
| *FLPKDCKG_03237* | *acoD* | + | 3528738 | 3530267 | 67.28 | 56.60 | 77.54 | 24.33 | Acetaldehyde dehydrogenase 2 |
| *FLPKDCKG_03238* | | + | 3530357 | 3530734 | 6.33 | 12.63 | 8.34 | 4.37 | Hypothetical protein |
| *FLPKDCKG_03239* | *nsaD-2* | - | 3531020 | 3531664 | 13.19 | 27.00 | 11.52 | 4.90 | 2-hydroxychromene-2-carboxylate isomerase |

| Locus name | Gene name | Strand | Gene start (bp) | Gene end (bp) | Average TPM Inoculum | Average TPM $t_6$ | Average TPM $t_{24}$ | Average TPM $t_{168}$ | Annotation description |
|---|---|---|---|---|---|---|---|---|---|
| *FLPKDCKG_03240* | | - | 3531661 | 3531870 | 42.71 | 49.71 | 41.01 | 13.82 | Hypothetical protein |
| *FLPKDCKG_03241* | *gcvA-8* | + | 3532073 | 3532993 | 2.95 | 3.23 | 2.47 | 0.29 | Glycine cleavage system transcriptional activator |
| *FLPKDCKG_03242* | *ltaS1* | + | 3533237 | 3535309 | 2.37 | 7.66 | 4.21 | 0.77 | Lipoteichoic acid synthase 1 |
| *FLPKDCKG_03243* | *selD* | - | 3535348 | 3536409 | 19.76 | 29.49 | 21.93 | 4.32 | Selenide, water dikinase |
| *FLPKDCKG_03244* | | + | 3536552 | 3537655 | 122.27 | 64.97 | 44.04 | 22.44 | Hypothetical protein |
| *FLPKDCKG_03245* | *resA-2* | + | 3537668 | 3538210 | 9.55 | 14.38 | 11.73 | 2.85 | Thiol-disulfide oxidoreductase ResA |
| *FLPKDCKG_03246* | *epsJ* | - | 3538228 | 3539262 | 1.60 | 7.36 | 8.30 | 1.44 | putative glycosyltransferase EpsJ |
| *FLPKDCKG_03247* | | - | 3539455 | 3539550 | 13.27 | 76.51 | 31.83 | 2.92 | tRNA |
| *FLPKDCKG_03248* | *selB* | - | 3539577 | 3541514 | 4.63 | 5.26 | 2.82 | 3.10 | Selenocysteine-specific elongation factor |
| *FLPKDCKG_03249* | *selA* | - | 3541511 | 3542971 | 7.64 | 3.29 | 2.46 | 0.71 | L-seryl-tRNA(Sec) selenium transferase |
| *FLPKDCKG_03250* | *fdhE* | - | 3542974 | 3543933 | 10.51 | 9.38 | 4.96 | 1.71 | Protein FdhE |
| *FLPKDCKG_03251* | *fdoI* | - | 3544036 | 3544683 | 17.00 | 17.27 | 4.22 | 1.64 | Formate dehydrogenase, cytochrome b556(fdo) subunit |
| *FLPKDCKG_03252* | *fdnH* | - | 3544676 | 3545575 | 37.26 | 16.90 | 14.14 | 2.47 | Formate dehydrogenase, nitrate-inducible, iron-sulfur subunit |
| *FLPKDCKG_03253* | *fdoG-1* | - | 3545586 | 3548015 | 47.68 | 29.00 | 17.25 | 4.56 | Formate dehydrogenase-O major subunit |
| *FLPKDCKG_03254* | *fdoG-2* | - | 3548064 | 3548654 | 123.93 | 103.24 | 50.49 | 16.86 | Formate dehydrogenase-O major subunit |
| *FLPKDCKG_03255* | | + | 3549215 | 3550471 | 1.76 | 5.38 | 3.58 | 0.96 | Hypothetical protein |
| *FLPKDCKG_03256* | | + | 3550549 | 3551046 | 17.47 | 24.19 | 19.47 | 9.20 | putative acyl-CoA thioester hydrolase |
| *FLPKDCKG_03257* | | - | 3551387 | 3552400 | 5.90 | 3.45 | 3.74 | 1.95 | Hypothetical protein |
| *FLPKDCKG_03258* | *pdaA* | - | 3552403 | 3553197 | 7.38 | 8.09 | 11.06 | 2.63 | Peptidoglycan-N-acetylmuramic acid deacetylase PdaA |
| *FLPKDCKG_03259* | *pgl-1* | - | 3553181 | 3554170 | 22.05 | 33.28 | 21.34 | 6.88 | 6-phosphogluconolactonase |
| *FLPKDCKG_03260* | | - | 3554336 | 3555229 | 43.23 | 17.63 | 9.29 | 5.73 | Hypothetical protein |
| *FLPKDCKG_03261* | | - | 3555235 | 3556533 | 137.99 | 39.42 | 29.13 | 6.66 | Hypothetical protein |
| *FLPKDCKG_03262* | *mopII* | - | 3556654 | 3556869 | 0.65 | 0.96 | 4.44 | 2.06 | Molybdenum-pterin-binding protein 2 |
| *FLPKDCKG_03263* | *ssuB-4* | - | 3556961 | 3557866 | 0.24 | 0.00 | 0.91 | 2.22 | Aliphatic sulfonates import ATP-binding protein SsuB |
| *FLPKDCKG_03264* | *metQ-3* | - | 3557892 | 3558725 | 0.17 | 0.74 | 6.40 | 1.77 | D-methionine-binding lipoprotein MetQ |
| *FLPKDCKG_03265* | *metI-2* | - | 3558742 | 3559404 | 0.20 | 0.00 | 3.05 | 1.89 | D-methionine transport system permease protein MetI |
| *FLPKDCKG_03266* | *metN-2* | - | 3559404 | 3560213 | 0.00 | 1.10 | 1.48 | 2.04 | Methionine import ATP-binding protein MetN |
| *FLPKDCKG_03267* | *soxC-1* | - | 3560242 | 3561471 | 0.19 | 0.00 | 2.83 | 1.97 | Dibenzothiophene desulfurization enzyme C |
| *FLPKDCKG_03268* | *ssuC-8* | - | 3561497 | 3562279 | 0.06 | 0.00 | 2.05 | 1.23 | Putative aliphatic sulfonates transport permease protein SsuC |
| *FLPKDCKG_03269* | *msuD* | - | 3562279 | 3563451 | 0.09 | 0.37 | 2.84 | 1.68 | Methanesulfonate monooxygenase |
| *FLPKDCKG_03270* | *ssuA-4* | - | 3563469 | 3564434 | 0.09 | 0.12 | 2.08 | 0.30 | Putative aliphatic sulfonates-binding protein |
| *FLPKDCKG_03271* | *ssuE* | - | 3564452 | 3565045 | 0.00 | 0.00 | 4.00 | 1.46 | FMN reductase (NADPH) |
| *FLPKDCKG_03272* | *metQ-4* | - | 3565095 | 3565901 | 0.23 | 1.77 | 6.86 | 3.62 | D-methionine-binding lipoprotein MetQ |
| *FLPKDCKG_03273* | *dmoA* | - | 3566821 | 3568413 | 0.23 | 0.94 | 4.44 | 1.28 | Dimethyl-sulfide monooxygenase |
| *FLPKDCKG_03274* | *soxC-2* | - | 3568434 | 3569648 | 0.12 | 0.67 | 0.62 | 0.63 | Dibenzothiophene desulfurization enzyme C |
| *FLPKDCKG_03275* | *soxC-3* | - | 3569652 | 3570893 | 0.17 | 0.42 | 0.54 | 0.90 | Dibenzothiophene desulfurization enzyme C |
| *FLPKDCKG_03276* | | + | 3571122 | 3571949 | 0.12 | 0.00 | 1.47 | 0.93 | Hypothetical protein |
| *FLPKDCKG_03277* | *gsiB-6* | + | 3571985 | 3573673 | 0.20 | 0.30 | 0.96 | 1.07 | Glutathione-binding protein GsiB |
| *FLPKDCKG_03278* | *dppB-6* | + | 3573686 | 3574663 | 0.00 | 0.38 | 0.96 | 0.02 | Dipeptide transport system permease protein DppB |
| *FLPKDCKG_03279* | *oppC* | + | 3574660 | 3575544 | 0.05 | 0.57 | 1.52 | 0.21 | Oligopeptide transport system permease protein OppC |
| *FLPKDCKG_03280* | *oppD-7* | + | 3575541 | 3575924 | 0.00 | 1.63 | 0.00 | 1.21 | Oligopeptide transport ATP-binding protein OppD |
| *FLPKDCKG_03281* | *gsiA-7* | + | 3575945 | 3577249 | 0.03 | 0.00 | 0.19 | 0.25 | Glutathione import ATP-binding protein GsiA |
| *FLPKDCKG_03282* | *teaD-5* | + | 3577369 | 3577806 | 57.02 | 67.24 | 37.17 | 11.17 | TRAP-T-associated universal stress protein TeaD |
| *FLPKDCKG_03283* | *ascD-2* | - | 3577828 | 3578844 | 2.87 | 7.16 | 6.08 | 0.74 | CDP-6-deoxy-L-threo-D-glycero-4-hexulose-3-dehydrase reductase |
| *FLPKDCKG_03284* | | - | 3578856 | 3579335 | 1.93 | 2.41 | 1.54 | 0.00 | Hypothetical protein |
| *FLPKDCKG_03285* | *aldHT* | - | 3579357 | 3580817 | 2.36 | 0.96 | 0.95 | 0.09 | Aldehyde dehydrogenase, thermostable |
| *FLPKDCKG_03286* | | - | 3580936 | 3582468 | 1.55 | 0.29 | 0.25 | 0.32 | Hypothetical protein |
| *FLPKDCKG_03287* | | + | 3582971 | 3583429 | 0.72 | 0.00 | 0.00 | 0.00 | Hypothetical protein |
| *FLPKDCKG_03288* | *rspR-8* | + | 3583504 | 3584409 | 25.24 | 13.50 | 8.58 | 4.50 | HTH-type transcriptional repressor RspR |
| *FLPKDCKG_03289* | *paaE-2* | - | 3584243 | 3584584 | 0.16 | 2.71 | 1.08 | 0.00 | 1,2-phenylacetyl-CoA epoxidase, subunit E |
| *FLPKDCKG_03290* | | - | 3584590 | 3584931 | 0.33 | 2.66 | 2.00 | 0.28 | Hypothetical protein |
| *FLPKDCKG_03291* | *nqrF* | - | 3585005 | 3585343 | 0.60 | 1.39 | 0.00 | 0.00 | Na(+)-translocating NADH-quinone reductase subunit F |
| *FLPKDCKG_03292* | | - | 3585423 | 3586154 | 5.17 | 4.32 | 1.79 | 2.97 | PqqC-like protein |
| *FLPKDCKG_03293* | | + | 3586377 | 3586586 | 5.89 | 4.05 | 2.66 | 9.30 | Hypothetical protein |
| *FLPKDCKG_03294* | | - | 3586674 | 3587078 | 1.30 | 6.37 | 3.59 | 0.98 | Hypothetical protein |
| *FLPKDCKG_03295* | *msrP-3* | + | 3587221 | 3588456 | 9.34 | 33.96 | 17.79 | 2.80 | Protein-methionine-sulfoxide reductase catalytic subunit MsrP |
| *FLPKDCKG_03296* | | + | 3588467 | 3588790 | 3.65 | 13.83 | 9.17 | 1.10 | Hypothetical protein |
| *FLPKDCKG_03297* | | + | 3588908 | 3590611 | 5.26 | 8.29 | 2.54 | 0.81 | putative multidrug-efflux transporter |
| *FLPKDCKG_03298* | | + | 3590628 | 3592688 | 4.96 | 9.88 | 5.57 | 1.16 | Hypothetical protein |
| *FLPKDCKG_03299* | | - | 3592678 | 3593673 | 2.74 | 12.13 | 4.43 | 0.99 | Hypothetical protein |
| *FLPKDCKG_03300* | *yehB* | - | 3593670 | 3596015 | 5.72 | 28.52 | 16.52 | 2.27 | Outer membrane usher protein YehB |
| *FLPKDCKG_03301* | | - | 3596073 | 3596798 | 30.97 | 115.15 | 53.49 | 10.49 | Hypothetical protein |
| *FLPKDCKG_03302* | | - | 3596798 | 3597313 | 229.85 | 302.17 | 195.45 | 43.97 | Hypothetical protein |
| *FLPKDCKG_03303* | *ribZ-2* | + | 3597783 | 3599192 | 1.45 | 4.90 | 3.64 | 1.37 | Riboflavin transporter RibZ |
| *FLPKDCKG_03304* | *tsr-5* | + | 3599333 | 3600955 | 2.78 | 4.19 | 1.88 | 0.37 | Methyl-accepting chemotaxis protein I |
| *FLPKDCKG_03305* | *hutI-1* | - | 3600997 | 3602214 | 8.83 | 2.26 | 2.60 | 1.37 | Imidazolonepropionase |
| *FLPKDCKG_03306* | *hutH-1* | - | 3602240 | 3603769 | 16.42 | 4.67 | 2.79 | 0.65 | Histidine ammonia-lyase |
| *FLPKDCKG_03307* | *ybbW* | - | 3603799 | 3605223 | 9.38 | 4.35 | 4.76 | 0.07 | Putative allantoin permease |
| *FLPKDCKG_03308* | *hutU-1* | - | 3605293 | 3606984 | 24.56 | 7.36 | 3.65 | 1.75 | Urocanate hydratase |

| Locus name | Gene name | Strand | Gene start (bp) | Gene end (bp) | Average TPM Inoculum | Average TPM $t_6$ | Average TPM $t_{24}$ | Average TPM $t_{168}$ | Annotation description |
|---|---|---|---|---|---|---|---|---|---|
| *FLPKDCKG_03309* | *ves* | - | 3607128 | 3607748 | 0.94 | 4.98 | 2.77 | 2.14 | Protein Ves |
| *FLPKDCKG_03310* | *yvoA* | - | 3607745 | 3608488 | 5.14 | 7.58 | 3.37 | 1.79 | HTH-type transcriptional repressor YvoA |
| *FLPKDCKG_03311* | *frmR* | + | 3608620 | 3608910 | 13.10 | 30.29 | 28.89 | 2.87 | Transcriptional repressor FrmR |
| *FLPKDCKG_03312* | *frmA* | + | 3608937 | 3610046 | 9.77 | 8.22 | 6.76 | 1.56 | S-(hydroxymethyl)glutathione dehydrogenase |
| *FLPKDCKG_03313* | | + | 3610215 | 3612140 | 0.09 | 0.27 | 0.64 | 0.06 | NAD 5'-nucleotidase |
| *FLPKDCKG_03314* | *bshA-2* | - | 3612267 | 3613469 | 0.54 | 0.84 | 1.15 | 0.00 | N-acetyl-alpha-D-glucosaminyl L-malate synthase |
| *FLPKDCKG_03315* | | - | 3613466 | 3614467 | 0.08 | 1.15 | 1.22 | 0.34 | Hypothetical protein |
| *FLPKDCKG_03316* | | - | 3614498 | 3614866 | 0.39 | 1.80 | 2.29 | 0.00 | Hypothetical protein |
| *FLPKDCKG_03317* | *cusS-2* | - | 3614963 | 3616303 | 1.16 | 2.78 | 2.28 | 1.24 | Sensor kinase CusS |
| *FLPKDCKG_03318* | *czcR-2* | - | 3616303 | 3616992 | 5.91 | 12.25 | 11.22 | 1.16 | Transcriptional activator protein CzcR |
| *FLPKDCKG_03319* | | - | 3617169 | 3617510 | 0.17 | 24.02 | 28.59 | 2.78 | Hypothetical protein |
| *FLPKDCKG_03320* | *czcC* | + | 3617763 | 3619016 | 0.07 | 4.24 | 1.49 | 0.00 | Cobalt-zinc-cadmium resistance protein CzcC |
| *FLPKDCKG_03321* | *czcB* | + | 3619031 | 3620245 | 0.07 | 2.43 | 2.32 | 0.59 | Cobalt-zinc-cadmium resistance protein CzcB |
| *FLPKDCKG_03322* | *czcA* | + | 3620262 | 3623447 | 0.08 | 1.47 | 0.33 | 0.29 | Cobalt-zinc-cadmium resistance protein CzcA |
| *FLPKDCKG_03323* | | + | 3623596 | 3624627 | 0.32 | 0.30 | 2.58 | 0.00 | putative glycosyltransferase |
| *FLPKDCKG_03324* | *arnT-3* | + | 3624611 | 3625993 | 0.41 | 1.96 | 1.05 | 0.00 | Undecaprenyl phosphate-alpha-4-amino-4-deoxy-L-arabinose arabinosyl transferase |
| *FLPKDCKG_03325* | *gpuA* | - | 3626068 | 3627114 | 0.18 | 0.66 | 1.30 | 0.46 | Guanidinopropionase |
| *FLPKDCKG_03326* | *putP* | - | 3627111 | 3627683 | 0.16 | 5.66 | 4.25 | 1.22 | High-affinity proline transporter PutP |
| *FLPKDCKG_03327* | *opuE* | - | 3627689 | 3628510 | 0.21 | 2.21 | 2.88 | 0.04 | Osmoregulated proline transporter OpuE |
| *FLPKDCKG_03328* | *cynR-10* | + | 3628761 | 3629678 | 1.69 | 2.81 | 0.64 | 0.37 | HTH-type transcriptional regulator CynR |
| *FLPKDCKG_03329* | *panB* | + | 3629741 | 3630571 | 3.81 | 5.33 | 3.56 | 0.78 | 3-methyl-2-oxobutanoate hydroxymethyltransferase |
| *FLPKDCKG_03330* | *engB-1* | + | 3630637 | 3631746 | 32.14 | 17.70 | 11.34 | 2.51 | GTP-binding protein EngB |
| *FLPKDCKG_03331* | *arcB* | - | 3631777 | 3632844 | 225.35 | 12.90 | 5.73 | 1.55 | Delta(1)-pyrroline-2-carboxylate reductase |
| *FLPKDCKG_03332* | | + | 3633289 | 3634248 | 16.95 | 40.24 | 20.09 | 5.54 | N-carbamoyl-D-amino acid hydrolase |
| *FLPKDCKG_03333* | *dmlR-18* | - | 3634267 | 3635214 | 2.83 | 11.18 | 5.91 | 1.07 | HTH-type transcriptional regulator DmlR |
| *FLPKDCKG_03334* | | + | 3635328 | 3636566 | 0.48 | 0.54 | 0.00 | 0.26 | putative transporter |
| *FLPKDCKG_03335* | *nadA* | + | 3636713 | 3637888 | 3.77 | 4.61 | 4.07 | 2.16 | Quinolinate synthase A |
| *FLPKDCKG_03336* | *nadB-1* | + | 3638209 | 3639204 | 1.28 | 0.34 | 1.30 | 0.07 | L-aspartate oxidase |
| *FLPKDCKG_03337* | *nadB-2* | + | 3639204 | 3639479 | 0.84 | 0.99 | 4.03 | 0.28 | L-aspartate oxidase |
| *FLPKDCKG_03338* | | - | 3639545 | 3640195 | 0.59 | 0.50 | 0.85 | 0.46 | Putative lipoprotein/NMB1162 |
| *FLPKDCKG_03339* | | - | 3640197 | 3640709 | 0.35 | 0.00 | 0.00 | 0.00 | Hypothetical protein |
| *FLPKDCKG_03340* | | - | 3640721 | 3641398 | 0.79 | 1.01 | 0.83 | 0.45 | Putative lipoprotein/NMB1164 |
| *FLPKDCKG_03341* | | - | 3641690 | 3642598 | 1.09 | 6.01 | 4.53 | 0.40 | Hypothetical protein |
| *FLPKDCKG_03342* | *speE-2* | - | 3642729 | 3643526 | 3.63 | 4.42 | 3.37 | 1.88 | Polyamine aminopropyltransferase |
| *FLPKDCKG_03343* | | + | 3643699 | 3644715 | 0.62 | 0.00 | 0.43 | 0.07 | Hypothetical protein |
| *FLPKDCKG_03344* | | + | 3644836 | 3645327 | 1.86 | 2.98 | 5.08 | 0.00 | Hypothetical protein |
| *FLPKDCKG_03345* | *moxC* | - | 3645324 | 3646661 | 0.10 | 0.72 | 0.30 | 0.00 | Putative monooxygenase MoxC |
| *FLPKDCKG_03346* | *fepC-2* | - | 3646688 | 3647476 | 0.16 | 0.73 | 0.71 | 0.00 | Ferric enterobactin transport ATP-binding protein FepC |
| *FLPKDCKG_03347* | *fepG* | - | 3647533 | 3648558 | 2.54 | 3.23 | 1.43 | 0.29 | Ferric enterobactin transport system permease protein FepG |
| *FLPKDCKG_03348* | *yfiZ-2* | - | 3648578 | 3649594 | 0.52 | 1.13 | 0.52 | 0.00 | putative siderophore transport system permease protein YfiZ |
| *FLPKDCKG_03349* | *fepB* | - | 3649600 | 3650553 | 0.44 | 0.52 | 0.36 | 0.00 | Ferrienterobactin-binding periplasmic protein |
| *FLPKDCKG_03350* | *fhuA-10* | - | 3650547 | 3652763 | 0.63 | 0.73 | 0.54 | 0.28 | Ferrichrome-iron receptor |
| *FLPKDCKG_03351* | *soxS* | - | 3652891 | 3653778 | 2.05 | 3.94 | 4.03 | 1.14 | Regulatory protein SoxS |
| *FLPKDCKG_03352* | *rapA* | - | 3654426 | 3657023 | 3.71 | 10.30 | 3.39 | 0.80 | RNA polymerase-associated protein RapA |
| *FLPKDCKG_03353* | | - | 3657020 | 3659017 | 3.71 | 8.98 | 3.23 | 1.13 | Hypothetical protein |
| *FLPKDCKG_03354* | | - | 3659014 | 3660351 | 2.78 | 5.28 | 2.43 | 0.00 | Hypothetical protein |
| *FLPKDCKG_03355* | | - | 3660471 | 3661421 | 6.80 | 8.58 | 3.47 | 1.34 | Hypothetical protein |
| *FLPKDCKG_03356* | | - | 3661517 | 3662464 | 3.21 | 2.16 | 3.77 | 0.31 | Hypothetical protein |
| *FLPKDCKG_03357* | *addA-1* | - | 3662457 | 3665660 | 2.55 | 1.71 | 0.87 | 0.51 | ATP-dependent helicase/nuclease subunit A |
| *FLPKDCKG_03358* | | - | 3665663 | 3668356 | 3.21 | 3.40 | 1.14 | 0.99 | Hypothetical protein |
| *FLPKDCKG_03359* | | - | 3668429 | 3670084 | 7.28 | 6.24 | 5.07 | 1.46 | Hypothetical protein |
| *FLPKDCKG_03360* | | - | 3670864 | 3671889 | 2.31 | 8.25 | 3.45 | 0.94 | Hypothetical protein |
| *FLPKDCKG_03361* | | - | 3672018 | 3672746 | 1.05 | 3.96 | 1.76 | 0.18 | Hypothetical protein |
| *FLPKDCKG_03362* | *glmS-1* | - | 3674039 | 3675871 | 24.57 | 63.89 | 50.17 | 14.86 | Glutamine--fructose-6-phosphate aminotransferase [isomerizing] |
| *FLPKDCKG_03363* | *lrp-8* | + | 3676023 | 3676511 | 1.91 | 6.02 | 2.10 | 0.79 | Leucine-responsive regulatory protein |
| *FLPKDCKG_03364* | *kanF* | - | 3676539 | 3677600 | 6.32 | 10.38 | 9.52 | 2.25 | 2-deoxystreptamine glucosyltransferase |
| *FLPKDCKG_03365* | *rkpK-1* | - | 3677597 | 3678919 | 11.07 | 14.73 | 13.44 | 2.88 | UDP-glucose 6-dehydrogenase |
| *FLPKDCKG_03366* | *ribZ-3* | - | 3679040 | 3680431 | 3.19 | 10.95 | 7.64 | 2.45 | Riboflavin transporter RibZ |
| *FLPKDCKG_03367* | *glmU* | - | 3680428 | 3681801 | 19.87 | 46.90 | 36.70 | 9.78 | Bifunctional protein GlmU |
| *FLPKDCKG_03368* | | - | 3681865 | 3682656 | 54.15 | 68.91 | 42.60 | 21.00 | Hypothetical protein |
| *FLPKDCKG_03369* | | - | 3682700 | 3683377 | 73.54 | 97.33 | 83.37 | 25.64 | Hypothetical protein |
| *FLPKDCKG_03370* | | + | 3683509 | 3683730 | 39.18 | 132.33 | 50.02 | 17.36 | Hypothetical protein |
| *FLPKDCKG_03371* | *ypmQ-2* | - | 3683814 | 3684419 | 8.15 | 33.87 | 19.56 | 6.82 | SCO1 protein |
| *FLPKDCKG_03372* | *ctaB* | - | 3684436 | 3685326 | 10.92 | 26.05 | 19.21 | 3.99 | Protoheme IX farnesyltransferase |
| *FLPKDCKG_03373* | *ctaA* | - | 3685340 | 3686374 | 2.03 | 6.29 | 2.81 | 1.81 | Heme A synthase |
| *FLPKDCKG_03374* | | - | 3686396 | 3687031 | 1.92 | 8.56 | 6.72 | 2.15 | Hypothetical protein |
| *FLPKDCKG_03375* | | - | 3687056 | 3687853 | 7.56 | 19.39 | 6.46 | 2.44 | putative SURF1-like protein |
| *FLPKDCKG_03376* | | + | 3687890 | 3688090 | 18.11 | 40.33 | 29.39 | 5.94 | Hypothetical protein |
| *FLPKDCKG_03377* | *ctaE* | - | 3688245 | 3689120 | 12.80 | 18.24 | 13.74 | 6.04 | Cytochrome c oxidase subunit 3 |

| Locus name | Gene name | Strand | Gene start (bp) | Gene end (bp) | Average TPM Inoculum | Average TPM $t_6$ | Average TPM $t_{24}$ | Average TPM $t_{168}$ | Annotation description |
|---|---|---|---|---|---|---|---|---|---|
| *FLPKDCKG_03378* | | - | 3689159 | 3689383 | 19.91 | 44.25 | 41.08 | 7.40 | Hypothetical protein |
| *FLPKDCKG_03379* | | - | 3689395 | 3689523 | 31.14 | 40.49 | 30.23 | 17.01 | Hypothetical protein |
| *FLPKDCKG_03380* | *ctaD* | - | 3689559 | 3691166 | 27.17 | 27.70 | 29.87 | 9.38 | Cytochrome c oxidase subunit 1 |
| *FLPKDCKG_03381* | *ctaC* | - | 3691267 | 3692424 | 14.02 | 26.58 | 23.52 | 8.58 | Cytochrome c oxidase subunit 2 |
| *FLPKDCKG_03382* | *idi* | + | 3692717 | 3693523 | 5.28 | 12.78 | 5.59 | 2.46 | Isopentenyl-diphosphate Delta-isomerase |
| *FLPKDCKG_03383* | *apnhaP* | - | 3693544 | 3696315 | 6.06 | 10.18 | 5.80 | 2.87 | Na(+)/H(+) antiporter ApNhaP |
| *FLPKDCKG_03384* | *cpdA-3* | + | 3696504 | 3697346 | 10.71 | 52.88 | 35.52 | 7.69 | 3',5'-cyclic adenosine monophosphate phosphodiesterase CpdA |
| *FLPKDCKG_03385* | *rpoH* | - | 3697732 | 3698616 | 59.99 | 113.73 | 127.46 | 64.25 | RNA polymerase sigma factor RpoH |
| *FLPKDCKG_03386* | | - | 3698820 | 3699509 | 7.03 | 65.40 | 43.30 | 8.48 | Hypothetical protein |
| *FLPKDCKG_03387* | *yeiG* | - | 3699606 | 3700469 | 9.38 | 22.35 | 10.64 | 1.84 | S-formylglutathione hydrolase YeiG |
| *FLPKDCKG_03388* | | + | 3700705 | 3701508 | 62.28 | 90.92 | 57.17 | 17.04 | Hypothetical protein |
| *FLPKDCKG_03389* | *gltB* | + | 3701866 | 3706605 | 91.87 | 56.40 | 62.18 | 17.19 | Ferredoxin-dependent glutamate synthase 1 |
| *FLPKDCKG_03390* | *gltD* | + | 3706614 | 3708080 | 21.31 | 41.33 | 35.29 | 10.66 | Glutamate synthase [NADPH] small chain |
| *FLPKDCKG_03391* | | - | 3708211 | 3709548 | 0.15 | 1.26 | 4.31 | 3.28 | Hypothetical protein |
| *FLPKDCKG_03392* | *kipA-2* | - | 3709702 | 3710691 | 0.74 | 5.38 | 3.70 | 2.24 | KipI antagonist |
| *FLPKDCKG_03393* | *kipI-2* | - | 3710681 | 3711556 | 1.09 | 5.42 | 9.88 | 2.25 | Kinase A inhibitor |
| *FLPKDCKG_03394* | *accC-2* | - | 3711553 | 3712941 | 0.86 | 4.19 | 4.39 | 3.66 | Biotin carboxylase |
| *FLPKDCKG_03395* | *accB-2* | - | 3712945 | 3713187 | 2.64 | 5.49 | 5.69 | 3.09 | Biotin carboxyl carrier protein of acetyl-CoA carboxylase |
| *FLPKDCKG_03396* | | - | 3713227 | 3713997 | 1.96 | 9.58 | 14.78 | 4.55 | Hypothetical protein |
| *FLPKDCKG_03397* | *cmpR-12* | + | 3714256 | 3715173 | 2.94 | 6.67 | 8.08 | 0.89 | HTH-type transcriptional activator CmpR |
| *FLPKDCKG_03398* | *ompW* | + | 3715302 | 3715943 | 68.82 | 5.57 | 2.14 | 1.37 | Outer membrane protein W |
| *FLPKDCKG_03399* | *gltR-5* | - | 3715961 | 3716833 | 3.09 | 5.66 | 3.35 | 1.34 | HTH-type transcriptional regulator GltR |
| *FLPKDCKG_03400* | | + | 3716899 | 3718125 | 0.55 | 0.12 | 0.34 | 0.07 | Hypothetical protein |
| *FLPKDCKG_03401* | | + | 3718139 | 3719116 | 8.44 | 7.23 | 7.56 | 4.22 | Hypothetical protein |
| *FLPKDCKG_03402* | | + | 3719136 | 3720197 | 3.64 | 4.39 | 2.92 | 2.02 | Nitronate monooxygenase |
| *FLPKDCKG_03403* | *mlaF* | + | 3720483 | 3721319 | 4.08 | 15.57 | 9.11 | 2.51 | putative phospholipid import ATP-binding protein MlaF |
| *FLPKDCKG_03404* | *mlaE-1* | + | 3721316 | 3722104 | 3.36 | 3.27 | 3.83 | 1.28 | putative phospholipid ABC transporter permease protein MlaE |
| *FLPKDCKG_03405* | *mlaD* | + | 3722118 | 3722618 | 21.97 | 36.71 | 27.48 | 4.78 | putative phospholipid ABC transporter-binding protein MlaD |
| *FLPKDCKG_03406* | *mlaA* | + | 3722703 | 3723512 | 22.31 | 26.31 | 30.82 | 9.75 | putative phospholipid-binding lipoprotein MlaA |
| *FLPKDCKG_03407* | *mlaC* | + | 3723551 | 3724180 | 15.04 | 45.07 | 25.70 | 9.08 | putative phospholipid-binding protein MlaC |
| *FLPKDCKG_03408* | *ybhF-2* | + | 3724356 | 3725153 | 33.33 | 84.65 | 40.88 | 13.69 | putative ABC transporter ATP-binding protein YbhF |
| *FLPKDCKG_03409* | *yadH-1* | + | 3725150 | 3725962 | 8.63 | 17.39 | 10.05 | 5.26 | Inner membrane transport permease YadH |
| *FLPKDCKG_03410* | *ibaG* | + | 3725968 | 3726213 | 20.24 | 55.62 | 23.78 | 14.12 | Acid stress protein IbaG |
| *FLPKDCKG_03411* | *murA-2* | + | 3726216 | 3727496 | 4.18 | 13.28 | 12.06 | 7.97 | UDP-N-acetylglucosamine 1-carboxyvinyltransferase |
| *FLPKDCKG_03412* | *hisG* | + | 3727493 | 3728167 | 6.51 | 28.39 | 20.92 | 8.04 | ATP phosphoribosyltransferase |
| *FLPKDCKG_03413* | *hisD* | + | 3728213 | 3729517 | 1.45 | 6.97 | 5.05 | 3.30 | Histidinol dehydrogenase |
| *FLPKDCKG_03414* | *hisC2* | + | 3729514 | 3730605 | 32.21 | 76.95 | 39.69 | 10.84 | Histidinol-phosphate aminotransferase 2 |
| *FLPKDCKG_03415* | *hisB* | + | 3730619 | 3731206 | 27.75 | 62.89 | 51.79 | 17.71 | Histidine biosynthesis bifunctional protein HisB |
| *FLPKDCKG_03416* | *hisH-1* | + | 3731244 | 3731942 | 16.36 | 59.89 | 47.87 | 11.19 | Imidazole glycerol phosphate synthase subunit HisH |
| *FLPKDCKG_03417* | *hisA* | + | 3732011 | 3732751 | 33.02 | 71.75 | 72.62 | 30.90 | 1-(5-phosphoribosyl)-5-[(5-phosphoribosylamino)methylideneamino] imidazole-4-carboxamide isomerase |
| *FLPKDCKG_03418* | *hisF-1* | + | 3732748 | 3733566 | 8.86 | 12.96 | 14.96 | 5.02 | Imidazole glycerol phosphate synthase subunit HisF |
| *FLPKDCKG_03419* | *hisI* | + | 3733569 | 3733973 | 9.90 | 8.69 | 18.09 | 13.59 | Phosphoribosyl-AMP cyclohydrolase |
| *FLPKDCKG_03420* | *hisE* | + | 3733970 | 3734320 | 8.38 | 30.12 | 36.33 | 5.48 | Phosphoribosyl-ATP pyrophosphatase |
| *FLPKDCKG_03421* | | + | 3734336 | 3734704 | 28.77 | 73.51 | 48.17 | 12.87 | Purine nucleoside phosphoramidase |
| *FLPKDCKG_03422* | *tatA* | + | 3734719 | 3734946 | 28.95 | 58.57 | 47.27 | 11.48 | Sec-independent protein translocase protein TatA |
| *FLPKDCKG_03423* | *tatB* | + | 3735018 | 3735629 | 10.45 | 7.82 | 11.89 | 4.34 | Sec-independent protein translocase protein TatB |
| *FLPKDCKG_03424* | *tatC* | + | 3735626 | 3736417 | 2.37 | 9.49 | 9.32 | 3.18 | Sec-independent protein translocase protein TatC |
| *FLPKDCKG_03425* | | + | 3736522 | 3736911 | 1.00 | 6.96 | 3.44 | 0.81 | Hypothetical protein |
| *FLPKDCKG_03426* | *degP-1* | - | 3736947 | 3738107 | 3.97 | 6.21 | 6.41 | 1.34 | Periplasmic serine endoprotease DegP |
| *FLPKDCKG_03427* | | + | 3738127 | 3738903 | 21.19 | 24.41 | 16.03 | 7.26 | GTP cyclohydrolase 1 type 2 |
| *FLPKDCKG_03428* | *mscL* | - | 3739215 | 3739673 | 36.66 | 60.24 | 53.51 | 13.82 | Large-conductance mechanosensitive channel |
| *FLPKDCKG_03429* | *petA* | + | 3739814 | 3740455 | 40.65 | 98.02 | 63.36 | 26.98 | Ubiquinol-cytochrome c reductase iron-sulfur subunit |
| *FLPKDCKG_03430* | *fbcH* | + | 3740530 | 3741918 | 18.79 | 35.43 | 32.86 | 10.34 | Cytochrome b/c1 |
| *FLPKDCKG_03431* | *petC* | + | 3741943 | 3742797 | 31.19 | 64.42 | 59.03 | 20.27 | Cytochrome c1 |
| *FLPKDCKG_03432* | *sspA-1* | + | 3742951 | 3743562 | 88.24 | 107.45 | 81.17 | 30.25 | Stringent starvation protein A |
| *FLPKDCKG_03433* | *sspB* | + | 3743575 | 3744018 | 9.71 | 24.09 | 20.77 | 7.82 | Stringent starvation protein B |
| *FLPKDCKG_03434* | | + | 3744293 | 3744973 | 1.71 | 0.61 | 1.02 | 0.25 | Hypothetical protein |
| *FLPKDCKG_03435* | | + | 3745070 | 3745145 | 4.50 | 10.79 | 1.66 | 1.93 | tRNA |
| *FLPKDCKG_03436* | *gntT* | - | 3745268 | 3746656 | 3.46 | 3.81 | 1.78 | 2.14 | High-affinity gluconate transporter |
| *FLPKDCKG_03437* | | + | 3746810 | 3746885 | 8.50 | 9.13 | 30.24 | 7.13 | tRNA |
| *FLPKDCKG_03438* | | + | 3747067 | 3747480 | 2.72 | 1.52 | 3.12 | 0.07 | Hypothetical protein |
| *FLPKDCKG_03439* | *gcvA-9* | + | 3747541 | 3748443 | 3.13 | 1.41 | 0.87 | 0.74 | Glycine cleavage system transcriptional activator |
| *FLPKDCKG_03440* | | + | 3748552 | 3749025 | 9.61 | 18.01 | 8.43 | 3.97 | Hypothetical protein |
| *FLPKDCKG_03441* | *sufS* | + | 3749073 | 3750281 | 2.33 | 4.03 | 4.07 | 1.41 | Cysteine desulfurase |
| *FLPKDCKG_03442* | *eamA-3* | - | 3750309 | 3751202 | 0.61 | 0.42 | 0.54 | 0.21 | putative amino-acid metabolite efflux pump |
| *FLPKDCKG_03443* | *marA-1* | + | 3751372 | 3752259 | 9.48 | 7.04 | 3.52 | 0.25 | Multiple antibiotic resistance protein MarA |
| *FLPKDCKG_03444* | | + | 3752326 | 3753756 | 2.60 | 2.86 | 1.99 | 0.52 | putative membrane protein |
| *FLPKDCKG_03445* | *der-2* | + | 3753749 | 3755119 | 2.93 | 5.13 | 3.97 | 0.75 | GTPase Der |
| *FLPKDCKG_03446* | *gsiB-7* | + | 3755330 | 3756937 | 41.09 | 21.15 | 18.88 | 12.03 | Glutathione-binding protein GsiB |

| Locus name | Gene name | Strand | Gene start (bp) | Gene end (bp) | Average TPM Inoculum | Average TPM $t_6$ | Average TPM $t_{24}$ | Average TPM $t_{168}$ | Annotation description |
|---|---|---|---|---|---|---|---|---|---|
| *FLPKDCKG_03447* | *dppB-7* | + | 3757094 | 3758065 | 3.31 | 4.13 | 4.43 | 1.76 | Dipeptide transport system permease protein DppB |
| *FLPKDCKG_03448* | *dppC-4* | + | 3758078 | 3759004 | 1.82 | 3.01 | 1.53 | 0.53 | Dipeptide transport system permease protein DppC |
| *FLPKDCKG_03449* | *gsiA-8* | + | 3759012 | 3760772 | 4.34 | 1.75 | 2.70 | 2.19 | Glutathione import ATP-binding protein GsiA |
| *FLPKDCKG_03450* | *hutI-2* | + | 3760769 | 3761329 | 4.35 | 1.82 | 1.61 | 1.29 | Imidazolonepropionase |
| *FLPKDCKG_03451* | *hutI-3* | + | 3761308 | 3761979 | 1.50 | 1.87 | 1.93 | 0.52 | Imidazolonepropionase |
| *FLPKDCKG_03452* | | - | 3762012 | 3763448 | 3.49 | 8.32 | 2.16 | 0.54 | Hypothetical protein |
| *FLPKDCKG_03453* | | - | 3763603 | 3765075 | 41.73 | 38.84 | 28.99 | 8.35 | Hypothetical protein |
| *FLPKDCKG_03454* | *rhlE-2* | + | 3765377 | 3766891 | 4.41 | 29.46 | 40.27 | 13.65 | ATP-dependent RNA helicase RhlE |
| *FLPKDCKG_03455* | *rhtB-3* | + | 3766937 | 3767566 | 4.23 | 9.81 | 8.05 | 7.42 | Homoserine/homoserine lactone efflux protein |
| *FLPKDCKG_03456* | *prfC* | + | 3767678 | 3769285 | 33.56 | 51.93 | 35.84 | 12.41 | Peptide chain release factor 3 |
| *FLPKDCKG_03457* | | + | 3769323 | 3770009 | 39.60 | 89.90 | 37.22 | 20.03 | Hypothetical protein |
| *FLPKDCKG_03458* | *bfr-1* | - | 3770094 | 3770570 | 72.49 | 312.07 | 151.42 | 26.09 | Bacterioferritin |
| *FLPKDCKG_03459* | *yijE-7* | - | 3770995 | 3771909 | 1.77 | 3.25 | 3.66 | 1.47 | putative cystine transporter YijE |
| *FLPKDCKG_03460* | *gltC-12* | + | 3772023 | 3772943 | 8.85 | 26.02 | 15.95 | 4.73 | HTH-type transcriptional regulator GltC |
| *FLPKDCKG_03461* | | - | 3772913 | 3773194 | 82.24 | 93.15 | 108.66 | 80.90 | Hypothetical protein |
| *FLPKDCKG_03462* | *relE2-1* | - | 3773391 | 3773669 | 3.38 | 9.47 | 4.74 | 0.23 | Toxin RelE2 |
| *FLPKDCKG_03463* | | - | 3773738 | 3773944 | 52.93 | 51.95 | 24.70 | 13.04 | Hypothetical protein |
| *FLPKDCKG_03464* | *rpoN-2* | - | 3774038 | 3775465 | 8.27 | 10.62 | 8.57 | 0.66 | RNA polymerase sigma-54 factor |
| *FLPKDCKG_03465* | | + | 3775482 | 3776132 | 81.23 | 76.91 | 48.69 | 9.02 | Hypothetical protein |
| *FLPKDCKG_03466* | | + | 3776325 | 3777302 | 8.01 | 59.03 | 36.75 | 11.97 | Hypothetical protein |
| *FLPKDCKG_03467* | *leuA-1* | + | 3777696 | 3779402 | 63.86 | 43.62 | 50.40 | 15.57 | 2-isopropylmalate synthase |
| *FLPKDCKG_03468* | *amiD-4* | - | 3779565 | 3780893 | 14.15 | 13.08 | 9.72 | 1.79 | Putative amidase AmiD |
| *FLPKDCKG_03469* | *cytR-1* | + | 3781174 | 3782223 | 7.78 | 5.32 | 6.00 | 0.90 | HTH-type transcriptional repressor CytR |
| *FLPKDCKG_03470* | *cpdA-4* | + | 3782261 | 3783058 | 6.36 | 9.01 | 6.78 | 0.94 | 3',5'-cyclic adenosine monophosphate phosphodiesterase CpdA |
| *FLPKDCKG_03471* | *ugpC-4* | + | 3783181 | 3784236 | 2.49 | 2.92 | 2.28 | 0.40 | sn-glycerol-3-phosphate import ATP-binding protein UgpC |
| *FLPKDCKG_03472* | *ugpB-3* | + | 3784270 | 3785565 | 4.77 | 7.45 | 4.86 | 1.50 | sn-glycerol-3-phosphate-binding periplasmic protein UgpB |
| *FLPKDCKG_03473* | *ugpA-3* | + | 3785579 | 3786454 | 1.65 | 3.61 | 0.82 | 1.41 | sn-glycerol-3-phosphate transport system permease protein UgpA |
| *FLPKDCKG_03474* | *lacG* | + | 3786451 | 3787251 | 0.76 | 2.76 | 1.77 | 0.81 | Lactose transport system permease protein LacG |
| *FLPKDCKG_03475* | | - | 3787316 | 3787633 | 28.28 | 34.25 | 19.18 | 9.26 | Hypothetical protein |
| *FLPKDCKG_03476* | | - | 3787714 | 3788058 | 167.41 | 152.69 | 79.39 | 20.36 | Hypothetical protein |
| *FLPKDCKG_03477* | *argS* | + | 3788375 | 3790075 | 8.26 | 18.04 | 16.92 | 7.27 | Arginine--tRNA ligase |
| *FLPKDCKG_03478* | *ftsN* | + | 3790085 | 3790801 | 17.30 | 16.51 | 6.38 | 4.47 | Cell division protein FtsN |
| *FLPKDCKG_03479* | *dsbA* | + | 3790888 | 3791520 | 18.32 | 31.79 | 39.32 | 11.70 | Thiol:disulfide interchange protein DsbA |
| *FLPKDCKG_03480* | *amiC-7* | + | 3791683 | 3792894 | 2.73 | 3.09 | 1.93 | 0.00 | Aliphatic amidase expression-regulating protein |
| *FLPKDCKG_03481* | | + | 3792873 | 3793832 | 2.31 | 0.60 | 2.63 | 1.06 | Hypothetical protein |
| *FLPKDCKG_03482* | *metC-3* | - | 3793837 | 3795030 | 10.50 | 6.55 | 5.55 | 1.52 | Cystathionine beta-lyase |
| *FLPKDCKG_03483* | *nfnB* | - | 3795114 | 3795812 | 11.65 | 6.67 | 3.70 | 0.92 | Nitroreductase NfnB |
| *FLPKDCKG_03484* | *leuE-2* | + | 3795890 | 3796516 | 5.57 | 5.89 | 4.06 | 2.87 | Leucine efflux protein |
| *FLPKDCKG_03485* | *gloB-2* | - | 3796619 | 3797686 | 8.56 | 15.21 | 8.06 | 3.45 | Hydroxyacylglutathione hydrolase |
| *FLPKDCKG_03486* | *merR* | + | 3797760 | 3798143 | 3.62 | 10.48 | 5.19 | 2.25 | Mercuric resistance operon regulatory protein |
| *FLPKDCKG_03487* | | + | 3798154 | 3798486 | 1.06 | 4.72 | 2.39 | 0.00 | Hypothetical protein |
| *FLPKDCKG_03488* | | - | 3798654 | 3799625 | 1.73 | 1.04 | 0.95 | 0.82 | Hypothetical protein |
| *FLPKDCKG_03489* | *yxeP-3* | - | 3799672 | 3800850 | 1.80 | 0.73 | 3.18 | 0.46 | putative hydrolase YxeP |
| *FLPKDCKG_03490* | *acdA* | + | 3801096 | 3802274 | 23.84 | 16.54 | 9.39 | 4.97 | Acyl-CoA dehydrogenase |
| *FLPKDCKG_03491* | *aceK* | + | 3802298 | 3804163 | 7.20 | 3.23 | 3.00 | 1.86 | Isocitrate dehydrogenase kinase/phosphatase |
| *FLPKDCKG_03492* | *thlA-1* | + | 3804197 | 3805378 | 40.10 | 23.99 | 22.43 | 5.80 | Acetyl-CoA acetyltransferase |
| *FLPKDCKG_03493* | | + | 3805666 | 3807273 | 4.04 | 4.28 | 2.69 | 0.61 | Methylmalonyl-CoA carboxyltransferase 12S subunit |
| *FLPKDCKG_03494* | *accA1-1* | + | 3807328 | 3809343 | 4.62 | 3.52 | 1.88 | 2.68 | Acetyl-/propionyl-coenzyme A carboxylase alpha chain |
| *FLPKDCKG_03495* | | + | 3809402 | 3810199 | 0.77 | 0.42 | 4.68 | 1.89 | Hypothetical protein |
| *FLPKDCKG_03496* | | + | 3810270 | 3810680 | 6.87 | 28.85 | 17.52 | 3.41 | Hypothetical protein |
| *FLPKDCKG_03497* | *nphR-2* | - | 3810735 | 3811646 | 7.15 | 17.13 | 12.58 | 2.86 | Transcriptional activator NphR |
| *FLPKDCKG_03498* | *apc3-2* | - | 3811719 | 3813803 | 0.48 | 19.24 | 27.01 | 3.13 | Acetophenone carboxylase gamma subunit |
| *FLPKDCKG_03499* | *apc4-1* | - | 3813814 | 3815586 | 2.54 | 60.28 | 47.71 | 11.81 | Acetophenone carboxylase delta subunit |
| *FLPKDCKG_03500* | *entS-1* | + | 3815803 | 3817035 | 0.21 | 1.70 | 1.17 | 0.29 | Enterobactin exporter EntS |
| *FLPKDCKG_03501* | *allR-1* | + | 3817161 | 3817931 | 2.49 | 1.31 | 0.09 | 0.24 | HTH-type transcriptional repressor AllR |
| *FLPKDCKG_03502* | | + | 3817924 | 3819072 | 3.67 | 4.01 | 2.92 | 0.30 | Hypothetical protein |
| *FLPKDCKG_03503* | | + | 3819069 | 3819410 | 4.17 | 4.98 | 4.21 | 0.04 | Hypothetical protein |
| *FLPKDCKG_03504* | | + | 3819472 | 3820443 | 1.16 | 2.31 | 0.66 | 0.02 | Hypothetical protein |
| *FLPKDCKG_03505* | | + | 3820480 | 3822627 | 2.29 | 4.02 | 3.85 | 1.25 | Hypothetical protein |
| *FLPKDCKG_03506* | *gno-2* | + | 3822661 | 3823422 | 9.74 | 13.48 | 7.70 | 0.31 | Gluconate 5-dehydrogenase |
| *FLPKDCKG_03507* | *bbsG* | + | 3823451 | 3825739 | 8.30 | 7.37 | 4.29 | 1.98 | (R)-benzylsuccinyl-CoA dehydrogenase |
| *FLPKDCKG_03508* | | + | 3825749 | 3827401 | 3.56 | 3.01 | 2.86 | 0.58 | Long-chain-fatty-acid--CoA ligase |
| *FLPKDCKG_03509* | *fecI-9* | + | 3827572 | 3828117 | 3.15 | 5.38 | 2.88 | 0.00 | putative RNA polymerase sigma factor FecI |
| *FLPKDCKG_03510* | | + | 3828168 | 3829139 | 1.46 | 3.24 | 0.86 | 0.14 | Hypothetical protein |
| *FLPKDCKG_03511* | *fpvA-4* | + | 3829237 | 3831666 | 0.36 | 0.19 | 0.33 | 0.07 | Ferripyoverdine receptor |
| *FLPKDCKG_03512* | | - | 3831724 | 3832785 | 2.39 | 2.41 | 3.57 | 0.07 | Hypothetical protein |
| *FLPKDCKG_03513* | | - | 3832856 | 3833623 | 0.63 | 1.34 | 1.13 | 0.73 | Hypothetical protein |
| *FLPKDCKG_03514* | *lysN-2* | + | 3833738 | 3835069 | 2.17 | 3.82 | 1.99 | 0.76 | 2-aminoadipate transaminase |
| *FLPKDCKG_03515* | | + | 3835270 | 3836220 | 93.85 | 89.81 | 66.43 | 15.72 | Hypothetical protein |

| Locus name | Gene name | Strand | Gene start (bp) | Gene end (bp) | Average TPM Inoculum | Average TPM $t_6$ | Average TPM $t_{24}$ | Average TPM $t_{168}$ | Annotation description |
|---|---|---|---|---|---|---|---|---|---|
| *FLPKDCKG_03516* | *nicF-2* | + | 3836338 | 3836928 | 27.67 | 20.21 | 17.53 | 5.74 | Maleamate amidohydrolase |
| *FLPKDCKG_03517* | | + | 3836937 | 3837461 | 16.82 | 24.78 | 14.71 | 4.84 | Hypothetical protein |
| *FLPKDCKG_03518* | *rhtB-4* | + | 3837494 | 3838126 | 4.47 | 4.03 | 7.78 | 1.49 | Homoserine/homoserine lactone efflux protein |
| *FLPKDCKG_03519* | | + | 3838139 | 3838840 | 4.36 | 20.45 | 5.90 | 1.11 | Hypothetical protein |
| *FLPKDCKG_03520* | | - | 3838854 | 3839240 | 1.61 | 7.50 | 3.85 | 0.71 | Hypothetical protein |
| *FLPKDCKG_03521* | | + | 3839361 | 3839699 | 2.77 | 3.71 | 7.40 | 0.47 | Hypothetical protein |
| *FLPKDCKG_03522* | *higB-2-1* | - | 3839730 | 3840155 | 0.70 | 4.28 | 2.52 | 0.00 | Toxin HigB-2 |
| *FLPKDCKG_03523* | *higA-2-1* | - | 3840293 | 3840610 | 0.78 | 1.03 | 0.00 | 0.00 | Antitoxin HigA-2 |
| *FLPKDCKG_03524* | | - | 3840914 | 3841342 | 0.55 | 2.95 | 0.66 | 0.39 | Hypothetical protein |
| *FLPKDCKG_03525* | | - | 3841346 | 3841657 | 0.00 | 1.62 | 0.00 | 0.92 | Hypothetical protein |
| *FLPKDCKG_03526* | *cdhR-6* | + | 3841796 | 3842752 | 0.98 | 1.58 | 1.48 | 0.29 | HTH-type transcriptional regulator CdhR |
| *FLPKDCKG_03527* | | - | 3842847 | 3843179 | 0.95 | 1.57 | 0.00 | 0.87 | Hypothetical protein |
| *FLPKDCKG_03528* | *bspRIM* | + | 3843833 | 3845371 | 13.63 | 25.91 | 14.75 | 4.08 | Modification methylase BspRI |
| *FLPKDCKG_03529* | *vsr* | - | 3845500 | 3845832 | 19.21 | 55.64 | 23.80 | 6.10 | Very short patch repair protein |
| *FLPKDCKG_03530* | | - | 3846030 | 3847814 | 10.39 | 23.91 | 16.55 | 3.01 | Hypothetical protein |
| *FLPKDCKG_03531* | | - | 3847811 | 3848662 | 3.76 | 14.21 | 4.50 | 1.49 | Hypothetical protein |
| *FLPKDCKG_03532* | | - | 3848799 | 3851687 | 6.75 | 10.17 | 6.77 | 1.35 | Hypothetical protein |
| *FLPKDCKG_03533* | *mutL-3* | - | 3851716 | 3853173 | 6.90 | 11.24 | 6.65 | 1.96 | DNA mismatch repair protein MutL |
| *FLPKDCKG_03534* | *rpoD-2* | - | 3853212 | 3854252 | 4.92 | 6.30 | 4.51 | 0.15 | RNA polymerase sigma factor RpoD |
| *FLPKDCKG_03535* | *argD-2* | - | 3857087 | 3858280 | 31.79 | 39.39 | 31.84 | 12.61 | Acetylornithine aminotransferase |
| *FLPKDCKG_03536* | | + | 3858494 | 3859459 | 1.18 | 4.35 | 5.99 | 3.98 | Hypothetical protein |
| *FLPKDCKG_03537* | | - | 3859617 | 3859946 | 110.11 | 33.76 | 27.06 | 12.59 | Hypothetical protein |
| *FLPKDCKG_03538* | *fliY-4* | + | 3860332 | 3861114 | 0.00 | 32.15 | 58.92 | 27.77 | L-cystine-binding protein FliY |
| *FLPKDCKG_03539* | *glnP-6* | + | 3861143 | 3861811 | 0.31 | 10.09 | 10.69 | 4.04 | Glutamine transport system permease protein GlnP |
| *FLPKDCKG_03540* | *yecS-4* | + | 3861808 | 3862464 | 1.67 | 15.55 | 15.34 | 3.81 | L-cystine transport system permease protein YecS |
| *FLPKDCKG_03541* | *glnQ-10* | + | 3862445 | 3863182 | 0.20 | 22.49 | 25.35 | 12.94 | Glutamine transport ATP-binding protein GlnQ |
| *FLPKDCKG_03542* | | - | 3863226 | 3864617 | 1.38 | 6.68 | 11.24 | 3.07 | Hypothetical protein |
| *FLPKDCKG_03543* | | - | 3864617 | 3865756 | 5.89 | 7.68 | 5.34 | 2.13 | D-threonine aldolase |
| *FLPKDCKG_03544* | | + | 3865955 | 3866914 | 87.97 | 27.67 | 18.36 | 11.53 | Hypothetical protein |
| *FLPKDCKG_03545* | *siaT-2* | + | 3867018 | 3869042 | 5.02 | 3.22 | 1.89 | 1.77 | Sialic acid TRAP transporter permease protein SiaT |
| *FLPKDCKG_03546* | *smg* | + | 3869278 | 3869736 | 244.05 | 289.28 | 221.47 | 79.98 | Protein Smg |
| *FLPKDCKG_03547* | *gluQ* | + | 3869836 | 3870717 | 2.52 | 11.10 | 5.63 | 2.84 | Glutamyl-Q tRNA(Asp) synthetase |
| *FLPKDCKG_03548* | | - | 3870733 | 3873363 | 7.84 | 14.09 | 7.40 | 3.67 | Hypothetical protein |
| *FLPKDCKG_03549* | *topB* | + | 3873426 | 3876077 | 8.76 | 15.00 | 11.05 | 5.37 | DNA topoisomerase 3 |
| *FLPKDCKG_03550* | | + | 3876211 | 3877269 | 12.18 | 6.44 | 5.59 | 1.41 | Hypothetical protein |
| *FLPKDCKG_03551* | *ssuB-5* | + | 3877269 | 3878039 | 3.11 | 1.42 | 1.37 | 0.54 | Aliphatic sulfonates import ATP-binding protein SsuB |
| *FLPKDCKG_03552* | *ribX-2* | + | 3878047 | 3878901 | 2.21 | 2.35 | 3.34 | 0.35 | Riboflavin transport system permease protein RibX |
| *FLPKDCKG_03553* | *gno-3* | + | 3878916 | 3879698 | 3.45 | 3.41 | 5.50 | 3.38 | Gluconate 5-dehydrogenase |
| *FLPKDCKG_03554* | *ilvI-3* | - | 3879804 | 3881624 | 27.83 | 48.46 | 20.99 | 6.77 | Acetolactate synthase isozyme 3 large subunit |
| *FLPKDCKG_03555* | *lrp-9* | + | 3881718 | 3882188 | 21.93 | 44.95 | 22.06 | 4.68 | Leucine-responsive regulatory protein |
| *FLPKDCKG_03556* | | - | 3882220 | 3883140 | 2.28 | 9.23 | 7.09 | 2.11 | Hypothetical protein |
| *FLPKDCKG_03557* | | + | 3883249 | 3883773 | 25.25 | 41.40 | 18.19 | 5.88 | Hypothetical protein |
| *FLPKDCKG_03558* | *ribB-1* | + | 3884540 | 3885259 | 3.44 | 5.09 | 3.43 | 2.07 | 3,4-dihydroxy-2-butanone 4-phosphate synthase |
| *FLPKDCKG_03559* | *yajR-1* | - | 3885318 | 3886526 | 0.33 | 0.68 | 0.13 | 0.00 | Inner membrane transport protein YajR |
| *FLPKDCKG_03560* | *fadR-3* | + | 3886678 | 3887355 | 3.84 | 6.08 | 3.41 | 0.00 | Fatty acid metabolism regulator protein |
| *FLPKDCKG_03561* | | - | 3887399 | 3887896 | 1.18 | 1.96 | 0.77 | 0.18 | Hypothetical protein |
| *FLPKDCKG_03562* | *flr-1* | - | 3887901 | 3888470 | 1.77 | 3.52 | 3.50 | 1.16 | Flavoredoxin |
| *FLPKDCKG_03563* | | - | 3888483 | 3888935 | 14.42 | 24.16 | 14.73 | 5.08 | Hypothetical protein |
| *FLPKDCKG_03564* | | - | 3889093 | 3889704 | 4.47 | 9.23 | 3.89 | 1.57 | Hypothetical protein |
| *FLPKDCKG_03565* | | - | 3889871 | 3890797 | 7.21 | 11.77 | 7.36 | 2.97 | Hypothetical protein |
| *FLPKDCKG_03566* | | - | 3890923 | 3892008 | 1.33 | 3.17 | 3.91 | 1.54 | putative membrane protein |
| *FLPKDCKG_03567* | | + | 3892287 | 3893318 | 0.99 | 6.92 | 10.99 | 5.53 | Purine-binding protein |
| *FLPKDCKG_03568* | *mglA* | + | 3893315 | 3894844 | 0.19 | 1.61 | 4.82 | 1.79 | Galactose/methyl galactoside import ATP-binding protein MglA |
| *FLPKDCKG_03569* | | + | 3894876 | 3895949 | 0.13 | 0.23 | 1.38 | 0.97 | Hypothetical protein |
| *FLPKDCKG_03570* | | + | 3895960 | 3896910 | 0.42 | 0.40 | 1.62 | 0.50 | Hypothetical protein |
| *FLPKDCKG_03571* | *codA* | + | 3896927 | 3898204 | 1.28 | 3.23 | 2.35 | 1.69 | Cytosine deaminase |
| *FLPKDCKG_03572* | | - | 3898544 | 3899002 | 8.78 | 10.19 | 10.54 | 2.51 | Hypothetical protein |
| *FLPKDCKG_03573* | *cdiA-3* | - | 3898999 | 3909087 | 0.42 | 1.33 | 0.90 | 0.30 | 16S rRNA endonuclease CdiA |
| *FLPKDCKG_03574* | *shlB-2* | - | 3909159 | 3910508 | 0.10 | 1.50 | 1.44 | 0.11 | Hemolysin transporter protein ShlB |
| *FLPKDCKG_03575* | *rutC-2* | - | 3911193 | 3911579 | 0.74 | 0.27 | 6.15 | 0.19 | Putative aminoacrylate peracid reductase RutC |
| *FLPKDCKG_03576* | *hcnC-1* | - | 3911572 | 3912648 | 1.03 | 0.90 | 3.41 | 0.39 | Hydrogen cyanide synthase subunit HcnC |
| *FLPKDCKG_03577* | *hcnB-1* | - | 3912746 | 3914191 | 0.33 | 2.16 | 1.49 | 1.00 | Hydrogen cyanide synthase subunit HcnB |
| *FLPKDCKG_03578* | *hcnA-1* | - | 3914188 | 3914496 | 0.76 | 2.33 | 2.70 | 3.28 | Hydrogen cyanide synthase subunit HcnA |
| *FLPKDCKG_03579* | *hcnC-2* | - | 3914480 | 3915637 | 0.22 | 1.08 | 1.83 | 2.53 | Hydrogen cyanide synthase subunit HcnC |
| *FLPKDCKG_03580* | *ydcV-2* | - | 3915637 | 3916437 | 0.58 | 0.82 | 2.04 | 0.46 | Inner membrane ABC transporter permease protein YdcV |
| *FLPKDCKG_03581* | *potB-1* | - | 3916439 | 3917305 | 0.25 | 0.56 | 1.34 | 0.13 | Spermidine/putrescine transport system permease protein PotB |
| *FLPKDCKG_03582* | *potA-4* | - | 3917302 | 3918357 | 0.21 | 1.01 | 0.96 | 1.09 | Spermidine/putrescine import ATP-binding protein PotA |
| *FLPKDCKG_03583* | *potD-1* | - | 3918414 | 3919445 | 1.07 | 2.75 | 7.52 | 2.22 | Spermidine/putrescine-binding periplasmic protein |
| *FLPKDCKG_03584* | *treR* | + | 3919677 | 3920474 | 5.17 | 14.47 | 6.99 | 1.48 | HTH-type transcriptional regulator TreR |

| Locus name | Gene name | Strand | Gene start (bp) | Gene end (bp) | Average TPM Inoculum | Average TPM $t_6$ | Average TPM $t_{24}$ | Average TPM $t_{168}$ | Annotation description |
|---|---|---|---|---|---|---|---|---|---|
| *FLPKDCKG_03585* | *apc4-2* | - | 3920461 | 3922191 | 1.84 | 1.88 | 0.99 | 0.58 | Acetophenone carboxylase delta subunit |
| *FLPKDCKG_03586* | *apc3-3* | - | 3922184 | 3924241 | 0.91 | 0.24 | 0.29 | 0.00 | Acetophenone carboxylase gamma subunit |
| *FLPKDCKG_03587* | | - | 3924282 | 3925262 | 0.99 | 0.67 | 0.62 | 0.32 | Hypothetical protein |
| *FLPKDCKG_03588* | *pcpR-7* | + | 3925432 | 3926328 | 3.14 | 1.40 | 0.10 | 1.22 | PCP degradation transcriptional activation protein |
| *FLPKDCKG_03589* | *tetD-2* | - | 3926402 | 3927235 | 2.49 | 5.62 | 2.16 | 1.26 | Transposon Tn10 TetD protein |
| *FLPKDCKG_03590* | *ghrA-3* | - | 3927512 | 3928546 | 7.75 | 10.51 | 6.95 | 1.48 | Glyoxylate/hydroxypyruvate reductase A |
| *FLPKDCKG_03591* | *phnS-2* | - | 3928637 | 3929704 | 2.64 | 2.51 | 2.08 | 0.38 | Putative 2-aminoethylphosphonate-binding periplasmic protein |
| *FLPKDCKG_03592* | *abo-2* | - | 3929701 | 3930873 | 0.57 | 2.07 | 0.44 | 0.28 | 4-methylaminobutanoate oxidase (formaldehyde-forming) |
| *FLPKDCKG_03593* | *hcnB-2* | - | 3930870 | 3932249 | 0.12 | 0.24 | 0.68 | 0.14 | Hydrogen cyanide synthase subunit HcnB |
| *FLPKDCKG_03594* | *hcnA-2* | - | 3932246 | 3932575 | 0.96 | 0.00 | 1.10 | 0.00 | Hydrogen cyanide synthase subunit HcnA |
| *FLPKDCKG_03595* | *modB-1* | - | 3932613 | 3933407 | 1.44 | 0.00 | 0.00 | 0.44 | Molybdenum transport system permease protein ModB |
| *FLPKDCKG_03596* | *potB-2* | - | 3933423 | 3934310 | 0.58 | 1.25 | 0.42 | 1.02 | Spermidine/putrescine transport system permease protein PotB |
| *FLPKDCKG_03597* | *ugpC-5* | - | 3934312 | 3935382 | 0.78 | 0.86 | 0.07 | 1.44 | sn-glycerol-3-phosphate import ATP-binding protein UgpC |
| *FLPKDCKG_03598* | *kdgR-5* | + | 3935581 | 3936423 | 7.14 | 8.48 | 10.69 | 3.15 | Pectin degradation repressor protein KdgR |
| *FLPKDCKG_03599* | *lldD* | - | 3936841 | 3938004 | 6.78 | 9.73 | 6.38 | 2.72 | L-lactate dehydrogenase |
| *FLPKDCKG_03600* | *ectD-2* | - | 3938214 | 3939110 | 1.21 | 10.26 | 5.47 | 0.64 | Ectoine dioxygenase |
| *FLPKDCKG_03601* | *acyP* | + | 3939272 | 3939556 | 2.72 | 14.31 | 22.22 | 4.03 | Acylphosphatase |
| *FLPKDCKG_03602* | *eamA-4* | - | 3939650 | 3940555 | 2.99 | 9.74 | 6.46 | 0.51 | putative amino-acid metabolite efflux pump |
| *FLPKDCKG_03603* | *ltaA* | - | 3940596 | 3941654 | 0.51 | 0.31 | 1.77 | 0.35 | L-allo-threonine aldolase |
| *FLPKDCKG_03604* | *glnQ-11* | - | 3941660 | 3942418 | 1.92 | 1.12 | 1.01 | 1.91 | Glutamine transport ATP-binding protein GlnQ |
| *FLPKDCKG_03605* | *gltK* | - | 3942446 | 3943135 | 0.72 | 1.16 | 0.89 | 0.56 | Glutamate/aspartate import permease protein GltK |
| *FLPKDCKG_03606* | *yecS-5* | - | 3943144 | 3943896 | 1.54 | 0.00 | 0.98 | 0.42 | L-cystine transport system permease protein YecS |
| *FLPKDCKG_03607* | *gltI* | - | 3944001 | 3944909 | 15.97 | 21.25 | 16.43 | 6.63 | Glutamate/aspartate import solute-binding protein |
| *FLPKDCKG_03608* | *cynR-11* | + | 3945081 | 3945578 | 16.49 | 5.84 | 5.93 | 5.30 | HTH-type transcriptional regulator CynR |
| *FLPKDCKG_03609* | | + | 3945533 | 3946012 | 3.38 | 1.31 | 3.22 | 0.72 | Hypothetical protein |
| *FLPKDCKG_03610* | | - | 3946086 | 3946541 | 2.79 | 1.52 | 0.00 | 0.67 | Hypothetical protein |
| *FLPKDCKG_03611* | *nfdA-2* | - | 3946553 | 3948448 | 3.10 | 4.80 | 1.84 | 0.94 | N-substituted formamide deformylase |
| *FLPKDCKG_03612* | | - | 3948578 | 3948838 | 0.71 | 0.42 | 0.00 | 0.14 | Hypothetical protein |
| *FLPKDCKG_03613* | *yhhW-3* | - | 3948865 | 3949569 | 0.67 | 0.75 | 0.00 | 1.09 | Quercetin 2,3-dioxygenase |
| *FLPKDCKG_03614* | *entS-2* | - | 3949609 | 3951201 | 0.29 | 0.52 | 0.00 | 0.00 | Enterobactin exporter EntS |
| *FLPKDCKG_03615* | | - | 3951319 | 3952005 | 5.04 | 5.82 | 3.18 | 1.36 | Hypothetical protein |
| *FLPKDCKG_03616* | *dmlR-19* | + | 3952321 | 3953244 | 2.38 | 4.75 | 2.51 | 1.02 | HTH-type transcriptional regulator DmlR |
| *FLPKDCKG_03617* | *gyrB* | - | 3953366 | 3955813 | 8.08 | 23.11 | 17.11 | 6.62 | DNA gyrase subunit B |
| *FLPKDCKG_03618* | *dnaN* | - | 3955909 | 3957018 | 30.57 | 57.66 | 42.87 | 12.53 | DNA polymerase III subunit beta |
| *FLPKDCKG_03619* | *dnaA-1* | - | 3957021 | 3958133 | 14.02 | 20.89 | 14.79 | 5.45 | Chromosomal replication initiator protein DnaA |
| *FLPKDCKG_03620* | *dnaA-2* | - | 3958073 | 3958480 | 42.09 | 83.17 | 44.23 | 15.58 | Chromosomal replication initiator protein DnaA |
| *FLPKDCKG_03621* | *rpmH* | + | 3958894 | 3959028 | 28.22 | 238.26 | 152.28 | 84.16 | 50S ribosomal protein L34 |
| *FLPKDCKG_03622* | *rnpA* | + | 3959073 | 3959444 | 1.28 | 16.18 | 9.91 | 3.91 | Ribonuclease P protein component |
| *FLPKDCKG_03623* | *yidD* | + | 3959441 | 3959764 | 3.01 | 11.23 | 16.89 | 6.70 | Putative membrane protein insertion efficiency factor |
| *FLPKDCKG_03624* | *yidC* | + | 3959813 | 3961498 | 10.73 | 19.91 | 19.48 | 7.22 | Membrane protein insertase YidC |
| *FLPKDCKG_03625* | *katA* | + | 3961768 | 3963213 | 69.90 | 231.64 | 245.69 | 42.36 | Catalase |
| *FLPKDCKG_03626* | *cysK* | - | 3963507 | 3964433 | 2.00 | 4.26 | 5.71 | 2.67 | Cysteine synthase |
| *FLPKDCKG_03627* | | - | 3964546 | 3964731 | 0.52 | 0.00 | 15.09 | 1.60 | Hypothetical protein |
| *FLPKDCKG_03628* | *tauD-2* | - | 3964763 | 3965677 | 0.86 | 13.64 | 63.56 | 19.84 | Alpha-ketoglutarate-dependent taurine dioxygenase |
| *FLPKDCKG_03629* | *yxeP-4* | - | 3965812 | 3967104 | 4.35 | 7.00 | 8.04 | 1.63 | putative hydrolase YxeP |
| *FLPKDCKG_03630* | *oppF-6* | - | 3967237 | 3968253 | 4.13 | 5.17 | 6.64 | 2.13 | Oligopeptide transport ATP-binding protein OppF |
| *FLPKDCKG_03631* | *oppD-8* | - | 3968250 | 3969245 | 6.32 | 7.13 | 2.85 | 3.19 | Oligopeptide transport ATP-binding protein OppD |
| *FLPKDCKG_03632* | *ddpC-4* | - | 3969257 | 3970174 | 2.95 | 5.47 | 4.12 | 2.00 | putative D,D-dipeptide transport system permease protein DdpC |
| *FLPKDCKG_03633* | *gsiC-7* | - | 3970171 | 3971100 | 5.45 | 4.96 | 7.63 | 1.56 | Glutathione transport system permease protein GsiC |
| *FLPKDCKG_03634* | *gsiB-8* | - | 3971168 | 3972754 | 48.21 | 29.28 | 26.65 | 9.58 | Glutathione-binding protein GsiB |
| *FLPKDCKG_03635* | *mnmE* | + | 3972870 | 3974222 | 5.95 | 4.42 | 5.29 | 1.89 | tRNA modification GTPase MnmE |
| *FLPKDCKG_03636* | *mshD-1* | + | 3974228 | 3974698 | 1.28 | 3.49 | 2.82 | 1.06 | Mycothiol acetyltransferase |
| *FLPKDCKG_03637* | *clcB* | - | 3974869 | 3976260 | 2.66 | 6.10 | 6.17 | 1.29 | Voltage-gated ClC-type chloride channel ClcB |
| *FLPKDCKG_03638* | *tctD-5* | + | 3976752 | 3977429 | 4.66 | 10.00 | 5.88 | 3.55 | Transcriptional regulatory protein tctD |
| *FLPKDCKG_03639* | *qseC-6* | + | 3977419 | 3978903 | 1.16 | 3.89 | 1.76 | 0.93 | Sensor protein QseC |
| *FLPKDCKG_03640* | | + | 3979016 | 3980014 | 1.04 | 13.78 | 9.78 | 8.00 | Hypothetical protein |
| *FLPKDCKG_03641* | | + | 3980134 | 3980571 | 0.29 | 4.45 | 1.76 | 3.16 | Hypothetical protein |
| *FLPKDCKG_03642* | | + | 3980568 | 3982094 | 0.25 | 1.17 | 2.19 | 3.08 | Hypothetical protein |
| *FLPKDCKG_03643* | *ybdL* | - | 3982258 | 3983424 | 30.92 | 35.87 | 21.70 | 7.36 | Methionine aminotransferase |
| *FLPKDCKG_03644* | | + | 3983539 | 3984573 | 0.89 | 2.37 | 1.34 | 0.53 | Hypothetical protein |
| *FLPKDCKG_03645* | *cspA-3* | + | 3984805 | 3985011 | 21.90 | 35.28 | 41.87 | 12.47 | Cold shock protein CspA |
| *FLPKDCKG_03646* | *mnmG* | + | 3987527 | 3989446 | 17.12 | 28.20 | 18.67 | 8.85 | tRNA uridine 5-carboxymethylaminomethyl modification enzyme MnmG |
| *FLPKDCKG_03647* | *rsmG* | + | 3989443 | 3990132 | 11.99 | 24.21 | 17.39 | 6.11 | Ribosomal RNA small subunit methyltransferase G |
| *FLPKDCKG_03648* | *parA* | + | 3990129 | 3990926 | 9.03 | 19.13 | 14.60 | 8.39 | Chromosome partitioning protein ParA |
| *FLPKDCKG_03649* | *rimI* | + | 3990935 | 3991747 | 5.14 | 14.60 | 12.13 | 4.39 | Ribosomal-protein-alanine acetyltransferase |
| *FLPKDCKG_03650* | *parB* | + | 3991792 | 3992709 | 6.80 | 11.00 | 9.90 | 5.99 | putative chromosome-partitioning protein ParB |
| *FLPKDCKG_03651* | *ansA* | + | 3992702 | 3993706 | 3.26 | 7.33 | 6.74 | 0.71 | L-asparaginase |
| *FLPKDCKG_03652* | *gltC-13* | - | 3993754 | 3994635 | 2.58 | 4.10 | 0.92 | 1.02 | HTH-type transcriptional regulator GltC |
| *FLPKDCKG_03653* | | + | 3994758 | 3995345 | 8.69 | 5.30 | 5.64 | 1.61 | Hypothetical protein |

| Locus name | Gene name | Strand | Gene start (bp) | Gene end (bp) | Average TPM Inoculum | Average TPM $t_6$ | Average TPM $t_{24}$ | Average TPM $t_{168}$ | Annotation description |
|---|---|---|---|---|---|---|---|---|---|
| *FLPKDCKG_03654* | *metC-4* | + | 3995351 | 3996562 | 0.79 | 1.20 | 1.67 | 0.72 | Cystathionine beta-lyase |
| *FLPKDCKG_03655* | *glpE-2* | + | 3996565 | 3998160 | 0.39 | 1.51 | 1.01 | 0.36 | Thiosulfate sulfurtransferase GlpE |
| *FLPKDCKG_03656* | | + | 3998313 | 3999314 | 0.28 | 1.68 | 1.03 | 0.00 | Hypothetical protein |
| *FLPKDCKG_03657* | *glcR* | - | 3999323 | 4000084 | 2.42 | 4.71 | 3.59 | 1.57 | HTH-type transcriptional repressor GlcR |
| *FLPKDCKG_03658* | *cysW-3* | - | 4000104 | 4001786 | 1.55 | 2.33 | 2.67 | 1.03 | Sulfate transport system permease protein CysW |
| *FLPKDCKG_03659* | *potA-5* | - | 4001824 | 4002831 | 1.60 | 1.99 | 1.48 | 1.19 | Spermidine/putrescine import ATP-binding protein PotA |
| *FLPKDCKG_03660* | *idiA* | - | 4002883 | 4003893 | 4.59 | 7.12 | 3.44 | 2.40 | Iron deficiency-induced protein A |
| *FLPKDCKG_03661* | *hisN* | + | 4004108 | 4004914 | 4.09 | 7.87 | 7.66 | 3.30 | Histidinol-phosphatase |
| *FLPKDCKG_03662* | *ipuC* | + | 4004979 | 4006364 | 5.11 | 9.01 | 6.89 | 1.19 | Glutamate--isopropylamine ligase |
| *FLPKDCKG_03663* | *sad* | + | 4006408 | 4007799 | 2.72 | 2.04 | 3.60 | 2.49 | Succinate semialdehyde dehydrogenase [NAD(P)+] Sad |
| *FLPKDCKG_03664* | *mdh-3* | + | 4007820 | 4008983 | 3.96 | 6.35 | 6.31 | 3.00 | NAD-dependent methanol dehydrogenase |
| *FLPKDCKG_03665* | | + | 4009097 | 4009183 | 14.19 | 79.89 | 52.28 | 12.42 | tRNA |
| *FLPKDCKG_03666* | | + | 4009334 | 4009407 | 25.57 | 158.98 | 143.56 | 60.86 | tRNA |
| *FLPKDCKG_03667* | | + | 4009420 | 4009494 | 8.75 | 85.38 | 74.52 | 19.23 | tRNA |
| *FLPKDCKG_03668* | *tuf1-1* | + | 4009563 | 4010753 | 47.81 | 168.45 | 179.12 | 77.15 | Elongation factor Tu |
| *FLPKDCKG_03669* | | + | 4010918 | 4010993 | 111.50 | 235.25 | 402.44 | 75.75 | tRNA |
| *FLPKDCKG_03670* | *secE* | + | 4011175 | 4011438 | 41.31 | 72.30 | 72.73 | 24.55 | Protein translocase subunit SecE |
| *FLPKDCKG_03671* | *nusG* | + | 4011448 | 4011981 | 39.21 | 106.59 | 94.53 | 36.09 | Transcription termination/antitermination protein NusG |
| *FLPKDCKG_03672* | *rplK* | + | 4012043 | 4012474 | 35.17 | 91.41 | 85.14 | 51.61 | 50S ribosomal protein L11 |
| *FLPKDCKG_03673* | *rplA* | + | 4012477 | 4013178 | 11.59 | 46.01 | 39.54 | 20.27 | 50S ribosomal protein L1 |
| *FLPKDCKG_03674* | *rplJ* | + | 4013428 | 4013952 | 26.74 | 275.62 | 266.13 | 176.68 | 50S ribosomal protein L10 |
| *FLPKDCKG_03675* | *rplL* | + | 4014034 | 4014414 | 15.32 | 121.42 | 131.59 | 93.33 | 50S ribosomal protein L7/L12 |
| *FLPKDCKG_03676* | *rpoB* | + | 4014578 | 4018690 | 8.96 | 33.00 | 37.63 | 19.37 | DNA-directed RNA polymerase subunit beta |
| *FLPKDCKG_03677* | *rpoC* | + | 4018690 | 4022934 | 7.75 | 25.86 | 27.79 | 16.19 | DNA-directed RNA polymerase subunit beta' |
| *FLPKDCKG_03678* | | - | 4023045 | 4023398 | 17.91 | 18.01 | 22.24 | 6.02 | Hypothetical protein |
| *FLPKDCKG_03679* | | + | 4023597 | 4024916 | 15.52 | 28.45 | 14.33 | 6.55 | Hypothetical protein |
| *FLPKDCKG_03680* | *phoR-2* | - | 4024943 | 4027267 | 3.54 | 6.48 | 4.59 | 1.88 | Alkaline phosphatase synthesis sensor protein PhoR |
| *FLPKDCKG_03681* | | - | 4027273 | 4028382 | 5.67 | 6.63 | 9.78 | 2.33 | Hypothetical protein |
| *FLPKDCKG_03682* | *phoP-2* | - | 4028416 | 4029132 | 29.86 | 44.40 | 21.76 | 6.05 | Alkaline phosphatase synthesis transcriptional regulatory protein PhoP |
| *FLPKDCKG_03683* | *ompR-4* | + | 4029392 | 4030093 | 188.26 | 43.66 | 12.73 | 1.92 | Transcriptional regulatory protein OmpR |
| *FLPKDCKG_03684* | *rpsL* | + | 4030876 | 4031091 | 21.53 | 253.46 | 200.38 | 52.72 | 30S ribosomal protein S12 |
| *FLPKDCKG_03685* | *rpsG* | + | 4031267 | 4031737 | 17.44 | 144.59 | 153.23 | 61.16 | 30S ribosomal protein S7 |
| *FLPKDCKG_03686* | *fusA-2* | + | 4031756 | 4033858 | 9.28 | 68.47 | 66.41 | 31.59 | Elongation factor G |
| *FLPKDCKG_03687* | *tuf1-2* | + | 4033935 | 4035125 | 34.46 | 179.51 | 169.59 | 85.15 | Elongation factor Tu |
| *FLPKDCKG_03688* | *rpsJ* | + | 4035225 | 4035509 | 3.83 | 36.51 | 69.25 | 24.96 | 30S ribosomal protein S10 |
| *FLPKDCKG_03689* | *rplC* | + | 4035841 | 4036530 | 55.85 | 186.96 | 156.51 | 65.93 | 50S ribosomal protein L3 |
| *FLPKDCKG_03690* | *rplD* | + | 4036535 | 4037152 | 29.48 | 88.81 | 98.79 | 44.35 | 50S ribosomal protein L4 |
| *FLPKDCKG_03691* | *rplW* | + | 4037149 | 4037445 | 33.43 | 103.54 | 91.33 | 37.71 | 50S ribosomal protein L23 |
| *FLPKDCKG_03692* | *rplB* | + | 4037446 | 4038273 | 31.92 | 111.08 | 105.60 | 56.20 | 50S ribosomal protein L2 |
| *FLPKDCKG_03693* | *rpsS* | + | 4038286 | 4038561 | 30.04 | 101.00 | 67.36 | 46.16 | 30S ribosomal protein S19 |
| *FLPKDCKG_03694* | *rplV* | + | 4038565 | 4038894 | 18.10 | 76.77 | 87.81 | 31.40 | 50S ribosomal protein L22 |
| *FLPKDCKG_03695* | *rpsC* | + | 4038904 | 4039695 | 26.51 | 117.08 | 103.41 | 57.25 | 30S ribosomal protein S3 |
| *FLPKDCKG_03696* | *rplP* | + | 4039698 | 4040114 | 15.25 | 75.57 | 75.39 | 41.03 | 50S ribosomal protein L16 |
| *FLPKDCKG_03697* | *rpmC* | + | 4040128 | 4040319 | 38.38 | 172.62 | 173.51 | 84.81 | 50S ribosomal protein L29 |
| *FLPKDCKG_03698* | *rpsQ* | + | 4040322 | 4040603 | 3.42 | 29.44 | 51.72 | 13.97 | 30S ribosomal protein S17 |
| *FLPKDCKG_03699* | | + | 4040715 | 4041281 | 1.80 | 1.64 | 0.99 | 0.00 | Hypothetical protein |
| *FLPKDCKG_03700* | | - | 4041330 | 4042397 | 2.24 | 3.07 | 0.73 | 0.39 | Hypothetical protein |
| *FLPKDCKG_03701* | *livH-15* | - | 4042408 | 4043418 | 2.53 | 0.26 | 0.00 | 0.09 | High-affinity branched-chain amino acid transport system permease protein LivH |
| *FLPKDCKG_03702* | *livF-15* | - | 4043415 | 4044167 | 3.70 | 0.50 | 0.00 | 0.00 | High-affinity branched-chain amino acid transport ATP-binding protein LivF |
| *FLPKDCKG_03703* | *artM-2* | - | 4044164 | 4044934 | 3.03 | 0.60 | 0.73 | 0.71 | Arginine transport ATP-binding protein ArtM |
| *FLPKDCKG_03704* | | - | 4044946 | 4046118 | 13.81 | 1.65 | 0.98 | 0.97 | Leu/Ile/Val-binding protein |
| *FLPKDCKG_03705* | *lip1* | - | 4046135 | 4047031 | 8.17 | 2.63 | 0.14 | 0.00 | Lipase 1 |
| *FLPKDCKG_03706* | *feaR-2* | + | 4047195 | 4048154 | 6.07 | 3.82 | 3.24 | 0.69 | Transcriptional activator FeaR |
| *FLPKDCKG_03707* | *megL* | - | 4048163 | 4049344 | 3.96 | 5.75 | 2.94 | 0.03 | L-methionine gamma-lyase |
| *FLPKDCKG_03708* | *araC-3* | + | 4049350 | 4050267 | 3.76 | 3.32 | 5.09 | 0.00 | Arabinose operon regulatory protein |
| *FLPKDCKG_03709* | *araE* | - | 4050330 | 4051757 | 52.03 | 79.28 | 62.72 | 23.04 | Alpha-ketoglutaric semialdehyde dehydrogenase |
| *FLPKDCKG_03710* | | + | 4051943 | 4052707 | 13.32 | 24.58 | 14.35 | 5.92 | Hypothetical protein |
| *FLPKDCKG_03711* | *rplN* | + | 4053060 | 4053428 | 43.49 | 149.95 | 126.30 | 70.78 | 50S ribosomal protein L14 |
| *FLPKDCKG_03712* | *rplX* | + | 4053440 | 4053760 | 30.08 | 124.80 | 115.56 | 48.48 | 50S ribosomal protein L24 |
| *FLPKDCKG_03713* | *rplE* | + | 4053773 | 4054312 | 39.64 | 143.11 | 141.31 | 60.61 | 50S ribosomal protein L5 |
| *FLPKDCKG_03714* | *rpsN* | + | 4054325 | 4054630 | 51.59 | 157.87 | 151.91 | 72.99 | 30S ribosomal protein S14 |
| *FLPKDCKG_03715* | *rpsH* | + | 4054641 | 4055036 | 31.43 | 96.44 | 113.45 | 46.30 | 30S ribosomal protein S8 |
| *FLPKDCKG_03716* | *rplF* | + | 4055049 | 4055582 | 36.61 | 113.48 | 132.69 | 59.37 | 50S ribosomal protein L6 |
| *FLPKDCKG_03717* | *rplR* | + | 4055604 | 4055969 | 23.67 | 101.85 | 88.52 | 38.14 | 50S ribosomal protein L18 |
| *FLPKDCKG_03718* | *rpsE* | + | 4055985 | 4056506 | 24.77 | 73.16 | 91.31 | 56.05 | 30S ribosomal protein S5 |
| *FLPKDCKG_03719* | *rpmD* | + | 4056510 | 4056695 | 19.73 | 46.80 | 85.40 | 36.55 | 50S ribosomal protein L30 |
| *FLPKDCKG_03720* | *rplO* | + | 4056705 | 4057145 | 7.54 | 45.82 | 57.37 | 31.53 | 50S ribosomal protein L15 |
| *FLPKDCKG_03721* | *secY* | + | 4057160 | 4058485 | 9.40 | 57.63 | 45.67 | 23.64 | Protein translocase subunit SecY |
| *FLPKDCKG_03722* | *infA-1* | + | 4058492 | 4058710 | 21.37 | 120.53 | 112.15 | 42.17 | Translation initiation factor IF-1 |

| Locus name | Gene name | Strand | Gene start (bp) | Gene end (bp) | Average TPM Inoculum | Average TPM $t_6$ | Average TPM $t_{24}$ | Average TPM $t_{168}$ | Annotation description |
|---|---|---|---|---|---|---|---|---|---|
| *FLPKDCKG_03723* | *rpmJ* | + | 4058745 | 4058858 | 20.02 | 95.52 | 174.24 | 31.33 | 50S ribosomal protein L36 |
| *FLPKDCKG_03724* | *rpsM* | + | 4058902 | 4059267 | 17.39 | 119.74 | 114.81 | 35.02 | 30S ribosomal protein S13 |
| *FLPKDCKG_03725* | *rpsK* | + | 4059283 | 4059684 | 19.76 | 136.33 | 137.39 | 78.00 | 30S ribosomal protein S11 |
| *FLPKDCKG_03726* | *rpsD* | + | 4059696 | 4060319 | 6.19 | 75.12 | 66.30 | 42.77 | 30S ribosomal protein S4 |
| *FLPKDCKG_03727* | *rpoA* | + | 4060532 | 4061518 | 1.79 | 41.00 | 35.33 | 21.79 | DNA-directed RNA polymerase subunit alpha |
| *FLPKDCKG_03728* | *rplQ* | + | 4061706 | 4062098 | 3.65 | 29.87 | 40.99 | 24.70 | 50S ribosomal protein L17 |
| *FLPKDCKG_03729* | | - | 4062201 | 4062638 | 58.84 | 33.77 | 27.53 | 12.53 | putative HIT-like protein |
| *FLPKDCKG_03730* | *cutA* | + | 4062759 | 4063106 | 35.06 | 71.15 | 34.08 | 8.26 | Divalent-cation tolerance protein CutA |
| *FLPKDCKG_03731* | *dsbD-1* | + | 4063106 | 4064980 | 3.65 | 6.76 | 5.46 | 3.16 | Thiol:disulfide interchange protein DsbD |
| *FLPKDCKG_03732* | *eamA-5* | + | 4065052 | 4065975 | 8.76 | 11.21 | 9.32 | 1.32 | putative amino-acid metabolite efflux pump |
| *FLPKDCKG_03733* | *hemB* | - | 4066104 | 4067120 | 12.03 | 24.71 | 18.57 | 5.81 | Delta-aminolevulinic acid dehydratase |
| *FLPKDCKG_03734* | *engB-2* | - | 4067117 | 4067740 | 25.82 | 32.37 | 21.90 | 4.29 | putative GTP-binding protein EngB |
| *FLPKDCKG_03735* | *cycA-2* | + | 4067996 | 4068697 | 28.02 | 51.23 | 41.28 | 14.86 | Cytochrome c4 |
| *FLPKDCKG_03736* | *ccsB* | + | 4068818 | 4070896 | 3.48 | 8.49 | 6.44 | 1.76 | Cytochrome c biogenesis protein CcsB |
| *FLPKDCKG_03737* | *ccsA* | + | 4070905 | 4072251 | 5.18 | 7.78 | 8.12 | 2.60 | Cytochrome c biogenesis protein CcsA |
| *FLPKDCKG_03738* | *lysA-1* | - | 4072312 | 4073604 | 8.84 | 11.38 | 9.99 | 2.04 | Diaminopimelate decarboxylase |
| *FLPKDCKG_03739* | *cyaY* | + | 4073824 | 4074153 | 31.40 | 43.67 | 26.37 | 9.31 | Protein CyaY |
| *FLPKDCKG_03740* | *mrcA-2* | - | 4074203 | 4076650 | 8.92 | 21.65 | 19.44 | 6.36 | Penicillin-binding protein 1A |
| *FLPKDCKG_03741* | *aroK-1* | + | 4076871 | 4077407 | 30.27 | 50.84 | 27.16 | 9.96 | Shikimate kinase 1 |
| *FLPKDCKG_03742* | *aroB* | + | 4077404 | 4078480 | 10.88 | 21.20 | 14.72 | 3.94 | 3-dehydroquinate synthase |
| *FLPKDCKG_03743* | *dgt* | + | 4078498 | 4079634 | 4.89 | 20.54 | 9.35 | 3.16 | Deoxyguanosinetriphosphate triphosphohydrolase-like protein |
| *FLPKDCKG_03744* | *sspA-2* | - | 4079721 | 4080383 | 4.34 | 10.88 | 10.85 | 3.18 | Stringent starvation protein A |
| *FLPKDCKG_03745* | *uvrA-2* | - | 4080413 | 4083268 | 10.34 | 19.24 | 12.27 | 3.58 | UvrABC system protein A |
| *FLPKDCKG_03746* | *yajR-2* | + | 4083403 | 4084587 | 1.67 | 6.17 | 3.95 | 0.49 | Inner membrane transport protein YajR |
| *FLPKDCKG_03747* | *ssb* | + | 4084870 | 4085373 | 42.49 | 40.70 | 36.52 | 15.38 | Single-stranded DNA-binding protein |
| *FLPKDCKG_03748* | *bioC-1* | + | 4085858 | 4086586 | 2.68 | 7.33 | 5.90 | 0.96 | Malonyl-[acyl-carrier protein] O-methyltransferase |
| *FLPKDCKG_03749* | | + | 4086851 | 4087156 | 188.09 | 422.46 | 264.41 | 85.31 | Hypothetical protein |
| *FLPKDCKG_03750* | | + | 4087237 | 4087542 | 101.18 | 189.18 | 137.45 | 68.48 | Hypothetical protein |
| *FLPKDCKG_03751* | *acuR* | + | 4087768 | 4088262 | 26.50 | 46.04 | 34.09 | 10.51 | Transcriptional regulator AcuR |
| *FLPKDCKG_03752* | | - | 4088398 | 4089255 | 8.80 | 11.23 | 8.67 | 5.62 | Hypothetical protein |
| *FLPKDCKG_03753* | | - | 4089359 | 4089475 | 59.93 | 34.14 | 14.02 | 0.68 | Hypothetical protein |
| *FLPKDCKG_03754* | | - | 4089561 | 4090178 | 78.27 | 26.35 | 14.46 | 5.22 | Hypothetical protein |
| *FLPKDCKG_03755* | *glcB* | - | 4090297 | 4092471 | 48.31 | 44.15 | 37.79 | 12.07 | Malate synthase G |
| *FLPKDCKG_03756* | *glcC-2* | + | 4092708 | 4093466 | 10.26 | 25.89 | 15.47 | 3.01 | Glc operon transcriptional activator |
| *FLPKDCKG_03757* | *lysDH* | - | 4093546 | 4094664 | 116.34 | 12.04 | 7.10 | 1.24 | Lysine 6-dehydrogenase |
| *FLPKDCKG_03758* | | - | 4094870 | 4095259 | 129.75 | 39.48 | 33.14 | 13.61 | Hypothetical protein |
| *FLPKDCKG_03759* | | - | 4095641 | 4096051 | 23.30 | 13.52 | 16.63 | 3.22 | Hypothetical protein |
| *FLPKDCKG_03760* | *ubiA* | - | 4096122 | 4097048 | 3.87 | 7.71 | 8.28 | 3.72 | 4-hydroxybenzoate octaprenyltransferase |
| *FLPKDCKG_03761* | | + | 4097734 | 4097830 | 69.31 | 92.67 | 68.75 | 12.37 | tRNA |
| *FLPKDCKG_03762* | *msrC* | + | 4098519 | 4099004 | 11.46 | 14.72 | 10.42 | 2.09 | Free methionine-R-sulfoxide reductase |
| *FLPKDCKG_03763* | *dapA-3* | + | 4099312 | 4100205 | 0.67 | 2.99 | 0.81 | 0.31 | 4-hydroxy-tetrahydrodipicolinate synthase |
| *FLPKDCKG_03764* | | + | 4100329 | 4101105 | 8.08 | 8.25 | 7.99 | 1.41 | Hypothetical protein |
| *FLPKDCKG_03765* | | + | 4101145 | 4101996 | 4.68 | 10.50 | 8.80 | 1.78 | Hypothetical protein |
| *FLPKDCKG_03766* | *fabH2* | + | 4102032 | 4103963 | 3.85 | 10.12 | 5.98 | 1.70 | 3-oxoacyl-[acyl-carrier-protein] synthase 3 protein 2 |
| *FLPKDCKG_03767* | | + | 4103960 | 4104856 | 1.23 | 7.01 | 5.01 | 0.76 | Hypothetical protein |
| *FLPKDCKG_03768* | | + | 4104984 | 4106267 | 0.63 | 2.52 | 0.68 | 0.50 | Hypothetical protein |
| *FLPKDCKG_03769* | | + | 4106515 | 4106829 | 3.05 | 3.59 | 4.21 | 1.77 | Hypothetical protein |
| *FLPKDCKG_03770* | | + | 4106841 | 4107509 | 4.25 | 11.87 | 6.94 | 3.79 | Hypothetical protein |
| *FLPKDCKG_03771* | *aspS* | + | 4107550 | 4109337 | 6.10 | 21.16 | 16.13 | 8.41 | Aspartate--tRNA(Asp/Asn) ligase |
| *FLPKDCKG_03772* | | + | 4109693 | 4110559 | 4.49 | 11.05 | 5.84 | 1.28 | Hypothetical protein |
| *FLPKDCKG_03773* | *clsB-1* | + | 4110556 | 4111767 | 2.13 | 5.71 | 6.23 | 0.85 | Cardiolipin synthase B |
| *FLPKDCKG_03774* | *yijE-8* | - | 4112315 | 4113214 | 6.44 | 23.77 | 12.20 | 3.19 | putative cystine transporter YijE |
| *FLPKDCKG_03775* | *bmr3-1* | - | 4113216 | 4114274 | 3.95 | 13.54 | 8.21 | 2.90 | Multidrug resistance protein 3 |
| *FLPKDCKG_03776* | | - | 4114678 | 4115247 | 287.77 | 337.48 | 152.83 | 59.24 | Hypothetical protein |
| *FLPKDCKG_03777* | *rkpK-2* | + | 4115507 | 4116871 | 25.38 | 68.65 | 48.68 | 15.13 | UDP-glucose 6-dehydrogenase |
| *FLPKDCKG_03778* | | + | 4116893 | 4117837 | 15.11 | 45.84 | 37.96 | 10.53 | UDP-glucose 4-epimerase |
| *FLPKDCKG_03779* | *pseB* | + | 4117857 | 4118867 | 9.36 | 23.91 | 18.67 | 5.34 | UDP-N-acetylglucosamine 4,6-dehydratase (inverting) |
| *FLPKDCKG_03780* | *pseC* | + | 4118878 | 4120032 | 13.63 | 48.91 | 34.98 | 9.46 | UDP-4-amino-4,6-dideoxy-N-acetyl-beta-L-altrosamine transaminase |
| *FLPKDCKG_03781* | *kdsB-1* | + | 4120029 | 4121408 | 12.38 | 61.26 | 36.47 | 15.72 | 3-deoxy-manno-octulosonate cytidylyltransferase |
| *FLPKDCKG_03782* | | + | 4121410 | 4122561 | 5.06 | 39.31 | 36.13 | 8.80 | Hypothetical protein |
| *FLPKDCKG_03783* | *hisH-2* | + | 4122672 | 4123205 | 5.75 | 40.70 | 36.03 | 7.05 | Imidazole glycerol phosphate synthase subunit HisH |
| *FLPKDCKG_03784* | *hisF-2* | + | 4123199 | 4123981 | 12.61 | 55.16 | 38.87 | 7.62 | Imidazole glycerol phosphate synthase subunit HisF |
| *FLPKDCKG_03785* | *pseG* | + | 4124095 | 4125606 | 2.88 | 17.75 | 17.74 | 4.60 | UDP-2,4-diacetamido-2,4,6-trideoxy-beta-L-altropyranose hydrolase |
| *FLPKDCKG_03786* | *pseI* | + | 4125603 | 4126658 | 14.81 | 48.83 | 40.47 | 11.58 | Pseudaminic acid synthase |
| *FLPKDCKG_03787* | | + | 4126751 | 4128238 | 10.97 | 41.94 | 35.98 | 5.54 | Hypothetical protein |
| *FLPKDCKG_03788* | | + | 4128485 | 4129186 | 8.61 | 34.80 | 25.40 | 4.85 | Hypothetical protein |
| *FLPKDCKG_03789* | | + | 4129214 | 4130467 | 4.34 | 22.50 | 13.35 | 6.31 | Hypothetical protein |
| *FLPKDCKG_03790* | *lacA-1* | - | 4130476 | 4131501 | 4.26 | 13.37 | 7.24 | 0.64 | Galactoside O-acetyltransferase |
| *FLPKDCKG_03791* | | - | 4131491 | 4131742 | 2.71 | 11.02 | 2.74 | 2.80 | Hypothetical protein |

| Locus name | Gene name | Strand | Gene start (bp) | Gene end (bp) | Average TPM Inoculum | Average TPM $t_6$ | Average TPM $t_{24}$ | Average TPM $t_{168}$ | Annotation description |
|---|---|---|---|---|---|---|---|---|---|
| *FLPKDCKG_03792* | | - | 4131882 | 4132826 | 4.25 | 8.55 | 7.71 | 0.97 | Hypothetical protein |
| *FLPKDCKG_03793* | | - | 4132789 | 4133736 | 14.20 | 23.04 | 18.05 | 2.94 | Hypothetical protein |
| *FLPKDCKG_03794* | *tuaG* | - | 4133772 | 4134530 | 12.40 | 26.52 | 19.52 | 5.28 | Putative teichuronic acid biosynthesis glycosyltransferase TuaG |
| *FLPKDCKG_03795* | *tagO-1* | + | 4134774 | 4135784 | 3.90 | 12.25 | 6.26 | 3.10 | putative undecaprenyl-phosphate N-acetylglucosaminyl 1-phosphate transferase |
| *FLPKDCKG_03796* | *pglF* | + | 4135884 | 4137779 | 12.30 | 24.83 | 21.68 | 6.51 | UDP-N-acetyl-alpha-D-glucosamine C6 dehydratase |
| *FLPKDCKG_03797* | *rfaC* | + | 4137841 | 4138905 | 11.81 | 39.90 | 26.55 | 5.29 | Lipopolysaccharide heptosyltransferase 1 |
| *FLPKDCKG_03798* | *waaA* | + | 4138909 | 4140183 | 5.29 | 20.86 | 9.58 | 4.32 | 3-deoxy-D-manno-octulosonic acid transferase |
| *FLPKDCKG_03799* | | - | 4140206 | 4140430 | 1.77 | 3.63 | 1.93 | 2.73 | Hypothetical protein |
| *FLPKDCKG_03800* | | - | 4141184 | 4141342 | 3.55 | 2.20 | 5.24 | 0.18 | Hypothetical protein |
| *FLPKDCKG_03801* | *coaX* | - | 4141361 | 4142167 | 3.75 | 7.40 | 5.83 | 0.63 | Type III pantothenate kinase |
| *FLPKDCKG_03802* | *birA* | - | 4142164 | 4143018 | 4.64 | 4.76 | 4.87 | 0.64 | Bifunctional ligase/repressor BirA |
| *FLPKDCKG_03803* | *mlaE-2* | + | 4143113 | 4144246 | 6.16 | 6.18 | 3.38 | 1.82 | putative phospholipid ABC transporter permease protein MlaE |
| *FLPKDCKG_03804* | *mkl* | + | 4144246 | 4145085 | 4.47 | 3.65 | 2.94 | 0.02 | putative ribonucleotide transport ATP-binding protein mkl |
| *FLPKDCKG_03805* | | + | 4145072 | 4146028 | 4.21 | 6.82 | 4.95 | 1.15 | Hypothetical protein |
| *FLPKDCKG_03806* | | + | 4146025 | 4146636 | 3.92 | 2.57 | 1.19 | 0.71 | Hypothetical protein |
| *FLPKDCKG_03807* | | + | 4146669 | 4147322 | 3.26 | 4.91 | 4.39 | 1.32 | Hypothetical protein |
| *FLPKDCKG_03808* | | - | 4147566 | 4147646 | 3.41 | 0.87 | 4.90 | 2.52 | tRNA |
| *FLPKDCKG_03809* | *dat* | + | 4148871 | 4149737 | 10.84 | 19.59 | 9.72 | 7.11 | D-alanine aminotransferase |
| *FLPKDCKG_03810* | | + | 4149839 | 4150111 | 15.44 | 28.36 | 23.97 | 11.89 | Hypothetical protein |
| *FLPKDCKG_03811* | *lipB* | + | 4150123 | 4150767 | 2.06 | 6.64 | 6.34 | 2.91 | Octanoyltransferase |
| *FLPKDCKG_03812* | *esiB-2* | + | 4150764 | 4151522 | 2.83 | 8.39 | 9.92 | 1.04 | Secretory immunoglobulin A-binding protein EsiB |
| *FLPKDCKG_03813* | | - | 4151556 | 4153409 | 10.03 | 21.83 | 15.85 | 2.92 | Hypothetical protein |
| *FLPKDCKG_03814* | *rfbA* | - | 4153414 | 4154298 | 7.00 | 36.96 | 17.51 | 2.71 | Glucose-1-phosphate thymidylyltransferase 1 |
| *FLPKDCKG_03815* | | - | 4154295 | 4154546 | 1.95 | 12.81 | 8.67 | 1.92 | Hypothetical protein |
| *FLPKDCKG_03816* | | + | 4154811 | 4155974 | 10.77 | 35.80 | 12.43 | 2.94 | Hypothetical protein |
| *FLPKDCKG_03817* | *acpP-4* | - | 4155934 | 4156161 | 7.00 | 21.90 | 13.58 | 4.69 | Acyl carrier protein |
| *FLPKDCKG_03818* | *fabH-3* | - | 4156158 | 4157093 | 4.92 | 33.45 | 22.92 | 5.46 | 3-oxoacyl-[acyl-carrier-protein] synthase 3 |
| *FLPKDCKG_03819* | | - | 4157090 | 4157713 | 3.57 | 20.68 | 14.74 | 2.06 | Hypothetical protein |
| *FLPKDCKG_03820* | *lpxA-2* | - | 4157710 | 4158621 | 6.81 | 42.14 | 21.28 | 6.77 | Acyl-[acyl-carrier-protein]--UDP-N-acetylglucosamine O-acyltransferase |
| *FLPKDCKG_03821* | | - | 4158621 | 4159706 | 13.85 | 47.51 | 23.43 | 6.03 | putative glycosyltransferase |
| *FLPKDCKG_03822* | *cytR-2* | - | 4159763 | 4160605 | 1.83 | 7.61 | 6.42 | 3.35 | HTH-type transcriptional repressor CytR |
| *FLPKDCKG_03823* | *lipA* | + | 4160717 | 4161721 | 20.48 | 56.58 | 28.70 | 9.35 | Lipoyl synthase |
| *FLPKDCKG_03824* | *rfbB* | - | 4161782 | 4162843 | 10.63 | 43.83 | 34.74 | 3.01 | dTDP-glucose 4,6-dehydratase |
| *FLPKDCKG_03825* | | + | 4163233 | 4163436 | 25.98 | 97.08 | 42.67 | 10.32 | Hypothetical protein |
| *FLPKDCKG_03826* | *cusF* | + | 4163715 | 4164020 | 28.31 | 61.86 | 55.28 | 16.28 | Cation efflux system protein CusF |
| *FLPKDCKG_03827* | *hslU* | - | 4164175 | 4165509 | 10.38 | 17.23 | 31.51 | 11.73 | ATP-dependent protease ATPase subunit HslU |
| *FLPKDCKG_03828* | *hslV* | - | 4165583 | 4166122 | 15.01 | 20.49 | 37.09 | 13.82 | ATP-dependent protease subunit HslV |
| *FLPKDCKG_03829* | *dksA* | - | 4166330 | 4166794 | 80.64 | 135.21 | 129.90 | 60.15 | RNA polymerase-binding transcription factor DksA |
| *FLPKDCKG_03830* | *yjiA-2* | - | 4166922 | 4168052 | 23.84 | 43.75 | 45.62 | 23.48 | putative GTP-binding protein YjiA |
| *FLPKDCKG_03831* | *zur* | - | 4168414 | 4168917 | 40.33 | 47.35 | 25.89 | 18.24 | Zinc uptake regulation protein |
| *FLPKDCKG_03832* | *znuC* | + | 4169232 | 4169954 | 0.67 | 0.83 | 2.11 | 1.99 | High-affinity zinc uptake system ATP-binding protein ZnuC |
| *FLPKDCKG_03833* | *mntB* | + | 4169951 | 4170817 | 0.21 | 0.66 | 1.68 | 3.82 | Manganese transport system membrane protein MntB |
| *FLPKDCKG_03834* | *fimA* | + | 4170837 | 4171835 | 1.09 | 2.53 | 7.67 | 5.90 | Manganese ABC transporter substrate-binding lipoprotein |
| *FLPKDCKG_03835* | | + | 4171796 | 4172479 | 4.01 | 8.15 | 6.97 | 3.78 | Hypothetical protein |
| *FLPKDCKG_03836* | | + | 4172470 | 4173618 | 1.60 | 4.48 | 1.80 | 1.61 | Hypothetical protein |
| *FLPKDCKG_03837* | *xerC-2* | - | 4173632 | 4174606 | 4.14 | 6.15 | 4.62 | 1.02 | Tyrosine recombinase XerC |
| *FLPKDCKG_03838* | | - | 4174641 | 4175318 | 27.15 | 41.24 | 25.86 | 8.25 | Hypothetical protein |
| *FLPKDCKG_03839* | *dapF* | - | 4175323 | 4176234 | 40.85 | 65.11 | 40.01 | 5.93 | Diaminopimelate epimerase |
| *FLPKDCKG_03840* | *lpxL* | - | 4176283 | 4177173 | 3.10 | 7.69 | 5.81 | 1.91 | Lipid A biosynthesis lauroyltransferase |
| *FLPKDCKG_03841* | *lpxP* | - | 4177223 | 4178086 | 5.62 | 9.57 | 6.27 | 1.98 | Lipid A biosynthesis palmitoleoyltransferase |
| *FLPKDCKG_03842* | *metK* | + | 4178362 | 4179525 | 67.02 | 88.98 | 75.03 | 28.68 | S-adenosylmethionine synthase |
| *FLPKDCKG_03843* | *gatA-1* | - | 4179588 | 4181015 | 0.19 | 0.85 | 0.91 | 0.34 | Glutamyl-tRNA(Gln) amidotransferase subunit A |
| *FLPKDCKG_03844* | *kgtP-1* | - | 4181109 | 4182413 | 0.57 | 2.08 | 0.76 | 0.95 | Alpha-ketoglutarate permease |
| *FLPKDCKG_03845* | | + | 4182489 | 4183499 | 0.77 | 1.68 | 0.97 | 0.50 | Hypothetical protein |
| *FLPKDCKG_03846* | *livF-16* | - | 4183541 | 4186030 | 0.18 | 3.04 | 3.75 | 0.94 | High-affinity branched-chain amino acid transport ATP-binding protein LivF |
| *FLPKDCKG_03847* | *livH-16* | - | 4186030 | 4186908 | 0.10 | 1.18 | 2.66 | 0.23 | High-affinity branched-chain amino acid transport system permease protein LivH |
| *FLPKDCKG_03848* | | - | 4186927 | 4188117 | 0.54 | 8.78 | 11.68 | 3.39 | Leu/Ile/Val-binding protein |
| *FLPKDCKG_03849* | *pgdA-2* | + | 4188315 | 4189274 | 0.70 | 5.28 | 5.19 | 2.55 | Peptidoglycan deacetylase |
| *FLPKDCKG_03850* | *lutR-2* | + | 4189309 | 4190073 | 1.20 | 1.75 | 4.50 | 2.26 | HTH-type transcriptional regulator LutR |
| *FLPKDCKG_03851* | *flr-2* | + | 4190075 | 4190698 | 1.08 | 4.83 | 2.71 | 2.16 | Flavoredoxin |
| *FLPKDCKG_03852* | *dmlR-20* | - | 4190717 | 4191640 | 0.84 | 1.97 | 3.07 | 0.87 | HTH-type transcriptional regulator DmlR |
| *FLPKDCKG_03853* | | + | 4191771 | 4192757 | 0.87 | 1.58 | 0.50 | 0.00 | Hypothetical protein |
| *FLPKDCKG_03854* | *leuC-2* | + | 4192819 | 4194240 | 0.63 | 0.98 | 0.18 | 0.13 | 3-isopropylmalate dehydratase large subunit |
| *FLPKDCKG_03855* | *leuD-2* | + | 4194237 | 4194875 | 1.54 | 2.55 | 1.02 | 0.76 | 3-isopropylmalate dehydratase small subunit |
| *FLPKDCKG_03856* | *dehH1* | + | 4194905 | 4195798 | 1.95 | 0.80 | 3.53 | 0.49 | Haloacetate dehalogenase H-1 |
| *FLPKDCKG_03857* | | + | 4195997 | 4196233 | 41.93 | 91.49 | 58.59 | 13.62 | Hypothetical protein |
| *FLPKDCKG_03858* | *mgtA-2* | + | 4196313 | 4197353 | 0.97 | 2.32 | 4.50 | 0.75 | GDP-mannose-dependent alpha-mannosyltransferase |
| *FLPKDCKG_03859* | *ahcY* | + | 4197657 | 4199075 | 18.36 | 55.73 | 57.56 | 23.19 | Adenosylhomocysteinase |
| *FLPKDCKG_03860* | | + | 4199196 | 4199531 | 62.11 | 117.21 | 82.10 | 32.09 | Hypothetical protein |

| Locus name | Gene name | Strand | Gene start (bp) | Gene end (bp) | Average TPM Inoculum | Average TPM $t_6$ | Average TPM $t_{24}$ | Average TPM $t_{168}$ | Annotation description |
|---|---|---|---|---|---|---|---|---|---|
| *FLPKDCKG_03861* | *metF* | + | 4199667 | 4200509 | 5.52 | 28.64 | 33.55 | 10.43 | 5,10-methylenetetrahydrofolate reductase |
| *FLPKDCKG_03862* | | - | 4200678 | 4201676 | 0.60 | 1.40 | 1.82 | 0.73 | Hypothetical protein |
| *FLPKDCKG_03863* | | - | 4201830 | 4202693 | 12.54 | 27.90 | 16.39 | 4.42 | Hypothetical protein |
| *FLPKDCKG_03864* | | - | 4202749 | 4203552 | 4.94 | 25.64 | 28.87 | 5.60 | Hypothetical protein |
| *FLPKDCKG_03865* | | - | 4203566 | 4204552 | 3.94 | 25.84 | 16.42 | 4.25 | Hypothetical protein |
| *FLPKDCKG_03866* | | - | 4204555 | 4205985 | 6.16 | 36.37 | 32.81 | 7.46 | Hypothetical protein |
| *FLPKDCKG_03867* | | - | 4206389 | 4206955 | 13.11 | 22.55 | 14.57 | 2.52 | 5-formyltetrahydrofolate cyclo-ligase |
| *FLPKDCKG_03868* | | + | 4207089 | 4207250 | 162.36 | 190.91 | 108.74 | 29.67 | Hypothetical protein |
| *FLPKDCKG_03869* | *slt-2* | + | 4207247 | 4209394 | 22.45 | 17.56 | 14.62 | 6.18 | Soluble lytic murein transglycosylase |
| *FLPKDCKG_03870* | *cca* | + | 4209391 | 4210494 | 3.61 | 5.05 | 6.69 | 1.96 | Multifunctional CCA protein |
| *FLPKDCKG_03871* | | - | 4210507 | 4211688 | 3.08 | 3.58 | 5.01 | 2.82 | Hypothetical protein |
| *FLPKDCKG_03872* | *rhlB* | - | 4211774 | 4212721 | 2.81 | 17.97 | 30.67 | 13.69 | ATP-dependent RNA helicase RhlB |
| *FLPKDCKG_03873* | *rhlE-3* | - | 4212718 | 4213554 | 4.44 | 51.08 | 72.79 | 27.06 | ATP-dependent RNA helicase RhlE |
| *FLPKDCKG_03874* | *ung* | - | 4213832 | 4214593 | 2.99 | 8.05 | 4.13 | 1.72 | Uracil-DNA glycosylase |
| *FLPKDCKG_03875* | *shlB-3* | + | 4215161 | 4215457 | 9.04 | 37.17 | 17.53 | 2.19 | Hemolysin transporter protein ShlB |
| *FLPKDCKG_03876* | | + | 4215793 | 4216140 | 19.12 | 21.72 | 16.21 | 2.21 | Hypothetical protein |
| *FLPKDCKG_03877* | *tenA* | - | 4216180 | 4216842 | 8.29 | 19.80 | 23.09 | 2.78 | Aminopyrimidine aminohydrolase |
| *FLPKDCKG_03878* | *bdhA-2* | - | 4216919 | 4217704 | 12.96 | 108.62 | 51.00 | 8.02 | D-beta-hydroxybutyrate dehydrogenase |
| *FLPKDCKG_03879* | | + | 4218088 | 4218774 | 4.17 | 8.09 | 2.86 | 1.06 | Hypothetical protein |
| *FLPKDCKG_03880* | | + | 4218956 | 4220161 | 3.83 | 7.61 | 5.12 | 1.15 | Hypothetical protein |
| *FLPKDCKG_03881* | | + | 4220192 | 4221040 | 3.90 | 6.38 | 4.30 | 1.02 | Hypothetical protein |
| *FLPKDCKG_03882* | | - | 4221107 | 4221508 | 18.30 | 19.61 | 13.43 | 2.22 | Hypothetical protein |
| *FLPKDCKG_03883* | *nhaA* | - | 4221537 | 4222781 | 1.89 | 6.61 | 2.87 | 1.00 | Na(+)/H(+) antiporter NhaA |
| *FLPKDCKG_03884* | | - | 4222940 | 4223404 | 18.40 | 14.31 | 15.79 | 2.89 | Hypothetical protein |
| *FLPKDCKG_03885* | | - | 4223401 | 4223850 | 51.53 | 19.94 | 22.19 | 5.51 | Hypothetical protein |
| *FLPKDCKG_03886* | *tmk-2* | - | 4223871 | 4225337 | 13.28 | 14.64 | 10.08 | 1.71 | Thymidylate kinase |
| *FLPKDCKG_03887* | | - | 4225438 | 4226301 | 89.45 | 77.45 | 34.17 | 10.44 | putative metallo-hydrolase |
| *FLPKDCKG_03888* | *bigR-1* | + | 4226425 | 4226763 | 21.82 | 31.40 | 24.23 | 6.57 | Biofilm growth-associated repressor |
| *FLPKDCKG_03889* | | + | 4226760 | 4227188 | 10.20 | 14.69 | 3.54 | 2.14 | Hypothetical protein |
| *FLPKDCKG_03890* | | + | 4227190 | 4227627 | 14.43 | 14.93 | 13.31 | 3.82 | Hypothetical protein |
| *FLPKDCKG_03891* | *fccB* | + | 4227641 | 4228879 | 16.85 | 10.96 | 10.10 | 4.17 | Sulfide dehydrogenase [flavocytochrome c] flavoprotein chain |
| *FLPKDCKG_03892* | | - | 4228957 | 4230225 | 2.18 | 2.00 | 3.64 | 2.54 | Phenylacetate-coenzyme A ligase |
| *FLPKDCKG_03893* | *livF-17* | - | 4230308 | 4231150 | 4.07 | 4.44 | 7.44 | 3.68 | High-affinity branched-chain amino acid transport ATP-binding protein LivF |
| *FLPKDCKG_03894* | | - | 4231209 | 4232543 | 13.19 | 7.99 | 10.83 | 8.34 | Hypothetical protein |
| *FLPKDCKG_03895* | | - | 4232665 | 4232775 | 1.72 | 2.70 | 1.66 | 0.62 | Hypothetical protein |
| *FLPKDCKG_03896* | | - | 4232775 | 4233839 | 1.51 | 3.49 | 1.58 | 1.04 | Hypothetical protein |
| *FLPKDCKG_03897* | *cysA-5* | - | 4233864 | 4235699 | 2.65 | 3.56 | 3.23 | 2.98 | Sulfate/thiosulfate import ATP-binding protein CysA |
| *FLPKDCKG_03898* | | - | 4235705 | 4237651 | 5.77 | 5.29 | 5.92 | 3.02 | Long-chain-fatty-acid--CoA ligase FadD15 |
| *FLPKDCKG_03899* | *crp* | + | 4237851 | 4238558 | 6.80 | 20.06 | 14.79 | 3.12 | cAMP-activated global transcriptional regulator CRP |
| *FLPKDCKG_03900* | *ohrR-1* | + | 4238638 | 4239099 | 7.94 | 21.91 | 16.32 | 7.11 | Organic hydroperoxide resistance transcriptional regulator |
| *FLPKDCKG_03901* | *ohrB-2* | + | 4239252 | 4239680 | 9.59 | 45.47 | 39.90 | 11.20 | Organic hydroperoxide resistance protein OhrB |
| *FLPKDCKG_03902* | *livF-18* | - | 4240158 | 4240880 | 6.11 | 19.83 | 9.22 | 7.25 | High-affinity branched-chain amino acid transport ATP-binding protein LivF |
| *FLPKDCKG_03903* | *metN2* | - | 4240897 | 4242819 | 1.25 | 5.26 | 5.55 | 2.64 | Methionine import ATP-binding protein MetN 2 |
| *FLPKDCKG_03904* | *livH-17* | - | 4242823 | 4243695 | 1.04 | 5.84 | 4.13 | 1.71 | High-affinity branched-chain amino acid transport system permease protein LivH |
| *FLPKDCKG_03905* | *braC-5* | - | 4243838 | 4245001 | 21.19 | 43.39 | 52.43 | 23.45 | Leucine-, isoleucine-, valine-, threonine-, and alanine-binding protein |
| *FLPKDCKG_03906* | *gcvA-10* | - | 4245332 | 4246273 | 3.53 | 5.42 | 5.95 | 3.45 | Glycine cleavage system transcriptional activator |
| *FLPKDCKG_03907* | | - | 4246326 | 4246688 | 2.93 | 3.20 | 0.00 | 0.00 | Hypothetical protein |
| *FLPKDCKG_03908* | *argO-1* | - | 4246694 | 4247326 | 1.42 | 0.01 | 0.88 | 0.60 | Arginine exporter protein ArgO |
| *FLPKDCKG_03909* | | + | 4247475 | 4248374 | 4.49 | 5.14 | 3.97 | 0.98 | putative HTH-type transcriptional regulator |
| *FLPKDCKG_03910* | | + | 4248393 | 4248884 | 20.06 | 20.00 | 20.56 | 3.97 | Hypothetical protein |
| *FLPKDCKG_03911* | *gdhA-2* | - | 4248991 | 4250334 | 2.78 | 6.69 | 7.01 | 0.43 | NADP-specific glutamate dehydrogenase |
| *FLPKDCKG_03912* | *exoA* | - | 4250789 | 4251577 | 3.29 | 7.66 | 6.62 | 1.48 | Exodeoxyribonuclease |
| *FLPKDCKG_03913* | *pyrE* | + | 4251712 | 4252404 | 12.90 | 23.84 | 18.68 | 6.29 | Orotate phosphoribosyltransferase |
| *FLPKDCKG_03914* | | - | 4252894 | 4253145 | 30.88 | 23.97 | 12.97 | 6.67 | Hypothetical protein |
| *FLPKDCKG_03915* | *gatB* | - | 4253273 | 4254730 | 5.17 | 8.52 | 7.48 | 4.18 | Aspartyl/glutamyl-tRNA(Asn/Gln) amidotransferase subunit B |
| *FLPKDCKG_03916* | *gatA-2* | - | 4254733 | 4256271 | 5.92 | 11.12 | 7.28 | 3.61 | Glutamyl-tRNA(Gln) amidotransferase subunit A |
| *FLPKDCKG_03917* | *gatC* | - | 4256275 | 4256583 | 38.88 | 55.54 | 50.33 | 10.49 | Glutamyl-tRNA(Gln) amidotransferase subunit C |
| *FLPKDCKG_03918* | *mreB* | + | 4256812 | 4257855 | 25.99 | 49.30 | 47.40 | 18.90 | Rod shape-determining protein MreB |
| *FLPKDCKG_03919* | *mreC* | + | 4257940 | 4258824 | 3.58 | 9.70 | 5.98 | 4.71 | Cell shape-determining protein MreC |
| *FLPKDCKG_03920* | *mreD* | + | 4258811 | 4259380 | 1.11 | 2.16 | 3.01 | 2.36 | Rod shape-determining protein MreD |
| *FLPKDCKG_03921* | *mrdA* | + | 4259398 | 4261326 | 0.94 | 3.81 | 2.96 | 1.13 | Peptidoglycan D,D-transpeptidase MrdA |
| *FLPKDCKG_03922* | *mrdB* | + | 4261323 | 4262459 | 1.41 | 3.75 | 4.24 | 1.11 | Peptidoglycan glycosyltransferase MrdB |
| *FLPKDCKG_03923* | *ldh* | - | 4263284 | 4264333 | 29.82 | 10.97 | 7.11 | 2.90 | L-lactate dehydrogenase |
| *FLPKDCKG_03924* | *ybhI-1* | - | 4264346 | 4265839 | 44.22 | 19.66 | 12.42 | 5.95 | Inner membrane protein YbhI |
| *FLPKDCKG_03925* | *slmA-2* | - | 4266126 | 4266701 | 4.13 | 10.00 | 10.34 | 4.24 | Nucleoid occlusion factor SlmA |
| *FLPKDCKG_03926* | *gph-1* | - | 4266757 | 4267503 | 5.48 | 13.05 | 7.31 | 2.80 | Phosphoglycolate phosphatase |
| *FLPKDCKG_03927* | *argB* | - | 4267503 | 4268405 | 28.97 | 28.17 | 34.07 | 15.74 | Acetylglutamate kinase |
| *FLPKDCKG_03928* | *metQ-5* | + | 4268741 | 4269529 | 38.64 | 67.04 | 73.24 | 23.81 | D-methionine-binding lipoprotein MetQ |
| *FLPKDCKG_03929* | *resA-3* | - | 4269729 | 4270223 | 33.08 | 40.46 | 31.52 | 13.86 | Thiol-disulfide oxidoreductase ResA |

| Locus name | Gene name | Strand | Gene start (bp) | Gene end (bp) | Average TPM Inoculum | Average TPM $t_6$ | Average TPM $t_{24}$ | Average TPM $t_{168}$ | Annotation description |
|---|---|---|---|---|---|---|---|---|---|
| *FLPKDCKG_03930* | | - | 4270220 | 4270966 | 80.61 | 93.81 | 62.61 | 19.67 | Hypothetical protein |
| *FLPKDCKG_03931* | *gmhA* | - | 4270963 | 4271556 | 35.95 | 47.25 | 32.68 | 9.02 | Phosphoheptose isomerase |
| *FLPKDCKG_03932* | | - | 4271587 | 4272066 | 0.00 | 0.00 | 0.00 | 0.48 | Hypothetical protein |
| *FLPKDCKG_03933* | *rsmI-2* | + | 4272133 | 4273062 | 9.20 | 13.60 | 8.28 | 1.24 | Ribosomal RNA small subunit methyltransferase I |
| *FLPKDCKG_03934* | *ogt-1* | + | 4273066 | 4273683 | 6.54 | 12.46 | 16.55 | 5.93 | Methylated-DNA--protein-cysteine methyltransferase |
| *FLPKDCKG_03935* | *rlpA* | - | 4273687 | 4274643 | 8.54 | 7.92 | 11.01 | 1.57 | Endolytic peptidoglycan transglycosylase RlpA |
| *FLPKDCKG_03936* | *rapZ* | - | 4274640 | 4275518 | 47.05 | 45.96 | 28.35 | 9.11 | RNase adapter protein RapZ |
| *FLPKDCKG_03937* | *cc4-3* | + | 4275790 | 4276467 | 5.04 | 11.54 | 8.06 | 1.67 | Cytochrome c4 |
| *FLPKDCKG_03938* | *adhB-1* | + | 4276479 | 4277762 | 4.45 | 7.25 | 3.92 | 1.77 | Alcohol dehydrogenase cytochrome c subunit |
| *FLPKDCKG_03939* | *lysN-3* | - | 4277763 | 4279196 | 1.94 | 5.29 | 3.74 | 1.74 | 2-aminoadipate transaminase |
| *FLPKDCKG_03940* | *hprK* | - | 4279218 | 4280144 | 8.15 | 15.24 | 16.06 | 3.24 | HPr kinase/phosphorylase |
| *FLPKDCKG_03941* | *ptsN* | - | 4280164 | 4280619 | 32.21 | 104.72 | 56.08 | 14.62 | Nitrogen regulatory protein |
| *FLPKDCKG_03942* | *hpf* | - | 4281006 | 4281341 | 654.84 | 588.73 | 410.22 | 146.40 | Ribosome hibernation promoting factor |
| *FLPKDCKG_03943* | *lptB-7* | - | 4281562 | 4282356 | 4.35 | 15.61 | 12.26 | 4.99 | Lipopolysaccharide export system ATP-binding protein LptB |
| *FLPKDCKG_03944* | *lptA* | - | 4282389 | 4283030 | 24.90 | 67.92 | 43.10 | 16.73 | Lipopolysaccharide export system protein LptA |
| *FLPKDCKG_03945* | *lptC* | - | 4283027 | 4283659 | 22.74 | 66.08 | 47.01 | 16.63 | Lipopolysaccharide export system protein LptC |
| *FLPKDCKG_03946* | | - | 4283691 | 4284299 | 14.43 | 27.47 | 23.50 | 7.95 | 3-deoxy-D-manno-octulosonate 8-phosphate phosphatase KdsC |
| *FLPKDCKG_03947* | *kdsD-2* | - | 4284296 | 4285285 | 21.06 | 28.50 | 14.99 | 5.48 | Arabinose 5-phosphate isomerase KdsD |
| *FLPKDCKG_03948* | *purT* | + | 4285391 | 4286608 | 10.75 | 10.06 | 8.08 | 5.32 | Formate-dependent phosphoribosylglycinamide formyltransferase |
| *FLPKDCKG_03949* | | + | 4286879 | 4287424 | 8.36 | 6.53 | 1.08 | 1.34 | Hypothetical protein |
| *FLPKDCKG_03950* | *cydA* | + | 4287554 | 4289128 | 18.15 | 6.53 | 6.82 | 2.53 | Cytochrome bd-I ubiquinol oxidase subunit 1 |
| *FLPKDCKG_03951* | *cydB-2* | + | 4289142 | 4290293 | 16.81 | 7.02 | 8.70 | 2.93 | Cytochrome bd-I ubiquinol oxidase subunit 2 |
| *FLPKDCKG_03952* | *cydX* | + | 4290309 | 4290434 | 12.37 | 7.37 | 0.00 | 0.92 | Cytochrome bd ubiquinol oxidase subunit X |
| *FLPKDCKG_03953* | *cydD* | + | 4290431 | 4292176 | 3.76 | 4.19 | 1.72 | 1.71 | ATP-binding/permease protein CydD |
| *FLPKDCKG_03954* | | + | 4292173 | 4293870 | 1.96 | 3.25 | 3.53 | 0.14 | putative ABC transporter ATP-binding protein |
| *FLPKDCKG_03955* | *apt* | + | 4293925 | 4294476 | 3.50 | 9.55 | 6.38 | 0.23 | Adenine phosphoribosyltransferase |
| *FLPKDCKG_03956* | *dmlR-21* | - | 4294507 | 4295397 | 7.21 | 11.70 | 7.20 | 3.29 | HTH-type transcriptional regulator DmlR |
| *FLPKDCKG_03957* | | + | 4295772 | 4296155 | 41.65 | 66.06 | 56.60 | 9.42 | Hypothetical protein |
| *FLPKDCKG_03958* | | + | 4296586 | 4297710 | 1.35 | 4.47 | 6.95 | 2.48 | Outer membrane porin protein |
| *FLPKDCKG_03959* | *ilvA-2* | - | 4297871 | 4299379 | 2.38 | 4.09 | 3.53 | 1.59 | L-threonine dehydratase biosynthetic IlvA |
| *FLPKDCKG_03960* | *yqaA* | + | 4299545 | 4300069 | 4.78 | 8.29 | 5.57 | 0.92 | Inner membrane protein YqaA |
| *FLPKDCKG_03961* | | + | 4300202 | 4304164 | 11.87 | 19.17 | 10.25 | 3.33 | putative FAD-linked oxidoreductase |
| *FLPKDCKG_03962* | | + | 4304328 | 4306199 | 0.41 | 0.14 | 1.22 | 0.19 | Hypothetical protein |
| *FLPKDCKG_03963* | | - | 4306283 | 4307251 | 5.90 | 6.42 | 8.16 | 2.92 | Hypothetical protein |
| *FLPKDCKG_03964* | | - | 4307337 | 4307684 | 12.05 | 13.52 | 12.94 | 2.18 | Hypothetical protein |
| *FLPKDCKG_03965* | | - | 4307687 | 4308055 | 24.97 | 40.38 | 20.26 | 13.50 | Hypothetical protein |
| *FLPKDCKG_03966* | *lysN-4* | + | 4308269 | 4309510 | 2.84 | 11.19 | 5.03 | 3.02 | 2-aminoadipate transaminase |
| *FLPKDCKG_03967* | *yhdW-3* | + | 4309611 | 4310627 | 7.84 | 21.80 | 22.70 | 13.54 | Putative amino-acid ABC transporter-binding protein YhdW |
| *FLPKDCKG_03968* | | - | 4310739 | 4311113 | 5.25 | 11.45 | 11.08 | 5.04 | Hypothetical protein |
| *FLPKDCKG_03969* | *liuE* | - | 4311308 | 4312216 | 20.21 | 29.43 | 20.92 | 7.31 | 3-hydroxy-3-isohexenylglutaryl-CoA/hydroxy-methylglutaryl-CoA lyase |
| *FLPKDCKG_03970* | *dprA* | - | 4312365 | 4313492 | 1.55 | 3.15 | 3.04 | 1.81 | DNA processing protein DprA |
| *FLPKDCKG_03971* | *def1* | + | 4313749 | 4314261 | 42.90 | 76.44 | 105.91 | 18.65 | Peptide deformylase 1 |
| *FLPKDCKG_03972* | *fmt* | + | 4314293 | 4315234 | 1.83 | 12.30 | 5.26 | 1.61 | Methionyl-tRNA formyltransferase |
| *FLPKDCKG_03973* | | - | 4315282 | 4315797 | 3.78 | 22.43 | 22.18 | 2.10 | Hypothetical protein |
| *FLPKDCKG_03974* | *gph-2* | - | 4315823 | 4316449 | 8.19 | 28.62 | 16.42 | 3.61 | Phosphoglycolate phosphatase |
| *FLPKDCKG_03975* | | - | 4316470 | 4317030 | 18.85 | 27.71 | 14.66 | 2.44 | Hypothetical protein |
| *FLPKDCKG_03976* | *yvdD* | - | 4317053 | 4317655 | 25.72 | 30.86 | 15.72 | 6.04 | LOG family protein YvdD |
| *FLPKDCKG_03977* | *bfr-2* | + | 4317883 | 4318443 | 49.91 | 347.52 | 170.31 | 20.93 | Bacterioferritin |
| *FLPKDCKG_03978* | *yybR* | - | 4318499 | 4318885 | 2.64 | 23.99 | 18.52 | 1.71 | putative HTH-type transcriptional regulator YybR |
| *FLPKDCKG_03979* | *sugE-3* | + | 4318981 | 4319298 | 0.35 | 3.13 | 0.62 | 1.45 | Quaternary ammonium compound-resistance protein SugE |
| *FLPKDCKG_03980* | *rimO* | - | 4319353 | 4320669 | 9.51 | 15.61 | 13.05 | 6.60 | Ribosomal protein S12 methylthiotransferase RimO |
| *FLPKDCKG_03981* | *rhtB-5* | - | 4320821 | 4321453 | 9.42 | 12.29 | 10.54 | 3.20 | Homoserine/homoserine lactone efflux protein |
| *FLPKDCKG_03982* | *rsmB-1* | + | 4321583 | 4322920 | 4.16 | 8.59 | 4.91 | 0.69 | Ribosomal RNA small subunit methyltransferase B |
| *FLPKDCKG_03983* | | + | 4322962 | 4323552 | 28.12 | 23.64 | 21.38 | 7.71 | Hypothetical protein |
| *FLPKDCKG_03984* | *gchK-1* | + | 4323549 | 4325870 | 13.04 | 17.67 | 9.28 | 3.02 | Globin-coupled histidine kinase |
| *FLPKDCKG_03985* | *walR* | + | 4325882 | 4326571 | 60.13 | 59.02 | 52.58 | 18.83 | Transcriptional regulatory protein WalR |
| *FLPKDCKG_03986* | *trkA* | + | 4326568 | 4327947 | 11.73 | 19.99 | 13.29 | 5.01 | Trk system potassium uptake protein TrkA |
| *FLPKDCKG_03987* | *trkH* | + | 4328003 | 4329472 | 3.89 | 10.41 | 6.92 | 2.46 | Trk system potassium uptake protein TrkH |
| *FLPKDCKG_03988* | *ybdK* | + | 4329860 | 4331104 | 71.48 | 248.54 | 195.33 | 67.99 | Putative glutamate--cysteine ligase 2 |
| *FLPKDCKG_03989* | *bioC-2* | - | 4331172 | 4332104 | 2.10 | 10.93 | 9.31 | 1.55 | Malonyl-[acyl-carrier protein] O-methyltransferase |
| *FLPKDCKG_03990* | | + | 4332210 | 4332905 | 0.21 | 0.84 | 0.07 | 0.00 | Hypothetical protein |
| *FLPKDCKG_03991* | *trmL* | + | 4333037 | 4333507 | 6.82 | 6.85 | 6.38 | 0.83 | tRNA (cytidine(34)-2'-O)-methyltransferase |
| *FLPKDCKG_03992* | | - | 4333593 | 4334588 | 5.82 | 12.18 | 13.28 | 5.11 | Hypothetical protein |
| *FLPKDCKG_03993* | *gpsA* | - | 4334661 | 4335710 | 3.94 | 11.02 | 16.06 | 3.09 | Glycerol-3-phosphate dehydrogenase [NAD(P)+] |
| *FLPKDCKG_03994* | *secB* | - | 4335729 | 4336271 | 60.56 | 126.78 | 180.10 | 59.02 | Protein-export protein SecB |
| *FLPKDCKG_03995* | *grxC* | - | 4336321 | 4336590 | 86.93 | 143.30 | 145.38 | 46.73 | Glutaredoxin 3 |
| *FLPKDCKG_03996* | *glpE-3* | - | 4336641 | 4337063 | 14.77 | 74.58 | 61.16 | 14.91 | Thiosulfate sulfurtransferase GlpE |
| *FLPKDCKG_03997* | *gpmA* | + | 4337225 | 4337977 | 12.54 | 29.41 | 12.58 | 6.40 | 2,3-bisphosphoglycerate-dependent phosphoglycerate mutase |
| *FLPKDCKG_03998* | *envC* | + | 4337983 | 4339572 | 9.07 | 20.20 | 12.78 | 3.08 | Murein hydrolase activator EnvC |

| Locus name | Gene name | Strand | Gene start (bp) | Gene end (bp) | Average TPM Inoculum | Average TPM $t_6$ | Average TPM $t_{24}$ | Average TPM $t_{168}$ | Annotation description |
|---|---|---|---|---|---|---|---|---|---|
| *FLPKDCKG_03999* | *ctpB* | + | 4339634 | 4341079 | 46.87 | 77.93 | 49.69 | 24.04 | Carboxy-terminal processing protease CtpB |
| *FLPKDCKG_04000* | *moeB* | + | 4341076 | 4341831 | 22.55 | 73.99 | 56.65 | 12.59 | Molybdopterin-synthase adenylyltransferase |
| *FLPKDCKG_04001* | *adrA-2* | + | 4341971 | 4343161 | 2.67 | 3.33 | 2.61 | 1.90 | putative diguanylate cyclase AdrA |
| *FLPKDCKG_04002* | *lutR-3* | + | 4343250 | 4343969 | 4.89 | 13.49 | 15.69 | 2.92 | HTH-type transcriptional regulator LutR |
| *FLPKDCKG_04003* | | + | 4343966 | 4345378 | 3.05 | 4.30 | 3.44 | 2.16 | putative FAD-linked oxidoreductase |
| *FLPKDCKG_04004* | *thiC* | - | 4345577 | 4347490 | 21.73 | 24.22 | 23.76 | 18.79 | Phosphomethylpyrimidine synthase |
| *FLPKDCKG_04005* | | + | 4347850 | 4348422 | 226.43 | 459.38 | 248.28 | 54.36 | Hypothetical protein |
| *FLPKDCKG_04006* | | - | 4348502 | 4348879 | 7.34 | 15.80 | 9.54 | 3.80 | Hypothetical protein |
| *FLPKDCKG_04007* | | + | 4349030 | 4349719 | 1.20 | 5.10 | 2.51 | 0.00 | Hypothetical protein |
| *FLPKDCKG_04008* | *proB* | - | 4349888 | 4351024 | 3.99 | 5.84 | 3.57 | 1.90 | Glutamate 5-kinase |
| *FLPKDCKG_04009* | *obg* | - | 4351083 | 4352216 | 3.35 | 5.91 | 8.99 | 5.59 | GTPase Obg |
| *FLPKDCKG_04010* | *rpmA* | - | 4352392 | 4352652 | 33.48 | 77.58 | 73.78 | 35.33 | 50S ribosomal protein L27 |
| *FLPKDCKG_04011* | *rplU* | - | 4352688 | 4352999 | 191.32 | 509.85 | 393.54 | 201.52 | 50S ribosomal protein L21 |
| *FLPKDCKG_04012* | *ispB* | + | 4353483 | 4354409 | 7.26 | 22.01 | 10.86 | 4.92 | Octaprenyl diphosphate synthase |
| *FLPKDCKG_04013* | | + | 4354571 | 4354647 | 9.60 | 29.32 | 1.69 | 0.00 | tRNA |
| *FLPKDCKG_04014* | | + | 4354763 | 4355893 | 0.17 | 47.04 | 33.95 | 2.20 | Hypothetical protein |
| *FLPKDCKG_04015* | *menH-2* | + | 4355880 | 4356638 | 0.97 | 48.63 | 29.71 | 1.14 | 2-succinyl-6-hydroxy-2,4-cyclohexadiene-1-carboxylate synthase |
| *FLPKDCKG_04016* | | - | 4356714 | 4357529 | 0.77 | 21.59 | 7.23 | 1.10 | 2-oxoglutaramate amidase |
| *FLPKDCKG_04017* | | - | 4357542 | 4358405 | 0.15 | 1.24 | 0.72 | 0.00 | Beta-lactamase |
| *FLPKDCKG_04018* | | - | 4358417 | 4358899 | 0.86 | 3.33 | 0.00 | 0.00 | Hypothetical protein |
| *FLPKDCKG_04019* | | - | 4358848 | 4359861 | 0.32 | 0.06 | 0.00 | 0.00 | Hypothetical protein |
| *FLPKDCKG_04020* | | - | 4360101 | 4361096 | 1.73 | 3.36 | 3.45 | 0.16 | Hypothetical protein |
| *FLPKDCKG_04021* | *cimH* | - | 4361086 | 4362450 | 2.69 | 7.84 | 6.79 | 1.93 | Citrate/malate transporter |
| *FLPKDCKG_04022* | *cynR-12* | + | 4362546 | 4363484 | 2.30 | 4.51 | 1.47 | 1.62 | HTH-type transcriptional regulator CynR |
| *FLPKDCKG_04023* | *qorA-2* | - | 4363501 | 4364532 | 0.65 | 1.10 | 1.71 | 0.67 | Quinone oxidoreductase 1 |
| *FLPKDCKG_04024* | *besA* | + | 4364842 | 4365813 | 0.67 | 0.00 | 0.00 | 0.04 | Ferri-bacillibactin esterase BesA |
| *FLPKDCKG_04025* | *gcvA-11* | + | 4365927 | 4366790 | 1.76 | 5.31 | 2.45 | 1.07 | Glycine cleavage system transcriptional activator |
| *FLPKDCKG_04026* | *nolA-2* | - | 4366806 | 4367192 | 6.01 | 7.05 | 15.63 | 7.35 | Nodulation protein NolA |
| *FLPKDCKG_04027* | | + | 4367283 | 4368188 | 0.17 | 2.97 | 3.51 | 0.62 | Arylesterase |
| *FLPKDCKG_04028* | | - | 4368227 | 4368661 | 14.18 | 29.48 | 13.67 | 6.68 | Hypothetical protein |
| *FLPKDCKG_04029* | | + | 4368851 | 4369297 | 4.25 | 3.34 | 4.86 | 1.74 | Hypothetical protein |
| *FLPKDCKG_04030* | *mtnB* | + | 4369484 | 4370122 | 1.27 | 18.83 | 22.22 | 1.67 | Methylthioribulose-1-phosphate dehydratase |
| *FLPKDCKG_04031* | *ygbM* | + | 4370174 | 4370965 | 0.33 | 5.84 | 5.95 | 2.52 | Putative hydroxypyruvate isomerase YgbM |
| *FLPKDCKG_04032* | | + | 4370962 | 4371939 | 0.70 | 9.42 | 12.56 | 3.83 | putative epimerase/dehydratase |
| *FLPKDCKG_04033* | *Hgd* | + | 4371936 | 4372841 | 0.33 | 6.04 | 5.32 | 0.80 | 2-(hydroxymethyl)glutarate dehydrogenase |
| *FLPKDCKG_04034* | | + | 4372838 | 4373758 | 1.12 | 5.39 | 6.54 | 0.98 | Hypothetical protein |
| *FLPKDCKG_04035* | | + | 4373704 | 4374096 | 1.23 | 16.19 | 12.78 | 2.96 | Hypothetical protein |
| *FLPKDCKG_04036* | *lutR-4* | - | 4374143 | 4374850 | 1.95 | 8.26 | 4.96 | 1.01 | HTH-type transcriptional regulator LutR |
| *FLPKDCKG_04037* | *tsaR* | - | 4374919 | 4375809 | 5.62 | 11.21 | 7.62 | 3.78 | HTH-type transcriptional regulator TsaR |
| *FLPKDCKG_04038* | *proP-5* | + | 4375955 | 4377286 | 0.74 | 1.18 | 1.64 | 0.33 | Proline/betaine transporter |
| *FLPKDCKG_04039* | | + | 4377286 | 4378377 | 1.15 | 1.05 | 3.48 | 1.03 | Hypothetical protein |
| *FLPKDCKG_04040* | | + | 4378482 | 4380611 | 0.48 | 1.51 | 12.59 | 14.77 | putative TonB-dependent receptor |
| *FLPKDCKG_04041* | *hyi* | + | 4380686 | 4381471 | 6.41 | 4.68 | 7.18 | 2.01 | Hydroxypyruvate isomerase |
| *FLPKDCKG_04042* | | + | 4381591 | 4382574 | 2.54 | 2.69 | 0.35 | 0.50 | Hypothetical protein |
| *FLPKDCKG_04043* | *rhtA* | - | 4382625 | 4383524 | 2.47 | 1.33 | 0.32 | 0.61 | Threonine/homoserine exporter RhtA |
| *FLPKDCKG_04044* | *rhtB-6* | - | 4383547 | 4384191 | 15.13 | 3.11 | 4.19 | 2.04 | Homoserine/homoserine lactone efflux protein |
| *FLPKDCKG_04045* | *decR-4* | + | 4384385 | 4384861 | 4.89 | 8.22 | 5.41 | 0.24 | DNA-binding transcriptional activator DecR |
| *FLPKDCKG_04046* | *rhtB-7* | + | 4384939 | 4385547 | 7.71 | 15.22 | 6.32 | 1.62 | Homoserine/homoserine lactone efflux protein |
| *FLPKDCKG_04047* | *wecE* | - | 4385593 | 4386744 | 24.44 | 39.71 | 23.52 | 5.13 | dTDP-4-amino-4,6-dideoxygalactose transaminase |
| *FLPKDCKG_04048* | | - | 4386984 | 4387059 | 2.26 | 3.57 | 5.20 | 3.86 | tRNA |
| *FLPKDCKG_04049* | *hemA* | + | 4387343 | 4388617 | 27.45 | 30.39 | 18.82 | 6.91 | Glutamyl-tRNA reductase |
| *FLPKDCKG_04050* | *prfA* | + | 4388712 | 4389782 | 2.10 | 5.36 | 3.88 | 1.62 | Peptide chain release factor RF1 |
| *FLPKDCKG_04051* | *prmC* | + | 4389820 | 4390647 | 0.55 | 4.41 | 3.20 | 0.53 | Release factor glutamine methyltransferase |
| *FLPKDCKG_04052* | *grxD* | + | 4390787 | 4391113 | 57.94 | 48.16 | 33.73 | 17.88 | Glutaredoxin 4 |
| *FLPKDCKG_04053* | *ubiX* | + | 4391118 | 4391681 | 9.92 | 29.50 | 7.97 | 2.07 | Flavin prenyltransferase UbiX |
| *FLPKDCKG_04054* | | - | 4391793 | 4392866 | 6.08 | 6.47 | 7.25 | 1.88 | Hypothetical protein |
| *FLPKDCKG_04055* | *metQ-6* | + | 4393047 | 4393844 | 1.68 | 8.17 | 24.26 | 9.52 | D-methionine-binding lipoprotein MetQ |
| *FLPKDCKG_04056* | *mtnD* | + | 4393848 | 4394387 | 0.87 | 4.87 | 8.13 | 5.20 | Acireductone dioxygenase |
| *FLPKDCKG_04057* | *cpnA* | - | 4394497 | 4395312 | 0.21 | 1.43 | 0.94 | 0.61 | Cyclopentanol dehydrogenase |
| *FLPKDCKG_04058* | *citA-2* | - | 4395309 | 4396097 | 0.42 | 1.99 | 0.20 | 0.55 | Putative citrate synthase 2 |
| *FLPKDCKG_04059* | *fadD3-1* | - | 4396094 | 4397599 | 0.30 | 0.65 | 1.06 | 0.53 | 3-[(3aS,4S,7aS)-7a-methyl-1,5-dioxo-octahydro-1H-inden-4-yl]propanoyl:CoA ligase |
| *FLPKDCKG_04060* | | - | 4397611 | 4398600 | 0.48 | 0.00 | 0.43 | 0.16 | Hypothetical protein |
| *FLPKDCKG_04061* | *allR-2* | + | 4398703 | 4399506 | 1.14 | 4.27 | 6.60 | 2.27 | HTH-type transcriptional repressor AllR |
| *FLPKDCKG_04062* | | - | 4399557 | 4400102 | 1.53 | 20.94 | 38.72 | 13.79 | Hypothetical protein |
| *FLPKDCKG_04063* | *sir* | - | 4400095 | 4401831 | 3.11 | 41.73 | 83.24 | 23.72 | Sulfite reductase [ferredoxin] |
| *FLPKDCKG_04064* | *stp-3* | - | 4401991 | 4403520 | 0.29 | 0.46 | 1.47 | 0.32 | Multidrug resistance protein Stp |
| *FLPKDCKG_04065* | *gltC-14* | + | 4403645 | 4404508 | 3.87 | 9.41 | 3.65 | 0.96 | HTH-type transcriptional regulator GltC |
| *FLPKDCKG_04066* | | - | 4404586 | 4404777 | 13.75 | 10.67 | 13.38 | 5.09 | Hypothetical protein |
| *FLPKDCKG_04067* | *gltC-15* | - | 4404862 | 4405782 | 0.80 | 2.92 | 0.50 | 0.85 | HTH-type transcriptional regulator GltC |

| Locus name | Gene name | Strand | Gene start (bp) | Gene end (bp) | Average TPM Inoculum | Average TPM $t_6$ | Average TPM $t_{24}$ | Average TPM $t_{168}$ | Annotation description |
|---|---|---|---|---|---|---|---|---|---|
| *FLPKDCKG_04068* | | + | 4406104 | 4407438 | 1.29 | 19.14 | 11.32 | 1.79 | Hypothetical protein |
| *FLPKDCKG_04069* | | - | 4407601 | 4408185 | 221.16 | 448.37 | 247.72 | 57.09 | Hypothetical protein |
| *FLPKDCKG_04070* | *ubiG-2* | - | 4408274 | 4409026 | 4.29 | 15.46 | 11.02 | 3.45 | Ubiquinone biosynthesis O-methyltransferase |
| *FLPKDCKG_04071* | *glmS-2* | + | 4409525 | 4410292 | 1.28 | 7.67 | 4.85 | 0.93 | Glutamine--fructose-6-phosphate aminotransferase [isomerizing] |
| *FLPKDCKG_04072* | | + | 4410829 | 4411854 | 9.54 | 28.12 | 21.64 | 5.05 | putative glycosyltransferase |
| *FLPKDCKG_04073* | | + | 4411906 | 4412982 | 3.78 | 18.29 | 12.28 | 2.62 | Hypothetical protein |
| *FLPKDCKG_04074* | | + | 4412960 | 4414540 | 3.95 | 9.87 | 5.72 | 1.44 | Hypothetical protein |
| *FLPKDCKG_04075* | | + | 4414542 | 4415336 | 2.43 | 8.89 | 5.46 | 1.37 | Hypothetical protein |
| *FLPKDCKG_04076* | *iolE* | + | 4415333 | 4416169 | 0.99 | 3.17 | 5.46 | 0.84 | Inosose dehydratase |
| *FLPKDCKG_04077* | *arnE-2* | + | 4416166 | 4416522 | 1.87 | 8.83 | 7.55 | 2.64 | 4-amino-4-deoxy-L-arabinose-phosphoundecaprenol flippase subunit ArnE |
| *FLPKDCKG_04078* | | + | 4416533 | 4417342 | 1.82 | 6.90 | 3.63 | 1.54 | Undecaprenyl-phosphate mannosyltransferase |
| *FLPKDCKG_04079* | | - | 4417753 | 4417962 | 0.00 | 0.00 | 0.00 | 0.00 | Hypothetical protein |
| *FLPKDCKG_04080* | | - | 4418317 | 4419135 | 0.49 | 1.43 | 0.43 | 0.71 | Hypothetical protein |
| *FLPKDCKG_04081* | *yibF-2* | - | 4419489 | 4420094 | 10.39 | 3.80 | 2.26 | 0.41 | putative GST-like protein YibF |
| *FLPKDCKG_04082* | *kstD* | - | 4420120 | 4421826 | 1.28 | 0.92 | 0.63 | 0.09 | 3-oxosteroid 1-dehydrogenase |
| *FLPKDCKG_04083* | *butA* | - | 4421832 | 4422572 | 0.74 | 0.44 | 0.13 | 0.00 | Diacetyl reductase [(S)-acetoin forming] |
| *FLPKDCKG_04084* | *gno-4* | - | 4422581 | 4423336 | 0.73 | 0.00 | 0.00 | 0.03 | Gluconate 5-dehydrogenase |
| *FLPKDCKG_04085* | *livF-19* | - | 4423346 | 4424065 | 0.93 | 0.37 | 0.00 | 0.81 | High-affinity branched-chain amino acid transport ATP-binding protein LivF |
| *FLPKDCKG_04086* | *araG* | - | 4424058 | 4424822 | 1.54 | 0.97 | 0.71 | 0.00 | Arabinose import ATP-binding protein AraG |
| *FLPKDCKG_04087* | *livH-18* | - | 4424819 | 4425823 | 1.35 | 0.41 | 0.00 | 0.00 | High-affinity branched-chain amino acid transport system permease protein LivH |
| *FLPKDCKG_04088* | *livH-19* | - | 4425820 | 4426722 | 1.33 | 0.38 | 0.74 | 0.06 | High-affinity branched-chain amino acid transport system permease protein LivH |
| *FLPKDCKG_04089* | *braC-6* | - | 4426784 | 4427995 | 6.44 | 1.99 | 2.18 | 0.60 | Leucine-, isoleucine-, valine-, threonine-, and alanine-binding protein |
| *FLPKDCKG_04090* | *fabG-11* | - | 4428088 | 4428879 | 5.91 | 5.58 | 0.00 | 0.00 | 3-oxoacyl-[acyl-carrier-protein] reductase FabG |
| *FLPKDCKG_04091* | *gntR-2* | + | 4429085 | 4429762 | 26.34 | 12.28 | 13.15 | 4.79 | putative D-xylose utilization operon transcriptional repressor |
| *FLPKDCKG_04092* | *cynT* | - | 4430272 | 4430922 | 17.15 | 58.12 | 31.78 | 11.04 | Carbonic anhydrase 1 |
| *FLPKDCKG_04093* | | - | 4431439 | 4431774 | 34.89 | 70.79 | 29.23 | 17.76 | Hypothetical protein |
| *FLPKDCKG_04094* | *psiF* | + | 4432153 | 4432521 | 444.03 | 456.07 | 269.32 | 62.96 | Phosphate starvation-inducible protein PsiF |
| *FLPKDCKG_04095* | | - | 4432702 | 4433742 | 0.89 | 3.91 | 1.48 | 1.19 | Hypothetical protein |
| *FLPKDCKG_04096* | *catD-3* | - | 4433746 | 4434525 | 2.94 | 6.62 | 5.89 | 2.55 | 3-oxoadipate enol-lactonase 2 |
| *FLPKDCKG_04097* | | - | 4434581 | 4434877 | 3.43 | 9.00 | 7.93 | 0.00 | Hypothetical protein |
| *FLPKDCKG_04098* | *catC* | - | 4434890 | 4435165 | 14.07 | 18.20 | 13.23 | 4.86 | Muconolactone Delta-isomerase |
| *FLPKDCKG_04099* | *pcaJ-2* | - | 4435218 | 4435862 | 3.64 | 4.43 | 2.68 | 1.68 | 3-oxoadipate CoA-transferase subunit B |
| *FLPKDCKG_04100* | *pcaI-2* | - | 4435859 | 4436548 | 6.25 | 8.36 | 9.33 | 1.64 | 3-oxoadipate CoA-transferase subunit A |
| *FLPKDCKG_04101* | *pcaR-1* | + | 4436718 | 4437500 | 2.01 | 4.80 | 2.29 | 0.59 | Pca regulon regulatory protein |
| *FLPKDCKG_04102* | *fabG-12* | - | 4437551 | 4438279 | 1.59 | 1.37 | 3.32 | 1.36 | 3-oxoacyl-[acyl-carrier-protein] reductase FabG |
| *FLPKDCKG_04103* | *atoE* | - | 4438431 | 4439771 | 0.58 | 2.06 | 1.53 | 0.18 | Putative short-chain fatty acid transporter |
| *FLPKDCKG_04104* | *hbd-1* | - | 4439957 | 4440892 | 3.87 | 4.01 | 2.31 | 1.14 | 3-hydroxybutyryl-CoA dehydrogenase |
| *FLPKDCKG_04105* | *ppc* | - | 4441238 | 4444084 | 4.59 | 7.84 | 7.44 | 2.87 | Phosphoenolpyruvate carboxylase |
| *FLPKDCKG_04106* | *queC* | + | 4444256 | 4444969 | 18.35 | 22.36 | 11.54 | 4.12 | 7-cyano-7-deazaguanine synthase |
| *FLPKDCKG_04107* | *cph2-4* | + | 4445268 | 4446926 | 0.65 | 0.50 | 1.61 | 0.38 | Phytochrome-like protein cph2 |
| *FLPKDCKG_04108* | *mdtL-2* | - | 4446904 | 4448085 | 0.66 | 1.10 | 4.01 | 1.34 | Multidrug resistance protein MdtL |
| *FLPKDCKG_04109* | *bar* | - | 4448082 | 4448627 | 4.87 | 8.81 | 6.52 | 3.13 | Phosphinothricin N-acetyltransferase |
| *FLPKDCKG_04110* | *cmpR-13* | + | 4448753 | 4449643 | 2.51 | 6.59 | 3.76 | 2.40 | HTH-type transcriptional activator CmpR |
| *FLPKDCKG_04111* | *leuA-2* | - | 4449591 | 4450628 | 0.80 | 2.94 | 1.44 | 0.00 | 2-isopropylmalate synthase |
| *FLPKDCKG_04112* | *nimR-8* | + | 4450871 | 4451899 | 1.97 | 5.28 | 0.60 | 0.17 | HTH-type transcriptional regulator NimR |
| *FLPKDCKG_04113* | *ilvD-2* | + | 4452013 | 4453878 | 13.49 | 12.34 | 15.15 | 6.17 | Dihydroxy-acid dehydratase |
| *FLPKDCKG_04114* | *dmlR-22* | + | 4454035 | 4454916 | 2.61 | 4.48 | 1.69 | 0.60 | HTH-type transcriptional regulator DmlR |
| *FLPKDCKG_04115* | *lpd-2* | - | 4454980 | 4456377 | 0.21 | 0.30 | 0.00 | 0.33 | Dihydrolipoyl dehydrogenase |
| *FLPKDCKG_04116* | *sucB-2* | - | 4456386 | 4457039 | 0.27 | 1.04 | 0.00 | 0.62 | Dihydrolipoyllysine-residue succinyltransferase component of 2-oxoglutarate dehydrogenase complex |
| *FLPKDCKG_04117* | | - | 4457665 | 4458156 | 0.68 | 0.85 | 0.00 | 0.00 | Hypothetical protein |
| *FLPKDCKG_04118* | *acoB* | - | 4458153 | 4460336 | 0.49 | 0.00 | 0.41 | 0.20 | Acetoin:2,6-dichlorophenolindophenol oxidoreductase subunit beta |
| *FLPKDCKG_04119* | *bacC-5* | - | 4460366 | 4461154 | 0.88 | 0.00 | 1.53 | 0.24 | Dihydroanticapsin 7-dehydrogenase |
| *FLPKDCKG_04120* | *livF-20* | - | 4461183 | 4461902 | 0.37 | 1.20 | 0.00 | 0.00 | High-affinity branched-chain amino acid transport ATP-binding protein LivF |
| *FLPKDCKG_04121* | *lptB-8* | - | 4461899 | 4462651 | 0.65 | 0.00 | 0.00 | 0.33 | Lipopolysaccharide export system ATP-binding protein LptB |
| *FLPKDCKG_04122* | | - | 4462648 | 4463922 | 0.33 | 0.00 | 0.00 | 0.00 | Hypothetical protein |
| *FLPKDCKG_04123* | *livH-20* | - | 4463919 | 4464860 | 0.42 | 1.06 | 0.78 | 0.00 | High-affinity branched-chain amino acid transport system permease protein LivH |
| *FLPKDCKG_04124* | *braC-7* | - | 4464999 | 4466234 | 1.11 | 2.11 | 2.52 | 0.71 | Leucine-, isoleucine-, valine-, threonine-, and alanine-binding protein |
| *FLPKDCKG_04125* | *aroE-2* | - | 4466275 | 4467096 | 0.80 | 0.73 | 0.70 | 0.59 | Shikimate dehydrogenase (NADP(+)) |
| *FLPKDCKG_04126* | *ephD* | + | 4467335 | 4467664 | 4.46 | 2.15 | 8.98 | 1.09 | putative oxidoreductase EphD |
| *FLPKDCKG_04127* | *fabG-13* | + | 4467664 | 4468071 | 3.70 | 5.50 | 3.44 | 2.92 | 3-oxoacyl-[acyl-carrier-protein] reductase FabG |
| *FLPKDCKG_04128* | *lutR-5* | + | 4468224 | 4468976 | 5.17 | 6.42 | 4.24 | 0.69 | HTH-type transcriptional regulator LutR |
| *FLPKDCKG_04129* | *iolG* | - | 4469036 | 4470373 | 31.30 | 24.25 | 19.85 | 6.49 | Inositol 2-dehydrogenase/D-chiro-inositol 3-dehydrogenase |
| *FLPKDCKG_04130* | | - | 4470478 | 4471038 | 4.31 | 5.31 | 6.29 | 0.89 | Putative 5'(3')-deoxyribonucleotidase |
| *FLPKDCKG_04131* | *thpR* | - | 4471095 | 4471625 | 11.51 | 18.74 | 11.67 | 1.14 | RNA 2',3'-cyclic phosphodiesterase |
| *FLPKDCKG_04132* | *srpC-3* | - | 4472530 | 4474017 | 0.69 | 1.41 | 1.20 | 0.55 | Solvent efflux pump outer membrane protein SrpC |
| *FLPKDCKG_04133* | *bepE-2* | - | 4474087 | 4477302 | 0.70 | 1.12 | 1.08 | 0.36 | Efflux pump membrane transporter BepE |
| *FLPKDCKG_04134* | *bepF* | - | 4477339 | 4478529 | 1.15 | 3.01 | 2.64 | 0.93 | Efflux pump periplasmic linker BepF |
| *FLPKDCKG_04135* | *pgrR-7* | + | 4478925 | 4479989 | 46.02 | 60.06 | 47.79 | 12.38 | HTH-type transcriptional regulator PgrR |
| *FLPKDCKG_04136* | | + | 4480168 | 4481289 | 4.80 | 6.90 | 7.52 | 2.15 | Hypothetical protein |

| Locus name | Gene name | Strand | Gene start (bp) | Gene end (bp) | Average TPM Inoculum | Average TPM $t_6$ | Average TPM $t_{24}$ | Average TPM $t_{168}$ | Annotation description |
|---|---|---|---|---|---|---|---|---|---|
| *FLPKDCKG_04137* | | + | 4481402 | 4482169 | 13.33 | 6.60 | 3.77 | 3.71 | Hypothetical protein |
| *FLPKDCKG_04138* | *pcaR-2* | + | 4482374 | 4483201 | 7.28 | 4.77 | 0.22 | 0.60 | Pca regulon regulatory protein |
| *FLPKDCKG_04139* | *dapA-4* | - | 4483253 | 4484155 | 0.94 | 1.16 | 1.42 | 0.39 | 4-hydroxy-tetrahydrodipicolinate synthase |
| *FLPKDCKG_04140* | *potH-4* | - | 4484192 | 4485967 | 0.42 | 0.22 | 0.39 | 0.39 | Putrescine transport system permease protein PotH |
| *FLPKDCKG_04141* | *potA-6* | - | 4485972 | 4487060 | 0.42 | 0.50 | 0.78 | 0.26 | Spermidine/putrescine import ATP-binding protein PotA |
| *FLPKDCKG_04142* | *potD-2* | - | 4487083 | 4488165 | 0.97 | 1.70 | 0.44 | 1.00 | Spermidine/putrescine-binding periplasmic protein |
| *FLPKDCKG_04143* | | - | 4488205 | 4489632 | 1.00 | 2.60 | 2.17 | 0.57 | putative FAD-linked oxidoreductase |
| *FLPKDCKG_04144* | *mdlB-4* | + | 4489853 | 4491001 | 2.12 | 0.84 | 1.03 | 0.90 | (S)-mandelate dehydrogenase |
| *FLPKDCKG_04145* | *proC-2* | + | 4491020 | 4491826 | 1.28 | 1.55 | 0.32 | 0.83 | Pyrroline-5-carboxylate reductase |
| *FLPKDCKG_04146* | *stp-4* | - | 4491841 | 4493286 | 0.63 | 1.36 | 0.00 | 0.14 | Multidrug resistance protein Stp |
| *FLPKDCKG_04147* | | + | 4493381 | 4494319 | 1.45 | 2.67 | 3.06 | 0.19 | Hypothetical protein |
| *FLPKDCKG_04148* | *yijE-9* | + | 4494398 | 4495294 | 0.17 | 0.45 | 2.11 | 0.52 | putative cystine transporter YijE |
| *FLPKDCKG_04149* | *yqjI* | + | 4495402 | 4496262 | 5.69 | 7.00 | 6.26 | 1.66 | Transcriptional regulator YqjI |
| *FLPKDCKG_04150* | *yqjH-2* | + | 4496289 | 4497104 | 2.94 | 5.92 | 3.06 | 2.66 | NADPH-dependent ferric-chelate reductase |
| *FLPKDCKG_04151* | *benM-3* | - | 4497189 | 4498115 | 1.20 | 0.67 | 0.36 | 0.27 | HTH-type transcriptional regulator BenM |
| *FLPKDCKG_04152* | *Dml* | + | 4498172 | 4499032 | 1.37 | 4.09 | 2.81 | 0.26 | 2,3-dimethylmalate lyase |
| *FLPKDCKG_04153* | | - | 4499077 | 4501953 | 1.74 | 6.34 | 7.30 | 2.52 | Hypothetical protein |
| *FLPKDCKG_04154* | | - | 4501950 | 4505051 | 2.82 | 9.42 | 7.93 | 2.64 | Hypothetical protein |
| *FLPKDCKG_04155* | *rpoE-2* | - | 4505254 | 4505832 | 0.58 | 2.95 | 0.72 | 0.00 | ECF RNA polymerase sigma-E factor |
| *FLPKDCKG_04156* | *uvrY-2* | - | 4506108 | 4506746 | 9.24 | 9.15 | 4.35 | 1.51 | Response regulator UvrY |
| *FLPKDCKG_04157* | *tmoS-1* | + | 4506812 | 4508233 | 2.93 | 5.11 | 2.66 | 0.61 | Sensor histidine kinase TmoS |
| *FLPKDCKG_04158* | *yhjX-1* | + | 4508318 | 4509568 | 3.60 | 3.60 | 4.05 | 1.00 | putative MFS-type transporter YhjX |
| *FLPKDCKG_04159* | | - | 4509624 | 4510004 | 9.90 | 6.35 | 4.47 | 2.63 | Hypothetical protein |
| *FLPKDCKG_04160* | | + | 4510150 | 4510389 | 6.71 | 16.66 | 7.94 | 1.46 | Hypothetical protein |
| *FLPKDCKG_04161* | *allA* | + | 4510386 | 4510934 | 2.00 | 5.00 | 3.67 | 0.99 | Ureidoglycolate lyase |
| *FLPKDCKG_04162* | | + | 4511122 | 4512579 | 6.22 | 3.95 | 5.23 | 1.63 | Hypothetical protein |
| *FLPKDCKG_04163* | | - | 4512632 | 4513327 | 5.56 | 11.93 | 6.45 | 1.93 | Hypothetical protein |
| *FLPKDCKG_04164* | | + | 4513600 | 4514193 | 7.41 | 5.99 | 2.25 | 1.26 | Hypothetical protein |
| *FLPKDCKG_04165* | | + | 4514441 | 4514908 | 45.69 | 15.66 | 16.27 | 2.54 | Hypothetical protein |
| *FLPKDCKG_04166* | | + | 4514914 | 4515387 | 11.75 | 15.05 | 7.58 | 1.49 | Hypothetical protein |
| *FLPKDCKG_04167* | | + | 4515387 | 4515677 | 33.45 | 8.57 | 5.98 | 1.99 | Hypothetical protein |
| *FLPKDCKG_04168* | | + | 4515735 | 4517033 | 20.68 | 10.80 | 6.39 | 0.47 | Hypothetical protein |
| *FLPKDCKG_04169* | | + | 4517030 | 4517374 | 35.11 | 12.27 | 8.62 | 3.56 | Hypothetical protein |
| *FLPKDCKG_04170* | | + | 4517371 | 4519332 | 25.01 | 8.54 | 5.99 | 0.53 | Hypothetical protein |
| *FLPKDCKG_04171* | | + | 4519335 | 4520063 | 15.00 | 5.23 | 4.58 | 1.02 | Hypothetical protein |
| *FLPKDCKG_04172* | | + | 4520066 | 4520845 | 7.51 | 1.25 | 4.03 | 1.98 | Hypothetical protein |
| *FLPKDCKG_04173* | | + | 4520842 | 4521447 | 25.41 | 12.47 | 8.66 | 1.71 | Hypothetical protein |
| *FLPKDCKG_04174* | | + | 4521444 | 4525076 | 15.66 | 4.61 | 3.05 | 0.47 | Hypothetical protein |
| *FLPKDCKG_04175* | | + | 4525081 | 4525446 | 36.32 | 4.79 | 8.65 | 1.91 | Hypothetical protein |
| *FLPKDCKG_04176* | | + | 4525427 | 4525663 | 52.31 | 14.19 | 20.30 | 5.25 | Hypothetical protein |
| *FLPKDCKG_04177* | | + | 4525660 | 4526256 | 16.38 | 6.70 | 5.42 | 1.47 | Hypothetical protein |
| *FLPKDCKG_04178* | *cynR-13* | + | 4526398 | 4527327 | 5.86 | 3.96 | 2.73 | 0.83 | HTH-type transcriptional regulator CynR |
| *FLPKDCKG_04179* | *bdhA-3* | - | 4527346 | 4528164 | 8.21 | 9.27 | 2.52 | 0.91 | D-beta-hydroxybutyrate dehydrogenase |
| *FLPKDCKG_04180* | | - | 4528195 | 4529394 | 8.47 | 1.73 | 1.46 | 0.15 | Hypothetical protein |
| *FLPKDCKG_04181* | *kce-2* | - | 4529447 | 4530382 | 8.13 | 5.15 | 1.48 | 1.04 | 3-keto-5-aminohexanoate cleavage enzyme |
| *FLPKDCKG_04182* | *ppiC-2* | - | 4530732 | 4531013 | 27.51 | 90.16 | 50.94 | 18.84 | Peptidyl-prolyl cis-trans isomerase C |
| *FLPKDCKG_04183* | *yhdW-4* | - | 4531213 | 4532229 | 6.06 | 4.36 | 4.30 | 0.89 | Putative amino-acid ABC transporter-binding protein YhdW |
| *FLPKDCKG_04184* | | + | 4532521 | 4532844 | 3.03 | 60.08 | 17.78 | 12.10 | Hypothetical protein |
| *FLPKDCKG_04185* | | - | 4532916 | 4533518 | 0.37 | 3.96 | 1.85 | 0.94 | Hypothetical protein |
| *FLPKDCKG_04186* | | + | 4533884 | 4534192 | 9.80 | 23.44 | 13.73 | 3.77 | Hypothetical protein |
| *FLPKDCKG_04187* | | + | 4534189 | 4534491 | 7.67 | 13.13 | 9.53 | 6.98 | Hypothetical protein |
| *FLPKDCKG_04188* | | - | 4534547 | 4534631 | 42.05 | 17.04 | 18.71 | 4.68 | tRNA |
| *FLPKDCKG_04189* | *xanP* | - | 4534750 | 4536141 | 2.03 | 3.29 | 1.33 | 1.53 | Xanthine permease XanP |
| *FLPKDCKG_04190* | | + | 4536292 | 4537695 | 2.34 | 2.77 | 2.48 | 0.19 | Hypothetical protein |
| *FLPKDCKG_04191* | *htpG* | + | 4537897 | 4539786 | 7.26 | 10.26 | 22.18 | 12.55 | Chaperone protein HtpG |
| *FLPKDCKG_04192* | *hipA* | + | 4540050 | 4541564 | 1.57 | 2.70 | 2.60 | 0.85 | Serine/threonine-protein kinase HipA |
| *FLPKDCKG_04193* | | - | 4542058 | 4542261 | 1.10 | 0.00 | 0.00 | 2.55 | Hypothetical protein |
| *FLPKDCKG_04194* | *fhuA-11* | - | 4542784 | 4545147 | 0.34 | 0.53 | 0.83 | 2.62 | Ferrichrome-iron receptor |
| *FLPKDCKG_04195* | | - | 4545189 | 4546649 | 0.00 | 0.79 | 1.46 | 1.26 | Hypothetical protein |
| *FLPKDCKG_04196* | | - | 4546660 | 4548048 | 0.23 | 1.05 | 0.88 | 1.73 | Hypothetical protein |
| *FLPKDCKG_04197* | *ybhF-3* | - | 4548045 | 4548755 | 0.18 | 2.20 | 1.31 | 3.57 | putative ABC transporter ATP-binding protein YbhF |
| *FLPKDCKG_04198* | *dgoT* | - | 4548988 | 4550298 | 2.12 | 4.00 | 2.50 | 0.79 | D-galactonate transporter |
| *FLPKDCKG_04199* | *dgoD-1* | - | 4550331 | 4551479 | 6.79 | 0.13 | 0.54 | 1.76 | D-galactonate dehydratase |
| *FLPKDCKG_04200* | *dgoA* | - | 4551476 | 4552129 | 2.98 | 0.79 | 0.00 | 1.98 | 2-dehydro-3-deoxy-6-phosphogalactonate aldolase |
| *FLPKDCKG_04201* | *dgoK1* | - | 4552126 | 4553142 | 3.19 | 0.00 | 0.00 | 0.29 | putative 2-dehydro-3-deoxygalactonokinase DgoK1 |
| *FLPKDCKG_04202* | *kdgR-6* | + | 4553400 | 4554239 | 5.50 | 0.69 | 1.59 | 0.88 | Transcriptional regulator KdgR |
| *FLPKDCKG_04203* | *sigM* | + | 4554432 | 4554938 | 1.38 | 2.83 | 2.92 | 0.87 | ECF RNA polymerase sigma factor SigM |
| *FLPKDCKG_04204* | | + | 4555011 | 4556030 | 0.99 | 3.35 | 2.34 | 0.54 | Hypothetical protein |
| *FLPKDCKG_04205* | *tdhA* | + | 4556496 | 4559504 | 0.65 | 1.59 | 0.43 | 0.20 | TonB-dependent heme receptor A |

| Locus name | Gene name | Strand | Gene start (bp) | Gene end (bp) | Average TPM Inoculum | Average TPM $t_6$ | Average TPM $t_{24}$ | Average TPM $t_{168}$ | Annotation description |
|---|---|---|---|---|---|---|---|---|---|
| *FLPKDCKG_04206* | *hasA-1* | + | 4559647 | 4560249 | 1.04 | 0.43 | 1.80 | 0.70 | Hemophore HasA |
| *FLPKDCKG_04207* | *hasA-2* | + | 4560488 | 4561099 | 1.70 | 3.68 | 3.73 | 0.71 | Hemophore HasA |
| *FLPKDCKG_04208* | *prsD* | + | 4561231 | 4563153 | 0.30 | 0.23 | 0.76 | 0.32 | Type I secretion system ATP-binding protein PrsD |
| *FLPKDCKG_04209* | *prsE-2* | + | 4563150 | 4564496 | 0.18 | 1.06 | 0.00 | 0.00 | Type I secretion system membrane fusion protein PrsE |
| *FLPKDCKG_04210* | *tolC-2* | + | 4564493 | 4565851 | 0.59 | 0.88 | 0.36 | 0.00 | Outer membrane protein TolC |
| *FLPKDCKG_04211* | *oxyR-3* | - | 4565982 | 4566917 | 2.76 | 6.13 | 4.36 | 1.62 | Hydrogen peroxide-inducible genes activator |
| *FLPKDCKG_04212* | *aldA* | + | 4567016 | 4568491 | 0.11 | 0.27 | 0.42 | 0.15 | Putative aldehyde dehydrogenase AldA |
| *FLPKDCKG_04213* | *strE* | + | 4568497 | 4569528 | 0.55 | 0.00 | 0.69 | 0.19 | dTDP-glucose 4,6-dehydratase |
| *FLPKDCKG_04214* | | + | 4569636 | 4570595 | 0.14 | 0.79 | 0.12 | 0.00 | Hypothetical protein |
| *FLPKDCKG_04215* | | + | 4570784 | 4572802 | 1.90 | 3.19 | 1.90 | 0.82 | Hypothetical protein |
| *FLPKDCKG_04216* | | + | 4572841 | 4573377 | 1.52 | 10.73 | 2.07 | 0.04 | Hypothetical protein |
| *FLPKDCKG_04217* | | + | 4573421 | 4573972 | 2.81 | 10.18 | 7.12 | 0.20 | Hypothetical protein |
| *FLPKDCKG_04218* | *gstB-6* | - | 4574051 | 4574671 | 10.53 | 25.67 | 22.23 | 6.28 | Glutathione S-transferase GstB |
| *FLPKDCKG_04219* | *ampD* | - | 4574689 | 4575267 | 12.63 | 21.02 | 22.97 | 2.85 | 1,6-anhydro-N-acetylmuramyl-L-alanine amidase AmpD |
| *FLPKDCKG_04220* | | - | 4575475 | 4575741 | 3.37 | 1.42 | 2.79 | 3.38 | Hypothetical protein |
| *FLPKDCKG_04221* | *ypjD* | - | 4575743 | 4576231 | 3.03 | 3.62 | 4.45 | 3.89 | Inner membrane protein YpjD |
| *FLPKDCKG_04222* | *ffh* | + | 4576690 | 4578096 | 19.50 | 26.56 | 18.38 | 5.93 | Signal recognition particle protein |
| *FLPKDCKG_04223* | | - | 4578280 | 4578726 | 110.88 | 265.22 | 206.37 | 98.34 | Hypothetical protein |
| *FLPKDCKG_04224* | | - | 4578773 | 4579090 | 99.49 | 234.21 | 121.71 | 54.50 | Hypothetical protein |
| *FLPKDCKG_04225* | *thiS* | - | 4579341 | 4579541 | 4.41 | 9.85 | 11.33 | 0.00 | Sulfur carrier protein ThiS |
| *FLPKDCKG_04226* | *pip* | - | 4579552 | 4580493 | 13.89 | 17.71 | 12.95 | 5.35 | Proline iminopeptidase |
| *FLPKDCKG_04227* | *trmB* | - | 4580493 | 4581260 | 32.85 | 26.65 | 24.74 | 8.00 | tRNA (guanine-N(7)-)-methyltransferase |
| *FLPKDCKG_04228* | *gltC-16* | + | 4581442 | 4582353 | 0.57 | 1.39 | 2.00 | 1.86 | HTH-type transcriptional regulator GltC |
| *FLPKDCKG_04229* | *smtA* | + | 4582498 | 4583691 | 5.89 | 9.40 | 5.27 | 0.26 | Succinyl-CoA--L-malate CoA-transferase alpha subunit |
| *FLPKDCKG_04230* | *yngG-1* | + | 4583732 | 4584694 | 6.31 | 1.04 | 3.45 | 0.43 | Hydroxymethylglutaryl-CoA lyase YngG |
| *FLPKDCKG_04231* | | + | 4584756 | 4585742 | 11.26 | 7.35 | 4.59 | 0.38 | Hypothetical protein |
| *FLPKDCKG_04232* | | + | 4585822 | 4585895 | 29.60 | 47.76 | 41.94 | 14.21 | tRNA |
| *FLPKDCKG_04233* | | + | 4586150 | 4587103 | 2.67 | 2.68 | 3.94 | 1.00 | Hypothetical protein |
| *FLPKDCKG_04234* | | - | 4587135 | 4588109 | 1.43 | 4.62 | 3.18 | 2.15 | Hypothetical protein |
| *FLPKDCKG_04235* | *dgoD-2* | - | 4588138 | 4589343 | 1.48 | 1.50 | 0.87 | 1.29 | D-galactonate dehydratase |
| *FLPKDCKG_04236* | *citZ* | + | 4589465 | 4590658 | 6.89 | 9.72 | 5.64 | 1.69 | Citrate synthase 2 |
| *FLPKDCKG_04237* | *gatA-3* | - | 4590693 | 4592237 | 1.15 | 1.48 | 2.59 | 0.34 | Glutamyl-tRNA(Gln) amidotransferase subunit A |
| *FLPKDCKG_04238* | *proA-5* | + | 4592432 | 4593133 | 1.74 | 3.75 | 0.91 | 0.61 | 4-hydroxy-4-methyl-2-oxoglutarate aldolase/4-carboxy-4-hydroxy-2-oxoadipate aldolase |
| *FLPKDCKG_04239* | *rlmN-2* | + | 4593244 | 4594290 | 1.88 | 4.24 | 3.17 | 0.21 | putative dual-specificity RNA methyltransferase RlmN |
| *FLPKDCKG_04240* | *fnr-2* | - | 4594343 | 4595041 | 4.45 | 5.42 | 5.36 | 2.66 | Anaerobic regulatory protein |
| *FLPKDCKG_04241* | | + | 4595257 | 4596480 | 14.83 | 1.21 | 0.84 | 0.44 | Hypothetical protein |
| *FLPKDCKG_04242* | *kce-3* | + | 4596491 | 4597405 | 12.69 | 0.44 | 0.52 | 1.19 | 3-keto-5-aminohexanoate cleavage enzyme |
| *FLPKDCKG_04243* | *fabG-14* | + | 4597440 | 4598216 | 7.13 | 2.48 | 2.40 | 0.45 | 3-oxoacyl-[acyl-carrier-protein] reductase FabG |
| *FLPKDCKG_04244* | *fumB-2* | - | 4598241 | 4599797 | 0.64 | 0.29 | 0.28 | 0.22 | Fumarate hydratase class I, anaerobic |
| *FLPKDCKG_04245* | *yhjE-4* | - | 4599861 | 4601219 | 0.25 | 0.33 | 0.00 | 0.00 | Inner membrane metabolite transport protein YhjE |
| *FLPKDCKG_04246* | *nagL-3* | - | 4601310 | 4601981 | 0.19 | 0.73 | 0.00 | 0.00 | Maleylpyruvate isomerase |
| *FLPKDCKG_04247* | | - | 4601995 | 4603017 | 0.39 | 0.84 | 1.19 | 0.42 | Hypothetical protein |
| *FLPKDCKG_04248* | *ccrA* | - | 4603077 | 4604036 | 0.43 | 1.01 | 0.43 | 0.88 | Metallo-beta-lactamase type 2 |
| *FLPKDCKG_04249* | | - | 4604073 | 4604534 | 0.87 | 0.56 | 0.00 | 0.00 | Hypothetical protein |
| *FLPKDCKG_04250* | | - | 4604685 | 4604891 | 1.12 | 1.49 | 0.00 | 0.00 | Hypothetical protein |
| *FLPKDCKG_04251* | *mhpA-2* | - | 4604866 | 4606566 | 0.26 | 0.21 | 0.65 | 0.00 | 3-(3-hydroxy-phenyl)propionate/3-hydroxycinnamic acid hydroxylase |
| *FLPKDCKG_04252* | *uvrY-3* | + | 4606724 | 4607509 | 6.19 | 6.63 | 4.10 | 2.43 | Response regulator UvrY |
| *FLPKDCKG_04253* | *rstB-4* | + | 4607752 | 4609053 | 0.37 | 0.43 | 0.82 | 0.00 | Sensor protein RstB |
| *FLPKDCKG_04254* | *argH-4* | + | 4609216 | 4610751 | 3.91 | 13.78 | 13.66 | 2.01 | Argininosuccinate lyase |
| *FLPKDCKG_04255* | *fpvA-5* | - | 4610838 | 4613213 | 1.33 | 3.32 | 2.31 | 0.50 | Ferripyoverdine receptor |
| *FLPKDCKG_04256* | | - | 4613390 | 4614358 | 1.13 | 0.00 | 0.57 | 0.35 | Hypothetical protein |
| *FLPKDCKG_04257* | *fecI-10* | - | 4614352 | 4614852 | 1.78 | 1.62 | 2.91 | 0.25 | putative RNA polymerase sigma factor FecI |
| *FLPKDCKG_04258* | *prpF* | - | 4614983 | 4616167 | 3.97 | 10.43 | 10.37 | 5.58 | 2-methyl-aconitate isomerase |
| *FLPKDCKG_04259* | *acnD* | - | 4616232 | 4618520 | 7.90 | 18.93 | 10.24 | 5.67 | 2-methylcitrate dehydratase (2-methyl-trans-aconitate forming) |
| *FLPKDCKG_04260* | *acnM* | - | 4618502 | 4618849 | 4.66 | 13.41 | 4.12 | 2.28 | Aconitate hydratase A |
| *FLPKDCKG_04261* | *gntR-3* | - | 4618846 | 4619556 | 9.45 | 17.17 | 12.74 | 3.98 | putative D-xylose utilization operon transcriptional repressor |
| *FLPKDCKG_04262* | *ykgO* | + | 4619772 | 4619912 | 1.58 | 0.60 | 8.35 | 3.36 | 50S ribosomal protein L36 2 |
| *FLPKDCKG_04263* | | + | 4619919 | 4620029 | 0.78 | 4.98 | 6.13 | 3.61 | Hypothetical protein |
| *FLPKDCKG_04264* | | + | 4620041 | 4620271 | 3.80 | 6.37 | 4.12 | 0.20 | Hypothetical protein |
| *FLPKDCKG_04265* | | - | 4620343 | 4620510 | 39.86 | 64.45 | 32.87 | 19.43 | Hypothetical protein |
| *FLPKDCKG_04266* | | + | 4620711 | 4621151 | 0.95 | 0.66 | 0.00 | 0.00 | Hypothetical protein |
| *FLPKDCKG_04267* | *amaB-4* | - | 4621148 | 4622410 | 2.27 | 1.64 | 2.11 | 0.81 | N-carbamoyl-L-amino acid hydrolase |
| *FLPKDCKG_04268* | *allB* | - | 4622412 | 4623767 | 6.13 | 1.79 | 0.69 | 1.33 | Allantoinase |
| *FLPKDCKG_04269* | *cmpR-14* | + | 4623869 | 4624165 | 21.45 | 21.43 | 8.72 | 6.15 | HTH-type transcriptional activator CmpR |
| *FLPKDCKG_04270* | *hcaR-6* | + | 4624239 | 4624763 | 3.18 | 10.01 | 2.41 | 1.25 | Hca operon transcriptional activator HcaR |
| *FLPKDCKG_04271* | | + | 4624979 | 4626082 | 1.13 | 2.10 | 1.93 | 1.22 | Hypothetical protein |
| *FLPKDCKG_04272* | *hutU-2* | + | 4626093 | 4627748 | 1.08 | 1.03 | 0.96 | 1.59 | Urocanate hydratase |
| *FLPKDCKG_04273* | *proP-6* | + | 4627799 | 4629139 | 0.25 | 0.53 | 0.68 | 0.86 | Proline/betaine transporter |
| *FLPKDCKG_04274* | *rnz* | + | 4629147 | 4630280 | 0.28 | 0.00 | 0.00 | 0.57 | Ribonuclease Z |

| Locus name | Gene name | Strand | Gene start (bp) | Gene end (bp) | Average TPM Inoculum | Average TPM $t_6$ | Average TPM $t_{24}$ | Average TPM $t_{168}$ | Annotation description |
|---|---|---|---|---|---|---|---|---|---|
| *FLPKDCKG_04275* | *hutH-2* | + | 4630298 | 4632025 | 0.09 | 0.16 | 0.97 | 0.91 | Histidine ammonia-lyase |
| *FLPKDCKG_04276* | *yfcG-4* | + | 4632258 | 4632956 | 12.49 | 13.48 | 14.64 | 5.94 | Disulfide-bond oxidoreductase YfcG |
| *FLPKDCKG_04277* | *adhB-2* | + | 4633211 | 4634563 | 3.41 | 2.48 | 1.50 | 0.19 | Alcohol dehydrogenase cytochrome c subunit |
| *FLPKDCKG_04278* | *tsdA-2* | + | 4634556 | 4635455 | 1.34 | 2.41 | 1.79 | 1.66 | Thiosulfate dehydrogenase |
| *FLPKDCKG_04279* | *cynR-14* | + | 4635576 | 4636502 | 3.91 | 2.77 | 3.32 | 0.12 | HTH-type transcriptional regulator CynR |
| *FLPKDCKG_04280* | *ifcA* | + | 4636649 | 4638082 | 0.21 | 0.81 | 0.70 | 0.14 | Fumarate reductase flavoprotein subunit |
| *FLPKDCKG_04281* | | + | 4638069 | 4639274 | 0.07 | 0.27 | 0.00 | 0.21 | Hypothetical protein |
| *FLPKDCKG_04282* | | + | 4639365 | 4640342 | 0.27 | 2.88 | 3.61 | 0.55 | Hypothetical protein |
| *FLPKDCKG_04283* | | - | 4640382 | 4640858 | 1.49 | 8.48 | 5.59 | 0.54 | Hypothetical protein |
| *FLPKDCKG_04284* | | - | 4640913 | 4641632 | 4.63 | 23.77 | 9.58 | 3.39 | Hypothetical protein |
| *FLPKDCKG_04285* | *lysN-5* | - | 4641736 | 4643145 | 2.66 | 3.65 | 2.69 | 1.47 | 2-aminoadipate transaminase |
| *FLPKDCKG_04286* | | - | 4643171 | 4643311 | 1.11 | 2.35 | 0.00 | 1.35 | Hypothetical protein |
| *FLPKDCKG_04287* | *appB* | - | 4643311 | 4644318 | 1.27 | 4.52 | 2.00 | 1.16 | Cytochrome bd-II ubiquinol oxidase subunit 2 |
| *FLPKDCKG_04288* | *appC-2* | - | 4644323 | 4645759 | 1.15 | 2.13 | 0.85 | 0.12 | Cytochrome bd-II ubiquinol oxidase subunit 1 |
| *FLPKDCKG_04289* | | - | 4646071 | 4646757 | 39.83 | 59.41 | 39.99 | 21.27 | Hypothetical protein |
| *FLPKDCKG_04290* | *yedA* | - | 4646889 | 4647752 | 0.93 | 0.81 | 0.13 | 0.65 | putative inner membrane transporter YedA |
| *FLPKDCKG_04291* | *dmlR-23* | + | 4647896 | 4648792 | 1.71 | 7.22 | 6.31 | 0.30 | HTH-type transcriptional regulator DmlR |
| *FLPKDCKG_04292* | *phaB-3* | - | 4648825 | 4649571 | 3.04 | 5.47 | 0.70 | 0.49 | Acetoacetyl-CoA reductase |
| *FLPKDCKG_04293* | | - | 4649561 | 4650043 | 11.20 | 3.07 | 8.47 | 3.40 | Hypothetical protein |
| *FLPKDCKG_04294* | *yhhJ* | - | 4650091 | 4651218 | 2.85 | 7.00 | 8.56 | 0.44 | Inner membrane transport permease YhhJ |
| *FLPKDCKG_04295* | *rbbA* | - | 4651222 | 4653975 | 1.97 | 1.00 | 0.77 | 0.45 | Ribosome-associated ATPase |
| *FLPKDCKG_04296* | *oprM-6* | - | 4653975 | 4656545 | 2.42 | 1.11 | 0.52 | 0.25 | Outer membrane protein OprM |
| *FLPKDCKG_04297* | *phaC-3* | - | 4656548 | 4658068 | 5.26 | 1.20 | 1.10 | 0.19 | Poly(3-hydroxyalkanoate) polymerase subunit PhaC |
| *FLPKDCKG_04298* | *srpR* | + | 4658305 | 4658907 | 2.96 | 2.10 | 0.42 | 0.35 | HTH-type transcriptional regulator SrpR |
| *FLPKDCKG_04299* | *adrB* | + | 4659009 | 4660592 | 3.92 | 4.66 | 3.22 | 0.81 | Putative cyclic-di-GMP phosphodiesterase AdrB |
| *FLPKDCKG_04300* | | - | 4660604 | 4660951 | 2.46 | 4.91 | 1.10 | 0.36 | Hypothetical protein |
| *FLPKDCKG_04301* | | + | 4661049 | 4661711 | 1.54 | 4.09 | 1.08 | 0.70 | Hypothetical protein |
| *FLPKDCKG_04302* | | + | 4661711 | 4661944 | 1.61 | 1.16 | 6.75 | 2.22 | Hypothetical protein |
| *FLPKDCKG_04303* | | + | 4662029 | 4662397 | 1.79 | 0.89 | 4.96 | 0.21 | Hypothetical protein |
| *FLPKDCKG_04304* | *kch* | - | 4662394 | 4663596 | 4.23 | 7.55 | 3.54 | 2.42 | Voltage-gated potassium channel Kch |
| *FLPKDCKG_04305* | *ompR-5* | - | 4664024 | 4664839 | 6.94 | 6.34 | 1.71 | 1.19 | Transcriptional regulatory protein OmpR |
| *FLPKDCKG_04306* | | + | 4665572 | 4665913 | 6.56 | 9.18 | 9.41 | 2.18 | Hypothetical protein |
| *FLPKDCKG_04307* | *oatA-2* | - | 4665894 | 4667099 | 0.80 | 2.17 | 1.19 | 0.64 | O-acetyltransferase OatA |
| *FLPKDCKG_04308* | | + | 4667359 | 4667889 | 6.62 | 12.94 | 6.18 | 2.31 | Hypothetical protein |
| *FLPKDCKG_04309* | | + | 4668028 | 4668822 | 7.09 | 10.33 | 11.81 | 3.76 | Hypothetical protein |
| *FLPKDCKG_04310* | *glpE-4* | + | 4668918 | 4669655 | 1.75 | 2.58 | 3.79 | 1.73 | Thiosulfate sulfurtransferase GlpE |
| *FLPKDCKG_04311* | *metG* | - | 4669742 | 4671808 | 10.38 | 21.84 | 19.10 | 4.11 | Methionine--tRNA ligase |
| *FLPKDCKG_04312* | | - | 4672020 | 4672295 | 241.77 | 984.80 | 746.61 | 161.98 | Hypothetical protein |
| *FLPKDCKG_04313* | *apbC* | + | 4672397 | 4673485 | 10.02 | 25.74 | 18.21 | 6.30 | Iron-sulfur cluster carrier protein |
| *FLPKDCKG_04314* | *tamA* | + | 4673489 | 4675381 | 1.33 | 8.52 | 6.18 | 1.06 | Translocation and assembly module TamA |
| *FLPKDCKG_04315* | *tamB* | + | 4675378 | 4679121 | 1.00 | 6.29 | 4.65 | 0.98 | Translocation and assembly module TamB |
| *FLPKDCKG_04316* | *dcd* | + | 4679213 | 4679776 | 49.71 | 69.15 | 39.55 | 17.36 | dCTP deaminase |
| *FLPKDCKG_04317* | | - | 4679820 | 4680671 | 5.53 | 5.70 | 8.62 | 1.52 | Hypothetical protein |
| *FLPKDCKG_04318* | *vdcA* | + | 4680832 | 4682070 | 1.37 | 3.92 | 1.68 | 0.54 | Diguanylate cyclase VdcA |
| *FLPKDCKG_04319* | *speG-3* | - | 4682040 | 4682612 | 4.79 | 4.72 | 3.18 | 1.47 | Spermidine N(1)-acetyltransferase |
| *FLPKDCKG_04320* | | + | 4682729 | 4683181 | 161.57 | 263.05 | 168.47 | 40.19 | Hypothetical protein |
| *FLPKDCKG_04321* | *adiA* | + | 4683436 | 4685697 | 27.55 | 38.41 | 27.50 | 8.59 | Biodegradative arginine decarboxylase |
| *FLPKDCKG_04322* | *solA* | + | 4685715 | 4686917 | 2.58 | 6.59 | 3.06 | 1.05 | N-methyl-L-tryptophan oxidase |
| *FLPKDCKG_04323* | *sodB-1* | + | 4687114 | 4687497 | 10.81 | 18.12 | 28.83 | 4.22 | Superoxide dismutase [Mn/Fe] |
| *FLPKDCKG_04324* | *sodB-2* | + | 4687530 | 4687739 | 5.56 | 18.31 | 11.99 | 1.90 | Superoxide dismutase [Mn/Fe] |
| *FLPKDCKG_04325* | *zupT-2* | + | 4687767 | 4688660 | 4.49 | 13.00 | 5.46 | 2.21 | Zinc transporter ZupT |
| *FLPKDCKG_04326* | *gcvT* | + | 4689066 | 4690166 | 7.10 | 4.43 | 5.63 | 6.05 | Aminomethyltransferase |
| *FLPKDCKG_04327* | *gcvH* | + | 4690265 | 4690642 | 77.80 | 61.92 | 54.83 | 27.87 | Glycine cleavage system H protein |
| *FLPKDCKG_04328* | *gcvP* | + | 4690734 | 4693607 | 6.12 | 4.17 | 3.59 | 2.68 | Glycine dehydrogenase (decarboxylating) |
| *FLPKDCKG_04329* | | - | 4695323 | 4696237 | 2.51 | 12.13 | 7.19 | 0.93 | Hypothetical protein |
| *FLPKDCKG_04330* | | + | 4696464 | 4696709 | 0.34 | 2.15 | 0.14 | 0.00 | Hypothetical protein |
| *FLPKDCKG_04331* | | - | 4696839 | 4698809 | 24.24 | 46.03 | 40.09 | 12.08 | Hypothetical protein |
| *FLPKDCKG_04332* | | - | 4699019 | 4700494 | 1.06 | 4.82 | 2.11 | 2.17 | Hypothetical protein |
| *FLPKDCKG_04333* | *gatA-4* | - | 4700547 | 4701956 | 1.33 | 2.04 | 4.08 | 2.44 | Glutamyl-tRNA(Gln) amidotransferase subunit A |
| *FLPKDCKG_04334* | *gsiA-9* | - | 4702001 | 4703686 | 1.73 | 3.41 | 3.72 | 1.74 | Glutathione import ATP-binding protein GsiA |
| *FLPKDCKG_04335* | *gsiD-6* | - | 4703692 | 4704597 | 0.70 | 2.92 | 0.41 | 1.52 | Glutathione transport system permease protein GsiD |
| *FLPKDCKG_04336* | *gsiC-8* | - | 4704607 | 4705548 | 0.85 | 4.20 | 2.32 | 2.56 | Glutathione transport system permease protein GsiC |
| *FLPKDCKG_04337* | *gsiB-9* | - | 4705575 | 4707140 | 7.43 | 22.27 | 17.63 | 7.90 | Glutathione-binding protein GsiB |
| *FLPKDCKG_04338* | | - | 4707225 | 4708730 | 0.54 | 3.95 | 4.30 | 0.76 | Hypothetical protein |
| *FLPKDCKG_04339* | *bauC* | - | 4708809 | 4710305 | 1.64 | 7.32 | 5.04 | 2.17 | Putative 3-oxopropanoate dehydrogenase |
| *FLPKDCKG_04340* | *bauA-2* | - | 4710403 | 4711728 | 5.65 | 15.33 | 10.57 | 2.52 | Beta-alanine--pyruvate aminotransferase |
| *FLPKDCKG_04341* | *lysN-6* | + | 4711901 | 4713319 | 6.89 | 4.94 | 3.81 | 3.48 | 2-aminoadipate transaminase |
| *FLPKDCKG_04342* | *dhfrIII* | - | 4713390 | 4713875 | 7.92 | 17.60 | 13.05 | 4.48 | Dihydrofolate reductase type 3 |
| *FLPKDCKG_04343* | | - | 4713990 | 4716053 | 0.80 | 3.73 | 1.97 | 0.73 | putative signaling protein |

| Locus name | Gene name | Strand | Gene start (bp) | Gene end (bp) | Average TPM Inoculum | Average TPM $t_6$ | Average TPM $t_{24}$ | Average TPM $t_{168}$ | Annotation description |
|---|---|---|---|---|---|---|---|---|---|
| *FLPKDCKG_04344* | *thyA* | - | 4716452 | 4717423 | 23.51 | 36.59 | 20.91 | 10.32 | Thymidylate synthase |
| *FLPKDCKG_04345* | | - | 4717474 | 4717914 | 44.11 | 65.52 | 49.44 | 10.50 | Hypothetical protein |
| *FLPKDCKG_04346* | *coq7* | + | 4718031 | 4718708 | 12.30 | 14.57 | 15.76 | 3.06 | 2-nonaprenyl-3-methyl-6-methoxy-1,4-benzoquinol hydroxylase |
| *FLPKDCKG_04347* | *tagO-2* | + | 4719516 | 4720619 | 41.82 | 141.76 | 90.15 | 20.02 | putative undecaprenyl-phosphate N-acetylglucosaminyl 1-phosphate transferase |
| *FLPKDCKG_04348* | *wbpA-2* | + | 4720716 | 4721993 | 15.66 | 79.48 | 57.31 | 17.11 | UDP-N-acetyl-D-glucosamine 6-dehydrogenase |
| *FLPKDCKG_04349* | *wbgU* | + | 4722006 | 4723031 | 6.89 | 37.72 | 36.29 | 8.94 | UDP-N-acetylglucosamine 4-epimerase |
| *FLPKDCKG_04350* | *murJ-1* | + | 4723033 | 4724382 | 0.56 | 10.44 | 9.29 | 2.88 | Lipid II flippase MurJ |
| *FLPKDCKG_04351* | *pglH-1* | + | 4724383 | 4725528 | 2.56 | 20.17 | 15.38 | 3.59 | GalNAc-alpha-(1-4)-GalNAc-alpha-(1-3)-diNAcBac-PP-undecaprenol alpha-1,4-N-acetyl-D-galactosaminyltransferase |
| *FLPKDCKG_04352* | *asnB* | + | 4725575 | 4727506 | 4.08 | 23.84 | 22.48 | 4.03 | Asparagine synthetase [glutamine-hydrolyzing] 1 |
| *FLPKDCKG_04353* | *pglH-2* | + | 4727510 | 4728625 | 0.58 | 8.92 | 10.66 | 0.86 | GalNAc-alpha-(1-4)-GalNAc-alpha-(1-3)-diNAcBac-PP-undecaprenol alpha-1,4-N-acetyl-D-galactosaminyltransferase |
| *FLPKDCKG_04354* | *epsF-3* | + | 4728622 | 4729779 | 0.99 | 6.85 | 6.11 | 2.72 | Putative glycosyltransferase EpsF |
| *FLPKDCKG_04355* | *pglA* | + | 4729776 | 4730909 | 1.33 | 8.43 | 7.28 | 1.22 | N,N'-diacetylbacillosaminyl-diphospho-undecaprenol alpha-1,3-N-acetylgalactosaminyltransferase |
| *FLPKDCKG_04356* | *ggt-2* | - | 4730917 | 4732590 | 5.70 | 7.37 | 7.22 | 2.94 | Gamma-glutamyltranspeptidase |
| *FLPKDCKG_04357* | *pgi-1* | - | 4732673 | 4734232 | 8.41 | 13.96 | 10.15 | 2.19 | Glucose-6-phosphate isomerase |
| *FLPKDCKG_04358* | *algC-1* | - | 4734240 | 4735631 | 30.48 | 57.00 | 23.78 | 10.89 | Phosphomannomutase/phosphoglucomutase |
| *FLPKDCKG_04359* | | + | 4735940 | 4736584 | 25.86 | 31.40 | 27.46 | 3.83 | Hypothetical protein |
| *FLPKDCKG_04360* | | + | 4736679 | 4738094 | 21.59 | 35.98 | 14.71 | 6.80 | putative FAD-linked oxidoreductase |
| *FLPKDCKG_04361* | *qseB-4* | + | 4738133 | 4738846 | 4.81 | 4.06 | 4.29 | 1.30 | Transcriptional regulatory protein QseB |
| *FLPKDCKG_04362* | *tcrY* | + | 4738843 | 4740171 | 3.60 | 6.44 | 3.98 | 0.55 | putative sensor histidine kinase TcrY |
| *FLPKDCKG_04363* | | + | 4740284 | 4741069 | 2.35 | 23.78 | 7.01 | 2.49 | Hypothetical protein |
| *FLPKDCKG_04364* | *dctM-5* | - | 4741021 | 4742304 | 0.51 | 6.01 | 4.83 | 2.33 | C4-dicarboxylate TRAP transporter large permease protein DctM |
| *FLPKDCKG_04365* | *yiaM-3* | - | 4742295 | 4742858 | 0.25 | 1.84 | 1.37 | 1.36 | 2,3-diketo-L-gulonate TRAP transporter small permease protein YiaM |
| *FLPKDCKG_04366* | | - | 4742868 | 4743845 | 0.80 | 10.00 | 5.76 | 6.43 | Solute-binding protein |
| *FLPKDCKG_04367* | | - | 4743966 | 4744847 | 1.15 | 9.41 | 5.77 | 4.84 | 6-deoxy-6-sulfogluconolactonase |
| *FLPKDCKG_04368* | *udh-1* | - | 4744866 | 4745705 | 13.34 | 14.72 | 13.20 | 3.86 | Uronate dehydrogenase |
| *FLPKDCKG_04369* | *lutR-6* | + | 4745823 | 4746551 | 14.02 | 21.04 | 13.30 | 2.84 | HTH-type transcriptional regulator LutR |
| *FLPKDCKG_04370* | *ltaS2-2* | + | 4746781 | 4748676 | 1.42 | 2.68 | 1.56 | 0.52 | Lipoteichoic acid synthase 2 |
| *FLPKDCKG_04371* | *yihG* | + | 4748754 | 4749770 | 1.15 | 0.00 | 0.90 | 0.81 | putative acyltransferase YihG |
| *FLPKDCKG_04372* | | - | 4749781 | 4751100 | 12.65 | 6.36 | 5.92 | 4.56 | Hypothetical protein |
| *FLPKDCKG_04373* | *hmgA* | - | 4751219 | 4752517 | 53.89 | 15.15 | 12.25 | 5.10 | Homogentisate 1,2-dioxygenase |
| *FLPKDCKG_04374* | *pcpR-8* | + | 4752683 | 4753600 | 1.04 | 4.59 | 0.37 | 0.23 | PCP degradation transcriptional activation protein |
| *FLPKDCKG_04375* | | - | 4753566 | 4753784 | 3.16 | 0.00 | 0.00 | 0.00 | Hypothetical protein |
| *FLPKDCKG_04376* | *otcC* | - | 4753805 | 4755484 | 3.38 | 1.19 | 0.62 | 1.48 | Anhydrotetracycline monooxygenase |
| *FLPKDCKG_04377* | | - | 4755613 | 4756566 | 16.21 | 2.74 | 4.58 | 3.22 | Metallo-beta-lactamase type 2 |
| *FLPKDCKG_04378* | *mutM* | - | 4756795 | 4757526 | 15.45 | 27.22 | 21.28 | 2.84 | Formamidopyrimidine-DNA glycosylase |
| *FLPKDCKG_04379* | *bepA-5* | + | 4757621 | 4759507 | 26.12 | 38.39 | 22.58 | 6.08 | Beta-barrel assembly-enhancing protease |
| *FLPKDCKG_04380* | *lolB* | + | 4759504 | 4760100 | 5.84 | 13.99 | 12.20 | 5.17 | Outer-membrane lipoprotein LolB |
| *FLPKDCKG_04381* | *tar-5* | + | 4760337 | 4761935 | 2.10 | 0.93 | 0.25 | 0.00 | Methyl-accepting chemotaxis protein II |
| *FLPKDCKG_04382* | *ispE* | + | 4762128 | 4763069 | 28.65 | 78.43 | 53.93 | 17.47 | 4-diphosphocytidyl-2-C-methyl-D-erythritol kinase |
| *FLPKDCKG_04383* | | + | 4763071 | 4763147 | 32.98 | 19.41 | 52.99 | 28.57 | tRNA |
| *FLPKDCKG_04384* | *prs* | + | 4763283 | 4764233 | 12.11 | 26.39 | 26.60 | 11.29 | Ribose-phosphate pyrophosphokinase |
| *FLPKDCKG_04385* | *rplY* | + | 4764361 | 4764975 | 4.73 | 34.18 | 45.40 | 21.04 | 50S ribosomal protein L25 |
| *FLPKDCKG_04386* | *pth* | + | 4765073 | 4765669 | 3.76 | 3.52 | 9.99 | 2.03 | Peptidyl-tRNA hydrolase |
| *FLPKDCKG_04387* | | + | 4765669 | 4766448 | 0.64 | 2.20 | 2.44 | 0.69 | Hypothetical protein |
| *FLPKDCKG_04388* | *ychF* | + | 4766544 | 4767635 | 12.20 | 29.52 | 22.35 | 8.89 | Ribosome-binding ATPase YchF |
| *FLPKDCKG_04389* | *intA-1* | + | 4767832 | 4769043 | 1.46 | 5.01 | 0.96 | 0.22 | Prophage integrase IntA |
| *FLPKDCKG_04390* | *fic* | + | 4769392 | 4770438 | 15.73 | 4.50 | 3.91 | 1.87 | Adenosine monophosphate-protein transferase SoFic |
| *FLPKDCKG_04391* | | + | 4770446 | 4770589 | 18.34 | 5.02 | 4.54 | 1.51 | Hypothetical protein |
| *FLPKDCKG_04392* | | + | 4770586 | 4773672 | 6.65 | 3.56 | 2.70 | 1.29 | Hypothetical protein |
| *FLPKDCKG_04393* | *dpnA* | + | 4773674 | 4776580 | 7.60 | 6.17 | 3.03 | 1.12 | Modification methylase DpnIIB |
| *FLPKDCKG_04394* | | + | 4776596 | 4778146 | 4.69 | 6.98 | 2.79 | 0.92 | Hypothetical protein |
| *FLPKDCKG_04395* | *yjcD* | + | 4778146 | 4780233 | 3.12 | 6.98 | 4.40 | 0.89 | Putative ATP-dependent DNA helicase YjcD |
| *FLPKDCKG_04396* | *xerC-3* | - | 4780312 | 4781175 | 0.34 | 1.77 | 0.86 | 0.00 | Tyrosine recombinase XerC |
| *FLPKDCKG_04397* | | - | 4781777 | 4782934 | 0.43 | 0.00 | 0.61 | 0.00 | Hypothetical protein |
| *FLPKDCKG_04398* | *fhuE-2* | - | 4783297 | 4785588 | 0.41 | 2.72 | 1.25 | 0.69 | FhuE receptor |
| *FLPKDCKG_04399* | *ettA-2* | - | 4785832 | 4787445 | 2.28 | 2.83 | 2.22 | 0.72 | Energy-dependent translational throttle protein EttA |
| *FLPKDCKG_04400* | | - | 4787653 | 4788150 | 1.15 | 1.10 | 2.23 | 0.00 | Hypothetical protein |
| *FLPKDCKG_04401* | *sarZ* | - | 4788455 | 4788901 | 3.48 | 10.96 | 8.81 | 1.17 | HTH-type transcriptional regulator SarZ |
| *FLPKDCKG_04402* | *bmr3-2* | - | 4788905 | 4790503 | 4.76 | 9.00 | 5.26 | 0.03 | Multidrug resistance protein 3 |
| *FLPKDCKG_04403* | | + | 4790841 | 4793339 | 8.82 | 15.62 | 15.31 | 4.42 | Hypothetical protein |
| *FLPKDCKG_04404* | | + | 4793336 | 4793722 | 4.33 | 10.84 | 8.56 | 2.24 | Hypothetical protein |
| *FLPKDCKG_04405* | *ydcO-2* | - | 4793787 | 4795007 | 3.74 | 8.10 | 3.42 | 1.20 | Inner membrane protein YdcO |
| *FLPKDCKG_04406* | *fdx-3* | - | 4795004 | 4795255 | 130.94 | 119.32 | 59.26 | 14.23 | Ferredoxin |
| *FLPKDCKG_04407* | *coaD* | - | 4795433 | 4795936 | 22.04 | 26.80 | 30.99 | 9.32 | Phosphopantetheine adenylyltransferase |
| *FLPKDCKG_04408* | *rsmD* | - | 4795995 | 4796561 | 5.46 | 27.32 | 15.70 | 3.37 | Ribosomal RNA small subunit methyltransferase D |
| *FLPKDCKG_04409* | *ftsY* | + | 4796634 | 4797977 | 8.14 | 16.60 | 10.62 | 6.14 | Signal recognition particle receptor FtsY |
| *FLPKDCKG_04410* | *ygiF* | - | 4798111 | 4799769 | 6.13 | 6.68 | 4.63 | 4.32 | Inorganic triphosphatase |
| *FLPKDCKG_04411* | | + | 4800095 | 4804318 | 0.44 | 1.50 | 0.24 | 0.26 | Extracellular serine protease |
| *FLPKDCKG_04412* | *phnE-1* | - | 4804496 | 4805320 | 1.93 | 4.91 | 1.62 | 1.07 | Phosphate-import permease protein PhnE |

| Locus name | Gene name | Strand | Gene start (bp) | Gene end (bp) | Average TPM Inoculum | Average TPM $t_6$ | Average TPM $t_{24}$ | Average TPM $t_{168}$ | Annotation description |
|---|---|---|---|---|---|---|---|---|---|
| *FLPKDCKG_04413* | *phnE-2* | - | 4805317 | 4806216 | 2.16 | 2.80 | 1.95 | 1.28 | Phosphate-import permease protein PhnE |
| *FLPKDCKG_04414* | *tcyC-3* | - | 4806216 | 4806986 | 3.91 | 8.07 | 2.01 | 1.02 | L-cystine import ATP-binding protein TcyC |
| *FLPKDCKG_04415* | *phnD-1* | - | 4806992 | 4807999 | 21.29 | 23.47 | 16.81 | 2.87 | Phosphate-import protein PhnD |
| *FLPKDCKG_04416* | *gylR-1* | - | 4808213 | 4809028 | 5.82 | 4.62 | 4.16 | 1.42 | Glycerol operon regulatory protein |
| *FLPKDCKG_04417* | *mngB* | - | 4809039 | 4809845 | 15.64 | 12.75 | 13.75 | 5.81 | Mannosyl-3-phosphoglycerate phosphatase |
| *FLPKDCKG_04418* | *ribB-2* | + | 4809993 | 4811168 | 45.13 | 62.60 | 39.55 | 13.07 | 3,4-dihydroxy-2-butanone 4-phosphate synthase |
| *FLPKDCKG_04419* | *ribH* | + | 4811179 | 4811709 | 125.29 | 144.60 | 112.05 | 64.95 | 6,7-dimethyl-8-ribityllumazine synthase |
| *FLPKDCKG_04420* | *nusB* | + | 4811678 | 4812169 | 6.68 | 9.60 | 8.16 | 5.91 | N utilization substance protein B |
| *FLPKDCKG_04421* | *thiL* | + | 4812399 | 4813364 | 1.71 | 2.11 | 5.64 | 2.11 | Thiamine-monophosphate kinase |
| *FLPKDCKG_04422* | *pgpA* | + | 4813514 | 4814020 | 39.75 | 32.85 | 30.67 | 11.03 | Phosphatidylglycerophosphatase A |
| *FLPKDCKG_04423* | *pncC-1* | + | 4814017 | 4814583 | 18.32 | 38.80 | 20.97 | 5.64 | Nicotinamide-nucleotide amidohydrolase PncC |
| *FLPKDCKG_04424* | *pyrF* | - | 4814653 | 4815471 | 16.13 | 27.19 | 16.20 | 2.55 | Orotidine 5'-phosphate decarboxylase |
| *FLPKDCKG_04425* | *ndh* | - | 4815479 | 4816795 | 46.14 | 50.92 | 31.43 | 6.32 | NADH dehydrogenase |
| *FLPKDCKG_04426* | *dgkA* | - | 4816851 | 4817249 | 1.76 | 5.44 | 7.55 | 1.17 | Diacylglycerol kinase |
| *FLPKDCKG_04427* | *por-2* | + | 4817518 | 4819146 | 0.92 | 2.00 | 1.73 | 0.61 | Polyol:NADP oxidoreductase |
| *FLPKDCKG_04428* | *livF-21* | - | 4819177 | 4819887 | 0.00 | 0.00 | 0.00 | 0.00 | High-affinity branched-chain amino acid transport ATP-binding protein LivF |
| *FLPKDCKG_04429* | *lptB-9* | - | 4819884 | 4820663 | 0.05 | 0.00 | 0.00 | 0.00 | Lipopolysaccharide export system ATP-binding protein LptB |
| *FLPKDCKG_04430* | | - | 4820660 | 4821646 | 0.04 | 0.00 | 0.00 | 0.00 | Hypothetical protein |
| *FLPKDCKG_04431* | *livH-21* | - | 4821666 | 4822586 | 0.21 | 0.00 | 0.00 | 0.00 | High-affinity branched-chain amino acid transport system permease protein LivH |
| *FLPKDCKG_04432* | | - | 4822622 | 4823797 | 0.23 | 2.11 | 0.33 | 0.55 | Leu/Ile/Val-binding protein |
| *FLPKDCKG_04433* | | - | 4823848 | 4824849 | 0.37 | 1.25 | 0.03 | 0.33 | Hypothetical protein |
| *FLPKDCKG_04434* | *nbaC* | - | 4824880 | 4825431 | 6.40 | 10.82 | 6.53 | 3.01 | 3-hydroxyanthranilate 3,4-dioxygenase |
| *FLPKDCKG_04435* | *gbpR-3* | - | 4825584 | 4826549 | 2.55 | 4.74 | 3.96 | 0.70 | HTH-type transcriptional regulator GbpR |
| *FLPKDCKG_04436* | *amnD-1* | - | 4826634 | 4827065 | 7.45 | 21.72 | 10.41 | 4.72 | 2-aminomuconate deaminase |
| *FLPKDCKG_04437* | *amnC-2* | - | 4827091 | 4828548 | 1.82 | 0.10 | 0.81 | 0.72 | 2-aminomuconic 6-semialdehyde dehydrogenase |
| *FLPKDCKG_04438* | *hosA* | + | 4828763 | 4829257 | 5.34 | 12.08 | 15.37 | 0.19 | Transcriptional regulator HosA |
| *FLPKDCKG_04439* | | - | 4829336 | 4830283 | 6.58 | 16.39 | 6.70 | 1.02 | Hypothetical protein |
| *FLPKDCKG_04440* | *groL-2* | - | 4830487 | 4832127 | 40.49 | 42.37 | 87.14 | 66.37 | 60 kDa chaperonin |
| *FLPKDCKG_04441* | *groS-2* | - | 4832165 | 4832452 | 47.80 | 70.36 | 109.43 | 81.04 | 10 kDa chaperonin |
| *FLPKDCKG_04442* | | - | 4833220 | 4833981 | 6.67 | 25.55 | 11.27 | 4.51 | Hypothetical protein |
| *FLPKDCKG_04443* | | - | 4834131 | 4834460 | 9.44 | 9.25 | 5.05 | 0.00 | Insertion element IS6110 uncharacterized 12.0 kDa protein |
| *FLPKDCKG_04444* | | + | 4834745 | 4835218 | 1.70 | 2.13 | 1.14 | 1.05 | Hypothetical protein |
| *FLPKDCKG_04445* | | - | 4835739 | 4836410 | 13.35 | 20.04 | 14.22 | 5.65 | Hypothetical protein |
| *FLPKDCKG_04446* | | - | 4837185 | 4837562 | 6.66 | 18.95 | 16.11 | 1.64 | putative peptidase |
| *FLPKDCKG_04447* | *der-3* | + | 4838093 | 4840348 | 5.49 | 6.57 | 5.28 | 0.88 | GTPase Der |
| *FLPKDCKG_04448* | *der-4* | + | 4840368 | 4842218 | 5.90 | 3.21 | 2.21 | 1.55 | GTPase Der |
| *FLPKDCKG_04449* | | - | 4842697 | 4843641 | 15.60 | 19.68 | 10.07 | 4.73 | Hypothetical protein |
| *FLPKDCKG_04450* | *oprM-7* | - | 4843927 | 4845060 | 7.77 | 10.44 | 10.97 | 2.89 | Outer membrane protein OprM |
| *FLPKDCKG_04451* | *ltxD* | - | 4845404 | 4846843 | 3.57 | 9.18 | 4.22 | 0.10 | Leukotoxin export protein LtxD |
| *FLPKDCKG_04452* | *apxIB-2* | - | 4846840 | 4848966 | 3.64 | 6.58 | 5.69 | 0.98 | Toxin RTX-I translocation ATP-binding protein |
| *FLPKDCKG_04453* | *cya* | - | 4849074 | 4855778 | 32.56 | 21.26 | 10.66 | 3.28 | Bifunctional hemolysin/adenylate cyclase |
| *FLPKDCKG_04454* | | - | 4855780 | 4856289 | 17.67 | 15.93 | 4.94 | 1.12 | Hypothetical protein |
| *FLPKDCKG_04455* | | + | 4856872 | 4857354 | 3.73 | 8.43 | 9.70 | 2.73 | Hypothetical protein |
| *FLPKDCKG_04456* | | - | 4857366 | 4857779 | 1.18 | 1.65 | 1.48 | 0.54 | Hypothetical protein |
| *FLPKDCKG_04457* | *betA* | - | 4857779 | 4859392 | 0.84 | 0.80 | 1.18 | 1.12 | Oxygen-dependent choline dehydrogenase |
| *FLPKDCKG_04458* | *phdK* | - | 4859389 | 4860885 | 1.31 | 0.65 | 0.00 | 0.23 | 2-formylbenzoate dehydrogenase |
| *FLPKDCKG_04459* | *ribY* | - | 4860895 | 4861908 | 3.30 | 0.87 | 2.23 | 1.39 | Riboflavin-binding protein RibY |
| *FLPKDCKG_04460* | *ribX-3* | - | 4861932 | 4862738 | 1.36 | 1.52 | 0.50 | 0.73 | Riboflavin transport system permease protein RibX |
| *FLPKDCKG_04461* | *cmpD-2* | - | 4862753 | 4863568 | 1.15 | 2.63 | 2.08 | 0.02 | Bicarbonate transport ATP-binding protein CmpD |
| *FLPKDCKG_04462* | *gstB-7* | - | 4863576 | 4864178 | 3.59 | 2.57 | 0.39 | 1.20 | Glutathione S-transferase GstB |
| *FLPKDCKG_04463* | *amnD-2* | - | 4864215 | 4864592 | 4.19 | 0.57 | 2.49 | 0.76 | 2-aminomuconate deaminase |
| *FLPKDCKG_04464* | | - | 4864594 | 4865730 | 3.16 | 3.75 | 3.07 | 1.66 | Hypothetical protein |
| *FLPKDCKG_04465* | *nagI* | - | 4865742 | 4866869 | 8.93 | 8.47 | 3.75 | 3.00 | Gentisate 1,2-dioxygenase |
| *FLPKDCKG_04466* | *lldR-3* | + | 4867135 | 4867842 | 5.27 | 6.05 | 6.73 | 0.06 | Putative L-lactate dehydrogenase operon regulatory protein |
| *FLPKDCKG_04467* | *apc4-3* | - | 4867902 | 4868801 | 1.07 | 4.82 | 2.07 | 0.74 | Acetophenone carboxylase delta subunit |
| *FLPKDCKG_04468* | *apc4-4* | - | 4868832 | 4869158 | 0.67 | 0.46 | 0.53 | 1.15 | Acetophenone carboxylase delta subunit |
| *FLPKDCKG_04469* | *lprI-2* | + | 4869783 | 4870436 | 15.40 | 33.40 | 15.78 | 2.69 | Lipoprotein LprI |
| *FLPKDCKG_04470* | | + | 4870426 | 4870557 | 14.65 | 20.81 | 26.56 | 2.65 | Hypothetical protein |
| *FLPKDCKG_04471* | *bluF* | + | 4870754 | 4871980 | 18.83 | 35.22 | 14.57 | 2.00 | Blue light- and temperature-regulated antirepressor BluF |
| *FLPKDCKG_04472* | | + | 4871986 | 4872183 | 4.61 | 80.22 | 32.67 | 5.30 | Hypothetical protein |
| *FLPKDCKG_04473* | *blc-2* | - | 4872224 | 4872739 | 3.49 | 170.83 | 100.06 | 9.70 | Outer membrane lipoprotein Blc |
| *FLPKDCKG_04474* | *cfa-4* | - | 4872750 | 4873832 | 4.62 | 238.99 | 131.07 | 20.14 | Cyclopropane-fatty-acyl-phospholipid synthase |
| *FLPKDCKG_04475* | | - | 4873844 | 4874611 | 3.58 | 182.48 | 95.96 | 11.39 | Hypothetical protein |
| *FLPKDCKG_04476* | | - | 4874608 | 4875120 | 0.68 | 28.89 | 20.18 | 6.95 | Hypothetical protein |
| *FLPKDCKG_04477* | *cfa-5* | - | 4875120 | 4876394 | 6.39 | 304.04 | 162.59 | 17.12 | Cyclopropane-fatty-acyl-phospholipid synthase |
| *FLPKDCKG_04478* | | - | 4876391 | 4877167 | 5.61 | 354.92 | 166.53 | 18.05 | Hypothetical protein |
| *FLPKDCKG_04479* | | - | 4877164 | 4878429 | 3.19 | 179.55 | 94.88 | 9.44 | Hypothetical protein |
| *FLPKDCKG_04480* | | - | 4878432 | 4879361 | 7.94 | 497.08 | 250.03 | 21.63 | Hypothetical protein |
| *FLPKDCKG_04481* | | - | 4879433 | 4880185 | 1.04 | 62.14 | 31.39 | 3.48 | Hypothetical protein |

| Locus name | Gene name | Strand | Gene start (bp) | Gene end (bp) | Average TPM Inoculum | Average TPM $t_6$ | Average TPM $t_{24}$ | Average TPM $t_{168}$ | Annotation description |
|---|---|---|---|---|---|---|---|---|---|
| *FLPKDCKG_04482* | *nlhH-2* | - | 4880187 | 4881083 | 0.22 | 27.12 | 14.56 | 1.61 | Carboxylesterase NlhH |
| *FLPKDCKG_04483* | | - | 4881143 | 4881814 | 3.86 | 35.27 | 15.23 | 5.00 | Hypothetical protein |
| *FLPKDCKG_04484* | *sigK-3* | - | 4881811 | 4882356 | 7.48 | 69.90 | 31.15 | 5.65 | ECF RNA polymerase sigma factor SigK |
| *FLPKDCKG_04485* | *bphP-2* | - | 4882514 | 4883650 | 0.72 | 3.45 | 2.24 | 0.42 | Bacteriophytochrome |
| *FLPKDCKG_04486* | *cheB3* | - | 4883647 | 4884282 | 1.49 | 2.23 | 1.30 | 0.59 | Chemotaxis response regulator protein-glutamate methylesterase of group 3 operon |
| *FLPKDCKG_04487* | *cheR-2* | - | 4884279 | 4885124 | 1.65 | 3.97 | 1.37 | 0.86 | Chemotaxis protein methyltransferase |
| *FLPKDCKG_04488* | *barA* | - | 4885140 | 4888544 | 2.51 | 3.42 | 5.20 | 1.06 | Signal transduction histidine-protein kinase BarA |
| *FLPKDCKG_04489* | *phoP-3* | + | 4888713 | 4889147 | 17.68 | 110.52 | 102.95 | 22.01 | Alkaline phosphatase synthesis transcriptional regulatory protein PhoP |
| *FLPKDCKG_04490* | | + | 4889287 | 4889769 | 1.89 | 29.40 | 24.96 | 5.00 | Hypothetical protein |
| *FLPKDCKG_04491* | | - | 4889809 | 4890228 | 4.76 | 22.86 | 9.23 | 3.17 | Hypothetical protein |
| *FLPKDCKG_04492* | *pupA-2* | - | 4890531 | 4893008 | 0.78 | 2.00 | 2.82 | 0.31 | Ferric-pseudobactin 358 receptor |
| *FLPKDCKG_04493* | | - | 4893141 | 4894169 | 0.18 | 0.43 | 0.00 | 0.00 | Hypothetical protein |
| *FLPKDCKG_04494* | *fecI-11* | - | 4894174 | 4894668 | 0.65 | 1.81 | 0.00 | 0.12 | putative RNA polymerase sigma factor FecI |
| *FLPKDCKG_04495* | | - | 4894920 | 4895765 | 3.68 | 2.71 | 4.83 | 1.60 | Hypothetical protein |
| *FLPKDCKG_04496* | | - | 4895877 | 4897952 | 4.72 | 7.56 | 5.34 | 0.95 | Hypothetical protein |
| *FLPKDCKG_04497* | | - | 4898090 | 4898764 | 4.04 | 4.99 | 5.91 | 0.93 | Hypothetical protein |
| *FLPKDCKG_04498* | *dmlR-24* | + | 4898966 | 4899874 | 0.61 | 1.82 | 1.57 | 0.17 | HTH-type transcriptional regulator DmlR |
| *FLPKDCKG_04499* | | + | 4899967 | 4900668 | 0.49 | 2.03 | 0.12 | 0.04 | Hypothetical protein |
| *FLPKDCKG_04500* | *kynB-3* | + | 4900754 | 4901767 | 0.14 | 1.22 | 0.31 | 0.19 | Kynurenine formamidase |
| *FLPKDCKG_04501* | *vraR-1* | + | 4901886 | 4902662 | 0.90 | 1.25 | 1.64 | 0.52 | Response regulator protein VraR |
| *FLPKDCKG_04502* | | + | 4902805 | 4904331 | 1.58 | 1.66 | 0.66 | 0.69 | Hypothetical protein |
| *FLPKDCKG_04503* | *dmdB* | + | 4904448 | 4906097 | 0.96 | 1.10 | 0.00 | 0.39 | 3-methylmercaptopropionyl-CoA ligase |
| *FLPKDCKG_04504* | | + | 4906135 | 4907274 | 1.15 | 4.91 | 4.23 | 1.48 | Outer membrane porin protein 32 |
| *FLPKDCKG_04505* | | - | 4907264 | 4908355 | 0.12 | 1.31 | 0.40 | 0.17 | Hydroxypyruvate reductase |
| *FLPKDCKG_04506* | *tarD-1* | - | 4908371 | 4909546 | 0.46 | 1.38 | 0.37 | 0.43 | D(-)-tartrate dehydratase |
| *FLPKDCKG_04507* | | - | 4909576 | 4910580 | 0.53 | 7.47 | 2.37 | 0.17 | Hypothetical protein |
| *FLPKDCKG_04508* | *dapA-5* | - | 4910658 | 4911581 | 1.74 | 5.66 | 3.97 | 0.45 | 4-hydroxy-tetrahydrodipicolinate synthase |
| *FLPKDCKG_04509* | *yjiE-3* | + | 4911759 | 4912718 | 4.80 | 5.28 | 6.04 | 2.08 | HTH-type transcriptional regulator YjiE |
| *FLPKDCKG_04510* | | + | 4912845 | 4913432 | 6.32 | 36.80 | 12.32 | 8.23 | Hypothetical protein |
| *FLPKDCKG_04511* | | + | 4913523 | 4914041 | 2.25 | 3.00 | 1.85 | 0.74 | Hypothetical protein |
| *FLPKDCKG_04512* | | + | 4914038 | 4914649 | 1.31 | 2.40 | 1.08 | 0.00 | Hypothetical protein |
| *FLPKDCKG_04513* | *cynR-15* | + | 4915244 | 4916155 | 1.12 | 1.25 | 1.96 | 0.07 | HTH-type transcriptional regulator CynR |
| *FLPKDCKG_04514* | *cmpR-15* | - | 4916163 | 4917077 | 1.52 | 2.14 | 1.70 | 0.87 | HTH-type transcriptional activator CmpR |
| *FLPKDCKG_04515* | *aspC-2* | + | 4917151 | 4918350 | 0.25 | 1.25 | 1.13 | 0.00 | Aspartate aminotransferase |
| *FLPKDCKG_04516* | *alsS* | + | 4918457 | 4920064 | 0.15 | 1.53 | 0.78 | 0.35 | Acetolactate synthase |
| *FLPKDCKG_04517* | | + | 4920128 | 4921090 | 0.09 | 0.13 | 0.53 | 0.00 | Hypothetical protein |
| *FLPKDCKG_04518* | *argE-4* | + | 4921090 | 4922250 | 0.40 | 2.33 | 0.74 | 0.26 | Acetylornithine deacetylase |
| *FLPKDCKG_04519* | *eamA-6* | - | 4922267 | 4923238 | 3.13 | 4.51 | 4.82 | 1.61 | putative amino-acid metabolite efflux pump |
| *FLPKDCKG_04520* | *uctC-5* | + | 4923410 | 4924606 | 1.18 | 2.66 | 2.00 | 0.18 | Acetyl-CoA:oxalate CoA-transferase |
| *FLPKDCKG_04521* | | + | 4924687 | 4925646 | 1.59 | 1.30 | 0.00 | 0.00 | Hypothetical protein |
| *FLPKDCKG_04522* | *bcd* | + | 4925692 | 4926897 | 1.12 | 2.75 | 1.09 | 0.93 | Acyl-CoA dehydrogenase, short-chain specific |
| *FLPKDCKG_04523* | | + | 4927015 | 4927512 | 6.41 | 18.40 | 13.39 | 2.08 | Hypothetical protein |
| *FLPKDCKG_04524* | *dsbG* | + | 4927784 | 4928551 | 2.48 | 7.11 | 4.01 | 1.07 | Thiol:disulfide interchange protein DsbG |
| *FLPKDCKG_04525* | | - | 4928568 | 4929044 | 3.32 | 12.38 | 6.59 | 1.89 | Hypothetical protein |
| *FLPKDCKG_04526* | *tmoS-2* | + | 4929214 | 4932579 | 6.68 | 49.39 | 23.41 | 2.95 | Sensor histidine kinase TmoS |
| *FLPKDCKG_04527* | *fecI-12* | + | 4932616 | 4933152 | 2.60 | 9.54 | 8.91 | 0.00 | putative RNA polymerase sigma factor FecI |
| *FLPKDCKG_04528* | | + | 4933158 | 4934141 | 0.18 | 0.67 | 0.26 | 1.06 | Hypothetical protein |
| *FLPKDCKG_04529* | *pupB-2* | + | 4934276 | 4936780 | 0.45 | 1.48 | 0.76 | 0.42 | Ferric-pseudobactin BN7/BN8 receptor |
| *FLPKDCKG_04530* | *ligB* | + | 4936876 | 4939383 | 5.12 | 37.04 | 22.65 | 2.05 | DNA ligase B |
| *FLPKDCKG_04531* | *deaD* | + | 4939380 | 4941836 | 1.59 | 16.16 | 9.82 | 0.42 | ATP-dependent RNA helicase DeaD |
| *FLPKDCKG_04532* | | + | 4941820 | 4942485 | 0.70 | 12.55 | 6.51 | 1.00 | Hypothetical protein |
| *FLPKDCKG_04533* | *msbA-4* | - | 4942569 | 4944311 | 0.06 | 2.86 | 3.59 | 0.23 | Lipid A export ATP-binding/permease protein MsbA |
| *FLPKDCKG_04534* | *irtA-2* | - | 4944308 | 4946044 | 0.45 | 1.17 | 0.65 | 0.00 | Iron import ATP-binding/permease protein IrtA |
| *FLPKDCKG_04535* | | - | 4946041 | 4947231 | 0.59 | 2.66 | 0.85 | 0.00 | Hypothetical protein |
| *FLPKDCKG_04536* | *rcnR* | + | 4947564 | 4947836 | 1.60 | 4.95 | 20.41 | 19.95 | Transcriptional repressor RcnR |
| *FLPKDCKG_04537* | *zitB-2* | + | 4947893 | 4948939 | 0.64 | 1.06 | 0.00 | 3.23 | Zinc transporter ZitB |
| *FLPKDCKG_04538* | | - | 4948958 | 4949386 | 0.37 | 1.57 | 0.60 | 4.23 | Hypothetical protein |
| *FLPKDCKG_04539* | | - | 4949393 | 4949488 | 0.00 | 0.26 | 0.00 | 0.00 | Hypothetical protein |
| *FLPKDCKG_04540* | | + | 4949618 | 4950097 | 9.19 | 50.56 | 37.70 | 12.87 | Hypothetical protein |
| *FLPKDCKG_04541* | *ydaP* | + | 4950322 | 4952091 | 0.38 | 38.91 | 27.61 | 1.88 | Putative thiamine pyrophosphate-containing protein YdaP |
| *FLPKDCKG_04542* | | + | 4952203 | 4952508 | 0.47 | 764.95 | 481.15 | 42.75 | Hypothetical protein |
| *FLPKDCKG_04543* | | + | 4952566 | 4952964 | 2.22 | 874.90 | 634.42 | 53.35 | Hypothetical protein |
| *FLPKDCKG_04544* | | - | 4953041 | 4953784 | 4.66 | 50.28 | 47.69 | 4.42 | Hypothetical protein |
| *FLPKDCKG_04545* | | - | 4953781 | 4954656 | 26.59 | 12.78 | 9.11 | 3.55 | Hypothetical protein |
| *FLPKDCKG_04546* | *novR-3* | + | 4954896 | 4955663 | 2.70 | 3.91 | 8.83 | 2.16 | Decarboxylase NovR |
| *FLPKDCKG_04547* | | - | 4955736 | 4956524 | 15.30 | 29.63 | 17.55 | 2.60 | Hypothetical protein |
| *FLPKDCKG_04548* | *cynR-16* | - | 4956648 | 4957547 | 7.33 | 8.29 | 8.57 | 2.40 | HTH-type transcriptional regulator CynR |
| *FLPKDCKG_04549* | | + | 4957637 | 4958608 | 4.17 | 5.90 | 7.30 | 1.14 | Hydroxypyruvate reductase |
| *FLPKDCKG_04550* | *glxR-3* | + | 4958709 | 4959584 | 19.69 | 26.16 | 14.68 | 1.28 | 2-hydroxy-3-oxopropionate reductase |

| Locus name | Gene name | Strand | Gene start (bp) | Gene end (bp) | Average TPM Inoculum | Average TPM $t_6$ | Average TPM $t_{24}$ | Average TPM $t_{168}$ | Annotation description |
|---|---|---|---|---|---|---|---|---|---|
| *FLPKDCKG_04551* | *yjaB-1* | + | 4959621 | 4960070 | 5.49 | 6.77 | 5.03 | 0.63 | putative N-acetyltransferase YjaB |
| *FLPKDCKG_04552* | *nap* | + | 4960067 | 4960918 | 1.89 | 3.99 | 0.48 | 1.81 | putative carboxylesterase nap |
| *FLPKDCKG_04553* | *dmsA* | - | 4960974 | 4963292 | 6.41 | 3.60 | 4.99 | 1.52 | Dimethyl sulfoxide/trimethylamine N-oxide reductase |
| *FLPKDCKG_04554* | *dapA-6* | + | 4963527 | 4964456 | 45.33 | 26.01 | 35.75 | 17.87 | 4-hydroxy-tetrahydrodipicolinate synthase |
| *FLPKDCKG_04555* | | + | 4964482 | 4965468 | 0.76 | 2.17 | 4.35 | 1.19 | Hypothetical protein |
| *FLPKDCKG_04556* | *cdhR-7* | - | 4965458 | 4966438 | 2.15 | 3.46 | 1.23 | 1.58 | HTH-type transcriptional regulator CdhR |
| *FLPKDCKG_04557* | *inhA-3* | + | 4966580 | 4967716 | 0.36 | 0.00 | 0.49 | 0.18 | Isonitrile hydratase |
| *FLPKDCKG_04558* | *tetA-2* | - | 4967754 | 4969004 | 0.12 | 0.09 | 0.42 | 0.11 | Tetracycline resistance protein, class C |
| *FLPKDCKG_04559* | *pchR-6* | + | 4969065 | 4970018 | 1.73 | 0.94 | 1.02 | 0.42 | Regulatory protein PchR |
| *FLPKDCKG_04560* | *yjhB* | + | 4970155 | 4971417 | 3.54 | 5.05 | 5.14 | 1.40 | Putative metabolite transport protein YjhB |
| *FLPKDCKG_04561* | *clpP-2* | - | 4971481 | 4972086 | 2.07 | 6.57 | 0.00 | 0.43 | ATP-dependent Clp protease proteolytic subunit |
| *FLPKDCKG_04562* | | - | 4972125 | 4972484 | 10.96 | 5.55 | 4.74 | 1.84 | Hypothetical protein |
| *FLPKDCKG_04563* | *sdpR-2* | - | 4972481 | 4972795 | 14.08 | 8.10 | 8.80 | 4.75 | Transcriptional repressor SdpR |
| *FLPKDCKG_04564* | *yeaP* | - | 4972864 | 4974018 | 8.36 | 9.24 | 6.27 | 1.96 | putative diguanylate cyclase YeaP |
| *FLPKDCKG_04565* | *trg-4* | + | 4974044 | 4976035 | 0.50 | 0.79 | 0.85 | 0.00 | Methyl-accepting chemotaxis protein III |
| *FLPKDCKG_04566* | *hpxO-3* | - | 4976082 | 4977293 | 0.11 | 0.00 | 0.49 | 0.20 | FAD-dependent urate hydroxylase |
| *FLPKDCKG_04567* | *nimR-9* | + | 4977384 | 4978220 | 2.44 | 2.99 | 2.85 | 0.66 | HTH-type transcriptional regulator NimR |
| *FLPKDCKG_04568* | *tarD-2* | - | 4978262 | 4979428 | 1.29 | 3.68 | 1.71 | 0.74 | D(-)-tartrate dehydratase |
| *FLPKDCKG_04569* | *dmlR-25* | + | 4979587 | 4980477 | 1.90 | 2.84 | 1.25 | 0.69 | HTH-type transcriptional regulator DmlR |
| *FLPKDCKG_04570* | | - | 4980531 | 4980983 | 2.51 | 2.86 | 0.51 | 0.56 | Hypothetical protein |
| *FLPKDCKG_04571* | | - | 4981045 | 4981395 | 28.75 | 39.89 | 33.13 | 5.57 | Hypothetical protein |
| *FLPKDCKG_04572* | | - | 4981401 | 4981760 | 7.62 | 4.35 | 3.06 | 1.76 | Hypothetical protein |
| *FLPKDCKG_04573* | *dmlR-26* | - | 4981834 | 4982736 | 15.69 | 29.24 | 19.54 | 3.92 | HTH-type transcriptional regulator DmlR |
| *FLPKDCKG_04574* | *dkgB-3* | + | 4982838 | 4983641 | 0.90 | 2.20 | 0.00 | 0.22 | 2,5-diketo-D-gluconic acid reductase B |
| *FLPKDCKG_04575* | *luxA* | + | 4983650 | 4984609 | 0.37 | 1.76 | 0.00 | 0.00 | Alkanal monooxygenase alpha chain |
| *FLPKDCKG_04576* | | + | 4984646 | 4985536 | 0.34 | 0.06 | 2.09 | 0.11 | Hypothetical protein |
| *FLPKDCKG_04577* | *gpr-2* | + | 4985657 | 4986706 | 6.80 | 22.68 | 27.23 | 4.75 | L-glyceraldehyde 3-phosphate reductase |
| *FLPKDCKG_04578* | *narB* | - | 4986775 | 4990992 | 0.12 | 4.54 | 9.52 | 8.04 | Nitrate reductase |
| *FLPKDCKG_04579* | *nasD* | + | 4991399 | 4993951 | 0.08 | 11.52 | 22.23 | 21.87 | Nitrite reductase [NAD(P)H] |
| *FLPKDCKG_04580* | *nirD* | + | 4993951 | 4994355 | 0.14 | 4.43 | 38.32 | 35.53 | Nitrite reductase (NADH) small subunit |
| *FLPKDCKG_04581* | *nasR-1* | + | 4994367 | 4995209 | 0.00 | 3.52 | 15.80 | 11.11 | Nitrate regulatory protein |
| *FLPKDCKG_04582* | *nasR-2* | + | 4995173 | 4995634 | 0.83 | 9.03 | 40.14 | 29.48 | Nitrate regulatory protein |
| *FLPKDCKG_04583* | | - | 4995659 | 4996471 | 4.28 | 15.01 | 15.93 | 10.76 | Hypothetical protein |
| *FLPKDCKG_04584* | *mexR* | + | 4996544 | 4997521 | 2.29 | 4.15 | 3.63 | 2.56 | Multidrug resistance operon repressor |
| *FLPKDCKG_04585* | *plcN-4* | + | 4997886 | 5000129 | 0.53 | 1.09 | 0.68 | 0.96 | Non-hemolytic phospholipase C |
| *FLPKDCKG_04586* | | - | 5000143 | 5001555 | 7.45 | 3.29 | 2.73 | 1.10 | Hypothetical protein |
| *FLPKDCKG_04587* | | - | 5001665 | 5002360 | 3.23 | 11.17 | 3.19 | 0.44 | Pentapeptide repeat protein |
| *FLPKDCKG_04588* | *rcsB-2* | - | 5002392 | 5003090 | 9.81 | 15.30 | 9.01 | 4.09 | Transcriptional regulatory protein RcsB |
| *FLPKDCKG_04589* | | + | 5003466 | 5004263 | 0.96 | 3.16 | 1.48 | 0.18 | Hypothetical protein |
| *FLPKDCKG_04590* | | - | 5004324 | 5004767 | 6.97 | 1.38 | 2.03 | 1.13 | Hypothetical protein |
| *FLPKDCKG_04591* | | + | 5004944 | 5005309 | 4.76 | 0.00 | 0.00 | 0.00 | Hypothetical protein |
| *FLPKDCKG_04592* | | - | 5005329 | 5006333 | 1.28 | 7.35 | 3.32 | 2.06 | Hypothetical protein |
| *FLPKDCKG_04593* | *yfcG-5* | + | 5006450 | 5007058 | 2.00 | 2.13 | 3.90 | 0.45 | Disulfide-bond oxidoreductase YfcG |
| *FLPKDCKG_04594* | | - | 5007072 | 5008394 | 1.71 | 1.97 | 2.04 | 1.85 | Hypothetical protein |
| *FLPKDCKG_04595* | *bigR-2* | + | 5008534 | 5009223 | 2.04 | 5.02 | 7.56 | 1.94 | Biofilm growth-associated repressor |
| *FLPKDCKG_04596* | *mdtL-3* | - | 5009272 | 5010438 | 0.13 | 0.49 | 0.21 | 0.00 | Multidrug resistance protein MdtL |
| *FLPKDCKG_04597* | *nimR-10* | + | 5010529 | 5011317 | 0.94 | 2.37 | 3.39 | 1.11 | HTH-type transcriptional regulator NimR |
| *FLPKDCKG_04598* | | - | 5011314 | 5012420 | 2.53 | 31.42 | 14.14 | 1.53 | Outer membrane porin protein |
| *FLPKDCKG_04599* | | - | 5012689 | 5013885 | 0.71 | 0.29 | 1.54 | 0.31 | Outer membrane porin protein |
| *FLPKDCKG_04600* | | - | 5014177 | 5014485 | 2.60 | 8.13 | 3.41 | 3.01 | Hypothetical protein |
| *FLPKDCKG_04601* | *cyoD-1* | - | 5014972 | 5015370 | 72.13 | 79.49 | 33.25 | 5.41 | Cytochrome bo(3) ubiquinol oxidase subunit 4 |
| *FLPKDCKG_04602* | *cyoC-1* | - | 5015367 | 5016008 | 33.08 | 23.02 | 11.78 | 3.00 | Cytochrome bo(3) ubiquinol oxidase subunit 3 |
| *FLPKDCKG_04603* | *cyoB-1* | - | 5016001 | 5018004 | 36.84 | 31.74 | 19.17 | 4.98 | Cytochrome bo(3) ubiquinol oxidase subunit 1 |
| *FLPKDCKG_04604* | *cyoA-1* | - | 5018079 | 5019122 | 58.57 | 52.19 | 33.19 | 7.15 | Cytochrome bo(3) ubiquinol oxidase subunit 2 |
| *FLPKDCKG_04605* | *yhjE-5* | + | 5019347 | 5020681 | 40.55 | 34.43 | 17.55 | 11.13 | Inner membrane metabolite transport protein YhjE |
| *FLPKDCKG_04606* | *yhdW-5* | - | 5020737 | 5021753 | 5.66 | 6.56 | 3.17 | 0.69 | Putative amino-acid ABC transporter-binding protein YhdW |
| *FLPKDCKG_04607* | *dgdA-1* | - | 5021798 | 5023126 | 0.58 | 1.91 | 0.00 | 0.00 | 2,2-dialkylglycine decarboxylase |
| *FLPKDCKG_04608* | *yjiE-4* | + | 5023286 | 5024185 | 1.91 | 0.85 | 0.97 | 0.00 | HTH-type transcriptional regulator YjiE |
| *FLPKDCKG_04609* | *aphA-4* | - | 5024173 | 5025216 | 3.41 | 6.97 | 3.71 | 1.28 | Acetylpolyamine aminohydrolase |
| *FLPKDCKG_04610* | | - | 5025247 | 5025849 | 3.74 | 4.24 | 0.93 | 0.70 | Hypothetical protein |
| *FLPKDCKG_04611* | | - | 5025863 | 5027209 | 8.29 | 6.49 | 6.88 | 2.22 | Hypothetical protein |
| *FLPKDCKG_04612* | | + | 5027326 | 5027727 | 11.33 | 27.36 | 15.86 | 1.95 | Hypothetical protein |
| *FLPKDCKG_04613* | | - | 5027830 | 5028432 | 8.24 | 26.22 | 45.94 | 18.87 | Hypothetical protein |
| *FLPKDCKG_04614* | | - | 5028485 | 5028886 | 3.41 | 137.68 | 229.61 | 124.93 | Hypothetical protein |
| *FLPKDCKG_04615* | *nhaR* | - | 5028973 | 5029866 | 2.17 | 4.04 | 2.84 | 0.89 | Transcriptional activator protein NhaR |
| *FLPKDCKG_04616* | *alx* | + | 5029973 | 5030971 | 0.60 | 70.49 | 130.04 | 68.92 | Inner membrane protein alx |
| *FLPKDCKG_04617* | | - | 5031091 | 5031804 | 2.09 | 2.58 | 3.46 | 3.62 | Hypothetical protein |
| *FLPKDCKG_04618* | *tam-2* | + | 5032055 | 5032720 | 3.21 | 13.60 | 8.12 | 4.31 | Trans-aconitate 2-methyltransferase |
| *FLPKDCKG_04619* | *nimR-11* | - | 5032769 | 5033554 | 1.33 | 5.39 | 0.39 | 0.72 | HTH-type transcriptional regulator NimR |

| Locus name | Gene name | Strand | Gene start (bp) | Gene end (bp) | Average TPM Inoculum | Average TPM $t_6$ | Average TPM $t_{24}$ | Average TPM $t_{168}$ | Annotation description |
|---|---|---|---|---|---|---|---|---|---|
| *FLPKDCKG_04620* | *nimT* | + | 5033642 | 5034874 | 1.48 | 11.07 | 9.57 | 0.60 | 2-nitroimidazole transporter |
| *FLPKDCKG_04621* | | - | 5034909 | 5035244 | 5.02 | 40.22 | 29.38 | 6.91 | Hypothetical protein |
| *FLPKDCKG_04622* | *prtS* | - | 5035241 | 5036305 | 16.63 | 94.98 | 51.14 | 9.88 | Protease PrtS |
| *FLPKDCKG_04623* | | + | 5036516 | 5037145 | 4.54 | 20.36 | 14.24 | 4.20 | Hypothetical protein |
| *FLPKDCKG_04624* | | - | 5037168 | 5037644 | 156.54 | 122.35 | 89.32 | 30.74 | Hypothetical protein |
| *FLPKDCKG_04625* | *hflC-2* | + | 5037915 | 5038808 | 3.28 | 8.54 | 8.56 | 3.55 | Modulator of FtsH protease HflC |
| *FLPKDCKG_04626* | *chbC* | - | 5038801 | 5040879 | 1.46 | 2.85 | 1.58 | 1.24 | PTS system N,N'-diacetylchitobiose-specific EIIC component |
| *FLPKDCKG_04627* | | + | 5041183 | 5041425 | 213.62 | 168.06 | 116.13 | 40.74 | Hypothetical protein |
| *FLPKDCKG_04628* | *sigE-2* | + | 5041534 | 5042037 | 6.72 | 18.51 | 8.46 | 2.57 | ECF RNA polymerase sigma factor SigE |
| *FLPKDCKG_04629* | | + | 5042090 | 5043085 | 2.45 | 5.25 | 8.75 | 3.43 | Hypothetical protein |
| *FLPKDCKG_04630* | *fpvA-6* | + | 5043277 | 5045607 | 0.50 | 1.11 | 0.84 | 0.11 | Ferripyoverdine receptor |
| *FLPKDCKG_04631* | *fecI-13* | + | 5045693 | 5046214 | 3.15 | 0.23 | 0.00 | 0.25 | putative RNA polymerase sigma factor FecI |
| *FLPKDCKG_04632* | | + | 5046211 | 5047200 | 0.92 | 0.76 | 0.90 | 0.00 | Hypothetical protein |
| *FLPKDCKG_04633* | *fcuA-2* | + | 5047294 | 5049747 | 0.65 | 0.65 | 0.82 | 0.42 | Ferrichrome receptor FcuA |
| *FLPKDCKG_04634* | *nanR* | - | 5049764 | 5050657 | 1.07 | 2.36 | 0.95 | 0.30 | Transcriptional regulator NanR |
| *FLPKDCKG_04635* | *citB-1* | - | 5050654 | 5053314 | 0.72 | 1.01 | 2.35 | 0.73 | Aconitate hydratase A |
| *FLPKDCKG_04636* | | - | 5053408 | 5054058 | 0.88 | 1.05 | 0.00 | 0.22 | Hypothetical protein |
| *FLPKDCKG_04637* | *alkA-1* | - | 5054055 | 5054759 | 1.13 | 1.68 | 1.64 | 0.53 | DNA-3-methyladenine glycosylase 2 |
| *FLPKDCKG_04638* | *yieH-2* | + | 5054929 | 5055618 | 4.22 | 9.36 | 8.64 | 2.63 | 6-phosphogluconate phosphatase |
| *FLPKDCKG_04639* | *fabG-15* | - | 5055635 | 5056411 | 11.74 | 12.04 | 5.85 | 3.51 | 3-oxoacyl-[acyl-carrier-protein] reductase FabG |
| *FLPKDCKG_04640* | | + | 5056530 | 5057048 | 17.60 | 14.62 | 12.90 | 6.70 | Hypothetical protein |
| *FLPKDCKG_04641* | | - | 5057056 | 5057424 | 6.78 | 4.87 | 3.27 | 1.08 | Hypothetical protein |
| *FLPKDCKG_04642* | *marA-2* | - | 5057506 | 5058327 | 3.70 | 3.15 | 3.15 | 0.73 | Multiple antibiotic resistance protein MarA |
| *FLPKDCKG_04643* | | - | 5058340 | 5058723 | 9.01 | 10.48 | 11.26 | 4.33 | Hypothetical protein |
| *FLPKDCKG_04644* | *pkn1-3* | - | 5058770 | 5059765 | 0.90 | 2.05 | 2.69 | 1.90 | Serine/threonine-protein kinase pkn1 |
| *FLPKDCKG_04645* | *rclR-4* | - | 5059858 | 5060715 | 3.27 | 2.73 | 1.76 | 0.31 | RCS-specific HTH-type transcriptional activator RclR |
| *FLPKDCKG_04646* | | + | 5060851 | 5061393 | 1.30 | 2.50 | 1.04 | 0.81 | Hypothetical protein |
| *FLPKDCKG_04647* | *alkB* | - | 5061405 | 5062064 | 2.51 | 4.67 | 5.96 | 1.16 | Alpha-ketoglutarate-dependent dioxygenase AlkB |
| *FLPKDCKG_04648* | | - | 5062066 | 5062761 | 8.75 | 5.81 | 0.87 | 0.57 | Hypothetical protein |
| *FLPKDCKG_04649* | *clsC* | - | 5062903 | 5064447 | 2.67 | 3.56 | 1.95 | 1.52 | Cardiolipin synthase C |
| *FLPKDCKG_04650* | *sutR-1* | + | 5064578 | 5065150 | 1.36 | 0.23 | 0.96 | 1.20 | HTH-type transcriptional regulator SutR |
| *FLPKDCKG_04651* | *ttr* | + | 5065158 | 5065694 | 1.52 | 1.56 | 1.36 | 3.55 | Acetyltransferase |
| *FLPKDCKG_04652* | | - | 5065729 | 5066055 | 5.23 | 0.94 | 2.76 | 0.49 | Hypothetical protein |
| *FLPKDCKG_04653* | *ephA* | - | 5066172 | 5067074 | 0.39 | 1.34 | 0.00 | 0.00 | Epoxide hydrolase A |
| *FLPKDCKG_04654* | *oxyR-4* | + | 5067182 | 5068069 | 1.79 | 3.64 | 0.03 | 1.65 | Hydrogen peroxide-inducible genes activator |
| *FLPKDCKG_04655* | | + | 5068214 | 5068366 | 2.07 | 22.52 | 29.95 | 10.18 | Hypothetical protein |
| *FLPKDCKG_04656* | *fabG-16* | + | 5068586 | 5069395 | 0.44 | 2.77 | 4.89 | 1.00 | 3-oxoacyl-[acyl-carrier-protein] reductase FabG |
| *FLPKDCKG_04657* | | - | 5069412 | 5070182 | 0.51 | 2.99 | 3.67 | 1.28 | Hypothetical protein |
| *FLPKDCKG_04658* | | - | 5070179 | 5070589 | 0.45 | 0.00 | 1.62 | 0.58 | Hypothetical protein |
| *FLPKDCKG_04659* | *gltC-17* | + | 5070715 | 5071605 | 3.38 | 3.34 | 3.69 | 0.54 | HTH-type transcriptional regulator GltC |
| *FLPKDCKG_04660* | | - | 5071616 | 5071861 | 1.47 | 2.53 | 2.03 | 0.09 | Hypothetical protein |
| *FLPKDCKG_04661* | *mepA* | - | 5071828 | 5073024 | 0.95 | 0.48 | 0.48 | 0.89 | Multidrug export protein MepA |
| *FLPKDCKG_04662* | *badR-3* | + | 5073213 | 5073683 | 4.74 | 6.32 | 2.34 | 1.75 | Transcriptional activatory protein BadR |
| *FLPKDCKG_04663* | | - | 5073706 | 5074407 | 2.18 | 4.23 | 3.68 | 4.25 | Hypothetical protein |
| *FLPKDCKG_04664* | *ytfF-2* | - | 5074411 | 5075298 | 1.03 | 3.89 | 0.99 | 1.24 | Inner membrane protein YtfF |
| *FLPKDCKG_04665* | *nphR-3* | - | 5075395 | 5076189 | 3.15 | 6.72 | 3.85 | 0.76 | Transcriptional activator NphR |
| *FLPKDCKG_04666* | *gbpR-4* | - | 5076229 | 5077191 | 5.34 | 11.50 | 8.95 | 2.22 | HTH-type transcriptional regulator GbpR |
| *FLPKDCKG_04667* | *maeB-2* | + | 5077298 | 5079592 | 5.19 | 17.87 | 9.46 | 4.40 | NADP-dependent malic enzyme |
| *FLPKDCKG_04668* | | - | 5079697 | 5080074 | 97.12 | 61.20 | 43.12 | 10.03 | Hypothetical protein |
| *FLPKDCKG_04669* | | - | 5080266 | 5080625 | 9.18 | 11.10 | 15.93 | 5.29 | Hypothetical protein |
| *FLPKDCKG_04670* | *slyX* | - | 5080854 | 5081117 | 8.85 | 31.22 | 12.56 | 1.81 | Protein SlyX |
| *FLPKDCKG_04671* | *fhuA-12* | + | 5081234 | 5083288 | 3.39 | 9.11 | 3.17 | 0.29 | Ferrichrome-iron receptor |
| *FLPKDCKG_04672* | | - | 5083308 | 5083925 | 9.25 | 5.91 | 7.15 | 0.94 | Hypothetical protein |
| *FLPKDCKG_04673* | | - | 5084075 | 5084497 | 2.29 | 1.53 | 2.01 | 0.00 | Hypothetical protein |
| *FLPKDCKG_04674* | *cyoD-2* | - | 5084671 | 5085021 | 0.98 | 1.79 | 1.71 | 6.33 | Cytochrome bo(3) ubiquinol oxidase subunit 4 |
| *FLPKDCKG_04675* | *cyoC-2* | - | 5085021 | 5085635 | 0.21 | 1.55 | 4.84 | 0.27 | Cytochrome bo(3) ubiquinol oxidase subunit 3 |
| *FLPKDCKG_04676* | *cyoB-2* | - | 5085639 | 5087615 | 0.73 | 2.64 | 7.02 | 3.42 | Cytochrome bo(3) ubiquinol oxidase subunit 1 |
| *FLPKDCKG_04677* | *cyoA-2* | - | 5087615 | 5088472 | 1.34 | 9.04 | 14.50 | 6.81 | Cytochrome bo(3) ubiquinol oxidase subunit 2 |
| *FLPKDCKG_04678* | *regA* | - | 5088785 | 5089339 | 53.94 | 145.96 | 86.00 | 40.45 | Photosynthetic apparatus regulatory protein RegA |
| *FLPKDCKG_04679* | *regB* | - | 5089341 | 5090543 | 5.10 | 22.58 | 17.56 | 4.13 | Sensor histidine kinase RegB |
| *FLPKDCKG_04680* | | - | 5090760 | 5091251 | 24.31 | 36.26 | 34.88 | 14.20 | Hypothetical protein |
| *FLPKDCKG_04681* | *ypmQ-3* | - | 5091280 | 5091897 | 46.42 | 51.84 | 46.88 | 21.90 | SCO1 protein |
| *FLPKDCKG_04682* | | - | 5091957 | 5092391 | 27.12 | 38.35 | 26.42 | 8.05 | Hypothetical protein |
| *FLPKDCKG_04683* | *est-2* | - | 5092424 | 5093941 | 3.31 | 2.07 | 3.95 | 1.49 | Carboxylesterase |
| *FLPKDCKG_04684* | | - | 5093931 | 5095217 | 4.84 | 2.34 | 8.14 | 3.12 | Hypothetical protein |
| *FLPKDCKG_04685* | *nlpD-2* | + | 5095386 | 5096174 | 33.56 | 55.61 | 46.34 | 12.72 | Murein hydrolase activator NlpD |
| *FLPKDCKG_04686* | *tcyC-4* | - | 5096254 | 5097015 | 9.93 | 28.92 | 18.09 | 5.79 | L-cystine import ATP-binding protein TcyC |
| *FLPKDCKG_04687* | *tcyB* | - | 5097012 | 5097761 | 10.73 | 17.56 | 11.90 | 4.40 | L-cystine transport system permease protein TcyB |
| *FLPKDCKG_04688* | | - | 5097774 | 5098559 | 63.80 | 65.58 | 70.12 | 22.12 | putative amino-acid ABC transporter-binding protein |

| Locus name | Gene name | Strand | Gene start (bp) | Gene end (bp) | Average TPM Inoculum | Average TPM $t_6$ | Average TPM $t_{24}$ | Average TPM $t_{168}$ | Annotation description |
|---|---|---|---|---|---|---|---|---|---|
| *FLPKDCKG_04689* | | + | 5098794 | 5099840 | 5.21 | 11.30 | 7.70 | 2.06 | putative aminopeptidase |
| *FLPKDCKG_04690* | | + | 5099989 | 5100612 | 0.69 | 9.27 | 9.55 | 3.76 | Hypothetical protein |
| *FLPKDCKG_04691* | *lrp-10* | - | 5100623 | 5101096 | 4.17 | 6.18 | 6.08 | 2.39 | Leucine-responsive regulatory protein |
| *FLPKDCKG_04692* | *kynU* | + | 5101235 | 5102485 | 10.87 | 184.17 | 120.93 | 17.31 | Kynureninase |
| *FLPKDCKG_04693* | *rocC* | + | 5102544 | 5103950 | 1.74 | 14.20 | 10.31 | 2.19 | Amino-acid permease RocC |
| *FLPKDCKG_04694* | *dmlR-27* | - | 5103994 | 5104905 | 1.50 | 5.04 | 3.26 | 0.97 | HTH-type transcriptional regulator DmlR |
| *FLPKDCKG_04695* | *thrC-3* | + | 5105007 | 5106140 | 0.44 | 1.04 | 0.08 | 0.29 | Threonine synthase |
| *FLPKDCKG_04696* | *cmpR-16* | - | 5106112 | 5107032 | 1.58 | 2.77 | 1.26 | 0.89 | HTH-type transcriptional activator CmpR |
| *FLPKDCKG_04697* | *ybhI-2* | + | 5107142 | 5108452 | 2.30 | 6.64 | 1.65 | 0.52 | Inner membrane protein YbhI |
| *FLPKDCKG_04698* | *ehpR* | - | 5108424 | 5108846 | 20.92 | 39.34 | 22.70 | 4.61 | Phenazine antibiotic resistance protein EhpR |
| *FLPKDCKG_04699* | *niaR* | - | 5108943 | 5109635 | 3.93 | 4.57 | 6.49 | 0.41 | putative transcription repressor NiaR |
| *FLPKDCKG_04700* | | - | 5109877 | 5110740 | 3.95 | 6.50 | 5.44 | 1.02 | Hypothetical protein |
| *FLPKDCKG_04701* | | - | 5110905 | 5111453 | 5.11 | 6.28 | 2.36 | 0.46 | Hypothetical protein |
| *FLPKDCKG_04702* | *fecI-14* | + | 5111556 | 5112065 | 4.51 | 0.11 | 1.34 | 0.00 | putative RNA polymerase sigma factor FecI |
| *FLPKDCKG_04703* | | + | 5112351 | 5113043 | 4.14 | 0.36 | 2.30 | 0.98 | Hypothetical protein |
| *FLPKDCKG_04704* | *hemR* | + | 5113536 | 5115770 | 4.06 | 1.18 | 1.81 | 0.68 | Hemin receptor |
| *FLPKDCKG_04705* | *hemS* | + | 5115836 | 5116894 | 5.17 | 1.18 | 2.44 | 1.26 | Hemin transport protein HemS |
| *FLPKDCKG_04706* | *hmuT* | + | 5116905 | 5117738 | 1.86 | 2.00 | 2.90 | 0.39 | Hemin-binding periplasmic protein HmuT |
| *FLPKDCKG_04707* | *hmuU-5* | + | 5117722 | 5118720 | 0.84 | 0.87 | 0.12 | 1.99 | Hemin transport system permease protein HmuU |
| *FLPKDCKG_04708* | *hmuV* | + | 5118720 | 5119505 | 0.59 | 0.98 | 0.33 | 0.64 | Hemin import ATP-binding protein HmuV |
| *FLPKDCKG_04709* | | + | 5119586 | 5120068 | 49.74 | 32.58 | 28.09 | 10.90 | Hypothetical protein |
| *FLPKDCKG_04710* | *alkA-2* | - | 5120529 | 5120852 | 38.70 | 18.50 | 14.42 | 5.45 | putative bifunctional transcriptional activator/DNA repair enzyme AlkA |
| *FLPKDCKG_04711* | *alkA-3* | - | 5120909 | 5121985 | 19.63 | 5.05 | 2.18 | 2.23 | putative bifunctional transcriptional activator/DNA repair enzyme AlkA |
| *FLPKDCKG_04712* | *ogt-2* | + | 5122122 | 5122673 | 19.47 | 2.52 | 2.56 | 0.97 | Methylated-DNA--protein-cysteine methyltransferase |
| *FLPKDCKG_04713* | *ybjG* | - | 5122736 | 5123341 | 3.55 | 8.96 | 3.76 | 0.99 | Putative undecaprenyl-diphosphatase YbjG |
| *FLPKDCKG_04714* | | - | 5123434 | 5123868 | 3.23 | 2.31 | 2.64 | 0.14 | Hypothetical protein |
| *FLPKDCKG_04715* | *liaS* | - | 5124022 | 5125818 | 0.28 | 0.33 | 0.47 | 0.05 | Sensor histidine kinase LiaS |
| *FLPKDCKG_04716* | *nreC-2* | - | 5125893 | 5126525 | 1.91 | 2.19 | 1.44 | 2.12 | Oxygen regulatory protein NreC |
| *FLPKDCKG_04717* | *arsC-2* | - | 5126682 | 5127110 | 25.26 | 20.89 | 20.13 | 3.94 | Arsenate reductase |
| *FLPKDCKG_04718* | | - | 5127188 | 5128720 | 0.54 | 1.61 | 0.51 | 0.85 | Hypothetical protein |
| *FLPKDCKG_04719* | *bfrD-4* | - | 5128729 | 5130930 | 0.53 | 3.88 | 2.71 | 0.94 | putative TonB-dependent receptor BfrD |
| *FLPKDCKG_04720* | | - | 5131099 | 5131497 | 7.08 | 11.60 | 25.39 | 7.99 | Hypothetical protein |
| *FLPKDCKG_04721* | *livF-22* | - | 5131523 | 5132221 | 0.30 | 10.71 | 13.93 | 6.36 | High-affinity branched-chain amino acid transport ATP-binding protein LivF |
| *FLPKDCKG_04722* | *rbsA-3* | - | 5132223 | 5133062 | 0.23 | 15.97 | 15.77 | 5.46 | Ribose import ATP-binding protein RbsA |
| *FLPKDCKG_04723* | | - | 5133059 | 5134186 | 0.13 | 17.61 | 20.50 | 8.74 | Hypothetical protein |
| *FLPKDCKG_04724* | *livH-22* | - | 5134183 | 5135814 | 0.08 | 6.00 | 17.48 | 6.26 | High-affinity branched-chain amino acid transport system permease protein LivH |
| *FLPKDCKG_04725* | | - | 5135835 | 5136035 | 0.00 | 27.89 | 35.04 | 35.45 | Limonene hydroxylase |
| *FLPKDCKG_04726* | *amiC-8* | - | 5136044 | 5137087 | 0.25 | 147.01 | 197.57 | 78.81 | Aliphatic amidase expression-regulating protein |
| *FLPKDCKG_04727* | *atzF* | + | 5137434 | 5139263 | 0.07 | 24.36 | 34.46 | 16.02 | Allophanate hydrolase |
| *FLPKDCKG_04728* | *kipA-3* | + | 5140263 | 5142923 | 0.13 | 16.75 | 24.80 | 12.21 | KipI antagonist |
| *FLPKDCKG_04729* | *lgoR* | + | 5142908 | 5143645 | 0.61 | 68.65 | 79.99 | 34.22 | putative HTH-type transcriptional regulator LgoR |
| *FLPKDCKG_04730* | | - | 5144112 | 5144927 | 7.05 | 16.25 | 28.51 | 10.93 | Hypothetical protein |
| *FLPKDCKG_04731* | *ecfG* | - | 5144929 | 5145441 | 17.88 | 20.03 | 22.90 | 9.34 | ECF RNA polymerase sigma factor EcfG |
| *FLPKDCKG_04732* | | - | 5145473 | 5145841 | 99.72 | 92.75 | 88.78 | 26.12 | Hypothetical protein |
| *FLPKDCKG_04733* | *mshD-2* | - | 5145946 | 5146431 | 53.15 | 45.15 | 27.52 | 26.70 | Mycothiol acetyltransferase |
| *FLPKDCKG_04734* | *oxyR-5* | - | 5146441 | 5147367 | 20.40 | 12.38 | 2.29 | 3.07 | Hydrogen peroxide-inducible genes activator |
| *FLPKDCKG_04735* | *galD* | + | 5147420 | 5148568 | 0.86 | 0.21 | 0.00 | 0.43 | 4-oxalomesaconate tautomerase |
| *FLPKDCKG_04736* | | + | 5148707 | 5149675 | 0.83 | 1.21 | 0.00 | 0.57 | Hypothetical protein |
| *FLPKDCKG_04737* | | - | 5149632 | 5151032 | 0.27 | 0.00 | 0.28 | 0.36 | Hypothetical protein |
| *FLPKDCKG_04738* | | - | 5151029 | 5152591 | 1.16 | 0.26 | 0.53 | 0.00 | Hypothetical protein |
| *FLPKDCKG_04739* | *fabG-17* | - | 5152744 | 5153448 | 10.27 | 17.92 | 11.33 | 1.66 | 3-oxoacyl-[acyl-carrier-protein] reductase FabG |
| *FLPKDCKG_04740* | | - | 5153536 | 5153988 | 166.66 | 215.49 | 130.22 | 37.13 | Azurin |
| *FLPKDCKG_04741* | | - | 5154153 | 5155478 | 12.42 | 27.99 | 14.99 | 3.44 | Hypothetical protein |
| *FLPKDCKG_04742* | *nagAa* | - | 5155581 | 5156906 | 0.98 | 10.55 | 33.72 | 13.08 | Ferredoxin--NAD(P)(+) reductase (naphthalene dioxygenase/salicylate 5-hydroxylase ferredoxin-specific) |
| *FLPKDCKG_04743* | | - | 5156917 | 5157252 | 12.16 | 97.17 | 200.72 | 94.56 | Hypothetical protein |
| *FLPKDCKG_04744* | | + | 5157450 | 5157797 | 24.52 | 46.20 | 56.15 | 24.27 | Hypothetical protein |
| *FLPKDCKG_04745* | *phoP-4* | + | 5157797 | 5158459 | 5.49 | 9.61 | 16.92 | 0.91 | Virulence transcriptional regulatory protein PhoP |
| *FLPKDCKG_04746* | *phoQ* | + | 5158456 | 5159838 | 3.92 | 8.26 | 9.75 | 4.62 | Sensor protein PhoQ |
| *FLPKDCKG_04747* | *uspG* | - | 5159879 | 5160307 | 37.35 | 27.77 | 17.52 | 5.16 | Universal stress protein UP12 |
| *FLPKDCKG_04748* | | - | 5160317 | 5160742 | 36.15 | 20.90 | 17.94 | 2.21 | Hypothetical protein |
| *FLPKDCKG_04749* | *murI* | - | 5160877 | 5161680 | 1.64 | 1.77 | 1.93 | 2.96 | Glutamate racemase |
| *FLPKDCKG_04750* | *benM-4* | - | 5161718 | 5162644 | 2.54 | 1.66 | 3.34 | 2.46 | HTH-type transcriptional regulator BenM |
| *FLPKDCKG_04751* | *srpC-4* | + | 5162859 | 5163356 | 1.41 | 8.01 | 5.60 | 2.44 | putative chromate transport protein |
| *FLPKDCKG_04752* | *srpC-5* | + | 5163353 | 5163877 | 2.27 | 2.21 | 4.69 | 1.11 | putative chromate transport protein |
| *FLPKDCKG_04753* | | + | 5163910 | 5164599 | 7.53 | 9.47 | 7.69 | 0.94 | Hypothetical protein |
| *FLPKDCKG_04754* | *prmA-2* | - | 5164649 | 5165779 | 1.53 | 4.78 | 3.44 | 2.71 | Ribosomal protein L11 methyltransferase |
| *FLPKDCKG_04755* | *yhjX-2* | - | 5165934 | 5167601 | 1.00 | 3.89 | 5.42 | 5.33 | putative MFS-type transporter YhjX |
| *FLPKDCKG_04756* | | - | 5167748 | 5168005 | 2.70 | 31.42 | 18.58 | 16.41 | Hypothetical protein |
| *FLPKDCKG_04757* | *fdhD* | - | 5168011 | 5168847 | 2.84 | 12.36 | 6.31 | 3.59 | Sulfurtransferase FdhD |

| Locus name | Gene name | Strand | Gene start (bp) | Gene end (bp) | Average TPM Inoculum | Average TPM $t_6$ | Average TPM $t_{24}$ | Average TPM $t_{168}$ | Annotation description |
|---|---|---|---|---|---|---|---|---|---|
| *FLPKDCKG_04758* | | - | 5168872 | 5171748 | 3.79 | 16.53 | 14.87 | 9.85 | Putative formate dehydrogenase |
| *FLPKDCKG_04759* | *hndC* | - | 5171762 | 5173333 | 3.01 | 14.94 | 10.09 | 7.47 | NADP-reducing hydrogenase subunit HndC |
| *FLPKDCKG_04760* | *hndA* | - | 5173330 | 5173887 | 7.22 | 35.35 | 28.72 | 13.38 | NADP-reducing hydrogenase subunit HndA |
| *FLPKDCKG_04761* | | + | 5174046 | 5175122 | 3.58 | 30.00 | 16.03 | 3.30 | Hypothetical protein |
| *FLPKDCKG_04762* | *moaA* | + | 5175158 | 5176252 | 25.54 | 29.57 | 23.09 | 10.48 | GTP 3',8-cyclase |
| *FLPKDCKG_04763* | *mobA* | + | 5176245 | 5176868 | 6.59 | 16.76 | 15.85 | 3.84 | Molybdenum cofactor guanylyltransferase |
| *FLPKDCKG_04764* | *moeA* | - | 5176853 | 5178058 | 7.20 | 12.39 | 15.42 | 2.11 | Molybdopterin molybdenumtransferase |
| *FLPKDCKG_04765* | *moaB* | - | 5178064 | 5179065 | 42.36 | 55.55 | 35.91 | 11.62 | Molybdenum cofactor biosynthesis protein B |
| *FLPKDCKG_04766* | *moaD* | - | 5179071 | 5179322 | 15.91 | 22.75 | 9.80 | 6.09 | Molybdopterin synthase sulfur carrier subunit |
| *FLPKDCKG_04767* | *moaC* | - | 5179303 | 5179791 | 10.61 | 16.50 | 11.12 | 1.07 | Cyclic pyranopterin monophosphate synthase |
| *FLPKDCKG_04768* | | + | 5180107 | 5183829 | 21.60 | 18.69 | 12.20 | 3.65 | Hypothetical protein |
| *FLPKDCKG_04769* | *clcD-3* | + | 5183892 | 5184776 | 35.88 | 38.15 | 23.32 | 8.99 | Carboxymethylenebutenolidase |
| *FLPKDCKG_04770* | | - | 5185404 | 5185829 | 57.37 | 166.97 | 91.73 | 11.20 | Hypothetical protein |
| *FLPKDCKG_04771* | | + | 5186059 | 5186562 | 33.72 | 35.79 | 42.50 | 7.79 | Putative peroxiredoxin |
| *FLPKDCKG_04772* | *ycaD* | + | 5186651 | 5188045 | 2.38 | 5.14 | 3.62 | 1.14 | putative MFS-type transporter YcaD |
| *FLPKDCKG_04773* | | - | 5188059 | 5188946 | 0.85 | 3.58 | 0.00 | 0.26 | Hypothetical protein |
| *FLPKDCKG_04774* | *bioB* | - | 5188943 | 5189968 | 3.19 | 9.28 | 5.08 | 2.73 | Biotin synthase |
| *FLPKDCKG_04775* | *kinA* | - | 5190219 | 5191709 | 5.98 | 13.81 | 9.94 | 1.67 | Sporulation kinase A |
| *FLPKDCKG_04776* | | - | 5191939 | 5192015 | 56.47 | 50.47 | 88.61 | 18.68 | tRNA |
| *FLPKDCKG_04777* | | - | 5192091 | 5192166 | 347.40 | 280.15 | 105.91 | 46.96 | tRNA |
| *FLPKDCKG_04778* | | - | 5192212 | 5192287 | 86.39 | 170.12 | 186.54 | 60.46 | tRNA |
| *FLPKDCKG_04779* | | - | 5192391 | 5192467 | 103.41 | 141.69 | 150.39 | 44.15 | tRNA |
| *FLPKDCKG_04780* | | - | 5192529 | 5192604 | 208.73 | 206.38 | 179.84 | 52.88 | tRNA |
| *FLPKDCKG_04781* | | - | 5192673 | 5192748 | 184.13 | 358.98 | 302.83 | 107.34 | tRNA |
| *FLPKDCKG_04782* | *yhdN-2* | + | 5193061 | 5194116 | 10.48 | 30.54 | 21.09 | 6.63 | General stress protein 69 |
| *FLPKDCKG_04783* | *lnt* | - | 5194254 | 5195885 | 1.14 | 7.95 | 8.32 | 3.64 | Apolipoprotein N-acyltransferase |
| *FLPKDCKG_04784* | *corC-2* | - | 5195882 | 5196772 | 8.71 | 39.10 | 40.30 | 12.02 | Magnesium and cobalt efflux protein CorC |
| *FLPKDCKG_04785* | *ybeY* | - | 5196927 | 5197400 | 1.87 | 5.95 | 10.74 | 3.23 | Endoribonuclease YbeY |
| *FLPKDCKG_04786* | | - | 5197390 | 5198415 | 2.58 | 12.06 | 9.72 | 3.87 | PhoH-like protein |
| *FLPKDCKG_04787* | *miaB* | - | 5198412 | 5199845 | 13.65 | 30.79 | 16.64 | 8.36 | tRNA-2-methylthio-N(6)-dimethylallyladenosine synthase |
| *FLPKDCKG_04788* | | + | 5200011 | 5200772 | 4.76 | 13.15 | 8.99 | 7.38 | Hypothetical protein |
| *FLPKDCKG_04789* | *alaS-2* | - | 5200851 | 5201564 | 4.98 | 9.76 | 2.49 | 0.00 | Alanine--tRNA ligase |
| *FLPKDCKG_04790* | | - | 5201593 | 5202270 | 10.48 | 13.24 | 10.83 | 4.61 | Hypothetical protein |
| *FLPKDCKG_04791* | | - | 5202361 | 5202672 | 9.24 | 15.80 | 6.11 | 1.85 | Hypothetical protein |
| *FLPKDCKG_04792* | | - | 5202669 | 5203037 | 11.93 | 19.75 | 16.19 | 6.34 | Hypothetical protein |
| *FLPKDCKG_04793* | | - | 5203207 | 5204142 | 5.04 | 11.76 | 9.47 | 3.58 | Hypothetical protein |
| *FLPKDCKG_04794* | | - | 5204194 | 5206074 | 13.06 | 31.71 | 29.21 | 10.86 | Hypothetical protein |
| *FLPKDCKG_04795* | | - | 5206190 | 5206534 | 122.54 | 147.73 | 140.87 | 52.92 | Hypothetical protein |
| *FLPKDCKG_04796* | *dinJ* | + | 5206907 | 5207203 | 45.05 | 44.64 | 51.80 | 9.13 | Antitoxin DinJ |
| *FLPKDCKG_04797* | *relE2-2* | + | 5207190 | 5207471 | 4.30 | 22.63 | 7.44 | 3.84 | Toxin RelE2 |
| *FLPKDCKG_04798* | *tgt* | - | 5207521 | 5208657 | 2.79 | 6.24 | 6.37 | 2.51 | Queuine tRNA-ribosyltransferase |
| *FLPKDCKG_04799* | *queA* | - | 5208654 | 5209778 | 4.83 | 12.96 | 7.88 | 4.81 | S-adenosylmethionine:tRNA ribosyltransferase-isomerase |
| *FLPKDCKG_04800* | *gylR-2* | - | 5209948 | 5210781 | 0.95 | 2.58 | 2.02 | 0.31 | Glycerol operon regulatory protein |
| *FLPKDCKG_04801* | *limB-2* | + | 5210903 | 5211916 | 2.59 | 4.14 | 3.46 | 3.12 | Limonene 1,2-monooxygenase |
| *FLPKDCKG_04802* | *C1-hpah-2* | + | 5211940 | 5212443 | 1.62 | 3.55 | 1.83 | 1.58 | p-hydroxyphenylacetate 3-hydroxylase, reductase component |
| *FLPKDCKG_04803* | *braC-8* | + | 5212510 | 5213664 | 2.41 | 2.97 | 2.58 | 2.13 | Leucine-, isoleucine-, valine-, threonine-, and alanine-binding protein |
| *FLPKDCKG_04804* | *dacB* | + | 5213734 | 5215224 | 4.54 | 6.54 | 11.16 | 2.63 | D-alanyl-D-alanine carboxypeptidase DacB |
| *FLPKDCKG_04805* | *upp* | + | 5215349 | 5215987 | 8.50 | 22.74 | 22.71 | 7.92 | Uracil phosphoribosyltransferase |
| *FLPKDCKG_04806* | *cysG* | - | 5216048 | 5217463 | 0.75 | 19.07 | 38.34 | 15.43 | Siroheme synthase |
| *FLPKDCKG_04807* | *nrtA-2* | + | 5218071 | 5219369 | 0.03 | 15.62 | 44.82 | 51.23 | Nitrate transport protein NrtA |
| *FLPKDCKG_04808* | *cmpB-4* | + | 5219520 | 5220269 | 0.25 | 9.97 | 23.23 | 17.56 | Bicarbonate transport system permease protein CmpB |
| *FLPKDCKG_04809* | *cmpD-3* | + | 5220274 | 5221152 | 0.15 | 20.40 | 41.18 | 30.64 | Bicarbonate transport ATP-binding protein CmpD |
| *FLPKDCKG_04810* | | + | 5221376 | 5222107 | 55.02 | 111.76 | 100.07 | 25.76 | Hypothetical protein |
| *FLPKDCKG_04811* | | + | 5222315 | 5222506 | 166.69 | 319.67 | 322.66 | 63.73 | Hypothetical protein |
| *FLPKDCKG_04812* | | + | 5222575 | 5222985 | 13.72 | 58.29 | 38.74 | 10.67 | Hypothetical protein |
| *FLPKDCKG_04813* | | + | 5222989 | 5223348 | 5.61 | 20.02 | 14.26 | 1.42 | Hypothetical protein |
| *FLPKDCKG_04814* | *ubiD* | - | 5223447 | 5224991 | 5.52 | 7.73 | 5.36 | 1.39 | 3-octaprenyl-4-hydroxybenzoate carboxy-lyase |
| *FLPKDCKG_04815* | *mltC* | + | 5225075 | 5225893 | 17.17 | 112.66 | 80.38 | 23.05 | Membrane-bound lytic murein transglycosylase C |
| *FLPKDCKG_04816* | | - | 5225924 | 5227030 | 1.23 | 29.17 | 17.29 | 4.80 | Hypothetical protein |
| *FLPKDCKG_04817* | | - | 5227349 | 5228458 | 1.51 | 7.31 | 2.70 | 2.16 | Hypothetical protein |
| *FLPKDCKG_04818* | *aspC-3* | - | 5228839 | 5229999 | 6.27 | 8.40 | 6.23 | 2.76 | Aspartate aminotransferase |
| *FLPKDCKG_04819* | | + | 5230044 | 5230940 | 2.86 | 4.02 | 2.08 | 1.39 | Hypothetical protein |
| *FLPKDCKG_04820* | | - | 5231257 | 5231604 | 9.97 | 10.13 | 13.51 | 4.45 | Hypothetical protein |
| *FLPKDCKG_04821* | *maeB-3* | + | 5232456 | 5233370 | 4.57 | 2.13 | 5.68 | 1.97 | NADP-dependent malic enzyme |
| *FLPKDCKG_04822* | | + | 5233351 | 5234586 | 6.78 | 9.94 | 5.36 | 1.58 | Hypothetical protein |
| *FLPKDCKG_04823* | | + | 5234583 | 5235050 | 53.36 | 28.06 | 25.31 | 11.02 | Hypothetical protein |
| *FLPKDCKG_04824* | *rsmE* | - | 5235125 | 5235856 | 5.86 | 9.43 | 3.92 | 2.12 | Ribosomal RNA small subunit methyltransferase E |
| *FLPKDCKG_04825* | | - | 5235961 | 5236467 | 6.22 | 4.48 | 9.09 | 2.24 | Hypothetical protein |
| *FLPKDCKG_04826* | *tktA* | + | 5236726 | 5238762 | 23.10 | 51.89 | 33.70 | 16.27 | Transketolase 1 |

| Locus name | Gene name | Strand | Gene start (bp) | Gene end (bp) | Average TPM Inoculum | Average TPM $t_6$ | Average TPM $t_{24}$ | Average TPM $t_{168}$ | Annotation description |
|---|---|---|---|---|---|---|---|---|---|
| *FLPKDCKG_04827* | *gap1* | + | 5238778 | 5239788 | 50.68 | 81.59 | 68.36 | 30.09 | Glyceraldehyde-3-phosphate dehydrogenase 1 |
| *FLPKDCKG_04828* | *pgk* | + | 5239886 | 5241079 | 2.47 | 5.21 | 8.80 | 1.92 | Phosphoglycerate kinase |
| *FLPKDCKG_04829* | | - | 5241168 | 5242151 | 0.48 | 1.17 | 0.64 | 0.78 | Hypothetical protein |
| *FLPKDCKG_04830* | | + | 5242449 | 5244140 | 1.48 | 2.27 | 2.23 | 1.24 | putative sulfate transporter |
| *FLPKDCKG_04831* | *fabI-2* | - | 5244170 | 5244943 | 1.40 | 7.42 | 6.27 | 1.16 | Enoyl-[acyl-carrier-protein] reductase [NADH] FabI |
| *FLPKDCKG_04832* | *ackA* | - | 5244940 | 5246124 | 2.71 | 8.73 | 16.05 | 6.48 | Acetate kinase |
| *FLPKDCKG_04833* | *pta-2* | - | 5246206 | 5247150 | 10.32 | 28.85 | 39.65 | 8.43 | Phosphate acetyltransferase |
| *FLPKDCKG_04834* | *phaC-4* | - | 5247147 | 5249549 | 19.61 | 43.39 | 42.88 | 14.30 | Poly(3-hydroxyalkanoate) polymerase subunit PhaC |
| *FLPKDCKG_04835* | | - | 5249734 | 5250135 | 19.84 | 6.02 | 4.16 | 2.38 | Hypothetical protein |
| *FLPKDCKG_04836* | *yfcF* | - | 5250155 | 5250790 | 41.52 | 28.44 | 25.92 | 7.34 | Glutathione S-transferase YfcF |
| *FLPKDCKG_04837* | | - | 5250949 | 5251278 | 2.54 | 6.80 | 6.14 | 1.84 | Hypothetical protein |
| *FLPKDCKG_04838* | *ypeA-2* | - | 5251328 | 5251876 | 13.87 | 28.65 | 17.11 | 4.49 | Acetyltransferase YpeA |
| *FLPKDCKG_04839* | *cbpA* | + | 5251992 | 5252954 | 24.70 | 24.39 | 22.95 | 14.95 | Curved DNA-binding protein |
| *FLPKDCKG_04840* | *cbpM* | + | 5252957 | 5253274 | 7.49 | 12.05 | 10.97 | 9.94 | Chaperone modulatory protein CbpM |
| *FLPKDCKG_04841* | *yfcG-6* | + | 5253399 | 5254013 | 93.11 | 62.78 | 23.09 | 15.10 | Disulfide-bond oxidoreductase YfcG |
| *FLPKDCKG_04842* | | - | 5254014 | 5254661 | 4.31 | 8.01 | 4.14 | 2.77 | Hypothetical protein |
| *FLPKDCKG_04843* | *pitA* | - | 5254761 | 5256236 | 2.53 | 8.70 | 7.43 | 3.70 | Low-affinity inorganic phosphate transporter 1 |
| *FLPKDCKG_04844* | | - | 5256392 | 5256517 | 1.84 | 0.00 | 0.00 | 0.00 | Hypothetical protein |
| *FLPKDCKG_04845* | | - | 5256561 | 5257685 | 4.53 | 1.20 | 0.60 | 1.08 | Hypothetical protein |
| *FLPKDCKG_04846* | *mog* | + | 5257950 | 5258597 | 7.71 | 10.56 | 15.68 | 4.09 | Molybdopterin adenylyltransferase |
| *FLPKDCKG_04847* | | + | 5258637 | 5258984 | 15.30 | 32.53 | 20.37 | 7.26 | Hypothetical protein |
| *FLPKDCKG_04848* | *mepM-3* | + | 5258984 | 5259868 | 3.10 | 5.84 | 6.49 | 2.12 | Murein DD-endopeptidase MepM |
| *FLPKDCKG_04849* | | - | 5259883 | 5260209 | 6.46 | 20.99 | 10.28 | 4.80 | Hypothetical protein |
| *FLPKDCKG_04850* | *infA-2* | - | 5260372 | 5260635 | 25.32 | 201.60 | 170.37 | 30.69 | Translation initiation factor IF-1 |
| *FLPKDCKG_04851* | | - | 5261152 | 5261691 | 7.84 | 19.19 | 7.98 | 4.29 | Hypothetical protein |
| *FLPKDCKG_04852* | *sdpR-3* | - | 5261688 | 5262035 | 12.62 | 46.38 | 24.67 | 11.06 | Transcriptional repressor SdpR |
| *FLPKDCKG_04853* | *gfa-1* | - | 5262100 | 5262531 | 50.96 | 150.23 | 50.57 | 12.49 | Glutathione-dependent formaldehyde-activating enzyme |
| *FLPKDCKG_04854* | *phnC* | + | 5262730 | 5263572 | 0.17 | 0.45 | 0.67 | 0.13 | Phosphate-import ATP-binding protein PhnC |
| *FLPKDCKG_04855* | *phnD-2* | + | 5263602 | 5264582 | 0.26 | 1.10 | 1.09 | 1.79 | Phosphate-import protein PhnD |
| *FLPKDCKG_04856* | *phnE-3* | + | 5264660 | 5265454 | 0.06 | 0.62 | 0.00 | 0.00 | Phosphate-import permease protein PhnE |
| *FLPKDCKG_04857* | | - | 5265487 | 5265978 | 0.28 | 0.00 | 0.59 | 0.00 | Hypothetical protein |
| *FLPKDCKG_04858* | | - | 5265975 | 5266946 | 0.10 | 0.00 | 0.45 | 0.54 | Hypothetical protein |
| *FLPKDCKG_04859* | *ywrD-4* | - | 5267003 | 5267659 | 0.06 | 0.97 | 0.00 | 0.00 | Putative gamma-glutamyltransferase YwrD |
| *FLPKDCKG_04860* | | + | 5267688 | 5268884 | 0.16 | 0.00 | 0.03 | 0.00 | Hypothetical protein |
| *FLPKDCKG_04861* | *gbpR-5* | + | 5268941 | 5269900 | 1.16 | 4.57 | 4.19 | 0.99 | HTH-type transcriptional regulator GbpR |
| *FLPKDCKG_04862* | *phnN* | - | 5269942 | 5270520 | 0.99 | 9.14 | 4.52 | 0.22 | Ribose 1,5-bisphosphate phosphokinase PhnN |
| *FLPKDCKG_04863* | | - | 5270517 | 5271260 | 0.27 | 0.50 | 1.64 | 0.78 | Hypothetical protein |
| *FLPKDCKG_04864* | *phnM-1* | - | 5271263 | 5272459 | 0.90 | 1.86 | 0.88 | 0.29 | Alpha-D-ribose 1-methylphosphonate 5-triphosphate diphosphatase |
| *FLPKDCKG_04865* | *phnF* | - | 5272557 | 5273279 | 7.55 | 9.00 | 10.94 | 2.56 | putative transcriptional regulator PhnF |
| *FLPKDCKG_04866* | *phnG* | + | 5273464 | 5273934 | 0.25 | 0.00 | 0.00 | 0.00 | Alpha-D-ribose 1-methylphosphonate 5-triphosphate synthase subunit PhnG |
| *FLPKDCKG_04867* | *phnH* | + | 5273936 | 5274553 | 0.23 | 1.41 | 0.00 | 0.00 | Alpha-D-ribose 1-methylphosphonate 5-triphosphate synthase subunit PhnH |
| *FLPKDCKG_04868* | *phnI* | + | 5274555 | 5275670 | 0.08 | 0.34 | 0.05 | 0.00 | Alpha-D-ribose 1-methylphosphonate 5-triphosphate synthase subunit PhnI |
| *FLPKDCKG_04869* | *phnJ* | + | 5275667 | 5276557 | 0.56 | 2.20 | 1.39 | 0.78 | Alpha-D-ribose 1-methylphosphonate 5-phosphate C-P lyase |
| *FLPKDCKG_04870* | *phnK* | + | 5276831 | 5277328 | 0.00 | 0.37 | 0.00 | 0.80 | Putative phosphonates utilization ATP-binding protein PhnK |
| *FLPKDCKG_04871* | *phnL* | + | 5277331 | 5278047 | 0.00 | 0.77 | 0.00 | 0.00 | Alpha-D-ribose 1-methylphosphonate 5-triphosphate synthase subunit PhnL |
| *FLPKDCKG_04872* | *phnM-2* | + | 5278265 | 5279254 | 0.18 | 0.91 | 0.00 | 0.00 | Alpha-D-ribose 1-methylphosphonate 5-triphosphate diphosphatase |
| *FLPKDCKG_04873* | | + | 5279372 | 5279806 | 29.10 | 47.64 | 25.52 | 11.63 | Hypothetical protein |
| *FLPKDCKG_04874* | *dctA-2* | - | 5279920 | 5281281 | 2.65 | 7.58 | 5.24 | 1.43 | Aerobic C4-dicarboxylate transport protein |
| *FLPKDCKG_04875* | | + | 5281557 | 5282456 | 14.84 | 6.37 | 5.22 | 0.62 | 6-deoxy-6-sulfogluconolactonase |
| *FLPKDCKG_04876* | *fda* | + | 5282630 | 5283694 | 130.79 | 151.51 | 111.53 | 39.40 | Fructose-bisphosphate aldolase |
| *FLPKDCKG_04877* | | - | 5283780 | 5284235 | 10.77 | 19.24 | 18.14 | 4.94 | Hypothetical protein |
| *FLPKDCKG_04878* | *purC* | + | 5284406 | 5285287 | 34.33 | 43.65 | 32.73 | 12.09 | Phosphoribosylaminoimidazole-succinocarboxamide synthase |
| *FLPKDCKG_04879* | | + | 5285534 | 5285692 | 7.05 | 0.00 | 2.44 | 1.40 | Hypothetical protein |
| *FLPKDCKG_04880* | *dsbB-2* | + | 5285704 | 5286276 | 1.57 | 0.36 | 0.54 | 0.43 | Disulfide bond formation protein B |
| *FLPKDCKG_04881* | *purE* | + | 5286351 | 5286872 | 10.02 | 11.98 | 11.28 | 3.60 | N5-carboxyaminoimidazole ribonucleotide mutase |
| *FLPKDCKG_04882* | *purK* | + | 5286878 | 5288068 | 5.06 | 11.04 | 8.82 | 2.72 | N5-carboxyaminoimidazole ribonucleotide synthase |
| *FLPKDCKG_04883* | *ywlC* | + | 5288073 | 5289119 | 2.05 | 2.72 | 5.06 | 0.52 | Threonylcarbamoyl-AMP synthase |
| *FLPKDCKG_04884* | *pcaR-3* | - | 5289136 | 5289963 | 2.13 | 3.66 | 4.07 | 1.71 | Pca regulon regulatory protein |
| *FLPKDCKG_04885* | *pcaJ-3* | - | 5289963 | 5290604 | 1.52 | 2.40 | 0.00 | 1.06 | 3-oxoadipate CoA-transferase subunit B |
| *FLPKDCKG_04886* | *pcaI-3* | - | 5290614 | 5291294 | 1.00 | 2.04 | 0.41 | 1.01 | 3-oxoadipate CoA-transferase subunit A |
| *FLPKDCKG_04887* | | - | 5291339 | 5292355 | 3.33 | 6.53 | 4.74 | 2.10 | Hypothetical protein |
| *FLPKDCKG_04888* | *yhdW-6* | + | 5292711 | 5293733 | 39.37 | 21.70 | 28.59 | 17.43 | Putative amino-acid ABC transporter-binding protein YhdW |
| *FLPKDCKG_04889* | *ilvX* | + | 5293962 | 5295518 | 8.88 | 8.86 | 3.97 | 2.27 | Putative acetolactate synthase large subunit IlvX |
| *FLPKDCKG_04890* | *livF-23* | - | 5295558 | 5296268 | 0.46 | 1.92 | 0.00 | 1.32 | High-affinity branched-chain amino acid transport ATP-binding protein LivF |
| *FLPKDCKG_04891* | *ecfA3* | - | 5296256 | 5296684 | 0.16 | 1.92 | 0.80 | 2.76 | Energy-coupling factor transporter ATP-binding protein EcfA3 |
| *FLPKDCKG_04892* | | - | 5297021 | 5298061 | 0.15 | 1.05 | 0.38 | 0.86 | Hypothetical protein |
| *FLPKDCKG_04893* | *livH-23* | - | 5298064 | 5298945 | 0.36 | 0.27 | 0.00 | 0.14 | High-affinity branched-chain amino acid transport system permease protein LivH |
| *FLPKDCKG_04894* | | - | 5298961 | 5300121 | 0.22 | 1.09 | 1.79 | 3.22 | Leu/Ile/Val-binding protein |
| *FLPKDCKG_04895* | *menH-3* | - | 5300141 | 5300986 | 1.01 | 0.22 | 0.57 | 3.30 | 2-succinyl-6-hydroxy-2,4-cyclohexadiene-1-carboxylate synthase |

| Locus name | Gene name | Strand | Gene start (bp) | Gene end (bp) | Average TPM Inoculum | Average TPM $t_6$ | Average TPM $t_{24}$ | Average TPM $t_{168}$ | Annotation description |
|---|---|---|---|---|---|---|---|---|---|
| *FLPKDCKG_04896* | *bclA-2* | - | 5300983 | 5302542 | 2.11 | 3.25 | 0.58 | 1.59 | Benzoate--CoA ligase |
| *FLPKDCKG_04897* | | - | 5302702 | 5303172 | 13.31 | 3.38 | 1.85 | 11.63 | Hypothetical protein |
| *FLPKDCKG_04898* | *aroK-2* | + | 5303351 | 5304262 | 26.76 | 5.45 | 3.72 | 6.14 | Shikimate kinase |
| *FLPKDCKG_04899* | *boxC* | + | 5304501 | 5306162 | 32.83 | 5.39 | 4.36 | 7.01 | Benzoyl-CoA-dihydrodiol lyase |
| *FLPKDCKG_04900* | *boxB* | + | 5306272 | 5307699 | 12.05 | 3.97 | 5.79 | 7.07 | Benzoyl-CoA oxygenase component B |
| *FLPKDCKG_04901* | *boxA* | + | 5307751 | 5308998 | 7.04 | 8.39 | 2.57 | 4.42 | Benzoyl-CoA oxygenase component A |
| *FLPKDCKG_04902* | *paaZ-1* | + | 5309059 | 5309499 | 6.39 | 5.11 | 5.41 | 1.49 | Bifunctional protein PaaZ |
| *FLPKDCKG_04903* | | + | 5309804 | 5311513 | 14.54 | 9.42 | 6.56 | 1.07 | Hypothetical protein |
| *FLPKDCKG_04904* | | - | 5311834 | 5312502 | 11.25 | 6.05 | 4.09 | 1.55 | Hypothetical protein |
| *FLPKDCKG_04905* | | - | 5312767 | 5313753 | 8.48 | 7.82 | 12.20 | 3.85 | Hypothetical protein |
| *FLPKDCKG_04906* | *mmgC-3* | - | 5314005 | 5315198 | 26.55 | 18.26 | 11.38 | 9.04 | Acyl-CoA dehydrogenase |
| *FLPKDCKG_04907* | *ilvG-2* | - | 5315331 | 5317013 | 34.13 | 10.80 | 10.48 | 4.11 | Acetolactate synthase isozyme 2 large subunit |
| *FLPKDCKG_04908* | *perR* | + | 5317193 | 5318095 | 2.92 | 5.80 | 3.42 | 0.23 | HTH-type transcriptional regulator PerR |
| *FLPKDCKG_04909* | *xerD* | + | 5318226 | 5319209 | 3.02 | 2.43 | 3.42 | 1.46 | Tyrosine recombinase XerD |
| *FLPKDCKG_04910* | *ybaK-2* | + | 5319276 | 5319764 | 2.52 | 3.23 | 3.49 | 0.69 | Cys-tRNA(Pro)/Cys-tRNA(Cys) deacylase YbaK |
| *FLPKDCKG_04911* | *yraA-2* | + | 5319912 | 5320490 | 152.23 | 102.18 | 47.37 | 19.31 | Putative cysteine protease YraA |
| *FLPKDCKG_04912* | *adhT-2* | - | 5320641 | 5321669 | 94.95 | 22.91 | 12.57 | 5.54 | Alcohol dehydrogenase |
| *FLPKDCKG_04913* | | + | 5321881 | 5322270 | 39.30 | 25.64 | 24.39 | 8.86 | Hypothetical protein |
| *FLPKDCKG_04914* | *pbpG* | - | 5322406 | 5322621 | 9.39 | 8.65 | 2.38 | 0.00 | D-alanyl-D-alanine endopeptidase |
| *FLPKDCKG_04915* | | + | 5322924 | 5323142 | 23.17 | 27.03 | 17.97 | 6.92 | Hypothetical protein |
| *FLPKDCKG_04916* | | - | 5324000 | 5324074 | 4.18 | 7.39 | 4.69 | 1.11 | tRNA |
| *FLPKDCKG_04917* | *crt-3* | + | 5324293 | 5325057 | 1.21 | 1.62 | 0.65 | 0.99 | Short-chain-enoyl-CoA hydratase |
| *FLPKDCKG_04918* | *fadD3-2* | + | 5325054 | 5326676 | 0.70 | 1.16 | 1.13 | 0.30 | 3-[(3aS,4S,7aS)-7a-methyl-1,5-dioxo-octahydro-1H-inden-4-yl]propanoyl:CoA ligase |
| *FLPKDCKG_04919* | *yngG-2* | + | 5326673 | 5327623 | 0.88 | 2.83 | 0.93 | 0.33 | Hydroxymethylglutaryl-CoA lyase YngG |
| *FLPKDCKG_04920* | *mhqR-2* | + | 5327711 | 5328247 | 3.59 | 3.14 | 2.46 | 0.36 | HTH-type transcriptional regulator MhqR |
| *FLPKDCKG_04921* | *paaF-3* | + | 5328339 | 5329073 | 0.40 | 0.00 | 0.98 | 0.51 | 2,3-dehydroadipyl-CoA hydratase |
| *FLPKDCKG_04922* | *uctC-6* | + | 5329070 | 5330266 | 0.46 | 0.78 | 0.62 | 0.00 | Acetyl-CoA:oxalate CoA-transferase |
| *FLPKDCKG_04923* | | + | 5330338 | 5331309 | 0.72 | 0.43 | 0.82 | 0.00 | Hypothetical protein |
| *FLPKDCKG_04924* | | + | 5331461 | 5331715 | 66.58 | 19.05 | 16.44 | 4.21 | Hypothetical protein |
| *FLPKDCKG_04925* | *fitB* | + | 5331715 | 5332140 | 3.91 | 5.69 | 4.82 | 0.87 | Toxin FitB |
| *FLPKDCKG_04926* | | - | 5332216 | 5333169 | 0.69 | 1.25 | 1.14 | 0.00 | Nitronate monooxygenase |
| *FLPKDCKG_04927* | *yhjX-3* | + | 5333385 | 5334584 | 5.36 | 7.01 | 3.97 | 2.19 | putative MFS-type transporter YhjX |
| *FLPKDCKG_04928* | | + | 5334601 | 5334762 | 3.71 | 1.67 | 6.59 | 0.21 | Hypothetical protein |
| *FLPKDCKG_04929* | | + | 5334822 | 5336609 | 3.24 | 4.15 | 3.31 | 1.33 | Hypothetical protein |
| *FLPKDCKG_04930* | | + | 5336548 | 5337480 | 3.99 | 3.65 | 3.61 | 1.87 | Hypothetical protein |
| *FLPKDCKG_04931* | *addA-2* | + | 5337484 | 5341056 | 2.93 | 4.34 | 2.11 | 1.39 | ATP-dependent helicase/nuclease subunit A |
| *FLPKDCKG_04932* | *dnaX* | + | 5341645 | 5343936 | 11.57 | 17.09 | 9.92 | 4.64 | DNA polymerase III subunit tau |
| *FLPKDCKG_04933* | *ybaB* | + | 5343982 | 5344308 | 23.02 | 40.13 | 35.18 | 15.04 | Nucleoid-associated protein YbaB |
| *FLPKDCKG_04934* | *recR* | + | 5344376 | 5344984 | 5.52 | 16.95 | 12.50 | 4.29 | Recombination protein RecR |
| *FLPKDCKG_04935* | *smtB* | + | 5345083 | 5346372 | 5.98 | 10.25 | 11.35 | 2.18 | Succinyl-CoA--L-malate CoA-transferase beta subunit |
| *FLPKDCKG_04936* | *ssuA-5* | + | 5346353 | 5347393 | 42.81 | 27.81 | 24.47 | 7.26 | Putative aliphatic sulfonates-binding protein |
| *FLPKDCKG_04937* | *tauB-6* | + | 5347386 | 5348198 | 4.02 | 9.22 | 6.71 | 2.56 | Taurine import ATP-binding protein TauB |
| *FLPKDCKG_04938* | *ssuC-9* | + | 5348250 | 5349044 | 3.91 | 8.28 | 2.29 | 1.43 | Putative aliphatic sulfonates transport permease protein SsuC |
| *FLPKDCKG_04939* | | - | 5349085 | 5349660 | 3.61 | 2.19 | 1.93 | 1.20 | Hypothetical protein |
| *FLPKDCKG_04940* | | - | 5349657 | 5350307 | 6.83 | 4.17 | 0.89 | 1.18 | Hypothetical protein |
| *FLPKDCKG_04941* | *exaE* | - | 5350315 | 5350680 | 12.28 | 4.83 | 6.57 | 2.17 | Transcriptional activator protein ExaE |
| *FLPKDCKG_04942* | *bphP-3* | - | 5350677 | 5352884 | 28.79 | 15.78 | 6.99 | 2.41 | Bacteriophytochrome |
| *FLPKDCKG_04943* | | + | 5353051 | 5353578 | 5.80 | 14.76 | 12.10 | 2.61 | Hypothetical protein |
| *FLPKDCKG_04944* | *tal* | - | 5353620 | 5354573 | 36.02 | 57.41 | 27.52 | 15.24 | Transaldolase |
| *FLPKDCKG_04945* | *carA-1* | + | 5354797 | 5355951 | 58.24 | 83.74 | 58.44 | 22.57 | Carbamoyl-phosphate synthase small chain |
| *FLPKDCKG_04946* | *carB* | + | 5355962 | 5359204 | 18.03 | 25.57 | 21.96 | 6.79 | Carbamoyl-phosphate synthase large chain |
| *FLPKDCKG_04947* | | + | 5359253 | 5360170 | 7.19 | 15.11 | 12.32 | 3.20 | Hypothetical protein |
| *FLPKDCKG_04948* | | + | 5360235 | 5361008 | 2.07 | 2.66 | 3.73 | 0.42 | putative oxidoreductase |
| *FLPKDCKG_04949* | *pspA* | - | 5361061 | 5361705 | 2.25 | 5.59 | 3.11 | 5.02 | Phosphoserine phosphatase 1 |
| *FLPKDCKG_04950* | *cmpR-17* | - | 5361755 | 5362663 | 2.81 | 3.64 | 5.76 | 0.38 | HTH-type transcriptional activator CmpR |
| *FLPKDCKG_04951* | *potA-7* | - | 5362688 | 5363773 | 3.94 | 2.48 | 2.73 | 2.34 | Spermidine/putrescine import ATP-binding protein PotA |
| *FLPKDCKG_04952* | *phnU* | - | 5363790 | 5365499 | 1.50 | 1.26 | 1.11 | 0.22 | Putative 2-aminoethylphosphonate transport system permease protein PhnU |
| *FLPKDCKG_04953* | *pgtC* | - | 5365587 | 5366912 | 23.21 | 11.03 | 11.16 | 4.85 | Phosphoglycerate transport regulatory protein PgtC |
| *FLPKDCKG_04954* | | - | 5367056 | 5367487 | 13.42 | 2.86 | 3.00 | 0.71 | Hypothetical protein |
| *FLPKDCKG_04955* | *rutG* | + | 5367619 | 5368929 | 5.25 | 4.33 | 1.83 | 2.86 | Putative pyrimidine permease RutG |
| *FLPKDCKG_04956* | *tsr-6* | - | 5369044 | 5370663 | 41.37 | 6.41 | 5.70 | 5.32 | Methyl-accepting chemotaxis protein I |
| *FLPKDCKG_04957* | *greA* | + | 5371361 | 5371837 | 32.17 | 13.68 | 24.04 | 9.55 | Transcription elongation factor GreA |
| *FLPKDCKG_04958* | *yhbY* | - | 5371962 | 5372564 | 85.65 | 74.75 | 65.60 | 25.59 | RNA-binding protein YhbY |
| *FLPKDCKG_04959* | *rlmE* | + | 5372583 | 5373215 | 18.14 | 48.15 | 75.30 | 15.51 | Ribosomal RNA large subunit methyltransferase E |
| *FLPKDCKG_04960* | *ftsH* | + | 5373444 | 5375342 | 76.25 | 135.32 | 90.30 | 29.92 | ATP-dependent zinc metalloprotease FtsH |
| *FLPKDCKG_04961* | *folP* | + | 5375418 | 5376206 | 10.68 | 35.67 | 27.01 | 8.96 | Dihydropteroate synthase |
| *FLPKDCKG_04962* | *glmM* | + | 5376203 | 5377546 | 8.63 | 29.97 | 29.13 | 5.87 | Phosphoglucosamine mutase |
| *FLPKDCKG_04963* | | + | 5377765 | 5378568 | 51.90 | 45.49 | 36.33 | 4.45 | Hypothetical protein |
| *FLPKDCKG_04964* | | + | 5378571 | 5378681 | 24.41 | 44.66 | 20.97 | 5.70 | Hypothetical protein |

| Locus name | Gene name | Strand | Gene start (bp) | Gene end (bp) | Average TPM Inoculum | Average TPM $t_6$ | Average TPM $t_{24}$ | Average TPM $t_{168}$ | Annotation description |
|---|---|---|---|---|---|---|---|---|---|
| *FLPKDCKG_04965* | *gppA* | - | 5378739 | 5379062 | 3.66 | 6.07 | 1.26 | 1.21 | Guanosine-5'-triphosphate,3'-diphosphate pyrophosphatase |
| *FLPKDCKG_04966* | *ppx* | - | 5379073 | 5380224 | 10.41 | 19.56 | 16.31 | 6.21 | Exopolyphosphatase |
| *FLPKDCKG_04967* | *ppk* | + | 5380444 | 5382519 | 17.92 | 16.19 | 12.15 | 3.21 | Polyphosphate kinase |
| *FLPKDCKG_04968* | *pstS* | + | 5383046 | 5384086 | 18.40 | 47.63 | 76.79 | 34.57 | Phosphate-binding protein PstS |
| *FLPKDCKG_04969* | *pstC* | + | 5384224 | 5385240 | 5.58 | 9.48 | 8.70 | 5.17 | Phosphate transport system permease protein PstC |
| *FLPKDCKG_04970* | *pstA* | + | 5385265 | 5386065 | 5.12 | 12.19 | 8.46 | 2.59 | Phosphate transport system permease protein PstA |
| *FLPKDCKG_04971* | *pstB* | + | 5386169 | 5386945 | 6.98 | 10.49 | 15.10 | 7.76 | Phosphate import ATP-binding protein PstB |
| *FLPKDCKG_04972* | *genK* | - | 5387090 | 5388445 | 0.26 | 2.04 | 2.33 | 2.14 | Gentisate transporter |
| *FLPKDCKG_04973* | | + | 5388613 | 5388689 | 2.26 | 0.38 | 8.91 | 0.00 | tRNA |
| *FLPKDCKG_04974* | | + | 5391111 | 5394041 | 12.49 | 15.91 | 7.80 | 2.28 | Hypothetical protein |
| *FLPKDCKG_04975* | *dmlR-28* | - | 5394986 | 5395885 | 15.31 | 80.27 | 38.64 | 6.56 | HTH-type transcriptional regulator DmlR |
| *FLPKDCKG_04976* | *mmgC-4* | + | 5395989 | 5397149 | 0.42 | 1.11 | 0.47 | 0.00 | Acyl-CoA dehydrogenase |
| *FLPKDCKG_04977* | *fadB-2* | + | 5397158 | 5397577 | 1.72 | 12.07 | 2.43 | 0.00 | putative enoyl-CoA hydratase |
| *FLPKDCKG_04978* | *mmgB* | + | 5397604 | 5399205 | 3.58 | 15.37 | 7.19 | 1.34 | putative 3-hydroxybutyryl-CoA dehydrogenase |
| *FLPKDCKG_04979* | *pcaF-2* | + | 5399219 | 5400427 | 0.82 | 3.40 | 1.96 | 0.16 | Beta-ketoadipyl-CoA thiolase |
| *FLPKDCKG_04980* | | + | 5400533 | 5402206 | 0.45 | 1.16 | 1.48 | 0.24 | Electron transfer flavoprotein-ubiquinone oxidoreductase |
| *FLPKDCKG_04981* | | - | 5402500 | 5403108 | 0.31 | 0.71 | 0.80 | 0.00 | Hypothetical protein |
| *FLPKDCKG_04982* | | - | 5403188 | 5403601 | 0.23 | 1.77 | 0.98 | 0.96 | Hypothetical protein |
| *FLPKDCKG_04983* | *ilvA-3* | + | 5403855 | 5404109 | 1.44 | 0.00 | 2.79 | 1.19 | L-threonine dehydratase biosynthetic IlvA |
| *FLPKDCKG_04984* | | - | 5404031 | 5405137 | 12.19 | 33.54 | 16.58 | 5.84 | Hypothetical protein |
| *FLPKDCKG_04985* | *qorA-3* | - | 5405950 | 5406939 | 1.28 | 2.58 | 1.67 | 0.64 | Quinone oxidoreductase 1 |
| *FLPKDCKG_04986* | *pcaK* | + | 5407287 | 5408636 | 1.73 | 3.05 | 1.36 | 0.03 | 4-hydroxybenzoate transporter PcaK |
| *FLPKDCKG_04987* | *fadH-2* | + | 5408962 | 5410992 | 1.94 | 1.85 | 0.68 | 0.14 | 2,4-dienoyl-CoA reductase [NADPH] |
| *FLPKDCKG_04988* | | + | 5411021 | 5412670 | 1.30 | 0.95 | 1.82 | 0.00 | Long-chain-fatty-acid--CoA ligase |
| *FLPKDCKG_04989* | *fabG-18* | + | 5412698 | 5413465 | 3.91 | 2.70 | 2.01 | 0.05 | 3-oxoacyl-[acyl-carrier-protein] reductase FabG |
| *FLPKDCKG_04990* | *menB* | + | 5413484 | 5414284 | 3.03 | 1.79 | 1.32 | 0.00 | 1,4-dihydroxy-2-naphthoyl-CoA synthase |
| *FLPKDCKG_04991* | *crt-4* | + | 5414281 | 5415054 | 0.80 | 4.15 | 1.95 | 0.83 | Short-chain-enoyl-CoA hydratase |
| *FLPKDCKG_04992* | | + | 5415123 | 5416025 | 1.06 | 2.01 | 1.41 | 1.01 | Hypothetical protein |
| *FLPKDCKG_04993* | *paaG-3* | + | 5416049 | 5416843 | 1.78 | 4.12 | 1.16 | 0.56 | 1,2-epoxyphenylacetyl-CoA isomerase |
| *FLPKDCKG_04994* | *dmdC-3* | + | 5416976 | 5418769 | 1.79 | 3.50 | 2.44 | 0.56 | 3-methylmercaptopropionyl-CoA dehydrogenase |
| *FLPKDCKG_04995* | | + | 5419848 | 5420075 | 0.28 | 2.15 | 0.00 | 0.00 | Hypothetical protein |
| *FLPKDCKG_04996* | | + | 5420293 | 5420472 | 0.00 | 0.00 | 0.00 | 0.00 | Hypothetical protein |
| *FLPKDCKG_04997* | *gcvA-12* | + | 5420523 | 5422637 | 0.20 | 0.78 | 0.50 | 0.00 | Glycine cleavage system transcriptional activator |
| *FLPKDCKG_04998* | | + | 5423292 | 5424668 | 2.30 | 6.86 | 2.93 | 1.69 | Hypothetical protein |
| *FLPKDCKG_04999* | | + | 5424680 | 5424997 | 1.14 | 0.00 | 0.00 | 0.00 | Hypothetical protein |
| *FLPKDCKG_05000* | | + | 5425034 | 5426638 | 0.60 | 1.70 | 0.71 | 1.06 | Methylmalonyl-CoA carboxyltransferase 12S subunit |
| *FLPKDCKG_05001* | *accA1-2* | + | 5426642 | 5428657 | 0.82 | 1.83 | 0.39 | 0.25 | Acetyl-/propionyl-coenzyme A carboxylase alpha chain |
| *FLPKDCKG_05002* | *yfdE-3* | + | 5428671 | 5429891 | 0.88 | 0.95 | 0.93 | 0.34 | Acetyl-CoA:oxalate CoA-transferase |
| *FLPKDCKG_05003* | *paaZ-2* | + | 5429932 | 5430420 | 1.27 | 3.79 | 4.35 | 1.89 | Bifunctional protein PaaZ |
| *FLPKDCKG_05004* | | + | 5430485 | 5431450 | 0.53 | 4.79 | 0.24 | 0.79 | Hypothetical protein |
| *FLPKDCKG_05005* | | + | 5431509 | 5432471 | 0.80 | 3.90 | 2.06 | 1.14 | Putative monooxygenase |
| *FLPKDCKG_05006* | *dpgD* | + | 5432512 | 5433300 | 2.64 | 5.54 | 2.24 | 0.16 | Enoyl-CoA-hydratase |
| *FLPKDCKG_05007* | *paaF-4* | + | 5433303 | 5434103 | 1.35 | 6.63 | 4.32 | 0.51 | 2,3-dehydroadipyl-CoA hydratase |
| *FLPKDCKG_05008* | *tsaQ1-2* | - | 5434100 | 5434888 | 7.64 | 9.63 | 7.64 | 1.34 | HTH-type transcriptional regulator TsaQ1/TsaQ2 |
| *FLPKDCKG_05009* | *tsaQ1-3* | - | 5434891 | 5435697 | 5.55 | 14.04 | 5.02 | 1.52 | HTH-type transcriptional regulator TsaQ1/TsaQ2 |
| *FLPKDCKG_05010* | *namA-1* | - | 5435803 | 5436933 | 0.80 | 1.75 | 2.02 | 0.00 | NADPH dehydrogenase |
| *FLPKDCKG_05011* | | - | 5436972 | 5437262 | 2.55 | 4.44 | 1.06 | 0.00 | Hypothetical protein |
| *FLPKDCKG_05012* | *bacC-6* | - | 5437309 | 5438070 | 0.68 | 2.80 | 2.55 | 0.00 | Dihydroanticapsin 7-dehydrogenase |
| *FLPKDCKG_05013* | *livF-24* | - | 5438117 | 5438839 | 0.19 | 0.00 | 0.28 | 0.00 | High-affinity branched-chain amino acid transport ATP-binding protein LivF |
| *FLPKDCKG_05014* | *cysA-6* | - | 5438829 | 5439563 | 0.55 | 1.32 | 1.13 | 0.00 | Sulfate/thiosulfate import ATP-binding protein CysA |
| *FLPKDCKG_05015* | | - | 5439571 | 5440515 | 0.28 | 1.84 | 0.81 | 0.00 | Hypothetical protein |
| *FLPKDCKG_05016* | *livH-24* | - | 5440508 | 5441371 | 0.46 | 0.35 | 0.00 | 0.00 | High-affinity branched-chain amino acid transport system permease protein LivH |
| *FLPKDCKG_05017* | *braC-9* | - | 5441421 | 5442629 | 0.63 | 3.24 | 0.93 | 0.50 | Leucine-, isoleucine-, valine-, threonine-, and alanine-binding protein |
| *FLPKDCKG_05018* | *mmsB-2* | - | 5442705 | 5443595 | 0.42 | 0.63 | 0.00 | 0.45 | 3-hydroxyisobutyrate dehydrogenase |
| *FLPKDCKG_05019* | *crt-5* | - | 5443610 | 5444425 | 1.22 | 3.67 | 0.90 | 0.93 | Short-chain-enoyl-CoA hydratase |
| *FLPKDCKG_05020* | *gcvA-13* | + | 5444739 | 5445509 | 2.99 | 2.49 | 3.60 | 2.81 | Glycine cleavage system transcriptional activator |
| *FLPKDCKG_05021* | *scoA-2* | + | 5445758 | 5446330 | 1.72 | 3.87 | 0.47 | 0.91 | putative succinyl-CoA:3-ketoacid coenzyme A transferase subunit A |
| *FLPKDCKG_05022* | *scoB-2* | + | 5446343 | 5446981 | 2.82 | 4.76 | 3.42 | 1.36 | putative succinyl-CoA:3-ketoacid coenzyme A transferase subunit B |
| *FLPKDCKG_05023* | *proC-3* | - | 5447082 | 5447858 | 4.93 | 9.22 | 1.54 | 2.24 | Pyrroline-5-carboxylate reductase |
| *FLPKDCKG_05024* | *rutR-2* | - | 5447996 | 5448667 | 9.42 | 11.76 | 5.65 | 0.67 | HTH-type transcriptional regulator RutR |
| *FLPKDCKG_05025* | | + | 5448774 | 5449736 | 2.24 | 2.26 | 3.14 | 0.71 | Putative monooxygenase |
| *FLPKDCKG_05026* | *paaG-4* | + | 5449817 | 5450623 | 9.19 | 15.47 | 9.12 | 3.28 | 1,2-epoxyphenylacetyl-CoA isomerase |
| *FLPKDCKG_05027* | *iscS-2* | - | 5450728 | 5451945 | 17.11 | 17.17 | 7.03 | 1.76 | Cysteine desulfurase IscS |
| *FLPKDCKG_05028* | *lysN-7* | + | 5452042 | 5453481 | 5.81 | 12.10 | 8.59 | 1.99 | 2-aminoadipate transaminase |
| *FLPKDCKG_05029* | *intA-2* | - | 5454249 | 5455679 | 16.87 | 19.31 | 13.23 | 2.96 | Prophage integrase IntA |
| *FLPKDCKG_05030* | | + | 5456265 | 5456621 | 1.53 | 0.00 | 1.54 | 2.57 | Hypothetical protein |
| *FLPKDCKG_05031* | *bshA-3* | + | 5456621 | 5457724 | 0.33 | 0.64 | 0.19 | 0.39 | N-acetyl-alpha-D-glucosaminyl L-malate synthase |
| *FLPKDCKG_05032* | *ahpD-3* | + | 5457830 | 5458297 | 3.62 | 6.37 | 5.39 | 1.30 | Alkyl hydroperoxide reductase AhpD |
| *FLPKDCKG_05033* | *argO-2* | - | 5458317 | 5458919 | 4.06 | 7.89 | 1.54 | 1.19 | Arginine exporter protein ArgO |

| Locus name | Gene name | Strand | Gene start (bp) | Gene end (bp) | Average TPM Inoculum | Average TPM $t_6$ | Average TPM $t_{24}$ | Average TPM $t_{168}$ | Annotation description |
|---|---|---|---|---|---|---|---|---|---|
| *FLPKDCKG_05034* | | - | 5459025 | 5459276 | 10.85 | 10.43 | 9.90 | 3.46 | Hypothetical protein |
| *FLPKDCKG_05035* | | - | 5459521 | 5459745 | 267.11 | 491.71 | 263.61 | 82.83 | Hypothetical protein |
| *FLPKDCKG_05036* | | + | 5459947 | 5461149 | 7.66 | 15.61 | 15.27 | 4.35 | Hypothetical protein |
| *FLPKDCKG_05037* | | - | 5461179 | 5461802 | 69.21 | 93.07 | 67.04 | 15.76 | Hypothetical protein |
| *FLPKDCKG_05038* | *dmlR-29* | - | 5461932 | 5462903 | 8.34 | 15.68 | 4.97 | 3.57 | HTH-type transcriptional regulator DmlR |
| *FLPKDCKG_05039* | *exsA-1* | - | 5463030 | 5463977 | 2.81 | 3.98 | 5.09 | 1.50 | Exoenzyme S synthesis regulatory protein ExsA |
| *FLPKDCKG_05040* | *gpr-3* | + | 5464297 | 5465343 | 1.37 | 3.92 | 1.11 | 1.22 | L-glyceraldehyde 3-phosphate reductase |
| *FLPKDCKG_05041* | | + | 5465474 | 5465587 | 18.45 | 14.57 | 11.68 | 5.08 | Hypothetical protein |
| *FLPKDCKG_05042* | | + | 5465602 | 5466360 | 2.32 | 3.69 | 4.99 | 0.50 | Hypothetical protein |
| *FLPKDCKG_05043* | | - | 5466396 | 5467154 | 8.97 | 52.46 | 45.21 | 6.24 | Hypothetical protein |
| *FLPKDCKG_05044* | | - | 5467275 | 5467820 | 28.81 | 1583.12 | 802.74 | 74.88 | Immunogenic protein MPB70 |
| *FLPKDCKG_05045* | *tpm* | + | 5468587 | 5469246 | 9.54 | 13.33 | 9.63 | 1.54 | Thiopurine S-methyltransferase |
| *FLPKDCKG_05046* | | - | 5469335 | 5469934 | 4.02 | 4.11 | 4.66 | 3.20 | Hypothetical protein |
| *FLPKDCKG_05047* | *acnB* | - | 5470162 | 5472753 | 10.68 | 45.46 | 37.93 | 19.24 | Aconitate hydratase B |
| *FLPKDCKG_05048* | | - | 5473074 | 5473973 | 2.32 | 13.07 | 6.54 | 1.92 | Hypothetical protein |
| *FLPKDCKG_05049* | | - | 5473963 | 5474847 | 2.42 | 7.78 | 9.11 | 3.61 | Hypothetical protein |
| *FLPKDCKG_05050* | | + | 5475140 | 5475904 | 3.94 | 12.17 | 7.24 | 2.65 | Hypothetical protein |
| *FLPKDCKG_05051* | *hrp1-2* | - | 5475938 | 5476378 | 99.08 | 100.11 | 65.67 | 10.12 | Hypoxic response protein 1 |
| *FLPKDCKG_05052* | | - | 5476362 | 5476835 | 62.53 | 61.09 | 26.92 | 7.38 | Hypothetical protein |
| *FLPKDCKG_05053* | | - | 5477221 | 5478234 | 2.57 | 8.22 | 2.51 | 0.09 | 4-hydroxyproline 2-epimerase |
| *FLPKDCKG_05054* | *ydcV-3* | - | 5478274 | 5479065 | 0.64 | 3.40 | 0.70 | 0.16 | Inner membrane ABC transporter permease protein YdcV |
| *FLPKDCKG_05055* | *potB-3* | - | 5479062 | 5479910 | 0.33 | 0.00 | 0.00 | 0.59 | Spermidine/putrescine transport system permease protein PotB |
| *FLPKDCKG_05056* | *potA-8* | - | 5479907 | 5480956 | 0.47 | 0.00 | 0.60 | 0.18 | Spermidine/putrescine import ATP-binding protein PotA |
| *FLPKDCKG_05057* | *potD-3* | - | 5480973 | 5482028 | 1.89 | 2.40 | 2.49 | 0.05 | Spermidine/putrescine-binding periplasmic protein |
| *FLPKDCKG_05058* | *cdhR-8* | + | 5482153 | 5483235 | 11.46 | 18.96 | 12.74 | 1.77 | HTH-type transcriptional regulator CdhR |
| *FLPKDCKG_05059* | | + | 5483260 | 5483970 | 12.53 | 39.88 | 16.07 | 5.79 | Hypothetical protein |
| *FLPKDCKG_05060* | *bdlA-2* | + | 5484347 | 5485816 | 13.86 | 5.80 | 3.88 | 2.93 | Biofilm dispersion protein BdlA |
| *FLPKDCKG_05061* | | + | 5485937 | 5486161 | 2.02 | 2.83 | 1.49 | 1.76 | Hypothetical protein |
| *FLPKDCKG_05062* | *polC* | + | 5486165 | 5486794 | 0.72 | 1.87 | 2.86 | 0.00 | DNA polymerase III PolC-type |
| *FLPKDCKG_05063* | | - | 5486876 | 5487166 | 8.66 | 6.85 | 5.87 | 2.56 | Hypothetical protein |
| *FLPKDCKG_05064* | | + | 5487692 | 5488171 | 1.22 | 0.79 | 0.64 | 1.27 | Hypothetical protein |
| *FLPKDCKG_05065* | | + | 5488219 | 5488920 | 0.13 | 2.53 | 1.21 | 0.57 | Hypothetical protein |
| *FLPKDCKG_05066* | *mscM* | - | 5488953 | 5491367 | 1.91 | 4.61 | 5.34 | 2.12 | Miniconductance mechanosensitive channel MscM |
| *FLPKDCKG_05067* | *cynR-17* | - | 5491478 | 5492383 | 2.13 | 3.37 | 2.15 | 0.54 | HTH-type transcriptional regulator CynR |
| *FLPKDCKG_05068* | *cat1* | + | 5492622 | 5494139 | 1.05 | 1.89 | 2.88 | 0.11 | Succinyl-CoA:coenzyme A transferase |
| *FLPKDCKG_05069* | *gcvA-14* | - | 5494169 | 5495116 | 4.70 | 8.31 | 9.41 | 2.87 | Glycine cleavage system transcriptional activator |
| *FLPKDCKG_05070* | | + | 5495283 | 5495624 | 3.00 | 8.19 | 5.35 | 5.35 | Hypothetical protein |
| *FLPKDCKG_05071* | | + | 5496932 | 5498452 | 186307.66 | 155145.78 | 143339.12 | 135907.70 | rRNA |
| *FLPKDCKG_05072* | | + | 5498588 | 5498664 | 34.01 | 67.22 | 67.45 | 26.03 | tRNA |
| *FLPKDCKG_05073* | | + | 5498695 | 5498770 | 27.99 | 5.10 | 12.78 | 2.91 | tRNA |
| *FLPKDCKG_05074* | | + | 5499069 | 5501947 | 125852.86 | 123299.07 | 161841.23 | 218766.54 | rRNA |
| *FLPKDCKG_05075* | | + | 5502150 | 5502261 | 153.45 | 280.09 | 308.65 | 253.48 | rRNA |
| *FLPKDCKG_05076* | | + | 5502439 | 5502550 | 27.67 | 78.68 | 97.67 | 69.07 | rRNA |
| *FLPKDCKG_05077* | *novR-4* | - | 5502685 | 5503449 | 10.66 | 11.52 | 7.71 | 2.86 | Decarboxylase NovR |
| *FLPKDCKG_05078* | | - | 5503573 | 5504286 | 13.02 | 17.62 | 7.47 | 3.38 | (S)-2-haloacid dehalogenase |
| *FLPKDCKG_05079* | *yofA-6* | + | 5504482 | 5505369 | 2.83 | 5.82 | 10.28 | 3.62 | HTH-type transcriptional regulator YofA |
| *FLPKDCKG_05080* | | - | 5505361 | 5505756 | 7.29 | 12.94 | 9.88 | 4.58 | Hypothetical protein |
| *FLPKDCKG_05081* | | - | 5505765 | 5506037 | 38.84 | 32.31 | 39.22 | 6.98 | Hypothetical protein |
| *FLPKDCKG_05082* | *lcdH-1* | - | 5506133 | 5507110 | 30.47 | 23.28 | 14.94 | 3.35 | L-carnitine dehydrogenase |
| *FLPKDCKG_05083* | | + | 5507223 | 5507855 | 3.62 | 4.67 | 2.65 | 0.81 | Hypothetical protein |
| *FLPKDCKG_05084* | *stp-5* | - | 5507941 | 5509344 | 1.46 | 4.84 | 2.98 | 1.47 | Multidrug resistance protein Stp |
| *FLPKDCKG_05085* | *dmlR-30* | + | 5509493 | 5510359 | 2.87 | 3.47 | 3.52 | 0.28 | HTH-type transcriptional regulator DmlR |
| *FLPKDCKG_05086* | *aam-3* | + | 5510474 | 5511877 | 5.09 | 7.84 | 6.66 | 1.47 | Acylamidase |
| *FLPKDCKG_05087* | | - | 5511892 | 5512431 | 2.88 | 2.56 | 2.63 | 2.15 | Acetyltransferase |
| *FLPKDCKG_05088* | *hpcG-1* | - | 5512463 | 5513245 | 3.13 | 4.35 | 4.13 | 1.29 | 2-oxo-hept-4-ene-1,7-dioate hydratase |
| *FLPKDCKG_05089* | | - | 5513251 | 5513631 | 3.14 | 4.35 | 3.71 | 1.44 | Hypothetical protein |
| *FLPKDCKG_05090* | *proA-6* | - | 5513682 | 5514962 | 15.05 | 11.47 | 10.32 | 2.67 | Gamma-glutamyl phosphate reductase |
| *FLPKDCKG_05091* | *holA* | - | 5515050 | 5516108 | 2.74 | 6.22 | 5.22 | 2.27 | DNA polymerase III subunit delta |
| *FLPKDCKG_05092* | *lptE* | - | 5516112 | 5516774 | 14.81 | 17.65 | 14.43 | 5.00 | LPS-assembly lipoprotein LptE |
| *FLPKDCKG_05093* | *leuS* | - | 5516786 | 5519443 | 11.55 | 18.76 | 11.56 | 5.13 | Leucine--tRNA ligase |
| *FLPKDCKG_05094* | | - | 5519530 | 5520744 | 2.55 | 2.74 | 3.31 | 1.09 | putative transporter |
| *FLPKDCKG_05095* | *mhqR-3* | - | 5520797 | 5521309 | 17.50 | 19.38 | 10.19 | 2.13 | HTH-type transcriptional regulator MhqR |
| *FLPKDCKG_05096* | | - | 5521473 | 5521931 | 85.16 | 82.19 | 73.56 | 41.00 | Hypothetical protein |
| *FLPKDCKG_05097* | *dacC* | + | 5522289 | 5523464 | 5.76 | 8.72 | 6.83 | 1.72 | D-alanyl-D-alanine carboxypeptidase DacC |
| *FLPKDCKG_05098* | *scdA* | + | 5523480 | 5524028 | 7.08 | 11.49 | 10.97 | 2.41 | Iron-sulfur cluster repair protein ScdA |
| *FLPKDCKG_05099* | *pphA-2* | + | 5524112 | 5524984 | 15.56 | 10.59 | 8.96 | 5.04 | Phosphonopyruvate hydrolase |
| *FLPKDCKG_05100* | | + | 5525022 | 5526806 | 2.07 | 3.49 | 2.52 | 0.25 | putative sulfate transporter |
| *FLPKDCKG_05101* | *lptB-10* | - | 5526814 | 5527578 | 0.86 | 3.73 | 2.04 | 1.47 | Lipopolysaccharide export system ATP-binding protein LptB |
| *FLPKDCKG_05102* | *livH-25* | - | 5527575 | 5529452 | 1.01 | 1.85 | 3.07 | 0.70 | High-affinity branched-chain amino acid transport system permease protein LivH |

| Locus name | Gene name | Strand | Gene start (bp) | Gene end (bp) | Average TPM Inoculum | Average TPM $t_6$ | Average TPM $t_{24}$ | Average TPM $t_{168}$ | Annotation description |
|---|---|---|---|---|---|---|---|---|---|
| *FLPKDCKG_05103* | *amiC-9* | - | 5529452 | 5530672 | 9.51 | 8.63 | 7.83 | 1.66 | Aliphatic amidase expression-regulating protein |
| *FLPKDCKG_05104* | *rpmG* | - | 5530871 | 5531038 | 15.93 | 25.66 | 14.27 | 18.47 | 50S ribosomal protein L33 |
| *FLPKDCKG_05105* | *rpmB* | - | 5531076 | 5531312 | 170.50 | 485.31 | 369.25 | 142.40 | 50S ribosomal protein L28 |
| *FLPKDCKG_05106* | *bcr-2* | - | 5531766 | 5532935 | 2.60 | 8.07 | 6.13 | 2.47 | Bicyclomycin resistance protein |
| *FLPKDCKG_05107* | *cmaA1* | + | 5533124 | 5534398 | 28.83 | 37.09 | 15.64 | 3.87 | Cyclopropane mycolic acid synthase 1 |
| *FLPKDCKG_05108* | *proX-2* | + | 5534815 | 5535816 | 2.37 | 91.40 | 82.19 | 22.84 | Glycine betaine/proline betaine-binding periplasmic protein |
| *FLPKDCKG_05109* | *proW* | + | 5536102 | 5536959 | 0.61 | 6.13 | 8.96 | 3.07 | Glycine betaine/proline betaine transport system permease protein ProW |
| *FLPKDCKG_05110* | *proV* | + | 5536956 | 5538254 | 4.18 | 18.31 | 15.62 | 2.25 | Glycine betaine/proline betaine transport system ATP-binding protein ProV |
| *FLPKDCKG_05111* | *xylH-1* | + | 5538390 | 5538764 | 10.52 | 13.18 | 14.57 | 9.03 | 2-hydroxymuconate tautomerase |
| *FLPKDCKG_05112* | *ccpA* | + | 5538774 | 5539793 | 3.53 | 8.93 | 7.40 | 2.96 | Catabolite control protein A |
| *FLPKDCKG_05113* | *udh-2* | - | 5539806 | 5540717 | 1.51 | 15.93 | 9.73 | 2.89 | Uronate dehydrogenase |
| *FLPKDCKG_05114* | *pgl-2* | - | 5540734 | 5541816 | 2.82 | 34.17 | 15.04 | 3.72 | 6-phosphogluconolactonase |
| *FLPKDCKG_05115* | *xylH-2* | - | 5541809 | 5542996 | 2.08 | 16.31 | 7.28 | 2.30 | Xylose transport system permease protein XylH |
| *FLPKDCKG_05116* | *xylG* | - | 5543002 | 5545578 | 4.17 | 35.22 | 18.47 | 4.84 | Xylose import ATP-binding protein XylG |
| *FLPKDCKG_05117* | *oprB* | - | 5545605 | 5546984 | 2.79 | 51.93 | 18.21 | 3.76 | Porin B |
| *FLPKDCKG_05118* | | - | 5547236 | 5548210 | 0.96 | 0.35 | 0.00 | 0.00 | Hypothetical protein |
| *FLPKDCKG_05119* | *paaZ-3* | - | 5548297 | 5548770 | 5.75 | 6.39 | 0.78 | 0.00 | Bifunctional protein PaaZ |
| *FLPKDCKG_05120* | *hcaR-7* | + | 5550081 | 5550977 | 3.46 | 8.75 | 3.88 | 2.59 | Hca operon transcriptional activator HcaR |
| *FLPKDCKG_05121* | | + | 5551861 | 5551958 | 4.85 | 0.00 | 0.00 | 0.00 | tRNA |
| *FLPKDCKG_05122* | | - | 5552012 | 5553163 | 0.92 | 0.58 | 0.36 | 0.00 | Acyl-CoA dehydrogenase FadE26 |
| *FLPKDCKG_05123* | *bktB-1* | + | 5553344 | 5553772 | 129.75 | 107.56 | 66.64 | 20.51 | Beta-ketothiolase BktB |
| *FLPKDCKG_05124* | *bktB-2* | + | 5553745 | 5554524 | 34.65 | 54.13 | 37.36 | 12.38 | Beta-ketothiolase BktB |
| *FLPKDCKG_05125* | *speE-3* | + | 5554686 | 5555456 | 4.66 | 8.70 | 6.06 | 1.95 | Polyamine aminopropyltransferase |
| *FLPKDCKG_05126* | *rspR-9* | + | 5555568 | 5556317 | 56.05 | 18.23 | 11.15 | 5.15 | HTH-type transcriptional repressor RspR |
| *FLPKDCKG_05127* | | + | 5556268 | 5557581 | 37.41 | 10.02 | 9.62 | 2.55 | Hypothetical protein |
| *FLPKDCKG_05128* | *echA8-5* | + | 5557630 | 5558421 | 22.71 | 8.63 | 4.57 | 2.77 | putative enoyl-CoA hydratase echA8 |
| *FLPKDCKG_05129* | *lcfB-4* | + | 5558427 | 5559968 | 12.92 | 2.76 | 5.38 | 2.64 | Long-chain-fatty-acid--CoA ligase |
| *FLPKDCKG_05130* | | + | 5560130 | 5561104 | 14.36 | 6.83 | 8.97 | 4.56 | Hypothetical protein |
| *FLPKDCKG_05131* | *gpr-4* | - | 5561132 | 5562088 | 23.36 | 28.07 | 16.95 | 7.06 | L-glyceraldehyde 3-phosphate reductase |
| *FLPKDCKG_05132* | | + | 5562330 | 5563301 | 53.00 | 50.28 | 47.92 | 25.29 | Hypothetical protein |
| *FLPKDCKG_05133* | | + | 5563693 | 5564583 | 2.04 | 4.19 | 3.90 | 2.43 | Hypothetical protein |
| *FLPKDCKG_05134* | *ecfA2* | + | 5564585 | 5565379 | 1.94 | 6.15 | 3.24 | 4.14 | Energy-coupling factor transporter ATP-binding protein EcfA2 |
| *FLPKDCKG_05135* | *dadA-3* | - | 5565482 | 5566732 | 2.25 | 2.09 | 3.13 | 0.09 | D-amino acid dehydrogenase |
| *FLPKDCKG_05136* | *mnmC-3* | - | 5566892 | 5568529 | 0.61 | 2.07 | 1.06 | 0.72 | tRNA 5-methylaminomethyl-2-thiouridine biosynthesis bifunctional protein MnmC |
| *FLPKDCKG_05137* | *speE-4* | - | 5568529 | 5570040 | 2.10 | 4.00 | 3.56 | 1.07 | Polyamine aminopropyltransferase |
| *FLPKDCKG_05138* | | + | 5570136 | 5570522 | 46.66 | 104.43 | 57.36 | 12.62 | Hypothetical protein |
| *FLPKDCKG_05139* | | + | 5570567 | 5571604 | 10.57 | 22.60 | 8.39 | 6.23 | Hypothetical protein |
| *FLPKDCKG_05140* | | + | 5571618 | 5573087 | 5.37 | 15.11 | 9.85 | 4.12 | Hypothetical protein |
| *FLPKDCKG_05141* | | + | 5573091 | 5573249 | 2.24 | 17.26 | 11.63 | 3.09 | Hypothetical protein |
| *FLPKDCKG_05142* | *speH* | + | 5573246 | 5573596 | 0.61 | 3.51 | 4.94 | 1.28 | S-adenosylmethionine decarboxylase proenzyme |
| *FLPKDCKG_05143* | *modA* | + | 5573722 | 5574480 | 21.68 | 2.89 | 5.33 | 0.80 | Molybdate-binding periplasmic protein |
| *FLPKDCKG_05144* | *modB-2* | + | 5574511 | 5575194 | 2.90 | 0.40 | 3.91 | 0.76 | Molybdenum transport system permease protein ModB |
| *FLPKDCKG_05145* | *modC* | + | 5575191 | 5575877 | 2.99 | 5.31 | 3.67 | 2.61 | Molybdenum import ATP-binding protein ModC |
| *FLPKDCKG_05146* | *mopA* | - | 5575937 | 5576731 | 12.06 | 21.86 | 15.04 | 2.27 | Molybdenum-pterin-binding protein MopA |
| *FLPKDCKG_05147* | | + | 5576930 | 5577529 | 140.08 | 25.13 | 23.78 | 11.31 | Hypothetical protein |
| *FLPKDCKG_05148* | | + | 5577553 | 5578749 | 17.16 | 10.04 | 8.80 | 2.43 | Hypothetical protein |
| *FLPKDCKG_05149* | *mprA-2* | + | 5578905 | 5579348 | 1.05 | 0.77 | 1.39 | 0.00 | Response regulator MprA |
| *FLPKDCKG_05150* | *ttgA* | + | 5579722 | 5580864 | 25.70 | 30.08 | 29.18 | 11.36 | Toluene efflux pump periplasmic linker protein TtgA |
| *FLPKDCKG_05151* | *acrB-2* | + | 5580928 | 5584149 | 12.97 | 15.20 | 13.39 | 6.57 | Multidrug efflux pump subunit AcrB |
| *FLPKDCKG_05152* | *oprM-8* | + | 5584146 | 5585633 | 12.04 | 14.97 | 15.92 | 4.83 | Outer membrane protein OprM |
| *FLPKDCKG_05153* | *pdxJ* | + | 5585630 | 5586370 | 14.22 | 22.51 | 13.35 | 6.23 | Pyridoxine 5'-phosphate synthase |
| *FLPKDCKG_05154* | *acpS* | + | 5586406 | 5586843 | 8.86 | 11.50 | 12.14 | 0.98 | Holo-[acyl-carrier-protein] synthase |
| *FLPKDCKG_05155* | *nagZ* | + | 5586881 | 5587936 | 3.87 | 13.23 | 8.86 | 2.91 | Beta-hexosaminidase |
| *FLPKDCKG_05156* | *uvrC* | + | 5587899 | 5589818 | 1.47 | 5.14 | 4.60 | 1.22 | UvrABC system protein C |
| *FLPKDCKG_05157* | *pgsA* | + | 5589808 | 5590380 | 47.24 | 55.75 | 36.37 | 6.35 | CDP-diacylglycerol--glycerol-3-phosphate 3-phosphatidyltransferase |
| *FLPKDCKG_05158* | *fsr-2* | + | 5590577 | 5591863 | 1.91 | 2.45 | 2.08 | 2.97 | Fosmidomycin resistance protein |
| *FLPKDCKG_05159* | *dmlA-2* | - | 5591973 | 5593049 | 0.26 | 0.56 | 2.03 | 0.66 | D-malate dehydrogenase [decarboxylating] |
| *FLPKDCKG_05160* | *dmlR-31* | + | 5593189 | 5594097 | 1.97 | 2.25 | 2.12 | 0.02 | HTH-type transcriptional regulator DmlR |
| *FLPKDCKG_05161* | *ccp-1* | + | 5594359 | 5595363 | 6.57 | 1.75 | 3.21 | 0.00 | Cytochrome c551 peroxidase |
| *FLPKDCKG_05162* | *queF* | - | 5595437 | 5596261 | 3.15 | 11.70 | 7.99 | 2.89 | NADPH-dependent 7-cyano-7-deazaguanine reductase |
| *FLPKDCKG_05163* | *aceA* | + | 5596585 | 5597904 | 1340.02 | 103.02 | 214.22 | 54.89 | Isocitrate lyase |
| *FLPKDCKG_05164* | *nimR-12* | - | 5598019 | 5598399 | 4.22 | 2.87 | 3.77 | 0.57 | HTH-type transcriptional regulator NimR |
| *FLPKDCKG_05165* | *nimR-13* | - | 5598399 | 5598902 | 1.70 | 0.73 | 2.60 | 1.75 | HTH-type transcriptional regulator NimR |
| *FLPKDCKG_05166* | | - | 5599070 | 5599282 | 0.00 | 0.00 | 0.00 | 0.00 | Hypothetical protein |
| *FLPKDCKG_05167* | *menH-4* | + | 5599384 | 5599680 | 0.15 | 1.44 | 0.00 | 0.00 | 2-succinyl-6-hydroxy-2,4-cyclohexadiene-1-carboxylate synthase |
| *FLPKDCKG_05168* | *bioD1* | - | 5599702 | 5600364 | 1.03 | 3.23 | 3.55 | 0.26 | ATP-dependent dethiobiotin synthetase BioD 1 |
| *FLPKDCKG_05169* | *bioC-3* | - | 5600357 | 5601130 | 1.12 | 1.01 | 0.00 | 0.40 | Malonyl-[acyl-carrier protein] O-methyltransferase |
| *FLPKDCKG_05170* | *bioH* | - | 5601127 | 5601792 | 0.25 | 1.85 | 0.38 | 0.00 | Pimeloyl-[acyl-carrier protein] methyl ester esterase |
| *FLPKDCKG_05171* | *bioF* | - | 5601792 | 5602976 | 0.31 | 2.75 | 1.67 | 0.15 | 8-amino-7-oxononanoate synthase |

| Locus name | Gene name | Strand | Gene start (bp) | Gene end (bp) | Average TPM Inoculum | Average TPM $t_6$ | Average TPM $t_{24}$ | Average TPM $t_{168}$ | Annotation description |
|---|---|---|---|---|---|---|---|---|---|
| *FLPKDCKG_05172* | *bioA* | + | 5603089 | 5604357 | 2.75 | 3.90 | 1.86 | 0.84 | Adenosylmethionine-8-amino-7-oxononanoate aminotransferase |
| *FLPKDCKG_05173* | *ctpV-1* | + | 5604447 | 5605421 | 13.59 | 0.46 | 0.78 | 0.57 | putative copper-exporting P-type ATPase V |
| *FLPKDCKG_05174* | | + | 5605418 | 5605579 | 5.32 | 2.68 | 0.00 | 0.78 | Hypothetical protein |
| *FLPKDCKG_05175* | *ccoN1-1* | + | 5605727 | 5607211 | 16.88 | 0.58 | 0.61 | 0.05 | Cbb3-type cytochrome c oxidase subunit CcoN1 |
| *FLPKDCKG_05176* | | + | 5607222 | 5607947 | 18.53 | 3.14 | 0.00 | 0.09 | Hypothetical protein |
| *FLPKDCKG_05177* | | + | 5607950 | 5608123 | 40.57 | 5.44 | 0.00 | 1.83 | Hypothetical protein |
| *FLPKDCKG_05178* | *ccoP-1* | + | 5608120 | 5609070 | 27.64 | 2.12 | 2.02 | 0.60 | Cbb3-type cytochrome c oxidase subunit CcoP |
| *FLPKDCKG_05179* | | + | 5609139 | 5610629 | 3.42 | 0.61 | 0.23 | 0.00 | Hypothetical protein |
| *FLPKDCKG_05180* | | + | 5610652 | 5610888 | 32.68 | 1.74 | 0.00 | 0.00 | Hypothetical protein |
| *FLPKDCKG_05181* | | + | 5610908 | 5611144 | 19.50 | 3.41 | 0.00 | 0.00 | Hypothetical protein |
| *FLPKDCKG_05182* | *gbpR-6* | - | 5611541 | 5612467 | 5.81 | 4.65 | 0.56 | 0.09 | HTH-type transcriptional regulator GbpR |
| *FLPKDCKG_05183* | | + | 5612588 | 5613562 | 1.04 | 16.87 | 24.99 | 7.77 | Hypothetical protein |
| *FLPKDCKG_05184* | *pdxA2-3* | + | 5613570 | 5614613 | 0.68 | 13.42 | 14.89 | 3.22 | 4-hydroxythreonine-4-phosphate dehydrogenase 2 |
| *FLPKDCKG_05185* | *rhmD* | + | 5614671 | 5615843 | 0.48 | 8.71 | 13.86 | 3.78 | L-rhamnonate dehydratase |
| *FLPKDCKG_05186* | *araD* | + | 5615849 | 5616784 | 1.06 | 8.30 | 11.87 | 3.31 | L-2-keto-3-deoxyarabonate dehydratase |
| *FLPKDCKG_05187* | | - | 5616812 | 5618095 | 6.20 | 25.00 | 16.15 | 5.50 | Hypothetical protein |
| *FLPKDCKG_05188* | *gabD* | - | 5618341 | 5619843 | 43.96 | 18.18 | 11.71 | 3.38 | Succinate-semialdehyde dehydrogenase [NADP(+)] |
| *FLPKDCKG_05189* | *glnQ-12* | + | 5619968 | 5620723 | 21.90 | 19.21 | 13.05 | 0.85 | Glutamine transport ATP-binding protein GlnQ |
| *FLPKDCKG_05190* | *artP-2* | + | 5620918 | 5621736 | 96.10 | 28.65 | 15.71 | 3.06 | Arginine-binding extracellular protein ArtP |
| *FLPKDCKG_05191* | *occM-2* | + | 5621774 | 5622466 | 13.85 | 4.09 | 5.18 | 1.50 | Octopine transport system permease protein OccM |
| *FLPKDCKG_05192* | *yecS-6* | + | 5622469 | 5623158 | 15.11 | 5.21 | 4.58 | 0.00 | L-cystine transport system permease protein YecS |
| *FLPKDCKG_05193* | *puuB-3* | + | 5623168 | 5624460 | 11.87 | 5.40 | 3.43 | 0.46 | Gamma-glutamylputrescine oxidoreductase |
| *FLPKDCKG_05194* | *sutR-2* | + | 5624473 | 5625057 | 6.44 | 8.39 | 2.40 | 0.00 | HTH-type transcriptional regulator SutR |
| *FLPKDCKG_05195* | | + | 5625166 | 5625242 | 20.49 | 46.93 | 27.68 | 10.88 | tRNA |
| *FLPKDCKG_05196* | | - | 5625338 | 5625799 | 21.39 | 29.12 | 23.11 | 3.80 | Hypothetical protein |
| *FLPKDCKG_05197* | | + | 5625936 | 5626823 | 1.02 | 89.70 | 81.95 | 6.37 | Manganese catalase |
| *FLPKDCKG_05198* | *yegE-2* | - | 5626958 | 5628418 | 3.97 | 4.59 | 3.09 | 0.47 | putative diguanylate cyclase YegE |
| *FLPKDCKG_05199* | | + | 5628964 | 5629215 | 0.38 | 51.22 | 45.15 | 1.84 | Hypothetical protein |
| *FLPKDCKG_05200* | | - | 5629227 | 5630057 | 0.26 | 23.44 | 23.73 | 2.47 | Hypothetical protein |
| *FLPKDCKG_05201* | | - | 5630069 | 5630350 | 1.23 | 281.03 | 138.64 | 17.30 | Hypothetical protein |
| *FLPKDCKG_05202* | | + | 5630492 | 5631100 | 2.04 | 528.54 | 240.72 | 28.46 | Hypothetical protein |
| *FLPKDCKG_05203* | | - | 5631160 | 5631396 | 77.94 | 106.86 | 86.82 | 13.86 | Hypothetical protein |
| *FLPKDCKG_05204* | | + | 5631569 | 5631865 | 12.25 | 63.42 | 55.40 | 10.53 | Hypothetical protein |
| *FLPKDCKG_05205* | | - | 5631892 | 5632083 | 133.26 | 850.28 | 466.07 | 292.65 | Hypothetical protein |
| *FLPKDCKG_05206* | | + | 5632302 | 5632691 | 0.44 | 11.02 | 9.54 | 0.53 | Hypothetical protein |
| *FLPKDCKG_05207* | | + | 5632739 | 5633746 | 1.50 | 4.32 | 1.51 | 2.04 | Hypothetical protein |
| *FLPKDCKG_05208* | *coaE-2* | + | 5633806 | 5634393 | 3.14 | 2.61 | 2.94 | 1.01 | Dephospho-CoA kinase |
| *FLPKDCKG_05209* | *lysN-8* | + | 5634510 | 5635895 | 1.84 | 3.17 | 4.77 | 0.31 | 2-aminoadipate transaminase |
| *FLPKDCKG_05210* | *yabI-1* | + | 5635944 | 5636471 | 10.79 | 11.35 | 16.14 | 5.15 | Inner membrane protein YabI |
| *FLPKDCKG_05211* | | - | 5636542 | 5637558 | 0.42 | 1.47 | 0.27 | 0.84 | Outer membrane porin protein |
| *FLPKDCKG_05212* | *cocE* | - | 5637667 | 5639565 | 0.23 | 0.84 | 0.99 | 0.34 | Cocaine esterase |
| *FLPKDCKG_05213* | *gsiB-10* | - | 5639562 | 5641157 | 0.34 | 0.00 | 0.39 | 0.43 | Glutathione-binding protein GsiB |
| *FLPKDCKG_05214* | *ddpF* | - | 5641154 | 5642188 | 0.12 | 0.80 | 0.31 | 0.00 | putative D,D-dipeptide transport ATP-binding protein DdpF |
| *FLPKDCKG_05215* | *oppD-9* | - | 5642185 | 5643168 | 0.04 | 0.51 | 0.00 | 0.00 | Oligopeptide transport ATP-binding protein OppD |
| *FLPKDCKG_05216* | *gsiD-7* | - | 5643177 | 5644082 | 0.05 | 0.00 | 0.00 | 0.27 | Glutathione transport system permease protein GsiD |
| *FLPKDCKG_05217* | *dppB-8* | - | 5644079 | 5645053 | 0.28 | 1.37 | 0.27 | 0.00 | Dipeptide transport system permease protein DppB |
| *FLPKDCKG_05218* | *gltC-18* | + | 5645322 | 5646275 | 2.04 | 3.95 | 0.29 | 1.45 | HTH-type transcriptional regulator GltC |
| *FLPKDCKG_05219* | *ohrB-3* | + | 5646399 | 5646824 | 0.48 | 3.74 | 4.92 | 1.91 | Organic hydroperoxide resistance protein OhrB |
| *FLPKDCKG_05220* | *cpxA-2* | - | 5646913 | 5648238 | 0.62 | 2.36 | 3.31 | 0.52 | Sensor histidine kinase CpxA |
| *FLPKDCKG_05221* | *ompR-6* | - | 5648239 | 5648976 | 1.70 | 3.94 | 4.27 | 0.00 | Transcriptional regulatory protein OmpR |
| *FLPKDCKG_05222* | | - | 5648985 | 5649149 | 5.97 | 10.78 | 11.96 | 1.72 | Hypothetical protein |
| *FLPKDCKG_05223* | *dsbD-2* | + | 5649332 | 5650024 | 0.15 | 0.43 | 0.04 | 0.00 | Thiol:disulfide interchange protein DsbD |
| *FLPKDCKG_05224* | *ykuV* | + | 5650115 | 5650660 | 0.32 | 0.60 | 0.00 | 0.04 | Thiol-disulfide oxidoreductase YkuV |
| *FLPKDCKG_05225* | *arsB* | - | 5650759 | 5652021 | 0.56 | 1.37 | 1.47 | 0.50 | Arsenical pump membrane protein |
| *FLPKDCKG_05226* | *degP-2* | - | 5652153 | 5653493 | 0.52 | 1.73 | 0.46 | 0.14 | Periplasmic serine endoprotease DegP |
| *FLPKDCKG_05227* | *ywrD-5* | - | 5653600 | 5654475 | 1.35 | 2.41 | 4.12 | 2.18 | Putative gamma-glutamyltransferase YwrD |
| *FLPKDCKG_05228* | *acyI* | - | 5654472 | 5655194 | 1.16 | 1.19 | 1.96 | 0.31 | Acylase ACY 1 |
| *FLPKDCKG_05229* | | - | 5655220 | 5656194 | 2.50 | 5.75 | 1.89 | 1.11 | Hypothetical protein |
| *FLPKDCKG_05230* | *gntR-4* | + | 5656410 | 5657066 | 1.86 | 1.30 | 1.93 | 1.10 | putative D-xylose utilization operon transcriptional repressor |
| *FLPKDCKG_05231* | | - | 5657092 | 5657973 | 1.11 | 1.66 | 2.16 | 0.04 | Hypothetical protein |
| *FLPKDCKG_05232* | *oxlT* | + | 5658268 | 5659539 | 3.06 | 4.16 | 1.84 | 0.58 | Oxalate:formate antiporter |
| *FLPKDCKG_05233* | *garR* | - | 5659564 | 5660436 | 24.53 | 23.55 | 13.04 | 5.05 | 2-hydroxy-3-oxopropionate reductase |
| *FLPKDCKG_05234* | *nagK-3* | - | 5660633 | 5661340 | 1.45 | 2.01 | 1.76 | 0.34 | Fumarylpyruvate hydrolase |
| *FLPKDCKG_05235* | *btr-2* | - | 5661413 | 5662420 | 6.83 | 7.65 | 2.98 | 3.70 | HTH-type transcriptional activator Btr |
| *FLPKDCKG_05236* | | + | 5662591 | 5663019 | 9.75 | 18.97 | 7.09 | 2.51 | Hypothetical protein |
| *FLPKDCKG_05237* | *clsA* | - | 5663098 | 5664513 | 5.06 | 6.67 | 5.41 | 1.21 | Major cardiolipin synthase ClsA |
| *FLPKDCKG_05238* | | + | 5664787 | 5664862 | 10.79 | 21.83 | 41.49 | 3.98 | tRNA |
| *FLPKDCKG_05239* | | + | 5665670 | 5666272 | 0.68 | 1.17 | 3.82 | 2.19 | Hypothetical protein |
| *FLPKDCKG_05240* | | + | 5666235 | 5666414 | 3.24 | 10.22 | 1.79 | 0.00 | Hypothetical protein |

| Locus name | Gene name | Strand | Gene start (bp) | Gene end (bp) | Average TPM Inoculum | Average TPM $t_6$ | Average TPM $t_{24}$ | Average TPM $t_{168}$ | Annotation description |
|---|---|---|---|---|---|---|---|---|---|
| *FLPKDCKG_05241* | | - | 5666420 | 5666659 | 38.65 | 72.44 | 42.01 | 9.24 | Hypothetical protein |
| *FLPKDCKG_05242* | | + | 5666677 | 5666925 | 1.59 | 8.93 | 10.52 | 0.73 | Hypothetical protein |
| *FLPKDCKG_05243* | | + | 5667145 | 5667324 | 197.52 | 111.00 | 69.88 | 14.18 | Hypothetical protein |
| *FLPKDCKG_05244* | | + | 5667614 | 5667853 | 3.40 | 25.29 | 11.82 | 1.85 | Hypothetical protein |
| *FLPKDCKG_05245* | | - | 5667934 | 5668167 | 7.50 | 26.55 | 13.55 | 4.06 | Hypothetical protein |
| *FLPKDCKG_05246* | | + | 5668690 | 5668881 | 1.01 | 12.14 | 4.94 | 0.00 | Hypothetical protein |
| *FLPKDCKG_05247* | | - | 5669713 | 5669907 | 25.52 | 59.17 | 17.39 | 9.29 | Hypothetical protein |
| *FLPKDCKG_05248* | | - | 5671617 | 5673377 | 16.29 | 19.99 | 10.48 | 2.01 | Hypothetical protein |
| *FLPKDCKG_05249* | | + | 5674388 | 5674483 | 3.92 | 6.51 | 4.05 | 2.89 | Hypothetical protein |
| *FLPKDCKG_05250* | | + | 5674486 | 5674863 | 1.38 | 6.84 | 0.92 | 0.81 | Hypothetical protein |
| *FLPKDCKG_05251* | *inhA-4* | - | 5674959 | 5675582 | 0.52 | 0.85 | 1.76 | 0.77 | Isonitrile hydratase |
| *FLPKDCKG_05252* | | + | 5676053 | 5676205 | 16.24 | 34.51 | 35.59 | 17.83 | Hypothetical protein |
| *FLPKDCKG_05253* | *leuE-3* | - | 5676314 | 5676934 | 2.37 | 15.43 | 15.74 | 5.23 | Leucine efflux protein |
| *FLPKDCKG_05254* | *lrp-11* | + | 5677016 | 5677468 | 8.41 | 24.90 | 12.09 | 4.27 | Leucine-responsive regulatory protein |
| *FLPKDCKG_05255* | *leuE-4* | - | 5677590 | 5678207 | 26.51 | 32.76 | 14.73 | 8.20 | Leucine efflux protein |
| *FLPKDCKG_05256* | | - | 5678362 | 5679327 | 0.42 | 4.08 | 1.14 | 0.07 | Hypothetical protein |
| *FLPKDCKG_05257* | *tar-6* | + | 5679693 | 5681183 | 4.53 | 1.29 | 1.05 | 0.04 | Methyl-accepting chemotaxis protein II |
| *FLPKDCKG_05258* | *fliC-3* | - | 5681302 | 5682987 | 0.94 | 1.97 | 1.43 | 0.37 | Flagellin |
| *FLPKDCKG_05259* | *mprA-3* | - | 5683341 | 5684048 | 1.00 | 6.91 | 1.85 | 1.09 | Response regulator MprA |
| *FLPKDCKG_05260* | | - | 5684201 | 5686018 | 1.34 | 7.06 | 3.79 | 1.25 | Trehalase |
| *FLPKDCKG_05261* | *fcuA-3* | + | 5686476 | 5688635 | 3.95 | 1.28 | 1.20 | 0.52 | Ferrichrome receptor FcuA |
| *FLPKDCKG_05262* | | + | 5688718 | 5689005 | 0.68 | 0.00 | 0.00 | 0.00 | Hypothetical protein |
| *FLPKDCKG_05263* | | + | 5688992 | 5689282 | 2.16 | 0.00 | 0.00 | 0.00 | Hypothetical protein |
| *FLPKDCKG_05264* | | + | 5689297 | 5690934 | 0.54 | 0.71 | 0.72 | 0.16 | Hypothetical protein |
| *FLPKDCKG_05265* | | - | 5690973 | 5694071 | 0.28 | 1.18 | 0.40 | 0.02 | Extracellular serine protease |
| *FLPKDCKG_05266* | *tar-7* | + | 5694468 | 5696111 | 0.10 | 0.88 | 0.00 | 0.05 | Methyl-accepting chemotaxis protein II |
| *FLPKDCKG_05267* | *caf1A* | - | 5696014 | 5698455 | 1.49 | 3.25 | 1.29 | 0.97 | F1 capsule-anchoring protein |
| *FLPKDCKG_05268* | *yraI* | - | 5698452 | 5699165 | 0.07 | 0.84 | 3.82 | 0.00 | putative fimbrial chaperone YraI |
| *FLPKDCKG_05269* | | - | 5699235 | 5699870 | 0.00 | 2.37 | 0.00 | 0.45 | Hypothetical protein |
| *FLPKDCKG_05270* | | - | 5699909 | 5700580 | 0.16 | 1.57 | 0.74 | 0.50 | Hypothetical protein |
| *FLPKDCKG_05271* | | - | 5700631 | 5701383 | 0.40 | 4.68 | 2.56 | 3.86 | Hypothetical protein |
| *FLPKDCKG_05272* | | - | 5701430 | 5702167 | 1.20 | 3.38 | 1.97 | 9.65 | Hypothetical protein |
| *FLPKDCKG_05273* | *phnS-3* | - | 5702340 | 5703425 | 0.94 | 5.02 | 2.32 | 2.00 | Putative 2-aminoethylphosphonate-binding periplasmic protein |
| *FLPKDCKG_05274* | *rcsB-3* | - | 5703479 | 5704123 | 2.73 | 4.03 | 7.51 | 3.03 | Transcriptional regulatory protein RcsB |
| *FLPKDCKG_05275* | *rcsC-1* | - | 5704188 | 5707466 | 6.14 | 24.30 | 14.30 | 4.79 | Sensor histidine kinase RcsC |
| *FLPKDCKG_05276* | *dosP-2* | - | 5707463 | 5708653 | 13.97 | 31.14 | 17.33 | 5.06 | Oxygen sensor protein DosP |
| *FLPKDCKG_05277* | | + | 5708900 | 5709316 | 0.17 | 1.60 | 1.21 | 0.46 | Hypothetical protein |
| *FLPKDCKG_05278* | *papD* | - | 5709687 | 5710433 | 0.06 | 0.00 | 0.84 | 0.45 | Chaperone protein PapD |
| *FLPKDCKG_05279* | *papC* | - | 5710475 | 5712964 | 0.24 | 0.75 | 0.01 | 0.19 | Outer membrane usher protein PapC |
| *FLPKDCKG_05280* | *ybgD* | - | 5713129 | 5713683 | 0.16 | 3.33 | 0.58 | 0.00 | putative fimbrial-like protein YbgD |
| *FLPKDCKG_05281* | | - | 5714181 | 5714978 | 12.25 | 22.05 | 5.30 | 1.01 | Hypothetical protein |
| *FLPKDCKG_05282* | *qseB-5* | + | 5715045 | 5715710 | 7.71 | 15.11 | 0.75 | 0.56 | Transcriptional regulatory protein QseB |
| *FLPKDCKG_05283* | *qseC-7* | + | 5715707 | 5716990 | 0.33 | 1.04 | 2.07 | 0.34 | Sensor protein QseC |
| *FLPKDCKG_05284* | | - | 5716926 | 5717561 | 0.41 | 2.12 | 0.46 | 0.73 | Hypothetical protein |
| *FLPKDCKG_05285* | *cysD-2* | - | 5717991 | 5719214 | 0.47 | 3.15 | 0.55 | 0.14 | Sulfate adenylyltransferase subunit 2 |
| *FLPKDCKG_05286* | *csaA* | - | 5719454 | 5719825 | 12.10 | 18.35 | 11.40 | 5.34 | putative chaperone CsaA |
| *FLPKDCKG_05287* | | - | 5719884 | 5721512 | 2.67 | 9.34 | 8.02 | 1.35 | Hypothetical protein |
| *FLPKDCKG_05288* | | - | 5721694 | 5721864 | 68.14 | 104.22 | 61.20 | 12.00 | Hypothetical protein |
| *FLPKDCKG_05289* | | + | 5722344 | 5722538 | 28.13 | 28.64 | 16.09 | 1.71 | Hypothetical protein |
| *FLPKDCKG_05290* | | + | 5722535 | 5723053 | 1.94 | 5.13 | 4.22 | 1.98 | Hypothetical protein |
| *FLPKDCKG_05291* | *paiB* | + | 5723177 | 5723809 | 1.92 | 1.06 | 3.71 | 2.08 | Protease synthase and sporulation protein PAI 2 |
| *FLPKDCKG_05292* | | + | 5724630 | 5724965 | 0.32 | 2.73 | 0.00 | 0.00 | Hypothetical protein |
| *FLPKDCKG_05293* | *cmtR* | + | 5725136 | 5725876 | 34.23 | 27.35 | 15.36 | 4.89 | HTH-type transcriptional regulator CmtR |
| *FLPKDCKG_05294* | | + | 5725982 | 5726632 | 12.54 | 7.12 | 6.93 | 1.41 | Hypothetical protein |
| *FLPKDCKG_05295* | | + | 5726858 | 5727130 | 31.10 | 96.53 | 65.21 | 32.29 | Hypothetical protein |
| *FLPKDCKG_05296* | | - | 5727173 | 5727940 | 1.29 | 4.75 | 6.24 | 4.22 | Hypothetical protein |
| *FLPKDCKG_05297* | | - | 5727937 | 5728473 | 3.56 | 5.85 | 3.67 | 2.63 | Hypothetical protein |
| *FLPKDCKG_05298* | | - | 5728654 | 5729508 | 22.93 | 37.91 | 23.22 | 11.06 | Hypothetical protein |
| *FLPKDCKG_05299* | | + | 5729598 | 5730224 | 60.99 | 85.15 | 62.77 | 15.72 | Putative threonylcarbamoyl-AMP synthase |
| *FLPKDCKG_05300* | | + | 5730285 | 5730932 | 7.18 | 14.16 | 7.67 | 4.29 | Hypothetical protein |
| *FLPKDCKG_05301* | *dapA-7* | + | 5731020 | 5731925 | 19.90 | 41.68 | 40.21 | 16.73 | 4-hydroxy-tetrahydrodipicolinate synthase |
| *FLPKDCKG_05302* | *bamC* | + | 5731943 | 5733067 | 8.62 | 37.92 | 42.02 | 15.51 | Outer membrane protein assembly factor BamC |
| *FLPKDCKG_05303* | | - | 5733274 | 5733501 | 206.15 | 622.10 | 491.11 | 91.73 | Hypothetical protein |
| *FLPKDCKG_05304* | *roxA* | - | 5733719 | 5734912 | 4.31 | 6.99 | 4.34 | 1.29 | 50S ribosomal protein L16 3-hydroxylase |
| *FLPKDCKG_05305* | *mutS* | - | 5734909 | 5737563 | 6.49 | 7.21 | 4.65 | 2.54 | DNA mismatch repair protein MutS |
| *FLPKDCKG_05306* | *hpxO-4* | + | 5737869 | 5739125 | 13.65 | 14.42 | 9.34 | 1.62 | FAD-dependent urate hydroxylase |
| *FLPKDCKG_05307* | | - | 5739196 | 5739822 | 18.87 | 70.31 | 45.36 | 9.18 | Hypothetical protein |
| *FLPKDCKG_05308* | | - | 5739899 | 5740081 | 57.94 | 231.63 | 126.17 | 23.46 | Hypothetical protein |
| *FLPKDCKG_05309* | | - | 5740323 | 5740586 | 28.70 | 82.99 | 49.46 | 5.32 | Hypothetical protein |

| Locus name | Gene name | Strand | Gene start (bp) | Gene end (bp) | Average TPM Inoculum | Average TPM $t_6$ | Average TPM $t_{24}$ | Average TPM $t_{168}$ | Annotation description |
|---|---|---|---|---|---|---|---|---|---|
| *FLPKDCKG_05310* | *galM* | + | 5740780 | 5741688 | 1.44 | 2.00 | 2.54 | 0.04 | Aldose 1-epimerase |
| *FLPKDCKG_05311* | | + | 5741766 | 5742122 | 22.39 | 28.32 | 24.24 | 4.75 | Hypothetical protein |
| *FLPKDCKG_05312* | | - | 5742201 | 5742464 | 6.72 | 10.18 | 8.62 | 3.19 | Hypothetical protein |
| *FLPKDCKG_05313* | | - | 5742625 | 5743638 | 7.60 | 25.23 | 11.46 | 3.93 | Zinc-type alcohol dehydrogenase-like protein |
| *FLPKDCKG_05314* | *fabG-19* | - | 5743665 | 5744459 | 31.28 | 62.93 | 33.16 | 2.79 | 3-oxoacyl-[acyl-carrier-protein] reductase FabG |
| *FLPKDCKG_05315* | *dmlR-32* | + | 5744569 | 5745486 | 2.42 | 29.96 | 14.57 | 3.42 | HTH-type transcriptional regulator DmlR |
| *FLPKDCKG_05316* | | - | 5745499 | 5746002 | 18.95 | 144.94 | 92.29 | 20.24 | Hypothetical protein |
| *FLPKDCKG_05317* | | + | 5746193 | 5746411 | 148.24 | 423.86 | 252.88 | 42.82 | Dodecin |
| *FLPKDCKG_05318* | *chrR* | - | 5747366 | 5747920 | 233.18 | 274.20 | 150.20 | 51.83 | Chromate reductase |
| *FLPKDCKG_05319* | | - | 5748056 | 5748469 | 0.47 | 19.27 | 29.28 | 5.87 | Hypothetical protein |
| *FLPKDCKG_05320* | | + | 5748642 | 5749187 | 0.12 | 25.85 | 22.69 | 2.40 | Hypothetical protein |
| *FLPKDCKG_05321* | *lutR-7* | - | 5749214 | 5749921 | 0.00 | 18.99 | 22.15 | 6.72 | HTH-type transcriptional regulator LutR |
| *FLPKDCKG_05322* | *atzE* | - | 5749942 | 5751330 | 0.17 | 16.76 | 23.64 | 8.04 | Biuret hydrolase |
| *FLPKDCKG_05323* | | - | 5751327 | 5751509 | 0.00 | 8.95 | 21.20 | 3.71 | Hypothetical protein |
| *FLPKDCKG_05324* | *tauB-7* | - | 5751538 | 5752359 | 0.34 | 42.69 | 39.40 | 13.90 | Taurine import ATP-binding protein TauB |
| *FLPKDCKG_05325* | *ssuC-10* | - | 5752362 | 5753249 | 0.21 | 16.99 | 16.46 | 3.88 | Putative aliphatic sulfonates transport permease protein SsuC |
| *FLPKDCKG_05326* | *ssuA-6* | - | 5753253 | 5754260 | 0.31 | 27.34 | 38.73 | 16.18 | Putative aliphatic sulfonates-binding protein |
| *FLPKDCKG_05327* | *gltR-6* | + | 5754541 | 5755452 | 4.35 | 8.81 | 2.93 | 0.98 | HTH-type transcriptional regulator GltR |
| *FLPKDCKG_05328* | | + | 5755575 | 5756243 | 0.75 | 11.32 | 4.41 | 2.44 | Hypothetical protein |
| *FLPKDCKG_05329* | *yhjE-6* | + | 5756292 | 5757617 | 0.23 | 1.34 | 2.31 | 0.00 | Inner membrane metabolite transport protein YhjE |
| *FLPKDCKG_05330* | *irpA* | + | 5757835 | 5759133 | 18.92 | 4.55 | 3.72 | 3.61 | Iron-regulated protein A |
| *FLPKDCKG_05331* | | + | 5759130 | 5760647 | 3.66 | 0.23 | 0.11 | 0.94 | Hypothetical protein |
| *FLPKDCKG_05332* | | + | 5760647 | 5761279 | 1.81 | 0.87 | 0.00 | 0.00 | Hypothetical protein |
| *FLPKDCKG_05333* | | + | 5761279 | 5762373 | 1.82 | 1.36 | 1.29 | 0.00 | Hypothetical protein |
| *FLPKDCKG_05334* | | + | 5762390 | 5763538 | 1.19 | 0.68 | 0.45 | 0.72 | Hypothetical protein |
| *FLPKDCKG_05335* | *mscS-1* | + | 5763684 | 5764532 | 24.08 | 16.62 | 18.77 | 7.19 | Small-conductance mechanosensitive channel |
| *FLPKDCKG_05336* | *iclR-5* | - | 5764616 | 5765377 | 8.95 | 13.21 | 15.28 | 4.44 | Transcriptional repressor IclR |
| *FLPKDCKG_05337* | | + | 5765539 | 5766225 | 16.92 | 13.95 | 16.33 | 7.50 | Hypothetical protein |
| *FLPKDCKG_05338* | | + | 5766222 | 5766527 | 8.73 | 12.63 | 4.65 | 4.98 | Hypothetical protein |
| *FLPKDCKG_05339* | *trmJ* | - | 5766597 | 5767355 | 6.11 | 17.83 | 6.62 | 2.41 | tRNA (cytidine/uridine-2'-O-)-methyltransferase TrmJ |
| *FLPKDCKG_05340* | *suhB-2* | + | 5767509 | 5768294 | 9.47 | 15.96 | 11.78 | 5.47 | Inositol-1-monophosphatase |
| *FLPKDCKG_05341* | | + | 5768550 | 5769650 | 0.54 | 2.38 | 0.45 | 0.23 | Hypothetical protein |
| *FLPKDCKG_05342* | *apbE* | - | 5769699 | 5770709 | 0.73 | 0.86 | 0.00 | 0.00 | FAD:protein FMN transferase |
| *FLPKDCKG_05343* | | - | 5770746 | 5771246 | 1.30 | 0.00 | 0.00 | 0.00 | Hypothetical protein |
| *FLPKDCKG_05344* | | - | 5771243 | 5772073 | 0.27 | 1.50 | 0.00 | 0.00 | Hypothetical protein |
| *FLPKDCKG_05345* | *yxlF* | - | 5772070 | 5772975 | 0.27 | 0.10 | 0.00 | 0.46 | putative ABC transporter ATP-binding protein YxlF |
| *FLPKDCKG_05346* | | - | 5772972 | 5774285 | 1.53 | 1.03 | 1.20 | 0.05 | Hypothetical protein |
| *FLPKDCKG_05347* | *nosZ* | - | 5774291 | 5776207 | 3.25 | 1.50 | 0.58 | 0.17 | Nitrous-oxide reductase |
| *FLPKDCKG_05348* | *yccM* | - | 5776254 | 5778593 | 0.89 | 0.20 | 0.22 | 0.07 | Putative electron transport protein YccM |
| *FLPKDCKG_05349* | *azu* | + | 5778847 | 5779296 | 4.60 | 1.70 | 1.79 | 0.88 | Azurin |
| *FLPKDCKG_05350* | *corC-3* | + | 5779486 | 5781069 | 12.03 | 19.43 | 13.49 | 3.07 | Magnesium and cobalt efflux protein CorC |
| *FLPKDCKG_05351* | *uppP* | + | 5781103 | 5781966 | 3.51 | 6.64 | 8.57 | 5.33 | Undecaprenyl-diphosphatase |
| *FLPKDCKG_05352* | *lpxH-2* | - | 5782076 | 5782849 | 5.46 | 12.64 | 16.05 | 3.88 | UDP-2,3-diacylglucosamine hydrolase |
| *FLPKDCKG_05353* | *ppiB* | - | 5782842 | 5783486 | 70.29 | 96.42 | 81.30 | 26.22 | Peptidyl-prolyl cis-trans isomerase B |
| *FLPKDCKG_05354* | | - | 5783483 | 5784166 | 39.01 | 46.55 | 30.53 | 7.15 | Hypothetical protein |
| *FLPKDCKG_05355* | *cysS* | + | 5784492 | 5785946 | 7.61 | 19.44 | 14.59 | 6.17 | Cysteine--tRNA ligase |
| *FLPKDCKG_05356* | *alkA-4* | + | 5785967 | 5786611 | 23.63 | 26.17 | 31.48 | 9.62 | DNA-3-methyladenine glycosylase |
| *FLPKDCKG_05357* | *accA* | + | 5786648 | 5787613 | 5.75 | 10.18 | 10.49 | 3.20 | Acetyl-coenzyme A carboxylase carboxyl transferase subunit alpha |
| *FLPKDCKG_05358* | | - | 5788232 | 5788322 | 2.29 | 3.54 | 8.01 | 2.08 | tRNA |
| *FLPKDCKG_05359* | | + | 5788974 | 5790239 | 26.41 | 41.34 | 27.48 | 9.88 | Aspartate kinase |
| *FLPKDCKG_05360* | | + | 5790327 | 5790419 | 11.81 | 13.74 | 13.15 | 1.36 | tRNA |
| *FLPKDCKG_05361* | | - | 5790588 | 5790821 | 0.33 | 4.04 | 0.00 | 0.49 | Hypothetical protein |
| *FLPKDCKG_05362* | | - | 5790843 | 5792333 | 1.38 | 0.66 | 0.21 | 0.00 | Hypothetical protein |
| *FLPKDCKG_05363* | *ccoP-2* | - | 5792344 | 5793297 | 0.09 | 0.48 | 0.00 | 0.00 | Cbb3-type cytochrome c oxidase subunit CcoP |
| *FLPKDCKG_05364* | | - | 5793294 | 5793467 | 0.50 | 0.00 | 0.00 | 0.00 | Hypothetical protein |
| *FLPKDCKG_05365* | | - | 5793470 | 5794159 | 4.84 | 0.64 | 0.00 | 0.00 | Hypothetical protein |
| *FLPKDCKG_05366* | *ccoN1-2* | - | 5794170 | 5795654 | 1.74 | 1.40 | 0.13 | 0.47 | Cbb3-type cytochrome c oxidase subunit CcoN1 |
| *FLPKDCKG_05367* | | - | 5795713 | 5795874 | 0.54 | 8.08 | 0.00 | 1.06 | Hypothetical protein |
| *FLPKDCKG_05368* | *ctpV-2* | - | 5795871 | 5796845 | 0.40 | 0.36 | 0.40 | 0.00 | putative copper-exporting P-type ATPase V |
| *FLPKDCKG_05369* | *tsdA-3* | - | 5796897 | 5797934 | 0.64 | 0.80 | 0.00 | 0.03 | Thiosulfate dehydrogenase |
| *FLPKDCKG_05370* | *cc4-4* | - | 5797931 | 5798656 | 1.81 | 2.34 | 2.44 | 0.80 | Cytochrome c4 |
| *FLPKDCKG_05371* | | - | 5798944 | 5799282 | 43.66 | 42.31 | 42.47 | 14.91 | Hypothetical protein |
| *FLPKDCKG_05372* | | - | 5799356 | 5799763 | 36.78 | 43.89 | 43.83 | 16.39 | Hypothetical protein |
| *FLPKDCKG_05373* | *rcsC-2* | + | 5800314 | 5803091 | 0.27 | 1.57 | 1.18 | 0.15 | Sensor histidine kinase RcsC |
| *FLPKDCKG_05374* | *cph2-5* | + | 5803088 | 5804362 | 5.72 | 11.02 | 5.33 | 1.49 | Phytochrome-like protein cph2 |
| *FLPKDCKG_05375* | *rcsC-3* | + | 5804456 | 5805967 | 1.39 | 1.67 | 1.42 | 1.11 | Sensor histidine kinase RcsC |
| *FLPKDCKG_05376* | *rcsC-4* | + | 5805964 | 5807694 | 2.21 | 1.91 | 2.25 | 1.48 | Sensor histidine kinase RcsC |
| *FLPKDCKG_05377* | *icsA* | - | 5807717 | 5811064 | 0.26 | 0.36 | 0.37 | 0.62 | Outer membrane protein IcsA autotransporter |
| *FLPKDCKG_05378* | *selU* | - | 5811376 | 5812419 | 8.26 | 10.25 | 9.64 | 2.08 | tRNA 2-selenouridine synthase |

| Locus name | Gene name | Strand | Gene start (bp) | Gene end (bp) | Average TPM Inoculum | Average TPM $t_6$ | Average TPM $t_{24}$ | Average TPM $t_{168}$ | Annotation description |
|---|---|---|---|---|---|---|---|---|---|
| *FLPKDCKG_05379* | *nudC-2* | - | 5812483 | 5813100 | 48.86 | 24.44 | 19.06 | 7.47 | NADH pyrophosphatase |
| *FLPKDCKG_05380* | *plcB* | - | 5813181 | 5814779 | 0.61 | 2.65 | 1.99 | 0.57 | Phospholipase C 2 |
| *FLPKDCKG_05381* | *mtrR-1* | + | 5815129 | 5815761 | 36.31 | 49.06 | 26.07 | 8.10 | HTH-type transcriptional regulator MtrR |
| *FLPKDCKG_05382* | *cysA-7* | - | 5815787 | 5816476 | 1.63 | 8.93 | 6.39 | 0.87 | Sulfate/thiosulfate import ATP-binding protein CysA |
| *FLPKDCKG_05383* | | + | 5816727 | 5816803 | 83.13 | 366.66 | 149.65 | 75.48 | tRNA |
| *FLPKDCKG_05384* | | + | 5816925 | 5817001 | 56.05 | 220.71 | 75.18 | 38.00 | tRNA |
| *FLPKDCKG_05385* | | - | 5817159 | 5817371 | 32.31 | 2.90 | 4.51 | 3.25 | Hypothetical protein |
| *FLPKDCKG_05386* | | + | 5817683 | 5817759 | 52.04 | 214.01 | 76.93 | 43.50 | tRNA |
| *FLPKDCKG_05387* | | - | 5817967 | 5818173 | 7.97 | 6.82 | 13.72 | 0.20 | Hypothetical protein |
| *FLPKDCKG_05388* | *mshD-3* | + | 5818228 | 5818689 | 0.39 | 0.98 | 3.20 | 0.00 | Mycothiol acetyltransferase |
| *FLPKDCKG_05389* | | - | 5818735 | 5819331 | 0.91 | 1.48 | 2.27 | 0.00 | Hypothetical protein |
| *FLPKDCKG_05390* | *dmlR-33* | + | 5819460 | 5820380 | 5.19 | 6.37 | 2.82 | 0.58 | HTH-type transcriptional regulator DmlR |
| *FLPKDCKG_05391* | | - | 5820417 | 5820665 | 83.28 | 90.29 | 61.16 | 10.86 | Hypothetical protein |
| *FLPKDCKG_05392* | | - | 5820756 | 5821355 | 85.71 | 129.39 | 70.62 | 18.55 | Hypothetical protein |
| *FLPKDCKG_05393* | *cobB-1* | - | 5822373 | 5823155 | 2.77 | 4.57 | 8.14 | 2.13 | NAD-dependent protein deacylase |
| *FLPKDCKG_05394* | | + | 5823354 | 5823980 | 6.36 | 1.68 | 1.79 | 0.92 | 3-mercaptopropionate dioxygenase |
| *FLPKDCKG_05395* | *kdgR-7* | + | 5824043 | 5824807 | 4.39 | 2.61 | 1.96 | 0.89 | Transcriptional regulator KdgR |
| *FLPKDCKG_05396* | *uctC-7* | + | 5824940 | 5826103 | 1.84 | 2.48 | 0.58 | 0.40 | Acetyl-CoA:oxalate CoA-transferase |
| *FLPKDCKG_05397* | *echA6* | + | 5826100 | 5826822 | 1.17 | 1.76 | 1.77 | 0.65 | putative enoyl-CoA hydratase echA6 |
| *FLPKDCKG_05398* | *catI* | + | 5826875 | 5827729 | 0.75 | 0.83 | 1.48 | 1.11 | 3-oxoadipate CoA-transferase subunit A |
| *FLPKDCKG_05399* | *catJ* | + | 5827722 | 5828492 | 0.59 | 0.42 | 0.81 | 0.52 | 3-oxoadipate CoA-transferase subunit B |
| *FLPKDCKG_05400* | *dasR-2* | - | 5828503 | 5829396 | 0.79 | 1.25 | 2.01 | 0.54 | HTH-type transcriptional repressor DasR |
| *FLPKDCKG_05401* | *pdxA2-4* | - | 5829401 | 5830420 | 3.77 | 5.35 | 3.36 | 1.54 | 4-hydroxythreonine-4-phosphate dehydrogenase 2 |
| *FLPKDCKG_05402* | | - | 5830448 | 5831668 | 0.74 | 0.71 | 0.80 | 0.37 | Hypothetical protein |
| *FLPKDCKG_05403* | *kdgT* | - | 5831665 | 5832792 | 1.11 | 2.89 | 1.34 | 0.22 | 2-keto-3-deoxygluconate permease |
| *FLPKDCKG_05404* | | - | 5832844 | 5834697 | 2.95 | 3.22 | 1.23 | 0.51 | Putative multidrug export ATP-binding/permease protein |
| *FLPKDCKG_05405* | *marR-2* | - | 5834769 | 5835284 | 1.77 | 4.53 | 2.63 | 0.00 | Multiple antibiotic resistance protein MarR |
| *FLPKDCKG_05406* | | + | 5835414 | 5836388 | 0.39 | 1.41 | 2.16 | 0.11 | Hypothetical protein |
| *FLPKDCKG_05407* | | + | 5836400 | 5837968 | 0.33 | 1.35 | 0.75 | 1.24 | Arylsulfatase |
| *FLPKDCKG_05408* | *fadD-2* | - | 5838012 | 5839706 | 5.97 | 8.16 | 5.56 | 1.46 | Long-chain-fatty-acid--CoA ligase |
| *FLPKDCKG_05409* | | - | 5839709 | 5840305 | 16.28 | 16.51 | 6.58 | 2.85 | Hypothetical protein |
| *FLPKDCKG_05410* | | - | 5840266 | 5840814 | 5.93 | 6.31 | 3.71 | 3.33 | Hypothetical protein |
| *FLPKDCKG_05411* | *fadA-3* | - | 5840833 | 5842032 | 9.47 | 6.39 | 3.32 | 2.58 | 3-ketoacyl-CoA thiolase |
| *FLPKDCKG_05412* | *fadN* | - | 5842238 | 5844628 | 22.46 | 8.17 | 10.10 | 3.55 | putative 3-hydroxyacyl-CoA dehydrogenase |
| *FLPKDCKG_05413* | | - | 5844689 | 5845999 | 84.06 | 22.63 | 18.67 | 5.89 | Putative outer membrane protein |
| *FLPKDCKG_05414* | *fadJ-2* | - | 5846097 | 5848031 | 15.24 | 3.92 | 8.14 | 2.44 | Fatty acid oxidation complex subunit alpha |
| *FLPKDCKG_05415* | *phaA* | - | 5848031 | 5849320 | 23.17 | 6.06 | 5.45 | 5.28 | Acetyl-CoA acetyltransferase |
| *FLPKDCKG_05416* | *fadE-2* | - | 5849348 | 5851696 | 22.92 | 5.01 | 4.72 | 1.92 | Acyl-coenzyme A dehydrogenase |
| *FLPKDCKG_05417* | *icaR* | - | 5851756 | 5852427 | 42.47 | 5.01 | 12.97 | 3.69 | Biofilm operon icaADBC HTH-type negative transcriptional regulator IcaR |
| *FLPKDCKG_05418* | | + | 5852785 | 5854386 | 0.72 | 3.19 | 1.77 | 0.00 | Hypothetical protein |
| *FLPKDCKG_05419* | *vraR-2* | + | 5854952 | 5855698 | 71.46 | 36.91 | 33.67 | 8.37 | Response regulator protein VraR |
| *FLPKDCKG_05420* | *degU-3* | + | 5856007 | 5856669 | 0.41 | 2.99 | 1.52 | 0.00 | Transcriptional regulatory protein DegU |
| *FLPKDCKG_05421* | *mshA-2* | - | 5856682 | 5857956 | 0.21 | 2.14 | 0.40 | 1.05 | D-inositol 3-phosphate glycosyltransferase |
| *FLPKDCKG_05422* | *lacA-2* | - | 5857953 | 5858513 | 0.02 | 0.05 | 0.00 | 0.00 | Galactoside O-acetyltransferase |
| *FLPKDCKG_05423* | | - | 5858510 | 5859397 | 1.34 | 1.02 | 0.74 | 0.04 | Hypothetical protein |
| *FLPKDCKG_05424* | *pglJ-2* | - | 5859411 | 5860541 | 0.14 | 0.14 | 0.00 | 0.00 | N-acetylgalactosamine-N,N'-diacetylbacillosaminyl-diphospho-undecaprenol 4-alpha-N-acetylgalactosaminyltransferase |
| *FLPKDCKG_05425* | | - | 5860534 | 5861826 | 0.24 | 0.09 | 0.36 | 0.00 | Hypothetical protein |
| *FLPKDCKG_05426* | | - | 5861906 | 5862808 | 0.44 | 0.32 | 0.00 | 0.10 | putative glycosyltransferase |
| *FLPKDCKG_05427* | | - | 5862829 | 5864145 | 0.18 | 0.00 | 0.40 | 0.43 | Hypothetical protein |
| *FLPKDCKG_05428* | *fcl* | - | 5864175 | 5865128 | 0.32 | 0.36 | 0.10 | 0.00 | GDP-L-fucose synthase |
| *FLPKDCKG_05429* | *gmd* | - | 5865135 | 5866244 | 1.15 | 2.07 | 0.34 | 1.28 | GDP-mannose 4,6-dehydratase |
| *FLPKDCKG_05430* | | - | 5866255 | 5866662 | 0.72 | 0.28 | 1.15 | 0.00 | Hypothetical protein |
| *FLPKDCKG_05431* | *algA* | - | 5866655 | 5868085 | 0.33 | 0.47 | 0.67 | 0.51 | Alginate biosynthesis protein AlgA |
| *FLPKDCKG_05432* | *gmm* | - | 5868082 | 5868627 | 0.13 | 0.90 | 0.00 | 0.00 | GDP-mannose mannosyl hydrolase |
| *FLPKDCKG_05433* | *wcaJ* | - | 5868617 | 5870026 | 0.13 | 0.37 | 0.40 | 0.00 | UDP-glucose:undecaprenyl-phosphate glucose-1-phosphate transferase |
| *FLPKDCKG_05434* | | + | 5870274 | 5871410 | 0.56 | 2.69 | 0.82 | 0.63 | Hypothetical protein |
| *FLPKDCKG_05435* | *ptk* | + | 5871516 | 5873777 | 0.37 | 0.24 | 0.34 | 0.26 | Tyrosine-protein kinase ptk |
| *FLPKDCKG_05436* | *algC-2* | + | 5873796 | 5875199 | 0.26 | 1.00 | 0.59 | 0.00 | Phosphomannomutase/phosphoglucomutase |
| *FLPKDCKG_05437* | *pgi-2* | + | 5875213 | 5876886 | 0.31 | 0.51 | 1.04 | 0.00 | Glucose-6-phosphate isomerase |
| *FLPKDCKG_05438* | *galE-2* | + | 5876899 | 5877951 | 0.34 | 0.00 | 0.44 | 0.47 | UDP-glucose 4-epimerase |
| *FLPKDCKG_05439* | *cysE* | + | 5878135 | 5879082 | 13.22 | 26.52 | 14.37 | 7.06 | Serine acetyltransferase |
| *FLPKDCKG_05440* | *cobB-2* | + | 5879174 | 5880001 | 2.55 | 7.15 | 5.26 | 3.15 | NAD-dependent protein deacetylase |
| *FLPKDCKG_05441* | *suhB-3* | - | 5880173 | 5881027 | 3.40 | 5.97 | 4.46 | 1.54 | Inositol-1-monophosphatase |
| *FLPKDCKG_05442* | *davD* | - | 5881227 | 5882696 | 8.82 | 55.21 | 38.48 | 7.36 | Glutarate-semialdehyde dehydrogenase DavD |
| *FLPKDCKG_05443* | *puuE* | - | 5882821 | 5884083 | 10.62 | 25.70 | 18.10 | 6.31 | 4-aminobutyrate aminotransferase PuuE |
| *FLPKDCKG_05444* | *gabR-4* | + | 5884179 | 5885708 | 4.87 | 6.03 | 6.01 | 1.78 | HTH-type transcriptional regulatory protein GabR |
| *FLPKDCKG_05445* | | - | 5885811 | 5886722 | 3.62 | 10.88 | 3.82 | 2.57 | Hypothetical protein |
| *FLPKDCKG_05446* | *pcrA* | + | 5886833 | 5888923 | 2.87 | 10.18 | 5.46 | 1.72 | ATP-dependent DNA helicase PcrA |
| *FLPKDCKG_05447* | *corA* | + | 5889049 | 5890059 | 10.09 | 10.11 | 6.54 | 2.95 | Cobalt/magnesium transport protein CorA |

| Locus name | Gene name | Strand | Gene start (bp) | Gene end (bp) | Average TPM Inoculum | Average TPM $t_6$ | Average TPM $t_{24}$ | Average TPM $t_{168}$ | Annotation description |
|---|---|---|---|---|---|---|---|---|---|
| *FLPKDCKG_05448* | *nucH* | + | 5890162 | 5890827 | 1.68 | 0.95 | 1.16 | 1.48 | Thermonuclease |
| *FLPKDCKG_05449* | *hbpA-5* | + | 5890967 | 5893204 | 3.75 | 5.39 | 4.55 | 1.45 | Heme-binding protein A |
| *FLPKDCKG_05450* | *gsiC-9* | + | 5893201 | 5894178 | 2.31 | 2.24 | 1.88 | 1.47 | Glutathione transport system permease protein GsiC |
| *FLPKDCKG_05451* | | + | 5894184 | 5895692 | 3.04 | 3.83 | 3.18 | 0.33 | Putative peptide transport permease protein |
| *FLPKDCKG_05452* | *gsiA-10* | + | 5895689 | 5897386 | 2.65 | 3.73 | 2.89 | 1.37 | Glutathione import ATP-binding protein GsiA |
| *FLPKDCKG_05453* | | - | 5897444 | 5897983 | 8.99 | 136.05 | 86.41 | 32.71 | Hypothetical protein |
| *FLPKDCKG_05454* | | - | 5898012 | 5899706 | 37.61 | 286.51 | 178.91 | 58.38 | Hypothetical protein |
| *FLPKDCKG_05455* | *hxuB-2* | - | 5900063 | 5901724 | 16.19 | 15.43 | 11.18 | 3.69 | Heme/hemopexin transporter protein HuxB |
| *FLPKDCKG_05456* | | + | 5902215 | 5903096 | 122.87 | 133.58 | 58.73 | 29.17 | putative signaling protein |
| *FLPKDCKG_05457* | | + | 5903242 | 5903418 | 244.21 | 125.57 | 103.96 | 73.94 | Hypothetical protein |
| *FLPKDCKG_05458* | | + | 5903637 | 5904158 | 20.12 | 20.51 | 11.49 | 6.37 | Hypothetical protein |
| *FLPKDCKG_05459* | | + | 5904125 | 5904565 | 35.43 | 20.15 | 11.99 | 6.36 | Hypothetical protein |
| *FLPKDCKG_05460* | | + | 5904567 | 5905058 | 34.24 | 11.87 | 10.51 | 6.22 | Hypothetical protein |
| *FLPKDCKG_05461* | | + | 5905055 | 5906008 | 33.72 | 12.56 | 7.17 | 4.58 | Hypothetical protein |
| *FLPKDCKG_05462* | *outD-2* | + | 5906076 | 5907452 | 31.68 | 16.90 | 14.16 | 6.68 | Type II secretion system protein D |
| *FLPKDCKG_05463* | | + | 5907488 | 5908789 | 48.16 | 15.60 | 14.70 | 6.52 | Hypothetical protein |
| *FLPKDCKG_05464* | | + | 5908810 | 5910165 | 31.42 | 14.64 | 7.18 | 5.00 | Putative conjugal transfer protein |
| *FLPKDCKG_05465* | | + | 5910165 | 5911094 | 11.79 | 8.24 | 4.78 | 1.11 | Hypothetical protein |
| *FLPKDCKG_05466* | | + | 5911106 | 5912107 | 38.97 | 19.78 | 12.03 | 6.62 | Hypothetical protein |
| *FLPKDCKG_05467* | *bepA-6* | + | 5912107 | 5912997 | 35.04 | 12.36 | 13.69 | 6.64 | Beta-barrel assembly-enhancing protease |
| *FLPKDCKG_05468* | | + | 5913020 | 5913469 | 12.45 | 3.16 | 1.44 | 2.14 | Hypothetical protein |
| *FLPKDCKG_05469* | | + | 5913502 | 5915601 | 12.16 | 6.22 | 4.80 | 2.50 | Hypothetical protein |
| *FLPKDCKG_05470* | *hyfR* | + | 5915651 | 5917057 | 9.95 | 5.80 | 6.43 | 4.76 | DNA-binding transcriptional activator HyfR |
| *FLPKDCKG_05471* | | + | 5917173 | 5917880 | 17.62 | 13.85 | 17.26 | 7.67 | Hypothetical protein |
| *FLPKDCKG_05472* | | + | 5917968 | 5918426 | 26.57 | 28.21 | 25.41 | 17.65 | Hypothetical protein |
| *FLPKDCKG_05473* | *efp* | - | 5918515 | 5919075 | 18.21 | 65.13 | 51.58 | 25.46 | Elongation factor P |
| *FLPKDCKG_05474* | | - | 5919283 | 5920398 | 2.68 | 6.18 | 3.85 | 1.28 | Hypothetical protein |
| *FLPKDCKG_05475* | | - | 5920408 | 5921610 | 15.24 | 30.36 | 22.49 | 5.60 | Hypothetical protein |
| *FLPKDCKG_05476* | *ygiD* | + | 5921855 | 5922628 | 2.49 | 3.82 | 3.26 | 2.06 | 4,5-DOPA dioxygenase extradiol |
| *FLPKDCKG_05477* | | + | 5922782 | 5923747 | 2.46 | 7.16 | 8.62 | 2.91 | Outer membrane protein Omp38 |
| *FLPKDCKG_05478* | *citB-2* | - | 5923851 | 5926556 | 39.71 | 63.83 | 44.34 | 21.81 | Aconitate hydratase A |
| *FLPKDCKG_05479* | *dmlR-34* | - | 5926861 | 5927790 | 1.36 | 2.09 | 2.05 | 0.43 | HTH-type transcriptional regulator DmlR |
| *FLPKDCKG_05480* | *fadH-3* | + | 5927896 | 5928795 | 2.52 | 1.83 | 2.12 | 1.55 | putative 2,4-dienoyl-CoA reductase |
| *FLPKDCKG_05481* | | + | 5928792 | 5930303 | 0.88 | 2.74 | 1.22 | 0.73 | Long-chain-fatty-acid--CoA ligase FadD13 |
| *FLPKDCKG_05482* | | + | 5930361 | 5931344 | 0.57 | 1.16 | 2.32 | 0.09 | Hypothetical protein |
| *FLPKDCKG_05483* | *dmlR-35* | - | 5931278 | 5932264 | 0.98 | 2.28 | 1.71 | 0.21 | HTH-type transcriptional regulator DmlR |
| *FLPKDCKG_05484* | | + | 5932383 | 5933357 | 0.45 | 0.77 | 0.00 | 0.00 | Hypothetical protein |
| *FLPKDCKG_05485* | *fcuA-4* | + | 5933665 | 5935842 | 0.91 | 0.59 | 2.58 | 2.81 | Ferrichrome receptor FcuA |
| *FLPKDCKG_05486* | *fabG-20* | - | 5935904 | 5936671 | 2.82 | 2.58 | 0.00 | 0.29 | 3-oxoacyl-[acyl-carrier-protein] reductase FabG |
| *FLPKDCKG_05487* | *ppsC* | - | 5936668 | 5937663 | 4.65 | 0.38 | 1.40 | 0.24 | Phthiocerol/phenolphthiocerol synthesis polyketide synthase type I PpsC |
| *FLPKDCKG_05488* | *livF-25* | - | 5937683 | 5938387 | 7.05 | 1.45 | 0.00 | 2.14 | High-affinity branched-chain amino acid transport ATP-binding protein LivF |
| *FLPKDCKG_05489* | *lptB-11* | - | 5938380 | 5940161 | 3.07 | 0.37 | 0.74 | 0.23 | Lipopolysaccharide export system ATP-binding protein LptB |
| *FLPKDCKG_05490* | *livH-26* | - | 5940184 | 5941050 | 4.16 | 1.35 | 1.46 | 0.87 | High-affinity branched-chain amino acid transport system permease protein LivH |
| *FLPKDCKG_05491* | *braC-10* | - | 5941200 | 5942399 | 29.42 | 9.39 | 4.71 | 4.20 | Leucine-, isoleucine-, valine-, threonine-, and alanine-binding protein |
| *FLPKDCKG_05492* | *fabG-21* | + | 5942732 | 5943514 | 4.89 | 1.17 | 3.62 | 0.63 | 3-oxoacyl-[acyl-carrier-protein] reductase FabG |
| *FLPKDCKG_05493* | *lcdH-2* | + | 5943514 | 5944476 | 4.53 | 1.33 | 0.97 | 1.15 | L-carnitine dehydrogenase |
| *FLPKDCKG_05494* | *kce-4* | + | 5944469 | 5945404 | 9.73 | 1.72 | 2.63 | 0.35 | 3-keto-5-aminohexanoate cleavage enzyme |
| *FLPKDCKG_05495* | *gldA* | - | 5945529 | 5946596 | 2.80 | 1.62 | 1.33 | 1.60 | Glycerol dehydrogenase |
| *FLPKDCKG_05496* | *mcbR* | - | 5946606 | 5947310 | 10.23 | 9.94 | 8.51 | 1.82 | HTH-type transcriptional regulator McbR |
| *FLPKDCKG_05497* | *tktB* | + | 5947446 | 5948303 | 4.71 | 2.75 | 4.90 | 1.47 | Transketolase 2 |
| *FLPKDCKG_05498* | *dxs-2* | + | 5948304 | 5949290 | 3.65 | 1.90 | 1.84 | 2.08 | 1-deoxy-D-xylulose-5-phosphate synthase |
| *FLPKDCKG_05499* | *aroE-3* | + | 5949347 | 5950156 | 2.38 | 0.37 | 1.71 | 0.42 | Shikimate dehydrogenase (NADP(+)) |
| *FLPKDCKG_05500* | *alkJ-3* | + | 5950175 | 5951773 | 1.10 | 0.67 | 1.12 | 0.11 | Alcohol dehydrogenase [acceptor] |
| *FLPKDCKG_05501* | | - | 5951858 | 5952961 | 0.66 | 1.74 | 3.25 | 1.14 | L-talarate/galactarate dehydratase |
| *FLPKDCKG_05502* | *glnH-5* | - | 5952975 | 5953793 | 1.82 | 6.66 | 1.38 | 1.94 | ABC transporter glutamine-binding protein GlnH |
| *FLPKDCKG_05503* | *glnQ-13* | - | 5953847 | 5954578 | 0.96 | 5.11 | 1.06 | 2.40 | Glutamine transport ATP-binding protein GlnQ |
| *FLPKDCKG_05504* | *yecS-7* | - | 5954562 | 5955221 | 0.50 | 3.65 | 0.97 | 0.89 | L-cystine transport system permease protein YecS |
| *FLPKDCKG_05505* | *occM-3* | - | 5955232 | 5955915 | 0.43 | 1.95 | 1.99 | 0.31 | Octopine transport system permease protein OccM |
| *FLPKDCKG_05506* | *suyB-2* | - | 5955938 | 5957170 | 0.74 | 2.86 | 4.23 | 1.49 | (2R)-sulfolactate sulfo-lyase subunit beta |
| *FLPKDCKG_05507* | *uxaA* | - | 5957181 | 5957507 | 1.34 | 12.79 | 16.27 | 3.04 | Altronate dehydratase |
| *FLPKDCKG_05508* | | + | 5957756 | 5958526 | 0.53 | 0.93 | 5.76 | 0.01 | 2-keto-3-deoxy-L-fuconate dehydrogenase |
| *FLPKDCKG_05509* | | + | 5958564 | 5959418 | 4.22 | 6.69 | 4.34 | 2.75 | Ureidoglycolate lyase |
| *FLPKDCKG_05510* | *kdgR-8* | + | 5959447 | 5960295 | 4.32 | 4.96 | 6.35 | 2.50 | Transcriptional regulator KdgR |
| *FLPKDCKG_05511* | | - | 5960354 | 5961490 | 0.26 | 1.92 | 1.59 | 1.28 | Hypothetical protein |
| *FLPKDCKG_05512* | | - | 5961487 | 5962356 | 0.57 | 0.77 | 0.85 | 0.50 | Hypothetical protein |
| *FLPKDCKG_05513* | | - | 5962722 | 5963768 | 0.84 | 6.78 | 3.15 | 1.73 | Epoxide hydrolase B |
| *FLPKDCKG_05514* | *nylB'* | + | 5963962 | 5965161 | 1.64 | 0.57 | 0.04 | 0.18 | 6-aminohexanoate-dimer hydrolase |
| *FLPKDCKG_05515* | | - | 5965201 | 5966646 | 0.38 | 0.82 | 1.27 | 0.32 | 3'3'-cGAMP-specific phosphodiesterase 3 |
| *FLPKDCKG_05516* | *pytH-2* | + | 5966795 | 5967568 | 0.49 | 0.00 | 0.15 | 0.00 | Pyrethroid hydrolase |

| Locus name | Gene name | Strand | Gene start (bp) | Gene end (bp) | Average TPM Inoculum | Average TPM $t_6$ | Average TPM $t_{24}$ | Average TPM $t_{168}$ | Annotation description |
|---|---|---|---|---|---|---|---|---|---|
| *FLPKDCKG_05517* | *proP-7* | - | 5967674 | 5968315 | 0.29 | 0.32 | 0.00 | 0.00 | Proline/betaine transporter |
| *FLPKDCKG_05518* | | + | 5968519 | 5968914 | 0.00 | 0.00 | 0.00 | 0.00 | Hypothetical protein |
| *FLPKDCKG_05519* | *gabR-5* | + | 5969124 | 5970626 | 3.28 | 5.13 | 3.82 | 0.80 | HTH-type transcriptional regulatory protein GabR |
| *FLPKDCKG_05520* | *dgdA-2* | - | 5970780 | 5972108 | 0.42 | 0.62 | 1.43 | 0.30 | 2,2-dialkylglycine decarboxylase |
| *FLPKDCKG_05521* | *thrB-2* | - | 5972105 | 5973184 | 0.17 | 0.00 | 0.86 | 0.00 | Homoserine kinase |
| *FLPKDCKG_05522* | *ylmB-2* | - | 5973181 | 5974479 | 0.56 | 0.94 | 0.33 | 0.44 | N-formyl-4-amino-5-aminomethyl-2-methylpyrimidine deformylase |
| *FLPKDCKG_05523* | | - | 5974573 | 5975052 | 0.18 | 0.00 | 0.72 | 0.23 | Hypothetical protein |
| *FLPKDCKG_05524* | | - | 5975112 | 5976626 | 0.26 | 0.92 | 1.08 | 0.00 | Hypothetical protein |
| *FLPKDCKG_05525* | | - | 5976630 | 5977115 | 0.73 | 2.57 | 0.00 | 0.00 | Hypothetical protein |
| *FLPKDCKG_05526* | *gchK-2* | + | 5977460 | 5978845 | 1.15 | 3.52 | 1.15 | 0.36 | Globin-coupled histidine kinase |
| *FLPKDCKG_05527* | *glnG-2* | + | 5978842 | 5980137 | 1.61 | 7.62 | 4.62 | 0.02 | Nitrogen regulation protein NR(I) |
| *FLPKDCKG_05528* | *pncC-2* | - | 5980165 | 5980683 | 0.46 | 44.65 | 26.81 | 0.75 | Nicotinamide-nucleotide amidohydrolase PncC |
| *FLPKDCKG_05529* | *ku* | + | 5980894 | 5981847 | 14.69 | 112.65 | 55.88 | 6.04 | Non-homologous end joining protein Ku |
| *FLPKDCKG_05530* | *ligD* | + | 5981850 | 5984474 | 5.66 | 54.60 | 30.03 | 3.09 | Multifunctional non-homologous end joining protein LigD |
| *FLPKDCKG_05531* | *higB-2-2* | + | 5984646 | 5984957 | 34.65 | 74.08 | 35.86 | 9.75 | Toxin HigB-2 |
| *FLPKDCKG_05532* | *higA-2-2* | + | 5984959 | 5985279 | 28.73 | 77.20 | 66.12 | 10.15 | Antitoxin HigA-2 |
| *FLPKDCKG_05533* | *gabP* | - | 5985358 | 5986803 | 2.15 | 11.58 | 8.46 | 1.82 | GABA permease |
| *FLPKDCKG_05534* | *treY* | - | 5986985 | 5989681 | 0.32 | 33.50 | 21.66 | 1.84 | Maltooligosyl trehalose synthase |
| *FLPKDCKG_05535* | *malQ* | - | 5989678 | 5991795 | 0.13 | 33.76 | 17.77 | 2.42 | 4-alpha-glucanotransferase |
| *FLPKDCKG_05536* | *treZ* | - | 5991792 | 5993585 | 0.33 | 60.27 | 35.61 | 3.94 | Malto-oligosyltrehalose trehalohydrolase |
| *FLPKDCKG_05537* | *glgX* | - | 5993582 | 5995693 | 0.13 | 84.93 | 52.79 | 5.58 | Glycogen operon protein GlgX |
| *FLPKDCKG_05538* | *glgB* | - | 5995696 | 5997912 | 0.12 | 62.02 | 37.05 | 2.84 | 1,4-alpha-glucan branching enzyme GlgB |
| *FLPKDCKG_05539* | *treS* | - | 5997909 | 6001274 | 0.22 | 79.08 | 42.37 | 2.90 | Trehalose synthase/amylase TreS |
| *FLPKDCKG_05540* | *glgE2* | - | 6001297 | 6004494 | 0.10 | 72.39 | 45.23 | 4.30 | Alpha-1,4-glucan:maltose-1-phosphate maltosyltransferase 2 |
| *FLPKDCKG_05541* | *glgA* | - | 6004504 | 6006075 | 0.05 | 147.94 | 91.01 | 9.51 | Glycogen synthase |
| *FLPKDCKG_05542* | *ybhN* | - | 6006268 | 6007251 | 0.27 | 49.47 | 25.39 | 2.62 | Inner membrane protein YbhN |
| *FLPKDCKG_05543* | *clsB-2* | - | 6007248 | 6008450 | 0.22 | 19.56 | 13.32 | 1.52 | Cardiolipin synthase B |
| *FLPKDCKG_05544* | | - | 6008447 | 6009202 | 0.31 | 50.83 | 33.74 | 1.31 | Hypothetical protein |
| *FLPKDCKG_05545* | *chaA* | - | 6009264 | 6010349 | 4.38 | 15.24 | 9.05 | 3.80 | Sodium-potassium/proton antiporter ChaA |
| *FLPKDCKG_05546* | | + | 6010462 | 6010932 | 0.29 | 197.53 | 91.51 | 5.93 | Hypothetical protein |
| *FLPKDCKG_05547* | | + | 6010929 | 6012011 | 0.08 | 84.16 | 40.92 | 4.26 | Hypothetical protein |
| *FLPKDCKG_05548* | | + | 6012069 | 6012560 | 0.26 | 42.16 | 26.28 | 3.05 | Hypothetical protein |
| *FLPKDCKG_05549* | | + | 6012715 | 6013221 | 0.29 | 419.87 | 356.84 | 36.15 | Hypothetical protein |
| *FLPKDCKG_05550* | | + | 6013253 | 6013507 | 0.33 | 209.33 | 159.15 | 10.75 | Hypothetical protein |
| *FLPKDCKG_05551* | *pgi-3* | + | 6013516 | 6015453 | 0.05 | 69.14 | 58.17 | 7.67 | Glucose-6-phosphate isomerase |
| *FLPKDCKG_05552* | | + | 6015450 | 6015836 | 0.11 | 25.96 | 29.51 | 6.80 | Hypothetical protein |
| *FLPKDCKG_05553* | *mscS-2* | - | 6015837 | 6016928 | 0.08 | 33.68 | 23.07 | 2.16 | Small-conductance mechanosensitive channel |
| *FLPKDCKG_05554* | *neoL* | + | 6017046 | 6017756 | 0.00 | 33.58 | 28.11 | 1.15 | 2'-N-acetylparomamine deacetylase |
| *FLPKDCKG_05555* | *wbbL* | + | 6017753 | 6018640 | 0.05 | 16.87 | 10.77 | 0.49 | N-acetylglucosaminyl-diphospho-decaprenol L-rhamnosyltransferase |
| *FLPKDCKG_05556* | | + | 6018669 | 6019148 | 0.18 | 5.69 | 11.58 | 1.49 | Hypothetical protein |
| *FLPKDCKG_05557* | *atm1-3* | + | 6019279 | 6021993 | 0.09 | 5.06 | 4.26 | 0.53 | ATM1-type heavy metal exporter |
| *FLPKDCKG_05558* | | + | 6022210 | 6022878 | 0.00 | 59.84 | 57.75 | 4.47 | Hypothetical protein |
| *FLPKDCKG_05559* | *mshA-3* | + | 6022875 | 6024146 | 0.15 | 37.58 | 34.89 | 3.06 | D-inositol 3-phosphate glycosyltransferase |
| *FLPKDCKG_05560* | *bacC-7* | + | 6024162 | 6024890 | 0.40 | 50.55 | 48.93 | 4.83 | Dihydroanticapsin 7-dehydrogenase |
| *FLPKDCKG_05561* | *gmhB* | + | 6024881 | 6025504 | 0.10 | 34.58 | 38.27 | 3.54 | D-glycero-beta-D-manno-heptose-1,7-bisphosphate 7-phosphatase |
| *FLPKDCKG_05562* | *rfaF-2* | + | 6025504 | 6026613 | 0.57 | 29.82 | 24.65 | 2.45 | ADP-heptose--LPS heptosyltransferase 2 |
| *FLPKDCKG_05563* | *novN* | + | 6026601 | 6028340 | 0.42 | 32.70 | 34.20 | 3.51 | Decarbamoylnovobiocin carbamoyltransferase |
| *FLPKDCKG_05564* | *pglI-2* | + | 6028337 | 6029323 | 0.04 | 19.01 | 17.12 | 1.41 | GalNAc(5)-diNAcBac-PP-undecaprenol beta-1,3-glucosyltransferase |
| *FLPKDCKG_05565* | *rfaF-3* | + | 6029320 | 6030381 | 0.06 | 14.43 | 15.05 | 0.42 | ADP-heptose--LPS heptosyltransferase 2 |
| *FLPKDCKG_05566* | *bshA-4* | + | 6030378 | 6031346 | 0.11 | 29.05 | 22.61 | 1.16 | N-acetyl-alpha-D-glucosaminyl L-malate synthase |
| *FLPKDCKG_05567* | *galE1* | + | 6031343 | 6032362 | 0.25 | 23.10 | 17.27 | 1.67 | UDP-glucose 4-epimerase |
| *FLPKDCKG_05568* | | + | 6032483 | 6032662 | 0.76 | 254.55 | 196.35 | 18.51 | Hypothetical protein |
| *FLPKDCKG_05569* | | + | 6032742 | 6033119 | 0.12 | 25.79 | 30.68 | 4.93 | Hypothetical protein |
| *FLPKDCKG_05570* | | - | 6033205 | 6033471 | 0.00 | 34.37 | 36.70 | 4.07 | Hypothetical protein |
| *FLPKDCKG_05571* | | - | 6033468 | 6033698 | 0.00 | 95.91 | 60.24 | 2.43 | Hypothetical protein |
| *FLPKDCKG_05572* | | - | 6033736 | 6034284 | 0.31 | 324.13 | 235.88 | 31.21 | Hypothetical protein |
| *FLPKDCKG_05573* | | - | 6034508 | 6034804 | 0.10 | 49.51 | 37.18 | 5.02 | Hypothetical protein |
| *FLPKDCKG_05574* | | - | 6034874 | 6034984 | 2.17 | 105.14 | 109.72 | 9.60 | Hypothetical protein |
| *FLPKDCKG_05575* | *osmY-3* | - | 6035027 | 6035350 | 0.00 | 137.33 | 108.65 | 10.39 | Osmotically-inducible protein Y |
| *FLPKDCKG_05576* | | - | 6035437 | 6035745 | 0.29 | 8.50 | 8.18 | 1.61 | Hypothetical protein |
| *FLPKDCKG_05577* | | - | 6035950 | 6036216 | 58.02 | 76.50 | 44.43 | 6.35 | Hypothetical protein |
| *FLPKDCKG_05578* | *phzF-3* | - | 6036286 | 6037194 | 1.59 | 4.94 | 4.04 | 0.31 | Trans-2,3-dihydro-3-hydroxyanthranilate isomerase |
| *FLPKDCKG_05579* | *rhtC-2* | - | 6037201 | 6037830 | 0.67 | 0.99 | 1.78 | 0.58 | Threonine efflux protein |
| *FLPKDCKG_05580* | | - | 6037867 | 6038376 | 4.58 | 8.01 | 7.15 | 1.46 | Hypothetical protein |
| *FLPKDCKG_05581* | *gabR-6* | + | 6038405 | 6039751 | 3.01 | 4.20 | 2.16 | 1.20 | HTH-type transcriptional regulatory protein GabR |
| *FLPKDCKG_05582* | *cdhR-9* | - | 6039794 | 6040762 | 7.67 | 19.83 | 20.81 | 4.62 | HTH-type transcriptional regulator CdhR |
| *FLPKDCKG_05583* | *iorA* | + | 6040964 | 6041434 | 10.00 | 30.65 | 51.64 | 13.44 | Isoquinoline 1-oxidoreductase subunit alpha |
| *FLPKDCKG_05584* | *iorB* | + | 6041434 | 6043722 | 8.63 | 17.71 | 35.18 | 12.66 | Isoquinoline 1-oxidoreductase subunit beta |
| *FLPKDCKG_05585* | *pucB* | + | 6043766 | 6044347 | 4.07 | 11.40 | 16.86 | 3.24 | Purine catabolism protein PucB |

| Locus name | Gene name | Strand | Gene start (bp) | Gene end (bp) | Average TPM Inoculum | Average TPM $t_6$ | Average TPM $t_{24}$ | Average TPM $t_{168}$ | Annotation description |
|---|---|---|---|---|---|---|---|---|---|
| *FLPKDCKG_05586* | *pucA* | + | 6044348 | 6045361 | 2.06 | 2.74 | 12.48 | 3.48 | putative xanthine dehydrogenase subunit A |
| *FLPKDCKG_05587* | | - | 6045398 | 6045471 | 0.48 | 3.63 | 5.70 | 0.19 | tRNA |
| *FLPKDCKG_05588* | *cadA-2* | + | 6045639 | 6047903 | 1.35 | 6.45 | 2.47 | 0.82 | Inducible lysine decarboxylase |
| *FLPKDCKG_05589* | *lldR-4* | + | 6048014 | 6048757 | 6.04 | 8.17 | 5.56 | 0.00 | Putative L-lactate dehydrogenase operon regulatory protein |
| *FLPKDCKG_05590* | *hisC-3* | + | 6048757 | 6049308 | 1.33 | 2.72 | 3.28 | 2.18 | Histidinol-phosphate aminotransferase |
| *FLPKDCKG_05591* | *carA-2* | + | 6049497 | 6050216 | 0.33 | 1.59 | 2.06 | 0.63 | Carbamoyl-phosphate synthase arginine-specific small chain |
| *FLPKDCKG_05592* | | + | 6050195 | 6050431 | 0.18 | 0.00 | 2.97 | 0.00 | Hypothetical protein |
| *FLPKDCKG_05593* | | + | 6050645 | 6050854 | 0.62 | 144.21 | 95.71 | 8.75 | Hypothetical protein |
| *FLPKDCKG_05594* | | + | 6050910 | 6051542 | 0.36 | 31.79 | 31.30 | 1.49 | Hypothetical protein |
| *FLPKDCKG_05595* | | + | 6051741 | 6052865 | 57.50 | 5.21 | 5.66 | 2.83 | Monocarboxylate 2-oxoacid-binding periplasmic protein |
| *FLPKDCKG_05596* | | + | 6053015 | 6053638 | 5.02 | 2.66 | 1.92 | 1.67 | Hypothetical protein |
| *FLPKDCKG_05597* | *teaC* | + | 6053635 | 6055695 | 2.86 | 0.39 | 0.45 | 0.45 | Ectoine TRAP transporter large permease protein TeaC |
| *FLPKDCKG_05598* | | + | 6055830 | 6056363 | 1.11 | 1.02 | 1.43 | 0.55 | Hypothetical protein |
| *FLPKDCKG_05599* | *oprM-9* | - | 6056418 | 6057911 | 0.24 | 0.32 | 0.00 | 0.29 | Outer membrane protein OprM |
| *FLPKDCKG_05600* | *bepE-3* | - | 6057932 | 6061108 | 0.18 | 0.14 | 0.55 | 0.21 | Efflux pump membrane transporter BepE |
| *FLPKDCKG_05601* | *acrA-2* | - | 6061105 | 6062301 | 0.05 | 0.00 | 1.03 | 0.00 | Multidrug efflux pump subunit AcrA |
| *FLPKDCKG_05602* | *katE* | + | 6062594 | 6064978 | 0.46 | 114.39 | 79.25 | 7.56 | Catalase HPII |
| *FLPKDCKG_05603* | | - | 6065019 | 6065942 | 0.20 | 33.71 | 22.10 | 0.27 | Hypothetical protein |
| *FLPKDCKG_05604* | *hpxO-5* | - | 6066026 | 6067183 | 0.03 | 0.53 | 0.50 | 0.31 | FAD-dependent urate hydroxylase |
| *FLPKDCKG_05605* | *mhpB* | - | 6067254 | 6068246 | 0.09 | 0.32 | 0.44 | 0.00 | 2,3-dihydroxyphenylpropionate/2,3-dihydroxicinnamic acid 1,2-dioxygenase |
| *FLPKDCKG_05606* | | + | 6068495 | 6069622 | 2.68 | 6.10 | 3.26 | 1.60 | Hypothetical protein |
| *FLPKDCKG_05607* | *nlhH-3* | - | 6069668 | 6070552 | 0.95 | 3.78 | 5.35 | 0.29 | Carboxylesterase NlhH |
| *FLPKDCKG_05608* | *oatA-3* | - | 6070769 | 6071878 | 0.12 | 0.00 | 0.31 | 0.13 | O-acetyltransferase OatA |
| *FLPKDCKG_05609* | *bla* | - | 6071896 | 6072723 | 0.06 | 0.63 | 0.61 | 0.00 | Beta-lactamase OXA-18 |
| *FLPKDCKG_05610* | | + | 6072977 | 6073069 | 0.63 | 0.00 | 0.00 | 0.00 | tRNA |
| *FLPKDCKG_05611* | *blaI* | - | 6074462 | 6074851 | 4.06 | 4.25 | 2.40 | 2.25 | Transcriptional regulator BlaI |
| *FLPKDCKG_05612* | *bmaC-1* | - | 6074933 | 6075349 | 3.27 | 0.00 | 0.60 | 1.40 | Adhesin BmaC autotransporter |
| *FLPKDCKG_05613* | *bmaC-2* | - | 6075409 | 6077640 | 12.39 | 7.59 | 3.62 | 1.78 | Adhesin BmaC autotransporter |
| *FLPKDCKG_05614* | *ydaM-3* | - | 6077714 | 6078928 | 14.97 | 1.78 | 1.66 | 0.78 | putative diguanylate cyclase YdaM |
| *FLPKDCKG_05615* | *gst-2* | - | 6079151 | 6079876 | 1.05 | 1.61 | 3.57 | 0.00 | Glutathione S-transferase |
| *FLPKDCKG_05616* | *dmlR-36* | + | 6079977 | 6080882 | 3.07 | 6.25 | 5.86 | 2.12 | HTH-type transcriptional regulator DmlR |
| *FLPKDCKG_05617* | *yhjQ* | + | 6080975 | 6081304 | 1.74 | 209.99 | 128.14 | 12.76 | putative cysteine-rich protein YhjQ |
| *FLPKDCKG_05618* | | - | 6081359 | 6081727 | 4.89 | 69.37 | 34.50 | 4.76 | Hypothetical protein |
| *FLPKDCKG_05619* | *zraS* | + | 6081896 | 6083266 | 4.22 | 10.97 | 9.57 | 2.87 | Sensor protein ZraS |
| *FLPKDCKG_05620* | *zraR-3* | + | 6083251 | 6084582 | 1.56 | 5.57 | 1.47 | 1.15 | Transcriptional regulatory protein ZraR |
| *FLPKDCKG_05621* | | + | 6084734 | 6085390 | 0.69 | 19.35 | 4.06 | 0.90 | Hypothetical protein |
| *FLPKDCKG_05622* | | + | 6085457 | 6085807 | 0.36 | 1.22 | 0.58 | 1.47 | Hypothetical protein |
| *FLPKDCKG_05623* | | + | 6085817 | 6087055 | 0.24 | 1.28 | 0.30 | 0.13 | putative MFS-type transporter |
| *FLPKDCKG_05624* | | - | 6087262 | 6088218 | 3.38 | 4.68 | 1.90 | 2.93 | Hypothetical protein |
| *FLPKDCKG_05625* | | + | 6088857 | 6089438 | 2.64 | 2.52 | 1.94 | 1.69 | Hypothetical protein |
| *FLPKDCKG_05626* | *torA* | - | 6089458 | 6089757 | 0.52 | 2.34 | 2.98 | 0.80 | Trimethylamine-N-oxide reductase 1 |
| *FLPKDCKG_05627* | | - | 6089840 | 6090493 | 5.34 | 15.30 | 12.60 | 3.25 | Hypothetical protein |
| *FLPKDCKG_05628* | | - | 6090493 | 6090855 | 29.27 | 33.60 | 17.64 | 7.44 | Hypothetical protein |
| *FLPKDCKG_05629* | *tsaQ1-4* | - | 6091409 | 6092233 | 6.28 | 12.06 | 4.02 | 1.70 | HTH-type transcriptional regulator TsaQ1/TsaQ2 |
| *FLPKDCKG_05630* | *crt-6* | + | 6092360 | 6093166 | 0.88 | 2.06 | 2.21 | 0.78 | Short-chain-enoyl-CoA hydratase |
| *FLPKDCKG_05631* | | + | 6093163 | 6095268 | 1.27 | 5.37 | 4.81 | 0.96 | Hypothetical protein |
| *FLPKDCKG_05632* | | + | 6095372 | 6096364 | 2.26 | 3.93 | 1.10 | 0.94 | Hypothetical protein |
| *FLPKDCKG_05633* | | - | 6096436 | 6097404 | 0.12 | 0.98 | 0.58 | 0.00 | Hypothetical protein |
| *FLPKDCKG_05634* | *gnl-2* | - | 6097433 | 6098371 | 0.25 | 0.56 | 0.18 | 0.00 | Gluconolactonase |
| *FLPKDCKG_05635* | *ligI-1* | - | 6098401 | 6099210 | 0.07 | 1.07 | 0.02 | 0.00 | 2-pyrone-4,6-dicarbaxylate hydrolase |
| *FLPKDCKG_05636* | | - | 6099224 | 6100228 | 0.43 | 2.42 | 0.47 | 0.43 | Hypothetical protein |
| *FLPKDCKG_05637* | *gdhI* | - | 6100280 | 6101080 | 0.60 | 2.68 | 1.36 | 0.00 | Glucose 1-dehydrogenase 1 |
| *FLPKDCKG_05638* | | - | 6101115 | 6101684 | 0.01 | 0.00 | 0.00 | 0.00 | Hypothetical protein |
| *FLPKDCKG_05639* | | - | 6101698 | 6102669 | 0.18 | 1.07 | 0.48 | 0.02 | Hypothetical protein |
| *FLPKDCKG_05640* | *catM* | + | 6102839 | 6103702 | 1.93 | 2.62 | 1.15 | 0.70 | HTH-type transcriptional regulator CatM |
| *FLPKDCKG_05641* | *prpD-3* | + | 6103859 | 6105193 | 0.86 | 1.63 | 1.13 | 0.91 | 2-methylcitrate dehydratase |
| *FLPKDCKG_05642* | | + | 6105364 | 6106371 | 0.25 | 0.10 | 0.43 | 0.58 | Hypothetical protein |
| *FLPKDCKG_05643* | | + | 6106384 | 6107784 | 0.62 | 1.73 | 0.00 | 0.00 | Hypothetical protein |
| *FLPKDCKG_05644* | *hcaR-8* | - | 6107791 | 6108675 | 5.79 | 9.32 | 6.33 | 1.52 | Hca operon transcriptional activator HcaR |
| *FLPKDCKG_05645* | *bpoC* | + | 6108798 | 6109673 | 1.29 | 8.43 | 3.40 | 1.63 | Putative non-heme bromoperoxidase BpoC |
| *FLPKDCKG_05646* | | + | 6109920 | 6110144 | 0.60 | 6.59 | 2.47 | 0.00 | Hypothetical protein |
| *FLPKDCKG_05647* | | + | 6110450 | 6111412 | 1.35 | 6.95 | 2.10 | 0.49 | Hypothetical protein |
| *FLPKDCKG_05648* | *nagL-4* | + | 6111415 | 6112050 | 0.95 | 4.24 | 1.51 | 0.74 | Maleylpyruvate isomerase |
| *FLPKDCKG_05649* | *novR-5* | + | 6112064 | 6112849 | 0.89 | 2.56 | 0.26 | 2.70 | Decarboxylase NovR |
| *FLPKDCKG_05650* | *puuR-2* | + | 6112968 | 6113522 | 5.62 | 8.04 | 4.68 | 2.75 | HTH-type transcriptional regulator PuuR |
| *FLPKDCKG_05651* | | + | 6113540 | 6113905 | 1.96 | 6.80 | 6.29 | 1.43 | Hypothetical protein |
| *FLPKDCKG_05652* | | - | 6113927 | 6114307 | 4.85 | 9.07 | 7.51 | 0.00 | Hypothetical protein |
| *FLPKDCKG_05653* | *fecA-2* | + | 6114535 | 6115218 | 0.12 | 0.00 | 0.00 | 0.00 | Fe(3+) dicitrate transport protein FecA |
| *FLPKDCKG_05654* | *fecI-15* | + | 6115208 | 6115756 | 0.23 | 0.00 | 0.74 | 0.26 | putative RNA polymerase sigma factor FecI |

| Locus name | Gene name | Strand | Gene start (bp) | Gene end (bp) | Average TPM Inoculum | Average TPM $t_6$ | Average TPM $t_{24}$ | Average TPM $t_{168}$ | Annotation description |
|---|---|---|---|---|---|---|---|---|---|
| *FLPKDCKG_05655* | | + | 6115753 | 6116748 | 0.19 | 0.00 | 0.71 | 0.09 | Hypothetical protein |
| *FLPKDCKG_05656* | *hxuA-2* | + | 6116874 | 6129464 | 0.17 | 0.24 | 0.17 | 0.32 | Heme/hemopexin-binding protein |
| *FLPKDCKG_05657* | *fecA-3* | + | 6129638 | 6130333 | 0.00 | 0.86 | 0.00 | 0.00 | Fe(3+) dicitrate transport protein FecA |
| *FLPKDCKG_05658* | *fecI-16* | + | 6130339 | 6130881 | 0.11 | 0.00 | 0.00 | 0.00 | putative RNA polymerase sigma factor FecI |
| *FLPKDCKG_05659* | | + | 6130886 | 6131851 | 0.56 | 0.83 | 0.32 | 0.54 | Hypothetical protein |
| *FLPKDCKG_05660* | *hxuA-3* | + | 6132045 | 6144737 | 0.11 | 0.30 | 0.18 | 0.05 | Heme/hemopexin-binding protein |
| *FLPKDCKG_05661* | | + | 6145057 | 6146673 | 0.05 | 0.00 | 0.39 | 0.00 | Hypothetical protein |
| *FLPKDCKG_05662* | *exbB-3* | + | 6146711 | 6148633 | 0.05 | 0.80 | 0.05 | 0.13 | Biopolymer transport protein ExbB |
| *FLPKDCKG_05663* | *exbD-3* | + | 6148647 | 6149066 | 0.00 | 0.00 | 0.00 | 0.00 | Biopolymer transport protein ExbD |
| *FLPKDCKG_05664* | | + | 6149084 | 6149800 | 0.09 | 1.26 | 0.13 | 0.00 | Hypothetical protein |
| *FLPKDCKG_05665* | | + | 6149837 | 6151540 | 0.14 | 0.29 | 0.00 | 0.00 | Hypothetical protein |
| *FLPKDCKG_05666* | | + | 6151551 | 6152246 | 0.07 | 0.00 | 0.00 | 0.00 | Hypothetical protein |
| *FLPKDCKG_05667* | | + | 6152354 | 6152839 | 0.69 | 0.00 | 1.07 | 0.62 | Hypothetical protein |
| *FLPKDCKG_05668* | *prsA2* | + | 6152944 | 6154014 | 0.14 | 0.05 | 0.00 | 0.00 | Foldase protein PrsA 2 |
| *FLPKDCKG_05669* | | + | 6154109 | 6154429 | 0.13 | 0.00 | 0.00 | 0.00 | Hypothetical protein |
| *FLPKDCKG_05670* | | + | 6154426 | 6154908 | 0.09 | 0.00 | 0.43 | 0.00 | Hypothetical protein |
| *FLPKDCKG_05671* | *pdxK* | + | 6155016 | 6155879 | 5.86 | 12.63 | 5.80 | 1.29 | Pyridoxine/pyridoxal/pyridoxamine kinase |
| *FLPKDCKG_05672* | *cynS* | - | 6155913 | 6156359 | 1.99 | 3.12 | 3.31 | 1.09 | Cyanate hydratase |
| *FLPKDCKG_05673* | *cmpD-4* | - | 6156365 | 6157159 | 0.22 | 0.38 | 1.14 | 0.73 | Bicarbonate transport ATP-binding protein CmpD |
| *FLPKDCKG_05674* | *cmpB-5* | - | 6157165 | 6158007 | 0.45 | 1.42 | 0.87 | 0.56 | Bicarbonate transport system permease protein CmpB |
| *FLPKDCKG_05675* | *cmpC-2* | - | 6158007 | 6159392 | 0.26 | 1.97 | 0.33 | 1.36 | Bicarbonate transport ATP-binding protein CmpC |
| *FLPKDCKG_05676* | *uvrY-4* | - | 6159525 | 6160163 | 1.08 | 1.76 | 0.78 | 0.96 | Response regulator UvrY |
| *FLPKDCKG_05677* | *degS-2* | - | 6160176 | 6161318 | 2.21 | 4.17 | 4.57 | 1.01 | Signal transduction histidine-protein kinase/phosphatase DegS |
| *FLPKDCKG_05678* | *ybhF-4* | + | 6161511 | 6162434 | 9.83 | 18.97 | 16.98 | 2.61 | putative ABC transporter ATP-binding protein YbhF |
| *FLPKDCKG_05679* | *yadH-2* | + | 6162431 | 6163192 | 3.00 | 7.78 | 6.34 | 2.75 | Inner membrane transport permease YadH |
| *FLPKDCKG_05680* | *ligE* | - | 6163278 | 6163976 | 13.76 | 25.23 | 21.54 | 9.00 | Beta-etherase |
| *FLPKDCKG_05681* | | - | 6164058 | 6164450 | 10.79 | 425.08 | 297.26 | 95.29 | Hypothetical protein |
| *FLPKDCKG_05682* | *thlA-2* | - | 6164646 | 6165827 | 43.39 | 34.55 | 29.59 | 10.29 | Acetyl-CoA acetyltransferase |
| *FLPKDCKG_05683* | *npt* | - | 6165873 | 6166532 | 1.43 | 4.62 | 2.47 | 1.40 | 4'-phosphopantetheinyl transferase Npt |
| *FLPKDCKG_05684* | *ttrR* | + | 6166770 | 6167330 | 4.47 | 6.16 | 2.47 | 0.97 | Tetrathionate response regulatory protein TtrR |
| *FLPKDCKG_05685* | *dctM-6* | - | 6167351 | 6168667 | 1.24 | 1.33 | 0.38 | 0.48 | C4-dicarboxylate TRAP transporter large permease protein DctM |
| *FLPKDCKG_05686* | | - | 6168690 | 6169220 | 1.76 | 0.72 | 0.78 | 0.00 | Hypothetical protein |
| *FLPKDCKG_05687* | | - | 6169276 | 6170370 | 10.41 | 3.21 | 0.20 | 0.24 | Solute-binding protein |
| *FLPKDCKG_05688* | *lcfB-5* | - | 6170411 | 6171970 | 9.16 | 2.20 | 0.48 | 0.00 | Long-chain-fatty-acid--CoA ligase |
| *FLPKDCKG_05689* | *pcaF-3* | - | 6171975 | 6173156 | 26.91 | 6.83 | 2.29 | 1.75 | Beta-ketoadipyl-CoA thiolase |
| *FLPKDCKG_05690* | *arnA* | + | 6173635 | 6174231 | 32.05 | 4.82 | 0.80 | 0.00 | Bifunctional polymyxin resistance protein ArnA |
| *FLPKDCKG_05691* | | + | 6174555 | 6185762 | 14.44 | 2.87 | 2.70 | 0.66 | Phenolphthiocerol synthesis polyketide synthase type I Pks15/1 |
| *FLPKDCKG_05692* | | + | 6186158 | 6187186 | 31.91 | 5.55 | 6.71 | 1.04 | Hypothetical protein |
| *FLPKDCKG_05693* | *menH-5* | + | 6187189 | 6187995 | 8.49 | 5.22 | 1.84 | 1.07 | 2-succinyl-6-hydroxy-2,4-cyclohexadiene-1-carboxylate synthase |
| *FLPKDCKG_05694* | *bedC1* | + | 6188071 | 6189216 | 15.63 | 3.03 | 3.88 | 0.72 | Benzene 1,2-dioxygenase subunit alpha |
| *FLPKDCKG_05695* | | + | 6189229 | 6189840 | 8.45 | 8.84 | 4.30 | 0.38 | Hypothetical protein |
| *FLPKDCKG_05696* | *acoC-2* | + | 6189849 | 6190745 | 35.72 | 30.53 | 21.79 | 7.30 | Dihydrolipoyllysine-residue acetyltransferase component of acetoin cleaving system |
| *FLPKDCKG_05697* | | - | 6190822 | 6191637 | 1.32 | 4.25 | 0.43 | 0.80 | Hypothetical protein |
| *FLPKDCKG_05698* | *nimR-14* | + | 6191738 | 6192508 | 3.11 | 3.86 | 1.05 | 1.47 | HTH-type transcriptional regulator NimR |
| *FLPKDCKG_05699* | | - | 6192525 | 6192962 | 4.38 | 16.62 | 12.90 | 1.76 | Hypothetical protein |
| *FLPKDCKG_05700* | *cspA-4* | + | 6193377 | 6193580 | 41.43 | 142.93 | 73.44 | 21.72 | Cold shock protein CspA |
| *FLPKDCKG_05701* | | + | 6193621 | 6193902 | 25.25 | 77.31 | 58.32 | 22.71 | Hypothetical protein |
| *FLPKDCKG_05702* | | - | 6193959 | 6194648 | 8.34 | 20.76 | 18.93 | 5.02 | Hypothetical protein |
| *FLPKDCKG_05703* | | + | 6194881 | 6195063 | 4.63 | 10.37 | 9.55 | 2.83 | Hypothetical protein |
| *FLPKDCKG_05704* | | + | 6195084 | 6196151 | 1.32 | 4.19 | 3.68 | 1.04 | Hypothetical protein |
| *FLPKDCKG_05705* | | + | 6196306 | 6196818 | 5.19 | 7.20 | 3.65 | 3.58 | Hypothetical protein |
| *FLPKDCKG_05706* | | - | 6196866 | 6197369 | 8.25 | 9.85 | 9.05 | 7.73 | Hypothetical protein |
| *FLPKDCKG_05707* | | - | 6197495 | 6197866 | 19.44 | 35.80 | 17.66 | 27.55 | Hypothetical protein |
| *FLPKDCKG_05708* | *cdhR-10* | + | 6198036 | 6199010 | 9.49 | 10.06 | 10.44 | 4.91 | HTH-type transcriptional regulator CdhR |
| *FLPKDCKG_05709* | *ribZ-4* | - | 6199086 | 6200414 | 1.05 | 4.00 | 2.58 | 0.50 | Riboflavin transporter RibZ |
| *FLPKDCKG_05710* | *dmlR-37* | + | 6200641 | 6201576 | 5.14 | 13.87 | 4.01 | 2.43 | HTH-type transcriptional regulator DmlR |
| *FLPKDCKG_05711* | | + | 6201697 | 6202506 | 4.07 | 5.36 | 3.56 | 2.35 | Hypothetical protein |
| *FLPKDCKG_05712* | *yfeX* | + | 6202615 | 6203562 | 2.85 | 2.44 | 6.81 | 2.06 | putative deferrochelatase/peroxidase YfeX |
| *FLPKDCKG_05713* | | - | 6203569 | 6204015 | 12.23 | 2.53 | 3.25 | 1.30 | Hypothetical protein |
| *FLPKDCKG_05714* | *ascD-3* | - | 6204113 | 6205828 | 0.35 | 1.57 | 0.85 | 0.05 | CDP-6-deoxy-L-threo-D-glycero-4-hexulose-3-dehydrase reductase |
| *FLPKDCKG_05715* | *gabR-7* | + | 6205910 | 6207412 | 2.14 | 2.48 | 1.42 | 1.08 | HTH-type transcriptional regulatory protein GabR |
| *FLPKDCKG_05716* | *yddE* | + | 6207472 | 6208260 | 10.26 | 13.81 | 7.35 | 2.06 | putative isomerase YddE |
| *FLPKDCKG_05717* | | + | 6208405 | 6209148 | 1.82 | 4.62 | 4.40 | 1.28 | Hypothetical protein |
| *FLPKDCKG_05718* | *dadA-4* | - | 6209200 | 6210429 | 5.92 | 2.36 | 3.10 | 3.51 | D-amino acid dehydrogenase |
| *FLPKDCKG_05719* | *ampC* | + | 6210792 | 6212084 | 4.83 | 4.56 | 3.43 | 0.59 | Beta-lactamase |
| *FLPKDCKG_05720* | *hbd-2* | - | 6212112 | 6213125 | 0.61 | 2.50 | 0.00 | 1.08 | 3-hydroxybutyryl-CoA dehydrogenase |
| *FLPKDCKG_05721* | *odh* | - | 6213128 | 6214225 | 0.32 | 0.82 | 1.10 | 0.00 | Opine dehydrogenase |
| *FLPKDCKG_05722* | *exsA-2* | - | 6214416 | 6215408 | 3.67 | 6.65 | 6.36 | 1.62 | Exoenzyme S synthesis regulatory protein ExsA |
| *FLPKDCKG_05723* | *lgoT* | + | 6215571 | 6216974 | 1.13 | 9.13 | 4.00 | 1.67 | putative L-galactonate transporter |

| Locus name | Gene name | Strand | Gene start (bp) | Gene end (bp) | Average TPM Inoculum | Average TPM $t_6$ | Average TPM $t_{24}$ | Average TPM $t_{168}$ | Annotation description |
|---|---|---|---|---|---|---|---|---|---|
| *FLPKDCKG_05724* | | + | 6217144 | 6217569 | 0.42 | 1.43 | 3.68 | 0.17 | Hypothetical protein |
| *FLPKDCKG_05725* | *metQ-7* | + | 6217771 | 6218610 | 0.19 | 1.74 | 25.41 | 10.32 | D-methionine-binding lipoprotein MetQ |
| *FLPKDCKG_05726* | *metN-3* | + | 6218703 | 6219821 | 0.27 | 0.85 | 4.23 | 0.67 | Methionine import ATP-binding protein MetN |
| *FLPKDCKG_05727* | *metI-3* | + | 6219799 | 6220455 | 0.43 | 0.00 | 9.64 | 2.87 | D-methionine transport system permease protein MetI |
| *FLPKDCKG_05728* | *mdeA-3* | + | 6220502 | 6221698 | 0.28 | 0.84 | 10.89 | 4.46 | L-methionine gamma-lyase |
| *FLPKDCKG_05729* | | - | 6221741 | 6222817 | 5.62 | 10.14 | 8.19 | 2.85 | Hypothetical protein |
| *FLPKDCKG_05730* | | - | 6222854 | 6223297 | 13.21 | 30.86 | 18.93 | 9.00 | Hypothetical protein |
| *FLPKDCKG_05731* | | + | 6223497 | 6224756 | 7.10 | 6.43 | 3.70 | 1.85 | Hypothetical protein |
| *FLPKDCKG_05732* | *hslR* | - | 6224779 | 6225159 | 17.23 | 17.55 | 11.19 | 9.65 | Heat shock protein 15 |
| *FLPKDCKG_05733* | | - | 6225210 | 6226808 | 4.01 | 6.06 | 3.61 | 1.28 | putative signaling protein |
| *FLPKDCKG_05734* | | - | 6227055 | 6228044 | 0.27 | 1.38 | 1.02 | 0.13 | Hypothetical protein |
| *FLPKDCKG_05735* | | - | 6228041 | 6228874 | 0.14 | 3.18 | 0.72 | 0.36 | 4-sulfomuconolactone hydrolase |
| *FLPKDCKG_05736* | *benM-5* | + | 6229005 | 6229892 | 2.81 | 6.85 | 6.19 | 2.13 | HTH-type transcriptional regulator BenM |
| *FLPKDCKG_05737* | *bacA* | + | 6230037 | 6231869 | 7.13 | 19.09 | 10.79 | 2.40 | Vitamin B12 transport ATP-binding protein BacA |
| *FLPKDCKG_05738* | *nadX* | - | 6231901 | 6232704 | 3.34 | 5.46 | 4.03 | 1.91 | L-aspartate dehydrogenase |
| *FLPKDCKG_05739* | *mshD-4* | + | 6232888 | 6233397 | 2.20 | 0.07 | 2.82 | 1.44 | Mycothiol acetyltransferase |
| *FLPKDCKG_05740* | *yxeP-5* | - | 6233493 | 6234881 | 5.11 | 2.74 | 1.30 | 1.77 | putative hydrolase YxeP |
| *FLPKDCKG_05741* | *stp-6* | - | 6234967 | 6236628 | 0.25 | 0.36 | 0.00 | 0.00 | Multidrug resistance protein Stp |
| *FLPKDCKG_05742* | *mtrR-2* | + | 6236765 | 6237427 | 8.10 | 10.06 | 8.95 | 2.37 | HTH-type transcriptional regulator MtrR |
| *FLPKDCKG_05743* | | - | 6237414 | 6238364 | 0.48 | 2.17 | 3.16 | 0.46 | Hypothetical protein |
| *FLPKDCKG_05744* | *pcd* | - | 6238420 | 6238635 | 0.00 | 1.85 | 4.19 | 0.00 | Phenmedipham hydrolase |
| *FLPKDCKG_05745* | *pnbA* | - | 6238657 | 6239685 | 0.26 | 0.76 | 1.85 | 0.41 | Para-nitrobenzyl esterase |
| *FLPKDCKG_05746* | | - | 6239843 | 6240793 | 6.88 | 3.11 | 2.37 | 0.76 | Adenine deaminase |
| *FLPKDCKG_05747* | *gbpR-7* | - | 6240802 | 6241722 | 9.73 | 4.97 | 3.72 | 1.30 | HTH-type transcriptional regulator GbpR |
| *FLPKDCKG_05748* | | + | 6241829 | 6242554 | 1.63 | 1.77 | 3.66 | 1.15 | Hypothetical protein |
| *FLPKDCKG_05749* | *mltB-2* | + | 6242657 | 6244207 | 9.14 | 6.51 | 5.04 | 2.96 | Membrane-bound lytic murein transglycosylase B |
| *FLPKDCKG_05750* | | + | 6244204 | 6244839 | 3.28 | 1.96 | 1.92 | 1.61 | putative methyltransferase |
| *FLPKDCKG_05751* | | + | 6245015 | 6245590 | 7.86 | 8.02 | 3.48 | 0.74 | Hypothetical protein |
| *FLPKDCKG_05752* | | + | 6245587 | 6245793 | 4.91 | 20.94 | 1.71 | 4.51 | Hypothetical protein |
| *FLPKDCKG_05753* | *fcuA-5* | - | 6245846 | 6248272 | 0.94 | 2.80 | 0.47 | 0.18 | Ferrichrome receptor FcuA |
| *FLPKDCKG_05754* | | - | 6248532 | 6249581 | 0.45 | 1.88 | 0.16 | 0.27 | Hypothetical protein |
| *FLPKDCKG_05755* | *fecI-17* | - | 6249574 | 6250101 | 0.30 | 0.95 | 0.00 | 0.47 | putative RNA polymerase sigma factor FecI |
| *FLPKDCKG_05756* | | + | 6250317 | 6250613 | 19.94 | 35.57 | 12.40 | 10.93 | Hypothetical protein |
| *FLPKDCKG_05757* | | + | 6250610 | 6250888 | 14.13 | 21.56 | 13.53 | 2.34 | Hypothetical protein |
| *FLPKDCKG_05758* | *yjaB-2* | - | 6250894 | 6251331 | 7.46 | 15.74 | 11.40 | 1.80 | putative N-acetyltransferase YjaB |
| *FLPKDCKG_05759* | | - | 6251344 | 6253083 | 3.02 | 5.51 | 4.99 | 0.97 | Hypothetical protein |
| *FLPKDCKG_05760* | *gsiD-8* | - | 6253142 | 6254008 | 1.87 | 1.22 | 0.52 | 0.10 | Glutathione transport system permease protein GsiD |
| *FLPKDCKG_05761* | *nikB* | - | 6254014 | 6254937 | 0.79 | 0.57 | 0.65 | 0.00 | Nickel transport system permease protein NikB |
| *FLPKDCKG_05762* | *dppA-6* | - | 6255119 | 6256651 | 0.99 | 0.81 | 1.08 | 0.56 | Periplasmic dipeptide transport protein |
| *FLPKDCKG_05763* | *gsiA-11* | - | 6256706 | 6258559 | 0.44 | 2.01 | 0.34 | 0.45 | Glutathione import ATP-binding protein GsiA |
| *FLPKDCKG_05764* | | - | 6258569 | 6259081 | 1.52 | 3.88 | 8.69 | 4.65 | Hypothetical protein |
| *FLPKDCKG_05765* | *decR-5* | - | 6259254 | 6259742 | 4.05 | 7.75 | 0.70 | 1.86 | DNA-binding transcriptional activator DecR |
| *FLPKDCKG_05766* | *csd-2* | + | 6259866 | 6261347 | 9.82 | 15.69 | 9.27 | 3.38 | putative cysteine desulfurase |
| *FLPKDCKG_05767* | | - | 6261404 | 6261865 | 6.12 | 17.57 | 8.35 | 3.75 | Hypothetical protein |
| *FLPKDCKG_05768* | | + | 6261897 | 6262514 | 1.17 | 8.22 | 11.13 | 0.32 | Hypothetical protein |
| *FLPKDCKG_05769* | *fmdA* | + | 6262713 | 6263942 | 109.88 | 64.82 | 54.70 | 12.56 | Formamidase |
| *FLPKDCKG_05770* | | + | 6263965 | 6264297 | 18.65 | 18.86 | 10.05 | 6.46 | Hypothetical protein |
| *FLPKDCKG_05771* | | - | 6264449 | 6265195 | 11.75 | 20.40 | 15.26 | 6.26 | Hypothetical protein |
| *FLPKDCKG_05772* | *nlpE* | - | 6265354 | 6265779 | 41.30 | 79.59 | 53.92 | 11.47 | Lipoprotein NlpE |
| *FLPKDCKG_05773* | | - | 6265913 | 6266257 | 0.41 | 8.29 | 0.60 | 0.80 | Hypothetical protein |
| *FLPKDCKG_05774* | | - | 6266845 | 6267843 | 2.27 | 5.35 | 6.30 | 0.39 | Hypothetical protein |
| *FLPKDCKG_05775* | | - | 6267903 | 6268853 | 0.43 | 1.41 | 0.72 | 0.11 | Hypothetical protein |
| *FLPKDCKG_05776* | | - | 6268924 | 6269994 | 0.38 | 0.91 | 0.00 | 0.00 | Hypothetical protein |
| *FLPKDCKG_05777* | | - | 6270006 | 6270953 | 0.13 | 0.57 | 1.06 | 0.00 | Hydroxypyruvate reductase |
| *FLPKDCKG_05778* | *proA-7* | - | 6270950 | 6271540 | 0.78 | 1.15 | 0.57 | 0.73 | 4-hydroxy-4-methyl-2-oxoglutarate aldolase/4-carboxy-4-hydroxy-2-oxoadipate aldolase |
| *FLPKDCKG_05779* | | - | 6271483 | 6271629 | 0.00 | 0.01 | 0.00 | 2.35 | Hypothetical protein |
| *FLPKDCKG_05780* | | - | 6271669 | 6272664 | 0.21 | 0.99 | 0.90 | 0.46 | Hypothetical protein |
| *FLPKDCKG_05781* | *ligI-2* | - | 6272691 | 6273557 | 0.43 | 0.79 | 0.45 | 0.32 | 2-pyrone-4,6-dicarbaxylate hydrolase |
| *FLPKDCKG_05782* | | - | 6273565 | 6273744 | 0.81 | 2.30 | 0.00 | 0.00 | Hypothetical protein |
| *FLPKDCKG_05783* | | - | 6273732 | 6274529 | 0.73 | 8.53 | 2.84 | 1.11 | Hypothetical protein |
| *FLPKDCKG_05784* | *kipR-3* | + | 6274575 | 6275363 | 3.76 | 8.79 | 4.63 | 2.40 | HTH-type transcriptional regulator KipR |
| *FLPKDCKG_05785* | *gfa-2* | + | 6275520 | 6275915 | 1.64 | 3.22 | 0.00 | 0.28 | Glutathione-dependent formaldehyde-activating enzyme |
| *FLPKDCKG_05786* | | + | 6276006 | 6276479 | 7.00 | 10.72 | 5.33 | 3.98 | Hypothetical protein |
| *FLPKDCKG_05787* | *gstB-8* | + | 6276496 | 6277149 | 0.67 | 2.46 | 0.86 | 0.00 | Glutathione S-transferase GstB |
| *FLPKDCKG_05788* | *speG-4* | + | 6277227 | 6277802 | 8.16 | 14.48 | 11.89 | 2.30 | Spermidine N(1)-acetyltransferase |
| *FLPKDCKG_05789* | | + | 6277867 | 6278001 | 6.00 | 18.99 | 6.55 | 1.21 | Hypothetical protein |
| *FLPKDCKG_05790* | *sttH* | + | 6277998 | 6278411 | 4.90 | 23.40 | 7.64 | 1.11 | Streptothricin hydrolase |
| *FLPKDCKG_05791* | | + | 6278418 | 6278642 | 1.32 | 13.55 | 6.42 | 3.92 | Hypothetical protein |
| *FLPKDCKG_05792* | | + | 6278789 | 6279121 | 44.96 | 153.82 | 117.43 | 46.58 | Hypothetical protein |

| Locus name | Gene name | Strand | Gene start (bp) | Gene end (bp) | Average TPM Inoculum | Average TPM $t_6$ | Average TPM $t_{24}$ | Average TPM $t_{168}$ | Annotation description |
|---|---|---|---|---|---|---|---|---|---|
| *FLPKDCKG_05793* | *rfnT* | - | 6279204 | 6280220 | 1.37 | 14.19 | 9.20 | 3.25 | Riboflavin transporter RfnT |
| *FLPKDCKG_05794* | *inhA-5* | + | 6280530 | 6281495 | 7.88 | 24.93 | 18.37 | 1.26 | Isonitrile hydratase |
| *FLPKDCKG_05795* | *pchR-7* | + | 6281616 | 6282440 | 1.45 | 1.13 | 1.68 | 0.41 | Regulatory protein PchR |
| *FLPKDCKG_05796* | *fptA-4* | + | 6282534 | 6284681 | 0.34 | 0.73 | 1.07 | 0.21 | Fe(3+)-pyochelin receptor |
| *FLPKDCKG_05797* | *fptA-5* | + | 6284861 | 6286993 | 0.05 | 0.64 | 0.13 | 0.00 | Fe(3+)-pyochelin receptor |
| *FLPKDCKG_05798* | | + | 6287053 | 6287460 | 0.10 | 0.00 | 0.00 | 0.10 | Hypothetical protein |
| *FLPKDCKG_05799* | | + | 6287472 | 6288677 | 0.18 | 0.00 | 0.37 | 0.10 | Hypothetical protein |
| *FLPKDCKG_05800* | *dnaK-2* | + | 6288742 | 6289995 | 0.79 | 1.97 | 1.04 | 0.68 | Chaperone protein DnaK |
| *FLPKDCKG_05801* | *dmlR-38* | - | 6290051 | 6290749 | 1.54 | 2.07 | 2.08 | 0.09 | HTH-type transcriptional regulator DmlR |
| *FLPKDCKG_05802* | *nudF* | + | 6291528 | 6292115 | 5.91 | 13.79 | 18.12 | 4.13 | ADP-ribose pyrophosphatase |
| *FLPKDCKG_05803* | | + | 6292134 | 6292469 | 4.67 | 6.38 | 16.01 | 2.60 | Ferredoxin |
| *FLPKDCKG_05804* | | - | 6292509 | 6292955 | 13.43 | 24.51 | 21.24 | 4.26 | Hypothetical protein |
| *FLPKDCKG_05805* | *kgtP-2* | - | 6293427 | 6294728 | 21.36 | 58.16 | 23.60 | 6.16 | Alpha-ketoglutarate permease |
| *FLPKDCKG_05806* | | - | 6294956 | 6296158 | 11.40 | 33.25 | 18.17 | 3.07 | Hypothetical protein |
| *FLPKDCKG_05807* | *anr-2* | - | 6296173 | 6296850 | 44.30 | 64.30 | 33.25 | 12.44 | Transcriptional activator protein Anr |
| *FLPKDCKG_05808* | *btrG* | - | 6297026 | 6297460 | 3.84 | 2.38 | 1.92 | 0.00 | Gamma-L-glutamyl-butirosin B gamma-glutamyl cyclotransferase |
| *FLPKDCKG_05809* | *clpB-2* | + | 6297613 | 6300219 | 40.26 | 26.88 | 55.13 | 26.41 | Chaperone protein ClpB |
| *FLPKDCKG_05810* | | - | 6300332 | 6300772 | 54.84 | 41.49 | 15.00 | 2.20 | Hypothetical protein |
| *FLPKDCKG_05811* | *sqr* | - | 6300769 | 6301959 | 251.09 | 151.56 | 67.77 | 7.01 | Sulfide-quinone reductase |
| *FLPKDCKG_05812* | | - | 6302132 | 6303778 | 0.19 | 2.09 | 1.33 | 0.40 | Hypothetical protein |
| *FLPKDCKG_05813* | | - | 6303927 | 6304676 | 0.68 | 2.47 | 0.00 | 1.59 | Hypothetical protein |
| *FLPKDCKG_05814* | | + | 6304753 | 6305964 | 3.13 | 22.68 | 6.09 | 1.64 | Hypothetical protein |
| *FLPKDCKG_05815* | | + | 6305976 | 6306806 | 2.34 | 20.92 | 3.16 | 0.86 | Hypothetical protein |
| *FLPKDCKG_05816* | | + | 6306861 | 6307304 | 0.21 | 4.01 | 0.35 | 0.09 | Hypothetical protein |
| *FLPKDCKG_05817* | *cynR-18* | - | 6307374 | 6308342 | 2.47 | 8.12 | 3.53 | 0.59 | HTH-type transcriptional regulator CynR |
| *FLPKDCKG_05818* | *dag* | + | 6308493 | 6309959 | 0.89 | 1.15 | 1.80 | 0.56 | N-acyl-D-glutamate deacylase |
| *FLPKDCKG_05819* | | + | 6309993 | 6310952 | 0.75 | 2.10 | 0.80 | 0.12 | Hypothetical protein |
| *FLPKDCKG_05820* | | + | 6311079 | 6311633 | 50.63 | 30.98 | 20.89 | 4.51 | Hypothetical protein |
| *FLPKDCKG_05821* | *crcB-2* | + | 6311638 | 6312030 | 8.82 | 9.92 | 8.22 | 2.51 | Putative fluoride ion transporter CrcB |
| *FLPKDCKG_05822* | *pds* | - | 6312109 | 6313434 | 2.01 | 6.33 | 4.40 | 1.63 | 15-cis-phytoene desaturase |
| *FLPKDCKG_05823* | *crtB-1* | - | 6313431 | 6314294 | 4.93 | 10.47 | 4.67 | 2.91 | All-trans-phytoene synthase |
| *FLPKDCKG_05824* | *crtB-2* | - | 6314314 | 6315144 | 14.43 | 29.21 | 9.53 | 5.10 | All-trans-phytoene synthase |
| *FLPKDCKG_05825* | *dadX-2* | + | 6315309 | 6316439 | 17.47 | 15.24 | 12.09 | 4.14 | Alanine racemase, catabolic |
| *FLPKDCKG_05826* | *fabG-22* | + | 6316703 | 6317422 | 73.05 | 49.54 | 37.07 | 24.74 | 3-oxoacyl-[acyl-carrier-protein] reductase FabG |
| *FLPKDCKG_05827* | *hpcH* | - | 6317534 | 6318334 | 5.89 | 11.70 | 11.06 | 2.02 | 4-hydroxy-2-oxo-heptane-1,7-dioate aldolase |
| *FLPKDCKG_05828* | *hpcG-2* | - | 6318349 | 6319152 | 6.54 | 6.68 | 9.07 | 1.31 | 2-oxo-hept-4-ene-1,7-dioate hydratase |
| *FLPKDCKG_05829* | *hpcE-1* | - | 6319178 | 6319927 | 1.03 | 0.40 | 0.00 | 1.01 | Homoprotocatechuate catabolism bifunctional isomerase/decarboxylase |
| *FLPKDCKG_05830* | *hpcE-2* | - | 6319949 | 6320977 | 0.98 | 0.04 | 0.59 | 0.42 | Homoprotocatechuate catabolism bifunctional isomerase/decarboxylase |
| *FLPKDCKG_05831* | *betB* | + | 6321188 | 6322654 | 0.76 | 0.88 | 0.00 | 0.38 | NAD/NADP-dependent betaine aldehyde dehydrogenase |
| *FLPKDCKG_05832* | *hpcB* | + | 6322694 | 6323545 | 1.41 | 2.22 | 6.79 | 0.71 | 3,4-dihydroxyphenylacetate 2,3-dioxygenase |
| *FLPKDCKG_05833* | *ohrR-2* | - | 6323547 | 6323993 | 51.60 | 127.08 | 78.13 | 21.96 | Organic hydroperoxide resistance transcriptional regulator |
| *FLPKDCKG_05834* | | - | 6324106 | 6324750 | 58.08 | 48.43 | 51.88 | 27.75 | Hypothetical protein |
| *FLPKDCKG_05835* | *radA* | + | 6325182 | 6326546 | 2.79 | 6.76 | 5.29 | 3.34 | DNA repair protein RadA |
| *FLPKDCKG_05836* | | + | 6326559 | 6327332 | 0.61 | 4.23 | 0.53 | 0.00 | Hypothetical protein |
| *FLPKDCKG_05837* | | + | 6327534 | 6327773 | 124.91 | 1946.91 | 1822.60 | 361.56 | Hypothetical protein |
| *FLPKDCKG_05838* | | - | 6327976 | 6328932 | 34.43 | 67.14 | 34.65 | 13.09 | 2-ketogluconate reductase |
| *FLPKDCKG_05839* | | - | 6329043 | 6330014 | 2.45 | 8.41 | 7.53 | 4.70 | Hypothetical protein |
| *FLPKDCKG_05840* | | - | 6330088 | 6331089 | 1.19 | 9.45 | 5.66 | 2.97 | Hypothetical protein |
| *FLPKDCKG_05841* | | + | 6331319 | 6332248 | 1.63 | 16.47 | 6.49 | 4.39 | 5-dehydro-4-deoxyglucarate dehydratase |
| *FLPKDCKG_05842* | *pdhR* | + | 6332309 | 6333082 | 6.27 | 13.05 | 7.80 | 3.49 | Pyruvate dehydrogenase complex repressor |
| *FLPKDCKG_05843* | *yjmC-2* | + | 6333137 | 6334243 | 2.11 | 6.35 | 6.27 | 1.33 | putative oxidoreductase YjmC |
| *FLPKDCKG_05844* | | + | 6334359 | 6335354 | 2.05 | 8.63 | 8.45 | 1.36 | Hypothetical protein |
| *FLPKDCKG_05845* | *garL* | + | 6335424 | 6336191 | 5.55 | 14.59 | 8.20 | 3.52 | 5-keto-4-deoxy-D-glucarate aldolase |
| *FLPKDCKG_05846* | *lysA-2* | - | 6336240 | 6337469 | 0.66 | 10.48 | 4.70 | 1.83 | Diaminopimelate decarboxylase |
| *FLPKDCKG_05847* | *cmpR-18* | + | 6337588 | 6338502 | 2.99 | 10.43 | 5.80 | 0.61 | HTH-type transcriptional activator CmpR |
| *FLPKDCKG_05848* | *rpsT* | - | 6338704 | 6339003 | 52.97 | 109.83 | 118.00 | 38.50 | 30S ribosomal protein S20 |
| *FLPKDCKG_05849* | | - | 6339346 | 6340317 | 41.68 | 4.51 | 4.23 | 2.56 | Hypothetical protein |
| *FLPKDCKG_05850* | *murJ-2* | - | 6340517 | 6342076 | 0.50 | 1.92 | 1.72 | 0.45 | Lipid II flippase MurJ |
| *FLPKDCKG_05851* | | - | 6342254 | 6343012 | 16.80 | 19.16 | 33.68 | 12.16 | putative oxidoreductase |
| *FLPKDCKG_05852* | *adk* | - | 6343145 | 6343801 | 22.59 | 46.66 | 43.92 | 12.07 | Adenylate kinase |
| *FLPKDCKG_05853* | *kdsB-2* | - | 6343894 | 6344658 | 5.86 | 20.68 | 18.30 | 7.04 | 3-deoxy-manno-octulosonate cytidylyltransferase |
| *FLPKDCKG_05854* | | - | 6344671 | 6344853 | 85.17 | 159.29 | 102.78 | 34.64 | Hypothetical protein |
| *FLPKDCKG_05855* | *igaA* | + | 6345049 | 6347286 | 1.43 | 3.47 | 1.53 | 0.74 | Intracellular growth attenuator protein igaA |
| *FLPKDCKG_05856* | *lpxK* | - | 6347283 | 6348332 | 0.98 | 4.43 | 3.17 | 0.97 | Tetraacyldisaccharide 4'-kinase |
| *FLPKDCKG_05857* | *exbD-4* | - | 6348329 | 6348742 | 8.68 | 43.04 | 24.68 | 4.72 | Biopolymer transport protein ExbD |
| *FLPKDCKG_05858* | *exbB-4* | - | 6348739 | 6349356 | 12.86 | 48.09 | 40.61 | 14.55 | Biopolymer transport protein ExbB |
| *FLPKDCKG_05859* | *xseA* | + | 6349712 | 6351076 | 2.06 | 5.48 | 5.68 | 1.43 | Exodeoxyribonuclease 7 large subunit |
| *FLPKDCKG_05860* | | + | 6351144 | 6354419 | 0.88 | 1.41 | 1.83 | 0.30 | Hypothetical protein |
| *FLPKDCKG_05861* | *sodB-3* | + | 6354549 | 6355127 | 286.18 | 549.79 | 332.28 | 76.88 | Superoxide dismutase [Fe] |

| Locus name | Gene name | Strand | Gene start (bp) | Gene end (bp) | Average TPM Inoculum | Average TPM $t_6$ | Average TPM $t_{24}$ | Average TPM $t_{168}$ | Annotation description |
|---|---|---|---|---|---|---|---|---|---|
| *FLPKDCKG_05862* | *trg-5* | + | 6355414 | 6357141 | 0.87 | 0.87 | 0.07 | 0.00 | Methyl-accepting chemotaxis protein III |
| *FLPKDCKG_05863* | *clcA-2* | + | 6357238 | 6358647 | 1.74 | 5.99 | 7.64 | 4.20 | H(+)/Cl(-) exchange transporter ClcA |
| *FLPKDCKG_05864* | *pcaF-4* | + | 6358768 | 6359973 | 12.71 | 24.35 | 13.32 | 5.66 | Beta-ketoadipyl-CoA thiolase |
| *FLPKDCKG_05865* | | - | 6360090 | 6360608 | 82.37 | 161.91 | 93.47 | 31.42 | Hypothetical protein |
| *FLPKDCKG_05866* | *cspE* | - | 6360701 | 6360946 | 57.51 | 104.58 | 67.51 | 37.50 | Cold shock-like protein CspE |
| *FLPKDCKG_05867* | *clpS* | + | 6361173 | 6361487 | 20.24 | 71.13 | 55.15 | 9.41 | ATP-dependent Clp protease adapter protein ClpS |
| *FLPKDCKG_05868* | *yfhM* | + | 6361640 | 6366865 | 9.58 | 8.66 | 10.95 | 2.90 | putative lipoprotein YfhM |
| *FLPKDCKG_05869* | *pbpC-1* | + | 6366817 | 6368694 | 17.10 | 76.85 | 39.29 | 18.51 | Penicillin-binding protein 1C |
| *FLPKDCKG_05870* | *pbpC-2* | + | 6368658 | 6369083 | 10.92 | 60.62 | 48.14 | 14.42 | Penicillin-binding protein 1C |
| *FLPKDCKG_05871* | *clpA* | + | 6369139 | 6371448 | 151.29 | 181.03 | 110.38 | 44.94 | ATP-dependent Clp protease ATP-binding subunit ClpA |
| *FLPKDCKG_05872* | *pqiA* | + | 6371453 | 6372769 | 7.48 | 19.61 | 14.86 | 3.87 | Paraquat-inducible protein A |
| *FLPKDCKG_05873* | *pqiB* | + | 6372825 | 6374489 | 9.29 | 21.35 | 14.67 | 6.06 | Paraquat-inducible protein B |
| *FLPKDCKG_05874* | | + | 6374494 | 6375186 | 3.67 | 13.41 | 8.65 | 5.59 | Hypothetical protein |
| *FLPKDCKG_05875* | *putA* | + | 6375355 | 6379176 | 5.31 | 1.93 | 2.05 | 1.79 | Bifunctional protein PutA |
| *FLPKDCKG_05876* | *braC-11* | + | 6379364 | 6380509 | 38.15 | 25.15 | 23.66 | 15.54 | Leucine-, isoleucine-, valine-, threonine-, and alanine-binding protein |
| *FLPKDCKG_05877* | *livH-27* | + | 6380672 | 6381601 | 1.25 | 4.16 | 1.84 | 1.21 | High-affinity branched-chain amino acid transport system permease protein LivH |
| *FLPKDCKG_05878* | *livH-28* | + | 6381598 | 6382686 | 1.96 | 0.38 | 1.70 | 1.17 | High-affinity branched-chain amino acid transport system permease protein LivH |
| *FLPKDCKG_05879* | *lptB-12* | + | 6382683 | 6383483 | 2.64 | 1.77 | 1.69 | 0.63 | Lipopolysaccharide export system ATP-binding protein LptB |
| *FLPKDCKG_05880* | *livF-26* | + | 6383649 | 6384215 | 0.99 | 0.67 | 0.10 | 0.92 | High-affinity branched-chain amino acid transport ATP-binding protein LivF |
| *FLPKDCKG_05881* | *yhjX-4* | - | 6384288 | 6385571 | 0.36 | 9.65 | 0.67 | 2.27 | putative MFS-type transporter YhjX |
| *FLPKDCKG_05882* | *kstR2-5* | + | 6385704 | 6386333 | 4.46 | 27.56 | 9.06 | 4.38 | HTH-type transcriptional repressor KstR2 |
| *FLPKDCKG_05883* | | + | 6386510 | 6386731 | 216.98 | 255.18 | 204.13 | 142.70 | Hypothetical protein |
| *FLPKDCKG_05884* | *gcvA-15* | - | 6386797 | 6387681 | 1.84 | 5.62 | 4.56 | 2.84 | Glycine cleavage system transcriptional activator |
| *FLPKDCKG_05885* | *yfcG-7* | + | 6387859 | 6388482 | 0.30 | 1.76 | 0.00 | 1.04 | Disulfide-bond oxidoreductase YfcG |
| *FLPKDCKG_05886* | *luxB* | + | 6388508 | 6389554 | 1.09 | 2.70 | 3.79 | 1.95 | Alkanal monooxygenase beta chain |
| *FLPKDCKG_05887* | *mntH-2* | - | 6389558 | 6390847 | 2.51 | 16.96 | 21.16 | 9.49 | Divalent metal cation transporter MntH |
| *FLPKDCKG_05888* | *osmC* | - | 6390972 | 6391397 | 36.18 | 157.92 | 121.35 | 32.84 | Peroxiredoxin OsmC |
| *FLPKDCKG_05889* | *lhpI* | + | 6391647 | 6392591 | 2.69 | 7.86 | 3.74 | 2.34 | Delta(1)-pyrroline-2-carboxylate reductase |
| *FLPKDCKG_05890* | | + | 6392629 | 6393801 | 4.76 | 6.82 | 5.90 | 1.51 | Leu/Ile/Val-binding protein |
| *FLPKDCKG_05891* | | + | 6393979 | 6394890 | 2.16 | 69.66 | 47.26 | 4.68 | putative amino-acid ABC transporter-binding protein |
| *FLPKDCKG_05892* | *yecS-8* | + | 6394900 | 6395553 | 0.46 | 25.48 | 19.70 | 1.28 | L-cystine transport system permease protein YecS |
| *FLPKDCKG_05893* | *yecS-9* | + | 6395550 | 6396221 | 0.58 | 45.03 | 26.20 | 1.17 | L-cystine transport system permease protein YecS |
| *FLPKDCKG_05894* | *glnQ-14* | + | 6396218 | 6396988 | 0.79 | 46.96 | 31.38 | 1.03 | Glutamine transport ATP-binding protein GlnQ |
| *FLPKDCKG_05895* | *namA-2* | + | 6397118 | 6398230 | 6.45 | 19.03 | 12.95 | 4.23 | NADPH dehydrogenase |
| *FLPKDCKG_05896* | | - | 6398299 | 6398967 | 4.46 | 5.11 | 5.76 | 0.74 | Hypothetical protein |
| *FLPKDCKG_05897* | | - | 6399045 | 6399581 | 18.74 | 7.42 | 7.76 | 6.11 | Hypothetical protein |
| *FLPKDCKG_05898* | *yabI-2* | - | 6399593 | 6400231 | 17.50 | 9.06 | 8.21 | 1.94 | Inner membrane protein YabI |
| *FLPKDCKG_05899* | | + | 6400337 | 6401377 | 0.67 | 1.98 | 0.17 | 0.30 | Hypothetical protein |
| *FLPKDCKG_05900* | *kdgR-9* | + | 6401582 | 6402349 | 6.59 | 1.34 | 1.97 | 1.64 | Transcriptional regulator KdgR |
| *FLPKDCKG_05901* | *pepE-2* | + | 6402440 | 6403144 | 4.65 | 1.47 | 1.73 | 0.95 | Peptidase E |
| *FLPKDCKG_05902* | | + | 6403141 | 6404070 | 0.67 | 0.82 | 1.17 | 1.44 | Hypothetical protein |
| *FLPKDCKG_05903* | | + | 6404067 | 6404756 | 0.58 | 0.78 | 0.29 | 0.64 | Hypothetical protein |
| *FLPKDCKG_05904* | | + | 6404770 | 6405759 | 0.62 | 0.44 | 0.00 | 0.48 | Hypothetical protein |
| *FLPKDCKG_05905* | *coaBC* | - | 6405867 | 6407063 | 28.73 | 21.99 | 15.61 | 7.75 | Coenzyme A biosynthesis bifunctional protein CoaBC |
| *FLPKDCKG_05906* | *lspA* | - | 6407221 | 6407745 | 1.35 | 4.96 | 5.99 | 0.17 | Lipoprotein signal peptidase |
| *FLPKDCKG_05907* | *ileS* | - | 6407748 | 6410609 | 7.73 | 19.99 | 15.21 | 5.42 | Isoleucine--tRNA ligase |
| *FLPKDCKG_05908* | *ribF* | - | 6410661 | 6411455 | 14.99 | 38.45 | 29.83 | 9.03 | Riboflavin biosynthesis protein RibF |
| *FLPKDCKG_05909* | *purN* | + | 6411690 | 6412292 | 17.78 | 57.05 | 32.49 | 9.96 | Phosphoribosylglycinamide formyltransferase |
| *FLPKDCKG_05910* | *rsmB-2* | + | 6412378 | 6414075 | 3.11 | 11.43 | 6.39 | 1.02 | Ribosomal RNA small subunit methyltransferase B |
| *FLPKDCKG_05911* | *rsmB-3* | + | 6414363 | 6414725 | 10.65 | 49.14 | 30.46 | 6.20 | Ribosomal RNA small subunit methyltransferase B |
| *FLPKDCKG_05912* | | - | 6414902 | 6415297 | 7.53 | 13.01 | 7.34 | 4.78 | Hypothetical protein |
| *FLPKDCKG_05913* | | + | 6415622 | 6416830 | 193.92 | 330.82 | 290.20 | 93.14 | Hypothetical protein |
| *FLPKDCKG_05914* | *uvrD* | + | 6416827 | 6419118 | 7.67 | 24.87 | 19.92 | 5.37 | DNA helicase II |
| *FLPKDCKG_05915* | *tyrA-2* | + | 6419124 | 6419429 | 3.89 | 7.97 | 7.23 | 1.37 | T-protein |
| *FLPKDCKG_05916* | *ccp-2* | - | 6419548 | 6420870 | 0.03 | 1.52 | 0.58 | 0.22 | Cytochrome c551 peroxidase |
| *FLPKDCKG_05917* | | - | 6421060 | 6421722 | 0.24 | 2.16 | 1.00 | 0.10 | Hypothetical protein |
| *FLPKDCKG_05918* | *fpvA-7* | - | 6421850 | 6424471 | 0.27 | 0.79 | 1.70 | 0.12 | Ferripyoverdine receptor |
| *FLPKDCKG_05919* | *fpvA-8* | - | 6424574 | 6427210 | 1.29 | 5.43 | 1.73 | 0.71 | Ferripyoverdine receptor |
| *FLPKDCKG_05920* | *fpvA-9* | - | 6427402 | 6430032 | 0.18 | 0.51 | 0.25 | 0.00 | Ferripyoverdine receptor |
| *FLPKDCKG_05921* | | - | 6430206 | 6431138 | 1.19 | 2.41 | 0.06 | 1.11 | Hypothetical protein |
| *FLPKDCKG_05922* | *ylaC* | - | 6431135 | 6431623 | 2.12 | 1.74 | 0.00 | 0.00 | RNA polymerase sigma factor YlaC |
| *FLPKDCKG_05923* | *yheS-2* | - | 6431747 | 6433681 | 1.41 | 3.96 | 4.69 | 0.44 | putative ABC transporter ATP-binding protein YheS |
| *FLPKDCKG_05924* | *prmB* | - | 6433678 | 6434574 | 2.56 | 5.58 | 5.23 | 3.32 | 50S ribosomal protein L3 glutamine methyltransferase |
| *FLPKDCKG_05925* | *dapE* | - | 6434585 | 6435802 | 6.81 | 9.29 | 7.17 | 4.26 | Succinyl-diaminopimelate desuccinylase |
| *FLPKDCKG_05926* | *dapD-2* | - | 6435802 | 6436623 | 12.98 | 15.03 | 15.56 | 6.37 | 2,3,4,5-tetrahydropyridine-2,6-dicarboxylate N-succinyltransferase |
| *FLPKDCKG_05927* | *dapL* | - | 6436652 | 6437848 | 10.61 | 25.93 | 11.98 | 4.19 | LL-diaminopimelate aminotransferase |
| *FLPKDCKG_05928* | | + | 6437896 | 6437982 | 32.24 | 65.87 | 26.45 | 6.20 | tRNA |
| *FLPKDCKG_05929* | | + | 6438223 | 6438498 | 47.53 | 128.50 | 63.60 | 38.38 | Hypothetical protein |
| *FLPKDCKG_05930* | | - | 6438561 | 6439073 | 5.99 | 31.57 | 26.96 | 5.62 | Hypothetical protein |

| Locus name | Gene name | Strand | Gene start (bp) | Gene end (bp) | Average TPM Inoculum | Average TPM $t_6$ | Average TPM $t_{24}$ | Average TPM $t_{168}$ | Annotation description |
|---|---|---|---|---|---|---|---|---|---|
| *FLPKDCKG_05931* | | - | 6439084 | 6439443 | 15.37 | 12.12 | 13.99 | 6.80 | Hypothetical protein |
| *FLPKDCKG_05932* | | - | 6439548 | 6439937 | 4.91 | 9.03 | 3.11 | 1.47 | Hypothetical protein |
| *FLPKDCKG_05933* | | + | 6440884 | 6441615 | 0.61 | 0.50 | 0.39 | 0.08 | Hypothetical protein |
| *FLPKDCKG_05934* | *glpK-4* | + | 6441630 | 6443168 | 0.31 | 0.45 | 0.71 | 0.08 | Glycerol kinase |
| *FLPKDCKG_05935* | *glpD2-4* | + | 6443165 | 6444844 | 0.42 | 0.73 | 0.25 | 0.24 | Glycerol-3-phosphate dehydrogenase 2 |
| *FLPKDCKG_05936* | | + | 6444841 | 6445275 | 0.60 | 1.21 | 1.62 | 0.32 | Hypothetical protein |
| *FLPKDCKG_05937* | | + | 6445254 | 6446396 | 0.87 | 0.84 | 0.19 | 0.12 | putative FAD-linked oxidoreductase |
| *FLPKDCKG_05938* | *glpP-2* | + | 6446398 | 6446979 | 0.99 | 3.03 | 3.32 | 1.13 | Glycerol uptake operon antiterminator regulatory protein |
| *FLPKDCKG_05939* | *aqpZ* | + | 6447122 | 6447832 | 32.81 | 12.30 | 4.24 | 2.34 | Aquaporin Z |
| *FLPKDCKG_05940* | | - | 6447876 | 6449093 | 4.21 | 6.42 | 4.26 | 1.16 | Hypothetical protein |
| *FLPKDCKG_05941* | | - | 6449117 | 6450076 | 10.42 | 10.51 | 7.17 | 2.17 | Hypothetical protein |
| *FLPKDCKG_05942* | *kdsA-2* | + | 6450313 | 6451170 | 5.63 | 6.03 | 7.19 | 2.67 | 2-dehydro-3-deoxyphosphooctonate aldolase |
| *FLPKDCKG_05943* | | + | 6451237 | 6451545 | 4.57 | 8.00 | 6.31 | 3.74 | Hypothetical protein |
| *FLPKDCKG_05944* | | - | 6451569 | 6451871 | 7.10 | 6.94 | 2.87 | 1.24 | Hypothetical protein |
| *FLPKDCKG_05945* | | - | 6451942 | 6452469 | 6.43 | 5.79 | 4.65 | 2.87 | Hypothetical protein |
| *FLPKDCKG_05946* | *tetC* | - | 6452593 | 6453174 | 1.49 | 0.89 | 3.00 | 0.98 | Transposon Tn10 TetC protein |
| *FLPKDCKG_05947* | | + | 6453309 | 6454499 | 0.20 | 0.16 | 0.00 | 0.14 | Hypothetical protein |
| *FLPKDCKG_05948* | *rutC-3* | + | 6454567 | 6454965 | 5.49 | 2.72 | 1.96 | 0.72 | Putative aminoacrylate peracid reductase RutC |
| *FLPKDCKG_05949* | *dmlR-39* | - | 6455148 | 6456065 | 4.76 | 8.06 | 5.45 | 2.28 | HTH-type transcriptional regulator DmlR |

**Supplementary Table S4. Average gene expression in bacterial endophytes. (C)** Gene expression in *Pandoraea* sp. Col-0-28.

Gene expression was calculated as Transcripts per million (TPM) using StringTie in the inoculum, 6 ($t_6$), 24 ($t_{24}$) and 168 ($t_{168}$) hours post-inoculation. Each treatment had four biological replicates. When available, gene name for each locus is shown. DNA template strand, coordinates for the start and end of each gene, and a brief annotation are also shown.

| Locus name | Gene name | Strand | Gene start (bp) | Gene end (bp) | Average TPM Inoculum | Average TPM $t_6$ | Average TPM $t_{24}$ | Average TPM $t_{168}$ | Annotation description |
|---|---|---|---|---|---|---|---|---|---|
| *FEPFGCPE_00001* | *amnD-1* | - | 409 | 843 | 2.49 | 13.43 | 10.79 | 3.70 | 2-aminomuconate deaminase |
| *FEPFGCPE_00002* | *amnC* | - | 932 | 2404 | 1.18 | 6.73 | 3.39 | 1.23 | 2-aminomuconic 6-semialdehyde dehydrogenase |
| *FEPFGCPE_00003* | | - | 2597 | 3163 | 12.67 | 43.08 | 50.62 | 9.76 | Hypothetical protein |
| *FEPFGCPE_00004* | *degA* | + | 3478 | 4521 | 2.13 | 9.89 | 5.09 | 2.41 | HTH-type transcriptional regulator DegA |
| *FEPFGCPE_00005* | *phnS-1* | + | 4708 | 5721 | 2.00 | 9.16 | 8.82 | 2.59 | Putative 2-aminoethylphosphonate-binding periplasmic protein |
| *FEPFGCPE_00006* | | + | 5777 | 6073 | 1.36 | 2.54 | 5.99 | 2.67 | Hypothetical protein |
| *FEPFGCPE_00007* | *phnU-1* | + | 6067 | 6600 | 0.31 | 0.29 | 1.86 | 0.90 | Putative 2-aminoethylphosphonate transport system permease protein PhnU |
| *FEPFGCPE_00008* | *phnV-1* | + | 6606 | 7433 | 0.74 | 3.85 | 0.69 | 1.23 | Putative 2-aminoethylphosphonate transport system permease protein PhnV |
| *FEPFGCPE_00009* | *potA-1* | + | 7436 | 8479 | 1.32 | 1.29 | 2.18 | 0.32 | Spermidine/putrescine import ATP-binding protein PotA |
| *FEPFGCPE_00010* | *cpdA-1* | + | 8487 | 9314 | 0.91 | 5.31 | 3.27 | 1.17 | 3',5'-cyclic adenosine monophosphate phosphodiesterase CpdA |
| *FEPFGCPE_00011* | *hmp-1* | - | 9421 | 10623 | 5.22 | 59.70 | 47.02 | 48.29 | Flavohemoprotein |
| *FEPFGCPE_00012* | | - | 10947 | 12518 | 0.55 | 0.67 | 1.18 | 0.26 | Hypothetical protein |
| *FEPFGCPE_00013* | | - | 12804 | 13880 | 1.70 | 3.86 | 3.99 | 0.80 | Hypothetical protein |
| *FEPFGCPE_00014* | | + | 14042 | 14182 | 4.66 | 11.34 | 4.22 | 4.60 | Hypothetical protein |
| *FEPFGCPE_00015* | | + | 14179 | 14850 | 8.91 | 32.29 | 22.75 | 8.97 | Hypothetical protein |
| *FEPFGCPE_00016* | | - | 15044 | 15119 | 83.60 | 24.58 | 36.95 | 17.14 | tRNA |
| *FEPFGCPE_00017* | *fbp-1* | - | 15323 | 16345 | 14.20 | 51.91 | 43.06 | 28.63 | Fructose-1,6-bisphosphatase class 1 |
| *FEPFGCPE_00018* | *pyrB-1* | - | 16497 | 17780 | 15.78 | 37.25 | 38.56 | 22.40 | Aspartate carbamoyltransferase catalytic subunit |
| *FEPFGCPE_00019* | *gltC-1* | + | 17987 | 18835 | 1.77 | 6.71 | 3.12 | 1.95 | HTH-type transcriptional regulator GltC |
| *FEPFGCPE_00020* | | - | 18679 | 19299 | 7.94 | 36.37 | 23.39 | 6.38 | Hypothetical protein |
| *FEPFGCPE_00021* | *ydjE-1* | + | 19327 | 20766 | 6.94 | 15.08 | 11.21 | 4.60 | Inner membrane metabolite transport protein YdjE |
| *FEPFGCPE_00022* | *tehB-1* | + | 20824 | 21417 | 4.09 | 9.66 | 7.64 | 3.34 | Tellurite methyltransferase |
| *FEPFGCPE_00023* | *metR-1* | - | 21452 | 22375 | 4.76 | 9.33 | 2.56 | 7.02 | HTH-type transcriptional regulator MetR |
| *FEPFGCPE_00024* | *metE-1* | + | 22511 | 24817 | 7.96 | 62.87 | 94.17 | 103.91 | 5-methyltetrahydropteroyltriglutamate--homocysteine methyltransferase |
| *FEPFGCPE_00025* | *pepN* | - | 24923 | 27634 | 4.28 | 33.64 | 30.63 | 12.83 | Aminopeptidase N |
| *FEPFGCPE_00026* | | + | 27835 | 28443 | 41.53 | 72.79 | 75.56 | 28.09 | Hypothetical protein |
| *FEPFGCPE_00027* | | - | 28613 | 29245 | 10.52 | 17.94 | 21.99 | 14.68 | Hypothetical protein |
| *FEPFGCPE_00028* | *guaD* | - | 29751 | 31088 | 4.67 | 8.05 | 5.23 | 3.85 | Guanine deaminase |
| *FEPFGCPE_00029* | *adeP-1* | - | 31108 | 32532 | 3.38 | 5.48 | 5.56 | 1.24 | Adenine permease AdeP |
| *FEPFGCPE_00030* | | - | 32809 | 33849 | 3.75 | 19.23 | 16.83 | 4.79 | Adenine deaminase |
| *FEPFGCPE_00031* | | - | 33936 | 35015 | 5.39 | 12.15 | 10.90 | 2.37 | Hypothetical protein |
| *FEPFGCPE_00032* | *xdhA-1* | - | 35015 | 37363 | 11.45 | 18.42 | 13.74 | 4.66 | Putative xanthine dehydrogenase molybdenum-binding subunit XdhA |
| *FEPFGCPE_00033* | *cutS-1* | - | 37376 | 38848 | 15.38 | 28.84 | 27.86 | 6.74 | Carbon monoxide dehydrogenase small chain |
| *FEPFGCPE_00034* | *nodD2* | + | 39039 | 40004 | 14.83 | 36.48 | 21.14 | 7.29 | Nodulation protein D 2 |
| *FEPFGCPE_00035* | *dsbB* | - | 40100 | 40612 | 6.56 | 27.63 | 19.36 | 6.62 | Disulfide bond formation protein B |
| *FEPFGCPE_00036* | *atzE* | - | 40834 | 42219 | 2.34 | 12.06 | 4.16 | 1.35 | Biuret hydrolase |
| *FEPFGCPE_00037* | *rspR-1* | - | 42222 | 42935 | 1.47 | 4.83 | 1.54 | 1.18 | HTH-type transcriptional repressor RspR |
| *FEPFGCPE_00038* | | - | 43007 | 43693 | 7.24 | 42.66 | 25.35 | 8.56 | Hypothetical protein |
| *FEPFGCPE_00039* | *ybaL* | + | 44154 | 45935 | 3.52 | 9.76 | 9.67 | 4.22 | Inner membrane protein YbaL |
| *FEPFGCPE_00040* | | - | 46042 | 46608 | 3.70 | 6.39 | 5.65 | 2.20 | Hypothetical protein |
| *FEPFGCPE_00041* | *ribB* | - | 46797 | 47540 | 2.44 | 12.01 | 9.85 | 4.27 | 3,4-dihydroxy-2-butanone 4-phosphate synthase |
| *FEPFGCPE_00042* | *copZ* | - | 48046 | 48246 | 9.05 | 38.26 | 38.63 | 11.45 | Copper chaperone CopZ |
| *FEPFGCPE_00043* | *copA-1* | + | 48518 | 51064 | 9.27 | 17.72 | 18.88 | 6.29 | Copper-exporting P-type ATPase A |
| *FEPFGCPE_00044* | | + | 51239 | 52057 | 48.41 | 110.15 | 104.59 | 37.38 | Hypothetical protein |
| *FEPFGCPE_00045* | *hmrR* | + | 52054 | 52617 | 32.90 | 90.19 | 92.11 | 36.03 | HTH-type transcriptional regulator HmrR |
| *FEPFGCPE_00046* | | + | 52928 | 53068 | 45.26 | 89.37 | 95.91 | 63.14 | Hypothetical protein |
| *FEPFGCPE_00047* | | - | 53312 | 53776 | 45.30 | 61.19 | 57.99 | 15.07 | Hypothetical protein |
| *FEPFGCPE_00048* | | + | 54177 | 54653 | 12.33 | 43.50 | 27.78 | 17.46 | Hypothetical protein |
| *FEPFGCPE_00049* | *mntH-1* | - | 54712 | 56022 | 2.50 | 15.62 | 9.42 | 3.92 | Divalent metal cation transporter MntH |
| *FEPFGCPE_00050* | | + | 56042 | 57040 | 2.38 | 7.31 | 6.96 | 2.55 | Hypothetical protein |
| *FEPFGCPE_00051* | | + | 57530 | 58507 | 0.81 | 1.52 | 2.45 | 0.72 | Hypothetical protein |
| *FEPFGCPE_00052* | *dnaE2* | + | 58529 | 61837 | 1.27 | 2.66 | 2.71 | 0.25 | Error-prone DNA polymerase |
| *FEPFGCPE_00053* | *higB-1* | + | 61853 | 62068 | 0.51 | 0.00 | 3.40 | 0.21 | Toxin HigB-1 |
| *FEPFGCPE_00054* | *higA* | + | 62132 | 62443 | 3.68 | 8.67 | 4.80 | 1.27 | Antitoxin HigA |
| *FEPFGCPE_00055* | *uacT-1* | - | 62505 | 63875 | 5.57 | 7.28 | 2.71 | 2.58 | Uric acid transporter UacT |
| *FEPFGCPE_00056* | | - | 64087 | 64206 | 1.00 | 0.00 | 0.00 | 2.37 | Hypothetical protein |
| *FEPFGCPE_00057* | | - | 64307 | 65815 | 2.50 | 5.13 | 2.88 | 0.56 | Hypothetical protein |
| *FEPFGCPE_00058* | | - | 65834 | 67651 | 3.06 | 14.89 | 9.51 | 1.55 | putative signaling protein |
| *FEPFGCPE_00059* | | - | 67910 | 68662 | 10.49 | 2.18 | 5.15 | 2.05 | Aspartate racemase |
| *FEPFGCPE_00060* | *gltI-1* | - | 68740 | 69642 | 62.56 | 20.09 | 22.76 | 11.27 | Glutamate/aspartate import solute-binding protein |
| *FEPFGCPE_00061* | *dadA1-1* | - | 69834 | 71090 | 30.90 | 11.65 | 11.35 | 5.49 | D-amino acid dehydrogenase 1 |
| *FEPFGCPE_00062* | *cmpR-1* | + | 71311 | 72270 | 4.42 | 26.21 | 11.29 | 2.17 | HTH-type transcriptional activator CmpR |
| *FEPFGCPE_00063* | | - | 72286 | 72897 | 9.89 | 31.19 | 23.74 | 6.38 | Hypothetical protein |
| *FEPFGCPE_00064* | *dbpA* | - | 72973 | 74385 | 1.92 | 14.45 | 10.02 | 3.09 | ATP-dependent RNA helicase DbpA |
| *FEPFGCPE_00065* | *gdhB* | + | 74527 | 79587 | 6.96 | 35.40 | 18.97 | 8.07 | NAD-specific glutamate dehydrogenase |
| *FEPFGCPE_00066* | | + | 79824 | 81044 | 1.99 | 10.22 | 5.50 | 2.24 | Hypothetical protein |
| *FEPFGCPE_00067* | *gabR-1* | - | 81060 | 82697 | 5.56 | 13.50 | 17.51 | 3.22 | HTH-type transcriptional regulatory protein GabR |

| Locus name | Gene name | Strand | Gene start (bp) | Gene end (bp) | Average TPM Inoculum | Average TPM $t_6$ | Average TPM $t_{24}$ | Average TPM $t_{168}$ | Annotation description |
|---|---|---|---|---|---|---|---|---|---|
| *FEPFGCPE_00068* | | + | 82794 | 83519 | 4.88 | 13.89 | 11.64 | 5.76 | Hypothetical protein |
| *FEPFGCPE_00069* | *adeP-2* | - | 83592 | 84884 | 2.80 | 10.16 | 8.30 | 4.90 | Adenine permease AdeP |
| *FEPFGCPE_00070* | | - | 84976 | 86016 | 6.05 | 10.19 | 2.86 | 0.49 | Outer membrane porin protein 32 |
| *FEPFGCPE_00071* | | - | 87137 | 87976 | 1.86 | 4.32 | 7.42 | 2.02 | Hypothetical protein |
| *FEPFGCPE_00072* | *ethR* | + | 88134 | 88784 | 1.77 | 7.46 | 7.72 | 3.92 | HTH-type transcriptional regulator EthR |
| *FEPFGCPE_00073* | *icd-1* | - | 88906 | 91146 | 4.95 | 34.37 | 41.85 | 38.84 | Isocitrate dehydrogenase [NADP] |
| *FEPFGCPE_00074* | | - | 91746 | 91820 | 1.98 | 0.00 | 0.14 | 5.66 | tRNA |
| *FEPFGCPE_00075* | *dnaQ* | - | 92078 | 92818 | 2.65 | 25.99 | 16.61 | 8.23 | DNA polymerase III subunit epsilon |
| *FEPFGCPE_00076* | *rnhA* | - | 92931 | 93380 | 3.53 | 20.91 | 22.33 | 1.51 | Ribonuclease HI |
| *FEPFGCPE_00077* | | - | 93392 | 94252 | 5.49 | 23.36 | 21.42 | 3.88 | Hypothetical protein |
| *FEPFGCPE_00078* | *gloB-1* | + | 94267 | 95094 | 7.92 | 29.12 | 21.09 | 10.79 | Hydroxyacylglutathione hydrolase |
| *FEPFGCPE_00079* | *mltD-1* | + | 95420 | 97096 | 10.83 | 86.73 | 51.03 | 30.28 | Membrane-bound lytic murein transglycosylase D |
| *FEPFGCPE_00080* | | - | 97172 | 98449 | 1.74 | 19.31 | 9.74 | 3.90 | Hypothetical protein |
| *FEPFGCPE_00081* | | - | 98436 | 99131 | 1.21 | 5.05 | 0.97 | 0.55 | Hypothetical protein |
| *FEPFGCPE_00082* | *tehB-2* | - | 99138 | 100016 | 0.50 | 2.67 | 1.85 | 0.22 | Tellurite methyltransferase |
| *FEPFGCPE_00083* | | - | 100046 | 101218 | 1.34 | 9.15 | 3.12 | 0.23 | Hypothetical protein |
| *FEPFGCPE_00084* | | - | 101218 | 102408 | 0.44 | 6.04 | 3.98 | 2.10 | Hypothetical protein |
| *FEPFGCPE_00085* | | - | 102509 | 102691 | 1.25 | 11.75 | 6.81 | 2.31 | Hypothetical protein |
| *FEPFGCPE_00086* | *acsA-1* | + | 103447 | 105348 | 0.86 | 24.21 | 14.99 | 15.61 | Acetyl-coenzyme A synthetase |
| *FEPFGCPE_00087* | *gltC-2* | - | 105432 | 106349 | 1.19 | 9.73 | 2.40 | 2.64 | HTH-type transcriptional regulator GltC |
| *FEPFGCPE_00088* | *dapE-1* | + | 106567 | 107964 | 0.72 | 1.42 | 1.76 | 2.18 | Succinyl-diaminopimelate desuccinylase |
| *FEPFGCPE_00089* | *proP-1* | + | 107961 | 109316 | 1.41 | 4.14 | 4.14 | 2.26 | Proline/betaine transporter |
| *FEPFGCPE_00090* | *sdgC-1* | + | 109884 | 112226 | 0.81 | 2.84 | 1.92 | 1.30 | Salicyloyl-CoA 5-hydroxylase |
| *FEPFGCPE_00091* | *actIII* | + | 112236 | 113048 | 2.22 | 18.14 | 8.52 | 2.88 | Putative ketoacyl reductase |
| *FEPFGCPE_00092* | *paaG-1* | + | 113084 | 113947 | 1.74 | 7.94 | 7.16 | 4.26 | 1,2-epoxyphenylacetyl-CoA isomerase |
| *FEPFGCPE_00093* | | + | 113961 | 115145 | 1.51 | 11.59 | 11.25 | 5.52 | putative acyl-CoA dehydrogenase fadE25 |
| *FEPFGCPE_00094* | *bclA-1* | + | 115205 | 116848 | 2.50 | 19.66 | 16.55 | 3.07 | Benzoate--CoA ligase |
| *FEPFGCPE_00095* | | + | 116936 | 117406 | 1.26 | 11.73 | 6.30 | 2.97 | 1,4-dihydroxy-2-naphthoyl-CoA hydrolase |
| *FEPFGCPE_00096* | *ridA-1* | + | 117406 | 117816 | 7.50 | 29.43 | 19.90 | 13.88 | 2-iminobutanoate/2-iminopropanoate deaminase |
| *FEPFGCPE_00097* | | + | 118025 | 118255 | 6.46 | 74.05 | 29.46 | 4.77 | Hypothetical protein |
| *FEPFGCPE_00098* | *mctP-1* | + | 118252 | 119802 | 2.82 | 15.78 | 7.81 | 1.72 | Monocarboxylate transport permease protein |
| *FEPFGCPE_00099* | *allA-1* | - | 120229 | 120762 | 60.73 | 49.01 | 35.67 | 11.82 | Ureidoglycolate lyase |
| *FEPFGCPE_00100* | *alc-1* | - | 120836 | 121858 | 27.00 | 57.96 | 39.74 | 16.73 | allantoicase |
| *FEPFGCPE_00101* | *dctA-1* | - | 121937 | 123268 | 1.36 | 16.09 | 16.71 | 10.55 | Aerobic C4-dicarboxylate transport protein |
| *FEPFGCPE_00102* | *gdhI* | + | 123703 | 124470 | 12.16 | 16.07 | 24.68 | 14.07 | Glucose 1-dehydrogenase 1 |
| *FEPFGCPE_00103* | *treF* | + | 124473 | 126110 | 3.82 | 3.36 | 8.16 | 3.63 | Cytoplasmic trehalase |
| *FEPFGCPE_00104* | *lutR-1* | + | 126255 | 127016 | 8.65 | 33.70 | 27.03 | 14.94 | HTH-type transcriptional regulator LutR |
| *FEPFGCPE_00105* | *carA-1* | + | 127212 | 128357 | 11.10 | 40.49 | 34.07 | 18.43 | Carbamoyl-phosphate synthase small chain |
| *FEPFGCPE_00106* | *leuE* | + | 128397 | 129065 | 8.16 | 14.34 | 23.32 | 8.03 | Leucine efflux protein |
| *FEPFGCPE_00107* | *carB* | + | 129084 | 132326 | 5.88 | 18.92 | 16.38 | 12.19 | Carbamoyl-phosphate synthase large chain |
| *FEPFGCPE_00108* | *greA* | + | 132573 | 133049 | 1.71 | 14.60 | 25.92 | 16.42 | Transcription elongation factor GreA |
| *FEPFGCPE_00109* | | + | 133033 | 133548 | 2.16 | 10.29 | 10.44 | 3.20 | Hypothetical protein |
| *FEPFGCPE_00110* | *yhbY* | - | 133667 | 134167 | 8.93 | 45.41 | 43.74 | 18.23 | RNA-binding protein YhbY |
| *FEPFGCPE_00111* | *rlmE* | + | 134307 | 134957 | 9.06 | 13.86 | 28.89 | 12.14 | Ribosomal RNA large subunit methyltransferase E |
| *FEPFGCPE_00112* | *ftsH* | + | 135077 | 136951 | 41.94 | 133.19 | 120.38 | 73.86 | ATP-dependent zinc metalloprotease FtsH |
| *FEPFGCPE_00113* | *folP* | + | 137224 | 138153 | 3.80 | 25.13 | 17.56 | 9.42 | Dihydropteroate synthase |
| *FEPFGCPE_00114* | *glmM* | + | 138345 | 139688 | 6.14 | 27.46 | 24.62 | 11.43 | Phosphoglucosamine mutase |
| *FEPFGCPE_00115* | *pstS* | + | 140179 | 141213 | 17.25 | 72.40 | 65.26 | 39.93 | Phosphate-binding protein PstS |
| *FEPFGCPE_00116* | *pstC* | + | 141350 | 142324 | 6.07 | 24.62 | 16.96 | 3.33 | Phosphate transport system permease protein PstC |
| *FEPFGCPE_00117* | *pstA* | + | 142324 | 143220 | 11.98 | 32.82 | 29.99 | 10.42 | Phosphate transport system permease protein PstA |
| *FEPFGCPE_00118* | *pstB-1* | + | 143281 | 143691 | 6.14 | 24.21 | 17.32 | 6.08 | Phosphate import ATP-binding protein PstB |
| *FEPFGCPE_00119* | *pstB-2* | + | 143664 | 144062 | 14.25 | 51.69 | 51.69 | 24.28 | Phosphate import ATP-binding protein PstB |
| *FEPFGCPE_00120* | *phoU* | + | 144083 | 144790 | 6.59 | 34.14 | 24.51 | 9.37 | Phosphate-specific transport system accessory protein PhoU |
| *FEPFGCPE_00121* | *phoB* | + | 144827 | 145528 | 7.62 | 36.80 | 26.18 | 8.65 | Phosphate regulon transcriptional regulatory protein PhoB |
| *FEPFGCPE_00122* | *phoR-1* | + | 145620 | 146963 | 2.98 | 20.49 | 14.64 | 4.49 | Phosphate regulon sensor protein PhoR |
| *FEPFGCPE_00123* | *ppk* | - | 147016 | 149094 | 5.49 | 28.04 | 19.79 | 6.31 | Polyphosphate kinase |
| *FEPFGCPE_00124* | *ppx* | + | 149289 | 150824 | 11.00 | 28.52 | 25.00 | 6.38 | Exopolyphosphatase |
| *FEPFGCPE_00125* | | - | 150944 | 151819 | 13.15 | 64.57 | 52.60 | 21.06 | Hypothetical protein |
| *FEPFGCPE_00126* | *sixA* | - | 151959 | 152453 | 13.68 | 35.27 | 33.09 | 7.23 | Phosphohistidine phosphatase SixA |
| *FEPFGCPE_00127* | | + | 152613 | 152689 | 15.37 | 115.80 | 33.98 | 23.28 | tRNA |
| *FEPFGCPE_00128* | *yjcD* | + | 153183 | 155012 | 0.35 | 2.11 | 0.11 | 0.04 | Putative ATP-dependent DNA helicase YjcD |
| *FEPFGCPE_00129* | | + | 155210 | 155566 | 11.49 | 45.05 | 20.23 | 9.11 | Hypothetical protein |
| *FEPFGCPE_00130* | *rpsU* | + | 155934 | 156146 | 17.65 | 39.66 | 39.90 | 49.08 | 30S ribosomal protein S21 |
| *FEPFGCPE_00131* | | + | 156308 | 156781 | 1.49 | 8.25 | 5.52 | 4.21 | Hypothetical protein |
| *FEPFGCPE_00132* | | - | 156830 | 157831 | 0.97 | 6.73 | 5.65 | 0.42 | Hypothetical protein |
| *FEPFGCPE_00133* | | - | 158217 | 158291 | 11.63 | 62.98 | 30.48 | 11.77 | tRNA |
| *FEPFGCPE_00134* | *yfdE-1* | - | 158450 | 159673 | 1.73 | 7.41 | 9.82 | 3.76 | Acetyl-CoA:oxalate CoA-transferase |
| *FEPFGCPE_00135* | *alaS-1* | + | 159917 | 162541 | 21.09 | 54.30 | 56.49 | 29.72 | Alanine--tRNA ligase |
| *FEPFGCPE_00136* | | + | 162915 | 164180 | 2.14 | 5.14 | 4.85 | 2.53 | Hypothetical protein |
| *FEPFGCPE_00137* | *novR* | + | 164235 | 165005 | 3.10 | 6.47 | 5.43 | 3.01 | Decarboxylase NovR |
| *FEPFGCPE_00138* | *ywhB* | + | 165113 | 165313 | 6.48 | 28.27 | 18.51 | 12.98 | 2-hydroxymuconate tautomerase |

| Locus name | Gene name | Strand | Gene start (bp) | Gene end (bp) | Average TPM Inoculum | Average TPM $t_6$ | Average TPM $t_{24}$ | Average TPM $t_{168}$ | Annotation description |
|---|---|---|---|---|---|---|---|---|---|
| *FEPFGCPE_00139* | *ltaE* | + | 165400 | 166533 | 2.09 | 12.53 | 10.01 | 5.56 | Low specificity L-threonine aldolase |
| *FEPFGCPE_00140* | *clcA-1* | - | 166576 | 167802 | 2.07 | 15.11 | 13.83 | 6.34 | H(+)/Cl(-) exchange transporter ClcA |
| *FEPFGCPE_00141* | *nudC-1* | + | 168105 | 168635 | 3.06 | 11.77 | 12.41 | 3.43 | NADH pyrophosphatase |
| *FEPFGCPE_00142* | *tusA* | + | 168735 | 168962 | 14.98 | 155.63 | 90.94 | 43.90 | Sulfurtransferase TusA |
| *FEPFGCPE_00143* | *gtaB* | - | 169079 | 169963 | 15.19 | 24.30 | 16.52 | 19.45 | UTP--glucose-1-phosphate uridylyltransferase |
| *FEPFGCPE_00144* | *valS* | - | 169993 | 172854 | 5.41 | 22.64 | 22.17 | 12.11 | Valine--tRNA ligase |
| *FEPFGCPE_00145* | *uvrD* | + | 172966 | 175287 | 2.04 | 12.12 | 11.53 | 7.48 | DNA helicase II |
| *FEPFGCPE_00146* | | - | 175376 | 176185 | 4.32 | 11.58 | 4.98 | 1.21 | Hypothetical protein |
| *FEPFGCPE_00147* | *thrB* | - | 176217 | 177188 | 4.03 | 16.08 | 8.76 | 3.06 | Homoserine kinase |
| *FEPFGCPE_00148* | | - | 177317 | 177685 | 86.01 | 89.82 | 90.46 | 29.72 | Hypothetical protein |
| *FEPFGCPE_00149* | | + | 178041 | 179276 | 5.01 | 16.84 | 14.58 | 4.58 | Hypothetical protein |
| *FEPFGCPE_00150* | | - | 179392 | 180090 | 33.89 | 42.34 | 42.22 | 25.20 | Cytokinin riboside 5'-monophosphate phosphoribohydrolase |
| *FEPFGCPE_00151* | *polA* | + | 180127 | 182865 | 3.72 | 18.66 | 18.92 | 6.81 | DNA polymerase I |
| *FEPFGCPE_00152* | *ndh-1* | + | 183075 | 184472 | 3.68 | 22.61 | 17.19 | 8.80 | NADH dehydrogenase |
| *FEPFGCPE_00153* | *clcD-1* | + | 184737 | 185612 | 5.49 | 34.73 | 28.38 | 9.48 | Carboxymethylenebutenolidase |
| *FEPFGCPE_00154* | *zupT* | + | 185800 | 186669 | 8.10 | 30.57 | 23.48 | 9.51 | Zinc transporter ZupT |
| *FEPFGCPE_00155* | *sseB* | - | 186600 | 187484 | 20.58 | 67.48 | 43.50 | 15.93 | Putative thiosulfate sulfurtransferase SseB |
| *FEPFGCPE_00156* | *ribN* | - | 187536 | 188390 | 50.79 | 121.12 | 51.26 | 23.12 | Riboflavin transporter |
| *FEPFGCPE_00157* | *bphA1* | - | 188661 | 189776 | 21.50 | 18.77 | 17.68 | 10.36 | Biphenyl 2,3-dioxygenase subunit alpha |
| *FEPFGCPE_00158* | *xseB* | + | 190005 | 190274 | 43.70 | 110.42 | 111.08 | 45.97 | Exodeoxyribonuclease 7 small subunit |
| *FEPFGCPE_00159* | | + | 190454 | 191350 | 25.92 | 64.00 | 63.84 | 30.32 | Farnesyl diphosphate synthase |
| *FEPFGCPE_00160* | *dxs-1* | + | 191549 | 193465 | 15.63 | 38.21 | 39.36 | 22.83 | 1-deoxy-D-xylulose-5-phosphate synthase |
| *FEPFGCPE_00161* | *folE2* | + | 193665 | 194474 | 0.94 | 5.79 | 4.19 | 1.08 | GTP cyclohydrolase FolE2 |
| *FEPFGCPE_00162* | *tsaD* | - | 194557 | 195588 | 4.98 | 8.15 | 7.53 | 5.04 | tRNA N6-adenosine threonylcarbamoyltransferase |
| *FEPFGCPE_00163* | | + | 195671 | 196930 | 1.58 | 6.67 | 6.12 | 2.55 | Ferredoxin--NADP reductase |
| *FEPFGCPE_00164* | | + | 197192 | 197638 | 74.40 | 392.50 | 333.11 | 107.22 | Hypothetical protein |
| *FEPFGCPE_00165* | *dnaG* | + | 197679 | 199589 | 7.64 | 80.19 | 84.85 | 27.55 | DNA primase |
| *FEPFGCPE_00166* | *rpoD* | + | 199653 | 202283 | 19.64 | 83.80 | 87.20 | 49.25 | RNA polymerase sigma factor RpoD |
| *FEPFGCPE_00167* | | + | 202312 | 202384 | 4.19 | 3.84 | 0.44 | 0.93 | tRNA |
| *FEPFGCPE_00168* | *xerC-1* | + | 202708 | 203700 | 1.29 | 9.33 | 5.46 | 1.91 | Tyrosine recombinase XerC |
| *FEPFGCPE_00169* | | - | 203779 | 204615 | 4.75 | 23.06 | 14.14 | 4.43 | Hypothetical protein |
| *FEPFGCPE_00170* | | - | 204612 | 204878 | 2.02 | 6.85 | 11.29 | 2.19 | Hypothetical protein |
| *FEPFGCPE_00171* | *thcD* | + | 205729 | 206955 | 1.84 | 4.43 | 3.75 | 0.89 | Rhodocoxin reductase |
| *FEPFGCPE_00172* | | + | 207155 | 207415 | 0.76 | 2.98 | 5.17 | 0.00 | Hypothetical protein |
| *FEPFGCPE_00173* | *hpcG-1* | + | 207448 | 208251 | 0.95 | 10.73 | 5.75 | 2.60 | 2-oxo-hept-4-ene-1,7-dioate hydratase |
| *FEPFGCPE_00174* | *hpcH-1* | + | 208248 | 209066 | 2.80 | 28.51 | 13.07 | 4.42 | 4-hydroxy-2-oxo-heptane-1,7-dioate aldolase |
| *FEPFGCPE_00175* | *dmlR-1* | + | 209280 | 210224 | 4.68 | 26.91 | 13.54 | 3.10 | HTH-type transcriptional regulator DmlR |
| *FEPFGCPE_00176* | *glnQ-1* | - | 211099 | 211848 | 0.63 | 10.49 | 5.75 | 4.53 | Glutamine transport ATP-binding protein GlnQ |
| *FEPFGCPE_00177* | *yxeN* | - | 211849 | 212982 | 3.07 | 9.95 | 8.96 | 2.80 | putative amino-acid permease protein YxeN |
| *FEPFGCPE_00178* | *fliY-1* | - | 212984 | 213796 | 2.52 | 10.23 | 11.25 | 1.33 | L-cystine-binding protein FliY |
| *FEPFGCPE_00179* | *adc* | - | 213890 | 214639 | 0.76 | 8.98 | 6.29 | 3.48 | putative acetoacetate decarboxylase |
| *FEPFGCPE_00180* | | + | 214738 | 214983 | 0.74 | 21.82 | 12.55 | 4.67 | Hypothetical protein |
| *FEPFGCPE_00181* | | - | 215252 | 216103 | 1.32 | 14.61 | 7.40 | 1.25 | Hypothetical protein |
| *FEPFGCPE_00182* | | - | 216263 | 216397 | 4.89 | 21.82 | 8.01 | 3.24 | Hypothetical protein |
| *FEPFGCPE_00183* | | - | 216787 | 217407 | 0.65 | 3.81 | 5.14 | 0.00 | Hypothetical protein |
| *FEPFGCPE_00184* | | - | ` | 217806 | 3.50 | 12.37 | 7.48 | 0.00 | Hypothetical protein |
| *FEPFGCPE_00185* | *nagAb-1* | + | 218075 | 218398 | 0.00 | 6.01 | 1.73 | 1.06 | Naphthalene 1,2-dioxygenase/salicylate 5-hydroxylase systems, ferredoxin component |
| *FEPFGCPE_00186* | | + | 218513 | 219397 | 8.67 | 68.28 | 51.48 | 10.44 | Ureidoglycolate lyase |
| *FEPFGCPE_00187* | | + | 219390 | 220388 | 6.40 | 41.27 | 36.76 | 13.10 | Delta(1)-pyrroline-2-carboxylate reductase |
| *FEPFGCPE_00188* | *phdI* | + | 220435 | 221532 | 2.74 | 22.66 | 12.34 | 4.27 | 1-hydroxy-2-naphthoate 1,2-dioxygenasee |
| *FEPFGCPE_00189* | *andAd* | + | 221539 | 222057 | 3.37 | 32.13 | 23.75 | 10.47 | Anthranilate 1,2-dioxygenase small subunit |
| *FEPFGCPE_00190* | *nagG-1* | + | 222077 | 223330 | 5.15 | 41.67 | 36.92 | 11.14 | Salicylate 5-hydroxylase, large oxygenase component |
| *FEPFGCPE_00191* | | + | 223414 | 224637 | 3.31 | 43.49 | 26.04 | 6.24 | Leu/Ile/Val-binding protein |
| *FEPFGCPE_00192* | *livH-1* | + | 224714 | 225577 | 1.62 | 19.72 | 12.24 | 1.94 | High-affinity branched-chain amino acid transport system permease protein LivH |
| *FEPFGCPE_00193* | | + | 225581 | 226588 | 3.66 | 32.70 | 23.79 | 8.79 | Hypothetical protein |
| *FEPFGCPE_00194* | *lptB-1* | + | 226575 | 227318 | 3.23 | 38.06 | 31.70 | 7.29 | Lipopolysaccharide export system ATP-binding protein LptB |
| *FEPFGCPE_00195* | *livF-1* | + | 227318 | 228013 | 1.44 | 23.46 | 19.15 | 2.89 | High-affinity branched-chain amino acid transport ATP-binding protein LivF |
| *FEPFGCPE_00196* | | - | 228248 | 228556 | 0.76 | 8.31 | 0.11 | 0.71 | Hypothetical protein |
| *FEPFGCPE_00197* | *cynR-1* | - | 229035 | 229955 | 1.90 | 14.47 | 11.44 | 2.96 | HTH-type transcriptional regulator CynR |
| *FEPFGCPE_00198* | *citA* | + | 230116 | 230931 | 0.96 | 183.11 | 87.20 | 3.40 | Citrate synthase 1 |
| *FEPFGCPE_00199* | *uctC-1* | + | 230982 | 232160 | 1.55 | 131.50 | 62.84 | 5.31 | Acetyl-CoA:oxalate CoA-transferase |
| *FEPFGCPE_00200* | *nicT-1* | + | 232408 | 233769 | 2.50 | 106.49 | 49.71 | 3.83 | Putative metabolite transport protein NicT |
| *FEPFGCPE_00201* | | - | 234266 | 234778 | 1.24 | 26.69 | 10.12 | 0.53 | Bacteriohemerythrin |
| *FEPFGCPE_00202* | *ophA1-1* | - | 234964 | 235932 | 0.17 | 2.05 | 1.85 | 0.00 | Phthalate dioxygenase reductase |
| *FEPFGCPE_00203* | *ttuB-1* | + | 236169 | 237512 | 0.32 | 2.18 | 1.03 | 0.58 | Putative tartrate transporter |
| *FEPFGCPE_00204* | *pht5-1* | + | 237559 | 238551 | 0.16 | 1.91 | 0.00 | 0.97 | 4,5-dihydroxyphthalate decarboxylase |
| *FEPFGCPE_00205* | | + | 239123 | 239710 | 1.04 | 9.86 | 9.38 | 3.32 | Hypothetical protein |
| *FEPFGCPE_00206* | *nadC-1* | + | 239852 | 240697 | 0.49 | 9.23 | 2.57 | 0.56 | Nicotinate-nucleotide pyrophosphorylase [carboxylating] |
| *FEPFGCPE_00207* | *pht4-1* | - | 240866 | 242041 | 0.44 | 2.18 | 1.83 | 0.80 | Putative 4,5-dihydroxyphthalate dehydrogenase |
| *FEPFGCPE_00208* | *pht3-1* | - | 242058 | 243389 | 1.27 | 6.97 | 3.47 | 0.61 | Phthalate 4,5-dioxygenase oxygenase subunit |
| *FEPFGCPE_00209* | *trg-1* | + | 244891 | 246435 | 0.30 | 1.18 | 2.01 | 0.00 | Methyl-accepting chemotaxis protein III |

| Locus name | Gene name | Strand | Gene start (bp) | Gene end (bp) | Average TPM Inoculum | Average TPM $t_6$ | Average TPM $t_{24}$ | Average TPM $t_{168}$ | Annotation description |
|---|---|---|---|---|---|---|---|---|---|
| *FEPFGCPE_00210* | | + | 246432 | 246668 | 1.85 | 13.30 | 0.00 | 3.68 | Hypothetical protein |
| *FEPFGCPE_00211* | | + | 246887 | 247117 | 2.89 | 13.35 | 6.17 | 1.49 | Hypothetical protein |
| *FEPFGCPE_00212* | | + | 247114 | 247950 | 3.60 | 22.68 | 13.31 | 3.96 | Hypothetical protein |
| *FEPFGCPE_00213* | | - | 247891 | 248805 | 1.24 | 5.50 | 2.07 | 0.67 | Hypothetical protein |
| *FEPFGCPE_00214* | *ssuB* | - | 249012 | 249791 | 1.22 | 7.30 | 2.24 | 1.33 | Aliphatic sulfonates import ATP-binding protein SsuB |
| *FEPFGCPE_00215* | *ssuC-1* | - | 249801 | 250613 | 3.25 | 36.26 | 15.23 | 3.40 | Putative aliphatic sulfonates transport permease protein SsuC |
| *FEPFGCPE_00216* | | - | 250624 | 251622 | 1.22 | 4.19 | 7.02 | 0.62 | Hypothetical protein |
| *FEPFGCPE_00217* | *mdlC-1* | - | 251756 | 253564 | 0.64 | 6.96 | 2.82 | 1.29 | Benzoylformate decarboxylase |
| *FEPFGCPE_00218* | *phdK* | - | 253587 | 255074 | 0.38 | 6.72 | 2.43 | 0.29 | 2-formylbenzoate dehydrogenase |
| *FEPFGCPE_00219* | | + | 255821 | 256060 | 0.68 | 4.59 | 1.69 | 2.36 | Hypothetical protein |
| *FEPFGCPE_00220* | *kdgR-1* | - | 256195 | 256959 | 0.68 | 3.72 | 3.38 | 2.24 | Transcriptional regulator KdgR |
| *FEPFGCPE_00221* | | + | 257308 | 257616 | 1.91 | 11.98 | 5.07 | 0.65 | Hypothetical protein |
| *FEPFGCPE_00222* | *actP-1* | + | 257618 | 259162 | 0.77 | 8.37 | 8.38 | 1.38 | Cation/acetate symporter ActP |
| *FEPFGCPE_00223* | | + | 259163 | 259888 | 1.42 | 14.93 | 8.55 | 3.20 | Arylmalonate decarboxylase |
| *FEPFGCPE_00224* | | + | 259957 | 260478 | 0.64 | 7.46 | 7.05 | 7.65 | Hypothetical protein |
| *FEPFGCPE_00225* | | + | 260967 | 261443 | 2.61 | 47.05 | 33.68 | 12.96 | 4-sulfomuconolactone hydrolase |
| *FEPFGCPE_00226* | | + | 261811 | 261933 | 2.76 | 8.62 | 4.75 | 1.31 | Hypothetical protein |
| *FEPFGCPE_00227* | *ompR-1* | - | 262564 | 263283 | 0.80 | 7.43 | 5.79 | 0.60 | Transcriptional regulatory protein OmpR |
| *FEPFGCPE_00228* | | + | 263518 | 264156 | 0.54 | 2.76 | 1.53 | 1.10 | Hypothetical protein |
| *FEPFGCPE_00229* | *exbB-1* | + | 264210 | 265031 | 0.15 | 1.70 | 2.35 | 0.00 | Biopolymer transport protein ExbB |
| *FEPFGCPE_00230* | *exbD-1* | + | 265036 | 265470 | 0.12 | 0.46 | 0.00 | 0.00 | Biopolymer transport protein ExbD |
| *FEPFGCPE_00231* | *exbD-2* | + | 265502 | 265930 | 0.16 | 0.59 | 0.00 | 0.00 | Biopolymer transport protein ExbD |
| *FEPFGCPE_00232* | *lgoT-1* | + | 265987 | 267339 | 0.07 | 1.86 | 0.93 | 0.94 | putative L-galactonate transporter |
| *FEPFGCPE_00233* | *ycjG* | + | 267346 | 268362 | 0.29 | 3.23 | 1.85 | 0.00 | L-Ala-D/L-Glu epimerase |
| *FEPFGCPE_00234* | *dap-1* | + | 268349 | 270061 | 0.41 | 2.98 | 1.46 | 0.54 | D-aminopeptidase |
| *FEPFGCPE_00235* | *dap-2* | - | 270139 | 271770 | 1.36 | 3.62 | 3.54 | 1.19 | D-aminopeptidase |
| *FEPFGCPE_00236* | *fyuA* | - | 271801 | 274158 | 1.41 | 4.93 | 3.52 | 1.30 | Pesticin receptor |
| *FEPFGCPE_00237* | *envZ-1* | + | 274447 | 275478 | 0.81 | 6.56 | 6.32 | 1.72 | Osmolarity sensor protein EnvZ |
| *FEPFGCPE_00238* | | + | 275603 | 277060 | 1.10 | 6.58 | 3.32 | 1.19 | Hypothetical protein |
| *FEPFGCPE_00239* | *dadA1-2* | - | 277238 | 278470 | 0.75 | 1.87 | 1.45 | 0.35 | D-amino acid dehydrogenase 1 |
| *FEPFGCPE_00240* | *yabJ-1* | - | 278618 | 278971 | 1.49 | 7.69 | 3.26 | 0.18 | 2-iminobutanoate/2-iminopropanoate deaminase |
| *FEPFGCPE_00241* | | + | 279157 | 279813 | 2.14 | 7.89 | 6.15 | 1.96 | Hypothetical protein |
| *FEPFGCPE_00242* | | + | 279810 | 279941 | 0.53 | 4.92 | 1.73 | 1.59 | Hypothetical protein |
| *FEPFGCPE_00243* | | - | 280046 | 280276 | 4.41 | 35.37 | 20.82 | 4.99 | Hypothetical protein |
| *FEPFGCPE_00244* | *cspG* | - | 280479 | 280682 | 4.26 | 14.56 | 22.74 | 6.59 | Cold shock-like protein CspG |
| *FEPFGCPE_00245* | *kipR-1* | - | 281001 | 281711 | 3.22 | 18.00 | 5.65 | 1.36 | HTH-type transcriptional regulator KipR |
| *FEPFGCPE_00246* | *proA-1* | + | 281873 | 282571 | 1.29 | 1.74 | 6.69 | 2.53 | 4-hydroxy-4-methyl-2-oxoglutarate aldolase/4-carboxy-4-hydroxy-2-oxoadipate aldolase |
| *FEPFGCPE_00247* | | + | 282571 | 283518 | 1.09 | 2.42 | 1.43 | 0.47 | Hydroxypyruvate reductase |
| *FEPFGCPE_00248* | | + | 283542 | 284471 | 1.04 | 3.37 | 1.80 | 0.04 | Lactonase drp35 |
| *FEPFGCPE_00249* | *ttuB-2* | + | 284549 | 285967 | 0.95 | 2.11 | 2.36 | 0.15 | Putative tartrate transporter |
| *FEPFGCPE_00250* | | + | 285981 | 287027 | 0.98 | 6.11 | 2.40 | 0.55 | Hypothetical protein |
| *FEPFGCPE_00251* | *naiP-1* | + | 287024 | 288304 | 0.58 | 1.65 | 4.48 | 0.15 | Putative niacin/nicotinamide transporter NaiP |
| *FEPFGCPE_00252* | *ttuB-3* | + | 288453 | 289778 | 1.01 | 2.42 | 1.95 | 0.77 | Putative tartrate transporter |
| *FEPFGCPE_00253* | | + | 289897 | 290046 | 9.43 | 95.81 | 87.22 | 27.89 | Hypothetical protein |
| *FEPFGCPE_00254* | *ansA-1* | + | 290310 | 291320 | 2.02 | 7.81 | 3.45 | 2.25 | L-asparaginase |
| *FEPFGCPE_00255* | *mcbR* | + | 291369 | 292082 | 1.27 | 12.63 | 6.61 | 1.80 | HTH-type transcriptional regulator McbR |
| *FEPFGCPE_00256* | *cat1* | - | 292417 | 293913 | 0.46 | 1.97 | 2.70 | 0.00 | Succinyl-CoA:coenzyme A transferase |
| *FEPFGCPE_00257* | *hcaR-1* | + | 294109 | 295026 | 1.61 | 11.84 | 3.61 | 0.17 | Hca operon transcriptional activator HcaR |
| *FEPFGCPE_00258* | | - | 294988 | 295473 | 5.86 | 65.28 | 29.88 | 6.05 | Hypothetical protein |
| *FEPFGCPE_00259* | | + | 295913 | 296176 | 15.45 | 57.62 | 30.50 | 17.19 | Hypothetical protein |
| *FEPFGCPE_00260* | *pcaK-1* | + | 296764 | 298119 | 0.57 | 1.36 | 0.69 | 0.54 | 4-hydroxybenzoate transporter PcaK |
| *FEPFGCPE_00261* | *bcr-1* | + | 298295 | 299509 | 0.45 | 5.12 | 4.18 | 2.05 | Bicyclomycin resistance protein |
| *FEPFGCPE_00262* | *thcB* | - | 299544 | 301898 | 1.75 | 9.24 | 10.07 | 5.25 | Cytochrome P450 116 |
| *FEPFGCPE_00263* | *iclR-1* | + | 302090 | 303052 | 3.02 | 16.30 | 15.28 | 6.59 | Transcriptional repressor IclR |
| *FEPFGCPE_00264* | *dppA-1* | + | 303203 | 304807 | 3.32 | 15.90 | 11.21 | 6.01 | Periplasmic dipeptide transport protein |
| *FEPFGCPE_00265* | | + | 305590 | 305985 | 0.50 | 3.12 | 2.68 | 1.52 | Hypothetical protein |
| *FEPFGCPE_00266* | *metQ-1* | + | 306402 | 307211 | 0.62 | 84.73 | 27.63 | 16.75 | D-methionine-binding lipoprotein MetQ |
| *FEPFGCPE_00267* | | + | 307379 | 307540 | 146.50 | 689.07 | 475.10 | 80.29 | Hypothetical protein |
| *FEPFGCPE_00268* | *prpR* | - | 307528 | 309699 | 7.70 | 49.06 | 42.75 | 11.08 | Propionate catabolism operon regulatory protein |
| *FEPFGCPE_00269* | *prpB-1* | + | 310053 | 310943 | 6.23 | 18.72 | 15.34 | 7.77 | 2-methylisocitrate lyase |
| *FEPFGCPE_00270* | *prpC* | + | 310972 | 312123 | 7.53 | 21.66 | 18.25 | 9.81 | 2-methylcitrate synthase |
| *FEPFGCPE_00271* | *acnD* | + | 312267 | 314867 | 3.42 | 14.42 | 6.73 | 4.87 | 2-methylcitrate dehydratase (2-methyl-trans-aconitate forming) |
| *FEPFGCPE_00272* | *prpF* | + | 314888 | 316078 | 7.34 | 24.36 | 20.72 | 9.03 | 2-methyl-aconitate isomerase |
| *FEPFGCPE_00273* | *prpD* | + | 316233 | 317681 | 5.01 | 13.72 | 11.03 | 4.09 | 2-methylcitrate dehydratase |
| *FEPFGCPE_00274* | *glxK* | - | 317804 | 318955 | 2.37 | 19.15 | 7.62 | 4.59 | Glycerate 3-kinase |
| *FEPFGCPE_00275* | | + | 319340 | 319852 | 8.60 | 28.17 | 14.61 | 7.52 | Hypothetical protein |
| *FEPFGCPE_00276* | *glpR-1* | + | 319983 | 320807 | 3.93 | 12.41 | 15.05 | 0.64 | Glycerol-3-phosphate regulon repressor |
| *FEPFGCPE_00277* | *glpD* | + | 320985 | 322523 | 1.93 | 8.13 | 6.81 | 3.30 | Aerobic glycerol-3-phosphate dehydrogenase |
| *FEPFGCPE_00278* | *glpK* | + | 322554 | 324140 | 2.02 | 10.21 | 4.20 | 3.38 | Glycerol kinase |
| *FEPFGCPE_00279* | *nagK-1* | - | 324758 | 325462 | 2.32 | 8.85 | 5.04 | 3.64 | Fumarylpyruvate hydrolase |
| *FEPFGCPE_00280* | | - | 325573 | 325992 | 2.50 | 24.02 | 8.22 | 2.40 | Hypothetical protein |

| Locus name | Gene name | Strand | Gene start (bp) | Gene end (bp) | Average TPM Inoculum | Average TPM $t_6$ | Average TPM $t_{24}$ | Average TPM $t_{168}$ | Annotation description |
|---|---|---|---|---|---|---|---|---|---|
| *FEPFGCPE_00281* | *oxyR-1* | + | 326172 | 327089 | 1.94 | 10.05 | 4.25 | 1.45 | Hydrogen peroxide-inducible genes activator |
| *FEPFGCPE_00282* | *ttgC* | - | 327086 | 328594 | 2.54 | 12.28 | 8.41 | 3.93 | Toluene efflux pump outer membrane protein TtgC |
| *FEPFGCPE_00283* | *ybhS* | - | 328602 | 329744 | 2.85 | 19.82 | 9.24 | 2.21 | Inner membrane transport permease YbhS |
| *FEPFGCPE_00284* | *ybhF-1* | - | 329741 | 330724 | 2.13 | 10.28 | 9.60 | 0.40 | putative ABC transporter ATP-binding protein YbhF |
| *FEPFGCPE_00285* | *mdtA-1* | - | 330721 | 331686 | 7.36 | 31.59 | 21.12 | 5.29 | Multidrug resistance protein MdtA |
| *FEPFGCPE_00286* | *folE-1* | + | 331869 | 332435 | 5.80 | 19.85 | 23.81 | 15.60 | GTP cyclohydrolase 1 |
| *FEPFGCPE_00287* | *lppC* | - | 332514 | 333092 | 23.01 | 53.87 | 69.92 | 45.64 | Putative lipoprotein LppC |
| *FEPFGCPE_00288* | | - | 333104 | 333289 | 8.33 | 10.50 | 15.07 | 12.12 | Hypothetical protein |
| *FEPFGCPE_00289* | *cydB-1* | - | 333273 | 334271 | 4.66 | 3.58 | 10.85 | 8.15 | Cytochrome bd-I ubiquinol oxidase subunit 2 |
| *FEPFGCPE_00290* | *appC* | - | 334312 | 335757 | 5.39 | 7.27 | 20.75 | 21.56 | Cytochrome bd-II ubiquinol oxidase subunit 1 |
| *FEPFGCPE_00291* | | - | 335988 | 336659 | 2.82 | 8.68 | 12.33 | 4.09 | Hypothetical protein |
| *FEPFGCPE_00292* | *nasA-1* | + | 337132 | 338520 | 0.82 | 21.78 | 58.73 | 51.38 | Nitrate transporter |
| *FEPFGCPE_00293* | *prpB-2* | - | 338694 | 339518 | 7.48 | 34.14 | 31.41 | 10.43 | 2-methylisocitrate lyase |
| *FEPFGCPE_00294* | *yceJ-1* | + | 339780 | 340412 | 7.15 | 9.78 | 12.31 | 1.97 | Cytochrome b561 |
| *FEPFGCPE_00295* | *icaA* | + | 340409 | 341914 | 6.09 | 13.26 | 9.89 | 5.09 | Poly-beta-1,6-N-acetyl-D-glucosamine synthase |
| *FEPFGCPE_00296* | | + | 341911 | 342582 | 4.20 | 14.88 | 14.83 | 3.18 | Hypothetical protein |
| *FEPFGCPE_00297* | *selU* | + | 342579 | 343634 | 1.57 | 8.44 | 7.71 | 1.23 | tRNA 2-selenouridine synthase |
| *FEPFGCPE_00298* | *arsH-1* | - | 343665 | 344429 | 1.00 | 6.56 | 2.17 | 1.52 | NADPH-dependent FMN reductase ArsH |
| *FEPFGCPE_00299* | *arsC-1* | - | 344426 | 344848 | 0.41 | 3.43 | 1.27 | 1.86 | Arsenate reductase |
| *FEPFGCPE_00300* | *acr3-1* | - | 344858 | 345922 | 1.38 | 4.09 | 2.35 | 0.45 | Arsenical-resistance protein Acr3 |
| *FEPFGCPE_00301* | *ygaV* | - | 345919 | 346254 | 2.41 | 7.46 | 0.00 | 0.21 | putative HTH-type transcriptional regulator YgaV |
| *FEPFGCPE_00302* | *gltC-3* | - | 346342 | 347232 | 1.82 | 8.55 | 2.14 | 0.60 | HTH-type transcriptional regulator GltC |
| *FEPFGCPE_00303* | *potD-1* | + | 347407 | 348474 | 1.01 | 5.12 | 5.17 | 1.96 | Spermidine/putrescine-binding periplasmic protein |
| *FEPFGCPE_00304* | *potA-2* | + | 348498 | 349577 | 0.81 | 3.47 | 2.96 | 1.04 | Spermidine/putrescine import ATP-binding protein PotA |
| *FEPFGCPE_00305* | *potB-1* | + | 349613 | 350512 | 0.28 | 2.47 | 1.33 | 0.00 | Spermidine/putrescine transport system permease protein PotB |
| *FEPFGCPE_00306* | *ydcV-1* | + | 350524 | 351339 | 0.22 | 2.02 | 1.26 | 0.00 | Inner membrane ABC transporter permease protein YdcV |
| *FEPFGCPE_00307* | *hcnC* | + | 351343 | 352443 | 0.24 | 3.68 | 0.48 | 0.28 | Hydrogen cyanide synthase subunit HcnC |
| *FEPFGCPE_00308* | *hcnA-1* | + | 352421 | 352729 | 0.73 | 0.50 | 0.00 | 0.86 | Hydrogen cyanide synthase subunit HcnA |
| *FEPFGCPE_00309* | *hcnB-1* | + | 352726 | 354159 | 0.49 | 4.25 | 2.21 | 0.39 | Hydrogen cyanide synthase subunit HcnB |
| *FEPFGCPE_00310* | *yabJ-2* | + | 354218 | 354574 | 1.51 | 12.08 | 12.21 | 3.51 | 2-iminobutanoate/2-iminopropanoate deaminase |
| *FEPFGCPE_00311* | *crp* | - | 354682 | 355353 | 1.31 | 14.11 | 7.27 | 2.66 | cAMP-activated global transcriptional regulator CRP |
| *FEPFGCPE_00312* | | + | 355688 | 357661 | 0.70 | 6.58 | 2.76 | 0.34 | Long-chain-fatty-acid--CoA ligase FadD15 |
| *FEPFGCPE_00313* | *lptB-2* | + | 357728 | 358534 | 0.21 | 5.56 | 3.96 | 0.04 | Lipopolysaccharide export system ATP-binding protein LptB |
| *FEPFGCPE_00314* | *livH-2* | + | 358587 | 359501 | 0.20 | 4.43 | 1.45 | 0.90 | High-affinity branched-chain amino acid transport system permease protein LivH |
| *FEPFGCPE_00315* | | + | 359557 | 360633 | 1.41 | 5.94 | 1.54 | 0.47 | Hypothetical protein |
| *FEPFGCPE_00316* | | + | 360716 | 362041 | 0.55 | 12.31 | 4.49 | 1.14 | Hypothetical protein |
| *FEPFGCPE_00317* | *livF-2* | + | 362122 | 362934 | 0.82 | 13.35 | 6.20 | 2.64 | High-affinity branched-chain amino acid transport ATP-binding protein LivF |
| *FEPFGCPE_00318* | | + | 363009 | 364247 | 0.72 | 17.13 | 5.93 | 1.94 | Phenylacetate-coenzyme A ligase |
| *FEPFGCPE_00319* | *yhjE-1* | - | 364313 | 365629 | 0.22 | 5.73 | 3.22 | 0.73 | Inner membrane metabolite transport protein YhjE |
| *FEPFGCPE_00320* | *yfdE-2* | - | 365723 | 366913 | 0.30 | 2.49 | 0.00 | 0.00 | Acetyl-CoA:oxalate CoA-transferase |
| *FEPFGCPE_00321* | *aidB-1* | - | 366910 | 368664 | 0.22 | 0.24 | 0.90 | 0.04 | Putative acyl-CoA dehydrogenase AidB |
| *FEPFGCPE_00322* | *benM-1* | + | 368807 | 369700 | 1.03 | 1.59 | 2.27 | 0.13 | HTH-type transcriptional regulator BenM |
| *FEPFGCPE_00323* | *mch* | + | 369887 | 370396 | 0.39 | 0.99 | 1.05 | 0.00 | Mesaconyl-CoA hydratase |
| *FEPFGCPE_00324* | | + | 370509 | 370832 | 0.17 | 0.00 | 0.00 | 1.79 | Hypothetical protein |
| *FEPFGCPE_00325* | *gatA-1* | + | 370813 | 372324 | 0.58 | 0.95 | 0.91 | 0.26 | Glutamyl-tRNA(Gln) amidotransferase subunit A |
| *FEPFGCPE_00326* | *ydjE-2* | + | 372430 | 373827 | 0.75 | 1.86 | 1.08 | 0.41 | Inner membrane metabolite transport protein YdjE |
| *FEPFGCPE_00327* | | + | 373864 | 375297 | 0.91 | 5.17 | 2.87 | 2.09 | Hypothetical protein |
| *FEPFGCPE_00328* | *mmgC-1* | + | 375327 | 376499 | 1.22 | 12.17 | 6.47 | 1.97 | Acyl-CoA dehydrogenase |
| *FEPFGCPE_00329* | *uctC-2* | + | 376496 | 377686 | 1.09 | 5.03 | 3.25 | 1.53 | Acetyl-CoA:oxalate CoA-transferase |
| *FEPFGCPE_00330* | *yvoA-1* | + | 377723 | 378472 | 3.71 | 15.61 | 12.80 | 4.82 | HTH-type transcriptional repressor YvoA |
| *FEPFGCPE_00331* | | + | 378519 | 379328 | 2.58 | 18.88 | 7.40 | 1.33 | Citrate lyase subunit beta-like protein |
| *FEPFGCPE_00332* | *cmpR-2* | - | 379362 | 380279 | 1.40 | 6.18 | 5.14 | 0.20 | HTH-type transcriptional activator CmpR |
| *FEPFGCPE_00333* | *nicT-2* | + | 380399 | 380839 | 0.99 | 3.29 | 4.92 | 1.71 | Putative metabolite transport protein NicT |
| *FEPFGCPE_00334* | *ttuB-4* | + | 380843 | 381685 | 0.19 | 0.00 | 1.27 | 0.28 | Putative tartrate transporter |
| *FEPFGCPE_00335* | | + | 381682 | 383238 | 0.40 | 0.47 | 3.01 | 1.24 | Hypothetical protein |
| *FEPFGCPE_00336* | | - | 383379 | 383657 | 0.45 | 7.32 | 12.08 | 1.96 | Hypothetical protein |
| *FEPFGCPE_00337* | *guaB-1* | - | 383686 | 384930 | 0.76 | 12.42 | 5.95 | 2.31 | Inosine-5'-monophosphate dehydrogenase |
| *FEPFGCPE_00338* | *cynR-2* | + | 385011 | 385967 | 0.83 | 2.90 | 2.55 | 1.29 | HTH-type transcriptional regulator CynR |
| *FEPFGCPE_00339* | | - | 385948 | 386640 | 1.25 | 11.55 | 8.68 | 2.75 | Hypothetical protein |
| *FEPFGCPE_00340* | | - | 386792 | 387763 | 1.44 | 8.02 | 2.95 | 0.33 | Hypothetical protein |
| *FEPFGCPE_00341* | | - | 387787 | 388764 | 0.54 | 1.69 | 1.54 | 0.59 | Hypothetical protein |
| *FEPFGCPE_00342* | | - | 388807 | 390174 | 0.48 | 3.28 | 1.43 | 0.60 | Putative conjugal transfer protein |
| *FEPFGCPE_00343* | | - | 390205 | 391413 | 0.21 | 3.17 | 1.16 | 0.32 | Hypothetical protein |
| *FEPFGCPE_00344* | | - | 391442 | 391882 | 0.49 | 6.66 | 0.29 | 0.18 | Hypothetical protein |
| *FEPFGCPE_00345* | | - | 391879 | 392400 | 0.35 | 2.13 | 1.61 | 0.00 | Hypothetical protein |
| *FEPFGCPE_00346* | | - | 392410 | 393681 | 0.25 | 0.82 | 0.00 | 0.00 | Hypothetical protein |
| *FEPFGCPE_00347* | | - | 393797 | 394129 | 0.00 | 0.18 | 1.22 | 0.00 | Hypothetical protein |
| *FEPFGCPE_00348* | *xcpQ* | - | 394154 | 395929 | 0.56 | 3.37 | 1.24 | 0.00 | Type II secretion system protein D |
| *FEPFGCPE_00349* | | - | 396012 | 396854 | 0.06 | 2.44 | 0.04 | 0.00 | Hypothetical protein |
| *FEPFGCPE_00350* | | - | 396867 | 398300 | 0.28 | 0.67 | 0.50 | 0.00 | Hypothetical protein |
| *FEPFGCPE_00351* | | - | 398332 | 398871 | 0.46 | 4.52 | 1.57 | 1.48 | Hypothetical protein |

| Locus name | Gene name | Strand | Gene start (bp) | Gene end (bp) | Average TPM Inoculum | Average TPM $t_6$ | Average TPM $t_{24}$ | Average TPM $t_{168}$ | Annotation description |
|---|---|---|---|---|---|---|---|---|---|
| *FEPFGCPE_00352* | | - | 398992 | 399177 | 0.41 | 0.85 | 0.00 | 0.00 | Hypothetical protein |
| *FEPFGCPE_00353* | | + | 400383 | 401678 | 0.30 | 0.04 | 0.43 | 0.00 | Hypothetical protein |
| *FEPFGCPE_00354* | | + | 401675 | 402622 | 0.32 | 2.79 | 0.61 | 0.42 | Hypothetical protein |
| *FEPFGCPE_00355* | | + | 402603 | 403664 | 0.57 | 4.72 | 0.63 | 0.14 | putative glycosyltransferase |
| *FEPFGCPE_00356* | | - | 403669 | 404043 | 1.10 | 11.36 | 2.40 | 0.00 | Hypothetical protein |
| *FEPFGCPE_00357* | | - | 404047 | 404784 | 3.44 | 12.20 | 9.17 | 1.41 | putative S-adenosylmethionine-dependent methyltransferase/MSMEI_2290 |
| *FEPFGCPE_00358* | *rnz* | - | 405029 | 406018 | 0.98 | 4.69 | 2.33 | 0.59 | Ribonuclease Z |
| *FEPFGCPE_00359* | *gbpR-1* | + | 406122 | 407054 | 2.68 | 16.02 | 7.29 | 1.96 | HTH-type transcriptional regulator GbpR |
| *FEPFGCPE_00360* | *gabR-2* | - | 407520 | 409007 | 2.12 | 9.93 | 8.23 | 1.76 | HTH-type transcriptional regulatory protein GabR |
| *FEPFGCPE_00361* | | + | 409103 | 410668 | 0.45 | 1.17 | 0.58 | 0.21 | Hypothetical protein |
| *FEPFGCPE_00362* | *ydhC-1* | + | 410709 | 411911 | 0.25 | 3.32 | 0.55 | 0.16 | Inner membrane transport protein YdhC |
| *FEPFGCPE_00363* | *rhlE-1* | + | 412229 | 413566 | 0.90 | 11.24 | 12.28 | 3.18 | ATP-dependent RNA helicase RhlE |
| *FEPFGCPE_00364* | *mhbT-1* | - | 413766 | 414890 | 0.11 | 1.72 | 0.74 | 0.29 | 3-hydroxybenzoate transporter MhbT |
| *FEPFGCPE_00365* | *pcaK-2* | - | 414857 | 415141 | 0.00 | 0.00 | 3.71 | 0.00 | 4-hydroxybenzoate transporter PcaK |
| *FEPFGCPE_00366* | *sdgC-2* | - | 415214 | 416365 | 0.15 | 2.50 | 1.03 | 1.15 | Salicyloyl-CoA 5-hydroxylase |
| *FEPFGCPE_00367* | *maiA-1* | - | 416408 | 417187 | 0.44 | 3.00 | 2.71 | 0.40 | Maleate isomerase |
| *FEPFGCPE_00368* | | - | 417184 | 417672 | 0.34 | 2.68 | 0.00 | 1.42 | 1,4-dihydroxy-2-naphthoyl-CoA hydrolase |
| *FEPFGCPE_00369* | *dhbE* | - | 417669 | 419378 | 0.39 | 1.93 | 1.89 | 1.47 | 2,3-dihydroxybenzoate-AMP ligase |
| *FEPFGCPE_00370* | | - | 419408 | 419686 | 0.27 | 0.30 | 2.29 | 0.00 | Hypothetical protein |
| *FEPFGCPE_00371* | *fosB* | - | 419683 | 420330 | 0.46 | 1.48 | 2.11 | 0.00 | Metallothiol transferase FosB |
| *FEPFGCPE_00372* | | - | 420382 | 421665 | 0.71 | 1.14 | 3.22 | 0.64 | Hypothetical protein |
| *FEPFGCPE_00373* | *kstR2-1* | + | 421850 | 422533 | 3.61 | 19.57 | 13.50 | 4.31 | HTH-type transcriptional repressor KstR2 |
| *FEPFGCPE_00374* | *czcO-1* | - | 422893 | 423945 | 0.69 | 4.74 | 5.67 | 1.95 | putative oxidoreductase CzcO |
| *FEPFGCPE_00375* | *arsA* | - | 424022 | 425785 | 2.20 | 7.31 | 5.77 | 3.89 | Arsenical pump-driving ATPase |
| *FEPFGCPE_00376* | *arsD* | - | 425795 | 426640 | 4.81 | 17.34 | 20.40 | 2.20 | Arsenical resistance operon trans-acting repressor ArsD |
| *FEPFGCPE_00377* | *thlA-1* | + | 427144 | 428328 | 5.02 | 11.25 | 16.73 | 6.22 | Acetyl-CoA acetyltransferase |
| *FEPFGCPE_00378* | | + | 428373 | 428789 | 5.83 | 7.40 | 4.46 | 2.51 | Hypothetical protein |
| *FEPFGCPE_00379* | | - | 428835 | 429176 | 6.36 | 7.30 | 5.28 | 1.22 | Hypothetical protein |
| *FEPFGCPE_00380* | | - | 429163 | 429435 | 17.44 | 16.03 | 12.82 | 1.72 | Hypothetical protein |
| *FEPFGCPE_00381* | | + | 429506 | 430075 | 1.16 | 3.58 | 0.81 | 0.16 | Hypothetical protein |
| *FEPFGCPE_00382* | | - | 430459 | 431226 | 3.26 | 31.68 | 22.23 | 5.78 | Hypothetical protein |
| *FEPFGCPE_00383* | *dmlR-2* | - | 431763 | 432719 | 1.86 | 7.11 | 7.05 | 2.79 | HTH-type transcriptional regulator DmlR |
| *FEPFGCPE_00384* | *araE-1* | + | 433060 | 434496 | 0.22 | 1.26 | 0.90 | 0.00 | Alpha-ketoglutaric semialdehyde dehydrogenase |
| *FEPFGCPE_00385* | | + | 434510 | 434839 | 0.52 | 2.00 | 5.42 | 3.70 | Hypothetical protein |
| *FEPFGCPE_00386* | *ppsC-1* | + | 434854 | 435840 | 0.72 | 3.58 | 2.32 | 1.02 | Phthiocerol/phenolphthiocerol synthesis polyketide synthase type I PpsC |
| *FEPFGCPE_00387* | | - | 436100 | 436189 | 2.59 | 17.06 | 16.05 | 0.00 | tRNA |
| *FEPFGCPE_00388* | *acsA-2* | + | 436503 | 438227 | 8.49 | 23.51 | 13.78 | 3.70 | Acetyl-coenzyme A synthetase |
| *FEPFGCPE_00389* | *hemN-1* | - | 438246 | 439466 | 1.94 | 18.73 | 12.25 | 2.48 | Oxygen-independent coproporphyrinogen-III oxidase-like protein |
| *FEPFGCPE_00390* | *rdgB* | - | 439472 | 440089 | 1.79 | 9.57 | 10.14 | 4.49 | dITP/XTP pyrophosphatase |
| *FEPFGCPE_00391* | *rph* | - | 440092 | 440820 | 3.85 | 32.38 | 18.73 | 10.96 | Ribonuclease PH |
| *FEPFGCPE_00392* | | + | 441032 | 441961 | 3.88 | 33.53 | 34.88 | 15.18 | Hypothetical protein |
| *FEPFGCPE_00393* | *gmk* | + | 441967 | 442644 | 4.64 | 42.42 | 43.59 | 15.59 | Guanylate kinase |
| *FEPFGCPE_00394* | *rpoZ* | + | 442735 | 442941 | 13.76 | 152.10 | 142.88 | 37.31 | DNA-directed RNA polymerase subunit omega |
| *FEPFGCPE_00395* | *spoT* | + | 443217 | 445424 | 2.58 | 26.67 | 23.15 | 8.52 | Guanosine-3',5'-bis(diphosphate) 3'-pyrophosphohydrolase |
| *FEPFGCPE_00396* | *greB* | - | 445491 | 446054 | 4.32 | 29.64 | 20.42 | 9.99 | Transcription elongation factor GreB |
| *FEPFGCPE_00397* | | + | 446254 | 447153 | 5.20 | 25.28 | 19.06 | 6.56 | Putative quercetin 2,3-dioxygenase |
| *FEPFGCPE_00398* | *gerN* | - | 447254 | 449773 | 1.66 | 9.34 | 6.36 | 2.19 | Na(+)/H(+)-K(+) antiporter GerN |
| *FEPFGCPE_00399* | *trxA-1* | + | 449962 | 450816 | 4.72 | 26.10 | 8.50 | 9.29 | Thioredoxin 1 |
| *FEPFGCPE_00400* | | + | 450849 | 451226 | 4.71 | 22.55 | 19.64 | 2.24 | Hypothetical protein |
| *FEPFGCPE_00401* | | + | 451223 | 451810 | 3.06 | 14.18 | 12.65 | 1.43 | Hypothetical protein |
| *FEPFGCPE_00402* | *gltI-2* | + | 451921 | 453066 | 2.39 | 18.30 | 14.93 | 2.50 | Glutamate/aspartate import solute-binding protein |
| *FEPFGCPE_00403* | | + | 453503 | 453595 | 0.58 | 0.00 | 0.00 | 1.94 | Hypothetical protein |
| *FEPFGCPE_00404* | | + | 453592 | 453684 | 1.31 | 0.00 | 3.36 | 6.90 | Hypothetical protein |
| *FEPFGCPE_00405* | *kdpA* | + | 453681 | 455486 | 0.79 | 1.42 | 2.59 | 5.70 | Potassium-transporting ATPase potassium-binding subunit |
| *FEPFGCPE_00406* | *kdpB* | + | 455515 | 457641 | 0.26 | 0.96 | 2.04 | 5.33 | Potassium-transporting ATPase ATP-binding subunit |
| *FEPFGCPE_00407* | *kdpC* | + | 457719 | 458303 | 0.61 | 0.44 | 1.31 | 2.78 | Potassium-transporting ATPase KdpC subunit |
| *FEPFGCPE_00408* | *kdpD* | + | 458478 | 460808 | 1.77 | 5.56 | 4.27 | 3.30 | Sensor protein KdpD |
| *FEPFGCPE_00409* | *kdpE* | + | 461294 | 462004 | 2.55 | 14.40 | 6.02 | 1.27 | KDP operon transcriptional regulatory protein KdpE |
| *FEPFGCPE_00410* | *sugE-1* | + | 462291 | 462608 | 3.78 | 20.82 | 9.23 | 2.88 | Quaternary ammonium compound-resistance protein SugE |
| *FEPFGCPE_00411* | | + | 462889 | 463485 | 8.99 | 29.30 | 14.69 | 5.61 | Hypothetical protein |
| *FEPFGCPE_00412* | | + | 463472 | 466021 | 6.48 | 12.20 | 11.09 | 2.99 | Hypothetical protein |
| *FEPFGCPE_00413* | *yjbI* | + | 466047 | 466544 | 12.85 | 53.42 | 37.02 | 7.84 | Group 2 truncated hemoglobin YjbI |
| *FEPFGCPE_00414* | | - | 466619 | 467539 | 5.94 | 10.58 | 5.51 | 4.41 | Hypothetical protein |
| *FEPFGCPE_00415* | | - | 467562 | 468107 | 5.01 | 10.01 | 18.93 | 2.39 | Hypothetical protein |
| *FEPFGCPE_00416* | | - | 468264 | 469289 | 0.39 | 1.20 | 0.78 | 0.98 | Hypothetical protein |
| *FEPFGCPE_00417* | | - | 469316 | 469753 | 1.02 | 6.77 | 1.59 | 1.68 | Hypothetical protein |
| *FEPFGCPE_00418* | *alaS-2* | + | 470123 | 470854 | 8.87 | 17.76 | 6.71 | 4.12 | Alanine--tRNA ligase |
| *FEPFGCPE_00419* | | + | 470910 | 471521 | 5.35 | 14.92 | 8.99 | 1.37 | Hypothetical protein |
| *FEPFGCPE_00420* | | - | 471542 | 472087 | 16.74 | 26.02 | 30.86 | 10.34 | Hypothetical protein |
| *FEPFGCPE_00421* | *yvdD* | - | 472343 | 472939 | 5.47 | 27.20 | 18.25 | 5.57 | LOG family protein YvdD |
| *FEPFGCPE_00422* | *acrR-1* | - | 472943 | 473611 | 14.49 | 65.34 | 42.79 | 18.10 | HTH-type transcriptional regulator AcrR |

| Locus name | Gene name | Strand | Gene start (bp) | Gene end (bp) | Average TPM Inoculum | Average TPM $t_6$ | Average TPM $t_{24}$ | Average TPM $t_{168}$ | Annotation description |
|---|---|---|---|---|---|---|---|---|---|
| *FEPFGCPE_00423* | *dgkA* | - | 473754 | 474188 | 4.59 | 34.22 | 24.46 | 5.18 | Diacylglycerol kinase |
| *FEPFGCPE_00424* | *mgtA-1* | - | 474227 | 475261 | 3.44 | 26.45 | 15.39 | 2.52 | GDP-mannose-dependent alpha-mannosyltransferase |
| *FEPFGCPE_00425* | *lpxH-1* | - | 475467 | 476411 | 1.17 | 4.62 | 4.28 | 0.25 | UDP-2,3-diacylglucosamine hydrolase |
| *FEPFGCPE_00426* | | - | 476558 | 477514 | 2.15 | 5.28 | 6.67 | 4.05 | Hypothetical protein |
| *FEPFGCPE_00427* | | - | 477511 | 478083 | 5.77 | 25.01 | 23.53 | 5.76 | Hypothetical protein |
| *FEPFGCPE_00428* | | - | 478301 | 478741 | 110.44 | 160.79 | 75.09 | 31.75 | Hypothetical protein |
| *FEPFGCPE_00429* | | - | 479160 | 479588 | 396.41 | 388.13 | 216.96 | 84.09 | Hypothetical protein |
| *FEPFGCPE_00430* | *sigE* | - | 479585 | 480157 | 262.23 | 383.58 | 197.56 | 61.78 | RNA polymerase sigma-E factor |
| *FEPFGCPE_00431* | *ilvI* | + | 480607 | 482376 | 14.77 | 21.99 | 32.88 | 23.02 | Acetolactate synthase isozyme 3 large subunit |
| *FEPFGCPE_00432* | *ilvH* | + | 482459 | 482950 | 55.96 | 165.81 | 167.17 | 79.02 | Acetolactate synthase isozyme 3 small subunit |
| *FEPFGCPE_00433* | *ilvC* | + | 483029 | 484045 | 17.48 | 42.82 | 46.10 | 48.38 | Ketol-acid reductoisomerase (NADP(+)) |
| *FEPFGCPE_00434* | *psd* | + | 484423 | 485058 | 4.44 | 38.18 | 20.05 | 6.81 | Phosphatidylserine decarboxylase proenzyme |
| *FEPFGCPE_00435* | | + | 485216 | 486094 | 3.23 | 19.07 | 9.08 | 3.66 | Hypothetical protein |
| *FEPFGCPE_00436* | | + | 486289 | 486774 | 8.21 | 60.69 | 47.15 | 25.65 | Hypothetical protein |
| *FEPFGCPE_00437* | | - | 486929 | 487081 | 39.56 | 164.62 | 157.08 | 37.62 | Hypothetical protein |
| *FEPFGCPE_00438* | *leuA-1* | + | 487200 | 488744 | 13.99 | 80.60 | 72.55 | 29.20 | 2-isopropylmalate synthase |
| *FEPFGCPE_00439* | | - | 488850 | 489431 | 12.10 | 56.73 | 33.05 | 11.63 | Hypothetical protein |
| *FEPFGCPE_00440* | | + | 489574 | 490848 | 2.30 | 14.49 | 8.74 | 3.67 | Hypothetical protein |
| *FEPFGCPE_00441* | *rpsO* | + | 491144 | 491413 | 11.72 | 25.82 | 52.82 | 37.97 | 30S ribosomal protein S15 |
| *FEPFGCPE_00442* | *pnp* | + | 491761 | 493911 | 6.37 | 31.73 | 37.80 | 26.76 | Polyribonucleotide nucleotidyltransferase |
| *FEPFGCPE_00443* | | + | 494095 | 494170 | 80.37 | 109.55 | 59.67 | 30.60 | tRNA |
| *FEPFGCPE_00444* | | + | 494334 | 494409 | 15.96 | 129.07 | 83.67 | 22.80 | tRNA |
| *FEPFGCPE_00445* | | + | 494507 | 494583 | 18.41 | 55.97 | 47.99 | 26.15 | tRNA |
| *FEPFGCPE_00446* | | + | 494683 | 494759 | 12.79 | 27.62 | 24.08 | 18.93 | tRNA |
| *FEPFGCPE_00447* | *norM* | - | 495014 | 496363 | 1.71 | 8.46 | 7.20 | 0.91 | Multidrug resistance protein NorM |
| *FEPFGCPE_00448* | | - | 496401 | 498131 | 1.37 | 6.33 | 5.30 | 2.13 | Hypothetical protein |
| *FEPFGCPE_00449* | *rpmE2* | - | 498438 | 498704 | 19.02 | 47.66 | 72.13 | 127.96 | 50S ribosomal protein L31 type B |
| *FEPFGCPE_00450* | *mtfA* | - | 498892 | 499743 | 1.52 | 6.05 | 7.81 | 3.82 | Protein MtfA |
| *FEPFGCPE_00451* | | - | 499882 | 501303 | 2.71 | 14.65 | 6.89 | 1.08 | putative FAD-linked oxidoreductase |
| *FEPFGCPE_00452* | | - | 501369 | 501752 | 4.14 | 9.83 | 18.90 | 5.24 | Hypothetical protein |
| *FEPFGCPE_00453* | | - | 501915 | 502502 | 1.91 | 9.36 | 10.24 | 8.20 | NAD(P)H dehydrogenase (quinone) |
| *FEPFGCPE_00454* | | + | 502841 | 504157 | 3.13 | 18.38 | 13.72 | 3.82 | Hypothetical protein |
| *FEPFGCPE_00455* | | - | 504147 | 504470 | 5.09 | 35.05 | 30.85 | 7.18 | Hypothetical protein |
| *FEPFGCPE_00456* | *ybfF* | - | 504503 | 505318 | 1.26 | 8.40 | 9.43 | 3.41 | Esterase YbfF |
| *FEPFGCPE_00457* | *megL* | - | 505315 | 506622 | 2.74 | 13.38 | 11.78 | 4.19 | L-methionine gamma-lyase |
| *FEPFGCPE_00458* | *hrp1* | + | 507034 | 507489 | 16.13 | 72.80 | 38.74 | 11.99 | Hypoxic response protein 1 |
| *FEPFGCPE_00459* | *lplT-1* | + | 507486 | 509405 | 3.37 | 13.44 | 9.52 | 2.20 | Lysophospholipid transporter LplT |
| *FEPFGCPE_00460* | *aroC* | + | 509586 | 510689 | 5.02 | 36.88 | 30.67 | 12.70 | Chorismate synthase |
| *FEPFGCPE_00461* | | - | 510763 | 512433 | 10.13 | 55.82 | 44.80 | 15.83 | Electron transfer flavoprotein-ubiquinone oxidoreductase |
| *FEPFGCPE_00462* | *gno-1* | + | 512612 | 513388 | 11.95 | 54.26 | 37.40 | 14.17 | Gluconate 5-dehydrogenase |
| *FEPFGCPE_00463* | *dctP* | + | 513568 | 514632 | 2.83 | 10.86 | 8.95 | 0.97 | C4-dicarboxylate-binding periplasmic protein DctP |
| *FEPFGCPE_00464* | | + | 514851 | 515228 | 4.06 | 38.01 | 27.10 | 9.22 | Hypothetical protein |
| *FEPFGCPE_00465* | *uhpA* | - | 515280 | 515825 | 6.29 | 40.48 | 24.27 | 12.52 | Transcriptional regulatory protein UhpA |
| *FEPFGCPE_00466* | *scoA* | + | 516054 | 516752 | 8.96 | 41.62 | 23.45 | 26.76 | putative succinyl-CoA:3-ketoacid coenzyme A transferase subunit A |
| *FEPFGCPE_00467* | *scoB* | + | 516776 | 517414 | 7.82 | 25.23 | 28.43 | 17.69 | putative succinyl-CoA:3-ketoacid coenzyme A transferase subunit B |
| *FEPFGCPE_00468* | | + | 517690 | 518568 | 4.86 | 43.27 | 36.03 | 17.65 | Tropinesterase |
| *FEPFGCPE_00469* | *yabJ-3* | + | 518635 | 518991 | 4.46 | 68.55 | 40.64 | 13.93 | 2-iminobutanoate/2-iminopropanoate deaminase |
| *FEPFGCPE_00470* | *relA* | + | 519040 | 521337 | 3.00 | 24.93 | 17.71 | 8.55 | GTP pyrophosphokinase |
| *FEPFGCPE_00471* | | + | 521412 | 521488 | 19.72 | 114.48 | 116.90 | 50.77 | tRNA |
| *FEPFGCPE_00472* | *thrS* | + | 521726 | 523633 | 20.91 | 72.75 | 62.60 | 25.48 | Threonine--tRNA ligase |
| *FEPFGCPE_00473* | *infC* | + | 523736 | 524203 | 26.52 | 85.78 | 99.95 | 88.46 | Translation initiation factor IF-3 |
| *FEPFGCPE_00474* | *rpmI* | + | 524380 | 524577 | 21.63 | 143.92 | 156.60 | 127.27 | 50S ribosomal protein L35 |
| *FEPFGCPE_00475* | *rplT* | + | 524603 | 524962 | 8.43 | 62.01 | 55.21 | 43.10 | 50S ribosomal protein L20 |
| *FEPFGCPE_00476* | *pheS* | + | 525114 | 526127 | 1.56 | 16.24 | 12.85 | 7.70 | Phenylalanine--tRNA ligase alpha subunit |
| *FEPFGCPE_00477* | *pheT* | + | 526214 | 528652 | 3.41 | 27.18 | 21.30 | 8.82 | Phenylalanine--tRNA ligase beta subunit |
| *FEPFGCPE_00478* | *ihfA* | + | 528724 | 529107 | 13.58 | 134.65 | 119.24 | 43.03 | Integration host factor subunit alpha |
| *FEPFGCPE_00479* | *carA-2* | + | 529187 | 529573 | 2.94 | 8.57 | 17.21 | 4.24 | HTH-type transcriptional repressor CarA |
| *FEPFGCPE_00480* | *gmr-1* | + | 529985 | 531571 | 2.29 | 12.91 | 6.60 | 1.66 | Cyclic di-GMP phosphodiesterase Gmr |
| *FEPFGCPE_00481* | | + | 531581 | 532255 | 3.58 | 18.21 | 12.18 | 1.46 | putative signaling protein |
| *FEPFGCPE_00482* | *pleD-1* | + | 532720 | 534303 | 4.38 | 19.18 | 11.74 | 1.84 | Response regulator PleD |
| *FEPFGCPE_00483* | | + | 534482 | 534724 | 15.51 | 118.51 | 90.40 | 51.75 | Hypothetical protein |
| *FEPFGCPE_00484* | *rubA-1* | - | 534756 | 534950 | 13.59 | 86.91 | 62.88 | 19.25 | Rubredoxin |
| *FEPFGCPE_00485* | | - | 535141 | 535452 | 11.73 | 68.75 | 69.03 | 25.91 | Hypothetical protein |
| *FEPFGCPE_00486* | | - | 535900 | 536226 | 5.84 | 26.20 | 45.15 | 15.46 | Hypothetical protein |
| *FEPFGCPE_00487* | | - | 536305 | 536769 | 2.05 | 13.98 | 11.43 | 0.13 | Hypothetical protein |
| *FEPFGCPE_00488* | | + | 537097 | 537330 | 14.74 | 74.00 | 60.76 | 22.04 | Hypothetical protein |
| *FEPFGCPE_00489* | *dosP* | - | 537342 | 539972 | 3.84 | 17.36 | 12.63 | 6.43 | Oxygen sensor protein DosP |
| *FEPFGCPE_00490* | | - | 540093 | 540311 | 7.61 | 47.73 | 24.06 | 5.02 | Hypothetical protein |
| *FEPFGCPE_00491* | | - | 540379 | 540960 | 4.08 | 24.95 | 24.90 | 3.22 | Hypothetical protein |
| *FEPFGCPE_00492* | *adrA* | - | 540995 | 542347 | 13.66 | 84.40 | 47.28 | 12.92 | putative diguanylate cyclase AdrA |
| *FEPFGCPE_00493* | | + | 542650 | 543657 | 3.77 | 20.83 | 9.27 | 4.40 | Hypothetical protein |

| Locus name | Gene name | Strand | Gene start (bp) | Gene end (bp) | Average TPM Inoculum | Average TPM $t_6$ | Average TPM $t_{24}$ | Average TPM $t_{168}$ | Annotation description |
|---|---|---|---|---|---|---|---|---|---|
| *FEPFGCPE_00494* | | + | 543788 | 544093 | 20.95 | 233.23 | 123.07 | 22.80 | Hypothetical protein |
| *FEPFGCPE_00495* | | + | 544133 | 544300 | 7.42 | 66.67 | 29.01 | 3.64 | Hypothetical protein |
| *FEPFGCPE_00496* | | + | 544297 | 544788 | 6.89 | 48.99 | 38.20 | 8.03 | Hypothetical protein |
| *FEPFGCPE_00497* | | - | 544866 | 545993 | 8.51 | 46.88 | 32.70 | 17.32 | Soluble hydrogenase 42 kDa subunit |
| *FEPFGCPE_00498* | *yfcG-1* | + | 546279 | 546917 | 2.61 | 28.43 | 15.13 | 5.30 | Disulfide-bond oxidoreductase YfcG |
| *FEPFGCPE_00499* | | - | 546982 | 547458 | 2.10 | 20.68 | 8.31 | 4.93 | Hypothetical protein |
| *FEPFGCPE_00500* | *kipR-2* | - | 547488 | 548357 | 2.27 | 20.61 | 6.06 | 2.43 | HTH-type transcriptional regulator KipR |
| *FEPFGCPE_00501* | *potA-3* | + | 548655 | 549755 | 0.98 | 2.05 | 0.81 | 1.63 | Spermidine/putrescine import ATP-binding protein PotA |
| *FEPFGCPE_00502* | *potB-2* | + | 549748 | 550653 | 1.03 | 3.79 | 4.36 | 1.21 | Spermidine/putrescine transport system permease protein PotB |
| *FEPFGCPE_00503* | *ydcV-2* | + | 550697 | 551491 | 0.94 | 3.23 | 1.80 | 0.83 | Inner membrane ABC transporter permease protein YdcV |
| *FEPFGCPE_00504* | *hcnA-2* | + | 551515 | 551910 | 0.28 | 1.01 | 0.00 | 2.24 | Hydrogen cyanide synthase subunit HcnA |
| *FEPFGCPE_00505* | *hcnB-2* | + | 551903 | 553327 | 0.35 | 2.44 | 1.98 | 0.39 | Hydrogen cyanide synthase subunit HcnB |
| *FEPFGCPE_00506* | *thiO* | + | 553320 | 554456 | 0.51 | 5.24 | 3.25 | 1.33 | Glycine oxidase |
| *FEPFGCPE_00507* | *phnS-2* | + | 554536 | 555582 | 0.68 | 9.47 | 4.80 | 4.51 | Putative 2-aminoethylphosphonate-binding periplasmic protein |
| *FEPFGCPE_00508* | | - | 555736 | 556131 | 4.22 | 29.89 | 11.87 | 9.63 | Hypothetical protein |
| *FEPFGCPE_00509* | | + | 556242 | 556991 | 7.72 | 23.36 | 18.19 | 4.73 | Hypothetical protein |
| *FEPFGCPE_00510* | | - | 557027 | 557272 | 8.95 | 54.64 | 47.81 | 5.66 | Hypothetical protein |
| *FEPFGCPE_00511* | *sir* | + | 557556 | 559562 | 3.34 | 200.28 | 91.03 | 51.41 | Sulfite reductase [ferredoxin] |
| *FEPFGCPE_00512* | | + | 559552 | 560028 | 3.84 | 310.34 | 138.49 | 57.95 | Hypothetical protein |
| *FEPFGCPE_00513* | | - | 560250 | 560326 | 8.23 | 18.93 | 24.24 | 17.98 | tRNA |
| *FEPFGCPE_00514* | *acsA-3* | + | 560561 | 562249 | 10.94 | 52.34 | 15.81 | 5.76 | Acetyl-coenzyme A synthetase |
| *FEPFGCPE_00515* | | - | 562305 | 563078 | 1.34 | 16.57 | 8.55 | 1.95 | Hypothetical protein |
| *FEPFGCPE_00516* | *gshA* | + | 563351 | 564913 | 4.15 | 22.22 | 18.12 | 4.85 | Glutamate--cysteine ligase |
| *FEPFGCPE_00517* | | - | 565002 | 565325 | 0.63 | 0.00 | 1.48 | 0.00 | Hypothetical protein |
| *FEPFGCPE_00518* | *ygaZ* | - | 565322 | 566035 | 0.81 | 4.26 | 1.57 | 3.87 | Inner membrane protein YgaZ |
| *FEPFGCPE_00519* | *mrcA-1* | - | 566245 | 568752 | 1.80 | 10.75 | 9.81 | 5.96 | Penicillin-binding protein 1A |
| *FEPFGCPE_00520* | | - | 569021 | 569323 | 2.35 | 16.57 | 8.99 | 3.07 | Hypothetical protein |
| *FEPFGCPE_00521* | *tsr-1* | + | 569535 | 571157 | 2.55 | 13.43 | 5.41 | 2.11 | Methyl-accepting chemotaxis protein I |
| *FEPFGCPE_00522* | | + | 571289 | 571696 | 15.87 | 61.62 | 39.53 | 5.73 | Hypothetical protein |
| *FEPFGCPE_00523* | *cysI* | - | 571823 | 573547 | 3.06 | 104.28 | 52.31 | 22.87 | Sulfite reductase [NADPH] hemoprotein beta-component |
| *FEPFGCPE_00524* | *cysJ-1* | - | 573637 | 575469 | 2.45 | 168.24 | 89.78 | 48.49 | Sulfite reductase [NADPH] flavoprotein alpha-component |
| *FEPFGCPE_00525* | *sucA* | + | 575882 | 578743 | 12.74 | 71.79 | 67.85 | 57.39 | 2-oxoglutarate dehydrogenase E1 component |
| *FEPFGCPE_00526* | *sucB* | + | 578834 | 580096 | 6.60 | 32.58 | 27.76 | 26.12 | Dihydrolipoyllysine-residue succinyltransferase component of 2-oxoglutarate dehydrogenase complex |
| *FEPFGCPE_00527* | *lpdG* | + | 580194 | 581621 | 2.93 | 21.94 | 28.41 | 18.50 | Dihydrolipoyl dehydrogenase |
| *FEPFGCPE_00528* | *zapE-1* | + | 581715 | 582812 | 1.47 | 11.08 | 11.93 | 4.89 | Cell division protein ZapE |
| *FEPFGCPE_00529* | | - | 582852 | 583178 | 10.78 | 108.53 | 84.48 | 29.02 | Hypothetical protein |
| *FEPFGCPE_00530* | | + | 583398 | 584048 | 0.96 | 4.15 | 12.99 | 1.63 | Hypothetical protein |
| *FEPFGCPE_00531* | | + | 584225 | 584707 | 25.00 | 63.11 | 26.27 | 20.68 | Hypothetical protein |
| *FEPFGCPE_00532* | | - | 585136 | 586368 | 0.59 | 71.72 | 23.68 | 13.52 | Hypothetical protein |
| *FEPFGCPE_00533* | | - | 586409 | 587140 | 0.41 | 50.53 | 18.38 | 9.96 | Hypothetical protein |
| *FEPFGCPE_00534* | | - | 587208 | 588371 | 0.11 | 69.78 | 24.53 | 14.11 | Hypothetical protein |
| *FEPFGCPE_00535* | | - | 588411 | 590120 | 0.15 | 83.62 | 26.07 | 15.63 | Hypothetical protein |
| *FEPFGCPE_00536* | | - | 590149 | 590469 | 1.19 | 155.19 | 36.70 | 31.11 | Hypothetical protein |
| *FEPFGCPE_00537* | | + | 591394 | 591579 | 151.70 | 528.29 | 369.88 | 113.09 | Hypothetical protein |
| *FEPFGCPE_00538* | | + | 591725 | 591925 | 54.50 | 179.16 | 71.08 | 21.24 | Hypothetical protein |
| *FEPFGCPE_00539* | | + | 591939 | 592418 | 85.46 | 305.91 | 190.36 | 58.52 | Hypothetical protein |
| *FEPFGCPE_00540* | | + | 592630 | 593106 | 17.71 | 47.09 | 18.28 | 3.58 | Hypothetical protein |
| *FEPFGCPE_00541* | | + | 593262 | 594284 | 14.66 | 54.03 | 28.99 | 5.24 | Hypothetical protein |
| *FEPFGCPE_00542* | *outD* | + | 594359 | 595747 | 9.17 | 45.66 | 19.59 | 1.78 | Type II secretion system protein D |
| *FEPFGCPE_00543* | | + | 595824 | 597164 | 4.06 | 34.68 | 12.97 | 4.00 | Hypothetical protein |
| *FEPFGCPE_00544* | | + | 597166 | 598542 | 2.25 | 30.11 | 12.82 | 1.98 | Putative conjugal transfer protein |
| *FEPFGCPE_00545* | | + | 598654 | 599460 | 1.23 | 19.08 | 4.67 | 1.35 | Hypothetical protein |
| *FEPFGCPE_00546* | | + | 599480 | 600499 | 1.34 | 26.35 | 11.12 | 1.24 | Hypothetical protein |
| *FEPFGCPE_00547* | *bepA-1* | + | 600496 | 601473 | 1.57 | 31.86 | 14.19 | 4.04 | Beta-barrel assembly-enhancing protease |
| *FEPFGCPE_00548* | | + | 601547 | 602083 | 1.16 | 52.36 | 16.31 | 6.32 | Hypothetical protein |
| *FEPFGCPE_00549* | | + | 602126 | 604174 | 0.48 | 22.78 | 7.35 | 2.52 | Hypothetical protein |
| *FEPFGCPE_00550* | *hyfR-1* | + | 604305 | 605720 | 0.70 | 20.62 | 7.89 | 2.39 | DNA-binding transcriptional activator HyfR |
| *FEPFGCPE_00551* | | + | 606162 | 606893 | 6.45 | 38.11 | 23.36 | 8.29 | Hypothetical protein |
| *FEPFGCPE_00552* | *kup* | + | 607308 | 609200 | 2.88 | 12.27 | 6.60 | 2.83 | Low affinity potassium transport system protein kup |
| *FEPFGCPE_00553* | *ypfD* | - | 609442 | 611793 | 1.46 | 15.88 | 8.34 | 7.18 | 30S ribosomal protein S1 |
| *FEPFGCPE_00554* | | + | 611866 | 612777 | 14.05 | 74.62 | 57.01 | 19.86 | Hypothetical protein |
| *FEPFGCPE_00555* | | + | 612823 | 613026 | 19.46 | 70.43 | 66.32 | 17.05 | Hypothetical protein |
| *FEPFGCPE_00556* | *dinG* | + | 613112 | 615370 | 1.64 | 9.55 | 10.48 | 2.72 | putative ATP-dependent helicase DinG |
| *FEPFGCPE_00557* | *bamD* | - | 615485 | 616309 | 31.75 | 173.50 | 143.56 | 53.93 | Outer membrane protein assembly factor BamD |
| *FEPFGCPE_00558* | *rluD* | + | 616331 | 617491 | 3.12 | 14.59 | 13.66 | 5.03 | Ribosomal large subunit pseudouridine synthase D |
| *FEPFGCPE_00559* | *yfiH* | + | 617502 | 618293 | 3.75 | 17.37 | 19.93 | 5.87 | Laccase domain protein YfiH |
| *FEPFGCPE_00560* | | - | 618332 | 618577 | 13.01 | 48.73 | 41.92 | 8.40 | Hypothetical protein |
| *FEPFGCPE_00561* | *phaC-1* | + | 618772 | 620622 | 9.79 | 40.62 | 27.77 | 15.42 | Poly(3-hydroxyalkanoate) polymerase subunit PhaC |
| *FEPFGCPE_00562* | *phaA* | + | 620684 | 621865 | 22.51 | 156.12 | 107.96 | 61.98 | Acetyl-CoA acetyltransferase |
| *FEPFGCPE_00563* | *phaB-1* | + | 622047 | 622787 | 45.14 | 243.04 | 175.63 | 73.84 | Acetoacetyl-CoA reductase |
| *FEPFGCPE_00564* | | + | 622952 | 623527 | 33.59 | 179.87 | 111.64 | 45.18 | Hypothetical protein |

| Locus name | Gene name | Strand | Gene start (bp) | Gene end (bp) | Average TPM Inoculum | Average TPM $t_6$ | Average TPM $t_{24}$ | Average TPM $t_{168}$ | Annotation description |
|---|---|---|---|---|---|---|---|---|---|
| *FEPFGCPE_00565* | *rimO* | + | 623784 | 625151 | 5.74 | 14.32 | 12.61 | 7.18 | Ribosomal protein S12 methylthiotransferase RimO |
| *FEPFGCPE_00566* | *kdgK* | + | 625151 | 626089 | 2.14 | 2.33 | 5.22 | 4.41 | 2-dehydro-3-deoxygluconokinase |
| *FEPFGCPE_00567* | *bktB* | + | 626356 | 627537 | 15.44 | 44.20 | 21.14 | 13.41 | Beta-ketothiolase BktB |
| *FEPFGCPE_00568* | *metC-1* | + | 627590 | 628807 | 6.70 | 34.50 | 19.80 | 7.65 | Cystathionine beta-lyase MetC |
| *FEPFGCPE_00569* | *serB* | - | 628866 | 629759 | 2.12 | 14.46 | 8.19 | 2.16 | Phosphoserine phosphatase |
| *FEPFGCPE_00570* | *cdhR-1* | - | 629894 | 630910 | 4.37 | 18.40 | 9.34 | 2.18 | HTH-type transcriptional regulator CdhR |
| *FEPFGCPE_00571* | | - | 630969 | 631535 | 4.13 | 15.99 | 8.85 | 2.53 | Hypothetical protein |
| *FEPFGCPE_00572* | | - | 631695 | 632720 | 1.81 | 16.61 | 12.12 | 4.73 | Delta(1)-pyrroline-2-carboxylate reductase |
| *FEPFGCPE_00573* | *psdht* | - | 632846 | 633934 | 3.66 | 21.83 | 19.81 | 7.80 | Phenylserine dehydratase |
| *FEPFGCPE_00574* | *proP-2* | - | 633976 | 635262 | 3.49 | 38.94 | 27.42 | 6.94 | Proline/betaine transporter |
| *FEPFGCPE_00575* | *argE* | - | 635453 | 636670 | 2.37 | 30.30 | 17.37 | 8.41 | Acetylornithine deacetylase |
| *FEPFGCPE_00576* | *mfd* | - | 636828 | 640316 | 3.30 | 17.40 | 12.94 | 5.87 | Transcription-repair-coupling factor |
| *FEPFGCPE_00577* | *ispD* | + | 640500 | 641123 | 3.82 | 36.28 | 21.05 | 12.24 | 2-C-methyl-D-erythritol 4-phosphate cytidylyltransferase |
| *FEPFGCPE_00578* | *ispF* | + | 641224 | 641709 | 1.86 | 19.18 | 8.31 | 3.17 | 2-C-methyl-D-erythritol 2,4-cyclodiphosphate synthase |
| *FEPFGCPE_00579* | *ahpD-1* | - | 641861 | 642388 | 1.97 | 43.50 | 66.86 | 36.35 | Alkyl hydroperoxide reductase AhpD |
| *FEPFGCPE_00580* | | - | 642448 | 642996 | 46.66 | 180.60 | 221.68 | 238.81 | Alkyl hydroperoxide reductase subunit C |
| *FEPFGCPE_00581* | | - | 643102 | 643206 | 47.94 | 367.87 | 419.36 | 86.87 | Hypothetical protein |
| *FEPFGCPE_00582* | *envZ-2* | - | 643227 | 644567 | 1.60 | 12.97 | 4.73 | 2.46 | Osmolarity sensor protein EnvZ |
| *FEPFGCPE_00583* | *ompR-2* | - | 644748 | 645473 | 24.36 | 93.10 | 67.59 | 22.73 | Transcriptional regulatory protein OmpR |
| *FEPFGCPE_00584* | | + | 645643 | 645972 | 12.78 | 46.77 | 53.56 | 16.38 | Hypothetical protein |
| *FEPFGCPE_00585* | | + | 646084 | 647832 | 6.96 | 26.15 | 13.48 | 6.10 | Hypothetical protein |
| *FEPFGCPE_00586* | | + | 647964 | 649190 | 26.92 | 70.56 | 37.37 | 13.40 | Hypothetical protein |
| *FEPFGCPE_00587* | *crtB-1* | - | 649329 | 650204 | 15.70 | 63.78 | 42.86 | 11.89 | All-trans-phytoene synthase |
| *FEPFGCPE_00588* | | + | 650521 | 650607 | 0.00 | 6.43 | 5.71 | 0.00 | tRNA |
| *FEPFGCPE_00589* | | - | 650712 | 651014 | 1.98 | 5.40 | 6.34 | 0.69 | Hypothetical protein |
| *FEPFGCPE_00590* | *yheS-1* | - | 651356 | 653035 | 0.54 | 3.92 | 1.06 | 1.88 | putative ABC transporter ATP-binding protein YheS |
| *FEPFGCPE_00591* | | + | 653441 | 654802 | 6.70 | 6.62 | 3.06 | 2.47 | Hypothetical protein |
| *FEPFGCPE_00592* | | + | 654962 | 655216 | 8.62 | 54.32 | 49.57 | 14.67 | Hypothetical protein |
| *FEPFGCPE_00593* | *map-1* | + | 655216 | 656046 | 6.02 | 27.19 | 16.39 | 9.97 | Methionine aminopeptidase |
| *FEPFGCPE_00594* | | - | 656084 | 656911 | 4.88 | 16.16 | 12.40 | 8.14 | Hypothetical protein |
| *FEPFGCPE_00595* | *guaB-2* | - | 657180 | 658301 | 0.77 | 3.35 | 4.18 | 0.74 | Inosine-5'-monophosphate dehydrogenase |
| *FEPFGCPE_00596* | *mepC-1* | - | 658670 | 660250 | 2.02 | 6.95 | 5.59 | 2.55 | Multidrug/solvent efflux pump outer membrane protein MepC |
| *FEPFGCPE_00597* | *bepE* | - | 660263 | 663448 | 1.38 | 5.73 | 3.48 | 1.83 | Efflux pump membrane transporter BepE |
| *FEPFGCPE_00598* | *bepF* | - | 663513 | 664826 | 2.26 | 10.34 | 3.48 | 7.87 | Efflux pump periplasmic linker BepF |
| *FEPFGCPE_00599* | *nlhH-1* | - | 664856 | 665761 | 1.72 | 17.49 | 12.62 | 4.51 | Carboxylesterase NlhH |
| *FEPFGCPE_00600* | *pgrR-1* | + | 666003 | 667016 | 3.93 | 18.47 | 14.89 | 11.05 | HTH-type transcriptional regulator PgrR |
| *FEPFGCPE_00601* | | + | 667522 | 668520 | 1.12 | 2.20 | 2.21 | 0.88 | Hypothetical protein |
| *FEPFGCPE_00602* | *nadE-1* | + | 668672 | 669733 | 0.40 | 2.18 | 1.31 | 1.15 | NH(3)-dependent NAD(+) synthetase |
| *FEPFGCPE_00603* | | + | 669739 | 670296 | 0.72 | 1.68 | 3.15 | 1.33 | Hypothetical protein |
| *FEPFGCPE_00604* | | + | 670329 | 670691 | 2.68 | 6.73 | 3.61 | 2.91 | Hypothetical protein |
| *FEPFGCPE_00605* | *chuR* | + | 670717 | 671892 | 5.45 | 28.58 | 8.50 | 3.37 | Anaerobic sulfatase-maturating enzyme |
| *FEPFGCPE_00606* | *cphA-1* | - | 671946 | 674522 | 31.40 | 81.24 | 29.37 | 10.55 | Cyanophycin synthetase |
| *FEPFGCPE_00607* | *cphA-2* | - | 674675 | 677329 | 77.19 | 94.71 | 39.47 | 10.43 | Cyanophycin synthetase |
| *FEPFGCPE_00608* | | + | 677606 | 680008 | 3.32 | 12.93 | 6.17 | 2.55 | putative ABC transporter ATP-binding protein |
| *FEPFGCPE_00609* | | + | 680027 | 680494 | 4.46 | 13.11 | 7.53 | 0.30 | Hypothetical protein |
| *FEPFGCPE_00610* | *rdgC* | + | 680567 | 681490 | 1.40 | 20.47 | 17.66 | 15.89 | Recombination-associated protein RdgC |
| *FEPFGCPE_00611* | | - | 681587 | 682291 | 1.73 | 6.10 | 11.10 | 3.64 | putative oxidoreductase |
| *FEPFGCPE_00612* | *bepR* | + | 682516 | 683118 | 2.67 | 2.30 | 6.67 | 5.01 | HTH-type transcriptional repressor BepR |
| *FEPFGCPE_00613* | *dmlR-3* | - | 683179 | 684084 | 1.39 | 2.73 | 3.03 | 2.80 | HTH-type transcriptional regulator DmlR |
| *FEPFGCPE_00614* | *entA* | + | 684269 | 684982 | 0.82 | 7.30 | 5.34 | 1.74 | 2,3-dihydro-2,3-dihydroxybenzoate dehydrogenase |
| *FEPFGCPE_00615* | | + | 685205 | 685354 | 70.86 | 280.80 | 145.76 | 71.50 | Hypothetical protein |
| *FEPFGCPE_00616* | | - | 685454 | 685642 | 184.56 | 1040.31 | 847.58 | 177.21 | Hypothetical protein |
| *FEPFGCPE_00617* | *minC* | + | 685992 | 686744 | 6.63 | 47.23 | 53.74 | 22.90 | Septum site-determining protein MinC |
| *FEPFGCPE_00618* | *minD* | + | 686788 | 687603 | 5.60 | 52.64 | 55.00 | 32.50 | Septum site-determining protein MinD |
| *FEPFGCPE_00619* | *minE* | + | 687613 | 687867 | 3.33 | 29.93 | 15.86 | 16.09 | Cell division topological specificity factor |
| *FEPFGCPE_00620* | | - | 688052 | 689050 | 1.22 | 8.60 | 11.18 | 4.83 | Hypothetical protein |
| *FEPFGCPE_00621* | *fadD-1* | - | 689065 | 690780 | 2.96 | 8.04 | 6.97 | 3.61 | Long-chain-fatty-acid--CoA ligase |
| *FEPFGCPE_00622* | *fadI* | - | 690898 | 692217 | 11.07 | 14.44 | 17.10 | 5.95 | 3-ketoacyl-CoA thiolase |
| *FEPFGCPE_00623* | | - | 692214 | 693167 | 10.59 | 11.97 | 12.28 | 5.15 | Hypothetical protein |
| *FEPFGCPE_00624* | *fabG-1* | - | 693164 | 694615 | 17.79 | 9.95 | 7.43 | 1.14 | 3-oxoacyl-[acyl-carrier-protein] reductase FabG |
| *FEPFGCPE_00625* | *fadE* | - | 694661 | 697162 | 52.60 | 23.24 | 21.95 | 10.38 | Acyl-coenzyme A dehydrogenase |
| *FEPFGCPE_00626* | *ttgW* | - | 697342 | 698172 | 150.03 | 52.85 | 35.15 | 16.95 | putative HTH-type transcriptional regulator TtgW |
| *FEPFGCPE_00627* | *tig* | + | 698616 | 699956 | 11.73 | 43.42 | 42.42 | 36.64 | Trigger factor |
| *FEPFGCPE_00628* | *clpP* | + | 700080 | 700736 | 16.52 | 124.34 | 76.18 | 38.21 | ATP-dependent Clp protease proteolytic subunit |
| *FEPFGCPE_00629* | *clpX* | + | 700973 | 702253 | 26.62 | 122.94 | 85.47 | 35.06 | ATP-dependent Clp protease ATP-binding subunit ClpX |
| *FEPFGCPE_00630* | *lon-1* | + | 702450 | 704870 | 119.62 | 223.10 | 118.85 | 56.68 | Lon protease |
| *FEPFGCPE_00631* | *hupB-1* | + | 705172 | 705444 | 99.00 | 501.92 | 323.48 | 148.04 | DNA-binding protein HU-beta |
| *FEPFGCPE_00632* | | + | 705501 | 705576 | 73.88 | 357.29 | 239.84 | 106.35 | tRNA |
| *FEPFGCPE_00633* | | + | 705591 | 705667 | 15.62 | 31.55 | 25.57 | 18.28 | tRNA |
| *FEPFGCPE_00634* | *ppiD-1* | + | 705750 | 707636 | 4.40 | 14.69 | 10.13 | 6.09 | Peptidyl-prolyl cis-trans isomerase D |
| *FEPFGCPE_00635* | *tesA* | - | 707731 | 708378 | 3.97 | 13.76 | 8.37 | 3.37 | Esterase TesA |

| Locus name | Gene name | Strand | Gene start (bp) | Gene end (bp) | Average TPM Inoculum | Average TPM $t_6$ | Average TPM $t_{24}$ | Average TPM $t_{168}$ | Annotation description |
|---|---|---|---|---|---|---|---|---|---|
| *FEPFGCPE_00636* | *ytrE* | + | 708431 | 709153 | 10.83 | 21.38 | 15.41 | 4.94 | ABC transporter ATP-binding protein YtrE |
| *FEPFGCPE_00637* | *pgi* | - | 709177 | 710844 | 6.09 | 29.37 | 26.61 | 8.98 | Glucose-6-phosphate isomerase |
| *FEPFGCPE_00638* | *nnr* | - | 710965 | 712695 | 5.32 | 25.32 | 14.19 | 6.75 | Bifunctional NAD(P)H-hydrate repair enzyme Nnr |
| *FEPFGCPE_00639* | *dadA1-3* | - | 712991 | 714265 | 1.13 | 11.71 | 3.55 | 0.68 | D-amino acid dehydrogenase 1 |
| *FEPFGCPE_00640* | *purL* | + | 714519 | 718550 | 3.81 | 15.60 | 19.62 | 12.86 | Phosphoribosylformylglycinamidine synthase |
| *FEPFGCPE_00641* | | - | 718658 | 719356 | 2.22 | 19.65 | 13.37 | 11.27 | putative parvulin-type peptidyl-prolyl cis-trans isomerase |
| *FEPFGCPE_00642* | | - | 719353 | 719451 | 7.80 | 38.90 | 55.40 | 26.84 | Hypothetical protein |
| *FEPFGCPE_00643* | *bolA* | - | 719522 | 719785 | 14.30 | 93.39 | 62.17 | 27.63 | DNA-binding transcriptional regulator BolA |
| *FEPFGCPE_00644* | *yciB* | - | 719808 | 720341 | 5.42 | 26.52 | 21.16 | 15.38 | Intracellular septation protein |
| *FEPFGCPE_00645* | | - | 720545 | 720964 | 2.62 | 13.53 | 15.30 | 7.60 | Hypothetical protein |
| *FEPFGCPE_00646* | *msrB-1* | - | 721144 | 721566 | 4.87 | 26.87 | 22.51 | 6.89 | Peptide methionine sulfoxide reductase MsrB |
| *FEPFGCPE_00647* | | - | 721730 | 723247 | 5.88 | 29.32 | 20.49 | 9.25 | Hypothetical protein |
| *FEPFGCPE_00648* | *dmdB* | + | 723526 | 725154 | 7.90 | 28.93 | 21.53 | 7.99 | 3-methylmercaptopropionyl-CoA ligase |
| *FEPFGCPE_00649* | *livH-3* | + | 725376 | 726359 | 1.91 | 11.59 | 8.09 | 1.74 | High-affinity branched-chain amino acid transport system permease protein LivH |
| *FEPFGCPE_00650* | | + | 726359 | 727639 | 2.67 | 10.80 | 8.06 | 2.11 | Hypothetical protein |
| *FEPFGCPE_00651* | *lptB-3* | + | 727636 | 728469 | 2.45 | 20.36 | 6.93 | 2.20 | Lipopolysaccharide export system ATP-binding protein LptB |
| *FEPFGCPE_00652* | *livF-3* | + | 728462 | 729184 | 1.72 | 12.51 | 9.07 | 0.71 | High-affinity branched-chain amino acid transport ATP-binding protein LivF |
| *FEPFGCPE_00653* | *fadB* | + | 729298 | 731394 | 2.59 | 19.95 | 8.87 | 3.46 | Fatty acid oxidation complex subunit alpha |
| *FEPFGCPE_00654* | *pncA* | + | 731456 | 732061 | 3.37 | 20.73 | 12.52 | 6.83 | Nicotinamidase |
| *FEPFGCPE_00655* | | + | 732175 | 732615 | 4.43 | 30.76 | 21.09 | 3.96 | Hypothetical protein |
| *FEPFGCPE_00656* | *curA* | + | 732737 | 733732 | 2.53 | 22.51 | 13.93 | 5.91 | NADPH-dependent curcumin reductase |
| *FEPFGCPE_00657* | | + | 733869 | 734636 | 0.77 | 8.49 | 2.74 | 0.51 | putative S-adenosylmethionine-dependent methyltransferase/MSMEI_2290 |
| *FEPFGCPE_00658* | *fabB* | - | 734712 | 735959 | 0.99 | 2.27 | 2.07 | 1.46 | 3-oxoacyl-[acyl-carrier-protein] synthase 1 |
| *FEPFGCPE_00659* | | - | 735959 | 736438 | 0.50 | 9.47 | 11.10 | 6.82 | Hypothetical protein |
| *FEPFGCPE_00660* | *fabF-1* | - | 736487 | 737716 | 0.43 | 0.45 | 1.47 | 0.37 | 3-oxoacyl-[acyl-carrier-protein] synthase 2 |
| *FEPFGCPE_00661* | *fabG-2* | - | 737716 | 738441 | 0.08 | 4.26 | 2.67 | 0.83 | 3-oxoacyl-[acyl-carrier-protein] reductase FabG |
| *FEPFGCPE_00662* | *fabZ-1* | - | 738438 | 738905 | 0.25 | 1.30 | 0.00 | 0.00 | 3-hydroxyacyl-[acyl-carrier-protein] dehydratase FabZ |
| *FEPFGCPE_00663* | *fabF-2* | - | 738905 | 739948 | 0.11 | 1.44 | 0.74 | 0.00 | 3-oxoacyl-[acyl-carrier-protein] synthase 2 |
| *FEPFGCPE_00664* | | - | 739939 | 740109 | 0.59 | 3.20 | 0.32 | 0.00 | Hypothetical protein |
| *FEPFGCPE_00665* | | - | 740106 | 742532 | 0.39 | 0.83 | 0.30 | 0.96 | Hypothetical protein |
| *FEPFGCPE_00666* | *lolA-1* | - | 742529 | 743218 | 0.17 | 1.69 | 0.00 | 0.84 | Outer-membrane lipoprotein carrier protein |
| *FEPFGCPE_00667* | | - | 743203 | 743568 | 0.15 | 3.19 | 0.00 | 0.54 | Hypothetical protein |
| *FEPFGCPE_00668* | *hutH-1* | - | 743649 | 745289 | 0.56 | 2.37 | 1.85 | 1.05 | Histidine ammonia-lyase |
| *FEPFGCPE_00669* | *lpxL-1* | - | 745286 | 747082 | 0.44 | 1.18 | 2.11 | 0.93 | Lipid A biosynthesis lauroyltransferase |
| *FEPFGCPE_00670* | *bclA-2* | - | 747079 | 749010 | 0.71 | 2.58 | 1.16 | 0.05 | Benzoate--CoA ligase |
| *FEPFGCPE_00671* | | - | 749007 | 749657 | 0.77 | 0.96 | 2.48 | 0.00 | Hypothetical protein |
| *FEPFGCPE_00672* | *acpP-1* | - | 749667 | 749933 | 1.71 | 19.25 | 7.70 | 6.53 | Acyl carrier protein |
| *FEPFGCPE_00673* | *acpP-2* | - | 749946 | 750209 | 1.62 | 7.69 | 0.24 | 0.00 | Acyl carrier protein |
| *FEPFGCPE_00674* | | - | 750190 | 751029 | 2.69 | 7.42 | 2.79 | 0.07 | Hypothetical protein |
| *FEPFGCPE_00675* | | - | 751016 | 751855 | 1.91 | 6.09 | 6.63 | 1.58 | Hypothetical protein |
| *FEPFGCPE_00676* | | - | 752179 | 753012 | 2.75 | 15.60 | 8.73 | 4.07 | Hypothetical protein |
| *FEPFGCPE_00677* | | - | 753003 | 753308 | 3.86 | 7.95 | 11.99 | 7.40 | Hypothetical protein |
| *FEPFGCPE_00678* | *ygfZ* | - | 753332 | 754447 | 11.66 | 34.49 | 24.87 | 15.50 | tRNA-modifying protein YgfZ |
| *FEPFGCPE_00679* | *mltG* | + | 754630 | 755625 | 2.07 | 11.51 | 9.53 | 4.78 | Endolytic murein transglycosylase |
| *FEPFGCPE_00680* | *tmk-1* | + | 755622 | 755828 | 3.46 | 9.14 | 6.79 | 2.81 | Thymidylate kinase |
| *FEPFGCPE_00681* | *tmk-2* | + | 755833 | 756291 | 4.66 | 15.66 | 18.03 | 4.35 | Thymidylate kinase |
| *FEPFGCPE_00682* | *holB* | + | 756401 | 757441 | 3.43 | 15.69 | 14.73 | 4.43 | DNA polymerase III subunit delta' |
| *FEPFGCPE_00683* | *ycfH* | + | 757471 | 758253 | 2.53 | 11.56 | 12.16 | 2.62 | putative metal-dependent hydrolase YcfH |
| *FEPFGCPE_00684* | | + | 758327 | 759001 | 2.26 | 22.54 | 18.27 | 10.40 | Hypothetical protein |
| *FEPFGCPE_00685* | *otsA* | - | 759080 | 760510 | 4.89 | 24.26 | 70.19 | 13.04 | Trehalose-6-phosphate synthase |
| *FEPFGCPE_00686* | *otsB* | - | 760507 | 761328 | 7.30 | 40.61 | 89.10 | 15.72 | Trehalose-6-phosphate phosphatase |
| *FEPFGCPE_00687* | *cysJ-2* | + | 761683 | 764271 | 1.93 | 7.90 | 15.26 | 15.46 | Sulfite reductase [NADPH] flavoprotein alpha-component |
| *FEPFGCPE_00688* | *acyP-1* | + | 764381 | 764665 | 12.09 | 26.96 | 21.17 | 13.78 | Acylphosphatase |
| *FEPFGCPE_00689* | *tsr-2* | - | 764682 | 766325 | 1.56 | 4.67 | 3.48 | 0.88 | Methyl-accepting chemotaxis protein I |
| *FEPFGCPE_00690* | *rpfC-1* | + | 766598 | 768289 | 1.14 | 3.64 | 2.57 | 1.09 | Sensory/regulatory protein RpfC |
| *FEPFGCPE_00691* | | - | 768280 | 768924 | 4.17 | 27.27 | 20.04 | 5.07 | Hypothetical protein |
| *FEPFGCPE_00692* | *bcp-1* | - | 768953 | 769510 | 16.99 | 63.00 | 66.49 | 19.35 | Putative peroxiredoxin bcp |
| *FEPFGCPE_00693* | *decR-1* | - | 769785 | 770261 | 0.49 | 2.45 | 4.99 | 1.47 | DNA-binding transcriptional activator DecR |
| *FEPFGCPE_00694* | *ilvG* | + | 770434 | 772275 | 16.42 | 5.72 | 6.28 | 3.35 | Acetolactate synthase isozyme 2 large subunit |
| *FEPFGCPE_00695* | *artJ-1* | + | 772480 | 773259 | 12.49 | 24.74 | 20.95 | 6.56 | ABC transporter arginine-binding protein 1 |
| *FEPFGCPE_00696* | *yecS-1* | + | 773410 | 774162 | 3.02 | 8.60 | 3.91 | 1.32 | L-cystine transport system permease protein YecS |
| *FEPFGCPE_00697* | *glnQ-2* | + | 774146 | 774868 | 1.83 | 8.71 | 3.45 | 1.32 | Glutamine transport ATP-binding protein GlnQ |
| *FEPFGCPE_00698* | *astC* | + | 775076 | 776317 | 8.52 | 38.60 | 25.02 | 12.12 | Succinylornithine transaminase |
| *FEPFGCPE_00699* | *astA* | + | 776422 | 777501 | 3.35 | 12.50 | 7.23 | 3.51 | Arginine N-succinyltransferase |
| *FEPFGCPE_00700* | | - | 777590 | 778567 | 0.96 | 4.57 | 3.11 | 0.73 | Hypothetical protein |
| *FEPFGCPE_00701* | *caf1A* | - | 778564 | 781167 | 0.34 | 2.44 | 1.90 | 0.37 | F1 capsule-anchoring protein |
| *FEPFGCPE_00702* | | - | 781154 | 781921 | 0.29 | 1.29 | 0.87 | 0.55 | Hypothetical protein |
| *FEPFGCPE_00703* | | - | 781937 | 782464 | 0.41 | 6.80 | 1.06 | 0.92 | Hypothetical protein |
| *FEPFGCPE_00704* | *dps* | + | 783212 | 783697 | 30.99 | 109.56 | 68.73 | 60.69 | DNA protection during starvation protein |
| *FEPFGCPE_00705* | *corA* | + | 783841 | 787593 | 14.67 | 53.41 | 34.19 | 16.26 | Cobalt/magnesium transport protein CorA |
| *FEPFGCPE_00706* | *uctC-3* | + | 787842 | 789140 | 14.93 | 27.95 | 18.26 | 6.35 | Acetyl-CoA:oxalate CoA-transferase |

| Locus name | Gene name | Strand | Gene start (bp) | Gene end (bp) | Average TPM Inoculum | Average TPM $t_6$ | Average TPM $t_{24}$ | Average TPM $t_{168}$ | Annotation description |
|---|---|---|---|---|---|---|---|---|---|
| *FEPFGCPE_00707* | | + | 789387 | 790208 | 7.88 | 7.15 | 12.53 | 8.12 | Hypothetical protein |
| *FEPFGCPE_00708* | | + | 790683 | 791192 | 23.07 | 84.89 | 76.14 | 34.03 | Hypothetical protein |
| *FEPFGCPE_00709* | *rcsB-1* | + | 791403 | 792077 | 10.48 | 59.34 | 52.00 | 27.06 | Transcriptional regulatory protein RcsB |
| *FEPFGCPE_00710* | *rcsC* | + | 792128 | 795496 | 2.10 | 11.43 | 10.51 | 3.93 | Sensor histidine kinase RcsC |
| *FEPFGCPE_00711* | *pcpR-1* | - | 795519 | 796427 | 1.91 | 9.33 | 8.15 | 1.21 | PCP degradation transcriptional activation protein |
| *FEPFGCPE_00712* | *hmgA* | + | 796541 | 797860 | 4.72 | 17.91 | 20.47 | 10.44 | Homogentisate 1,2-dioxygenase |
| *FEPFGCPE_00713* | | + | 797961 | 799244 | 1.32 | 9.18 | 12.55 | 4.25 | Hypothetical protein |
| *FEPFGCPE_00714* | | - | 799724 | 800902 | 4.23 | 6.64 | 3.82 | 2.36 | Hypothetical protein |
| *FEPFGCPE_00715* | | - | 801033 | 802964 | 0.78 | 5.99 | 2.43 | 0.54 | Hypothetical protein |
| *FEPFGCPE_00716* | *astE* | - | 803170 | 804225 | 1.87 | 8.32 | 6.23 | 0.78 | Succinylglutamate desuccinylase |
| *FEPFGCPE_00717* | *astD* | + | 804432 | 805913 | 9.68 | 30.36 | 20.59 | 4.62 | N-succinylglutamate 5-semialdehyde dehydrogenase |
| *FEPFGCPE_00718* | *astB* | + | 805956 | 807317 | 4.25 | 22.62 | 18.48 | 7.53 | N-succinylarginine dihydrolase |
| *FEPFGCPE_00719* | | + | 807339 | 807908 | 9.60 | 82.55 | 49.29 | 11.85 | NAD(P)H-dependent FMN reductase |
| *FEPFGCPE_00720* | | + | 808106 | 808408 | 93.11 | 393.37 | 335.43 | 76.51 | Hypothetical protein |
| *FEPFGCPE_00721* | | + | 808602 | 808820 | 18.17 | 82.64 | 89.33 | 17.10 | Hypothetical protein |
| *FEPFGCPE_00722* | | + | 808984 | 809175 | 15.30 | 47.64 | 29.13 | 12.74 | Hypothetical protein |
| *FEPFGCPE_00723* | *bioC-1* | - | 809211 | 809960 | 4.59 | 6.77 | 6.72 | 1.50 | Malonyl-[acyl-carrier protein] O-methyltransferase |
| *FEPFGCPE_00724* | | + | 810197 | 811417 | 1.64 | 10.04 | 4.84 | 2.72 | Hypothetical protein |
| *FEPFGCPE_00725* | *prmA-1* | - | 811495 | 812658 | 1.61 | 9.63 | 9.04 | 3.57 | Ribosomal protein L11 methyltransferase |
| *FEPFGCPE_00726* | *yccS-1* | - | 812944 | 814869 | 1.97 | 11.35 | 10.18 | 2.22 | Inner membrane protein YccS |
| *FEPFGCPE_00727* | | - | 814890 | 815540 | 6.81 | 28.73 | 19.43 | 7.31 | Hypothetical protein |
| *FEPFGCPE_00728* | | - | 815547 | 815849 | 21.16 | 75.80 | 44.45 | 14.28 | Hypothetical protein |
| *FEPFGCPE_00729* | *bcsB* | + | 816137 | 818506 | 10.12 | 35.83 | 16.76 | 2.43 | Cyclic di-GMP-binding protein |
| *FEPFGCPE_00730* | *bcsZ* | + | 818507 | 819730 | 10.34 | 46.58 | 22.44 | 6.92 | Endoglucanase |
| *FEPFGCPE_00731* | *bcsC* | + | 819703 | 823569 | 2.58 | 21.16 | 12.36 | 2.06 | Cellulose synthase operon protein C |
| *FEPFGCPE_00732* | | + | 823650 | 824420 | 2.25 | 12.34 | 7.96 | 1.78 | Iron-sulfur cluster carrier protein |
| *FEPFGCPE_00733* | *bcsA* | + | 824417 | 827074 | 1.73 | 12.03 | 6.05 | 0.75 | Cellulose synthase catalytic subunit [UDP-forming] |
| *FEPFGCPE_00734* | | + | 827108 | 828613 | 1.16 | 20.04 | 12.46 | 2.83 | Hypothetical protein |
| *FEPFGCPE_00735* | *gpr* | + | 828903 | 829901 | 4.24 | 28.69 | 22.33 | 16.86 | L-glyceraldehyde 3-phosphate reductase |
| *FEPFGCPE_00736* | *nhaX-1* | + | 830149 | 831105 | 8.96 | 27.01 | 19.63 | 6.60 | Stress response protein NhaX |
| *FEPFGCPE_00737* | *camA* | - | 831185 | 832444 | 6.19 | 19.98 | 13.71 | 6.37 | Putidaredoxin reductase |
| *FEPFGCPE_00738* | *nicS-1* | - | 832561 | 833214 | 3.67 | 10.99 | 8.74 | 2.41 | HTH-type transcriptional repressor NicS |
| *FEPFGCPE_00739* | *mepC-2* | - | 833211 | 834719 | 2.06 | 6.30 | 2.11 | 1.72 | Multidrug/solvent efflux pump outer membrane protein MepC |
| *FEPFGCPE_00740* | *czcA-1* | - | 834716 | 837916 | 1.54 | 5.12 | 2.41 | 1.52 | Cobalt-zinc-cadmium resistance protein CzcA |
| *FEPFGCPE_00741* | *mdtA-2* | - | 837913 | 839154 | 4.68 | 4.97 | 3.39 | 3.46 | Multidrug resistance protein MdtA |
| *FEPFGCPE_00742* | *phaB-2* | + | 839510 | 840253 | 44.27 | 196.33 | 167.47 | 66.84 | Acetoacetyl-CoA reductase |
| *FEPFGCPE_00743* | | + | 840295 | 840885 | 43.52 | 174.55 | 139.87 | 78.99 | Hypothetical protein |
| *FEPFGCPE_00744* | *phaC-2* | - | 840867 | 842864 | 12.18 | 27.63 | 16.10 | 8.06 | Poly(3-hydroxyalkanoate) polymerase subunit PhaC |
| *FEPFGCPE_00745* | | - | 843064 | 843771 | 1.14 | 5.35 | 7.30 | 0.00 | Hypothetical protein |
| *FEPFGCPE_00746* | | + | 844063 | 844869 | 0.20 | 2.43 | 0.82 | 0.35 | Glycyl-glycine endopeptidase ALE-1 |
| *FEPFGCPE_00747* | *cpdA-2* | + | 844995 | 845990 | 1.51 | 4.05 | 1.95 | 0.26 | 3',5'-cyclic adenosine monophosphate phosphodiesterase CpdA |
| *FEPFGCPE_00748* | *mauC-1* | + | 846016 | 846396 | 2.62 | 5.11 | 1.53 | 1.03 | Amicyanin-alpha |
| *FEPFGCPE_00749* | *sigR-1* | + | 846512 | 847186 | 5.72 | 19.20 | 7.35 | 4.82 | ECF RNA polymerase sigma factor SigR |
| *FEPFGCPE_00750* | | + | 847173 | 848030 | 4.73 | 6.42 | 6.74 | 3.49 | Hypothetical protein |
| *FEPFGCPE_00751* | *tam* | + | 848070 | 848861 | 2.90 | 16.97 | 12.52 | 3.66 | Trans-aconitate 2-methyltransferase |
| *FEPFGCPE_00752* | *rhaS-1* | + | 848886 | 849890 | 7.31 | 24.84 | 17.34 | 4.60 | HTH-type transcriptional activator RhaS |
| *FEPFGCPE_00753* | | - | 849770 | 851011 | 1.64 | 6.66 | 6.75 | 2.30 | Hypothetical protein |
| *FEPFGCPE_00754* | *dmlR-4* | - | 851191 | 852117 | 3.60 | 10.64 | 6.35 | 2.05 | HTH-type transcriptional regulator DmlR |
| *FEPFGCPE_00755* | *gstB-1* | + | 852240 | 852911 | 1.99 | 8.06 | 4.31 | 1.67 | Glutathione S-transferase GST-6.0 |
| *FEPFGCPE_00756* | *tsr-3* | + | 853141 | 854802 | 2.72 | 8.19 | 3.41 | 0.83 | Methyl-accepting chemotaxis protein I |
| *FEPFGCPE_00757* | *dapA-1* | - | 854840 | 855793 | 3.59 | 10.68 | 3.12 | 0.90 | 4-hydroxy-tetrahydrodipicolinate synthase |
| *FEPFGCPE_00758* | | + | 856423 | 856671 | 68.07 | 300.22 | 175.01 | 82.02 | Hypothetical protein |
| *FEPFGCPE_00759* | *dosC-1* | + | 856794 | 858194 | 11.42 | 37.60 | 16.22 | 7.00 | Diguanylate cyclase DosC |
| *FEPFGCPE_00760* | | - | 858223 | 859008 | 4.60 | 13.16 | 7.47 | 2.87 | Hypothetical protein |
| *FEPFGCPE_00761* | *aroK-1* | - | 859170 | 860087 | 1.67 | 7.01 | 4.06 | 3.12 | Shikimate kinase |
| *FEPFGCPE_00762* | *dmdD* | + | 860308 | 861066 | 0.98 | 8.31 | 5.51 | 2.64 | Methylthioacryloyl-CoA hydratase |
| *FEPFGCPE_00763* | *lcdH* | + | 861099 | 862025 | 1.67 | 22.32 | 9.36 | 3.41 | L-carnitine dehydrogenase |
| *FEPFGCPE_00764* | *kce-1* | + | 862125 | 863045 | 0.99 | 15.15 | 7.18 | 1.96 | 3-keto-5-aminohexanoate cleavage enzyme |
| *FEPFGCPE_00765* | | + | 863108 | 864289 | 0.48 | 6.92 | 5.07 | 0.78 | Leu/Ile/Val-binding protein |
| *FEPFGCPE_00766* | *fadD3-1* | + | 864274 | 866127 | 2.39 | 4.75 | 6.28 | 1.78 | 3-[(3aS,4S,7aS)-7a-methyl-1,5-dioxo-octahydro-1H-inden-4-yl]propanoyl:CoA ligase |
| *FEPFGCPE_00767* | | - | 866140 | 867174 | 0.67 | 3.57 | 3.76 | 0.98 | Hypothetical protein |
| *FEPFGCPE_00768* | *cysL-1* | + | 867274 | 868161 | 3.74 | 12.63 | 15.28 | 2.88 | HTH-type transcriptional regulator CysL |
| *FEPFGCPE_00769* | | + | 868257 | 868988 | 5.27 | 2.69 | 6.53 | 1.52 | Hypothetical protein |
| *FEPFGCPE_00770* | *ftsH2* | + | 869050 | 870675 | 0.90 | 3.58 | 2.22 | 0.07 | ATP-dependent zinc metalloprotease FtsH 2 |
| *FEPFGCPE_00771* | *nreB-1* | + | 870881 | 872065 | 3.01 | 9.83 | 3.78 | 1.48 | Oxygen sensor histidine kinase NreB |
| *FEPFGCPE_00772* | *nreC-1* | + | 872065 | 872709 | 2.49 | 6.47 | 3.02 | 1.41 | Oxygen regulatory protein NreC |
| *FEPFGCPE_00773* | *codA* | + | 872966 | 874234 | 0.27 | 2.39 | 0.95 | 0.38 | Cytosine deaminase |
| *FEPFGCPE_00774* | | + | 874266 | 874802 | 0.42 | 3.11 | 1.95 | 0.00 | Hypothetical protein |
| *FEPFGCPE_00775* | *ywrD-1* | + | 874952 | 876721 | 0.50 | 4.28 | 2.07 | 0.28 | Putative gamma-glutamyltransferase YwrD |
| *FEPFGCPE_00776* | *metR-2* | + | 876781 | 877668 | 2.55 | 9.84 | 6.06 | 2.56 | HTH-type transcriptional regulator MetR |
| *FEPFGCPE_00777* | *allE* | - | 877777 | 878616 | 1.13 | 3.12 | 3.78 | 0.53 | (S)-ureidoglycine aminohydrolase |

| Locus name | Gene name | Strand | Gene start (bp) | Gene end (bp) | Average TPM Inoculum | Average TPM $t_6$ | Average TPM $t_{24}$ | Average TPM $t_{168}$ | Annotation description |
|---|---|---|---|---|---|---|---|---|---|
| *FEPFGCPE_00778* | | - | 878763 | 879287 | 0.50 | 0.51 | 2.74 | 0.00 | Hypothetical protein |
| *FEPFGCPE_00779* | | - | 879367 | 879870 | 4.42 | 5.01 | 4.70 | 0.42 | Hypothetical protein |
| *FEPFGCPE_00780* | *gntR-1* | - | 880226 | 880879 | 4.62 | 4.00 | 5.30 | 3.05 | putative D-xylose utilization operon transcriptional repressor |
| *FEPFGCPE_00781* | | + | 881132 | 881599 | 0.41 | 3.50 | 5.66 | 0.00 | Hypothetical protein |
| *FEPFGCPE_00782* | | + | 881876 | 882925 | 1.91 | 2.48 | 1.79 | 0.15 | Outer membrane porin protein 32 |
| *FEPFGCPE_00783* | | + | 883007 | 883897 | 1.03 | 5.99 | 2.31 | 1.15 | Hypothetical protein |
| *FEPFGCPE_00784* | | + | 883894 | 884808 | 0.51 | 1.38 | 2.58 | 0.46 | Hypothetical protein |
| *FEPFGCPE_00785* | | - | 884944 | 885927 | 0.87 | 1.58 | 1.93 | 0.83 | 2-ketogluconate reductase |
| *FEPFGCPE_00786* | | + | 886088 | 887044 | 1.42 | 6.82 | 1.58 | 0.98 | Ureidoglycolate lyase |
| *FEPFGCPE_00787* | *ribY-1* | + | 887082 | 888143 | 0.79 | 2.37 | 1.19 | 1.38 | Riboflavin-binding protein RibY |
| *FEPFGCPE_00788* | *dbfB-1* | + | 888178 | 888735 | 1.98 | 6.46 | 2.01 | 1.38 | 2,2',3-trihydroxybiphenyl dioxygenase |
| *FEPFGCPE_00789* | *ribX-1* | + | 888764 | 889507 | 0.59 | 2.61 | 1.04 | 1.09 | Riboflavin transport system permease protein RibX |
| *FEPFGCPE_00790* | *cmpD-1* | + | 889504 | 890367 | 1.89 | 3.97 | 1.97 | 0.00 | Bicarbonate transport ATP-binding protein CmpD |
| *FEPFGCPE_00791* | *nlhH-2* | + | 890367 | 891311 | 0.65 | 4.51 | 1.74 | 1.24 | Carboxylesterase NlhH |
| *FEPFGCPE_00792* | | - | 891198 | 891458 | 0.91 | 2.68 | 0.00 | 0.00 | Hypothetical protein |
| *FEPFGCPE_00793* | *yagE* | + | 891483 | 892388 | 0.50 | 0.44 | 0.00 | 0.00 | putative 2-keto-3-deoxy-galactonate aldolase YagE |
| *FEPFGCPE_00794* | *porA* | + | 892395 | 893750 | 0.12 | 0.86 | 1.01 | 0.00 | Major outer membrane protein P.IA |
| *FEPFGCPE_00795* | *pheP* | + | 893781 | 894626 | 0.24 | 0.87 | 0.96 | 0.02 | Phenylalanine-specific permease |
| *FEPFGCPE_00796* | *yifK* | + | 894626 | 895174 | 0.09 | 4.25 | 1.72 | 0.02 | putative transport protein YifK |
| *FEPFGCPE_00797* | *puuB-1* | - | 895200 | 896528 | 1.03 | 2.62 | 2.50 | 0.85 | Gamma-glutamylputrescine oxidoreductase |
| *FEPFGCPE_00798* | *tauB-1* | - | 896559 | 897401 | 0.58 | 1.43 | 0.71 | 1.24 | Taurine import ATP-binding protein TauB |
| *FEPFGCPE_00799* | *ssuC-2* | - | 897398 | 898165 | 1.08 | 4.64 | 1.45 | 0.76 | Putative aliphatic sulfonates transport permease protein SsuC |
| *FEPFGCPE_00800* | *tauA-1* | - | 898236 | 899240 | 0.72 | 4.03 | 1.70 | 0.00 | Taurine-binding periplasmic protein |
| *FEPFGCPE_00801* | *gltC-4* | - | 899352 | 900314 | 1.68 | 10.06 | 6.65 | 0.97 | HTH-type transcriptional regulator GltC |
| *FEPFGCPE_00802* | *hutG-1* | - | 900538 | 901701 | 0.52 | 3.10 | 4.55 | 1.03 | Formimidoylglutamase |
| *FEPFGCPE_00803* | *hutI* | - | 901728 | 902945 | 0.66 | 5.22 | 2.25 | 1.85 | Imidazolonepropionase |
| *FEPFGCPE_00804* | *proY* | - | 902953 | 904332 | 1.44 | 6.09 | 6.66 | 2.25 | Proline-specific permease ProY |
| *FEPFGCPE_00805* | *hutH-2* | - | 904474 | 906015 | 2.14 | 21.28 | 10.45 | 3.92 | Histidine ammonia-lyase |
| *FEPFGCPE_00806* | *hutU* | - | 906057 | 907736 | 2.98 | 23.05 | 12.01 | 8.75 | Urocanate hydratase |
| *FEPFGCPE_00807* | *yvoA-2* | - | 907899 | 908675 | 6.42 | 24.07 | 14.73 | 3.33 | HTH-type transcriptional repressor YvoA |
| *FEPFGCPE_00808* | *dthadh* | - | 908820 | 910034 | 1.34 | 9.15 | 7.90 | 1.30 | D-threo-3-hydroxyaspartate dehydratase |
| *FEPFGCPE_00809* | *yjhB-1* | - | 910213 | 911508 | 1.52 | 4.69 | 2.39 | 0.33 | Putative metabolite transport protein YjhB |
| *FEPFGCPE_00810* | *pgrR-2* | - | 911569 | 912543 | 1.62 | 2.67 | 0.55 | 0.43 | HTH-type transcriptional regulator PgrR |
| *FEPFGCPE_00811* | *bacC-1* | + | 912686 | 913420 | 2.57 | 6.27 | 4.65 | 0.83 | Dihydroanticapsin 7-dehydrogenase |
| *FEPFGCPE_00812* | | + | 913502 | 913711 | 6.27 | 24.11 | 4.69 | 3.61 | putative tautomerase |
| *FEPFGCPE_00813* | *dgdA* | - | 913738 | 915039 | 4.92 | 15.72 | 3.68 | 2.39 | 2,2-dialkylglycine decarboxylase |
| *FEPFGCPE_00814* | *gltR-1* | + | 915160 | 915975 | 6.99 | 16.28 | 11.51 | 3.97 | HTH-type transcriptional regulator GltR |
| *FEPFGCPE_00815* | | + | 916207 | 916335 | 12.76 | 26.48 | 33.40 | 6.28 | Hypothetical protein |
| *FEPFGCPE_00816* | *cysE-1* | + | 916864 | 917805 | 13.42 | 46.09 | 36.30 | 11.46 | Serine acetyltransferase |
| *FEPFGCPE_00817* | *srpC-1* | - | 917877 | 919112 | 1.51 | 12.22 | 6.65 | 4.18 | putative chromate transport protein |
| *FEPFGCPE_00818* | | + | 919460 | 920143 | 0.72 | 1.97 | 0.32 | 0.44 | Hypothetical protein |
| *FEPFGCPE_00819* | | + | 920146 | 922272 | 0.58 | 1.79 | 3.33 | 0.00 | Hypothetical protein |
| *FEPFGCPE_00820* | *msbA-1* | + | 922439 | 924178 | 0.88 | 3.41 | 3.28 | 1.71 | Lipid A export ATP-binding/permease protein MsbA |
| *FEPFGCPE_00821* | *narG* | + | 924496 | 928257 | 15.15 | 52.85 | 34.43 | 11.77 | Respiratory nitrate reductase 1 alpha chain |
| *FEPFGCPE_00822* | *narH* | + | 928254 | 929780 | 7.92 | 70.37 | 55.08 | 12.90 | Respiratory nitrate reductase 1 beta chain |
| *FEPFGCPE_00823* | *narJ* | + | 929782 | 930507 | 2.17 | 32.39 | 15.09 | 5.93 | Nitrate reductase molybdenum cofactor assembly chaperone NarJ |
| *FEPFGCPE_00824* | *narI* | + | 930532 | 931215 | 2.39 | 45.94 | 22.80 | 4.38 | Respiratory nitrate reductase 1 gamma chain |
| *FEPFGCPE_00825* | *ppiD-2* | + | 931289 | 932092 | 1.92 | 40.83 | 29.52 | 9.87 | Peptidyl-prolyl cis-trans isomerase D |
| *FEPFGCPE_00826* | *narK2* | + | 932103 | 933371 | 3.09 | 73.16 | 45.92 | 9.47 | putative nitrate/nitrite transporter NarK2 |
| *FEPFGCPE_00827* | *dmlR-5* | + | 933454 | 934365 | 1.68 | 15.98 | 8.64 | 1.65 | HTH-type transcriptional regulator DmlR |
| *FEPFGCPE_00828* | | + | 934553 | 935695 | 1.51 | 2.95 | 3.16 | 0.72 | Outer membrane porin protein |
| *FEPFGCPE_00829* | *pcaK-3* | + | 935793 | 937091 | 0.57 | 2.54 | 2.10 | 0.36 | 4-hydroxybenzoate transporter PcaK |
| *FEPFGCPE_00830* | | + | 937130 | 937810 | 0.67 | 2.54 | 1.53 | 0.00 | Hypothetical protein |
| *FEPFGCPE_00831* | *tdh* | + | 937857 | 938885 | 0.90 | 2.20 | 2.85 | 2.92 | L-threonine 3-dehydrogenase |
| *FEPFGCPE_00832* | | - | 938942 | 939241 | 0.88 | 6.75 | 9.39 | 5.13 | Hypothetical protein |
| *FEPFGCPE_00833* | | - | 939231 | 940688 | 1.56 | 4.99 | 5.90 | 4.79 | Ribonuclease |
| *FEPFGCPE_00834* | *nhaX-2* | + | 940974 | 941477 | 2.10 | 7.28 | 5.23 | 4.08 | Stress response protein NhaX |
| *FEPFGCPE_00835* | | + | 941482 | 941949 | 3.31 | 5.17 | 5.93 | 1.95 | Putative universal stress protein |
| *FEPFGCPE_00836* | *nreC-2* | + | 942118 | 942762 | 13.49 | 39.13 | 29.41 | 6.78 | Oxygen regulatory protein NreC |
| *FEPFGCPE_00837* | *fnr-1* | + | 942914 | 943702 | 5.66 | 22.47 | 28.19 | 13.07 | Fumarate and nitrate reduction regulatory protein |
| *FEPFGCPE_00838* | *adhA* | + | 943749 | 944732 | 2.29 | 11.25 | 13.61 | 4.07 | putative alcohol dehydrogenase AdhA |
| *FEPFGCPE_00839* | *nreB-2* | - | 944739 | 946724 | 6.25 | 26.99 | 11.07 | 5.89 | Oxygen sensor histidine kinase NreB |
| *FEPFGCPE_00840* | | + | 946954 | 947796 | 3.26 | 15.97 | 15.31 | 9.59 | Hypothetical protein |
| *FEPFGCPE_00841* | *adh* | + | 947821 | 948855 | 2.91 | 11.46 | 8.05 | 7.08 | Alcohol dehydrogenase |
| *FEPFGCPE_00842* | *prs-1* | - | 948906 | 949814 | 4.15 | 6.12 | 8.10 | 5.48 | Ribose-phosphate pyrophosphokinase |
| *FEPFGCPE_00843* | *deoA* | - | 949811 | 951415 | 0.90 | 2.62 | 5.28 | 3.96 | Thymidine phosphorylase |
| *FEPFGCPE_00844* | | + | 951839 | 952348 | 3.44 | 1.38 | 4.83 | 2.81 | Hypothetical protein |
| *FEPFGCPE_00845* | *yknY* | + | 952435 | 953142 | 2.77 | 0.34 | 0.99 | 0.61 | putative ABC transporter ATP-binding protein YknY |
| *FEPFGCPE_00846* | *aaeA-1* | + | 953139 | 954224 | 1.05 | 0.16 | 2.52 | 0.36 | p-hydroxybenzoic acid efflux pump subunit AaeA |
| *FEPFGCPE_00847* | | + | 954256 | 955398 | 1.73 | 4.52 | 8.56 | 1.53 | Hypothetical protein |
| *FEPFGCPE_00848* | | + | 955602 | 956639 | 4.81 | 4.63 | 6.15 | 2.71 | Putative NAD(P)H nitroreductase |

| Locus name | Gene name | Strand | Gene start (bp) | Gene end (bp) | Average TPM Inoculum | Average TPM $t_6$ | Average TPM $t_{24}$ | Average TPM $t_{168}$ | Annotation description |
|---|---|---|---|---|---|---|---|---|---|
| *FEPFGCPE_00849* | *mntH-2* | + | 956714 | 958027 | 5.94 | 5.96 | 8.54 | 0.99 | Divalent metal cation transporter MntH |
| *FEPFGCPE_00850* | | - | 958090 | 958917 | 2.12 | 6.13 | 13.20 | 4.38 | Hypothetical protein |
| *FEPFGCPE_00851* | *osmY-1* | + | 959232 | 959885 | 1.00 | 1.13 | 13.20 | 6.38 | Osmotically-inducible protein Y |
| *FEPFGCPE_00852* | | + | 959952 | 960425 | 2.11 | 8.19 | 18.90 | 14.78 | Hypothetical protein |
| *FEPFGCPE_00853* | *nhaX-3* | + | 960481 | 960942 | 2.93 | 8.74 | 9.09 | 5.90 | Stress response protein NhaX |
| *FEPFGCPE_00854* | *nhaX-4* | - | 961004 | 961963 | 13.35 | 30.93 | 19.52 | 8.47 | Stress response protein NhaX |
| *FEPFGCPE_00855* | | + | 962092 | 963381 | 4.05 | 8.79 | 6.71 | 1.15 | Hypothetical protein |
| *FEPFGCPE_00856* | | + | 963507 | 963935 | 4.17 | 46.77 | 73.38 | 48.41 | 18 kDa heat shock protein |
| *FEPFGCPE_00857* | | - | 963987 | 964331 | 1.92 | 7.07 | 8.91 | 11.92 | Hypothetical protein |
| *FEPFGCPE_00858* | *nreC-3* | + | 965026 | 965673 | 6.74 | 46.87 | 26.35 | 5.97 | Oxygen regulatory protein NreC |
| *FEPFGCPE_00859* | *anr* | - | 965738 | 966523 | 0.90 | 5.62 | 12.37 | 3.73 | Transcriptional activator protein Anr |
| *FEPFGCPE_00860* | | + | 966890 | 967516 | 0.85 | 4.68 | 11.55 | 2.25 | Putative phosphoribosyl transferase |
| *FEPFGCPE_00861* | *polX* | + | 967587 | 969377 | 0.88 | 1.75 | 1.93 | 0.75 | DNA polymerase/3'-5' exonuclease PolX |
| *FEPFGCPE_00862* | | + | 969527 | 969901 | 7.41 | 20.32 | 15.81 | 5.73 | Hypothetical protein |
| *FEPFGCPE_00863* | *yocK* | + | 969915 | 970292 | 3.83 | 6.60 | 8.18 | 5.25 | General stress protein 16O |
| *FEPFGCPE_00864* | *icaB* | + | 970298 | 971098 | 2.25 | 3.26 | 1.59 | 0.38 | Poly-beta-1,6-N-acetyl-D-glucosamine N-deacetylase |
| *FEPFGCPE_00865* | | + | 971091 | 972359 | 0.77 | 2.10 | 2.64 | 0.19 | Hypothetical protein |
| *FEPFGCPE_00866* | | + | 972606 | 973109 | 4.58 | 8.68 | 10.35 | 6.57 | Hypothetical protein |
| *FEPFGCPE_00867* | *nrdZ* | + | 973126 | 974931 | 4.48 | 2.94 | 3.80 | 1.53 | Vitamin B12-dependent ribonucleoside-diphosphate reductase |
| *FEPFGCPE_00868* | | + | 975096 | 975749 | 1.13 | 7.62 | 10.12 | 6.55 | Hypothetical protein |
| *FEPFGCPE_00869* | | + | 975915 | 976121 | 2.89 | 10.28 | 18.02 | 4.86 | Dodecin |
| *FEPFGCPE_00870* | *tar-1* | - | 976156 | 977772 | 0.97 | 5.73 | 2.16 | 1.06 | Methyl-accepting chemotaxis protein II |
| *FEPFGCPE_00871* | | - | 977974 | 978411 | 1.34 | 1.22 | 0.39 | 1.28 | Hypothetical protein |
| *FEPFGCPE_00872* | *ftsH4* | - | 978751 | 980697 | 8.82 | 6.41 | 7.84 | 2.33 | ATP-dependent zinc metalloprotease FtsH 4 |
| *FEPFGCPE_00873* | | + | 980920 | 981765 | 6.16 | 6.09 | 4.00 | 2.36 | Hypothetical protein |
| *FEPFGCPE_00874* | | + | 981765 | 982265 | 2.20 | 4.60 | 3.33 | 0.80 | Hypothetical protein |
| *FEPFGCPE_00875* | *czcS-1* | - | 982285 | 983685 | 0.93 | 3.27 | 3.47 | 0.26 | Sensor protein CzcS |
| *FEPFGCPE_00876* | *copR* | - | 983682 | 984320 | 1.96 | 5.37 | 4.37 | 1.79 | Transcriptional activator protein CopR |
| *FEPFGCPE_00877* | *copD-1* | - | 984474 | 985397 | 0.44 | 3.30 | 2.71 | 1.14 | Copper resistance protein D |
| *FEPFGCPE_00878* | *pcoC-1* | - | 985403 | 985627 | 0.07 | 5.71 | 4.16 | 0.13 | Copper resistance protein C |
| *FEPFGCPE_00879* | | - | 985627 | 985788 | 1.85 | 11.84 | 16.03 | 0.00 | Hypothetical protein |
| *FEPFGCPE_00880* | *copB-1* | - | 985864 | 986775 | 1.00 | 9.27 | 8.19 | 1.71 | Copper resistance protein B |
| *FEPFGCPE_00881* | *copA-2* | - | 986772 | 988688 | 1.06 | 7.09 | 3.18 | 1.67 | Copper resistance protein A |
| *FEPFGCPE_00882* | *steT* | - | 988935 | 990305 | 1.96 | 8.56 | 5.36 | 3.19 | Serine/threonine exchanger SteT |
| *FEPFGCPE_00883* | *dmlR-6* | - | 990534 | 991454 | 1.11 | 9.36 | 6.87 | 2.77 | HTH-type transcriptional regulator DmlR |
| *FEPFGCPE_00884* | *ald-1* | + | 991679 | 993139 | 1.50 | 7.03 | 7.19 | 1.21 | 3-succinoylsemialdehyde-pyridine dehydrogenase |
| *FEPFGCPE_00885* | *soxB* | + | 993260 | 994588 | 1.21 | 10.08 | 5.50 | 1.97 | Sarcosine oxidase subunit beta |
| *FEPFGCPE_00886* | *cynR-3* | + | 994765 | 995718 | 1.59 | 20.77 | 10.88 | 3.56 | HTH-type transcriptional regulator CynR |
| *FEPFGCPE_00887* | *ifcA* | + | 995782 | 997191 | 0.45 | 6.37 | 2.73 | 0.30 | Fumarate reductase flavoprotein subunit |
| *FEPFGCPE_00888* | | + | 997178 | 998401 | 0.54 | 2.56 | 0.15 | 0.92 | Hypothetical protein |
| *FEPFGCPE_00889* | *proP-3* | + | 998460 | 999779 | 0.53 | 4.41 | 0.33 | 0.00 | Proline/betaine transporter |
| *FEPFGCPE_00890* | | + | 1000360 | 1001634 | 1.93 | 8.82 | 4.95 | 1.75 | NADPH oxidoreductase |
| *FEPFGCPE_00891* | | + | 1001631 | 1002020 | 7.52 | 33.36 | 22.93 | 2.55 | Soluble cytochrome b558 |
| *FEPFGCPE_00892* | | + | 1002134 | 1002592 | 8.54 | 44.67 | 27.87 | 8.15 | Hypothetical protein |
| *FEPFGCPE_00893* | | + | 1003005 | 1004198 | 2.65 | 19.47 | 14.59 | 2.72 | Hypothetical protein |
| *FEPFGCPE_00894* | *xerD-1* | + | 1004719 | 1005009 | 1.06 | 1.57 | 5.33 | 0.00 | Tyrosine recombinase XerD |
| *FEPFGCPE_00895* | | + | 1005063 | 1005317 | 47.44 | 93.54 | 53.40 | 16.75 | Hypothetical protein |
| *FEPFGCPE_00896* | *mazF* | + | 1005317 | 1005679 | 24.32 | 49.71 | 23.12 | 8.39 | Endoribonuclease MazF |
| *FEPFGCPE_00897* | | + | 1005747 | 1005941 | 26.07 | 48.88 | 21.71 | 10.03 | Hypothetical protein |
| *FEPFGCPE_00898* | | + | 1005953 | 1006372 | 24.07 | 51.41 | 27.29 | 11.44 | Hypothetical protein |
| *FEPFGCPE_00899* | | + | 1006392 | 1006892 | 12.50 | 28.81 | 8.12 | 3.84 | Hypothetical protein |
| *FEPFGCPE_00900* | | + | 1006919 | 1007176 | 21.03 | 51.94 | 30.88 | 12.73 | Hypothetical protein |
| *FEPFGCPE_00901* | *mprA* | + | 1007299 | 1007997 | 4.98 | 10.88 | 7.81 | 4.12 | Response regulator MprA |
| *FEPFGCPE_00902* | *phoR-2* | + | 1007978 | 1009249 | 8.01 | 23.22 | 16.76 | 2.76 | Phosphate regulon sensor protein PhoR |
| *FEPFGCPE_00903* | *czcC-1* | + | 1009369 | 1010766 | 1.31 | 2.04 | 3.06 | 0.81 | Cobalt-zinc-cadmium resistance protein CzcC |
| *FEPFGCPE_00904* | | + | 1010763 | 1011827 | 1.42 | 4.19 | 1.92 | 1.05 | Hypothetical protein |
| *FEPFGCPE_00905* | *czcA-2* | + | 1011867 | 1014986 | 1.53 | 12.18 | 7.45 | 0.86 | Cobalt-zinc-cadmium resistance protein CzcA |
| *FEPFGCPE_00906* | *yajR-1* | - | 1015063 | 1016307 | 4.30 | 26.27 | 16.23 | 6.24 | Inner membrane transport protein YajR |
| *FEPFGCPE_00907* | | + | 1017162 | 1017530 | 47.38 | 169.91 | 118.14 | 32.63 | Hypothetical protein |
| *FEPFGCPE_00908* | *czcC-2* | + | 1017618 | 1018862 | 0.92 | 5.76 | 3.13 | 1.44 | Cobalt-zinc-cadmium resistance protein CzcC |
| *FEPFGCPE_00909* | *cusB* | + | 1018859 | 1020382 | 1.96 | 10.69 | 8.96 | 1.57 | Cation efflux system protein CusB |
| *FEPFGCPE_00910* | *cusA* | + | 1020379 | 1023513 | 3.15 | 19.39 | 11.33 | 2.88 | Cation efflux system protein CusA |
| *FEPFGCPE_00911* | *cusF* | + | 1023565 | 1023933 | 1.85 | 19.84 | 23.92 | 4.50 | Cation efflux system protein CusF |
| *FEPFGCPE_00912* | *glnB-1* | - | 1024682 | 1024984 | 7.32 | 50.48 | 37.72 | 15.18 | Nitrogen regulatory protein P-II |
| *FEPFGCPE_00913* | *mco* | - | 1025024 | 1026640 | 4.34 | 14.11 | 10.89 | 3.70 | Multicopper oxidase mco |
| *FEPFGCPE_00914* | | - | 1026693 | 1026857 | 4.62 | 19.57 | 23.96 | 7.61 | Hypothetical protein |
| *FEPFGCPE_00915* | | - | 1026892 | 1027857 | 2.36 | 12.71 | 7.58 | 2.72 | Hypothetical protein |
| *FEPFGCPE_00916* | | - | 1027923 | 1028174 | 9.32 | 36.33 | 41.51 | 12.10 | Hypothetical protein |
| *FEPFGCPE_00917* | | - | 1028334 | 1028636 | 5.38 | 26.12 | 23.57 | 5.87 | Hypothetical protein |
| *FEPFGCPE_00918* | | - | 1028873 | 1029274 | 6.44 | 11.51 | 5.41 | 1.47 | Hypothetical protein |
| *FEPFGCPE_00919* | *actP-2* | + | 1029553 | 1029921 | 15.62 | 19.27 | 19.46 | 14.72 | Copper-transporting P-type ATPase |

| Locus name | Gene name | Strand | Gene start (bp) | Gene end (bp) | Average TPM Inoculum | Average TPM $t_6$ | Average TPM $t_{24}$ | Average TPM $t_{168}$ | Annotation description |
|---|---|---|---|---|---|---|---|---|---|
| *FEPFGCPE_00920* | *silP* | + | 1029921 | 1030121 | 53.43 | 45.70 | 61.74 | 28.48 | Silver exporting P-type ATPase |
| *FEPFGCPE_00921* | *frmR-1* | + | 1030221 | 1030493 | 2.64 | 46.32 | 43.17 | 17.96 | Transcriptional repressor FrmR |
| *FEPFGCPE_00922* | *zitB* | + | 1030500 | 1031852 | 0.94 | 15.39 | 9.94 | 4.88 | Zinc transporter ZitB |
| *FEPFGCPE_00923* | *crnA* | + | 1031973 | 1032788 | 1.30 | 9.77 | 10.87 | 3.64 | Creatinine amidohydrolase |
| *FEPFGCPE_00924* | | + | 1032845 | 1034017 | 3.62 | 19.53 | 11.86 | 7.25 | PE-PGRS family protein PE_PGRS18 |
| *FEPFGCPE_00925* | | - | 1034104 | 1034313 | 4.29 | 25.63 | 15.24 | 5.23 | Hypothetical protein |
| *FEPFGCPE_00926* | | - | 1034658 | 1035095 | 2.44 | 9.50 | 6.41 | 4.67 | Hypothetical protein |
| *FEPFGCPE_00927* | | + | 1035369 | 1035878 | 6.59 | 1.87 | 8.62 | 7.40 | Hypothetical protein |
| *FEPFGCPE_00928* | *copA-3* | + | 1036011 | 1038143 | 3.90 | 3.05 | 4.77 | 1.63 | Copper-exporting P-type ATPase A |
| *FEPFGCPE_00929* | | + | 1038140 | 1038358 | 1.99 | 3.25 | 2.76 | 1.90 | Hypothetical protein |
| *FEPFGCPE_00930* | | - | 1038305 | 1038475 | 1.66 | 5.30 | 1.61 | 0.00 | Hypothetical protein |
| *FEPFGCPE_00931* | | + | 1038552 | 1038815 | 4.21 | 5.01 | 10.72 | 5.76 | Hypothetical protein |
| *FEPFGCPE_00932* | | + | 1038818 | 1039480 | 3.97 | 12.66 | 16.45 | 2.79 | Hypothetical protein |
| *FEPFGCPE_00933* | | + | 1039533 | 1039754 | 1.74 | 10.39 | 20.72 | 8.69 | Hypothetical protein |
| *FEPFGCPE_00934* | | + | 1039943 | 1040737 | 1.06 | 3.42 | 0.66 | 0.69 | Hypothetical protein |
| *FEPFGCPE_00935* | | + | 1040737 | 1041237 | 0.32 | 4.09 | 0.98 | 2.56 | Hypothetical protein |
| *FEPFGCPE_00936* | *czcS-2* | - | 1041256 | 1042656 | 0.88 | 4.27 | 3.15 | 1.21 | Sensor protein CzcS |
| *FEPFGCPE_00937* | *czcR-1* | - | 1042653 | 1043291 | 0.44 | 4.28 | 2.67 | 2.26 | Transcriptional activator protein CzcR |
| *FEPFGCPE_00938* | *copD-2* | - | 1043449 | 1044372 | 0.39 | 2.41 | 3.05 | 3.63 | Copper resistance protein D |
| *FEPFGCPE_00939* | *copC* | - | 1044378 | 1044764 | 0.75 | 13.25 | 12.32 | 4.21 | Copper resistance protein C |
| *FEPFGCPE_00940* | *copB-2* | - | 1044837 | 1045745 | 1.20 | 13.09 | 13.01 | 3.33 | Copper resistance protein B |
| *FEPFGCPE_00941* | *copA-4* | - | 1045745 | 1047670 | 1.30 | 10.20 | 6.62 | 3.12 | Copper resistance protein A |
| *FEPFGCPE_00942* | | - | 1047952 | 1048191 | 3.06 | 17.68 | 12.14 | 2.56 | Hypothetical protein |
| *FEPFGCPE_00943* | | + | 1048716 | 1048862 | 0.27 | 6.77 | 0.64 | 3.13 | Hypothetical protein |
| *FEPFGCPE_00944* | *czcD* | - | 1049328 | 1050383 | 2.77 | 14.54 | 24.50 | 15.76 | Cadmium, cobalt and zinc/H(+)-K(+) antiporter |
| *FEPFGCPE_00945* | *frmR-2* | - | 1050594 | 1050764 | 4.04 | 14.83 | 28.82 | 20.46 | Transcriptional repressor FrmR |
| *FEPFGCPE_00946* | | + | 1051055 | 1051342 | 0.75 | 1.13 | 2.27 | 1.25 | Hypothetical protein |
| *FEPFGCPE_00947* | | - | 1051485 | 1051769 | 6.31 | 32.81 | 22.00 | 6.26 | Hypothetical protein |
| *FEPFGCPE_00948* | *merR* | - | 1051889 | 1052284 | 3.67 | 11.61 | 5.38 | 1.57 | Mercuric resistance operon regulatory protein |
| *FEPFGCPE_00949* | | + | 1052361 | 1052708 | 0.47 | 0.97 | 1.28 | 2.77 | Hypothetical protein |
| *FEPFGCPE_00950* | *merP* | + | 1052726 | 1053001 | 2.44 | 10.28 | 2.30 | 11.01 | Mercuric transport protein periplasmic component |
| *FEPFGCPE_00951* | | + | 1053009 | 1053221 | 0.79 | 3.63 | 4.49 | 0.00 | Hypothetical protein |
| *FEPFGCPE_00952* | | + | 1053602 | 1053802 | 5.21 | 17.65 | 7.96 | 3.27 | Hypothetical protein |
| *FEPFGCPE_00953* | | + | 1054646 | 1054783 | 7.00 | 52.07 | 33.47 | 13.23 | Hypothetical protein |
| *FEPFGCPE_00954* | | + | 1054874 | 1055284 | 4.73 | 30.18 | 17.74 | 6.18 | Hypothetical protein |
| *FEPFGCPE_00955* | *xerC-2* | + | 1055655 | 1057634 | 1.61 | 14.30 | 11.44 | 3.51 | Tyrosine recombinase XerC |
| *FEPFGCPE_00956* | | + | 1057986 | 1058684 | 1.14 | 26.96 | 21.22 | 8.19 | Hypothetical protein |
| *FEPFGCPE_00957* | | + | 1058681 | 1060612 | 1.46 | 42.22 | 20.70 | 6.71 | Hypothetical protein |
| *FEPFGCPE_00958* | | + | 1060623 | 1061639 | 1.03 | 55.45 | 28.91 | 8.84 | Hypothetical protein |
| *FEPFGCPE_00959* | | + | 1061636 | 1064326 | 3.32 | 43.51 | 33.65 | 6.66 | Hypothetical protein |
| *FEPFGCPE_00960* | *hdl IVa-1* | + | 1064475 | 1065116 | 2.57 | 30.92 | 22.40 | 9.19 | (S)-2-haloacid dehalogenase 4A |
| *FEPFGCPE_00961* | *gltR-2* | + | 1065667 | 1066629 | 4.40 | 22.33 | 16.63 | 6.10 | HTH-type transcriptional regulator GltR |
| *FEPFGCPE_00962* | *caiD* | - | 1066696 | 1067502 | 1.12 | 8.59 | 4.01 | 2.54 | Carnitinyl-CoA dehydratase |
| *FEPFGCPE_00963* | *ubiX* | - | 1067578 | 1068186 | 1.35 | 0.37 | 3.38 | 0.35 | Flavin prenyltransferase UbiX |
| *FEPFGCPE_00964* | *shdC* | - | 1068183 | 1069640 | 1.02 | 1.46 | 2.07 | 0.52 | Phenolic acid decarboxylase subunit C |
| *FEPFGCPE_00965* | *fcbA1* | - | 1069661 | 1071214 | 0.71 | 2.74 | 2.90 | 2.43 | 4-chlorobenzoate--CoA ligase |
| *FEPFGCPE_00966* | *cutS-2* | - | 1071226 | 1071738 | 1.24 | 4.54 | 1.18 | 1.57 | Carbon monoxide dehydrogenase small chain |
| *FEPFGCPE_00967* | *kdhA* | - | 1071735 | 1072553 | 0.74 | 2.35 | 2.91 | 1.15 | 6-hydroxypseudooxynicotine dehydrogenase complex subunit alpha |
| *FEPFGCPE_00968* | *cdhA* | - | 1072550 | 1075594 | 1.00 | 1.90 | 0.84 | 0.65 | Caffeine dehydrogenase subunit alpha |
| *FEPFGCPE_00969* | | - | 1076033 | 1076827 | 2.70 | 8.51 | 12.17 | 2.07 | Hypothetical protein |
| *FEPFGCPE_00970* | | + | 1076951 | 1077640 | 2.76 | 15.18 | 9.26 | 2.83 | Hypothetical protein |
| *FEPFGCPE_00971* | *gbsA-1* | - | 1077721 | 1079178 | 0.60 | 2.42 | 0.69 | 0.15 | Betaine aldehyde dehydrogenase |
| *FEPFGCPE_00972* | *proP-4* | - | 1079335 | 1080690 | 0.67 | 3.20 | 2.84 | 0.60 | Proline/betaine transporter |
| *FEPFGCPE_00973* | | - | 1080805 | 1082538 | 0.92 | 5.74 | 2.77 | 1.65 | 5-(hydroxymethyl)furfural oxidase |
| *FEPFGCPE_00974* | *pucA-1* | + | 1082930 | 1083970 | 2.09 | 8.35 | 4.98 | 1.17 | putative xanthine dehydrogenase subunit A |
| *FEPFGCPE_00975* | | + | 1084355 | 1085203 | 4.85 | 24.23 | 30.11 | 10.15 | Hypothetical protein |
| *FEPFGCPE_00976* | | - | 1085260 | 1085427 | 22.53 | 11.22 | 23.45 | 8.16 | Hypothetical protein |
| *FEPFGCPE_00977* | | - | 1085451 | 1085738 | 9.14 | 17.15 | 30.09 | 12.34 | Hypothetical protein |
| *FEPFGCPE_00978* | | + | 1086076 | 1086549 | 5.98 | 2.58 | 12.10 | 4.64 | Hypothetical protein |
| *FEPFGCPE_00979* | | + | 1086747 | 1089143 | 1.69 | 2.96 | 2.67 | 1.14 | Calcium-transporting ATPase 1 |
| *FEPFGCPE_00980* | *sigK-1* | + | 1089350 | 1089967 | 12.03 | 72.39 | 31.67 | 12.75 | ECF RNA polymerase sigma factor SigK |
| *FEPFGCPE_00981* | | + | 1089964 | 1090698 | 3.02 | 32.09 | 12.20 | 3.07 | Hypothetical protein |
| *FEPFGCPE_00982* | | + | 1090886 | 1091677 | 0.88 | 7.58 | 6.48 | 1.55 | Hypothetical protein |
| *FEPFGCPE_00983* | *cfa* | + | 1091709 | 1092812 | 1.03 | 5.66 | 4.95 | 1.48 | Cyclopropane-fatty-acyl-phospholipid synthase |
| *FEPFGCPE_00984* | *dadA-1* | + | 1092964 | 1094268 | 0.89 | 7.46 | 8.07 | 0.42 | D-amino acid dehydrogenase |
| *FEPFGCPE_00985* | | + | 1094265 | 1095092 | 0.49 | 7.14 | 5.61 | 0.99 | Hypothetical protein |
| *FEPFGCPE_00986* | *ufaA1* | + | 1095089 | 1096300 | 0.38 | 9.95 | 4.17 | 1.38 | Tuberculostearic acid methyltransferase UfaA1 |
| *FEPFGCPE_00987* | *dosC-2* | - | 1096363 | 1098138 | 1.25 | 6.40 | 2.93 | 3.38 | Diguanylate cyclase DosC |
| *FEPFGCPE_00988* | *fadH* | - | 1098365 | 1100386 | 3.16 | 5.84 | 5.53 | 1.73 | 2,4-dienoyl-CoA reductase [NADPH] |
| *FEPFGCPE_00989* | | + | 1100575 | 1101144 | 2.99 | 7.48 | 7.98 | 2.31 | Hypothetical protein |
| *FEPFGCPE_00990* | | + | 1101405 | 1102427 | 136.89 | 198.42 | 80.87 | 32.79 | PE-PGRS family protein PE_PGRS18 |

| Locus name | Gene name | Strand | Gene start (bp) | Gene end (bp) | Average TPM Inoculum | Average TPM $t_6$ | Average TPM $t_{24}$ | Average TPM $t_{168}$ | Annotation description |
|---|---|---|---|---|---|---|---|---|---|
| *FEPFGCPE_00991* | | + | 1102439 | 1103446 | 41.75 | 73.74 | 29.39 | 7.65 | Hypothetical protein |
| *FEPFGCPE_00992* | | - | 1103478 | 1104071 | 11.60 | 47.71 | 19.91 | 7.96 | Hypothetical protein |
| *FEPFGCPE_00993* | *cadI-1* | - | 1104253 | 1104732 | 7.20 | 24.79 | 14.46 | 6.17 | Cadmium-induced protein CadI |
| *FEPFGCPE_00994* | | - | 1104795 | 1105370 | 20.94 | 28.52 | 18.27 | 7.04 | Hypothetical protein |
| *FEPFGCPE_00995* | | - | 1105423 | 1106220 | 21.09 | 6.40 | 12.05 | 5.39 | Sulfide-quinone reductase |
| *FEPFGCPE_00996* | *metE-2* | + | 1106447 | 1108735 | 10.70 | 104.83 | 137.34 | 182.43 | 5-methyltetrahydropteroyltriglutamate--homocysteine methyltransferase |
| *FEPFGCPE_00997* | *rssB* | + | 1109118 | 1109777 | 2.42 | 4.75 | 8.99 | 7.47 | Swarming motility regulation protein RssB |
| *FEPFGCPE_00998* | *qseC-1* | + | 1109774 | 1111141 | 2.62 | 10.05 | 13.04 | 5.09 | Sensor protein QseC |
| *FEPFGCPE_00999* | *arnT-1* | + | 1111202 | 1112917 | 1.54 | 8.96 | 8.41 | 3.47 | Undecaprenyl phosphate-alpha-4-amino-4-deoxy-L-arabinose arabinosyl transferase |
| *FEPFGCPE_01000* | *czcA-3* | + | 1113235 | 1116408 | 0.59 | 2.43 | 0.94 | 0.55 | Cobalt-zinc-cadmium resistance protein CzcA |
| *FEPFGCPE_01001* | *mdtA-3* | + | 1116421 | 1117674 | 1.53 | 4.24 | 2.39 | 2.58 | Multidrug resistance protein MdtA |
| *FEPFGCPE_01002* | *ttgF* | + | 1117671 | 1119137 | 0.48 | 0.71 | 0.69 | 0.25 | Toluene efflux pump outer membrane protein TtgF |
| *FEPFGCPE_01003* | *czcR-2* | + | 1119147 | 1119821 | 3.81 | 26.47 | 13.02 | 2.53 | Transcriptional activator protein CzcR |
| *FEPFGCPE_01004* | *cusS-1* | + | 1119887 | 1121308 | 1.72 | 8.49 | 5.75 | 1.04 | Sensor kinase CusS |
| *FEPFGCPE_01005* | *arfB* | + | 1121427 | 1121843 | 5.78 | 36.57 | 36.99 | 7.32 | Peptidyl-tRNA hydrolase ArfB |
| *FEPFGCPE_01006* | | + | 1121939 | 1122253 | 4.56 | 11.22 | 9.61 | 1.70 | Hypothetical protein |
| *FEPFGCPE_01007* | *hipA* | + | 1122280 | 1123575 | 2.29 | 6.06 | 10.85 | 1.42 | Serine/threonine-protein kinase HipA |
| *FEPFGCPE_01008* | *ydhP-1* | - | 1123608 | 1124822 | 1.37 | 2.95 | 11.82 | 3.54 | Inner membrane transport protein YdhP |
| *FEPFGCPE_01009* | *chaB* | - | 1125018 | 1125260 | 4.11 | 17.09 | 88.21 | 44.83 | Putative cation transport regulator ChaB |
| *FEPFGCPE_01010* | | - | 1125326 | 1125556 | 11.56 | 81.72 | 393.43 | 78.94 | Hypothetical protein |
| *FEPFGCPE_01011* | | - | 1125576 | 1126007 | 9.79 | 59.62 | 392.18 | 125.32 | Hypothetical protein |
| *FEPFGCPE_01012* | *osmY-2* | - | 1126098 | 1126460 | 14.10 | 99.71 | 699.47 | 243.63 | Osmotically-inducible protein Y |
| *FEPFGCPE_01013* | | - | 1126685 | 1127122 | 11.59 | 115.80 | 749.10 | 107.15 | Hypothetical protein |
| *FEPFGCPE_01014* | | + | 1127277 | 1127546 | 0.61 | 3.22 | 2.08 | 0.00 | Hypothetical protein |
| *FEPFGCPE_01015* | | + | 1127832 | 1127984 | 2.71 | 3.25 | 4.05 | 1.16 | Hypothetical protein |
| *FEPFGCPE_01016* | | - | 1128000 | 1131083 | 2.39 | 11.40 | 10.40 | 1.89 | Hypothetical protein |
| *FEPFGCPE_01017* | | - | 1131237 | 1132736 | 4.97 | 19.95 | 9.98 | 3.73 | Hypothetical protein |
| *FEPFGCPE_01018* | | + | 1133256 | 1135376 | 1.79 | 4.83 | 2.77 | 1.87 | putative signaling protein |
| *FEPFGCPE_01019* | | - | 1135471 | 1135875 | 7.08 | 2.94 | 7.55 | 3.75 | Hypothetical protein |
| *FEPFGCPE_01020* | | - | 1135914 | 1136666 | 3.19 | 9.00 | 5.92 | 2.91 | Hypothetical protein |
| *FEPFGCPE_01021* | *gabR-3* | + | 1136853 | 1138394 | 3.64 | 10.24 | 6.43 | 2.71 | HTH-type transcriptional regulatory protein GabR |
| *FEPFGCPE_01022* | *smc-1* | - | 1138453 | 1139478 | 4.82 | 8.22 | 8.06 | 4.83 | Chromosome partition protein Smc |
| *FEPFGCPE_01023* | *xerC-3* | + | 1139656 | 1140777 | 0.46 | 1.33 | 1.35 | 0.38 | Tyrosine recombinase XerC |
| *FEPFGCPE_01024* | | + | 1140943 | 1142076 | 1.77 | 24.22 | 15.74 | 7.60 | Putative monooxygenase |
| *FEPFGCPE_01025* | *livF-4* | - | 1142097 | 1142819 | 2.39 | 30.64 | 19.03 | 10.01 | High-affinity branched-chain amino acid transport ATP-binding protein LivF |
| *FEPFGCPE_01026* | *xylG* | - | 1142893 | 1144881 | 1.75 | 18.80 | 7.75 | 3.66 | Xylose import ATP-binding protein XylG |
| *FEPFGCPE_01027* | *livH-4* | - | 1144884 | 1145762 | 1.70 | 14.71 | 5.38 | 6.48 | High-affinity branched-chain amino acid transport system permease protein LivH |
| *FEPFGCPE_01028* | *braC-1* | - | 1145991 | 1147205 | 11.15 | 172.79 | 64.03 | 35.35 | Leucine-, isoleucine-, valine-, threonine-, and alanine-binding protein |
| *FEPFGCPE_01029* | *ftsK-1* | - | 1147596 | 1151189 | 6.78 | 44.49 | 33.22 | 9.85 | DNA translocase FtsK |
| *FEPFGCPE_01030* | | - | 1151309 | 1152316 | 1.39 | 8.45 | 7.43 | 0.73 | Hypothetical protein |
| *FEPFGCPE_01031* | *pdeB* | - | 1152808 | 1155825 | 3.84 | 17.66 | 8.61 | 2.18 | Cyclic di-GMP phosphodiesterase PdeB |
| *FEPFGCPE_01032* | | - | 1156044 | 1156916 | 4.27 | 46.63 | 29.57 | 14.16 | Hypothetical protein |
| *FEPFGCPE_01033* | *slyA-1* | + | 1157280 | 1157774 | 3.39 | 6.27 | 5.65 | 2.35 | Transcriptional regulator SlyA |
| *FEPFGCPE_01034* | *emrB-1* | + | 1157771 | 1159324 | 1.47 | 4.02 | 0.57 | 0.34 | Multidrug export protein EmrB |
| *FEPFGCPE_01035* | *pcaR-1* | - | 1159332 | 1160063 | 1.90 | 3.87 | 4.92 | 2.19 | Pca regulon regulatory protein |
| *FEPFGCPE_01036* | *caiC* | - | 1160307 | 1161941 | 0.91 | 3.15 | 0.94 | 0.81 | putative crotonobetaine/carnitine-CoA ligase |
| *FEPFGCPE_01037* | *potH-1* | - | 1161959 | 1163707 | 0.38 | 1.77 | 0.70 | 0.00 | Putrescine transport system permease protein PotH |
| *FEPFGCPE_01038* | *potA-4* | - | 1163754 | 1164827 | 2.09 | 4.38 | 1.71 | 1.68 | Spermidine/putrescine import ATP-binding protein PotA |
| *FEPFGCPE_01039* | *potD-2* | - | 1164862 | 1165944 | 1.44 | 4.12 | 2.05 | 3.52 | Spermidine/putrescine-binding periplasmic protein |
| *FEPFGCPE_01040* | *crt-1* | - | 1166063 | 1166575 | 1.56 | 4.77 | 3.58 | 1.21 | Short-chain-enoyl-CoA hydratase |
| *FEPFGCPE_01041* | *crt-2* | - | 1166572 | 1166862 | 1.66 | 0.00 | 0.00 | 0.59 | Short-chain-enoyl-CoA hydratase |
| *FEPFGCPE_01042* | *atpC-1* | - | 1167138 | 1167563 | 7.46 | 42.35 | 37.59 | 9.67 | ATP synthase epsilon chain |
| *FEPFGCPE_01043* | | - | 1168053 | 1168565 | 0.79 | 2.22 | 3.58 | 0.32 | Hypothetical protein |
| *FEPFGCPE_01044* | *garB* | + | 1168953 | 1170314 | 1.87 | 10.74 | 5.11 | 3.21 | Glutathione amide reductase |
| *FEPFGCPE_01045* | | - | 1170377 | 1171771 | 1.24 | 9.35 | 7.08 | 2.81 | Hypothetical protein |
| *FEPFGCPE_01046* | *ectD* | - | 1172033 | 1172794 | 2.09 | 9.82 | 10.19 | 5.75 | Ectoine dioxygenase |
| *FEPFGCPE_01047* | | - | 1172899 | 1173267 | 13.98 | 53.39 | 41.64 | 11.82 | Hypothetical protein |
| *FEPFGCPE_01048* | | - | 1173361 | 1173564 | 35.35 | 224.77 | 153.98 | 61.82 | Hypothetical protein |
| *FEPFGCPE_01049* | | - | 1173651 | 1173884 | 5.47 | 35.57 | 43.86 | 8.39 | Hypothetical protein |
| *FEPFGCPE_01050* | *fdhD* | - | 1173912 | 1174871 | 4.54 | 15.85 | 8.45 | 4.21 | Sulfurtransferase FdhD |
| *FEPFGCPE_01051* | | - | 1174911 | 1177796 | 10.47 | 21.47 | 19.50 | 5.80 | Putative formate dehydrogenase |
| *FEPFGCPE_01052* | *hndC-1* | - | 1177827 | 1179467 | 8.95 | 18.84 | 15.18 | 4.26 | NADP-reducing hydrogenase subunit HndC |
| *FEPFGCPE_01053* | *nqo2-1* | - | 1179460 | 1179933 | 4.43 | 23.57 | 18.56 | 3.68 | NADH-quinone oxidoreductase subunit 2 |
| *FEPFGCPE_01054* | | - | 1180199 | 1181272 | 0.50 | 2.46 | 7.27 | 0.82 | Hypothetical protein |
| *FEPFGCPE_01055* | | + | 1181508 | 1183820 | 1.20 | 4.63 | 4.24 | 1.29 | putative oxidoreductase |
| *FEPFGCPE_01056* | | + | 1184184 | 1184462 | 1.19 | 7.06 | 16.06 | 12.32 | Hypothetical protein |
| *FEPFGCPE_01057* | | + | 1184452 | 1184706 | 1.74 | 11.84 | 6.56 | 9.31 | Hypothetical protein |
| *FEPFGCPE_01058* | | + | 1184717 | 1186303 | 1.53 | 4.36 | 8.39 | 9.12 | Hypothetical protein |
| *FEPFGCPE_01059* | | + | 1186816 | 1188396 | 2.96 | 20.69 | 61.39 | 101.81 | Isocitrate lyase |
| *FEPFGCPE_01060* | *fpr-1* | - | 1188489 | 1189421 | 26.32 | 54.79 | 56.43 | 31.78 | Ferredoxin--NADP reductase |
| *FEPFGCPE_01061* | *hupB-2* | + | 1189995 | 1190273 | 143.11 | 659.97 | 335.60 | 187.78 | DNA-binding protein HU-beta |

| Locus name | Gene name | Strand | Gene start (bp) | Gene end (bp) | Average TPM Inoculum | Average TPM $t_6$ | Average TPM $t_{24}$ | Average TPM $t_{168}$ | Annotation description |
|---|---|---|---|---|---|---|---|---|---|
| *FEPFGCPE_01062* | | + | 1190644 | 1191678 | 4.74 | 19.08 | 12.31 | 6.56 | Hypothetical protein |
| *FEPFGCPE_01063* | *cadA* | - | 1191683 | 1194121 | 12.82 | 42.75 | 22.90 | 26.90 | putative cadmium-transporting ATPase |
| *FEPFGCPE_01064* | *zntR-1* | + | 1194190 | 1194660 | 14.48 | 104.91 | 107.14 | 37.85 | HTH-type transcriptional regulator ZntR |
| *FEPFGCPE_01065* | *fccB-1* | + | 1195002 | 1196267 | 11.13 | 30.15 | 25.73 | 7.34 | Sulfide dehydrogenase [flavocytochrome c] flavoprotein chain |
| *FEPFGCPE_01066* | *cph2-1* | + | 1196584 | 1197918 | 4.49 | 18.90 | 12.49 | 5.20 | Phytochrome-like protein cph2 |
| *FEPFGCPE_01067* | | + | 1198429 | 1199370 | 1.10 | 11.21 | 8.10 | 3.57 | Putative phospholipase A1 |
| *FEPFGCPE_01068* | *arnE-1* | - | 1199451 | 1199873 | 6.25 | 25.11 | 16.89 | 5.02 | 4-amino-4-deoxy-L-arabinose-phosphoundecaprenol flippase subunit ArnE |
| *FEPFGCPE_01069* | | + | 1200043 | 1200258 | 9.93 | 38.33 | 38.00 | 17.71 | Hypothetical protein |
| *FEPFGCPE_01070* | *ald-2* | - | 1200239 | 1201672 | 5.29 | 19.26 | 16.33 | 6.61 | 3-succinoylsemialdehyde-pyridine dehydrogenase |
| *FEPFGCPE_01071* | | + | 1202008 | 1203207 | 6.07 | 10.80 | 9.77 | 3.65 | Hypothetical protein |
| *FEPFGCPE_01072* | *ybiT-1* | - | 1203305 | 1203565 | 2.59 | 3.44 | 4.97 | 1.25 | putative ABC transporter ATP-binding protein YbiT |
| *FEPFGCPE_01073* | *ybiT-2* | - | 1203562 | 1204899 | 12.81 | 17.57 | 31.26 | 14.83 | putative ABC transporter ATP-binding protein YbiT |
| *FEPFGCPE_01074* | *syrM1-1* | + | 1205370 | 1206308 | 16.36 | 18.31 | 15.08 | 6.93 | HTH-type transcriptional regulator SyrM 1 |
| *FEPFGCPE_01075* | | - | 1206528 | 1207886 | 24.14 | 10.74 | 10.26 | 1.52 | 8-oxoguanine deaminase |
| *FEPFGCPE_01076* | *uacT-2* | - | 1207886 | 1209253 | 24.30 | 11.14 | 5.07 | 2.95 | Uric acid transporter UacT |
| *FEPFGCPE_01077* | *pucM* | + | 1209690 | 1210040 | 40.55 | 16.06 | 6.80 | 6.65 | 5-hydroxyisourate hydrolase |
| *FEPFGCPE_01078* | | + | 1210083 | 1211288 | 14.16 | 13.59 | 7.58 | 3.56 | Hypothetical protein |
| *FEPFGCPE_01079* | *mepA* | + | 1211844 | 1213223 | 4.53 | 18.01 | 13.37 | 3.23 | Multidrug export protein MepA |
| *FEPFGCPE_01080* | | - | 1213301 | 1214146 | 2.42 | 24.99 | 21.00 | 5.30 | Hypothetical protein |
| *FEPFGCPE_01081* | *tgpA* | - | 1214196 | 1215002 | 6.89 | 40.43 | 44.19 | 11.11 | Protein-glutamine gamma-glutamyltransferase |
| *FEPFGCPE_01082* | | - | 1215007 | 1216005 | 3.25 | 40.46 | 73.24 | 19.34 | Hypothetical protein |
| *FEPFGCPE_01083* | | - | 1216051 | 1217475 | 2.79 | 42.78 | 53.54 | 17.21 | Hypothetical protein |
| *FEPFGCPE_01084* | *allA-2* | - | 1217934 | 1218494 | 75.42 | 62.66 | 40.58 | 13.64 | Ureidoglycolate lyase |
| *FEPFGCPE_01085* | *alc-2* | - | 1218544 | 1219566 | 53.68 | 71.30 | 46.49 | 13.49 | allantoicase |
| *FEPFGCPE_01086* | *uao* | - | 1219862 | 1220200 | 22.67 | 23.91 | 10.53 | 3.29 | Uric acid degradation bifunctional protein |
| *FEPFGCPE_01087* | *pgdA* | - | 1220335 | 1221279 | 16.75 | 21.03 | 15.46 | 7.37 | Peptidoglycan deacetylase |
| *FEPFGCPE_01088* | *ldcA* | - | 1221647 | 1222579 | 4.24 | 9.67 | 8.39 | 2.98 | Murein tetrapeptide carboxypeptidase |
| *FEPFGCPE_01089* | *tadA* | - | 1222576 | 1223085 | 11.36 | 36.49 | 24.39 | 11.85 | tRNA-specific adenosine deaminase |
| *FEPFGCPE_01090* | *dkgB* | - | 1223299 | 1224102 | 3.10 | 18.27 | 17.37 | 5.26 | 2,5-diketo-D-gluconic acid reductase B |
| *FEPFGCPE_01091* | *rspR-2* | - | 1224289 | 1224897 | 3.14 | 13.84 | 6.34 | 1.66 | HTH-type transcriptional repressor RspR |
| *FEPFGCPE_01092* | | - | 1224963 | 1225898 | 3.62 | 24.89 | 24.53 | 4.07 | Hypothetical protein |
| *FEPFGCPE_01093* | *lon2* | - | 1225980 | 1226651 | 10.79 | 22.64 | 9.04 | 3.41 | Lon protease 2 |
| *FEPFGCPE_01094* | *fabG-3* | + | 1226849 | 1227553 | 2.70 | 18.25 | 14.64 | 3.73 | 3-oxoacyl-[acyl-carrier-protein] reductase FabG |
| *FEPFGCPE_01095* | *oxyR-2* | - | 1227747 | 1228619 | 14.00 | 36.07 | 15.29 | 4.82 | Hydrogen peroxide-inducible genes activator |
| *FEPFGCPE_01096* | *proP-5* | + | 1228789 | 1230114 | 2.60 | 11.85 | 10.92 | 3.30 | Proline/betaine transporter |
| *FEPFGCPE_01097* | | - | 1230172 | 1230588 | 1.25 | 3.03 | 3.68 | 5.10 | Hypothetical protein |
| *FEPFGCPE_01098* | *yxeP-1* | + | 1230648 | 1231457 | 1.86 | 5.55 | 2.98 | 1.45 | putative hydrolase YxeP |
| *FEPFGCPE_01099* | | + | 1231464 | 1232558 | 2.21 | 6.32 | 4.68 | 0.84 | Hypothetical protein |
| *FEPFGCPE_01100* | *mtnN-1* | - | 1233497 | 1235473 | 45.58 | 50.99 | 39.65 | 14.66 | 5'-methylthioadenosine/S-adenosylhomocysteine nucleosidase |
| *FEPFGCPE_01101* | | - | 1236076 | 1236855 | 1.55 | 5.39 | 3.74 | 1.59 | Hypothetical protein |
| *FEPFGCPE_01102* | | + | 1236878 | 1237471 | 31.80 | 50.25 | 30.99 | 9.33 | Hypothetical protein |
| *FEPFGCPE_01103* | | + | 1237464 | 1238501 | 19.73 | 23.36 | 15.65 | 5.88 | Hypothetical protein |
| *FEPFGCPE_01104* | | + | 1238601 | 1239404 | 4.91 | 7.28 | 8.57 | 2.69 | Hypothetical protein |
| *FEPFGCPE_01105* | | + | 1239416 | 1240195 | 9.83 | 29.59 | 24.25 | 5.01 | Hypothetical protein |
| *FEPFGCPE_01106* | *helY* | + | 1240203 | 1243700 | 3.04 | 9.78 | 3.94 | 1.98 | putative helicase HelY |
| *FEPFGCPE_01107* | | + | 1243708 | 1244874 | 5.69 | 24.33 | 11.22 | 3.14 | Hypothetical protein |
| *FEPFGCPE_01108* | | - | 1245015 | 1248308 | 3.34 | 21.97 | 12.20 | 3.95 | Hypothetical protein |
| *FEPFGCPE_01109* | | + | 1248929 | 1249603 | 5.39 | 48.43 | 36.36 | 6.44 | Hypothetical protein |
| *FEPFGCPE_01110* | | - | 1249574 | 1250356 | 1.14 | 11.58 | 7.41 | 1.09 | Insertion sequence IS5376 putative ATP-binding protein |
| *FEPFGCPE_01111* | | - | 1250366 | 1251910 | 0.16 | 0.37 | 0.42 | 0.00 | Hypothetical protein |
| *FEPFGCPE_01112* | | + | 1252186 | 1252929 | 0.53 | 6.67 | 0.72 | 1.09 | Hypothetical protein |
| *FEPFGCPE_01113* | | + | 1252988 | 1253563 | 0.31 | 3.71 | 1.27 | 0.00 | Hypothetical protein |
| *FEPFGCPE_01114* | | + | 1253610 | 1254314 | 1.66 | 3.18 | 2.87 | 0.51 | Hypothetical protein |
| *FEPFGCPE_01115* | | + | 1254627 | 1255565 | 1.95 | 8.87 | 7.24 | 2.32 | Hypothetical protein |
| *FEPFGCPE_01116* | | - | 1255750 | 1257945 | 1.73 | 14.65 | 8.70 | 2.63 | Hypothetical protein |
| *FEPFGCPE_01117* | | - | 1257942 | 1259090 | 4.96 | 21.70 | 26.22 | 5.20 | Hypothetical protein |
| *FEPFGCPE_01118* | | - | 1259137 | 1259577 | 7.79 | 51.87 | 23.43 | 8.04 | Hypothetical protein |
| *FEPFGCPE_01119* | | + | 1260910 | 1261836 | 13.78 | 23.26 | 14.44 | 4.16 | Hypothetical protein |
| *FEPFGCPE_01120* | | + | 1262366 | 1264018 | 3.80 | 20.13 | 10.60 | 4.26 | Hypothetical protein |
| *FEPFGCPE_01121* | *amnD-2* | - | 1264184 | 1264543 | 5.70 | 21.85 | 21.69 | 2.82 | 2-aminomuconate deaminase |
| *FEPFGCPE_01122* | | + | 1264765 | 1265037 | 4.04 | 18.93 | 10.04 | 2.24 | Hypothetical protein |
| *FEPFGCPE_01123* | | + | 1265055 | 1265906 | 1.36 | 15.10 | 7.41 | 1.26 | Hypothetical protein |
| *FEPFGCPE_01124* | *intA* | - | 1266802 | 1268106 | 3.96 | 10.95 | 12.35 | 3.43 | Prophage integrase IntA |
| *FEPFGCPE_01125* | *guaA* | - | 1268329 | 1269912 | 7.50 | 30.78 | 32.89 | 19.80 | GMP synthase [glutamine-hydrolyzing] |
| *FEPFGCPE_01126* | | - | 1269887 | 1270579 | 7.00 | 51.02 | 48.20 | 22.19 | Hypothetical protein |
| *FEPFGCPE_01127* | *guaB-3* | - | 1270586 | 1272046 | 7.01 | 54.23 | 56.02 | 30.89 | Inosine-5'-monophosphate dehydrogenase |
| *FEPFGCPE_01128* | *ytfF* | + | 1272333 | 1273262 | 0.73 | 11.04 | 9.28 | 2.00 | Inner membrane protein YtfF |
| *FEPFGCPE_01129* | *pasI* | - | 1273402 | 1273737 | 11.04 | 106.49 | 99.46 | 27.57 | Persistence and stress-resistance antitoxin PasI |
| *FEPFGCPE_01130* | *pasT* | - | 1273727 | 1274209 | 2.80 | 28.96 | 31.03 | 12.79 | Persistence and stress-resistance toxin PasT |
| *FEPFGCPE_01131* | *smpB* | + | 1274319 | 1274765 | 8.80 | 71.34 | 52.10 | 23.61 | SsrA-binding protein |
| *FEPFGCPE_01132* | *hflK-1* | - | 1274904 | 1275890 | 10.65 | 47.60 | 41.68 | 18.23 | Modulator of FtsH protease HflK |

| Locus name | Gene name | Strand | Gene start (bp) | Gene end (bp) | Average TPM Inoculum | Average TPM $t_6$ | Average TPM $t_{24}$ | Average TPM $t_{168}$ | Annotation description |
|---|---|---|---|---|---|---|---|---|---|
| *FEPFGCPE_01133* | | - | 1276029 | 1276463 | 7.58 | 54.62 | 39.62 | 14.69 | Hypothetical protein |
| *FEPFGCPE_01134* | *ppsA* | - | 1276630 | 1279020 | 13.83 | 45.42 | 46.12 | 37.04 | Phosphoenolpyruvate synthase |
| *FEPFGCPE_01135* | *ppsR* | + | 1279286 | 1280164 | 7.98 | 32.18 | 32.48 | 15.28 | Phosphoenolpyruvate synthase regulatory protein |
| *FEPFGCPE_01136* | *rlmB-1* | - | 1280274 | 1281053 | 1.87 | 9.82 | 9.26 | 5.49 | 23S rRNA (guanosine-2'-O-)-methyltransferase RlmB |
| *FEPFGCPE_01137* | *rnhB* | - | 1281050 | 1281793 | 1.87 | 5.92 | 6.60 | 2.49 | Ribonuclease HII |
| *FEPFGCPE_01138* | *lpxB* | - | 1281790 | 1282965 | 2.03 | 8.36 | 7.51 | 3.63 | Lipid-A-disaccharide synthase |
| *FEPFGCPE_01139* | *lpxA* | - | 1282969 | 1283760 | 6.76 | 39.56 | 31.97 | 14.61 | Acyl-[acyl-carrier-protein]--UDP-N-acetylglucosamine O-acyltransferase |
| *FEPFGCPE_01140* | *fabZ-2* | - | 1283778 | 1284236 | 9.56 | 36.18 | 33.01 | 21.91 | 3-hydroxyacyl-[acyl-carrier-protein] dehydratase FabZ |
| *FEPFGCPE_01141* | *lpxD* | - | 1284233 | 1285303 | 2.91 | 30.57 | 29.84 | 17.93 | UDP-3-O-acylglucosamine N-acyltransferase |
| *FEPFGCPE_01142* | *skp* | - | 1285318 | 1285860 | 2.74 | 18.97 | 22.93 | 9.95 | Chaperone protein Skp |
| *FEPFGCPE_01143* | *bamA* | - | 1285908 | 1288211 | 3.61 | 41.01 | 35.30 | 18.52 | Outer membrane protein assembly factor BamA |
| *FEPFGCPE_01144* | *rseP* | - | 1288283 | 1289674 | 3.47 | 15.26 | 9.59 | 2.98 | Regulator of sigma-E protease RseP |
| *FEPFGCPE_01145* | *dxr* | - | 1289718 | 1290917 | 1.68 | 6.43 | 6.86 | 3.88 | 1-deoxy-D-xylulose 5-phosphate reductoisomerase |
| *FEPFGCPE_01146* | *cdsA* | - | 1290920 | 1291729 | 4.26 | 13.76 | 23.34 | 4.04 | Phosphatidate cytidylyltransferase |
| *FEPFGCPE_01147* | *uppS* | - | 1291729 | 1292442 | 19.03 | 49.88 | 35.17 | 13.57 | Ditrans,polycis-undecaprenyl-diphosphate synthase ((2E,6E)-farnesyl-diphosphate specific) |
| *FEPFGCPE_01148* | *frr* | - | 1292576 | 1293136 | 21.52 | 45.02 | 37.61 | 25.54 | Ribosome-recycling factor |
| *FEPFGCPE_01149* | *pyrH* | - | 1293222 | 1293935 | 8.95 | 42.34 | 24.99 | 16.12 | Uridylate kinase |
| *FEPFGCPE_01150* | *tsf* | - | 1294033 | 1294914 | 13.42 | 43.19 | 59.37 | 51.49 | Elongation factor Ts |
| *FEPFGCPE_01151* | *rpsB* | - | 1295070 | 1295822 | 21.77 | 54.80 | 58.52 | 37.77 | 30S ribosomal protein S2 |
| *FEPFGCPE_01152* | *map-2* | + | 1296343 | 1297170 | 10.22 | 48.05 | 29.78 | 16.17 | Methionine aminopeptidase |
| *FEPFGCPE_01153* | *glnD* | + | 1297247 | 1299811 | 4.96 | 25.02 | 18.29 | 9.89 | Bifunctional uridylyltransferase/uridylyl-removing enzyme |
| *FEPFGCPE_01154* | *rluF* | + | 1299808 | 1301679 | 2.52 | 11.34 | 14.67 | 5.58 | 23S rRNA pseudouridine(2604) synthase |
| *FEPFGCPE_01155* | *def* | - | 1301877 | 1302410 | 22.96 | 72.54 | 32.41 | 19.51 | Peptide deformylase |
| *FEPFGCPE_01156* | *ligA* | - | 1302427 | 1304520 | 9.04 | 33.45 | 18.61 | 7.00 | DNA ligase |
| *FEPFGCPE_01157* | *zipA* | - | 1304698 | 1306215 | 4.01 | 19.16 | 18.31 | 11.64 | Cell division protein ZipA |
| *FEPFGCPE_01158* | *smc-2* | - | 1306255 | 1309770 | 8.06 | 34.45 | 24.65 | 11.86 | Chromosome partition protein Smc |
| *FEPFGCPE_01159* | | - | 1309894 | 1310307 | 6.55 | 32.08 | 7.90 | 9.67 | Hypothetical protein |
| *FEPFGCPE_01160* | | - | 1310300 | 1310878 | 3.93 | 5.82 | 10.91 | 3.90 | Hypothetical protein |
| *FEPFGCPE_01161* | *dapL* | + | 1311233 | 1312435 | 7.34 | 30.02 | 31.82 | 12.98 | LL-diaminopimelate aminotransferase |
| *FEPFGCPE_01162* | *dapD* | + | 1312460 | 1313287 | 7.89 | 47.93 | 29.35 | 26.24 | 2,3,4,5-tetrahydropyridine-2,6-dicarboxylate N-succinyltransferase |
| *FEPFGCPE_01163* | *spxA* | + | 1313375 | 1313722 | 5.39 | 16.02 | 23.63 | 9.82 | Regulatory protein Spx |
| *FEPFGCPE_01164* | *dapE-2* | + | 1313727 | 1314878 | 3.71 | 20.28 | 21.87 | 10.68 | Succinyl-diaminopimelate desuccinylase |
| *FEPFGCPE_01165* | *prmB* | + | 1314899 | 1315792 | 2.50 | 13.61 | 7.77 | 6.50 | 50S ribosomal protein L3 glutamine methyltransferase |
| *FEPFGCPE_01166* | *clsA* | + | 1315904 | 1317331 | 2.43 | 21.77 | 9.92 | 3.30 | Major cardiolipin synthase ClsA |
| *FEPFGCPE_01167* | *yheS-2* | + | 1317504 | 1319444 | 1.91 | 9.78 | 9.07 | 4.91 | putative ABC transporter ATP-binding protein YheS |
| *FEPFGCPE_01168* | *radA* | - | 1319526 | 1320899 | 5.04 | 17.53 | 17.25 | 6.10 | DNA repair protein RadA |
| *FEPFGCPE_01169* | *dadX* | - | 1320923 | 1321993 | 5.23 | 13.07 | 15.43 | 5.71 | Alanine racemase, catabolic |
| *FEPFGCPE_01170* | *lplT-2* | + | 1322331 | 1323623 | 2.33 | 19.63 | 12.17 | 5.87 | Lysophospholipid transporter LplT |
| *FEPFGCPE_01171* | *thiD-1* | + | 1323625 | 1324449 | 1.94 | 15.56 | 17.04 | 6.00 | Hydroxymethylpyrimidine/phosphomethylpyrimidine kinase |
| *FEPFGCPE_01172* | | - | 1324494 | 1325636 | 1.90 | 13.50 | 12.81 | 4.62 | Hypothetical protein |
| *FEPFGCPE_01173* | | - | 1325597 | 1326667 | 4.78 | 24.47 | 21.06 | 6.57 | Hypothetical protein |
| *FEPFGCPE_01174* | *rimI* | - | 1326654 | 1327133 | 5.10 | 35.10 | 37.20 | 14.58 | Ribosomal-protein-alanine acetyltransferase |
| *FEPFGCPE_01175* | *tsaB* | - | 1327133 | 1327903 | 2.97 | 12.80 | 13.63 | 4.31 | tRNA threonylcarbamoyladenosine biosynthesis protein TsaB |
| *FEPFGCPE_01176* | | - | 1328089 | 1328244 | 80.24 | 401.03 | 292.98 | 99.17 | Hypothetical protein |
| *FEPFGCPE_01177* | *rhlE-2* | + | 1328737 | 1329942 | 2.10 | 14.04 | 30.53 | 27.58 | ATP-dependent RNA helicase RhlE |
| *FEPFGCPE_01178* | | + | 1329905 | 1330129 | 3.88 | 4.20 | 17.06 | 5.37 | Hypothetical protein |
| *FEPFGCPE_01179* | *pip* | - | 1330306 | 1331364 | 16.22 | 27.87 | 22.88 | 8.49 | Proline iminopeptidase |
| *FEPFGCPE_01180* | *aceB* | - | 1331646 | 1333244 | 0.38 | 2.80 | 1.84 | 0.71 | Malate synthase A |
| *FEPFGCPE_01181* | | - | 1333338 | 1334045 | 1.42 | 4.85 | 4.75 | 0.83 | 2-haloalkanoic acid dehalogenase |
| *FEPFGCPE_01182* | *dmlR-7* | + | 1334165 | 1335106 | 1.87 | 7.69 | 3.58 | 1.67 | HTH-type transcriptional regulator DmlR |
| *FEPFGCPE_01183* | *teaD* | + | 1335420 | 1335857 | 25.19 | 70.87 | 38.83 | 7.10 | TRAP-T-associated universal stress protein TeaD |
| *FEPFGCPE_01184* | *aceA* | - | 1336092 | 1337414 | 46.91 | 25.84 | 63.67 | 139.97 | Isocitrate lyase |
| *FEPFGCPE_01185* | *yijE* | - | 1337748 | 1338752 | 2.54 | 8.27 | 6.14 | 1.54 | putative cystine transporter YijE |
| *FEPFGCPE_01186* | *queF-1* | + | 1339081 | 1339767 | 1.47 | 12.13 | 8.72 | 3.20 | NADPH-dependent 7-cyano-7-deazaguanine reductase |
| *FEPFGCPE_01187* | | + | 1339764 | 1340210 | 4.51 | 42.20 | 27.90 | 11.91 | Hypothetical protein |
| *FEPFGCPE_01188* | *btrG* | - | 1340338 | 1340832 | 5.83 | 17.87 | 22.76 | 13.17 | Gamma-L-glutamyl-butirosin B gamma-glutamyl cyclotransferase |
| *FEPFGCPE_01189* | *clpB* | - | 1341056 | 1343692 | 24.56 | 66.28 | 72.15 | 32.65 | Chaperone protein ClpB |
| *FEPFGCPE_01190* | *ctb* | + | 1343977 | 1344462 | 3.55 | 9.24 | 9.07 | 2.93 | Group 3 truncated hemoglobin ctb |
| *FEPFGCPE_01191* | *moaE* | - | 1344520 | 1345002 | 3.73 | 23.32 | 29.94 | 7.31 | Molybdopterin synthase catalytic subunit |
| *FEPFGCPE_01192* | *moaD* | - | 1345055 | 1345333 | 7.23 | 77.06 | 68.13 | 12.95 | Molybdopterin synthase sulfur carrier subunit |
| *FEPFGCPE_01193* | *moeA-1* | - | 1345352 | 1346563 | 2.06 | 8.02 | 7.96 | 7.92 | Molybdopterin molybdenumtransferase |
| *FEPFGCPE_01194* | *thrC-1* | - | 1346617 | 1348083 | 2.56 | 13.38 | 9.70 | 10.25 | Threonine synthase |
| *FEPFGCPE_01195* | *hom* | - | 1348124 | 1349434 | 5.47 | 18.63 | 14.11 | 11.87 | Homoserine dehydrogenase |
| *FEPFGCPE_01196* | *alaA* | - | 1349472 | 1350809 | 10.95 | 34.18 | 28.06 | 12.83 | Glutamate-pyruvate aminotransferase AlaA |
| *FEPFGCPE_01197* | | + | 1350837 | 1351406 | 13.23 | 52.12 | 54.94 | 23.31 | Hypothetical protein |
| *FEPFGCPE_01198* | *arnT-2* | + | 1351403 | 1352389 | 1.34 | 4.06 | 5.56 | 1.34 | Undecaprenyl phosphate-alpha-4-amino-4-deoxy-L-arabinose arabinosyl transferase |
| *FEPFGCPE_01199* | | + | 1352480 | 1352767 | 1.95 | 6.78 | 3.68 | 0.80 | Hypothetical protein |
| *FEPFGCPE_01200* | | + | 1352752 | 1353165 | 3.31 | 13.34 | 24.79 | 11.40 | Hypothetical protein |
| *FEPFGCPE_01201* | *arnF* | + | 1353309 | 1353680 | 4.87 | 56.82 | 38.30 | 18.59 | putative 4-amino-4-deoxy-L-arabinose-phosphoundecaprenol flippase subunit ArnF |
| *FEPFGCPE_01202* | *arnB-1* | + | 1353769 | 1354935 | 4.99 | 29.68 | 20.43 | 11.44 | UDP-4-amino-4-deoxy-L-arabinose--oxoglutarate aminotransferase |
| *FEPFGCPE_01203* | *arnC-1* | + | 1354932 | 1355954 | 8.05 | 39.97 | 29.37 | 8.07 | Undecaprenyl-phosphate 4-deoxy-4-formamido-L-arabinose transferase |

| Locus name | Gene name | Strand | Gene start (bp) | Gene end (bp) | Average TPM Inoculum | Average TPM $t_6$ | Average TPM $t_{24}$ | Average TPM $t_{168}$ | Annotation description |
|---|---|---|---|---|---|---|---|---|---|
| FEPFGCPE_01204 | arnA-1 | + | 1355951 | 1356874 | 3.64 | 28.82 | 17.62 | 9.62 | Bifunctional polymyxin resistance protein ArnA |
| FEPFGCPE_01205 | arnA-2 | + | 1356882 | 1357931 | 11.19 | 62.99 | 53.17 | 27.20 | Bifunctional polymyxin resistance protein ArnA |
| FEPFGCPE_01206 | arnD | + | 1358142 | 1359047 | 7.24 | 25.81 | 20.07 | 4.03 | putative 4-deoxy-4-formamido-L-arabinose-phosphoundecaprenol deformylase ArnD |
| FEPFGCPE_01207 | bcp-2 | + | 1359106 | 1359567 | 10.53 | 40.70 | 26.73 | 21.34 | Putative peroxiredoxin bcp |
| FEPFGCPE_01208 | | + | 1359900 | 1361642 | 29.25 | 53.89 | 40.31 | 18.01 | PhoH-like protein |
| FEPFGCPE_01209 | mepH | + | 1361874 | 1362692 | 26.02 | 66.76 | 38.04 | 16.04 | Murein DD-endopeptidase MepH |
| FEPFGCPE_01210 | | - | 1362736 | 1362843 | 24.21 | 207.77 | 223.59 | 44.34 | Hypothetical protein |
| FEPFGCPE_01211 | pitA-1 | - | 1363059 | 1364069 | 1.79 | 13.63 | 7.50 | 2.86 | Low-affinity inorganic phosphate transporter 1 |
| FEPFGCPE_01212 | | - | 1364083 | 1364709 | 5.06 | 30.63 | 35.78 | 11.34 | Hypothetical protein |
| FEPFGCPE_01213 | dnaB | - | 1364824 | 1366299 | 5.27 | 35.39 | 31.23 | 12.03 | Replicative DNA helicase |
| FEPFGCPE_01214 | rplI | - | 1366493 | 1366945 | 7.56 | 30.90 | 22.61 | 39.56 | 50S ribosomal protein L9 |
| FEPFGCPE_01215 | rpsR | - | 1366973 | 1367254 | 15.56 | 81.71 | 72.09 | 89.86 | 30S ribosomal protein S18 |
| FEPFGCPE_01216 | priB | - | 1367257 | 1367559 | 18.60 | 104.51 | 98.77 | 90.40 | Primosomal replication protein N |
| FEPFGCPE_01217 | rpsF | - | 1367606 | 1367983 | 54.67 | 193.43 | 245.53 | 172.02 | 30S ribosomal protein S6 |
| FEPFGCPE_01218 | dmlR-8 | - | 1368102 | 1369013 | 3.11 | 10.19 | 12.29 | 5.44 | HTH-type transcriptional regulator DmlR |
| FEPFGCPE_01219 | gcl | + | 1369135 | 1370901 | 0.53 | 2.56 | 1.59 | 0.53 | Glyoxylate carboligase |
| FEPFGCPE_01220 | hyi | + | 1371058 | 1371849 | 1.84 | 5.68 | 3.65 | 0.54 | Hydroxypyruvate isomerase |
| FEPFGCPE_01221 | glxR-1 | + | 1372089 | 1372970 | 1.40 | 9.21 | 4.44 | 1.69 | 2-hydroxy-3-oxopropionate reductase |
| FEPFGCPE_01222 | ansA-2 | - | 1373097 | 1374137 | 5.57 | 35.36 | 30.43 | 9.94 | L-asparaginase |
| FEPFGCPE_01223 | cbl | - | 1374324 | 1375262 | 0.34 | 93.95 | 34.11 | 18.43 | HTH-type transcriptional regulator cbl |
| FEPFGCPE_01224 | cysA-1 | - | 1375518 | 1376606 | 0.49 | 193.75 | 92.45 | 35.61 | Sulfate/thiosulfate import ATP-binding protein CysA |
| FEPFGCPE_01225 | cysW | - | 1376634 | 1377539 | 1.08 | 177.12 | 74.24 | 31.72 | Sulfate transport system permease protein CysW |
| FEPFGCPE_01226 | cysT | - | 1377560 | 1378360 | 0.27 | 61.10 | 24.33 | 15.01 | Sulfate transport system permease protein CysT |
| FEPFGCPE_01227 | sbp | - | 1378458 | 1379489 | 1.33 | 449.29 | 169.76 | 83.36 | Sulfate-binding protein |
| FEPFGCPE_01228 | lexA-1 | + | 1379747 | 1380400 | 5.87 | 46.86 | 25.55 | 11.52 | LexA repressor |
| FEPFGCPE_01229 | | + | 1380575 | 1380985 | 3.80 | 12.08 | 13.98 | 2.24 | Hypothetical protein |
| FEPFGCPE_01230 | crt-3 | - | 1381024 | 1382136 | 2.09 | 9.35 | 6.50 | 2.57 | Short-chain-enoyl-CoA hydratase |
| FEPFGCPE_01231 | typA | - | 1382273 | 1384093 | 7.33 | 18.24 | 23.02 | 17.21 | GTP-binding protein TypA/BipA |
| FEPFGCPE_01232 | | + | 1384388 | 1384897 | 10.18 | 60.59 | 70.25 | 13.27 | putative HTH-type transcriptional regulator/GBAA_1941/BAS1801 |
| FEPFGCPE_01233 | mepC-3 | + | 1385015 | 1386532 | 5.74 | 53.69 | 62.61 | 17.31 | Multidrug/solvent efflux pump outer membrane protein MepC |
| FEPFGCPE_01234 | emrA-1 | + | 1386606 | 1387841 | 3.40 | 37.89 | 41.78 | 22.81 | Multidrug export protein EmrA |
| FEPFGCPE_01235 | emrB-2 | + | 1387879 | 1389429 | 1.64 | 33.57 | 35.54 | 10.37 | Multidrug export protein EmrB |
| FEPFGCPE_01236 | | + | 1389674 | 1390168 | 3.55 | 17.37 | 8.48 | 2.21 | putative acyl-CoA thioester hydrolase |
| FEPFGCPE_01237 | truB | - | 1390310 | 1391296 | 0.62 | 6.31 | 3.16 | 1.31 | tRNA pseudouridine synthase B |
| FEPFGCPE_01238 | rbfA | - | 1391414 | 1391848 | 2.36 | 22.29 | 21.63 | 10.66 | Ribosome-binding factor A |
| FEPFGCPE_01239 | infB | - | 1391932 | 1394880 | 8.90 | 47.48 | 53.04 | 34.16 | Translation initiation factor IF-2 |
| FEPFGCPE_01240 | nusA | - | 1394961 | 1396436 | 7.25 | 36.07 | 43.41 | 24.96 | Transcription termination/antitermination protein NusA |
| FEPFGCPE_01241 | rimP | - | 1396433 | 1396891 | 6.90 | 33.12 | 35.07 | 27.04 | Ribosome maturation factor RimP |
| FEPFGCPE_01242 | rluB | - | 1397352 | 1399172 | 10.62 | 52.31 | 53.70 | 28.37 | Ribosomal large subunit pseudouridine synthase B |
| FEPFGCPE_01243 | scpB | - | 1399326 | 1400546 | 42.56 | 206.44 | 185.84 | 42.14 | Segregation and condensation protein B |
| FEPFGCPE_01244 | fnr-2 | + | 1400874 | 1401602 | 21.53 | 60.96 | 61.44 | 22.81 | Fumarate and nitrate reduction regulatory protein |
| FEPFGCPE_01245 | | + | 1401862 | 1402296 | 5.14 | 20.21 | 11.41 | 6.96 | Hypothetical protein |
| FEPFGCPE_01246 | dmlR-9 | - | 1402309 | 1403241 | 2.18 | 7.78 | 7.72 | 2.02 | HTH-type transcriptional regulator DmlR |
| FEPFGCPE_01247 | | + | 1403379 | 1404395 | 0.49 | 1.50 | 0.93 | 0.07 | Zinc-type alcohol dehydrogenase-like protein |
| FEPFGCPE_01248 | cinA | - | 1404511 | 1405350 | 2.35 | 15.76 | 14.62 | 6.11 | CinA-like protein |
| FEPFGCPE_01249 | | - | 1405438 | 1406358 | 1.06 | 7.39 | 4.76 | 2.47 | Hypothetical protein |
| FEPFGCPE_01250 | | - | 1406477 | 1407472 | 5.53 | 7.94 | 7.16 | 3.92 | PE-PGRS family protein PE_PGRS18 |
| FEPFGCPE_01251 | | - | 1407840 | 1411931 | 0.99 | 3.62 | 2.53 | 0.75 | Hypothetical protein |
| FEPFGCPE_01252 | argA-1 | + | 1412027 | 1413376 | 8.81 | 29.12 | 21.69 | 7.35 | Amino-acid acetyltransferase |
| FEPFGCPE_01253 | | + | 1413554 | 1413829 | 15.23 | 81.32 | 61.09 | 17.53 | putative Fe(2+)-trafficking protein |
| FEPFGCPE_01254 | | - | 1414016 | 1415083 | 3.13 | 20.96 | 20.95 | 8.77 | Hypothetical protein |
| FEPFGCPE_01255 | rpiA | - | 1415099 | 1415791 | 2.15 | 22.53 | 22.68 | 9.80 | Ribose-5-phosphate isomerase A |
| FEPFGCPE_01256 | talB | + | 1415919 | 1416869 | 6.85 | 18.39 | 18.31 | 9.36 | Transaldolase B |
| FEPFGCPE_01257 | rlmB-2 | - | 1416988 | 1417761 | 2.76 | 9.97 | 7.70 | 7.08 | 23S rRNA (guanosine-2'-O-)-methyltransferase RlmB |
| FEPFGCPE_01258 | rnr | - | 1417942 | 1420623 | 15.50 | 59.97 | 45.66 | 17.81 | Ribonuclease R |
| FEPFGCPE_01259 | | + | 1420787 | 1420871 | 4.45 | 11.16 | 6.51 | 1.16 | tRNA |
| FEPFGCPE_01260 | pgrR-3 | + | 1421222 | 1422136 | 2.49 | 7.26 | 5.58 | 1.72 | HTH-type transcriptional regulator PgrR |
| FEPFGCPE_01261 | | + | 1422271 | 1422355 | 5.95 | 22.24 | 3.85 | 7.74 | tRNA |
| FEPFGCPE_01262 | rsuA | + | 1422474 | 1423145 | 0.83 | 6.25 | 5.46 | 1.73 | Ribosomal small subunit pseudouridine synthase A |
| FEPFGCPE_01263 | rlmJ | - | 1423226 | 1424098 | 4.88 | 38.93 | 30.94 | 8.45 | Ribosomal RNA large subunit methyltransferase J |
| FEPFGCPE_01264 | tag | - | 1424098 | 1424760 | 5.54 | 24.68 | 15.09 | 7.64 | DNA-3-methyladenine glycosylase 1 |
| FEPFGCPE_01265 | | - | 1424973 | 1425833 | 6.53 | 21.11 | 10.60 | 6.63 | Ureidoglycolate lyase |
| FEPFGCPE_01266 | chaA | + | 1426246 | 1427328 | 2.16 | 12.44 | 9.54 | 3.52 | Sodium-potassium/proton antiporter ChaA |
| FEPFGCPE_01267 | | - | 1427351 | 1427581 | 2.49 | 44.58 | 28.76 | 18.55 | Hypothetical protein |
| FEPFGCPE_01268 | | - | 1427636 | 1427890 | 10.43 | 65.96 | 55.54 | 19.94 | Hypothetical protein |
| FEPFGCPE_01269 | ribC | + | 1428005 | 1428742 | 4.01 | 27.27 | 12.41 | 7.52 | Riboflavin synthase |
| FEPFGCPE_01270 | xthA | + | 1428771 | 1429550 | 2.92 | 21.11 | 10.12 | 6.37 | Exodeoxyribonuclease III |
| FEPFGCPE_01271 | | - | 1429656 | 1430639 | 1.58 | 9.47 | 7.51 | 3.34 | Hypothetical protein |
| FEPFGCPE_01272 | pipB2 | - | 1430915 | 1433764 | 0.78 | 3.56 | 2.59 | 1.54 | Secreted effector protein PipB2 |
| FEPFGCPE_01273 | rmlD | - | 1433890 | 1434771 | 6.34 | 34.45 | 22.00 | 10.01 | dTDP-4-dehydrorhamnose reductase |
| FEPFGCPE_01274 | glnG-1 | - | 1434827 | 1436413 | 2.27 | 106.30 | 111.68 | 39.12 | Nitrogen regulation protein NR(I) |

| Locus name | Gene name | Strand | Gene start (bp) | Gene end (bp) | Average TPM Inoculum | Average TPM $t_6$ | Average TPM $t_{24}$ | Average TPM $t_{168}$ | Annotation description |
|---|---|---|---|---|---|---|---|---|---|
| *FEPFGCPE_01275* | *glnL* | - | 1436476 | 1437690 | 1.94 | 66.64 | 52.93 | 15.02 | Nitrogen regulation protein NR(II) |
| *FEPFGCPE_01276* | | - | 1437848 | 1439263 | 13.58 | 424.13 | 428.16 | 248.95 | 3-hydroxylaminophenol mutase |
| *FEPFGCPE_01277* | | + | 1439727 | 1440194 | 24.78 | 117.80 | 77.61 | 32.20 | Hypothetical protein |
| *FEPFGCPE_01278* | *rhtB-1* | + | 1440302 | 1440919 | 5.80 | 36.96 | 37.76 | 5.99 | Homoserine/homoserine lactone efflux protein |
| *FEPFGCPE_01279* | | + | 1441170 | 1441502 | 0.86 | 7.05 | 19.05 | 15.71 | Hypothetical protein |
| *FEPFGCPE_01280* | | + | 1441936 | 1442175 | 6.33 | 46.47 | 32.27 | 9.48 | Hypothetical protein |
| *FEPFGCPE_01281* | | + | 1442359 | 1442544 | 12.82 | 158.86 | 82.10 | 23.77 | Hypothetical protein |
| *FEPFGCPE_01282* | | + | 1442616 | 1442691 | 24.25 | 122.02 | 83.76 | 23.39 | tRNA |
| *FEPFGCPE_01283* | | + | 1442749 | 1442824 | 24.53 | 118.00 | 71.12 | 24.64 | tRNA |
| *FEPFGCPE_01284* | | + | 1442880 | 1442955 | 18.85 | 103.61 | 70.12 | 19.60 | tRNA |
| *FEPFGCPE_01285* | | + | 1443155 | 1443343 | 8.64 | 103.92 | 372.01 | 69.45 | Hypothetical protein |
| *FEPFGCPE_01286* | | - | 1443371 | 1443733 | 3.95 | 25.93 | 54.29 | 5.61 | Hypothetical protein |
| *FEPFGCPE_01287* | | + | 1444003 | 1444362 | 8.27 | 90.46 | 81.11 | 10.92 | Hypothetical protein |
| *FEPFGCPE_01288* | *hcf136* | + | 1444435 | 1445550 | 4.04 | 28.08 | 34.34 | 7.23 | Ycf48-like protein |
| *FEPFGCPE_01289* | *tar-2* | + | 1445865 | 1447511 | 0.59 | 3.91 | 3.75 | 1.32 | Methyl-accepting chemotaxis protein II |
| *FEPFGCPE_01290* | *fusA-1* | + | 1447665 | 1449827 | 24.57 | 122.48 | 99.33 | 31.41 | Elongation factor G |
| *FEPFGCPE_01291* | *fghA* | - | 1449991 | 1450845 | 3.14 | 30.94 | 18.82 | 7.20 | S-formylglutathione hydrolase |
| *FEPFGCPE_01292* | *norG-1* | - | 1451317 | 1452054 | 1.60 | 7.99 | 6.50 | 1.77 | HTH-type transcriptional regulator NorG |
| *FEPFGCPE_01293* | | + | 1452118 | 1452564 | 0.35 | 1.14 | 0.85 | 0.09 | Hypothetical protein |
| *FEPFGCPE_01294* | *yjcC-1* | - | 1452855 | 1454411 | 3.60 | 14.05 | 8.65 | 2.38 | Putative cyclic-di-GMP phosphodiesterase YjcC |
| *FEPFGCPE_01295* | *norG-2* | + | 1454660 | 1456111 | 2.14 | 9.65 | 3.33 | 2.02 | HTH-type transcriptional regulator NorG |
| *FEPFGCPE_01296* | *gmr-2* | - | 1456126 | 1457349 | 1.05 | 1.47 | 2.74 | 0.82 | Cyclic di-GMP phosphodiesterase Gmr |
| *FEPFGCPE_01297* | | + | 1457477 | 1457662 | 2.53 | 7.20 | 9.52 | 3.16 | Hypothetical protein |
| *FEPFGCPE_01298* | *adrB-1* | - | 1457676 | 1458662 | 5.45 | 5.92 | 3.88 | 1.87 | Putative cyclic-di-GMP phosphodiesterase AdrB |
| *FEPFGCPE_01299* | | - | 1458798 | 1459169 | 1.57 | 4.37 | 3.55 | 1.37 | S-formylglutathione hydrolase |
| *FEPFGCPE_01300* | *sdhB-1* | + | 1459324 | 1460031 | 3.46 | 6.60 | 8.54 | 1.38 | Succinate dehydrogenase iron-sulfur subunit |
| *FEPFGCPE_01301* | | + | 1460531 | 1460737 | 4.44 | 7.42 | 5.28 | 1.22 | Hypothetical protein |
| *FEPFGCPE_01302* | | + | 1460850 | 1461038 | 59.83 | 475.18 | 237.19 | 39.78 | Hypothetical protein |
| *FEPFGCPE_01303* | | + | 1461431 | 1461724 | 25.23 | 51.25 | 52.49 | 7.10 | Hypothetical protein |
| *FEPFGCPE_01304* | *lutR-2* | + | 1461955 | 1462716 | 4.98 | 14.28 | 5.50 | 3.88 | HTH-type transcriptional regulator LutR |
| *FEPFGCPE_01305* | | + | 1462719 | 1463666 | 4.82 | 31.62 | 10.48 | 3.82 | Hypothetical protein |
| *FEPFGCPE_01306* | *cph2-2* | - | 1463716 | 1464798 | 8.15 | 29.34 | 19.21 | 3.01 | Phytochrome-like protein cph2 |
| *FEPFGCPE_01307* | *cspA* | + | 1465680 | 1465883 | 49.98 | 254.14 | 199.81 | 158.41 | Cold shock-like protein CspA |
| *FEPFGCPE_01308* | *purT* | - | 1466021 | 1467223 | 12.00 | 30.86 | 20.46 | 17.56 | Formate-dependent phosphoribosylglycinamide formyltransferase |
| *FEPFGCPE_01309* | *betI* | - | 1467320 | 1468030 | 15.63 | 23.94 | 14.67 | 2.93 | HTH-type transcriptional regulator BetI |
| *FEPFGCPE_01310* | *sadH* | + | 1468137 | 1469033 | 42.77 | 13.93 | 13.31 | 2.75 | Putative oxidoreductase SadH |
| *FEPFGCPE_01311* | *ethA* | + | 1469059 | 1470633 | 21.84 | 9.63 | 8.47 | 1.40 | FAD-containing monooxygenase EthA |
| *FEPFGCPE_01312* | *acoC* | + | 1470746 | 1471741 | 5.58 | 6.55 | 11.07 | 3.32 | Dihydrolipoyllysine-residue acetyltransferase component of acetoin cleaving system |
| *FEPFGCPE_01313* | *gloC-1* | + | 1471826 | 1473475 | 4.54 | 17.92 | 10.87 | 2.91 | Hydroxyacylglutathione hydrolase GloC |
| *FEPFGCPE_01314* | | - | 1473498 | 1473746 | 26.93 | 55.05 | 34.84 | 21.96 | Hypothetical protein |
| *FEPFGCPE_01315* | *yedQ* | + | 1474347 | 1475558 | 6.03 | 38.53 | 22.60 | 6.74 | putative diguanylate cyclase YedQ |
| *FEPFGCPE_01316* | | - | 1475638 | 1475713 | 65.73 | 112.88 | 82.19 | 47.86 | tRNA |
| *FEPFGCPE_01317* | *dusA* | - | 1475902 | 1476915 | 2.34 | 23.15 | 16.81 | 5.05 | tRNA-dihydrouridine(20/20a) synthase |
| *FEPFGCPE_01318* | *eamA-1* | + | 1477159 | 1478043 | 1.61 | 8.77 | 7.90 | 1.39 | putative amino-acid metabolite efflux pump |
| *FEPFGCPE_01319* | | - | 1478191 | 1478267 | 5.40 | 3.31 | 10.88 | 12.84 | tRNA |
| *FEPFGCPE_01320* | *nudC-2* | + | 1478450 | 1479085 | 8.04 | 53.18 | 34.33 | 16.78 | NADH pyrophosphatase |
| *FEPFGCPE_01321* | *aat* | + | 1479145 | 1479909 | 12.99 | 55.64 | 46.34 | 10.74 | Leucyl/phenylalanyl-tRNA--protein transferase |
| *FEPFGCPE_01322* | *ate* | + | 1479906 | 1480682 | 8.71 | 27.78 | 23.16 | 4.42 | Putative arginyl-tRNA--protein transferase |
| *FEPFGCPE_01323* | *pyrD* | + | 1480869 | 1481897 | 2.27 | 16.45 | 10.88 | 6.50 | Dihydroorotate dehydrogenase (quinone) |
| *FEPFGCPE_01324* | *fliY-2* | + | 1481870 | 1482745 | 18.94 | 114.93 | 60.02 | 28.68 | L-cystine-binding protein FliY |
| *FEPFGCPE_01325* | *yecS-2* | + | 1482857 | 1483516 | 1.01 | 7.60 | 5.32 | 0.75 | L-cystine transport system permease protein YecS |
| *FEPFGCPE_01326* | *lutB-1* | - | 1483640 | 1485049 | 4.00 | 16.04 | 9.54 | 3.15 | Lactate utilization protein B |
| *FEPFGCPE_01327* | *lutA-1* | - | 1485138 | 1485863 | 13.51 | 65.71 | 48.69 | 17.85 | Lactate utilization protein A |
| *FEPFGCPE_01328* | | - | 1485954 | 1487318 | 5.08 | 20.11 | 19.70 | 4.86 | Hypothetical protein |
| *FEPFGCPE_01329* | *yiaJ* | + | 1487559 | 1488386 | 13.57 | 23.76 | 32.74 | 12.07 | DNA-binding transcriptional repressor YiaJ |
| *FEPFGCPE_01330* | *pbpG-1* | - | 1488479 | 1489732 | 16.53 | 73.02 | 57.10 | 37.88 | D-alanyl-D-alanine endopeptidase |
| *FEPFGCPE_01331* | *nfnB* | - | 1489921 | 1490742 | 4.72 | 33.66 | 33.65 | 12.97 | Nitroreductase NfnB |
| *FEPFGCPE_01332* | *yedA* | - | 1490790 | 1491800 | 4.50 | 32.72 | 31.06 | 16.46 | putative inner membrane transporter YedA |
| *FEPFGCPE_01333* | *hdaH-1* | - | 1491797 | 1492738 | 2.60 | 18.98 | 15.66 | 5.65 | Histone deacetylase-like amidohydrolase |
| *FEPFGCPE_01334* | | + | 1493225 | 1493788 | 17.24 | 165.66 | 379.39 | 651.50 | Hypothetical protein |
| *FEPFGCPE_01335* | *lpdA* | - | 1494270 | 1495802 | 4.16 | 20.88 | 18.63 | 23.43 | Dihydrolipoyl dehydrogenase |
| *FEPFGCPE_01336* | *hslV-1* | - | 1495823 | 1496404 | 6.77 | 25.43 | 25.86 | 12.47 | ATP-dependent protease subunit HslV |
| *FEPFGCPE_01337* | *aceF* | - | 1496487 | 1498151 | 7.12 | 34.64 | 25.43 | 20.97 | Dihydrolipoyllysine-residue acetyltransferase component of pyruvate dehydrogenase complex |
| *FEPFGCPE_01338* | *aceE* | - | 1498247 | 1500934 | 10.62 | 53.80 | 51.28 | 42.34 | Pyruvate dehydrogenase E1 component |
| *FEPFGCPE_01339* | *fixL-1* | + | 1501277 | 1503760 | 5.24 | 23.77 | 15.68 | 5.31 | Sensor protein FixL |
| *FEPFGCPE_01340* | *tdiR* | + | 1503757 | 1504401 | 27.66 | 137.62 | 92.45 | 30.72 | Transcriptional regulatory protein TdiR |
| *FEPFGCPE_01341* | *folD* | + | 1504707 | 1505561 | 5.82 | 31.03 | 27.36 | 8.50 | Bifunctional protein FolD protein |
| *FEPFGCPE_01342* | *prlC* | + | 1505797 | 1507875 | 14.11 | 51.81 | 36.47 | 20.50 | Oligopeptidase A |
| *FEPFGCPE_01343* | | - | 1507957 | 1508592 | 1.21 | 2.72 | 5.88 | 0.58 | Hypothetical protein |
| *FEPFGCPE_01344* | *gpt* | - | 1508869 | 1509405 | 9.92 | 40.99 | 45.85 | 14.95 | Xanthine phosphoribosyltransferase |
| *FEPFGCPE_01345* | *purA-1* | - | 1509433 | 1510773 | 15.99 | 78.45 | 67.49 | 31.56 | Adenylosuccinate synthetase |

| Locus name | Gene name | Strand | Gene start (bp) | Gene end (bp) | Average TPM Inoculum | Average TPM $t_6$ | Average TPM $t_{24}$ | Average TPM $t_{168}$ | Annotation description |
|---|---|---|---|---|---|---|---|---|---|
| *FEPFGCPE_01346* | *hisZ* | - | 1510848 | 1512008 | 4.09 | 29.70 | 23.13 | 11.31 | ATP phosphoribosyltransferase regulatory subunit |
| *FEPFGCPE_01347* | | - | 1512047 | 1512202 | 40.95 | 152.33 | 123.67 | 53.13 | Hypothetical protein |
| *FEPFGCPE_01348* | *hflC* | - | 1512310 | 1513212 | 17.65 | 36.61 | 49.18 | 26.78 | Modulator of FtsH protease HflC |
| *FEPFGCPE_01349* | *hflK-2* | - | 1513225 | 1514559 | 12.26 | 27.06 | 24.89 | 15.52 | Modulator of FtsH protease HflK |
| *FEPFGCPE_01350* | | + | 1514643 | 1514876 | 39.50 | 162.60 | 109.59 | 40.29 | Hypothetical protein |
| *FEPFGCPE_01351* | *hflX* | - | 1514799 | 1516004 | 17.24 | 49.04 | 44.16 | 16.89 | GTPase HflX |
| *FEPFGCPE_01352* | *hfq* | - | 1516068 | 1516304 | 176.38 | 1106.73 | 872.85 | 248.82 | RNA-binding protein Hfq |
| *FEPFGCPE_01353* | *der* | - | 1516431 | 1517768 | 19.50 | 48.92 | 45.19 | 24.77 | GTPase Der |
| *FEPFGCPE_01354* | *bamB* | - | 1517846 | 1519063 | 12.08 | 52.98 | 29.61 | 17.65 | Outer membrane protein assembly factor BamB |
| *FEPFGCPE_01355* | | - | 1519120 | 1519749 | 8.64 | 25.56 | 22.36 | 15.33 | Hypothetical protein |
| *FEPFGCPE_01356* | *hisS* | - | 1519782 | 1521158 | 15.42 | 26.18 | 18.43 | 12.43 | Histidine--tRNA ligase |
| *FEPFGCPE_01357* | *ispG* | - | 1521180 | 1522460 | 23.27 | 34.11 | 36.40 | 15.98 | 4-hydroxy-3-methylbut-2-en-1-yl diphosphate synthase (flavodoxin) |
| *FEPFGCPE_01358* | *rodZ* | - | 1522479 | 1523576 | 10.12 | 22.48 | 21.59 | 14.71 | Cytoskeleton protein RodZ |
| *FEPFGCPE_01359* | *rlmN* | - | 1523769 | 1524890 | 10.49 | 17.41 | 20.16 | 15.94 | Dual-specificity RNA methyltransferase RlmN |
| *FEPFGCPE_01360* | *ndk* | - | 1524950 | 1525375 | 26.24 | 70.05 | 65.10 | 63.58 | Nucleoside diphosphate kinase |
| *FEPFGCPE_01361* | *yccA* | + | 1525784 | 1526485 | 134.13 | 128.13 | 159.94 | 93.77 | Modulator of FtsH protease YccA |
| *FEPFGCPE_01362* | *rlmD* | - | 1526717 | 1528177 | 4.15 | 13.71 | 11.24 | 3.07 | 23S rRNA (uracil(1939)-C(5))-methyltransferase RlmD |
| *FEPFGCPE_01363* | | - | 1528366 | 1529142 | 12.16 | 54.90 | 36.66 | 13.97 | Hypothetical protein |
| *FEPFGCPE_01364* | *rpoS* | - | 1529257 | 1530342 | 41.31 | 84.37 | 60.06 | 22.33 | RNA polymerase sigma factor RpoS |
| *FEPFGCPE_01365* | *nlpD-1* | - | 1530358 | 1531185 | 38.75 | 121.08 | 74.83 | 36.33 | Murein hydrolase activator NlpD |
| *FEPFGCPE_01366* | *pcm-1* | - | 1531218 | 1532366 | 14.76 | 44.15 | 31.75 | 11.93 | Protein-L-isoaspartate O-methyltransferase |
| *FEPFGCPE_01367* | *surE* | - | 1532363 | 1533124 | 8.51 | 45.40 | 29.24 | 15.82 | 5'-nucleotidase SurE |
| *FEPFGCPE_01368* | *ssuC-3* | - | 1533289 | 1534107 | 6.86 | 26.10 | 18.13 | 3.76 | Putative aliphatic sulfonates transport permease protein SsuC |
| *FEPFGCPE_01369* | | - | 1534129 | 1534263 | 2.57 | 14.15 | 9.25 | 3.56 | Hypothetical protein |
| *FEPFGCPE_01370* | *tauB-2* | - | 1534270 | 1535085 | 2.18 | 22.00 | 15.31 | 4.58 | Taurine import ATP-binding protein TauB |
| *FEPFGCPE_01371* | *cmpA* | - | 1535082 | 1536113 | 4.65 | 41.79 | 24.99 | 6.70 | Bicarbonate-binding protein CmpA |
| *FEPFGCPE_01372* | *smtA* | - | 1536218 | 1537543 | 4.05 | 21.04 | 12.81 | 6.20 | Succinyl-CoA--L-malate CoA-transferase alpha subunit |
| *FEPFGCPE_01373* | *yhjB* | - | 1537746 | 1538471 | 4.17 | 7.87 | 3.86 | 0.00 | Putative HTH-type transcriptional regulator YhjB |
| *FEPFGCPE_01374* | | + | 1538843 | 1539304 | 243.39 | 762.22 | 447.09 | 98.37 | Hypothetical protein |
| *FEPFGCPE_01375* | | - | 1539443 | 1540888 | 8.91 | 43.32 | 24.79 | 2.82 | Hypothetical protein |
| *FEPFGCPE_01376* | | - | 1541137 | 1541898 | 1.95 | 4.64 | 5.32 | 0.00 | Hypothetical protein |
| *FEPFGCPE_01377* | *lagD* | - | 1541911 | 1542621 | 3.01 | 3.37 | 3.73 | 0.00 | Lactococcin-G-processing and transport ATP-binding protein LagD |
| *FEPFGCPE_01378* | | - | 1542674 | 1544056 | 3.14 | 9.52 | 4.29 | 1.53 | Hypothetical protein |
| *FEPFGCPE_01379* | | - | 1544087 | 1544737 | 8.15 | 14.34 | 8.43 | 1.67 | Hypothetical protein |
| *FEPFGCPE_01380* | | - | 1544734 | 1545246 | 2.10 | 2.15 | 2.27 | 0.00 | Hypothetical protein |
| *FEPFGCPE_01381* | | - | 1545252 | 1545824 | 2.56 | 0.00 | 0.28 | 0.41 | Hypothetical protein |
| *FEPFGCPE_01382* | *zraR-1* | - | 1546577 | 1547935 | 15.34 | 56.48 | 15.77 | 4.54 | Transcriptional regulatory protein ZraR |
| *FEPFGCPE_01383* | *recR* | - | 1548185 | 1548781 | 5.73 | 25.65 | 20.75 | 6.28 | Recombination protein RecR |
| *FEPFGCPE_01384* | *ybaB* | - | 1548817 | 1549143 | 31.31 | 45.51 | 36.89 | 34.32 | Nucleoid-associated protein YbaB |
| *FEPFGCPE_01385* | *dnaX* | - | 1549302 | 1551263 | 7.05 | 25.15 | 26.74 | 9.19 | DNA polymerase III subunit tau |
| *FEPFGCPE_01386* | *trxA-2* | + | 1551978 | 1552304 | 53.83 | 279.76 | 265.11 | 150.22 | Thioredoxin 1 |
| *FEPFGCPE_01387* | *rho* | + | 1552498 | 1553760 | 8.50 | 42.13 | 32.79 | 25.68 | Transcription termination factor Rho |
| *FEPFGCPE_01388* | | + | 1553933 | 1554934 | 7.65 | 26.05 | 17.90 | 11.80 | Hypothetical protein |
| *FEPFGCPE_01389* | *idiA* | - | 1555123 | 1556178 | 17.69 | 68.90 | 110.37 | 134.79 | Iron deficiency-induced protein A |
| *FEPFGCPE_01390* | | + | 1556825 | 1557724 | 2.75 | 6.39 | 6.62 | 4.89 | putative NTE family protein |
| *FEPFGCPE_01391* | *pgdS* | - | 1557863 | 1558570 | 14.79 | 134.94 | 112.93 | 70.39 | Gamma-DL-glutamyl hydrolase |
| *FEPFGCPE_01392* | *gsiA-1* | - | 1558882 | 1560495 | 3.20 | 10.16 | 3.48 | 1.53 | Glutathione import ATP-binding protein GsiA |
| *FEPFGCPE_01393* | *yejE* | - | 1560492 | 1561469 | 6.66 | 17.83 | 11.32 | 2.63 | Inner membrane ABC transporter permease protein YejE |
| *FEPFGCPE_01394* | *yejB* | - | 1561588 | 1562643 | 2.16 | 8.01 | 3.30 | 2.48 | Inner membrane ABC transporter permease protein YejB |
| *FEPFGCPE_01395* | *oppA* | - | 1562685 | 1564709 | 4.57 | 11.44 | 6.94 | 2.16 | Periplasmic oligopeptide-binding protein |
| *FEPFGCPE_01396* | | + | 1564829 | 1565782 | 10.54 | 33.86 | 21.62 | 5.80 | Hypothetical protein |
| *FEPFGCPE_01397* | *mdfA* | - | 1565843 | 1567075 | 2.38 | 10.25 | 9.46 | 3.68 | Multidrug transporter MdfA |
| *FEPFGCPE_01398* | *fbp-2* | - | 1567209 | 1567550 | 3.35 | 21.36 | 7.26 | 18.38 | FK506-binding protein |
| *FEPFGCPE_01399* | | - | 1567672 | 1567765 | 5.09 | 18.38 | 4.55 | 9.13 | tRNA |
| *FEPFGCPE_01400* | | + | 1567843 | 1568097 | 8.64 | 14.59 | 12.71 | 7.61 | Hypothetical protein |
| *FEPFGCPE_01401* | | - | 1567989 | 1569239 | 12.09 | 24.94 | 27.35 | 19.16 | Aspartate kinase |
| *FEPFGCPE_01402* | *tilS* | - | 1569437 | 1570951 | 1.17 | 18.05 | 23.72 | 8.39 | tRNA(Ile)-lysidine synthase |
| *FEPFGCPE_01403* | *accA* | - | 1571001 | 1572044 | 19.56 | 96.44 | 76.67 | 34.34 | Acetyl-coenzyme A carboxylase carboxyl transferase subunit alpha |
| *FEPFGCPE_01404* | *alkA-1* | - | 1572125 | 1573327 | 1.63 | 17.37 | 16.30 | 8.78 | DNA-3-methyladenine glycosylase |
| *FEPFGCPE_01405* | *cysS* | - | 1573352 | 1574755 | 1.40 | 16.85 | 12.61 | 7.44 | Cysteine--tRNA ligase |
| *FEPFGCPE_01406* | *bepA-2* | + | 1575002 | 1575730 | 5.03 | 50.20 | 40.85 | 14.15 | Beta-barrel assembly-enhancing protease |
| *FEPFGCPE_01407* | | + | 1575784 | 1576020 | 3.44 | 60.42 | 34.84 | 39.27 | Hypothetical protein |
| *FEPFGCPE_01408* | | + | 1576032 | 1576379 | 20.12 | 251.01 | 167.17 | 61.37 | Peptidyl-prolyl cis-trans isomerase cyp18 |
| *FEPFGCPE_01409* | *ppiB* | + | 1576424 | 1576918 | 20.40 | 183.11 | 154.46 | 131.91 | Peptidyl-prolyl cis-trans isomerase B |
| *FEPFGCPE_01410* | *lpxH-2* | + | 1577146 | 1577796 | 0.88 | 2.97 | 5.43 | 0.48 | UDP-2,3-diacylglucosamine hydrolase |
| *FEPFGCPE_01411* | *cysE-2* | - | 1577878 | 1578657 | 0.75 | 11.77 | 7.07 | 4.72 | Serine acetyltransferase |
| *FEPFGCPE_01412* | *trmJ* | - | 1578805 | 1579623 | 3.75 | 13.11 | 12.95 | 6.16 | tRNA (cytidine/uridine-2'-O-)-methyltransferase TrmJ |
| *FEPFGCPE_01413* | *suhB* | + | 1580056 | 1580922 | 5.36 | 11.65 | 18.16 | 12.87 | Inositol-1-monophosphatase |
| *FEPFGCPE_01414* | *gltX* | - | 1581053 | 1582444 | 2.36 | 14.80 | 12.72 | 8.07 | Glutamate--tRNA ligase |
| *FEPFGCPE_01415* | *rhaR* | + | 1582667 | 1583443 | 3.42 | 11.99 | 3.45 | 0.00 | HTH-type transcriptional activator RhaR |
| *FEPFGCPE_01416* | *cptA* | + | 1583672 | 1585336 | 8.44 | 10.82 | 5.87 | 4.18 | Phosphoethanolamine transferase CptA |

| Locus name | Gene name | Strand | Gene start (bp) | Gene end (bp) | Average TPM Inoculum | Average TPM $t_6$ | Average TPM $t_{24}$ | Average TPM $t_{168}$ | Annotation description |
|---|---|---|---|---|---|---|---|---|---|
| *FEPFGCPE_01417* | *araE-2* | - | 1585403 | 1586836 | 4.68 | 37.43 | 37.92 | 13.87 | Alpha-ketoglutaric semialdehyde dehydrogenase |
| *FEPFGCPE_01418* | *hisC* | - | 1587073 | 1588143 | 1.93 | 15.86 | 18.03 | 6.11 | Histidinol-phosphate aminotransferase |
| *FEPFGCPE_01419* | *metZ* | - | 1588158 | 1589345 | 3.45 | 29.91 | 11.19 | 11.34 | O-succinylhomoserine sulfhydrylase |
| *FEPFGCPE_01420* | *purF* | - | 1589570 | 1591090 | 9.94 | 46.02 | 40.04 | 18.80 | Amidophosphoribosyltransferase |
| *FEPFGCPE_01421* | *cvpA* | - | 1591146 | 1592027 | 6.07 | 27.67 | 28.03 | 12.95 | Colicin V production protein |
| *FEPFGCPE_01422* | *dedD* | - | 1592029 | 1593054 | 6.95 | 47.27 | 48.47 | 17.61 | Cell division protein DedD |
| *FEPFGCPE_01423* | *folC* | - | 1593100 | 1594398 | 3.36 | 27.29 | 19.57 | 8.46 | Dihydrofolate synthase/folylpolyglutamate synthase |
| *FEPFGCPE_01424* | *accD* | - | 1594588 | 1595463 | 8.97 | 49.70 | 26.73 | 20.43 | Acetyl-coenzyme A carboxylase carboxyl transferase subunit beta |
| *FEPFGCPE_01425* | *trpA* | - | 1595593 | 1596417 | 2.17 | 12.75 | 10.36 | 4.25 | Tryptophan synthase alpha chain |
| *FEPFGCPE_01426* | *rsrIM* | - | 1596371 | 1597300 | 2.06 | 17.32 | 10.56 | 4.98 | Modification methylase RsrI |
| *FEPFGCPE_01427* | *trpB* | - | 1597433 | 1598635 | 6.61 | 31.15 | 32.51 | 9.53 | Tryptophan synthase beta chain |
| *FEPFGCPE_01428* | *trpF* | - | 1598693 | 1599349 | 6.97 | 32.86 | 33.56 | 9.89 | N-(5'-phosphoribosyl)anthranilate isomerase |
| *FEPFGCPE_01429* | *truA* | - | 1599346 | 1600146 | 1.07 | 6.43 | 6.24 | 3.45 | tRNA pseudouridine synthase A |
| *FEPFGCPE_01430* | | - | 1600244 | 1603288 | 6.04 | 26.38 | 23.93 | 11.45 | Hypothetical protein |
| *FEPFGCPE_01431* | *asd* | - | 1603414 | 1604544 | 19.06 | 61.22 | 64.15 | 32.88 | Aspartate-semialdehyde dehydrogenase |
| *FEPFGCPE_01432* | *leuB* | - | 1604605 | 1605672 | 24.13 | 126.83 | 104.22 | 43.94 | 3-isopropylmalate dehydrogenase |
| *FEPFGCPE_01433* | *leuD1* | - | 1605697 | 1606347 | 14.17 | 48.27 | 34.12 | 21.34 | 3-isopropylmalate dehydratase small subunit 1 |
| *FEPFGCPE_01434* | | - | 1606469 | 1606597 | 41.37 | 50.15 | 44.28 | 24.45 | Hypothetical protein |
| *FEPFGCPE_01435* | *leuC-1* | - | 1606653 | 1608062 | 11.37 | 31.46 | 27.72 | 15.73 | 3-isopropylmalate dehydratase large subunit |
| *FEPFGCPE_01436* | *gltA-1* | - | 1608344 | 1609645 | 6.12 | 46.34 | 47.39 | 40.38 | Citrate synthase |
| *FEPFGCPE_01437* | *sdhE* | - | 1609675 | 1609947 | 6.10 | 36.98 | 73.58 | 26.57 | FAD assembly factor SdhE |
| *FEPFGCPE_01438* | *sdhB-2* | - | 1609958 | 1610659 | 9.20 | 62.73 | 69.90 | 32.06 | Succinate dehydrogenase iron-sulfur subunit |
| *FEPFGCPE_01439* | *sdhA* | - | 1610675 | 1612450 | 5.58 | 38.11 | 49.36 | 34.14 | Succinate dehydrogenase flavoprotein subunit |
| *FEPFGCPE_01440* | *sdhD* | - | 1612454 | 1612819 | 6.21 | 47.73 | 30.59 | 28.79 | Succinate dehydrogenase hydrophobic membrane anchor subunit |
| *FEPFGCPE_01441* | *sdhC* | - | 1612821 | 1613234 | 10.23 | 35.82 | 30.84 | 25.97 | Succinate dehydrogenase cytochrome b556 subunit |
| *FEPFGCPE_01442* | *yurK* | - | 1613425 | 1614267 | 5.72 | 49.31 | 54.41 | 24.30 | putative HTH-type transcriptional regulator YurK |
| *FEPFGCPE_01443* | *mdh-1* | + | 1614608 | 1615591 | 4.19 | 56.61 | 88.25 | 104.35 | Malate dehydrogenase |
| *FEPFGCPE_01444* | | + | 1616164 | 1616661 | 10.91 | 87.88 | 69.59 | 43.58 | Hypothetical protein |
| *FEPFGCPE_01445* | *acn* | + | 1616872 | 1619580 | 7.56 | 55.56 | 59.20 | 37.35 | Aconitate hydratase A |
| *FEPFGCPE_01446* | *alkA-2* | + | 1619787 | 1620848 | 4.69 | 10.34 | 8.08 | 3.32 | putative bifunctional transcriptional activator/DNA repair enzyme AlkA |
| *FEPFGCPE_01447* | *alkA-3* | + | 1620959 | 1621273 | 9.53 | 20.12 | 13.31 | 6.97 | putative bifunctional transcriptional activator/DNA repair enzyme AlkA |
| *FEPFGCPE_01448* | *ogt* | + | 1621270 | 1621833 | 4.05 | 11.49 | 6.59 | 1.34 | Methylated-DNA--protein-cysteine methyltransferase, constitutive |
| *FEPFGCPE_01449* | | + | 1621969 | 1623147 | 6.08 | 30.68 | 13.57 | 9.39 | Hypothetical protein |
| *FEPFGCPE_01450* | *trmB-1* | - | 1623249 | 1623974 | 0.81 | 4.29 | 0.77 | 0.00 | tRNA (guanine-N(7)-)-methyltransferase |
| *FEPFGCPE_01451* | | + | 1624328 | 1624951 | 2.12 | 24.78 | 27.54 | 8.47 | Hypothetical protein |
| *FEPFGCPE_01452* | *scpD* | - | 1624996 | 1626144 | 3.02 | 18.39 | 23.93 | 8.88 | Zinc carboxypeptidase |
| *FEPFGCPE_01453* | | - | 1626210 | 1626989 | 4.39 | 33.12 | 20.76 | 8.53 | Hypothetical protein |
| *FEPFGCPE_01454* | | + | 1627037 | 1627549 | 2.01 | 7.38 | 3.29 | 3.96 | Hypothetical protein |
| *FEPFGCPE_01455* | *ybdL* | + | 1627689 | 1628885 | 2.06 | 11.16 | 9.91 | 6.66 | Methionine aminotransferase |
| *FEPFGCPE_01456* | | - | 1628994 | 1629650 | 3.31 | 28.40 | 30.58 | 11.23 | Glutathione S-transferase |
| *FEPFGCPE_01457* | *paaH* | + | 1630251 | 1631807 | 4.51 | 13.26 | 12.90 | 7.11 | 3-hydroxyadipyl-CoA dehydrogenase |
| *FEPFGCPE_01458* | *paaF-1* | + | 1631824 | 1632603 | 1.97 | 8.02 | 6.71 | 0.76 | 2,3-dehydroadipyl-CoA hydratase |
| *FEPFGCPE_01459* | *gpmA-1* | + | 1632707 | 1633465 | 7.18 | 28.59 | 18.90 | 6.85 | 2,3-bisphosphoglycerate-dependent phosphoglycerate mutase |
| *FEPFGCPE_01460* | | + | 1633590 | 1634738 | 6.70 | 27.29 | 19.52 | 9.36 | Putative aminoglycoside phosphotransferase |
| *FEPFGCPE_01461* | *bbsG* | + | 1634735 | 1636009 | 6.51 | 30.14 | 27.46 | 7.18 | (R)-benzylsuccinyl-CoA dehydrogenase |
| *FEPFGCPE_01462* | *yfcG-2* | + | 1636006 | 1636698 | 7.85 | 46.95 | 34.26 | 13.79 | Disulfide-bond oxidoreductase YfcG |
| *FEPFGCPE_01463* | | + | 1636952 | 1637287 | 19.78 | 76.27 | 41.56 | 7.04 | Hypothetical protein |
| *FEPFGCPE_01464* | *paaZ* | - | 1637340 | 1637795 | 9.60 | 53.22 | 40.26 | 5.66 | Bifunctional protein PaaZ |
| *FEPFGCPE_01465* | | - | 1637848 | 1638321 | 4.91 | 57.06 | 25.10 | 6.50 | Hypothetical protein |
| *FEPFGCPE_01466* | *act* | - | 1638387 | 1638980 | 2.80 | 18.64 | 10.97 | 7.97 | Methanol dehydrogenase activator |
| *FEPFGCPE_01467* | | - | 1639057 | 1639377 | 18.21 | 66.17 | 83.74 | 15.76 | Hypothetical protein |
| *FEPFGCPE_01468* | *nuoN* | - | 1639462 | 1640931 | 1.39 | 10.59 | 10.40 | 3.52 | NADH-quinone oxidoreductase subunit N |
| *FEPFGCPE_01469* | *nuoM* | - | 1640952 | 1642433 | 1.62 | 19.91 | 13.08 | 10.70 | NADH-quinone oxidoreductase subunit M |
| *FEPFGCPE_01470* | *nuoL* | - | 1642452 | 1644497 | 1.13 | 16.16 | 11.58 | 5.90 | NADH-quinone oxidoreductase subunit L |
| *FEPFGCPE_01471* | | - | 1644515 | 1644826 | 0.86 | 8.52 | 8.68 | 6.39 | NADH-quinone oxidoreductase subunit 11 |
| *FEPFGCPE_01472* | *nuoJ* | - | 1644836 | 1645471 | 2.00 | 20.71 | 19.98 | 7.80 | NADH-quinone oxidoreductase subunit J |
| *FEPFGCPE_01473* | *nuoI* | - | 1645551 | 1646042 | 2.81 | 19.85 | 13.30 | 10.73 | NADH-quinone oxidoreductase subunit I |
| *FEPFGCPE_01474* | *nuoH* | - | 1646066 | 1647130 | 4.80 | 25.63 | 24.82 | 15.02 | NADH-quinone oxidoreductase subunit H |
| *FEPFGCPE_01475* | *nqo3* | - | 1647131 | 1649443 | 2.83 | 16.13 | 12.31 | 6.16 | NADH-quinone oxidoreductase chain 3 |
| *FEPFGCPE_01476* | *nuoF* | - | 1649473 | 1650765 | 5.57 | 27.32 | 22.56 | 13.86 | NADH-quinone oxidoreductase subunit F |
| *FEPFGCPE_01477* | *nqo2-2* | - | 1650762 | 1651262 | 13.59 | 47.65 | 38.28 | 30.99 | NADH-quinone oxidoreductase subunit 2 |
| *FEPFGCPE_01478* | *nuoD* | - | 1651434 | 1652687 | 6.74 | 27.71 | 15.72 | 10.63 | NADH-quinone oxidoreductase subunit D |
| *FEPFGCPE_01479* | *nqo5* | - | 1652697 | 1653296 | 9.70 | 30.06 | 26.06 | 20.05 | NADH-quinone oxidoreductase chain 5 |
| *FEPFGCPE_01480* | *nuoB* | - | 1653309 | 1653788 | 13.88 | 38.22 | 37.00 | 27.75 | NADH-quinone oxidoreductase subunit B |
| *FEPFGCPE_01481* | *ndhC* | - | 1653836 | 1654297 | 17.89 | 59.86 | 36.60 | 12.56 | NAD(P)H-quinone oxidoreductase subunit 3 |
| *FEPFGCPE_01482* | | - | 1654325 | 1654409 | 9.02 | 20.51 | 8.23 | 3.28 | tRNA |
| *FEPFGCPE_01483* | *secG* | - | 1654553 | 1654984 | 21.69 | 51.15 | 54.23 | 37.59 | Protein-export membrane protein SecG |
| *FEPFGCPE_01484* | *tpiA* | - | 1655015 | 1655824 | 21.61 | 38.33 | 50.86 | 27.38 | Triosephosphate isomerase |
| *FEPFGCPE_01485* | *ppsC-2* | - | 1655973 | 1656977 | 3.92 | 20.81 | 21.87 | 10.99 | Phthiocerol/phenolphthiocerol synthesis polyketide synthase type I PpsC |
| *FEPFGCPE_01486* | | + | 1657106 | 1657336 | 1.07 | 3.76 | 3.86 | 1.99 | Hypothetical protein |
| *FEPFGCPE_01487* | *vraR-1* | - | 1657400 | 1657771 | 18.99 | 132.63 | 161.45 | 69.02 | Response regulator protein VraR |

| Locus name | Gene name | Strand | Gene start (bp) | Gene end (bp) | Average TPM Inoculum | Average TPM $t_6$ | Average TPM $t_{24}$ | Average TPM $t_{168}$ | Annotation description |
|---|---|---|---|---|---|---|---|---|---|
| *FEPFGCPE_01488* | *yegE* | - | 1657918 | 1659888 | 2.17 | 5.00 | 2.33 | 0.79 | putative diguanylate cyclase YegE |
| *FEPFGCPE_01489* | | - | 1659942 | 1661270 | 4.06 | 16.48 | 8.94 | 4.80 | Hypothetical protein |
| *FEPFGCPE_01490* | | - | 1661331 | 1661657 | 1.39 | 11.97 | 5.36 | 2.18 | Hypothetical protein |
| *FEPFGCPE_01491* | *mutS* | + | 1661839 | 1662921 | 3.17 | 16.27 | 10.21 | 7.26 | DNA mismatch repair protein MutS |
| *FEPFGCPE_01492* | | + | 1662933 | 1664309 | 2.99 | 11.60 | 9.50 | 1.89 | Hypothetical protein |
| *FEPFGCPE_01493* | *slyD* | - | 1664461 | 1664994 | 20.25 | 162.37 | 112.04 | 58.05 | FKBP-type peptidyl-prolyl cis-trans isomerase SlyD |
| *FEPFGCPE_01494* | *roxA* | + | 1665125 | 1666348 | 13.09 | 33.44 | 31.10 | 8.39 | 50S ribosomal protein L16 3-hydroxylase |
| *FEPFGCPE_01495* | *yycJ* | - | 1666881 | 1667672 | 1.79 | 7.09 | 2.79 | 3.36 | Putative metallo-hydrolase YycJ |
| *FEPFGCPE_01496* | *bamC* | - | 1667681 | 1668823 | 9.57 | 58.80 | 48.92 | 26.24 | Outer membrane protein assembly factor BamC |
| *FEPFGCPE_01497* | *dapA-2* | - | 1668875 | 1669792 | 12.37 | 84.87 | 50.53 | 30.43 | 4-hydroxy-tetrahydrodipicolinate synthase |
| *FEPFGCPE_01498* | | - | 1669789 | 1670445 | 5.91 | 18.19 | 22.48 | 6.27 | Hypothetical protein |
| *FEPFGCPE_01499* | *trpS* | - | 1670442 | 1671644 | 3.27 | 17.18 | 11.86 | 5.84 | Tryptophan--tRNA ligase |
| *FEPFGCPE_01500* | | - | 1671670 | 1672335 | 7.05 | 27.36 | 14.02 | 10.06 | Hypothetical protein |
| *FEPFGCPE_01501* | *tsaC* | - | 1672409 | 1673035 | 4.98 | 21.96 | 15.03 | 4.69 | Threonylcarbamoyl-AMP synthase |
| *FEPFGCPE_01502* | | - | 1673048 | 1673881 | 28.53 | 61.31 | 42.89 | 18.98 | 3',5'-nucleoside bisphosphate phosphatase |
| *FEPFGCPE_01503* | | - | 1673994 | 1674884 | 1.49 | 4.37 | 5.91 | 1.93 | Tropinesterase |
| *FEPFGCPE_01504* | | - | 1674966 | 1675910 | 2.61 | 12.07 | 4.35 | 2.66 | Hypothetical protein |
| *FEPFGCPE_01505* | | + | 1675960 | 1676484 | 6.48 | 33.17 | 36.61 | 12.41 | Putative acetyltransferase |
| *FEPFGCPE_01506* | *hslO* | + | 1676653 | 1677588 | 3.03 | 14.66 | 13.35 | 8.07 | 33 kDa chaperonin |
| *FEPFGCPE_01507* | | + | 1677609 | 1678145 | 5.87 | 15.61 | 17.70 | 6.13 | Hypothetical protein |
| *FEPFGCPE_01508* | *ftsB* | - | 1678235 | 1678666 | 4.46 | 23.47 | 27.30 | 12.66 | Cell division protein FtsB |
| *FEPFGCPE_01509* | *eno* | - | 1678739 | 1680022 | 4.41 | 37.38 | 33.74 | 27.21 | Enolase |
| *FEPFGCPE_01510* | *kdsA* | - | 1680413 | 1681267 | 5.62 | 15.33 | 18.07 | 10.58 | 2-dehydro-3-deoxyphosphooctonate aldolase |
| *FEPFGCPE_01511* | *pyrG* | - | 1681305 | 1682963 | 7.08 | 48.89 | 39.34 | 17.78 | CTP synthase |
| *FEPFGCPE_01512* | | - | 1683150 | 1683980 | 5.42 | 9.37 | 9.48 | 2.12 | 2-(acetamidomethylene)succinate hydrolase |
| *FEPFGCPE_01513* | *comEC* | - | 1684284 | 1687220 | 1.48 | 3.54 | 3.84 | 1.48 | ComE operon protein 3 |
| *FEPFGCPE_01514* | *yjjV* | - | 1687265 | 1688131 | 3.90 | 5.62 | 5.28 | 1.02 | putative metal-dependent hydrolase YjjV |
| *FEPFGCPE_01515* | *lolD* | - | 1688339 | 1689076 | 4.40 | 4.65 | 6.86 | 2.44 | Lipoprotein-releasing system ATP-binding protein LolD |
| *FEPFGCPE_01516* | *lolE* | - | 1689069 | 1690322 | 4.05 | 6.68 | 8.62 | 2.11 | Lipoprotein-releasing system transmembrane protein LolE |
| *FEPFGCPE_01517* | | + | 1690539 | 1690868 | 38.19 | 36.27 | 30.98 | 24.33 | Hypothetical protein |
| *FEPFGCPE_01518* | | + | 1690862 | 1691617 | 1.20 | 3.56 | 6.90 | 3.64 | Hypothetical protein |
| *FEPFGCPE_01519* | *recJ* | + | 1691614 | 1693311 | 1.27 | 6.07 | 4.27 | 1.04 | Single-stranded-DNA-specific exonuclease RecJ |
| *FEPFGCPE_01520* | *prfB* | + | 1693518 | 1694564 | 3.89 | 27.13 | 22.33 | 8.94 | Peptide chain release factor RF2 |
| *FEPFGCPE_01521* | *ivy* | + | 1694568 | 1695047 | 2.53 | 19.72 | 21.09 | 5.52 | Inhibitor of vertebrate lysozyme |
| *FEPFGCPE_01522* | *lysS* | + | 1695064 | 1696608 | 2.87 | 11.51 | 13.48 | 8.47 | Lysine--tRNA ligase |
| *FEPFGCPE_01523* | | - | 1696632 | 1697234 | 5.52 | 26.95 | 12.32 | 7.48 | Hypothetical protein |
| *FEPFGCPE_01524* | *slyB-1* | + | 1697563 | 1698270 | 7.43 | 78.62 | 56.00 | 28.29 | Outer membrane lipoprotein SlyB |
| *FEPFGCPE_01525* | | - | 1698392 | 1698583 | 12.37 | 33.69 | 29.61 | 17.48 | Hypothetical protein |
| *FEPFGCPE_01526* | *fdx-1* | - | 1698637 | 1698978 | 24.64 | 45.67 | 43.13 | 15.28 | 2Fe-2S ferredoxin |
| *FEPFGCPE_01527* | *hscA* | - | 1699002 | 1700879 | 8.66 | 15.68 | 14.19 | 7.31 | Chaperone protein HscA |
| *FEPFGCPE_01528* | *hscB* | - | 1700966 | 1701496 | 6.62 | 18.21 | 13.25 | 3.07 | Co-chaperone protein HscB |
| *FEPFGCPE_01529* | *iscA* | - | 1701906 | 1702229 | 56.12 | 87.46 | 82.38 | 55.41 | Iron-binding protein IscA |
| *FEPFGCPE_01530* | *iscU* | - | 1702261 | 1702668 | 93.43 | 153.18 | 146.31 | 88.62 | Iron-sulfur cluster assembly scaffold protein IscU |
| *FEPFGCPE_01531* | *iscS-1* | - | 1702687 | 1703910 | 76.81 | 124.52 | 115.49 | 78.31 | Cysteine desulfurase IscS |
| *FEPFGCPE_01532* | *iscR* | - | 1703941 | 1704468 | 186.08 | 229.39 | 205.36 | 97.79 | HTH-type transcriptional regulator IscR |
| *FEPFGCPE_01533* | | - | 1704585 | 1706252 | 1.62 | 3.58 | 4.65 | 0.85 | Hypothetical protein |
| *FEPFGCPE_01534* | | - | 1706300 | 1706692 | 1.30 | 4.59 | 4.90 | 0.70 | Hypothetical protein |
| *FEPFGCPE_01535* | *yfkJ* | - | 1706771 | 1707217 | 0.36 | 4.78 | 4.06 | 0.00 | Low molecular weight protein-tyrosine-phosphatase YfkJ |
| *FEPFGCPE_01536* | | - | 1707651 | 1708094 | 24.72 | 96.18 | 67.80 | 21.88 | Hypothetical protein |
| *FEPFGCPE_01537* | | - | 1708512 | 1708955 | 43.95 | 140.96 | 86.20 | 24.63 | Hypothetical protein |
| *FEPFGCPE_01538* | | + | 1709211 | 1709468 | 0.00 | 6.25 | 1.36 | 0.00 | Hypothetical protein |
| *FEPFGCPE_01539* | | - | 1709837 | 1710949 | 5.73 | 21.42 | 13.02 | 4.99 | Hypothetical protein |
| *FEPFGCPE_01540* | | - | 1711137 | 1711892 | 22.45 | 33.65 | 11.42 | 5.84 | Hypothetical protein |
| *FEPFGCPE_01541* | | + | 1712115 | 1712249 | 0.80 | 0.00 | 0.00 | 0.00 | Hypothetical protein |
| *FEPFGCPE_01542* | | + | 1712246 | 1712503 | 0.17 | 2.52 | 0.62 | 0.00 | Hypothetical protein |
| *FEPFGCPE_01543* | | + | 1712500 | 1713009 | 0.21 | 2.38 | 1.46 | 0.00 | Hypothetical protein |
| *FEPFGCPE_01544* | | + | 1713019 | 1713414 | 0.14 | 3.10 | 0.83 | 0.00 | Hypothetical protein |
| *FEPFGCPE_01545* | | + | 1713416 | 1713577 | 1.25 | 11.09 | 14.13 | 0.00 | Hypothetical protein |
| *FEPFGCPE_01546* | | + | 1714548 | 1714877 | 0.21 | 4.21 | 2.13 | 0.00 | Hypothetical protein |
| *FEPFGCPE_01547* | | + | 1714874 | 1715140 | 0.81 | 0.65 | 2.85 | 0.00 | Hypothetical protein |
| *FEPFGCPE_01548* | | + | 1715133 | 1716230 | 0.46 | 4.44 | 1.36 | 1.04 | Hypothetical protein |
| *FEPFGCPE_01549* | | + | 1716230 | 1716586 | 0.78 | 8.33 | 7.99 | 3.53 | Hypothetical protein |
| *FEPFGCPE_01550* | | - | 1716876 | 1717505 | 1.56 | 15.92 | 7.41 | 2.68 | Hypothetical protein |
| *FEPFGCPE_01551* | *bspRIM* | + | 1717538 | 1718821 | 0.50 | 1.79 | 0.39 | 0.33 | Modification methylase BspRI |
| *FEPFGCPE_01552* | | + | 1718874 | 1719323 | 0.36 | 0.00 | 0.00 | 0.00 | Hypothetical protein |
| *FEPFGCPE_01553* | | + | 1719320 | 1719520 | 0.27 | 8.65 | 1.11 | 1.17 | Hypothetical protein |
| *FEPFGCPE_01554* | | + | 1719517 | 1719612 | 0.00 | 2.49 | 4.64 | 0.00 | Hypothetical protein |
| *FEPFGCPE_01555* | | + | 1719593 | 1719790 | 1.11 | 5.21 | 4.18 | 2.28 | Hypothetical protein |
| *FEPFGCPE_01556* | | + | 1719796 | 1720881 | 0.58 | 6.33 | 3.50 | 1.47 | Hypothetical protein |
| *FEPFGCPE_01557* | *uvrB* | - | 1720978 | 1723065 | 3.36 | 18.36 | 10.76 | 4.42 | UvrABC system protein B |
| *FEPFGCPE_01558* | *tyrB-1* | + | 1723385 | 1724584 | 9.14 | 24.08 | 28.03 | 26.73 | Aromatic-amino-acid aminotransferase |

| Locus name | Gene name | Strand | Gene start (bp) | Gene end (bp) | Average TPM Inoculum | Average TPM $t_6$ | Average TPM $t_{24}$ | Average TPM $t_{168}$ | Annotation description |
|---|---|---|---|---|---|---|---|---|---|
| *FEPFGCPE_01559* | | + | 1724743 | 1726044 | 6.02 | 27.85 | 20.43 | 11.54 | Hypothetical protein |
| *FEPFGCPE_01560* | *nicS-2* | - | 1726152 | 1726790 | 6.57 | 26.44 | 19.96 | 6.56 | HTH-type transcriptional repressor NicS |
| *FEPFGCPE_01561* | *rsxB* | + | 1726895 | 1727788 | 1.64 | 11.28 | 11.94 | 4.20 | Electron transport complex subunit RsxB |
| *FEPFGCPE_01562* | *nth* | + | 1727854 | 1728498 | 0.59 | 11.52 | 8.60 | 0.80 | Endonuclease III |
| *FEPFGCPE_01563* | *rutE* | - | 1728638 | 1729231 | 4.78 | 33.76 | 15.70 | 17.22 | putative malonic semialdehyde reductase RutE |
| *FEPFGCPE_01564* | | - | 1729323 | 1730267 | 1.14 | 6.34 | 5.89 | 1.57 | Hypothetical protein |
| *FEPFGCPE_01565* | | + | 1730423 | 1730854 | 7.21 | 24.81 | 17.10 | 7.69 | Hypothetical protein |
| *FEPFGCPE_01566* | | - | 1730929 | 1731342 | 12.80 | 55.21 | 33.48 | 13.54 | Cytochrome c4 |
| *FEPFGCPE_01567* | | - | 1731368 | 1731733 | 12.01 | 37.48 | 31.33 | 11.76 | Cytochrome c-554(548) |
| *FEPFGCPE_01568* | | + | 1732015 | 1732857 | 8.86 | 42.97 | 39.93 | 16.82 | Hypothetical protein |
| *FEPFGCPE_01569* | | + | 1732871 | 1734046 | 9.54 | 36.62 | 35.01 | 13.81 | Hypothetical protein |
| *FEPFGCPE_01570* | *ydcO* | + | 1734126 | 1735322 | 1.71 | 12.26 | 9.88 | 3.90 | Inner membrane protein YdcO |
| *FEPFGCPE_01571* | | + | 1735411 | 1735755 | 4.84 | 27.17 | 16.49 | 13.93 | Hypothetical protein |
| *FEPFGCPE_01572* | *speE-1* | - | 1735868 | 1736710 | 1.63 | 3.51 | 5.61 | 3.44 | Polyamine aminopropyltransferase |
| *FEPFGCPE_01573* | *mcpA* | + | 1737156 | 1737800 | 0.53 | 8.13 | 17.58 | 5.50 | Methyl-accepting chemotaxis protein McpA |
| *FEPFGCPE_01574* | | - | 1737904 | 1738407 | 0.67 | 7.21 | 12.28 | 3.92 | Hypothetical protein |
| *FEPFGCPE_01575* | *ubiC* | - | 1738404 | 1739012 | 0.73 | 4.84 | 7.60 | 2.24 | Chorismate pyruvate-lyase |
| *FEPFGCPE_01576* | *htpG* | - | 1739223 | 1741109 | 5.48 | 14.22 | 11.70 | 8.56 | Chaperone protein HtpG |
| *FEPFGCPE_01577* | *lysN-1* | - | 1741364 | 1742809 | 7.18 | 8.24 | 4.26 | 2.71 | 2-aminoadipate transaminase |
| *FEPFGCPE_01578* | | + | 1743021 | 1743374 | 40.80 | 8.81 | 13.50 | 4.89 | Hypothetical protein |
| *FEPFGCPE_01579* | | + | 1743429 | 1744115 | 15.70 | 25.75 | 24.04 | 5.96 | Hypothetical protein |
| *FEPFGCPE_01580* | | + | 1744189 | 1745085 | 5.79 | 6.31 | 5.79 | 3.28 | Hypothetical protein |
| *FEPFGCPE_01581* | *lysN-2* | + | 1745227 | 1746417 | 7.50 | 41.67 | 22.60 | 8.84 | 2-aminoadipate transaminase |
| *FEPFGCPE_01582* | | + | 1746550 | 1747005 | 4.33 | 45.03 | 36.75 | 17.99 | Hypothetical protein |
| *FEPFGCPE_01583* | *fabG-4* | - | 1747156 | 1747947 | 2.68 | 29.05 | 39.21 | 9.09 | 3-oxoacyl-[acyl-carrier-protein] reductase FabG |
| *FEPFGCPE_01584* | | - | 1747962 | 1748162 | 3.58 | 24.10 | 35.22 | 5.12 | Sulfoquinovose 1-dehydrogenase |
| *FEPFGCPE_01585* | *dmlR-10* | + | 1748179 | 1748613 | 0.07 | 5.36 | 13.78 | 1.46 | HTH-type transcriptional regulator DmlR |
| *FEPFGCPE_01586* | *ynfM* | - | 1748708 | 1749976 | 0.58 | 24.78 | 65.30 | 4.64 | Inner membrane transport protein YnfM |
| *FEPFGCPE_01587* | *benM-2* | + | 1750109 | 1751032 | 1.18 | 9.41 | 3.65 | 1.97 | HTH-type transcriptional regulator BenM |
| *FEPFGCPE_01588* | | - | 1751054 | 1751506 | 2.93 | 15.56 | 14.85 | 2.90 | Hypothetical protein |
| *FEPFGCPE_01589* | *parC* | - | 1751519 | 1753855 | 1.91 | 10.74 | 8.89 | 3.35 | DNA topoisomerase 4 subunit A |
| *FEPFGCPE_01590* | *mltC-1* | - | 1754118 | 1755080 | 2.24 | 13.83 | 11.77 | 4.19 | Membrane-bound lytic murein transglycosylase C |
| *FEPFGCPE_01591* | | - | 1755064 | 1755480 | 7.31 | 52.12 | 40.76 | 15.60 | Hypothetical protein |
| *FEPFGCPE_01592* | *parE* | - | 1755589 | 1757568 | 4.02 | 16.15 | 18.18 | 5.36 | DNA topoisomerase 4 subunit B |
| *FEPFGCPE_01593* | *uup* | - | 1757874 | 1759805 | 2.96 | 14.06 | 7.09 | 4.18 | ABC transporter ATP-binding protein uup |
| *FEPFGCPE_01594* | | - | 1759892 | 1760380 | 7.03 | 19.67 | 21.14 | 11.80 | Acetyltransferase |
| *FEPFGCPE_01595* | | - | 1760482 | 1760715 | 17.25 | 19.24 | 37.50 | 34.51 | Hypothetical protein |
| *FEPFGCPE_01596* | | - | 1761274 | 1761789 | 26.49 | 94.51 | 121.56 | 27.67 | Hypothetical protein |
| *FEPFGCPE_01597* | | + | 1761944 | 1762141 | 11.45 | 46.21 | 36.28 | 9.52 | Hypothetical protein |
| *FEPFGCPE_01598* | *glnS* | + | 1762251 | 1763969 | 9.58 | 48.32 | 39.49 | 21.83 | Glutamine--tRNA ligase |
| *FEPFGCPE_01599* | | + | 1764254 | 1765192 | 47.51 | 64.30 | 16.91 | 5.91 | Urea transporter |
| *FEPFGCPE_01600* | *phrA* | - | 1765483 | 1766979 | 4.44 | 20.16 | 13.67 | 3.25 | Deoxyribodipyrimidine photo-lyase |
| *FEPFGCPE_01601* | | + | 1767077 | 1767532 | 6.48 | 26.09 | 11.65 | 9.40 | Hypothetical protein |
| *FEPFGCPE_01602* | | - | 1767784 | 1768293 | 3.75 | 16.19 | 13.66 | 3.45 | Hypothetical protein |
| *FEPFGCPE_01603* | *dinB* | - | 1768364 | 1769494 | 1.59 | 7.93 | 4.05 | 2.18 | DNA polymerase IV |
| *FEPFGCPE_01604* | | - | 1769574 | 1770287 | 2.61 | 13.11 | 15.94 | 2.19 | Hypothetical protein |
| *FEPFGCPE_01605* | *uctC-4* | - | 1770484 | 1771566 | 5.39 | 31.56 | 24.49 | 3.32 | Acetyl-CoA:oxalate CoA-transferase |
| *FEPFGCPE_01606* | *uctC-5* | - | 1771669 | 1772952 | 6.21 | 24.42 | 10.42 | 1.30 | Acetyl-CoA:oxalate CoA-transferase |
| *FEPFGCPE_01607* | *dmdA* | - | 1772965 | 1774983 | 23.41 | 40.55 | 22.76 | 5.05 | 2,3-dimethylmalate dehydratase large subunit |
| *FEPFGCPE_01608* | *ssuA-1* | - | 1775111 | 1776187 | 36.69 | 58.17 | 35.43 | 11.96 | Putative aliphatic sulfonates-binding protein |
| *FEPFGCPE_01609* | *cysL-2* | - | 1776368 | 1777279 | 7.94 | 27.09 | 19.00 | 1.36 | HTH-type transcriptional regulator CysL |
| *FEPFGCPE_01610* | | + | 1777325 | 1777912 | 102.62 | 321.56 | 204.18 | 48.45 | Hypothetical protein |
| *FEPFGCPE_01611* | *phnU-2* | - | 1777868 | 1779589 | 1.40 | 2.94 | 3.51 | 0.76 | Putative 2-aminoethylphosphonate transport system permease protein PhnU |
| *FEPFGCPE_01612* | *fbpC2* | + | 1779889 | 1780998 | 6.81 | 9.49 | 7.93 | 6.26 | Fe(3+) ions import ATP-binding protein FbpC 2 |
| *FEPFGCPE_01613* | *thyA* | + | 1781058 | 1781852 | 5.76 | 24.64 | 19.81 | 13.96 | Thymidylate synthase |
| *FEPFGCPE_01614* | *dhfrIII* | + | 1781849 | 1782337 | 6.46 | 33.10 | 34.18 | 13.92 | Dihydrofolate reductase type 3 |
| *FEPFGCPE_01615* | *cph2-3* | - | 1782384 | 1783823 | 4.08 | 25.65 | 12.23 | 4.90 | Phytochrome-like protein cph2 |
| *FEPFGCPE_01616* | *livF-5* | - | 1784028 | 1784741 | 4.12 | 13.50 | 7.84 | 1.73 | High-affinity branched-chain amino acid transport ATP-binding protein LivF |
| *FEPFGCPE_01617* | *lptB-4* | - | 1784738 | 1785541 | 2.67 | 5.88 | 13.15 | 4.86 | Lipopolysaccharide export system ATP-binding protein LptB |
| *FEPFGCPE_01618* | *livH-5* | - | 1785560 | 1786435 | 1.30 | 5.77 | 2.90 | 1.25 | High-affinity branched-chain amino acid transport system permease protein LivH |
| *FEPFGCPE_01619* | *livH-6* | - | 1786449 | 1787330 | 1.21 | 7.91 | 2.70 | 2.51 | High-affinity branched-chain amino acid transport system permease protein LivH |
| *FEPFGCPE_01620* | *braC-2* | - | 1787461 | 1788612 | 12.65 | 76.01 | 53.13 | 32.15 | Leucine-, isoleucine-, valine-, threonine-, and alanine-binding protein |
| *FEPFGCPE_01621* | *pmbA* | - | 1788831 | 1790189 | 5.10 | 30.48 | 23.77 | 7.55 | Metalloprotease PmbA |
| *FEPFGCPE_01622* | | + | 1790305 | 1790910 | 14.46 | 61.97 | 62.55 | 22.84 | Hypothetical protein |
| *FEPFGCPE_01623* | *mog* | + | 1790910 | 1791530 | 7.94 | 51.60 | 47.80 | 16.05 | Molybdopterin adenylyltransferase |
| *FEPFGCPE_01624* | *orn* | - | 1791627 | 1792265 | 7.02 | 17.89 | 18.44 | 10.75 | Oligoribonuclease |
| *FEPFGCPE_01625* | *htpX-1* | + | 1792369 | 1793616 | 5.55 | 27.24 | 17.05 | 8.64 | Protease HtpX |
| *FEPFGCPE_01626* | *rsgA* | + | 1793613 | 1794554 | 2.49 | 9.50 | 9.08 | 1.97 | Small ribosomal subunit biogenesis GTPase RsgA |
| *FEPFGCPE_01627* | *cobD-1* | - | 1794643 | 1795659 | 4.87 | 29.69 | 21.51 | 7.17 | Cobalamin biosynthesis protein CobD |
| *FEPFGCPE_01628* | *nudL* | - | 1795717 | 1796403 | 1.67 | 16.07 | 12.94 | 2.67 | putative Nudix hydrolase NudL |
| *FEPFGCPE_01629* | *rplS* | - | 1796636 | 1797022 | 9.13 | 19.27 | 70.77 | 47.48 | 50S ribosomal protein L19 |

| Locus name | Gene name | Strand | Gene start (bp) | Gene end (bp) | Average TPM Inoculum | Average TPM $t_6$ | Average TPM $t_{24}$ | Average TPM $t_{168}$ | Annotation description |
|---|---|---|---|---|---|---|---|---|---|
| *FEPFGCPE_01630* | *trmD* | - | 1797126 | 1797896 | 46.66 | 146.44 | 171.61 | 83.69 | tRNA (guanine-N(1)-)-methyltransferase |
| *FEPFGCPE_01631* | *rimM* | - | 1797957 | 1798607 | 13.47 | 40.28 | 36.15 | 25.34 | Ribosome maturation factor RimM |
| *FEPFGCPE_01632* | *rpsP* | - | 1798737 | 1798988 | 18.83 | 88.17 | 78.94 | 59.39 | 30S ribosomal protein S16 |
| *FEPFGCPE_01633* | | + | 1799188 | 1800393 | 5.10 | 24.37 | 22.96 | 7.78 | Hypothetical protein |
| *FEPFGCPE_01634* | | + | 1800584 | 1801390 | 19.67 | 42.02 | 65.20 | 22.82 | Hypothetical protein |
| *FEPFGCPE_01635* | | + | 1801400 | 1801849 | 9.65 | 40.61 | 38.94 | 19.48 | Hypothetical protein |
| *FEPFGCPE_01636* | *proC-1* | + | 1801859 | 1802791 | 6.30 | 17.39 | 19.57 | 7.77 | Pyrroline-5-carboxylate reductase |
| *FEPFGCPE_01637* | | + | 1802979 | 1804133 | 2.83 | 12.47 | 8.43 | 3.73 | Hypothetical protein |
| *FEPFGCPE_01638* | *lrp-1* | - | 1804200 | 1804688 | 4.42 | 7.74 | 4.24 | 2.86 | Leucine-responsive regulatory protein |
| *FEPFGCPE_01639* | *dadA-2* | + | 1804793 | 1806079 | 14.08 | 8.12 | 20.34 | 15.87 | D-amino acid dehydrogenase |
| *FEPFGCPE_01640* | *ald-3* | + | 1806185 | 1807318 | 13.00 | 3.56 | 6.86 | 5.56 | Alanine dehydrogenase |
| *FEPFGCPE_01641* | *ubiG* | + | 1807539 | 1808234 | 10.67 | 36.11 | 32.11 | 17.96 | Ubiquinone biosynthesis O-methyltransferase |
| *FEPFGCPE_01642* | *gph-1* | + | 1808227 | 1808958 | 5.18 | 37.21 | 21.45 | 5.47 | Phosphoglycolate phosphatase |
| *FEPFGCPE_01643* | | + | 1809070 | 1813074 | 4.85 | 38.38 | 23.36 | 10.00 | Hypothetical protein |
| *FEPFGCPE_01644* | *ssrA* | + | 1813105 | 1813463 | 1938.26 | 13528.55 | 13583.92 | 5850.60 | tmRNA |
| *FEPFGCPE_01645* | | - | 1813635 | 1814141 | 3.84 | 2.99 | 4.14 | 2.86 | Hypothetical protein |
| *FEPFGCPE_01646* | *lcfB-1* | - | 1814235 | 1815857 | 1.87 | 7.13 | 5.13 | 1.20 | Long-chain-fatty-acid--CoA ligase |
| *FEPFGCPE_01647* | *fadA-1* | - | 1815854 | 1817023 | 1.47 | 4.07 | 2.73 | 0.70 | 3-ketoacyl-CoA thiolase |
| *FEPFGCPE_01648* | *fabG-5* | - | 1817058 | 1817813 | 0.27 | 3.32 | 0.88 | 0.00 | 3-oxoacyl-[acyl-carrier-protein] reductase FabG |
| *FEPFGCPE_01649* | *bacC-2* | - | 1817928 | 1818749 | 0.82 | 5.42 | 2.17 | 1.41 | Dihydroanticapsin 7-dehydrogenase |
| *FEPFGCPE_01650* | *kipR-3* | + | 1819003 | 1819896 | 0.54 | 1.65 | 2.66 | 0.00 | HTH-type transcriptional regulator KipR |
| *FEPFGCPE_01651* | | + | 1819968 | 1820870 | 0.31 | 0.95 | 0.75 | 0.43 | Hypothetical protein |
| *FEPFGCPE_01652* | *dorA* | + | 1820867 | 1823209 | 0.19 | 1.06 | 0.66 | 0.28 | Dimethyl sulfoxide/trimethylamine N-oxide reductase |
| *FEPFGCPE_01653* | *siaP-1* | + | 1823265 | 1824284 | 0.28 | 0.69 | 1.12 | 0.75 | Sialic acid-binding periplasmic protein SiaP |
| *FEPFGCPE_01654* | *siaT-1* | + | 1824316 | 1826205 | 0.54 | 1.45 | 1.56 | 0.12 | Sialic acid TRAP transporter permease protein SiaT |
| *FEPFGCPE_01655* | *tktA-1* | + | 1826315 | 1827172 | 0.93 | 6.19 | 5.16 | 1.74 | Transketolase 1 |
| *FEPFGCPE_01656* | *dxs-2* | + | 1827169 | 1828197 | 0.56 | 4.45 | 1.63 | 0.62 | 1-deoxy-D-xylulose-5-phosphate synthase |
| *FEPFGCPE_01657* | *cbh* | - | 1828325 | 1829437 | 2.04 | 13.14 | 7.73 | 3.27 | Choloylglycine hydrolase |
| *FEPFGCPE_01658* | | - | 1829606 | 1831009 | 1.29 | 8.09 | 4.56 | 1.18 | Gluconate 2-dehydrogenase cytochrome c subunit |
| *FEPFGCPE_01659* | | - | 1831021 | 1832796 | 0.74 | 9.35 | 6.25 | 2.90 | Gluconate 2-dehydrogenase flavoprotein |
| *FEPFGCPE_01660* | | - | 1832800 | 1833522 | 1.80 | 6.64 | 9.72 | 2.44 | Gluconate 2-dehydrogenase subunit 3 |
| *FEPFGCPE_01661* | | + | 1833556 | 1833720 | 2.08 | 15.51 | 30.99 | 7.55 | Hypothetical protein |
| *FEPFGCPE_01662* | | - | 1833846 | 1835507 | 0.61 | 1.74 | 0.70 | 0.32 | Hypothetical protein |
| *FEPFGCPE_01663* | | + | 1835534 | 1835803 | 1.36 | 5.74 | 3.00 | 1.33 | Hypothetical protein |
| *FEPFGCPE_01664* | | - | 1835693 | 1836607 | 0.83 | 5.80 | 4.14 | 0.56 | Hypothetical protein |
| *FEPFGCPE_01665* | | + | 1836954 | 1838003 | 3.67 | 16.40 | 10.97 | 0.97 | Outer membrane porin protein 32 |
| *FEPFGCPE_01666* | | + | 1838175 | 1838498 | 3.60 | 6.82 | 8.35 | 0.00 | Hypothetical protein |
| *FEPFGCPE_01667* | *naiP-2* | + | 1838648 | 1839910 | 1.08 | 4.74 | 3.91 | 1.82 | Putative niacin/nicotinamide transporter NaiP |
| *FEPFGCPE_01668* | *pyrK* | + | 1839960 | 1840892 | 1.56 | 3.21 | 0.82 | 1.03 | Dihydroorotate dehydrogenase B (NAD(+)), electron transfer subunit |
| *FEPFGCPE_01669* | *pyrDB* | + | 1840885 | 1841835 | 0.61 | 2.60 | 1.29 | 1.80 | Dihydroorotate dehydrogenase B (NAD(+)), catalytic subunit |
| *FEPFGCPE_01670* | | + | 1841892 | 1843145 | 0.79 | 4.08 | 2.12 | 0.76 | putative hydrolase |
| *FEPFGCPE_01671* | *dapE-3* | - | 1843263 | 1844723 | 7.34 | 21.77 | 15.28 | 3.63 | Succinyl-diaminopimelate desuccinylase |
| *FEPFGCPE_01672* | | - | 1844965 | 1845330 | 38.06 | 79.74 | 30.83 | 18.03 | Hypothetical protein |
| *FEPFGCPE_01673* | | - | 1845330 | 1845497 | 96.82 | 338.68 | 176.53 | 46.78 | Hypothetical protein |
| *FEPFGCPE_01674* | | - | 1846045 | 1846230 | 179.76 | 1603.13 | 562.50 | 179.07 | Hypothetical protein |
| *FEPFGCPE_01675* | | - | 1846716 | 1846997 | 1.08 | 5.22 | 7.27 | 3.26 | Hypothetical protein |
| *FEPFGCPE_01676* | *nucA* | - | 1847104 | 1848036 | 0.83 | 2.29 | 2.96 | 0.00 | Nuclease |
| *FEPFGCPE_01677* | | - | 1848088 | 1848246 | 1.64 | 11.90 | 5.89 | 4.16 | Hypothetical protein |
| *FEPFGCPE_01678* | *paiB* | - | 1848365 | 1849009 | 3.19 | 20.25 | 8.40 | 3.20 | Protease synthase and sporulation protein PAI 2 |
| *FEPFGCPE_01679* | | - | 1849119 | 1850741 | 2.39 | 8.47 | 5.65 | 1.20 | Hypothetical protein |
| *FEPFGCPE_01680* | | - | 1850785 | 1850985 | 26.66 | 72.46 | 67.15 | 19.69 | Hypothetical protein |
| *FEPFGCPE_01681* | *bcrC* | + | 1851315 | 1852004 | 0.94 | 6.88 | 2.96 | 1.55 | Undecaprenyl-diphosphatase BcrC |
| *FEPFGCPE_01682* | *gcvR* | + | 1852142 | 1852672 | 20.50 | 87.70 | 43.52 | 29.25 | Glycine cleavage system transcriptional repressor |
| *FEPFGCPE_01683* | | + | 1852683 | 1853270 | 4.49 | 19.13 | 16.53 | 6.16 | Hypothetical protein |
| *FEPFGCPE_01684* | *iclR-2* | - | 1853376 | 1854134 | 5.44 | 39.81 | 20.40 | 5.19 | Transcriptional repressor IclR |
| *FEPFGCPE_01685* | *glxR-2* | - | 1854485 | 1855393 | 7.56 | 42.97 | 27.69 | 20.42 | 2-hydroxy-3-oxopropionate reductase |
| *FEPFGCPE_01686* | *ybgC* | + | 1855681 | 1856148 | 30.18 | 117.72 | 140.99 | 34.98 | Acyl-CoA thioesterase YbgC |
| *FEPFGCPE_01687* | *exbB-2* | + | 1856188 | 1856862 | 12.67 | 67.62 | 58.42 | 31.73 | Biopolymer transport protein ExbB |
| *FEPFGCPE_01688* | *exbD-3* | + | 1856884 | 1857318 | 6.68 | 30.22 | 35.07 | 18.20 | Biopolymer transport protein ExbD |
| *FEPFGCPE_01689* | | + | 1857315 | 1858361 | 8.23 | 29.10 | 34.44 | 14.26 | Hypothetical protein |
| *FEPFGCPE_01690* | *dpp5* | + | 1858463 | 1859755 | 18.05 | 99.11 | 94.15 | 45.15 | Dipeptidyl-peptidase 5 |
| *FEPFGCPE_01691* | *pal* | + | 1859786 | 1860283 | 26.54 | 109.52 | 102.74 | 72.52 | Outer membrane protein P6 |
| *FEPFGCPE_01692* | *cpoB* | + | 1860294 | 1861037 | 6.40 | 29.45 | 35.64 | 25.29 | Cell division coordinator CpoB |
| *FEPFGCPE_01693* | | + | 1861443 | 1862558 | 1.75 | 14.49 | 17.51 | 5.38 | Hypothetical protein |
| *FEPFGCPE_01694* | | + | 1862665 | 1862740 | 0.71 | 0.26 | 9.46 | 0.94 | tRNA |
| *FEPFGCPE_01695* | | + | 1862880 | 1863500 | 9.72 | 51.95 | 30.44 | 12.61 | Hypothetical protein |
| *FEPFGCPE_01696* | | + | 1863656 | 1864381 | 1.08 | 4.02 | 1.22 | 0.70 | Hypothetical protein |
| *FEPFGCPE_01697* | | + | 1864384 | 1864518 | 0.40 | 1.13 | 0.00 | 0.00 | Hypothetical protein |
| *FEPFGCPE_01698* | | + | 1864626 | 1866404 | 0.37 | 1.26 | 0.99 | 0.18 | Hypothetical protein |
| *FEPFGCPE_01699* | | + | 1866508 | 1867398 | 0.41 | 2.81 | 2.01 | 0.00 | Hypothetical protein |
| *FEPFGCPE_01700* | | + | 1867395 | 1867571 | 0.63 | 3.06 | 1.26 | 0.00 | Hypothetical protein |

| Locus name | Gene name | Strand | Gene start (bp) | Gene end (bp) | Average TPM Inoculum | Average TPM $t_6$ | Average TPM $t_{24}$ | Average TPM $t_{168}$ | Annotation description |
|---|---|---|---|---|---|---|---|---|---|
| *FEPFGCPE_01701* | | + | 1867575 | 1867832 | 0.99 | 0.00 | 0.00 | 0.00 | Hypothetical protein |
| *FEPFGCPE_01702* | | + | 1867822 | 1868079 | 0.70 | 0.88 | 0.00 | 0.00 | Hypothetical protein |
| *FEPFGCPE_01703* | *xerC-4* | + | 1868076 | 1868798 | 2.38 | 6.71 | 1.59 | 1.34 | Tyrosine recombinase XerC |
| *FEPFGCPE_01704* | | + | 1869347 | 1870285 | 3.05 | 15.00 | 9.31 | 4.23 | Hypothetical protein |
| *FEPFGCPE_01705* | | - | 1870794 | 1871165 | 37.07 | 50.95 | 25.72 | 20.89 | Single-stranded DNA-binding protein |
| *FEPFGCPE_01706* | | - | 1871569 | 1871922 | 45.38 | 30.74 | 30.30 | 12.63 | Hypothetical protein |
| *FEPFGCPE_01707* | | - | 1871909 | 1872175 | 152.47 | 63.94 | 56.59 | 20.51 | Hypothetical protein |
| *FEPFGCPE_01708* | *yecD* | - | 1872356 | 1872778 | 1.40 | 7.96 | 7.71 | 0.64 | Isochorismatase family protein YecD |
| *FEPFGCPE_01709* | *kipR-4* | - | 1872982 | 1873794 | 2.71 | 20.71 | 20.19 | 4.85 | HTH-type transcriptional regulator KipR |
| *FEPFGCPE_01710* | *bphC2-1* | - | 1873969 | 1874535 | 0.46 | 2.60 | 1.75 | 0.14 | Biphenyl-2,3-diol 1,2-dioxygenase 2 |
| *FEPFGCPE_01711* | | + | 1874681 | 1875532 | 0.27 | 0.33 | 0.05 | 0.00 | Ureidoglycolate lyase |
| *FEPFGCPE_01712* | *mhpA-1* | + | 1875589 | 1877118 | 0.41 | 1.54 | 2.48 | 0.00 | 3-(3-hydroxy-phenyl)propionate/3-hydroxycinnamic acid hydroxylase |
| *FEPFGCPE_01713* | *proP-6* | + | 1877183 | 1878529 | 1.23 | 3.00 | 2.30 | 0.29 | Proline/betaine transporter |
| *FEPFGCPE_01714* | *Dml* | - | 1878599 | 1879471 | 1.13 | 5.15 | 8.40 | 1.08 | 2,3-dimethylmalate lyase |
| *FEPFGCPE_01715* | *ydjE-3* | - | 1879567 | 1880964 | 1.88 | 6.63 | 1.89 | 0.30 | Inner membrane metabolite transport protein YdjE |
| *FEPFGCPE_01716* | *lutR-3* | - | 1881022 | 1881726 | 5.20 | 4.82 | 3.03 | 0.00 | HTH-type transcriptional regulator LutR |
| *FEPFGCPE_01717* | | - | 1881924 | 1882544 | 1.62 | 9.66 | 2.99 | 3.42 | Hypothetical protein |
| *FEPFGCPE_01718* | | + | 1882839 | 1883492 | 3.91 | 26.32 | 19.94 | 8.38 | Hypothetical protein |
| *FEPFGCPE_01719* | *ribZ-1* | + | 1883499 | 1884905 | 2.74 | 16.12 | 21.12 | 6.58 | Riboflavin transporter RibZ |
| *FEPFGCPE_01720* | *nsrR* | - | 1884957 | 1885454 | 4.66 | 28.57 | 29.80 | 6.26 | HTH-type transcriptional repressor NsrR |
| *FEPFGCPE_01721* | | - | 1885604 | 1886338 | 0.86 | 1.40 | 2.54 | 0.38 | Hypothetical protein |
| *FEPFGCPE_01722* | *mgtB* | - | 1886493 | 1889246 | 0.23 | 0.44 | 0.72 | 0.53 | Magnesium-transporting ATPase, P-type 1 |
| *FEPFGCPE_01723* | | - | 1889555 | 1889911 | 0.89 | 10.22 | 6.34 | 3.89 | Hypothetical protein |
| *FEPFGCPE_01724* | | + | 1891082 | 1891384 | 17.40 | 86.76 | 76.88 | 28.81 | Hypothetical protein |
| *FEPFGCPE_01725* | *uvrA-1* | + | 1891415 | 1894066 | 2.83 | 15.86 | 16.36 | 4.31 | UvrABC system protein A |
| *FEPFGCPE_01726* | *ghrB-1* | + | 1895049 | 1895639 | 0.96 | 5.90 | 6.02 | 1.40 | Glyoxylate/hydroxypyruvate reductase B |
| *FEPFGCPE_01727* | *cmpR-3* | + | 1895632 | 1896576 | 0.52 | 8.50 | 3.21 | 0.34 | HTH-type transcriptional activator CmpR |
| *FEPFGCPE_01728* | *naiP-3* | + | 1896783 | 1898084 | 0.52 | 7.31 | 3.60 | 1.25 | Putative niacin/nicotinamide transporter NaiP |
| *FEPFGCPE_01729* | *maiA-2* | + | 1898112 | 1898864 | 0.22 | 5.31 | 5.89 | 1.86 | Maleate isomerase |
| *FEPFGCPE_01730* | *ribBA-1* | - | 1899428 | 1900075 | 3.45 | 28.86 | 40.23 | 11.98 | Riboflavin biosynthesis protein RibBA |
| *FEPFGCPE_01731* | *rhtB-2* | + | 1900457 | 1901098 | 0.37 | 2.94 | 2.11 | 1.19 | Homoserine/homoserine lactone efflux protein |
| *FEPFGCPE_01732* | | + | 1901314 | 1901610 | 0.43 | 7.04 | 1.29 | 2.57 | Hypothetical protein |
| *FEPFGCPE_01733* | *czcR-3* | + | 1901868 | 1902545 | 1.01 | 7.01 | 1.88 | 1.29 | Transcriptional activator protein CzcR |
| *FEPFGCPE_01734* | *cusS-2* | + | 1902542 | 1904005 | 0.56 | 5.07 | 2.39 | 0.45 | Sensor kinase CusS |
| *FEPFGCPE_01735* | | - | 1904034 | 1904792 | 0.06 | 1.96 | 1.31 | 0.00 | Hypothetical protein |
| *FEPFGCPE_01736* | | - | 1904823 | 1906238 | 0.40 | 1.13 | 3.23 | 0.58 | D-hydantoinase |
| *FEPFGCPE_01737* | | - | 1906243 | 1906977 | 0.22 | 2.45 | 0.55 | 0.32 | Arylmalonate decarboxylase |
| *FEPFGCPE_01738* | *garP* | - | 1907051 | 1908349 | 0.80 | 3.85 | 2.79 | 1.06 | putative galactarate transporter |
| *FEPFGCPE_01739* | *gntR-2* | + | 1908496 | 1909245 | 19.13 | 16.70 | 15.73 | 3.42 | putative D-xylose utilization operon transcriptional repressor |
| *FEPFGCPE_01740* | | + | 1909430 | 1910161 | 5.32 | 2.08 | 11.45 | 3.37 | Hypothetical protein |
| *FEPFGCPE_01741* | *proP-7* | - | 1910439 | 1911791 | 1.24 | 4.51 | 3.96 | 1.93 | Proline/betaine transporter |
| *FEPFGCPE_01742* | *echA8-1* | - | 1911852 | 1912679 | 0.49 | 3.84 | 1.57 | 1.13 | putative enoyl-CoA hydratase echA8 |
| *FEPFGCPE_01743* | *uctC-6* | - | 1913015 | 1914208 | 0.52 | 3.37 | 1.53 | 0.00 | Acetyl-CoA:oxalate CoA-transferase |
| *FEPFGCPE_01744* | *gcvA-1* | - | 1914391 | 1915275 | 0.66 | 4.99 | 6.16 | 0.72 | Glycine cleavage system transcriptional activator |
| *FEPFGCPE_01745* | *btr* | - | 1915303 | 1916253 | 0.76 | 4.29 | 8.16 | 0.31 | HTH-type transcriptional activator Btr |
| *FEPFGCPE_01746* | *gabD-1* | + | 1916520 | 1917908 | 0.66 | 4.54 | 2.34 | 1.79 | Succinate-semialdehyde dehydrogenase [NADP(+)] GabD |
| *FEPFGCPE_01747* | *vnfA* | + | 1918308 | 1919837 | 2.65 | 7.35 | 7.58 | 2.33 | Nitrogen fixation protein VnfA |
| *FEPFGCPE_01748* | | + | 1920056 | 1920595 | 0.27 | 23.56 | 148.59 | 160.73 | Hypothetical protein |
| *FEPFGCPE_01749* | | + | 1920640 | 1921779 | 0.93 | 5.64 | 40.75 | 24.64 | Hypothetical protein |
| *FEPFGCPE_01750* | *norW* | + | 1921776 | 1923131 | 0.29 | 3.90 | 32.46 | 28.09 | Nitric oxide reductase FlRd-NAD(+) reductase |
| *FEPFGCPE_01751* | *cmpC-1* | + | 1923191 | 1924585 | 0.12 | 6.63 | 44.85 | 33.84 | Bicarbonate transport ATP-binding protein CmpC |
| *FEPFGCPE_01752* | *cmpB-1* | + | 1924601 | 1925470 | 0.68 | 3.98 | 54.29 | 34.23 | Bicarbonate transport system permease protein CmpB |
| *FEPFGCPE_01753* | *cmpD-2* | + | 1925460 | 1926389 | 1.87 | 8.55 | 72.06 | 62.66 | Bicarbonate transport ATP-binding protein CmpD |
| *FEPFGCPE_01754* | *cynS* | + | 1926423 | 1926872 | 0.77 | 17.47 | 136.42 | 168.03 | Cyanate hydratase |
| *FEPFGCPE_01755* | | - | 1927077 | 1927173 | 8.46 | 32.39 | 19.37 | 24.72 | tRNA |
| *FEPFGCPE_01756* | *selB* | - | 1927259 | 1929169 | 4.68 | 28.50 | 20.96 | 8.17 | Selenocysteine-specific elongation factor |
| *FEPFGCPE_01757* | *selA* | - | 1929166 | 1930596 | 3.26 | 14.44 | 10.83 | 4.96 | L-seryl-tRNA(Sec) selenium transferase |
| *FEPFGCPE_01758* | *fdhE* | - | 1930615 | 1931652 | 10.70 | 18.51 | 21.13 | 6.95 | Protein FdhE |
| *FEPFGCPE_01759* | *fdoI* | - | 1931695 | 1932342 | 12.87 | 20.72 | 12.68 | 4.22 | Formate dehydrogenase, cytochrome b556(fdo) subunit |
| *FEPFGCPE_01760* | *fdoH* | - | 1932345 | 1933295 | 15.77 | 8.70 | 13.46 | 4.26 | Formate dehydrogenase-O iron-sulfur subunit |
| *FEPFGCPE_01761* | *fdoG-1* | - | 1933305 | 1935734 | 25.31 | 20.87 | 17.25 | 4.50 | Formate dehydrogenase-O major subunit |
| *FEPFGCPE_01762* | *fdoG-2* | - | 1935783 | 1936373 | 62.17 | 276.91 | 194.27 | 51.00 | Formate dehydrogenase-O major subunit |
| *FEPFGCPE_01763* | *yebS* | + | 1936553 | 1937221 | 8.69 | 9.98 | 7.69 | 2.57 | Inner membrane protein YebS |
| *FEPFGCPE_01764* | *pqiA* | + | 1937218 | 1937862 | 7.31 | 23.49 | 16.20 | 4.15 | Paraquat-inducible protein A |
| *FEPFGCPE_01765* | *pqiB* | + | 1937855 | 1939519 | 5.56 | 16.84 | 10.42 | 5.00 | Paraquat-inducible protein B |
| *FEPFGCPE_01766* | | + | 1939516 | 1940190 | 5.44 | 16.96 | 16.34 | 1.17 | Hypothetical protein |
| *FEPFGCPE_01767* | | + | 1940371 | 1941630 | 1.70 | 5.81 | 6.22 | 1.10 | Hypothetical protein |
| *FEPFGCPE_01768* | *hsrA-1* | - | 1941640 | 1943082 | 6.18 | 13.25 | 14.79 | 5.73 | putative transport protein HsrA |
| *FEPFGCPE_01769* | | - | 1943195 | 1945066 | 14.66 | 22.05 | 21.38 | 23.71 | Trehalase |
| *FEPFGCPE_01770* | | - | 1945050 | 1945217 | 8.80 | 35.74 | 23.29 | 21.12 | Hypothetical protein |
| *FEPFGCPE_01771* | *yfdE-3* | - | 1945373 | 1946602 | 1.96 | 5.53 | 4.14 | 2.06 | Acetyl-CoA:oxalate CoA-transferase |

| Locus name | Gene name | Strand | Gene start (bp) | Gene end (bp) | Average TPM Inoculum | Average TPM $t_6$ | Average TPM $t_{24}$ | Average TPM $t_{168}$ | Annotation description |
|---|---|---|---|---|---|---|---|---|---|
| *FEPFGCPE_01772* | *crt-4* | - | 1946604 | 1947530 | 1.87 | 4.14 | 1.23 | 0.00 | Short-chain-enoyl-CoA hydratase |
| *FEPFGCPE_01773* | *rspR-3* | + | 1947594 | 1948268 | 1.94 | 5.03 | 2.77 | 1.71 | HTH-type transcriptional repressor RspR |
| *FEPFGCPE_01774* | *rspR-4* | - | 1948341 | 1949060 | 16.50 | 55.54 | 29.37 | 19.56 | HTH-type transcriptional repressor RspR |
| *FEPFGCPE_01775* | *tesE* | + | 1949352 | 1950110 | 2.28 | 18.37 | 11.43 | 6.44 | 2-hydroxyhexa-2,4-dienoate hydratase |
| *FEPFGCPE_01776* | *cynR-4* | - | 1950380 | 1951330 | 1.75 | 5.67 | 1.86 | 0.30 | HTH-type transcriptional regulator CynR |
| *FEPFGCPE_01777* | *pntB-1* | - | 1951437 | 1951958 | 1.10 | 5.57 | 2.47 | 0.63 | NAD(P) transhydrogenase subunit beta |
| *FEPFGCPE_01778* | *pntB-2* | - | 1951955 | 1952878 | 1.18 | 0.94 | 3.51 | 0.83 | NAD(P) transhydrogenase subunit beta |
| *FEPFGCPE_01779* | *pntA-1* | - | 1952880 | 1953224 | 1.39 | 13.41 | 4.33 | 5.86 | NAD(P) transhydrogenase subunit alpha |
| *FEPFGCPE_01780* | *pntAA-1* | - | 1953240 | 1954370 | 2.10 | 7.17 | 9.82 | 4.62 | NAD(P) transhydrogenase subunit alpha part 1 |
| *FEPFGCPE_01781* | | - | 1954413 | 1954787 | 2.08 | 4.25 | 21.29 | 2.78 | Hypothetical protein |
| *FEPFGCPE_01782* | *cmpR-4* | + | 1955038 | 1956033 | 4.19 | 3.18 | 1.78 | 2.33 | HTH-type transcriptional activator CmpR |
| *FEPFGCPE_01783* | | - | 1956096 | 1956869 | 0.85 | 0.44 | 5.22 | 0.00 | Ureidoglycolate lyase |
| *FEPFGCPE_01784* | *frc-1* | - | 1957007 | 1958254 | 0.79 | 2.49 | 2.59 | 0.39 | Formyl-CoA:oxalate CoA-transferase |
| *FEPFGCPE_01785* | *fixL-2* | - | 1958955 | 1959461 | 4.81 | 44.65 | 21.96 | 7.52 | Sensor protein FixL |
| *FEPFGCPE_01786* | *frc-2* | - | 1959635 | 1960885 | 2.09 | 21.22 | 18.72 | 4.36 | Formyl-CoA:oxalate CoA-transferase |
| *FEPFGCPE_01787* | *oxc* | - | 1960911 | 1962701 | 2.17 | 13.26 | 12.38 | 2.88 | Oxalyl-CoA decarboxylase |
| *FEPFGCPE_01788* | *oxlT-1* | - | 1962893 | 1964236 | 2.64 | 14.27 | 17.70 | 4.57 | Oxalate:formate antiporter |
| *FEPFGCPE_01789* | *panE* | + | 1964978 | 1965991 | 0.95 | 4.27 | 2.21 | 1.87 | 2-dehydropantoate 2-reductase |
| *FEPFGCPE_01790* | | + | 1966558 | 1966776 | 27.39 | 138.27 | 101.40 | 51.48 | Hypothetical protein |
| *FEPFGCPE_01791* | *rspR-5* | + | 1966806 | 1967519 | 11.70 | 51.91 | 47.09 | 20.01 | HTH-type transcriptional repressor RspR |
| *FEPFGCPE_01792* | *hndA* | + | 1968000 | 1968467 | 4.55 | 6.06 | 10.01 | 0.23 | NADP-reducing hydrogenase subunit HndA |
| *FEPFGCPE_01793* | *hndC-2* | + | 1968464 | 1970065 | 1.34 | 4.68 | 4.18 | 1.43 | NADP-reducing hydrogenase subunit HndC |
| *FEPFGCPE_01794* | | + | 1970079 | 1972943 | 0.84 | 2.45 | 4.51 | 0.44 | Putative formate dehydrogenase |
| *FEPFGCPE_01795* | | + | 1973154 | 1973387 | 2.22 | 10.43 | 12.24 | 1.83 | Hypothetical protein |
| *FEPFGCPE_01796* | | - | 1973506 | 1974354 | 0.86 | 6.77 | 5.41 | 1.64 | Hypothetical protein |
| *FEPFGCPE_01797* | | - | 1974777 | 1975271 | 3.66 | 17.41 | 18.18 | 7.72 | Hypothetical protein |
| *FEPFGCPE_01798* | | + | 1975626 | 1976174 | 1.25 | 3.99 | 2.85 | 3.96 | Hypothetical protein |
| *FEPFGCPE_01799* | | - | 1976158 | 1977525 | 2.72 | 12.88 | 9.29 | 3.05 | Hypothetical protein |
| *FEPFGCPE_01800* | | - | 1977850 | 1978281 | 2.22 | 14.88 | 12.60 | 4.35 | Hypothetical protein |
| *FEPFGCPE_01801* | *dinF* | + | 1978517 | 1979848 | 1.29 | 13.62 | 4.94 | 1.78 | DNA damage-inducible protein F |
| *FEPFGCPE_01802* | *hisN* | - | 1979907 | 1980707 | 1.31 | 14.06 | 11.09 | 10.51 | Histidinol-phosphatase |
| *FEPFGCPE_01803* | | + | 1980906 | 1981826 | 1.07 | 5.27 | 2.85 | 1.58 | Hypothetical protein |
| *FEPFGCPE_01804* | | + | 1981920 | 1982174 | 11.20 | 48.22 | 21.75 | 11.85 | Hypothetical protein |
| *FEPFGCPE_01805* | | + | 1982441 | 1982902 | 17.35 | 51.23 | 44.75 | 16.99 | Universal stress protein/MSMEI_3859 |
| *FEPFGCPE_01806* | *araQ* | - | 1983007 | 1983891 | 1.41 | 2.59 | 1.58 | 0.91 | L-arabinose transport system permease protein AraQ |
| *FEPFGCPE_01807* | *lacF* | - | 1983888 | 1984763 | 1.15 | 4.46 | 4.78 | 1.95 | Lactose transport system permease protein LacF |
| *FEPFGCPE_01808* | *ugpB-1* | - | 1984872 | 1986176 | 1.09 | 1.31 | 1.29 | 0.29 | sn-glycerol-3-phosphate-binding periplasmic protein UgpB |
| *FEPFGCPE_01809* | *ugpC-1* | - | 1986286 | 1987473 | 0.37 | 1.36 | 0.30 | 0.23 | sn-glycerol-3-phosphate import ATP-binding protein UgpC |
| *FEPFGCPE_01810* | | - | 1987594 | 1988463 | 2.76 | 3.15 | 6.43 | 0.95 | Thermostable monoacylglycerol lipase |
| *FEPFGCPE_01811* | | - | 1988466 | 1989347 | 8.12 | 22.22 | 10.05 | 2.11 | Hypothetical protein |
| *FEPFGCPE_01812* | *dppA-2* | - | 1989527 | 1990345 | 0.86 | 10.91 | 14.43 | 4.52 | D-aminopeptidase |
| *FEPFGCPE_01813* | | - | 1990370 | 1991479 | 1.32 | 5.73 | 5.52 | 0.80 | Beta-peptidyl aminopeptidase BapA |
| *FEPFGCPE_01814* | *gsiD-1* | - | 1991500 | 1992408 | 1.13 | 3.95 | 3.17 | 3.04 | Glutathione transport system permease protein GsiD |
| *FEPFGCPE_01815* | *gsiC-1* | - | 1992419 | 1993417 | 0.91 | 5.30 | 3.69 | 2.22 | Glutathione transport system permease protein GsiC |
| *FEPFGCPE_01816* | *gsiB-1* | - | 1993438 | 1994997 | 2.01 | 26.08 | 29.14 | 22.58 | Glutathione-binding protein GsiB |
| *FEPFGCPE_01817* | *gsiA-2* | - | 1995121 | 1997007 | 1.14 | 5.42 | 11.45 | 3.30 | Glutathione import ATP-binding protein GsiA |
| *FEPFGCPE_01818* | *iaaA* | - | 1997134 | 1998090 | 1.87 | 10.27 | 5.76 | 2.81 | Isoaspartyl peptidase |
| *FEPFGCPE_01819* | *hexR-1* | - | 1998087 | 1999193 | 2.23 | 15.76 | 11.82 | 2.97 | HTH-type transcriptional regulator HexR |
| *FEPFGCPE_01820* | | + | 1999612 | 1999833 | 148.06 | 44.44 | 106.79 | 76.64 | Hypothetical protein |
| *FEPFGCPE_01821* | | - | 2000032 | 2000589 | 38.41 | 33.45 | 28.67 | 11.85 | Hypothetical protein |
| *FEPFGCPE_01822* | *crt-5* | - | 2000654 | 2001958 | 7.51 | 19.77 | 19.35 | 6.12 | Short-chain-enoyl-CoA hydratase |
| *FEPFGCPE_01823* | *mmgC-2* | - | 2002001 | 2003134 | 16.30 | 50.26 | 28.15 | 18.21 | Acyl-CoA dehydrogenase |
| *FEPFGCPE_01824* | *acsA-4* | + | 2003576 | 2005237 | 10.13 | 11.70 | 6.71 | 3.45 | Acetyl-coenzyme A synthetase |
| *FEPFGCPE_01825* | | + | 2005273 | 2006040 | 6.26 | 9.99 | 4.91 | 3.56 | putative oxidoreductase |
| *FEPFGCPE_01826* | | + | 2006184 | 2007194 | 3.54 | 14.50 | 13.00 | 4.87 | putative HTH-type transcriptional regulator |
| *FEPFGCPE_01827* | *gudP-1* | - | 2007210 | 2008589 | 1.10 | 6.21 | 2.63 | 0.40 | putative glucarate transporter |
| *FEPFGCPE_01828* | *pcaR-2* | - | 2008822 | 2009637 | 2.91 | 9.70 | 8.44 | 3.97 | Pca regulon regulatory protein |
| *FEPFGCPE_01829* | | + | 2010365 | 2010760 | 34.58 | 227.48 | 275.05 | 133.07 | Hypothetical protein |
| *FEPFGCPE_01830* | *lhgO* | - | 2010931 | 2012034 | 1.75 | 15.16 | 11.12 | 2.80 | L-2-hydroxyglutarate oxidase LhgO |
| *FEPFGCPE_01831* | *etfA-1* | - | 2012149 | 2013081 | 9.52 | 33.15 | 28.91 | 19.42 | Electron transfer flavoprotein subunit alpha |
| *FEPFGCPE_01832* | *etfB-1* | - | 2013096 | 2013845 | 23.01 | 56.80 | 58.27 | 32.75 | Electron transfer flavoprotein subunit beta |
| *FEPFGCPE_01833* | *naiP-4* | - | 2013922 | 2015151 | 3.92 | 7.74 | 6.58 | 2.70 | Putative niacin/nicotinamide transporter NaiP |
| *FEPFGCPE_01834* | | - | 2015242 | 2016132 | 12.18 | 10.08 | 12.45 | 2.44 | 4-sulfomuconolactone hydrolase |
| *FEPFGCPE_01835* | | - | 2016240 | 2017652 | 16.80 | 8.41 | 2.42 | 1.44 | putative FAD-linked oxidoreductase |
| *FEPFGCPE_01836* | *lutR-4* | - | 2017703 | 2018434 | 107.51 | 35.92 | 25.54 | 4.79 | HTH-type transcriptional regulator LutR |
| *FEPFGCPE_01837* | | - | 2018605 | 2018904 | 9.00 | 63.85 | 45.05 | 24.97 | Hypothetical protein |
| *FEPFGCPE_01838* | *amtB-1* | + | 2019288 | 2020490 | 0.67 | 2.47 | 4.79 | 1.27 | Ammonia channel |
| *FEPFGCPE_01839* | | - | 2021007 | 2021600 | 1.16 | 9.05 | 3.41 | 1.38 | Hypothetical protein |
| *FEPFGCPE_01840* | | - | 2021578 | 2022462 | 2.66 | 11.07 | 11.67 | 3.55 | Hypothetical protein |
| *FEPFGCPE_01841* | | - | 2022531 | 2022722 | 17.46 | 40.59 | 27.47 | 19.46 | Hypothetical protein |
| *FEPFGCPE_01842* | | + | 2022872 | 2023606 | 2.32 | 9.18 | 3.83 | 1.25 | Hypothetical protein |

| Locus name | Gene name | Strand | Gene start (bp) | Gene end (bp) | Average TPM Inoculum | Average TPM $t_6$ | Average TPM $t_{24}$ | Average TPM $t_{168}$ | Annotation description |
|---|---|---|---|---|---|---|---|---|---|
| *FEPFGCPE_01843* | *argA-2* | - | 2023651 | 2024109 | 1.46 | 21.05 | 4.70 | 4.40 | Amino-acid acetyltransferase |
| *FEPFGCPE_01844* | | - | 2024116 | 2025417 | 0.63 | 1.93 | 1.40 | 0.00 | Hypothetical protein |
| *FEPFGCPE_01845* | *arsH-2* | + | 2025563 | 2026333 | 1.63 | 9.39 | 1.80 | 2.59 | NADPH-dependent FMN reductase ArsH |
| *FEPFGCPE_01846* | *sdpR* | + | 2026337 | 2026666 | 1.22 | 15.69 | 5.45 | 13.71 | Transcriptional repressor SdpR |
| *FEPFGCPE_01847* | *cadI-2* | + | 2026667 | 2027131 | 0.70 | 5.07 | 2.39 | 2.34 | Cadmium-induced protein CadI |
| *FEPFGCPE_01848* | *hlyU* | - | 2027211 | 2027552 | 4.99 | 39.65 | 33.07 | 14.88 | Transcriptional activator HlyU |
| *FEPFGCPE_01849* | *arsC-2* | + | 2027761 | 2028300 | 5.32 | 6.07 | 8.83 | 6.20 | Protein ArsC |
| *FEPFGCPE_01850* | *acr3-2* | + | 2028303 | 2029376 | 1.65 | 5.27 | 2.46 | 0.15 | Arsenical-resistance protein Acr3 |
| *FEPFGCPE_01851* | *glrR* | - | 2029411 | 2030772 | 2.28 | 5.86 | 3.54 | 0.31 | Transcriptional regulatory protein GlrR |
| *FEPFGCPE_01852* | *dctB-1* | - | 2030769 | 2032232 | 1.68 | 2.51 | 1.66 | 0.21 | C4-dicarboxylate transport sensor protein DctB |
| *FEPFGCPE_01853* | *phnD-1* | - | 2032240 | 2033130 | 7.97 | 8.18 | 3.66 | 1.04 | Phosphate-import protein PhnD |
| *FEPFGCPE_01854* | *aioB* | + | 2033297 | 2033821 | 0.57 | 1.99 | 2.79 | 0.00 | Arsenite oxidase subunit AioB |
| *FEPFGCPE_01855* | *aioA* | + | 2033832 | 2036309 | 0.35 | 0.97 | 1.62 | 0.25 | Arsenite oxidase subunit AioA |
| *FEPFGCPE_01856* | *nirM-1* | + | 2036375 | 2036659 | 0.33 | 4.15 | 0.00 | 0.00 | Cytochrome c-551 |
| *FEPFGCPE_01857* | *arsC-3* | - | 2036722 | 2037150 | 1.67 | 14.57 | 1.11 | 2.44 | Arsenate reductase |
| *FEPFGCPE_01858* | *czcO-2* | - | 2037147 | 2038229 | 2.36 | 2.84 | 1.42 | 0.92 | putative oxidoreductase CzcO |
| *FEPFGCPE_01859* | | - | 2038601 | 2038840 | 1.84 | 10.51 | 8.45 | 1.78 | Hypothetical protein |
| *FEPFGCPE_01860* | | - | 2039012 | 2040490 | 0.77 | 2.99 | 1.18 | 1.87 | Hypothetical protein |
| *FEPFGCPE_01861* | | - | 2040497 | 2041456 | 1.34 | 4.39 | 3.97 | 1.05 | Hypothetical protein |
| *FEPFGCPE_01862* | | + | 2042112 | 2043101 | 0.70 | 5.40 | 3.25 | 0.00 | Hypothetical protein |
| *FEPFGCPE_01863* | *tnsB* | + | 2043095 | 2045116 | 0.96 | 7.42 | 7.18 | 1.08 | Transposon Tn7 transposition protein TnsB |
| *FEPFGCPE_01864* | | + | 2045113 | 2046441 | 2.55 | 22.00 | 11.50 | 1.86 | Hypothetical protein |
| *FEPFGCPE_01865* | *gltC-5* | - | 2046862 | 2047755 | 1.82 | 10.55 | 2.58 | 2.59 | HTH-type transcriptional regulator GltC |
| *FEPFGCPE_01866* | *citN* | + | 2047855 | 2049159 | 1.68 | 6.36 | 2.57 | 0.00 | Citrate transporter |
| *FEPFGCPE_01867* | | + | 2049185 | 2050552 | 0.80 | 3.86 | 0.52 | 0.64 | Hypothetical protein |
| *FEPFGCPE_01868* | | + | 2050554 | 2050865 | 0.40 | 0.00 | 0.73 | 0.00 | Hypothetical protein |
| *FEPFGCPE_01869* | | - | 2050885 | 2052504 | 0.27 | 1.80 | 0.85 | 0.45 | Hypothetical protein |
| *FEPFGCPE_01870* | | + | 2052992 | 2053939 | 2.86 | 18.87 | 4.68 | 2.63 | Hypothetical protein |
| *FEPFGCPE_01871* | *kdgR-2* | + | 2053939 | 2054730 | 1.04 | 8.68 | 6.64 | 2.10 | Transcriptional regulator KdgR |
| *FEPFGCPE_01872* | *mhpE* | - | 2054853 | 2055872 | 1.06 | 4.93 | 1.86 | 0.91 | 4-hydroxy-2-oxovalerate aldolase |
| *FEPFGCPE_01873* | *dmpF* | - | 2055907 | 2056851 | 0.84 | 2.74 | 3.82 | 0.98 | Acetaldehyde dehydrogenase |
| *FEPFGCPE_01874* | *pagP* | + | 2057572 | 2058234 | 4.67 | 7.42 | 10.98 | 6.56 | Lipid A palmitoyltransferase PagP |
| *FEPFGCPE_01875* | *fucP* | - | 2058210 | 2059562 | 4.07 | 6.83 | 3.33 | 2.09 | L-fucose-proton symporter |
| *FEPFGCPE_01876* | | + | 2059810 | 2060709 | 2.11 | 9.42 | 4.97 | 1.44 | Hypothetical protein |
| *FEPFGCPE_01877* | | + | 2061022 | 2061492 | 54.66 | 476.31 | 239.51 | 159.50 | Hypothetical protein |
| *FEPFGCPE_01878* | *fliY-3* | + | 2061831 | 2062769 | 7.67 | 61.13 | 27.48 | 10.20 | L-cystine-binding protein FliY |
| *FEPFGCPE_01879* | *yqjI* | + | 2063058 | 2063705 | 1.91 | 5.37 | 7.41 | 5.83 | Transcriptional regulator YqjI |
| *FEPFGCPE_01880* | *yqjH* | + | 2063705 | 2064544 | 2.16 | 3.97 | 4.44 | 2.36 | NADPH-dependent ferric-chelate reductase |
| *FEPFGCPE_01881* | | + | 2064603 | 2064953 | 0.66 | 0.80 | 6.30 | 3.17 | Hypothetical protein |
| *FEPFGCPE_01882* | | + | 2065111 | 2065458 | 0.62 | 2.89 | 2.25 | 0.59 | Hypothetical protein |
| *FEPFGCPE_01883* | *yueD* | - | 2065583 | 2066371 | 2.19 | 13.65 | 13.54 | 5.83 | Benzil reductase ((S)-benzoin forming) |
| *FEPFGCPE_01884* | *spsI-1* | + | 2066809 | 2068194 | 6.07 | 15.82 | 14.09 | 3.39 | Bifunctional IPC transferase and DIPP synthase |
| *FEPFGCPE_01885* | | + | 2068195 | 2069217 | 1.99 | 5.09 | 9.60 | 3.88 | Hypothetical protein |
| *FEPFGCPE_01886* | *spsI-2* | + | 2069233 | 2069997 | 1.56 | 7.44 | 9.10 | 4.22 | Bifunctional IPC transferase and DIPP synthase |
| *FEPFGCPE_01887* | *pphA-1* | + | 2069994 | 2071703 | 3.08 | 12.20 | 6.88 | 3.01 | Phosphonopyruvate hydrolase |
| *FEPFGCPE_01888* | | + | 2071696 | 2072883 | 2.32 | 9.34 | 8.64 | 6.44 | Hypothetical protein |
| *FEPFGCPE_01889* | *phnW-1* | + | 2072922 | 2073995 | 2.54 | 7.48 | 10.41 | 3.95 | 2-aminoethylphosphonate--pyruvate transaminase |
| *FEPFGCPE_01890* | *dxs-3* | + | 2074018 | 2075934 | 0.83 | 5.47 | 5.46 | 3.01 | 1-deoxy-D-xylulose-5-phosphate synthase |
| *FEPFGCPE_01891* | | - | 2076020 | 2078656 | 0.43 | 2.98 | 2.53 | 1.80 | Acyltrehalose exporter MmpL10 |
| *FEPFGCPE_01892* | | + | 2078833 | 2079525 | 21.72 | 22.91 | 25.77 | 18.43 | Hypothetical protein |
| *FEPFGCPE_01893* | *ispH-1* | + | 2079865 | 2080824 | 8.71 | 62.32 | 53.75 | 16.33 | 4-hydroxy-3-methylbut-2-enyl diphosphate reductase |
| *FEPFGCPE_01894* | *moaA1* | + | 2080848 | 2081969 | 5.19 | 35.45 | 28.73 | 16.68 | GTP 3',8-cyclase 1 |
| *FEPFGCPE_01895* | | + | 2081970 | 2082965 | 2.37 | 16.12 | 14.52 | 5.61 | 3 beta-hydroxysteroid dehydrogenase/Delta 5--4-isomerase |
| *FEPFGCPE_01896* | *crtQ* | + | 2082962 | 2084164 | 1.10 | 5.17 | 4.88 | 1.13 | 4,4'-diaponeurosporenoate glycosyltransferase |
| *FEPFGCPE_01897* | | - | 2084281 | 2084976 | 0.50 | 5.74 | 4.04 | 1.25 | Hypothetical protein |
| *FEPFGCPE_01898* | *shc* | - | 2084988 | 2087060 | 1.49 | 11.36 | 10.68 | 4.71 | Squalene--hopene cyclase |
| *FEPFGCPE_01899* | *mnmC-1* | - | 2087182 | 2088435 | 2.20 | 17.83 | 17.59 | 6.40 | tRNA 5-methylaminomethyl-2-thiouridine biosynthesis bifunctional protein MnmC |
| *FEPFGCPE_01900* | *crtB-2* | - | 2088435 | 2089406 | 2.63 | 8.92 | 14.36 | 5.07 | All-trans-phytoene synthase |
| *FEPFGCPE_01901* | *crtB-3* | + | 2089460 | 2090323 | 1.62 | 4.36 | 7.74 | 1.10 | All-trans-phytoene synthase |
| *FEPFGCPE_01902* | | + | 2090522 | 2091700 | 0.92 | 0.85 | 1.57 | 0.48 | Hypothetical protein |
| *FEPFGCPE_01903* | | - | 2091707 | 2092345 | 4.01 | 23.49 | 19.84 | 3.87 | Hypothetical protein |
| *FEPFGCPE_01904* | *csgA-1* | - | 2092527 | 2093258 | 1.82 | 2.28 | 4.84 | 3.40 | C-factor |
| *FEPFGCPE_01905* | *syrM1-2* | + | 2093424 | 2094377 | 5.00 | 18.84 | 7.26 | 2.86 | HTH-type transcriptional regulator SyrM 1 |
| *FEPFGCPE_01906* | | - | 2094399 | 2094839 | 4.15 | 18.75 | 11.15 | 3.21 | Hypothetical protein |
| *FEPFGCPE_01907* | *atoE-1* | - | 2094948 | 2096396 | 2.52 | 1.44 | 1.44 | 0.77 | Putative short-chain fatty acid transporter |
| *FEPFGCPE_01908* | | - | 2096523 | 2097467 | 0.39 | 2.38 | 1.20 | 0.57 | Delta(1)-pyrroline-2-carboxylate reductase |
| *FEPFGCPE_01909* | *hmp-2* | - | 2097502 | 2100018 | 0.80 | 0.87 | 0.92 | 0.00 | Flavohemoprotein |
| *FEPFGCPE_01910* | | - | 2100046 | 2100963 | 0.18 | 3.35 | 4.19 | 0.12 | Acetylene hydratase |
| *FEPFGCPE_01911* | *oxyR-3* | + | 2101062 | 2101988 | 2.61 | 14.65 | 9.77 | 2.43 | Hydrogen peroxide-inducible genes activator |
| *FEPFGCPE_01912* | *pdxA2* | - | 2102030 | 2103052 | 1.34 | 5.67 | 3.46 | 0.90 | 4-hydroxythreonine-4-phosphate dehydrogenase 2 |
| *FEPFGCPE_01913* | | - | 2103123 | 2104424 | 0.36 | 3.12 | 3.04 | 0.85 | Hypothetical protein |

| Locus name | Gene name | Strand | Gene start (bp) | Gene end (bp) | Average TPM Inoculum | Average TPM $t_6$ | Average TPM $t_{24}$ | Average TPM $t_{168}$ | Annotation description |
|---|---|---|---|---|---|---|---|---|---|
| *FEPFGCPE_01914* | *kdgT* | - | 2104421 | 2105410 | 1.30 | 4.90 | 5.37 | 1.15 | 2-keto-3-deoxygluconate permease |
| *FEPFGCPE_01915* | *srlR* | + | 2105748 | 2106512 | 0.70 | 7.87 | 8.58 | 3.23 | Glucitol operon repressor |
| *FEPFGCPE_01916* | *dctA-2* | - | 2106567 | 2107892 | 1.32 | 3.23 | 2.84 | 0.36 | C4-dicarboxylate transport protein |
| *FEPFGCPE_01917* | *arcB* | - | 2107937 | 2108851 | 0.58 | 3.83 | 0.54 | 0.66 | Delta(1)-pyrroline-2-carboxylate reductase |
| *FEPFGCPE_01918* | *yddE* | - | 2108848 | 2109774 | 0.06 | 3.43 | 1.79 | 0.00 | putative isomerase YddE |
| *FEPFGCPE_01919* | *soxS* | + | 2110223 | 2111008 | 2.14 | 13.59 | 5.31 | 2.29 | Regulatory protein SoxS |
| *FEPFGCPE_01920* | | - | 2111030 | 2111782 | 1.61 | 12.49 | 6.75 | 1.47 | Hypothetical protein |
| *FEPFGCPE_01921* | *gabD-2* | - | 2111872 | 2113344 | 1.19 | 8.73 | 4.82 | 2.92 | Succinate-semialdehyde dehydrogenase [NADP(+)] GabD |
| *FEPFGCPE_01922* | *lrp-2* | - | 2113432 | 2113926 | 2.95 | 24.09 | 11.52 | 3.00 | Leucine-responsive regulatory protein |
| *FEPFGCPE_01923* | | - | 2114046 | 2115449 | 1.53 | 6.98 | 7.29 | 3.77 | putative aminotransferase |
| *FEPFGCPE_01924* | *ghrA-1* | - | 2115613 | 2116539 | 0.31 | 11.89 | 5.15 | 5.18 | Glyoxylate/hydroxypyruvate reductase A |
| *FEPFGCPE_01925* | | - | 2116553 | 2117437 | 0.65 | 18.78 | 12.90 | 9.17 | Hypothetical protein |
| *FEPFGCPE_01926* | *puuC-1* | + | 2117771 | 2119249 | 0.42 | 3.64 | 0.42 | 0.18 | Aldehyde dehydrogenase PuuC |
| *FEPFGCPE_01927* | *hdl IVa-2* | + | 2119560 | 2120228 | 4.40 | 16.96 | 9.26 | 8.13 | (S)-2-haloacid dehalogenase 4A |
| *FEPFGCPE_01928* | *puuB-2* | + | 2120277 | 2121554 | 3.49 | 8.78 | 6.02 | 1.62 | Gamma-glutamylputrescine oxidoreductase |
| *FEPFGCPE_01929* | *gsiB-2* | + | 2121629 | 2123242 | 5.09 | 16.27 | 12.06 | 4.99 | Glutathione-binding protein GsiB |
| *FEPFGCPE_01930* | *gsiC-2* | + | 2123359 | 2124315 | 1.92 | 8.79 | 2.72 | 1.58 | Glutathione transport system permease protein GsiC |
| *FEPFGCPE_01931* | *gsiD-2* | + | 2124312 | 2125100 | 0.65 | 4.86 | 3.16 | 0.25 | Glutathione transport system permease protein GsiD |
| *FEPFGCPE_01932* | *gsiA-3* | + | 2125253 | 2127139 | 1.34 | 6.10 | 4.15 | 1.48 | Glutathione import ATP-binding protein GsiA |
| *FEPFGCPE_01933* | *hcnA-3* | + | 2127136 | 2127510 | 1.04 | 8.56 | 2.27 | 1.47 | Hydrogen cyanide synthase subunit HcnA |
| *FEPFGCPE_01934* | *hcnB-3* | + | 2127495 | 2128934 | 0.66 | 5.66 | 4.73 | 0.84 | Hydrogen cyanide synthase subunit HcnB |
| *FEPFGCPE_01935* | *abo-1* | + | 2128931 | 2130148 | 0.80 | 4.48 | 4.08 | 1.51 | 4-methylaminobutanoate oxidase (formaldehyde-forming) |
| *FEPFGCPE_01936* | *araC-1* | + | 2130145 | 2130582 | 0.79 | 7.38 | 2.02 | 0.76 | Arabinose operon regulatory protein |
| *FEPFGCPE_01937* | *sauU-1* | - | 2130609 | 2131985 | 0.74 | 2.68 | 0.00 | 0.39 | putative sulfoacetate transporter SauU |
| *FEPFGCPE_01938* | *sauT* | - | 2132113 | 2133732 | 0.54 | 1.32 | 0.56 | 0.00 | putative sulfoacetate--CoA ligase |
| *FEPFGCPE_01939* | *sauS* | - | 2133729 | 2135192 | 0.98 | 1.08 | 0.44 | 0.12 | Sulfoacetaldehyde dehydrogenase (acylating) |
| *FEPFGCPE_01940* | *iclR-3* | + | 2135480 | 2136244 | 2.23 | 8.69 | 4.61 | 0.66 | Transcriptional repressor IclR |
| *FEPFGCPE_01941* | | - | 2136234 | 2137622 | 1.08 | 12.90 | 13.96 | 1.52 | Hypothetical protein |
| *FEPFGCPE_01942* | *rlp2* | - | 2137658 | 2138938 | 0.28 | 10.56 | 8.77 | 0.00 | Ribulose bisphosphate carboxylase-like protein 2 |
| *FEPFGCPE_01943* | | - | 2138935 | 2139489 | 0.54 | 10.40 | 9.27 | 0.07 | Hypothetical protein |
| *FEPFGCPE_01944* | *siaT-2* | - | 2139653 | 2141572 | 0.35 | 9.23 | 8.94 | 1.01 | Sialic acid TRAP transporter permease protein SiaT |
| *FEPFGCPE_01945* | *siaP-2* | - | 2141574 | 2142626 | 0.24 | 12.87 | 12.67 | 2.13 | Sialic acid-binding periplasmic protein SiaP |
| *FEPFGCPE_01946* | *xecD* | - | 2142738 | 2143553 | 1.59 | 24.54 | 29.38 | 3.70 | 2-(R)-hydroxypropyl-CoM dehydrogenase |
| *FEPFGCPE_01947* | *alsB* | + | 2143757 | 2144791 | 0.92 | 9.27 | 4.96 | 0.79 | D-allose-binding periplasmic protein |
| *FEPFGCPE_01948* | *lnpD* | - | 2144918 | 2145940 | 2.61 | 29.65 | 18.55 | 6.22 | UDP-glucose 4-epimerase |
| *FEPFGCPE_01949* | *glmS-1* | - | 2145971 | 2147803 | 0.77 | 13.69 | 7.36 | 3.47 | Glutamine--fructose-6-phosphate aminotransferase [isomerizing] |
| *FEPFGCPE_01950* | | - | 2147966 | 2148805 | 0.70 | 32.84 | 12.94 | 4.01 | Hypothetical protein |
| *FEPFGCPE_01951* | | - | 2148892 | 2150271 | 0.74 | 23.81 | 9.06 | 3.44 | Hypothetical protein |
| *FEPFGCPE_01952* | *mshA* | - | 2150271 | 2151818 | 1.09 | 47.61 | 18.91 | 7.15 | D-inositol 3-phosphate glycosyltransferase |
| *FEPFGCPE_01953* | | - | 2151818 | 2152681 | 0.66 | 18.79 | 5.72 | 2.75 | Hypothetical protein |
| *FEPFGCPE_01954* | | - | 2152811 | 2154223 | 1.78 | 34.80 | 13.10 | 6.58 | Hypothetical protein |
| *FEPFGCPE_01955* | | - | 2154235 | 2154780 | 2.79 | 55.39 | 36.46 | 18.56 | Hypothetical protein |
| *FEPFGCPE_01956* | | - | 2154811 | 2158929 | 2.52 | 28.85 | 14.05 | 6.12 | Hypothetical protein |
| *FEPFGCPE_01957* | | - | 2158967 | 2159602 | 6.94 | 45.34 | 24.61 | 8.64 | Hypothetical protein |
| *FEPFGCPE_01958* | | - | 2159613 | 2161856 | 5.47 | 40.36 | 18.86 | 7.90 | Hypothetical protein |
| *FEPFGCPE_01959* | | - | 2162347 | 2162619 | 21.73 | 117.01 | 47.57 | 9.29 | Hypothetical protein |
| *FEPFGCPE_01960* | | - | 2162879 | 2164153 | 0.56 | 1.58 | 2.30 | 0.00 | putative transporter |
| *FEPFGCPE_01961* | *oxyR-4* | + | 2164284 | 2165216 | 3.64 | 5.32 | 2.18 | 1.86 | Hydrogen peroxide-inducible genes activator |
| *FEPFGCPE_01962* | | + | 2165380 | 2165796 | 4.59 | 125.65 | 94.29 | 26.17 | Hypothetical protein |
| *FEPFGCPE_01963* | *pfeA* | + | 2165879 | 2168179 | 0.70 | 21.99 | 20.40 | 16.40 | Ferric enterobactin receptor |
| *FEPFGCPE_01964* | | + | 2168206 | 2169765 | 1.80 | 12.91 | 12.27 | 8.36 | Hypothetical protein |
| *FEPFGCPE_01965* | | + | 2170019 | 2171494 | 5.16 | 12.68 | 5.23 | 2.40 | Hypothetical protein |
| *FEPFGCPE_01966* | | + | 2171759 | 2172604 | 22.36 | 5.40 | 5.82 | 2.11 | Hypothetical protein |
| *FEPFGCPE_01967* | | + | 2172662 | 2173651 | 49.55 | 6.28 | 11.06 | 6.62 | Hypothetical protein |
| *FEPFGCPE_01968* | | + | 2173857 | 2174354 | 22.66 | 16.44 | 12.49 | 13.93 | Hypothetical protein |
| *FEPFGCPE_01969* | | - | 2174523 | 2174972 | 11.28 | 29.10 | 31.42 | 16.67 | Hypothetical protein |
| *FEPFGCPE_01970* | *cpo-1* | - | 2175017 | 2175838 | 1.85 | 10.34 | 11.26 | 6.87 | Non-heme chloroperoxidase |
| *FEPFGCPE_01971* | | + | 2176292 | 2176843 | 0.53 | 3.41 | 6.12 | 0.39 | Hypothetical protein |
| *FEPFGCPE_01972* | | + | 2177091 | 2178116 | 0.80 | 2.72 | 0.50 | 0.61 | Hypothetical protein |
| *FEPFGCPE_01973* | | - | 2178178 | 2180139 | 1.27 | 5.19 | 4.55 | 0.72 | Hypothetical protein |
| *FEPFGCPE_01974* | | - | 2180136 | 2184284 | 1.66 | 10.35 | 4.62 | 2.37 | Hypothetical protein |
| *FEPFGCPE_01975* | *vraR-2* | + | 2184614 | 2185336 | 5.32 | 25.05 | 14.66 | 5.01 | Response regulator protein VraR |
| *FEPFGCPE_01976* | *liaS-1* | + | 2185435 | 2187297 | 1.14 | 7.42 | 5.43 | 0.73 | Sensor histidine kinase LiaS |
| *FEPFGCPE_01977* | *pcaR-3* | - | 2187449 | 2188294 | 1.58 | 14.34 | 7.38 | 2.10 | Pca regulon regulatory protein |
| *FEPFGCPE_01978* | *mhpA-2* | + | 2188552 | 2190405 | 0.10 | 1.29 | 0.79 | 0.13 | 3-(3-hydroxy-phenyl)propionate/3-hydroxycinnamic acid hydroxylase |
| *FEPFGCPE_01979* | *mhpB-1* | + | 2190409 | 2191368 | 0.17 | 1.25 | 0.05 | 0.00 | 2,3-dihydroxyphenylpropionate/2,3-dihydroxicinnamic acid 1,2-dioxygenase |
| *FEPFGCPE_01980* | *mhpD-1* | + | 2191384 | 2192169 | 0.23 | 2.42 | 1.80 | 0.08 | 2-keto-4-pentenoate hydratase |
| *FEPFGCPE_01981* | *mhpT-1* | + | 2192455 | 2193741 | 0.11 | 0.53 | 1.32 | 0.00 | 3-(3-hydroxy-phenyl)propionate transporter |
| *FEPFGCPE_01982* | *mhpC-1* | + | 2193853 | 2194731 | 0.29 | 3.24 | 1.44 | 0.23 | 2-hydroxy-6-oxononadienedioate/2-hydroxy-6-oxononatrienedioate hydrolase |
| *FEPFGCPE_01983* | | - | 2194937 | 2195788 | 1.30 | 14.49 | 7.33 | 1.25 | Hypothetical protein |
| *FEPFGCPE_01984* | | - | 2195806 | 2196075 | 3.83 | 16.66 | 9.41 | 2.25 | Hypothetical protein |

| Locus name | Gene name | Strand | Gene start (bp) | Gene end (bp) | Average TPM Inoculum | Average TPM $t_6$ | Average TPM $t_{24}$ | Average TPM $t_{168}$ | Annotation description |
|---|---|---|---|---|---|---|---|---|---|
| *FEPFGCPE_01985* | *yhjE-2* | - | 2196398 | 2197735 | 1.65 | 11.24 | 7.15 | 1.23 | Inner membrane metabolite transport protein YhjE |
| *FEPFGCPE_01986* | *yabJ-4* | - | 2197854 | 2198216 | 0.00 | 0.00 | 2.10 | 0.00 | 2-iminobutanoate/2-iminopropanoate deaminase |
| *FEPFGCPE_01987* | *lhpI* | - | 2198333 | 2199274 | 1.74 | 5.09 | 9.28 | 1.42 | Delta(1)-pyrroline-2-carboxylate reductase |
| *FEPFGCPE_01988* | | - | 2199358 | 2200335 | 1.47 | 11.66 | 5.72 | 3.04 | Hypothetical protein |
| *FEPFGCPE_01989* | *ade* | - | 2200406 | 2201641 | 2.54 | 10.48 | 9.43 | 3.73 | Adenine deaminase |
| *FEPFGCPE_01990* | | + | 2202543 | 2202887 | 7.36 | 47.89 | 37.21 | 12.57 | Hypothetical protein |
| *FEPFGCPE_01991* | | + | 2203034 | 2203432 | 2.96 | 20.79 | 14.41 | 3.35 | Hypothetical protein |
| *FEPFGCPE_01992* | | - | 2204000 | 2204587 | 0.39 | 2.59 | 3.69 | 0.55 | Hypothetical protein |
| *FEPFGCPE_01993* | | + | 2204880 | 2205551 | 0.91 | 6.91 | 3.89 | 0.57 | Hypothetical protein |
| *FEPFGCPE_01994* | | - | 2205708 | 2205798 | 17.94 | 45.59 | 24.21 | 25.00 | tRNA |
| *FEPFGCPE_01995* | *serS* | - | 2205901 | 2207232 | 3.33 | 29.66 | 25.30 | 14.49 | Serine--tRNA ligase |
| *FEPFGCPE_01996* | *rarA* | - | 2207370 | 2208836 | 0.77 | 9.31 | 8.23 | 2.67 | Replication-associated recombination protein A |
| *FEPFGCPE_01997* | *lolA-2* | - | 2208919 | 2209569 | 9.57 | 40.18 | 35.45 | 26.70 | Outer-membrane lipoprotein carrier protein |
| *FEPFGCPE_01998* | *ftsK-2* | - | 2209575 | 2211905 | 6.66 | 33.71 | 27.47 | 14.18 | DNA translocase FtsK |
| *FEPFGCPE_01999* | *trxB-1* | + | 2211985 | 2213181 | 5.74 | 33.55 | 29.00 | 18.66 | Thioredoxin reductase |
| *FEPFGCPE_02000* | *smrA* | + | 2213280 | 2213957 | 1.03 | 6.07 | 7.52 | 3.82 | putative DNA endonuclease SmrA |
| *FEPFGCPE_02001* | *glnB-2* | - | 2214179 | 2214517 | 21.47 | 192.63 | 157.17 | 92.24 | Nitrogen regulatory protein P-II |
| *FEPFGCPE_02002* | *nadE-2* | - | 2214605 | 2216707 | 3.43 | 23.73 | 18.32 | 5.79 | Glutamine-dependent NAD(+) synthetase |
| *FEPFGCPE_02003* | | - | 2216863 | 2217999 | 1.83 | 11.34 | 7.12 | 2.14 | Hypothetical protein |
| *FEPFGCPE_02004* | *ppa* | - | 2218172 | 2218699 | 8.47 | 54.34 | 48.10 | 37.63 | Inorganic pyrophosphatase |
| *FEPFGCPE_02005* | *mdlB* | - | 2218802 | 2219986 | 0.44 | 3.10 | 2.18 | 0.65 | (S)-mandelate dehydrogenase |
| *FEPFGCPE_02006* | *pcaR-4* | + | 2220137 | 2221012 | 1.29 | 4.48 | 7.03 | 1.55 | Pca regulon regulatory protein |
| *FEPFGCPE_02007* | *dapA-3* | - | 2221051 | 2221866 | 0.55 | 6.58 | 2.14 | 0.00 | 4-hydroxy-tetrahydrodipicolinate synthase |
| *FEPFGCPE_02008* | *potH-2* | - | 2222005 | 2223711 | 0.46 | 2.02 | 1.47 | 0.30 | Putrescine transport system permease protein PotH |
| *FEPFGCPE_02009* | *potA-5* | - | 2223811 | 2224956 | 0.64 | 3.77 | 1.62 | 2.22 | Spermidine/putrescine import ATP-binding protein PotA |
| *FEPFGCPE_02010* | *potD-3* | - | 2225002 | 2226150 | 1.01 | 2.83 | 3.04 | 0.76 | Spermidine/putrescine-binding periplasmic protein |
| *FEPFGCPE_02011* | | - | 2226260 | 2227696 | 0.66 | 2.74 | 2.75 | 0.98 | putative FAD-linked oxidoreductase |
| *FEPFGCPE_02012* | *puuC-2* | + | 2227957 | 2229396 | 6.37 | 28.71 | 17.81 | 9.35 | Aldehyde dehydrogenase PuuC |
| *FEPFGCPE_02013* | *tsaC1-1* | + | 2229503 | 2230261 | 5.04 | 35.39 | 29.71 | 13.87 | 4-formylbenzenesulfonate dehydrogenase TsaC1/TsaC2 |
| *FEPFGCPE_02014* | *nicT-3* | + | 2230355 | 2231671 | 3.81 | 44.76 | 21.96 | 10.77 | Putative metabolite transport protein NicT |
| *FEPFGCPE_02015* | *bmr3-1* | + | 2231840 | 2233348 | 1.87 | 6.43 | 5.44 | 2.41 | Multidrug resistance protein 3 |
| *FEPFGCPE_02016* | | - | 2233382 | 2234554 | 4.32 | 30.32 | 14.63 | 9.22 | Hypothetical protein |
| *FEPFGCPE_02017* | *hemX* | - | 2234565 | 2235728 | 5.28 | 28.36 | 28.11 | 13.22 | Putative uroporphyrinogen-III C-methyltransferase |
| *FEPFGCPE_02018* | *hemD* | - | 2235725 | 2236642 | 1.54 | 16.05 | 6.86 | 3.69 | Uroporphyrinogen-III synthase |
| *FEPFGCPE_02019* | *hemC* | - | 2236720 | 2237688 | 5.24 | 30.82 | 18.21 | 7.95 | Porphobilinogen deaminase |
| *FEPFGCPE_02020* | *ppc* | + | 2237768 | 2240734 | 3.13 | 9.20 | 15.07 | 8.40 | Phosphoenolpyruvate carboxylase |
| *FEPFGCPE_02021* | *aroQ-1* | + | 2240961 | 2241737 | 4.17 | 11.70 | 10.81 | 1.84 | Monofunctional chorismate mutase |
| *FEPFGCPE_02022* | *argH* | + | 2241951 | 2243348 | 2.93 | 17.80 | 15.36 | 12.33 | Argininosuccinate lyase |
| *FEPFGCPE_02023* | *gltS* | - | 2243462 | 2244673 | 0.28 | 3.32 | 2.16 | 1.28 | Sodium/glutamate symporter |
| *FEPFGCPE_02024* | *adiA* | - | 2245027 | 2247312 | 2.24 | 18.64 | 12.28 | 5.39 | Biodegradative arginine decarboxylase |
| *FEPFGCPE_02025* | *dcd* | - | 2247418 | 2247987 | 6.45 | 42.75 | 32.88 | 17.27 | dCTP deaminase |
| *FEPFGCPE_02026* | *apbC* | - | 2248111 | 2249193 | 7.69 | 23.29 | 16.04 | 7.88 | Iron-sulfur cluster carrier protein |
| *FEPFGCPE_02027* | *yiaD-1* | + | 2249469 | 2250113 | 41.34 | 223.09 | 123.10 | 66.39 | putative lipoprotein YiaD |
| *FEPFGCPE_02028* | *metG* | + | 2250575 | 2252674 | 5.27 | 18.67 | 22.28 | 13.57 | Methionine--tRNA ligase |
| *FEPFGCPE_02029* | *decR-2* | - | 2252839 | 2253300 | 1.41 | 4.62 | 0.99 | 0.33 | DNA-binding transcriptional activator DecR |
| *FEPFGCPE_02030* | | + | 2253418 | 2253681 | 265.16 | 36.43 | 44.77 | 18.14 | Hypothetical protein |
| *FEPFGCPE_02031* | *bamE-1* | - | 2253729 | 2254334 | 30.21 | 3.15 | 8.39 | 6.79 | Outer membrane protein assembly factor BamE |
| *FEPFGCPE_02032* | *tamB* | - | 2254400 | 2256838 | 1.11 | 6.72 | 3.71 | 2.12 | Translocation and assembly module TamB |
| *FEPFGCPE_02033* | | - | 2256786 | 2258543 | 0.78 | 2.95 | 3.37 | 1.07 | Hypothetical protein |
| *FEPFGCPE_02034* | *tamA* | - | 2258543 | 2260351 | 1.11 | 4.02 | 3.49 | 1.42 | Translocation and assembly module TamA |
| *FEPFGCPE_02035* | | + | 2260628 | 2260810 | 15.48 | 144.88 | 106.67 | 62.51 | Hypothetical protein |
| *FEPFGCPE_02036* | *scpA* | + | 2260838 | 2261860 | 3.06 | 36.18 | 18.91 | 6.03 | Segregation and condensation protein A |
| *FEPFGCPE_02037* | *panC* | + | 2262045 | 2262881 | 6.54 | 30.96 | 16.80 | 5.24 | Pantothenate synthetase |
| *FEPFGCPE_02038* | *panD* | + | 2263009 | 2263431 | 2.38 | 13.34 | 11.97 | 6.24 | Aspartate 1-decarboxylase |
| *FEPFGCPE_02039* | *soj-1* | - | 2263533 | 2264156 | 5.40 | 34.39 | 28.84 | 11.67 | Chromosome-partitioning ATPase Soj |
| *FEPFGCPE_02040* | *cobQ* | - | 2264217 | 2265773 | 1.63 | 9.34 | 6.28 | 3.03 | Cobyric acid synthase |
| *FEPFGCPE_02041* | *cobP* | + | 2265814 | 2266365 | 6.74 | 25.34 | 21.98 | 6.21 | Bifunctional adenosylcobalamin biosynthesis protein CobP |
| *FEPFGCPE_02042* | *cobD-2* | + | 2266372 | 2267352 | 3.49 | 11.35 | 5.33 | 0.37 | Cobalamin biosynthesis protein CobD |
| *FEPFGCPE_02043* | *cobD-3* | + | 2267349 | 2268407 | 3.04 | 13.35 | 11.86 | 5.49 | Threonine-phosphate decarboxylase |
| *FEPFGCPE_02044* | | + | 2268631 | 2268849 | 42.06 | 214.57 | 175.81 | 127.54 | Hypothetical protein |
| *FEPFGCPE_02045* | | - | 2269109 | 2269273 | 44.42 | 80.62 | 103.70 | 58.19 | Hypothetical protein |
| *FEPFGCPE_02046* | *btuF* | + | 2269522 | 2270475 | 1.46 | 3.91 | 5.35 | 1.63 | Vitamin B12-binding protein |
| *FEPFGCPE_02047* | *pspA-1* | - | 2270503 | 2271078 | 0.45 | 3.19 | 3.65 | 1.23 | Phosphoserine phosphatase 1 |
| *FEPFGCPE_02048* | *cobS-1* | - | 2271084 | 2271431 | 0.17 | 6.17 | 3.65 | 0.90 | Adenosylcobinamide-GDP ribazoletransferase |
| *FEPFGCPE_02049* | *cobS-2* | - | 2271407 | 2271961 | 0.69 | 4.78 | 2.02 | 0.83 | Adenosylcobinamide-GDP ribazoletransferase |
| *FEPFGCPE_02050* | *cobT-1* | - | 2271958 | 2273043 | 0.84 | 7.82 | 12.70 | 3.29 | Nicotinate-nucleotide--dimethylbenzimidazole phosphoribosyltransferase |
| *FEPFGCPE_02051* | *cobO* | - | 2273075 | 2273719 | 1.85 | 20.47 | 9.97 | 4.11 | Cob(I)yrinic acid a,c-diamide adenosyltransferase |
| *FEPFGCPE_02052* | *yusV* | - | 2273812 | 2274636 | 0.88 | 5.77 | 3.79 | 1.66 | putative siderophore transport system ATP-binding protein YusV |
| *FEPFGCPE_02053* | *hmuU-1* | - | 2274633 | 2275628 | 0.95 | 4.69 | 3.52 | 0.98 | Hemin transport system permease protein HmuU |
| *FEPFGCPE_02054* | *btuB* | - | 2275656 | 2277680 | 2.78 | 19.22 | 10.96 | 7.21 | Vitamin B12 transporter BtuB |
| *FEPFGCPE_02055* | | + | 2278176 | 2278487 | 24.97 | 90.92 | 87.11 | 41.41 | Hypothetical protein |

| Locus name | Gene name | Strand | Gene start (bp) | Gene end (bp) | Average TPM Inoculum | Average TPM $t_6$ | Average TPM $t_{24}$ | Average TPM $t_{168}$ | Annotation description |
|---|---|---|---|---|---|---|---|---|---|
| *FEPFGCPE_02056* | *zapA* | + | 2278484 | 2278795 | 12.32 | 58.20 | 40.60 | 18.35 | Cell division protein ZapA |
| *FEPFGCPE_02057* | *dmdC-1* | - | 2279156 | 2280937 | 10.80 | 34.39 | 32.21 | 15.37 | 3-methylmercaptopropionyl-CoA dehydrogenase |
| *FEPFGCPE_02058* | *etfA-2* | - | 2281114 | 2282046 | 24.61 | 65.75 | 53.69 | 42.39 | Electron transfer flavoprotein subunit alpha |
| *FEPFGCPE_02059* | *etfB-2* | - | 2282061 | 2282810 | 42.92 | 103.16 | 108.97 | 61.48 | Electron transfer flavoprotein subunit beta |
| *FEPFGCPE_02060* | | + | 2283180 | 2284085 | 7.79 | 19.75 | 19.45 | 3.79 | Monoacylglycerol lipase |
| *FEPFGCPE_02061* | *fpr-2* | + | 2284208 | 2284990 | 32.03 | 48.31 | 32.70 | 10.54 | Ferredoxin--NADP reductase |
| *FEPFGCPE_02062* | *menB-1* | - | 2285069 | 2285893 | 7.46 | 23.40 | 10.58 | 3.91 | 1,4-dihydroxy-2-naphthoyl-CoA synthase |
| *FEPFGCPE_02063* | *hdaH-2* | - | 2285954 | 2286877 | 6.94 | 35.20 | 21.71 | 8.63 | Histone deacetylase-like amidohydrolase |
| *FEPFGCPE_02064* | *mltB* | + | 2287326 | 2288537 | 2.12 | 18.70 | 11.60 | 7.54 | Membrane-bound lytic murein transglycosylase B |
| *FEPFGCPE_02065* | *cysM* | - | 2288644 | 2289546 | 50.67 | 83.58 | 54.80 | 23.24 | Cysteine synthase B |
| *FEPFGCPE_02066* | *comEA* | - | 2289664 | 2290347 | 148.48 | 111.96 | 60.12 | 23.54 | ComE operon protein 1 |
| *FEPFGCPE_02067* | *hldD* | - | 2290523 | 2291527 | 2.85 | 17.09 | 10.56 | 8.05 | ADP-L-glycero-D-manno-heptose-6-epimerase |
| *FEPFGCPE_02068* | *rfaE* | - | 2291524 | 2292513 | 3.99 | 31.84 | 32.39 | 10.05 | D-beta-D-heptose 7-phosphate kinase |
| *FEPFGCPE_02069* | *rkpK* | - | 2292531 | 2293904 | 1.90 | 26.88 | 22.35 | 16.69 | UDP-glucose 6-dehydrogenase |
| *FEPFGCPE_02070* | *lapB-1* | - | 2293925 | 2295100 | 1.57 | 21.25 | 18.14 | 5.05 | Lipopolysaccharide assembly protein B |
| *FEPFGCPE_02071* | *lapA* | - | 2295165 | 2295458 | 5.74 | 34.62 | 54.09 | 24.49 | Lipopolysaccharide assembly protein A |
| *FEPFGCPE_02072* | *ihfB* | - | 2295699 | 2295992 | 15.32 | 80.90 | 66.09 | 25.42 | Integration host factor subunit beta |
| *FEPFGCPE_02073* | *rpsA* | - | 2296005 | 2297708 | 19.82 | 94.24 | 103.11 | 86.50 | 30S ribosomal protein S1 |
| *FEPFGCPE_02074* | *cmk* | - | 2297823 | 2298518 | 21.05 | 101.78 | 100.72 | 43.94 | Cytidylate kinase |
| *FEPFGCPE_02075* | *dctD-1* | - | 2299026 | 2300366 | 1.02 | 5.22 | 1.97 | 1.57 | C4-dicarboxylate transport transcriptional regulatory protein DctD |
| *FEPFGCPE_02076* | *dctB-2* | - | 2300363 | 2302258 | 0.93 | 10.70 | 5.17 | 1.08 | C4-dicarboxylate transport sensor protein DctB |
| *FEPFGCPE_02077* | *livF-6* | - | 2302512 | 2303216 | 0.33 | 6.02 | 1.53 | 2.75 | High-affinity branched-chain amino acid transport ATP-binding protein LivF |
| *FEPFGCPE_02078* | *lptB-5* | - | 2303216 | 2303923 | 0.41 | 1.65 | 0.61 | 1.01 | Lipopolysaccharide export system ATP-binding protein LptB |
| *FEPFGCPE_02079* | | - | 2303913 | 2304869 | 0.33 | 5.82 | 1.86 | 1.23 | Hypothetical protein |
| *FEPFGCPE_02080* | *livH-7* | - | 2304862 | 2305869 | 0.60 | 5.81 | 4.61 | 1.35 | High-affinity branched-chain amino acid transport system permease protein LivH |
| *FEPFGCPE_02081* | | - | 2305829 | 2307088 | 1.55 | 25.22 | 15.97 | 6.59 | Leu/Ile/Val-binding protein |
| *FEPFGCPE_02082* | *bag* | + | 2307308 | 2307811 | 0.55 | 1.39 | 3.83 | 1.46 | IgA FC receptor |
| *FEPFGCPE_02083* | *aroA* | - | 2307971 | 2309293 | 2.86 | 10.88 | 7.13 | 5.80 | 3-phosphoshikimate 1-carboxyvinyltransferase |
| *FEPFGCPE_02084* | *tyrA* | - | 2309293 | 2310165 | 3.04 | 12.25 | 12.52 | 7.21 | Prephenate dehydrogenase |
| *FEPFGCPE_02085* | *pheA* | - | 2310187 | 2311269 | 2.43 | 10.98 | 5.87 | 2.25 | P-protein |
| *FEPFGCPE_02086* | *serC* | - | 2311380 | 2312492 | 8.05 | 33.85 | 26.25 | 11.19 | Phosphoserine aminotransferase |
| *FEPFGCPE_02087* | | - | 2312775 | 2313362 | 27.75 | 187.97 | 145.05 | 113.07 | Hypothetical protein |
| *FEPFGCPE_02088* | *gyrA* | - | 2313502 | 2316129 | 11.47 | 29.66 | 24.90 | 11.58 | DNA gyrase subunit A |
| *FEPFGCPE_02089* | *ompA* | + | 2316541 | 2317206 | 153.47 | 869.94 | 573.66 | 391.05 | Outer membrane protein A |
| *FEPFGCPE_02090* | | - | 2317316 | 2318104 | 21.59 | 79.84 | 72.27 | 36.43 | Hypothetical protein |
| *FEPFGCPE_02091* | | + | 2318391 | 2318861 | 19.55 | 30.12 | 22.72 | 13.02 | Hypothetical protein |
| *FEPFGCPE_02092* | | + | 2319048 | 2319755 | 23.50 | 41.68 | 37.76 | 22.88 | Hypothetical protein |
| *FEPFGCPE_02093* | *lcfB-2* | - | 2319863 | 2321395 | 3.89 | 8.12 | 8.02 | 6.41 | Long-chain-fatty-acid--CoA ligase |
| *FEPFGCPE_02094* | *echA8-2* | - | 2321463 | 2322251 | 10.05 | 23.20 | 19.36 | 3.96 | putative enoyl-CoA hydratase echA8 |
| *FEPFGCPE_02095* | | - | 2322313 | 2323890 | 6.51 | 8.14 | 3.60 | 3.48 | Hypothetical protein |
| *FEPFGCPE_02096* | *dctM-1* | - | 2324004 | 2324672 | 5.01 | 6.36 | 3.90 | 0.52 | C4-dicarboxylate TRAP transporter large permease protein DctM |
| *FEPFGCPE_02097* | *dctM-2* | - | 2324627 | 2325346 | 5.23 | 0.86 | 2.02 | 0.73 | C4-dicarboxylate TRAP transporter large permease protein DctM |
| *FEPFGCPE_02098* | | - | 2325343 | 2325933 | 8.97 | 4.75 | 2.11 | 4.55 | Hypothetical protein |
| *FEPFGCPE_02099* | | - | 2325930 | 2326976 | 14.40 | 7.59 | 5.25 | 1.84 | Solute-binding protein |
| *FEPFGCPE_02100* | *rspR-6* | + | 2327317 | 2327988 | 5.81 | 0.90 | 3.43 | 0.92 | HTH-type transcriptional repressor RspR |
| *FEPFGCPE_02101* | *lgt* | - | 2328078 | 2328968 | 7.11 | 3.58 | 6.07 | 9.03 | Prolipoprotein diacylglyceryl transferase |
| *FEPFGCPE_02102* | *benM-3* | - | 2329170 | 2330117 | 1.43 | 7.31 | 4.84 | 1.06 | HTH-type transcriptional regulator BenM |
| *FEPFGCPE_02103* | *ilvD-1* | + | 2330338 | 2332206 | 1.91 | 17.14 | 10.79 | 6.30 | Dihydroxy-acid dehydratase |
| *FEPFGCPE_02104* | *ilvD-2* | + | 2332464 | 2334137 | 11.51 | 61.67 | 40.75 | 19.50 | Dihydroxy-acid dehydratase |
| *FEPFGCPE_02105* | *nirM-2* | + | 2334338 | 2334724 | 3.96 | 21.19 | 15.63 | 7.57 | Cytochrome c-551 |
| *FEPFGCPE_02106* | | - | 2334814 | 2335167 | 1.60 | 15.49 | 15.51 | 0.60 | Hypothetical protein |
| *FEPFGCPE_02107* | | - | 2335164 | 2335586 | 7.30 | 40.57 | 37.22 | 16.62 | Hypothetical protein |
| *FEPFGCPE_02108* | *pepA* | - | 2335583 | 2337094 | 4.55 | 29.85 | 24.80 | 17.09 | Cytosol aminopeptidase |
| *FEPFGCPE_02109* | *lptF* | + | 2337186 | 2338313 | 1.65 | 10.98 | 6.78 | 1.23 | Lipopolysaccharide export system permease protein LptF |
| *FEPFGCPE_02110* | *lptG* | + | 2338313 | 2339452 | 1.64 | 12.36 | 6.03 | 2.07 | Lipopolysaccharide export system permease protein LptG |
| *FEPFGCPE_02111* | *cbiX* | + | 2339463 | 2339843 | 2.14 | 34.27 | 17.33 | 10.41 | Sirohydrochlorin cobaltochelatase |
| *FEPFGCPE_02112* | *sumT-1* | - | 2339965 | 2340792 | 3.49 | 60.31 | 28.70 | 14.60 | Uroporphyrinogen-III C-methyltransferase |
| *FEPFGCPE_02113* | *cysN* | - | 2340838 | 2342139 | 3.96 | 117.10 | 61.82 | 29.73 | Sulfate adenylyltransferase subunit 1 |
| *FEPFGCPE_02114* | *cysD-1* | - | 2342239 | 2343186 | 3.51 | 179.74 | 107.18 | 52.44 | Sulfate adenylyltransferase subunit 2 |
| *FEPFGCPE_02115* | *cysH* | - | 2343316 | 2344107 | 3.97 | 211.32 | 100.30 | 62.91 | Thioredoxin-dependent 5'-adenylylsulfate reductase |
| *FEPFGCPE_02116* | | - | 2344136 | 2344912 | 3.21 | 106.75 | 60.97 | 23.43 | Hypothetical protein |
| *FEPFGCPE_02117* | *cysB-1* | + | 2345045 | 2345986 | 3.07 | 13.18 | 10.74 | 5.51 | HTH-type transcriptional regulator CysB |
| *FEPFGCPE_02118* | | + | 2346142 | 2347311 | 6.21 | 51.72 | 39.31 | 18.14 | Leu/Ile/Val-binding protein |
| *FEPFGCPE_02119* | *garR* | + | 2347319 | 2348203 | 3.56 | 26.23 | 24.08 | 8.88 | 2-hydroxy-3-oxopropionate reductase |
| *FEPFGCPE_02120* | | + | 2348324 | 2348408 | 26.07 | 21.95 | 30.12 | 5.34 | tRNA |
| *FEPFGCPE_02121* | | - | 2349261 | 2349383 | 1.51 | 16.20 | 4.49 | 6.17 | Hypothetical protein |
| *FEPFGCPE_02122* | | - | 2349380 | 2349958 | 3.46 | 45.05 | 15.72 | 8.41 | Hypothetical protein |
| *FEPFGCPE_02123* | | - | 2349960 | 2350511 | 0.59 | 4.68 | 1.67 | 0.84 | Hypothetical protein |
| *FEPFGCPE_02124* | | - | 2350508 | 2350765 | 1.23 | 2.64 | 0.00 | 0.23 | Hypothetical protein |
| *FEPFGCPE_02125* | | - | 2350790 | 2351764 | 2.75 | 9.92 | 3.57 | 4.03 | Hypothetical protein |
| *FEPFGCPE_02126* | | - | 2351768 | 2354296 | 1.21 | 4.96 | 4.34 | 1.18 | Hypothetical protein |

| Locus name | Gene name | Strand | Gene start (bp) | Gene end (bp) | Average TPM Inoculum | Average TPM $t_6$ | Average TPM $t_{24}$ | Average TPM $t_{168}$ | Annotation description |
|---|---|---|---|---|---|---|---|---|---|
| *FEPFGCPE_02127* | | - | 2354374 | 2360049 | 0.60 | 3.54 | 1.77 | 0.38 | Hypothetical protein |
| *FEPFGCPE_02128* | | - | 2360049 | 2362349 | 0.70 | 5.91 | 2.13 | 0.52 | Hypothetical protein |
| *FEPFGCPE_02129* | | - | 2362359 | 2362886 | 0.52 | 0.19 | 3.42 | 2.04 | Hypothetical protein |
| *FEPFGCPE_02130* | | - | 2362888 | 2363346 | 0.36 | 1.98 | 2.99 | 0.00 | Hypothetical protein |
| *FEPFGCPE_02131* | | - | 2363343 | 2366165 | 0.56 | 2.72 | 2.44 | 0.53 | Hypothetical protein |
| *FEPFGCPE_02132* | | - | 2366167 | 2366907 | 0.15 | 0.27 | 0.00 | 0.00 | Hypothetical protein |
| *FEPFGCPE_02133* | | - | 2366911 | 2367258 | 0.00 | 0.00 | 0.00 | 0.00 | Hypothetical protein |
| *FEPFGCPE_02134* | | - | 2367255 | 2367614 | 0.30 | 1.24 | 0.00 | 0.00 | Hypothetical protein |
| *FEPFGCPE_02135* | | - | 2367625 | 2367852 | 0.68 | 0.00 | 0.00 | 0.00 | Hypothetical protein |
| *FEPFGCPE_02136* | | - | 2367910 | 2368386 | 0.23 | 2.92 | 4.41 | 0.33 | Hypothetical protein |
| *FEPFGCPE_02137* | | - | 2368398 | 2369444 | 0.33 | 1.23 | 0.58 | 0.78 | Hypothetical protein |
| *FEPFGCPE_02138* | | - | 2369544 | 2370272 | 0.15 | 2.40 | 3.64 | 0.00 | Hypothetical protein |
| *FEPFGCPE_02139* | | - | 2370262 | 2370588 | 0.00 | 4.30 | 0.55 | 0.00 | Hypothetical protein |
| *FEPFGCPE_02140* | | - | 2370572 | 2371255 | 0.20 | 2.32 | 1.25 | 0.89 | Hypothetical protein |
| *FEPFGCPE_02141* | | - | 2371252 | 2371485 | 0.20 | 3.05 | 8.53 | 0.42 | Hypothetical protein |
| *FEPFGCPE_02142* | | - | 2371488 | 2371736 | 0.93 | 1.73 | 4.71 | 1.34 | Hypothetical protein |
| *FEPFGCPE_02143* | | - | 2371739 | 2372077 | 0.81 | 0.00 | 2.81 | 0.74 | Hypothetical protein |
| *FEPFGCPE_02144* | | - | 2372077 | 2372445 | 0.28 | 0.54 | 0.00 | 0.00 | Hypothetical protein |
| *FEPFGCPE_02145* | | - | 2372442 | 2374112 | 0.27 | 0.67 | 1.04 | 0.18 | Hypothetical protein |
| *FEPFGCPE_02146* | | - | 2374123 | 2374440 | 0.00 | 0.00 | 0.00 | 0.00 | Hypothetical protein |
| *FEPFGCPE_02147* | | - | 2374443 | 2374649 | 0.00 | 0.00 | 0.00 | 0.00 | Hypothetical protein |
| *FEPFGCPE_02148* | | - | 2374661 | 2376160 | 0.12 | 0.59 | 0.54 | 0.00 | Hypothetical protein |
| *FEPFGCPE_02149* | | - | 2376326 | 2376955 | 0.09 | 0.63 | 0.47 | 0.31 | Hypothetical protein |
| *FEPFGCPE_02150* | | - | 2376958 | 2377497 | 0.10 | 2.12 | 2.00 | 2.06 | Hypothetical protein |
| *FEPFGCPE_02151* | | - | 2377584 | 2377781 | 0.27 | 0.17 | 1.65 | 0.14 | Hypothetical protein |
| *FEPFGCPE_02152* | | - | 2377861 | 2378016 | 1.81 | 12.90 | 4.98 | 1.74 | Hypothetical protein |
| *FEPFGCPE_02153* | | - | 2378016 | 2378522 | 0.33 | 1.29 | 2.67 | 1.03 | Hypothetical protein |
| *FEPFGCPE_02154* | *rusA* | - | 2378532 | 2378909 | 0.70 | 3.93 | 0.60 | 0.00 | Crossover junction endodeoxyribonuclease RusA |
| *FEPFGCPE_02155* | *dnaC* | - | 2378906 | 2379703 | 0.33 | 0.00 | 1.14 | 0.00 | DNA replication protein DnaC |
| *FEPFGCPE_02156* | | - | 2379645 | 2380463 | 2.42 | 3.76 | 1.70 | 0.80 | Hypothetical protein |
| *FEPFGCPE_02157* | | - | 2380464 | 2380829 | 1.40 | 0.81 | 0.74 | 1.36 | Hypothetical protein |
| *FEPFGCPE_02158* | | - | 2380826 | 2381239 | 1.30 | 1.37 | 2.48 | 0.56 | Hypothetical protein |
| *FEPFGCPE_02159* | | - | 2381316 | 2381720 | 0.53 | 0.18 | 0.00 | 0.00 | Hypothetical protein |
| *FEPFGCPE_02160* | | + | 2382193 | 2382903 | 17.76 | 82.85 | 44.48 | 14.67 | putative HTH-type transcriptional regulator |
| *FEPFGCPE_02161* | | + | 2382900 | 2383262 | 12.08 | 50.75 | 27.08 | 12.92 | Hypothetical protein |
| *FEPFGCPE_02162* | | + | 2383612 | 2384034 | 2.57 | 1.61 | 0.00 | 0.97 | Hypothetical protein |
| *FEPFGCPE_02163* | | + | 2384031 | 2384759 | 1.15 | 3.79 | 0.85 | 0.98 | Hypothetical protein |
| *FEPFGCPE_02164* | | + | 2384756 | 2384878 | 1.18 | 2.53 | 0.00 | 0.29 | Hypothetical protein |
| *FEPFGCPE_02165* | | + | 2385060 | 2385260 | 2.17 | 13.06 | 7.68 | 0.00 | Hypothetical protein |
| *FEPFGCPE_02166* | | + | 2385301 | 2385798 | 1.79 | 6.94 | 8.84 | 0.67 | Hypothetical protein |
| *FEPFGCPE_02167* | | + | 2385791 | 2386420 | 1.04 | 4.20 | 3.99 | 2.32 | Hypothetical protein |
| *FEPFGCPE_02168* | | + | 2386488 | 2387777 | 1.10 | 3.51 | 3.15 | 0.78 | Hypothetical protein |
| *FEPFGCPE_02169* | | + | 2387774 | 2388481 | 1.15 | 4.64 | 3.73 | 1.51 | Hypothetical protein |
| *FEPFGCPE_02170* | *pphA-2* | + | 2388609 | 2389373 | 0.38 | 2.27 | 1.79 | 0.00 | Serine/threonine-protein phosphatase 1 |
| *FEPFGCPE_02171* | | + | 2389370 | 2389837 | 0.38 | 5.47 | 2.25 | 2.44 | Hypothetical protein |
| *FEPFGCPE_02172* | | + | 2390131 | 2391225 | 1.44 | 5.55 | 2.21 | 1.96 | Hypothetical protein |
| *FEPFGCPE_02173* | | - | 2391828 | 2392136 | 1.27 | 0.94 | 0.55 | 1.65 | Hypothetical protein |
| *FEPFGCPE_02174* | | + | 2392551 | 2393060 | 0.45 | 3.02 | 1.89 | 0.43 | Hypothetical protein |
| *FEPFGCPE_02175* | | - | 2393345 | 2394307 | 0.44 | 4.77 | 2.60 | 0.37 | Glycerate dehydrogenase |
| *FEPFGCPE_02176* | *gno-2* | + | 2394582 | 2395340 | 0.22 | 1.50 | 2.13 | 0.51 | Gluconate 5-dehydrogenase |
| *FEPFGCPE_02177* | *dgoT-1* | + | 2395398 | 2396720 | 0.49 | 2.29 | 0.67 | 0.00 | D-galactonate transporter |
| *FEPFGCPE_02178* | | + | 2396747 | 2397139 | 0.17 | 1.08 | 0.54 | 0.54 | Hypothetical protein |
| *FEPFGCPE_02179* | *citZ* | + | 2397239 | 2398483 | 1.52 | 3.69 | 3.89 | 1.15 | Citrate synthase 2 |
| *FEPFGCPE_02180* | *fhuA* | + | 2398672 | 2400846 | 0.43 | 5.98 | 6.45 | 8.42 | Ferrichrome-iron receptor |
| *FEPFGCPE_02181* | | + | 2401019 | 2401339 | 0.20 | 5.00 | 3.96 | 2.91 | Hypothetical protein |
| *FEPFGCPE_02182* | *azoB* | + | 2401448 | 2402311 | 4.70 | 21.92 | 14.12 | 8.66 | NAD(P)H azoreductase |
| *FEPFGCPE_02183* | | - | 2402346 | 2403215 | 2.28 | 12.70 | 6.09 | 0.77 | Hypothetical protein |
| *FEPFGCPE_02184* | *bepA-3* | - | 2403238 | 2405340 | 0.91 | 5.41 | 3.17 | 0.88 | Beta-barrel assembly-enhancing protease |
| *FEPFGCPE_02185* | *ompB* | - | 2405481 | 2408216 | 3.60 | 13.34 | 8.76 | 3.22 | Outer membrane protein B |
| *FEPFGCPE_02186* | *btuE* | + | 2408652 | 2409197 | 9.42 | 92.75 | 59.21 | 31.57 | Thioredoxin/glutathione peroxidase BtuE |
| *FEPFGCPE_02187* | *dbfB-2* | - | 2409426 | 2410397 | 1.92 | 27.81 | 18.80 | 3.96 | 2,2',3-trihydroxybiphenyl dioxygenase |
| *FEPFGCPE_02188* | *pobB* | - | 2410432 | 2411397 | 0.55 | 16.19 | 10.02 | 2.87 | Phenoxybenzoate dioxygenase subunit beta |
| *FEPFGCPE_02189* | *ridA-2* | - | 2411442 | 2411888 | 1.16 | 18.47 | 12.08 | 1.08 | 2-iminobutanoate/2-iminopropanoate deaminase |
| *FEPFGCPE_02190* | *kshA* | - | 2411906 | 2412937 | 1.08 | 22.69 | 8.62 | 2.26 | 3-ketosteroid-9-alpha-monooxygenase, oxygenase component |
| *FEPFGCPE_02191* | | - | 2413003 | 2413221 | 3.53 | 55.38 | 18.21 | 7.21 | Hypothetical protein |
| *FEPFGCPE_02192* | | - | 2413221 | 2413574 | 1.68 | 53.35 | 19.10 | 10.03 | Hypothetical protein |
| *FEPFGCPE_02193* | *ribY-2* | - | 2413588 | 2414598 | 1.43 | 32.34 | 21.94 | 3.13 | Riboflavin-binding protein RibY |
| *FEPFGCPE_02194* | *gbsA-2* | - | 2414673 | 2416169 | 0.27 | 15.11 | 7.65 | 0.22 | Betaine aldehyde dehydrogenase |
| *FEPFGCPE_02195* | *nadX* | - | 2416183 | 2417001 | 0.63 | 6.74 | 4.30 | 0.61 | L-aspartate dehydrogenase |
| *FEPFGCPE_02196* | *ilvB* | - | 2416994 | 2418646 | 0.19 | 11.97 | 5.04 | 1.28 | Acetolactate synthase isozyme 1 large subunit |
| *FEPFGCPE_02197* | *bacC-3* | - | 2418639 | 2419445 | 1.01 | 19.24 | 7.63 | 2.58 | Dihydroanticapsin 7-dehydrogenase |

| Locus name | Gene name | Strand | Gene start (bp) | Gene end (bp) | Average TPM Inoculum | Average TPM $t_6$ | Average TPM $t_{24}$ | Average TPM $t_{168}$ | Annotation description |
|---|---|---|---|---|---|---|---|---|---|
| *FEPFGCPE_02198* | *pcaR-5* | - | 2419442 | 2420389 | 0.58 | 14.82 | 9.93 | 1.39 | Pca regulon regulatory protein |
| *FEPFGCPE_02199* | *hsaD-1* | - | 2420391 | 2421299 | 0.62 | 10.22 | 5.59 | 0.39 | 4,5:9,10-diseco-3-hydroxy-5,9,17-trioxoandrosta-1(10),2-diene-4-oate hydrolase |
| *FEPFGCPE_02200* | | - | 2421401 | 2421958 | 2.91 | 35.12 | 23.95 | 2.00 | Hypothetical protein |
| *FEPFGCPE_02201* | *bacC-4* | - | 2421999 | 2422790 | 1.40 | 19.71 | 7.64 | 0.42 | Dihydroanticapsin 7-dehydrogenase |
| *FEPFGCPE_02202* | *ribX-2* | - | 2422809 | 2423591 | 0.84 | 8.83 | 4.97 | 1.22 | Riboflavin transport system permease protein RibX |
| *FEPFGCPE_02203* | *tauB-3* | - | 2423631 | 2424500 | 2.46 | 18.67 | 11.37 | 3.59 | Taurine import ATP-binding protein TauB |
| *FEPFGCPE_02204* | | - | 2424814 | 2425923 | 0.92 | 9.31 | 5.34 | 1.69 | Outer membrane porin protein |
| *FEPFGCPE_02205* | | + | 2426383 | 2427540 | 1.19 | 2.82 | 3.57 | 0.96 | 6-hydroxynicotinate 3-monooxygenase |
| *FEPFGCPE_02206* | *iclR-4* | + | 2427623 | 2428468 | 1.60 | 4.71 | 8.42 | 0.68 | Transcriptional repressor IclR |
| *FEPFGCPE_02207* | | - | 2429085 | 2429150 | 4.73 | 5.72 | 11.85 | 4.57 | tRNA |
| *FEPFGCPE_02208* | | + | 2429346 | 2430575 | 1.39 | 7.01 | 4.02 | 3.78 | Cytochrome P450 107B1 |
| *FEPFGCPE_02209* | *ribZ-2* | + | 2430789 | 2432177 | 1.10 | 14.68 | 10.79 | 4.68 | Riboflavin transporter RibZ |
| *FEPFGCPE_02210* | *dnaK-1* | + | 2432278 | 2433531 | 1.57 | 20.16 | 16.69 | 6.13 | Chaperone protein DnaK |
| *FEPFGCPE_02211* | *amn* | - | 2433549 | 2435048 | 6.20 | 32.80 | 20.69 | 5.56 | AMP nucleosidase |
| *FEPFGCPE_02212* | *glk* | - | 2435071 | 2436969 | 6.03 | 16.38 | 13.21 | 3.79 | Glucokinase |
| *FEPFGCPE_02213* | *pgl* | - | 2436947 | 2437624 | 5.52 | 23.53 | 15.17 | 6.54 | 6-phosphogluconolactonase |
| *FEPFGCPE_02214* | *zwf* | - | 2437689 | 2439176 | 9.20 | 31.97 | 23.09 | 9.14 | Glucose-6-phosphate 1-dehydrogenase |
| *FEPFGCPE_02215* | *edd-1* | + | 2439409 | 2441256 | 8.98 | 30.72 | 29.14 | 5.88 | Phosphogluconate dehydratase |
| *FEPFGCPE_02216* | | - | 2441279 | 2441491 | 10.44 | 53.48 | 33.55 | 12.08 | Hypothetical protein |
| *FEPFGCPE_02217* | | + | 2441801 | 2442151 | 6.44 | 21.03 | 18.88 | 21.06 | Hypothetical protein |
| *FEPFGCPE_02218* | *dapA-4* | + | 2442207 | 2443088 | 1.95 | 27.66 | 7.44 | 5.63 | 4-hydroxy-tetrahydrodipicolinate synthase |
| *FEPFGCPE_02219* | | + | 2443138 | 2443665 | 0.84 | 28.63 | 0.91 | 4.67 | Hypothetical protein |
| *FEPFGCPE_02220* | *mgtA-2* | - | 2443882 | 2446623 | 8.59 | 23.09 | 10.72 | 9.51 | Magnesium-transporting ATPase, P-type 1 |
| *FEPFGCPE_02221* | | - | 2446703 | 2446885 | 36.52 | 101.11 | 41.71 | 36.31 | Hypothetical protein |
| *FEPFGCPE_02222* | *ybjG-1* | + | 2447503 | 2448105 | 2.85 | 10.47 | 7.40 | 4.57 | Putative undecaprenyl-diphosphatase YbjG |
| *FEPFGCPE_02223* | *rnk* | - | 2448253 | 2448678 | 13.16 | 55.42 | 20.76 | 24.53 | Regulator of nucleoside diphosphate kinase |
| *FEPFGCPE_02224* | *hemL* | + | 2448810 | 2450114 | 6.89 | 28.80 | 21.74 | 11.73 | Glutamate-1-semialdehyde 2,1-aminomutase |
| *FEPFGCPE_02225* | | + | 2450199 | 2450813 | 7.47 | 38.43 | 31.22 | 15.71 | Hypothetical protein |
| *FEPFGCPE_02226* | | + | 2450824 | 2451798 | 6.63 | 30.39 | 27.11 | 12.51 | Hypothetical protein |
| *FEPFGCPE_02227* | | + | 2451798 | 2452343 | 3.83 | 10.28 | 10.65 | 3.38 | Hypothetical protein |
| *FEPFGCPE_02228* | *ribD* | + | 2452377 | 2453447 | 2.62 | 14.93 | 9.57 | 4.77 | Riboflavin biosynthesis protein RibD |
| *FEPFGCPE_02229* | *amnE* | - | 2453477 | 2454238 | 0.78 | 0.52 | 1.65 | 0.67 | 4-oxalocrotonate decarboxylase |
| *FEPFGCPE_02230* | *amnF* | - | 2454235 | 2455065 | 0.50 | 3.39 | 1.03 | 0.00 | 2-oxopent-4-enoate hydratase |
| *FEPFGCPE_02231* | *dmpI* | - | 2455062 | 2455274 | 2.16 | 1.09 | 2.98 | 0.44 | 2-hydroxymuconate tautomerase |
| *FEPFGCPE_02232* | *tfdS* | + | 2455417 | 2456322 | 0.85 | 1.28 | 1.01 | 1.93 | HTH-type transcriptional regulator TfdS |
| *FEPFGCPE_02233* | *braC-3* | + | 2456748 | 2457860 | 1.51 | 14.83 | 7.71 | 3.35 | Leucine-, isoleucine-, valine-, threonine-, and alanine-binding protein |
| *FEPFGCPE_02234* | *livH-8* | + | 2457964 | 2458887 | 0.56 | 3.03 | 3.42 | 0.70 | High-affinity branched-chain amino acid transport system permease protein LivH |
| *FEPFGCPE_02235* | | + | 2458896 | 2460170 | 0.51 | 2.18 | 0.95 | 0.60 | Hypothetical protein |
| *FEPFGCPE_02236* | *lptB-6* | + | 2460167 | 2460934 | 0.69 | 2.56 | 2.63 | 1.67 | Lipopolysaccharide export system ATP-binding protein LptB |
| *FEPFGCPE_02237* | *livF-7* | + | 2460934 | 2461635 | 0.48 | 4.83 | 1.12 | 0.00 | High-affinity branched-chain amino acid transport ATP-binding protein LivF |
| *FEPFGCPE_02238* | *ribBA-2* | + | 2461888 | 2463009 | 8.14 | 75.43 | 61.90 | 19.50 | Riboflavin biosynthesis protein RibBA |
| *FEPFGCPE_02239* | *ribH* | + | 2463141 | 2463656 | 13.65 | 108.73 | 70.90 | 75.34 | 6,7-dimethyl-8-ribityllumazine synthase |
| *FEPFGCPE_02240* | *nusB* | + | 2463656 | 2464108 | 4.70 | 46.92 | 35.64 | 22.59 | N utilization substance protein B |
| *FEPFGCPE_02241* | *aspC* | + | 2464257 | 2465462 | 0.92 | 8.90 | 14.24 | 5.91 | Aspartate aminotransferase |
| *FEPFGCPE_02242* | *mltC-2* | - | 2465632 | 2466675 | 3.97 | 61.52 | 110.16 | 65.00 | Membrane-bound lytic murein transglycosylase C |
| *FEPFGCPE_02243* | *ubiD* | + | 2467246 | 2468796 | 2.73 | 10.29 | 11.28 | 3.71 | 3-octaprenyl-4-hydroxybenzoate carboxy-lyase |
| *FEPFGCPE_02244* | *yveA* | + | 2469330 | 2470946 | 1.11 | 4.80 | 5.40 | 1.54 | Aspartate-proton symporter |
| *FEPFGCPE_02245* | *ylaB* | - | 2471041 | 2472933 | 0.90 | 0.54 | 0.36 | 0.30 | Putative cyclic-di-GMP phosphodiesterase YlaB |
| *FEPFGCPE_02246* | *metQ-2* | - | 2473354 | 2474166 | 3.46 | 48.98 | 36.06 | 11.47 | D-methionine-binding lipoprotein MetQ |
| *FEPFGCPE_02247* | *metI* | - | 2474224 | 2474877 | 3.41 | 32.25 | 16.88 | 5.90 | D-methionine transport system permease protein MetI |
| *FEPFGCPE_02248* | *metN* | - | 2474867 | 2475994 | 1.59 | 18.81 | 18.04 | 6.77 | Methionine import ATP-binding protein MetN |
| *FEPFGCPE_02249* | *pdtaR-1* | - | 2476060 | 2476482 | 8.93 | 64.98 | 154.76 | 31.10 | putative transcriptional regulatory protein pdtaR |
| *FEPFGCPE_02250* | *uvrY-1* | - | 2476496 | 2477140 | 13.41 | 98.66 | 380.86 | 43.03 | Response regulator UvrY |
| *FEPFGCPE_02251* | | - | 2477280 | 2477441 | 18.28 | 165.58 | 500.25 | 89.40 | Hypothetical protein |
| *FEPFGCPE_02252* | | - | 2477539 | 2477715 | 9.12 | 115.95 | 309.47 | 74.97 | Hypothetical protein |
| *FEPFGCPE_02253* | *liaS-2* | + | 2477916 | 2479466 | 3.96 | 30.79 | 27.79 | 7.31 | Sensor histidine kinase LiaS |
| *FEPFGCPE_02254* | *csbD* | + | 2479720 | 2479926 | 7.59 | 99.54 | 889.72 | 215.11 | Stress response protein CsbD |
| *FEPFGCPE_02255* | | + | 2480033 | 2480275 | 5.59 | 52.18 | 518.21 | 129.18 | Hypothetical protein |
| *FEPFGCPE_02256* | | + | 2480290 | 2480739 | 3.98 | 34.54 | 246.55 | 59.03 | Hypothetical protein |
| *FEPFGCPE_02257* | | + | 2480747 | 2480881 | 0.92 | 46.63 | 149.95 | 28.35 | Hypothetical protein |
| *FEPFGCPE_02258* | | + | 2481419 | 2481721 | 42.53 | 169.96 | 192.99 | 87.42 | Hypothetical protein |
| *FEPFGCPE_02259* | *mcp4* | - | 2481846 | 2483258 | 2.64 | 28.15 | 21.28 | 6.30 | Methyl-accepting chemotaxis protein 4 |
| *FEPFGCPE_02260* | *cmpR-5* | + | 2483567 | 2484475 | 2.14 | 10.00 | 3.59 | 5.46 | HTH-type transcriptional activator CmpR |
| *FEPFGCPE_02261* | *ahpF* | - | 2484905 | 2486503 | 0.34 | 24.99 | 26.54 | 26.91 | Alkyl hydroperoxide reductase subunit F |
| *FEPFGCPE_02262* | *ahpC* | - | 2486710 | 2487273 | 7.28 | 263.41 | 270.01 | 171.41 | Alkyl hydroperoxide reductase subunit C |
| *FEPFGCPE_02263* | | - | 2487465 | 2488142 | 2.59 | 13.17 | 5.92 | 2.44 | Hypothetical protein |
| *FEPFGCPE_02264* | | - | 2488187 | 2489419 | 1.46 | 6.34 | 5.04 | 1.99 | Hypothetical protein |
| *FEPFGCPE_02265* | | - | 2489563 | 2489805 | 5.57 | 56.44 | 34.94 | 10.69 | Hypothetical protein |
| *FEPFGCPE_02266* | *yusO-1* | + | 2490017 | 2490481 | 12.52 | 124.29 | 52.25 | 16.31 | putative HTH-type transcriptional regulator YusO |
| *FEPFGCPE_02267* | *clcA-2* | + | 2490534 | 2492288 | 1.27 | 5.43 | 1.47 | 1.11 | H(+)/Cl(-) exchange transporter ClcA |
| *FEPFGCPE_02268* | *pcaK-4* | - | 2492438 | 2493814 | 0.82 | 4.99 | 3.15 | 0.45 | 4-hydroxybenzoate transporter PcaK |

| Locus name | Gene name | Strand | Gene start (bp) | Gene end (bp) | Average TPM Inoculum | Average TPM $t_6$ | Average TPM $t_{24}$ | Average TPM $t_{168}$ | Annotation description |
|---|---|---|---|---|---|---|---|---|---|
| *FEPFGCPE_02269* | *praI* | - | 2494083 | 2495261 | 0.79 | 2.02 | 1.66 | 0.23 | 4-hydroxybenzoate 3-monooxygenase (NAD(P)H) |
| *FEPFGCPE_02270* | *araC-2* | + | 2495481 | 2496470 | 6.49 | 13.66 | 6.56 | 1.12 | Arabinose operon regulatory protein |
| *FEPFGCPE_02271* | *catD-1* | - | 2496564 | 2497370 | 1.21 | 7.35 | 3.44 | 3.45 | 3-oxoadipate enol-lactonase 2 |
| *FEPFGCPE_02272* | | - | 2497384 | 2497809 | 1.15 | 7.57 | 9.12 | 3.68 | Hypothetical protein |
| *FEPFGCPE_02273* | *pcaB* | - | 2497806 | 2499182 | 2.54 | 9.58 | 11.30 | 4.28 | 3-carboxy-cis,cis-muconate cycloisomerase |
| *FEPFGCPE_02274* | *pcaG* | - | 2499323 | 2499913 | 1.77 | 10.86 | 7.46 | 2.15 | Protocatechuate 3,4-dioxygenase alpha chain |
| *FEPFGCPE_02275* | *pcaH* | - | 2499917 | 2500627 | 0.23 | 4.89 | 8.94 | 0.43 | Protocatechuate 3,4-dioxygenase beta chain |
| *FEPFGCPE_02276* | *gbpR-2* | + | 2500747 | 2501694 | 1.23 | 6.16 | 4.40 | 0.79 | HTH-type transcriptional regulator GbpR |
| *FEPFGCPE_02277* | | - | 2501731 | 2502636 | 0.68 | 3.63 | 8.07 | 0.44 | Hypothetical protein |
| *FEPFGCPE_02278* | *dmdC-2* | - | 2502633 | 2504375 | 0.40 | 4.72 | 5.35 | 0.29 | 3-methylmercaptopropionyl-CoA dehydrogenase |
| *FEPFGCPE_02279* | *fadD-2* | - | 2504377 | 2506251 | 0.50 | 6.08 | 7.85 | 0.49 | Long-chain-fatty-acid--CoA ligase |
| *FEPFGCPE_02280* | *vdh-1* | - | 2506333 | 2507784 | 1.03 | 15.06 | 14.69 | 2.47 | Vanillin dehydrogenase |
| *FEPFGCPE_02281* | | - | 2507825 | 2508655 | 2.16 | 22.60 | 39.34 | 5.37 | Hydroxycinnamoyl-CoA hydratase-lyase |
| *FEPFGCPE_02282* | | - | 2508797 | 2509939 | 1.31 | 13.61 | 12.80 | 2.43 | Outer membrane porin protein |
| *FEPFGCPE_02283* | *mhpT-2* | - | 2510086 | 2511342 | 0.25 | 8.92 | 12.01 | 0.73 | 3-(3-hydroxy-phenyl)propionate transporter |
| *FEPFGCPE_02284* | | - | 2511624 | 2512166 | 1.23 | 14.87 | 8.42 | 4.35 | putative HTH-type transcriptional regulator |
| *FEPFGCPE_02285* | *cheY-1* | - | 2512316 | 2512681 | 0.45 | 5.43 | 1.09 | 0.00 | Chemotaxis protein CheY |
| *FEPFGCPE_02286* | | - | 2512729 | 2513370 | 0.56 | 3.84 | 2.92 | 0.47 | Hypothetical protein |
| *FEPFGCPE_02287* | | - | 2513410 | 2513979 | 0.00 | 0.34 | 1.59 | 0.00 | Hypothetical protein |
| *FEPFGCPE_02288* | *mcpQ* | - | 2513996 | 2514724 | 0.53 | 0.00 | 0.49 | 0.80 | Methyl-accepting chemotaxis protein McpQ |
| *FEPFGCPE_02289* | *frzE-1* | - | 2514721 | 2517126 | 0.15 | 0.94 | 0.72 | 0.00 | Gliding motility regulatory protein |
| *FEPFGCPE_02290* | *ywrD-2* | - | 2517443 | 2519080 | 0.56 | 4.82 | 4.59 | 2.50 | Putative gamma-glutamyltransferase YwrD |
| *FEPFGCPE_02291* | *gsiD-3* | - | 2519158 | 2520081 | 0.34 | 6.71 | 3.86 | 0.59 | Glutathione transport system permease protein GsiD |
| *FEPFGCPE_02292* | *dppB-1* | - | 2520095 | 2521039 | 0.29 | 2.78 | 2.12 | 0.94 | Dipeptide transport system permease protein DppB |
| *FEPFGCPE_02293* | *gsiB-3* | - | 2521080 | 2522582 | 0.68 | 6.45 | 3.70 | 1.25 | Glutathione-binding protein GsiB |
| *FEPFGCPE_02294* | *oppF-1* | - | 2522624 | 2523721 | 0.71 | 3.00 | 2.91 | 0.29 | Oligopeptide transport ATP-binding protein OppF |
| *FEPFGCPE_02295* | *oppD-1* | - | 2523718 | 2524749 | 1.74 | 4.81 | 3.99 | 0.69 | Oligopeptide transport ATP-binding protein OppD |
| *FEPFGCPE_02296* | *gcvA-2* | + | 2524896 | 2525765 | 3.53 | 19.10 | 6.77 | 5.29 | Glycine cleavage system transcriptional activator |
| *FEPFGCPE_02297* | *yhjE-3* | - | 2525848 | 2527182 | 26.62 | 99.58 | 59.89 | 18.83 | Inner membrane metabolite transport protein YhjE |
| *FEPFGCPE_02298* | *cyoA-1* | + | 2527301 | 2528377 | 10.73 | 140.73 | 96.97 | 28.95 | Cytochrome bo(3) ubiquinol oxidase subunit 2 |
| *FEPFGCPE_02299* | *cyoB-1* | + | 2528443 | 2530434 | 4.96 | 83.45 | 46.91 | 14.25 | Cytochrome bo(3) ubiquinol oxidase subunit 1 |
| *FEPFGCPE_02300* | *cyoC-1* | + | 2530427 | 2531095 | 5.27 | 62.96 | 49.40 | 9.82 | Cytochrome bo(3) ubiquinol oxidase subunit 3 |
| *FEPFGCPE_02301* | *cyoD-1* | + | 2531092 | 2531499 | 4.43 | 79.27 | 45.22 | 9.69 | Cytochrome bo(3) ubiquinol oxidase subunit 4 |
| *FEPFGCPE_02302* | | + | 2531626 | 2532357 | 1.21 | 9.14 | 4.46 | 2.11 | putative SURF1-like protein |
| *FEPFGCPE_02303* | *ggt* | + | 2532836 | 2534563 | 14.74 | 87.66 | 42.60 | 19.16 | Gamma-glutamyltranspeptidase |
| *FEPFGCPE_02304* | *pbpG-2* | + | 2534857 | 2536143 | 1.77 | 10.60 | 7.03 | 2.76 | D-alanyl-D-alanine endopeptidase |
| *FEPFGCPE_02305* | | - | 2536835 | 2536946 | 450.65 | 63.31 | 73.96 | 75.46 | rRNA |
| *FEPFGCPE_02306* | | - | 2537081 | 2539953 | 108852.82 | 93420.25 | 105035.38 | 141071.01 | rRNA |
| *FEPFGCPE_02307* | | - | 2540246 | 2540320 | 11.74 | 0.00 | 0.00 | 0.00 | tRNA |
| *FEPFGCPE_02308* | | - | 2540431 | 2540507 | 186.24 | 760.31 | 558.17 | 257.94 | tRNA |
| *FEPFGCPE_02309* | | - | 2540577 | 2542090 | 119324.36 | 98430.24 | 99115.74 | 87233.57 | rRNA |
| *FEPFGCPE_02310* | *todT* | - | 2542932 | 2543630 | 2.43 | 16.73 | 6.88 | 6.40 | Response regulator protein TodT |
| *FEPFGCPE_02311* | | + | 2543863 | 2544549 | 1.57 | 21.68 | 15.06 | 4.08 | Hypothetical protein |
| *FEPFGCPE_02312* | *nfdA-1* | + | 2544563 | 2546422 | 1.99 | 14.33 | 10.56 | 4.34 | N-substituted formamide deformylase |
| *FEPFGCPE_02313* | | + | 2546449 | 2546730 | 8.30 | 81.17 | 34.58 | 2.20 | Hypothetical protein |
| *FEPFGCPE_02314* | | - | 2546841 | 2547443 | 0.56 | 9.39 | 6.63 | 1.28 | Hypothetical protein |
| *FEPFGCPE_02315* | *tauA-2* | - | 2547733 | 2548767 | 0.58 | 5.49 | 2.98 | 1.37 | Taurine-binding periplasmic protein |
| *FEPFGCPE_02316* | *gpuA* | - | 2548847 | 2549821 | 0.57 | 7.21 | 2.25 | 0.00 | Guanidinopropionase |
| *FEPFGCPE_02317* | *gltR-3* | + | 2550031 | 2550936 | 1.92 | 7.86 | 7.92 | 2.23 | HTH-type transcriptional regulator GltR |
| *FEPFGCPE_02318* | | + | 2551594 | 2552307 | 4.61 | 17.06 | 16.15 | 4.92 | Hypothetical protein |
| *FEPFGCPE_02319* | | + | 2552481 | 2553542 | 2.49 | 18.68 | 9.69 | 4.58 | Hypothetical protein |
| *FEPFGCPE_02320* | | - | 2553834 | 2554505 | 4.63 | 36.64 | 27.10 | 6.65 | Hypothetical protein |
| *FEPFGCPE_02321* | | + | 2554943 | 2555215 | 4.04 | 18.93 | 10.04 | 2.24 | Hypothetical protein |
| *FEPFGCPE_02322* | | + | 2555233 | 2556084 | 1.38 | 15.31 | 7.49 | 1.17 | Hypothetical protein |
| *FEPFGCPE_02323* | | - | 2556363 | 2556887 | 5.57 | 33.35 | 25.21 | 5.93 | Hypothetical protein |
| *FEPFGCPE_02324* | | - | 2556909 | 2558150 | 9.59 | 53.38 | 31.34 | 10.36 | Hypothetical protein |
| *FEPFGCPE_02325* | | + | 2558224 | 2558472 | 13.10 | 40.27 | 47.36 | 4.64 | Hypothetical protein |
| *FEPFGCPE_02326* | | + | 2558532 | 2558840 | 4.38 | 23.67 | 17.19 | 3.72 | Hypothetical protein |
| *FEPFGCPE_02327* | | + | 2559023 | 2559694 | 1.11 | 6.91 | 3.68 | 0.57 | Hypothetical protein |
| *FEPFGCPE_02328* | | + | 2559749 | 2560549 | 1.66 | 10.95 | 6.07 | 1.18 | Hypothetical protein |
| *FEPFGCPE_02329* | | + | 2561225 | 2561662 | 6.12 | 19.70 | 9.06 | 3.23 | Hypothetical protein |
| *FEPFGCPE_02330* | | + | 2561659 | 2562123 | 2.32 | 10.05 | 5.29 | 1.63 | Hypothetical protein |
| *FEPFGCPE_02331* | | - | 2562417 | 2562507 | 18.85 | 14.15 | 14.91 | 13.48 | tRNA |
| *FEPFGCPE_02332* | *actP-3* | - | 2562594 | 2564612 | 4.63 | 72.71 | 45.59 | 14.76 | Cation/acetate symporter ActP |
| *FEPFGCPE_02333* | | - | 2564609 | 2565106 | 3.61 | 59.66 | 37.30 | 11.03 | Hypothetical protein |
| *FEPFGCPE_02334* | *acs* | - | 2565129 | 2567111 | 11.03 | 181.11 | 134.14 | 71.05 | Acetyl-coenzyme A synthetase |
| *FEPFGCPE_02335* | *rhtA* | + | 2567348 | 2568340 | 1.42 | 4.97 | 3.34 | 1.03 | Threonine/homoserine exporter RhtA |
| *FEPFGCPE_02336* | | + | 2568527 | 2569099 | 3.59 | 26.41 | 15.84 | 2.88 | Hypothetical protein |
| *FEPFGCPE_02337* | *fumB* | + | 2569115 | 2570638 | 1.79 | 16.13 | 11.02 | 8.79 | Fumarate hydratase class I, anaerobic |
| *FEPFGCPE_02338* | *bfr* | + | 2570927 | 2571406 | 14.61 | 62.67 | 39.36 | 19.30 | Bacterioferritin |
| *FEPFGCPE_02339* | *yrpC* | + | 2571519 | 2572397 | 6.59 | 25.54 | 23.07 | 10.34 | Glutamate racemase 2 |

| Locus name | Gene name | Strand | Gene start (bp) | Gene end (bp) | Average TPM Inoculum | Average TPM $t_6$ | Average TPM $t_{24}$ | Average TPM $t_{168}$ | Annotation description |
|---|---|---|---|---|---|---|---|---|---|
| *FEPFGCPE_02340* | | + | 2572511 | 2572768 | 32.92 | 142.14 | 222.09 | 189.33 | Hypothetical protein |
| *FEPFGCPE_02341* | | + | 2573203 | 2573907 | 11.16 | 52.80 | 65.26 | 63.08 | Hypothetical protein |
| *FEPFGCPE_02342* | *exbB-3* | + | 2573947 | 2574684 | 4.27 | 22.60 | 38.03 | 73.11 | Biopolymer transport protein ExbB |
| *FEPFGCPE_02343* | *exbD-4* | + | 2574712 | 2575137 | 1.24 | 17.79 | 21.01 | 28.59 | Biopolymer transport protein ExbD |
| *FEPFGCPE_02344* | | + | 2575371 | 2576267 | 1.47 | 10.46 | 10.72 | 3.90 | Putative quercetin 2,3-dioxygenase |
| *FEPFGCPE_02345* | | + | 2576439 | 2576849 | 20.95 | 141.35 | 76.44 | 39.78 | Hypothetical protein |
| *FEPFGCPE_02346* | | + | 2576898 | 2577386 | 4.07 | 19.04 | 18.48 | 6.72 | Hypothetical protein |
| *FEPFGCPE_02347* | | + | 2577551 | 2577778 | 13.37 | 135.83 | 283.29 | 720.32 | Hypothetical protein |
| *FEPFGCPE_02348* | | - | 2578187 | 2578594 | 3.86 | 27.60 | 28.07 | 20.49 | Hypothetical protein |
| *FEPFGCPE_02349* | | - | 2578713 | 2579501 | 6.26 | 41.90 | 29.95 | 13.45 | Hypothetical protein |
| *FEPFGCPE_02350* | *dps2* | - | 2579587 | 2580075 | 2.22 | 27.02 | 25.11 | 13.89 | DNA protection during starvation protein 2 |
| *FEPFGCPE_02351* | *thiE-1* | + | 2580318 | 2581019 | 5.44 | 44.68 | 51.90 | 21.71 | Thiamine-phosphate synthase |
| *FEPFGCPE_02352* | *oxyR-5* | - | 2581195 | 2582139 | 7.00 | 25.47 | 19.81 | 9.52 | Hydrogen peroxide-inducible genes activator |
| *FEPFGCPE_02353* | *recG* | - | 2582306 | 2584582 | 1.37 | 7.56 | 9.97 | 4.60 | ATP-dependent DNA helicase RecG |
| *FEPFGCPE_02354* | *queA* | + | 2584700 | 2585782 | 2.42 | 15.85 | 15.57 | 6.41 | S-adenosylmethionine:tRNA ribosyltransferase-isomerase |
| *FEPFGCPE_02355* | *tgt* | + | 2585920 | 2587050 | 20.21 | 50.16 | 35.14 | 12.59 | Queuine tRNA-ribosyltransferase |
| *FEPFGCPE_02356* | | + | 2587215 | 2587541 | 21.84 | 67.26 | 66.76 | 45.54 | Hypothetical protein |
| *FEPFGCPE_02357* | | + | 2587677 | 2589569 | 4.52 | 12.70 | 14.84 | 10.75 | Hypothetical protein |
| *FEPFGCPE_02358* | | + | 2589594 | 2590571 | 6.61 | 14.62 | 11.63 | 7.11 | Hypothetical protein |
| *FEPFGCPE_02359* | *purB* | - | 2590756 | 2592129 | 3.92 | 28.13 | 23.82 | 14.36 | Adenylosuccinate lyase |
| *FEPFGCPE_02360* | *yibF-1* | + | 2592353 | 2592964 | 4.60 | 20.79 | 17.15 | 11.46 | putative GST-like protein YibF |
| *FEPFGCPE_02361* | *mltD-2* | - | 2593305 | 2594003 | 3.73 | 31.40 | 22.29 | 10.88 | Membrane-bound lytic murein transglycosylase D |
| *FEPFGCPE_02362* | *yscU* | - | 2594000 | 2595151 | 0.49 | 3.84 | 0.43 | 0.53 | Yop proteins translocation protein U |
| *FEPFGCPE_02363* | | - | 2595159 | 2596001 | 0.39 | 0.94 | 0.00 | 0.00 | Hypothetical protein |
| *FEPFGCPE_02364* | | - | 2596007 | 2596273 | 0.20 | 0.00 | 0.00 | 0.68 | Hypothetical protein |
| *FEPFGCPE_02365* | *fliP-1* | - | 2596293 | 2596943 | 0.72 | 3.31 | 0.41 | 0.24 | Flagellar biosynthetic protein FliP |
| *FEPFGCPE_02366* | | - | 2596992 | 2598161 | 0.16 | 2.02 | 0.27 | 0.00 | Hypothetical protein |
| *FEPFGCPE_02367* | | - | 2598151 | 2598597 | 0.00 | 0.01 | 1.94 | 0.00 | Hypothetical protein |
| *FEPFGCPE_02368* | | - | 2598716 | 2599126 | 0.31 | 2.64 | 0.31 | 0.11 | Hypothetical protein |
| *FEPFGCPE_02369* | *yscN* | - | 2599102 | 2600376 | 0.27 | 0.96 | 0.71 | 0.24 | putative ATP synthase YscN |
| *FEPFGCPE_02370* | *ssaV* | - | 2600465 | 2602540 | 0.52 | 1.58 | 0.09 | 0.04 | Secretion system apparatus protein SsaV |
| *FEPFGCPE_02371* | | - | 2602537 | 2602926 | 0.14 | 0.00 | 0.57 | 0.00 | Hypothetical protein |
| *FEPFGCPE_02372* | *nreC-4* | - | 2603083 | 2603742 | 0.21 | 2.08 | 0.20 | 0.35 | Oxygen regulatory protein NreC |
| *FEPFGCPE_02373* | | + | 2603988 | 2604584 | 0.60 | 3.53 | 1.43 | 0.00 | Hypothetical protein |
| *FEPFGCPE_02374* | *spiA* | + | 2604587 | 2606203 | 0.32 | 0.79 | 1.95 | 0.14 | Type III secretion system outer membrane protein SpiA |
| *FEPFGCPE_02375* | | + | 2606200 | 2607531 | 0.27 | 1.32 | 0.00 | 0.00 | Hypothetical protein |
| *FEPFGCPE_02376* | | + | 2607506 | 2607745 | 0.98 | 0.00 | 5.23 | 0.13 | Hypothetical protein |
| *FEPFGCPE_02377* | *spaK* | + | 2607947 | 2608360 | 1.49 | 4.55 | 4.86 | 0.55 | Surface presentation of antigens protein SpaK |
| *FEPFGCPE_02378* | | - | 2608437 | 2609573 | 0.25 | 0.54 | 1.72 | 0.00 | Hypothetical protein |
| *FEPFGCPE_02379* | | - | 2609566 | 2610255 | 0.15 | 0.00 | 0.63 | 0.00 | Hypothetical protein |
| *FEPFGCPE_02380* | | - | 2610231 | 2610935 | 0.24 | 1.29 | 0.33 | 0.56 | Hypothetical protein |
| *FEPFGCPE_02381* | *yscJ* | - | 2610875 | 2611717 | 0.40 | 1.15 | 0.00 | 0.00 | Yop proteins translocation lipoprotein J |
| *FEPFGCPE_02382* | | - | 2611714 | 2612070 | 0.16 | 0.00 | 0.00 | 0.00 | Hypothetical protein |
| *FEPFGCPE_02383* | | - | 2612067 | 2612459 | 0.13 | 0.22 | 0.00 | 0.99 | Hypothetical protein |
| *FEPFGCPE_02384* | | - | 2612456 | 2612674 | 0.00 | 2.75 | 0.00 | 0.00 | Hypothetical protein |
| *FEPFGCPE_02385* | *virF* | - | 2612671 | 2613471 | 0.57 | 3.32 | 0.85 | 0.87 | Virulence regulon transcriptional activator VirF |
| *FEPFGCPE_02386* | *gltA-2* | - | 2613696 | 2615297 | 0.83 | 6.63 | 2.26 | 0.96 | Glutamate synthase [NADPH] large chain |
| *FEPFGCPE_02387* | *mnmA* | - | 2615366 | 2616520 | 1.59 | 6.63 | 8.59 | 2.18 | tRNA-specific 2-thiouridylase MnmA |
| *FEPFGCPE_02388* | *nudJ* | - | 2616517 | 2616990 | 3.58 | 18.43 | 22.71 | 6.25 | Phosphatase NudJ |
| *FEPFGCPE_02389* | *pntAA-2* | + | 2617450 | 2618580 | 4.41 | 29.83 | 20.40 | 10.75 | NAD(P) transhydrogenase subunit alpha part 1 |
| *FEPFGCPE_02390* | *pntA-2* | + | 2618596 | 2618916 | 3.27 | 25.95 | 15.97 | 15.03 | NAD(P) transhydrogenase subunit alpha |
| *FEPFGCPE_02391* | *pntB-3* | + | 2618913 | 2620355 | 3.17 | 24.12 | 20.57 | 8.12 | NAD(P) transhydrogenase subunit beta |
| *FEPFGCPE_02392* | *mtnN-2* | + | 2620458 | 2621219 | 0.44 | 2.13 | 1.22 | 0.51 | 5'-methylthioadenosine/S-adenosylhomocysteine nucleosidase |
| *FEPFGCPE_02393* | | + | 2621375 | 2622490 | 0.57 | 1.24 | 2.97 | 0.46 | Hypothetical protein |
| *FEPFGCPE_02394* | *rlmL* | - | 2622570 | 2623901 | 4.41 | 28.72 | 22.19 | 10.00 | Ribosomal RNA large subunit methyltransferase L |
| *FEPFGCPE_02395* | *gntK* | - | 2623951 | 2624484 | 5.40 | 24.25 | 18.89 | 3.66 | Thermoresistant gluconokinase |
| *FEPFGCPE_02396* | *gntT* | - | 2624555 | 2625910 | 5.43 | 26.79 | 20.25 | 6.79 | High-affinity gluconate transporter |
| *FEPFGCPE_02397* | *eda* | - | 2626015 | 2626641 | 6.85 | 41.31 | 26.78 | 2.65 | KHG/KDPG aldolase |
| *FEPFGCPE_02398* | *edd-2* | - | 2626678 | 2628513 | 3.78 | 19.33 | 12.29 | 2.06 | Phosphogluconate dehydratase |
| *FEPFGCPE_02399* | *hexR-2* | + | 2628812 | 2629657 | 2.21 | 18.50 | 11.31 | 3.46 | HTH-type transcriptional regulator HexR |
| *FEPFGCPE_02400* | *ghrB-2* | - | 2629719 | 2630699 | 5.88 | 19.49 | 11.30 | 8.89 | Glyoxylate/hydroxypyruvate reductase B |
| *FEPFGCPE_02401* | *nsaD-1* | - | 2630771 | 2631391 | 3.02 | 13.51 | 15.03 | 6.65 | 2-hydroxychromene-2-carboxylate isomerase |
| *FEPFGCPE_02402* | | - | 2631498 | 2632010 | 6.52 | 20.26 | 14.94 | 13.38 | Hypothetical protein |
| *FEPFGCPE_02403* | *ruvC* | + | 2632216 | 2632758 | 2.00 | 17.63 | 10.95 | 0.44 | Crossover junction endodeoxyribonuclease RuvC |
| *FEPFGCPE_02404* | *ruvB* | + | 2632901 | 2633971 | 6.36 | 45.80 | 41.62 | 10.51 | Holliday junction ATP-dependent DNA helicase RuvB |
| *FEPFGCPE_02405* | | + | 2634008 | 2634994 | 2.85 | 12.53 | 13.40 | 3.29 | Hypothetical protein |
| *FEPFGCPE_02406* | *pspA-2* | - | 2635098 | 2635808 | 2.12 | 17.73 | 12.52 | 3.70 | Phosphoserine phosphatase 1 |
| *FEPFGCPE_02407* | | - | 2635870 | 2636496 | 5.74 | 22.48 | 24.68 | 9.46 | Hypothetical protein |
| *FEPFGCPE_02408* | *dtd* | - | 2636680 | 2637147 | 3.26 | 17.66 | 8.88 | 3.53 | D-aminoacyl-tRNA deacylase |
| *FEPFGCPE_02409* | *tyrS* | - | 2637200 | 2638444 | 6.05 | 33.50 | 18.47 | 13.10 | Tyrosine--tRNA ligase |
| *FEPFGCPE_02410* | *mepM* | + | 2638644 | 2640014 | 1.19 | 15.26 | 11.69 | 8.01 | Murein DD-endopeptidase MepM |

| Locus name | Gene name | Strand | Gene start (bp) | Gene end (bp) | Average TPM Inoculum | Average TPM $t_6$ | Average TPM $t_{24}$ | Average TPM $t_{168}$ | Annotation description |
|---|---|---|---|---|---|---|---|---|---|
| *FEPFGCPE_02411* | *anmK* | + | 2640108 | 2641256 | 2.81 | 22.68 | 14.43 | 6.41 | Anhydro-N-acetylmuramic acid kinase |
| *FEPFGCPE_02412* | *erpA* | - | 2641356 | 2641721 | 44.65 | 125.71 | 161.46 | 92.21 | Iron-sulfur cluster insertion protein ErpA |
| *FEPFGCPE_02413* | *argC-1* | - | 2641819 | 2642862 | 6.81 | 40.89 | 36.73 | 22.25 | N-acetyl-gamma-glutamyl-phosphate reductase |
| *FEPFGCPE_02414* | *rpsI* | - | 2643046 | 2643441 | 14.27 | 28.42 | 43.94 | 34.06 | 30S ribosomal protein S9 |
| *FEPFGCPE_02415* | *rplM* | - | 2643452 | 2643880 | 38.94 | 161.58 | 197.31 | 180.00 | 50S ribosomal protein L13 |
| *FEPFGCPE_02416* | | + | 2644518 | 2644940 | 8.09 | 40.10 | 36.49 | 12.64 | Hypothetical protein |
| *FEPFGCPE_02417* | | + | 2644959 | 2645516 | 6322.34 | 11088.82 | 10505.45 | 6491.30 | Hypothetical protein |
| *FEPFGCPE_02418* | | - | 2645562 | 2646383 | 3.33 | 11.07 | 8.37 | 8.43 | Hypothetical protein |
| *FEPFGCPE_02419* | *pyrC-1* | - | 2646389 | 2647426 | 10.94 | 19.10 | 16.95 | 17.06 | Dihydroorotase |
| *FEPFGCPE_02420* | *yafJ* | + | 2647621 | 2648451 | 6.24 | 14.48 | 5.20 | 4.81 | Putative glutamine amidotransferase YafJ |
| *FEPFGCPE_02421* | *glnQ-3* | - | 2648527 | 2649252 | 6.34 | 37.43 | 29.38 | 24.28 | Glutamine transport ATP-binding protein GlnQ |
| *FEPFGCPE_02422* | *gltK* | - | 2649249 | 2649929 | 4.72 | 44.97 | 29.51 | 16.85 | Glutamate/aspartate import permease protein GltK |
| *FEPFGCPE_02423* | *yecS-3* | - | 2649929 | 2650669 | 2.54 | 23.40 | 24.83 | 19.60 | L-cystine transport system permease protein YecS |
| *FEPFGCPE_02424* | *gltI-3* | - | 2650779 | 2651687 | 34.42 | 284.97 | 254.52 | 168.92 | Glutamate/aspartate import solute-binding protein |
| *FEPFGCPE_02425* | *gdhA* | - | 2651852 | 2653138 | 4.30 | 22.74 | 22.68 | 17.97 | Glutamate dehydrogenase |
| *FEPFGCPE_02426* | *yjiE-1* | - | 2653348 | 2654304 | 1.08 | 6.32 | 5.78 | 2.18 | HTH-type transcriptional regulator YjiE |
| *FEPFGCPE_02427* | *atm1* | - | 2654450 | 2656354 | 1.46 | 8.91 | 4.86 | 3.10 | ATM1-type heavy metal exporter |
| *FEPFGCPE_02428* | | + | 2656567 | 2656986 | 0.87 | 18.40 | 16.28 | 7.45 | putative acyl-CoA thioester hydrolase |
| *FEPFGCPE_02429* | *paaF-2* | - | 2657087 | 2657854 | 2.94 | 17.02 | 16.57 | 1.12 | 2,3-dehydroadipyl-CoA hydratase |
| *FEPFGCPE_02430* | *fadA-2* | - | 2658025 | 2659224 | 4.03 | 20.23 | 15.55 | 5.03 | 3-ketoacyl-CoA thiolase |
| *FEPFGCPE_02431* | *fadN* | - | 2659242 | 2661686 | 8.81 | 24.24 | 18.63 | 5.91 | putative 3-hydroxyacyl-CoA dehydrogenase |
| *FEPFGCPE_02432* | *dmdC-3* | - | 2661704 | 2663491 | 12.18 | 21.64 | 22.43 | 7.58 | 3-methylmercaptopropionyl-CoA dehydrogenase |
| *FEPFGCPE_02433* | | - | 2663540 | 2664133 | 26.29 | 29.37 | 23.14 | 5.35 | Hypothetical protein |
| *FEPFGCPE_02434* | *clsB* | - | 2664345 | 2665502 | 1.82 | 10.79 | 3.39 | 2.13 | Cardiolipin synthase B |
| *FEPFGCPE_02435* | *nudB* | - | 2665521 | 2665985 | 1.90 | 19.22 | 10.79 | 1.30 | Dihydroneopterin triphosphate diphosphatase |
| *FEPFGCPE_02436* | *aspS* | - | 2666159 | 2667961 | 4.45 | 14.80 | 11.94 | 9.78 | Aspartate--tRNA(Asp/Asn) ligase |
| *FEPFGCPE_02437* | | - | 2668009 | 2668653 | 6.12 | 31.84 | 25.84 | 8.58 | Hypothetical protein |
| *FEPFGCPE_02438* | | - | 2668756 | 2669097 | 17.02 | 30.29 | 47.78 | 26.37 | Hypothetical protein |
| *FEPFGCPE_02439* | *opuE* | - | 2669256 | 2670737 | 1.45 | 7.55 | 3.85 | 2.99 | Osmoregulated proline transporter OpuE |
| *FEPFGCPE_02440* | *tpm* | - | 2671076 | 2671771 | 0.93 | 8.15 | 5.88 | 3.93 | Thiopurine S-methyltransferase |
| *FEPFGCPE_02441* | *ubiB* | - | 2671789 | 2673354 | 1.43 | 8.07 | 8.09 | 3.22 | putative protein kinase UbiB |
| *FEPFGCPE_02442* | | - | 2673358 | 2673993 | 2.09 | 8.34 | 11.54 | 3.91 | Hypothetical protein |
| *FEPFGCPE_02443* | | - | 2674137 | 2675129 | 238.84 | 190.70 | 198.61 | 147.62 | Hypothetical protein |
| *FEPFGCPE_02444* | *ubiE-1* | - | 2675338 | 2676072 | 2.06 | 16.31 | 7.46 | 7.50 | Ubiquinone/menaquinone biosynthesis C-methyltransferase UbiE |
| *FEPFGCPE_02445* | | - | 2676133 | 2676546 | 5.81 | 30.39 | 31.23 | 12.72 | Hypothetical protein |
| *FEPFGCPE_02446* | | - | 2676626 | 2677066 | 1.39 | 8.48 | 12.80 | 2.03 | AP-4-A phosphorylase |
| *FEPFGCPE_02447* | *nhaK* | - | 2677138 | 2678829 | 1.57 | 9.91 | 7.92 | 4.04 | Sodium, potassium, lithium and rubidium/H(+) antiporter |
| *FEPFGCPE_02448* | | - | 2678882 | 2682907 | 1.39 | 13.95 | 12.80 | 7.57 | putative FAD-linked oxidoreductase |
| *FEPFGCPE_02449* | *iorA-1* | + | 2683452 | 2683910 | 7.76 | 30.53 | 30.43 | 18.67 | Isoquinoline 1-oxidoreductase subunit alpha |
| *FEPFGCPE_02450* | *iorB-1* | + | 2683926 | 2686148 | 7.33 | 31.71 | 32.24 | 10.96 | Isoquinoline 1-oxidoreductase subunit beta |
| *FEPFGCPE_02451* | *yqaA* | - | 2686364 | 2686963 | 4.13 | 33.73 | 21.47 | 9.86 | Inner membrane protein YqaA |
| *FEPFGCPE_02452* | *pucB* | + | 2687106 | 2687726 | 11.99 | 49.63 | 44.57 | 11.82 | Purine catabolism protein PucB |
| *FEPFGCPE_02453* | *ilvA* | + | 2687848 | 2689377 | 4.91 | 23.51 | 14.24 | 8.80 | L-threonine dehydratase biosynthetic IlvA |
| *FEPFGCPE_02454* | | + | 2689434 | 2690366 | 6.47 | 23.11 | 24.49 | 4.09 | Hypothetical protein |
| *FEPFGCPE_02455* | *queF-2* | + | 2690353 | 2691180 | 4.39 | 34.16 | 18.92 | 9.07 | NADPH-dependent 7-cyano-7-deazaguanine reductase |
| *FEPFGCPE_02456* | | - | 2691248 | 2691769 | 2.04 | 14.35 | 9.16 | 3.46 | Hypothetical protein |
| *FEPFGCPE_02457* | | - | 2691766 | 2692230 | 2.23 | 17.63 | 11.82 | 3.33 | Hypothetical protein |
| *FEPFGCPE_02458* | *yabJ-5* | - | 2692388 | 2692780 | 1.40 | 15.79 | 8.92 | 3.12 | 2-iminobutanoate/2-iminopropanoate deaminase |
| *FEPFGCPE_02459* | *dan* | - | 2692820 | 2694310 | 0.98 | 4.31 | 4.02 | 1.13 | D-aminoacylase |
| *FEPFGCPE_02460* | *hexR-3* | - | 2694312 | 2695193 | 1.25 | 6.37 | 4.50 | 2.67 | HTH-type transcriptional regulator HexR |
| *FEPFGCPE_02461* | | + | 2695359 | 2696630 | 4.13 | 13.12 | 15.43 | 4.98 | D-threonine aldolase |
| *FEPFGCPE_02462* | *mexR-1* | - | 2696820 | 2697371 | 1.38 | 33.54 | 34.80 | 12.31 | Multidrug resistance operon repressor |
| *FEPFGCPE_02463* | *bmr3-2* | - | 2697376 | 2698971 | 1.16 | 35.25 | 28.26 | 8.09 | Multidrug resistance protein 3 |
| *FEPFGCPE_02464* | *nlhH-3* | - | 2699172 | 2700029 | 1.79 | 10.29 | 6.34 | 2.20 | Carboxylesterase NlhH |
| *FEPFGCPE_02465* | *nimR-1* | + | 2700155 | 2700928 | 0.86 | 6.00 | 5.11 | 0.58 | HTH-type transcriptional regulator NimR |
| *FEPFGCPE_02466* | | + | 2701049 | 2701513 | 0.23 | 0.31 | 2.60 | 0.49 | Hypothetical protein |
| *FEPFGCPE_02467* | *purA-2* | + | 2701510 | 2702901 | 0.44 | 2.79 | 3.21 | 0.07 | Adenylosuccinate synthetase |
| *FEPFGCPE_02468* | | + | 2702891 | 2703511 | 0.65 | 7.41 | 4.39 | 2.73 | Hypothetical protein |
| *FEPFGCPE_02469* | *naiP-5* | + | 2703718 | 2705091 | 0.13 | 3.93 | 2.10 | 0.96 | Putative niacin/nicotinamide transporter NaiP |
| *FEPFGCPE_02470* | | - | 2705258 | 2705827 | 0.17 | 3.68 | 2.20 | 0.17 | Hypothetical protein |
| *FEPFGCPE_02471* | | - | 2706158 | 2707048 | 0.20 | 2.42 | 0.50 | 2.20 | Putative 2-dehydropantoate 2-reductase |
| *FEPFGCPE_02472* | *sauU-2* | - | 2707095 | 2708309 | 0.15 | 0.34 | 2.39 | 0.16 | putative sulfoacetate transporter SauU |
| *FEPFGCPE_02473* | *gcvA-3* | + | 2708439 | 2709389 | 3.17 | 14.88 | 10.12 | 1.35 | Glycine cleavage system transcriptional activator |
| *FEPFGCPE_02474* | *dhmA1* | - | 2709745 | 2710665 | 0.71 | 5.60 | 2.93 | 3.54 | Haloalkane dehalogenase 1 |
| *FEPFGCPE_02475* | *zntR-2* | + | 2710770 | 2711138 | 2.10 | 19.40 | 12.76 | 4.07 | HTH-type transcriptional regulator ZntR |
| *FEPFGCPE_02476* | *argC-2* | - | 2711445 | 2712374 | 0.25 | 6.94 | 3.54 | 1.41 | N-acetyl-gamma-glutamyl-phosphate reductase |
| *FEPFGCPE_02477* | *oxyR-6* | + | 2712511 | 2713395 | 1.12 | 4.48 | 0.77 | 0.78 | Hydrogen peroxide-inducible genes activator |
| *FEPFGCPE_02478* | | + | 2714118 | 2715014 | 0.63 | 3.15 | 0.61 | 0.67 | N-carbamoyl-D-amino acid hydrolase |
| *FEPFGCPE_02479* | *amiC-1* | + | 2715076 | 2716239 | 0.30 | 3.92 | 1.23 | 1.14 | Aliphatic amidase expression-regulating protein |
| *FEPFGCPE_02480* | *livH-9* | + | 2716316 | 2717149 | 0.91 | 5.35 | 3.73 | 1.66 | High-affinity branched-chain amino acid transport system permease protein LivH |
| *FEPFGCPE_02481* | *glnQ-4* | + | 2717142 | 2719004 | 1.38 | 9.48 | 8.41 | 3.67 | Glutamine transport ATP-binding protein GlnQ |

| Locus name | Gene name | Strand | Gene start (bp) | Gene end (bp) | Average TPM Inoculum | Average TPM $t_6$ | Average TPM $t_{24}$ | Average TPM $t_{168}$ | Annotation description |
|---|---|---|---|---|---|---|---|---|---|
| *FEPFGCPE_02482* | *livF-8* | + | 2719001 | 2719708 | 0.63 | 7.88 | 6.23 | 1.61 | High-affinity branched-chain amino acid transport ATP-binding protein LivF |
| *FEPFGCPE_02483* | *ansA-3* | + | 2719866 | 2720801 | 3.49 | 17.36 | 14.51 | 4.96 | L-asparaginase |
| *FEPFGCPE_02484* | *amiC-2* | + | 2720812 | 2721972 | 3.74 | 25.22 | 11.28 | 2.37 | Aliphatic amidase expression-regulating protein |
| *FEPFGCPE_02485* | *amiR* | + | 2721969 | 2722589 | 2.23 | 18.48 | 6.71 | 2.27 | Aliphatic amidase regulator |
| *FEPFGCPE_02486* | *ywnA* | - | 2722958 | 2723410 | 4.91 | 12.78 | 3.99 | 3.06 | Putative HTH-type transcriptional regulator YwnA |
| *FEPFGCPE_02487* | *trxB-2* | + | 2723512 | 2724417 | 1.54 | 2.74 | 0.86 | 0.77 | Thioredoxin reductase |
| *FEPFGCPE_02488* | *dnrC* | - | 2724818 | 2725681 | 1.31 | 4.28 | 1.87 | 0.00 | Aklanonic acid methyltransferase DnrC |
| *FEPFGCPE_02489* | *trxB-3* | - | 2725778 | 2726671 | 1.79 | 5.71 | 3.62 | 1.99 | Thioredoxin reductase |
| *FEPFGCPE_02490* | *srpR* | + | 2726820 | 2727452 | 1.94 | 3.09 | 5.28 | 3.35 | HTH-type transcriptional regulator SrpR |
| *FEPFGCPE_02491* | | + | 2728025 | 2728369 | 7.25 | 57.99 | 45.85 | 13.72 | HTH-type transcriptional regulator |
| *FEPFGCPE_02492* | | + | 2728359 | 2728877 | 4.24 | 27.87 | 14.88 | 5.48 | Hypothetical protein |
| *FEPFGCPE_02493* | | - | 2728995 | 2729885 | 1.12 | 7.78 | 8.96 | 2.44 | Soluble epoxide hydrolase |
| *FEPFGCPE_02494* | *comR* | - | 2730062 | 2730667 | 1.02 | 1.92 | 3.67 | 1.52 | HTH-type transcriptional repressor ComR |
| *FEPFGCPE_02495* | | + | 2731092 | 2732666 | 0.67 | 6.57 | 3.69 | 1.13 | putative sulfate transporter |
| *FEPFGCPE_02496* | | - | 2732744 | 2733172 | 1.31 | 12.85 | 8.84 | 0.99 | Hypothetical protein |
| *FEPFGCPE_02497* | | - | 2733722 | 2734495 | 1.55 | 16.77 | 17.20 | 3.11 | Hypothetical protein |
| *FEPFGCPE_02498* | | - | 2734988 | 2735149 | 0.69 | 7.21 | 0.00 | 1.03 | Hypothetical protein |
| *FEPFGCPE_02499* | *ahpD-2* | - | 2735149 | 2735442 | 0.63 | 3.76 | 4.95 | 1.84 | Alkyl hydroperoxide reductase AhpD |
| *FEPFGCPE_02500* | | - | 2735466 | 2735882 | 2.21 | 2.62 | 1.17 | 2.50 | Hypothetical protein |
| *FEPFGCPE_02501* | *gabR-4* | - | 2736002 | 2737411 | 0.82 | 6.15 | 6.27 | 1.77 | HTH-type transcriptional regulatory protein GabR |
| *FEPFGCPE_02502* | | + | 2737594 | 2738238 | 2.33 | 17.50 | 10.69 | 4.27 | Hypothetical protein |
| *FEPFGCPE_02503* | *fabG-6* | + | 2738308 | 2739030 | 4.11 | 13.88 | 13.07 | 7.26 | 3-oxoacyl-[acyl-carrier-protein] reductase FabG |
| *FEPFGCPE_02504* | | + | 2739307 | 2739705 | 2.96 | 20.82 | 14.45 | 3.35 | Hypothetical protein |
| *FEPFGCPE_02505* | | - | 2739820 | 2740863 | 0.92 | 5.18 | 3.73 | 0.91 | Hypothetical protein |
| *FEPFGCPE_02506* | | + | 2741156 | 2741827 | 0.91 | 6.91 | 3.63 | 0.57 | Hypothetical protein |
| *FEPFGCPE_02507* | *gbpR-3* | - | 2742128 | 2743111 | 0.37 | 2.87 | 0.45 | 0.87 | HTH-type transcriptional regulator GbpR |
| *FEPFGCPE_02508* | *proP-8* | + | 2743301 | 2744641 | 0.61 | 0.94 | 0.98 | 0.46 | Proline/betaine transporter |
| *FEPFGCPE_02509* | | + | 2744676 | 2746241 | 0.54 | 1.19 | 0.39 | 0.71 | Arylsulfatase |
| *FEPFGCPE_02510* | *pcaR-6* | - | 2746355 | 2747140 | 0.40 | 2.14 | 0.24 | 0.00 | Pca regulon regulatory protein |
| *FEPFGCPE_02511* | *lcfB-3* | + | 2747439 | 2749073 | 1.55 | 5.55 | 2.87 | 0.57 | Long-chain-fatty-acid--CoA ligase |
| *FEPFGCPE_02512* | *echA6* | + | 2749066 | 2749845 | 1.12 | 1.99 | 2.49 | 0.54 | putative enoyl-CoA hydratase echA6 |
| *FEPFGCPE_02513* | *potD-4* | + | 2749899 | 2750963 | 1.21 | 4.73 | 3.28 | 2.24 | Spermidine/putrescine-binding periplasmic protein |
| *FEPFGCPE_02514* | *cynR-5* | - | 2750980 | 2751882 | 0.28 | 0.87 | 3.37 | 0.00 | HTH-type transcriptional regulator CynR |
| *FEPFGCPE_02515* | *madA* | + | 2752033 | 2753694 | 1.69 | 1.26 | 1.58 | 0.46 | Acetyl-S-ACP:malonate ACP transferase |
| *FEPFGCPE_02516* | *citG* | + | 2753694 | 2754602 | 0.75 | 1.53 | 0.93 | 0.27 | 2-(5''-triphosphoribosyl)-3'-dephosphocoenzyme-A synthase |
| *FEPFGCPE_02517* | *mdcC* | + | 2754614 | 2754916 | 2.39 | 0.00 | 1.91 | 0.00 | Malonate decarboxylase acyl carrier protein |
| *FEPFGCPE_02518* | *madC* | + | 2754929 | 2755786 | 1.23 | 1.20 | 0.00 | 0.05 | Malonyl-S-ACP:biotin-protein carboxyltransferase MADC |
| *FEPFGCPE_02519* | *madD* | + | 2755783 | 2756604 | 2.20 | 1.74 | 0.03 | 1.52 | Malonyl-S-ACP:biotin-protein carboxyltransferase MADD |
| *FEPFGCPE_02520* | | + | 2756783 | 2757178 | 1.58 | 2.89 | 2.44 | 0.59 | Hypothetical protein |
| *FEPFGCPE_02521* | | + | 2757192 | 2757956 | 1.22 | 0.92 | 1.48 | 0.67 | Hypothetical protein |
| *FEPFGCPE_02522* | *mdcG* | + | 2758043 | 2758612 | 1.08 | 2.44 | 2.47 | 0.34 | Phosphoribosyl-dephospho-CoA transferase |
| *FEPFGCPE_02523* | *fabD-1* | + | 2758609 | 2759547 | 1.80 | 2.90 | 2.27 | 0.45 | Malonyl CoA-acyl carrier protein transacylase |
| *FEPFGCPE_02524* | *yddG* | - | 2759624 | 2760556 | 1.07 | 14.36 | 5.77 | 0.71 | Aromatic amino acid exporter YddG |
| *FEPFGCPE_02525* | *tauR* | - | 2760644 | 2762179 | 3.65 | 24.01 | 13.90 | 4.25 | HTH-type transcriptional regulator TauR |
| *FEPFGCPE_02526* | *tauA-3* | + | 2762477 | 2763535 | 5.10 | 23.41 | 7.42 | 3.23 | Taurine-binding periplasmic protein |
| *FEPFGCPE_02527* | *tauB-4* | + | 2763615 | 2764439 | 4.92 | 15.40 | 10.32 | 2.07 | Taurine import ATP-binding protein TauB |
| *FEPFGCPE_02528* | *ssuC-4* | + | 2764460 | 2765395 | 0.91 | 4.45 | 5.16 | 2.28 | Putative aliphatic sulfonates transport permease protein SsuC |
| *FEPFGCPE_02529* | *gabP* | + | 2765639 | 2767042 | 0.45 | 3.29 | 2.36 | 0.47 | GABA permease |
| *FEPFGCPE_02530* | *ydcV-3* | - | 2767188 | 2768075 | 2.72 | 10.98 | 7.87 | 2.10 | Inner membrane ABC transporter permease protein YdcV |
| *FEPFGCPE_02531* | *potH-3* | - | 2768079 | 2769341 | 1.22 | 6.17 | 6.86 | 1.07 | Putrescine transport system permease protein PotH |
| *FEPFGCPE_02532* | *potD-5* | - | 2769446 | 2770480 | 1.39 | 16.98 | 11.50 | 5.25 | Spermidine/putrescine-binding periplasmic protein |
| *FEPFGCPE_02533* | *potA-6* | - | 2770538 | 2771674 | 1.77 | 7.83 | 4.87 | 2.45 | Spermidine/putrescine import ATP-binding protein PotA |
| *FEPFGCPE_02534* | *gabR-5* | + | 2771843 | 2773351 | 2.42 | 5.18 | 4.41 | 1.89 | HTH-type transcriptional regulatory protein GabR |
| *FEPFGCPE_02535* | *puuE* | + | 2773585 | 2774868 | 1.70 | 8.28 | 8.19 | 2.56 | 4-aminobutyrate aminotransferase PuuE |
| *FEPFGCPE_02536* | *yciC-1* | - | 2774905 | 2775849 | 1.14 | 10.45 | 4.36 | 1.35 | Putative metal chaperone YciC |
| *FEPFGCPE_02537* | *nfdA-2* | - | 2775918 | 2777585 | 0.78 | 2.95 | 1.67 | 0.59 | N-substituted formamide deformylase |
| *FEPFGCPE_02538* | *mctP-2* | - | 2777585 | 2779006 | 1.80 | 4.33 | 2.89 | 1.62 | Monocarboxylate transport permease protein |
| *FEPFGCPE_02539* | | - | 2779115 | 2779315 | 1.39 | 0.00 | 4.08 | 0.00 | Hypothetical protein |
| *FEPFGCPE_02540* | *puuB-3* | + | 2779616 | 2781049 | 2.87 | 3.68 | 2.66 | 1.38 | Gamma-glutamylputrescine oxidoreductase |
| *FEPFGCPE_02541* | | + | 2781084 | 2781473 | 3.70 | 6.10 | 1.69 | 0.00 | Hypothetical protein |
| *FEPFGCPE_02542* | *xsc* | + | 2781650 | 2783482 | 1.84 | 7.75 | 3.37 | 1.41 | Sulfoacetaldehyde acetyltransferase |
| *FEPFGCPE_02543* | *eutD* | + | 2783563 | 2784603 | 0.71 | 2.64 | 1.49 | 0.02 | Ethanolamine utilization protein EutD |
| *FEPFGCPE_02544* | *tsaS* | + | 2784600 | 2785367 | 0.26 | 0.33 | 3.33 | 0.38 | putative inner membrane transporter protein TsaS |
| *FEPFGCPE_02545* | *cobT-2* | - | 2785389 | 2787131 | 0.44 | 4.72 | 1.89 | 0.33 | Aerobic cobaltochelatase subunit CobT |
| *FEPFGCPE_02546* | *cobS-3* | - | 2787128 | 2788114 | 0.67 | 4.60 | 1.11 | 0.57 | Aerobic cobaltochelatase subunit CobS |
| *FEPFGCPE_02547* | *iclR-5* | + | 2788266 | 2789123 | 3.35 | 20.13 | 12.20 | 6.28 | Transcriptional repressor IclR |
| *FEPFGCPE_02548* | *kdgR-3* | - | 2789128 | 2790000 | 1.26 | 4.13 | 1.81 | 1.33 | Transcriptional regulator KdgR |
| *FEPFGCPE_02549* | *alkJ-1* | + | 2790342 | 2792012 | 0.46 | 1.40 | 1.52 | 0.07 | Alcohol dehydrogenase [acceptor] |
| *FEPFGCPE_02550* | *proP-9* | + | 2792056 | 2793384 | 0.39 | 3.05 | 1.09 | 0.00 | Proline/betaine transporter |
| *FEPFGCPE_02551* | *yjcC-2* | + | 2793622 | 2794872 | 4.12 | 25.13 | 20.28 | 11.84 | Putative cyclic-di-GMP phosphodiesterase YjcC |
| *FEPFGCPE_02552* | *namA* | - | 2794961 | 2796067 | 0.41 | 1.06 | 2.03 | 1.24 | NADPH dehydrogenase |

| Locus name | Gene name | Strand | Gene start (bp) | Gene end (bp) | Average TPM Inoculum | Average TPM $t_6$ | Average TPM $t_{24}$ | Average TPM $t_{168}$ | Annotation description |
|---|---|---|---|---|---|---|---|---|---|
| *FEPFGCPE_02553* | *cynR-6* | + | 2796244 | 2797167 | 11.38 | 12.45 | 8.45 | 5.45 | HTH-type transcriptional regulator CynR |
| *FEPFGCPE_02554* | *comC* | - | 2797258 | 2798289 | 2.91 | 8.92 | 5.72 | 3.23 | (2R)-3-sulfolactate dehydrogenase (NADP(+)) |
| *FEPFGCPE_02555* | *dctA-3* | - | 2798286 | 2799608 | 4.19 | 9.06 | 6.43 | 2.73 | Aerobic C4-dicarboxylate transport protein |
| *FEPFGCPE_02556* | *suyB* | - | 2799792 | 2800967 | 2.60 | 6.76 | 4.30 | 2.97 | (2R)-sulfolactate sulfo-lyase subunit beta |
| *FEPFGCPE_02557* | *suyA* | - | 2801017 | 2801358 | 3.33 | 13.24 | 9.35 | 3.12 | (2R)-sulfolactate sulfo-lyase subunit alpha |
| *FEPFGCPE_02558* | *mngR-1* | + | 2801529 | 2802296 | 5.32 | 17.58 | 12.27 | 3.82 | Mannosyl-D-glycerate transport/metabolism system repressor MngR |
| *FEPFGCPE_02559* | | - | 2802404 | 2803513 | 1.91 | 9.75 | 9.46 | 5.23 | Tartrate dehydrogenase/decarboxylase |
| *FEPFGCPE_02560* | *dmlR-11* | + | 2803540 | 2804520 | 0.59 | 1.09 | 2.84 | 0.79 | HTH-type transcriptional regulator DmlR |
| *FEPFGCPE_02561* | | - | 2804563 | 2806899 | 0.64 | 6.47 | 3.47 | 2.12 | Beta-glucosidase BoGH3B |
| *FEPFGCPE_02562* | *mdoG-1* | + | 2806914 | 2808734 | 7.68 | 10.01 | 6.92 | 7.21 | Glucans biosynthesis protein G |
| *FEPFGCPE_02563* | *pcaK-5* | + | 2809079 | 2810458 | 7.35 | 24.83 | 13.28 | 3.29 | 4-hydroxybenzoate transporter PcaK |
| *FEPFGCPE_02564* | | + | 2810716 | 2811141 | 6.72 | 49.17 | 19.18 | 6.42 | Hypothetical protein |
| *FEPFGCPE_02565* | *gylR-1* | - | 2811239 | 2812117 | 1.06 | 6.50 | 5.79 | 1.69 | Glycerol operon regulatory protein |
| *FEPFGCPE_02566* | *gylR-2* | - | 2812114 | 2812920 | 0.88 | 9.70 | 3.35 | 2.90 | Glycerol operon regulatory protein |
| *FEPFGCPE_02567* | *ccrA* | + | 2813187 | 2814146 | 3.05 | 18.96 | 9.22 | 2.93 | Metallo-beta-lactamase type 2 |
| *FEPFGCPE_02568* | *mhpA-3* | + | 2814233 | 2815969 | 2.21 | 12.05 | 5.28 | 1.95 | 3-(3-hydroxy-phenyl)propionate/3-hydroxycinnamic acid hydroxylase |
| *FEPFGCPE_02569* | | + | 2816036 | 2816269 | 2.31 | 14.93 | 7.08 | 2.61 | Hypothetical protein |
| *FEPFGCPE_02570* | *pcaR-7* | - | 2816352 | 2817152 | 1.90 | 6.69 | 3.50 | 1.99 | Pca regulon regulatory protein |
| *FEPFGCPE_02571* | *pcaI* | + | 2817455 | 2818132 | 4.95 | 7.53 | 10.91 | 2.72 | 3-oxoadipate CoA-transferase subunit A |
| *FEPFGCPE_02572* | *pcaJ* | + | 2818158 | 2818808 | 5.50 | 12.69 | 4.99 | 3.15 | 3-oxoadipate CoA-transferase subunit B |
| *FEPFGCPE_02573* | *pcaF-1* | + | 2818907 | 2820109 | 4.31 | 12.04 | 7.54 | 2.79 | Beta-ketoadipyl-CoA thiolase |
| *FEPFGCPE_02574* | *leuA-2* | - | 2820265 | 2821977 | 4.44 | 11.64 | 9.02 | 3.13 | 2-isopropylmalate synthase |
| *FEPFGCPE_02575* | *lrp-3* | - | 2822329 | 2822802 | 4.02 | 10.13 | 6.75 | 0.47 | Leucine-responsive regulatory protein |
| *FEPFGCPE_02576* | *hpd-1* | + | 2823093 | 2824166 | 3.59 | 16.39 | 16.02 | 6.75 | 4-hydroxyphenylpyruvate dioxygenase |
| *FEPFGCPE_02577* | *aroP* | + | 2824330 | 2825718 | 0.49 | 5.82 | 4.44 | 2.99 | Aromatic amino acid transport protein AroP |
| *FEPFGCPE_02578* | *tyrB-2* | + | 2825933 | 2827132 | 4.32 | 15.46 | 12.01 | 5.94 | Aromatic-amino-acid aminotransferase |
| *FEPFGCPE_02579* | | - | 2827033 | 2827248 | 0.00 | 0.09 | 0.00 | 1.89 | Hypothetical protein |
| *FEPFGCPE_02580* | *ytnP* | - | 2827245 | 2828135 | 0.14 | 5.90 | 3.20 | 2.88 | putative quorum-quenching lactonase YtnP |
| *FEPFGCPE_02581* | *lgoT-2* | - | 2828193 | 2829551 | 1.09 | 6.19 | 4.18 | 0.68 | putative L-galactonate transporter |
| *FEPFGCPE_02582* | *doeB* | - | 2829616 | 2830632 | 0.08 | 1.52 | 1.42 | 0.00 | N-alpha-acetyl-L-2,4-diaminobutyric acid deacetylase |
| *FEPFGCPE_02583* | *hcaR-2* | - | 2830789 | 2831682 | 0.75 | 6.73 | 4.78 | 0.82 | Hca operon transcriptional activator HcaR |
| *FEPFGCPE_02584* | *yghU* | + | 2831866 | 2832585 | 4.80 | 21.22 | 20.08 | 11.92 | Disulfide-bond oxidoreductase YghU |
| *FEPFGCPE_02585* | | - | 2832751 | 2833956 | 0.95 | 3.63 | 1.51 | 0.19 | Formate dehydrogenase |
| *FEPFGCPE_02586* | *oxyR-7* | - | 2834132 | 2835043 | 1.76 | 1.48 | 2.06 | 0.38 | Hydrogen peroxide-inducible genes activator |
| *FEPFGCPE_02587* | *mrcA-2* | - | 2835214 | 2837772 | 0.77 | 3.92 | 2.82 | 1.36 | Penicillin-binding protein 1A |
| *FEPFGCPE_02588* | *pdhR-1* | - | 2837834 | 2838550 | 2.57 | 6.91 | 7.37 | 0.25 | Pyruvate dehydrogenase complex repressor |
| *FEPFGCPE_02589* | *glcA* | + | 2838874 | 2840580 | 0.57 | 4.31 | 0.89 | 0.77 | Glycolate permease GlcA |
| *FEPFGCPE_02590* | *lutA-2* | + | 2840660 | 2841442 | 0.35 | 0.91 | 1.18 | 1.90 | Lactate utilization protein A |
| *FEPFGCPE_02591* | *lutB-2* | + | 2841439 | 2842848 | 0.53 | 0.86 | 1.35 | 0.39 | Lactate utilization protein B |
| *FEPFGCPE_02592* | | + | 2842845 | 2843474 | 0.41 | 1.91 | 2.54 | 0.89 | Hypothetical protein |
| *FEPFGCPE_02593* | *ompW-1* | + | 2843615 | 2844322 | 3.84 | 9.99 | 11.51 | 3.29 | Outer membrane protein W |
| *FEPFGCPE_02594* | *yodB* | - | 2844499 | 2845071 | 35.63 | 196.47 | 155.31 | 51.68 | Cytochrome b561 |
| *FEPFGCPE_02595* | | + | 2845665 | 2846498 | 53.41 | 99.85 | 70.67 | 31.01 | putative oxidoreductase |
| *FEPFGCPE_02596* | *dadA1-4* | + | 2846584 | 2847852 | 11.28 | 31.64 | 20.32 | 6.21 | D-amino acid dehydrogenase 1 |
| *FEPFGCPE_02597* | *limB-1* | - | 2847934 | 2848935 | 2.50 | 8.39 | 10.06 | 1.19 | Limonene 1,2-monooxygenase |
| *FEPFGCPE_02598* | *hxlR* | + | 2849008 | 2849505 | 2.00 | 15.19 | 8.10 | 0.81 | HTH-type transcriptional activator HxlR |
| *FEPFGCPE_02599* | | - | 2849508 | 2850491 | 0.97 | 10.60 | 7.39 | 1.27 | aspartoacylase |
| *FEPFGCPE_02600* | *proP-10* | - | 2850633 | 2851952 | 1.32 | 5.09 | 3.67 | 1.54 | Proline/betaine transporter |
| *FEPFGCPE_02601* | | - | 2852127 | 2853047 | 3.57 | 21.42 | 11.28 | 7.53 | Soluble epoxide hydrolase |
| *FEPFGCPE_02602* | *hosA* | - | 2853279 | 2853806 | 1.82 | 10.25 | 15.58 | 3.40 | Transcriptional regulator HosA |
| *FEPFGCPE_02603* | *yxeP-2* | + | 2854042 | 2855196 | 1.09 | 2.61 | 2.22 | 0.17 | putative hydrolase YxeP |
| *FEPFGCPE_02604* | *proP-11* | + | 2855237 | 2856553 | 2.79 | 14.43 | 9.09 | 1.47 | Proline/betaine transporter |
| *FEPFGCPE_02605* | | - | 2856612 | 2856929 | 18.03 | 32.98 | 27.41 | 7.24 | Hypothetical protein |
| *FEPFGCPE_02606* | | + | 2857351 | 2857914 | 8.38 | 45.66 | 19.30 | 2.35 | Hypothetical protein |
| *FEPFGCPE_02607* | | - | 2857898 | 2858497 | 0.27 | 7.38 | 5.91 | 1.25 | Hypothetical protein |
| *FEPFGCPE_02608* | *glnQ-5* | - | 2858556 | 2859302 | 0.29 | 2.67 | 4.56 | 0.70 | Glutamine transport ATP-binding protein GlnQ |
| *FEPFGCPE_02609* | | - | 2859299 | 2859628 | 0.61 | 1.21 | 0.99 | 1.53 | N(1)-aminopropylagmatine ureohydrolase |
| *FEPFGCPE_02610* | *gbuA* | - | 2859625 | 2860257 | 2.73 | 13.73 | 12.55 | 2.10 | Guanidinobutyrase |
| *FEPFGCPE_02611* | *yecS-4* | - | 2860289 | 2860951 | 0.69 | 5.33 | 6.34 | 0.78 | L-cystine transport system permease protein YecS |
| *FEPFGCPE_02612* | *yecS-5* | - | 2860941 | 2861636 | 0.52 | 2.22 | 0.04 | 2.19 | L-cystine transport system permease protein YecS |
| *FEPFGCPE_02613* | *tcyA* | - | 2861633 | 2862493 | 0.56 | 3.44 | 2.87 | 0.00 | L-cystine-binding protein TcyA |
| *FEPFGCPE_02614* | *gltC-6* | + | 2862694 | 2863566 | 1.09 | 2.51 | 3.13 | 0.77 | HTH-type transcriptional regulator GltC |
| *FEPFGCPE_02615* | *kgtP* | + | 2863960 | 2865249 | 2.84 | 24.76 | 26.27 | 12.44 | Alpha-ketoglutarate permease |
| *FEPFGCPE_02616* | *yjiE-2* | - | 2865278 | 2866186 | 1.69 | 8.66 | 7.11 | 3.75 | HTH-type transcriptional regulator YjiE |
| *FEPFGCPE_02617* | *ruvA* | - | 2866240 | 2866821 | 1.60 | 10.29 | 11.32 | 9.68 | Holliday junction ATP-dependent DNA helicase RuvA |
| *FEPFGCPE_02618* | *gbpR-4* | - | 2866895 | 2867845 | 1.14 | 4.99 | 4.23 | 1.11 | HTH-type transcriptional regulator GbpR |
| *FEPFGCPE_02619* | *sdgD* | + | 2868039 | 2869076 | 1.89 | 139.44 | 476.06 | 141.09 | Gentisate 1,2-dioxygenase |
| *FEPFGCPE_02620* | *nagK-2* | + | 2869176 | 2869874 | 1.23 | 117.34 | 518.76 | 202.65 | Fumarylpyruvate hydrolase |
| *FEPFGCPE_02621* | *nagL-1* | + | 2869884 | 2870525 | 0.50 | 36.55 | 216.58 | 95.78 | Maleylpyruvate isomerase |
| *FEPFGCPE_02622* | *mhbM* | + | 2870566 | 2871783 | 0.29 | 54.32 | 230.34 | 80.68 | 3-hydroxybenzoate 6-hydroxylase |
| *FEPFGCPE_02623* | *mhbT-2* | + | 2871991 | 2873334 | 0.22 | 26.47 | 89.22 | 35.35 | 3-hydroxybenzoate transporter MhbT |

| Locus name | Gene name | Strand | Gene start (bp) | Gene end (bp) | Average TPM Inoculum | Average TPM $t_6$ | Average TPM $t_{24}$ | Average TPM $t_{168}$ | Annotation description |
|---|---|---|---|---|---|---|---|---|---|
| *FEPFGCPE_02624* | | + | 2873470 | 2874546 | 3.09 | 10.04 | 14.12 | 4.37 | 2-ketogluconate reductase |
| *FEPFGCPE_02625* | | - | 2874560 | 2874811 | 29.72 | 96.71 | 63.54 | 25.00 | Hypothetical protein |
| *FEPFGCPE_02626* | *rhaS-2* | - | 2875175 | 2876152 | 1.35 | 8.64 | 7.30 | 2.58 | HTH-type transcriptional activator RhaS |
| *FEPFGCPE_02627* | *yhdN* | + | 2876393 | 2877442 | 1.78 | 7.39 | 5.12 | 3.02 | General stress protein 69 |
| *FEPFGCPE_02628* | *yfcF* | + | 2877667 | 2878341 | 2.32 | 11.88 | 5.01 | 2.45 | Glutathione S-transferase YfcF |
| *FEPFGCPE_02629* | *ilvE* | + | 2878666 | 2879763 | 3.77 | 21.44 | 21.47 | 12.17 | Branched-chain-amino-acid aminotransferase |
| *FEPFGCPE_02630* | | + | 2879824 | 2880561 | 2.11 | 6.85 | 3.14 | 6.38 | Hypothetical protein |
| *FEPFGCPE_02631* | *gcvA-4* | - | 2880568 | 2881476 | 1.49 | 4.22 | 3.90 | 3.08 | Glycine cleavage system transcriptional activator |
| *FEPFGCPE_02632* | | + | 2881643 | 2882668 | 67.05 | 11.85 | 6.60 | 1.18 | Hypothetical protein |
| *FEPFGCPE_02633* | *aldA-1* | + | 2882785 | 2883681 | 35.71 | 9.44 | 4.35 | 2.00 | Lactaldehyde dehydrogenase |
| *FEPFGCPE_02634* | *aldA-2* | + | 2883674 | 2884282 | 40.28 | 11.33 | 5.82 | 5.50 | Putative aldehyde dehydrogenase AldA |
| *FEPFGCPE_02635* | *lysN-3* | + | 2884444 | 2885646 | 2.38 | 1.68 | 1.21 | 0.87 | 2-aminoadipate transaminase |
| *FEPFGCPE_02636* | *yxeP-3* | - | 2885647 | 2886813 | 1.19 | 1.26 | 3.85 | 1.08 | putative hydrolase YxeP |
| *FEPFGCPE_02637* | *ygeX* | - | 2886851 | 2888083 | 0.25 | 0.94 | 1.42 | 0.22 | Diaminopropionate ammonia-lyase |
| *FEPFGCPE_02638* | *amaB-1* | - | 2888135 | 2889415 | 0.75 | 2.60 | 0.73 | 0.00 | N-carbamoyl-L-amino acid hydrolase |
| *FEPFGCPE_02639* | *decR-3* | + | 2889596 | 2890069 | 0.93 | 9.56 | 2.67 | 0.62 | DNA-binding transcriptional activator DecR |
| *FEPFGCPE_02640* | | - | 2890140 | 2890724 | 2.52 | 21.21 | 10.39 | 9.07 | Immunogenic protein MPB70 |
| *FEPFGCPE_02641* | *nlhH-4* | - | 2890771 | 2891664 | 0.12 | 1.31 | 1.40 | 0.00 | Carboxylesterase NlhH |
| *FEPFGCPE_02642* | *blc-1* | + | 2891860 | 2892546 | 0.16 | 4.96 | 3.41 | 0.96 | Outer membrane lipoprotein Blc |
| *FEPFGCPE_02643* | | + | 2892543 | 2893073 | 0.34 | 5.08 | 6.43 | 2.01 | Hypothetical protein |
| *FEPFGCPE_02644* | | + | 2893080 | 2893559 | 0.78 | 5.35 | 4.97 | 2.58 | Hypothetical protein |
| *FEPFGCPE_02645* | *pcaK-6* | - | 2893696 | 2895087 | 0.54 | 4.07 | 1.52 | 0.00 | 4-hydroxybenzoate transporter PcaK |
| *FEPFGCPE_02646* | *mhpA-4* | - | 2895197 | 2896765 | 0.29 | 2.27 | 1.62 | 0.86 | 3-(3-hydroxy-phenyl)propionate/3-hydroxycinnamic acid hydroxylase |
| *FEPFGCPE_02647* | | - | 2896762 | 2897634 | 0.53 | 5.61 | 2.33 | 0.55 | Ureidoglycolate lyase |
| *FEPFGCPE_02648* | *bphC* | - | 2897677 | 2898573 | 0.28 | 1.26 | 1.66 | 1.11 | Manganese-dependent 2,3-dihydroxybiphenyl 1,2-dioxygenase |
| *FEPFGCPE_02649* | | - | 2898570 | 2899622 | 0.30 | 1.96 | 1.05 | 0.60 | Hypothetical protein |
| *FEPFGCPE_02650* | | - | 2899615 | 2900043 | 0.13 | 2.42 | 0.00 | 0.00 | Hypothetical protein |
| *FEPFGCPE_02651* | *rspR-7* | + | 2900259 | 2900981 | 3.80 | 36.75 | 20.91 | 10.78 | HTH-type transcriptional repressor RspR |
| *FEPFGCPE_02652* | | + | 2901333 | 2902373 | 1.25 | 6.94 | 2.88 | 2.37 | Outer membrane porin protein |
| *FEPFGCPE_02653* | *cimH* | - | 2902482 | 2903837 | 1.31 | 6.49 | 5.98 | 4.28 | Citrate/malate transporter |
| *FEPFGCPE_02654* | *atoE-2* | - | 2903869 | 2905233 | 0.59 | 5.20 | 3.30 | 1.67 | Putative short-chain fatty acid transporter |
| *FEPFGCPE_02655* | | - | 2905337 | 2906170 | 0.45 | 3.53 | 3.40 | 2.50 | 4-sulfomuconolactone hydrolase |
| *FEPFGCPE_02656* | *uctC-7* | - | 2906239 | 2907432 | 0.15 | 1.75 | 2.36 | 1.59 | Acetyl-CoA:oxalate CoA-transferase |
| *FEPFGCPE_02657* | *citE* | - | 2907512 | 2908411 | 0.41 | 7.65 | 12.26 | 1.82 | Citrate lyase subunit beta |
| *FEPFGCPE_02658* | *cmpR-6* | + | 2908523 | 2909422 | 1.22 | 2.27 | 3.24 | 0.39 | HTH-type transcriptional activator CmpR |
| *FEPFGCPE_02659* | *rhdA* | - | 2909470 | 2910330 | 1.83 | 7.58 | 10.55 | 3.30 | Thiosulfate sulfurtransferase |
| *FEPFGCPE_02660* | *ssb* | - | 2910696 | 2911217 | 15.73 | 76.25 | 58.73 | 36.76 | Single-stranded DNA-binding protein |
| *FEPFGCPE_02661* | *yajR-2* | - | 2911248 | 2912411 | 13.00 | 35.07 | 39.79 | 20.30 | Inner membrane transport protein YajR |
| *FEPFGCPE_02662* | *uvrA-2* | + | 2912901 | 2915765 | 2.30 | 7.29 | 8.90 | 5.33 | UvrABC system protein A |
| *FEPFGCPE_02663* | | + | 2915869 | 2916897 | 2.54 | 7.39 | 5.86 | 6.35 | Hypothetical protein |
| *FEPFGCPE_02664* | *dauR* | + | 2917039 | 2917683 | 2.01 | 14.10 | 12.23 | 1.61 | Transcriptional regulator DauR |
| *FEPFGCPE_02665* | *tdcB* | + | 2917680 | 2918660 | 1.28 | 8.46 | 9.49 | 5.68 | L-threonine dehydratase catabolic TdcB |
| *FEPFGCPE_02666* | *yabJ-6* | + | 2918758 | 2919054 | 1.87 | 14.11 | 7.24 | 15.05 | 2-iminobutanoate/2-iminopropanoate deaminase |
| *FEPFGCPE_02667* | | - | 2919009 | 2919281 | 5.18 | 26.54 | 7.10 | 3.26 | Hypothetical protein |
| *FEPFGCPE_02668* | | + | 2919375 | 2920283 | 9.88 | 34.53 | 28.19 | 9.96 | Hypothetical protein |
| *FEPFGCPE_02669* | | - | 2920277 | 2920825 | 7.08 | 34.28 | 22.79 | 4.97 | Hypothetical protein |
| *FEPFGCPE_02670* | | - | 2920785 | 2922602 | 11.28 | 23.96 | 13.70 | 5.88 | Hypothetical protein |
| *FEPFGCPE_02671* | *safD* | - | 2922941 | 2924395 | 0.59 | 0.97 | 1.52 | 0.08 | Sulfoacetaldehyde dehydrogenase |
| *FEPFGCPE_02672* | *phnA* | - | 2924496 | 2925749 | 0.77 | 2.50 | 2.57 | 0.82 | Phosphonoacetate hydrolase |
| *FEPFGCPE_02673* | *rspR-8* | - | 2925856 | 2926572 | 0.62 | 5.05 | 2.19 | 0.16 | HTH-type transcriptional repressor RspR |
| *FEPFGCPE_02674* | *phnW-2* | + | 2926965 | 2928098 | 0.39 | 1.37 | 0.95 | 0.19 | 2-aminoethylphosphonate--pyruvate transaminase |
| *FEPFGCPE_02675* | | + | 2928265 | 2928711 | 2.57 | 14.91 | 17.40 | 4.61 | Hypothetical protein |
| *FEPFGCPE_02676* | *hsrA-2* | + | 2929080 | 2930450 | 0.81 | 6.16 | 7.24 | 0.30 | putative transport protein HsrA |
| *FEPFGCPE_02677* | *purU* | - | 2930459 | 2931331 | 0.79 | 18.89 | 14.57 | 8.89 | Formyltetrahydrofolate deformylase |
| *FEPFGCPE_02678* | | - | 2931411 | 2932262 | 0.44 | 4.65 | 5.32 | 3.87 | Hypothetical protein |
| *FEPFGCPE_02679* | *abo-2* | - | 2932409 | 2934262 | 0.17 | 0.88 | 0.70 | 0.49 | 4-methylaminobutanoate oxidase (formaldehyde-forming) |
| *FEPFGCPE_02680* | *abo-3* | - | 2934300 | 2934893 | 0.19 | 0.91 | 0.34 | 0.00 | 4-methylaminobutanoate oxidase (formaldehyde-forming) |
| *FEPFGCPE_02681* | *phnE* | - | 2935060 | 2935866 | 0.37 | 0.00 | 0.44 | 0.53 | Phosphate-import permease protein PhnE |
| *FEPFGCPE_02682* | *phnD-2* | - | 2936040 | 2937008 | 0.07 | 1.56 | 0.00 | 0.65 | Phosphate-import protein PhnD |
| *FEPFGCPE_02683* | *phnC* | - | 2937025 | 2937945 | 0.31 | 2.28 | 1.57 | 0.00 | Phosphate-import ATP-binding protein PhnC |
| *FEPFGCPE_02684* | *glpR-2* | - | 2938268 | 2939023 | 1.42 | 12.06 | 6.40 | 1.49 | Glycerol-3-phosphate regulon repressor |
| *FEPFGCPE_02685* | *rhtB-3* | - | 2939107 | 2939721 | 0.94 | 6.57 | 7.38 | 2.77 | Homoserine/homoserine lactone efflux protein |
| *FEPFGCPE_02686* | *apt* | - | 2939797 | 2940348 | 3.55 | 28.22 | 23.93 | 16.43 | Adenine phosphoribosyltransferase |
| *FEPFGCPE_02687* | *kefC-1* | - | 2940527 | 2942509 | 2.25 | 13.36 | 10.85 | 3.93 | Glutathione-regulated potassium-efflux system protein KefC |
| *FEPFGCPE_02688* | *kdsD* | + | 2942715 | 2943695 | 9.68 | 75.76 | 60.25 | 17.17 | Arabinose 5-phosphate isomerase KdsD |
| *FEPFGCPE_02689* | *kdsC* | + | 2943707 | 2944288 | 1.56 | 16.01 | 7.40 | 3.20 | 3-deoxy-D-manno-octulosonate 8-phosphate phosphatase KdsC |
| *FEPFGCPE_02690* | *lptC* | + | 2944312 | 2944971 | 4.25 | 22.00 | 27.39 | 8.17 | Lipopolysaccharide export system protein LptC |
| *FEPFGCPE_02691* | *lptA* | + | 2944968 | 2945666 | 3.17 | 13.66 | 14.03 | 4.36 | Lipopolysaccharide export system protein LptA |
| *FEPFGCPE_02692* | *lptB-7* | + | 2945659 | 2946462 | 3.18 | 18.97 | 11.67 | 3.41 | Lipopolysaccharide export system ATP-binding protein LptB |
| *FEPFGCPE_02693* | *rpoN* | + | 2946557 | 2948044 | 49.43 | 149.35 | 104.59 | 42.83 | RNA polymerase sigma-54 factor |
| *FEPFGCPE_02694* | *hpf* | + | 2948187 | 2948540 | 76.60 | 266.95 | 243.77 | 214.28 | Ribosome hibernation promoting factor |

| Locus name | Gene name | Strand | Gene start (bp) | Gene end (bp) | Average TPM Inoculum | Average TPM $t_6$ | Average TPM $t_{24}$ | Average TPM $t_{168}$ | Annotation description |
|---|---|---|---|---|---|---|---|---|---|
| *FEPFGCPE_02695* | *ptsN* | + | 2948770 | 2949225 | 7.95 | 75.81 | 62.60 | 34.04 | Nitrogen regulatory protein |
| *FEPFGCPE_02696* | *hprK* | + | 2949323 | 2950288 | 3.06 | 15.44 | 14.89 | 5.83 | HPr kinase/phosphorylase |
| *FEPFGCPE_02697* | *psiF* | + | 2950435 | 2950746 | 412.85 | 1470.02 | 594.00 | 269.52 | Phosphate starvation-inducible protein PsiF |
| *FEPFGCPE_02698* | *rapZ* | + | 2950977 | 2951852 | 17.55 | 45.40 | 35.92 | 12.78 | RNase adapter protein RapZ |
| *FEPFGCPE_02699* | *mutY* | - | 2951868 | 2953025 | 4.33 | 10.12 | 7.75 | 1.73 | Adenine DNA glycosylase |
| *FEPFGCPE_02700* | *mutM* | - | 2953108 | 2953938 | 2.36 | 12.88 | 14.26 | 2.97 | Formamidopyrimidine-DNA glycosylase |
| *FEPFGCPE_02701* | *bepA-4* | + | 2954018 | 2955943 | 23.84 | 44.17 | 27.87 | 16.57 | Beta-barrel assembly-enhancing protease |
| *FEPFGCPE_02702* | *lolB* | + | 2955943 | 2956566 | 18.52 | 39.98 | 27.15 | 7.26 | Outer-membrane lipoprotein LolB |
| *FEPFGCPE_02703* | *ispE* | + | 2956808 | 2957677 | 41.20 | 121.83 | 90.09 | 38.85 | 4-diphosphocytidyl-2-C-methyl-D-erythritol kinase |
| *FEPFGCPE_02704* | | + | 2957731 | 2957807 | 28.58 | 34.19 | 36.49 | 23.16 | tRNA |
| *FEPFGCPE_02705* | *prs-2* | + | 2957857 | 2958804 | 27.00 | 53.10 | 45.49 | 22.86 | Ribose-phosphate pyrophosphokinase |
| *FEPFGCPE_02706* | *rplY* | + | 2958959 | 2959579 | 15.54 | 67.17 | 76.87 | 42.22 | 50S ribosomal protein L25 |
| *FEPFGCPE_02707* | | + | 2959830 | 2961035 | 1.13 | 2.60 | 4.30 | 3.04 | Putative prophage phiRv2 integrase |
| *FEPFGCPE_02708* | | + | 2961032 | 2961178 | 0.74 | 5.74 | 10.07 | 0.00 | Hypothetical protein |
| *FEPFGCPE_02709* | | + | 2961175 | 2961558 | 114.73 | 122.30 | 63.11 | 45.32 | Hypothetical protein |
| *FEPFGCPE_02710* | *higA-2* | + | 2961539 | 2961844 | 42.17 | 37.03 | 34.48 | 15.05 | Antitoxin HigA-2 |
| *FEPFGCPE_02711* | | + | 2961870 | 2963153 | 11.66 | 12.40 | 9.05 | 3.80 | Hypothetical protein |
| *FEPFGCPE_02712* | | - | 2963115 | 2963261 | 4.31 | 7.72 | 8.03 | 5.28 | Hypothetical protein |
| *FEPFGCPE_02713* | | - | 2963547 | 2965229 | 10.90 | 37.77 | 28.84 | 11.86 | Hypothetical protein |
| *FEPFGCPE_02714* | | - | 2965195 | 2966322 | 10.94 | 36.70 | 20.57 | 11.96 | Hypothetical protein |
| *FEPFGCPE_02715* | | + | 2966981 | 2967805 | 1.31 | 3.06 | 1.29 | 0.11 | Hypothetical protein |
| *FEPFGCPE_02716* | *spo0C* | + | 2967871 | 2969955 | 0.53 | 1.60 | 0.21 | 0.92 | Chromosome-partitioning protein Spo0J |
| *FEPFGCPE_02717* | | + | 2970018 | 2970227 | 1.90 | 12.97 | 3.54 | 0.72 | Hypothetical protein |
| *FEPFGCPE_02718* | | - | 2970320 | 2971132 | 2.22 | 11.82 | 7.74 | 2.41 | Hypothetical protein |
| *FEPFGCPE_02719* | | + | 2971988 | 2972302 | 3.52 | 8.04 | 4.59 | 2.08 | Hypothetical protein |
| *FEPFGCPE_02720* | | - | 2972583 | 2973719 | 5.52 | 18.72 | 14.25 | 4.18 | Hypothetical protein |
| *FEPFGCPE_02721* | *sutR-1* | - | 2973754 | 2974044 | 6.51 | 42.74 | 30.26 | 6.12 | HTH-type transcriptional regulator SutR |
| *FEPFGCPE_02722* | | + | 2974303 | 2974653 | 6.33 | 9.11 | 6.65 | 0.09 | Hypothetical protein |
| *FEPFGCPE_02723* | | + | 2974979 | 2975734 | 0.18 | 2.25 | 0.94 | 0.00 | Hypothetical protein |
| *FEPFGCPE_02724* | | + | 2975818 | 2976108 | 0.43 | 0.09 | 0.00 | 0.92 | Hypothetical protein |
| *FEPFGCPE_02725* | | + | 2976135 | 2976992 | 0.13 | 0.56 | 0.00 | 0.00 | Hypothetical protein |
| *FEPFGCPE_02726* | *soj-2* | + | 2977246 | 2977884 | 0.26 | 0.00 | 2.43 | 0.00 | Chromosome-partitioning ATPase Soj |
| *FEPFGCPE_02727* | | + | 2977881 | 2978165 | 0.98 | 4.02 | 0.00 | 1.65 | Hypothetical protein |
| *FEPFGCPE_02728* | | + | 2978162 | 2978707 | 0.12 | 0.00 | 0.00 | 0.71 | Hypothetical protein |
| *FEPFGCPE_02729* | | + | 2978704 | 2979291 | 0.28 | 0.00 | 0.46 | 0.00 | Hypothetical protein |
| *FEPFGCPE_02730* | | + | 2979288 | 2979743 | 0.13 | 0.00 | 0.32 | 1.86 | Hypothetical protein |
| *FEPFGCPE_02731* | | + | 2979740 | 2981707 | 0.41 | 3.12 | 1.91 | 1.76 | Hypothetical protein |
| *FEPFGCPE_02732* | *paaG-2* | - | 2981754 | 2982542 | 3.80 | 40.97 | 39.84 | 24.32 | 1,2-epoxyphenylacetyl-CoA isomerase |
| *FEPFGCPE_02733* | | - | 2982574 | 2983356 | 0.84 | 38.12 | 70.25 | 101.70 | putative oxidoreductase |
| *FEPFGCPE_02734* | *paaF-3* | - | 2983387 | 2984172 | 0.58 | 59.33 | 87.50 | 126.17 | 2,3-dehydroadipyl-CoA hydratase |
| *FEPFGCPE_02735* | *fadA-3* | - | 2984184 | 2985365 | 1.26 | 57.49 | 94.78 | 162.53 | 3-ketoacyl-CoA thiolase |
| *FEPFGCPE_02736* | | - | 2985393 | 2987069 | 1.47 | 76.45 | 116.16 | 153.01 | Long-chain-fatty-acid--CoA ligase |
| *FEPFGCPE_02737* | *uctC-8* | - | 2987081 | 2988139 | 0.97 | 56.96 | 99.64 | 155.88 | Acetyl-CoA:oxalate CoA-transferase |
| *FEPFGCPE_02738* | *mmgC-3* | - | 2988142 | 2989287 | 1.63 | 57.12 | 96.92 | 163.17 | Acyl-CoA dehydrogenase |
| *FEPFGCPE_02739* | | + | 2989481 | 2990521 | 6.80 | 16.18 | 11.83 | 4.74 | putative HTH-type transcriptional regulator |
| *FEPFGCPE_02740* | | - | 2990482 | 2990721 | 6.64 | 17.73 | 19.32 | 3.15 | Hypothetical protein |
| *FEPFGCPE_02741* | *gloB-2* | + | 2990827 | 2991876 | 4.23 | 18.16 | 24.76 | 49.62 | Hydroxyacylglutathione hydrolase |
| *FEPFGCPE_02742* | *pcaK-7* | + | 2992276 | 2993643 | 1.59 | 11.84 | 19.35 | 36.62 | 4-hydroxybenzoate transporter PcaK |
| *FEPFGCPE_02743* | *dmdC-4* | + | 2993668 | 2995452 | 2.47 | 20.79 | 38.76 | 87.05 | 3-methylmercaptopropionyl-CoA dehydrogenase |
| *FEPFGCPE_02744* | *phaJ* | + | 2995465 | 2995935 | 2.32 | 21.13 | 28.79 | 58.48 | (R)-specific enoyl-CoA hydratase |
| *FEPFGCPE_02745* | *etfB-3* | + | 2995957 | 2996706 | 3.13 | 38.97 | 57.67 | 164.41 | Electron transfer flavoprotein subunit beta |
| *FEPFGCPE_02746* | *etfA-3* | + | 2996741 | 2997673 | 4.71 | 28.23 | 59.64 | 93.07 | Electron transfer flavoprotein subunit alpha |
| *FEPFGCPE_02747* | | + | 2997688 | 2997888 | 37.78 | 234.66 | 165.17 | 192.87 | Hypothetical protein |
| *FEPFGCPE_02748* | *hcaR-3* | + | 2998048 | 2998959 | 4.95 | 27.31 | 25.81 | 30.07 | Hca operon transcriptional activator HcaR |
| *FEPFGCPE_02749* | | + | 2998985 | 2999215 | 6.44 | 69.73 | 52.27 | 80.33 | Hypothetical protein |
| *FEPFGCPE_02750* | *traG* | + | 2999442 | 3001472 | 0.29 | 1.30 | 0.83 | 1.12 | Conjugal transfer protein TraG |
| *FEPFGCPE_02751* | | + | 3001469 | 3001924 | 0.12 | 0.00 | 0.00 | 3.16 | Hypothetical protein |
| *FEPFGCPE_02752* | *ptlH* | + | 3001921 | 3002958 | 0.00 | 0.00 | 0.00 | 5.15 | Type IV secretion system protein PtlH |
| *FEPFGCPE_02753* | | + | 3002955 | 3003338 | 0.29 | 0.00 | 0.00 | 5.77 | Hypothetical protein |
| *FEPFGCPE_02754* | | + | 3003335 | 3003619 | 0.26 | 0.00 | 0.48 | 0.41 | Hypothetical protein |
| *FEPFGCPE_02755* | *virB4* | + | 3003630 | 3006077 | 0.32 | 1.65 | 1.24 | 2.03 | Type IV secretion system protein virB4 |
| *FEPFGCPE_02756* | | + | 3006074 | 3006811 | 0.15 | 0.00 | 2.16 | 5.76 | Hypothetical protein |
| *FEPFGCPE_02757* | | + | 3006824 | 3008131 | 0.34 | 2.40 | 0.51 | 1.62 | Hypothetical protein |
| *FEPFGCPE_02758* | | + | 3008143 | 3008847 | 0.40 | 1.37 | 2.33 | 4.40 | Hypothetical protein |
| *FEPFGCPE_02759* | *ptlF* | + | 3008844 | 3009842 | 0.24 | 1.20 | 0.36 | 2.50 | Type IV secretion system protein PtlF |
| *FEPFGCPE_02760* | | + | 3009845 | 3011107 | 0.90 | 0.93 | 2.29 | 2.68 | Type IV secretion system protein virB10 |
| *FEPFGCPE_02761* | | + | 3011104 | 3011352 | 0.45 | 1.37 | 0.31 | 0.00 | Hypothetical protein |
| *FEPFGCPE_02762* | *pth* | + | 3011680 | 3012276 | 5.93 | 11.86 | 10.89 | 4.11 | Peptidyl-tRNA hydrolase |
| *FEPFGCPE_02763* | *fdx-2* | - | 3012324 | 3012581 | 12.11 | 58.42 | 33.86 | 10.16 | Ferredoxin |
| *FEPFGCPE_02764* | *coaD* | - | 3012646 | 3013140 | 20.31 | 76.61 | 40.61 | 12.77 | Phosphopantetheine adenylyltransferase |
| *FEPFGCPE_02765* | *rsmD* | - | 3013196 | 3013789 | 5.71 | 72.08 | 48.30 | 25.98 | Ribosomal RNA small subunit methyltransferase D |

| Locus name | Gene name | Strand | Gene start (bp) | Gene end (bp) | Average TPM Inoculum | Average TPM $t_6$ | Average TPM $t_{24}$ | Average TPM $t_{168}$ | Annotation description |
|---|---|---|---|---|---|---|---|---|---|
| *FEPFGCPE_02766* | *drrA-1* | + | 3013957 | 3014979 | 4.20 | 15.25 | 12.61 | 2.17 | Daunorubicin/doxorubicin resistance ATP-binding protein DrrA |
| *FEPFGCPE_02767* | *yadH-1* | + | 3014972 | 3015883 | 1.76 | 3.99 | 7.98 | 2.51 | Inner membrane transport permease YadH |
| *FEPFGCPE_02768* | | - | 3016014 | 3017462 | 2.83 | 21.00 | 11.40 | 5.10 | putative zinc protease |
| *FEPFGCPE_02769* | | - | 3017475 | 3018893 | 3.70 | 17.54 | 11.77 | 4.56 | putative zinc protease |
| *FEPFGCPE_02770* | *ftsY* | + | 3019065 | 3020498 | 2.21 | 22.86 | 19.12 | 11.50 | Signal recognition particle receptor FtsY |
| *FEPFGCPE_02771* | *argO* | - | 3020616 | 3021242 | 0.36 | 3.28 | 1.93 | 0.34 | Arginine exporter protein ArgO |
| *FEPFGCPE_02772* | *nagL-2* | - | 3021251 | 3021898 | 2.27 | 8.36 | 13.35 | 4.84 | Maleylpyruvate isomerase |
| *FEPFGCPE_02773* | *trpD-1* | + | 3022097 | 3023053 | 17.82 | 41.27 | 25.36 | 4.90 | Anthranilate phosphoribosyltransferase |
| *FEPFGCPE_02774* | | - | 3023253 | 3023636 | 47.59 | 153.75 | 101.01 | 29.27 | Hypothetical protein |
| *FEPFGCPE_02775* | *rpoH* | + | 3023857 | 3024795 | 49.52 | 46.72 | 54.36 | 36.30 | RNA polymerase sigma factor RpoH |
| *FEPFGCPE_02776* | | + | 3025180 | 3025500 | 35.55 | 88.27 | 52.01 | 15.55 | Hypothetical protein |
| *FEPFGCPE_02777* | *cydA* | + | 3025490 | 3027094 | 12.30 | 39.24 | 20.32 | 7.82 | Cytochrome bd ubiquinol oxidase subunit 1 |
| *FEPFGCPE_02778* | *cydB-2* | + | 3027116 | 3028258 | 9.28 | 31.67 | 25.76 | 9.01 | Cytochrome bd-I ubiquinol oxidase subunit 2 |
| *FEPFGCPE_02779* | *cydX* | + | 3028274 | 3028396 | 4.09 | 22.87 | 16.55 | 2.93 | Cytochrome bd ubiquinol oxidase subunit X |
| *FEPFGCPE_02780* | *ypmQ* | - | 3028483 | 3029085 | 3.66 | 20.29 | 16.67 | 8.71 | SCO1 protein |
| *FEPFGCPE_02781* | *ctaB* | - | 3029230 | 3030135 | 7.52 | 20.19 | 28.23 | 7.97 | Protoheme IX farnesyltransferase |
| *FEPFGCPE_02782* | *ctaA* | - | 3030154 | 3031341 | 10.98 | 67.72 | 61.08 | 15.23 | Heme A synthase |
| *FEPFGCPE_02783* | | - | 3031418 | 3032050 | 10.52 | 20.05 | 32.41 | 20.95 | Hypothetical protein |
| *FEPFGCPE_02784* | | - | 3032058 | 3032816 | 14.96 | 36.45 | 55.67 | 37.71 | Hypothetical protein |
| *FEPFGCPE_02785* | | + | 3032924 | 3033127 | 17.68 | 117.56 | 106.19 | 37.79 | Hypothetical protein |
| *FEPFGCPE_02786* | *ctaE-1* | - | 3033453 | 3034313 | 9.68 | 67.61 | 46.57 | 16.63 | Cytochrome c oxidase subunit 3 |
| *FEPFGCPE_02787* | | - | 3034353 | 3034574 | 32.67 | 221.15 | 219.44 | 41.76 | Hypothetical protein |
| *FEPFGCPE_02788* | *ctaG* | - | 3034567 | 3035184 | 14.82 | 110.02 | 72.91 | 12.95 | Cytochrome c oxidase assembly protein CtaG |
| *FEPFGCPE_02789* | | - | 3035315 | 3035434 | 23.92 | 207.07 | 166.68 | 53.83 | Hypothetical protein |
| *FEPFGCPE_02790* | *ctaD* | - | 3035437 | 3037029 | 7.60 | 71.18 | 43.38 | 16.16 | Cytochrome c oxidase subunit 1 |
| *FEPFGCPE_02791* | *ctaC* | - | 3037065 | 3038294 | 18.33 | 118.01 | 82.44 | 26.01 | Cytochrome c oxidase subunit 2 |
| *FEPFGCPE_02792* | | - | 3038335 | 3038808 | 23.51 | 129.67 | 84.29 | 26.89 | Hypothetical protein |
| *FEPFGCPE_02793* | *lasR-1* | + | 3039388 | 3040215 | 0.51 | 5.06 | 2.86 | 0.77 | Transcriptional activator protein LasR |
| *FEPFGCPE_02794* | *flr* | - | 3040163 | 3040810 | 0.45 | 2.57 | 0.68 | 0.71 | Flavoredoxin |
| *FEPFGCPE_02795* | *aao* | - | 3040859 | 3041815 | 1.57 | 5.05 | 3.61 | 0.75 | putative D-amino-acid oxidase |
| *FEPFGCPE_02796* | *thrC-2* | - | 3041812 | 3043083 | 0.35 | 2.48 | 2.40 | 0.18 | Threonine synthase |
| *FEPFGCPE_02797* | *argF-1* | - | 3043071 | 3044117 | 0.11 | 0.48 | 1.20 | 0.31 | Ornithine carbamoyltransferase |
| *FEPFGCPE_02798* | | - | 3044239 | 3045165 | 0.20 | 2.35 | 1.13 | 0.70 | Hypothetical protein |
| *FEPFGCPE_02799* | *purA-3* | - | 3045197 | 3046162 | 0.22 | 0.47 | 1.11 | 0.00 | Adenylosuccinate synthetase |
| *FEPFGCPE_02800* | *purA-4* | - | 3046162 | 3046665 | 0.56 | 1.31 | 0.00 | 0.78 | Adenylosuccinate synthetase |
| *FEPFGCPE_02801* | | - | 3046662 | 3047717 | 0.32 | 0.77 | 0.45 | 0.00 | Delta(1)-pyrroline-2-carboxylate reductase |
| *FEPFGCPE_02802* | *sbnA* | - | 3047714 | 3048799 | 0.47 | 0.94 | 1.78 | 0.00 | putative siderophore biosynthesis protein SbnA |
| *FEPFGCPE_02803* | *bioC-2* | - | 3049009 | 3049929 | 1.33 | 11.03 | 4.33 | 6.69 | Malonyl-[acyl-carrier protein] O-methyltransferase |
| *FEPFGCPE_02804* | *pyrE-1* | + | 3050033 | 3050881 | 4.92 | 16.27 | 14.70 | 4.28 | Orotate phosphoribosyltransferase |
| *FEPFGCPE_02805* | *trmL* | + | 3050898 | 3051368 | 3.22 | 21.93 | 16.58 | 3.60 | tRNA (cytidine(34)-2'-O)-methyltransferase |
| *FEPFGCPE_02806* | *ymdB* | - | 3051350 | 3051880 | 9.15 | 37.89 | 45.21 | 11.70 | O-acetyl-ADP-ribose deacetylase |
| *FEPFGCPE_02807* | *gpsA* | - | 3052035 | 3053054 | 5.43 | 27.48 | 18.98 | 4.96 | Glycerol-3-phosphate dehydrogenase [NAD(P)+] |
| *FEPFGCPE_02808* | *secB* | - | 3053118 | 3053624 | 10.34 | 63.20 | 41.46 | 25.85 | Protein-export protein SecB |
| *FEPFGCPE_02809* | *grxC* | - | 3053690 | 3053950 | 23.51 | 89.57 | 98.05 | 34.85 | Glutaredoxin 3 |
| *FEPFGCPE_02810* | *ygaP* | - | 3053968 | 3054375 | 14.04 | 53.00 | 40.50 | 21.92 | Inner membrane protein YgaP |
| *FEPFGCPE_02811* | *gpmA-2* | + | 3054610 | 3055356 | 2.75 | 25.48 | 32.25 | 13.53 | 2,3-bisphosphoglycerate-dependent phosphoglycerate mutase |
| *FEPFGCPE_02812* | | + | 3055786 | 3057309 | 23.64 | 70.33 | 67.73 | 33.69 | putative CtpA-like serine protease |
| *FEPFGCPE_02813* | *moeB* | + | 3057380 | 3058132 | 7.82 | 43.05 | 29.32 | 10.91 | Molybdopterin-synthase adenylyltransferase |
| *FEPFGCPE_02814* | *ptsI* | - | 3058286 | 3060022 | 4.70 | 18.87 | 15.73 | 6.32 | Phosphoenolpyruvate-protein phosphotransferase |
| *FEPFGCPE_02815* | *ptsH* | - | 3060167 | 3060436 | 23.67 | 94.75 | 92.23 | 28.36 | Phosphocarrier protein HPr |
| *FEPFGCPE_02816* | *manX* | - | 3060440 | 3060853 | 8.17 | 44.92 | 39.29 | 24.06 | PTS system mannose-specific EIIAB component |
| *FEPFGCPE_02817* | *gshB* | - | 3061023 | 3061961 | 3.82 | 24.36 | 27.93 | 11.32 | Glutathione synthetase |
| *FEPFGCPE_02818* | | - | 3062022 | 3063308 | 0.96 | 2.59 | 3.41 | 0.54 | Hypothetical protein |
| *FEPFGCPE_02819* | | - | 3063750 | 3065039 | 5.59 | 29.93 | 34.65 | 10.01 | Hypothetical protein |
| *FEPFGCPE_02820* | *amtB-2* | - | 3065397 | 3066926 | 0.94 | 72.89 | 96.35 | 56.96 | Ammonia channel |
| *FEPFGCPE_02821* | *glnK* | - | 3066953 | 3067291 | 2.11 | 179.60 | 210.88 | 110.04 | Nitrogen regulatory protein P-II 2 |
| *FEPFGCPE_02822* | | - | 3067305 | 3068126 | 1.71 | 271.88 | 292.89 | 106.13 | Hypothetical protein |
| *FEPFGCPE_02823* | | + | 3068520 | 3068768 | 13.90 | 35.92 | 34.21 | 29.18 | Hypothetical protein |
| *FEPFGCPE_02824* | *comM* | + | 3069055 | 3070593 | 0.42 | 2.38 | 2.55 | 0.95 | Competence protein ComM |
| *FEPFGCPE_02825* | | - | 3070616 | 3071203 | 4.01 | 30.80 | 18.80 | 10.92 | Hypothetical protein |
| *FEPFGCPE_02826* | | + | 3071390 | 3072001 | 5.04 | 23.42 | 19.23 | 1.33 | Hypothetical protein |
| *FEPFGCPE_02827* | *resA-1* | + | 3072126 | 3072650 | 9.00 | 58.65 | 39.66 | 10.80 | Thiol-disulfide oxidoreductase ResA |
| *FEPFGCPE_02828* | *dctD-2* | - | 3072768 | 3074123 | 2.18 | 17.10 | 9.15 | 6.32 | C4-dicarboxylate transport transcriptional regulatory protein DctD |
| *FEPFGCPE_02829* | *dctB-3* | - | 3074120 | 3075973 | 0.81 | 16.06 | 5.29 | 1.06 | C4-dicarboxylate transport sensor protein DctB |
| *FEPFGCPE_02830* | *dctA-4* | - | 3076353 | 3077639 | 1.74 | 86.51 | 19.91 | 10.26 | C4-dicarboxylate transport protein |
| *FEPFGCPE_02831* | *cmpD-3* | - | 3077684 | 3078478 | 7.02 | 233.92 | 56.28 | 26.74 | Bicarbonate transport ATP-binding protein CmpD |
| *FEPFGCPE_02832* | *ssuC-5* | - | 3078489 | 3079325 | 2.40 | 64.31 | 24.03 | 6.32 | Putative aliphatic sulfonates transport permease protein SsuC |
| *FEPFGCPE_02833* | *ssuA-2* | - | 3079340 | 3080308 | 4.81 | 153.57 | 59.69 | 24.85 | Putative aliphatic sulfonates-binding protein |
| *FEPFGCPE_02834* | *kdgR-4* | - | 3080598 | 3081377 | 3.92 | 10.61 | 8.08 | 2.09 | Transcriptional regulator KdgR |
| *FEPFGCPE_02835* | | - | 3081377 | 3082261 | 4.65 | 15.90 | 8.45 | 1.31 | Hypothetical protein |
| *FEPFGCPE_02836* | *lipA* | - | 3082631 | 3083641 | 5.54 | 30.04 | 26.24 | 14.05 | Lipoyl synthase |

| Locus name | Gene name | Strand | Gene start (bp) | Gene end (bp) | Average TPM Inoculum | Average TPM $t_6$ | Average TPM $t_{24}$ | Average TPM $t_{168}$ | Annotation description |
|---|---|---|---|---|---|---|---|---|---|
| *FEPFGCPE_02837* | *lipB* | - | 3083666 | 3084373 | 10.99 | 85.63 | 70.05 | 25.92 | Octanoyltransferase |
| *FEPFGCPE_02838* | | - | 3084635 | 3084967 | 19.37 | 48.90 | 41.92 | 14.15 | Hypothetical protein |
| *FEPFGCPE_02839* | *gcvA-5* | + | 3085068 | 3086108 | 5.82 | 20.12 | 17.13 | 5.78 | Glycine cleavage system transcriptional activator |
| *FEPFGCPE_02840* | | - | 3086177 | 3086449 | 4.96 | 15.02 | 9.10 | 4.12 | Hypothetical protein |
| *FEPFGCPE_02841* | *dat* | - | 3086446 | 3087294 | 4.56 | 27.11 | 10.03 | 4.54 | D-alanine aminotransferase |
| *FEPFGCPE_02842* | *dacC-1* | - | 3087297 | 3088484 | 5.09 | 16.24 | 17.69 | 12.34 | D-alanyl-D-alanine carboxypeptidase DacC |
| *FEPFGCPE_02843* | | - | 3088737 | 3089366 | 2.75 | 24.81 | 22.70 | 13.33 | Hypothetical protein |
| *FEPFGCPE_02844* | *fdx4* | - | 3089429 | 3089857 | 6.76 | 77.80 | 47.86 | 19.65 | Ferredoxin, 2Fe-2S |
| *FEPFGCPE_02845* | *emrB-3* | - | 3089925 | 3091466 | 0.48 | 3.33 | 2.03 | 0.00 | Multidrug export protein EmrB |
| *FEPFGCPE_02846* | | - | 3091664 | 3092836 | 1.26 | 4.52 | 5.50 | 0.49 | Hypothetical protein |
| *FEPFGCPE_02847* | | - | 3092850 | 3093719 | 2.29 | 11.93 | 6.18 | 1.15 | Hypothetical protein |
| *FEPFGCPE_02848* | | - | 3093719 | 3094678 | 2.89 | 13.51 | 8.54 | 3.84 | Hypothetical protein |
| *FEPFGCPE_02849* | *mlaF-1* | - | 3094668 | 3095624 | 2.02 | 10.27 | 3.27 | 1.92 | putative phospholipid import ATP-binding protein MlaF |
| *FEPFGCPE_02850* | *mlaE-1* | - | 3095624 | 3096745 | 1.77 | 6.33 | 5.85 | 2.32 | putative phospholipid ABC transporter permease protein MlaE |
| *FEPFGCPE_02851* | *birA-1* | + | 3096951 | 3097949 | 3.79 | 18.20 | 8.54 | 4.37 | Bifunctional ligase/repressor BirA |
| *FEPFGCPE_02852* | *coaX* | + | 3097946 | 3099067 | 2.41 | 12.19 | 7.00 | 3.42 | Type III pantothenate kinase |
| *FEPFGCPE_02853* | | + | 3099039 | 3099983 | 2.22 | 15.54 | 16.58 | 2.42 | Hypothetical protein |
| *FEPFGCPE_02854* | | - | 3100005 | 3101012 | 2.37 | 19.02 | 12.03 | 5.15 | Hypothetical protein |
| *FEPFGCPE_02855* | | - | 3101009 | 3101512 | 3.00 | 21.27 | 15.90 | 5.76 | D-beta-D-heptose 1-phosphate adenylyltransferase |
| *FEPFGCPE_02856* | | - | 3101619 | 3102494 | 13.49 | 34.52 | 35.97 | 23.57 | Hypothetical protein |
| *FEPFGCPE_02857* | | + | 3103208 | 3103768 | 4.59 | 12.74 | 10.53 | 3.53 | Hypothetical protein |
| *FEPFGCPE_02858* | | + | 3103893 | 3104408 | 19.75 | 30.72 | 13.46 | 4.81 | Hypothetical protein |
| *FEPFGCPE_02859* | *eamB* | + | 3104522 | 3105109 | 2.34 | 9.64 | 1.55 | 2.03 | Cysteine/O-acetylserine efflux protein |
| *FEPFGCPE_02860* | *araC-3* | - | 3105137 | 3105979 | 2.29 | 17.46 | 5.61 | 1.85 | Arabinose operon regulatory protein |
| *FEPFGCPE_02861* | | - | 3106007 | 3107296 | 4.39 | 15.52 | 7.69 | 2.83 | Hypothetical protein |
| *FEPFGCPE_02862* | | - | 3107413 | 3108333 | 17.57 | 80.61 | 50.06 | 25.15 | Hypothetical protein |
| *FEPFGCPE_02863* | *camK* | - | 3108478 | 3109299 | 0.63 | 7.31 | 2.75 | 1.99 | 6-oxocamphor hydrolase |
| *FEPFGCPE_02864* | | - | 3109463 | 3110614 | 7.27 | 58.69 | 25.60 | 10.27 | Hypothetical protein |
| *FEPFGCPE_02865* | | - | 3110891 | 3111877 | 1.10 | 14.29 | 14.14 | 5.73 | Hypothetical protein |
| *FEPFGCPE_02866* | | + | 3112247 | 3113284 | 3.67 | 6.35 | 3.18 | 2.59 | Hypothetical protein |
| *FEPFGCPE_02867* | *catD-2* | + | 3113504 | 3114298 | 50.14 | 115.33 | 72.14 | 29.97 | 3-oxoadipate enol-lactonase 2 |
| *FEPFGCPE_02868* | | + | 3114471 | 3114728 | 124.87 | 358.48 | 225.73 | 84.14 | Hypothetical protein |
| *FEPFGCPE_02869* | *glpE-1* | - | 3114861 | 3115253 | 4.58 | 9.14 | 11.65 | 1.26 | Thiosulfate sulfurtransferase GlpE |
| *FEPFGCPE_02870* | *metH-1* | + | 3115400 | 3116542 | 6.25 | 13.79 | 11.79 | 10.73 | Methionine synthase |
| *FEPFGCPE_02871* | *metH-2* | + | 3116659 | 3119418 | 5.42 | 9.64 | 8.31 | 6.65 | Methionine synthase |
| *FEPFGCPE_02872* | | + | 3119593 | 3120312 | 7.10 | 10.60 | 13.66 | 2.84 | Hypothetical protein |
| *FEPFGCPE_02873* | | - | 3120527 | 3120853 | 75.86 | 146.08 | 101.97 | 37.10 | Hypothetical protein |
| *FEPFGCPE_02874* | *argS* | + | 3121021 | 3122802 | 3.41 | 19.77 | 12.56 | 9.58 | Arginine--tRNA ligase |
| *FEPFGCPE_02875* | *cwlC* | + | 3122869 | 3123540 | 2.35 | 18.11 | 16.83 | 11.58 | Sporulation-specific N-acetylmuramoyl-L-alanine amidase |
| *FEPFGCPE_02876* | *dsbA* | + | 3123790 | 3124473 | 36.60 | 49.84 | 49.21 | 35.41 | Thiol:disulfide interchange protein DsbA |
| *FEPFGCPE_02877* | | + | 3124520 | 3125374 | 4.16 | 11.08 | 6.38 | 7.99 | putative oxidoreductase |
| *FEPFGCPE_02878* | | + | 3125474 | 3125986 | 2.87 | 7.04 | 6.82 | 2.93 | Hypothetical protein |
| *FEPFGCPE_02879* | *proP-12* | + | 3126212 | 3127531 | 4.91 | 16.59 | 22.33 | 6.02 | Proline/betaine transporter |
| *FEPFGCPE_02880* | *gloB-3* | - | 3127725 | 3128807 | 1.75 | 5.72 | 8.02 | 3.07 | Hydroxyacylglutathione hydrolase |
| *FEPFGCPE_02881* | *cueR* | + | 3128936 | 3129373 | 3.80 | 14.25 | 15.84 | 15.33 | HTH-type transcriptional regulator CueR |
| *FEPFGCPE_02882* | | + | 3129617 | 3130102 | 11.02 | 34.33 | 18.59 | 8.03 | Putative esterase |
| *FEPFGCPE_02883* | *acdA-1* | + | 3130201 | 3131382 | 7.45 | 19.16 | 13.52 | 8.76 | Acyl-CoA dehydrogenase |
| *FEPFGCPE_02884* | *aceK* | + | 3131467 | 3133272 | 6.37 | 34.62 | 22.55 | 8.97 | Isocitrate dehydrogenase kinase/phosphatase |
| *FEPFGCPE_02885* | *can* | + | 3133321 | 3134001 | 4.18 | 24.61 | 20.90 | 14.78 | Carbonic anhydrase 2 |
| *FEPFGCPE_02886* | *thlA-2* | + | 3134146 | 3135330 | 8.78 | 46.54 | 31.16 | 16.38 | Acetyl-CoA acetyltransferase |
| *FEPFGCPE_02887* | *csgA-2* | + | 3135371 | 3136048 | 7.93 | 44.17 | 25.50 | 14.49 | C-factor |
| *FEPFGCPE_02888* | *nsaD-2* | + | 3136176 | 3136781 | 5.74 | 42.85 | 25.28 | 7.11 | 2-hydroxychromene-2-carboxylate isomerase |
| *FEPFGCPE_02889* | | + | 3136918 | 3137223 | 2.01 | 12.61 | 14.64 | 5.91 | Hypothetical protein |
| *FEPFGCPE_02890* | | + | 3137291 | 3138898 | 1.20 | 13.25 | 8.92 | 4.21 | Methylmalonyl-CoA carboxyltransferase 12S subunit |
| *FEPFGCPE_02891* | *dpgD* | + | 3138933 | 3139724 | 0.83 | 11.49 | 14.57 | 3.91 | Enoyl-CoA-hydratase |
| *FEPFGCPE_02892* | *bioA* | + | 3140038 | 3141414 | 0.74 | 9.05 | 6.11 | 2.82 | Adenosylmethionine-8-amino-7-oxononanoate aminotransferase |
| *FEPFGCPE_02893* | *bioF* | + | 3141411 | 3142619 | 0.53 | 13.96 | 9.20 | 3.27 | 8-amino-7-oxononanoate synthase |
| *FEPFGCPE_02894* | *bioD1* | + | 3142612 | 3143382 | 0.41 | 12.69 | 6.48 | 1.94 | ATP-dependent dethiobiotin synthetase BioD 1 |
| *FEPFGCPE_02895* | *bioB* | + | 3143442 | 3144500 | 0.43 | 14.55 | 9.28 | 6.02 | Biotin synthase |
| *FEPFGCPE_02896* | *accA1* | + | 3144705 | 3146057 | 0.41 | 16.14 | 9.76 | 3.96 | Acetyl-/propionyl-coenzyme A carboxylase alpha chain |
| *FEPFGCPE_02897* | *gcdC* | + | 3146153 | 3146725 | 0.28 | 27.14 | 15.52 | 5.26 | Glutaconyl-CoA decarboxylase subunit gamma |
| *FEPFGCPE_02898* | | + | 3147004 | 3148200 | 0.50 | 2.25 | 0.83 | 0.09 | Outer membrane porin protein |
| *FEPFGCPE_02899* | | + | 3149175 | 3149420 | 0.45 | 2.45 | 1.51 | 1.55 | Hypothetical protein |
| *FEPFGCPE_02900* | | + | 3149683 | 3150495 | 1.87 | 9.26 | 5.78 | 0.47 | Hypothetical protein |
| *FEPFGCPE_02901* | | + | 3151074 | 3151451 | 0.33 | 0.42 | 0.72 | 2.99 | Hypothetical protein |
| *FEPFGCPE_02902* | | - | 3151549 | 3152385 | 0.87 | 9.77 | 5.91 | 2.33 | Tropinesterase |
| *FEPFGCPE_02903* | | + | 3152538 | 3153089 | 2.57 | 5.41 | 11.55 | 0.77 | Hypothetical protein |
| *FEPFGCPE_02904* | *ghrA-2* | + | 3153325 | 3154377 | 8.91 | 15.24 | 13.89 | 8.33 | Glyoxylate/hydroxypyruvate reductase A |
| *FEPFGCPE_02905* | *liuE* | + | 3154425 | 3155366 | 6.94 | 22.82 | 18.30 | 14.62 | 3-hydroxy-3-isohexenylglutaryl-CoA/hydroxy-methylglutaryl-CoA lyase |
| *FEPFGCPE_02906* | *ybaK-1* | + | 3155425 | 3155943 | 4.26 | 18.02 | 10.15 | 8.18 | Cys-tRNA(Pro)/Cys-tRNA(Cys) deacylase YbaK |
| *FEPFGCPE_02907* | | + | 3155977 | 3156219 | 3.71 | 21.04 | 21.06 | 8.74 | Hypothetical protein |

| Locus name | Gene name | Strand | Gene start (bp) | Gene end (bp) | Average TPM Inoculum | Average TPM $t_6$ | Average TPM $t_{24}$ | Average TPM $t_{168}$ | Annotation description |
|---|---|---|---|---|---|---|---|---|---|
| *FEPFGCPE_02908* | | + | 3156254 | 3156685 | 3.08 | 12.26 | 5.28 | 1.28 | Hypothetical protein |
| *FEPFGCPE_02909* | *gloC-2* | + | 3156729 | 3157652 | 2.18 | 7.22 | 5.84 | 3.91 | Hydroxyacylglutathione hydrolase GloC |
| *FEPFGCPE_02910* | | + | 3158023 | 3158970 | 7.51 | 24.41 | 17.95 | 7.41 | Nitronate monooxygenase |
| *FEPFGCPE_02911* | *hbd* | - | 3159309 | 3160157 | 3.22 | 8.93 | 10.89 | 6.20 | 3-hydroxybutyryl-CoA dehydrogenase |
| *FEPFGCPE_02912* | | - | 3160444 | 3161484 | 7.03 | 4.35 | 5.15 | 1.91 | Outer membrane porin protein 32 |
| *FEPFGCPE_02913* | | - | 3161948 | 3162223 | 10.49 | 41.08 | 32.40 | 26.07 | Hypothetical protein |
| *FEPFGCPE_02914* | | - | 3162512 | 3163156 | 8.57 | 43.50 | 37.83 | 10.63 | Hypothetical protein |
| *FEPFGCPE_02915* | | + | 3163682 | 3164143 | 10.78 | 9.19 | 13.34 | 1.89 | Hypothetical protein |
| *FEPFGCPE_02916* | | + | 3164312 | 3164716 | 2.06 | 13.16 | 9.59 | 2.67 | Hypothetical protein |
| *FEPFGCPE_02917* | *yigZ* | - | 3164793 | 3165374 | 3.85 | 19.06 | 15.72 | 8.68 | IMPACT family member YigZ |
| *FEPFGCPE_02918* | *phzF* | - | 3165397 | 3166266 | 1.75 | 5.42 | 7.16 | 3.86 | Trans-2,3-dihydro-3-hydroxyanthranilate isomerase |
| *FEPFGCPE_02919* | *yhhW-1* | + | 3166644 | 3167363 | 0.67 | 1.98 | 3.07 | 0.00 | Quercetin 2,3-dioxygenase |
| *FEPFGCPE_02920* | *atpD* | + | 3167527 | 3168999 | 2.14 | 5.82 | 4.01 | 2.69 | ATP synthase subunit beta |
| *FEPFGCPE_02921* | *atpC-2* | + | 3169014 | 3169445 | 1.44 | 2.73 | 4.72 | 5.53 | ATP synthase epsilon chain |
| *FEPFGCPE_02922* | | + | 3169442 | 3169765 | 0.66 | 0.00 | 2.67 | 4.62 | Hypothetical protein |
| *FEPFGCPE_02923* | | + | 3169758 | 3170051 | 1.92 | 6.78 | 9.00 | 7.47 | Hypothetical protein |
| *FEPFGCPE_02924* | *atpB-1* | + | 3170048 | 3170746 | 0.42 | 1.06 | 2.31 | 0.69 | ATP synthase subunit a |
| *FEPFGCPE_02925* | *atpE-1* | + | 3170743 | 3170988 | 0.14 | 3.53 | 0.00 | 2.36 | ATP synthase subunit c |
| *FEPFGCPE_02926* | *atpF-1* | + | 3170997 | 3171743 | 0.12 | 0.98 | 2.25 | 2.25 | ATP synthase subunit b |
| *FEPFGCPE_02927* | *atpA-1* | + | 3171706 | 3173283 | 0.42 | 4.15 | 3.23 | 1.72 | ATP synthase subunit alpha |
| *FEPFGCPE_02928* | *atpG-1* | + | 3173264 | 3174169 | 0.81 | 2.14 | 8.25 | 3.77 | ATP synthase gamma chain, sodium ion specific |
| *FEPFGCPE_02929* | *yciC-2* | - | 3174287 | 3175537 | 0.31 | 3.86 | 23.84 | 23.46 | Putative metal chaperone YciC |
| *FEPFGCPE_02930* | *pcoC-2* | + | 3175973 | 3176383 | 1.34 | 5.16 | 1.63 | 3.39 | Copper resistance protein C |
| *FEPFGCPE_02931* | | + | 3176383 | 3177372 | 0.70 | 1.09 | 0.00 | 0.00 | Hypothetical protein |
| *FEPFGCPE_02932* | *ada* | + | 3177461 | 3178666 | 1.23 | 4.74 | 3.33 | 1.85 | Bifunctional transcriptional activator/DNA repair enzyme Ada |
| *FEPFGCPE_02933* | | + | 3178663 | 3179394 | 1.54 | 10.32 | 9.33 | 1.42 | Hypothetical protein |
| *FEPFGCPE_02934* | *alkB* | + | 3179391 | 3180050 | 1.13 | 3.28 | 8.04 | 1.26 | Alpha-ketoglutarate-dependent dioxygenase AlkB |
| *FEPFGCPE_02935* | | - | 3180090 | 3180710 | 3.22 | 18.65 | 9.75 | 4.55 | Hypothetical protein |
| *FEPFGCPE_02936* | | - | 3180760 | 3181476 | 2.67 | 23.45 | 15.37 | 5.30 | Hypothetical protein |
| *FEPFGCPE_02937* | *dmlR-12* | + | 3181602 | 3182531 | 2.61 | 5.27 | 6.29 | 2.50 | HTH-type transcriptional regulator DmlR |
| *FEPFGCPE_02938* | *plcC* | + | 3182770 | 3184488 | 0.94 | 1.37 | 1.29 | 1.09 | Phospholipase C 3 |
| *FEPFGCPE_02939* | *yusO-2* | - | 3184683 | 3185138 | 3.74 | 26.64 | 16.98 | 7.44 | putative HTH-type transcriptional regulator YusO |
| *FEPFGCPE_02940* | *mhpA-5* | + | 3185330 | 3187018 | 1.06 | 1.15 | 0.67 | 0.00 | 3-(3-hydroxy-phenyl)propionate/3-hydroxycinnamic acid hydroxylase |
| *FEPFGCPE_02941* | | + | 3187048 | 3187242 | 0.98 | 4.61 | 5.25 | 2.31 | Hypothetical protein |
| *FEPFGCPE_02942* | *ttuB-5* | + | 3187323 | 3188750 | 0.61 | 1.29 | 1.30 | 0.24 | Putative tartrate transporter |
| *FEPFGCPE_02943* | | + | 3188823 | 3189611 | 4.44 | 26.45 | 15.35 | 6.01 | Hypothetical protein |
| *FEPFGCPE_02944* | | + | 3189693 | 3189769 | 18.47 | 103.59 | 91.82 | 48.62 | tRNA |
| *FEPFGCPE_02945* | | + | 3190076 | 3190378 | 9.01 | 176.11 | 127.21 | 49.67 | Hypothetical protein |
| *FEPFGCPE_02946* | *dmlR-13* | - | 3190466 | 3191368 | 1.66 | 6.70 | 4.42 | 7.15 | HTH-type transcriptional regulator DmlR |
| *FEPFGCPE_02947* | *yfcG-3* | + | 3191506 | 3192195 | 0.36 | 1.36 | 1.32 | 0.00 | Disulfide-bond oxidoreductase YfcG |
| *FEPFGCPE_02948* | | + | 3192212 | 3193186 | 0.26 | 3.38 | 4.21 | 0.16 | Hypothetical protein |
| *FEPFGCPE_02949* | | + | 3193726 | 3194574 | 0.90 | 6.61 | 4.13 | 0.04 | Hypothetical protein |
| *FEPFGCPE_02950* | *dmlR-14* | + | 3194680 | 3195576 | 1.37 | 5.20 | 4.89 | 0.36 | HTH-type transcriptional regulator DmlR |
| *FEPFGCPE_02951* | *argG* | + | 3195689 | 3197023 | 25.13 | 59.29 | 61.56 | 39.57 | Argininosuccinate synthase |
| *FEPFGCPE_02952* | | + | 3197240 | 3198634 | 1.57 | 8.70 | 8.21 | 4.26 | Hypothetical protein |
| *FEPFGCPE_02953* | | + | 3198792 | 3199964 | 3.99 | 16.23 | 13.94 | 3.58 | Hypothetical protein |
| *FEPFGCPE_02954* | | - | 3199874 | 3200548 | 6.55 | 21.11 | 14.91 | 7.48 | Hypothetical protein |
| *FEPFGCPE_02955* | | - | 3200632 | 3200913 | 14.52 | 44.37 | 15.14 | 25.95 | Hypothetical protein |
| *FEPFGCPE_02956* | | - | 3200980 | 3201771 | 5.29 | 18.54 | 13.45 | 8.40 | Hypothetical protein |
| *FEPFGCPE_02957* | *pcrA* | - | 3201981 | 3204065 | 6.11 | 6.17 | 3.89 | 1.93 | ATP-dependent DNA helicase PcrA |
| *FEPFGCPE_02958* | | + | 3204232 | 3205182 | 1.37 | 4.90 | 1.27 | 0.95 | Hypothetical protein |
| *FEPFGCPE_02959* | *eriC* | - | 3205179 | 3206597 | 1.26 | 3.90 | 2.89 | 0.80 | Chloride/fluoride channel protein |
| *FEPFGCPE_02960* | | + | 3206857 | 3207417 | 5.35 | 3.05 | 3.56 | 3.34 | Hypothetical protein |
| *FEPFGCPE_02961* | | + | 3207430 | 3208023 | 15.30 | 4.94 | 3.57 | 1.74 | Hypothetical protein |
| *FEPFGCPE_02962* | *iscS-2* | - | 3208071 | 3209231 | 46.05 | 4.36 | 6.41 | 4.73 | Cysteine desulfurase IscS |
| *FEPFGCPE_02963* | *qseC-2* | - | 3209347 | 3210789 | 0.58 | 1.47 | 3.06 | 0.83 | Sensor protein QseC |
| *FEPFGCPE_02964* | *tctD-1* | - | 3210814 | 3211626 | 0.83 | 7.16 | 4.59 | 1.20 | Transcriptional regulatory protein tctD |
| *FEPFGCPE_02965* | | + | 3211647 | 3212750 | 0.95 | 27.43 | 45.91 | 53.28 | Hypothetical protein |
| *FEPFGCPE_02966* | *phnU-3* | + | 3212920 | 3214683 | 0.82 | 4.49 | 8.08 | 6.35 | Putative 2-aminoethylphosphonate transport system permease protein PhnU |
| *FEPFGCPE_02967* | *potA-7* | + | 3214676 | 3215746 | 0.54 | 5.67 | 17.34 | 15.12 | Spermidine/putrescine import ATP-binding protein PotA |
| *FEPFGCPE_02968* | | + | 3215846 | 3216922 | 17.44 | 14.55 | 32.07 | 30.93 | Outer membrane porin protein 32 |
| *FEPFGCPE_02969* | *lysDH* | - | 3217012 | 3218109 | 57.61 | 18.25 | 5.62 | 3.80 | Lysine 6-dehydrogenase |
| *FEPFGCPE_02970* | *rcnA* | - | 3218422 | 3219540 | 1.02 | 4.22 | 5.01 | 3.80 | Nickel/cobalt efflux system RcnA |
| *FEPFGCPE_02971* | *leuC-2* | - | 3219662 | 3221692 | 1.18 | 3.91 | 1.51 | 0.37 | 3-isopropylmalate dehydratase large subunit |
| *FEPFGCPE_02972* | *tehA* | + | 3222003 | 3222989 | 11.85 | 10.60 | 9.67 | 3.37 | Tellurite resistance protein TehA |
| *FEPFGCPE_02973* | *sugE-2* | + | 3223071 | 3223397 | 2.01 | 7.22 | 8.61 | 0.00 | Quaternary ammonium compound-resistance protein SugE |
| *FEPFGCPE_02974* | *cdhR-2* | - | 3223477 | 3224436 | 3.26 | 12.04 | 9.80 | 3.25 | HTH-type transcriptional regulator CdhR |
| *FEPFGCPE_02975* | *htrC* | - | 3224901 | 3225482 | 2.22 | 2.83 | 4.98 | 0.36 | Heat shock protein C |
| *FEPFGCPE_02976* | *glmS-2* | - | 3225851 | 3227680 | 9.12 | 23.29 | 21.30 | 13.98 | Glutamine--fructose-6-phosphate aminotransferase [isomerizing] |
| *FEPFGCPE_02977* | *glmU* | - | 3227799 | 3229160 | 10.68 | 18.00 | 21.62 | 9.51 | Bifunctional protein GlmU |
| *FEPFGCPE_02978* | *ttcA* | - | 3229272 | 3230228 | 10.05 | 25.81 | 16.63 | 13.18 | tRNA 2-thiocytidine biosynthesis protein TtcA |

| Locus name | Gene name | Strand | Gene start (bp) | Gene end (bp) | Average TPM Inoculum | Average TPM $t_6$ | Average TPM $t_{24}$ | Average TPM $t_{168}$ | Annotation description |
|---|---|---|---|---|---|---|---|---|---|
| *FEPFGCPE_02979* | *folB* | - | 3230230 | 3230580 | 8.80 | 20.37 | 15.75 | 6.33 | Dihydroneopterin aldolase |
| *FEPFGCPE_02980* | *fabL* | - | 3230647 | 3231498 | 7.38 | 13.21 | 6.34 | 3.16 | Enoyl-[acyl-carrier-protein] reductase [NADPH] FabL |
| *FEPFGCPE_02981* | | + | 3231757 | 3232971 | 2.15 | 13.00 | 20.17 | 5.53 | Hypothetical protein |
| *FEPFGCPE_02982* | | + | 3233000 | 3233194 | 1.20 | 9.33 | 14.34 | 3.26 | Hypothetical protein |
| *FEPFGCPE_02983* | *gltI-4* | + | 3233482 | 3234384 | 0.66 | 1.01 | 1.55 | 0.22 | Glutamate/aspartate import solute-binding protein |
| *FEPFGCPE_02984* | *cca* | - | 3234444 | 3235694 | 2.32 | 18.70 | 12.08 | 4.16 | Multifunctional CCA protein |
| *FEPFGCPE_02985* | *sspA-1* | - | 3235691 | 3236335 | 4.66 | 26.10 | 19.74 | 3.61 | Stringent starvation protein A |
| *FEPFGCPE_02986* | | - | 3236400 | 3237350 | 5.41 | 21.54 | 11.74 | 3.56 | 3 beta-hydroxysteroid dehydrogenase/Delta 5--4-isomerase |
| *FEPFGCPE_02987* | *slt* | - | 3237402 | 3239408 | 6.39 | 31.01 | 22.18 | 9.56 | Soluble lytic murein transglycosylase |
| *FEPFGCPE_02988* | | + | 3239520 | 3240104 | 5.18 | 17.22 | 11.95 | 4.72 | 5-formyltetrahydrofolate cyclo-ligase |
| *FEPFGCPE_02989* | *kipI* | + | 3240101 | 3240754 | 3.75 | 15.17 | 6.36 | 2.30 | Kinase A inhibitor |
| *FEPFGCPE_02990* | *kipA* | + | 3240751 | 3241761 | 3.69 | 23.42 | 12.83 | 2.41 | KipI antagonist |
| *FEPFGCPE_02991* | | + | 3241804 | 3242550 | 3.07 | 21.44 | 13.85 | 7.75 | Hypothetical protein |
| *FEPFGCPE_02992* | *bauC-1* | - | 3242713 | 3244239 | 1.57 | 8.94 | 4.39 | 2.54 | Putative 3-oxopropanoate dehydrogenase |
| *FEPFGCPE_02993* | *bauA* | - | 3244362 | 3245690 | 2.19 | 19.19 | 10.41 | 2.60 | Beta-alanine--pyruvate aminotransferase |
| *FEPFGCPE_02994* | *lysN-4* | + | 3245879 | 3247327 | 9.66 | 5.86 | 5.64 | 1.36 | 2-aminoadipate transaminase |
| *FEPFGCPE_02995* | | - | 3247424 | 3248443 | 1.96 | 12.11 | 9.15 | 1.27 | Hypothetical protein |
| *FEPFGCPE_02996* | *basS* | - | 3248517 | 3249893 | 1.38 | 4.77 | 5.58 | 0.22 | Sensor protein BasS |
| *FEPFGCPE_02997* | *tctD-2* | - | 3249890 | 3250606 | 3.23 | 32.89 | 13.22 | 7.22 | Transcriptional regulatory protein tctD |
| *FEPFGCPE_02998* | | - | 3250983 | 3252032 | 0.66 | 11.25 | 5.96 | 2.57 | Hypothetical protein |
| *FEPFGCPE_02999* | *ssuC-6* | - | 3252118 | 3253008 | 0.34 | 1.11 | 1.74 | 0.00 | Putative aliphatic sulfonates transport permease protein SsuC |
| *FEPFGCPE_03000* | *tauB-5* | - | 3253005 | 3253847 | 0.00 | 1.07 | 0.00 | 0.00 | Taurine import ATP-binding protein TauB |
| *FEPFGCPE_03001* | | - | 3254216 | 3255706 | 0.54 | 2.12 | 1.92 | 0.99 | Hypothetical protein |
| *FEPFGCPE_03002* | | - | 3255877 | 3256041 | 0.14 | 0.88 | 0.08 | 1.88 | Hypothetical protein |
| *FEPFGCPE_03003* | *cdhR-3* | - | 3256351 | 3257337 | 2.34 | 14.55 | 11.94 | 2.69 | HTH-type transcriptional regulator CdhR |
| *FEPFGCPE_03004* | *rfnT* | - | 3257359 | 3258738 | 1.53 | 7.47 | 4.47 | 2.14 | Riboflavin transporter RfnT |
| *FEPFGCPE_03005* | | - | 3258731 | 3258931 | 0.00 | 0.00 | 0.00 | 0.00 | Hypothetical protein |
| *FEPFGCPE_03006* | *amiC-3* | + | 3259127 | 3260383 | 0.14 | 97.15 | 155.89 | 75.62 | Aliphatic amidase expression-regulating protein |
| *FEPFGCPE_03007* | *livH-10* | + | 3260446 | 3262068 | 0.03 | 8.53 | 17.64 | 10.04 | High-affinity branched-chain amino acid transport system permease protein LivH |
| *FEPFGCPE_03008* | | + | 3262069 | 3263271 | 0.13 | 18.50 | 31.37 | 12.61 | Hypothetical protein |
| *FEPFGCPE_03009* | *lptB-8* | + | 3263268 | 3264134 | 0.26 | 11.77 | 35.23 | 17.55 | Lipopolysaccharide export system ATP-binding protein LptB |
| *FEPFGCPE_03010* | *livF-9* | + | 3264137 | 3264838 | 0.24 | 27.30 | 49.84 | 16.76 | High-affinity branched-chain amino acid transport ATP-binding protein LivF |
| *FEPFGCPE_03011* | *ureD* | + | 3264916 | 3265767 | 0.33 | 8.23 | 20.67 | 3.58 | Urease accessory protein UreD |
| *FEPFGCPE_03012* | *fliY-4* | + | 3266111 | 3266974 | 0.90 | 34.92 | 76.52 | 57.77 | L-cystine-binding protein FliY |
| *FEPFGCPE_03013* | *artQ* | + | 3267015 | 3267701 | 1.19 | 8.39 | 25.09 | 12.98 | Arginine transport system permease protein ArtQ |
| *FEPFGCPE_03014* | *yecS-6* | + | 3267698 | 3268348 | 0.67 | 6.71 | 14.33 | 9.29 | L-cystine transport system permease protein YecS |
| *FEPFGCPE_03015* | *glnQ-6* | + | 3268335 | 3269057 | 0.45 | 21.34 | 32.08 | 15.60 | Glutamine transport ATP-binding protein GlnQ |
| *FEPFGCPE_03016* | *ureA* | + | 3269222 | 3269524 | 3.61 | 42.32 | 35.82 | 26.43 | Urease subunit gamma |
| *FEPFGCPE_03017* | | + | 3269650 | 3270333 | 0.61 | 19.33 | 28.84 | 16.47 | Hypothetical protein |
| *FEPFGCPE_03018* | *ureB* | + | 3270379 | 3270684 | 0.41 | 30.80 | 34.98 | 15.63 | Urease subunit beta |
| *FEPFGCPE_03019* | *ureC* | + | 3270681 | 3272381 | 3.19 | 35.59 | 41.15 | 24.01 | Urease subunit alpha |
| *FEPFGCPE_03020* | *ureE* | + | 3272385 | 3273260 | 1.20 | 6.81 | 12.54 | 7.13 | Urease accessory protein UreE |
| *FEPFGCPE_03021* | *ureF* | + | 3273229 | 3273954 | 0.68 | 9.83 | 13.57 | 3.04 | Urease accessory protein UreF |
| *FEPFGCPE_03022* | *ureG* | + | 3274061 | 3274699 | 0.84 | 20.52 | 23.01 | 18.09 | Urease accessory protein UreG |
| *FEPFGCPE_03023* | | - | 3274732 | 3275061 | 2.57 | 10.08 | 12.81 | 5.68 | Hypothetical protein |
| *FEPFGCPE_03024* | | - | 3275031 | 3275441 | 4.29 | 38.98 | 46.05 | 16.86 | Hypothetical protein |
| *FEPFGCPE_03025* | | - | 3275588 | 3276589 | 4.97 | 15.63 | 9.15 | 2.86 | Hypothetical protein |
| *FEPFGCPE_03026* | *oppF-2* | - | 3276586 | 3277632 | 10.21 | 38.59 | 20.45 | 4.59 | Oligopeptide transport ATP-binding protein OppF |
| *FEPFGCPE_03027* | *oppD-2* | - | 3277629 | 3278633 | 1.01 | 5.72 | 2.34 | 2.75 | Oligopeptide transport ATP-binding protein OppD |
| *FEPFGCPE_03028* | *dppC* | - | 3278630 | 3279550 | 0.67 | 2.63 | 0.93 | 1.67 | Dipeptide transport system permease protein DppC |
| *FEPFGCPE_03029* | *dppB-2* | - | 3279563 | 3280552 | 1.21 | 3.00 | 4.40 | 2.89 | Dipeptide transport system permease protein DppB |
| *FEPFGCPE_03030* | *dppA-3* | - | 3280797 | 3282437 | 8.25 | 49.50 | 29.30 | 19.00 | Periplasmic dipeptide transport protein |
| *FEPFGCPE_03031* | *metF* | - | 3282642 | 3283502 | 5.82 | 35.59 | 20.60 | 10.45 | 5,10-methylenetetrahydrofolate reductase |
| *FEPFGCPE_03032* | *tlyA* | - | 3283499 | 3284338 | 16.89 | 82.13 | 62.94 | 45.34 | 16S/23S rRNA (cytidine-2'-O)-methyltransferase TlyA |
| *FEPFGCPE_03033* | | - | 3284386 | 3284745 | 5.77 | 29.76 | 35.58 | 16.26 | Hypothetical protein |
| *FEPFGCPE_03034* | *ahcY* | - | 3284919 | 3286334 | 9.97 | 81.48 | 49.54 | 58.12 | Adenosylhomocysteinase |
| *FEPFGCPE_03035* | *cidB* | - | 3286601 | 3287284 | 0.94 | 1.62 | 1.46 | 2.17 | Holin-like protein CidB |
| *FEPFGCPE_03036* | *cidA* | - | 3287355 | 3287798 | 0.52 | 4.99 | 8.60 | 5.77 | Holin-like protein CidA |
| *FEPFGCPE_03037* | *gltC-7* | + | 3287995 | 3288939 | 0.96 | 4.05 | 0.65 | 0.50 | HTH-type transcriptional regulator GltC |
| *FEPFGCPE_03038* | *exoA* | + | 3289033 | 3289806 | 3.87 | 23.30 | 18.38 | 7.82 | Exodeoxyribonuclease |
| *FEPFGCPE_03039* | | - | 3289895 | 3291190 | 2.10 | 6.39 | 4.64 | 3.70 | Hypothetical protein |
| *FEPFGCPE_03040* | | - | 3291297 | 3291959 | 0.72 | 6.43 | 5.98 | 4.37 | Hypothetical protein |
| *FEPFGCPE_03041* | *metX* | - | 3291956 | 3293137 | 1.71 | 9.03 | 8.70 | 2.85 | Homoserine O-acetyltransferase |
| *FEPFGCPE_03042* | *slmA* | - | 3293446 | 3294102 | 1.91 | 19.94 | 7.64 | 2.13 | Nucleoid occlusion factor SlmA |
| *FEPFGCPE_03043* | *gph-2* | - | 3294113 | 3294916 | 3.15 | 19.37 | 5.24 | 6.96 | Phosphoglycolate phosphatase |
| *FEPFGCPE_03044* | | + | 3294952 | 3295107 | 2.64 | 53.77 | 9.25 | 7.50 | Hypothetical protein |
| *FEPFGCPE_03045* | | - | 3295187 | 3296266 | 0.10 | 0.92 | 3.52 | 0.73 | Outer membrane porin protein 32 |
| *FEPFGCPE_03046* | | - | 3296462 | 3296944 | 0.22 | 0.22 | 4.56 | 0.96 | Hypothetical protein |
| *FEPFGCPE_03047* | *mhpC-2* | - | 3296941 | 3297696 | 0.00 | 0.64 | 0.00 | 0.00 | 2-hydroxy-6-oxononadienedioate/2-hydroxy-6-oxononatrienedioate hydrolase |
| *FEPFGCPE_03048* | *galD* | - | 3297693 | 3298811 | 0.05 | 1.44 | 0.00 | 0.21 | 4-oxalomesaconate tautomerase |
| *FEPFGCPE_03049* | *proA-2* | - | 3298822 | 3299526 | 0.00 | 1.71 | 0.32 | 0.60 | 4-hydroxy-4-methyl-2-oxoglutarate aldolase/4-carboxy-4-hydroxy-2-oxoadipate aldolase |

| Locus name | Gene name | Strand | Gene start (bp) | Gene end (bp) | Average TPM Inoculum | Average TPM $t_6$ | Average TPM $t_{24}$ | Average TPM $t_{168}$ | Annotation description |
|---|---|---|---|---|---|---|---|---|---|
| *FEPFGCPE_03050* | *galB* | - | 3299523 | 3300278 | 0.00 | 2.04 | 0.00 | 0.00 | 4-oxalmesaconate hydratase |
| *FEPFGCPE_03051* | *livF-10* | - | 3300314 | 3301033 | 0.29 | 4.40 | 4.28 | 0.41 | High-affinity branched-chain amino acid transport ATP-binding protein LivF |
| *FEPFGCPE_03052* | *cysA-2* | - | 3301026 | 3301778 | 0.47 | 2.22 | 1.66 | 0.48 | Sulfate/thiosulfate import ATP-binding protein CysA |
| *FEPFGCPE_03053* | | - | 3301775 | 3302860 | 0.42 | 1.52 | 2.06 | 0.14 | Hypothetical protein |
| *FEPFGCPE_03054* | *livH-11* | - | 3302874 | 3303746 | 0.42 | 2.75 | 0.90 | 0.00 | High-affinity branched-chain amino acid transport system permease protein LivH |
| *FEPFGCPE_03055* | *amiC-4* | - | 3303784 | 3304980 | 0.15 | 0.71 | 0.40 | 0.32 | Aliphatic amidase expression-regulating protein |
| *FEPFGCPE_03056* | *tsaR-1* | - | 3305370 | 3306710 | 0.20 | 0.17 | 3.39 | 0.81 | HTH-type transcriptional regulator TsaR |
| *FEPFGCPE_03057* | *galA* | - | 3306730 | 3307089 | 0.89 | 5.08 | 1.78 | 0.69 | Gallate dioxygenase |
| *FEPFGCPE_03058* | *ligB* | - | 3307086 | 3307928 | 0.34 | 3.46 | 3.73 | 2.07 | Protocatechuate 4,5-dioxygenase beta chain |
| *FEPFGCPE_03059* | *yvoA-3* | + | 3308139 | 3308807 | 1.18 | 8.00 | 9.53 | 1.29 | HTH-type transcriptional repressor YvoA |
| *FEPFGCPE_03060* | *tsaM1* | + | 3308892 | 3309965 | 0.69 | 5.58 | 14.09 | 1.66 | Toluene-4-sulfonate monooxygenase system iron-sulfur subunit TsaM1 |
| *FEPFGCPE_03061* | *ophA1-2* | + | 3309962 | 3310906 | 0.19 | 3.22 | 4.04 | 0.00 | Phthalate dioxygenase reductase |
| *FEPFGCPE_03062* | *yhjE-4* | - | 3311022 | 3312389 | 0.72 | 2.11 | 2.28 | 0.29 | Inner membrane metabolite transport protein YhjE |
| *FEPFGCPE_03063* | *vdh-2* | - | 3312484 | 3313953 | 1.99 | 9.31 | 3.38 | 1.92 | Vanillin dehydrogenase |
| *FEPFGCPE_03064* | *mdlC-2* | - | 3314007 | 3315635 | 2.73 | 5.51 | 2.23 | 0.56 | Benzoylformate decarboxylase |
| *FEPFGCPE_03065* | *tsaR-2* | + | 3315761 | 3316792 | 1.61 | 4.56 | 1.83 | 1.63 | HTH-type transcriptional regulator TsaR |
| *FEPFGCPE_03066* | | - | 3316808 | 3317506 | 0.82 | 1.85 | 1.08 | 0.00 | Hypothetical protein |
| *FEPFGCPE_03067* | *argB* | - | 3317620 | 3318525 | 14.74 | 58.50 | 49.60 | 36.87 | Acetylglutamate kinase |
| *FEPFGCPE_03068* | | - | 3318891 | 3319169 | 20.04 | 81.88 | 69.17 | 41.11 | Hypothetical protein |
| *FEPFGCPE_03069* | | - | 3319189 | 3319716 | 3.00 | 16.47 | 22.62 | 9.23 | Hypothetical protein |
| *FEPFGCPE_03070* | *regB* | + | 3319958 | 3321226 | 11.01 | 8.59 | 21.51 | 9.02 | Sensor histidine kinase RegB |
| *FEPFGCPE_03071* | *regA* | + | 3321223 | 3321765 | 9.34 | 8.13 | 12.98 | 9.59 | Photosynthetic apparatus regulatory protein RegA |
| *FEPFGCPE_03072* | *hslU* | - | 3321841 | 3323199 | 5.14 | 21.44 | 23.12 | 21.15 | ATP-dependent protease ATPase subunit HslU |
| *FEPFGCPE_03073* | *hslV-2* | - | 3323307 | 3323858 | 18.55 | 61.07 | 66.08 | 41.12 | ATP-dependent protease subunit HslV |
| *FEPFGCPE_03074* | | - | 3324148 | 3326337 | 0.66 | 2.39 | 4.17 | 7.81 | putative TonB-dependent receptor |
| *FEPFGCPE_03075* | | - | 3326563 | 3327045 | 2.21 | 21.88 | 142.47 | 99.61 | Hypothetical protein |
| *FEPFGCPE_03076* | *dksA* | - | 3327282 | 3327956 | 102.41 | 456.39 | 350.82 | 101.90 | RNA polymerase-binding transcription factor DksA |
| *FEPFGCPE_03077* | *yjiA* | - | 3328092 | 3329159 | 19.45 | 60.38 | 50.22 | 22.10 | putative GTP-binding protein YjiA |
| *FEPFGCPE_03078* | | - | 3329251 | 3329874 | 16.70 | 73.03 | 52.29 | 20.60 | Hypothetical protein |
| *FEPFGCPE_03079* | | + | 3329890 | 3330765 | 2.20 | 9.83 | 9.01 | 0.70 | Hypothetical protein |
| *FEPFGCPE_03080* | *rlmI* | - | 3330785 | 3331999 | 1.91 | 7.89 | 3.87 | 4.33 | Ribosomal RNA large subunit methyltransferase I |
| *FEPFGCPE_03081* | *xerC-5* | - | 3332120 | 3333166 | 1.17 | 5.42 | 9.24 | 2.74 | Tyrosine recombinase XerC |
| *FEPFGCPE_03082* | | - | 3333225 | 3333893 | 2.87 | 25.89 | 38.90 | 17.75 | Hypothetical protein |
| *FEPFGCPE_03083* | *dapF* | - | 3334062 | 3334946 | 14.26 | 57.67 | 50.68 | 11.32 | Diaminopimelate epimerase |
| *FEPFGCPE_03084* | *lpxL-2* | - | 3334957 | 3335853 | 11.30 | 16.70 | 9.71 | 3.49 | Lipid A biosynthesis lauroyltransferase |
| *FEPFGCPE_03085* | *metK* | + | 3336042 | 3337220 | 10.76 | 91.96 | 51.74 | 28.67 | S-adenosylmethionine synthase |
| *FEPFGCPE_03086* | *calB* | - | 3337349 | 3338827 | 3.91 | 18.31 | 13.28 | 3.09 | Coniferyl aldehyde dehydrogenase |
| *FEPFGCPE_03087* | *aidB-2* | - | 3338898 | 3340562 | 15.51 | 49.70 | 23.71 | 9.60 | Putative acyl-CoA dehydrogenase AidB |
| *FEPFGCPE_03088* | *alkJ-2* | + | 3340853 | 3342532 | 5.49 | 15.09 | 6.38 | 1.73 | Alcohol dehydrogenase [acceptor] |
| *FEPFGCPE_03089* | *outO-1* | - | 3342503 | 3343396 | 0.89 | 10.57 | 1.21 | 0.60 | Type 4 prepilin-like proteins leader peptide-processing enzyme |
| *FEPFGCPE_03090* | *tcpE* | - | 3343396 | 3344487 | 0.44 | 6.48 | 5.04 | 0.93 | Toxin coregulated pilus biosynthesis protein E |
| *FEPFGCPE_03091* | *tcpT* | - | 3344530 | 3346248 | 0.26 | 3.71 | 1.68 | 1.08 | Toxin coregulated pilus biosynthesis protein T |
| *FEPFGCPE_03092* | | - | 3346232 | 3346693 | 0.61 | 2.75 | 1.04 | 1.02 | Hypothetical protein |
| *FEPFGCPE_03093* | | - | 3346683 | 3348047 | 0.31 | 2.77 | 0.47 | 0.55 | Hypothetical protein |
| *FEPFGCPE_03094* | *bfpB* | - | 3348049 | 3349830 | 0.35 | 2.01 | 1.96 | 0.77 | Outer membrane lipoprotein BfpB |
| *FEPFGCPE_03095* | | - | 3349883 | 3350440 | 0.84 | 1.66 | 0.67 | 0.00 | Hypothetical protein |
| *FEPFGCPE_03096* | | - | 3350496 | 3351137 | 1.11 | 9.07 | 1.43 | 0.44 | Hypothetical protein |
| *FEPFGCPE_03097* | | - | 3351635 | 3352441 | 1.79 | 12.80 | 14.99 | 1.60 | Hypothetical protein |
| *FEPFGCPE_03098* | *gatB* | - | 3352567 | 3354039 | 1.98 | 10.86 | 13.40 | 6.50 | Aspartyl/glutamyl-tRNA(Asn/Gln) amidotransferase subunit B |
| *FEPFGCPE_03099* | | - | 3354167 | 3354658 | 6.24 | 42.87 | 24.24 | 28.03 | Hypothetical protein |
| *FEPFGCPE_03100* | *gatA-2* | - | 3354661 | 3356151 | 2.01 | 12.99 | 10.20 | 6.69 | Glutamyl-tRNA(Gln) amidotransferase subunit A |
| *FEPFGCPE_03101* | *gatC* | - | 3356274 | 3356573 | 4.08 | 20.52 | 28.23 | 14.81 | Glutamyl-tRNA(Gln) amidotransferase subunit C |
| *FEPFGCPE_03102* | *mreB* | + | 3357124 | 3358167 | 8.34 | 26.51 | 25.52 | 16.40 | Rod shape-determining protein MreB |
| *FEPFGCPE_03103* | *mreC* | + | 3358352 | 3359320 | 0.97 | 8.95 | 2.57 | 0.85 | Cell shape-determining protein MreC |
| *FEPFGCPE_03104* | *mreD* | + | 3359330 | 3359842 | 1.06 | 5.51 | 3.97 | 3.09 | Rod shape-determining protein MreD |
| *FEPFGCPE_03105* | *mrdA* | + | 3359886 | 3362099 | 0.47 | 3.42 | 3.28 | 1.73 | Peptidoglycan D,D-transpeptidase MrdA |
| *FEPFGCPE_03106* | *mrdB* | + | 3362096 | 3363247 | 3.76 | 23.85 | 15.19 | 6.12 | Peptidoglycan glycosyltransferase MrdB |
| *FEPFGCPE_03107* | *esiB* | - | 3363244 | 3363957 | 9.76 | 37.28 | 37.35 | 11.20 | Secretory immunoglobulin A-binding protein EsiB |
| *FEPFGCPE_03108* | | - | 3364128 | 3364550 | 5.29 | 38.89 | 28.29 | 9.40 | Hypothetical protein |
| *FEPFGCPE_03109* | | - | 3364589 | 3364975 | 1.47 | 11.04 | 6.11 | 2.33 | Hypothetical protein |
| *FEPFGCPE_03110* | *queD* | - | 3365159 | 3365602 | 1.71 | 4.26 | 5.70 | 6.82 | 6-carboxy-5,6,7,8-tetrahydropterin synthase |
| *FEPFGCPE_03111* | *queE* | - | 3365612 | 3366247 | 5.42 | 22.71 | 15.69 | 5.96 | 7-carboxy-7-deazaguanine synthase |
| *FEPFGCPE_03112* | *queC* | - | 3366270 | 3367007 | 6.73 | 27.53 | 19.71 | 8.88 | 7-cyano-7-deazaguanine synthase |
| *FEPFGCPE_03113* | | - | 3367224 | 3367299 | 1.45 | 6.01 | 0.00 | 0.00 | tRNA |
| *FEPFGCPE_03114* | *glnG-2* | - | 3367405 | 3368145 | 10.48 | 61.15 | 34.18 | 22.51 | Nitrogen regulation protein NR(I) |
| *FEPFGCPE_03115* | *resE-1* | - | 3368146 | 3370497 | 4.89 | 24.63 | 15.69 | 4.97 | Sensor histidine kinase ResE |
| *FEPFGCPE_03116* | | - | 3370487 | 3371230 | 2.82 | 9.46 | 4.87 | 3.14 | Hypothetical protein |
| *FEPFGCPE_03117* | *rsmB-1* | - | 3371227 | 3372669 | 2.68 | 10.35 | 6.36 | 2.14 | Ribosomal RNA small subunit methyltransferase B |
| *FEPFGCPE_03118* | *htpX-2* | - | 3372713 | 3373573 | 38.73 | 32.08 | 35.98 | 20.18 | Protease HtpX |
| *FEPFGCPE_03119* | *fmt* | - | 3373785 | 3374768 | 1.94 | 9.29 | 12.11 | 3.32 | Methionyl-tRNA formyltransferase |
| *FEPFGCPE_03120* | | - | 3374798 | 3376546 | 2.47 | 8.42 | 4.09 | 3.44 | Hypothetical protein |

| Locus name | Gene name | Strand | Gene start (bp) | Gene end (bp) | Average TPM Inoculum | Average TPM $t_6$ | Average TPM $t_{24}$ | Average TPM $t_{168}$ | Annotation description |
|---|---|---|---|---|---|---|---|---|---|
| *FEPFGCPE_03121* | *def1* | - | 3376693 | 3377196 | 8.85 | 35.72 | 20.67 | 14.75 | Peptide deformylase 1 |
| *FEPFGCPE_03122* | *dprA* | + | 3377434 | 3378774 | 2.67 | 22.04 | 16.66 | 5.26 | DNA processing protein DprA |
| *FEPFGCPE_03123* | *trxC* | + | 3378866 | 3379255 | 2.88 | 23.16 | 18.55 | 2.41 | Putative thioredoxin 2 |
| *FEPFGCPE_03124* | *topB* | + | 3379481 | 3382168 | 6.11 | 48.37 | 34.64 | 13.95 | DNA topoisomerase 3 |
| *FEPFGCPE_03125* | *dmlR-15* | - | 3382267 | 3383235 | 2.34 | 5.98 | 5.79 | 2.84 | HTH-type transcriptional regulator DmlR |
| *FEPFGCPE_03126* | *hemN-2* | + | 3383659 | 3385200 | 1.54 | 1.14 | 0.93 | 0.25 | Oxygen-independent coproporphyrinogen III oxidase |
| *FEPFGCPE_03127* | | - | 3385307 | 3385504 | 0.63 | 0.00 | 3.55 | 4.15 | Hypothetical protein |
| *FEPFGCPE_03128* | *cstA* | - | 3385521 | 3387587 | 0.59 | 4.17 | 5.51 | 4.03 | Carbon starvation protein A |
| *FEPFGCPE_03129* | *nreB-3* | + | 3387952 | 3389337 | 0.88 | 11.38 | 5.36 | 0.58 | Oxygen sensor histidine kinase NreB |
| *FEPFGCPE_03130* | *nreC-5* | + | 3389334 | 3389966 | 0.83 | 10.86 | 4.10 | 4.55 | Oxygen regulatory protein NreC |
| *FEPFGCPE_03131* | *fitB* | - | 3390041 | 3390460 | 0.15 | 1.95 | 4.14 | 3.32 | Toxin FitB |
| *FEPFGCPE_03132* | | - | 3390476 | 3390682 | 1.73 | 6.61 | 11.53 | 1.82 | Hypothetical protein |
| *FEPFGCPE_03133* | *pckG* | - | 3390928 | 3392817 | 5.94 | 59.49 | 57.48 | 29.61 | Phosphoenolpyruvate carboxykinase [GTP] |
| *FEPFGCPE_03134* | | + | 3393159 | 3393383 | 23.64 | 103.41 | 76.21 | 12.30 | Hypothetical protein |
| *FEPFGCPE_03135* | *ghrB-3* | + | 3393380 | 3394390 | 9.59 | 28.02 | 22.71 | 6.37 | Glyoxylate/hydroxypyruvate reductase B |
| *FEPFGCPE_03136* | | - | 3394414 | 3395262 | 2.88 | 14.93 | 17.20 | 2.77 | Hypothetical protein |
| *FEPFGCPE_03137* | | - | 3395318 | 3395719 | 3.15 | 39.51 | 31.48 | 6.03 | Putative esterase |
| *FEPFGCPE_03138* | | - | 3395857 | 3396444 | 28.41 | 257.05 | 140.67 | 37.08 | Hypothetical protein |
| *FEPFGCPE_03139* | *catD-3* | + | 3396828 | 3397235 | 2.73 | 12.14 | 8.19 | 8.69 | Putative oxidoreductase CatD |
| *FEPFGCPE_03140* | | + | 3397471 | 3398031 | 1.37 | 47.56 | 57.31 | 36.86 | Hypothetical protein |
| *FEPFGCPE_03141* | | - | 3398089 | 3398775 | 0.96 | 14.88 | 11.35 | 13.65 | PKHD-type hydroxylase |
| *FEPFGCPE_03142* | *bfrD* | - | 3398791 | 3401025 | 0.62 | 7.66 | 9.27 | 11.11 | putative TonB-dependent receptor BfrD |
| *FEPFGCPE_03143* | | + | 3401553 | 3405311 | 3.10 | 9.63 | 6.33 | 4.18 | Hypothetical protein |
| *FEPFGCPE_03144* | | + | 3405436 | 3406017 | 2.51 | 6.56 | 1.56 | 3.21 | Hypothetical protein |
| *FEPFGCPE_03145* | *aguA* | + | 3406081 | 3407151 | 1.35 | 5.02 | 4.26 | 0.39 | Agmatine deiminase |
| *FEPFGCPE_03146* | *lutR-5* | - | 3407194 | 3408210 | 4.27 | 3.58 | 1.89 | 1.11 | HTH-type transcriptional regulator LutR |
| *FEPFGCPE_03147* | | + | 3408502 | 3409707 | 0.49 | 2.67 | 4.59 | 1.33 | Leu/Ile/Val-binding protein |
| *FEPFGCPE_03148* | *livH-12* | + | 3409860 | 3410717 | 0.46 | 2.48 | 1.55 | 1.43 | High-affinity branched-chain amino acid transport system permease protein LivH |
| *FEPFGCPE_03149* | | + | 3410717 | 3411706 | 0.87 | 2.49 | 1.71 | 0.47 | Hypothetical protein |
| *FEPFGCPE_03150* | *lptB-9* | + | 3411703 | 3412470 | 0.07 | 3.50 | 0.66 | 0.89 | Lipopolysaccharide export system ATP-binding protein LptB |
| *FEPFGCPE_03151* | *livF-11* | + | 3412467 | 3413168 | 0.00 | 1.37 | 0.77 | 0.88 | High-affinity branched-chain amino acid transport ATP-binding protein LivF |
| *FEPFGCPE_03152* | *amaB-2* | + | 3413295 | 3414557 | 0.28 | 2.09 | 1.61 | 1.05 | N-carbamoyl-L-amino acid hydrolase |
| *FEPFGCPE_03153* | | + | 3414554 | 3416053 | 0.34 | 4.59 | 3.54 | 0.63 | Hypothetical protein |
| *FEPFGCPE_03154* | *amiE* | - | 3416072 | 3417118 | 3.58 | 18.82 | 16.13 | 5.36 | Aliphatic amidase |
| *FEPFGCPE_03155* | | - | 3417198 | 3417551 | 3.48 | 26.20 | 23.75 | 4.09 | Hypothetical protein |
| *FEPFGCPE_03156* | *fmdA* | - | 3417604 | 3418842 | 2.90 | 14.13 | 10.42 | 2.73 | Formamidase |
| *FEPFGCPE_03157* | *livF-12* | - | 3418911 | 3419600 | 2.10 | 3.80 | 1.11 | 0.41 | High-affinity branched-chain amino acid transport ATP-binding protein LivF |
| *FEPFGCPE_03158* | *lptB-10* | - | 3419615 | 3420358 | 2.91 | 9.02 | 3.30 | 1.61 | Lipopolysaccharide export system ATP-binding protein LptB |
| *FEPFGCPE_03159* | | - | 3420355 | 3421539 | 0.23 | 0.67 | 0.30 | 0.80 | Hypothetical protein |
| *FEPFGCPE_03160* | *livH-13* | - | 3421598 | 3422515 | 0.20 | 1.41 | 0.39 | 1.10 | High-affinity branched-chain amino acid transport system permease protein LivH |
| *FEPFGCPE_03161* | *amiC-5* | - | 3422571 | 3423845 | 0.41 | 3.58 | 1.27 | 1.69 | Aliphatic amidase expression-regulating protein |
| *FEPFGCPE_03162* | *rpfC-2* | + | 3424043 | 3427477 | 0.16 | 1.46 | 2.57 | 1.02 | Sensory/regulatory protein RpfC |
| *FEPFGCPE_03163* | *pleD-2* | + | 3427440 | 3428426 | 2.39 | 14.23 | 7.34 | 2.93 | Response regulator PleD |
| *FEPFGCPE_03164* | *proP-13* | - | 3428491 | 3429801 | 7.00 | 7.77 | 10.40 | 5.46 | Proline/betaine transporter |
| *FEPFGCPE_03165* | *yxeP-4* | - | 3429798 | 3430988 | 12.23 | 16.20 | 9.72 | 4.32 | putative hydrolase YxeP |
| *FEPFGCPE_03166* | *gltC-8* | + | 3431256 | 3432176 | 1.95 | 4.38 | 2.54 | 0.37 | HTH-type transcriptional regulator GltC |
| *FEPFGCPE_03167* | | + | 3432262 | 3433689 | 2.78 | 15.33 | 9.02 | 3.62 | Hypothetical protein |
| *FEPFGCPE_03168* | *pitA-2* | + | 3433823 | 3434473 | 1.58 | 19.59 | 7.68 | 3.97 | L-methionine sulfoximine/L-methionine sulfone acetyltransferase |
| *FEPFGCPE_03169* | | - | 3434440 | 3435177 | 4.23 | 29.08 | 9.37 | 4.46 | Hypothetical protein |
| *FEPFGCPE_03170* | *cph2-4* | - | 3435196 | 3436239 | 5.03 | 21.58 | 15.08 | 5.70 | Phytochrome-like protein cph2 |
| *FEPFGCPE_03171* | *cheB-1* | - | 3436253 | 3437263 | 7.86 | 25.10 | 20.17 | 10.03 | Chemotaxis response regulator protein-glutamate methylesterase |
| *FEPFGCPE_03172* | *frzE-2* | - | 3437260 | 3439683 | 1.86 | 15.36 | 5.41 | 3.07 | Gliding motility regulatory protein |
| *FEPFGCPE_03173* | | - | 3439680 | 3440333 | 1.29 | 12.45 | 8.99 | 0.55 | Hypothetical protein |
| *FEPFGCPE_03174* | *wspC* | - | 3440330 | 3441808 | 3.25 | 16.89 | 6.32 | 2.79 | putative biofilm formation methyltransferase WspC |
| *FEPFGCPE_03175* | *cheW-1* | - | 3441805 | 3442293 | 2.14 | 13.48 | 9.30 | 1.91 | Chemotaxis protein CheW |
| *FEPFGCPE_03176* | *yoaH* | - | 3442329 | 3444002 | 9.88 | 39.69 | 22.81 | 7.05 | Putative methyl-accepting chemotaxis protein YoaH |
| *FEPFGCPE_03177* | *kce-2* | - | 3444158 | 3445009 | 1.55 | 6.00 | 3.29 | 2.00 | 3-keto-5-aminohexanoate cleavage enzyme |
| *FEPFGCPE_03178* | | - | 3445009 | 3446349 | 0.40 | 2.80 | 2.65 | 0.09 | putative aminotransferase |
| *FEPFGCPE_03179* | *hexR-4* | + | 3446615 | 3447547 | 1.93 | 4.28 | 7.17 | 2.44 | HTH-type transcriptional regulator HexR |
| *FEPFGCPE_03180* | | - | 3447531 | 3448748 | 0.87 | 4.85 | 5.84 | 0.86 | Hypothetical protein |
| *FEPFGCPE_03181* | *lysN-5* | + | 3448879 | 3450243 | 2.23 | 0.44 | 1.51 | 0.74 | 2-aminoadipate transaminase |
| *FEPFGCPE_03182* | | + | 3450374 | 3450787 | 16.96 | 77.55 | 40.33 | 26.96 | Hypothetical protein |
| *FEPFGCPE_03183* | *fsr* | - | 3450901 | 3452136 | 1.96 | 9.64 | 6.47 | 4.55 | Fosmidomycin resistance protein |
| *FEPFGCPE_03184* | *adrB-2* | + | 3452548 | 3454152 | 6.32 | 19.58 | 11.34 | 3.16 | Putative cyclic-di-GMP phosphodiesterase AdrB |
| *FEPFGCPE_03185* | | - | 3454235 | 3454471 | 28.61 | 82.29 | 36.02 | 17.61 | Hypothetical protein |
| *FEPFGCPE_03186* | *mopA* | + | 3454761 | 3455612 | 3.22 | 14.12 | 7.13 | 1.72 | Molybdenum-pterin-binding protein MopA |
| *FEPFGCPE_03187* | *ydhP-2* | - | 3455602 | 3456831 | 1.00 | 4.77 | 1.45 | 1.13 | Inner membrane transport protein YdhP |
| *FEPFGCPE_03188* | *dmlR-16* | + | 3457012 | 3457920 | 1.10 | 8.50 | 9.29 | 2.35 | HTH-type transcriptional regulator DmlR |
| *FEPFGCPE_03189* | | - | 3458020 | 3459324 | 0.46 | 2.79 | 0.78 | 1.95 | putative MFS-type transporter |
| *FEPFGCPE_03190* | *allS-1* | + | 3459557 | 3460450 | 1.31 | 3.75 | 3.05 | 3.11 | HTH-type transcriptional activator AllS |
| *FEPFGCPE_03191* | | + | 3460500 | 3460877 | 1.78 | 14.11 | 12.97 | 7.42 | Hypothetical protein |

| Locus name | Gene name | Strand | Gene start (bp) | Gene end (bp) | Average TPM Inoculum | Average TPM $t_6$ | Average TPM $t_{24}$ | Average TPM $t_{168}$ | Annotation description |
|---|---|---|---|---|---|---|---|---|---|
| *FEPFGCPE_03192* | | - | 3460940 | 3461293 | 1.47 | 8.97 | 5.47 | 2.15 | Hypothetical protein |
| *FEPFGCPE_03193* | | + | 3461485 | 3461907 | 5.62 | 16.14 | 13.62 | 5.46 | Hypothetical protein |
| *FEPFGCPE_03194* | | - | 3462042 | 3462944 | 0.42 | 7.08 | 4.69 | 0.00 | Hypothetical protein |
| *FEPFGCPE_03195* | *pepE-1* | - | 3462973 | 3463686 | 0.56 | 6.05 | 4.35 | 5.29 | Peptidase E |
| *FEPFGCPE_03196* | *glcB* | + | 3464149 | 3466329 | 1.86 | 10.73 | 12.52 | 19.42 | Malate synthase G |
| *FEPFGCPE_03197* | | + | 3466425 | 3467702 | 1.09 | 2.91 | 5.05 | 1.68 | Hypothetical protein |
| *FEPFGCPE_03198* | | + | 3468042 | 3468236 | 96.53 | 93.53 | 63.63 | 32.59 | Hypothetical protein |
| *FEPFGCPE_03199* | | + | 3468454 | 3468699 | 41.28 | 53.72 | 209.20 | 25.63 | Hypothetical protein |
| *FEPFGCPE_03200* | *yhjE-5* | - | 3468799 | 3470106 | 9.24 | 8.01 | 11.74 | 2.41 | Inner membrane metabolite transport protein YhjE |
| *FEPFGCPE_03201* | *mdh-2* | - | 3470307 | 3471308 | 0.76 | 0.85 | 1.44 | 1.46 | NAD-dependent methanol dehydrogenase |
| *FEPFGCPE_03202* | *cmpR-7* | + | 3471553 | 3472416 | 2.40 | 1.86 | 5.67 | 2.42 | HTH-type transcriptional activator CmpR |
| *FEPFGCPE_03203* | *uppP-1* | - | 3472480 | 3473331 | 2.81 | 13.37 | 11.93 | 3.90 | Undecaprenyl-diphosphatase |
| *FEPFGCPE_03204* | | - | 3473747 | 3474049 | 0.18 | 0.40 | 6.03 | 0.50 | Hypothetical protein |
| *FEPFGCPE_03205* | *uvrA-3* | - | 3474320 | 3480250 | 1.18 | 6.51 | 5.92 | 2.67 | UvrABC system protein A |
| *FEPFGCPE_03206* | | + | 3480523 | 3482190 | 1.97 | 16.30 | 14.26 | 3.65 | Hypothetical protein |
| *FEPFGCPE_03207* | *rutR* | - | 3482254 | 3482913 | 2.40 | 12.94 | 12.71 | 5.69 | HTH-type transcriptional regulator RutR |
| *FEPFGCPE_03208* | *ndh-2* | + | 3483069 | 3484361 | 6.32 | 5.45 | 5.95 | 7.23 | NADH dehydrogenase |
| *FEPFGCPE_03209* | | + | 3484508 | 3484840 | 2.38 | 5.11 | 2.57 | 4.70 | Hypothetical protein |
| *FEPFGCPE_03210* | *shlB* | + | 3485442 | 3487208 | 0.25 | 0.45 | 0.38 | 0.00 | Hemolysin transporter protein ShlB |
| *FEPFGCPE_03211* | *cdiA2-1* | + | 3487269 | 3496871 | 0.87 | 4.17 | 2.80 | 0.74 | tRNA nuclease CdiA-2 |
| *FEPFGCPE_03212* | | + | 3496871 | 3497149 | 6.30 | 30.84 | 32.50 | 8.50 | Hypothetical protein |
| *FEPFGCPE_03213* | *cdiA2-2* | + | 3497253 | 3499916 | 1.29 | 6.10 | 4.30 | 1.04 | tRNA nuclease CdiA-2 |
| *FEPFGCPE_03214* | | + | 3501458 | 3501709 | 0.49 | 5.61 | 5.76 | 3.27 | Hypothetical protein |
| *FEPFGCPE_03215* | *ampC* | + | 3502213 | 3503439 | 2.58 | 6.73 | 4.95 | 1.98 | Beta-lactamase |
| *FEPFGCPE_03216* | *metC-2* | - | 3503515 | 3504705 | 1.22 | 8.28 | 5.70 | 3.69 | Cystathionine beta-lyase |
| *FEPFGCPE_03217* | *glpE-2* | - | 3504801 | 3506402 | 0.29 | 1.43 | 0.87 | 0.88 | Thiosulfate sulfurtransferase GlpE |
| *FEPFGCPE_03218* | | - | 3506560 | 3507156 | 0.74 | 11.88 | 6.11 | 4.44 | Hypothetical protein |
| *FEPFGCPE_03219* | *glnQ-7* | - | 3507266 | 3508039 | 0.92 | 8.92 | 3.99 | 3.99 | Glutamine transport ATP-binding protein GlnQ |
| *FEPFGCPE_03220* | *glnP-1* | - | 3508069 | 3508761 | 0.42 | 2.10 | 0.93 | 0.91 | putative glutamine ABC transporter permease protein GlnP |
| *FEPFGCPE_03221* | *glnM-1* | - | 3508766 | 3509458 | 0.31 | 3.01 | 0.62 | 0.36 | putative glutamine ABC transporter permease protein GlnM |
| *FEPFGCPE_03222* | *glnH-1* | - | 3509474 | 3510298 | 0.51 | 7.22 | 3.47 | 2.02 | ABC transporter glutamine-binding protein GlnH |
| *FEPFGCPE_03223* | *gltC-9* | + | 3510487 | 3511365 | 0.27 | 6.03 | 1.88 | 0.86 | HTH-type transcriptional regulator GltC |
| *FEPFGCPE_03224* | *rcnR* | + | 3511480 | 3511749 | 3.60 | 23.65 | 24.02 | 11.16 | Transcriptional repressor RcnR |
| *FEPFGCPE_03225* | *natA* | + | 3511763 | 3512392 | 1.22 | 7.37 | 8.53 | 2.37 | ABC transporter ATP-binding protein NatA |
| *FEPFGCPE_03226* | *trg-2* | + | 3512590 | 3514458 | 1.01 | 3.33 | 2.57 | 0.41 | Methyl-accepting chemotaxis protein III |
| *FEPFGCPE_03227* | | + | 3514515 | 3514599 | 0.64 | 7.14 | 0.00 | 0.00 | tRNA |
| *FEPFGCPE_03228* | | - | 3514678 | 3515193 | 1.51 | 7.15 | 4.47 | 0.83 | Hypothetical protein |
| *FEPFGCPE_03229* | *qseC-3* | - | 3515193 | 3516590 | 2.06 | 12.29 | 8.20 | 1.26 | Sensor protein QseC |
| *FEPFGCPE_03230* | *qseB-1* | - | 3516566 | 3517231 | 3.33 | 20.05 | 16.06 | 4.01 | Transcriptional regulatory protein QseB |
| *FEPFGCPE_03231* | | - | 3517255 | 3518013 | 0.93 | 9.56 | 4.53 | 1.19 | Hypothetical protein |
| *FEPFGCPE_03232* | | - | 3518036 | 3518812 | 3.35 | 2.45 | 2.22 | 0.00 | Hypothetical protein |
| *FEPFGCPE_03233* | *ybjG-2* | - | 3518897 | 3519493 | 0.36 | 3.69 | 0.02 | 1.81 | Putative undecaprenyl-diphosphatase YbjG |
| *FEPFGCPE_03234* | *yceJ-2* | - | 3519764 | 3520324 | 4.23 | 16.52 | 7.74 | 2.43 | Cytochrome b561 |
| *FEPFGCPE_03235* | | - | 3520486 | 3520896 | 0.97 | 4.77 | 3.19 | 0.00 | Hypothetical protein |
| *FEPFGCPE_03236* | | - | 3521074 | 3523467 | 0.79 | 3.43 | 1.80 | 0.34 | Calcium-transporting ATPase 1 |
| *FEPFGCPE_03237* | | + | 3523812 | 3524360 | 4.97 | 14.01 | 6.32 | 3.34 | Hypothetical protein |
| *FEPFGCPE_03238* | | - | 3524652 | 3524927 | 0.74 | 2.52 | 6.75 | 1.09 | Hypothetical protein |
| *FEPFGCPE_03239* | *aziB2* | + | 3525235 | 3526233 | 4.05 | 20.80 | 25.16 | 3.19 | 3-hydroxy-5-methyl-1-naphthoate 3-O-methyltransferase |
| *FEPFGCPE_03240* | *mauC-2* | - | 3526367 | 3526699 | 1.16 | 6.25 | 6.50 | 5.03 | Amicyanin |
| *FEPFGCPE_03241* | | - | 3526703 | 3527239 | 2.63 | 5.16 | 7.84 | 4.34 | Hypothetical protein |
| *FEPFGCPE_03242* | *rpoE-1* | - | 3527290 | 3527994 | 2.07 | 6.95 | 7.75 | 2.81 | ECF RNA polymerase sigma-E factor |
| *FEPFGCPE_03243* | *acnB* | + | 3528288 | 3530873 | 0.61 | 11.46 | 14.73 | 20.71 | Aconitate hydratase B |
| *FEPFGCPE_03244* | *glpE-3* | - | 3531032 | 3534292 | 0.44 | 3.79 | 2.58 | 1.94 | Thiosulfate sulfurtransferase GlpE |
| *FEPFGCPE_03245* | *glnQ-8* | - | 3534337 | 3535122 | 0.07 | 1.62 | 1.14 | 0.78 | Glutamine transport ATP-binding protein GlnQ |
| *FEPFGCPE_03246* | *glnM-2* | - | 3535155 | 3535844 | 0.16 | 1.25 | 3.19 | 1.90 | putative glutamine ABC transporter permease protein GlnM |
| *FEPFGCPE_03247* | *glnM-3* | - | 3535850 | 3536545 | 0.42 | 4.52 | 3.19 | 0.67 | putative glutamine ABC transporter permease protein GlnM |
| *FEPFGCPE_03248* | *fliY-5* | - | 3536563 | 3537399 | 0.20 | 6.43 | 6.27 | 2.21 | L-cystine-binding protein FliY |
| *FEPFGCPE_03249* | *metC-3* | - | 3537432 | 3538598 | 0.55 | 6.72 | 3.59 | 0.59 | Cystathionine beta-lyase MetC |
| *FEPFGCPE_03250* | *gstB-2* | - | 3538752 | 3539360 | 0.45 | 2.09 | 4.02 | 1.91 | Glutathione S-transferase GstB |
| *FEPFGCPE_03251* | | - | 3539434 | 3540045 | 0.69 | 2.97 | 2.91 | 1.49 | Hypothetical protein |
| *FEPFGCPE_03252* | *cysB-2* | + | 3540157 | 3541083 | 3.93 | 14.22 | 12.63 | 5.48 | HTH-type transcriptional regulator CysB |
| *FEPFGCPE_03253* | | + | 3541489 | 3542613 | 0.67 | 4.12 | 4.89 | 0.74 | Outer membrane porin protein |
| *FEPFGCPE_03254* | *gltC-10* | - | 3542876 | 3543739 | 0.69 | 0.75 | 1.11 | 0.62 | HTH-type transcriptional regulator GltC |
| *FEPFGCPE_03255* | | + | 3543850 | 3544713 | 5.87 | 2.96 | 3.90 | 3.02 | Soluble epoxide hydrolase |
| *FEPFGCPE_03256* | | - | 3545064 | 3545408 | 80.64 | 5.28 | 19.85 | 5.98 | Hypothetical protein |
| *FEPFGCPE_03257* | *metC-4* | - | 3545535 | 3546722 | 58.92 | 6.59 | 7.82 | 6.47 | Cystathionine beta-lyase MetC |
| *FEPFGCPE_03258* | *decR-4* | + | 3546891 | 3547358 | 3.50 | 14.23 | 11.12 | 0.39 | DNA-binding transcriptional activator DecR |
| *FEPFGCPE_03259* | *dcyD* | - | 3547452 | 3548468 | 16.14 | 9.50 | 5.17 | 1.64 | D-cysteine desulfhydrase |
| *FEPFGCPE_03260* | *glnQ-9* | - | 3548527 | 3549252 | 55.59 | 9.94 | 7.65 | 4.19 | Glutamine transport ATP-binding protein GlnQ |
| *FEPFGCPE_03261* | *yecS-7* | - | 3549288 | 3549947 | 70.02 | 4.18 | 10.57 | 4.07 | L-cystine transport system permease protein YecS |
| *FEPFGCPE_03262* | *fliY-6* | - | 3550018 | 3550821 | 136.08 | 13.97 | 14.53 | 7.67 | L-cystine-binding protein FliY |

| Locus name | Gene name | Strand | Gene start (bp) | Gene end (bp) | Average TPM Inoculum | Average TPM $t_6$ | Average TPM $t_{24}$ | Average TPM $t_{168}$ | Annotation description |
|---|---|---|---|---|---|---|---|---|---|
| *FEPFGCPE_03263* | *pht3-2* | + | 3551757 | 3553088 | 0.43 | 0.37 | 2.59 | 0.00 | Phthalate 4,5-dioxygenase oxygenase subunit |
| *FEPFGCPE_03264* | *pht4-2* | + | 3553105 | 3554280 | 0.58 | 2.09 | 0.61 | 0.86 | Putative 4,5-dihydroxyphthalate dehydrogenase |
| *FEPFGCPE_03265* | *nadC-2* | - | 3554450 | 3555295 | 0.90 | 4.36 | 2.07 | 1.65 | Nicotinate-nucleotide pyrophosphorylase [carboxylating] |
| *FEPFGCPE_03266* | | - | 3555428 | 3556018 | 2.68 | 18.67 | 6.90 | 5.06 | Hypothetical protein |
| *FEPFGCPE_03267* | *pht5-2* | - | 3556144 | 3557136 | 0.28 | 1.54 | 0.00 | 0.04 | 4,5-dihydroxyphthalate decarboxylase |
| *FEPFGCPE_03268* | *ttuB-6* | - | 3557190 | 3558524 | 0.84 | 2.35 | 1.91 | 1.36 | Putative tartrate transporter |
| *FEPFGCPE_03269* | *ophA1-3* | + | 3558763 | 3559728 | 0.06 | 1.07 | 0.60 | 0.00 | Phthalate dioxygenase reductase |
| *FEPFGCPE_03270* | *dmlR-17* | - | 3559863 | 3560747 | 0.59 | 5.56 | 3.47 | 0.36 | HTH-type transcriptional regulator DmlR |
| *FEPFGCPE_03271* | *cpnA* | + | 3560940 | 3561704 | 0.87 | 0.59 | 1.02 | 1.14 | Cyclopentanol dehydrogenase |
| *FEPFGCPE_03272* | *yhhW-2* | + | 3561773 | 3562480 | 1.14 | 2.97 | 1.61 | 0.44 | Quercetin 2,3-dioxygenase |
| *FEPFGCPE_03273* | | - | 3562584 | 3563027 | 6.94 | 6.51 | 5.71 | 3.91 | putative HTH-type transcriptional regulator |
| *FEPFGCPE_03274* | *bpoC* | + | 3563132 | 3564049 | 0.19 | 1.59 | 2.16 | 0.43 | Putative non-heme bromoperoxidase BpoC |
| *FEPFGCPE_03275* | *pikAII* | + | 3564079 | 3565092 | 1.00 | 3.30 | 1.34 | 1.04 | Narbonolide/10-deoxymethynolide synthase PikA2, modules 3 and 4 |
| *FEPFGCPE_03276* | *isfD* | + | 3565149 | 3565937 | 0.46 | 2.91 | 4.09 | 1.48 | Sulfoacetaldehyde reductase |
| *FEPFGCPE_03277* | *nemA-1* | + | 3565988 | 3567055 | 1.27 | 6.98 | 6.33 | 1.28 | N-ethylmaleimide reductase |
| *FEPFGCPE_03278* | *gyrB* | - | 3567617 | 3570094 | 2.93 | 22.67 | 21.28 | 12.18 | DNA gyrase subunit B |
| *FEPFGCPE_03279* | *dnaN* | - | 3570273 | 3571379 | 6.11 | 35.47 | 32.59 | 16.97 | DNA polymerase III subunit beta |
| *FEPFGCPE_03280* | *dnaA* | - | 3571619 | 3572806 | 6.66 | 21.86 | 17.80 | 9.55 | Chromosomal replication initiator protein DnaA |
| *FEPFGCPE_03281* | | + | 3572959 | 3573783 | 7.35 | 56.77 | 44.48 | 14.72 | Hypothetical protein |
| *FEPFGCPE_03282* | *rpmH* | + | 3573967 | 3574101 | 8.25 | 22.02 | 73.06 | 27.92 | 50S ribosomal protein L34 |
| *FEPFGCPE_03283* | *rnpA* | + | 3574163 | 3574669 | 7.16 | 17.54 | 36.89 | 20.50 | Ribonuclease P protein component |
| *FEPFGCPE_03284* | *yidD* | + | 3574684 | 3575043 | 14.13 | 33.74 | 28.20 | 19.08 | Putative membrane protein insertion efficiency factor |
| *FEPFGCPE_03285* | *yidC* | + | 3575068 | 3576723 | 4.90 | 9.89 | 11.38 | 14.17 | Membrane protein insertase YidC |
| *FEPFGCPE_03286* | *mnmE* | + | 3576832 | 3578226 | 3.43 | 15.18 | 20.63 | 7.54 | tRNA modification GTPase MnmE |
| *FEPFGCPE_03287* | | + | 3578382 | 3578573 | 3.80 | 28.52 | 6.00 | 1.09 | Hypothetical protein |
| *FEPFGCPE_03288* | | + | 3578722 | 3579078 | 8.07 | 36.41 | 23.90 | 10.40 | Hypothetical protein |
| *FEPFGCPE_03289* | | + | 3579093 | 3579707 | 8.22 | 58.20 | 29.82 | 10.96 | Hypothetical protein |
| *FEPFGCPE_03290* | | + | 3579788 | 3580171 | 6.14 | 43.24 | 35.87 | 11.43 | Hypothetical protein |
| *FEPFGCPE_03291* | *thpR* | + | 3580329 | 3580913 | 9.19 | 69.00 | 39.60 | 10.30 | RNA 2',3'-cyclic phosphodiesterase |
| *FEPFGCPE_03292* | | + | 3580971 | 3581186 | 23.18 | 149.20 | 98.73 | 41.22 | Hypothetical protein |
| *FEPFGCPE_03293* | | - | 3581284 | 3582078 | 2.37 | 15.30 | 9.82 | 2.67 | putative signaling protein |
| *FEPFGCPE_03294* | *emrB-4* | + | 3582501 | 3584066 | 3.22 | 4.32 | 4.04 | 0.71 | Multidrug export protein EmrB |
| *FEPFGCPE_03295* | | + | 3584222 | 3585169 | 1.07 | 9.71 | 2.35 | 1.03 | putative membrane protein |
| *FEPFGCPE_03296* | *ohrR-1* | - | 3585251 | 3585706 | 3.19 | 17.26 | 13.24 | 3.65 | Organic hydroperoxide resistance transcriptional regulator |
| *FEPFGCPE_03297* | | + | 3585928 | 3586164 | 2.15 | 15.24 | 10.54 | 1.90 | Hypothetical protein |
| *FEPFGCPE_03298* | *clcB* | + | 3586237 | 3588012 | 2.52 | 15.15 | 9.85 | 4.27 | Voltage-gated ClC-type chloride channel ClcB |
| *FEPFGCPE_03299* | *mnmC-2* | + | 3588009 | 3590015 | 1.40 | 6.36 | 4.28 | 1.74 | tRNA 5-methylaminomethyl-2-thiouridine biosynthesis bifunctional protein MnmC |
| *FEPFGCPE_03300* | | + | 3590530 | 3591735 | 2.67 | 19.28 | 19.80 | 13.73 | Hypothetical protein |
| *FEPFGCPE_03301* | *ybdK* | + | 3591739 | 3592854 | 3.01 | 15.97 | 18.70 | 11.81 | Putative glutamate--cysteine ligase 2 |
| *FEPFGCPE_03302* | *lexA-2* | + | 3594393 | 3594929 | 230.05 | 3641.60 | 2947.16 | 460.56 | LexA repressor |
| *FEPFGCPE_03303* | | + | 3595091 | 3595543 | 7.80 | 48.15 | 27.61 | 6.12 | Hypothetical protein |
| *FEPFGCPE_03304* | *phhB* | - | 3595587 | 3595892 | 16.40 | 97.45 | 52.79 | 27.97 | Putative pterin-4-alpha-carbinolamine dehydratase |
| *FEPFGCPE_03305* | *phhA* | - | 3596182 | 3597027 | 25.39 | 81.17 | 67.41 | 29.75 | Phenylalanine-4-hydroxylase |
| *FEPFGCPE_03306* | *lrp-4* | + | 3597169 | 3597660 | 8.92 | 32.47 | 35.01 | 14.97 | Leucine-responsive regulatory protein |
| *FEPFGCPE_03307* | | - | 3597810 | 3598325 | 2.42 | 31.05 | 18.57 | 7.55 | Hypothetical protein |
| *FEPFGCPE_03308* | *yfhM* | + | 3598522 | 3604554 | 3.14 | 21.01 | 13.68 | 4.12 | putative lipoprotein YfhM |
| *FEPFGCPE_03309* | | + | 3604577 | 3605302 | 1.36 | 20.24 | 7.47 | 1.76 | putative membrane protein |
| *FEPFGCPE_03310* | *pbpC* | + | 3605286 | 3607493 | 2.34 | 11.06 | 7.26 | 2.02 | Penicillin-binding protein 1C |
| *FEPFGCPE_03311* | | + | 3607724 | 3607858 | 13.20 | 59.49 | 10.22 | 2.23 | Hypothetical protein |
| *FEPFGCPE_03312* | *ocd* | + | 3607855 | 3608697 | 8.68 | 29.07 | 9.69 | 8.95 | Delta(1)-pyrroline-2-carboxylate reductase 1 |
| *FEPFGCPE_03313* | *hmuV* | - | 3608768 | 3609628 | 0.98 | 8.71 | 11.57 | 20.95 | Hemin import ATP-binding protein HmuV |
| *FEPFGCPE_03314* | *hmuU-2* | - | 3609666 | 3610682 | 0.56 | 3.40 | 7.48 | 12.52 | Hemin transport system permease protein HmuU |
| *FEPFGCPE_03315* | *hmuT* | - | 3610862 | 3611791 | 0.92 | 8.69 | 24.20 | 58.67 | Hemin-binding periplasmic protein HmuT |
| *FEPFGCPE_03316* | *hemS* | - | 3611788 | 3612915 | 0.84 | 10.12 | 24.32 | 77.37 | Hemin transport protein HemS |
| *FEPFGCPE_03317* | *hemR* | - | 3612991 | 3615414 | 1.76 | 10.59 | 19.34 | 50.97 | Hemin receptor |
| *FEPFGCPE_03318* | *lrp-5* | - | 3615679 | 3616140 | 4.04 | 4.24 | 7.09 | 0.59 | Leucine-responsive regulatory protein |
| *FEPFGCPE_03319* | | + | 3616366 | 3619983 | 2.88 | 7.33 | 5.70 | 2.07 | Hypothetical protein |
| *FEPFGCPE_03320* | *mdtL* | - | 3620112 | 3621170 | 2.10 | 6.82 | 6.26 | 1.01 | Multidrug resistance protein MdtL |
| *FEPFGCPE_03321* | *trpS2* | + | 3621744 | 3622772 | 1.78 | 19.70 | 10.22 | 7.00 | Tryptophan--tRNA ligase 2 |
| *FEPFGCPE_03322* | | - | 3622874 | 3623236 | 1.41 | 3.04 | 3.06 | 0.54 | Hypothetical protein |
| *FEPFGCPE_03323* | | - | 3623291 | 3624463 | 0.83 | 5.53 | 1.69 | 1.49 | Hypothetical protein |
| *FEPFGCPE_03324* | *lhyD* | - | 3624692 | 3625924 | 2.61 | 27.40 | 21.89 | 8.53 | L-hydantoinase |
| *FEPFGCPE_03325* | *proP-14* | - | 3625989 | 3627269 | 0.79 | 6.56 | 3.28 | 2.23 | Proline/betaine transporter |
| *FEPFGCPE_03326* | *gcvA-6* | - | 3627495 | 3628397 | 2.32 | 12.57 | 9.69 | 2.88 | Glycine cleavage system transcriptional activator |
| *FEPFGCPE_03327* | *fumC* | - | 3628436 | 3629824 | 7.37 | 28.03 | 24.11 | 11.19 | Fumarate hydratase class II |
| *FEPFGCPE_03328* | *cynR-7* | - | 3630060 | 3630935 | 1.27 | 2.96 | 3.09 | 1.10 | HTH-type transcriptional regulator CynR |
| *FEPFGCPE_03329* | *bauC-2* | + | 3631067 | 3632659 | 2.24 | 17.55 | 13.30 | 14.49 | Putative 3-oxopropanoate dehydrogenase |
| *FEPFGCPE_03330* | *alkJ-3* | + | 3632827 | 3634461 | 4.19 | 7.54 | 6.30 | 6.80 | Alcohol dehydrogenase [acceptor] |
| *FEPFGCPE_03331* | *ydhC-2* | - | 3634403 | 3635578 | 2.41 | 16.16 | 5.65 | 1.64 | Inner membrane transport protein YdhC |
| *FEPFGCPE_03332* | *opuBA* | + | 3636100 | 3636879 | 13.67 | 34.48 | 25.90 | 11.89 | Choline transport ATP-binding protein OpuBA |
| *FEPFGCPE_03333* | *livF-13* | + | 3636888 | 3637649 | 8.53 | 26.70 | 12.16 | 8.05 | High-affinity branched-chain amino acid transport ATP-binding protein LivF |

| Locus name | Gene name | Strand | Gene start (bp) | Gene end (bp) | Average TPM Inoculum | Average TPM $t_6$ | Average TPM $t_{24}$ | Average TPM $t_{168}$ | Annotation description |
|---|---|---|---|---|---|---|---|---|---|
| *FEPFGCPE_03334* | | + | 3637720 | 3638925 | 17.52 | 59.05 | 32.01 | 24.58 | Hypothetical protein |
| *FEPFGCPE_03335* | *livH-14* | + | 3639002 | 3639886 | 5.36 | 23.83 | 11.47 | 4.50 | High-affinity branched-chain amino acid transport system permease protein LivH |
| *FEPFGCPE_03336* | | + | 3639898 | 3640905 | 5.99 | 17.36 | 10.10 | 6.46 | Hypothetical protein |
| *FEPFGCPE_03337* | | + | 3641334 | 3642116 | 0.79 | 4.24 | 2.36 | 0.85 | Hypothetical protein |
| *FEPFGCPE_03338* | | + | 3642360 | 3643505 | 8.78 | 36.39 | 25.13 | 23.61 | Leu/Ile/Val-binding protein |
| *FEPFGCPE_03339* | *ohrR-2* | + | 3643763 | 3644263 | 5.43 | 11.64 | 4.78 | 2.92 | Organic hydroperoxide resistance transcriptional regulator |
| *FEPFGCPE_03340* | *livH-15* | + | 3644556 | 3645611 | 2.17 | 5.79 | 3.28 | 2.62 | High-affinity branched-chain amino acid transport system permease protein LivH |
| *FEPFGCPE_03341* | *lptB-11* | + | 3645630 | 3647426 | 6.03 | 10.02 | 5.36 | 4.04 | Lipopolysaccharide export system ATP-binding protein LptB |
| *FEPFGCPE_03342* | *livF-14* | + | 3647426 | 3648172 | 5.99 | 7.67 | 3.52 | 4.46 | High-affinity branched-chain amino acid transport ATP-binding protein LivF |
| *FEPFGCPE_03343* | *mnmG* | + | 3649484 | 3651454 | 3.77 | 17.06 | 21.17 | 8.88 | tRNA uridine 5-carboxymethylaminomethyl modification enzyme MnmG |
| *FEPFGCPE_03344* | *rsmG* | + | 3651467 | 3652147 | 1.20 | 6.19 | 5.05 | 6.62 | Ribosomal RNA small subunit methyltransferase G |
| *FEPFGCPE_03345* | *blc-2* | + | 3652144 | 3652770 | 6.36 | 41.20 | 28.31 | 14.08 | Outer membrane lipoprotein Blc |
| *FEPFGCPE_03346* | *soj-3* | + | 3652940 | 3653710 | 6.31 | 22.86 | 27.83 | 12.81 | Sporulation initiation inhibitor protein Soj |
| *FEPFGCPE_03347* | *parB* | + | 3653738 | 3654637 | 1.65 | 10.11 | 7.61 | 3.11 | putative chromosome-partitioning protein ParB |
| *FEPFGCPE_03348* | *ybiR* | + | 3654662 | 3655909 | 1.60 | 12.72 | 6.27 | 1.52 | Inner membrane protein YbiR |
| *FEPFGCPE_03349* | | + | 3656154 | 3656657 | 21.86 | 63.28 | 47.90 | 22.40 | Hypothetical protein |
| *FEPFGCPE_03350* | *atpB-2* | + | 3656727 | 3657575 | 12.42 | 45.42 | 54.98 | 31.87 | ATP synthase subunit a |
| *FEPFGCPE_03351* | *atpE-2* | + | 3657661 | 3657927 | 17.68 | 67.44 | 91.71 | 77.96 | ATP synthase subunit c |
| *FEPFGCPE_03352* | *atpF-2* | + | 3658053 | 3658523 | 14.83 | 112.33 | 87.86 | 115.31 | ATP synthase subunit b |
| *FEPFGCPE_03353* | *atpH* | + | 3658526 | 3659056 | 16.53 | 120.92 | 127.77 | 96.53 | ATP synthase subunit delta |
| *FEPFGCPE_03354* | *atpA-2* | + | 3659107 | 3660648 | 12.77 | 65.95 | 69.38 | 70.79 | ATP synthase subunit alpha |
| *FEPFGCPE_03355* | *atpG-2* | + | 3660737 | 3661624 | 8.07 | 51.12 | 44.97 | 43.21 | ATP synthase gamma chain |
| *FEPFGCPE_03356* | *atpD1* | + | 3661686 | 3663077 | 9.25 | 70.13 | 66.36 | 56.90 | ATP synthase subunit beta 1 |
| *FEPFGCPE_03357* | *atpC-3* | + | 3663225 | 3663641 | 3.99 | 38.18 | 29.88 | 30.49 | ATP synthase epsilon chain |
| *FEPFGCPE_03358* | *fadD3-2* | + | 3664035 | 3665753 | 8.99 | 25.90 | 23.15 | 11.91 | 3-[(3aS,4S,7aS)-7a-methyl-1,5-dioxo-octahydro-1H-inden-4-yl]propanoyl:CoA ligase |
| *FEPFGCPE_03359* | *bcr-2* | + | 3665858 | 3667069 | 3.08 | 4.51 | 14.41 | 2.47 | Bicyclomycin resistance protein |
| *FEPFGCPE_03360* | | - | 3667116 | 3667307 | 121.96 | 68.88 | 118.34 | 40.76 | Hypothetical protein |
| *FEPFGCPE_03361* | *flhD* | + | 3667627 | 3667944 | 4.15 | 22.90 | 6.51 | 0.74 | Flagellar transcriptional regulator FlhD |
| *FEPFGCPE_03362* | *flhC* | + | 3668010 | 3668585 | 2.30 | 8.92 | 5.71 | 1.48 | Flagellar transcriptional regulator FlhC |
| *FEPFGCPE_03363* | *motA* | + | 3668702 | 3669562 | 2.77 | 12.63 | 3.37 | 0.58 | Motility protein A |
| *FEPFGCPE_03364* | *motB* | + | 3669575 | 3670552 | 3.97 | 7.53 | 3.52 | 0.34 | Motility protein B |
| *FEPFGCPE_03365* | *cheY-2* | + | 3670570 | 3670953 | 1.79 | 10.16 | 6.38 | 0.00 | Chemotaxis protein CheY |
| *FEPFGCPE_03366* | | - | 3671301 | 3671915 | 1.07 | 2.54 | 1.19 | 0.00 | Hypothetical protein |
| *FEPFGCPE_03367* | *cheA* | + | 3672617 | 3673213 | 1.94 | 1.94 | 3.18 | 0.00 | Chemotaxis protein CheA |
| *FEPFGCPE_03368* | *cheW-2* | + | 3673231 | 3673746 | 2.09 | 10.34 | 5.59 | 0.35 | Chemotaxis protein CheW |
| *FEPFGCPE_03369* | *tar-3* | + | 3673819 | 3676158 | 1.28 | 4.43 | 1.24 | 0.18 | Methyl-accepting chemotaxis protein II |
| *FEPFGCPE_03370* | *tsr-4* | + | 3676230 | 3678005 | 1.28 | 4.87 | 1.64 | 0.51 | Methyl-accepting chemotaxis protein I |
| *FEPFGCPE_03371* | *tar-4* | + | 3678098 | 3679720 | 1.75 | 7.09 | 3.56 | 0.97 | Methyl-accepting chemotaxis protein II |
| *FEPFGCPE_03372* | *cheR* | + | 3679756 | 3680697 | 0.71 | 3.46 | 2.89 | 0.78 | Chemotaxis protein methyltransferase |
| *FEPFGCPE_03373* | *cheD-1* | + | 3680699 | 3681373 | 1.86 | 11.24 | 8.19 | 4.58 | Chemoreceptor glutamine deamidase CheD |
| *FEPFGCPE_03374* | *cheB-2* | + | 3681370 | 3682449 | 1.11 | 13.15 | 3.82 | 1.65 | Chemotaxis response regulator protein-glutamate methylesterase |
| *FEPFGCPE_03375* | *cheY-3* | + | 3682585 | 3682983 | 3.02 | 42.87 | 17.93 | 13.24 | Chemotaxis protein CheY |
| *FEPFGCPE_03376* | | + | 3682986 | 3683660 | 2.94 | 15.99 | 7.64 | 3.56 | Hypothetical protein |
| *FEPFGCPE_03377* | *flhB-1* | + | 3683847 | 3684998 | 1.14 | 4.66 | 3.23 | 0.31 | Flagellar biosynthetic protein FlhB |
| *FEPFGCPE_03378* | *flhA-1* | + | 3685002 | 3687107 | 0.55 | 3.05 | 1.39 | 0.42 | Flagellar biosynthesis protein FlhA |
| *FEPFGCPE_03379* | *flhF* | + | 3687104 | 3688378 | 0.80 | 4.36 | 2.05 | 0.22 | Flagellar biosynthesis protein FlhF |
| *FEPFGCPE_03380* | *ylxH* | + | 3688371 | 3689243 | 3.08 | 11.25 | 1.53 | 0.00 | Flagellum site-determining protein YlxH |
| *FEPFGCPE_03381* | *fliA* | + | 3689281 | 3690009 | 2.26 | 10.02 | 2.19 | 2.82 | RNA polymerase sigma factor FliA |
| *FEPFGCPE_03382* | | - | 3690084 | 3690347 | 2.78 | 8.09 | 3.19 | 1.46 | Hypothetical protein |
| *FEPFGCPE_03383* | | + | 3690856 | 3691182 | 1.44 | 10.13 | 6.15 | 0.98 | Hypothetical protein |
| *FEPFGCPE_03384* | *flgN* | - | 3691200 | 3691658 | 4.75 | 11.57 | 12.87 | 4.54 | Flagella synthesis protein FlgN |
| *FEPFGCPE_03385* | *flgM* | - | 3691655 | 3691993 | 9.94 | 37.80 | 21.10 | 10.74 | Negative regulator of flagellin synthesis |
| *FEPFGCPE_03386* | *flgA* | - | 3692120 | 3693148 | 1.30 | 3.32 | 1.50 | 0.30 | Flagella basal body P-ring formation protein FlgA |
| *FEPFGCPE_03387* | *flgB* | + | 3693551 | 3693988 | 2.96 | 8.26 | 0.41 | 0.00 | Flagellar basal body rod protein FlgB |
| *FEPFGCPE_03388* | *flgC* | + | 3693994 | 3694398 | 2.09 | 10.20 | 0.00 | 0.00 | Flagellar basal-body rod protein FlgC |
| *FEPFGCPE_03389* | *flgD* | + | 3694421 | 3695146 | 1.88 | 6.27 | 1.22 | 0.00 | Basal-body rod modification protein FlgD |
| *FEPFGCPE_03390* | *flgE* | + | 3695235 | 3696491 | 2.88 | 17.75 | 1.72 | 1.95 | Flagellar hook protein FlgE |
| *FEPFGCPE_03391* | *flgF* | + | 3696533 | 3697276 | 1.93 | 13.48 | 2.71 | 0.78 | Flagellar basal-body rod protein FlgF |
| *FEPFGCPE_03392* | *flgG* | + | 3697344 | 3698126 | 1.22 | 7.13 | 1.34 | 0.75 | Flagellar basal-body rod protein FlgG |
| *FEPFGCPE_03393* | *flgH-1* | + | 3698154 | 3698879 | 2.21 | 6.81 | 0.63 | 0.70 | Flagellar L-ring protein |
| *FEPFGCPE_03394* | *flgI* | + | 3698879 | 3700054 | 0.38 | 2.88 | 0.70 | 0.00 | Flagellar P-ring protein |
| *FEPFGCPE_03395* | *flgJ* | + | 3700056 | 3700940 | 0.88 | 4.28 | 0.65 | 0.93 | Peptidoglycan hydrolase FlgJ |
| *FEPFGCPE_03396* | *flgK* | + | 3701077 | 3703065 | 3.71 | 8.58 | 3.55 | 0.39 | Flagellar hook-associated protein 1 |
| *FEPFGCPE_03397* | *flgL* | + | 3703082 | 3704317 | 3.75 | 6.32 | 3.71 | 0.10 | Flagellar hook-associated protein 3 |
| *FEPFGCPE_03398* | | - | 3704445 | 3705236 | 0.66 | 5.01 | 0.42 | 0.15 | Hypothetical protein |
| *FEPFGCPE_03399* | | - | 3705249 | 3705518 | 1.08 | 2.64 | 0.67 | 0.00 | Hypothetical protein |
| *FEPFGCPE_03400* | *fliP-2* | - | 3705533 | 3706327 | 2.56 | 9.61 | 1.78 | 0.04 | Flagellar biosynthetic protein FliP |
| *FEPFGCPE_03401* | *fliO* | - | 3706327 | 3706818 | 3.35 | 15.47 | 9.07 | 3.00 | Flagellar protein FliO |
| *FEPFGCPE_03402* | *fliN* | - | 3706815 | 3707243 | 3.50 | 3.73 | 1.92 | 2.50 | Flagellar motor switch protein FliN |
| *FEPFGCPE_03403* | *fliM* | - | 3707243 | 3708241 | 2.41 | 4.70 | 1.07 | 0.14 | Flagellar motor switch protein FliM |
| *FEPFGCPE_03404* | | - | 3708249 | 3708683 | 3.61 | 10.83 | 0.81 | 0.00 | Hypothetical protein |

| Locus name | Gene name | Strand | Gene start (bp) | Gene end (bp) | Average TPM Inoculum | Average TPM $t_6$ | Average TPM $t_{24}$ | Average TPM $t_{168}$ | Annotation description |
|---|---|---|---|---|---|---|---|---|---|
| *FEPFGCPE_03405* | *fliK* | - | 3708905 | 3710341 | 0.90 | 6.94 | 1.63 | 1.00 | Flagellar hook-length control protein |
| *FEPFGCPE_03406* | *fliJ* | - | 3710361 | 3710813 | 1.20 | 5.11 | 3.34 | 1.26 | Flagellar FliJ protein |
| *FEPFGCPE_03407* | *fliI* | - | 3710826 | 3712601 | 0.73 | 5.40 | 1.40 | 0.26 | Flagellum-specific ATP synthase |
| *FEPFGCPE_03408* | *fliH* | - | 3712595 | 3713254 | 1.94 | 7.75 | 0.00 | 0.11 | Flagellar assembly protein FliH |
| *FEPFGCPE_03409* | *fliG* | - | 3713276 | 3714274 | 0.80 | 4.13 | 1.03 | 0.72 | Flagellar motor switch protein FliG |
| *FEPFGCPE_03410* | *fliF* | - | 3714261 | 3715946 | 1.88 | 6.51 | 4.72 | 0.45 | Flagellar M-ring protein |
| *FEPFGCPE_03411* | *fliE* | + | 3716297 | 3716632 | 3.42 | 10.14 | 9.77 | 3.53 | Flagellar hook-basal body complex protein FliE |
| *FEPFGCPE_03412* | | - | 3716797 | 3718059 | 0.77 | 3.20 | 1.91 | 0.11 | Hypothetical protein |
| *FEPFGCPE_03413* | *flhB-2* | - | 3718236 | 3718628 | 2.54 | 6.34 | 4.25 | 0.00 | Flagellar biosynthetic protein FlhB |
| *FEPFGCPE_03414* | | - | 3718625 | 3720058 | 1.39 | 7.63 | 1.09 | 0.72 | Hypothetical protein |
| *FEPFGCPE_03415* | *fliT* | - | 3720103 | 3720441 | 2.63 | 15.53 | 9.85 | 2.52 | Flagellar protein FliT |
| *FEPFGCPE_03416* | *fliS* | - | 3720434 | 3720868 | 1.04 | 3.81 | 4.31 | 2.71 | Flagellar protein FliS |
| *FEPFGCPE_03417* | *fliD* | - | 3720908 | 3723013 | 4.31 | 13.65 | 5.42 | 1.60 | Flagellar hook-associated protein 2 |
| *FEPFGCPE_03418* | *fliC* | - | 3723170 | 3724582 | 13.40 | 77.16 | 20.18 | 7.62 | Flagellin |
| *FEPFGCPE_03419* | *uvrY-2* | - | 3725075 | 3725734 | 6.46 | 26.74 | 7.69 | 3.91 | Response regulator UvrY |
| *FEPFGCPE_03420* | *degS* | - | 3725731 | 3726612 | 18.15 | 41.98 | 27.65 | 7.72 | Signal transduction histidine-protein kinase/phosphatase DegS |
| *FEPFGCPE_03421* | *rssA* | - | 3726983 | 3728377 | 2.31 | 12.83 | 7.22 | 1.19 | Swarming motility regulation sensor protein RssA |
| *FEPFGCPE_03422* | *tctD-3* | - | 3728370 | 3729161 | 2.96 | 12.73 | 7.60 | 2.36 | Transcriptional regulatory protein tctD |
| *FEPFGCPE_03423* | | + | 3729326 | 3730375 | 59.55 | 55.61 | 54.53 | 31.72 | Hypothetical protein |
| *FEPFGCPE_03424* | *cmpC-2* | + | 3730545 | 3731378 | 8.53 | 16.99 | 13.09 | 8.36 | Bicarbonate transport ATP-binding protein CmpC |
| *FEPFGCPE_03425* | *ribX-3* | + | 3731467 | 3732330 | 4.31 | 5.57 | 9.70 | 5.44 | Riboflavin transport system permease protein RibX |
| *FEPFGCPE_03426* | | - | 3732460 | 3733062 | 130.04 | 509.30 | 512.62 | 188.22 | Hypothetical protein |
| *FEPFGCPE_03427* | | - | 3733304 | 3733837 | 34.62 | 113.30 | 82.68 | 33.32 | Hypothetical protein |
| *FEPFGCPE_03428* | *baeR* | - | 3733880 | 3734791 | 0.55 | 6.05 | 2.46 | 0.00 | Transcriptional regulatory protein BaeR |
| *FEPFGCPE_03429* | *baeS* | - | 3734791 | 3735978 | 0.52 | 3.34 | 3.02 | 0.14 | Signal transduction histidine-protein kinase BaeS |
| *FEPFGCPE_03430* | *flgH-2* | + | 3736198 | 3736896 | 1.64 | 1.50 | 1.79 | 0.00 | Flagellar L-ring protein |
| *FEPFGCPE_03431* | | + | 3736896 | 3737429 | 1.29 | 7.57 | 5.76 | 1.03 | Hypothetical protein |
| *FEPFGCPE_03432* | *bmaC* | - | 3737472 | 3741146 | 7.80 | 34.14 | 22.14 | 8.27 | Adhesin BmaC autotransporter |
| *FEPFGCPE_03433* | | - | 3741783 | 3741858 | 10.72 | 13.65 | 9.20 | 4.33 | tRNA |
| *FEPFGCPE_03434* | | + | 3742240 | 3742893 | 6.13 | 28.89 | 14.18 | 6.84 | Cytochrome c-555 |
| *FEPFGCPE_03435* | | + | 3742844 | 3743137 | 6.61 | 28.41 | 11.50 | 8.87 | Cytochrome c-555 |
| *FEPFGCPE_03436* | *cpdA-3* | - | 3743284 | 3744060 | 9.12 | 11.12 | 15.45 | 1.36 | 3',5'-cyclic adenosine monophosphate phosphodiesterase CpdA |
| *FEPFGCPE_03437* | *rep* | - | 3744540 | 3746645 | 4.48 | 17.74 | 14.29 | 8.98 | ATP-dependent DNA helicase Rep |
| *FEPFGCPE_03438* | | + | 3746913 | 3747728 | 25.32 | 50.26 | 24.39 | 9.37 | Hypothetical protein |
| *FEPFGCPE_03439* | *gcvT* | + | 3748259 | 3749365 | 21.74 | 38.73 | 33.75 | 16.54 | Aminomethyltransferase |
| *FEPFGCPE_03440* | *gcvH* | + | 3749443 | 3749826 | 46.59 | 118.06 | 92.89 | 41.12 | Glycine cleavage system H protein |
| *FEPFGCPE_03441* | *gcvP* | + | 3749924 | 3752851 | 11.43 | 13.53 | 12.61 | 6.00 | Glycine dehydrogenase (decarboxylating) |
| *FEPFGCPE_03442* | *sdaA* | + | 3752991 | 3754400 | 8.31 | 13.38 | 6.38 | 4.17 | L-serine dehydratase 1 |
| *FEPFGCPE_03443* | | + | 3754651 | 3756273 | 4.79 | 20.91 | 10.22 | 3.51 | Hypothetical protein |
| *FEPFGCPE_03444* | *alsT* | + | 3756557 | 3757972 | 0.62 | 1.82 | 1.68 | 0.64 | Amino-acid carrier protein AlsT |
| *FEPFGCPE_03445* | *braC-4* | - | 3758177 | 3759319 | 8.18 | 68.32 | 51.28 | 39.27 | Leucine-, isoleucine-, valine-, threonine-, and alanine-binding protein |
| *FEPFGCPE_03446* | *putP* | - | 3759528 | 3761072 | 0.15 | 1.79 | 1.51 | 0.39 | Sodium/proline symporter |
| *FEPFGCPE_03447* | *putA* | - | 3761573 | 3765496 | 1.30 | 4.87 | 2.66 | 1.27 | Bifunctional protein PutA |
| *FEPFGCPE_03448* | | - | 3765671 | 3766009 | 5.07 | 34.26 | 19.06 | 6.37 | Hypothetical protein |
| *FEPFGCPE_03449* | *msrP-1* | - | 3766024 | 3767337 | 6.01 | 34.83 | 25.89 | 10.98 | Protein-methionine-sulfoxide reductase catalytic subunit MsrP |
| *FEPFGCPE_03450* | *priA* | - | 3767468 | 3769975 | 1.30 | 6.16 | 4.96 | 1.83 | Primosomal protein N' |
| *FEPFGCPE_03451* | *hemE* | - | 3770370 | 3771479 | 17.32 | 50.07 | 50.17 | 27.32 | Uroporphyrinogen decarboxylase |
| *FEPFGCPE_03452* | *pheC* | + | 3771686 | 3772507 | 5.94 | 35.54 | 16.06 | 8.90 | Cyclohexadienyl dehydratase |
| *FEPFGCPE_03453* | *nlhH-5* | - | 3772528 | 3773466 | 2.35 | 8.37 | 12.66 | 3.75 | Carboxylesterase NlhH |
| *FEPFGCPE_03454* | *ddpX* | - | 3773503 | 3773997 | 2.90 | 29.98 | 20.67 | 10.42 | D-alanyl-D-alanine dipeptidase |
| *FEPFGCPE_03455* | *ygbM* | - | 3774726 | 3775514 | 3.12 | 18.39 | 11.75 | 2.62 | Putative hydroxypyruvate isomerase YgbM |
| *FEPFGCPE_03456* | | - | 3775572 | 3776078 | 8.35 | 30.18 | 12.06 | 2.80 | Hypothetical protein |
| *FEPFGCPE_03457* | *arsC-4* | + | 3776377 | 3776733 | 3.66 | 20.46 | 17.53 | 5.22 | Arsenate reductase |
| *FEPFGCPE_03458* | | - | 3776807 | 3777568 | 2.53 | 15.32 | 18.84 | 5.29 | Hypothetical protein |
| *FEPFGCPE_03459* | *gloC-3* | - | 3777565 | 3778221 | 5.49 | 37.75 | 26.24 | 15.71 | Hydroxyacylglutathione hydrolase GloC |
| *FEPFGCPE_03460* | *rlpA* | + | 3778456 | 3779073 | 4.02 | 38.96 | 37.85 | 14.88 | Endolytic peptidoglycan transglycosylase RlpA |
| *FEPFGCPE_03461* | | + | 3779519 | 3780505 | 5.57 | 49.58 | 50.92 | 20.81 | Hypothetical protein |
| *FEPFGCPE_03462* | | + | 3780740 | 3781645 | 1.73 | 8.93 | 2.81 | 2.78 | Hypothetical protein |
| *FEPFGCPE_03463* | *ecfA2* | + | 3781648 | 3782442 | 1.51 | 9.12 | 5.86 | 3.19 | Energy-coupling factor transporter ATP-binding protein EcfA2 |
| *FEPFGCPE_03464* | *rsmI-1* | - | 3782577 | 3783461 | 2.63 | 13.33 | 11.30 | 2.95 | Ribosomal RNA small subunit methyltransferase I |
| *FEPFGCPE_03465* | *nucS* | + | 3783557 | 3783910 | 3.10 | 8.47 | 10.32 | 3.95 | Endonuclease NucS |
| *FEPFGCPE_03466* | *gmhA* | + | 3784065 | 3784652 | 4.73 | 31.66 | 25.38 | 11.76 | Phosphoheptose isomerase |
| *FEPFGCPE_03467* | | + | 3784771 | 3785538 | 12.17 | 16.89 | 25.31 | 25.87 | Hypothetical protein |
| *FEPFGCPE_03468* | *fccA* | + | 3785643 | 3786086 | 1.90 | 7.29 | 4.27 | 1.14 | Cytochrome subunit of sulfide dehydrogenase |
| *FEPFGCPE_03469* | *fccB-2* | + | 3786097 | 3787392 | 3.45 | 7.76 | 9.56 | 1.63 | Sulfide dehydrogenase [flavocytochrome c] flavoprotein chain |
| *FEPFGCPE_03470* | | - | 3787405 | 3788727 | 3.14 | 16.69 | 12.08 | 4.70 | Hypothetical protein |
| *FEPFGCPE_03471* | *hxuB* | + | 3789144 | 3790907 | 7.08 | 43.81 | 17.41 | 5.44 | Heme/hemopexin transporter protein HuxB |
| *FEPFGCPE_03472* | | - | 3790890 | 3791045 | 2.63 | 15.00 | 1.43 | 0.00 | Hypothetical protein |
| *FEPFGCPE_03473* | | + | 3791300 | 3791647 | 2.71 | 25.19 | 28.82 | 10.02 | putative HTH-type transcriptional regulator |
| *FEPFGCPE_03474* | *yedY* | + | 3791805 | 3793172 | 0.91 | 3.77 | 10.00 | 3.78 | Sulfoxide reductase catalytic subunit YedY |
| *FEPFGCPE_03475* | | + | 3793156 | 3794349 | 5.39 | 23.48 | 17.77 | 9.22 | Cytochrome c-551 |

| Locus name | Gene name | Strand | Gene start (bp) | Gene end (bp) | Average TPM Inoculum | Average TPM $t_6$ | Average TPM $t_{24}$ | Average TPM $t_{168}$ | Annotation description |
|---|---|---|---|---|---|---|---|---|---|
| *FEPFGCPE_03476* | *cyt* | + | 3794433 | 3794762 | 13.82 | 79.39 | 53.06 | 23.59 | Cytochrome c-552 |
| *FEPFGCPE_03477* | | + | 3794899 | 3795366 | 14.55 | 92.51 | 51.10 | 56.69 | Hypothetical protein |
| *FEPFGCPE_03478* | | + | 3795398 | 3795709 | 19.04 | 139.54 | 78.69 | 39.11 | Hypothetical protein |
| *FEPFGCPE_03479* | *soxA* | + | 3795875 | 3796750 | 5.73 | 34.98 | 22.40 | 16.12 | SoxAX cytochrome complex subunit A |
| *FEPFGCPE_03480* | | + | 3796770 | 3797420 | 4.93 | 28.09 | 13.68 | 5.86 | Hypothetical protein |
| *FEPFGCPE_03481* | *resA-2* | + | 3797721 | 3798224 | 7.02 | 39.61 | 19.73 | 7.61 | Thiol-disulfide oxidoreductase ResA |
| *FEPFGCPE_03482* | *mggB* | + | 3798521 | 3800230 | 5.86 | 42.42 | 30.94 | 8.81 | Mannosylglucosyl-3-phosphoglycerate phosphatase |
| *FEPFGCPE_03483* | | + | 3800355 | 3800430 | 35.81 | 52.02 | 45.53 | 33.71 | tRNA |
| *FEPFGCPE_03484* | *speG* | + | 3800468 | 3801007 | 7.25 | 43.48 | 27.36 | 4.88 | Spermidine N(1)-acetyltransferase |
| *FEPFGCPE_03485* | *pglF* | - | 3801085 | 3803055 | 3.59 | 28.35 | 17.44 | 5.02 | UDP-N-acetyl-alpha-D-glucosamine C6 dehydratase |
| *FEPFGCPE_03486* | *wcaJ* | - | 3803132 | 3803722 | 2.35 | 17.54 | 14.13 | 7.18 | UDP-glucose:undecaprenyl-phosphate glucose-1-phosphate transferase |
| *FEPFGCPE_03487* | *arnB-2* | - | 3803722 | 3804876 | 3.03 | 27.65 | 22.12 | 11.70 | UDP-4-amino-4-deoxy-L-arabinose--oxoglutarate aminotransferase |
| *FEPFGCPE_03488* | | - | 3804937 | 3805599 | 2.49 | 18.47 | 16.00 | 4.18 | Putative acetyltransferase |
| *FEPFGCPE_03489* | | - | 3805609 | 3806844 | 0.69 | 8.64 | 5.01 | 3.29 | Hypothetical protein |
| *FEPFGCPE_03490* | | - | 3806841 | 3808049 | 1.14 | 9.29 | 6.72 | 1.98 | Hypothetical protein |
| *FEPFGCPE_03491* | | - | 3808046 | 3808669 | 0.68 | 9.07 | 10.33 | 3.53 | Putative acetyltransferase |
| *FEPFGCPE_03492* | | - | 3808673 | 3809911 | 1.39 | 7.91 | 9.35 | 2.91 | Hypothetical protein |
| *FEPFGCPE_03493* | | - | 3809990 | 3810925 | 0.69 | 17.39 | 12.77 | 6.48 | Hypothetical protein |
| *FEPFGCPE_03494* | | - | 3810978 | 3812540 | 1.13 | 18.54 | 22.41 | 5.96 | Hypothetical protein |
| *FEPFGCPE_03495* | *pseI* | - | 3812571 | 3813620 | 1.18 | 28.53 | 19.96 | 10.70 | Pseudaminic acid synthase |
| *FEPFGCPE_03496* | *pseG* | - | 3813765 | 3814880 | 1.89 | 22.09 | 17.46 | 4.83 | UDP-2,4-diacetamido-2,4,6-trideoxy-beta-L-altropyranose hydrolase |
| *FEPFGCPE_03497* | *kat* | - | 3814885 | 3816174 | 1.37 | 15.58 | 12.28 | 4.79 | 3-aminobutyryl-CoA aminotransferase |
| *FEPFGCPE_03498* | *kdsB-1* | - | 3816178 | 3816948 | 1.23 | 17.68 | 13.17 | 3.86 | 8-amino-3,8-dideoxy-manno-octulosonate cytidylyltransferase |
| *FEPFGCPE_03499* | | - | 3816945 | 3817835 | 2.22 | 9.74 | 12.68 | 3.80 | putative oxidoreductase |
| *FEPFGCPE_03500* | *pseC* | - | 3817832 | 3818980 | 2.57 | 24.11 | 19.58 | 5.24 | UDP-4-amino-4,6-dideoxy-N-acetyl-beta-L-altrosamine transaminase |
| *FEPFGCPE_03501* | *pseB* | - | 3818977 | 3819978 | 5.80 | 41.36 | 28.65 | 15.48 | UDP-N-acetylglucosamine 4,6-dehydratase (inverting) |
| *FEPFGCPE_03502* | *galE1* | - | 3820030 | 3820980 | 10.13 | 45.98 | 39.69 | 24.24 | UDP-glucose 4-epimerase |
| *FEPFGCPE_03503* | | + | 3821875 | 3822705 | 9.91 | 57.97 | 35.26 | 21.50 | Hypothetical protein |
| *FEPFGCPE_03504* | *coxM* | + | 3823075 | 3824556 | 9.81 | 62.75 | 45.74 | 10.56 | Alternative cytochrome c oxidase subunit 2 |
| *FEPFGCPE_03505* | *coxN* | + | 3824849 | 3826696 | 2.32 | 36.64 | 25.58 | 3.75 | Alternative cytochrome c oxidase subunit 1 |
| *FEPFGCPE_03506* | *ctaE-2* | + | 3826693 | 3827328 | 1.39 | 18.27 | 17.71 | 3.40 | putative cytochrome c oxidase subunit 3 |
| *FEPFGCPE_03507* | *cyoC-2* | + | 3827348 | 3828040 | 1.35 | 32.04 | 20.82 | 5.69 | Cytochrome bo(3) ubiquinol oxidase subunit 3 |
| *FEPFGCPE_03508* | | + | 3828049 | 3828405 | 2.01 | 37.72 | 22.07 | 4.04 | Hypothetical protein |
| *FEPFGCPE_03509* | *oxyR-8* | - | 3828433 | 3829365 | 1.03 | 10.07 | 9.27 | 1.27 | Hydrogen peroxide-inducible genes activator |
| *FEPFGCPE_03510* | | + | 3829570 | 3830391 | 2.34 | 14.21 | 5.97 | 1.59 | Hypothetical protein |
| *FEPFGCPE_03511* | *yjhB-2* | + | 3830658 | 3831989 | 0.79 | 7.43 | 4.47 | 2.03 | Putative metabolite transport protein YjhB |
| *FEPFGCPE_03512* | *apc3* | + | 3832220 | 3835891 | 0.51 | 0.34 | 1.11 | 0.84 | Acetophenone carboxylase gamma subunit |
| *FEPFGCPE_03513* | | + | 3836048 | 3836449 | 0.13 | 1.04 | 0.00 | 0.00 | Hypothetical protein |
| *FEPFGCPE_03514* | | + | 3836463 | 3836993 | 0.00 | 0.05 | 0.36 | 0.00 | Hypothetical protein |
| *FEPFGCPE_03515* | | + | 3837097 | 3837723 | 0.60 | 1.03 | 0.81 | 0.74 | Hypothetical protein |
| *FEPFGCPE_03516* | | + | 3837720 | 3838235 | 0.10 | 0.00 | 0.00 | 0.00 | Hypothetical protein |
| *FEPFGCPE_03517* | *sseC* | + | 3838232 | 3839764 | 0.42 | 0.58 | 0.00 | 0.00 | Secreted effector protein SseC |
| *FEPFGCPE_03518* | | + | 3839825 | 3840499 | 0.52 | 0.00 | 0.57 | 1.33 | Hypothetical protein |
| *FEPFGCPE_03519* | | + | 3840504 | 3840932 | 0.52 | 3.79 | 3.21 | 0.42 | Hypothetical protein |
| *FEPFGCPE_03520* | | - | 3840939 | 3842414 | 1.47 | 4.14 | 3.41 | 0.70 | Hypothetical protein |
| *FEPFGCPE_03521* | | + | 3842653 | 3842982 | 1.21 | 19.89 | 4.80 | 0.67 | Hypothetical protein |
| *FEPFGCPE_03522* | *pbpE* | - | 3842954 | 3844267 | 3.31 | 31.77 | 26.20 | 5.34 | Penicillin-binding protein 4* |
| *FEPFGCPE_03523* | *mprF* | + | 3844639 | 3847362 | 2.26 | 9.39 | 7.96 | 2.76 | Phosphatidylglycerol lysyltransferase |
| *FEPFGCPE_03524* | | + | 3847359 | 3848960 | 1.54 | 15.91 | 8.00 | 4.23 | Hypothetical protein |
| *FEPFGCPE_03525* | *rutG* | + | 3849110 | 3850417 | 1.31 | 9.56 | 5.86 | 3.24 | Putative pyrimidine permease RutG |
| *FEPFGCPE_03526* | *gltC-11* | - | 3850517 | 3851647 | 4.77 | 33.29 | 20.68 | 5.26 | HTH-type transcriptional regulator GltC |
| *FEPFGCPE_03527* | *rhlE-3* | + | 3851799 | 3853448 | 0.46 | 3.23 | 6.48 | 3.30 | ATP-dependent RNA helicase RhlE |
| *FEPFGCPE_03528* | *panS* | + | 3853601 | 3854554 | 0.51 | 2.77 | 0.19 | 0.00 | Pantothenate precursors transporter PanS |
| *FEPFGCPE_03529* | *ppiC* | + | 3854726 | 3855487 | 56.56 | 159.62 | 79.88 | 32.79 | Peptidyl-prolyl cis-trans isomerase C |
| *FEPFGCPE_03530* | *hemY* | + | 3855899 | 3857935 | 1.82 | 8.50 | 4.24 | 1.88 | Protoporphyrinogen oxidase |
| *FEPFGCPE_03531* | | + | 3857978 | 3858784 | 1.80 | 4.17 | 4.99 | 1.47 | Hypothetical protein |
| *FEPFGCPE_03532* | | + | 3858798 | 3860129 | 0.53 | 6.33 | 3.60 | 1.50 | Putative epoxidase LasC |
| *FEPFGCPE_03533* | *ycgJ* | - | 3860363 | 3861202 | 1.27 | 12.16 | 7.26 | 7.15 | putative methyltransferase YcgJ |
| *FEPFGCPE_03534* | | + | 3861335 | 3862171 | 1.68 | 6.24 | 4.99 | 1.40 | Hypothetical protein |
| *FEPFGCPE_03535* | *crcB* | - | 3862181 | 3862579 | 1.09 | 10.88 | 9.70 | 2.04 | Putative fluoride ion transporter CrcB |
| *FEPFGCPE_03536* | *pyrE-2* | - | 3862683 | 3863372 | 4.47 | 25.60 | 28.14 | 16.25 | Orotate phosphoribosyltransferase |
| *FEPFGCPE_03537* | *yjhB-3* | + | 3863729 | 3864967 | 0.84 | 4.53 | 2.80 | 2.31 | Putative metabolite transport protein YjhB |
| *FEPFGCPE_03538* | *maeB-1* | + | 3865283 | 3867562 | 1.27 | 20.68 | 14.75 | 10.22 | NADP-dependent malic enzyme |
| *FEPFGCPE_03539* | | - | 3867708 | 3871274 | 14.44 | 36.04 | 27.22 | 9.67 | Hypothetical protein |
| *FEPFGCPE_03540* | | + | 3871872 | 3872111 | 29.07 | 184.36 | 217.10 | 135.36 | Hypothetical protein |
| *FEPFGCPE_03541* | *lrp-6* | + | 3872313 | 3872801 | 0.88 | 12.07 | 4.13 | 5.26 | Leucine-responsive regulatory protein |
| *FEPFGCPE_03542* | | - | 3872888 | 3873544 | 23.35 | 25.27 | 27.85 | 28.60 | Hypothetical protein |
| *FEPFGCPE_03543* | *kstR2-2* | - | 3873786 | 3874442 | 2.77 | 11.13 | 13.38 | 4.61 | HTH-type transcriptional repressor KstR2 |
| *FEPFGCPE_03544* | | - | 3874452 | 3875324 | 5.33 | 45.05 | 32.22 | 9.26 | Hypothetical protein |
| *FEPFGCPE_03545* | *paaE* | - | 3875454 | 3876542 | 1.30 | 2.98 | 4.89 | 3.74 | 1,2-phenylacetyl-CoA epoxidase, subunit E |
| *FEPFGCPE_03546* | *paaD* | - | 3876628 | 3877260 | 2.83 | 6.89 | 4.26 | 4.37 | Putative 1,2-phenylacetyl-CoA epoxidase, subunit D |

| Locus name | Gene name | Strand | Gene start (bp) | Gene end (bp) | Average TPM Inoculum | Average TPM $t_6$ | Average TPM $t_{24}$ | Average TPM $t_{168}$ | Annotation description |
|---|---|---|---|---|---|---|---|---|---|
| *FEPFGCPE_03547* | *paaC* | - | 3877283 | 3878086 | 2.85 | 10.55 | 5.25 | 10.12 | 1,2-phenylacetyl-CoA epoxidase, subunit C |
| *FEPFGCPE_03548* | *paaB* | - | 3878136 | 3878432 | 10.47 | 42.14 | 24.02 | 34.02 | 1,2-phenylacetyl-CoA epoxidase, subunit B |
| *FEPFGCPE_03549* | *paaA* | - | 3878485 | 3879492 | 7.11 | 28.69 | 21.03 | 31.93 | 1,2-phenylacetyl-CoA epoxidase, subunit A |
| *FEPFGCPE_03550* | *gbpR-5* | - | 3879707 | 3880648 | 2.42 | 9.75 | 9.32 | 5.48 | HTH-type transcriptional regulator GbpR |
| *FEPFGCPE_03551* | | + | 3881059 | 3882159 | 0.46 | 1.59 | 1.29 | 0.23 | Outer membrane porin protein 32 |
| *FEPFGCPE_03552* | *atzA* | + | 3882188 | 3883648 | 1.02 | 5.36 | 0.57 | 0.73 | Atrazine chlorohydrolase |
| *FEPFGCPE_03553* | *clsC* | - | 3883953 | 3885596 | 3.68 | 12.16 | 11.19 | 2.28 | Cardiolipin synthase C |
| *FEPFGCPE_03554* | | + | 3886309 | 3887829 | 133082.47 | 108538.31 | 108916.68 | 103597.18 | rRNA |
| *FEPFGCPE_03555* | | + | 3887899 | 3887975 | 191.09 | 779.63 | 568.80 | 264.70 | tRNA |
| *FEPFGCPE_03556* | | + | 3888088 | 3888163 | 44.55 | 4.74 | 8.45 | 2.87 | tRNA |
| *FEPFGCPE_03557* | | + | 3888457 | 3891330 | 109967.65 | 94761.38 | 106707.85 | 143977.60 | rRNA |
| *FEPFGCPE_03558* | | + | 3891465 | 3891576 | 533.99 | 65.44 | 76.93 | 82.92 | rRNA |
| *FEPFGCPE_03559* | *arnC-2* | + | 3892109 | 3893038 | 0.91 | 24.52 | 18.76 | 9.73 | Undecaprenyl-phosphate 4-deoxy-4-formamido-L-arabinose transferase |
| *FEPFGCPE_03560* | | + | 3893035 | 3893811 | 0.61 | 23.59 | 15.83 | 7.72 | Hypothetical protein |
| *FEPFGCPE_03561* | | + | 3893808 | 3894755 | 0.88 | 9.47 | 5.14 | 3.71 | Hypothetical protein |
| *FEPFGCPE_03562* | | + | 3894755 | 3895297 | 2.27 | 16.83 | 9.29 | 4.99 | Hypothetical protein |
| *FEPFGCPE_03563* | *per* | + | 3895333 | 3896595 | 1.70 | 14.01 | 8.24 | 4.82 | GDP-perosamine synthase |
| *FEPFGCPE_03564* | | + | 3896579 | 3898213 | 5.52 | 26.96 | 21.35 | 6.65 | Hypothetical protein |
| *FEPFGCPE_03565* | *wecE* | + | 3898210 | 3899361 | 4.43 | 19.07 | 18.43 | 10.45 | dTDP-4-amino-4,6-dideoxygalactose transaminase |
| *FEPFGCPE_03566* | *cysNC* | - | 3899383 | 3901326 | 5.23 | 48.74 | 18.70 | 10.77 | Bifunctional enzyme CysN/CysC |
| *FEPFGCPE_03567* | *cysD-2* | - | 3901326 | 3902759 | 9.60 | 84.76 | 37.30 | 17.87 | Sulfate adenylyltransferase subunit 2 |
| *FEPFGCPE_03568* | | + | 3902912 | 3903691 | 4.00 | 50.93 | 23.88 | 17.74 | Hypothetical protein |
| *FEPFGCPE_03569* | | + | 3903691 | 3905661 | 4.23 | 20.71 | 16.77 | 5.68 | Hypothetical protein |
| *FEPFGCPE_03570* | | + | 3905726 | 3905811 | 56.12 | 271.60 | 222.18 | 106.88 | tRNA |
| *FEPFGCPE_03571* | | + | 3905885 | 3905958 | 140.75 | 254.33 | 288.92 | 168.06 | tRNA |
| *FEPFGCPE_03572* | | + | 3905993 | 3906067 | 51.76 | 123.34 | 113.94 | 127.73 | tRNA |
| *FEPFGCPE_03573* | *tufA-1* | + | 3906140 | 3907330 | 57.75 | 148.18 | 157.30 | 137.52 | Elongation factor Tu |
| *FEPFGCPE_03574* | | + | 3907395 | 3907470 | 28.83 | 111.17 | 171.54 | 56.24 | tRNA |
| *FEPFGCPE_03575* | *secE* | + | 3907493 | 3907873 | 23.94 | 77.41 | 71.08 | 59.78 | Protein translocase subunit SecE |
| *FEPFGCPE_03576* | *nusG* | + | 3907875 | 3908432 | 28.32 | 67.56 | 68.76 | 46.86 | Transcription termination/antitermination protein NusG |
| *FEPFGCPE_03577* | *rplK* | + | 3908575 | 3909006 | 24.72 | 51.26 | 55.88 | 47.04 | 50S ribosomal protein L11 |
| *FEPFGCPE_03578* | *rplA* | + | 3909006 | 3909701 | 22.50 | 66.76 | 52.26 | 52.23 | 50S ribosomal protein L1 |
| *FEPFGCPE_03579* | *rplJ* | + | 3910025 | 3910522 | 24.84 | 111.87 | 130.71 | 102.72 | 50S ribosomal protein L10 |
| *FEPFGCPE_03580* | *rplL* | + | 3910583 | 3910957 | 27.17 | 94.55 | 94.70 | 106.50 | 50S ribosomal protein L7/L12 |
| *FEPFGCPE_03581* | *rpoB* | + | 3911466 | 3915572 | 9.30 | 47.00 | 52.41 | 39.20 | DNA-directed RNA polymerase subunit beta |
| *FEPFGCPE_03582* | *rpoC* | + | 3915599 | 3919810 | 6.28 | 52.15 | 52.87 | 40.25 | DNA-directed RNA polymerase subunit beta' |
| *FEPFGCPE_03583* | *fecA* | - | 3919945 | 3922122 | 0.55 | 4.64 | 4.02 | 8.12 | Fe(3+) dicitrate transport protein FecA |
| *FEPFGCPE_03584* | *hpcG-2* | + | 3922441 | 3923250 | 1.31 | 3.58 | 0.27 | 0.17 | 2-oxo-hept-4-ene-1,7-dioate hydratase |
| *FEPFGCPE_03585* | *hpcH-2* | + | 3923298 | 3924083 | 1.17 | 2.40 | 0.78 | 0.41 | 4-hydroxy-2-oxo-heptane-1,7-dioate aldolase |
| *FEPFGCPE_03586* | *aldA-3* | + | 3924161 | 3925636 | 1.03 | 1.88 | 4.49 | 0.00 | Putative aldehyde dehydrogenase AldA |
| *FEPFGCPE_03587* | *hpcB* | + | 3925681 | 3926544 | 0.49 | 3.36 | 2.22 | 0.36 | 3,4-dihydroxyphenylacetate 2,3-dioxygenase |
| *FEPFGCPE_03588* | *hpcE* | + | 3926557 | 3927507 | 4.67 | 21.99 | 15.36 | 7.05 | Homoprotocatechuate catabolism bifunctional isomerase/decarboxylase |
| *FEPFGCPE_03589* | | + | 3927512 | 3928501 | 3.52 | 13.50 | 10.53 | 4.36 | Hypothetical protein |
| *FEPFGCPE_03590* | *gltC-12* | + | 3928649 | 3929581 | 0.73 | 6.99 | 6.76 | 1.01 | HTH-type transcriptional regulator GltC |
| *FEPFGCPE_03591* | | - | 3929606 | 3930103 | 1.02 | 3.78 | 5.88 | 0.00 | Hypothetical protein |
| *FEPFGCPE_03592* | *clcD-2* | + | 3930561 | 3931271 | 5.20 | 27.11 | 17.75 | 13.54 | Carboxymethylenebutenolidase |
| *FEPFGCPE_03593* | | - | 3931563 | 3931871 | 0.15 | 7.40 | 20.57 | 9.15 | Hypothetical protein |
| *FEPFGCPE_03594* | | + | 3932023 | 3932421 | 1.60 | 9.33 | 17.14 | 1.93 | Hypothetical protein |
| *FEPFGCPE_03595* | *nimR-2* | + | 3932545 | 3933336 | 5.31 | 26.91 | 15.13 | 7.77 | HTH-type transcriptional regulator NimR |
| *FEPFGCPE_03596* | | - | 3933284 | 3934006 | 3.62 | 26.01 | 17.67 | 8.86 | Hypothetical protein |
| *FEPFGCPE_03597* | | - | 3934236 | 3935447 | 1.06 | 5.41 | 2.66 | 0.84 | Hypothetical protein |
| *FEPFGCPE_03598* | *rpsL* | + | 3935860 | 3936234 | 26.32 | 116.79 | 104.33 | 119.50 | 30S ribosomal protein S12 |
| *FEPFGCPE_03599* | *rpsG* | + | 3936361 | 3936831 | 31.83 | 127.13 | 128.53 | 128.71 | 30S ribosomal protein S7 |
| *FEPFGCPE_03600* | *fusA-2* | + | 3936949 | 3939051 | 16.15 | 77.96 | 83.57 | 84.94 | Elongation factor G |
| *FEPFGCPE_03601* | *tufA-2* | + | 3939134 | 3940342 | 27.09 | 83.35 | 100.27 | 112.84 | Elongation factor Tu |
| *FEPFGCPE_03602* | *rpsJ* | + | 3940457 | 3940768 | 11.36 | 26.48 | 28.07 | 31.17 | 30S ribosomal protein S10 |
| *FEPFGCPE_03603* | *rplC* | + | 3940974 | 3941630 | 34.21 | 92.43 | 97.76 | 95.41 | 50S ribosomal protein L3 |
| *FEPFGCPE_03604* | *rplD* | + | 3941633 | 3942253 | 25.57 | 83.33 | 61.64 | 71.83 | 50S ribosomal protein L4 |
| *FEPFGCPE_03605* | *rplW* | + | 3942250 | 3942564 | 25.10 | 72.12 | 101.18 | 115.88 | 50S ribosomal protein L23 |
| *FEPFGCPE_03606* | *rplB* | + | 3942567 | 3943394 | 31.91 | 85.90 | 83.24 | 78.19 | 50S ribosomal protein L2 |
| *FEPFGCPE_03607* | *rpsS* | + | 3943408 | 3943683 | 55.91 | 133.94 | 137.40 | 137.11 | 30S ribosomal protein S19 |
| *FEPFGCPE_03608* | *rplV* | + | 3943694 | 3944023 | 9.05 | 22.98 | 18.10 | 23.98 | 50S ribosomal protein L22 |
| *FEPFGCPE_03609* | *rpsC* | + | 3944033 | 3944839 | 53.55 | 146.90 | 171.48 | 151.28 | 30S ribosomal protein S3 |
| *FEPFGCPE_03610* | *rplP* | + | 3944842 | 3945258 | 64.70 | 217.68 | 215.88 | 212.72 | 50S ribosomal protein L16 |
| *FEPFGCPE_03611* | *rpmC* | + | 3945273 | 3945467 | 64.49 | 145.78 | 195.12 | 253.82 | 50S ribosomal protein L29 |
| *FEPFGCPE_03612* | *rpsQ* | + | 3945464 | 3945736 | 16.50 | 41.18 | 44.30 | 37.66 | 30S ribosomal protein S17 |
| *FEPFGCPE_03613* | *rplN* | + | 3946002 | 3946370 | 22.67 | 87.68 | 98.46 | 97.86 | 50S ribosomal protein L14 |
| *FEPFGCPE_03614* | *rplX* | + | 3946382 | 3946690 | 15.03 | 58.69 | 51.08 | 57.85 | 50S ribosomal protein L24 |
| *FEPFGCPE_03615* | *rplE* | + | 3946710 | 3947249 | 21.15 | 116.34 | 129.72 | 90.02 | 50S ribosomal protein L5 |
| *FEPFGCPE_03616* | *rpsN* | + | 3947257 | 3947562 | 36.54 | 167.03 | 180.43 | 147.94 | 30S ribosomal protein S14 |
| *FEPFGCPE_03617* | *rpsH* | + | 3947576 | 3947971 | 30.00 | 120.11 | 125.01 | 120.41 | 30S ribosomal protein S8 |

| Locus name | Gene name | Strand | Gene start (bp) | Gene end (bp) | Average TPM Inoculum | Average TPM $t_6$ | Average TPM $t_{24}$ | Average TPM $t_{168}$ | Annotation description |
|---|---|---|---|---|---|---|---|---|---|
| *FEPFGCPE_03618* | *rplF* | + | 3947987 | 3948520 | 22.05 | 126.22 | 147.31 | 160.48 | 50S ribosomal protein L6 |
| *FEPFGCPE_03619* | *rplR* | + | 3948532 | 3948897 | 15.07 | 87.92 | 127.85 | 94.06 | 50S ribosomal protein L18 |
| *FEPFGCPE_03620* | *rpsE* | + | 3948911 | 3949429 | 19.29 | 128.15 | 142.08 | 115.62 | 30S ribosomal protein S5 |
| *FEPFGCPE_03621* | *rpmD* | + | 3949441 | 3949623 | 53.44 | 328.89 | 329.43 | 220.82 | 50S ribosomal protein L30 |
| *FEPFGCPE_03622* | *rplO* | + | 3949635 | 3950069 | 9.29 | 61.23 | 54.81 | 61.31 | 50S ribosomal protein L15 |
| *FEPFGCPE_03623* | *secY* | + | 3950093 | 3951430 | 12.14 | 96.03 | 89.89 | 49.12 | Protein translocase subunit SecY |
| *FEPFGCPE_03624* | *infA-1* | + | 3951440 | 3951658 | 52.60 | 310.38 | 285.46 | 147.70 | Translation initiation factor IF-1 |
| *FEPFGCPE_03625* | *rpmJ* | + | 3951678 | 3951794 | 18.78 | 270.46 | 391.66 | 197.16 | 50S ribosomal protein L36 |
| *FEPFGCPE_03626* | *rpsM* | + | 3951830 | 3952195 | 11.38 | 111.85 | 107.97 | 108.99 | 30S ribosomal protein S13 |
| *FEPFGCPE_03627* | *rpsK* | + | 3952213 | 3952617 | 7.88 | 107.57 | 105.43 | 82.19 | 30S ribosomal protein S11 |
| *FEPFGCPE_03628* | *rpsD* | + | 3952770 | 3953393 | 5.08 | 52.46 | 54.07 | 36.80 | 30S ribosomal protein S4 |
| *FEPFGCPE_03629* | *rpoA* | + | 3953499 | 3954476 | 7.34 | 68.07 | 69.42 | 57.07 | DNA-directed RNA polymerase subunit alpha |
| *FEPFGCPE_03630* | *rplQ* | + | 3954642 | 3955034 | 2.47 | 34.08 | 50.60 | 47.55 | 50S ribosomal protein L17 |
| *FEPFGCPE_03631* | *cutA* | + | 3955229 | 3955561 | 71.58 | 114.64 | 125.04 | 31.37 | Divalent-cation tolerance protein CutA |
| *FEPFGCPE_03632* | *dsbD-1* | + | 3955554 | 3957545 | 7.56 | 22.32 | 13.98 | 8.06 | Thiol:disulfide interchange protein DsbD |
| *FEPFGCPE_03633* | *hemB* | - | 3957631 | 3958629 | 11.82 | 41.56 | 29.32 | 15.65 | Delta-aminolevulinic acid dehydratase |
| *FEPFGCPE_03634* | *engB* | - | 3959116 | 3959784 | 2.44 | 10.62 | 15.19 | 2.95 | putative GTP-binding protein EngB |
| *FEPFGCPE_03635* | *cycA* | + | 3960083 | 3960739 | 23.11 | 46.35 | 43.98 | 16.73 | Cytochrome c4 |
| *FEPFGCPE_03636* | *ccsB* | + | 3960906 | 3963041 | 4.10 | 13.25 | 9.78 | 4.78 | Cytochrome c biogenesis protein CcsB |
| *FEPFGCPE_03637* | *ccsA* | + | 3963053 | 3964237 | 6.61 | 30.72 | 24.33 | 4.95 | Cytochrome c biogenesis protein CcsA |
| *FEPFGCPE_03638* | *msrP-2* | + | 3964441 | 3965439 | 20.75 | 23.19 | 17.59 | 7.89 | Protein-methionine-sulfoxide reductase catalytic subunit MsrP |
| *FEPFGCPE_03639* | *msrQ* | + | 3965467 | 3966267 | 8.67 | 6.94 | 6.60 | 3.51 | Protein-methionine-sulfoxide reductase heme-binding subunit MsrQ |
| *FEPFGCPE_03640* | *lysA* | - | 3966195 | 3967463 | 5.31 | 17.03 | 14.73 | 6.56 | Diaminopimelate decarboxylase |
| *FEPFGCPE_03641* | *cyaY* | + | 3967837 | 3968157 | 5.90 | 21.63 | 26.57 | 18.92 | Protein CyaY |
| *FEPFGCPE_03642* | *mrcA-3* | - | 3968252 | 3970693 | 3.02 | 19.43 | 14.33 | 4.66 | Penicillin-binding protein 1A |
| *FEPFGCPE_03643* | | + | 3970938 | 3971987 | 4.28 | 2.36 | 3.71 | 2.45 | Hypothetical protein |
| *FEPFGCPE_03644* | | + | 3971978 | 3972649 | 0.67 | 0.00 | 1.26 | 0.86 | Hypothetical protein |
| *FEPFGCPE_03645* | *pilQ* | + | 3972639 | 3975248 | 2.62 | 15.22 | 12.64 | 3.19 | Type IV pilus biogenesis and competence protein PilQ |
| *FEPFGCPE_03646* | *aroK-2* | + | 3975217 | 3975870 | 8.65 | 33.51 | 45.03 | 13.69 | Shikimate kinase 1 |
| *FEPFGCPE_03647* | *aroB* | + | 3975891 | 3976982 | 7.40 | 40.73 | 31.08 | 17.34 | 3-dehydroquinate synthase |
| *FEPFGCPE_03648* | *dgt* | + | 3976999 | 3978135 | 5.24 | 19.75 | 15.76 | 7.08 | Deoxyguanosinetriphosphate triphosphohydrolase-like protein |
| *FEPFGCPE_03649* | | - | 3978280 | 3978657 | 31.53 | 88.82 | 132.36 | 74.87 | Hypothetical protein |
| *FEPFGCPE_03650* | *ompW-2* | - | 3978975 | 3979724 | 77.90 | 120.76 | 85.37 | 35.81 | Outer membrane protein W |
| *FEPFGCPE_03651* | *ompW-3* | - | 3979992 | 3980840 | 19.31 | 65.06 | 33.11 | 7.39 | Outer membrane protein W |
| *FEPFGCPE_03652* | | + | 3981038 | 3981739 | 3.11 | 12.99 | 5.66 | 0.76 | Hypothetical protein |
| *FEPFGCPE_03653* | *gltB* | + | 3982118 | 3986851 | 42.86 | 54.85 | 54.32 | 30.35 | Ferredoxin-dependent glutamate synthase 1 |
| *FEPFGCPE_03654* | *gltD* | + | 3986948 | 3988414 | 42.43 | 171.90 | 152.16 | 47.28 | Glutamate synthase [NADPH] small chain |
| *FEPFGCPE_03655* | *udh* | + | 3988907 | 3989731 | 0.85 | 11.43 | 5.43 | 3.65 | Uronate dehydrogenase |
| *FEPFGCPE_03656* | | + | 3989728 | 3990630 | 0.92 | 11.28 | 2.10 | 0.21 | 6-deoxy-6-sulfogluconolactonase |
| *FEPFGCPE_03657* | *yddA* | - | 3990627 | 3992426 | 3.20 | 10.19 | 7.01 | 3.59 | Inner membrane ABC transporter ATP-binding protein YddA |
| *FEPFGCPE_03658* | *mnmC-3* | + | 3992989 | 3994119 | 5.93 | 13.81 | 13.58 | 5.57 | tRNA 5-methylaminomethyl-2-thiouridine biosynthesis bifunctional protein MnmC |
| *FEPFGCPE_03659* | *thiS* | + | 3994135 | 3994332 | 2.49 | 9.32 | 11.15 | 5.33 | Sulfur carrier protein ThiS |
| *FEPFGCPE_03660* | *thiG* | + | 3994334 | 3995179 | 2.13 | 16.46 | 15.47 | 5.57 | Thiazole synthase |
| *FEPFGCPE_03661* | *thiE-2* | + | 3995244 | 3996437 | 2.11 | 10.45 | 7.96 | 2.73 | Thiamine-phosphate synthase |
| *FEPFGCPE_03662* | | - | 3996458 | 3997042 | 1.21 | 8.26 | 8.81 | 2.16 | Hypothetical protein |
| *FEPFGCPE_03663* | *mlaF-2* | + | 3997177 | 3998124 | 8.97 | 57.77 | 45.95 | 14.46 | putative phospholipid import ATP-binding protein MlaF |
| *FEPFGCPE_03664* | *mlaE-2* | + | 3998121 | 3998888 | 4.08 | 13.14 | 17.23 | 16.57 | putative phospholipid ABC transporter permease protein MlaE |
| *FEPFGCPE_03665* | *mlaD* | + | 3998938 | 3999492 | 8.61 | 62.26 | 39.07 | 16.53 | putative phospholipid ABC transporter-binding protein MlaD |
| *FEPFGCPE_03666* | *mlaA* | + | 3999498 | 4000364 | 3.81 | 12.04 | 8.79 | 5.05 | putative phospholipid-binding lipoprotein MlaA |
| *FEPFGCPE_03667* | *mlaC* | + | 4000553 | 4001173 | 19.98 | 104.58 | 107.32 | 67.00 | putative phospholipid-binding protein MlaC |
| *FEPFGCPE_03668* | | + | 4001178 | 4001435 | 11.43 | 44.04 | 52.17 | 34.37 | Hypothetical protein |
| *FEPFGCPE_03669* | *drrA-2* | + | 4001790 | 4002716 | 1.62 | 1.99 | 3.71 | 5.41 | Daunorubicin/doxorubicin resistance ATP-binding protein DrrA |
| *FEPFGCPE_03670* | *yadH-2* | + | 4002713 | 4003480 | 1.95 | 9.23 | 5.80 | 2.67 | Inner membrane transport permease YadH |
| *FEPFGCPE_03671* | *ibaG* | + | 4003499 | 4003738 | 2.69 | 25.02 | 8.84 | 2.10 | Acid stress protein IbaG |
| *FEPFGCPE_03672* | *murA-1* | + | 4003735 | 4005075 | 1.69 | 4.71 | 6.00 | 6.16 | UDP-N-acetylglucosamine 1-carboxyvinyltransferase |
| *FEPFGCPE_03673* | *hisG* | + | 4005072 | 4005752 | 8.94 | 31.28 | 23.35 | 11.05 | ATP phosphoribosyltransferase |
| *FEPFGCPE_03674* | *hisD* | + | 4005749 | 4007071 | 8.34 | 19.60 | 16.58 | 16.01 | Histidinol dehydrogenase |
| *FEPFGCPE_03675* | | + | 4007073 | 4007636 | 3.77 | 2.68 | 6.20 | 3.36 | Hypothetical protein |
| *FEPFGCPE_03676* | *hisC2* | + | 4007647 | 4008750 | 6.56 | 36.77 | 23.93 | 8.75 | Histidinol-phosphate aminotransferase 2 |
| *FEPFGCPE_03677* | *hisB* | + | 4008778 | 4009365 | 4.11 | 33.95 | 21.86 | 10.56 | Histidine biosynthesis bifunctional protein HisB |
| *FEPFGCPE_03678* | | + | 4009478 | 4010104 | 8.43 | 45.89 | 36.46 | 12.13 | Hypothetical protein |
| *FEPFGCPE_03679* | *hisH* | + | 4010101 | 4010883 | 4.87 | 30.54 | 21.49 | 5.39 | Imidazole glycerol phosphate synthase subunit HisH |
| *FEPFGCPE_03680* | *hisA* | + | 4010931 | 4011680 | 9.04 | 95.90 | 61.88 | 26.88 | 1-(5-phosphoribosyl)-5-[(5-phosphoribosylamino)methylideneamino] imidazole-4-carboxamide isomerase |
| *FEPFGCPE_03681* | *hisF* | + | 4011719 | 4012492 | 5.24 | 30.38 | 25.26 | 10.13 | Imidazole glycerol phosphate synthase subunit HisF |
| *FEPFGCPE_03682* | *hisI* | + | 4012510 | 4012908 | 4.74 | 17.83 | 31.61 | 11.32 | Phosphoribosyl-AMP cyclohydrolase |
| *FEPFGCPE_03683* | *hisE* | + | 4012914 | 4013270 | 4.68 | 34.06 | 39.23 | 5.96 | Phosphoribosyl-ATP pyrophosphatase |
| *FEPFGCPE_03684* | | + | 4013336 | 4013758 | 8.12 | 53.79 | 33.80 | 9.96 | Hypothetical protein |
| *FEPFGCPE_03685* | | + | 4013814 | 4014188 | 4.80 | 56.16 | 47.53 | 15.68 | Purine nucleoside phosphoramidase |
| *FEPFGCPE_03686* | *tatA* | + | 4014468 | 4014713 | 23.02 | 129.54 | 124.40 | 52.57 | Sec-independent protein translocase protein TatA |
| *FEPFGCPE_03687* | | + | 4014884 | 4015267 | 16.86 | 91.90 | 77.30 | 37.61 | Hypothetical protein |
| *FEPFGCPE_03688* | *tatC* | + | 4015333 | 4016124 | 2.58 | 8.91 | 13.54 | 2.75 | Sec-independent protein translocase protein TatC |

| Locus name | Gene name | Strand | Gene start (bp) | Gene end (bp) | Average TPM Inoculum | Average TPM $t_6$ | Average TPM $t_{24}$ | Average TPM $t_{168}$ | Annotation description |
|---|---|---|---|---|---|---|---|---|---|
| *FEPFGCPE_03689* | *degQ* | - | 4016351 | 4017595 | 4.67 | 27.07 | 16.20 | 8.82 | Periplasmic pH-dependent serine endoprotease DegQ |
| *FEPFGCPE_03690* | *ybgI* | + | 4017614 | 4018363 | 3.92 | 12.53 | 13.35 | 5.18 | GTP cyclohydrolase 1 type 2 |
| *FEPFGCPE_03691* | *mscL* | - | 4018409 | 4018855 | 7.50 | 63.09 | 36.69 | 16.33 | Large-conductance mechanosensitive channel |
| *FEPFGCPE_03692* | *petA* | + | 4019273 | 4019887 | 14.51 | 62.13 | 33.18 | 21.05 | Ubiquinol-cytochrome c reductase iron-sulfur subunit |
| *FEPFGCPE_03693* | *fbcH* | + | 4019890 | 4021275 | 10.29 | 27.70 | 15.08 | 8.14 | Cytochrome b/c1 |
| *FEPFGCPE_03694* | *petC* | + | 4021307 | 4022026 | 23.31 | 74.99 | 59.25 | 20.83 | Cytochrome c1 |
| *FEPFGCPE_03695* | *sspA-2* | + | 4022110 | 4022721 | 22.08 | 106.63 | 88.69 | 39.77 | Stringent starvation protein A |
| *FEPFGCPE_03696* | *sspB* | + | 4022798 | 4023256 | 13.14 | 99.06 | 70.50 | 37.13 | Stringent starvation protein B |
| *FEPFGCPE_03697* | | + | 4023383 | 4023458 | 0.71 | 0.00 | 2.74 | 0.00 | tRNA |
| *FEPFGCPE_03698* | *kipR-5* | - | 4023678 | 4024418 | 3.31 | 31.07 | 22.84 | 4.95 | HTH-type transcriptional regulator KipR |
| *FEPFGCPE_03699* | *bphC2-2* | - | 4024940 | 4025503 | 0.33 | 0.75 | 1.95 | 0.49 | Biphenyl-2,3-diol 1,2-dioxygenase 2 |
| *FEPFGCPE_03700* | *hapE* | + | 4026027 | 4028006 | 0.72 | 3.84 | 2.15 | 1.87 | 4-hydroxyacetophenone monooxygenase |
| *FEPFGCPE_03701* | *nlhH-6* | + | 4028279 | 4028977 | 0.73 | 4.20 | 4.16 | 2.59 | Carboxylesterase NlhH |
| *FEPFGCPE_03702* | | + | 4029423 | 4030274 | 0.93 | 1.42 | 1.22 | 0.00 | Ureidoglycolate lyase |
| *FEPFGCPE_03703* | *mhpA-6* | + | 4030550 | 4032016 | 0.52 | 0.60 | 1.27 | 0.13 | 3-(3-hydroxy-phenyl)propionate/3-hydroxycinnamic acid hydroxylase |
| *FEPFGCPE_03704* | *fabG-7* | - | 4032138 | 4032878 | 0.46 | 2.18 | 0.43 | 0.00 | 3-oxoacyl-[acyl-carrier-protein] reductase FabG |
| *FEPFGCPE_03705* | *pcaK-8* | + | 4033005 | 4034390 | 0.75 | 2.98 | 2.68 | 0.22 | 4-hydroxybenzoate transporter PcaK |
| *FEPFGCPE_03706* | *mhpB-2* | + | 4034432 | 4035367 | 0.21 | 1.45 | 0.00 | 0.00 | 2,3-dihydroxyphenylpropionate/2,3-dihydroxicinnamic acid 1,2-dioxygenase |
| *FEPFGCPE_03707* | *mhpC-3* | + | 4035364 | 4036239 | 0.33 | 1.51 | 1.01 | 0.19 | 2-hydroxy-6-oxononadienedioate/2-hydroxy-6-oxononatrienedioate hydrolase |
| *FEPFGCPE_03708* | *mhpD-2* | + | 4036247 | 4037050 | 0.22 | 1.36 | 0.00 | 0.91 | 2-keto-4-pentenoate hydratase |
| *FEPFGCPE_03709* | | + | 4037072 | 4037533 | 0.64 | 0.55 | 0.10 | 1.04 | Hypothetical protein |
| *FEPFGCPE_03710* | | + | 4037530 | 4038300 | 0.31 | 3.63 | 0.92 | 0.58 | 2-dehydro-3,6-dideoxy-6-sulfogluconate aldolase |
| *FEPFGCPE_03711* | *mhpA-7* | + | 4038297 | 4039922 | 0.48 | 1.48 | 1.28 | 0.68 | 3-(3-hydroxy-phenyl)propionate/3-hydroxycinnamic acid hydroxylase |
| *FEPFGCPE_03712* | | + | 4040270 | 4041379 | 0.50 | 1.69 | 2.06 | 0.17 | Outer membrane porin protein |
| *FEPFGCPE_03713* | | + | 4041523 | 4041687 | 0.47 | 0.00 | 2.72 | 0.00 | Hypothetical protein |
| *FEPFGCPE_03714* | *tsr-5* | + | 4041882 | 4043891 | 1.33 | 11.49 | 7.13 | 3.57 | Methyl-accepting chemotaxis protein I |
| *FEPFGCPE_03715* | *cph2-5* | - | 4044144 | 4045196 | 2.87 | 9.85 | 9.26 | 3.09 | Phytochrome-like protein cph2 |
| *FEPFGCPE_03716* | | - | 4045473 | 4045775 | 4.66 | 7.35 | 2.99 | 1.78 | Hypothetical protein |
| *FEPFGCPE_03717* | *zraR-2* | - | 4046074 | 4047384 | 3.05 | 5.64 | 9.12 | 6.84 | Transcriptional regulatory protein ZraR |
| *FEPFGCPE_03718* | | + | 4047601 | 4048476 | 114.37 | 3.60 | 3.58 | 4.72 | putative metallo-hydrolase |
| *FEPFGCPE_03719* | | + | 4048494 | 4048688 | 213.52 | 9.68 | 3.20 | 2.54 | Hypothetical protein |
| *FEPFGCPE_03720* | *blh* | + | 4048772 | 4050439 | 94.19 | 3.53 | 3.46 | 2.43 | Beta-lactamase hydrolase-like protein |
| *FEPFGCPE_03721* | | + | 4050480 | 4051268 | 23.53 | 5.29 | 2.83 | 1.34 | Hypothetical protein |
| *FEPFGCPE_03722* | | - | 4051463 | 4052782 | 4.84 | 2.95 | 3.57 | 1.08 | Hypothetical protein |
| *FEPFGCPE_03723* | | - | 4052779 | 4053453 | 6.88 | 5.27 | 5.61 | 3.58 | Putative phosphoribosyl transferase |
| *FEPFGCPE_03724* | *trg-3* | + | 4054352 | 4055512 | 1.04 | 4.92 | 1.91 | 0.41 | Methyl-accepting chemotaxis protein III |
| *FEPFGCPE_03725* | | - | 4055530 | 4055946 | 0.70 | 3.94 | 3.85 | 0.83 | Hypothetical protein |
| *FEPFGCPE_03726* | *bdhA-1* | - | 4056074 | 4056859 | 0.66 | 3.19 | 1.54 | 0.94 | D-beta-hydroxybutyrate dehydrogenase |
| *FEPFGCPE_03727* | | - | 4056863 | 4058263 | 1.07 | 4.09 | 7.68 | 2.58 | Hypothetical protein |
| *FEPFGCPE_03728* | *hyfR-2* | + | 4058575 | 4060101 | 3.70 | 6.29 | 7.22 | 4.06 | DNA-binding transcriptional activator HyfR |
| *FEPFGCPE_03729* | | - | 4060211 | 4061389 | 1.08 | 5.28 | 3.52 | 0.40 | Outer membrane porin protein |
| *FEPFGCPE_03730* | *hpd-2* | - | 4061464 | 4063353 | 0.28 | 0.78 | 0.97 | 0.00 | 4-hydroxyphenylpyruvate dioxygenase |
| *FEPFGCPE_03731* | *aroQ-2* | + | 4063676 | 4064137 | 0.43 | 0.18 | 0.00 | 0.25 | 3-dehydroquinate dehydratase |
| *FEPFGCPE_03732* | *aroE-1* | + | 4064140 | 4065015 | 0.14 | 1.24 | 1.22 | 0.67 | Quinate/shikimate dehydrogenase (NAD(+)) |
| *FEPFGCPE_03733* | *dgoT-2* | + | 4065038 | 4066390 | 1.06 | 5.42 | 0.41 | 1.03 | D-galactonate transporter |
| *FEPFGCPE_03734* | *nicS-3* | - | 4066548 | 4067228 | 2.73 | 45.49 | 24.99 | 11.98 | HTH-type transcriptional repressor NicS |
| *FEPFGCPE_03735* | *ald1* | - | 4067422 | 4068942 | 7.63 | 315.11 | 119.16 | 24.35 | Long-chain-aldehyde dehydrogenase |
| *FEPFGCPE_03736* | *acoR-1* | + | 4069237 | 4071141 | 4.09 | 26.74 | 11.36 | 7.44 | Acetoin catabolism regulatory protein |
| *FEPFGCPE_03737* | | - | 4071172 | 4071636 | 4.01 | 30.12 | 18.45 | 5.50 | Hypothetical protein |
| *FEPFGCPE_03738* | | + | 4071827 | 4071902 | 80.57 | 102.85 | 48.79 | 30.60 | tRNA |
| *FEPFGCPE_03739* | | + | 4071920 | 4071995 | 4.23 | 25.36 | 2.42 | 4.75 | tRNA |
| *FEPFGCPE_03740* | | + | 4072260 | 4072478 | 1.09 | 25.15 | 10.64 | 1.88 | Hypothetical protein |
| *FEPFGCPE_03741* | | + | 4072540 | 4073088 | 0.40 | 6.13 | 10.74 | 2.99 | Hypothetical protein |
| *FEPFGCPE_03742* | *slyB-2* | - | 4073180 | 4073719 | 21.78 | 103.92 | 71.78 | 44.53 | Outer membrane lipoprotein SlyB |
| *FEPFGCPE_03743* | *guaB-4* | - | 4074208 | 4074684 | 39.94 | 181.57 | 142.49 | 47.80 | Inosine-5'-monophosphate dehydrogenase |
| *FEPFGCPE_03744* | *infA-2* | + | 4075271 | 4075492 | 7.54 | 38.35 | 39.62 | 35.22 | Translation initiation factor IF-1 |
| *FEPFGCPE_03745* | *acyP-2* | - | 4075823 | 4076119 | 4.99 | 35.80 | 17.29 | 3.79 | Acylphosphatase |
| *FEPFGCPE_03746* | *cpdA-4* | - | 4076234 | 4077346 | 1.58 | 14.14 | 7.48 | 2.84 | 3',5'-cyclic adenosine monophosphate phosphodiesterase CpdA |
| *FEPFGCPE_03747* | | - | 4077534 | 4077914 | 6.69 | 6.35 | 11.35 | 4.74 | Hypothetical protein |
| *FEPFGCPE_03748* | *mphP* | + | 4078083 | 4078430 | 0.94 | 7.37 | 0.09 | 0.61 | Phenol hydroxylase P5 protein |
| *FEPFGCPE_03749* | *ettA* | + | 4078494 | 4080161 | 8.35 | 40.51 | 43.88 | 23.37 | Energy-dependent translational throttle protein EttA |
| *FEPFGCPE_03750* | *bdhA-2* | + | 4080329 | 4081126 | 5.67 | 24.86 | 32.03 | 30.12 | D-beta-hydroxybutyrate dehydrogenase |
| *FEPFGCPE_03751* | | + | 4081226 | 4082407 | 4.17 | 14.96 | 17.29 | 7.54 | Hypothetical protein |
| *FEPFGCPE_03752* | *cheW-3* | - | 4082533 | 4084158 | 1.19 | 7.26 | 1.76 | 1.30 | Chemotaxis protein CheW |
| *FEPFGCPE_03753* | *cheD-2* | - | 4084176 | 4084682 | 1.60 | 8.14 | 0.55 | 0.00 | Chemoreceptor glutamine deamidase CheD |
| *FEPFGCPE_03754* | *cheR2* | - | 4084666 | 4085508 | 0.58 | 5.01 | 3.83 | 0.30 | Chemotaxis protein methyltransferase Cher2 |
| *FEPFGCPE_03755* | *bdlA* | - | 4085617 | 4087314 | 1.58 | 9.07 | 4.98 | 1.22 | Biofilm dispersion protein BdlA |
| *FEPFGCPE_03756* | | + | 4088011 | 4088124 | 14.05 | 60.11 | 35.81 | 24.85 | Hypothetical protein |
| *FEPFGCPE_03757* | | - | 4088189 | 4088482 | 12.09 | 53.48 | 44.37 | 24.16 | Hypothetical protein |
| *FEPFGCPE_03758* | *bsdB* | - | 4088571 | 4089185 | 5.16 | 28.17 | 18.30 | 8.45 | putative UbiX-like flavin prenyltransferase |
| *FEPFGCPE_03759* | *grxD* | - | 4089203 | 4089517 | 8.39 | 81.23 | 51.33 | 35.50 | Glutaredoxin 4 |

| Locus name | Gene name | Strand | Gene start (bp) | Gene end (bp) | Average TPM Inoculum | Average TPM $t_6$ | Average TPM $t_{24}$ | Average TPM $t_{168}$ | Annotation description |
|---|---|---|---|---|---|---|---|---|---|
| *FEPFGCPE_03760* | *prmC* | - | 4089598 | 4090455 | 1.98 | 17.92 | 12.20 | 5.49 | Release factor glutamine methyltransferase |
| *FEPFGCPE_03761* | *prfA* | - | 4090455 | 4091525 | 10.13 | 31.52 | 23.59 | 5.58 | Peptide chain release factor RF1 |
| *FEPFGCPE_03762* | *hemA* | - | 4091538 | 4092818 | 20.05 | 66.95 | 52.59 | 17.17 | Glutamyl-tRNA reductase |
| *FEPFGCPE_03763* | | + | 4093128 | 4093382 | 30.26 | 136.62 | 107.81 | 25.59 | Hypothetical protein |
| *FEPFGCPE_03764* | | + | 4093404 | 4093643 | 10.86 | 44.09 | 27.15 | 6.52 | Hypothetical protein |
| *FEPFGCPE_03765* | *bmr3-3* | + | 4093874 | 4095352 | 2.27 | 11.93 | 8.96 | 1.72 | Multidrug resistance protein 3 |
| *FEPFGCPE_03766* | | - | 4095372 | 4096487 | 3.33 | 15.33 | 10.88 | 0.66 | putative glycosyltransferase |
| *FEPFGCPE_03767* | | - | 4096493 | 4097824 | 3.39 | 9.34 | 7.38 | 2.51 | Hypothetical protein |
| *FEPFGCPE_03768* | *ychF* | - | 4098206 | 4099300 | 3.05 | 18.64 | 15.66 | 10.00 | Ribosome-binding ATPase YchF |
| *FEPFGCPE_03769* | | - | 4099490 | 4100359 | 3.83 | 28.39 | 17.02 | 6.04 | Hypothetical protein |
| *FEPFGCPE_03770* | *ubiI* | + | 4100758 | 4101735 | 2.60 | 10.52 | 8.09 | 4.05 | 2-octaprenylphenol hydroxylase |
| *FEPFGCPE_03771* | *dsbC* | + | 4101835 | 4102599 | 3.41 | 18.30 | 17.48 | 8.37 | putative thiol:disulfide interchange protein DsbC |
| *FEPFGCPE_03772* | | + | 4102660 | 4104474 | 1.63 | 9.15 | 7.62 | 1.57 | Hypothetical protein |
| *FEPFGCPE_03773* | *dmlR-18* | - | 4104488 | 4105426 | 3.82 | 4.40 | 2.17 | 2.55 | HTH-type transcriptional regulator DmlR |
| *FEPFGCPE_03774* | *azoR* | + | 4105638 | 4106237 | 1.89 | 10.17 | 11.61 | 12.37 | FMN-dependent NADH-azoreductase |
| *FEPFGCPE_03775* | | + | 4106302 | 4107222 | 0.11 | 1.67 | 4.01 | 1.10 | Hypothetical protein |
| *FEPFGCPE_03776* | *ung* | - | 4107208 | 4108233 | 1.33 | 12.47 | 10.54 | 3.05 | Uracil-DNA glycosylase |
| *FEPFGCPE_03777* | *ygiF* | - | 4108301 | 4108996 | 1.10 | 10.69 | 11.44 | 5.49 | Inorganic triphosphatase |
| *FEPFGCPE_03778* | *trpC* | - | 4109062 | 4109874 | 3.24 | 11.67 | 13.37 | 7.44 | Indole-3-glycerol phosphate synthase |
| *FEPFGCPE_03779* | *trpD-2* | - | 4109991 | 4111019 | 10.37 | 22.83 | 24.40 | 16.80 | Anthranilate phosphoribosyltransferase |
| *FEPFGCPE_03780* | *trpG* | - | 4111082 | 4111669 | 25.51 | 48.20 | 62.07 | 38.57 | Anthranilate synthase component 2 |
| *FEPFGCPE_03781* | *puuB-4* | - | 4112463 | 4113758 | 0.68 | 7.26 | 0.87 | 0.44 | Gamma-glutamylputrescine oxidoreductase |
| *FEPFGCPE_03782* | *glnQ-10* | - | 4113820 | 4114551 | 0.56 | 1.03 | 4.26 | 1.32 | Glutamine transport ATP-binding protein GlnQ |
| *FEPFGCPE_03783* | *yecS-8* | - | 4114548 | 4115207 | 0.84 | 7.66 | 3.08 | 1.07 | L-cystine transport system permease protein YecS |
| *FEPFGCPE_03784* | *artJ-2* | - | 4115299 | 4116051 | 0.36 | 0.96 | 1.11 | 1.24 | ABC transporter arginine-binding protein 1 |
| *FEPFGCPE_03785* | *gntR-3* | + | 4116456 | 4117130 | 7.17 | 27.49 | 20.20 | 4.45 | putative D-xylose utilization operon transcriptional repressor |
| *FEPFGCPE_03786* | | + | 4117678 | 4118706 | 3.29 | 8.32 | 9.79 | 1.75 | Hypothetical protein |
| *FEPFGCPE_03787* | *glpC-1* | + | 4118708 | 4121683 | 0.81 | 4.98 | 3.73 | 1.40 | Anaerobic glycerol-3-phosphate dehydrogenase subunit C |
| *FEPFGCPE_03788* | *araE-3* | - | 4121825 | 4123258 | 0.62 | 6.26 | 3.78 | 1.39 | Alpha-ketoglutaric semialdehyde dehydrogenase |
| *FEPFGCPE_03789* | *abo-4* | - | 4123377 | 4124480 | 0.37 | 1.04 | 1.08 | 0.00 | 4-methylaminobutanoate oxidase (formaldehyde-forming) |
| *FEPFGCPE_03790* | | - | 4125036 | 4125416 | 0.54 | 2.37 | 4.83 | 1.14 | Hypothetical protein |
| *FEPFGCPE_03791* | *proP-15* | - | 4125442 | 4126722 | 4.43 | 28.39 | 14.05 | 3.04 | Proline/betaine transporter |
| *FEPFGCPE_03792* | *gcvA-7* | + | 4127022 | 4127936 | 3.35 | 14.24 | 8.87 | 2.09 | Glycine cleavage system transcriptional activator |
| *FEPFGCPE_03793* | *resA-3* | - | 4128093 | 4128932 | 1.75 | 15.79 | 3.68 | 3.91 | Thiol-disulfide oxidoreductase ResA |
| *FEPFGCPE_03794* | *dsbD-2* | - | 4128929 | 4130773 | 0.37 | 0.27 | 1.44 | 0.31 | Thiol:disulfide interchange protein DsbD |
| *FEPFGCPE_03795* | *qseB-2* | + | 4130977 | 4131741 | 2.57 | 4.22 | 5.02 | 1.53 | Transcriptional regulatory protein QseB |
| *FEPFGCPE_03796* | *qseC-4* | + | 4131767 | 4133047 | 2.47 | 6.24 | 9.63 | 2.43 | Sensor protein QseC |
| *FEPFGCPE_03797* | *yhhJ* | - | 4133086 | 4134276 | 1.22 | 7.86 | 3.32 | 1.39 | Inner membrane transport permease YhhJ |
| *FEPFGCPE_03798* | *rbbA* | - | 4134276 | 4137071 | 1.76 | 6.70 | 3.35 | 0.40 | Ribosome-associated ATPase |
| *FEPFGCPE_03799* | *mdtN* | - | 4137082 | 4138155 | 3.25 | 9.79 | 5.83 | 0.51 | Multidrug resistance protein MdtN |
| *FEPFGCPE_03800* | *oprM-1* | - | 4138152 | 4139753 | 4.37 | 8.91 | 5.81 | 2.34 | Outer membrane protein OprM |
| *FEPFGCPE_03801* | *trpE* | - | 4140001 | 4141506 | 5.29 | 14.25 | 6.79 | 6.11 | Anthranilate synthase component 1 |
| *FEPFGCPE_03802* | *gph-3* | - | 4141892 | 4142608 | 3.54 | 28.63 | 20.73 | 8.09 | Phosphoglycolate phosphatase |
| *FEPFGCPE_03803* | *rpe* | - | 4142605 | 4143285 | 12.57 | 53.65 | 47.18 | 22.86 | Ribulose-phosphate 3-epimerase |
| *FEPFGCPE_03804* | *apaG* | + | 4143488 | 4143862 | 16.29 | 17.82 | 19.70 | 7.59 | Protein ApaG |
| *FEPFGCPE_03805* | | - | 4143890 | 4144141 | 12.53 | 33.01 | 32.28 | 4.25 | Hypothetical protein |
| *FEPFGCPE_03806* | *mltA* | + | 4144308 | 4145282 | 7.10 | 19.50 | 10.12 | 5.37 | Membrane-bound lytic murein transglycosylase A |
| *FEPFGCPE_03807* | *paaK* | - | 4145357 | 4146652 | 1.41 | 4.57 | 4.87 | 1.99 | Phenylacetate-coenzyme A ligase |
| *FEPFGCPE_03808* | *paaI* | - | 4146746 | 4147204 | 1.46 | 3.45 | 3.71 | 3.41 | Acyl-coenzyme A thioesterase PaaI |
| *FEPFGCPE_03809* | *paaG-3* | - | 4147237 | 4148019 | 3.16 | 11.59 | 11.12 | 7.82 | 1,2-epoxyphenylacetyl-CoA isomerase |
| *FEPFGCPE_03810* | *pcaF-2* | - | 4148073 | 4149275 | 2.86 | 8.47 | 7.76 | 6.50 | Beta-ketoadipyl-CoA thiolase |
| *FEPFGCPE_03811* | *xylC* | - | 4149387 | 4151081 | 2.86 | 9.00 | 7.70 | 4.95 | Benzaldehyde dehydrogenase [NAD(+)] |
| *FEPFGCPE_03812* | *echA8-3* | + | 4151614 | 4152390 | 8.21 | 51.17 | 39.69 | 31.35 | putative enoyl-CoA hydratase echA8 |
| *FEPFGCPE_03813* | *artI* | + | 4152475 | 4153278 | 8.27 | 55.77 | 36.61 | 19.70 | Putative ABC transporter arginine-binding protein 2 |
| *FEPFGCPE_03814* | | + | 4153860 | 4155378 | 122920.04 | 101911.32 | 102054.11 | 90383.14 | rRNA |
| *FEPFGCPE_03815* | | + | 4155448 | 4155524 | 191.30 | 779.63 | 568.80 | 264.81 | tRNA |
| *FEPFGCPE_03816* | | + | 4155633 | 4155705 | 0.00 | 0.00 | 0.89 | 0.00 | tRNA |
| *FEPFGCPE_03817* | | + | 4155999 | 4158871 | 112993.75 | 97957.32 | 109933.60 | 146291.51 | rRNA |
| *FEPFGCPE_03818* | | + | 4159006 | 4159117 | 534.16 | 65.36 | 77.09 | 82.97 | rRNA |
| *FEPFGCPE_03819* | | + | 4159316 | 4160089 | 1.75 | 20.06 | 13.70 | 2.40 | Hypothetical protein |
| *FEPFGCPE_03820* | *yxeP-5* | - | 4160180 | 4161376 | 17.90 | 80.61 | 66.92 | 29.60 | putative hydrolase YxeP |
| *FEPFGCPE_03821* | *dmsA* | + | 4161591 | 4163666 | 5.97 | 23.85 | 19.09 | 8.67 | Dimethyl sulfoxide reductase DmsA |
| *FEPFGCPE_03822* | | + | 4163716 | 4164879 | 2.07 | 10.90 | 9.29 | 1.18 | Hypothetical protein |
| *FEPFGCPE_03823* | *badR-1* | - | 4165063 | 4165542 | 6.39 | 21.33 | 17.21 | 5.42 | Transcriptional activatory protein BadR |
| *FEPFGCPE_03824* | | - | 4165745 | 4166899 | 263.53 | 1159.20 | 736.82 | 772.40 | Outer membrane porin protein |
| *FEPFGCPE_03825* | *coq7* | + | 4167218 | 4167874 | 7.54 | 39.85 | 32.95 | 12.42 | 2-nonaprenyl-3-methyl-6-methoxy-1,4-benzoquinol hydroxylase |
| *FEPFGCPE_03826* | *mraZ* | + | 4168337 | 4168765 | 23.03 | 39.03 | 18.05 | 12.24 | Transcriptional regulator MraZ |
| *FEPFGCPE_03827* | *rsmH* | + | 4168788 | 4169741 | 32.62 | 74.89 | 48.46 | 21.84 | Ribosomal RNA small subunit methyltransferase H |
| *FEPFGCPE_03828* | *ftsL* | + | 4169738 | 4170082 | 8.28 | 20.44 | 3.95 | 4.96 | Cell division protein FtsL |
| *FEPFGCPE_03829* | *penA-1* | + | 4170079 | 4171854 | 7.77 | 29.37 | 12.40 | 8.78 | putative peptidoglycan D,D-transpeptidase PenA |
| *FEPFGCPE_03830* | *murE* | + | 4171842 | 4173341 | 4.80 | 19.49 | 5.58 | 2.59 | UDP-N-acetylmuramoyl-L-alanyl-D-glutamate--2,6-diaminopimelate ligase |

| Locus name | Gene name | Strand | Gene start (bp) | Gene end (bp) | Average TPM Inoculum | Average TPM $t_6$ | Average TPM $t_{24}$ | Average TPM $t_{168}$ | Annotation description |
|---|---|---|---|---|---|---|---|---|---|
| *FEPFGCPE_03831* | *murF* | + | 4173420 | 4174829 | 3.77 | 12.42 | 6.76 | 4.77 | UDP-N-acetylmuramoyl-tripeptide--D-alanyl-D-alanine ligase |
| *FEPFGCPE_03832* | *mraY* | + | 4174845 | 4176014 | 2.38 | 14.18 | 6.14 | 2.84 | Phospho-N-acetylmuramoyl-pentapeptide-transferase |
| *FEPFGCPE_03833* | *murD* | + | 4176177 | 4177703 | 4.41 | 21.91 | 7.93 | 7.49 | UDP-N-acetylmuramoylalanine--D-glutamate ligase |
| *FEPFGCPE_03834* | *ftsW* | + | 4177700 | 4178968 | 4.12 | 15.50 | 4.50 | 4.57 | putative peptidoglycan glycosyltransferase FtsW |
| *FEPFGCPE_03835* | *murG* | + | 4178965 | 4180068 | 2.18 | 18.13 | 4.85 | 3.97 | UDP-N-acetylglucosamine--N-acetylmuramyl-(pentapeptide) pyrophosphoryl-undecaprenol N-acetylglucosamine transferase |
| *FEPFGCPE_03836* | *murC* | + | 4180070 | 4181479 | 1.80 | 11.00 | 8.75 | 5.22 | UDP-N-acetylmuramate--L-alanine ligase |
| *FEPFGCPE_03837* | *ddl* | + | 4181481 | 4182464 | 3.59 | 45.56 | 22.05 | 9.46 | D-alanine--D-alanine ligase |
| *FEPFGCPE_03838* | *ftsQ* | + | 4182512 | 4183348 | 4.10 | 39.39 | 16.11 | 6.23 | Cell division protein FtsQ |
| *FEPFGCPE_03839* | *ftsA* | + | 4183371 | 4184603 | 4.67 | 46.89 | 26.06 | 12.02 | Cell division protein FtsA |
| *FEPFGCPE_03840* | *ftsZ* | + | 4184710 | 4185900 | 20.82 | 144.05 | 75.72 | 78.94 | Cell division protein FtsZ |
| *FEPFGCPE_03841* | *lpxC* | + | 4186439 | 4187353 | 38.48 | 41.42 | 50.07 | 23.29 | UDP-3-O-acyl-N-acetylglucosamine deacetylase |
| *FEPFGCPE_03842* | | - | 4187497 | 4187964 | 3.43 | 35.52 | 22.96 | 10.14 | Hypothetical protein |
| *FEPFGCPE_03843* | *secA* | + | 4188270 | 4191065 | 4.82 | 17.95 | 17.38 | 9.52 | Protein translocase subunit SecA |
| *FEPFGCPE_03844* | *argJ* | + | 4191229 | 4192458 | 1.84 | 11.09 | 11.77 | 5.08 | Arginine biosynthesis bifunctional protein ArgJ |
| *FEPFGCPE_03845* | | + | 4192665 | 4193546 | 2.50 | 15.54 | 11.94 | 5.36 | Hypothetical protein |
| *FEPFGCPE_03846* | *mutT* | + | 4193657 | 4194130 | 3.65 | 4.68 | 15.25 | 4.77 | 8-oxo-dGTP diphosphatase |
| *FEPFGCPE_03847* | *yacG* | - | 4194254 | 4194448 | 5.52 | 21.76 | 10.09 | 11.16 | DNA gyrase inhibitor YacG |
| *FEPFGCPE_03848* | *zapD* | - | 4194453 | 4195208 | 7.59 | 39.32 | 31.16 | 8.10 | Cell division protein ZapD |
| *FEPFGCPE_03849* | *coaE* | - | 4195349 | 4195963 | 2.51 | 12.17 | 5.32 | 5.34 | Dephospho-CoA kinase |
| *FEPFGCPE_03850* | *outO-2* | - | 4195960 | 4196802 | 3.78 | 17.02 | 7.91 | 2.48 | Type 4 prepilin-like proteins leader peptide-processing enzyme |
| *FEPFGCPE_03851* | *gspF* | - | 4196799 | 4198019 | 1.71 | 6.61 | 3.92 | 0.85 | Putative type II secretion system protein F |
| *FEPFGCPE_03852* | *hxcR* | - | 4198137 | 4199333 | 0.32 | 1.61 | 0.04 | 1.79 | putative type II secretion system protein HxcR |
| *FEPFGCPE_03853* | *corC-1* | - | 4199447 | 4200826 | 1.62 | 15.07 | 15.10 | 3.54 | Magnesium and cobalt efflux protein CorC |
| *FEPFGCPE_03854* | | + | 4201338 | 4202132 | 10.03 | 26.31 | 7.94 | 3.91 | Hypothetical protein |
| *FEPFGCPE_03855* | | + | 4202143 | 4202778 | 9.40 | 16.24 | 17.17 | 6.10 | Hypothetical protein |
| *FEPFGCPE_03856* | | + | 4202788 | 4203621 | 5.69 | 11.76 | 10.32 | 4.32 | Hypothetical protein |
| *FEPFGCPE_03857* | | + | 4203789 | 4204247 | 3.74 | 12.26 | 8.79 | 1.10 | Hypothetical protein |
| *FEPFGCPE_03858* | *hin* | - | 4204305 | 4204874 | 2.82 | 5.25 | 3.45 | 0.81 | DNA-invertase hin |
| *FEPFGCPE_03859* | | - | 4205089 | 4205430 | 0.36 | 1.92 | 0.00 | 0.00 | Hypothetical protein |
| *FEPFGCPE_03860* | | + | 4205555 | 4206016 | 3.05 | 15.10 | 11.20 | 5.35 | Hypothetical protein |
| *FEPFGCPE_03861* | | + | 4206029 | 4206364 | 1.98 | 11.00 | 7.41 | 2.14 | Hypothetical protein |
| *FEPFGCPE_03862* | | + | 4206401 | 4206802 | 16.72 | 30.69 | 34.33 | 5.61 | Hypothetical protein |
| *FEPFGCPE_03863* | | + | 4206799 | 4207635 | 4.43 | 13.04 | 11.71 | 1.44 | Hypothetical protein |
| *FEPFGCPE_03864* | | + | 4207685 | 4208704 | 3.42 | 15.00 | 10.70 | 2.82 | Hypothetical protein |
| *FEPFGCPE_03865* | | - | 4208903 | 4209100 | 16.36 | 38.99 | 21.41 | 10.26 | Hypothetical protein |
| *FEPFGCPE_03866* | | + | 4209410 | 4209868 | 1.91 | 5.69 | 3.93 | 0.00 | Hypothetical protein |
| *FEPFGCPE_03867* | | + | 4209884 | 4209985 | 0.84 | 8.27 | 8.46 | 0.00 | Hypothetical protein |
| *FEPFGCPE_03868* | | + | 4210023 | 4210250 | 2.94 | 16.26 | 12.88 | 0.00 | Hypothetical protein |
| *FEPFGCPE_03869* | | + | 4210745 | 4211179 | 0.36 | 6.56 | 5.23 | 2.21 | Hypothetical protein |
| *FEPFGCPE_03870* | | + | 4211569 | 4213242 | 6.65 | 12.72 | 11.94 | 3.82 | Hypothetical protein |
| *FEPFGCPE_03871* | | + | 4213239 | 4215905 | 2.59 | 10.78 | 8.70 | 3.29 | Hypothetical protein |
| *FEPFGCPE_03872* | | + | 4216779 | 4217570 | 1.36 | 6.43 | 4.05 | 0.04 | Hypothetical protein |
| *FEPFGCPE_03873* | | - | 4217734 | 4219371 | 1.35 | 6.88 | 4.05 | 0.67 | Hypothetical protein |
| *FEPFGCPE_03874* | | - | 4219587 | 4219663 | 8.75 | 6.47 | 6.55 | 3.44 | tRNA |
| *FEPFGCPE_03875* | *ispB* | - | 4219687 | 4220667 | 12.90 | 25.86 | 20.99 | 9.42 | Octaprenyl diphosphate synthase |
| *FEPFGCPE_03876* | *rplU* | + | 4220945 | 4221256 | 63.45 | 254.16 | 280.38 | 223.31 | 50S ribosomal protein L21 |
| *FEPFGCPE_03877* | *rpmA* | + | 4221285 | 4221545 | 8.24 | 25.52 | 51.89 | 48.97 | 50S ribosomal protein L27 |
| *FEPFGCPE_03878* | *obg-1* | + | 4221694 | 4222161 | 3.99 | 17.12 | 17.13 | 20.26 | GTPase Obg |
| *FEPFGCPE_03879* | *obg-2* | + | 4222143 | 4222823 | 2.40 | 6.07 | 2.11 | 4.76 | GTPase Obg |
| *FEPFGCPE_03880* | *proB* | + | 4222936 | 4224069 | 2.76 | 7.99 | 11.02 | 7.10 | Glutamate 5-kinase |
| *FEPFGCPE_03881* | | - | 4224188 | 4224766 | 2.06 | 23.08 | 12.26 | 9.16 | Hypothetical protein |
| *FEPFGCPE_03882* | *rppH* | - | 4224908 | 4225558 | 36.32 | 242.16 | 139.41 | 33.03 | RNA pyrophosphohydrolase |
| *FEPFGCPE_03883* | *proS* | + | 4225772 | 4227511 | 4.14 | 15.88 | 12.39 | 7.84 | Proline--tRNA ligase |
| *FEPFGCPE_03884* | *hpt* | - | 4227631 | 4228179 | 11.47 | 47.92 | 30.23 | 17.36 | Hypoxanthine-guanine phosphoribosyltransferase |
| *FEPFGCPE_03885* | *ffh* | - | 4228290 | 4229654 | 11.00 | 39.92 | 36.72 | 14.45 | Signal recognition particle protein |
| *FEPFGCPE_03886* | *ypjD* | + | 4229768 | 4230652 | 3.23 | 22.37 | 22.76 | 7.73 | Inner membrane protein YpjD |
| *FEPFGCPE_03887* | | + | 4230659 | 4230937 | 6.59 | 46.98 | 36.66 | 5.66 | Hypothetical protein |
| *FEPFGCPE_03888* | | + | 4231110 | 4231628 | 97.43 | 2567.33 | 2836.56 | 963.98 | Hypothetical protein |
| *FEPFGCPE_03889* | *ampD* | + | 4232089 | 4232688 | 2.87 | 10.44 | 23.95 | 4.86 | 1,6-anhydro-N-acetylmuramyl-L-alanine amidase AmpD |
| *FEPFGCPE_03890* | *nrdA* | + | 4233063 | 4236017 | 13.44 | 52.34 | 42.46 | 34.66 | Ribonucleoside-diphosphate reductase 1 subunit alpha |
| *FEPFGCPE_03891* | | - | 4236129 | 4237091 | 2.96 | 25.08 | 18.08 | 5.27 | Hypothetical protein |
| *FEPFGCPE_03892* | *nrdB* | + | 4237636 | 4238778 | 12.94 | 43.64 | 29.08 | 22.15 | Ribonucleoside-diphosphate reductase subunit beta |
| *FEPFGCPE_03893* | *hctB* | + | 4239485 | 4240096 | 182.56 | 2177.72 | 1094.51 | 360.50 | Histone H1-like protein HC2 |
| *FEPFGCPE_03894* | *pcp* | - | 4240211 | 4240690 | 60.62 | 66.37 | 59.78 | 32.30 | Outer membrane lipoprotein pcp |
| *FEPFGCPE_03895* | *adoK* | - | 4240710 | 4241645 | 35.70 | 38.40 | 34.80 | 14.18 | Adenosine kinase |
| *FEPFGCPE_03896* | *tpx* | - | 4241824 | 4242324 | 6.74 | 78.17 | 50.32 | 30.09 | putative thiol peroxidase |
| *FEPFGCPE_03897* | | - | 4242357 | 4243310 | 3.68 | 27.95 | 31.30 | 6.08 | Hypothetical protein |
| *FEPFGCPE_03898* | *prmA-2* | - | 4243374 | 4244267 | 2.06 | 20.98 | 17.75 | 10.33 | Ribosomal protein L11 methyltransferase |
| *FEPFGCPE_03899* | *accC* | - | 4244307 | 4245668 | 2.32 | 26.18 | 25.87 | 14.89 | Biotin carboxylase |
| *FEPFGCPE_03900* | *accB* | - | 4245768 | 4246229 | 2.15 | 12.52 | 21.70 | 13.75 | Biotin carboxyl carrier protein of acetyl-CoA carboxylase |
| *FEPFGCPE_03901* | *aroQ-3* | - | 4246299 | 4246745 | 4.70 | 52.14 | 42.67 | 12.89 | 3-dehydroquinate dehydratase |

| Locus name | Gene name | Strand | Gene start (bp) | Gene end (bp) | Average TPM Inoculum | Average TPM $t_6$ | Average TPM $t_{24}$ | Average TPM $t_{168}$ | Annotation description |
|---|---|---|---|---|---|---|---|---|---|
| *FEPFGCPE_03902* | *tlpA* | - | 4246878 | 4247402 | 1.64 | 16.05 | 9.44 | 2.43 | Thiol:disulfide interchange protein TlpA |
| *FEPFGCPE_03903* | | - | 4247504 | 4248238 | 8.01 | 60.68 | 45.43 | 14.38 | Hypothetical protein |
| *FEPFGCPE_03904* | *mpl* | + | 4248281 | 4249663 | 4.26 | 25.49 | 29.08 | 12.74 | UDP-N-acetylmuramate--L-alanyl-gamma-D-glutamyl-meso-2,6-diaminoheptandioate ligase |
| *FEPFGCPE_03905* | | + | 4249713 | 4250324 | 3.74 | 7.65 | 5.52 | 5.46 | Hypothetical protein |
| *FEPFGCPE_03906* | *rnb* | + | 4250372 | 4252621 | 5.12 | 30.04 | 32.37 | 10.67 | Exoribonuclease 2 |
| *FEPFGCPE_03907* | | + | 4252678 | 4253574 | 0.92 | 8.30 | 7.98 | 3.54 | Hypothetical protein |
| *FEPFGCPE_03908* | *aroE-2* | + | 4253787 | 4254659 | 1.55 | 7.39 | 7.02 | 3.16 | Shikimate dehydrogenase (NADP(+)) |
| *FEPFGCPE_03909* | *mtgA* | + | 4254668 | 4255408 | 2.98 | 10.40 | 14.11 | 2.32 | Biosynthetic peptidoglycan transglycosylase |
| *FEPFGCPE_03910* | *mscS-1* | - | 4255468 | 4256343 | 22.13 | 18.10 | 20.05 | 18.44 | Small-conductance mechanosensitive channel |
| *FEPFGCPE_03911* | *pyrF* | + | 4256441 | 4257334 | 2.36 | 27.27 | 22.68 | 4.38 | Orotidine 5'-phosphate decarboxylase |
| *FEPFGCPE_03912* | *pncC* | - | 4257391 | 4257894 | 3.89 | 10.04 | 8.23 | 7.34 | Nicotinamide-nucleotide amidohydrolase PncC |
| *FEPFGCPE_03913* | *pgpA* | - | 4258027 | 4258548 | 2.45 | 14.24 | 11.37 | 3.46 | Phosphatidylglycerophosphatase A |
| *FEPFGCPE_03914* | *thiL* | - | 4258545 | 4259546 | 2.35 | 15.21 | 15.62 | 3.67 | Thiamine-monophosphate kinase |
| *FEPFGCPE_03915* | *maeB-2* | + | 4259978 | 4262281 | 13.15 | 61.93 | 51.46 | 19.67 | NADP-dependent malic enzyme |
| *FEPFGCPE_03916* | *rnaSA* | + | 4262568 | 4262969 | 60.47 | 255.01 | 215.27 | 46.96 | Guanyl-specific ribonuclease Sa |
| *FEPFGCPE_03917* | | + | 4262977 | 4263525 | 27.95 | 104.17 | 75.83 | 35.38 | Barstar |
| *FEPFGCPE_03918* | | - | 4263693 | 4264337 | 16.79 | 94.24 | 96.96 | 55.68 | Hypothetical protein |
| *FEPFGCPE_03919* | *rsmE* | - | 4264386 | 4265123 | 15.26 | 108.47 | 82.98 | 24.76 | Ribosomal RNA small subunit methyltransferase E |
| *FEPFGCPE_03920* | | - | 4265203 | 4265625 | 11.12 | 23.00 | 28.37 | 5.93 | Hypothetical protein |
| *FEPFGCPE_03921* | *speE-2* | - | 4265787 | 4266893 | 6.24 | 22.28 | 15.67 | 6.90 | Polyamine aminopropyltransferase |
| *FEPFGCPE_03922* | *tktA-2* | + | 4266925 | 4268904 | 6.62 | 28.66 | 31.41 | 25.60 | Transketolase 1 |
| *FEPFGCPE_03923* | *gap* | + | 4268933 | 4269943 | 12.87 | 48.63 | 58.53 | 41.45 | Glyceraldehyde-3-phosphate dehydrogenase |
| *FEPFGCPE_03924* | *fur* | - | 4270242 | 4270679 | 29.15 | 182.71 | 116.76 | 53.25 | Ferric uptake regulation protein |
| *FEPFGCPE_03925* | *bamE-2* | + | 4270768 | 4271514 | 4.88 | 41.60 | 38.29 | 32.08 | Outer membrane protein assembly factor BamE |
| *FEPFGCPE_03926* | *dapB* | + | 4271539 | 4272339 | 3.79 | 45.44 | 33.00 | 18.61 | 4-hydroxy-tetrahydrodipicolinate reductase |
| *FEPFGCPE_03927* | | + | 4272389 | 4272550 | 4.42 | 29.29 | 36.85 | 15.65 | Hypothetical protein |
| *FEPFGCPE_03928* | *leuS* | + | 4272655 | 4275249 | 6.00 | 28.88 | 25.70 | 13.06 | Leucine--tRNA ligase |
| *FEPFGCPE_03929* | *lptE* | + | 4275258 | 4275860 | 4.73 | 19.72 | 15.17 | 11.01 | LPS-assembly lipoprotein LptE |
| *FEPFGCPE_03930* | *holA* | + | 4275872 | 4276951 | 6.56 | 14.76 | 15.76 | 4.85 | DNA polymerase III subunit delta |
| *FEPFGCPE_03931* | *proA-3* | + | 4277158 | 4278426 | 2.53 | 12.21 | 8.94 | 7.92 | Gamma-glutamyl phosphate reductase |
| *FEPFGCPE_03932* | *ves* | + | 4278449 | 4279057 | 1.59 | 4.96 | 1.27 | 1.62 | Protein Ves |
| *FEPFGCPE_03933* | *rclR* | + | 4279283 | 4280224 | 0.40 | 0.94 | 2.67 | 0.26 | RCS-specific HTH-type transcriptional activator RclR |
| *FEPFGCPE_03934* | *iorA-2* | + | 4280396 | 4280839 | 0.64 | 0.00 | 0.00 | 0.95 | Isoquinoline 1-oxidoreductase subunit alpha |
| *FEPFGCPE_03935* | *iorB-2* | + | 4280820 | 4282709 | 0.23 | 0.38 | 0.53 | 0.00 | Isoquinoline 1-oxidoreductase subunit beta |
| *FEPFGCPE_03936* | *iorB-3* | + | 4282691 | 4283110 | 0.13 | 0.00 | 0.00 | 0.00 | Isoquinoline 1-oxidoreductase subunit beta |
| *FEPFGCPE_03937* | *pucA-2* | + | 4283110 | 4284120 | 0.29 | 0.90 | 0.66 | 0.46 | putative xanthine dehydrogenase subunit A |
| *FEPFGCPE_03938* | | + | 4284164 | 4285219 | 0.42 | 2.51 | 1.48 | 0.41 | Outer membrane porin protein |
| *FEPFGCPE_03939* | *purH* | - | 4285308 | 4286873 | 2.05 | 14.55 | 15.92 | 16.48 | Bifunctional purine biosynthesis protein PurH |
| *FEPFGCPE_03940* | *fis* | - | 4287017 | 4287250 | 7.57 | 49.73 | 33.25 | 20.46 | DNA-binding protein Fis |
| *FEPFGCPE_03941* | *dus* | - | 4287247 | 4288227 | 9.79 | 48.60 | 51.55 | 15.85 | putative tRNA-dihydrouridine synthase |
| *FEPFGCPE_03942* | *ubiH* | - | 4288384 | 4289718 | 1.61 | 14.30 | 9.04 | 4.70 | 2-octaprenyl-6-methoxyphenol hydroxylase |
| *FEPFGCPE_03943* | *pepP* | - | 4289718 | 4291049 | 3.45 | 22.57 | 16.12 | 10.72 | Xaa-Pro aminopeptidase |
| *FEPFGCPE_03944* | *cugP* | - | 4291092 | 4291841 | 3.59 | 14.01 | 7.95 | 5.25 | UTP--glucose-1-phosphate uridylyltransferase |
| *FEPFGCPE_03945* | | - | 4291852 | 4292937 | 4.27 | 22.59 | 12.84 | 9.20 | Hypothetical protein |
| *FEPFGCPE_03946* | *lptD* | + | 4293047 | 4295461 | 3.49 | 21.33 | 16.71 | 7.47 | LPS-assembly protein LptD |
| *FEPFGCPE_03947* | *surA* | + | 4295454 | 4296821 | 4.76 | 21.47 | 13.77 | 11.85 | Chaperone SurA |
| *FEPFGCPE_03948* | *pdxA* | + | 4296873 | 4297952 | 4.51 | 20.45 | 15.55 | 7.62 | 4-hydroxythreonine-4-phosphate dehydrogenase |
| *FEPFGCPE_03949* | *rsmA* | + | 4297963 | 4298796 | 2.21 | 12.37 | 11.19 | 3.85 | Ribosomal RNA small subunit methyltransferase A |
| *FEPFGCPE_03950* | *eamA-2* | - | 4298771 | 4299661 | 3.25 | 20.10 | 10.34 | 4.42 | putative amino-acid metabolite efflux pump |
| *FEPFGCPE_03951* | *rubA2* | + | 4299902 | 4300339 | 2.70 | 25.53 | 12.27 | 4.16 | Rubredoxin-2 |
| *FEPFGCPE_03952* | *gloA* | - | 4300448 | 4300915 | 6.23 | 61.57 | 59.55 | 20.38 | Lactoylglutathione lyase |
| *FEPFGCPE_03953* | | - | 4300961 | 4301902 | 2.66 | 7.07 | 5.77 | 3.42 | Hypothetical protein |
| *FEPFGCPE_03954* | | - | 4301925 | 4302125 | 8.95 | 20.71 | 29.48 | 22.94 | Hypothetical protein |
| *FEPFGCPE_03955* | | - | 4302818 | 4303411 | 4.65 | 19.59 | 11.07 | 9.52 | D-glycero-beta-D-manno-heptose-1,7-bisphosphate 7-phosphatase |
| *FEPFGCPE_03956* | *glyS* | - | 4303444 | 4305573 | 3.74 | 12.54 | 7.98 | 6.92 | Glycine--tRNA ligase beta subunit |
| *FEPFGCPE_03957* | *glyQ* | - | 4305570 | 4306508 | 6.06 | 25.26 | 21.33 | 24.34 | Glycine--tRNA ligase alpha subunit |
| *FEPFGCPE_03958* | *lnt* | - | 4306714 | 4308279 | 0.79 | 9.44 | 3.77 | 2.66 | Apolipoprotein N-acyltransferase |
| *FEPFGCPE_03959* | *corC-2* | - | 4308319 | 4309194 | 9.76 | 93.34 | 76.20 | 23.84 | Magnesium and cobalt efflux protein CorC |
| *FEPFGCPE_03960* | | - | 4309347 | 4310033 | 1.81 | 16.78 | 7.97 | 1.65 | Hypothetical protein |
| *FEPFGCPE_03961* | *ybeY* | - | 4310045 | 4310572 | 4.05 | 21.41 | 12.68 | 8.67 | Endoribonuclease YbeY |
| *FEPFGCPE_03962* | | - | 4310569 | 4311384 | 2.50 | 17.22 | 18.68 | 4.08 | PhoH-like protein |
| *FEPFGCPE_03963* | *miaB* | - | 4311750 | 4313093 | 6.42 | 35.10 | 24.61 | 12.35 | tRNA-2-methylthio-N(6)-dimethylallyladenosine synthase |
| *FEPFGCPE_03964* | *mntA* | - | 4313306 | 4314235 | 1.87 | 7.71 | 21.57 | 15.31 | Manganese-binding lipoprotein MntA |
| *FEPFGCPE_03965* | *mntB* | - | 4314247 | 4315110 | 2.13 | 9.52 | 23.11 | 8.82 | Manganese transport system membrane protein MntB |
| *FEPFGCPE_03966* | *fecE* | - | 4315107 | 4315922 | 1.78 | 4.77 | 17.05 | 11.00 | Fe(3+) dicitrate transport ATP-binding protein FecE |
| *FEPFGCPE_03967* | | - | 4315919 | 4316041 | 0.55 | 1.39 | 51.46 | 0.83 | Hypothetical protein |
| *FEPFGCPE_03968* | *lon-2* | + | 4317070 | 4319022 | 206.86 | 758.04 | 301.22 | 110.19 | Lon protease |
| *FEPFGCPE_03969* | | + | 4319038 | 4320300 | 57.16 | 419.00 | 153.78 | 57.22 | Hypothetical protein |
| *FEPFGCPE_03970* | | + | 4320338 | 4321882 | 16.33 | 174.47 | 66.04 | 20.41 | Hypothetical protein |
| *FEPFGCPE_03971* | | + | 4321882 | 4322301 | 5.38 | 108.34 | 32.87 | 12.86 | Hypothetical protein |
| *FEPFGCPE_03972* | *ycdT* | - | 4322315 | 4323682 | 2.80 | 31.73 | 12.93 | 4.29 | putative diguanylate cyclase YcdT |

| Locus name | Gene name | Strand | Gene start (bp) | Gene end (bp) | Average TPM Inoculum | Average TPM $t_6$ | Average TPM $t_{24}$ | Average TPM $t_{168}$ | Annotation description |
|---|---|---|---|---|---|---|---|---|---|
| *FEPFGCPE_03973* | *osmY-3* | + | 4324573 | 4325301 | 0.81 | 2.77 | 5.09 | 1.27 | Osmoprotectant import permease protein OsmY |
| *FEPFGCPE_03974* | *osmX* | + | 4325322 | 4326263 | 0.59 | 2.70 | 4.21 | 3.45 | Osmoprotectant-binding protein OsmX |
| *FEPFGCPE_03975* | *osmW* | + | 4326312 | 4326959 | 1.26 | 2.63 | 7.65 | 4.08 | Osmoprotectant import permease protein OsmW |
| *FEPFGCPE_03976* | *osmV* | + | 4326956 | 4328155 | 1.13 | 4.16 | 4.22 | 5.44 | Osmoprotectant import ATP-binding protein OsmV |
| *FEPFGCPE_03977* | *ybdG* | + | 4328322 | 4329569 | 1.18 | 1.25 | 5.83 | 2.28 | Miniconductance mechanosensitive channel YbdG |
| *FEPFGCPE_03978* | | + | 4329606 | 4329749 | 1.56 | 2.34 | 1.75 | 1.04 | Hypothetical protein |
| *FEPFGCPE_03979* | *nemA-2* | - | 4329965 | 4331089 | 3.41 | 17.20 | 15.84 | 12.88 | N-ethylmaleimide reductase |
| *FEPFGCPE_03980* | *oxlT-2* | - | 4331267 | 4332553 | 1.40 | 7.80 | 4.11 | 1.77 | Oxalate:formate antiporter |
| *FEPFGCPE_03981* | *acuR-1* | + | 4332764 | 4333531 | 4.01 | 17.97 | 15.73 | 4.60 | Transcriptional regulator AcuR |
| *FEPFGCPE_03982* | *sutR-2* | - | 4333592 | 4334182 | 4.42 | 16.76 | 9.86 | 2.50 | HTH-type transcriptional regulator SutR |
| *FEPFGCPE_03983* | *kbl* | + | 4334392 | 4335609 | 73.66 | 67.99 | 33.52 | 16.47 | 2-amino-3-ketobutyrate coenzyme A ligase |
| *FEPFGCPE_03984* | | + | 4335612 | 4336577 | 117.15 | 227.55 | 89.13 | 48.57 | putative epimerase/dehydratase |
| *FEPFGCPE_03985* | | - | 4336735 | 4336878 | 83.12 | 350.83 | 96.47 | 18.06 | Hypothetical protein |
| *FEPFGCPE_03986* | | + | 4337320 | 4338204 | 7.38 | 22.75 | 10.15 | 5.07 | Hypothetical protein |
| *FEPFGCPE_03987* | *cmpD-4* | - | 4338276 | 4339592 | 3.06 | 15.47 | 5.88 | 5.56 | Bicarbonate transport ATP-binding protein CmpD |
| *FEPFGCPE_03988* | *cmpB-2* | - | 4339603 | 4341342 | 1.85 | 11.10 | 5.28 | 1.69 | Bicarbonate transport system permease protein CmpB |
| *FEPFGCPE_03989* | *lapB-2* | + | 4341809 | 4343410 | 12.38 | 8.33 | 9.24 | 7.15 | Lipopolysaccharide assembly protein B |
| *FEPFGCPE_03990* | | - | 4343392 | 4344465 | 5.98 | 6.26 | 6.47 | 3.66 | Hypothetical protein |
| *FEPFGCPE_03991* | *nhaR* | - | 4344452 | 4345417 | 3.63 | 7.40 | 8.20 | 2.58 | Transcriptional activator protein NhaR |
| *FEPFGCPE_03992* | | + | 4345603 | 4346382 | 5.17 | 81.19 | 204.29 | 319.11 | Hypothetical protein |
| *FEPFGCPE_03993* | | + | 4346556 | 4346876 | 11.57 | 275.11 | 542.03 | 535.93 | Hypothetical protein |
| *FEPFGCPE_03994* | | - | 4346989 | 4347065 | 5.11 | 14.24 | 23.87 | 31.24 | tRNA |
| *FEPFGCPE_03995* | *proP-16* | - | 4347246 | 4348904 | 8.74 | 623.39 | 279.60 | 133.86 | Proline/betaine transporter |
| *FEPFGCPE_03996* | *qseC-5* | - | 4349076 | 4350719 | 3.78 | 15.16 | 6.09 | 3.04 | Sensor protein QseC |
| *FEPFGCPE_03997* | *tctD-4* | - | 4350770 | 4351459 | 22.72 | 76.00 | 61.26 | 19.10 | Transcriptional regulatory protein tctD |
| *FEPFGCPE_03998* | *recA* | + | 4351662 | 4352726 | 16.21 | 79.02 | 69.11 | 33.42 | Protein RecA |
| *FEPFGCPE_03999* | *recX* | + | 4352827 | 4353372 | 3.34 | 19.38 | 4.26 | 2.36 | Regulatory protein RecX |
| *FEPFGCPE_04000* | | + | 4353682 | 4354368 | 23.50 | 171.81 | 170.50 | 79.22 | Hypothetical protein |
| *FEPFGCPE_04001* | *sucC* | + | 4354365 | 4355531 | 7.01 | 69.58 | 90.27 | 93.35 | Succinate--CoA ligase [ADP-forming] subunit beta |
| *FEPFGCPE_04002* | *sucD* | + | 4355614 | 4356495 | 6.52 | 38.09 | 47.33 | 43.73 | Succinate--CoA ligase [ADP-forming] subunit alpha |
| *FEPFGCPE_04003* | | + | 4356651 | 4357379 | 2.68 | 3.68 | 3.75 | 3.43 | Hypothetical protein |
| *FEPFGCPE_04004* | *pilE* | + | 4357609 | 4358133 | 4.11 | 4.38 | 14.04 | 8.14 | Fimbrial protein |
| *FEPFGCPE_04005* | | + | 4358452 | 4360224 | 1.56 | 6.71 | 3.55 | 1.41 | Hypothetical protein |
| *FEPFGCPE_04006* | *thiD-2* | - | 4360308 | 4361150 | 4.92 | 31.35 | 14.41 | 5.47 | Hydroxymethylpyrimidine/phosphomethylpyrimidine kinase |
| *FEPFGCPE_04007* | *rubA-2* | + | 4361368 | 4361538 | 8.04 | 99.14 | 60.35 | 39.49 | Rubredoxin |
| *FEPFGCPE_04008* | | + | 4361779 | 4363167 | 1.50 | 13.28 | 13.45 | 5.08 | Hypothetical protein |
| *FEPFGCPE_04009* | | + | 4363312 | 4363881 | 10.20 | 37.55 | 34.33 | 12.42 | Hypothetical protein |
| *FEPFGCPE_04010* | *yqgF* | + | 4363878 | 4364306 | 2.72 | 15.82 | 14.35 | 5.08 | Putative pre-16S rRNA nuclease |
| *FEPFGCPE_04011* | *pyrB-2* | + | 4364381 | 4365358 | 1.99 | 10.12 | 10.68 | 3.65 | Aspartate carbamoyltransferase |
| *FEPFGCPE_04012* | *pyrC-2* | + | 4365366 | 4366634 | 1.88 | 16.62 | 11.39 | 3.50 | Dihydroorotase |
| *FEPFGCPE_04013* | *plsC* | + | 4366647 | 4367477 | 0.71 | 6.77 | 3.33 | 0.97 | 1-acyl-sn-glycerol-3-phosphate acyltransferase |
| *FEPFGCPE_04014* | *apaH* | - | 4367602 | 4368609 | 2.44 | 16.53 | 9.02 | 2.74 | Bis(5'-nucleosyl)-tetraphosphatase, symmetrical |
| *FEPFGCPE_04015* | *tagO* | - | 4368706 | 4369797 | 4.86 | 18.96 | 9.24 | 1.03 | putative undecaprenyl-phosphate N-acetylglucosaminyl 1-phosphate transferase |
| *FEPFGCPE_04016* | *pglA* | - | 4369791 | 4370933 | 1.65 | 4.56 | 5.60 | 1.30 | N,N'-diacetylbacillosaminyl-diphospho-undecaprenol alpha-1,3-N-acetylgalactosaminyltransferase |
| *FEPFGCPE_04017* | *epsF* | - | 4370953 | 4372083 | 1.31 | 6.08 | 1.97 | 1.87 | Putative glycosyltransferase EpsF |
| *FEPFGCPE_04018* | *pglH* | - | 4372080 | 4373216 | 2.53 | 9.30 | 4.02 | 1.54 | GalNAc-alpha-(1-4)-GalNAc-alpha-(1-3)-diNAcBac-PP-undecaprenol alpha-1,4-N-acetyl-D-galactosaminyltransferase |
| *FEPFGCPE_04019* | | - | 4373221 | 4374438 | 0.46 | 3.40 | 2.18 | 0.78 | Hypothetical protein |
| *FEPFGCPE_04020* | *algC* | + | 4374796 | 4376193 | 3.14 | 17.08 | 16.64 | 14.42 | Phosphomannomutase/phosphoglucomutase |
| *FEPFGCPE_04021* | *rfaC* | + | 4376389 | 4377414 | 3.35 | 13.47 | 14.61 | 6.40 | Lipopolysaccharide heptosyltransferase 1 |
| *FEPFGCPE_04022* | | + | 4377411 | 4378307 | 3.15 | 12.10 | 11.60 | 3.23 | Hypothetical protein |
| *FEPFGCPE_04023* | *waaA* | + | 4378308 | 4379618 | 3.35 | 12.99 | 11.14 | 3.03 | 3-deoxy-D-manno-octulosonic acid transferase |
| *FEPFGCPE_04024* | *moeZ* | - | 4379631 | 4379954 | 11.05 | 53.76 | 32.25 | 20.73 | putative adenylyltransferase/sulfurtransferase MoeZ |
| *FEPFGCPE_04025* | *pcm-2* | - | 4379988 | 4380695 | 19.68 | 63.57 | 54.57 | 33.52 | Protein-L-isoaspartate O-methyltransferase |
| *FEPFGCPE_04026* | *kstR* | - | 4380717 | 4381403 | 6.88 | 21.12 | 18.60 | 8.28 | HTH-type transcriptional repressor KstR |
| *FEPFGCPE_04027* | *fabG-8* | + | 4381502 | 4382062 | 7.57 | 49.09 | 38.49 | 21.45 | 3-oxoacyl-[acyl-carrier-protein] reductase FabG |
| *FEPFGCPE_04028* | | + | 4382065 | 4382283 | 3.64 | 10.40 | 32.93 | 5.54 | Hypothetical protein |
| *FEPFGCPE_04029* | | + | 4382355 | 4382552 | 1.55 | 4.53 | 13.59 | 0.00 | Hypothetical protein |
| *FEPFGCPE_04030* | | + | 4382621 | 4383262 | 0.84 | 5.28 | 1.56 | 2.23 | Hypothetical protein |
| *FEPFGCPE_04031* | *uppP-2* | + | 4383350 | 4384192 | 0.75 | 4.07 | 1.31 | 0.04 | Undecaprenyl-diphosphatase |
| *FEPFGCPE_04032* | *trmB-2* | - | 4384241 | 4385107 | 0.88 | 22.55 | 12.03 | 3.93 | tRNA (guanine-N(7)-)-methyltransferase |
| *FEPFGCPE_04033* | | - | 4385328 | 4385846 | 0.46 | 4.95 | 2.89 | 2.83 | Hypothetical protein |
| *FEPFGCPE_04034* | *yiaD-2* | - | 4386186 | 4386854 | 0.82 | 1.36 | 5.35 | 1.71 | putative lipoprotein YiaD |
| *FEPFGCPE_04035* | | - | 4386851 | 4389877 | 0.80 | 5.50 | 4.41 | 2.07 | Hypothetical protein |
| *FEPFGCPE_04036* | | - | 4389858 | 4390655 | 0.96 | 3.56 | 9.63 | 4.88 | Hypothetical protein |
| *FEPFGCPE_04037* | | - | 4390729 | 4391685 | 0.41 | 2.90 | 0.56 | 2.39 | Hypothetical protein |
| *FEPFGCPE_04038* | | - | 4391822 | 4391895 | 21.89 | 17.76 | 3.48 | 3.66 | tRNA |
| *FEPFGCPE_04039* | | - | 4392007 | 4392693 | 5.72 | 17.77 | 10.55 | 3.47 | Hypothetical protein |
| *FEPFGCPE_04040* | | - | 4392660 | 4393523 | 10.18 | 29.61 | 17.27 | 3.64 | Hypothetical protein |
| *FEPFGCPE_04041* | | - | 4393636 | 4393878 | 8.33 | 25.56 | 14.74 | 2.30 | Hypothetical protein |
| *FEPFGCPE_04042* | *dpnA* | + | 4394352 | 4395398 | 13.05 | 44.64 | 25.03 | 7.23 | Modification methylase DpnIIB |
| *FEPFGCPE_04043* | *mboIIM* | + | 4395395 | 4396267 | 19.90 | 91.86 | 50.32 | 22.46 | Modification methylase MboII |

| Locus name | Gene name | Strand | Gene start (bp) | Gene end (bp) | Average TPM Inoculum | Average TPM $t_6$ | Average TPM $t_{24}$ | Average TPM $t_{168}$ | Annotation description |
|---|---|---|---|---|---|---|---|---|---|
| *FEPFGCPE_04044* | *ndhI* | - | 4396221 | 4397282 | 11.55 | 48.20 | 29.01 | 7.34 | NAD(P)H-quinone oxidoreductase subunit I, chloroplastic |
| *FEPFGCPE_04045* | | - | 4397260 | 4398912 | 22.78 | 69.58 | 42.79 | 17.90 | Hypothetical protein |
| *FEPFGCPE_04046* | | - | 4399501 | 4399755 | 6.22 | 42.11 | 33.21 | 14.15 | Hypothetical protein |
| *FEPFGCPE_04047* | | + | 4399892 | 4400140 | 5.86 | 13.42 | 3.98 | 3.15 | Hypothetical protein |
| *FEPFGCPE_04048* | | + | 4400137 | 4400415 | 5.24 | 34.37 | 17.18 | 4.92 | Hypothetical protein |
| *FEPFGCPE_04049* | | - | 4400752 | 4401459 | 2.28 | 12.86 | 7.54 | 3.56 | Hypothetical protein |
| *FEPFGCPE_04050* | | + | 4402184 | 4402972 | 5.06 | 12.94 | 6.22 | 4.86 | Hydrolase |
| *FEPFGCPE_04051* | | - | 4403063 | 4404874 | 4.35 | 20.11 | 6.71 | 1.40 | putative signaling protein |
| *FEPFGCPE_04052* | | - | 4405339 | 4405530 | 78.22 | 147.38 | 107.88 | 25.00 | Hypothetical protein |
| *FEPFGCPE_04053* | *kefF-1* | - | 4405911 | 4406687 | 1.04 | 4.15 | 0.94 | 0.60 | Glutathione-regulated potassium-efflux system ancillary protein KefF |
| *FEPFGCPE_04054* | *syrM1-3* | + | 4406794 | 4407687 | 0.37 | 1.64 | 1.70 | 0.17 | HTH-type transcriptional regulator SyrM 1 |
| *FEPFGCPE_04055* | *zapE-2* | - | 4407691 | 4408725 | 1.18 | 5.96 | 6.14 | 1.07 | Cell division protein ZapE |
| *FEPFGCPE_04056* | *nhoA* | + | 4408958 | 4409815 | 1.86 | 10.28 | 9.04 | 2.22 | N-hydroxyarylamine O-acetyltransferase |
| *FEPFGCPE_04057* | | - | 4409921 | 4411159 | 1.62 | 6.44 | 4.45 | 1.87 | putative transporter |
| *FEPFGCPE_04058* | *dmlR-19* | + | 4411258 | 4412238 | 1.16 | 6.22 | 6.12 | 2.67 | HTH-type transcriptional regulator DmlR |
| *FEPFGCPE_04059* | *gcvA-8* | - | 4412245 | 4413222 | 0.81 | 2.88 | 3.16 | 1.05 | Glycine cleavage system transcriptional activator |
| *FEPFGCPE_04060* | | + | 4413364 | 4414602 | 17.92 | 12.32 | 8.43 | 4.00 | Hypothetical protein |
| *FEPFGCPE_04061* | *mmsA* | + | 4414782 | 4416281 | 31.76 | 25.86 | 20.72 | 11.41 | Methylmalonate-semialdehyde dehydrogenase [acylating] |
| *FEPFGCPE_04062* | *mmsB* | + | 4416317 | 4417246 | 28.10 | 35.95 | 19.73 | 4.73 | 3-hydroxyisobutyrate dehydrogenase |
| *FEPFGCPE_04063* | | - | 4417333 | 4417755 | 6.18 | 65.09 | 53.08 | 19.23 | Hypothetical protein |
| *FEPFGCPE_04064* | | + | 4417794 | 4418858 | 3.80 | 11.82 | 10.52 | 2.50 | Hypothetical protein |
| *FEPFGCPE_04065* | | + | 4418914 | 4419270 | 4.73 | 23.16 | 36.23 | 5.06 | HTH-type transcriptional regulator |
| *FEPFGCPE_04066* | | + | 4419254 | 4420201 | 1.90 | 14.46 | 11.91 | 3.16 | Hypothetical protein |
| *FEPFGCPE_04067* | | + | 4420437 | 4420844 | 2.20 | 58.31 | 307.93 | 57.52 | Hypothetical protein |
| *FEPFGCPE_04068* | *kefC-2* | - | 4420939 | 4422858 | 1.18 | 12.41 | 15.65 | 2.57 | Glutathione-regulated potassium-efflux system protein KefC |
| *FEPFGCPE_04069* | *kefF-2* | - | 4422858 | 4423592 | 2.44 | 13.90 | 6.01 | 2.09 | Glutathione-regulated potassium-efflux system ancillary protein KefF |
| *FEPFGCPE_04070* | | + | 4423877 | 4424170 | 0.86 | 3.92 | 1.30 | 0.00 | Hypothetical protein |
| *FEPFGCPE_04071* | *yhdG* | - | 4424228 | 4425709 | 1.07 | 1.18 | 2.04 | 0.26 | putative amino acid permease YhdG |
| *FEPFGCPE_04072* | | - | 4426256 | 4427026 | 8.87 | 41.36 | 17.30 | 11.34 | Hypothetical protein |
| *FEPFGCPE_04073* | *flhA-2* | - | 4427105 | 4428211 | 13.78 | 46.06 | 29.52 | 8.97 | S-(hydroxymethyl)glutathione dehydrogenase |
| *FEPFGCPE_04074* | *cph2-6* | - | 4428357 | 4430078 | 3.19 | 7.47 | 5.36 | 3.22 | Phytochrome-like protein cph2 |
| *FEPFGCPE_04075* | | + | 4430575 | 4431129 | 4.98 | 29.98 | 11.56 | 2.45 | Hypothetical protein |
| *FEPFGCPE_04076* | *yfiN* | + | 4431126 | 4432496 | 5.47 | 22.66 | 10.54 | 2.68 | putative diguanylate cyclase YfiN |
| *FEPFGCPE_04077* | *yiaD-3* | + | 4432641 | 4433168 | 5.84 | 12.08 | 15.77 | 3.89 | putative lipoprotein YiaD |
| *FEPFGCPE_04078* | | + | 4433313 | 4434632 | 1.95 | 8.60 | 5.31 | 1.08 | Hypothetical protein |
| *FEPFGCPE_04079* | *nsaD-3* | - | 4434651 | 4435313 | 4.14 | 25.89 | 18.24 | 10.26 | 2-hydroxychromene-2-carboxylate isomerase |
| *FEPFGCPE_04080* | | + | 4435572 | 4436624 | 1.90 | 5.10 | 5.28 | 2.51 | Hypothetical protein |
| *FEPFGCPE_04081* | | + | 4436747 | 4436917 | 10.69 | 16.99 | 49.74 | 13.16 | Hypothetical protein |
| *FEPFGCPE_04082* | | - | 4437011 | 4437904 | 4.57 | 24.50 | 17.43 | 11.74 | Hypothetical protein |
| *FEPFGCPE_04083* | | - | 4437898 | 4438092 | 7.45 | 37.79 | 40.98 | 10.29 | Hypothetical protein |
| *FEPFGCPE_04084* | | - | 4438083 | 4438640 | 5.56 | 44.97 | 36.42 | 16.28 | Hypothetical protein |
| *FEPFGCPE_04085* | | - | 4438767 | 4439114 | 2.02 | 20.48 | 11.44 | 1.21 | Hypothetical protein |
| *FEPFGCPE_04086* | | - | 4439177 | 4439389 | 2.30 | 66.50 | 22.40 | 13.93 | Hypothetical protein |
| *FEPFGCPE_04087* | *yofA* | + | 4439675 | 4440628 | 1.71 | 11.35 | 10.98 | 3.41 | HTH-type transcriptional regulator YofA |
| *FEPFGCPE_04088* | | + | 4440843 | 4441496 | 0.56 | 4.21 | 5.17 | 1.04 | Hypothetical protein |
| *FEPFGCPE_04089* | *yhjE-6* | + | 4441562 | 4442911 | 0.65 | 1.69 | 1.53 | 1.31 | Inner membrane metabolite transport protein YhjE |
| *FEPFGCPE_04090* | *torS* | - | 4442980 | 4445742 | 0.46 | 2.01 | 0.85 | 0.23 | Sensor protein TorS |
| *FEPFGCPE_04091* | *pleD-3* | - | 4445739 | 4446644 | 1.86 | 8.67 | 5.86 | 2.54 | Response regulator PleD |
| *FEPFGCPE_04092* | *murA-2* | - | 4446927 | 4448228 | 4.41 | 29.30 | 16.84 | 16.27 | UDP-N-acetylglucosamine 1-carboxyvinyltransferase |
| *FEPFGCPE_04093* | *rspR-9* | - | 4448436 | 4449236 | 0.57 | 11.21 | 7.16 | 3.37 | HTH-type transcriptional repressor RspR |
| *FEPFGCPE_04094* | | + | 4449551 | 4449775 | 0.79 | 0.12 | 2.13 | 0.00 | Hypothetical protein |
| *FEPFGCPE_04095* | *yodF* | + | 4449772 | 4451247 | 1.83 | 10.18 | 3.24 | 2.70 | putative symporter YodF |
| *FEPFGCPE_04096* | *atzC* | + | 4451268 | 4452458 | 1.04 | 5.01 | 7.15 | 1.57 | N-isopropylammelide isopropyl amidohydrolase |
| *FEPFGCPE_04097* | *yycB* | + | 4452455 | 4453726 | 1.21 | 7.44 | 4.63 | 1.16 | putative transporter YycB |
| *FEPFGCPE_04098* | | + | 4453974 | 4454960 | 13.58 | 33.84 | 17.46 | 8.96 | Hypothetical protein |
| *FEPFGCPE_04099* | *mdoG-2* | + | 4455286 | 4456845 | 11.76 | 41.63 | 36.65 | 17.95 | Glucans biosynthesis protein G |
| *FEPFGCPE_04100* | *opgH* | + | 4456842 | 4459484 | 4.50 | 14.28 | 13.33 | 6.87 | Glucans biosynthesis glucosyltransferase H |
| *FEPFGCPE_04101* | *nlpD-2* | - | 4459597 | 4460361 | 4.88 | 10.62 | 11.72 | 7.34 | Murein hydrolase activator NlpD |
| *FEPFGCPE_04102* | *cspD-1* | + | 4460700 | 4460903 | 9.90 | 26.81 | 74.22 | 46.25 | Cold shock-like protein CspD |
| *FEPFGCPE_04103* | *rhtB-4* | + | 4461126 | 4461761 | 12.03 | 45.71 | 42.58 | 20.26 | Homoserine/homoserine lactone efflux protein |
| *FEPFGCPE_04104* | *ycgR* | + | 4462037 | 4462798 | 6.63 | 15.45 | 12.96 | 6.18 | Flagellar brake protein YcgR |
| *FEPFGCPE_04105* | *corC-3* | - | 4462877 | 4463611 | 1.04 | 14.01 | 9.87 | 5.44 | Magnesium and cobalt efflux protein CorC |
| *FEPFGCPE_04106* | | + | 4463923 | 4464216 | 4.04 | 27.42 | 34.45 | 15.90 | Hypothetical protein |
| *FEPFGCPE_04107* | *dmlR-20* | - | 4464373 | 4465284 | 2.09 | 10.21 | 9.58 | 2.92 | HTH-type transcriptional regulator DmlR |
| *FEPFGCPE_04108* | *ygiD* | - | 4465409 | 4466236 | 2.21 | 6.76 | 7.37 | 4.91 | 4,5-DOPA dioxygenase extradiol |
| *FEPFGCPE_04109* | *catD-4* | - | 4466284 | 4466688 | 0.95 | 9.02 | 4.64 | 1.75 | Putative oxidoreductase CatD |
| *FEPFGCPE_04110* | *lutA-3* | - | 4466872 | 4468122 | 1.74 | 10.49 | 5.31 | 3.18 | Lactate utilization protein A |
| *FEPFGCPE_04111* | | - | 4468131 | 4469255 | 2.78 | 14.62 | 13.57 | 0.44 | putative FAD-linked oxidoreductase |
| *FEPFGCPE_04112* | | - | 4469255 | 4470841 | 0.90 | 5.56 | 4.33 | 2.15 | putative FAD-linked oxidoreductase |
| *FEPFGCPE_04113* | *pdhR-2* | + | 4471120 | 4471926 | 1.43 | 8.57 | 5.08 | 3.41 | Pyruvate dehydrogenase complex repressor |
| *FEPFGCPE_04114* | *oxlT-3* | + | 4472922 | 4474247 | 1.46 | 6.31 | 4.58 | 1.26 | Oxalate:formate antiporter |

| Locus name | Gene name | Strand | Gene start (bp) | Gene end (bp) | Average TPM Inoculum | Average TPM $t_6$ | Average TPM $t_{24}$ | Average TPM $t_{168}$ | Annotation description |
|---|---|---|---|---|---|---|---|---|---|
| *FEPFGCPE_04115* | | - | 4474396 | 4474471 | 24.33 | 16.63 | 16.28 | 6.25 | tRNA |
| *FEPFGCPE_04116* | | - | 4474558 | 4474633 | 37.96 | 38.15 | 32.38 | 10.34 | tRNA |
| *FEPFGCPE_04117* | | - | 4474737 | 4475060 | 11.71 | 42.17 | 43.57 | 13.89 | Ferredoxin 1 |
| *FEPFGCPE_04118* | *pncB2* | + | 4475323 | 4476516 | 2.58 | 12.01 | 13.01 | 5.09 | Nicotinate phosphoribosyltransferase 2 |
| *FEPFGCPE_04119* | *lutC* | + | 4476707 | 4477369 | 3.87 | 25.03 | 14.75 | 9.20 | Lactate utilization protein C |
| *FEPFGCPE_04120* | | + | 4477616 | 4479034 | 2.18 | 9.63 | 8.42 | 3.26 | Hypothetical protein |
| *FEPFGCPE_04121* | *ghrB-4* | + | 4479046 | 4480065 | 7.22 | 19.22 | 15.42 | 6.85 | Glyoxylate/hydroxypyruvate reductase B |
| *FEPFGCPE_04122* | | + | 4480144 | 4481607 | 5.51 | 25.22 | 12.90 | 5.31 | Hypothetical protein |
| *FEPFGCPE_04123* | | - | 4481728 | 4482234 | 3.38 | 21.37 | 10.67 | 1.91 | Hypothetical protein |
| *FEPFGCPE_04124* | | - | 4482349 | 4482633 | 11.63 | 40.05 | 16.49 | 13.33 | Hypothetical protein |
| *FEPFGCPE_04125* | *moeA-2* | - | 4482737 | 4484020 | 14.31 | 19.12 | 11.71 | 3.47 | Molybdopterin molybdenumtransferase |
| *FEPFGCPE_04126* | *mobA* | - | 4484107 | 4484685 | 42.08 | 75.13 | 26.57 | 9.11 | Molybdenum cofactor guanylyltransferase |
| *FEPFGCPE_04127* | | - | 4484795 | 4485679 | 4.52 | 17.57 | 17.38 | 2.65 | Hypothetical protein |
| *FEPFGCPE_04128* | *moaA* | - | 4485747 | 4486874 | 10.85 | 26.81 | 16.75 | 7.67 | GTP 3',8-cyclase |
| *FEPFGCPE_04129* | *rne* | - | 4487172 | 4490387 | 3.48 | 25.49 | 25.05 | 22.12 | Ribonuclease E |
| *FEPFGCPE_04130* | *rluC* | + | 4490964 | 4492013 | 18.54 | 56.31 | 41.91 | 15.58 | Ribosomal large subunit pseudouridine synthase C |
| *FEPFGCPE_04131* | | + | 4492000 | 4492662 | 7.64 | 19.08 | 13.05 | 6.61 | 5'-nucleotidase |
| *FEPFGCPE_04132* | | + | 4492659 | 4493075 | 19.52 | 71.91 | 63.12 | 20.90 | Hypothetical protein |
| *FEPFGCPE_04133* | *sppA* | + | 4493139 | 4494194 | 6.85 | 31.74 | 22.65 | 11.45 | Putative signal peptide peptidase SppA |
| *FEPFGCPE_04134* | *rsmI-2* | - | 4494285 | 4495001 | 1.83 | 13.86 | 13.87 | 2.31 | Ribosomal RNA small subunit methyltransferase I |
| *FEPFGCPE_04135* | *yceF* | - | 4494998 | 4495600 | 1.51 | 4.91 | 8.28 | 6.98 | Maf-like protein YceF |
| *FEPFGCPE_04136* | *yceD* | + | 4495813 | 4496406 | 12.50 | 79.43 | 51.16 | 28.83 | Large ribosomal RNA subunit accumulation protein YceD |
| *FEPFGCPE_04137* | *rpmF* | + | 4496535 | 4496714 | 56.75 | 315.79 | 319.52 | 138.01 | 50S ribosomal protein L32 |
| *FEPFGCPE_04138* | *plsX* | + | 4496833 | 4497885 | 6.22 | 28.08 | 16.85 | 9.39 | Phosphate acyltransferase |
| *FEPFGCPE_04139* | *fabH* | + | 4497925 | 4498914 | 5.35 | 29.06 | 25.20 | 12.76 | 3-oxoacyl-[acyl-carrier-protein] synthase 3 |
| *FEPFGCPE_04140* | *fabD-2* | + | 4499022 | 4499984 | 5.12 | 24.73 | 27.44 | 14.79 | Malonyl CoA-acyl carrier protein transacylase |
| *FEPFGCPE_04141* | *fabG-9* | + | 4500009 | 4500758 | 4.88 | 40.64 | 32.83 | 19.77 | 3-oxoacyl-[acyl-carrier-protein] reductase FabG |
| *FEPFGCPE_04142* | *acpP-3* | + | 4500895 | 4501134 | 38.31 | 441.06 | 352.91 | 173.74 | Acyl carrier protein |
| *FEPFGCPE_04143* | *fabF-3* | + | 4501282 | 4502520 | 9.14 | 48.16 | 49.16 | 32.14 | 3-oxoacyl-[acyl-carrier-protein] synthase 2 |
| *FEPFGCPE_04144* | *rpoE-2* | + | 4502688 | 4503287 | 120.50 | 138.17 | 149.22 | 95.63 | ECF RNA polymerase sigma-E factor |
| *FEPFGCPE_04145* | *rseA* | + | 4503326 | 4504021 | 105.70 | 81.00 | 139.47 | 104.38 | Anti-sigma-E factor RseA |
| *FEPFGCPE_04146* | *mucB* | + | 4504022 | 4505080 | 31.72 | 67.42 | 45.59 | 26.92 | Sigma factor AlgU regulatory protein MucB |
| *FEPFGCPE_04147* | *htrA* | + | 4505211 | 4506698 | 18.27 | 30.51 | 37.03 | 22.96 | Putative serine protease HtrA |
| *FEPFGCPE_04148* | | + | 4506711 | 4506956 | 19.24 | 34.15 | 53.97 | 20.88 | Hypothetical protein |
| *FEPFGCPE_04149* | *lepA* | + | 4507156 | 4508961 | 6.47 | 25.61 | 20.55 | 8.58 | Elongation factor 4 |
| *FEPFGCPE_04150* | *lepB* | + | 4508980 | 4509894 | 4.71 | 34.62 | 25.16 | 8.91 | Signal peptidase I |
| *FEPFGCPE_04151* | *rnc* | + | 4510003 | 4511124 | 10.82 | 46.31 | 38.61 | 17.61 | Ribonuclease 3 |
| *FEPFGCPE_04152* | *era* | + | 4511181 | 4512074 | 11.81 | 60.25 | 44.91 | 18.58 | GTPase Era |
| *FEPFGCPE_04153* | *recO* | + | 4512061 | 4513074 | 3.71 | 22.64 | 23.73 | 10.59 | DNA repair protein RecO |
| *FEPFGCPE_04154* | *pdxJ* | + | 4513071 | 4513832 | 2.51 | 6.09 | 7.09 | 7.31 | Pyridoxine 5'-phosphate synthase |
| *FEPFGCPE_04155* | *acpS* | + | 4513872 | 4514390 | 6.01 | 14.41 | 12.63 | 14.31 | Holo-[acyl-carrier-protein] synthase |
| *FEPFGCPE_04156* | *nagZ* | + | 4514387 | 4515421 | 1.30 | 9.33 | 6.04 | 6.19 | Beta-hexosaminidase |
| *FEPFGCPE_04157* | *efp* | - | 4515545 | 4516102 | 11.28 | 29.42 | 28.09 | 20.35 | Elongation factor P |
| *FEPFGCPE_04158* | | - | 4516242 | 4517483 | 1.56 | 13.39 | 10.05 | 4.50 | Hypothetical protein |
| *FEPFGCPE_04159* | *uvrC* | + | 4517617 | 4519866 | 2.31 | 15.21 | 13.88 | 3.18 | UvrABC system protein C |
| *FEPFGCPE_04160* | *kch* | - | 4519880 | 4521163 | 2.61 | 19.97 | 11.40 | 4.30 | Voltage-gated potassium channel Kch |
| *FEPFGCPE_04161* | *pgsA* | + | 4521283 | 4521876 | 9.38 | 45.98 | 30.72 | 11.71 | CDP-diacylglycerol--glycerol-3-phosphate 3-phosphatidyltransferase |
| *FEPFGCPE_04162* | | + | 4522054 | 4522129 | 7.07 | 11.83 | 9.27 | 26.43 | tRNA |
| *FEPFGCPE_04163* | | + | 4522179 | 4522254 | 7.08 | 14.40 | 8.48 | 13.16 | tRNA |
| *FEPFGCPE_04164* | | + | 4522366 | 4522439 | 0.28 | 0.00 | 0.00 | 0.39 | tRNA |
| *FEPFGCPE_04165* | | - | 4522704 | 4523465 | 1.92 | 4.78 | 0.69 | 1.55 | Metallo-beta-lactamase L1 type 3 |
| *FEPFGCPE_04166* | | - | 4523507 | 4524409 | 1.44 | 6.26 | 3.22 | 2.00 | Arylesterase |
| *FEPFGCPE_04167* | *pytH* | - | 4524462 | 4525157 | 1.23 | 2.86 | 2.91 | 0.45 | Pyrethroid hydrolase |
| *FEPFGCPE_04168* | | - | 4525196 | 4525855 | 2.18 | 16.23 | 4.38 | 0.41 | Hypothetical protein |
| *FEPFGCPE_04169* | *cdhR-4* | + | 4526129 | 4527094 | 1.30 | 7.79 | 8.90 | 0.66 | HTH-type transcriptional regulator CdhR |
| *FEPFGCPE_04170* | *dmlR-21* | - | 4527191 | 4527514 | 2.14 | 4.78 | 7.68 | 0.00 | HTH-type transcriptional regulator DmlR |
| *FEPFGCPE_04171* | *yajR-3* | - | 4527637 | 4528842 | 2.88 | 12.47 | 7.78 | 1.28 | Inner membrane transport protein YajR |
| *FEPFGCPE_04172* | *yxeP-6* | + | 4529161 | 4530429 | 12.24 | 31.12 | 16.84 | 6.03 | putative hydrolase YxeP |
| *FEPFGCPE_04173* | | + | 4530426 | 4531070 | 6.03 | 54.24 | 30.48 | 6.91 | Hypothetical protein |
| *FEPFGCPE_04174* | *lutR-6* | - | 4531161 | 4531907 | 5.11 | 45.96 | 17.95 | 11.13 | HTH-type transcriptional regulator LutR |
| *FEPFGCPE_04175* | | - | 4532145 | 4533320 | 4.12 | 288.38 | 54.39 | 8.00 | Outer membrane porin protein |
| *FEPFGCPE_04176* | *gudD-1* | - | 4533387 | 4534670 | 0.54 | 32.02 | 1.91 | 0.99 | Glucarate dehydratase |
| *FEPFGCPE_04177* | | + | 4534982 | 4536157 | 0.73 | 110.72 | 18.04 | 4.78 | Hypothetical protein |
| *FEPFGCPE_04178* | *potA-8* | + | 4536179 | 4537261 | 0.47 | 41.74 | 17.12 | 2.36 | Spermidine/putrescine import ATP-binding protein PotA |
| *FEPFGCPE_04179* | *sugA* | + | 4537258 | 4538142 | 3.11 | 51.13 | 16.45 | 5.55 | Trehalose transport system permease protein SugA |
| *FEPFGCPE_04180* | *sugB* | + | 4538139 | 4539095 | 0.71 | 23.70 | 9.52 | 1.16 | Trehalose transport system permease protein SugB |
| *FEPFGCPE_04181* | | + | 4539313 | 4539705 | 8.04 | 197.66 | 82.23 | 36.98 | Hypothetical protein |
| *FEPFGCPE_04182* | *sigR-2* | + | 4539882 | 4540445 | 3.24 | 30.75 | 15.30 | 2.88 | ECF RNA polymerase sigma factor SigR |
| *FEPFGCPE_04183* | | + | 4540445 | 4541434 | 3.43 | 21.85 | 10.36 | 4.46 | Hypothetical protein |
| *FEPFGCPE_04184* | | - | 4541537 | 4543111 | 0.62 | 26.17 | 6.94 | 4.12 | Alpha-ketoglutaric semialdehyde dehydrogenase |
| *FEPFGCPE_04185* | | - | 4543123 | 4544034 | 1.01 | 32.68 | 16.02 | 2.73 | putative 5-dehydro-4-deoxyglucarate dehydratase |

| Locus name | Gene name | Strand | Gene start (bp) | Gene end (bp) | Average TPM Inoculum | Average TPM $t_6$ | Average TPM $t_{24}$ | Average TPM $t_{168}$ | Annotation description |
|---|---|---|---|---|---|---|---|---|---|
| *FEPFGCPE_04186* | *garD* | + | 4544412 | 4546028 | 0.99 | 10.65 | 5.02 | 3.05 | Galactarate dehydratase (L-threo-forming) |
| *FEPFGCPE_04187* | | - | 4546043 | 4546519 | 7.56 | 19.69 | 13.12 | 7.21 | Hypothetical protein |
| *FEPFGCPE_04188* | | + | 4546991 | 4547305 | 0.69 | 0.06 | 1.41 | 2.70 | Hypothetical protein |
| *FEPFGCPE_04189* | | + | 4547302 | 4548282 | 0.86 | 1.05 | 2.70 | 0.28 | Hypothetical protein |
| *FEPFGCPE_04190* | | + | 4548299 | 4549279 | 3.02 | 5.12 | 3.71 | 1.35 | Hypothetical protein |
| *FEPFGCPE_04191* | | - | 4549328 | 4550155 | 2.52 | 8.28 | 3.83 | 1.21 | Hypothetical protein |
| *FEPFGCPE_04192* | *egtD* | - | 4550588 | 4551571 | 6.47 | 32.95 | 22.36 | 4.67 | Histidine N-alpha-methyltransferase |
| *FEPFGCPE_04193* | *egtB* | - | 4551571 | 4552794 | 19.01 | 54.00 | 48.24 | 12.36 | Hercynine oxygenase |
| *FEPFGCPE_04194* | | - | 4552881 | 4553084 | 35.77 | 55.74 | 42.63 | 7.08 | Hypothetical protein |
| *FEPFGCPE_04195* | | - | 4554064 | 4554930 | 1.34 | 3.02 | 2.02 | 0.41 | 4-sulfomuconolactone hydrolase |
| *FEPFGCPE_04196* | *naiP-6* | - | 4555036 | 4556313 | 2.83 | 15.87 | 10.70 | 3.77 | Putative niacin/nicotinamide transporter NaiP |
| *FEPFGCPE_04197* | *mngR-2* | - | 4556348 | 4557070 | 2.50 | 7.47 | 7.70 | 1.26 | Mannosyl-D-glycerate transport/metabolism system repressor MngR |
| *FEPFGCPE_04198* | | - | 4557464 | 4558093 | 0.90 | 2.88 | 10.42 | 0.53 | Hypothetical protein |
| *FEPFGCPE_04199* | *paaF-4* | + | 4558581 | 4559447 | 2.54 | 8.54 | 3.12 | 0.93 | 2,3-dehydroadipyl-CoA hydratase |
| *FEPFGCPE_04200* | *gmr-3* | - | 4559488 | 4561491 | 10.73 | 28.86 | 20.24 | 4.42 | Cyclic di-GMP phosphodiesterase Gmr |
| *FEPFGCPE_04201* | *katG* | + | 4562017 | 4564191 | 7.32 | 45.32 | 28.45 | 21.30 | Catalase-peroxidase |
| *FEPFGCPE_04202* | *chrA1-1* | - | 4564240 | 4565424 | 4.25 | 20.91 | 13.67 | 2.36 | Chromate transport protein |
| *FEPFGCPE_04203* | | - | 4565421 | 4566401 | 0.38 | 0.90 | 1.81 | 1.16 | Hypothetical protein |
| *FEPFGCPE_04204* | *arnE-2* | + | 4566532 | 4567593 | 2.40 | 8.46 | 9.08 | 2.19 | 4-amino-4-deoxy-L-arabinose-phosphoundecaprenol flippase subunit ArnE |
| *FEPFGCPE_04205* | | + | 4567798 | 4568487 | 1.47 | 2.57 | 4.83 | 3.58 | Hypothetical protein |
| *FEPFGCPE_04206* | | - | 4568529 | 4568921 | 2.75 | 8.22 | 9.87 | 4.12 | Hypothetical protein |
| *FEPFGCPE_04207* | *gntR-4* | - | 4569049 | 4570086 | 3.84 | 11.57 | 6.95 | 2.61 | HTH-type transcriptional regulator GntR |
| *FEPFGCPE_04208* | *gudP-2* | + | 4570255 | 4571676 | 0.88 | 23.98 | 6.98 | 1.43 | putative glucarate transporter |
| *FEPFGCPE_04209* | *gnl* | + | 4571716 | 4572798 | 1.32 | 24.26 | 3.74 | 0.49 | Gluconolactonase |
| *FEPFGCPE_04210* | *gudD-2* | + | 4572845 | 4574194 | 0.77 | 16.20 | 2.91 | 0.68 | Glucarate dehydratase |
| *FEPFGCPE_04211* | *bicA* | + | 4574268 | 4575755 | 1.42 | 14.42 | 4.18 | 2.13 | Bicarbonate transporter BicA |
| *FEPFGCPE_04212* | | - | 4575786 | 4576262 | 2.41 | 15.15 | 8.40 | 6.24 | Cytochrome c2 |
| *FEPFGCPE_04213* | *fliY-7* | - | 4576422 | 4577219 | 0.59 | 41.34 | 13.71 | 3.96 | L-cystine-binding protein FliY |
| *FEPFGCPE_04214* | *glnQ-11* | - | 4577263 | 4577991 | 0.95 | 63.39 | 9.21 | 5.15 | Glutamine transport ATP-binding protein GlnQ |
| *FEPFGCPE_04215* | *yecS-9* | - | 4577975 | 4578628 | 1.05 | 14.61 | 5.76 | 2.98 | L-cystine transport system permease protein YecS |
| *FEPFGCPE_04216* | *tcyB* | - | 4578625 | 4579440 | 1.85 | 12.92 | 10.13 | 1.06 | L-cystine transport system permease protein TcyB |
| *FEPFGCPE_04217* | *lrp-7* | + | 4579865 | 4580326 | 1.83 | 17.36 | 6.31 | 1.20 | Leucine-responsive regulatory protein |
| *FEPFGCPE_04218* | *tpd* | + | 4580516 | 4581139 | 38.42 | 82.09 | 126.73 | 118.06 | 34 kDa membrane antigen |
| *FEPFGCPE_04219* | | + | 4581207 | 4581563 | 25.16 | 72.11 | 96.43 | 88.26 | Hypothetical protein |
| *FEPFGCPE_04220* | *efeU* | + | 4581622 | 4582458 | 8.68 | 25.00 | 38.11 | 35.55 | Ferrous iron permease EfeU |
| *FEPFGCPE_04221* | | + | 4582604 | 4583998 | 4.71 | 11.35 | 15.27 | 10.76 | Hypothetical protein |
| *FEPFGCPE_04222* | *ubiA* | + | 4584075 | 4584947 | 3.41 | 17.39 | 14.96 | 5.30 | 4-hydroxybenzoate octaprenyltransferase |
| *FEPFGCPE_04223* | *proC-2* | - | 4585061 | 4585873 | 2.42 | 7.94 | 6.17 | 4.69 | Pyrroline-5-carboxylate reductase |
| *FEPFGCPE_04224* | | - | 4585965 | 4586678 | 1.74 | 4.37 | 3.42 | 0.44 | Hypothetical protein |
| *FEPFGCPE_04225* | *gpx1* | + | 4586906 | 4587403 | 2.79 | 16.49 | 9.84 | 1.25 | Hydroperoxy fatty acid reductase gpx1 |
| *FEPFGCPE_04226* | | - | 4587457 | 4587948 | 1.39 | 3.86 | 4.05 | 0.60 | Hypothetical protein |
| *FEPFGCPE_04227* | *norG-3* | + | 4588107 | 4589663 | 2.15 | 11.95 | 12.53 | 2.97 | HTH-type transcriptional regulator NorG |
| *FEPFGCPE_04228* | | - | 4589713 | 4590717 | 1.77 | 13.47 | 10.61 | 2.19 | putative FAD-linked oxidoreductase |
| *FEPFGCPE_04229* | | - | 4590794 | 4591663 | 4.95 | 27.75 | 18.67 | 6.05 | Cyclic nucleotide-gated potassium channel |
| *FEPFGCPE_04230* | | - | 4591665 | 4593200 | 3.80 | 17.38 | 10.30 | 3.79 | putative FAD-linked oxidoreductase |
| *FEPFGCPE_04231* | | - | 4593208 | 4594629 | 11.16 | 43.71 | 26.51 | 10.08 | putative FAD-linked oxidoreductase |
| *FEPFGCPE_04232* | *yvqK* | + | 4594825 | 4595523 | 10.92 | 62.04 | 36.57 | 16.76 | Cob(I)yrinic acid a,c-diamide adenosyltransferase |
| *FEPFGCPE_04233* | *aroG* | - | 4595638 | 4596711 | 8.15 | 21.86 | 26.69 | 21.04 | Phospho-2-dehydro-3-deoxyheptonate aldolase, Phe-sensitive |
| *FEPFGCPE_04234* | *tldD* | - | 4597165 | 4598625 | 9.37 | 53.53 | 46.29 | 16.80 | Metalloprotease TldD |
| *FEPFGCPE_04235* | | - | 4599046 | 4600374 | 12.40 | 71.09 | 55.63 | 13.60 | Hypothetical protein |
| *FEPFGCPE_04236* | | - | 4600371 | 4603406 | 2.27 | 13.05 | 6.72 | 2.19 | Hypothetical protein |
| *FEPFGCPE_04237* | *glnE* | + | 4603580 | 4606444 | 3.46 | 21.01 | 13.72 | 4.47 | Bifunctional glutamine synthetase adenylyltransferase/adenylyl-removing enzyme |
| *FEPFGCPE_04238* | *recN* | - | 4606526 | 4608190 | 3.23 | 15.10 | 8.05 | 2.24 | DNA repair protein RecN |
| *FEPFGCPE_04239* | *nadK* | - | 4608214 | 4609122 | 10.88 | 32.46 | 21.69 | 7.02 | NAD kinase |
| *FEPFGCPE_04240* | *hrcA* | + | 4609235 | 4610263 | 10.04 | 34.82 | 21.66 | 11.49 | Heat-inducible transcription repressor HrcA |
| *FEPFGCPE_04241* | | + | 4610524 | 4612938 | 2.84 | 34.56 | 35.75 | 18.07 | Hypothetical protein |
| *FEPFGCPE_04242* | *hemH* | + | 4613044 | 4614228 | 3.89 | 24.05 | 16.35 | 10.91 | Ferrochelatase |
| *FEPFGCPE_04243* | *hslR* | + | 4614225 | 4614626 | 5.57 | 83.90 | 42.89 | 19.41 | Heat shock protein 15 |
| *FEPFGCPE_04244* | | - | 4614681 | 4614869 | 16.82 | 100.11 | 67.07 | 46.03 | Hypothetical protein |
| *FEPFGCPE_04245* | *grpE* | + | 4615168 | 4615713 | 7.68 | 38.89 | 46.02 | 22.18 | Protein GrpE |
| *FEPFGCPE_04246* | *dnaK-2* | + | 4615965 | 4617890 | 20.83 | 61.69 | 65.92 | 56.72 | Chaperone protein DnaK |
| *FEPFGCPE_04247* | *dnaJ* | + | 4618018 | 4619169 | 3.33 | 18.36 | 17.08 | 12.34 | Chaperone protein DnaJ |
| *FEPFGCPE_04248* | *pabB* | + | 4619181 | 4621184 | 0.73 | 3.52 | 4.18 | 1.73 | Aminodeoxychorismate synthase component 1 |
| *FEPFGCPE_04249* | | - | 4621225 | 4622253 | 1.63 | 8.24 | 5.34 | 1.93 | Hypothetical protein |
| *FEPFGCPE_04250* | *ssuC-7* | - | 4622343 | 4623263 | 0.81 | 0.62 | 5.10 | 0.94 | Putative aliphatic sulfonates transport permease protein SsuC |
| *FEPFGCPE_04251* | *cmpD-5* | - | 4623274 | 4624074 | 0.58 | 6.74 | 3.00 | 2.77 | Bicarbonate transport ATP-binding protein CmpD |
| *FEPFGCPE_04252* | | - | 4624116 | 4624268 | 1.85 | 19.16 | 13.94 | 4.66 | Hypothetical protein |
| *FEPFGCPE_04253* | *qseB-3* | + | 4624288 | 4624950 | 1.01 | 5.27 | 5.36 | 2.96 | Transcriptional regulatory protein QseB |
| *FEPFGCPE_04254* | *prrB* | + | 4624947 | 4626314 | 1.67 | 11.50 | 7.12 | 1.60 | Sensor-type histidine kinase PrrB |
| *FEPFGCPE_04255* | *panB* | - | 4626458 | 4627276 | 2.89 | 13.53 | 4.68 | 3.48 | 3-methyl-2-oxobutanoate hydroxymethyltransferase |
| *FEPFGCPE_04256* | *dgk* | - | 4627331 | 4627987 | 5.90 | 42.25 | 29.13 | 9.27 | Deoxyguanosine kinase |

| Locus name | Gene name | Strand | Gene start (bp) | Gene end (bp) | Average TPM Inoculum | Average TPM $t_6$ | Average TPM $t_{24}$ | Average TPM $t_{168}$ | Annotation description |
|---|---|---|---|---|---|---|---|---|---|
| *FEPFGCPE_04257* | *folK* | - | 4627984 | 4628496 | 12.66 | 73.19 | 59.23 | 14.73 | 2-amino-4-hydroxy-6-hydroxymethyldihydropteridine pyrophosphokinase |
| *FEPFGCPE_04258* | *pcnB* | - | 4628493 | 4630007 | 2.77 | 19.39 | 18.07 | 8.75 | Poly(A) polymerase I |
| *FEPFGCPE_04259* | | - | 4630004 | 4630693 | 4.76 | 51.28 | 42.14 | 17.94 | putative phosphatase |
| *FEPFGCPE_04260* | *hda* | - | 4630749 | 4631459 | 2.22 | 10.58 | 7.25 | 3.63 | DnaA regulatory inactivator Hda |
| *FEPFGCPE_04261* | *purM* | + | 4631606 | 4632652 | 11.42 | 81.49 | 66.19 | 38.25 | Phosphoribosylformylglycinamidine cyclo-ligase |
| *FEPFGCPE_04262* | *gltC-13* | - | 4632790 | 4633704 | 0.51 | 1.78 | 2.72 | 1.25 | HTH-type transcriptional regulator GltC |
| *FEPFGCPE_04263* | *pyrC-3* | - | 4633752 | 4634993 | 0.75 | 2.99 | 2.80 | 1.79 | Dihydroorotase |
| *FEPFGCPE_04264* | *pepE-2* | - | 4635068 | 4636264 | 0.52 | 5.17 | 1.83 | 1.71 | putative dipeptidase PepE |
| *FEPFGCPE_04265* | *potB-3* | - | 4636381 | 4637301 | 1.03 | 5.42 | 4.76 | 3.43 | Spermidine/putrescine transport system permease protein PotB |
| *FEPFGCPE_04266* | *potD-6* | - | 4637328 | 4638392 | 1.90 | 7.43 | 4.86 | 4.36 | Spermidine/putrescine-binding periplasmic protein |
| *FEPFGCPE_04267* | *ugpC-2* | - | 4638469 | 4639521 | 1.01 | 3.34 | 1.08 | 1.19 | sn-glycerol-3-phosphate import ATP-binding protein UgpC |
| *FEPFGCPE_04268* | *phnV-2* | - | 4639523 | 4640398 | 2.08 | 4.95 | 0.15 | 0.66 | Putative 2-aminoethylphosphonate transport system permease protein PhnV |
| *FEPFGCPE_04269* | *miaA* | - | 4640599 | 4641579 | 1.57 | 5.84 | 5.59 | 1.82 | tRNA dimethylallyltransferase |
| *FEPFGCPE_04270* | *mutL* | - | 4641587 | 4643650 | 3.39 | 18.19 | 16.66 | 4.99 | DNA mismatch repair protein MutL |
| *FEPFGCPE_04271* | | + | 4643900 | 4644568 | 15.47 | 34.95 | 41.28 | 24.75 | putative membrane protein |
| *FEPFGCPE_04272* | *amiC-6* | - | 4644684 | 4646138 | 12.72 | 22.98 | 15.03 | 10.36 | N-acetylmuramoyl-L-alanine amidase AmiC |
| *FEPFGCPE_04273* | *tsaE* | - | 4646132 | 4646656 | 20.79 | 67.94 | 27.58 | 8.93 | tRNA threonylcarbamoyladenosine biosynthesis protein TsaE |
| *FEPFGCPE_04274* | *queG* | + | 4646665 | 4647936 | 2.73 | 16.54 | 13.63 | 1.53 | Epoxyqueuosine reductase |
| *FEPFGCPE_04275* | *xerD-2* | + | 4647933 | 4648922 | 1.08 | 17.93 | 9.09 | 1.90 | Tyrosine recombinase XerD |
| *FEPFGCPE_04276* | *lcfB-4* | - | 4649014 | 4650579 | 1.16 | 4.52 | 4.17 | 2.08 | Long-chain-fatty-acid--CoA ligase |
| *FEPFGCPE_04277* | *ybaK-2* | + | 4650694 | 4651188 | 1.37 | 9.01 | 5.02 | 1.39 | Cys-tRNA(Pro)/Cys-tRNA(Cys) deacylase YbaK |
| *FEPFGCPE_04278* | *plsY* | + | 4651322 | 4651930 | 3.34 | 17.17 | 11.78 | 3.20 | putative glycerol-3-phosphate acyltransferase |
| *FEPFGCPE_04279* | | - | 4652030 | 4652515 | 8.22 | 27.30 | 23.55 | 15.49 | Hypothetical protein |
| *FEPFGCPE_04280* | *murB* | + | 4652768 | 4653784 | 2.12 | 14.06 | 9.74 | 3.64 | UDP-N-acetylenolpyruvoylglucosamine reductase |
| *FEPFGCPE_04281* | | - | 4653774 | 4655222 | 4.30 | 27.97 | 20.79 | 11.42 | Hypothetical protein |
| *FEPFGCPE_04282* | | - | 4655313 | 4655636 | 4.82 | 58.15 | 77.47 | 49.05 | Hypothetical protein |
| *FEPFGCPE_04283* | *argF-2* | - | 4656053 | 4657042 | 4.00 | 40.77 | 31.42 | 22.91 | Ornithine carbamoyltransferase |
| *FEPFGCPE_04284* | | + | 4657331 | 4657645 | 4.22 | 14.06 | 10.28 | 14.75 | Hypothetical protein |
| *FEPFGCPE_04285* | *rpsT* | - | 4657940 | 4658212 | 33.07 | 64.04 | 73.98 | 64.86 | 30S ribosomal protein S20 |
| *FEPFGCPE_04286* | *murJ* | + | 4658341 | 4659942 | 2.97 | 13.84 | 16.01 | 3.71 | Lipid II flippase MurJ |
| *FEPFGCPE_04287* | | + | 4659989 | 4660855 | 5.89 | 37.45 | 25.81 | 6.68 | Hypothetical protein |
| *FEPFGCPE_04288* | | - | 4660969 | 4661463 | 11.51 | 75.29 | 62.32 | 32.25 | Hypothetical protein |
| *FEPFGCPE_04289* | *adk* | - | 4661759 | 4662424 | 17.14 | 82.46 | 76.88 | 42.75 | Adenylate kinase |
| *FEPFGCPE_04290* | *kdsB-2* | - | 4662730 | 4663509 | 1.81 | 21.99 | 13.92 | 7.32 | 3-deoxy-manno-octulosonate cytidylyltransferase |
| *FEPFGCPE_04291* | | - | 4663518 | 4663724 | 0.97 | 17.17 | 10.62 | 2.09 | Hypothetical protein |
| *FEPFGCPE_04292* | *lpxK* | - | 4663705 | 4664778 | 1.84 | 11.84 | 9.54 | 3.12 | Tetraacyldisaccharide 4'-kinase |
| *FEPFGCPE_04293* | *exbD-5* | - | 4664775 | 4665236 | 1.48 | 6.47 | 5.43 | 2.83 | Biopolymer transport protein ExbD |
| *FEPFGCPE_04294* | *exbB-4* | - | 4665252 | 4665866 | 1.22 | 5.16 | 5.18 | 2.65 | Biopolymer transport protein ExbB |
| *FEPFGCPE_04295* | *xseA* | + | 4666316 | 4667689 | 1.39 | 7.62 | 7.01 | 2.90 | Exodeoxyribonuclease 7 large subunit |
| *FEPFGCPE_04296* | *sodB* | + | 4667893 | 4668471 | 56.77 | 168.57 | 191.89 | 118.03 | Superoxide dismutase [Fe] |
| *FEPFGCPE_04297* | *rpfG* | + | 4668850 | 4669443 | 4.82 | 8.23 | 2.83 | 1.30 | Cyclic di-GMP phosphodiesterase response regulator RpfG |
| *FEPFGCPE_04298* | | - | 4669459 | 4669938 | 22.37 | 62.66 | 42.63 | 10.25 | Hypothetical protein |
| *FEPFGCPE_04299* | | - | 4670075 | 4670284 | 5609.95 | 16905.27 | 7691.10 | 2610.93 | Hypothetical protein |
| *FEPFGCPE_04300* | | - | 4670412 | 4670678 | 5040.86 | 10611.08 | 4174.50 | 1096.44 | Hypothetical protein |
| *FEPFGCPE_04301* | *rluE* | - | 4670729 | 4671391 | 1.48 | 15.14 | 10.86 | 6.54 | Ribosomal large subunit pseudouridine synthase E |
| *FEPFGCPE_04302* | *icd-2* | + | 4671603 | 4672856 | 7.21 | 47.04 | 51.17 | 31.91 | Isocitrate dehydrogenase [NADP] |
| *FEPFGCPE_04303* | | - | 4672931 | 4673359 | 11.02 | 60.82 | 36.23 | 15.90 | Hypothetical protein |
| *FEPFGCPE_04304* | *pgrR-4* | - | 4673580 | 4674503 | 0.78 | 4.68 | 5.06 | 1.31 | HTH-type transcriptional regulator PgrR |
| *FEPFGCPE_04305* | *bdcA* | + | 4674612 | 4675355 | 2.02 | 6.29 | 2.31 | 1.97 | Cyclic-di-GMP-binding biofilm dispersal mediator protein |
| *FEPFGCPE_04306* | *pcpR-2* | - | 4675383 | 4676294 | 1.77 | 4.75 | 3.77 | 0.00 | PCP degradation transcriptional activation protein |
| *FEPFGCPE_04307* | *ndoR* | + | 4676461 | 4677540 | 2.18 | 120.70 | 361.63 | 230.96 | Ferredoxin--NAD(P)(+) reductase (naphthalene dioxygenase ferredoxin-specific) |
| *FEPFGCPE_04308* | *nagG-2* | + | 4677615 | 4678904 | 1.69 | 112.89 | 507.65 | 290.11 | Salicylate 5-hydroxylase, large oxygenase component |
| *FEPFGCPE_04309* | *nagH* | + | 4678917 | 4679390 | 1.74 | 102.73 | 557.77 | 369.75 | Salicylate 5-hydroxylase, small oxygenase component |
| *FEPFGCPE_04310* | *nagAb-2* | + | 4679404 | 4679718 | 0.92 | 36.52 | 250.24 | 159.87 | Naphthalene 1,2-dioxygenase/salicylate 5-hydroxylase systems, ferredoxin component |
| *FEPFGCPE_04311* | *cspD-2* | - | 4680312 | 4680518 | 108.07 | 572.54 | 343.96 | 148.53 | Cold shock-like protein CspD |
| *FEPFGCPE_04312* | *clpS* | + | 4680957 | 4681271 | 210.18 | 345.85 | 308.94 | 140.29 | ATP-dependent Clp protease adapter protein ClpS |
| *FEPFGCPE_04313* | *clpA* | + | 4681268 | 4683565 | 33.31 | 97.54 | 72.62 | 44.61 | ATP-dependent Clp protease ATP-binding subunit ClpA |
| *FEPFGCPE_04314* | *dut* | - | 4683657 | 4684103 | 5.08 | 11.63 | 7.13 | 2.04 | Deoxyuridine 5'-triphosphate nucleotidohydrolase |
| *FEPFGCPE_04315* | *limB-2* | - | 4684190 | 4685224 | 3.61 | 12.30 | 5.31 | 5.02 | Limonene 1,2-monooxygenase |
| *FEPFGCPE_04316* | | - | 4685281 | 4686444 | 0.55 | 5.12 | 2.48 | 0.48 | Hypothetical protein |
| *FEPFGCPE_04317* | *coaBC* | - | 4686682 | 4687893 | 6.92 | 29.66 | 17.19 | 10.76 | Coenzyme A biosynthesis bifunctional protein CoaBC |
| *FEPFGCPE_04318* | *lspA* | - | 4687970 | 4688515 | 9.43 | 37.44 | 37.20 | 12.93 | Lipoprotein signal peptidase |
| *FEPFGCPE_04319* | *ileS* | - | 4688515 | 4691352 | 4.26 | 21.02 | 18.06 | 11.72 | Isoleucine--tRNA ligase |
| *FEPFGCPE_04320* | *ribF* | - | 4691512 | 4692504 | 5.61 | 20.48 | 22.70 | 8.74 | Riboflavin biosynthesis protein RibF |
| *FEPFGCPE_04321* | *purN* | + | 4692785 | 4693393 | 8.82 | 29.97 | 27.03 | 14.53 | Phosphoribosylglycinamide formyltransferase |
| *FEPFGCPE_04322* | *rsmB-2* | + | 4693390 | 4694433 | 1.97 | 11.01 | 10.86 | 5.83 | Ribosomal RNA small subunit methyltransferase B |
| *FEPFGCPE_04323* | *rsmF* | + | 4694430 | 4694855 | 4.41 | 36.42 | 24.35 | 11.29 | Ribosomal RNA small subunit methyltransferase F |
| *FEPFGCPE_04324* | | + | 4694902 | 4696296 | 2.76 | 11.05 | 13.33 | 2.00 | putative MscS family protein.1 |
| *FEPFGCPE_04325* | | + | 4696609 | 4697805 | 15.44 | 40.34 | 53.97 | 53.71 | Hypothetical protein |
| *FEPFGCPE_04326* | *nadA* | + | 4697900 | 4699033 | 7.36 | 35.10 | 26.16 | 15.31 | Quinolinate synthase A |
| *FEPFGCPE_04327* | *nadC-3* | + | 4699030 | 4699935 | 6.79 | 20.39 | 24.81 | 11.72 | Nicotinate-nucleotide pyrophosphorylase [carboxylating] |

| Locus name | Gene name | Strand | Gene start (bp) | Gene end (bp) | Average TPM Inoculum | Average TPM $t_6$ | Average TPM $t_{24}$ | Average TPM $t_{168}$ | Annotation description |
|---|---|---|---|---|---|---|---|---|---|
| *FEPFGCPE_04328* | *nadB* | - | 4700039 | 4701739 | 8.40 | 19.58 | 16.94 | 9.86 | L-aspartate oxidase |
| *FEPFGCPE_04329* | *rpmG* | - | 4701886 | 4702053 | 7.45 | 12.19 | 11.98 | 21.75 | 50S ribosomal protein L33 |
| *FEPFGCPE_04330* | *rpmB* | - | 4702070 | 4702303 | 75.04 | 360.72 | 299.86 | 212.87 | 50S ribosomal protein L28 |
| *FEPFGCPE_04331* | | - | 4702529 | 4703203 | 5.97 | 12.61 | 6.63 | 5.35 | Hypothetical protein |
| *FEPFGCPE_04332* | *fkpB* | + | 4703366 | 4703821 | 34.10 | 93.62 | 93.73 | 40.18 | FKBP-type 16 kDa peptidyl-prolyl cis-trans isomerase |
| *FEPFGCPE_04333* | *ispH-2* | + | 4703830 | 4704804 | 15.31 | 42.22 | 38.98 | 22.13 | 4-hydroxy-3-methylbut-2-enyl diphosphate reductase |
| *FEPFGCPE_04334* | *braC-5* | + | 4705525 | 4706733 | 48.74 | 346.50 | 174.74 | 98.60 | Leucine-, isoleucine-, valine-, threonine-, and alanine-binding protein |
| *FEPFGCPE_04335* | *livH-16* | + | 4706946 | 4707896 | 4.96 | 43.09 | 26.59 | 17.40 | High-affinity branched-chain amino acid transport system permease protein LivH |
| *FEPFGCPE_04336* | | + | 4707915 | 4709054 | 3.12 | 23.51 | 12.72 | 7.13 | Hypothetical protein |
| *FEPFGCPE_04337* | *lptB-12* | + | 4709071 | 4709844 | 4.72 | 38.42 | 26.18 | 12.89 | Lipopolysaccharide export system ATP-binding protein LptB |
| *FEPFGCPE_04338* | *livF-15* | + | 4709844 | 4710557 | 2.69 | 31.69 | 13.26 | 6.98 | High-affinity branched-chain amino acid transport ATP-binding protein LivF |
| *FEPFGCPE_04339* | *ypeA* | - | 4710676 | 4711155 | 2.73 | 1.94 | 4.31 | 1.05 | Acetyltransferase YpeA |
| *FEPFGCPE_04340* | | - | 4711152 | 4711649 | 8.67 | 26.13 | 25.16 | 9.41 | Hypothetical protein |
| *FEPFGCPE_04341* | *argD* | - | 4711733 | 4713001 | 14.26 | 35.78 | 22.02 | 8.57 | Acetylornithine aminotransferase |
| *FEPFGCPE_04342* | *ascD* | - | 4713286 | 4714320 | 2.41 | 17.30 | 6.80 | 6.14 | CDP-6-deoxy-L-threo-D-glycero-4-hexulose-3-dehydrase reductase |
| *FEPFGCPE_04343* | | + | 4714336 | 4715406 | 4.51 | 31.41 | 28.46 | 5.70 | Hypothetical protein |
| *FEPFGCPE_04344* | | + | 4715977 | 4716198 | 5.22 | 27.23 | 8.68 | 1.51 | Hypothetical protein |
| *FEPFGCPE_04345* | *ompR-3* | + | 4716216 | 4717058 | 1.05 | 4.65 | 5.07 | 1.45 | Transcriptional regulatory protein OmpR |
| *FEPFGCPE_04346* | *envZ-3* | + | 4717055 | 4718386 | 1.01 | 7.33 | 4.36 | 0.49 | Osmolarity sensor protein EnvZ |
| *FEPFGCPE_04347* | | - | 4718441 | 4719361 | 3.38 | 12.39 | 8.35 | 3.32 | Hypothetical protein |
| *FEPFGCPE_04348* | *zntB* | + | 4719694 | 4720710 | 1.99 | 7.76 | 4.16 | 2.94 | Zinc transport protein ZntB |
| *FEPFGCPE_04349* | | - | 4720748 | 4721164 | 11.36 | 65.57 | 55.84 | 11.11 | Hypothetical protein |
| *FEPFGCPE_04350* | *exsA* | - | 4721161 | 4721718 | 5.84 | 42.06 | 21.25 | 6.02 | Exoenzyme S synthesis regulatory protein ExsA |
| *FEPFGCPE_04351* | | - | 4721836 | 4722879 | 0.86 | 4.47 | 2.94 | 0.03 | Outer membrane porin protein |
| *FEPFGCPE_04352* | *betC* | + | 4723152 | 4724969 | 1.50 | 8.50 | 5.29 | 1.08 | Choline-sulfatase |
| *FEPFGCPE_04353* | *tibC* | - | 4725012 | 4726274 | 0.89 | 2.78 | 2.05 | 0.64 | Glycosyltransferase TibC |
| *FEPFGCPE_04354* | *gbpR-6* | - | 4726481 | 4727452 | 3.54 | 12.55 | 8.22 | 3.16 | HTH-type transcriptional regulator GbpR |
| *FEPFGCPE_04355* | *maeB-3* | + | 4727649 | 4729937 | 1.72 | 11.87 | 12.85 | 9.87 | NADP-dependent malic enzyme |
| *FEPFGCPE_04356* | | - | 4730079 | 4730483 | 9.58 | 272.82 | 115.74 | 30.80 | Hypothetical protein |
| *FEPFGCPE_04357* | | - | 4730693 | 4731094 | 1.21 | 4.03 | 4.60 | 0.00 | Hypothetical protein |
| *FEPFGCPE_04358* | *allS-2* | + | 4731224 | 4732111 | 1.81 | 8.41 | 4.53 | 1.76 | HTH-type transcriptional activator AllS |
| *FEPFGCPE_04359* | | - | 4732098 | 4732967 | 7.52 | 7.99 | 5.98 | 2.78 | putative metallo-hydrolase |
| *FEPFGCPE_04360* | *bigR* | + | 4733077 | 4733439 | 24.13 | 10.56 | 10.79 | 4.13 | Biofilm growth-associated repressor |
| *FEPFGCPE_04361* | | + | 4733463 | 4733909 | 7.57 | 6.06 | 7.41 | 3.24 | Hypothetical protein |
| *FEPFGCPE_04362* | | + | 4733906 | 4734328 | 10.12 | 24.23 | 12.61 | 4.62 | Hypothetical protein |
| *FEPFGCPE_04363* | *hsrA-3* | - | 4734339 | 4735721 | 3.90 | 15.85 | 8.76 | 3.49 | putative transport protein HsrA |
| *FEPFGCPE_04364* | | - | 4735797 | 4736405 | 11.37 | 18.00 | 9.89 | 4.15 | Hypothetical protein |
| *FEPFGCPE_04365* | | + | 4737180 | 4737470 | 42.17 | 123.21 | 69.35 | 41.68 | DNA-binding protein Bv3F |
| *FEPFGCPE_04366* | *envZ-4* | - | 4737756 | 4739030 | 1.34 | 8.28 | 3.07 | 4.22 | Osmolarity sensor protein EnvZ |
| *FEPFGCPE_04367* | *ompR-4* | - | 4739152 | 4739898 | 2.16 | 20.41 | 11.19 | 2.95 | Transcriptional regulatory protein OmpR |
| *FEPFGCPE_04368* | *mdtA-4* | + | 4740401 | 4741753 | 4.23 | 12.25 | 8.21 | 2.28 | Multidrug resistance protein MdtA |
| *FEPFGCPE_04369* | *mdtB* | + | 4741846 | 4745028 | 1.04 | 4.91 | 4.40 | 1.47 | Multidrug resistance protein MdtB |
| *FEPFGCPE_04370* | *mdtC* | + | 4745025 | 4748165 | 0.85 | 5.00 | 3.60 | 1.11 | Multidrug resistance protein MdtC |
| *FEPFGCPE_04371* | *oprM-2* | + | 4748365 | 4749864 | 0.97 | 4.88 | 5.79 | 2.02 | Outer membrane protein OprM |
| *FEPFGCPE_04372* | *rcsB-2* | + | 4750398 | 4751036 | 0.65 | 3.34 | 0.95 | 0.66 | Transcriptional regulatory protein RcsB |
| *FEPFGCPE_04373* | *yccS-2* | - | 4751095 | 4753317 | 0.90 | 4.03 | 3.19 | 0.35 | Inner membrane protein YccS |
| *FEPFGCPE_04374* | *recC* | + | 4753574 | 4756990 | 1.60 | 8.25 | 5.47 | 2.22 | RecBCD enzyme subunit RecC |
| *FEPFGCPE_04375* | *recB* | + | 4756987 | 4760808 | 1.22 | 11.31 | 5.02 | 1.86 | RecBCD enzyme subunit RecB |
| *FEPFGCPE_04376* | *recD* | + | 4760805 | 4762949 | 0.91 | 13.68 | 10.32 | 3.13 | RecBCD enzyme subunit RecD |
| *FEPFGCPE_04377* | *tlyC* | - | 4762900 | 4764258 | 2.00 | 14.06 | 12.28 | 3.37 | Hemolysin C |
| *FEPFGCPE_04378* | | + | 4764728 | 4765831 | 0.71 | 2.85 | 1.39 | 0.50 | Hypothetical protein |
| *FEPFGCPE_04379* | *thiC* | - | 4765971 | 4767887 | 3.04 | 33.50 | 32.01 | 19.91 | Phosphomethylpyrimidine synthase |
| *FEPFGCPE_04380* | | + | 4768452 | 4768907 | 2.76 | 20.73 | 7.48 | 3.84 | Hypothetical protein |
| *FEPFGCPE_04381* | | - | 4768945 | 4770630 | 1.77 | 8.74 | 5.71 | 0.98 | Hypothetical protein |
| *FEPFGCPE_04382* | *bla* | + | 4770864 | 4771730 | 9.40 | 32.70 | 15.82 | 8.04 | Beta-lactamase OXA-10 |
| *FEPFGCPE_04383* | *pnpB-1* | - | 4771804 | 4772436 | 27.98 | 181.65 | 155.80 | 64.52 | p-benzoquinone reductase |
| *FEPFGCPE_04384* | | - | 4772558 | 4772851 | 2.24 | 13.88 | 5.32 | 1.76 | Hypothetical protein |
| *FEPFGCPE_04385* | *cynR-8* | - | 4773091 | 4774053 | 2.99 | 13.01 | 8.52 | 1.33 | HTH-type transcriptional regulator CynR |
| *FEPFGCPE_04386* | | + | 4774267 | 4775334 | 3.97 | 11.23 | 7.55 | 3.53 | Hypothetical protein |
| *FEPFGCPE_04387* | *rhlE-4* | + | 4775595 | 4777106 | 5.91 | 25.02 | 25.17 | 15.05 | ATP-dependent RNA helicase RhlE |
| *FEPFGCPE_04388* | *gluQ* | - | 4777219 | 4778169 | 1.86 | 10.17 | 5.69 | 0.26 | Glutamyl-Q tRNA(Asp) synthetase |
| *FEPFGCPE_04389* | | - | 4778178 | 4779227 | 8.49 | 20.42 | 25.01 | 6.41 | Hypothetical protein |
| *FEPFGCPE_04390* | *dnaE* | + | 4779304 | 4782894 | 3.58 | 18.82 | 12.29 | 6.45 | DNA polymerase III subunit alpha |
| *FEPFGCPE_04391* | | + | 4782969 | 4783820 | 1.66 | 6.78 | 6.79 | 1.66 | Hypothetical protein |
| *FEPFGCPE_04392* | *sunS* | - | 4783837 | 4784637 | 0.91 | 6.89 | 9.75 | 2.53 | SPBc2 prophage-derived glycosyltransferase SunS |
| *FEPFGCPE_04393* | *recQ* | - | 4784685 | 4786565 | 2.11 | 18.75 | 15.55 | 6.61 | ATP-dependent DNA helicase RecQ |
| *FEPFGCPE_04394* | *msbA-2* | + | 4786667 | 4788418 | 1.69 | 10.57 | 8.82 | 3.58 | Lipid A export ATP-binding/permease protein MsbA |
| *FEPFGCPE_04395* | | + | 4788476 | 4789057 | 3.70 | 19.80 | 23.29 | 6.82 | Hypothetical protein |
| *FEPFGCPE_04396* | *rfbB* | + | 4789087 | 4790151 | 2.29 | 21.95 | 21.61 | 7.66 | dTDP-glucose 4,6-dehydratase |
| *FEPFGCPE_04397* | *rfbA* | + | 4790159 | 4791046 | 2.72 | 14.99 | 11.07 | 5.08 | Glucose-1-phosphate thymidylyltransferase 1 |
| *FEPFGCPE_04398* | *rmlC* | + | 4791090 | 4791638 | 2.14 | 27.34 | 12.52 | 7.96 | dTDP-4-dehydrorhamnose 3,5-epimerase |

| Locus name | Gene name | Strand | Gene start (bp) | Gene end (bp) | Average TPM Inoculum | Average TPM $t_6$ | Average TPM $t_{24}$ | Average TPM $t_{168}$ | Annotation description |
|---|---|---|---|---|---|---|---|---|---|
| *FEPFGCPE_04399* | | - | 4791695 | 4792981 | 1.83 | 11.48 | 7.84 | 2.47 | putative cation efflux system protein |
| *FEPFGCPE_04400* | *rng* | - | 4793012 | 4794481 | 3.87 | 13.83 | 11.91 | 5.93 | Ribonuclease G |
| *FEPFGCPE_04401* | *yhdE* | - | 4794527 | 4795126 | 4.15 | 16.84 | 9.82 | 5.29 | Maf-like protein YhdE |
| *FEPFGCPE_04402* | *ugpQ* | - | 4795341 | 4796120 | 0.66 | 2.73 | 2.41 | 0.85 | Glycerophosphodiester phosphodiesterase, cytoplasmic |
| *FEPFGCPE_04403* | *ugpC-3* | - | 4796117 | 4797214 | 0.51 | 10.45 | 8.67 | 2.21 | sn-glycerol-3-phosphate import ATP-binding protein UgpC |
| *FEPFGCPE_04404* | *lacG* | - | 4797217 | 4798062 | 0.60 | 7.86 | 5.85 | 3.07 | Lactose transport system permease protein LacG |
| *FEPFGCPE_04405* | *ugpA* | - | 4798059 | 4798955 | 0.73 | 9.53 | 4.42 | 3.49 | sn-glycerol-3-phosphate transport system permease protein UgpA |
| *FEPFGCPE_04406* | *ugpB-2* | - | 4799113 | 4800444 | 2.77 | 33.00 | 26.58 | 12.18 | sn-glycerol-3-phosphate-binding periplasmic protein UgpB |
| *FEPFGCPE_04407* | *rlmH* | - | 4800708 | 4801178 | 8.28 | 17.94 | 10.95 | 5.74 | Ribosomal RNA large subunit methyltransferase H |
| *FEPFGCPE_04408* | *rsfS* | - | 4801206 | 4801646 | 3.96 | 53.77 | 39.86 | 26.95 | Ribosomal silencing factor RsfS |
| *FEPFGCPE_04409* | *nadD* | - | 4801709 | 4802440 | 5.70 | 77.48 | 51.12 | 13.36 | putative nicotinate-nucleotide adenylyltransferase |
| *FEPFGCPE_04410* | *hemF* | - | 4802454 | 4803389 | 7.49 | 44.59 | 26.04 | 8.58 | Oxygen-dependent coproporphyrinogen-III oxidase |
| *FEPFGCPE_04411* | *mscS-2* | - | 4803515 | 4804399 | 12.97 | 49.14 | 32.25 | 8.19 | Small-conductance mechanosensitive channel |
| *FEPFGCPE_04412* | *purD* | - | 4804703 | 4805980 | 5.99 | 41.17 | 23.92 | 13.95 | Phosphoribosylamine--glycine ligase |
| *FEPFGCPE_04413* | *pmpR* | - | 4805982 | 4806707 | 5.38 | 46.31 | 35.31 | 28.56 | Transcriptional regulatory protein PmpR |
| *FEPFGCPE_04414* | | + | 4807016 | 4808671 | 1.95 | 14.79 | 8.64 | 2.75 | Hypothetical protein |
| *FEPFGCPE_04415* | *eamA-3* | + | 4808790 | 4809725 | 2.07 | 13.34 | 12.51 | 4.60 | putative amino-acid metabolite efflux pump |
| *FEPFGCPE_04416* | *upp* | - | 4809779 | 4810429 | 2.06 | 22.77 | 12.96 | 2.32 | Uracil phosphoribosyltransferase |
| *FEPFGCPE_04417* | *fabG-10* | + | 4810641 | 4811420 | 5.46 | 26.71 | 16.07 | 7.98 | 3-oxoacyl-[acyl-carrier-protein] reductase FabG |
| *FEPFGCPE_04418* | *mgsA* | + | 4811440 | 4811853 | 6.88 | 43.32 | 24.98 | 10.32 | Methylglyoxal synthase |
| *FEPFGCPE_04419* | *qorA* | + | 4811906 | 4812880 | 6.35 | 52.21 | 31.43 | 18.65 | Quinone oxidoreductase 1 |
| *FEPFGCPE_04420* | | - | 4813014 | 4813304 | 12.22 | 37.65 | 28.67 | 24.58 | Hypothetical protein |
| *FEPFGCPE_04421* | | + | 4813654 | 4815063 | 9.69 | 46.84 | 44.27 | 15.04 | Putative glutamine amidotransferase |
| *FEPFGCPE_04422* | | - | 4815175 | 4815444 | 6.42 | 54.09 | 91.85 | 108.79 | Outer membrane porin protein 32 |
| *FEPFGCPE_04423* | | - | 4815535 | 4816335 | 23.12 | 225.65 | 363.59 | 251.61 | Outer membrane porin protein |
| *FEPFGCPE_04424* | *acoR-2* | - | 4817168 | 4818400 | 1.44 | 17.82 | 6.61 | 0.90 | Acetoin catabolism regulatory protein |
| *FEPFGCPE_04425* | | - | 4818779 | 4818949 | 3.51 | 0.00 | 0.59 | 0.00 | Hypothetical protein |
| *FEPFGCPE_04426* | *fabG-11* | - | 4819551 | 4820306 | 1.89 | 11.28 | 4.64 | 0.96 | 3-oxoacyl-[acyl-carrier-protein] reductase FabG |
| *FEPFGCPE_04427* | *siaT-3* | - | 4820409 | 4822271 | 0.28 | 5.85 | 2.70 | 0.35 | Sialic acid TRAP transporter permease protein SiaT |
| *FEPFGCPE_04428* | *siaP-3* | - | 4822287 | 4823306 | 1.69 | 36.70 | 11.92 | 8.04 | Sialic acid-binding periplasmic protein SiaP |
| *FEPFGCPE_04429* | *fadA-4* | + | 4823892 | 4825058 | 3.04 | 32.21 | 17.74 | 9.88 | 3-ketoacyl-CoA thiolase |
| *FEPFGCPE_04430* | | + | 4825141 | 4826610 | 1.96 | 34.65 | 11.30 | 5.71 | Long-chain-fatty-acid--CoA ligase FadD13 |
| *FEPFGCPE_04431* | | + | 4826715 | 4827251 | 1.80 | 51.68 | 19.45 | 2.85 | Hypothetical protein |
| *FEPFGCPE_04432* | | + | 4827248 | 4827661 | 1.67 | 40.01 | 16.20 | 8.15 | Hypothetical protein |
| *FEPFGCPE_04433* | *alaC* | - | 4827742 | 4828959 | 4.14 | 8.71 | 3.43 | 2.44 | Glutamate-pyruvate aminotransferase AlaC |
| *FEPFGCPE_04434* | *emrE* | + | 4829135 | 4829464 | 2.01 | 5.47 | 6.81 | 1.53 | Multidrug transporter EmrE |
| *FEPFGCPE_04435* | *rocD2* | - | 4829525 | 4830742 | 10.56 | 21.42 | 13.87 | 2.57 | Ornithine aminotransferase 2 |
| *FEPFGCPE_04436* | | + | 4830930 | 4831006 | 3.13 | 14.69 | 3.98 | 1.37 | tRNA |
| *FEPFGCPE_04437* | *gltC-14* | - | 4831155 | 4832021 | 1.42 | 6.08 | 1.14 | 5.13 | HTH-type transcriptional regulator GltC |
| *FEPFGCPE_04438* | *stp* | + | 4832122 | 4833546 | 1.84 | 12.10 | 4.11 | 0.50 | Multidrug resistance protein Stp |
| *FEPFGCPE_04439* | | + | 4833622 | 4833918 | 1.53 | 16.45 | 5.75 | 3.60 | Hypothetical protein |
| *FEPFGCPE_04440* | | - | 4834077 | 4834886 | 0.40 | 4.48 | 4.27 | 2.09 | Hypothetical protein |
| *FEPFGCPE_04441* | *naiP-7* | - | 4834994 | 4836376 | 0.85 | 5.66 | 2.08 | 0.50 | Putative niacin/nicotinamide transporter NaiP |
| *FEPFGCPE_04442* | *adh1-1* | - | 4836507 | 4837772 | 1.26 | 7.05 | 4.75 | 0.53 | Long-chain-alcohol dehydrogenase 1 |
| *FEPFGCPE_04443* | *menH* | - | 4837769 | 4838530 | 0.78 | 0.30 | 1.69 | 0.63 | 2-succinyl-6-hydroxy-2,4-cyclohexadiene-1-carboxylate synthase |
| *FEPFGCPE_04444* | *rspR-10* | + | 4838845 | 4839480 | 8.81 | 16.27 | 17.66 | 8.00 | HTH-type transcriptional repressor RspR |
| *FEPFGCPE_04445* | *catC* | - | 4839539 | 4839814 | 3.04 | 15.11 | 10.38 | 7.43 | Muconolactone Delta-isomerase |
| *FEPFGCPE_04446* | *catD-5* | - | 4839931 | 4840767 | 0.51 | 1.25 | 4.50 | 0.64 | 3-oxoadipate enol-lactonase 2 |
| *FEPFGCPE_04447* | *catB* | - | 4840922 | 4842067 | 0.25 | 3.29 | 1.22 | 0.49 | Muconate cycloisomerase 1 |
| *FEPFGCPE_04448* | *benM-4* | - | 4842173 | 4842676 | 1.22 | 5.93 | 4.51 | 3.30 | HTH-type transcriptional regulator BenM |
| *FEPFGCPE_04449* | *catA1* | + | 4843280 | 4844215 | 0.80 | 3.72 | 3.56 | 4.04 | Catechol 1,2-dioxygenase 1 |
| *FEPFGCPE_04450* | *cbdA* | + | 4844323 | 4845681 | 0.26 | 0.58 | 0.41 | 4.12 | 2-halobenzoate 1,2-dioxygenase large subunit |
| *FEPFGCPE_04451* | *cbdB* | + | 4845678 | 4846166 | 0.34 | 2.97 | 1.19 | 0.67 | 2-halobenzoate 1,2-dioxygenase small subunit |
| *FEPFGCPE_04452* | *benC* | + | 4846207 | 4847229 | 0.41 | 0.46 | 0.02 | 1.67 | Benzoate 1,2-dioxygenase electron transfer component |
| *FEPFGCPE_04453* | *lvr* | + | 4847226 | 4847999 | 0.42 | 2.96 | 0.15 | 1.33 | Levodione reductase |
| *FEPFGCPE_04454* | | + | 4848151 | 4848609 | 2.75 | 5.59 | 5.47 | 2.45 | Hypothetical protein |
| *FEPFGCPE_04455* | *metR-3* | - | 4848641 | 4849513 | 8.71 | 14.31 | 10.19 | 3.75 | HTH-type transcriptional regulator MetR |
| *FEPFGCPE_04456* | *mexA-1* | + | 4849816 | 4851063 | 0.43 | 2.41 | 1.30 | 1.52 | Multidrug resistance protein MexA |
| *FEPFGCPE_04457* | *acrB* | + | 4851066 | 4854203 | 0.60 | 1.66 | 0.87 | 0.18 | Multidrug efflux pump subunit AcrB |
| *FEPFGCPE_04458* | *oprM-3* | + | 4854200 | 4855660 | 0.57 | 3.39 | 3.87 | 0.74 | Outer membrane protein OprM |
| *FEPFGCPE_04459* | *cysL-3* | + | 4855771 | 4856664 | 1.77 | 3.60 | 4.50 | 1.48 | HTH-type transcriptional regulator CysL |
| *FEPFGCPE_04460* | *lasR-2* | + | 4856813 | 4857580 | 7.54 | 26.93 | 21.77 | 8.92 | Transcriptional activator protein LasR |
| *FEPFGCPE_04461* | | + | 4857850 | 4858110 | 5.49 | 9.12 | 22.19 | 5.57 | Hypothetical protein |
| *FEPFGCPE_04462* | *oprM-4* | - | 4858323 | 4859855 | 0.41 | 2.87 | 2.00 | 0.51 | Outer membrane protein OprM |
| *FEPFGCPE_04463* | *aaeA-2* | - | 4859852 | 4860817 | 0.32 | 1.23 | 0.34 | 0.74 | p-hydroxybenzoic acid efflux pump subunit AaeA |
| *FEPFGCPE_04464* | *aaeX* | - | 4860814 | 4861023 | 1.48 | 3.42 | 1.27 | 0.00 | Protein AaeX |
| *FEPFGCPE_04465* | *aaeB* | - | 4861020 | 4863029 | 0.27 | 1.12 | 0.89 | 0.12 | p-hydroxybenzoic acid efflux pump subunit AaeB |
| *FEPFGCPE_04466* | *slyA-2* | - | 4863033 | 4863506 | 0.57 | 5.21 | 2.32 | 0.00 | Transcriptional regulator SlyA |
| *FEPFGCPE_04467* | *badR-2* | - | 4863687 | 4864238 | 1.75 | 4.65 | 8.71 | 2.00 | Transcriptional activatory protein BadR |
| *FEPFGCPE_04468* | *tsaC1-2* | + | 4864383 | 4865150 | 0.51 | 0.18 | 1.48 | 0.51 | 4-formylbenzenesulfonate dehydrogenase TsaC1/TsaC2 |
| *FEPFGCPE_04469* | *menB-2* | + | 4865259 | 4866041 | 0.17 | 0.33 | 1.96 | 0.15 | 1,4-dihydroxy-2-naphthoyl-CoA synthase |

| Locus name | Gene name | Strand | Gene start (bp) | Gene end (bp) | Average TPM Inoculum | Average TPM $t_6$ | Average TPM $t_{24}$ | Average TPM $t_{168}$ | Annotation description |
|---|---|---|---|---|---|---|---|---|---|
| *FEPFGCPE_04470* | *acdA-2* | + | 4866056 | 4867210 | 0.40 | 1.68 | 0.69 | 0.98 | Acyl-CoA dehydrogenase |
| *FEPFGCPE_04471* | *fadK* | + | 4867258 | 4868901 | 0.77 | 4.27 | 3.41 | 1.53 | Short-chain-fatty-acid--CoA ligase |
| *FEPFGCPE_04472* | *carC* | + | 4868944 | 4870113 | 1.11 | 9.05 | 10.06 | 5.30 | Caffeyl-CoA reductase-Etf complex subunit CarC |
| *FEPFGCPE_04473* | *nasD* | + | 4870492 | 4872951 | 0.21 | 58.97 | 133.14 | 97.89 | Nitrite reductase [NAD(P)H] |
| *FEPFGCPE_04474* | *nasE* | + | 4872962 | 4873327 | 0.30 | 97.39 | 279.02 | 163.51 | Assimilatory nitrite reductase [NAD(P)H] small subunit |
| *FEPFGCPE_04475* | *nasA-2* | + | 4873375 | 4876092 | 0.22 | 33.65 | 104.44 | 49.96 | Nitrate reductase |
| *FEPFGCPE_04476* | *sumT-2* | + | 4876183 | 4876986 | 1.59 | 49.30 | 133.65 | 77.01 | Uroporphyrinogen-III C-methyltransferase |
| *FEPFGCPE_04477* | *pdtaR-2* | + | 4877235 | 4877810 | 42.44 | 63.90 | 48.73 | 28.01 | putative transcriptional regulatory protein pdtaR |
| *FEPFGCPE_04478* | *nrtA* | + | 4877820 | 4878998 | 25.25 | 55.87 | 44.23 | 19.86 | Nitrate transport protein NrtA |
| *FEPFGCPE_04479* | | - | 4879022 | 4879363 | 12.75 | 66.13 | 38.06 | 22.74 | Hypothetical protein |
| *FEPFGCPE_04480* | | + | 4879660 | 4879995 | 1.68 | 19.57 | 12.47 | 4.20 | Hypothetical protein |
| *FEPFGCPE_04481* | *mexR-2* | + | 4880132 | 4881112 | 8.97 | 33.54 | 30.16 | 10.25 | Multidrug resistance operon repressor |
| *FEPFGCPE_04482* | *ubiE-2* | - | 4881194 | 4882006 | 5.44 | 26.29 | 16.19 | 4.08 | Ubiquinone/menaquinone biosynthesis C-methyltransferase UbiE |
| *FEPFGCPE_04483* | | - | 4882164 | 4882240 | 5.40 | 1.13 | 6.45 | 11.43 | tRNA |
| *FEPFGCPE_04484* | | - | 4882394 | 4883122 | 1.55 | 4.58 | 3.02 | 2.46 | Hypothetical protein |
| *FEPFGCPE_04485* | *ydjM* | + | 4883455 | 4883913 | 0.29 | 1.71 | 1.70 | 0.00 | Inner membrane protein YdjM |
| *FEPFGCPE_04486* | *srkA* | - | 4884011 | 4885132 | 3.09 | 19.10 | 19.77 | 4.22 | Stress response kinase A |
| *FEPFGCPE_04487* | | + | 4885635 | 4887155 | 132963.68 | 108422.29 | 108785.94 | 103469.52 | rRNA |
| *FEPFGCPE_04488* | | + | 4887225 | 4887301 | 191.09 | 779.51 | 568.84 | 264.21 | tRNA |
| *FEPFGCPE_04489* | | + | 4887414 | 4887489 | 44.58 | 4.20 | 7.29 | 2.50 | tRNA |
| *FEPFGCPE_04490* | | + | 4887774 | 4890635 | 100467.91 | 85379.65 | 91554.03 | 103690.87 | rRNA |
| *FEPFGCPE_04491* | | + | 4890769 | 4890880 | 533.83 | 65.44 | 77.10 | 82.70 | rRNA |
| *FEPFGCPE_04492* | *cho* | - | 4891069 | 4892820 | 1.37 | 9.71 | 9.48 | 2.06 | Excinuclease cho |
| *FEPFGCPE_04493* | | - | 4892808 | 4893332 | 5.82 | 39.03 | 29.87 | 8.40 | Hypothetical protein |
| *FEPFGCPE_04494* | | + | 4893477 | 4894196 | 1.87 | 14.81 | 6.41 | 2.39 | Hypothetical protein |
| *FEPFGCPE_04495* | *ydaM* | - | 4894294 | 4895472 | 9.10 | 40.38 | 30.69 | 7.02 | putative diguanylate cyclase YdaM |
| *FEPFGCPE_04496* | | + | 4895601 | 4896152 | 7.93 | 39.15 | 34.62 | 16.61 | Hypothetical protein |
| *FEPFGCPE_04497* | *ttr* | - | 4896279 | 4896845 | 6.77 | 16.63 | 18.01 | 5.69 | Acetyltransferase |
| *FEPFGCPE_04498* | *sinR* | - | 4896845 | 4897435 | 2.31 | 10.70 | 13.04 | 1.81 | HTH-type transcriptional regulator SinR |
| *FEPFGCPE_04499* | | + | 4897527 | 4898039 | 1.20 | 5.47 | 4.59 | 0.76 | L-methionine sulfoximine/L-methionine sulfone acetyltransferase |
| *FEPFGCPE_04500* | *xdhA-2* | + | 4898464 | 4901415 | 4.81 | 18.80 | 10.82 | 3.46 | Putative xanthine dehydrogenase molybdenum-binding subunit XdhA |
| *FEPFGCPE_04501* | *hcrC* | + | 4901412 | 4902107 | 6.21 | 24.67 | 8.14 | 2.16 | 4-hydroxybenzoyl-CoA reductase subunit gamma |
| *FEPFGCPE_04502* | *adhB* | + | 4902119 | 4903498 | 2.43 | 16.08 | 10.03 | 1.38 | Alcohol dehydrogenase cytochrome c subunit |
| *FEPFGCPE_04503* | *birA-2* | + | 4903575 | 4904228 | 2.62 | 5.80 | 8.86 | 0.00 | Bifunctional ligase/repressor BirA |
| *FEPFGCPE_04504* | *ehpR* | + | 4904391 | 4904768 | 0.43 | 7.44 | 10.65 | 6.26 | Phenazine antibiotic resistance protein EhpR |
| *FEPFGCPE_04505* | *msrAB* | - | 4904855 | 4905643 | 2.14 | 12.43 | 7.64 | 1.62 | Peptide methionine sulfoxide reductase MsrA/MsrB |
| *FEPFGCPE_04506* | *ykuV* | - | 4905757 | 4907580 | 0.90 | 5.92 | 1.98 | 1.26 | Thiol-disulfide oxidoreductase YkuV |
| *FEPFGCPE_04507* | *msrB-2* | - | 4907657 | 4908241 | 5.69 | 14.94 | 6.83 | 1.58 | Peptide methionine sulfoxide reductase MsrB |
| *FEPFGCPE_04508* | *regX3* | + | 4908489 | 4909232 | 18.18 | 38.22 | 30.58 | 6.23 | Sensory transduction protein regX3 |
| *FEPFGCPE_04509* | *resE-2* | + | 4909232 | 4910719 | 3.44 | 10.18 | 4.33 | 2.95 | Sensor histidine kinase ResE |
| *FEPFGCPE_04510* | | - | 4910703 | 4911566 | 2.47 | 36.72 | 21.00 | 9.36 | Hypothetical protein |
| *FEPFGCPE_04511* | *blc-3* | - | 4912009 | 4912590 | 1.46 | 6.88 | 5.82 | 0.80 | Outer membrane lipoprotein Blc |
| *FEPFGCPE_04512* | *sigK-2* | - | 4912699 | 4913376 | 1.37 | 6.59 | 8.11 | 2.55 | ECF RNA polymerase sigma factor SigK |
| *FEPFGCPE_04513* | *folE-2* | - | 4913416 | 4914147 | 0.99 | 6.60 | 3.98 | 1.01 | GTP cyclohydrolase 1 |
| *FEPFGCPE_04514* | *mlrA* | - | 4914189 | 4915148 | 2.38 | 15.47 | 11.60 | 3.77 | HTH-type transcriptional regulator MlrA |
| *FEPFGCPE_04515* | *xerS* | + | 4915856 | 4916548 | 2.92 | 5.85 | 8.74 | 2.39 | Tyrosine recombinase XerS |
| *FEPFGCPE_04516* | | + | 4916918 | 4917148 | 14.54 | 63.46 | 37.44 | 7.85 | Hypothetical protein |
| *FEPFGCPE_04517* | | + | 4917379 | 4917684 | 3.12 | 12.31 | 10.14 | 4.02 | Hypothetical protein |
| *FEPFGCPE_04518* | | + | 4917698 | 4918258 | 3.31 | 27.31 | 20.11 | 4.29 | Hypothetical protein |
| *FEPFGCPE_04519* | | + | 4918277 | 4918468 | 3.86 | 38.40 | 16.10 | 10.21 | Hypothetical protein |
| *FEPFGCPE_04520* | | + | 4918575 | 4919015 | 1.63 | 8.87 | 8.90 | 0.84 | Hypothetical protein |
| *FEPFGCPE_04521* | | + | 4919127 | 4919648 | 2.00 | 14.92 | 12.83 | 3.38 | Hypothetical protein |
| *FEPFGCPE_04522* | | + | 4919706 | 4920302 | 1.14 | 8.73 | 10.30 | 1.12 | Hypothetical protein |
| *FEPFGCPE_04523* | | + | 4920374 | 4920874 | 0.97 | 8.25 | 4.00 | 2.02 | Hypothetical protein |
| *FEPFGCPE_04524* | | + | 4920932 | 4921255 | 1.11 | 14.83 | 9.68 | 2.70 | Hypothetical protein |
| *FEPFGCPE_04525* | | + | 4921272 | 4921910 | 0.58 | 5.44 | 7.02 | 2.12 | Hypothetical protein |
| *FEPFGCPE_04526* | | + | 4922211 | 4922429 | 2.34 | 4.23 | 12.12 | 1.57 | Hypothetical protein |
| *FEPFGCPE_04527* | | + | 4922398 | 4922655 | 3.91 | 25.10 | 15.63 | 4.19 | Hypothetical protein |
| *FEPFGCPE_04528* | | + | 4923094 | 4923558 | 0.91 | 7.55 | 2.80 | 0.43 | Hypothetical protein |
| *FEPFGCPE_04529* | | + | 4923909 | 4924517 | 1.82 | 16.84 | 9.05 | 3.96 | Hypothetical protein |
| *FEPFGCPE_04530* | | + | 4924650 | 4924973 | 1.66 | 5.20 | 8.50 | 3.42 | Hypothetical protein |
| *FEPFGCPE_04531* | | + | 4925717 | 4925956 | 2.48 | 24.95 | 35.81 | 8.47 | Hypothetical protein |
| *FEPFGCPE_04532* | | + | 4925972 | 4926391 | 2.14 | 23.42 | 11.75 | 4.57 | Hypothetical protein |
| *FEPFGCPE_04533* | | + | 4926414 | 4926713 | 4.71 | 26.95 | 17.13 | 5.94 | Hypothetical protein |
| *FEPFGCPE_04534* | | + | 4926781 | 4927557 | 4.25 | 24.57 | 27.87 | 6.17 | Hypothetical protein |
| *FEPFGCPE_04535* | | + | 4927639 | 4927932 | 1.59 | 10.38 | 12.16 | 1.94 | Hypothetical protein |
| *FEPFGCPE_04536* | | + | 4927947 | 4928315 | 1.16 | 14.70 | 7.56 | 4.70 | Hypothetical protein |
| *FEPFGCPE_04537* | | + | 4928326 | 4928490 | 2.27 | 55.14 | 12.17 | 8.51 | Hypothetical protein |
| *FEPFGCPE_04538* | | + | 4928553 | 4929173 | 1.83 | 12.28 | 11.92 | 7.61 | Hypothetical protein |
| *FEPFGCPE_04539* | | + | 4929204 | 4929932 | 2.07 | 15.95 | 9.28 | 4.75 | Hypothetical protein |
| *FEPFGCPE_04540* | | + | 4929971 | 4930222 | 4.49 | 41.55 | 28.47 | 7.94 | Hypothetical protein |

| Locus name | Gene name | Strand | Gene start (bp) | Gene end (bp) | Average TPM Inoculum | Average TPM $t_6$ | Average TPM $t_{24}$ | Average TPM $t_{168}$ | Annotation description |
|---|---|---|---|---|---|---|---|---|---|
| *FEPFGCPE_04541* | | + | 4930241 | 4930822 | 2.43 | 28.99 | 20.23 | 5.12 | Hypothetical protein |
| *FEPFGCPE_04542* | | + | 4931584 | 4931847 | 1.41 | 3.83 | 9.13 | 0.00 | Hypothetical protein |
| *FEPFGCPE_04543* | | + | 4931863 | 4932177 | 0.93 | 9.79 | 10.36 | 2.53 | Hypothetical protein |
| *FEPFGCPE_04544* | | + | 4932354 | 4932584 | 4.80 | 29.59 | 18.48 | 7.27 | Hypothetical protein |
| *FEPFGCPE_04545* | | + | 4932605 | 4932769 | 3.14 | 6.96 | 10.32 | 3.66 | Hypothetical protein |
| *FEPFGCPE_04546* | *groS3* | + | 4933612 | 4933902 | 17.61 | 96.73 | 86.58 | 49.72 | 10 kDa chaperonin 3 |
| *FEPFGCPE_04547* | *groL* | + | 4933952 | 4935598 | 12.75 | 46.07 | 42.26 | 43.09 | 60 kDa chaperonin |
| *FEPFGCPE_04548* | *gcvA-9* | - | 4935741 | 4936712 | 0.90 | 6.42 | 3.05 | 0.04 | Glycine cleavage system transcriptional activator |
| *FEPFGCPE_04549* | *sauU-3* | + | 4936816 | 4938027 | 1.87 | 5.35 | 4.37 | 1.94 | putative sulfoacetate transporter SauU |
| *FEPFGCPE_04550* | *moaC* | - | 4938112 | 4938591 | 6.74 | 49.60 | 42.45 | 11.56 | Cyclic pyranopterin monophosphate synthase |
| *FEPFGCPE_04551* | *bepA-5* | + | 4938981 | 4940492 | 8.19 | 15.58 | 9.11 | 7.10 | Beta-barrel assembly-enhancing protease |
| *FEPFGCPE_04552* | | + | 4940814 | 4941176 | 11.34 | 59.52 | 39.04 | 20.29 | Hypothetical protein |
| *FEPFGCPE_04553* | | + | 4941306 | 4942529 | 7.94 | 29.44 | 25.89 | 6.10 | Hypothetical protein |
| *FEPFGCPE_04554* | | - | 4942542 | 4943207 | 2.67 | 18.49 | 8.93 | 4.12 | Hypothetical protein |
| *FEPFGCPE_04555* | | - | 4943217 | 4944248 | 3.24 | 11.98 | 7.35 | 3.84 | Hypothetical protein |
| *FEPFGCPE_04556* | | - | 4944265 | 4944720 | 8.72 | 15.74 | 12.93 | 10.40 | Hypothetical protein |
| *FEPFGCPE_04557* | *rfaF* | - | 4944781 | 4945806 | 7.59 | 14.81 | 10.03 | 5.86 | ADP-heptose--LPS heptosyltransferase 2 |
| *FEPFGCPE_04558* | | - | 4945914 | 4946111 | 5.21 | 54.45 | 25.92 | 20.75 | Hypothetical protein |
| *FEPFGCPE_04559* | | + | 4946567 | 4947352 | 1.81 | 5.38 | 5.58 | 1.33 | Hypothetical protein |
| *FEPFGCPE_04560* | | + | 4947349 | 4947672 | 1.06 | 8.73 | 6.13 | 0.60 | Hypothetical protein |
| *FEPFGCPE_04561* | *pgk* | + | 4947759 | 4948964 | 5.71 | 24.08 | 28.77 | 12.84 | Phosphoglycerate kinase |
| *FEPFGCPE_04562* | *pykA* | + | 4949189 | 4950631 | 6.78 | 23.53 | 21.59 | 13.34 | Pyruvate kinase II |
| *FEPFGCPE_04563* | *fda* | + | 4950754 | 4951818 | 21.63 | 82.09 | 78.79 | 36.56 | Fructose-bisphosphate aldolase |
| *FEPFGCPE_04564* | | - | 4951828 | 4952523 | 4.19 | 11.03 | 12.61 | 3.33 | Hypothetical protein |
| *FEPFGCPE_04565* | | + | 4952723 | 4954297 | 0.43 | 0.81 | 0.42 | 0.42 | Hypothetical protein |
| *FEPFGCPE_04566* | *purC* | + | 4954442 | 4955347 | 10.81 | 52.31 | 57.15 | 33.26 | Phosphoribosylaminoimidazole-succinocarboxamide synthase |
| *FEPFGCPE_04567* | *purE* | + | 4955364 | 4955870 | 5.77 | 28.68 | 40.02 | 16.48 | N5-carboxyaminoimidazole ribonucleotide mutase |
| *FEPFGCPE_04568* | *purK* | + | 4955960 | 4957105 | 1.92 | 10.23 | 11.51 | 2.34 | N5-carboxyaminoimidazole ribonucleotide synthase |
| *FEPFGCPE_04569* | *ywlC* | + | 4957109 | 4958170 | 3.38 | 15.08 | 11.06 | 2.66 | Threonylcarbamoyl-AMP synthase |
| *FEPFGCPE_04570* | | - | 4958214 | 4959152 | 10.48 | 24.09 | 20.25 | 7.03 | Hypothetical protein |
| *FEPFGCPE_04571* | *dacC-2* | - | 4959329 | 4960846 | 6.87 | 14.70 | 4.90 | 6.11 | D-alanyl-D-alanine carboxypeptidase DacC |
| *FEPFGCPE_04572* | *qseB-4* | + | 4961241 | 4961948 | 4.31 | 7.13 | 7.37 | 3.58 | Transcriptional regulatory protein QseB |
| *FEPFGCPE_04573* | *qseC-6* | + | 4961948 | 4963357 | 1.57 | 6.76 | 3.78 | 1.40 | Sensor protein QseC |
| *FEPFGCPE_04574* | | - | 4963388 | 4963786 | 2.00 | 8.51 | 6.07 | 2.24 | Hypothetical protein |
| *FEPFGCPE_04575* | *degP* | + | 4964040 | 4965575 | 141.15 | 92.37 | 132.81 | 56.66 | Periplasmic serine endoprotease DegP |
| *FEPFGCPE_04576* | | + | 4965704 | 4966147 | 215.30 | 181.21 | 200.27 | 92.61 | Hypothetical protein |
| *FEPFGCPE_04577* | *pucA-3* | - | 4966307 | 4967326 | 6.80 | 21.35 | 19.09 | 6.82 | putative xanthine dehydrogenase subunit A |
| *FEPFGCPE_04578* | | + | 4967629 | 4968246 | 23.52 | 58.75 | 28.62 | 13.47 | Hypothetical protein |
| *FEPFGCPE_04579* | *mopB* | + | 4968236 | 4968760 | 13.11 | 29.49 | 37.54 | 4.77 | Molybdenum-pterin-binding protein MopB |
| *FEPFGCPE_04580* | | + | 4968870 | 4969505 | 9.29 | 30.95 | 17.69 | 4.59 | Hypothetical protein |
| *FEPFGCPE_04581* | *msrP-3* | + | 4969535 | 4970353 | 8.82 | 150.27 | 74.29 | 11.71 | Protein-methionine-sulfoxide reductase catalytic subunit MsrP |
| *FEPFGCPE_04582* | | + | 4970435 | 4970728 | 18.87 | 328.95 | 207.71 | 58.74 | Hypothetical protein |
| *FEPFGCPE_04583* | *cysA-3* | - | 4970842 | 4971951 | 1.02 | 4.46 | 3.34 | 0.97 | Sulfate/thiosulfate import ATP-binding protein CysA |
| *FEPFGCPE_04584* | *modB* | - | 4971941 | 4972639 | 2.54 | 3.92 | 6.72 | 0.39 | Molybdenum transport system permease protein ModB |
| *FEPFGCPE_04585* | *modA* | - | 4972659 | 4973483 | 6.47 | 12.24 | 9.43 | 9.02 | Molybdate-binding periplasmic protein |
| *FEPFGCPE_04586* | *acrR-2* | - | 4973802 | 4974464 | 12.99 | 15.95 | 14.59 | 4.62 | HTH-type transcriptional regulator AcrR |
| *FEPFGCPE_04587* | *mexA-2* | + | 4974949 | 4976094 | 3.37 | 27.37 | 11.89 | 10.25 | Multidrug resistance protein MexA |
| *FEPFGCPE_04588* | *mexB* | + | 4976174 | 4979353 | 3.46 | 23.08 | 16.50 | 9.92 | Multidrug resistance protein MexB |
| *FEPFGCPE_04589* | *oprM-5* | + | 4979357 | 4980847 | 4.54 | 18.94 | 13.55 | 9.76 | Outer membrane protein OprM |
| *FEPFGCPE_04590* | | + | 4981163 | 4981420 | 25.78 | 56.17 | 42.61 | 12.07 | Hypothetical protein |
| *FEPFGCPE_04591* | | + | 4981849 | 4982106 | 0.53 | 2.88 | 0.00 | 0.00 | Hypothetical protein |
| *FEPFGCPE_04592* | | + | 4982103 | 4983056 | 1.25 | 1.23 | 3.05 | 1.92 | Hypothetical protein |
| *FEPFGCPE_04593* | | + | 4983035 | 4983688 | 0.96 | 3.04 | 2.69 | 0.00 | Hypothetical protein |
| *FEPFGCPE_04594* | | + | 4983678 | 4984205 | 4.40 | 14.79 | 7.22 | 4.14 | Hypothetical protein |
| *FEPFGCPE_04595* | | - | 4984306 | 4984824 | 11.67 | 74.37 | 60.89 | 14.13 | Hypothetical protein |
| *FEPFGCPE_04596* | | - | 4984838 | 4985221 | 9.32 | 26.49 | 24.79 | 25.03 | Hypothetical protein |
| *FEPFGCPE_04597* | | - | 4985317 | 4985592 | 125.80 | 333.56 | 248.56 | 144.46 | Hypothetical protein |
| *FEPFGCPE_04598* | *prxU* | + | 4985831 | 4986469 | 24.61 | 1577.84 | 661.46 | 336.52 | Selenocysteine-containing peroxiredoxin PrxU |
| *FEPFGCPE_04599* | *mmgC-4* | - | 4986728 | 4987921 | 3.21 | 19.73 | 13.92 | 7.09 | Acyl-CoA dehydrogenase |
| *FEPFGCPE_04600* | *tsaQ1* | + | 4988153 | 4989001 | 1.96 | 8.00 | 9.16 | 2.92 | HTH-type transcriptional regulator TsaQ1/TsaQ2 |
| *FEPFGCPE_04601* | *kynA* | - | 4989389 | 4990357 | 2.28 | 10.93 | 10.01 | 5.57 | Tryptophan 2,3-dioxygenase |
| *FEPFGCPE_04602* | *kynU* | - | 4990374 | 4991627 | 3.08 | 21.66 | 18.52 | 4.75 | Kynureninase |
| *FEPFGCPE_04603* | *kynB* | - | 4991703 | 4992368 | 2.62 | 13.72 | 17.95 | 3.05 | Kynurenine formamidase |
| *FEPFGCPE_04604* | *mhqR* | - | 4992624 | 4993088 | 1.63 | 4.07 | 1.46 | 0.68 | HTH-type transcriptional regulator MhqR |
| *FEPFGCPE_04605* | *C1-hpah* | - | 4993097 | 4993615 | 4.63 | 19.42 | 16.00 | 6.40 | p-hydroxyphenylacetate 3-hydroxylase, reductase component |
| *FEPFGCPE_04606* | *nicX* | + | 4993776 | 4994825 | 2.05 | 12.12 | 9.52 | 1.17 | 2,5-dihydroxypyridine 5,6-dioxygenase |
| *FEPFGCPE_04607* | *cpo-2* | + | 4994822 | 4995619 | 1.38 | 12.60 | 2.02 | 2.17 | Non-heme chloroperoxidase |
| *FEPFGCPE_04608* | *ydbM* | + | 4995629 | 4996873 | 3.63 | 34.48 | 15.33 | 2.23 | Putative acyl-CoA dehydrogenase YdbM |
| *FEPFGCPE_04609* | | + | 4996919 | 4998469 | 1.17 | 14.56 | 3.36 | 0.98 | Long-chain-fatty-acid--CoA ligase FadD13 |
| *FEPFGCPE_04610* | *kipR-6* | + | 4998715 | 4999461 | 3.74 | 16.40 | 10.33 | 2.57 | HTH-type transcriptional regulator KipR |
| *FEPFGCPE_04611* | *glnH-2* | + | 4999499 | 5000236 | 4.55 | 20.83 | 9.36 | 5.65 | Glutamine-binding periplasmic protein |

| Locus name | Gene name | Strand | Gene start (bp) | Gene end (bp) | Average TPM Inoculum | Average TPM $t_6$ | Average TPM $t_{24}$ | Average TPM $t_{168}$ | Annotation description |
|---|---|---|---|---|---|---|---|---|---|
| *FEPFGCPE_04612* | *glnP-2* | + | 5000349 | 5001017 | 1.05 | 7.66 | 5.85 | 2.66 | Glutamine transport system permease protein GlnP |
| *FEPFGCPE_04613* | *glnQ-12* | + | 5001014 | 5001790 | 2.87 | 13.83 | 11.47 | 4.01 | Glutamine transport ATP-binding protein GlnQ |
| *FEPFGCPE_04614* | *hutG-2* | + | 5001810 | 5002757 | 2.40 | 12.14 | 8.84 | 2.11 | Formimidoylglutamase |
| *FEPFGCPE_04615* | *msrA2* | + | 5002919 | 5003491 | 46.35 | 139.47 | 55.47 | 34.53 | Peptide methionine sulfoxide reductase MsrA 2 |
| *FEPFGCPE_04616* | | - | 5003658 | 5004008 | 13.08 | 27.35 | 16.93 | 7.33 | Hypothetical protein |
| *FEPFGCPE_04617* | *bmr3-4* | + | 5004393 | 5005967 | 1.30 | 2.57 | 0.69 | 1.02 | Multidrug resistance protein 3 |
| *FEPFGCPE_04618* | *fabV* | + | 5006137 | 5007330 | 1.15 | 9.35 | 9.58 | 6.77 | Enoyl-[acyl-carrier-protein] reductase [NADH] |
| *FEPFGCPE_04619* | | - | 5007534 | 5008667 | 6.39 | 23.92 | 22.87 | 7.26 | Hypothetical protein |
| *FEPFGCPE_04620* | | - | 5008664 | 5009875 | 17.84 | 101.43 | 62.66 | 26.88 | putative fatty acid methyltransferase |
| *FEPFGCPE_04621* | *pdxH* | - | 5009989 | 5010735 | 11.47 | 46.74 | 31.28 | 14.60 | Pyridoxine/pyridoxamine 5'-phosphate oxidase |
| *FEPFGCPE_04622* | | - | 5010919 | 5012955 | 3.72 | 11.28 | 10.70 | 2.85 | Hypothetical protein |
| *FEPFGCPE_04623* | *tcdA* | + | 5012957 | 5013832 | 9.98 | 73.10 | 69.92 | 13.41 | tRNA threonylcarbamoyladenosine dehydratase |
| *FEPFGCPE_04624* | *fadD-3* | - | 5014043 | 5015695 | 7.58 | 19.89 | 12.65 | 5.84 | Long-chain-fatty-acid--CoA ligase |
| *FEPFGCPE_04625* | | + | 5016105 | 5016181 | 10.77 | 56.16 | 37.65 | 7.73 | tRNA |
| *FEPFGCPE_04626* | | + | 5016336 | 5016412 | 5.93 | 25.25 | 26.28 | 5.97 | tRNA |
| *FEPFGCPE_04627* | *oprM-6* | - | 5016582 | 5018054 | 1.03 | 1.31 | 1.85 | 0.62 | Outer membrane protein OprM |
| *FEPFGCPE_04628* | *emrB-5* | - | 5018061 | 5019776 | 1.46 | 4.14 | 1.06 | 0.00 | Multidrug export protein EmrB |
| *FEPFGCPE_04629* | *emrA-2* | - | 5019787 | 5021127 | 0.60 | 2.68 | 1.04 | 1.68 | Multidrug export protein EmrA |
| *FEPFGCPE_04630* | *ybiH* | + | 5021230 | 5021979 | 7.79 | 36.01 | 18.52 | 13.27 | putative HTH-type transcriptional regulator YbiH |
| *FEPFGCPE_04631* | *lrp-8* | + | 5022181 | 5022654 | 2.03 | 12.24 | 6.52 | 5.34 | Leucine-responsive regulatory protein |
| *FEPFGCPE_04632* | *lldD* | + | 5022720 | 5023907 | 2.66 | 5.84 | 9.62 | 2.49 | L-lactate dehydrogenase |
| *FEPFGCPE_04633* | | + | 5024183 | 5024302 | 0.52 | 8.40 | 0.00 | 3.05 | Hypothetical protein |
| *FEPFGCPE_04634* | | - | 5024392 | 5024826 | 4.69 | 30.89 | 36.36 | 20.91 | Hypothetical protein |
| *FEPFGCPE_04635* | *hcaR-4* | - | 5024962 | 5025999 | 2.43 | 6.49 | 3.71 | 1.92 | Hca operon transcriptional activator HcaR |
| *FEPFGCPE_04636* | *proP-17* | + | 5026194 | 5027513 | 1.78 | 10.71 | 21.22 | 21.44 | Proline/betaine transporter |
| *FEPFGCPE_04637* | *ais* | + | 5027621 | 5028436 | 0.89 | 29.07 | 31.21 | 29.25 | Aconitate isomerase |
| *FEPFGCPE_04638* | *aam* | + | 5028585 | 5030135 | 3.01 | 17.60 | 12.22 | 6.19 | Acylamidase |
| *FEPFGCPE_04639* | | - | 5030165 | 5030413 | 2.94 | 23.28 | 16.76 | 6.07 | Hypothetical protein |
| *FEPFGCPE_04640* | | + | 5030522 | 5031145 | 5.82 | 14.14 | 17.75 | 5.16 | 3-mercaptopropionate dioxygenase |
| *FEPFGCPE_04641* | *glpE-4* | + | 5031289 | 5032947 | 3.48 | 17.52 | 14.10 | 5.83 | Thiosulfate sulfurtransferase GlpE |
| *FEPFGCPE_04642* | | + | 5033116 | 5034240 | 4.52 | 30.39 | 28.45 | 6.16 | Hypothetical protein |
| *FEPFGCPE_04643* | *adh1-2* | - | 5034438 | 5035598 | 4.09 | 18.44 | 16.12 | 4.30 | Long-chain-alcohol dehydrogenase 1 |
| *FEPFGCPE_04644* | *davD* | - | 5035748 | 5037196 | 3.13 | 6.00 | 5.71 | 4.85 | Glutarate-semialdehyde dehydrogenase DavD |
| *FEPFGCPE_04645* | *iclR-6* | + | 5037442 | 5038263 | 0.45 | 2.04 | 0.49 | 0.20 | Transcriptional repressor IclR |
| *FEPFGCPE_04646* | *pnpB-2* | + | 5038801 | 5039442 | 15.65 | 44.27 | 22.07 | 8.40 | p-benzoquinone reductase |
| *FEPFGCPE_04647* | *rspR-11* | - | 5039497 | 5040213 | 8.72 | 42.11 | 15.85 | 8.38 | HTH-type transcriptional repressor RspR |
| *FEPFGCPE_04648* | | + | 5040390 | 5041610 | 0.91 | 9.72 | 2.45 | 1.08 | Soluble hydrogenase 42 kDa subunit |
| *FEPFGCPE_04649* | *sauU-4* | + | 5041798 | 5043075 | 1.71 | 7.73 | 7.37 | 1.91 | putative sulfoacetate transporter SauU |
| *FEPFGCPE_04650* | *glpC-2* | + | 5043175 | 5046237 | 2.05 | 16.05 | 8.92 | 3.02 | Anaerobic glycerol-3-phosphate dehydrogenase subunit C |
| *FEPFGCPE_04651* | | + | 5046275 | 5047477 | 3.40 | 19.53 | 8.85 | 5.17 | D-threonine aldolase |
| *FEPFGCPE_04652* | | + | 5047599 | 5047674 | 94.84 | 111.01 | 55.39 | 34.09 | tRNA |
| *FEPFGCPE_04653* | | + | 5047691 | 5047766 | 5.05 | 22.43 | 5.70 | 6.50 | tRNA |
| *FEPFGCPE_04654* | | - | 5047948 | 5048244 | 4.56 | 57.52 | 25.95 | 12.45 | Hypothetical protein |
| *FEPFGCPE_04655* | | - | 5048316 | 5048633 | 1.34 | 5.90 | 6.39 | 4.06 | Hypothetical protein |
| *FEPFGCPE_04656* | | - | 5048847 | 5049359 | 1.68 | 8.15 | 9.59 | 4.60 | Hypothetical protein |
| *FEPFGCPE_04657* | | - | 5049510 | 5050244 | 2.53 | 14.98 | 13.62 | 2.30 | Hypothetical protein |
| *FEPFGCPE_04658* | | + | 5050520 | 5051041 | 12.07 | 67.08 | 58.79 | 16.49 | Hypothetical protein |
| *FEPFGCPE_04659* | | - | 5051149 | 5051448 | 8.15 | 24.00 | 41.26 | 21.91 | Hypothetical protein |
| *FEPFGCPE_04660* | | - | 5051505 | 5051858 | 18.24 | 119.61 | 121.29 | 47.75 | Hypothetical protein |
| *FEPFGCPE_04661* | *yibF-2* | + | 5052098 | 5052712 | 6.60 | 27.89 | 19.74 | 5.94 | putative GST-like protein YibF |
| *FEPFGCPE_04662* | *yadH-3* | - | 5052770 | 5053543 | 5.72 | 18.84 | 13.57 | 7.04 | Inner membrane transport permease YadH |
| *FEPFGCPE_04663* | *ybhF-2* | - | 5053540 | 5054469 | 4.78 | 20.86 | 17.32 | 4.44 | putative ABC transporter ATP-binding protein YbhF |
| *FEPFGCPE_04664* | | - | 5054617 | 5055045 | 6.07 | 10.15 | 17.94 | 9.58 | Hypothetical protein |
| *FEPFGCPE_04665* | | - | 5055107 | 5055514 | 12.13 | 47.27 | 45.76 | 15.21 | Hypothetical protein |
| *FEPFGCPE_04666* | | + | 5055680 | 5056351 | 2.30 | 8.36 | 13.53 | 6.32 | Hypothetical protein |
| *FEPFGCPE_04667* | *dmlR-22* | - | 5056422 | 5057327 | 1.74 | 1.70 | 4.26 | 3.54 | HTH-type transcriptional regulator DmlR |
| *FEPFGCPE_04668* | *fabG-12* | + | 5057546 | 5058304 | 0.82 | 4.41 | 1.67 | 1.27 | 3-oxoacyl-[acyl-carrier-protein] reductase FabG |
| *FEPFGCPE_04669* | | - | 5058350 | 5059306 | 1.34 | 2.22 | 1.05 | 0.97 | Hypothetical protein |
| *FEPFGCPE_04670* | *lrp-9* | - | 5059299 | 5059778 | 2.07 | 1.65 | 5.47 | 2.94 | Leucine-responsive regulatory protein |
| *FEPFGCPE_04671* | *fcuA* | + | 5060065 | 5062347 | 2.13 | 16.71 | 13.10 | 20.11 | Ferrichrome receptor FcuA |
| *FEPFGCPE_04672* | *acuR-2* | + | 5062540 | 5063124 | 1.50 | 11.39 | 8.04 | 12.00 | Transcriptional regulator AcuR |
| *FEPFGCPE_04673* | *ohrB* | + | 5063260 | 5063685 | 2.14 | 16.11 | 18.78 | 27.00 | Organic hydroperoxide resistance protein OhrB |
| *FEPFGCPE_04674* | | - | 5063800 | 5064435 | 34.25 | 31.98 | 31.51 | 16.17 | Hypothetical protein |
| *FEPFGCPE_04675* | *slyX* | - | 5064631 | 5064843 | 4.33 | 30.21 | 24.66 | 12.09 | Protein SlyX |
| *FEPFGCPE_04676* | | + | 5064910 | 5065263 | 2.62 | 36.51 | 15.36 | 5.27 | Hypothetical protein |
| *FEPFGCPE_04677* | | - | 5065269 | 5066834 | 3.67 | 16.28 | 12.29 | 6.05 | putative signaling protein |
| *FEPFGCPE_04678* | *lrp-10* | - | 5066912 | 5067376 | 1.10 | 7.15 | 6.98 | 4.71 | Leucine-responsive regulatory protein |
| *FEPFGCPE_04679* | *rhtB-5* | + | 5067644 | 5068273 | 0.74 | 3.54 | 5.35 | 0.00 | Homoserine/homoserine lactone efflux protein |
| *FEPFGCPE_04680* | *cyoA-2* | + | 5068742 | 5069833 | 3.11 | 19.64 | 43.09 | 46.91 | Cytochrome bo(3) ubiquinol oxidase subunit 2 |
| *FEPFGCPE_04681* | *cyoB-2* | + | 5069840 | 5071828 | 4.20 | 11.87 | 22.87 | 30.69 | Cytochrome bo(3) ubiquinol oxidase subunit 1 |
| *FEPFGCPE_04682* | *cyoC-3* | + | 5071831 | 5072457 | 6.82 | 10.31 | 21.84 | 32.14 | Cytochrome bo(3) ubiquinol oxidase subunit 3 |

| Locus name | Gene name | Strand | Gene start (bp) | Gene end (bp) | Average TPM Inoculum | Average TPM $t_6$ | Average TPM $t_{24}$ | Average TPM $t_{168}$ | Annotation description |
|---|---|---|---|---|---|---|---|---|---|
| *FEPFGCPE_04683* | *cyoD-2* | + | 5072458 | 5072823 | 6.38 | 9.00 | 22.46 | 63.89 | Cytochrome bo(3) ubiquinol oxidase subunit 4 |
| *FEPFGCPE_04684* | *penA-2* | - | 5073550 | 5075337 | 22.20 | 24.59 | 10.70 | 4.84 | putative peptidoglycan D,D-transpeptidase PenA |
| *FEPFGCPE_04685* | | - | 5075579 | 5076430 | 2.66 | 11.12 | 6.91 | 1.69 | Hypothetical protein |
| *FEPFGCPE_04686* | | + | 5076582 | 5077931 | 0.23 | 1.46 | 1.37 | 1.20 | Hypothetical protein |
| *FEPFGCPE_04687* | *srpC-2* | - | 5077959 | 5078489 | 0.62 | 4.58 | 4.74 | 1.29 | putative chromate transport protein |
| *FEPFGCPE_04688* | *chrA1-2* | - | 5078486 | 5079088 | 0.73 | 5.09 | 5.95 | 2.51 | Chromate transport protein |
| *FEPFGCPE_04689* | *hcaR-5* | + | 5079247 | 5080170 | 2.73 | 10.51 | 8.64 | 2.48 | Hca operon transcriptional activator HcaR |
| *FEPFGCPE_04690* | *resA-4* | - | 5080230 | 5080829 | 15.19 | 70.34 | 40.11 | 8.10 | Thiol-disulfide oxidoreductase ResA |
| *FEPFGCPE_04691* | | - | 5080856 | 5082013 | 28.90 | 117.02 | 61.83 | 22.87 | Hypothetical protein |
| *FEPFGCPE_04692* | *selD* | + | 5082308 | 5083387 | 5.56 | 29.78 | 17.87 | 8.68 | Selenide, water dikinase |
| *FEPFGCPE_04693* | *hsaD-2* | - | 5083442 | 5084272 | 5.00 | 16.18 | 10.05 | 2.48 | 4,5:9,10-diseco-3-hydroxy-5,9,17-trioxoandrosta-1(10),2-diene-4-oate hydrolase |
| *FEPFGCPE_04694* | *bclA-3* | - | 5084287 | 5085654 | 5.84 | 12.86 | 9.37 | 2.48 | Benzoate--CoA ligase |
| *FEPFGCPE_04695* | | - | 5085674 | 5085856 | 5.46 | 13.84 | 11.13 | 4.43 | Hypothetical protein |
| *FEPFGCPE_04696* | *boxD* | - | 5085993 | 5087579 | 3.90 | 6.50 | 5.64 | 2.48 | 3,4-dehydroadipyl-CoA semialdehyde dehydrogenase |
| *FEPFGCPE_04697* | | - | 5087615 | 5088085 | 15.31 | 30.65 | 22.33 | 11.18 | Hypothetical protein |
| *FEPFGCPE_04698* | *aroK-3* | + | 5088246 | 5089160 | 6.79 | 30.02 | 12.21 | 6.06 | Shikimate kinase |
| *FEPFGCPE_04699* | *boxC* | + | 5089273 | 5090928 | 29.44 | 89.47 | 52.33 | 51.67 | Benzoyl-CoA-dihydrodiol lyase |
| *FEPFGCPE_04700* | *boxB* | + | 5091028 | 5092455 | 9.59 | 33.26 | 22.75 | 16.96 | Benzoyl-CoA oxygenase component B |
| *FEPFGCPE_04701* | *boxA* | + | 5092503 | 5093759 | 8.66 | 46.98 | 29.16 | 12.95 | Benzoyl-CoA oxygenase component A |
| *FEPFGCPE_04702* | *yfkM* | + | 5093994 | 5094575 | 21.22 | 92.16 | 73.51 | 36.09 | General stress protein 18 |
| *FEPFGCPE_04703* | | + | 5094717 | 5096057 | 4.57 | 21.04 | 12.47 | 3.65 | Hypothetical protein |
| *FEPFGCPE_04704* | *glyA1* | + | 5096398 | 5097645 | 9.50 | 44.66 | 41.44 | 30.50 | Serine hydroxymethyltransferase 1 |
| *FEPFGCPE_04705* | *nrdR* | + | 5097664 | 5098146 | 2.75 | 31.24 | 19.14 | 16.35 | Transcriptional repressor NrdR |
| *FEPFGCPE_04706* | | - | 5100181 | 5101119 | 0.21 | 2.67 | 1.41 | 0.63 | Hypothetical protein |
| *FEPFGCPE_04707* | | - | 5101101 | 5101664 | 1.52 | 2.08 | 2.23 | 1.05 | Hypothetical protein |

**Supplementary Table S5. Bacterial strains used in this study.**

| Strains | Comments | References |
|---|---|---|
| *Achromobacter xylosoxidans* Col-0–50 | Non-pathogenic Betaproteobacteria isolated from Arabidopsis Col-0 leaves. Kanamycin-resistant strain. | Chen *et al.* (2020) |
| *Escherichia coli* DH5α | *E. coli* strain used for cloning of most constructs. | Hanahan (1985) |
| *E. coli* DH5α pUC18-mini-Tn7T-Gm | Strain used for integration of genes into the attTn7 site of *Pseudomonas*. | Choi & Schweizer (2006) |
| *E. coli* MaH1 | Strain is a derivative of *E. coli* DH5α with attTn7::*pir116* integrated into the genome for expression of the π protein, which is necessary for R6K plasmid replication. | Kvitko *et al.* (2012) |
| *E. coli* PIR2 pBG42 | Strain used for integration of *msfGFP* into the attTn7 site of *Pseudomonas*. | Zobel *et al.* (2015) |
| *E. coli* MaH1 pBG42_mod | Strain carrying pBG42_mod, a derivative of pBG42 that lacks the *msfGFP* gene and has an EcoRI restriction site immediately downstream of the BCD2 translational coupler. | This study |
| *E. coli* RHO3 pTNS3 | Tn7 transposase helper strain, site-specific TnsD pathway only. | Choi *et al.* (2008) |
| *E. coli* RHO5 | *E. coli* SM10 derivative carrying the *pir116* gene for replication of R6K plasmids. Used for conjugation of plasmids into *Pseudomonas*. | Kvitko *et al.* (2012) |
| *E. coli* RHO5 pUC18-mini-Tn7T-Gm::*tetR(BD)-*$P_{tet}$*-mCitrine_mCerulean3-BCD2-*$P_{14g}$ | Strain used for integration of a dual fluorescent reporter for cell division into the attTn7 site of *Pseudomonas*. *mCitrine* is under the control of a tetracycline inducible promoter, while *mCerulean3* is under the control of the constitutive promoter $P_{14g}$ and the translational enhancer BCD2. | This study |
| *Pandoraea* sp. Col-0–28 | Non-pathogenic Betaproteobacteria isolated from Arabidopsis Col-0 leaves. Ampicillin-resistant strain. | Chen *et al.* (2020) |
| *Pseudomonas fluorescens* 0-1 | Non-pathogenic strain isolated from soil. Ampicillin-resistant strain. | Silby *et al.* (2009) |
| *Pseudomonas syringae* pv. *tomato* DC3000 | Rifampicin-resistant derivative from *Pseudomonas syringae* pv. *tomato* strain DC52. | Cuppels (1986) |
| *Pst* DC3000 attTn7::*tetR(BD)-*$P_{tet}$*-mCitrine_mCerulean-BCD2-*$P_{14g}$ | *Pst* DC3000 strain carrying the cell division reporter: *mCitrine* expressed under the control of a tetracycline inducible promoter and *mCerulean3* under the control of the constitutive promoter $P_{14g}$ and the translational enhancer BCD2. | This study |
| *Pst* DC3000 pDSK600::*avrPphB* | *Pst* DC3000 expressing *avrPphB* effector from plasmid pPPY3031 (pDSK600::*avrPphB*). AvrPphB is expressed under the control of a triple lacUV5 promoter. | Puri *et al.* (1997) |
| *Pst* DC3000 pDSK600::*avrRpt2* | *P*st DC3000 expressing *avrRpt2* effector from plasmid pDSK600. | Mudgett & Staskawicz (1999) |
| *Pst* DC3000 pVSP61::*avrRps4* | *P*st DC3000 expressing *avrRps4* effector from plasmid pDSK600. | Hinsch & Staskawicz (1996) |
| *Pst* Δ*hrcC*Δ*CFA* | Non-pathogenic *Pseudomonas syringae* strain with the coronafacic acid cluster (*CFA*; 11 genes are deleted) and a gene necessary for type III secretion system formation (*hrcC*) deleted. | Worley *et al.* (2013) |
| *Pst* Δ*hrcC*Δ*CFA* attTn7T::*tetR(BD)-*$P_{tet}$*-mCitrine_mCerulean-BCD2-*$P_{14g}$ | Non-pathogenic *Pst* Δ*hrcC*Δ*CFA* strain carrying the cell division reporter: *mCitrine* expressed under the control of a tetracycline inducible promoter and *mCerulean3* under the control of the constitutive promoter $P_{14g}$ and the translational enhancer BCD2. | This study |
| *Rhodococcus* sp. 964 | Non-pathogenic Actinobacteria strain isolated from Arabidopsis leaves. Rifampicin-resistant strain. | Bodenhausen *et al.* (2014) |

**Supplementary Table S6. Plant genotypes used in this study.**

| Plant genotypes | Comments | References |
|---|---|---|
| *A. thaliana* Bu-22 | Carries *RPS7*, a resistance locus that recognizes the AvrPto effector. | Velásquez *et al.* (2017) |
| *Arabidopsis thaliana* Col-0 | | |
| *A. thaliana* Col-0 *bak1-5 bkk1-1 cerk1* | Three genes involved in pattern recognition receptor signaling are mutated: *bak1*, *bkk1*, and *cerk1*. | Xin *et al.* (2016) |

**Supplementary Table S7. Primer sequences used in this study.**

| Primer name | Primer sequence (5' to 3') | Comments | References |
|---|---|---|---|
| AVL001 | GCCAGGTATAATTGCACGAC | Used with primer AVL002 to create plasmid pBG42_mod, which lacks the fluorescent *msfGFP* gene. | This study |
| AVL002 | CCGGGTACCGAGCTCGaattCATTAGAAAACCTCCTTAGCATG | Used with primer AVL001 to create plasmid pBG42_mod. In lowercase, the sequence used to restore an EcoRI site. | This study |
| AVL003 | **GCCTGCAAGGGAGCTCACTAG**TTAATTAAGCCCATTGACAAGG | Used with primer AVL004 to amplify the 14g promoter and BCD2 translational enhancer from pBG42 for Gibson assembly of pUC18-mini-Tn7T-Gm::*tetR(BD)-$P_{tet}$-mCitrine_ mCerulean3-BCD2-$P_{14g}$*. In bold, sequence complementary to pUC18-mini-Tn7T-Gm. | This study |
| AVL004 | **CCTCGCCCTTGCTCACCAT**TAGAAAACCTCCTTAGCATG | Used with primer AVL003 to amplify the 14g promoter and BCD2 translational enhancer from pBG42 for Gibson assembly of pUC18-mini-Tn7T-Gm::*tetR(BD)-$P_{tet}$-mCitrine_ mCerulean3-BCD2-$P_{14g}$*. In bold, sequence complementary to *mCerulean3*. | This study |
| AVL005 | **CATGCTAAGGAGGTTTTCTA**ATGGTGAGCAAGGGCGAGG | Used with primer AVL005 to amplify *mCerulean3* for Gibson assembly *of* pUC18-mini-Tn7T-Gm::*tetR(BD)-$P_{tet}$-mCitrine_ mCerulean3-BCD2-$P_{14g}$* In bold, sequence complementary to the BCD2 translational enhancer. | This study |
| AVL006 | CCGAGAcCGAAGAGGCACTA | Used with primer AVL006 for Gibson assembly of pUC18-mini-Tn7T-Gm::*tetR(BD)-$P_{tet}$-mCitrine_ mCerulean3-BCD2-$P_{14g}$*. | This study |
| AVL007 | GTTTAGCGGACGACAATCCT | Used with primer AVL008 to check for transposants while with primer AVL009, it is used to check for bacteria in which transposition has not occurred. Primer hybridizes near the 5' end of the CDS of *PSPTO_5594*. | Kvitko lab (University of Georgia) |
| AVL008 | ATTAGCTTACGACGCTACACCC | Used with primer AVL007 to check for transposants. | Kvitko lab (University of Georgia) |
| AVL009 | GCAGCTGCTTTCGTATTATGTG | Used with primer AVL010 to check for transposants while with primer AVL007, it is used to check for bacteria in which transposition has not occurred. Primer hybridizes near the 3' end of *glmS* (*PSPTO_5595*). | Kvitko lab (University of Georgia) |
| AVL010 | CACAGCATAACTGGACTGATTTC | Used with primer AVL009 to check for transposants. | Kvitko lab (University of Georgia) |
| AVL011 | TTGACTcatATGGTCGCATTTGCAGGAT | Used with primer AVL012 to amplify *hrpA* (*PSPTO_1381*) from *Pseudomonas syringae* pv. *tomato* (*Pst*) DC3000 to check for genomic DNA contamination in RNA samples. In lowercase, an added NdeI site. | This study |

| Primer name | Primer sequence (5' to 3') | Comments | References |
|---|---|---|---|
| AVL012 | GCCgaattcTTAGTAACTGATACCTTTAGCG | Used with primer AVL011 to amplify *hrpA* (*PSPTO_1381*) from *Pst* DC3000 to check for genomic DNA contamination in RNA samples. In lowercase, an added EcoRI site. | This study |
| AVL013 | GAAACTGATCGACTTGCAGG | Used with primer AVL014 to amplify *rpoD* (*FEPFGCPE_00166*) of *Pandoraea* sp. Col-0-28 to check for genomic DNA contamination in RNA samples. | This study |
| AVL014 | ATGTGCACCGGAATACGGAT | Used with primer AVL013 to amplify *rpoD* (*FEPFGCPE_00166*) of *Pandoraea* sp. Col-0-28 to check for genomic DNA contamination in RNA samples. | This study |
| AVL015 | TGCAGTGGGTCGTCGACGAA | Used with primer AVL016 to amplify *rpoD1* (*FLPKDCKG_01693*) of *Achromobacter xylosoxidans* Col-0-50 to check for genomic DNA contamination in RNA samples. | This study |
| AVL016 | CGACGGTACTCGAACTTGTC | Used with primer AVL015 to amplify *rpoD1* (*FLPKDCKG_01693*) of *A. xylosoxidans* Col-0-50 to check for genomic DNA contamination in RNA samples. | This study |
| AVL017 | ATCTCATACTGACCCATATGTTCC | Used with primer AVL018 to amplify microsatellite marker *nga151a* from *Arabidopsis thaliana* to check for genomic DNA contamination in RNA samples. | Berendzen *et al.* (2005) |
| AVL018 | ATTGTACAGTCTAAAAGCGAGAG | Used with primer AVL017 to amplify microsatellite marker *nga151a* from *A. thaliana* to check for genomic DNA contamination in RNA samples. | Berendzen *et al.* (2005) |
| AVL019 | CTGACCGGTGGCATCTATTT | Used with primer AVL020 for quantitative real-time PCR (qRT-PCR) of reference *Pst* DC3000 gene *leuB* (*PSPTO_2175*). | This study |
| AVL020 | ATTTCGCTCTCGCTGTACG | Used with primer AVL019 for qRT-PCR of reference *Pst* DC3000 gene *leuB* (*PSPTO_2175*). | This study |
| AVL021 | GTCTCCAGAGCGATTTGTTG | Used with primer AVL022 for qRT-PCR of *Pst* DC3000 effector gene *avrPto* (*PSPTO_4001*). | González-Lamothe *et al.* (2012) |
| AVL022 | GGGCTAGGAGAAAGTGTTGA | Used with primer AVL021 for qRT-PCR of *Pst* DC3000 effector gene *avrPto* (*PSPTO_4001*). | González-Lamothe *et al.* (2012) |
| AVL023 | TGCTCGTCTCGTCGCCAAG | Used with primer AVL024 for qRT-PCR of *Pst* DC3000 *cfl* (*PSPTO_4680*); a coronafacic acid ligase involved in the biosynthesis of the toxin coronatine. | Worley *et al.* (2013) |
| AVL024 | CGATACCCTTAGTTAGTCCTGTGG | Used with primer AVL023 for qRT-PCR of *Pst* DC3000 *cfl* (*PSPTO_4680*); a coronafacic acid ligase involved in the biosynthesis of the toxin coronatine. | Worley *et al.* (2013) |
| AVL025 | CAACATGACATCCCGGTCA | Used with primer AVL026 for qRT-PCR of *Pst* DC3000 *fmt* (*PSPTO_0178*), a $tRNA^{fMet}$-formyl transferase involved in formylating the methionine of the initiator tRNA. | This study |
| AVL026 | CGGAATATCCAGCACCACTT | Used with primer AVL025 for qRT-PCR of *Pst* DC3000 *fmt* (*PSPTO_0178*), a $tRNA^{fMet}$-formyl transferase involved in formylating the methionine of the initiator tRNA. | This study |

| Primer name | Primer sequence (5' to 3') | Comments | References |
|---|---|---|---|
| AVL027 | TCACCATTCCCAACGAGAAG | Used with primer AVL028 for qRT-PCR of *Pst* DC3000 cell division gene *ftsZ* (*PSPTO_4403*). | This study |
| AVL028 | CGCTTGATGATGTCGGAGATA | Used with primer AVL027 for qRT-PCR of *Pst* DC3000 cell division gene *ftsZ* (*PSPTO_4403*). | This study |
| AVL029 | AGATGCTCAGGGCGTTTATC | Used with primer AVL030 for qRT-PCR of *Pst* DC3000 *hrpL* RNA polymerase sigma factor (*PSPTO_1404*). | Kvitko lab (University of Georgia) |
| AVL030 | ACGCAGGGCTTCAAGAAA | Used with primer AVL029 for qRT-PCR of *Pst* DC3000 *hrpL* RNA polymerase sigma factor (*PSPTO_1404*). | Kvitko lab (University of Georgia) |
| AVL031 | TGAAGGTCGGTGACAAGTTC | Used with primer AVL032 for qRT-PCR of *Pst* DC3000 gene *PSPTO_4337* that encodes a pyruvate kinase involved in the Entner-Doudoroff pathway. | This study |
| AVL032 | CCGCAATCCTTGACCAGAT | Used with primer AVL031 for qRT-PCR of *Pst* DC3000 gene *PSPTO_4337* that encodes a pyruvate kinase involved in the Entner-Doudoroff pathway. | This study |
| AVL033 | CAACTCGGGTGACGAAGATAAC | Used with primer AVL034 for qRT-PCR of *Pst* DC3000 housekeeping RNA polymerase sigma factor *rpoD* (*PSPTO_0537*). | This study |
| AVL034 | GCTTTGAGGTGCTTGGTTACTA | Used with primer AVL033 for qRT-PCR of *Pst* DC3000 housekeeping RNA polymerase sigma factor *rpoD* (*PSPTO_0537*). | This study |
| AVL035 | GTATAATTGCACGACCTAGGG | Used for sequencing genes cloned into pBG42_mod. | This study |
| AVL036 | GATCTATCAACAGGAGTCCAAG | Used for sequencing genes cloned into pBG42_mod. | This study |
| AVL037 | TACAACGCCATCCACGGCAA | Primer specific for *mCerulean3* that will not hybridize with the sequence of *mCitrine*. Used for sequencing. | This study |
| AVL038 | GAACTTCAAGATCCGCCACA | Primer specific for *mCitrine* that will not hybridize with the sequence of *mCerulean3*. Used for sequencing. | This study |
